# Notch signaling in the embryonic ectoderm promotes periderm cell fate and represses mineralization of vibrissa hair follicles

**DOI:** 10.64898/2026.01.27.702034

**Authors:** Dianzheng Zhao, Yunus Ozekin, Erin Binne, Irene Choi, Aftab Taiyab, Trevor Williams, Hong Li

## Abstract

The Notch signaling pathway is a critical means to regulate cell fate choice in animals. Appropriate regulation of this pathway is also required for human face formation as both loss and gain of function mutations of Notch signaling can cause syndromes with craniofacial abnormalities. Here we examine the consequences of manipulation of Notch signaling in the early mouse embryonic ectoderm by either removing the transcriptional effector *Rbpj* or expressing a constitutively active form of the Notch1 intracellular domain. Loss of *Rbpj* resulted in cleft secondary palate but strikingly was also associated with the ectopic mineralization of vibrissa follicles. In contrast, activation of Notch signaling resulted in multiple embryonic defects including a fully penetrant bilateral cleft lip and palate. Further, single cell RNA-seq data indicated a switch from a basal epithelial cell identity towards periderm when Notch signal transduction was elevated. These cell fate changes were accompanied by misregulation of genes and pathways known to impact human and mouse orofacial clefting including *Grhl3*, *Irf6*, and Wnt pathway. Together, these findings provide insight into human craniofacial conditions caused by misregulated Notch activity.

**SUMMARY STATEMENT:** Our studies demonstrate that Notch signaling in the embryonic ectoderm stimulates periderm cell fate while also repressing transformation of the inner root sheath of whisker follicles into mineralized tissue.

## INTRODUCTION

The Notch signaling pathway is a one of the fundamental conserved metazoan cell:cell communication systems (Bray and Bigas, 2025, Gozlan and Sprinzak, 2023, Siebel and Lendahl, 2017, Yao Meng, 2024, Zhou et al., 2022a). In mammals, the pathway consists of four receptors (Notch1–4) and five canonical ligands (Jag1, Jag2, Dll1, Dll3, and Dll4). In this canonical pathway, signaling is initiated when a transmembrane ligand binds to a Notch receptor on a neighboring cell, triggering a series of proteolytic cleavages that release the Notch intracellular domain (ICD). The ICD then translocates to the nucleus where it forms a transcriptional activation complex with Rbpjκ and coactivators of the Maml family to regulate downstream target gene expression (Bray and Bigas, 2025, Gozlan and Sprinzak, 2023, Meng et al., 2024, Siebel and Lendahl, 2017, Zhou et al., 2022b). All four Notch receptors contain an ICD, with a nuclear localization sequence, an Rbpj interaction motif, and a PEST sequence to direct rapid protein turnover (Meng et al., 2024, Siebel and Lendahl, 2017, Zhou et al., 2022b). Notch1 and Notch2 ICDs also have a well-defined transcriptional activation domain that is not equivalently defined in the other two family members, while further analysis has shown that exchanging the ICDs between Notch1 and Notch2 does not significantly affect developmental outcomes, indicating a significant degree of functional overlap (Liu et al., 2015, Meng et al., 2024). Downstream transcriptional targets of the Notch ICD activation complex include multiple Hes and Hey basic helix-loop-helix transcription repressors which are major effectors of canonical Notch signaling dynamics (Kobayashi and Kageyama, 2014, Weber et al., 2014). The Notch receptors can also function via non-canonical mechanisms in which other ligands, such as the Delta-like kinases, or different downstream effector molecules are involved, but these systems are less well understood (Andersen et al., 2012, Huang et al., 2025, Siebel and Lendahl, 2017).

The Notch pathway is critical for regulating numerous cell fate decisions during mammalian development, and is involved in diverse processes including somitogenesis, skeletogenesis, neurogenesis, hematopoiesis and tissue homeostasis (Bray and Bigas, 2025, Gozlan and Sprinzak, 2023, Pakvasa et al., 2021, Siebel and Lendahl, 2017, Zieba et al., 2020). Regarding cell fate, Notch signaling is involved in stem cell maintenance, and can also influence cell differentiation by processes such as lateral inhibition or manipulation of the expression and stability of various pathway components (Bray and Bigas, 2025, Gozlan and Sprinzak, 2023, Siebel and Lendahl, 2017). The importance of appropriately regulated Notch signaling is also apparent from mutations in this pathway that occur in human cancer or inherited disease conditions (Bray and Bigas, 2025, Huang et al., 2025, Meng et al., 2024, Siebel and Lendahl, 2017, Zhou et al., 2022b). With respect to the current study, the Notch pathway is involved in four genetically inherited birth defects impacting the skin and craniofacial complex. The first, Alagille Syndrome, is caused by heterozygous mutations in *JAG1*, or less frequently *NOTCH2* (Spinner et al., 1993). This condition has a combination of liver, bone, cardiovascular and eye defects along with facial features typified by a broad forehead and pointed chin. Hajdu-Cheney Syndrome is also caused by genetic modification of *NOTCH2*, but in this instance the mutation leads to loss of the PEST domain and a gain of function in Notch signaling (Canalis and Zanotti, 2014). This condition is associated with multiple skeletal defects including short stature, bowed long bones, open skull sutures, dental anomalies, facial defects, and osteoporosis. The third, Lateral Meningocele Syndrome, is caused by mutations in *NOTCH3* that are also thought to be activating changes that disrupt the PEST sequence (Ejaz et al., 1993). Alongside the spinal meningoceles and facial defects, patients may have skin, joint, neurologic, and cardiovascular abnormalities. For both Hajdu-Cheney Syndrome and Lateral Meningocele Syndrome patients faces are described as exhibiting hypertelorism, micrognathia, midfacial flattening, and high arched palate, sometimes with cleft secondary palate only (CPO) (Canalis and Zanotti, 2014, Ejaz et al., 1993). Finally, Adams-Oliver Syndrome (AOS) is typified by aplasia cutis of the scalp as well as by limb defects, usually terminal truncations of the fingers and toes (Hassed et al., 2017). Loss of function mutations in several Notch pathway genes can underlie this Syndrome including *RBPJ* in AOS3, *NOTCH1* in AOS5, and *DLL4* in AOS6.

Multiple studies in mouse have also explored the role of the Notch pathway in skin and face development. In seminal studies, homozygous knockout of *Jag2* was shown to result in CPO by promoting intraoral fusions between the tongue and palatal shelves. Such fusions would prevent the normal elevation of the palatal shelves above the tongue to allow appropriate fusion at the midline (Jiang et al., 1998). Further analyses indicated that *Jag2* was necessary for the activation of Notch1, particularly in the periderm layer covering the tongue and mandible (Casey et al., 2006). Detailed analysis of control and mutant embryonic tongue epithelia showed that while the former had a standard of cuboidal basal cells covered with a flattened layer of periderm, in the absence of Jag2 the basal cells were more disorganized and the periderm abnormal and discontinuous (Casey et al., 2006). The periderm is an embryo-specific surface epithelial cell layer that is thought to have a dual role, first in providing a barrier function to protect the embryonic surface from the surrounding amniotic fluid, and second to prevent inappropriate fusions between juxtaposed skin layers (Eckhart et al., 2024, Hammond et al., 2019, Jacob et al., 2023, McGowan and Coulombe, 1998, Richardson et al., 2014). The periderm forms soon after the appearance of the basal epithelial cell layer, beginning around E9.5, and eventually comes to cover the entire embryo before being lost late in embryogenesis coincident with the appearance of a cornified stratified epithelium (McGowan and Coulombe, 1998, Richardson et al., 2014). In contrast to our understanding of the post-natal skin development, much less is known about the signals and mechanisms driving periderm formation from basal epithelia. However, it is thought that some basal cells are induced to detach, migrate to the surface, and flatten out to form a superficial protective layer. Defects in periderm formation and function are referred to as peridermopathies and in humans can be caused by mutations in *IRF6*, *CHUK*, and *RIPK4* (Hammond et al., 2019). Such conditions include various types of popliteal pterygium syndrome and Cocoon Syndrome, in which inappropriate epithelial fusions involve structures including the limbs, genitalia, and face accompanied by cleft lip and/or palate. Notably, intraoral adhesions have also been observed in mouse models with altered *Irf6*, *Chuk*, and *Ripk4*, as well as in additional models involving periderm-expressed genes, including *Grhl3* and *Sfn*. Critically, such adhesions were also observed when the periderm was ablated using a diphtheria toxin expression cassette (Hammond et al., 2019, Rountree et al., 2010). Together, these findings indicate the importance of periderm for normal embryogenesis, as well as a potential role for Notch signaling in its regulatory circuitry.

The Notch signaling pathway also has critical roles in development of the more mature stratified epithelium, as well as skin appendages, notably hair follicles (Siebel and Lendahl, 2017, Zhou et al., 2022a). Conditional deletion of the Notch canonical transcriptional effector *Rbpj* from E14.5 using a *K14-Cre* transgene resulted in a thinner epidermis with reductions in the spinous and granular layers (Blanpain et al., 2006). In contrast, the ectopic expression of a floxed Notch1 ICD (N1-ICD) construct using a *K14-Cre* transgene caused an expansion of the spinous cell layer (Blanpain et al., 2006). Loss of *Rbpj* also caused defects in hair formation, with the follicles reported to form degenerative cystic structures lacking a normal hair shaft (Yamamoto et al., 2003, Blanpain et al., 2006), whereas N1-ICD overexpression resulted in wider and shallower vibrissa (Logan et al., 2018). These findings indicate critical roles for Notch signaling in the ectoderm and the secondary palate, but the timing and tissue-specificity of the Cre recombinase transgenes utilized were unable to address whether Notch pathway affected early ectoderm stratification into periderm. Further, although manipulation of the pathway can result in CPO, its potential roles in upper lip and primary palate formation remains largely unknown.

Here, we investigated the function of Notch signaling in early facial development using the ectoderm specific *Crect* transgene which begins expression at E8.5, prior to periderm formation. Conditional deletion of Rbpj caused intraoral epithelial fusions, CPO, and skin stratification defects. Further, this genetic manipulation caused ossification of hair follicles which may provide a mechanistic insight into human condition osteoma cutis (Limaiem and Sergent, 2025). Activation of the pathway by ectopic expression of the N1-ICD instead resulted in fully penetrant cleft lip and palate. Single-cell transcriptomic and bioinformatic analyses further revealed a shift from a basal to periderm cell fate accompanied by disrupted epithelial–mesenchymal signaling crosstalk.

## RESULTS

### Notch pathway gene expression in the developing mouse upper face

Previous studies have shown that Notch signaling is critical for mouse secondary palate development by maintaining periderm integrity and preventing premature adhesion within the oral cavity (Casey et al., 2006, Jiang et al., 1998). However, whether Notch signaling in the ectoderm can impact the earlier development of the facial ectoderm and its function in upper lip and primary palate formation remains unknown. To begin addressing this question, we first examined the expression of the four Notch receptors by mining a previously published bulk RNA-seq dataset spanning the critical stages of upper lip and primary palate development, which included separated facial prominences and tissue layers between E10.5-E12.5 (Hooper et al., 2020). Among the four Notch receptors, *Notch1* and *Notch3* were the most abundantly expressed in the ectoderm across all facial prominences during development as well as in the olfactory epithelia, while *Notch2* and *Notch4* had much lower levels of expression (Fig. 1A). Next, we refined and extended these data by mapping the expression of various components of the Notch signaling pathway with respect to their position in the ectoderm of the fusion sites of the lip and primary palate using a previously published single-cell RNA-seq dataset derived from the E11.5 mouse lambdoid (λ) junction (Li et al., 2019) (Fig. 1B, Fig. S1). *Notch1* showed most prominent expression in the periderm, where *Notch3* was also detected (Fig. 1C). *Notch3* expression was also enriched in the epithelial tissues of the palate and nasal cavity, whereas only sporadic expression of the four Notch receptors were present in the surface ectoderm. *Jag1*, *Jag2*, *Dlk1* and *Dlk2* were the most significantly expressed Notch ligands in the ectoderm, with *Jag1* having the greatest presence in the periderm. With respect to Notch targets, the transcription factor *Hey1* displayed periderm enrichment with transcripts present in the nasal epithelium while *Nrarp*, a feedback inhibitor of Notch signaling, was enriched specifically in the periderm. Finally, other components of Notch signaling, including genes for the transcription factors *Rbpj*, *Hes1*, and *Hes6*, were broadly expressed in the various ectodermal compartments (Fig. 1C). Together, these finding suggest that major components of the Notch signaling pathway are present in early embryonic mouse facial ectoderm, with a concentration of *Notch1* in the periderm.

**Figure 1.**
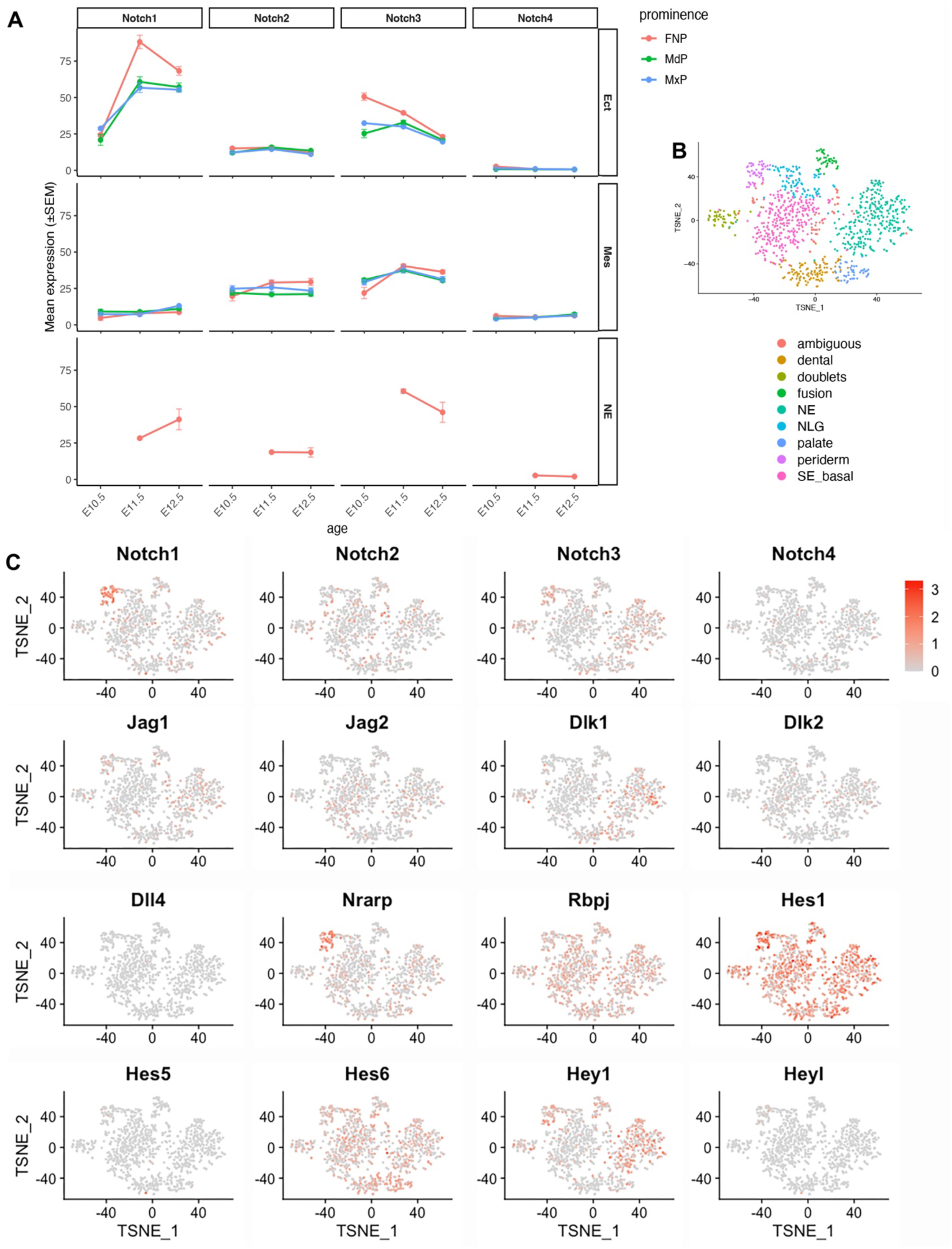
Expression of Notch signaling genes during facial development. (A) *Notch1* is the most abundantly expressed Notch receptor in the facial ectoderm. The plot shows the average expression (FPKM) ± standard error of the mean (SEM) of Notch receptors during facial development at embryonic stages E10.5, E11.5, and E12.5, stratified by tissue layer (Ect, Mes, NE) and facial prominence (FNP, MdP, and MxP), obtained from our previously published RNA-seq dataset (Hooper et al., 2020). (B) TSNE plot of the reclustered ectoderm derived cells at E11.5 lambdoidal junction with colors annotating the clusters, obtained from our previously published single-cell RNA-seq dataset (Li et al., 2019). (C) Expression of genes in the Notch signaling based on our previously published single-cell RNA-seq dataset (Li et al., 2019). In each feature plot, the name of the gene is shown on the top, and each dot represents a single cell, with the intensity of red indicating expression level and gray indicating no expression. Note that all the plots share the same color scale for expression as shown in the top right corner. Ect, ectoderm; FNP, frontonasal process; MdP, mandibular prominence; Mes, mesenchyme; MxP, maxillary prominence; NLG, nasolacrimal groove; NE, nasal epithelium; SE_basal, surface ectoderm basal cells.

### Loss of the Notch effector Rbpj in the ectoderm results in oral adhesions, cleft palate and mineralization of vibrissa follicles

Previous studies aimed at targeting all canonical Notch signaling in skin have focused on targeting *Rbpj*, a transcriptional cofactor that is required by the Notch ICD of all four paralogs (Yamamoto et al., 2003, Blanpain et al., 2006). Therefore, we took a similar approach using *Crect* Cre recombinase transgenic mice in conjunction with the *Rbpj* conditional allele to determine how loss of the canonical Notch pathway in the embryonic ectoderm beginning at E8.5 would impact craniofacial development. All *Crect;Rbpj^flox/flox^*mutant mice had shinier skin, open eyelids, and a lack of visible whiskers at birth (Fig. 2A-B). They also died shortly after birth and were found to have full penetrant CPO (Fig. 2D, F). Alizarin red and alcian blue staining analysis of P0 animals was used to determine how loss of *Rbpj* in the ectoderm altered the gross morphology of the craniofacial skeleton. As expected, the mutants exhibited defects in the formation of the palatine bones, consistent with CPO but otherwise the arrangement of the skeleton looked similar between controls and mutants (Fig. 2C-D, Fig. S2A-D). Analysis of the secondary palate phenotype using histological sections revealed that at E13.5 the mutant mice had a fusion between the tongue and the nasal septum, the tongue and palatal shelves, and between the palatal shelves and the mandible (Fig. 2E-J, Fig. S2H). At this stage, the control epithelia covering the tongue, palatal shelves and mandible generally has the appearance of a bilayer with cuboidal basal cells covered by flattened cells corresponding to the periderm (Fig. 2G, I). The same arrangement can be observed on the surface epithelia of the facial prominences (Fig. 2K). In the mutant, a similar arrangement of basal cells and periderm appeared to persist on the facial ectoderm, as well as on the palatal shelves and mandible (Fig. 2H, L). However, the epithelium of the tongue was notably more disorganized and a continuous periderm layer was not always observed (Fig. 2H, asterisk). In areas of fusion between the tongue, palatal shelves, nasal septum, and mandible the epithelial was intermingled such that the structural origin of the epithelial was not apparent (Fig. 2H, J). Such abnormal fusions of the oral epithelia and defects in tongue periderm were also observed previously when the Notch ligand *Jag2* was deleted in the ectoderm and this prevented normal elevation of the palatal shelves above the tongue (Jiang et al., 1998, Casey et al., 2006).

**Figure 2.**
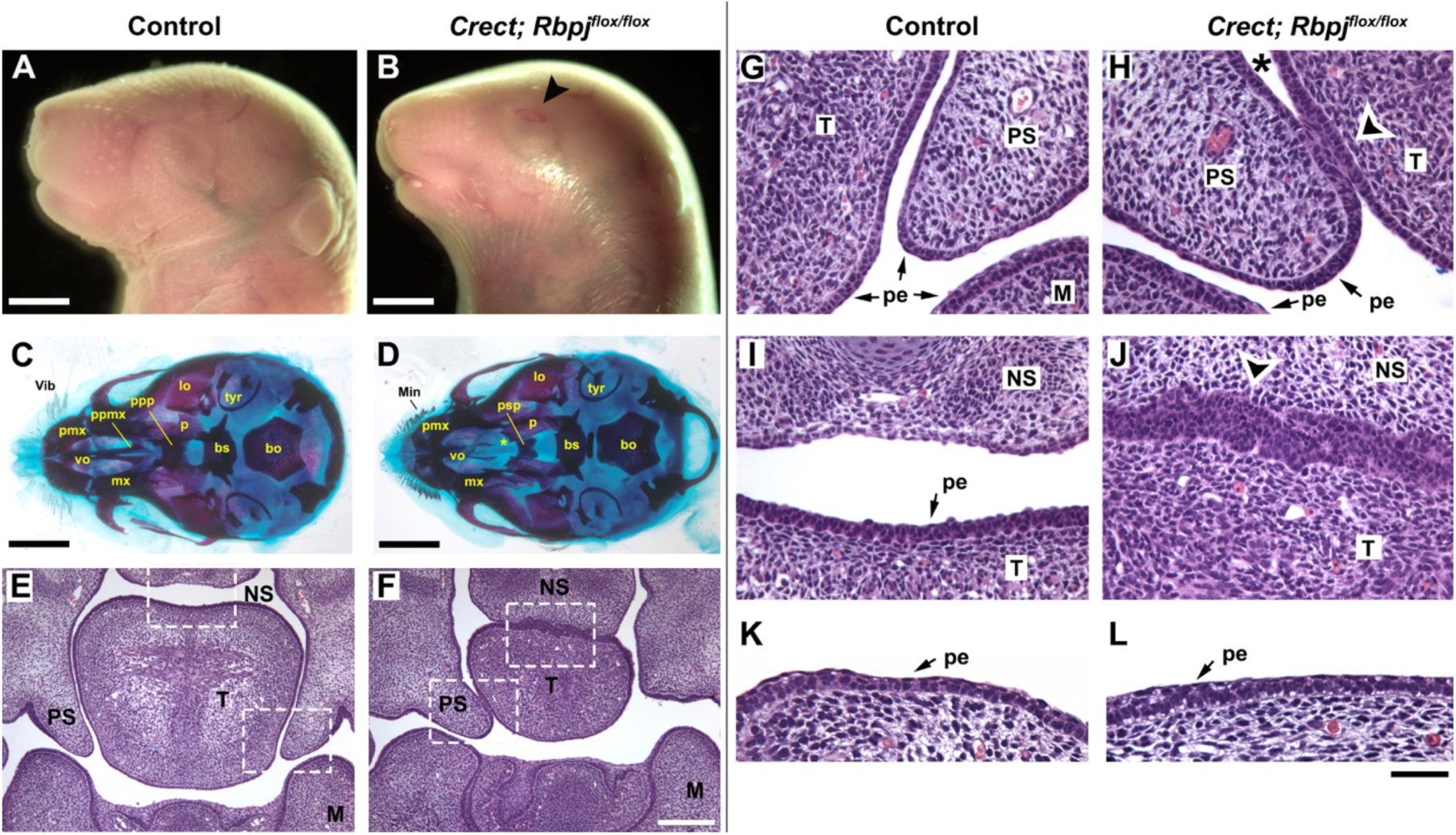
Loss of Rbpj in the ectoderm results in defects in the skin, secondary palate, and vibrissae. (A, B) Side view of E18.5 control (A) or mutant (B) mouse face showing external differences in skin, eyelids, and vibrissae between the two samples. Arrowhead in B shows open eyelid. (C, D) E18.5 craniofacial skeletal preparations of either control (C) or mutant embryos (D) shown in ventral views with the mandible removed. Yellow “*” denotes region of hypoplastic development of palatal processes (maxillary and palatine bones). (E - L) H&E stained coronal sections of E13.5 control (E, G, I, K) or mutant (F, H, J, L) head. White dotted outlines in E and F are shown in more detail in G-J. Arrowheads in H and J show regions of fusion between tongue and palatal shelf or tongue and nasal septum in mutant, respectively. Black asterisk in H shows region of tongue epithelium that lacks apparent flattened periderm layer. (K, L) show presence of a flattened periderm layer on surface of the external facial epithelium of both control (K) and mutant (L). bs, basisphenoid; bo, basioccipital; lo, lamina obturans; M, mandible; mx, maxilla: min, mineralization; NS, nasal septum; p, palatine; pe, periderm; pmx, premaxilla; ppmx, palatal process of maxilla; ppp, palatal process of palatine; PS, palatal shelf; psp, presphenoid; T, tongue; tyr, tympanic ring; Vib, vibrissae; vo, vomer. Scale bars: 2 mm in A-D; 200 μm in E-F; 50 μm in G-L.

The phenotype of shiny skin in the perinatal mutant mice (Fig. 2B) also indicated a defect in later skin development. Therefore, we also examined the histology of the facial skin at E18.5 using Masson’s trichrome staining. This analysis indicated a much thinner epidermis in mutants, with a reduction in the spinous and granular layers compared with controls (Fig. 3A-B). Surprisingly, further defects in the skin were evident from the skeletal preparations in which we saw new areas of alizarin red staining in the mutants corresponding to the location of vibrissae – structures that had not been removed to keep the nasal capsule intact for skeletal preparation (Fig. 2D, Fig. S2B, D, G). Such staining is suggestive of bone formation and this possibility was examined in more detail using histology focusing on the large mystacial vibrissae. A normal vibrissa consists of a dermal papilla surrounded by an epithelial matrix that produces the keratinized hair shaft (Fig. 3C). The inner root sheath (IRS) surrounds the vibrissa as it grows, and this is in turn encircled by an outer root sheath that eventually merges with the surface epithelia. While the E18.5 *Rbpj* mutant mice also had a vibrissa follicle with a dermal papilla, there was no clear vibrissa shaft or IRS layer. Instead, the follicle was wider and shorter with a cyst like appearance (Fig. 3D, Fig. S2F). This aberrant cystic morphology has previously been reported for hair follicles derived from *nes-Cre* or *K14-Cre Rbpj* conditional knockout mice at later timepoints (Yamamoto et al., 2003, Blanpain et al., 2006). We further analyzed the mutant vibrissae by Von Kossa and Alizarin red staining of histological sections, techniques which in combination can detect calcium-based mineralization. Both techniques detected staining in mutant vibrissae beginning after E16.5, whereas no specific staining was observed in the control vibrissa follicles (Fig. 3E-G). Intriguingly, the mineralized tissue staining in mutants was not in the position normally corresponding to the shaft of the vibrissa, but instead occupied the location of the IRS. In combination, these findings confirm a fundamental role of *Rbpj* in maintaining the normal development of the surface epithelia, as well as revealing a novel role in repressing the mineralization of vibrissa hair follicles.

**Figure 3.**
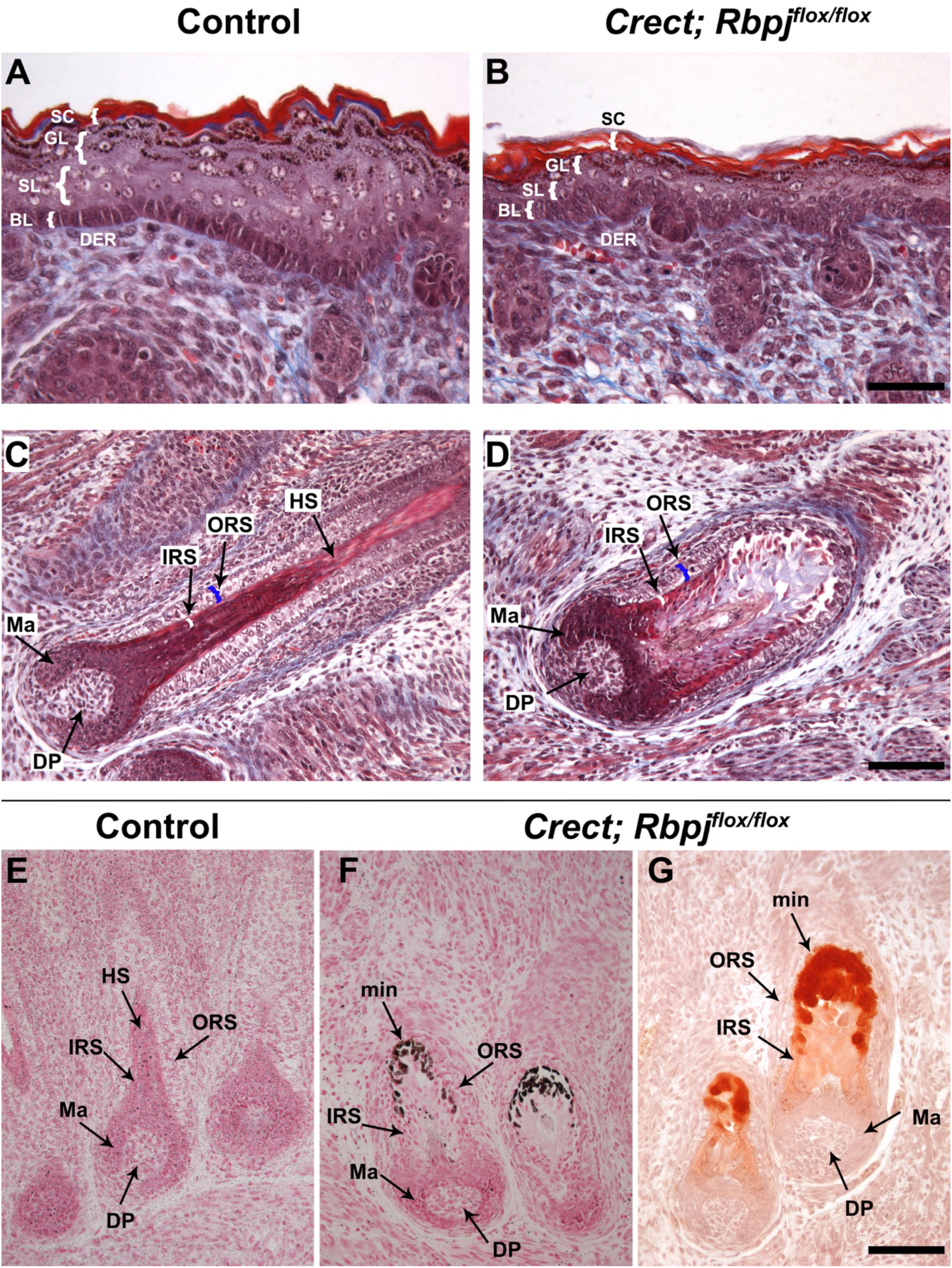
Loss of Rbpj results in skin defects and mineralization of vibrissae. (A-D) Masson’s trichrome stain of E18.5 facial skin (A, B) and mystacial vibrissa follicles (C, D) from control (A, C) or Rbpj mutant (B, D). (E-G) Mystacial vibrissa follicles of control (E) or mutant (F, G) stained using Von Kossa (E, F), Alizarin red (G). BL, basal layer; DER, dermis: DP, Dermal papilla; GL, granular layer; HS, hair shaft; IRS, Inner root sheath; Ma, Matrix; min, mineralization; ORS, outer root sheath; SC, stratum corneum; SL, spinous layer. Scale bars: 50 μm in A-B; 100 μm in C-G.

### Activation of Notch signaling in the embryonic ectoderm causes multiple defects including cleft lip and palate

We next sought to determine how constitutive activation of the Notch pathway impacted craniofacial development. In this gain-of-function model (NotchGOF) the *Crect* driver was used in combination with the *Rosa^N1-IC^* floxed allele (Murtaugh et al., 2003) to express the N1-ICD in the early embryonic ectoderm beginning around E8.5. NotchGOF mice did not survive into the postnatal period and we therefore began by examining relevant litters in late embryogenesis. Compared to control animals, all E18.5 NotchGOF embryos displayed structural defects including bilateral cleft lip and palate, severe limb hypoplasia (meromelia), kinked tail, and skin lesions across the face and anterior cervicothoracic region (Fig. 4A; 5/5 mutants). Partial penetrance was also observed for ventral body wall closure defects (3/5 mutants) and exencephaly (1/5 mutants) (Fig. 4A, Fig. S3B). These findings illustrate that normal regulation of Notch signaling in the ectoderm is critical for numerous major developmental processes. In the remainder of this study, we focus on the changes caused by the forced expression of the N1-ICD on development of the midface.

**Figure 4.**
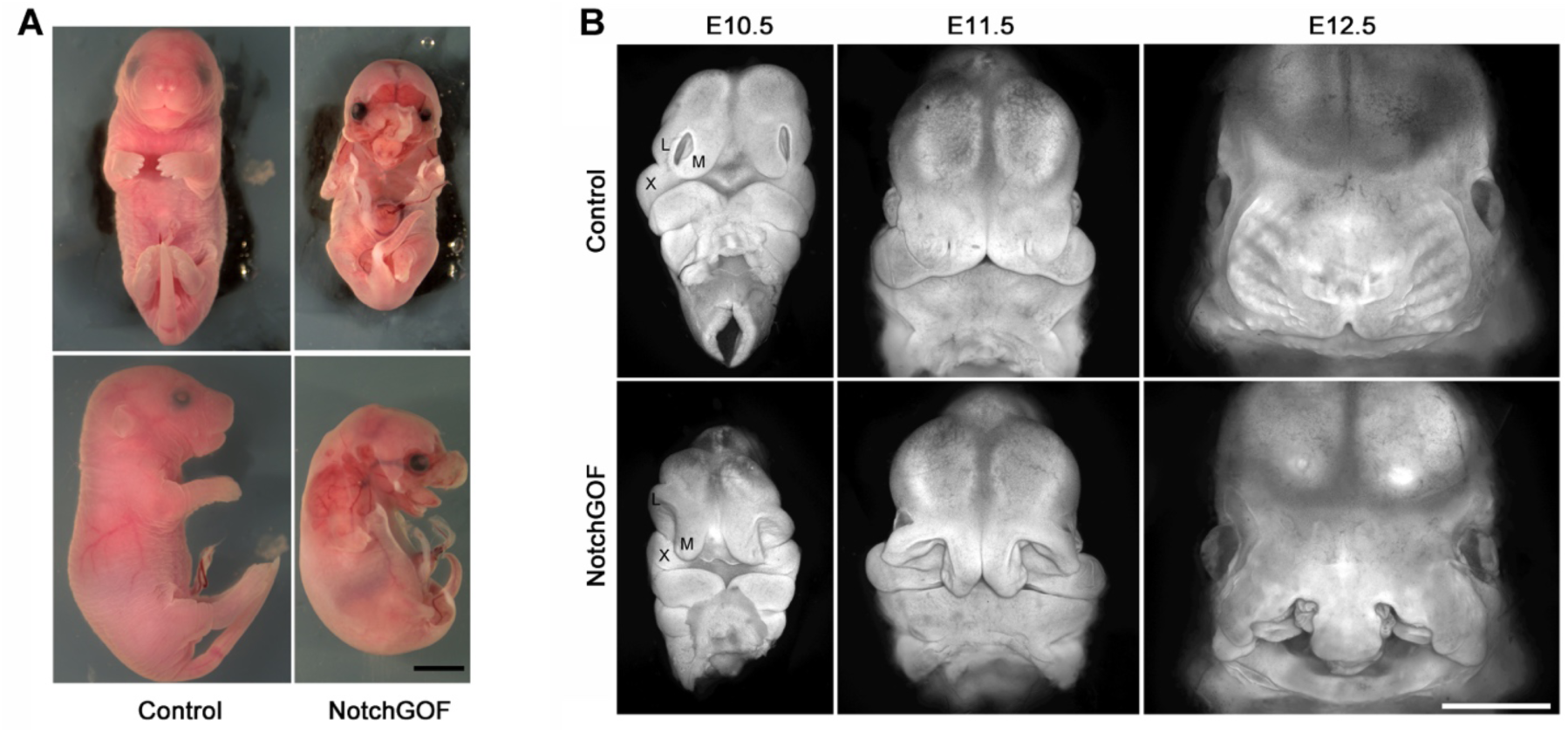
Gross morphology and fully penetrant cleft lip and palate phenotype in NotchGOF embryos. (A) Frontal (top panels) and lateral (bottom panels) views showing the gross morphology of E18.5 control (left panels) and NotchGOF mutant (right panels) embryos. (B) Fully penetrant cleft lip and palate in NotchGOF mutants. Shown are frontal views of control and NotchGOF embryos at E10.5, E11.5, and E12.5 following whole-mount DAPI staining. Note that the bodies were removed prior to imaging. L, lateral nasal process; M, medial nasal process; X, maxillary prominence. Scale bars: 4 mm in A; 1 mm in B.

To understand the development of the orofacial clefting phenotype we compared the external morphology of control and NotchGOF embryos from E10.5-E12.5 using whole-mount DAPI staining (Fig. 4B, Fig. S3). At E10.5, NotchGOF mutants displayed distinct facial morphology, particularly in the maxillary prominences and frontonasal process, with widened separation between the lateral and medial nasal processes (Fig. 4B). By E11.5, these morphological abnormalities were exacerbated, and by E12.5 NotchGOF embryos exhibited fully penetrant bilateral cleft lip and palate (CL/P) (Fig. 4B, Fig. S3). A subset of NotchGOF mutants also showed exencephaly, alongside bilateral CL/P (Fig. S3B). In addition, the mandibles were smaller by E12.5 (Fig. S3A).

The craniofacial developmental defects were further examined by comparing Alizarin red and alcian blue stained head skeletons of E18.5 control and mutant mice (Fig. S4). NotchGOF mutant animals had severe disruptions in the morphology of the skeletal elements comprising the midface, including an absence of the palatal bones derived from the maxilla and palatine structures, indicative of cleft secondary palate. The premaxilla was also found to protrude from the center of the face with no significant attachment to the maxilla, a phenotype consistent with bilateral cleft lip and primary palate. All mutant skeletons were also found to have reduced size of tympanic rings and mandibles (Fig. S4D, F). The presence of a cleft secondary palate and hypoplastic development of the palatal shelves was also confirmed using histological sections at E16.5 (Fig. S5). These results demonstrate that ectodermal activation of Notch signaling prevented the appropriate juxtaposition of the lateral and medial nasal prominences with the maxillary prominence at the λ junction, with the severity of the defect resulting in bilateral CL/P.

### Disrupted cell proliferation in facial prominences of NotchGOF mutants

To determine whether the abnormal morphology of facial prominences in the NotchGOF model results from altered cell proliferation, we performed EdU incorporation followed by flow cytometry at E10.5 on dissected facial prominences, including both ectoderm and mesenchyme, from control and NotchGOF embryos. This analysis revealed a modest but significant reduction in the overall proliferation when cells from the three prominences (frontonasal, maxillary, and mandibular) were combined (Fig. 5A). When analyzed separately, a statistically significant decrease was detected only in the frontonasal process (Fig. 5B). Given the relatively small proportion of ectoderm within facial prominences, these changes most likely reflect alterations in the mesenchymal cells.

**Figure 5.**
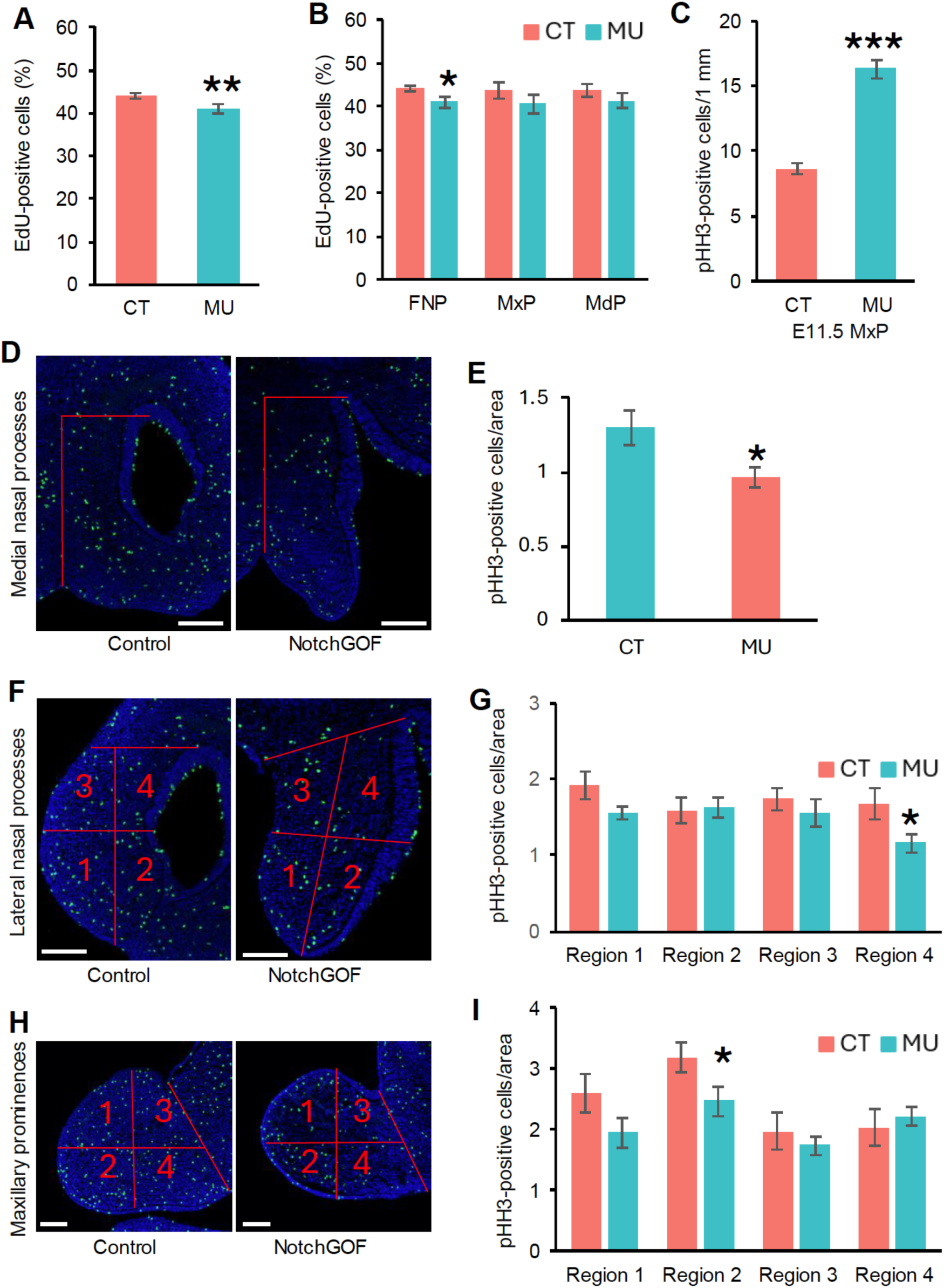
Cell proliferation analysis in NotchGOF facial prominences. (A, B) Bar charts showing the percentage of EdU-positive cells combined from all three facial prominences (A) or within each individual prominence (B) of E10.5 control and NotchGOF embryos, measured by flow cytometry following EdU labeling. Data are presented as mean ± SEM (*n = 5*; each sample pooled from 2–3 embryos). (C) Quantification of Phospho-Histone H3 (pHH3)-positive cells in the maxillary process (MxP) ectoderm, normalized to epithelial length (pHH3-positive cells per 1 mm ectoderm). Two sections were analyzed per embryo (three embryos total; *n = 6*). Data are presented as mean ± SEM. (D, F, H) Representative images of pHH3 staining in the medial nasal (D), lateral nasal (F), and maxillary (H) prominences of control and NotchGOF embryos. Red lines and numbers indicate the quantified regions. Scale bars, 100 μm. (E, G, I) Bar charts showing quantification of pHH3-positive cells per area in the medial nasal (E) as well as in subregions of the lateral nasal (G) and maxillary (I) prominences. Two sections were quantified per embryo (three embryos total; *n = 6*). Data are presented as mean ± SEM. Statistical significance was determined using a two-tailed *t*-test ( *, p < 0.05; **, p < 0.01; *** p < 0.001). CT, control; FNP, frontonasal process; MdP, mandibular prominence; MxP, maxillary prominence; MU, NotchGOF mutant.

Because spatial information is lost during cell dissociation for flow cytometry, we next performed immunostaining for phosphorylated histone H3 (pHH3) on E11.5 facial sections, focusing on the two most affected prominences (frontonasal and maxillary), which fuse to form the upper lip and primary palate. As Notch activation occurred specifically in ectodermal cells, we first quantified pHH3-positive cells in the facial ectoderm. A significant increase in pHH3-positive cells was observed in the NotchGOF maxillary ectoderm (Fig. 5C), whereas no change was detected in the surface ectoderm of the medial or lateral nasal prominences, or in the nasal epithelium (data not shown). We then quantified pHH3-positive cells within the mesenchyme of each prominence. The mesenchyme of the medial nasal process of NotchGOF embryos showed a significant reduction in proliferating cells (Fig. 5D-E), while region-specific decreases were also observed in the lateral nasal and maxillary prominences (Fig. 5F–I). Together, these results reveal spatially and temporally specific disruptions in cell proliferation within NotchGOF embryos and suggest non-cell-autonomous effects of ectodermal Notch activation on the underlying mesenchyme that likely contribute to prominence hypoplasia and abnormal facial morphogenesis.

### Prominent cell composition changes in λ junction of the NotchGOF model

To investigate the molecular basis underlying the CL/P phenotype in NotchGOF embryos, we performed single cell RNA-seq (scRNA-seq) analysis on micro-dissected tissues from the upper face of E11.5 control and NotchGOF somite-matched embryos. The region purified corresponded to the convergence between the maxillary, lateral nasal and medial nasal prominences, equivalent to the λ junction in a control sample. After quality assessment and filtering, approximately 10,000 cells per sample were retained for downstream analysis. Initial clustering of the combined control and NotchGOF datasets identified 24 clusters which could be generally grouped into seven well-defined cell populations (Fig. 6A, Fig. S6, Table S1). These seven groups were identified as mesenchymal cells, ectodermal cells, olfactory neurons (OFNs), Schwann cell progenitors, endothelial cells, red blood cells (RBCs), and other blood-derived cells (Fig. 6A) based on canonical marker gene expression (Table S2) and with reference to previously annotated λ-junction cell types (Fig. S6B-H, Fig. S7) (Li et al., 2019). Next, we examined how sample identity was arranged in the merged UMAP plot and discovered notable differences in cell distribution between control and NotchGOF embryos, particularly for the ectoderm, mesenchyme, and OFNs (Fig. 6B). The ectoderm comprised a higher proportion of total cells in the NotchGOF, whereas mesenchymal cells were somewhat reduced (Fig. 6C). OFNs were also markedly decreased in NotchGOF (Fig. 6B-C, F), consistent with the established role of Notch signaling in neural differentiation (McLaren and Butts, 2025), whereas blood and endothelial populations were comparable between samples (Fig. 6B-C).

**Figure 6.**
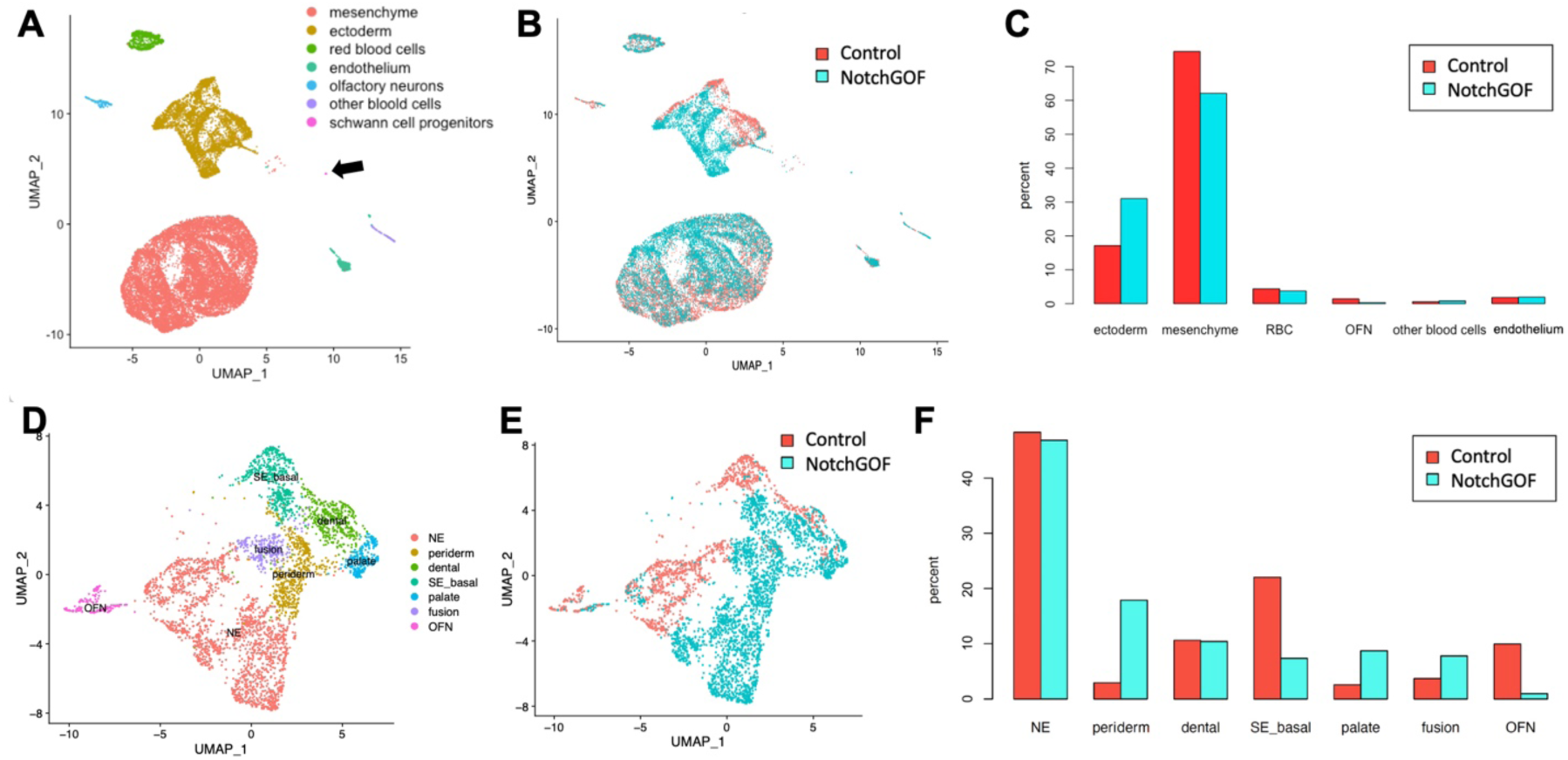
Single-cell transcriptomic analysis of control and NotchGOF λ-junction regions. (A) UMAP plot of combined E11.5 λ junction cells isolated from control and somite-matched NotchGOF embryos. Major cell types are annotated by colors. Arrow, Schwann cell progenitors. (B) Sample information overlaid on the UMAP plot shown in (A), with red cells representing control and blue cells representing NotchGOF samples. (C) Bar chart showing the relative proportions of major cell types by sample. Note that Schwann progenitor cells were not included. (D) UMAP plot of reclustered ectoderm derived cells with cell type annotations. (E) Sample information overlaid on the UMAP of reclustered ectoderm derived cells shown in (D). (F) Bar chart showing the composition of ectoderm-derived cells by sample. NE, nasal epithelial cells; SE_basal, surface ectoderm basal cells; OFN, olfactory neurons; RBC, red blood cells.

Out of all seven cell types, ectoderm cells were separated most by sample on the UMAP plot (Fig. 6B), suggesting there were major transcriptomic differences between the control and mutant cells in the ectoderm. Therefore, we reclustered all ectoderm-derived cells, including OFNs, to gain more insight into this phenomenon. This resulted in 12 ectodermal clusters (Fig. S8A, Table S3) that again could be classified into seven general cell populations (Fig. 6D) based on marker gene expression and prior annotations (Fig. S8B-I, Fig. S9, Table S4) (Li et al., 2019). These seven groups were annotated as nasal epithelial cells (NE), surface ectoderm basal cells (SE_basal), periderm, dental cells, palatal cells, fusion cells, and OFNs. Sample identity was again mapped on the reclustered ectodermal UMAP plot and confirmed that there was a significant difference in the distribution of control and mutant cells for most cell populations, suggesting that the ectopic expression of the N1-ICD had significantly changed the ectodermal transcriptome (Fig. 6D-F).

To assess how the phenotype and gene expression alterations correlated with altered Notch1 signaling in the NotchGOF embryos we examined the distribution of *Notch1* transcripts between control and mutant samples in the scRNA-seq datasets. This analysis revealed that the levels and distribution of *Notch1* transcripts was similar in endothelial and mesenchymal tissues of the mutant but significantly increased in ectodermal derivatives consistent with the ability of the *Crect* transgene to activate the floxed NICD allele specifically in embryonic epithelia (Fig. S10A-B). We next mined the data to determine if there were any additional changes associated in components of the Notch signaling pathway in the mutant ectoderm (Fig. S10C, Table S5). This examination showed that there were significant increases in the expression of the Notch receptors *Notch2* and especially *Notch3* RNA in addition to *Notch1* throughout the ectoderm. Differential expression of Notch ligands was observed, with increased *Jag1* transcript levels and broader distribution in ectodermal populations, accompanied by decreased expression and distribution of *Jag2*. *Dlk1* and *Dlk2*, which have been considered as non-canonical inhibitory ligands of Notch signaling, also show similar dynamics as the expression of the former gene becomes more widespread, but the latter is less widely expressed. Multiple downstream effectors and inhibitors of Notch signaling, including *Nrarp*, *Hes1*, *Hes5*, *Hey1*, *Hey2*, and *Heyl* exhibit increased expression. The significant alteration in *Hes1* expression in the ectoderm was verified by WMISH (Fig. S10D). Together, these findings indicate that the expansion of N1-ICD expression was confined to ectoderm and, further, that this change resulted in significant alterations of the Notch pathway genes in this tissue layer.

### Altered ratio between periderm and basal epithelial cells in NotchGOF embryos

Analysis of cell-type contributions by sample revealed significant differences between control and mutant ectodermal populations (Fig. 6F). We considered that some of these changes might be caused by altered morphology, for example additional fusion zone or palatal tissue available due to the failure of the prominences to align appropriately. However, the notable reciprocal shift between surface basal epithelial and periderm populations for the two genotypes was striking and we explored these changes further using both gene expression and histological analyses. We first examined the expression of three periderm marker genes (*Gabrp*, *Irf6*, and *Grhl3)* in the reclustered ectodermal cells. Analysis of the scRNA-seq data demonstrated that a higher percentage of cells expressed all three markers in regions corresponding to the periderm in NotchGOF embryos, while *Irf6* was also expressed at higher levels compared with controls (Fig. 7A, C, Fig. S11, Table S5). Whole mount in situ hybridization (WMISH) for these genes confirmed their upregulation in the facial region of NotchGOF mutants (Fig. 7B, D and Fig. S11B). In addition, RNAscope analysis of facial sections from E11.5 embryos revealed increased *Gabrp* expression in the lateral nasal process (Fig. 8C). Examination of additional periderm markers revealed significant increases in other genes associated with this layer including *Zfp750*, *Rhov*, *Ppl*, *Nectin4* and *Sfn,* consistent with the alteration and expansion of this tissue layer (Fig. S12B, Table S5). Nevertheless, such changes were not uniform as in some instances expression differences were confined to the presumptive periderm such as for *Gabrp*, *Ppl*, *Lypd3*, and *Cldn8*, whereas for genes such as *Nectin4*, *Dynlt5* (previously termed *Tctex1d1*), and *Rhov* the domain of expression was significantly expanded to other cell types (Fig 7A-B, Fig. S12B). Notably, the expression of *Irf6* and *Sfn* was also upregulated in the surface basal cells at the single-cell level (Fig. 7A, Fig. S12B).

**Figure 7.**
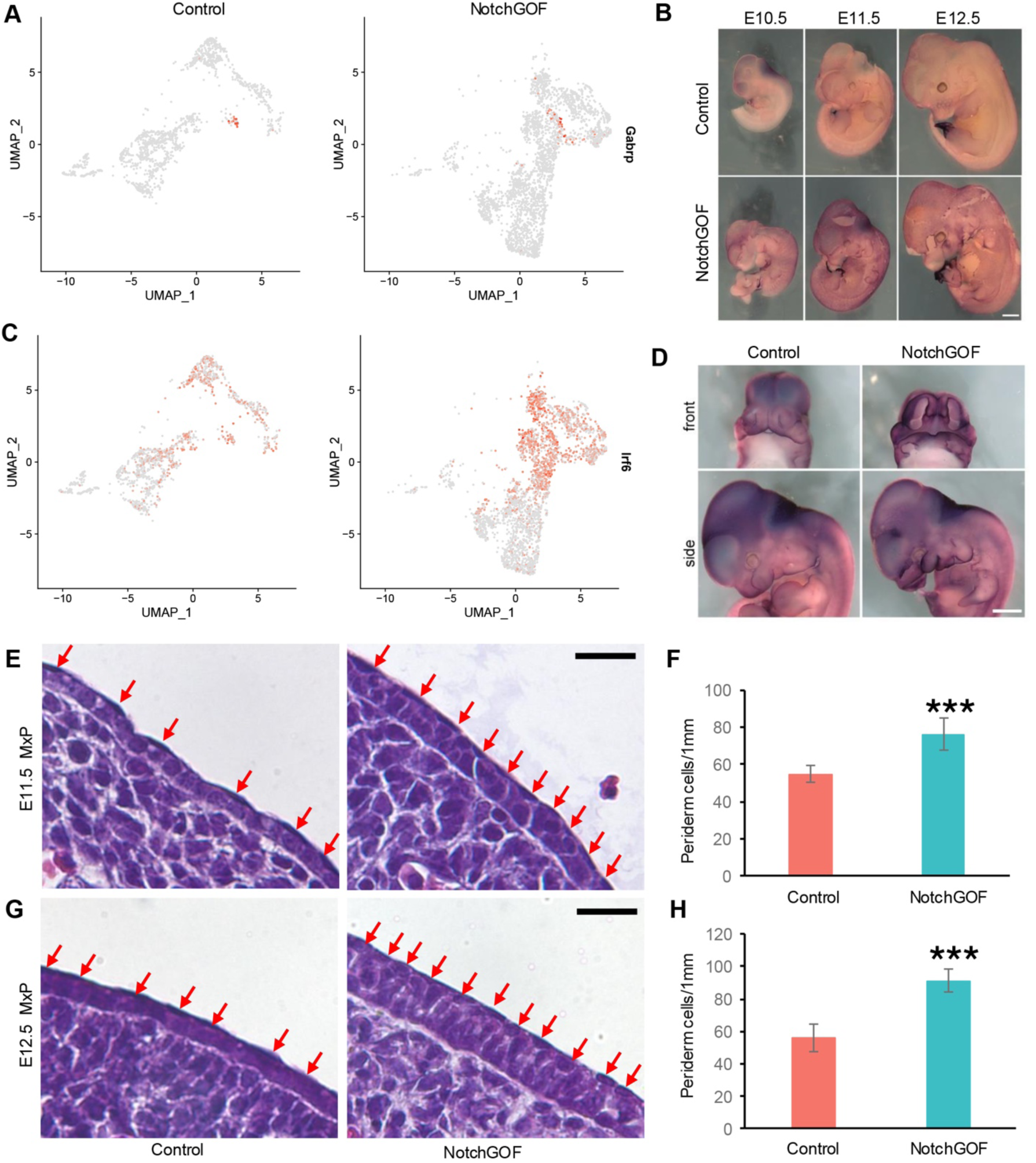
Expression of periderm marker genes and histological analysis of periderm in control and NotchGOF ectoderm. (A) Feature plots split by sample showing expression of the periderm marker gene *Gabrp* in ectoderm-derived cells, with control ectodermal cells on the left and mutant ectodermal cells on the right. (B) WMISH of *Gabrp* in E10.5, E11.5, and E12.5 embryos (side views shown). Scale bar, 1 mm. (C) Feature plot showing expression of the periderm marker gene *Irf6* in ectodermal cells by sample. (D) WMISH of *Irf6* in E11.5 embryos (both frontal with body removed and side views shown). Scale bar, 1 mm. (E, G) H&E staining of the maxillary process (MxP) shows an increased number of periderm cells in NotchGOF embryos at E11.5 (E) and E12.5 (G). Red arrows indicate periderm cells. Scale bars, 20 μm. (F, H) Quantification of periderm cells in the MxP at E11.5 (F) and E12.5 (H). The number of periderm cells was normalized to epithelial length (cells per 1 mm of ectoderm). Two sections were quantified per embryo (three embryos total; *n = 6*). Data are presented as mean ± SD. Statistical significance was determined using a two-tailed *t*-test. ***, p < 0.001.

**Figure 8.**
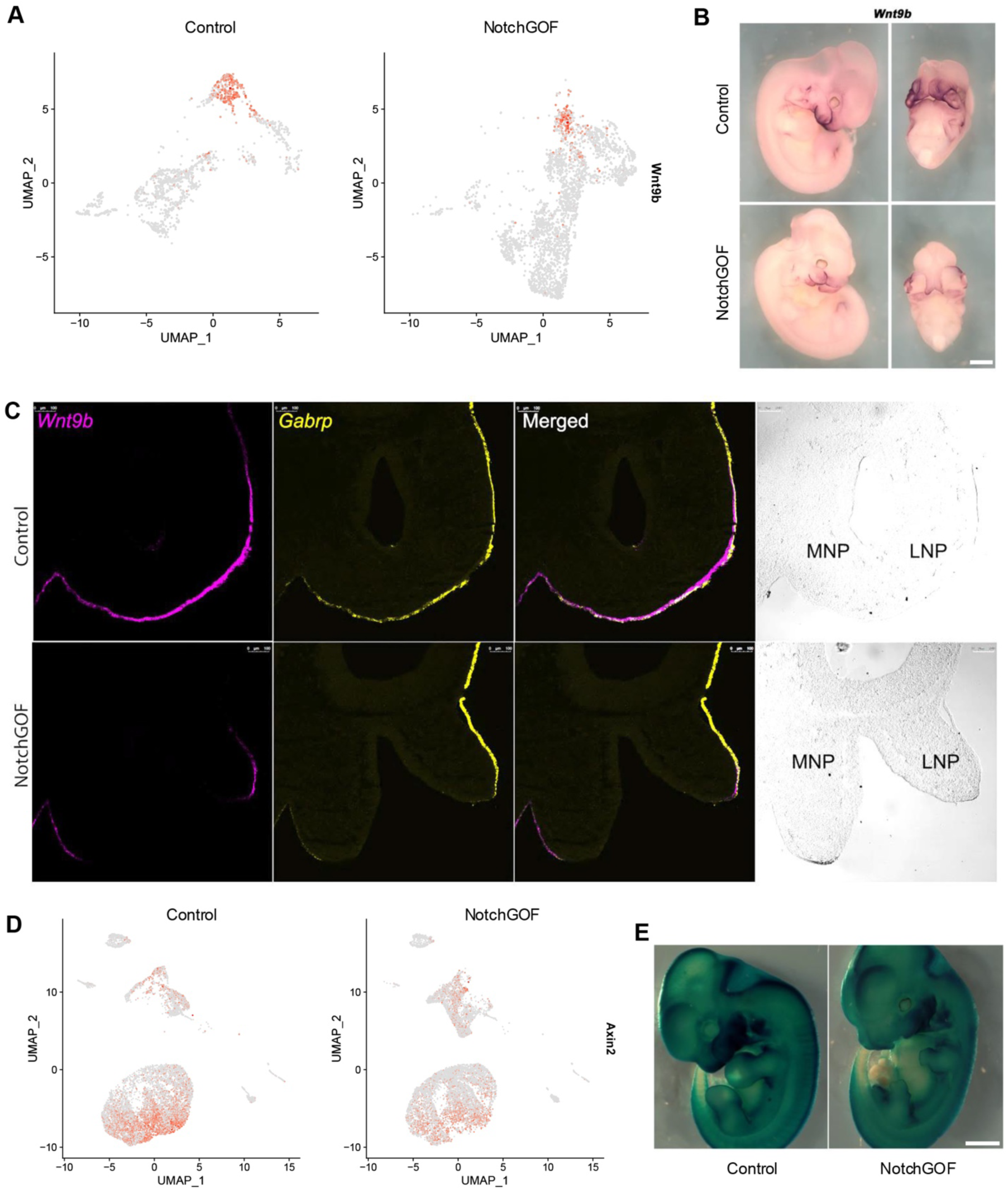
Disrupted Wnt signaling activity in NotchGOF mutants. (A) Feature plot showing expression of the Wnt ligand *Wnt9b* in ectodermal cells by sample. (B) WMISH of *Wnt9b* in E11.5 control and NotchGOF embryos. Both side views (left panels) and frontal views (right panels, body removed) are shown. Scale bar, 1 mm. (C) RNAscope analysis of *Wnt9b* (magenta) and *Gabrp* (yellow) in E11.5 control (top panels) and NotchGOF mutant (bottom panels) facial sections. Bright-field images are included to visualize tissue morphology. Scale bars, 100 μm. (D) Feature plot showing expression of *Axin2* in all cells within the λ-junction region by sample. (E) Axin2-lacZ staining of E11.5 control and NotchGOF embryos. Scale bar, 1 mm. LNP, lateral nasal process; MNP, medial nasal process.

To assess potential changes in periderm morphology, we performed H&E staining on facial prominences from E11.5 and E12.5 embryos. In control embryos, the ectoderm consisted of a single layer of cuboidal basal cells covered by flattened periderm cells. In contrast, NotchGOF mutants displayed disorganized basal cells and an increased number of periderm cells, which appeared rounder and smaller at both stages (Fig. 7E, G). Quantification confirmed a significant (p < 0.001) increase in periderm cell number in NotchGOF mutants at both E11.5 and E12.5 when normalized to the length of the ectoderm (Fig. 7F, H). Together, these results demonstrate that activation of Notch signaling expands the peridermal cell population, suggesting that excessive periderm formation may contribute to cleft lip and palate development.

### Notch activation in the ectoderm alters basal cell gene expression

The alterations in the morphology of the basal cells, and the switch in its relative abundance compared to the periderm prompted an examination of gene expression in additional cell types associated with the surface ectoderm of the face. First, an examination of keratin gene expression indicated that *Krt8* and *Krt18*, two genes expressed in the periderm and nasal epithelia, but less so in the basal surface epithelia and fusion zone, remained relatively unchanged between control and mutant samples (Fig. S13B). *Krt5* and *Krt14*, which are normally expressed in the basal cells of stratified epithelia also displayed relatively similar expression and distribution between control and NotchGOF in basal, dental and fusion clusters. In contrast, *Krt15*, whose expression is concentrated in the basal surface epithelium and is thought to maintain keratinocyte stemness and inhibit differentiation, is significantly down-regulated in the NotchGOF mutants (Fig. S13B-C, Table S5). Examination of additional markers of basal cells, including *Trp63*, *Perp*, and *Cxcl14* indicated that these were also expressed similarly between control and mutant samples (Fig. S13B-C). However, additional genes marking this cluster were either expressed at significantly lower levels, such as *Lmo1* and *Rprml*, or shifted expression to alternative epithelial populations like *Gjb2* and *Gjb6*, the latter two alterations suggesting aberrant basal epithelial cell:cell communication. Down-regulation of *Kitl* and *Edaradd* further indicated potential misregulation of cell signaling within the basal epithelia. Next, the analysis was extended to genes expressed in the fusion zone epithelia. Surprisingly, in this instance, genes such as *Arhgap29*, *Adamts9*, and the cell cycle regulators *Cdkn1a*, *Cdkn2c*, and *Gadd45g* were still strongly expressed in this population for both control and mutant samples, even though the prominences in the latter embryos do not become juxtaposed (Fig. S13B).

### Altered signaling crosstalk between ectoderm and mesenchyme

To gain additional insight into the cellular and molecular changes that occurred in both the ectoderm and mesenchyme, we used Enrichr (Kuleshov et al., 2016) to examine terms highlighted by gene expression changes in these two layers between the control and NotchGOF samples (Tables S5-7). As might be expected, the term Notch signaling pathway was highlighted in the ectoderm and orofacial cleft was enriched in both the ectoderm and mesenchyme gene sets (Table S7). Other terms enriched in the ectoderm included several associated with neurogenesis, as well as cell junctions and cadherin binding indicating an alteration in epithelial function and or identity. Notably, the gene set up-regulated in the mutant ectoderm included the term “Wnt receptor activity”, while the down-regulated group contained “Ligand Receptor Activity” and “Frizzled Binding” in which five Wnt ligands were present: *Wnt3*, *Wnt6*, *Wnt7b*, *Wnt9b*, and *Wnt10a*. Examination of the terms enriched in the mesenchyme also revealed terms associated with the Wnt signaling pathway, such as “Regulation of Canonical Wnt Signaling Pathway”, “Negative Regulation of Wnt Signaling Pathway”, and “beta-catenin-TCF Complex”, were highlighted when either all genes in Table S6, or only the genes down-regulated in the mutant, were used for the analysis. These findings suggested that disruptions in signal transduction mediated by the Wnt signaling pathway occur in both the ectoderm and mesenchyme of NotchGOF embryos. Since previous studies in several mouse models have shown that CL/P is often associated with defective Wnt signaling between the ectoderm and mesenchyme, and because mutations in Wnt pathway components are associated with cleft lip and palate in humans we examined how the ectopic activation of Notch impacted Wnt pathway gene expression in more detail (Fig. 8, Fig. S14-15).

Expression analysis of the Wnt ligand *Wnt9b*, a marker of surface ectodermal basal cells whose loss causes cleft lip and palate in mice, showed reduced numbers of *Wnt9b* expressing cells in NotchGOF embryos (Fig. 8A). This downregulation was validated by both WMISH and RNAscope (Fig. 8B-C). There was also a decrease in the distribution and expression of other Wnt ligands and Wnt inhibitors in the basal epithelial cells of NotchGOF mutants (Table S5, Fig. S14). Specifically, *Wnt3*, *Wnt6*, *Sostdc1*, and *Kremen2* were expressed in fewer cells, while *Wnt7b* and *Wnt10a* were detected in a smaller number of cells at significantly decreased levels. We next examined if there were corresponding changes to the Wnt signaling pathway in the mesenchyme (Fig. 8D, Fig. S15, Table S6). Here there was a significant reduction in the expression and distribution of several Wnt pathway genes, including repressors *Axin2*, *Wif1*, *Nkd1*, *Nkd2*, and *Apcdd1*. Consistent with these findings, when the *Axin2-lacZ* reporter allele was introduced into the appropriate mouse genetic background, reduced *LacZ* staining indicative of diminished Wnt signaling activity in the facial region of NotchGOF embryos was apparent (Fig. 8E). The location of expression of these Wnt pathway genes in a control sample largely overlapped with *Hey2*, a marker of the immediate subsurface mesenchyme (Li et al., 2019). *Hey2* was less expressed in this region of the NotchGOF mutant, and fewer mutant cells appeared to map to this location in the mutant suggesting a defect in this particular mesenchymal population (Fig. S15). In support of this hypothesis, ∼60 genes down-regulated in the NotchGOF mesenchyme (Table S6) were previously identified as being enriched in the subsurface mesenchyme (Table S2 from (Li et al., 2019)) including *Stmn2*, *Pax1*, and *Smoc2*. Together, these findings indicate that activation of Notch signaling in the ectoderm caused a significant reduction of Wnt ligand expression in basal epithelial cells of the face and disrupted the normal gene expression program in the adjacent subsurface mesenchyme consistent with the appearance of CL/P.

## DISCUSSION

Previous studies have established the importance of ectodermal Notch signaling in secondary palate development (Casey et al., 2006, Jiang et al., 1998), but its role during early facial morphogenesis, particularly in upper lip and primary palate formation, has remained unclear. Here, we addressed this question by conditionally activating or ablating Notch signaling in the embryonic ectoderm starting from E8.5. Ectodermal deletion of *Rbpj*, the key transcriptional factor of the canonical Notch pathway, produced CPO, consistent with the phenotype of *Jag2* mutants (Casey et al., 2006, Jiang et al., 1998). Surprisingly, using this model we also uncovered a previously unrecognized role for canonical Notch signaling in suppressing ossification of vibrissa follicles. In contrast, ectodermal overactivation of Notch signaling through expression of the N1-ICD promoted differentiation of basal cells into periderm, disrupted ectoderm–mesenchyme crosstalk, and led to fully penetrant bilateral cleft lip and palate. Together, our findings highlight a dual requirement for precise Notch dosage control and reveal spatiotemporal context-dependent roles for Notch signaling in the embryonic ectoderm during craniofacial development.

### Functions of Notch signaling in craniofacial development

Here we have shown that ectodermal overactivation of Notch signaling causes fully penetrant cleft lip and palate providing a further mouse model which can be used to study this critical human pathology. Our histological analysis of the cleft secondary palate phenotype revealed severely hypoplastic palatal shelves in the NotchGOF model. This stands in contrast to the CPO phenotype observed with the *Rbpj* conditional knockout. Previous work has primarily focused on Jag2–Notch1 signaling in the oral epithelium (Casey et al., 2006, Jiang et al., 1998). In this context, loss of *Jag2* disrupts proper periderm differentiation, leads to aberrant epithelial adhesions within the oral cavity, and results in CPO. Loss of *Rbpj* in the ectoderm similarly produces CPO associated with periderm defects and intraoral adhesions, mirroring the phenotype of *Jag2* knockout embryos and supporting the conclusion that Jag2 is the major ligand mediating canonical Notch signaling during secondary palate formation. However, our studies show that extending deletion of *Rbpj* to the early facial ectoderm did not result in any cleft lip and primary palate. Furthermore, there was no obvious change to the periderm of the surface epithelia of the external facial prominences. However, note that *Jag2* mutants also have syndactyly affecting both forelimbs and hindlimbs (Jiang et al., 1998), a condition thought to involve defective periderm function (Kashgari et al., 2020) and this phenotype was also observed in the absence of *Rbpj*. The *Crect*;*Rbpj* model also had a defect in in skin stratification, with thinner spinous and granular cell layers, a phenotype also observed when *Rbpj* is removed later in embryogenesis (Blanpain et al., 2006). Together, these findings indicate that the secondary palate is more sensitive to loss of canonical Notch signaling in the ectoderm than the upper lip and primary palate. Further, these findings suggest that either endogenous canonical Notch signaling has a greater influence over the periderm of the oral cavity and limbs than the rest of the embryo surface, or that these former regions have fundamentally different cellular and molecular properties.

### Canonical Notch signaling suppresses mineralization of vibrissa follicles

Previous studies had shown that loss of *Rbpj* in the ectoderm also resulted in aberrant hair follicles with a cystic appearance (Yamamoto et al., 2003, Blanpain et al., 2006, Turkoz et al., 2016). A similar phenotype of follicle cysts also occurred when presenilin or Notch ligands were removed from the ectoderm (Pan et al., 2004). This phenotype was also apparent in the vibrissa follicles of the *Crect*;*Rbpj* mutant mice and further analysis indicated that the IRS of the follicle had been replaced by ossified tissue. Note, the type of ossification associated with the whisker is distinct from that caused by loss of *Abcc6* which instead involved the connective tissue capsule surrounding the vibrissa follicles (Klement et al., 2005). The ossified tissue in the *Rbpj* mutant raises two important questions. The first issue concerns development and evolution since most calcified tissue in mammals is derived from neural crest and mesoderm, and not directly from ectoderm. One possibility is that the hair follicles have switched to forming a different epithelial appendage that will stain for ossification, namely towards a dental fate. In this hypothesis, Notch signaling in the hair follicles of the surface epithelia would be required to suppress an ossification program that can occur in the oral cavity to form teeth. The second consideration involves a rare type of human pathology termed osteoma cutis in which bony nodules form in the dermis (Limaiem and Sergent, 2025). The number of these nodules can vary between individuals, and in many cases are triggered by skin conditions such as chromic acne. However, about 15% of cases occur spontaneously with no documented pre-existing condition. At present, little is known concerning the cellular and molecular origin of such lesions, but it is possible that they are related to the type of aberrant Notch signaling creating the ossified structures seen in the absence of *Rbpj*. Further analysis will be required to test these hypotheses.

### Roles of Notch in periderm formation and function

Our analysis, coupled with previous studies on Jag2, demonstrate that a functioning canonical Notch signaling pathway is critical for aspects of periderm development and function. This conclusion seems especially true for the oral cavity and limbs, where periderm related defects are observed. Nevertheless, the periderm still forms on the facial prominences of *Rbpj* mutants indicating that canonical Notch signaling is not required for its formation and separation from the basal epithelial layer. The NotchGOF phenotype we describe provides further evidence that this pathway is critical for periderm biology. The activation of Notch signaling was found to drive a shift from basal cells towards a periderm gene expression signature. Periderm markers including *Gabrp*, *Irf6, Grhl3* and *Sfn* were upregulated in NotchGOF ectoderm based on single-cell transcriptomic analysis. Further analysis revealed that while the mean expression level of *Gabrp* and *Grhl3* per cell remains similar, a larger proportion of cells express these genes. In contrast, *Irf6* and *Sfn* were upregulated at the single-cell level, particularly in the surface ectoderm basal cells, and across a larger fraction of cells, supporting the notion that Notch signaling acts upstream of *Irf6* and *Sfn* during early periderm differentiation in the basal cells.

A previous study of Notch function at the later stages of skin development demonstrated that canonical Notch signaling functions as a molecular switch controlling the commitment of basal cells to a spinous cell fate during epithelial stratification (Blanpain et al., 2006). Further, while active Notch1 protein is largely restricted to periderm cells in the oral epithelium, a small fraction of basal cells exhibited active nuclear Notch1 (Casey et al., 2006). Our scRNA-seq data are also consistent with the presence of Notch1 mainly in periderm cells with some sporadic expression in surface ectoderm basal cells. Therefore, it is possible that Notch has a similar function in the early embryonic ectoderm as in the later ectoderm: in driving select cells in which it is expressed away from a basal cell fate. Histologically, periderm cells in *NotchGOF* mutants appear rounder rather than squamous as in wild-type embryos (Fig. 7), indicating that although Notch promotes the initiation of periderm differentiation, additional signaling pathways may be required for full periderm maturation. This hypothesis is supported by our transcriptomic analysis which revealed disruptions in multiple cell-junction related pathways, including cadherin and desmosomal genes, in the *NotchGOF* mutants (Table S7). Further, these data demonstrated that NotchGOF mutants have a subset of cells which co-express select basal and periderm markers unlike the control population. Based on these observations, we propose that Notch signaling plays dual roles during periderm formation. First, at early fate-decision stages, it is involved in periderm initiation from basal cells. This would need to occur via a non-canonical pathway since periderm can still form in the absence of Rbpj. Second, the pathway is involved in maintaining periderm integrity and/or normal function through Rbpj-dependent canonical signaling to prevent inappropriate epithelial adhesion.

### Notch–Wnt crosstalk

The Wnt signaling pathway is evolutionarily conserved and serves as a major regulator of embryonic development by controlling cell proliferation, differentiation, and tissue patterning. During craniofacial morphogenesis, Wnt ligands produced by the ectoderm provide essential cues for mesenchymal growth and morphogenesis (Goodnough et al., 2014, Brugmann et al., 2007, Reynolds et al., 2019, Van Otterloo et al., 2022, Ferretti et al., 2011). Mutations in multiple components of Wnt signaling pathway have been associated with both syndromic and nonsyndromic orofacial clefts (Reynolds et al., 2019). Crosstalk between the Wnt and Notch pathways is well established, with multiple studies demonstrating their opposing roles in balancing progenitor cell maintenance and differentiation. Indeed, some of the non-canonical Notch pathway also involves components of the Wnt pathway (Andersson et al., 2011, Collu et al., 2014, Siebel and Lendahl, 2017).

In our NotchGOF model, we observed reduced expression of five Wnt ligands in the ectoderm. For *Wnt3*, *Wnt6,* and *Wnt9b,* this reduction mainly reflected a decrease in the proportion of cells expressing these transcripts, whereas for *Wnt7b* and *Wnt10a* both the number of expressing cells and the per-cell expression levels were reduced. These findings suggest that activation of Notch signaling suppresses Wnt signal in the ectoderm through both indirect mechanisms—via depletion of the basal cell population—and potentially direct transcriptional repression. Reduced ectodermal Wnt ligand expression was accompanied by downregulation of Wnt signaling output in the mesenchyme, reduced mesenchymal cell proliferation, and a possible decrease in the mesenchymal cell population adjacent to the surface ectoderm. Supporting a role for reduced Wnt signaling in the clefting phenotype, all five Wnt ligands have been implicated in nonsyndromic orofacial clefting in humans (Reynolds et al., 2019), and disruption of *Wnt9b* in mice results in cleft lip and palate (Jin et al., 2012, Juriloff et al., 2006). Furthermore, models of mouse facial clefting based on changes in Pbx and AP-2 transcription factors show reduced Wnt signaling and such defects can be at least partially rescued by ectopic expression of Wnt1 in the ectoderm (Ferretti et al., 2011, Van Otterloo et al., 2022).

In addition to Wnt signaling, genes involved in other signaling pathways are also perturbed in the *NotchGOF* mutant including Fgf, Pdgf, Bmp, Kit, and Edar. We propose that disruption of the balance between basal and peridermal cell populations leads to reduced population of basal cells capable of producing autocrine and paracrine signals. Although changes in Wnt signaling are a clear consequence of this genetic manipulation, likely impairing mesenchymal proliferation and morphogenesis resulting in cleft lip and palate in the NotchGOF model, it is likely that the overall phenotype is caused by a combination of all these signaling pathway alterations.

### Clinical implications of the NotchGOF model

Mutations in components of the Notch signaling pathway give rise to a spectrum of congenital disorders affecting multiple organ systems. Craniofacial anomalies are a feature of several of these conditions including Alagille, Hajdu-Cheney, Lateral Meningomyocele and AOS. Multiple tissues are required for face formation (Dixon et al., 2011, Hammond and Dixon, 2022, Chai and Maxson, 2006) and tissue-specific loss and gain of function studies assist in determining the critical sites of action of specific genes and signaling pathways. Alagille Syndrome and AOS are caused by heterozygous mutations in various components of the Notch pathway and so would be predicted to cause a partial loss of function rather than a completely null phenotype (Hassed et al., 2017, Spinner et al., 1993). With respect to the craniofacial complex, Alagille Syndrome patients have a broad forehead, pointed chin, and broad nasal tip, while those with AOS have aplasia cutis of the scalp. None of these phenotypes are apparent in the *Crect;Rbpj* conditional knockout mice, which are instead typified by cleft palate. Conversely, loss of *Rbpj* in the neural crest does cause defects in the shape of the craniofacial skeleton as well as delayed ossification of the cranial vault (Mead and Yutzey, 2012). Further, a *Wnt1-Cre* knockout of *Jag1* caused severe midfacial retrusion with potential relevance to Alagille Syndrome, indicating that the non-canonical pathway must also operate in the neural crest to impact facial morphology (Humphreys et al., 2012). Together, the findings suggest that Alagille Syndrome and AOS probably result more from disruption of Notch signaling in the neural crest than in the ectoderm.

The two human gain of function syndromes, Hajdu-Cheney and Lateral Meningocele, caused by mutations in NOTCH2 and NOTCH3 respectively, have similar facial phenotypes including hypertelorism, micrognathia, midfacial flattening, and high arched palate, and sometimes CPO (Canalis and Zanotti, 2014, Ejaz et al., 1993). Mouse models have been generated expressing knock-ins of these *Notch2* and *Notch3* mutations (Canalis et al., 2016, Canalis et al., 2018). Models of both conditions caused reduced bone density, but no impact on craniofacial development was reported. In other studies, activation of the N1-ICD was driven in the mouse neural crest (Mead and Yutzey, 2012). This paradigm resulted in lethality prior to E14.5 accompanied by exencephaly preventing an analysis of the later craniofacial skeleton. Even so, defects in face formation were apparent earlier in embryogenesis with defects in formation of the midface and mandible. Similarly, we observe severe craniofacial phenotypes in our ectodermal NotchGOF model, including full-penetrance cleft lip and palate, micrognathia, and occasional exencephaly. The potential micrognathia phenotype shared across GOF models supports both a cell-autonomous role for Notch signaling within neural crest cells and a non-cell-autonomous influence from the ectoderm during lower jaw development and potentially palate formation. Together, these findings indicate that both ectodermal and neural crest cell populations are highly sensitive to Notch overactivation and suggest that misregulated Notch signaling in both tissue layers can underlie the defects seen in Hajdu-Cheney and Lateral Meningocele Syndromes.

In summary, our study demonstrates that precise regulation of Notch signaling is critical for periderm formation and epithelial–mesenchymal communication during facial development. Disruption of this balance, either through loss or gain of function, leads to orofacial clefting. These findings clarify the developmental roles of Notch signaling in the facial ectoderm and provide a framework for understanding craniofacial anomalies associated with human mutations in the Notch pathway.

## MATERIALS AND METHODS

### Mouse strains and animal husbandry

All animal experiments were conducted in accordance with protocols approved by the Institutional Animal Care and Use Committee (IACUC) of the University of Colorado Anschutz Medical Campus. The Rosa^N1-IC^ strain (*Gt(ROSA)26Sor^tm1(Notch1)Dam^/J*; stock no. 008159), which allows Cre-dependent overexpression of the N1-ICD (Murtaugh et al., 2003), and the Axin2LacZ reporter strain (*B6N.129P2-Axin2^tm1Wbm^/J*; stock no. 009120) (Lustig et al., 2002) were obtained from The Jackson Laboratory. The floxed *Rbpj^tm1Hon^* conditional knockout mice and the ectoderm-specific Cre recombinase driver line *Crect* have been described previously (Tanigaki et al., 2002, Reid et al., 2011). Note that the Crect transgene was always introduced from the male to generate animals for experimental analysis to avoid more widespread mosaic recombination observed when females were used for the final mating. To generate NotchGOF embryos, Rosa^N1-IC^ homozygous females were bred with heterozygous Crect males. This breeding scheme produced approximately 50% NotchGOF mutant embryos (*Crect^+/−^; Rosa^N1-IC^ ^+/−^*) and 50% wild-type littermates (*Rosa^N1-IC^ ^+/−^*) used as controls. To generate NotchGOF with Axin2LacZ reporter embryos, *RosaN1-IC* homozygous females were crossed with *Crect^+/−^;Axin2LacZ^+/−^*double-heterozygous males, yielding an expected 25% mutant (*Crect^+/−^;Axin2LacZ^+/−^*; *Rosa^N1-IC^ ^+/−^*) ratio. Crect-Rbpj mice were generated by breeding a male *Crect^+/-^;Rbpj^flox/+^*mouse with *Rbpj^flox/flox^*females to give an expected 25% of *Crect^+/-^;Rbpj^flox/flox^* mutants. For embryo staging, noon on the day a copulatory plug was detected was designated as embryonic day (E) 0.5. Embryos were collected at the desired developmental stages for analysis. Yolk sacs were collected and digested in 100 µL DirectPCR Lysis Reagent (Mouse Tail) (Viagen Biotech) supplemented with 10 µg/mL Proteinase K overnight at 56 °C, followed by enzyme inactivation at 85 °C for 45 min. The resulting lysates were used directly for PCR genotyping with DreamTaq DNA Polymerase (Thermo Fisher Scientific).

### Skeletal staining

Skeletons were prepared by Alizarin Red and Alcian Blue staining as previously described (McLeod, 1980). Briefly, E18.5 embryos were euthanized, skinned, and eviscerated, then dehydrated in 95% ethanol for 3 days followed by 100% acetone for 2 days. Specimens were stained for 5 days at 37 °C with agitation in a solution containing Alcian Blue 8GX (Sigma) and Alizarin Red S (Sigma). After staining, skeletons were rinsed in distilled water and cleared in 2% potassium hydroxide (KOH) for 2 days at room temperature. Samples were then transferred through a graded glycerol/1% KOH series (20:80, 50:50, and 80:20) for 1–2 days each and finally stored in 100% glycerol at room temperature. The skeletons were imaged using a SPOT RT3 camera mounted on a Leica M420 microscope.

### Histological analyses

Sectioning and histological analyses were performed at the Research Histology Core at the University of Colorado Anschutz Medical Campus. Briefly, embryo heads were fixed overnight in 4% paraformaldehyde (PFA) in PBS at 4 °C, dehydrated, and embedded in paraffin according to a previously published protocol (Fischer et al., 2008). Paraffin-embedded samples were sectioned at 10 µm. Sections were deparaffinized in xylene and rehydrated through a graded ethanol series, followed by hematoxylin and eosin staining as previously described (Cardiff et al., 2014). Masson’s trichrome, von Kossa, and Alizarin Red staining were also performed on control and Rbpj mutant samples as previously described (Clark, 1981). Stained sections were imaged using a SPOT RT3 camera mounted on a Nikon Eclipse E600 microscope.

### Whole mount DAPI staining

NotchGOF and control embryos at E10.5, E11.5, and E12.5 were processed for whole mount DAPI staining following a previously published protocol (Sandell et al., 2012) with a slight modification. Briefly, dissected embryos were fixed in 4% paraformaldehyde (PFA) in phosphate-buffered saline (PBS) at 4 °C overnight with gentle rocking, then washed three times in PBS. Embryos were stained with 10 µg/mL DAPI (Sigma) in PBS at room temperature for 1 h, followed by three additional PBS washes. The craniofacial regions of stained embryos were imaged using an Axiocam 506 mono digital camera (Carl Zeiss) mounted on an Axio Observer 7 fluorescence microscope (Carl Zeiss), as previously described (Dennison et al., 2021).

### EdU labeling followed by flow cytometry

Pregnant females were administered 150 µL of Click-iT™ EdU (Invitrogen/Life Technologies) solution (0.5 mg/mL in PBS) via intraperitoneal injection. Twenty minutes after injection, the females were sacrificed, and embryonic facial prominences were dissected in cold PBS. Following microdissection, the frontonasal process (FNP), maxillary prominence (MxP), and mandibular prominence (MdP) from two to three embryos were pooled and incubated in 1.2 mL of 0.25% trypsin-EDTA (Invitrogen/Life Technologies) at 37 °C for 10 min with gentle agitation. Enzymatic digestion was terminated by adding 300 µL of fetal bovine serum (Invitrogen/Life Technologies). The suspensions were pipetted up and down for about 20 times with 1 mL tips to aid cell dissociation. Cells were collected by centrifugation at 300 × *g* for 5 min and washed with PBS containing 1% bovine serum albumin (BSA). EdU labeling and detection were performed according to the manufacturer’s instructions using the Click-iT™ Plus EdU Alexa Fluor™ 594 Flow Cytometry Kit (Invitrogen/Life Technologies). Briefly, 100 µL of Click-iT™ fixative (Component D) was added to the cells, followed by incubation for 15 min at room temperature in the dark. After washing with PBS containing 1% BSA, cells were resuspended in 100 µL of Click-iT™ permeabilization and wash reagent and incubated for 15 min. Subsequently, 0.5 mL of Click-iT™ Plus reaction cocktail was added, and samples were incubated for 30 min. Cells were then washed again with the permeabilization and wash reagent and resuspended in PBS containing 2% FBS and DAPI (0.5 µg/mL). Unlabeled and single-color controls were used to establish compensation settings for flow cytometry. Data acquisition was performed on a BD LSRFortessa™ Flow Cytometer (BD Biosciences), with 100,000 cells collected per sample to ensure robust statistical representation. All flow cytometry analyses were conducted at the University of Colorado Allergy and Clinical Immunology Flow Cytometry Facility.

### Cell proliferation analysis by immunofluorescence

Immunofluorescence for phospho-Histone H3 (pHH3) on frozen sections was performed as previously described (Van Otterloo et al., 2022). Stained sections were imaged using a Leica SP8 confocal microscope. For quantification, each facial prominence was outlined in ImageJ, and pHH3-positive cells within the defined regions were quantified using the thresholding and particle analysis functions. The number of pHH3-positive cells per unit area was calculated from sections of three control and three NotchGOF embryos. The number of pHH3-positive cells per millimeter of ectodermal length was determined by automated counting in ImageJ. Statistical significance between groups was assessed using an unpaired Student’s *t*-test.

### Single cell RNA sequencing and data analysis

The λ junction regions were dissected and processed for single cell preparation as previously described (Li et al., 2019). Both control and NotchGOF samples were obtained from two E11.5 littermate embryos, each with 43 somites. The sex of the embryos was not determined, as no sex-specific phenotypic differences were observed. Cell suspensions were processed for droplet generation and library construction using the Chromium Next GEM 3′ v3.1 kit (10x Genomics) following the manufacturer’s protocol. Sequencing was performed on an Illumina NovaSeq 6000 platform with 2×150 bp reads, generating approximately 440 million raw reads per sample. Library preparation and sequencing were carried out by the University of Colorado Genomics and Microarray Shared Resource.

Reads were aligned to the mm10-2020-A reference genome using Cell Ranger v4.0.0 (10x Genomics), and gene expression matrices were generated. This initial processing yielded 11,618 control cells (average 38,111 reads per cell) and 12,220 NotchGOF mutant cells (average 36,112 reads per cell). Secondary analysis was conducted in R using Seurat v4.0.0 (Hao et al., 2021) for data preprocessing, normalization, identification of highly variable genes, cell-cycle regression, scaling, dimensional reduction, clustering, and differential expression analysis under default parameters unless stated otherwise. Cells expressing fewer than 200 or more than 8,000 genes or with >15% mitochondrial transcripts were excluded. Clustering was performed using the first 30 principal components with a resolution of 0.8. Clusters exhibiting low UMI counts or doublet signatures were removed. After quality control, 9,482 control and 10,378 NotchGOF cells remained in the dataset. Reclustering of ectoderm-derived cells was performed with the same parameters, resulting in an ectodermal dataset containing 1,299 control and 2,660 NotchGOF mutant cells.

### In situ hybridization

WMISH was performed on control and NotchGOF embryos at the desired embryonic stages using probes specific for *Gabrp*, *Irf6*, *Grhl3*, *Wnt9b*, and *Hes1*, as previously described (Li et al., 2019). The plasmid used to generate the *Irf6* probe was kindly provided by Dr. Brian C. Schutte, while probes for the other genes were generated in-house and verified by sequencing. Images of stained embryos were captured using a SPOT RT3 camera mounted on a Leica M420 microscope.

RNAscope was performed on fixed frozen sections to detect *Wnt9b* and *Gabrp* using the Multiplex Fluorescent Reagent Kit v2 (Advanced Cell Diagnostics) following the manufacturer’s protocol and as described previously (Li et al., 2019). Images were acquired with a Leica SP8 confocal microscope.

### β-Galactosidase staining

Whole mount β-galactosidase staining was performed as previously described (Van Otterloo et al., 2022). Briefly, E11.5 embryos were fixed in 0.25% glutaraldehyde in PBS for 30 minutes at room temperature, washed three times in lacZ rinse buffer (2 mM magnesium chloride, 0.2 M sodium phosphate pH 7.3, 0.01% sodium deoxycholate, 0.02% NP-40), and incubated in rinse buffer containing 1 mg/ml X-gal at 37°C until the desired color was achieved. Embryos were post-fixed in 4% paraformaldehyde (PFA) in PBS overnight following brief rinses in PBS. Stained samples were imaged using a SPOT RT3 camera mounted on a Leica M420 microscope.

## Acknowledgements

Special thanks to Dr. Katie Fantauzzo for assistance with imaging embryos prepared by whole-mount DAPI staining; to Dr. Raphael Kopan for generously providing the *Rbpj^flox/flox^* mouse line used in this study; to the University of Colorado Genomics and Microarray Core Facility for performing the library preparation and single cell sequencing; to E. Erin Smith, HTL (ASCP)CM QIHC, Allison Quador, HTL(ASCP)CM, and Jessica Arnold HTL(ASCP)CM of the University of Colorado Cancer Center Pathology Shared Resource for histology analyses; to the University of Colorado Allergy and Clinical Immunology Flow Cytometry Facility for flow cytometry analyses; and to all our colleagues in the Department of Craniofacial Biology for their insightful discussions and feedback on this work.

## Data and code availability

The scRNA-seq datasets have been deposited in GEO under accession number GSE315754.

## Author contributions

Conceptualization: A.T., T.W., H.L.; Methodology: D.Z., Y.O., E.B., I.C., A.T., T.W., H.L.; Validation: D.Z., Y.O., E.B., A.T., T.W., H.L.; Formal analysis: D.Z., Y.O., A.T., T.W., H.L.; Investigation: D.Z., Y.O., A.T., T.W., H.L.; Resources: T.W., H.L.; Data curation: T.W., H.L.; Writing - original draft: D.Z., E.B., T.W., H.L.; Writing - review & editing: D.Z., Y.O., E.B., I.C., A.T., T.W., H.L.; Visualization: D.Z., Y.O., E.B., A.T., T.W., H.L.; Supervision: T.W., H.L.; Project administration: T.W., H.L.; Funding acquisition: T.W., H.L.

