## supplementary materials for "Notch signaling in the embryonic ectoderm promotes periderm cell fate and represses mineralization of vibrissa hair follicles"

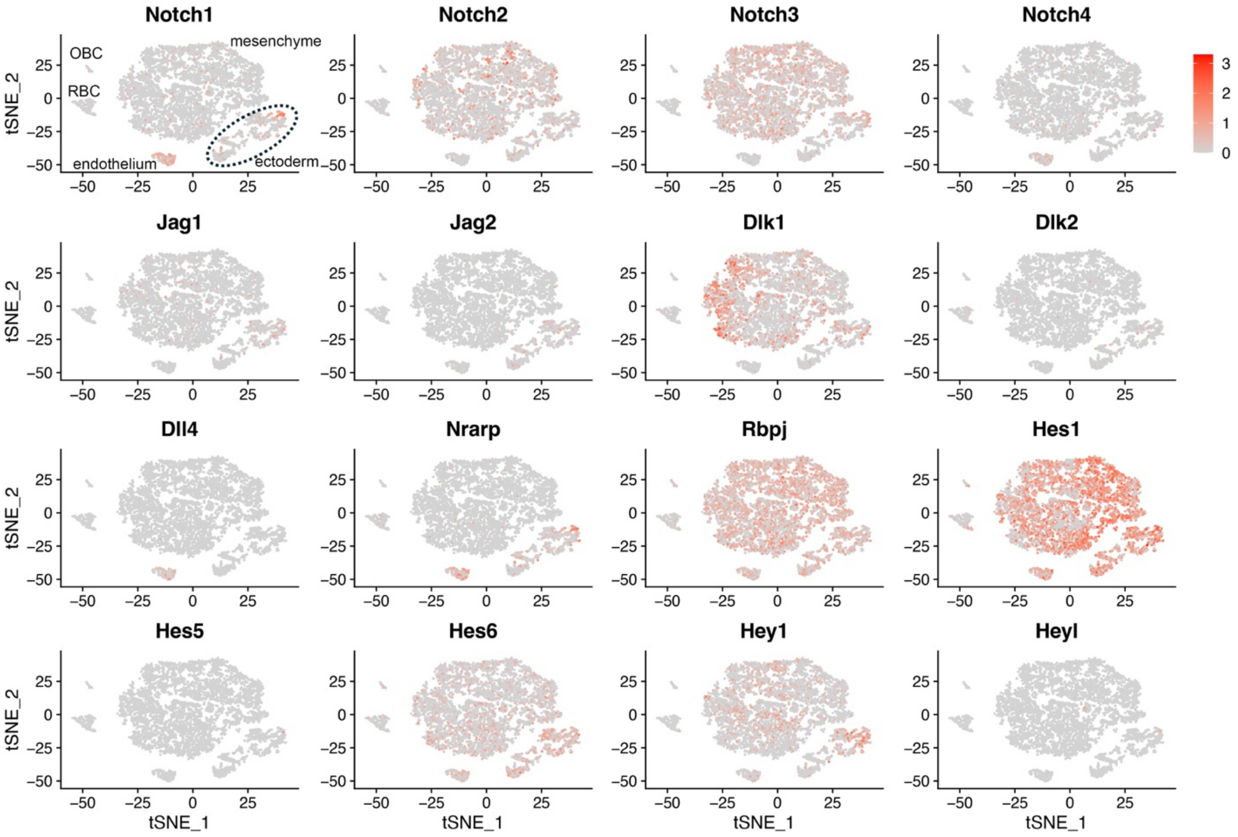


**Supplementary Figure 1. Expression of Notch signaling genes in the E11.5 mouse λ junction**

Expression of genes in the Notch signaling pathway based on a previously published single-cell RNA-seq dataset of E11.5 λ junction (Li et al., 2019). All the plots share the same color scale for expression as shown in the top right corner. The major cell type annotations are shown in the feature plot of *Notch1*. OBC, other blood cells; RBC, red blood cells.

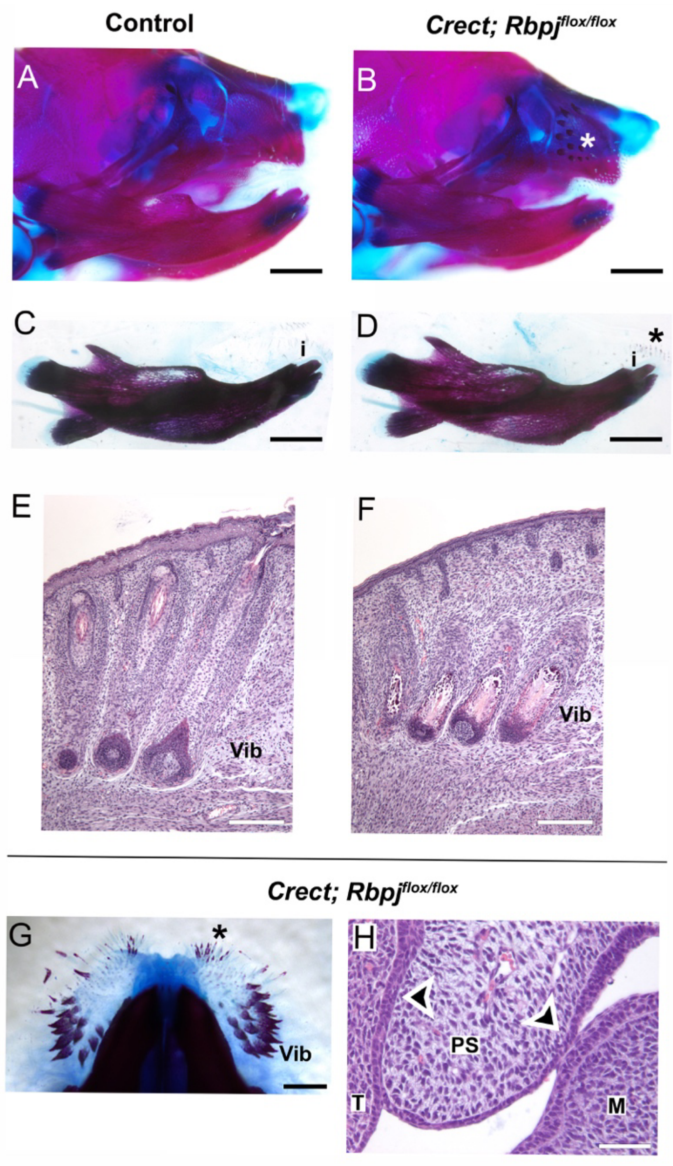


**Supplementary Figure 2. Rbpj mutant skeletons.** Side view of E18.5 control (A, C) or mutant (B, D) facial skeleton (A, B) and isolated mandible (C, D). (E, F) E18.5 mystacial vibrissa follicles of control (E) or mutant (F) stained with H&E. (G) Ventral view of snout skeleton of P1 mutant showing mineralization of mystacial vibrissae (Vib) as well as smaller more anterior vibrissae (*). (H) Coronal section of E13.5 mutant oral cavity stained with H&E showing adhesion between palatal shelf and mandible as well as between palatal shelf and tongue (arrowheads). "*" ectopic mineralization of the vibrissae: i, lower incisor; M, mandible; PS, palatal shelf; T, tongue; Vib, mystacial vibrissae.

Scale bars: 1 mm in A-D, G; 200 μm in E-F; 50 μm in H.

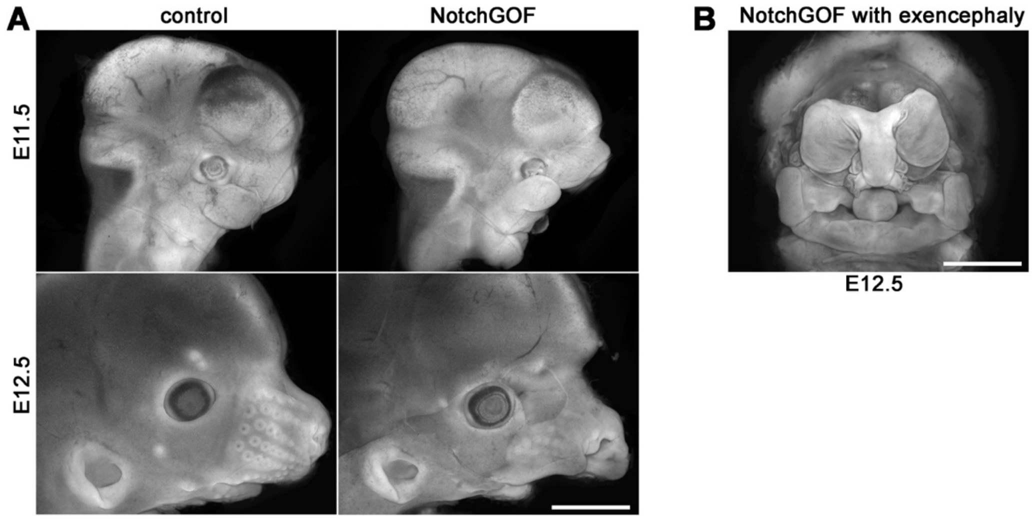


**Supplementary Figure 3. Morphological features of NotchGOF embryos.**
(A) Side views of control and NotchGOF embryos at E11.5 and E12.5 following whole-mount DAPI staining.
(B) Frontal view of an E12.5 NotchGOF embryo displaying exencephaly.

Note that the bodies were removed prior to imaging. Scale bars, 1 mm (all panels).

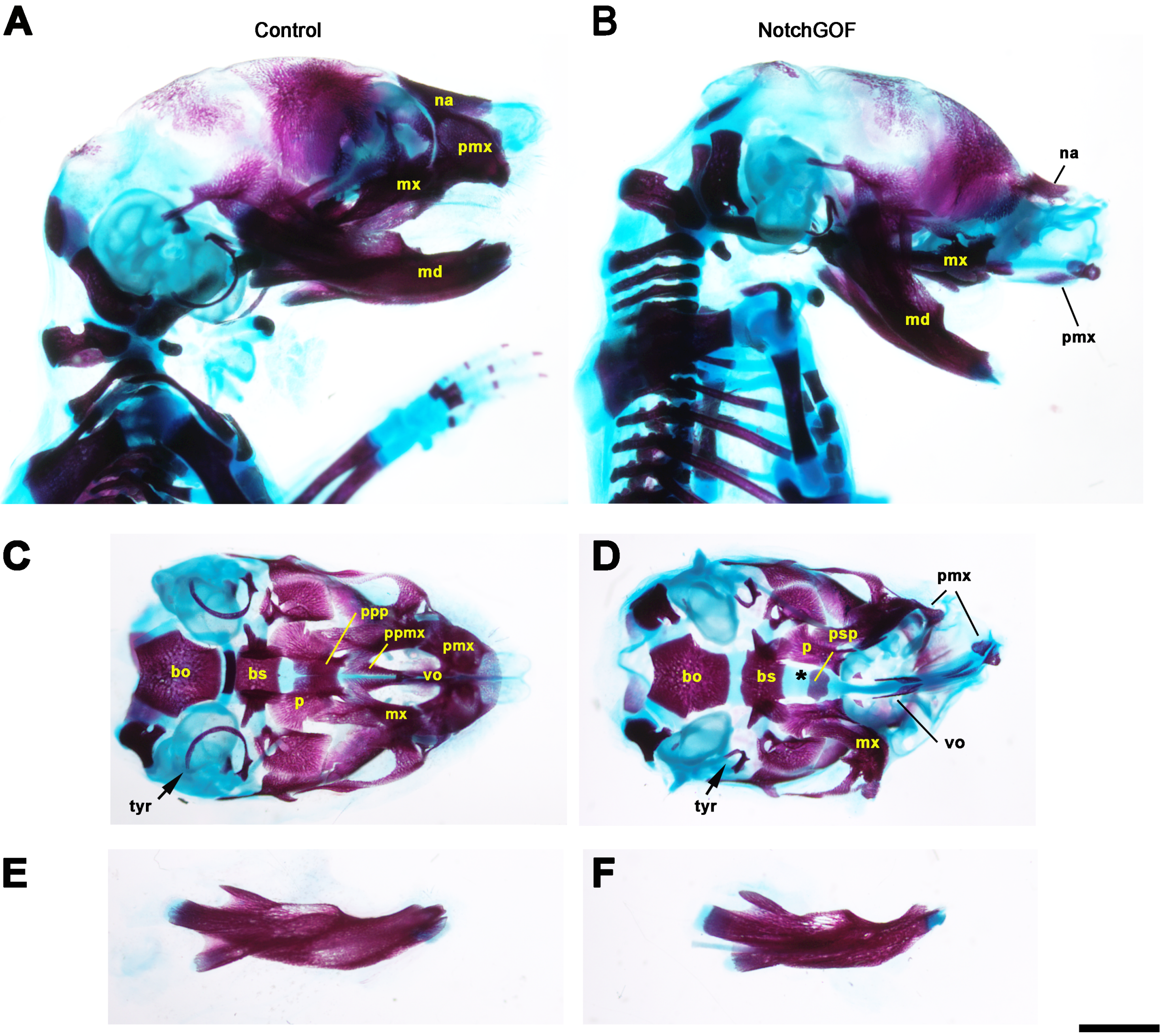


**Supplementary Figure 4. NotchGOF skeleton.**

Skeletons of E18.5 control (A, C, E) and NotchGOF (B, D, F) embryos prepared by Alizarin Red (bone) and Alcian Blue (cartilage) staining. Shown are lateral views of the craniofacial region (A, B), cranial base views with the mandible removed (C, D), and isolated mandibles (E, F). Major skeletal phenotypes in NotchGOF embryos include abnormalities of the premaxilla, maxilla, and palatine bones, consistent with complete cleft lip and palate. Asterisk indicates region of hypoplastic development of the palatal processes derived from the maxillary and palatine bones.

bo, basioccipital; bs, basisphenoid; md, mandible; mx, maxilla; na, nasal; p, palatine; pmx, premaxilla; ppmx, palatal process of maxilla; ppp, palatal process of palatine; psp, presphenoid; tyr, tympanic ring; vo, vomer.

Scale bar, 2 mm (all panels).

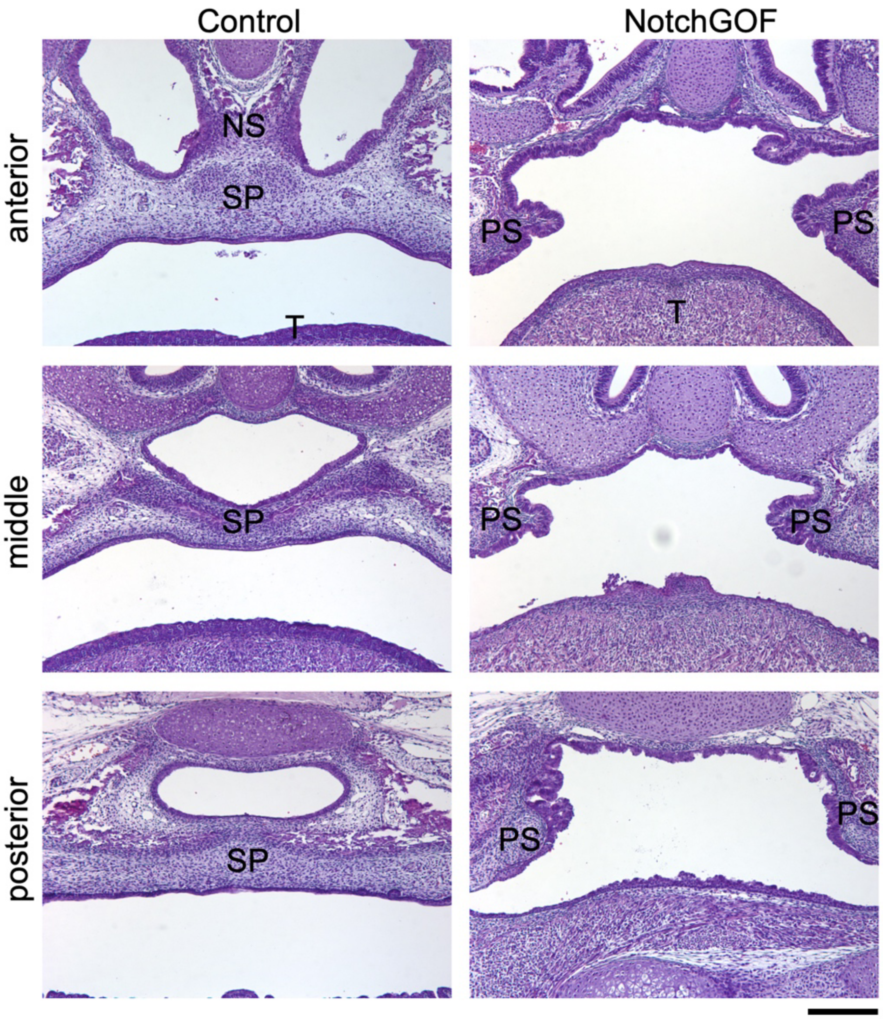


**Supplementary Figure 5. Histological comparison of E16.5 secondary palate morphology.** Hematoxylin and eosin (H&E) staining of coronal sections through the E16.5 mouse secondary palate comparing control (left panels) and NotchGOF mutant (right panels) embryos. Sections are oriented from anterior to posterior. NS, nasal septum; PS, palatal shelf; SP, secondary palate; T, tongue. Scale bar: 200 µm (all panels).

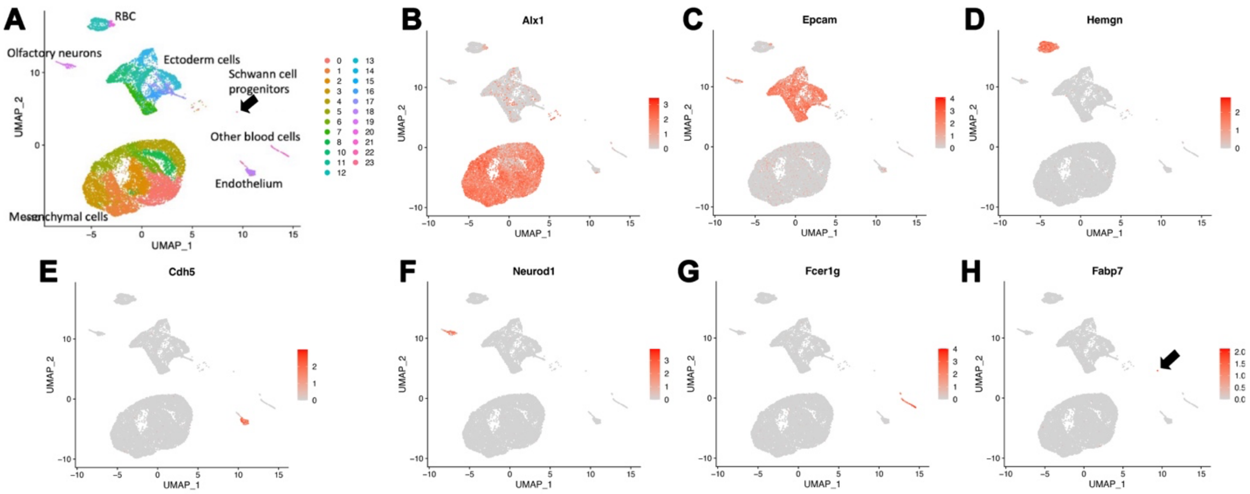


**Supplementary Figure 6. Integrated clustering of E11.5 control and NotchGOF λ-junction cells.**

(A) UMAP plot of cells from E11.5 control and NotchGOF λ-junction regions, showing original clusters (numbered) and annotations of major cell types. Cluster 9 was excluded from the analyses due to low UMI counts.

(B–H) Feature plots showing the expression of representative marker genes for each major cell type: *Alx1* (B), mesenchymal cells; *Epcam* (C), ectodermal cells; *Hemgn* (D), red blood cells; *Cdh5* (E), endothelial cells; *Neurod1* (F), olfactory neurons; *Fcer1g* (G), other blood cells; and *Fabp7* (H), Schwann progenitor cells.

Arrows indicate the smallest cluster, corresponding to Schwann progenitor cells.

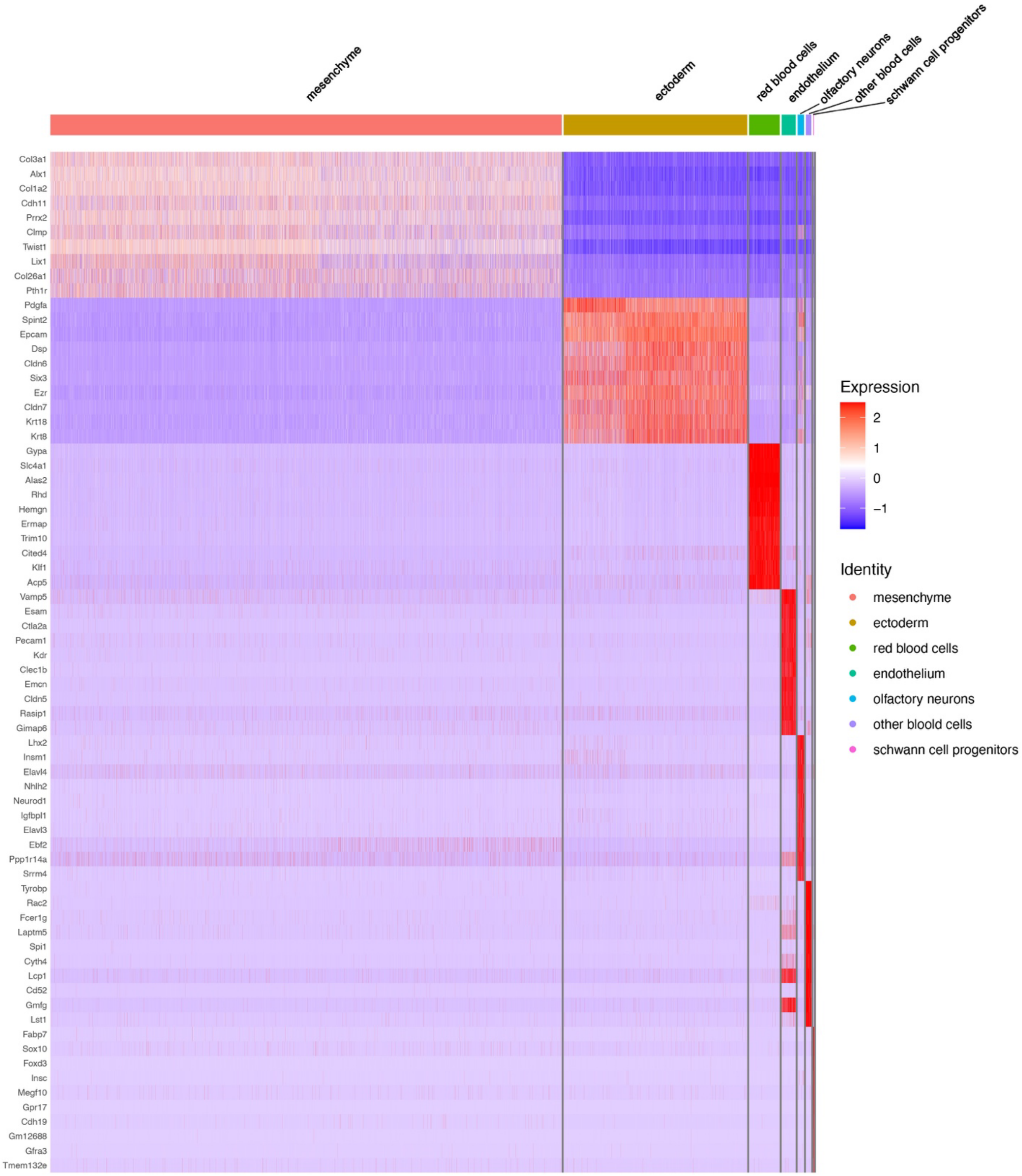


**Supplementary Figure 7. Heatmap of the top 10 marker genes for major cell types in integrated control and NotchGOF λ-junction cells.**

Each row represents the scaled expression of a gene, and each column represents an individual cell. Red indicates higher expression and blue indicates lower expression. The annotation bar above the heatmap denotes cell-type identities.

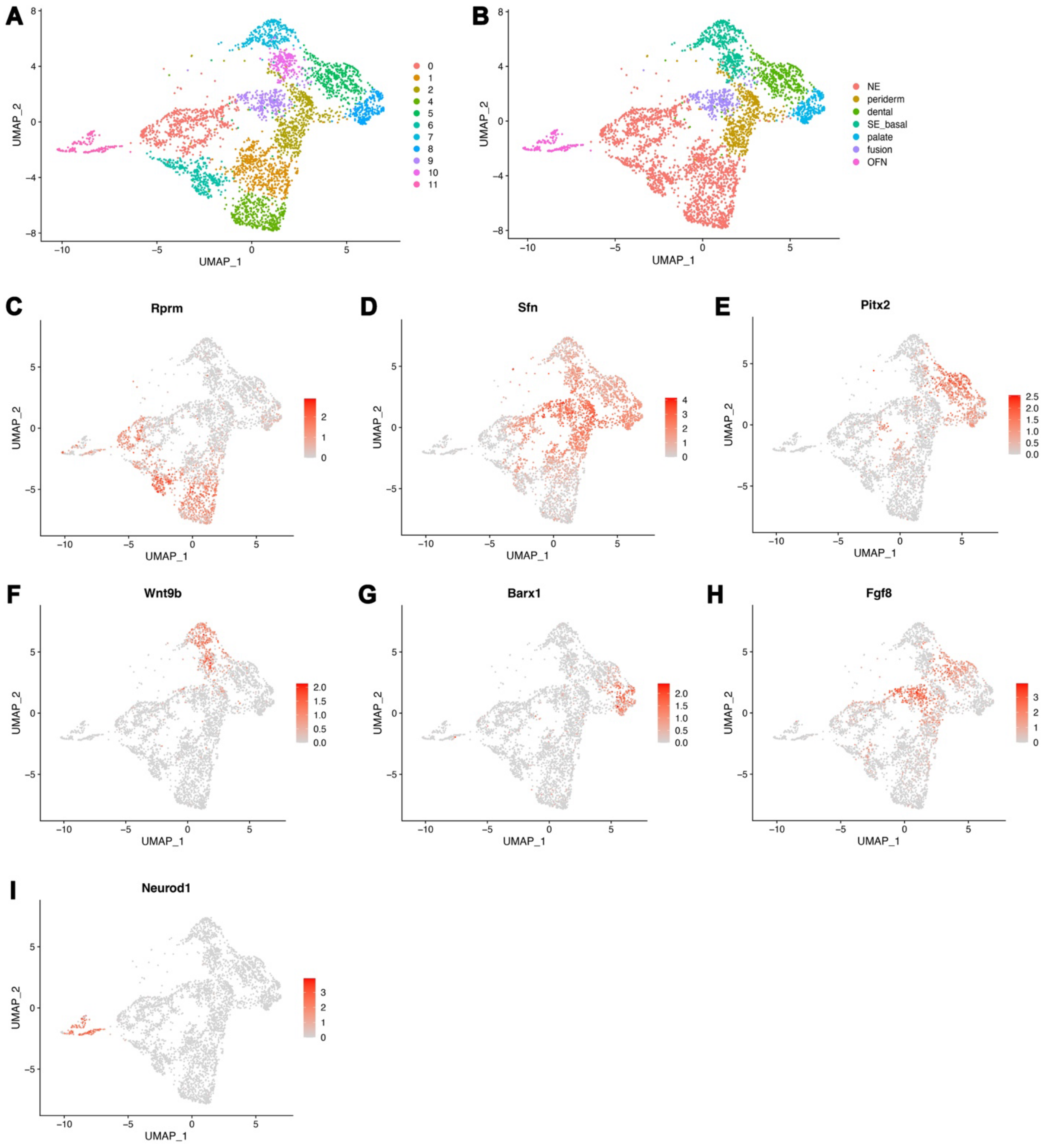


**Supplementary Figure 8. Integrated reclustering of E11.5 control and NotchGOF λ-junction ectoderm cells.**

(A, B) UMAP plot of ectodermal derived cells from E11.5 control and NotchGOF λ-junction regions, showing original clusters (A) and annotations of cell populations (B). Cluster 3 was excluded from the analyses due to low UMI counts.

(C–I) Feature plots showing the expression of representative marker genes for each cell population: *Rprm* (C), nasal epithelial cells; *Sfn* (D), periderm; *Pitx2* (E), dental cells; *Wnt9b* (F), surface ectoderm basal cells; *Barx1* (G), palatal cells; *Fgf8* (H), fusion zone cells; and *Neurod1* (I), olfactory neurons.

NE, nasal epithelial cells; SE_basal, surface ectoderm basal cells; OFN, olfactory neurons.

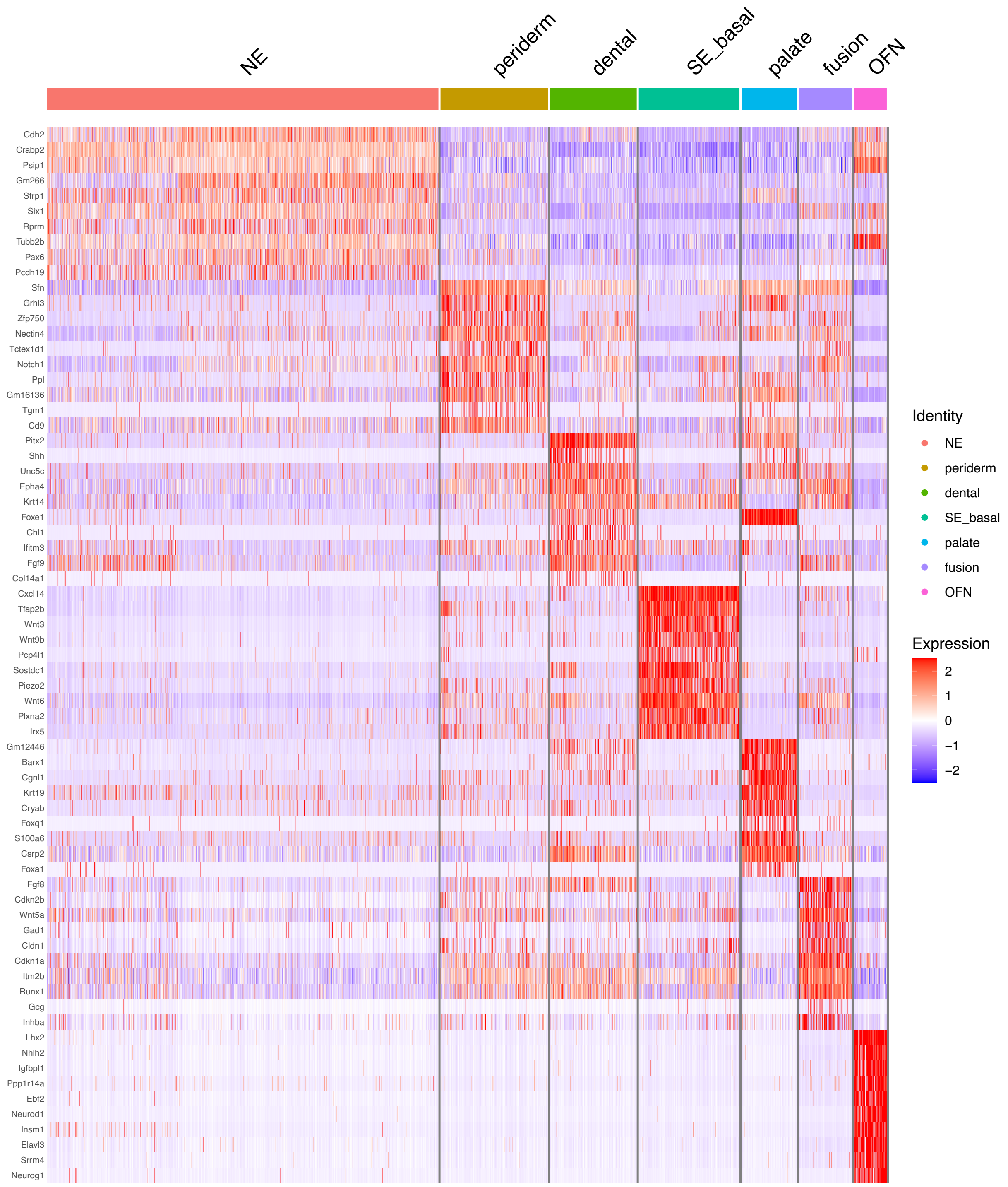


**Supplementary Figure 9. Heatmap of the top 10 marker genes for cell populations in integrated control and NotchGOF λ-junction ectodermal cells.**

Each row represents the scaled expression of a gene, and each column represents an individual cell. Red indicates higher expression and blue indicates lower expression. The annotation bar above the heatmap denotes cell-type identities.

NE, nasal epithelial cells; SE_basal, surface ectoderm basal cells; OFN, olfactory neurons.

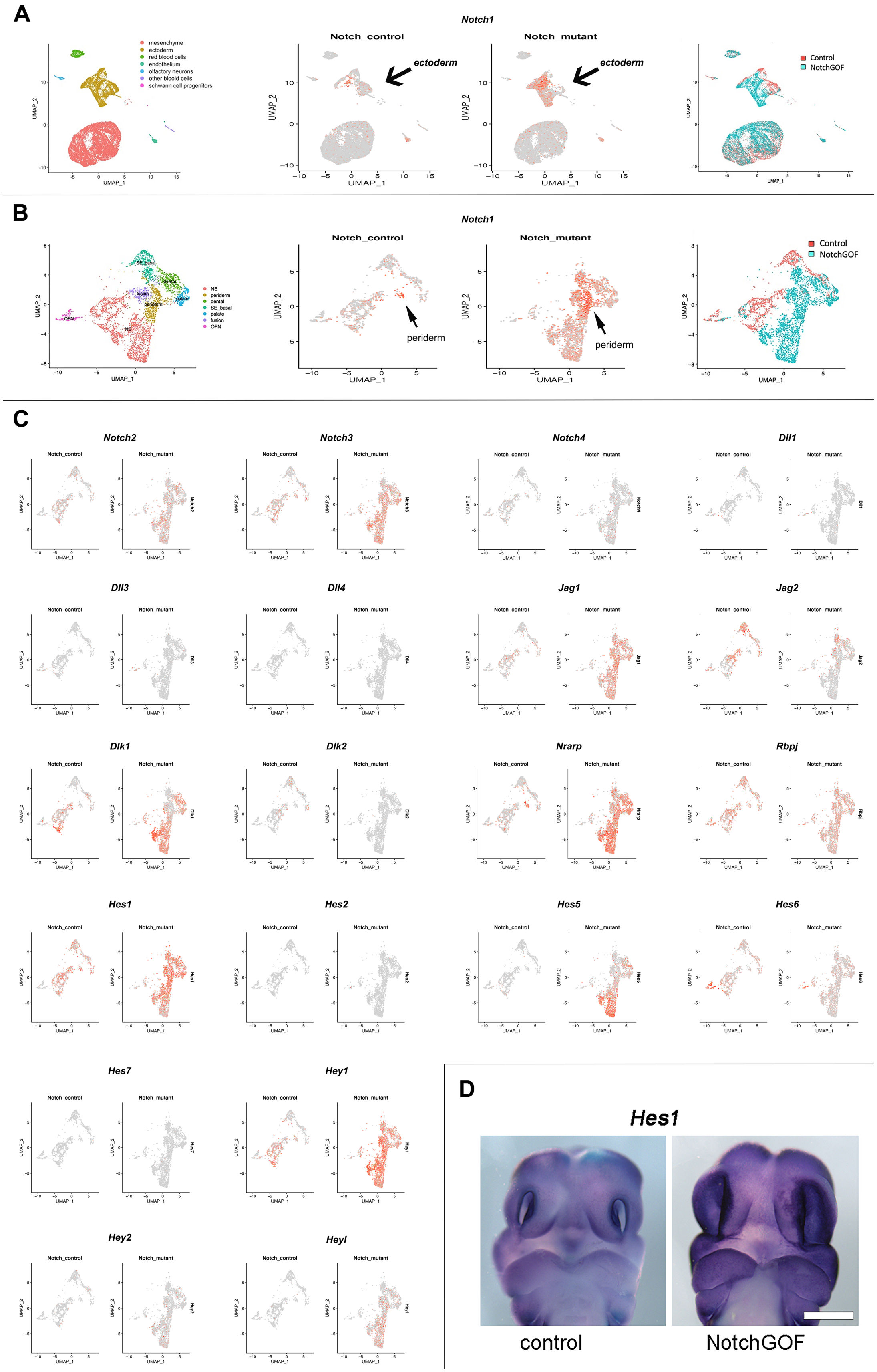
A
Notch1
Notch_control
Notch_mutant
ectoderm
mesenchyme
ectoderm
red blood cells
endothelium
olfactory neurons
other blood cells
schwann cell progenitors
Control
NotchGOF
B
periderm
NE
periderm
dental
SE_basal
palate
fusion
OFN
C
Notch2
Notch3
Notch4
Dll1
Dll3
Dll4
Jag1
Jag2
Dlk1
Dlk2
Nrarp
Rbpj
Hes1
Hes2
Hes5
Hes6
Hes7
Hey1
Hey2
Heyl
D
Hes1
control
NotchGOF

**Supplementary Figure 10. Notch pathway gene expression in the control and NotchGOF ectoderm**. (A) Left panel: UMAP plot of cells from E11.5 control and NotchGOF λ-junction regions showing annotations of major cell types. Middle panel: Feature plots of *Notch1* expression in control and mutant samples. Right panel: Sample identity overlaid on the UMAP plot. Arrow highlights the ectoderm. (B) An equivalent set of panels for the reclustered ectoderm shown in (A). Arrow highlights the periderm. (C). Feature plots of selected Notch pathway genes in control and mutant samples. (D) WMISH of *Hes1* in E10.5 embryos. Frontal views of the embryos are shown, with bodies removed prior to imaging. Scale bar, 1 mm.

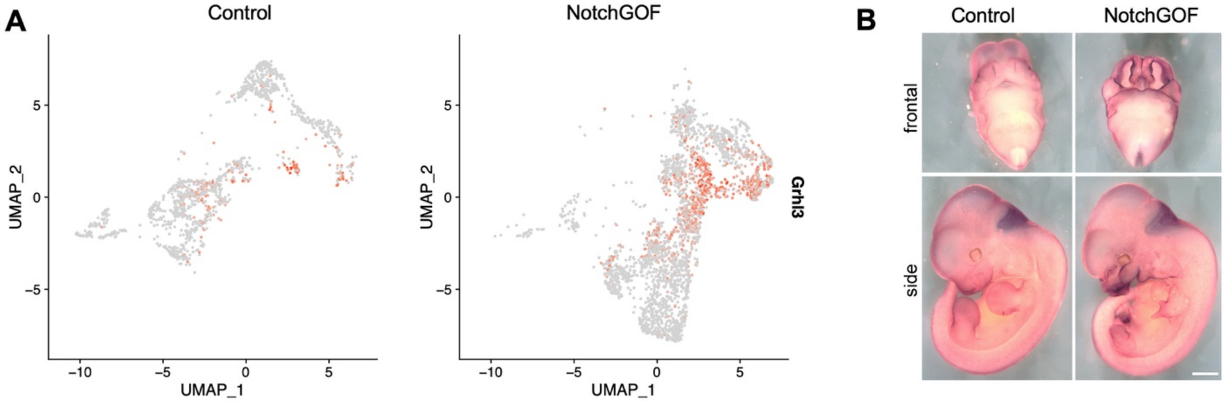


**Supplementary Figure 11. Expression of the periderm marker gene *Grhl3* in the ectoderm**.
(A) Feature plot showing expression of the periderm marker gene *Grhl3* in ectodermal cells by sample.
(B) WMISH of *Grhl3* in E11.5 embryos. Both frontal (top panels) and lateral (bottom panels) views of the embryos are shown, with bodies removed prior to frontal-view imaging. Scale bar, 1 mm (all panels).

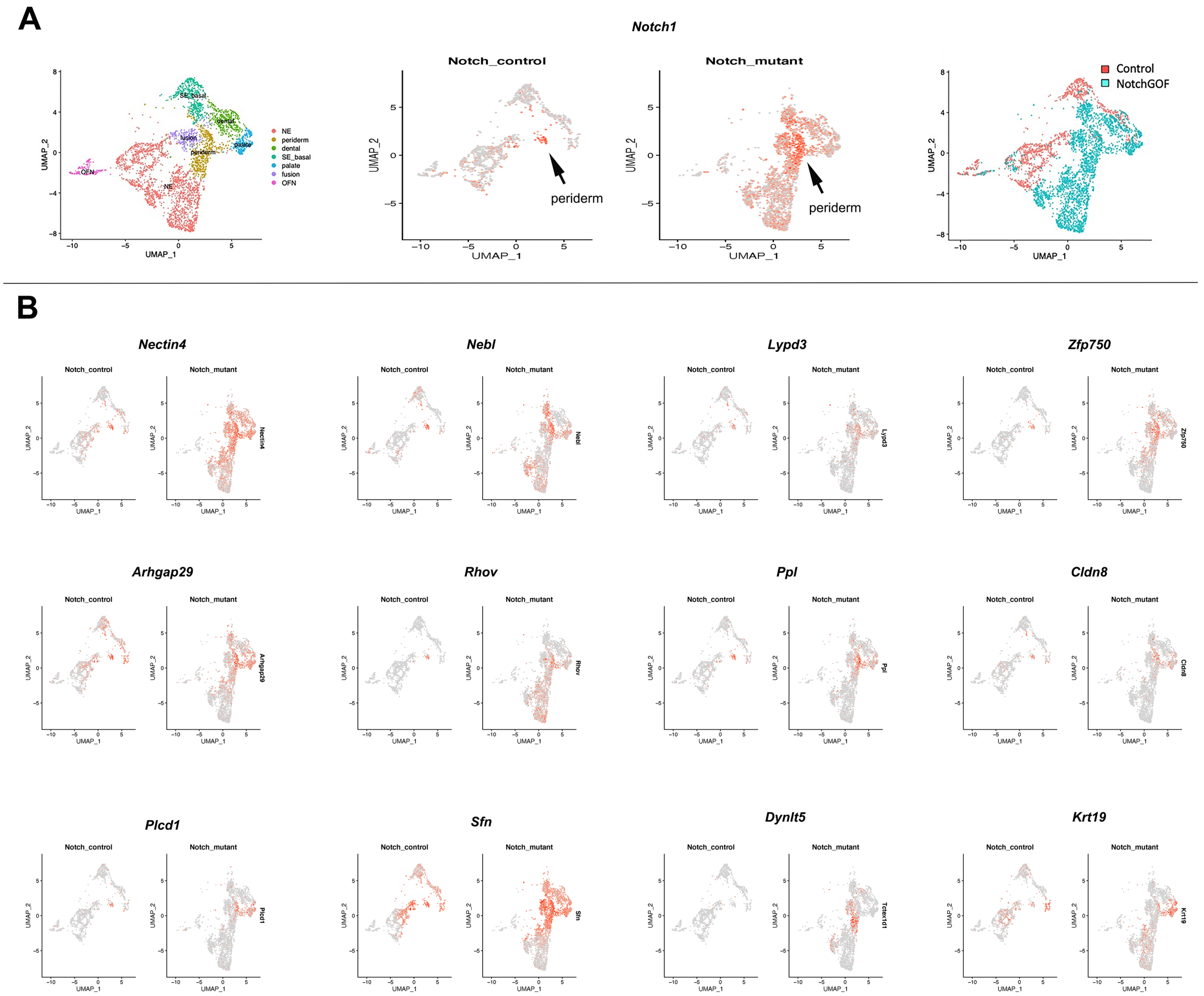


**Supplementary Figure 12. Periderm gene expression in the control and NotchGOF ectoderm**. (A) Left panel: UMAP plot of reclustered ectoderm cells from E11.5 control and NotchGOF λ-junction regions, showing annotations of major cell types. Middle panel: Feature plots of *Notch1* expression in control and mutant samples. Arrows highlight the periderm. Right panel: Sample identity overlaid on the UMAP plot. (B). Feature plots of select examples of periderm gene expression in control and mutant samples.

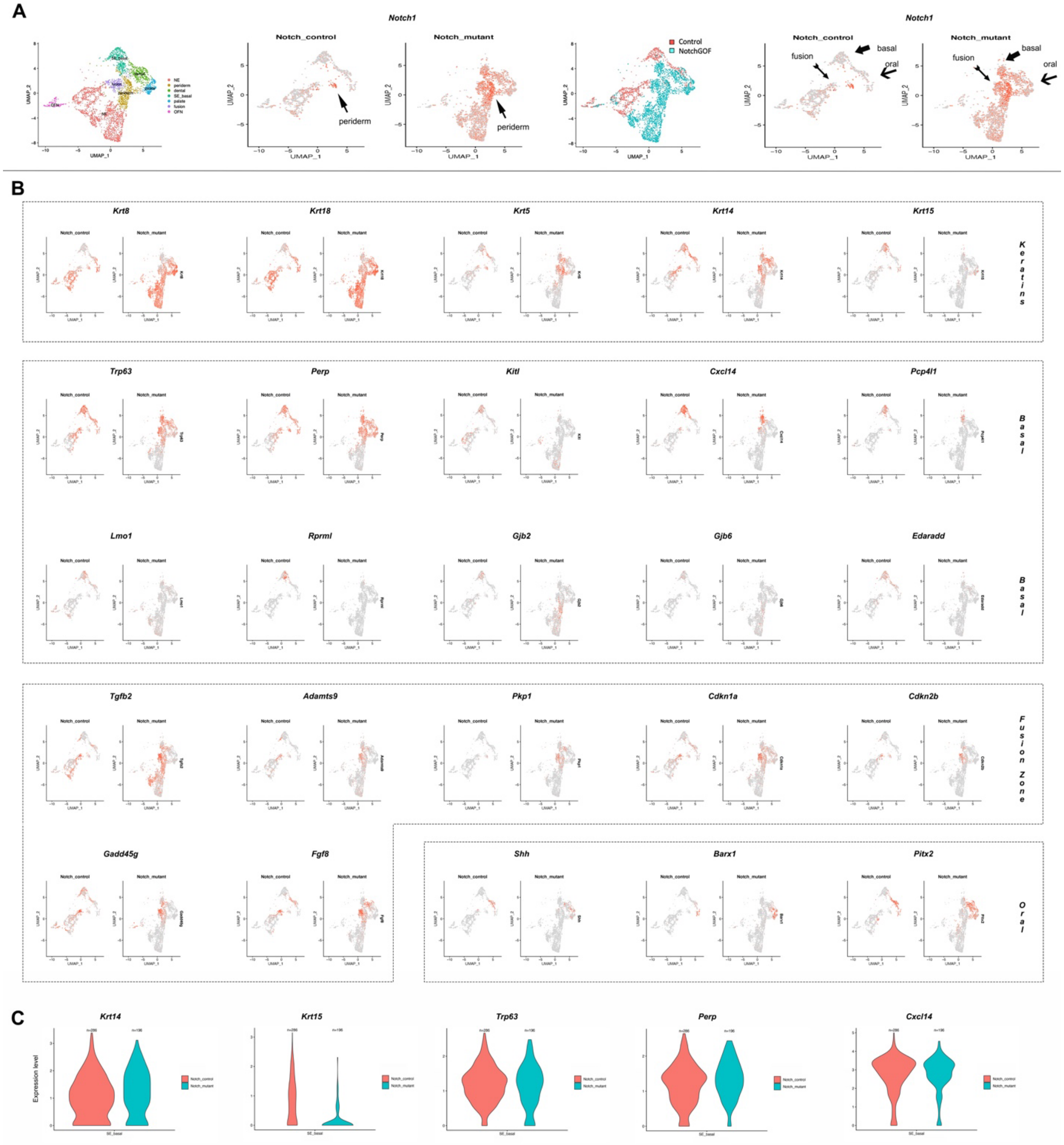


**Supplementary Figure 13. Basal, fusion zone, and oral gene expression in the control and NotchGOF ectoderm**. (A) First panel: UMAP plot of reclustered ectoderm cells from E11.5 control and NotchGOF λ-junction regions, showing annotations of major cell types. Second panel: Feature plots of *Notch1* expression in control and mutant samples. Arrows highlight the periderm. Third panel: Sample identity overlaid on the UMAP plot. Fourth panel: Feature plots of *Notch1* expression in control and mutant samples with different arrows illustrating major cell groupings including fusion, surface ectoderm basal cells (basal), and oral cells. (B). Feature plots of select examples of gene expression in control and mutant samples. (C). Violin plots showing gene expression of *Krt14*, *Krt5*, *Trp63*, *Perp*, and *Cxcl14* in the control and mutant surface ectoderm basal cells (SE_basal), with violin width scaled to the number of cells in each sample.

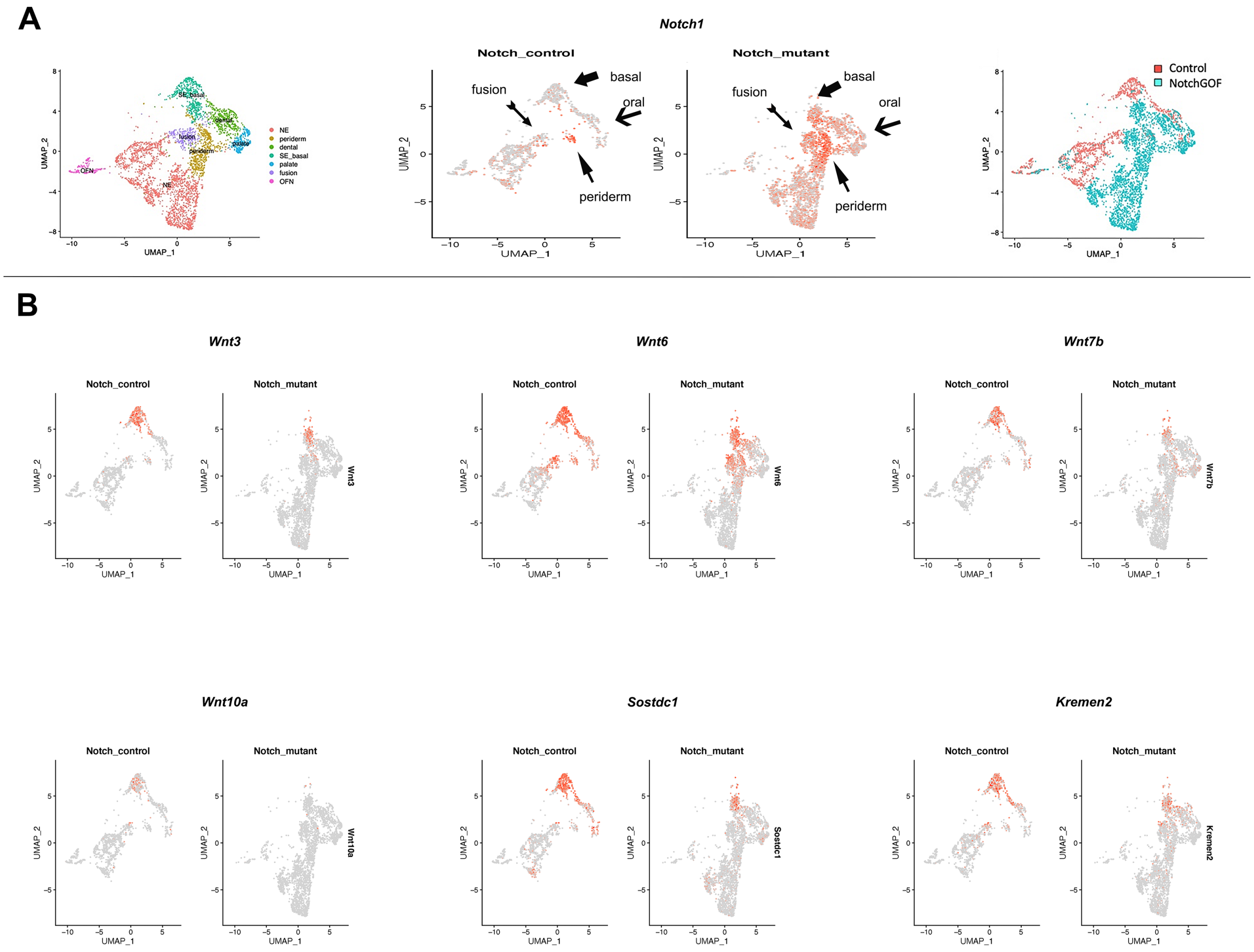


**Supplementary Figure 14. Differentially expressed Wnt ligands and Wnt pathway genes in NotchGOF λ-junction ectodermal cells.**

(A) Left panel: UMAP plot of reclustered ectoderm cells from E11.5 control and NotchGOF λ-junction regions, showing annotations of major cell types. Middle panel: Feature plots of *Notch1* expression in control and mutant samples. Arrows highlight the relevant cluster identities. Right panel: Sample identity overlaid on the UMAP plot. (B). Feature plots of select examples of Wnt pathway gene expression in control and mutant samples.

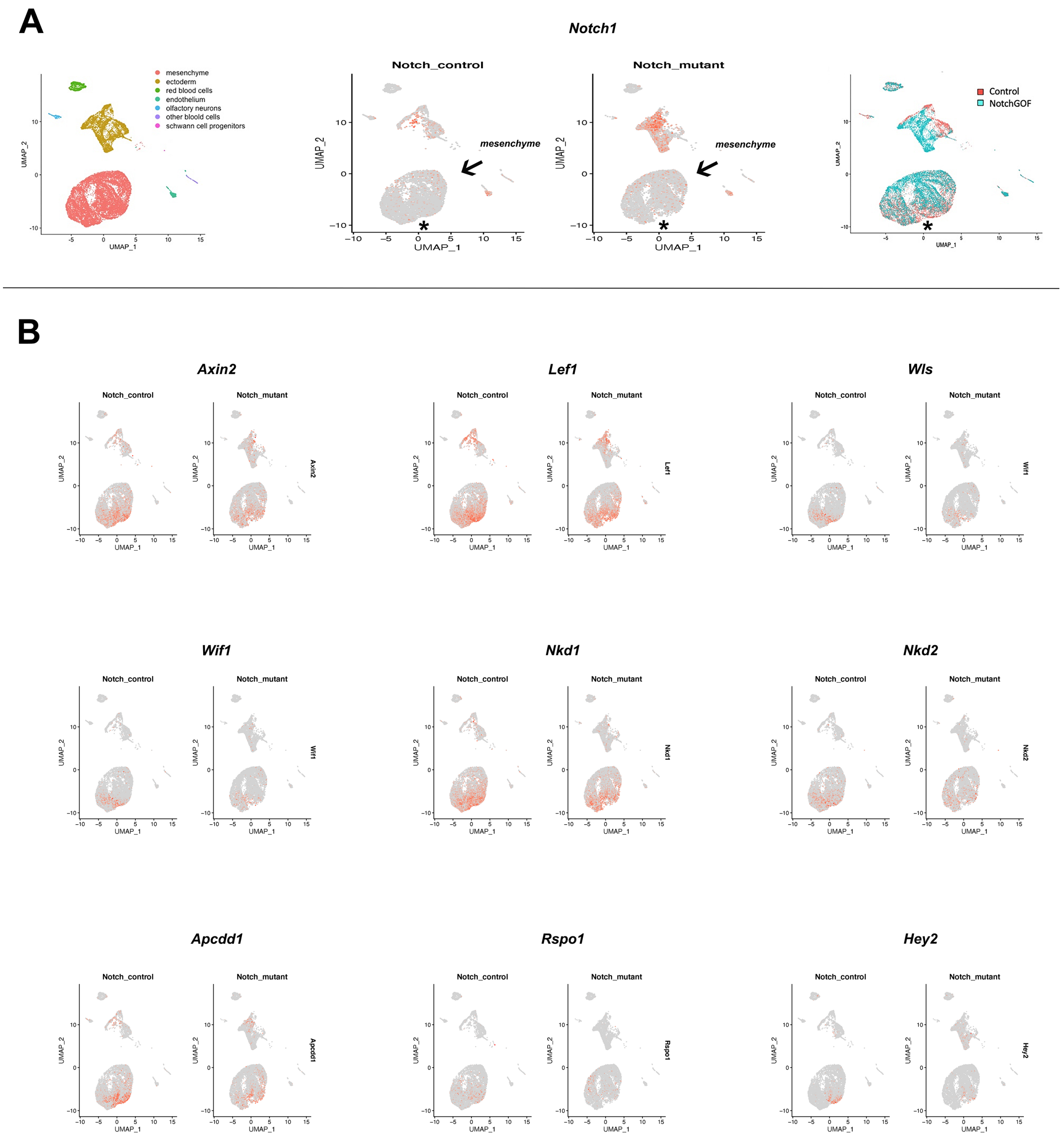


**Supplementary Figure 15. Wnt pathway gene expression in the control and NotchGOF mesenchyme**. (A) Left panel: UMAP plot of cells from E11.5 control and NotchGOF λ-junction regions, showing annotations of major cell types. Middle panel: Feature plots of *Notch1* expression in control and mutant samples with mesenchyme cluster shown by an arrow. Right panel: Sample identity overlaid on the UMAP plot. Asterisk shows area of the mesenchyme in the UMAP that is depleted for mutant cells. (C). Feature plots of select examples of Wnt pathway gene expression in control and mutant samples, as well as *Hey2*, a marker of the cells adjacent to the surface ectoderm.

**Supplementary Tables:**

**Supplementary Table 1. Marker genes for the original clusters of integrated clustering of E11.5 control and NotchGOF λ-junction cells.**

This table was generated using the FindAllMarkers function in the Seurat R package with default parameters, except that min.pct (minimal percent of expression) was set to 0.10, only.pos (only return positive markers) was set to TRUE, and logfc.threshold was set to 0.25. Marker genes were further filtered based on an adjusted *p*-value **(**p_val_adj**)** < 0.05.

avg_logFC, average log fold change in expression; pct.1, percentage of cells within the cluster expressing the gene; pct.2, percentage of cells outside the cluster expressing the gene; p_val, *p*-value; p_val_adj, adjusted *p*-value. #NA indicates a value of pct1/pct2 where the pct2 value was 0 and produced a division error.

Note that Cluster 9 has been omitted from Table S1 due to low UMI counts.

Tables S1–S4 were generated using the same Seurat workflow and parameter settings as described above, and abbreviations are identical across all tables.

**Supplementary Table 2. Marker genes for the major cell types of integrated clustering of E11.5 control and NotchGOF λ-junction cells.**

**Supplementary Table 3. Marker genes for the original clusters of integrated clustering of E11.5 control and NotchGOF λ-junction ectoderm cells.**

Note that Cluster 3 has been omitted from Table S3 due to low UMI counts.

**Supplementary Table 4. Marker genes for the annotated cell populations of integrated clustering of E11.5 control and NotchGOF λ-junction ectoderm cells.**

**Supplementary Table 5. Differentially expressed genes in the mutant λ-junction ectoderm cells.**

Differentially expressed genes were identified using the FindMarkers function in the Seurat R package with default parameters, except that min.pct (minimum percentage of cells expressing the gene) was set to 0.05 and logfc.threshold was set to 0.25, comparing NotchGOF ectodermal cells with control ectodermal cells. Genes were further filtered based on an adjusted p-value (p_val_adj) < 0.05. The presence of sex-related genes in the differential expression analysis likely reflects differences in embryo sex between control and mutant samples, as embryos were not sexed due to the absence of sex-specific phenotypic differences.

avg_log2FC, average log fold change in expression, positive values indicate higher expression in the mutant than control, and negative values indicate lower expression in mutant than control; pct.1, percentage of NotchGOF cells expressing the gene; pct.2 , percentage of control cells expressing the gene; p_val, p-value; p_val_adj, adjusted p-value.

**Supplementary Table 6. Differentially expressed genes in the mutant λ-junction mesenchymal cells.**

This table, referring to mesenchymal cells, was generated using the same analysis pipeline and parameter settings as described for Table S5.

**Supplementary Table 7. Enriched terms detected using Enrichr for differentially expressed genes in the mutant λ-junction ectoderm or mesenchymal cells.** Separate Tabs show the top 10 terms based on p-value with an adjusted p value < 0.05. Terms are derived from all differentially expressed genes detected between the control and mutant ectoderm (All Ect), the up-regulated genes in the mutant ectoderm (Ect Up), or the down-regulated genes in the ectoderm (Ect Down) derived from Supplementary Table 5. Equivalent data are presented for the mesenchyme genes shown in Supplementary Table 6. Results are shown for GO Biological Process 2025 (GO BP 2025), GO Cellular Component 2025 (GO CC 2025), GO Molecular Function 2025 (GO MF 2025) and Jensen DISEASES Experimental 2025 (Diseases 2025).

**Supplementary Table 1. Marker genes for the original clusters of integrated clustering of E11.5 control and NotchGOF λ-junction cells.**

| cluster | gene | avg_log2FC | p_val | p_val_adj | pct.1 | pct.2 | pct1-pct2 | pct1/pct2 |
|---|---|---|---|---|---|---|---|---|
| 0 | Mab21l2 | 1.98 | 0 | 0 | 0.95 | 0.43 | 0.52 | 2.20 |
| 0 | Lhx8 | 1.97 | 0 | 0 | 0.68 | 0.21 | 0.46 | 3.15 |
| 0 | Kctd12 | 2.28 | 0 | 0 | 0.69 | 0.24 | 0.45 | 2.85 |
| 0 | Cdh6 | 2.35 | 0 | 0 | 0.67 | 0.23 | 0.44 | 2.90 |
| 0 | Ndnf | 2.59 | 0 | 0 | 0.59 | 0.16 | 0.43 | 3.75 |
| 0 | Col1a1 | 2.22 | 0 | 0 | 0.67 | 0.25 | 0.42 | 2.69 |
| 0 | Tbx3 | 1.49 | 0 | 0 | 0.80 | 0.39 | 0.41 | 2.05 |
| 0 | Lix1 | 1.26 | 0 | 0 | 0.98 | 0.57 | 0.40 | 1.70 |
| 0 | Msx2 | 1.49 | 0 | 0 | 0.78 | 0.38 | 0.40 | 2.04 |
| 0 | Gm49708 | 1.65 | 0 | 0 | 0.93 | 0.57 | 0.36 | 1.62 |
| 0 | Slitrk6 | 1.80 | 0 | 0 | 0.63 | 0.28 | 0.34 | 2.22 |
| 0 | Lef1 | 1.63 | 0 | 0 | 0.74 | 0.42 | 0.32 | 1.76 |
| 0 | Serpinf1 | 1.15 | 0 | 0 | 0.92 | 0.60 | 0.31 | 1.52 |
| 0 | Col26a1 | 1.21 | 0 | 0 | 0.89 | 0.57 | 0.31 | 1.54 |
| 0 | Cdh11 | 1.08 | 0 | 0 | 0.96 | 0.66 | 0.30 | 1.45 |
| 0 | Msx1 | 1.37 | 0 | 0 | 0.98 | 0.68 | 0.30 | 1.44 |
| 0 | Col1a2 | 0.96 | 0 | 0 | 0.99 | 0.71 | 0.28 | 1.40 |
| 0 | Prrx2 | 1.02 | 0 | 0 | 0.99 | 0.71 | 0.28 | 1.40 |
| 0 | Col6a1 | 2.34 | 0 | 0 | 0.38 | 0.11 | 0.27 | 3.38 |
| 0 | Serpine2 | 1.14 | 0 | 0 | 0.92 | 0.68 | 0.25 | 1.37 |
| 0 | Lgals1 | 1.09 | 0 | 0 | 1.00 | 0.81 | 0.19 | 1.24 |
| 0 | Mpped2 | 0.94 | 0 | 0 | 0.99 | 0.83 | 0.16 | 1.19 |
| 0 | Tcf12 | 0.86 | 0 | 0 | 0.99 | 0.86 | 0.13 | 1.15 |
| 0 | Cald1 | 0.84 | 0 | 0 | 1.00 | 0.88 | 0.12 | 1.14 |
| 0 | Ptn | 1.32 | 0 | 0 | 1.00 | 0.89 | 0.11 | 1.13 |
| 0 | Serpinh1 | 0.68 | 0 | 0 | 1.00 | 0.94 | 0.06 | 1.07 |
| 0 | Tuba1a | 0.55 | 0 | 0 | 1.00 | 0.98 | 0.02 | 1.02 |
| 0 | Tmsb10 | 0.40 | 6.29E-306 | 1.31E-301 | 1.00 | 1.00 | 0.00 | 1.00 |
| 0 | Twist1 | 0.86 | 3.03E-303 | 6.30E-299 | 1.00 | 0.74 | 0.26 | 1.36 |
| 0 | Nkd1 | 1.49 | 3.53E-296 | 7.35E-292 | 0.65 | 0.32 | 0.32 | 2.00 |
| 0 | H2afy2 | 0.60 | 9.17E-294 | 1.91E-289 | 1.00 | 0.93 | 0.07 | 1.07 |
| 0 | Tril | 1.20 | 4.86E-292 | 1.01E-287 | 0.80 | 0.51 | 0.29 | 1.56 |
| 0 | Wnt5a | 1.00 | 9.96E-292 | 2.07E-287 | 0.88 | 0.60 | 0.28 | 1.46 |
| 0 | Col5a2 | 1.21 | 1.18E-287 | 2.45E-283 | 0.78 | 0.49 | 0.29 | 1.58 |
| 0 | Epha7 | 1.25 | 1.06E-284 | 2.21E-280 | 0.80 | 0.53 | 0.27 | 1.51 |
| 0 | Id2 | 0.77 | 2.69E-281 | 5.60E-277 | 0.99 | 0.85 | 0.15 | 1.17 |
| 0 | Vim | 0.62 | 5.28E-275 | 1.10E-270 | 0.96 | 0.82 | 0.14 | 1.17 |
| 0 | Ptms | 0.60 | 6.73E-273 | 1.40E-268 | 1.00 | 0.95 | 0.05 | 1.05 |
| 0 | Zeb2 | 1.57 | 2.20E-272 | 4.57E-268 | 0.65 | 0.36 | 0.29 | 1.80 |
| 0 | Postn | 2.50 | 1.38E-269 | 2.86E-265 | 0.27 | 0.06 | 0.21 | 4.32 |
| 0 | Fzd10os | 1.84 | 3.40E-267 | 7.07E-263 | 0.44 | 0.17 | 0.28 | 2.65 |
| 0 | Col3a1 | 0.73 | 1.73E-264 | 3.59E-260 | 0.97 | 0.69 | 0.28 | 1.41 |
| 0 | Col6a2 | 2.29 | 1.34E-261 | 2.79E-257 | 0.36 | 0.12 | 0.25 | 3.08 |
| 0 | Pax3 | 1.18 | 3.13E-258 | 6.51E-254 | 0.76 | 0.41 | 0.35 | 1.85 |
| 0 | Phldb2 | 1.31 | 1.37E-253 | 2.84E-249 | 0.76 | 0.50 | 0.26 | 1.51 |
| 0 | Zfp503 | 1.34 | 9.66E-253 | 2.01E-248 | 0.64 | 0.34 | 0.30 | 1.89 |
| 0 | Adamts20 | 2.04 | 6.32E-252 | 1.31E-247 | 0.36 | 0.12 | 0.24 | 3.10 |
| 0 | Gata3 | 1.00 | 6.09E-248 | 1.27E-243 | 0.80 | 0.48 | 0.32 | 1.66 |
| 0 | Kif26b | 1.24 | 1.13E-245 | 2.35E-241 | 0.67 | 0.37 | 0.30 | 1.81 |
| 0 | Angpt1 | 2.06 | 2.21E-241 | 4.59E-237 | 0.35 | 0.12 | 0.24 | 3.03 |
| 0 | Nbl1 | 1.06 | 1.60E-237 | 3.32E-233 | 0.78 | 0.45 | 0.33 | 1.72 |
| 0 | Mfap2 | 0.59 | 2.07E-233 | 4.30E-229 | 0.99 | 0.91 | 0.08 | 1.09 |
| 0 | Sgk1 | 1.67 | 1.75E-230 | 3.64E-226 | 0.44 | 0.18 | 0.26 | 2.48 |
| 0 | Dnm3os | 1.07 | 6.33E-230 | 1.32E-225 | 0.77 | 0.49 | 0.29 | 1.59 |
| 0 | Tmem26 | 1.59 | 1.03E-226 | 2.15E-222 | 0.49 | 0.22 | 0.27 | 2.23 |

**Supplementary Table 1. Marker genes for the original clusters of integrated clustering of E11.5 control and NotchGOF λ-junction cells.**

| | | | | | | | | |
|---|---|---|---|---|---|---|---|---|
| 0 | Alx1 | 0.69 | 2.94E-225 | 6.11E-221 | 0.98 | 0.69 | 0.29 | 1.41 |
| 0 | Ankrd12 | 1.21 | 1.06E-223 | 2.21E-219 | 0.73 | 0.50 | 0.23 | 1.45 |
| 0 | Fscn1 | 0.57 | 4.44E-218 | 9.24E-214 | 0.98 | 0.90 | 0.08 | 1.09 |
| 0 | Bambi | 0.75 | 9.14E-218 | 1.90E-213 | 0.85 | 0.61 | 0.23 | 1.38 |
| 0 | Sox11 | 0.52 | 7.51E-217 | 1.56E-212 | 1.00 | 0.96 | 0.04 | 1.04 |
| 0 | Bdnf | 1.64 | 2.83E-214 | 5.88E-210 | 0.52 | 0.26 | 0.26 | 2.03 |
| 0 | Tpm4 | 0.50 | 8.67E-202 | 1.80E-197 | 0.99 | 0.93 | 0.06 | 1.07 |
| 0 | AI606473 | 2.14 | 3.17E-201 | 6.60E-197 | 0.25 | 0.07 | 0.18 | 3.69 |
| 0 | Pth1r | 0.89 | 3.71E-201 | 7.72E-197 | 0.80 | 0.54 | 0.26 | 1.48 |
| 0 | Snai2 | 1.11 | 1.59E-196 | 3.30E-192 | 0.69 | 0.44 | 0.25 | 1.57 |
| 0 | Bmp4 | 1.05 | 1.76E-196 | 3.66E-192 | 0.73 | 0.48 | 0.25 | 1.51 |
| 0 | Clmp | 0.78 | 4.14E-193 | 8.61E-189 | 0.87 | 0.61 | 0.26 | 1.42 |
| 0 | Marcks | 0.42 | 1.50E-191 | 3.12E-187 | 1.00 | 0.97 | 0.03 | 1.03 |
| 0 | Apcdd1 | 1.84 | 5.87E-191 | 1.22E-186 | 0.42 | 0.19 | 0.23 | 2.23 |
| 0 | Hey2 | 2.80 | 4.69E-190 | 9.75E-186 | 0.21 | 0.05 | 0.16 | 4.06 |
| 0 | Igfbp7 | 1.26 | 2.05E-189 | 4.26E-185 | 0.49 | 0.23 | 0.26 | 2.13 |
| 0 | 9130024F11Rik | 1.47 | 7.45E-186 | 1.55E-181 | 0.43 | 0.19 | 0.25 | 2.30 |
| 0 | Basp1 | 0.41 | 1.12E-185 | 2.33E-181 | 1.00 | 0.95 | 0.05 | 1.05 |
| 0 | Frmd6 | 1.03 | 8.81E-178 | 1.83E-173 | 0.69 | 0.46 | 0.23 | 1.50 |
| 0 | Satb2 | 1.24 | 2.99E-177 | 6.22E-173 | 0.48 | 0.23 | 0.25 | 2.12 |
| 0 | Tead2 | 0.50 | 1.82E-169 | 3.78E-165 | 0.97 | 0.89 | 0.08 | 1.09 |
| 0 | Lpar1 | 0.67 | 2.07E-167 | 4.31E-163 | 0.90 | 0.68 | 0.23 | 1.34 |
| 0 | Ednra | 1.18 | 3.24E-165 | 6.73E-161 | 0.55 | 0.31 | 0.24 | 1.78 |
| 0 | Cxcl14 | 0.58 | 3.40E-165 | 7.07E-161 | 0.34 | 0.13 | 0.21 | 2.67 |
| 0 | Akr7a5 | 0.69 | 9.74E-163 | 2.03E-158 | 0.90 | 0.78 | 0.12 | 1.16 |
| 0 | Rai14 | 0.79 | 8.81E-162 | 1.83E-157 | 0.82 | 0.62 | 0.20 | 1.31 |
| 0 | Irx3 | 1.25 | 4.68E-161 | 9.73E-157 | 0.48 | 0.24 | 0.24 | 1.98 |
| 0 | Axin2 | 1.30 | 1.58E-156 | 3.29E-152 | 0.53 | 0.31 | 0.22 | 1.69 |
| 0 | Cachd1 | 1.11 | 5.86E-156 | 1.22E-151 | 0.68 | 0.50 | 0.18 | 1.37 |
| 0 | Hmgb3 | 0.36 | 4.95E-154 | 1.03E-149 | 1.00 | 0.99 | 0.01 | 1.01 |
| 0 | Rcn3 | 0.69 | 7.69E-149 | 1.60E-144 | 0.86 | 0.71 | 0.14 | 1.20 |
| 0 | Actg1 | 0.28 | 1.56E-147 | 3.24E-143 | 1.00 | 1.00 | 0.00 | 1.00 |
| 0 | Rgs4 | 2.60 | 4.85E-146 | 1.01E-141 | 0.13 | 0.02 | 0.10 | 5.29 |
| 0 | Crabp1 | 0.45 | 5.69E-146 | 1.18E-141 | 0.99 | 0.84 | 0.15 | 1.18 |
| 0 | Thy1 | 2.98 | 1.85E-145 | 3.86E-141 | 0.12 | 0.02 | 0.10 | 5.62 |
| 0 | Tbx2 | 0.61 | 4.31E-142 | 8.96E-138 | 0.87 | 0.61 | 0.26 | 1.42 |
| 0 | Alx4 | 0.97 | 1.14E-141 | 2.37E-137 | 0.62 | 0.40 | 0.22 | 1.55 |
| 0 | Snhg18 | 0.97 | 2.35E-141 | 4.89E-137 | 0.68 | 0.50 | 0.18 | 1.37 |
| 0 | Atpif1 | 0.28 | 8.41E-141 | 1.75E-136 | 1.00 | 0.99 | 0.01 | 1.01 |
| 0 | Gas1 | 0.62 | 9.18E-141 | 1.91E-136 | 0.89 | 0.68 | 0.21 | 1.31 |
| 0 | Bzw2 | 0.47 | 8.46E-139 | 1.76E-134 | 0.98 | 0.93 | 0.04 | 1.05 |
| 0 | Hddc2 | 0.69 | 4.88E-138 | 1.01E-133 | 0.85 | 0.75 | 0.11 | 1.14 |
| 0 | Pax1 | 2.97 | 6.56E-137 | 1.37E-132 | 0.11 | 0.02 | 0.09 | 5.43 |
| 0 | Dkk1 | 0.28 | 9.91E-134 | 2.06E-129 | 0.50 | 0.25 | 0.25 | 2.00 |
| 0 | Pcdh18 | 0.68 | 1.38E-133 | 2.88E-129 | 0.81 | 0.61 | 0.20 | 1.32 |
| 0 | 6330403K07Rik | 0.52 | 8.38E-131 | 1.74E-126 | 0.94 | 0.84 | 0.10 | 1.12 |
| 0 | Runx2 | 1.62 | 1.09E-130 | 2.26E-126 | 0.32 | 0.14 | 0.18 | 2.27 |
| 0 | Limd2 | 0.43 | 3.34E-128 | 6.95E-124 | 0.98 | 0.89 | 0.09 | 1.10 |
| 0 | Sox4 | 0.50 | 3.43E-128 | 7.14E-124 | 0.99 | 0.93 | 0.06 | 1.06 |
| 0 | Micos10 | 0.40 | 4.39E-128 | 9.13E-124 | 0.99 | 0.96 | 0.03 | 1.03 |
| 0 | Stx18 | 0.86 | 1.86E-127 | 3.86E-123 | 0.67 | 0.50 | 0.18 | 1.35 |
| 0 | Pde3a | 1.64 | 8.58E-126 | 1.78E-121 | 0.28 | 0.11 | 0.16 | 2.46 |
| 0 | Msrb2 | 1.18 | 1.05E-125 | 2.19E-121 | 0.49 | 0.30 | 0.20 | 1.66 |
| 0 | Sertad4 | 0.97 | 1.49E-122 | 3.11E-118 | 0.55 | 0.34 | 0.21 | 1.61 |
| 0 | Tnfaip8 | 0.68 | 4.67E-121 | 9.71E-117 | 0.86 | 0.74 | 0.12 | 1.16 |
| 0 | Serf1 | 0.42 | 5.62E-121 | 1.17E-116 | 0.97 | 0.92 | 0.06 | 1.06 |

**Supplementary Table 1. Marker genes for the original clusters of integrated clustering of E11.5 control and NotchGOF λ-junction cells.**

| | | | | | | | | |
|---|---|---|---|---|---|---|---|---|
| 0 | Lima1 | 0.56 | 6.00E-121 | 1.25E-116 | 0.88 | 0.72 | 0.16 | 1.22 |
| 0 | Mir100hg | 0.65 | 1.42E-119 | 2.95E-115 | 0.88 | 0.74 | 0.14 | 1.19 |
| 0 | Npc2 | 0.43 | 2.51E-119 | 5.23E-115 | 0.98 | 0.92 | 0.06 | 1.06 |
| 0 | Nradd | 0.81 | 1.08E-117 | 2.25E-113 | 0.68 | 0.51 | 0.18 | 1.35 |
| 0 | Cacna1g | 1.06 | 4.52E-117 | 9.41E-113 | 0.50 | 0.31 | 0.20 | 1.64 |
| 0 | Crlf1 | 0.53 | 1.13E-116 | 2.36E-112 | 0.85 | 0.61 | 0.24 | 1.39 |
| 0 | Tmem119 | 0.81 | 1.22E-115 | 2.53E-111 | 0.64 | 0.43 | 0.21 | 1.48 |
| 0 | Sdc1 | 0.60 | 3.76E-115 | 7.82E-111 | 0.82 | 0.69 | 0.14 | 1.20 |
| 0 | Rerg | 1.38 | 3.84E-115 | 7.98E-111 | 0.41 | 0.23 | 0.18 | 1.81 |
| 0 | Tmem132c | 1.18 | 6.74E-114 | 1.40E-109 | 0.49 | 0.30 | 0.19 | 1.65 |
| 0 | Ifitm2 | 0.40 | 2.17E-113 | 4.52E-109 | 0.99 | 0.94 | 0.05 | 1.06 |
| 0 | Ppfibp2 | 0.86 | 2.90E-113 | 6.04E-109 | 0.60 | 0.41 | 0.20 | 1.48 |
| 0 | Nrp2 | 0.75 | 3.81E-110 | 7.93E-106 | 0.56 | 0.35 | 0.21 | 1.62 |
| 0 | Sema5a | 1.09 | 3.57E-109 | 7.43E-105 | 0.53 | 0.35 | 0.18 | 1.50 |
| 0 | Rtn4 | 0.40 | 2.51E-107 | 5.21E-103 | 0.98 | 0.93 | 0.04 | 1.05 |
| 0 | Cnn3 | 0.37 | 4.79E-107 | 9.95E-103 | 0.98 | 0.93 | 0.05 | 1.05 |
| 0 | Arl2bp | 0.53 | 8.03E-106 | 1.67E-101 | 0.87 | 0.78 | 0.09 | 1.11 |
| 0 | Nfatc4 | 0.72 | 1.61E-105 | 3.34E-101 | 0.70 | 0.53 | 0.17 | 1.32 |
| 0 | Six1 | 0.62 | 3.27E-105 | 6.80E-101 | 0.78 | 0.59 | 0.19 | 1.33 |
| 0 | AW551984 | 1.65 | 3.55E-105 | 7.39E-101 | 0.25 | 0.11 | 0.15 | 2.38 |
| 0 | Ntf3 | 1.13 | 4.88E-105 | 1.01E-100 | 0.45 | 0.27 | 0.18 | 1.66 |
| 0 | Map1lc3a | 0.54 | 7.41E-104 | 1.54E-99 | 0.89 | 0.80 | 0.09 | 1.11 |
| 0 | Aopep | 0.66 | 1.51E-103 | 3.14E-99 | 0.75 | 0.60 | 0.15 | 1.24 |
| 0 | Dbn1 | 0.45 | 3.22E-103 | 6.69E-99 | 0.95 | 0.87 | 0.08 | 1.10 |
| 0 | Ywhaq | 0.28 | 7.14E-103 | 1.49E-98 | 0.99 | 0.98 | 0.01 | 1.01 |
| 0 | Lix1l | 0.87 | 8.96E-103 | 1.86E-98 | 0.54 | 0.36 | 0.19 | 1.52 |
| 0 | Fbln1 | 0.48 | 9.10E-103 | 1.89E-98 | 0.90 | 0.78 | 0.12 | 1.15 |
| 0 | Snai1 | 0.86 | 3.28E-102 | 6.82E-98 | 0.58 | 0.39 | 0.19 | 1.48 |
| 0 | Ctsk | 1.50 | 1.35E-100 | 2.82E-96 | 0.26 | 0.11 | 0.15 | 2.32 |
| 0 | Irx5 | 1.10 | 1.32E-97 | 2.75E-93 | 0.44 | 0.26 | 0.18 | 1.71 |
| 0 | Bcl11b | 0.59 | 2.74E-97 | 5.71E-93 | 0.74 | 0.57 | 0.17 | 1.29 |
| 0 | Bcl11a | 0.53 | 2.61E-96 | 5.43E-92 | 0.85 | 0.72 | 0.12 | 1.17 |
| 0 | Rmst | 1.96 | 4.76E-96 | 9.89E-92 | 0.18 | 0.06 | 0.11 | 2.87 |
| 0 | Prickle1 | 1.16 | 5.38E-95 | 1.12E-90 | 0.37 | 0.20 | 0.17 | 1.82 |
| 0 | Ssbp3 | 0.35 | 1.61E-94 | 3.35E-90 | 0.98 | 0.93 | 0.05 | 1.05 |
| 0 | Tmem59 | 0.44 | 1.77E-94 | 3.69E-90 | 0.93 | 0.88 | 0.05 | 1.06 |
| 0 | Ccbe1 | 2.09 | 2.39E-94 | 4.97E-90 | 0.15 | 0.05 | 0.10 | 3.20 |
| 0 | Lum | 1.20 | 3.06E-93 | 6.37E-89 | 0.30 | 0.14 | 0.16 | 2.12 |
| 0 | Ppp1r14b | 0.25 | 9.61E-92 | 2.00E-87 | 1.00 | 0.99 | 0.01 | 1.01 |
| 0 | Pmaip1 | 1.05 | 1.12E-91 | 2.34E-87 | 0.40 | 0.22 | 0.17 | 1.77 |
| 0 | Gja1 | 0.43 | 1.41E-91 | 2.93E-87 | 0.88 | 0.76 | 0.12 | 1.16 |
| 0 | Gpx8 | 0.53 | 1.02E-89 | 2.13E-85 | 0.82 | 0.69 | 0.12 | 1.18 |
| 0 | Pcolce | 0.67 | 1.38E-88 | 2.86E-84 | 0.62 | 0.44 | 0.18 | 1.41 |
| 0 | Gli3 | 0.67 | 5.01E-88 | 1.04E-83 | 0.70 | 0.57 | 0.14 | 1.24 |
| 0 | Rspo1 | 1.80 | 2.01E-87 | 4.17E-83 | 0.16 | 0.06 | 0.11 | 2.88 |
| 0 | Timp2 | 0.54 | 2.74E-87 | 5.70E-83 | 0.79 | 0.66 | 0.14 | 1.21 |
| 0 | Pja1 | 0.58 | 6.66E-87 | 1.38E-82 | 0.77 | 0.66 | 0.10 | 1.16 |
| 0 | Tsc22d1 | 0.34 | 1.02E-86 | 2.12E-82 | 0.98 | 0.93 | 0.05 | 1.05 |
| 0 | Capzb | 0.37 | 6.39E-86 | 1.33E-81 | 0.96 | 0.91 | 0.05 | 1.06 |
| 0 | Rest | 0.44 | 5.61E-85 | 1.17E-80 | 0.90 | 0.80 | 0.10 | 1.12 |
| 0 | Cyp26b1 | 1.79 | 3.82E-84 | 7.94E-80 | 0.24 | 0.11 | 0.13 | 2.16 |
| 0 | Txnrd1 | 0.42 | 1.36E-83 | 2.83E-79 | 0.91 | 0.86 | 0.05 | 1.06 |
| 0 | Mdk | 0.27 | 4.27E-83 | 8.88E-79 | 1.00 | 0.99 | 0.01 | 1.01 |
| 0 | Ddah2 | 0.32 | 4.31E-83 | 8.97E-79 | 0.99 | 0.94 | 0.05 | 1.05 |
| 0 | Myo1b | 0.49 | 6.46E-83 | 1.34E-78 | 0.83 | 0.72 | 0.11 | 1.16 |
| 0 | Zfp462 | 0.53 | 7.22E-83 | 1.50E-78 | 0.82 | 0.72 | 0.10 | 1.14 |

**Supplementary Table 1. Marker genes for the original clusters of integrated clustering of E11.5 control and NotchGOF λ-junction cells.**

| | | | | | | | | |
|---|---|---|---|---|---|---|---|---|
| 0 | Trps1 | 1.27 | 7.52E-83 | 1.56E-78 | 0.59 | 0.47 | 0.12 | 1.24 |
| 0 | Crabp2 | 0.35 | 6.10E-81 | 1.27E-76 | 0.99 | 0.91 | 0.09 | 1.10 |
| 0 | Dlc1 | 0.82 | 7.30E-81 | 1.52E-76 | 0.51 | 0.34 | 0.17 | 1.49 |
| 0 | Rgs5 | 1.64 | 1.71E-80 | 3.56E-76 | 0.18 | 0.07 | 0.11 | 2.57 |
| 0 | Daam2 | 0.90 | 2.51E-80 | 5.23E-76 | 0.44 | 0.28 | 0.16 | 1.59 |
| 0 | Crnde | 1.38 | 3.77E-80 | 7.83E-76 | 0.27 | 0.14 | 0.13 | 1.99 |
| 0 | Atxn7l3b | 0.25 | 5.68E-80 | 1.18E-75 | 1.00 | 0.98 | 0.02 | 1.02 |
| 0 | Prex2 | 0.74 | 1.90E-79 | 3.95E-75 | 0.55 | 0.38 | 0.17 | 1.43 |
| 0 | Fat4 | 1.23 | 2.58E-79 | 5.36E-75 | 0.41 | 0.26 | 0.15 | 1.57 |
| 0 | S1pr3 | 1.01 | 7.52E-79 | 1.56E-74 | 0.42 | 0.26 | 0.16 | 1.61 |
| 0 | Hmgcs1 | 0.51 | 1.36E-78 | 2.84E-74 | 0.84 | 0.76 | 0.08 | 1.11 |
| 0 | Fam89b | 0.45 | 6.01E-78 | 1.25E-73 | 0.86 | 0.79 | 0.07 | 1.09 |
| 0 | Pde4d | 0.99 | 1.08E-77 | 2.25E-73 | 0.40 | 0.25 | 0.15 | 1.62 |
| 0 | Bmper | 1.86 | 3.06E-77 | 6.36E-73 | 0.17 | 0.07 | 0.10 | 2.51 |
| 0 | Robo2 | 1.38 | 3.48E-77 | 7.24E-73 | 0.32 | 0.18 | 0.14 | 1.79 |
| 0 | Robo1 | 0.75 | 8.11E-77 | 1.69E-72 | 0.59 | 0.45 | 0.14 | 1.30 |
| 0 | Khdrbs1 | 0.30 | 1.46E-76 | 3.04E-72 | 0.99 | 0.95 | 0.03 | 1.04 |
| 0 | Golm1 | 0.64 | 4.78E-76 | 9.93E-72 | 0.67 | 0.54 | 0.13 | 1.24 |
| 0 | Setbp1 | 0.87 | 7.00E-76 | 1.46E-71 | 0.52 | 0.37 | 0.15 | 1.41 |
| 0 | Mmp14 | 0.58 | 7.84E-76 | 1.63E-71 | 0.74 | 0.62 | 0.12 | 1.19 |
| 0 | Asph | 0.57 | 1.76E-75 | 3.66E-71 | 0.76 | 0.64 | 0.12 | 1.19 |
| 0 | Unc119 | 0.44 | 2.74E-75 | 5.70E-71 | 0.85 | 0.77 | 0.08 | 1.10 |
| 0 | Rhoq | 0.74 | 1.04E-74 | 2.15E-70 | 0.57 | 0.43 | 0.14 | 1.33 |
| 0 | Fam107b | 0.91 | 1.09E-74 | 2.27E-70 | 0.48 | 0.34 | 0.14 | 1.42 |
| 0 | Eid1 | 0.25 | 1.34E-74 | 2.78E-70 | 0.99 | 0.97 | 0.02 | 1.02 |
| 0 | Mmp2 | 0.59 | 1.55E-74 | 3.23E-70 | 0.66 | 0.50 | 0.16 | 1.33 |
| 0 | Gap43 | 0.64 | 2.03E-74 | 4.22E-70 | 0.56 | 0.40 | 0.16 | 1.41 |
| 0 | Fzd10 | 1.27 | 4.01E-73 | 8.33E-69 | 0.25 | 0.12 | 0.13 | 2.02 |
| 0 | Ptov1 | 0.26 | 3.23E-72 | 6.73E-68 | 0.99 | 0.97 | 0.03 | 1.03 |
| 0 | Dync1i2 | 0.37 | 3.53E-72 | 7.33E-68 | 0.93 | 0.88 | 0.05 | 1.05 |
| 0 | Nbea | 0.81 | 8.93E-72 | 1.86E-67 | 0.56 | 0.43 | 0.13 | 1.31 |
| 0 | Eml4 | 0.38 | 5.01E-71 | 1.04E-66 | 0.90 | 0.83 | 0.08 | 1.09 |
| 0 | Aldh1a2 | 1.65 | 6.42E-71 | 1.33E-66 | 0.15 | 0.06 | 0.09 | 2.65 |
| 0 | Samd5 | 0.33 | 6.98E-71 | 1.45E-66 | 0.54 | 0.34 | 0.20 | 1.60 |
| 0 | Cxxc4 | 1.03 | 1.13E-70 | 2.35E-66 | 0.43 | 0.29 | 0.14 | 1.49 |
| 0 | A830018L16Rik | 1.41 | 1.46E-70 | 3.05E-66 | 0.23 | 0.11 | 0.12 | 2.07 |
| 0 | Tpd52l1 | 0.93 | 2.14E-70 | 4.45E-66 | 0.46 | 0.32 | 0.14 | 1.45 |
| 0 | Tshz2 | 0.83 | 2.44E-70 | 5.08E-66 | 0.48 | 0.32 | 0.16 | 1.49 |
| 0 | Cyp51 | 0.50 | 5.48E-70 | 1.14E-65 | 0.78 | 0.68 | 0.10 | 1.15 |
| 0 | C430049B03Rik | 0.83 | 9.63E-70 | 2.00E-65 | 0.48 | 0.34 | 0.15 | 1.44 |
| 0 | Emilin2 | 1.01 | 2.03E-69 | 4.22E-65 | 0.38 | 0.24 | 0.14 | 1.59 |
| 0 | Atat1 | 0.63 | 5.82E-69 | 1.21E-64 | 0.67 | 0.54 | 0.13 | 1.23 |
| 0 | Gnai3 | 0.38 | 6.08E-69 | 1.26E-64 | 0.89 | 0.83 | 0.06 | 1.07 |
| 0 | Sulf2 | 0.47 | 9.58E-69 | 1.99E-64 | 0.79 | 0.65 | 0.14 | 1.21 |
| 0 | Ahnak | 0.75 | 1.01E-68 | 2.10E-64 | 0.58 | 0.44 | 0.14 | 1.31 |
| 0 | Rnf122 | 0.61 | 1.39E-68 | 2.89E-64 | 0.65 | 0.52 | 0.13 | 1.26 |
| 0 | Flna | 0.41 | 1.65E-68 | 3.43E-64 | 0.88 | 0.78 | 0.10 | 1.12 |
| 0 | 2610318N02Rik | 0.59 | 2.22E-68 | 4.62E-64 | 0.69 | 0.59 | 0.11 | 1.18 |
| 0 | Ube2r2 | 0.29 | 2.51E-68 | 5.22E-64 | 0.98 | 0.96 | 0.02 | 1.02 |
| 0 | Gm8797 | 0.29 | 5.08E-68 | 1.06E-63 | 0.97 | 0.94 | 0.03 | 1.03 |
| 0 | Sh3bgrl | 0.35 | 5.37E-68 | 1.12E-63 | 0.92 | 0.85 | 0.08 | 1.09 |
| 0 | Cdo1 | 0.70 | 1.76E-67 | 3.67E-63 | 0.62 | 0.49 | 0.13 | 1.26 |
| 0 | Marveld1 | 1.04 | 1.79E-67 | 3.71E-63 | 0.36 | 0.22 | 0.14 | 1.63 |
| 0 | Fst | 0.88 | 3.09E-67 | 6.43E-63 | 0.23 | 0.11 | 0.12 | 2.09 |
| 0 | Paip2 | 0.28 | 1.67E-66 | 3.48E-62 | 0.99 | 0.97 | 0.01 | 1.01 |
| 0 | Dynlrb1 | 0.29 | 1.85E-66 | 3.85E-62 | 0.98 | 0.94 | 0.04 | 1.04 |

**Supplementary Table 1. Marker genes for the original clusters of integrated clustering of E11.5 control and NotchGOF λ-junction cells.**

| | | | | | | | | |
|---|---|---|---|---|---|---|---|---|
| 0 | Ppdpf | 0.26 | 2.05E-66 | 4.27E-62 | 0.98 | 0.96 | 0.02 | 1.02 |
| 0 | Arid5b | 0.55 | 7.48E-66 | 1.56E-61 | 0.76 | 0.66 | 0.10 | 1.16 |
| 0 | a | 1.32 | 1.67E-65 | 3.48E-61 | 0.24 | 0.12 | 0.12 | 1.97 |
| 0 | Zfhx3 | 0.64 | 1.71E-65 | 3.57E-61 | 0.88 | 0.81 | 0.08 | 1.09 |
| 0 | Hsd17b10 | 0.34 | 1.81E-65 | 3.77E-61 | 0.95 | 0.90 | 0.05 | 1.05 |
| 0 | Otor | 1.56 | 4.24E-65 | 8.81E-61 | 0.16 | 0.07 | 0.09 | 2.45 |
| 0 | Sgcd | 1.02 | 5.23E-65 | 1.09E-60 | 0.34 | 0.21 | 0.14 | 1.66 |
| 0 | Ctxn1 | 0.41 | 5.81E-65 | 1.21E-60 | 0.86 | 0.77 | 0.09 | 1.12 |
| 0 | Unc5b | 0.64 | 1.82E-64 | 3.80E-60 | 0.53 | 0.37 | 0.16 | 1.41 |
| 0 | Eci2 | 0.48 | 6.08E-64 | 1.26E-59 | 0.77 | 0.67 | 0.09 | 1.14 |
| 0 | Tnfaip6 | 0.58 | 6.16E-64 | 1.28E-59 | 0.62 | 0.45 | 0.17 | 1.38 |
| 0 | Tcf4 | 0.37 | 7.33E-64 | 1.52E-59 | 0.96 | 0.89 | 0.06 | 1.07 |
| 0 | Ilf2 | 0.28 | 9.56E-64 | 1.99E-59 | 0.98 | 0.94 | 0.04 | 1.05 |
| 0 | Spock3 | 1.41 | 1.12E-63 | 2.32E-59 | 0.23 | 0.12 | 0.11 | 1.97 |
| 0 | Kdm2b | 0.61 | 2.04E-62 | 4.25E-58 | 0.66 | 0.55 | 0.10 | 1.19 |
| 0 | Ubr5 | 0.56 | 2.89E-62 | 6.02E-58 | 0.74 | 0.66 | 0.08 | 1.12 |
| 0 | Ostc | 0.29 | 3.08E-62 | 6.41E-58 | 0.98 | 0.94 | 0.03 | 1.03 |
| 0 | Idh1 | 0.53 | 4.69E-62 | 9.76E-58 | 0.71 | 0.62 | 0.09 | 1.15 |
| 0 | Ankrd44 | 0.93 | 8.06E-62 | 1.68E-57 | 0.40 | 0.26 | 0.13 | 1.50 |
| 0 | Rnf182 | 0.88 | 8.63E-62 | 1.80E-57 | 0.33 | 0.20 | 0.13 | 1.66 |
| 0 | Rasl11b | 0.61 | 1.98E-61 | 4.11E-57 | 0.61 | 0.47 | 0.14 | 1.30 |
| 0 | Crispld1 | 0.74 | 7.18E-61 | 1.49E-56 | 0.45 | 0.31 | 0.14 | 1.46 |
| 0 | Igfbp4 | 0.28 | 8.00E-61 | 1.66E-56 | 0.99 | 0.89 | 0.10 | 1.12 |
| 0 | Ddr2 | 0.53 | 1.24E-60 | 2.58E-56 | 0.63 | 0.47 | 0.16 | 1.34 |
| 0 | Tbc1d16 | 0.67 | 1.65E-60 | 3.44E-56 | 0.56 | 0.43 | 0.13 | 1.29 |
| 0 | Zfp428 | 0.45 | 2.32E-60 | 4.82E-56 | 0.78 | 0.68 | 0.10 | 1.14 |
| 0 | Nsg1 | 0.49 | 2.94E-60 | 6.11E-56 | 0.76 | 0.66 | 0.11 | 1.16 |
| 0 | Psma4 | 0.25 | 3.04E-60 | 6.32E-56 | 0.98 | 0.96 | 0.01 | 1.01 |
| 0 | Arl3 | 0.35 | 8.42E-60 | 1.75E-55 | 0.89 | 0.84 | 0.05 | 1.06 |
| 0 | Jarid2 | 0.55 | 1.10E-59 | 2.29E-55 | 0.69 | 0.59 | 0.09 | 1.16 |
| 0 | Pxdc1 | 1.39 | 3.15E-59 | 6.55E-55 | 0.20 | 0.09 | 0.10 | 2.10 |
| 0 | Mfap4 | 0.29 | 4.10E-59 | 8.52E-55 | 0.96 | 0.83 | 0.13 | 1.16 |
| 0 | Pgm2l1 | 0.80 | 7.56E-59 | 1.57E-54 | 0.47 | 0.34 | 0.13 | 1.38 |
| 0 | Efna2 | 0.42 | 9.18E-59 | 1.91E-54 | 0.79 | 0.70 | 0.09 | 1.13 |
| 0 | Hdac2 | 0.31 | 9.66E-59 | 2.01E-54 | 0.94 | 0.90 | 0.04 | 1.05 |
| 0 | Rora | 1.07 | 2.25E-58 | 4.68E-54 | 0.29 | 0.17 | 0.12 | 1.72 |
| 0 | Hnrnph3 | 0.34 | 2.37E-58 | 4.93E-54 | 0.90 | 0.85 | 0.06 | 1.06 |
| 0 | Fnbp1l | 0.36 | 3.18E-58 | 6.62E-54 | 0.91 | 0.84 | 0.07 | 1.08 |
| 0 | Arhgdib | 0.43 | 5.44E-58 | 1.13E-53 | 0.63 | 0.48 | 0.14 | 1.30 |
| 0 | Glipr2 | 0.72 | 9.09E-58 | 1.89E-53 | 0.50 | 0.37 | 0.13 | 1.35 |
| 0 | Pafah1b3 | 0.28 | 2.19E-57 | 4.56E-53 | 0.96 | 0.92 | 0.04 | 1.04 |
| 0 | Fblim1 | 0.59 | 2.41E-57 | 5.02E-53 | 0.58 | 0.45 | 0.13 | 1.29 |
| 0 | Igsf10 | 1.17 | 2.83E-57 | 5.89E-53 | 0.34 | 0.22 | 0.12 | 1.54 |
| 0 | Cacna1d | 0.87 | 4.20E-57 | 8.73E-53 | 0.35 | 0.22 | 0.13 | 1.60 |
| 0 | Tbcb | 0.32 | 5.05E-57 | 1.05E-52 | 0.93 | 0.88 | 0.05 | 1.05 |
| 0 | Bgn | 0.75 | 8.80E-57 | 1.83E-52 | 0.39 | 0.25 | 0.14 | 1.58 |
| 0 | Galnt5 | 1.82 | 9.82E-57 | 2.04E-52 | 0.11 | 0.04 | 0.07 | 2.87 |
| 0 | Fbln2 | 0.53 | 1.96E-56 | 4.08E-52 | 0.64 | 0.50 | 0.14 | 1.28 |
| 0 | Acat2 | 0.37 | 3.87E-56 | 8.05E-52 | 0.88 | 0.79 | 0.08 | 1.10 |
| 0 | Tgfb1i1 | 1.07 | 5.03E-56 | 1.05E-51 | 0.23 | 0.12 | 0.11 | 1.89 |
| 0 | Commd3 | 0.38 | 1.58E-55 | 3.29E-51 | 0.85 | 0.80 | 0.05 | 1.06 |
| 0 | Snrpa | 0.29 | 1.59E-55 | 3.30E-51 | 0.95 | 0.92 | 0.03 | 1.04 |
| 0 | Rftn2 | 0.89 | 1.82E-55 | 3.79E-51 | 0.34 | 0.21 | 0.13 | 1.58 |
| 0 | Lrrc17 | 1.56 | 2.90E-55 | 6.02E-51 | 0.16 | 0.07 | 0.09 | 2.28 |
| 0 | Fzd2 | 0.30 | 1.63E-54 | 3.40E-50 | 0.96 | 0.86 | 0.09 | 1.11 |
| 0 | Tax1bp1 | 0.28 | 1.91E-54 | 3.98E-50 | 0.95 | 0.93 | 0.02 | 1.02 |

**Supplementary Table 1. Marker genes for the original clusters of integrated clustering of E11.5 control and NotchGOF λ-junction cells.**

| | | | | | | | | |
|---|---|---|---|---|---|---|---|---|
| 0 | Chd4 | 0.27 | 1.69E-53 | 3.52E-49 | 0.99 | 0.96 | 0.03 | 1.03 |
| 0 | Rbfox2 | 0.38 | 2.02E-53 | 4.20E-49 | 0.83 | 0.76 | 0.07 | 1.10 |
| 0 | Rspo3 | 0.48 | 3.15E-53 | 6.56E-49 | 0.48 | 0.33 | 0.15 | 1.47 |
| 0 | Pgrmc1 | 0.31 | 4.14E-53 | 8.60E-49 | 0.96 | 0.91 | 0.05 | 1.05 |
| 0 | Dock4 | 0.79 | 4.46E-53 | 9.28E-49 | 0.42 | 0.30 | 0.12 | 1.41 |
| 0 | Efnb1 | 0.51 | 4.85E-53 | 1.01E-48 | 0.64 | 0.52 | 0.13 | 1.24 |
| 0 | Zmat2 | 0.27 | 7.66E-53 | 1.59E-48 | 0.97 | 0.95 | 0.02 | 1.03 |
| 0 | Igdcc4 | 1.21 | 1.30E-52 | 2.71E-48 | 0.22 | 0.12 | 0.10 | 1.88 |
| 0 | Ahsa2 | 0.43 | 5.75E-52 | 1.20E-47 | 0.76 | 0.69 | 0.07 | 1.10 |
| 0 | Tbx3os1 | 1.14 | 6.94E-52 | 1.44E-47 | 0.23 | 0.12 | 0.11 | 1.85 |
| 0 | Dpysl2 | 0.41 | 9.26E-52 | 1.93E-47 | 0.78 | 0.69 | 0.09 | 1.13 |
| 0 | Mxd4 | 0.54 | 1.29E-51 | 2.69E-47 | 0.58 | 0.45 | 0.13 | 1.28 |
| 0 | Mab21l1 | 0.84 | 2.52E-51 | 5.24E-47 | 0.27 | 0.15 | 0.12 | 1.77 |
| 0 | Gtf3c6 | 0.33 | 2.67E-51 | 5.55E-47 | 0.91 | 0.86 | 0.05 | 1.06 |
| 0 | Plekha2 | 1.69 | 5.38E-51 | 1.12E-46 | 0.11 | 0.04 | 0.07 | 2.72 |
| 0 | Top2b | 0.28 | 7.36E-51 | 1.53E-46 | 0.95 | 0.92 | 0.03 | 1.04 |
| 0 | Map4k4 | 0.29 | 7.70E-51 | 1.60E-46 | 0.96 | 0.90 | 0.06 | 1.07 |
| 0 | Frem1 | 0.54 | 1.97E-50 | 4.10E-46 | 0.53 | 0.39 | 0.14 | 1.35 |
| 0 | Rcn2 | 0.26 | 2.22E-50 | 4.62E-46 | 0.96 | 0.91 | 0.06 | 1.06 |
| 0 | Jam3 | 0.49 | 5.42E-50 | 1.13E-45 | 0.67 | 0.56 | 0.11 | 1.20 |
| 0 | Gng10 | 0.35 | 1.33E-49 | 2.76E-45 | 0.87 | 0.81 | 0.06 | 1.08 |
| 0 | Cmtm3 | 0.59 | 1.45E-49 | 3.02E-45 | 0.53 | 0.41 | 0.12 | 1.30 |
| 0 | Hp1bp3 | 0.31 | 1.64E-49 | 3.40E-45 | 0.92 | 0.89 | 0.04 | 1.04 |
| 0 | Peg10 | 0.57 | 1.70E-49 | 3.53E-45 | 0.55 | 0.42 | 0.13 | 1.31 |
| 0 | Selenok | 0.28 | 2.20E-49 | 4.58E-45 | 0.94 | 0.91 | 0.03 | 1.03 |
| 0 | Septin3 | 0.59 | 3.16E-49 | 6.57E-45 | 0.55 | 0.43 | 0.12 | 1.28 |
| 0 | Mthfd2l | 0.47 | 6.04E-49 | 1.26E-44 | 0.72 | 0.63 | 0.09 | 1.14 |
| 0 | Trappc4 | 0.33 | 9.84E-49 | 2.05E-44 | 0.87 | 0.82 | 0.05 | 1.06 |
| 0 | Rspo2 | 0.50 | 1.12E-48 | 2.34E-44 | 0.50 | 0.35 | 0.15 | 1.42 |
| 0 | Elovl6 | 0.44 | 1.28E-48 | 2.65E-44 | 0.73 | 0.64 | 0.09 | 1.14 |
| 0 | Pdlim4 | 0.39 | 1.98E-48 | 4.11E-44 | 0.73 | 0.61 | 0.12 | 1.19 |
| 0 | Pygo1 | 0.57 | 2.80E-48 | 5.83E-44 | 0.57 | 0.45 | 0.11 | 1.25 |
| 0 | Foxl2 | 0.99 | 3.23E-48 | 6.72E-44 | 0.22 | 0.12 | 0.10 | 1.87 |
| 0 | Tagln3 | 0.65 | 4.41E-48 | 9.18E-44 | 0.37 | 0.25 | 0.12 | 1.47 |
| 0 | Cd1d1 | 1.07 | 1.85E-47 | 3.86E-43 | 0.26 | 0.15 | 0.10 | 1.68 |
| 0 | Tcf7 | 1.04 | 3.59E-47 | 7.47E-43 | 0.24 | 0.14 | 0.10 | 1.75 |
| 0 | Sncaip | 0.70 | 4.57E-47 | 9.51E-43 | 0.41 | 0.29 | 0.12 | 1.42 |
| 0 | Sall4 | 0.97 | 7.99E-47 | 1.66E-42 | 0.29 | 0.18 | 0.11 | 1.62 |
| 0 | Hmgcr | 0.48 | 1.82E-46 | 3.78E-42 | 0.74 | 0.66 | 0.08 | 1.11 |
| 0 | Pttg1 | 0.48 | 1.83E-46 | 3.80E-42 | 0.76 | 0.71 | 0.05 | 1.07 |
| 0 | Septin6 | 0.47 | 2.20E-46 | 4.59E-42 | 0.66 | 0.56 | 0.10 | 1.18 |
| 0 | Strap | 0.30 | 2.70E-46 | 5.61E-42 | 0.93 | 0.89 | 0.04 | 1.05 |
| 0 | Creb5 | 0.84 | 8.49E-46 | 1.77E-41 | 0.43 | 0.31 | 0.12 | 1.37 |
| 0 | Trim24 | 0.38 | 1.04E-45 | 2.17E-41 | 0.77 | 0.70 | 0.07 | 1.10 |
| 0 | Dclk2 | 1.26 | 1.48E-45 | 3.07E-41 | 0.19 | 0.10 | 0.09 | 1.87 |
| 0 | Ldb2 | 0.79 | 1.85E-45 | 3.84E-41 | 0.38 | 0.26 | 0.12 | 1.45 |
| 0 | Chn1 | 0.64 | 2.39E-45 | 4.97E-41 | 0.50 | 0.39 | 0.11 | 1.28 |
| 0 | Nkd2 | 1.26 | 3.39E-45 | 7.05E-41 | 0.21 | 0.12 | 0.09 | 1.78 |
| 0 | Cbfb | 0.48 | 7.99E-45 | 1.66E-40 | 0.65 | 0.56 | 0.09 | 1.15 |
| 0 | Mdga2 | 1.45 | 1.14E-44 | 2.37E-40 | 0.14 | 0.06 | 0.08 | 2.21 |
| 0 | Sdk2 | 1.53 | 2.42E-44 | 5.03E-40 | 0.11 | 0.04 | 0.07 | 2.48 |
| 0 | Cenpa | 0.48 | 2.77E-44 | 5.76E-40 | 0.79 | 0.79 | -0.01 | 0.99 |
| 0 | Psmd6 | 0.29 | 2.94E-44 | 6.11E-40 | 0.92 | 0.89 | 0.03 | 1.03 |
| 0 | Ick | 0.61 | 3.45E-44 | 7.19E-40 | 0.53 | 0.43 | 0.10 | 1.23 |
| 0 | Ptbp2 | 0.34 | 4.02E-44 | 8.35E-40 | 0.84 | 0.76 | 0.08 | 1.11 |
| 0 | Wdr86 | 0.49 | 9.94E-44 | 2.07E-39 | 0.61 | 0.51 | 0.11 | 1.21 |

**Supplementary Table 1. Marker genes for the original clusters of integrated clustering of E11.5 control and NotchGOF λ-junction cells.**

| | | | | | | | | |
|---|---|---|---|---|---|---|---|---|
| 0 | Vegfd | 1.55 | 1.12E-43 | 2.32E-39 | 0.11 | 0.04 | 0.06 | 2.45 |
| 0 | Dact3 | 0.47 | 1.13E-43 | 2.35E-39 | 0.60 | 0.48 | 0.12 | 1.25 |
| 0 | Eva1b | 0.36 | 1.61E-43 | 3.35E-39 | 0.66 | 0.52 | 0.14 | 1.27 |
| 0 | Map2k2 | 0.26 | 1.74E-43 | 3.62E-39 | 0.94 | 0.92 | 0.02 | 1.02 |
| 0 | Pdlim2 | 1.11 | 4.31E-43 | 8.96E-39 | 0.25 | 0.16 | 0.10 | 1.62 |
| 0 | Lhfpl2 | 0.73 | 4.33E-43 | 9.01E-39 | 0.43 | 0.32 | 0.11 | 1.34 |
| 0 | Rhobtb3 | 0.32 | 5.26E-43 | 1.09E-38 | 0.88 | 0.81 | 0.07 | 1.09 |
| 0 | Pcdh10 | 1.15 | 7.33E-43 | 1.52E-38 | 0.18 | 0.10 | 0.09 | 1.91 |
| 0 | Casp6 | 0.47 | 9.93E-43 | 2.07E-38 | 0.63 | 0.55 | 0.08 | 1.15 |
| 0 | Adamts19 | 0.87 | 1.47E-42 | 3.05E-38 | 0.34 | 0.23 | 0.11 | 1.45 |
| 0 | Dact1 | 0.65 | 1.69E-42 | 3.51E-38 | 0.42 | 0.30 | 0.12 | 1.39 |
| 0 | Tlnrd1 | 0.50 | 1.73E-42 | 3.59E-38 | 0.59 | 0.49 | 0.10 | 1.20 |
| 0 | Stard8 | 1.31 | 2.35E-42 | 4.88E-38 | 0.16 | 0.08 | 0.08 | 2.00 |
| 0 | Baiap2l2 | 1.18 | 4.54E-42 | 9.43E-38 | 0.19 | 0.11 | 0.09 | 1.84 |
| 0 | Fbn2 | 0.28 | 8.18E-42 | 1.70E-37 | 0.84 | 0.72 | 0.12 | 1.16 |
| 0 | Lzts2 | 0.47 | 1.10E-41 | 2.28E-37 | 0.67 | 0.60 | 0.08 | 1.13 |
| 0 | Gpx7 | 0.55 | 1.14E-41 | 2.38E-37 | 0.54 | 0.44 | 0.10 | 1.24 |
| 0 | Gm1673 | 0.25 | 1.49E-41 | 3.10E-37 | 0.98 | 0.92 | 0.05 | 1.06 |
| 0 | Gm11627 | 1.06 | 5.55E-41 | 1.16E-36 | 0.22 | 0.13 | 0.09 | 1.74 |
| 0 | Rab34 | 0.34 | 8.36E-41 | 1.74E-36 | 0.79 | 0.71 | 0.08 | 1.11 |
| 0 | Fbxl7 | 0.59 | 1.10E-40 | 2.29E-36 | 0.56 | 0.45 | 0.10 | 1.22 |
| 0 | Pdgfrb | 0.47 | 1.58E-40 | 3.29E-36 | 0.57 | 0.44 | 0.13 | 1.29 |
| 0 | Eya1 | 0.45 | 1.72E-40 | 3.58E-36 | 0.60 | 0.49 | 0.12 | 1.25 |
| 0 | Rbm4b | 0.41 | 1.73E-40 | 3.60E-36 | 0.73 | 0.66 | 0.07 | 1.11 |
| 0 | Bin1 | 0.42 | 1.08E-39 | 2.24E-35 | 0.69 | 0.61 | 0.08 | 1.14 |
| 0 | Trim35 | 0.36 | 1.29E-39 | 2.68E-35 | 0.77 | 0.70 | 0.07 | 1.10 |
| 0 | Nrep | 0.29 | 1.45E-39 | 3.01E-35 | 0.88 | 0.81 | 0.06 | 1.07 |
| 0 | As3mt | 0.66 | 2.20E-39 | 4.57E-35 | 0.45 | 0.36 | 0.10 | 1.28 |
| 0 | Slc38a2 | 0.27 | 2.72E-39 | 5.65E-35 | 0.94 | 0.90 | 0.05 | 1.05 |
| 0 | Zfp771 | 0.39 | 3.51E-39 | 7.30E-35 | 0.74 | 0.68 | 0.05 | 1.08 |
| 0 | Fn1 | 0.33 | 4.45E-39 | 9.25E-35 | 0.86 | 0.77 | 0.08 | 1.11 |
| 0 | Nenf | 0.32 | 4.49E-39 | 9.34E-35 | 0.82 | 0.78 | 0.04 | 1.06 |
| 0 | Etv4 | 0.66 | 4.96E-39 | 1.03E-34 | 0.43 | 0.32 | 0.11 | 1.33 |
| 0 | Gtf2e2 | 0.33 | 6.22E-39 | 1.29E-34 | 0.80 | 0.76 | 0.04 | 1.06 |
| 0 | Gmps | 0.26 | 6.23E-39 | 1.30E-34 | 0.93 | 0.89 | 0.04 | 1.04 |
| 0 | Snrpb2 | 0.27 | 9.56E-39 | 1.99E-34 | 0.90 | 0.85 | 0.05 | 1.06 |
| 0 | Cdca7 | 0.35 | 1.26E-38 | 2.61E-34 | 0.79 | 0.71 | 0.07 | 1.10 |
| 0 | Myh10 | 0.28 | 1.39E-38 | 2.90E-34 | 0.93 | 0.86 | 0.07 | 1.08 |
| 0 | Pex5l | 0.92 | 1.65E-38 | 3.43E-34 | 0.16 | 0.08 | 0.08 | 1.98 |
| 0 | Casp8ap2 | 0.33 | 1.89E-38 | 3.92E-34 | 0.84 | 0.79 | 0.05 | 1.07 |
| 0 | Usp2 | 1.17 | 1.89E-38 | 3.93E-34 | 0.22 | 0.13 | 0.09 | 1.66 |
| 0 | Tmem50a | 0.33 | 2.77E-38 | 5.76E-34 | 0.84 | 0.80 | 0.04 | 1.05 |
| 0 | Emilin1 | 0.36 | 2.98E-38 | 6.19E-34 | 0.64 | 0.52 | 0.13 | 1.25 |
| 0 | Dancr | 0.45 | 3.15E-38 | 6.55E-34 | 0.66 | 0.58 | 0.08 | 1.14 |
| 0 | Acads | 0.42 | 3.24E-38 | 6.73E-34 | 0.66 | 0.60 | 0.07 | 1.11 |
| 0 | Pdgfra | 0.38 | 4.41E-38 | 9.17E-34 | 0.66 | 0.53 | 0.14 | 1.26 |
| 0 | Dchs1 | 0.56 | 6.00E-38 | 1.25E-33 | 0.51 | 0.41 | 0.10 | 1.24 |
| 0 | Rab18 | 0.28 | 6.39E-38 | 1.33E-33 | 0.85 | 0.82 | 0.04 | 1.05 |
| 0 | Tmsb15b2 | 0.44 | 6.54E-38 | 1.36E-33 | 0.63 | 0.54 | 0.09 | 1.18 |
| 0 | Zcrb1 | 0.26 | 8.59E-38 | 1.79E-33 | 0.94 | 0.93 | 0.02 | 1.02 |
| 0 | Ndufa1 | 0.29 | 9.46E-38 | 1.97E-33 | 0.89 | 0.86 | 0.03 | 1.03 |
| 0 | Actn1 | 0.32 | 1.15E-37 | 2.39E-33 | 0.79 | 0.71 | 0.08 | 1.11 |
| 0 | Vegfb | 0.38 | 1.24E-37 | 2.57E-33 | 0.72 | 0.65 | 0.07 | 1.11 |
| 0 | Plekho1 | 0.36 | 1.98E-37 | 4.11E-33 | 0.70 | 0.61 | 0.10 | 1.16 |
| 0 | Nisch | 0.25 | 2.79E-37 | 5.80E-33 | 0.96 | 0.91 | 0.04 | 1.05 |
| 0 | Hacd1 | 0.59 | 2.83E-37 | 5.88E-33 | 0.47 | 0.38 | 0.09 | 1.23 |

**Supplementary Table 1. Marker genes for the original clusters of integrated clustering of E11.5 control and NotchGOF λ-junction cells.**

| | | | | | | | | |
|---|---|---|---|---|---|---|---|---|
| 0 | Rrbp1 | 0.26 | 2.97E-37 | 6.17E-33 | 0.94 | 0.89 | 0.05 | 1.06 |
| 0 | Plppr3 | 0.58 | 5.25E-37 | 1.09E-32 | 0.47 | 0.37 | 0.10 | 1.26 |
| 0 | Dnm1 | 0.64 | 5.48E-37 | 1.14E-32 | 0.42 | 0.32 | 0.10 | 1.33 |
| 0 | Plin2 | 0.42 | 9.27E-37 | 1.93E-32 | 0.60 | 0.52 | 0.08 | 1.15 |
| 0 | Cables1 | 1.23 | 9.65E-37 | 2.01E-32 | 0.15 | 0.08 | 0.07 | 1.95 |
| 0 | Frmd4a | 0.35 | 1.39E-36 | 2.89E-32 | 0.75 | 0.66 | 0.08 | 1.13 |
| 0 | Usp3 | 0.35 | 1.63E-36 | 3.40E-32 | 0.79 | 0.73 | 0.06 | 1.08 |
| 0 | Ap3s1 | 0.34 | 2.53E-36 | 5.26E-32 | 0.76 | 0.71 | 0.05 | 1.07 |
| 0 | Mapre2 | 0.53 | 2.84E-36 | 5.91E-32 | 0.49 | 0.39 | 0.10 | 1.25 |
| 0 | Clip3 | 0.46 | 3.19E-36 | 6.63E-32 | 0.59 | 0.51 | 0.08 | 1.17 |
| 0 | Adgrl3 | 0.63 | 3.64E-36 | 7.58E-32 | 0.52 | 0.43 | 0.09 | 1.21 |
| 0 | Zcchc24 | 0.67 | 5.40E-36 | 1.12E-31 | 0.38 | 0.28 | 0.10 | 1.36 |
| 0 | Tmem223 | 0.29 | 6.54E-36 | 1.36E-31 | 0.87 | 0.84 | 0.03 | 1.04 |
| 0 | Rcn1 | 0.27 | 1.48E-35 | 3.08E-31 | 0.88 | 0.82 | 0.06 | 1.08 |
| 0 | Tmem160 | 0.34 | 1.59E-35 | 3.32E-31 | 0.78 | 0.73 | 0.05 | 1.07 |
| 0 | Il11ra1 | 0.28 | 1.64E-35 | 3.40E-31 | 0.74 | 0.60 | 0.13 | 1.22 |
| 0 | Prrx1 | 0.29 | 2.13E-35 | 4.43E-31 | 0.80 | 0.63 | 0.17 | 1.27 |
| 0 | Hook3 | 0.39 | 2.70E-35 | 5.63E-31 | 0.72 | 0.65 | 0.06 | 1.10 |
| 0 | Sox12 | 0.37 | 3.24E-35 | 6.75E-31 | 0.72 | 0.66 | 0.06 | 1.10 |
| 0 | Fgfr1 | 0.37 | 3.92E-35 | 8.14E-31 | 0.73 | 0.64 | 0.09 | 1.14 |
| 0 | Asb4 | 0.34 | 6.33E-35 | 1.32E-30 | 0.41 | 0.29 | 0.12 | 1.42 |
| 0 | Arid4b | 0.27 | 9.14E-35 | 1.90E-30 | 0.90 | 0.86 | 0.04 | 1.04 |
| 0 | Septin9 | 0.40 | 1.35E-34 | 2.80E-30 | 0.70 | 0.63 | 0.06 | 1.10 |
| 0 | Anapc16 | 0.29 | 1.48E-34 | 3.08E-30 | 0.81 | 0.75 | 0.05 | 1.07 |
| 0 | Plekha5 | 0.53 | 1.66E-34 | 3.45E-30 | 0.54 | 0.45 | 0.09 | 1.19 |
| 0 | Smad7 | 0.30 | 2.44E-34 | 5.07E-30 | 0.78 | 0.69 | 0.09 | 1.14 |
| 0 | Six4 | 0.56 | 2.63E-34 | 5.48E-30 | 0.53 | 0.45 | 0.08 | 1.18 |
| 0 | Smo | 0.34 | 3.42E-34 | 7.11E-30 | 0.78 | 0.73 | 0.05 | 1.07 |
| 0 | Bmi1 | 0.59 | 3.57E-34 | 7.43E-30 | 0.49 | 0.40 | 0.09 | 1.22 |
| 0 | Cep170 | 0.47 | 3.62E-34 | 7.53E-30 | 0.58 | 0.50 | 0.08 | 1.16 |
| 0 | Igf2bp1 | 0.26 | 4.34E-34 | 9.03E-30 | 0.93 | 0.88 | 0.05 | 1.05 |
| 0 | Cacna2d3 | 1.45 | 6.67E-34 | 1.39E-29 | 0.14 | 0.07 | 0.07 | 1.93 |
| 0 | Smad9 | 1.34 | 9.02E-34 | 1.88E-29 | 0.11 | 0.05 | 0.06 | 2.16 |
| 0 | Pdlim7 | 0.33 | 1.07E-33 | 2.23E-29 | 0.81 | 0.75 | 0.07 | 1.09 |
| 0 | Rala | 0.26 | 1.07E-33 | 2.23E-29 | 0.88 | 0.85 | 0.03 | 1.03 |
| 0 | Cnst | 1.01 | 1.87E-33 | 3.89E-29 | 0.26 | 0.17 | 0.08 | 1.48 |
| 0 | Pin4 | 0.27 | 2.12E-33 | 4.41E-29 | 0.86 | 0.83 | 0.03 | 1.03 |
| 0 | Cdk5r1 | 1.12 | 2.20E-33 | 4.57E-29 | 0.18 | 0.10 | 0.08 | 1.75 |
| 0 | Cntrl | 0.53 | 3.23E-33 | 6.73E-29 | 0.53 | 0.45 | 0.08 | 1.17 |
| 0 | Mylk | 0.93 | 3.50E-33 | 7.28E-29 | 0.25 | 0.16 | 0.09 | 1.53 |
| 0 | Cnrip1 | 0.90 | 4.90E-33 | 1.02E-28 | 0.21 | 0.13 | 0.08 | 1.65 |
| 0 | Sav1 | 0.52 | 5.38E-33 | 1.12E-28 | 0.52 | 0.44 | 0.08 | 1.18 |
| 0 | Ppp3ca | 0.38 | 5.97E-33 | 1.24E-28 | 0.78 | 0.72 | 0.06 | 1.08 |
| 0 | Fkbp7 | 0.65 | 6.95E-33 | 1.45E-28 | 0.37 | 0.27 | 0.10 | 1.37 |
| 0 | Kansl1l | 0.75 | 7.25E-33 | 1.51E-28 | 0.34 | 0.25 | 0.09 | 1.37 |
| 0 | Itgb5 | 0.39 | 7.95E-33 | 1.65E-28 | 0.67 | 0.57 | 0.09 | 1.16 |
| 0 | Dgcr6 | 0.33 | 9.62E-33 | 2.00E-28 | 0.77 | 0.73 | 0.04 | 1.06 |
| 0 | Pik3r3 | 0.71 | 1.04E-32 | 2.16E-28 | 0.36 | 0.27 | 0.09 | 1.34 |
| 0 | Gm4258 | 0.48 | 1.22E-32 | 2.54E-28 | 0.57 | 0.49 | 0.09 | 1.18 |
| 0 | Rangrf | 0.45 | 1.65E-32 | 3.43E-28 | 0.60 | 0.53 | 0.07 | 1.13 |
| 0 | Cops8 | 0.27 | 1.70E-32 | 3.54E-28 | 0.85 | 0.81 | 0.04 | 1.05 |
| 0 | Gpc2 | 0.60 | 1.89E-32 | 3.93E-28 | 0.47 | 0.39 | 0.09 | 1.22 |
| 0 | Map7d1 | 0.39 | 2.18E-32 | 4.54E-28 | 0.67 | 0.61 | 0.06 | 1.10 |
| 0 | Gli2 | 0.70 | 2.87E-32 | 5.97E-28 | 0.34 | 0.25 | 0.09 | 1.36 |
| 0 | Dixdc1 | 0.52 | 3.13E-32 | 6.51E-28 | 0.48 | 0.39 | 0.10 | 1.25 |
| 0 | Eid2 | 0.34 | 3.50E-32 | 7.29E-28 | 0.72 | 0.64 | 0.07 | 1.11 |

**Supplementary Table 1. Marker genes for the original clusters of integrated clustering of E11.5 control and NotchGOF λ-junction cells.**

| | | | | | | | | |
|---|---|---|---|---|---|---|---|---|
| 0 | Mex3a | 0.26 | 3.57E-32 | 7.42E-28 | 0.91 | 0.85 | 0.06 | 1.07 |
| 0 | Dld | 0.32 | 3.74E-32 | 7.78E-28 | 0.80 | 0.77 | 0.04 | 1.05 |
| 0 | Echs1 | 0.25 | 3.83E-32 | 7.96E-28 | 0.89 | 0.85 | 0.03 | 1.04 |
| 0 | Cpne8 | 1.19 | 1.41E-31 | 2.92E-27 | 0.14 | 0.07 | 0.07 | 1.92 |
| 0 | Zfp397 | 0.36 | 1.45E-31 | 3.02E-27 | 0.74 | 0.68 | 0.06 | 1.09 |
| 0 | Vcam1 | 0.43 | 1.85E-31 | 3.84E-27 | 0.53 | 0.42 | 0.11 | 1.26 |
| 0 | Tns3 | 0.44 | 2.31E-31 | 4.81E-27 | 0.57 | 0.48 | 0.09 | 1.18 |
| 0 | Limch1 | 0.26 | 2.42E-31 | 5.04E-27 | 0.78 | 0.66 | 0.12 | 1.17 |
| 0 | Slco3a1 | 0.90 | 2.61E-31 | 5.43E-27 | 0.20 | 0.12 | 0.08 | 1.66 |
| 0 | Vps72 | 0.28 | 3.02E-31 | 6.29E-27 | 0.83 | 0.79 | 0.04 | 1.05 |
| 0 | Spin2c | 0.49 | 4.96E-31 | 1.03E-26 | 0.51 | 0.41 | 0.10 | 1.24 |
| 0 | Ppm1h | 1.44 | 1.01E-30 | 2.11E-26 | 0.14 | 0.08 | 0.06 | 1.84 |
| 0 | Phlda1 | 0.49 | 1.03E-30 | 2.14E-26 | 0.50 | 0.41 | 0.09 | 1.21 |
| 0 | Itm2c | 0.28 | 1.04E-30 | 2.15E-26 | 0.80 | 0.72 | 0.07 | 1.10 |
| 0 | Epha4 | 0.34 | 1.85E-30 | 3.86E-26 | 0.74 | 0.64 | 0.10 | 1.16 |
| 0 | Rbpms2 | 0.67 | 2.02E-30 | 4.21E-26 | 0.30 | 0.22 | 0.09 | 1.41 |
| 0 | Fez1 | 0.93 | 2.31E-30 | 4.80E-26 | 0.22 | 0.14 | 0.08 | 1.58 |
| 0 | Tle6 | 1.38 | 2.33E-30 | 4.84E-26 | 0.10 | 0.05 | 0.05 | 2.13 |
| 0 | Tnrc6c | 0.31 | 2.54E-30 | 5.28E-26 | 0.83 | 0.80 | 0.03 | 1.04 |
| 0 | Mylip | 0.56 | 3.45E-30 | 7.17E-26 | 0.40 | 0.30 | 0.09 | 1.31 |
| 0 | Dlx5 | 0.44 | 3.70E-30 | 7.70E-26 | 0.30 | 0.21 | 0.10 | 1.46 |
| 0 | Plscr3 | 0.39 | 5.28E-30 | 1.10E-25 | 0.62 | 0.55 | 0.06 | 1.12 |
| 0 | Twistnb | 0.29 | 5.80E-30 | 1.21E-25 | 0.81 | 0.76 | 0.05 | 1.06 |
| 0 | Mex3b | 0.29 | 7.46E-30 | 1.55E-25 | 0.78 | 0.68 | 0.10 | 1.14 |
| 0 | Pakap.1 | 0.49 | 9.31E-30 | 1.94E-25 | 0.49 | 0.40 | 0.09 | 1.22 |
| 0 | Chmp5 | 0.27 | 1.12E-29 | 2.33E-25 | 0.83 | 0.80 | 0.03 | 1.04 |
| 0 | Akr1b8 | 1.23 | 1.36E-29 | 2.84E-25 | 0.13 | 0.07 | 0.06 | 1.89 |
| 0 | Mmp23 | 1.31 | 1.49E-29 | 3.09E-25 | 0.10 | 0.05 | 0.05 | 2.08 |
| 0 | Kdm4a | 0.50 | 1.83E-29 | 3.80E-25 | 0.48 | 0.40 | 0.08 | 1.20 |
| 0 | Mme | 0.37 | 2.04E-29 | 4.24E-25 | 0.56 | 0.46 | 0.10 | 1.23 |
| 0 | Hsd17b7 | 0.70 | 2.81E-29 | 5.84E-25 | 0.35 | 0.26 | 0.09 | 1.32 |
| 0 | Impact | 0.61 | 3.48E-29 | 7.23E-25 | 0.40 | 0.32 | 0.09 | 1.27 |
| 0 | Kcp | 1.06 | 3.54E-29 | 7.37E-25 | 0.17 | 0.10 | 0.07 | 1.69 |
| 0 | Zmiz1 | 0.34 | 3.95E-29 | 8.21E-25 | 0.74 | 0.69 | 0.06 | 1.08 |
| 0 | Ubtf | 0.27 | 4.38E-29 | 9.12E-25 | 0.85 | 0.83 | 0.02 | 1.02 |
| 0 | Pnpla8 | 0.47 | 4.49E-29 | 9.35E-25 | 0.56 | 0.49 | 0.07 | 1.13 |
| 0 | Tfap2c | 0.72 | 5.36E-29 | 1.11E-24 | 0.35 | 0.27 | 0.09 | 1.32 |
| 0 | Cux2 | 1.14 | 6.54E-29 | 1.36E-24 | 0.16 | 0.09 | 0.07 | 1.74 |
| 0 | Reck | 0.71 | 7.08E-29 | 1.47E-24 | 0.32 | 0.24 | 0.09 | 1.36 |
| 0 | Tle2 | 0.99 | 7.49E-29 | 1.56E-24 | 0.18 | 0.11 | 0.07 | 1.64 |
| 0 | Naa35 | 0.29 | 7.67E-29 | 1.59E-24 | 0.77 | 0.72 | 0.04 | 1.06 |
| 0 | Tiam2 | 0.96 | 8.03E-29 | 1.67E-24 | 0.15 | 0.09 | 0.07 | 1.79 |
| 0 | Ctnnbip1 | 0.26 | 9.12E-29 | 1.90E-24 | 0.85 | 0.81 | 0.04 | 1.04 |
| 0 | Tsc22d3 | 0.40 | 2.53E-28 | 5.25E-24 | 0.59 | 0.52 | 0.07 | 1.13 |
| 0 | Msrb1 | 0.42 | 2.63E-28 | 5.47E-24 | 0.55 | 0.48 | 0.07 | 1.14 |
| 0 | Bad | 0.44 | 3.40E-28 | 7.08E-24 | 0.58 | 0.52 | 0.06 | 1.12 |
| 0 | P4ha2 | 0.59 | 4.03E-28 | 8.37E-24 | 0.35 | 0.26 | 0.09 | 1.33 |
| 0 | Ift74 | 0.31 | 4.47E-28 | 9.30E-24 | 0.79 | 0.73 | 0.06 | 1.08 |
| 0 | Wipi1 | 0.54 | 6.16E-28 | 1.28E-23 | 0.47 | 0.39 | 0.08 | 1.21 |
| 0 | Inka1 | 0.35 | 6.27E-28 | 1.30E-23 | 0.59 | 0.50 | 0.09 | 1.18 |
| 0 | Mpc1 | 0.36 | 7.83E-28 | 1.63E-23 | 0.67 | 0.62 | 0.04 | 1.07 |
| 0 | Adprh | 0.28 | 9.40E-28 | 1.95E-23 | 0.82 | 0.78 | 0.04 | 1.05 |
| 0 | Ccni | 0.27 | 1.00E-27 | 2.08E-23 | 0.83 | 0.80 | 0.03 | 1.04 |
| 0 | Lactb | 0.54 | 1.06E-27 | 2.20E-23 | 0.42 | 0.33 | 0.08 | 1.25 |
| 0 | Mex3d | 0.50 | 1.21E-27 | 2.51E-23 | 0.52 | 0.45 | 0.07 | 1.15 |
| 0 | 2700081O15Rik | 0.39 | 1.22E-27 | 2.53E-23 | 0.61 | 0.55 | 0.06 | 1.11 |

**Supplementary Table 1. Marker genes for the original clusters of integrated clustering of E11.5 control and NotchGOF λ-junction cells.**

| | | | | | | | | |
|---|---|---|---|---|---|---|---|---|
| 0 | Serinc1 | 0.32 | 1.22E-27 | 2.54E-23 | 0.74 | 0.69 | 0.05 | 1.06 |
| 0 | Epb41l2 | 0.30 | 1.25E-27 | 2.61E-23 | 0.73 | 0.66 | 0.07 | 1.10 |
| 0 | Kank2 | 0.58 | 1.37E-27 | 2.85E-23 | 0.39 | 0.30 | 0.09 | 1.30 |
| 0 | Cdc40 | 0.31 | 1.59E-27 | 3.31E-23 | 0.74 | 0.70 | 0.04 | 1.06 |
| 0 | Tmem100 | 0.84 | 2.15E-27 | 4.46E-23 | 0.21 | 0.14 | 0.08 | 1.56 |
| 0 | L1cam | 1.00 | 2.37E-27 | 4.93E-23 | 0.14 | 0.07 | 0.06 | 1.85 |
| 0 | Nfia | 0.53 | 3.34E-27 | 6.96E-23 | 0.54 | 0.46 | 0.07 | 1.16 |
| 0 | Pcsk5 | 0.49 | 3.52E-27 | 7.33E-23 | 0.45 | 0.36 | 0.10 | 1.27 |
| 0 | Kcnj2 | 0.92 | 3.56E-27 | 7.40E-23 | 0.25 | 0.17 | 0.08 | 1.46 |
| 0 | Scd1 | 0.78 | 4.36E-27 | 9.07E-23 | 0.26 | 0.18 | 0.08 | 1.46 |
| 0 | Runx1t1 | 0.34 | 4.72E-27 | 9.82E-23 | 0.62 | 0.52 | 0.11 | 1.20 |
| 0 | Kif3a | 0.34 | 6.28E-27 | 1.31E-22 | 0.67 | 0.61 | 0.06 | 1.10 |
| 0 | Cnr1 | 1.11 | 6.86E-27 | 1.43E-22 | 0.14 | 0.08 | 0.06 | 1.74 |
| 0 | Zfand5 | 0.25 | 9.02E-27 | 1.88E-22 | 0.85 | 0.83 | 0.02 | 1.02 |
| 0 | Dhx29 | 0.41 | 1.03E-26 | 2.14E-22 | 0.58 | 0.51 | 0.07 | 1.13 |
| 0 | Rbis | 0.26 | 1.17E-26 | 2.43E-22 | 0.84 | 0.82 | 0.02 | 1.03 |
| 0 | Glul | 0.33 | 1.37E-26 | 2.85E-22 | 0.66 | 0.61 | 0.05 | 1.08 |
| 0 | Arhgef40 | 0.35 | 1.63E-26 | 3.40E-22 | 0.60 | 0.52 | 0.09 | 1.17 |
| 0 | Gnb4 | 0.57 | 2.20E-26 | 4.58E-22 | 0.38 | 0.29 | 0.09 | 1.29 |
| 0 | 4930402H24Rik | 0.67 | 4.48E-26 | 9.32E-22 | 0.32 | 0.24 | 0.08 | 1.33 |
| 0 | Chst2 | 0.81 | 4.95E-26 | 1.03E-21 | 0.27 | 0.20 | 0.08 | 1.40 |
| 0 | Psd3 | 0.71 | 5.52E-26 | 1.15E-21 | 0.30 | 0.22 | 0.08 | 1.37 |
| 0 | Asphd2 | 0.90 | 6.52E-26 | 1.36E-21 | 0.18 | 0.12 | 0.07 | 1.59 |
| 0 | Tpmt | 0.53 | 1.16E-25 | 2.40E-21 | 0.44 | 0.36 | 0.07 | 1.20 |
| 0 | Caml | 0.43 | 1.78E-25 | 3.71E-21 | 0.55 | 0.49 | 0.06 | 1.12 |
| 0 | Trnau1ap | 0.33 | 1.94E-25 | 4.04E-21 | 0.67 | 0.63 | 0.05 | 1.07 |
| 0 | Tnrc18 | 0.38 | 2.18E-25 | 4.53E-21 | 0.62 | 0.55 | 0.06 | 1.12 |
| 0 | Adgrl2 | 0.37 | 2.22E-25 | 4.61E-21 | 0.73 | 0.68 | 0.04 | 1.06 |
| 0 | Ackr3 | 0.48 | 2.50E-25 | 5.19E-21 | 0.49 | 0.41 | 0.08 | 1.18 |
| 0 | Lrp1 | 0.33 | 2.72E-25 | 5.66E-21 | 0.64 | 0.57 | 0.07 | 1.12 |
| 0 | Gm3830 | 1.19 | 2.97E-25 | 6.18E-21 | 0.11 | 0.06 | 0.05 | 1.91 |
| 0 | Nid1 | 0.44 | 4.07E-25 | 8.47E-21 | 0.43 | 0.34 | 0.09 | 1.27 |
| 0 | Tiprl | 0.29 | 5.40E-25 | 1.12E-20 | 0.73 | 0.69 | 0.04 | 1.06 |
| 0 | B230217C12Rik | 0.74 | 9.49E-25 | 1.97E-20 | 0.24 | 0.17 | 0.08 | 1.44 |
| 0 | Asns | 0.48 | 9.55E-25 | 1.99E-20 | 0.50 | 0.43 | 0.06 | 1.15 |
| 0 | Tmeff1 | 0.29 | 1.38E-24 | 2.88E-20 | 0.74 | 0.68 | 0.05 | 1.08 |
| 0 | Meis1 | 0.49 | 1.61E-24 | 3.36E-20 | 0.53 | 0.45 | 0.08 | 1.17 |
| 0 | Tlcd2 | 0.85 | 1.92E-24 | 4.00E-20 | 0.20 | 0.13 | 0.07 | 1.54 |
| 0 | Guk1 | 0.26 | 2.45E-24 | 5.09E-20 | 0.80 | 0.78 | 0.02 | 1.03 |
| 0 | Sbk1 | 0.36 | 2.53E-24 | 5.26E-20 | 0.59 | 0.52 | 0.07 | 1.14 |
| 0 | Tanc1 | 0.45 | 2.90E-24 | 6.02E-20 | 0.51 | 0.44 | 0.07 | 1.16 |
| 0 | Srek1ip1 | 0.28 | 4.91E-24 | 1.02E-19 | 0.74 | 0.71 | 0.03 | 1.05 |
| 0 | Gsto1 | 0.28 | 5.33E-24 | 1.11E-19 | 0.70 | 0.64 | 0.06 | 1.10 |
| 0 | Tes | 0.30 | 5.45E-24 | 1.13E-19 | 0.74 | 0.69 | 0.05 | 1.08 |
| 0 | Hdgfl3 | 0.27 | 5.80E-24 | 1.21E-19 | 0.77 | 0.70 | 0.07 | 1.10 |
| 0 | Thyn1 | 0.27 | 6.40E-24 | 1.33E-19 | 0.77 | 0.73 | 0.05 | 1.06 |
| 0 | Rbmx | 0.26 | 7.48E-24 | 1.56E-19 | 0.79 | 0.74 | 0.05 | 1.06 |
| 0 | Ogfr | 0.29 | 9.43E-24 | 1.96E-19 | 0.71 | 0.66 | 0.05 | 1.07 |
| 0 | Zkscan3 | 0.32 | 9.43E-24 | 1.96E-19 | 0.71 | 0.67 | 0.04 | 1.06 |
| 0 | Mocs2 | 0.34 | 1.17E-23 | 2.44E-19 | 0.64 | 0.60 | 0.04 | 1.07 |
| 0 | Cdr1os | 1.18 | 1.28E-23 | 2.67E-19 | 0.10 | 0.06 | 0.05 | 1.89 |
| 0 | Kifap3 | 0.28 | 1.55E-23 | 3.23E-19 | 0.82 | 0.77 | 0.05 | 1.06 |
| 0 | Spats2 | 0.27 | 2.27E-23 | 4.73E-19 | 0.73 | 0.68 | 0.05 | 1.07 |
| 0 | Sra1 | 0.30 | 2.28E-23 | 4.74E-19 | 0.73 | 0.70 | 0.03 | 1.05 |
| 0 | Dlg2 | 1.15 | 3.07E-23 | 6.38E-19 | 0.12 | 0.06 | 0.05 | 1.81 |
| 0 | Lmna | 0.59 | 3.30E-23 | 6.86E-19 | 0.34 | 0.27 | 0.07 | 1.27 |

**Supplementary Table 1. Marker genes for the original clusters of integrated clustering of E11.5 control and NotchGOF λ-junction cells.**

| | | | | | | | | |
|---|---|---|---|---|---|---|---|---|
| 0 | Glrx2 | 0.26 | 3.31E-23 | 6.88E-19 | 0.77 | 0.75 | 0.02 | 1.03 |
| 0 | Ywhag | 0.30 | 3.52E-23 | 7.32E-19 | 0.67 | 0.63 | 0.04 | 1.07 |
| 0 | Zfp553 | 0.36 | 3.54E-23 | 7.37E-19 | 0.65 | 0.60 | 0.05 | 1.09 |
| 0 | Nav1 | 0.38 | 3.74E-23 | 7.78E-19 | 0.60 | 0.54 | 0.06 | 1.12 |
| 0 | Sh3gl3 | 0.62 | 4.01E-23 | 8.33E-19 | 0.40 | 0.32 | 0.08 | 1.23 |
| 0 | Supt20 | 0.35 | 4.22E-23 | 8.79E-19 | 0.63 | 0.59 | 0.05 | 1.08 |
| 0 | Yaf2 | 0.28 | 4.78E-23 | 9.94E-19 | 0.73 | 0.70 | 0.04 | 1.05 |
| 0 | Gpr45 | 0.76 | 6.97E-23 | 1.45E-18 | 0.25 | 0.18 | 0.07 | 1.40 |
| 0 | Psmd9 | 0.31 | 7.98E-23 | 1.66E-18 | 0.69 | 0.65 | 0.04 | 1.06 |
| 0 | Nfkbia | 0.38 | 8.03E-23 | 1.67E-18 | 0.57 | 0.51 | 0.06 | 1.12 |
| 0 | Dusp7 | 0.39 | 9.42E-23 | 1.96E-18 | 0.56 | 0.49 | 0.07 | 1.14 |
| 0 | 3110021N24Rik | 0.97 | 9.59E-23 | 2.00E-18 | 0.17 | 0.11 | 0.06 | 1.58 |
| 0 | Brms1l | 0.52 | 1.22E-22 | 2.54E-18 | 0.43 | 0.36 | 0.07 | 1.19 |
| 0 | Pla2g7 | 1.07 | 1.67E-22 | 3.48E-18 | 0.11 | 0.06 | 0.05 | 1.85 |
| 0 | Tax1bp3 | 0.30 | 1.79E-22 | 3.73E-18 | 0.69 | 0.64 | 0.05 | 1.07 |
| 0 | Loxl2 | 0.41 | 2.00E-22 | 4.17E-18 | 0.50 | 0.43 | 0.07 | 1.17 |
| 0 | Phf6 | 0.27 | 2.44E-22 | 5.08E-18 | 0.77 | 0.72 | 0.05 | 1.07 |
| 0 | Msantd4 | 0.39 | 4.11E-22 | 8.56E-18 | 0.57 | 0.52 | 0.05 | 1.11 |
| 0 | Dzip1 | 0.51 | 5.37E-22 | 1.12E-17 | 0.40 | 0.33 | 0.07 | 1.23 |
| 0 | Cd248 | 0.57 | 6.26E-22 | 1.30E-17 | 0.32 | 0.24 | 0.08 | 1.32 |
| 0 | Tspan11 | 0.79 | 6.87E-22 | 1.43E-17 | 0.23 | 0.16 | 0.07 | 1.43 |
| 0 | Acot1 | 0.45 | 8.86E-22 | 1.84E-17 | 0.42 | 0.34 | 0.08 | 1.22 |
| 0 | Hvcn1 | 0.96 | 8.88E-22 | 1.85E-17 | 0.13 | 0.07 | 0.05 | 1.73 |
| 0 | Fndc5 | 0.75 | 9.09E-22 | 1.89E-17 | 0.23 | 0.16 | 0.07 | 1.42 |
| 0 | Mllt3 | 0.38 | 1.02E-21 | 2.12E-17 | 0.68 | 0.66 | 0.02 | 1.03 |
| 0 | Iah1 | 0.36 | 1.11E-21 | 2.31E-17 | 0.57 | 0.52 | 0.05 | 1.11 |
| 0 | Adamts4 | 1.21 | 1.18E-21 | 2.45E-17 | 0.10 | 0.05 | 0.05 | 1.85 |
| 0 | Lrrc4b | 0.84 | 1.53E-21 | 3.18E-17 | 0.19 | 0.12 | 0.06 | 1.51 |
| 0 | Copb1 | 0.26 | 1.76E-21 | 3.67E-17 | 0.79 | 0.77 | 0.02 | 1.03 |
| 0 | Klhl24 | 0.29 | 2.07E-21 | 4.30E-17 | 0.65 | 0.59 | 0.06 | 1.10 |
| 0 | Oaf | 0.37 | 2.45E-21 | 5.11E-17 | 0.52 | 0.45 | 0.07 | 1.16 |
| 0 | Rnd3 | 0.25 | 3.90E-21 | 8.11E-17 | 0.70 | 0.63 | 0.07 | 1.11 |
| 0 | Zfp334 | 0.54 | 6.36E-21 | 1.32E-16 | 0.39 | 0.33 | 0.06 | 1.20 |
| 0 | Prtn3 | 0.58 | 6.41E-21 | 1.33E-16 | 0.19 | 0.13 | 0.06 | 1.48 |
| 0 | Maco1 | 0.29 | 7.09E-21 | 1.47E-16 | 0.72 | 0.69 | 0.04 | 1.06 |
| 0 | Ptpn9 | 0.39 | 7.54E-21 | 1.57E-16 | 0.54 | 0.48 | 0.05 | 1.11 |
| 0 | Ift43 | 0.30 | 7.63E-21 | 1.59E-16 | 0.60 | 0.54 | 0.06 | 1.11 |
| 0 | Hsd17b4 | 0.40 | 9.21E-21 | 1.92E-16 | 0.59 | 0.55 | 0.04 | 1.08 |
| 0 | Pam | 1.03 | 9.48E-21 | 1.97E-16 | 0.21 | 0.14 | 0.06 | 1.44 |
| 0 | Tmem131l | 0.36 | 1.02E-20 | 2.13E-16 | 0.59 | 0.54 | 0.05 | 1.09 |
| 0 | Ccnb2 | 0.39 | 1.06E-20 | 2.21E-16 | 0.62 | 0.59 | 0.02 | 1.04 |
| 0 | Gga2 | 0.31 | 1.19E-20 | 2.46E-16 | 0.68 | 0.64 | 0.04 | 1.06 |
| 0 | Cdc14b | 0.45 | 1.26E-20 | 2.62E-16 | 0.46 | 0.39 | 0.06 | 1.16 |
| 0 | Hdhd2 | 0.40 | 1.93E-20 | 4.02E-16 | 0.55 | 0.50 | 0.05 | 1.10 |
| 0 | C1qtnf6 | 0.75 | 2.22E-20 | 4.62E-16 | 0.22 | 0.15 | 0.07 | 1.42 |
| 0 | Serinc2 | 0.45 | 2.26E-20 | 4.71E-16 | 0.45 | 0.38 | 0.07 | 1.19 |
| 0 | Shd | 0.65 | 2.54E-20 | 5.28E-16 | 0.23 | 0.16 | 0.07 | 1.41 |
| 0 | Fdft1 | 0.33 | 2.83E-20 | 5.88E-16 | 0.64 | 0.60 | 0.04 | 1.07 |
| 0 | Dync2li1 | 0.52 | 3.31E-20 | 6.88E-16 | 0.41 | 0.35 | 0.06 | 1.18 |
| 0 | Efemp2 | 0.34 | 3.36E-20 | 6.99E-16 | 0.53 | 0.47 | 0.06 | 1.13 |
| 0 | Cop1 | 0.28 | 4.31E-20 | 8.97E-16 | 0.70 | 0.67 | 0.03 | 1.05 |
| 0 | Samd14 | 0.43 | 4.45E-20 | 9.25E-16 | 0.46 | 0.40 | 0.06 | 1.15 |
| 0 | Tmem30b | 0.71 | 4.70E-20 | 9.77E-16 | 0.21 | 0.14 | 0.06 | 1.45 |
| 0 | Tmem44 | 0.64 | 6.29E-20 | 1.31E-15 | 0.28 | 0.21 | 0.07 | 1.33 |
| 0 | Fam110b | 0.49 | 6.52E-20 | 1.36E-15 | 0.37 | 0.30 | 0.07 | 1.23 |
| 0 | Fibp | 0.31 | 6.88E-20 | 1.43E-15 | 0.65 | 0.61 | 0.04 | 1.07 |

**Supplementary Table 1. Marker genes for the original clusters of integrated clustering of E11.5 control and NotchGOF λ-junction cells.**

| | | | | | | | | |
|---|---|---|---|---|---|---|---|---|
| 0 | Dpep1 | 0.53 | 6.94E-20 | 1.44E-15 | 0.29 | 0.22 | 0.07 | 1.33 |
| 0 | Tro | 0.43 | 7.53E-20 | 1.57E-15 | 0.48 | 0.41 | 0.07 | 1.16 |
| 0 | Rbm4 | 0.26 | 7.93E-20 | 1.65E-15 | 0.74 | 0.70 | 0.04 | 1.06 |
| 0 | Spg20 | 0.35 | 8.88E-20 | 1.85E-15 | 0.58 | 0.53 | 0.05 | 1.09 |
| 0 | Angpt2 | 1.15 | 9.04E-20 | 1.88E-15 | 0.12 | 0.07 | 0.05 | 1.70 |
| 0 | Lsp1 | 0.54 | 9.94E-20 | 2.07E-15 | 0.22 | 0.15 | 0.06 | 1.42 |
| 0 | Papss1 | 0.29 | 1.19E-19 | 2.47E-15 | 0.69 | 0.66 | 0.03 | 1.04 |
| 0 | Zdhhc20 | 0.30 | 1.43E-19 | 2.98E-15 | 0.63 | 0.58 | 0.05 | 1.09 |
| 0 | Tns1 | 0.51 | 1.44E-19 | 3.00E-15 | 0.37 | 0.30 | 0.07 | 1.24 |
| 0 | Thap12 | 0.27 | 1.47E-19 | 3.07E-15 | 0.72 | 0.69 | 0.03 | 1.05 |
| 0 | Chd9 | 0.34 | 1.52E-19 | 3.15E-15 | 0.58 | 0.52 | 0.05 | 1.10 |
| 0 | Hpse2 | 0.75 | 1.81E-19 | 3.77E-15 | 0.23 | 0.16 | 0.06 | 1.38 |
| 0 | Ugdh | 0.37 | 1.81E-19 | 3.77E-15 | 0.52 | 0.46 | 0.06 | 1.13 |
| 0 | Tm2d2 | 0.26 | 1.95E-19 | 4.06E-15 | 0.75 | 0.72 | 0.03 | 1.04 |
| 0 | Gcc2 | 0.29 | 2.04E-19 | 4.24E-15 | 0.70 | 0.66 | 0.04 | 1.06 |
| 0 | Mafb | 0.50 | 2.19E-19 | 4.56E-15 | 0.29 | 0.21 | 0.07 | 1.34 |
| 0 | Kcnk2 | 0.61 | 2.46E-19 | 5.11E-15 | 0.36 | 0.29 | 0.07 | 1.23 |
| 0 | Gm45716 | 0.45 | 2.49E-19 | 5.17E-15 | 0.43 | 0.37 | 0.07 | 1.18 |
| 0 | Angptl4 | 0.46 | 2.84E-19 | 5.91E-15 | 0.40 | 0.33 | 0.07 | 1.21 |
| 0 | Oaz2 | 0.36 | 2.94E-19 | 6.12E-15 | 0.59 | 0.55 | 0.04 | 1.08 |
| 0 | Itih5 | 0.63 | 3.16E-19 | 6.56E-15 | 0.18 | 0.12 | 0.06 | 1.51 |
| 0 | Gpsm3 | 0.80 | 3.47E-19 | 7.23E-15 | 0.13 | 0.08 | 0.05 | 1.64 |
| 0 | Hsf2 | 0.37 | 3.58E-19 | 7.44E-15 | 0.53 | 0.47 | 0.06 | 1.12 |
| 0 | Zfp68 | 0.43 | 3.78E-19 | 7.86E-15 | 0.48 | 0.43 | 0.05 | 1.13 |
| 0 | Rxylt1 | 0.40 | 3.79E-19 | 7.89E-15 | 0.52 | 0.47 | 0.05 | 1.10 |
| 0 | Rnf157 | 0.79 | 4.27E-19 | 8.87E-15 | 0.21 | 0.15 | 0.06 | 1.42 |
| 0 | Smad1 | 0.41 | 4.39E-19 | 9.13E-15 | 0.52 | 0.47 | 0.05 | 1.11 |
| 0 | Hax1 | 0.29 | 4.40E-19 | 9.15E-15 | 0.69 | 0.67 | 0.02 | 1.03 |
| 0 | Got1 | 0.38 | 7.62E-19 | 1.58E-14 | 0.50 | 0.45 | 0.05 | 1.12 |
| 0 | Lrrc58 | 0.28 | 8.88E-19 | 1.85E-14 | 0.67 | 0.62 | 0.04 | 1.07 |
| 0 | Rassf3 | 0.40 | 9.14E-19 | 1.90E-14 | 0.43 | 0.36 | 0.07 | 1.20 |
| 0 | Svep1 | 0.96 | 9.15E-19 | 1.90E-14 | 0.13 | 0.08 | 0.05 | 1.64 |
| 0 | Wdr6 | 0.28 | 9.34E-19 | 1.94E-14 | 0.70 | 0.66 | 0.04 | 1.06 |
| 0 | Rac3 | 0.49 | 1.10E-18 | 2.29E-14 | 0.39 | 0.32 | 0.06 | 1.20 |
| 0 | Rarg | 0.30 | 1.13E-18 | 2.35E-14 | 0.58 | 0.51 | 0.07 | 1.13 |
| 0 | Pdxp | 0.46 | 1.23E-18 | 2.56E-14 | 0.44 | 0.39 | 0.06 | 1.15 |
| 0 | Snhg4 | 0.43 | 1.29E-18 | 2.68E-14 | 0.51 | 0.46 | 0.05 | 1.11 |
| 0 | Camk1 | 0.38 | 1.67E-18 | 3.48E-14 | 0.48 | 0.42 | 0.06 | 1.14 |
| 0 | Aff4 | 0.26 | 1.74E-18 | 3.62E-14 | 0.67 | 0.63 | 0.04 | 1.06 |
| 0 | Abcd2 | 0.90 | 1.78E-18 | 3.70E-14 | 0.12 | 0.07 | 0.05 | 1.71 |
| 0 | Acsl3 | 0.29 | 1.81E-18 | 3.76E-14 | 0.77 | 0.73 | 0.03 | 1.04 |
| 0 | Dctn4 | 0.30 | 1.82E-18 | 3.78E-14 | 0.67 | 0.64 | 0.03 | 1.05 |
| 0 | Zfp516 | 0.34 | 1.87E-18 | 3.89E-14 | 0.54 | 0.49 | 0.05 | 1.11 |
| 0 | Pabpc4l | 0.46 | 2.05E-18 | 4.27E-14 | 0.44 | 0.38 | 0.06 | 1.16 |
| 0 | Aff3 | 0.69 | 2.32E-18 | 4.83E-14 | 0.28 | 0.21 | 0.06 | 1.30 |
| 0 | Net1 | 0.43 | 2.35E-18 | 4.88E-14 | 0.62 | 0.59 | 0.03 | 1.05 |
| 0 | St3gal2 | 0.63 | 2.67E-18 | 5.56E-14 | 0.31 | 0.25 | 0.06 | 1.26 |
| 0 | Zfp322a | 0.30 | 3.08E-18 | 6.40E-14 | 0.67 | 0.63 | 0.04 | 1.06 |
| 0 | Mrpl4 | 0.29 | 3.34E-18 | 6.95E-14 | 0.68 | 0.65 | 0.03 | 1.05 |
| 0 | Slc39a6 | 0.31 | 3.64E-18 | 7.57E-14 | 0.61 | 0.56 | 0.05 | 1.09 |
| 0 | Mycn | 0.39 | 4.07E-18 | 8.46E-14 | 0.44 | 0.37 | 0.06 | 1.17 |
| 0 | Rnf150 | 0.58 | 4.16E-18 | 8.66E-14 | 0.36 | 0.30 | 0.06 | 1.20 |
| 0 | Sash1 | 0.37 | 4.73E-18 | 9.84E-14 | 0.46 | 0.39 | 0.07 | 1.17 |
| 0 | Zfp30 | 0.42 | 4.99E-18 | 1.04E-13 | 0.48 | 0.42 | 0.06 | 1.14 |
| 0 | Dcp2 | 0.28 | 5.28E-18 | 1.10E-13 | 0.66 | 0.62 | 0.03 | 1.05 |
| 0 | Twist2 | 0.51 | 5.36E-18 | 1.12E-13 | 0.29 | 0.22 | 0.07 | 1.31 |

**Supplementary Table 1. Marker genes for the original clusters of integrated clustering of E11.5 control and NotchGOF λ-junction cells.**

| | | | | | | | | |
|---|---|---|---|---|---|---|---|---|
| 0 | Plgrkt | 0.38 | 6.12E-18 | 1.27E-13 | 0.52 | 0.47 | 0.05 | 1.10 |
| 0 | Peg12 | 0.56 | 7.03E-18 | 1.46E-13 | 0.31 | 0.25 | 0.07 | 1.27 |
| 0 | Wif1 | 1.12 | 7.63E-18 | 1.59E-13 | 0.14 | 0.09 | 0.05 | 1.56 |
| 0 | Zfp9 | 0.73 | 7.84E-18 | 1.63E-13 | 0.22 | 0.16 | 0.06 | 1.38 |
| 0 | Tmem164 | 0.56 | 9.33E-18 | 1.94E-13 | 0.37 | 0.31 | 0.06 | 1.19 |
| 0 | Dnm3 | 0.78 | 9.67E-18 | 2.01E-13 | 0.22 | 0.16 | 0.06 | 1.38 |
| 0 | Map2k6 | 0.40 | 9.98E-18 | 2.08E-13 | 0.49 | 0.43 | 0.06 | 1.13 |
| 0 | Asxl3 | 0.34 | 1.00E-17 | 2.09E-13 | 0.51 | 0.46 | 0.05 | 1.12 |
| 0 | Mtmr11 | 0.84 | 1.40E-17 | 2.92E-13 | 0.15 | 0.10 | 0.05 | 1.53 |
| 0 | Efna4 | 0.28 | 1.44E-17 | 2.99E-13 | 0.66 | 0.62 | 0.05 | 1.07 |
| 0 | Ripk3 | 0.65 | 1.85E-17 | 3.85E-13 | 0.22 | 0.16 | 0.06 | 1.38 |
| 0 | Dkk3 | 0.72 | 2.22E-17 | 4.62E-13 | 0.17 | 0.11 | 0.06 | 1.49 |
| 0 | G430095P16Rik | 0.94 | 2.25E-17 | 4.69E-13 | 0.15 | 0.10 | 0.05 | 1.52 |
| 0 | Impa1 | 0.30 | 2.59E-17 | 5.39E-13 | 0.62 | 0.58 | 0.04 | 1.06 |
| 0 | Leprot | 0.33 | 3.03E-17 | 6.29E-13 | 0.56 | 0.52 | 0.04 | 1.09 |
| 0 | Brd9 | 0.32 | 3.31E-17 | 6.88E-13 | 0.59 | 0.55 | 0.04 | 1.08 |
| 0 | Zfp260 | 0.26 | 3.65E-17 | 7.59E-13 | 0.68 | 0.65 | 0.04 | 1.06 |
| 0 | Gpx3 | 0.53 | 3.96E-17 | 8.23E-13 | 0.28 | 0.21 | 0.07 | 1.31 |
| 0 | Ap1m1 | 0.43 | 4.49E-17 | 9.34E-13 | 0.44 | 0.38 | 0.05 | 1.13 |
| 0 | Eno3 | 0.45 | 5.16E-17 | 1.07E-12 | 0.43 | 0.37 | 0.06 | 1.17 |
| 0 | Vegfc | 0.52 | 6.12E-17 | 1.27E-12 | 0.33 | 0.26 | 0.07 | 1.25 |
| 0 | Atoh8 | 0.89 | 6.31E-17 | 1.31E-12 | 0.14 | 0.09 | 0.05 | 1.57 |
| 0 | Bnip3 | 0.41 | 7.22E-17 | 1.50E-12 | 0.46 | 0.41 | 0.05 | 1.13 |
| 0 | Sh3bp5 | 0.26 | 8.90E-17 | 1.85E-12 | 0.56 | 0.50 | 0.06 | 1.11 |
| 0 | 4833420G17Rik | 0.28 | 9.50E-17 | 1.98E-12 | 0.67 | 0.65 | 0.02 | 1.04 |
| 0 | Arhgef25 | 0.41 | 1.04E-16 | 2.17E-12 | 0.43 | 0.36 | 0.06 | 1.17 |
| 0 | Plin3 | 0.29 | 1.14E-16 | 2.38E-12 | 0.59 | 0.55 | 0.04 | 1.08 |
| 0 | Dpf2 | 0.27 | 1.15E-16 | 2.39E-12 | 0.68 | 0.65 | 0.03 | 1.05 |
| 0 | Edil3 | 0.82 | 1.37E-16 | 2.84E-12 | 0.22 | 0.17 | 0.06 | 1.34 |
| 0 | Lpgat1 | 0.32 | 1.50E-16 | 3.11E-12 | 0.60 | 0.56 | 0.04 | 1.07 |
| 0 | Osbpl1a | 0.41 | 1.53E-16 | 3.19E-12 | 0.44 | 0.38 | 0.06 | 1.15 |
| 0 | Cfap36 | 0.29 | 1.66E-16 | 3.45E-12 | 0.59 | 0.55 | 0.04 | 1.08 |
| 0 | Smarcc2 | 0.28 | 1.99E-16 | 4.14E-12 | 0.68 | 0.65 | 0.03 | 1.04 |
| 0 | Mcfd2 | 0.40 | 2.27E-16 | 4.72E-12 | 0.50 | 0.46 | 0.05 | 1.10 |
| 0 | Kctd15 | 0.38 | 2.51E-16 | 5.21E-12 | 0.47 | 0.41 | 0.06 | 1.14 |
| 0 | Lrba | 0.60 | 2.55E-16 | 5.30E-12 | 0.28 | 0.22 | 0.06 | 1.27 |
| 0 | Slc16a3 | 0.54 | 3.30E-16 | 6.85E-12 | 0.38 | 0.32 | 0.05 | 1.17 |
| 0 | Scarf2 | 0.56 | 3.40E-16 | 7.07E-12 | 0.31 | 0.25 | 0.06 | 1.23 |
| 0 | Evi5 | 0.29 | 3.53E-16 | 7.33E-12 | 0.63 | 0.61 | 0.02 | 1.04 |
| 0 | Cacnb3 | 0.47 | 3.72E-16 | 7.73E-12 | 0.33 | 0.27 | 0.06 | 1.23 |
| 0 | Map3k7 | 0.38 | 4.48E-16 | 9.31E-12 | 0.52 | 0.47 | 0.05 | 1.10 |
| 0 | Slc50a1 | 0.39 | 5.24E-16 | 1.09E-11 | 0.43 | 0.37 | 0.06 | 1.17 |
| 0 | Gse1 | 0.57 | 8.10E-16 | 1.69E-11 | 0.35 | 0.30 | 0.05 | 1.17 |
| 0 | Ephb4 | 0.40 | 9.94E-16 | 2.07E-11 | 0.45 | 0.39 | 0.06 | 1.14 |
| 0 | Cul5 | 0.26 | 1.21E-15 | 2.53E-11 | 0.70 | 0.68 | 0.02 | 1.03 |
| 0 | Fcgrt | 0.61 | 1.27E-15 | 2.64E-11 | 0.19 | 0.14 | 0.05 | 1.39 |
| 0 | Syne1 | 0.79 | 1.56E-15 | 3.25E-11 | 0.18 | 0.13 | 0.05 | 1.40 |
| 0 | Ip6k2 | 0.29 | 1.59E-15 | 3.30E-11 | 0.59 | 0.55 | 0.04 | 1.07 |
| 0 | Cacfd1 | 0.40 | 1.63E-15 | 3.40E-11 | 0.48 | 0.45 | 0.04 | 1.09 |
| 0 | Ubqln2 | 0.27 | 1.91E-15 | 3.98E-11 | 0.68 | 0.65 | 0.03 | 1.04 |
| 0 | Pitpnc1 | 0.30 | 2.09E-15 | 4.36E-11 | 0.63 | 0.58 | 0.04 | 1.07 |
| 0 | Zfp57 | 0.43 | 2.27E-15 | 4.73E-11 | 0.39 | 0.33 | 0.06 | 1.17 |
| 0 | Rps6ka3 | 0.40 | 2.35E-15 | 4.90E-11 | 0.49 | 0.45 | 0.04 | 1.08 |
| 0 | D10Wsu102e | 0.32 | 2.43E-15 | 5.06E-11 | 0.62 | 0.59 | 0.03 | 1.05 |
| 0 | Ppp2r3a | 0.48 | 2.59E-15 | 5.39E-11 | 0.40 | 0.35 | 0.05 | 1.15 |
| 0 | Spag9 | 0.29 | 2.63E-15 | 5.47E-11 | 0.66 | 0.62 | 0.04 | 1.06 |

**Supplementary Table 1. Marker genes for the original clusters of integrated clustering of E11.5 control and NotchGOF λ-junction cells.**

| | | | | | | | | |
|---|---|---|---|---|---|---|---|---|
| 0 | Shf | 0.50 | 2.85E-15 | 5.93E-11 | 0.33 | 0.27 | 0.06 | 1.21 |
| 0 | Apip | 0.26 | 3.04E-15 | 6.32E-11 | 0.63 | 0.61 | 0.03 | 1.04 |
| 0 | Rnmt | 0.31 | 3.08E-15 | 6.40E-11 | 0.56 | 0.53 | 0.04 | 1.07 |
| 0 | Unc5c | 0.32 | 3.37E-15 | 7.00E-11 | 0.41 | 0.33 | 0.07 | 1.22 |
| 0 | Raph1 | 0.38 | 3.94E-15 | 8.19E-11 | 0.47 | 0.42 | 0.05 | 1.12 |
| 0 | Fkbp11 | 0.46 | 4.16E-15 | 8.65E-11 | 0.35 | 0.29 | 0.06 | 1.20 |
| 0 | 2610008E11Rik | 0.44 | 4.29E-15 | 8.93E-11 | 0.44 | 0.39 | 0.05 | 1.13 |
| 0 | Trim26 | 0.36 | 4.49E-15 | 9.34E-11 | 0.47 | 0.42 | 0.05 | 1.11 |
| 0 | Apopt1 | 0.29 | 5.08E-15 | 1.06E-10 | 0.59 | 0.56 | 0.03 | 1.06 |
| 0 | Sgtb | 0.78 | 5.22E-15 | 1.08E-10 | 0.15 | 0.10 | 0.05 | 1.47 |
| 0 | Zbed3 | 0.30 | 5.51E-15 | 1.15E-10 | 0.59 | 0.56 | 0.03 | 1.06 |
| 0 | Mvk | 0.36 | 6.05E-15 | 1.26E-10 | 0.47 | 0.42 | 0.05 | 1.11 |
| 0 | Aplp1 | 0.49 | 6.19E-15 | 1.29E-10 | 0.26 | 0.21 | 0.06 | 1.28 |
| 0 | Adam19 | 0.67 | 6.33E-15 | 1.32E-10 | 0.20 | 0.15 | 0.05 | 1.36 |
| 0 | Arl8a | 0.37 | 6.33E-15 | 1.32E-10 | 0.47 | 0.42 | 0.05 | 1.11 |
| 0 | Stmn2 | 0.39 | 6.77E-15 | 1.41E-10 | 0.14 | 0.10 | 0.05 | 1.48 |
| 0 | Nlgn2 | 0.55 | 8.92E-15 | 1.86E-10 | 0.29 | 0.23 | 0.06 | 1.25 |
| 0 | Rassf8 | 0.63 | 9.35E-15 | 1.94E-10 | 0.23 | 0.18 | 0.06 | 1.31 |
| 0 | Tnks | 0.30 | 9.49E-15 | 1.97E-10 | 0.58 | 0.54 | 0.04 | 1.07 |
| 0 | Erc1 | 0.31 | 9.80E-15 | 2.04E-10 | 0.59 | 0.55 | 0.04 | 1.07 |
| 0 | Selenos | 0.26 | 1.12E-14 | 2.33E-10 | 0.64 | 0.61 | 0.03 | 1.05 |
| 0 | Kat6b | 0.47 | 1.15E-14 | 2.39E-10 | 0.51 | 0.47 | 0.04 | 1.09 |
| 0 | Fbln5 | 0.61 | 1.33E-14 | 2.76E-10 | 0.19 | 0.14 | 0.05 | 1.39 |
| 0 | Rcbtb1 | 0.34 | 1.36E-14 | 2.82E-10 | 0.51 | 0.47 | 0.04 | 1.09 |
| 0 | Itch | 0.26 | 1.40E-14 | 2.90E-10 | 0.57 | 0.54 | 0.03 | 1.06 |
| 0 | 4833439L19Rik | 0.31 | 1.40E-14 | 2.92E-10 | 0.54 | 0.50 | 0.03 | 1.07 |
| 0 | Nfe2l2 | 0.40 | 1.48E-14 | 3.07E-10 | 0.48 | 0.44 | 0.04 | 1.09 |
| 0 | Ech1 | 0.31 | 2.57E-14 | 5.35E-10 | 0.55 | 0.51 | 0.04 | 1.07 |
| 0 | Pex19 | 0.29 | 2.65E-14 | 5.51E-10 | 0.56 | 0.52 | 0.04 | 1.07 |
| 0 | Taf7 | 0.25 | 2.74E-14 | 5.69E-10 | 0.70 | 0.69 | 0.01 | 1.01 |
| 0 | Tpm2 | 0.29 | 3.00E-14 | 6.24E-10 | 0.49 | 0.43 | 0.06 | 1.13 |
| 0 | Wnk3 | 0.44 | 3.45E-14 | 7.18E-10 | 0.37 | 0.32 | 0.05 | 1.17 |
| 0 | Kcnmb4 | 0.39 | 3.48E-14 | 7.23E-10 | 0.37 | 0.32 | 0.06 | 1.18 |
| 0 | Col6a3 | 0.77 | 4.45E-14 | 9.25E-10 | 0.21 | 0.16 | 0.05 | 1.31 |
| 0 | Jakmip2 | 0.56 | 4.55E-14 | 9.47E-10 | 0.19 | 0.14 | 0.05 | 1.38 |
| 0 | Samd1 | 0.41 | 4.79E-14 | 9.96E-10 | 0.45 | 0.40 | 0.05 | 1.12 |
| 0 | Col13a1 | 0.59 | 4.90E-14 | 1.02E-09 | 0.15 | 0.10 | 0.05 | 1.48 |
| 0 | Trim71 | 0.57 | 4.98E-14 | 1.04E-09 | 0.26 | 0.21 | 0.05 | 1.26 |
| 0 | Ift57 | 0.43 | 5.10E-14 | 1.06E-09 | 0.36 | 0.31 | 0.05 | 1.18 |
| 0 | Samd4 | 0.43 | 5.24E-14 | 1.09E-09 | 0.42 | 0.36 | 0.06 | 1.15 |
| 0 | Capn6 | 0.33 | 6.07E-14 | 1.26E-09 | 0.59 | 0.54 | 0.04 | 1.08 |
| 0 | Dusp9 | 0.36 | 7.13E-14 | 1.48E-09 | 0.46 | 0.41 | 0.05 | 1.13 |
| 0 | Snhg3 | 0.32 | 7.45E-14 | 1.55E-09 | 0.57 | 0.53 | 0.04 | 1.07 |
| 0 | Scpep1 | 0.35 | 7.51E-14 | 1.56E-09 | 0.45 | 0.41 | 0.05 | 1.12 |
| 0 | Cacna1h | 0.91 | 7.83E-14 | 1.63E-09 | 0.11 | 0.07 | 0.04 | 1.56 |
| 0 | Cnih2 | 0.39 | 7.95E-14 | 1.65E-09 | 0.38 | 0.32 | 0.06 | 1.18 |
| 0 | Cdc42ep4 | 0.34 | 8.16E-14 | 1.70E-09 | 0.49 | 0.44 | 0.05 | 1.10 |
| 0 | 2310009A05Rik | 0.25 | 9.14E-14 | 1.90E-09 | 0.61 | 0.58 | 0.03 | 1.05 |
| 0 | Tmem216 | 0.33 | 1.04E-13 | 2.16E-09 | 0.48 | 0.43 | 0.05 | 1.11 |
| 0 | B3gnt5 | 0.40 | 1.43E-13 | 2.97E-09 | 0.40 | 0.35 | 0.05 | 1.14 |
| 0 | Wnt5b | 0.46 | 1.52E-13 | 3.17E-09 | 0.31 | 0.25 | 0.06 | 1.22 |
| 0 | Filip1l | 0.51 | 1.75E-13 | 3.64E-09 | 0.31 | 0.25 | 0.06 | 1.22 |
| 0 | Efnb3 | 0.45 | 1.78E-13 | 3.70E-09 | 0.36 | 0.31 | 0.05 | 1.17 |
| 0 | Vash2 | 0.49 | 1.82E-13 | 3.79E-09 | 0.34 | 0.28 | 0.05 | 1.18 |
| 0 | Bloc1s1 | 0.26 | 1.87E-13 | 3.88E-09 | 0.62 | 0.59 | 0.03 | 1.05 |
| 0 | Meis3 | 0.59 | 1.87E-13 | 3.89E-09 | 0.25 | 0.19 | 0.05 | 1.26 |

**Supplementary Table 1. Marker genes for the original clusters of integrated clustering of E11.5 control and NotchGOF λ-junction cells.**

| | | | | | | | | |
|---|---|---|---|---|---|---|---|---|
| 0 | Kctd12b | 0.51 | 2.00E-13 | 4.16E-09 | 0.23 | 0.18 | 0.06 | 1.32 |
| 0 | Aida | 0.32 | 2.13E-13 | 4.43E-09 | 0.53 | 0.50 | 0.03 | 1.07 |
| 0 | Jagn1 | 0.27 | 2.13E-13 | 4.43E-09 | 0.60 | 0.58 | 0.02 | 1.04 |
| 0 | Pex13 | 0.26 | 2.35E-13 | 4.89E-09 | 0.61 | 0.58 | 0.03 | 1.06 |
| 0 | Cby1 | 0.30 | 2.63E-13 | 5.48E-09 | 0.55 | 0.52 | 0.03 | 1.06 |
| 0 | Shkbp1 | 0.42 | 2.68E-13 | 5.57E-09 | 0.38 | 0.33 | 0.05 | 1.14 |
| 0 | 2810001G20Rik | 0.37 | 2.74E-13 | 5.70E-09 | 0.43 | 0.38 | 0.05 | 1.13 |
| 0 | Psrc1 | 0.44 | 3.02E-13 | 6.29E-09 | 0.45 | 0.42 | 0.03 | 1.08 |
| 0 | Ube2q2 | 0.31 | 3.14E-13 | 6.52E-09 | 0.53 | 0.50 | 0.03 | 1.06 |
| 0 | Sccpdh | 0.43 | 4.33E-13 | 9.01E-09 | 0.37 | 0.32 | 0.05 | 1.15 |
| 0 | 4931406C07Rik | 0.51 | 4.40E-13 | 9.16E-09 | 0.31 | 0.26 | 0.05 | 1.21 |
| 0 | Zfp354c | 0.39 | 5.07E-13 | 1.05E-08 | 0.43 | 0.39 | 0.04 | 1.11 |
| 0 | Sqle | 0.29 | 5.82E-13 | 1.21E-08 | 0.60 | 0.58 | 0.02 | 1.04 |
| 0 | Pls3 | 0.39 | 5.94E-13 | 1.24E-08 | 0.35 | 0.29 | 0.06 | 1.19 |
| 0 | Slc35a1 | 0.32 | 6.73E-13 | 1.40E-08 | 0.51 | 0.47 | 0.03 | 1.07 |
| 0 | Rcor2 | 0.30 | 7.99E-13 | 1.66E-08 | 0.53 | 0.49 | 0.04 | 1.08 |
| 0 | Enho | 0.31 | 8.07E-13 | 1.68E-08 | 0.50 | 0.45 | 0.05 | 1.11 |
| 0 | Triqk | 0.69 | 8.85E-13 | 1.84E-08 | 0.17 | 0.12 | 0.05 | 1.38 |
| 0 | Arf2 | 0.39 | 9.02E-13 | 1.88E-08 | 0.42 | 0.38 | 0.04 | 1.10 |
| 0 | Ino80e | 0.32 | 9.50E-13 | 1.98E-08 | 0.55 | 0.52 | 0.03 | 1.05 |
| 0 | Mvd | 0.46 | 1.01E-12 | 2.11E-08 | 0.34 | 0.30 | 0.05 | 1.16 |
| 0 | Prrg1 | 0.57 | 1.03E-12 | 2.15E-08 | 0.24 | 0.19 | 0.05 | 1.27 |
| 0 | Fkbp10 | 0.37 | 1.20E-12 | 2.50E-08 | 0.39 | 0.33 | 0.05 | 1.16 |
| 0 | Rwdd4a | 0.26 | 1.25E-12 | 2.61E-08 | 0.60 | 0.58 | 0.02 | 1.04 |
| 0 | Hspb11 | 0.35 | 1.30E-12 | 2.70E-08 | 0.48 | 0.44 | 0.04 | 1.09 |
| 0 | Cblb | 0.40 | 1.31E-12 | 2.72E-08 | 0.39 | 0.34 | 0.05 | 1.14 |
| 0 | 5730409E04Rik | 0.81 | 1.37E-12 | 2.86E-08 | 0.12 | 0.08 | 0.04 | 1.49 |
| 0 | Phf20 | 0.26 | 1.55E-12 | 3.22E-08 | 0.63 | 0.60 | 0.02 | 1.04 |
| 0 | Acadsb | 0.42 | 1.69E-12 | 3.51E-08 | 0.40 | 0.36 | 0.04 | 1.11 |
| 0 | Cacna1c | 0.84 | 1.87E-12 | 3.89E-08 | 0.13 | 0.09 | 0.04 | 1.47 |
| 0 | Mmp11 | 0.40 | 2.03E-12 | 4.22E-08 | 0.36 | 0.31 | 0.05 | 1.16 |
| 0 | Rab31 | 0.55 | 2.35E-12 | 4.88E-08 | 0.27 | 0.22 | 0.05 | 1.22 |
| 0 | Col5a1 | 0.52 | 2.38E-12 | 4.96E-08 | 0.31 | 0.26 | 0.05 | 1.18 |
| 0 | Tmem59l | 0.52 | 2.80E-12 | 5.83E-08 | 0.25 | 0.20 | 0.05 | 1.26 |
| 0 | Palm | 0.32 | 3.11E-12 | 6.47E-08 | 0.49 | 0.46 | 0.04 | 1.08 |
| 0 | Bach1 | 0.36 | 3.55E-12 | 7.38E-08 | 0.50 | 0.48 | 0.03 | 1.05 |
| 0 | P4ha1 | 0.32 | 3.62E-12 | 7.54E-08 | 0.55 | 0.51 | 0.03 | 1.06 |
| 0 | Arpin | 0.59 | 3.90E-12 | 8.10E-08 | 0.22 | 0.17 | 0.05 | 1.28 |
| 0 | Chst15 | 0.59 | 3.92E-12 | 8.14E-08 | 0.21 | 0.17 | 0.05 | 1.30 |
| 0 | Uba3 | 0.29 | 4.08E-12 | 8.48E-08 | 0.54 | 0.52 | 0.02 | 1.05 |
| 0 | Abl1 | 0.30 | 4.10E-12 | 8.52E-08 | 0.53 | 0.50 | 0.03 | 1.07 |
| 0 | Ncs1 | 0.39 | 4.19E-12 | 8.70E-08 | 0.36 | 0.31 | 0.05 | 1.17 |
| 0 | Zfp329 | 0.27 | 4.25E-12 | 8.84E-08 | 0.58 | 0.55 | 0.03 | 1.05 |
| 0 | 2610307P16Rik | 0.68 | 4.63E-12 | 9.64E-08 | 0.22 | 0.17 | 0.05 | 1.26 |
| 0 | Cep85l | 0.67 | 5.20E-12 | 1.08E-07 | 0.18 | 0.13 | 0.04 | 1.33 |
| 0 | Zc3hc1 | 0.37 | 5.35E-12 | 1.11E-07 | 0.44 | 0.41 | 0.03 | 1.08 |
| 0 | Gng8 | 0.40 | 5.56E-12 | 1.16E-07 | 0.18 | 0.13 | 0.05 | 1.36 |
| 0 | Ppil3 | 0.29 | 5.57E-12 | 1.16E-07 | 0.56 | 0.53 | 0.03 | 1.06 |
| 0 | Tmco3 | 0.46 | 5.66E-12 | 1.18E-07 | 0.34 | 0.29 | 0.05 | 1.17 |
| 0 | Dhx8 | 0.28 | 5.92E-12 | 1.23E-07 | 0.60 | 0.59 | 0.02 | 1.03 |
| 0 | Slain2 | 0.31 | 6.33E-12 | 1.32E-07 | 0.48 | 0.44 | 0.04 | 1.09 |
| 0 | Mterf3 | 0.34 | 6.97E-12 | 1.45E-07 | 0.46 | 0.43 | 0.03 | 1.07 |
| 0 | Itga8 | 0.41 | 7.58E-12 | 1.58E-07 | 0.24 | 0.19 | 0.05 | 1.29 |
| 0 | Inpp4b | 0.66 | 8.03E-12 | 1.67E-07 | 0.20 | 0.15 | 0.05 | 1.31 |
| 0 | Twsg1 | 0.27 | 8.53E-12 | 1.77E-07 | 0.52 | 0.49 | 0.03 | 1.07 |
| 0 | Smim14 | 0.29 | 9.19E-12 | 1.91E-07 | 0.45 | 0.41 | 0.04 | 1.11 |

**Supplementary Table 1. Marker genes for the original clusters of integrated clustering of E11.5 control and NotchGOF λ-junction cells.**

| | | | | | | | | |
|---|---|---|---|---|---|---|---|---|
| 0 | Nkiras2 | 0.30 | 9.43E-12 | 1.96E-07 | 0.47 | 0.43 | 0.04 | 1.10 |
| 0 | St6galnac3 | 0.35 | 9.88E-12 | 2.06E-07 | 0.44 | 0.39 | 0.05 | 1.12 |
| 0 | Gm15222 | 0.74 | 1.26E-11 | 2.62E-07 | 0.16 | 0.11 | 0.04 | 1.37 |
| 0 | Tmem98 | 0.25 | 1.42E-11 | 2.95E-07 | 0.53 | 0.50 | 0.03 | 1.07 |
| 0 | 2300009A05Rik | 0.41 | 1.42E-11 | 2.95E-07 | 0.42 | 0.39 | 0.03 | 1.09 |
| 0 | Map4 | 0.34 | 1.42E-11 | 2.95E-07 | 0.43 | 0.39 | 0.04 | 1.10 |
| 0 | Rap2b | 0.38 | 1.50E-11 | 3.11E-07 | 0.51 | 0.48 | 0.03 | 1.05 |
| 0 | Antxr1 | 0.38 | 1.52E-11 | 3.16E-07 | 0.36 | 0.31 | 0.05 | 1.16 |
| 0 | Sdk1 | 0.75 | 1.58E-11 | 3.28E-07 | 0.12 | 0.08 | 0.04 | 1.48 |
| 0 | Kctd20 | 0.34 | 1.61E-11 | 3.34E-07 | 0.52 | 0.49 | 0.02 | 1.05 |
| 0 | Kctd10 | 0.43 | 1.64E-11 | 3.40E-07 | 0.34 | 0.29 | 0.04 | 1.15 |
| 0 | Abhd8 | 0.31 | 1.67E-11 | 3.48E-07 | 0.51 | 0.48 | 0.03 | 1.06 |
| 0 | Gtf3c1 | 0.25 | 1.68E-11 | 3.50E-07 | 0.61 | 0.57 | 0.03 | 1.06 |
| 0 | Bbx | 0.31 | 1.95E-11 | 4.06E-07 | 0.60 | 0.59 | 0.02 | 1.03 |
| 0 | Gfra4 | 0.70 | 1.96E-11 | 4.08E-07 | 0.14 | 0.10 | 0.04 | 1.40 |
| 0 | Bak1 | 0.36 | 2.04E-11 | 4.25E-07 | 0.44 | 0.41 | 0.04 | 1.09 |
| 0 | Ccdc90b | 0.25 | 2.14E-11 | 4.45E-07 | 0.59 | 0.55 | 0.04 | 1.07 |
| 0 | Rilpl1 | 0.39 | 2.50E-11 | 5.20E-07 | 0.33 | 0.29 | 0.05 | 1.17 |
| 0 | Nxpe3 | 0.76 | 2.70E-11 | 5.61E-07 | 0.14 | 0.10 | 0.04 | 1.39 |
| 0 | Ush1c | 0.69 | 2.86E-11 | 5.94E-07 | 0.14 | 0.10 | 0.04 | 1.40 |
| 0 | Mad2l2 | 0.34 | 3.00E-11 | 6.24E-07 | 0.45 | 0.42 | 0.03 | 1.08 |
| 0 | Zfp131 | 0.26 | 3.33E-11 | 6.92E-07 | 0.59 | 0.57 | 0.02 | 1.03 |
| 0 | Lrrc75a | 0.51 | 3.40E-11 | 7.07E-07 | 0.25 | 0.20 | 0.05 | 1.23 |
| 0 | Casc4 | 0.31 | 3.61E-11 | 7.50E-07 | 0.50 | 0.47 | 0.03 | 1.06 |
| 0 | Grem1 | 0.69 | 4.32E-11 | 8.98E-07 | 0.13 | 0.09 | 0.04 | 1.43 |
| 0 | Mmd | 0.37 | 4.39E-11 | 9.13E-07 | 0.39 | 0.35 | 0.05 | 1.13 |
| 0 | Dnajc12 | 0.58 | 5.04E-11 | 1.05E-06 | 0.21 | 0.16 | 0.04 | 1.27 |
| 0 | Ror2 | 0.38 | 5.45E-11 | 1.13E-06 | 0.35 | 0.30 | 0.05 | 1.17 |
| 0 | Sirt2 | 0.27 | 7.28E-11 | 1.51E-06 | 0.53 | 0.51 | 0.02 | 1.05 |
| 0 | Cstf3 | 0.28 | 7.46E-11 | 1.55E-06 | 0.57 | 0.56 | 0.02 | 1.03 |
| 0 | Mmgt2 | 0.32 | 7.89E-11 | 1.64E-06 | 0.47 | 0.43 | 0.04 | 1.09 |
| 0 | Usp5 | 0.28 | 8.54E-11 | 1.78E-06 | 0.52 | 0.50 | 0.02 | 1.04 |
| 0 | Armcx1 | 0.39 | 8.88E-11 | 1.85E-06 | 0.38 | 0.34 | 0.04 | 1.13 |
| 0 | Mtfr1 | 0.30 | 9.07E-11 | 1.89E-06 | 0.54 | 0.52 | 0.02 | 1.04 |
| 0 | Ehd1 | 0.27 | 9.16E-11 | 1.90E-06 | 0.55 | 0.52 | 0.03 | 1.05 |
| 0 | Dohh | 0.26 | 9.56E-11 | 1.99E-06 | 0.54 | 0.52 | 0.02 | 1.04 |
| 0 | Rnf14 | 0.40 | 9.96E-11 | 2.07E-06 | 0.39 | 0.35 | 0.04 | 1.11 |
| 0 | Ubtd2 | 0.40 | 1.01E-10 | 2.10E-06 | 0.36 | 0.32 | 0.04 | 1.12 |
| 0 | Tasor2 | 0.27 | 1.10E-10 | 2.28E-06 | 0.54 | 0.51 | 0.03 | 1.06 |
| 0 | Fam181b | 0.75 | 1.12E-10 | 2.33E-06 | 0.15 | 0.11 | 0.04 | 1.37 |
| 0 | Gmcl1 | 0.31 | 1.12E-10 | 2.33E-06 | 0.48 | 0.44 | 0.03 | 1.08 |
| 0 | Klhdc10 | 0.32 | 1.22E-10 | 2.53E-06 | 0.47 | 0.44 | 0.03 | 1.07 |
| 0 | Mapk8 | 0.32 | 1.30E-10 | 2.71E-06 | 0.46 | 0.42 | 0.03 | 1.08 |
| 0 | Camk1g | 0.77 | 1.34E-10 | 2.79E-06 | 0.11 | 0.08 | 0.04 | 1.44 |
| 0 | Snx4 | 0.32 | 1.85E-10 | 3.85E-06 | 0.45 | 0.42 | 0.03 | 1.08 |
| 0 | Pmp22 | 0.45 | 1.99E-10 | 4.15E-06 | 0.23 | 0.18 | 0.05 | 1.26 |
| 0 | Tnfrsf19 | 0.38 | 2.00E-10 | 4.15E-06 | 0.19 | 0.15 | 0.05 | 1.31 |
| 0 | Tmem184c | 0.40 | 2.01E-10 | 4.18E-06 | 0.38 | 0.34 | 0.04 | 1.11 |
| 0 | Polb | 0.26 | 2.01E-10 | 4.19E-06 | 0.61 | 0.59 | 0.02 | 1.04 |
| 0 | Snx18 | 0.36 | 2.04E-10 | 4.24E-06 | 0.41 | 0.37 | 0.04 | 1.09 |
| 0 | Rnf115 | 0.27 | 2.18E-10 | 4.54E-06 | 0.54 | 0.52 | 0.02 | 1.04 |
| 0 | Egfl6 | 0.71 | 2.27E-10 | 4.72E-06 | 0.14 | 0.10 | 0.04 | 1.39 |
| 0 | Sgce | 0.36 | 2.32E-10 | 4.82E-06 | 0.37 | 0.32 | 0.05 | 1.14 |
| 0 | Selenon | 0.37 | 2.73E-10 | 5.68E-06 | 0.40 | 0.37 | 0.04 | 1.10 |
| 0 | Ubxn6 | 0.26 | 2.74E-10 | 5.70E-06 | 0.54 | 0.52 | 0.02 | 1.04 |
| 0 | Zc3h7a | 0.37 | 3.10E-10 | 6.45E-06 | 0.38 | 0.34 | 0.04 | 1.11 |

**Supplementary Table 1. Marker genes for the original clusters of integrated clustering of E11.5 control and NotchGOF λ-junction cells.**

| | | | | | | | | |
|---|---|---|---|---|---|---|---|---|
| 0 | Amer2 | 0.64 | 3.12E-10 | 6.49E-06 | 0.14 | 0.10 | 0.04 | 1.37 |
| 0 | Taf13 | 0.28 | 3.41E-10 | 7.10E-06 | 0.53 | 0.52 | 0.02 | 1.03 |
| 0 | Plcg1 | 0.35 | 3.48E-10 | 7.24E-06 | 0.43 | 0.40 | 0.03 | 1.08 |
| 0 | Ccdc28b | 0.34 | 3.73E-10 | 7.75E-06 | 0.41 | 0.37 | 0.04 | 1.11 |
| 0 | Nuak1 | 0.41 | 3.98E-10 | 8.28E-06 | 0.25 | 0.20 | 0.05 | 1.23 |
| 0 | Twf2 | 0.26 | 4.40E-10 | 9.15E-06 | 0.46 | 0.42 | 0.04 | 1.09 |
| 0 | Dpf1 | 0.58 | 4.55E-10 | 9.46E-06 | 0.20 | 0.16 | 0.04 | 1.26 |
| 0 | Phactr3 | 0.74 | 4.77E-10 | 9.92E-06 | 0.11 | 0.07 | 0.03 | 1.47 |
| 0 | Agps | 0.35 | 5.33E-10 | 1.11E-05 | 0.43 | 0.40 | 0.03 | 1.07 |
| 0 | Pus7l | 0.60 | 5.42E-10 | 1.13E-05 | 0.19 | 0.15 | 0.04 | 1.27 |
| 0 | Mcc | 0.62 | 5.68E-10 | 1.18E-05 | 0.17 | 0.13 | 0.04 | 1.32 |
| 0 | Hacd3 | 0.31 | 5.80E-10 | 1.21E-05 | 0.46 | 0.43 | 0.03 | 1.08 |
| 0 | Pcca | 0.37 | 6.31E-10 | 1.31E-05 | 0.33 | 0.29 | 0.04 | 1.15 |
| 0 | Klhl22 | 0.40 | 6.79E-10 | 1.41E-05 | 0.37 | 0.34 | 0.03 | 1.10 |
| 0 | Bcl9l | 0.39 | 6.91E-10 | 1.44E-05 | 0.37 | 0.33 | 0.04 | 1.12 |
| 0 | Nicn1 | 0.38 | 6.97E-10 | 1.45E-05 | 0.37 | 0.33 | 0.04 | 1.12 |
| 0 | Shox2 | 0.34 | 7.03E-10 | 1.46E-05 | 0.17 | 0.12 | 0.04 | 1.35 |
| 0 | Ercc5 | 0.39 | 7.44E-10 | 1.55E-05 | 0.35 | 0.31 | 0.04 | 1.12 |
| 0 | Tvp23a | 0.60 | 9.87E-10 | 2.05E-05 | 0.17 | 0.13 | 0.04 | 1.30 |
| 0 | Tspan2 | 0.60 | 1.09E-09 | 2.27E-05 | 0.13 | 0.10 | 0.04 | 1.39 |
| 0 | Podxl2 | 0.25 | 1.10E-09 | 2.29E-05 | 0.46 | 0.42 | 0.04 | 1.10 |
| 0 | Dnajb4 | 0.40 | 1.37E-09 | 2.86E-05 | 0.34 | 0.30 | 0.04 | 1.13 |
| 0 | Aebp1 | 0.45 | 1.51E-09 | 3.14E-05 | 0.25 | 0.20 | 0.04 | 1.22 |
| 0 | BC028528 | 0.54 | 1.54E-09 | 3.20E-05 | 0.11 | 0.07 | 0.03 | 1.45 |
| 0 | Kcne1l | 0.49 | 1.67E-09 | 3.47E-05 | 0.16 | 0.12 | 0.04 | 1.33 |
| 0 | Ndst1 | 0.28 | 1.78E-09 | 3.70E-05 | 0.50 | 0.47 | 0.03 | 1.05 |
| 0 | Atn1 | 0.29 | 1.82E-09 | 3.79E-05 | 0.47 | 0.44 | 0.03 | 1.06 |
| 0 | Sema6c | 0.58 | 1.88E-09 | 3.92E-05 | 0.20 | 0.16 | 0.04 | 1.26 |
| 0 | Bmp1 | 0.32 | 1.92E-09 | 3.98E-05 | 0.45 | 0.42 | 0.03 | 1.07 |
| 0 | Zdhhc17 | 0.48 | 1.94E-09 | 4.03E-05 | 0.29 | 0.25 | 0.04 | 1.16 |
| 0 | Ccar2 | 0.30 | 2.11E-09 | 4.40E-05 | 0.49 | 0.46 | 0.03 | 1.05 |
| 0 | Zfand4 | 0.78 | 2.19E-09 | 4.55E-05 | 0.11 | 0.08 | 0.03 | 1.42 |
| 0 | Trafd1 | 0.44 | 2.29E-09 | 4.77E-05 | 0.33 | 0.30 | 0.04 | 1.13 |
| 0 | Cdc27 | 0.25 | 2.33E-09 | 4.84E-05 | 0.54 | 0.53 | 0.02 | 1.03 |
| 0 | Ralgds | 0.46 | 2.98E-09 | 6.19E-05 | 0.26 | 0.22 | 0.04 | 1.19 |
| 0 | Ptpdc1 | 0.51 | 3.16E-09 | 6.58E-05 | 0.20 | 0.16 | 0.04 | 1.25 |
| 0 | Dipk1b | 0.28 | 3.18E-09 | 6.62E-05 | 0.51 | 0.48 | 0.03 | 1.06 |
| 0 | Thra | 0.33 | 3.30E-09 | 6.86E-05 | 0.45 | 0.42 | 0.03 | 1.08 |
| 0 | Phf21a | 0.28 | 3.39E-09 | 7.04E-05 | 0.50 | 0.47 | 0.03 | 1.06 |
| 0 | Ttyh3 | 0.26 | 3.47E-09 | 7.22E-05 | 0.55 | 0.54 | 0.02 | 1.03 |
| 0 | Tmem45a | 0.68 | 3.55E-09 | 7.38E-05 | 0.11 | 0.08 | 0.03 | 1.43 |
| 0 | St6galnac4 | 0.35 | 3.65E-09 | 7.58E-05 | 0.24 | 0.19 | 0.05 | 1.23 |
| 0 | Sh3pxd2b | 0.45 | 3.76E-09 | 7.81E-05 | 0.32 | 0.29 | 0.04 | 1.12 |
| 0 | Dusp18 | 0.53 | 3.90E-09 | 8.11E-05 | 0.20 | 0.16 | 0.04 | 1.24 |
| 0 | Gm26648 | 0.80 | 3.95E-09 | 8.21E-05 | 0.12 | 0.09 | 0.03 | 1.39 |
| 0 | Sec23a | 0.36 | 4.09E-09 | 8.50E-05 | 0.37 | 0.34 | 0.03 | 1.10 |
| 0 | Ypel1 | 0.26 | 4.12E-09 | 8.57E-05 | 0.53 | 0.50 | 0.03 | 1.06 |
| 0 | Trio | 0.26 | 4.36E-09 | 9.08E-05 | 0.54 | 0.51 | 0.03 | 1.05 |
| 0 | Lss | 0.55 | 4.50E-09 | 9.36E-05 | 0.19 | 0.16 | 0.04 | 1.25 |
| 0 | Agk | 0.37 | 5.09E-09 | 1.06E-04 | 0.35 | 0.32 | 0.04 | 1.11 |
| 0 | Vps4a | 0.28 | 5.14E-09 | 1.07E-04 | 0.52 | 0.51 | 0.01 | 1.02 |
| 0 | Sh2b2 | 0.53 | 5.34E-09 | 1.11E-04 | 0.20 | 0.16 | 0.04 | 1.25 |
| 0 | Tmem198b | 0.62 | 5.63E-09 | 1.17E-04 | 0.19 | 0.15 | 0.04 | 1.26 |
| 0 | Soat1 | 0.26 | 5.95E-09 | 1.24E-04 | 0.47 | 0.44 | 0.03 | 1.07 |
| 0 | Snx15 | 0.43 | 6.08E-09 | 1.26E-04 | 0.33 | 0.30 | 0.03 | 1.10 |
| 0 | Clip2 | 0.55 | 6.17E-09 | 1.28E-04 | 0.22 | 0.19 | 0.04 | 1.21 |

**Supplementary Table 1. Marker genes for the original clusters of integrated clustering of E11.5 control and NotchGOF λ-junction cells.**

| | | | | | | | | |
|---|---|---|---|---|---|---|---|---|
| 0 | Gm10762 | 0.36 | 6.57E-09 | 1.37E-04 | 0.42 | 0.39 | 0.03 | 1.07 |
| 0 | Zcchc9 | 0.26 | 6.64E-09 | 1.38E-04 | 0.50 | 0.48 | 0.02 | 1.05 |
| 0 | Mogat2 | 0.60 | 6.68E-09 | 1.39E-04 | 0.15 | 0.11 | 0.04 | 1.32 |
| 0 | Cnpy4 | 0.32 | 6.87E-09 | 1.43E-04 | 0.38 | 0.35 | 0.03 | 1.09 |
| 0 | Nif3l1 | 0.33 | 7.96E-09 | 1.66E-04 | 0.39 | 0.36 | 0.03 | 1.10 |
| 0 | Pxylp1 | 0.65 | 8.02E-09 | 1.67E-04 | 0.14 | 0.11 | 0.04 | 1.33 |
| 0 | 4933439C10Rik | 0.68 | 9.40E-09 | 1.96E-04 | 0.13 | 0.10 | 0.03 | 1.34 |
| 0 | Ncoa7 | 0.32 | 1.08E-08 | 2.24E-04 | 0.43 | 0.40 | 0.02 | 1.06 |
| 0 | Zfp518b | 0.34 | 1.25E-08 | 2.60E-04 | 0.39 | 0.36 | 0.03 | 1.09 |
| 0 | Hip1 | 0.27 | 1.32E-08 | 2.76E-04 | 0.47 | 0.45 | 0.03 | 1.06 |
| 0 | Camsap1 | 0.26 | 1.33E-08 | 2.76E-04 | 0.50 | 0.48 | 0.03 | 1.05 |
| 0 | Ccdc66 | 0.33 | 1.55E-08 | 3.22E-04 | 0.41 | 0.39 | 0.03 | 1.07 |
| 0 | Emp3 | 0.36 | 1.62E-08 | 3.36E-04 | 0.17 | 0.13 | 0.04 | 1.29 |
| 0 | Hacd4 | 0.33 | 1.62E-08 | 3.37E-04 | 0.29 | 0.25 | 0.04 | 1.17 |
| 0 | Lrrc20 | 0.59 | 1.63E-08 | 3.39E-04 | 0.18 | 0.15 | 0.04 | 1.24 |
| 0 | Xbp1 | 0.28 | 1.63E-08 | 3.40E-04 | 0.50 | 0.48 | 0.03 | 1.05 |
| 0 | Trp53inp2 | 0.59 | 1.71E-08 | 3.56E-04 | 0.13 | 0.09 | 0.03 | 1.35 |
| 0 | Akap8l | 0.35 | 1.74E-08 | 3.62E-04 | 0.40 | 0.37 | 0.03 | 1.08 |
| 0 | Limk1 | 0.66 | 1.78E-08 | 3.69E-04 | 0.18 | 0.15 | 0.04 | 1.24 |
| 0 | Arid1b | 0.29 | 2.34E-08 | 4.86E-04 | 0.50 | 0.48 | 0.01 | 1.03 |
| 0 | Inhba | 0.27 | 2.37E-08 | 4.93E-04 | 0.21 | 0.16 | 0.04 | 1.26 |
| 0 | Zfp846 | 0.45 | 2.51E-08 | 5.22E-04 | 0.28 | 0.25 | 0.04 | 1.14 |
| 0 | Gas8 | 0.29 | 2.54E-08 | 5.28E-04 | 0.42 | 0.39 | 0.03 | 1.08 |
| 0 | Tmem230 | 0.27 | 2.67E-08 | 5.55E-04 | 0.44 | 0.42 | 0.02 | 1.06 |
| 0 | Adamts10 | 0.40 | 2.73E-08 | 5.68E-04 | 0.35 | 0.32 | 0.03 | 1.10 |
| 0 | Lockd | 0.37 | 3.04E-08 | 6.32E-04 | 0.39 | 0.36 | 0.03 | 1.09 |
| 0 | Kirrel | 0.47 | 3.11E-08 | 6.46E-04 | 0.26 | 0.22 | 0.04 | 1.17 |
| 0 | Tmem222 | 0.30 | 3.23E-08 | 6.72E-04 | 0.47 | 0.45 | 0.02 | 1.04 |
| 0 | Zc2hc1a | 0.29 | 3.62E-08 | 7.52E-04 | 0.44 | 0.42 | 0.03 | 1.06 |
| 0 | Ddx3y | 0.71 | 3.91E-08 | 8.13E-04 | 0.10 | 0.07 | 0.03 | 1.41 |
| 0 | Smyd5 | 0.34 | 4.17E-08 | 8.67E-04 | 0.36 | 0.32 | 0.03 | 1.10 |
| 0 | Simc1 | 0.56 | 4.23E-08 | 8.79E-04 | 0.17 | 0.13 | 0.04 | 1.26 |
| 0 | Cbx2 | 0.28 | 4.27E-08 | 8.88E-04 | 0.45 | 0.43 | 0.03 | 1.06 |
| 0 | Oxr1 | 0.36 | 4.81E-08 | 1.00E-03 | 0.36 | 0.33 | 0.03 | 1.09 |
| 0 | Fez2 | 0.39 | 4.86E-08 | 1.01E-03 | 0.34 | 0.31 | 0.03 | 1.09 |
| 0 | Gadd45b | 0.41 | 4.95E-08 | 1.03E-03 | 0.18 | 0.14 | 0.04 | 1.26 |
| 0 | Rufy3 | 0.33 | 5.16E-08 | 1.07E-03 | 0.26 | 0.23 | 0.04 | 1.17 |
| 0 | Slc38a6 | 0.55 | 5.65E-08 | 1.18E-03 | 0.18 | 0.15 | 0.04 | 1.24 |
| 0 | Tmem218 | 0.39 | 5.77E-08 | 1.20E-03 | 0.33 | 0.30 | 0.03 | 1.11 |
| 0 | Rasl11a | 0.37 | 5.83E-08 | 1.21E-03 | 0.31 | 0.27 | 0.04 | 1.14 |
| 0 | Pheta1 | 0.34 | 5.91E-08 | 1.23E-03 | 0.35 | 0.32 | 0.03 | 1.10 |
| 0 | Ppp1r18 | 0.40 | 6.03E-08 | 1.25E-03 | 0.29 | 0.25 | 0.04 | 1.15 |
| 0 | Zyg11b | 0.34 | 6.27E-08 | 1.30E-03 | 0.37 | 0.34 | 0.03 | 1.09 |
| 0 | Gskip | 0.36 | 6.29E-08 | 1.31E-03 | 0.38 | 0.35 | 0.03 | 1.08 |
| 0 | Insig1 | 0.39 | 6.43E-08 | 1.34E-03 | 0.34 | 0.31 | 0.04 | 1.11 |
| 0 | Ggps1 | 0.26 | 6.89E-08 | 1.43E-03 | 0.45 | 0.42 | 0.03 | 1.06 |
| 0 | 2900076A07Rik | 0.37 | 6.98E-08 | 1.45E-03 | 0.34 | 0.30 | 0.03 | 1.11 |
| 0 | Camkk2 | 0.44 | 8.31E-08 | 1.73E-03 | 0.27 | 0.23 | 0.04 | 1.16 |
| 0 | S1pr2 | 0.34 | 8.61E-08 | 1.79E-03 | 0.35 | 0.32 | 0.03 | 1.10 |
| 0 | P3h1 | 0.51 | 8.77E-08 | 1.82E-03 | 0.18 | 0.14 | 0.04 | 1.24 |
| 0 | 2810013P06Rik | 0.39 | 1.03E-07 | 2.15E-03 | 0.33 | 0.29 | 0.03 | 1.11 |
| 0 | Pfkfb4 | 0.71 | 1.05E-07 | 2.18E-03 | 0.11 | 0.08 | 0.03 | 1.36 |
| 0 | Dennd5a | 0.34 | 1.13E-07 | 2.35E-03 | 0.37 | 0.35 | 0.03 | 1.08 |
| 0 | Cib2 | 0.39 | 1.19E-07 | 2.47E-03 | 0.29 | 0.25 | 0.03 | 1.13 |
| 0 | Usp11 | 0.34 | 1.19E-07 | 2.48E-03 | 0.37 | 0.34 | 0.03 | 1.08 |
| 0 | Ankrd50 | 0.46 | 1.24E-07 | 2.58E-03 | 0.26 | 0.23 | 0.03 | 1.15 |

**Supplementary Table 1. Marker genes for the original clusters of integrated clustering of E11.5 control and NotchGOF λ-junction cells.**

| | | | | | | | | |
|---|---|---|---|---|---|---|---|---|
| 0 | Ethe1 | 0.26 | 1.30E-07 | 2.70E-03 | 0.47 | 0.45 | 0.02 | 1.05 |
| 0 | Ppp2r5d | 0.28 | 1.32E-07 | 2.75E-03 | 0.48 | 0.47 | 0.02 | 1.03 |
| 0 | Mbtps2 | 0.45 | 1.45E-07 | 3.01E-03 | 0.25 | 0.22 | 0.03 | 1.15 |
| 0 | 5430416N02Rik | 0.31 | 1.47E-07 | 3.06E-03 | 0.48 | 0.46 | 0.02 | 1.04 |
| 0 | Ints8 | 0.26 | 1.48E-07 | 3.09E-03 | 0.45 | 0.43 | 0.02 | 1.05 |
| 0 | Smad6 | 0.39 | 1.51E-07 | 3.14E-03 | 0.28 | 0.25 | 0.03 | 1.14 |
| 0 | Rab2b | 0.36 | 1.53E-07 | 3.17E-03 | 0.31 | 0.27 | 0.04 | 1.13 |
| 0 | Slit3 | 0.30 | 1.61E-07 | 3.34E-03 | 0.23 | 0.19 | 0.04 | 1.21 |
| 0 | 3010003L21Rik | 0.36 | 1.64E-07 | 3.40E-03 | 0.34 | 0.31 | 0.03 | 1.10 |
| 0 | Gstm7 | 0.38 | 1.91E-07 | 3.98E-03 | 0.31 | 0.28 | 0.04 | 1.13 |
| 0 | Nln | 0.37 | 1.99E-07 | 4.13E-03 | 0.32 | 0.29 | 0.03 | 1.12 |
| 0 | Rbms2 | 0.26 | 2.10E-07 | 4.37E-03 | 0.48 | 0.47 | 0.02 | 1.03 |
| 0 | Timp1 | 0.50 | 2.16E-07 | 4.50E-03 | 0.20 | 0.17 | 0.03 | 1.20 |
| 0 | Ptges2 | 0.36 | 2.23E-07 | 4.63E-03 | 0.34 | 0.31 | 0.03 | 1.09 |
| 0 | Polr3gl | 0.32 | 2.24E-07 | 4.66E-03 | 0.34 | 0.31 | 0.03 | 1.10 |
| 0 | Coq10b | 0.41 | 2.41E-07 | 5.01E-03 | 0.28 | 0.25 | 0.03 | 1.12 |
| 0 | Sipa1l2 | 0.56 | 2.54E-07 | 5.29E-03 | 0.16 | 0.13 | 0.03 | 1.24 |
| 0 | Dyrk2 | 0.29 | 2.62E-07 | 5.46E-03 | 0.40 | 0.38 | 0.03 | 1.07 |
| 0 | Crmp1 | 0.26 | 2.64E-07 | 5.48E-03 | 0.15 | 0.12 | 0.03 | 1.29 |
| 0 | Tnks1bp1 | 0.51 | 2.77E-07 | 5.76E-03 | 0.20 | 0.17 | 0.03 | 1.19 |
| 0 | Chp1 | 0.32 | 2.90E-07 | 6.03E-03 | 0.41 | 0.39 | 0.02 | 1.05 |
| 0 | Prxl2b | 0.27 | 2.94E-07 | 6.11E-03 | 0.44 | 0.41 | 0.03 | 1.06 |
| 0 | Zadh2 | 0.60 | 2.97E-07 | 6.18E-03 | 0.14 | 0.10 | 0.03 | 1.30 |
| 0 | Gbf1 | 0.28 | 3.15E-07 | 6.54E-03 | 0.45 | 0.44 | 0.02 | 1.04 |
| 0 | Pcgf1 | 0.51 | 3.27E-07 | 6.80E-03 | 0.20 | 0.17 | 0.03 | 1.19 |
| 0 | Sh3kbp1 | 0.42 | 3.32E-07 | 6.90E-03 | 0.26 | 0.22 | 0.04 | 1.16 |
| 0 | Ric1 | 0.39 | 3.49E-07 | 7.27E-03 | 0.36 | 0.33 | 0.03 | 1.08 |
| 0 | Micu1 | 0.36 | 3.50E-07 | 7.27E-03 | 0.36 | 0.33 | 0.02 | 1.07 |
| 0 | Med22 | 0.37 | 3.68E-07 | 7.65E-03 | 0.36 | 0.34 | 0.02 | 1.07 |
| 0 | Nsdhl | 0.32 | 3.78E-07 | 7.87E-03 | 0.40 | 0.37 | 0.03 | 1.08 |
| 0 | Sgip1 | 0.58 | 3.84E-07 | 7.99E-03 | 0.14 | 0.11 | 0.03 | 1.29 |
| 0 | Ankrd29 | 0.62 | 3.90E-07 | 8.11E-03 | 0.12 | 0.09 | 0.03 | 1.32 |
| 0 | Pcyt1b | 0.51 | 4.00E-07 | 8.32E-03 | 0.15 | 0.12 | 0.03 | 1.27 |
| 0 | Gm49700 | 0.51 | 4.17E-07 | 8.67E-03 | 0.19 | 0.16 | 0.03 | 1.21 |
| 0 | Epop | 0.28 | 4.29E-07 | 8.92E-03 | 0.41 | 0.38 | 0.03 | 1.07 |
| 0 | Rhou | 0.25 | 4.40E-07 | 9.15E-03 | 0.44 | 0.41 | 0.03 | 1.07 |
| 0 | Eif4e3 | 0.35 | 4.70E-07 | 9.77E-03 | 0.31 | 0.28 | 0.03 | 1.12 |
| 0 | Zbtb1 | 0.31 | 4.71E-07 | 9.80E-03 | 0.38 | 0.35 | 0.03 | 1.08 |
| 0 | Plekhg2 | 0.28 | 4.84E-07 | 1.01E-02 | 0.40 | 0.38 | 0.03 | 1.07 |
| 0 | Adcy3 | 0.63 | 4.85E-07 | 1.01E-02 | 0.11 | 0.08 | 0.03 | 1.35 |
| 0 | Adgre5 | 0.55 | 5.29E-07 | 1.10E-02 | 0.16 | 0.12 | 0.03 | 1.26 |
| 0 | Mettl7a1 | 0.58 | 6.39E-07 | 1.33E-02 | 0.16 | 0.13 | 0.03 | 1.23 |
| 0 | Phyh | 0.30 | 6.45E-07 | 1.34E-02 | 0.32 | 0.29 | 0.03 | 1.11 |
| 0 | Ebp | 0.30 | 6.53E-07 | 1.36E-02 | 0.42 | 0.40 | 0.02 | 1.04 |
| 0 | Alkbh8 | 0.30 | 7.27E-07 | 1.51E-02 | 0.44 | 0.42 | 0.02 | 1.04 |
| 0 | Actr1a | 0.26 | 8.40E-07 | 1.75E-02 | 0.48 | 0.46 | 0.01 | 1.03 |
| 0 | B230354K17Rik | 0.34 | 8.51E-07 | 1.77E-02 | 0.38 | 0.36 | 0.03 | 1.07 |
| 0 | Lrp3 | 0.45 | 8.63E-07 | 1.79E-02 | 0.25 | 0.21 | 0.03 | 1.15 |
| 0 | Spire1 | 0.34 | 9.61E-07 | 2.00E-02 | 0.37 | 0.35 | 0.02 | 1.06 |
| 0 | Leng1 | 0.41 | 1.07E-06 | 2.24E-02 | 0.26 | 0.23 | 0.03 | 1.13 |
| 0 | Cand2 | 0.26 | 1.12E-06 | 2.32E-02 | 0.50 | 0.49 | 0.01 | 1.02 |
| 0 | Snhg20 | 0.37 | 1.21E-06 | 2.52E-02 | 0.39 | 0.38 | 0.01 | 1.04 |
| 0 | Ahcy | 0.39 | 1.22E-06 | 2.54E-02 | 0.30 | 0.27 | 0.03 | 1.10 |
| 0 | Unc119b | 0.35 | 1.26E-06 | 2.62E-02 | 0.37 | 0.34 | 0.02 | 1.06 |
| 0 | G0s2 | 0.50 | 1.31E-06 | 2.73E-02 | 0.18 | 0.15 | 0.03 | 1.21 |
| 0 | Ccdc30 | 0.55 | 1.34E-06 | 2.79E-02 | 0.13 | 0.10 | 0.03 | 1.29 |

**Supplementary Table 1. Marker genes for the original clusters of integrated clustering of E11.5 control and NotchGOF λ-junction cells.**

| | | | | | | | | |
|---|---|---|---|---|---|---|---|---|
| 0 | Zbed5 | 0.52 | 1.36E-06 | 2.83E-02 | 0.18 | 0.15 | 0.03 | 1.20 |
| 0 | Ppp2r1b | 0.27 | 1.41E-06 | 2.93E-02 | 0.41 | 0.40 | 0.02 | 1.05 |
| 0 | Osbpl9 | 0.25 | 1.42E-06 | 2.96E-02 | 0.45 | 0.43 | 0.02 | 1.05 |
| 0 | Trim8 | 0.28 | 1.50E-06 | 3.11E-02 | 0.33 | 0.30 | 0.03 | 1.11 |
| 0 | Afap1 | 0.34 | 1.51E-06 | 3.15E-02 | 0.29 | 0.25 | 0.03 | 1.13 |
| 0 | Pik3ca | 0.29 | 1.62E-06 | 3.36E-02 | 0.45 | 0.43 | 0.02 | 1.04 |
| 0 | Zfp945 | 0.31 | 1.62E-06 | 3.37E-02 | 0.38 | 0.36 | 0.02 | 1.06 |
| 0 | 4921531C22Rik | 0.49 | 1.67E-06 | 3.48E-02 | 0.21 | 0.18 | 0.03 | 1.18 |
| 0 | Ccl25 | 0.34 | 1.75E-06 | 3.63E-02 | 0.30 | 0.27 | 0.03 | 1.11 |
| 0 | Tmtc2 | 0.41 | 1.79E-06 | 3.73E-02 | 0.23 | 0.20 | 0.03 | 1.17 |
| 0 | Ccl27a | 0.34 | 1.82E-06 | 3.80E-02 | 0.33 | 0.30 | 0.03 | 1.09 |
| 0 | Tet1 | 0.32 | 1.83E-06 | 3.81E-02 | 0.37 | 0.35 | 0.03 | 1.07 |
| 0 | 2610027K06Rik | 0.49 | 1.83E-06 | 3.81E-02 | 0.16 | 0.13 | 0.03 | 1.24 |
| 0 | Xpc | 0.34 | 1.85E-06 | 3.84E-02 | 0.36 | 0.33 | 0.02 | 1.07 |
| 0 | Cdkl3 | 0.53 | 1.89E-06 | 3.92E-02 | 0.16 | 0.14 | 0.03 | 1.21 |
| 0 | Ormdl3 | 0.27 | 1.96E-06 | 4.08E-02 | 0.47 | 0.46 | 0.01 | 1.02 |
| 0 | Foxf2 | 0.25 | 2.02E-06 | 4.20E-02 | 0.12 | 0.09 | 0.03 | 1.33 |
| 0 | Myof | 0.50 | 2.21E-06 | 4.60E-02 | 0.12 | 0.09 | 0.03 | 1.32 |
| 0 | Prex1 | 0.38 | 2.24E-06 | 4.66E-02 | 0.22 | 0.19 | 0.03 | 1.16 |
| 0 | Ttc23 | 0.32 | 2.40E-06 | 4.99E-02 | 0.33 | 0.30 | 0.03 | 1.09 |
| 0 | Mras | 0.41 | 2.40E-06 | 4.99E-02 | 0.24 | 0.21 | 0.03 | 1.15 |
| 1 | Pax7 | 2.48 | 0 | 0 | 0.83 | 0.17 | 0.66 | 5.00 |
| 1 | Nbl1 | 1.69 | 0 | 0 | 0.95 | 0.44 | 0.51 | 2.17 |
| 1 | Igf1 | 1.96 | 0 | 0 | 0.93 | 0.42 | 0.51 | 2.20 |
| 1 | Pax3 | 1.34 | 0 | 0 | 0.88 | 0.41 | 0.47 | 2.17 |
| 1 | Hpgd | 1.77 | 0 | 0 | 0.69 | 0.31 | 0.39 | 2.25 |
| 1 | Rarb | 1.44 | 0 | 0 | 0.69 | 0.31 | 0.38 | 2.21 |
| 1 | Jun | 1.48 | 0 | 0 | 0.97 | 0.64 | 0.33 | 1.52 |
| 1 | Lix1 | 1.30 | 0 | 0 | 0.92 | 0.59 | 0.33 | 1.55 |
| 1 | Gas1 | 1.02 | 0 | 0 | 0.95 | 0.67 | 0.28 | 1.41 |
| 1 | Gm30564 | 2.67 | 0 | 0 | 0.34 | 0.07 | 0.27 | 5.09 |
| 1 | Twist1 | 0.94 | 0 | 0 | 1.00 | 0.74 | 0.26 | 1.34 |
| 1 | Lgals1 | 0.80 | 0 | 0 | 1.00 | 0.81 | 0.19 | 1.24 |
| 1 | Crabp1 | 1.28 | 0 | 0 | 1.00 | 0.84 | 0.16 | 1.19 |
| 1 | Micos10 | 0.86 | 0 | 0 | 1.00 | 0.96 | 0.04 | 1.04 |
| 1 | Nme2 | 0.44 | 0 | 0 | 1.00 | 0.98 | 0.02 | 1.02 |
| 1 | Txn1 | 0.50 | 0 | 0 | 1.00 | 0.99 | 0.01 | 1.01 |
| 1 | Tcf15 | 1.53 | 2.13E-298 | 4.42E-294 | 0.61 | 0.24 | 0.36 | 2.48 |
| 1 | Ptn | 0.76 | 2.53E-292 | 5.27E-288 | 1.00 | 0.89 | 0.11 | 1.12 |
| 1 | Gm9866 | 2.79 | 8.45E-283 | 1.76E-278 | 0.25 | 0.05 | 0.21 | 5.29 |
| 1 | Septin11 | 0.78 | 2.09E-281 | 4.35E-277 | 0.98 | 0.86 | 0.12 | 1.14 |
| 1 | Alx1 | 0.80 | 1.22E-274 | 2.54E-270 | 0.99 | 0.70 | 0.29 | 1.42 |
| 1 | Tmsb10 | 0.37 | 1.99E-262 | 4.14E-258 | 1.00 | 1.00 | 0.00 | 1.00 |
| 1 | Prdm16os | 2.18 | 4.12E-258 | 8.56E-254 | 0.34 | 0.09 | 0.25 | 3.67 |
| 1 | Rnf128 | 2.21 | 5.07E-257 | 1.05E-252 | 0.38 | 0.12 | 0.26 | 3.24 |
| 1 | Dkk2 | 1.47 | 1.88E-252 | 3.90E-248 | 0.56 | 0.23 | 0.32 | 2.40 |
| 1 | Vcam1 | 1.15 | 1.67E-246 | 3.47E-242 | 0.74 | 0.40 | 0.34 | 1.86 |
| 1 | Tnfaip6 | 1.07 | 4.51E-240 | 9.38E-236 | 0.78 | 0.43 | 0.34 | 1.79 |
| 1 | Crabp2 | 0.59 | 8.99E-233 | 1.87E-228 | 1.00 | 0.91 | 0.09 | 1.10 |
| 1 | Cntfr | 1.56 | 1.10E-229 | 2.29E-225 | 0.48 | 0.19 | 0.29 | 2.52 |
| 1 | Crlf1 | 1.03 | 2.77E-226 | 5.77E-222 | 0.88 | 0.61 | 0.27 | 1.44 |
| 1 | Rspo2 | 1.30 | 6.65E-224 | 1.38E-219 | 0.66 | 0.34 | 0.32 | 1.95 |
| 1 | Fap | 1.59 | 1.17E-223 | 2.43E-219 | 0.45 | 0.17 | 0.28 | 2.65 |
| 1 | Hmgb1 | 0.32 | 1.29E-222 | 2.69E-218 | 1.00 | 1.00 | 0.00 | 1.00 |
| 1 | Tcf12 | 0.67 | 2.46E-222 | 5.12E-218 | 0.98 | 0.86 | 0.12 | 1.14 |
| 1 | Igfbp4 | 0.60 | 1.14E-219 | 2.37E-215 | 1.00 | 0.89 | 0.11 | 1.12 |

**Supplementary Table 1. Marker genes for the original clusters of integrated clustering of E11.5 control and NotchGOF λ-junction cells.**

| | | | | | | | | |
|---|---|---|---|---|---|---|---|---|
| 1 | Anxa7 | 1.40 | 1.62E-218 | 3.38E-214 | 0.62 | 0.33 | 0.29 | 1.86 |
| 1 | Lpar1 | 0.80 | 2.14E-218 | 4.44E-214 | 0.94 | 0.68 | 0.27 | 1.39 |
| 1 | Gch1 | 2.52 | 8.63E-208 | 1.79E-203 | 0.24 | 0.06 | 0.18 | 4.14 |
| 1 | Gata2 | 1.05 | 5.99E-202 | 1.25E-197 | 0.73 | 0.41 | 0.32 | 1.79 |
| 1 | H2afz | 0.34 | 6.52E-195 | 1.36E-190 | 1.00 | 1.00 | 0.00 | 1.00 |
| 1 | Akr7a5 | 0.72 | 2.01E-187 | 4.18E-183 | 0.93 | 0.77 | 0.16 | 1.20 |
| 1 | Capzb | 0.55 | 2.15E-186 | 4.47E-182 | 0.99 | 0.91 | 0.08 | 1.08 |
| 1 | Sox11 | 0.44 | 6.70E-186 | 1.39E-181 | 1.00 | 0.96 | 0.04 | 1.04 |
| 1 | H2afy | 0.41 | 2.02E-184 | 4.19E-180 | 1.00 | 0.96 | 0.04 | 1.04 |
| 1 | Hspe1 | 0.41 | 4.86E-183 | 1.01E-178 | 1.00 | 0.99 | 0.01 | 1.01 |
| 1 | Ybx1 | 0.30 | 1.23E-181 | 2.55E-177 | 1.00 | 1.00 | 0.00 | 1.00 |
| 1 | Il11ra1 | 0.78 | 5.56E-181 | 1.16E-176 | 0.86 | 0.59 | 0.28 | 1.47 |
| 1 | Ran | 0.33 | 4.58E-174 | 9.53E-170 | 1.00 | 1.00 | 0.00 | 1.00 |
| 1 | Prdm16 | 1.19 | 7.85E-167 | 1.63E-162 | 0.57 | 0.30 | 0.27 | 1.90 |
| 1 | Gsc | 1.18 | 3.07E-164 | 6.39E-160 | 0.54 | 0.27 | 0.27 | 2.00 |
| 1 | Sh3gl3 | 1.14 | 2.14E-163 | 4.44E-159 | 0.58 | 0.30 | 0.27 | 1.90 |
| 1 | Arhgdib | 0.87 | 1.09E-162 | 2.26E-158 | 0.74 | 0.47 | 0.27 | 1.57 |
| 1 | Hmgb3 | 0.38 | 5.67E-162 | 1.18E-157 | 1.00 | 0.99 | 0.01 | 1.01 |
| 1 | Bzw1 | 0.40 | 1.32E-161 | 2.76E-157 | 1.00 | 0.98 | 0.02 | 1.02 |
| 1 | Prrx2 | 0.62 | 1.56E-160 | 3.24E-156 | 0.97 | 0.72 | 0.26 | 1.36 |
| 1 | Jaml | 3.04 | 1.70E-160 | 3.54E-156 | 0.12 | 0.02 | 0.10 | 6.72 |
| 1 | Limd2 | 0.49 | 1.09E-157 | 2.26E-153 | 0.99 | 0.89 | 0.11 | 1.12 |
| 1 | Reep5 | 0.68 | 6.99E-154 | 1.45E-149 | 0.92 | 0.77 | 0.15 | 1.20 |
| 1 | Csrp1 | 1.08 | 9.50E-152 | 1.98E-147 | 0.57 | 0.31 | 0.26 | 1.82 |
| 1 | Sec61b | 0.39 | 1.75E-150 | 3.64E-146 | 1.00 | 0.98 | 0.01 | 1.01 |
| 1 | Nme1 | 0.37 | 1.42E-145 | 2.96E-141 | 1.00 | 0.99 | 0.01 | 1.01 |
| 1 | Hmgn1 | 0.26 | 2.89E-145 | 6.01E-141 | 1.00 | 0.99 | 0.01 | 1.01 |
| 1 | Lsm6 | 0.45 | 1.65E-144 | 3.43E-140 | 0.98 | 0.93 | 0.05 | 1.05 |
| 1 | Rflnb | 0.80 | 1.30E-141 | 2.70E-137 | 0.76 | 0.53 | 0.23 | 1.44 |
| 1 | Gap43 | 0.73 | 4.76E-135 | 9.91E-131 | 0.66 | 0.39 | 0.27 | 1.70 |
| 1 | Ctsb | 0.52 | 1.16E-134 | 2.40E-130 | 0.89 | 0.73 | 0.16 | 1.21 |
| 1 | Enho | 0.90 | 3.47E-134 | 7.22E-130 | 0.67 | 0.42 | 0.24 | 1.58 |
| 1 | Hmga2 | 0.51 | 2.32E-133 | 4.83E-129 | 0.99 | 0.92 | 0.07 | 1.07 |
| 1 | Cenpa | 0.62 | 8.05E-132 | 1.67E-127 | 0.91 | 0.78 | 0.13 | 1.16 |
| 1 | Fscn1 | 0.44 | 1.51E-131 | 3.14E-127 | 0.99 | 0.90 | 0.09 | 1.09 |
| 1 | Hsp90b1 | 0.39 | 6.98E-130 | 1.45E-125 | 1.00 | 0.97 | 0.03 | 1.03 |
| 1 | Cox7a2 | 0.29 | 4.48E-128 | 9.31E-124 | 1.00 | 0.99 | 0.01 | 1.01 |
| 1 | Spin2c | 0.91 | 2.68E-127 | 5.57E-123 | 0.64 | 0.39 | 0.24 | 1.61 |
| 1 | Vcan | 0.44 | 4.67E-127 | 9.72E-123 | 0.98 | 0.85 | 0.13 | 1.16 |
| 1 | Selenoh | 0.36 | 6.45E-126 | 1.34E-121 | 1.00 | 0.96 | 0.04 | 1.04 |
| 1 | Mex3b | 0.65 | 3.26E-125 | 6.78E-121 | 0.86 | 0.68 | 0.19 | 1.27 |
| 1 | Mogat2 | 1.65 | 3.63E-125 | 7.56E-121 | 0.27 | 0.10 | 0.17 | 2.71 |
| 1 | 1110017D15Rik | 1.81 | 1.31E-124 | 2.72E-120 | 0.21 | 0.07 | 0.15 | 3.19 |
| 1 | Tbx2 | 0.56 | 5.77E-124 | 1.20E-119 | 0.89 | 0.62 | 0.27 | 1.44 |
| 1 | D030045P18Rik | 2.21 | 1.04E-123 | 2.17E-119 | 0.15 | 0.03 | 0.11 | 4.29 |
| 1 | Cct2 | 0.32 | 3.47E-122 | 7.22E-118 | 1.00 | 0.98 | 0.02 | 1.02 |
| 1 | Eif5a | 0.28 | 6.04E-122 | 1.26E-117 | 1.00 | 1.00 | 0.00 | 1.00 |
| 1 | Pcsk5 | 1.01 | 9.63E-122 | 2.00E-117 | 0.58 | 0.34 | 0.24 | 1.69 |
| 1 | Eif2s2 | 0.33 | 1.64E-120 | 3.41E-116 | 1.00 | 0.99 | 0.01 | 1.01 |
| 1 | Manf | 0.52 | 2.28E-120 | 4.75E-116 | 0.97 | 0.89 | 0.08 | 1.09 |
| 1 | Cycs | 0.39 | 1.15E-119 | 2.39E-115 | 1.00 | 0.96 | 0.04 | 1.04 |
| 1 | Apod | 2.60 | 1.96E-119 | 4.09E-115 | 0.11 | 0.02 | 0.09 | 5.58 |
| 1 | Rassf3 | 1.03 | 4.75E-118 | 9.88E-114 | 0.57 | 0.35 | 0.22 | 1.64 |
| 1 | Gxylt2 | 1.10 | 4.88E-116 | 1.01E-111 | 0.45 | 0.23 | 0.22 | 1.96 |
| 1 | Cped1 | 0.72 | 4.00E-114 | 8.32E-110 | 0.70 | 0.44 | 0.26 | 1.58 |
| 1 | Adgre5 | 1.48 | 1.28E-113 | 2.67E-109 | 0.28 | 0.11 | 0.17 | 2.54 |

**Supplementary Table 1. Marker genes for the original clusters of integrated clustering of E11.5 control and NotchGOF λ-junction cells.**

| | | | | | | | | |
|---|---|---|---|---|---|---|---|---|
| 1 | Hpse2 | 1.18 | 6.00E-112 | 1.25E-107 | 0.35 | 0.15 | 0.19 | 2.28 |
| 1 | Hsd17b10 | 0.43 | 6.52E-112 | 1.36E-107 | 0.97 | 0.90 | 0.07 | 1.08 |
| 1 | Pdia6 | 0.37 | 1.11E-111 | 2.31E-107 | 1.00 | 0.95 | 0.05 | 1.05 |
| 1 | Car14 | 0.82 | 8.14E-111 | 1.69E-106 | 0.63 | 0.40 | 0.23 | 1.58 |
| 1 | Rps27l | 0.29 | 1.01E-110 | 2.10E-106 | 1.00 | 0.99 | 0.01 | 1.01 |
| 1 | Wnt5a | 0.66 | 1.82E-110 | 3.78E-106 | 0.83 | 0.62 | 0.21 | 1.34 |
| 1 | Dixdc1 | 0.89 | 1.19E-109 | 2.48E-105 | 0.60 | 0.37 | 0.23 | 1.62 |
| 1 | Rassf9 | 1.47 | 5.23E-109 | 1.09E-104 | 0.32 | 0.15 | 0.18 | 2.21 |
| 1 | Dnm3 | 1.32 | 5.30E-109 | 1.10E-104 | 0.33 | 0.14 | 0.18 | 2.26 |
| 1 | Cox7b | 0.30 | 2.56E-107 | 5.31E-103 | 1.00 | 0.98 | 0.02 | 1.02 |
| 1 | Twistnb | 0.64 | 1.06E-106 | 2.21E-102 | 0.88 | 0.76 | 0.12 | 1.16 |
| 1 | Bmerb1 | 1.68 | 2.43E-106 | 5.06E-102 | 0.21 | 0.07 | 0.14 | 2.89 |
| 1 | Cd34 | 0.47 | 2.97E-106 | 6.18E-102 | 0.28 | 0.12 | 0.17 | 2.47 |
| 1 | Emilin2 | 1.04 | 7.13E-106 | 1.48E-101 | 0.44 | 0.24 | 0.21 | 1.89 |
| 1 | Tpm4 | 0.38 | 1.26E-105 | 2.61E-101 | 0.99 | 0.93 | 0.07 | 1.07 |
| 1 | Atp5f1 | 0.25 | 2.78E-105 | 5.79E-101 | 1.00 | 0.99 | 0.01 | 1.01 |
| 1 | Dnaja1 | 0.30 | 7.11E-105 | 1.48E-100 | 1.00 | 0.99 | 0.01 | 1.01 |
| 1 | Dpep1 | 0.99 | 2.22E-104 | 4.63E-100 | 0.41 | 0.21 | 0.21 | 2.00 |
| 1 | Ccne1 | 0.78 | 7.11E-104 | 1.48E-99 | 0.69 | 0.48 | 0.21 | 1.43 |
| 1 | Sulf2 | 0.60 | 1.84E-102 | 3.84E-98 | 0.84 | 0.65 | 0.19 | 1.30 |
| 1 | Tmem119 | 0.70 | 7.18E-102 | 1.49E-97 | 0.67 | 0.43 | 0.24 | 1.56 |
| 1 | Nucks1 | 0.33 | 1.71E-101 | 3.55E-97 | 1.00 | 0.98 | 0.02 | 1.02 |
| 1 | Lsm5 | 0.38 | 2.69E-101 | 5.60E-97 | 0.98 | 0.94 | 0.05 | 1.05 |
| 1 | Atp5k | 0.32 | 1.90E-100 | 3.95E-96 | 1.00 | 0.97 | 0.02 | 1.02 |
| 1 | Ccnb2 | 0.70 | 8.78E-99 | 1.83E-94 | 0.75 | 0.58 | 0.17 | 1.29 |
| 1 | Cox5a | 0.28 | 1.09E-98 | 2.27E-94 | 1.00 | 0.99 | 0.01 | 1.01 |
| 1 | C630043F03Rik | 1.39 | 2.84E-98 | 5.91E-94 | 0.30 | 0.13 | 0.16 | 2.25 |
| 1 | Uqcc2 | 0.34 | 1.15E-97 | 2.39E-93 | 0.99 | 0.95 | 0.04 | 1.04 |
| 1 | Samd4 | 0.86 | 1.66E-97 | 3.44E-93 | 0.56 | 0.35 | 0.21 | 1.61 |
| 1 | Cyp51 | 0.63 | 3.33E-96 | 6.93E-92 | 0.83 | 0.67 | 0.16 | 1.24 |
| 1 | Pdcd5 | 0.33 | 1.17E-95 | 2.44E-91 | 0.99 | 0.97 | 0.02 | 1.02 |
| 1 | Eif3e | 0.30 | 1.27E-95 | 2.64E-91 | 0.99 | 0.98 | 0.01 | 1.01 |
| 1 | Scd1 | 1.17 | 2.53E-95 | 5.26E-91 | 0.35 | 0.17 | 0.18 | 2.05 |
| 1 | Traf3ip2 | 1.05 | 8.30E-95 | 1.73E-90 | 0.39 | 0.20 | 0.19 | 1.94 |
| 1 | Ranbp1 | 0.27 | 9.29E-95 | 1.93E-90 | 1.00 | 0.99 | 0.01 | 1.01 |
| 1 | Gm49708 | 0.67 | 2.46E-94 | 5.13E-90 | 0.78 | 0.60 | 0.18 | 1.30 |
| 1 | Asph | 0.58 | 2.16E-93 | 4.50E-89 | 0.81 | 0.63 | 0.18 | 1.29 |
| 1 | Arl2bp | 0.47 | 4.63E-93 | 9.62E-89 | 0.91 | 0.78 | 0.13 | 1.17 |
| 1 | Dusp9 | 0.81 | 5.37E-93 | 1.12E-88 | 0.60 | 0.39 | 0.20 | 1.52 |
| 1 | Ret | 2.04 | 1.26E-92 | 2.62E-88 | 0.16 | 0.05 | 0.11 | 3.16 |
| 1 | Zmat2 | 0.36 | 3.81E-91 | 7.92E-87 | 0.98 | 0.94 | 0.04 | 1.04 |
| 1 | Rangrf | 0.69 | 1.43E-90 | 2.97E-86 | 0.70 | 0.52 | 0.18 | 1.35 |
| 1 | Calm3 | 0.37 | 5.98E-90 | 1.24E-85 | 0.97 | 0.91 | 0.06 | 1.07 |
| 1 | Cct6a | 0.29 | 9.41E-90 | 1.96E-85 | 1.00 | 0.98 | 0.02 | 1.02 |
| 1 | Glipr2 | 0.79 | 1.64E-89 | 3.41E-85 | 0.57 | 0.37 | 0.21 | 1.56 |
| 1 | Rtraf | 0.28 | 2.40E-89 | 5.00E-85 | 1.00 | 0.98 | 0.02 | 1.02 |
| 1 | Mif | 0.26 | 1.47E-88 | 3.06E-84 | 1.00 | 0.99 | 0.01 | 1.01 |
| 1 | Ppdpf | 0.32 | 2.20E-88 | 4.57E-84 | 1.00 | 0.96 | 0.04 | 1.04 |
| 1 | Marcks | 0.29 | 8.02E-88 | 1.67E-83 | 1.00 | 0.97 | 0.03 | 1.03 |
| 1 | Pdgfra | 0.52 | 1.14E-87 | 2.37E-83 | 0.76 | 0.52 | 0.24 | 1.47 |
| 1 | Wif1 | 1.60 | 4.52E-87 | 9.39E-83 | 0.21 | 0.08 | 0.13 | 2.60 |
| 1 | Rai14 | 0.51 | 1.05E-86 | 2.19E-82 | 0.83 | 0.63 | 0.21 | 1.33 |
| 1 | Ppp1r14b | 0.26 | 1.31E-86 | 2.73E-82 | 1.00 | 0.99 | 0.01 | 1.01 |
| 1 | Pttg1 | 0.54 | 1.50E-86 | 3.13E-82 | 0.83 | 0.70 | 0.13 | 1.18 |
| 1 | Pth1r | 0.52 | 3.12E-86 | 6.49E-82 | 0.79 | 0.55 | 0.24 | 1.45 |
| 1 | Nkd1 | 0.83 | 5.16E-86 | 1.07E-81 | 0.55 | 0.34 | 0.20 | 1.59 |

**Supplementary Table 1. Marker genes for the original clusters of integrated clustering of E11.5 control and NotchGOF λ-junction cells.**

| | | | | | | | | |
|---|---|---|---|---|---|---|---|---|
| 1 | Asns | 0.77 | 8.34E-86 | 1.73E-81 | 0.61 | 0.42 | 0.19 | 1.44 |
| 1 | Rev3l | 0.61 | 4.42E-85 | 9.20E-81 | 0.75 | 0.57 | 0.18 | 1.31 |
| 1 | Sfpq | 0.28 | 1.01E-84 | 2.11E-80 | 1.00 | 0.98 | 0.02 | 1.02 |
| 1 | Col23a1 | 0.83 | 2.16E-84 | 4.49E-80 | 0.49 | 0.29 | 0.20 | 1.68 |
| 1 | Prdx6 | 0.33 | 3.04E-84 | 6.32E-80 | 0.99 | 0.96 | 0.03 | 1.03 |
| 1 | Ostc | 0.33 | 3.08E-84 | 6.40E-80 | 0.99 | 0.94 | 0.04 | 1.04 |
| 1 | Mpped2 | 0.53 | 9.71E-83 | 2.02E-78 | 0.92 | 0.84 | 0.08 | 1.10 |
| 1 | Gsto1 | 0.56 | 1.24E-82 | 2.58E-78 | 0.80 | 0.63 | 0.16 | 1.26 |
| 1 | Ptgr1 | 0.98 | 8.61E-82 | 1.79E-77 | 0.43 | 0.25 | 0.18 | 1.73 |
| 1 | St6galnac3 | 0.81 | 1.67E-81 | 3.47E-77 | 0.56 | 0.38 | 0.19 | 1.50 |
| 1 | Tmem132c | 0.76 | 1.22E-80 | 2.55E-76 | 0.50 | 0.30 | 0.20 | 1.66 |
| 1 | Psma4 | 0.29 | 1.27E-80 | 2.64E-76 | 0.99 | 0.96 | 0.03 | 1.03 |
| 1 | Snai1 | 0.68 | 1.48E-80 | 3.08E-76 | 0.60 | 0.39 | 0.21 | 1.53 |
| 1 | Psma3 | 0.27 | 2.79E-80 | 5.79E-76 | 1.00 | 0.98 | 0.02 | 1.02 |
| 1 | Pmaip1 | 1.07 | 3.57E-80 | 7.43E-76 | 0.40 | 0.23 | 0.17 | 1.77 |
| 1 | Fbln1 | 0.40 | 5.16E-80 | 1.07E-75 | 0.93 | 0.78 | 0.14 | 1.18 |
| 1 | Irs2 | 0.76 | 1.07E-79 | 2.23E-75 | 0.60 | 0.42 | 0.18 | 1.44 |
| 1 | Btg1 | 0.53 | 1.18E-79 | 2.45E-75 | 0.89 | 0.78 | 0.11 | 1.14 |
| 1 | Cct8 | 0.26 | 1.20E-79 | 2.51E-75 | 1.00 | 0.98 | 0.02 | 1.02 |
| 1 | Hnrnpdl | 0.31 | 1.64E-79 | 3.42E-75 | 1.00 | 0.96 | 0.03 | 1.03 |
| 1 | Ssbp3 | 0.34 | 1.84E-79 | 3.82E-75 | 0.98 | 0.93 | 0.05 | 1.05 |
| 1 | Ifngr2 | 0.77 | 1.85E-79 | 3.84E-75 | 0.54 | 0.35 | 0.19 | 1.53 |
| 1 | Timm8b | 0.31 | 1.11E-78 | 2.31E-74 | 0.98 | 0.95 | 0.03 | 1.04 |
| 1 | Ndufc1 | 0.28 | 2.70E-78 | 5.61E-74 | 0.99 | 0.97 | 0.02 | 1.02 |
| 1 | Ruvbl1 | 0.42 | 2.89E-78 | 6.02E-74 | 0.93 | 0.84 | 0.09 | 1.11 |
| 1 | Cbx1 | 0.25 | 3.88E-78 | 8.06E-74 | 1.00 | 0.98 | 0.02 | 1.02 |
| 1 | Sox8 | 1.83 | 5.23E-78 | 1.09E-73 | 0.15 | 0.05 | 0.10 | 3.06 |
| 1 | Cnn2 | 0.41 | 8.35E-77 | 1.74E-72 | 0.81 | 0.61 | 0.21 | 1.34 |
| 1 | Ifitm1 | 0.70 | 9.14E-77 | 1.90E-72 | 0.33 | 0.17 | 0.16 | 1.92 |
| 1 | Clmp | 0.44 | 1.38E-76 | 2.87E-72 | 0.85 | 0.62 | 0.24 | 1.38 |
| 1 | Eif1b | 0.34 | 1.40E-76 | 2.92E-72 | 0.98 | 0.92 | 0.05 | 1.06 |
| 1 | Tlnrd1 | 0.66 | 2.10E-76 | 4.36E-72 | 0.66 | 0.48 | 0.18 | 1.36 |
| 1 | Dbi | 0.27 | 2.77E-76 | 5.76E-72 | 1.00 | 0.97 | 0.02 | 1.02 |
| 1 | Pcdh18 | 0.44 | 1.55E-75 | 3.22E-71 | 0.84 | 0.61 | 0.23 | 1.37 |
| 1 | Ppa1 | 0.34 | 1.82E-75 | 3.79E-71 | 0.97 | 0.89 | 0.08 | 1.09 |
| 1 | Runx1t1 | 0.48 | 5.71E-75 | 1.19E-70 | 0.73 | 0.51 | 0.22 | 1.43 |
| 1 | Col25a1 | 0.98 | 3.72E-74 | 7.74E-70 | 0.34 | 0.18 | 0.16 | 1.89 |
| 1 | Gng2 | 0.58 | 4.01E-74 | 8.34E-70 | 0.73 | 0.55 | 0.18 | 1.32 |
| 1 | Ndufs5 | 0.31 | 8.58E-74 | 1.78E-69 | 0.98 | 0.95 | 0.04 | 1.04 |
| 1 | Nasp | 0.26 | 8.71E-74 | 1.81E-69 | 1.00 | 0.98 | 0.02 | 1.02 |
| 1 | Rbpj | 0.51 | 1.71E-73 | 3.55E-69 | 0.83 | 0.69 | 0.14 | 1.20 |
| 1 | Ube2s | 0.34 | 1.12E-72 | 2.33E-68 | 0.99 | 0.97 | 0.02 | 1.02 |
| 1 | Lima1 | 0.43 | 1.52E-72 | 3.17E-68 | 0.89 | 0.72 | 0.17 | 1.23 |
| 1 | 2810001G20Rik | 0.74 | 2.65E-72 | 5.51E-68 | 0.55 | 0.37 | 0.18 | 1.49 |
| 1 | Snhg4 | 0.68 | 4.09E-72 | 8.51E-68 | 0.62 | 0.45 | 0.17 | 1.38 |
| 1 | Pid1 | 0.96 | 5.26E-72 | 1.09E-67 | 0.34 | 0.18 | 0.16 | 1.89 |
| 1 | Ddt | 0.39 | 8.44E-72 | 1.75E-67 | 0.93 | 0.84 | 0.09 | 1.11 |
| 1 | Atp1b3 | 0.50 | 1.15E-71 | 2.39E-67 | 0.84 | 0.70 | 0.14 | 1.19 |
| 1 | Gng10 | 0.41 | 1.24E-71 | 2.58E-67 | 0.92 | 0.80 | 0.11 | 1.14 |
| 1 | Rwdd1 | 0.33 | 4.72E-71 | 9.81E-67 | 0.97 | 0.93 | 0.04 | 1.05 |
| 1 | Col26a1 | 0.31 | 8.15E-71 | 1.70E-66 | 0.85 | 0.58 | 0.26 | 1.45 |
| 1 | Mrpl11 | 0.34 | 1.41E-70 | 2.94E-66 | 0.96 | 0.90 | 0.06 | 1.07 |
| 1 | Ddx39b | 0.27 | 1.70E-69 | 3.54E-65 | 1.00 | 0.97 | 0.03 | 1.03 |
| 1 | Dynlrb1 | 0.29 | 1.99E-69 | 4.15E-65 | 0.98 | 0.94 | 0.04 | 1.04 |
| 1 | S1pr3 | 0.81 | 2.52E-69 | 5.24E-65 | 0.43 | 0.26 | 0.18 | 1.68 |
| 1 | Hmgcr | 0.53 | 2.87E-69 | 5.97E-65 | 0.79 | 0.66 | 0.14 | 1.21 |

**Supplementary Table 1. Marker genes for the original clusters of integrated clustering of E11.5 control and NotchGOF λ-junction cells.**

| | | | | | | | | |
|---|---|---|---|---|---|---|---|---|
| 1 | Hnrnph3 | 0.38 | 4.27E-69 | 8.88E-65 | 0.93 | 0.85 | 0.09 | 1.10 |
| 1 | Efna2 | 0.44 | 4.88E-69 | 1.01E-64 | 0.84 | 0.69 | 0.15 | 1.22 |
| 1 | Ifitm2 | 0.29 | 5.57E-69 | 1.16E-64 | 1.00 | 0.94 | 0.06 | 1.06 |
| 1 | Pitx2 | 0.66 | 1.03E-68 | 2.14E-64 | 0.27 | 0.13 | 0.14 | 2.10 |
| 1 | Alx4 | 0.59 | 1.09E-68 | 2.27E-64 | 0.61 | 0.40 | 0.20 | 1.50 |
| 1 | Gtf2h5 | 0.29 | 1.32E-68 | 2.75E-64 | 0.98 | 0.94 | 0.04 | 1.04 |
| 1 | Dnm3os | 0.50 | 1.67E-68 | 3.47E-64 | 0.72 | 0.50 | 0.22 | 1.44 |
| 1 | Laptm4b | 0.64 | 2.76E-68 | 5.75E-64 | 0.70 | 0.53 | 0.17 | 1.31 |
| 1 | Mtap | 0.48 | 4.17E-68 | 8.67E-64 | 0.82 | 0.68 | 0.14 | 1.21 |
| 1 | Hnrnpm | 0.26 | 4.06E-67 | 8.45E-63 | 1.00 | 0.98 | 0.02 | 1.02 |
| 1 | Map1b | 0.47 | 9.33E-67 | 1.94E-62 | 0.76 | 0.61 | 0.15 | 1.24 |
| 1 | Pdlim7 | 0.42 | 1.00E-66 | 2.09E-62 | 0.88 | 0.74 | 0.14 | 1.19 |
| 1 | Arxes2 | 0.67 | 1.38E-66 | 2.87E-62 | 0.55 | 0.38 | 0.17 | 1.46 |
| 1 | Serpine2 | 0.52 | 1.41E-66 | 2.93E-62 | 0.86 | 0.69 | 0.17 | 1.25 |
| 1 | Ptbp2 | 0.41 | 4.33E-66 | 9.00E-62 | 0.88 | 0.76 | 0.12 | 1.16 |
| 1 | Cacna1g | 0.67 | 1.13E-65 | 2.34E-61 | 0.50 | 0.31 | 0.18 | 1.59 |
| 1 | Farsb | 0.56 | 2.41E-65 | 5.01E-61 | 0.70 | 0.55 | 0.15 | 1.27 |
| 1 | Hspa5 | 0.29 | 2.89E-65 | 6.02E-61 | 1.00 | 0.97 | 0.03 | 1.03 |
| 1 | Rarg | 0.55 | 5.05E-65 | 1.05E-60 | 0.69 | 0.50 | 0.18 | 1.37 |
| 1 | Rcn3 | 0.41 | 1.03E-64 | 2.14E-60 | 0.86 | 0.72 | 0.15 | 1.20 |
| 1 | Rerg | 0.68 | 1.53E-64 | 3.18E-60 | 0.40 | 0.23 | 0.17 | 1.73 |
| 1 | Cacybp | 0.29 | 1.92E-64 | 3.99E-60 | 0.99 | 0.96 | 0.03 | 1.03 |
| 1 | Zcchc12 | 1.65 | 2.45E-64 | 5.10E-60 | 0.14 | 0.05 | 0.09 | 2.89 |
| 1 | Denr | 0.31 | 2.67E-64 | 5.54E-60 | 0.97 | 0.92 | 0.04 | 1.05 |
| 1 | Scube1 | 0.57 | 5.19E-64 | 1.08E-59 | 0.55 | 0.36 | 0.19 | 1.54 |
| 1 | Fzd1 | 0.71 | 5.49E-64 | 1.14E-59 | 0.50 | 0.32 | 0.17 | 1.53 |
| 1 | Lsm3 | 0.31 | 1.15E-63 | 2.38E-59 | 0.97 | 0.92 | 0.05 | 1.06 |
| 1 | Dnah7a | 1.21 | 1.61E-63 | 3.36E-59 | 0.23 | 0.11 | 0.12 | 2.11 |
| 1 | Ndufc2 | 0.25 | 2.02E-63 | 4.19E-59 | 1.00 | 0.97 | 0.03 | 1.03 |
| 1 | Pja1 | 0.45 | 2.79E-63 | 5.81E-59 | 0.81 | 0.66 | 0.15 | 1.22 |
| 1 | Dab2 | 0.57 | 4.52E-63 | 9.40E-59 | 0.51 | 0.34 | 0.18 | 1.52 |
| 1 | Id3 | 0.25 | 4.67E-63 | 9.71E-59 | 0.99 | 0.95 | 0.04 | 1.04 |
| 1 | Hdgfl3 | 0.44 | 8.32E-63 | 1.73E-58 | 0.84 | 0.69 | 0.16 | 1.23 |
| 1 | Gcsh | 0.41 | 1.20E-62 | 2.49E-58 | 0.84 | 0.72 | 0.12 | 1.16 |
| 1 | Gtf3c6 | 0.36 | 1.49E-62 | 3.10E-58 | 0.94 | 0.85 | 0.08 | 1.10 |
| 1 | Pdgfrb | 0.54 | 5.03E-62 | 1.05E-57 | 0.63 | 0.44 | 0.20 | 1.45 |
| 1 | Car12 | 1.32 | 2.00E-61 | 4.16E-57 | 0.19 | 0.08 | 0.11 | 2.32 |
| 1 | Scd2 | 0.30 | 2.67E-61 | 5.54E-57 | 0.97 | 0.91 | 0.07 | 1.07 |
| 1 | Zfhx3 | 0.47 | 1.19E-60 | 2.48E-56 | 0.91 | 0.80 | 0.10 | 1.13 |
| 1 | Syngr3 | 1.56 | 2.52E-60 | 5.25E-56 | 0.11 | 0.04 | 0.08 | 3.17 |
| 1 | Smim11 | 0.42 | 5.38E-60 | 1.12E-55 | 0.85 | 0.75 | 0.10 | 1.14 |
| 1 | Tes | 0.43 | 1.45E-59 | 3.01E-55 | 0.81 | 0.68 | 0.13 | 1.20 |
| 1 | Cpxm2 | 1.55 | 1.83E-59 | 3.81E-55 | 0.12 | 0.04 | 0.08 | 3.03 |
| 1 | Epha8 | 1.78 | 2.04E-59 | 4.24E-55 | 0.10 | 0.03 | 0.07 | 3.25 |
| 1 | Fibin | 1.07 | 3.20E-59 | 6.66E-55 | 0.23 | 0.11 | 0.12 | 2.09 |
| 1 | Qprt | 1.75 | 4.92E-59 | 1.02E-54 | 0.11 | 0.04 | 0.08 | 3.08 |
| 1 | Phf5a | 0.27 | 5.83E-59 | 1.21E-54 | 0.99 | 0.95 | 0.04 | 1.04 |
| 1 | Cct4 | 0.25 | 5.87E-59 | 1.22E-54 | 0.99 | 0.96 | 0.03 | 1.03 |
| 1 | Flrt3 | 0.66 | 1.06E-58 | 2.20E-54 | 0.59 | 0.43 | 0.17 | 1.38 |
| 1 | Fgfr1 | 0.40 | 1.77E-58 | 3.68E-54 | 0.80 | 0.64 | 0.17 | 1.26 |
| 1 | Ror2 | 0.73 | 3.19E-58 | 6.64E-54 | 0.45 | 0.28 | 0.16 | 1.58 |
| 1 | Sh3bp5 | 0.47 | 3.28E-58 | 6.83E-54 | 0.66 | 0.49 | 0.18 | 1.36 |
| 1 | Fcgrt | 0.97 | 4.46E-58 | 9.28E-54 | 0.26 | 0.13 | 0.13 | 1.96 |
| 1 | Hmgcs1 | 0.43 | 5.85E-58 | 1.22E-53 | 0.87 | 0.76 | 0.11 | 1.14 |
| 1 | Efnb1 | 0.50 | 7.40E-58 | 1.54E-53 | 0.69 | 0.51 | 0.18 | 1.35 |
| 1 | Ddx25 | 1.07 | 1.40E-57 | 2.90E-53 | 0.25 | 0.13 | 0.12 | 1.97 |

**Supplementary Table 1. Marker genes for the original clusters of integrated clustering of E11.5 control and NotchGOF λ-junction cells.**

| | | | | | | | | |
|---|---|---|---|---|---|---|---|---|
| 1 | Mrpl20 | 0.28 | 2.83E-57 | 5.89E-53 | 0.97 | 0.92 | 0.05 | 1.05 |
| 1 | Dancr | 0.51 | 4.28E-57 | 8.91E-53 | 0.72 | 0.58 | 0.15 | 1.25 |
| 1 | Ccdc124 | 0.35 | 5.28E-57 | 1.10E-52 | 0.89 | 0.81 | 0.08 | 1.10 |
| 1 | Psmc6 | 0.27 | 6.37E-57 | 1.33E-52 | 0.97 | 0.94 | 0.04 | 1.04 |
| 1 | Phf6 | 0.44 | 1.08E-56 | 2.25E-52 | 0.82 | 0.71 | 0.11 | 1.15 |
| 1 | Pygo1 | 0.56 | 4.29E-56 | 8.93E-52 | 0.61 | 0.45 | 0.16 | 1.35 |
| 1 | a | 1.12 | 1.33E-55 | 2.76E-51 | 0.24 | 0.12 | 0.12 | 1.98 |
| 1 | Khdrbs1 | 0.25 | 1.60E-55 | 3.32E-51 | 0.99 | 0.95 | 0.04 | 1.04 |
| 1 | Cdca7 | 0.37 | 5.44E-55 | 1.13E-50 | 0.84 | 0.71 | 0.13 | 1.19 |
| 1 | Hspa9 | 0.32 | 7.95E-55 | 1.65E-50 | 0.95 | 0.89 | 0.06 | 1.07 |
| 1 | Atoh8 | 1.20 | 1.13E-54 | 2.34E-50 | 0.18 | 0.08 | 0.10 | 2.25 |
| 1 | Svep1 | 1.37 | 1.52E-54 | 3.16E-50 | 0.17 | 0.07 | 0.10 | 2.30 |
| 1 | Tmem167 | 0.26 | 3.24E-54 | 6.75E-50 | 0.99 | 0.95 | 0.04 | 1.04 |
| 1 | Adamtsl1 | 1.01 | 3.49E-54 | 7.26E-50 | 0.20 | 0.09 | 0.11 | 2.16 |
| 1 | Gulo | 1.34 | 3.49E-54 | 7.27E-50 | 0.19 | 0.09 | 0.10 | 2.19 |
| 1 | Arf4 | 0.26 | 4.20E-54 | 8.74E-50 | 0.98 | 0.95 | 0.03 | 1.04 |
| 1 | Tle1 | 0.56 | 5.10E-54 | 1.06E-49 | 0.66 | 0.52 | 0.14 | 1.27 |
| 1 | Fn1 | 0.26 | 1.27E-53 | 2.65E-49 | 0.91 | 0.77 | 0.14 | 1.19 |
| 1 | Sqle | 0.51 | 1.37E-53 | 2.85E-49 | 0.71 | 0.57 | 0.14 | 1.24 |
| 1 | Knstrn | 0.51 | 2.81E-53 | 5.85E-49 | 0.71 | 0.61 | 0.10 | 1.17 |
| 1 | Rrm2 | 0.33 | 3.07E-53 | 6.39E-49 | 0.93 | 0.79 | 0.14 | 1.17 |
| 1 | Cdc20 | 0.63 | 3.41E-53 | 7.09E-49 | 0.66 | 0.57 | 0.10 | 1.17 |
| 1 | Ddx39 | 0.37 | 5.88E-53 | 1.22E-48 | 0.88 | 0.78 | 0.10 | 1.12 |
| 1 | Anp32e | 0.26 | 6.36E-53 | 1.32E-48 | 0.98 | 0.94 | 0.03 | 1.04 |
| 1 | Nkd2 | 1.17 | 3.52E-52 | 7.33E-48 | 0.23 | 0.12 | 0.11 | 1.96 |
| 1 | Tcf4 | 0.34 | 6.17E-52 | 1.28E-47 | 0.96 | 0.89 | 0.07 | 1.08 |
| 1 | Cfl2 | 0.35 | 7.70E-52 | 1.60E-47 | 0.89 | 0.80 | 0.10 | 1.12 |
| 1 | C1qbp | 0.29 | 1.05E-51 | 2.18E-47 | 0.96 | 0.92 | 0.05 | 1.05 |
| 1 | 2700069I18Rik | 1.06 | 1.21E-51 | 2.52E-47 | 0.23 | 0.12 | 0.12 | 1.97 |
| 1 | Snrpc | 0.26 | 1.49E-51 | 3.11E-47 | 0.98 | 0.95 | 0.04 | 1.04 |
| 1 | Smad7 | 0.41 | 1.90E-51 | 3.94E-47 | 0.83 | 0.68 | 0.15 | 1.22 |
| 1 | Mettl9 | 0.27 | 2.01E-51 | 4.19E-47 | 0.98 | 0.93 | 0.05 | 1.05 |
| 1 | Mrpl42 | 0.26 | 3.09E-51 | 6.44E-47 | 0.97 | 0.93 | 0.04 | 1.04 |
| 1 | Mmp2 | 0.44 | 4.92E-51 | 1.02E-46 | 0.68 | 0.50 | 0.18 | 1.35 |
| 1 | Ctxn1 | 0.37 | 6.10E-51 | 1.27E-46 | 0.88 | 0.77 | 0.12 | 1.15 |
| 1 | Cdkn3 | 0.68 | 8.17E-51 | 1.70E-46 | 0.43 | 0.28 | 0.15 | 1.52 |
| 1 | Hsph1 | 0.43 | 2.77E-50 | 5.76E-46 | 0.79 | 0.67 | 0.12 | 1.18 |
| 1 | Sh3bgrl | 0.31 | 5.88E-50 | 1.22E-45 | 0.94 | 0.84 | 0.10 | 1.11 |
| 1 | Mme | 0.43 | 1.56E-49 | 3.24E-45 | 0.62 | 0.45 | 0.17 | 1.38 |
| 1 | Ddit4l | 0.63 | 2.09E-49 | 4.35E-45 | 0.34 | 0.20 | 0.14 | 1.71 |
| 1 | Myh10 | 0.32 | 2.67E-49 | 5.55E-45 | 0.96 | 0.86 | 0.10 | 1.11 |
| 1 | Ppid | 0.30 | 4.63E-49 | 9.64E-45 | 0.94 | 0.87 | 0.07 | 1.08 |
| 1 | Loxl2 | 0.54 | 5.90E-49 | 1.23E-44 | 0.58 | 0.42 | 0.15 | 1.36 |
| 1 | Rasl11a | 0.72 | 1.59E-48 | 3.31E-44 | 0.41 | 0.26 | 0.14 | 1.54 |
| 1 | Tbcb | 0.29 | 1.84E-48 | 3.82E-44 | 0.95 | 0.88 | 0.07 | 1.08 |
| 1 | Eif4ebp1 | 0.39 | 1.87E-48 | 3.89E-44 | 0.81 | 0.68 | 0.12 | 1.18 |
| 1 | Dgcr6 | 0.38 | 2.77E-48 | 5.76E-44 | 0.83 | 0.72 | 0.11 | 1.15 |
| 1 | Tle5 | 0.25 | 3.11E-48 | 6.47E-44 | 0.99 | 0.93 | 0.06 | 1.06 |
| 1 | Timp2 | 0.38 | 5.25E-48 | 1.09E-43 | 0.80 | 0.66 | 0.14 | 1.21 |
| 1 | Calb2 | 2.05 | 5.50E-48 | 1.14E-43 | 0.12 | 0.05 | 0.07 | 2.55 |
| 1 | Hbegf | 1.21 | 5.69E-48 | 1.18E-43 | 0.16 | 0.07 | 0.09 | 2.22 |
| 1 | Pcolce | 0.39 | 6.64E-48 | 1.38E-43 | 0.63 | 0.44 | 0.18 | 1.42 |
| 1 | Epop | 0.60 | 7.27E-48 | 1.51E-43 | 0.51 | 0.37 | 0.14 | 1.38 |
| 1 | Tmem59l | 0.91 | 9.80E-48 | 2.04E-43 | 0.32 | 0.19 | 0.13 | 1.66 |
| 1 | Myorg | 1.37 | 1.70E-47 | 3.53E-43 | 0.16 | 0.07 | 0.09 | 2.26 |
| 1 | Fez1 | 0.85 | 2.01E-47 | 4.18E-43 | 0.25 | 0.14 | 0.12 | 1.87 |

**Supplementary Table 1. Marker genes for the original clusters of integrated clustering of E11.5 control and NotchGOF λ-junction cells.**

| | | | | | | | | |
|---|---|---|---|---|---|---|---|---|
| 1 | Kctd15 | 0.57 | 3.32E-47 | 6.91E-43 | 0.56 | 0.40 | 0.15 | 1.38 |
| 1 | Rbmx | 0.37 | 4.69E-47 | 9.75E-43 | 0.84 | 0.74 | 0.10 | 1.14 |
| 1 | Alkal2 | 1.33 | 9.57E-47 | 1.99E-42 | 0.16 | 0.07 | 0.09 | 2.19 |
| 1 | Mrpl13 | 0.27 | 1.01E-46 | 2.09E-42 | 0.95 | 0.90 | 0.05 | 1.06 |
| 1 | Psmd6 | 0.28 | 1.37E-46 | 2.86E-42 | 0.95 | 0.89 | 0.06 | 1.07 |
| 1 | Bmp7 | 0.60 | 1.45E-46 | 3.01E-42 | 0.52 | 0.37 | 0.15 | 1.41 |
| 1 | Pinx1 | 0.49 | 1.57E-46 | 3.26E-42 | 0.66 | 0.53 | 0.13 | 1.24 |
| 1 | Emc2 | 0.47 | 1.72E-46 | 3.57E-42 | 0.70 | 0.58 | 0.12 | 1.21 |
| 1 | Mesd | 0.34 | 2.01E-46 | 4.18E-42 | 0.88 | 0.77 | 0.11 | 1.15 |
| 1 | Olfml3 | 0.56 | 2.39E-46 | 4.96E-42 | 0.43 | 0.27 | 0.15 | 1.57 |
| 1 | Vstm2b | 1.22 | 3.05E-46 | 6.34E-42 | 0.18 | 0.09 | 0.09 | 2.09 |
| 1 | Lef1 | 0.25 | 5.56E-46 | 1.16E-41 | 0.62 | 0.44 | 0.18 | 1.41 |
| 1 | Pdlim2 | 0.87 | 8.96E-46 | 1.86E-41 | 0.27 | 0.16 | 0.12 | 1.76 |
| 1 | Gpx8 | 0.37 | 1.72E-45 | 3.57E-41 | 0.83 | 0.69 | 0.13 | 1.19 |
| 1 | Dph3 | 0.36 | 2.99E-45 | 6.22E-41 | 0.84 | 0.75 | 0.09 | 1.12 |
| 1 | Dcun1d5 | 0.30 | 3.11E-45 | 6.47E-41 | 0.93 | 0.85 | 0.08 | 1.09 |
| 1 | Chchd1 | 0.26 | 4.05E-45 | 8.43E-41 | 0.97 | 0.94 | 0.03 | 1.03 |
| 1 | Psmc1 | 0.32 | 4.11E-45 | 8.56E-41 | 0.87 | 0.80 | 0.08 | 1.10 |
| 1 | Mir100hg | 0.28 | 5.41E-45 | 1.13E-40 | 0.89 | 0.75 | 0.14 | 1.19 |
| 1 | Plaat3 | 0.86 | 5.92E-45 | 1.23E-40 | 0.33 | 0.21 | 0.12 | 1.59 |
| 1 | Map2k2 | 0.26 | 1.72E-44 | 3.57E-40 | 0.96 | 0.92 | 0.04 | 1.04 |
| 1 | Psrc1 | 0.59 | 1.87E-44 | 3.88E-40 | 0.53 | 0.41 | 0.13 | 1.32 |
| 1 | Gm42372 | 1.44 | 4.89E-44 | 1.02E-39 | 0.11 | 0.04 | 0.07 | 2.60 |
| 1 | Plk2 | 0.36 | 9.73E-44 | 2.02E-39 | 0.52 | 0.36 | 0.16 | 1.46 |
| 1 | Map4k4 | 0.26 | 5.55E-43 | 1.15E-38 | 0.97 | 0.89 | 0.08 | 1.09 |
| 1 | Bbip1 | 0.37 | 9.30E-43 | 1.93E-38 | 0.79 | 0.69 | 0.10 | 1.14 |
| 1 | Slc35b1 | 0.44 | 1.19E-42 | 2.47E-38 | 0.71 | 0.58 | 0.13 | 1.22 |
| 1 | Lactb | 0.58 | 1.41E-42 | 2.92E-38 | 0.47 | 0.33 | 0.14 | 1.43 |
| 1 | Homer3 | 0.49 | 1.53E-42 | 3.19E-38 | 0.60 | 0.47 | 0.14 | 1.29 |
| 1 | 2610027K06Rik | 0.92 | 1.66E-42 | 3.46E-38 | 0.23 | 0.12 | 0.11 | 1.85 |
| 1 | Igfbp7 | 0.61 | 2.38E-42 | 4.96E-38 | 0.39 | 0.25 | 0.14 | 1.56 |
| 1 | 1110038B12Rik | 0.26 | 4.83E-42 | 1.00E-37 | 0.97 | 0.92 | 0.05 | 1.05 |
| 1 | Nradd | 0.46 | 5.90E-42 | 1.23E-37 | 0.65 | 0.51 | 0.14 | 1.26 |
| 1 | Shisa4 | 0.59 | 1.11E-41 | 2.31E-37 | 0.46 | 0.32 | 0.14 | 1.44 |
| 1 | Lockd | 0.59 | 1.14E-41 | 2.37E-37 | 0.49 | 0.35 | 0.14 | 1.39 |
| 1 | Nde1 | 0.48 | 1.15E-41 | 2.40E-37 | 0.70 | 0.61 | 0.09 | 1.15 |
| 1 | Stra6 | 0.54 | 1.21E-41 | 2.51E-37 | 0.47 | 0.33 | 0.14 | 1.44 |
| 1 | Arxes1 | 0.59 | 1.26E-41 | 2.63E-37 | 0.49 | 0.36 | 0.14 | 1.38 |
| 1 | Nme4 | 0.34 | 1.48E-41 | 3.07E-37 | 0.83 | 0.75 | 0.09 | 1.11 |
| 1 | Id2 | 0.28 | 1.62E-41 | 3.38E-37 | 0.94 | 0.86 | 0.09 | 1.10 |
| 1 | Polr2i | 0.27 | 1.74E-41 | 3.62E-37 | 0.92 | 0.87 | 0.06 | 1.06 |
| 1 | Wnk3 | 0.64 | 2.06E-41 | 4.29E-37 | 0.44 | 0.31 | 0.13 | 1.44 |
| 1 | Rftn2 | 0.63 | 2.07E-41 | 4.31E-37 | 0.35 | 0.22 | 0.13 | 1.61 |
| 1 | Yaf2 | 0.36 | 3.37E-41 | 7.00E-37 | 0.80 | 0.69 | 0.11 | 1.16 |
| 1 | Msn | 0.27 | 4.06E-41 | 8.45E-37 | 0.65 | 0.50 | 0.15 | 1.31 |
| 1 | Prex2 | 0.48 | 4.73E-41 | 9.84E-37 | 0.55 | 0.39 | 0.16 | 1.40 |
| 1 | Bend5 | 0.50 | 5.29E-41 | 1.10E-36 | 0.56 | 0.42 | 0.14 | 1.34 |
| 1 | Csrnp3 | 1.03 | 5.73E-41 | 1.19E-36 | 0.18 | 0.09 | 0.09 | 1.98 |
| 1 | Bcat1 | 0.62 | 6.00E-41 | 1.25E-36 | 0.43 | 0.30 | 0.13 | 1.43 |
| 1 | Rbmxl1 | 0.27 | 6.23E-41 | 1.30E-36 | 0.95 | 0.88 | 0.07 | 1.08 |
| 1 | Slirp | 0.27 | 8.21E-41 | 1.71E-36 | 0.93 | 0.88 | 0.05 | 1.06 |
| 1 | B3gnt5 | 0.57 | 9.19E-41 | 1.91E-36 | 0.48 | 0.35 | 0.13 | 1.39 |
| 1 | 0610010K14Rik | 0.30 | 1.10E-40 | 2.28E-36 | 0.89 | 0.81 | 0.08 | 1.10 |
| 1 | Stx18 | 0.54 | 1.14E-40 | 2.37E-36 | 0.63 | 0.51 | 0.12 | 1.24 |
| 1 | Cyp27a1 | 1.16 | 2.03E-40 | 4.21E-36 | 0.16 | 0.08 | 0.08 | 2.09 |
| 1 | Casp6 | 0.43 | 2.24E-40 | 4.65E-36 | 0.68 | 0.55 | 0.13 | 1.24 |

**Supplementary Table 1. Marker genes for the original clusters of integrated clustering of E11.5 control and NotchGOF λ-junction cells.**

| | | | | | | | | |
|---|---|---|---|---|---|---|---|---|
| 1 | Psmg1 | 0.41 | 2.43E-40 | 5.05E-36 | 0.74 | 0.62 | 0.12 | 1.19 |
| 1 | P3h2 | 1.19 | 2.67E-40 | 5.55E-36 | 0.15 | 0.07 | 0.08 | 2.16 |
| 1 | Fam89b | 0.30 | 3.02E-40 | 6.28E-36 | 0.88 | 0.79 | 0.10 | 1.12 |
| 1 | Tceal8 | 0.26 | 3.05E-40 | 6.33E-36 | 0.94 | 0.88 | 0.06 | 1.06 |
| 1 | Bud31 | 0.31 | 7.05E-40 | 1.47E-35 | 0.88 | 0.82 | 0.06 | 1.08 |
| 1 | Jam3 | 0.42 | 1.08E-39 | 2.24E-35 | 0.70 | 0.56 | 0.14 | 1.25 |
| 1 | Utrn | 0.36 | 1.25E-39 | 2.59E-35 | 0.73 | 0.60 | 0.13 | 1.22 |
| 1 | Gdf6 | 0.77 | 1.27E-39 | 2.64E-35 | 0.19 | 0.10 | 0.09 | 1.94 |
| 1 | Etv4 | 0.66 | 1.42E-39 | 2.95E-35 | 0.45 | 0.32 | 0.13 | 1.41 |
| 1 | Cep89 | 0.64 | 2.03E-39 | 4.22E-35 | 0.44 | 0.31 | 0.13 | 1.40 |
| 1 | Srp9 | 0.25 | 2.97E-39 | 6.17E-35 | 0.96 | 0.91 | 0.04 | 1.05 |
| 1 | Ddx21 | 0.35 | 3.27E-39 | 6.80E-35 | 0.83 | 0.72 | 0.11 | 1.15 |
| 1 | Limch1 | 0.32 | 3.40E-39 | 7.06E-35 | 0.81 | 0.66 | 0.15 | 1.23 |
| 1 | Adamts1 | 0.47 | 3.46E-39 | 7.20E-35 | 0.40 | 0.26 | 0.13 | 1.51 |
| 1 | Ctbp2 | 0.38 | 3.85E-39 | 8.01E-35 | 0.78 | 0.68 | 0.11 | 1.16 |
| 1 | Smn1 | 0.35 | 5.13E-39 | 1.07E-34 | 0.81 | 0.71 | 0.09 | 1.13 |
| 1 | 1190005I06Rik | 0.87 | 6.89E-39 | 1.43E-34 | 0.22 | 0.12 | 0.10 | 1.84 |
| 1 | Unc119 | 0.29 | 8.93E-39 | 1.86E-34 | 0.88 | 0.77 | 0.11 | 1.14 |
| 1 | Ctnnbip1 | 0.29 | 1.77E-38 | 3.68E-34 | 0.90 | 0.80 | 0.09 | 1.12 |
| 1 | Irx5 | 0.75 | 1.84E-38 | 3.82E-34 | 0.39 | 0.27 | 0.12 | 1.45 |
| 1 | Tspan11 | 0.71 | 1.91E-38 | 3.98E-34 | 0.27 | 0.16 | 0.11 | 1.71 |
| 1 | Parvb | 0.85 | 3.16E-38 | 6.56E-34 | 0.22 | 0.12 | 0.10 | 1.81 |
| 1 | Nfib | 0.43 | 4.33E-38 | 9.00E-34 | 0.84 | 0.73 | 0.11 | 1.15 |
| 1 | Cul4b | 0.39 | 6.13E-38 | 1.27E-33 | 0.72 | 0.60 | 0.12 | 1.19 |
| 1 | Pcnp | 0.27 | 8.12E-38 | 1.69E-33 | 0.93 | 0.88 | 0.06 | 1.06 |
| 1 | Ets1 | 0.28 | 1.25E-37 | 2.60E-33 | 0.28 | 0.17 | 0.12 | 1.69 |
| 1 | Tmem160 | 0.33 | 1.90E-37 | 3.95E-33 | 0.82 | 0.72 | 0.10 | 1.13 |
| 1 | Kcnk2 | 0.53 | 2.00E-37 | 4.16E-33 | 0.42 | 0.29 | 0.13 | 1.45 |
| 1 | Kcp | 0.96 | 4.82E-37 | 1.00E-32 | 0.19 | 0.10 | 0.09 | 1.88 |
| 1 | Chordc1 | 0.34 | 5.66E-37 | 1.18E-32 | 0.82 | 0.74 | 0.09 | 1.12 |
| 1 | Boc | 0.76 | 8.48E-37 | 1.76E-32 | 0.30 | 0.19 | 0.11 | 1.59 |
| 1 | Fam92a | 0.30 | 9.25E-37 | 1.92E-32 | 0.87 | 0.74 | 0.13 | 1.17 |
| 1 | Cby1 | 0.43 | 1.43E-36 | 2.97E-32 | 0.63 | 0.51 | 0.12 | 1.23 |
| 1 | 2610318N02Rik | 0.35 | 1.84E-36 | 3.84E-32 | 0.72 | 0.58 | 0.13 | 1.22 |
| 1 | Fli1 | 0.39 | 1.97E-36 | 4.10E-32 | 0.51 | 0.36 | 0.15 | 1.42 |
| 1 | Rab4a | 0.54 | 2.43E-36 | 5.05E-32 | 0.46 | 0.34 | 0.12 | 1.36 |
| 1 | Rpl7l1 | 0.27 | 3.78E-36 | 7.85E-32 | 0.92 | 0.85 | 0.07 | 1.08 |
| 1 | Snrpb2 | 0.27 | 4.98E-36 | 1.04E-31 | 0.91 | 0.85 | 0.06 | 1.07 |
| 1 | Casp8ap2 | 0.32 | 8.47E-36 | 1.76E-31 | 0.86 | 0.79 | 0.08 | 1.10 |
| 1 | Nfil3 | 0.83 | 8.89E-36 | 1.85E-31 | 0.22 | 0.13 | 0.10 | 1.76 |
| 1 | Snhg18 | 0.38 | 1.13E-35 | 2.35E-31 | 0.65 | 0.51 | 0.15 | 1.29 |
| 1 | Hypk | 0.27 | 1.62E-35 | 3.37E-31 | 0.90 | 0.84 | 0.06 | 1.08 |
| 1 | Adamts19 | 0.63 | 2.55E-35 | 5.31E-31 | 0.35 | 0.24 | 0.12 | 1.50 |
| 1 | Sigmar1 | 0.41 | 2.80E-35 | 5.82E-31 | 0.66 | 0.54 | 0.12 | 1.22 |
| 1 | Plekho1 | 0.34 | 7.18E-35 | 1.49E-30 | 0.74 | 0.60 | 0.13 | 1.22 |
| 1 | Cend1 | 1.14 | 9.05E-35 | 1.88E-30 | 0.12 | 0.05 | 0.07 | 2.25 |
| 1 | 2810468N07Rik | 1.19 | 9.35E-35 | 1.95E-30 | 0.15 | 0.07 | 0.07 | 2.04 |
| 1 | Tmem126a | 0.31 | 9.49E-35 | 1.97E-30 | 0.82 | 0.71 | 0.11 | 1.15 |
| 1 | Ttpa | 1.20 | 1.02E-34 | 2.12E-30 | 0.14 | 0.07 | 0.07 | 2.07 |
| 1 | 1500009L16Rik | 0.36 | 1.08E-34 | 2.24E-30 | 0.69 | 0.57 | 0.13 | 1.22 |
| 1 | Dhx29 | 0.44 | 1.17E-34 | 2.43E-30 | 0.62 | 0.51 | 0.11 | 1.23 |
| 1 | Tcim | 0.49 | 1.28E-34 | 2.66E-30 | 0.48 | 0.36 | 0.13 | 1.36 |
| 1 | Scpep1 | 0.50 | 1.31E-34 | 2.72E-30 | 0.53 | 0.40 | 0.13 | 1.32 |
| 1 | Fdft1 | 0.44 | 1.49E-34 | 3.10E-30 | 0.69 | 0.59 | 0.09 | 1.16 |
| 1 | Ciao2a | 0.26 | 1.65E-34 | 3.43E-30 | 0.91 | 0.84 | 0.06 | 1.08 |
| 1 | Col6a3 | 0.75 | 2.42E-34 | 5.03E-30 | 0.26 | 0.16 | 0.10 | 1.67 |

**Supplementary Table 1. Marker genes for the original clusters of integrated clustering of E11.5 control and NotchGOF λ-junction cells.**

| | | | | | | | | |
|---|---|---|---|---|---|---|---|---|
| 1 | Akr1b3 | 0.35 | 2.77E-34 | 5.76E-30 | 0.76 | 0.67 | 0.10 | 1.14 |
| 1 | Dld | 0.29 | 2.93E-34 | 6.09E-30 | 0.85 | 0.76 | 0.09 | 1.12 |
| 1 | Cdc6 | 0.73 | 3.06E-34 | 6.36E-30 | 0.38 | 0.27 | 0.11 | 1.41 |
| 1 | Rpp25l | 0.32 | 3.52E-34 | 7.31E-30 | 0.81 | 0.73 | 0.08 | 1.11 |
| 1 | Cldnd1 | 0.34 | 3.95E-34 | 8.21E-30 | 0.78 | 0.68 | 0.10 | 1.14 |
| 1 | Fkbp5 | 0.47 | 7.58E-34 | 1.58E-29 | 0.53 | 0.41 | 0.12 | 1.30 |
| 1 | Zfas1 | 0.27 | 7.63E-34 | 1.59E-29 | 0.89 | 0.82 | 0.08 | 1.09 |
| 1 | Cetn2 | 0.37 | 8.58E-34 | 1.78E-29 | 0.74 | 0.63 | 0.11 | 1.17 |
| 1 | Hells | 0.26 | 1.13E-33 | 2.36E-29 | 0.88 | 0.77 | 0.12 | 1.15 |
| 1 | Dctd | 0.45 | 1.20E-33 | 2.49E-29 | 0.58 | 0.46 | 0.12 | 1.26 |
| 1 | Ddr2 | 0.29 | 1.42E-33 | 2.96E-29 | 0.64 | 0.47 | 0.17 | 1.36 |
| 1 | Mrps15 | 0.27 | 1.78E-33 | 3.71E-29 | 0.88 | 0.81 | 0.07 | 1.08 |
| 1 | Gtpbp4 | 0.26 | 2.21E-33 | 4.60E-29 | 0.92 | 0.85 | 0.07 | 1.08 |
| 1 | Fdps | 0.26 | 2.26E-33 | 4.71E-29 | 0.89 | 0.81 | 0.08 | 1.10 |
| 1 | Hacd4 | 0.56 | 2.51E-33 | 5.22E-29 | 0.35 | 0.24 | 0.12 | 1.49 |
| 1 | Gm45716 | 0.53 | 3.04E-33 | 6.33E-29 | 0.48 | 0.36 | 0.12 | 1.33 |
| 1 | Pbdc1 | 0.25 | 6.36E-33 | 1.32E-28 | 0.92 | 0.85 | 0.06 | 1.08 |
| 1 | Rbm28 | 0.30 | 7.25E-33 | 1.51E-28 | 0.83 | 0.74 | 0.09 | 1.12 |
| 1 | Idi1 | 0.40 | 8.06E-33 | 1.68E-28 | 0.69 | 0.58 | 0.11 | 1.18 |
| 1 | Rab3il1 | 0.50 | 1.08E-32 | 2.24E-28 | 0.41 | 0.29 | 0.12 | 1.42 |
| 1 | Twist2 | 0.78 | 1.16E-32 | 2.41E-28 | 0.33 | 0.22 | 0.11 | 1.49 |
| 1 | Ak6 | 0.33 | 1.18E-32 | 2.46E-28 | 0.77 | 0.68 | 0.09 | 1.13 |
| 1 | Cdt1 | 0.46 | 1.44E-32 | 3.00E-28 | 0.50 | 0.38 | 0.12 | 1.33 |
| 1 | Rab30 | 0.80 | 2.64E-32 | 5.50E-28 | 0.23 | 0.13 | 0.09 | 1.69 |
| 1 | Chek1 | 0.40 | 3.09E-32 | 6.43E-28 | 0.66 | 0.54 | 0.12 | 1.22 |
| 1 | Mpc1 | 0.35 | 3.83E-32 | 7.97E-28 | 0.71 | 0.62 | 0.10 | 1.16 |
| 1 | Sdf2l1 | 0.42 | 5.88E-32 | 1.22E-27 | 0.58 | 0.46 | 0.12 | 1.25 |
| 1 | Col5a2 | 0.28 | 6.79E-32 | 1.41E-27 | 0.67 | 0.51 | 0.15 | 1.30 |
| 1 | Msmo1 | 0.47 | 9.10E-32 | 1.89E-27 | 0.60 | 0.49 | 0.11 | 1.22 |
| 1 | Dnajc15 | 0.28 | 1.72E-31 | 3.58E-27 | 0.85 | 0.77 | 0.08 | 1.11 |
| 1 | Ackr3 | 0.43 | 1.96E-31 | 4.08E-27 | 0.53 | 0.41 | 0.12 | 1.30 |
| 1 | Smoc1 | 1.04 | 2.27E-31 | 4.73E-27 | 0.14 | 0.07 | 0.07 | 2.01 |
| 1 | Psmd14 | 0.29 | 2.46E-31 | 5.12E-27 | 0.85 | 0.79 | 0.06 | 1.08 |
| 1 | Soat1 | 0.44 | 5.95E-31 | 1.24E-26 | 0.55 | 0.43 | 0.12 | 1.28 |
| 1 | Septin9 | 0.31 | 7.64E-31 | 1.59E-26 | 0.74 | 0.63 | 0.12 | 1.19 |
| 1 | Creb5 | 0.41 | 7.76E-31 | 1.61E-26 | 0.44 | 0.32 | 0.13 | 1.41 |
| 1 | Cdon | 0.63 | 8.63E-31 | 1.80E-26 | 0.32 | 0.21 | 0.10 | 1.49 |
| 1 | Sertad4 | 0.57 | 1.01E-30 | 2.11E-26 | 0.47 | 0.36 | 0.11 | 1.31 |
| 1 | Ppfibp2 | 0.31 | 1.23E-30 | 2.55E-26 | 0.57 | 0.42 | 0.15 | 1.37 |
| 1 | Lrp1 | 0.29 | 2.01E-30 | 4.17E-26 | 0.70 | 0.57 | 0.13 | 1.23 |
| 1 | Ttc39c | 0.65 | 2.22E-30 | 4.62E-26 | 0.29 | 0.19 | 0.10 | 1.53 |
| 1 | Elovl6 | 0.32 | 2.26E-30 | 4.70E-26 | 0.75 | 0.64 | 0.11 | 1.17 |
| 1 | Eva1b | 0.27 | 2.87E-30 | 5.97E-26 | 0.67 | 0.53 | 0.15 | 1.28 |
| 1 | Kmt5a | 0.26 | 2.87E-30 | 5.97E-26 | 0.89 | 0.83 | 0.05 | 1.06 |
| 1 | Magt1 | 0.28 | 3.26E-30 | 6.79E-26 | 0.84 | 0.74 | 0.10 | 1.13 |
| 1 | Tagln3 | 0.37 | 3.27E-30 | 6.81E-26 | 0.37 | 0.25 | 0.12 | 1.46 |
| 1 | Ung | 0.50 | 3.31E-30 | 6.89E-26 | 0.44 | 0.33 | 0.11 | 1.35 |
| 1 | Bloc1s2 | 0.33 | 3.76E-30 | 7.82E-26 | 0.75 | 0.66 | 0.09 | 1.14 |
| 1 | Sema5a | 0.27 | 4.27E-30 | 8.88E-26 | 0.50 | 0.36 | 0.14 | 1.39 |
| 1 | Cacng8 | 1.04 | 4.60E-30 | 9.57E-26 | 0.12 | 0.06 | 0.06 | 2.08 |
| 1 | Anxa11 | 0.90 | 5.05E-30 | 1.05E-25 | 0.23 | 0.15 | 0.09 | 1.60 |
| 1 | Ncs1 | 0.51 | 5.41E-30 | 1.13E-25 | 0.41 | 0.30 | 0.11 | 1.38 |
| 1 | Gmfb | 0.40 | 6.14E-30 | 1.28E-25 | 0.64 | 0.54 | 0.10 | 1.19 |
| 1 | Snhg3 | 0.42 | 6.35E-30 | 1.32E-25 | 0.63 | 0.53 | 0.10 | 1.19 |
| 1 | Utp14a | 0.37 | 6.60E-30 | 1.37E-25 | 0.67 | 0.56 | 0.10 | 1.18 |
| 1 | Mns1 | 0.41 | 8.88E-30 | 1.85E-25 | 0.63 | 0.53 | 0.10 | 1.18 |

**Supplementary Table 1. Marker genes for the original clusters of integrated clustering of E11.5 control and NotchGOF λ-junction cells.**

| | | | | | | | | |
|---|---|---|---|---|---|---|---|---|
| 1 | Tpgs2 | 0.36 | 9.45E-30 | 1.97E-25 | 0.68 | 0.57 | 0.11 | 1.19 |
| 1 | Gyg | 0.41 | 1.27E-29 | 2.64E-25 | 0.53 | 0.41 | 0.12 | 1.29 |
| 1 | Rfc3 | 0.28 | 1.40E-29 | 2.91E-25 | 0.84 | 0.76 | 0.08 | 1.10 |
| 1 | Pim1 | 0.40 | 2.08E-29 | 4.32E-25 | 0.58 | 0.46 | 0.12 | 1.26 |
| 1 | Tm2d2 | 0.28 | 2.27E-29 | 4.73E-25 | 0.81 | 0.72 | 0.09 | 1.13 |
| 1 | Rnf150 | 0.53 | 2.78E-29 | 5.78E-25 | 0.41 | 0.30 | 0.11 | 1.38 |
| 1 | Fkbp11 | 0.53 | 4.33E-29 | 9.01E-25 | 0.40 | 0.29 | 0.11 | 1.39 |
| 1 | Nfe2l2 | 0.46 | 4.81E-29 | 1.00E-24 | 0.55 | 0.44 | 0.11 | 1.25 |
| 1 | Bccip | 0.26 | 7.17E-29 | 1.49E-24 | 0.86 | 0.79 | 0.06 | 1.08 |
| 1 | Gfpt2 | 0.61 | 9.66E-29 | 2.01E-24 | 0.27 | 0.17 | 0.10 | 1.56 |
| 1 | Idh1 | 0.31 | 1.48E-28 | 3.07E-24 | 0.73 | 0.62 | 0.11 | 1.17 |
| 1 | Arf2 | 0.44 | 1.59E-28 | 3.31E-24 | 0.49 | 0.37 | 0.11 | 1.30 |
| 1 | Glud1 | 0.30 | 1.80E-28 | 3.74E-24 | 0.74 | 0.64 | 0.10 | 1.16 |
| 1 | Guk1 | 0.27 | 2.46E-28 | 5.11E-24 | 0.85 | 0.77 | 0.08 | 1.10 |
| 1 | Ethe1 | 0.41 | 3.49E-28 | 7.26E-24 | 0.56 | 0.44 | 0.12 | 1.27 |
| 1 | Prpsap1 | 0.30 | 3.65E-28 | 7.59E-24 | 0.77 | 0.68 | 0.10 | 1.14 |
| 1 | Tmem29 | 0.37 | 4.47E-28 | 9.30E-24 | 0.61 | 0.51 | 0.10 | 1.19 |
| 1 | Dhcr7 | 0.52 | 5.70E-28 | 1.19E-23 | 0.43 | 0.32 | 0.11 | 1.35 |
| 1 | Insig1 | 0.59 | 5.72E-28 | 1.19E-23 | 0.40 | 0.30 | 0.10 | 1.34 |
| 1 | Klhdc8b | 0.71 | 5.79E-28 | 1.20E-23 | 0.24 | 0.15 | 0.09 | 1.61 |
| 1 | 1110008P14Rik | 0.41 | 5.99E-28 | 1.25E-23 | 0.57 | 0.46 | 0.11 | 1.24 |
| 1 | Zfhx4 | 0.34 | 6.01E-28 | 1.25E-23 | 0.81 | 0.73 | 0.08 | 1.11 |
| 1 | Igsf3 | 0.32 | 1.01E-27 | 2.10E-23 | 0.70 | 0.57 | 0.13 | 1.22 |
| 1 | Pam16 | 0.27 | 1.06E-27 | 2.20E-23 | 0.85 | 0.79 | 0.06 | 1.08 |
| 1 | Pmpcb | 0.29 | 1.80E-27 | 3.74E-23 | 0.78 | 0.69 | 0.09 | 1.12 |
| 1 | Bak1 | 0.41 | 2.18E-27 | 4.53E-23 | 0.51 | 0.40 | 0.11 | 1.28 |
| 1 | Apip | 0.29 | 2.34E-27 | 4.87E-23 | 0.71 | 0.60 | 0.11 | 1.18 |
| 1 | Col27a1 | 0.64 | 2.42E-27 | 5.02E-23 | 0.27 | 0.18 | 0.09 | 1.53 |
| 1 | Zbed3 | 0.36 | 2.54E-27 | 5.29E-23 | 0.65 | 0.55 | 0.10 | 1.18 |
| 1 | Tfdp2 | 0.26 | 2.59E-27 | 5.38E-23 | 0.82 | 0.74 | 0.08 | 1.11 |
| 1 | Dlg2 | 1.07 | 3.12E-27 | 6.48E-23 | 0.13 | 0.06 | 0.06 | 1.97 |
| 1 | Mrps6 | 0.31 | 3.59E-27 | 7.46E-23 | 0.72 | 0.62 | 0.10 | 1.16 |
| 1 | Ncald | 0.95 | 5.44E-27 | 1.13E-22 | 0.16 | 0.09 | 0.07 | 1.81 |
| 1 | Usp14 | 0.26 | 5.45E-27 | 1.13E-22 | 0.85 | 0.79 | 0.06 | 1.08 |
| 1 | Sdc1 | 0.26 | 6.08E-27 | 1.26E-22 | 0.81 | 0.69 | 0.12 | 1.18 |
| 1 | Gpx7 | 0.34 | 7.39E-27 | 1.54E-22 | 0.56 | 0.44 | 0.13 | 1.29 |
| 1 | Raet1e | 0.54 | 8.12E-27 | 1.69E-22 | 0.33 | 0.22 | 0.11 | 1.48 |
| 1 | Mphosph10 | 0.27 | 8.65E-27 | 1.80E-22 | 0.83 | 0.75 | 0.09 | 1.12 |
| 1 | Actn1 | 0.28 | 9.19E-27 | 1.91E-22 | 0.82 | 0.71 | 0.11 | 1.16 |
| 1 | Tmem100 | 0.74 | 9.95E-27 | 2.07E-22 | 0.22 | 0.14 | 0.08 | 1.62 |
| 1 | Ldb2 | 0.45 | 1.12E-26 | 2.33E-22 | 0.37 | 0.26 | 0.11 | 1.41 |
| 1 | Emilin1 | 0.26 | 1.26E-26 | 2.62E-22 | 0.67 | 0.52 | 0.15 | 1.29 |
| 1 | Abca5 | 0.75 | 1.30E-26 | 2.70E-22 | 0.20 | 0.12 | 0.08 | 1.66 |
| 1 | Ube2f | 0.35 | 1.54E-26 | 3.20E-22 | 0.65 | 0.56 | 0.10 | 1.17 |
| 1 | Efnb3 | 0.46 | 1.61E-26 | 3.35E-22 | 0.42 | 0.31 | 0.11 | 1.37 |
| 1 | Clic4 | 0.29 | 1.75E-26 | 3.64E-22 | 0.76 | 0.67 | 0.10 | 1.14 |
| 1 | Coq7 | 0.31 | 2.05E-26 | 4.27E-22 | 0.75 | 0.66 | 0.09 | 1.13 |
| 1 | Prkar2b | 0.31 | 2.15E-26 | 4.47E-22 | 0.59 | 0.48 | 0.11 | 1.23 |
| 1 | Axin2 | 0.48 | 2.22E-26 | 4.61E-22 | 0.44 | 0.33 | 0.11 | 1.33 |
| 1 | Crcp | 0.44 | 2.25E-26 | 4.68E-22 | 0.47 | 0.36 | 0.10 | 1.28 |
| 1 | Sc5d | 0.49 | 2.38E-26 | 4.95E-22 | 0.47 | 0.37 | 0.10 | 1.27 |
| 1 | Adora2b | 0.79 | 3.43E-26 | 7.14E-22 | 0.17 | 0.10 | 0.07 | 1.74 |
| 1 | Fndc5 | 0.57 | 3.45E-26 | 7.18E-22 | 0.25 | 0.16 | 0.09 | 1.58 |
| 1 | Slc9a3r2 | 0.33 | 5.34E-26 | 1.11E-21 | 0.36 | 0.26 | 0.11 | 1.43 |
| 1 | S1pr2 | 0.46 | 7.16E-26 | 1.49E-21 | 0.42 | 0.31 | 0.11 | 1.35 |
| 1 | Hadha | 0.25 | 7.76E-26 | 1.61E-21 | 0.86 | 0.80 | 0.06 | 1.08 |

**Supplementary Table 1. Marker genes for the original clusters of integrated clustering of E11.5 control and NotchGOF λ-junction cells.**

| | | | | | | | | |
|---|---|---|---|---|---|---|---|---|
| 1 | Septin3 | 0.36 | 8.56E-26 | 1.78E-21 | 0.54 | 0.43 | 0.11 | 1.26 |
| 1 | Ick | 0.37 | 9.73E-26 | 2.02E-21 | 0.54 | 0.43 | 0.11 | 1.26 |
| 1 | Fblim1 | 0.33 | 1.11E-25 | 2.31E-21 | 0.57 | 0.45 | 0.12 | 1.26 |
| 1 | Zfp57 | 0.44 | 1.12E-25 | 2.33E-21 | 0.44 | 0.33 | 0.11 | 1.34 |
| 1 | Cth | 0.64 | 1.24E-25 | 2.58E-21 | 0.29 | 0.20 | 0.09 | 1.46 |
| 1 | Ccn1 | 0.39 | 1.41E-25 | 2.93E-21 | 0.41 | 0.30 | 0.11 | 1.37 |
| 1 | Gusb | 0.43 | 1.53E-25 | 3.18E-21 | 0.40 | 0.30 | 0.10 | 1.35 |
| 1 | Sec13 | 0.26 | 1.54E-25 | 3.21E-21 | 0.80 | 0.72 | 0.09 | 1.12 |
| 1 | Bmp2 | 0.93 | 1.59E-25 | 3.30E-21 | 0.12 | 0.06 | 0.06 | 1.97 |
| 1 | As3mt | 0.45 | 3.41E-25 | 7.10E-21 | 0.46 | 0.36 | 0.11 | 1.29 |
| 1 | Mrpl22 | 0.30 | 5.00E-25 | 1.04E-20 | 0.73 | 0.65 | 0.08 | 1.13 |
| 1 | Lmcd1 | 0.83 | 5.05E-25 | 1.05E-20 | 0.15 | 0.08 | 0.07 | 1.80 |
| 1 | Dtl | 0.33 | 5.55E-25 | 1.15E-20 | 0.71 | 0.61 | 0.10 | 1.16 |
| 1 | Net1 | 0.32 | 5.80E-25 | 1.21E-20 | 0.67 | 0.58 | 0.09 | 1.15 |
| 1 | Abhd8 | 0.36 | 6.95E-25 | 1.45E-20 | 0.58 | 0.47 | 0.11 | 1.23 |
| 1 | Tctex1d2 | 0.39 | 7.70E-25 | 1.60E-20 | 0.53 | 0.43 | 0.10 | 1.24 |
| 1 | Kyat3 | 0.75 | 7.87E-25 | 1.64E-20 | 0.21 | 0.13 | 0.08 | 1.61 |
| 1 | Lpin2 | 0.35 | 8.00E-25 | 1.66E-20 | 0.63 | 0.54 | 0.10 | 1.18 |
| 1 | Sorcs2 | 0.79 | 8.24E-25 | 1.71E-20 | 0.17 | 0.10 | 0.07 | 1.72 |
| 1 | Rcl1 | 0.41 | 8.81E-25 | 1.83E-20 | 0.54 | 0.44 | 0.10 | 1.22 |
| 1 | Anapc16 | 0.27 | 1.04E-24 | 2.16E-20 | 0.83 | 0.75 | 0.08 | 1.11 |
| 1 | Cnih2 | 0.38 | 1.09E-24 | 2.27E-20 | 0.43 | 0.32 | 0.11 | 1.35 |
| 1 | Bcat2 | 0.53 | 1.30E-24 | 2.71E-20 | 0.38 | 0.28 | 0.10 | 1.34 |
| 1 | Rangap1 | 0.25 | 1.69E-24 | 3.52E-20 | 0.79 | 0.71 | 0.07 | 1.10 |
| 1 | Vegfb | 0.26 | 1.71E-24 | 3.56E-20 | 0.75 | 0.65 | 0.10 | 1.15 |
| 1 | Got1 | 0.39 | 1.79E-24 | 3.72E-20 | 0.55 | 0.45 | 0.11 | 1.24 |
| 1 | Dcp2 | 0.30 | 2.69E-24 | 5.58E-20 | 0.71 | 0.62 | 0.10 | 1.16 |
| 1 | Ric1 | 0.43 | 3.25E-24 | 6.76E-20 | 0.43 | 0.32 | 0.10 | 1.32 |
| 1 | Ube2d1 | 0.29 | 3.29E-24 | 6.84E-20 | 0.72 | 0.62 | 0.10 | 1.15 |
| 1 | Iglon5 | 0.63 | 3.99E-24 | 8.29E-20 | 0.25 | 0.16 | 0.09 | 1.52 |
| 1 | Elmod2 | 0.57 | 4.31E-24 | 8.96E-20 | 0.31 | 0.22 | 0.09 | 1.42 |
| 1 | Clmn | 0.52 | 4.73E-24 | 9.83E-20 | 0.36 | 0.26 | 0.10 | 1.38 |
| 1 | Syde1 | 0.61 | 6.27E-24 | 1.30E-19 | 0.25 | 0.17 | 0.09 | 1.51 |
| 1 | Ap4m1 | 0.43 | 7.99E-24 | 1.66E-19 | 0.51 | 0.41 | 0.10 | 1.24 |
| 1 | Gypc | 0.57 | 8.89E-24 | 1.85E-19 | 0.30 | 0.21 | 0.09 | 1.43 |
| 1 | Tns1 | 0.44 | 9.60E-24 | 2.00E-19 | 0.40 | 0.29 | 0.11 | 1.36 |
| 1 | Faap20 | 0.30 | 1.13E-23 | 2.35E-19 | 0.69 | 0.59 | 0.10 | 1.16 |
| 1 | Gm11627 | 0.66 | 1.71E-23 | 3.55E-19 | 0.21 | 0.13 | 0.08 | 1.61 |
| 1 | Dusp18 | 0.62 | 1.84E-23 | 3.82E-19 | 0.24 | 0.16 | 0.08 | 1.52 |
| 1 | Fat4 | 0.44 | 1.89E-23 | 3.94E-19 | 0.37 | 0.27 | 0.10 | 1.39 |
| 1 | Ptprg | 0.43 | 1.99E-23 | 4.13E-19 | 0.46 | 0.36 | 0.10 | 1.28 |
| 1 | Spa17 | 0.64 | 2.78E-23 | 5.78E-19 | 0.22 | 0.15 | 0.08 | 1.54 |
| 1 | Scarf2 | 0.41 | 4.54E-23 | 9.44E-19 | 0.35 | 0.25 | 0.10 | 1.41 |
| 1 | Acads | 0.29 | 4.75E-23 | 9.88E-19 | 0.69 | 0.60 | 0.09 | 1.16 |
| 1 | Zmat5 | 0.38 | 5.11E-23 | 1.06E-18 | 0.57 | 0.47 | 0.10 | 1.20 |
| 1 | Cdc25a | 0.30 | 5.79E-23 | 1.20E-18 | 0.71 | 0.61 | 0.09 | 1.15 |
| 1 | 9130401M01Rik | 0.32 | 5.98E-23 | 1.24E-18 | 0.67 | 0.58 | 0.10 | 1.17 |
| 1 | Hadh | 0.26 | 6.99E-23 | 1.45E-18 | 0.84 | 0.77 | 0.07 | 1.09 |
| 1 | Dnph1 | 0.37 | 7.55E-23 | 1.57E-18 | 0.60 | 0.51 | 0.09 | 1.17 |
| 1 | Notch2 | 0.44 | 8.06E-23 | 1.68E-18 | 0.46 | 0.36 | 0.10 | 1.29 |
| 1 | Pdxp | 0.39 | 8.92E-23 | 1.85E-18 | 0.48 | 0.38 | 0.10 | 1.27 |
| 1 | Eif4e3 | 0.43 | 9.42E-23 | 1.96E-18 | 0.37 | 0.27 | 0.10 | 1.36 |
| 1 | B9d1 | 0.32 | 1.11E-22 | 2.30E-18 | 0.66 | 0.56 | 0.10 | 1.18 |
| 1 | Tro | 0.34 | 1.33E-22 | 2.77E-18 | 0.52 | 0.41 | 0.11 | 1.26 |
| 1 | Bri3bp | 0.30 | 1.61E-22 | 3.34E-18 | 0.68 | 0.58 | 0.09 | 1.16 |
| 1 | Letm1 | 0.40 | 1.64E-22 | 3.42E-18 | 0.53 | 0.43 | 0.09 | 1.21 |

**Supplementary Table 1. Marker genes for the original clusters of integrated clustering of E11.5 control and NotchGOF λ-junction cells.**

| | | | | | | | | |
|---|---|---|---|---|---|---|---|---|
| 1 | Ror1 | 0.60 | 1.92E-22 | 3.99E-18 | 0.19 | 0.12 | 0.07 | 1.63 |
| 1 | Gm44257 | 0.92 | 1.94E-22 | 4.03E-18 | 0.11 | 0.06 | 0.05 | 1.95 |
| 1 | Alpl | 0.76 | 1.99E-22 | 4.15E-18 | 0.16 | 0.10 | 0.07 | 1.68 |
| 1 | Ei24 | 0.26 | 2.15E-22 | 4.47E-18 | 0.78 | 0.69 | 0.09 | 1.13 |
| 1 | Crnde | 0.76 | 2.25E-22 | 4.68E-18 | 0.22 | 0.14 | 0.08 | 1.53 |
| 1 | Thyn1 | 0.25 | 2.46E-22 | 5.11E-18 | 0.82 | 0.72 | 0.09 | 1.13 |
| 1 | Ankrd29 | 0.87 | 2.52E-22 | 5.23E-18 | 0.15 | 0.09 | 0.06 | 1.74 |
| 1 | Palm | 0.33 | 2.70E-22 | 5.63E-18 | 0.56 | 0.45 | 0.11 | 1.24 |
| 1 | Cers2 | 0.33 | 3.35E-22 | 6.97E-18 | 0.62 | 0.53 | 0.09 | 1.17 |
| 1 | Mcm4 | 0.27 | 3.82E-22 | 7.94E-18 | 0.81 | 0.73 | 0.08 | 1.11 |
| 1 | Xbp1 | 0.35 | 5.12E-22 | 1.06E-17 | 0.56 | 0.47 | 0.09 | 1.20 |
| 1 | Atp1b1 | 0.70 | 5.83E-22 | 1.21E-17 | 0.14 | 0.08 | 0.06 | 1.74 |
| 1 | Fam118b | 0.40 | 6.08E-22 | 1.27E-17 | 0.46 | 0.36 | 0.10 | 1.27 |
| 1 | Tmem216 | 0.37 | 8.35E-22 | 1.74E-17 | 0.53 | 0.43 | 0.10 | 1.23 |
| 1 | Marveld1 | 0.41 | 1.06E-21 | 2.20E-17 | 0.33 | 0.23 | 0.10 | 1.42 |
| 1 | Cenpv | 0.35 | 1.20E-21 | 2.50E-17 | 0.59 | 0.49 | 0.10 | 1.20 |
| 1 | Gnpnat1 | 0.36 | 1.25E-21 | 2.61E-17 | 0.52 | 0.42 | 0.10 | 1.24 |
| 1 | Tsc22d3 | 0.30 | 1.33E-21 | 2.76E-17 | 0.62 | 0.52 | 0.10 | 1.20 |
| 1 | Dnm1 | 0.38 | 1.60E-21 | 3.33E-17 | 0.42 | 0.32 | 0.11 | 1.33 |
| 1 | Gm7324 | 0.30 | 1.86E-21 | 3.88E-17 | 0.70 | 0.61 | 0.09 | 1.14 |
| 1 | Txndc15 | 0.27 | 2.01E-21 | 4.18E-17 | 0.68 | 0.59 | 0.09 | 1.16 |
| 1 | Dchs1 | 0.33 | 2.54E-21 | 5.29E-17 | 0.51 | 0.41 | 0.11 | 1.26 |
| 1 | Antxr2 | 0.59 | 2.61E-21 | 5.42E-17 | 0.26 | 0.18 | 0.08 | 1.46 |
| 1 | Thoc1 | 0.26 | 2.61E-21 | 5.43E-17 | 0.75 | 0.66 | 0.08 | 1.13 |
| 1 | Fbxo17 | 0.63 | 2.82E-21 | 5.87E-17 | 0.23 | 0.16 | 0.08 | 1.50 |
| 1 | Ppp2r3c | 0.33 | 2.87E-21 | 5.97E-17 | 0.56 | 0.46 | 0.10 | 1.21 |
| 1 | Tmem183a | 0.27 | 3.57E-21 | 7.42E-17 | 0.73 | 0.65 | 0.08 | 1.13 |
| 1 | Gfer | 0.32 | 3.75E-21 | 7.79E-17 | 0.64 | 0.56 | 0.08 | 1.15 |
| 1 | Cenps | 0.35 | 3.95E-21 | 8.22E-17 | 0.59 | 0.50 | 0.09 | 1.19 |
| 1 | Nadk | 0.36 | 4.11E-21 | 8.56E-17 | 0.51 | 0.42 | 0.09 | 1.22 |
| 1 | Cacna1d | 0.48 | 4.86E-21 | 1.01E-16 | 0.32 | 0.23 | 0.09 | 1.41 |
| 1 | Cox20 | 0.27 | 5.41E-21 | 1.12E-16 | 0.73 | 0.66 | 0.07 | 1.11 |
| 1 | Map2k6 | 0.33 | 5.46E-21 | 1.14E-16 | 0.53 | 0.43 | 0.10 | 1.23 |
| 1 | Mvk | 0.35 | 5.83E-21 | 1.21E-16 | 0.51 | 0.42 | 0.10 | 1.24 |
| 1 | Umps | 0.27 | 7.15E-21 | 1.49E-16 | 0.69 | 0.60 | 0.09 | 1.15 |
| 1 | Cntrl | 0.35 | 7.63E-21 | 1.59E-16 | 0.55 | 0.45 | 0.09 | 1.20 |
| 1 | Galt | 0.48 | 9.04E-21 | 1.88E-16 | 0.36 | 0.27 | 0.09 | 1.33 |
| 1 | Gpn1 | 0.29 | 9.49E-21 | 1.97E-16 | 0.67 | 0.58 | 0.09 | 1.16 |
| 1 | Bach2 | 0.44 | 1.30E-20 | 2.70E-16 | 0.46 | 0.37 | 0.09 | 1.26 |
| 1 | Hax1 | 0.26 | 1.37E-20 | 2.85E-16 | 0.74 | 0.66 | 0.08 | 1.12 |
| 1 | Nipsnap1 | 0.28 | 1.38E-20 | 2.88E-16 | 0.66 | 0.55 | 0.11 | 1.19 |
| 1 | Scn3a | 0.95 | 1.41E-20 | 2.93E-16 | 0.10 | 0.05 | 0.05 | 1.94 |
| 1 | Lrrc49 | 0.49 | 1.42E-20 | 2.96E-16 | 0.33 | 0.24 | 0.09 | 1.36 |
| 1 | Atat1 | 0.29 | 1.54E-20 | 3.20E-16 | 0.64 | 0.55 | 0.10 | 1.18 |
| 1 | Chn1 | 0.36 | 1.79E-20 | 3.73E-16 | 0.49 | 0.39 | 0.10 | 1.25 |
| 1 | Fbxo16 | 0.78 | 2.08E-20 | 4.34E-16 | 0.17 | 0.10 | 0.06 | 1.62 |
| 1 | Ccdc58 | 0.25 | 2.59E-20 | 5.38E-16 | 0.76 | 0.68 | 0.09 | 1.13 |
| 1 | Jakmip2 | 0.58 | 2.76E-20 | 5.73E-16 | 0.21 | 0.14 | 0.07 | 1.53 |
| 1 | Fbxo22 | 0.28 | 3.25E-20 | 6.76E-16 | 0.68 | 0.59 | 0.09 | 1.16 |
| 1 | Gdf11 | 0.40 | 3.77E-20 | 7.84E-16 | 0.41 | 0.31 | 0.10 | 1.31 |
| 1 | B230217C12Rik | 0.57 | 3.91E-20 | 8.13E-16 | 0.25 | 0.17 | 0.08 | 1.46 |
| 1 | Anapc15 | 0.34 | 4.09E-20 | 8.51E-16 | 0.67 | 0.60 | 0.08 | 1.13 |
| 1 | Slc39a6 | 0.30 | 4.46E-20 | 9.28E-16 | 0.66 | 0.56 | 0.10 | 1.18 |
| 1 | Gstp1 | 0.35 | 4.80E-20 | 9.98E-16 | 0.59 | 0.51 | 0.08 | 1.16 |
| 1 | Cep19 | 0.36 | 7.09E-20 | 1.48E-15 | 0.44 | 0.34 | 0.10 | 1.29 |
| 1 | E2f3 | 0.29 | 9.62E-20 | 2.00E-15 | 0.66 | 0.57 | 0.08 | 1.14 |

**Supplementary Table 1. Marker genes for the original clusters of integrated clustering of E11.5 control and NotchGOF λ-junction cells.**

| | | | | | | | | |
|---|---|---|---|---|---|---|---|---|
| 1 | Ivd | 0.37 | 9.81E-20 | 2.04E-15 | 0.49 | 0.40 | 0.09 | 1.23 |
| 1 | Mir17hg | 0.45 | 1.33E-19 | 2.77E-15 | 0.35 | 0.26 | 0.09 | 1.34 |
| 1 | Lix1l | 0.29 | 2.48E-19 | 5.16E-15 | 0.48 | 0.37 | 0.11 | 1.29 |
| 1 | Kif21b | 0.65 | 2.89E-19 | 6.01E-15 | 0.23 | 0.16 | 0.07 | 1.46 |
| 1 | Rhou | 0.33 | 2.92E-19 | 6.06E-15 | 0.51 | 0.41 | 0.10 | 1.25 |
| 1 | Plekhh2 | 0.62 | 2.99E-19 | 6.22E-15 | 0.22 | 0.15 | 0.07 | 1.49 |
| 1 | Carhsp1 | 0.25 | 3.73E-19 | 7.76E-15 | 0.68 | 0.58 | 0.09 | 1.16 |
| 1 | Wdr35 | 0.45 | 3.93E-19 | 8.18E-15 | 0.38 | 0.29 | 0.09 | 1.31 |
| 1 | Scn2b | 0.85 | 4.48E-19 | 9.31E-15 | 0.13 | 0.08 | 0.06 | 1.71 |
| 1 | Tcf7 | 0.81 | 4.69E-19 | 9.75E-15 | 0.21 | 0.14 | 0.07 | 1.46 |
| 1 | Fgfr1op | 0.36 | 5.93E-19 | 1.23E-14 | 0.47 | 0.39 | 0.08 | 1.22 |
| 1 | Cenpw | 0.26 | 5.97E-19 | 1.24E-14 | 0.79 | 0.73 | 0.06 | 1.09 |
| 1 | Snrnp25 | 0.30 | 7.34E-19 | 1.53E-14 | 0.69 | 0.61 | 0.08 | 1.13 |
| 1 | Coq10b | 0.49 | 8.31E-19 | 1.73E-14 | 0.33 | 0.25 | 0.08 | 1.32 |
| 1 | Dlgap5 | 0.41 | 9.06E-19 | 1.88E-14 | 0.54 | 0.48 | 0.06 | 1.13 |
| 1 | Wdyhv1 | 0.49 | 1.00E-18 | 2.08E-14 | 0.34 | 0.26 | 0.08 | 1.31 |
| 1 | Tbc1d7 | 0.33 | 1.41E-18 | 2.92E-14 | 0.48 | 0.38 | 0.09 | 1.24 |
| 1 | Mertk | 0.51 | 1.73E-18 | 3.60E-14 | 0.23 | 0.16 | 0.07 | 1.47 |
| 1 | Praf2 | 0.29 | 1.79E-18 | 3.72E-14 | 0.62 | 0.52 | 0.10 | 1.18 |
| 1 | 5430416N02Rik | 0.34 | 2.26E-18 | 4.70E-14 | 0.54 | 0.45 | 0.09 | 1.19 |
| 1 | Tmem97 | 0.34 | 2.78E-18 | 5.79E-14 | 0.54 | 0.46 | 0.09 | 1.19 |
| 1 | Mrps22 | 0.29 | 3.47E-18 | 7.21E-14 | 0.65 | 0.57 | 0.08 | 1.14 |
| 1 | Il17rd | 0.35 | 3.68E-18 | 7.65E-14 | 0.59 | 0.49 | 0.09 | 1.19 |
| 1 | Selenos | 0.27 | 4.22E-18 | 8.78E-14 | 0.68 | 0.60 | 0.08 | 1.13 |
| 1 | Fads2 | 0.35 | 4.31E-18 | 8.97E-14 | 0.55 | 0.46 | 0.08 | 1.18 |
| 1 | Srxn1 | 0.74 | 9.50E-18 | 1.97E-13 | 0.17 | 0.11 | 0.06 | 1.56 |
| 1 | Eepd1 | 0.66 | 1.01E-17 | 2.09E-13 | 0.17 | 0.11 | 0.06 | 1.55 |
| 1 | Papss1 | 0.26 | 1.01E-17 | 2.11E-13 | 0.73 | 0.66 | 0.08 | 1.12 |
| 1 | Loxl1 | 0.70 | 1.08E-17 | 2.25E-13 | 0.12 | 0.07 | 0.05 | 1.74 |
| 1 | Tiam2 | 0.74 | 1.11E-17 | 2.31E-13 | 0.14 | 0.09 | 0.06 | 1.64 |
| 1 | Ptpmt1 | 0.30 | 1.52E-17 | 3.17E-13 | 0.63 | 0.55 | 0.08 | 1.14 |
| 1 | Spg20 | 0.29 | 1.71E-17 | 3.56E-13 | 0.61 | 0.53 | 0.08 | 1.15 |
| 1 | Arhgef40 | 0.27 | 1.98E-17 | 4.11E-13 | 0.61 | 0.52 | 0.10 | 1.19 |
| 1 | Exosc9 | 0.30 | 1.98E-17 | 4.13E-13 | 0.58 | 0.50 | 0.08 | 1.16 |
| 1 | Nab2 | 0.36 | 3.99E-17 | 8.30E-13 | 0.39 | 0.31 | 0.09 | 1.28 |
| 1 | Cib2 | 0.41 | 5.90E-17 | 1.23E-12 | 0.33 | 0.25 | 0.08 | 1.34 |
| 1 | Ccdc137 | 0.46 | 6.11E-17 | 1.27E-12 | 0.29 | 0.21 | 0.08 | 1.36 |
| 1 | Pwp1 | 0.27 | 6.40E-17 | 1.33E-12 | 0.64 | 0.57 | 0.07 | 1.13 |
| 1 | Por | 0.33 | 7.61E-17 | 1.58E-12 | 0.55 | 0.46 | 0.08 | 1.18 |
| 1 | Atg5 | 0.28 | 7.76E-17 | 1.61E-12 | 0.64 | 0.56 | 0.08 | 1.14 |
| 1 | Smarca1 | 0.29 | 8.40E-17 | 1.75E-12 | 0.55 | 0.46 | 0.10 | 1.21 |
| 1 | R3hcc1 | 0.29 | 8.99E-17 | 1.87E-12 | 0.63 | 0.55 | 0.08 | 1.14 |
| 1 | Tmem164 | 0.41 | 9.08E-17 | 1.89E-12 | 0.38 | 0.31 | 0.08 | 1.25 |
| 1 | Tex9 | 0.29 | 1.10E-16 | 2.29E-12 | 0.53 | 0.45 | 0.09 | 1.19 |
| 1 | Chchd6 | 0.30 | 1.14E-16 | 2.37E-12 | 0.60 | 0.53 | 0.07 | 1.14 |
| 1 | Dusp7 | 0.27 | 1.22E-16 | 2.53E-12 | 0.58 | 0.49 | 0.09 | 1.19 |
| 1 | Nln | 0.43 | 1.32E-16 | 2.74E-12 | 0.36 | 0.28 | 0.08 | 1.28 |
| 1 | Slc18b1 | 0.63 | 1.37E-16 | 2.84E-12 | 0.18 | 0.12 | 0.06 | 1.50 |
| 1 | Nudt2 | 0.26 | 1.40E-16 | 2.91E-12 | 0.66 | 0.58 | 0.08 | 1.13 |
| 1 | Tpmt | 0.36 | 1.42E-16 | 2.96E-12 | 0.45 | 0.36 | 0.09 | 1.23 |
| 1 | Eln | 0.80 | 1.48E-16 | 3.08E-12 | 0.12 | 0.07 | 0.05 | 1.68 |
| 1 | Taf13 | 0.27 | 1.49E-16 | 3.10E-12 | 0.60 | 0.51 | 0.09 | 1.18 |
| 1 | Samd14 | 0.28 | 1.55E-16 | 3.22E-12 | 0.49 | 0.40 | 0.09 | 1.23 |
| 1 | Pycr2 | 0.25 | 1.65E-16 | 3.43E-12 | 0.69 | 0.60 | 0.08 | 1.14 |
| 1 | Tfap2c | 0.45 | 1.96E-16 | 4.09E-12 | 0.35 | 0.27 | 0.08 | 1.29 |
| 1 | Angptl4 | 0.33 | 2.21E-16 | 4.60E-12 | 0.42 | 0.33 | 0.09 | 1.28 |

**Supplementary Table 1. Marker genes for the original clusters of integrated clustering of E11.5 control and NotchGOF λ-junction cells.**

| | | | | | | | | |
|---|---|---|---|---|---|---|---|---|
| 1 | Tpx2 | 0.31 | 3.07E-16 | 6.39E-12 | 0.68 | 0.65 | 0.03 | 1.05 |
| 1 | Ankrd44 | 0.40 | 3.18E-16 | 6.61E-12 | 0.35 | 0.27 | 0.08 | 1.30 |
| 1 | Ehd1 | 0.26 | 3.90E-16 | 8.11E-12 | 0.60 | 0.52 | 0.09 | 1.16 |
| 1 | Ptges2 | 0.41 | 4.46E-16 | 9.28E-12 | 0.39 | 0.31 | 0.08 | 1.25 |
| 1 | Sv2a | 0.86 | 4.96E-16 | 1.03E-11 | 0.12 | 0.07 | 0.05 | 1.66 |
| 1 | Tmem138 | 0.33 | 5.51E-16 | 1.15E-11 | 0.46 | 0.38 | 0.08 | 1.22 |
| 1 | Kdm4a | 0.29 | 5.62E-16 | 1.17E-11 | 0.49 | 0.40 | 0.09 | 1.22 |
| 1 | Pmp22 | 0.39 | 8.11E-16 | 1.69E-11 | 0.25 | 0.18 | 0.07 | 1.41 |
| 1 | Gpr45 | 0.49 | 8.40E-16 | 1.75E-11 | 0.25 | 0.18 | 0.07 | 1.39 |
| 1 | Ptcd3 | 0.26 | 8.52E-16 | 1.77E-11 | 0.57 | 0.49 | 0.08 | 1.17 |
| 1 | Sgcd | 0.38 | 1.07E-15 | 2.22E-11 | 0.29 | 0.21 | 0.08 | 1.37 |
| 1 | Srprb | 0.27 | 1.11E-15 | 2.31E-11 | 0.57 | 0.49 | 0.08 | 1.16 |
| 1 | Sdad1 | 0.29 | 1.20E-15 | 2.50E-11 | 0.55 | 0.47 | 0.08 | 1.18 |
| 1 | Sema6c | 0.51 | 1.34E-15 | 2.79E-11 | 0.22 | 0.15 | 0.07 | 1.42 |
| 1 | Ccdc167 | 0.28 | 1.37E-15 | 2.86E-11 | 0.53 | 0.45 | 0.08 | 1.19 |
| 1 | 4921531C22Rik | 0.54 | 1.39E-15 | 2.89E-11 | 0.24 | 0.17 | 0.07 | 1.38 |
| 1 | Prps1 | 0.30 | 1.40E-15 | 2.91E-11 | 0.52 | 0.44 | 0.08 | 1.18 |
| 1 | Wdr55 | 0.34 | 1.43E-15 | 2.98E-11 | 0.44 | 0.36 | 0.08 | 1.22 |
| 1 | Arhgap45 | 0.77 | 1.53E-15 | 3.18E-11 | 0.10 | 0.06 | 0.04 | 1.73 |
| 1 | Armcx6 | 0.50 | 1.66E-15 | 3.45E-11 | 0.27 | 0.20 | 0.07 | 1.36 |
| 1 | Tmem44 | 0.45 | 1.67E-15 | 3.47E-11 | 0.28 | 0.21 | 0.07 | 1.34 |
| 1 | Tsr2 | 0.31 | 1.73E-15 | 3.61E-11 | 0.47 | 0.39 | 0.09 | 1.22 |
| 1 | Nf2 | 0.40 | 1.78E-15 | 3.71E-11 | 0.36 | 0.28 | 0.08 | 1.27 |
| 1 | Hsf2 | 0.29 | 1.83E-15 | 3.81E-11 | 0.55 | 0.47 | 0.08 | 1.18 |
| 1 | Czib | 0.28 | 2.08E-15 | 4.32E-11 | 0.60 | 0.53 | 0.07 | 1.14 |
| 1 | Zc3h8 | 0.49 | 2.44E-15 | 5.08E-11 | 0.27 | 0.20 | 0.07 | 1.33 |
| 1 | Pstk | 0.26 | 2.90E-15 | 6.02E-11 | 0.63 | 0.56 | 0.08 | 1.14 |
| 1 | Ahcy | 0.44 | 2.95E-15 | 6.14E-11 | 0.34 | 0.27 | 0.07 | 1.27 |
| 1 | Timm21 | 0.26 | 3.18E-15 | 6.62E-11 | 0.61 | 0.53 | 0.08 | 1.15 |
| 1 | Aifm1 | 0.25 | 4.33E-15 | 9.01E-11 | 0.67 | 0.60 | 0.07 | 1.12 |
| 1 | Tmem121 | 0.36 | 4.37E-15 | 9.08E-11 | 0.30 | 0.22 | 0.08 | 1.34 |
| 1 | Cdca3 | 0.27 | 4.71E-15 | 9.80E-11 | 0.70 | 0.64 | 0.06 | 1.09 |
| 1 | Nqo2 | 0.66 | 4.79E-15 | 9.97E-11 | 0.18 | 0.12 | 0.06 | 1.47 |
| 1 | Bckdhb | 0.32 | 5.69E-15 | 1.18E-10 | 0.44 | 0.36 | 0.08 | 1.23 |
| 1 | Cdc27 | 0.27 | 5.72E-15 | 1.19E-10 | 0.59 | 0.52 | 0.07 | 1.14 |
| 1 | Tmem98 | 0.25 | 5.77E-15 | 1.20E-10 | 0.58 | 0.49 | 0.08 | 1.16 |
| 1 | Stradb | 0.56 | 6.22E-15 | 1.29E-10 | 0.25 | 0.19 | 0.06 | 1.34 |
| 1 | Ripk3 | 0.50 | 6.30E-15 | 1.31E-10 | 0.23 | 0.16 | 0.07 | 1.42 |
| 1 | Nsdhl | 0.35 | 6.96E-15 | 1.45E-10 | 0.44 | 0.37 | 0.07 | 1.20 |
| 1 | Parm1 | 0.35 | 6.98E-15 | 1.45E-10 | 0.22 | 0.16 | 0.07 | 1.43 |
| 1 | Ppil3 | 0.26 | 7.79E-15 | 1.62E-10 | 0.60 | 0.53 | 0.08 | 1.14 |
| 1 | Socs1 | 0.50 | 7.82E-15 | 1.63E-10 | 0.22 | 0.15 | 0.06 | 1.42 |
| 1 | Rusc1 | 0.57 | 8.22E-15 | 1.71E-10 | 0.20 | 0.14 | 0.06 | 1.43 |
| 1 | Tgfbr3 | 0.75 | 1.04E-14 | 2.17E-10 | 0.12 | 0.08 | 0.05 | 1.63 |
| 1 | Rpf2 | 0.27 | 1.07E-14 | 2.23E-10 | 0.60 | 0.53 | 0.07 | 1.14 |
| 1 | Ebp | 0.35 | 1.09E-14 | 2.27E-10 | 0.47 | 0.40 | 0.07 | 1.19 |
| 1 | Msrb3 | 0.58 | 1.14E-14 | 2.37E-10 | 0.16 | 0.11 | 0.06 | 1.52 |
| 1 | Pcca | 0.38 | 1.28E-14 | 2.67E-10 | 0.37 | 0.29 | 0.08 | 1.27 |
| 1 | Ptgis | 0.39 | 1.35E-14 | 2.81E-10 | 0.17 | 0.11 | 0.06 | 1.54 |
| 1 | Hccs | 0.38 | 1.46E-14 | 3.03E-10 | 0.38 | 0.31 | 0.07 | 1.24 |
| 1 | Avpi1 | 0.32 | 1.71E-14 | 3.55E-10 | 0.39 | 0.31 | 0.08 | 1.25 |
| 1 | Msantd3 | 0.28 | 1.98E-14 | 4.12E-10 | 0.58 | 0.50 | 0.08 | 1.15 |
| 1 | Cd2bp2 | 0.29 | 2.01E-14 | 4.19E-10 | 0.56 | 0.49 | 0.07 | 1.14 |
| 1 | Armcx3 | 0.28 | 2.16E-14 | 4.49E-10 | 0.50 | 0.42 | 0.08 | 1.19 |
| 1 | Irx3 | 0.56 | 2.21E-14 | 4.59E-10 | 0.33 | 0.27 | 0.07 | 1.26 |
| 1 | Nudt1 | 0.27 | 2.29E-14 | 4.75E-10 | 0.53 | 0.45 | 0.08 | 1.18 |

**Supplementary Table 1. Marker genes for the original clusters of integrated clustering of E11.5 control and NotchGOF λ-junction cells.**

| | | | | | | | | |
|---|---|---|---|---|---|---|---|---|
| 1 | Ppa2 | 0.25 | 2.34E-14 | 4.87E-10 | 0.63 | 0.55 | 0.08 | 1.14 |
| 1 | Gm10762 | 0.33 | 2.50E-14 | 5.21E-10 | 0.46 | 0.39 | 0.08 | 1.20 |
| 1 | Kank4 | 0.39 | 2.68E-14 | 5.58E-10 | 0.27 | 0.19 | 0.07 | 1.37 |
| 1 | Wars | 0.35 | 3.07E-14 | 6.38E-10 | 0.48 | 0.41 | 0.07 | 1.18 |
| 1 | Plpp1 | 0.46 | 3.36E-14 | 6.98E-10 | 0.20 | 0.14 | 0.06 | 1.43 |
| 1 | Sox5 | 0.28 | 3.46E-14 | 7.19E-10 | 0.47 | 0.39 | 0.08 | 1.20 |
| 1 | G0s2 | 0.56 | 3.71E-14 | 7.71E-10 | 0.21 | 0.15 | 0.06 | 1.42 |
| 1 | Rtca | 0.25 | 4.29E-14 | 8.92E-10 | 0.55 | 0.47 | 0.08 | 1.17 |
| 1 | Scamp3 | 0.26 | 4.50E-14 | 9.35E-10 | 0.60 | 0.53 | 0.07 | 1.14 |
| 1 | Lama2 | 0.44 | 4.62E-14 | 9.60E-10 | 0.20 | 0.14 | 0.06 | 1.44 |
| 1 | Zc3hc1 | 0.29 | 5.26E-14 | 1.09E-09 | 0.48 | 0.41 | 0.08 | 1.19 |
| 1 | Prmt6 | 0.36 | 6.13E-14 | 1.28E-09 | 0.34 | 0.26 | 0.07 | 1.28 |
| 1 | 2700062C07Rik | 0.33 | 6.26E-14 | 1.30E-09 | 0.45 | 0.38 | 0.08 | 1.20 |
| 1 | Pex13 | 0.26 | 6.47E-14 | 1.35E-09 | 0.65 | 0.58 | 0.07 | 1.12 |
| 1 | AW551984 | 0.57 | 6.60E-14 | 1.37E-09 | 0.17 | 0.12 | 0.06 | 1.47 |
| 1 | Prmt7 | 0.26 | 6.67E-14 | 1.39E-09 | 0.58 | 0.50 | 0.07 | 1.15 |
| 1 | Prr7 | 0.39 | 7.69E-14 | 1.60E-09 | 0.30 | 0.23 | 0.07 | 1.32 |
| 1 | Npl | 0.32 | 8.26E-14 | 1.72E-09 | 0.15 | 0.09 | 0.05 | 1.54 |
| 1 | Vldlr | 0.59 | 8.50E-14 | 1.77E-09 | 0.17 | 0.11 | 0.05 | 1.48 |
| 1 | Spata24 | 0.37 | 9.05E-14 | 1.88E-09 | 0.35 | 0.28 | 0.07 | 1.25 |
| 1 | Dhrs3 | 0.37 | 9.23E-14 | 1.92E-09 | 0.36 | 0.29 | 0.07 | 1.25 |
| 1 | Abat | 0.51 | 9.32E-14 | 1.94E-09 | 0.21 | 0.15 | 0.06 | 1.41 |
| 1 | Zfp68 | 0.25 | 9.87E-14 | 2.05E-09 | 0.50 | 0.43 | 0.08 | 1.18 |
| 1 | Itgb1bp1 | 0.29 | 1.02E-13 | 2.12E-09 | 0.47 | 0.39 | 0.08 | 1.20 |
| 1 | Osr1 | 0.55 | 1.11E-13 | 2.30E-09 | 0.11 | 0.07 | 0.04 | 1.67 |
| 1 | Clip2 | 0.46 | 1.12E-13 | 2.33E-09 | 0.25 | 0.18 | 0.06 | 1.34 |
| 1 | Slc25a38 | 0.40 | 1.36E-13 | 2.83E-09 | 0.34 | 0.27 | 0.07 | 1.26 |
| 1 | 4933404O12Rik | 0.41 | 1.54E-13 | 3.20E-09 | 0.31 | 0.24 | 0.07 | 1.28 |
| 1 | Pakap.1 | 0.25 | 1.63E-13 | 3.38E-09 | 0.49 | 0.41 | 0.09 | 1.21 |
| 1 | Nup37 | 0.26 | 1.71E-13 | 3.56E-09 | 0.59 | 0.53 | 0.06 | 1.12 |
| 1 | Iah1 | 0.26 | 1.77E-13 | 3.67E-09 | 0.59 | 0.52 | 0.07 | 1.14 |
| 1 | Eef1aknmt | 0.40 | 1.85E-13 | 3.85E-09 | 0.29 | 0.22 | 0.07 | 1.30 |
| 1 | Hspb11 | 0.25 | 1.89E-13 | 3.93E-09 | 0.52 | 0.44 | 0.08 | 1.18 |
| 1 | Tacc3 | 0.25 | 2.12E-13 | 4.41E-09 | 0.64 | 0.60 | 0.05 | 1.08 |
| 1 | Ndufaf6 | 0.31 | 3.02E-13 | 6.29E-09 | 0.41 | 0.34 | 0.07 | 1.21 |
| 1 | Ice2 | 0.40 | 3.85E-13 | 8.01E-09 | 0.31 | 0.25 | 0.07 | 1.28 |
| 1 | Orc1 | 0.55 | 4.04E-13 | 8.41E-09 | 0.18 | 0.13 | 0.05 | 1.42 |
| 1 | Bcas3 | 0.46 | 4.61E-13 | 9.58E-09 | 0.27 | 0.21 | 0.06 | 1.31 |
| 1 | Arhgap6 | 0.79 | 4.70E-13 | 9.77E-09 | 0.10 | 0.06 | 0.04 | 1.67 |
| 1 | Ccdc102a | 0.33 | 4.75E-13 | 9.88E-09 | 0.34 | 0.27 | 0.07 | 1.26 |
| 1 | Rhobtb2 | 0.54 | 4.77E-13 | 9.92E-09 | 0.17 | 0.12 | 0.05 | 1.45 |
| 1 | Olfml1 | 0.46 | 5.04E-13 | 1.05E-08 | 0.17 | 0.11 | 0.05 | 1.47 |
| 1 | Ldah | 0.27 | 5.33E-13 | 1.11E-08 | 0.51 | 0.44 | 0.07 | 1.16 |
| 1 | 2610008E11Rik | 0.30 | 5.37E-13 | 1.12E-08 | 0.46 | 0.39 | 0.07 | 1.19 |
| 1 | Ino80e | 0.27 | 5.59E-13 | 1.16E-08 | 0.58 | 0.52 | 0.06 | 1.12 |
| 1 | Prex1 | 0.39 | 5.81E-13 | 1.21E-08 | 0.25 | 0.19 | 0.06 | 1.33 |
| 1 | Pheta1 | 0.29 | 6.04E-13 | 1.26E-08 | 0.39 | 0.31 | 0.08 | 1.24 |
| 1 | Smpd2 | 0.32 | 6.67E-13 | 1.39E-08 | 0.38 | 0.31 | 0.07 | 1.23 |
| 1 | Ccdc66 | 0.31 | 8.07E-13 | 1.68E-08 | 0.45 | 0.38 | 0.07 | 1.19 |
| 1 | Sgcb | 0.44 | 8.97E-13 | 1.87E-08 | 0.27 | 0.20 | 0.06 | 1.31 |
| 1 | Dlc1 | 0.26 | 9.69E-13 | 2.02E-08 | 0.44 | 0.35 | 0.08 | 1.24 |
| 1 | Dse | 0.39 | 1.01E-12 | 2.09E-08 | 0.33 | 0.26 | 0.07 | 1.26 |
| 1 | Cdr2l | 0.46 | 1.18E-12 | 2.46E-08 | 0.25 | 0.19 | 0.06 | 1.33 |
| 1 | Actr1a | 0.26 | 1.28E-12 | 2.67E-08 | 0.53 | 0.46 | 0.07 | 1.15 |
| 1 | Gm26648 | 0.67 | 1.41E-12 | 2.94E-08 | 0.13 | 0.08 | 0.05 | 1.55 |
| 1 | Brms1l | 0.31 | 1.47E-12 | 3.06E-08 | 0.44 | 0.37 | 0.07 | 1.19 |

**Supplementary Table 1. Marker genes for the original clusters of integrated clustering of E11.5 control and NotchGOF λ-junction cells.**

| | | | | | | | | |
|---|---|---|---|---|---|---|---|---|
| 1 | Acyp1 | 0.27 | 1.57E-12 | 3.26E-08 | 0.49 | 0.42 | 0.07 | 1.16 |
| 1 | 2310010J17Rik | 0.66 | 1.63E-12 | 3.38E-08 | 0.13 | 0.08 | 0.04 | 1.54 |
| 1 | Rpap3 | 0.27 | 1.67E-12 | 3.47E-08 | 0.53 | 0.46 | 0.07 | 1.15 |
| 1 | Aen | 0.29 | 1.75E-12 | 3.63E-08 | 0.46 | 0.39 | 0.07 | 1.18 |
| 1 | Mmd | 0.30 | 2.21E-12 | 4.60E-08 | 0.42 | 0.34 | 0.08 | 1.22 |
| 1 | Dclk1 | 0.34 | 2.56E-12 | 5.33E-08 | 0.32 | 0.25 | 0.07 | 1.28 |
| 1 | Alkbh7 | 0.26 | 2.66E-12 | 5.53E-08 | 0.46 | 0.40 | 0.07 | 1.17 |
| 1 | Ptx3 | 0.34 | 3.10E-12 | 6.45E-08 | 0.16 | 0.11 | 0.05 | 1.49 |
| 1 | Gmpr2 | 0.27 | 3.15E-12 | 6.55E-08 | 0.48 | 0.41 | 0.07 | 1.17 |
| 1 | Peg12 | 0.33 | 3.65E-12 | 7.58E-08 | 0.31 | 0.25 | 0.07 | 1.28 |
| 1 | Dcun1d4 | 0.43 | 3.82E-12 | 7.94E-08 | 0.24 | 0.18 | 0.06 | 1.33 |
| 1 | Uxs1 | 0.35 | 4.39E-12 | 9.13E-08 | 0.37 | 0.30 | 0.07 | 1.22 |
| 1 | Vcpkmt | 0.43 | 4.76E-12 | 9.90E-08 | 0.27 | 0.21 | 0.06 | 1.29 |
| 1 | Zfp354c | 0.26 | 5.06E-12 | 1.05E-07 | 0.46 | 0.38 | 0.07 | 1.19 |
| 1 | 1500015A07Rik | 0.51 | 5.18E-12 | 1.08E-07 | 0.17 | 0.12 | 0.05 | 1.42 |
| 1 | Col6a2 | 0.34 | 5.76E-12 | 1.20E-07 | 0.20 | 0.14 | 0.06 | 1.40 |
| 1 | Cdc42ep4 | 0.26 | 5.98E-12 | 1.24E-07 | 0.51 | 0.44 | 0.07 | 1.16 |
| 1 | Grem1 | 0.38 | 6.01E-12 | 1.25E-07 | 0.14 | 0.09 | 0.05 | 1.51 |
| 1 | Elp6 | 0.28 | 7.29E-12 | 1.52E-07 | 0.45 | 0.38 | 0.07 | 1.19 |
| 1 | Fhod3 | 0.66 | 9.09E-12 | 1.89E-07 | 0.13 | 0.09 | 0.04 | 1.51 |
| 1 | Elovl2 | 0.60 | 1.06E-11 | 2.20E-07 | 0.15 | 0.11 | 0.05 | 1.45 |
| 1 | Il6st | 0.32 | 1.07E-11 | 2.22E-07 | 0.35 | 0.28 | 0.07 | 1.24 |
| 1 | Castor2 | 0.53 | 1.24E-11 | 2.59E-07 | 0.18 | 0.13 | 0.05 | 1.39 |
| 1 | Baiap2l2 | 0.46 | 1.49E-11 | 3.10E-07 | 0.16 | 0.11 | 0.05 | 1.44 |
| 1 | Gpr180 | 0.33 | 1.65E-11 | 3.44E-07 | 0.36 | 0.29 | 0.07 | 1.23 |
| 1 | Atp23 | 0.37 | 1.77E-11 | 3.68E-07 | 0.29 | 0.23 | 0.06 | 1.26 |
| 1 | Acot1 | 0.27 | 2.10E-11 | 4.36E-07 | 0.42 | 0.34 | 0.07 | 1.21 |
| 1 | Adgra2 | 0.31 | 2.20E-11 | 4.57E-07 | 0.30 | 0.23 | 0.07 | 1.29 |
| 1 | Hsd17b7 | 0.35 | 2.36E-11 | 4.91E-07 | 0.33 | 0.27 | 0.06 | 1.24 |
| 1 | Unc119b | 0.31 | 2.68E-11 | 5.57E-07 | 0.40 | 0.34 | 0.06 | 1.19 |
| 1 | Rgl1 | 0.34 | 2.72E-11 | 5.66E-07 | 0.20 | 0.15 | 0.06 | 1.38 |
| 1 | Ppp2r1b | 0.31 | 3.44E-11 | 7.15E-07 | 0.45 | 0.39 | 0.06 | 1.16 |
| 1 | Ush1c | 0.61 | 3.84E-11 | 7.98E-07 | 0.15 | 0.10 | 0.05 | 1.45 |
| 1 | 5330438D12Rik | 0.67 | 3.85E-11 | 8.01E-07 | 0.14 | 0.10 | 0.04 | 1.44 |
| 1 | Hgsnat | 0.39 | 3.91E-11 | 8.14E-07 | 0.25 | 0.19 | 0.06 | 1.30 |
| 1 | Eif2s3y | 0.53 | 4.03E-11 | 8.38E-07 | 0.12 | 0.08 | 0.04 | 1.53 |
| 1 | Nhsl2 | 0.49 | 4.25E-11 | 8.83E-07 | 0.21 | 0.16 | 0.05 | 1.34 |
| 1 | Dnajc17 | 0.29 | 4.32E-11 | 8.98E-07 | 0.39 | 0.33 | 0.07 | 1.21 |
| 1 | Tmem184c | 0.28 | 4.72E-11 | 9.82E-07 | 0.41 | 0.34 | 0.07 | 1.20 |
| 1 | Crat | 0.39 | 4.74E-11 | 9.86E-07 | 0.26 | 0.20 | 0.06 | 1.29 |
| 1 | Magee1 | 0.46 | 5.19E-11 | 1.08E-06 | 0.18 | 0.13 | 0.05 | 1.39 |
| 1 | Antxr1 | 0.26 | 5.75E-11 | 1.20E-06 | 0.38 | 0.31 | 0.07 | 1.23 |
| 1 | Rab39b | 0.46 | 5.86E-11 | 1.22E-06 | 0.21 | 0.16 | 0.05 | 1.34 |
| 1 | Mvd | 0.31 | 5.98E-11 | 1.24E-06 | 0.36 | 0.30 | 0.06 | 1.22 |
| 1 | Fam174b | 0.51 | 6.37E-11 | 1.33E-06 | 0.16 | 0.11 | 0.05 | 1.43 |
| 1 | Nid2 | 0.31 | 7.35E-11 | 1.53E-06 | 0.30 | 0.23 | 0.07 | 1.28 |
| 1 | Zfp239 | 0.31 | 7.56E-11 | 1.57E-06 | 0.30 | 0.23 | 0.06 | 1.27 |
| 1 | Poglut2 | 0.28 | 8.16E-11 | 1.70E-06 | 0.45 | 0.39 | 0.06 | 1.16 |
| 1 | Tmem41a | 0.36 | 8.40E-11 | 1.75E-06 | 0.27 | 0.22 | 0.06 | 1.27 |
| 1 | 3110056K07Rik | 0.35 | 8.59E-11 | 1.79E-06 | 0.30 | 0.24 | 0.06 | 1.26 |
| 1 | Nphp1 | 0.30 | 9.87E-11 | 2.05E-06 | 0.36 | 0.30 | 0.06 | 1.21 |
| 1 | Gm42067 | 0.55 | 1.00E-10 | 2.08E-06 | 0.17 | 0.12 | 0.05 | 1.37 |
| 1 | Hilpda | 0.27 | 1.05E-10 | 2.19E-06 | 0.42 | 0.35 | 0.07 | 1.19 |
| 1 | Psph | 0.33 | 1.06E-10 | 2.20E-06 | 0.33 | 0.28 | 0.06 | 1.21 |
| 1 | Impdh1 | 0.38 | 1.06E-10 | 2.20E-06 | 0.27 | 0.21 | 0.06 | 1.27 |
| 1 | Gm36988 | 0.42 | 1.16E-10 | 2.41E-06 | 0.19 | 0.14 | 0.05 | 1.37 |

**Supplementary Table 1. Marker genes for the original clusters of integrated clustering of E11.5 control and NotchGOF λ-junction cells.**

| | | | | | | | | |
|---|---|---|---|---|---|---|---|---|
| 1 | Rita1 | 0.41 | 1.17E-10 | 2.44E-06 | 0.25 | 0.20 | 0.06 | 1.29 |
| 1 | Stat5b | 0.51 | 1.21E-10 | 2.52E-06 | 0.17 | 0.12 | 0.05 | 1.38 |
| 1 | Reck | 0.28 | 1.39E-10 | 2.90E-06 | 0.31 | 0.24 | 0.07 | 1.27 |
| 1 | 2810013P06Rik | 0.35 | 1.41E-10 | 2.92E-06 | 0.35 | 0.29 | 0.06 | 1.21 |
| 1 | Ints8 | 0.27 | 1.49E-10 | 3.10E-06 | 0.48 | 0.43 | 0.06 | 1.14 |
| 1 | Dpysl4 | 0.36 | 1.51E-10 | 3.14E-06 | 0.23 | 0.18 | 0.06 | 1.32 |
| 1 | 1110002L01Rik | 0.43 | 1.80E-10 | 3.74E-06 | 0.24 | 0.18 | 0.05 | 1.29 |
| 1 | Lrrc20 | 0.45 | 1.81E-10 | 3.77E-06 | 0.20 | 0.15 | 0.05 | 1.35 |
| 1 | Aars | 0.26 | 1.96E-10 | 4.08E-06 | 0.52 | 0.46 | 0.05 | 1.12 |
| 1 | Cdc25b | 0.37 | 2.06E-10 | 4.28E-06 | 0.26 | 0.20 | 0.06 | 1.28 |
| 1 | Ptges | 0.41 | 2.07E-10 | 4.30E-06 | 0.19 | 0.14 | 0.05 | 1.37 |
| 1 | Ykt6 | 0.29 | 2.14E-10 | 4.45E-06 | 0.47 | 0.40 | 0.06 | 1.15 |
| 1 | Ube2o | 0.29 | 2.18E-10 | 4.53E-06 | 0.45 | 0.38 | 0.06 | 1.16 |
| 1 | Gosr2 | 0.27 | 2.19E-10 | 4.56E-06 | 0.41 | 0.35 | 0.06 | 1.16 |
| 1 | Aspm | 0.48 | 2.19E-10 | 4.56E-06 | 0.44 | 0.41 | 0.03 | 1.07 |
| 1 | Bub1b | 0.27 | 2.24E-10 | 4.67E-06 | 0.59 | 0.55 | 0.04 | 1.07 |
| 1 | Cryz | 0.26 | 2.29E-10 | 4.77E-06 | 0.43 | 0.37 | 0.06 | 1.17 |
| 1 | Smim26 | 0.27 | 3.04E-10 | 6.32E-06 | 0.49 | 0.43 | 0.06 | 1.14 |
| 1 | Spock3 | 0.34 | 3.24E-10 | 6.74E-06 | 0.18 | 0.13 | 0.05 | 1.40 |
| 1 | Ube2g2 | 0.31 | 3.38E-10 | 7.03E-06 | 0.38 | 0.32 | 0.06 | 1.18 |
| 1 | Vasn | 0.27 | 3.57E-10 | 7.43E-06 | 0.28 | 0.21 | 0.06 | 1.30 |
| 1 | Hyou1 | 0.36 | 3.91E-10 | 8.13E-06 | 0.34 | 0.29 | 0.06 | 1.20 |
| 1 | Vhl | 0.34 | 4.52E-10 | 9.41E-06 | 0.29 | 0.24 | 0.06 | 1.24 |
| 1 | Zfp286 | 0.31 | 5.61E-10 | 1.17E-05 | 0.30 | 0.24 | 0.06 | 1.25 |
| 1 | Ccnb1 | 0.35 | 5.65E-10 | 1.18E-05 | 0.61 | 0.58 | 0.03 | 1.06 |
| 1 | Fbxo21 | 0.25 | 6.50E-10 | 1.35E-05 | 0.44 | 0.38 | 0.06 | 1.17 |
| 1 | Mettl2 | 0.26 | 7.54E-10 | 1.57E-05 | 0.42 | 0.35 | 0.06 | 1.18 |
| 1 | Insyn1 | 0.63 | 7.63E-10 | 1.59E-05 | 0.10 | 0.07 | 0.04 | 1.53 |
| 1 | Snx24 | 0.28 | 7.77E-10 | 1.62E-05 | 0.42 | 0.35 | 0.07 | 1.18 |
| 1 | Dpf1 | 0.42 | 8.03E-10 | 1.67E-05 | 0.21 | 0.16 | 0.05 | 1.32 |
| 1 | Cenpe | 0.29 | 9.35E-10 | 1.94E-05 | 0.69 | 0.65 | 0.03 | 1.05 |
| 1 | Cdkl3 | 0.47 | 1.03E-09 | 2.14E-05 | 0.18 | 0.13 | 0.05 | 1.36 |
| 1 | Kbtbd4 | 0.29 | 1.03E-09 | 2.15E-05 | 0.40 | 0.34 | 0.06 | 1.18 |
| 1 | Gm49700 | 0.41 | 1.05E-09 | 2.19E-05 | 0.21 | 0.16 | 0.05 | 1.32 |
| 1 | Dhcr24 | 0.31 | 1.12E-09 | 2.33E-05 | 0.41 | 0.35 | 0.06 | 1.16 |
| 1 | Zfp768 | 0.27 | 1.12E-09 | 2.34E-05 | 0.39 | 0.33 | 0.06 | 1.19 |
| 1 | Chst10 | 0.64 | 1.28E-09 | 2.66E-05 | 0.12 | 0.08 | 0.04 | 1.47 |
| 1 | Ppcs | 0.29 | 1.30E-09 | 2.71E-05 | 0.35 | 0.29 | 0.06 | 1.21 |
| 1 | Dnajc25 | 0.36 | 1.36E-09 | 2.84E-05 | 0.32 | 0.27 | 0.06 | 1.21 |
| 1 | Zfp668 | 0.33 | 1.58E-09 | 3.29E-05 | 0.29 | 0.24 | 0.06 | 1.23 |
| 1 | Pik3r3 | 0.28 | 1.59E-09 | 3.30E-05 | 0.33 | 0.27 | 0.06 | 1.22 |
| 1 | Syne1 | 0.44 | 1.61E-09 | 3.34E-05 | 0.18 | 0.13 | 0.05 | 1.35 |
| 1 | Vash2 | 0.27 | 1.67E-09 | 3.46E-05 | 0.35 | 0.28 | 0.06 | 1.22 |
| 1 | Ccdc136 | 0.26 | 1.94E-09 | 4.03E-05 | 0.34 | 0.28 | 0.06 | 1.23 |
| 1 | Nif3l1 | 0.27 | 1.97E-09 | 4.10E-05 | 0.41 | 0.35 | 0.06 | 1.17 |
| 1 | Slc4a8 | 0.55 | 2.07E-09 | 4.31E-05 | 0.14 | 0.10 | 0.04 | 1.42 |
| 1 | Ckap2 | 0.30 | 2.45E-09 | 5.09E-05 | 0.44 | 0.39 | 0.05 | 1.12 |
| 1 | 3010003L21Rik | 0.39 | 2.58E-09 | 5.37E-05 | 0.36 | 0.31 | 0.05 | 1.17 |
| 1 | Mcm3ap | 0.26 | 2.59E-09 | 5.39E-05 | 0.39 | 0.33 | 0.06 | 1.18 |
| 1 | Sapcd2 | 0.38 | 2.66E-09 | 5.53E-05 | 0.30 | 0.25 | 0.05 | 1.20 |
| 1 | 2300009A05Rik | 0.27 | 2.73E-09 | 5.67E-05 | 0.44 | 0.39 | 0.05 | 1.14 |
| 1 | Mras | 0.32 | 2.87E-09 | 5.96E-05 | 0.26 | 0.20 | 0.06 | 1.27 |
| 1 | Sgip1 | 0.40 | 2.90E-09 | 6.02E-05 | 0.15 | 0.11 | 0.04 | 1.41 |
| 1 | Fam229b | 0.47 | 3.12E-09 | 6.49E-05 | 0.15 | 0.11 | 0.04 | 1.40 |
| 1 | Aifm2 | 0.49 | 3.72E-09 | 7.74E-05 | 0.14 | 0.10 | 0.04 | 1.40 |
| 1 | Gfod2 | 0.31 | 4.84E-09 | 1.01E-04 | 0.36 | 0.30 | 0.06 | 1.19 |

**Supplementary Table 1. Marker genes for the original clusters of integrated clustering of E11.5 control and NotchGOF λ-junction cells.**

| | | | | | | | | |
|---|---|---|---|---|---|---|---|---|
| 1 | Shkbp1 | 0.26 | 5.12E-09 | 1.06E-04 | 0.39 | 0.33 | 0.06 | 1.18 |
| 1 | Gt(ROSA)26Sor | 0.32 | 5.48E-09 | 1.14E-04 | 0.29 | 0.23 | 0.05 | 1.22 |
| 1 | Gstk1 | 0.32 | 5.57E-09 | 1.16E-04 | 0.22 | 0.17 | 0.05 | 1.29 |
| 1 | Micu1 | 0.27 | 5.71E-09 | 1.19E-04 | 0.39 | 0.33 | 0.06 | 1.18 |
| 1 | Zfp334 | 0.25 | 6.27E-09 | 1.30E-04 | 0.39 | 0.33 | 0.06 | 1.18 |
| 1 | Zfp599 | 0.38 | 6.40E-09 | 1.33E-04 | 0.23 | 0.18 | 0.05 | 1.27 |
| 1 | Tlcd2 | 0.46 | 6.53E-09 | 1.36E-04 | 0.18 | 0.13 | 0.04 | 1.33 |
| 1 | Sirt5 | 0.27 | 7.90E-09 | 1.64E-04 | 0.32 | 0.26 | 0.06 | 1.21 |
| 1 | Gm8251 | 0.64 | 8.33E-09 | 1.73E-04 | 0.10 | 0.07 | 0.03 | 1.47 |
| 1 | 3110040N11Rik | 0.36 | 8.64E-09 | 1.80E-04 | 0.24 | 0.20 | 0.05 | 1.25 |
| 1 | Lrrc4b | 0.36 | 8.68E-09 | 1.80E-04 | 0.17 | 0.13 | 0.05 | 1.35 |
| 1 | Ccl25 | 0.31 | 8.76E-09 | 1.82E-04 | 0.32 | 0.27 | 0.05 | 1.20 |
| 1 | Raet1d | 0.58 | 9.23E-09 | 1.92E-04 | 0.12 | 0.08 | 0.04 | 1.43 |
| 1 | Ttl | 0.38 | 9.30E-09 | 1.93E-04 | 0.27 | 0.22 | 0.05 | 1.23 |
| 1 | Fam241b | 0.31 | 9.88E-09 | 2.06E-04 | 0.21 | 0.16 | 0.05 | 1.30 |
| 1 | Bnc2 | 0.26 | 1.07E-08 | 2.24E-04 | 0.23 | 0.18 | 0.05 | 1.30 |
| 1 | Slc5a3 | 0.33 | 1.25E-08 | 2.59E-04 | 0.25 | 0.20 | 0.05 | 1.25 |
| 1 | D430019H16Rik | 0.32 | 1.51E-08 | 3.14E-04 | 0.24 | 0.19 | 0.05 | 1.27 |
| 1 | Adam19 | 0.31 | 1.61E-08 | 3.34E-04 | 0.20 | 0.15 | 0.05 | 1.32 |
| 1 | Slc25a20 | 0.30 | 1.61E-08 | 3.35E-04 | 0.28 | 0.23 | 0.05 | 1.23 |
| 1 | Cep41 | 0.30 | 1.66E-08 | 3.44E-04 | 0.27 | 0.22 | 0.05 | 1.23 |
| 1 | E2f1 | 0.30 | 1.85E-08 | 3.84E-04 | 0.32 | 0.27 | 0.05 | 1.20 |
| 1 | Trip4 | 0.27 | 2.01E-08 | 4.18E-04 | 0.34 | 0.28 | 0.06 | 1.20 |
| 1 | Sacs | 0.25 | 2.28E-08 | 4.73E-04 | 0.22 | 0.17 | 0.05 | 1.29 |
| 1 | Tmem267 | 0.44 | 2.28E-08 | 4.74E-04 | 0.21 | 0.17 | 0.04 | 1.27 |
| 1 | Rilpl2 | 0.25 | 2.31E-08 | 4.81E-04 | 0.33 | 0.27 | 0.06 | 1.21 |
| 1 | Mxra8 | 0.26 | 2.52E-08 | 5.24E-04 | 0.19 | 0.14 | 0.05 | 1.32 |
| 1 | 2810410L24Rik | 0.51 | 2.58E-08 | 5.36E-04 | 0.11 | 0.08 | 0.03 | 1.45 |
| 1 | Vat1 | 0.40 | 2.66E-08 | 5.54E-04 | 0.18 | 0.13 | 0.04 | 1.32 |
| 1 | Sipa1l2 | 0.44 | 3.40E-08 | 7.07E-04 | 0.17 | 0.13 | 0.04 | 1.31 |
| 1 | Zfp932 | 0.29 | 3.48E-08 | 7.24E-04 | 0.29 | 0.23 | 0.05 | 1.22 |
| 1 | Alg14 | 0.26 | 3.49E-08 | 7.25E-04 | 0.36 | 0.31 | 0.05 | 1.18 |
| 1 | Spata6 | 0.33 | 3.81E-08 | 7.93E-04 | 0.25 | 0.20 | 0.05 | 1.24 |
| 1 | Heg1 | 0.32 | 3.93E-08 | 8.17E-04 | 0.19 | 0.15 | 0.05 | 1.30 |
| 1 | Rab36 | 0.39 | 4.13E-08 | 8.59E-04 | 0.16 | 0.12 | 0.04 | 1.34 |
| 1 | Fam171b | 0.37 | 4.22E-08 | 8.78E-04 | 0.29 | 0.24 | 0.05 | 1.21 |
| 1 | Rpusd4 | 0.29 | 4.54E-08 | 9.45E-04 | 0.36 | 0.30 | 0.05 | 1.17 |
| 1 | Pcolce2 | 0.37 | 5.23E-08 | 1.09E-03 | 0.18 | 0.13 | 0.04 | 1.31 |
| 1 | Pacsin1 | 0.46 | 5.98E-08 | 1.24E-03 | 0.14 | 0.10 | 0.04 | 1.37 |
| 1 | Pdxk | 0.43 | 6.16E-08 | 1.28E-03 | 0.15 | 0.11 | 0.04 | 1.35 |
| 1 | Rnf26 | 0.26 | 6.36E-08 | 1.32E-03 | 0.41 | 0.36 | 0.05 | 1.14 |
| 1 | Mppe1 | 0.39 | 6.48E-08 | 1.35E-03 | 0.17 | 0.13 | 0.04 | 1.32 |
| 1 | Rcc1l | 0.30 | 7.48E-08 | 1.56E-03 | 0.28 | 0.24 | 0.05 | 1.21 |
| 1 | Ddx3y | 0.44 | 7.89E-08 | 1.64E-03 | 0.11 | 0.07 | 0.03 | 1.45 |
| 1 | Lnpep | 0.30 | 8.19E-08 | 1.70E-03 | 0.33 | 0.29 | 0.05 | 1.17 |
| 1 | Endog | 0.35 | 8.40E-08 | 1.75E-03 | 0.26 | 0.22 | 0.05 | 1.21 |
| 1 | Plcl2 | 0.55 | 8.84E-08 | 1.84E-03 | 0.15 | 0.11 | 0.04 | 1.33 |
| 1 | 9330020H09Rik | 0.46 | 9.21E-08 | 1.92E-03 | 0.15 | 0.11 | 0.04 | 1.34 |
| 1 | Ndufaf1 | 0.26 | 9.69E-08 | 2.01E-03 | 0.34 | 0.29 | 0.05 | 1.18 |
| 1 | 2010204K13Rik | 0.33 | 9.74E-08 | 2.03E-03 | 0.30 | 0.25 | 0.05 | 1.19 |
| 1 | Lmf1 | 0.33 | 1.16E-07 | 2.41E-03 | 0.27 | 0.22 | 0.05 | 1.22 |
| 1 | Mgat5 | 0.32 | 1.62E-07 | 3.36E-03 | 0.31 | 0.26 | 0.05 | 1.19 |
| 1 | Agbl5 | 0.38 | 1.67E-07 | 3.46E-03 | 0.24 | 0.19 | 0.04 | 1.23 |
| 1 | Def8 | 0.33 | 1.73E-07 | 3.59E-03 | 0.23 | 0.18 | 0.04 | 1.24 |
| 1 | Abitram | 0.35 | 1.76E-07 | 3.67E-03 | 0.27 | 0.23 | 0.05 | 1.20 |
| 1 | Ccdc117 | 0.33 | 1.79E-07 | 3.72E-03 | 0.24 | 0.20 | 0.04 | 1.22 |

**Supplementary Table 1. Marker genes for the original clusters of integrated clustering of E11.5 control and NotchGOF λ-junction cells.**

| | | | | | | | | |
|---|---|---|---|---|---|---|---|---|
| 1 | Ndufaf5 | 0.30 | 1.84E-07 | 3.83E-03 | 0.24 | 0.20 | 0.05 | 1.23 |
| 1 | Fastkd2 | 0.26 | 1.84E-07 | 3.83E-03 | 0.36 | 0.31 | 0.05 | 1.16 |
| 1 | Atg4c | 0.41 | 1.92E-07 | 3.98E-03 | 0.11 | 0.08 | 0.03 | 1.42 |
| 1 | Lrrc8a | 0.31 | 2.00E-07 | 4.15E-03 | 0.31 | 0.27 | 0.04 | 1.16 |
| 1 | Suox | 0.47 | 2.04E-07 | 4.25E-03 | 0.11 | 0.08 | 0.03 | 1.41 |
| 1 | Camkk2 | 0.27 | 2.28E-07 | 4.75E-03 | 0.28 | 0.23 | 0.05 | 1.21 |
| 1 | Sh2b2 | 0.28 | 2.45E-07 | 5.10E-03 | 0.20 | 0.16 | 0.05 | 1.29 |
| 1 | Slc25a25 | 0.32 | 2.46E-07 | 5.12E-03 | 0.22 | 0.18 | 0.04 | 1.25 |
| 1 | Nmb | 0.52 | 2.50E-07 | 5.19E-03 | 0.11 | 0.08 | 0.03 | 1.41 |
| 1 | Rabl3 | 0.29 | 2.54E-07 | 5.28E-03 | 0.23 | 0.18 | 0.05 | 1.24 |
| 1 | Zc4h2 | 0.35 | 2.56E-07 | 5.33E-03 | 0.23 | 0.19 | 0.05 | 1.24 |
| 1 | Simc1 | 0.36 | 2.58E-07 | 5.37E-03 | 0.17 | 0.13 | 0.04 | 1.29 |
| 1 | Gm5784 | 0.56 | 2.63E-07 | 5.47E-03 | 0.10 | 0.07 | 0.03 | 1.43 |
| 1 | Coq3 | 0.34 | 2.64E-07 | 5.48E-03 | 0.22 | 0.18 | 0.04 | 1.23 |
| 1 | 4930581F22Rik | 0.47 | 2.66E-07 | 5.52E-03 | 0.12 | 0.09 | 0.03 | 1.38 |
| 1 | Cip2a | 0.29 | 3.03E-07 | 6.30E-03 | 0.43 | 0.39 | 0.04 | 1.10 |
| 1 | Edn3 | 0.29 | 3.05E-07 | 6.34E-03 | 0.18 | 0.14 | 0.04 | 1.30 |
| 1 | Bdh2 | 0.41 | 3.09E-07 | 6.43E-03 | 0.13 | 0.09 | 0.03 | 1.37 |
| 1 | Mtmr11 | 0.36 | 3.53E-07 | 7.34E-03 | 0.14 | 0.10 | 0.04 | 1.36 |
| 1 | Tstd2 | 0.29 | 4.30E-07 | 8.94E-03 | 0.31 | 0.27 | 0.05 | 1.17 |
| 1 | Dcun1d3 | 0.41 | 4.43E-07 | 9.21E-03 | 0.19 | 0.15 | 0.04 | 1.27 |
| 1 | Glce | 0.33 | 4.58E-07 | 9.53E-03 | 0.22 | 0.18 | 0.04 | 1.23 |
| 1 | Slc11a2 | 0.46 | 4.73E-07 | 9.83E-03 | 0.21 | 0.17 | 0.04 | 1.22 |
| 1 | C230038L03Rik | 0.38 | 5.00E-07 | 1.04E-02 | 0.16 | 0.12 | 0.04 | 1.32 |
| 1 | Zfp454 | 0.40 | 5.17E-07 | 1.07E-02 | 0.16 | 0.12 | 0.04 | 1.31 |
| 1 | Jmjd8 | 0.27 | 5.40E-07 | 1.12E-02 | 0.33 | 0.28 | 0.05 | 1.17 |
| 1 | Rufy1 | 0.28 | 5.56E-07 | 1.16E-02 | 0.29 | 0.25 | 0.05 | 1.19 |
| 1 | Pdp2 | 0.41 | 5.57E-07 | 1.16E-02 | 0.21 | 0.17 | 0.04 | 1.23 |
| 1 | Ccnj | 0.36 | 5.59E-07 | 1.16E-02 | 0.21 | 0.17 | 0.04 | 1.24 |
| 1 | Fam149a | 0.37 | 5.69E-07 | 1.18E-02 | 0.18 | 0.14 | 0.04 | 1.27 |
| 1 | Lhpp | 0.44 | 7.23E-07 | 1.50E-02 | 0.13 | 0.10 | 0.03 | 1.34 |
| 1 | Stard4 | 0.25 | 7.25E-07 | 1.51E-02 | 0.32 | 0.28 | 0.05 | 1.17 |
| 1 | Slc2a4 | 0.32 | 8.32E-07 | 1.73E-02 | 0.21 | 0.17 | 0.04 | 1.24 |
| 1 | Lipe | 0.48 | 8.53E-07 | 1.77E-02 | 0.11 | 0.08 | 0.03 | 1.37 |
| 1 | St3gal1 | 0.55 | 8.63E-07 | 1.80E-02 | 0.13 | 0.10 | 0.03 | 1.31 |
| 1 | Oscp1 | 0.30 | 8.85E-07 | 1.84E-02 | 0.24 | 0.19 | 0.04 | 1.22 |
| 1 | Lpar2 | 0.57 | 9.10E-07 | 1.89E-02 | 0.10 | 0.07 | 0.03 | 1.39 |
| 1 | Phactr3 | 0.42 | 9.88E-07 | 2.05E-02 | 0.10 | 0.07 | 0.03 | 1.41 |
| 1 | Hist1h2ag | 0.28 | 1.00E-06 | 2.08E-02 | 0.35 | 0.31 | 0.04 | 1.14 |
| 1 | Tchp | 0.27 | 1.34E-06 | 2.78E-02 | 0.34 | 0.29 | 0.04 | 1.15 |
| 1 | Zmat3 | 0.36 | 1.35E-06 | 2.80E-02 | 0.23 | 0.19 | 0.04 | 1.21 |
| 1 | Pxdc1 | 0.63 | 1.37E-06 | 2.85E-02 | 0.13 | 0.10 | 0.03 | 1.31 |
| 1 | Hsd11b2 | 0.44 | 1.39E-06 | 2.89E-02 | 0.13 | 0.10 | 0.03 | 1.33 |
| 1 | Cdr2 | 0.28 | 1.43E-06 | 2.98E-02 | 0.26 | 0.22 | 0.04 | 1.19 |
| 1 | Tvp23a | 0.36 | 1.48E-06 | 3.08E-02 | 0.17 | 0.13 | 0.04 | 1.28 |
| 1 | Uck1 | 0.26 | 1.54E-06 | 3.20E-02 | 0.32 | 0.27 | 0.05 | 1.17 |
| 1 | Spsb1 | 0.33 | 1.67E-06 | 3.48E-02 | 0.19 | 0.15 | 0.04 | 1.26 |
| 1 | Trmt61a | 0.36 | 1.76E-06 | 3.66E-02 | 0.18 | 0.15 | 0.04 | 1.26 |
| 1 | Sergef | 0.37 | 1.77E-06 | 3.68E-02 | 0.17 | 0.13 | 0.04 | 1.27 |
| 1 | Lin28b | 0.37 | 1.95E-06 | 4.06E-02 | 0.24 | 0.20 | 0.04 | 1.19 |
| 1 | 3110021N24Rik | 0.40 | 2.10E-06 | 4.37E-02 | 0.14 | 0.11 | 0.03 | 1.30 |
| 1 | Nqo1 | 0.52 | 2.38E-06 | 4.95E-02 | 0.13 | 0.10 | 0.03 | 1.31 |
| 1 | 5930430L01Rik | 0.38 | 2.40E-06 | 5.00E-02 | 0.13 | 0.10 | 0.03 | 1.33 |
| 2 | Lhx8 | 2.37 | 0 | 0 | 0.82 | 0.21 | 0.61 | 3.85 |
| 2 | Mab21l2 | 1.63 | 0 | 0 | 0.97 | 0.45 | 0.53 | 2.18 |
| 2 | Hist1h1a | 1.70 | 0 | 0 | 0.90 | 0.37 | 0.53 | 2.41 |

**Supplementary Table 1. Marker genes for the original clusters of integrated clustering of E11.5 control and NotchGOF λ-junction cells.**

| | | | | | | | | |
|---|---|---|---|---|---|---|---|---|
| 2 | Hist1h2ab | 1.83 | 0 | 0 | 0.84 | 0.32 | 0.52 | 2.60 |
| 2 | Hist2h2ac | 1.81 | 0 | 0 | 0.78 | 0.28 | 0.50 | 2.82 |
| 2 | Satb2 | 2.06 | 0 | 0 | 0.71 | 0.21 | 0.50 | 3.42 |
| 2 | Hist1h4d | 1.55 | 0 | 0 | 0.89 | 0.41 | 0.48 | 2.16 |
| 2 | Tbx3 | 1.50 | 0 | 0 | 0.87 | 0.39 | 0.48 | 2.21 |
| 2 | Hist1h2an | 1.73 | 0 | 0 | 0.71 | 0.26 | 0.45 | 2.70 |
| 2 | Hist1h3e | 1.44 | 0 | 0 | 0.83 | 0.38 | 0.44 | 2.16 |
| 2 | Hist1h1d | 1.44 | 0 | 0 | 0.73 | 0.31 | 0.42 | 2.33 |
| 2 | Hist1h3c | 1.52 | 0 | 0 | 0.97 | 0.56 | 0.41 | 1.74 |
| 2 | Hist1h4h | 1.40 | 0 | 0 | 0.78 | 0.37 | 0.41 | 2.09 |
| 2 | 9130024F11Rik | 1.56 | 0 | 0 | 0.59 | 0.18 | 0.41 | 3.26 |
| 2 | Ndnf | 1.68 | 0 | 0 | 0.57 | 0.17 | 0.40 | 3.34 |
| 2 | Hist1h1b | 1.91 | 0 | 0 | 1.00 | 0.61 | 0.39 | 1.65 |
| 2 | Fbxo5 | 1.06 | 0 | 0 | 0.94 | 0.55 | 0.38 | 1.69 |
| 2 | Hist1h1e | 1.42 | 0 | 0 | 0.99 | 0.61 | 0.38 | 1.63 |
| 2 | Hist1h2ae | 1.79 | 0 | 0 | 1.00 | 0.67 | 0.33 | 1.49 |
| 2 | Hist1h2ap | 1.75 | 0 | 0 | 1.00 | 0.73 | 0.27 | 1.38 |
| 2 | AI606473 | 2.33 | 0 | 0 | 0.34 | 0.06 | 0.27 | 5.33 |
| 2 | Top2a | 1.08 | 0 | 0 | 1.00 | 0.77 | 0.23 | 1.30 |
| 2 | Pclaf | 0.84 | 0 | 0 | 1.00 | 0.88 | 0.12 | 1.14 |
| 2 | Hist1h3f | 1.59 | 3.60E-303 | 7.48E-299 | 0.61 | 0.24 | 0.38 | 2.59 |
| 2 | Hist1h2bm | 1.52 | 3.87E-295 | 8.05E-291 | 0.66 | 0.27 | 0.38 | 2.41 |
| 2 | Hist2h2bb | 1.56 | 8.75E-281 | 1.82E-276 | 0.58 | 0.22 | 0.36 | 2.68 |
| 2 | Hist1h2be | 1.56 | 1.82E-277 | 3.78E-273 | 0.60 | 0.24 | 0.36 | 2.52 |
| 2 | Hist1h1c | 1.32 | 2.71E-275 | 5.64E-271 | 0.76 | 0.38 | 0.38 | 1.98 |
| 2 | Postn | 2.07 | 4.18E-275 | 8.70E-271 | 0.31 | 0.07 | 0.24 | 4.65 |
| 2 | Hist1h3b | 1.64 | 3.40E-272 | 7.07E-268 | 0.54 | 0.20 | 0.34 | 2.73 |
| 2 | Epha7 | 1.11 | 4.75E-271 | 9.87E-267 | 0.89 | 0.53 | 0.35 | 1.66 |
| 2 | Dek | 0.56 | 5.95E-264 | 1.24E-259 | 1.00 | 0.96 | 0.04 | 1.04 |
| 2 | Cdh11 | 0.82 | 8.85E-255 | 1.84E-250 | 0.98 | 0.67 | 0.31 | 1.46 |
| 2 | Col26a1 | 1.05 | 5.18E-243 | 1.08E-238 | 0.91 | 0.58 | 0.33 | 1.57 |
| 2 | Atad2 | 0.93 | 3.90E-241 | 8.10E-237 | 0.95 | 0.72 | 0.24 | 1.33 |
| 2 | Basp1 | 0.50 | 4.49E-237 | 9.33E-233 | 1.00 | 0.95 | 0.05 | 1.05 |
| 2 | Tk1 | 1.04 | 1.14E-236 | 2.36E-232 | 0.85 | 0.52 | 0.34 | 1.66 |
| 2 | Smc2 | 0.68 | 6.75E-231 | 1.40E-226 | 0.99 | 0.85 | 0.14 | 1.16 |
| 2 | Rrm2 | 0.80 | 1.55E-230 | 3.22E-226 | 0.99 | 0.79 | 0.20 | 1.26 |
| 2 | Msx1 | 1.10 | 2.73E-230 | 5.68E-226 | 0.90 | 0.70 | 0.20 | 1.28 |
| 2 | Asb4 | 1.51 | 3.50E-230 | 7.28E-226 | 0.60 | 0.27 | 0.33 | 2.25 |
| 2 | Hist1h3i | 1.50 | 4.67E-230 | 9.72E-226 | 0.53 | 0.21 | 0.32 | 2.54 |
| 2 | Hist1h2bb | 1.59 | 2.34E-214 | 4.86E-210 | 0.47 | 0.18 | 0.30 | 2.65 |
| 2 | Rrm1 | 0.70 | 4.82E-211 | 1.00E-206 | 0.98 | 0.80 | 0.18 | 1.23 |
| 2 | Esco2 | 0.95 | 2.12E-206 | 4.41E-202 | 0.78 | 0.42 | 0.36 | 1.87 |
| 2 | Hist1h3d | 1.60 | 1.50E-205 | 3.12E-201 | 0.43 | 0.16 | 0.28 | 2.76 |
| 2 | Lhx6 | 2.31 | 3.35E-205 | 6.97E-201 | 0.22 | 0.05 | 0.18 | 4.83 |
| 2 | Serpinf1 | 0.77 | 1.21E-204 | 2.52E-200 | 0.93 | 0.61 | 0.31 | 1.51 |
| 2 | Tubb5 | 0.36 | 3.29E-201 | 6.85E-197 | 1.00 | 1.00 | 0.00 | 1.00 |
| 2 | Hist1h2bn | 1.69 | 8.55E-200 | 1.78E-195 | 0.38 | 0.13 | 0.25 | 3.00 |
| 2 | Nasp | 0.45 | 1.43E-198 | 2.97E-194 | 1.00 | 0.98 | 0.02 | 1.02 |
| 2 | Incenp | 0.83 | 1.64E-198 | 3.41E-194 | 0.89 | 0.56 | 0.33 | 1.58 |
| 2 | Mab21l1 | 1.40 | 5.75E-198 | 1.20E-193 | 0.41 | 0.14 | 0.27 | 2.96 |
| 2 | Clspn | 0.87 | 9.00E-197 | 1.87E-192 | 0.89 | 0.59 | 0.30 | 1.50 |
| 2 | Col1a1 | 1.15 | 1.42E-196 | 2.95E-192 | 0.60 | 0.27 | 0.33 | 2.22 |
| 2 | Col1a2 | 0.67 | 2.76E-196 | 5.74E-192 | 0.99 | 0.72 | 0.27 | 1.38 |
| 2 | Tyms | 0.66 | 1.06E-194 | 2.20E-190 | 0.99 | 0.82 | 0.17 | 1.21 |
| 2 | Gmnn | 0.67 | 7.08E-190 | 1.47E-185 | 0.98 | 0.78 | 0.19 | 1.25 |
| 2 | Six1 | 0.78 | 2.48E-189 | 5.16E-185 | 0.90 | 0.58 | 0.32 | 1.55 |

**Supplementary Table 1. Marker genes for the original clusters of integrated clustering of E11.5 control and NotchGOF λ-junction cells.**

| | | | | | | | | |
|---|---|---|---|---|---|---|---|---|
| 2 | Smc4 | 0.65 | 3.93E-189 | 8.17E-185 | 0.99 | 0.78 | 0.21 | 1.27 |
| 2 | Rad51ap1 | 0.94 | 1.08E-187 | 2.24E-183 | 0.78 | 0.46 | 0.33 | 1.72 |
| 2 | Foxl2 | 1.52 | 6.54E-187 | 1.36E-182 | 0.34 | 0.11 | 0.24 | 3.21 |
| 2 | Hnrnpab | 0.39 | 1.24E-186 | 2.58E-182 | 1.00 | 0.99 | 0.01 | 1.01 |
| 2 | Kctd12 | 1.25 | 1.48E-186 | 3.09E-182 | 0.57 | 0.27 | 0.30 | 2.13 |
| 2 | Kif11 | 0.85 | 1.66E-185 | 3.45E-181 | 0.85 | 0.51 | 0.35 | 1.69 |
| 2 | Adgrl3 | 0.97 | 3.58E-182 | 7.44E-178 | 0.74 | 0.41 | 0.33 | 1.79 |
| 2 | Cdk1 | 0.66 | 4.46E-181 | 9.27E-177 | 0.95 | 0.64 | 0.31 | 1.49 |
| 2 | Mis18bp1 | 0.85 | 2.13E-177 | 4.43E-173 | 0.82 | 0.48 | 0.34 | 1.70 |
| 2 | Pbk | 0.83 | 2.69E-177 | 5.60E-173 | 0.85 | 0.51 | 0.34 | 1.66 |
| 2 | Dnmt1 | 0.69 | 2.97E-176 | 6.19E-172 | 0.95 | 0.77 | 0.18 | 1.24 |
| 2 | Kif15 | 0.79 | 5.32E-176 | 1.11E-171 | 0.90 | 0.60 | 0.29 | 1.49 |
| 2 | Tcf12 | 0.61 | 9.06E-175 | 1.88E-170 | 0.98 | 0.87 | 0.12 | 1.14 |
| 2 | Fst | 1.35 | 3.36E-172 | 6.98E-168 | 0.32 | 0.10 | 0.22 | 3.17 |
| 2 | Aurkb | 0.81 | 3.52E-171 | 7.33E-167 | 0.80 | 0.46 | 0.34 | 1.75 |
| 2 | Prrx2 | 0.67 | 1.00E-170 | 2.08E-166 | 0.99 | 0.72 | 0.27 | 1.38 |
| 2 | Cdh6 | 1.37 | 5.51E-169 | 1.15E-164 | 0.53 | 0.26 | 0.28 | 2.06 |
| 2 | Sgk1 | 1.24 | 7.17E-169 | 1.49E-164 | 0.45 | 0.18 | 0.27 | 2.48 |
| 2 | Slitrk6 | 1.21 | 1.62E-168 | 3.36E-164 | 0.59 | 0.30 | 0.29 | 1.98 |
| 2 | Foxf2 | 1.53 | 1.11E-167 | 2.30E-163 | 0.27 | 0.08 | 0.20 | 3.61 |
| 2 | Id2 | 0.64 | 9.30E-167 | 1.93E-162 | 0.99 | 0.85 | 0.14 | 1.16 |
| 2 | Lsm2 | 0.49 | 2.17E-166 | 4.50E-162 | 1.00 | 0.92 | 0.08 | 1.08 |
| 2 | Tbx3os1 | 1.45 | 5.38E-166 | 1.12E-161 | 0.34 | 0.12 | 0.23 | 2.99 |
| 2 | Spc25 | 0.75 | 1.00E-163 | 2.08E-159 | 0.85 | 0.52 | 0.33 | 1.63 |
| 2 | Stmn1 | 0.32 | 7.28E-163 | 1.51E-158 | 1.00 | 0.99 | 0.01 | 1.01 |
| 2 | H3f3b | 0.30 | 2.35E-161 | 4.88E-157 | 1.00 | 1.00 | 0.00 | 1.00 |
| 2 | Cdca2 | 0.88 | 1.10E-160 | 2.30E-156 | 0.73 | 0.40 | 0.33 | 1.83 |
| 2 | Mpped2 | 0.56 | 1.56E-160 | 3.25E-156 | 1.00 | 0.84 | 0.16 | 1.19 |
| 2 | Hist2h2aa1 | 1.48 | 2.34E-158 | 4.87E-154 | 0.39 | 0.15 | 0.24 | 2.57 |
| 2 | Ptn | 0.47 | 6.49E-156 | 1.35E-151 | 1.00 | 0.89 | 0.11 | 1.12 |
| 2 | Exo1 | 0.89 | 2.29E-154 | 4.77E-150 | 0.73 | 0.42 | 0.31 | 1.72 |
| 2 | Sema5a | 1.31 | 4.48E-154 | 9.31E-150 | 0.62 | 0.35 | 0.27 | 1.78 |
| 2 | Hmgb2 | 0.47 | 1.07E-152 | 2.22E-148 | 1.00 | 0.98 | 0.02 | 1.02 |
| 2 | Pcna | 0.63 | 5.27E-151 | 1.10E-146 | 0.97 | 0.82 | 0.15 | 1.18 |
| 2 | Peg10 | 0.96 | 1.34E-150 | 2.79E-146 | 0.71 | 0.41 | 0.30 | 1.74 |
| 2 | Tbx2 | 0.65 | 2.31E-150 | 4.80E-146 | 0.92 | 0.62 | 0.30 | 1.49 |
| 2 | Hist1h2af | 1.55 | 2.60E-150 | 5.41E-146 | 0.34 | 0.12 | 0.22 | 2.73 |
| 2 | Baz1b | 0.55 | 1.56E-149 | 3.25E-145 | 0.98 | 0.88 | 0.10 | 1.11 |
| 2 | Mdga2 | 1.96 | 6.49E-148 | 1.35E-143 | 0.21 | 0.06 | 0.16 | 3.82 |
| 2 | Tril | 0.90 | 3.15E-147 | 6.55E-143 | 0.79 | 0.52 | 0.27 | 1.52 |
| 2 | Lig1 | 0.63 | 3.70E-147 | 7.70E-143 | 0.96 | 0.77 | 0.19 | 1.25 |
| 2 | Hist1h2ak | 1.44 | 9.94E-147 | 2.07E-142 | 0.37 | 0.14 | 0.22 | 2.57 |
| 2 | Htr1b | 2.20 | 6.60E-145 | 1.37E-140 | 0.16 | 0.03 | 0.13 | 5.03 |
| 2 | Cdca5 | 0.99 | 1.75E-141 | 3.65E-137 | 0.58 | 0.31 | 0.28 | 1.90 |
| 2 | Pgm2l1 | 1.09 | 3.66E-141 | 7.62E-137 | 0.60 | 0.33 | 0.27 | 1.81 |
| 2 | Dnajc9 | 0.52 | 8.07E-141 | 1.68E-136 | 0.98 | 0.85 | 0.13 | 1.15 |
| 2 | Dnm3os | 0.68 | 2.64E-139 | 5.49E-135 | 0.81 | 0.49 | 0.32 | 1.65 |
| 2 | Angpt1 | 1.33 | 8.76E-138 | 1.82E-133 | 0.34 | 0.13 | 0.21 | 2.70 |
| 2 | Ranbp1 | 0.33 | 1.22E-137 | 2.53E-133 | 1.00 | 0.99 | 0.01 | 1.01 |
| 2 | Zfp503 | 0.99 | 2.13E-137 | 4.42E-133 | 0.63 | 0.35 | 0.28 | 1.81 |
| 2 | Fbln1 | 0.59 | 9.41E-137 | 1.96E-132 | 0.96 | 0.78 | 0.18 | 1.23 |
| 2 | Tpm4 | 0.42 | 2.59E-136 | 5.39E-132 | 1.00 | 0.93 | 0.07 | 1.07 |
| 2 | Mki67 | 0.59 | 2.97E-136 | 6.18E-132 | 0.96 | 0.70 | 0.26 | 1.38 |
| 2 | Ndc80 | 0.84 | 3.56E-135 | 7.40E-131 | 0.68 | 0.39 | 0.29 | 1.73 |
| 2 | Ezh2 | 0.43 | 2.07E-134 | 4.31E-130 | 1.00 | 0.94 | 0.06 | 1.06 |
| 2 | Lsm3 | 0.43 | 4.50E-134 | 9.36E-130 | 0.99 | 0.91 | 0.08 | 1.09 |

**Supplementary Table 1. Marker genes for the original clusters of integrated clustering of E11.5 control and NotchGOF λ-junction cells.**

| | | | | | | | | |
|---|---|---|---|---|---|---|---|---|
| 2 | Rad51 | 0.75 | 1.50E-132 | 3.13E-128 | 0.80 | 0.52 | 0.28 | 1.54 |
| 2 | Pth1r | 0.67 | 2.35E-129 | 4.90E-125 | 0.85 | 0.54 | 0.31 | 1.56 |
| 2 | Nuf2 | 0.78 | 3.65E-129 | 7.59E-125 | 0.74 | 0.45 | 0.29 | 1.63 |
| 2 | Twist1 | 0.51 | 3.18E-128 | 6.62E-124 | 1.00 | 0.74 | 0.26 | 1.34 |
| 2 | Hist1h2bh | 1.61 | 4.79E-128 | 9.95E-124 | 0.28 | 0.10 | 0.18 | 2.80 |
| 2 | Plk4 | 0.66 | 2.37E-127 | 4.92E-123 | 0.80 | 0.53 | 0.28 | 1.53 |
| 2 | Gli3 | 0.74 | 2.82E-127 | 5.87E-123 | 0.81 | 0.56 | 0.25 | 1.45 |
| 2 | Fam107b | 1.04 | 3.45E-127 | 7.18E-123 | 0.59 | 0.33 | 0.26 | 1.77 |
| 2 | Fen1 | 0.67 | 5.57E-127 | 1.16E-122 | 0.87 | 0.63 | 0.24 | 1.37 |
| 2 | Ncapg | 0.76 | 5.58E-127 | 1.16E-122 | 0.74 | 0.45 | 0.29 | 1.64 |
| 2 | Orc6 | 0.56 | 9.85E-127 | 2.05E-122 | 0.95 | 0.78 | 0.17 | 1.21 |
| 2 | Melk | 0.78 | 3.48E-126 | 7.25E-122 | 0.73 | 0.45 | 0.29 | 1.64 |
| 2 | Cenph | 0.66 | 3.00E-125 | 6.25E-121 | 0.86 | 0.61 | 0.25 | 1.41 |
| 2 | Wnt11 | 1.91 | 3.91E-125 | 8.13E-121 | 0.16 | 0.04 | 0.13 | 4.21 |
| 2 | Cycs | 0.39 | 1.01E-124 | 2.09E-120 | 1.00 | 0.96 | 0.04 | 1.04 |
| 2 | Srsf3 | 0.30 | 2.35E-124 | 4.89E-120 | 1.00 | 1.00 | 0.00 | 1.00 |
| 2 | Hist1h2ad | 1.62 | 1.08E-123 | 2.26E-119 | 0.25 | 0.08 | 0.17 | 3.00 |
| 2 | Dut | 0.42 | 1.89E-123 | 3.92E-119 | 1.00 | 0.94 | 0.06 | 1.07 |
| 2 | Lix1 | 0.53 | 2.32E-123 | 4.82E-119 | 0.90 | 0.60 | 0.30 | 1.50 |
| 2 | Cenpq | 0.70 | 2.67E-123 | 5.56E-119 | 0.81 | 0.53 | 0.27 | 1.51 |
| 2 | Spdl1 | 0.80 | 3.99E-123 | 8.31E-119 | 0.69 | 0.41 | 0.28 | 1.68 |
| 2 | Adamts20 | 1.31 | 1.26E-122 | 2.61E-118 | 0.33 | 0.13 | 0.20 | 2.57 |
| 2 | Ncaph | 0.77 | 1.68E-122 | 3.49E-118 | 0.73 | 0.47 | 0.27 | 1.57 |
| 2 | Mxd3 | 1.13 | 4.24E-122 | 8.82E-118 | 0.46 | 0.22 | 0.23 | 2.06 |
| 2 | Serpine2 | 0.58 | 6.85E-122 | 1.43E-117 | 0.90 | 0.69 | 0.22 | 1.32 |
| 2 | Prim2 | 0.89 | 7.42E-122 | 1.54E-117 | 0.63 | 0.36 | 0.27 | 1.74 |
| 2 | Col3a1 | 0.44 | 1.54E-121 | 3.20E-117 | 0.98 | 0.70 | 0.29 | 1.41 |
| 2 | Shox2 | 1.45 | 1.62E-121 | 3.37E-117 | 0.30 | 0.11 | 0.19 | 2.70 |
| 2 | Bub3 | 0.51 | 1.63E-121 | 3.39E-117 | 0.96 | 0.81 | 0.15 | 1.18 |
| 2 | Skp2 | 0.63 | 2.65E-121 | 5.51E-117 | 0.90 | 0.68 | 0.22 | 1.33 |
| 2 | Ednra | 0.88 | 9.02E-121 | 1.88E-116 | 0.59 | 0.31 | 0.28 | 1.89 |
| 2 | Hjurp | 0.67 | 1.16E-120 | 2.41E-116 | 0.84 | 0.58 | 0.26 | 1.44 |
| 2 | Ntf3 | 1.00 | 4.27E-120 | 8.89E-116 | 0.52 | 0.27 | 0.25 | 1.94 |
| 2 | Spc24 | 0.56 | 8.45E-120 | 1.76E-115 | 0.94 | 0.71 | 0.23 | 1.32 |
| 2 | Blm | 0.74 | 1.41E-119 | 2.93E-115 | 0.76 | 0.49 | 0.27 | 1.55 |
| 2 | Anp32b | 0.27 | 5.83E-119 | 1.21E-114 | 1.00 | 0.99 | 0.01 | 1.01 |
| 2 | Smarcc1 | 0.38 | 2.10E-118 | 4.38E-114 | 1.00 | 0.96 | 0.04 | 1.04 |
| 2 | Snai2 | 0.80 | 3.61E-118 | 7.51E-114 | 0.71 | 0.44 | 0.27 | 1.61 |
| 2 | Cks1b | 0.37 | 3.64E-118 | 7.58E-114 | 1.00 | 0.95 | 0.04 | 1.05 |
| 2 | Pde3a | 1.44 | 5.17E-118 | 1.07E-113 | 0.30 | 0.11 | 0.19 | 2.64 |
| 2 | Ppfibp2 | 0.77 | 5.46E-118 | 1.14E-113 | 0.69 | 0.40 | 0.29 | 1.71 |
| 2 | Smchd1 | 0.48 | 6.88E-118 | 1.43E-113 | 0.97 | 0.85 | 0.12 | 1.14 |
| 2 | Mlf1 | 1.17 | 3.92E-117 | 8.15E-113 | 0.41 | 0.19 | 0.22 | 2.18 |
| 2 | Epha3 | 0.71 | 7.88E-117 | 1.64E-112 | 0.62 | 0.33 | 0.28 | 1.85 |
| 2 | Ccdc34 | 0.43 | 2.30E-116 | 4.79E-112 | 0.99 | 0.91 | 0.09 | 1.09 |
| 2 | Clmp | 0.56 | 1.35E-115 | 2.82E-111 | 0.91 | 0.61 | 0.30 | 1.49 |
| 2 | Nkd1 | 0.97 | 2.68E-115 | 5.57E-111 | 0.59 | 0.34 | 0.25 | 1.75 |
| 2 | Tmpo | 0.39 | 4.91E-115 | 1.02E-110 | 1.00 | 0.95 | 0.05 | 1.05 |
| 2 | Hist1h4c | 1.39 | 2.09E-114 | 4.34E-110 | 0.32 | 0.13 | 0.19 | 2.45 |
| 2 | Col13a1 | 1.30 | 2.92E-114 | 6.07E-110 | 0.27 | 0.09 | 0.17 | 2.89 |
| 2 | Mcm10 | 0.89 | 1.20E-113 | 2.50E-109 | 0.58 | 0.33 | 0.25 | 1.77 |
| 2 | H2afz | 0.28 | 1.21E-113 | 2.52E-109 | 1.00 | 1.00 | 0.00 | 1.00 |
| 2 | Smc6 | 0.49 | 1.35E-113 | 2.82E-109 | 0.98 | 0.86 | 0.12 | 1.14 |
| 2 | Chaf1a | 0.62 | 2.35E-113 | 4.89E-109 | 0.86 | 0.63 | 0.24 | 1.37 |
| 2 | Ckap2l | 0.66 | 5.01E-113 | 1.04E-108 | 0.74 | 0.46 | 0.28 | 1.61 |
| 2 | Phldb2 | 0.70 | 1.19E-112 | 2.48E-108 | 0.77 | 0.51 | 0.26 | 1.50 |

**Supplementary Table 1. Marker genes for the original clusters of integrated clustering of E11.5 control and NotchGOF λ-junction cells.**

| | | | | | | | | |
|---|---|---|---|---|---|---|---|---|
| 2 | Gpc3 | 0.55 | 1.37E-112 | 2.85E-108 | 0.96 | 0.82 | 0.15 | 1.18 |
| 2 | Aff2 | 1.57 | 1.55E-110 | 3.23E-106 | 0.21 | 0.06 | 0.15 | 3.27 |
| 2 | Diaph3 | 0.86 | 2.42E-110 | 5.03E-106 | 0.60 | 0.35 | 0.25 | 1.73 |
| 2 | Hint1 | 0.27 | 3.05E-110 | 6.35E-106 | 1.00 | 0.99 | 0.01 | 1.01 |
| 2 | Kif2c | 0.58 | 3.74E-110 | 7.78E-106 | 0.74 | 0.44 | 0.30 | 1.68 |
| 2 | Frzb | 0.45 | 6.16E-110 | 1.28E-105 | 0.62 | 0.33 | 0.29 | 1.89 |
| 2 | Ube2e3 | 0.41 | 5.93E-109 | 1.23E-104 | 0.98 | 0.91 | 0.08 | 1.08 |
| 2 | Hist1h2br | 1.42 | 5.99E-109 | 1.24E-104 | 0.32 | 0.14 | 0.19 | 2.36 |
| 2 | Mis18a | 0.74 | 1.50E-108 | 3.11E-104 | 0.70 | 0.44 | 0.26 | 1.60 |
| 2 | Kif18a | 0.89 | 2.16E-108 | 4.50E-104 | 0.57 | 0.32 | 0.25 | 1.77 |
| 2 | Pde4d | 0.94 | 3.33E-108 | 6.92E-104 | 0.48 | 0.24 | 0.24 | 2.00 |
| 2 | Sox4 | 0.44 | 1.91E-107 | 3.96E-103 | 1.00 | 0.94 | 0.06 | 1.07 |
| 2 | Sdc1 | 0.54 | 6.00E-107 | 1.25E-102 | 0.90 | 0.68 | 0.22 | 1.32 |
| 2 | Col6a1 | 1.19 | 1.09E-106 | 2.27E-102 | 0.32 | 0.13 | 0.19 | 2.49 |
| 2 | Axin2 | 0.91 | 3.58E-106 | 7.45E-102 | 0.56 | 0.32 | 0.24 | 1.76 |
| 2 | Sgo2a | 0.61 | 3.67E-106 | 7.63E-102 | 0.74 | 0.45 | 0.29 | 1.64 |
| 2 | Pola1 | 0.73 | 5.69E-106 | 1.18E-101 | 0.72 | 0.47 | 0.25 | 1.53 |
| 2 | Cenpu | 0.94 | 8.64E-106 | 1.80E-101 | 0.53 | 0.29 | 0.24 | 1.82 |
| 2 | Hist1h2bl | 1.58 | 1.26E-105 | 2.62E-101 | 0.25 | 0.09 | 0.16 | 2.74 |
| 2 | Hspd1 | 0.39 | 1.80E-105 | 3.74E-101 | 1.00 | 0.98 | 0.02 | 1.02 |
| 2 | Supt16 | 0.36 | 2.48E-105 | 5.16E-101 | 1.00 | 0.96 | 0.04 | 1.04 |
| 2 | Ttk | 0.86 | 2.94E-105 | 6.11E-101 | 0.58 | 0.34 | 0.24 | 1.71 |
| 2 | Haspin | 1.02 | 5.74E-105 | 1.19E-100 | 0.49 | 0.26 | 0.23 | 1.86 |
| 2 | Msx2 | 0.84 | 9.24E-105 | 1.92E-100 | 0.65 | 0.41 | 0.25 | 1.61 |
| 2 | Rbm39 | 0.27 | 2.09E-104 | 4.35E-100 | 1.00 | 0.99 | 0.01 | 1.01 |
| 2 | Kif20b | 0.60 | 7.10E-104 | 1.48E-99 | 0.80 | 0.53 | 0.27 | 1.50 |
| 2 | Nfia | 0.77 | 2.19E-103 | 4.56E-99 | 0.70 | 0.45 | 0.26 | 1.57 |
| 2 | Anln | 0.77 | 5.72E-103 | 1.19E-98 | 0.62 | 0.37 | 0.25 | 1.68 |
| 2 | Oxct1 | 0.47 | 7.10E-103 | 1.48E-98 | 0.96 | 0.81 | 0.16 | 1.20 |
| 2 | Zeb2 | 0.80 | 7.27E-103 | 1.51E-98 | 0.62 | 0.38 | 0.24 | 1.65 |
| 2 | Snrpd1 | 0.28 | 7.60E-103 | 1.58E-98 | 1.00 | 0.99 | 0.01 | 1.01 |
| 2 | Setbp1 | 0.76 | 1.56E-102 | 3.25E-98 | 0.62 | 0.36 | 0.25 | 1.69 |
| 2 | Ska1 | 0.75 | 3.04E-102 | 6.32E-98 | 0.65 | 0.39 | 0.25 | 1.64 |
| 2 | Prr16 | 1.84 | 4.37E-102 | 9.08E-98 | 0.14 | 0.03 | 0.10 | 4.15 |
| 2 | Lyar | 0.44 | 5.16E-102 | 1.07E-97 | 0.96 | 0.86 | 0.11 | 1.13 |
| 2 | Gjc1 | 0.49 | 6.41E-102 | 1.33E-97 | 0.96 | 0.79 | 0.16 | 1.20 |
| 2 | Shcbp1 | 0.71 | 6.61E-102 | 1.38E-97 | 0.70 | 0.44 | 0.26 | 1.58 |
| 2 | Sgo1 | 0.70 | 2.33E-101 | 4.85E-97 | 0.69 | 0.42 | 0.27 | 1.63 |
| 2 | Sgcd | 0.99 | 7.85E-101 | 1.63E-96 | 0.42 | 0.20 | 0.22 | 2.08 |
| 2 | Runx2 | 1.11 | 3.80E-100 | 7.91E-96 | 0.34 | 0.14 | 0.19 | 2.33 |
| 2 | Brca1 | 0.84 | 4.10E-100 | 8.52E-96 | 0.60 | 0.36 | 0.24 | 1.68 |
| 2 | Cdca4 | 0.67 | 7.70E-100 | 1.60E-95 | 0.77 | 0.53 | 0.24 | 1.44 |
| 2 | Marcks | 0.31 | 4.16E-99 | 8.66E-95 | 1.00 | 0.97 | 0.03 | 1.03 |
| 2 | Tnfaip6 | 0.59 | 2.06E-98 | 4.29E-94 | 0.74 | 0.44 | 0.30 | 1.68 |
| 2 | Hnrnpd | 0.28 | 9.43E-98 | 1.96E-93 | 1.00 | 0.99 | 0.01 | 1.01 |
| 2 | Prex2 | 0.75 | 1.22E-97 | 2.54E-93 | 0.64 | 0.38 | 0.26 | 1.68 |
| 2 | Prrx1 | 0.61 | 8.73E-97 | 1.82E-92 | 0.89 | 0.63 | 0.26 | 1.41 |
| 2 | Lpar1 | 0.45 | 1.01E-96 | 2.11E-92 | 0.93 | 0.68 | 0.25 | 1.37 |
| 2 | Vegfd | 1.98 | 1.49E-96 | 3.10E-92 | 0.15 | 0.04 | 0.11 | 3.71 |
| 2 | Mns1 | 0.62 | 1.68E-96 | 3.49E-92 | 0.78 | 0.52 | 0.26 | 1.49 |
| 2 | Cdc45 | 0.83 | 1.86E-96 | 3.88E-92 | 0.58 | 0.35 | 0.24 | 1.69 |
| 2 | Prc1 | 0.49 | 3.52E-96 | 7.33E-92 | 0.79 | 0.50 | 0.29 | 1.58 |
| 2 | Pbx3 | 0.65 | 8.53E-96 | 1.77E-91 | 0.75 | 0.51 | 0.24 | 1.48 |
| 2 | Dctpp1 | 0.37 | 8.58E-96 | 1.78E-91 | 0.99 | 0.93 | 0.06 | 1.07 |
| 2 | Eif4a3 | 0.42 | 2.98E-95 | 6.19E-91 | 0.96 | 0.86 | 0.10 | 1.12 |
| 2 | H1f0 | 0.53 | 5.27E-95 | 1.10E-90 | 0.95 | 0.84 | 0.11 | 1.13 |

**Supplementary Table 1. Marker genes for the original clusters of integrated clustering of E11.5 control and NotchGOF λ-junction cells.**

| | | | | | | | | |
|---|---|---|---|---|---|---|---|---|
| 2 | Hist1h2bj | 1.02 | 4.15E-94 | 8.64E-90 | 0.43 | 0.23 | 0.21 | 1.91 |
| 2 | Mthfd2 | 0.73 | 4.98E-94 | 1.04E-89 | 0.69 | 0.45 | 0.24 | 1.53 |
| 2 | Otor | 1.43 | 8.85E-94 | 1.84E-89 | 0.20 | 0.06 | 0.13 | 3.06 |
| 2 | Bdnf | 0.79 | 1.24E-93 | 2.59E-89 | 0.50 | 0.27 | 0.23 | 1.87 |
| 2 | Nme1 | 0.31 | 2.87E-93 | 5.97E-89 | 1.00 | 0.99 | 0.01 | 1.01 |
| 2 | Hirip3 | 0.60 | 3.15E-93 | 6.54E-89 | 0.86 | 0.67 | 0.19 | 1.29 |
| 2 | Nbea | 0.70 | 3.68E-93 | 7.66E-89 | 0.66 | 0.42 | 0.24 | 1.57 |
| 2 | Neil3 | 1.03 | 1.32E-92 | 2.75E-88 | 0.41 | 0.21 | 0.20 | 1.95 |
| 2 | Slbp | 0.46 | 1.46E-92 | 3.04E-88 | 0.97 | 0.86 | 0.11 | 1.13 |
| 2 | E130114P18Rik | 1.02 | 9.48E-92 | 1.97E-87 | 0.41 | 0.21 | 0.21 | 1.99 |
| 2 | Slc38a2 | 0.40 | 1.98E-91 | 4.13E-87 | 0.98 | 0.90 | 0.08 | 1.09 |
| 2 | Eme1 | 0.79 | 2.01E-91 | 4.17E-87 | 0.57 | 0.34 | 0.23 | 1.67 |
| 2 | Cenpj | 0.80 | 2.72E-91 | 5.66E-87 | 0.58 | 0.35 | 0.23 | 1.65 |
| 2 | Smc3 | 0.34 | 5.93E-91 | 1.23E-86 | 0.99 | 0.94 | 0.05 | 1.05 |
| 2 | Cbx5 | 0.34 | 1.19E-90 | 2.47E-86 | 1.00 | 0.95 | 0.05 | 1.05 |
| 2 | Inpp4b | 1.03 | 1.93E-90 | 4.01E-86 | 0.32 | 0.14 | 0.18 | 2.30 |
| 2 | Psat1 | 0.71 | 2.44E-90 | 5.07E-86 | 0.67 | 0.43 | 0.24 | 1.55 |
| 2 | Rnps1 | 0.33 | 4.79E-90 | 9.97E-86 | 0.99 | 0.94 | 0.05 | 1.05 |
| 2 | Ncapg2 | 0.73 | 6.42E-90 | 1.34E-85 | 0.62 | 0.39 | 0.23 | 1.60 |
| 2 | Gm47283 | 0.40 | 6.80E-90 | 1.41E-85 | 0.98 | 0.91 | 0.08 | 1.09 |
| 2 | Cenpm | 0.58 | 8.97E-90 | 1.86E-85 | 0.81 | 0.58 | 0.23 | 1.40 |
| 2 | Bzw2 | 0.38 | 2.08E-89 | 4.33E-85 | 0.99 | 0.93 | 0.05 | 1.06 |
| 2 | Ddr2 | 0.59 | 2.41E-89 | 5.01E-85 | 0.73 | 0.46 | 0.27 | 1.58 |
| 2 | Smarca5 | 0.31 | 2.53E-89 | 5.26E-85 | 1.00 | 0.97 | 0.03 | 1.03 |
| 2 | Srsf10 | 0.35 | 3.47E-89 | 7.21E-85 | 0.99 | 0.94 | 0.05 | 1.05 |
| 2 | Rnf227 | 0.70 | 1.93E-88 | 4.02E-84 | 0.64 | 0.39 | 0.25 | 1.63 |
| 2 | Frem1 | 0.60 | 2.51E-88 | 5.22E-84 | 0.64 | 0.38 | 0.26 | 1.68 |
| 2 | Trip13 | 0.60 | 7.01E-88 | 1.46E-83 | 0.76 | 0.52 | 0.25 | 1.48 |
| 2 | Dcakd | 0.37 | 2.84E-87 | 5.91E-83 | 0.99 | 0.90 | 0.09 | 1.10 |
| 2 | Rad54l | 0.84 | 3.00E-87 | 6.24E-83 | 0.51 | 0.29 | 0.22 | 1.75 |
| 2 | G2e3 | 0.68 | 8.36E-87 | 1.74E-82 | 0.68 | 0.45 | 0.24 | 1.53 |
| 2 | U2af1 | 0.31 | 1.16E-86 | 2.40E-82 | 1.00 | 0.97 | 0.03 | 1.03 |
| 2 | Prdx4 | 0.38 | 1.22E-86 | 2.53E-82 | 0.98 | 0.88 | 0.10 | 1.11 |
| 2 | Kif26b | 0.59 | 2.54E-86 | 5.28E-82 | 0.64 | 0.38 | 0.26 | 1.67 |
| 2 | Cenpp | 0.70 | 6.85E-86 | 1.43E-81 | 0.61 | 0.38 | 0.23 | 1.60 |
| 2 | Rfc4 | 0.46 | 8.90E-86 | 1.85E-81 | 0.94 | 0.78 | 0.16 | 1.21 |
| 2 | Dhfr | 0.53 | 1.11E-85 | 2.31E-81 | 0.87 | 0.66 | 0.21 | 1.33 |
| 2 | Lrr1 | 0.94 | 1.35E-85 | 2.80E-81 | 0.45 | 0.25 | 0.20 | 1.82 |
| 2 | Rnf122 | 0.61 | 1.79E-85 | 3.73E-81 | 0.75 | 0.51 | 0.24 | 1.46 |
| 2 | Nfatc4 | 0.59 | 2.65E-85 | 5.52E-81 | 0.77 | 0.53 | 0.24 | 1.45 |
| 2 | Hist1h3a | 1.05 | 4.07E-85 | 8.47E-81 | 0.35 | 0.17 | 0.18 | 2.05 |
| 2 | Hist1h2ac | 1.37 | 8.39E-85 | 1.74E-80 | 0.23 | 0.09 | 0.14 | 2.52 |
| 2 | Exosc8 | 0.45 | 2.47E-84 | 5.13E-80 | 0.93 | 0.78 | 0.15 | 1.20 |
| 2 | Haus3 | 0.64 | 5.98E-84 | 1.24E-79 | 0.71 | 0.48 | 0.23 | 1.48 |
| 2 | Pole | 0.87 | 8.47E-84 | 1.76E-79 | 0.48 | 0.28 | 0.21 | 1.74 |
| 2 | Anp32e | 0.32 | 8.55E-84 | 1.78E-79 | 1.00 | 0.94 | 0.05 | 1.06 |
| 2 | Bub1 | 0.54 | 1.12E-83 | 2.33E-79 | 0.79 | 0.54 | 0.25 | 1.46 |
| 2 | Hist1h4i | 0.67 | 2.45E-83 | 5.11E-79 | 0.67 | 0.44 | 0.23 | 1.53 |
| 2 | H2afx | 0.35 | 4.82E-83 | 1.00E-78 | 1.00 | 0.92 | 0.08 | 1.08 |
| 2 | Apcdd1 | 1.04 | 9.45E-83 | 1.96E-78 | 0.39 | 0.20 | 0.19 | 1.94 |
| 2 | Bambi | 0.40 | 1.31E-82 | 2.73E-78 | 0.85 | 0.62 | 0.23 | 1.37 |
| 2 | Eya1 | 0.52 | 5.49E-82 | 1.14E-77 | 0.72 | 0.48 | 0.24 | 1.51 |
| 2 | Ncl | 0.27 | 1.06E-81 | 2.21E-77 | 1.00 | 1.00 | 0.00 | 1.00 |
| 2 | Auts2 | 0.37 | 2.09E-81 | 4.34E-77 | 0.98 | 0.86 | 0.13 | 1.15 |
| 2 | Ptch1 | 0.67 | 2.21E-81 | 4.59E-77 | 0.66 | 0.43 | 0.23 | 1.52 |
| 2 | Pold3 | 0.58 | 3.25E-81 | 6.76E-77 | 0.77 | 0.55 | 0.22 | 1.40 |

**Supplementary Table 1. Marker genes for the original clusters of integrated clustering of E11.5 control and NotchGOF λ-junction cells.**

| | | | | | | | | |
|---|---|---|---|---|---|---|---|---|
| 2 | Aurka | 0.61 | 4.52E-81 | 9.40E-77 | 0.67 | 0.43 | 0.24 | 1.57 |
| 2 | Rai14 | 0.51 | 1.36E-80 | 2.82E-76 | 0.85 | 0.63 | 0.22 | 1.35 |
| 2 | Smc1a | 0.34 | 2.18E-80 | 4.54E-76 | 0.99 | 0.94 | 0.05 | 1.06 |
| 2 | Cxcl14 | 0.40 | 4.16E-80 | 8.65E-76 | 0.30 | 0.14 | 0.16 | 2.20 |
| 2 | Ctcf | 0.38 | 9.82E-80 | 2.04E-75 | 0.97 | 0.88 | 0.09 | 1.10 |
| 2 | Ncapd3 | 0.63 | 1.08E-79 | 2.24E-75 | 0.72 | 0.51 | 0.21 | 1.41 |
| 2 | Arhgef40 | 0.55 | 1.24E-79 | 2.58E-75 | 0.74 | 0.50 | 0.24 | 1.47 |
| 2 | Rfc5 | 0.56 | 2.43E-79 | 5.06E-75 | 0.80 | 0.57 | 0.23 | 1.41 |
| 2 | Dbf4 | 0.49 | 5.05E-79 | 1.05E-74 | 0.87 | 0.64 | 0.23 | 1.37 |
| 2 | Siah1b | 0.81 | 6.13E-79 | 1.27E-74 | 0.48 | 0.27 | 0.21 | 1.75 |
| 2 | Sass6 | 0.60 | 9.54E-79 | 1.98E-74 | 0.70 | 0.48 | 0.23 | 1.47 |
| 2 | Pcdh18 | 0.44 | 1.27E-78 | 2.64E-74 | 0.87 | 0.61 | 0.25 | 1.41 |
| 2 | Eml4 | 0.39 | 1.42E-78 | 2.96E-74 | 0.96 | 0.82 | 0.14 | 1.16 |
| 2 | Phip | 0.38 | 1.76E-78 | 3.67E-74 | 0.96 | 0.87 | 0.09 | 1.11 |
| 2 | Mad2l1 | 0.55 | 2.77E-78 | 5.75E-74 | 0.77 | 0.54 | 0.23 | 1.41 |
| 2 | Brd3 | 0.34 | 3.02E-78 | 6.29E-74 | 0.99 | 0.92 | 0.07 | 1.07 |
| 2 | Slco3a1 | 1.10 | 1.01E-77 | 2.09E-73 | 0.27 | 0.12 | 0.15 | 2.32 |
| 2 | Serpinh1 | 0.32 | 1.10E-77 | 2.28E-73 | 1.00 | 0.94 | 0.06 | 1.06 |
| 2 | C430049B03Rik | 0.70 | 2.78E-77 | 5.78E-73 | 0.56 | 0.33 | 0.23 | 1.68 |
| 2 | Ercc6l | 0.74 | 4.13E-77 | 8.59E-73 | 0.55 | 0.34 | 0.21 | 1.61 |
| 2 | Hist1h2bg | 1.40 | 4.84E-77 | 1.01E-72 | 0.22 | 0.09 | 0.13 | 2.51 |
| 2 | Cks2 | 0.35 | 4.95E-77 | 1.03E-72 | 0.98 | 0.82 | 0.16 | 1.19 |
| 2 | Dtymk | 0.37 | 1.06E-76 | 2.20E-72 | 0.97 | 0.88 | 0.09 | 1.10 |
| 2 | Mex3a | 0.38 | 2.95E-76 | 6.14E-72 | 0.96 | 0.84 | 0.12 | 1.14 |
| 2 | Hmmr | 0.44 | 8.29E-76 | 1.72E-71 | 0.78 | 0.51 | 0.27 | 1.54 |
| 2 | Ccdc88a | 0.40 | 1.22E-75 | 2.53E-71 | 0.94 | 0.80 | 0.14 | 1.18 |
| 2 | Spock3 | 1.26 | 1.39E-75 | 2.90E-71 | 0.26 | 0.12 | 0.15 | 2.28 |
| 2 | Lef1 | 0.61 | 1.43E-75 | 2.98E-71 | 0.66 | 0.44 | 0.22 | 1.49 |
| 2 | Ptges3 | 0.28 | 2.05E-75 | 4.25E-71 | 1.00 | 0.97 | 0.03 | 1.03 |
| 2 | Irak1bp1 | 0.63 | 2.76E-75 | 5.73E-71 | 0.74 | 0.53 | 0.21 | 1.40 |
| 2 | Aff3 | 0.94 | 3.37E-75 | 7.02E-71 | 0.39 | 0.20 | 0.18 | 1.91 |
| 2 | Rfc1 | 0.39 | 5.62E-75 | 1.17E-70 | 0.96 | 0.86 | 0.11 | 1.12 |
| 2 | Net1 | 0.55 | 6.68E-75 | 1.39E-70 | 0.77 | 0.57 | 0.20 | 1.36 |
| 2 | Pax3 | 0.54 | 9.71E-75 | 2.02E-70 | 0.68 | 0.43 | 0.25 | 1.57 |
| 2 | Cald1 | 0.35 | 1.68E-74 | 3.49E-70 | 1.00 | 0.88 | 0.12 | 1.13 |
| 2 | Ctps | 0.58 | 2.55E-74 | 5.31E-70 | 0.77 | 0.57 | 0.20 | 1.36 |
| 2 | Frmd6 | 0.57 | 2.57E-74 | 5.34E-70 | 0.70 | 0.47 | 0.23 | 1.49 |
| 2 | Thrap3 | 0.35 | 2.66E-74 | 5.54E-70 | 0.98 | 0.92 | 0.06 | 1.07 |
| 2 | Usp1 | 0.42 | 4.52E-74 | 9.40E-70 | 0.95 | 0.80 | 0.15 | 1.19 |
| 2 | Kif4 | 0.55 | 8.43E-74 | 1.75E-69 | 0.68 | 0.45 | 0.24 | 1.52 |
| 2 | Daam2 | 0.65 | 1.01E-73 | 2.09E-69 | 0.49 | 0.28 | 0.22 | 1.79 |
| 2 | Prickle1 | 0.82 | 1.43E-73 | 2.96E-69 | 0.40 | 0.21 | 0.19 | 1.92 |
| 2 | Lgals1 | 0.34 | 1.84E-73 | 3.82E-69 | 0.99 | 0.81 | 0.18 | 1.22 |
| 2 | Col5a2 | 0.53 | 1.94E-73 | 4.03E-69 | 0.74 | 0.51 | 0.23 | 1.46 |
| 2 | Prpf4 | 0.56 | 2.61E-73 | 5.43E-69 | 0.77 | 0.57 | 0.20 | 1.36 |
| 2 | Nap1l4 | 0.33 | 3.03E-73 | 6.30E-69 | 0.99 | 0.92 | 0.06 | 1.07 |
| 2 | Actn1 | 0.46 | 3.14E-73 | 6.54E-69 | 0.89 | 0.71 | 0.19 | 1.27 |
| 2 | Fbn2 | 0.40 | 6.12E-73 | 1.27E-68 | 0.92 | 0.72 | 0.20 | 1.28 |
| 2 | Nrp2 | 0.44 | 6.65E-73 | 1.38E-68 | 0.58 | 0.35 | 0.23 | 1.65 |
| 2 | Iqgap2 | 0.78 | 7.05E-73 | 1.47E-68 | 0.55 | 0.35 | 0.20 | 1.58 |
| 2 | Sh3kbp1 | 0.89 | 1.07E-72 | 2.23E-68 | 0.39 | 0.21 | 0.18 | 1.88 |
| 2 | Bmi1 | 0.65 | 1.21E-72 | 2.51E-68 | 0.60 | 0.39 | 0.21 | 1.55 |
| 2 | Rad21 | 0.40 | 1.30E-72 | 2.70E-68 | 0.94 | 0.79 | 0.15 | 1.19 |
| 2 | Jarid2 | 0.54 | 1.42E-72 | 2.95E-68 | 0.79 | 0.59 | 0.20 | 1.34 |
| 2 | Cenpw | 0.44 | 2.73E-72 | 5.67E-68 | 0.90 | 0.72 | 0.19 | 1.26 |
| 2 | Tpr | 0.27 | 3.13E-72 | 6.51E-68 | 1.00 | 0.98 | 0.02 | 1.02 |

**Supplementary Table 1. Marker genes for the original clusters of integrated clustering of E11.5 control and NotchGOF λ-junction cells.**

| | | | | | | | | |
|---|---|---|---|---|---|---|---|---|
| 2 | Rbbp4 | 0.29 | 4.19E-72 | 8.71E-68 | 0.99 | 0.96 | 0.03 | 1.04 |
| 2 | Tpd52l1 | 0.70 | 4.82E-72 | 1.00E-67 | 0.53 | 0.31 | 0.22 | 1.69 |
| 2 | Traip | 0.84 | 4.88E-72 | 1.02E-67 | 0.45 | 0.26 | 0.19 | 1.75 |
| 2 | H2afy2 | 0.29 | 6.28E-72 | 1.31E-67 | 0.99 | 0.93 | 0.06 | 1.07 |
| 2 | Mybl2 | 0.90 | 6.44E-72 | 1.34E-67 | 0.41 | 0.23 | 0.18 | 1.79 |
| 2 | Gap43 | 0.48 | 1.02E-71 | 2.13E-67 | 0.63 | 0.40 | 0.23 | 1.58 |
| 2 | Fzd2 | 0.36 | 1.08E-71 | 2.25E-67 | 0.98 | 0.86 | 0.12 | 1.14 |
| 2 | Kif20a | 0.52 | 1.15E-71 | 2.39E-67 | 0.67 | 0.45 | 0.23 | 1.51 |
| 2 | Stx18 | 0.62 | 2.07E-71 | 4.30E-67 | 0.69 | 0.50 | 0.19 | 1.38 |
| 2 | Limd2 | 0.32 | 4.04E-71 | 8.41E-67 | 0.99 | 0.89 | 0.10 | 1.11 |
| 2 | Nudc | 0.28 | 4.30E-71 | 8.95E-67 | 0.99 | 0.96 | 0.04 | 1.04 |
| 2 | Birc5 | 0.35 | 4.93E-71 | 1.03E-66 | 0.98 | 0.79 | 0.19 | 1.24 |
| 2 | Lepr | 1.56 | 5.80E-71 | 1.21E-66 | 0.13 | 0.04 | 0.09 | 3.33 |
| 2 | Bora | 0.60 | 6.24E-71 | 1.30E-66 | 0.64 | 0.43 | 0.22 | 1.51 |
| 2 | AW551984 | 1.10 | 8.35E-71 | 1.74E-66 | 0.25 | 0.11 | 0.14 | 2.30 |
| 2 | Rnf182 | 0.72 | 9.15E-71 | 1.90E-66 | 0.38 | 0.20 | 0.18 | 1.91 |
| 2 | Fxyd6 | 0.48 | 1.33E-70 | 2.76E-66 | 0.87 | 0.66 | 0.21 | 1.31 |
| 2 | Ccng2 | 0.64 | 1.99E-70 | 4.14E-66 | 0.62 | 0.41 | 0.21 | 1.51 |
| 2 | Cdca8 | 0.36 | 2.76E-70 | 5.73E-66 | 0.96 | 0.73 | 0.23 | 1.31 |
| 2 | Cenpn | 0.67 | 3.32E-70 | 6.91E-66 | 0.57 | 0.37 | 0.21 | 1.57 |
| 2 | Msn | 0.37 | 5.35E-70 | 1.11E-65 | 0.73 | 0.49 | 0.24 | 1.49 |
| 2 | Cenpl | 0.79 | 8.08E-70 | 1.68E-65 | 0.45 | 0.26 | 0.19 | 1.71 |
| 2 | Iws1 | 0.48 | 4.04E-69 | 8.40E-65 | 0.86 | 0.69 | 0.18 | 1.26 |
| 2 | Scaf11 | 0.32 | 5.81E-69 | 1.21E-64 | 0.98 | 0.92 | 0.06 | 1.06 |
| 2 | Dscc1 | 0.80 | 7.72E-69 | 1.60E-64 | 0.48 | 0.28 | 0.19 | 1.68 |
| 2 | Ier5l | 0.51 | 1.72E-68 | 3.57E-64 | 0.82 | 0.65 | 0.17 | 1.27 |
| 2 | Rhobtb3 | 0.38 | 3.61E-68 | 7.51E-64 | 0.94 | 0.80 | 0.14 | 1.17 |
| 2 | Cep192 | 0.65 | 5.71E-68 | 1.19E-63 | 0.63 | 0.42 | 0.21 | 1.49 |
| 2 | Tmem26 | 0.69 | 6.45E-68 | 1.34E-63 | 0.43 | 0.24 | 0.20 | 1.84 |
| 2 | Gen1 | 0.84 | 7.83E-68 | 1.63E-63 | 0.44 | 0.26 | 0.18 | 1.68 |
| 2 | Srek1 | 0.38 | 9.12E-68 | 1.90E-63 | 0.95 | 0.82 | 0.13 | 1.16 |
| 2 | Cachd1 | 0.59 | 2.92E-67 | 6.07E-63 | 0.70 | 0.50 | 0.20 | 1.40 |
| 2 | Alx1 | 0.37 | 5.05E-67 | 1.05E-62 | 0.96 | 0.71 | 0.26 | 1.36 |
| 2 | Cd248 | 0.68 | 6.00E-67 | 1.25E-62 | 0.43 | 0.23 | 0.20 | 1.83 |
| 2 | Bcl11a | 0.43 | 9.24E-67 | 1.92E-62 | 0.90 | 0.72 | 0.18 | 1.25 |
| 2 | 2810408I11Rik | 0.77 | 1.86E-66 | 3.87E-62 | 0.47 | 0.28 | 0.19 | 1.65 |
| 2 | Rhno1 | 0.47 | 3.06E-66 | 6.36E-62 | 0.84 | 0.65 | 0.19 | 1.29 |
| 2 | Alx4 | 0.60 | 5.15E-66 | 1.07E-61 | 0.62 | 0.40 | 0.22 | 1.54 |
| 2 | Sae1 | 0.32 | 9.87E-66 | 2.05E-61 | 0.98 | 0.90 | 0.09 | 1.09 |
| 2 | Nolc1 | 0.41 | 1.05E-65 | 2.19E-61 | 0.92 | 0.79 | 0.13 | 1.17 |
| 2 | A830018L16Rik | 0.95 | 1.24E-65 | 2.58E-61 | 0.25 | 0.11 | 0.14 | 2.25 |
| 2 | Tipin | 0.41 | 1.29E-65 | 2.67E-61 | 0.91 | 0.76 | 0.16 | 1.21 |
| 2 | Pcdh10 | 1.11 | 1.36E-65 | 2.83E-61 | 0.22 | 0.09 | 0.13 | 2.36 |
| 2 | Ppp3ca | 0.39 | 2.10E-65 | 4.36E-61 | 0.89 | 0.71 | 0.18 | 1.25 |
| 2 | Gli1 | 1.00 | 2.77E-65 | 5.76E-61 | 0.26 | 0.12 | 0.14 | 2.18 |
| 2 | Trir | 0.29 | 2.92E-65 | 6.07E-61 | 0.99 | 0.94 | 0.05 | 1.05 |
| 2 | Ticrr | 0.90 | 6.88E-65 | 1.43E-60 | 0.38 | 0.21 | 0.17 | 1.78 |
| 2 | Tra2a | 0.31 | 8.57E-65 | 1.78E-60 | 0.99 | 0.93 | 0.06 | 1.06 |
| 2 | Cenpe | 0.31 | 1.42E-64 | 2.95E-60 | 0.88 | 0.63 | 0.24 | 1.39 |
| 2 | Trim59 | 0.41 | 1.73E-64 | 3.60E-60 | 0.88 | 0.70 | 0.18 | 1.26 |
| 2 | Asph | 0.43 | 2.00E-64 | 4.15E-60 | 0.83 | 0.63 | 0.20 | 1.32 |
| 2 | Snai1 | 0.57 | 2.97E-64 | 6.18E-60 | 0.62 | 0.39 | 0.23 | 1.58 |
| 2 | Wnt5a | 0.35 | 5.33E-64 | 1.11E-59 | 0.85 | 0.62 | 0.23 | 1.37 |
| 2 | Zc3h18 | 0.45 | 5.42E-64 | 1.13E-59 | 0.86 | 0.70 | 0.16 | 1.23 |
| 2 | Gata3 | 0.48 | 6.55E-64 | 1.36E-59 | 0.73 | 0.50 | 0.22 | 1.45 |
| 2 | Foxm1 | 0.83 | 6.59E-64 | 1.37E-59 | 0.42 | 0.25 | 0.17 | 1.70 |

**Supplementary Table 1. Marker genes for the original clusters of integrated clustering of E11.5 control and NotchGOF λ-junction cells.**

| | | | | | | | | |
|---|---|---|---|---|---|---|---|---|
| 2 | Nup85 | 0.67 | 8.61E-64 | 1.79E-59 | 0.54 | 0.35 | 0.19 | 1.55 |
| 2 | Knl1 | 0.42 | 1.02E-63 | 2.13E-59 | 0.71 | 0.47 | 0.24 | 1.50 |
| 2 | Igfbp7 | 0.65 | 1.20E-63 | 2.50E-59 | 0.43 | 0.25 | 0.19 | 1.76 |
| 2 | Fndc3c1 | 0.59 | 1.38E-63 | 2.87E-59 | 0.65 | 0.45 | 0.20 | 1.45 |
| 2 | Mif | 0.27 | 1.77E-63 | 3.68E-59 | 1.00 | 0.99 | 0.01 | 1.01 |
| 2 | Lsm8 | 0.30 | 1.95E-63 | 4.06E-59 | 0.98 | 0.90 | 0.08 | 1.09 |
| 2 | Brip1 | 0.90 | 2.56E-63 | 5.33E-59 | 0.38 | 0.21 | 0.16 | 1.77 |
| 2 | Ska3 | 0.74 | 3.15E-63 | 6.56E-59 | 0.45 | 0.27 | 0.18 | 1.68 |
| 2 | E2f7 | 0.82 | 4.39E-63 | 9.14E-59 | 0.41 | 0.24 | 0.17 | 1.73 |
| 2 | Creb5 | 0.62 | 5.15E-63 | 1.07E-58 | 0.51 | 0.31 | 0.20 | 1.65 |
| 2 | Adamts19 | 0.73 | 1.34E-62 | 2.79E-58 | 0.41 | 0.23 | 0.18 | 1.79 |
| 2 | Topbp1 | 0.41 | 2.24E-62 | 4.66E-58 | 0.89 | 0.72 | 0.17 | 1.24 |
| 2 | Zwilch | 0.72 | 2.41E-62 | 5.00E-58 | 0.48 | 0.30 | 0.18 | 1.61 |
| 2 | Greb1l | 1.31 | 2.81E-62 | 5.85E-58 | 0.16 | 0.06 | 0.10 | 2.75 |
| 2 | Xist | 0.34 | 4.04E-62 | 8.40E-58 | 0.90 | 0.86 | 0.03 | 1.04 |
| 2 | Eno3 | 0.61 | 4.29E-62 | 8.92E-58 | 0.56 | 0.36 | 0.21 | 1.58 |
| 2 | Hist1h2bp | 1.32 | 5.75E-62 | 1.20E-57 | 0.18 | 0.07 | 0.11 | 2.48 |
| 2 | Arhgap11a | 0.55 | 8.74E-62 | 1.82E-57 | 0.59 | 0.39 | 0.20 | 1.53 |
| 2 | Mfap2 | 0.27 | 8.89E-62 | 1.85E-57 | 1.00 | 0.92 | 0.08 | 1.09 |
| 2 | Atad5 | 0.56 | 9.71E-62 | 2.02E-57 | 0.71 | 0.51 | 0.20 | 1.39 |
| 2 | Ywhah | 0.29 | 1.04E-61 | 2.17E-57 | 0.98 | 0.90 | 0.08 | 1.09 |
| 2 | Arhgap28 | 0.44 | 1.11E-61 | 2.30E-57 | 0.74 | 0.51 | 0.23 | 1.44 |
| 2 | Wdhd1 | 0.67 | 1.51E-61 | 3.14E-57 | 0.50 | 0.31 | 0.19 | 1.61 |
| 2 | Eif4g1 | 0.31 | 1.75E-61 | 3.65E-57 | 0.98 | 0.91 | 0.07 | 1.08 |
| 2 | Asf1b | 0.56 | 2.56E-61 | 5.32E-57 | 0.62 | 0.42 | 0.20 | 1.47 |
| 2 | Tcof1 | 0.40 | 2.89E-61 | 6.02E-57 | 0.89 | 0.74 | 0.15 | 1.21 |
| 2 | Adgrl2 | 0.46 | 4.71E-61 | 9.79E-57 | 0.85 | 0.67 | 0.17 | 1.26 |
| 2 | Nup35 | 0.56 | 6.51E-61 | 1.35E-56 | 0.65 | 0.46 | 0.20 | 1.43 |
| 2 | Plk1 | 0.48 | 8.34E-61 | 1.73E-56 | 0.66 | 0.44 | 0.22 | 1.49 |
| 2 | Shmt1 | 0.63 | 1.32E-60 | 2.74E-56 | 0.54 | 0.34 | 0.19 | 1.57 |
| 2 | Myg1 | 0.47 | 1.32E-60 | 2.75E-56 | 0.79 | 0.60 | 0.19 | 1.31 |
| 2 | Bcat1 | 0.70 | 1.43E-60 | 2.98E-56 | 0.48 | 0.29 | 0.19 | 1.64 |
| 2 | Srrm1 | 0.26 | 1.45E-60 | 3.02E-56 | 0.99 | 0.96 | 0.03 | 1.03 |
| 2 | Galnt5 | 1.58 | 1.75E-60 | 3.64E-56 | 0.12 | 0.04 | 0.08 | 3.13 |
| 2 | Wee1 | 0.57 | 2.06E-60 | 4.28E-56 | 0.63 | 0.44 | 0.19 | 1.44 |
| 2 | Snrnp70 | 0.25 | 2.46E-60 | 5.13E-56 | 0.99 | 0.96 | 0.03 | 1.03 |
| 2 | Hist1h2bf | 1.30 | 3.38E-60 | 7.03E-56 | 0.19 | 0.08 | 0.11 | 2.41 |
| 2 | Kif22 | 0.46 | 4.99E-60 | 1.04E-55 | 0.70 | 0.48 | 0.22 | 1.46 |
| 2 | Tmem119 | 0.47 | 6.58E-60 | 1.37E-55 | 0.67 | 0.44 | 0.23 | 1.54 |
| 2 | Pin1 | 0.32 | 6.66E-60 | 1.39E-55 | 0.96 | 0.87 | 0.08 | 1.10 |
| 2 | Ndufaf2 | 0.40 | 7.74E-60 | 1.61E-55 | 0.88 | 0.73 | 0.16 | 1.22 |
| 2 | Cpne8 | 1.26 | 8.39E-60 | 1.74E-55 | 0.17 | 0.07 | 0.11 | 2.52 |
| 2 | Dlc1 | 0.58 | 1.44E-59 | 2.99E-55 | 0.55 | 0.34 | 0.20 | 1.60 |
| 2 | Ilf2 | 0.27 | 2.08E-59 | 4.33E-55 | 0.99 | 0.94 | 0.05 | 1.05 |
| 2 | Lrba | 0.75 | 3.44E-59 | 7.16E-55 | 0.38 | 0.21 | 0.17 | 1.80 |
| 2 | Ppp1cc | 0.28 | 3.91E-59 | 8.13E-55 | 0.99 | 0.94 | 0.05 | 1.06 |
| 2 | Alyref | 0.27 | 8.15E-59 | 1.69E-54 | 0.99 | 0.94 | 0.05 | 1.05 |
| 2 | Trim24 | 0.42 | 8.34E-59 | 1.74E-54 | 0.85 | 0.70 | 0.16 | 1.23 |
| 2 | Cacna1g | 0.61 | 9.18E-59 | 1.91E-54 | 0.51 | 0.31 | 0.20 | 1.65 |
| 2 | Gna11 | 0.44 | 1.15E-58 | 2.39E-54 | 0.83 | 0.66 | 0.17 | 1.26 |
| 2 | Aldh1a2 | 1.02 | 1.32E-58 | 2.75E-54 | 0.16 | 0.06 | 0.10 | 2.68 |
| 2 | Uba2 | 0.33 | 1.37E-58 | 2.86E-54 | 0.95 | 0.86 | 0.10 | 1.11 |
| 2 | Suv39h2 | 0.60 | 1.54E-58 | 3.20E-54 | 0.59 | 0.40 | 0.19 | 1.49 |
| 2 | Sall4 | 0.84 | 1.64E-58 | 3.41E-54 | 0.33 | 0.18 | 0.16 | 1.89 |
| 2 | Rad54b | 0.84 | 1.88E-58 | 3.91E-54 | 0.34 | 0.19 | 0.16 | 1.82 |
| 2 | Rtf1 | 0.30 | 3.71E-58 | 7.73E-54 | 0.98 | 0.92 | 0.06 | 1.06 |

**Supplementary Table 1. Marker genes for the original clusters of integrated clustering of E11.5 control and NotchGOF λ-junction cells.**

| | | | | | | | | |
|---|---|---|---|---|---|---|---|---|
| 2 | Hist1h4a | 0.97 | 3.85E-58 | 8.02E-54 | 0.32 | 0.17 | 0.15 | 1.86 |
| 2 | Fbn1 | 0.65 | 4.74E-58 | 9.87E-54 | 0.51 | 0.32 | 0.19 | 1.60 |
| 2 | Hist1h2ai | 1.54 | 6.78E-58 | 1.41E-53 | 0.14 | 0.05 | 0.09 | 2.80 |
| 2 | Atp11c | 0.74 | 1.06E-57 | 2.20E-53 | 0.40 | 0.23 | 0.17 | 1.72 |
| 2 | Ccne1 | 0.46 | 1.53E-57 | 3.18E-53 | 0.70 | 0.49 | 0.21 | 1.44 |
| 2 | Fam111a | 0.58 | 1.59E-57 | 3.31E-53 | 0.49 | 0.31 | 0.19 | 1.60 |
| 2 | Pimreg | 0.36 | 2.16E-57 | 4.50E-53 | 0.74 | 0.50 | 0.24 | 1.49 |
| 2 | Stag1 | 0.39 | 2.19E-57 | 4.55E-53 | 0.89 | 0.74 | 0.15 | 1.20 |
| 2 | 4930503L19Rik | 0.64 | 2.39E-57 | 4.97E-53 | 0.51 | 0.32 | 0.19 | 1.60 |
| 2 | Fscn1 | 0.27 | 2.97E-57 | 6.18E-53 | 0.99 | 0.90 | 0.09 | 1.09 |
| 2 | Ccp110 | 0.54 | 4.45E-57 | 9.26E-53 | 0.61 | 0.41 | 0.20 | 1.48 |
| 2 | Rspo3 | 0.28 | 8.00E-57 | 1.66E-52 | 0.53 | 0.32 | 0.21 | 1.64 |
| 2 | Nusap1 | 0.29 | 1.03E-56 | 2.14E-52 | 0.71 | 0.47 | 0.24 | 1.52 |
| 2 | Ubr5 | 0.47 | 1.16E-56 | 2.42E-52 | 0.81 | 0.65 | 0.16 | 1.24 |
| 2 | Prr11 | 0.90 | 1.39E-56 | 2.88E-52 | 0.33 | 0.18 | 0.15 | 1.81 |
| 2 | Ppig | 0.27 | 1.39E-56 | 2.90E-52 | 0.99 | 0.93 | 0.06 | 1.06 |
| 2 | Taf1 | 0.49 | 1.40E-56 | 2.91E-52 | 0.76 | 0.56 | 0.19 | 1.34 |
| 2 | Eif4e | 0.30 | 1.64E-56 | 3.41E-52 | 0.96 | 0.88 | 0.08 | 1.09 |
| 2 | Ssrp1 | 0.28 | 2.46E-56 | 5.11E-52 | 0.99 | 0.92 | 0.07 | 1.07 |
| 2 | Chtf18 | 0.78 | 2.50E-56 | 5.20E-52 | 0.38 | 0.22 | 0.16 | 1.74 |
| 2 | Zwint | 0.31 | 2.63E-56 | 5.47E-52 | 0.97 | 0.86 | 0.11 | 1.12 |
| 2 | Rps6ka3 | 0.67 | 3.77E-56 | 7.84E-52 | 0.62 | 0.44 | 0.18 | 1.41 |
| 2 | Prim1 | 0.41 | 4.58E-56 | 9.53E-52 | 0.83 | 0.66 | 0.18 | 1.27 |
| 2 | Dpyd | 1.50 | 6.06E-56 | 1.26E-51 | 0.12 | 0.04 | 0.08 | 3.03 |
| 2 | Cacna1c | 1.10 | 8.02E-56 | 1.67E-51 | 0.19 | 0.08 | 0.11 | 2.35 |
| 2 | Cbfb | 0.48 | 8.78E-56 | 1.83E-51 | 0.74 | 0.55 | 0.19 | 1.34 |
| 2 | Wdr86 | 0.48 | 1.44E-55 | 3.00E-51 | 0.71 | 0.50 | 0.21 | 1.42 |
| 2 | Sms | 0.32 | 1.83E-55 | 3.80E-51 | 0.96 | 0.85 | 0.11 | 1.13 |
| 2 | Plekho1 | 0.40 | 3.34E-55 | 6.95E-51 | 0.81 | 0.60 | 0.21 | 1.35 |
| 2 | Bmp4 | 0.52 | 3.67E-55 | 7.64E-51 | 0.68 | 0.50 | 0.19 | 1.37 |
| 2 | Kdm2b | 0.56 | 4.87E-55 | 1.01E-50 | 0.72 | 0.55 | 0.17 | 1.31 |
| 2 | Dock4 | 0.62 | 5.14E-55 | 1.07E-50 | 0.48 | 0.30 | 0.19 | 1.63 |
| 2 | Car14 | 0.55 | 6.09E-55 | 1.27E-50 | 0.61 | 0.40 | 0.20 | 1.50 |
| 2 | Tfdp1 | 0.44 | 6.41E-55 | 1.33E-50 | 0.82 | 0.64 | 0.18 | 1.28 |
| 2 | Bin1 | 0.40 | 7.85E-55 | 1.63E-50 | 0.79 | 0.60 | 0.19 | 1.31 |
| 2 | Mastl | 0.80 | 8.74E-55 | 1.82E-50 | 0.36 | 0.21 | 0.15 | 1.72 |
| 2 | Nxt1 | 0.42 | 9.46E-55 | 1.97E-50 | 0.83 | 0.66 | 0.17 | 1.26 |
| 2 | Ccna2 | 0.31 | 1.03E-54 | 2.14E-50 | 0.86 | 0.61 | 0.25 | 1.41 |
| 2 | Zeb1 | 0.55 | 1.29E-54 | 2.68E-50 | 0.53 | 0.33 | 0.20 | 1.60 |
| 2 | Trps1 | 0.60 | 1.38E-54 | 2.86E-50 | 0.65 | 0.47 | 0.18 | 1.37 |
| 2 | Dnm1 | 0.56 | 1.65E-54 | 3.44E-50 | 0.51 | 0.31 | 0.20 | 1.63 |
| 2 | Nup160 | 0.63 | 1.87E-54 | 3.89E-50 | 0.47 | 0.29 | 0.18 | 1.60 |
| 2 | Gins3 | 0.72 | 3.93E-54 | 8.18E-50 | 0.41 | 0.24 | 0.17 | 1.69 |
| 2 | Nsg1 | 0.37 | 4.86E-54 | 1.01E-49 | 0.84 | 0.65 | 0.18 | 1.28 |
| 2 | Gga2 | 0.42 | 5.63E-54 | 1.17E-49 | 0.81 | 0.63 | 0.19 | 1.30 |
| 2 | Kpna2 | 0.42 | 6.29E-54 | 1.31E-49 | 0.60 | 0.40 | 0.20 | 1.51 |
| 2 | Ndc1 | 0.69 | 6.32E-54 | 1.31E-49 | 0.46 | 0.29 | 0.17 | 1.59 |
| 2 | Depdc1b | 1.11 | 6.53E-54 | 1.36E-49 | 0.22 | 0.11 | 0.12 | 2.07 |
| 2 | Spag5 | 0.68 | 7.70E-54 | 1.60E-49 | 0.46 | 0.29 | 0.17 | 1.57 |
| 2 | Dpf1 | 0.88 | 7.85E-54 | 1.63E-49 | 0.29 | 0.15 | 0.14 | 1.93 |
| 2 | Acat2 | 0.33 | 9.06E-54 | 1.88E-49 | 0.93 | 0.79 | 0.14 | 1.17 |
| 2 | Tmem132c | 0.61 | 9.19E-54 | 1.91E-49 | 0.49 | 0.31 | 0.18 | 1.59 |
| 2 | Kansl1l | 0.73 | 1.01E-53 | 2.11E-49 | 0.41 | 0.25 | 0.17 | 1.67 |
| 2 | Bclaf1 | 0.28 | 1.05E-53 | 2.19E-49 | 0.99 | 0.93 | 0.06 | 1.06 |
| 2 | Nsmce4a | 0.30 | 1.30E-53 | 2.71E-49 | 0.97 | 0.87 | 0.10 | 1.11 |
| 2 | Mpp6 | 0.38 | 1.59E-53 | 3.30E-49 | 0.88 | 0.73 | 0.15 | 1.20 |

**Supplementary Table 1. Marker genes for the original clusters of integrated clustering of E11.5 control and NotchGOF λ-junction cells.**

| | | | | | | | | |
|---|---|---|---|---|---|---|---|---|
| 2 | Fnbp1l | 0.34 | 2.05E-53 | 4.26E-49 | 0.95 | 0.84 | 0.11 | 1.13 |
| 2 | Sart3 | 0.49 | 2.57E-53 | 5.34E-49 | 0.73 | 0.54 | 0.18 | 1.34 |
| 2 | Hist1h2ag | 0.65 | 2.61E-53 | 5.43E-49 | 0.47 | 0.30 | 0.18 | 1.59 |
| 2 | Pask | 0.92 | 2.61E-53 | 5.43E-49 | 0.28 | 0.14 | 0.14 | 1.94 |
| 2 | Tead2 | 0.28 | 2.90E-53 | 6.04E-49 | 0.99 | 0.89 | 0.09 | 1.11 |
| 2 | Osbpl1a | 0.53 | 2.93E-53 | 6.10E-49 | 0.56 | 0.37 | 0.19 | 1.52 |
| 2 | Pitpnc1 | 0.40 | 2.95E-53 | 6.14E-49 | 0.77 | 0.57 | 0.20 | 1.35 |
| 2 | St3gal4 | 0.67 | 3.43E-53 | 7.14E-49 | 0.41 | 0.24 | 0.17 | 1.72 |
| 2 | Nsl1 | 0.87 | 3.77E-53 | 7.83E-49 | 0.32 | 0.18 | 0.14 | 1.81 |
| 2 | Dbn1 | 0.30 | 4.10E-53 | 8.53E-49 | 0.98 | 0.87 | 0.11 | 1.13 |
| 2 | Brd2 | 0.34 | 6.45E-53 | 1.34E-48 | 0.92 | 0.81 | 0.11 | 1.14 |
| 2 | Tns3 | 0.50 | 6.62E-53 | 1.38E-48 | 0.67 | 0.48 | 0.19 | 1.40 |
| 2 | Brd8 | 0.40 | 7.44E-53 | 1.55E-48 | 0.88 | 0.74 | 0.14 | 1.19 |
| 2 | Tube1 | 0.84 | 7.48E-53 | 1.56E-48 | 0.31 | 0.16 | 0.14 | 1.86 |
| 2 | Ifrd1 | 0.56 | 7.84E-53 | 1.63E-48 | 0.66 | 0.49 | 0.17 | 1.34 |
| 2 | Fn1 | 0.32 | 9.87E-53 | 2.05E-48 | 0.94 | 0.77 | 0.17 | 1.22 |
| 2 | Taf5 | 0.59 | 1.19E-52 | 2.48E-48 | 0.60 | 0.42 | 0.18 | 1.43 |
| 2 | Dchs1 | 0.52 | 1.46E-52 | 3.04E-48 | 0.60 | 0.40 | 0.20 | 1.51 |
| 2 | Hpf1 | 0.34 | 4.49E-52 | 9.33E-48 | 0.94 | 0.83 | 0.10 | 1.12 |
| 2 | Dact3 | 0.39 | 5.08E-52 | 1.06E-47 | 0.70 | 0.47 | 0.23 | 1.48 |
| 2 | Safb | 0.36 | 6.89E-52 | 1.43E-47 | 0.89 | 0.77 | 0.12 | 1.15 |
| 2 | Pdgfra | 0.35 | 1.33E-51 | 2.76E-47 | 0.76 | 0.52 | 0.24 | 1.46 |
| 2 | Ctsk | 0.84 | 1.43E-51 | 2.97E-47 | 0.24 | 0.12 | 0.13 | 2.07 |
| 2 | Lix1l | 0.47 | 2.56E-51 | 5.31E-47 | 0.57 | 0.36 | 0.21 | 1.57 |
| 2 | Gspt1 | 0.27 | 2.61E-51 | 5.44E-47 | 0.98 | 0.92 | 0.06 | 1.07 |
| 2 | Emsy | 0.46 | 3.62E-51 | 7.53E-47 | 0.78 | 0.60 | 0.18 | 1.30 |
| 2 | Plcb1 | 1.09 | 4.00E-51 | 8.31E-47 | 0.19 | 0.09 | 0.11 | 2.22 |
| 2 | Rfc2 | 0.34 | 5.69E-51 | 1.18E-46 | 0.93 | 0.79 | 0.14 | 1.18 |
| 2 | Abracl | 0.27 | 8.50E-51 | 1.77E-46 | 0.98 | 0.92 | 0.06 | 1.06 |
| 2 | Raph1 | 0.56 | 8.88E-51 | 1.85E-46 | 0.59 | 0.41 | 0.18 | 1.44 |
| 2 | Cenps | 0.49 | 9.50E-51 | 1.98E-46 | 0.68 | 0.49 | 0.19 | 1.39 |
| 2 | Tnrc18 | 0.45 | 1.07E-50 | 2.22E-46 | 0.73 | 0.54 | 0.19 | 1.36 |
| 2 | Matr3 | 0.28 | 1.08E-50 | 2.25E-46 | 0.96 | 0.89 | 0.07 | 1.08 |
| 2 | Naa50 | 0.35 | 1.51E-50 | 3.14E-46 | 0.91 | 0.78 | 0.13 | 1.17 |
| 2 | Pcolce | 0.40 | 1.84E-50 | 3.83E-46 | 0.66 | 0.44 | 0.22 | 1.50 |
| 2 | Gm13429 | 1.27 | 3.01E-50 | 6.26E-46 | 0.13 | 0.05 | 0.08 | 2.71 |
| 2 | Cep295 | 0.51 | 3.66E-50 | 7.62E-46 | 0.66 | 0.47 | 0.19 | 1.40 |
| 2 | Cblb | 0.56 | 3.95E-50 | 8.23E-46 | 0.51 | 0.33 | 0.18 | 1.56 |
| 2 | Gnb4 | 0.55 | 4.22E-50 | 8.78E-46 | 0.47 | 0.28 | 0.18 | 1.64 |
| 2 | Ogdh | 0.49 | 4.45E-50 | 9.26E-46 | 0.69 | 0.51 | 0.18 | 1.36 |
| 2 | Gpc6 | 0.41 | 4.97E-50 | 1.03E-45 | 0.77 | 0.57 | 0.20 | 1.35 |
| 2 | Cep57 | 0.46 | 5.97E-50 | 1.24E-45 | 0.76 | 0.58 | 0.18 | 1.30 |
| 2 | Pbrm1 | 0.27 | 6.92E-50 | 1.44E-45 | 0.98 | 0.94 | 0.05 | 1.05 |
| 2 | Hey2 | 1.12 | 8.03E-50 | 1.67E-45 | 0.15 | 0.06 | 0.09 | 2.48 |
| 2 | Klf6 | 0.59 | 9.51E-50 | 1.98E-45 | 0.38 | 0.22 | 0.15 | 1.69 |
| 2 | Suv39h1 | 0.54 | 1.12E-49 | 2.33E-45 | 0.63 | 0.45 | 0.18 | 1.39 |
| 2 | Sf3a3 | 0.31 | 1.40E-49 | 2.91E-45 | 0.95 | 0.85 | 0.11 | 1.13 |
| 2 | Ssbp2 | 0.35 | 1.95E-49 | 4.06E-45 | 0.86 | 0.69 | 0.17 | 1.24 |
| 2 | Mtx2 | 0.32 | 2.14E-49 | 4.45E-45 | 0.93 | 0.83 | 0.10 | 1.12 |
| 2 | Cdk5rap2 | 0.42 | 3.23E-49 | 6.72E-45 | 0.77 | 0.59 | 0.18 | 1.31 |
| 2 | Tln2 | 0.43 | 5.55E-49 | 1.15E-44 | 0.75 | 0.57 | 0.18 | 1.32 |
| 2 | Nsd2 | 0.36 | 5.72E-49 | 1.19E-44 | 0.90 | 0.74 | 0.16 | 1.21 |
| 2 | Asap1 | 0.34 | 6.28E-49 | 1.31E-44 | 0.88 | 0.72 | 0.16 | 1.22 |
| 2 | Akap12 | 0.51 | 8.17E-49 | 1.70E-44 | 0.33 | 0.18 | 0.15 | 1.80 |
| 2 | Hddc2 | 0.37 | 1.03E-48 | 2.13E-44 | 0.88 | 0.75 | 0.13 | 1.17 |
| 2 | Ccnf | 0.54 | 1.08E-48 | 2.24E-44 | 0.50 | 0.32 | 0.17 | 1.54 |

**Supplementary Table 1. Marker genes for the original clusters of integrated clustering of E11.5 control and NotchGOF λ-junction cells.**

| | | | | | | | | |
|---|---|---|---|---|---|---|---|---|
| 2 | Psd3 | 0.67 | 1.17E-48 | 2.44E-44 | 0.36 | 0.21 | 0.15 | 1.73 |
| 2 | Dusp7 | 0.45 | 1.55E-48 | 3.22E-44 | 0.68 | 0.48 | 0.20 | 1.41 |
| 2 | Mrpl18 | 0.26 | 1.66E-48 | 3.46E-44 | 0.98 | 0.93 | 0.06 | 1.06 |
| 2 | Cdc7 | 0.57 | 2.09E-48 | 4.34E-44 | 0.52 | 0.35 | 0.17 | 1.50 |
| 2 | Trim28 | 0.26 | 2.22E-48 | 4.62E-44 | 0.99 | 0.93 | 0.06 | 1.06 |
| 2 | Gm3550 | 1.08 | 2.38E-48 | 4.94E-44 | 0.20 | 0.10 | 0.11 | 2.08 |
| 2 | Sfrp2 | 0.37 | 2.95E-48 | 6.13E-44 | 0.77 | 0.60 | 0.18 | 1.30 |
| 2 | Ncapd2 | 0.45 | 3.02E-48 | 6.27E-44 | 0.72 | 0.54 | 0.19 | 1.35 |
| 2 | Chek1 | 0.42 | 5.09E-48 | 1.06E-43 | 0.73 | 0.53 | 0.19 | 1.36 |
| 2 | Zfp462 | 0.37 | 6.09E-48 | 1.27E-43 | 0.87 | 0.72 | 0.15 | 1.21 |
| 2 | Tcf19 | 0.77 | 6.96E-48 | 1.45E-43 | 0.32 | 0.18 | 0.14 | 1.77 |
| 2 | Nup107 | 0.57 | 9.23E-48 | 1.92E-43 | 0.54 | 0.36 | 0.18 | 1.48 |
| 2 | Nfya | 0.61 | 1.13E-47 | 2.34E-43 | 0.47 | 0.30 | 0.16 | 1.54 |
| 2 | Capn6 | 0.43 | 2.08E-47 | 4.32E-43 | 0.72 | 0.53 | 0.19 | 1.35 |
| 2 | Peg12 | 0.60 | 2.25E-47 | 4.68E-43 | 0.40 | 0.24 | 0.16 | 1.69 |
| 2 | Pum1 | 0.28 | 2.53E-47 | 5.26E-43 | 0.96 | 0.87 | 0.09 | 1.11 |
| 2 | Pbdc1 | 0.31 | 2.88E-47 | 5.99E-43 | 0.95 | 0.85 | 0.10 | 1.12 |
| 2 | Smo | 0.35 | 3.30E-47 | 6.86E-43 | 0.88 | 0.72 | 0.16 | 1.22 |
| 2 | Irx5 | 0.32 | 3.38E-47 | 7.04E-43 | 0.45 | 0.27 | 0.18 | 1.68 |
| 2 | Sf3a1 | 0.35 | 3.64E-47 | 7.57E-43 | 0.88 | 0.73 | 0.16 | 1.22 |
| 2 | Hist1h2bk | 0.68 | 3.85E-47 | 8.00E-43 | 0.36 | 0.21 | 0.15 | 1.69 |
| 2 | Lima1 | 0.30 | 4.78E-47 | 9.94E-43 | 0.90 | 0.72 | 0.17 | 1.24 |
| 2 | Tcf3 | 0.28 | 5.78E-47 | 1.20E-42 | 0.97 | 0.89 | 0.08 | 1.09 |
| 2 | Aarsd1 | 0.34 | 6.02E-47 | 1.25E-42 | 0.89 | 0.76 | 0.14 | 1.18 |
| 2 | Depdc1a | 0.48 | 6.21E-47 | 1.29E-42 | 0.52 | 0.34 | 0.18 | 1.51 |
| 2 | Cmc2 | 0.44 | 6.32E-47 | 1.31E-42 | 0.76 | 0.59 | 0.17 | 1.29 |
| 2 | Arid1a | 0.32 | 8.00E-47 | 1.66E-42 | 0.92 | 0.80 | 0.12 | 1.15 |
| 2 | Rasl11b | 0.40 | 1.04E-46 | 2.15E-42 | 0.67 | 0.47 | 0.20 | 1.43 |
| 2 | Ythdc2 | 0.66 | 1.09E-46 | 2.26E-42 | 0.41 | 0.25 | 0.16 | 1.61 |
| 2 | Kpnb1 | 0.31 | 1.18E-46 | 2.45E-42 | 0.93 | 0.83 | 0.11 | 1.13 |
| 2 | Ppp2r2a | 0.33 | 1.31E-46 | 2.72E-42 | 0.91 | 0.78 | 0.13 | 1.17 |
| 2 | Fbxl7 | 0.48 | 1.38E-46 | 2.86E-42 | 0.63 | 0.45 | 0.19 | 1.41 |
| 2 | Iqgap3 | 0.79 | 1.44E-46 | 3.00E-42 | 0.35 | 0.21 | 0.14 | 1.67 |
| 2 | Gfra4 | 0.90 | 1.55E-46 | 3.22E-42 | 0.20 | 0.09 | 0.11 | 2.15 |
| 2 | Mre11a | 0.55 | 1.80E-46 | 3.74E-42 | 0.54 | 0.37 | 0.17 | 1.47 |
| 2 | Pold1 | 0.56 | 1.99E-46 | 4.13E-42 | 0.49 | 0.33 | 0.17 | 1.52 |
| 2 | Elovl6 | 0.40 | 2.67E-46 | 5.55E-42 | 0.81 | 0.63 | 0.18 | 1.28 |
| 2 | Kcnj2 | 0.63 | 3.22E-46 | 6.70E-42 | 0.30 | 0.16 | 0.14 | 1.84 |
| 2 | Cenpk | 0.42 | 3.83E-46 | 7.97E-42 | 0.74 | 0.55 | 0.18 | 1.33 |
| 2 | Bcl9l | 0.58 | 4.28E-46 | 8.90E-42 | 0.49 | 0.32 | 0.17 | 1.53 |
| 2 | Pla2g4a | 0.57 | 4.70E-46 | 9.77E-42 | 0.40 | 0.24 | 0.16 | 1.68 |
| 2 | Ppp1r8 | 0.48 | 5.33E-46 | 1.11E-41 | 0.61 | 0.43 | 0.18 | 1.42 |
| 2 | Greb1 | 0.60 | 7.75E-46 | 1.61E-41 | 0.45 | 0.28 | 0.16 | 1.58 |
| 2 | Lpl | 0.67 | 8.09E-46 | 1.68E-41 | 0.28 | 0.15 | 0.13 | 1.85 |
| 2 | BC030867 | 0.96 | 9.04E-46 | 1.88E-41 | 0.25 | 0.13 | 0.12 | 1.88 |
| 2 | Hacd1 | 0.53 | 1.27E-45 | 2.64E-41 | 0.55 | 0.38 | 0.18 | 1.47 |
| 2 | Pmf1 | 0.39 | 1.32E-45 | 2.74E-41 | 0.83 | 0.66 | 0.17 | 1.26 |
| 2 | Pcnt | 0.50 | 1.69E-45 | 3.51E-41 | 0.63 | 0.46 | 0.17 | 1.37 |
| 2 | Map2 | 0.63 | 1.70E-45 | 3.54E-41 | 0.35 | 0.20 | 0.14 | 1.70 |
| 2 | Tlnrd1 | 0.44 | 2.12E-45 | 4.40E-41 | 0.67 | 0.49 | 0.18 | 1.37 |
| 2 | Cse1l | 0.39 | 2.40E-45 | 5.00E-41 | 0.78 | 0.61 | 0.17 | 1.28 |
| 2 | Samd1 | 0.51 | 2.67E-45 | 5.56E-41 | 0.56 | 0.39 | 0.17 | 1.44 |
| 2 | Rasl11a | 0.60 | 4.49E-45 | 9.33E-41 | 0.43 | 0.26 | 0.16 | 1.62 |
| 2 | Gmps | 0.27 | 4.70E-45 | 9.77E-41 | 0.97 | 0.89 | 0.08 | 1.09 |
| 2 | Meis2 | 0.30 | 5.27E-45 | 1.10E-40 | 0.88 | 0.65 | 0.22 | 1.34 |
| 2 | Ankrd44 | 0.57 | 7.14E-45 | 1.49E-40 | 0.43 | 0.27 | 0.16 | 1.62 |

**Supplementary Table 1. Marker genes for the original clusters of integrated clustering of E11.5 control and NotchGOF λ-junction cells.**

| | | | | | | | | |
|---|---|---|---|---|---|---|---|---|
| 2 | Mms22l | 0.61 | 7.19E-45 | 1.49E-40 | 0.51 | 0.34 | 0.16 | 1.47 |
| 2 | AI314278 | 1.26 | 8.10E-45 | 1.68E-40 | 0.12 | 0.04 | 0.07 | 2.70 |
| 2 | Suz12 | 0.36 | 1.06E-44 | 2.20E-40 | 0.85 | 0.71 | 0.14 | 1.19 |
| 2 | Meaf6 | 0.37 | 1.07E-44 | 2.23E-40 | 0.82 | 0.65 | 0.16 | 1.25 |
| 2 | Ciapin1 | 0.35 | 1.13E-44 | 2.36E-40 | 0.87 | 0.72 | 0.15 | 1.20 |
| 2 | Pds5b | 0.39 | 1.19E-44 | 2.48E-40 | 0.81 | 0.64 | 0.18 | 1.28 |
| 2 | Six4 | 0.46 | 1.37E-44 | 2.86E-40 | 0.63 | 0.45 | 0.18 | 1.41 |
| 2 | Rpa1 | 0.53 | 1.49E-44 | 3.09E-40 | 0.58 | 0.40 | 0.17 | 1.43 |
| 2 | Cntln | 0.48 | 2.92E-44 | 6.07E-40 | 0.65 | 0.47 | 0.18 | 1.37 |
| 2 | Asphd2 | 0.84 | 3.06E-44 | 6.37E-40 | 0.22 | 0.11 | 0.11 | 2.00 |
| 2 | Robo1 | 0.45 | 3.27E-44 | 6.79E-40 | 0.63 | 0.45 | 0.18 | 1.39 |
| 2 | Arid1b | 0.47 | 6.25E-44 | 1.30E-39 | 0.64 | 0.47 | 0.17 | 1.36 |
| 2 | Sncaip | 0.51 | 7.94E-44 | 1.65E-39 | 0.46 | 0.29 | 0.17 | 1.58 |
| 2 | Timm8a1 | 0.37 | 7.94E-44 | 1.65E-39 | 0.83 | 0.68 | 0.15 | 1.22 |
| 2 | Vegfc | 0.50 | 9.42E-44 | 1.96E-39 | 0.41 | 0.25 | 0.16 | 1.63 |
| 2 | Cenpi | 0.81 | 9.42E-44 | 1.96E-39 | 0.28 | 0.16 | 0.12 | 1.79 |
| 2 | Stag2 | 0.39 | 1.01E-43 | 2.10E-39 | 0.79 | 0.64 | 0.16 | 1.25 |
| 2 | Tcf7 | 0.80 | 1.05E-43 | 2.18E-39 | 0.26 | 0.14 | 0.12 | 1.88 |
| 2 | Robo2 | 0.65 | 1.17E-43 | 2.44E-39 | 0.32 | 0.18 | 0.14 | 1.75 |
| 2 | Slf1 | 0.43 | 1.36E-43 | 2.82E-39 | 0.70 | 0.52 | 0.18 | 1.34 |
| 2 | Pa2g4 | 0.25 | 1.42E-43 | 2.95E-39 | 0.97 | 0.91 | 0.07 | 1.07 |
| 2 | Cdk14 | 0.42 | 1.56E-43 | 3.24E-39 | 0.66 | 0.48 | 0.19 | 1.39 |
| 2 | Tut4 | 0.26 | 1.74E-43 | 3.62E-39 | 0.99 | 0.92 | 0.07 | 1.07 |
| 2 | Dock7 | 0.40 | 1.79E-43 | 3.72E-39 | 0.77 | 0.60 | 0.17 | 1.29 |
| 2 | Hspa9 | 0.28 | 1.82E-43 | 3.78E-39 | 0.97 | 0.89 | 0.08 | 1.09 |
| 2 | Tmem30b | 0.59 | 2.59E-43 | 5.39E-39 | 0.26 | 0.14 | 0.13 | 1.91 |
| 2 | Ankrd12 | 0.47 | 2.75E-43 | 5.73E-39 | 0.67 | 0.52 | 0.16 | 1.31 |
| 2 | Thsd7a | 0.59 | 2.88E-43 | 5.99E-39 | 0.26 | 0.14 | 0.13 | 1.91 |
| 2 | Wdr75 | 0.44 | 3.08E-43 | 6.40E-39 | 0.73 | 0.57 | 0.16 | 1.29 |
| 2 | Etv1 | 0.44 | 5.10E-43 | 1.06E-38 | 0.45 | 0.28 | 0.17 | 1.60 |
| 2 | Spin2c | 0.42 | 5.36E-43 | 1.11E-38 | 0.60 | 0.40 | 0.19 | 1.48 |
| 2 | Tkt | 0.26 | 5.43E-43 | 1.13E-38 | 0.98 | 0.91 | 0.06 | 1.07 |
| 2 | Lin54 | 0.71 | 5.62E-43 | 1.17E-38 | 0.36 | 0.22 | 0.14 | 1.61 |
| 2 | Rrp1b | 0.48 | 5.73E-43 | 1.19E-38 | 0.69 | 0.52 | 0.17 | 1.34 |
| 2 | Ppm1g | 0.30 | 6.71E-43 | 1.40E-38 | 0.92 | 0.80 | 0.13 | 1.16 |
| 2 | Mycn | 0.50 | 7.33E-43 | 1.52E-38 | 0.53 | 0.36 | 0.17 | 1.46 |
| 2 | Nop56 | 0.28 | 7.96E-43 | 1.66E-38 | 0.95 | 0.86 | 0.09 | 1.11 |
| 2 | Zcchc24 | 0.52 | 8.30E-43 | 1.73E-38 | 0.44 | 0.28 | 0.17 | 1.61 |
| 2 | Haus5 | 0.67 | 9.56E-43 | 1.99E-38 | 0.38 | 0.24 | 0.14 | 1.60 |
| 2 | Kntc1 | 0.80 | 1.22E-42 | 2.54E-38 | 0.31 | 0.18 | 0.13 | 1.70 |
| 2 | Adnp | 0.29 | 1.38E-42 | 2.88E-38 | 0.94 | 0.83 | 0.11 | 1.13 |
| 2 | Fbln2 | 0.27 | 1.47E-42 | 3.05E-38 | 0.71 | 0.50 | 0.21 | 1.42 |
| 2 | S1pr3 | 0.50 | 1.53E-42 | 3.19E-38 | 0.42 | 0.26 | 0.16 | 1.62 |
| 2 | Paics | 0.25 | 1.67E-42 | 3.46E-38 | 0.98 | 0.92 | 0.07 | 1.07 |
| 2 | Epb41l2 | 0.35 | 1.86E-42 | 3.87E-38 | 0.82 | 0.65 | 0.16 | 1.25 |
| 2 | Pcsk5 | 0.44 | 2.23E-42 | 4.63E-38 | 0.53 | 0.35 | 0.18 | 1.52 |
| 2 | Erc1 | 0.44 | 2.65E-42 | 5.52E-38 | 0.71 | 0.54 | 0.17 | 1.31 |
| 2 | Snrnp40 | 0.27 | 2.65E-42 | 5.52E-38 | 0.94 | 0.85 | 0.09 | 1.11 |
| 2 | Ntng1 | 0.71 | 3.23E-42 | 6.72E-38 | 0.22 | 0.11 | 0.11 | 1.98 |
| 2 | Sertad4 | 0.48 | 3.26E-42 | 6.78E-38 | 0.53 | 0.35 | 0.18 | 1.50 |
| 2 | Rprd1b | 0.43 | 3.48E-42 | 7.24E-38 | 0.69 | 0.52 | 0.17 | 1.33 |
| 2 | Espl1 | 0.72 | 3.93E-42 | 8.18E-38 | 0.34 | 0.21 | 0.13 | 1.64 |
| 2 | Mme | 0.38 | 4.18E-42 | 8.70E-38 | 0.64 | 0.45 | 0.19 | 1.43 |
| 2 | Pdgfrb | 0.32 | 4.56E-42 | 9.47E-38 | 0.65 | 0.44 | 0.22 | 1.49 |
| 2 | Sirt1 | 0.51 | 5.01E-42 | 1.04E-37 | 0.55 | 0.38 | 0.17 | 1.45 |
| 2 | Nup133 | 0.50 | 5.16E-42 | 1.07E-37 | 0.53 | 0.36 | 0.16 | 1.45 |

**Supplementary Table 1. Marker genes for the original clusters of integrated clustering of E11.5 control and NotchGOF λ-junction cells.**

| | | | | | | | | |
|---|---|---|---|---|---|---|---|---|
| 2 | Magohb | 0.43 | 5.54E-42 | 1.15E-37 | 0.73 | 0.56 | 0.17 | 1.31 |
| 2 | Fignl1 | 0.42 | 5.98E-42 | 1.24E-37 | 0.67 | 0.50 | 0.17 | 1.35 |
| 2 | Dhx9 | 0.27 | 6.17E-42 | 1.28E-37 | 0.97 | 0.89 | 0.08 | 1.08 |
| 2 | Gm4258 | 0.40 | 6.62E-42 | 1.38E-37 | 0.66 | 0.48 | 0.18 | 1.38 |
| 2 | Sf3b3 | 0.28 | 7.18E-42 | 1.49E-37 | 0.95 | 0.85 | 0.10 | 1.12 |
| 2 | Vrk1 | 0.40 | 7.30E-42 | 1.52E-37 | 0.77 | 0.60 | 0.17 | 1.29 |
| 2 | Pitpnb | 0.37 | 1.02E-41 | 2.13E-37 | 0.82 | 0.66 | 0.16 | 1.24 |
| 2 | Pwwp3a | 0.41 | 1.17E-41 | 2.43E-37 | 0.71 | 0.54 | 0.17 | 1.33 |
| 2 | Map3k7 | 0.49 | 1.19E-41 | 2.48E-37 | 0.63 | 0.46 | 0.17 | 1.36 |
| 2 | Brd7 | 0.30 | 1.22E-41 | 2.54E-37 | 0.92 | 0.82 | 0.10 | 1.12 |
| 2 | Taf3 | 0.44 | 1.33E-41 | 2.76E-37 | 0.66 | 0.49 | 0.16 | 1.33 |
| 2 | Hspa14 | 0.37 | 1.36E-41 | 2.82E-37 | 0.80 | 0.65 | 0.15 | 1.24 |
| 2 | Sephs1 | 0.38 | 1.41E-41 | 2.94E-37 | 0.77 | 0.60 | 0.17 | 1.29 |
| 2 | Sap30 | 0.38 | 1.46E-41 | 3.03E-37 | 0.80 | 0.64 | 0.16 | 1.25 |
| 2 | Itih5 | 0.76 | 1.66E-41 | 3.46E-37 | 0.23 | 0.12 | 0.11 | 1.94 |
| 2 | Smarca4 | 0.26 | 2.64E-41 | 5.49E-37 | 0.97 | 0.89 | 0.08 | 1.09 |
| 2 | Mis12 | 0.45 | 3.58E-41 | 7.44E-37 | 0.64 | 0.47 | 0.17 | 1.36 |
| 2 | Srsf4 | 0.32 | 6.88E-41 | 1.43E-36 | 0.86 | 0.71 | 0.15 | 1.21 |
| 2 | Vars | 0.40 | 1.03E-40 | 2.14E-36 | 0.75 | 0.59 | 0.16 | 1.26 |
| 2 | Wbp11 | 0.28 | 1.19E-40 | 2.48E-36 | 0.95 | 0.85 | 0.10 | 1.12 |
| 2 | Paxip1 | 0.57 | 1.26E-40 | 2.62E-36 | 0.47 | 0.32 | 0.15 | 1.46 |
| 2 | Ubap2 | 0.32 | 1.32E-40 | 2.74E-36 | 0.89 | 0.76 | 0.13 | 1.17 |
| 2 | Ubr7 | 0.46 | 1.37E-40 | 2.84E-36 | 0.63 | 0.46 | 0.17 | 1.37 |
| 2 | Pdzrn3 | 0.67 | 1.39E-40 | 2.89E-36 | 0.29 | 0.16 | 0.13 | 1.78 |
| 2 | Myo1b | 0.30 | 1.42E-40 | 2.96E-36 | 0.87 | 0.72 | 0.15 | 1.21 |
| 2 | Rsrc1 | 0.35 | 1.56E-40 | 3.24E-36 | 0.87 | 0.73 | 0.13 | 1.18 |
| 2 | Sin3a | 0.46 | 1.59E-40 | 3.31E-36 | 0.67 | 0.51 | 0.17 | 1.33 |
| 2 | Actl6a | 0.26 | 1.73E-40 | 3.61E-36 | 0.96 | 0.87 | 0.09 | 1.10 |
| 2 | Numa1 | 0.35 | 1.96E-40 | 4.08E-36 | 0.86 | 0.71 | 0.15 | 1.22 |
| 2 | Cdk11b | 0.31 | 2.10E-40 | 4.38E-36 | 0.89 | 0.76 | 0.13 | 1.17 |
| 2 | Cep128 | 0.75 | 2.20E-40 | 4.57E-36 | 0.28 | 0.16 | 0.12 | 1.73 |
| 2 | Ift27 | 0.39 | 2.51E-40 | 5.22E-36 | 0.73 | 0.55 | 0.18 | 1.32 |
| 2 | Haus6 | 0.55 | 2.63E-40 | 5.47E-36 | 0.50 | 0.35 | 0.16 | 1.45 |
| 2 | Ipo9 | 0.41 | 3.17E-40 | 6.58E-36 | 0.69 | 0.52 | 0.17 | 1.33 |
| 2 | Gemin6 | 0.45 | 3.80E-40 | 7.91E-36 | 0.61 | 0.44 | 0.17 | 1.38 |
| 2 | Angptl4 | 0.47 | 4.14E-40 | 8.60E-36 | 0.49 | 0.32 | 0.17 | 1.52 |
| 2 | Etv4 | 0.43 | 5.76E-40 | 1.20E-35 | 0.49 | 0.32 | 0.17 | 1.53 |
| 2 | Col6a2 | 0.73 | 6.10E-40 | 1.27E-35 | 0.26 | 0.14 | 0.12 | 1.85 |
| 2 | Prr12 | 0.50 | 6.81E-40 | 1.42E-35 | 0.58 | 0.42 | 0.16 | 1.38 |
| 2 | Igf2bp1 | 0.26 | 6.94E-40 | 1.44E-35 | 0.97 | 0.88 | 0.09 | 1.10 |
| 2 | Tns1 | 0.47 | 8.22E-40 | 1.71E-35 | 0.46 | 0.29 | 0.17 | 1.57 |
| 2 | Crybg3 | 0.67 | 9.01E-40 | 1.87E-35 | 0.33 | 0.20 | 0.13 | 1.66 |
| 2 | Syne2 | 0.32 | 1.36E-39 | 2.82E-35 | 0.87 | 0.71 | 0.16 | 1.22 |
| 2 | Emilin1 | 0.27 | 1.36E-39 | 2.82E-35 | 0.73 | 0.51 | 0.22 | 1.43 |
| 2 | Aunip | 0.65 | 1.38E-39 | 2.88E-35 | 0.38 | 0.25 | 0.14 | 1.55 |
| 2 | Zfp101 | 0.54 | 1.57E-39 | 3.26E-35 | 0.51 | 0.35 | 0.16 | 1.46 |
| 2 | Tsen34 | 0.27 | 1.60E-39 | 3.33E-35 | 0.95 | 0.83 | 0.12 | 1.14 |
| 2 | Sulf2 | 0.26 | 2.60E-39 | 5.41E-35 | 0.84 | 0.65 | 0.19 | 1.30 |
| 2 | Osr2 | 1.21 | 3.14E-39 | 6.53E-35 | 0.17 | 0.08 | 0.09 | 2.10 |
| 2 | Rgs5 | 0.84 | 3.51E-39 | 7.29E-35 | 0.16 | 0.08 | 0.09 | 2.19 |
| 2 | Glul | 0.37 | 4.03E-39 | 8.38E-35 | 0.77 | 0.60 | 0.16 | 1.27 |
| 2 | Rspo1 | 1.10 | 5.58E-39 | 1.16E-34 | 0.14 | 0.06 | 0.08 | 2.29 |
| 2 | Gins4 | 0.36 | 8.28E-39 | 1.72E-34 | 0.78 | 0.63 | 0.15 | 1.24 |
| 2 | Hist1h4j | 1.33 | 9.18E-39 | 1.91E-34 | 0.12 | 0.05 | 0.07 | 2.47 |
| 2 | Mrpl28 | 0.26 | 9.64E-39 | 2.00E-34 | 0.95 | 0.88 | 0.07 | 1.08 |
| 2 | Tmem131l | 0.40 | 1.08E-38 | 2.26E-34 | 0.70 | 0.53 | 0.17 | 1.32 |

**Supplementary Table 1. Marker genes for the original clusters of integrated clustering of E11.5 control and NotchGOF λ-junction cells.**

| | | | | | | | | |
|---|---|---|---|---|---|---|---|---|
| 2 | Mir17hg | 0.54 | 1.78E-38 | 3.71E-34 | 0.40 | 0.25 | 0.15 | 1.58 |
| 2 | Lhfpl2 | 0.44 | 1.81E-38 | 3.76E-34 | 0.48 | 0.32 | 0.17 | 1.53 |
| 2 | Frmd4a | 0.30 | 2.42E-38 | 5.04E-34 | 0.82 | 0.66 | 0.17 | 1.26 |
| 2 | Pcmt1 | 0.35 | 2.64E-38 | 5.50E-34 | 0.86 | 0.73 | 0.13 | 1.17 |
| 2 | Rcc1 | 0.39 | 3.04E-38 | 6.33E-34 | 0.67 | 0.50 | 0.18 | 1.35 |
| 2 | Dusp9 | 0.43 | 3.37E-38 | 7.00E-34 | 0.57 | 0.40 | 0.17 | 1.43 |
| 2 | Nup155 | 0.49 | 3.45E-38 | 7.17E-34 | 0.55 | 0.39 | 0.16 | 1.40 |
| 2 | Meis1 | 0.55 | 4.80E-38 | 9.97E-34 | 0.59 | 0.44 | 0.15 | 1.34 |
| 2 | Timm23 | 0.26 | 5.24E-38 | 1.09E-33 | 0.98 | 0.91 | 0.06 | 1.07 |
| 2 | Aif1l | 0.47 | 6.46E-38 | 1.34E-33 | 0.53 | 0.37 | 0.16 | 1.44 |
| 2 | Chaf1b | 0.42 | 6.66E-38 | 1.39E-33 | 0.66 | 0.50 | 0.16 | 1.33 |
| 2 | BC055324 | 0.67 | 7.60E-38 | 1.58E-33 | 0.35 | 0.22 | 0.13 | 1.59 |
| 2 | Rad50 | 0.37 | 7.89E-38 | 1.64E-33 | 0.81 | 0.67 | 0.15 | 1.22 |
| 2 | Mdc1 | 0.49 | 8.29E-38 | 1.72E-33 | 0.53 | 0.37 | 0.16 | 1.43 |
| 2 | Slc2a4 | 0.75 | 9.26E-38 | 1.93E-33 | 0.28 | 0.16 | 0.12 | 1.73 |
| 2 | Spidr | 0.58 | 9.46E-38 | 1.97E-33 | 0.44 | 0.30 | 0.14 | 1.48 |
| 2 | Wdr82 | 0.36 | 1.20E-37 | 2.49E-33 | 0.73 | 0.57 | 0.16 | 1.29 |
| 2 | Polr2b | 0.34 | 1.42E-37 | 2.95E-33 | 0.81 | 0.66 | 0.15 | 1.22 |
| 2 | Cand2 | 0.42 | 1.66E-37 | 3.46E-33 | 0.64 | 0.48 | 0.17 | 1.35 |
| 2 | Tex30 | 0.45 | 2.47E-37 | 5.14E-33 | 0.62 | 0.46 | 0.16 | 1.35 |
| 2 | Cables1 | 0.82 | 2.50E-37 | 5.20E-33 | 0.17 | 0.08 | 0.09 | 2.12 |
| 2 | Mcm7 | 0.27 | 2.53E-37 | 5.26E-33 | 0.95 | 0.86 | 0.09 | 1.11 |
| 2 | Bicc1 | 0.60 | 2.70E-37 | 5.62E-33 | 0.36 | 0.22 | 0.14 | 1.62 |
| 2 | Nudt5 | 0.31 | 2.94E-37 | 6.12E-33 | 0.90 | 0.78 | 0.12 | 1.15 |
| 2 | Bms1 | 0.31 | 3.05E-37 | 6.34E-33 | 0.83 | 0.67 | 0.16 | 1.24 |
| 2 | Stil | 0.66 | 3.86E-37 | 8.03E-33 | 0.30 | 0.18 | 0.12 | 1.66 |
| 2 | Bbx | 0.41 | 4.09E-37 | 8.51E-33 | 0.73 | 0.57 | 0.16 | 1.28 |
| 2 | Dnaaf2 | 0.40 | 4.16E-37 | 8.66E-33 | 0.67 | 0.50 | 0.17 | 1.34 |
| 2 | Fgf13 | 0.89 | 4.77E-37 | 9.93E-33 | 0.20 | 0.10 | 0.10 | 1.93 |
| 2 | St3gal2 | 0.61 | 5.06E-37 | 1.05E-32 | 0.38 | 0.24 | 0.14 | 1.57 |
| 2 | Polr2a | 0.29 | 6.17E-37 | 1.28E-32 | 0.90 | 0.79 | 0.12 | 1.15 |
| 2 | Fancd2 | 0.89 | 6.31E-37 | 1.31E-32 | 0.22 | 0.12 | 0.10 | 1.85 |
| 2 | Cit | 0.49 | 9.45E-37 | 1.97E-32 | 0.51 | 0.36 | 0.15 | 1.42 |
| 2 | Rbmx2 | 0.43 | 1.02E-36 | 2.13E-32 | 0.58 | 0.42 | 0.16 | 1.38 |
| 2 | Kif23 | 0.36 | 1.03E-36 | 2.14E-32 | 0.60 | 0.43 | 0.17 | 1.41 |
| 2 | Tagln3 | 0.34 | 1.13E-36 | 2.34E-32 | 0.40 | 0.25 | 0.15 | 1.58 |
| 2 | Mapre2 | 0.41 | 1.14E-36 | 2.38E-32 | 0.56 | 0.39 | 0.17 | 1.44 |
| 2 | Col6a3 | 0.68 | 1.17E-36 | 2.44E-32 | 0.27 | 0.16 | 0.12 | 1.75 |
| 2 | Hist1h4b | 1.31 | 1.50E-36 | 3.12E-32 | 0.11 | 0.04 | 0.06 | 2.56 |
| 2 | Cnst | 0.64 | 1.52E-36 | 3.17E-32 | 0.29 | 0.17 | 0.12 | 1.71 |
| 2 | Jakmip2 | 0.61 | 1.55E-36 | 3.23E-32 | 0.25 | 0.14 | 0.11 | 1.83 |
| 2 | 4930579G24Rik | 0.59 | 1.67E-36 | 3.47E-32 | 0.38 | 0.24 | 0.14 | 1.57 |
| 2 | Kansl1 | 0.37 | 1.92E-36 | 4.00E-32 | 0.77 | 0.63 | 0.14 | 1.23 |
| 2 | Pdlim7 | 0.29 | 2.22E-36 | 4.62E-32 | 0.88 | 0.74 | 0.14 | 1.19 |
| 2 | Ahctf1 | 0.43 | 2.53E-36 | 5.27E-32 | 0.68 | 0.53 | 0.15 | 1.29 |
| 2 | Irx3 | 0.34 | 2.58E-36 | 5.36E-32 | 0.41 | 0.26 | 0.15 | 1.59 |
| 2 | Gtse1 | 0.75 | 2.76E-36 | 5.74E-32 | 0.29 | 0.17 | 0.12 | 1.66 |
| 2 | Itpr1 | 0.59 | 3.09E-36 | 6.43E-32 | 0.33 | 0.20 | 0.13 | 1.63 |
| 2 | Rab31 | 0.58 | 3.89E-36 | 8.08E-32 | 0.35 | 0.21 | 0.13 | 1.62 |
| 2 | Fzd10 | 0.66 | 4.00E-36 | 8.31E-32 | 0.24 | 0.13 | 0.11 | 1.83 |
| 2 | Septin3 | 0.41 | 5.53E-36 | 1.15E-31 | 0.60 | 0.43 | 0.18 | 1.41 |
| 2 | Ilf3 | 0.28 | 6.88E-36 | 1.43E-31 | 0.91 | 0.79 | 0.12 | 1.16 |
| 2 | Fam193a | 0.33 | 8.33E-36 | 1.73E-31 | 0.76 | 0.60 | 0.16 | 1.27 |
| 2 | E2f3 | 0.33 | 8.88E-36 | 1.85E-31 | 0.74 | 0.57 | 0.18 | 1.31 |
| 2 | Itga8 | 0.42 | 1.11E-35 | 2.30E-31 | 0.31 | 0.18 | 0.13 | 1.73 |
| 2 | Lrrc17 | 0.87 | 1.41E-35 | 2.94E-31 | 0.15 | 0.07 | 0.08 | 2.15 |

**Supplementary Table 1. Marker genes for the original clusters of integrated clustering of E11.5 control and NotchGOF λ-junction cells.**

| | | | | | | | | |
|---|---|---|---|---|---|---|---|---|
| 2 | Ivns1abp | 0.29 | 1.99E-35 | 4.13E-31 | 0.90 | 0.76 | 0.13 | 1.17 |
| 2 | Bub1b | 0.27 | 2.12E-35 | 4.41E-31 | 0.73 | 0.54 | 0.19 | 1.35 |
| 2 | Ect2 | 0.35 | 2.42E-35 | 5.03E-31 | 0.56 | 0.39 | 0.17 | 1.43 |
| 2 | Cnih2 | 0.39 | 2.58E-35 | 5.36E-31 | 0.47 | 0.31 | 0.16 | 1.51 |
| 2 | Myh10 | 0.25 | 2.61E-35 | 5.43E-31 | 0.97 | 0.86 | 0.11 | 1.13 |
| 2 | Syce2 | 0.39 | 2.72E-35 | 5.66E-31 | 0.73 | 0.58 | 0.15 | 1.26 |
| 2 | Txlna | 0.33 | 3.00E-35 | 6.23E-31 | 0.79 | 0.64 | 0.16 | 1.24 |
| 2 | Pcolce2 | 0.67 | 3.42E-35 | 7.11E-31 | 0.23 | 0.13 | 0.10 | 1.81 |
| 2 | Mmp2 | 0.32 | 3.73E-35 | 7.75E-31 | 0.70 | 0.50 | 0.20 | 1.40 |
| 2 | Pwwp3b | 0.48 | 4.79E-35 | 9.97E-31 | 0.45 | 0.30 | 0.15 | 1.50 |
| 2 | Wnt5b | 0.54 | 5.18E-35 | 1.08E-30 | 0.39 | 0.25 | 0.14 | 1.57 |
| 2 | Dsn1 | 0.62 | 5.79E-35 | 1.20E-30 | 0.36 | 0.23 | 0.13 | 1.55 |
| 2 | Timm17a | 0.26 | 5.82E-35 | 1.21E-30 | 0.91 | 0.81 | 0.10 | 1.12 |
| 2 | Phrf1 | 0.39 | 6.16E-35 | 1.28E-30 | 0.68 | 0.52 | 0.16 | 1.32 |
| 2 | Cep290 | 0.39 | 6.34E-35 | 1.32E-30 | 0.66 | 0.50 | 0.16 | 1.31 |
| 2 | Brca2 | 0.42 | 6.35E-35 | 1.32E-30 | 0.62 | 0.46 | 0.16 | 1.34 |
| 2 | Pdlim4 | 0.26 | 6.74E-35 | 1.40E-30 | 0.79 | 0.61 | 0.18 | 1.30 |
| 2 | Rbm28 | 0.29 | 7.21E-35 | 1.50E-30 | 0.87 | 0.74 | 0.13 | 1.18 |
| 2 | Kif18b | 0.78 | 7.47E-35 | 1.55E-30 | 0.26 | 0.15 | 0.11 | 1.71 |
| 2 | Gtf2e2 | 0.31 | 7.53E-35 | 1.57E-30 | 0.87 | 0.75 | 0.12 | 1.16 |
| 2 | Gli2 | 0.50 | 7.71E-35 | 1.60E-30 | 0.39 | 0.25 | 0.14 | 1.56 |
| 2 | Arhgap19 | 0.52 | 8.78E-35 | 1.83E-30 | 0.41 | 0.28 | 0.14 | 1.49 |
| 2 | Chn1 | 0.41 | 9.07E-35 | 1.89E-30 | 0.55 | 0.39 | 0.16 | 1.41 |
| 2 | Map7d1 | 0.32 | 1.00E-34 | 2.09E-30 | 0.77 | 0.60 | 0.17 | 1.27 |
| 2 | Dcp2 | 0.33 | 1.19E-34 | 2.47E-30 | 0.77 | 0.61 | 0.16 | 1.26 |
| 2 | Cops3 | 0.27 | 1.26E-34 | 2.63E-30 | 0.91 | 0.79 | 0.12 | 1.15 |
| 2 | Gpatch8 | 0.27 | 1.30E-34 | 2.70E-30 | 0.89 | 0.78 | 0.12 | 1.15 |
| 2 | Tmem45a | 0.87 | 1.33E-34 | 2.77E-30 | 0.16 | 0.08 | 0.08 | 2.11 |
| 2 | Rif1 | 0.27 | 1.36E-34 | 2.83E-30 | 0.91 | 0.79 | 0.12 | 1.15 |
| 2 | Cdkn2d | 0.44 | 1.45E-34 | 3.02E-30 | 0.52 | 0.37 | 0.15 | 1.40 |
| 2 | Tle4 | 0.31 | 1.47E-34 | 3.05E-30 | 0.74 | 0.57 | 0.17 | 1.30 |
| 2 | Emc8 | 0.28 | 1.48E-34 | 3.09E-30 | 0.91 | 0.78 | 0.13 | 1.17 |
| 2 | Clic4 | 0.32 | 1.63E-34 | 3.38E-30 | 0.82 | 0.66 | 0.15 | 1.23 |
| 2 | Cbx2 | 0.42 | 1.65E-34 | 3.44E-30 | 0.57 | 0.42 | 0.16 | 1.38 |
| 2 | Kifc5b | 0.81 | 1.67E-34 | 3.48E-30 | 0.25 | 0.15 | 0.10 | 1.70 |
| 2 | Med4 | 0.30 | 1.71E-34 | 3.57E-30 | 0.84 | 0.70 | 0.14 | 1.20 |
| 2 | Nkap | 0.37 | 2.50E-34 | 5.19E-30 | 0.73 | 0.58 | 0.15 | 1.26 |
| 2 | Supt20 | 0.34 | 2.62E-34 | 5.46E-30 | 0.73 | 0.58 | 0.15 | 1.26 |
| 2 | Uhrf1 | 0.31 | 2.66E-34 | 5.52E-30 | 0.83 | 0.67 | 0.16 | 1.24 |
| 2 | Bard1 | 0.73 | 2.69E-34 | 5.59E-30 | 0.29 | 0.18 | 0.11 | 1.62 |
| 2 | Nsd3 | 0.35 | 3.00E-34 | 6.24E-30 | 0.78 | 0.62 | 0.16 | 1.25 |
| 2 | Cdr1os | 0.86 | 3.58E-34 | 7.44E-30 | 0.13 | 0.05 | 0.07 | 2.31 |
| 2 | Tmem100 | 0.72 | 3.83E-34 | 7.96E-30 | 0.24 | 0.13 | 0.11 | 1.79 |
| 2 | Fmr1 | 0.33 | 4.24E-34 | 8.81E-30 | 0.79 | 0.65 | 0.14 | 1.21 |
| 2 | Ube2t | 0.49 | 4.28E-34 | 8.91E-30 | 0.45 | 0.32 | 0.14 | 1.44 |
| 2 | Zik1 | 0.50 | 4.62E-34 | 9.61E-30 | 0.47 | 0.33 | 0.15 | 1.45 |
| 2 | Dixdc1 | 0.36 | 5.32E-34 | 1.11E-29 | 0.56 | 0.38 | 0.18 | 1.46 |
| 2 | Ppp2r2d | 0.30 | 5.97E-34 | 1.24E-29 | 0.81 | 0.67 | 0.14 | 1.21 |
| 2 | Vezf1 | 0.27 | 7.09E-34 | 1.47E-29 | 0.91 | 0.79 | 0.12 | 1.15 |
| 2 | Cdk2 | 0.45 | 8.16E-34 | 1.70E-29 | 0.55 | 0.40 | 0.15 | 1.37 |
| 2 | E2f6 | 0.36 | 9.20E-34 | 1.91E-29 | 0.68 | 0.51 | 0.17 | 1.33 |
| 2 | Mtrex | 0.34 | 1.03E-33 | 2.15E-29 | 0.76 | 0.61 | 0.15 | 1.25 |
| 2 | Rbm5 | 0.25 | 1.20E-33 | 2.49E-29 | 0.92 | 0.81 | 0.10 | 1.13 |
| 2 | Hmgxb4 | 0.35 | 1.60E-33 | 3.32E-29 | 0.77 | 0.61 | 0.16 | 1.26 |
| 2 | Kif2a | 0.27 | 1.66E-33 | 3.44E-29 | 0.90 | 0.78 | 0.12 | 1.15 |
| 2 | Rfwd3 | 0.50 | 1.67E-33 | 3.47E-29 | 0.50 | 0.36 | 0.14 | 1.39 |

**Supplementary Table 1. Marker genes for the original clusters of integrated clustering of E11.5 control and NotchGOF λ-junction cells.**

| | | | | | | | | |
|---|---|---|---|---|---|---|---|---|
| 2 | Kank2 | 0.40 | 1.68E-33 | 3.50E-29 | 0.45 | 0.30 | 0.15 | 1.52 |
| 2 | Sp3 | 0.36 | 1.79E-33 | 3.72E-29 | 0.76 | 0.62 | 0.15 | 1.24 |
| 2 | Slit2 | 0.40 | 1.86E-33 | 3.86E-29 | 0.51 | 0.35 | 0.15 | 1.44 |
| 2 | Jam3 | 0.30 | 1.98E-33 | 4.12E-29 | 0.73 | 0.56 | 0.18 | 1.31 |
| 2 | Mtf2 | 0.26 | 2.08E-33 | 4.32E-29 | 0.92 | 0.80 | 0.13 | 1.16 |
| 2 | Peli1 | 0.30 | 2.15E-33 | 4.47E-29 | 0.80 | 0.66 | 0.14 | 1.21 |
| 2 | Timm50 | 0.29 | 2.37E-33 | 4.92E-29 | 0.86 | 0.73 | 0.14 | 1.19 |
| 2 | Ipo5 | 0.30 | 2.38E-33 | 4.96E-29 | 0.83 | 0.69 | 0.15 | 1.22 |
| 2 | Timp1 | 0.62 | 2.46E-33 | 5.12E-29 | 0.27 | 0.16 | 0.11 | 1.71 |
| 2 | Timeless | 0.37 | 2.59E-33 | 5.39E-29 | 0.69 | 0.52 | 0.17 | 1.32 |
| 2 | Mybbp1a | 0.30 | 2.63E-33 | 5.48E-29 | 0.89 | 0.76 | 0.12 | 1.16 |
| 2 | Rgs2 | 0.29 | 2.98E-33 | 6.21E-29 | 0.49 | 0.33 | 0.16 | 1.48 |
| 2 | Prps1 | 0.41 | 3.17E-33 | 6.60E-29 | 0.58 | 0.43 | 0.15 | 1.35 |
| 2 | Lum | 0.37 | 3.43E-33 | 7.12E-29 | 0.26 | 0.15 | 0.11 | 1.76 |
| 2 | Msantd2 | 0.40 | 3.67E-33 | 7.63E-29 | 0.60 | 0.43 | 0.16 | 1.38 |
| 2 | Rad18 | 0.44 | 3.85E-33 | 8.00E-29 | 0.50 | 0.36 | 0.15 | 1.42 |
| 2 | Racgap1 | 0.28 | 4.34E-33 | 9.04E-29 | 0.69 | 0.51 | 0.18 | 1.35 |
| 2 | Adpgk | 0.48 | 4.90E-33 | 1.02E-28 | 0.44 | 0.30 | 0.14 | 1.48 |
| 2 | Plpp3 | 0.32 | 5.09E-33 | 1.06E-28 | 0.81 | 0.65 | 0.16 | 1.25 |
| 2 | Ttc28 | 0.32 | 8.89E-33 | 1.85E-28 | 0.75 | 0.58 | 0.17 | 1.29 |
| 2 | Eif3b | 0.26 | 1.07E-32 | 2.23E-28 | 0.91 | 0.81 | 0.10 | 1.12 |
| 2 | Setdb1 | 0.33 | 1.21E-32 | 2.52E-28 | 0.75 | 0.60 | 0.15 | 1.25 |
| 2 | Dpysl2 | 0.29 | 1.45E-32 | 3.03E-28 | 0.83 | 0.68 | 0.14 | 1.21 |
| 2 | Adgra3 | 0.34 | 1.63E-32 | 3.39E-28 | 0.71 | 0.54 | 0.16 | 1.30 |
| 2 | Cep55 | 0.46 | 2.27E-32 | 4.73E-28 | 0.36 | 0.23 | 0.13 | 1.56 |
| 2 | Impa2 | 0.51 | 2.33E-32 | 4.84E-28 | 0.42 | 0.28 | 0.14 | 1.48 |
| 2 | Rapgef6 | 0.38 | 2.45E-32 | 5.09E-28 | 0.59 | 0.42 | 0.16 | 1.38 |
| 2 | Gpt2 | 0.61 | 2.49E-32 | 5.18E-28 | 0.32 | 0.20 | 0.12 | 1.59 |
| 2 | Mmd | 0.40 | 2.51E-32 | 5.22E-28 | 0.49 | 0.34 | 0.16 | 1.46 |
| 2 | Smc5 | 0.34 | 2.86E-32 | 5.95E-28 | 0.77 | 0.63 | 0.14 | 1.23 |
| 2 | Snhg5 | 0.41 | 2.89E-32 | 6.01E-28 | 0.58 | 0.43 | 0.16 | 1.36 |
| 2 | Prrg1 | 0.62 | 3.53E-32 | 7.34E-28 | 0.30 | 0.18 | 0.12 | 1.64 |
| 2 | Cdk12 | 0.30 | 3.68E-32 | 7.65E-28 | 0.84 | 0.72 | 0.13 | 1.17 |
| 2 | Golm1 | 0.33 | 3.84E-32 | 7.99E-28 | 0.70 | 0.54 | 0.16 | 1.30 |
| 2 | Ccn1 | 0.48 | 4.23E-32 | 8.79E-28 | 0.44 | 0.29 | 0.14 | 1.48 |
| 2 | Oaf | 0.34 | 4.48E-32 | 9.32E-28 | 0.61 | 0.44 | 0.17 | 1.37 |
| 2 | Lnpk | 0.40 | 5.10E-32 | 1.06E-27 | 0.62 | 0.47 | 0.15 | 1.32 |
| 2 | Igsf10 | 0.50 | 5.23E-32 | 1.09E-27 | 0.35 | 0.23 | 0.13 | 1.57 |
| 2 | Bmper | 1.05 | 5.25E-32 | 1.09E-27 | 0.15 | 0.07 | 0.08 | 2.04 |
| 2 | Osbpl6 | 0.74 | 5.51E-32 | 1.15E-27 | 0.24 | 0.14 | 0.10 | 1.73 |
| 2 | Rbp4 | 1.11 | 6.25E-32 | 1.30E-27 | 0.12 | 0.05 | 0.07 | 2.26 |
| 2 | Septin6 | 0.31 | 7.22E-32 | 1.50E-27 | 0.71 | 0.55 | 0.16 | 1.29 |
| 2 | Tnfaip8 | 0.34 | 7.52E-32 | 1.56E-27 | 0.87 | 0.75 | 0.12 | 1.16 |
| 2 | Nrm | 0.41 | 7.97E-32 | 1.66E-27 | 0.49 | 0.34 | 0.15 | 1.44 |
| 2 | Rpa3 | 0.25 | 8.31E-32 | 1.73E-27 | 0.92 | 0.80 | 0.11 | 1.14 |
| 2 | Lmnb2 | 0.44 | 8.34E-32 | 1.74E-27 | 0.51 | 0.36 | 0.15 | 1.40 |
| 2 | Nelfe | 0.32 | 8.90E-32 | 1.85E-27 | 0.79 | 0.64 | 0.14 | 1.22 |
| 2 | Hist2h3b | 0.99 | 9.77E-32 | 2.03E-27 | 0.14 | 0.07 | 0.07 | 2.11 |
| 2 | Ehmt1 | 0.26 | 9.96E-32 | 2.07E-27 | 0.93 | 0.81 | 0.12 | 1.14 |
| 2 | Pom121 | 0.33 | 1.07E-31 | 2.22E-27 | 0.73 | 0.58 | 0.15 | 1.26 |
| 2 | Prkd1 | 0.68 | 1.17E-31 | 2.44E-27 | 0.22 | 0.13 | 0.10 | 1.79 |
| 2 | Ddx21 | 0.29 | 1.27E-31 | 2.65E-27 | 0.84 | 0.72 | 0.12 | 1.17 |
| 2 | Rad17 | 0.39 | 1.41E-31 | 2.93E-27 | 0.62 | 0.46 | 0.16 | 1.35 |
| 2 | Kifc1 | 0.46 | 1.68E-31 | 3.50E-27 | 0.40 | 0.27 | 0.13 | 1.49 |
| 2 | Ubash3b | 0.44 | 2.34E-31 | 4.86E-27 | 0.35 | 0.23 | 0.13 | 1.56 |
| 2 | Yaf2 | 0.27 | 2.45E-31 | 5.10E-27 | 0.84 | 0.69 | 0.15 | 1.22 |

**Supplementary Table 1. Marker genes for the original clusters of integrated clustering of E11.5 control and NotchGOF λ-junction cells.**

| | | | | | | | | |
|---|---|---|---|---|---|---|---|---|
| 2 | Cdyl | 0.46 | 2.49E-31 | 5.18E-27 | 0.51 | 0.37 | 0.14 | 1.39 |
| 2 | Tbc1d10b | 0.43 | 2.67E-31 | 5.55E-27 | 0.48 | 0.34 | 0.15 | 1.43 |
| 2 | Prkar2a | 0.38 | 3.08E-31 | 6.41E-27 | 0.65 | 0.50 | 0.15 | 1.29 |
| 2 | Dleu2 | 0.57 | 3.24E-31 | 6.74E-27 | 0.36 | 0.24 | 0.12 | 1.49 |
| 2 | Zranb3 | 0.60 | 3.41E-31 | 7.09E-27 | 0.32 | 0.20 | 0.12 | 1.58 |
| 2 | Tshz1 | 0.33 | 3.78E-31 | 7.86E-27 | 0.46 | 0.30 | 0.15 | 1.50 |
| 2 | Gmeb1 | 0.43 | 3.89E-31 | 8.10E-27 | 0.55 | 0.40 | 0.15 | 1.38 |
| 2 | Msrb2 | 0.52 | 3.92E-31 | 8.15E-27 | 0.44 | 0.31 | 0.14 | 1.45 |
| 2 | Fancm | 0.47 | 4.65E-31 | 9.68E-27 | 0.49 | 0.35 | 0.14 | 1.41 |
| 2 | Fam181b | 0.71 | 4.98E-31 | 1.04E-26 | 0.19 | 0.10 | 0.09 | 1.87 |
| 2 | Hcfc1 | 0.31 | 5.48E-31 | 1.14E-26 | 0.80 | 0.66 | 0.15 | 1.22 |
| 2 | Stxbp6 | 0.69 | 6.33E-31 | 1.32E-26 | 0.22 | 0.13 | 0.10 | 1.76 |
| 2 | Exosc10 | 0.41 | 6.37E-31 | 1.32E-26 | 0.57 | 0.42 | 0.15 | 1.36 |
| 2 | Zw10 | 0.46 | 6.37E-31 | 1.33E-26 | 0.45 | 0.31 | 0.14 | 1.45 |
| 2 | Mapk8 | 0.40 | 7.31E-31 | 1.52E-26 | 0.57 | 0.41 | 0.16 | 1.38 |
| 2 | Scoc | 0.29 | 7.58E-31 | 1.58E-26 | 0.77 | 0.62 | 0.15 | 1.25 |
| 2 | Dkc1 | 0.26 | 8.12E-31 | 1.69E-26 | 0.89 | 0.78 | 0.10 | 1.13 |
| 2 | Zfp280d | 0.32 | 8.20E-31 | 1.70E-26 | 0.79 | 0.65 | 0.14 | 1.21 |
| 2 | Med30 | 0.29 | 8.99E-31 | 1.87E-26 | 0.83 | 0.69 | 0.14 | 1.20 |
| 2 | Dact1 | 0.37 | 1.01E-30 | 2.10E-26 | 0.45 | 0.30 | 0.15 | 1.50 |
| 2 | Med22 | 0.43 | 1.11E-30 | 2.31E-26 | 0.47 | 0.33 | 0.14 | 1.43 |
| 2 | BC003965 | 0.33 | 1.14E-30 | 2.37E-26 | 0.78 | 0.62 | 0.16 | 1.25 |
| 2 | Idh3a | 0.33 | 1.16E-30 | 2.40E-26 | 0.71 | 0.55 | 0.16 | 1.29 |
| 2 | Rbms3 | 0.32 | 1.17E-30 | 2.44E-26 | 0.56 | 0.40 | 0.16 | 1.41 |
| 2 | Tmem132a | 0.30 | 1.20E-30 | 2.50E-26 | 0.78 | 0.62 | 0.16 | 1.25 |
| 2 | Rora | 0.57 | 1.24E-30 | 2.58E-26 | 0.28 | 0.17 | 0.11 | 1.65 |
| 2 | Ldlr | 0.57 | 1.31E-30 | 2.73E-26 | 0.35 | 0.23 | 0.12 | 1.51 |
| 2 | Atf1 | 0.38 | 1.34E-30 | 2.80E-26 | 0.64 | 0.49 | 0.15 | 1.30 |
| 2 | Hist4h4 | 0.94 | 1.38E-30 | 2.86E-26 | 0.17 | 0.09 | 0.08 | 1.88 |
| 2 | Uhrf2 | 0.27 | 1.40E-30 | 2.91E-26 | 0.85 | 0.72 | 0.13 | 1.18 |
| 2 | H2-Q7 | 0.70 | 1.61E-30 | 3.35E-26 | 0.19 | 0.10 | 0.09 | 1.88 |
| 2 | 2700081O15Rik | 0.30 | 1.66E-30 | 3.45E-26 | 0.70 | 0.55 | 0.16 | 1.29 |
| 2 | Donson | 0.59 | 1.74E-30 | 3.63E-26 | 0.35 | 0.23 | 0.12 | 1.50 |
| 2 | Nup205 | 0.47 | 1.75E-30 | 3.65E-26 | 0.47 | 0.33 | 0.14 | 1.41 |
| 2 | Zcchc3 | 0.28 | 2.28E-30 | 4.75E-26 | 0.84 | 0.70 | 0.14 | 1.19 |
| 2 | Nfyb | 0.26 | 2.34E-30 | 4.88E-26 | 0.90 | 0.78 | 0.12 | 1.15 |
| 2 | Mta1 | 0.31 | 2.51E-30 | 5.21E-26 | 0.83 | 0.68 | 0.16 | 1.23 |
| 2 | Fzd10os | 0.47 | 3.05E-30 | 6.35E-26 | 0.31 | 0.19 | 0.12 | 1.62 |
| 2 | Clasp1 | 0.35 | 3.23E-30 | 6.71E-26 | 0.69 | 0.54 | 0.15 | 1.28 |
| 2 | Haus8 | 0.47 | 3.30E-30 | 6.87E-26 | 0.47 | 0.34 | 0.13 | 1.39 |
| 2 | Lzts2 | 0.31 | 3.67E-30 | 7.63E-26 | 0.74 | 0.59 | 0.15 | 1.25 |
| 2 | Arhgap10 | 0.37 | 3.81E-30 | 7.93E-26 | 0.56 | 0.41 | 0.16 | 1.38 |
| 2 | Ipo7 | 0.34 | 5.18E-30 | 1.08E-25 | 0.75 | 0.60 | 0.14 | 1.24 |
| 2 | Eftud2 | 0.30 | 5.26E-30 | 1.09E-25 | 0.82 | 0.68 | 0.14 | 1.21 |
| 2 | Col5a1 | 0.45 | 7.04E-30 | 1.46E-25 | 0.39 | 0.26 | 0.14 | 1.53 |
| 2 | Ddx27 | 0.31 | 7.60E-30 | 1.58E-25 | 0.82 | 0.69 | 0.13 | 1.19 |
| 2 | Aifm1 | 0.32 | 7.74E-30 | 1.61E-25 | 0.74 | 0.59 | 0.15 | 1.26 |
| 2 | Gtf3c1 | 0.33 | 7.76E-30 | 1.61E-25 | 0.72 | 0.56 | 0.16 | 1.28 |
| 2 | Sdk2 | 0.99 | 9.11E-30 | 1.89E-25 | 0.11 | 0.05 | 0.06 | 2.33 |
| 2 | Phlda1 | 0.31 | 9.19E-30 | 1.91E-25 | 0.55 | 0.41 | 0.15 | 1.36 |
| 2 | Rnf144a | 0.38 | 1.08E-29 | 2.24E-25 | 0.58 | 0.43 | 0.15 | 1.35 |
| 2 | Tmeff1 | 0.28 | 1.17E-29 | 2.43E-25 | 0.82 | 0.67 | 0.14 | 1.21 |
| 2 | Dock11 | 0.52 | 1.25E-29 | 2.60E-25 | 0.34 | 0.22 | 0.12 | 1.55 |
| 2 | Dna2 | 0.54 | 1.38E-29 | 2.87E-25 | 0.37 | 0.25 | 0.12 | 1.49 |
| 2 | Ppp3cb | 0.31 | 1.41E-29 | 2.92E-25 | 0.74 | 0.61 | 0.14 | 1.22 |
| 2 | Gps1 | 0.27 | 1.66E-29 | 3.46E-25 | 0.85 | 0.73 | 0.12 | 1.17 |

**Supplementary Table 1. Marker genes for the original clusters of integrated clustering of E11.5 control and NotchGOF λ-junction cells.**

| | | | | | | | | |
|---|---|---|---|---|---|---|---|---|
| 2 | Neto2 | 0.54 | 1.89E-29 | 3.93E-25 | 0.34 | 0.22 | 0.12 | 1.54 |
| 2 | Pja1 | 0.25 | 2.28E-29 | 4.73E-25 | 0.81 | 0.66 | 0.15 | 1.22 |
| 2 | Pcyt1a | 0.47 | 2.39E-29 | 4.97E-25 | 0.44 | 0.31 | 0.13 | 1.43 |
| 2 | Gm42047 | 0.86 | 2.52E-29 | 5.24E-25 | 0.23 | 0.13 | 0.09 | 1.69 |
| 2 | Magi1 | 0.43 | 2.73E-29 | 5.67E-25 | 0.43 | 0.30 | 0.14 | 1.46 |
| 2 | Adprh | 0.26 | 2.84E-29 | 5.91E-25 | 0.89 | 0.77 | 0.12 | 1.15 |
| 2 | Chd1 | 0.26 | 2.88E-29 | 5.98E-25 | 0.89 | 0.78 | 0.11 | 1.14 |
| 2 | Acaca | 0.47 | 2.91E-29 | 6.05E-25 | 0.45 | 0.32 | 0.13 | 1.41 |
| 2 | Cnksr2 | 0.35 | 3.38E-29 | 7.02E-25 | 0.46 | 0.31 | 0.15 | 1.47 |
| 2 | Naa20 | 0.27 | 3.49E-29 | 7.25E-25 | 0.84 | 0.71 | 0.13 | 1.19 |
| 2 | Rbpms2 | 0.39 | 3.77E-29 | 7.84E-25 | 0.34 | 0.21 | 0.13 | 1.58 |
| 2 | Hyls1 | 0.65 | 4.17E-29 | 8.68E-25 | 0.27 | 0.17 | 0.10 | 1.61 |
| 2 | Nup153 | 0.40 | 4.31E-29 | 8.97E-25 | 0.57 | 0.42 | 0.15 | 1.35 |
| 2 | Mcph1 | 0.49 | 4.32E-29 | 8.98E-25 | 0.41 | 0.29 | 0.12 | 1.43 |
| 2 | Tinf2 | 0.65 | 5.06E-29 | 1.05E-24 | 0.30 | 0.20 | 0.11 | 1.55 |
| 2 | Senp2 | 0.34 | 5.46E-29 | 1.13E-24 | 0.70 | 0.56 | 0.14 | 1.26 |
| 2 | Lap3 | 0.30 | 5.60E-29 | 1.17E-24 | 0.79 | 0.66 | 0.13 | 1.20 |
| 2 | Col23a1 | 0.40 | 5.90E-29 | 1.23E-24 | 0.45 | 0.30 | 0.15 | 1.48 |
| 2 | Smco4 | 0.37 | 6.26E-29 | 1.30E-24 | 0.62 | 0.48 | 0.15 | 1.31 |
| 2 | Nup62 | 0.32 | 6.43E-29 | 1.34E-24 | 0.69 | 0.54 | 0.15 | 1.27 |
| 2 | Dlst | 0.33 | 6.96E-29 | 1.45E-24 | 0.70 | 0.55 | 0.15 | 1.27 |
| 2 | Tanc1 | 0.36 | 9.44E-29 | 1.96E-24 | 0.58 | 0.43 | 0.15 | 1.34 |
| 2 | Rab23 | 0.42 | 1.03E-28 | 2.14E-24 | 0.45 | 0.31 | 0.14 | 1.44 |
| 2 | Smad7 | 0.25 | 1.06E-28 | 2.21E-24 | 0.84 | 0.68 | 0.16 | 1.23 |
| 2 | Sf1 | 0.25 | 1.07E-28 | 2.23E-24 | 0.90 | 0.78 | 0.12 | 1.16 |
| 2 | Slc43a3 | 0.44 | 1.30E-28 | 2.70E-24 | 0.41 | 0.27 | 0.13 | 1.48 |
| 2 | Ranbp3 | 0.34 | 1.32E-28 | 2.74E-24 | 0.62 | 0.48 | 0.14 | 1.30 |
| 2 | Abi2 | 0.37 | 1.48E-28 | 3.08E-24 | 0.57 | 0.43 | 0.15 | 1.34 |
| 2 | E2f8 | 0.58 | 1.68E-28 | 3.49E-24 | 0.30 | 0.20 | 0.11 | 1.55 |
| 2 | Usp2 | 0.72 | 1.74E-28 | 3.61E-24 | 0.23 | 0.13 | 0.09 | 1.70 |
| 2 | Cad | 0.40 | 2.03E-28 | 4.22E-24 | 0.51 | 0.37 | 0.14 | 1.37 |
| 2 | Cdk5r1 | 0.67 | 3.07E-28 | 6.38E-24 | 0.19 | 0.10 | 0.08 | 1.83 |
| 2 | Kirrel | 0.54 | 3.61E-28 | 7.51E-24 | 0.33 | 0.21 | 0.12 | 1.55 |
| 2 | Dcaf15 | 0.42 | 3.69E-28 | 7.68E-24 | 0.47 | 0.33 | 0.14 | 1.41 |
| 2 | Specc1 | 0.38 | 3.84E-28 | 7.98E-24 | 0.56 | 0.41 | 0.15 | 1.37 |
| 2 | Srm | 0.29 | 4.25E-28 | 8.84E-24 | 0.80 | 0.67 | 0.14 | 1.20 |
| 2 | Xpo1 | 0.25 | 4.48E-28 | 9.31E-24 | 0.90 | 0.77 | 0.13 | 1.16 |
| 2 | Pola2 | 0.39 | 4.62E-28 | 9.61E-24 | 0.55 | 0.41 | 0.14 | 1.33 |
| 2 | Arhgap24 | 0.60 | 4.88E-28 | 1.01E-23 | 0.20 | 0.11 | 0.09 | 1.79 |
| 2 | Klhdc10 | 0.40 | 4.90E-28 | 1.02E-23 | 0.56 | 0.43 | 0.13 | 1.31 |
| 2 | Pknox1 | 0.51 | 5.02E-28 | 1.04E-23 | 0.34 | 0.22 | 0.12 | 1.52 |
| 2 | Rnaseh2a | 0.32 | 5.57E-28 | 1.16E-23 | 0.70 | 0.56 | 0.15 | 1.27 |
| 2 | Arhgap42 | 0.32 | 8.49E-28 | 1.76E-23 | 0.51 | 0.36 | 0.15 | 1.43 |
| 2 | Ppp5c | 0.31 | 8.50E-28 | 1.77E-23 | 0.68 | 0.52 | 0.16 | 1.30 |
| 2 | Gpsm3 | 0.64 | 8.52E-28 | 1.77E-23 | 0.16 | 0.08 | 0.08 | 1.95 |
| 2 | Pde3b | 0.45 | 9.33E-28 | 1.94E-23 | 0.36 | 0.23 | 0.12 | 1.52 |
| 2 | Rbl1 | 0.51 | 9.66E-28 | 2.01E-23 | 0.40 | 0.28 | 0.12 | 1.43 |
| 2 | Slc39a6 | 0.28 | 9.94E-28 | 2.07E-23 | 0.71 | 0.55 | 0.16 | 1.29 |
| 2 | Gm17092 | 0.53 | 1.05E-27 | 2.19E-23 | 0.35 | 0.24 | 0.12 | 1.50 |
| 2 | Lin9 | 0.45 | 1.10E-27 | 2.29E-23 | 0.45 | 0.32 | 0.13 | 1.41 |
| 2 | Tasor2 | 0.34 | 1.21E-27 | 2.52E-23 | 0.64 | 0.50 | 0.14 | 1.29 |
| 2 | Prrc2a | 0.25 | 1.27E-27 | 2.63E-23 | 0.87 | 0.76 | 0.12 | 1.15 |
| 2 | Cdc14b | 0.38 | 1.28E-27 | 2.66E-23 | 0.53 | 0.39 | 0.15 | 1.38 |
| 2 | Ddias | 0.70 | 1.47E-27 | 3.05E-23 | 0.24 | 0.15 | 0.09 | 1.63 |
| 2 | Bnc2 | 0.52 | 1.49E-27 | 3.09E-23 | 0.28 | 0.17 | 0.11 | 1.63 |
| 2 | Ptprd | 0.34 | 1.68E-27 | 3.49E-23 | 0.58 | 0.43 | 0.15 | 1.36 |

**Supplementary Table 1. Marker genes for the original clusters of integrated clustering of E11.5 control and NotchGOF λ-junction cells.**

| | | | | | | | | |
|---|---|---|---|---|---|---|---|---|
| 2 | Fgf10 | 0.79 | 1.73E-27 | 3.60E-23 | 0.18 | 0.10 | 0.08 | 1.81 |
| 2 | Twist2 | 0.30 | 1.88E-27 | 3.92E-23 | 0.35 | 0.22 | 0.13 | 1.58 |
| 2 | Colec12 | 0.41 | 2.09E-27 | 4.35E-23 | 0.33 | 0.21 | 0.12 | 1.56 |
| 2 | Mcm5 | 0.25 | 2.63E-27 | 5.46E-23 | 0.86 | 0.73 | 0.14 | 1.19 |
| 2 | Ogfr | 0.26 | 2.83E-27 | 5.88E-23 | 0.79 | 0.65 | 0.14 | 1.21 |
| 2 | Umps | 0.30 | 2.88E-27 | 6.00E-23 | 0.74 | 0.59 | 0.15 | 1.26 |
| 2 | Nhlrc2 | 0.41 | 2.93E-27 | 6.08E-23 | 0.52 | 0.38 | 0.14 | 1.36 |
| 2 | Ror2 | 0.38 | 2.95E-27 | 6.14E-23 | 0.42 | 0.29 | 0.14 | 1.47 |
| 2 | Nedd1 | 0.44 | 3.34E-27 | 6.94E-23 | 0.46 | 0.33 | 0.13 | 1.39 |
| 2 | Eea1 | 0.38 | 3.77E-27 | 7.84E-23 | 0.55 | 0.42 | 0.13 | 1.31 |
| 2 | Utp15 | 0.38 | 3.92E-27 | 8.15E-23 | 0.53 | 0.38 | 0.14 | 1.38 |
| 2 | Abcd2 | 0.77 | 4.38E-27 | 9.10E-23 | 0.14 | 0.07 | 0.07 | 2.00 |
| 2 | Zcchc8 | 0.37 | 4.64E-27 | 9.66E-23 | 0.61 | 0.46 | 0.15 | 1.31 |
| 2 | Oip5 | 0.58 | 5.92E-27 | 1.23E-22 | 0.29 | 0.19 | 0.10 | 1.54 |
| 2 | Nfatc2ip | 0.62 | 5.92E-27 | 1.23E-22 | 0.25 | 0.15 | 0.10 | 1.64 |
| 2 | Nup188 | 0.42 | 7.72E-27 | 1.61E-22 | 0.52 | 0.38 | 0.13 | 1.34 |
| 2 | Aebp1 | 0.49 | 7.74E-27 | 1.61E-22 | 0.31 | 0.20 | 0.11 | 1.56 |
| 2 | Phf20 | 0.31 | 8.28E-27 | 1.72E-22 | 0.74 | 0.59 | 0.15 | 1.24 |
| 2 | Tdp2 | 0.31 | 8.43E-27 | 1.75E-22 | 0.70 | 0.56 | 0.14 | 1.26 |
| 2 | Pif1 | 0.76 | 1.07E-26 | 2.22E-22 | 0.22 | 0.13 | 0.09 | 1.67 |
| 2 | Eed | 0.31 | 1.15E-26 | 2.40E-22 | 0.68 | 0.53 | 0.15 | 1.28 |
| 2 | Tubb6 | 0.27 | 1.17E-26 | 2.43E-22 | 0.19 | 0.11 | 0.09 | 1.80 |
| 2 | Tnks | 0.33 | 1.32E-26 | 2.75E-22 | 0.67 | 0.53 | 0.14 | 1.27 |
| 2 | Abce1 | 0.26 | 1.39E-26 | 2.90E-22 | 0.86 | 0.75 | 0.11 | 1.15 |
| 2 | Spata7 | 0.45 | 1.51E-26 | 3.14E-22 | 0.51 | 0.37 | 0.14 | 1.36 |
| 2 | Gnai1 | 0.63 | 1.61E-26 | 3.34E-22 | 0.24 | 0.15 | 0.10 | 1.66 |
| 2 | 1810026B05Rik | 0.30 | 1.86E-26 | 3.87E-22 | 0.73 | 0.58 | 0.15 | 1.26 |
| 2 | Ppp1r18 | 0.40 | 1.93E-26 | 4.02E-22 | 0.36 | 0.24 | 0.12 | 1.51 |
| 2 | Cdc25c | 0.38 | 2.03E-26 | 4.23E-22 | 0.47 | 0.34 | 0.13 | 1.39 |
| 2 | Midn | 0.27 | 2.10E-26 | 4.36E-22 | 0.84 | 0.72 | 0.12 | 1.17 |
| 2 | Zfp553 | 0.28 | 2.19E-26 | 4.56E-22 | 0.74 | 0.59 | 0.15 | 1.25 |
| 2 | Grem1 | 0.89 | 2.21E-26 | 4.60E-22 | 0.17 | 0.09 | 0.08 | 1.83 |
| 2 | Ep400 | 0.29 | 2.27E-26 | 4.73E-22 | 0.80 | 0.66 | 0.13 | 1.20 |
| 2 | Ska2 | 0.28 | 2.30E-26 | 4.78E-22 | 0.68 | 0.53 | 0.16 | 1.29 |
| 2 | Dclk2 | 0.58 | 2.36E-26 | 4.91E-22 | 0.19 | 0.11 | 0.08 | 1.79 |
| 2 | Dmwd | 0.33 | 2.49E-26 | 5.18E-22 | 0.58 | 0.43 | 0.15 | 1.34 |
| 2 | Poc1b | 0.40 | 2.82E-26 | 5.86E-22 | 0.45 | 0.32 | 0.13 | 1.40 |
| 2 | Pum2 | 0.27 | 2.94E-26 | 6.12E-22 | 0.87 | 0.74 | 0.13 | 1.17 |
| 2 | Cdc27 | 0.31 | 3.07E-26 | 6.38E-22 | 0.65 | 0.51 | 0.14 | 1.27 |
| 2 | Eri2 | 0.58 | 3.12E-26 | 6.48E-22 | 0.29 | 0.19 | 0.10 | 1.55 |
| 2 | Crispld1 | 0.29 | 3.85E-26 | 8.01E-22 | 0.46 | 0.32 | 0.14 | 1.45 |
| 2 | Akap13 | 0.29 | 3.89E-26 | 8.10E-22 | 0.64 | 0.49 | 0.15 | 1.31 |
| 2 | Akap11 | 0.31 | 3.94E-26 | 8.20E-22 | 0.62 | 0.47 | 0.15 | 1.32 |
| 2 | D10Wsu102e | 0.33 | 3.98E-26 | 8.28E-22 | 0.71 | 0.58 | 0.13 | 1.23 |
| 2 | Pfas | 0.46 | 4.06E-26 | 8.44E-22 | 0.40 | 0.28 | 0.12 | 1.44 |
| 2 | Qrsl1 | 0.56 | 5.36E-26 | 1.11E-21 | 0.31 | 0.20 | 0.11 | 1.52 |
| 2 | Riok1 | 0.29 | 5.90E-26 | 1.23E-21 | 0.72 | 0.57 | 0.15 | 1.25 |
| 2 | Rfc3 | 0.25 | 6.00E-26 | 1.25E-21 | 0.88 | 0.76 | 0.13 | 1.17 |
| 2 | Rtel1 | 0.65 | 6.01E-26 | 1.25E-21 | 0.28 | 0.18 | 0.10 | 1.56 |
| 2 | Naf1 | 0.38 | 6.41E-26 | 1.33E-21 | 0.53 | 0.39 | 0.14 | 1.37 |
| 2 | Adgrl1 | 0.32 | 6.87E-26 | 1.43E-21 | 0.58 | 0.43 | 0.15 | 1.36 |
| 2 | Sdk1 | 0.82 | 7.64E-26 | 1.59E-21 | 0.15 | 0.08 | 0.07 | 1.90 |
| 2 | Cep83 | 0.32 | 7.70E-26 | 1.60E-21 | 0.67 | 0.54 | 0.13 | 1.24 |
| 2 | Pde5a | 0.46 | 7.83E-26 | 1.63E-21 | 0.28 | 0.17 | 0.10 | 1.59 |
| 2 | Smad1 | 0.32 | 8.39E-26 | 1.75E-21 | 0.61 | 0.46 | 0.15 | 1.32 |
| 2 | Ddx20 | 0.42 | 8.49E-26 | 1.76E-21 | 0.45 | 0.33 | 0.12 | 1.38 |

**Supplementary Table 1. Marker genes for the original clusters of integrated clustering of E11.5 control and NotchGOF λ-junction cells.**

| | | | | | | | | |
|---|---|---|---|---|---|---|---|---|
| 2 | Reck | 0.40 | 9.17E-26 | 1.91E-21 | 0.36 | 0.24 | 0.12 | 1.52 |
| 2 | St6galnac3 | 0.31 | 9.29E-26 | 1.93E-21 | 0.53 | 0.38 | 0.15 | 1.40 |
| 2 | Drosha | 0.31 | 1.01E-25 | 2.10E-21 | 0.65 | 0.49 | 0.16 | 1.32 |
| 2 | Atxn2l | 0.30 | 1.03E-25 | 2.14E-21 | 0.71 | 0.57 | 0.14 | 1.24 |
| 2 | Mthfd1 | 0.37 | 1.05E-25 | 2.18E-21 | 0.51 | 0.37 | 0.14 | 1.37 |
| 2 | Cnot9 | 0.37 | 1.11E-25 | 2.31E-21 | 0.52 | 0.38 | 0.14 | 1.37 |
| 2 | Cenpc1 | 0.32 | 1.23E-25 | 2.57E-21 | 0.59 | 0.44 | 0.15 | 1.33 |
| 2 | Hunk | 0.31 | 1.30E-25 | 2.70E-21 | 0.64 | 0.49 | 0.15 | 1.31 |
| 2 | Dusp1 | 0.44 | 1.30E-25 | 2.71E-21 | 0.40 | 0.28 | 0.12 | 1.42 |
| 2 | Eri1 | 0.33 | 1.38E-25 | 2.86E-21 | 0.63 | 0.49 | 0.14 | 1.28 |
| 2 | Ankrd26 | 0.36 | 1.41E-25 | 2.93E-21 | 0.56 | 0.42 | 0.13 | 1.32 |
| 2 | Arhgef26 | 0.63 | 1.44E-25 | 2.99E-21 | 0.20 | 0.12 | 0.09 | 1.73 |
| 2 | B3gnt5 | 0.36 | 1.66E-25 | 3.45E-21 | 0.48 | 0.35 | 0.14 | 1.39 |
| 2 | Gria3 | 0.51 | 1.74E-25 | 3.61E-21 | 0.25 | 0.16 | 0.10 | 1.62 |
| 2 | Tpst1 | 0.39 | 1.74E-25 | 3.62E-21 | 0.46 | 0.33 | 0.13 | 1.41 |
| 2 | AI506816 | 0.34 | 1.80E-25 | 3.75E-21 | 0.66 | 0.53 | 0.14 | 1.26 |
| 2 | Gse1 | 0.38 | 1.85E-25 | 3.85E-21 | 0.42 | 0.29 | 0.12 | 1.41 |
| 2 | Ppat | 0.30 | 1.91E-25 | 3.97E-21 | 0.63 | 0.49 | 0.15 | 1.30 |
| 2 | L1cam | 0.59 | 2.00E-25 | 4.15E-21 | 0.15 | 0.07 | 0.07 | 1.96 |
| 2 | Fbxo30 | 0.35 | 2.01E-25 | 4.19E-21 | 0.54 | 0.40 | 0.14 | 1.34 |
| 2 | Cand1 | 0.27 | 2.14E-25 | 4.45E-21 | 0.78 | 0.63 | 0.15 | 1.23 |
| 2 | Dot1l | 0.33 | 2.15E-25 | 4.47E-21 | 0.56 | 0.42 | 0.14 | 1.33 |
| 2 | Ppp4r1 | 0.38 | 2.18E-25 | 4.54E-21 | 0.52 | 0.39 | 0.13 | 1.34 |
| 2 | P4ha2 | 0.42 | 2.25E-25 | 4.68E-21 | 0.39 | 0.26 | 0.12 | 1.46 |
| 2 | Fzr1 | 0.34 | 2.41E-25 | 5.01E-21 | 0.57 | 0.43 | 0.14 | 1.33 |
| 2 | Dyrk1a | 0.35 | 2.62E-25 | 5.45E-21 | 0.51 | 0.37 | 0.14 | 1.37 |
| 2 | Fam72a | 0.93 | 3.02E-25 | 6.27E-21 | 0.13 | 0.07 | 0.07 | 1.94 |
| 2 | Tyw3 | 0.47 | 3.02E-25 | 6.29E-21 | 0.31 | 0.20 | 0.11 | 1.55 |
| 2 | Mcm8 | 0.45 | 3.19E-25 | 6.63E-21 | 0.37 | 0.25 | 0.12 | 1.45 |
| 2 | Pitrm1 | 0.31 | 3.19E-25 | 6.64E-21 | 0.67 | 0.53 | 0.14 | 1.27 |
| 2 | Klhl22 | 0.38 | 3.40E-25 | 7.07E-21 | 0.46 | 0.33 | 0.14 | 1.41 |
| 2 | Aaas | 0.41 | 3.59E-25 | 7.46E-21 | 0.45 | 0.32 | 0.13 | 1.40 |
| 2 | Rttn | 0.62 | 3.60E-25 | 7.50E-21 | 0.25 | 0.16 | 0.09 | 1.60 |
| 2 | Agps | 0.34 | 3.61E-25 | 7.52E-21 | 0.53 | 0.39 | 0.14 | 1.36 |
| 2 | Gxylt1 | 0.34 | 3.96E-25 | 8.24E-21 | 0.59 | 0.45 | 0.14 | 1.31 |
| 2 | Get4 | 0.30 | 3.99E-25 | 8.31E-21 | 0.68 | 0.54 | 0.15 | 1.27 |
| 2 | Pold2 | 0.35 | 4.02E-25 | 8.36E-21 | 0.56 | 0.42 | 0.14 | 1.34 |
| 2 | Sfxn1 | 0.25 | 4.16E-25 | 8.66E-21 | 0.86 | 0.73 | 0.13 | 1.18 |
| 2 | Abcf2 | 0.27 | 4.21E-25 | 8.75E-21 | 0.76 | 0.63 | 0.13 | 1.20 |
| 2 | A430005L14Rik | 0.33 | 5.31E-25 | 1.10E-20 | 0.63 | 0.49 | 0.13 | 1.27 |
| 2 | Edil3 | 0.53 | 5.74E-25 | 1.19E-20 | 0.26 | 0.16 | 0.10 | 1.60 |
| 2 | Phf23 | 0.27 | 5.76E-25 | 1.20E-20 | 0.75 | 0.61 | 0.14 | 1.23 |
| 2 | Cth | 0.47 | 5.86E-25 | 1.22E-20 | 0.31 | 0.20 | 0.11 | 1.54 |
| 2 | Poc5 | 0.55 | 6.58E-25 | 1.37E-20 | 0.29 | 0.19 | 0.10 | 1.53 |
| 2 | Mtbp | 0.52 | 6.76E-25 | 1.41E-20 | 0.32 | 0.21 | 0.11 | 1.50 |
| 2 | Efnb1 | 0.28 | 7.01E-25 | 1.46E-20 | 0.67 | 0.52 | 0.15 | 1.29 |
| 2 | Cinp | 0.35 | 7.05E-25 | 1.47E-20 | 0.61 | 0.47 | 0.14 | 1.29 |
| 2 | Kctd15 | 0.33 | 7.22E-25 | 1.50E-20 | 0.55 | 0.41 | 0.15 | 1.36 |
| 2 | Gabpb2 | 0.34 | 7.36E-25 | 1.53E-20 | 0.55 | 0.41 | 0.14 | 1.34 |
| 2 | Ccdc25 | 0.26 | 7.86E-25 | 1.63E-20 | 0.77 | 0.63 | 0.14 | 1.21 |
| 2 | Nol10 | 0.37 | 7.87E-25 | 1.64E-20 | 0.45 | 0.32 | 0.13 | 1.40 |
| 2 | Gnl3 | 0.25 | 8.23E-25 | 1.71E-20 | 0.86 | 0.74 | 0.12 | 1.16 |
| 2 | Gnaq | 0.29 | 8.80E-25 | 1.83E-20 | 0.66 | 0.51 | 0.15 | 1.29 |
| 2 | Cep72 | 0.55 | 9.71E-25 | 2.02E-20 | 0.23 | 0.14 | 0.09 | 1.63 |
| 2 | Cactin | 0.41 | 1.06E-24 | 2.20E-20 | 0.43 | 0.31 | 0.12 | 1.39 |
| 2 | Stk3 | 0.35 | 1.07E-24 | 2.23E-20 | 0.55 | 0.41 | 0.14 | 1.33 |

**Supplementary Table 1. Marker genes for the original clusters of integrated clustering of E11.5 control and NotchGOF λ-junction cells.**

| | | | | | | | | |
|---|---|---|---|---|---|---|---|---|
| 2 | Mrps22 | 0.29 | 1.13E-24 | 2.35E-20 | 0.70 | 0.56 | 0.14 | 1.24 |
| 2 | Nampt | 0.39 | 1.30E-24 | 2.71E-20 | 0.49 | 0.36 | 0.13 | 1.35 |
| 2 | Plcg1 | 0.35 | 1.32E-24 | 2.75E-20 | 0.53 | 0.39 | 0.14 | 1.37 |
| 2 | Tle2 | 0.54 | 1.41E-24 | 2.94E-20 | 0.20 | 0.11 | 0.08 | 1.75 |
| 2 | 9430015G10Rik | 0.48 | 1.45E-24 | 3.02E-20 | 0.33 | 0.22 | 0.11 | 1.50 |
| 2 | Cnep1r1 | 0.30 | 1.47E-24 | 3.05E-20 | 0.67 | 0.53 | 0.14 | 1.27 |
| 2 | Nufip2 | 0.29 | 1.56E-24 | 3.25E-20 | 0.68 | 0.54 | 0.15 | 1.28 |
| 2 | Dnajc11 | 0.33 | 1.81E-24 | 3.76E-20 | 0.61 | 0.48 | 0.14 | 1.29 |
| 2 | Ptprg | 0.37 | 1.92E-24 | 3.99E-20 | 0.49 | 0.36 | 0.13 | 1.37 |
| 2 | Lpgat1 | 0.28 | 2.29E-24 | 4.76E-20 | 0.70 | 0.54 | 0.16 | 1.29 |
| 2 | Trim71 | 0.46 | 2.36E-24 | 4.90E-20 | 0.31 | 0.20 | 0.11 | 1.53 |
| 2 | Stard8 | 0.74 | 2.55E-24 | 5.31E-20 | 0.15 | 0.08 | 0.07 | 1.85 |
| 2 | Gtf2a1 | 0.32 | 2.77E-24 | 5.75E-20 | 0.62 | 0.48 | 0.14 | 1.28 |
| 2 | Polq | 0.84 | 2.90E-24 | 6.04E-20 | 0.16 | 0.09 | 0.07 | 1.76 |
| 2 | Bud13 | 0.42 | 3.49E-24 | 7.26E-20 | 0.39 | 0.28 | 0.12 | 1.43 |
| 2 | 9930021J03Rik | 0.29 | 3.56E-24 | 7.41E-20 | 0.69 | 0.54 | 0.15 | 1.28 |
| 2 | Eif1ad | 0.27 | 3.64E-24 | 7.57E-20 | 0.77 | 0.64 | 0.13 | 1.20 |
| 2 | Wwp1 | 0.37 | 3.74E-24 | 7.79E-20 | 0.54 | 0.40 | 0.13 | 1.33 |
| 2 | Dab2ip | 0.26 | 3.99E-24 | 8.30E-20 | 0.73 | 0.59 | 0.14 | 1.24 |
| 2 | Nup50 | 0.29 | 4.05E-24 | 8.42E-20 | 0.73 | 0.60 | 0.13 | 1.21 |
| 2 | Smad9 | 0.84 | 4.11E-24 | 8.54E-20 | 0.11 | 0.05 | 0.06 | 2.08 |
| 2 | Rbm6 | 0.26 | 4.94E-24 | 1.03E-19 | 0.83 | 0.71 | 0.12 | 1.16 |
| 2 | Mthfd1l | 0.62 | 5.50E-24 | 1.14E-19 | 0.24 | 0.15 | 0.09 | 1.58 |
| 2 | Anapc7 | 0.30 | 5.65E-24 | 1.18E-19 | 0.68 | 0.54 | 0.14 | 1.26 |
| 2 | Ush1c | 0.68 | 5.70E-24 | 1.19E-19 | 0.17 | 0.10 | 0.08 | 1.78 |
| 2 | Cnrip1 | 0.48 | 6.27E-24 | 1.30E-19 | 0.22 | 0.13 | 0.09 | 1.68 |
| 2 | Prpf3 | 0.37 | 6.70E-24 | 1.39E-19 | 0.51 | 0.38 | 0.13 | 1.33 |
| 2 | Pms2 | 0.38 | 7.36E-24 | 1.53E-19 | 0.43 | 0.31 | 0.12 | 1.40 |
| 2 | Dcaf1 | 0.34 | 8.14E-24 | 1.69E-19 | 0.56 | 0.43 | 0.13 | 1.30 |
| 2 | Cep78 | 0.34 | 9.15E-24 | 1.90E-19 | 0.52 | 0.38 | 0.13 | 1.35 |
| 2 | Tmem109 | 0.34 | 1.09E-23 | 2.28E-19 | 0.49 | 0.35 | 0.13 | 1.37 |
| 2 | Cmtr1 | 0.37 | 1.21E-23 | 2.51E-19 | 0.45 | 0.33 | 0.13 | 1.39 |
| 2 | Orc3 | 0.33 | 1.39E-23 | 2.89E-19 | 0.51 | 0.38 | 0.13 | 1.36 |
| 2 | Senp1 | 0.29 | 1.46E-23 | 3.04E-19 | 0.59 | 0.45 | 0.14 | 1.32 |
| 2 | Mmp23 | 0.92 | 1.55E-23 | 3.22E-19 | 0.11 | 0.05 | 0.06 | 2.10 |
| 2 | Camsap2 | 0.36 | 1.55E-23 | 3.22E-19 | 0.44 | 0.31 | 0.13 | 1.40 |
| 2 | Zfp131 | 0.27 | 1.76E-23 | 3.66E-19 | 0.70 | 0.56 | 0.14 | 1.25 |
| 2 | Me2 | 0.34 | 1.89E-23 | 3.93E-19 | 0.58 | 0.44 | 0.13 | 1.30 |
| 2 | Adamts10 | 0.35 | 2.03E-23 | 4.21E-19 | 0.44 | 0.31 | 0.13 | 1.41 |
| 2 | Enkd1 | 0.43 | 2.23E-23 | 4.63E-19 | 0.39 | 0.27 | 0.12 | 1.43 |
| 2 | Tbc1d16 | 0.30 | 2.31E-23 | 4.80E-19 | 0.57 | 0.44 | 0.14 | 1.32 |
| 2 | Snx30 | 0.56 | 2.34E-23 | 4.86E-19 | 0.23 | 0.14 | 0.09 | 1.62 |
| 2 | Slfn9 | 0.36 | 2.35E-23 | 4.88E-19 | 0.35 | 0.24 | 0.11 | 1.46 |
| 2 | Tm9sf1 | 0.34 | 2.58E-23 | 5.37E-19 | 0.57 | 0.43 | 0.14 | 1.32 |
| 2 | Hdgfl2 | 0.26 | 2.62E-23 | 5.45E-19 | 0.80 | 0.67 | 0.12 | 1.18 |
| 2 | Orc2 | 0.27 | 2.63E-23 | 5.47E-19 | 0.71 | 0.56 | 0.15 | 1.27 |
| 2 | A330076H08Rik | 0.79 | 2.83E-23 | 5.88E-19 | 0.13 | 0.07 | 0.06 | 1.91 |
| 2 | Myo5a | 0.36 | 3.10E-23 | 6.44E-19 | 0.43 | 0.31 | 0.12 | 1.39 |
| 2 | Ttf2 | 0.38 | 3.11E-23 | 6.46E-19 | 0.43 | 0.31 | 0.12 | 1.38 |
| 2 | Mpdz | 0.29 | 3.12E-23 | 6.48E-19 | 0.66 | 0.51 | 0.15 | 1.29 |
| 2 | Abhd10 | 0.40 | 3.14E-23 | 6.53E-19 | 0.43 | 0.31 | 0.12 | 1.38 |
| 2 | Ap1m1 | 0.33 | 3.18E-23 | 6.61E-19 | 0.51 | 0.38 | 0.13 | 1.34 |
| 2 | Rnf8 | 0.26 | 3.19E-23 | 6.62E-19 | 0.75 | 0.61 | 0.14 | 1.23 |
| 2 | Lcorl | 0.26 | 3.27E-23 | 6.80E-19 | 0.61 | 0.47 | 0.14 | 1.29 |
| 2 | Eif2b1 | 0.33 | 3.48E-23 | 7.23E-19 | 0.61 | 0.48 | 0.13 | 1.28 |
| 2 | Spata5 | 0.40 | 3.58E-23 | 7.45E-19 | 0.44 | 0.32 | 0.12 | 1.36 |

**Supplementary Table 1. Marker genes for the original clusters of integrated clustering of E11.5 control and NotchGOF λ-junction cells.**

| | | | | | | | | |
|---|---|---|---|---|---|---|---|---|
| 2 | Ncbp1 | 0.28 | 3.72E-23 | 7.74E-19 | 0.72 | 0.59 | 0.14 | 1.23 |
| 2 | Atm | 0.36 | 3.74E-23 | 7.77E-19 | 0.43 | 0.31 | 0.12 | 1.40 |
| 2 | Ldb2 | 0.37 | 3.85E-23 | 8.00E-19 | 0.38 | 0.26 | 0.12 | 1.46 |
| 2 | Tspan2 | 0.69 | 4.14E-23 | 8.61E-19 | 0.17 | 0.09 | 0.07 | 1.78 |
| 2 | Dlg4 | 0.36 | 4.32E-23 | 8.98E-19 | 0.42 | 0.30 | 0.12 | 1.41 |
| 2 | Pias4 | 0.33 | 4.34E-23 | 9.03E-19 | 0.57 | 0.43 | 0.13 | 1.30 |
| 2 | Lca5 | 0.44 | 4.57E-23 | 9.51E-19 | 0.31 | 0.21 | 0.10 | 1.50 |
| 2 | Map1s | 0.32 | 4.65E-23 | 9.67E-19 | 0.49 | 0.36 | 0.13 | 1.37 |
| 2 | Odf2l | 0.37 | 4.83E-23 | 1.00E-18 | 0.47 | 0.34 | 0.12 | 1.36 |
| 2 | Nup214 | 0.33 | 4.89E-23 | 1.02E-18 | 0.59 | 0.46 | 0.13 | 1.28 |
| 2 | Rassf8 | 0.49 | 4.89E-23 | 1.02E-18 | 0.27 | 0.18 | 0.10 | 1.56 |
| 2 | Rngtt | 0.37 | 5.12E-23 | 1.06E-18 | 0.51 | 0.39 | 0.13 | 1.33 |
| 2 | Uba6 | 0.34 | 5.91E-23 | 1.23E-18 | 0.52 | 0.39 | 0.13 | 1.34 |
| 2 | Hist1h3g | 0.97 | 6.54E-23 | 1.36E-18 | 0.12 | 0.06 | 0.06 | 1.97 |
| 2 | Wdcp | 0.43 | 7.63E-23 | 1.59E-18 | 0.48 | 0.36 | 0.13 | 1.35 |
| 2 | Strn4 | 0.29 | 7.95E-23 | 1.65E-18 | 0.62 | 0.48 | 0.14 | 1.29 |
| 2 | Usp39 | 0.27 | 8.41E-23 | 1.75E-18 | 0.71 | 0.57 | 0.13 | 1.23 |
| 2 | Eif4enif1 | 0.30 | 8.82E-23 | 1.83E-18 | 0.66 | 0.53 | 0.14 | 1.26 |
| 2 | Ppm1h | 0.84 | 8.94E-23 | 1.86E-18 | 0.14 | 0.08 | 0.07 | 1.83 |
| 2 | Trib2 | 0.43 | 9.07E-23 | 1.89E-18 | 0.29 | 0.19 | 0.10 | 1.54 |
| 2 | Dimt1 | 0.36 | 9.49E-23 | 1.97E-18 | 0.52 | 0.40 | 0.12 | 1.31 |
| 2 | Nfx1 | 0.26 | 9.68E-23 | 2.01E-18 | 0.70 | 0.56 | 0.14 | 1.25 |
| 2 | Dda1 | 0.26 | 1.06E-22 | 2.20E-18 | 0.69 | 0.56 | 0.13 | 1.23 |
| 2 | Ehmt2 | 0.29 | 1.18E-22 | 2.46E-18 | 0.68 | 0.56 | 0.13 | 1.23 |
| 2 | Phf2 | 0.31 | 1.23E-22 | 2.56E-18 | 0.42 | 0.29 | 0.13 | 1.43 |
| 2 | Nfix | 0.64 | 1.25E-22 | 2.61E-18 | 0.23 | 0.15 | 0.08 | 1.57 |
| 2 | Xpo5 | 0.33 | 1.33E-22 | 2.78E-18 | 0.55 | 0.42 | 0.13 | 1.30 |
| 2 | Setd1a | 0.37 | 1.82E-22 | 3.79E-18 | 0.41 | 0.29 | 0.12 | 1.40 |
| 2 | Fem1c | 0.31 | 1.83E-22 | 3.80E-18 | 0.50 | 0.37 | 0.13 | 1.34 |
| 2 | Dis3 | 0.29 | 1.90E-22 | 3.95E-18 | 0.68 | 0.54 | 0.14 | 1.25 |
| 2 | Tfap2c | 0.43 | 1.90E-22 | 3.95E-18 | 0.38 | 0.27 | 0.11 | 1.42 |
| 2 | Ptpn9 | 0.27 | 1.94E-22 | 4.04E-18 | 0.62 | 0.48 | 0.14 | 1.30 |
| 2 | Ireb2 | 0.30 | 1.96E-22 | 4.07E-18 | 0.67 | 0.54 | 0.14 | 1.25 |
| 2 | Impa1 | 0.29 | 1.99E-22 | 4.14E-18 | 0.70 | 0.57 | 0.13 | 1.22 |
| 2 | Ephb4 | 0.33 | 2.00E-22 | 4.17E-18 | 0.52 | 0.38 | 0.13 | 1.34 |
| 2 | Anapc15 | 0.27 | 2.00E-22 | 4.17E-18 | 0.72 | 0.59 | 0.13 | 1.22 |
| 2 | Cstf1 | 0.38 | 2.02E-22 | 4.20E-18 | 0.37 | 0.26 | 0.11 | 1.43 |
| 2 | Zfp143 | 0.47 | 2.12E-22 | 4.41E-18 | 0.36 | 0.26 | 0.11 | 1.41 |
| 2 | Mphosph9 | 0.43 | 2.25E-22 | 4.68E-18 | 0.35 | 0.24 | 0.11 | 1.44 |
| 2 | Phf12 | 0.31 | 2.34E-22 | 4.86E-18 | 0.56 | 0.42 | 0.13 | 1.32 |
| 2 | E2f5 | 0.34 | 2.39E-22 | 4.96E-18 | 0.60 | 0.48 | 0.13 | 1.27 |
| 2 | Fndc4 | 0.32 | 2.50E-22 | 5.21E-18 | 0.53 | 0.40 | 0.13 | 1.33 |
| 2 | Gtf3c2 | 0.27 | 2.52E-22 | 5.25E-18 | 0.72 | 0.58 | 0.14 | 1.24 |
| 2 | Ada | 0.73 | 2.62E-22 | 5.45E-18 | 0.14 | 0.08 | 0.06 | 1.85 |
| 2 | Gabra4 | 1.00 | 2.74E-22 | 5.69E-18 | 0.10 | 0.05 | 0.05 | 2.08 |
| 2 | Akap8 | 0.27 | 3.49E-22 | 7.26E-18 | 0.77 | 0.65 | 0.12 | 1.18 |
| 2 | Asns | 0.34 | 3.55E-22 | 7.38E-18 | 0.56 | 0.43 | 0.13 | 1.30 |
| 2 | Tubgcp2 | 0.27 | 3.66E-22 | 7.61E-18 | 0.65 | 0.51 | 0.14 | 1.27 |
| 2 | Rnf157 | 0.57 | 3.72E-22 | 7.73E-18 | 0.23 | 0.14 | 0.09 | 1.60 |
| 2 | Mosmo | 0.30 | 4.07E-22 | 8.47E-18 | 0.61 | 0.48 | 0.13 | 1.28 |
| 2 | Cep170 | 0.28 | 4.22E-22 | 8.77E-18 | 0.63 | 0.50 | 0.14 | 1.28 |
| 2 | Fasn | 0.30 | 4.40E-22 | 9.15E-18 | 0.63 | 0.49 | 0.14 | 1.28 |
| 2 | Ttyh3 | 0.28 | 4.63E-22 | 9.63E-18 | 0.66 | 0.52 | 0.14 | 1.26 |
| 2 | Mrc2 | 0.40 | 4.74E-22 | 9.87E-18 | 0.32 | 0.21 | 0.11 | 1.51 |
| 2 | Rbm4b | 0.25 | 4.89E-22 | 1.02E-17 | 0.79 | 0.66 | 0.13 | 1.20 |
| 2 | Pxdc1 | 0.73 | 4.95E-22 | 1.03E-17 | 0.17 | 0.10 | 0.07 | 1.72 |

**Supplementary Table 1. Marker genes for the original clusters of integrated clustering of E11.5 control and NotchGOF λ-junction cells.**

| | | | | | | | | |
|---|---|---|---|---|---|---|---|---|
| 2 | 9430038I01Rik | 0.34 | 5.07E-22 | 1.05E-17 | 0.45 | 0.33 | 0.13 | 1.38 |
| 2 | Dse | 0.34 | 5.08E-22 | 1.06E-17 | 0.38 | 0.26 | 0.12 | 1.45 |
| 2 | Dpy19l1 | 0.32 | 5.44E-22 | 1.13E-17 | 0.48 | 0.35 | 0.13 | 1.37 |
| 2 | Pkmyt1 | 0.60 | 5.53E-22 | 1.15E-17 | 0.25 | 0.16 | 0.09 | 1.53 |
| 2 | Skida1 | 0.50 | 5.57E-22 | 1.16E-17 | 0.23 | 0.15 | 0.09 | 1.58 |
| 2 | Leo1 | 0.25 | 6.25E-22 | 1.30E-17 | 0.75 | 0.62 | 0.13 | 1.20 |
| 2 | Cacna2d1 | 0.36 | 6.44E-22 | 1.34E-17 | 0.19 | 0.11 | 0.08 | 1.70 |
| 2 | Slitrk4 | 0.74 | 6.55E-22 | 1.36E-17 | 0.10 | 0.05 | 0.05 | 2.06 |
| 2 | Lrch3 | 0.31 | 7.38E-22 | 1.53E-17 | 0.48 | 0.36 | 0.13 | 1.36 |
| 2 | S1pr2 | 0.32 | 7.88E-22 | 1.64E-17 | 0.44 | 0.31 | 0.13 | 1.41 |
| 2 | Rpp30 | 0.30 | 8.58E-22 | 1.78E-17 | 0.68 | 0.55 | 0.13 | 1.24 |
| 2 | Unc5b | 0.27 | 9.00E-22 | 1.87E-17 | 0.52 | 0.38 | 0.14 | 1.35 |
| 2 | Rerg | 0.60 | 9.79E-22 | 2.04E-17 | 0.34 | 0.24 | 0.10 | 1.41 |
| 2 | Uck2 | 0.28 | 1.00E-21 | 2.08E-17 | 0.65 | 0.51 | 0.14 | 1.26 |
| 2 | Elmo2 | 0.43 | 1.00E-21 | 2.09E-17 | 0.35 | 0.24 | 0.11 | 1.44 |
| 2 | Ccm2 | 0.32 | 1.03E-21 | 2.14E-17 | 0.52 | 0.39 | 0.13 | 1.35 |
| 2 | Pus7l | 0.55 | 1.10E-21 | 2.29E-17 | 0.23 | 0.14 | 0.08 | 1.58 |
| 2 | Mad1l1 | 0.39 | 1.12E-21 | 2.33E-17 | 0.46 | 0.35 | 0.12 | 1.34 |
| 2 | Mark1 | 0.36 | 1.12E-21 | 2.33E-17 | 0.35 | 0.24 | 0.11 | 1.45 |
| 2 | Mllt3 | 0.29 | 1.14E-21 | 2.37E-17 | 0.77 | 0.65 | 0.12 | 1.18 |
| 2 | Terf1 | 0.26 | 1.14E-21 | 2.38E-17 | 0.65 | 0.51 | 0.14 | 1.27 |
| 2 | Tmem97 | 0.28 | 1.27E-21 | 2.65E-17 | 0.59 | 0.45 | 0.14 | 1.31 |
| 2 | Slit3 | 0.31 | 1.42E-21 | 2.96E-17 | 0.28 | 0.18 | 0.10 | 1.54 |
| 2 | Pds5a | 0.25 | 1.49E-21 | 3.10E-17 | 0.79 | 0.66 | 0.12 | 1.19 |
| 2 | Vrk2 | 0.44 | 1.67E-21 | 3.48E-17 | 0.32 | 0.22 | 0.10 | 1.46 |
| 2 | Recql | 0.49 | 1.68E-21 | 3.50E-17 | 0.28 | 0.19 | 0.09 | 1.50 |
| 2 | Zfp61 | 0.37 | 1.89E-21 | 3.93E-17 | 0.34 | 0.23 | 0.11 | 1.47 |
| 2 | Fam110b | 0.32 | 1.92E-21 | 3.98E-17 | 0.42 | 0.30 | 0.12 | 1.42 |
| 2 | Cdkn2aipnl | 0.28 | 1.94E-21 | 4.04E-17 | 0.63 | 0.50 | 0.13 | 1.27 |
| 2 | Map3k12 | 0.40 | 2.05E-21 | 4.27E-17 | 0.30 | 0.20 | 0.10 | 1.51 |
| 2 | Exoc5 | 0.28 | 2.14E-21 | 4.44E-17 | 0.69 | 0.56 | 0.13 | 1.24 |
| 2 | Ppp2r3a | 0.35 | 2.22E-21 | 4.61E-17 | 0.47 | 0.34 | 0.12 | 1.36 |
| 2 | Pphln1 | 0.32 | 2.29E-21 | 4.77E-17 | 0.51 | 0.39 | 0.12 | 1.32 |
| 2 | Nup98 | 0.29 | 2.74E-21 | 5.70E-17 | 0.60 | 0.47 | 0.13 | 1.28 |
| 2 | Gtpbp10 | 0.34 | 2.88E-21 | 5.99E-17 | 0.42 | 0.30 | 0.12 | 1.39 |
| 2 | Tnks1bp1 | 0.49 | 3.32E-21 | 6.91E-17 | 0.25 | 0.17 | 0.09 | 1.53 |
| 2 | Eepd1 | 0.61 | 3.41E-21 | 7.10E-17 | 0.19 | 0.11 | 0.08 | 1.67 |
| 2 | Ppp2r1b | 0.31 | 3.55E-21 | 7.38E-17 | 0.52 | 0.39 | 0.13 | 1.34 |
| 2 | Dock1 | 0.27 | 3.56E-21 | 7.40E-17 | 0.63 | 0.49 | 0.14 | 1.29 |
| 2 | Myo9a | 0.32 | 3.77E-21 | 7.85E-17 | 0.48 | 0.36 | 0.13 | 1.35 |
| 2 | Itsn1 | 0.32 | 3.88E-21 | 8.07E-17 | 0.55 | 0.43 | 0.13 | 1.30 |
| 2 | Gle1 | 0.31 | 4.00E-21 | 8.31E-17 | 0.59 | 0.47 | 0.12 | 1.26 |
| 2 | Fam83d | 0.49 | 4.37E-21 | 9.09E-17 | 0.28 | 0.19 | 0.09 | 1.47 |
| 2 | Pls3 | 0.29 | 4.81E-21 | 1.00E-16 | 0.41 | 0.29 | 0.12 | 1.41 |
| 2 | Ctdspl2 | 0.27 | 5.23E-21 | 1.09E-16 | 0.70 | 0.57 | 0.14 | 1.24 |
| 2 | Virma | 0.26 | 5.28E-21 | 1.10E-16 | 0.71 | 0.58 | 0.13 | 1.23 |
| 2 | Bola1 | 0.26 | 5.48E-21 | 1.14E-16 | 0.74 | 0.61 | 0.13 | 1.22 |
| 2 | Hsd17b7 | 0.38 | 5.92E-21 | 1.23E-16 | 0.37 | 0.26 | 0.11 | 1.42 |
| 2 | Rad9a | 0.43 | 6.03E-21 | 1.25E-16 | 0.30 | 0.21 | 0.10 | 1.47 |
| 2 | Spsb4 | 0.43 | 6.10E-21 | 1.27E-16 | 0.30 | 0.20 | 0.10 | 1.50 |
| 2 | Phactr4 | 0.28 | 6.16E-21 | 1.28E-16 | 0.61 | 0.48 | 0.13 | 1.28 |
| 2 | Utp18 | 0.28 | 6.41E-21 | 1.33E-16 | 0.69 | 0.57 | 0.12 | 1.21 |
| 2 | Herc2 | 0.33 | 6.63E-21 | 1.38E-16 | 0.52 | 0.40 | 0.12 | 1.30 |
| 2 | Irf2bpl | 0.26 | 6.68E-21 | 1.39E-16 | 0.50 | 0.37 | 0.13 | 1.35 |
| 2 | Kpna3 | 0.29 | 7.30E-21 | 1.52E-16 | 0.72 | 0.60 | 0.12 | 1.20 |
| 2 | Dab2 | 0.26 | 8.05E-21 | 1.67E-16 | 0.47 | 0.34 | 0.13 | 1.36 |

**Supplementary Table 1. Marker genes for the original clusters of integrated clustering of E11.5 control and NotchGOF λ-junction cells.**

| | | | | | | | | |
|---|---|---|---|---|---|---|---|---|
| 2 | Rbm10 | 0.26 | 8.46E-21 | 1.76E-16 | 0.72 | 0.58 | 0.14 | 1.24 |
| 2 | Aars | 0.30 | 8.86E-21 | 1.84E-16 | 0.58 | 0.46 | 0.13 | 1.28 |
| 2 | Larp4 | 0.26 | 9.24E-21 | 1.92E-16 | 0.72 | 0.59 | 0.13 | 1.23 |
| 2 | Rbm15 | 0.33 | 1.01E-20 | 2.11E-16 | 0.48 | 0.36 | 0.12 | 1.34 |
| 2 | Gins1 | 0.34 | 1.05E-20 | 2.18E-16 | 0.50 | 0.38 | 0.12 | 1.31 |
| 2 | Gtpbp1 | 0.49 | 1.08E-20 | 2.25E-16 | 0.29 | 0.20 | 0.09 | 1.46 |
| 2 | Safb2 | 0.27 | 1.12E-20 | 2.32E-16 | 0.68 | 0.55 | 0.13 | 1.24 |
| 2 | Gps2 | 0.29 | 1.12E-20 | 2.33E-16 | 0.67 | 0.54 | 0.13 | 1.24 |
| 2 | Cntrob | 0.44 | 1.15E-20 | 2.39E-16 | 0.30 | 0.20 | 0.10 | 1.48 |
| 2 | Cacng4 | 0.34 | 1.18E-20 | 2.45E-16 | 0.36 | 0.25 | 0.11 | 1.43 |
| 2 | Fam13a | 0.66 | 1.18E-20 | 2.46E-16 | 0.16 | 0.09 | 0.07 | 1.74 |
| 2 | Fign | 0.39 | 1.26E-20 | 2.62E-16 | 0.40 | 0.28 | 0.11 | 1.39 |
| 2 | Isy1 | 0.27 | 1.47E-20 | 3.05E-16 | 0.66 | 0.53 | 0.13 | 1.24 |
| 2 | Mdn1 | 0.34 | 1.51E-20 | 3.15E-16 | 0.52 | 0.40 | 0.12 | 1.30 |
| 2 | Abl1 | 0.30 | 1.59E-20 | 3.30E-16 | 0.62 | 0.49 | 0.13 | 1.27 |
| 2 | Xrcc6 | 0.30 | 1.59E-20 | 3.31E-16 | 0.56 | 0.43 | 0.13 | 1.31 |
| 2 | Kbtbd4 | 0.33 | 1.73E-20 | 3.61E-16 | 0.46 | 0.34 | 0.12 | 1.35 |
| 2 | Foxk2 | 0.35 | 1.86E-20 | 3.86E-16 | 0.48 | 0.36 | 0.12 | 1.33 |
| 2 | Megf9 | 0.51 | 2.10E-20 | 4.37E-16 | 0.27 | 0.18 | 0.09 | 1.49 |
| 2 | Cox10 | 0.30 | 2.11E-20 | 4.39E-16 | 0.44 | 0.32 | 0.12 | 1.36 |
| 2 | Rnf214 | 0.26 | 2.12E-20 | 4.41E-16 | 0.64 | 0.51 | 0.13 | 1.26 |
| 2 | Ccnk | 0.31 | 2.21E-20 | 4.61E-16 | 0.55 | 0.41 | 0.13 | 1.32 |
| 2 | 4930453N24Rik | 0.26 | 2.32E-20 | 4.83E-16 | 0.70 | 0.58 | 0.12 | 1.20 |
| 2 | Anapc10 | 0.29 | 2.57E-20 | 5.34E-16 | 0.63 | 0.51 | 0.12 | 1.24 |
| 2 | Tbc1d31 | 0.44 | 2.70E-20 | 5.62E-16 | 0.31 | 0.21 | 0.10 | 1.45 |
| 2 | Ccne2 | 0.56 | 2.86E-20 | 5.95E-16 | 0.29 | 0.20 | 0.09 | 1.45 |
| 2 | Yeats2 | 0.25 | 3.26E-20 | 6.79E-16 | 0.69 | 0.57 | 0.13 | 1.22 |
| 2 | Sap130 | 0.36 | 3.29E-20 | 6.85E-16 | 0.36 | 0.25 | 0.11 | 1.43 |
| 2 | Tfap4 | 0.30 | 3.46E-20 | 7.20E-16 | 0.44 | 0.32 | 0.12 | 1.38 |
| 2 | Agk | 0.34 | 3.46E-20 | 7.20E-16 | 0.43 | 0.31 | 0.12 | 1.38 |
| 2 | Axin1 | 0.30 | 3.51E-20 | 7.31E-16 | 0.47 | 0.35 | 0.12 | 1.35 |
| 2 | Caml | 0.30 | 3.78E-20 | 7.85E-16 | 0.62 | 0.48 | 0.13 | 1.27 |
| 2 | Usp48 | 0.26 | 4.31E-20 | 8.96E-16 | 0.64 | 0.51 | 0.13 | 1.25 |
| 2 | Mllt1 | 0.36 | 4.64E-20 | 9.65E-16 | 0.40 | 0.29 | 0.11 | 1.39 |
| 2 | Elp6 | 0.33 | 4.74E-20 | 9.87E-16 | 0.50 | 0.38 | 0.12 | 1.32 |
| 2 | Zbtb10 | 0.42 | 4.84E-20 | 1.01E-15 | 0.29 | 0.19 | 0.10 | 1.50 |
| 2 | Syne1 | 0.50 | 5.38E-20 | 1.12E-15 | 0.21 | 0.13 | 0.08 | 1.61 |
| 2 | Mtfr2 | 0.77 | 5.58E-20 | 1.16E-15 | 0.17 | 0.10 | 0.07 | 1.65 |
| 2 | Utp6 | 0.29 | 6.09E-20 | 1.27E-15 | 0.55 | 0.44 | 0.12 | 1.27 |
| 2 | Ankrd50 | 0.38 | 6.21E-20 | 1.29E-15 | 0.32 | 0.22 | 0.10 | 1.45 |
| 2 | Ankrd45 | 0.84 | 6.34E-20 | 1.32E-15 | 0.11 | 0.06 | 0.05 | 1.95 |
| 2 | Slc25a13 | 0.32 | 8.45E-20 | 1.76E-15 | 0.56 | 0.43 | 0.12 | 1.28 |
| 2 | Daxx | 0.30 | 8.52E-20 | 1.77E-15 | 0.52 | 0.40 | 0.12 | 1.31 |
| 2 | Casc3 | 0.26 | 8.55E-20 | 1.78E-15 | 0.66 | 0.52 | 0.13 | 1.25 |
| 2 | Cacna1d | 0.33 | 8.60E-20 | 1.79E-15 | 0.33 | 0.23 | 0.10 | 1.46 |
| 2 | Dhps | 0.30 | 8.70E-20 | 1.81E-15 | 0.61 | 0.49 | 0.12 | 1.25 |
| 2 | Cdc42se2 | 0.28 | 9.39E-20 | 1.95E-15 | 0.62 | 0.49 | 0.14 | 1.28 |
| 2 | Snx7 | 0.68 | 9.57E-20 | 1.99E-15 | 0.16 | 0.10 | 0.07 | 1.69 |
| 2 | Zfp423 | 0.39 | 9.66E-20 | 2.01E-15 | 0.30 | 0.20 | 0.10 | 1.49 |
| 2 | Dck | 0.34 | 9.81E-20 | 2.04E-15 | 0.56 | 0.45 | 0.12 | 1.26 |
| 2 | Coa7 | 0.33 | 1.02E-19 | 2.11E-15 | 0.49 | 0.37 | 0.12 | 1.32 |
| 2 | Plppr3 | 0.27 | 1.09E-19 | 2.26E-15 | 0.50 | 0.37 | 0.13 | 1.35 |
| 2 | Bag2 | 0.44 | 1.09E-19 | 2.27E-15 | 0.50 | 0.39 | 0.11 | 1.28 |
| 2 | Shf | 0.35 | 1.14E-19 | 2.38E-15 | 0.38 | 0.27 | 0.11 | 1.41 |
| 2 | Vopp1 | 0.40 | 1.32E-19 | 2.74E-15 | 0.37 | 0.27 | 0.10 | 1.38 |
| 2 | Cast | 0.42 | 1.32E-19 | 2.74E-15 | 0.33 | 0.24 | 0.10 | 1.41 |

**Supplementary Table 1. Marker genes for the original clusters of integrated clustering of E11.5 control and NotchGOF λ-junction cells.**

| | | | | | | | | |
|---|---|---|---|---|---|---|---|---|
| 2 | Cep162 | 0.37 | 1.39E-19 | 2.88E-15 | 0.35 | 0.25 | 0.10 | 1.41 |
| 2 | Ptpdc1 | 0.39 | 1.47E-19 | 3.06E-15 | 0.25 | 0.16 | 0.09 | 1.56 |
| 2 | Cars | 0.37 | 1.53E-19 | 3.18E-15 | 0.40 | 0.29 | 0.11 | 1.37 |
| 2 | Pms1 | 0.47 | 1.57E-19 | 3.26E-15 | 0.30 | 0.20 | 0.09 | 1.45 |
| 2 | Prkce | 0.53 | 1.61E-19 | 3.36E-15 | 0.23 | 0.15 | 0.08 | 1.55 |
| 2 | Rufy3 | 0.32 | 1.63E-19 | 3.38E-15 | 0.32 | 0.22 | 0.10 | 1.44 |
| 2 | Med1 | 0.28 | 1.65E-19 | 3.44E-15 | 0.69 | 0.57 | 0.13 | 1.22 |
| 2 | Smpd4 | 0.30 | 1.87E-19 | 3.89E-15 | 0.50 | 0.38 | 0.12 | 1.31 |
| 2 | Arl6 | 0.29 | 1.88E-19 | 3.91E-15 | 0.54 | 0.41 | 0.13 | 1.32 |
| 2 | Vps16 | 0.34 | 1.91E-19 | 3.98E-15 | 0.39 | 0.28 | 0.11 | 1.38 |
| 2 | Mad2l2 | 0.27 | 1.95E-19 | 4.05E-15 | 0.53 | 0.41 | 0.12 | 1.30 |
| 2 | Cep135 | 0.37 | 2.30E-19 | 4.79E-15 | 0.39 | 0.29 | 0.10 | 1.36 |
| 2 | Exosc2 | 0.28 | 2.46E-19 | 5.12E-15 | 0.56 | 0.44 | 0.13 | 1.29 |
| 2 | D16Ertd472e | 0.31 | 2.55E-19 | 5.30E-15 | 0.37 | 0.26 | 0.11 | 1.41 |
| 2 | Nol9 | 0.35 | 2.62E-19 | 5.44E-15 | 0.42 | 0.31 | 0.11 | 1.35 |
| 2 | Lrrc4b | 0.44 | 2.78E-19 | 5.77E-15 | 0.20 | 0.12 | 0.08 | 1.62 |
| 2 | Pdzd8 | 0.29 | 2.98E-19 | 6.19E-15 | 0.56 | 0.44 | 0.12 | 1.28 |
| 2 | Cpsf1 | 0.33 | 3.10E-19 | 6.45E-15 | 0.43 | 0.31 | 0.11 | 1.37 |
| 2 | Eloa | 0.27 | 3.38E-19 | 7.03E-15 | 0.58 | 0.46 | 0.13 | 1.27 |
| 2 | Ctu2 | 0.36 | 3.41E-19 | 7.08E-15 | 0.43 | 0.33 | 0.11 | 1.34 |
| 2 | Gfm1 | 0.32 | 3.56E-19 | 7.40E-15 | 0.45 | 0.34 | 0.11 | 1.33 |
| 2 | Poc1a | 0.47 | 3.72E-19 | 7.73E-15 | 0.30 | 0.21 | 0.09 | 1.43 |
| 2 | Nkd2 | 0.41 | 3.74E-19 | 7.77E-15 | 0.20 | 0.12 | 0.08 | 1.62 |
| 2 | Med13 | 0.28 | 4.08E-19 | 8.48E-15 | 0.67 | 0.54 | 0.13 | 1.23 |
| 2 | Pelp1 | 0.29 | 4.13E-19 | 8.58E-15 | 0.63 | 0.50 | 0.13 | 1.25 |
| 2 | Morc4 | 0.58 | 4.34E-19 | 9.02E-15 | 0.22 | 0.14 | 0.08 | 1.55 |
| 2 | Bcl7b | 0.28 | 4.39E-19 | 9.12E-15 | 0.51 | 0.39 | 0.12 | 1.30 |
| 2 | Map2k6 | 0.25 | 4.59E-19 | 9.56E-15 | 0.56 | 0.42 | 0.14 | 1.32 |
| 2 | Ddx19a | 0.30 | 5.59E-19 | 1.16E-14 | 0.56 | 0.44 | 0.12 | 1.27 |
| 2 | Fmnl3 | 0.34 | 6.63E-19 | 1.38E-14 | 0.30 | 0.20 | 0.10 | 1.49 |
| 2 | Gtf2f2 | 0.32 | 7.31E-19 | 1.52E-14 | 0.50 | 0.38 | 0.12 | 1.31 |
| 2 | Pidd1 | 0.63 | 7.84E-19 | 1.63E-14 | 0.19 | 0.12 | 0.07 | 1.60 |
| 2 | Bnip3 | 0.25 | 7.95E-19 | 1.65E-14 | 0.53 | 0.41 | 0.12 | 1.30 |
| 2 | Nop9 | 0.30 | 8.06E-19 | 1.68E-14 | 0.49 | 0.37 | 0.12 | 1.33 |
| 2 | Zfp647 | 0.38 | 8.29E-19 | 1.72E-14 | 0.36 | 0.25 | 0.10 | 1.41 |
| 2 | Fgfrl1 | 0.35 | 8.60E-19 | 1.79E-14 | 0.44 | 0.33 | 0.11 | 1.32 |
| 2 | Prkrip1 | 0.27 | 9.15E-19 | 1.90E-14 | 0.60 | 0.48 | 0.12 | 1.25 |
| 2 | Stn1 | 0.28 | 9.60E-19 | 2.00E-14 | 0.47 | 0.35 | 0.12 | 1.35 |
| 2 | Sp1 | 0.32 | 9.70E-19 | 2.02E-14 | 0.52 | 0.40 | 0.12 | 1.29 |
| 2 | Dennd5a | 0.29 | 1.05E-18 | 2.19E-14 | 0.45 | 0.34 | 0.11 | 1.34 |
| 2 | Nrf1 | 0.29 | 1.07E-18 | 2.23E-14 | 0.56 | 0.44 | 0.12 | 1.28 |
| 2 | Cdc23 | 0.26 | 1.20E-18 | 2.49E-14 | 0.68 | 0.55 | 0.13 | 1.23 |
| 2 | Haus4 | 0.38 | 1.23E-18 | 2.55E-14 | 0.41 | 0.31 | 0.10 | 1.33 |
| 2 | Zgrf1 | 0.44 | 1.36E-18 | 2.82E-14 | 0.29 | 0.20 | 0.09 | 1.45 |
| 2 | Haus7 | 0.27 | 1.43E-18 | 2.97E-14 | 0.58 | 0.45 | 0.13 | 1.28 |
| 2 | Tmem39b | 0.27 | 1.66E-18 | 3.46E-14 | 0.47 | 0.35 | 0.12 | 1.35 |
| 2 | Jcad | 0.77 | 1.67E-18 | 3.48E-14 | 0.10 | 0.05 | 0.05 | 1.92 |
| 2 | Rdh11 | 0.40 | 1.68E-18 | 3.50E-14 | 0.30 | 0.21 | 0.09 | 1.44 |
| 2 | Nop2 | 0.32 | 1.69E-18 | 3.51E-14 | 0.43 | 0.32 | 0.11 | 1.35 |
| 2 | Mdm1 | 0.27 | 1.71E-18 | 3.55E-14 | 0.40 | 0.29 | 0.11 | 1.37 |
| 2 | Fsd1l | 0.36 | 1.78E-18 | 3.71E-14 | 0.37 | 0.27 | 0.10 | 1.39 |
| 2 | Gcn1 | 0.28 | 1.84E-18 | 3.82E-14 | 0.62 | 0.50 | 0.12 | 1.25 |
| 2 | Kdm3b | 0.25 | 1.84E-18 | 3.84E-14 | 0.56 | 0.44 | 0.12 | 1.29 |
| 2 | Xrcc3 | 0.55 | 1.89E-18 | 3.92E-14 | 0.16 | 0.10 | 0.07 | 1.68 |
| 2 | Wipf1 | 0.34 | 1.97E-18 | 4.10E-14 | 0.32 | 0.22 | 0.10 | 1.44 |
| 2 | Srfbp1 | 0.32 | 2.04E-18 | 4.25E-14 | 0.50 | 0.38 | 0.12 | 1.31 |

**Supplementary Table 1. Marker genes for the original clusters of integrated clustering of E11.5 control and NotchGOF λ-junction cells.**

| | | | | | | | | |
|---|---|---|---|---|---|---|---|---|
| 2 | Btaf1 | 0.34 | 2.14E-18 | 4.45E-14 | 0.46 | 0.35 | 0.11 | 1.30 |
| 2 | Clock | 0.28 | 2.20E-18 | 4.58E-14 | 0.42 | 0.30 | 0.11 | 1.38 |
| 2 | Troap | 0.46 | 2.43E-18 | 5.06E-14 | 0.26 | 0.18 | 0.08 | 1.47 |
| 2 | Tmem107 | 0.36 | 2.48E-18 | 5.16E-14 | 0.38 | 0.28 | 0.10 | 1.35 |
| 2 | Cpsf4 | 0.33 | 2.71E-18 | 5.64E-14 | 0.49 | 0.38 | 0.11 | 1.29 |
| 2 | Evc | 0.44 | 2.72E-18 | 5.66E-14 | 0.22 | 0.15 | 0.08 | 1.54 |
| 2 | Camk2n2 | 0.69 | 2.74E-18 | 5.70E-14 | 0.15 | 0.09 | 0.06 | 1.69 |
| 2 | Kif24 | 0.68 | 2.79E-18 | 5.80E-14 | 0.16 | 0.09 | 0.06 | 1.68 |
| 2 | Taf6 | 0.26 | 2.90E-18 | 6.03E-14 | 0.62 | 0.50 | 0.12 | 1.25 |
| 2 | Surf6 | 0.33 | 3.17E-18 | 6.60E-14 | 0.37 | 0.27 | 0.10 | 1.39 |
| 2 | Ahi1 | 0.27 | 3.30E-18 | 6.86E-14 | 0.53 | 0.41 | 0.12 | 1.30 |
| 2 | Gm49797 | 0.35 | 3.37E-18 | 7.00E-14 | 0.33 | 0.23 | 0.10 | 1.42 |
| 2 | Dipk1a | 0.37 | 3.39E-18 | 7.05E-14 | 0.29 | 0.20 | 0.09 | 1.46 |
| 2 | Gsdme | 0.38 | 3.56E-18 | 7.40E-14 | 0.33 | 0.23 | 0.10 | 1.41 |
| 2 | Sclt1 | 0.29 | 3.97E-18 | 8.26E-14 | 0.53 | 0.41 | 0.11 | 1.27 |
| 2 | Dhodh | 0.32 | 4.32E-18 | 8.99E-14 | 0.39 | 0.28 | 0.11 | 1.38 |
| 2 | Slc9a5 | 0.83 | 4.45E-18 | 9.25E-14 | 0.10 | 0.05 | 0.05 | 1.91 |
| 2 | Ccdc138 | 0.47 | 4.64E-18 | 9.64E-14 | 0.27 | 0.19 | 0.09 | 1.46 |
| 2 | Scaf1 | 0.31 | 4.67E-18 | 9.70E-14 | 0.51 | 0.40 | 0.11 | 1.28 |
| 2 | Rps6ka5 | 0.52 | 4.93E-18 | 1.02E-13 | 0.20 | 0.13 | 0.07 | 1.58 |
| 2 | Zfp566 | 0.32 | 5.06E-18 | 1.05E-13 | 0.29 | 0.19 | 0.09 | 1.48 |
| 2 | Plekhg2 | 0.30 | 5.21E-18 | 1.08E-13 | 0.49 | 0.37 | 0.12 | 1.31 |
| 2 | Apex2 | 0.35 | 5.28E-18 | 1.10E-13 | 0.32 | 0.23 | 0.09 | 1.41 |
| 2 | Nup93 | 0.26 | 5.89E-18 | 1.23E-13 | 0.59 | 0.46 | 0.13 | 1.27 |
| 2 | Pias2 | 0.26 | 5.95E-18 | 1.24E-13 | 0.51 | 0.39 | 0.12 | 1.32 |
| 2 | Entpd6 | 0.50 | 6.39E-18 | 1.33E-13 | 0.22 | 0.14 | 0.08 | 1.54 |
| 2 | Raet1e | 0.35 | 6.53E-18 | 1.36E-13 | 0.32 | 0.22 | 0.10 | 1.44 |
| 2 | Clcf1 | 0.64 | 6.98E-18 | 1.45E-13 | 0.10 | 0.05 | 0.05 | 1.91 |
| 2 | Asrgl1 | 0.31 | 7.15E-18 | 1.49E-13 | 0.42 | 0.31 | 0.11 | 1.34 |
| 2 | Atad3a | 0.26 | 7.17E-18 | 1.49E-13 | 0.69 | 0.56 | 0.13 | 1.23 |
| 2 | Snx27 | 0.25 | 7.23E-18 | 1.50E-13 | 0.59 | 0.47 | 0.13 | 1.27 |
| 2 | Zfp521 | 0.32 | 7.23E-18 | 1.50E-13 | 0.39 | 0.28 | 0.11 | 1.38 |
| 2 | S100pbp | 0.33 | 7.90E-18 | 1.64E-13 | 0.40 | 0.30 | 0.10 | 1.35 |
| 2 | Tsix | 0.34 | 8.34E-18 | 1.74E-13 | 0.48 | 0.36 | 0.11 | 1.31 |
| 2 | Unk | 0.40 | 8.68E-18 | 1.80E-13 | 0.29 | 0.20 | 0.09 | 1.44 |
| 2 | Sertad3 | 0.36 | 9.58E-18 | 1.99E-13 | 0.33 | 0.24 | 0.09 | 1.40 |
| 2 | Selenoi | 0.30 | 9.67E-18 | 2.01E-13 | 0.44 | 0.33 | 0.11 | 1.34 |
| 2 | Cnpy4 | 0.30 | 9.71E-18 | 2.02E-13 | 0.45 | 0.34 | 0.12 | 1.34 |
| 2 | Pcgf2 | 0.27 | 9.94E-18 | 2.07E-13 | 0.50 | 0.38 | 0.12 | 1.33 |
| 2 | Sec23b | 0.31 | 1.02E-17 | 2.12E-13 | 0.47 | 0.36 | 0.11 | 1.30 |
| 2 | Ado | 0.26 | 1.13E-17 | 2.34E-13 | 0.50 | 0.38 | 0.12 | 1.30 |
| 2 | Pmp22 | 0.36 | 1.15E-17 | 2.39E-13 | 0.26 | 0.18 | 0.09 | 1.49 |
| 2 | Tedc2 | 0.43 | 1.20E-17 | 2.50E-13 | 0.25 | 0.17 | 0.08 | 1.47 |
| 2 | Itfg2 | 0.30 | 1.20E-17 | 2.50E-13 | 0.44 | 0.33 | 0.11 | 1.33 |
| 2 | Slc35e1 | 0.48 | 1.28E-17 | 2.66E-13 | 0.24 | 0.16 | 0.08 | 1.49 |
| 2 | Recql4 | 0.71 | 1.47E-17 | 3.06E-13 | 0.14 | 0.08 | 0.06 | 1.69 |
| 2 | Aco1 | 0.28 | 1.48E-17 | 3.08E-13 | 0.44 | 0.32 | 0.11 | 1.35 |
| 2 | Morc3 | 0.27 | 1.49E-17 | 3.10E-13 | 0.53 | 0.41 | 0.12 | 1.28 |
| 2 | Tpmt | 0.28 | 1.54E-17 | 3.20E-13 | 0.48 | 0.36 | 0.12 | 1.32 |
| 2 | Etv6 | 0.32 | 1.54E-17 | 3.21E-13 | 0.38 | 0.27 | 0.10 | 1.38 |
| 2 | Stambpl1 | 0.64 | 1.59E-17 | 3.30E-13 | 0.12 | 0.06 | 0.05 | 1.83 |
| 2 | Telo2 | 0.33 | 1.66E-17 | 3.45E-13 | 0.39 | 0.29 | 0.10 | 1.36 |
| 2 | Pdzd4 | 0.48 | 1.74E-17 | 3.62E-13 | 0.22 | 0.14 | 0.08 | 1.54 |
| 2 | Mast2 | 0.38 | 1.81E-17 | 3.77E-13 | 0.39 | 0.28 | 0.10 | 1.36 |
| 2 | Clpx | 0.27 | 1.84E-17 | 3.84E-13 | 0.59 | 0.46 | 0.12 | 1.27 |
| 2 | Plag1 | 0.36 | 1.88E-17 | 3.90E-13 | 0.27 | 0.18 | 0.09 | 1.46 |

**Supplementary Table 1. Marker genes for the original clusters of integrated clustering of E11.5 control and NotchGOF λ-junction cells.**

| | | | | | | | | |
|---|---|---|---|---|---|---|---|---|
| 2 | Poglut2 | 0.29 | 1.93E-17 | 4.02E-13 | 0.50 | 0.38 | 0.12 | 1.31 |
| 2 | Cops7b | 0.28 | 2.03E-17 | 4.21E-13 | 0.46 | 0.35 | 0.11 | 1.32 |
| 2 | Ltn1 | 0.27 | 2.21E-17 | 4.60E-13 | 0.43 | 0.32 | 0.11 | 1.34 |
| 2 | Actr6 | 0.28 | 2.22E-17 | 4.62E-13 | 0.50 | 0.38 | 0.12 | 1.30 |
| 2 | Pdss1 | 0.50 | 2.23E-17 | 4.63E-13 | 0.23 | 0.16 | 0.08 | 1.48 |
| 2 | Ccbe1 | 0.82 | 2.23E-17 | 4.65E-13 | 0.10 | 0.05 | 0.05 | 1.89 |
| 2 | Nek2 | 0.30 | 2.27E-17 | 4.72E-13 | 0.32 | 0.23 | 0.09 | 1.39 |
| 2 | Fgfr1op | 0.29 | 2.49E-17 | 5.19E-13 | 0.50 | 0.39 | 0.11 | 1.29 |
| 2 | 2610307P16Rik | 0.39 | 2.65E-17 | 5.52E-13 | 0.25 | 0.17 | 0.08 | 1.49 |
| 2 | Marveld1 | 0.29 | 2.72E-17 | 5.65E-13 | 0.33 | 0.23 | 0.10 | 1.44 |
| 2 | Samd4 | 0.26 | 2.84E-17 | 5.91E-13 | 0.48 | 0.36 | 0.12 | 1.34 |
| 2 | Decr1 | 0.26 | 3.05E-17 | 6.35E-13 | 0.52 | 0.40 | 0.12 | 1.30 |
| 2 | Ankrd29 | 0.54 | 3.29E-17 | 6.84E-13 | 0.15 | 0.09 | 0.06 | 1.70 |
| 2 | Mras | 0.35 | 3.32E-17 | 6.90E-13 | 0.29 | 0.20 | 0.09 | 1.45 |
| 2 | Usp42 | 0.30 | 3.45E-17 | 7.18E-13 | 0.46 | 0.35 | 0.11 | 1.32 |
| 2 | Map4 | 0.25 | 3.50E-17 | 7.28E-13 | 0.51 | 0.39 | 0.12 | 1.31 |
| 2 | Tada2a | 0.27 | 4.03E-17 | 8.37E-13 | 0.53 | 0.41 | 0.12 | 1.30 |
| 2 | Mmp16 | 0.27 | 4.10E-17 | 8.52E-13 | 0.33 | 0.23 | 0.10 | 1.42 |
| 2 | Mapk7 | 0.30 | 4.15E-17 | 8.64E-13 | 0.44 | 0.33 | 0.11 | 1.32 |
| 2 | Dync1i1 | 0.30 | 4.18E-17 | 8.69E-13 | 0.11 | 0.06 | 0.05 | 1.85 |
| 2 | Capn15 | 0.31 | 4.38E-17 | 9.12E-13 | 0.48 | 0.37 | 0.11 | 1.30 |
| 2 | Usp13 | 0.68 | 4.73E-17 | 9.84E-13 | 0.12 | 0.07 | 0.05 | 1.79 |
| 2 | Grwd1 | 0.29 | 4.80E-17 | 9.99E-13 | 0.54 | 0.42 | 0.12 | 1.28 |
| 2 | Zfp410 | 0.27 | 4.94E-17 | 1.03E-12 | 0.53 | 0.41 | 0.12 | 1.29 |
| 2 | Mcc | 0.44 | 4.94E-17 | 1.03E-12 | 0.20 | 0.12 | 0.07 | 1.58 |
| 2 | Agap1 | 0.25 | 4.95E-17 | 1.03E-12 | 0.53 | 0.41 | 0.12 | 1.29 |
| 2 | Clasp2 | 0.40 | 5.22E-17 | 1.09E-12 | 0.32 | 0.23 | 0.09 | 1.38 |
| 2 | Lrrc40 | 0.31 | 5.46E-17 | 1.14E-12 | 0.43 | 0.32 | 0.10 | 1.32 |
| 2 | Kctd12b | 0.28 | 5.54E-17 | 1.15E-12 | 0.26 | 0.17 | 0.09 | 1.50 |
| 2 | Stam | 0.31 | 5.71E-17 | 1.19E-12 | 0.39 | 0.29 | 0.10 | 1.36 |
| 2 | Dgcr8 | 0.29 | 6.14E-17 | 1.28E-12 | 0.47 | 0.36 | 0.11 | 1.30 |
| 2 | Mepce | 0.33 | 6.16E-17 | 1.28E-12 | 0.43 | 0.33 | 0.10 | 1.31 |
| 2 | Slc12a7 | 0.40 | 6.56E-17 | 1.37E-12 | 0.31 | 0.22 | 0.09 | 1.39 |
| 2 | Ntn3 | 0.41 | 6.57E-17 | 1.37E-12 | 0.23 | 0.15 | 0.08 | 1.51 |
| 2 | Ncoa7 | 0.25 | 6.71E-17 | 1.40E-12 | 0.52 | 0.39 | 0.12 | 1.31 |
| 2 | Zmym1 | 0.31 | 7.09E-17 | 1.48E-12 | 0.43 | 0.33 | 0.11 | 1.32 |
| 2 | Necap1 | 0.35 | 7.71E-17 | 1.60E-12 | 0.33 | 0.23 | 0.09 | 1.40 |
| 2 | Cited1 | 0.49 | 7.81E-17 | 1.62E-12 | 0.18 | 0.11 | 0.07 | 1.60 |
| 2 | Naa40 | 0.28 | 7.88E-17 | 1.64E-12 | 0.47 | 0.36 | 0.11 | 1.31 |
| 2 | Cep85l | 0.43 | 7.90E-17 | 1.64E-12 | 0.20 | 0.13 | 0.07 | 1.55 |
| 2 | Saal1 | 0.29 | 8.13E-17 | 1.69E-12 | 0.57 | 0.45 | 0.12 | 1.26 |
| 2 | Hsf1 | 0.30 | 8.25E-17 | 1.72E-12 | 0.44 | 0.34 | 0.11 | 1.31 |
| 2 | Ankmy2 | 0.31 | 8.55E-17 | 1.78E-12 | 0.38 | 0.28 | 0.10 | 1.38 |
| 2 | Phf13 | 0.36 | 9.38E-17 | 1.95E-12 | 0.36 | 0.26 | 0.10 | 1.37 |
| 2 | Lrch2 | 0.33 | 9.53E-17 | 1.98E-12 | 0.37 | 0.27 | 0.10 | 1.37 |
| 2 | Nr4a2 | 0.59 | 9.64E-17 | 2.00E-12 | 0.16 | 0.10 | 0.06 | 1.63 |
| 2 | Smyd3 | 0.37 | 9.90E-17 | 2.06E-12 | 0.32 | 0.23 | 0.09 | 1.40 |
| 2 | Aard | 0.55 | 1.04E-16 | 2.16E-12 | 0.12 | 0.07 | 0.05 | 1.79 |
| 2 | Large1 | 0.40 | 1.12E-16 | 2.33E-12 | 0.27 | 0.19 | 0.08 | 1.44 |
| 2 | Alms1 | 0.37 | 1.16E-16 | 2.41E-12 | 0.35 | 0.25 | 0.09 | 1.37 |
| 2 | Mypop | 0.44 | 1.20E-16 | 2.50E-12 | 0.17 | 0.11 | 0.07 | 1.61 |
| 2 | Vps4a | 0.27 | 1.24E-16 | 2.57E-12 | 0.61 | 0.50 | 0.11 | 1.23 |
| 2 | Dusp6 | 0.31 | 1.29E-16 | 2.68E-12 | 0.46 | 0.35 | 0.10 | 1.29 |
| 2 | Ciz1 | 0.31 | 1.30E-16 | 2.71E-12 | 0.44 | 0.33 | 0.10 | 1.31 |
| 2 | St8sia4 | 0.52 | 1.35E-16 | 2.81E-12 | 0.15 | 0.09 | 0.06 | 1.68 |
| 2 | Qtrt2 | 0.31 | 1.37E-16 | 2.85E-12 | 0.34 | 0.24 | 0.10 | 1.40 |

**Supplementary Table 1. Marker genes for the original clusters of integrated clustering of E11.5 control and NotchGOF λ-junction cells.**

| | | | | | | | | |
|---|---|---|---|---|---|---|---|---|
| 2 | Ints6l | 0.42 | 1.46E-16 | 3.03E-12 | 0.24 | 0.17 | 0.08 | 1.47 |
| 2 | Fto | 0.27 | 1.53E-16 | 3.17E-12 | 0.58 | 0.46 | 0.12 | 1.26 |
| 2 | Slc4a1ap | 0.31 | 1.55E-16 | 3.23E-12 | 0.48 | 0.37 | 0.11 | 1.29 |
| 2 | Nbn | 0.33 | 1.56E-16 | 3.24E-12 | 0.43 | 0.33 | 0.10 | 1.32 |
| 2 | Nid2 | 0.29 | 1.73E-16 | 3.61E-12 | 0.33 | 0.23 | 0.10 | 1.43 |
| 2 | Zfp41 | 0.32 | 1.80E-16 | 3.75E-12 | 0.44 | 0.34 | 0.10 | 1.30 |
| 2 | Jpt2 | 0.27 | 1.97E-16 | 4.11E-12 | 0.55 | 0.44 | 0.12 | 1.26 |
| 2 | Lrp12 | 0.35 | 2.12E-16 | 4.42E-12 | 0.28 | 0.19 | 0.09 | 1.44 |
| 2 | Edrf1 | 0.27 | 2.26E-16 | 4.70E-12 | 0.38 | 0.28 | 0.10 | 1.36 |
| 2 | Nacc1 | 0.26 | 2.37E-16 | 4.93E-12 | 0.50 | 0.38 | 0.11 | 1.30 |
| 2 | Ppm1m | 0.34 | 2.37E-16 | 4.93E-12 | 0.33 | 0.24 | 0.10 | 1.40 |
| 2 | Xpo4 | 0.29 | 2.66E-16 | 5.53E-12 | 0.54 | 0.43 | 0.11 | 1.26 |
| 2 | Gk | 0.74 | 3.03E-16 | 6.30E-12 | 0.13 | 0.07 | 0.05 | 1.71 |
| 2 | Naa16 | 0.30 | 3.12E-16 | 6.49E-12 | 0.48 | 0.37 | 0.11 | 1.29 |
| 2 | Tex10 | 0.26 | 3.38E-16 | 7.03E-12 | 0.56 | 0.44 | 0.12 | 1.26 |
| 2 | Mars | 0.29 | 3.43E-16 | 7.14E-12 | 0.46 | 0.35 | 0.11 | 1.30 |
| 2 | Setd1b | 0.29 | 3.52E-16 | 7.32E-12 | 0.53 | 0.42 | 0.11 | 1.27 |
| 2 | Smyd2 | 0.28 | 3.54E-16 | 7.37E-12 | 0.51 | 0.40 | 0.11 | 1.27 |
| 2 | Nelfa | 0.26 | 4.20E-16 | 8.74E-12 | 0.65 | 0.53 | 0.12 | 1.22 |
| 2 | Ccdc9 | 0.29 | 4.74E-16 | 9.86E-12 | 0.39 | 0.29 | 0.10 | 1.34 |
| 2 | St3gal3 | 0.50 | 5.14E-16 | 1.07E-11 | 0.17 | 0.11 | 0.06 | 1.58 |
| 2 | Sugp2 | 0.35 | 5.22E-16 | 1.08E-11 | 0.27 | 0.19 | 0.08 | 1.43 |
| 2 | Kat7 | 0.28 | 5.40E-16 | 1.12E-11 | 0.50 | 0.39 | 0.11 | 1.29 |
| 2 | Cherp | 0.27 | 5.43E-16 | 1.13E-11 | 0.54 | 0.42 | 0.12 | 1.27 |
| 2 | Smoc1 | 0.71 | 5.54E-16 | 1.15E-11 | 0.13 | 0.07 | 0.05 | 1.71 |
| 2 | Cep250 | 0.32 | 5.68E-16 | 1.18E-11 | 0.35 | 0.26 | 0.09 | 1.37 |
| 2 | Tmem43 | 0.41 | 5.98E-16 | 1.24E-11 | 0.29 | 0.20 | 0.08 | 1.40 |
| 2 | 1700021F05Rik | 0.26 | 6.71E-16 | 1.40E-11 | 0.57 | 0.45 | 0.12 | 1.26 |
| 2 | Arpin | 0.43 | 7.10E-16 | 1.48E-11 | 0.25 | 0.17 | 0.08 | 1.46 |
| 2 | Tubgcp3 | 0.28 | 7.24E-16 | 1.51E-11 | 0.42 | 0.31 | 0.11 | 1.34 |
| 2 | Tbp | 0.31 | 8.63E-16 | 1.79E-11 | 0.33 | 0.24 | 0.09 | 1.38 |
| 2 | Foxn2 | 0.49 | 8.68E-16 | 1.80E-11 | 0.23 | 0.16 | 0.07 | 1.46 |
| 2 | Nudcd1 | 0.25 | 9.23E-16 | 1.92E-11 | 0.51 | 0.40 | 0.11 | 1.28 |
| 2 | Trerf1 | 0.30 | 9.63E-16 | 2.00E-11 | 0.36 | 0.26 | 0.10 | 1.38 |
| 2 | Zfp609 | 0.25 | 1.00E-15 | 2.09E-11 | 0.53 | 0.42 | 0.12 | 1.28 |
| 2 | Cluh | 0.37 | 1.06E-15 | 2.20E-11 | 0.30 | 0.22 | 0.08 | 1.38 |
| 2 | Simc1 | 0.47 | 1.15E-15 | 2.40E-11 | 0.20 | 0.13 | 0.07 | 1.53 |
| 2 | Srgap2 | 0.26 | 1.35E-15 | 2.80E-11 | 0.46 | 0.35 | 0.11 | 1.30 |
| 2 | Gm3830 | 0.69 | 1.41E-15 | 2.93E-11 | 0.11 | 0.06 | 0.05 | 1.80 |
| 2 | Pias3 | 0.34 | 1.43E-15 | 2.98E-11 | 0.28 | 0.20 | 0.08 | 1.42 |
| 2 | Klhl12 | 0.32 | 1.50E-15 | 3.12E-11 | 0.36 | 0.27 | 0.10 | 1.36 |
| 2 | Car13 | 0.54 | 1.52E-15 | 3.15E-11 | 0.15 | 0.09 | 0.06 | 1.63 |
| 2 | Col4a3bp | 0.28 | 1.57E-15 | 3.27E-11 | 0.41 | 0.31 | 0.10 | 1.32 |
| 2 | Gas2 | 0.36 | 1.60E-15 | 3.33E-11 | 0.34 | 0.25 | 0.09 | 1.35 |
| 2 | Rnf165 | 0.29 | 2.22E-15 | 4.61E-11 | 0.33 | 0.24 | 0.09 | 1.38 |
| 2 | Yes1 | 0.25 | 2.28E-15 | 4.75E-11 | 0.45 | 0.34 | 0.11 | 1.32 |
| 2 | Polr3f | 0.25 | 2.36E-15 | 4.92E-11 | 0.44 | 0.33 | 0.11 | 1.33 |
| 2 | Zbtb45 | 0.31 | 2.37E-15 | 4.92E-11 | 0.30 | 0.21 | 0.09 | 1.40 |
| 2 | Vash2 | 0.26 | 2.45E-15 | 5.09E-11 | 0.38 | 0.28 | 0.10 | 1.36 |
| 2 | Sufu | 0.35 | 2.53E-15 | 5.26E-11 | 0.30 | 0.22 | 0.09 | 1.39 |
| 2 | Epdr1 | 0.57 | 2.64E-15 | 5.48E-11 | 0.16 | 0.10 | 0.06 | 1.58 |
| 2 | Dnajc14 | 0.28 | 2.68E-15 | 5.58E-11 | 0.40 | 0.30 | 0.10 | 1.33 |
| 2 | Mboat7 | 0.25 | 2.86E-15 | 5.95E-11 | 0.43 | 0.32 | 0.11 | 1.33 |
| 2 | Tbc1d1 | 0.29 | 3.06E-15 | 6.37E-11 | 0.47 | 0.36 | 0.11 | 1.29 |
| 2 | Ghr | 0.33 | 3.28E-15 | 6.83E-11 | 0.21 | 0.13 | 0.07 | 1.53 |
| 2 | Rnf150 | 0.26 | 3.51E-15 | 7.29E-11 | 0.40 | 0.30 | 0.10 | 1.34 |

**Supplementary Table 1. Marker genes for the original clusters of integrated clustering of E11.5 control and NotchGOF λ-junction cells.**

| | | | | | | | | |
|---|---|---|---|---|---|---|---|---|
| 2 | Zfp628 | 0.52 | 3.57E-15 | 7.43E-11 | 0.16 | 0.10 | 0.06 | 1.61 |
| 2 | Fbxl5 | 0.26 | 4.17E-15 | 8.67E-11 | 0.46 | 0.36 | 0.10 | 1.29 |
| 2 | Bmpr1b | 0.52 | 4.19E-15 | 8.72E-11 | 0.14 | 0.08 | 0.05 | 1.65 |
| 2 | Vkorc1l1 | 0.27 | 4.35E-15 | 9.05E-11 | 0.50 | 0.39 | 0.11 | 1.27 |
| 2 | Cchcr1 | 0.43 | 4.54E-15 | 9.44E-11 | 0.21 | 0.14 | 0.07 | 1.49 |
| 2 | Wdr20 | 0.33 | 4.61E-15 | 9.58E-11 | 0.37 | 0.28 | 0.09 | 1.33 |
| 2 | Elac2 | 0.32 | 4.70E-15 | 9.78E-11 | 0.31 | 0.22 | 0.09 | 1.38 |
| 2 | Rmdn3 | 0.33 | 4.71E-15 | 9.79E-11 | 0.35 | 0.26 | 0.09 | 1.34 |
| 2 | Polr1a | 0.29 | 4.79E-15 | 9.97E-11 | 0.45 | 0.35 | 0.10 | 1.29 |
| 2 | Nfatc3 | 0.27 | 5.11E-15 | 1.06E-10 | 0.46 | 0.35 | 0.11 | 1.30 |
| 2 | Tbc1d19 | 0.29 | 5.21E-15 | 1.08E-10 | 0.29 | 0.21 | 0.08 | 1.41 |
| 2 | Arhgap33 | 0.53 | 5.44E-15 | 1.13E-10 | 0.19 | 0.13 | 0.07 | 1.50 |
| 2 | Fat3 | 0.46 | 6.28E-15 | 1.31E-10 | 0.14 | 0.09 | 0.06 | 1.64 |
| 2 | 4930452B06Rik | 0.47 | 6.50E-15 | 1.35E-10 | 0.18 | 0.12 | 0.06 | 1.53 |
| 2 | Npat | 0.30 | 6.64E-15 | 1.38E-10 | 0.43 | 0.34 | 0.10 | 1.28 |
| 2 | Med23 | 0.28 | 6.68E-15 | 1.39E-10 | 0.36 | 0.26 | 0.09 | 1.36 |
| 2 | Dtnbp1 | 0.26 | 6.87E-15 | 1.43E-10 | 0.52 | 0.41 | 0.11 | 1.27 |
| 2 | Usp51 | 0.67 | 6.87E-15 | 1.43E-10 | 0.13 | 0.08 | 0.05 | 1.65 |
| 2 | Cenpo | 0.54 | 6.90E-15 | 1.43E-10 | 0.19 | 0.13 | 0.06 | 1.50 |
| 2 | Zfp574 | 0.29 | 7.34E-15 | 1.53E-10 | 0.28 | 0.20 | 0.08 | 1.41 |
| 2 | Nploc4 | 0.33 | 7.59E-15 | 1.58E-10 | 0.32 | 0.23 | 0.09 | 1.37 |
| 2 | Brpf1 | 0.28 | 7.63E-15 | 1.59E-10 | 0.41 | 0.31 | 0.10 | 1.31 |
| 2 | Zbtb14 | 0.36 | 8.90E-15 | 1.85E-10 | 0.31 | 0.22 | 0.08 | 1.37 |
| 2 | Bbs7 | 0.39 | 9.29E-15 | 1.93E-10 | 0.23 | 0.16 | 0.07 | 1.46 |
| 2 | Crem | 0.33 | 9.39E-15 | 1.95E-10 | 0.25 | 0.17 | 0.08 | 1.45 |
| 2 | Zfp711 | 0.32 | 9.63E-15 | 2.00E-10 | 0.29 | 0.21 | 0.08 | 1.40 |
| 2 | Ints1 | 0.31 | 9.90E-15 | 2.06E-10 | 0.37 | 0.28 | 0.09 | 1.33 |
| 2 | Zfp689 | 0.50 | 9.92E-15 | 2.06E-10 | 0.15 | 0.09 | 0.06 | 1.62 |
| 2 | Shroom4 | 0.57 | 1.03E-14 | 2.15E-10 | 0.14 | 0.08 | 0.05 | 1.63 |
| 2 | Pank3 | 0.26 | 1.04E-14 | 2.16E-10 | 0.48 | 0.37 | 0.10 | 1.28 |
| 2 | Gm16536 | 0.39 | 1.08E-14 | 2.24E-10 | 0.22 | 0.15 | 0.07 | 1.47 |
| 2 | Tnfrsf21 | 0.50 | 1.10E-14 | 2.30E-10 | 0.20 | 0.14 | 0.07 | 1.47 |
| 2 | Rab35 | 0.26 | 1.12E-14 | 2.33E-10 | 0.46 | 0.36 | 0.11 | 1.29 |
| 2 | 5031439G07Rik | 0.31 | 1.15E-14 | 2.40E-10 | 0.33 | 0.24 | 0.09 | 1.36 |
| 2 | Emp2 | 0.63 | 1.20E-14 | 2.50E-10 | 0.12 | 0.07 | 0.05 | 1.70 |
| 2 | Ddx11 | 0.34 | 1.27E-14 | 2.63E-10 | 0.33 | 0.24 | 0.09 | 1.35 |
| 2 | Dcp1a | 0.33 | 1.28E-14 | 2.65E-10 | 0.26 | 0.19 | 0.08 | 1.42 |
| 2 | Cep120 | 0.29 | 1.29E-14 | 2.68E-10 | 0.32 | 0.23 | 0.09 | 1.37 |
| 2 | Sgf29 | 0.31 | 1.35E-14 | 2.81E-10 | 0.41 | 0.32 | 0.09 | 1.27 |
| 2 | Tmem129 | 0.38 | 1.45E-14 | 3.02E-10 | 0.24 | 0.17 | 0.07 | 1.44 |
| 2 | Kif14 | 0.40 | 1.47E-14 | 3.06E-10 | 0.26 | 0.18 | 0.07 | 1.40 |
| 2 | Scmh1 | 0.26 | 1.64E-14 | 3.42E-10 | 0.41 | 0.31 | 0.10 | 1.32 |
| 2 | 2900052L18Rik | 0.38 | 1.68E-14 | 3.49E-10 | 0.22 | 0.15 | 0.07 | 1.47 |
| 2 | Zscan2 | 0.28 | 1.79E-14 | 3.72E-10 | 0.26 | 0.18 | 0.08 | 1.44 |
| 2 | 9930104L06Rik | 0.43 | 1.83E-14 | 3.80E-10 | 0.22 | 0.15 | 0.07 | 1.47 |
| 2 | Tsga10 | 0.51 | 2.05E-14 | 4.26E-10 | 0.15 | 0.10 | 0.06 | 1.60 |
| 2 | Trub2 | 0.26 | 2.40E-14 | 4.98E-10 | 0.40 | 0.30 | 0.10 | 1.34 |
| 2 | Cep131 | 0.30 | 2.51E-14 | 5.22E-10 | 0.33 | 0.24 | 0.09 | 1.36 |
| 2 | Hic2 | 0.27 | 2.73E-14 | 5.68E-10 | 0.38 | 0.29 | 0.09 | 1.32 |
| 2 | Ttbk2 | 0.35 | 2.74E-14 | 5.69E-10 | 0.25 | 0.17 | 0.08 | 1.43 |
| 2 | Pank4 | 0.42 | 3.17E-14 | 6.59E-10 | 0.21 | 0.14 | 0.07 | 1.48 |
| 2 | Smyd4 | 0.36 | 3.47E-14 | 7.22E-10 | 0.24 | 0.17 | 0.07 | 1.43 |
| 2 | Ttll5 | 0.43 | 3.51E-14 | 7.31E-10 | 0.20 | 0.14 | 0.07 | 1.48 |
| 2 | Mms19 | 0.25 | 3.60E-14 | 7.49E-10 | 0.35 | 0.26 | 0.09 | 1.35 |
| 2 | Ahcy | 0.28 | 3.61E-14 | 7.52E-10 | 0.36 | 0.26 | 0.09 | 1.34 |
| 2 | Sp2 | 0.44 | 3.87E-14 | 8.06E-10 | 0.23 | 0.17 | 0.07 | 1.42 |

**Supplementary Table 1. Marker genes for the original clusters of integrated clustering of E11.5 control and NotchGOF λ-junction cells.**

| | | | | | | | | |
|---|---|---|---|---|---|---|---|---|
| 2 | Xrcc2 | 0.42 | 4.02E-14 | 8.35E-10 | 0.21 | 0.14 | 0.07 | 1.46 |
| 2 | Tchp | 0.30 | 4.18E-14 | 8.69E-10 | 0.38 | 0.29 | 0.09 | 1.32 |
| 2 | Tmtc2 | 0.37 | 4.51E-14 | 9.37E-10 | 0.27 | 0.19 | 0.08 | 1.40 |
| 2 | Trim12c | 0.35 | 4.81E-14 | 1.00E-09 | 0.25 | 0.17 | 0.07 | 1.43 |
| 2 | Antxr2 | 0.35 | 4.99E-14 | 1.04E-09 | 0.25 | 0.18 | 0.07 | 1.41 |
| 2 | B4galt1 | 0.34 | 5.02E-14 | 1.04E-09 | 0.30 | 0.22 | 0.08 | 1.36 |
| 2 | Micu1 | 0.27 | 5.08E-14 | 1.06E-09 | 0.42 | 0.32 | 0.10 | 1.30 |
| 2 | Dxo | 0.25 | 5.46E-14 | 1.14E-09 | 0.50 | 0.40 | 0.10 | 1.25 |
| 2 | Lrrc49 | 0.29 | 5.80E-14 | 1.21E-09 | 0.33 | 0.24 | 0.09 | 1.36 |
| 2 | Crocc | 0.46 | 5.96E-14 | 1.24E-09 | 0.21 | 0.15 | 0.07 | 1.45 |
| 2 | Fhl3 | 0.53 | 6.05E-14 | 1.26E-09 | 0.16 | 0.10 | 0.06 | 1.56 |
| 2 | Ranbp6 | 0.37 | 6.46E-14 | 1.34E-09 | 0.26 | 0.19 | 0.08 | 1.40 |
| 2 | Tent4a | 0.32 | 6.50E-14 | 1.35E-09 | 0.34 | 0.26 | 0.08 | 1.32 |
| 2 | Esyt2 | 0.35 | 6.51E-14 | 1.35E-09 | 0.29 | 0.21 | 0.08 | 1.38 |
| 2 | Osgin2 | 0.31 | 6.63E-14 | 1.38E-09 | 0.26 | 0.18 | 0.08 | 1.42 |
| 2 | Kat14 | 0.30 | 8.27E-14 | 1.72E-09 | 0.36 | 0.27 | 0.09 | 1.34 |
| 2 | Tubgcp6 | 0.46 | 8.46E-14 | 1.76E-09 | 0.19 | 0.13 | 0.06 | 1.48 |
| 2 | Nepro | 0.37 | 9.03E-14 | 1.88E-09 | 0.21 | 0.14 | 0.07 | 1.48 |
| 2 | Top3a | 0.38 | 9.17E-14 | 1.91E-09 | 0.23 | 0.16 | 0.07 | 1.44 |
| 2 | Bap1 | 0.30 | 1.11E-13 | 2.30E-09 | 0.42 | 0.32 | 0.09 | 1.29 |
| 2 | Snrnp35 | 0.29 | 1.16E-13 | 2.40E-09 | 0.36 | 0.27 | 0.09 | 1.32 |
| 2 | Epb41l4a | 0.50 | 1.25E-13 | 2.60E-09 | 0.16 | 0.10 | 0.06 | 1.55 |
| 2 | Gm10184 | 0.50 | 1.42E-13 | 2.96E-09 | 0.20 | 0.14 | 0.06 | 1.45 |
| 2 | Ttc37 | 0.25 | 1.44E-13 | 3.00E-09 | 0.41 | 0.31 | 0.10 | 1.30 |
| 2 | AU020206 | 0.37 | 1.45E-13 | 3.01E-09 | 0.29 | 0.22 | 0.08 | 1.35 |
| 2 | Map3k3 | 0.39 | 1.81E-13 | 3.77E-09 | 0.22 | 0.15 | 0.07 | 1.46 |
| 2 | Scx | 0.43 | 1.95E-13 | 4.05E-09 | 0.23 | 0.16 | 0.07 | 1.44 |
| 2 | Setdb2 | 0.56 | 2.00E-13 | 4.15E-09 | 0.14 | 0.09 | 0.05 | 1.59 |
| 2 | Coq6 | 0.28 | 2.10E-13 | 4.37E-09 | 0.38 | 0.29 | 0.09 | 1.30 |
| 2 | Ncoa1 | 0.33 | 2.15E-13 | 4.47E-09 | 0.24 | 0.17 | 0.07 | 1.43 |
| 2 | Nsf | 0.26 | 2.17E-13 | 4.50E-09 | 0.43 | 0.33 | 0.10 | 1.29 |
| 2 | Mid1 | 0.31 | 2.23E-13 | 4.64E-09 | 0.32 | 0.24 | 0.08 | 1.35 |
| 2 | E130307A14Rik | 0.32 | 2.27E-13 | 4.73E-09 | 0.27 | 0.19 | 0.08 | 1.41 |
| 2 | Tbl3 | 0.27 | 2.36E-13 | 4.92E-09 | 0.43 | 0.34 | 0.09 | 1.27 |
| 2 | Wrap53 | 0.33 | 2.44E-13 | 5.07E-09 | 0.30 | 0.22 | 0.08 | 1.35 |
| 2 | Ift80 | 0.34 | 2.45E-13 | 5.09E-09 | 0.29 | 0.21 | 0.08 | 1.37 |
| 2 | Zfp28 | 0.52 | 3.00E-13 | 6.25E-09 | 0.13 | 0.08 | 0.05 | 1.64 |
| 2 | Pkd1 | 0.33 | 3.01E-13 | 6.26E-09 | 0.23 | 0.16 | 0.07 | 1.43 |
| 2 | Ppp1r12c | 0.26 | 3.15E-13 | 6.55E-09 | 0.40 | 0.31 | 0.09 | 1.29 |
| 2 | Cnr1 | 0.46 | 3.29E-13 | 6.85E-09 | 0.14 | 0.08 | 0.05 | 1.62 |
| 2 | Spin4 | 0.37 | 3.30E-13 | 6.87E-09 | 0.18 | 0.12 | 0.06 | 1.51 |
| 2 | Panx1 | 0.28 | 3.33E-13 | 6.92E-09 | 0.40 | 0.31 | 0.09 | 1.30 |
| 2 | Rptor | 0.34 | 3.57E-13 | 7.43E-09 | 0.31 | 0.23 | 0.08 | 1.35 |
| 2 | Rmi2 | 0.32 | 3.58E-13 | 7.44E-09 | 0.27 | 0.20 | 0.08 | 1.38 |
| 2 | Spry4 | 0.36 | 4.03E-13 | 8.38E-09 | 0.21 | 0.14 | 0.07 | 1.46 |
| 2 | Ifit2 | 0.56 | 4.06E-13 | 8.44E-09 | 0.14 | 0.09 | 0.05 | 1.60 |
| 2 | Clcn7 | 0.48 | 4.76E-13 | 9.90E-09 | 0.14 | 0.09 | 0.05 | 1.59 |
| 2 | Maml1 | 0.35 | 4.79E-13 | 9.96E-09 | 0.29 | 0.21 | 0.08 | 1.36 |
| 2 | Ankle2 | 0.26 | 4.92E-13 | 1.02E-08 | 0.45 | 0.35 | 0.09 | 1.26 |
| 2 | Manea | 0.26 | 4.99E-13 | 1.04E-08 | 0.31 | 0.23 | 0.08 | 1.36 |
| 2 | Hcfc2 | 0.39 | 5.26E-13 | 1.09E-08 | 0.21 | 0.14 | 0.06 | 1.45 |
| 2 | Lmf2 | 0.37 | 5.59E-13 | 1.16E-08 | 0.25 | 0.18 | 0.07 | 1.39 |
| 2 | Gtf3c5 | 0.31 | 5.82E-13 | 1.21E-08 | 0.28 | 0.21 | 0.08 | 1.37 |
| 2 | Nle1 | 0.29 | 5.91E-13 | 1.23E-08 | 0.31 | 0.23 | 0.08 | 1.35 |
| 2 | Eldr | 0.33 | 5.96E-13 | 1.24E-08 | 0.25 | 0.18 | 0.07 | 1.40 |
| 2 | Pvt1 | 0.51 | 6.12E-13 | 1.27E-08 | 0.12 | 0.07 | 0.05 | 1.64 |

**Supplementary Table 1. Marker genes for the original clusters of integrated clustering of E11.5 control and NotchGOF λ-junction cells.**

| | | | | | | | | |
|---|---|---|---|---|---|---|---|---|
| 2 | Slc25a25 | 0.31 | 6.66E-13 | 1.39E-08 | 0.25 | 0.18 | 0.07 | 1.41 |
| 2 | Atr | 0.27 | 6.71E-13 | 1.39E-08 | 0.42 | 0.33 | 0.09 | 1.29 |
| 2 | Snip1 | 0.26 | 7.05E-13 | 1.47E-08 | 0.42 | 0.33 | 0.09 | 1.29 |
| 2 | Adamts7 | 0.46 | 7.05E-13 | 1.47E-08 | 0.14 | 0.09 | 0.05 | 1.57 |
| 2 | Mbtps2 | 0.29 | 7.44E-13 | 1.55E-08 | 0.29 | 0.22 | 0.08 | 1.36 |
| 2 | Cdkal1 | 0.30 | 7.99E-13 | 1.66E-08 | 0.32 | 0.24 | 0.08 | 1.33 |
| 2 | Cd1d1 | 0.33 | 8.04E-13 | 1.67E-08 | 0.23 | 0.16 | 0.07 | 1.43 |
| 2 | Plscr1 | 0.28 | 8.06E-13 | 1.68E-08 | 0.31 | 0.23 | 0.08 | 1.36 |
| 2 | Msh3 | 0.26 | 9.83E-13 | 2.04E-08 | 0.51 | 0.41 | 0.10 | 1.25 |
| 2 | E130309D02Rik | 0.26 | 1.02E-12 | 2.12E-08 | 0.44 | 0.35 | 0.10 | 1.27 |
| 2 | Tiam2 | 0.38 | 1.02E-12 | 2.13E-08 | 0.14 | 0.09 | 0.05 | 1.59 |
| 2 | Soga1 | 0.46 | 1.03E-12 | 2.14E-08 | 0.17 | 0.11 | 0.06 | 1.49 |
| 2 | Pnpt1 | 0.26 | 1.09E-12 | 2.27E-08 | 0.37 | 0.28 | 0.09 | 1.32 |
| 2 | Fancf | 0.45 | 1.10E-12 | 2.29E-08 | 0.19 | 0.13 | 0.06 | 1.45 |
| 2 | Nqo1 | 0.52 | 1.11E-12 | 2.31E-08 | 0.15 | 0.09 | 0.05 | 1.54 |
| 2 | Zfp948 | 0.54 | 1.18E-12 | 2.45E-08 | 0.14 | 0.09 | 0.05 | 1.57 |
| 2 | Utp20 | 0.29 | 1.38E-12 | 2.86E-08 | 0.42 | 0.33 | 0.09 | 1.27 |
| 2 | Tob1 | 0.26 | 1.49E-12 | 3.09E-08 | 0.36 | 0.27 | 0.09 | 1.32 |
| 2 | Pus7 | 0.26 | 1.51E-12 | 3.14E-08 | 0.41 | 0.32 | 0.09 | 1.28 |
| 2 | Tsc1 | 0.28 | 1.67E-12 | 3.48E-08 | 0.29 | 0.21 | 0.08 | 1.36 |
| 2 | Slc38a6 | 0.29 | 1.82E-12 | 3.78E-08 | 0.21 | 0.15 | 0.07 | 1.45 |
| 2 | Rmi1 | 0.25 | 1.88E-12 | 3.90E-08 | 0.38 | 0.29 | 0.09 | 1.30 |
| 2 | Dgkd | 0.30 | 2.24E-12 | 4.66E-08 | 0.31 | 0.23 | 0.08 | 1.35 |
| 2 | Pxylp1 | 0.39 | 2.31E-12 | 4.81E-08 | 0.16 | 0.10 | 0.06 | 1.53 |
| 2 | Zfand4 | 0.53 | 2.66E-12 | 5.54E-08 | 0.12 | 0.08 | 0.05 | 1.62 |
| 2 | Ap1s2 | 0.32 | 2.78E-12 | 5.78E-08 | 0.19 | 0.13 | 0.06 | 1.47 |
| 2 | Zbtb6 | 0.28 | 2.94E-12 | 6.10E-08 | 0.27 | 0.20 | 0.07 | 1.37 |
| 2 | Rbm19 | 0.26 | 2.99E-12 | 6.22E-08 | 0.35 | 0.27 | 0.08 | 1.30 |
| 2 | Mcm9 | 0.36 | 3.09E-12 | 6.42E-08 | 0.22 | 0.16 | 0.07 | 1.41 |
| 2 | Nemp1 | 0.39 | 3.34E-12 | 6.94E-08 | 0.18 | 0.12 | 0.06 | 1.47 |
| 2 | Chek2 | 0.34 | 3.38E-12 | 7.03E-08 | 0.30 | 0.23 | 0.07 | 1.32 |
| 2 | Cebpd | 0.41 | 3.48E-12 | 7.24E-08 | 0.15 | 0.10 | 0.05 | 1.56 |
| 2 | AW554918 | 0.30 | 3.66E-12 | 7.61E-08 | 0.27 | 0.20 | 0.07 | 1.37 |
| 2 | Csrnp3 | 0.39 | 3.82E-12 | 7.94E-08 | 0.15 | 0.10 | 0.05 | 1.54 |
| 2 | Cep85 | 0.32 | 3.92E-12 | 8.15E-08 | 0.22 | 0.16 | 0.07 | 1.41 |
| 2 | Nhsl2 | 0.30 | 4.30E-12 | 8.95E-08 | 0.22 | 0.16 | 0.07 | 1.41 |
| 2 | Ttc7b | 0.34 | 4.47E-12 | 9.30E-08 | 0.22 | 0.15 | 0.06 | 1.42 |
| 2 | Depdc5 | 0.31 | 5.13E-12 | 1.07E-07 | 0.27 | 0.20 | 0.07 | 1.36 |
| 2 | Atrnl1 | 0.33 | 5.37E-12 | 1.12E-07 | 0.26 | 0.19 | 0.07 | 1.37 |
| 2 | Gnasas1 | 0.28 | 5.39E-12 | 1.12E-07 | 0.25 | 0.18 | 0.07 | 1.40 |
| 2 | Ercc8 | 0.28 | 5.59E-12 | 1.16E-07 | 0.30 | 0.22 | 0.08 | 1.34 |
| 2 | Bcas3 | 0.28 | 5.81E-12 | 1.21E-07 | 0.28 | 0.21 | 0.07 | 1.36 |
| 2 | Cep126 | 0.29 | 6.18E-12 | 1.28E-07 | 0.22 | 0.15 | 0.07 | 1.43 |
| 2 | Ctif | 0.43 | 6.58E-12 | 1.37E-07 | 0.15 | 0.10 | 0.05 | 1.51 |
| 2 | 1700066M21Rik | 0.54 | 7.08E-12 | 1.47E-07 | 0.13 | 0.08 | 0.05 | 1.58 |
| 2 | Slc7a2 | 0.53 | 7.54E-12 | 1.57E-07 | 0.11 | 0.06 | 0.04 | 1.66 |
| 2 | Pprc1 | 0.26 | 7.97E-12 | 1.66E-07 | 0.33 | 0.25 | 0.08 | 1.32 |
| 2 | Fbxo42 | 0.29 | 8.04E-12 | 1.67E-07 | 0.25 | 0.18 | 0.07 | 1.38 |
| 2 | Armh4 | 0.46 | 8.16E-12 | 1.70E-07 | 0.20 | 0.15 | 0.06 | 1.41 |
| 2 | Mylk | 0.31 | 8.22E-12 | 1.71E-07 | 0.23 | 0.17 | 0.06 | 1.38 |
| 2 | Tonsl | 0.46 | 8.44E-12 | 1.76E-07 | 0.19 | 0.13 | 0.06 | 1.44 |
| 2 | Slc25a28 | 0.26 | 9.31E-12 | 1.94E-07 | 0.34 | 0.26 | 0.08 | 1.31 |
| 2 | Lrrc14 | 0.26 | 9.63E-12 | 2.00E-07 | 0.32 | 0.25 | 0.08 | 1.31 |
| 2 | Pxk | 0.34 | 1.05E-11 | 2.19E-07 | 0.23 | 0.16 | 0.06 | 1.40 |
| 2 | Bahcc1 | 0.28 | 1.18E-11 | 2.46E-07 | 0.35 | 0.27 | 0.08 | 1.30 |
| 2 | Mrm2 | 0.32 | 1.29E-11 | 2.68E-07 | 0.27 | 0.20 | 0.07 | 1.34 |

**Supplementary Table 1. Marker genes for the original clusters of integrated clustering of E11.5 control and NotchGOF λ-junction cells.**

| | | | | | | | | |
|---|---|---|---|---|---|---|---|---|
| 2 | Tspan9 | 0.26 | 1.32E-11 | 2.74E-07 | 0.26 | 0.19 | 0.07 | 1.37 |
| 2 | Batf3 | 0.37 | 1.39E-11 | 2.89E-07 | 0.21 | 0.15 | 0.06 | 1.42 |
| 2 | Lrrc75a | 0.27 | 1.42E-11 | 2.95E-07 | 0.28 | 0.20 | 0.07 | 1.36 |
| 2 | Pbld1 | 0.91 | 1.47E-11 | 3.06E-07 | 0.12 | 0.07 | 0.04 | 1.60 |
| 2 | Adam19 | 0.32 | 1.51E-11 | 3.14E-07 | 0.21 | 0.15 | 0.06 | 1.42 |
| 2 | Slc36a4 | 0.30 | 1.52E-11 | 3.16E-07 | 0.22 | 0.16 | 0.07 | 1.41 |
| 2 | Evc2 | 0.28 | 1.52E-11 | 3.17E-07 | 0.23 | 0.17 | 0.07 | 1.39 |
| 2 | Heatr5b | 0.33 | 1.73E-11 | 3.60E-07 | 0.18 | 0.12 | 0.06 | 1.46 |
| 2 | Pcx | 0.38 | 1.83E-11 | 3.80E-07 | 0.18 | 0.13 | 0.06 | 1.45 |
| 2 | Agtpbp1 | 0.28 | 2.09E-11 | 4.34E-07 | 0.24 | 0.17 | 0.07 | 1.38 |
| 2 | Zfp449 | 0.35 | 2.13E-11 | 4.44E-07 | 0.17 | 0.12 | 0.06 | 1.47 |
| 2 | Cep76 | 0.42 | 2.27E-11 | 4.71E-07 | 0.22 | 0.16 | 0.06 | 1.37 |
| 2 | Focad | 0.51 | 2.58E-11 | 5.36E-07 | 0.13 | 0.08 | 0.05 | 1.55 |
| 2 | Rab11fip3 | 0.37 | 2.67E-11 | 5.56E-07 | 0.20 | 0.14 | 0.06 | 1.41 |
| 2 | Zfp946 | 0.29 | 2.79E-11 | 5.81E-07 | 0.27 | 0.20 | 0.07 | 1.36 |
| 2 | Nxpe3 | 0.33 | 2.81E-11 | 5.84E-07 | 0.15 | 0.10 | 0.05 | 1.52 |
| 2 | Itga5 | 0.26 | 3.00E-11 | 6.23E-07 | 0.28 | 0.21 | 0.07 | 1.35 |
| 2 | Nars2 | 0.30 | 3.10E-11 | 6.45E-07 | 0.17 | 0.11 | 0.06 | 1.48 |
| 2 | Rpp40 | 0.30 | 3.51E-11 | 7.29E-07 | 0.30 | 0.23 | 0.07 | 1.31 |
| 2 | Lyst | 0.39 | 3.73E-11 | 7.77E-07 | 0.17 | 0.12 | 0.05 | 1.45 |
| 2 | Cerk | 0.28 | 3.76E-11 | 7.81E-07 | 0.21 | 0.14 | 0.06 | 1.42 |
| 2 | Exoc2 | 0.28 | 3.81E-11 | 7.93E-07 | 0.31 | 0.24 | 0.07 | 1.31 |
| 2 | Eaf1 | 0.32 | 4.25E-11 | 8.83E-07 | 0.25 | 0.19 | 0.07 | 1.35 |
| 2 | 2810429I04Rik | 0.36 | 4.36E-11 | 9.06E-07 | 0.21 | 0.15 | 0.06 | 1.40 |
| 2 | 1810055G02Rik | 0.28 | 4.52E-11 | 9.40E-07 | 0.28 | 0.21 | 0.07 | 1.35 |
| 2 | Dpagt1 | 0.26 | 5.48E-11 | 1.14E-06 | 0.36 | 0.28 | 0.08 | 1.29 |
| 2 | Wasf3 | 0.29 | 5.95E-11 | 1.24E-06 | 0.23 | 0.16 | 0.06 | 1.40 |
| 2 | Plekhn1 | 0.29 | 6.41E-11 | 1.33E-06 | 0.26 | 0.20 | 0.07 | 1.35 |
| 2 | Mier2 | 0.28 | 6.41E-11 | 1.33E-06 | 0.26 | 0.19 | 0.07 | 1.35 |
| 2 | Apaf1 | 0.29 | 6.55E-11 | 1.36E-06 | 0.29 | 0.22 | 0.07 | 1.33 |
| 2 | Inpp5a | 0.33 | 7.16E-11 | 1.49E-06 | 0.21 | 0.15 | 0.06 | 1.40 |
| 2 | Acap3 | 0.45 | 7.17E-11 | 1.49E-06 | 0.11 | 0.07 | 0.04 | 1.58 |
| 2 | Efr3b | 0.35 | 7.74E-11 | 1.61E-06 | 0.14 | 0.09 | 0.05 | 1.52 |
| 2 | Ehd3 | 0.50 | 7.92E-11 | 1.65E-06 | 0.16 | 0.11 | 0.05 | 1.46 |
| 2 | Atf7 | 0.29 | 7.93E-11 | 1.65E-06 | 0.26 | 0.19 | 0.07 | 1.35 |
| 2 | Tmem243 | 0.27 | 8.00E-11 | 1.66E-06 | 0.28 | 0.21 | 0.07 | 1.34 |
| 2 | Sbf1 | 0.27 | 8.28E-11 | 1.72E-06 | 0.34 | 0.27 | 0.08 | 1.29 |
| 2 | Chml | 0.29 | 8.49E-11 | 1.77E-06 | 0.20 | 0.14 | 0.06 | 1.41 |
| 2 | Crnde | 0.27 | 8.50E-11 | 1.77E-06 | 0.21 | 0.15 | 0.06 | 1.40 |
| 2 | Cux2 | 0.52 | 8.59E-11 | 1.79E-06 | 0.14 | 0.10 | 0.05 | 1.48 |
| 2 | Kcne1l | 0.26 | 9.11E-11 | 1.90E-06 | 0.18 | 0.12 | 0.06 | 1.47 |
| 2 | Coq3 | 0.33 | 9.27E-11 | 1.93E-06 | 0.24 | 0.18 | 0.06 | 1.36 |
| 2 | Mb21d2 | 0.27 | 9.28E-11 | 1.93E-06 | 0.21 | 0.15 | 0.06 | 1.41 |
| 2 | Neurl4 | 0.29 | 9.95E-11 | 2.07E-06 | 0.28 | 0.21 | 0.07 | 1.33 |
| 2 | Gm26648 | 0.42 | 1.14E-10 | 2.36E-06 | 0.13 | 0.09 | 0.05 | 1.53 |
| 2 | Ccdc91 | 0.25 | 1.22E-10 | 2.54E-06 | 0.34 | 0.27 | 0.08 | 1.29 |
| 2 | Pot1b | 0.34 | 1.31E-10 | 2.72E-06 | 0.19 | 0.13 | 0.06 | 1.41 |
| 2 | Bicra | 0.32 | 1.34E-10 | 2.78E-06 | 0.20 | 0.14 | 0.06 | 1.40 |
| 2 | Zcchc4 | 0.30 | 1.68E-10 | 3.50E-06 | 0.24 | 0.18 | 0.06 | 1.36 |
| 2 | Pip4k2a | 0.33 | 1.71E-10 | 3.56E-06 | 0.16 | 0.11 | 0.05 | 1.45 |
| 2 | Dlg2 | 0.54 | 1.92E-10 | 4.00E-06 | 0.11 | 0.07 | 0.04 | 1.60 |
| 2 | Fancc | 0.49 | 1.94E-10 | 4.04E-06 | 0.14 | 0.10 | 0.05 | 1.47 |
| 2 | Svep1 | 0.43 | 2.05E-10 | 4.27E-06 | 0.12 | 0.08 | 0.04 | 1.55 |
| 2 | Plcl2 | 0.36 | 2.20E-10 | 4.57E-06 | 0.16 | 0.11 | 0.05 | 1.46 |
| 2 | Akr1b8 | 0.43 | 2.21E-10 | 4.59E-06 | 0.12 | 0.07 | 0.04 | 1.57 |
| 2 | Mtr | 0.25 | 2.36E-10 | 4.90E-06 | 0.30 | 0.23 | 0.07 | 1.31 |

**Supplementary Table 1. Marker genes for the original clusters of integrated clustering of E11.5 control and NotchGOF λ-junction cells.**

| | | | | | | | | |
|---|---|---|---|---|---|---|---|---|
| 2 | Zfp882 | 0.30 | 2.79E-10 | 5.81E-06 | 0.19 | 0.13 | 0.06 | 1.42 |
| 2 | Polh | 0.43 | 3.04E-10 | 6.31E-06 | 0.13 | 0.09 | 0.04 | 1.52 |
| 2 | Zfp335 | 0.45 | 3.11E-10 | 6.46E-06 | 0.14 | 0.09 | 0.05 | 1.50 |
| 2 | Adnp2 | 0.29 | 3.12E-10 | 6.50E-06 | 0.23 | 0.17 | 0.06 | 1.36 |
| 2 | Pcgf1 | 0.26 | 3.18E-10 | 6.61E-06 | 0.23 | 0.17 | 0.06 | 1.37 |
| 2 | Eml3 | 0.36 | 3.40E-10 | 7.07E-06 | 0.17 | 0.12 | 0.05 | 1.43 |
| 2 | Slx4 | 0.42 | 3.48E-10 | 7.23E-06 | 0.18 | 0.12 | 0.05 | 1.41 |
| 2 | Lhpp | 0.36 | 3.66E-10 | 7.61E-06 | 0.15 | 0.10 | 0.05 | 1.47 |
| 2 | Gtpbp3 | 0.28 | 3.68E-10 | 7.66E-06 | 0.25 | 0.19 | 0.07 | 1.34 |
| 2 | Srf | 0.36 | 3.70E-10 | 7.69E-06 | 0.18 | 0.13 | 0.05 | 1.42 |
| 2 | Sgtb | 0.34 | 3.72E-10 | 7.73E-06 | 0.15 | 0.10 | 0.05 | 1.47 |
| 2 | Dkk3 | 0.31 | 3.77E-10 | 7.84E-06 | 0.17 | 0.11 | 0.05 | 1.46 |
| 2 | Alkbh4 | 0.36 | 3.94E-10 | 8.20E-06 | 0.18 | 0.12 | 0.05 | 1.43 |
| 2 | Tvp23a | 0.28 | 4.03E-10 | 8.39E-06 | 0.18 | 0.13 | 0.06 | 1.44 |
| 2 | Ephb6 | 0.38 | 4.83E-10 | 1.00E-05 | 0.13 | 0.08 | 0.04 | 1.53 |
| 2 | Gucy1a2 | 0.51 | 5.21E-10 | 1.08E-05 | 0.11 | 0.07 | 0.04 | 1.59 |
| 2 | Tecpr2 | 0.42 | 6.04E-10 | 1.26E-05 | 0.14 | 0.09 | 0.04 | 1.48 |
| 2 | Rragb | 0.37 | 7.70E-10 | 1.60E-05 | 0.13 | 0.08 | 0.04 | 1.52 |
| 2 | Slc8a1 | 0.40 | 8.07E-10 | 1.68E-05 | 0.17 | 0.12 | 0.05 | 1.41 |
| 2 | Ambra1 | 0.28 | 8.23E-10 | 1.71E-05 | 0.21 | 0.15 | 0.06 | 1.38 |
| 2 | Fbxl15 | 0.28 | 8.36E-10 | 1.74E-05 | 0.25 | 0.19 | 0.06 | 1.32 |
| 2 | Tbc1d12 | 0.27 | 8.58E-10 | 1.78E-05 | 0.22 | 0.16 | 0.06 | 1.36 |
| 2 | Mlst8 | 0.30 | 8.68E-10 | 1.80E-05 | 0.18 | 0.13 | 0.05 | 1.41 |
| 2 | Tulp3 | 0.27 | 9.06E-10 | 1.88E-05 | 0.31 | 0.23 | 0.07 | 1.31 |
| 2 | Gphn | 0.30 | 1.18E-09 | 2.45E-05 | 0.24 | 0.18 | 0.06 | 1.33 |
| 2 | Zfp472 | 0.33 | 1.36E-09 | 2.84E-05 | 0.20 | 0.14 | 0.05 | 1.38 |
| 2 | Trmt5 | 0.34 | 1.46E-09 | 3.04E-05 | 0.21 | 0.16 | 0.06 | 1.35 |
| 2 | Fanca | 0.34 | 1.51E-09 | 3.15E-05 | 0.16 | 0.11 | 0.05 | 1.42 |
| 2 | Cyld | 0.26 | 1.55E-09 | 3.23E-05 | 0.29 | 0.22 | 0.07 | 1.30 |
| 2 | Nprl3 | 0.34 | 1.61E-09 | 3.34E-05 | 0.17 | 0.12 | 0.05 | 1.40 |
| 2 | Polr1b | 0.29 | 1.64E-09 | 3.42E-05 | 0.23 | 0.17 | 0.06 | 1.35 |
| 2 | Zfp280b | 0.28 | 1.91E-09 | 3.97E-05 | 0.21 | 0.16 | 0.06 | 1.35 |
| 2 | Fanci | 0.58 | 2.04E-09 | 4.24E-05 | 0.14 | 0.09 | 0.04 | 1.45 |
| 2 | Ptcd1 | 0.29 | 2.04E-09 | 4.24E-05 | 0.19 | 0.13 | 0.05 | 1.39 |
| 2 | Ctdp1 | 0.28 | 2.10E-09 | 4.37E-05 | 0.26 | 0.20 | 0.06 | 1.30 |
| 2 | Gpd2 | 0.27 | 2.25E-09 | 4.67E-05 | 0.29 | 0.22 | 0.07 | 1.29 |
| 2 | Kri1 | 0.27 | 2.44E-09 | 5.08E-05 | 0.23 | 0.17 | 0.06 | 1.33 |
| 2 | Parpbp | 0.39 | 2.52E-09 | 5.24E-05 | 0.22 | 0.17 | 0.05 | 1.32 |
| 2 | Smg8 | 0.27 | 2.63E-09 | 5.46E-05 | 0.23 | 0.17 | 0.06 | 1.35 |
| 2 | 6030442K20Rik | 0.46 | 2.93E-09 | 6.10E-05 | 0.11 | 0.07 | 0.04 | 1.53 |
| 2 | Fcsk | 0.32 | 3.21E-09 | 6.67E-05 | 0.13 | 0.09 | 0.04 | 1.50 |
| 2 | Rnf19a | 0.26 | 3.33E-09 | 6.93E-05 | 0.28 | 0.22 | 0.06 | 1.29 |
| 2 | Bcs1l | 0.29 | 3.39E-09 | 7.06E-05 | 0.23 | 0.18 | 0.06 | 1.33 |
| 2 | Eif4ebp3 | 0.30 | 3.48E-09 | 7.24E-05 | 0.24 | 0.19 | 0.06 | 1.31 |
| 2 | Smg9 | 0.33 | 3.52E-09 | 7.32E-05 | 0.19 | 0.14 | 0.05 | 1.37 |
| 2 | Gm4737 | 0.34 | 4.16E-09 | 8.65E-05 | 0.17 | 0.12 | 0.05 | 1.41 |
| 2 | Eya4 | 0.25 | 4.64E-09 | 9.64E-05 | 0.35 | 0.28 | 0.07 | 1.24 |
| 2 | Fancb | 0.52 | 4.79E-09 | 9.95E-05 | 0.12 | 0.08 | 0.04 | 1.46 |
| 2 | Ccdc14 | 0.27 | 4.83E-09 | 1.00E-04 | 0.25 | 0.19 | 0.06 | 1.32 |
| 2 | Ell | 0.34 | 4.90E-09 | 1.02E-04 | 0.16 | 0.11 | 0.05 | 1.42 |
| 2 | Eml1 | 0.28 | 4.98E-09 | 1.04E-04 | 0.22 | 0.16 | 0.06 | 1.34 |
| 2 | Vps11 | 0.28 | 5.26E-09 | 1.09E-04 | 0.21 | 0.15 | 0.05 | 1.35 |
| 2 | Micall2 | 0.47 | 5.67E-09 | 1.18E-04 | 0.10 | 0.06 | 0.04 | 1.56 |
| 2 | Sdccag8 | 0.29 | 6.01E-09 | 1.25E-04 | 0.17 | 0.12 | 0.05 | 1.40 |
| 2 | Raet1d | 0.35 | 6.07E-09 | 1.26E-04 | 0.12 | 0.08 | 0.04 | 1.49 |
| 2 | Gpr161 | 0.40 | 7.66E-09 | 1.59E-04 | 0.15 | 0.10 | 0.04 | 1.42 |

**Supplementary Table 1. Marker genes for the original clusters of integrated clustering of E11.5 control and NotchGOF λ-junction cells.**

| | | | | | | | | |
|---|---|---|---|---|---|---|---|---|
| 2 | Ror1 | 0.25 | 7.69E-09 | 1.60E-04 | 0.17 | 0.12 | 0.05 | 1.41 |
| 2 | Dusp4 | 0.35 | 1.09E-08 | 2.26E-04 | 0.12 | 0.08 | 0.04 | 1.49 |
| 2 | Zfp658 | 0.36 | 1.28E-08 | 2.65E-04 | 0.10 | 0.07 | 0.04 | 1.55 |
| 2 | Fam118a | 0.28 | 1.33E-08 | 2.76E-04 | 0.22 | 0.17 | 0.05 | 1.33 |
| 2 | Zfp954 | 0.41 | 1.37E-08 | 2.85E-04 | 0.12 | 0.08 | 0.04 | 1.48 |
| 2 | Usp54 | 0.30 | 1.42E-08 | 2.94E-04 | 0.17 | 0.13 | 0.05 | 1.38 |
| 2 | Iffo1 | 0.37 | 1.47E-08 | 3.06E-04 | 0.13 | 0.09 | 0.04 | 1.45 |
| 2 | Samd8 | 0.26 | 1.52E-08 | 3.17E-04 | 0.26 | 0.21 | 0.06 | 1.29 |
| 2 | Adcy3 | 0.28 | 1.88E-08 | 3.91E-04 | 0.12 | 0.08 | 0.04 | 1.49 |
| 2 | 3110021N24Rik | 0.30 | 2.00E-08 | 4.17E-04 | 0.15 | 0.11 | 0.05 | 1.42 |
| 2 | Gm12184 | 0.32 | 2.01E-08 | 4.18E-04 | 0.15 | 0.11 | 0.04 | 1.40 |
| 2 | Mettl15 | 0.48 | 2.29E-08 | 4.76E-04 | 0.10 | 0.07 | 0.04 | 1.51 |
| 2 | Sh3bgrl2 | 0.44 | 2.38E-08 | 4.95E-04 | 0.12 | 0.08 | 0.04 | 1.47 |
| 2 | Pck2 | 0.29 | 2.41E-08 | 5.02E-04 | 0.26 | 0.20 | 0.06 | 1.27 |
| 2 | Zbtb48 | 0.27 | 2.43E-08 | 5.05E-04 | 0.17 | 0.13 | 0.05 | 1.38 |
| 2 | Kctd9 | 0.39 | 2.43E-08 | 5.05E-04 | 0.15 | 0.11 | 0.04 | 1.38 |
| 2 | Piga | 0.27 | 2.69E-08 | 5.59E-04 | 0.21 | 0.16 | 0.05 | 1.33 |
| 2 | Cdh24 | 0.27 | 2.81E-08 | 5.84E-04 | 0.18 | 0.13 | 0.05 | 1.36 |
| 2 | Mtrr | 0.37 | 2.85E-08 | 5.93E-04 | 0.16 | 0.11 | 0.04 | 1.39 |
| 2 | Zdhhc15 | 0.35 | 2.87E-08 | 5.97E-04 | 0.17 | 0.13 | 0.05 | 1.36 |
| 2 | Dnajc27 | 0.35 | 2.95E-08 | 6.14E-04 | 0.13 | 0.09 | 0.04 | 1.44 |
| 2 | Thada | 0.33 | 3.41E-08 | 7.08E-04 | 0.16 | 0.12 | 0.04 | 1.37 |
| 2 | Faxc | 0.32 | 3.74E-08 | 7.78E-04 | 0.14 | 0.10 | 0.04 | 1.44 |
| 2 | Vac14 | 0.25 | 3.83E-08 | 7.96E-04 | 0.27 | 0.22 | 0.06 | 1.27 |
| 2 | Gm21887 | 0.28 | 3.90E-08 | 8.11E-04 | 0.18 | 0.14 | 0.05 | 1.35 |
| 2 | Ap4e1 | 0.35 | 4.59E-08 | 9.56E-04 | 0.15 | 0.10 | 0.04 | 1.39 |
| 2 | Cpq | 0.28 | 4.72E-08 | 9.82E-04 | 0.21 | 0.16 | 0.05 | 1.32 |
| 2 | Cbx8 | 0.30 | 4.74E-08 | 9.87E-04 | 0.16 | 0.12 | 0.04 | 1.37 |
| 2 | Ufsp1 | 0.34 | 4.75E-08 | 9.88E-04 | 0.16 | 0.11 | 0.04 | 1.38 |
| 2 | Mcoln1 | 0.30 | 4.80E-08 | 9.99E-04 | 0.14 | 0.10 | 0.04 | 1.42 |
| 2 | Atg2a | 0.31 | 6.59E-08 | 1.37E-03 | 0.15 | 0.11 | 0.04 | 1.40 |
| 2 | Nkiras1 | 0.25 | 6.93E-08 | 1.44E-03 | 0.18 | 0.14 | 0.05 | 1.35 |
| 2 | Ttc26 | 0.30 | 7.06E-08 | 1.47E-03 | 0.15 | 0.11 | 0.04 | 1.40 |
| 2 | Tbc1d2b | 0.40 | 7.69E-08 | 1.60E-03 | 0.12 | 0.08 | 0.04 | 1.44 |
| 2 | Mutyh | 0.35 | 9.52E-08 | 1.98E-03 | 0.15 | 0.11 | 0.04 | 1.38 |
| 2 | Gm43566 | 0.34 | 1.03E-07 | 2.14E-03 | 0.10 | 0.07 | 0.03 | 1.51 |
| 2 | Kif3c | 0.29 | 1.07E-07 | 2.24E-03 | 0.16 | 0.12 | 0.04 | 1.37 |
| 2 | Nupl2 | 0.27 | 1.16E-07 | 2.41E-03 | 0.20 | 0.15 | 0.05 | 1.32 |
| 2 | Lss | 0.27 | 1.40E-07 | 2.92E-03 | 0.20 | 0.16 | 0.05 | 1.32 |
| 2 | Nr1h5 | 0.30 | 1.42E-07 | 2.96E-03 | 0.18 | 0.13 | 0.05 | 1.35 |
| 2 | Ogg1 | 0.28 | 1.56E-07 | 3.24E-03 | 0.15 | 0.11 | 0.04 | 1.39 |
| 2 | Palb2 | 0.33 | 1.77E-07 | 3.67E-03 | 0.17 | 0.13 | 0.04 | 1.34 |
| 2 | Inpp5b | 0.26 | 1.85E-07 | 3.86E-03 | 0.16 | 0.11 | 0.04 | 1.37 |
| 2 | Clec2l | 0.35 | 1.96E-07 | 4.07E-03 | 0.10 | 0.07 | 0.03 | 1.49 |
| 2 | Fmn2 | 0.31 | 2.19E-07 | 4.55E-03 | 0.10 | 0.07 | 0.03 | 1.49 |
| 2 | Gm17494 | 0.49 | 3.54E-07 | 7.36E-03 | 0.11 | 0.08 | 0.03 | 1.43 |
| 2 | Intu | 0.27 | 3.58E-07 | 7.45E-03 | 0.14 | 0.11 | 0.04 | 1.37 |
| 2 | Lama1 | 0.35 | 3.98E-07 | 8.29E-03 | 0.14 | 0.10 | 0.04 | 1.37 |
| 2 | Pcdhgc4 | 0.44 | 4.19E-07 | 8.72E-03 | 0.10 | 0.07 | 0.03 | 1.44 |
| 2 | Tceanc | 0.26 | 4.88E-07 | 1.01E-02 | 0.13 | 0.09 | 0.04 | 1.41 |
| 2 | Gmeb2 | 0.26 | 5.36E-07 | 1.11E-02 | 0.18 | 0.14 | 0.04 | 1.31 |
| 2 | Atoh8 | 0.29 | 5.55E-07 | 1.16E-02 | 0.13 | 0.09 | 0.04 | 1.40 |
| 2 | G430095P16Rik | 0.25 | 5.73E-07 | 1.19E-02 | 0.14 | 0.10 | 0.04 | 1.37 |
| 2 | Bbs5 | 0.27 | 5.74E-07 | 1.19E-02 | 0.14 | 0.10 | 0.04 | 1.37 |
| 2 | Ica1 | 0.26 | 6.04E-07 | 1.26E-02 | 0.16 | 0.12 | 0.04 | 1.35 |
| 2 | Proser3 | 0.26 | 6.14E-07 | 1.28E-02 | 0.22 | 0.17 | 0.05 | 1.28 |

**Supplementary Table 1. Marker genes for the original clusters of integrated clustering of E11.5 control and NotchGOF λ-junction cells.**

| | | | | | | | | |
|---|---|---|---|---|---|---|---|---|
| 2 | Rexo5 | 0.29 | 6.61E-07 | 1.37E-02 | 0.16 | 0.12 | 0.04 | 1.34 |
| 2 | Kif7 | 0.26 | 7.23E-07 | 1.50E-02 | 0.16 | 0.12 | 0.04 | 1.34 |
| 2 | Cog7 | 0.26 | 7.29E-07 | 1.52E-02 | 0.18 | 0.13 | 0.04 | 1.33 |
| 2 | Ccsap | 0.32 | 8.12E-07 | 1.69E-02 | 0.15 | 0.11 | 0.04 | 1.34 |
| 2 | Setd6 | 0.29 | 8.19E-07 | 1.70E-02 | 0.16 | 0.12 | 0.04 | 1.32 |
| 2 | Spata2 | 0.27 | 9.44E-07 | 1.96E-02 | 0.16 | 0.12 | 0.04 | 1.34 |
| 2 | Zfp763 | 0.25 | 9.81E-07 | 2.04E-02 | 0.22 | 0.17 | 0.05 | 1.28 |
| 2 | Zfand2a | 0.36 | 9.96E-07 | 2.07E-02 | 0.10 | 0.07 | 0.03 | 1.42 |
| 2 | Snhg11 | 0.26 | 1.02E-06 | 2.12E-02 | 0.14 | 0.10 | 0.04 | 1.38 |
| 2 | Aplf | 0.31 | 1.04E-06 | 2.16E-02 | 0.14 | 0.11 | 0.04 | 1.35 |
| 2 | Cacna2d3 | 0.34 | 1.04E-06 | 2.16E-02 | 0.11 | 0.08 | 0.03 | 1.41 |
| 2 | Cbx4 | 0.30 | 1.21E-06 | 2.52E-02 | 0.14 | 0.10 | 0.04 | 1.36 |
| 2 | Cdan1 | 0.28 | 1.31E-06 | 2.73E-02 | 0.13 | 0.09 | 0.04 | 1.39 |
| 2 | Pir | 0.29 | 1.36E-06 | 2.84E-02 | 0.16 | 0.12 | 0.04 | 1.32 |
| 2 | St6gal2 | 0.28 | 1.48E-06 | 3.08E-02 | 0.14 | 0.10 | 0.04 | 1.37 |
| 2 | Hexdc | 0.26 | 1.49E-06 | 3.10E-02 | 0.14 | 0.10 | 0.04 | 1.38 |
| 2 | Eif5a2 | 0.26 | 1.57E-06 | 3.26E-02 | 0.15 | 0.11 | 0.04 | 1.34 |
| 2 | Gm38037 | 0.43 | 1.61E-06 | 3.34E-02 | 0.12 | 0.09 | 0.03 | 1.37 |
| 2 | Nkrf | 0.42 | 1.83E-06 | 3.81E-02 | 0.12 | 0.09 | 0.03 | 1.38 |
| 2 | Dclre1b | 0.30 | 1.94E-06 | 4.03E-02 | 0.15 | 0.12 | 0.04 | 1.32 |
| 2 | Gba2 | 0.26 | 2.00E-06 | 4.15E-02 | 0.16 | 0.12 | 0.04 | 1.32 |
| 2 | Arl10 | 0.31 | 2.00E-06 | 4.16E-02 | 0.10 | 0.07 | 0.03 | 1.43 |
| 2 | Btbd8 | 0.37 | 2.26E-06 | 4.71E-02 | 0.10 | 0.07 | 0.03 | 1.41 |
| 3 | Pax7 | 2.79 | 0 | 0 | 0.88 | 0.17 | 0.71 | 5.06 |
| 3 | Hist1h2ab | 1.79 | 0 | 0 | 0.86 | 0.33 | 0.53 | 2.62 |
| 3 | Hist1h1a | 1.61 | 0 | 0 | 0.90 | 0.38 | 0.52 | 2.39 |
| 3 | Igf1 | 1.86 | 0 | 0 | 0.95 | 0.43 | 0.52 | 2.20 |
| 3 | Pax3 | 1.53 | 0 | 0 | 0.92 | 0.41 | 0.51 | 2.23 |
| 3 | Hist2h2ac | 1.80 | 0 | 0 | 0.78 | 0.28 | 0.50 | 2.77 |
| 3 | Nbl1 | 1.28 | 0 | 0 | 0.95 | 0.45 | 0.50 | 2.10 |
| 3 | Hist1h4d | 1.44 | 0 | 0 | 0.89 | 0.42 | 0.47 | 2.13 |
| 3 | Hist1h2an | 1.68 | 0 | 0 | 0.74 | 0.27 | 0.47 | 2.78 |
| 3 | Tnfaip6 | 1.44 | 0 | 0 | 0.89 | 0.43 | 0.46 | 2.08 |
| 3 | Hist1h3e | 1.38 | 0 | 0 | 0.83 | 0.39 | 0.45 | 2.15 |
| 3 | Hist1h1d | 1.38 | 0 | 0 | 0.76 | 0.32 | 0.44 | 2.40 |
| 3 | Prdm16 | 1.53 | 0 | 0 | 0.73 | 0.29 | 0.44 | 2.51 |
| 3 | Tcf15 | 1.60 | 0 | 0 | 0.68 | 0.24 | 0.44 | 2.79 |
| 3 | Rarb | 1.53 | 0 | 0 | 0.75 | 0.31 | 0.43 | 2.38 |
| 3 | Cntfr | 2.02 | 0 | 0 | 0.61 | 0.19 | 0.43 | 3.30 |
| 3 | Hist1h2bm | 1.61 | 0 | 0 | 0.69 | 0.27 | 0.42 | 2.54 |
| 3 | Hist1h3c | 1.51 | 0 | 0 | 0.96 | 0.56 | 0.40 | 1.72 |
| 3 | Hpgd | 1.74 | 0 | 0 | 0.71 | 0.32 | 0.40 | 2.26 |
| 3 | Hist1h1b | 1.88 | 0 | 0 | 1.00 | 0.61 | 0.39 | 1.64 |
| 3 | Hist1h1e | 1.40 | 0 | 0 | 0.99 | 0.61 | 0.38 | 1.61 |
| 3 | Jun | 1.32 | 0 | 0 | 0.97 | 0.65 | 0.33 | 1.51 |
| 3 | Hist1h2ae | 1.68 | 0 | 0 | 1.00 | 0.67 | 0.33 | 1.49 |
| 3 | Hist1h2ap | 1.56 | 0 | 0 | 1.00 | 0.73 | 0.27 | 1.38 |
| 3 | Crabp1 | 1.28 | 0 | 0 | 1.00 | 0.84 | 0.16 | 1.19 |
| 3 | Pclaf | 0.92 | 0 | 0 | 1.00 | 0.88 | 0.12 | 1.14 |
| 3 | Micos10 | 0.84 | 0 | 0 | 1.00 | 0.96 | 0.04 | 1.04 |
| 3 | Top2a | 0.97 | 1.69E-307 | 3.52E-303 | 1.00 | 0.77 | 0.23 | 1.30 |
| 3 | Gm30564 | 2.15 | 2.26E-294 | 4.69E-290 | 0.35 | 0.07 | 0.27 | 4.73 |
| 3 | Nme2 | 0.45 | 1.59E-292 | 3.30E-288 | 1.00 | 0.98 | 0.02 | 1.02 |
| 3 | Hist1h3f | 1.53 | 7.54E-292 | 1.57E-287 | 0.63 | 0.24 | 0.40 | 2.67 |
| 3 | Fbxo5 | 1.04 | 7.02E-288 | 1.46E-283 | 0.94 | 0.56 | 0.39 | 1.69 |
| 3 | Hist2h2bb | 1.55 | 9.70E-280 | 2.02E-275 | 0.60 | 0.22 | 0.38 | 2.75 |

**Supplementary Table 1. Marker genes for the original clusters of integrated clustering of E11.5 control and NotchGOF λ-junction cells.**

| | | | | | | | | |
|---|---|---|---|---|---|---|---|---|
| 3 | Gas1 | 1.04 | 3.14E-279 | 6.54E-275 | 0.95 | 0.68 | 0.28 | 1.41 |
| 3 | Hist1h2be | 1.57 | 4.60E-278 | 9.56E-274 | 0.62 | 0.24 | 0.38 | 2.56 |
| 3 | Hist1h3i | 1.59 | 1.49E-276 | 3.10E-272 | 0.58 | 0.21 | 0.37 | 2.79 |
| 3 | H2afz | 0.41 | 3.79E-273 | 7.88E-269 | 1.00 | 1.00 | 0.00 | 1.00 |
| 3 | Twist1 | 0.99 | 3.80E-270 | 7.90E-266 | 1.00 | 0.75 | 0.25 | 1.33 |
| 3 | Hist1h4h | 1.32 | 4.72E-265 | 9.82E-261 | 0.77 | 0.38 | 0.40 | 2.06 |
| 3 | Ranbp1 | 0.47 | 6.52E-259 | 1.36E-254 | 1.00 | 0.99 | 0.01 | 1.01 |
| 3 | Sh3gl3 | 1.40 | 2.27E-258 | 4.72E-254 | 0.69 | 0.30 | 0.39 | 2.31 |
| 3 | Ybx1 | 0.37 | 1.40E-255 | 2.90E-251 | 1.00 | 1.00 | 0.00 | 1.00 |
| 3 | Rrm2 | 0.89 | 1.62E-255 | 3.36E-251 | 0.99 | 0.79 | 0.20 | 1.26 |
| 3 | Alx1 | 0.83 | 3.54E-252 | 7.36E-248 | 0.99 | 0.71 | 0.29 | 1.41 |
| 3 | Hist1h1c | 1.15 | 2.92E-250 | 6.06E-246 | 0.78 | 0.39 | 0.40 | 2.03 |
| 3 | Hist1h3d | 1.78 | 1.23E-247 | 2.57E-243 | 0.47 | 0.16 | 0.32 | 3.03 |
| 3 | Txn1 | 0.45 | 2.38E-246 | 4.94E-242 | 1.00 | 0.99 | 0.01 | 1.01 |
| 3 | Dut | 0.63 | 3.76E-242 | 7.81E-238 | 1.00 | 0.94 | 0.06 | 1.07 |
| 3 | Pbk | 1.06 | 4.96E-238 | 1.03E-233 | 0.87 | 0.51 | 0.36 | 1.71 |
| 3 | Hist1h2bb | 1.58 | 2.90E-237 | 6.03E-233 | 0.51 | 0.18 | 0.33 | 2.82 |
| 3 | Esco2 | 1.10 | 1.08E-233 | 2.26E-229 | 0.80 | 0.42 | 0.38 | 1.92 |
| 3 | Tyms | 0.71 | 2.60E-225 | 5.40E-221 | 0.99 | 0.82 | 0.17 | 1.20 |
| 3 | Lsm2 | 0.59 | 1.10E-224 | 2.29E-220 | 1.00 | 0.92 | 0.08 | 1.08 |
| 3 | H2afy | 0.49 | 4.87E-220 | 1.01E-215 | 1.00 | 0.96 | 0.04 | 1.04 |
| 3 | Lix1 | 0.99 | 2.16E-219 | 4.49E-215 | 0.92 | 0.60 | 0.32 | 1.54 |
| 3 | Cycs | 0.55 | 1.48E-217 | 3.08E-213 | 1.00 | 0.96 | 0.04 | 1.04 |
| 3 | Hmgb1 | 0.33 | 3.30E-216 | 6.87E-212 | 1.00 | 1.00 | 0.00 | 1.00 |
| 3 | Dek | 0.50 | 2.28E-212 | 4.75E-208 | 1.00 | 0.96 | 0.04 | 1.04 |
| 3 | Gsc | 1.68 | 1.80E-207 | 3.74E-203 | 0.59 | 0.27 | 0.32 | 2.21 |
| 3 | D030045P18Rik | 2.80 | 2.52E-207 | 5.24E-203 | 0.19 | 0.03 | 0.16 | 6.00 |
| 3 | Hist1h3b | 1.44 | 2.66E-207 | 5.54E-203 | 0.52 | 0.20 | 0.32 | 2.57 |
| 3 | Ptn | 0.64 | 5.94E-204 | 1.23E-199 | 1.00 | 0.89 | 0.11 | 1.12 |
| 3 | Hist2h2aa1 | 1.64 | 1.37E-195 | 2.85E-191 | 0.43 | 0.15 | 0.28 | 2.83 |
| 3 | Dkk2 | 1.24 | 1.72E-194 | 3.58E-190 | 0.57 | 0.24 | 0.33 | 2.38 |
| 3 | Gmnn | 0.68 | 3.58E-191 | 7.45E-187 | 0.98 | 0.78 | 0.20 | 1.26 |
| 3 | Tcf12 | 0.66 | 3.99E-191 | 8.30E-187 | 0.99 | 0.87 | 0.13 | 1.14 |
| 3 | Septin11 | 0.63 | 1.69E-186 | 3.52E-182 | 0.99 | 0.86 | 0.13 | 1.15 |
| 3 | Tubb5 | 0.36 | 1.80E-186 | 3.75E-182 | 1.00 | 1.00 | 0.00 | 1.00 |
| 3 | Tk1 | 0.92 | 1.07E-180 | 2.23E-176 | 0.84 | 0.52 | 0.32 | 1.61 |
| 3 | Pcna | 0.72 | 2.44E-180 | 5.07E-176 | 0.98 | 0.83 | 0.15 | 1.18 |
| 3 | Car14 | 1.02 | 4.93E-180 | 1.03E-175 | 0.74 | 0.39 | 0.34 | 1.88 |
| 3 | Atad2 | 0.75 | 5.15E-180 | 1.07E-175 | 0.95 | 0.72 | 0.23 | 1.32 |
| 3 | Aurkb | 0.85 | 7.27E-180 | 1.51E-175 | 0.83 | 0.46 | 0.37 | 1.81 |
| 3 | Nasp | 0.43 | 9.12E-180 | 1.90E-175 | 1.00 | 0.98 | 0.02 | 1.02 |
| 3 | Fap | 1.31 | 9.49E-179 | 1.97E-174 | 0.47 | 0.17 | 0.29 | 2.69 |
| 3 | St6galnac3 | 1.06 | 5.59E-178 | 1.16E-173 | 0.71 | 0.37 | 0.34 | 1.94 |
| 3 | Lig1 | 0.67 | 2.81E-177 | 5.85E-173 | 0.97 | 0.77 | 0.20 | 1.26 |
| 3 | Eif5a | 0.38 | 1.89E-176 | 3.93E-172 | 1.00 | 1.00 | 0.00 | 1.00 |
| 3 | Rad51ap1 | 0.90 | 2.15E-174 | 4.46E-170 | 0.80 | 0.46 | 0.34 | 1.75 |
| 3 | Rspo2 | 1.08 | 4.35E-174 | 9.06E-170 | 0.68 | 0.34 | 0.34 | 1.99 |
| 3 | Rnf128 | 1.74 | 7.11E-174 | 1.48E-169 | 0.37 | 0.12 | 0.24 | 2.97 |
| 3 | 1110017D15Rik | 1.94 | 1.46E-173 | 3.04E-169 | 0.26 | 0.07 | 0.19 | 3.92 |
| 3 | Hist1h2ak | 1.54 | 1.50E-173 | 3.12E-169 | 0.40 | 0.14 | 0.26 | 2.80 |
| 3 | Vstm2b | 1.87 | 3.26E-173 | 6.78E-169 | 0.29 | 0.08 | 0.21 | 3.65 |
| 3 | Anxa7 | 1.22 | 4.62E-173 | 9.61E-169 | 0.65 | 0.34 | 0.31 | 1.92 |
| 3 | Hist1h2bn | 1.60 | 2.58E-171 | 5.37E-167 | 0.38 | 0.13 | 0.25 | 2.90 |
| 3 | Hnrnpab | 0.37 | 1.34E-170 | 2.79E-166 | 1.00 | 0.99 | 0.01 | 1.01 |
| 3 | Vcam1 | 0.91 | 5.55E-168 | 1.15E-163 | 0.76 | 0.41 | 0.35 | 1.87 |
| 3 | Mns1 | 0.85 | 4.93E-162 | 1.03E-157 | 0.84 | 0.52 | 0.33 | 1.63 |

**Supplementary Table 1. Marker genes for the original clusters of integrated clustering of E11.5 control and NotchGOF λ-junction cells.**

| | | | | | | | | |
|---|---|---|---|---|---|---|---|---|
| 3 | Gap43 | 0.79 | 1.32E-160 | 2.74E-156 | 0.74 | 0.39 | 0.35 | 1.91 |
| 3 | Hspe1 | 0.41 | 5.14E-160 | 1.07E-155 | 1.00 | 0.99 | 0.01 | 1.01 |
| 3 | Hist1h2af | 1.54 | 2.08E-159 | 4.32E-155 | 0.36 | 0.12 | 0.24 | 2.90 |
| 3 | Nme1 | 0.41 | 5.47E-159 | 1.14E-154 | 1.00 | 0.99 | 0.01 | 1.01 |
| 3 | Il11ra1 | 0.82 | 1.93E-158 | 4.01E-154 | 0.89 | 0.59 | 0.30 | 1.51 |
| 3 | Bub3 | 0.60 | 2.41E-158 | 5.01E-154 | 0.98 | 0.81 | 0.16 | 1.20 |
| 3 | Rad51 | 0.86 | 1.06E-157 | 2.21E-153 | 0.82 | 0.52 | 0.31 | 1.59 |
| 3 | Ran | 0.33 | 3.64E-156 | 7.57E-152 | 1.00 | 1.00 | 0.00 | 1.00 |
| 3 | Tmem132c | 1.09 | 5.62E-156 | 1.17E-151 | 0.60 | 0.30 | 0.31 | 2.04 |
| 3 | Lgals1 | 0.54 | 2.45E-155 | 5.10E-151 | 1.00 | 0.81 | 0.19 | 1.23 |
| 3 | Srsf3 | 0.34 | 2.71E-153 | 5.64E-149 | 1.00 | 1.00 | 0.00 | 1.00 |
| 3 | Spc25 | 0.76 | 1.07E-151 | 2.23E-147 | 0.85 | 0.52 | 0.33 | 1.63 |
| 3 | Tbx2 | 0.72 | 3.74E-151 | 7.78E-147 | 0.91 | 0.62 | 0.29 | 1.47 |
| 3 | Clspn | 0.75 | 5.08E-151 | 1.06E-146 | 0.90 | 0.59 | 0.30 | 1.51 |
| 3 | Ccne1 | 0.86 | 5.61E-149 | 1.17E-144 | 0.79 | 0.48 | 0.31 | 1.65 |
| 3 | Pcsk5 | 0.97 | 8.33E-149 | 1.73E-144 | 0.66 | 0.34 | 0.32 | 1.96 |
| 3 | Capzb | 0.50 | 1.89E-144 | 3.92E-140 | 0.99 | 0.91 | 0.08 | 1.09 |
| 3 | Lpar1 | 0.64 | 2.94E-143 | 6.11E-139 | 0.95 | 0.68 | 0.27 | 1.40 |
| 3 | Gata2 | 0.88 | 8.37E-143 | 1.74E-138 | 0.73 | 0.42 | 0.32 | 1.76 |
| 3 | Incenp | 0.68 | 1.03E-142 | 2.14E-138 | 0.89 | 0.57 | 0.32 | 1.57 |
| 3 | Cdk1 | 0.56 | 2.86E-141 | 5.95E-137 | 0.95 | 0.64 | 0.31 | 1.49 |
| 3 | Dnajc9 | 0.52 | 1.94E-140 | 4.03E-136 | 0.99 | 0.85 | 0.13 | 1.16 |
| 3 | Mif | 0.35 | 6.19E-140 | 1.29E-135 | 1.00 | 0.99 | 0.01 | 1.01 |
| 3 | Dnmt1 | 0.63 | 5.35E-138 | 1.11E-133 | 0.96 | 0.77 | 0.19 | 1.24 |
| 3 | Prrx2 | 0.63 | 1.97E-137 | 4.11E-133 | 0.98 | 0.72 | 0.26 | 1.36 |
| 3 | Col23a1 | 0.92 | 3.08E-137 | 6.41E-133 | 0.59 | 0.29 | 0.31 | 2.07 |
| 3 | Exo1 | 0.86 | 1.03E-136 | 2.14E-132 | 0.73 | 0.43 | 0.31 | 1.72 |
| 3 | Hist1h2ad | 1.69 | 1.40E-136 | 2.91E-132 | 0.27 | 0.08 | 0.19 | 3.21 |
| 3 | Cd34 | 0.49 | 4.21E-135 | 8.77E-131 | 0.33 | 0.11 | 0.22 | 2.89 |
| 3 | Rrm1 | 0.57 | 1.77E-134 | 3.67E-130 | 0.97 | 0.80 | 0.17 | 1.21 |
| 3 | Pitx2 | 1.05 | 3.04E-130 | 6.32E-126 | 0.34 | 0.13 | 0.22 | 2.74 |
| 3 | U2af1 | 0.38 | 2.44E-128 | 5.08E-124 | 1.00 | 0.97 | 0.03 | 1.03 |
| 3 | Sumo2 | 0.25 | 4.86E-128 | 1.01E-123 | 1.00 | 1.00 | 0.00 | 1.00 |
| 3 | Cenph | 0.68 | 5.96E-128 | 1.24E-123 | 0.88 | 0.61 | 0.27 | 1.44 |
| 3 | Spin2c | 0.86 | 4.54E-127 | 9.44E-123 | 0.69 | 0.39 | 0.30 | 1.77 |
| 3 | Hist1h2bj | 1.17 | 1.92E-126 | 3.99E-122 | 0.48 | 0.22 | 0.25 | 2.13 |
| 3 | Mthfd2 | 0.83 | 4.13E-126 | 8.59E-122 | 0.74 | 0.45 | 0.29 | 1.65 |
| 3 | Tmsb10 | 0.28 | 5.52E-126 | 1.15E-121 | 1.00 | 1.00 | 0.00 | 1.00 |
| 3 | Hist1h2br | 1.30 | 6.60E-126 | 1.37E-121 | 0.35 | 0.14 | 0.22 | 2.60 |
| 3 | Gm9866 | 1.83 | 1.43E-124 | 2.98E-120 | 0.21 | 0.06 | 0.15 | 3.67 |
| 3 | Gxylt2 | 1.02 | 1.49E-124 | 3.10E-120 | 0.50 | 0.23 | 0.27 | 2.17 |
| 3 | Cdca2 | 0.78 | 2.07E-124 | 4.31E-120 | 0.71 | 0.40 | 0.31 | 1.76 |
| 3 | Melk | 0.80 | 3.53E-124 | 7.35E-120 | 0.74 | 0.45 | 0.29 | 1.65 |
| 3 | Fen1 | 0.68 | 1.14E-122 | 2.37E-118 | 0.89 | 0.63 | 0.26 | 1.41 |
| 3 | Snrpg | 0.25 | 7.82E-122 | 1.63E-117 | 1.00 | 1.00 | 0.00 | 1.00 |
| 3 | Lsm3 | 0.43 | 1.14E-121 | 2.36E-117 | 0.99 | 0.91 | 0.08 | 1.09 |
| 3 | Prdm16os | 1.30 | 2.68E-121 | 5.58E-117 | 0.30 | 0.10 | 0.20 | 2.89 |
| 3 | Csrp1 | 0.84 | 4.70E-121 | 9.77E-117 | 0.61 | 0.32 | 0.29 | 1.93 |
| 3 | Hpse2 | 1.07 | 3.93E-119 | 8.17E-115 | 0.38 | 0.15 | 0.23 | 2.50 |
| 3 | Hist1h2ac | 1.55 | 1.78E-118 | 3.70E-114 | 0.27 | 0.09 | 0.18 | 2.95 |
| 3 | Smc4 | 0.51 | 3.37E-118 | 7.02E-114 | 0.99 | 0.78 | 0.21 | 1.27 |
| 3 | Ube2i | 0.33 | 4.17E-117 | 8.66E-113 | 1.00 | 0.98 | 0.02 | 1.02 |
| 3 | Hmga2 | 0.46 | 1.11E-116 | 2.30E-112 | 0.99 | 0.93 | 0.07 | 1.07 |
| 3 | Tuba1b | 0.32 | 1.82E-116 | 3.79E-112 | 1.00 | 0.99 | 0.01 | 1.01 |
| 3 | Hsp90aa1 | 0.30 | 1.91E-116 | 3.97E-112 | 1.00 | 1.00 | 0.00 | 1.00 |
| 3 | Raet1e | 1.09 | 2.32E-115 | 4.83E-111 | 0.46 | 0.21 | 0.25 | 2.18 |

**Supplementary Table 1. Marker genes for the original clusters of integrated clustering of E11.5 control and NotchGOF λ-junction cells.**

| | | | | | | | | |
|---|---|---|---|---|---|---|---|---|
| 3 | Dusp9 | 0.83 | 2.37E-115 | 4.93E-111 | 0.68 | 0.39 | 0.29 | 1.74 |
| 3 | Dhfr | 0.64 | 9.05E-115 | 1.88E-110 | 0.89 | 0.66 | 0.23 | 1.35 |
| 3 | Fbln1 | 0.54 | 9.74E-115 | 2.03E-110 | 0.97 | 0.78 | 0.18 | 1.24 |
| 3 | Ercc6l | 0.89 | 2.06E-114 | 4.29E-110 | 0.61 | 0.33 | 0.28 | 1.82 |
| 3 | Tipin | 0.54 | 2.82E-113 | 5.86E-109 | 0.95 | 0.75 | 0.20 | 1.27 |
| 3 | Hist1h4c | 1.37 | 1.42E-112 | 2.95E-108 | 0.33 | 0.13 | 0.20 | 2.52 |
| 3 | Cenpq | 0.69 | 2.16E-112 | 4.50E-108 | 0.81 | 0.54 | 0.28 | 1.52 |
| 3 | Wif1 | 1.51 | 4.38E-112 | 9.11E-108 | 0.24 | 0.08 | 0.17 | 3.09 |
| 3 | Mybl2 | 1.12 | 5.67E-112 | 1.18E-107 | 0.46 | 0.23 | 0.24 | 2.05 |
| 3 | Fxyd6 | 0.60 | 6.53E-112 | 1.36E-107 | 0.90 | 0.66 | 0.24 | 1.36 |
| 3 | Alyref | 0.38 | 2.44E-111 | 5.07E-107 | 1.00 | 0.94 | 0.06 | 1.06 |
| 3 | Samd4 | 0.83 | 1.03E-110 | 2.14E-106 | 0.62 | 0.34 | 0.28 | 1.81 |
| 3 | Id3 | 0.46 | 1.26E-110 | 2.61E-106 | 1.00 | 0.95 | 0.04 | 1.05 |
| 3 | Hist1h2bp | 1.70 | 1.65E-110 | 3.44E-106 | 0.22 | 0.07 | 0.15 | 3.17 |
| 3 | Ccn1 | 1.01 | 2.42E-110 | 5.03E-106 | 0.54 | 0.29 | 0.26 | 1.90 |
| 3 | Crabp2 | 0.42 | 3.32E-110 | 6.91E-106 | 1.00 | 0.91 | 0.09 | 1.10 |
| 3 | Phgdh | 0.52 | 6.39E-109 | 1.33E-104 | 0.97 | 0.81 | 0.16 | 1.20 |
| 3 | Pola1 | 0.74 | 6.58E-109 | 1.37E-104 | 0.74 | 0.47 | 0.27 | 1.59 |
| 3 | Rassf3 | 0.91 | 7.12E-108 | 1.48E-103 | 0.62 | 0.35 | 0.27 | 1.78 |
| 3 | Crlf1 | 0.69 | 9.07E-108 | 1.89E-103 | 0.87 | 0.62 | 0.25 | 1.41 |
| 3 | Brca1 | 0.80 | 1.38E-107 | 2.87E-103 | 0.64 | 0.36 | 0.28 | 1.78 |
| 3 | Hmgb2 | 0.41 | 2.58E-107 | 5.36E-103 | 1.00 | 0.98 | 0.02 | 1.02 |
| 3 | Twistnb | 0.60 | 2.96E-107 | 6.15E-103 | 0.92 | 0.76 | 0.16 | 1.21 |
| 3 | Hist1h2bh | 1.43 | 7.22E-107 | 1.50E-102 | 0.28 | 0.10 | 0.18 | 2.71 |
| 3 | Prim1 | 0.61 | 7.58E-107 | 1.58E-102 | 0.89 | 0.65 | 0.23 | 1.36 |
| 3 | Cox5a | 0.29 | 2.06E-106 | 4.28E-102 | 1.00 | 0.99 | 0.01 | 1.01 |
| 3 | Hist1h2bg | 1.51 | 2.20E-106 | 4.58E-102 | 0.25 | 0.09 | 0.16 | 2.92 |
| 3 | Akr7a5 | 0.53 | 2.90E-106 | 6.03E-102 | 0.94 | 0.78 | 0.16 | 1.21 |
| 3 | Pdap1 | 0.30 | 4.76E-106 | 9.91E-102 | 1.00 | 0.99 | 0.01 | 1.01 |
| 3 | Kif11 | 0.57 | 8.46E-106 | 1.76E-101 | 0.83 | 0.51 | 0.31 | 1.61 |
| 3 | Col25a1 | 0.98 | 1.19E-105 | 2.46E-101 | 0.40 | 0.18 | 0.22 | 2.25 |
| 3 | Igfbp4 | 0.43 | 1.30E-105 | 2.70E-101 | 1.00 | 0.89 | 0.10 | 1.12 |
| 3 | Srsf7 | 0.30 | 1.64E-105 | 3.41E-101 | 1.00 | 0.99 | 0.01 | 1.01 |
| 3 | Dctpp1 | 0.40 | 3.82E-105 | 7.95E-101 | 0.99 | 0.93 | 0.06 | 1.07 |
| 3 | Dpep1 | 0.86 | 4.80E-105 | 9.98E-101 | 0.45 | 0.21 | 0.24 | 2.18 |
| 3 | Hspd1 | 0.37 | 7.97E-105 | 1.66E-100 | 1.00 | 0.98 | 0.02 | 1.02 |
| 3 | Cped1 | 0.71 | 1.67E-104 | 3.47E-100 | 0.73 | 0.44 | 0.29 | 1.65 |
| 3 | Slbp | 0.49 | 3.93E-104 | 8.17E-100 | 0.98 | 0.86 | 0.12 | 1.13 |
| 3 | Prim2 | 0.88 | 4.07E-104 | 8.46E-100 | 0.62 | 0.36 | 0.26 | 1.71 |
| 3 | Pth1r | 0.57 | 6.01E-104 | 1.25E-99 | 0.86 | 0.55 | 0.32 | 1.58 |
| 3 | Epha8 | 2.14 | 7.03E-104 | 1.46E-99 | 0.14 | 0.03 | 0.11 | 4.39 |
| 3 | Irx5 | 1.20 | 2.51E-103 | 5.23E-99 | 0.49 | 0.26 | 0.23 | 1.86 |
| 3 | Ncl | 0.33 | 5.80E-103 | 1.21E-98 | 1.00 | 1.00 | 0.01 | 1.01 |
| 3 | Supt16 | 0.36 | 2.38E-102 | 4.95E-98 | 1.00 | 0.96 | 0.04 | 1.04 |
| 3 | Hist1h3a | 1.18 | 2.49E-102 | 5.17E-98 | 0.38 | 0.17 | 0.21 | 2.21 |
| 3 | Diaph3 | 0.85 | 2.80E-102 | 5.82E-98 | 0.60 | 0.35 | 0.25 | 1.72 |
| 3 | Cdc45 | 0.84 | 7.54E-102 | 1.57E-97 | 0.61 | 0.35 | 0.26 | 1.76 |
| 3 | Alx4 | 0.74 | 7.77E-102 | 1.62E-97 | 0.68 | 0.40 | 0.28 | 1.69 |
| 3 | Trip13 | 0.66 | 1.15E-101 | 2.39E-97 | 0.78 | 0.52 | 0.27 | 1.52 |
| 3 | Kcnb2 | 1.82 | 3.52E-101 | 7.33E-97 | 0.16 | 0.04 | 0.12 | 3.77 |
| 3 | Myorg | 1.70 | 1.87E-100 | 3.89E-96 | 0.21 | 0.07 | 0.14 | 3.15 |
| 3 | Hint1 | 0.26 | 7.16E-100 | 1.49E-95 | 1.00 | 0.99 | 0.01 | 1.01 |
| 3 | Phf5a | 0.35 | 3.57E-99 | 7.43E-95 | 1.00 | 0.95 | 0.04 | 1.05 |
| 3 | Smc6 | 0.46 | 4.05E-99 | 8.41E-95 | 0.97 | 0.86 | 0.11 | 1.13 |
| 3 | Mis18bp1 | 0.52 | 1.97E-98 | 4.09E-94 | 0.80 | 0.49 | 0.31 | 1.63 |
| 3 | Gm48898 | 2.25 | 3.05E-98 | 6.35E-94 | 0.11 | 0.02 | 0.09 | 4.91 |

**Supplementary Table 1. Marker genes for the original clusters of integrated clustering of E11.5 control and NotchGOF λ-junction cells.**

| | | | | | | | | |
|---|---|---|---|---|---|---|---|---|
| 3 | Fn1 | 0.42 | 3.35E-98 | 6.97E-94 | 0.96 | 0.77 | 0.19 | 1.24 |
| 3 | Haspin | 0.92 | 6.10E-98 | 1.27E-93 | 0.50 | 0.26 | 0.24 | 1.90 |
| 3 | Lsm8 | 0.41 | 1.86E-97 | 3.86E-93 | 0.98 | 0.90 | 0.08 | 1.09 |
| 3 | Fzd1 | 0.82 | 2.64E-97 | 5.48E-93 | 0.58 | 0.32 | 0.26 | 1.82 |
| 3 | Snrpd1 | 0.28 | 6.68E-97 | 1.39E-92 | 1.00 | 0.99 | 0.01 | 1.01 |
| 3 | Dnm3 | 1.12 | 1.05E-96 | 2.19E-92 | 0.35 | 0.15 | 0.20 | 2.37 |
| 3 | Hspa9 | 0.43 | 3.06E-96 | 6.36E-92 | 0.98 | 0.89 | 0.09 | 1.10 |
| 3 | Mpp6 | 0.53 | 5.30E-96 | 1.10E-91 | 0.92 | 0.73 | 0.20 | 1.27 |
| 3 | Cdca5 | 0.85 | 7.53E-96 | 1.57E-91 | 0.56 | 0.31 | 0.24 | 1.77 |
| 3 | Sox11 | 0.33 | 1.38E-95 | 2.87E-91 | 1.00 | 0.96 | 0.04 | 1.04 |
| 3 | Selenoh | 0.32 | 2.17E-95 | 4.52E-91 | 1.00 | 0.96 | 0.04 | 1.04 |
| 3 | Bicc1 | 0.91 | 4.76E-95 | 9.90E-91 | 0.44 | 0.21 | 0.23 | 2.07 |
| 3 | Mki67 | 0.47 | 1.23E-94 | 2.55E-90 | 0.96 | 0.70 | 0.26 | 1.37 |
| 3 | Sgo1 | 0.65 | 1.41E-94 | 2.93E-90 | 0.70 | 0.43 | 0.28 | 1.65 |
| 3 | Rbbp4 | 0.34 | 2.24E-94 | 4.66E-90 | 1.00 | 0.96 | 0.04 | 1.04 |
| 3 | Rfc4 | 0.49 | 3.14E-94 | 6.54E-90 | 0.94 | 0.78 | 0.16 | 1.21 |
| 3 | Net1 | 0.69 | 3.16E-94 | 6.57E-90 | 0.80 | 0.57 | 0.23 | 1.41 |
| 3 | Gch1 | 1.58 | 3.30E-94 | 6.87E-90 | 0.20 | 0.07 | 0.14 | 3.06 |
| 3 | Kif15 | 0.58 | 3.95E-94 | 8.22E-90 | 0.87 | 0.61 | 0.26 | 1.43 |
| 3 | Srsf10 | 0.37 | 1.08E-93 | 2.24E-89 | 1.00 | 0.94 | 0.06 | 1.06 |
| 3 | Ptges3 | 0.33 | 1.31E-93 | 2.73E-89 | 1.00 | 0.97 | 0.03 | 1.03 |
| 3 | Eepd1 | 1.29 | 2.18E-93 | 4.54E-89 | 0.27 | 0.10 | 0.17 | 2.62 |
| 3 | Cenpu | 0.92 | 3.91E-93 | 8.13E-89 | 0.52 | 0.29 | 0.23 | 1.79 |
| 3 | Nudc | 0.34 | 5.19E-93 | 1.08E-88 | 1.00 | 0.96 | 0.04 | 1.04 |
| 3 | Dcakd | 0.38 | 1.54E-92 | 3.20E-88 | 0.99 | 0.90 | 0.09 | 1.10 |
| 3 | Parm1 | 1.00 | 3.82E-92 | 7.95E-88 | 0.34 | 0.15 | 0.19 | 2.32 |
| 3 | Ppa1 | 0.39 | 1.80E-91 | 3.75E-87 | 0.99 | 0.89 | 0.10 | 1.11 |
| 3 | Fkbp3 | 0.28 | 2.64E-91 | 5.48E-87 | 1.00 | 0.98 | 0.02 | 1.02 |
| 3 | Nkd1 | 0.86 | 5.11E-90 | 1.06E-85 | 0.59 | 0.34 | 0.25 | 1.72 |
| 3 | Prps1 | 0.76 | 7.43E-90 | 1.55E-85 | 0.67 | 0.42 | 0.24 | 1.57 |
| 3 | Lsm6 | 0.36 | 1.02E-89 | 2.12E-85 | 0.99 | 0.93 | 0.06 | 1.07 |
| 3 | 2700069I18Rik | 1.28 | 1.18E-89 | 2.44E-85 | 0.29 | 0.12 | 0.17 | 2.49 |
| 3 | Hist1h2bl | 1.36 | 1.18E-88 | 2.45E-84 | 0.25 | 0.09 | 0.15 | 2.66 |
| 3 | Asns | 0.74 | 1.21E-88 | 2.51E-84 | 0.68 | 0.42 | 0.26 | 1.62 |
| 3 | Rai14 | 0.50 | 1.36E-88 | 2.83E-84 | 0.89 | 0.62 | 0.27 | 1.42 |
| 3 | Prkcq | 1.86 | 1.50E-88 | 3.12E-84 | 0.12 | 0.03 | 0.09 | 4.32 |
| 3 | Fam111a | 0.77 | 7.19E-88 | 1.50E-83 | 0.55 | 0.31 | 0.24 | 1.80 |
| 3 | Sms | 0.42 | 9.16E-88 | 1.90E-83 | 0.97 | 0.85 | 0.12 | 1.14 |
| 3 | H1f0 | 0.55 | 1.07E-87 | 2.22E-83 | 0.95 | 0.84 | 0.11 | 1.13 |
| 3 | Eif4a3 | 0.42 | 2.56E-87 | 5.32E-83 | 0.98 | 0.86 | 0.11 | 1.13 |
| 3 | Nuf2 | 0.63 | 2.81E-87 | 5.85E-83 | 0.72 | 0.46 | 0.27 | 1.58 |
| 3 | Dnah7a | 1.15 | 2.95E-87 | 6.14E-83 | 0.28 | 0.11 | 0.17 | 2.52 |
| 3 | Ska1 | 0.70 | 3.26E-87 | 6.78E-83 | 0.65 | 0.40 | 0.25 | 1.63 |
| 3 | Lpl | 0.93 | 1.26E-86 | 2.62E-82 | 0.33 | 0.15 | 0.18 | 2.26 |
| 3 | P3h2 | 1.42 | 7.73E-86 | 1.61E-81 | 0.20 | 0.07 | 0.13 | 3.02 |
| 3 | Hnrnpd | 0.28 | 9.16E-86 | 1.91E-81 | 1.00 | 0.99 | 0.01 | 1.01 |
| 3 | Cdon | 0.98 | 9.17E-86 | 1.91E-81 | 0.41 | 0.20 | 0.21 | 2.02 |
| 3 | Sgo2a | 0.51 | 1.65E-85 | 3.43E-81 | 0.75 | 0.46 | 0.29 | 1.64 |
| 3 | Mcm10 | 0.77 | 2.11E-85 | 4.39E-81 | 0.58 | 0.33 | 0.24 | 1.73 |
| 3 | Id2 | 0.49 | 7.16E-85 | 1.49E-80 | 0.97 | 0.85 | 0.12 | 1.14 |
| 3 | Ywhah | 0.34 | 7.77E-85 | 1.62E-80 | 0.98 | 0.90 | 0.08 | 1.09 |
| 3 | Snai1 | 0.71 | 3.40E-84 | 7.06E-80 | 0.65 | 0.39 | 0.25 | 1.65 |
| 3 | Dab2 | 0.58 | 5.60E-84 | 1.17E-79 | 0.59 | 0.33 | 0.25 | 1.76 |
| 3 | Plk4 | 0.54 | 7.76E-84 | 1.61E-79 | 0.79 | 0.53 | 0.26 | 1.49 |
| 3 | Scube1 | 0.63 | 1.27E-83 | 2.65E-79 | 0.62 | 0.35 | 0.26 | 1.75 |
| 3 | Lyar | 0.41 | 1.44E-83 | 2.99E-79 | 0.98 | 0.86 | 0.12 | 1.14 |

**Supplementary Table 1. Marker genes for the original clusters of integrated clustering of E11.5 control and NotchGOF λ-junction cells.**

| | | | | | | | | |
|---|---|---|---|---|---|---|---|---|
| 3 | Farsb | 0.63 | 2.82E-83 | 5.87E-79 | 0.77 | 0.54 | 0.23 | 1.43 |
| 3 | Mrpl12 | 0.38 | 3.25E-83 | 6.77E-79 | 0.99 | 0.89 | 0.09 | 1.10 |
| 3 | Dixdc1 | 0.69 | 7.49E-83 | 1.56E-78 | 0.63 | 0.38 | 0.25 | 1.67 |
| 3 | Dtymk | 0.39 | 9.84E-83 | 2.05E-78 | 0.98 | 0.88 | 0.10 | 1.12 |
| 3 | Paics | 0.36 | 1.02E-82 | 2.12E-78 | 0.99 | 0.92 | 0.08 | 1.08 |
| 3 | Jaml | 2.19 | 1.30E-82 | 2.70E-78 | 0.10 | 0.02 | 0.08 | 4.68 |
| 3 | Mtap | 0.51 | 1.59E-82 | 3.32E-78 | 0.89 | 0.68 | 0.21 | 1.31 |
| 3 | Ctsb | 0.36 | 1.95E-82 | 4.06E-78 | 0.92 | 0.74 | 0.19 | 1.26 |
| 3 | Ifitm1 | 0.55 | 3.76E-82 | 7.83E-78 | 0.37 | 0.17 | 0.20 | 2.16 |
| 3 | Rev3l | 0.55 | 6.82E-82 | 1.42E-77 | 0.81 | 0.57 | 0.24 | 1.41 |
| 3 | Sec61b | 0.29 | 1.42E-81 | 2.96E-77 | 1.00 | 0.98 | 0.02 | 1.02 |
| 3 | Hjurp | 0.53 | 1.44E-81 | 2.99E-77 | 0.84 | 0.58 | 0.26 | 1.44 |
| 3 | Bcat1 | 0.73 | 1.14E-80 | 2.37E-76 | 0.53 | 0.29 | 0.24 | 1.81 |
| 3 | Ndc80 | 0.65 | 1.21E-80 | 2.52E-76 | 0.65 | 0.40 | 0.25 | 1.62 |
| 3 | Snrnp40 | 0.39 | 1.88E-80 | 3.91E-76 | 0.97 | 0.84 | 0.13 | 1.15 |
| 3 | Anp32e | 0.34 | 1.96E-80 | 4.08E-76 | 1.00 | 0.94 | 0.05 | 1.06 |
| 3 | Tpm4 | 0.32 | 8.01E-80 | 1.67E-75 | 1.00 | 0.93 | 0.07 | 1.07 |
| 3 | Orc6 | 0.43 | 8.54E-80 | 1.78E-75 | 0.95 | 0.78 | 0.16 | 1.21 |
| 3 | Trir | 0.32 | 1.09E-79 | 2.27E-75 | 0.99 | 0.94 | 0.05 | 1.05 |
| 3 | Rps27l | 0.26 | 1.49E-79 | 3.10E-75 | 1.00 | 0.99 | 0.01 | 1.01 |
| 3 | Smc2 | 0.35 | 1.74E-79 | 3.62E-75 | 0.99 | 0.85 | 0.14 | 1.16 |
| 3 | Arhgdib | 0.43 | 2.19E-79 | 4.55E-75 | 0.74 | 0.48 | 0.27 | 1.56 |
| 3 | Ccdc34 | 0.34 | 2.48E-79 | 5.15E-75 | 0.99 | 0.91 | 0.08 | 1.09 |
| 3 | Ctps | 0.58 | 2.55E-79 | 5.31E-75 | 0.79 | 0.57 | 0.23 | 1.40 |
| 3 | Ncapg2 | 0.73 | 3.55E-79 | 7.39E-75 | 0.62 | 0.39 | 0.23 | 1.60 |
| 3 | Spdl1 | 0.62 | 6.60E-79 | 1.37E-74 | 0.66 | 0.42 | 0.25 | 1.60 |
| 3 | Nkd2 | 1.04 | 7.74E-79 | 1.61E-74 | 0.28 | 0.12 | 0.16 | 2.40 |
| 3 | Cpxm2 | 1.66 | 1.46E-78 | 3.03E-74 | 0.14 | 0.04 | 0.10 | 3.58 |
| 3 | Rflnb | 0.51 | 1.81E-78 | 3.76E-74 | 0.78 | 0.53 | 0.25 | 1.47 |
| 3 | Traf3ip2 | 0.75 | 2.24E-78 | 4.66E-74 | 0.41 | 0.20 | 0.21 | 2.01 |
| 3 | Diaph2 | 1.15 | 2.64E-78 | 5.49E-74 | 0.31 | 0.14 | 0.17 | 2.19 |
| 3 | Slfn9 | 0.83 | 2.69E-78 | 5.59E-74 | 0.45 | 0.23 | 0.21 | 1.91 |
| 3 | Tmpo | 0.32 | 3.63E-78 | 7.54E-74 | 1.00 | 0.95 | 0.05 | 1.05 |
| 3 | Haus3 | 0.62 | 5.48E-78 | 1.14E-73 | 0.72 | 0.48 | 0.25 | 1.52 |
| 3 | Sema5a | 0.53 | 8.93E-78 | 1.86E-73 | 0.61 | 0.35 | 0.26 | 1.73 |
| 3 | 2810408I11Rik | 0.80 | 2.92E-77 | 6.07E-73 | 0.50 | 0.28 | 0.22 | 1.76 |
| 3 | Srsf2 | 0.26 | 3.40E-77 | 7.08E-73 | 1.00 | 0.99 | 0.01 | 1.01 |
| 3 | Rnf227 | 0.65 | 4.32E-77 | 8.98E-73 | 0.65 | 0.40 | 0.26 | 1.64 |
| 3 | Sgpp2 | 1.98 | 1.22E-76 | 2.53E-72 | 0.11 | 0.03 | 0.08 | 4.28 |
| 3 | Blm | 0.61 | 2.64E-76 | 5.49E-72 | 0.74 | 0.50 | 0.24 | 1.48 |
| 3 | Limd2 | 0.35 | 3.58E-76 | 7.44E-72 | 0.99 | 0.89 | 0.10 | 1.12 |
| 3 | Ckap2l | 0.49 | 7.33E-76 | 1.52E-71 | 0.74 | 0.46 | 0.28 | 1.59 |
| 3 | Chaf1a | 0.50 | 9.21E-76 | 1.92E-71 | 0.85 | 0.63 | 0.22 | 1.35 |
| 3 | Cbx5 | 0.32 | 1.57E-75 | 3.26E-71 | 0.99 | 0.95 | 0.04 | 1.05 |
| 3 | Rpp25l | 0.48 | 1.98E-75 | 4.11E-71 | 0.91 | 0.73 | 0.18 | 1.25 |
| 3 | Rerg | 0.58 | 3.04E-75 | 6.31E-71 | 0.45 | 0.23 | 0.22 | 1.95 |
| 3 | Cacna1g | 0.67 | 3.48E-75 | 7.24E-71 | 0.55 | 0.31 | 0.24 | 1.77 |
| 3 | Hist1h2bf | 1.27 | 8.05E-75 | 1.67E-70 | 0.21 | 0.08 | 0.13 | 2.68 |
| 3 | Usp1 | 0.41 | 8.83E-75 | 1.84E-70 | 0.96 | 0.80 | 0.16 | 1.20 |
| 3 | Cenpk | 0.53 | 1.56E-74 | 3.25E-70 | 0.79 | 0.55 | 0.24 | 1.43 |
| 3 | Gm1673 | 0.34 | 2.14E-74 | 4.45E-70 | 1.00 | 0.92 | 0.08 | 1.08 |
| 3 | Hist1h4a | 1.11 | 3.78E-74 | 7.85E-70 | 0.34 | 0.17 | 0.17 | 2.02 |
| 3 | Flrt3 | 0.76 | 4.00E-74 | 8.32E-70 | 0.65 | 0.43 | 0.22 | 1.51 |
| 3 | Pdlim7 | 0.45 | 2.90E-73 | 6.02E-69 | 0.92 | 0.74 | 0.18 | 1.24 |
| 3 | Dbi | 0.27 | 4.38E-73 | 9.12E-69 | 1.00 | 0.98 | 0.02 | 1.02 |
| 3 | Atp1b3 | 0.53 | 5.27E-73 | 1.10E-68 | 0.88 | 0.70 | 0.17 | 1.24 |

**Supplementary Table 1. Marker genes for the original clusters of integrated clustering of E11.5 control and NotchGOF λ-junction cells.**

| | | | | | | | | |
|---|---|---|---|---|---|---|---|---|
| 3 | Bzw1 | 0.27 | 5.55E-73 | 1.15E-68 | 1.00 | 0.98 | 0.02 | 1.02 |
| 3 | Lsm5 | 0.30 | 6.68E-73 | 1.39E-68 | 0.99 | 0.94 | 0.05 | 1.06 |
| 3 | Fli1 | 0.57 | 1.08E-72 | 2.24E-68 | 0.61 | 0.35 | 0.26 | 1.73 |
| 3 | Mex3b | 0.47 | 1.54E-72 | 3.20E-68 | 0.88 | 0.68 | 0.20 | 1.29 |
| 3 | Ttk | 0.66 | 1.81E-72 | 3.76E-68 | 0.57 | 0.34 | 0.23 | 1.66 |
| 3 | Ror2 | 0.71 | 3.17E-72 | 6.60E-68 | 0.51 | 0.28 | 0.23 | 1.81 |
| 3 | Prc1 | 0.32 | 4.94E-72 | 1.03E-67 | 0.79 | 0.50 | 0.29 | 1.57 |
| 3 | Psat1 | 0.60 | 8.27E-72 | 1.72E-67 | 0.67 | 0.43 | 0.24 | 1.56 |
| 3 | Cmc2 | 0.54 | 1.40E-71 | 2.90E-67 | 0.81 | 0.58 | 0.22 | 1.38 |
| 3 | Tmem119 | 0.49 | 3.03E-71 | 6.31E-67 | 0.71 | 0.43 | 0.28 | 1.63 |
| 3 | Asph | 0.48 | 6.42E-71 | 1.33E-66 | 0.86 | 0.63 | 0.23 | 1.37 |
| 3 | Siah1b | 0.71 | 7.09E-71 | 1.47E-66 | 0.49 | 0.27 | 0.21 | 1.78 |
| 3 | Zfhx3 | 0.62 | 1.04E-70 | 2.16E-66 | 0.94 | 0.80 | 0.14 | 1.17 |
| 3 | Snu13 | 0.27 | 1.12E-70 | 2.32E-66 | 1.00 | 0.97 | 0.02 | 1.02 |
| 3 | Bub1 | 0.48 | 1.62E-70 | 3.37E-66 | 0.81 | 0.54 | 0.27 | 1.49 |
| 3 | Aurka | 0.51 | 2.60E-70 | 5.41E-66 | 0.69 | 0.43 | 0.26 | 1.59 |
| 3 | Clgn | 1.57 | 2.66E-70 | 5.53E-66 | 0.14 | 0.04 | 0.10 | 3.31 |
| 3 | Adamts19 | 0.81 | 8.65E-70 | 1.80E-65 | 0.43 | 0.23 | 0.20 | 1.88 |
| 3 | Mrpl18 | 0.32 | 1.10E-69 | 2.30E-65 | 0.99 | 0.93 | 0.06 | 1.06 |
| 3 | Eln | 1.25 | 1.30E-69 | 2.70E-65 | 0.19 | 0.07 | 0.12 | 2.79 |
| 3 | Mis18a | 0.57 | 1.59E-69 | 3.30E-65 | 0.68 | 0.44 | 0.24 | 1.55 |
| 3 | Cfl2 | 0.40 | 2.09E-69 | 4.36E-65 | 0.95 | 0.79 | 0.15 | 1.19 |
| 3 | Ipo5 | 0.47 | 3.52E-69 | 7.33E-65 | 0.88 | 0.68 | 0.20 | 1.29 |
| 3 | H2-Q7 | 1.19 | 5.85E-69 | 1.22E-64 | 0.24 | 0.10 | 0.14 | 2.43 |
| 3 | Nolc1 | 0.43 | 6.13E-69 | 1.27E-64 | 0.94 | 0.79 | 0.15 | 1.18 |
| 3 | Hirip3 | 0.52 | 7.77E-69 | 1.62E-64 | 0.85 | 0.67 | 0.18 | 1.27 |
| 3 | Ret | 1.59 | 8.17E-69 | 1.70E-64 | 0.16 | 0.05 | 0.11 | 2.96 |
| 3 | Fibin | 0.94 | 1.07E-68 | 2.23E-64 | 0.26 | 0.11 | 0.15 | 2.35 |
| 3 | Cct6a | 0.27 | 1.20E-68 | 2.50E-64 | 1.00 | 0.98 | 0.02 | 1.02 |
| 3 | Hist1h2bk | 0.92 | 2.75E-68 | 5.73E-64 | 0.39 | 0.21 | 0.18 | 1.87 |
| 3 | Hist4h4 | 1.29 | 3.19E-68 | 6.64E-64 | 0.22 | 0.09 | 0.13 | 2.48 |
| 3 | Lrr1 | 0.81 | 4.16E-68 | 8.65E-64 | 0.44 | 0.25 | 0.19 | 1.79 |
| 3 | Hdgfl3 | 0.43 | 5.19E-68 | 1.08E-63 | 0.89 | 0.69 | 0.20 | 1.30 |
| 3 | Zeb1 | 0.57 | 5.27E-68 | 1.10E-63 | 0.57 | 0.33 | 0.24 | 1.74 |
| 3 | Rnps1 | 0.30 | 5.68E-68 | 1.18E-63 | 0.99 | 0.94 | 0.05 | 1.05 |
| 3 | Atp1b1 | 1.21 | 1.26E-67 | 2.63E-63 | 0.20 | 0.08 | 0.12 | 2.56 |
| 3 | Wdhd1 | 0.70 | 1.50E-67 | 3.12E-63 | 0.52 | 0.31 | 0.21 | 1.69 |
| 3 | Tubg1 | 0.55 | 4.89E-67 | 1.02E-62 | 0.77 | 0.55 | 0.22 | 1.39 |
| 3 | Brip1 | 0.87 | 6.37E-67 | 1.33E-62 | 0.40 | 0.21 | 0.18 | 1.86 |
| 3 | Pcolce2 | 0.91 | 7.54E-67 | 1.57E-62 | 0.28 | 0.12 | 0.15 | 2.24 |
| 3 | Ncapg | 0.51 | 1.35E-66 | 2.80E-62 | 0.70 | 0.46 | 0.24 | 1.53 |
| 3 | Nxt1 | 0.47 | 1.61E-66 | 3.35E-62 | 0.87 | 0.66 | 0.21 | 1.32 |
| 3 | Gm47283 | 0.35 | 1.62E-66 | 3.37E-62 | 0.99 | 0.91 | 0.08 | 1.09 |
| 3 | Srm | 0.48 | 1.81E-66 | 3.76E-62 | 0.86 | 0.66 | 0.20 | 1.30 |
| 3 | Dscc1 | 0.75 | 2.48E-66 | 5.16E-62 | 0.49 | 0.29 | 0.20 | 1.71 |
| 3 | Eme1 | 0.64 | 2.58E-66 | 5.37E-62 | 0.57 | 0.35 | 0.23 | 1.65 |
| 3 | Abracl | 0.32 | 5.95E-66 | 1.24E-61 | 0.99 | 0.92 | 0.07 | 1.07 |
| 3 | Asf1b | 0.58 | 6.15E-66 | 1.28E-61 | 0.65 | 0.42 | 0.23 | 1.55 |
| 3 | Bach2 | 0.66 | 6.73E-66 | 1.40E-61 | 0.58 | 0.36 | 0.22 | 1.60 |
| 3 | Ppm1g | 0.42 | 6.82E-66 | 1.42E-61 | 0.94 | 0.80 | 0.14 | 1.17 |
| 3 | Ncaph | 0.54 | 7.76E-66 | 1.61E-61 | 0.71 | 0.47 | 0.24 | 1.50 |
| 3 | Eif4ebp1 | 0.49 | 8.02E-66 | 1.67E-61 | 0.86 | 0.68 | 0.18 | 1.27 |
| 3 | Emilin2 | 0.72 | 8.62E-66 | 1.79E-61 | 0.44 | 0.24 | 0.20 | 1.84 |
| 3 | Zcchc12 | 1.51 | 1.45E-65 | 3.01E-61 | 0.15 | 0.05 | 0.10 | 3.06 |
| 3 | Umps | 0.52 | 2.07E-65 | 4.32E-61 | 0.80 | 0.59 | 0.21 | 1.35 |
| 3 | Sox8 | 1.33 | 4.69E-65 | 9.76E-61 | 0.15 | 0.05 | 0.10 | 3.02 |

**Supplementary Table 1. Marker genes for the original clusters of integrated clustering of E11.5 control and NotchGOF λ-junction cells.**

| | | | | | | | | |
|---|---|---|---|---|---|---|---|---|
| 3 | Pole | 0.72 | 6.37E-65 | 1.32E-60 | 0.48 | 0.28 | 0.20 | 1.73 |
| 3 | Skp2 | 0.44 | 8.85E-65 | 1.84E-60 | 0.88 | 0.68 | 0.21 | 1.30 |
| 3 | Shcbp1 | 0.55 | 1.05E-64 | 2.19E-60 | 0.68 | 0.45 | 0.23 | 1.51 |
| 3 | Lmcd1 | 1.01 | 2.18E-64 | 4.53E-60 | 0.20 | 0.08 | 0.12 | 2.56 |
| 3 | Ndufaf2 | 0.44 | 2.42E-64 | 5.03E-60 | 0.90 | 0.73 | 0.17 | 1.24 |
| 3 | Rfc2 | 0.39 | 2.58E-64 | 5.36E-60 | 0.94 | 0.79 | 0.16 | 1.20 |
| 3 | Tlnrd1 | 0.54 | 3.34E-64 | 6.95E-60 | 0.71 | 0.48 | 0.23 | 1.47 |
| 3 | Scarf2 | 0.68 | 4.18E-64 | 8.70E-60 | 0.45 | 0.24 | 0.21 | 1.85 |
| 3 | Clmp | 0.38 | 7.87E-64 | 1.64E-59 | 0.90 | 0.62 | 0.28 | 1.46 |
| 3 | Cenpw | 0.42 | 7.97E-64 | 1.66E-59 | 0.92 | 0.72 | 0.20 | 1.28 |
| 3 | Shmt1 | 0.71 | 8.29E-64 | 1.72E-59 | 0.54 | 0.34 | 0.20 | 1.59 |
| 3 | Timm8a1 | 0.47 | 8.82E-64 | 1.84E-59 | 0.87 | 0.68 | 0.19 | 1.28 |
| 3 | Cks1b | 0.28 | 9.37E-64 | 1.95E-59 | 1.00 | 0.95 | 0.05 | 1.05 |
| 3 | Ank3 | 0.49 | 5.77E-63 | 1.20E-58 | 0.70 | 0.47 | 0.24 | 1.50 |
| 3 | Hist1h2ag | 0.68 | 9.13E-63 | 1.90E-58 | 0.50 | 0.30 | 0.20 | 1.69 |
| 3 | Pid1 | 0.70 | 9.92E-63 | 2.06E-58 | 0.36 | 0.18 | 0.18 | 1.98 |
| 3 | Nmral1 | 0.45 | 1.25E-62 | 2.60E-58 | 0.88 | 0.70 | 0.18 | 1.26 |
| 3 | Metrn | 0.49 | 2.04E-62 | 4.23E-58 | 0.72 | 0.50 | 0.23 | 1.46 |
| 3 | Alkal2 | 1.25 | 2.49E-62 | 5.18E-58 | 0.19 | 0.07 | 0.12 | 2.59 |
| 3 | Meaf6 | 0.46 | 3.69E-62 | 7.67E-58 | 0.85 | 0.65 | 0.19 | 1.30 |
| 3 | Ezh2 | 0.30 | 4.75E-62 | 9.87E-58 | 0.99 | 0.94 | 0.05 | 1.05 |
| 3 | Cenpm | 0.49 | 6.09E-62 | 1.27E-57 | 0.79 | 0.58 | 0.21 | 1.36 |
| 3 | Me2 | 0.60 | 6.50E-62 | 1.35E-57 | 0.65 | 0.44 | 0.22 | 1.49 |
| 3 | Uba2 | 0.33 | 8.72E-62 | 1.81E-57 | 0.96 | 0.86 | 0.10 | 1.12 |
| 3 | Rfc5 | 0.47 | 1.38E-61 | 2.88E-57 | 0.79 | 0.57 | 0.22 | 1.39 |
| 3 | Spc24 | 0.37 | 2.28E-61 | 4.75E-57 | 0.93 | 0.72 | 0.21 | 1.30 |
| 3 | Pbdc1 | 0.35 | 3.77E-61 | 7.85E-57 | 0.96 | 0.85 | 0.11 | 1.13 |
| 3 | Srrm1 | 0.26 | 4.64E-61 | 9.65E-57 | 0.99 | 0.96 | 0.03 | 1.03 |
| 3 | Col27a1 | 0.78 | 7.54E-61 | 1.57E-56 | 0.34 | 0.17 | 0.17 | 1.98 |
| 3 | Pa2g4 | 0.32 | 1.21E-60 | 2.52E-56 | 0.98 | 0.91 | 0.08 | 1.08 |
| 3 | Cdca4 | 0.50 | 1.62E-60 | 3.36E-56 | 0.75 | 0.54 | 0.22 | 1.40 |
| 3 | Hist1h4j | 1.60 | 1.68E-60 | 3.49E-56 | 0.14 | 0.05 | 0.09 | 3.04 |
| 3 | Gsto1 | 0.44 | 1.78E-60 | 3.70E-56 | 0.83 | 0.63 | 0.20 | 1.31 |
| 3 | Anln | 0.51 | 2.93E-60 | 6.10E-56 | 0.59 | 0.37 | 0.22 | 1.60 |
| 3 | Pdgfrb | 0.48 | 3.75E-60 | 7.81E-56 | 0.68 | 0.43 | 0.25 | 1.57 |
| 3 | Dnajc8 | 0.27 | 6.34E-60 | 1.32E-55 | 0.99 | 0.94 | 0.05 | 1.05 |
| 3 | Rasl11a | 0.69 | 8.72E-60 | 1.81E-55 | 0.46 | 0.26 | 0.19 | 1.74 |
| 3 | Rnf150 | 0.63 | 1.96E-59 | 4.07E-55 | 0.50 | 0.29 | 0.21 | 1.70 |
| 3 | Reep5 | 0.43 | 1.99E-59 | 4.15E-55 | 0.91 | 0.77 | 0.14 | 1.17 |
| 3 | Prex2 | 0.54 | 2.01E-59 | 4.17E-55 | 0.61 | 0.39 | 0.23 | 1.59 |
| 3 | Ssbp3 | 0.29 | 2.07E-59 | 4.30E-55 | 0.99 | 0.93 | 0.06 | 1.06 |
| 3 | Ccna2 | 0.33 | 2.59E-59 | 5.39E-55 | 0.88 | 0.61 | 0.28 | 1.45 |
| 3 | Twist2 | 0.90 | 3.03E-59 | 6.30E-55 | 0.40 | 0.22 | 0.18 | 1.83 |
| 3 | Pold3 | 0.48 | 4.03E-59 | 8.38E-55 | 0.77 | 0.55 | 0.22 | 1.39 |
| 3 | C1qbp | 0.31 | 5.51E-59 | 1.15E-54 | 0.98 | 0.92 | 0.07 | 1.08 |
| 3 | Cdc7 | 0.65 | 7.81E-59 | 1.62E-54 | 0.55 | 0.35 | 0.21 | 1.60 |
| 3 | Spink2 | 1.57 | 1.33E-58 | 2.77E-54 | 0.11 | 0.03 | 0.08 | 3.53 |
| 3 | Irx3 | 1.02 | 2.10E-58 | 4.37E-54 | 0.42 | 0.26 | 0.17 | 1.64 |
| 3 | Msn | 0.34 | 2.12E-58 | 4.41E-54 | 0.73 | 0.49 | 0.24 | 1.48 |
| 3 | Kif2c | 0.39 | 2.26E-58 | 4.70E-54 | 0.69 | 0.45 | 0.25 | 1.55 |
| 3 | Sulf2 | 0.41 | 4.58E-58 | 9.53E-54 | 0.87 | 0.65 | 0.22 | 1.34 |
| 3 | Dna2 | 0.77 | 6.44E-58 | 1.34E-53 | 0.43 | 0.24 | 0.18 | 1.74 |
| 3 | Rassf9 | 0.95 | 6.50E-58 | 1.35E-53 | 0.30 | 0.15 | 0.15 | 1.99 |
| 3 | Grem1 | 0.91 | 8.18E-58 | 1.70E-53 | 0.21 | 0.09 | 0.12 | 2.42 |
| 3 | Mrpl20 | 0.28 | 1.07E-57 | 2.23E-53 | 0.98 | 0.92 | 0.06 | 1.07 |
| 3 | Ebf1 | 0.35 | 1.34E-57 | 2.78E-53 | 0.48 | 0.28 | 0.20 | 1.71 |

**Supplementary Table 1. Marker genes for the original clusters of integrated clustering of E11.5 control and NotchGOF λ-junction cells.**

| | | | | | | | | |
|---|---|---|---|---|---|---|---|---|
| 3 | Plxna2 | 0.70 | 1.44E-57 | 2.99E-53 | 0.39 | 0.22 | 0.17 | 1.79 |
| 3 | Plaat3 | 0.75 | 1.54E-57 | 3.21E-53 | 0.38 | 0.21 | 0.17 | 1.83 |
| 3 | Kpna2 | 0.34 | 1.72E-57 | 3.58E-53 | 0.63 | 0.40 | 0.24 | 1.59 |
| 3 | Mpped2 | 0.40 | 1.89E-57 | 3.94E-53 | 0.94 | 0.84 | 0.09 | 1.11 |
| 3 | Plk1 | 0.40 | 1.95E-57 | 4.06E-53 | 0.69 | 0.44 | 0.25 | 1.56 |
| 3 | Hat1 | 0.38 | 2.17E-57 | 4.51E-53 | 0.91 | 0.75 | 0.16 | 1.21 |
| 3 | Bmp7 | 0.62 | 2.56E-57 | 5.33E-53 | 0.57 | 0.37 | 0.21 | 1.57 |
| 3 | S1pr3 | 0.64 | 5.18E-57 | 1.08E-52 | 0.46 | 0.26 | 0.20 | 1.75 |
| 3 | Syce2 | 0.51 | 5.22E-57 | 1.08E-52 | 0.77 | 0.58 | 0.20 | 1.35 |
| 3 | Timm23 | 0.30 | 8.18E-57 | 1.70E-52 | 0.98 | 0.91 | 0.07 | 1.08 |
| 3 | Sgcd | 0.75 | 9.76E-57 | 2.03E-52 | 0.38 | 0.21 | 0.18 | 1.85 |
| 3 | Efnb1 | 0.47 | 1.07E-56 | 2.22E-52 | 0.74 | 0.51 | 0.23 | 1.44 |
| 3 | Ror1 | 0.87 | 1.09E-56 | 2.28E-52 | 0.25 | 0.11 | 0.13 | 2.20 |
| 3 | Ccp110 | 0.51 | 1.62E-56 | 3.37E-52 | 0.63 | 0.41 | 0.22 | 1.53 |
| 3 | Pin1 | 0.32 | 2.08E-56 | 4.32E-52 | 0.97 | 0.87 | 0.09 | 1.11 |
| 3 | Oxct1 | 0.34 | 3.30E-56 | 6.86E-52 | 0.96 | 0.81 | 0.15 | 1.18 |
| 3 | Txnl1 | 0.26 | 4.58E-56 | 9.52E-52 | 0.99 | 0.94 | 0.05 | 1.05 |
| 3 | Dbf4 | 0.38 | 5.33E-56 | 1.11E-51 | 0.86 | 0.64 | 0.23 | 1.36 |
| 3 | Kif20b | 0.35 | 5.86E-56 | 1.22E-51 | 0.80 | 0.54 | 0.26 | 1.49 |
| 3 | Edil3 | 0.74 | 8.20E-56 | 1.70E-51 | 0.32 | 0.16 | 0.16 | 1.99 |
| 3 | Naa50 | 0.38 | 1.09E-55 | 2.27E-51 | 0.93 | 0.78 | 0.15 | 1.19 |
| 3 | Utrn | 0.41 | 1.10E-55 | 2.28E-51 | 0.82 | 0.60 | 0.23 | 1.38 |
| 3 | Spock3 | 0.86 | 1.51E-55 | 3.13E-51 | 0.26 | 0.12 | 0.14 | 2.14 |
| 3 | Manf | 0.34 | 1.53E-55 | 3.19E-51 | 0.97 | 0.89 | 0.08 | 1.09 |
| 3 | Kntc1 | 0.94 | 1.56E-55 | 3.25E-51 | 0.33 | 0.18 | 0.15 | 1.84 |
| 3 | Banf1 | 0.26 | 2.00E-55 | 4.15E-51 | 0.99 | 0.95 | 0.05 | 1.05 |
| 3 | Smchd1 | 0.31 | 2.09E-55 | 4.34E-51 | 0.97 | 0.85 | 0.13 | 1.15 |
| 3 | Mlf1 | 0.75 | 2.48E-55 | 5.15E-51 | 0.36 | 0.20 | 0.16 | 1.84 |
| 3 | Rad54l | 0.69 | 4.01E-55 | 8.34E-51 | 0.48 | 0.30 | 0.19 | 1.64 |
| 3 | Exosc8 | 0.36 | 4.51E-55 | 9.38E-51 | 0.94 | 0.78 | 0.16 | 1.20 |
| 3 | Eif1b | 0.27 | 6.70E-55 | 1.39E-50 | 0.99 | 0.92 | 0.06 | 1.07 |
| 3 | Hist1h4i | 0.50 | 7.30E-55 | 1.52E-50 | 0.66 | 0.44 | 0.22 | 1.49 |
| 3 | Baz1b | 0.32 | 1.39E-54 | 2.90E-50 | 0.98 | 0.88 | 0.09 | 1.11 |
| 3 | 4930503L19Rik | 0.60 | 2.54E-54 | 5.29E-50 | 0.52 | 0.32 | 0.20 | 1.63 |
| 3 | Mrto4 | 0.36 | 3.46E-54 | 7.19E-50 | 0.93 | 0.81 | 0.13 | 1.16 |
| 3 | Batf3 | 1.02 | 4.10E-54 | 8.53E-50 | 0.28 | 0.14 | 0.14 | 1.98 |
| 3 | Spag5 | 0.67 | 8.62E-54 | 1.79E-49 | 0.48 | 0.29 | 0.18 | 1.63 |
| 3 | Ndc1 | 0.64 | 1.19E-53 | 2.47E-49 | 0.48 | 0.29 | 0.19 | 1.65 |
| 3 | Srpk1 | 0.33 | 1.74E-53 | 3.63E-49 | 0.96 | 0.84 | 0.12 | 1.14 |
| 3 | Mrps22 | 0.47 | 1.76E-53 | 3.65E-49 | 0.76 | 0.56 | 0.21 | 1.37 |
| 3 | Cks2 | 0.27 | 2.05E-53 | 4.26E-49 | 0.98 | 0.82 | 0.16 | 1.19 |
| 3 | Kif4 | 0.41 | 2.30E-53 | 4.79E-49 | 0.68 | 0.45 | 0.23 | 1.51 |
| 3 | Timp3 | 0.28 | 2.47E-53 | 5.14E-49 | 0.32 | 0.16 | 0.16 | 1.99 |
| 3 | Bard1 | 0.82 | 2.87E-53 | 5.97E-49 | 0.33 | 0.18 | 0.15 | 1.86 |
| 3 | Wnt5a | 0.45 | 3.59E-53 | 7.46E-49 | 0.83 | 0.62 | 0.21 | 1.33 |
| 3 | Ak2 | 0.33 | 3.66E-53 | 7.61E-49 | 0.95 | 0.84 | 0.11 | 1.13 |
| 3 | Gm3550 | 1.09 | 5.71E-53 | 1.19E-48 | 0.22 | 0.10 | 0.12 | 2.20 |
| 3 | Ssrp1 | 0.28 | 6.84E-53 | 1.42E-48 | 0.98 | 0.92 | 0.06 | 1.07 |
| 3 | Prdx4 | 0.29 | 1.26E-52 | 2.61E-48 | 0.98 | 0.89 | 0.09 | 1.10 |
| 3 | Ccdc25 | 0.43 | 2.42E-52 | 5.04E-48 | 0.82 | 0.63 | 0.19 | 1.30 |
| 3 | Sass6 | 0.46 | 2.43E-52 | 5.05E-48 | 0.69 | 0.48 | 0.21 | 1.44 |
| 3 | Psmd13 | 0.30 | 2.82E-52 | 5.87E-48 | 0.96 | 0.88 | 0.08 | 1.09 |
| 3 | Gyg | 0.52 | 7.77E-52 | 1.62E-47 | 0.61 | 0.40 | 0.20 | 1.50 |
| 3 | Mrpl13 | 0.29 | 9.12E-52 | 1.90E-47 | 0.98 | 0.90 | 0.08 | 1.09 |
| 3 | Calb2 | 2.04 | 1.02E-51 | 2.12E-47 | 0.13 | 0.05 | 0.09 | 2.81 |
| 3 | Col6a3 | 0.66 | 1.12E-51 | 2.34E-47 | 0.31 | 0.15 | 0.15 | 1.99 |

**Supplementary Table 1. Marker genes for the original clusters of integrated clustering of E11.5 control and NotchGOF λ-junction cells.**

| | | | | | | | | |
|---|---|---|---|---|---|---|---|---|
| 3 | 0610010K14Rik | 0.35 | 1.13E-51 | 2.34E-47 | 0.93 | 0.80 | 0.12 | 1.15 |
| 3 | Crnde | 0.92 | 1.41E-51 | 2.93E-47 | 0.28 | 0.14 | 0.14 | 1.99 |
| 3 | Arxes2 | 0.56 | 2.41E-51 | 5.01E-47 | 0.58 | 0.38 | 0.21 | 1.54 |
| 3 | Mms22l | 0.59 | 2.94E-51 | 6.11E-47 | 0.53 | 0.34 | 0.19 | 1.56 |
| 3 | Ethe1 | 0.52 | 3.82E-51 | 7.94E-47 | 0.64 | 0.43 | 0.20 | 1.47 |
| 3 | Irs2 | 0.53 | 5.35E-51 | 1.11E-46 | 0.62 | 0.42 | 0.20 | 1.47 |
| 3 | Gins3 | 0.72 | 8.05E-51 | 1.67E-46 | 0.41 | 0.24 | 0.17 | 1.71 |
| 3 | Kif22 | 0.39 | 9.00E-51 | 1.87E-46 | 0.71 | 0.48 | 0.23 | 1.49 |
| 3 | Cox10 | 0.59 | 1.46E-50 | 3.04E-46 | 0.51 | 0.32 | 0.19 | 1.60 |
| 3 | Enho | 0.51 | 1.64E-50 | 3.41E-46 | 0.64 | 0.43 | 0.21 | 1.49 |
| 3 | Bora | 0.49 | 1.92E-50 | 4.00E-46 | 0.64 | 0.43 | 0.21 | 1.48 |
| 3 | Pdgfra | 0.35 | 2.10E-50 | 4.38E-46 | 0.77 | 0.52 | 0.25 | 1.48 |
| 3 | Actn1 | 0.40 | 3.29E-50 | 6.84E-46 | 0.89 | 0.71 | 0.19 | 1.26 |
| 3 | Ptprg | 0.65 | 1.01E-49 | 2.10E-45 | 0.54 | 0.35 | 0.19 | 1.53 |
| 3 | Adgre5 | 0.89 | 1.12E-49 | 2.34E-45 | 0.24 | 0.12 | 0.13 | 2.09 |
| 3 | Iws1 | 0.40 | 1.17E-49 | 2.43E-45 | 0.87 | 0.69 | 0.18 | 1.26 |
| 3 | Zwilch | 0.62 | 1.36E-49 | 2.82E-45 | 0.48 | 0.30 | 0.18 | 1.60 |
| 3 | Mme | 0.37 | 1.60E-49 | 3.32E-45 | 0.67 | 0.45 | 0.22 | 1.49 |
| 3 | Sorcs2 | 0.94 | 2.20E-49 | 4.57E-45 | 0.22 | 0.10 | 0.12 | 2.18 |
| 3 | Chek1 | 0.45 | 2.28E-49 | 4.75E-45 | 0.74 | 0.53 | 0.20 | 1.38 |
| 3 | Prpf4 | 0.43 | 2.42E-49 | 5.03E-45 | 0.77 | 0.57 | 0.20 | 1.34 |
| 3 | Chaf1b | 0.49 | 2.54E-49 | 5.29E-45 | 0.69 | 0.50 | 0.20 | 1.40 |
| 3 | Chtf18 | 0.71 | 2.82E-49 | 5.87E-45 | 0.38 | 0.22 | 0.16 | 1.73 |
| 3 | Pgm5 | 1.45 | 3.31E-49 | 6.89E-45 | 0.11 | 0.04 | 0.07 | 3.09 |
| 3 | Ctcf | 0.30 | 9.87E-49 | 2.05E-44 | 0.97 | 0.88 | 0.09 | 1.10 |
| 3 | Mxd3 | 0.65 | 1.31E-48 | 2.73E-44 | 0.39 | 0.23 | 0.16 | 1.72 |
| 3 | Mad2l1 | 0.42 | 1.34E-48 | 2.79E-44 | 0.77 | 0.55 | 0.22 | 1.41 |
| 3 | Pcdh18 | 0.29 | 1.84E-48 | 3.84E-44 | 0.87 | 0.62 | 0.25 | 1.40 |
| 3 | Ankrd44 | 0.53 | 2.43E-48 | 5.05E-44 | 0.45 | 0.27 | 0.18 | 1.69 |
| 3 | Fignl1 | 0.46 | 2.79E-48 | 5.80E-44 | 0.70 | 0.50 | 0.20 | 1.40 |
| 3 | Cenpe | 0.28 | 3.03E-48 | 6.30E-44 | 0.87 | 0.64 | 0.24 | 1.37 |
| 3 | Etf1 | 0.33 | 3.59E-48 | 7.46E-44 | 0.92 | 0.79 | 0.13 | 1.16 |
| 3 | Mcph1 | 0.64 | 3.60E-48 | 7.49E-44 | 0.46 | 0.29 | 0.17 | 1.60 |
| 3 | Trmt6 | 0.42 | 4.33E-48 | 9.01E-44 | 0.80 | 0.63 | 0.17 | 1.28 |
| 3 | Arhgap10 | 0.53 | 5.97E-48 | 1.24E-43 | 0.60 | 0.40 | 0.20 | 1.49 |
| 3 | Gspt1 | 0.27 | 6.60E-48 | 1.37E-43 | 0.98 | 0.92 | 0.06 | 1.07 |
| 3 | Glipr2 | 0.46 | 6.89E-48 | 1.43E-43 | 0.58 | 0.37 | 0.21 | 1.56 |
| 3 | Calm3 | 0.26 | 9.88E-48 | 2.06E-43 | 0.98 | 0.91 | 0.08 | 1.09 |
| 3 | Gmps | 0.28 | 1.04E-47 | 2.15E-43 | 0.98 | 0.89 | 0.09 | 1.10 |
| 3 | Gen1 | 0.70 | 2.96E-47 | 6.15E-43 | 0.43 | 0.26 | 0.16 | 1.62 |
| 3 | Septin3 | 0.48 | 3.13E-47 | 6.50E-43 | 0.63 | 0.42 | 0.21 | 1.50 |
| 3 | Ticrr | 0.73 | 3.34E-47 | 6.94E-43 | 0.37 | 0.22 | 0.16 | 1.72 |
| 3 | Nsl1 | 0.79 | 3.43E-47 | 7.13E-43 | 0.32 | 0.18 | 0.14 | 1.82 |
| 3 | Olfml3 | 0.46 | 3.92E-47 | 8.14E-43 | 0.46 | 0.27 | 0.19 | 1.70 |
| 3 | Cep78 | 0.51 | 4.61E-47 | 9.59E-43 | 0.58 | 0.38 | 0.20 | 1.51 |
| 3 | Smad7 | 0.38 | 4.78E-47 | 9.95E-43 | 0.85 | 0.68 | 0.17 | 1.25 |
| 3 | Depdc1a | 0.46 | 4.79E-47 | 9.97E-43 | 0.54 | 0.34 | 0.20 | 1.57 |
| 3 | Fgfr1 | 0.34 | 5.03E-47 | 1.05E-42 | 0.85 | 0.64 | 0.21 | 1.34 |
| 3 | Traip | 0.68 | 6.22E-47 | 1.29E-42 | 0.42 | 0.26 | 0.16 | 1.61 |
| 3 | Gabpb1 | 0.49 | 6.32E-47 | 1.31E-42 | 0.64 | 0.45 | 0.19 | 1.42 |
| 3 | Dkc1 | 0.34 | 1.27E-46 | 2.65E-42 | 0.92 | 0.78 | 0.14 | 1.17 |
| 3 | B3gnt5 | 0.49 | 1.49E-46 | 3.09E-42 | 0.54 | 0.34 | 0.20 | 1.58 |
| 3 | Ppid | 0.29 | 1.93E-46 | 4.02E-42 | 0.97 | 0.87 | 0.09 | 1.11 |
| 3 | Pmf1 | 0.37 | 2.34E-46 | 4.87E-42 | 0.84 | 0.66 | 0.18 | 1.27 |
| 3 | Plekho1 | 0.34 | 2.36E-46 | 4.90E-42 | 0.81 | 0.60 | 0.21 | 1.35 |
| 3 | Svep1 | 1.02 | 2.70E-46 | 5.62E-42 | 0.18 | 0.08 | 0.10 | 2.33 |

**Supplementary Table 1. Marker genes for the original clusters of integrated clustering of E11.5 control and NotchGOF λ-junction cells.**

| | | | | | | | | |
|---|---|---|---|---|---|---|---|---|
| 3 | Boc | 0.67 | 4.10E-46 | 8.53E-42 | 0.34 | 0.19 | 0.15 | 1.81 |
| 3 | Rarg | 0.40 | 4.53E-46 | 9.43E-42 | 0.72 | 0.50 | 0.22 | 1.43 |
| 3 | Creb5 | 0.52 | 5.61E-46 | 1.17E-41 | 0.49 | 0.31 | 0.18 | 1.57 |
| 3 | Tenm4 | 0.38 | 6.85E-46 | 1.43E-41 | 0.79 | 0.60 | 0.20 | 1.33 |
| 3 | Aunip | 0.69 | 8.45E-46 | 1.76E-41 | 0.41 | 0.25 | 0.16 | 1.64 |
| 3 | Sfxn1 | 0.38 | 8.56E-46 | 1.78E-41 | 0.88 | 0.72 | 0.16 | 1.22 |
| 3 | E2f7 | 0.64 | 9.99E-46 | 2.08E-41 | 0.40 | 0.24 | 0.16 | 1.68 |
| 3 | Nap1l4 | 0.26 | 1.10E-45 | 2.28E-41 | 0.99 | 0.92 | 0.07 | 1.07 |
| 3 | Gtf2a2 | 0.29 | 1.60E-45 | 3.34E-41 | 0.97 | 0.88 | 0.09 | 1.11 |
| 3 | Ccdc88a | 0.31 | 2.68E-45 | 5.57E-41 | 0.95 | 0.80 | 0.15 | 1.19 |
| 3 | Mrpl28 | 0.29 | 4.48E-45 | 9.32E-41 | 0.97 | 0.88 | 0.09 | 1.10 |
| 3 | 1110038B12Rik | 0.27 | 5.94E-45 | 1.23E-40 | 0.99 | 0.92 | 0.06 | 1.07 |
| 3 | Cenpj | 0.58 | 6.53E-45 | 1.36E-40 | 0.54 | 0.36 | 0.18 | 1.49 |
| 3 | Cenps | 0.48 | 6.92E-45 | 1.44E-40 | 0.68 | 0.49 | 0.19 | 1.39 |
| 3 | Ctbp2 | 0.37 | 8.89E-45 | 1.85E-40 | 0.84 | 0.67 | 0.17 | 1.25 |
| 3 | Ier5l | 0.37 | 1.23E-44 | 2.55E-40 | 0.84 | 0.65 | 0.20 | 1.30 |
| 3 | Adamtsl1 | 0.80 | 1.25E-44 | 2.59E-40 | 0.21 | 0.10 | 0.11 | 2.16 |
| 3 | Neil3 | 0.64 | 1.50E-44 | 3.13E-40 | 0.37 | 0.22 | 0.15 | 1.70 |
| 3 | Raet1d | 1.04 | 2.21E-44 | 4.60E-40 | 0.17 | 0.08 | 0.10 | 2.29 |
| 3 | Adamts1 | 0.49 | 3.06E-44 | 6.37E-40 | 0.43 | 0.26 | 0.17 | 1.63 |
| 3 | Impa2 | 0.55 | 5.85E-44 | 1.22E-39 | 0.46 | 0.28 | 0.17 | 1.62 |
| 3 | Epop | 0.51 | 1.00E-43 | 2.08E-39 | 0.56 | 0.37 | 0.19 | 1.52 |
| 3 | Brca2 | 0.45 | 1.06E-43 | 2.20E-39 | 0.65 | 0.46 | 0.19 | 1.41 |
| 3 | Nup35 | 0.46 | 1.06E-43 | 2.21E-39 | 0.65 | 0.46 | 0.19 | 1.42 |
| 3 | Pwwp3a | 0.42 | 1.16E-43 | 2.42E-39 | 0.73 | 0.53 | 0.20 | 1.37 |
| 3 | Qprt | 1.28 | 1.31E-43 | 2.72E-39 | 0.11 | 0.04 | 0.07 | 2.87 |
| 3 | Rbpj | 0.38 | 1.44E-43 | 3.00E-39 | 0.85 | 0.69 | 0.15 | 1.22 |
| 3 | Topbp1 | 0.35 | 1.56E-43 | 3.24E-39 | 0.87 | 0.72 | 0.15 | 1.21 |
| 3 | Aarsd1 | 0.33 | 1.57E-43 | 3.26E-39 | 0.91 | 0.76 | 0.16 | 1.20 |
| 3 | Mdc1 | 0.52 | 1.91E-43 | 3.97E-39 | 0.55 | 0.37 | 0.18 | 1.49 |
| 3 | Ivns1abp | 0.32 | 2.16E-43 | 4.49E-39 | 0.91 | 0.76 | 0.14 | 1.19 |
| 3 | Rcc1 | 0.43 | 2.92E-43 | 6.08E-39 | 0.70 | 0.50 | 0.20 | 1.40 |
| 3 | Foxp2 | 0.38 | 5.04E-43 | 1.05E-38 | 0.51 | 0.33 | 0.17 | 1.52 |
| 3 | Cenpl | 0.60 | 5.05E-43 | 1.05E-38 | 0.42 | 0.27 | 0.16 | 1.60 |
| 3 | Odc1 | 0.28 | 5.12E-43 | 1.07E-38 | 0.92 | 0.80 | 0.12 | 1.15 |
| 3 | Pkig | 0.31 | 5.21E-43 | 1.08E-38 | 0.91 | 0.77 | 0.14 | 1.18 |
| 3 | Epb41l3 | 0.46 | 6.29E-43 | 1.31E-38 | 0.63 | 0.43 | 0.20 | 1.46 |
| 3 | Ddx25 | 0.83 | 6.86E-43 | 1.43E-38 | 0.25 | 0.13 | 0.12 | 1.93 |
| 3 | Knl1 | 0.31 | 8.33E-43 | 1.73E-38 | 0.70 | 0.47 | 0.23 | 1.48 |
| 3 | Fhod3 | 1.09 | 8.98E-43 | 1.87E-38 | 0.18 | 0.08 | 0.10 | 2.20 |
| 3 | Rab30 | 0.74 | 1.05E-42 | 2.18E-38 | 0.26 | 0.13 | 0.13 | 1.94 |
| 3 | Zfhx4 | 0.44 | 1.17E-42 | 2.43E-38 | 0.85 | 0.73 | 0.12 | 1.16 |
| 3 | Bmp2 | 1.07 | 1.55E-42 | 3.23E-38 | 0.15 | 0.06 | 0.09 | 2.46 |
| 3 | Syne1 | 0.76 | 1.60E-42 | 3.34E-38 | 0.25 | 0.13 | 0.12 | 1.97 |
| 3 | Cnih2 | 0.49 | 1.66E-42 | 3.45E-38 | 0.49 | 0.31 | 0.18 | 1.58 |
| 3 | Prpf19 | 0.31 | 2.25E-42 | 4.68E-38 | 0.92 | 0.79 | 0.13 | 1.17 |
| 3 | Stag1 | 0.34 | 3.05E-42 | 6.34E-38 | 0.89 | 0.74 | 0.15 | 1.20 |
| 3 | Stub1 | 0.25 | 3.24E-42 | 6.73E-38 | 0.98 | 0.92 | 0.07 | 1.07 |
| 3 | Rpa1 | 0.50 | 3.38E-42 | 7.04E-38 | 0.58 | 0.40 | 0.18 | 1.45 |
| 3 | Mcm5 | 0.36 | 3.73E-42 | 7.76E-38 | 0.89 | 0.72 | 0.16 | 1.22 |
| 3 | Cinp | 0.45 | 4.00E-42 | 8.32E-38 | 0.65 | 0.47 | 0.18 | 1.39 |
| 3 | Ptbp2 | 0.31 | 4.39E-42 | 9.13E-38 | 0.91 | 0.76 | 0.15 | 1.20 |
| 3 | Rhno1 | 0.36 | 5.29E-42 | 1.10E-37 | 0.83 | 0.65 | 0.18 | 1.27 |
| 3 | Farp1 | 0.42 | 5.30E-42 | 1.10E-37 | 0.68 | 0.48 | 0.20 | 1.42 |
| 3 | Myg1 | 0.38 | 6.22E-42 | 1.29E-37 | 0.79 | 0.60 | 0.19 | 1.31 |
| 3 | Cenpn | 0.48 | 6.27E-42 | 1.30E-37 | 0.55 | 0.37 | 0.18 | 1.50 |

**Supplementary Table 1. Marker genes for the original clusters of integrated clustering of E11.5 control and NotchGOF λ-junction cells.**

| | | | | | | | | |
|---|---|---|---|---|---|---|---|---|
| 3 | Elof1 | 0.27 | 8.80E-42 | 1.83E-37 | 0.97 | 0.90 | 0.08 | 1.09 |
| 3 | Tcf19 | 0.70 | 1.19E-41 | 2.47E-37 | 0.32 | 0.18 | 0.14 | 1.77 |
| 3 | Arxes1 | 0.49 | 1.58E-41 | 3.30E-37 | 0.54 | 0.36 | 0.18 | 1.52 |
| 3 | Tns1 | 0.52 | 1.88E-41 | 3.91E-37 | 0.47 | 0.29 | 0.18 | 1.62 |
| 3 | Cse1l | 0.39 | 2.26E-41 | 4.70E-37 | 0.79 | 0.61 | 0.18 | 1.29 |
| 3 | Haus5 | 0.62 | 2.36E-41 | 4.90E-37 | 0.39 | 0.24 | 0.15 | 1.64 |
| 3 | Kctd15 | 0.47 | 2.51E-41 | 5.23E-37 | 0.59 | 0.40 | 0.18 | 1.46 |
| 3 | Zik1 | 0.49 | 4.30E-41 | 8.94E-37 | 0.50 | 0.32 | 0.18 | 1.56 |
| 3 | Tle1 | 0.43 | 5.49E-41 | 1.14E-36 | 0.70 | 0.52 | 0.18 | 1.35 |
| 3 | Ppp2r2d | 0.38 | 6.67E-41 | 1.39E-36 | 0.84 | 0.66 | 0.17 | 1.26 |
| 3 | Nup85 | 0.50 | 7.27E-41 | 1.51E-36 | 0.53 | 0.35 | 0.18 | 1.51 |
| 3 | Nudt5 | 0.31 | 1.07E-40 | 2.23E-36 | 0.92 | 0.78 | 0.14 | 1.18 |
| 3 | Taf1 | 0.39 | 1.68E-40 | 3.49E-36 | 0.75 | 0.57 | 0.19 | 1.33 |
| 3 | Taf5 | 0.46 | 2.90E-40 | 6.03E-36 | 0.60 | 0.42 | 0.19 | 1.44 |
| 3 | Hist2h3b | 1.00 | 3.14E-40 | 6.52E-36 | 0.15 | 0.07 | 0.09 | 2.35 |
| 3 | Bcl7c | 0.29 | 4.09E-40 | 8.50E-36 | 0.93 | 0.82 | 0.12 | 1.14 |
| 3 | Ska3 | 0.52 | 4.81E-40 | 1.00E-35 | 0.43 | 0.27 | 0.16 | 1.59 |
| 3 | Ptgr1 | 0.58 | 7.40E-40 | 1.54E-35 | 0.41 | 0.25 | 0.16 | 1.63 |
| 3 | Agpat5 | 0.41 | 7.64E-40 | 1.59E-35 | 0.62 | 0.43 | 0.19 | 1.43 |
| 3 | Fez1 | 0.70 | 7.88E-40 | 1.64E-35 | 0.26 | 0.14 | 0.12 | 1.90 |
| 3 | Rfc1 | 0.26 | 8.84E-40 | 1.84E-35 | 0.96 | 0.86 | 0.11 | 1.12 |
| 3 | Ly6e | 0.38 | 8.93E-40 | 1.86E-35 | 0.63 | 0.44 | 0.19 | 1.44 |
| 3 | Kpnb1 | 0.30 | 1.01E-39 | 2.11E-35 | 0.95 | 0.83 | 0.12 | 1.15 |
| 3 | Rab3il1 | 0.45 | 1.34E-39 | 2.78E-35 | 0.45 | 0.28 | 0.17 | 1.60 |
| 3 | Map4k4 | 0.26 | 1.60E-39 | 3.32E-35 | 0.98 | 0.90 | 0.08 | 1.09 |
| 3 | Tfdp1 | 0.35 | 1.78E-39 | 3.71E-35 | 0.82 | 0.65 | 0.18 | 1.27 |
| 3 | Fxn | 0.41 | 1.88E-39 | 3.92E-35 | 0.71 | 0.53 | 0.18 | 1.33 |
| 3 | Ric1 | 0.48 | 1.93E-39 | 4.00E-35 | 0.49 | 0.32 | 0.17 | 1.54 |
| 3 | Plk2 | 0.30 | 2.31E-39 | 4.81E-35 | 0.55 | 0.36 | 0.19 | 1.54 |
| 3 | Phf6 | 0.36 | 2.95E-39 | 6.14E-35 | 0.86 | 0.71 | 0.15 | 1.22 |
| 3 | Eif2s1 | 0.25 | 3.76E-39 | 7.82E-35 | 0.97 | 0.90 | 0.07 | 1.08 |
| 3 | Axin2 | 0.46 | 4.99E-39 | 1.04E-34 | 0.50 | 0.33 | 0.18 | 1.54 |
| 3 | BC030867 | 0.83 | 5.78E-39 | 1.20E-34 | 0.25 | 0.13 | 0.11 | 1.87 |
| 3 | C530008M17Rik | 0.44 | 6.02E-39 | 1.25E-34 | 0.59 | 0.40 | 0.19 | 1.47 |
| 3 | S1pr2 | 0.45 | 6.55E-39 | 1.36E-34 | 0.48 | 0.31 | 0.18 | 1.58 |
| 3 | Tmem59l | 0.62 | 1.03E-38 | 2.13E-34 | 0.33 | 0.19 | 0.14 | 1.73 |
| 3 | Gins4 | 0.35 | 1.03E-38 | 2.13E-34 | 0.81 | 0.63 | 0.18 | 1.28 |
| 3 | Rfc3 | 0.32 | 1.29E-38 | 2.69E-34 | 0.90 | 0.76 | 0.15 | 1.19 |
| 3 | Ncapd3 | 0.37 | 1.35E-38 | 2.81E-34 | 0.71 | 0.52 | 0.19 | 1.37 |
| 3 | Yaf2 | 0.34 | 1.67E-38 | 3.47E-34 | 0.85 | 0.69 | 0.17 | 1.24 |
| 3 | Mcm7 | 0.29 | 1.80E-38 | 3.74E-34 | 0.96 | 0.86 | 0.10 | 1.12 |
| 3 | Uchl3 | 0.28 | 1.84E-38 | 3.82E-34 | 0.93 | 0.79 | 0.13 | 1.17 |
| 3 | Jakmip2 | 0.69 | 2.87E-38 | 5.96E-34 | 0.25 | 0.14 | 0.12 | 1.88 |
| 3 | Ddr2 | 0.32 | 3.52E-38 | 7.32E-34 | 0.70 | 0.47 | 0.23 | 1.48 |
| 3 | C630043F03Rik | 0.72 | 5.04E-38 | 1.05E-33 | 0.26 | 0.14 | 0.12 | 1.87 |
| 3 | Slc8a1 | 0.76 | 7.36E-38 | 1.53E-33 | 0.23 | 0.12 | 0.11 | 1.96 |
| 3 | Magohb | 0.37 | 9.58E-38 | 1.99E-33 | 0.74 | 0.56 | 0.19 | 1.34 |
| 3 | 1500009L16Rik | 0.36 | 9.62E-38 | 2.00E-33 | 0.76 | 0.56 | 0.19 | 1.35 |
| 3 | Prr11 | 0.69 | 1.02E-37 | 2.12E-33 | 0.31 | 0.18 | 0.13 | 1.70 |
| 3 | Donson | 0.64 | 1.36E-37 | 2.83E-33 | 0.37 | 0.23 | 0.14 | 1.61 |
| 3 | Ddx21 | 0.32 | 1.40E-37 | 2.91E-33 | 0.86 | 0.72 | 0.14 | 1.19 |
| 3 | Gpc6 | 0.27 | 1.65E-37 | 3.43E-33 | 0.77 | 0.57 | 0.20 | 1.36 |
| 3 | Fndc3c1 | 0.37 | 2.08E-37 | 4.32E-33 | 0.65 | 0.45 | 0.20 | 1.44 |
| 3 | Myh10 | 0.27 | 2.92E-37 | 6.07E-33 | 0.97 | 0.86 | 0.12 | 1.14 |
| 3 | Uhrf1 | 0.35 | 3.63E-37 | 7.55E-33 | 0.84 | 0.67 | 0.17 | 1.26 |
| 3 | Tpgs2 | 0.36 | 3.70E-37 | 7.71E-33 | 0.75 | 0.57 | 0.19 | 1.33 |

**Supplementary Table 1. Marker genes for the original clusters of integrated clustering of E11.5 control and NotchGOF λ-junction cells.**

| | | | | | | | | |
|---|---|---|---|---|---|---|---|---|
| 3 | Atad5 | 0.36 | 4.29E-37 | 8.93E-33 | 0.70 | 0.51 | 0.19 | 1.37 |
| 3 | Haus4 | 0.50 | 4.36E-37 | 9.07E-33 | 0.46 | 0.30 | 0.16 | 1.52 |
| 3 | Arhgap11a | 0.39 | 4.58E-37 | 9.53E-33 | 0.57 | 0.39 | 0.18 | 1.46 |
| 3 | Kif18a | 0.42 | 6.84E-37 | 1.42E-32 | 0.50 | 0.33 | 0.17 | 1.52 |
| 3 | Hccs | 0.51 | 8.16E-37 | 1.70E-32 | 0.46 | 0.30 | 0.16 | 1.52 |
| 3 | Cachd1 | 0.32 | 1.05E-36 | 2.17E-32 | 0.70 | 0.51 | 0.20 | 1.39 |
| 3 | Rnf115 | 0.38 | 1.05E-36 | 2.19E-32 | 0.69 | 0.50 | 0.19 | 1.37 |
| 3 | Tmem132a | 0.33 | 1.13E-36 | 2.36E-32 | 0.81 | 0.62 | 0.19 | 1.31 |
| 3 | Arhgap6 | 0.98 | 1.18E-36 | 2.45E-32 | 0.14 | 0.06 | 0.08 | 2.34 |
| 3 | Fkbp5 | 0.42 | 1.19E-36 | 2.47E-32 | 0.59 | 0.41 | 0.18 | 1.45 |
| 3 | Pold1 | 0.53 | 1.50E-36 | 3.13E-32 | 0.48 | 0.33 | 0.16 | 1.47 |
| 3 | G2e3 | 0.39 | 1.90E-36 | 3.95E-32 | 0.64 | 0.45 | 0.18 | 1.40 |
| 3 | Gm17167 | 1.02 | 2.15E-36 | 4.48E-32 | 0.12 | 0.05 | 0.07 | 2.50 |
| 3 | Prelid3b | 0.34 | 2.36E-36 | 4.91E-32 | 0.81 | 0.64 | 0.17 | 1.27 |
| 3 | Grwd1 | 0.42 | 3.49E-36 | 7.26E-32 | 0.59 | 0.41 | 0.18 | 1.44 |
| 3 | Tmem100 | 0.76 | 3.53E-36 | 7.34E-32 | 0.25 | 0.13 | 0.12 | 1.86 |
| 3 | Rad54b | 0.65 | 3.76E-36 | 7.83E-32 | 0.32 | 0.19 | 0.13 | 1.67 |
| 3 | Nup43 | 0.35 | 4.15E-36 | 8.64E-32 | 0.79 | 0.61 | 0.19 | 1.31 |
| 3 | Akr1b3 | 0.34 | 7.52E-36 | 1.56E-31 | 0.83 | 0.66 | 0.17 | 1.26 |
| 3 | Cacna1d | 0.45 | 9.01E-36 | 1.87E-31 | 0.37 | 0.22 | 0.15 | 1.67 |
| 3 | Tomm40 | 0.28 | 1.19E-35 | 2.47E-31 | 0.92 | 0.80 | 0.13 | 1.16 |
| 3 | U2af2 | 0.27 | 1.39E-35 | 2.89E-31 | 0.94 | 0.84 | 0.11 | 1.13 |
| 3 | Ift27 | 0.36 | 2.10E-35 | 4.37E-31 | 0.73 | 0.55 | 0.18 | 1.33 |
| 3 | Ttpa | 1.02 | 2.32E-35 | 4.83E-31 | 0.15 | 0.07 | 0.08 | 2.20 |
| 3 | Utp18 | 0.35 | 3.58E-35 | 7.45E-31 | 0.75 | 0.57 | 0.18 | 1.32 |
| 3 | Foxm1 | 0.59 | 4.39E-35 | 9.13E-31 | 0.39 | 0.25 | 0.14 | 1.55 |
| 3 | Bc1 | 0.79 | 4.45E-35 | 9.27E-31 | 0.25 | 0.14 | 0.11 | 1.78 |
| 3 | Srek1 | 0.27 | 5.30E-35 | 1.10E-30 | 0.94 | 0.82 | 0.12 | 1.15 |
| 3 | BC055324 | 0.63 | 5.43E-35 | 1.13E-30 | 0.35 | 0.22 | 0.14 | 1.62 |
| 3 | Usp39 | 0.34 | 5.49E-35 | 1.14E-30 | 0.75 | 0.57 | 0.18 | 1.32 |
| 3 | Fam92a | 0.27 | 8.22E-35 | 1.71E-30 | 0.90 | 0.74 | 0.15 | 1.21 |
| 3 | Dctd | 0.40 | 8.56E-35 | 1.78E-30 | 0.63 | 0.45 | 0.18 | 1.40 |
| 3 | St3gal1 | 0.96 | 9.03E-35 | 1.88E-30 | 0.19 | 0.09 | 0.09 | 1.98 |
| 3 | Gemin6 | 0.40 | 9.11E-35 | 1.89E-30 | 0.62 | 0.44 | 0.17 | 1.39 |
| 3 | Dock11 | 0.59 | 9.33E-35 | 1.94E-30 | 0.35 | 0.22 | 0.14 | 1.63 |
| 3 | Gfra1 | 1.05 | 1.06E-34 | 2.20E-30 | 0.11 | 0.04 | 0.07 | 2.59 |
| 3 | Ddx20 | 0.49 | 1.16E-34 | 2.41E-30 | 0.48 | 0.32 | 0.16 | 1.50 |
| 3 | Pinx1 | 0.37 | 1.54E-34 | 3.20E-30 | 0.71 | 0.53 | 0.18 | 1.34 |
| 3 | Plcl2 | 0.74 | 1.54E-34 | 3.21E-30 | 0.20 | 0.11 | 0.10 | 1.94 |
| 3 | Cenpp | 0.44 | 1.82E-34 | 3.78E-30 | 0.55 | 0.39 | 0.17 | 1.43 |
| 3 | Thoc3 | 0.27 | 1.92E-34 | 3.99E-30 | 0.91 | 0.78 | 0.13 | 1.17 |
| 3 | Gfer | 0.35 | 2.01E-34 | 4.17E-30 | 0.72 | 0.55 | 0.17 | 1.30 |
| 3 | Zfas1 | 0.28 | 2.11E-34 | 4.39E-30 | 0.93 | 0.81 | 0.11 | 1.14 |
| 3 | Aars | 0.42 | 2.25E-34 | 4.69E-30 | 0.62 | 0.45 | 0.17 | 1.37 |
| 3 | Mrps7 | 0.28 | 2.26E-34 | 4.69E-30 | 0.90 | 0.77 | 0.13 | 1.17 |
| 3 | Ifrd1 | 0.44 | 2.38E-34 | 4.95E-30 | 0.66 | 0.50 | 0.16 | 1.33 |
| 3 | Mad1l1 | 0.44 | 2.39E-34 | 4.96E-30 | 0.51 | 0.34 | 0.17 | 1.48 |
| 3 | Dcp2 | 0.35 | 3.49E-34 | 7.26E-30 | 0.79 | 0.61 | 0.18 | 1.29 |
| 3 | Amotl2 | 0.38 | 3.84E-34 | 7.98E-30 | 0.50 | 0.33 | 0.17 | 1.51 |
| 3 | Csrnp3 | 0.80 | 4.18E-34 | 8.69E-30 | 0.19 | 0.10 | 0.10 | 2.00 |
| 3 | Dis3 | 0.37 | 4.67E-34 | 9.71E-30 | 0.72 | 0.54 | 0.18 | 1.33 |
| 3 | Wee1 | 0.37 | 4.96E-34 | 1.03E-29 | 0.63 | 0.44 | 0.19 | 1.42 |
| 3 | Vasn | 0.50 | 5.01E-34 | 1.04E-29 | 0.35 | 0.21 | 0.14 | 1.67 |
| 3 | AW551984 | 0.71 | 5.47E-34 | 1.14E-29 | 0.22 | 0.11 | 0.10 | 1.90 |
| 3 | Tcof1 | 0.30 | 6.38E-34 | 1.33E-29 | 0.88 | 0.74 | 0.14 | 1.19 |
| 3 | a | 0.65 | 7.00E-34 | 1.45E-29 | 0.23 | 0.13 | 0.11 | 1.87 |

**Supplementary Table 1. Marker genes for the original clusters of integrated clustering of E11.5 control and NotchGOF λ-junction cells.**

| | | | | | | | | |
|---|---|---|---|---|---|---|---|---|
| 3 | Wdr18 | 0.31 | 1.02E-33 | 2.12E-29 | 0.87 | 0.72 | 0.15 | 1.21 |
| 3 | Adora2b | 0.69 | 1.04E-33 | 2.15E-29 | 0.20 | 0.10 | 0.10 | 2.00 |
| 3 | Lap3 | 0.32 | 1.12E-33 | 2.32E-29 | 0.82 | 0.66 | 0.16 | 1.25 |
| 3 | Cenpi | 0.70 | 1.14E-33 | 2.38E-29 | 0.27 | 0.16 | 0.12 | 1.73 |
| 3 | Nup133 | 0.41 | 1.27E-33 | 2.65E-29 | 0.53 | 0.36 | 0.17 | 1.47 |
| 3 | Tex30 | 0.39 | 1.31E-33 | 2.72E-29 | 0.64 | 0.46 | 0.18 | 1.39 |
| 3 | Dlc1 | 0.36 | 1.57E-33 | 3.27E-29 | 0.53 | 0.35 | 0.18 | 1.52 |
| 3 | Ddit4l | 0.35 | 1.95E-33 | 4.05E-29 | 0.34 | 0.20 | 0.14 | 1.69 |
| 3 | Rpa3 | 0.26 | 2.00E-33 | 4.17E-29 | 0.92 | 0.80 | 0.12 | 1.15 |
| 3 | Smoc1 | 0.90 | 2.29E-33 | 4.77E-29 | 0.15 | 0.07 | 0.08 | 2.15 |
| 3 | Nup50 | 0.34 | 2.30E-33 | 4.79E-29 | 0.77 | 0.60 | 0.17 | 1.28 |
| 3 | Gm45716 | 0.43 | 2.32E-33 | 4.83E-29 | 0.53 | 0.36 | 0.17 | 1.46 |
| 3 | Safb | 0.28 | 2.46E-33 | 5.12E-29 | 0.91 | 0.77 | 0.13 | 1.17 |
| 3 | Clec2l | 0.89 | 2.52E-33 | 5.24E-29 | 0.14 | 0.06 | 0.08 | 2.25 |
| 3 | Clmn | 0.48 | 2.56E-33 | 5.33E-29 | 0.41 | 0.26 | 0.15 | 1.58 |
| 3 | Cap2 | 1.13 | 2.91E-33 | 6.06E-29 | 0.11 | 0.04 | 0.07 | 2.48 |
| 3 | Cad | 0.43 | 3.26E-33 | 6.79E-29 | 0.54 | 0.37 | 0.17 | 1.45 |
| 3 | Ube2t | 0.46 | 3.51E-33 | 7.29E-29 | 0.47 | 0.32 | 0.15 | 1.47 |
| 3 | Lama2 | 0.58 | 3.51E-33 | 7.30E-29 | 0.25 | 0.14 | 0.11 | 1.81 |
| 3 | Jpt2 | 0.42 | 4.25E-33 | 8.83E-29 | 0.60 | 0.44 | 0.17 | 1.39 |
| 3 | Cdk2 | 0.43 | 4.29E-33 | 8.93E-29 | 0.56 | 0.40 | 0.16 | 1.41 |
| 3 | Mre11a | 0.44 | 4.33E-33 | 9.01E-29 | 0.53 | 0.37 | 0.16 | 1.44 |
| 3 | Anapc15 | 0.34 | 6.38E-33 | 1.33E-28 | 0.76 | 0.59 | 0.18 | 1.30 |
| 3 | Mir17hg | 0.50 | 8.14E-33 | 1.69E-28 | 0.40 | 0.26 | 0.14 | 1.55 |
| 3 | Dlgap1 | 1.29 | 1.07E-32 | 2.22E-28 | 0.11 | 0.05 | 0.07 | 2.47 |
| 3 | Bmerb1 | 0.77 | 1.21E-32 | 2.52E-28 | 0.17 | 0.08 | 0.09 | 2.06 |
| 3 | Car13 | 0.80 | 1.34E-32 | 2.79E-28 | 0.18 | 0.09 | 0.09 | 2.00 |
| 3 | Samd1 | 0.41 | 1.48E-32 | 3.08E-28 | 0.56 | 0.40 | 0.16 | 1.41 |
| 3 | Zranb3 | 0.64 | 1.75E-32 | 3.64E-28 | 0.33 | 0.20 | 0.13 | 1.63 |
| 3 | Vrk1 | 0.33 | 1.82E-32 | 3.78E-28 | 0.77 | 0.60 | 0.18 | 1.29 |
| 3 | Ppil1 | 0.25 | 1.92E-32 | 4.00E-28 | 0.93 | 0.80 | 0.13 | 1.16 |
| 3 | Loxl2 | 0.38 | 2.03E-32 | 4.22E-28 | 0.61 | 0.42 | 0.19 | 1.44 |
| 3 | Orc2 | 0.38 | 2.31E-32 | 4.81E-28 | 0.73 | 0.56 | 0.17 | 1.31 |
| 3 | Mgat5 | 0.58 | 2.32E-32 | 4.82E-28 | 0.39 | 0.25 | 0.14 | 1.55 |
| 3 | Cth | 0.55 | 2.37E-32 | 4.93E-28 | 0.33 | 0.20 | 0.13 | 1.64 |
| 3 | Dock10 | 0.89 | 3.23E-32 | 6.72E-28 | 0.16 | 0.08 | 0.08 | 2.06 |
| 3 | E2f3 | 0.32 | 4.73E-32 | 9.85E-28 | 0.75 | 0.57 | 0.18 | 1.32 |
| 3 | Nufip1 | 0.37 | 4.78E-32 | 9.95E-28 | 0.66 | 0.49 | 0.17 | 1.35 |
| 3 | Nf2 | 0.49 | 5.83E-32 | 1.21E-27 | 0.43 | 0.28 | 0.15 | 1.54 |
| 3 | Chchd6 | 0.34 | 5.84E-32 | 1.21E-27 | 0.70 | 0.52 | 0.18 | 1.34 |
| 3 | Suv39h2 | 0.41 | 6.27E-32 | 1.30E-27 | 0.57 | 0.40 | 0.17 | 1.43 |
| 3 | Pold2 | 0.43 | 6.34E-32 | 1.32E-27 | 0.58 | 0.42 | 0.17 | 1.40 |
| 3 | Ptx3 | 0.42 | 6.88E-32 | 1.43E-27 | 0.20 | 0.10 | 0.10 | 1.96 |
| 3 | Nedd1 | 0.46 | 7.33E-32 | 1.52E-27 | 0.48 | 0.33 | 0.15 | 1.47 |
| 3 | Ncald | 0.80 | 8.26E-32 | 1.72E-27 | 0.17 | 0.09 | 0.09 | 2.01 |
| 3 | 2610027K06Rik | 0.69 | 8.27E-32 | 1.72E-27 | 0.23 | 0.13 | 0.11 | 1.83 |
| 3 | Nup107 | 0.44 | 8.38E-32 | 1.74E-27 | 0.53 | 0.37 | 0.16 | 1.43 |
| 3 | Gcsh | 0.28 | 8.87E-32 | 1.85E-27 | 0.88 | 0.72 | 0.15 | 1.21 |
| 3 | Mcm8 | 0.53 | 9.87E-32 | 2.05E-27 | 0.39 | 0.25 | 0.14 | 1.54 |
| 3 | Dpf1 | 0.65 | 1.01E-31 | 2.11E-27 | 0.27 | 0.15 | 0.11 | 1.74 |
| 3 | Hist1h2ai | 1.18 | 1.05E-31 | 2.18E-27 | 0.12 | 0.05 | 0.07 | 2.31 |
| 3 | Enkd1 | 0.48 | 1.50E-31 | 3.11E-27 | 0.42 | 0.27 | 0.15 | 1.54 |
| 3 | Aif1l | 0.41 | 1.74E-31 | 3.62E-27 | 0.53 | 0.37 | 0.16 | 1.44 |
| 3 | Gfpt2 | 0.52 | 2.17E-31 | 4.51E-27 | 0.30 | 0.17 | 0.12 | 1.71 |
| 3 | Olfml1 | 0.55 | 2.78E-31 | 5.79E-27 | 0.21 | 0.11 | 0.10 | 1.91 |
| 3 | Mertk | 0.49 | 2.98E-31 | 6.20E-27 | 0.27 | 0.15 | 0.12 | 1.77 |

**Supplementary Table 1. Marker genes for the original clusters of integrated clustering of E11.5 control and NotchGOF λ-junction cells.**

| | | | | | | | | |
|---|---|---|---|---|---|---|---|---|
| 3 | Utp15 | 0.43 | 3.38E-31 | 7.02E-27 | 0.54 | 0.38 | 0.16 | 1.41 |
| 3 | Sart3 | 0.33 | 3.57E-31 | 7.43E-27 | 0.73 | 0.55 | 0.18 | 1.34 |
| 3 | Cenpv | 0.41 | 4.44E-31 | 9.23E-27 | 0.65 | 0.49 | 0.16 | 1.32 |
| 3 | Nol11 | 0.28 | 4.88E-31 | 1.01E-26 | 0.85 | 0.70 | 0.15 | 1.22 |
| 3 | AI506816 | 0.35 | 6.54E-31 | 1.36E-26 | 0.70 | 0.52 | 0.18 | 1.33 |
| 3 | Apip | 0.30 | 8.20E-31 | 1.70E-26 | 0.76 | 0.60 | 0.16 | 1.28 |
| 3 | Alpl | 0.74 | 8.85E-31 | 1.84E-26 | 0.19 | 0.10 | 0.09 | 1.93 |
| 3 | Zc3h18 | 0.31 | 1.14E-30 | 2.37E-26 | 0.85 | 0.70 | 0.15 | 1.22 |
| 3 | Clic4 | 0.29 | 1.16E-30 | 2.42E-26 | 0.82 | 0.67 | 0.15 | 1.23 |
| 3 | Usp14 | 0.26 | 1.18E-30 | 2.45E-26 | 0.92 | 0.78 | 0.14 | 1.18 |
| 3 | Ak6 | 0.30 | 1.23E-30 | 2.56E-26 | 0.84 | 0.68 | 0.16 | 1.24 |
| 3 | Mcm6 | 0.27 | 1.36E-30 | 2.83E-26 | 0.85 | 0.69 | 0.16 | 1.23 |
| 3 | Snai2 | 0.27 | 1.44E-30 | 3.00E-26 | 0.64 | 0.45 | 0.19 | 1.43 |
| 3 | Dicer1 | 0.26 | 1.56E-30 | 3.24E-26 | 0.92 | 0.80 | 0.12 | 1.14 |
| 3 | Med7 | 0.35 | 1.83E-30 | 3.81E-26 | 0.69 | 0.52 | 0.17 | 1.32 |
| 3 | Mrps27 | 0.35 | 1.88E-30 | 3.91E-26 | 0.60 | 0.42 | 0.18 | 1.42 |
| 3 | Dusp7 | 0.36 | 1.94E-30 | 4.03E-26 | 0.66 | 0.49 | 0.18 | 1.36 |
| 3 | Hspa14 | 0.29 | 2.15E-30 | 4.48E-26 | 0.82 | 0.65 | 0.18 | 1.27 |
| 3 | Kyat3 | 0.65 | 2.29E-30 | 4.76E-26 | 0.23 | 0.13 | 0.10 | 1.80 |
| 3 | Ttf2 | 0.45 | 2.46E-30 | 5.12E-26 | 0.45 | 0.31 | 0.15 | 1.48 |
| 3 | Timm50 | 0.28 | 3.33E-30 | 6.92E-26 | 0.88 | 0.73 | 0.15 | 1.21 |
| 3 | Spata5 | 0.44 | 3.71E-30 | 7.73E-26 | 0.47 | 0.32 | 0.15 | 1.48 |
| 3 | Prkrip1 | 0.35 | 3.98E-30 | 8.27E-26 | 0.64 | 0.48 | 0.16 | 1.34 |
| 3 | Hyls1 | 0.69 | 4.08E-30 | 8.48E-26 | 0.28 | 0.17 | 0.11 | 1.65 |
| 3 | Fcgrt | 0.39 | 4.41E-30 | 9.18E-26 | 0.24 | 0.14 | 0.11 | 1.81 |
| 3 | 1700021F05Rik | 0.39 | 5.61E-30 | 1.17E-25 | 0.61 | 0.45 | 0.16 | 1.35 |
| 3 | Nab2 | 0.43 | 5.66E-30 | 1.18E-25 | 0.45 | 0.30 | 0.15 | 1.50 |
| 3 | Zfp143 | 0.47 | 5.77E-30 | 1.20E-25 | 0.39 | 0.26 | 0.13 | 1.53 |
| 3 | Dhrs3 | 0.48 | 6.64E-30 | 1.38E-25 | 0.42 | 0.28 | 0.14 | 1.49 |
| 3 | BC003965 | 0.32 | 6.75E-30 | 1.40E-25 | 0.79 | 0.62 | 0.17 | 1.27 |
| 3 | Kif24 | 0.89 | 8.74E-30 | 1.82E-25 | 0.18 | 0.09 | 0.09 | 1.92 |
| 3 | Dxo | 0.41 | 9.84E-30 | 2.05E-25 | 0.55 | 0.39 | 0.16 | 1.41 |
| 3 | Lpin2 | 0.32 | 1.04E-29 | 2.17E-25 | 0.70 | 0.53 | 0.17 | 1.33 |
| 3 | Rrp1b | 0.35 | 1.06E-29 | 2.20E-25 | 0.70 | 0.52 | 0.18 | 1.35 |
| 3 | Haus8 | 0.42 | 1.08E-29 | 2.25E-25 | 0.49 | 0.34 | 0.15 | 1.45 |
| 3 | Gm7324 | 0.32 | 1.29E-29 | 2.69E-25 | 0.77 | 0.61 | 0.17 | 1.28 |
| 3 | Brd8 | 0.27 | 1.51E-29 | 3.14E-25 | 0.88 | 0.74 | 0.13 | 1.18 |
| 3 | Dynlt1f | 0.26 | 1.75E-29 | 3.63E-25 | 0.90 | 0.76 | 0.14 | 1.18 |
| 3 | Bcas3 | 0.53 | 1.86E-29 | 3.88E-25 | 0.32 | 0.20 | 0.12 | 1.60 |
| 3 | Ect2 | 0.27 | 1.91E-29 | 3.96E-25 | 0.57 | 0.40 | 0.18 | 1.45 |
| 3 | Rad18 | 0.43 | 1.95E-29 | 4.06E-25 | 0.51 | 0.36 | 0.15 | 1.42 |
| 3 | Ncbp1 | 0.32 | 2.14E-29 | 4.46E-25 | 0.76 | 0.59 | 0.17 | 1.29 |
| 3 | Gli3 | 0.33 | 2.16E-29 | 4.49E-25 | 0.74 | 0.57 | 0.18 | 1.31 |
| 3 | Foxp1 | 0.38 | 2.29E-29 | 4.77E-25 | 0.73 | 0.59 | 0.14 | 1.23 |
| 3 | Rcl1 | 0.40 | 4.45E-29 | 9.25E-25 | 0.60 | 0.44 | 0.16 | 1.36 |
| 3 | Scpep1 | 0.36 | 4.53E-29 | 9.42E-25 | 0.56 | 0.40 | 0.17 | 1.42 |
| 3 | Hsph1 | 0.29 | 5.16E-29 | 1.07E-24 | 0.82 | 0.67 | 0.15 | 1.23 |
| 3 | Arhgef40 | 0.28 | 6.16E-29 | 1.28E-24 | 0.70 | 0.51 | 0.19 | 1.38 |
| 3 | Dpy19l1 | 0.41 | 7.61E-29 | 1.58E-24 | 0.50 | 0.35 | 0.16 | 1.45 |
| 3 | Nfib | 0.31 | 8.04E-29 | 1.67E-24 | 0.88 | 0.73 | 0.15 | 1.20 |
| 3 | Mrpl32 | 0.26 | 8.83E-29 | 1.84E-24 | 0.88 | 0.76 | 0.12 | 1.15 |
| 3 | Mis12 | 0.36 | 9.79E-29 | 2.04E-24 | 0.64 | 0.48 | 0.17 | 1.35 |
| 3 | Emx2 | 0.62 | 1.24E-28 | 2.58E-24 | 0.11 | 0.05 | 0.06 | 2.33 |
| 3 | Prkar2b | 0.27 | 1.35E-28 | 2.81E-24 | 0.65 | 0.48 | 0.17 | 1.36 |
| 3 | Ada | 0.82 | 1.38E-28 | 2.88E-24 | 0.15 | 0.08 | 0.08 | 2.01 |
| 3 | Ipo9 | 0.34 | 1.56E-28 | 3.25E-24 | 0.69 | 0.53 | 0.16 | 1.31 |

**Supplementary Table 1. Marker genes for the original clusters of integrated clustering of E11.5 control and NotchGOF λ-junction cells.**

| | | | | | | | | |
|---|---|---|---|---|---|---|---|---|
| 3 | Eefsec | 0.34 | 1.60E-28 | 3.32E-24 | 0.53 | 0.37 | 0.16 | 1.43 |
| 3 | Ppfibp2 | 0.25 | 1.60E-28 | 3.33E-24 | 0.61 | 0.41 | 0.19 | 1.46 |
| 3 | Uxs1 | 0.45 | 1.86E-28 | 3.87E-24 | 0.44 | 0.30 | 0.14 | 1.46 |
| 3 | Xpo1 | 0.26 | 2.00E-28 | 4.16E-24 | 0.91 | 0.77 | 0.14 | 1.18 |
| 3 | Mthfd1 | 0.39 | 2.01E-28 | 4.19E-24 | 0.53 | 0.37 | 0.16 | 1.42 |
| 3 | Gulo | 1.12 | 2.11E-28 | 4.40E-24 | 0.17 | 0.09 | 0.08 | 1.90 |
| 3 | Snhg5 | 0.40 | 2.42E-28 | 5.03E-24 | 0.58 | 0.43 | 0.15 | 1.36 |
| 3 | Srprb | 0.36 | 2.51E-28 | 5.22E-24 | 0.65 | 0.49 | 0.16 | 1.33 |
| 3 | Cldnd1 | 0.27 | 2.69E-28 | 5.59E-24 | 0.84 | 0.68 | 0.16 | 1.24 |
| 3 | Adamts3 | 0.88 | 3.30E-28 | 6.87E-24 | 0.12 | 0.06 | 0.07 | 2.22 |
| 3 | Mc4r | 1.00 | 3.45E-28 | 7.18E-24 | 0.11 | 0.05 | 0.06 | 2.33 |
| 3 | B3glct | 0.45 | 3.80E-28 | 7.91E-24 | 0.45 | 0.30 | 0.14 | 1.46 |
| 3 | Ppif | 0.34 | 3.83E-28 | 7.97E-24 | 0.69 | 0.52 | 0.17 | 1.32 |
| 3 | Tpst1 | 0.41 | 4.57E-28 | 9.50E-24 | 0.48 | 0.33 | 0.15 | 1.46 |
| 3 | Hist1h3g | 1.01 | 4.61E-28 | 9.59E-24 | 0.13 | 0.06 | 0.07 | 2.12 |
| 3 | 4930579G24Rik | 0.47 | 5.55E-28 | 1.15E-23 | 0.37 | 0.24 | 0.13 | 1.53 |
| 3 | Rell1 | 0.34 | 5.68E-28 | 1.18E-23 | 0.49 | 0.34 | 0.15 | 1.45 |
| 3 | Idh3a | 0.33 | 5.69E-28 | 1.18E-23 | 0.72 | 0.55 | 0.16 | 1.30 |
| 3 | Spidr | 0.47 | 6.61E-28 | 1.37E-23 | 0.44 | 0.30 | 0.14 | 1.45 |
| 3 | Rrp8 | 0.35 | 6.69E-28 | 1.39E-23 | 0.68 | 0.52 | 0.16 | 1.30 |
| 3 | Msantd2 | 0.35 | 6.99E-28 | 1.45E-23 | 0.60 | 0.43 | 0.16 | 1.37 |
| 3 | Vars | 0.31 | 8.07E-28 | 1.68E-23 | 0.76 | 0.59 | 0.17 | 1.28 |
| 3 | Prkar2a | 0.31 | 1.09E-27 | 2.26E-23 | 0.67 | 0.50 | 0.17 | 1.34 |
| 3 | Ccnf | 0.32 | 1.27E-27 | 2.63E-23 | 0.48 | 0.33 | 0.15 | 1.46 |
| 3 | Sf3a1 | 0.27 | 1.46E-27 | 3.04E-23 | 0.87 | 0.73 | 0.14 | 1.19 |
| 3 | Ciapin1 | 0.27 | 1.79E-27 | 3.73E-23 | 0.86 | 0.72 | 0.14 | 1.19 |
| 3 | Xrcc2 | 0.64 | 2.40E-27 | 5.00E-23 | 0.24 | 0.14 | 0.10 | 1.71 |
| 3 | B3galt1 | 1.04 | 2.42E-27 | 5.02E-23 | 0.10 | 0.04 | 0.06 | 2.38 |
| 3 | Ppm1m | 0.47 | 3.18E-27 | 6.61E-23 | 0.37 | 0.24 | 0.13 | 1.54 |
| 3 | Mdn1 | 0.36 | 4.05E-27 | 8.43E-23 | 0.55 | 0.40 | 0.15 | 1.39 |
| 3 | Exosc10 | 0.36 | 4.65E-27 | 9.68E-23 | 0.58 | 0.42 | 0.16 | 1.38 |
| 3 | Dcaf15 | 0.42 | 5.02E-27 | 1.04E-22 | 0.47 | 0.33 | 0.14 | 1.42 |
| 3 | Iqgap3 | 0.52 | 5.07E-27 | 1.05E-22 | 0.33 | 0.21 | 0.12 | 1.55 |
| 3 | Exosc1 | 0.31 | 5.08E-27 | 1.06E-22 | 0.70 | 0.53 | 0.17 | 1.31 |
| 3 | Cisd1 | 0.27 | 5.14E-27 | 1.07E-22 | 0.87 | 0.74 | 0.13 | 1.18 |
| 3 | Ddias | 0.70 | 6.06E-27 | 1.26E-22 | 0.25 | 0.15 | 0.10 | 1.66 |
| 3 | Gtf2f2 | 0.41 | 6.24E-27 | 1.30E-22 | 0.52 | 0.38 | 0.14 | 1.38 |
| 3 | Lin54 | 0.51 | 7.05E-27 | 1.47E-22 | 0.34 | 0.23 | 0.12 | 1.52 |
| 3 | Gtse1 | 0.61 | 7.81E-27 | 1.63E-22 | 0.28 | 0.18 | 0.11 | 1.61 |
| 3 | Pola2 | 0.35 | 7.89E-27 | 1.64E-22 | 0.58 | 0.41 | 0.17 | 1.40 |
| 3 | Ifngr2 | 0.35 | 8.50E-27 | 1.77E-22 | 0.51 | 0.36 | 0.15 | 1.42 |
| 3 | Syde1 | 0.50 | 8.69E-27 | 1.81E-22 | 0.28 | 0.17 | 0.11 | 1.67 |
| 3 | Coprs | 0.29 | 8.89E-27 | 1.85E-22 | 0.65 | 0.48 | 0.18 | 1.37 |
| 3 | Cand2 | 0.31 | 9.49E-27 | 1.97E-22 | 0.64 | 0.48 | 0.16 | 1.34 |
| 3 | Dnajb11 | 0.29 | 1.02E-26 | 2.12E-22 | 0.82 | 0.67 | 0.15 | 1.22 |
| 3 | Lpp | 0.26 | 1.03E-26 | 2.14E-22 | 0.72 | 0.54 | 0.18 | 1.33 |
| 3 | Ranbp3 | 0.37 | 1.37E-26 | 2.86E-22 | 0.64 | 0.48 | 0.16 | 1.33 |
| 3 | Zfp521 | 0.38 | 1.38E-26 | 2.86E-22 | 0.42 | 0.28 | 0.14 | 1.51 |
| 3 | Emc2 | 0.30 | 1.38E-26 | 2.88E-22 | 0.74 | 0.58 | 0.16 | 1.27 |
| 3 | Gins1 | 0.39 | 1.51E-26 | 3.14E-22 | 0.53 | 0.38 | 0.15 | 1.38 |
| 3 | Tsen15 | 0.28 | 1.66E-26 | 3.46E-22 | 0.79 | 0.62 | 0.17 | 1.27 |
| 3 | Wdr86 | 0.26 | 1.74E-26 | 3.62E-22 | 0.69 | 0.50 | 0.19 | 1.37 |
| 3 | Sigmar1 | 0.35 | 1.90E-26 | 3.95E-22 | 0.69 | 0.54 | 0.15 | 1.29 |
| 3 | H2-Ab1 | 0.94 | 1.98E-26 | 4.11E-22 | 0.11 | 0.05 | 0.06 | 2.25 |
| 3 | Prep | 0.38 | 2.08E-26 | 4.34E-22 | 0.53 | 0.38 | 0.15 | 1.40 |
| 3 | Ankrd29 | 0.70 | 2.09E-26 | 4.36E-22 | 0.17 | 0.09 | 0.08 | 1.93 |

**Supplementary Table 1. Marker genes for the original clusters of integrated clustering of E11.5 control and NotchGOF λ-junction cells.**

| | | | | | | | | |
|---|---|---|---|---|---|---|---|---|
| 3 | Saal1 | 0.33 | 2.32E-26 | 4.82E-22 | 0.61 | 0.45 | 0.16 | 1.36 |
| 3 | Dnph1 | 0.31 | 2.34E-26 | 4.87E-22 | 0.68 | 0.51 | 0.17 | 1.34 |
| 3 | Qdpr | 0.29 | 2.35E-26 | 4.89E-22 | 0.69 | 0.52 | 0.16 | 1.31 |
| 3 | Lnpep | 0.53 | 3.71E-26 | 7.72E-22 | 0.40 | 0.28 | 0.12 | 1.44 |
| 3 | Mogat2 | 0.64 | 3.75E-26 | 7.80E-22 | 0.20 | 0.11 | 0.09 | 1.80 |
| 3 | Dnaaf2 | 0.30 | 3.88E-26 | 8.07E-22 | 0.66 | 0.50 | 0.16 | 1.33 |
| 3 | Nol9 | 0.42 | 3.98E-26 | 8.28E-22 | 0.45 | 0.31 | 0.14 | 1.44 |
| 3 | Rps6ka6 | 0.27 | 4.09E-26 | 8.51E-22 | 0.61 | 0.44 | 0.17 | 1.38 |
| 3 | Pask | 0.60 | 4.45E-26 | 9.25E-22 | 0.25 | 0.15 | 0.10 | 1.67 |
| 3 | Adgra2 | 0.35 | 5.31E-26 | 1.11E-21 | 0.36 | 0.23 | 0.13 | 1.58 |
| 3 | Galt | 0.41 | 5.45E-26 | 1.13E-21 | 0.40 | 0.27 | 0.13 | 1.51 |
| 3 | Nip7 | 0.25 | 6.13E-26 | 1.28E-21 | 0.87 | 0.74 | 0.14 | 1.18 |
| 3 | Polr2b | 0.28 | 6.16E-26 | 1.28E-21 | 0.82 | 0.66 | 0.16 | 1.24 |
| 3 | Pkia | 0.46 | 6.91E-26 | 1.44E-21 | 0.29 | 0.18 | 0.11 | 1.63 |
| 3 | Zfp101 | 0.37 | 8.38E-26 | 1.74E-21 | 0.50 | 0.35 | 0.15 | 1.41 |
| 3 | Lmnb2 | 0.38 | 8.46E-26 | 1.76E-21 | 0.51 | 0.36 | 0.15 | 1.41 |
| 3 | Dipk1a | 0.51 | 8.73E-26 | 1.81E-21 | 0.31 | 0.20 | 0.11 | 1.58 |
| 3 | Sergef | 0.65 | 9.46E-26 | 1.97E-21 | 0.22 | 0.13 | 0.09 | 1.73 |
| 3 | Dnajc25 | 0.40 | 1.13E-25 | 2.35E-21 | 0.39 | 0.26 | 0.13 | 1.50 |
| 3 | Mrpl16 | 0.35 | 1.56E-25 | 3.24E-21 | 0.65 | 0.50 | 0.15 | 1.30 |
| 3 | Mtfr2 | 0.75 | 1.57E-25 | 3.26E-21 | 0.18 | 0.10 | 0.08 | 1.79 |
| 3 | Rngtt | 0.38 | 1.61E-25 | 3.34E-21 | 0.53 | 0.39 | 0.15 | 1.38 |
| 3 | Foxk2 | 0.41 | 2.02E-25 | 4.19E-21 | 0.49 | 0.36 | 0.14 | 1.38 |
| 3 | Recql4 | 0.82 | 2.13E-25 | 4.43E-21 | 0.16 | 0.08 | 0.08 | 1.90 |
| 3 | Wdr75 | 0.28 | 2.27E-25 | 4.72E-21 | 0.73 | 0.57 | 0.16 | 1.28 |
| 3 | Eps8 | 0.55 | 2.42E-25 | 5.04E-21 | 0.26 | 0.15 | 0.10 | 1.66 |
| 3 | Ddx27 | 0.28 | 2.67E-25 | 5.55E-21 | 0.82 | 0.69 | 0.14 | 1.20 |
| 3 | E2f6 | 0.30 | 2.79E-25 | 5.81E-21 | 0.67 | 0.52 | 0.15 | 1.30 |
| 3 | Gnl3 | 0.25 | 3.39E-25 | 7.05E-21 | 0.87 | 0.74 | 0.13 | 1.18 |
| 3 | Notch2 | 0.35 | 3.43E-25 | 7.13E-21 | 0.50 | 0.35 | 0.15 | 1.43 |
| 3 | Senp7 | 0.33 | 3.43E-25 | 7.14E-21 | 0.68 | 0.54 | 0.14 | 1.25 |
| 3 | Mastl | 0.50 | 3.61E-25 | 7.50E-21 | 0.32 | 0.21 | 0.11 | 1.51 |
| 3 | Pmaip1 | 0.47 | 3.84E-25 | 7.99E-21 | 0.35 | 0.24 | 0.12 | 1.50 |
| 3 | Nutf2-ps1 | 0.27 | 4.68E-25 | 9.74E-21 | 0.68 | 0.51 | 0.17 | 1.33 |
| 3 | Peg12 | 0.43 | 5.22E-25 | 1.08E-20 | 0.37 | 0.24 | 0.13 | 1.52 |
| 3 | Emsy | 0.28 | 5.70E-25 | 1.19E-20 | 0.76 | 0.61 | 0.15 | 1.25 |
| 3 | Cyp27a1 | 0.73 | 7.13E-25 | 1.48E-20 | 0.15 | 0.08 | 0.07 | 1.92 |
| 3 | Cep295 | 0.31 | 7.82E-25 | 1.63E-20 | 0.64 | 0.47 | 0.16 | 1.34 |
| 3 | Zgrf1 | 0.48 | 8.47E-25 | 1.76E-20 | 0.30 | 0.20 | 0.11 | 1.56 |
| 3 | Id1 | 0.29 | 9.73E-25 | 2.02E-20 | 0.93 | 0.85 | 0.08 | 1.10 |
| 3 | Kpna3 | 0.29 | 9.97E-25 | 2.07E-20 | 0.76 | 0.60 | 0.16 | 1.27 |
| 3 | A430005L14Rik | 0.32 | 1.22E-24 | 2.55E-20 | 0.65 | 0.49 | 0.16 | 1.32 |
| 3 | 5330434G04Rik | 0.66 | 1.50E-24 | 3.12E-20 | 0.16 | 0.08 | 0.08 | 1.90 |
| 3 | Tssc4 | 0.28 | 1.57E-24 | 3.26E-20 | 0.73 | 0.57 | 0.16 | 1.27 |
| 3 | Uck2 | 0.30 | 1.78E-24 | 3.70E-20 | 0.67 | 0.51 | 0.16 | 1.31 |
| 3 | Nid2 | 0.33 | 1.79E-24 | 3.72E-20 | 0.35 | 0.23 | 0.13 | 1.55 |
| 3 | Hacd1 | 0.34 | 1.98E-24 | 4.12E-20 | 0.53 | 0.38 | 0.15 | 1.38 |
| 3 | Nop9 | 0.36 | 2.02E-24 | 4.19E-20 | 0.51 | 0.37 | 0.14 | 1.39 |
| 3 | Dhodh | 0.39 | 2.06E-24 | 4.29E-20 | 0.41 | 0.28 | 0.13 | 1.47 |
| 3 | Ccng2 | 0.31 | 2.33E-24 | 4.84E-20 | 0.57 | 0.42 | 0.15 | 1.36 |
| 3 | Kank4 | 0.36 | 2.81E-24 | 5.85E-20 | 0.31 | 0.19 | 0.12 | 1.60 |
| 3 | Magee1 | 0.49 | 2.93E-24 | 6.10E-20 | 0.22 | 0.13 | 0.09 | 1.73 |
| 3 | Osr1 | 0.59 | 3.23E-24 | 6.72E-20 | 0.13 | 0.07 | 0.07 | 2.02 |
| 3 | Pcbp3 | 0.48 | 3.31E-24 | 6.88E-20 | 0.27 | 0.17 | 0.10 | 1.61 |
| 3 | Lrrfip2 | 0.29 | 3.77E-24 | 7.85E-20 | 0.65 | 0.49 | 0.16 | 1.33 |
| 3 | Sgcb | 0.43 | 3.85E-24 | 8.01E-20 | 0.31 | 0.20 | 0.11 | 1.57 |

**Supplementary Table 1. Marker genes for the original clusters of integrated clustering of E11.5 control and NotchGOF λ-junction cells.**

| | | | | | | | | |
|---|---|---|---|---|---|---|---|---|
| 3 | Aff3 | 0.41 | 4.50E-24 | 9.35E-20 | 0.33 | 0.21 | 0.12 | 1.56 |
| 3 | Camk2n2 | 0.72 | 4.68E-24 | 9.74E-20 | 0.17 | 0.09 | 0.08 | 1.84 |
| 3 | Kifc5b | 0.66 | 4.78E-24 | 9.95E-20 | 0.24 | 0.15 | 0.09 | 1.63 |
| 3 | Thop1 | 0.26 | 5.39E-24 | 1.12E-19 | 0.67 | 0.50 | 0.17 | 1.33 |
| 3 | Ado | 0.33 | 5.66E-24 | 1.18E-19 | 0.53 | 0.38 | 0.14 | 1.38 |
| 3 | Nfil3 | 0.54 | 6.22E-24 | 1.29E-19 | 0.22 | 0.13 | 0.09 | 1.70 |
| 3 | Tinf2 | 0.52 | 6.95E-24 | 1.45E-19 | 0.30 | 0.20 | 0.11 | 1.53 |
| 3 | Tmem246 | 0.55 | 7.59E-24 | 1.58E-19 | 0.21 | 0.12 | 0.09 | 1.74 |
| 3 | Prkaca | 0.27 | 7.91E-24 | 1.65E-19 | 0.72 | 0.55 | 0.16 | 1.30 |
| 3 | Nudt2 | 0.30 | 8.53E-24 | 1.77E-19 | 0.73 | 0.58 | 0.15 | 1.26 |
| 3 | Mrps26 | 0.26 | 1.02E-23 | 2.13E-19 | 0.79 | 0.64 | 0.15 | 1.23 |
| 3 | Apoo | 0.29 | 1.14E-23 | 2.38E-19 | 0.64 | 0.48 | 0.16 | 1.33 |
| 3 | Ahcy | 0.41 | 1.47E-23 | 3.05E-19 | 0.39 | 0.26 | 0.12 | 1.47 |
| 3 | Snrnp25 | 0.26 | 1.50E-23 | 3.12E-19 | 0.76 | 0.60 | 0.16 | 1.27 |
| 3 | Haus6 | 0.36 | 1.99E-23 | 4.15E-19 | 0.49 | 0.35 | 0.14 | 1.40 |
| 3 | Cdc25c | 0.36 | 1.99E-23 | 4.15E-19 | 0.47 | 0.34 | 0.13 | 1.39 |
| 3 | Ncapd2 | 0.25 | 2.03E-23 | 4.22E-19 | 0.71 | 0.54 | 0.17 | 1.31 |
| 3 | 4933404O12Rik | 0.46 | 2.48E-23 | 5.16E-19 | 0.36 | 0.24 | 0.12 | 1.49 |
| 3 | St3gal4 | 0.38 | 2.51E-23 | 5.22E-19 | 0.36 | 0.24 | 0.12 | 1.51 |
| 3 | Slc20a1 | 0.39 | 3.06E-23 | 6.36E-19 | 0.44 | 0.31 | 0.12 | 1.40 |
| 3 | Hmga1b | 0.31 | 3.10E-23 | 6.46E-19 | 0.58 | 0.42 | 0.16 | 1.37 |
| 3 | Stra6 | 0.30 | 3.39E-23 | 7.06E-19 | 0.48 | 0.33 | 0.15 | 1.45 |
| 3 | Paxip1 | 0.38 | 3.57E-23 | 7.43E-19 | 0.46 | 0.33 | 0.13 | 1.41 |
| 3 | Ttc39c | 0.45 | 3.79E-23 | 7.89E-19 | 0.30 | 0.19 | 0.11 | 1.56 |
| 3 | Slf1 | 0.30 | 4.05E-23 | 8.42E-19 | 0.68 | 0.53 | 0.15 | 1.29 |
| 3 | Safb2 | 0.30 | 4.21E-23 | 8.76E-19 | 0.70 | 0.55 | 0.15 | 1.26 |
| 3 | Pum3 | 0.26 | 4.38E-23 | 9.12E-19 | 0.74 | 0.60 | 0.15 | 1.25 |
| 3 | Ubr7 | 0.30 | 6.29E-23 | 1.31E-18 | 0.61 | 0.46 | 0.15 | 1.32 |
| 3 | Slc2a4 | 0.53 | 6.66E-23 | 1.38E-18 | 0.26 | 0.17 | 0.10 | 1.59 |
| 3 | Oip5 | 0.55 | 6.71E-23 | 1.39E-18 | 0.29 | 0.19 | 0.10 | 1.52 |
| 3 | Acadl | 0.27 | 8.41E-23 | 1.75E-18 | 0.68 | 0.52 | 0.16 | 1.30 |
| 3 | Fam149a | 0.56 | 8.96E-23 | 1.86E-18 | 0.23 | 0.14 | 0.09 | 1.66 |
| 3 | Dhx29 | 0.27 | 1.02E-22 | 2.12E-18 | 0.66 | 0.50 | 0.15 | 1.30 |
| 3 | Erc1 | 0.28 | 1.02E-22 | 2.13E-18 | 0.70 | 0.54 | 0.16 | 1.30 |
| 3 | Atg5 | 0.27 | 1.07E-22 | 2.22E-18 | 0.72 | 0.56 | 0.16 | 1.29 |
| 3 | Map7d3 | 0.78 | 1.31E-22 | 2.73E-18 | 0.12 | 0.06 | 0.06 | 2.03 |
| 3 | Nelfe | 0.26 | 1.35E-22 | 2.80E-18 | 0.78 | 0.65 | 0.14 | 1.21 |
| 3 | Phc1 | 0.31 | 1.35E-22 | 2.82E-18 | 0.50 | 0.36 | 0.15 | 1.41 |
| 3 | Fam72a | 0.90 | 1.35E-22 | 2.82E-18 | 0.13 | 0.07 | 0.06 | 1.91 |
| 3 | Mrps30 | 0.29 | 1.39E-22 | 2.90E-18 | 0.74 | 0.59 | 0.14 | 1.24 |
| 3 | Ppp2r1b | 0.34 | 1.50E-22 | 3.11E-18 | 0.53 | 0.39 | 0.14 | 1.37 |
| 3 | Tmem45a | 0.62 | 1.77E-22 | 3.67E-18 | 0.15 | 0.08 | 0.07 | 1.91 |
| 3 | Zfp57 | 0.29 | 1.79E-22 | 3.72E-18 | 0.47 | 0.33 | 0.14 | 1.44 |
| 3 | Hmga1 | 0.27 | 1.96E-22 | 4.08E-18 | 0.64 | 0.48 | 0.16 | 1.33 |
| 3 | Strn4 | 0.29 | 2.20E-22 | 4.57E-18 | 0.63 | 0.48 | 0.15 | 1.32 |
| 3 | Slc7a2 | 0.83 | 2.41E-22 | 5.02E-18 | 0.13 | 0.06 | 0.06 | 1.98 |
| 3 | Serpine2 | 0.29 | 2.51E-22 | 5.22E-18 | 0.84 | 0.69 | 0.15 | 1.22 |
| 3 | Nubp1 | 0.28 | 2.52E-22 | 5.24E-18 | 0.69 | 0.54 | 0.16 | 1.29 |
| 3 | Wnk3 | 0.27 | 2.79E-22 | 5.79E-18 | 0.45 | 0.31 | 0.14 | 1.46 |
| 3 | Parvb | 0.62 | 2.90E-22 | 6.03E-18 | 0.21 | 0.12 | 0.08 | 1.68 |
| 3 | Pidd1 | 0.62 | 3.08E-22 | 6.40E-18 | 0.20 | 0.12 | 0.08 | 1.71 |
| 3 | Sf3a2 | 0.29 | 3.17E-22 | 6.59E-18 | 0.67 | 0.52 | 0.15 | 1.28 |
| 3 | Ptges | 0.47 | 3.18E-22 | 6.61E-18 | 0.23 | 0.14 | 0.09 | 1.67 |
| 3 | Map2k6 | 0.29 | 3.40E-22 | 7.06E-18 | 0.58 | 0.42 | 0.16 | 1.37 |
| 3 | Snx24 | 0.34 | 3.42E-22 | 7.10E-18 | 0.49 | 0.35 | 0.14 | 1.40 |
| 3 | Cenpo | 0.62 | 3.43E-22 | 7.14E-18 | 0.21 | 0.12 | 0.08 | 1.67 |

**Supplementary Table 1. Marker genes for the original clusters of integrated clustering of E11.5 control and NotchGOF λ-junction cells.**

| | | | | | | | | |
|---|---|---|---|---|---|---|---|---|
| 3 | Anxa11 | 0.65 | 3.59E-22 | 7.46E-18 | 0.24 | 0.15 | 0.09 | 1.61 |
| 3 | Zmym1 | 0.32 | 3.61E-22 | 7.52E-18 | 0.46 | 0.33 | 0.13 | 1.40 |
| 3 | Pigf | 0.32 | 3.97E-22 | 8.26E-18 | 0.53 | 0.39 | 0.14 | 1.35 |
| 3 | 5430416N02Rik | 0.28 | 4.54E-22 | 9.43E-18 | 0.60 | 0.45 | 0.15 | 1.34 |
| 3 | Tmem164 | 0.37 | 4.62E-22 | 9.61E-18 | 0.43 | 0.30 | 0.13 | 1.42 |
| 3 | Tbl3 | 0.36 | 4.64E-22 | 9.65E-18 | 0.47 | 0.34 | 0.13 | 1.39 |
| 3 | Homer2 | 0.39 | 5.03E-22 | 1.05E-17 | 0.31 | 0.20 | 0.11 | 1.53 |
| 3 | Stil | 0.46 | 5.47E-22 | 1.14E-17 | 0.29 | 0.19 | 0.10 | 1.53 |
| 3 | Arnt2 | 0.46 | 5.64E-22 | 1.17E-17 | 0.28 | 0.18 | 0.10 | 1.57 |
| 3 | Eed | 0.26 | 5.82E-22 | 1.21E-17 | 0.69 | 0.53 | 0.16 | 1.30 |
| 3 | Car12 | 0.55 | 6.68E-22 | 1.39E-17 | 0.16 | 0.09 | 0.07 | 1.83 |
| 3 | Psmc3ip | 0.28 | 6.97E-22 | 1.45E-17 | 0.70 | 0.55 | 0.16 | 1.29 |
| 3 | Ahctf1 | 0.27 | 7.28E-22 | 1.51E-17 | 0.69 | 0.53 | 0.16 | 1.30 |
| 3 | Ush1c | 0.59 | 7.77E-22 | 1.62E-17 | 0.18 | 0.10 | 0.08 | 1.79 |
| 3 | Stau2 | 0.33 | 9.32E-22 | 1.94E-17 | 0.51 | 0.37 | 0.14 | 1.37 |
| 3 | Arhgap19 | 0.38 | 1.04E-21 | 2.17E-17 | 0.39 | 0.28 | 0.12 | 1.41 |
| 3 | Ndufaf6 | 0.34 | 1.12E-21 | 2.33E-17 | 0.47 | 0.34 | 0.14 | 1.40 |
| 3 | Ackr3 | 0.27 | 1.19E-21 | 2.47E-17 | 0.55 | 0.41 | 0.14 | 1.34 |
| 3 | Lnpk | 0.28 | 1.21E-21 | 2.52E-17 | 0.62 | 0.47 | 0.15 | 1.32 |
| 3 | Prex1 | 0.39 | 1.42E-21 | 2.96E-17 | 0.29 | 0.19 | 0.10 | 1.55 |
| 3 | Tyw3 | 0.45 | 1.43E-21 | 2.96E-17 | 0.31 | 0.20 | 0.10 | 1.51 |
| 3 | Rab23 | 0.33 | 1.52E-21 | 3.16E-17 | 0.45 | 0.31 | 0.13 | 1.43 |
| 3 | B930095G15Rik | 0.47 | 1.92E-21 | 3.99E-17 | 0.27 | 0.17 | 0.10 | 1.56 |
| 3 | Stx18 | 0.25 | 1.95E-21 | 4.05E-17 | 0.66 | 0.51 | 0.16 | 1.31 |
| 3 | Tmem97 | 0.28 | 2.04E-21 | 4.25E-17 | 0.61 | 0.45 | 0.16 | 1.35 |
| 3 | Pphln1 | 0.33 | 2.05E-21 | 4.26E-17 | 0.52 | 0.39 | 0.14 | 1.35 |
| 3 | Snrnp35 | 0.36 | 2.10E-21 | 4.36E-17 | 0.39 | 0.27 | 0.12 | 1.44 |
| 3 | Zw10 | 0.36 | 2.25E-21 | 4.67E-17 | 0.44 | 0.31 | 0.13 | 1.40 |
| 3 | Ythdc2 | 0.36 | 2.65E-21 | 5.52E-17 | 0.38 | 0.26 | 0.12 | 1.46 |
| 3 | Slc1a5 | 0.48 | 2.75E-21 | 5.71E-17 | 0.29 | 0.20 | 0.10 | 1.51 |
| 3 | Dtnb | 0.39 | 2.77E-21 | 5.77E-17 | 0.38 | 0.27 | 0.12 | 1.44 |
| 3 | Noc4l | 0.30 | 2.97E-21 | 6.18E-17 | 0.56 | 0.42 | 0.14 | 1.34 |
| 3 | Tada2a | 0.30 | 3.13E-21 | 6.51E-17 | 0.56 | 0.41 | 0.15 | 1.36 |
| 3 | Ppp3cb | 0.25 | 3.14E-21 | 6.54E-17 | 0.76 | 0.61 | 0.15 | 1.25 |
| 3 | Polq | 0.75 | 3.20E-21 | 6.65E-17 | 0.16 | 0.09 | 0.07 | 1.76 |
| 3 | Trmt10a | 0.32 | 3.74E-21 | 7.79E-17 | 0.46 | 0.33 | 0.13 | 1.40 |
| 3 | C230038L03Rik | 0.50 | 4.46E-21 | 9.28E-17 | 0.20 | 0.12 | 0.08 | 1.70 |
| 3 | Rrp9 | 0.32 | 4.49E-21 | 9.35E-17 | 0.47 | 0.34 | 0.13 | 1.40 |
| 3 | Reck | 0.33 | 5.19E-21 | 1.08E-16 | 0.36 | 0.24 | 0.12 | 1.50 |
| 3 | Has2 | 0.39 | 5.31E-21 | 1.10E-16 | 0.29 | 0.19 | 0.10 | 1.54 |
| 3 | Gsdme | 0.41 | 7.05E-21 | 1.47E-16 | 0.34 | 0.23 | 0.11 | 1.48 |
| 3 | Tex10 | 0.29 | 8.07E-21 | 1.68E-16 | 0.58 | 0.44 | 0.15 | 1.33 |
| 3 | Rad17 | 0.28 | 8.21E-21 | 1.71E-16 | 0.61 | 0.46 | 0.15 | 1.33 |
| 3 | Tchp | 0.34 | 9.38E-21 | 1.95E-16 | 0.41 | 0.29 | 0.12 | 1.44 |
| 3 | Iglon5 | 0.42 | 1.15E-20 | 2.40E-16 | 0.26 | 0.17 | 0.10 | 1.58 |
| 3 | Wdr35 | 0.33 | 1.16E-20 | 2.41E-16 | 0.41 | 0.29 | 0.13 | 1.45 |
| 3 | Spata6 | 0.44 | 1.18E-20 | 2.44E-16 | 0.30 | 0.19 | 0.10 | 1.53 |
| 3 | Xrcc6 | 0.33 | 1.20E-20 | 2.49E-16 | 0.56 | 0.43 | 0.13 | 1.31 |
| 3 | Dlg2 | 0.80 | 1.28E-20 | 2.67E-16 | 0.13 | 0.07 | 0.06 | 1.94 |
| 3 | Mtrex | 0.26 | 1.35E-20 | 2.81E-16 | 0.75 | 0.61 | 0.14 | 1.23 |
| 3 | Cib2 | 0.29 | 1.45E-20 | 3.02E-16 | 0.37 | 0.25 | 0.12 | 1.49 |
| 3 | Hbegf | 0.64 | 2.07E-20 | 4.30E-16 | 0.14 | 0.08 | 0.07 | 1.83 |
| 3 | Cited1 | 0.39 | 2.43E-20 | 5.05E-16 | 0.19 | 0.11 | 0.08 | 1.72 |
| 3 | Cactin | 0.31 | 2.49E-20 | 5.17E-16 | 0.44 | 0.31 | 0.13 | 1.42 |
| 3 | Nup155 | 0.34 | 2.51E-20 | 5.22E-16 | 0.53 | 0.39 | 0.13 | 1.34 |
| 3 | Large1 | 0.44 | 2.98E-20 | 6.20E-16 | 0.29 | 0.19 | 0.10 | 1.52 |

**Supplementary Table 1. Marker genes for the original clusters of integrated clustering of E11.5 control and NotchGOF λ-junction cells.**

| | | | | | | | | |
|---|---|---|---|---|---|---|---|---|
| 3 | Zfp41 | 0.32 | 3.08E-20 | 6.41E-16 | 0.47 | 0.33 | 0.13 | 1.40 |
| 3 | Zbtb14 | 0.41 | 3.94E-20 | 8.19E-16 | 0.33 | 0.22 | 0.11 | 1.47 |
| 3 | Pif1 | 0.60 | 4.00E-20 | 8.33E-16 | 0.21 | 0.13 | 0.08 | 1.62 |
| 3 | Nup205 | 0.31 | 4.10E-20 | 8.53E-16 | 0.46 | 0.33 | 0.13 | 1.38 |
| 3 | Ppp1r8 | 0.28 | 4.75E-20 | 9.87E-16 | 0.58 | 0.44 | 0.14 | 1.33 |
| 3 | Nup160 | 0.38 | 4.82E-20 | 1.00E-15 | 0.42 | 0.30 | 0.12 | 1.40 |
| 3 | Stx12 | 0.25 | 4.82E-20 | 1.00E-15 | 0.66 | 0.51 | 0.15 | 1.30 |
| 3 | Elp6 | 0.28 | 4.94E-20 | 1.03E-15 | 0.52 | 0.38 | 0.14 | 1.38 |
| 3 | Zfp532 | 0.26 | 5.10E-20 | 1.06E-15 | 0.61 | 0.47 | 0.14 | 1.30 |
| 3 | Pde3b | 0.39 | 5.46E-20 | 1.14E-15 | 0.35 | 0.24 | 0.11 | 1.46 |
| 3 | Pmp22 | 0.33 | 5.62E-20 | 1.17E-15 | 0.28 | 0.18 | 0.10 | 1.57 |
| 3 | Acaca | 0.31 | 6.35E-20 | 1.32E-15 | 0.45 | 0.32 | 0.13 | 1.40 |
| 3 | Zcchc8 | 0.28 | 7.43E-20 | 1.55E-15 | 0.60 | 0.47 | 0.14 | 1.30 |
| 3 | Cpsf4 | 0.29 | 7.99E-20 | 1.66E-15 | 0.52 | 0.38 | 0.14 | 1.36 |
| 3 | Tspan11 | 0.38 | 8.00E-20 | 1.66E-15 | 0.26 | 0.16 | 0.10 | 1.60 |
| 3 | Rbmx2 | 0.29 | 8.48E-20 | 1.76E-15 | 0.56 | 0.42 | 0.14 | 1.32 |
| 3 | Sertad4 | 0.39 | 1.05E-19 | 2.18E-15 | 0.48 | 0.36 | 0.12 | 1.35 |
| 3 | Lrrc40 | 0.32 | 1.07E-19 | 2.22E-15 | 0.45 | 0.32 | 0.13 | 1.39 |
| 3 | Tmem121 | 0.39 | 1.32E-19 | 2.74E-15 | 0.33 | 0.22 | 0.11 | 1.48 |
| 3 | Rad9a | 0.35 | 1.40E-19 | 2.92E-15 | 0.31 | 0.21 | 0.10 | 1.49 |
| 3 | Tedc2 | 0.43 | 1.49E-19 | 3.11E-15 | 0.26 | 0.17 | 0.09 | 1.53 |
| 3 | Kif14 | 0.46 | 1.64E-19 | 3.40E-15 | 0.27 | 0.18 | 0.09 | 1.50 |
| 3 | Rbl1 | 0.37 | 1.94E-19 | 4.03E-15 | 0.40 | 0.28 | 0.12 | 1.40 |
| 3 | Ptcd3 | 0.27 | 1.94E-19 | 4.04E-15 | 0.64 | 0.48 | 0.15 | 1.32 |
| 3 | Atp11c | 0.39 | 2.09E-19 | 4.34E-15 | 0.35 | 0.24 | 0.11 | 1.44 |
| 3 | Nrm | 0.27 | 2.11E-19 | 4.38E-15 | 0.47 | 0.34 | 0.13 | 1.38 |
| 3 | Wrap53 | 0.38 | 2.21E-19 | 4.59E-15 | 0.32 | 0.22 | 0.10 | 1.47 |
| 3 | E130307A14Rik | 0.41 | 2.32E-19 | 4.83E-15 | 0.29 | 0.19 | 0.10 | 1.52 |
| 3 | Polr3f | 0.29 | 2.45E-19 | 5.10E-15 | 0.46 | 0.33 | 0.13 | 1.39 |
| 3 | 2810429I04Rik | 0.51 | 2.58E-19 | 5.37E-15 | 0.23 | 0.15 | 0.09 | 1.58 |
| 3 | Cep126 | 0.41 | 2.63E-19 | 5.48E-15 | 0.24 | 0.15 | 0.09 | 1.58 |
| 3 | Tcf7 | 0.62 | 2.74E-19 | 5.70E-15 | 0.22 | 0.14 | 0.08 | 1.56 |
| 3 | Utp6 | 0.30 | 2.84E-19 | 5.90E-15 | 0.57 | 0.44 | 0.14 | 1.31 |
| 3 | Znhit6 | 0.26 | 3.42E-19 | 7.11E-15 | 0.73 | 0.59 | 0.14 | 1.24 |
| 3 | Bysl | 0.33 | 4.11E-19 | 8.54E-15 | 0.45 | 0.33 | 0.12 | 1.38 |
| 3 | Mllt1 | 0.33 | 4.17E-19 | 8.67E-15 | 0.41 | 0.29 | 0.12 | 1.41 |
| 3 | Rpp40 | 0.38 | 4.25E-19 | 8.85E-15 | 0.33 | 0.23 | 0.10 | 1.45 |
| 3 | Gm4737 | 0.56 | 4.71E-19 | 9.81E-15 | 0.19 | 0.12 | 0.08 | 1.65 |
| 3 | Cars | 0.37 | 5.29E-19 | 1.10E-14 | 0.40 | 0.29 | 0.11 | 1.38 |
| 3 | Vkorc1l1 | 0.30 | 5.48E-19 | 1.14E-14 | 0.52 | 0.39 | 0.13 | 1.34 |
| 3 | Gemin5 | 0.30 | 5.62E-19 | 1.17E-14 | 0.48 | 0.36 | 0.13 | 1.36 |
| 3 | Suv39h1 | 0.29 | 6.82E-19 | 1.42E-14 | 0.59 | 0.46 | 0.14 | 1.30 |
| 3 | Myo5a | 0.30 | 7.69E-19 | 1.60E-14 | 0.43 | 0.31 | 0.12 | 1.39 |
| 3 | Abca5 | 0.42 | 8.11E-19 | 1.69E-14 | 0.20 | 0.12 | 0.08 | 1.65 |
| 3 | Phykpl | 0.37 | 9.25E-19 | 1.92E-14 | 0.35 | 0.24 | 0.11 | 1.44 |
| 3 | Smyd2 | 0.26 | 9.84E-19 | 2.05E-14 | 0.54 | 0.40 | 0.14 | 1.35 |
| 3 | Crem | 0.32 | 1.07E-18 | 2.22E-14 | 0.27 | 0.17 | 0.09 | 1.54 |
| 3 | Ercc8 | 0.37 | 1.10E-18 | 2.29E-14 | 0.32 | 0.22 | 0.10 | 1.46 |
| 3 | Tex9 | 0.28 | 1.15E-18 | 2.40E-14 | 0.58 | 0.44 | 0.14 | 1.31 |
| 3 | Laptm4b | 0.30 | 1.16E-18 | 2.41E-14 | 0.69 | 0.54 | 0.16 | 1.29 |
| 3 | Mdm1 | 0.25 | 1.36E-18 | 2.83E-14 | 0.41 | 0.29 | 0.12 | 1.41 |
| 3 | Troap | 0.47 | 1.41E-18 | 2.93E-14 | 0.26 | 0.18 | 0.09 | 1.50 |
| 3 | Antxr2 | 0.36 | 1.43E-18 | 2.97E-14 | 0.27 | 0.18 | 0.10 | 1.54 |
| 3 | Cep128 | 0.51 | 1.55E-18 | 3.22E-14 | 0.25 | 0.17 | 0.09 | 1.51 |
| 3 | Emp2 | 0.63 | 1.72E-18 | 3.58E-14 | 0.13 | 0.07 | 0.06 | 1.83 |
| 3 | Ptges2 | 0.31 | 1.85E-18 | 3.85E-14 | 0.43 | 0.31 | 0.12 | 1.39 |

**Supplementary Table 1. Marker genes for the original clusters of integrated clustering of E11.5 control and NotchGOF λ-junction cells.**

| | | | | | | | | |
|---|---|---|---|---|---|---|---|---|
| 3 | Pfas | 0.34 | 1.94E-18 | 4.04E-14 | 0.39 | 0.28 | 0.11 | 1.40 |
| 3 | Aven | 0.27 | 2.05E-18 | 4.26E-14 | 0.52 | 0.38 | 0.13 | 1.34 |
| 3 | Dcbld2 | 0.35 | 2.08E-18 | 4.32E-14 | 0.35 | 0.25 | 0.11 | 1.43 |
| 3 | Pkmyt1 | 0.55 | 2.24E-18 | 4.66E-14 | 0.25 | 0.17 | 0.08 | 1.51 |
| 3 | Pex14 | 0.27 | 2.33E-18 | 4.85E-14 | 0.46 | 0.33 | 0.13 | 1.38 |
| 3 | Dse | 0.29 | 2.38E-18 | 4.95E-14 | 0.38 | 0.26 | 0.12 | 1.44 |
| 3 | E2f5 | 0.25 | 2.44E-18 | 5.08E-14 | 0.61 | 0.48 | 0.14 | 1.29 |
| 3 | Gypc | 0.35 | 3.16E-18 | 6.57E-14 | 0.31 | 0.21 | 0.10 | 1.47 |
| 3 | Smg5 | 0.28 | 3.49E-18 | 7.26E-14 | 0.44 | 0.32 | 0.12 | 1.38 |
| 3 | Kbtbd4 | 0.29 | 3.77E-18 | 7.84E-14 | 0.46 | 0.34 | 0.12 | 1.37 |
| 3 | Nfatc2ip | 0.53 | 3.83E-18 | 7.96E-14 | 0.24 | 0.15 | 0.08 | 1.54 |
| 3 | Mars | 0.31 | 4.05E-18 | 8.42E-14 | 0.47 | 0.35 | 0.12 | 1.34 |
| 3 | Rad1 | 0.33 | 4.18E-18 | 8.70E-14 | 0.43 | 0.31 | 0.12 | 1.37 |
| 3 | Pdcd11 | 0.27 | 4.27E-18 | 8.89E-14 | 0.60 | 0.46 | 0.14 | 1.29 |
| 3 | Senp1 | 0.28 | 4.38E-18 | 9.11E-14 | 0.58 | 0.45 | 0.13 | 1.29 |
| 3 | Recql | 0.41 | 4.62E-18 | 9.60E-14 | 0.28 | 0.19 | 0.09 | 1.49 |
| 3 | Mmd | 0.27 | 4.93E-18 | 1.02E-13 | 0.46 | 0.34 | 0.12 | 1.36 |
| 3 | Ogfrl1 | 0.27 | 5.03E-18 | 1.05E-13 | 0.45 | 0.33 | 0.12 | 1.38 |
| 3 | Slc25a25 | 0.41 | 5.07E-18 | 1.05E-13 | 0.26 | 0.17 | 0.09 | 1.52 |
| 3 | Uap1 | 0.32 | 5.07E-18 | 1.05E-13 | 0.42 | 0.31 | 0.12 | 1.37 |
| 3 | Pnpt1 | 0.33 | 5.08E-18 | 1.06E-13 | 0.39 | 0.28 | 0.11 | 1.39 |
| 3 | 6720427I07Rik | 0.34 | 5.12E-18 | 1.07E-13 | 0.29 | 0.20 | 0.10 | 1.49 |
| 3 | Nup93 | 0.25 | 5.38E-18 | 1.12E-13 | 0.60 | 0.46 | 0.14 | 1.30 |
| 3 | Dcaf1 | 0.26 | 5.55E-18 | 1.15E-13 | 0.57 | 0.43 | 0.14 | 1.32 |
| 3 | Arl6 | 0.26 | 6.68E-18 | 1.39E-13 | 0.54 | 0.41 | 0.13 | 1.33 |
| 3 | Snip1 | 0.31 | 7.07E-18 | 1.47E-13 | 0.45 | 0.33 | 0.12 | 1.37 |
| 3 | Haus7 | 0.26 | 7.35E-18 | 1.53E-13 | 0.59 | 0.45 | 0.14 | 1.30 |
| 3 | Zfp946 | 0.36 | 7.55E-18 | 1.57E-13 | 0.29 | 0.20 | 0.10 | 1.48 |
| 3 | Pcdh19 | 0.28 | 7.91E-18 | 1.64E-13 | 0.25 | 0.16 | 0.09 | 1.55 |
| 3 | Lin28b | 0.55 | 8.38E-18 | 1.74E-13 | 0.29 | 0.20 | 0.09 | 1.44 |
| 3 | 2810468N07Rik | 0.51 | 8.38E-18 | 1.74E-13 | 0.13 | 0.07 | 0.06 | 1.81 |
| 3 | Lin9 | 0.31 | 9.05E-18 | 1.88E-13 | 0.44 | 0.32 | 0.12 | 1.36 |
| 3 | Dpysl4 | 0.31 | 1.05E-17 | 2.18E-13 | 0.27 | 0.18 | 0.09 | 1.53 |
| 3 | Ctu2 | 0.28 | 1.07E-17 | 2.23E-13 | 0.45 | 0.33 | 0.12 | 1.38 |
| 3 | Cbx2 | 0.26 | 1.11E-17 | 2.30E-13 | 0.55 | 0.42 | 0.13 | 1.32 |
| 3 | Letm1 | 0.27 | 1.15E-17 | 2.39E-13 | 0.56 | 0.43 | 0.13 | 1.29 |
| 3 | Nfya | 0.32 | 1.29E-17 | 2.69E-13 | 0.42 | 0.31 | 0.11 | 1.37 |
| 3 | Sipa1l2 | 0.55 | 1.39E-17 | 2.89E-13 | 0.20 | 0.13 | 0.08 | 1.59 |
| 3 | Adam19 | 0.41 | 1.48E-17 | 3.07E-13 | 0.23 | 0.15 | 0.08 | 1.56 |
| 3 | Usp51 | 0.68 | 1.54E-17 | 3.21E-13 | 0.14 | 0.08 | 0.06 | 1.78 |
| 3 | Zfp536 | 0.57 | 2.02E-17 | 4.20E-13 | 0.16 | 0.10 | 0.07 | 1.67 |
| 3 | Rbm15 | 0.28 | 2.02E-17 | 4.20E-13 | 0.48 | 0.36 | 0.12 | 1.34 |
| 3 | Fanca | 0.62 | 2.07E-17 | 4.30E-13 | 0.18 | 0.11 | 0.07 | 1.63 |
| 3 | Tasp1 | 0.43 | 2.16E-17 | 4.49E-13 | 0.24 | 0.15 | 0.08 | 1.54 |
| 3 | Qrsl1 | 0.33 | 2.32E-17 | 4.83E-13 | 0.31 | 0.21 | 0.10 | 1.48 |
| 3 | Tor1a | 0.25 | 2.51E-17 | 5.23E-13 | 0.65 | 0.51 | 0.14 | 1.28 |
| 3 | Akap11 | 0.25 | 2.63E-17 | 5.47E-13 | 0.61 | 0.47 | 0.14 | 1.29 |
| 3 | Zfp639 | 0.26 | 2.76E-17 | 5.73E-13 | 0.59 | 0.45 | 0.14 | 1.31 |
| 3 | Rbp4 | 0.77 | 2.94E-17 | 6.10E-13 | 0.11 | 0.06 | 0.05 | 1.91 |
| 3 | Gm10184 | 0.51 | 3.18E-17 | 6.62E-13 | 0.21 | 0.14 | 0.07 | 1.55 |
| 3 | Srrd | 0.29 | 3.38E-17 | 7.02E-13 | 0.38 | 0.27 | 0.11 | 1.39 |
| 3 | Dcun1d3 | 0.41 | 3.52E-17 | 7.31E-13 | 0.23 | 0.15 | 0.08 | 1.57 |
| 3 | Ice2 | 0.31 | 3.75E-17 | 7.79E-13 | 0.35 | 0.24 | 0.11 | 1.43 |
| 3 | Loxl1 | 0.53 | 3.97E-17 | 8.25E-13 | 0.13 | 0.07 | 0.06 | 1.83 |
| 3 | Xrcc3 | 0.49 | 4.29E-17 | 8.92E-13 | 0.16 | 0.10 | 0.07 | 1.68 |
| 3 | Coq5 | 0.27 | 4.34E-17 | 9.03E-13 | 0.59 | 0.46 | 0.13 | 1.28 |

**Supplementary Table 1. Marker genes for the original clusters of integrated clustering of E11.5 control and NotchGOF λ-junction cells.**

| | | | | | | | | |
|---|---|---|---|---|---|---|---|---|
| 3 | Nfix | 0.45 | 4.76E-17 | 9.90E-13 | 0.22 | 0.15 | 0.08 | 1.53 |
| 3 | Faf1 | 0.25 | 4.91E-17 | 1.02E-12 | 0.62 | 0.49 | 0.14 | 1.28 |
| 3 | Taco1 | 0.38 | 5.03E-17 | 1.05E-12 | 0.26 | 0.17 | 0.09 | 1.51 |
| 3 | Gli2 | 0.34 | 5.30E-17 | 1.10E-12 | 0.36 | 0.25 | 0.10 | 1.41 |
| 3 | Greb1 | 0.25 | 6.86E-17 | 1.43E-12 | 0.41 | 0.29 | 0.12 | 1.42 |
| 3 | Creld2 | 0.34 | 6.94E-17 | 1.44E-12 | 0.56 | 0.44 | 0.12 | 1.28 |
| 3 | Pcnt | 0.26 | 7.37E-17 | 1.53E-12 | 0.59 | 0.46 | 0.13 | 1.28 |
| 3 | Trmt2a | 0.29 | 7.77E-17 | 1.62E-12 | 0.40 | 0.29 | 0.11 | 1.39 |
| 3 | Snx7 | 0.48 | 9.09E-17 | 1.89E-12 | 0.16 | 0.10 | 0.07 | 1.67 |
| 3 | Coq3 | 0.37 | 9.18E-17 | 1.91E-12 | 0.26 | 0.18 | 0.09 | 1.48 |
| 3 | Naa25 | 0.29 | 9.31E-17 | 1.94E-12 | 0.47 | 0.35 | 0.12 | 1.34 |
| 3 | Cstf1 | 0.33 | 9.49E-17 | 1.97E-12 | 0.36 | 0.26 | 0.10 | 1.39 |
| 3 | Zfp599 | 0.45 | 9.73E-17 | 2.02E-12 | 0.27 | 0.18 | 0.09 | 1.48 |
| 3 | Ufsp1 | 0.51 | 9.94E-17 | 2.07E-12 | 0.18 | 0.11 | 0.07 | 1.62 |
| 3 | Scn3a | 0.56 | 1.18E-16 | 2.46E-12 | 0.10 | 0.05 | 0.05 | 1.92 |
| 3 | Pdss1 | 0.44 | 1.29E-16 | 2.68E-12 | 0.24 | 0.16 | 0.08 | 1.51 |
| 3 | Mios | 0.33 | 1.32E-16 | 2.75E-12 | 0.38 | 0.27 | 0.10 | 1.38 |
| 3 | Tube1 | 0.43 | 1.35E-16 | 2.80E-12 | 0.25 | 0.17 | 0.08 | 1.49 |
| 3 | Dnajc30 | 0.26 | 1.35E-16 | 2.81E-12 | 0.57 | 0.44 | 0.13 | 1.30 |
| 3 | Banp | 0.27 | 1.44E-16 | 2.99E-12 | 0.46 | 0.35 | 0.12 | 1.34 |
| 3 | Megf9 | 0.39 | 1.57E-16 | 3.27E-12 | 0.27 | 0.18 | 0.09 | 1.48 |
| 3 | Yars2 | 0.27 | 1.64E-16 | 3.40E-12 | 0.52 | 0.39 | 0.13 | 1.33 |
| 3 | Glce | 0.38 | 1.72E-16 | 3.58E-12 | 0.26 | 0.18 | 0.09 | 1.50 |
| 3 | Dip2c | 0.29 | 1.92E-16 | 3.99E-12 | 0.33 | 0.23 | 0.10 | 1.43 |
| 3 | Dclk1 | 0.25 | 2.40E-16 | 4.99E-12 | 0.35 | 0.25 | 0.10 | 1.42 |
| 3 | Aebp1 | 0.31 | 2.40E-16 | 5.00E-12 | 0.29 | 0.20 | 0.09 | 1.47 |
| 3 | Eif2s3y | 0.57 | 2.45E-16 | 5.09E-12 | 0.13 | 0.08 | 0.06 | 1.75 |
| 3 | Ddx51 | 0.36 | 2.84E-16 | 5.90E-12 | 0.25 | 0.17 | 0.08 | 1.50 |
| 3 | Mtbp | 0.39 | 2.85E-16 | 5.93E-12 | 0.31 | 0.21 | 0.09 | 1.43 |
| 3 | Tmem41a | 0.31 | 2.88E-16 | 5.99E-12 | 0.31 | 0.21 | 0.10 | 1.44 |
| 3 | Zfand4 | 0.58 | 3.05E-16 | 6.35E-12 | 0.13 | 0.08 | 0.06 | 1.74 |
| 3 | Mthfd1l | 0.47 | 3.23E-16 | 6.71E-12 | 0.23 | 0.15 | 0.08 | 1.51 |
| 3 | Serp2 | 0.30 | 3.48E-16 | 7.24E-12 | 0.28 | 0.19 | 0.09 | 1.48 |
| 3 | Spata7 | 0.33 | 3.53E-16 | 7.35E-12 | 0.50 | 0.37 | 0.12 | 1.33 |
| 3 | Cacna1c | 0.47 | 3.71E-16 | 7.72E-12 | 0.15 | 0.09 | 0.06 | 1.70 |
| 3 | Nle1 | 0.31 | 3.80E-16 | 7.91E-12 | 0.32 | 0.23 | 0.10 | 1.42 |
| 3 | Elmo2 | 0.33 | 4.22E-16 | 8.79E-12 | 0.34 | 0.24 | 0.10 | 1.41 |
| 3 | AU020206 | 0.38 | 4.63E-16 | 9.64E-12 | 0.31 | 0.22 | 0.09 | 1.43 |
| 3 | Ccdc138 | 0.35 | 5.46E-16 | 1.14E-11 | 0.28 | 0.19 | 0.09 | 1.48 |
| 3 | Uspl1 | 0.27 | 5.86E-16 | 1.22E-11 | 0.50 | 0.38 | 0.12 | 1.32 |
| 3 | Rab31 | 0.32 | 6.70E-16 | 1.39E-11 | 0.31 | 0.22 | 0.10 | 1.44 |
| 3 | Oscp1 | 0.30 | 7.16E-16 | 1.49E-11 | 0.28 | 0.19 | 0.09 | 1.47 |
| 3 | Endog | 0.32 | 8.01E-16 | 1.67E-11 | 0.30 | 0.21 | 0.09 | 1.44 |
| 3 | Poc1b | 0.26 | 1.06E-15 | 2.20E-11 | 0.44 | 0.32 | 0.12 | 1.35 |
| 3 | Bckdhb | 0.25 | 1.08E-15 | 2.25E-11 | 0.48 | 0.36 | 0.12 | 1.35 |
| 3 | Slc36a4 | 0.34 | 1.09E-15 | 2.28E-11 | 0.24 | 0.16 | 0.08 | 1.52 |
| 3 | Qtrt2 | 0.33 | 1.15E-15 | 2.38E-11 | 0.34 | 0.24 | 0.10 | 1.40 |
| 3 | Elovl5 | 0.30 | 1.22E-15 | 2.54E-11 | 0.40 | 0.29 | 0.11 | 1.37 |
| 3 | Klhdc4 | 0.26 | 1.27E-15 | 2.64E-11 | 0.46 | 0.34 | 0.12 | 1.34 |
| 3 | Fancb | 0.69 | 1.34E-15 | 2.78E-11 | 0.13 | 0.08 | 0.06 | 1.71 |
| 3 | Tbc1d31 | 0.33 | 1.35E-15 | 2.81E-11 | 0.30 | 0.21 | 0.09 | 1.43 |
| 3 | Vldlr | 0.44 | 1.53E-15 | 3.18E-11 | 0.18 | 0.11 | 0.07 | 1.59 |
| 3 | Kif18b | 0.45 | 1.64E-15 | 3.41E-11 | 0.23 | 0.15 | 0.08 | 1.49 |
| 3 | Gfm1 | 0.29 | 1.67E-15 | 3.48E-11 | 0.45 | 0.34 | 0.11 | 1.33 |
| 3 | Ddx3y | 0.57 | 1.90E-15 | 3.95E-11 | 0.13 | 0.07 | 0.05 | 1.74 |
| 3 | Scd1 | 0.33 | 2.01E-15 | 4.18E-11 | 0.27 | 0.18 | 0.09 | 1.47 |

**Supplementary Table 1. Marker genes for the original clusters of integrated clustering of E11.5 control and NotchGOF λ-junction cells.**

| | | | | | | | | |
|---|---|---|---|---|---|---|---|---|
| 3 | Slc4a4 | 0.37 | 2.19E-15 | 4.56E-11 | 0.17 | 0.11 | 0.07 | 1.61 |
| 3 | Nbn | 0.29 | 2.33E-15 | 4.85E-11 | 0.43 | 0.33 | 0.11 | 1.33 |
| 3 | Cep76 | 0.41 | 2.41E-15 | 5.01E-11 | 0.24 | 0.16 | 0.08 | 1.49 |
| 3 | Gm10282 | 0.36 | 2.46E-15 | 5.11E-11 | 0.23 | 0.15 | 0.08 | 1.51 |
| 3 | Atm | 0.29 | 2.57E-15 | 5.34E-11 | 0.42 | 0.31 | 0.11 | 1.35 |
| 3 | 2010204K13Rik | 0.31 | 3.28E-15 | 6.82E-11 | 0.34 | 0.25 | 0.10 | 1.40 |
| 3 | Magi2 | 0.35 | 3.35E-15 | 6.97E-11 | 0.27 | 0.18 | 0.09 | 1.47 |
| 3 | Mindy3 | 0.28 | 3.45E-15 | 7.18E-11 | 0.37 | 0.27 | 0.10 | 1.37 |
| 3 | Dsn1 | 0.34 | 4.78E-15 | 9.95E-11 | 0.32 | 0.23 | 0.09 | 1.39 |
| 3 | Rmi2 | 0.38 | 4.96E-15 | 1.03E-10 | 0.28 | 0.20 | 0.08 | 1.42 |
| 3 | Fbxl17 | 0.29 | 5.33E-15 | 1.11E-10 | 0.28 | 0.19 | 0.09 | 1.46 |
| 3 | Atoh8 | 0.56 | 5.98E-15 | 1.24E-10 | 0.15 | 0.09 | 0.06 | 1.67 |
| 3 | Neto2 | 0.29 | 6.33E-15 | 1.32E-10 | 0.32 | 0.22 | 0.09 | 1.42 |
| 3 | Espl1 | 0.38 | 6.35E-15 | 1.32E-10 | 0.30 | 0.22 | 0.08 | 1.38 |
| 3 | Lmf1 | 0.31 | 6.52E-15 | 1.36E-10 | 0.31 | 0.22 | 0.09 | 1.43 |
| 3 | B4galt6 | 0.36 | 7.14E-15 | 1.49E-10 | 0.31 | 0.22 | 0.09 | 1.41 |
| 3 | Sgf29 | 0.26 | 7.24E-15 | 1.51E-10 | 0.43 | 0.32 | 0.11 | 1.34 |
| 3 | Astn1 | 0.43 | 7.37E-15 | 1.53E-10 | 0.18 | 0.11 | 0.07 | 1.59 |
| 3 | Trp53rkb | 0.26 | 8.62E-15 | 1.79E-10 | 0.35 | 0.25 | 0.10 | 1.39 |
| 3 | Fancd2 | 0.57 | 9.08E-15 | 1.89E-10 | 0.19 | 0.12 | 0.07 | 1.53 |
| 3 | Aaas | 0.27 | 9.21E-15 | 1.92E-10 | 0.43 | 0.32 | 0.11 | 1.34 |
| 3 | Pole2 | 0.26 | 1.20E-14 | 2.50E-10 | 0.38 | 0.27 | 0.10 | 1.37 |
| 3 | Gm17092 | 0.32 | 1.26E-14 | 2.62E-10 | 0.33 | 0.24 | 0.09 | 1.39 |
| 3 | Pdp1 | 0.57 | 1.30E-14 | 2.71E-10 | 0.12 | 0.07 | 0.05 | 1.73 |
| 3 | Cacng8 | 0.55 | 1.31E-14 | 2.73E-10 | 0.11 | 0.06 | 0.05 | 1.82 |
| 3 | Plcb1 | 0.37 | 1.39E-14 | 2.90E-10 | 0.15 | 0.09 | 0.06 | 1.64 |
| 3 | Tmem209 | 0.28 | 1.51E-14 | 3.13E-10 | 0.36 | 0.26 | 0.10 | 1.39 |
| 3 | Gm21814 | 0.59 | 1.53E-14 | 3.19E-10 | 0.11 | 0.06 | 0.05 | 1.79 |
| 3 | Atp23 | 0.35 | 1.57E-14 | 3.26E-10 | 0.32 | 0.23 | 0.09 | 1.39 |
| 3 | Ddx11 | 0.32 | 1.60E-14 | 3.32E-10 | 0.34 | 0.24 | 0.09 | 1.38 |
| 3 | Bcdin3d | 0.28 | 1.84E-14 | 3.83E-10 | 0.34 | 0.25 | 0.10 | 1.39 |
| 3 | Rbm46 | 0.49 | 2.03E-14 | 4.23E-10 | 0.14 | 0.08 | 0.06 | 1.69 |
| 3 | Rspo1 | 0.64 | 2.11E-14 | 4.39E-10 | 0.12 | 0.07 | 0.05 | 1.77 |
| 3 | Usp6nl | 0.33 | 2.15E-14 | 4.46E-10 | 0.31 | 0.22 | 0.09 | 1.40 |
| 3 | Gpt2 | 0.28 | 2.25E-14 | 4.69E-10 | 0.29 | 0.20 | 0.09 | 1.45 |
| 3 | Tpmt | 0.26 | 2.42E-14 | 5.03E-10 | 0.47 | 0.36 | 0.11 | 1.31 |
| 3 | Cldn34c1 | 0.38 | 2.43E-14 | 5.06E-10 | 0.21 | 0.14 | 0.07 | 1.52 |
| 3 | Pde12 | 0.28 | 2.47E-14 | 5.13E-10 | 0.37 | 0.27 | 0.10 | 1.37 |
| 3 | Cntrob | 0.35 | 2.62E-14 | 5.45E-10 | 0.29 | 0.20 | 0.08 | 1.41 |
| 3 | Cend1 | 0.56 | 2.90E-14 | 6.04E-10 | 0.10 | 0.06 | 0.05 | 1.82 |
| 3 | Ropn1l | 0.60 | 3.00E-14 | 6.24E-10 | 0.11 | 0.06 | 0.05 | 1.80 |
| 3 | Inpp5a | 0.40 | 3.17E-14 | 6.59E-10 | 0.22 | 0.15 | 0.07 | 1.49 |
| 3 | Ctdp1 | 0.30 | 3.30E-14 | 6.87E-10 | 0.28 | 0.20 | 0.08 | 1.43 |
| 3 | Rfwd3 | 0.25 | 3.36E-14 | 6.98E-10 | 0.47 | 0.36 | 0.11 | 1.30 |
| 3 | Prpf3 | 0.26 | 3.74E-14 | 7.78E-10 | 0.50 | 0.39 | 0.12 | 1.30 |
| 3 | Aldh9a1 | 0.26 | 4.30E-14 | 8.94E-10 | 0.41 | 0.31 | 0.10 | 1.33 |
| 3 | Gfra4 | 0.41 | 4.97E-14 | 1.03E-09 | 0.16 | 0.10 | 0.06 | 1.62 |
| 3 | Nbas | 0.30 | 5.15E-14 | 1.07E-09 | 0.30 | 0.21 | 0.09 | 1.41 |
| 3 | Med14 | 0.27 | 5.53E-14 | 1.15E-09 | 0.35 | 0.26 | 0.10 | 1.37 |
| 3 | Cep72 | 0.42 | 5.81E-14 | 1.21E-09 | 0.22 | 0.15 | 0.07 | 1.48 |
| 3 | A030001D20Rik | 0.59 | 5.92E-14 | 1.23E-09 | 0.12 | 0.07 | 0.05 | 1.75 |
| 3 | Gm36988 | 0.33 | 5.95E-14 | 1.24E-09 | 0.21 | 0.14 | 0.07 | 1.53 |
| 3 | Gm15446 | 0.44 | 6.47E-14 | 1.35E-09 | 0.17 | 0.11 | 0.06 | 1.56 |
| 3 | Gm44257 | 0.54 | 6.64E-14 | 1.38E-09 | 0.11 | 0.06 | 0.05 | 1.80 |
| 3 | Dis3l | 0.43 | 7.62E-14 | 1.58E-09 | 0.21 | 0.14 | 0.07 | 1.49 |
| 3 | Usp13 | 0.57 | 8.32E-14 | 1.73E-09 | 0.12 | 0.07 | 0.05 | 1.72 |

**Supplementary Table 1. Marker genes for the original clusters of integrated clustering of E11.5 control and NotchGOF λ-junction cells.**

| | | | | | | | | |
|---|---|---|---|---|---|---|---|---|
| 3 | Cast | 0.26 | 8.84E-14 | 1.84E-09 | 0.33 | 0.24 | 0.09 | 1.38 |
| 3 | Sv2a | 0.60 | 9.22E-14 | 1.92E-09 | 0.13 | 0.08 | 0.05 | 1.68 |
| 3 | Mfsd5 | 0.26 | 9.42E-14 | 1.96E-09 | 0.33 | 0.24 | 0.09 | 1.39 |
| 3 | Stk17b | 0.49 | 1.07E-13 | 2.22E-09 | 0.19 | 0.13 | 0.06 | 1.48 |
| 3 | Mmaa | 0.44 | 1.22E-13 | 2.54E-09 | 0.20 | 0.14 | 0.07 | 1.50 |
| 3 | Tubgcp3 | 0.26 | 1.30E-13 | 2.70E-09 | 0.41 | 0.31 | 0.10 | 1.32 |
| 3 | Myb | 0.54 | 1.44E-13 | 3.00E-09 | 0.13 | 0.08 | 0.05 | 1.65 |
| 3 | Xkr5 | 0.70 | 1.68E-13 | 3.49E-09 | 0.12 | 0.07 | 0.05 | 1.70 |
| 3 | Alg3 | 0.27 | 1.96E-13 | 4.08E-09 | 0.37 | 0.27 | 0.10 | 1.36 |
| 3 | Rab36 | 0.30 | 2.04E-13 | 4.24E-09 | 0.18 | 0.11 | 0.07 | 1.57 |
| 3 | Clasp2 | 0.28 | 2.18E-13 | 4.53E-09 | 0.32 | 0.23 | 0.09 | 1.38 |
| 3 | Hyou1 | 0.26 | 2.24E-13 | 4.65E-09 | 0.38 | 0.28 | 0.10 | 1.35 |
| 3 | Mphosph9 | 0.31 | 2.49E-13 | 5.18E-09 | 0.33 | 0.25 | 0.09 | 1.36 |
| 3 | Hsd17b7 | 0.29 | 2.73E-13 | 5.68E-09 | 0.36 | 0.27 | 0.09 | 1.34 |
| 3 | Srxn1 | 0.45 | 2.99E-13 | 6.23E-09 | 0.17 | 0.11 | 0.06 | 1.56 |
| 3 | St5 | 0.29 | 3.00E-13 | 6.23E-09 | 0.31 | 0.22 | 0.09 | 1.39 |
| 3 | Plpp1 | 0.27 | 3.17E-13 | 6.58E-09 | 0.21 | 0.14 | 0.07 | 1.50 |
| 3 | Nr2f1 | 0.29 | 3.19E-13 | 6.64E-09 | 0.43 | 0.33 | 0.10 | 1.30 |
| 3 | Kif21b | 0.31 | 3.37E-13 | 7.00E-09 | 0.23 | 0.16 | 0.07 | 1.46 |
| 3 | Noa1 | 0.27 | 4.75E-13 | 9.88E-09 | 0.41 | 0.31 | 0.10 | 1.33 |
| 3 | B930036N10Rik | 0.46 | 4.90E-13 | 1.02E-08 | 0.12 | 0.07 | 0.05 | 1.68 |
| 3 | Helq | 0.37 | 4.99E-13 | 1.04E-08 | 0.20 | 0.13 | 0.07 | 1.49 |
| 3 | Zfp930 | 0.35 | 5.07E-13 | 1.05E-08 | 0.23 | 0.16 | 0.07 | 1.45 |
| 3 | Tshz3 | 0.42 | 5.41E-13 | 1.13E-08 | 0.16 | 0.10 | 0.06 | 1.57 |
| 3 | Pms1 | 0.28 | 5.60E-13 | 1.16E-08 | 0.29 | 0.21 | 0.08 | 1.40 |
| 3 | Edrf1 | 0.26 | 5.99E-13 | 1.25E-08 | 0.38 | 0.28 | 0.10 | 1.34 |
| 3 | Elmod2 | 0.28 | 6.68E-13 | 1.39E-08 | 0.31 | 0.22 | 0.09 | 1.38 |
| 3 | Rmi1 | 0.30 | 7.37E-13 | 1.53E-08 | 0.38 | 0.29 | 0.10 | 1.33 |
| 3 | Gm26648 | 0.40 | 7.74E-13 | 1.61E-08 | 0.14 | 0.09 | 0.05 | 1.62 |
| 3 | Zc3h8 | 0.26 | 8.65E-13 | 1.80E-08 | 0.29 | 0.20 | 0.08 | 1.40 |
| 3 | Timp1 | 0.34 | 9.21E-13 | 1.92E-08 | 0.24 | 0.16 | 0.07 | 1.44 |
| 3 | Nqo2 | 0.35 | 9.30E-13 | 1.93E-08 | 0.19 | 0.13 | 0.06 | 1.51 |
| 3 | Fam171b | 0.35 | 9.43E-13 | 1.96E-08 | 0.32 | 0.24 | 0.09 | 1.36 |
| 3 | Tmem43 | 0.27 | 9.92E-13 | 2.06E-08 | 0.29 | 0.20 | 0.08 | 1.41 |
| 3 | Slc25a10 | 0.26 | 1.10E-12 | 2.30E-08 | 0.32 | 0.23 | 0.09 | 1.38 |
| 3 | Rab39b | 0.27 | 1.13E-12 | 2.35E-08 | 0.23 | 0.15 | 0.07 | 1.47 |
| 3 | Pakap | 0.36 | 1.30E-12 | 2.70E-08 | 0.18 | 0.12 | 0.06 | 1.53 |
| 3 | Cluh | 0.33 | 1.47E-12 | 3.06E-08 | 0.30 | 0.22 | 0.08 | 1.38 |
| 3 | Blvra | 0.30 | 2.38E-12 | 4.95E-08 | 0.28 | 0.20 | 0.08 | 1.39 |
| 3 | Rtel1 | 0.37 | 2.46E-12 | 5.11E-08 | 0.25 | 0.18 | 0.07 | 1.39 |
| 3 | Mbnl3 | 0.43 | 2.49E-12 | 5.18E-08 | 0.18 | 0.12 | 0.06 | 1.50 |
| 3 | Tram1l1 | 0.25 | 2.59E-12 | 5.39E-08 | 0.25 | 0.17 | 0.08 | 1.44 |
| 3 | Smyd5 | 0.25 | 2.85E-12 | 5.93E-08 | 0.42 | 0.32 | 0.10 | 1.32 |
| 3 | Mpp2 | 0.32 | 3.51E-12 | 7.30E-08 | 0.24 | 0.16 | 0.07 | 1.43 |
| 3 | Eif5a2 | 0.39 | 4.87E-12 | 1.01E-07 | 0.17 | 0.11 | 0.06 | 1.51 |
| 3 | Ifit2 | 0.49 | 5.46E-12 | 1.14E-07 | 0.14 | 0.09 | 0.05 | 1.58 |
| 3 | Fastkd3 | 0.29 | 5.77E-12 | 1.20E-07 | 0.23 | 0.16 | 0.07 | 1.44 |
| 3 | Ldlr | 0.31 | 6.18E-12 | 1.29E-07 | 0.32 | 0.24 | 0.08 | 1.33 |
| 3 | Rapgef2 | 0.28 | 6.21E-12 | 1.29E-07 | 0.30 | 0.22 | 0.08 | 1.38 |
| 3 | Mypop | 0.36 | 6.23E-12 | 1.30E-07 | 0.17 | 0.11 | 0.06 | 1.54 |
| 3 | 2610020C07Rik | 0.28 | 6.87E-12 | 1.43E-07 | 0.25 | 0.18 | 0.07 | 1.42 |
| 3 | Efr3b | 0.41 | 7.34E-12 | 1.53E-07 | 0.14 | 0.09 | 0.05 | 1.58 |
| 3 | Ctif | 0.47 | 7.54E-12 | 1.57E-07 | 0.16 | 0.10 | 0.05 | 1.53 |
| 3 | Serpini1 | 0.44 | 7.64E-12 | 1.59E-07 | 0.10 | 0.06 | 0.04 | 1.72 |
| 3 | Stard4 | 0.30 | 8.26E-12 | 1.72E-07 | 0.36 | 0.27 | 0.08 | 1.31 |
| 3 | Pter | 0.41 | 1.03E-11 | 2.14E-07 | 0.17 | 0.11 | 0.06 | 1.50 |

**Supplementary Table 1. Marker genes for the original clusters of integrated clustering of E11.5 control and NotchGOF λ-junction cells.**

| | | | | | | | | |
|---|---|---|---|---|---|---|---|---|
| 3 | Ankrd28 | 0.32 | 1.10E-11 | 2.29E-07 | 0.25 | 0.18 | 0.07 | 1.40 |
| 3 | Thap1 | 0.28 | 1.20E-11 | 2.49E-07 | 0.33 | 0.25 | 0.08 | 1.33 |
| 3 | Epb41l4b | 0.48 | 1.37E-11 | 2.84E-07 | 0.14 | 0.09 | 0.05 | 1.55 |
| 3 | Inpp5b | 0.42 | 1.78E-11 | 3.69E-07 | 0.17 | 0.11 | 0.06 | 1.50 |
| 3 | Mutyh | 0.42 | 2.18E-11 | 4.54E-07 | 0.17 | 0.11 | 0.06 | 1.51 |
| 3 | Arl15 | 0.28 | 2.35E-11 | 4.88E-07 | 0.23 | 0.16 | 0.07 | 1.43 |
| 3 | Rttn | 0.33 | 2.48E-11 | 5.16E-07 | 0.23 | 0.16 | 0.07 | 1.41 |
| 3 | Tonsl | 0.40 | 2.86E-11 | 5.94E-07 | 0.19 | 0.13 | 0.06 | 1.47 |
| 3 | 2900009J06Rik | 0.45 | 2.95E-11 | 6.13E-07 | 0.13 | 0.08 | 0.05 | 1.59 |
| 3 | Nqo1 | 0.40 | 3.19E-11 | 6.63E-07 | 0.15 | 0.09 | 0.05 | 1.55 |
| 3 | Msra | 0.29 | 3.98E-11 | 8.27E-07 | 0.23 | 0.16 | 0.07 | 1.42 |
| 3 | Zfp772 | 0.38 | 4.55E-11 | 9.45E-07 | 0.11 | 0.07 | 0.04 | 1.66 |
| 3 | Ccne2 | 0.35 | 4.81E-11 | 1.00E-06 | 0.27 | 0.20 | 0.07 | 1.35 |
| 3 | Gtpbp3 | 0.26 | 4.87E-11 | 1.01E-06 | 0.26 | 0.19 | 0.07 | 1.38 |
| 3 | Simc1 | 0.30 | 5.01E-11 | 1.04E-06 | 0.19 | 0.13 | 0.06 | 1.47 |
| 3 | Ppm1k | 0.38 | 5.67E-11 | 1.18E-06 | 0.15 | 0.10 | 0.05 | 1.52 |
| 3 | Gpr137b | 0.31 | 5.75E-11 | 1.20E-06 | 0.22 | 0.15 | 0.07 | 1.43 |
| 3 | Pbld1 | 1.09 | 5.81E-11 | 1.21E-06 | 0.12 | 0.07 | 0.04 | 1.58 |
| 3 | Dym | 0.26 | 5.91E-11 | 1.23E-06 | 0.30 | 0.23 | 0.08 | 1.33 |
| 3 | Prkd1 | 0.27 | 6.05E-11 | 1.26E-06 | 0.19 | 0.13 | 0.06 | 1.46 |
| 3 | Grk3 | 0.36 | 6.19E-11 | 1.29E-06 | 0.17 | 0.11 | 0.06 | 1.50 |
| 3 | Bbs7 | 0.33 | 6.44E-11 | 1.34E-06 | 0.22 | 0.16 | 0.07 | 1.42 |
| 3 | Rhobtb2 | 0.29 | 6.82E-11 | 1.42E-06 | 0.17 | 0.12 | 0.06 | 1.49 |
| 3 | Gtpbp1 | 0.27 | 7.39E-11 | 1.54E-06 | 0.28 | 0.20 | 0.07 | 1.36 |
| 3 | Palb2 | 0.37 | 7.83E-11 | 1.63E-06 | 0.19 | 0.13 | 0.06 | 1.44 |
| 3 | Spsb1 | 0.25 | 8.12E-11 | 1.69E-06 | 0.21 | 0.15 | 0.07 | 1.45 |
| 3 | Mcu | 0.27 | 8.92E-11 | 1.86E-06 | 0.21 | 0.15 | 0.06 | 1.43 |
| 3 | Eldr | 0.28 | 9.78E-11 | 2.03E-06 | 0.24 | 0.18 | 0.07 | 1.39 |
| 3 | Eri2 | 0.36 | 1.19E-10 | 2.48E-06 | 0.26 | 0.19 | 0.07 | 1.35 |
| 3 | Med18 | 0.32 | 1.24E-10 | 2.57E-06 | 0.18 | 0.12 | 0.06 | 1.48 |
| 3 | Zfp142 | 0.31 | 1.27E-10 | 2.63E-06 | 0.22 | 0.16 | 0.06 | 1.41 |
| 3 | Phactr3 | 0.35 | 1.27E-10 | 2.64E-06 | 0.12 | 0.07 | 0.04 | 1.59 |
| 3 | Mfsd2a | 0.59 | 1.44E-10 | 2.99E-06 | 0.10 | 0.06 | 0.04 | 1.67 |
| 3 | Cbx8 | 0.35 | 1.48E-10 | 3.08E-06 | 0.17 | 0.12 | 0.06 | 1.47 |
| 3 | Poc5 | 0.29 | 1.53E-10 | 3.18E-06 | 0.26 | 0.20 | 0.07 | 1.35 |
| 3 | Usp28 | 0.27 | 1.58E-10 | 3.28E-06 | 0.30 | 0.22 | 0.08 | 1.35 |
| 3 | Zfp820 | 0.26 | 1.69E-10 | 3.52E-06 | 0.27 | 0.20 | 0.07 | 1.36 |
| 3 | 2810410L24Rik | 0.36 | 1.72E-10 | 3.57E-06 | 0.12 | 0.08 | 0.05 | 1.59 |
| 3 | Snhg11 | 0.25 | 1.98E-10 | 4.13E-06 | 0.15 | 0.10 | 0.05 | 1.52 |
| 3 | Tubgcp6 | 0.35 | 1.99E-10 | 4.15E-06 | 0.19 | 0.13 | 0.06 | 1.45 |
| 3 | Shq1 | 0.26 | 2.06E-10 | 4.28E-06 | 0.26 | 0.19 | 0.07 | 1.37 |
| 3 | Stradb | 0.30 | 2.57E-10 | 5.34E-06 | 0.25 | 0.19 | 0.07 | 1.36 |
| 3 | Dnajc27 | 0.38 | 3.06E-10 | 6.36E-06 | 0.14 | 0.09 | 0.05 | 1.52 |
| 3 | Nkrf | 0.44 | 3.11E-10 | 6.48E-06 | 0.13 | 0.09 | 0.05 | 1.53 |
| 3 | Zfp948 | 0.41 | 3.12E-10 | 6.48E-06 | 0.14 | 0.09 | 0.05 | 1.52 |
| 3 | Tceanc | 0.43 | 3.27E-10 | 6.79E-06 | 0.14 | 0.09 | 0.05 | 1.52 |
| 3 | Baiap2l2 | 0.26 | 3.38E-10 | 7.02E-06 | 0.17 | 0.11 | 0.05 | 1.49 |
| 3 | Frk | 0.27 | 4.39E-10 | 9.14E-06 | 0.20 | 0.14 | 0.06 | 1.42 |
| 3 | Sik2 | 0.29 | 4.52E-10 | 9.40E-06 | 0.22 | 0.16 | 0.06 | 1.40 |
| 3 | Zfp354a | 0.30 | 4.54E-10 | 9.44E-06 | 0.16 | 0.11 | 0.05 | 1.48 |
| 3 | Foxc2 | 0.28 | 5.49E-10 | 1.14E-05 | 0.17 | 0.12 | 0.05 | 1.46 |
| 3 | Pank4 | 0.31 | 5.74E-10 | 1.19E-05 | 0.20 | 0.14 | 0.06 | 1.41 |
| 3 | Cnksr3 | 0.28 | 5.98E-10 | 1.24E-05 | 0.16 | 0.10 | 0.05 | 1.49 |
| 3 | Pdp2 | 0.26 | 6.42E-10 | 1.34E-05 | 0.24 | 0.17 | 0.06 | 1.37 |
| 3 | Entpd6 | 0.36 | 6.55E-10 | 1.36E-05 | 0.20 | 0.14 | 0.06 | 1.39 |
| 3 | 2410131K14Rik | 0.28 | 8.91E-10 | 1.85E-05 | 0.21 | 0.15 | 0.06 | 1.40 |

**Supplementary Table 1. Marker genes for the original clusters of integrated clustering of E11.5 control and NotchGOF λ-junction cells.**

| | | | | | | | | |
|---|---|---|---|---|---|---|---|---|
| 3 | Zfp689 | 0.32 | 9.30E-10 | 1.93E-05 | 0.14 | 0.09 | 0.05 | 1.52 |
| 3 | Wdcp | 0.30 | 1.15E-09 | 2.38E-05 | 0.46 | 0.36 | 0.10 | 1.26 |
| 3 | Csrnp2 | 0.29 | 1.22E-09 | 2.54E-05 | 0.19 | 0.14 | 0.06 | 1.41 |
| 3 | Smg8 | 0.26 | 1.24E-09 | 2.57E-05 | 0.23 | 0.17 | 0.06 | 1.38 |
| 3 | Septin5 | 0.36 | 1.42E-09 | 2.96E-05 | 0.10 | 0.06 | 0.04 | 1.61 |
| 3 | Nsun5 | 0.29 | 1.78E-09 | 3.69E-05 | 0.24 | 0.18 | 0.06 | 1.36 |
| 3 | Zdhhc15 | 0.34 | 1.95E-09 | 4.05E-05 | 0.18 | 0.13 | 0.05 | 1.42 |
| 3 | Nepro | 0.29 | 1.95E-09 | 4.06E-05 | 0.20 | 0.14 | 0.06 | 1.39 |
| 3 | Ttc7b | 0.28 | 2.00E-09 | 4.17E-05 | 0.22 | 0.16 | 0.06 | 1.39 |
| 3 | Tsga10 | 0.29 | 2.06E-09 | 4.29E-05 | 0.14 | 0.10 | 0.05 | 1.50 |
| 3 | Lig4 | 0.27 | 2.23E-09 | 4.64E-05 | 0.22 | 0.16 | 0.06 | 1.38 |
| 3 | Hcfc2 | 0.27 | 2.93E-09 | 6.10E-05 | 0.20 | 0.14 | 0.06 | 1.40 |
| 3 | Fut10 | 0.32 | 2.99E-09 | 6.22E-05 | 0.18 | 0.13 | 0.05 | 1.41 |
| 3 | Top3a | 0.30 | 4.39E-09 | 9.14E-05 | 0.22 | 0.16 | 0.06 | 1.37 |
| 3 | Chml | 0.29 | 4.66E-09 | 9.70E-05 | 0.20 | 0.15 | 0.06 | 1.39 |
| 3 | Lrrc57 | 0.26 | 4.74E-09 | 9.85E-05 | 0.22 | 0.16 | 0.06 | 1.37 |
| 3 | Ddx59 | 0.25 | 4.95E-09 | 1.03E-04 | 0.17 | 0.12 | 0.05 | 1.44 |
| 3 | Kctd9 | 0.39 | 5.64E-09 | 1.17E-04 | 0.16 | 0.11 | 0.05 | 1.42 |
| 3 | Nsun6 | 0.29 | 5.86E-09 | 1.22E-04 | 0.21 | 0.15 | 0.06 | 1.37 |
| 3 | Mettl18 | 0.33 | 6.00E-09 | 1.25E-04 | 0.17 | 0.12 | 0.05 | 1.44 |
| 3 | Hexim2 | 0.36 | 6.14E-09 | 1.28E-04 | 0.11 | 0.07 | 0.04 | 1.56 |
| 3 | Lipe | 0.39 | 6.16E-09 | 1.28E-04 | 0.12 | 0.08 | 0.04 | 1.53 |
| 3 | Nutf2 | 0.56 | 6.39E-09 | 1.33E-04 | 0.11 | 0.07 | 0.04 | 1.54 |
| 3 | Insyn1 | 0.32 | 8.17E-09 | 1.70E-04 | 0.11 | 0.07 | 0.04 | 1.57 |
| 3 | Piga | 0.27 | 8.36E-09 | 1.74E-04 | 0.21 | 0.16 | 0.06 | 1.36 |
| 3 | Tbc1d4 | 0.32 | 9.44E-09 | 1.96E-04 | 0.16 | 0.11 | 0.05 | 1.43 |
| 3 | Zfp775 | 0.26 | 1.01E-08 | 2.10E-04 | 0.18 | 0.13 | 0.05 | 1.42 |
| 3 | Phka1 | 0.41 | 1.14E-08 | 2.36E-04 | 0.10 | 0.07 | 0.04 | 1.55 |
| 3 | Fig4 | 0.37 | 1.38E-08 | 2.86E-04 | 0.11 | 0.07 | 0.04 | 1.52 |
| 3 | Stat5b | 0.34 | 1.57E-08 | 3.26E-04 | 0.17 | 0.12 | 0.05 | 1.40 |
| 3 | Hspe1-rs1 | 0.25 | 2.36E-08 | 4.90E-04 | 0.18 | 0.13 | 0.05 | 1.38 |
| 3 | Nmb | 0.38 | 2.40E-08 | 4.99E-04 | 0.11 | 0.08 | 0.04 | 1.51 |
| 3 | Slc29a4 | 0.35 | 2.47E-08 | 5.14E-04 | 0.14 | 0.10 | 0.05 | 1.45 |
| 3 | Fhl3 | 0.31 | 2.58E-08 | 5.36E-04 | 0.15 | 0.10 | 0.05 | 1.43 |
| 3 | Nudt10 | 0.26 | 3.00E-08 | 6.23E-04 | 0.16 | 0.11 | 0.05 | 1.44 |
| 3 | Pcdh9 | 0.30 | 3.64E-08 | 7.57E-04 | 0.16 | 0.11 | 0.05 | 1.41 |
| 3 | Gpam | 0.36 | 3.99E-08 | 8.30E-04 | 0.14 | 0.10 | 0.04 | 1.44 |
| 3 | 9930104L06Rik | 0.27 | 4.62E-08 | 9.62E-04 | 0.20 | 0.15 | 0.05 | 1.36 |
| 3 | Ccdc15 | 0.30 | 4.69E-08 | 9.76E-04 | 0.16 | 0.11 | 0.05 | 1.42 |
| 3 | Tpm3-rs7 | 0.30 | 4.72E-08 | 9.82E-04 | 0.15 | 0.11 | 0.05 | 1.42 |
| 3 | Zfp982 | 0.34 | 5.14E-08 | 1.07E-03 | 0.14 | 0.10 | 0.04 | 1.44 |
| 3 | Tecpr2 | 0.43 | 5.15E-08 | 1.07E-03 | 0.13 | 0.09 | 0.04 | 1.45 |
| 3 | Klhl25 | 0.31 | 5.34E-08 | 1.11E-03 | 0.14 | 0.10 | 0.04 | 1.44 |
| 3 | Ap1s3 | 0.47 | 6.29E-08 | 1.31E-03 | 0.10 | 0.07 | 0.04 | 1.53 |
| 3 | Tbc1d2b | 0.37 | 9.57E-08 | 1.99E-03 | 0.12 | 0.08 | 0.04 | 1.48 |
| 3 | Dclre1b | 0.30 | 1.11E-07 | 2.32E-03 | 0.16 | 0.12 | 0.05 | 1.39 |
| 3 | Zswim1 | 0.28 | 1.32E-07 | 2.74E-03 | 0.16 | 0.12 | 0.05 | 1.39 |
| 3 | Polh | 0.41 | 1.44E-07 | 3.00E-03 | 0.12 | 0.09 | 0.04 | 1.44 |
| 3 | Trim68 | 0.34 | 1.77E-07 | 3.67E-03 | 0.13 | 0.09 | 0.04 | 1.42 |
| 3 | Zbtb48 | 0.28 | 1.96E-07 | 4.07E-03 | 0.17 | 0.13 | 0.05 | 1.38 |
| 3 | Trpt1 | 0.32 | 2.24E-07 | 4.65E-03 | 0.10 | 0.07 | 0.03 | 1.51 |
| 3 | Helb | 0.36 | 2.39E-07 | 4.96E-03 | 0.17 | 0.13 | 0.05 | 1.35 |
| 3 | Zfp472 | 0.29 | 2.75E-07 | 5.72E-03 | 0.19 | 0.15 | 0.05 | 1.34 |
| 3 | Dtwd2 | 0.27 | 2.89E-07 | 6.02E-03 | 0.16 | 0.12 | 0.04 | 1.37 |
| 3 | Setd6 | 0.29 | 3.02E-07 | 6.28E-03 | 0.17 | 0.12 | 0.04 | 1.35 |
| 3 | Gm9774 | 0.37 | 3.04E-07 | 6.32E-03 | 0.11 | 0.07 | 0.03 | 1.46 |

**Supplementary Table 1. Marker genes for the original clusters of integrated clustering of E11.5 control and NotchGOF λ-junction cells.**

| | | | | | | | | |
|---|---|---|---|---|---|---|---|---|
| 3 | Mtfp1 | 0.35 | 4.00E-07 | 8.32E-03 | 0.14 | 0.10 | 0.04 | 1.38 |
| 3 | Nap1l3 | 0.28 | 4.97E-07 | 1.03E-02 | 0.11 | 0.07 | 0.04 | 1.47 |
| 3 | Zfp287 | 0.27 | 5.22E-07 | 1.09E-02 | 0.11 | 0.07 | 0.04 | 1.49 |
| 3 | Pxdc1 | 0.31 | 5.36E-07 | 1.11E-02 | 0.14 | 0.10 | 0.04 | 1.38 |
| 3 | Fbxo7 | 0.28 | 5.59E-07 | 1.16E-02 | 0.18 | 0.13 | 0.04 | 1.33 |
| 3 | Slc25a15 | 0.32 | 5.67E-07 | 1.18E-02 | 0.13 | 0.09 | 0.04 | 1.40 |
| 3 | Fanci | 0.41 | 5.72E-07 | 1.19E-02 | 0.13 | 0.09 | 0.04 | 1.39 |
| 3 | Aard | 0.25 | 5.89E-07 | 1.23E-02 | 0.10 | 0.07 | 0.03 | 1.49 |
| 3 | Ttc30a1 | 0.27 | 6.51E-07 | 1.35E-02 | 0.15 | 0.11 | 0.04 | 1.38 |
| 3 | Hps3 | 0.28 | 9.83E-07 | 2.05E-02 | 0.12 | 0.08 | 0.04 | 1.44 |
| 3 | Slc1a4 | 0.34 | 1.04E-06 | 2.17E-02 | 0.10 | 0.07 | 0.03 | 1.48 |
| 3 | Nr1h5 | 0.27 | 1.13E-06 | 2.35E-02 | 0.18 | 0.13 | 0.04 | 1.34 |
| 3 | Cetn4 | 0.30 | 1.25E-06 | 2.61E-02 | 0.11 | 0.08 | 0.04 | 1.45 |
| 3 | Mars2 | 0.37 | 1.27E-06 | 2.64E-02 | 0.10 | 0.07 | 0.03 | 1.46 |
| 3 | Gk | 0.36 | 1.34E-06 | 2.78E-02 | 0.11 | 0.08 | 0.03 | 1.44 |
| 3 | Pitpnm2 | 0.28 | 1.61E-06 | 3.35E-02 | 0.12 | 0.08 | 0.04 | 1.42 |
| 3 | 2310010J17Rik | 0.32 | 1.62E-06 | 3.37E-02 | 0.12 | 0.08 | 0.04 | 1.43 |
| 4 | Pax9 | 2.77 | 0 | 0 | 0.95 | 0.31 | 0.64 | 3.08 |
| 4 | Flrt2 | 2.83 | 0 | 0 | 0.99 | 0.41 | 0.58 | 2.43 |
| 4 | Foxf1 | 2.91 | 0 | 0 | 0.68 | 0.14 | 0.54 | 4.94 |
| 4 | Frzb | 2.61 | 0 | 0 | 0.84 | 0.31 | 0.54 | 2.74 |
| 4 | Dkk1 | 3.11 | 0 | 0 | 0.73 | 0.24 | 0.49 | 3.09 |
| 4 | Plagl1 | 2.22 | 0 | 0 | 0.88 | 0.39 | 0.49 | 2.24 |
| 4 | Col9a1 | 2.20 | 0 | 0 | 0.71 | 0.25 | 0.46 | 2.81 |
| 4 | Epha3 | 1.95 | 0 | 0 | 0.77 | 0.32 | 0.45 | 2.40 |
| 4 | Nrp2 | 2.17 | 0 | 0 | 0.75 | 0.34 | 0.41 | 2.21 |
| 4 | Sox9 | 2.38 | 0 | 0 | 0.94 | 0.55 | 0.39 | 1.70 |
| 4 | Isl2 | 2.87 | 0 | 0 | 0.47 | 0.10 | 0.37 | 4.71 |
| 4 | Sox6 | 3.13 | 0 | 0 | 0.45 | 0.10 | 0.36 | 4.77 |
| 4 | Lgr5 | 2.66 | 0 | 0 | 0.52 | 0.18 | 0.34 | 2.89 |
| 4 | Pdlim3 | 3.07 | 0 | 0 | 0.45 | 0.12 | 0.33 | 3.70 |
| 4 | Fendrr | 2.95 | 0 | 0 | 0.37 | 0.07 | 0.30 | 5.23 |
| 4 | Col3a1 | 1.28 | 0 | 0 | 0.98 | 0.70 | 0.28 | 1.40 |
| 4 | Tenm3 | 1.51 | 0 | 0 | 0.88 | 0.62 | 0.26 | 1.42 |
| 4 | Fstl1 | 1.44 | 0 | 0 | 0.98 | 0.81 | 0.17 | 1.21 |
| 4 | Peg3 | 1.22 | 0 | 0 | 0.98 | 0.81 | 0.16 | 1.20 |
| 4 | Mfap4 | 0.98 | 2.13E-306 | 4.42E-302 | 0.99 | 0.84 | 0.15 | 1.18 |
| 4 | Unc5c | 1.83 | 8.33E-304 | 1.73E-299 | 0.67 | 0.31 | 0.36 | 2.16 |
| 4 | Fbln2 | 1.62 | 1.33E-300 | 2.77E-296 | 0.80 | 0.49 | 0.30 | 1.62 |
| 4 | Col2a1 | 1.33 | 1.03E-293 | 2.14E-289 | 0.88 | 0.62 | 0.26 | 1.41 |
| 4 | Mecom | 1.72 | 2.56E-289 | 5.33E-285 | 0.72 | 0.38 | 0.35 | 1.92 |
| 4 | Phactr2 | 1.63 | 3.75E-278 | 7.80E-274 | 0.73 | 0.43 | 0.31 | 1.71 |
| 4 | Gata2 | 1.55 | 2.12E-275 | 4.41E-271 | 0.76 | 0.41 | 0.35 | 1.84 |
| 4 | Csrp2 | 0.81 | 1.86E-267 | 3.87E-263 | 0.99 | 0.92 | 0.07 | 1.08 |
| 4 | Sfrp1 | 1.27 | 3.37E-261 | 7.02E-257 | 0.85 | 0.60 | 0.25 | 1.41 |
| 4 | Prrx1 | 1.07 | 1.79E-256 | 3.73E-252 | 0.92 | 0.63 | 0.29 | 1.47 |
| 4 | Cdh11 | 0.99 | 3.63E-249 | 7.55E-245 | 0.94 | 0.68 | 0.27 | 1.40 |
| 4 | Ddit4l | 2.29 | 2.11E-247 | 4.40E-243 | 0.49 | 0.19 | 0.30 | 2.60 |
| 4 | Rab32 | 2.77 | 7.02E-245 | 1.46E-240 | 0.31 | 0.07 | 0.23 | 4.18 |
| 4 | Sall1 | 2.73 | 8.40E-241 | 1.75E-236 | 0.30 | 0.07 | 0.23 | 4.14 |
| 4 | Inhba | 2.63 | 1.16E-237 | 2.42E-233 | 0.43 | 0.15 | 0.28 | 2.93 |
| 4 | Gpc3 | 0.79 | 2.55E-227 | 5.29E-223 | 0.97 | 0.82 | 0.15 | 1.19 |
| 4 | Col9a3 | 2.47 | 5.12E-226 | 1.07E-221 | 0.38 | 0.12 | 0.26 | 3.12 |
| 4 | Enpp2 | 2.77 | 7.67E-225 | 1.60E-220 | 0.33 | 0.09 | 0.24 | 3.52 |
| 4 | Arid5b | 1.09 | 9.52E-216 | 1.98E-211 | 0.86 | 0.65 | 0.21 | 1.31 |
| 4 | Cald1 | 0.65 | 1.97E-213 | 4.10E-209 | 1.00 | 0.88 | 0.12 | 1.13 |

**Supplementary Table 1. Marker genes for the original clusters of integrated clustering of E11.5 control and NotchGOF λ-junction cells.**

| | | | | | | | | |
|---|---|---|---|---|---|---|---|---|
| 4 | Foxc1 | 1.44 | 5.17E-212 | 1.08E-207 | 0.58 | 0.26 | 0.32 | 2.20 |
| 4 | Crym | 1.19 | 4.10E-211 | 8.53E-207 | 0.76 | 0.45 | 0.31 | 1.68 |
| 4 | Slitrk5 | 2.20 | 1.02E-208 | 2.13E-204 | 0.39 | 0.14 | 0.26 | 2.89 |
| 4 | Pex5l | 2.89 | 2.39E-207 | 4.96E-203 | 0.28 | 0.07 | 0.21 | 3.88 |
| 4 | Gdf6 | 2.67 | 3.96E-206 | 8.23E-202 | 0.31 | 0.09 | 0.22 | 3.49 |
| 4 | Col1a2 | 0.80 | 3.75E-201 | 7.80E-197 | 0.98 | 0.72 | 0.26 | 1.37 |
| 4 | Arhgap28 | 1.27 | 6.81E-196 | 1.42E-191 | 0.75 | 0.52 | 0.23 | 1.45 |
| 4 | Sned1 | 2.14 | 9.71E-196 | 2.02E-191 | 0.35 | 0.11 | 0.24 | 3.15 |
| 4 | Igfbp4 | 0.64 | 4.27E-194 | 8.88E-190 | 1.00 | 0.89 | 0.10 | 1.12 |
| 4 | Piezo2 | 1.80 | 4.97E-191 | 1.03E-186 | 0.48 | 0.21 | 0.27 | 2.33 |
| 4 | Rcn2 | 0.71 | 7.63E-191 | 1.59E-186 | 0.97 | 0.91 | 0.06 | 1.07 |
| 4 | Alx3 | 1.40 | 1.01E-188 | 2.11E-184 | 0.57 | 0.27 | 0.30 | 2.10 |
| 4 | Itprid2 | 2.40 | 1.56E-185 | 3.25E-181 | 0.37 | 0.13 | 0.23 | 2.77 |
| 4 | Rhoj | 1.98 | 4.94E-185 | 1.03E-180 | 0.33 | 0.10 | 0.23 | 3.22 |
| 4 | Ndfip1 | 0.84 | 8.42E-183 | 1.75E-178 | 0.91 | 0.83 | 0.08 | 1.10 |
| 4 | Slc25a4 | 0.45 | 8.22E-182 | 1.71E-177 | 1.00 | 0.97 | 0.03 | 1.03 |
| 4 | Ldhb | 0.68 | 5.76E-180 | 1.20E-175 | 0.98 | 0.90 | 0.07 | 1.08 |
| 4 | Kif26b | 1.37 | 8.01E-180 | 1.67E-175 | 0.65 | 0.38 | 0.26 | 1.69 |
| 4 | Reep5 | 0.84 | 7.32E-179 | 1.52E-174 | 0.91 | 0.77 | 0.14 | 1.18 |
| 4 | Sobp | 1.57 | 7.36E-179 | 1.53E-174 | 0.58 | 0.32 | 0.26 | 1.83 |
| 4 | Gata3 | 1.30 | 2.79E-178 | 5.80E-174 | 0.75 | 0.50 | 0.25 | 1.50 |
| 4 | Sem1 | 0.41 | 5.09E-176 | 1.06E-171 | 1.00 | 1.00 | 0.00 | 1.00 |
| 4 | Adamts5 | 2.75 | 6.67E-174 | 1.39E-169 | 0.30 | 0.09 | 0.21 | 3.20 |
| 4 | Oat | 1.03 | 5.73E-172 | 1.19E-167 | 0.82 | 0.67 | 0.15 | 1.23 |
| 4 | Ubb | 0.36 | 1.44E-171 | 2.99E-167 | 1.00 | 1.00 | 0.00 | 1.00 |
| 4 | Sfr1 | 0.57 | 6.22E-169 | 1.29E-164 | 1.00 | 0.98 | 0.02 | 1.02 |
| 4 | Angptl1 | 1.66 | 1.83E-167 | 3.81E-163 | 0.46 | 0.20 | 0.26 | 2.27 |
| 4 | App | 0.92 | 1.09E-166 | 2.27E-162 | 0.89 | 0.77 | 0.12 | 1.16 |
| 4 | Tril | 1.12 | 1.06E-163 | 2.20E-159 | 0.76 | 0.53 | 0.24 | 1.45 |
| 4 | Pcdh18 | 0.97 | 1.26E-163 | 2.63E-159 | 0.83 | 0.62 | 0.21 | 1.34 |
| 4 | Ccnd2 | 0.70 | 8.42E-158 | 1.75E-153 | 0.94 | 0.85 | 0.09 | 1.10 |
| 4 | Rspo3 | 1.87 | 1.07E-156 | 2.23E-152 | 0.57 | 0.32 | 0.24 | 1.76 |
| 4 | Tox | 1.63 | 2.20E-156 | 4.58E-152 | 0.48 | 0.24 | 0.24 | 2.01 |
| 4 | Emilin1 | 1.07 | 2.99E-155 | 6.22E-151 | 0.73 | 0.51 | 0.22 | 1.43 |
| 4 | Prss23 | 1.58 | 3.79E-155 | 7.89E-151 | 0.54 | 0.29 | 0.25 | 1.86 |
| 4 | Prrx2 | 0.67 | 4.53E-154 | 9.43E-150 | 0.98 | 0.72 | 0.26 | 1.36 |
| 4 | Serpinf1 | 0.89 | 2.03E-153 | 4.22E-149 | 0.84 | 0.63 | 0.22 | 1.35 |
| 4 | Mfap2 | 0.55 | 3.29E-151 | 6.84E-147 | 0.99 | 0.92 | 0.08 | 1.08 |
| 4 | Inmt | 3.30 | 1.07E-149 | 2.22E-145 | 0.12 | 0.02 | 0.11 | 6.89 |
| 4 | Ptch1 | 1.27 | 1.87E-149 | 3.90E-145 | 0.66 | 0.44 | 0.22 | 1.51 |
| 4 | Eid1 | 0.49 | 8.82E-149 | 1.83E-144 | 0.99 | 0.97 | 0.03 | 1.03 |
| 4 | Eva1b | 1.02 | 1.26E-148 | 2.63E-144 | 0.74 | 0.52 | 0.22 | 1.42 |
| 4 | mt-Co3 | 0.29 | 6.21E-148 | 1.29E-143 | 1.00 | 0.99 | 0.01 | 1.01 |
| 4 | Maged2 | 0.68 | 1.22E-146 | 2.54E-142 | 0.92 | 0.82 | 0.10 | 1.13 |
| 4 | Cped1 | 1.12 | 4.45E-145 | 9.26E-141 | 0.68 | 0.45 | 0.23 | 1.52 |
| 4 | Limch1 | 0.93 | 2.81E-144 | 5.84E-140 | 0.83 | 0.66 | 0.17 | 1.26 |
| 4 | Runx1t1 | 1.08 | 4.11E-140 | 8.54E-136 | 0.72 | 0.51 | 0.21 | 1.41 |
| 4 | Itga8 | 1.88 | 3.14E-139 | 6.52E-135 | 0.39 | 0.17 | 0.21 | 2.24 |
| 4 | Pdlim5 | 1.73 | 1.30E-137 | 2.70E-133 | 0.42 | 0.21 | 0.22 | 2.05 |
| 4 | Ppp3ca | 0.88 | 4.30E-137 | 8.95E-133 | 0.84 | 0.72 | 0.12 | 1.16 |
| 4 | Ptms | 0.49 | 1.33E-135 | 2.76E-131 | 0.99 | 0.95 | 0.05 | 1.05 |
| 4 | Arglu1 | 0.49 | 2.23E-135 | 4.64E-131 | 0.99 | 0.96 | 0.03 | 1.03 |
| 4 | Thsd7a | 1.79 | 1.22E-133 | 2.54E-129 | 0.33 | 0.13 | 0.20 | 2.53 |
| 4 | Mxd4 | 1.10 | 6.49E-133 | 1.35E-128 | 0.66 | 0.45 | 0.21 | 1.46 |
| 4 | Auts2 | 0.63 | 7.33E-132 | 1.53E-127 | 0.95 | 0.86 | 0.09 | 1.10 |
| 4 | Grin3a | 2.59 | 1.07E-131 | 2.22E-127 | 0.17 | 0.04 | 0.13 | 4.46 |

**Supplementary Table 1. Marker genes for the original clusters of integrated clustering of E11.5 control and NotchGOF λ-junction cells.**

| | | | | | | | | |
|---|---|---|---|---|---|---|---|---|
| 4 | Cxxc4 | 1.47 | 1.86E-130 | 3.87E-126 | 0.51 | 0.29 | 0.22 | 1.78 |
| 4 | Pdlim4 | 0.90 | 3.33E-130 | 6.92E-126 | 0.77 | 0.61 | 0.16 | 1.26 |
| 4 | Tenm4 | 0.94 | 4.79E-128 | 9.97E-124 | 0.77 | 0.60 | 0.17 | 1.28 |
| 4 | Plcl1 | 2.48 | 5.91E-126 | 1.23E-121 | 0.19 | 0.05 | 0.14 | 3.78 |
| 4 | Hs3st6 | 2.45 | 9.92E-126 | 2.06E-121 | 0.23 | 0.07 | 0.16 | 3.22 |
| 4 | Ppib | 0.41 | 5.11E-124 | 1.06E-119 | 1.00 | 0.98 | 0.02 | 1.02 |
| 4 | Sfrp2 | 0.84 | 1.61E-121 | 3.35E-117 | 0.76 | 0.60 | 0.16 | 1.26 |
| 4 | Tbx2 | 0.81 | 2.41E-121 | 5.01E-117 | 0.83 | 0.63 | 0.20 | 1.32 |
| 4 | Mdfi | 0.96 | 9.87E-121 | 2.05E-116 | 0.71 | 0.55 | 0.16 | 1.28 |
| 4 | Dact3 | 1.02 | 5.36E-120 | 1.12E-115 | 0.67 | 0.48 | 0.19 | 1.41 |
| 4 | Sox5 | 1.25 | 9.92E-120 | 2.06E-115 | 0.58 | 0.38 | 0.20 | 1.53 |
| 4 | Crispld1 | 1.36 | 1.84E-115 | 3.83E-111 | 0.51 | 0.31 | 0.20 | 1.65 |
| 4 | Twist1 | 0.49 | 1.36E-114 | 2.83E-110 | 0.99 | 0.75 | 0.24 | 1.32 |
| 4 | Col9a2 | 2.12 | 6.74E-113 | 1.40E-108 | 0.25 | 0.09 | 0.16 | 2.77 |
| 4 | Frem1 | 1.10 | 1.06E-112 | 2.20E-108 | 0.59 | 0.39 | 0.20 | 1.52 |
| 4 | Etv5 | 1.29 | 7.58E-112 | 1.58E-107 | 0.50 | 0.30 | 0.20 | 1.67 |
| 4 | Efnb2 | 1.22 | 3.10E-111 | 6.45E-107 | 0.61 | 0.44 | 0.18 | 1.40 |
| 4 | Ramp2 | 0.80 | 7.43E-111 | 1.55E-106 | 0.37 | 0.17 | 0.20 | 2.18 |
| 4 | Pcdh7 | 1.08 | 8.06E-110 | 1.68E-105 | 0.59 | 0.38 | 0.22 | 1.58 |
| 4 | Synpo2 | 2.10 | 1.13E-109 | 2.36E-105 | 0.20 | 0.06 | 0.14 | 3.29 |
| 4 | Asap1 | 0.76 | 1.46E-109 | 3.03E-105 | 0.81 | 0.73 | 0.09 | 1.12 |
| 4 | Slit3 | 1.87 | 1.74E-109 | 3.61E-105 | 0.36 | 0.18 | 0.19 | 2.05 |
| 4 | Tle4 | 0.90 | 4.45E-109 | 9.25E-105 | 0.72 | 0.58 | 0.15 | 1.25 |
| 4 | Zeb2 | 0.86 | 2.15E-108 | 4.47E-104 | 0.62 | 0.38 | 0.24 | 1.64 |
| 4 | Acvrl1 | 2.12 | 3.93E-108 | 8.18E-104 | 0.25 | 0.09 | 0.16 | 2.78 |
| 4 | Itga4 | 1.37 | 9.99E-108 | 2.08E-103 | 0.38 | 0.18 | 0.20 | 2.11 |
| 4 | Sulf1 | 0.89 | 9.18E-107 | 1.91E-102 | 0.70 | 0.53 | 0.18 | 1.34 |
| 4 | Zfp637 | 0.77 | 2.95E-105 | 6.13E-101 | 0.80 | 0.73 | 0.08 | 1.11 |
| 4 | Pdgfrb | 1.02 | 8.76E-105 | 1.82E-100 | 0.63 | 0.44 | 0.19 | 1.43 |
| 4 | mt-Cytb | 0.26 | 9.85E-104 | 2.05E-99 | 1.00 | 0.99 | 0.01 | 1.01 |
| 4 | Crlf1 | 0.71 | 1.56E-102 | 3.24E-98 | 0.81 | 0.63 | 0.19 | 1.30 |
| 4 | Bgn | 1.53 | 8.91E-102 | 1.85E-97 | 0.44 | 0.25 | 0.19 | 1.77 |
| 4 | Mindy4 | 1.93 | 2.64E-100 | 5.49E-96 | 0.28 | 0.12 | 0.16 | 2.40 |
| 4 | Myo6 | 1.59 | 1.62E-99 | 3.36E-95 | 0.35 | 0.17 | 0.18 | 2.05 |
| 4 | 6330403K07Rik | 0.60 | 1.16E-97 | 2.40E-93 | 0.92 | 0.85 | 0.08 | 1.09 |
| 4 | Cd63 | 0.49 | 1.40E-97 | 2.91E-93 | 0.97 | 0.90 | 0.06 | 1.07 |
| 4 | Synpo | 1.46 | 1.94E-97 | 4.04E-93 | 0.41 | 0.23 | 0.18 | 1.80 |
| 4 | Vegfc | 1.51 | 7.41E-97 | 1.54E-92 | 0.43 | 0.25 | 0.18 | 1.73 |
| 4 | Wipi1 | 1.30 | 1.64E-96 | 3.41E-92 | 0.55 | 0.38 | 0.16 | 1.42 |
| 4 | 0610040J01Rik | 2.98 | 3.59E-96 | 7.46E-92 | 0.11 | 0.02 | 0.09 | 4.83 |
| 4 | Entpd1 | 2.29 | 4.49E-94 | 9.34E-90 | 0.17 | 0.05 | 0.12 | 3.35 |
| 4 | Rhoc | 0.61 | 2.81E-93 | 5.85E-89 | 0.87 | 0.78 | 0.09 | 1.11 |
| 4 | Rcn3 | 0.66 | 3.87E-92 | 8.06E-88 | 0.82 | 0.72 | 0.10 | 1.13 |
| 4 | Plod2 | 1.00 | 4.26E-92 | 8.86E-88 | 0.62 | 0.49 | 0.14 | 1.29 |
| 4 | Gabarap | 0.38 | 7.35E-92 | 1.53E-87 | 0.98 | 0.98 | 0.01 | 1.01 |
| 4 | Mylip | 1.33 | 7.58E-92 | 1.58E-87 | 0.47 | 0.30 | 0.17 | 1.58 |
| 4 | Mettl24 | 2.93 | 3.30E-91 | 6.87E-87 | 0.10 | 0.02 | 0.08 | 4.95 |
| 4 | Rin2 | 1.52 | 1.11E-89 | 2.31E-85 | 0.36 | 0.19 | 0.17 | 1.91 |
| 4 | Foxd2os | 1.38 | 8.32E-89 | 1.73E-84 | 0.31 | 0.14 | 0.17 | 2.26 |
| 4 | Atxn7l3b | 0.31 | 1.28E-87 | 2.66E-83 | 1.00 | 0.98 | 0.02 | 1.02 |
| 4 | Pgm2l1 | 1.20 | 1.49E-86 | 3.09E-82 | 0.51 | 0.34 | 0.17 | 1.49 |
| 4 | Pid1 | 1.70 | 5.62E-86 | 1.17E-81 | 0.35 | 0.18 | 0.17 | 1.92 |
| 4 | H2afy2 | 0.39 | 1.62E-85 | 3.36E-81 | 0.99 | 0.93 | 0.05 | 1.06 |
| 4 | Gabra1 | 2.45 | 2.65E-85 | 5.52E-81 | 0.16 | 0.05 | 0.11 | 3.37 |
| 4 | P3h3 | 1.21 | 8.41E-85 | 1.75E-80 | 0.46 | 0.29 | 0.17 | 1.57 |
| 4 | Fn1 | 0.77 | 8.88E-85 | 1.85E-80 | 0.85 | 0.78 | 0.07 | 1.09 |

**Supplementary Table 1. Marker genes for the original clusters of integrated clustering of E11.5 control and NotchGOF λ-junction cells.**

| | | | | | | | | |
|---|---|---|---|---|---|---|---|---|
| 4 | Fli1 | 0.96 | 9.73E-85 | 2.02E-80 | 0.54 | 0.36 | 0.18 | 1.51 |
| 4 | Pdgfra | 0.72 | 5.26E-83 | 1.09E-78 | 0.71 | 0.53 | 0.18 | 1.35 |
| 4 | Septin7 | 0.37 | 5.47E-83 | 1.14E-78 | 0.98 | 0.97 | 0.02 | 1.02 |
| 4 | Zbtb20 | 1.42 | 7.26E-82 | 1.51E-77 | 0.45 | 0.28 | 0.17 | 1.59 |
| 4 | Eif3f | 0.29 | 7.44E-82 | 1.55E-77 | 1.00 | 0.99 | 0.00 | 1.00 |
| 4 | Nedd4 | 0.30 | 8.40E-82 | 1.75E-77 | 1.00 | 0.98 | 0.02 | 1.02 |
| 4 | Foxd1 | 1.91 | 2.30E-81 | 4.79E-77 | 0.24 | 0.10 | 0.14 | 2.43 |
| 4 | Creb3l2 | 1.21 | 2.99E-81 | 6.21E-77 | 0.47 | 0.31 | 0.16 | 1.53 |
| 4 | Tbx3 | 0.74 | 9.73E-81 | 2.02E-76 | 0.63 | 0.42 | 0.21 | 1.50 |
| 4 | Npnt | 1.16 | 6.11E-80 | 1.27E-75 | 0.40 | 0.22 | 0.18 | 1.79 |
| 4 | Cthrc1 | 1.35 | 1.19E-79 | 2.47E-75 | 0.35 | 0.18 | 0.17 | 1.92 |
| 4 | Grcc10 | 0.36 | 4.42E-79 | 9.20E-75 | 1.00 | 0.97 | 0.02 | 1.02 |
| 4 | Nfat5 | 0.92 | 8.04E-79 | 1.67E-74 | 0.67 | 0.57 | 0.11 | 1.19 |
| 4 | Grb10 | 0.41 | 2.70E-78 | 5.61E-74 | 0.97 | 0.93 | 0.03 | 1.04 |
| 4 | Nts | 3.06 | 3.71E-78 | 7.71E-74 | 0.15 | 0.05 | 0.11 | 3.19 |
| 4 | Six1 | 0.57 | 1.53E-77 | 3.18E-73 | 0.80 | 0.60 | 0.20 | 1.33 |
| 4 | Scx | 1.28 | 4.19E-77 | 8.71E-73 | 0.31 | 0.15 | 0.16 | 2.06 |
| 4 | Rest | 0.56 | 6.05E-77 | 1.26E-72 | 0.88 | 0.81 | 0.07 | 1.09 |
| 4 | Selenom | 0.71 | 9.53E-77 | 1.98E-72 | 0.64 | 0.51 | 0.13 | 1.25 |
| 4 | Tpr | 0.41 | 3.57E-76 | 7.43E-72 | 0.99 | 0.98 | 0.02 | 1.02 |
| 4 | Vcan | 0.39 | 5.31E-76 | 1.10E-71 | 0.97 | 0.86 | 0.11 | 1.13 |
| 4 | P4ha3 | 1.59 | 2.15E-75 | 4.48E-71 | 0.29 | 0.15 | 0.15 | 2.01 |
| 4 | Ypel3 | 0.80 | 3.39E-75 | 7.04E-71 | 0.67 | 0.56 | 0.11 | 1.19 |
| 4 | Rgs5 | 1.86 | 5.66E-75 | 1.18E-70 | 0.20 | 0.07 | 0.12 | 2.67 |
| 4 | Igfbp7 | 1.36 | 2.69E-74 | 5.59E-70 | 0.42 | 0.25 | 0.17 | 1.69 |
| 4 | Selenop | 1.04 | 3.08E-74 | 6.40E-70 | 0.33 | 0.18 | 0.16 | 1.89 |
| 4 | Ptov1 | 0.33 | 3.77E-74 | 7.85E-70 | 0.99 | 0.97 | 0.02 | 1.02 |
| 4 | Lima1 | 0.60 | 5.20E-74 | 1.08E-69 | 0.83 | 0.73 | 0.10 | 1.14 |
| 4 | Igsf3 | 0.74 | 1.59E-73 | 3.30E-69 | 0.68 | 0.58 | 0.11 | 1.19 |
| 4 | E130114P18Rik | 1.20 | 3.50E-73 | 7.27E-69 | 0.38 | 0.21 | 0.16 | 1.77 |
| 4 | Cpxm1 | 0.94 | 3.57E-73 | 7.42E-69 | 0.56 | 0.42 | 0.14 | 1.33 |
| 4 | Tshz2 | 1.09 | 4.22E-73 | 8.78E-69 | 0.49 | 0.33 | 0.16 | 1.50 |
| 4 | Ypel2 | 1.73 | 5.78E-73 | 1.20E-68 | 0.26 | 0.12 | 0.14 | 2.16 |
| 4 | Septin11 | 0.45 | 5.21E-72 | 1.08E-67 | 0.95 | 0.87 | 0.08 | 1.09 |
| 4 | Inka1 | 0.82 | 4.61E-71 | 9.59E-67 | 0.63 | 0.50 | 0.13 | 1.25 |
| 4 | Ntrk2 | 2.64 | 5.11E-71 | 1.06E-66 | 0.12 | 0.03 | 0.09 | 3.74 |
| 4 | Gpx3 | 1.35 | 5.55E-71 | 1.15E-66 | 0.36 | 0.21 | 0.15 | 1.74 |
| 4 | Etv1 | 1.12 | 8.55E-71 | 1.78E-66 | 0.44 | 0.29 | 0.16 | 1.54 |
| 4 | Efnb1 | 0.79 | 1.08E-70 | 2.25E-66 | 0.65 | 0.52 | 0.13 | 1.24 |
| 4 | Eml4 | 0.49 | 1.23E-70 | 2.57E-66 | 0.89 | 0.83 | 0.06 | 1.07 |
| 4 | Capns1 | 0.56 | 2.03E-70 | 4.22E-66 | 0.84 | 0.80 | 0.04 | 1.05 |
| 4 | Tut4 | 0.40 | 2.39E-70 | 4.98E-66 | 0.96 | 0.92 | 0.04 | 1.04 |
| 4 | Ednrb | 2.40 | 2.73E-70 | 5.69E-66 | 0.12 | 0.03 | 0.09 | 3.69 |
| 4 | Clmp | 0.56 | 3.74E-70 | 7.79E-66 | 0.81 | 0.63 | 0.19 | 1.30 |
| 4 | Cdo1 | 0.71 | 3.83E-70 | 7.96E-66 | 0.64 | 0.50 | 0.14 | 1.29 |
| 4 | Slc40a1 | 2.33 | 6.55E-70 | 1.36E-65 | 0.15 | 0.05 | 0.10 | 2.96 |
| 4 | Mmp2 | 0.77 | 5.32E-69 | 1.11E-64 | 0.64 | 0.51 | 0.13 | 1.26 |
| 4 | Asxl3 | 1.02 | 6.89E-68 | 1.43E-63 | 0.58 | 0.45 | 0.12 | 1.27 |
| 4 | Eif3e | 0.31 | 2.02E-67 | 4.20E-63 | 0.99 | 0.98 | 0.01 | 1.01 |
| 4 | Ndn | 0.55 | 5.67E-67 | 1.18E-62 | 0.81 | 0.74 | 0.08 | 1.10 |
| 4 | Fuca2 | 1.27 | 1.44E-66 | 3.00E-62 | 0.37 | 0.23 | 0.15 | 1.64 |
| 4 | Meis2 | 0.77 | 2.35E-66 | 4.89E-62 | 0.79 | 0.66 | 0.12 | 1.19 |
| 4 | Anxa11 | 1.51 | 2.47E-66 | 5.14E-62 | 0.28 | 0.14 | 0.14 | 1.97 |
| 4 | Hmcn1 | 0.71 | 4.15E-66 | 8.63E-62 | 0.66 | 0.53 | 0.14 | 1.26 |
| 4 | Rftn2 | 1.31 | 7.76E-66 | 1.61E-61 | 0.36 | 0.22 | 0.14 | 1.66 |
| 4 | Kank4 | 1.38 | 8.42E-66 | 1.75E-61 | 0.33 | 0.19 | 0.14 | 1.75 |

**Supplementary Table 1. Marker genes for the original clusters of integrated clustering of E11.5 control and NotchGOF λ-junction cells.**

| | | | | | | | | |
|---|---|---|---|---|---|---|---|---|
| 4 | Il11ra1 | 0.59 | 2.02E-65 | 4.21E-61 | 0.76 | 0.61 | 0.16 | 1.26 |
| 4 | Prdm16 | 0.98 | 2.70E-64 | 5.61E-60 | 0.47 | 0.32 | 0.16 | 1.50 |
| 4 | Nrp1 | 0.97 | 3.25E-64 | 6.75E-60 | 0.36 | 0.20 | 0.16 | 1.83 |
| 4 | Itm2c | 0.58 | 3.63E-64 | 7.55E-60 | 0.80 | 0.73 | 0.07 | 1.10 |
| 4 | AI314278 | 2.09 | 7.76E-64 | 1.61E-59 | 0.13 | 0.04 | 0.09 | 3.12 |
| 4 | Rab13 | 0.93 | 1.15E-63 | 2.39E-59 | 0.54 | 0.41 | 0.13 | 1.31 |
| 4 | Tpd52l1 | 1.07 | 1.47E-63 | 3.05E-59 | 0.46 | 0.32 | 0.14 | 1.43 |
| 4 | Sh3bgrl | 0.45 | 1.47E-63 | 3.07E-59 | 0.91 | 0.85 | 0.06 | 1.07 |
| 4 | Cygb | 1.94 | 1.55E-63 | 3.23E-59 | 0.19 | 0.08 | 0.11 | 2.45 |
| 4 | Morf4l1 | 0.28 | 2.16E-63 | 4.50E-59 | 1.00 | 0.99 | 0.01 | 1.01 |
| 4 | Gja1 | 0.47 | 3.03E-63 | 6.30E-59 | 0.84 | 0.77 | 0.08 | 1.10 |
| 4 | Basp1 | 0.29 | 1.14E-62 | 2.37E-58 | 0.99 | 0.95 | 0.04 | 1.05 |
| 4 | Nid1 | 1.08 | 2.17E-62 | 4.51E-58 | 0.48 | 0.33 | 0.14 | 1.43 |
| 4 | Fxyd1 | 2.03 | 2.49E-62 | 5.18E-58 | 0.15 | 0.05 | 0.10 | 2.80 |
| 4 | Sesn3 | 0.72 | 2.66E-62 | 5.54E-58 | 0.67 | 0.59 | 0.09 | 1.15 |
| 4 | 2610035D17Rik | 2.12 | 3.23E-62 | 6.73E-58 | 0.14 | 0.05 | 0.09 | 2.88 |
| 4 | Gpx4 | 0.27 | 9.95E-62 | 2.07E-57 | 1.00 | 0.99 | 0.01 | 1.01 |
| 4 | Eya2 | 0.82 | 1.34E-61 | 2.78E-57 | 0.38 | 0.22 | 0.16 | 1.71 |
| 4 | Tnik | 1.16 | 1.91E-61 | 3.97E-57 | 0.38 | 0.23 | 0.14 | 1.62 |
| 4 | Slc1a3 | 1.24 | 6.19E-61 | 1.29E-56 | 0.36 | 0.22 | 0.14 | 1.67 |
| 4 | Mex3b | 0.56 | 1.31E-60 | 2.72E-56 | 0.78 | 0.69 | 0.10 | 1.14 |
| 4 | Fzd2 | 0.40 | 1.52E-60 | 3.17E-56 | 0.95 | 0.87 | 0.08 | 1.09 |
| 4 | Tspan3 | 0.38 | 2.05E-60 | 4.27E-56 | 0.96 | 0.91 | 0.05 | 1.06 |
| 4 | Daam2 | 1.17 | 3.29E-60 | 6.85E-56 | 0.42 | 0.29 | 0.14 | 1.48 |
| 4 | Dnm3os | 0.67 | 3.68E-60 | 7.65E-56 | 0.67 | 0.51 | 0.16 | 1.31 |
| 4 | Nfic | 1.31 | 5.79E-60 | 1.20E-55 | 0.42 | 0.29 | 0.13 | 1.46 |
| 4 | Arhgef25 | 1.01 | 9.19E-60 | 1.91E-55 | 0.49 | 0.36 | 0.13 | 1.35 |
| 4 | Tmem50a | 0.47 | 1.21E-59 | 2.51E-55 | 0.84 | 0.81 | 0.04 | 1.05 |
| 4 | Ppp1r12a | 0.65 | 1.33E-59 | 2.77E-55 | 0.77 | 0.73 | 0.04 | 1.05 |
| 4 | Chst15 | 1.46 | 1.72E-59 | 3.58E-55 | 0.29 | 0.16 | 0.13 | 1.83 |
| 4 | Fam110b | 1.23 | 1.89E-59 | 3.94E-55 | 0.43 | 0.30 | 0.13 | 1.43 |
| 4 | Zfhx4 | 0.53 | 2.61E-59 | 5.43E-55 | 0.83 | 0.73 | 0.10 | 1.14 |
| 4 | Wwtr1 | 0.93 | 3.86E-59 | 8.03E-55 | 0.54 | 0.43 | 0.11 | 1.26 |
| 4 | Limd2 | 0.37 | 7.04E-59 | 1.46E-54 | 0.97 | 0.89 | 0.07 | 1.08 |
| 4 | Scube1 | 0.96 | 1.24E-58 | 2.58E-54 | 0.50 | 0.37 | 0.14 | 1.38 |
| 4 | Cdkn1c | 0.36 | 2.19E-58 | 4.55E-54 | 0.98 | 0.93 | 0.06 | 1.06 |
| 4 | Hmgn3 | 0.51 | 4.10E-58 | 8.52E-54 | 0.80 | 0.75 | 0.05 | 1.07 |
| 4 | Negr1 | 2.11 | 4.53E-58 | 9.42E-54 | 0.13 | 0.05 | 0.09 | 2.89 |
| 4 | Fbn2 | 0.51 | 1.39E-57 | 2.88E-53 | 0.83 | 0.73 | 0.10 | 1.14 |
| 4 | Ctsf | 1.60 | 2.29E-57 | 4.77E-53 | 0.21 | 0.10 | 0.11 | 2.18 |
| 4 | Olfm2 | 1.64 | 3.74E-57 | 7.78E-53 | 0.21 | 0.10 | 0.11 | 2.18 |
| 4 | Rcn1 | 0.47 | 5.10E-57 | 1.06E-52 | 0.86 | 0.82 | 0.03 | 1.04 |
| 4 | Pitpnc1 | 0.68 | 5.89E-57 | 1.23E-52 | 0.67 | 0.58 | 0.09 | 1.15 |
| 4 | Selenof | 0.35 | 8.89E-57 | 1.85E-52 | 0.97 | 0.96 | 0.02 | 1.02 |
| 4 | Sncaip | 1.11 | 9.84E-57 | 2.05E-52 | 0.42 | 0.29 | 0.13 | 1.44 |
| 4 | Maged1 | 0.34 | 1.20E-56 | 2.50E-52 | 0.97 | 0.94 | 0.04 | 1.04 |
| 4 | Ak1 | 1.79 | 1.87E-56 | 3.89E-52 | 0.18 | 0.08 | 0.10 | 2.36 |
| 4 | Osr2 | 1.46 | 2.03E-56 | 4.22E-52 | 0.19 | 0.08 | 0.11 | 2.38 |
| 4 | Kmt2e | 0.45 | 8.22E-55 | 1.71E-50 | 0.87 | 0.84 | 0.03 | 1.03 |
| 4 | Camk2n1 | 1.15 | 1.01E-54 | 2.11E-50 | 0.31 | 0.18 | 0.13 | 1.74 |
| 4 | Npc2 | 0.34 | 2.97E-54 | 6.17E-50 | 0.97 | 0.93 | 0.04 | 1.04 |
| 4 | Prkar2b | 0.84 | 3.89E-54 | 8.08E-50 | 0.57 | 0.49 | 0.08 | 1.17 |
| 4 | Selenok | 0.38 | 6.44E-54 | 1.34E-49 | 0.93 | 0.92 | 0.02 | 1.02 |
| 4 | Gulo | 1.57 | 8.39E-54 | 1.74E-49 | 0.20 | 0.09 | 0.11 | 2.24 |
| 4 | Dab2ip | 0.65 | 8.90E-53 | 1.85E-48 | 0.67 | 0.60 | 0.07 | 1.12 |
| 4 | Cacna1d | 1.22 | 1.36E-52 | 2.82E-48 | 0.36 | 0.23 | 0.13 | 1.58 |

**Supplementary Table 1. Marker genes for the original clusters of integrated clustering of E11.5 control and NotchGOF λ-junction cells.**

| | | | | | | | | |
|---|---|---|---|---|---|---|---|---|
| 4 | Map3k1 | 1.20 | 3.04E-52 | 6.32E-48 | 0.38 | 0.25 | 0.12 | 1.49 |
| 4 | Nrep | 0.44 | 4.43E-52 | 9.21E-48 | 0.87 | 0.82 | 0.05 | 1.06 |
| 4 | Sox12 | 0.58 | 2.10E-51 | 4.36E-47 | 0.71 | 0.66 | 0.05 | 1.08 |
| 4 | Pakap.1 | 0.84 | 2.78E-51 | 5.77E-47 | 0.52 | 0.40 | 0.11 | 1.28 |
| 4 | Ube2e3 | 0.35 | 4.43E-51 | 9.22E-47 | 0.93 | 0.91 | 0.02 | 1.02 |
| 4 | Olfml3 | 1.03 | 4.45E-51 | 9.26E-47 | 0.41 | 0.28 | 0.13 | 1.46 |
| 4 | Adra2a | 2.02 | 4.71E-51 | 9.79E-47 | 0.10 | 0.03 | 0.07 | 3.26 |
| 4 | Pth1r | 0.52 | 1.87E-50 | 3.88E-46 | 0.70 | 0.56 | 0.14 | 1.25 |
| 4 | Ndufb11 | 0.31 | 1.97E-50 | 4.09E-46 | 0.97 | 0.96 | 0.01 | 1.01 |
| 4 | Aopep | 0.61 | 5.29E-50 | 1.10E-45 | 0.69 | 0.62 | 0.07 | 1.12 |
| 4 | Arhgef40 | 0.70 | 5.73E-50 | 1.19E-45 | 0.61 | 0.52 | 0.09 | 1.18 |
| 4 | Rell1 | 1.11 | 8.14E-50 | 1.69E-45 | 0.45 | 0.34 | 0.11 | 1.31 |
| 4 | Efemp2 | 0.76 | 9.37E-50 | 1.95E-45 | 0.57 | 0.47 | 0.10 | 1.20 |
| 4 | Snai1 | 0.71 | 2.61E-49 | 5.43E-45 | 0.54 | 0.40 | 0.14 | 1.34 |
| 4 | Scaper | 1.23 | 7.45E-49 | 1.55E-44 | 0.40 | 0.29 | 0.11 | 1.40 |
| 4 | Tmtc4 | 1.18 | 8.56E-49 | 1.78E-44 | 0.39 | 0.28 | 0.12 | 1.42 |
| 4 | Mme | 0.74 | 1.34E-48 | 2.79E-44 | 0.58 | 0.46 | 0.12 | 1.25 |
| 4 | Tcaf1 | 0.59 | 1.39E-48 | 2.90E-44 | 0.72 | 0.67 | 0.06 | 1.08 |
| 4 | Ppic | 0.46 | 1.53E-48 | 3.18E-44 | 0.77 | 0.70 | 0.07 | 1.10 |
| 4 | Tsc22d1 | 0.35 | 3.05E-48 | 6.35E-44 | 0.97 | 0.93 | 0.03 | 1.04 |
| 4 | Ssbp4 | 0.56 | 4.96E-48 | 1.03E-43 | 0.70 | 0.65 | 0.05 | 1.08 |
| 4 | Ddr2 | 0.63 | 8.31E-48 | 1.73E-43 | 0.61 | 0.48 | 0.13 | 1.27 |
| 4 | Mir100hg | 0.49 | 2.48E-47 | 5.16E-43 | 0.83 | 0.76 | 0.07 | 1.09 |
| 4 | Slitrk4 | 2.13 | 2.77E-47 | 5.77E-43 | 0.13 | 0.05 | 0.08 | 2.65 |
| 4 | Cnn2 | 0.54 | 3.58E-47 | 7.45E-43 | 0.71 | 0.62 | 0.09 | 1.15 |
| 4 | Arhgdib | 0.94 | 4.27E-47 | 8.87E-43 | 0.59 | 0.49 | 0.10 | 1.19 |
| 4 | St6galnac4 | 1.22 | 4.58E-47 | 9.53E-43 | 0.31 | 0.19 | 0.12 | 1.63 |
| 4 | Fkbp7 | 1.02 | 5.04E-47 | 1.05E-42 | 0.39 | 0.27 | 0.12 | 1.44 |
| 4 | Wasf1 | 0.92 | 5.10E-47 | 1.06E-42 | 0.49 | 0.40 | 0.09 | 1.24 |
| 4 | Fgfr3 | 2.12 | 1.22E-46 | 2.53E-42 | 0.10 | 0.03 | 0.07 | 3.13 |
| 4 | Nlgn3 | 1.95 | 1.07E-45 | 2.23E-41 | 0.12 | 0.05 | 0.08 | 2.65 |
| 4 | Pla2g4a | 1.12 | 2.10E-45 | 4.36E-41 | 0.36 | 0.24 | 0.12 | 1.49 |
| 4 | Tgfbi | 0.80 | 3.64E-45 | 7.56E-41 | 0.41 | 0.28 | 0.13 | 1.46 |
| 4 | Dpep1 | 1.39 | 3.66E-45 | 7.62E-41 | 0.34 | 0.22 | 0.12 | 1.54 |
| 4 | Tmem26 | 1.01 | 5.91E-45 | 1.23E-40 | 0.37 | 0.24 | 0.12 | 1.51 |
| 4 | Slc16a10 | 1.74 | 6.75E-45 | 1.40E-40 | 0.20 | 0.10 | 0.10 | 1.97 |
| 4 | Dad1 | 0.31 | 6.95E-45 | 1.45E-40 | 0.96 | 0.94 | 0.02 | 1.02 |
| 4 | Ccnd1 | 0.47 | 7.59E-45 | 1.58E-40 | 0.84 | 0.80 | 0.04 | 1.05 |
| 4 | Penk | 0.81 | 1.15E-44 | 2.40E-40 | 0.19 | 0.09 | 0.10 | 2.16 |
| 4 | Bambi | 0.45 | 3.13E-44 | 6.52E-40 | 0.73 | 0.63 | 0.10 | 1.16 |
| 4 | Rab18 | 0.42 | 2.39E-43 | 4.96E-39 | 0.83 | 0.82 | 0.02 | 1.02 |
| 4 | Gpx7 | 0.78 | 3.00E-43 | 6.24E-39 | 0.53 | 0.44 | 0.09 | 1.20 |
| 4 | Gm8797 | 0.31 | 3.92E-43 | 8.15E-39 | 0.94 | 0.94 | 0.00 | 1.00 |
| 4 | Myl12b | 0.36 | 3.92E-43 | 8.15E-39 | 0.90 | 0.89 | 0.01 | 1.01 |
| 4 | Plod1 | 1.22 | 9.74E-43 | 2.03E-38 | 0.30 | 0.19 | 0.11 | 1.57 |
| 4 | Smad7 | 0.52 | 4.05E-42 | 8.43E-38 | 0.75 | 0.69 | 0.06 | 1.09 |
| 4 | Slc50a1 | 0.86 | 4.61E-42 | 9.59E-38 | 0.46 | 0.37 | 0.10 | 1.27 |
| 4 | Mageh1 | 0.61 | 4.86E-42 | 1.01E-37 | 0.62 | 0.56 | 0.06 | 1.11 |
| 4 | Frmd4a | 0.56 | 4.57E-41 | 9.50E-37 | 0.72 | 0.67 | 0.05 | 1.07 |
| 4 | Slc25a21 | 1.91 | 4.78E-41 | 9.94E-37 | 0.14 | 0.06 | 0.08 | 2.30 |
| 4 | Trim35 | 0.51 | 5.30E-41 | 1.10E-36 | 0.75 | 0.70 | 0.05 | 1.06 |
| 4 | Alx1 | 0.30 | 1.68E-40 | 3.49E-36 | 0.95 | 0.71 | 0.24 | 1.34 |
| 4 | Rarg | 0.63 | 1.86E-40 | 3.87E-36 | 0.59 | 0.52 | 0.08 | 1.15 |
| 4 | Kdelr1 | 0.40 | 2.30E-40 | 4.79E-36 | 0.84 | 0.83 | 0.01 | 1.01 |
| 4 | Egflam | 1.73 | 3.36E-40 | 6.98E-36 | 0.15 | 0.07 | 0.08 | 2.22 |
| 4 | Oaf | 0.68 | 8.82E-40 | 1.83E-35 | 0.54 | 0.45 | 0.09 | 1.20 |

**Supplementary Table 1. Marker genes for the original clusters of integrated clustering of E11.5 control and NotchGOF λ-junction cells.**

| | | | | | | | | |
|---|---|---|---|---|---|---|---|---|
| 4 | Slc35f2 | 1.85 | 1.79E-39 | 3.71E-35 | 0.13 | 0.06 | 0.08 | 2.36 |
| 4 | Pla2g7 | 1.81 | 2.78E-39 | 5.77E-35 | 0.14 | 0.06 | 0.08 | 2.31 |
| 4 | Fgfr1 | 0.48 | 2.90E-39 | 6.04E-35 | 0.72 | 0.65 | 0.06 | 1.10 |
| 4 | Hp1bp3 | 0.36 | 4.10E-39 | 8.53E-35 | 0.91 | 0.89 | 0.02 | 1.02 |
| 4 | Tmem59 | 0.36 | 4.25E-39 | 8.85E-35 | 0.89 | 0.88 | 0.01 | 1.01 |
| 4 | Gm16104 | 1.83 | 4.49E-39 | 9.35E-35 | 0.10 | 0.04 | 0.07 | 2.71 |
| 4 | Mef2c | 0.79 | 6.50E-39 | 1.35E-34 | 0.29 | 0.18 | 0.11 | 1.64 |
| 4 | Fkbp10 | 0.88 | 1.05E-38 | 2.18E-34 | 0.43 | 0.33 | 0.10 | 1.30 |
| 4 | Zfp36l2 | 0.73 | 1.23E-38 | 2.56E-34 | 0.51 | 0.43 | 0.08 | 1.19 |
| 4 | Vkorc1 | 0.79 | 1.77E-38 | 3.68E-34 | 0.50 | 0.43 | 0.08 | 1.18 |
| 4 | Rab21 | 1.07 | 2.48E-38 | 5.15E-34 | 0.52 | 0.46 | 0.05 | 1.11 |
| 4 | Lama2 | 1.28 | 2.90E-38 | 6.04E-34 | 0.24 | 0.14 | 0.10 | 1.73 |
| 4 | Sec24d | 1.66 | 3.00E-38 | 6.23E-34 | 0.16 | 0.08 | 0.08 | 2.09 |
| 4 | Itga9 | 1.05 | 3.50E-38 | 7.29E-34 | 0.31 | 0.21 | 0.11 | 1.52 |
| 4 | Adgra2 | 1.03 | 4.13E-38 | 8.59E-34 | 0.33 | 0.23 | 0.11 | 1.46 |
| 4 | Septin6 | 0.63 | 4.18E-38 | 8.70E-34 | 0.62 | 0.57 | 0.06 | 1.10 |
| 4 | Pde1c | 1.60 | 5.49E-38 | 1.14E-33 | 0.19 | 0.10 | 0.09 | 1.89 |
| 4 | Sox4 | 0.26 | 7.24E-38 | 1.51E-33 | 0.99 | 0.94 | 0.05 | 1.06 |
| 4 | Fam114a1 | 1.34 | 1.05E-37 | 2.18E-33 | 0.26 | 0.16 | 0.10 | 1.63 |
| 4 | Akr1a1 | 0.26 | 1.48E-37 | 3.08E-33 | 0.98 | 0.96 | 0.02 | 1.02 |
| 4 | Pmepa1 | 0.72 | 2.10E-37 | 4.37E-33 | 0.55 | 0.47 | 0.08 | 1.17 |
| 4 | Etv4 | 0.81 | 2.47E-37 | 5.13E-33 | 0.43 | 0.32 | 0.11 | 1.33 |
| 4 | Rbms3 | 0.73 | 2.77E-37 | 5.75E-33 | 0.50 | 0.41 | 0.10 | 1.23 |
| 4 | Pde7b | 1.91 | 2.79E-37 | 5.81E-33 | 0.10 | 0.04 | 0.06 | 2.62 |
| 4 | Commd8 | 0.52 | 3.41E-37 | 7.09E-33 | 0.70 | 0.67 | 0.03 | 1.04 |
| 4 | Lrp4 | 1.55 | 5.42E-37 | 1.13E-32 | 0.18 | 0.09 | 0.09 | 1.95 |
| 4 | Rdx | 0.25 | 5.99E-37 | 1.25E-32 | 0.98 | 0.96 | 0.02 | 1.02 |
| 4 | Map1lc3a | 0.39 | 6.44E-37 | 1.34E-32 | 0.83 | 0.81 | 0.03 | 1.03 |
| 4 | Skil | 0.56 | 7.28E-37 | 1.52E-32 | 0.62 | 0.55 | 0.07 | 1.12 |
| 4 | Islr | 1.32 | 1.41E-36 | 2.93E-32 | 0.16 | 0.08 | 0.08 | 2.09 |
| 4 | Lix1l | 0.76 | 1.68E-36 | 3.49E-32 | 0.47 | 0.37 | 0.10 | 1.25 |
| 4 | Rp9 | 0.43 | 2.16E-36 | 4.49E-32 | 0.79 | 0.79 | 0.00 | 1.00 |
| 4 | Rai14 | 0.44 | 2.51E-36 | 5.21E-32 | 0.73 | 0.64 | 0.09 | 1.14 |
| 4 | Pdcd4 | 0.35 | 2.57E-36 | 5.35E-32 | 0.71 | 0.64 | 0.08 | 1.12 |
| 4 | Nenf | 0.40 | 3.59E-36 | 7.47E-32 | 0.81 | 0.78 | 0.03 | 1.03 |
| 4 | Itgb1 | 0.33 | 6.04E-36 | 1.26E-31 | 0.89 | 0.85 | 0.04 | 1.04 |
| 4 | Pclo | 1.56 | 9.08E-36 | 1.89E-31 | 0.19 | 0.10 | 0.09 | 1.87 |
| 4 | Ghr | 1.27 | 9.78E-36 | 2.03E-31 | 0.23 | 0.13 | 0.10 | 1.72 |
| 4 | Wwp2 | 1.04 | 1.56E-35 | 3.24E-31 | 0.39 | 0.30 | 0.09 | 1.28 |
| 4 | Arf4 | 0.28 | 2.23E-35 | 4.64E-31 | 0.96 | 0.95 | 0.01 | 1.01 |
| 4 | Ahr | 1.65 | 2.26E-35 | 4.71E-31 | 0.16 | 0.08 | 0.08 | 2.00 |
| 4 | Nog | 1.70 | 2.36E-35 | 4.92E-31 | 0.17 | 0.09 | 0.08 | 1.94 |
| 4 | Smarca2 | 0.58 | 3.98E-35 | 8.27E-31 | 0.65 | 0.62 | 0.03 | 1.05 |
| 4 | Plxdc2 | 0.98 | 5.00E-35 | 1.04E-30 | 0.34 | 0.24 | 0.10 | 1.42 |
| 4 | Gfpt1 | 0.84 | 6.54E-35 | 1.36E-30 | 0.45 | 0.38 | 0.08 | 1.21 |
| 4 | Tiparp | 0.90 | 9.10E-35 | 1.89E-30 | 0.41 | 0.31 | 0.10 | 1.31 |
| 4 | Foxd2 | 1.05 | 1.32E-34 | 2.75E-30 | 0.20 | 0.11 | 0.09 | 1.90 |
| 4 | St6galnac6 | 1.26 | 1.57E-34 | 3.26E-30 | 0.25 | 0.15 | 0.09 | 1.61 |
| 4 | Acadl | 0.64 | 2.14E-34 | 4.44E-30 | 0.58 | 0.53 | 0.05 | 1.09 |
| 4 | Metrn | 0.73 | 2.76E-34 | 5.75E-30 | 0.56 | 0.51 | 0.05 | 1.10 |
| 4 | Spry1 | 1.02 | 3.08E-34 | 6.42E-30 | 0.38 | 0.28 | 0.10 | 1.34 |
| 4 | Tmx4 | 0.73 | 3.57E-34 | 7.43E-30 | 0.52 | 0.46 | 0.06 | 1.14 |
| 4 | Serinc1 | 0.47 | 3.58E-34 | 7.45E-30 | 0.72 | 0.70 | 0.02 | 1.03 |
| 4 | Map2k4 | 0.71 | 4.40E-34 | 9.15E-30 | 0.56 | 0.52 | 0.04 | 1.07 |
| 4 | Cdk14 | 0.64 | 4.73E-34 | 9.85E-30 | 0.56 | 0.49 | 0.07 | 1.14 |
| 4 | Epha5 | 1.74 | 5.69E-34 | 1.18E-29 | 0.14 | 0.06 | 0.07 | 2.16 |

**Supplementary Table 1. Marker genes for the original clusters of integrated clustering of E11.5 control and NotchGOF λ-junction cells.**

| | | | | | | | | |
|---|---|---|---|---|---|---|---|---|
| 4 | Fzd4 | 1.04 | 7.19E-34 | 1.50E-29 | 0.35 | 0.25 | 0.10 | 1.41 |
| 4 | Prdm16os | 1.24 | 8.08E-34 | 1.68E-29 | 0.20 | 0.11 | 0.09 | 1.80 |
| 4 | Spire1 | 0.90 | 1.45E-33 | 3.02E-29 | 0.42 | 0.34 | 0.08 | 1.23 |
| 4 | Morf4l2 | 0.27 | 1.59E-33 | 3.31E-29 | 0.95 | 0.94 | 0.01 | 1.02 |
| 4 | Drap1 | 0.29 | 2.21E-33 | 4.59E-29 | 0.94 | 0.93 | 0.01 | 1.01 |
| 4 | Lpar4 | 0.86 | 2.60E-33 | 5.42E-29 | 0.41 | 0.32 | 0.09 | 1.28 |
| 4 | Ebpl | 0.57 | 7.29E-33 | 1.52E-28 | 0.60 | 0.54 | 0.06 | 1.10 |
| 4 | Ptpdc1 | 1.23 | 7.46E-33 | 1.55E-28 | 0.25 | 0.16 | 0.09 | 1.58 |
| 4 | Ss18 | 0.53 | 2.11E-32 | 4.39E-28 | 0.67 | 0.65 | 0.02 | 1.03 |
| 4 | Bmpr2 | 0.64 | 2.12E-32 | 4.41E-28 | 0.60 | 0.55 | 0.04 | 1.08 |
| 4 | Prkg2 | 1.74 | 2.83E-32 | 5.88E-28 | 0.11 | 0.04 | 0.06 | 2.44 |
| 4 | Ssbp3 | 0.26 | 4.33E-32 | 9.00E-28 | 0.96 | 0.94 | 0.02 | 1.02 |
| 4 | Sh3glb1 | 0.35 | 4.97E-32 | 1.03E-27 | 0.84 | 0.84 | 0.00 | 1.00 |
| 4 | Gm16070 | 1.67 | 6.38E-32 | 1.33E-27 | 0.11 | 0.05 | 0.06 | 2.31 |
| 4 | Tcf7l1 | 0.53 | 8.18E-32 | 1.70E-27 | 0.63 | 0.58 | 0.05 | 1.09 |
| 4 | Rhoq | 0.67 | 9.24E-32 | 1.92E-27 | 0.51 | 0.44 | 0.07 | 1.15 |
| 4 | Ech1 | 0.63 | 9.54E-32 | 1.98E-27 | 0.56 | 0.51 | 0.05 | 1.09 |
| 4 | Hook3 | 0.49 | 1.11E-31 | 2.31E-27 | 0.69 | 0.66 | 0.03 | 1.04 |
| 4 | Ppfibp2 | 0.65 | 1.64E-31 | 3.40E-27 | 0.51 | 0.42 | 0.09 | 1.21 |
| 4 | Cttnbp2nl | 0.63 | 4.45E-31 | 9.26E-27 | 0.55 | 0.49 | 0.06 | 1.11 |
| 4 | Pcolce | 0.60 | 4.54E-31 | 9.45E-27 | 0.54 | 0.45 | 0.09 | 1.19 |
| 4 | Copz2 | 0.92 | 6.68E-31 | 1.39E-26 | 0.35 | 0.25 | 0.09 | 1.36 |
| 4 | Hdac9 | 1.58 | 7.98E-31 | 1.66E-26 | 0.14 | 0.07 | 0.07 | 2.03 |
| 4 | Zfand6 | 0.37 | 8.44E-31 | 1.76E-26 | 0.78 | 0.79 | -0.01 | 0.99 |
| 4 | Kcne1l | 1.33 | 9.67E-31 | 2.01E-26 | 0.20 | 0.12 | 0.08 | 1.71 |
| 4 | Snhg18 | 0.59 | 1.09E-30 | 2.26E-26 | 0.58 | 0.52 | 0.06 | 1.12 |
| 4 | Gulp1 | 0.96 | 1.30E-30 | 2.71E-26 | 0.33 | 0.24 | 0.09 | 1.37 |
| 4 | Dusp22 | 1.20 | 1.92E-30 | 3.99E-26 | 0.28 | 0.20 | 0.09 | 1.44 |
| 4 | Tm9sf3 | 0.34 | 2.21E-30 | 4.59E-26 | 0.88 | 0.86 | 0.01 | 1.01 |
| 4 | Kifap3 | 0.38 | 2.84E-30 | 5.90E-26 | 0.79 | 0.78 | 0.02 | 1.02 |
| 4 | Ttc28 | 0.51 | 3.17E-30 | 6.60E-26 | 0.64 | 0.59 | 0.04 | 1.07 |
| 4 | Kcnq1ot1 | 0.31 | 3.68E-30 | 7.65E-26 | 0.95 | 0.92 | 0.03 | 1.03 |
| 4 | Fibp | 0.50 | 4.15E-30 | 8.64E-26 | 0.64 | 0.61 | 0.03 | 1.05 |
| 4 | Erbb3 | 1.07 | 6.15E-30 | 1.28E-25 | 0.26 | 0.17 | 0.10 | 1.56 |
| 4 | Ssbp2 | 0.46 | 6.19E-30 | 1.29E-25 | 0.73 | 0.71 | 0.02 | 1.03 |
| 4 | Rab2a | 0.28 | 6.31E-30 | 1.31E-25 | 0.93 | 0.91 | 0.02 | 1.02 |
| 4 | Dst | 0.66 | 6.68E-30 | 1.39E-25 | 0.44 | 0.35 | 0.09 | 1.26 |
| 4 | Epb41l2 | 0.42 | 7.19E-30 | 1.50E-25 | 0.71 | 0.67 | 0.04 | 1.06 |
| 4 | Fndc5 | 1.12 | 8.20E-30 | 1.71E-25 | 0.25 | 0.16 | 0.09 | 1.54 |
| 4 | Pex3 | 0.91 | 8.38E-30 | 1.74E-25 | 0.39 | 0.31 | 0.08 | 1.26 |
| 4 | Apip | 0.54 | 1.12E-29 | 2.32E-25 | 0.63 | 0.61 | 0.02 | 1.04 |
| 4 | B2m | 0.45 | 1.26E-29 | 2.62E-25 | 0.59 | 0.54 | 0.05 | 1.09 |
| 4 | Slit2 | 0.68 | 1.28E-29 | 2.66E-25 | 0.45 | 0.36 | 0.09 | 1.25 |
| 4 | Lrp1 | 0.50 | 1.30E-29 | 2.71E-25 | 0.63 | 0.58 | 0.05 | 1.09 |
| 4 | Myof | 1.44 | 1.36E-29 | 2.82E-25 | 0.16 | 0.09 | 0.08 | 1.89 |
| 4 | Ckap4 | 0.36 | 1.39E-29 | 2.90E-25 | 0.82 | 0.80 | 0.02 | 1.03 |
| 4 | Cuedc2 | 0.42 | 1.85E-29 | 3.85E-25 | 0.76 | 0.77 | -0.01 | 0.98 |
| 4 | Inpp4b | 1.15 | 1.85E-29 | 3.85E-25 | 0.24 | 0.15 | 0.09 | 1.59 |
| 4 | Ccdc88a | 0.34 | 2.15E-29 | 4.46E-25 | 0.84 | 0.81 | 0.03 | 1.04 |
| 4 | Vegfb | 0.45 | 2.67E-29 | 5.56E-25 | 0.70 | 0.66 | 0.04 | 1.06 |
| 4 | Sdcbp | 0.43 | 2.96E-29 | 6.16E-25 | 0.70 | 0.69 | 0.01 | 1.01 |
| 4 | Gng10 | 0.33 | 3.06E-29 | 6.36E-25 | 0.83 | 0.81 | 0.02 | 1.02 |
| 4 | Apbb2 | 0.88 | 3.48E-29 | 7.23E-25 | 0.38 | 0.30 | 0.08 | 1.27 |
| 4 | Clip3 | 0.56 | 4.38E-29 | 9.12E-25 | 0.57 | 0.52 | 0.05 | 1.10 |
| 4 | Etfa | 0.41 | 5.75E-29 | 1.19E-24 | 0.74 | 0.74 | 0.00 | 1.00 |
| 4 | Eif5b | 0.25 | 6.40E-29 | 1.33E-24 | 0.97 | 0.97 | 0.01 | 1.01 |

**Supplementary Table 1. Marker genes for the original clusters of integrated clustering of E11.5 control and NotchGOF λ-junction cells.**

| | | | | | | | | |
|---|---|---|---|---|---|---|---|---|
| 4 | Ift74 | 0.39 | 1.23E-28 | 2.55E-24 | 0.76 | 0.74 | 0.03 | 1.04 |
| 4 | Hacd4 | 0.94 | 1.65E-28 | 3.43E-24 | 0.33 | 0.24 | 0.09 | 1.36 |
| 4 | Tbx3os1 | 1.26 | 1.93E-28 | 4.00E-24 | 0.21 | 0.13 | 0.08 | 1.65 |
| 4 | Foxf2 | 1.42 | 2.55E-28 | 5.30E-24 | 0.16 | 0.09 | 0.08 | 1.84 |
| 4 | Cited2 | 0.50 | 3.28E-28 | 6.83E-24 | 0.67 | 0.64 | 0.03 | 1.05 |
| 4 | Idh1 | 0.48 | 4.43E-28 | 9.21E-24 | 0.67 | 0.63 | 0.04 | 1.06 |
| 4 | Kcnj2 | 1.08 | 4.61E-28 | 9.58E-24 | 0.26 | 0.17 | 0.09 | 1.52 |
| 4 | Jam3 | 0.48 | 4.90E-28 | 1.02E-23 | 0.62 | 0.57 | 0.05 | 1.09 |
| 4 | Tmsb15b2 | 0.53 | 4.97E-28 | 1.03E-23 | 0.59 | 0.55 | 0.04 | 1.07 |
| 4 | Mmp14 | 0.44 | 5.52E-28 | 1.15E-23 | 0.67 | 0.63 | 0.04 | 1.06 |
| 4 | Hsd11b2 | 1.20 | 7.13E-28 | 1.48E-23 | 0.17 | 0.10 | 0.08 | 1.79 |
| 4 | Ddah1 | 1.27 | 8.04E-28 | 1.67E-23 | 0.22 | 0.13 | 0.08 | 1.61 |
| 4 | Col5a2 | 0.48 | 9.61E-28 | 2.00E-23 | 0.61 | 0.52 | 0.08 | 1.16 |
| 4 | Wnt5a | 0.41 | 2.38E-27 | 4.95E-23 | 0.72 | 0.63 | 0.09 | 1.14 |
| 4 | Timm17b | 0.32 | 2.38E-27 | 4.96E-23 | 0.81 | 0.82 | -0.01 | 0.99 |
| 4 | Glmp | 0.62 | 2.92E-27 | 6.07E-23 | 0.52 | 0.47 | 0.05 | 1.11 |
| 4 | Helz | 0.53 | 3.35E-27 | 6.96E-23 | 0.61 | 0.58 | 0.03 | 1.05 |
| 4 | Emp3 | 1.06 | 3.96E-27 | 8.24E-23 | 0.21 | 0.13 | 0.08 | 1.63 |
| 4 | Marveld1 | 0.93 | 4.51E-27 | 9.39E-23 | 0.32 | 0.23 | 0.08 | 1.36 |
| 4 | Syt12 | 1.58 | 5.49E-27 | 1.14E-22 | 0.10 | 0.05 | 0.06 | 2.22 |
| 4 | Pkn1 | 0.79 | 5.77E-27 | 1.20E-22 | 0.42 | 0.35 | 0.07 | 1.20 |
| 4 | Kank2 | 0.87 | 6.45E-27 | 1.34E-22 | 0.38 | 0.31 | 0.08 | 1.25 |
| 4 | Mab21l1 | 1.02 | 7.00E-27 | 1.46E-22 | 0.25 | 0.16 | 0.09 | 1.55 |
| 4 | Pbx3 | 0.50 | 8.82E-27 | 1.83E-22 | 0.58 | 0.53 | 0.05 | 1.10 |
| 4 | Dcaf8 | 0.40 | 1.48E-26 | 3.09E-22 | 0.74 | 0.75 | -0.01 | 0.99 |
| 4 | Hbp1 | 0.51 | 1.56E-26 | 3.24E-22 | 0.60 | 0.56 | 0.03 | 1.06 |
| 4 | Vasn | 0.99 | 1.60E-26 | 3.33E-22 | 0.29 | 0.21 | 0.08 | 1.39 |
| 4 | Ap3s1 | 0.40 | 2.00E-26 | 4.17E-22 | 0.71 | 0.71 | 0.00 | 1.00 |
| 4 | Dnajc10 | 0.47 | 2.47E-26 | 5.13E-22 | 0.69 | 0.69 | 0.00 | 1.00 |
| 4 | Smim10l1 | 0.49 | 3.96E-26 | 8.25E-22 | 0.63 | 0.61 | 0.02 | 1.04 |
| 4 | Papss1 | 0.40 | 4.03E-26 | 8.38E-22 | 0.68 | 0.66 | 0.02 | 1.03 |
| 4 | Erbin | 0.44 | 4.83E-26 | 1.00E-21 | 0.66 | 0.63 | 0.03 | 1.04 |
| 4 | Elk3 | 0.54 | 6.37E-26 | 1.32E-21 | 0.48 | 0.41 | 0.07 | 1.16 |
| 4 | Kctd12b | 1.03 | 6.59E-26 | 1.37E-21 | 0.26 | 0.18 | 0.08 | 1.47 |
| 4 | Copb1 | 0.34 | 6.86E-26 | 1.43E-21 | 0.79 | 0.77 | 0.02 | 1.02 |
| 4 | 2610301B20Rik | 0.96 | 9.00E-26 | 1.87E-21 | 0.35 | 0.28 | 0.07 | 1.26 |
| 4 | As3mt | 0.73 | 9.27E-26 | 1.93E-21 | 0.43 | 0.36 | 0.06 | 1.18 |
| 4 | Lman1 | 0.30 | 9.53E-26 | 1.98E-21 | 0.88 | 0.86 | 0.02 | 1.02 |
| 4 | Vps13a | 0.84 | 1.24E-25 | 2.58E-21 | 0.40 | 0.34 | 0.06 | 1.19 |
| 4 | Fam217b | 1.38 | 1.36E-25 | 2.83E-21 | 0.16 | 0.09 | 0.07 | 1.79 |
| 4 | Dap | 0.50 | 1.79E-25 | 3.73E-21 | 0.60 | 0.57 | 0.03 | 1.06 |
| 4 | L1cam | 1.44 | 1.79E-25 | 3.73E-21 | 0.14 | 0.08 | 0.07 | 1.89 |
| 4 | Pnisr | 0.26 | 1.87E-25 | 3.88E-21 | 0.93 | 0.92 | 0.02 | 1.02 |
| 4 | Cir1 | 0.45 | 2.58E-25 | 5.37E-21 | 0.64 | 0.63 | 0.01 | 1.01 |
| 4 | Elf2 | 0.43 | 2.93E-25 | 6.10E-21 | 0.67 | 0.67 | 0.01 | 1.01 |
| 4 | Gnb4 | 0.77 | 3.67E-25 | 7.63E-21 | 0.37 | 0.30 | 0.08 | 1.26 |
| 4 | Ednra | 0.68 | 3.82E-25 | 7.94E-21 | 0.42 | 0.33 | 0.08 | 1.25 |
| 4 | Ddx17 | 0.28 | 3.87E-25 | 8.04E-21 | 0.86 | 0.87 | -0.01 | 0.99 |
| 4 | Tnrc6a | 0.34 | 3.98E-25 | 8.29E-21 | 0.79 | 0.78 | 0.01 | 1.01 |
| 4 | Nbea | 0.60 | 5.27E-25 | 1.10E-20 | 0.50 | 0.44 | 0.06 | 1.13 |
| 4 | Sacs | 1.01 | 7.04E-25 | 1.46E-20 | 0.25 | 0.17 | 0.08 | 1.47 |
| 4 | Nrip1 | 0.28 | 7.95E-25 | 1.65E-20 | 0.79 | 0.75 | 0.03 | 1.04 |
| 4 | Laptm4b | 0.56 | 9.64E-25 | 2.00E-20 | 0.58 | 0.55 | 0.02 | 1.05 |
| 4 | Baiap2l2 | 1.25 | 1.38E-24 | 2.87E-20 | 0.18 | 0.11 | 0.07 | 1.66 |
| 4 | Cox20 | 0.46 | 1.65E-24 | 3.43E-20 | 0.67 | 0.67 | 0.00 | 1.01 |
| 4 | Ergic3 | 0.35 | 2.43E-24 | 5.05E-20 | 0.77 | 0.77 | 0.00 | 1.00 |

**Supplementary Table 1. Marker genes for the original clusters of integrated clustering of E11.5 control and NotchGOF λ-junction cells.**

| | | | | | | | | |
|---|---|---|---|---|---|---|---|---|
| 4 | Igf1r | 0.30 | 2.46E-24 | 5.12E-20 | 0.79 | 0.77 | 0.02 | 1.03 |
| 4 | Rab34 | 0.36 | 2.58E-24 | 5.36E-20 | 0.75 | 0.72 | 0.03 | 1.04 |
| 4 | Sulf2 | 0.34 | 2.65E-24 | 5.50E-20 | 0.72 | 0.66 | 0.06 | 1.09 |
| 4 | Rasl11b | 0.65 | 2.79E-24 | 5.81E-20 | 0.53 | 0.48 | 0.05 | 1.10 |
| 4 | Aldh2 | 0.34 | 3.00E-24 | 6.24E-20 | 0.78 | 0.78 | 0.00 | 1.00 |
| 4 | Sec62 | 0.26 | 3.04E-24 | 6.32E-20 | 0.93 | 0.94 | 0.00 | 1.00 |
| 4 | Spry2 | 0.87 | 3.14E-24 | 6.53E-20 | 0.36 | 0.29 | 0.08 | 1.26 |
| 4 | Tab2 | 0.47 | 3.40E-24 | 7.08E-20 | 0.67 | 0.68 | -0.01 | 0.99 |
| 4 | Bnip3l | 0.25 | 3.80E-24 | 7.91E-20 | 0.93 | 0.93 | 0.00 | 1.00 |
| 4 | Sdc2 | 0.76 | 4.03E-24 | 8.37E-20 | 0.42 | 0.36 | 0.06 | 1.16 |
| 4 | Psme2 | 0.34 | 4.68E-24 | 9.73E-20 | 0.76 | 0.76 | 0.00 | 1.00 |
| 4 | 2900097C17Rik | 0.35 | 4.98E-24 | 1.04E-19 | 0.75 | 0.74 | 0.01 | 1.01 |
| 4 | Cyp26b1 | 0.95 | 7.25E-24 | 1.51E-19 | 0.20 | 0.12 | 0.08 | 1.64 |
| 4 | Gstm5 | 0.26 | 8.47E-24 | 1.76E-19 | 0.89 | 0.88 | 0.00 | 1.00 |
| 4 | Arl1 | 0.28 | 9.65E-24 | 2.01E-19 | 0.88 | 0.89 | -0.01 | 0.99 |
| 4 | Rbm5 | 0.31 | 1.17E-23 | 2.43E-19 | 0.83 | 0.82 | 0.01 | 1.01 |
| 4 | H13 | 0.46 | 1.48E-23 | 3.07E-19 | 0.64 | 0.64 | 0.00 | 1.01 |
| 4 | Hsd17b11 | 0.95 | 1.49E-23 | 3.11E-19 | 0.28 | 0.20 | 0.08 | 1.39 |
| 4 | Zcchc3 | 0.35 | 1.64E-23 | 3.42E-19 | 0.73 | 0.71 | 0.01 | 1.02 |
| 4 | Znrd1 | 0.30 | 1.69E-23 | 3.52E-19 | 0.82 | 0.82 | 0.00 | 1.00 |
| 4 | Rbfox2 | 0.33 | 2.24E-23 | 4.66E-19 | 0.78 | 0.77 | 0.01 | 1.02 |
| 4 | Selenbp1 | 1.45 | 2.44E-23 | 5.08E-19 | 0.14 | 0.08 | 0.06 | 1.80 |
| 4 | Plin3 | 0.48 | 2.45E-23 | 5.09E-19 | 0.58 | 0.55 | 0.03 | 1.05 |
| 4 | Plpp3 | 0.47 | 4.40E-23 | 9.16E-19 | 0.70 | 0.66 | 0.04 | 1.06 |
| 4 | Itm2a | 0.35 | 5.87E-23 | 1.22E-18 | 0.61 | 0.54 | 0.06 | 1.12 |
| 4 | Pmaip1 | 0.98 | 6.75E-23 | 1.40E-18 | 0.32 | 0.24 | 0.08 | 1.35 |
| 4 | Ets2 | 0.85 | 6.82E-23 | 1.42E-18 | 0.32 | 0.25 | 0.07 | 1.30 |
| 4 | Praf2 | 0.48 | 6.90E-23 | 1.43E-18 | 0.56 | 0.53 | 0.03 | 1.06 |
| 4 | Ufc1 | 0.30 | 7.24E-23 | 1.51E-18 | 0.84 | 0.83 | 0.00 | 1.00 |
| 4 | Ccdc90b | 0.52 | 9.83E-23 | 2.04E-18 | 0.57 | 0.55 | 0.02 | 1.04 |
| 4 | Bin1 | 0.55 | 1.12E-22 | 2.32E-18 | 0.64 | 0.62 | 0.02 | 1.04 |
| 4 | Maml3 | 0.70 | 1.20E-22 | 2.51E-18 | 0.45 | 0.40 | 0.05 | 1.13 |
| 4 | Pja1 | 0.41 | 1.29E-22 | 2.69E-18 | 0.69 | 0.68 | 0.01 | 1.02 |
| 4 | Ptprs | 0.38 | 1.50E-22 | 3.12E-18 | 0.71 | 0.69 | 0.02 | 1.03 |
| 4 | Fam171b | 0.81 | 1.50E-22 | 3.13E-18 | 0.32 | 0.24 | 0.08 | 1.32 |
| 4 | Shd | 1.04 | 1.53E-22 | 3.17E-18 | 0.24 | 0.17 | 0.07 | 1.45 |
| 4 | Txndc15 | 0.45 | 1.80E-22 | 3.74E-18 | 0.62 | 0.60 | 0.02 | 1.03 |
| 4 | Golgb1 | 0.38 | 2.00E-22 | 4.16E-18 | 0.75 | 0.74 | 0.01 | 1.01 |
| 4 | Tsc22d3 | 0.50 | 2.09E-22 | 4.35E-18 | 0.56 | 0.53 | 0.03 | 1.06 |
| 4 | Pde4a | 1.49 | 2.20E-22 | 4.58E-18 | 0.11 | 0.06 | 0.06 | 1.98 |
| 4 | Ift20 | 0.35 | 2.88E-22 | 6.00E-18 | 0.75 | 0.77 | -0.03 | 0.97 |
| 4 | Fermt2 | 0.29 | 3.46E-22 | 7.20E-18 | 0.75 | 0.72 | 0.02 | 1.03 |
| 4 | Fxr1 | 0.31 | 3.61E-22 | 7.51E-18 | 0.84 | 0.85 | -0.02 | 0.98 |
| 4 | Nfatc4 | 0.50 | 4.03E-22 | 8.38E-18 | 0.59 | 0.55 | 0.04 | 1.07 |
| 4 | Cops6 | 0.26 | 4.34E-22 | 9.02E-18 | 0.87 | 0.88 | 0.00 | 1.00 |
| 4 | Eno3 | 0.62 | 5.19E-22 | 1.08E-17 | 0.44 | 0.37 | 0.07 | 1.18 |
| 4 | Washc2 | 0.53 | 5.57E-22 | 1.16E-17 | 0.55 | 0.52 | 0.03 | 1.07 |
| 4 | Traf4 | 0.40 | 5.68E-22 | 1.18E-17 | 0.65 | 0.63 | 0.02 | 1.03 |
| 4 | Cenpv | 0.46 | 6.27E-22 | 1.30E-17 | 0.56 | 0.50 | 0.06 | 1.12 |
| 4 | Tcea1 | 0.27 | 6.35E-22 | 1.32E-17 | 0.86 | 0.88 | -0.01 | 0.99 |
| 4 | Sri | 0.34 | 7.00E-22 | 1.46E-17 | 0.77 | 0.77 | 0.00 | 1.00 |
| 4 | Hivep2 | 1.40 | 1.19E-21 | 2.46E-17 | 0.13 | 0.07 | 0.06 | 1.82 |
| 4 | 4931406C07Rik | 0.85 | 1.37E-21 | 2.85E-17 | 0.32 | 0.26 | 0.07 | 1.27 |
| 4 | Gfpt2 | 1.06 | 1.69E-21 | 3.51E-17 | 0.25 | 0.18 | 0.07 | 1.40 |
| 4 | Efna2 | 0.35 | 1.69E-21 | 3.51E-17 | 0.73 | 0.71 | 0.02 | 1.03 |
| 4 | Calu | 0.38 | 1.70E-21 | 3.54E-17 | 0.70 | 0.71 | 0.00 | 0.99 |

**Supplementary Table 1. Marker genes for the original clusters of integrated clustering of E11.5 control and NotchGOF λ-junction cells.**

| | | | | | | | | |
|---|---|---|---|---|---|---|---|---|
| 4 | Bcl9 | 0.40 | 2.52E-21 | 5.25E-17 | 0.69 | 0.69 | 0.00 | 1.00 |
| 4 | Polr3gl | 0.76 | 2.76E-21 | 5.74E-17 | 0.37 | 0.31 | 0.06 | 1.19 |
| 4 | Ssr3 | 0.27 | 3.88E-21 | 8.07E-17 | 0.88 | 0.88 | 0.00 | 1.00 |
| 4 | Dnm1 | 0.71 | 4.23E-21 | 8.79E-17 | 0.39 | 0.32 | 0.07 | 1.20 |
| 4 | Jph1 | 1.55 | 5.43E-21 | 1.13E-16 | 0.12 | 0.07 | 0.06 | 1.85 |
| 4 | Hs2st1 | 0.51 | 7.25E-21 | 1.51E-16 | 0.58 | 0.57 | 0.02 | 1.03 |
| 4 | Gpr155 | 1.56 | 8.27E-21 | 1.72E-16 | 0.11 | 0.05 | 0.05 | 1.94 |
| 4 | Gtf3c6 | 0.26 | 8.82E-21 | 1.83E-16 | 0.85 | 0.86 | -0.02 | 0.98 |
| 4 | Ttc19 | 0.64 | 9.52E-21 | 1.98E-16 | 0.45 | 0.40 | 0.04 | 1.10 |
| 4 | Cnrip1 | 1.17 | 1.01E-20 | 2.10E-16 | 0.20 | 0.14 | 0.07 | 1.50 |
| 4 | Sgpp1 | 1.32 | 1.46E-20 | 3.03E-16 | 0.16 | 0.10 | 0.06 | 1.63 |
| 4 | Glg1 | 0.37 | 1.51E-20 | 3.14E-16 | 0.72 | 0.71 | 0.01 | 1.01 |
| 4 | Dhrs7 | 1.22 | 1.58E-20 | 3.29E-16 | 0.19 | 0.12 | 0.07 | 1.53 |
| 4 | Col25a1 | 0.80 | 1.61E-20 | 3.34E-16 | 0.27 | 0.19 | 0.08 | 1.40 |
| 4 | Gstm7 | 0.74 | 2.73E-20 | 5.68E-16 | 0.34 | 0.28 | 0.07 | 1.25 |
| 4 | Ptgfrn | 0.72 | 3.52E-20 | 7.31E-16 | 0.39 | 0.33 | 0.06 | 1.17 |
| 4 | Prrc1 | 0.63 | 4.87E-20 | 1.01E-15 | 0.47 | 0.44 | 0.04 | 1.08 |
| 4 | Capn6 | 0.47 | 5.43E-20 | 1.13E-15 | 0.58 | 0.55 | 0.04 | 1.07 |
| 4 | Lama4 | 0.95 | 5.57E-20 | 1.16E-15 | 0.20 | 0.13 | 0.07 | 1.54 |
| 4 | Zfp318 | 0.58 | 5.95E-20 | 1.24E-15 | 0.50 | 0.47 | 0.03 | 1.07 |
| 4 | Srpk2 | 0.36 | 9.29E-20 | 1.93E-15 | 0.71 | 0.71 | 0.00 | 1.00 |
| 4 | Trpm7 | 0.39 | 9.44E-20 | 1.96E-15 | 0.69 | 0.69 | 0.00 | 1.00 |
| 4 | Mir99ahg | 0.49 | 1.09E-19 | 2.27E-15 | 0.53 | 0.48 | 0.05 | 1.09 |
| 4 | Epc2 | 0.47 | 1.63E-19 | 3.40E-15 | 0.63 | 0.62 | 0.01 | 1.01 |
| 4 | 9130024F11Rik | 0.71 | 1.70E-19 | 3.53E-15 | 0.30 | 0.21 | 0.08 | 1.38 |
| 4 | Decr1 | 0.63 | 2.03E-19 | 4.22E-15 | 0.45 | 0.41 | 0.04 | 1.09 |
| 4 | Olfml1 | 1.10 | 2.47E-19 | 5.14E-15 | 0.18 | 0.11 | 0.06 | 1.56 |
| 4 | Zcchc24 | 0.70 | 2.55E-19 | 5.30E-15 | 0.35 | 0.29 | 0.07 | 1.23 |
| 4 | Tmbim6 | 0.25 | 2.78E-19 | 5.77E-15 | 0.84 | 0.84 | -0.01 | 0.99 |
| 4 | Iah1 | 0.52 | 2.85E-19 | 5.93E-15 | 0.54 | 0.52 | 0.02 | 1.03 |
| 4 | Msrb1 | 0.51 | 3.17E-19 | 6.59E-15 | 0.51 | 0.49 | 0.02 | 1.04 |
| 4 | Ubn1 | 0.32 | 4.32E-19 | 8.98E-15 | 0.76 | 0.77 | -0.01 | 0.99 |
| 4 | Snhg8 | 0.30 | 5.35E-19 | 1.11E-14 | 0.78 | 0.80 | -0.02 | 0.97 |
| 4 | Plin2 | 0.39 | 5.65E-19 | 1.17E-14 | 0.56 | 0.53 | 0.03 | 1.05 |
| 4 | Unc119 | 0.28 | 5.72E-19 | 1.19E-14 | 0.79 | 0.78 | 0.01 | 1.01 |
| 4 | Lmcd1 | 1.49 | 6.00E-19 | 1.25E-14 | 0.14 | 0.09 | 0.06 | 1.67 |
| 4 | Smap1 | 0.26 | 7.46E-19 | 1.55E-14 | 0.86 | 0.87 | -0.01 | 0.99 |
| 4 | Kdelr2 | 0.27 | 7.49E-19 | 1.56E-14 | 0.84 | 0.83 | 0.01 | 1.01 |
| 4 | Hspa13 | 0.55 | 7.70E-19 | 1.60E-14 | 0.52 | 0.50 | 0.02 | 1.04 |
| 4 | Tnks2 | 0.28 | 1.41E-18 | 2.92E-14 | 0.81 | 0.81 | 0.00 | 1.01 |
| 4 | Fam102b | 0.95 | 1.42E-18 | 2.95E-14 | 0.26 | 0.20 | 0.06 | 1.33 |
| 4 | Lox | 1.28 | 1.80E-18 | 3.75E-14 | 0.12 | 0.06 | 0.05 | 1.81 |
| 4 | Ankrd12 | 0.41 | 1.85E-18 | 3.86E-14 | 0.56 | 0.53 | 0.04 | 1.07 |
| 4 | Dhrs3 | 0.74 | 2.12E-18 | 4.40E-14 | 0.36 | 0.29 | 0.07 | 1.22 |
| 4 | Ccdc28b | 0.63 | 2.31E-18 | 4.81E-14 | 0.42 | 0.37 | 0.05 | 1.13 |
| 4 | Tmem263 | 0.56 | 2.59E-18 | 5.39E-14 | 0.50 | 0.47 | 0.03 | 1.06 |
| 4 | Tbc1d15 | 0.66 | 2.85E-18 | 5.93E-14 | 0.52 | 0.51 | 0.00 | 1.01 |
| 4 | Tcf25 | 0.27 | 2.95E-18 | 6.13E-14 | 0.82 | 0.83 | -0.01 | 0.98 |
| 4 | Golim4 | 0.43 | 3.38E-18 | 7.04E-14 | 0.62 | 0.61 | 0.01 | 1.02 |
| 4 | Tmem223 | 0.28 | 3.42E-18 | 7.12E-14 | 0.83 | 0.84 | -0.01 | 0.98 |
| 4 | Angptl4 | 0.64 | 3.54E-18 | 7.36E-14 | 0.39 | 0.33 | 0.06 | 1.18 |
| 4 | Ikbip | 0.78 | 3.92E-18 | 8.15E-14 | 0.35 | 0.30 | 0.05 | 1.18 |
| 4 | Atxn1 | 1.14 | 3.99E-18 | 8.30E-14 | 0.15 | 0.09 | 0.06 | 1.65 |
| 4 | Tns1 | 0.70 | 4.13E-18 | 8.60E-14 | 0.36 | 0.30 | 0.06 | 1.19 |
| 4 | Mnat1 | 0.35 | 4.18E-18 | 8.69E-14 | 0.68 | 0.69 | -0.01 | 0.98 |
| 4 | Psme1 | 0.28 | 4.40E-18 | 9.15E-14 | 0.76 | 0.76 | -0.01 | 0.99 |

**Supplementary Table 1. Marker genes for the original clusters of integrated clustering of E11.5 control and NotchGOF λ-junction cells.**

| | | | | | | | | |
|---|---|---|---|---|---|---|---|---|
| 4 | 2610318N02Rik | 0.42 | 5.28E-18 | 1.10E-13 | 0.61 | 0.60 | 0.01 | 1.02 |
| 4 | Ptpn9 | 0.50 | 5.44E-18 | 1.13E-13 | 0.52 | 0.49 | 0.03 | 1.06 |
| 4 | Sash1 | 0.69 | 6.50E-18 | 1.35E-13 | 0.43 | 0.40 | 0.04 | 1.09 |
| 4 | Camk2d | 0.84 | 6.63E-18 | 1.38E-13 | 0.26 | 0.19 | 0.07 | 1.35 |
| 4 | 4632427E13Rik | 0.39 | 7.81E-18 | 1.62E-13 | 0.65 | 0.65 | 0.00 | 1.01 |
| 4 | Yaf2 | 0.35 | 8.11E-18 | 1.69E-13 | 0.70 | 0.70 | -0.01 | 0.99 |
| 4 | Pcmtd2 | 0.63 | 8.75E-18 | 1.82E-13 | 0.42 | 0.37 | 0.04 | 1.11 |
| 4 | Rb1cc1 | 0.37 | 9.66E-18 | 2.01E-13 | 0.65 | 0.66 | 0.00 | 0.99 |
| 4 | Wbp1 | 0.25 | 9.93E-18 | 2.06E-13 | 0.76 | 0.77 | -0.01 | 0.98 |
| 4 | Trim44 | 0.31 | 1.06E-17 | 2.21E-13 | 0.72 | 0.71 | 0.01 | 1.01 |
| 4 | Lrpap1 | 0.48 | 1.08E-17 | 2.25E-13 | 0.52 | 0.49 | 0.02 | 1.05 |
| 4 | Tmem119 | 0.47 | 1.25E-17 | 2.60E-13 | 0.51 | 0.45 | 0.06 | 1.13 |
| 4 | Mpzl1 | 0.40 | 1.60E-17 | 3.33E-13 | 0.64 | 0.63 | 0.00 | 1.00 |
| 4 | 9530068E07Rik | 0.31 | 1.69E-17 | 3.51E-13 | 0.73 | 0.74 | -0.01 | 0.99 |
| 4 | Casd1 | 0.52 | 1.82E-17 | 3.78E-13 | 0.53 | 0.50 | 0.03 | 1.05 |
| 4 | Speg | 0.99 | 1.93E-17 | 4.02E-13 | 0.22 | 0.16 | 0.06 | 1.39 |
| 4 | Eid2 | 0.34 | 2.65E-17 | 5.51E-13 | 0.66 | 0.65 | 0.00 | 1.00 |
| 4 | Reck | 0.81 | 3.27E-17 | 6.80E-13 | 0.30 | 0.24 | 0.06 | 1.25 |
| 4 | Itgav | 0.76 | 3.73E-17 | 7.75E-13 | 0.33 | 0.27 | 0.06 | 1.21 |
| 4 | Setd5 | 0.33 | 3.85E-17 | 8.01E-13 | 0.70 | 0.72 | -0.01 | 0.98 |
| 4 | Mllt3 | 0.35 | 3.86E-17 | 8.03E-13 | 0.66 | 0.66 | 0.00 | 1.00 |
| 4 | Kdelr3 | 1.40 | 4.05E-17 | 8.41E-13 | 0.12 | 0.07 | 0.05 | 1.71 |
| 4 | Bcl2l11 | 0.64 | 4.48E-17 | 9.32E-13 | 0.40 | 0.35 | 0.05 | 1.14 |
| 4 | Pip4k2b | 0.92 | 5.15E-17 | 1.07E-12 | 0.27 | 0.22 | 0.06 | 1.26 |
| 4 | Ube3a | 0.27 | 5.46E-17 | 1.14E-12 | 0.84 | 0.84 | -0.01 | 0.99 |
| 4 | Tln1 | 0.27 | 5.93E-17 | 1.23E-12 | 0.74 | 0.74 | 0.00 | 1.00 |
| 4 | Naxe | 0.29 | 6.47E-17 | 1.35E-12 | 0.71 | 0.72 | -0.02 | 0.97 |
| 4 | Nucb1 | 0.53 | 7.57E-17 | 1.57E-12 | 0.53 | 0.52 | 0.01 | 1.02 |
| 4 | Foxp4 | 0.53 | 8.69E-17 | 1.81E-12 | 0.46 | 0.42 | 0.04 | 1.09 |
| 4 | Timm22 | 0.30 | 1.01E-16 | 2.09E-12 | 0.76 | 0.77 | -0.01 | 0.99 |
| 4 | Slc35a1 | 0.55 | 1.02E-16 | 2.13E-12 | 0.49 | 0.47 | 0.02 | 1.04 |
| 4 | Adat2 | 0.75 | 1.17E-16 | 2.42E-12 | 0.35 | 0.30 | 0.05 | 1.16 |
| 4 | Ing1 | 0.33 | 1.19E-16 | 2.47E-12 | 0.73 | 0.73 | 0.00 | 0.99 |
| 4 | Hadh | 0.29 | 1.25E-16 | 2.60E-12 | 0.76 | 0.78 | -0.02 | 0.98 |
| 4 | Myo1b | 0.30 | 1.55E-16 | 3.21E-12 | 0.74 | 0.73 | 0.01 | 1.02 |
| 4 | Sec31a | 0.45 | 1.64E-16 | 3.41E-12 | 0.56 | 0.55 | 0.00 | 1.00 |
| 4 | Gpx8 | 0.30 | 1.75E-16 | 3.63E-12 | 0.72 | 0.71 | 0.01 | 1.01 |
| 4 | Lypla1 | 0.53 | 1.75E-16 | 3.64E-12 | 0.48 | 0.46 | 0.02 | 1.05 |
| 4 | Clk1 | 0.38 | 2.00E-16 | 4.15E-12 | 0.67 | 0.68 | -0.02 | 0.98 |
| 4 | 4932438A13Rik | 0.70 | 2.03E-16 | 4.22E-12 | 0.39 | 0.35 | 0.04 | 1.11 |
| 4 | Xpa | 0.55 | 2.19E-16 | 4.55E-12 | 0.47 | 0.44 | 0.03 | 1.06 |
| 4 | Ttc39b | 1.38 | 2.24E-16 | 4.65E-12 | 0.11 | 0.06 | 0.05 | 1.75 |
| 4 | P3h4 | 0.62 | 2.79E-16 | 5.81E-12 | 0.39 | 0.35 | 0.04 | 1.12 |
| 4 | Mgp | 0.98 | 4.12E-16 | 8.57E-12 | 0.12 | 0.07 | 0.05 | 1.75 |
| 4 | P3h1 | 1.00 | 4.21E-16 | 8.76E-12 | 0.20 | 0.14 | 0.06 | 1.40 |
| 4 | Loxl3 | 1.30 | 4.52E-16 | 9.40E-12 | 0.12 | 0.07 | 0.05 | 1.72 |
| 4 | Tfpi | 0.88 | 4.76E-16 | 9.90E-12 | 0.17 | 0.11 | 0.06 | 1.55 |
| 4 | Arl2bp | 0.26 | 5.01E-16 | 1.04E-11 | 0.79 | 0.79 | 0.00 | 1.00 |
| 4 | Copa | 0.36 | 7.42E-16 | 1.54E-11 | 0.65 | 0.66 | -0.01 | 0.98 |
| 4 | Mafb | 0.97 | 7.43E-16 | 1.55E-11 | 0.28 | 0.22 | 0.06 | 1.29 |
| 4 | Trim8 | 0.69 | 7.49E-16 | 1.56E-11 | 0.35 | 0.30 | 0.05 | 1.17 |
| 4 | Rbm46 | 1.11 | 7.78E-16 | 1.62E-11 | 0.13 | 0.08 | 0.05 | 1.65 |
| 4 | Inpp1 | 1.06 | 7.84E-16 | 1.63E-11 | 0.22 | 0.16 | 0.06 | 1.36 |
| 4 | Gpbp1 | 0.27 | 8.47E-16 | 1.76E-11 | 0.77 | 0.79 | -0.02 | 0.97 |
| 4 | Uba7 | 1.11 | 8.87E-16 | 1.84E-11 | 0.16 | 0.10 | 0.05 | 1.52 |
| 4 | Prrg1 | 0.95 | 9.97E-16 | 2.07E-11 | 0.25 | 0.19 | 0.06 | 1.30 |

**Supplementary Table 1. Marker genes for the original clusters of integrated clustering of E11.5 control and NotchGOF λ-junction cells.**

| | | | | | | | | |
|---|---|---|---|---|---|---|---|---|
| 4 | Sdf2 | 0.41 | 1.02E-15 | 2.12E-11 | 0.57 | 0.56 | 0.01 | 1.03 |
| 4 | P4ha2 | 0.74 | 1.02E-15 | 2.13E-11 | 0.32 | 0.27 | 0.05 | 1.19 |
| 4 | Tm9sf4 | 0.42 | 1.07E-15 | 2.22E-11 | 0.58 | 0.58 | 0.00 | 1.00 |
| 4 | Slc16a2 | 0.97 | 1.13E-15 | 2.36E-11 | 0.21 | 0.15 | 0.06 | 1.40 |
| 4 | Slc18b1 | 1.04 | 1.15E-15 | 2.40E-11 | 0.18 | 0.12 | 0.06 | 1.48 |
| 4 | Acot1 | 0.58 | 1.32E-15 | 2.74E-11 | 0.40 | 0.35 | 0.05 | 1.14 |
| 4 | Klf4 | 0.83 | 1.41E-15 | 2.93E-11 | 0.27 | 0.21 | 0.06 | 1.30 |
| 4 | Dync2h1 | 0.64 | 1.52E-15 | 3.16E-11 | 0.42 | 0.38 | 0.04 | 1.11 |
| 4 | Tln2 | 0.46 | 1.80E-15 | 3.74E-11 | 0.59 | 0.59 | 0.01 | 1.02 |
| 4 | Ntn3 | 0.93 | 1.97E-15 | 4.10E-11 | 0.21 | 0.16 | 0.06 | 1.37 |
| 4 | Tnrc18 | 0.42 | 2.17E-15 | 4.52E-11 | 0.57 | 0.56 | 0.01 | 1.02 |
| 4 | Gsto1 | 0.34 | 2.18E-15 | 4.53E-11 | 0.65 | 0.65 | 0.00 | 1.00 |
| 4 | Lysmd2 | 0.83 | 2.68E-15 | 5.58E-11 | 0.27 | 0.21 | 0.06 | 1.26 |
| 4 | Chd6 | 0.32 | 2.72E-15 | 5.66E-11 | 0.71 | 0.73 | -0.02 | 0.97 |
| 4 | Tmem134 | 0.28 | 2.90E-15 | 6.02E-11 | 0.72 | 0.75 | -0.03 | 0.96 |
| 4 | Tm2d2 | 0.32 | 3.04E-15 | 6.33E-11 | 0.72 | 0.73 | -0.01 | 0.99 |
| 4 | Plekha5 | 0.53 | 3.43E-15 | 7.13E-11 | 0.48 | 0.46 | 0.02 | 1.05 |
| 4 | Mcc | 1.00 | 3.67E-15 | 7.63E-11 | 0.18 | 0.13 | 0.06 | 1.45 |
| 4 | Satb2 | 0.33 | 4.03E-15 | 8.39E-11 | 0.33 | 0.25 | 0.09 | 1.34 |
| 4 | Tox3 | 0.75 | 4.11E-15 | 8.54E-11 | 0.29 | 0.23 | 0.06 | 1.26 |
| 4 | Mocs2 | 0.40 | 4.23E-15 | 8.81E-11 | 0.59 | 0.61 | -0.01 | 0.98 |
| 4 | Adamts6 | 1.26 | 5.60E-15 | 1.16E-10 | 0.11 | 0.06 | 0.05 | 1.70 |
| 4 | Smim14 | 0.48 | 5.76E-15 | 1.20E-10 | 0.45 | 0.41 | 0.03 | 1.08 |
| 4 | Ralb | 0.41 | 6.26E-15 | 1.30E-10 | 0.46 | 0.43 | 0.03 | 1.08 |
| 4 | Eci2 | 0.30 | 6.32E-15 | 1.32E-10 | 0.68 | 0.69 | -0.01 | 0.99 |
| 4 | Arcn1 | 0.35 | 6.60E-15 | 1.37E-10 | 0.64 | 0.66 | -0.01 | 0.98 |
| 4 | Copb2 | 0.26 | 6.64E-15 | 1.38E-10 | 0.81 | 0.82 | -0.01 | 0.99 |
| 4 | Nynrin | 0.41 | 7.36E-15 | 1.53E-10 | 0.56 | 0.55 | 0.01 | 1.02 |
| 4 | Hmg20a | 0.56 | 7.50E-15 | 1.56E-10 | 0.45 | 0.42 | 0.03 | 1.06 |
| 4 | Wwc2 | 0.39 | 1.06E-14 | 2.21E-10 | 0.59 | 0.58 | 0.01 | 1.01 |
| 4 | Tbrg1 | 0.27 | 1.08E-14 | 2.25E-10 | 0.80 | 0.83 | -0.03 | 0.97 |
| 4 | Bad | 0.44 | 1.10E-14 | 2.28E-10 | 0.54 | 0.53 | 0.01 | 1.01 |
| 4 | Phactr3 | 1.24 | 1.27E-14 | 2.64E-10 | 0.12 | 0.07 | 0.05 | 1.66 |
| 4 | Srp72 | 0.25 | 1.33E-14 | 2.77E-10 | 0.80 | 0.83 | -0.03 | 0.96 |
| 4 | Fam92a | 0.27 | 1.79E-14 | 3.71E-10 | 0.75 | 0.76 | -0.01 | 0.98 |
| 4 | Pabpc4l | 0.57 | 1.92E-14 | 3.98E-10 | 0.42 | 0.38 | 0.03 | 1.09 |
| 4 | Podxl2 | 0.52 | 2.01E-14 | 4.17E-10 | 0.45 | 0.42 | 0.03 | 1.08 |
| 4 | Jak1 | 0.38 | 2.21E-14 | 4.60E-10 | 0.60 | 0.61 | 0.00 | 0.99 |
| 4 | Crot | 0.56 | 2.63E-14 | 5.47E-10 | 0.44 | 0.42 | 0.02 | 1.06 |
| 4 | Dennd5b | 0.55 | 2.66E-14 | 5.52E-10 | 0.43 | 0.39 | 0.04 | 1.10 |
| 4 | Vcl | 0.61 | 2.67E-14 | 5.56E-10 | 0.42 | 0.39 | 0.03 | 1.08 |
| 4 | Pkdcc | 0.37 | 3.07E-14 | 6.39E-10 | 0.39 | 0.33 | 0.06 | 1.20 |
| 4 | Zbed3 | 0.45 | 5.81E-14 | 1.21E-09 | 0.56 | 0.56 | -0.01 | 0.99 |
| 4 | Lpp | 0.41 | 5.95E-14 | 1.24E-09 | 0.57 | 0.56 | 0.01 | 1.02 |
| 4 | Slc25a36 | 0.51 | 6.14E-14 | 1.28E-09 | 0.46 | 0.43 | 0.02 | 1.05 |
| 4 | Smyd3 | 0.82 | 6.17E-14 | 1.28E-09 | 0.28 | 0.24 | 0.05 | 1.20 |
| 4 | Chd9 | 0.44 | 6.54E-14 | 1.36E-09 | 0.54 | 0.53 | 0.01 | 1.02 |
| 4 | Cul7 | 0.60 | 7.05E-14 | 1.47E-09 | 0.39 | 0.36 | 0.03 | 1.09 |
| 4 | Ttc14 | 0.29 | 7.15E-14 | 1.49E-09 | 0.72 | 0.74 | -0.02 | 0.97 |
| 4 | Fkbp9 | 0.55 | 7.41E-14 | 1.54E-09 | 0.46 | 0.44 | 0.01 | 1.03 |
| 4 | Ddhd2 | 0.60 | 7.60E-14 | 1.58E-09 | 0.41 | 0.38 | 0.03 | 1.08 |
| 4 | Ugp2 | 0.43 | 7.60E-14 | 1.58E-09 | 0.51 | 0.50 | 0.01 | 1.01 |
| 4 | Pxdn | 0.45 | 7.97E-14 | 1.66E-09 | 0.53 | 0.52 | 0.01 | 1.03 |
| 4 | Pcdh11x | 0.87 | 8.58E-14 | 1.78E-09 | 0.16 | 0.10 | 0.05 | 1.52 |
| 4 | Lactb | 0.64 | 9.60E-14 | 2.00E-09 | 0.38 | 0.34 | 0.04 | 1.12 |
| 4 | Rgl1 | 0.90 | 1.26E-13 | 2.62E-09 | 0.20 | 0.15 | 0.05 | 1.36 |

**Supplementary Table 1. Marker genes for the original clusters of integrated clustering of E11.5 control and NotchGOF λ-junction cells.**

| | | | | | | | | |
|---|---|---|---|---|---|---|---|---|
| 4 | Bend4 | 1.10 | 1.34E-13 | 2.80E-09 | 0.16 | 0.11 | 0.05 | 1.44 |
| 4 | Hexa | 0.52 | 1.61E-13 | 3.35E-09 | 0.36 | 0.31 | 0.05 | 1.14 |
| 4 | Pnpla8 | 0.45 | 1.76E-13 | 3.67E-09 | 0.51 | 0.50 | 0.01 | 1.02 |
| 4 | Snx17 | 0.31 | 2.06E-13 | 4.29E-09 | 0.65 | 0.68 | -0.03 | 0.96 |
| 4 | Raet1e | 0.75 | 2.28E-13 | 4.75E-09 | 0.28 | 0.23 | 0.05 | 1.22 |
| 4 | Aebp1 | 0.91 | 2.53E-13 | 5.26E-09 | 0.25 | 0.20 | 0.05 | 1.25 |
| 4 | Acads | 0.38 | 2.54E-13 | 5.28E-09 | 0.60 | 0.61 | 0.00 | 0.99 |
| 4 | Plscr3 | 0.41 | 2.58E-13 | 5.37E-09 | 0.56 | 0.56 | 0.00 | 0.99 |
| 4 | Cul5 | 0.30 | 3.28E-13 | 6.81E-09 | 0.67 | 0.69 | -0.01 | 0.98 |
| 4 | Ophn1 | 0.85 | 4.99E-13 | 1.04E-08 | 0.24 | 0.19 | 0.05 | 1.28 |
| 4 | Unc5b | 0.55 | 5.15E-13 | 1.07E-08 | 0.43 | 0.39 | 0.04 | 1.10 |
| 4 | Taf7 | 0.31 | 5.54E-13 | 1.15E-08 | 0.67 | 0.70 | -0.03 | 0.96 |
| 4 | Tcf7l2 | 0.30 | 5.95E-13 | 1.24E-08 | 0.62 | 0.61 | 0.01 | 1.01 |
| 4 | Rps6ka3 | 0.48 | 7.57E-13 | 1.57E-08 | 0.47 | 0.46 | 0.02 | 1.03 |
| 4 | Krit1 | 0.35 | 8.02E-13 | 1.67E-08 | 0.61 | 0.64 | -0.02 | 0.97 |
| 4 | Sirt2 | 0.46 | 8.05E-13 | 1.67E-08 | 0.50 | 0.51 | -0.01 | 0.98 |
| 4 | Adamts1 | 0.69 | 8.44E-13 | 1.75E-08 | 0.33 | 0.27 | 0.05 | 1.20 |
| 4 | Pkd2 | 0.57 | 8.47E-13 | 1.76E-08 | 0.41 | 0.39 | 0.03 | 1.06 |
| 4 | Dsel | 0.63 | 8.76E-13 | 1.82E-08 | 0.37 | 0.33 | 0.04 | 1.12 |
| 4 | Stx5a | 0.47 | 9.39E-13 | 1.95E-08 | 0.49 | 0.48 | 0.01 | 1.01 |
| 4 | Epb41l1 | 0.99 | 9.98E-13 | 2.07E-08 | 0.19 | 0.14 | 0.05 | 1.35 |
| 4 | Col13a1 | 0.96 | 1.03E-12 | 2.14E-08 | 0.15 | 0.11 | 0.05 | 1.47 |
| 4 | Sspn | 1.17 | 1.04E-12 | 2.17E-08 | 0.11 | 0.07 | 0.04 | 1.61 |
| 4 | Ei24 | 0.31 | 1.27E-12 | 2.63E-08 | 0.67 | 0.71 | -0.03 | 0.96 |
| 4 | Me1 | 0.74 | 1.33E-12 | 2.77E-08 | 0.29 | 0.25 | 0.05 | 1.19 |
| 4 | Dtx3 | 0.47 | 1.39E-12 | 2.90E-08 | 0.48 | 0.48 | 0.01 | 1.01 |
| 4 | Dtnbp1 | 0.50 | 1.55E-12 | 3.21E-08 | 0.44 | 0.42 | 0.02 | 1.05 |
| 4 | Eya4 | 0.52 | 1.59E-12 | 3.30E-08 | 0.34 | 0.28 | 0.06 | 1.20 |
| 4 | Tgfb1i1 | 0.86 | 1.62E-12 | 3.38E-08 | 0.18 | 0.13 | 0.05 | 1.38 |
| 4 | Neo1 | 0.46 | 1.66E-12 | 3.45E-08 | 0.51 | 0.50 | 0.01 | 1.02 |
| 4 | Wsb1 | 0.29 | 1.66E-12 | 3.45E-08 | 0.67 | 0.69 | -0.02 | 0.97 |
| 4 | Frmd6 | 0.42 | 1.73E-12 | 3.60E-08 | 0.52 | 0.49 | 0.03 | 1.06 |
| 4 | Zfc3h1 | 0.65 | 2.06E-12 | 4.28E-08 | 0.39 | 0.36 | 0.03 | 1.08 |
| 4 | Rhod | 0.60 | 2.37E-12 | 4.92E-08 | 0.37 | 0.34 | 0.04 | 1.10 |
| 4 | Gm36988 | 0.94 | 2.44E-12 | 5.07E-08 | 0.19 | 0.14 | 0.05 | 1.36 |
| 4 | Slco3a1 | 0.87 | 2.73E-12 | 5.67E-08 | 0.18 | 0.13 | 0.05 | 1.40 |
| 4 | Egln2 | 0.31 | 3.03E-12 | 6.31E-08 | 0.60 | 0.62 | -0.02 | 0.97 |
| 4 | Dzip1 | 0.57 | 3.58E-12 | 7.45E-08 | 0.37 | 0.33 | 0.04 | 1.11 |
| 4 | Fibin | 0.89 | 3.62E-12 | 7.53E-08 | 0.17 | 0.12 | 0.05 | 1.42 |
| 4 | Astn1 | 0.94 | 3.79E-12 | 7.88E-08 | 0.16 | 0.12 | 0.05 | 1.42 |
| 4 | Foxn3 | 0.26 | 4.29E-12 | 8.92E-08 | 0.68 | 0.69 | -0.01 | 0.99 |
| 4 | Kif3a | 0.31 | 4.66E-12 | 9.69E-08 | 0.61 | 0.62 | -0.01 | 0.99 |
| 4 | Pja2 | 0.40 | 4.76E-12 | 9.90E-08 | 0.55 | 0.55 | -0.01 | 0.99 |
| 4 | Cntln | 0.44 | 4.94E-12 | 1.03E-07 | 0.51 | 0.49 | 0.02 | 1.03 |
| 4 | Sema3a | 0.44 | 4.99E-12 | 1.04E-07 | 0.44 | 0.40 | 0.04 | 1.09 |
| 4 | Snhg6 | 0.25 | 5.47E-12 | 1.14E-07 | 0.73 | 0.75 | -0.03 | 0.97 |
| 4 | Ift81 | 0.47 | 5.75E-12 | 1.20E-07 | 0.47 | 0.46 | 0.01 | 1.03 |
| 4 | Emc2 | 0.35 | 6.38E-12 | 1.33E-07 | 0.58 | 0.60 | -0.01 | 0.98 |
| 4 | Chpf | 0.96 | 6.57E-12 | 1.37E-07 | 0.19 | 0.15 | 0.05 | 1.33 |
| 4 | Rnf130 | 0.37 | 6.85E-12 | 1.42E-07 | 0.56 | 0.57 | 0.00 | 0.99 |
| 4 | Tfg | 0.29 | 7.24E-12 | 1.51E-07 | 0.71 | 0.74 | -0.03 | 0.96 |
| 4 | Ubxn6 | 0.45 | 7.88E-12 | 1.64E-07 | 0.52 | 0.52 | -0.01 | 0.98 |
| 4 | Cyth3 | 0.58 | 8.41E-12 | 1.75E-07 | 0.41 | 0.39 | 0.02 | 1.05 |
| 4 | Cdc42ep4 | 0.45 | 8.50E-12 | 1.77E-07 | 0.46 | 0.45 | 0.01 | 1.03 |
| 4 | Rsu1 | 0.38 | 8.52E-12 | 1.77E-07 | 0.49 | 0.49 | 0.00 | 1.01 |
| 4 | Has2 | 0.88 | 8.65E-12 | 1.80E-07 | 0.24 | 0.19 | 0.05 | 1.25 |

**Supplementary Table 1. Marker genes for the original clusters of integrated clustering of E11.5 control and NotchGOF λ-junction cells.**

| | | | | | | | | |
|---|---|---|---|---|---|---|---|---|
| 4 | St3gal4 | 0.72 | 9.63E-12 | 2.00E-07 | 0.29 | 0.25 | 0.04 | 1.16 |
| 4 | Greb1 | 0.63 | 9.88E-12 | 2.06E-07 | 0.33 | 0.29 | 0.04 | 1.14 |
| 4 | Timp2 | 0.25 | 1.01E-11 | 2.10E-07 | 0.67 | 0.67 | 0.00 | 1.00 |
| 4 | Csk | 0.38 | 1.18E-11 | 2.44E-07 | 0.54 | 0.55 | 0.00 | 0.99 |
| 4 | Angptl2 | 0.83 | 1.34E-11 | 2.79E-07 | 0.23 | 0.18 | 0.05 | 1.25 |
| 4 | Galnt1 | 0.32 | 1.51E-11 | 3.14E-07 | 0.64 | 0.66 | -0.02 | 0.97 |
| 4 | Cd248 | 0.64 | 1.56E-11 | 3.24E-07 | 0.30 | 0.25 | 0.05 | 1.19 |
| 4 | Cd302 | 0.63 | 1.75E-11 | 3.64E-07 | 0.32 | 0.28 | 0.04 | 1.14 |
| 4 | Igdcc4 | 0.92 | 1.78E-11 | 3.71E-07 | 0.17 | 0.12 | 0.05 | 1.39 |
| 4 | Kctd1 | 0.39 | 1.89E-11 | 3.93E-07 | 0.49 | 0.47 | 0.02 | 1.05 |
| 4 | Dyrk2 | 0.54 | 2.56E-11 | 5.33E-07 | 0.40 | 0.38 | 0.02 | 1.06 |
| 4 | Fign | 0.64 | 2.71E-11 | 5.65E-07 | 0.33 | 0.29 | 0.04 | 1.14 |
| 4 | Desi2 | 0.47 | 2.76E-11 | 5.74E-07 | 0.46 | 0.45 | 0.01 | 1.02 |
| 4 | Usp47 | 0.29 | 2.86E-11 | 5.94E-07 | 0.68 | 0.70 | -0.02 | 0.97 |
| 4 | Oaz2 | 0.35 | 2.94E-11 | 6.12E-07 | 0.54 | 0.55 | -0.01 | 0.98 |
| 4 | Rap1gds1 | 0.67 | 3.26E-11 | 6.79E-07 | 0.33 | 0.29 | 0.04 | 1.12 |
| 4 | Use1 | 0.30 | 3.56E-11 | 7.40E-07 | 0.63 | 0.65 | -0.03 | 0.96 |
| 4 | Vcam1 | 0.41 | 4.07E-11 | 8.47E-07 | 0.47 | 0.43 | 0.04 | 1.09 |
| 4 | Klhl24 | 0.29 | 4.08E-11 | 8.48E-07 | 0.60 | 0.60 | 0.00 | 1.00 |
| 4 | R3hdm4 | 0.33 | 4.36E-11 | 9.08E-07 | 0.60 | 0.62 | -0.02 | 0.96 |
| 4 | Spsb1 | 0.89 | 4.50E-11 | 9.36E-07 | 0.20 | 0.15 | 0.05 | 1.32 |
| 4 | Igbp1 | 0.30 | 4.59E-11 | 9.54E-07 | 0.65 | 0.69 | -0.04 | 0.95 |
| 4 | Chpt1 | 0.57 | 4.94E-11 | 1.03E-06 | 0.41 | 0.39 | 0.02 | 1.04 |
| 4 | Yipf5 | 0.35 | 5.32E-11 | 1.11E-06 | 0.60 | 0.62 | -0.02 | 0.97 |
| 4 | Anxa5 | 0.27 | 5.34E-11 | 1.11E-06 | 0.61 | 0.61 | 0.00 | 1.00 |
| 4 | Eef2k | 0.76 | 5.41E-11 | 1.13E-06 | 0.28 | 0.24 | 0.04 | 1.16 |
| 4 | Akt3 | 0.71 | 5.66E-11 | 1.18E-06 | 0.30 | 0.27 | 0.04 | 1.13 |
| 4 | Ilrun | 0.46 | 5.82E-11 | 1.21E-06 | 0.47 | 0.46 | 0.00 | 1.01 |
| 4 | Loxl2 | 0.39 | 6.04E-11 | 1.26E-06 | 0.46 | 0.44 | 0.02 | 1.05 |
| 4 | 5730409E04Rik | 1.06 | 6.31E-11 | 1.31E-06 | 0.12 | 0.08 | 0.04 | 1.51 |
| 4 | Zfp950 | 0.54 | 6.33E-11 | 1.32E-06 | 0.40 | 0.38 | 0.02 | 1.06 |
| 4 | Ptpra | 0.33 | 6.41E-11 | 1.33E-06 | 0.58 | 0.61 | -0.03 | 0.96 |
| 4 | Rflnb | 0.41 | 7.45E-11 | 1.55E-06 | 0.57 | 0.55 | 0.02 | 1.03 |
| 4 | Sqstm1 | 0.50 | 7.78E-11 | 1.62E-06 | 0.43 | 0.41 | 0.01 | 1.03 |
| 4 | Cpt1c | 1.09 | 8.31E-11 | 1.73E-06 | 0.13 | 0.09 | 0.04 | 1.47 |
| 4 | Rtl8b | 0.51 | 9.12E-11 | 1.90E-06 | 0.36 | 0.33 | 0.03 | 1.09 |
| 4 | Zc3h14 | 0.25 | 9.25E-11 | 1.92E-06 | 0.74 | 0.77 | -0.03 | 0.96 |
| 4 | Gcc2 | 0.31 | 1.01E-10 | 2.09E-06 | 0.65 | 0.67 | -0.02 | 0.97 |
| 4 | Uqcc3 | 0.31 | 1.16E-10 | 2.42E-06 | 0.64 | 0.68 | -0.04 | 0.95 |
| 4 | Bloc1s1 | 0.37 | 1.23E-10 | 2.55E-06 | 0.57 | 0.60 | -0.03 | 0.96 |
| 4 | Gars | 0.30 | 1.27E-10 | 2.63E-06 | 0.65 | 0.67 | -0.02 | 0.96 |
| 4 | Rora | 0.77 | 1.36E-10 | 2.83E-06 | 0.23 | 0.18 | 0.05 | 1.27 |
| 4 | Tmem35a | 1.03 | 1.37E-10 | 2.84E-06 | 0.14 | 0.10 | 0.04 | 1.44 |
| 4 | Prkacb | 0.40 | 1.67E-10 | 3.48E-06 | 0.53 | 0.53 | 0.00 | 0.99 |
| 4 | Maf | 0.29 | 1.76E-10 | 3.66E-06 | 0.11 | 0.07 | 0.04 | 1.55 |
| 4 | Nbdy | 0.31 | 1.88E-10 | 3.92E-06 | 0.61 | 0.63 | -0.02 | 0.97 |
| 4 | Txndc5 | 0.32 | 1.97E-10 | 4.09E-06 | 0.60 | 0.62 | -0.02 | 0.98 |
| 4 | Uhrf2 | 0.26 | 2.15E-10 | 4.47E-06 | 0.71 | 0.74 | -0.03 | 0.96 |
| 4 | Narf | 0.46 | 2.37E-10 | 4.94E-06 | 0.43 | 0.42 | 0.01 | 1.03 |
| 4 | Itpr1 | 0.82 | 2.44E-10 | 5.07E-06 | 0.25 | 0.21 | 0.04 | 1.19 |
| 4 | Mycbp2 | 0.27 | 2.65E-10 | 5.51E-06 | 0.68 | 0.70 | -0.02 | 0.97 |
| 4 | Slc22a17 | 0.52 | 2.66E-10 | 5.53E-06 | 0.37 | 0.35 | 0.03 | 1.08 |
| 4 | Tsen15 | 0.28 | 2.70E-10 | 5.61E-06 | 0.63 | 0.64 | -0.01 | 0.99 |
| 4 | Snx24 | 0.56 | 2.75E-10 | 5.72E-06 | 0.38 | 0.36 | 0.02 | 1.06 |
| 4 | Btbd7 | 0.28 | 2.88E-10 | 6.00E-06 | 0.65 | 0.68 | -0.03 | 0.96 |
| 4 | Scara3 | 0.79 | 3.09E-10 | 6.42E-06 | 0.22 | 0.18 | 0.04 | 1.24 |

**Supplementary Table 1. Marker genes for the original clusters of integrated clustering of E11.5 control and NotchGOF λ-junction cells.**

| | | | | | | | | |
|---|---|---|---|---|---|---|---|---|
| 4 | Trpc4ap | 0.43 | 3.91E-10 | 8.14E-06 | 0.52 | 0.53 | -0.02 | 0.97 |
| 4 | Adam23 | 1.13 | 3.92E-10 | 8.16E-06 | 0.12 | 0.08 | 0.04 | 1.48 |
| 4 | Sgk3 | 0.85 | 3.98E-10 | 8.28E-06 | 0.19 | 0.15 | 0.04 | 1.30 |
| 4 | Dipk1b | 0.38 | 4.34E-10 | 9.02E-06 | 0.49 | 0.48 | 0.01 | 1.02 |
| 4 | Usp29 | 0.75 | 5.48E-10 | 1.14E-05 | 0.24 | 0.20 | 0.04 | 1.20 |
| 4 | C230038L03Rik | 0.90 | 5.85E-10 | 1.22E-05 | 0.16 | 0.12 | 0.04 | 1.35 |
| 4 | Six5 | 0.71 | 5.88E-10 | 1.22E-05 | 0.24 | 0.20 | 0.04 | 1.22 |
| 4 | Gsn | 0.79 | 6.33E-10 | 1.32E-05 | 0.19 | 0.14 | 0.04 | 1.30 |
| 4 | Klf12 | 0.79 | 6.46E-10 | 1.34E-05 | 0.20 | 0.16 | 0.04 | 1.28 |
| 4 | Igf2r | 0.41 | 6.71E-10 | 1.40E-05 | 0.50 | 0.50 | 0.00 | 1.00 |
| 4 | Txndc12 | 0.25 | 6.93E-10 | 1.44E-05 | 0.68 | 0.70 | -0.02 | 0.97 |
| 4 | Arhgap24 | 0.89 | 7.45E-10 | 1.55E-05 | 0.16 | 0.12 | 0.04 | 1.36 |
| 4 | Azin2 | 0.91 | 7.49E-10 | 1.56E-05 | 0.15 | 0.11 | 0.04 | 1.38 |
| 4 | Cnnm2 | 1.09 | 7.54E-10 | 1.57E-05 | 0.12 | 0.08 | 0.04 | 1.47 |
| 4 | Kctd2 | 0.54 | 7.85E-10 | 1.63E-05 | 0.38 | 0.37 | 0.02 | 1.05 |
| 4 | Gpr45 | 0.80 | 8.01E-10 | 1.67E-05 | 0.22 | 0.18 | 0.04 | 1.23 |
| 4 | Elovl6 | 0.26 | 8.57E-10 | 1.78E-05 | 0.65 | 0.65 | 0.00 | 1.00 |
| 4 | Ubr1 | 0.65 | 8.68E-10 | 1.81E-05 | 0.35 | 0.33 | 0.02 | 1.05 |
| 4 | Atf7ip | 0.31 | 8.95E-10 | 1.86E-05 | 0.60 | 0.63 | -0.02 | 0.96 |
| 4 | Tm2d1 | 0.38 | 9.83E-10 | 2.04E-05 | 0.50 | 0.51 | -0.01 | 0.98 |
| 4 | Rtkn | 0.76 | 9.99E-10 | 2.08E-05 | 0.25 | 0.21 | 0.04 | 1.18 |
| 4 | Akap8l | 0.51 | 1.09E-09 | 2.27E-05 | 0.39 | 0.37 | 0.02 | 1.05 |
| 4 | Ift46 | 0.39 | 1.09E-09 | 2.28E-05 | 0.49 | 0.49 | 0.00 | 0.99 |
| 4 | Anapc4 | 0.36 | 1.19E-09 | 2.46E-05 | 0.52 | 0.54 | -0.01 | 0.97 |
| 4 | Pgm2 | 0.64 | 1.21E-09 | 2.51E-05 | 0.30 | 0.27 | 0.03 | 1.11 |
| 4 | Pygo1 | 0.41 | 1.23E-09 | 2.56E-05 | 0.47 | 0.47 | 0.00 | 1.01 |
| 4 | Idh3g | 0.28 | 1.23E-09 | 2.56E-05 | 0.63 | 0.66 | -0.03 | 0.95 |
| 4 | Trappc3 | 0.29 | 1.24E-09 | 2.57E-05 | 0.65 | 0.68 | -0.03 | 0.95 |
| 4 | Rxylt1 | 0.40 | 1.24E-09 | 2.57E-05 | 0.48 | 0.47 | 0.00 | 1.01 |
| 4 | Aamdc | 0.59 | 1.31E-09 | 2.72E-05 | 0.34 | 0.32 | 0.03 | 1.08 |
| 4 | Cnot6l | 0.35 | 1.36E-09 | 2.82E-05 | 0.54 | 0.54 | 0.00 | 0.99 |
| 4 | Mn1 | 0.56 | 1.36E-09 | 2.84E-05 | 0.32 | 0.28 | 0.04 | 1.13 |
| 4 | Ulk2 | 0.67 | 1.38E-09 | 2.86E-05 | 0.28 | 0.25 | 0.04 | 1.14 |
| 4 | Ttll7 | 1.02 | 1.42E-09 | 2.96E-05 | 0.11 | 0.07 | 0.04 | 1.51 |
| 4 | Rras | 0.59 | 1.49E-09 | 3.09E-05 | 0.26 | 0.22 | 0.04 | 1.19 |
| 4 | Prkra | 0.27 | 1.50E-09 | 3.12E-05 | 0.66 | 0.68 | -0.02 | 0.96 |
| 4 | N4bp2 | 0.30 | 1.55E-09 | 3.23E-05 | 0.64 | 0.68 | -0.03 | 0.95 |
| 4 | Bmpr1a | 0.30 | 1.69E-09 | 3.52E-05 | 0.63 | 0.65 | -0.02 | 0.97 |
| 4 | Spsb4 | 0.68 | 1.79E-09 | 3.72E-05 | 0.25 | 0.20 | 0.04 | 1.20 |
| 4 | Gatd1 | 0.36 | 1.83E-09 | 3.81E-05 | 0.52 | 0.53 | -0.01 | 0.98 |
| 4 | Notch2 | 0.51 | 1.91E-09 | 3.97E-05 | 0.39 | 0.36 | 0.02 | 1.06 |
| 4 | Tnfsf12 | 1.06 | 1.93E-09 | 4.01E-05 | 0.10 | 0.07 | 0.04 | 1.53 |
| 4 | Cib2 | 0.62 | 1.97E-09 | 4.09E-05 | 0.29 | 0.26 | 0.04 | 1.14 |
| 4 | Dlc1 | 0.48 | 2.10E-09 | 4.36E-05 | 0.39 | 0.36 | 0.03 | 1.09 |
| 4 | Tes | 0.27 | 2.28E-09 | 4.73E-05 | 0.68 | 0.70 | -0.02 | 0.98 |
| 4 | Stx7 | 0.59 | 2.30E-09 | 4.78E-05 | 0.34 | 0.31 | 0.02 | 1.08 |
| 4 | Mpv17 | 0.42 | 2.35E-09 | 4.89E-05 | 0.48 | 0.48 | 0.00 | 1.00 |
| 4 | Dzip3 | 0.32 | 2.87E-09 | 5.97E-05 | 0.59 | 0.61 | -0.02 | 0.97 |
| 4 | Arhgap31 | 0.67 | 2.98E-09 | 6.19E-05 | 0.23 | 0.19 | 0.04 | 1.22 |
| 4 | Cspp1 | 0.27 | 3.05E-09 | 6.34E-05 | 0.66 | 0.69 | -0.03 | 0.96 |
| 4 | Stk38 | 0.26 | 3.26E-09 | 6.79E-05 | 0.64 | 0.66 | -0.03 | 0.96 |
| 4 | Cnksr2 | 0.58 | 3.62E-09 | 7.54E-05 | 0.35 | 0.32 | 0.03 | 1.08 |
| 4 | Zmym5 | 0.31 | 3.76E-09 | 7.82E-05 | 0.60 | 0.62 | -0.02 | 0.97 |
| 4 | Fndc3a | 0.27 | 3.78E-09 | 7.85E-05 | 0.67 | 0.69 | -0.02 | 0.97 |
| 4 | Zfpm1 | 0.56 | 3.82E-09 | 7.94E-05 | 0.34 | 0.31 | 0.03 | 1.10 |
| 4 | Ogfr | 0.30 | 3.84E-09 | 7.98E-05 | 0.63 | 0.67 | -0.04 | 0.94 |

**Supplementary Table 1. Marker genes for the original clusters of integrated clustering of E11.5 control and NotchGOF λ-junction cells.**

| | | | | | | | | |
|---|---|---|---|---|---|---|---|---|
| 4 | Smarca1 | 0.40 | 4.02E-09 | 8.35E-05 | 0.47 | 0.47 | 0.01 | 1.02 |
| 4 | Gpr27 | 0.91 | 4.48E-09 | 9.32E-05 | 0.18 | 0.14 | 0.04 | 1.28 |
| 4 | Pakap | 0.96 | 4.81E-09 | 1.00E-04 | 0.16 | 0.12 | 0.04 | 1.33 |
| 4 | Atad2b | 0.28 | 4.96E-09 | 1.03E-04 | 0.62 | 0.65 | -0.03 | 0.96 |
| 4 | Pdgfrl | 0.73 | 5.01E-09 | 1.04E-04 | 0.22 | 0.18 | 0.04 | 1.23 |
| 4 | Tmed1 | 0.26 | 5.24E-09 | 1.09E-04 | 0.65 | 0.67 | -0.02 | 0.97 |
| 4 | Snap47 | 0.26 | 5.37E-09 | 1.12E-04 | 0.63 | 0.65 | -0.02 | 0.96 |
| 4 | Agpat4 | 0.45 | 6.45E-09 | 1.34E-04 | 0.44 | 0.44 | 0.00 | 1.00 |
| 4 | Mxra7 | 0.67 | 6.75E-09 | 1.40E-04 | 0.27 | 0.23 | 0.04 | 1.15 |
| 4 | Pttg1 | 0.31 | 6.88E-09 | 1.43E-04 | 0.67 | 0.72 | -0.04 | 0.94 |
| 4 | Dlx5 | 0.34 | 7.80E-09 | 1.62E-04 | 0.26 | 0.22 | 0.05 | 1.23 |
| 4 | Srek1ip1 | 0.27 | 8.04E-09 | 1.67E-04 | 0.68 | 0.71 | -0.03 | 0.95 |
| 4 | Zeb1 | 0.46 | 8.10E-09 | 1.68E-04 | 0.38 | 0.35 | 0.03 | 1.08 |
| 4 | 2610027K06Rik | 0.89 | 8.26E-09 | 1.72E-04 | 0.17 | 0.13 | 0.04 | 1.30 |
| 4 | Fgfrl1 | 0.52 | 9.30E-09 | 1.93E-04 | 0.36 | 0.34 | 0.02 | 1.06 |
| 4 | Enc1 | 0.79 | 1.01E-08 | 2.10E-04 | 0.26 | 0.22 | 0.04 | 1.16 |
| 4 | Mmp16 | 0.70 | 1.03E-08 | 2.13E-04 | 0.27 | 0.24 | 0.04 | 1.16 |
| 4 | Smco4 | 0.40 | 1.10E-08 | 2.28E-04 | 0.49 | 0.49 | 0.00 | 1.00 |
| 4 | Nid2 | 0.63 | 1.17E-08 | 2.42E-04 | 0.27 | 0.24 | 0.04 | 1.15 |
| 4 | Ncoa7 | 0.52 | 1.18E-08 | 2.46E-04 | 0.41 | 0.41 | 0.00 | 1.01 |
| 4 | Zfp354c | 0.46 | 1.19E-08 | 2.48E-04 | 0.40 | 0.39 | 0.01 | 1.03 |
| 4 | Pgf | 0.91 | 1.23E-08 | 2.56E-04 | 0.10 | 0.07 | 0.03 | 1.49 |
| 4 | Eva1a | 0.65 | 1.33E-08 | 2.76E-04 | 0.27 | 0.24 | 0.03 | 1.13 |
| 4 | Hmgcl | 0.45 | 1.33E-08 | 2.77E-04 | 0.41 | 0.41 | 0.00 | 1.01 |
| 4 | Plat | 0.42 | 1.37E-08 | 2.84E-04 | 0.35 | 0.32 | 0.03 | 1.11 |
| 4 | Sdc3 | 0.68 | 1.39E-08 | 2.89E-04 | 0.24 | 0.20 | 0.04 | 1.19 |
| 4 | Lyrm2 | 0.30 | 1.39E-08 | 2.90E-04 | 0.60 | 0.63 | -0.03 | 0.95 |
| 4 | Trp53inp1 | 0.58 | 1.45E-08 | 3.02E-04 | 0.23 | 0.18 | 0.04 | 1.22 |
| 4 | Tmem230 | 0.47 | 1.55E-08 | 3.22E-04 | 0.42 | 0.42 | 0.00 | 1.01 |
| 4 | Clcn5 | 0.51 | 1.55E-08 | 3.22E-04 | 0.40 | 0.39 | 0.01 | 1.03 |
| 4 | Il17rd | 0.27 | 1.82E-08 | 3.78E-04 | 0.52 | 0.50 | 0.02 | 1.04 |
| 4 | Prxl2b | 0.47 | 1.90E-08 | 3.94E-04 | 0.42 | 0.41 | 0.00 | 1.00 |
| 4 | Usp9x | 0.26 | 2.11E-08 | 4.40E-04 | 0.66 | 0.69 | -0.03 | 0.96 |
| 4 | Cpq | 0.83 | 2.13E-08 | 4.43E-04 | 0.20 | 0.16 | 0.04 | 1.23 |
| 4 | Qdpr | 0.31 | 2.26E-08 | 4.70E-04 | 0.52 | 0.54 | -0.02 | 0.96 |
| 4 | Pofut2 | 0.40 | 2.26E-08 | 4.71E-04 | 0.48 | 0.49 | -0.01 | 0.99 |
| 4 | Hip1 | 0.43 | 2.31E-08 | 4.80E-04 | 0.45 | 0.45 | 0.00 | 0.99 |
| 4 | Selenos | 0.30 | 2.57E-08 | 5.34E-04 | 0.60 | 0.61 | -0.02 | 0.98 |
| 4 | Zfp30 | 0.43 | 2.64E-08 | 5.49E-04 | 0.43 | 0.43 | 0.00 | 1.01 |
| 4 | Fgd1 | 0.57 | 2.76E-08 | 5.74E-04 | 0.34 | 0.32 | 0.02 | 1.06 |
| 4 | Pkp4 | 0.54 | 2.78E-08 | 5.77E-04 | 0.39 | 0.38 | 0.02 | 1.04 |
| 4 | Tacc2 | 0.44 | 2.82E-08 | 5.86E-04 | 0.48 | 0.48 | 0.00 | 1.00 |
| 4 | Alkbh1 | 0.30 | 3.18E-08 | 6.61E-04 | 0.65 | 0.67 | -0.03 | 0.96 |
| 4 | Six4 | 0.39 | 3.26E-08 | 6.77E-04 | 0.47 | 0.46 | 0.01 | 1.02 |
| 4 | Oxa1l | 0.37 | 3.79E-08 | 7.88E-04 | 0.51 | 0.54 | -0.03 | 0.95 |
| 4 | Slc27a3 | 0.70 | 4.02E-08 | 8.36E-04 | 0.25 | 0.21 | 0.03 | 1.16 |
| 4 | Tmed4 | 0.36 | 4.06E-08 | 8.44E-04 | 0.54 | 0.56 | -0.02 | 0.96 |
| 4 | Antxr1 | 0.56 | 4.13E-08 | 8.59E-04 | 0.33 | 0.31 | 0.02 | 1.07 |
| 4 | Slc39a7 | 0.37 | 4.17E-08 | 8.67E-04 | 0.52 | 0.54 | -0.02 | 0.96 |
| 4 | Aff4 | 0.27 | 4.20E-08 | 8.73E-04 | 0.61 | 0.64 | -0.03 | 0.96 |
| 4 | Dnpep | 0.37 | 4.32E-08 | 8.99E-04 | 0.50 | 0.51 | -0.01 | 0.97 |
| 4 | C1qtnf6 | 0.80 | 4.34E-08 | 9.03E-04 | 0.20 | 0.16 | 0.04 | 1.23 |
| 4 | Stox2 | 0.29 | 4.36E-08 | 9.06E-04 | 0.56 | 0.56 | -0.01 | 0.99 |
| 4 | Micu2 | 0.36 | 4.50E-08 | 9.35E-04 | 0.50 | 0.51 | -0.01 | 0.97 |
| 4 | Ccs | 0.40 | 4.57E-08 | 9.50E-04 | 0.45 | 0.45 | 0.00 | 0.99 |
| 4 | Peak1 | 0.69 | 4.59E-08 | 9.55E-04 | 0.28 | 0.25 | 0.03 | 1.12 |

**Supplementary Table 1. Marker genes for the original clusters of integrated clustering of E11.5 control and NotchGOF λ-junction cells.**

| | | | | | | | | |
|---|---|---|---|---|---|---|---|---|
| 4 | Rnf217 | 0.75 | 4.63E-08 | 9.62E-04 | 0.22 | 0.19 | 0.03 | 1.18 |
| 4 | Tspan15 | 0.95 | 4.70E-08 | 9.78E-04 | 0.22 | 0.19 | 0.03 | 1.17 |
| 4 | Rilpl2 | 0.58 | 4.73E-08 | 9.83E-04 | 0.30 | 0.28 | 0.03 | 1.10 |
| 4 | Gprasp2 | 0.55 | 4.74E-08 | 9.87E-04 | 0.32 | 0.30 | 0.02 | 1.08 |
| 4 | Zfp532 | 0.43 | 5.00E-08 | 1.04E-03 | 0.47 | 0.48 | -0.01 | 0.98 |
| 4 | Sec22b | 0.29 | 5.09E-08 | 1.06E-03 | 0.57 | 0.61 | -0.03 | 0.95 |
| 4 | Cttnbp2 | 0.80 | 5.58E-08 | 1.16E-03 | 0.19 | 0.15 | 0.04 | 1.24 |
| 4 | Nsmf | 0.83 | 5.92E-08 | 1.23E-03 | 0.20 | 0.16 | 0.03 | 1.21 |
| 4 | Rtl8a | 0.45 | 6.04E-08 | 1.26E-03 | 0.40 | 0.39 | 0.01 | 1.03 |
| 4 | Arhgef9 | 0.72 | 6.11E-08 | 1.27E-03 | 0.22 | 0.18 | 0.04 | 1.19 |
| 4 | Sh3bp1 | 0.56 | 6.19E-08 | 1.29E-03 | 0.31 | 0.29 | 0.02 | 1.08 |
| 4 | Prxl2c | 0.60 | 6.52E-08 | 1.36E-03 | 0.31 | 0.29 | 0.02 | 1.08 |
| 4 | Evi5 | 0.32 | 6.95E-08 | 1.45E-03 | 0.57 | 0.61 | -0.04 | 0.93 |
| 4 | Pdzrn3 | 0.78 | 7.01E-08 | 1.46E-03 | 0.21 | 0.17 | 0.04 | 1.21 |
| 4 | Dctn4 | 0.29 | 7.41E-08 | 1.54E-03 | 0.61 | 0.64 | -0.04 | 0.94 |
| 4 | Crtc3 | 0.63 | 7.45E-08 | 1.55E-03 | 0.27 | 0.24 | 0.03 | 1.12 |
| 4 | Upf3a | 0.28 | 7.81E-08 | 1.63E-03 | 0.59 | 0.62 | -0.04 | 0.94 |
| 4 | Ift122 | 0.62 | 7.98E-08 | 1.66E-03 | 0.25 | 0.22 | 0.03 | 1.15 |
| 4 | Slc39a13 | 0.88 | 8.47E-08 | 1.76E-03 | 0.17 | 0.14 | 0.04 | 1.26 |
| 4 | Tbck | 0.60 | 8.74E-08 | 1.82E-03 | 0.31 | 0.29 | 0.02 | 1.08 |
| 4 | Bcl2 | 0.47 | 9.05E-08 | 1.88E-03 | 0.37 | 0.35 | 0.02 | 1.05 |
| 4 | Dchs1 | 0.37 | 9.57E-08 | 1.99E-03 | 0.43 | 0.42 | 0.01 | 1.02 |
| 4 | 1810026B05Rik | 0.28 | 9.89E-08 | 2.06E-03 | 0.57 | 0.60 | -0.03 | 0.96 |
| 4 | Dguok | 0.32 | 1.07E-07 | 2.22E-03 | 0.50 | 0.52 | -0.01 | 0.98 |
| 4 | Rbm18 | 0.32 | 1.12E-07 | 2.33E-03 | 0.53 | 0.56 | -0.03 | 0.95 |
| 4 | Eps8 | 0.74 | 1.14E-07 | 2.38E-03 | 0.20 | 0.16 | 0.04 | 1.23 |
| 4 | Rev3l | 0.32 | 1.22E-07 | 2.54E-03 | 0.58 | 0.59 | -0.02 | 0.97 |
| 4 | B4galt6 | 0.71 | 1.33E-07 | 2.76E-03 | 0.25 | 0.23 | 0.03 | 1.12 |
| 4 | Zfp260 | 0.27 | 1.42E-07 | 2.96E-03 | 0.63 | 0.65 | -0.02 | 0.97 |
| 4 | Pgrmc2 | 0.30 | 1.42E-07 | 2.96E-03 | 0.56 | 0.59 | -0.03 | 0.95 |
| 4 | Dnajb14 | 0.56 | 1.48E-07 | 3.08E-03 | 0.34 | 0.32 | 0.02 | 1.05 |
| 4 | Eif2a | 0.28 | 1.54E-07 | 3.21E-03 | 0.62 | 0.67 | -0.05 | 0.93 |
| 4 | Lhfpl2 | 0.52 | 1.55E-07 | 3.22E-03 | 0.35 | 0.33 | 0.02 | 1.06 |
| 4 | Hint2 | 0.27 | 1.67E-07 | 3.47E-03 | 0.59 | 0.62 | -0.03 | 0.95 |
| 4 | Colec12 | 0.54 | 1.76E-07 | 3.67E-03 | 0.26 | 0.22 | 0.04 | 1.16 |
| 4 | Selenon | 0.52 | 1.83E-07 | 3.80E-03 | 0.38 | 0.37 | 0.01 | 1.02 |
| 4 | Tlcd2 | 0.83 | 2.01E-07 | 4.18E-03 | 0.17 | 0.14 | 0.03 | 1.25 |
| 4 | Bcap29 | 0.53 | 2.08E-07 | 4.33E-03 | 0.36 | 0.34 | 0.01 | 1.03 |
| 4 | Atraid | 0.35 | 2.09E-07 | 4.34E-03 | 0.50 | 0.51 | -0.01 | 0.98 |
| 4 | Slc9a3r2 | 0.41 | 2.10E-07 | 4.38E-03 | 0.29 | 0.26 | 0.03 | 1.11 |
| 4 | Bmpr1b | 0.91 | 2.12E-07 | 4.41E-03 | 0.12 | 0.09 | 0.03 | 1.39 |
| 4 | Cldn11 | 0.46 | 2.30E-07 | 4.78E-03 | 0.17 | 0.13 | 0.04 | 1.30 |
| 4 | Tnks | 0.35 | 2.31E-07 | 4.80E-03 | 0.52 | 0.54 | -0.02 | 0.96 |
| 4 | Rara | 0.38 | 2.45E-07 | 5.10E-03 | 0.46 | 0.47 | -0.01 | 0.98 |
| 4 | Vps28 | 0.25 | 2.78E-07 | 5.78E-03 | 0.58 | 0.63 | -0.04 | 0.93 |
| 4 | Cdc37l1 | 0.28 | 2.88E-07 | 6.00E-03 | 0.57 | 0.60 | -0.03 | 0.94 |
| 4 | Atp6v0d1 | 0.36 | 3.03E-07 | 6.30E-03 | 0.51 | 0.54 | -0.03 | 0.94 |
| 4 | Phpt1 | 0.31 | 3.35E-07 | 6.97E-03 | 0.55 | 0.57 | -0.02 | 0.96 |
| 4 | Six2 | 0.34 | 3.63E-07 | 7.54E-03 | 0.47 | 0.45 | 0.02 | 1.04 |
| 4 | Zbed5 | 0.81 | 3.72E-07 | 7.73E-03 | 0.18 | 0.15 | 0.03 | 1.22 |
| 4 | Pex13 | 0.27 | 3.99E-07 | 8.29E-03 | 0.56 | 0.59 | -0.02 | 0.96 |
| 4 | Ugdh | 0.42 | 4.13E-07 | 8.58E-03 | 0.46 | 0.47 | -0.01 | 0.97 |
| 4 | Pds5b | 0.27 | 4.23E-07 | 8.80E-03 | 0.62 | 0.66 | -0.04 | 0.94 |
| 4 | 2700081O15Rik | 0.31 | 4.42E-07 | 9.19E-03 | 0.54 | 0.56 | -0.02 | 0.96 |
| 4 | Tmem101 | 0.31 | 4.55E-07 | 9.47E-03 | 0.51 | 0.52 | -0.01 | 0.98 |
| 4 | Wnt5b | 0.52 | 4.72E-07 | 9.82E-03 | 0.29 | 0.26 | 0.03 | 1.11 |

**Supplementary Table 1. Marker genes for the original clusters of integrated clustering of E11.5 control and NotchGOF λ-junction cells.**

| | | | | | | | | |
|---|---|---|---|---|---|---|---|---|
| 4 | Tmem29 | 0.36 | 4.86E-07 | 1.01E-02 | 0.50 | 0.52 | -0.02 | 0.96 |
| 4 | Ndfip2 | 0.32 | 5.65E-07 | 1.17E-02 | 0.50 | 0.52 | -0.02 | 0.97 |
| 4 | Fbn1 | 0.50 | 5.78E-07 | 1.20E-02 | 0.35 | 0.33 | 0.02 | 1.04 |
| 4 | Cfap97 | 0.39 | 6.11E-07 | 1.27E-02 | 0.45 | 0.46 | -0.01 | 0.98 |
| 4 | Nhs | 0.50 | 6.63E-07 | 1.38E-02 | 0.36 | 0.34 | 0.02 | 1.06 |
| 4 | Brd9 | 0.28 | 6.65E-07 | 1.38E-02 | 0.53 | 0.56 | -0.02 | 0.96 |
| 4 | Atp7a | 0.68 | 6.65E-07 | 1.38E-02 | 0.26 | 0.24 | 0.02 | 1.10 |
| 4 | Akap7 | 0.62 | 7.32E-07 | 1.52E-02 | 0.25 | 0.22 | 0.03 | 1.13 |
| 4 | Itpripl2 | 0.83 | 7.60E-07 | 1.58E-02 | 0.16 | 0.13 | 0.03 | 1.25 |
| 4 | Mbtps1 | 0.28 | 9.05E-07 | 1.88E-02 | 0.53 | 0.56 | -0.02 | 0.96 |
| 4 | Bfar | 0.37 | 9.08E-07 | 1.89E-02 | 0.48 | 0.49 | -0.01 | 0.97 |
| 4 | Hibch | 0.59 | 9.15E-07 | 1.90E-02 | 0.30 | 0.28 | 0.02 | 1.08 |
| 4 | Zdhhc20 | 0.30 | 9.26E-07 | 1.93E-02 | 0.57 | 0.59 | -0.02 | 0.96 |
| 4 | Slc44a1 | 0.71 | 9.63E-07 | 2.00E-02 | 0.26 | 0.24 | 0.02 | 1.09 |
| 4 | Anxa6 | 0.56 | 9.68E-07 | 2.01E-02 | 0.21 | 0.17 | 0.03 | 1.19 |
| 4 | Gli1 | 0.79 | 1.14E-06 | 2.38E-02 | 0.16 | 0.13 | 0.03 | 1.25 |
| 4 | Shisa5 | 0.59 | 1.17E-06 | 2.44E-02 | 0.27 | 0.24 | 0.03 | 1.11 |
| 4 | 2610008E11Rik | 0.41 | 1.18E-06 | 2.45E-02 | 0.40 | 0.40 | 0.00 | 1.01 |
| 4 | Lrp5 | 0.77 | 1.20E-06 | 2.49E-02 | 0.20 | 0.18 | 0.03 | 1.17 |
| 4 | Rgma | 0.62 | 1.24E-06 | 2.59E-02 | 0.27 | 0.24 | 0.03 | 1.10 |
| 4 | Map7d1 | 0.25 | 1.33E-06 | 2.76E-02 | 0.59 | 0.62 | -0.03 | 0.95 |
| 4 | Armcx3 | 0.37 | 1.33E-06 | 2.77E-02 | 0.43 | 0.43 | 0.00 | 1.00 |
| 4 | Dusp4 | 0.82 | 1.34E-06 | 2.79E-02 | 0.11 | 0.08 | 0.03 | 1.36 |
| 4 | 2810001G20Rik | 0.48 | 1.37E-06 | 2.84E-02 | 0.39 | 0.39 | 0.00 | 1.01 |
| 4 | Slc2a8 | 0.83 | 1.37E-06 | 2.85E-02 | 0.15 | 0.12 | 0.03 | 1.25 |
| 4 | Tmem98 | 0.36 | 1.39E-06 | 2.89E-02 | 0.49 | 0.51 | -0.02 | 0.96 |
| 4 | Micall1 | 0.82 | 1.49E-06 | 3.11E-02 | 0.15 | 0.12 | 0.03 | 1.26 |
| 4 | Washc3 | 0.39 | 1.49E-06 | 3.11E-02 | 0.48 | 0.51 | -0.03 | 0.95 |
| 4 | Siah2 | 1.00 | 1.65E-06 | 3.42E-02 | 0.10 | 0.07 | 0.03 | 1.38 |
| 4 | Zswim7 | 0.61 | 1.67E-06 | 3.48E-02 | 0.27 | 0.25 | 0.02 | 1.08 |
| 4 | 2310009B15Rik | 0.34 | 1.77E-06 | 3.68E-02 | 0.52 | 0.55 | -0.03 | 0.94 |
| 4 | Dnajb4 | 0.57 | 1.78E-06 | 3.71E-02 | 0.32 | 0.31 | 0.01 | 1.04 |
| 4 | Gpc6 | 0.26 | 1.84E-06 | 3.84E-02 | 0.59 | 0.59 | 0.00 | 1.00 |
| 4 | Zfp9 | 0.72 | 1.86E-06 | 3.86E-02 | 0.19 | 0.16 | 0.03 | 1.19 |
| 4 | Fbh1 | 0.51 | 1.86E-06 | 3.88E-02 | 0.36 | 0.36 | 0.00 | 1.01 |
| 4 | Serp2 | 0.68 | 1.96E-06 | 4.07E-02 | 0.23 | 0.20 | 0.03 | 1.15 |
| 4 | Armcx6 | 0.63 | 1.97E-06 | 4.10E-02 | 0.23 | 0.20 | 0.03 | 1.14 |
| 4 | Casp6 | 0.28 | 1.98E-06 | 4.12E-02 | 0.54 | 0.56 | -0.03 | 0.95 |
| 4 | Nav1 | 0.31 | 1.98E-06 | 4.12E-02 | 0.52 | 0.55 | -0.02 | 0.96 |
| 4 | Zfp811 | 0.77 | 2.09E-06 | 4.34E-02 | 0.15 | 0.12 | 0.03 | 1.26 |
| 4 | Aplp1 | 0.54 | 2.13E-06 | 4.43E-02 | 0.24 | 0.21 | 0.03 | 1.13 |
| 4 | Xrn1 | 0.31 | 2.13E-06 | 4.44E-02 | 0.49 | 0.52 | -0.03 | 0.95 |
| 4 | Lmbrd1 | 0.69 | 2.19E-06 | 4.56E-02 | 0.25 | 0.22 | 0.02 | 1.10 |
| 4 | Spred1 | 0.34 | 2.23E-06 | 4.64E-02 | 0.49 | 0.51 | -0.02 | 0.96 |
| 4 | Arid1b | 0.37 | 2.38E-06 | 4.95E-02 | 0.47 | 0.49 | -0.02 | 0.97 |
| 4 | Gprasp1 | 0.35 | 2.39E-06 | 4.96E-02 | 0.48 | 0.49 | -0.02 | 0.97 |
| 5 | Gsc | 2.61 | 0 | 0 | 0.88 | 0.25 | 0.62 | 3.45 |
| 5 | Ebf1 | 3.01 | 0 | 0 | 0.84 | 0.26 | 0.58 | 3.26 |
| 5 | Foxp2 | 2.09 | 0 | 0 | 0.88 | 0.31 | 0.57 | 2.84 |
| 5 | Foxc1 | 2.37 | 0 | 0 | 0.80 | 0.25 | 0.54 | 3.15 |
| 5 | Barx1 | 2.14 | 0 | 0 | 0.68 | 0.19 | 0.50 | 3.68 |
| 5 | Pkdcc | 2.54 | 0 | 0 | 0.79 | 0.30 | 0.49 | 2.62 |
| 5 | Foxc2 | 3.31 | 0 | 0 | 0.57 | 0.09 | 0.48 | 6.38 |
| 5 | Cldn11 | 3.14 | 0 | 0 | 0.54 | 0.10 | 0.44 | 5.41 |
| 5 | Itm2a | 2.52 | 0 | 0 | 0.96 | 0.52 | 0.44 | 1.85 |
| 5 | Tgfbi | 2.02 | 0 | 0 | 0.69 | 0.26 | 0.43 | 2.64 |

**Supplementary Table 1. Marker genes for the original clusters of integrated clustering of E11.5 control and NotchGOF λ-junction cells.**

| | | | | | | | | |
|---|---|---|---|---|---|---|---|---|
| 5 | Hmcn1 | 2.10 | 0 | 0 | 0.93 | 0.51 | 0.42 | 1.83 |
| 5 | Crym | 1.98 | 0 | 0 | 0.87 | 0.45 | 0.42 | 1.94 |
| 5 | Nr2f1 | 2.19 | 0 | 0 | 0.71 | 0.31 | 0.40 | 2.32 |
| 5 | Tbx22 | 2.75 | 0 | 0 | 0.49 | 0.09 | 0.40 | 5.39 |
| 5 | Foxd2os | 2.34 | 0 | 0 | 0.52 | 0.12 | 0.39 | 4.18 |
| 5 | Osr1 | 4.18 | 0 | 0 | 0.43 | 0.04 | 0.38 | 9.73 |
| 5 | Foxp1 | 1.59 | 0 | 0 | 0.96 | 0.58 | 0.38 | 1.66 |
| 5 | Shox2 | 2.83 | 0 | 0 | 0.46 | 0.11 | 0.36 | 4.38 |
| 5 | Map1b | 2.03 | 0 | 0 | 0.94 | 0.61 | 0.33 | 1.55 |
| 5 | Irx1 | 4.21 | 0 | 0 | 0.36 | 0.03 | 0.33 | 10.94 |
| 5 | Foxd2 | 2.45 | 0 | 0 | 0.42 | 0.09 | 0.33 | 4.57 |
| 5 | Vcan | 1.32 | 0 | 0 | 0.99 | 0.86 | 0.13 | 1.15 |
| 5 | Nefl | 3.60 | 1.87E-306 | 3.89E-302 | 0.20 | 0.02 | 0.18 | 10.63 |
| 5 | Alx3 | 1.82 | 5.36E-300 | 1.12E-295 | 0.69 | 0.27 | 0.43 | 2.60 |
| 5 | Hic1 | 3.75 | 1.29E-294 | 2.68E-290 | 0.24 | 0.03 | 0.21 | 7.87 |
| 5 | Rbp1 | 0.85 | 1.04E-279 | 2.17E-275 | 0.99 | 0.91 | 0.09 | 1.10 |
| 5 | Ptx3 | 2.79 | 4.02E-261 | 8.36E-257 | 0.38 | 0.09 | 0.29 | 4.15 |
| 5 | Elmod1 | 4.92 | 3.73E-257 | 7.75E-253 | 0.12 | 0.01 | 0.11 | 19.50 |
| 5 | Plpp3 | 1.33 | 5.47E-237 | 1.14E-232 | 0.90 | 0.65 | 0.25 | 1.39 |
| 5 | Ebf3 | 2.12 | 1.18E-225 | 2.44E-221 | 0.47 | 0.15 | 0.31 | 3.05 |
| 5 | Nr2f2 | 1.90 | 6.91E-225 | 1.44E-220 | 0.70 | 0.34 | 0.35 | 2.03 |
| 5 | Apoe | 0.48 | 2.51E-223 | 5.22E-219 | 0.76 | 0.41 | 0.35 | 1.85 |
| 5 | Plat | 1.69 | 7.84E-219 | 1.63E-214 | 0.64 | 0.30 | 0.35 | 2.18 |
| 5 | Traf3ip2 | 2.01 | 4.06E-189 | 8.44E-185 | 0.50 | 0.20 | 0.30 | 2.49 |
| 5 | Alx1 | 0.97 | 1.28E-187 | 2.66E-183 | 0.98 | 0.71 | 0.27 | 1.38 |
| 5 | Pfn1 | 0.41 | 3.36E-183 | 6.98E-179 | 1.00 | 1.00 | 0.00 | 1.00 |
| 5 | Tpm2 | 1.44 | 9.99E-183 | 2.08E-178 | 0.72 | 0.42 | 0.31 | 1.73 |
| 5 | Cxcl12 | 2.26 | 5.36E-173 | 1.11E-168 | 0.43 | 0.16 | 0.27 | 2.73 |
| 5 | Six2 | 1.72 | 1.76E-172 | 3.66E-168 | 0.70 | 0.44 | 0.27 | 1.61 |
| 5 | Rarb | 1.45 | 5.46E-168 | 1.14E-163 | 0.64 | 0.33 | 0.31 | 1.95 |
| 5 | Arg1 | 3.71 | 2.30E-165 | 4.79E-161 | 0.15 | 0.02 | 0.13 | 7.25 |
| 5 | Alkal2 | 2.16 | 2.88E-163 | 6.00E-159 | 0.28 | 0.07 | 0.21 | 4.00 |
| 5 | Nefm | 3.38 | 3.34E-162 | 6.95E-158 | 0.14 | 0.02 | 0.12 | 7.42 |
| 5 | Foxp4 | 1.34 | 1.76E-159 | 3.67E-155 | 0.68 | 0.40 | 0.28 | 1.68 |
| 5 | Ntng1 | 2.26 | 7.71E-157 | 1.60E-152 | 0.34 | 0.11 | 0.23 | 3.15 |
| 5 | Lrig3 | 1.81 | 3.25E-155 | 6.75E-151 | 0.49 | 0.21 | 0.28 | 2.30 |
| 5 | Mdk | 0.45 | 1.68E-149 | 3.50E-145 | 1.00 | 0.99 | 0.01 | 1.01 |
| 5 | Gmds | 1.48 | 2.66E-149 | 5.54E-145 | 0.61 | 0.34 | 0.26 | 1.77 |
| 5 | Cdkn1c | 0.67 | 7.72E-149 | 1.61E-144 | 1.00 | 0.93 | 0.07 | 1.08 |
| 5 | Flrt2 | 0.88 | 1.96E-146 | 4.09E-142 | 0.76 | 0.43 | 0.32 | 1.75 |
| 5 | Prrx1 | 0.78 | 3.50E-146 | 7.29E-142 | 0.93 | 0.63 | 0.30 | 1.47 |
| 5 | Crtac1 | 2.65 | 5.58E-144 | 1.16E-139 | 0.21 | 0.05 | 0.16 | 4.52 |
| 5 | Gm10000 | 2.90 | 8.15E-143 | 1.70E-138 | 0.16 | 0.03 | 0.13 | 5.89 |
| 5 | Islr | 2.30 | 8.99E-142 | 1.87E-137 | 0.26 | 0.07 | 0.19 | 3.73 |
| 5 | Adamts1 | 1.90 | 1.00E-135 | 2.09E-131 | 0.51 | 0.26 | 0.25 | 1.98 |
| 5 | Tcf15 | 1.60 | 7.03E-135 | 1.46E-130 | 0.54 | 0.27 | 0.27 | 2.03 |
| 5 | Ntan1 | 1.43 | 2.67E-133 | 5.56E-129 | 0.80 | 0.66 | 0.14 | 1.21 |
| 5 | Calcrl | 3.00 | 3.18E-132 | 6.62E-128 | 0.15 | 0.03 | 0.12 | 5.59 |
| 5 | Myl9 | 1.54 | 1.30E-131 | 2.71E-127 | 0.48 | 0.23 | 0.26 | 2.12 |
| 5 | Lypd6 | 2.00 | 6.66E-131 | 1.39E-126 | 0.33 | 0.11 | 0.22 | 2.89 |
| 5 | Mmp16 | 1.72 | 3.51E-129 | 7.30E-125 | 0.47 | 0.22 | 0.25 | 2.11 |
| 5 | Tgfb2 | 0.85 | 6.63E-129 | 1.38E-124 | 0.75 | 0.47 | 0.29 | 1.61 |
| 5 | Penk | 3.57 | 1.12E-128 | 2.32E-124 | 0.27 | 0.08 | 0.19 | 3.27 |
| 5 | Runx1t1 | 0.92 | 5.50E-124 | 1.14E-119 | 0.78 | 0.51 | 0.27 | 1.53 |
| 5 | Add3 | 0.96 | 3.97E-120 | 8.26E-116 | 0.82 | 0.65 | 0.17 | 1.26 |
| 5 | H2afv | 0.35 | 4.70E-120 | 9.77E-116 | 1.00 | 0.99 | 0.01 | 1.01 |

**Supplementary Table 1. Marker genes for the original clusters of integrated clustering of E11.5 control and NotchGOF λ-junction cells.**

| | | | | | | | | |
|---|---|---|---|---|---|---|---|---|
| 5 | Shisa2 | 2.00 | 3.67E-119 | 7.64E-115 | 0.29 | 0.10 | 0.20 | 2.99 |
| 5 | Bnc2 | 1.66 | 1.48E-117 | 3.07E-113 | 0.39 | 0.16 | 0.23 | 2.40 |
| 5 | Pdgfra | 0.99 | 2.90E-117 | 6.04E-113 | 0.77 | 0.53 | 0.24 | 1.46 |
| 5 | Tle5 | 0.50 | 1.10E-115 | 2.30E-111 | 0.99 | 0.94 | 0.05 | 1.06 |
| 5 | Tsc22d1 | 0.57 | 2.78E-113 | 5.77E-109 | 0.98 | 0.93 | 0.05 | 1.05 |
| 5 | Hpgd | 1.40 | 1.28E-112 | 2.66E-108 | 0.60 | 0.33 | 0.26 | 1.79 |
| 5 | Fli1 | 1.14 | 3.12E-112 | 6.48E-108 | 0.61 | 0.36 | 0.26 | 1.72 |
| 5 | Tmsb10 | 0.30 | 4.24E-111 | 8.81E-107 | 1.00 | 1.00 | 0.00 | 1.00 |
| 5 | Rab3il1 | 1.36 | 9.14E-111 | 1.90E-106 | 0.52 | 0.28 | 0.24 | 1.84 |
| 5 | Creb5 | 1.24 | 1.29E-110 | 2.69E-106 | 0.57 | 0.31 | 0.25 | 1.82 |
| 5 | Palld | 0.87 | 1.78E-110 | 3.71E-106 | 0.81 | 0.61 | 0.19 | 1.31 |
| 5 | Cthrc1 | 1.64 | 3.48E-108 | 7.23E-104 | 0.40 | 0.18 | 0.22 | 2.25 |
| 5 | Lpar1 | 0.73 | 6.43E-105 | 1.34E-100 | 0.89 | 0.69 | 0.20 | 1.29 |
| 5 | Emx2 | 1.95 | 2.08E-104 | 4.33E-100 | 0.18 | 0.04 | 0.14 | 4.07 |
| 5 | Epha4 | 0.95 | 2.58E-102 | 5.37E-98 | 0.81 | 0.64 | 0.17 | 1.26 |
| 5 | Mgp | 1.87 | 6.18E-98 | 1.29E-93 | 0.21 | 0.06 | 0.15 | 3.44 |
| 5 | Maged2 | 0.70 | 3.96E-97 | 8.25E-93 | 0.92 | 0.82 | 0.10 | 1.12 |
| 5 | Mn1 | 1.30 | 4.77E-97 | 9.92E-93 | 0.50 | 0.27 | 0.23 | 1.84 |
| 5 | Syt12 | 2.38 | 1.12E-96 | 2.32E-92 | 0.17 | 0.04 | 0.13 | 4.00 |
| 5 | Nrip1 | 0.73 | 1.29E-94 | 2.69E-90 | 0.88 | 0.75 | 0.13 | 1.17 |
| 5 | Zfp536 | 2.27 | 3.12E-93 | 6.50E-89 | 0.26 | 0.09 | 0.16 | 2.77 |
| 5 | Col2a1 | 0.75 | 1.27E-91 | 2.64E-87 | 0.83 | 0.63 | 0.20 | 1.32 |
| 5 | Scube1 | 1.08 | 3.20E-91 | 6.65E-87 | 0.59 | 0.36 | 0.23 | 1.63 |
| 5 | Tut4 | 0.48 | 3.20E-91 | 6.65E-87 | 0.98 | 0.92 | 0.06 | 1.07 |
| 5 | Fgfr1 | 0.75 | 2.51E-89 | 5.21E-85 | 0.82 | 0.64 | 0.17 | 1.27 |
| 5 | Cenpv | 0.96 | 3.02E-89 | 6.28E-85 | 0.69 | 0.49 | 0.20 | 1.42 |
| 5 | Cnn2 | 0.82 | 9.22E-89 | 1.92E-84 | 0.80 | 0.62 | 0.18 | 1.30 |
| 5 | Foxd1 | 1.81 | 2.22E-88 | 4.61E-84 | 0.27 | 0.10 | 0.17 | 2.70 |
| 5 | Fzd4 | 1.26 | 2.25E-88 | 4.68E-84 | 0.45 | 0.24 | 0.22 | 1.90 |
| 5 | Mgst1 | 1.00 | 3.48E-88 | 7.24E-84 | 0.59 | 0.36 | 0.23 | 1.65 |
| 5 | Ramp2 | 0.70 | 1.10E-87 | 2.28E-83 | 0.38 | 0.17 | 0.20 | 2.18 |
| 5 | Lum | 1.96 | 3.16E-87 | 6.57E-83 | 0.33 | 0.15 | 0.19 | 2.28 |
| 5 | Gpc6 | 0.90 | 7.78E-87 | 1.62E-82 | 0.75 | 0.58 | 0.17 | 1.29 |
| 5 | Fzd2 | 0.51 | 3.20E-86 | 6.65E-82 | 0.97 | 0.87 | 0.10 | 1.12 |
| 5 | Slc1a3 | 1.49 | 4.16E-85 | 8.65E-81 | 0.42 | 0.21 | 0.21 | 1.96 |
| 5 | Ppib | 0.38 | 1.18E-84 | 2.46E-80 | 1.00 | 0.98 | 0.02 | 1.02 |
| 5 | Thbs1 | 1.59 | 6.11E-84 | 1.27E-79 | 0.27 | 0.11 | 0.17 | 2.51 |
| 5 | Rev3l | 0.84 | 1.07E-83 | 2.23E-79 | 0.74 | 0.58 | 0.16 | 1.27 |
| 5 | Tfap2b | 0.91 | 1.21E-82 | 2.51E-78 | 0.24 | 0.08 | 0.16 | 2.95 |
| 5 | Tpm1 | 0.50 | 3.37E-82 | 7.02E-78 | 0.97 | 0.91 | 0.06 | 1.07 |
| 5 | Tmem167 | 0.41 | 2.72E-81 | 5.65E-77 | 0.98 | 0.95 | 0.03 | 1.04 |
| 5 | Ephb3 | 1.22 | 1.20E-80 | 2.50E-76 | 0.48 | 0.29 | 0.20 | 1.69 |
| 5 | Mdfic | 1.33 | 1.09E-79 | 2.26E-75 | 0.41 | 0.21 | 0.20 | 1.93 |
| 5 | Phlda1 | 1.23 | 1.12E-78 | 2.32E-74 | 0.59 | 0.41 | 0.19 | 1.46 |
| 5 | Nnat | 0.31 | 6.71E-78 | 1.39E-73 | 0.99 | 0.94 | 0.06 | 1.06 |
| 5 | Atp2b1 | 0.55 | 8.84E-78 | 1.84E-73 | 0.93 | 0.84 | 0.09 | 1.11 |
| 5 | Ogn | 2.68 | 3.62E-77 | 7.52E-73 | 0.11 | 0.02 | 0.09 | 4.63 |
| 5 | Scx | 1.21 | 1.03E-76 | 2.14E-72 | 0.34 | 0.15 | 0.19 | 2.22 |
| 5 | Tmpo | 0.40 | 5.83E-76 | 1.21E-71 | 0.99 | 0.95 | 0.04 | 1.04 |
| 5 | Angptl1 | 1.17 | 1.37E-75 | 2.84E-71 | 0.41 | 0.21 | 0.20 | 1.96 |
| 5 | Hspa5 | 0.43 | 2.08E-75 | 4.33E-71 | 1.00 | 0.97 | 0.03 | 1.03 |
| 5 | Col26a1 | 0.57 | 2.23E-75 | 4.64E-71 | 0.83 | 0.60 | 0.24 | 1.39 |
| 5 | Oaf | 0.92 | 2.46E-75 | 5.11E-71 | 0.63 | 0.45 | 0.19 | 1.42 |
| 5 | Ldhb | 0.44 | 4.15E-75 | 8.63E-71 | 0.98 | 0.91 | 0.08 | 1.08 |
| 5 | Pcolce | 0.89 | 6.87E-75 | 1.43E-70 | 0.65 | 0.45 | 0.20 | 1.45 |
| 5 | Nrp1 | 1.21 | 1.62E-74 | 3.38E-70 | 0.40 | 0.20 | 0.20 | 2.01 |

**Supplementary Table 1. Marker genes for the original clusters of integrated clustering of E11.5 control and NotchGOF λ-junction cells.**

| | | | | | | | | |
|---|---|---|---|---|---|---|---|---|
| 5 | Dnaja1 | 0.32 | 1.05E-73 | 2.18E-69 | 1.00 | 0.99 | 0.01 | 1.01 |
| 5 | Fermt2 | 0.81 | 7.75E-73 | 1.61E-68 | 0.82 | 0.72 | 0.10 | 1.14 |
| 5 | Zcchc3 | 0.63 | 3.39E-72 | 7.04E-68 | 0.83 | 0.71 | 0.12 | 1.17 |
| 5 | Loxl1 | 1.96 | 3.93E-72 | 8.18E-68 | 0.19 | 0.07 | 0.13 | 2.91 |
| 5 | Hmgn2 | 0.32 | 4.65E-72 | 9.68E-68 | 1.00 | 0.98 | 0.02 | 1.02 |
| 5 | H2afy2 | 0.38 | 9.36E-72 | 1.95E-67 | 0.99 | 0.93 | 0.06 | 1.06 |
| 5 | Lhfp | 1.25 | 1.22E-71 | 2.54E-67 | 0.41 | 0.22 | 0.19 | 1.85 |
| 5 | Colec12 | 1.27 | 3.99E-71 | 8.30E-67 | 0.40 | 0.21 | 0.19 | 1.90 |
| 5 | A830082K12Rik | 1.91 | 6.67E-71 | 1.39E-66 | 0.19 | 0.07 | 0.13 | 2.92 |
| 5 | Dicer1 | 0.54 | 8.26E-71 | 1.72E-66 | 0.90 | 0.81 | 0.09 | 1.11 |
| 5 | Ly6e | 0.87 | 4.94E-69 | 1.03E-64 | 0.62 | 0.44 | 0.18 | 1.40 |
| 5 | Ptgis | 1.74 | 1.45E-67 | 3.02E-63 | 0.25 | 0.10 | 0.14 | 2.38 |
| 5 | B4galt4 | 1.25 | 7.70E-67 | 1.60E-62 | 0.40 | 0.22 | 0.18 | 1.81 |
| 5 | Igfbp4 | 0.40 | 3.90E-66 | 8.11E-62 | 1.00 | 0.90 | 0.10 | 1.11 |
| 5 | Atp5j | 0.26 | 4.59E-66 | 9.55E-62 | 1.00 | 0.99 | 0.00 | 1.00 |
| 5 | Col9a1 | 0.87 | 1.77E-65 | 3.68E-61 | 0.48 | 0.28 | 0.21 | 1.74 |
| 5 | Slc25a4 | 0.28 | 4.16E-65 | 8.66E-61 | 1.00 | 0.97 | 0.03 | 1.03 |
| 5 | Pcdh11x | 1.71 | 1.53E-64 | 3.17E-60 | 0.24 | 0.10 | 0.14 | 2.43 |
| 5 | Ddr2 | 0.75 | 1.11E-63 | 2.31E-59 | 0.67 | 0.48 | 0.19 | 1.40 |
| 5 | Eln | 1.89 | 1.80E-63 | 3.75E-59 | 0.19 | 0.07 | 0.12 | 2.74 |
| 5 | Tmem176b | 0.73 | 2.09E-62 | 4.35E-58 | 0.71 | 0.56 | 0.15 | 1.26 |
| 5 | Slc9a3r2 | 0.97 | 7.18E-62 | 1.49E-57 | 0.44 | 0.25 | 0.18 | 1.72 |
| 5 | Pcp2 | 2.32 | 8.44E-62 | 1.75E-57 | 0.14 | 0.04 | 0.10 | 3.34 |
| 5 | Il11ra1 | 0.69 | 1.10E-61 | 2.28E-57 | 0.78 | 0.61 | 0.18 | 1.29 |
| 5 | Sox9 | 0.26 | 2.74E-61 | 5.69E-57 | 0.79 | 0.57 | 0.22 | 1.38 |
| 5 | Smoc1 | 1.87 | 6.69E-60 | 1.39E-55 | 0.19 | 0.07 | 0.12 | 2.67 |
| 5 | Fam43a | 1.98 | 6.97E-60 | 1.45E-55 | 0.15 | 0.05 | 0.10 | 3.15 |
| 5 | Rbms3 | 0.89 | 7.92E-60 | 1.65E-55 | 0.57 | 0.40 | 0.17 | 1.42 |
| 5 | Tnik | 1.04 | 9.34E-60 | 1.94E-55 | 0.41 | 0.23 | 0.18 | 1.78 |
| 5 | Lama4 | 1.26 | 1.41E-59 | 2.94E-55 | 0.27 | 0.12 | 0.15 | 2.20 |
| 5 | Cul4b | 0.77 | 4.16E-59 | 8.66E-55 | 0.72 | 0.61 | 0.11 | 1.18 |
| 5 | Mdfi | 0.73 | 8.05E-59 | 1.67E-54 | 0.70 | 0.56 | 0.14 | 1.26 |
| 5 | Cd248 | 1.07 | 2.06E-58 | 4.29E-54 | 0.42 | 0.24 | 0.17 | 1.72 |
| 5 | Sparc | 0.43 | 4.56E-58 | 9.48E-54 | 0.88 | 0.73 | 0.15 | 1.21 |
| 5 | Timp2 | 0.60 | 1.88E-57 | 3.91E-53 | 0.79 | 0.67 | 0.13 | 1.19 |
| 5 | Prss23 | 0.95 | 4.67E-57 | 9.72E-53 | 0.48 | 0.30 | 0.18 | 1.61 |
| 5 | Eva1b | 0.63 | 8.56E-57 | 1.78E-52 | 0.71 | 0.53 | 0.18 | 1.34 |
| 5 | Hsd11b2 | 1.39 | 9.81E-57 | 2.04E-52 | 0.23 | 0.09 | 0.13 | 2.40 |
| 5 | Mfap2 | 0.36 | 1.17E-56 | 2.44E-52 | 0.99 | 0.92 | 0.07 | 1.07 |
| 5 | Pbx1 | 0.40 | 1.19E-56 | 2.48E-52 | 0.97 | 0.92 | 0.05 | 1.05 |
| 5 | Hdgfl3 | 0.56 | 1.78E-55 | 3.70E-51 | 0.82 | 0.70 | 0.12 | 1.17 |
| 5 | Cdc42ep3 | 1.58 | 1.99E-55 | 4.14E-51 | 0.30 | 0.16 | 0.15 | 1.94 |
| 5 | Traf4 | 0.63 | 3.71E-55 | 7.72E-51 | 0.74 | 0.62 | 0.12 | 1.19 |
| 5 | Rhoj | 1.06 | 4.49E-55 | 9.33E-51 | 0.26 | 0.11 | 0.14 | 2.28 |
| 5 | Pfn2 | 0.61 | 7.17E-55 | 1.49E-50 | 0.78 | 0.66 | 0.13 | 1.19 |
| 5 | Epb41l3 | 0.79 | 9.29E-55 | 1.93E-50 | 0.60 | 0.44 | 0.16 | 1.37 |
| 5 | Calr | 0.33 | 9.51E-55 | 1.98E-50 | 1.00 | 0.97 | 0.03 | 1.03 |
| 5 | Ccnd1 | 0.47 | 1.07E-54 | 2.22E-50 | 0.91 | 0.79 | 0.12 | 1.15 |
| 5 | Tuba1b | 0.26 | 2.01E-54 | 4.18E-50 | 1.00 | 0.99 | 0.01 | 1.01 |
| 5 | Glipr2 | 0.89 | 2.92E-54 | 6.07E-50 | 0.54 | 0.38 | 0.16 | 1.43 |
| 5 | Etfb | 0.42 | 3.02E-54 | 6.28E-50 | 0.93 | 0.89 | 0.04 | 1.05 |
| 5 | Nrk | 1.30 | 8.25E-54 | 1.72E-49 | 0.29 | 0.15 | 0.15 | 2.01 |
| 5 | Fap | 1.21 | 9.81E-53 | 2.04E-48 | 0.34 | 0.19 | 0.16 | 1.82 |
| 5 | Trio | 0.82 | 1.48E-52 | 3.08E-48 | 0.64 | 0.51 | 0.13 | 1.25 |
| 5 | Fndc3c1 | 0.78 | 1.66E-52 | 3.45E-48 | 0.61 | 0.46 | 0.15 | 1.32 |
| 5 | Rsrc1 | 0.56 | 1.72E-52 | 3.59E-48 | 0.82 | 0.74 | 0.08 | 1.11 |

**Supplementary Table 1. Marker genes for the original clusters of integrated clustering of E11.5 control and NotchGOF λ-junction cells.**

| | | | | | | | | |
|---|---|---|---|---|---|---|---|---|
| 5 | Emilin1 | 0.62 | 3.70E-52 | 7.70E-48 | 0.70 | 0.52 | 0.18 | 1.34 |
| 5 | Sulf2 | 0.51 | 5.02E-52 | 1.04E-47 | 0.81 | 0.66 | 0.15 | 1.23 |
| 5 | Usp46 | 0.64 | 5.43E-52 | 1.13E-47 | 0.73 | 0.62 | 0.11 | 1.18 |
| 5 | Gng2 | 0.79 | 7.71E-52 | 1.60E-47 | 0.69 | 0.56 | 0.13 | 1.22 |
| 5 | Tns3 | 0.78 | 2.59E-51 | 5.39E-47 | 0.63 | 0.49 | 0.14 | 1.29 |
| 5 | Adgrl3 | 0.85 | 5.42E-51 | 1.13E-46 | 0.58 | 0.43 | 0.15 | 1.35 |
| 5 | Ssbp4 | 0.56 | 7.01E-51 | 1.46E-46 | 0.76 | 0.65 | 0.12 | 1.18 |
| 5 | Khdrbs2 | 2.12 | 9.16E-51 | 1.90E-46 | 0.11 | 0.03 | 0.08 | 3.52 |
| 5 | Homer2 | 1.20 | 1.28E-50 | 2.67E-46 | 0.35 | 0.20 | 0.15 | 1.73 |
| 5 | Tes | 0.56 | 3.14E-50 | 6.52E-46 | 0.80 | 0.69 | 0.11 | 1.16 |
| 5 | Fstl1 | 0.38 | 8.93E-50 | 1.86E-45 | 0.93 | 0.82 | 0.11 | 1.14 |
| 5 | Fbn2 | 0.50 | 1.26E-49 | 2.63E-45 | 0.85 | 0.73 | 0.12 | 1.17 |
| 5 | Nectin2 | 0.76 | 2.68E-49 | 5.57E-45 | 0.61 | 0.47 | 0.14 | 1.30 |
| 5 | Ndufb8 | 0.26 | 1.44E-48 | 3.00E-44 | 0.99 | 0.98 | 0.01 | 1.01 |
| 5 | Tnfaip8 | 0.46 | 2.88E-48 | 5.99E-44 | 0.84 | 0.75 | 0.09 | 1.12 |
| 5 | Zyx | 0.68 | 9.59E-48 | 1.99E-43 | 0.63 | 0.50 | 0.13 | 1.25 |
| 5 | Sned1 | 1.19 | 1.47E-47 | 3.05E-43 | 0.25 | 0.12 | 0.13 | 2.10 |
| 5 | Rex1bd | 0.42 | 2.41E-47 | 5.02E-43 | 0.90 | 0.86 | 0.04 | 1.05 |
| 5 | Bex3 | 0.29 | 3.50E-47 | 7.28E-43 | 0.99 | 0.98 | 0.02 | 1.02 |
| 5 | Marcks | 0.29 | 3.94E-46 | 8.20E-42 | 1.00 | 0.97 | 0.03 | 1.03 |
| 5 | Cped1 | 0.61 | 6.64E-46 | 1.38E-41 | 0.63 | 0.46 | 0.18 | 1.38 |
| 5 | Gjc1 | 0.43 | 1.22E-45 | 2.53E-41 | 0.90 | 0.80 | 0.10 | 1.12 |
| 5 | Pfdn1 | 0.33 | 1.74E-45 | 3.62E-41 | 0.97 | 0.94 | 0.03 | 1.04 |
| 5 | Gm1673 | 0.31 | 2.20E-45 | 4.58E-41 | 0.98 | 0.93 | 0.05 | 1.05 |
| 5 | Pdia6 | 0.33 | 3.62E-45 | 7.54E-41 | 0.98 | 0.95 | 0.03 | 1.03 |
| 5 | Chd3 | 0.34 | 1.08E-44 | 2.25E-40 | 0.95 | 0.89 | 0.06 | 1.07 |
| 5 | Maged1 | 0.32 | 1.48E-44 | 3.07E-40 | 0.98 | 0.94 | 0.05 | 1.05 |
| 5 | Arhgap31 | 1.02 | 2.11E-44 | 4.38E-40 | 0.32 | 0.18 | 0.14 | 1.78 |
| 5 | Pknox2 | 1.93 | 3.27E-44 | 6.80E-40 | 0.13 | 0.05 | 0.08 | 2.84 |
| 5 | Fxyd6 | 0.54 | 7.44E-43 | 1.55E-38 | 0.78 | 0.68 | 0.10 | 1.15 |
| 5 | Epha3 | 0.68 | 8.20E-43 | 1.71E-38 | 0.52 | 0.35 | 0.18 | 1.50 |
| 5 | Ddah2 | 0.30 | 2.14E-42 | 4.46E-38 | 0.99 | 0.95 | 0.04 | 1.04 |
| 5 | Serp2 | 1.13 | 2.21E-42 | 4.59E-38 | 0.32 | 0.19 | 0.13 | 1.70 |
| 5 | Ccne1 | 0.67 | 1.25E-41 | 2.60E-37 | 0.64 | 0.50 | 0.14 | 1.29 |
| 5 | Gm17167 | 1.83 | 3.11E-41 | 6.46E-37 | 0.13 | 0.05 | 0.08 | 2.73 |
| 5 | Rab34 | 0.47 | 3.32E-41 | 6.90E-37 | 0.81 | 0.71 | 0.10 | 1.14 |
| 5 | Spc24 | 0.49 | 6.59E-41 | 1.37E-36 | 0.80 | 0.73 | 0.07 | 1.09 |
| 5 | Adamtsl1 | 1.47 | 7.93E-41 | 1.65E-36 | 0.21 | 0.10 | 0.11 | 2.12 |
| 5 | Uqcc2 | 0.28 | 9.60E-41 | 2.00E-36 | 0.98 | 0.95 | 0.03 | 1.03 |
| 5 | Asxl3 | 0.91 | 1.25E-40 | 2.60E-36 | 0.57 | 0.46 | 0.11 | 1.24 |
| 5 | Ndufa13 | 0.27 | 2.07E-40 | 4.30E-36 | 1.00 | 0.98 | 0.02 | 1.02 |
| 5 | Serpinh1 | 0.29 | 4.04E-40 | 8.40E-36 | 1.00 | 0.94 | 0.06 | 1.06 |
| 5 | Selenop | 0.63 | 6.37E-40 | 1.32E-35 | 0.32 | 0.18 | 0.14 | 1.75 |
| 5 | Rgma | 0.93 | 7.38E-40 | 1.54E-35 | 0.37 | 0.23 | 0.14 | 1.59 |
| 5 | Clic1 | 0.31 | 1.31E-39 | 2.73E-35 | 0.95 | 0.92 | 0.03 | 1.03 |
| 5 | Dusp6 | 0.74 | 1.91E-39 | 3.97E-35 | 0.50 | 0.35 | 0.15 | 1.43 |
| 5 | Zfp608 | 0.63 | 4.85E-39 | 1.01E-34 | 0.65 | 0.53 | 0.12 | 1.22 |
| 5 | Lrp1 | 0.59 | 8.04E-39 | 1.67E-34 | 0.68 | 0.57 | 0.11 | 1.19 |
| 5 | Dab2 | 0.60 | 1.98E-38 | 4.11E-34 | 0.50 | 0.35 | 0.15 | 1.43 |
| 5 | Ppic | 0.41 | 5.24E-38 | 1.09E-33 | 0.81 | 0.70 | 0.10 | 1.15 |
| 5 | Cst3 | 0.26 | 1.05E-37 | 2.18E-33 | 0.95 | 0.92 | 0.04 | 1.04 |
| 5 | Clmp | 0.42 | 1.24E-37 | 2.57E-33 | 0.81 | 0.63 | 0.18 | 1.28 |
| 5 | Eya2 | 0.54 | 1.79E-37 | 3.73E-33 | 0.37 | 0.22 | 0.15 | 1.67 |
| 5 | Cald1 | 0.30 | 2.10E-37 | 4.37E-33 | 1.00 | 0.88 | 0.11 | 1.13 |
| 5 | Fat4 | 0.81 | 2.67E-37 | 5.55E-33 | 0.41 | 0.27 | 0.14 | 1.52 |
| 5 | Idh2 | 0.33 | 4.00E-37 | 8.31E-33 | 0.95 | 0.89 | 0.05 | 1.06 |

**Supplementary Table 1. Marker genes for the original clusters of integrated clustering of E11.5 control and NotchGOF λ-junction cells.**

| | | | | | | | | |
|---|---|---|---|---|---|---|---|---|
| 5 | Cited1 | 1.36 | 4.05E-37 | 8.41E-33 | 0.22 | 0.11 | 0.11 | 1.96 |
| 5 | Itm2c | 0.44 | 4.61E-37 | 9.58E-33 | 0.82 | 0.72 | 0.10 | 1.14 |
| 5 | Igsf3 | 0.55 | 7.62E-37 | 1.59E-32 | 0.68 | 0.58 | 0.10 | 1.18 |
| 5 | Ndufc1 | 0.26 | 2.16E-36 | 4.49E-32 | 0.98 | 0.97 | 0.01 | 1.01 |
| 5 | Zcchc24 | 0.84 | 1.20E-35 | 2.51E-31 | 0.42 | 0.28 | 0.13 | 1.47 |
| 5 | Col23a1 | 0.78 | 1.48E-35 | 3.07E-31 | 0.44 | 0.31 | 0.14 | 1.46 |
| 5 | Megf6 | 1.03 | 1.48E-35 | 3.07E-31 | 0.33 | 0.21 | 0.12 | 1.60 |
| 5 | Selenom | 0.47 | 1.49E-35 | 3.11E-31 | 0.63 | 0.52 | 0.12 | 1.22 |
| 5 | Sorbs2 | 0.96 | 1.54E-35 | 3.21E-31 | 0.35 | 0.21 | 0.13 | 1.63 |
| 5 | Ssr2 | 0.30 | 3.22E-35 | 6.69E-31 | 0.98 | 0.94 | 0.04 | 1.04 |
| 5 | Nid2 | 0.99 | 3.54E-35 | 7.36E-31 | 0.36 | 0.23 | 0.13 | 1.55 |
| 5 | Kifap3 | 0.42 | 3.81E-35 | 7.93E-31 | 0.84 | 0.77 | 0.07 | 1.09 |
| 5 | Mageh1 | 0.56 | 5.82E-35 | 1.21E-30 | 0.66 | 0.56 | 0.10 | 1.18 |
| 5 | Akap7 | 1.09 | 6.75E-35 | 1.40E-30 | 0.34 | 0.22 | 0.12 | 1.55 |
| 5 | Hsp90b1 | 0.26 | 2.07E-34 | 4.30E-30 | 1.00 | 0.98 | 0.02 | 1.02 |
| 5 | Reep5 | 0.34 | 3.41E-34 | 7.09E-30 | 0.87 | 0.78 | 0.10 | 1.12 |
| 5 | Hnrnpdl | 0.25 | 6.07E-34 | 1.26E-29 | 0.99 | 0.97 | 0.02 | 1.02 |
| 5 | Fibin | 1.24 | 7.07E-34 | 1.47E-29 | 0.23 | 0.12 | 0.11 | 1.91 |
| 5 | Flna | 0.37 | 1.77E-33 | 3.67E-29 | 0.87 | 0.79 | 0.08 | 1.10 |
| 5 | Cnpy2 | 0.28 | 1.92E-33 | 3.99E-29 | 0.97 | 0.93 | 0.03 | 1.04 |
| 5 | Zfp422 | 0.34 | 2.19E-33 | 4.55E-29 | 0.90 | 0.85 | 0.06 | 1.06 |
| 5 | Nsmce4a | 0.32 | 2.40E-33 | 5.00E-29 | 0.91 | 0.88 | 0.04 | 1.04 |
| 5 | Kdelr2 | 0.37 | 9.88E-33 | 2.06E-28 | 0.90 | 0.83 | 0.07 | 1.08 |
| 5 | Chchd7 | 0.51 | 1.25E-32 | 2.61E-28 | 0.77 | 0.72 | 0.05 | 1.06 |
| 5 | Fez1 | 1.24 | 1.89E-32 | 3.93E-28 | 0.25 | 0.14 | 0.11 | 1.77 |
| 5 | Slit2 | 0.83 | 2.91E-32 | 6.04E-28 | 0.49 | 0.36 | 0.13 | 1.35 |
| 5 | Cacna2d1 | 1.13 | 3.44E-32 | 7.15E-28 | 0.22 | 0.11 | 0.10 | 1.90 |
| 5 | Med13l | 0.60 | 4.81E-32 | 1.00E-27 | 0.59 | 0.50 | 0.09 | 1.19 |
| 5 | Polr2m | 0.26 | 9.20E-32 | 1.91E-27 | 0.98 | 0.95 | 0.03 | 1.03 |
| 5 | Lama2 | 1.02 | 9.29E-32 | 1.93E-27 | 0.25 | 0.14 | 0.11 | 1.79 |
| 5 | Txndc12 | 0.43 | 1.71E-31 | 3.55E-27 | 0.77 | 0.70 | 0.07 | 1.10 |
| 5 | Stra6 | 0.74 | 3.48E-31 | 7.24E-27 | 0.46 | 0.34 | 0.12 | 1.36 |
| 5 | Limch1 | 0.32 | 3.89E-31 | 8.09E-27 | 0.83 | 0.67 | 0.16 | 1.24 |
| 5 | Ebf2 | 0.70 | 4.64E-31 | 9.65E-27 | 0.10 | 0.04 | 0.06 | 2.73 |
| 5 | Gpc5 | 1.67 | 7.31E-31 | 1.52E-26 | 0.12 | 0.05 | 0.07 | 2.49 |
| 5 | Pcdh7 | 0.60 | 7.51E-31 | 1.56E-26 | 0.53 | 0.39 | 0.14 | 1.37 |
| 5 | Shisa4 | 0.73 | 1.15E-30 | 2.39E-26 | 0.45 | 0.32 | 0.12 | 1.38 |
| 5 | Kctd1 | 0.55 | 1.26E-30 | 2.61E-26 | 0.58 | 0.46 | 0.12 | 1.25 |
| 5 | Cmtm3 | 0.62 | 1.88E-30 | 3.91E-26 | 0.53 | 0.41 | 0.12 | 1.28 |
| 5 | Pax7 | 0.45 | 3.65E-30 | 7.60E-26 | 0.38 | 0.23 | 0.15 | 1.65 |
| 5 | Epha7 | 0.50 | 4.92E-30 | 1.02E-25 | 0.68 | 0.56 | 0.12 | 1.21 |
| 5 | Ruvbl1 | 0.33 | 8.42E-30 | 1.75E-25 | 0.90 | 0.84 | 0.06 | 1.07 |
| 5 | Hmgn3 | 0.39 | 8.79E-30 | 1.83E-25 | 0.82 | 0.75 | 0.07 | 1.10 |
| 5 | Metrn | 0.52 | 9.10E-30 | 1.89E-25 | 0.61 | 0.51 | 0.10 | 1.20 |
| 5 | Lsm2 | 0.26 | 1.49E-29 | 3.09E-25 | 0.95 | 0.92 | 0.02 | 1.02 |
| 5 | Eef2k | 0.85 | 1.72E-29 | 3.58E-25 | 0.35 | 0.24 | 0.12 | 1.48 |
| 5 | Ackr3 | 0.74 | 3.77E-29 | 7.83E-25 | 0.51 | 0.41 | 0.10 | 1.25 |
| 5 | Erbin | 0.47 | 4.09E-29 | 8.51E-25 | 0.71 | 0.63 | 0.08 | 1.13 |
| 5 | Gkap1 | 0.57 | 4.29E-29 | 8.93E-25 | 0.54 | 0.43 | 0.12 | 1.27 |
| 5 | Drap1 | 0.27 | 5.82E-29 | 1.21E-24 | 0.96 | 0.93 | 0.02 | 1.03 |
| 5 | P4hb | 0.32 | 6.67E-29 | 1.39E-24 | 0.93 | 0.88 | 0.05 | 1.05 |
| 5 | Slc38a1 | 0.37 | 8.83E-29 | 1.84E-24 | 0.83 | 0.76 | 0.07 | 1.09 |
| 5 | Gulp1 | 0.83 | 9.42E-29 | 1.96E-24 | 0.36 | 0.24 | 0.11 | 1.47 |
| 5 | Zeb1 | 0.62 | 9.46E-29 | 1.97E-24 | 0.47 | 0.34 | 0.13 | 1.37 |
| 5 | Pigx | 0.39 | 1.03E-28 | 2.14E-24 | 0.81 | 0.75 | 0.06 | 1.08 |
| 5 | Col12a1 | 1.29 | 1.27E-28 | 2.65E-24 | 0.13 | 0.06 | 0.07 | 2.28 |

**Supplementary Table 1. Marker genes for the original clusters of integrated clustering of E11.5 control and NotchGOF λ-junction cells.**

| | | | | | | | | |
|---|---|---|---|---|---|---|---|---|
| 5 | Ets1 | 0.57 | 1.59E-28 | 3.31E-24 | 0.28 | 0.17 | 0.11 | 1.65 |
| 5 | Sfrp2 | 0.57 | 1.66E-28 | 3.44E-24 | 0.69 | 0.61 | 0.08 | 1.13 |
| 5 | E2f6 | 0.55 | 2.91E-28 | 6.04E-24 | 0.62 | 0.52 | 0.10 | 1.19 |
| 5 | Tmem35a | 1.16 | 4.74E-28 | 9.85E-24 | 0.18 | 0.09 | 0.09 | 1.95 |
| 5 | Rrbp1 | 0.28 | 8.20E-28 | 1.70E-23 | 0.94 | 0.89 | 0.04 | 1.05 |
| 5 | Frem1 | 0.58 | 8.21E-28 | 1.71E-23 | 0.53 | 0.40 | 0.13 | 1.32 |
| 5 | Kcnq1ot1 | 0.31 | 9.06E-28 | 1.88E-23 | 0.97 | 0.92 | 0.05 | 1.06 |
| 5 | Cpxm1 | 0.61 | 9.16E-28 | 1.91E-23 | 0.54 | 0.43 | 0.11 | 1.26 |
| 5 | Sec62 | 0.29 | 9.75E-28 | 2.03E-23 | 0.96 | 0.93 | 0.02 | 1.03 |
| 5 | Eif1b | 0.26 | 1.31E-27 | 2.72E-23 | 0.96 | 0.93 | 0.03 | 1.04 |
| 5 | 2610002M06Rik | 0.84 | 2.61E-27 | 5.44E-23 | 0.39 | 0.29 | 0.11 | 1.37 |
| 5 | Tmem119 | 0.51 | 3.25E-27 | 6.76E-23 | 0.58 | 0.45 | 0.13 | 1.28 |
| 5 | Eya4 | 0.84 | 3.44E-27 | 7.15E-23 | 0.39 | 0.28 | 0.11 | 1.40 |
| 5 | Slc16a2 | 1.11 | 4.04E-27 | 8.40E-23 | 0.25 | 0.15 | 0.10 | 1.68 |
| 5 | Ndn | 0.34 | 4.59E-27 | 9.56E-23 | 0.81 | 0.74 | 0.07 | 1.09 |
| 5 | Sumo3 | 0.28 | 5.49E-27 | 1.14E-22 | 0.93 | 0.90 | 0.03 | 1.03 |
| 5 | Slc25a24 | 0.74 | 5.61E-27 | 1.17E-22 | 0.45 | 0.34 | 0.11 | 1.31 |
| 5 | Sh2d3c | 1.17 | 7.29E-27 | 1.52E-22 | 0.18 | 0.10 | 0.09 | 1.91 |
| 5 | P3h3 | 0.74 | 8.74E-27 | 1.82E-22 | 0.41 | 0.30 | 0.11 | 1.38 |
| 5 | Zfp36l1 | 0.30 | 1.36E-26 | 2.83E-22 | 0.90 | 0.85 | 0.06 | 1.07 |
| 5 | Plod2 | 0.59 | 1.80E-26 | 3.74E-22 | 0.59 | 0.49 | 0.10 | 1.20 |
| 5 | Dpysl4 | 0.89 | 1.81E-26 | 3.77E-22 | 0.28 | 0.18 | 0.11 | 1.60 |
| 5 | Adamts3 | 1.62 | 2.31E-26 | 4.80E-22 | 0.13 | 0.06 | 0.07 | 2.19 |
| 5 | Mpc2 | 0.37 | 2.64E-26 | 5.49E-22 | 0.83 | 0.79 | 0.04 | 1.05 |
| 5 | Car14 | 0.56 | 5.41E-26 | 1.12E-21 | 0.53 | 0.42 | 0.11 | 1.27 |
| 5 | AY036118 | 0.63 | 8.50E-26 | 1.77E-21 | 0.67 | 0.61 | 0.06 | 1.10 |
| 5 | Maml3 | 0.63 | 1.10E-25 | 2.29E-21 | 0.49 | 0.39 | 0.10 | 1.26 |
| 5 | Irf2bp2 | 0.62 | 1.43E-25 | 2.98E-21 | 0.45 | 0.34 | 0.11 | 1.32 |
| 5 | Flrt3 | 0.71 | 1.45E-25 | 3.03E-21 | 0.54 | 0.44 | 0.10 | 1.22 |
| 5 | Fat1 | 0.55 | 2.00E-25 | 4.17E-21 | 0.59 | 0.51 | 0.09 | 1.17 |
| 5 | Tceal8 | 0.29 | 2.26E-25 | 4.70E-21 | 0.92 | 0.89 | 0.04 | 1.04 |
| 5 | Lin28b | 0.92 | 2.91E-25 | 6.05E-21 | 0.30 | 0.20 | 0.10 | 1.50 |
| 5 | Lima1 | 0.32 | 3.79E-25 | 7.89E-21 | 0.83 | 0.73 | 0.10 | 1.13 |
| 5 | Nes | 0.47 | 3.82E-25 | 7.95E-21 | 0.53 | 0.42 | 0.12 | 1.28 |
| 5 | Rbms1 | 0.35 | 3.85E-25 | 8.01E-21 | 0.82 | 0.78 | 0.04 | 1.06 |
| 5 | Ccdc107 | 0.64 | 6.25E-25 | 1.30E-20 | 0.51 | 0.41 | 0.09 | 1.23 |
| 5 | Ogfrl1 | 0.76 | 6.48E-25 | 1.35E-20 | 0.43 | 0.33 | 0.10 | 1.29 |
| 5 | Spred3 | 1.16 | 7.22E-25 | 1.50E-20 | 0.17 | 0.09 | 0.08 | 1.90 |
| 5 | Eno3 | 0.60 | 9.08E-25 | 1.89E-20 | 0.48 | 0.37 | 0.11 | 1.30 |
| 5 | Itpr1 | 0.87 | 1.07E-24 | 2.23E-20 | 0.31 | 0.21 | 0.10 | 1.47 |
| 5 | Inppl1 | 0.66 | 1.39E-24 | 2.88E-20 | 0.48 | 0.39 | 0.09 | 1.24 |
| 5 | Tmed10 | 0.26 | 1.75E-24 | 3.63E-20 | 0.96 | 0.94 | 0.02 | 1.02 |
| 5 | Lmo4 | 0.37 | 2.13E-24 | 4.44E-20 | 0.87 | 0.81 | 0.06 | 1.07 |
| 5 | Col4a2 | 0.38 | 2.76E-24 | 5.73E-20 | 0.53 | 0.42 | 0.11 | 1.26 |
| 5 | Tmem165 | 0.31 | 3.31E-24 | 6.88E-20 | 0.85 | 0.80 | 0.04 | 1.05 |
| 5 | Plscr1 | 0.94 | 3.35E-24 | 6.96E-20 | 0.32 | 0.23 | 0.09 | 1.41 |
| 5 | Hmgn5 | 0.44 | 3.57E-24 | 7.43E-20 | 0.77 | 0.71 | 0.07 | 1.09 |
| 5 | Pclaf | 0.29 | 3.65E-24 | 7.59E-20 | 0.95 | 0.88 | 0.06 | 1.07 |
| 5 | Parm1 | 0.84 | 4.03E-24 | 8.38E-20 | 0.26 | 0.16 | 0.10 | 1.63 |
| 5 | Erp29 | 0.29 | 5.12E-24 | 1.06E-19 | 0.91 | 0.88 | 0.03 | 1.03 |
| 5 | Isyna1 | 0.41 | 1.59E-23 | 3.31E-19 | 0.74 | 0.68 | 0.06 | 1.09 |
| 5 | Scarf2 | 0.73 | 2.04E-23 | 4.24E-19 | 0.36 | 0.25 | 0.10 | 1.41 |
| 5 | Svbp | 0.29 | 2.14E-23 | 4.45E-19 | 0.89 | 0.87 | 0.03 | 1.03 |
| 5 | Prim1 | 0.41 | 2.46E-23 | 5.11E-19 | 0.73 | 0.67 | 0.06 | 1.09 |
| 5 | Ccdc85b | 0.36 | 3.27E-23 | 6.79E-19 | 0.65 | 0.58 | 0.06 | 1.11 |
| 5 | Tmem176a | 0.54 | 3.95E-23 | 8.22E-19 | 0.50 | 0.40 | 0.10 | 1.25 |

**Supplementary Table 1. Marker genes for the original clusters of integrated clustering of E11.5 control and NotchGOF λ-junction cells.**

| | | | | | | | | |
|---|---|---|---|---|---|---|---|---|
| 5 | Il6st | 0.77 | 5.84E-23 | 1.22E-18 | 0.37 | 0.28 | 0.10 | 1.35 |
| 5 | St3gal1 | 1.38 | 6.65E-23 | 1.38E-18 | 0.17 | 0.10 | 0.08 | 1.78 |
| 5 | Txndc5 | 0.42 | 8.06E-23 | 1.68E-18 | 0.69 | 0.61 | 0.08 | 1.13 |
| 5 | Wdr86 | 0.46 | 9.21E-23 | 1.91E-18 | 0.60 | 0.51 | 0.09 | 1.18 |
| 5 | Nid1 | 0.53 | 1.06E-22 | 2.20E-18 | 0.45 | 0.34 | 0.11 | 1.34 |
| 5 | Inka1 | 0.39 | 2.17E-22 | 4.52E-18 | 0.61 | 0.51 | 0.11 | 1.21 |
| 5 | Rhoc | 0.26 | 2.90E-22 | 6.03E-18 | 0.87 | 0.78 | 0.08 | 1.10 |
| 5 | Tln1 | 0.31 | 5.86E-22 | 1.22E-17 | 0.79 | 0.74 | 0.05 | 1.06 |
| 5 | Slbp | 0.30 | 7.29E-22 | 1.52E-17 | 0.90 | 0.87 | 0.03 | 1.03 |
| 5 | Cldn10 | 1.20 | 9.66E-22 | 2.01E-17 | 0.10 | 0.04 | 0.06 | 2.30 |
| 5 | Rhod | 0.70 | 1.05E-21 | 2.19E-17 | 0.42 | 0.34 | 0.09 | 1.26 |
| 5 | Tmod2 | 0.93 | 1.34E-21 | 2.79E-17 | 0.28 | 0.19 | 0.09 | 1.48 |
| 5 | Tspyl4 | 0.81 | 1.49E-21 | 3.09E-17 | 0.33 | 0.23 | 0.09 | 1.40 |
| 5 | Hmg20b | 0.30 | 1.90E-21 | 3.95E-17 | 0.84 | 0.80 | 0.04 | 1.06 |
| 5 | Arhgap28 | 0.38 | 2.11E-21 | 4.39E-17 | 0.63 | 0.53 | 0.10 | 1.19 |
| 5 | Dnajb11 | 0.36 | 2.80E-21 | 5.83E-17 | 0.74 | 0.68 | 0.05 | 1.08 |
| 5 | Sgip1 | 1.23 | 2.90E-21 | 6.03E-17 | 0.18 | 0.11 | 0.08 | 1.72 |
| 5 | Dact3 | 0.47 | 3.86E-21 | 8.02E-17 | 0.59 | 0.49 | 0.10 | 1.20 |
| 5 | Vav3 | 1.37 | 3.91E-21 | 8.14E-17 | 0.12 | 0.06 | 0.06 | 2.11 |
| 5 | Adora2b | 1.09 | 3.95E-21 | 8.21E-17 | 0.18 | 0.10 | 0.08 | 1.76 |
| 5 | Bicc1 | 0.81 | 4.88E-21 | 1.02E-16 | 0.32 | 0.23 | 0.09 | 1.41 |
| 5 | Tmem256 | 0.26 | 5.46E-21 | 1.13E-16 | 0.91 | 0.89 | 0.02 | 1.02 |
| 5 | Pmp22 | 0.77 | 6.07E-21 | 1.26E-16 | 0.27 | 0.18 | 0.09 | 1.53 |
| 5 | Sdc3 | 0.77 | 7.01E-21 | 1.46E-16 | 0.29 | 0.20 | 0.10 | 1.48 |
| 5 | Rragd | 0.57 | 7.25E-21 | 1.51E-16 | 0.32 | 0.22 | 0.10 | 1.45 |
| 5 | Fam92a | 0.34 | 1.14E-20 | 2.37E-16 | 0.81 | 0.75 | 0.06 | 1.07 |
| 5 | Fbxl7 | 0.51 | 1.35E-20 | 2.80E-16 | 0.55 | 0.46 | 0.09 | 1.20 |
| 5 | Gas2 | 0.72 | 2.47E-20 | 5.14E-16 | 0.34 | 0.25 | 0.09 | 1.35 |
| 5 | Fkbp10 | 0.63 | 3.79E-20 | 7.89E-16 | 0.43 | 0.33 | 0.10 | 1.29 |
| 5 | Armcx3 | 0.53 | 5.09E-20 | 1.06E-15 | 0.51 | 0.43 | 0.08 | 1.19 |
| 5 | Tyms | 0.33 | 5.45E-20 | 1.13E-15 | 0.86 | 0.83 | 0.03 | 1.03 |
| 5 | Rbfox2 | 0.28 | 7.76E-20 | 1.61E-15 | 0.81 | 0.77 | 0.05 | 1.06 |
| 5 | Tmed1 | 0.39 | 8.12E-20 | 1.69E-15 | 0.73 | 0.67 | 0.06 | 1.09 |
| 5 | Arhgef9 | 0.81 | 9.35E-20 | 1.94E-15 | 0.27 | 0.18 | 0.09 | 1.48 |
| 5 | Gipc1 | 0.37 | 1.16E-19 | 2.41E-15 | 0.74 | 0.69 | 0.05 | 1.08 |
| 5 | Pgf | 1.22 | 1.48E-19 | 3.07E-15 | 0.13 | 0.07 | 0.06 | 1.93 |
| 5 | Timm17b | 0.29 | 1.50E-19 | 3.12E-15 | 0.84 | 0.82 | 0.02 | 1.02 |
| 5 | Mpzl1 | 0.36 | 1.66E-19 | 3.44E-15 | 0.70 | 0.63 | 0.06 | 1.10 |
| 5 | Mcrip1 | 0.25 | 1.85E-19 | 3.85E-15 | 0.89 | 0.86 | 0.04 | 1.04 |
| 5 | Plekho1 | 0.37 | 2.22E-19 | 4.62E-15 | 0.69 | 0.61 | 0.07 | 1.12 |
| 5 | Wbp1 | 0.33 | 3.18E-19 | 6.61E-15 | 0.81 | 0.77 | 0.04 | 1.05 |
| 5 | Brd8 | 0.30 | 3.43E-19 | 7.14E-15 | 0.80 | 0.75 | 0.05 | 1.06 |
| 5 | Gng10 | 0.26 | 4.06E-19 | 8.45E-15 | 0.87 | 0.81 | 0.06 | 1.07 |
| 5 | 0610010K14Rik | 0.28 | 4.17E-19 | 8.68E-15 | 0.85 | 0.81 | 0.04 | 1.05 |
| 5 | Xpnpep1 | 0.49 | 4.41E-19 | 9.18E-15 | 0.58 | 0.52 | 0.06 | 1.11 |
| 5 | Foxo4 | 1.09 | 5.01E-19 | 1.04E-14 | 0.19 | 0.11 | 0.07 | 1.65 |
| 5 | Tmem108 | 0.69 | 5.29E-19 | 1.10E-14 | 0.31 | 0.21 | 0.09 | 1.43 |
| 5 | Pdgfrb | 0.39 | 6.17E-19 | 1.28E-14 | 0.56 | 0.45 | 0.11 | 1.24 |
| 5 | Sec24d | 1.02 | 1.15E-18 | 2.39E-14 | 0.15 | 0.08 | 0.07 | 1.84 |
| 5 | Fdx1 | 0.45 | 1.23E-18 | 2.56E-14 | 0.60 | 0.53 | 0.07 | 1.13 |
| 5 | Tcf7l1 | 0.46 | 1.24E-18 | 2.58E-14 | 0.64 | 0.58 | 0.06 | 1.10 |
| 5 | Frk | 1.06 | 1.25E-18 | 2.60E-14 | 0.22 | 0.14 | 0.08 | 1.54 |
| 5 | Ror1 | 0.98 | 1.75E-18 | 3.64E-14 | 0.19 | 0.12 | 0.08 | 1.64 |
| 5 | Cfl2 | 0.27 | 2.12E-18 | 4.40E-14 | 0.85 | 0.81 | 0.04 | 1.05 |
| 5 | Prkcsh | 0.34 | 2.96E-18 | 6.15E-14 | 0.76 | 0.72 | 0.04 | 1.06 |
| 5 | Pdia4 | 0.33 | 2.96E-18 | 6.15E-14 | 0.83 | 0.78 | 0.05 | 1.06 |

**Supplementary Table 1. Marker genes for the original clusters of integrated clustering of E11.5 control and NotchGOF λ-junction cells.**

| | | | | | | | | |
|---|---|---|---|---|---|---|---|---|
| 5 | Angptl4 | 0.56 | 3.15E-18 | 6.55E-14 | 0.42 | 0.33 | 0.09 | 1.28 |
| 5 | St6galnac3 | 0.53 | 5.62E-18 | 1.17E-13 | 0.48 | 0.39 | 0.09 | 1.23 |
| 5 | 5330434G04Rik | 1.20 | 5.70E-18 | 1.19E-13 | 0.15 | 0.09 | 0.07 | 1.75 |
| 5 | Fam111a | 0.64 | 5.94E-18 | 1.24E-13 | 0.41 | 0.32 | 0.09 | 1.27 |
| 5 | Bcl7c | 0.26 | 6.02E-18 | 1.25E-13 | 0.86 | 0.83 | 0.03 | 1.04 |
| 5 | Prkra | 0.35 | 6.52E-18 | 1.36E-13 | 0.72 | 0.68 | 0.04 | 1.05 |
| 5 | Galnt1 | 0.34 | 7.42E-18 | 1.54E-13 | 0.71 | 0.66 | 0.05 | 1.07 |
| 5 | Phgdh | 0.26 | 8.50E-18 | 1.77E-13 | 0.88 | 0.82 | 0.06 | 1.07 |
| 5 | Frmd4a | 0.32 | 9.51E-18 | 1.98E-13 | 0.73 | 0.67 | 0.06 | 1.09 |
| 5 | Hist1h3c | 0.47 | 9.75E-18 | 2.03E-13 | 0.66 | 0.59 | 0.07 | 1.12 |
| 5 | Klf12 | 0.83 | 1.08E-17 | 2.25E-13 | 0.23 | 0.15 | 0.08 | 1.52 |
| 5 | Wwp1 | 0.57 | 1.14E-17 | 2.38E-13 | 0.48 | 0.41 | 0.07 | 1.16 |
| 5 | Spry1 | 0.60 | 1.16E-17 | 2.42E-13 | 0.37 | 0.28 | 0.09 | 1.31 |
| 5 | Zfp637 | 0.32 | 1.17E-17 | 2.44E-13 | 0.76 | 0.73 | 0.03 | 1.03 |
| 5 | Col27a1 | 0.86 | 1.23E-17 | 2.56E-13 | 0.26 | 0.18 | 0.08 | 1.45 |
| 5 | Loxl2 | 0.43 | 1.24E-17 | 2.58E-13 | 0.52 | 0.43 | 0.09 | 1.20 |
| 5 | Sptbn1 | 0.27 | 1.48E-17 | 3.09E-13 | 0.82 | 0.76 | 0.06 | 1.08 |
| 5 | Tsen34 | 0.26 | 1.49E-17 | 3.09E-13 | 0.88 | 0.84 | 0.04 | 1.05 |
| 5 | Mesd | 0.29 | 1.58E-17 | 3.29E-13 | 0.81 | 0.78 | 0.04 | 1.05 |
| 5 | Sgcb | 0.82 | 2.52E-17 | 5.23E-13 | 0.28 | 0.20 | 0.08 | 1.39 |
| 5 | Cenpm | 0.42 | 3.02E-17 | 6.28E-13 | 0.65 | 0.60 | 0.06 | 1.09 |
| 5 | Timm22 | 0.28 | 3.90E-17 | 8.11E-13 | 0.80 | 0.77 | 0.03 | 1.04 |
| 5 | Galt | 0.77 | 5.04E-17 | 1.05E-12 | 0.35 | 0.27 | 0.08 | 1.28 |
| 5 | Calu | 0.33 | 5.40E-17 | 1.12E-12 | 0.75 | 0.70 | 0.05 | 1.07 |
| 5 | Pop4 | 0.50 | 7.84E-17 | 1.63E-12 | 0.52 | 0.46 | 0.07 | 1.14 |
| 5 | Bach2 | 0.51 | 9.55E-17 | 1.99E-12 | 0.46 | 0.37 | 0.09 | 1.23 |
| 5 | Gyg | 0.52 | 1.11E-16 | 2.31E-12 | 0.49 | 0.42 | 0.07 | 1.17 |
| 5 | Stc1 | 1.26 | 1.51E-16 | 3.14E-12 | 0.12 | 0.07 | 0.06 | 1.85 |
| 5 | Fbln5 | 0.92 | 1.73E-16 | 3.61E-12 | 0.21 | 0.14 | 0.07 | 1.53 |
| 5 | Cbr1 | 0.28 | 1.83E-16 | 3.81E-12 | 0.79 | 0.76 | 0.03 | 1.04 |
| 5 | Itga9 | 0.72 | 2.36E-16 | 4.91E-12 | 0.29 | 0.21 | 0.08 | 1.39 |
| 5 | Tsen15 | 0.36 | 3.75E-16 | 7.80E-12 | 0.68 | 0.63 | 0.05 | 1.07 |
| 5 | Zmym4 | 0.32 | 3.86E-16 | 8.03E-12 | 0.75 | 0.70 | 0.05 | 1.07 |
| 5 | Nmral1 | 0.31 | 4.06E-16 | 8.44E-12 | 0.75 | 0.71 | 0.04 | 1.06 |
| 5 | Fzd7 | 0.57 | 4.21E-16 | 8.76E-12 | 0.43 | 0.35 | 0.08 | 1.23 |
| 5 | Fbn1 | 0.59 | 5.16E-16 | 1.07E-11 | 0.41 | 0.33 | 0.08 | 1.25 |
| 5 | Gm36988 | 0.86 | 5.20E-16 | 1.08E-11 | 0.21 | 0.14 | 0.07 | 1.52 |
| 5 | Hirip3 | 0.36 | 1.02E-15 | 2.12E-11 | 0.74 | 0.69 | 0.06 | 1.08 |
| 5 | Csrp1 | 0.53 | 1.19E-15 | 2.48E-11 | 0.42 | 0.34 | 0.08 | 1.23 |
| 5 | Mtap | 0.29 | 1.21E-15 | 2.53E-11 | 0.75 | 0.69 | 0.05 | 1.07 |
| 5 | Ssr1 | 0.32 | 1.22E-15 | 2.54E-11 | 0.78 | 0.76 | 0.02 | 1.02 |
| 5 | Coprs | 0.44 | 1.47E-15 | 3.05E-11 | 0.55 | 0.49 | 0.06 | 1.13 |
| 5 | Ap1s3 | 1.16 | 1.80E-15 | 3.74E-11 | 0.12 | 0.07 | 0.05 | 1.83 |
| 5 | Rbm4b | 0.31 | 1.86E-15 | 3.87E-11 | 0.72 | 0.67 | 0.05 | 1.07 |
| 5 | Tmem147 | 0.34 | 1.88E-15 | 3.90E-11 | 0.71 | 0.68 | 0.03 | 1.04 |
| 5 | Vcl | 0.44 | 2.44E-15 | 5.08E-11 | 0.46 | 0.39 | 0.07 | 1.19 |
| 5 | Sh3gl3 | 0.34 | 2.72E-15 | 5.67E-11 | 0.42 | 0.33 | 0.10 | 1.30 |
| 5 | Rpn1 | 0.30 | 2.88E-15 | 5.99E-11 | 0.80 | 0.76 | 0.03 | 1.04 |
| 5 | Tln2 | 0.36 | 2.95E-15 | 6.13E-11 | 0.65 | 0.58 | 0.06 | 1.11 |
| 5 | BC028528 | 0.95 | 2.97E-15 | 6.17E-11 | 0.13 | 0.07 | 0.06 | 1.78 |
| 5 | Hist1h2ag | 0.62 | 3.09E-15 | 6.42E-11 | 0.38 | 0.31 | 0.07 | 1.23 |
| 5 | Mex3b | 0.29 | 3.40E-15 | 7.07E-11 | 0.76 | 0.69 | 0.07 | 1.10 |
| 5 | Ldb2 | 0.44 | 3.59E-15 | 7.47E-11 | 0.36 | 0.27 | 0.09 | 1.34 |
| 5 | Tigd2 | 0.55 | 3.64E-15 | 7.57E-11 | 0.46 | 0.39 | 0.07 | 1.17 |
| 5 | P3h2 | 1.16 | 4.14E-15 | 8.61E-11 | 0.13 | 0.07 | 0.06 | 1.76 |
| 5 | St6galnac6 | 0.80 | 4.23E-15 | 8.80E-11 | 0.23 | 0.16 | 0.07 | 1.47 |

**Supplementary Table 1. Marker genes for the original clusters of integrated clustering of E11.5 control and NotchGOF λ-junction cells.**

| | | | | | | | | |
|---|---|---|---|---|---|---|---|---|
| 5 | Flywch2 | 0.40 | 4.36E-15 | 9.07E-11 | 0.52 | 0.45 | 0.07 | 1.15 |
| 5 | Tspan15 | 0.73 | 4.47E-15 | 9.29E-11 | 0.26 | 0.19 | 0.08 | 1.40 |
| 5 | Enho | 0.41 | 5.44E-15 | 1.13E-10 | 0.52 | 0.45 | 0.08 | 1.17 |
| 5 | Rbmx | 0.29 | 6.18E-15 | 1.29E-10 | 0.78 | 0.74 | 0.04 | 1.05 |
| 5 | Pdlim7 | 0.27 | 7.32E-15 | 1.52E-10 | 0.80 | 0.75 | 0.05 | 1.06 |
| 5 | Fblim1 | 0.44 | 8.43E-15 | 1.75E-10 | 0.53 | 0.46 | 0.07 | 1.15 |
| 5 | Ccdc88a | 0.25 | 8.52E-15 | 1.77E-10 | 0.86 | 0.81 | 0.04 | 1.06 |
| 5 | Epb41l2 | 0.31 | 9.36E-15 | 1.95E-10 | 0.73 | 0.67 | 0.06 | 1.10 |
| 5 | Zmiz1 | 0.30 | 9.57E-15 | 1.99E-10 | 0.72 | 0.69 | 0.03 | 1.04 |
| 5 | Adcy4 | 1.16 | 1.05E-14 | 2.19E-10 | 0.11 | 0.06 | 0.05 | 1.84 |
| 5 | Top2a | 0.30 | 1.24E-14 | 2.59E-10 | 0.83 | 0.79 | 0.04 | 1.05 |
| 5 | Olfml3 | 0.56 | 1.27E-14 | 2.64E-10 | 0.36 | 0.28 | 0.08 | 1.29 |
| 5 | Zfp521 | 0.63 | 1.54E-14 | 3.21E-10 | 0.36 | 0.28 | 0.07 | 1.25 |
| 5 | Fyn | 0.44 | 1.68E-14 | 3.49E-10 | 0.51 | 0.44 | 0.07 | 1.15 |
| 5 | Fam184a | 0.96 | 1.74E-14 | 3.61E-10 | 0.18 | 0.11 | 0.06 | 1.56 |
| 5 | Boc | 0.78 | 1.83E-14 | 3.81E-10 | 0.27 | 0.20 | 0.07 | 1.36 |
| 5 | 1500009L16Rik | 0.34 | 1.88E-14 | 3.91E-10 | 0.63 | 0.58 | 0.05 | 1.09 |
| 5 | Wtip | 0.36 | 1.91E-14 | 3.96E-10 | 0.62 | 0.56 | 0.06 | 1.11 |
| 5 | Zfp580 | 0.48 | 2.00E-14 | 4.15E-10 | 0.50 | 0.44 | 0.06 | 1.14 |
| 5 | Desi2 | 0.44 | 2.11E-14 | 4.38E-10 | 0.51 | 0.44 | 0.06 | 1.14 |
| 5 | Spin2c | 0.42 | 3.01E-14 | 6.26E-10 | 0.50 | 0.42 | 0.08 | 1.19 |
| 5 | Tmem37 | 0.93 | 3.18E-14 | 6.61E-10 | 0.14 | 0.08 | 0.06 | 1.71 |
| 5 | Tk1 | 0.41 | 3.52E-14 | 7.31E-10 | 0.60 | 0.55 | 0.05 | 1.10 |
| 5 | Fam171b | 0.63 | 4.38E-14 | 9.11E-10 | 0.32 | 0.24 | 0.08 | 1.32 |
| 5 | Sox12 | 0.27 | 4.63E-14 | 9.63E-10 | 0.72 | 0.66 | 0.05 | 1.08 |
| 5 | Rraga | 0.28 | 6.68E-14 | 1.39E-09 | 0.74 | 0.73 | 0.01 | 1.02 |
| 5 | Tspan6 | 0.26 | 6.69E-14 | 1.39E-09 | 0.79 | 0.74 | 0.05 | 1.07 |
| 5 | 1810026B05Rik | 0.33 | 6.89E-14 | 1.43E-09 | 0.64 | 0.59 | 0.05 | 1.08 |
| 5 | Apbb2 | 0.54 | 7.33E-14 | 1.52E-09 | 0.38 | 0.30 | 0.07 | 1.25 |
| 5 | Gsto1 | 0.31 | 7.86E-14 | 1.63E-09 | 0.69 | 0.65 | 0.04 | 1.06 |
| 5 | Bmp1 | 0.52 | 9.02E-14 | 1.88E-09 | 0.48 | 0.42 | 0.06 | 1.14 |
| 5 | Ipo5 | 0.29 | 1.24E-13 | 2.57E-09 | 0.74 | 0.70 | 0.04 | 1.06 |
| 5 | Edn3 | 0.83 | 1.33E-13 | 2.77E-09 | 0.21 | 0.14 | 0.07 | 1.46 |
| 5 | B3gnt5 | 0.50 | 1.43E-13 | 2.98E-09 | 0.42 | 0.36 | 0.07 | 1.19 |
| 5 | Rimklb | 0.50 | 1.59E-13 | 3.32E-09 | 0.39 | 0.32 | 0.07 | 1.22 |
| 5 | Man1a2 | 0.29 | 2.55E-13 | 5.31E-09 | 0.72 | 0.69 | 0.03 | 1.05 |
| 5 | Timm50 | 0.29 | 3.36E-13 | 6.99E-09 | 0.76 | 0.74 | 0.02 | 1.03 |
| 5 | Adamts5 | 0.30 | 3.41E-13 | 7.10E-09 | 0.17 | 0.11 | 0.07 | 1.61 |
| 5 | Tcp11l1 | 0.56 | 3.50E-13 | 7.28E-09 | 0.36 | 0.30 | 0.07 | 1.23 |
| 5 | Sdf4 | 0.30 | 3.77E-13 | 7.84E-09 | 0.69 | 0.67 | 0.02 | 1.04 |
| 5 | Prom1 | 0.50 | 4.07E-13 | 8.47E-09 | 0.47 | 0.41 | 0.06 | 1.15 |
| 5 | Lox | 0.99 | 4.07E-13 | 8.47E-09 | 0.12 | 0.07 | 0.05 | 1.77 |
| 5 | Arxes1 | 0.51 | 5.47E-13 | 1.14E-08 | 0.44 | 0.37 | 0.07 | 1.18 |
| 5 | Hip1 | 0.41 | 6.08E-13 | 1.27E-08 | 0.51 | 0.44 | 0.06 | 1.14 |
| 5 | Lrrc8c | 0.80 | 7.03E-13 | 1.46E-08 | 0.18 | 0.12 | 0.06 | 1.50 |
| 5 | Skp2 | 0.30 | 7.54E-13 | 1.57E-08 | 0.73 | 0.69 | 0.04 | 1.05 |
| 5 | Lpl | 0.66 | 8.09E-13 | 1.68E-08 | 0.22 | 0.16 | 0.06 | 1.41 |
| 5 | Dpy19l1 | 0.55 | 8.49E-13 | 1.77E-08 | 0.42 | 0.36 | 0.06 | 1.17 |
| 5 | Sox8 | 1.13 | 8.67E-13 | 1.80E-08 | 0.10 | 0.06 | 0.05 | 1.82 |
| 5 | Nfib | 0.28 | 8.78E-13 | 1.83E-08 | 0.79 | 0.74 | 0.05 | 1.06 |
| 5 | Cfap20 | 0.26 | 9.51E-13 | 1.98E-08 | 0.79 | 0.74 | 0.04 | 1.06 |
| 5 | Neo1 | 0.40 | 1.05E-12 | 2.18E-08 | 0.55 | 0.50 | 0.05 | 1.11 |
| 5 | Hist1h2ae | 0.32 | 1.18E-12 | 2.46E-08 | 0.75 | 0.70 | 0.06 | 1.08 |
| 5 | Abtb2 | 1.03 | 1.25E-12 | 2.60E-08 | 0.12 | 0.07 | 0.05 | 1.71 |
| 5 | Cdk14 | 0.37 | 1.34E-12 | 2.78E-08 | 0.55 | 0.49 | 0.06 | 1.12 |
| 5 | Col6a3 | 0.72 | 1.39E-12 | 2.90E-08 | 0.23 | 0.16 | 0.07 | 1.41 |

**Supplementary Table 1. Marker genes for the original clusters of integrated clustering of E11.5 control and NotchGOF λ-junction cells.**

| | | | | | | | | |
|---|---|---|---|---|---|---|---|---|
| 5 | Ptgfrn | 0.51 | 1.43E-12 | 2.97E-08 | 0.40 | 0.33 | 0.07 | 1.20 |
| 5 | Olfml1 | 0.80 | 1.45E-12 | 3.01E-08 | 0.18 | 0.12 | 0.06 | 1.52 |
| 5 | Tipin | 0.26 | 1.67E-12 | 3.47E-08 | 0.81 | 0.77 | 0.04 | 1.05 |
| 5 | Sema3c | 0.65 | 1.78E-12 | 3.71E-08 | 0.15 | 0.09 | 0.06 | 1.62 |
| 5 | Mprip | 0.27 | 1.81E-12 | 3.76E-08 | 0.76 | 0.73 | 0.03 | 1.04 |
| 5 | Podxl2 | 0.40 | 2.57E-12 | 5.35E-08 | 0.48 | 0.42 | 0.06 | 1.15 |
| 5 | Qdpr | 0.33 | 3.20E-12 | 6.65E-08 | 0.58 | 0.54 | 0.04 | 1.07 |
| 5 | Mcc | 0.73 | 3.21E-12 | 6.68E-08 | 0.19 | 0.13 | 0.06 | 1.49 |
| 5 | Zfp57 | 0.55 | 3.31E-12 | 6.88E-08 | 0.40 | 0.33 | 0.06 | 1.19 |
| 5 | Lap3 | 0.28 | 3.78E-12 | 7.86E-08 | 0.71 | 0.67 | 0.04 | 1.05 |
| 5 | Sall2 | 0.40 | 4.35E-12 | 9.05E-08 | 0.46 | 0.39 | 0.07 | 1.17 |
| 5 | Camk2n1 | 0.39 | 4.36E-12 | 9.07E-08 | 0.26 | 0.19 | 0.07 | 1.39 |
| 5 | Nt5c3b | 0.27 | 4.65E-12 | 9.66E-08 | 0.71 | 0.68 | 0.03 | 1.04 |
| 5 | Sigmar1 | 0.44 | 4.73E-12 | 9.83E-08 | 0.59 | 0.55 | 0.04 | 1.08 |
| 5 | Cacna1h | 1.00 | 4.93E-12 | 1.03E-07 | 0.12 | 0.07 | 0.05 | 1.66 |
| 5 | Kmt2c | 0.44 | 5.46E-12 | 1.14E-07 | 0.53 | 0.49 | 0.04 | 1.08 |
| 5 | Bmp2 | 1.07 | 9.42E-12 | 1.96E-07 | 0.11 | 0.06 | 0.05 | 1.72 |
| 5 | Pwwp3a | 0.35 | 9.59E-12 | 2.00E-07 | 0.59 | 0.55 | 0.04 | 1.08 |
| 5 | Setbp1 | 0.44 | 1.23E-11 | 2.56E-07 | 0.45 | 0.38 | 0.07 | 1.17 |
| 5 | Tmem263 | 0.35 | 1.27E-11 | 2.64E-07 | 0.52 | 0.47 | 0.05 | 1.10 |
| 5 | Ctbp2 | 0.28 | 1.38E-11 | 2.87E-07 | 0.72 | 0.69 | 0.03 | 1.04 |
| 5 | Capn2 | 0.37 | 1.63E-11 | 3.40E-07 | 0.51 | 0.46 | 0.05 | 1.11 |
| 5 | Pxdn | 0.38 | 1.74E-11 | 3.61E-07 | 0.56 | 0.52 | 0.05 | 1.09 |
| 5 | Tspan4 | 0.87 | 1.83E-11 | 3.80E-07 | 0.18 | 0.12 | 0.06 | 1.44 |
| 5 | Fkbp7 | 0.62 | 1.96E-11 | 4.09E-07 | 0.34 | 0.28 | 0.06 | 1.22 |
| 5 | Hmox2 | 0.30 | 1.99E-11 | 4.14E-07 | 0.67 | 0.64 | 0.03 | 1.04 |
| 5 | Pds5b | 0.28 | 2.11E-11 | 4.40E-07 | 0.69 | 0.65 | 0.04 | 1.06 |
| 5 | Klhdc8b | 0.66 | 2.36E-11 | 4.91E-07 | 0.22 | 0.15 | 0.06 | 1.40 |
| 5 | Hpse2 | 0.73 | 2.55E-11 | 5.31E-07 | 0.23 | 0.17 | 0.06 | 1.36 |
| 5 | Tor1a | 0.33 | 3.08E-11 | 6.40E-07 | 0.56 | 0.52 | 0.04 | 1.08 |
| 5 | Col5a1 | 0.59 | 3.21E-11 | 6.68E-07 | 0.33 | 0.26 | 0.06 | 1.24 |
| 5 | Rspo2 | 0.64 | 3.40E-11 | 7.08E-07 | 0.43 | 0.37 | 0.07 | 1.18 |
| 5 | Lrpap1 | 0.44 | 3.45E-11 | 7.19E-07 | 0.53 | 0.49 | 0.03 | 1.07 |
| 5 | Srgap1 | 0.45 | 3.50E-11 | 7.29E-07 | 0.40 | 0.34 | 0.06 | 1.18 |
| 5 | Dchs1 | 0.34 | 3.76E-11 | 7.82E-07 | 0.49 | 0.42 | 0.08 | 1.18 |
| 5 | Rnd2 | 1.00 | 3.95E-11 | 8.21E-07 | 0.13 | 0.09 | 0.05 | 1.58 |
| 5 | Map7d1 | 0.30 | 5.43E-11 | 1.13E-06 | 0.66 | 0.62 | 0.04 | 1.07 |
| 5 | Jam3 | 0.31 | 6.60E-11 | 1.37E-06 | 0.62 | 0.57 | 0.05 | 1.09 |
| 5 | Eci1 | 0.36 | 7.47E-11 | 1.55E-06 | 0.56 | 0.52 | 0.04 | 1.07 |
| 5 | Aldh1a2 | 0.66 | 7.99E-11 | 1.66E-06 | 0.11 | 0.07 | 0.05 | 1.70 |
| 5 | Nsd2 | 0.26 | 8.48E-11 | 1.76E-06 | 0.76 | 0.75 | 0.01 | 1.01 |
| 5 | Kansl2 | 0.26 | 9.32E-11 | 1.94E-06 | 0.67 | 0.63 | 0.04 | 1.06 |
| 5 | Hddc3 | 0.30 | 9.33E-11 | 1.94E-06 | 0.61 | 0.58 | 0.04 | 1.06 |
| 5 | Txndc15 | 0.32 | 9.56E-11 | 1.99E-06 | 0.63 | 0.60 | 0.03 | 1.05 |
| 5 | Twist2 | 0.42 | 1.16E-10 | 2.40E-06 | 0.30 | 0.23 | 0.07 | 1.30 |
| 5 | Batf3 | 0.73 | 1.17E-10 | 2.44E-06 | 0.21 | 0.15 | 0.06 | 1.38 |
| 5 | Col25a1 | 0.53 | 1.23E-10 | 2.56E-06 | 0.26 | 0.19 | 0.07 | 1.35 |
| 5 | Mxra8 | 0.83 | 1.28E-10 | 2.67E-06 | 0.20 | 0.14 | 0.06 | 1.39 |
| 5 | Cdc42ep1 | 1.06 | 1.31E-10 | 2.72E-06 | 0.10 | 0.06 | 0.04 | 1.70 |
| 5 | St6gal1 | 0.67 | 1.36E-10 | 2.82E-06 | 0.24 | 0.18 | 0.06 | 1.33 |
| 5 | St6galnac4 | 0.63 | 1.47E-10 | 3.05E-06 | 0.26 | 0.20 | 0.06 | 1.31 |
| 5 | Pkn1 | 0.49 | 1.50E-10 | 3.11E-06 | 0.41 | 0.35 | 0.05 | 1.15 |
| 5 | Smarcd3 | 0.48 | 1.56E-10 | 3.25E-06 | 0.34 | 0.28 | 0.06 | 1.22 |
| 5 | Eefsec | 0.46 | 1.63E-10 | 3.39E-06 | 0.44 | 0.39 | 0.05 | 1.13 |
| 5 | Xrcc2 | 0.91 | 1.64E-10 | 3.41E-06 | 0.20 | 0.15 | 0.05 | 1.36 |
| 5 | Yif1b | 0.27 | 1.76E-10 | 3.66E-06 | 0.69 | 0.67 | 0.02 | 1.03 |

**Supplementary Table 1. Marker genes for the original clusters of integrated clustering of E11.5 control and NotchGOF λ-junction cells.**

| | | | | | | | | |
|---|---|---|---|---|---|---|---|---|
| 5 | Armcx2 | 0.34 | 1.88E-10 | 3.92E-06 | 0.57 | 0.53 | 0.04 | 1.08 |
| 5 | Ext2 | 0.40 | 1.88E-10 | 3.92E-06 | 0.46 | 0.42 | 0.05 | 1.11 |
| 5 | Rspo1 | 1.01 | 2.20E-10 | 4.57E-06 | 0.11 | 0.07 | 0.04 | 1.64 |
| 5 | Vldlr | 0.94 | 2.78E-10 | 5.79E-06 | 0.17 | 0.12 | 0.05 | 1.44 |
| 5 | Evi5 | 0.29 | 2.94E-10 | 6.11E-06 | 0.62 | 0.61 | 0.02 | 1.02 |
| 5 | Marchf2 | 0.29 | 3.04E-10 | 6.33E-06 | 0.56 | 0.51 | 0.04 | 1.09 |
| 5 | Ctdspl2 | 0.32 | 3.12E-10 | 6.49E-06 | 0.61 | 0.58 | 0.04 | 1.06 |
| 5 | Hist1h4h | 0.56 | 3.16E-10 | 6.58E-06 | 0.46 | 0.41 | 0.05 | 1.11 |
| 5 | Ccdc136 | 0.52 | 3.18E-10 | 6.61E-06 | 0.34 | 0.28 | 0.06 | 1.22 |
| 5 | Sumf2 | 0.40 | 4.04E-10 | 8.40E-06 | 0.46 | 0.41 | 0.05 | 1.12 |
| 5 | Slc35b1 | 0.32 | 4.20E-10 | 8.73E-06 | 0.61 | 0.59 | 0.02 | 1.03 |
| 5 | Hist1h2ap | 0.37 | 4.26E-10 | 8.85E-06 | 0.76 | 0.75 | 0.01 | 1.01 |
| 5 | Rbm18 | 0.31 | 4.26E-10 | 8.85E-06 | 0.59 | 0.55 | 0.03 | 1.06 |
| 5 | Mgmt | 0.64 | 4.29E-10 | 8.91E-06 | 0.26 | 0.21 | 0.06 | 1.27 |
| 5 | Gstp1 | 0.32 | 4.71E-10 | 9.79E-06 | 0.56 | 0.52 | 0.04 | 1.08 |
| 5 | Fst | 1.45 | 4.75E-10 | 9.89E-06 | 0.17 | 0.12 | 0.05 | 1.42 |
| 5 | Ltbp1 | 0.63 | 5.12E-10 | 1.07E-05 | 0.28 | 0.22 | 0.06 | 1.25 |
| 5 | Eva1a | 0.52 | 5.20E-10 | 1.08E-05 | 0.30 | 0.24 | 0.06 | 1.24 |
| 5 | Tmem131l | 0.34 | 5.48E-10 | 1.14E-05 | 0.58 | 0.55 | 0.03 | 1.06 |
| 5 | Nipsnap1 | 0.28 | 5.83E-10 | 1.21E-05 | 0.60 | 0.56 | 0.04 | 1.07 |
| 5 | Sp3 | 0.31 | 6.03E-10 | 1.25E-05 | 0.64 | 0.63 | 0.01 | 1.01 |
| 5 | Spats2 | 0.25 | 6.86E-10 | 1.43E-05 | 0.72 | 0.68 | 0.03 | 1.05 |
| 5 | Xbp1 | 0.40 | 6.94E-10 | 1.44E-05 | 0.52 | 0.48 | 0.05 | 1.09 |
| 5 | Ddrgk1 | 0.33 | 7.58E-10 | 1.58E-05 | 0.58 | 0.55 | 0.03 | 1.06 |
| 5 | Ncoa7 | 0.45 | 7.74E-10 | 1.61E-05 | 0.44 | 0.40 | 0.04 | 1.10 |
| 5 | Vcam1 | 0.25 | 7.75E-10 | 1.61E-05 | 0.51 | 0.43 | 0.08 | 1.19 |
| 5 | Tmem98 | 0.36 | 9.42E-10 | 1.96E-05 | 0.54 | 0.50 | 0.04 | 1.08 |
| 5 | Haus7 | 0.37 | 9.74E-10 | 2.03E-05 | 0.50 | 0.46 | 0.04 | 1.08 |
| 5 | Grn | 0.35 | 1.00E-09 | 2.08E-05 | 0.40 | 0.35 | 0.05 | 1.15 |
| 5 | Hist1h3i | 0.65 | 1.01E-09 | 2.11E-05 | 0.29 | 0.24 | 0.06 | 1.24 |
| 5 | Cdca7 | 0.25 | 1.08E-09 | 2.25E-05 | 0.76 | 0.72 | 0.04 | 1.06 |
| 5 | Nfu1 | 0.31 | 1.24E-09 | 2.58E-05 | 0.62 | 0.60 | 0.02 | 1.02 |
| 5 | Smad5 | 0.37 | 1.24E-09 | 2.58E-05 | 0.53 | 0.49 | 0.04 | 1.08 |
| 5 | Mmp2 | 0.28 | 1.26E-09 | 2.62E-05 | 0.58 | 0.52 | 0.06 | 1.12 |
| 5 | Mertk | 0.77 | 1.55E-09 | 3.23E-05 | 0.21 | 0.16 | 0.05 | 1.34 |
| 5 | Aamdc | 0.44 | 1.56E-09 | 3.25E-05 | 0.37 | 0.31 | 0.06 | 1.18 |
| 5 | Gdpd1 | 0.50 | 1.59E-09 | 3.31E-05 | 0.32 | 0.27 | 0.06 | 1.21 |
| 5 | Castor2 | 0.79 | 1.61E-09 | 3.35E-05 | 0.18 | 0.13 | 0.05 | 1.40 |
| 5 | Dzip1 | 0.49 | 1.65E-09 | 3.42E-05 | 0.38 | 0.33 | 0.05 | 1.16 |
| 5 | G0s2 | 0.78 | 1.69E-09 | 3.52E-05 | 0.20 | 0.15 | 0.05 | 1.36 |
| 5 | Vasn | 0.53 | 1.77E-09 | 3.68E-05 | 0.27 | 0.22 | 0.06 | 1.27 |
| 5 | Wnt5b | 0.57 | 1.78E-09 | 3.69E-05 | 0.31 | 0.26 | 0.06 | 1.22 |
| 5 | Cep19 | 0.45 | 2.08E-09 | 4.32E-05 | 0.40 | 0.35 | 0.05 | 1.15 |
| 5 | Mns1 | 0.27 | 2.25E-09 | 4.67E-05 | 0.59 | 0.54 | 0.04 | 1.08 |
| 5 | Pde3a | 0.64 | 2.28E-09 | 4.75E-05 | 0.18 | 0.13 | 0.05 | 1.41 |
| 5 | Tpd52l1 | 0.36 | 2.72E-09 | 5.66E-05 | 0.39 | 0.33 | 0.06 | 1.19 |
| 5 | Slc6a6 | 0.46 | 2.82E-09 | 5.86E-05 | 0.40 | 0.35 | 0.05 | 1.14 |
| 5 | Smarca1 | 0.30 | 2.89E-09 | 6.01E-05 | 0.51 | 0.46 | 0.05 | 1.11 |
| 5 | Dock1 | 0.33 | 2.92E-09 | 6.07E-05 | 0.54 | 0.50 | 0.04 | 1.08 |
| 5 | Hist1h1e | 0.31 | 3.27E-09 | 6.79E-05 | 0.68 | 0.65 | 0.03 | 1.05 |
| 5 | Tor2a | 0.39 | 3.32E-09 | 6.90E-05 | 0.46 | 0.42 | 0.04 | 1.10 |
| 5 | Coq7 | 0.26 | 3.44E-09 | 7.15E-05 | 0.70 | 0.67 | 0.03 | 1.04 |
| 5 | Adamts9 | 0.30 | 3.54E-09 | 7.37E-05 | 0.20 | 0.15 | 0.05 | 1.36 |
| 5 | Surf4 | 0.35 | 3.55E-09 | 7.38E-05 | 0.61 | 0.59 | 0.03 | 1.04 |
| 5 | Smyd2 | 0.40 | 4.04E-09 | 8.41E-05 | 0.45 | 0.41 | 0.04 | 1.11 |
| 5 | Ncam1 | 0.27 | 4.52E-09 | 9.41E-05 | 0.42 | 0.36 | 0.06 | 1.16 |

**Supplementary Table 1. Marker genes for the original clusters of integrated clustering of E11.5 control and NotchGOF λ-junction cells.**

| | | | | | | | | |
|---|---|---|---|---|---|---|---|---|
| 5 | Dnm1 | 0.39 | 4.61E-09 | 9.59E-05 | 0.38 | 0.33 | 0.06 | 1.17 |
| 5 | Psmc3ip | 0.29 | 5.20E-09 | 1.08E-04 | 0.58 | 0.56 | 0.02 | 1.04 |
| 5 | Map3k4 | 0.34 | 5.54E-09 | 1.15E-04 | 0.56 | 0.54 | 0.02 | 1.04 |
| 5 | Nfic | 0.29 | 6.00E-09 | 1.25E-04 | 0.36 | 0.30 | 0.06 | 1.21 |
| 5 | Mmp11 | 0.40 | 6.05E-09 | 1.26E-04 | 0.37 | 0.32 | 0.06 | 1.18 |
| 5 | Arhgap24 | 0.74 | 6.05E-09 | 1.26E-04 | 0.17 | 0.12 | 0.05 | 1.42 |
| 5 | Rnf168 | 0.35 | 6.15E-09 | 1.28E-04 | 0.55 | 0.53 | 0.03 | 1.05 |
| 5 | Cntln | 0.29 | 6.40E-09 | 1.33E-04 | 0.53 | 0.49 | 0.04 | 1.09 |
| 5 | Atad2 | 0.26 | 6.82E-09 | 1.42E-04 | 0.76 | 0.74 | 0.03 | 1.04 |
| 5 | Rab13 | 0.29 | 8.70E-09 | 1.81E-04 | 0.47 | 0.42 | 0.05 | 1.12 |
| 5 | Fat3 | 0.82 | 8.89E-09 | 1.85E-04 | 0.14 | 0.09 | 0.04 | 1.48 |
| 5 | P3h4 | 0.48 | 9.81E-09 | 2.04E-04 | 0.40 | 0.35 | 0.04 | 1.13 |
| 5 | Plekha8 | 0.36 | 9.93E-09 | 2.06E-04 | 0.50 | 0.47 | 0.04 | 1.07 |
| 5 | Kirrel | 0.58 | 1.03E-08 | 2.14E-04 | 0.27 | 0.22 | 0.05 | 1.23 |
| 5 | Nsmce1 | 0.26 | 1.07E-08 | 2.22E-04 | 0.71 | 0.70 | 0.01 | 1.01 |
| 5 | 1110008P14Rik | 0.41 | 1.11E-08 | 2.31E-04 | 0.49 | 0.47 | 0.02 | 1.05 |
| 5 | Pkd2 | 0.41 | 1.17E-08 | 2.43E-04 | 0.43 | 0.38 | 0.05 | 1.12 |
| 5 | Arxes2 | 0.40 | 1.22E-08 | 2.54E-04 | 0.44 | 0.39 | 0.05 | 1.12 |
| 5 | Adgra2 | 0.54 | 1.24E-08 | 2.59E-04 | 0.29 | 0.23 | 0.06 | 1.24 |
| 5 | Bcl2 | 0.41 | 1.39E-08 | 2.88E-04 | 0.40 | 0.35 | 0.05 | 1.14 |
| 5 | Spsb1 | 0.68 | 1.44E-08 | 3.00E-04 | 0.20 | 0.15 | 0.05 | 1.34 |
| 5 | Zfp281 | 0.38 | 1.53E-08 | 3.18E-04 | 0.49 | 0.46 | 0.03 | 1.07 |
| 5 | Ripk3 | 0.72 | 1.54E-08 | 3.20E-04 | 0.21 | 0.16 | 0.05 | 1.31 |
| 5 | Brwd1 | 0.26 | 1.63E-08 | 3.39E-04 | 0.65 | 0.64 | 0.02 | 1.02 |
| 5 | 2310009B15Rik | 0.35 | 1.87E-08 | 3.90E-04 | 0.56 | 0.54 | 0.02 | 1.04 |
| 5 | Slc25a36 | 0.33 | 1.90E-08 | 3.95E-04 | 0.48 | 0.43 | 0.04 | 1.10 |
| 5 | Ror2 | 0.43 | 1.94E-08 | 4.04E-04 | 0.35 | 0.30 | 0.05 | 1.17 |
| 5 | Smim10l1 | 0.27 | 1.96E-08 | 4.07E-04 | 0.64 | 0.61 | 0.03 | 1.04 |
| 5 | Slc35a1 | 0.34 | 1.98E-08 | 4.12E-04 | 0.51 | 0.47 | 0.03 | 1.07 |
| 5 | Praf2 | 0.26 | 2.10E-08 | 4.36E-04 | 0.57 | 0.53 | 0.04 | 1.07 |
| 5 | Ip6k2 | 0.31 | 2.16E-08 | 4.50E-04 | 0.58 | 0.56 | 0.02 | 1.04 |
| 5 | Arhgef10 | 0.69 | 2.21E-08 | 4.60E-04 | 0.18 | 0.13 | 0.05 | 1.36 |
| 5 | Ctdsp2 | 0.29 | 2.22E-08 | 4.61E-04 | 0.59 | 0.57 | 0.02 | 1.04 |
| 5 | 2610318N02Rik | 0.26 | 2.27E-08 | 4.72E-04 | 0.63 | 0.60 | 0.03 | 1.05 |
| 5 | Clk4 | 0.33 | 2.35E-08 | 4.90E-04 | 0.49 | 0.45 | 0.04 | 1.08 |
| 5 | Cacng8 | 0.88 | 2.39E-08 | 4.96E-04 | 0.10 | 0.06 | 0.04 | 1.60 |
| 5 | Cacna1a | 0.50 | 2.65E-08 | 5.52E-04 | 0.35 | 0.30 | 0.05 | 1.16 |
| 5 | Fkbp9 | 0.35 | 2.68E-08 | 5.58E-04 | 0.48 | 0.44 | 0.04 | 1.10 |
| 5 | St8sia2 | 0.56 | 2.74E-08 | 5.71E-04 | 0.24 | 0.19 | 0.05 | 1.26 |
| 5 | Syde1 | 0.65 | 2.88E-08 | 5.99E-04 | 0.22 | 0.17 | 0.05 | 1.28 |
| 5 | Obsl1 | 0.47 | 2.91E-08 | 6.06E-04 | 0.36 | 0.31 | 0.05 | 1.15 |
| 5 | Gm10320 | 0.42 | 3.28E-08 | 6.83E-04 | 0.44 | 0.40 | 0.05 | 1.11 |
| 5 | Csad | 0.46 | 3.67E-08 | 7.63E-04 | 0.36 | 0.32 | 0.05 | 1.15 |
| 5 | Tmem9 | 0.27 | 3.70E-08 | 7.70E-04 | 0.65 | 0.65 | 0.01 | 1.01 |
| 5 | Iqgap2 | 0.42 | 3.79E-08 | 7.88E-04 | 0.41 | 0.36 | 0.05 | 1.12 |
| 5 | Plaat3 | 0.66 | 4.08E-08 | 8.49E-04 | 0.27 | 0.22 | 0.05 | 1.21 |
| 5 | Chaf1b | 0.31 | 4.31E-08 | 8.97E-04 | 0.55 | 0.51 | 0.04 | 1.07 |
| 5 | S1pr3 | 0.38 | 4.88E-08 | 1.01E-03 | 0.33 | 0.27 | 0.06 | 1.20 |
| 5 | Gabpb1 | 0.36 | 5.96E-08 | 1.24E-03 | 0.49 | 0.47 | 0.03 | 1.06 |
| 5 | Slc49a4 | 0.53 | 6.06E-08 | 1.26E-03 | 0.31 | 0.27 | 0.05 | 1.18 |
| 5 | Egln2 | 0.26 | 7.40E-08 | 1.54E-03 | 0.63 | 0.61 | 0.02 | 1.03 |
| 5 | Bcl2l11 | 0.33 | 7.49E-08 | 1.56E-03 | 0.40 | 0.35 | 0.05 | 1.15 |
| 5 | Robo1 | 0.37 | 8.92E-08 | 1.86E-03 | 0.50 | 0.46 | 0.04 | 1.09 |
| 5 | Czib | 0.31 | 9.38E-08 | 1.95E-03 | 0.56 | 0.53 | 0.02 | 1.05 |
| 5 | Cnep1r1 | 0.28 | 1.04E-07 | 2.16E-03 | 0.56 | 0.54 | 0.02 | 1.03 |
| 5 | Fbxo45 | 0.36 | 1.10E-07 | 2.29E-03 | 0.40 | 0.36 | 0.04 | 1.12 |

**Supplementary Table 1. Marker genes for the original clusters of integrated clustering of E11.5 control and NotchGOF λ-junction cells.**

| | | | | | | | | |
|---|---|---|---|---|---|---|---|---|
| 5 | Wwtr1 | 0.30 | 1.16E-07 | 2.41E-03 | 0.47 | 0.43 | 0.04 | 1.09 |
| 5 | Hist1h3e | 0.40 | 1.20E-07 | 2.50E-03 | 0.46 | 0.42 | 0.04 | 1.09 |
| 5 | Gnpnat1 | 0.34 | 1.24E-07 | 2.59E-03 | 0.46 | 0.42 | 0.04 | 1.09 |
| 5 | Pofut2 | 0.32 | 1.45E-07 | 3.02E-03 | 0.51 | 0.48 | 0.03 | 1.06 |
| 5 | Aurkb | 0.30 | 1.47E-07 | 3.05E-03 | 0.53 | 0.49 | 0.05 | 1.09 |
| 5 | Adam11 | 0.86 | 1.58E-07 | 3.28E-03 | 0.12 | 0.08 | 0.04 | 1.46 |
| 5 | Slc22a17 | 0.40 | 1.79E-07 | 3.72E-03 | 0.39 | 0.34 | 0.04 | 1.13 |
| 5 | Vstm2b | 0.62 | 1.79E-07 | 3.73E-03 | 0.14 | 0.09 | 0.04 | 1.44 |
| 5 | Pbk | 0.30 | 1.92E-07 | 3.99E-03 | 0.57 | 0.54 | 0.02 | 1.04 |
| 5 | Fam216a | 0.29 | 2.02E-07 | 4.21E-03 | 0.55 | 0.53 | 0.02 | 1.04 |
| 5 | Gpat4 | 0.31 | 2.06E-07 | 4.28E-03 | 0.52 | 0.49 | 0.03 | 1.06 |
| 5 | Pcyox1l | 0.59 | 2.08E-07 | 4.32E-03 | 0.25 | 0.20 | 0.05 | 1.23 |
| 5 | Mgat3 | 0.66 | 2.12E-07 | 4.40E-03 | 0.20 | 0.15 | 0.05 | 1.30 |
| 5 | Abhd8 | 0.32 | 2.32E-07 | 4.82E-03 | 0.50 | 0.48 | 0.02 | 1.05 |
| 5 | Rab11fip2 | 0.33 | 2.36E-07 | 4.90E-03 | 0.41 | 0.37 | 0.04 | 1.12 |
| 5 | Dixdc1 | 0.33 | 2.44E-07 | 5.07E-03 | 0.44 | 0.39 | 0.05 | 1.11 |
| 5 | Dipk1b | 0.30 | 2.51E-07 | 5.22E-03 | 0.51 | 0.48 | 0.03 | 1.07 |
| 5 | Fmnl2 | 0.27 | 2.68E-07 | 5.57E-03 | 0.48 | 0.44 | 0.04 | 1.08 |
| 5 | Lysmd2 | 0.52 | 2.94E-07 | 6.11E-03 | 0.26 | 0.22 | 0.04 | 1.20 |
| 5 | Pxn | 0.61 | 2.98E-07 | 6.20E-03 | 0.23 | 0.18 | 0.04 | 1.24 |
| 5 | Creld2 | 0.36 | 3.12E-07 | 6.48E-03 | 0.47 | 0.45 | 0.03 | 1.07 |
| 5 | Phf20l1 | 0.25 | 3.17E-07 | 6.60E-03 | 0.66 | 0.65 | 0.01 | 1.01 |
| 5 | 1700096K18Rik | 0.84 | 3.24E-07 | 6.74E-03 | 0.13 | 0.09 | 0.04 | 1.40 |
| 5 | Gde1 | 0.27 | 3.43E-07 | 7.13E-03 | 0.55 | 0.52 | 0.03 | 1.05 |
| 5 | Hist1h2bn | 0.73 | 3.58E-07 | 7.44E-03 | 0.19 | 0.15 | 0.04 | 1.28 |
| 5 | Gstt2 | 0.88 | 3.77E-07 | 7.83E-03 | 0.11 | 0.07 | 0.04 | 1.47 |
| 5 | Arhgef25 | 0.35 | 3.83E-07 | 7.97E-03 | 0.41 | 0.37 | 0.04 | 1.11 |
| 5 | Spc25 | 0.29 | 4.03E-07 | 8.38E-03 | 0.59 | 0.55 | 0.03 | 1.06 |
| 5 | Adpgk | 0.44 | 4.60E-07 | 9.56E-03 | 0.35 | 0.31 | 0.04 | 1.14 |
| 5 | Lpar4 | 0.49 | 4.67E-07 | 9.71E-03 | 0.37 | 0.32 | 0.04 | 1.13 |
| 5 | Vangl2 | 0.30 | 5.23E-07 | 1.09E-02 | 0.59 | 0.57 | 0.02 | 1.03 |
| 5 | Wdr82 | 0.28 | 5.31E-07 | 1.10E-02 | 0.60 | 0.59 | 0.01 | 1.02 |
| 5 | Samd14 | 0.31 | 6.17E-07 | 1.28E-02 | 0.44 | 0.41 | 0.03 | 1.08 |
| 5 | Creb3l2 | 0.35 | 6.67E-07 | 1.39E-02 | 0.37 | 0.32 | 0.05 | 1.15 |
| 5 | Rad51 | 0.28 | 7.41E-07 | 1.54E-02 | 0.57 | 0.54 | 0.03 | 1.05 |
| 5 | Svil | 0.39 | 7.55E-07 | 1.57E-02 | 0.42 | 0.38 | 0.03 | 1.08 |
| 5 | Efnb3 | 0.40 | 7.90E-07 | 1.64E-02 | 0.36 | 0.31 | 0.04 | 1.13 |
| 5 | Gins2 | 0.26 | 8.10E-07 | 1.69E-02 | 0.63 | 0.61 | 0.03 | 1.04 |
| 5 | Slc27a3 | 0.50 | 8.34E-07 | 1.73E-02 | 0.26 | 0.21 | 0.05 | 1.21 |
| 5 | Dnph1 | 0.33 | 8.36E-07 | 1.74E-02 | 0.54 | 0.52 | 0.02 | 1.03 |
| 5 | Rgmb | 0.82 | 8.38E-07 | 1.74E-02 | 0.16 | 0.12 | 0.04 | 1.31 |
| 5 | Rarg | 0.26 | 8.49E-07 | 1.77E-02 | 0.55 | 0.52 | 0.03 | 1.06 |
| 5 | Ghr | 0.63 | 8.58E-07 | 1.78E-02 | 0.18 | 0.14 | 0.04 | 1.31 |
| 5 | Nog | 0.85 | 8.64E-07 | 1.80E-02 | 0.13 | 0.09 | 0.04 | 1.40 |
| 5 | Ctsk | 0.75 | 8.65E-07 | 1.80E-02 | 0.17 | 0.13 | 0.04 | 1.32 |
| 5 | Prps1 | 0.32 | 9.50E-07 | 1.98E-02 | 0.47 | 0.44 | 0.03 | 1.06 |
| 5 | Rnf115 | 0.31 | 9.54E-07 | 1.98E-02 | 0.54 | 0.52 | 0.02 | 1.03 |
| 5 | Dna2 | 0.46 | 1.09E-06 | 2.26E-02 | 0.30 | 0.26 | 0.04 | 1.16 |
| 5 | Ggh | 0.49 | 1.16E-06 | 2.40E-02 | 0.29 | 0.25 | 0.04 | 1.16 |
| 5 | Cadm4 | 0.75 | 1.19E-06 | 2.48E-02 | 0.12 | 0.09 | 0.04 | 1.42 |
| 5 | Cttnbp2nl | 0.28 | 1.23E-06 | 2.56E-02 | 0.52 | 0.50 | 0.03 | 1.05 |
| 5 | 4930503L19Rik | 0.40 | 1.24E-06 | 2.58E-02 | 0.37 | 0.33 | 0.04 | 1.11 |
| 5 | Slc50a1 | 0.35 | 1.32E-06 | 2.75E-02 | 0.41 | 0.37 | 0.04 | 1.09 |
| 5 | Igsf8 | 0.48 | 1.35E-06 | 2.81E-02 | 0.28 | 0.24 | 0.04 | 1.16 |
| 5 | Ophn1 | 0.53 | 1.36E-06 | 2.83E-02 | 0.23 | 0.19 | 0.04 | 1.23 |
| 5 | Nek6 | 0.43 | 1.58E-06 | 3.29E-02 | 0.35 | 0.31 | 0.04 | 1.12 |

**Supplementary Table 1. Marker genes for the original clusters of integrated clustering of E11.5 control and NotchGOF λ-junction cells.**

| | | | | | | | | |
|---|---|---|---|---|---|---|---|---|
| 5 | Zfyve16 | 0.38 | 1.64E-06 | 3.41E-02 | 0.39 | 0.35 | 0.04 | 1.11 |
| 5 | Rbm46 | 0.65 | 1.72E-06 | 3.57E-02 | 0.12 | 0.08 | 0.04 | 1.44 |
| 5 | Speg | 0.52 | 1.78E-06 | 3.69E-02 | 0.21 | 0.16 | 0.04 | 1.26 |
| 5 | Fxn | 0.26 | 1.80E-06 | 3.75E-02 | 0.55 | 0.55 | 0.00 | 1.01 |
| 5 | Gng3 | 0.40 | 1.87E-06 | 3.89E-02 | 0.10 | 0.07 | 0.03 | 1.47 |
| 5 | Lsp1 | 0.40 | 2.05E-06 | 4.26E-02 | 0.20 | 0.16 | 0.04 | 1.27 |
| 5 | Slitrk5 | 0.43 | 2.08E-06 | 4.32E-02 | 0.20 | 0.16 | 0.05 | 1.29 |
| 5 | Wnk3 | 0.37 | 2.08E-06 | 4.33E-02 | 0.36 | 0.32 | 0.04 | 1.13 |
| 5 | Ctps2 | 0.45 | 2.20E-06 | 4.58E-02 | 0.32 | 0.29 | 0.04 | 1.13 |
| 5 | Ikbip | 0.42 | 2.26E-06 | 4.70E-02 | 0.34 | 0.30 | 0.04 | 1.13 |
| 5 | H2-Q7 | 0.66 | 2.30E-06 | 4.79E-02 | 0.15 | 0.11 | 0.04 | 1.36 |
| 5 | Mfhas1 | 0.51 | 2.37E-06 | 4.94E-02 | 0.23 | 0.19 | 0.04 | 1.22 |
| 5 | Rhoq | 0.29 | 2.38E-06 | 4.94E-02 | 0.48 | 0.44 | 0.03 | 1.07 |
| 6 | Asb4 | 2.60 | 0 | 0 | 0.81 | 0.27 | 0.54 | 2.95 |
| 6 | Nrp1 | 2.83 | 0 | 0 | 0.67 | 0.19 | 0.48 | 3.56 |
| 6 | Sfrp2 | 3.02 | 0 | 0 | 0.96 | 0.60 | 0.37 | 1.61 |
| 6 | Prrx1 | 1.63 | 0 | 0 | 0.99 | 0.64 | 0.36 | 1.56 |
| 6 | Col9a1 | 2.41 | 8.03E-287 | 1.67E-282 | 0.74 | 0.27 | 0.47 | 2.76 |
| 6 | Meis2 | 1.46 | 1.28E-285 | 2.67E-281 | 0.99 | 0.66 | 0.33 | 1.51 |
| 6 | Foxd2os | 2.68 | 5.64E-281 | 1.17E-276 | 0.53 | 0.13 | 0.39 | 3.98 |
| 6 | Gpc3 | 1.24 | 1.20E-277 | 2.49E-273 | 0.99 | 0.83 | 0.17 | 1.20 |
| 6 | Mecom | 1.67 | 2.29E-241 | 4.77E-237 | 0.83 | 0.39 | 0.44 | 2.14 |
| 6 | Marcks | 0.74 | 1.83E-236 | 3.80E-232 | 1.00 | 0.97 | 0.03 | 1.03 |
| 6 | Lhx6 | 3.06 | 2.20E-233 | 4.57E-229 | 0.30 | 0.05 | 0.25 | 5.96 |
| 6 | Ebf2 | 2.64 | 4.75E-228 | 9.88E-224 | 0.24 | 0.03 | 0.21 | 7.68 |
| 6 | Osr2 | 2.84 | 1.16E-223 | 2.41E-219 | 0.36 | 0.08 | 0.29 | 4.84 |
| 6 | Itga4 | 2.37 | 2.26E-219 | 4.70E-215 | 0.55 | 0.18 | 0.37 | 3.09 |
| 6 | Foxd2 | 2.53 | 5.11E-216 | 1.06E-211 | 0.41 | 0.10 | 0.31 | 4.19 |
| 6 | Cdkn1c | 1.36 | 2.74E-215 | 5.70E-211 | 1.00 | 0.93 | 0.07 | 1.08 |
| 6 | Barx1 | 2.43 | 5.04E-194 | 1.05E-189 | 0.56 | 0.20 | 0.36 | 2.79 |
| 6 | Sulf1 | 1.84 | 1.26E-193 | 2.62E-189 | 0.84 | 0.53 | 0.31 | 1.59 |
| 6 | Foxp2 | 2.35 | 7.05E-189 | 1.47E-184 | 0.68 | 0.33 | 0.35 | 2.05 |
| 6 | Ccnd2 | 0.91 | 1.00E-188 | 2.08E-184 | 0.99 | 0.85 | 0.14 | 1.16 |
| 6 | Itm2a | 1.58 | 8.22E-184 | 1.71E-179 | 0.87 | 0.53 | 0.33 | 1.63 |
| 6 | 9130024F11Rik | 1.95 | 2.58E-166 | 5.37E-162 | 0.54 | 0.20 | 0.34 | 2.65 |
| 6 | Mab21l1 | 2.15 | 5.52E-165 | 1.15E-160 | 0.46 | 0.15 | 0.31 | 3.03 |
| 6 | Lgi1 | 3.91 | 4.18E-164 | 8.70E-160 | 0.13 | 0.01 | 0.12 | 11.45 |
| 6 | Slc1a3 | 2.00 | 1.34E-161 | 2.78E-157 | 0.54 | 0.21 | 0.33 | 2.56 |
| 6 | Plpp3 | 1.29 | 1.01E-152 | 2.09E-148 | 0.89 | 0.65 | 0.23 | 1.36 |
| 6 | Satb2 | 1.83 | 1.33E-150 | 2.77E-146 | 0.56 | 0.24 | 0.32 | 2.34 |
| 6 | Tbx22 | 2.28 | 3.63E-145 | 7.55E-141 | 0.36 | 0.11 | 0.26 | 3.45 |
| 6 | Foxf1 | 1.53 | 1.89E-144 | 3.93E-140 | 0.49 | 0.17 | 0.32 | 2.92 |
| 6 | Tgfbi | 1.84 | 4.76E-142 | 9.89E-138 | 0.59 | 0.28 | 0.31 | 2.14 |
| 6 | Scx | 1.96 | 7.90E-142 | 1.64E-137 | 0.44 | 0.15 | 0.29 | 2.93 |
| 6 | Eno3 | 1.57 | 2.93E-136 | 6.10E-132 | 0.67 | 0.36 | 0.31 | 1.85 |
| 6 | Tmem37 | 2.58 | 1.61E-129 | 3.35E-125 | 0.29 | 0.08 | 0.21 | 3.78 |
| 6 | Mfap2 | 0.73 | 2.63E-129 | 5.48E-125 | 1.00 | 0.92 | 0.08 | 1.08 |
| 6 | Aff2 | 2.59 | 4.24E-129 | 8.82E-125 | 0.27 | 0.07 | 0.20 | 4.00 |
| 6 | Nr2f1 | 1.40 | 8.22E-126 | 1.71E-121 | 0.64 | 0.32 | 0.32 | 1.99 |
| 6 | Foxf2 | 2.20 | 1.92E-124 | 3.99E-120 | 0.31 | 0.09 | 0.22 | 3.60 |
| 6 | Col1a2 | 0.81 | 2.96E-124 | 6.15E-120 | 0.98 | 0.73 | 0.25 | 1.34 |
| 6 | Angptl1 | 1.79 | 3.15E-124 | 6.55E-120 | 0.50 | 0.21 | 0.29 | 2.38 |
| 6 | Frzb | 1.10 | 1.61E-122 | 3.34E-118 | 0.67 | 0.34 | 0.33 | 1.99 |
| 6 | Mfap4 | 0.84 | 1.01E-120 | 2.11E-116 | 0.98 | 0.84 | 0.14 | 1.17 |
| 6 | Ldhb | 0.63 | 1.15E-116 | 2.39E-112 | 0.99 | 0.91 | 0.08 | 1.09 |
| 6 | Eya4 | 1.58 | 3.15E-116 | 6.56E-112 | 0.56 | 0.27 | 0.29 | 2.04 |

**Supplementary Table 1. Marker genes for the original clusters of integrated clustering of E11.5 control and NotchGOF λ-junction cells.**

| | | | | | | | | |
|---|---|---|---|---|---|---|---|---|
| 6 | Fzd2 | 0.72 | 1.14E-115 | 2.38E-111 | 0.98 | 0.87 | 0.12 | 1.13 |
| 6 | Cga | 3.95 | 1.21E-114 | 2.51E-110 | 0.11 | 0.01 | 0.10 | 8.46 |
| 6 | Frem1 | 1.34 | 2.45E-114 | 5.10E-110 | 0.68 | 0.39 | 0.29 | 1.73 |
| 6 | Peg3 | 0.82 | 7.34E-114 | 1.53E-109 | 0.96 | 0.82 | 0.14 | 1.16 |
| 6 | Maged2 | 0.94 | 4.03E-112 | 8.39E-108 | 0.93 | 0.82 | 0.11 | 1.13 |
| 6 | Flrt2 | 0.96 | 5.26E-112 | 1.09E-107 | 0.77 | 0.44 | 0.33 | 1.75 |
| 6 | Epha3 | 1.40 | 7.26E-112 | 1.51E-107 | 0.65 | 0.35 | 0.30 | 1.87 |
| 6 | Col3a1 | 1.03 | 8.31E-112 | 1.73E-107 | 0.97 | 0.71 | 0.25 | 1.36 |
| 6 | Nnat | 0.51 | 2.49E-111 | 5.19E-107 | 0.99 | 0.94 | 0.06 | 1.06 |
| 6 | Mafb | 2.05 | 7.30E-107 | 1.52E-102 | 0.47 | 0.21 | 0.26 | 2.23 |
| 6 | Cdh11 | 0.79 | 1.17E-103 | 2.43E-99 | 0.95 | 0.69 | 0.27 | 1.39 |
| 6 | Eva1b | 1.13 | 1.35E-102 | 2.80E-98 | 0.79 | 0.53 | 0.26 | 1.48 |
| 6 | Tsc22d1 | 0.63 | 3.86E-101 | 8.02E-97 | 0.99 | 0.93 | 0.05 | 1.06 |
| 6 | Mdk | 0.41 | 7.01E-97 | 1.46E-92 | 1.00 | 0.99 | 0.01 | 1.01 |
| 6 | Serpinf1 | 0.81 | 9.35E-96 | 1.94E-91 | 0.88 | 0.63 | 0.25 | 1.39 |
| 6 | Tut4 | 0.61 | 2.53E-94 | 5.26E-90 | 0.98 | 0.92 | 0.06 | 1.06 |
| 6 | Foxl2 | 1.97 | 1.50E-93 | 3.13E-89 | 0.33 | 0.12 | 0.21 | 2.78 |
| 6 | Epha4 | 0.79 | 4.90E-92 | 1.02E-87 | 0.88 | 0.64 | 0.24 | 1.37 |
| 6 | Cdh2 | 1.10 | 2.48E-90 | 5.15E-86 | 0.68 | 0.41 | 0.27 | 1.64 |
| 6 | Ptch1 | 1.12 | 5.12E-85 | 1.07E-80 | 0.68 | 0.44 | 0.23 | 1.53 |
| 6 | Gpc5 | 2.77 | 8.96E-85 | 1.86E-80 | 0.18 | 0.05 | 0.14 | 4.04 |
| 6 | Slit2 | 1.25 | 2.12E-82 | 4.42E-78 | 0.61 | 0.36 | 0.25 | 1.70 |
| 6 | Shisa2 | 2.07 | 4.83E-82 | 1.00E-77 | 0.29 | 0.10 | 0.19 | 2.83 |
| 6 | Sox4 | 0.52 | 1.55E-81 | 3.22E-77 | 0.99 | 0.94 | 0.05 | 1.06 |
| 6 | Gpc6 | 1.13 | 1.76E-81 | 3.66E-77 | 0.76 | 0.58 | 0.18 | 1.31 |
| 6 | Ndn | 0.84 | 5.97E-81 | 1.24E-76 | 0.87 | 0.74 | 0.13 | 1.17 |
| 6 | Cpxm1 | 1.32 | 1.65E-79 | 3.42E-75 | 0.64 | 0.42 | 0.22 | 1.51 |
| 6 | Peg10 | 1.17 | 5.57E-79 | 1.16E-74 | 0.64 | 0.43 | 0.22 | 1.50 |
| 6 | Amot | 0.97 | 8.97E-79 | 1.87E-74 | 0.68 | 0.47 | 0.21 | 1.43 |
| 6 | Sfrp1 | 0.96 | 9.79E-79 | 2.04E-74 | 0.81 | 0.62 | 0.20 | 1.32 |
| 6 | Meg3 | 0.71 | 1.00E-74 | 2.08E-70 | 0.89 | 0.68 | 0.20 | 1.30 |
| 6 | Ubb | 0.30 | 1.53E-74 | 3.17E-70 | 1.00 | 1.00 | 0.00 | 1.00 |
| 6 | Cldn11 | 2.05 | 2.69E-74 | 5.60E-70 | 0.31 | 0.12 | 0.19 | 2.52 |
| 6 | Col25a1 | 1.83 | 1.55E-73 | 3.23E-69 | 0.40 | 0.19 | 0.21 | 2.12 |
| 6 | Grb10 | 0.60 | 7.09E-73 | 1.47E-68 | 0.98 | 0.94 | 0.05 | 1.05 |
| 6 | Ube2e3 | 0.53 | 1.17E-72 | 2.44E-68 | 0.95 | 0.91 | 0.04 | 1.04 |
| 6 | AI314278 | 2.33 | 1.18E-72 | 2.46E-68 | 0.17 | 0.04 | 0.13 | 3.86 |
| 6 | Adra2a | 2.69 | 2.54E-72 | 5.29E-68 | 0.14 | 0.03 | 0.11 | 4.41 |
| 6 | Pkdcc | 1.23 | 5.13E-70 | 1.07E-65 | 0.55 | 0.32 | 0.23 | 1.71 |
| 6 | Mdfic | 1.74 | 2.85E-69 | 5.92E-65 | 0.41 | 0.22 | 0.20 | 1.92 |
| 6 | Wnt11 | 2.43 | 4.71E-69 | 9.80E-65 | 0.17 | 0.05 | 0.12 | 3.76 |
| 6 | Lhx8 | 1.05 | 1.91E-67 | 3.97E-63 | 0.50 | 0.26 | 0.24 | 1.92 |
| 6 | Colec12 | 1.87 | 5.71E-67 | 1.19E-62 | 0.41 | 0.21 | 0.19 | 1.91 |
| 6 | Kcnq1ot1 | 0.70 | 3.25E-66 | 6.76E-62 | 0.97 | 0.92 | 0.05 | 1.06 |
| 6 | Ccnd1 | 0.60 | 2.83E-65 | 5.88E-61 | 0.93 | 0.79 | 0.14 | 1.17 |
| 6 | Serpinh1 | 0.41 | 5.62E-65 | 1.17E-60 | 1.00 | 0.94 | 0.06 | 1.06 |
| 6 | Dlk1 | 0.60 | 7.95E-65 | 1.65E-60 | 0.64 | 0.41 | 0.23 | 1.55 |
| 6 | Arhgap24 | 1.86 | 1.38E-64 | 2.88E-60 | 0.28 | 0.11 | 0.17 | 2.51 |
| 6 | Sned1 | 1.89 | 1.49E-63 | 3.10E-59 | 0.29 | 0.12 | 0.17 | 2.39 |
| 6 | Aldh1a2 | 2.57 | 1.71E-63 | 3.55E-59 | 0.20 | 0.06 | 0.13 | 3.13 |
| 6 | Igfbp4 | 0.50 | 2.74E-62 | 5.69E-58 | 1.00 | 0.90 | 0.10 | 1.11 |
| 6 | Pcolce | 1.01 | 1.03E-60 | 2.15E-56 | 0.65 | 0.45 | 0.20 | 1.44 |
| 6 | Sostdc1 | 1.30 | 1.78E-60 | 3.71E-56 | 0.32 | 0.14 | 0.18 | 2.30 |
| 6 | Arid5b | 0.76 | 7.21E-60 | 1.50E-55 | 0.82 | 0.66 | 0.16 | 1.24 |
| 6 | Ddr2 | 0.89 | 1.10E-59 | 2.28E-55 | 0.68 | 0.48 | 0.20 | 1.42 |
| 6 | Tle4 | 0.96 | 1.36E-58 | 2.83E-54 | 0.72 | 0.58 | 0.14 | 1.24 |

**Supplementary Table 1. Marker genes for the original clusters of integrated clustering of E11.5 control and NotchGOF λ-junction cells.**

| | | | | | | | | |
|---|---|---|---|---|---|---|---|---|
| 6 | Mpped2 | 0.52 | 3.50E-58 | 7.28E-54 | 0.98 | 0.85 | 0.13 | 1.16 |
| 6 | Arhgap28 | 0.85 | 5.63E-57 | 1.17E-52 | 0.70 | 0.53 | 0.17 | 1.32 |
| 6 | Synpo2 | 2.22 | 9.62E-57 | 2.00E-52 | 0.20 | 0.07 | 0.13 | 2.91 |
| 6 | E130114P18Rik | 1.38 | 1.60E-56 | 3.32E-52 | 0.41 | 0.22 | 0.19 | 1.86 |
| 6 | Sulf2 | 0.69 | 2.40E-56 | 4.99E-52 | 0.80 | 0.66 | 0.14 | 1.21 |
| 6 | Cited1 | 1.77 | 3.01E-56 | 6.27E-52 | 0.27 | 0.11 | 0.16 | 2.38 |
| 6 | Mme | 1.20 | 4.47E-56 | 9.29E-52 | 0.64 | 0.46 | 0.18 | 1.38 |
| 6 | Ebf1 | 0.74 | 8.94E-56 | 1.86E-51 | 0.51 | 0.29 | 0.22 | 1.77 |
| 6 | Foxc1 | 1.30 | 3.08E-55 | 6.41E-51 | 0.47 | 0.28 | 0.19 | 1.69 |
| 6 | Hsd11b2 | 1.91 | 3.16E-55 | 6.57E-51 | 0.24 | 0.10 | 0.15 | 2.52 |
| 6 | Angptl4 | 1.22 | 4.88E-55 | 1.02E-50 | 0.52 | 0.33 | 0.19 | 1.59 |
| 6 | Fstl1 | 0.58 | 6.19E-55 | 1.29E-50 | 0.94 | 0.82 | 0.12 | 1.15 |
| 6 | Six1 | 0.61 | 5.32E-54 | 1.11E-49 | 0.81 | 0.60 | 0.20 | 1.34 |
| 6 | Nrk | 1.68 | 1.99E-53 | 4.14E-49 | 0.31 | 0.15 | 0.17 | 2.12 |
| 6 | Mest | 0.40 | 2.09E-53 | 4.34E-49 | 0.96 | 0.91 | 0.05 | 1.06 |
| 6 | Sobp | 1.02 | 2.31E-52 | 4.80E-48 | 0.52 | 0.33 | 0.19 | 1.57 |
| 6 | Basp1 | 0.35 | 2.73E-52 | 5.68E-48 | 1.00 | 0.95 | 0.05 | 1.05 |
| 6 | Col2a1 | 0.80 | 6.28E-52 | 1.31E-47 | 0.81 | 0.64 | 0.18 | 1.28 |
| 6 | Cd63 | 0.49 | 1.58E-51 | 3.30E-47 | 0.97 | 0.91 | 0.06 | 1.07 |
| 6 | Mir99ahg | 0.97 | 1.11E-49 | 2.31E-45 | 0.64 | 0.48 | 0.16 | 1.34 |
| 6 | Selenop | 1.02 | 7.62E-49 | 1.59E-44 | 0.35 | 0.18 | 0.17 | 1.95 |
| 6 | Idh2 | 0.47 | 1.43E-48 | 2.97E-44 | 0.95 | 0.90 | 0.06 | 1.07 |
| 6 | Capn6 | 0.79 | 1.49E-48 | 3.10E-44 | 0.68 | 0.54 | 0.14 | 1.26 |
| 6 | Cxcl12 | 1.44 | 2.19E-48 | 4.57E-44 | 0.33 | 0.17 | 0.16 | 1.98 |
| 6 | Smoc1 | 1.71 | 2.74E-48 | 5.69E-44 | 0.20 | 0.07 | 0.12 | 2.72 |
| 6 | Rhoc | 0.52 | 1.05E-47 | 2.19E-43 | 0.88 | 0.78 | 0.10 | 1.13 |
| 6 | Zcchc3 | 0.61 | 1.83E-47 | 3.80E-43 | 0.80 | 0.71 | 0.09 | 1.13 |
| 6 | Creb5 | 1.02 | 5.03E-47 | 1.05E-42 | 0.51 | 0.32 | 0.19 | 1.58 |
| 6 | App | 0.67 | 6.67E-47 | 1.39E-42 | 0.86 | 0.78 | 0.08 | 1.11 |
| 6 | Dpyd | 2.08 | 2.65E-46 | 5.52E-42 | 0.14 | 0.04 | 0.10 | 3.28 |
| 6 | Adgrl3 | 0.96 | 8.58E-46 | 1.78E-41 | 0.60 | 0.44 | 0.17 | 1.38 |
| 6 | Islr | 1.89 | 1.63E-45 | 3.39E-41 | 0.20 | 0.08 | 0.12 | 2.60 |
| 6 | Mxd4 | 0.87 | 9.43E-45 | 1.96E-40 | 0.63 | 0.46 | 0.17 | 1.36 |
| 6 | Hmcn1 | 0.70 | 1.21E-44 | 2.52E-40 | 0.70 | 0.53 | 0.17 | 1.33 |
| 6 | Rcn3 | 0.59 | 2.50E-44 | 5.20E-40 | 0.84 | 0.73 | 0.11 | 1.16 |
| 6 | Fzd4 | 1.19 | 2.93E-44 | 6.09E-40 | 0.41 | 0.25 | 0.17 | 1.68 |
| 6 | Rab34 | 0.59 | 1.19E-43 | 2.48E-39 | 0.81 | 0.71 | 0.10 | 1.14 |
| 6 | Col23a1 | 1.12 | 4.41E-43 | 9.17E-39 | 0.47 | 0.31 | 0.16 | 1.54 |
| 6 | Mab21l2 | 0.42 | 7.42E-43 | 1.54E-38 | 0.73 | 0.49 | 0.25 | 1.50 |
| 6 | Eml4 | 0.48 | 1.61E-42 | 3.35E-38 | 0.91 | 0.83 | 0.07 | 1.09 |
| 6 | Ppic | 0.60 | 1.06E-39 | 2.21E-35 | 0.81 | 0.71 | 0.10 | 1.14 |
| 6 | Cdk14 | 0.91 | 1.73E-39 | 3.60E-35 | 0.62 | 0.49 | 0.13 | 1.27 |
| 6 | Pdgfra | 0.69 | 2.19E-39 | 4.55E-35 | 0.70 | 0.54 | 0.17 | 1.32 |
| 6 | Dact3 | 0.71 | 3.27E-39 | 6.81E-35 | 0.64 | 0.49 | 0.16 | 1.32 |
| 6 | Ndfip1 | 0.46 | 4.55E-39 | 9.46E-35 | 0.89 | 0.83 | 0.06 | 1.07 |
| 6 | Dusp6 | 0.95 | 5.93E-39 | 1.23E-34 | 0.51 | 0.36 | 0.16 | 1.44 |
| 6 | Col26a1 | 0.45 | 7.43E-39 | 1.55E-34 | 0.80 | 0.60 | 0.19 | 1.32 |
| 6 | Ebf3 | 1.33 | 1.06E-38 | 2.21E-34 | 0.31 | 0.17 | 0.14 | 1.85 |
| 6 | Timp2 | 0.60 | 3.15E-38 | 6.56E-34 | 0.77 | 0.67 | 0.11 | 1.16 |
| 6 | Dnmt3a | 0.56 | 1.55E-37 | 3.23E-33 | 0.81 | 0.73 | 0.08 | 1.10 |
| 6 | Fbn2 | 0.54 | 8.43E-37 | 1.75E-32 | 0.86 | 0.73 | 0.13 | 1.18 |
| 6 | Pcp2 | 1.99 | 3.57E-36 | 7.43E-32 | 0.13 | 0.04 | 0.09 | 2.93 |
| 6 | Ptov1 | 0.27 | 4.66E-36 | 9.69E-32 | 0.99 | 0.97 | 0.02 | 1.02 |
| 6 | Dab2 | 0.91 | 5.08E-36 | 1.06E-31 | 0.50 | 0.35 | 0.15 | 1.43 |
| 6 | Fbn1 | 1.02 | 1.17E-35 | 2.43E-31 | 0.47 | 0.33 | 0.14 | 1.43 |
| 6 | Ier2 | 0.47 | 1.48E-35 | 3.07E-31 | 0.85 | 0.79 | 0.06 | 1.08 |

**Supplementary Table 1. Marker genes for the original clusters of integrated clustering of E11.5 control and NotchGOF λ-junction cells.**

| | | | | | | | | |
|---|---|---|---|---|---|---|---|---|
| 6 | 2610318N02Rik | 0.70 | 2.45E-35 | 5.10E-31 | 0.66 | 0.60 | 0.07 | 1.11 |
| 6 | Tox | 1.05 | 3.15E-35 | 6.56E-31 | 0.41 | 0.26 | 0.15 | 1.60 |
| 6 | Ssbp2 | 0.51 | 4.41E-35 | 9.18E-31 | 0.79 | 0.71 | 0.09 | 1.12 |
| 6 | Limd2 | 0.35 | 7.76E-35 | 1.61E-30 | 0.97 | 0.90 | 0.07 | 1.08 |
| 6 | Tiparp | 0.98 | 8.37E-35 | 1.74E-30 | 0.46 | 0.31 | 0.14 | 1.46 |
| 6 | Gas2 | 1.29 | 1.66E-34 | 3.45E-30 | 0.39 | 0.25 | 0.13 | 1.52 |
| 6 | Npc2 | 0.32 | 1.95E-34 | 4.05E-30 | 0.97 | 0.93 | 0.04 | 1.04 |
| 6 | Dnaja1 | 0.25 | 3.52E-34 | 7.33E-30 | 0.99 | 0.99 | 0.00 | 1.00 |
| 6 | Pbx3 | 0.67 | 4.18E-34 | 8.68E-30 | 0.65 | 0.53 | 0.12 | 1.24 |
| 6 | Slc38a1 | 0.53 | 6.44E-34 | 1.34E-29 | 0.84 | 0.77 | 0.08 | 1.10 |
| 6 | Tspan3 | 0.38 | 6.83E-34 | 1.42E-29 | 0.95 | 0.92 | 0.04 | 1.04 |
| 6 | Map2 | 1.11 | 9.86E-34 | 2.05E-29 | 0.35 | 0.21 | 0.14 | 1.65 |
| 6 | Twist2 | 0.99 | 1.35E-33 | 2.82E-29 | 0.37 | 0.23 | 0.15 | 1.66 |
| 6 | Clmp | 0.48 | 2.45E-33 | 5.10E-29 | 0.82 | 0.63 | 0.19 | 1.30 |
| 6 | Alkal2 | 1.65 | 6.30E-33 | 1.31E-28 | 0.18 | 0.08 | 0.10 | 2.33 |
| 6 | Sat1 | 1.64 | 1.10E-32 | 2.29E-28 | 0.42 | 0.30 | 0.12 | 1.39 |
| 6 | Tecr | 0.30 | 1.71E-32 | 3.55E-28 | 0.99 | 0.97 | 0.02 | 1.02 |
| 6 | Cd248 | 1.10 | 4.25E-32 | 8.83E-28 | 0.38 | 0.25 | 0.14 | 1.55 |
| 6 | Lpar1 | 0.42 | 5.40E-32 | 1.12E-27 | 0.86 | 0.70 | 0.16 | 1.23 |
| 6 | Oaf | 0.74 | 5.64E-32 | 1.17E-27 | 0.58 | 0.45 | 0.13 | 1.29 |
| 6 | Limch1 | 0.50 | 1.29E-31 | 2.68E-27 | 0.79 | 0.67 | 0.11 | 1.17 |
| 6 | Cdca7 | 0.52 | 1.32E-31 | 2.75E-27 | 0.79 | 0.72 | 0.07 | 1.09 |
| 6 | Mir100hg | 0.51 | 3.35E-31 | 6.97E-27 | 0.86 | 0.76 | 0.10 | 1.13 |
| 6 | Rbp1 | 0.29 | 4.62E-31 | 9.62E-27 | 0.97 | 0.91 | 0.07 | 1.07 |
| 6 | Sox9 | 0.36 | 7.37E-31 | 1.53E-26 | 0.73 | 0.58 | 0.15 | 1.26 |
| 6 | Mmp2 | 0.66 | 7.99E-31 | 1.66E-26 | 0.64 | 0.51 | 0.13 | 1.25 |
| 6 | Sesn3 | 0.63 | 8.80E-31 | 1.83E-26 | 0.68 | 0.59 | 0.10 | 1.16 |
| 6 | Tenm3 | 0.55 | 2.97E-30 | 6.18E-26 | 0.76 | 0.63 | 0.12 | 1.19 |
| 6 | Kcnj2 | 1.02 | 2.10E-29 | 4.36E-25 | 0.30 | 0.17 | 0.13 | 1.77 |
| 6 | Wdr86 | 0.68 | 2.37E-29 | 4.93E-25 | 0.62 | 0.51 | 0.10 | 1.20 |
| 6 | Tro | 0.81 | 4.63E-29 | 9.63E-25 | 0.53 | 0.42 | 0.12 | 1.28 |
| 6 | Efna2 | 0.48 | 5.44E-29 | 1.13E-24 | 0.79 | 0.71 | 0.09 | 1.12 |
| 6 | Shox2 | 1.15 | 8.28E-29 | 1.72E-24 | 0.24 | 0.12 | 0.12 | 1.95 |
| 6 | Inka1 | 0.64 | 8.96E-29 | 1.86E-24 | 0.62 | 0.51 | 0.12 | 1.23 |
| 6 | Foxp1 | 0.64 | 9.08E-29 | 1.89E-24 | 0.72 | 0.60 | 0.13 | 1.21 |
| 6 | Cmtm7 | 0.66 | 1.90E-28 | 3.95E-24 | 0.59 | 0.48 | 0.11 | 1.23 |
| 6 | Fendrr | 1.10 | 2.81E-28 | 5.85E-24 | 0.20 | 0.09 | 0.11 | 2.13 |
| 6 | Itm2c | 0.52 | 2.94E-28 | 6.10E-24 | 0.80 | 0.73 | 0.08 | 1.10 |
| 6 | Ldb2 | 1.07 | 3.31E-28 | 6.89E-24 | 0.39 | 0.27 | 0.13 | 1.47 |
| 6 | Slitrk5 | 1.10 | 4.29E-28 | 8.93E-24 | 0.28 | 0.15 | 0.12 | 1.80 |
| 6 | Mn1 | 0.90 | 4.57E-28 | 9.51E-24 | 0.41 | 0.28 | 0.13 | 1.48 |
| 6 | Cnpy2 | 0.34 | 4.93E-28 | 1.03E-23 | 0.96 | 0.93 | 0.03 | 1.03 |
| 6 | Igf1r | 0.47 | 9.76E-28 | 2.03E-23 | 0.82 | 0.77 | 0.05 | 1.07 |
| 6 | Ntng1 | 1.52 | 9.98E-28 | 2.08E-23 | 0.23 | 0.12 | 0.11 | 1.89 |
| 6 | Pcyt1a | 0.99 | 1.16E-27 | 2.41E-23 | 0.43 | 0.31 | 0.12 | 1.38 |
| 6 | Lgr5 | 1.16 | 1.40E-27 | 2.92E-23 | 0.33 | 0.20 | 0.13 | 1.64 |
| 6 | Tmem176b | 0.64 | 1.88E-27 | 3.91E-23 | 0.67 | 0.57 | 0.11 | 1.19 |
| 6 | Rbms3 | 1.02 | 2.43E-27 | 5.06E-23 | 0.52 | 0.41 | 0.11 | 1.28 |
| 6 | Reep5 | 0.44 | 1.03E-26 | 2.15E-22 | 0.85 | 0.78 | 0.07 | 1.09 |
| 6 | Hmgn3 | 0.44 | 2.50E-26 | 5.20E-22 | 0.80 | 0.75 | 0.05 | 1.07 |
| 6 | Igsf3 | 0.56 | 3.28E-26 | 6.83E-22 | 0.68 | 0.58 | 0.10 | 1.17 |
| 6 | Iqgap2 | 0.85 | 3.33E-26 | 6.93E-22 | 0.47 | 0.36 | 0.11 | 1.32 |
| 6 | Fbln1 | 0.40 | 7.80E-26 | 1.62E-21 | 0.85 | 0.79 | 0.06 | 1.07 |
| 6 | Foxc2 | 1.21 | 8.12E-26 | 1.69E-21 | 0.22 | 0.12 | 0.11 | 1.91 |
| 6 | Ptprd | 0.86 | 8.75E-26 | 1.82E-21 | 0.54 | 0.44 | 0.11 | 1.24 |
| 6 | Gm1673 | 0.29 | 1.16E-25 | 2.42E-21 | 0.98 | 0.93 | 0.05 | 1.05 |

**Supplementary Table 1. Marker genes for the original clusters of integrated clustering of E11.5 control and NotchGOF λ-junction cells.**

| | | | | | | | | |
|---|---|---|---|---|---|---|---|---|
| 6 | Fgf13 | 1.37 | 1.21E-25 | 2.52E-21 | 0.21 | 0.11 | 0.10 | 1.93 |
| 6 | Zeb1 | 0.75 | 1.23E-25 | 2.55E-21 | 0.47 | 0.34 | 0.13 | 1.37 |
| 6 | Setbp1 | 0.87 | 1.52E-25 | 3.17E-21 | 0.49 | 0.38 | 0.11 | 1.28 |
| 6 | Cacna2d1 | 1.28 | 2.33E-25 | 4.84E-21 | 0.22 | 0.12 | 0.10 | 1.90 |
| 6 | Traf4 | 0.55 | 2.44E-25 | 5.07E-21 | 0.70 | 0.63 | 0.07 | 1.11 |
| 6 | Tmem35a | 1.50 | 2.69E-25 | 5.60E-21 | 0.19 | 0.09 | 0.10 | 2.01 |
| 6 | Htr1b | 1.73 | 4.18E-25 | 8.68E-21 | 0.11 | 0.04 | 0.07 | 2.70 |
| 6 | Zeb2 | 0.52 | 9.11E-25 | 1.89E-20 | 0.55 | 0.39 | 0.15 | 1.39 |
| 6 | Sdc2 | 0.87 | 1.03E-24 | 2.14E-20 | 0.47 | 0.36 | 0.11 | 1.29 |
| 6 | Trim35 | 0.49 | 1.62E-24 | 3.36E-20 | 0.77 | 0.70 | 0.07 | 1.09 |
| 6 | Col13a1 | 1.41 | 2.04E-24 | 4.23E-20 | 0.20 | 0.11 | 0.10 | 1.92 |
| 6 | Cygb | 1.44 | 4.01E-24 | 8.35E-20 | 0.17 | 0.08 | 0.09 | 2.10 |
| 6 | Pbx1 | 0.26 | 4.78E-24 | 9.94E-20 | 0.97 | 0.92 | 0.05 | 1.06 |
| 6 | Asph | 0.47 | 7.49E-24 | 1.56E-19 | 0.74 | 0.65 | 0.09 | 1.14 |
| 6 | Spsb4 | 1.01 | 1.10E-23 | 2.28E-19 | 0.32 | 0.20 | 0.11 | 1.56 |
| 6 | Lrrc17 | 1.46 | 1.13E-23 | 2.34E-19 | 0.16 | 0.08 | 0.08 | 2.12 |
| 6 | Fndc3c1 | 0.70 | 1.22E-23 | 2.54E-19 | 0.56 | 0.47 | 0.09 | 1.20 |
| 6 | Gap43 | 0.51 | 2.03E-23 | 4.21E-19 | 0.54 | 0.41 | 0.13 | 1.30 |
| 6 | Maged1 | 0.28 | 2.43E-23 | 5.05E-19 | 0.98 | 0.94 | 0.04 | 1.04 |
| 6 | Zfp637 | 0.46 | 3.14E-23 | 6.53E-19 | 0.78 | 0.73 | 0.05 | 1.06 |
| 6 | Chd4 | 0.26 | 3.18E-23 | 6.62E-19 | 0.98 | 0.96 | 0.03 | 1.03 |
| 6 | Slc27a3 | 1.05 | 3.20E-23 | 6.66E-19 | 0.32 | 0.21 | 0.11 | 1.52 |
| 6 | Tead2 | 0.29 | 4.70E-23 | 9.78E-19 | 0.95 | 0.90 | 0.05 | 1.06 |
| 6 | Emilin1 | 0.52 | 4.89E-23 | 1.02E-18 | 0.65 | 0.53 | 0.12 | 1.23 |
| 6 | Dhx36 | 0.30 | 5.18E-23 | 1.08E-18 | 0.92 | 0.90 | 0.02 | 1.03 |
| 6 | Fkbp10 | 0.85 | 5.33E-23 | 1.11E-18 | 0.45 | 0.33 | 0.11 | 1.34 |
| 6 | Inpp4b | 1.26 | 5.34E-23 | 1.11E-18 | 0.25 | 0.15 | 0.10 | 1.69 |
| 6 | Irak1bp1 | 0.63 | 5.78E-23 | 1.20E-18 | 0.60 | 0.54 | 0.06 | 1.10 |
| 6 | Elovl6 | 0.52 | 6.39E-23 | 1.33E-18 | 0.71 | 0.65 | 0.06 | 1.09 |
| 6 | Rbp4 | 1.66 | 7.77E-23 | 1.62E-18 | 0.13 | 0.06 | 0.07 | 2.30 |
| 6 | 2700033N17Rik | 1.33 | 1.97E-22 | 4.10E-18 | 0.21 | 0.12 | 0.09 | 1.82 |
| 6 | Arf4 | 0.26 | 2.02E-22 | 4.20E-18 | 0.98 | 0.95 | 0.03 | 1.03 |
| 6 | Gulp1 | 1.04 | 2.93E-22 | 6.10E-18 | 0.35 | 0.25 | 0.11 | 1.43 |
| 6 | Dact1 | 0.90 | 5.18E-22 | 1.08E-17 | 0.41 | 0.31 | 0.11 | 1.34 |
| 6 | Tpr | 0.26 | 6.90E-22 | 1.43E-17 | 0.99 | 0.98 | 0.01 | 1.01 |
| 6 | Dbn1 | 0.31 | 8.28E-22 | 1.72E-17 | 0.93 | 0.88 | 0.05 | 1.06 |
| 6 | Zfp36l1 | 0.36 | 1.79E-21 | 3.73E-17 | 0.90 | 0.85 | 0.05 | 1.06 |
| 6 | Slc38a2 | 0.32 | 1.82E-21 | 3.79E-17 | 0.93 | 0.90 | 0.03 | 1.03 |
| 6 | Vegfb | 0.46 | 4.71E-21 | 9.79E-17 | 0.71 | 0.66 | 0.05 | 1.07 |
| 6 | Col12a1 | 1.88 | 5.80E-21 | 1.21E-16 | 0.13 | 0.06 | 0.07 | 2.19 |
| 6 | Mdfi | 0.52 | 5.95E-21 | 1.24E-16 | 0.63 | 0.56 | 0.07 | 1.13 |
| 6 | Hp1bp3 | 0.31 | 8.08E-21 | 1.68E-16 | 0.91 | 0.89 | 0.02 | 1.02 |
| 6 | Otor | 1.51 | 9.13E-21 | 1.90E-16 | 0.15 | 0.07 | 0.08 | 2.05 |
| 6 | Tnrc6a | 0.34 | 1.04E-20 | 2.17E-16 | 0.81 | 0.78 | 0.03 | 1.04 |
| 6 | Bnc2 | 1.01 | 1.06E-20 | 2.20E-16 | 0.28 | 0.18 | 0.10 | 1.59 |
| 6 | Nfia | 0.71 | 1.62E-20 | 3.37E-16 | 0.56 | 0.47 | 0.09 | 1.19 |
| 6 | Bzw2 | 0.26 | 3.04E-20 | 6.32E-16 | 0.96 | 0.94 | 0.02 | 1.02 |
| 6 | Gm13429 | 1.51 | 3.74E-20 | 7.77E-16 | 0.12 | 0.05 | 0.07 | 2.26 |
| 6 | Gjc1 | 0.31 | 4.45E-20 | 9.25E-16 | 0.87 | 0.81 | 0.07 | 1.08 |
| 6 | Slitrk4 | 1.29 | 4.63E-20 | 9.64E-16 | 0.12 | 0.05 | 0.07 | 2.29 |
| 6 | Asphd2 | 1.24 | 1.02E-19 | 2.11E-15 | 0.21 | 0.12 | 0.09 | 1.75 |
| 6 | Cdc26 | 0.28 | 1.16E-19 | 2.40E-15 | 0.94 | 0.93 | 0.01 | 1.01 |
| 6 | Wbp1 | 0.41 | 1.44E-19 | 3.00E-15 | 0.80 | 0.77 | 0.03 | 1.04 |
| 6 | Rbpms2 | 0.94 | 2.12E-19 | 4.42E-15 | 0.32 | 0.22 | 0.10 | 1.46 |
| 6 | Nisch | 0.29 | 2.13E-19 | 4.44E-15 | 0.94 | 0.92 | 0.02 | 1.03 |
| 6 | Rbm18 | 0.59 | 2.32E-19 | 4.83E-15 | 0.62 | 0.55 | 0.06 | 1.12 |

**Supplementary Table 1. Marker genes for the original clusters of integrated clustering of E11.5 control and NotchGOF λ-junction cells.**

| | | | | | | | | |
|---|---|---|---|---|---|---|---|---|
| 6 | Edn3 | 1.29 | 2.66E-19 | 5.53E-15 | 0.23 | 0.14 | 0.09 | 1.64 |
| 6 | Lix1l | 0.65 | 2.89E-19 | 6.01E-15 | 0.48 | 0.38 | 0.10 | 1.27 |
| 6 | Pnisr | 0.31 | 3.06E-19 | 6.36E-15 | 0.94 | 0.92 | 0.02 | 1.02 |
| 6 | Plce1 | 0.90 | 4.82E-19 | 1.00E-14 | 0.33 | 0.22 | 0.10 | 1.45 |
| 6 | Thsd7a | 1.31 | 5.91E-19 | 1.23E-14 | 0.24 | 0.15 | 0.09 | 1.63 |
| 6 | Plat | 0.90 | 6.45E-19 | 1.34E-14 | 0.41 | 0.32 | 0.10 | 1.31 |
| 6 | Polr2m | 0.26 | 7.12E-19 | 1.48E-14 | 0.97 | 0.95 | 0.02 | 1.02 |
| 6 | Ssr3 | 0.27 | 8.26E-19 | 1.72E-14 | 0.90 | 0.88 | 0.02 | 1.02 |
| 6 | Tmem223 | 0.36 | 1.00E-18 | 2.08E-14 | 0.86 | 0.84 | 0.02 | 1.03 |
| 6 | Epb41l2 | 0.46 | 1.24E-18 | 2.58E-14 | 0.73 | 0.67 | 0.06 | 1.09 |
| 6 | Phldb2 | 0.41 | 1.46E-18 | 3.03E-14 | 0.63 | 0.53 | 0.10 | 1.19 |
| 6 | Pdlim4 | 0.42 | 1.60E-18 | 3.33E-14 | 0.70 | 0.62 | 0.07 | 1.11 |
| 6 | Rnd3 | 0.49 | 1.74E-18 | 3.62E-14 | 0.70 | 0.64 | 0.06 | 1.09 |
| 6 | Jam3 | 0.48 | 2.46E-18 | 5.12E-14 | 0.65 | 0.57 | 0.07 | 1.13 |
| 6 | Echs1 | 0.31 | 5.45E-18 | 1.13E-13 | 0.87 | 0.86 | 0.01 | 1.01 |
| 6 | P3h3 | 0.83 | 5.46E-18 | 1.14E-13 | 0.40 | 0.30 | 0.09 | 1.31 |
| 6 | Dnm1 | 0.69 | 5.57E-18 | 1.16E-13 | 0.42 | 0.32 | 0.10 | 1.31 |
| 6 | Wdr6 | 0.44 | 6.43E-18 | 1.34E-13 | 0.70 | 0.66 | 0.04 | 1.06 |
| 6 | Cthrc1 | 1.44 | 6.99E-18 | 1.45E-13 | 0.28 | 0.19 | 0.09 | 1.48 |
| 6 | Zfp422 | 0.29 | 8.82E-18 | 1.84E-13 | 0.89 | 0.85 | 0.04 | 1.05 |
| 6 | Pcdh11x | 1.43 | 1.61E-17 | 3.36E-13 | 0.18 | 0.11 | 0.08 | 1.75 |
| 6 | Ssbp4 | 0.43 | 1.75E-17 | 3.63E-13 | 0.71 | 0.65 | 0.05 | 1.08 |
| 6 | Rcn2 | 0.26 | 1.96E-17 | 4.09E-13 | 0.97 | 0.91 | 0.06 | 1.06 |
| 6 | Etv1 | 0.82 | 2.25E-17 | 4.69E-13 | 0.40 | 0.30 | 0.10 | 1.34 |
| 6 | Sox12 | 0.40 | 2.45E-17 | 5.09E-13 | 0.71 | 0.66 | 0.05 | 1.07 |
| 6 | Six4 | 0.56 | 2.70E-17 | 5.61E-13 | 0.54 | 0.46 | 0.08 | 1.17 |
| 6 | Gsto1 | 0.40 | 3.12E-17 | 6.48E-13 | 0.70 | 0.65 | 0.05 | 1.07 |
| 6 | Zfhx4 | 0.49 | 3.83E-17 | 7.97E-13 | 0.74 | 0.74 | 0.01 | 1.01 |
| 6 | Gpx7 | 0.64 | 4.02E-17 | 8.35E-13 | 0.52 | 0.45 | 0.08 | 1.17 |
| 6 | Col11a1 | 0.60 | 4.78E-17 | 9.93E-13 | 0.52 | 0.42 | 0.09 | 1.22 |
| 6 | Mageh1 | 0.51 | 5.98E-17 | 1.24E-12 | 0.64 | 0.56 | 0.07 | 1.13 |
| 6 | Tbx3 | 0.33 | 7.19E-17 | 1.50E-12 | 0.57 | 0.43 | 0.13 | 1.30 |
| 6 | Auts2 | 0.30 | 9.16E-17 | 1.90E-12 | 0.92 | 0.87 | 0.06 | 1.06 |
| 6 | Palld | 0.47 | 1.00E-16 | 2.09E-12 | 0.69 | 0.62 | 0.06 | 1.10 |
| 6 | Ak1 | 1.31 | 1.15E-16 | 2.39E-12 | 0.15 | 0.08 | 0.07 | 1.88 |
| 6 | Greb1l | 1.44 | 1.38E-16 | 2.88E-12 | 0.13 | 0.06 | 0.06 | 2.00 |
| 6 | Arhgef25 | 0.70 | 2.00E-16 | 4.16E-12 | 0.45 | 0.37 | 0.08 | 1.22 |
| 6 | Pdlim7 | 0.32 | 4.63E-16 | 9.63E-12 | 0.80 | 0.75 | 0.04 | 1.06 |
| 6 | Adss | 0.32 | 5.08E-16 | 1.06E-11 | 0.74 | 0.73 | 0.02 | 1.03 |
| 6 | Pgf | 1.62 | 5.41E-16 | 1.13E-11 | 0.13 | 0.07 | 0.06 | 1.93 |
| 6 | AI606473 | 1.04 | 8.04E-16 | 1.67E-11 | 0.16 | 0.09 | 0.07 | 1.85 |
| 6 | Olfml3 | 0.80 | 1.33E-15 | 2.77E-11 | 0.37 | 0.29 | 0.09 | 1.31 |
| 6 | Morc4 | 1.12 | 1.64E-15 | 3.41E-11 | 0.23 | 0.14 | 0.08 | 1.57 |
| 6 | Fkbp7 | 0.74 | 1.86E-15 | 3.87E-11 | 0.37 | 0.28 | 0.09 | 1.33 |
| 6 | Ptgis | 1.29 | 2.27E-15 | 4.72E-11 | 0.18 | 0.11 | 0.07 | 1.67 |
| 6 | Rab13 | 0.56 | 3.40E-15 | 7.07E-11 | 0.50 | 0.42 | 0.08 | 1.18 |
| 6 | Runx1t1 | 0.48 | 3.44E-15 | 7.15E-11 | 0.62 | 0.53 | 0.10 | 1.18 |
| 6 | Gstm5 | 0.28 | 4.89E-15 | 1.02E-10 | 0.90 | 0.88 | 0.02 | 1.03 |
| 6 | Nynrin | 0.53 | 6.56E-15 | 1.37E-10 | 0.60 | 0.55 | 0.05 | 1.08 |
| 6 | Rnf122 | 0.46 | 7.19E-15 | 1.50E-10 | 0.60 | 0.53 | 0.06 | 1.12 |
| 6 | Rnf150 | 0.79 | 9.59E-15 | 1.99E-10 | 0.39 | 0.31 | 0.08 | 1.26 |
| 6 | Fli1 | 0.64 | 1.40E-14 | 2.91E-10 | 0.46 | 0.37 | 0.09 | 1.23 |
| 6 | Dnm3os | 0.45 | 1.53E-14 | 3.18E-10 | 0.61 | 0.52 | 0.09 | 1.18 |
| 6 | Il11ra1 | 0.30 | 1.74E-14 | 3.61E-10 | 0.72 | 0.61 | 0.11 | 1.18 |
| 6 | Foxp4 | 0.74 | 2.16E-14 | 4.48E-10 | 0.48 | 0.42 | 0.06 | 1.15 |
| 6 | Ddx50 | 0.32 | 4.19E-14 | 8.71E-10 | 0.81 | 0.79 | 0.01 | 1.02 |

**Supplementary Table 1. Marker genes for the original clusters of integrated clustering of E11.5 control and NotchGOF λ-junction cells.**

| | | | | | | | | |
|---|---|---|---|---|---|---|---|---|
| 6 | Add3 | 0.37 | 5.71E-14 | 1.19E-09 | 0.71 | 0.66 | 0.05 | 1.08 |
| 6 | Etfb | 0.27 | 8.17E-14 | 1.70E-09 | 0.90 | 0.89 | 0.01 | 1.01 |
| 6 | Itpr1 | 0.92 | 1.10E-13 | 2.29E-09 | 0.29 | 0.21 | 0.08 | 1.36 |
| 6 | Vcl | 0.65 | 1.18E-13 | 2.45E-09 | 0.46 | 0.39 | 0.07 | 1.17 |
| 6 | Tmem26 | 0.65 | 1.29E-13 | 2.68E-09 | 0.34 | 0.25 | 0.09 | 1.36 |
| 6 | Hook3 | 0.37 | 1.32E-13 | 2.74E-09 | 0.70 | 0.66 | 0.04 | 1.06 |
| 6 | Bmp1 | 0.67 | 1.34E-13 | 2.78E-09 | 0.49 | 0.42 | 0.06 | 1.15 |
| 6 | Pitpnc1 | 0.46 | 1.37E-13 | 2.84E-09 | 0.63 | 0.59 | 0.05 | 1.08 |
| 6 | Rest | 0.26 | 1.38E-13 | 2.86E-09 | 0.86 | 0.81 | 0.05 | 1.06 |
| 6 | Vegfc | 0.76 | 1.79E-13 | 3.72E-09 | 0.35 | 0.26 | 0.09 | 1.32 |
| 6 | Fign | 0.81 | 2.06E-13 | 4.29E-09 | 0.37 | 0.29 | 0.08 | 1.26 |
| 6 | Cdo1 | 0.32 | 2.15E-13 | 4.48E-09 | 0.59 | 0.50 | 0.09 | 1.18 |
| 6 | Mxra8 | 1.10 | 2.39E-13 | 4.97E-09 | 0.22 | 0.14 | 0.07 | 1.50 |
| 6 | Ube3a | 0.26 | 3.15E-13 | 6.56E-09 | 0.85 | 0.84 | 0.01 | 1.01 |
| 6 | Tmem256 | 0.25 | 3.90E-13 | 8.10E-09 | 0.89 | 0.89 | 0.00 | 1.00 |
| 6 | Lrig3 | 0.83 | 3.98E-13 | 8.27E-09 | 0.30 | 0.23 | 0.08 | 1.34 |
| 6 | Mmp14 | 0.38 | 4.00E-13 | 8.32E-09 | 0.68 | 0.63 | 0.05 | 1.08 |
| 6 | Bcl2l11 | 0.67 | 5.81E-13 | 1.21E-08 | 0.42 | 0.35 | 0.07 | 1.20 |
| 6 | Cobl | 1.64 | 6.46E-13 | 1.34E-08 | 0.12 | 0.07 | 0.06 | 1.82 |
| 6 | Ei24 | 0.39 | 6.59E-13 | 1.37E-08 | 0.72 | 0.70 | 0.01 | 1.02 |
| 6 | Aldh2 | 0.33 | 1.12E-12 | 2.32E-08 | 0.78 | 0.78 | 0.01 | 1.01 |
| 6 | Zfp292 | 0.28 | 1.12E-12 | 2.33E-08 | 0.88 | 0.86 | 0.02 | 1.03 |
| 6 | Cnn2 | 0.40 | 1.47E-12 | 3.06E-08 | 0.70 | 0.63 | 0.07 | 1.11 |
| 6 | Zfp9 | 0.90 | 1.49E-12 | 3.09E-08 | 0.24 | 0.16 | 0.07 | 1.46 |
| 6 | Praf2 | 0.47 | 2.09E-12 | 4.34E-08 | 0.57 | 0.53 | 0.04 | 1.08 |
| 6 | Itgb1 | 0.27 | 3.52E-12 | 7.33E-08 | 0.87 | 0.85 | 0.02 | 1.02 |
| 6 | Mdga2 | 1.08 | 4.06E-12 | 8.45E-08 | 0.13 | 0.07 | 0.06 | 1.81 |
| 6 | St6galnac6 | 1.02 | 4.31E-12 | 8.95E-08 | 0.23 | 0.16 | 0.07 | 1.45 |
| 6 | Synpo | 0.66 | 5.65E-12 | 1.17E-07 | 0.32 | 0.24 | 0.08 | 1.34 |
| 6 | Tmem9 | 0.39 | 6.18E-12 | 1.29E-07 | 0.67 | 0.64 | 0.03 | 1.04 |
| 6 | Rps6ka6 | 0.53 | 7.30E-12 | 1.52E-07 | 0.51 | 0.45 | 0.06 | 1.13 |
| 6 | Metrn | 0.49 | 7.68E-12 | 1.60E-07 | 0.56 | 0.51 | 0.05 | 1.09 |
| 6 | Ptx3 | 1.08 | 7.80E-12 | 1.62E-07 | 0.17 | 0.11 | 0.06 | 1.59 |
| 6 | B4galt4 | 0.95 | 9.19E-12 | 1.91E-07 | 0.29 | 0.23 | 0.06 | 1.28 |
| 6 | Septin6 | 0.39 | 9.83E-12 | 2.04E-07 | 0.62 | 0.57 | 0.05 | 1.10 |
| 6 | Snhg8 | 0.28 | 1.18E-11 | 2.45E-07 | 0.81 | 0.80 | 0.01 | 1.01 |
| 6 | Nktr | 0.25 | 1.24E-11 | 2.57E-07 | 0.86 | 0.85 | 0.02 | 1.02 |
| 6 | Rian | 0.60 | 1.85E-11 | 3.84E-07 | 0.39 | 0.32 | 0.07 | 1.23 |
| 6 | Gpx3 | 0.79 | 2.01E-11 | 4.17E-07 | 0.29 | 0.22 | 0.07 | 1.33 |
| 6 | Zfp266 | 0.26 | 2.22E-11 | 4.61E-07 | 0.82 | 0.80 | 0.02 | 1.03 |
| 6 | Gm26771 | 1.62 | 2.22E-11 | 4.63E-07 | 0.10 | 0.06 | 0.05 | 1.86 |
| 6 | Lrpap1 | 0.54 | 2.28E-11 | 4.75E-07 | 0.53 | 0.49 | 0.04 | 1.08 |
| 6 | Smo | 0.30 | 2.48E-11 | 5.16E-07 | 0.76 | 0.73 | 0.02 | 1.03 |
| 6 | Cdc42ep3 | 1.21 | 3.42E-11 | 7.11E-07 | 0.23 | 0.16 | 0.06 | 1.39 |
| 6 | Clk1 | 0.38 | 3.71E-11 | 7.72E-07 | 0.69 | 0.68 | 0.01 | 1.02 |
| 6 | Shisa4 | 0.67 | 3.90E-11 | 8.12E-07 | 0.39 | 0.33 | 0.07 | 1.20 |
| 6 | Zfp318 | 0.47 | 4.03E-11 | 8.38E-07 | 0.52 | 0.47 | 0.05 | 1.11 |
| 6 | Trpm7 | 0.31 | 4.13E-11 | 8.59E-07 | 0.71 | 0.69 | 0.03 | 1.04 |
| 6 | Mcm4 | 0.32 | 4.15E-11 | 8.63E-07 | 0.77 | 0.73 | 0.03 | 1.04 |
| 6 | Atraid | 0.52 | 4.46E-11 | 9.27E-07 | 0.54 | 0.51 | 0.03 | 1.07 |
| 6 | Emp3 | 0.72 | 5.61E-11 | 1.17E-06 | 0.20 | 0.13 | 0.07 | 1.51 |
| 6 | Nr4a2 | 1.09 | 5.88E-11 | 1.22E-06 | 0.16 | 0.10 | 0.06 | 1.58 |
| 6 | Dzip1 | 0.61 | 5.96E-11 | 1.24E-06 | 0.40 | 0.33 | 0.07 | 1.21 |
| 6 | Evi5 | 0.36 | 6.20E-11 | 1.29E-06 | 0.63 | 0.61 | 0.03 | 1.04 |
| 6 | Dchs1 | 0.51 | 7.45E-11 | 1.55E-06 | 0.48 | 0.42 | 0.06 | 1.15 |
| 6 | Pmepa1 | 0.48 | 7.58E-11 | 1.58E-06 | 0.54 | 0.48 | 0.06 | 1.13 |

**Supplementary Table 1. Marker genes for the original clusters of integrated clustering of E11.5 control and NotchGOF λ-junction cells.**

| | | | | | | | | |
|---|---|---|---|---|---|---|---|---|
| 6 | Tmem119 | 0.42 | 9.02E-11 | 1.88E-06 | 0.53 | 0.46 | 0.08 | 1.17 |
| 6 | Gmds | 0.64 | 9.87E-11 | 2.05E-06 | 0.42 | 0.36 | 0.06 | 1.17 |
| 6 | Nfatc4 | 0.36 | 9.91E-11 | 2.06E-06 | 0.61 | 0.55 | 0.06 | 1.11 |
| 6 | BC028528 | 1.11 | 1.14E-10 | 2.38E-06 | 0.13 | 0.07 | 0.05 | 1.72 |
| 6 | Six2 | 0.34 | 1.20E-10 | 2.49E-06 | 0.54 | 0.45 | 0.09 | 1.20 |
| 6 | Psme2 | 0.33 | 1.20E-10 | 2.49E-06 | 0.75 | 0.76 | -0.01 | 0.98 |
| 6 | Ephb3 | 0.80 | 1.24E-10 | 2.57E-06 | 0.36 | 0.30 | 0.06 | 1.20 |
| 6 | Ogfr | 0.34 | 1.27E-10 | 2.65E-06 | 0.69 | 0.67 | 0.02 | 1.03 |
| 6 | Kifap3 | 0.27 | 1.32E-10 | 2.75E-06 | 0.79 | 0.78 | 0.02 | 1.02 |
| 6 | Tspan11 | 0.91 | 1.39E-10 | 2.90E-06 | 0.23 | 0.17 | 0.07 | 1.40 |
| 6 | Rcor2 | 0.45 | 1.58E-10 | 3.28E-06 | 0.53 | 0.49 | 0.04 | 1.08 |
| 6 | Srpk2 | 0.31 | 1.60E-10 | 3.34E-06 | 0.73 | 0.71 | 0.02 | 1.03 |
| 6 | Neo1 | 0.50 | 1.85E-10 | 3.84E-06 | 0.53 | 0.50 | 0.03 | 1.07 |
| 6 | Mrc2 | 0.86 | 1.86E-10 | 3.88E-06 | 0.28 | 0.22 | 0.06 | 1.29 |
| 6 | Ung | 0.49 | 1.91E-10 | 3.97E-06 | 0.42 | 0.34 | 0.08 | 1.23 |
| 6 | Mycn | 0.49 | 2.04E-10 | 4.23E-06 | 0.44 | 0.38 | 0.07 | 1.18 |
| 6 | Map7d1 | 0.36 | 2.20E-10 | 4.57E-06 | 0.65 | 0.62 | 0.03 | 1.05 |
| 6 | Zfp703 | 0.85 | 2.25E-10 | 4.68E-06 | 0.31 | 0.25 | 0.06 | 1.24 |
| 6 | Lrp6 | 0.46 | 2.35E-10 | 4.89E-06 | 0.60 | 0.58 | 0.02 | 1.04 |
| 6 | Pth1r | 0.27 | 2.43E-10 | 5.06E-06 | 0.67 | 0.57 | 0.10 | 1.18 |
| 6 | Zfp354c | 0.55 | 2.52E-10 | 5.23E-06 | 0.45 | 0.39 | 0.06 | 1.15 |
| 6 | Pdcd4 | 0.26 | 3.01E-10 | 6.26E-06 | 0.69 | 0.64 | 0.05 | 1.07 |
| 6 | Efemp2 | 0.56 | 3.08E-10 | 6.41E-06 | 0.51 | 0.48 | 0.04 | 1.08 |
| 6 | B2m | 0.28 | 3.81E-10 | 7.93E-06 | 0.59 | 0.54 | 0.05 | 1.09 |
| 6 | Cyp51 | 0.35 | 3.85E-10 | 8.01E-06 | 0.72 | 0.69 | 0.03 | 1.04 |
| 6 | Klhl24 | 0.37 | 3.86E-10 | 8.04E-06 | 0.64 | 0.60 | 0.04 | 1.06 |
| 6 | Nadk2 | 0.74 | 3.87E-10 | 8.05E-06 | 0.40 | 0.35 | 0.05 | 1.14 |
| 6 | Fst | 0.41 | 4.16E-10 | 8.66E-06 | 0.19 | 0.12 | 0.07 | 1.55 |
| 6 | Fam92a | 0.29 | 4.22E-10 | 8.77E-06 | 0.76 | 0.76 | 0.00 | 1.00 |
| 6 | Dcaf8 | 0.28 | 4.31E-10 | 8.95E-06 | 0.76 | 0.75 | 0.01 | 1.02 |
| 6 | Ccdc8 | 0.57 | 4.65E-10 | 9.68E-06 | 0.38 | 0.33 | 0.06 | 1.17 |
| 6 | Ogt | 0.29 | 4.70E-10 | 9.77E-06 | 0.78 | 0.78 | 0.00 | 1.00 |
| 6 | Fmr1 | 0.32 | 4.89E-10 | 1.02E-05 | 0.68 | 0.66 | 0.02 | 1.02 |
| 6 | Casd1 | 0.44 | 5.49E-10 | 1.14E-05 | 0.54 | 0.50 | 0.04 | 1.08 |
| 6 | Gprasp1 | 0.46 | 6.12E-10 | 1.27E-05 | 0.53 | 0.49 | 0.04 | 1.08 |
| 6 | Epha7 | 0.30 | 6.29E-10 | 1.31E-05 | 0.62 | 0.56 | 0.06 | 1.10 |
| 6 | Dusp7 | 0.43 | 7.47E-10 | 1.55E-05 | 0.55 | 0.50 | 0.05 | 1.09 |
| 6 | Pbx2 | 0.26 | 7.49E-10 | 1.56E-05 | 0.83 | 0.81 | 0.02 | 1.02 |
| 6 | Flna | 0.27 | 7.88E-10 | 1.64E-05 | 0.82 | 0.79 | 0.02 | 1.03 |
| 6 | Erbin | 0.41 | 8.19E-10 | 1.70E-05 | 0.65 | 0.63 | 0.02 | 1.03 |
| 6 | Rbfox2 | 0.25 | 8.47E-10 | 1.76E-05 | 0.78 | 0.77 | 0.01 | 1.02 |
| 6 | Creb3l2 | 0.71 | 9.14E-10 | 1.90E-05 | 0.38 | 0.32 | 0.06 | 1.19 |
| 6 | Hspa12b | 0.94 | 9.33E-10 | 1.94E-05 | 0.19 | 0.13 | 0.06 | 1.46 |
| 6 | Adgra2 | 0.77 | 1.03E-09 | 2.15E-05 | 0.30 | 0.24 | 0.06 | 1.26 |
| 6 | Txndc12 | 0.31 | 1.09E-09 | 2.27E-05 | 0.72 | 0.70 | 0.02 | 1.03 |
| 6 | Lpp | 0.45 | 1.11E-09 | 2.32E-05 | 0.60 | 0.56 | 0.04 | 1.08 |
| 6 | Tmem165 | 0.31 | 1.16E-09 | 2.40E-05 | 0.82 | 0.80 | 0.02 | 1.02 |
| 6 | Arl6ip6 | 0.29 | 1.21E-09 | 2.52E-05 | 0.69 | 0.68 | 0.01 | 1.02 |
| 6 | Hexa | 0.49 | 1.23E-09 | 2.57E-05 | 0.38 | 0.31 | 0.06 | 1.20 |
| 6 | Pgm2 | 0.66 | 1.26E-09 | 2.62E-05 | 0.33 | 0.27 | 0.06 | 1.22 |
| 6 | Taf7 | 0.31 | 1.28E-09 | 2.66E-05 | 0.69 | 0.70 | 0.00 | 1.00 |
| 6 | Phf3 | 0.26 | 1.43E-09 | 2.98E-05 | 0.80 | 0.81 | -0.02 | 0.98 |
| 6 | Por | 0.52 | 1.46E-09 | 3.03E-05 | 0.50 | 0.47 | 0.03 | 1.07 |
| 6 | Ubtf | 0.26 | 1.60E-09 | 3.33E-05 | 0.82 | 0.83 | -0.01 | 0.99 |
| 6 | Ebpl | 0.37 | 1.69E-09 | 3.52E-05 | 0.59 | 0.55 | 0.04 | 1.08 |
| 6 | Golgb1 | 0.28 | 1.72E-09 | 3.59E-05 | 0.77 | 0.74 | 0.03 | 1.04 |

**Supplementary Table 1. Marker genes for the original clusters of integrated clustering of E11.5 control and NotchGOF λ-junction cells.**

| | | | | | | | | |
|---|---|---|---|---|---|---|---|---|
| 6 | Helz | 0.34 | 1.74E-09 | 3.62E-05 | 0.62 | 0.58 | 0.05 | 1.08 |
| 6 | Dse | 0.71 | 1.96E-09 | 4.08E-05 | 0.33 | 0.27 | 0.06 | 1.22 |
| 6 | Itga9 | 0.91 | 1.97E-09 | 4.11E-05 | 0.27 | 0.21 | 0.06 | 1.28 |
| 6 | Dmd | 1.25 | 2.03E-09 | 4.22E-05 | 0.14 | 0.09 | 0.05 | 1.56 |
| 6 | Foxd1 | 0.49 | 2.45E-09 | 5.10E-05 | 0.17 | 0.11 | 0.06 | 1.57 |
| 6 | Cbr1 | 0.26 | 2.83E-09 | 5.88E-05 | 0.77 | 0.76 | 0.01 | 1.01 |
| 6 | Slc50a1 | 0.60 | 2.83E-09 | 5.89E-05 | 0.43 | 0.37 | 0.05 | 1.14 |
| 6 | Inppl1 | 0.53 | 2.99E-09 | 6.23E-05 | 0.44 | 0.39 | 0.05 | 1.12 |
| 6 | Lrrc4b | 0.86 | 3.08E-09 | 6.40E-05 | 0.19 | 0.13 | 0.06 | 1.46 |
| 6 | Gli1 | 0.92 | 3.26E-09 | 6.78E-05 | 0.19 | 0.13 | 0.06 | 1.45 |
| 6 | Rin2 | 0.85 | 3.33E-09 | 6.92E-05 | 0.26 | 0.20 | 0.06 | 1.30 |
| 6 | Dusp23 | 0.82 | 4.02E-09 | 8.37E-05 | 0.23 | 0.17 | 0.06 | 1.36 |
| 6 | P4ha2 | 0.61 | 4.32E-09 | 8.98E-05 | 0.34 | 0.27 | 0.06 | 1.24 |
| 6 | Usp29 | 0.74 | 4.94E-09 | 1.03E-04 | 0.26 | 0.20 | 0.06 | 1.28 |
| 6 | Numa1 | 0.28 | 5.17E-09 | 1.08E-04 | 0.72 | 0.72 | 0.00 | 1.00 |
| 6 | Olfml1 | 0.94 | 5.27E-09 | 1.10E-04 | 0.17 | 0.12 | 0.06 | 1.48 |
| 6 | Tmem98 | 0.42 | 5.57E-09 | 1.16E-04 | 0.54 | 0.50 | 0.04 | 1.07 |
| 6 | Pnpla8 | 0.44 | 6.36E-09 | 1.32E-04 | 0.53 | 0.50 | 0.03 | 1.06 |
| 6 | Adamts9 | 0.57 | 6.55E-09 | 1.36E-04 | 0.21 | 0.15 | 0.06 | 1.41 |
| 6 | Rcbtb2 | 0.27 | 7.49E-09 | 1.56E-04 | 0.76 | 0.76 | 0.00 | 0.99 |
| 6 | Uhrf2 | 0.27 | 8.42E-09 | 1.75E-04 | 0.76 | 0.73 | 0.03 | 1.04 |
| 6 | Tcaf1 | 0.27 | 9.11E-09 | 1.89E-04 | 0.70 | 0.67 | 0.03 | 1.05 |
| 6 | S1pr3 | 0.66 | 9.73E-09 | 2.02E-04 | 0.33 | 0.28 | 0.06 | 1.21 |
| 6 | Plscr3 | 0.36 | 9.99E-09 | 2.08E-04 | 0.59 | 0.56 | 0.03 | 1.05 |
| 6 | Dld | 0.26 | 9.99E-09 | 2.08E-04 | 0.76 | 0.77 | -0.01 | 0.99 |
| 6 | Capns1 | 0.25 | 1.01E-08 | 2.10E-04 | 0.80 | 0.80 | 0.00 | 1.00 |
| 6 | Rxylt1 | 0.49 | 1.15E-08 | 2.39E-04 | 0.50 | 0.47 | 0.03 | 1.06 |
| 6 | Tln1 | 0.28 | 1.17E-08 | 2.44E-04 | 0.75 | 0.74 | 0.01 | 1.01 |
| 6 | Ccdc80 | 0.75 | 1.17E-08 | 2.44E-04 | 0.24 | 0.18 | 0.06 | 1.32 |
| 6 | 1810026B05Rik | 0.38 | 1.30E-08 | 2.70E-04 | 0.61 | 0.60 | 0.01 | 1.02 |
| 6 | Pmp22 | 0.85 | 1.34E-08 | 2.79E-04 | 0.24 | 0.18 | 0.06 | 1.31 |
| 6 | Eid2 | 0.30 | 1.35E-08 | 2.81E-04 | 0.67 | 0.65 | 0.01 | 1.02 |
| 6 | Cd320 | 0.38 | 1.42E-08 | 2.96E-04 | 0.58 | 0.56 | 0.02 | 1.04 |
| 6 | L3mbtl3 | 0.57 | 1.69E-08 | 3.51E-04 | 0.44 | 0.39 | 0.04 | 1.11 |
| 6 | Chpt1 | 0.56 | 1.87E-08 | 3.88E-04 | 0.43 | 0.39 | 0.04 | 1.10 |
| 6 | Tnrc18 | 0.39 | 1.94E-08 | 4.03E-04 | 0.59 | 0.56 | 0.03 | 1.05 |
| 6 | Glipr2 | 0.46 | 1.96E-08 | 4.08E-04 | 0.45 | 0.39 | 0.06 | 1.15 |
| 6 | Copz2 | 0.71 | 2.07E-08 | 4.31E-04 | 0.31 | 0.26 | 0.05 | 1.20 |
| 6 | Pxdn | 0.46 | 2.15E-08 | 4.47E-04 | 0.55 | 0.52 | 0.03 | 1.06 |
| 6 | Ttc28 | 0.34 | 2.21E-08 | 4.60E-04 | 0.63 | 0.60 | 0.03 | 1.05 |
| 6 | Ntn3 | 0.82 | 2.33E-08 | 4.85E-04 | 0.21 | 0.16 | 0.06 | 1.36 |
| 6 | Lap3 | 0.29 | 2.38E-08 | 4.95E-04 | 0.68 | 0.67 | 0.01 | 1.01 |
| 6 | Pygo1 | 0.42 | 2.40E-08 | 4.98E-04 | 0.52 | 0.47 | 0.05 | 1.11 |
| 6 | Asap1 | 0.28 | 2.46E-08 | 5.11E-04 | 0.75 | 0.73 | 0.01 | 1.02 |
| 6 | Tob1 | 0.63 | 2.47E-08 | 5.14E-04 | 0.33 | 0.28 | 0.06 | 1.20 |
| 6 | Cdk7 | 0.39 | 2.86E-08 | 5.96E-04 | 0.51 | 0.49 | 0.03 | 1.06 |
| 6 | Cntln | 0.48 | 3.14E-08 | 6.54E-04 | 0.52 | 0.49 | 0.03 | 1.06 |
| 6 | Fibp | 0.33 | 3.48E-08 | 7.24E-04 | 0.63 | 0.61 | 0.02 | 1.03 |
| 6 | Rsrc1 | 0.26 | 3.49E-08 | 7.26E-04 | 0.74 | 0.75 | -0.01 | 0.99 |
| 6 | Plod1 | 0.70 | 3.78E-08 | 7.85E-04 | 0.26 | 0.20 | 0.06 | 1.29 |
| 6 | Mydgf | 0.25 | 4.33E-08 | 9.01E-04 | 0.74 | 0.74 | 0.01 | 1.01 |
| 6 | Zfp771 | 0.27 | 4.52E-08 | 9.39E-04 | 0.68 | 0.69 | 0.00 | 0.99 |
| 6 | Dkk2 | 1.29 | 4.54E-08 | 9.45E-04 | 0.31 | 0.27 | 0.04 | 1.16 |
| 6 | Pdgfrb | 0.36 | 4.59E-08 | 9.54E-04 | 0.52 | 0.45 | 0.07 | 1.15 |
| 6 | L3hypdh | 0.76 | 5.30E-08 | 1.10E-03 | 0.24 | 0.19 | 0.06 | 1.30 |
| 6 | Adam11 | 1.06 | 5.31E-08 | 1.10E-03 | 0.13 | 0.08 | 0.05 | 1.55 |

**Supplementary Table 1. Marker genes for the original clusters of integrated clustering of E11.5 control and NotchGOF λ-junction cells.**

| | | | | | | | | |
|---|---|---|---|---|---|---|---|---|
| 6 | Sh3kbp1 | 0.71 | 6.00E-08 | 1.25E-03 | 0.28 | 0.22 | 0.05 | 1.24 |
| 6 | Uqcc3 | 0.32 | 6.29E-08 | 1.31E-03 | 0.67 | 0.67 | -0.01 | 0.99 |
| 6 | Rnf130 | 0.33 | 6.74E-08 | 1.40E-03 | 0.58 | 0.56 | 0.02 | 1.03 |
| 6 | Dipk1b | 0.42 | 7.29E-08 | 1.52E-03 | 0.51 | 0.48 | 0.04 | 1.07 |
| 6 | Pnrc2 | 0.33 | 7.70E-08 | 1.60E-03 | 0.62 | 0.61 | 0.00 | 1.01 |
| 6 | Dab2ip | 0.29 | 7.70E-08 | 1.60E-03 | 0.62 | 0.60 | 0.02 | 1.03 |
| 6 | Akap12 | 0.52 | 8.97E-08 | 1.87E-03 | 0.25 | 0.19 | 0.06 | 1.31 |
| 6 | Smim10l1 | 0.31 | 9.49E-08 | 1.97E-03 | 0.63 | 0.61 | 0.02 | 1.04 |
| 6 | Ccdc28b | 0.48 | 9.85E-08 | 2.05E-03 | 0.42 | 0.37 | 0.04 | 1.12 |
| 6 | Ift20 | 0.27 | 1.06E-07 | 2.20E-03 | 0.75 | 0.77 | -0.02 | 0.98 |
| 6 | Lox | 1.22 | 1.10E-07 | 2.30E-03 | 0.11 | 0.07 | 0.04 | 1.60 |
| 6 | Smarca2 | 0.32 | 1.16E-07 | 2.41E-03 | 0.64 | 0.62 | 0.02 | 1.03 |
| 6 | Nit2 | 0.42 | 1.18E-07 | 2.45E-03 | 0.52 | 0.51 | 0.02 | 1.03 |
| 6 | Nckap1 | 0.32 | 1.18E-07 | 2.46E-03 | 0.58 | 0.57 | 0.02 | 1.03 |
| 6 | Ehmt2 | 0.34 | 1.26E-07 | 2.62E-03 | 0.58 | 0.57 | 0.01 | 1.02 |
| 6 | Mthfd2l | 0.30 | 1.28E-07 | 2.66E-03 | 0.65 | 0.64 | 0.01 | 1.02 |
| 6 | Rarg | 0.32 | 1.44E-07 | 2.99E-03 | 0.56 | 0.52 | 0.03 | 1.07 |
| 6 | Sucla2 | 0.27 | 1.46E-07 | 3.04E-03 | 0.67 | 0.66 | 0.00 | 1.01 |
| 6 | St6gal2 | 1.01 | 1.46E-07 | 3.04E-03 | 0.15 | 0.10 | 0.05 | 1.47 |
| 6 | Tctex1d2 | 0.52 | 1.48E-07 | 3.08E-03 | 0.46 | 0.44 | 0.03 | 1.06 |
| 6 | Wnt5b | 0.65 | 1.50E-07 | 3.11E-03 | 0.31 | 0.26 | 0.05 | 1.20 |
| 6 | Mcfd2 | 0.42 | 1.52E-07 | 3.17E-03 | 0.49 | 0.46 | 0.03 | 1.07 |
| 6 | Zfp397 | 0.29 | 1.65E-07 | 3.42E-03 | 0.69 | 0.68 | 0.00 | 1.01 |
| 6 | Egflam | 1.05 | 1.67E-07 | 3.48E-03 | 0.12 | 0.07 | 0.04 | 1.57 |
| 6 | Rnf144a | 0.48 | 1.70E-07 | 3.54E-03 | 0.47 | 0.44 | 0.03 | 1.07 |
| 6 | Gatd1 | 0.39 | 1.75E-07 | 3.63E-03 | 0.54 | 0.53 | 0.01 | 1.02 |
| 6 | Hbp1 | 0.32 | 1.77E-07 | 3.68E-03 | 0.58 | 0.57 | 0.02 | 1.03 |
| 6 | Ptprs | 0.29 | 1.84E-07 | 3.84E-03 | 0.71 | 0.69 | 0.02 | 1.03 |
| 6 | Fzd1 | 0.70 | 1.88E-07 | 3.92E-03 | 0.38 | 0.34 | 0.04 | 1.12 |
| 6 | Ldb1 | 0.28 | 1.90E-07 | 3.95E-03 | 0.64 | 0.64 | 0.00 | 1.00 |
| 6 | Mrrf | 0.50 | 1.94E-07 | 4.03E-03 | 0.40 | 0.36 | 0.04 | 1.11 |
| 6 | Zmym4 | 0.27 | 2.12E-07 | 4.40E-03 | 0.70 | 0.70 | 0.00 | 1.00 |
| 6 | Prpsap1 | 0.29 | 2.20E-07 | 4.58E-03 | 0.68 | 0.69 | -0.01 | 0.99 |
| 6 | Tmed4 | 0.35 | 2.60E-07 | 5.41E-03 | 0.58 | 0.56 | 0.02 | 1.04 |
| 6 | Acads | 0.30 | 2.62E-07 | 5.45E-03 | 0.62 | 0.60 | 0.01 | 1.02 |
| 6 | Glg1 | 0.31 | 2.76E-07 | 5.75E-03 | 0.72 | 0.71 | 0.01 | 1.01 |
| 6 | Eci2 | 0.27 | 2.88E-07 | 6.00E-03 | 0.69 | 0.69 | 0.01 | 1.01 |
| 6 | A830082K12Rik | 1.09 | 2.94E-07 | 6.11E-03 | 0.11 | 0.07 | 0.04 | 1.56 |
| 6 | Sdc3 | 0.75 | 3.06E-07 | 6.37E-03 | 0.25 | 0.20 | 0.05 | 1.24 |
| 6 | P3h4 | 0.55 | 3.12E-07 | 6.49E-03 | 0.39 | 0.35 | 0.04 | 1.12 |
| 6 | Zbed3 | 0.32 | 3.30E-07 | 6.87E-03 | 0.58 | 0.56 | 0.01 | 1.02 |
| 6 | Fut8 | 0.66 | 3.45E-07 | 7.17E-03 | 0.34 | 0.29 | 0.04 | 1.14 |
| 6 | Vcam1 | 0.46 | 3.50E-07 | 7.27E-03 | 0.49 | 0.44 | 0.05 | 1.12 |
| 6 | Alkbh1 | 0.27 | 3.67E-07 | 7.64E-03 | 0.68 | 0.67 | 0.01 | 1.01 |
| 6 | Shd | 0.80 | 3.82E-07 | 7.95E-03 | 0.22 | 0.17 | 0.05 | 1.29 |
| 6 | Ubash3b | 0.59 | 3.94E-07 | 8.19E-03 | 0.29 | 0.24 | 0.05 | 1.22 |
| 6 | Mindy4 | 0.86 | 4.37E-07 | 9.09E-03 | 0.18 | 0.13 | 0.05 | 1.38 |
| 6 | Mapre2 | 0.44 | 4.49E-07 | 9.34E-03 | 0.44 | 0.40 | 0.04 | 1.10 |
| 6 | Zfp281 | 0.46 | 4.52E-07 | 9.41E-03 | 0.49 | 0.46 | 0.03 | 1.05 |
| 6 | Adamts10 | 0.56 | 4.74E-07 | 9.86E-03 | 0.37 | 0.32 | 0.05 | 1.14 |
| 6 | Col5a1 | 0.64 | 4.80E-07 | 9.99E-03 | 0.31 | 0.27 | 0.05 | 1.17 |
| 6 | Eci1 | 0.39 | 4.92E-07 | 1.02E-02 | 0.53 | 0.52 | 0.01 | 1.02 |
| 6 | Bmpr1b | 0.92 | 4.94E-07 | 1.03E-02 | 0.13 | 0.09 | 0.04 | 1.50 |
| 6 | Klf12 | 0.74 | 4.98E-07 | 1.04E-02 | 0.21 | 0.16 | 0.05 | 1.32 |
| 6 | Adgra3 | 0.33 | 5.80E-07 | 1.21E-02 | 0.57 | 0.56 | 0.01 | 1.02 |
| 6 | Sec24d | 1.01 | 5.81E-07 | 1.21E-02 | 0.12 | 0.08 | 0.04 | 1.52 |

**Supplementary Table 1. Marker genes for the original clusters of integrated clustering of E11.5 control and NotchGOF λ-junction cells.**

| | | | | | | | | |
|---|---|---|---|---|---|---|---|---|
| 6 | Lxn | 0.43 | 6.15E-07 | 1.28E-02 | 0.41 | 0.36 | 0.04 | 1.12 |
| 6 | Gpsm1 | 0.62 | 6.51E-07 | 1.35E-02 | 0.33 | 0.29 | 0.04 | 1.15 |
| 6 | Tsc22d4 | 0.29 | 7.49E-07 | 1.56E-02 | 0.62 | 0.61 | 0.01 | 1.01 |
| 6 | Yipf5 | 0.32 | 8.16E-07 | 1.70E-02 | 0.62 | 0.62 | 0.00 | 1.00 |
| 6 | Pja2 | 0.37 | 8.52E-07 | 1.77E-02 | 0.56 | 0.55 | 0.01 | 1.02 |
| 6 | Ror1 | 1.00 | 8.61E-07 | 1.79E-02 | 0.17 | 0.12 | 0.05 | 1.37 |
| 6 | St6galnac4 | 0.76 | 8.79E-07 | 1.83E-02 | 0.25 | 0.20 | 0.05 | 1.25 |
| 6 | Adcy4 | 1.05 | 8.82E-07 | 1.83E-02 | 0.10 | 0.06 | 0.04 | 1.59 |
| 6 | Prdm5 | 0.66 | 9.15E-07 | 1.90E-02 | 0.26 | 0.21 | 0.05 | 1.23 |
| 6 | Fgfr1 | 0.27 | 9.41E-07 | 1.96E-02 | 0.69 | 0.65 | 0.04 | 1.06 |
| 6 | 4632427E13Rik | 0.35 | 1.01E-06 | 2.09E-02 | 0.65 | 0.65 | 0.00 | 1.00 |
| 6 | Tsc22d3 | 0.35 | 1.02E-06 | 2.11E-02 | 0.55 | 0.53 | 0.02 | 1.04 |
| 6 | Hvcn1 | 1.06 | 1.02E-06 | 2.12E-02 | 0.12 | 0.08 | 0.04 | 1.50 |
| 6 | Gpt2 | 0.67 | 1.08E-06 | 2.24E-02 | 0.26 | 0.21 | 0.05 | 1.24 |
| 6 | Lama4 | 0.78 | 1.13E-06 | 2.35E-02 | 0.18 | 0.13 | 0.05 | 1.35 |
| 6 | Sox5 | 0.52 | 1.14E-06 | 2.37E-02 | 0.43 | 0.40 | 0.04 | 1.09 |
| 6 | Snap47 | 0.25 | 1.14E-06 | 2.38E-02 | 0.65 | 0.65 | -0.01 | 0.99 |
| 6 | Armcx2 | 0.38 | 1.15E-06 | 2.38E-02 | 0.55 | 0.53 | 0.02 | 1.03 |
| 6 | Selenos | 0.30 | 1.18E-06 | 2.46E-02 | 0.62 | 0.61 | 0.01 | 1.02 |
| 6 | Tfg | 0.26 | 1.23E-06 | 2.57E-02 | 0.72 | 0.74 | -0.02 | 0.97 |
| 6 | Zyx | 0.31 | 1.24E-06 | 2.58E-02 | 0.53 | 0.51 | 0.02 | 1.04 |
| 6 | Coprs | 0.41 | 1.26E-06 | 2.61E-02 | 0.51 | 0.49 | 0.02 | 1.03 |
| 6 | N4bp2 | 0.27 | 1.29E-06 | 2.69E-02 | 0.68 | 0.67 | 0.01 | 1.01 |
| 6 | Plppr3 | 0.48 | 1.38E-06 | 2.87E-02 | 0.42 | 0.38 | 0.04 | 1.10 |
| 6 | Myof | 0.92 | 1.43E-06 | 2.98E-02 | 0.13 | 0.09 | 0.04 | 1.47 |
| 6 | 2900076A07Rik | 0.55 | 1.58E-06 | 3.28E-02 | 0.35 | 0.31 | 0.04 | 1.14 |
| 6 | Lcorl | 0.37 | 1.66E-06 | 3.46E-02 | 0.52 | 0.48 | 0.03 | 1.07 |
| 6 | Trp53bp1 | 0.34 | 1.68E-06 | 3.50E-02 | 0.58 | 0.56 | 0.01 | 1.02 |
| 6 | Trio | 0.38 | 1.79E-06 | 3.72E-02 | 0.53 | 0.52 | 0.02 | 1.03 |
| 6 | Txndc15 | 0.33 | 1.96E-06 | 4.08E-02 | 0.60 | 0.60 | 0.01 | 1.01 |
| 6 | Ikbip | 0.62 | 1.96E-06 | 4.08E-02 | 0.34 | 0.30 | 0.04 | 1.13 |
| 6 | 2610002M06Rik | 0.60 | 2.39E-06 | 4.98E-02 | 0.33 | 0.29 | 0.04 | 1.14 |
| 7 | Tnfaip2 | 4.56 | 0 | 0 | 0.93 | 0.11 | 0.82 | 8.75 |
| 7 | Nrarp | 3.26 | 0 | 0 | 0.92 | 0.15 | 0.77 | 6.30 |
| 7 | Hes5 | 4.52 | 0 | 0 | 0.85 | 0.08 | 0.77 | 10.25 |
| 7 | Sox2 | 2.65 | 0 | 0 | 0.93 | 0.16 | 0.77 | 5.78 |
| 7 | Krt8 | 2.76 | 0 | 0 | 0.99 | 0.23 | 0.76 | 4.34 |
| 7 | Cldn6 | 2.55 | 0 | 0 | 0.95 | 0.20 | 0.75 | 4.78 |
| 7 | Dmrta2 | 3.54 | 0 | 0 | 0.86 | 0.11 | 0.75 | 7.90 |
| 7 | Cldn7 | 2.41 | 0 | 0 | 0.95 | 0.21 | 0.74 | 4.55 |
| 7 | Cldn3 | 2.87 | 0 | 0 | 0.87 | 0.13 | 0.74 | 6.60 |
| 7 | Pax6 | 3.53 | 0 | 0 | 0.84 | 0.10 | 0.74 | 8.61 |
| 7 | Pitx1 | 3.05 | 0 | 0 | 0.88 | 0.15 | 0.73 | 5.85 |
| 7 | Krt18 | 2.82 | 0 | 0 | 0.99 | 0.27 | 0.72 | 3.67 |
| 7 | Six3 | 2.43 | 0 | 0 | 0.93 | 0.21 | 0.72 | 4.44 |
| 7 | Cpm | 3.56 | 0 | 0 | 0.86 | 0.15 | 0.72 | 5.94 |
| 7 | Pdgfa | 1.80 | 0 | 0 | 0.96 | 0.25 | 0.71 | 3.79 |
| 7 | Hey1 | 2.65 | 0 | 0 | 0.97 | 0.27 | 0.70 | 3.58 |
| 7 | Spint2 | 2.16 | 0 | 0 | 0.98 | 0.28 | 0.70 | 3.48 |
| 7 | Ano1 | 3.65 | 0 | 0 | 0.80 | 0.10 | 0.70 | 8.12 |
| 7 | Slc9a3r1 | 2.77 | 0 | 0 | 0.91 | 0.21 | 0.70 | 4.29 |
| 7 | Dsp | 1.78 | 0 | 0 | 0.90 | 0.21 | 0.70 | 4.34 |
| 7 | Gm266 | 3.86 | 0 | 0 | 0.97 | 0.28 | 0.69 | 3.46 |
| 7 | Ezr | 2.44 | 0 | 0 | 0.98 | 0.29 | 0.69 | 3.35 |
| 7 | Itga6 | 2.89 | 0 | 0 | 0.96 | 0.28 | 0.68 | 3.40 |
| 7 | Epcam | 2.50 | 0 | 0 | 1.00 | 0.32 | 0.68 | 3.13 |

**Supplementary Table 1. Marker genes for the original clusters of integrated clustering of E11.5 control and NotchGOF λ-junction cells.**

| | | | | | | | | |
|---|---|---|---|---|---|---|---|---|
| 7 | Rab25 | 2.59 | 0 | 0 | 0.80 | 0.13 | 0.66 | 5.98 |
| 7 | Foxg1 | 2.88 | 0 | 0 | 0.77 | 0.12 | 0.65 | 6.56 |
| 7 | Oxtr | 4.48 | 0 | 0 | 0.68 | 0.04 | 0.64 | 16.93 |
| 7 | Mapk13 | 2.56 | 0 | 0 | 0.80 | 0.17 | 0.63 | 4.61 |
| 7 | Sp8 | 2.53 | 0 | 0 | 0.73 | 0.11 | 0.62 | 6.45 |
| 7 | Ephb1 | 3.50 | 0 | 0 | 0.69 | 0.07 | 0.61 | 9.38 |
| 7 | Jag1 | 2.47 | 0 | 0 | 0.76 | 0.15 | 0.61 | 5.04 |
| 7 | Socs2 | 2.55 | 0 | 0 | 0.93 | 0.32 | 0.61 | 2.90 |
| 7 | Igfbp5 | 1.10 | 0 | 0 | 0.99 | 0.38 | 0.61 | 2.58 |
| 7 | Cdh1 | 2.42 | 0 | 0 | 0.75 | 0.14 | 0.61 | 5.20 |
| 7 | Tpd52 | 1.77 | 0 | 0 | 0.83 | 0.24 | 0.60 | 3.51 |
| 7 | Six3os1 | 2.10 | 0 | 0 | 0.80 | 0.21 | 0.59 | 3.86 |
| 7 | Mpzl2 | 2.07 | 0 | 0 | 0.78 | 0.18 | 0.59 | 4.23 |
| 7 | Ikzf2 | 1.71 | 0 | 0 | 0.85 | 0.26 | 0.59 | 3.28 |
| 7 | Hist3h2ba | 1.68 | 0 | 0 | 0.84 | 0.25 | 0.59 | 3.35 |
| 7 | Cmtm8 | 2.37 | 0 | 0 | 0.74 | 0.15 | 0.58 | 4.81 |
| 7 | Slc22a23 | 3.66 | 0 | 0 | 0.65 | 0.07 | 0.58 | 9.29 |
| 7 | Otx2 | 3.03 | 0 | 0 | 0.65 | 0.08 | 0.58 | 8.47 |
| 7 | Col18a1 | 1.62 | 0 | 0 | 0.91 | 0.34 | 0.58 | 2.70 |
| 7 | Igfbp3 | 2.17 | 0 | 0 | 0.67 | 0.10 | 0.57 | 6.66 |
| 7 | Wwc1 | 2.31 | 0 | 0 | 0.72 | 0.16 | 0.57 | 4.61 |
| 7 | Crip2 | 1.92 | 0 | 0 | 0.71 | 0.15 | 0.56 | 4.77 |
| 7 | Rprm | 2.80 | 0 | 0 | 0.70 | 0.14 | 0.56 | 5.08 |
| 7 | Dlk1 | 1.70 | 0 | 0 | 0.94 | 0.40 | 0.55 | 2.37 |
| 7 | Cdh2 | 2.35 | 0 | 0 | 0.95 | 0.40 | 0.55 | 2.36 |
| 7 | Wfdc2 | 2.65 | 0 | 0 | 0.98 | 0.44 | 0.54 | 2.24 |
| 7 | Fjx1 | 1.86 | 0 | 0 | 0.91 | 0.37 | 0.54 | 2.47 |
| 7 | Sorbs2 | 2.36 | 0 | 0 | 0.74 | 0.20 | 0.54 | 3.74 |
| 7 | Cadm1 | 2.27 | 0 | 0 | 0.90 | 0.36 | 0.54 | 2.48 |
| 7 | Zdhhc2 | 3.46 | 0 | 0 | 0.64 | 0.11 | 0.54 | 6.01 |
| 7 | Ccdc88c | 2.37 | 0 | 0 | 0.67 | 0.14 | 0.54 | 4.96 |
| 7 | Celsr1 | 3.26 | 0 | 0 | 0.61 | 0.08 | 0.53 | 7.64 |
| 7 | Cldn9 | 3.18 | 0 | 0 | 0.60 | 0.07 | 0.53 | 9.05 |
| 7 | Ston2 | 2.24 | 0 | 0 | 0.69 | 0.16 | 0.53 | 4.33 |
| 7 | Espn | 2.76 | 0 | 0 | 0.61 | 0.09 | 0.52 | 7.02 |
| 7 | Fras1 | 2.14 | 0 | 0 | 0.77 | 0.25 | 0.52 | 3.14 |
| 7 | Plch1 | 3.17 | 0 | 0 | 0.60 | 0.08 | 0.52 | 7.69 |
| 7 | Lsr | 1.95 | 0 | 0 | 0.71 | 0.18 | 0.52 | 3.84 |
| 7 | Nectin4 | 2.06 | 0 | 0 | 0.62 | 0.11 | 0.52 | 5.72 |
| 7 | Aldh1a3 | 2.35 | 0 | 0 | 0.73 | 0.21 | 0.52 | 3.44 |
| 7 | Esrp1 | 1.90 | 0 | 0 | 0.66 | 0.15 | 0.51 | 4.48 |
| 7 | Galm | 2.36 | 0 | 0 | 0.70 | 0.19 | 0.51 | 3.66 |
| 7 | Tex15 | 3.27 | 0 | 0 | 0.59 | 0.09 | 0.51 | 6.74 |
| 7 | Kcnk1 | 2.00 | 0 | 0 | 0.67 | 0.17 | 0.50 | 4.05 |
| 7 | Erbb3 | 1.97 | 0 | 0 | 0.66 | 0.16 | 0.50 | 4.23 |
| 7 | Tspan13 | 1.86 | 0 | 0 | 0.89 | 0.39 | 0.50 | 2.27 |
| 7 | Mapk12 | 2.89 | 0 | 0 | 0.59 | 0.09 | 0.49 | 6.30 |
| 7 | Rhov | 3.74 | 0 | 0 | 0.52 | 0.04 | 0.49 | 14.50 |
| 7 | Tmem54 | 2.41 | 0 | 0 | 0.59 | 0.10 | 0.49 | 5.91 |
| 7 | Crb3 | 2.54 | 0 | 0 | 0.57 | 0.09 | 0.48 | 6.37 |
| 7 | Faah | 3.88 | 0 | 0 | 0.52 | 0.04 | 0.48 | 13.97 |
| 7 | Ap1m2 | 1.98 | 0 | 0 | 0.60 | 0.12 | 0.48 | 4.85 |
| 7 | AW011738 | 3.06 | 0 | 0 | 0.54 | 0.07 | 0.47 | 7.80 |
| 7 | Heyl | 3.79 | 0 | 0 | 0.51 | 0.04 | 0.47 | 13.65 |
| 7 | Ly6h | 1.86 | 0 | 0 | 0.60 | 0.13 | 0.47 | 4.57 |
| 7 | Prss8 | 2.74 | 0 | 0 | 0.54 | 0.08 | 0.47 | 6.97 |

**Supplementary Table 1. Marker genes for the original clusters of integrated clustering of E11.5 control and NotchGOF λ-junction cells.**

| | | | | | | | | |
|---|---|---|---|---|---|---|---|---|
| 7 | Cgn | 2.72 | 0 | 0 | 0.55 | 0.09 | 0.46 | 6.28 |
| 7 | Pls1 | 2.99 | 0 | 0 | 0.53 | 0.08 | 0.45 | 6.97 |
| 7 | St8sia6 | 4.58 | 0 | 0 | 0.48 | 0.03 | 0.45 | 16.66 |
| 7 | C130071C03Rik | 4.01 | 0 | 0 | 0.48 | 0.03 | 0.45 | 14.61 |
| 7 | Prkg1 | 2.49 | 0 | 0 | 0.59 | 0.14 | 0.45 | 4.23 |
| 7 | Ctnnd2 | 2.30 | 0 | 0 | 0.55 | 0.10 | 0.44 | 5.35 |
| 7 | 3110039M20Rik | 2.93 | 0 | 0 | 0.50 | 0.06 | 0.44 | 8.59 |
| 7 | Ppp2r2b | 2.56 | 0 | 0 | 0.52 | 0.08 | 0.44 | 6.63 |
| 7 | St14 | 2.19 | 0 | 0 | 0.53 | 0.10 | 0.43 | 5.29 |
| 7 | Bex1 | 1.88 | 0 | 0 | 0.99 | 0.57 | 0.42 | 1.73 |
| 7 | Cystm1 | 2.37 | 0 | 0 | 0.49 | 0.07 | 0.42 | 6.82 |
| 7 | Map7 | 2.38 | 0 | 0 | 0.51 | 0.09 | 0.41 | 5.55 |
| 7 | Rassf10 | 3.51 | 0 | 0 | 0.45 | 0.04 | 0.41 | 10.47 |
| 7 | Cbln2 | 4.32 | 0 | 0 | 0.43 | 0.02 | 0.41 | 18.61 |
| 7 | Spint1 | 2.36 | 0 | 0 | 0.48 | 0.08 | 0.40 | 5.81 |
| 7 | Jph4 | 4.14 | 0 | 0 | 0.43 | 0.04 | 0.39 | 11.34 |
| 7 | Nr2e1 | 3.86 | 0 | 0 | 0.42 | 0.03 | 0.39 | 16.00 |
| 7 | Dmrt3 | 3.44 | 0 | 0 | 0.42 | 0.04 | 0.38 | 11.03 |
| 7 | Sox3 | 4.06 | 0 | 0 | 0.40 | 0.02 | 0.38 | 16.63 |
| 7 | Exoc3l4 | 2.96 | 0 | 0 | 0.43 | 0.05 | 0.37 | 7.93 |
| 7 | Khdrbs3 | 1.85 | 0 | 0 | 0.97 | 0.60 | 0.37 | 1.62 |
| 7 | Six6 | 2.92 | 0 | 0 | 0.42 | 0.06 | 0.37 | 7.65 |
| 7 | Scg5 | 3.09 | 0 | 0 | 0.39 | 0.04 | 0.35 | 10.29 |
| 7 | Bex4 | 1.74 | 0 | 0 | 0.99 | 0.64 | 0.35 | 1.55 |
| 7 | Tenm2 | 3.02 | 0 | 0 | 0.41 | 0.06 | 0.35 | 7.05 |
| 7 | Cdkl1 | 4.75 | 0 | 0 | 0.37 | 0.02 | 0.35 | 18.40 |
| 7 | Bspry | 2.82 | 0 | 0 | 0.39 | 0.05 | 0.34 | 7.69 |
| 7 | Tubb2b | 2.25 | 0 | 0 | 0.99 | 0.65 | 0.34 | 1.52 |
| 7 | Nrcam | 3.54 | 0 | 0 | 0.37 | 0.04 | 0.34 | 10.08 |
| 7 | Dmbx1 | 3.61 | 0 | 0 | 0.35 | 0.03 | 0.32 | 11.80 |
| 7 | Uncx | 3.83 | 0 | 0 | 0.33 | 0.01 | 0.32 | 33.10 |
| 7 | Zmat4 | 3.16 | 0 | 0 | 0.36 | 0.04 | 0.32 | 8.57 |
| 7 | Prr15 | 4.32 | 0 | 0 | 0.34 | 0.02 | 0.32 | 16.80 |
| 7 | 4930426D05Rik | 4.17 | 0 | 0 | 0.33 | 0.02 | 0.31 | 17.26 |
| 7 | D230030E09Rik | 2.58 | 0 | 0 | 0.35 | 0.04 | 0.30 | 7.86 |
| 7 | Rims2 | 2.87 | 0 | 0 | 0.35 | 0.05 | 0.30 | 7.57 |
| 7 | Fgf12 | 3.20 | 0 | 0 | 0.33 | 0.04 | 0.30 | 9.28 |
| 7 | Ces1d | 3.47 | 0 | 0 | 0.33 | 0.03 | 0.30 | 10.55 |
| 7 | Scrn1 | 3.31 | 0 | 0 | 0.33 | 0.03 | 0.29 | 10.48 |
| 7 | Ctxn3 | 4.01 | 0 | 0 | 0.31 | 0.02 | 0.29 | 17.22 |
| 7 | Gjb2 | 2.63 | 0 | 0 | 0.31 | 0.03 | 0.28 | 9.12 |
| 7 | Fam107a | 3.05 | 0 | 0 | 0.31 | 0.03 | 0.27 | 9.27 |
| 7 | Rasef | 3.07 | 0 | 0 | 0.30 | 0.03 | 0.27 | 9.87 |
| 7 | Sox21 | 3.74 | 0 | 0 | 0.29 | 0.02 | 0.27 | 13.67 |
| 7 | Kazald1 | 3.97 | 0 | 0 | 0.28 | 0.02 | 0.27 | 16.65 |
| 7 | Gsap | 3.55 | 0 | 0 | 0.28 | 0.03 | 0.26 | 10.92 |
| 7 | Cyp4v3 | 3.68 | 0 | 0 | 0.28 | 0.02 | 0.26 | 11.75 |
| 7 | Myo5c | 3.31 | 0 | 0 | 0.29 | 0.03 | 0.26 | 10.18 |
| 7 | Crb2 | 4.00 | 0 | 0 | 0.25 | 0.02 | 0.24 | 15.88 |
| 7 | Fgf15 | 3.98 | 0 | 0 | 0.25 | 0.02 | 0.23 | 15.56 |
| 7 | Fam189a2 | 4.03 | 0 | 0 | 0.24 | 0.02 | 0.22 | 15.93 |
| 7 | Ttyh1 | 3.92 | 0 | 0 | 0.23 | 0.02 | 0.22 | 15.40 |
| 7 | Ap3b2 | 4.02 | 0 | 0 | 0.23 | 0.01 | 0.22 | 20.64 |
| 7 | Muc1 | 4.01 | 0 | 0 | 0.21 | 0.01 | 0.20 | 19.27 |
| 7 | Dmrta2os | 3.95 | 0 | 0 | 0.21 | 0.01 | 0.20 | 16.23 |
| 7 | Rims4 | 4.81 | 0 | 0 | 0.18 | 0.01 | 0.17 | 25.71 |

**Supplementary Table 1. Marker genes for the original clusters of integrated clustering of E11.5 control and NotchGOF λ-junction cells.**

| | | | | | | | | |
|---|---|---|---|---|---|---|---|---|
| 7 | Sox1 | 4.91 | 0 | 0 | 0.16 | 0.01 | 0.15 | 31.20 |
| 7 | Nipal1 | 5.40 | 0 | 0 | 0.14 | 0.00 | 0.14 | 34.75 |
| 7 | Psip1 | 1.04 | 0 | 0 | 1.00 | 0.98 | 0.02 | 1.02 |
| 7 | Isl1 | 1.92 | 1.04E-307 | 2.16E-303 | 0.72 | 0.20 | 0.52 | 3.64 |
| 7 | Syngr1 | 2.70 | 3.09E-306 | 6.42E-302 | 0.38 | 0.05 | 0.32 | 6.96 |
| 7 | Dab1 | 2.11 | 8.37E-304 | 1.74E-299 | 0.60 | 0.14 | 0.46 | 4.42 |
| 7 | Acer2 | 1.63 | 8.88E-304 | 1.85E-299 | 0.91 | 0.37 | 0.54 | 2.44 |
| 7 | Nuak2 | 3.17 | 1.16E-302 | 2.41E-298 | 0.27 | 0.03 | 0.24 | 9.96 |
| 7 | Ppp1r1a | 2.30 | 6.61E-302 | 1.38E-297 | 0.60 | 0.15 | 0.45 | 4.13 |
| 7 | Klhl14 | 4.35 | 1.38E-297 | 2.88E-293 | 0.15 | 0.01 | 0.15 | 21.86 |
| 7 | Ocln | 2.54 | 6.12E-296 | 1.27E-291 | 0.40 | 0.06 | 0.34 | 6.37 |
| 7 | Pcdh19 | 2.60 | 4.04E-295 | 8.40E-291 | 0.59 | 0.15 | 0.44 | 3.97 |
| 7 | Hook1 | 2.06 | 3.03E-294 | 6.30E-290 | 0.67 | 0.18 | 0.49 | 3.69 |
| 7 | 2900026A02Rik | 2.39 | 1.38E-289 | 2.86E-285 | 0.47 | 0.09 | 0.38 | 5.31 |
| 7 | Elavl2 | 2.02 | 1.04E-288 | 2.16E-284 | 0.64 | 0.17 | 0.47 | 3.80 |
| 7 | Marchf9 | 2.11 | 2.16E-288 | 4.49E-284 | 0.57 | 0.13 | 0.44 | 4.38 |
| 7 | Fgd2 | 2.23 | 4.25E-287 | 8.84E-283 | 0.46 | 0.09 | 0.38 | 5.42 |
| 7 | Notch1 | 1.45 | 1.99E-285 | 4.15E-281 | 0.75 | 0.23 | 0.53 | 3.34 |
| 7 | Cldn4 | 2.04 | 3.36E-284 | 6.98E-280 | 0.47 | 0.09 | 0.38 | 5.36 |
| 7 | Grb7 | 2.23 | 4.00E-283 | 8.32E-279 | 0.45 | 0.08 | 0.37 | 5.47 |
| 7 | Galnt14 | 3.58 | 1.99E-279 | 4.14E-275 | 0.21 | 0.02 | 0.19 | 12.81 |
| 7 | Kif21a | 1.69 | 3.13E-277 | 6.51E-273 | 0.81 | 0.29 | 0.52 | 2.79 |
| 7 | Trpm3 | 3.03 | 1.81E-273 | 3.77E-269 | 0.29 | 0.04 | 0.26 | 8.11 |
| 7 | Dpysl5 | 2.19 | 3.48E-273 | 7.24E-269 | 0.47 | 0.09 | 0.38 | 5.08 |
| 7 | Stard10 | 1.72 | 9.95E-272 | 2.07E-267 | 0.79 | 0.28 | 0.51 | 2.84 |
| 7 | Tspan12 | 2.14 | 5.79E-271 | 1.20E-266 | 0.66 | 0.19 | 0.46 | 3.41 |
| 7 | Apc | 1.77 | 1.13E-270 | 2.34E-266 | 0.93 | 0.60 | 0.33 | 1.55 |
| 7 | Gpm6a | 2.92 | 6.78E-270 | 1.41E-265 | 0.30 | 0.04 | 0.26 | 7.82 |
| 7 | Myh14 | 2.44 | 5.50E-266 | 1.14E-261 | 0.39 | 0.07 | 0.33 | 5.87 |
| 7 | Gm19938 | 3.30 | 6.78E-266 | 1.41E-261 | 0.20 | 0.02 | 0.18 | 12.44 |
| 7 | Tmem238 | 2.18 | 1.30E-265 | 2.70E-261 | 0.60 | 0.16 | 0.44 | 3.71 |
| 7 | Clu | 2.79 | 4.16E-265 | 8.66E-261 | 0.64 | 0.19 | 0.45 | 3.36 |
| 7 | Ptprf | 1.45 | 7.70E-264 | 1.60E-259 | 0.97 | 0.63 | 0.34 | 1.55 |
| 7 | Tom1l1 | 2.00 | 4.49E-263 | 9.34E-259 | 0.57 | 0.14 | 0.43 | 4.01 |
| 7 | Lypd2 | 3.06 | 5.32E-263 | 1.11E-258 | 0.26 | 0.03 | 0.23 | 9.14 |
| 7 | Nebl | 1.65 | 1.63E-261 | 3.39E-257 | 0.47 | 0.09 | 0.38 | 5.04 |
| 7 | Gpc1 | 1.80 | 1.96E-260 | 4.07E-256 | 0.84 | 0.38 | 0.46 | 2.22 |
| 7 | Krt19 | 1.96 | 3.12E-259 | 6.49E-255 | 0.37 | 0.06 | 0.31 | 6.11 |
| 7 | Trpc1 | 2.55 | 3.96E-258 | 8.23E-254 | 0.46 | 0.10 | 0.37 | 4.77 |
| 7 | Id4 | 2.23 | 2.31E-255 | 4.79E-251 | 0.75 | 0.29 | 0.46 | 2.61 |
| 7 | Hap1 | 2.75 | 1.08E-254 | 2.24E-250 | 0.30 | 0.04 | 0.26 | 7.45 |
| 7 | Slc12a2 | 1.79 | 5.20E-254 | 1.08E-249 | 0.87 | 0.42 | 0.45 | 2.09 |
| 7 | Dsg2 | 1.85 | 1.10E-253 | 2.28E-249 | 0.51 | 0.11 | 0.40 | 4.54 |
| 7 | Grhl1 | 3.44 | 1.43E-251 | 2.98E-247 | 0.19 | 0.02 | 0.17 | 12.60 |
| 7 | Grhl2 | 1.92 | 1.03E-250 | 2.15E-246 | 0.51 | 0.11 | 0.40 | 4.48 |
| 7 | Elf5 | 2.37 | 1.65E-250 | 3.44E-246 | 0.33 | 0.05 | 0.28 | 6.58 |
| 7 | Wipf3 | 2.61 | 9.32E-248 | 1.94E-243 | 0.32 | 0.05 | 0.27 | 6.59 |
| 7 | Tox2 | 2.96 | 1.76E-246 | 3.67E-242 | 0.25 | 0.03 | 0.22 | 8.55 |
| 7 | Stmn3 | 1.71 | 2.51E-242 | 5.21E-238 | 0.24 | 0.03 | 0.21 | 8.85 |
| 7 | Usp25 | 1.81 | 3.32E-241 | 6.91E-237 | 0.83 | 0.42 | 0.42 | 2.00 |
| 7 | Gm48435 | 3.29 | 1.23E-238 | 2.55E-234 | 0.20 | 0.02 | 0.18 | 11.00 |
| 7 | Gapdh | 0.55 | 5.48E-238 | 1.14E-233 | 1.00 | 1.00 | 0.00 | 1.00 |
| 7 | Arhgap8 | 2.78 | 9.69E-238 | 2.02E-233 | 0.26 | 0.03 | 0.23 | 7.91 |
| 7 | Slc27a2 | 4.32 | 1.81E-237 | 3.77E-233 | 0.14 | 0.01 | 0.14 | 16.00 |
| 7 | Arrdc1 | 2.27 | 5.58E-236 | 1.16E-231 | 0.45 | 0.10 | 0.35 | 4.60 |
| 7 | Prr36 | 2.33 | 2.86E-235 | 5.95E-231 | 0.39 | 0.07 | 0.32 | 5.31 |

**Supplementary Table 1. Marker genes for the original clusters of integrated clustering of E11.5 control and NotchGOF λ-junction cells.**

| | | | | | | | | |
|---|---|---|---|---|---|---|---|---|
| 7 | Hs6st2 | 1.83 | 4.96E-235 | 1.03E-230 | 0.73 | 0.27 | 0.46 | 2.74 |
| 7 | Bend7 | 2.33 | 2.03E-234 | 4.22E-230 | 0.38 | 0.07 | 0.31 | 5.40 |
| 7 | Gnao1 | 2.37 | 5.24E-233 | 1.09E-228 | 0.38 | 0.07 | 0.31 | 5.41 |
| 7 | Soga3 | 2.20 | 9.18E-233 | 1.91E-228 | 0.29 | 0.04 | 0.25 | 6.98 |
| 7 | Pawr | 2.14 | 1.19E-232 | 2.48E-228 | 0.49 | 0.11 | 0.37 | 4.25 |
| 7 | Cd9 | 1.41 | 2.87E-230 | 5.97E-226 | 0.76 | 0.28 | 0.49 | 2.74 |
| 7 | Fut9 | 3.56 | 4.17E-230 | 8.66E-226 | 0.21 | 0.02 | 0.18 | 9.76 |
| 7 | Pea15a | 1.91 | 4.90E-228 | 1.02E-223 | 0.65 | 0.21 | 0.44 | 3.03 |
| 7 | Gm16136 | 1.39 | 1.93E-227 | 4.01E-223 | 0.59 | 0.16 | 0.43 | 3.77 |
| 7 | Npdc1 | 2.19 | 2.68E-226 | 5.58E-222 | 0.43 | 0.09 | 0.34 | 4.74 |
| 7 | Rragd | 2.03 | 4.28E-226 | 8.90E-222 | 0.63 | 0.21 | 0.42 | 3.05 |
| 7 | Map7d2 | 2.37 | 7.41E-225 | 1.54E-220 | 0.37 | 0.07 | 0.30 | 5.27 |
| 7 | Stmn2 | 1.18 | 3.75E-223 | 7.81E-219 | 0.41 | 0.09 | 0.33 | 4.75 |
| 7 | Otx1 | 1.87 | 1.91E-222 | 3.98E-218 | 0.45 | 0.10 | 0.35 | 4.57 |
| 7 | Arhgef19 | 1.87 | 2.18E-222 | 4.53E-218 | 0.49 | 0.12 | 0.38 | 4.23 |
| 7 | Pkp2 | 2.54 | 3.83E-222 | 7.97E-218 | 0.29 | 0.04 | 0.25 | 6.64 |
| 7 | Tpi1 | 0.96 | 4.74E-221 | 9.86E-217 | 1.00 | 0.96 | 0.04 | 1.04 |
| 7 | Lrp2 | 2.47 | 2.45E-219 | 5.10E-215 | 0.29 | 0.04 | 0.25 | 6.61 |
| 7 | Nrg1 | 2.21 | 1.59E-218 | 3.30E-214 | 0.36 | 0.07 | 0.29 | 5.32 |
| 7 | Cyb5a | 0.89 | 3.72E-217 | 7.74E-213 | 0.99 | 0.92 | 0.07 | 1.08 |
| 7 | Arg2 | 2.48 | 1.84E-215 | 3.83E-211 | 0.29 | 0.04 | 0.24 | 6.63 |
| 7 | Gnai2 | 0.77 | 4.65E-215 | 9.67E-211 | 1.00 | 0.97 | 0.03 | 1.03 |
| 7 | Cdh4 | 3.00 | 5.69E-215 | 1.18E-210 | 0.27 | 0.04 | 0.23 | 6.68 |
| 7 | Kdf1 | 2.05 | 2.86E-213 | 5.96E-209 | 0.38 | 0.07 | 0.30 | 5.15 |
| 7 | Nup210 | 2.59 | 2.49E-212 | 5.18E-208 | 0.30 | 0.05 | 0.25 | 5.96 |
| 7 | Hsp90ab1 | 0.39 | 1.50E-211 | 3.12E-207 | 1.00 | 1.00 | 0.00 | 1.00 |
| 7 | Rps2 | 0.45 | 5.20E-211 | 1.08E-206 | 1.00 | 1.00 | 0.00 | 1.00 |
| 7 | Schip1 | 1.97 | 6.69E-210 | 1.39E-205 | 0.53 | 0.15 | 0.38 | 3.63 |
| 7 | Pla2g12a | 1.66 | 8.98E-210 | 1.87E-205 | 0.75 | 0.31 | 0.44 | 2.43 |
| 7 | Stox1 | 3.24 | 1.21E-209 | 2.52E-205 | 0.19 | 0.02 | 0.17 | 9.95 |
| 7 | Rbp1 | 1.28 | 4.79E-208 | 9.96E-204 | 0.99 | 0.91 | 0.08 | 1.09 |
| 7 | Dtna | 1.95 | 6.13E-208 | 1.27E-203 | 0.50 | 0.13 | 0.37 | 3.91 |
| 7 | Tst | 2.22 | 1.48E-207 | 3.07E-203 | 0.46 | 0.11 | 0.34 | 4.04 |
| 7 | Necab2 | 2.19 | 3.55E-207 | 7.38E-203 | 0.58 | 0.19 | 0.39 | 3.07 |
| 7 | Snn | 1.80 | 3.96E-207 | 8.24E-203 | 0.81 | 0.42 | 0.39 | 1.94 |
| 7 | Slc35f1 | 3.08 | 4.37E-206 | 9.09E-202 | 0.21 | 0.03 | 0.19 | 8.52 |
| 7 | Ermn | 2.51 | 5.70E-203 | 1.19E-198 | 0.25 | 0.04 | 0.21 | 7.11 |
| 7 | Svip | 2.39 | 9.48E-203 | 1.97E-198 | 0.38 | 0.08 | 0.30 | 4.75 |
| 7 | Wnt4 | 1.43 | 1.83E-201 | 3.81E-197 | 0.61 | 0.18 | 0.42 | 3.30 |
| 7 | 2610528J11Rik | 2.37 | 1.19E-199 | 2.48E-195 | 0.30 | 0.05 | 0.25 | 5.81 |
| 7 | Arhgap27os2 | 2.34 | 7.99E-199 | 1.66E-194 | 0.31 | 0.06 | 0.26 | 5.59 |
| 7 | Bnipl | 2.58 | 1.15E-198 | 2.39E-194 | 0.25 | 0.04 | 0.22 | 6.97 |
| 7 | Cdk6 | 1.67 | 1.97E-197 | 4.10E-193 | 0.74 | 0.32 | 0.42 | 2.30 |
| 7 | Eya1 | 1.38 | 8.44E-197 | 1.76E-192 | 0.88 | 0.48 | 0.40 | 1.83 |
| 7 | Igfbp2 | 0.75 | 7.93E-196 | 1.65E-191 | 0.99 | 0.75 | 0.25 | 1.33 |
| 7 | Chchd10 | 1.76 | 3.07E-195 | 6.38E-191 | 0.52 | 0.15 | 0.38 | 3.60 |
| 7 | Ildr1 | 2.81 | 3.35E-195 | 6.97E-191 | 0.23 | 0.03 | 0.20 | 7.50 |
| 7 | Myo5b | 2.64 | 1.29E-194 | 2.67E-190 | 0.25 | 0.04 | 0.21 | 7.00 |
| 7 | Rpl27a | 0.31 | 1.08E-193 | 2.25E-189 | 1.00 | 1.00 | 0.00 | 1.00 |
| 7 | Gsta4 | 1.09 | 3.08E-192 | 6.41E-188 | 0.96 | 0.75 | 0.22 | 1.29 |
| 7 | Lmo7 | 1.53 | 6.38E-191 | 1.33E-186 | 0.52 | 0.15 | 0.38 | 3.58 |
| 7 | Ckmt1 | 1.73 | 1.66E-190 | 3.45E-186 | 0.24 | 0.03 | 0.21 | 7.06 |
| 7 | Rab15 | 1.84 | 2.98E-190 | 6.20E-186 | 0.47 | 0.12 | 0.35 | 3.92 |
| 7 | Cadps2 | 2.62 | 2.57E-189 | 5.35E-185 | 0.22 | 0.03 | 0.19 | 7.48 |
| 7 | Hspa4l | 2.16 | 2.22E-188 | 4.61E-184 | 0.41 | 0.10 | 0.31 | 4.16 |
| 7 | Dsc2 | 2.24 | 6.32E-188 | 1.31E-183 | 0.28 | 0.05 | 0.23 | 5.88 |

**Supplementary Table 1. Marker genes for the original clusters of integrated clustering of E11.5 control and NotchGOF λ-junction cells.**

| | | | | | | | | |
|---|---|---|---|---|---|---|---|---|
| 7 | Smtnl2 | 2.22 | 1.19E-186 | 2.48E-182 | 0.33 | 0.06 | 0.26 | 5.13 |
| 7 | Cyb561 | 2.36 | 5.20E-186 | 1.08E-181 | 0.28 | 0.05 | 0.23 | 5.98 |
| 7 | Egfr | 2.35 | 4.30E-185 | 8.95E-181 | 0.39 | 0.09 | 0.30 | 4.33 |
| 7 | Cxcr4 | 2.31 | 9.34E-185 | 1.94E-180 | 0.26 | 0.04 | 0.22 | 6.29 |
| 7 | Cdcp1 | 2.38 | 1.50E-184 | 3.12E-180 | 0.28 | 0.05 | 0.23 | 5.78 |
| 7 | Muc4 | 3.69 | 5.64E-184 | 1.17E-179 | 0.12 | 0.01 | 0.11 | 15.13 |
| 7 | Cyb5r3 | 1.06 | 1.30E-183 | 2.70E-179 | 0.97 | 0.80 | 0.16 | 1.20 |
| 7 | Gad2 | 1.55 | 1.32E-183 | 2.75E-179 | 0.33 | 0.06 | 0.27 | 5.16 |
| 7 | Mgst1 | 1.63 | 2.49E-183 | 5.17E-179 | 0.78 | 0.36 | 0.42 | 2.18 |
| 7 | Misp | 3.38 | 2.49E-182 | 5.19E-178 | 0.15 | 0.01 | 0.13 | 11.15 |
| 7 | F11r | 1.49 | 1.29E-181 | 2.67E-177 | 0.56 | 0.17 | 0.39 | 3.38 |
| 7 | Mapk8ip2 | 2.60 | 4.49E-181 | 9.33E-177 | 0.22 | 0.03 | 0.19 | 7.13 |
| 7 | Camsap3 | 2.16 | 2.67E-180 | 5.56E-176 | 0.34 | 0.07 | 0.27 | 4.79 |
| 7 | Rps18 | 0.34 | 3.85E-180 | 8.01E-176 | 1.00 | 1.00 | 0.00 | 1.00 |
| 7 | Plekha7 | 2.06 | 8.42E-180 | 1.75E-175 | 0.37 | 0.08 | 0.29 | 4.58 |
| 7 | Slain1 | 2.27 | 9.75E-180 | 2.03E-175 | 0.28 | 0.05 | 0.23 | 5.68 |
| 7 | Synm | 3.37 | 4.71E-179 | 9.79E-175 | 0.15 | 0.02 | 0.14 | 10.20 |
| 7 | Upk3bl | 1.87 | 1.16E-178 | 2.41E-174 | 0.32 | 0.06 | 0.26 | 5.10 |
| 7 | Lad1 | 1.80 | 1.84E-178 | 3.83E-174 | 0.36 | 0.08 | 0.29 | 4.65 |
| 7 | Acvr2a | 1.61 | 2.28E-178 | 4.75E-174 | 0.66 | 0.25 | 0.41 | 2.62 |
| 7 | Rplp2 | 0.31 | 8.10E-178 | 1.68E-173 | 1.00 | 1.00 | 0.00 | 1.00 |
| 7 | Cep170b | 1.87 | 4.66E-177 | 9.70E-173 | 0.44 | 0.11 | 0.33 | 3.88 |
| 7 | Krtcap3 | 1.98 | 2.54E-176 | 5.27E-172 | 0.41 | 0.10 | 0.31 | 4.03 |
| 7 | Ppfia3 | 1.96 | 3.12E-176 | 6.48E-172 | 0.44 | 0.12 | 0.32 | 3.78 |
| 7 | Gstm1 | 2.55 | 9.09E-176 | 1.89E-171 | 0.59 | 0.23 | 0.36 | 2.55 |
| 7 | Pak3 | 1.89 | 1.29E-175 | 2.68E-171 | 0.46 | 0.12 | 0.33 | 3.67 |
| 7 | Rplp1 | 0.34 | 1.33E-175 | 2.76E-171 | 1.00 | 1.00 | 0.00 | 1.00 |
| 7 | Ormdl2 | 1.48 | 1.88E-175 | 3.91E-171 | 0.75 | 0.33 | 0.42 | 2.25 |
| 7 | Gm26793 | 3.32 | 1.07E-174 | 2.23E-170 | 0.15 | 0.01 | 0.13 | 10.43 |
| 7 | Rps19 | 0.29 | 6.86E-171 | 1.43E-166 | 1.00 | 1.00 | 0.00 | 1.00 |
| 7 | Bicd1 | 1.98 | 2.08E-169 | 4.33E-165 | 0.37 | 0.09 | 0.28 | 4.29 |
| 7 | Unc13b | 2.02 | 1.09E-168 | 2.27E-164 | 0.34 | 0.07 | 0.26 | 4.62 |
| 7 | Rpl12 | 0.32 | 1.94E-168 | 4.04E-164 | 1.00 | 1.00 | 0.00 | 1.00 |
| 7 | Meg3 | 1.11 | 4.52E-168 | 9.39E-164 | 0.95 | 0.68 | 0.26 | 1.39 |
| 7 | Samd5 | 1.73 | 2.41E-167 | 5.01E-163 | 0.73 | 0.35 | 0.38 | 2.11 |
| 7 | Prr15l | 2.48 | 2.08E-166 | 4.32E-162 | 0.20 | 0.03 | 0.18 | 7.03 |
| 7 | Cers4 | 1.74 | 2.50E-166 | 5.20E-162 | 0.49 | 0.14 | 0.34 | 3.37 |
| 7 | Rab11fip1 | 1.86 | 2.61E-166 | 5.43E-162 | 0.41 | 0.11 | 0.30 | 3.90 |
| 7 | Tjp3 | 2.29 | 6.62E-166 | 1.38E-161 | 0.24 | 0.04 | 0.20 | 6.10 |
| 7 | Ssx2ip | 1.42 | 1.01E-165 | 2.09E-161 | 0.75 | 0.34 | 0.41 | 2.18 |
| 7 | Igdcc3 | 1.65 | 2.12E-165 | 4.40E-161 | 0.61 | 0.22 | 0.39 | 2.74 |
| 7 | Nfe2l3 | 2.64 | 2.40E-165 | 4.98E-161 | 0.20 | 0.03 | 0.17 | 7.14 |
| 7 | Six1 | 1.03 | 2.71E-165 | 5.63E-161 | 0.96 | 0.60 | 0.36 | 1.60 |
| 7 | Emb | 1.22 | 5.22E-165 | 1.08E-160 | 0.80 | 0.39 | 0.41 | 2.06 |
| 7 | Tmem200c | 2.85 | 2.12E-162 | 4.40E-158 | 0.17 | 0.02 | 0.15 | 8.45 |
| 7 | Pcgf5 | 2.19 | 8.97E-162 | 1.87E-157 | 0.34 | 0.08 | 0.26 | 4.43 |
| 7 | Shisa3 | 3.02 | 2.19E-161 | 4.55E-157 | 0.13 | 0.01 | 0.12 | 11.45 |
| 7 | Fzd9 | 2.97 | 2.37E-161 | 4.93E-157 | 0.18 | 0.02 | 0.16 | 8.09 |
| 7 | Mbp | 1.86 | 3.04E-161 | 6.32E-157 | 0.36 | 0.08 | 0.27 | 4.25 |
| 7 | Lama5 | 1.85 | 7.82E-161 | 1.63E-156 | 0.40 | 0.10 | 0.30 | 3.88 |
| 7 | Il7 | 3.50 | 1.55E-160 | 3.23E-156 | 0.13 | 0.01 | 0.12 | 11.45 |
| 7 | Pkm | 0.55 | 3.88E-160 | 8.08E-156 | 1.00 | 0.98 | 0.02 | 1.02 |
| 7 | B3gnt7 | 2.51 | 5.10E-160 | 1.06E-155 | 0.22 | 0.04 | 0.19 | 6.37 |
| 7 | Stx3 | 1.60 | 1.06E-159 | 2.20E-155 | 0.43 | 0.11 | 0.31 | 3.75 |
| 7 | 2310030G06Rik | 2.00 | 2.97E-159 | 6.18E-155 | 0.30 | 0.06 | 0.24 | 4.90 |
| 7 | Adi1 | 1.63 | 6.65E-159 | 1.38E-154 | 0.68 | 0.30 | 0.38 | 2.24 |

**Supplementary Table 1. Marker genes for the original clusters of integrated clustering of E11.5 control and NotchGOF λ-junction cells.**

| | | | | | | | | |
|---|---|---|---|---|---|---|---|---|
| 7 | Rtn2 | 2.12 | 1.03E-157 | 2.13E-153 | 0.33 | 0.07 | 0.26 | 4.49 |
| 7 | Atox1 | 0.97 | 5.54E-157 | 1.15E-152 | 0.99 | 0.95 | 0.04 | 1.04 |
| 7 | Scube3 | 1.59 | 5.68E-157 | 1.18E-152 | 0.56 | 0.19 | 0.37 | 2.96 |
| 7 | Emid1 | 2.05 | 6.75E-157 | 1.40E-152 | 0.34 | 0.08 | 0.26 | 4.29 |
| 7 | Tnfrsf12a | 2.14 | 5.27E-156 | 1.10E-151 | 0.31 | 0.07 | 0.24 | 4.63 |
| 7 | Nectin3 | 1.30 | 5.82E-156 | 1.21E-151 | 0.85 | 0.53 | 0.32 | 1.60 |
| 7 | Srcin1 | 2.37 | 5.86E-156 | 1.22E-151 | 0.23 | 0.04 | 0.20 | 6.00 |
| 7 | Ide | 1.57 | 1.24E-153 | 2.59E-149 | 0.62 | 0.25 | 0.37 | 2.46 |
| 7 | Tbc1d30 | 2.60 | 1.03E-152 | 2.14E-148 | 0.18 | 0.03 | 0.16 | 7.24 |
| 7 | Rps12 | 0.29 | 1.05E-152 | 2.18E-148 | 1.00 | 1.00 | 0.00 | 1.00 |
| 7 | Frem2 | 1.33 | 1.38E-151 | 2.86E-147 | 0.55 | 0.18 | 0.37 | 3.06 |
| 7 | Epb41l5 | 1.49 | 1.39E-151 | 2.89E-147 | 0.61 | 0.24 | 0.38 | 2.58 |
| 7 | Rps10 | 0.26 | 8.11E-151 | 1.69E-146 | 1.00 | 1.00 | 0.00 | 1.00 |
| 7 | Plet1 | 3.60 | 1.08E-149 | 2.25E-145 | 0.11 | 0.01 | 0.10 | 13.25 |
| 7 | Rplp0 | 0.29 | 3.28E-149 | 6.82E-145 | 1.00 | 1.00 | 0.00 | 1.00 |
| 7 | Lypd6b | 2.02 | 3.73E-149 | 7.76E-145 | 0.29 | 0.06 | 0.23 | 4.85 |
| 7 | Sptbn2 | 1.82 | 5.15E-149 | 1.07E-144 | 0.38 | 0.10 | 0.28 | 3.76 |
| 7 | Ugt8a | 3.44 | 4.11E-147 | 8.55E-143 | 0.12 | 0.01 | 0.11 | 11.60 |
| 7 | Nckap5 | 1.77 | 5.03E-147 | 1.05E-142 | 0.34 | 0.08 | 0.26 | 4.19 |
| 7 | Rpl14 | 0.27 | 5.78E-147 | 1.20E-142 | 1.00 | 1.00 | 0.00 | 1.00 |
| 7 | Prps2 | 1.67 | 2.50E-145 | 5.19E-141 | 0.48 | 0.16 | 0.33 | 3.10 |
| 7 | Atp7b | 2.84 | 5.30E-145 | 1.10E-140 | 0.18 | 0.03 | 0.16 | 6.96 |
| 7 | Rpl17 | 0.28 | 8.87E-145 | 1.85E-140 | 1.00 | 1.00 | 0.00 | 1.00 |
| 7 | Gipc2 | 2.26 | 9.54E-144 | 1.98E-139 | 0.27 | 0.06 | 0.21 | 4.85 |
| 7 | Pgk1 | 0.69 | 1.46E-143 | 3.03E-139 | 1.00 | 0.95 | 0.05 | 1.05 |
| 7 | Kcnh8 | 2.30 | 2.13E-143 | 4.44E-139 | 0.19 | 0.03 | 0.16 | 6.71 |
| 7 | Kremen1 | 1.72 | 1.85E-142 | 3.85E-138 | 0.44 | 0.13 | 0.31 | 3.32 |
| 7 | Lgr4 | 1.40 | 2.82E-142 | 5.86E-138 | 0.57 | 0.21 | 0.36 | 2.67 |
| 7 | Nav2 | 0.98 | 3.80E-142 | 7.90E-138 | 0.62 | 0.22 | 0.40 | 2.77 |
| 7 | Slc2a1 | 1.32 | 6.85E-142 | 1.42E-137 | 0.77 | 0.41 | 0.36 | 1.87 |
| 7 | Slfn2 | 2.60 | 1.00E-141 | 2.08E-137 | 0.13 | 0.01 | 0.11 | 10.42 |
| 7 | Nuak1 | 1.70 | 1.55E-141 | 3.23E-137 | 0.53 | 0.19 | 0.34 | 2.78 |
| 7 | Atp6v0e2 | 2.09 | 8.71E-141 | 1.81E-136 | 0.24 | 0.05 | 0.20 | 5.33 |
| 7 | Maf | 1.07 | 1.09E-140 | 2.26E-136 | 0.29 | 0.06 | 0.23 | 4.59 |
| 7 | Ncam1 | 1.35 | 1.15E-140 | 2.40E-136 | 0.73 | 0.35 | 0.38 | 2.10 |
| 7 | Rbm47 | 2.59 | 1.68E-139 | 3.50E-135 | 0.19 | 0.03 | 0.16 | 6.64 |
| 7 | Pgam1 | 0.59 | 2.34E-139 | 4.86E-135 | 1.00 | 0.97 | 0.03 | 1.04 |
| 7 | Rian | 1.37 | 1.37E-138 | 2.86E-134 | 0.67 | 0.30 | 0.37 | 2.22 |
| 7 | B230118H07Rik | 1.31 | 1.39E-138 | 2.88E-134 | 0.80 | 0.49 | 0.30 | 1.62 |
| 7 | Nfasc | 2.55 | 1.54E-138 | 3.20E-134 | 0.16 | 0.02 | 0.14 | 7.85 |
| 7 | Ldha | 0.67 | 2.02E-138 | 4.21E-134 | 1.00 | 0.97 | 0.03 | 1.03 |
| 7 | Nap1l1 | 0.45 | 2.27E-138 | 4.72E-134 | 1.00 | 1.00 | 0.00 | 1.00 |
| 7 | Plp2 | 1.19 | 1.38E-137 | 2.87E-133 | 0.75 | 0.36 | 0.39 | 2.06 |
| 7 | Ppfibp1 | 1.34 | 1.69E-137 | 3.51E-133 | 0.76 | 0.42 | 0.35 | 1.83 |
| 7 | Stc2 | 3.50 | 1.87E-137 | 3.89E-133 | 0.11 | 0.01 | 0.10 | 11.00 |
| 7 | Pclo | 1.51 | 4.52E-137 | 9.41E-133 | 0.37 | 0.10 | 0.27 | 3.80 |
| 7 | Frmd3 | 2.91 | 5.52E-137 | 1.15E-132 | 0.14 | 0.02 | 0.12 | 9.13 |
| 7 | Rnh1 | 1.13 | 1.85E-136 | 3.84E-132 | 0.86 | 0.54 | 0.32 | 1.58 |
| 7 | Grtp1 | 1.85 | 5.32E-134 | 1.11E-129 | 0.41 | 0.13 | 0.29 | 3.26 |
| 7 | Lamc2 | 2.33 | 6.92E-134 | 1.44E-129 | 0.21 | 0.04 | 0.17 | 5.78 |
| 7 | Bcam | 1.04 | 1.19E-133 | 2.48E-129 | 0.56 | 0.19 | 0.37 | 2.94 |
| 7 | Fzd3 | 1.04 | 1.83E-133 | 3.81E-129 | 0.88 | 0.58 | 0.31 | 1.53 |
| 7 | Nhs | 1.35 | 1.57E-131 | 3.27E-127 | 0.68 | 0.32 | 0.36 | 2.11 |
| 7 | Rpl10a | 0.29 | 1.62E-131 | 3.38E-127 | 1.00 | 1.00 | 0.00 | 1.00 |
| 7 | Arhgdig | 0.90 | 2.85E-131 | 5.92E-127 | 0.30 | 0.07 | 0.23 | 4.18 |
| 7 | Frmd5 | 2.18 | 6.03E-131 | 1.25E-126 | 0.23 | 0.04 | 0.18 | 5.28 |

**Supplementary Table 1. Marker genes for the original clusters of integrated clustering of E11.5 control and NotchGOF λ-junction cells.**

| | | | | | | | | |
|---|---|---|---|---|---|---|---|---|
| 7 | Phlpp1 | 1.84 | 4.65E-130 | 9.67E-126 | 0.40 | 0.12 | 0.28 | 3.31 |
| 7 | Ust | 2.20 | 2.89E-129 | 6.02E-125 | 0.25 | 0.05 | 0.20 | 4.70 |
| 7 | Sparc | 0.87 | 6.61E-129 | 1.37E-124 | 0.95 | 0.73 | 0.22 | 1.30 |
| 7 | Notch3 | 1.28 | 1.50E-128 | 3.12E-124 | 0.75 | 0.39 | 0.36 | 1.91 |
| 7 | Cldn23 | 2.38 | 7.03E-128 | 1.46E-123 | 0.19 | 0.03 | 0.16 | 5.94 |
| 7 | Mdh1 | 0.64 | 2.09E-127 | 4.35E-123 | 0.99 | 0.95 | 0.05 | 1.05 |
| 7 | Rps6 | 0.31 | 3.64E-127 | 7.57E-123 | 1.00 | 1.00 | 0.00 | 1.00 |
| 7 | Gkap1 | 1.17 | 4.19E-127 | 8.71E-123 | 0.78 | 0.42 | 0.36 | 1.85 |
| 7 | Slc25a23 | 1.55 | 4.32E-127 | 8.99E-123 | 0.46 | 0.15 | 0.31 | 3.06 |
| 7 | Ndrg2 | 1.61 | 4.55E-127 | 9.46E-123 | 0.41 | 0.12 | 0.28 | 3.28 |
| 7 | Tmem229b | 2.89 | 5.01E-127 | 1.04E-122 | 0.15 | 0.02 | 0.13 | 7.40 |
| 7 | Cltb | 1.54 | 1.61E-126 | 3.35E-122 | 0.48 | 0.17 | 0.31 | 2.84 |
| 7 | Ccdc80 | 1.59 | 1.66E-126 | 3.45E-122 | 0.48 | 0.17 | 0.31 | 2.83 |
| 7 | Snx10 | 1.59 | 1.99E-126 | 4.14E-122 | 0.45 | 0.15 | 0.30 | 3.05 |
| 7 | H2-K1 | 1.73 | 6.10E-126 | 1.27E-121 | 0.40 | 0.12 | 0.28 | 3.26 |
| 7 | Tmem30b | 1.60 | 1.12E-125 | 2.32E-121 | 0.43 | 0.14 | 0.29 | 3.14 |
| 7 | Sh3gl2 | 2.35 | 5.51E-125 | 1.15E-120 | 0.19 | 0.03 | 0.16 | 5.79 |
| 7 | Cyp26a1 | 2.73 | 1.14E-124 | 2.37E-120 | 0.16 | 0.02 | 0.13 | 7.05 |
| 7 | Eno1 | 0.55 | 3.49E-124 | 7.27E-120 | 1.00 | 0.97 | 0.03 | 1.03 |
| 7 | A330048O09Rik | 2.98 | 7.38E-123 | 1.54E-118 | 0.12 | 0.01 | 0.11 | 9.08 |
| 7 | Aig1 | 1.29 | 1.59E-122 | 3.31E-118 | 0.68 | 0.32 | 0.36 | 2.13 |
| 7 | Klk8 | 1.22 | 2.13E-122 | 4.43E-118 | 0.54 | 0.20 | 0.34 | 2.70 |
| 7 | Kcna5 | 2.83 | 3.82E-122 | 7.95E-118 | 0.13 | 0.02 | 0.11 | 8.40 |
| 7 | D030055H07Rik | 2.84 | 6.99E-122 | 1.45E-117 | 0.12 | 0.01 | 0.11 | 8.79 |
| 7 | Npas3 | 2.29 | 8.67E-122 | 1.80E-117 | 0.24 | 0.05 | 0.19 | 4.67 |
| 7 | Arhgef16 | 1.85 | 1.59E-121 | 3.31E-117 | 0.24 | 0.05 | 0.19 | 4.76 |
| 7 | Ric3 | 2.24 | 3.07E-121 | 6.38E-117 | 0.22 | 0.04 | 0.18 | 5.02 |
| 7 | Nectin1 | 1.38 | 2.61E-120 | 5.44E-116 | 0.47 | 0.16 | 0.31 | 2.91 |
| 7 | Slc16a1 | 1.46 | 5.11E-119 | 1.06E-114 | 0.61 | 0.29 | 0.33 | 2.14 |
| 7 | Mtln | 1.23 | 9.11E-119 | 1.89E-114 | 0.66 | 0.32 | 0.35 | 2.10 |
| 7 | Fxyd7 | 2.71 | 1.93E-118 | 4.01E-114 | 0.19 | 0.03 | 0.16 | 5.59 |
| 7 | Filip1 | 2.40 | 2.15E-118 | 4.47E-114 | 0.18 | 0.03 | 0.15 | 5.97 |
| 7 | Pdgfc | 0.74 | 3.08E-118 | 6.41E-114 | 0.66 | 0.27 | 0.40 | 2.49 |
| 7 | Nes | 1.19 | 8.83E-118 | 1.84E-113 | 0.74 | 0.41 | 0.33 | 1.80 |
| 7 | 2010300C02Rik | 2.76 | 1.34E-117 | 2.78E-113 | 0.14 | 0.02 | 0.12 | 7.15 |
| 7 | Mfng | 1.00 | 1.50E-117 | 3.12E-113 | 0.21 | 0.04 | 0.17 | 5.31 |
| 7 | Mapk8ip1 | 1.33 | 1.91E-117 | 3.96E-113 | 0.58 | 0.24 | 0.33 | 2.36 |
| 7 | Smim22 | 2.22 | 4.12E-117 | 8.58E-113 | 0.19 | 0.03 | 0.15 | 5.78 |
| 7 | Mt2 | 0.37 | 5.07E-117 | 1.05E-112 | 0.40 | 0.12 | 0.28 | 3.35 |
| 7 | Ktn1 | 0.78 | 9.99E-117 | 2.08E-112 | 0.96 | 0.82 | 0.14 | 1.17 |
| 7 | Cdh3 | 1.16 | 1.08E-116 | 2.24E-112 | 0.53 | 0.20 | 0.34 | 2.71 |
| 7 | Eps8l1 | 2.26 | 1.28E-116 | 2.67E-112 | 0.19 | 0.03 | 0.15 | 5.67 |
| 7 | C330021F23Rik | 2.45 | 2.96E-116 | 6.17E-112 | 0.16 | 0.03 | 0.14 | 6.48 |
| 7 | Cdc14a | 1.64 | 3.18E-116 | 6.61E-112 | 0.56 | 0.24 | 0.31 | 2.29 |
| 7 | Hsp90aa1 | 0.43 | 6.18E-116 | 1.29E-111 | 1.00 | 1.00 | 0.00 | 1.00 |
| 7 | Rundc3a | 1.96 | 1.23E-115 | 2.56E-111 | 0.17 | 0.03 | 0.14 | 6.19 |
| 7 | Frat2 | 1.45 | 2.01E-115 | 4.19E-111 | 0.54 | 0.22 | 0.32 | 2.43 |
| 7 | Kcnb1 | 2.17 | 2.36E-113 | 4.90E-109 | 0.24 | 0.05 | 0.19 | 4.44 |
| 7 | Dag1 | 1.12 | 2.86E-113 | 5.94E-109 | 0.71 | 0.36 | 0.35 | 1.99 |
| 7 | Rpl38 | 0.28 | 1.13E-112 | 2.35E-108 | 1.00 | 1.00 | 0.00 | 1.00 |
| 7 | Marveld3 | 2.05 | 1.76E-112 | 3.66E-108 | 0.18 | 0.03 | 0.15 | 5.77 |
| 7 | Pard6b | 2.07 | 2.24E-112 | 4.65E-108 | 0.22 | 0.05 | 0.17 | 4.76 |
| 7 | Zfhx2 | 1.35 | 1.15E-111 | 2.38E-107 | 0.56 | 0.24 | 0.33 | 2.37 |
| 7 | Il17rd | 1.11 | 1.18E-110 | 2.45E-106 | 0.80 | 0.49 | 0.31 | 1.63 |
| 7 | Gucy1a1 | 1.85 | 1.51E-110 | 3.14E-106 | 0.27 | 0.07 | 0.20 | 3.99 |
| 7 | Lamc1 | 1.44 | 1.90E-110 | 3.94E-106 | 0.57 | 0.25 | 0.32 | 2.30 |

**Supplementary Table 1. Marker genes for the original clusters of integrated clustering of E11.5 control and NotchGOF λ-junction cells.**

| | | | | | | | | |
|---|---|---|---|---|---|---|---|---|
| 7 | Celf2 | 1.10 | 1.97E-110 | 4.10E-106 | 0.58 | 0.24 | 0.34 | 2.41 |
| 7 | Dusp5 | 1.75 | 2.78E-110 | 5.78E-106 | 0.35 | 0.11 | 0.25 | 3.29 |
| 7 | Lgi2 | 1.82 | 3.29E-110 | 6.84E-106 | 0.35 | 0.10 | 0.24 | 3.33 |
| 7 | Prr5l | 1.44 | 4.64E-110 | 9.65E-106 | 0.44 | 0.15 | 0.29 | 2.86 |
| 7 | Myl7 | 2.36 | 3.43E-109 | 7.14E-105 | 0.15 | 0.02 | 0.13 | 6.73 |
| 7 | Epn3 | 2.03 | 8.44E-109 | 1.75E-104 | 0.21 | 0.04 | 0.17 | 4.86 |
| 7 | Shc3 | 3.18 | 9.11E-109 | 1.89E-104 | 0.10 | 0.01 | 0.09 | 10.10 |
| 7 | Nr2f6 | 0.96 | 9.62E-109 | 2.00E-104 | 0.86 | 0.59 | 0.27 | 1.46 |
| 7 | Gjb6 | 2.89 | 1.15E-108 | 2.39E-104 | 0.11 | 0.01 | 0.10 | 8.54 |
| 7 | Pappa | 3.08 | 2.85E-108 | 5.94E-104 | 0.11 | 0.01 | 0.10 | 8.92 |
| 7 | Fam187b | 2.26 | 3.38E-108 | 7.03E-104 | 0.18 | 0.03 | 0.15 | 5.42 |
| 7 | Igsf9b | 2.15 | 3.59E-108 | 7.47E-104 | 0.22 | 0.05 | 0.18 | 4.57 |
| 7 | Jup | 1.26 | 7.93E-108 | 1.65E-103 | 0.50 | 0.19 | 0.31 | 2.66 |
| 7 | Scd2 | 0.71 | 3.79E-107 | 7.89E-103 | 0.99 | 0.91 | 0.08 | 1.09 |
| 7 | Rtn3 | 0.79 | 4.11E-107 | 8.54E-103 | 0.93 | 0.78 | 0.15 | 1.19 |
| 7 | Baiap2l1 | 1.95 | 6.24E-107 | 1.30E-102 | 0.24 | 0.06 | 0.18 | 4.33 |
| 7 | Atp2b1 | 0.77 | 1.95E-106 | 4.05E-102 | 0.97 | 0.84 | 0.13 | 1.15 |
| 7 | Fgf9 | 0.74 | 3.21E-106 | 6.68E-102 | 0.34 | 0.10 | 0.25 | 3.53 |
| 7 | Apoc1 | 1.28 | 3.88E-106 | 8.07E-102 | 0.29 | 0.08 | 0.21 | 3.77 |
| 7 | Nol7 | 0.63 | 6.21E-106 | 1.29E-101 | 0.95 | 0.88 | 0.08 | 1.09 |
| 7 | Pkp4 | 1.15 | 1.02E-105 | 2.12E-101 | 0.70 | 0.37 | 0.33 | 1.90 |
| 7 | Nt5dc2 | 0.59 | 1.57E-105 | 3.26E-101 | 0.98 | 0.92 | 0.06 | 1.07 |
| 7 | Ccdc171 | 1.73 | 1.80E-105 | 3.74E-101 | 0.31 | 0.09 | 0.22 | 3.57 |
| 7 | Llgl2 | 2.03 | 2.15E-105 | 4.48E-101 | 0.21 | 0.04 | 0.16 | 4.77 |
| 7 | Gm10076 | 0.27 | 3.41E-105 | 7.08E-101 | 1.00 | 1.00 | 0.00 | 1.00 |
| 7 | Dstn | 0.55 | 7.50E-105 | 1.56E-100 | 0.99 | 0.94 | 0.05 | 1.05 |
| 7 | Palm3 | 1.63 | 1.58E-104 | 3.30E-100 | 0.40 | 0.14 | 0.26 | 2.91 |
| 7 | Acsl6 | 2.71 | 6.54E-104 | 1.36E-99 | 0.12 | 0.02 | 0.11 | 7.56 |
| 7 | Nabp2 | 0.89 | 1.06E-103 | 2.21E-99 | 0.86 | 0.65 | 0.21 | 1.33 |
| 7 | Gsr | 1.17 | 1.12E-103 | 2.33E-99 | 0.66 | 0.32 | 0.34 | 2.04 |
| 7 | Srgap3 | 1.60 | 2.73E-103 | 5.68E-99 | 0.38 | 0.13 | 0.25 | 3.01 |
| 7 | Afdn | 0.85 | 4.60E-103 | 9.56E-99 | 0.86 | 0.56 | 0.30 | 1.53 |
| 7 | Cxadr | 1.09 | 6.41E-103 | 1.33E-98 | 0.78 | 0.45 | 0.33 | 1.73 |
| 7 | Zfp467 | 1.19 | 1.05E-102 | 2.19E-98 | 0.61 | 0.28 | 0.34 | 2.21 |
| 7 | Mal2 | 2.82 | 1.75E-102 | 3.65E-98 | 0.13 | 0.02 | 0.11 | 7.17 |
| 7 | Ago1 | 1.11 | 1.92E-102 | 3.98E-98 | 0.68 | 0.35 | 0.33 | 1.94 |
| 7 | Exph5 | 2.20 | 3.15E-102 | 6.54E-98 | 0.17 | 0.03 | 0.14 | 5.42 |
| 7 | Col11a1 | 1.18 | 5.59E-102 | 1.16E-97 | 0.73 | 0.41 | 0.31 | 1.76 |
| 7 | Celsr2 | 1.62 | 1.20E-101 | 2.49E-97 | 0.30 | 0.08 | 0.22 | 3.56 |
| 7 | Dhx32 | 1.35 | 2.23E-101 | 4.64E-97 | 0.51 | 0.21 | 0.30 | 2.46 |
| 7 | Pik3ap1 | 2.61 | 2.23E-101 | 4.64E-97 | 0.11 | 0.01 | 0.10 | 8.07 |
| 7 | Agrn | 1.09 | 3.70E-101 | 7.70E-97 | 0.70 | 0.38 | 0.32 | 1.86 |
| 7 | Nav3 | 1.28 | 4.03E-101 | 8.38E-97 | 0.37 | 0.12 | 0.25 | 3.09 |
| 7 | Lrrk1 | 1.48 | 1.24E-100 | 2.57E-96 | 0.40 | 0.14 | 0.26 | 2.88 |
| 7 | Mgat4a | 1.78 | 1.44E-100 | 2.99E-96 | 0.30 | 0.09 | 0.21 | 3.49 |
| 7 | Zfp219 | 1.05 | 1.90E-100 | 3.95E-96 | 0.72 | 0.39 | 0.33 | 1.86 |
| 7 | Rpl36 | 0.25 | 4.50E-100 | 9.35E-96 | 1.00 | 1.00 | 0.00 | 1.00 |
| 7 | Rgcc | 1.80 | 6.52E-100 | 1.36E-95 | 0.29 | 0.08 | 0.21 | 3.49 |
| 7 | Myc | 1.54 | 2.23E-99 | 4.64E-95 | 0.42 | 0.15 | 0.26 | 2.74 |
| 7 | Rnf208 | 1.86 | 3.64E-99 | 7.57E-95 | 0.25 | 0.06 | 0.18 | 4.02 |
| 7 | Pde1b | 2.19 | 5.36E-99 | 1.11E-94 | 0.17 | 0.03 | 0.14 | 5.42 |
| 7 | Zdbf2 | 1.14 | 2.07E-98 | 4.31E-94 | 0.71 | 0.40 | 0.32 | 1.80 |
| 7 | Dgka | 1.86 | 3.42E-98 | 7.11E-94 | 0.23 | 0.05 | 0.17 | 4.20 |
| 7 | Cdca7l | 1.18 | 4.01E-98 | 8.34E-94 | 0.61 | 0.29 | 0.32 | 2.08 |
| 7 | Ybx3 | 0.70 | 5.98E-98 | 1.24E-93 | 0.95 | 0.83 | 0.12 | 1.14 |
| 7 | Hmgn2 | 0.44 | 6.60E-98 | 1.37E-93 | 0.99 | 0.98 | 0.02 | 1.02 |

**Supplementary Table 1. Marker genes for the original clusters of integrated clustering of E11.5 control and NotchGOF λ-junction cells.**

| | | | | | | | | |
|---|---|---|---|---|---|---|---|---|
| 7 | Sorl1 | 1.36 | 7.19E-98 | 1.50E-93 | 0.40 | 0.14 | 0.26 | 2.85 |
| 7 | Uchl1 | 0.85 | 8.42E-98 | 1.75E-93 | 0.67 | 0.34 | 0.34 | 1.99 |
| 7 | Pard3b | 1.53 | 1.76E-97 | 3.65E-93 | 0.35 | 0.11 | 0.24 | 3.15 |
| 7 | Podn | 2.94 | 2.17E-97 | 4.50E-93 | 0.11 | 0.01 | 0.09 | 8.08 |
| 7 | Ckb | 1.86 | 2.35E-97 | 4.89E-93 | 0.59 | 0.32 | 0.27 | 1.84 |
| 7 | Mtus1 | 1.54 | 3.73E-97 | 7.76E-93 | 0.32 | 0.10 | 0.22 | 3.32 |
| 7 | Acot7 | 1.21 | 3.74E-97 | 7.78E-93 | 0.53 | 0.22 | 0.31 | 2.36 |
| 7 | Tkt | 0.58 | 4.26E-97 | 8.87E-93 | 0.99 | 0.92 | 0.07 | 1.08 |
| 7 | Abcg1 | 1.45 | 7.14E-97 | 1.48E-92 | 0.28 | 0.08 | 0.20 | 3.69 |
| 7 | Tmem108 | 1.71 | 7.71E-97 | 1.60E-92 | 0.48 | 0.21 | 0.27 | 2.31 |
| 7 | Cd83 | 2.19 | 8.08E-97 | 1.68E-92 | 0.12 | 0.02 | 0.11 | 6.83 |
| 7 | Abca3 | 1.85 | 1.54E-96 | 3.20E-92 | 0.23 | 0.06 | 0.18 | 4.11 |
| 7 | Jade1 | 1.45 | 3.68E-96 | 7.65E-92 | 0.43 | 0.16 | 0.27 | 2.65 |
| 7 | Hs6st1 | 1.39 | 5.11E-96 | 1.06E-91 | 0.43 | 0.16 | 0.27 | 2.69 |
| 7 | Rdh10 | 1.55 | 7.32E-96 | 1.52E-91 | 0.40 | 0.15 | 0.25 | 2.73 |
| 7 | Pcdh17 | 1.15 | 8.07E-96 | 1.68E-91 | 0.49 | 0.19 | 0.29 | 2.51 |
| 7 | Pla2r1 | 1.35 | 2.25E-95 | 4.68E-91 | 0.26 | 0.07 | 0.19 | 3.82 |
| 7 | Prom2 | 2.19 | 3.69E-95 | 7.68E-91 | 0.15 | 0.03 | 0.13 | 5.50 |
| 7 | Mapk11 | 1.83 | 6.42E-95 | 1.33E-90 | 0.25 | 0.07 | 0.18 | 3.82 |
| 7 | Rpl31 | 0.31 | 8.38E-95 | 1.74E-90 | 1.00 | 1.00 | 0.00 | 1.00 |
| 7 | Tyro3 | 1.54 | 1.36E-94 | 2.82E-90 | 0.42 | 0.16 | 0.26 | 2.63 |
| 7 | Pdk1 | 1.23 | 1.63E-94 | 3.38E-90 | 0.55 | 0.24 | 0.31 | 2.25 |
| 7 | Plekhh1 | 1.88 | 1.97E-94 | 4.10E-90 | 0.23 | 0.06 | 0.18 | 4.03 |
| 7 | Creld1 | 1.51 | 2.85E-94 | 5.93E-90 | 0.36 | 0.12 | 0.24 | 3.01 |
| 7 | Clstn1 | 1.12 | 2.89E-94 | 6.00E-90 | 0.70 | 0.38 | 0.32 | 1.85 |
| 7 | Kazn | 1.37 | 4.18E-94 | 8.69E-90 | 0.45 | 0.17 | 0.27 | 2.58 |
| 7 | Acaa2 | 0.87 | 1.01E-93 | 2.11E-89 | 0.78 | 0.47 | 0.31 | 1.66 |
| 7 | Mif | 0.40 | 1.70E-93 | 3.53E-89 | 1.00 | 0.99 | 0.01 | 1.01 |
| 7 | Synrg | 1.20 | 1.93E-93 | 4.02E-89 | 0.52 | 0.22 | 0.30 | 2.33 |
| 7 | Syngr2 | 1.45 | 4.86E-93 | 1.01E-88 | 0.38 | 0.13 | 0.25 | 2.85 |
| 7 | P2rx2 | 2.79 | 6.16E-93 | 1.28E-88 | 0.12 | 0.02 | 0.10 | 7.19 |
| 7 | Cgref1 | 2.57 | 9.61E-93 | 2.00E-88 | 0.12 | 0.02 | 0.10 | 7.19 |
| 7 | D930020B18Rik | 2.36 | 1.33E-92 | 2.77E-88 | 0.11 | 0.02 | 0.10 | 7.13 |
| 7 | Clic6 | 1.81 | 7.12E-92 | 1.48E-87 | 0.21 | 0.05 | 0.16 | 4.33 |
| 7 | Col2a1 | 0.68 | 7.89E-92 | 1.64E-87 | 0.91 | 0.63 | 0.28 | 1.44 |
| 7 | Npm3 | 0.62 | 3.05E-91 | 6.33E-87 | 0.94 | 0.79 | 0.15 | 1.19 |
| 7 | Sall2 | 1.13 | 4.39E-91 | 9.13E-87 | 0.69 | 0.38 | 0.31 | 1.81 |
| 7 | Lmo1 | 1.13 | 8.49E-91 | 1.77E-86 | 0.26 | 0.07 | 0.19 | 3.75 |
| 7 | Ak4 | 1.59 | 5.21E-90 | 1.08E-85 | 0.31 | 0.10 | 0.21 | 3.20 |
| 7 | Eml5 | 1.55 | 9.41E-90 | 1.96E-85 | 0.27 | 0.08 | 0.20 | 3.50 |
| 7 | Tnk1 | 2.05 | 1.20E-89 | 2.49E-85 | 0.16 | 0.03 | 0.13 | 5.09 |
| 7 | Rcsd1 | 1.52 | 2.81E-89 | 5.85E-85 | 0.29 | 0.09 | 0.20 | 3.40 |
| 7 | Ephb2 | 1.15 | 5.29E-89 | 1.10E-84 | 0.58 | 0.27 | 0.31 | 2.13 |
| 7 | Lrrc8b | 1.67 | 1.05E-88 | 2.18E-84 | 0.32 | 0.10 | 0.22 | 3.09 |
| 7 | Klc3 | 1.12 | 1.07E-88 | 2.22E-84 | 0.32 | 0.10 | 0.22 | 3.21 |
| 7 | Fam20c | 2.30 | 3.84E-88 | 8.00E-84 | 0.12 | 0.02 | 0.10 | 6.67 |
| 7 | Adgrg1 | 1.95 | 4.47E-88 | 9.29E-84 | 0.17 | 0.04 | 0.14 | 4.70 |
| 7 | Esrp2 | 1.64 | 5.52E-88 | 1.15E-83 | 0.23 | 0.06 | 0.17 | 3.90 |
| 7 | Olfm1 | 1.24 | 2.66E-87 | 5.54E-83 | 0.50 | 0.22 | 0.29 | 2.32 |
| 7 | Lrrn3 | 1.51 | 4.75E-87 | 9.89E-83 | 0.24 | 0.07 | 0.18 | 3.72 |
| 7 | Dock9 | 1.68 | 1.78E-86 | 3.69E-82 | 0.24 | 0.07 | 0.18 | 3.70 |
| 7 | Shroom3 | 1.37 | 3.51E-86 | 7.31E-82 | 0.39 | 0.14 | 0.25 | 2.71 |
| 7 | Mtcl1 | 1.50 | 3.63E-86 | 7.54E-82 | 0.34 | 0.11 | 0.22 | 2.98 |
| 7 | Gcat | 0.99 | 7.17E-86 | 1.49E-81 | 0.76 | 0.47 | 0.29 | 1.62 |
| 7 | Susd4 | 1.44 | 8.60E-86 | 1.79E-81 | 0.28 | 0.08 | 0.20 | 3.37 |
| 7 | Six2 | 0.75 | 9.74E-86 | 2.03E-81 | 0.77 | 0.44 | 0.33 | 1.76 |

**Supplementary Table 1. Marker genes for the original clusters of integrated clustering of E11.5 control and NotchGOF λ-junction cells.**

| | | | | | | | | |
|---|---|---|---|---|---|---|---|---|
| 7 | Aldoa | 0.52 | 1.47E-85 | 3.06E-81 | 0.99 | 0.95 | 0.04 | 1.04 |
| 7 | Stbd1 | 1.74 | 5.13E-85 | 1.07E-80 | 0.30 | 0.10 | 0.20 | 3.08 |
| 7 | Phyhipl | 2.07 | 7.77E-85 | 1.62E-80 | 0.16 | 0.03 | 0.13 | 4.94 |
| 7 | Cpz | 1.59 | 7.82E-85 | 1.63E-80 | 0.24 | 0.06 | 0.17 | 3.70 |
| 7 | Plekhg3 | 1.48 | 1.29E-84 | 2.68E-80 | 0.26 | 0.08 | 0.19 | 3.49 |
| 7 | Gcnt2 | 2.08 | 1.37E-84 | 2.86E-80 | 0.18 | 0.04 | 0.14 | 4.46 |
| 7 | Gdap1l1 | 1.81 | 1.62E-84 | 3.36E-80 | 0.10 | 0.01 | 0.09 | 7.36 |
| 7 | Prdx1 | 0.36 | 1.68E-84 | 3.49E-80 | 1.00 | 0.99 | 0.00 | 1.00 |
| 7 | Patj | 1.37 | 3.65E-84 | 7.59E-80 | 0.35 | 0.12 | 0.23 | 2.88 |
| 7 | Trpm4 | 1.47 | 2.20E-83 | 4.58E-79 | 0.29 | 0.09 | 0.20 | 3.23 |
| 7 | Arhgef5 | 1.11 | 5.44E-83 | 1.13E-78 | 0.46 | 0.19 | 0.27 | 2.44 |
| 7 | Psen2 | 1.79 | 5.62E-83 | 1.17E-78 | 0.25 | 0.07 | 0.18 | 3.51 |
| 7 | Macc1 | 1.71 | 5.91E-83 | 1.23E-78 | 0.20 | 0.05 | 0.15 | 4.08 |
| 7 | Dut | 0.57 | 8.38E-83 | 1.74E-78 | 0.98 | 0.94 | 0.04 | 1.04 |
| 7 | Zswim5 | 2.14 | 1.49E-82 | 3.11E-78 | 0.15 | 0.03 | 0.12 | 5.18 |
| 7 | Lrrfip1 | 1.10 | 4.82E-82 | 1.00E-77 | 0.63 | 0.33 | 0.30 | 1.92 |
| 7 | Pbx1 | 0.62 | 5.35E-82 | 1.11E-77 | 0.98 | 0.92 | 0.06 | 1.06 |
| 7 | Samd10 | 1.53 | 9.24E-82 | 1.92E-77 | 0.27 | 0.08 | 0.19 | 3.28 |
| 7 | Endod1 | 1.79 | 9.76E-82 | 2.03E-77 | 0.27 | 0.08 | 0.19 | 3.24 |
| 7 | Igsf9 | 1.00 | 2.40E-81 | 4.98E-77 | 0.70 | 0.39 | 0.31 | 1.78 |
| 7 | Tjp2 | 1.14 | 2.42E-81 | 5.03E-77 | 0.59 | 0.30 | 0.29 | 1.97 |
| 7 | Ggt7 | 2.21 | 2.44E-81 | 5.07E-77 | 0.15 | 0.03 | 0.12 | 5.18 |
| 7 | Metrnl | 1.66 | 3.56E-81 | 7.40E-77 | 0.22 | 0.06 | 0.16 | 3.76 |
| 7 | Vill | 1.65 | 8.26E-81 | 1.72E-76 | 0.26 | 0.08 | 0.18 | 3.41 |
| 7 | Ntn1 | 2.38 | 1.69E-80 | 3.52E-76 | 0.13 | 0.03 | 0.11 | 5.36 |
| 7 | Gpc4 | 1.36 | 3.40E-80 | 7.07E-76 | 0.36 | 0.13 | 0.23 | 2.75 |
| 7 | Ifi30 | 0.98 | 3.53E-80 | 7.34E-76 | 0.58 | 0.28 | 0.30 | 2.05 |
| 7 | Maneal | 2.43 | 4.04E-80 | 8.41E-76 | 0.12 | 0.02 | 0.10 | 6.16 |
| 7 | Gm15417 | 2.33 | 7.42E-80 | 1.54E-75 | 0.12 | 0.02 | 0.10 | 5.86 |
| 7 | Epha1 | 1.75 | 1.51E-79 | 3.15E-75 | 0.21 | 0.05 | 0.16 | 3.92 |
| 7 | Hpn | 1.94 | 1.88E-79 | 3.91E-75 | 0.16 | 0.03 | 0.13 | 4.79 |
| 7 | Pcbd1 | 1.34 | 2.38E-79 | 4.96E-75 | 0.33 | 0.11 | 0.22 | 2.89 |
| 7 | Ddr1 | 1.14 | 4.29E-79 | 8.91E-75 | 0.35 | 0.12 | 0.23 | 2.85 |
| 7 | mt-Nd6 | 1.15 | 6.58E-78 | 1.37E-73 | 0.62 | 0.34 | 0.28 | 1.81 |
| 7 | Cmtm4 | 1.14 | 7.51E-78 | 1.56E-73 | 0.52 | 0.25 | 0.28 | 2.11 |
| 7 | Acsl5 | 1.34 | 1.18E-77 | 2.45E-73 | 0.38 | 0.15 | 0.23 | 2.56 |
| 7 | Mettl26 | 0.89 | 1.55E-77 | 3.22E-73 | 0.76 | 0.48 | 0.28 | 1.57 |
| 7 | Marveld2 | 1.69 | 1.58E-77 | 3.30E-73 | 0.21 | 0.05 | 0.15 | 3.83 |
| 7 | Rpia | 1.16 | 1.76E-77 | 3.65E-73 | 0.56 | 0.28 | 0.28 | 1.98 |
| 7 | Cd82 | 1.72 | 7.17E-77 | 1.49E-72 | 0.22 | 0.06 | 0.16 | 3.67 |
| 7 | Ccnd1 | 0.84 | 7.26E-77 | 1.51E-72 | 0.89 | 0.80 | 0.10 | 1.12 |
| 7 | Dynll2 | 0.60 | 1.62E-76 | 3.38E-72 | 0.94 | 0.84 | 0.10 | 1.12 |
| 7 | Cox4i2 | 1.13 | 1.78E-76 | 3.71E-72 | 0.53 | 0.25 | 0.27 | 2.07 |
| 7 | Cobl | 1.48 | 1.81E-76 | 3.76E-72 | 0.23 | 0.06 | 0.16 | 3.56 |
| 7 | Kcnd2 | 2.71 | 2.71E-76 | 5.63E-72 | 0.10 | 0.02 | 0.09 | 6.50 |
| 7 | Ccdc181 | 1.65 | 3.00E-76 | 6.23E-72 | 0.22 | 0.06 | 0.16 | 3.66 |
| 7 | Gpi1 | 0.59 | 4.51E-76 | 9.38E-72 | 0.95 | 0.84 | 0.11 | 1.13 |
| 7 | Mcrip2 | 1.59 | 5.29E-76 | 1.10E-71 | 0.22 | 0.06 | 0.16 | 3.56 |
| 7 | Fgf13 | 1.48 | 5.49E-76 | 1.14E-71 | 0.30 | 0.10 | 0.20 | 2.94 |
| 7 | Rnf43 | 1.68 | 9.30E-76 | 1.94E-71 | 0.18 | 0.04 | 0.14 | 4.16 |
| 7 | Frmd4b | 0.88 | 1.14E-75 | 2.36E-71 | 0.41 | 0.16 | 0.25 | 2.58 |
| 7 | Hdgf | 0.39 | 1.86E-75 | 3.86E-71 | 1.00 | 0.97 | 0.02 | 1.02 |
| 7 | Sdc4 | 0.92 | 6.90E-75 | 1.43E-70 | 0.64 | 0.33 | 0.30 | 1.90 |
| 7 | Mid1ip1 | 0.92 | 8.83E-75 | 1.84E-70 | 0.47 | 0.21 | 0.26 | 2.27 |
| 7 | Psma7 | 0.36 | 3.12E-74 | 6.49E-70 | 1.00 | 0.99 | 0.01 | 1.01 |
| 7 | Fzd8 | 1.29 | 4.29E-74 | 8.93E-70 | 0.36 | 0.14 | 0.22 | 2.61 |

**Supplementary Table 1. Marker genes for the original clusters of integrated clustering of E11.5 control and NotchGOF λ-junction cells.**

| | | | | | | | | |
|---|---|---|---|---|---|---|---|---|
| 7 | Klhl13 | 1.09 | 4.34E-74 | 9.02E-70 | 0.53 | 0.26 | 0.27 | 2.06 |
| 7 | Cryab | 0.98 | 5.18E-74 | 1.08E-69 | 0.18 | 0.04 | 0.14 | 4.16 |
| 7 | Chd7 | 0.73 | 6.24E-74 | 1.30E-69 | 0.86 | 0.65 | 0.21 | 1.32 |
| 7 | Elf3 | 1.59 | 6.37E-74 | 1.33E-69 | 0.23 | 0.07 | 0.17 | 3.48 |
| 7 | Sox13 | 1.43 | 1.17E-73 | 2.43E-69 | 0.26 | 0.08 | 0.18 | 3.20 |
| 7 | Ccdc85c | 1.51 | 1.23E-73 | 2.55E-69 | 0.30 | 0.11 | 0.20 | 2.87 |
| 7 | Mkrn1 | 0.88 | 3.68E-73 | 7.66E-69 | 0.70 | 0.42 | 0.28 | 1.68 |
| 7 | Ppa1 | 0.59 | 1.23E-72 | 2.57E-68 | 0.97 | 0.89 | 0.07 | 1.08 |
| 7 | Kctd6 | 1.21 | 3.12E-72 | 6.48E-68 | 0.38 | 0.15 | 0.23 | 2.55 |
| 7 | Rad18 | 1.03 | 3.42E-72 | 7.12E-68 | 0.63 | 0.36 | 0.27 | 1.76 |
| 7 | Ntm | 1.37 | 3.63E-72 | 7.55E-68 | 0.29 | 0.10 | 0.19 | 2.94 |
| 7 | Vwa2 | 2.11 | 4.23E-72 | 8.80E-68 | 0.15 | 0.03 | 0.11 | 4.56 |
| 7 | Klf5 | 0.87 | 4.75E-72 | 9.89E-68 | 0.33 | 0.12 | 0.21 | 2.84 |
| 7 | Plxnb1 | 1.09 | 6.27E-72 | 1.30E-67 | 0.52 | 0.25 | 0.27 | 2.08 |
| 7 | Shtn1 | 1.42 | 6.75E-72 | 1.40E-67 | 0.19 | 0.05 | 0.14 | 3.94 |
| 7 | Gdap1 | 2.04 | 9.88E-71 | 2.05E-66 | 0.11 | 0.02 | 0.09 | 5.89 |
| 7 | Lratd2 | 1.11 | 1.17E-70 | 2.42E-66 | 0.30 | 0.10 | 0.20 | 2.92 |
| 7 | Il17re | 1.82 | 2.41E-70 | 5.02E-66 | 0.17 | 0.04 | 0.13 | 4.10 |
| 7 | Cnksr1 | 2.15 | 3.19E-70 | 6.63E-66 | 0.11 | 0.02 | 0.09 | 5.94 |
| 7 | Fam214a | 1.55 | 3.95E-70 | 8.22E-66 | 0.28 | 0.09 | 0.18 | 2.96 |
| 7 | Myo10 | 0.81 | 4.35E-70 | 9.04E-66 | 0.74 | 0.46 | 0.28 | 1.61 |
| 7 | Epb41l4a | 1.34 | 8.40E-70 | 1.75E-65 | 0.29 | 0.10 | 0.19 | 2.92 |
| 7 | Ubash3b | 1.07 | 1.20E-69 | 2.50E-65 | 0.48 | 0.23 | 0.25 | 2.12 |
| 7 | Dipk1c | 2.25 | 2.00E-69 | 4.16E-65 | 0.10 | 0.02 | 0.09 | 6.06 |
| 7 | Irs1 | 1.27 | 5.09E-69 | 1.06E-64 | 0.40 | 0.17 | 0.23 | 2.38 |
| 7 | Rassf9 | 1.12 | 8.55E-69 | 1.78E-64 | 0.38 | 0.16 | 0.23 | 2.47 |
| 7 | Fdps | 0.74 | 4.80E-68 | 9.98E-64 | 0.93 | 0.81 | 0.12 | 1.14 |
| 7 | Large2 | 1.45 | 5.38E-68 | 1.12E-63 | 0.25 | 0.08 | 0.17 | 3.11 |
| 7 | Ctnnal1 | 0.96 | 7.95E-68 | 1.65E-63 | 0.52 | 0.25 | 0.27 | 2.05 |
| 7 | Klhl23 | 0.97 | 9.15E-68 | 1.90E-63 | 0.64 | 0.36 | 0.28 | 1.76 |
| 7 | Sfrp1 | 0.75 | 9.74E-68 | 2.02E-63 | 0.83 | 0.62 | 0.22 | 1.35 |
| 7 | Mfsd13a | 2.03 | 1.04E-67 | 2.16E-63 | 0.15 | 0.03 | 0.12 | 4.38 |
| 7 | Cacnb3 | 0.95 | 1.14E-67 | 2.36E-63 | 0.54 | 0.26 | 0.28 | 2.05 |
| 7 | Gldc | 1.91 | 1.24E-67 | 2.59E-63 | 0.16 | 0.04 | 0.12 | 4.22 |
| 7 | B830012L14Rik | 1.62 | 2.23E-67 | 4.65E-63 | 0.19 | 0.05 | 0.14 | 3.80 |
| 7 | Ly75 | 1.29 | 4.45E-67 | 9.25E-63 | 0.36 | 0.14 | 0.22 | 2.50 |
| 7 | Cndp2 | 0.96 | 5.86E-67 | 1.22E-62 | 0.63 | 0.36 | 0.26 | 1.73 |
| 7 | Wscd1 | 2.14 | 9.23E-67 | 1.92E-62 | 0.12 | 0.02 | 0.09 | 5.27 |
| 7 | Pnck | 1.53 | 1.14E-66 | 2.37E-62 | 0.22 | 0.07 | 0.16 | 3.38 |
| 7 | Dennd1c | 1.88 | 2.01E-66 | 4.18E-62 | 0.13 | 0.03 | 0.11 | 4.62 |
| 7 | Cacna1b | 1.25 | 3.79E-66 | 7.89E-62 | 0.24 | 0.08 | 0.17 | 3.20 |
| 7 | Mtmr7 | 1.64 | 5.99E-66 | 1.25E-61 | 0.18 | 0.05 | 0.13 | 3.85 |
| 7 | Meis2 | 0.40 | 1.07E-65 | 2.22E-61 | 0.98 | 0.66 | 0.32 | 1.49 |
| 7 | Thbs3 | 1.37 | 1.68E-65 | 3.50E-61 | 0.27 | 0.09 | 0.18 | 2.91 |
| 7 | Tm7sf2 | 1.34 | 2.10E-65 | 4.37E-61 | 0.34 | 0.14 | 0.21 | 2.49 |
| 7 | Isyna1 | 0.68 | 3.82E-65 | 7.96E-61 | 0.86 | 0.67 | 0.19 | 1.28 |
| 7 | Tmsb4x | 0.73 | 4.31E-65 | 8.95E-61 | 1.00 | 0.99 | 0.01 | 1.01 |
| 7 | Cybrd1 | 1.70 | 4.31E-65 | 8.96E-61 | 0.17 | 0.05 | 0.13 | 3.87 |
| 7 | Aldoc | 2.37 | 4.37E-65 | 9.09E-61 | 0.11 | 0.02 | 0.09 | 5.38 |
| 7 | Pfkl | 0.72 | 6.49E-65 | 1.35E-60 | 0.85 | 0.64 | 0.22 | 1.34 |
| 7 | Pcbp4 | 0.50 | 7.15E-65 | 1.49E-60 | 0.97 | 0.86 | 0.10 | 1.12 |
| 7 | Fzd6 | 1.24 | 8.79E-65 | 1.83E-60 | 0.25 | 0.08 | 0.17 | 3.13 |
| 7 | B3gnt2 | 1.18 | 9.45E-65 | 1.97E-60 | 0.27 | 0.09 | 0.18 | 2.91 |
| 7 | Tubb4b | 0.63 | 2.22E-64 | 4.62E-60 | 0.95 | 0.85 | 0.11 | 1.12 |
| 7 | Retreg1 | 1.14 | 2.46E-64 | 5.11E-60 | 0.26 | 0.09 | 0.17 | 3.02 |
| 7 | Gnptg | 1.13 | 1.18E-63 | 2.46E-59 | 0.43 | 0.19 | 0.23 | 2.21 |

**Supplementary Table 1. Marker genes for the original clusters of integrated clustering of E11.5 control and NotchGOF λ-junction cells.**

| | | | | | | | | |
|---|---|---|---|---|---|---|---|---|
| 7 | Tacstd2 | 0.57 | 1.19E-63 | 2.47E-59 | 0.22 | 0.06 | 0.15 | 3.36 |
| 7 | Hes1 | 0.87 | 1.34E-63 | 2.79E-59 | 0.79 | 0.54 | 0.25 | 1.47 |
| 7 | Slc7a5 | 2.05 | 2.30E-63 | 4.78E-59 | 0.17 | 0.04 | 0.12 | 3.84 |
| 7 | Phf21b | 2.00 | 2.77E-63 | 5.75E-59 | 0.15 | 0.04 | 0.12 | 4.11 |
| 7 | Tgfa | 2.30 | 3.80E-63 | 7.90E-59 | 0.11 | 0.02 | 0.09 | 5.53 |
| 7 | Cblc | 1.90 | 4.29E-63 | 8.93E-59 | 0.13 | 0.03 | 0.10 | 4.55 |
| 7 | Ehbp1 | 0.90 | 5.71E-63 | 1.19E-58 | 0.48 | 0.23 | 0.26 | 2.11 |
| 7 | Mpst | 0.86 | 5.85E-63 | 1.22E-58 | 0.67 | 0.41 | 0.26 | 1.64 |
| 7 | Tceal3 | 1.29 | 6.20E-63 | 1.29E-58 | 0.30 | 0.11 | 0.19 | 2.71 |
| 7 | Phgdh | 0.50 | 6.42E-63 | 1.33E-58 | 0.95 | 0.82 | 0.14 | 1.17 |
| 7 | Wdsub1 | 1.10 | 1.14E-62 | 2.37E-58 | 0.43 | 0.20 | 0.24 | 2.19 |
| 7 | Myh9 | 0.69 | 1.31E-62 | 2.72E-58 | 0.76 | 0.50 | 0.26 | 1.53 |
| 7 | Ptpn3 | 2.13 | 2.16E-62 | 4.49E-58 | 0.14 | 0.03 | 0.11 | 4.24 |
| 7 | Lhfpl4 | 1.94 | 2.45E-62 | 5.09E-58 | 0.13 | 0.03 | 0.10 | 4.70 |
| 7 | Sgms2 | 2.31 | 3.12E-62 | 6.49E-58 | 0.11 | 0.02 | 0.09 | 5.53 |
| 7 | Tshz3 | 1.54 | 7.05E-62 | 1.47E-57 | 0.27 | 0.10 | 0.18 | 2.79 |
| 7 | Itga3 | 1.42 | 7.25E-62 | 1.51E-57 | 0.21 | 0.06 | 0.15 | 3.33 |
| 7 | Gstt3 | 1.61 | 8.67E-62 | 1.80E-57 | 0.17 | 0.04 | 0.12 | 3.82 |
| 7 | Frrs1 | 1.01 | 8.96E-62 | 1.86E-57 | 0.46 | 0.21 | 0.24 | 2.15 |
| 7 | Slc35g1 | 1.59 | 1.11E-61 | 2.32E-57 | 0.19 | 0.05 | 0.13 | 3.56 |
| 7 | Dnmbp | 1.25 | 1.25E-61 | 2.59E-57 | 0.31 | 0.12 | 0.19 | 2.61 |
| 7 | Plppr1 | 1.86 | 1.33E-61 | 2.77E-57 | 0.16 | 0.04 | 0.12 | 3.95 |
| 7 | Nr2f2 | 0.49 | 1.42E-61 | 2.95E-57 | 0.66 | 0.35 | 0.31 | 1.87 |
| 7 | Pmm1 | 1.04 | 1.66E-61 | 3.44E-57 | 0.51 | 0.26 | 0.25 | 1.94 |
| 7 | Crabp2 | 0.41 | 1.72E-61 | 3.58E-57 | 0.99 | 0.91 | 0.08 | 1.09 |
| 7 | Dtx3l | 1.89 | 1.86E-61 | 3.86E-57 | 0.15 | 0.04 | 0.11 | 4.05 |
| 7 | Lrp11 | 1.86 | 2.04E-61 | 4.24E-57 | 0.14 | 0.03 | 0.11 | 4.42 |
| 7 | Bphl | 0.89 | 2.26E-61 | 4.71E-57 | 0.64 | 0.37 | 0.27 | 1.73 |
| 7 | Stxbp2 | 1.22 | 3.64E-61 | 7.58E-57 | 0.33 | 0.13 | 0.20 | 2.53 |
| 7 | Exosc7 | 0.69 | 4.29E-61 | 8.92E-57 | 0.82 | 0.61 | 0.21 | 1.34 |
| 7 | Tspan33 | 0.51 | 4.97E-61 | 1.03E-56 | 0.24 | 0.08 | 0.17 | 3.14 |
| 7 | Rab27b | 1.01 | 6.17E-61 | 1.28E-56 | 0.26 | 0.09 | 0.17 | 2.89 |
| 7 | Egfl6 | 1.32 | 6.93E-61 | 1.44E-56 | 0.27 | 0.10 | 0.18 | 2.80 |
| 7 | Tedc1 | 1.22 | 7.15E-61 | 1.49E-56 | 0.32 | 0.12 | 0.19 | 2.58 |
| 7 | Grb14 | 1.51 | 8.22E-61 | 1.71E-56 | 0.16 | 0.04 | 0.12 | 3.88 |
| 7 | Tmem189 | 1.05 | 1.23E-60 | 2.56E-56 | 0.54 | 0.29 | 0.25 | 1.84 |
| 7 | Smim10l2a | 1.42 | 2.34E-60 | 4.87E-56 | 0.21 | 0.07 | 0.15 | 3.26 |
| 7 | Echdc1 | 1.02 | 7.69E-60 | 1.60E-55 | 0.47 | 0.23 | 0.24 | 2.04 |
| 7 | Sort1 | 1.09 | 1.16E-59 | 2.42E-55 | 0.42 | 0.20 | 0.23 | 2.17 |
| 7 | Lypd6 | 1.25 | 1.81E-59 | 3.76E-55 | 0.31 | 0.12 | 0.19 | 2.56 |
| 7 | Ddah1 | 1.04 | 3.11E-59 | 6.47E-55 | 0.33 | 0.13 | 0.20 | 2.51 |
| 7 | Acvr2b | 0.68 | 3.36E-59 | 6.98E-55 | 0.84 | 0.64 | 0.21 | 1.32 |
| 7 | Nrip1 | 0.56 | 4.52E-59 | 9.39E-55 | 0.91 | 0.75 | 0.16 | 1.21 |
| 7 | Dyrk3 | 1.46 | 7.17E-59 | 1.49E-54 | 0.26 | 0.09 | 0.17 | 2.80 |
| 7 | Srebf1 | 1.23 | 9.32E-59 | 1.94E-54 | 0.40 | 0.18 | 0.22 | 2.20 |
| 7 | Msi1 | 0.92 | 2.36E-58 | 4.90E-54 | 0.61 | 0.36 | 0.25 | 1.69 |
| 7 | Spata13 | 1.57 | 4.52E-58 | 9.40E-54 | 0.22 | 0.07 | 0.15 | 3.04 |
| 7 | Hist3h2a | 0.88 | 4.90E-58 | 1.02E-53 | 0.50 | 0.26 | 0.25 | 1.97 |
| 7 | Daam1 | 0.99 | 5.13E-58 | 1.07E-53 | 0.64 | 0.40 | 0.24 | 1.61 |
| 7 | Pdpn | 1.14 | 5.32E-58 | 1.11E-53 | 0.39 | 0.18 | 0.21 | 2.20 |
| 7 | Pcdh7 | 0.65 | 6.78E-58 | 1.41E-53 | 0.69 | 0.38 | 0.31 | 1.80 |
| 7 | Prkcz | 1.23 | 7.49E-58 | 1.56E-53 | 0.27 | 0.10 | 0.17 | 2.75 |
| 7 | Ablim1 | 0.86 | 7.50E-58 | 1.56E-53 | 0.46 | 0.22 | 0.24 | 2.12 |
| 7 | Tmx2 | 0.80 | 8.92E-58 | 1.85E-53 | 0.74 | 0.48 | 0.26 | 1.54 |
| 7 | Hip1r | 1.35 | 1.10E-57 | 2.28E-53 | 0.22 | 0.07 | 0.15 | 3.08 |
| 7 | Zbtb18 | 1.21 | 2.01E-57 | 4.19E-53 | 0.28 | 0.10 | 0.18 | 2.70 |

**Supplementary Table 1. Marker genes for the original clusters of integrated clustering of E11.5 control and NotchGOF λ-junction cells.**

| | | | | | | | | |
|---|---|---|---|---|---|---|---|---|
| 7 | Dsel | 0.97 | 4.31E-57 | 8.96E-53 | 0.58 | 0.32 | 0.25 | 1.79 |
| 7 | Prss22 | 1.61 | 4.54E-57 | 9.44E-53 | 0.19 | 0.06 | 0.13 | 3.30 |
| 7 | Slc2a3 | 0.37 | 6.08E-57 | 1.27E-52 | 0.33 | 0.13 | 0.20 | 2.50 |
| 7 | Dhrs4 | 0.69 | 7.30E-57 | 1.52E-52 | 0.83 | 0.63 | 0.20 | 1.31 |
| 7 | Col9a2 | 0.78 | 8.31E-57 | 1.73E-52 | 0.27 | 0.10 | 0.17 | 2.79 |
| 7 | Fam126a | 1.11 | 1.11E-56 | 2.32E-52 | 0.47 | 0.24 | 0.23 | 1.95 |
| 7 | Fgf11 | 2.10 | 1.82E-56 | 3.80E-52 | 0.10 | 0.02 | 0.08 | 5.32 |
| 7 | Bcar3 | 1.61 | 3.35E-56 | 6.96E-52 | 0.17 | 0.05 | 0.12 | 3.61 |
| 7 | Eif1ax | 0.40 | 3.94E-56 | 8.20E-52 | 0.97 | 0.94 | 0.04 | 1.04 |
| 7 | Gltp | 0.83 | 4.19E-56 | 8.71E-52 | 0.63 | 0.39 | 0.24 | 1.63 |
| 7 | Garem1 | 1.38 | 6.48E-56 | 1.35E-51 | 0.23 | 0.08 | 0.15 | 2.94 |
| 7 | Pradc1 | 1.11 | 7.95E-56 | 1.65E-51 | 0.43 | 0.21 | 0.22 | 2.07 |
| 7 | Abca2 | 1.96 | 9.05E-56 | 1.88E-51 | 0.12 | 0.03 | 0.09 | 4.58 |
| 7 | Aplp2 | 0.52 | 1.88E-55 | 3.91E-51 | 0.94 | 0.82 | 0.12 | 1.15 |
| 7 | Med19 | 0.57 | 1.97E-55 | 4.10E-51 | 0.89 | 0.74 | 0.15 | 1.20 |
| 7 | Wdr31 | 1.41 | 2.02E-55 | 4.19E-51 | 0.24 | 0.08 | 0.15 | 2.83 |
| 7 | Egln3 | 1.56 | 2.83E-55 | 5.89E-51 | 0.21 | 0.07 | 0.14 | 3.04 |
| 7 | Tmem159 | 0.98 | 4.70E-55 | 9.77E-51 | 0.49 | 0.25 | 0.24 | 1.94 |
| 7 | Zdhhc21 | 0.94 | 4.77E-55 | 9.92E-51 | 0.52 | 0.28 | 0.24 | 1.85 |
| 7 | Cyp39a1 | 1.12 | 1.55E-54 | 3.23E-50 | 0.34 | 0.14 | 0.20 | 2.37 |
| 7 | Cd151 | 0.75 | 1.69E-54 | 3.51E-50 | 0.58 | 0.32 | 0.26 | 1.82 |
| 7 | Abhd11 | 0.92 | 1.80E-54 | 3.73E-50 | 0.50 | 0.26 | 0.24 | 1.93 |
| 7 | Tcf7l2 | 0.68 | 2.35E-54 | 4.88E-50 | 0.81 | 0.60 | 0.21 | 1.35 |
| 7 | Lmln | 1.58 | 2.56E-54 | 5.33E-50 | 0.17 | 0.05 | 0.12 | 3.41 |
| 7 | Slc8a2 | 2.04 | 2.61E-54 | 5.43E-50 | 0.12 | 0.03 | 0.09 | 4.46 |
| 7 | Syt5 | 1.38 | 2.96E-54 | 6.15E-50 | 0.18 | 0.05 | 0.13 | 3.33 |
| 7 | Gm13648 | 1.77 | 3.88E-54 | 8.07E-50 | 0.11 | 0.02 | 0.08 | 4.82 |
| 7 | Irf2 | 0.87 | 4.03E-54 | 8.39E-50 | 0.49 | 0.25 | 0.24 | 1.98 |
| 7 | Pkdcc | 0.52 | 5.30E-54 | 1.10E-49 | 0.59 | 0.32 | 0.27 | 1.84 |
| 7 | Atp6v1f | 0.41 | 6.01E-54 | 1.25E-49 | 0.96 | 0.90 | 0.06 | 1.07 |
| 7 | Cbarp | 1.26 | 6.73E-54 | 1.40E-49 | 0.27 | 0.10 | 0.17 | 2.65 |
| 7 | Tubb4a | 1.69 | 6.81E-54 | 1.42E-49 | 0.13 | 0.03 | 0.10 | 4.06 |
| 7 | Galnt16 | 1.98 | 7.74E-54 | 1.61E-49 | 0.12 | 0.03 | 0.09 | 4.37 |
| 7 | Rrp12 | 1.14 | 8.24E-54 | 1.71E-49 | 0.40 | 0.19 | 0.21 | 2.11 |
| 7 | Lurap1l | 1.24 | 1.92E-53 | 3.99E-49 | 0.21 | 0.07 | 0.14 | 2.97 |
| 7 | Thrb | 1.51 | 1.95E-53 | 4.06E-49 | 0.20 | 0.07 | 0.14 | 3.03 |
| 7 | Msmo1 | 0.84 | 2.18E-53 | 4.54E-49 | 0.73 | 0.49 | 0.24 | 1.48 |
| 7 | Gpd2 | 0.98 | 4.32E-53 | 8.99E-49 | 0.44 | 0.22 | 0.22 | 2.02 |
| 7 | Dnajc1 | 0.78 | 5.02E-53 | 1.04E-48 | 0.76 | 0.55 | 0.21 | 1.39 |
| 7 | Ripk4 | 1.19 | 5.63E-53 | 1.17E-48 | 0.21 | 0.07 | 0.14 | 3.04 |
| 7 | Alad | 0.37 | 5.64E-53 | 1.17E-48 | 0.83 | 0.63 | 0.20 | 1.32 |
| 7 | Tmem200a | 1.68 | 5.73E-53 | 1.19E-48 | 0.17 | 0.05 | 0.12 | 3.39 |
| 7 | Tubb2a | 0.81 | 6.62E-53 | 1.38E-48 | 0.52 | 0.29 | 0.23 | 1.81 |
| 7 | Pomp | 0.32 | 1.40E-52 | 2.91E-48 | 1.00 | 0.98 | 0.02 | 1.02 |
| 7 | Efna1 | 0.89 | 1.84E-52 | 3.84E-48 | 0.33 | 0.13 | 0.19 | 2.44 |
| 7 | Pacsin3 | 1.03 | 2.61E-52 | 5.42E-48 | 0.43 | 0.21 | 0.22 | 2.03 |
| 7 | Fhl1 | 1.16 | 4.19E-52 | 8.71E-48 | 0.32 | 0.14 | 0.18 | 2.34 |
| 7 | Trak1 | 1.25 | 4.34E-52 | 9.03E-48 | 0.28 | 0.11 | 0.17 | 2.51 |
| 7 | Cpne5 | 1.46 | 4.36E-52 | 9.07E-48 | 0.17 | 0.05 | 0.12 | 3.46 |
| 7 | Tspan3 | 0.42 | 6.32E-52 | 1.32E-47 | 0.98 | 0.92 | 0.07 | 1.08 |
| 7 | Agpat3 | 0.91 | 7.35E-52 | 1.53E-47 | 0.49 | 0.26 | 0.24 | 1.93 |
| 7 | Anxa2 | 0.79 | 1.28E-51 | 2.66E-47 | 0.65 | 0.41 | 0.25 | 1.61 |
| 7 | Cdc42ep5 | 0.77 | 1.36E-51 | 2.83E-47 | 0.70 | 0.46 | 0.24 | 1.52 |
| 7 | Idi1 | 0.73 | 1.46E-51 | 3.04E-47 | 0.81 | 0.59 | 0.23 | 1.39 |
| 7 | Nrtn | 1.30 | 2.09E-51 | 4.34E-47 | 0.33 | 0.15 | 0.18 | 2.23 |
| 7 | D030056L22Rik | 0.81 | 2.23E-51 | 4.63E-47 | 0.69 | 0.47 | 0.22 | 1.46 |

**Supplementary Table 1. Marker genes for the original clusters of integrated clustering of E11.5 control and NotchGOF λ-junction cells.**

| | | | | | | | | |
|---|---|---|---|---|---|---|---|---|
| 7 | Uqcr11 | 0.32 | 2.96E-51 | 6.15E-47 | 1.00 | 0.97 | 0.02 | 1.02 |
| 7 | Taf10 | 0.39 | 4.15E-51 | 8.62E-47 | 0.97 | 0.93 | 0.05 | 1.05 |
| 7 | Tshz1 | 0.83 | 4.88E-51 | 1.02E-46 | 0.56 | 0.31 | 0.25 | 1.81 |
| 7 | Dbndd2 | 1.32 | 5.91E-51 | 1.23E-46 | 0.22 | 0.08 | 0.14 | 2.87 |
| 7 | Glrx | 0.96 | 7.54E-51 | 1.57E-46 | 0.45 | 0.23 | 0.22 | 1.96 |
| 7 | H2-D1 | 0.85 | 8.47E-51 | 1.76E-46 | 0.46 | 0.23 | 0.23 | 1.98 |
| 7 | Retreg2 | 0.92 | 1.05E-50 | 2.18E-46 | 0.51 | 0.28 | 0.23 | 1.82 |
| 7 | Ovol2 | 1.68 | 1.18E-50 | 2.45E-46 | 0.11 | 0.03 | 0.09 | 4.35 |
| 7 | Zfp750 | 1.34 | 1.79E-50 | 3.72E-46 | 0.16 | 0.05 | 0.11 | 3.43 |
| 7 | Rhbdf1 | 1.35 | 2.01E-50 | 4.18E-46 | 0.25 | 0.10 | 0.16 | 2.63 |
| 7 | Lin28b | 1.02 | 2.16E-50 | 4.49E-46 | 0.41 | 0.20 | 0.21 | 2.07 |
| 7 | Kank1 | 0.99 | 4.62E-50 | 9.61E-46 | 0.35 | 0.15 | 0.19 | 2.28 |
| 7 | Nudt14 | 0.81 | 7.16E-50 | 1.49E-45 | 0.64 | 0.39 | 0.24 | 1.61 |
| 7 | Rgs9 | 1.38 | 7.31E-50 | 1.52E-45 | 0.22 | 0.08 | 0.14 | 2.79 |
| 7 | Eef2kmt | 1.03 | 7.45E-50 | 1.55E-45 | 0.38 | 0.18 | 0.20 | 2.15 |
| 7 | Vim | 0.37 | 9.80E-50 | 2.04E-45 | 0.94 | 0.83 | 0.10 | 1.12 |
| 7 | Snhg4 | 0.92 | 1.84E-49 | 3.83E-45 | 0.67 | 0.46 | 0.22 | 1.47 |
| 7 | Ripply3 | 1.31 | 2.04E-49 | 4.25E-45 | 0.18 | 0.06 | 0.12 | 3.14 |
| 7 | Psmg4 | 0.56 | 3.30E-49 | 6.86E-45 | 0.87 | 0.74 | 0.14 | 1.18 |
| 7 | Paqr4 | 1.75 | 6.52E-49 | 1.36E-44 | 0.13 | 0.03 | 0.09 | 3.91 |
| 7 | Plekhf2 | 0.88 | 7.59E-49 | 1.58E-44 | 0.44 | 0.22 | 0.22 | 1.99 |
| 7 | Cadm4 | 1.24 | 7.87E-49 | 1.64E-44 | 0.23 | 0.08 | 0.14 | 2.78 |
| 7 | Gca | 1.79 | 1.52E-48 | 3.16E-44 | 0.10 | 0.02 | 0.08 | 4.59 |
| 7 | Uchl3 | 0.49 | 1.55E-48 | 3.23E-44 | 0.92 | 0.80 | 0.12 | 1.15 |
| 7 | Gm26917 | 0.88 | 1.63E-48 | 3.38E-44 | 0.75 | 0.55 | 0.20 | 1.36 |
| 7 | Oard1 | 0.60 | 2.11E-48 | 4.39E-44 | 0.84 | 0.67 | 0.17 | 1.25 |
| 7 | Fam162a | 0.40 | 2.24E-48 | 4.66E-44 | 0.97 | 0.89 | 0.07 | 1.08 |
| 7 | Ppm1l | 0.96 | 2.38E-48 | 4.94E-44 | 0.43 | 0.21 | 0.21 | 2.00 |
| 7 | Gucy1b1 | 1.64 | 2.48E-48 | 5.16E-44 | 0.13 | 0.04 | 0.10 | 3.72 |
| 7 | Atp8b1 | 1.77 | 2.50E-48 | 5.19E-44 | 0.14 | 0.04 | 0.10 | 3.66 |
| 7 | Firre | 1.34 | 2.78E-48 | 5.77E-44 | 0.23 | 0.08 | 0.14 | 2.71 |
| 7 | Tceal9 | 0.31 | 3.00E-48 | 6.25E-44 | 1.00 | 0.98 | 0.02 | 1.02 |
| 7 | Usp53 | 1.25 | 3.54E-48 | 7.37E-44 | 0.21 | 0.07 | 0.14 | 2.92 |
| 7 | Dnajc6 | 2.03 | 3.56E-48 | 7.41E-44 | 0.10 | 0.02 | 0.08 | 4.59 |
| 7 | Pbx4 | 1.29 | 3.97E-48 | 8.26E-44 | 0.23 | 0.08 | 0.14 | 2.76 |
| 7 | Gm10260 | 0.45 | 4.11E-48 | 8.55E-44 | 0.94 | 0.85 | 0.10 | 1.12 |
| 7 | Stxbp6 | 1.09 | 5.78E-48 | 1.20E-43 | 0.30 | 0.13 | 0.18 | 2.36 |
| 7 | Gm9922 | 1.50 | 1.15E-47 | 2.39E-43 | 0.14 | 0.04 | 0.10 | 3.68 |
| 7 | Aprt | 0.38 | 1.73E-47 | 3.59E-43 | 0.97 | 0.92 | 0.05 | 1.05 |
| 7 | Sypl | 0.70 | 2.22E-47 | 4.61E-43 | 0.71 | 0.47 | 0.23 | 1.49 |
| 7 | Tuba1c | 0.88 | 2.25E-47 | 4.69E-43 | 0.65 | 0.43 | 0.22 | 1.51 |
| 7 | Lrrc1 | 1.05 | 2.67E-47 | 5.55E-43 | 0.32 | 0.14 | 0.18 | 2.25 |
| 7 | Pak4 | 0.81 | 2.80E-47 | 5.82E-43 | 0.61 | 0.38 | 0.24 | 1.62 |
| 7 | Ptpru | 0.98 | 2.99E-47 | 6.23E-43 | 0.34 | 0.15 | 0.19 | 2.25 |
| 7 | Pnpla2 | 1.22 | 3.45E-47 | 7.17E-43 | 0.23 | 0.08 | 0.14 | 2.69 |
| 7 | Tmem184a | 1.61 | 3.91E-47 | 8.14E-43 | 0.16 | 0.05 | 0.11 | 3.27 |
| 7 | Slc35d1 | 1.14 | 3.99E-47 | 8.30E-43 | 0.33 | 0.15 | 0.18 | 2.23 |
| 7 | Mrps28 | 0.65 | 4.81E-47 | 1.00E-42 | 0.80 | 0.61 | 0.19 | 1.30 |
| 7 | Strbp | 0.55 | 6.11E-47 | 1.27E-42 | 0.87 | 0.70 | 0.17 | 1.24 |
| 7 | Capsl | 1.65 | 7.16E-47 | 1.49E-42 | 0.10 | 0.02 | 0.08 | 4.48 |
| 7 | Syt11 | 1.02 | 7.62E-47 | 1.58E-42 | 0.50 | 0.30 | 0.21 | 1.69 |
| 7 | Lnx1 | 1.73 | 8.55E-47 | 1.78E-42 | 0.12 | 0.03 | 0.09 | 4.03 |
| 7 | Hspb8 | 0.72 | 9.50E-47 | 1.98E-42 | 0.31 | 0.13 | 0.18 | 2.34 |
| 7 | Vps36 | 0.48 | 1.34E-46 | 2.78E-42 | 0.91 | 0.79 | 0.12 | 1.15 |
| 7 | Tmem51 | 1.28 | 1.58E-46 | 3.28E-42 | 0.24 | 0.09 | 0.15 | 2.59 |
| 7 | Cdk2ap2 | 0.74 | 2.26E-46 | 4.69E-42 | 0.68 | 0.45 | 0.23 | 1.51 |

**Supplementary Table 1. Marker genes for the original clusters of integrated clustering of E11.5 control and NotchGOF λ-junction cells.**

| | | | | | | | | |
|---|---|---|---|---|---|---|---|---|
| 7 | Dmd | 1.21 | 5.12E-46 | 1.06E-41 | 0.23 | 0.09 | 0.14 | 2.65 |
| 7 | Tafa5 | 1.08 | 7.78E-46 | 1.62E-41 | 0.28 | 0.12 | 0.16 | 2.40 |
| 7 | Tfrc | 0.75 | 8.96E-46 | 1.86E-41 | 0.62 | 0.39 | 0.24 | 1.60 |
| 7 | Ell2 | 0.98 | 9.80E-46 | 2.04E-41 | 0.23 | 0.08 | 0.14 | 2.69 |
| 7 | Sod1 | 0.52 | 9.98E-46 | 2.08E-41 | 0.91 | 0.79 | 0.12 | 1.15 |
| 7 | F3 | 1.12 | 1.67E-45 | 3.48E-41 | 0.25 | 0.10 | 0.15 | 2.53 |
| 7 | Tdrp | 1.78 | 2.05E-45 | 4.27E-41 | 0.11 | 0.03 | 0.08 | 4.12 |
| 7 | Gatm | 1.07 | 2.08E-45 | 4.32E-41 | 0.13 | 0.04 | 0.10 | 3.59 |
| 7 | Rassf7 | 0.91 | 2.64E-45 | 5.49E-41 | 0.39 | 0.19 | 0.20 | 2.04 |
| 7 | Aktip | 0.83 | 3.29E-45 | 6.84E-41 | 0.50 | 0.28 | 0.22 | 1.79 |
| 7 | Cds1 | 1.60 | 3.61E-45 | 7.51E-41 | 0.13 | 0.04 | 0.10 | 3.64 |
| 7 | Anxa5 | 0.71 | 7.23E-45 | 1.50E-40 | 0.79 | 0.60 | 0.19 | 1.31 |
| 7 | Mpp7 | 0.75 | 8.07E-45 | 1.68E-40 | 0.60 | 0.36 | 0.23 | 1.64 |
| 7 | Dnase2a | 0.81 | 8.50E-45 | 1.77E-40 | 0.42 | 0.21 | 0.21 | 1.99 |
| 7 | Dlx6 | 0.87 | 9.70E-45 | 2.02E-40 | 0.22 | 0.08 | 0.14 | 2.71 |
| 7 | Sgsm2 | 1.43 | 1.03E-44 | 2.14E-40 | 0.17 | 0.05 | 0.11 | 3.09 |
| 7 | Dennd1b | 1.02 | 1.04E-44 | 2.17E-40 | 0.33 | 0.15 | 0.18 | 2.20 |
| 7 | Xylt1 | 1.46 | 1.05E-44 | 2.19E-40 | 0.17 | 0.06 | 0.12 | 3.07 |
| 7 | Igf2 | 0.61 | 1.10E-44 | 2.29E-40 | 0.80 | 0.57 | 0.23 | 1.40 |
| 7 | Erap1 | 1.90 | 1.84E-44 | 3.83E-40 | 0.13 | 0.03 | 0.09 | 3.68 |
| 7 | Plekhg1 | 0.93 | 1.89E-44 | 3.93E-40 | 0.30 | 0.13 | 0.17 | 2.32 |
| 7 | Prr5 | 1.02 | 2.10E-44 | 4.37E-40 | 0.27 | 0.11 | 0.16 | 2.42 |
| 7 | Plagl2 | 0.81 | 2.99E-44 | 6.22E-40 | 0.57 | 0.35 | 0.22 | 1.62 |
| 7 | Trim37 | 0.73 | 3.51E-44 | 7.30E-40 | 0.61 | 0.38 | 0.23 | 1.60 |
| 7 | Cebpb | 0.72 | 5.04E-44 | 1.05E-39 | 0.27 | 0.11 | 0.16 | 2.40 |
| 7 | Krt7 | 1.18 | 5.49E-44 | 1.14E-39 | 0.16 | 0.05 | 0.11 | 3.20 |
| 7 | Hmgn5 | 0.50 | 7.10E-44 | 1.48E-39 | 0.87 | 0.71 | 0.17 | 1.24 |
| 7 | Tuba1b | 0.28 | 7.85E-44 | 1.63E-39 | 1.00 | 0.99 | 0.01 | 1.01 |
| 7 | Mirg | 1.48 | 8.40E-44 | 1.75E-39 | 0.15 | 0.04 | 0.10 | 3.32 |
| 7 | Irf2bpl | 0.79 | 1.23E-43 | 2.56E-39 | 0.60 | 0.38 | 0.22 | 1.59 |
| 7 | Arl6ip5 | 0.78 | 1.29E-43 | 2.68E-39 | 0.54 | 0.31 | 0.23 | 1.74 |
| 7 | Pacsin1 | 1.30 | 1.44E-43 | 3.00E-39 | 0.24 | 0.10 | 0.15 | 2.49 |
| 7 | Ccdc92 | 1.33 | 1.46E-43 | 3.04E-39 | 0.21 | 0.08 | 0.13 | 2.72 |
| 7 | Armc10 | 0.57 | 1.50E-43 | 3.11E-39 | 0.76 | 0.52 | 0.24 | 1.47 |
| 7 | Pdgfrl | 0.89 | 3.60E-43 | 7.50E-39 | 0.37 | 0.18 | 0.19 | 2.09 |
| 7 | Iqgap1 | 0.60 | 4.23E-43 | 8.80E-39 | 0.70 | 0.46 | 0.24 | 1.51 |
| 7 | Fmc1 | 0.61 | 4.86E-43 | 1.01E-38 | 0.78 | 0.59 | 0.20 | 1.33 |
| 7 | Rhpn2 | 1.54 | 5.55E-43 | 1.16E-38 | 0.10 | 0.02 | 0.08 | 4.21 |
| 7 | Rtkn2 | 1.39 | 6.21E-43 | 1.29E-38 | 0.20 | 0.07 | 0.13 | 2.74 |
| 7 | Trim25 | 1.43 | 7.01E-43 | 1.46E-38 | 0.17 | 0.06 | 0.11 | 3.04 |
| 7 | Reep3 | 0.64 | 7.46E-43 | 1.55E-38 | 0.73 | 0.51 | 0.22 | 1.42 |
| 7 | Sft2d2 | 0.94 | 7.70E-43 | 1.60E-38 | 0.32 | 0.14 | 0.17 | 2.22 |
| 7 | Aftph | 0.85 | 8.90E-43 | 1.85E-38 | 0.45 | 0.24 | 0.21 | 1.86 |
| 7 | Trmt9b | 1.30 | 9.75E-43 | 2.03E-38 | 0.19 | 0.07 | 0.12 | 2.86 |
| 7 | 1700023F06Rik | 1.63 | 1.10E-42 | 2.28E-38 | 0.14 | 0.04 | 0.10 | 3.43 |
| 7 | Bok | 0.78 | 1.24E-42 | 2.58E-38 | 0.39 | 0.19 | 0.20 | 2.03 |
| 7 | H2afj | 0.42 | 1.42E-42 | 2.95E-38 | 0.92 | 0.81 | 0.12 | 1.14 |
| 7 | Rasal2 | 0.85 | 1.93E-42 | 4.02E-38 | 0.45 | 0.24 | 0.21 | 1.86 |
| 7 | Sapcd1 | 1.75 | 2.05E-42 | 4.26E-38 | 0.13 | 0.04 | 0.09 | 3.57 |
| 7 | Fam173a | 0.66 | 2.15E-42 | 4.48E-38 | 0.72 | 0.50 | 0.22 | 1.45 |
| 7 | Rab3ip | 0.76 | 2.26E-42 | 4.69E-38 | 0.52 | 0.30 | 0.22 | 1.75 |
| 7 | Atl1 | 1.47 | 2.67E-42 | 5.55E-38 | 0.15 | 0.04 | 0.10 | 3.30 |
| 7 | Lgmn | 0.54 | 4.03E-42 | 8.38E-38 | 0.49 | 0.28 | 0.21 | 1.76 |
| 7 | Pcdh11x | 0.79 | 5.55E-42 | 1.15E-37 | 0.25 | 0.10 | 0.15 | 2.47 |
| 7 | Slc48a1 | 1.03 | 6.18E-42 | 1.28E-37 | 0.34 | 0.16 | 0.18 | 2.09 |
| 7 | E2f8 | 0.95 | 6.46E-42 | 1.34E-37 | 0.39 | 0.20 | 0.19 | 1.96 |

**Supplementary Table 1. Marker genes for the original clusters of integrated clustering of E11.5 control and NotchGOF λ-junction cells.**

| | | | | | | | | |
|---|---|---|---|---|---|---|---|---|
| 7 | Lrig3 | 0.75 | 7.82E-42 | 1.63E-37 | 0.43 | 0.22 | 0.21 | 1.95 |
| 7 | Arfip2 | 0.68 | 8.21E-42 | 1.71E-37 | 0.64 | 0.41 | 0.23 | 1.56 |
| 7 | Zfp36l1 | 0.44 | 8.32E-42 | 1.73E-37 | 0.94 | 0.85 | 0.09 | 1.11 |
| 7 | Ccn2 | 1.73 | 9.28E-42 | 1.93E-37 | 0.12 | 0.03 | 0.09 | 3.56 |
| 7 | Carmil3 | 1.30 | 1.01E-41 | 2.10E-37 | 0.16 | 0.05 | 0.11 | 3.10 |
| 7 | Ttc9 | 1.42 | 1.10E-41 | 2.29E-37 | 0.13 | 0.04 | 0.09 | 3.49 |
| 7 | Ndufa12 | 0.32 | 1.45E-41 | 3.01E-37 | 0.99 | 0.95 | 0.03 | 1.03 |
| 7 | Cdc42se1 | 0.63 | 1.61E-41 | 3.34E-37 | 0.66 | 0.42 | 0.24 | 1.58 |
| 7 | Grk5 | 1.10 | 1.62E-41 | 3.37E-37 | 0.27 | 0.11 | 0.16 | 2.36 |
| 7 | Vsig10 | 1.43 | 1.74E-41 | 3.63E-37 | 0.16 | 0.05 | 0.11 | 3.10 |
| 7 | Lipg | 1.50 | 2.20E-41 | 4.57E-37 | 0.12 | 0.03 | 0.09 | 3.64 |
| 7 | Iqgap2 | 0.70 | 2.86E-41 | 5.94E-37 | 0.59 | 0.36 | 0.23 | 1.66 |
| 7 | Gtf3a | 0.54 | 3.24E-41 | 6.73E-37 | 0.82 | 0.64 | 0.18 | 1.28 |
| 7 | Sult4a1 | 1.44 | 3.98E-41 | 8.28E-37 | 0.12 | 0.04 | 0.09 | 3.51 |
| 7 | Prss12 | 1.47 | 4.98E-41 | 1.04E-36 | 0.15 | 0.05 | 0.10 | 3.19 |
| 7 | Fam181b | 1.14 | 5.00E-41 | 1.04E-36 | 0.25 | 0.10 | 0.15 | 2.40 |
| 7 | Cyba | 0.51 | 6.14E-41 | 1.28E-36 | 0.68 | 0.44 | 0.23 | 1.52 |
| 7 | Rab11fip4 | 1.00 | 6.79E-41 | 1.41E-36 | 0.29 | 0.13 | 0.16 | 2.26 |
| 7 | Bex2 | 0.41 | 9.62E-41 | 2.00E-36 | 0.95 | 0.84 | 0.11 | 1.13 |
| 7 | Rif1 | 0.46 | 1.83E-40 | 3.81E-36 | 0.91 | 0.80 | 0.11 | 1.13 |
| 7 | Sec22a | 0.76 | 1.96E-40 | 4.08E-36 | 0.50 | 0.29 | 0.22 | 1.76 |
| 7 | Knop1 | 0.36 | 2.14E-40 | 4.44E-36 | 0.97 | 0.90 | 0.06 | 1.07 |
| 7 | Acss2 | 1.46 | 3.17E-40 | 6.60E-36 | 0.19 | 0.07 | 0.12 | 2.70 |
| 7 | Slc3a2 | 0.63 | 3.22E-40 | 6.70E-36 | 0.79 | 0.62 | 0.17 | 1.27 |
| 7 | Mboat1 | 1.52 | 3.59E-40 | 7.46E-36 | 0.15 | 0.05 | 0.10 | 3.15 |
| 7 | Tacc2 | 0.71 | 4.03E-40 | 8.39E-36 | 0.68 | 0.48 | 0.20 | 1.43 |
| 7 | Nhp2 | 0.40 | 4.85E-40 | 1.01E-35 | 0.95 | 0.88 | 0.07 | 1.08 |
| 7 | Akap1 | 0.96 | 7.71E-40 | 1.60E-35 | 0.36 | 0.18 | 0.18 | 1.98 |
| 7 | Xpr1 | 0.71 | 1.11E-39 | 2.31E-35 | 0.47 | 0.27 | 0.21 | 1.78 |
| 7 | Vgll3 | 1.83 | 1.22E-39 | 2.55E-35 | 0.10 | 0.03 | 0.08 | 4.04 |
| 7 | Apba2 | 1.27 | 1.22E-39 | 2.55E-35 | 0.17 | 0.06 | 0.11 | 2.91 |
| 7 | Nucb2 | 1.00 | 2.19E-39 | 4.55E-35 | 0.32 | 0.15 | 0.17 | 2.12 |
| 7 | Dach1 | 0.62 | 2.20E-39 | 4.57E-35 | 0.62 | 0.38 | 0.24 | 1.65 |
| 7 | Grhpr | 0.71 | 3.32E-39 | 6.91E-35 | 0.64 | 0.43 | 0.21 | 1.48 |
| 7 | Tjp1 | 0.58 | 3.84E-39 | 7.99E-35 | 0.70 | 0.49 | 0.21 | 1.44 |
| 7 | Snrpn | 0.58 | 5.62E-39 | 1.17E-34 | 0.69 | 0.46 | 0.23 | 1.49 |
| 7 | Arl4c | 0.56 | 7.14E-39 | 1.48E-34 | 0.54 | 0.32 | 0.22 | 1.69 |
| 7 | Syne4 | 0.79 | 8.63E-39 | 1.79E-34 | 0.38 | 0.19 | 0.19 | 2.01 |
| 7 | Cux2 | 1.03 | 9.04E-39 | 1.88E-34 | 0.23 | 0.09 | 0.14 | 2.48 |
| 7 | Plch2 | 1.77 | 9.98E-39 | 2.08E-34 | 0.10 | 0.03 | 0.08 | 3.92 |
| 7 | Nme3 | 1.12 | 1.34E-38 | 2.79E-34 | 0.19 | 0.07 | 0.12 | 2.67 |
| 7 | Tpbg | 0.67 | 1.43E-38 | 2.97E-34 | 0.52 | 0.30 | 0.22 | 1.74 |
| 7 | Npepl1 | 0.68 | 1.80E-38 | 3.74E-34 | 0.54 | 0.32 | 0.22 | 1.68 |
| 7 | Bmyc | 0.56 | 2.53E-38 | 5.26E-34 | 0.80 | 0.62 | 0.18 | 1.29 |
| 7 | Ydjc | 1.32 | 3.36E-38 | 6.99E-34 | 0.20 | 0.08 | 0.12 | 2.56 |
| 7 | Cd47 | 0.66 | 3.68E-38 | 7.66E-34 | 0.65 | 0.43 | 0.22 | 1.50 |
| 7 | Cntnap2 | 1.37 | 6.77E-38 | 1.41E-33 | 0.15 | 0.05 | 0.10 | 3.04 |
| 7 | Rimklb | 0.67 | 8.60E-38 | 1.79E-33 | 0.54 | 0.32 | 0.22 | 1.70 |
| 7 | Ralgps1 | 1.35 | 1.17E-37 | 2.42E-33 | 0.13 | 0.04 | 0.09 | 3.28 |
| 7 | Entpd5 | 1.15 | 1.27E-37 | 2.64E-33 | 0.20 | 0.07 | 0.12 | 2.64 |
| 7 | Ece1 | 0.85 | 2.23E-37 | 4.63E-33 | 0.41 | 0.22 | 0.19 | 1.84 |
| 7 | Acsl3 | 0.45 | 2.25E-37 | 4.69E-33 | 0.89 | 0.73 | 0.16 | 1.22 |
| 7 | Hist2h2be | 1.49 | 2.38E-37 | 4.95E-33 | 0.12 | 0.03 | 0.08 | 3.44 |
| 7 | Slc38a1 | 0.42 | 4.00E-37 | 8.33E-33 | 0.92 | 0.76 | 0.16 | 1.20 |
| 7 | Gpm6b | 1.17 | 5.07E-37 | 1.05E-32 | 0.17 | 0.06 | 0.11 | 2.76 |
| 7 | Etl4 | 0.67 | 5.39E-37 | 1.12E-32 | 0.52 | 0.30 | 0.21 | 1.71 |

**Supplementary Table 1. Marker genes for the original clusters of integrated clustering of E11.5 control and NotchGOF λ-junction cells.**

| | | | | | | | | |
|---|---|---|---|---|---|---|---|---|
| 7 | Dtx4 | 1.64 | 6.50E-37 | 1.35E-32 | 0.12 | 0.04 | 0.09 | 3.32 |
| 7 | Pick1 | 0.77 | 7.95E-37 | 1.65E-32 | 0.45 | 0.25 | 0.20 | 1.82 |
| 7 | Itgb6 | 1.36 | 1.09E-36 | 2.26E-32 | 0.13 | 0.04 | 0.09 | 3.26 |
| 7 | Mest | 0.41 | 1.12E-36 | 2.32E-32 | 0.98 | 0.91 | 0.08 | 1.08 |
| 7 | Gdpd1 | 0.67 | 1.47E-36 | 3.05E-32 | 0.47 | 0.26 | 0.21 | 1.78 |
| 7 | Naf1 | 0.67 | 1.48E-36 | 3.07E-32 | 0.60 | 0.39 | 0.21 | 1.54 |
| 7 | B4galt5 | 1.13 | 4.38E-36 | 9.12E-32 | 0.24 | 0.10 | 0.13 | 2.33 |
| 7 | Atp11b | 0.68 | 4.75E-36 | 9.88E-32 | 0.63 | 0.41 | 0.21 | 1.52 |
| 7 | Pcm1 | 0.38 | 5.28E-36 | 1.10E-31 | 0.96 | 0.89 | 0.07 | 1.08 |
| 7 | Dynlt3 | 0.76 | 8.40E-36 | 1.75E-31 | 0.44 | 0.25 | 0.19 | 1.77 |
| 7 | Larp1b | 1.44 | 8.84E-36 | 1.84E-31 | 0.19 | 0.07 | 0.11 | 2.55 |
| 7 | Chmp2b | 0.53 | 9.53E-36 | 1.98E-31 | 0.76 | 0.56 | 0.21 | 1.37 |
| 7 | Hells | 0.50 | 1.14E-35 | 2.36E-31 | 0.89 | 0.77 | 0.12 | 1.15 |
| 7 | Tekt1 | 1.06 | 1.17E-35 | 2.43E-31 | 0.21 | 0.09 | 0.13 | 2.47 |
| 7 | Satb1 | 0.61 | 1.21E-35 | 2.51E-31 | 0.60 | 0.38 | 0.22 | 1.59 |
| 7 | Pon2 | 0.58 | 1.21E-35 | 2.51E-31 | 0.35 | 0.17 | 0.18 | 2.02 |
| 7 | Trim2 | 0.61 | 1.71E-35 | 3.56E-31 | 0.62 | 0.41 | 0.21 | 1.52 |
| 7 | Ptprm | 1.29 | 2.31E-35 | 4.80E-31 | 0.14 | 0.05 | 0.09 | 3.04 |
| 7 | Nedd9 | 0.80 | 2.40E-35 | 4.99E-31 | 0.33 | 0.16 | 0.17 | 2.03 |
| 7 | Crim1 | 0.85 | 2.70E-35 | 5.61E-31 | 0.37 | 0.19 | 0.17 | 1.90 |
| 7 | Ung | 0.82 | 3.01E-35 | 6.25E-31 | 0.53 | 0.33 | 0.20 | 1.59 |
| 7 | Hey2 | 0.78 | 3.22E-35 | 6.70E-31 | 0.18 | 0.07 | 0.11 | 2.70 |
| 7 | Gm20324 | 1.66 | 3.73E-35 | 7.75E-31 | 0.10 | 0.03 | 0.07 | 3.61 |
| 7 | Airn | 0.91 | 3.86E-35 | 8.03E-31 | 0.35 | 0.18 | 0.17 | 1.92 |
| 7 | Morn2 | 0.69 | 4.53E-35 | 9.42E-31 | 0.53 | 0.32 | 0.21 | 1.65 |
| 7 | Lama1 | 1.04 | 4.63E-35 | 9.64E-31 | 0.23 | 0.10 | 0.13 | 2.33 |
| 7 | Prkag2 | 0.77 | 5.08E-35 | 1.06E-30 | 0.40 | 0.22 | 0.18 | 1.85 |
| 7 | Map4k5 | 0.77 | 5.85E-35 | 1.22E-30 | 0.50 | 0.30 | 0.19 | 1.64 |
| 7 | Mfsd6 | 1.61 | 7.44E-35 | 1.55E-30 | 0.11 | 0.03 | 0.08 | 3.41 |
| 7 | Zmiz2 | 0.67 | 8.90E-35 | 1.85E-30 | 0.51 | 0.30 | 0.20 | 1.68 |
| 7 | Fhdc1 | 1.34 | 9.10E-35 | 1.89E-30 | 0.12 | 0.04 | 0.08 | 3.15 |
| 7 | Sapcd2 | 0.79 | 9.89E-35 | 2.06E-30 | 0.43 | 0.25 | 0.18 | 1.75 |
| 7 | Fgf17 | 0.80 | 1.48E-34 | 3.08E-30 | 0.13 | 0.04 | 0.09 | 3.18 |
| 7 | Hk2 | 0.77 | 1.64E-34 | 3.41E-30 | 0.44 | 0.25 | 0.19 | 1.77 |
| 7 | Golga4 | 0.53 | 1.68E-34 | 3.50E-30 | 0.80 | 0.64 | 0.16 | 1.25 |
| 7 | Cnbp | 0.26 | 1.83E-34 | 3.81E-30 | 1.00 | 0.98 | 0.01 | 1.01 |
| 7 | Cecr2 | 0.75 | 1.87E-34 | 3.88E-30 | 0.40 | 0.22 | 0.18 | 1.84 |
| 7 | Jpt1 | 0.37 | 1.93E-34 | 4.01E-30 | 0.98 | 0.94 | 0.03 | 1.03 |
| 7 | Cux1 | 0.45 | 2.11E-34 | 4.39E-30 | 0.85 | 0.69 | 0.16 | 1.23 |
| 7 | Gas2l3 | 0.78 | 2.24E-34 | 4.66E-30 | 0.34 | 0.18 | 0.17 | 1.95 |
| 7 | 3110082I17Rik | 0.74 | 2.28E-34 | 4.74E-30 | 0.49 | 0.29 | 0.20 | 1.69 |
| 7 | Igf2r | 0.71 | 2.62E-34 | 5.45E-30 | 0.68 | 0.49 | 0.19 | 1.38 |
| 7 | Pwwp3b | 0.81 | 2.67E-34 | 5.56E-30 | 0.50 | 0.31 | 0.19 | 1.62 |
| 7 | Dhcr7 | 0.78 | 3.71E-34 | 7.71E-30 | 0.52 | 0.32 | 0.20 | 1.62 |
| 7 | Cpd | 0.76 | 4.41E-34 | 9.17E-30 | 0.53 | 0.33 | 0.19 | 1.59 |
| 7 | Ddx39 | 0.41 | 5.20E-34 | 1.08E-29 | 0.90 | 0.79 | 0.12 | 1.15 |
| 7 | Snx3 | 0.32 | 5.48E-34 | 1.14E-29 | 0.94 | 0.89 | 0.05 | 1.06 |
| 7 | Sh3rf1 | 0.72 | 5.90E-34 | 1.23E-29 | 0.41 | 0.22 | 0.19 | 1.86 |
| 7 | Magoh | 0.30 | 7.20E-34 | 1.50E-29 | 0.97 | 0.94 | 0.04 | 1.04 |
| 7 | Fgf10 | 1.24 | 8.32E-34 | 1.73E-29 | 0.23 | 0.10 | 0.13 | 2.23 |
| 7 | Grip1 | 0.97 | 9.67E-34 | 2.01E-29 | 0.23 | 0.10 | 0.13 | 2.30 |
| 7 | Adamts17 | 1.34 | 1.01E-33 | 2.10E-29 | 0.11 | 0.03 | 0.08 | 3.32 |
| 7 | Cyfip2 | 0.83 | 1.18E-33 | 2.45E-29 | 0.21 | 0.09 | 0.12 | 2.43 |
| 7 | Nomo1 | 0.84 | 1.57E-33 | 3.27E-29 | 0.36 | 0.19 | 0.17 | 1.90 |
| 7 | Plek2 | 1.14 | 1.89E-33 | 3.94E-29 | 0.16 | 0.06 | 0.10 | 2.72 |
| 7 | H2-Q4 | 1.06 | 2.00E-33 | 4.17E-29 | 0.23 | 0.10 | 0.13 | 2.29 |

**Supplementary Table 1. Marker genes for the original clusters of integrated clustering of E11.5 control and NotchGOF λ-junction cells.**

| | | | | | | | | |
|---|---|---|---|---|---|---|---|---|
| 7 | Fbxo27 | 1.13 | 2.02E-33 | 4.20E-29 | 0.17 | 0.06 | 0.11 | 2.67 |
| 7 | 1810037I17Rik | 0.38 | 2.26E-33 | 4.71E-29 | 0.93 | 0.83 | 0.10 | 1.12 |
| 7 | Slc16a3 | 0.77 | 2.48E-33 | 5.16E-29 | 0.52 | 0.32 | 0.20 | 1.61 |
| 7 | Capg | 0.81 | 3.00E-33 | 6.24E-29 | 0.30 | 0.15 | 0.15 | 2.06 |
| 7 | Ormdl1 | 0.66 | 3.15E-33 | 6.56E-29 | 0.63 | 0.44 | 0.20 | 1.45 |
| 7 | Slc35a3 | 0.96 | 3.21E-33 | 6.67E-29 | 0.26 | 0.12 | 0.14 | 2.19 |
| 7 | Scarb1 | 0.82 | 3.62E-33 | 7.53E-29 | 0.31 | 0.15 | 0.16 | 2.03 |
| 7 | Hlcs | 0.99 | 4.12E-33 | 8.57E-29 | 0.27 | 0.13 | 0.14 | 2.14 |
| 7 | Macrod2 | 0.68 | 4.20E-33 | 8.74E-29 | 0.40 | 0.22 | 0.18 | 1.84 |
| 7 | Parp9 | 1.75 | 4.24E-33 | 8.82E-29 | 0.10 | 0.03 | 0.07 | 3.48 |
| 7 | Dvl1 | 0.99 | 6.01E-33 | 1.25E-28 | 0.27 | 0.13 | 0.14 | 2.12 |
| 7 | Hmga2 | 0.29 | 8.41E-33 | 1.75E-28 | 0.99 | 0.93 | 0.06 | 1.07 |
| 7 | Hspd1 | 0.30 | 9.19E-33 | 1.91E-28 | 1.00 | 0.98 | 0.02 | 1.02 |
| 7 | mt-Nd5 | 0.41 | 9.28E-33 | 1.93E-28 | 0.99 | 0.95 | 0.04 | 1.04 |
| 7 | Gpr19 | 1.45 | 1.08E-32 | 2.24E-28 | 0.14 | 0.05 | 0.09 | 2.88 |
| 7 | Cryl1 | 0.79 | 1.26E-32 | 2.62E-28 | 0.30 | 0.14 | 0.15 | 2.06 |
| 7 | Gart | 0.55 | 1.35E-32 | 2.81E-28 | 0.72 | 0.56 | 0.16 | 1.29 |
| 7 | Ttc12 | 1.55 | 1.54E-32 | 3.20E-28 | 0.12 | 0.04 | 0.08 | 3.11 |
| 7 | Cenpb | 0.50 | 1.64E-32 | 3.40E-28 | 0.81 | 0.66 | 0.15 | 1.23 |
| 7 | Tdrkh | 1.28 | 1.66E-32 | 3.46E-28 | 0.15 | 0.06 | 0.10 | 2.70 |
| 7 | E330009J07Rik | 0.99 | 1.77E-32 | 3.69E-28 | 0.25 | 0.11 | 0.13 | 2.18 |
| 7 | Dedd2 | 1.09 | 1.96E-32 | 4.07E-28 | 0.18 | 0.07 | 0.11 | 2.52 |
| 7 | Hist1h2bc | 0.70 | 2.15E-32 | 4.46E-28 | 0.44 | 0.26 | 0.19 | 1.72 |
| 7 | Hspa8 | 0.26 | 2.23E-32 | 4.65E-28 | 1.00 | 1.00 | 0.00 | 1.00 |
| 7 | Noc2l | 0.55 | 2.72E-32 | 5.66E-28 | 0.77 | 0.60 | 0.17 | 1.28 |
| 7 | Nectin2 | 0.60 | 3.31E-32 | 6.88E-28 | 0.67 | 0.47 | 0.20 | 1.42 |
| 7 | Kitl | 0.80 | 3.68E-32 | 7.65E-28 | 0.18 | 0.07 | 0.11 | 2.51 |
| 7 | Med12l | 1.19 | 3.84E-32 | 7.99E-28 | 0.13 | 0.04 | 0.09 | 3.02 |
| 7 | Zdhhc12 | 0.77 | 4.92E-32 | 1.02E-27 | 0.44 | 0.25 | 0.18 | 1.72 |
| 7 | Clip4 | 1.38 | 5.68E-32 | 1.18E-27 | 0.13 | 0.04 | 0.09 | 3.02 |
| 7 | Gprc5c | 0.72 | 5.87E-32 | 1.22E-27 | 0.41 | 0.23 | 0.18 | 1.80 |
| 7 | Carmil1 | 0.96 | 6.05E-32 | 1.26E-27 | 0.29 | 0.14 | 0.15 | 2.04 |
| 7 | P4ha3 | 0.80 | 6.46E-32 | 1.34E-27 | 0.31 | 0.15 | 0.15 | 2.01 |
| 7 | Hif1a | 0.50 | 6.62E-32 | 1.38E-27 | 0.86 | 0.75 | 0.11 | 1.15 |
| 7 | Ccdc115 | 0.58 | 7.09E-32 | 1.47E-27 | 0.68 | 0.48 | 0.20 | 1.42 |
| 7 | Nudt18 | 1.08 | 9.28E-32 | 1.93E-27 | 0.16 | 0.06 | 0.10 | 2.69 |
| 7 | Aldh5a1 | 0.69 | 9.70E-32 | 2.02E-27 | 0.42 | 0.24 | 0.19 | 1.79 |
| 7 | Nlk | 0.69 | 9.88E-32 | 2.06E-27 | 0.45 | 0.27 | 0.18 | 1.69 |
| 7 | Fam3c | 0.61 | 1.28E-31 | 2.66E-27 | 0.57 | 0.36 | 0.21 | 1.59 |
| 7 | Rragb | 1.40 | 1.42E-31 | 2.94E-27 | 0.19 | 0.08 | 0.11 | 2.35 |
| 7 | Snhg14 | 0.85 | 1.56E-31 | 3.24E-27 | 0.33 | 0.17 | 0.16 | 1.91 |
| 7 | Myo6 | 0.66 | 1.61E-31 | 3.34E-27 | 0.35 | 0.18 | 0.17 | 1.93 |
| 7 | Kcnk2 | 0.85 | 1.93E-31 | 4.01E-27 | 0.47 | 0.29 | 0.18 | 1.61 |
| 7 | Vgll4 | 0.39 | 2.29E-31 | 4.77E-27 | 0.90 | 0.78 | 0.12 | 1.16 |
| 7 | Mycl | 0.46 | 2.58E-31 | 5.37E-27 | 0.50 | 0.30 | 0.21 | 1.71 |
| 7 | Gar1 | 0.43 | 2.60E-31 | 5.40E-27 | 0.87 | 0.74 | 0.13 | 1.18 |
| 7 | Cdt1 | 0.67 | 2.62E-31 | 5.45E-27 | 0.57 | 0.38 | 0.19 | 1.49 |
| 7 | Polr2f | 0.28 | 3.31E-31 | 6.89E-27 | 0.98 | 0.95 | 0.03 | 1.03 |
| 7 | Slc1a4 | 1.45 | 3.36E-31 | 6.99E-27 | 0.17 | 0.07 | 0.10 | 2.49 |
| 7 | Fbl | 0.31 | 3.43E-31 | 7.14E-27 | 0.96 | 0.91 | 0.04 | 1.05 |
| 7 | Igsf8 | 0.77 | 3.79E-31 | 7.88E-27 | 0.42 | 0.24 | 0.18 | 1.76 |
| 7 | Abhd14b | 1.23 | 4.50E-31 | 9.35E-27 | 0.15 | 0.06 | 0.10 | 2.67 |
| 7 | Shroom2 | 1.09 | 4.87E-31 | 1.01E-26 | 0.18 | 0.07 | 0.11 | 2.49 |
| 7 | Camta1 | 0.50 | 6.77E-31 | 1.41E-26 | 0.78 | 0.61 | 0.16 | 1.27 |
| 7 | Exoc6 | 1.16 | 7.13E-31 | 1.48E-26 | 0.18 | 0.07 | 0.10 | 2.46 |
| 7 | Pkp3 | 0.92 | 7.65E-31 | 1.59E-26 | 0.17 | 0.06 | 0.10 | 2.62 |

**Supplementary Table 1. Marker genes for the original clusters of integrated clustering of E11.5 control and NotchGOF λ-junction cells.**

| | | | | | | | | |
|---|---|---|---|---|---|---|---|---|
| 7 | AC149090.1 | 0.77 | 7.73E-31 | 1.61E-26 | 0.46 | 0.28 | 0.18 | 1.66 |
| 7 | Map3k5 | 1.69 | 7.79E-31 | 1.62E-26 | 0.11 | 0.03 | 0.07 | 3.28 |
| 7 | Magi3 | 0.72 | 9.11E-31 | 1.90E-26 | 0.35 | 0.18 | 0.17 | 1.90 |
| 7 | Ptprz1 | 0.79 | 1.02E-30 | 2.12E-26 | 0.23 | 0.10 | 0.13 | 2.23 |
| 7 | Cpne3 | 0.56 | 1.04E-30 | 2.17E-26 | 0.70 | 0.52 | 0.18 | 1.35 |
| 7 | Ccdc8 | 0.63 | 1.10E-30 | 2.30E-26 | 0.52 | 0.32 | 0.20 | 1.62 |
| 7 | Epha5 | 0.75 | 1.18E-30 | 2.45E-26 | 0.17 | 0.07 | 0.10 | 2.56 |
| 7 | Wrb | 0.70 | 1.28E-30 | 2.66E-26 | 0.41 | 0.23 | 0.18 | 1.77 |
| 7 | Rapgef5 | 0.63 | 1.35E-30 | 2.81E-26 | 0.25 | 0.11 | 0.14 | 2.19 |
| 7 | Parva | 0.51 | 1.54E-30 | 3.21E-26 | 0.73 | 0.54 | 0.18 | 1.34 |
| 7 | Phb2 | 0.29 | 1.71E-30 | 3.56E-26 | 0.98 | 0.96 | 0.02 | 1.02 |
| 7 | Zfp395 | 0.73 | 1.75E-30 | 3.64E-26 | 0.41 | 0.23 | 0.18 | 1.78 |
| 7 | Alpl | 1.03 | 2.38E-30 | 4.94E-26 | 0.22 | 0.10 | 0.12 | 2.23 |
| 7 | Adam22 | 1.27 | 2.69E-30 | 5.59E-26 | 0.18 | 0.07 | 0.10 | 2.46 |
| 7 | Clcn3 | 0.68 | 2.82E-30 | 5.86E-26 | 0.56 | 0.37 | 0.18 | 1.49 |
| 7 | Raver2 | 0.86 | 3.17E-30 | 6.59E-26 | 0.31 | 0.16 | 0.15 | 1.94 |
| 7 | Kpnb1 | 0.39 | 3.52E-30 | 7.32E-26 | 0.91 | 0.84 | 0.07 | 1.09 |
| 7 | Dnajc21 | 0.52 | 3.74E-30 | 7.79E-26 | 0.75 | 0.59 | 0.16 | 1.27 |
| 7 | Patz1 | 0.50 | 4.22E-30 | 8.77E-26 | 0.70 | 0.51 | 0.18 | 1.35 |
| 7 | Fndc3b | 0.70 | 5.22E-30 | 1.09E-25 | 0.41 | 0.24 | 0.18 | 1.73 |
| 7 | Ostf1 | 0.51 | 5.63E-30 | 1.17E-25 | 0.46 | 0.27 | 0.19 | 1.70 |
| 7 | Rreb1 | 0.75 | 9.84E-30 | 2.05E-25 | 0.37 | 0.20 | 0.16 | 1.81 |
| 7 | Lbr | 0.58 | 1.01E-29 | 2.10E-25 | 0.68 | 0.50 | 0.18 | 1.37 |
| 7 | A930004D18Rik | 1.29 | 1.35E-29 | 2.81E-25 | 0.13 | 0.05 | 0.08 | 2.79 |
| 7 | Scp2 | 0.39 | 1.53E-29 | 3.18E-25 | 0.87 | 0.77 | 0.10 | 1.14 |
| 7 | Ahr | 0.85 | 1.66E-29 | 3.44E-25 | 0.20 | 0.09 | 0.11 | 2.33 |
| 7 | Pole3 | 0.41 | 1.70E-29 | 3.53E-25 | 0.88 | 0.76 | 0.13 | 1.17 |
| 7 | Dusp28 | 0.92 | 1.74E-29 | 3.62E-25 | 0.24 | 0.11 | 0.13 | 2.15 |
| 7 | Clip1 | 0.59 | 1.85E-29 | 3.85E-25 | 0.67 | 0.49 | 0.18 | 1.36 |
| 7 | St7 | 0.68 | 2.29E-29 | 4.76E-25 | 0.39 | 0.22 | 0.17 | 1.77 |
| 7 | Iqcg | 1.32 | 2.87E-29 | 5.97E-25 | 0.11 | 0.03 | 0.07 | 3.15 |
| 7 | Ino80b | 0.59 | 2.94E-29 | 6.12E-25 | 0.63 | 0.45 | 0.18 | 1.40 |
| 7 | G3bp1 | 0.25 | 4.13E-29 | 8.59E-25 | 0.99 | 0.97 | 0.02 | 1.02 |
| 7 | Adgrb3 | 1.14 | 4.24E-29 | 8.82E-25 | 0.16 | 0.06 | 0.10 | 2.57 |
| 7 | Tusc2 | 0.62 | 5.03E-29 | 1.05E-24 | 0.58 | 0.38 | 0.20 | 1.51 |
| 7 | Emg1 | 0.29 | 5.86E-29 | 1.22E-24 | 0.96 | 0.92 | 0.04 | 1.05 |
| 7 | 1700096K18Rik | 0.97 | 5.94E-29 | 1.24E-24 | 0.21 | 0.09 | 0.11 | 2.25 |
| 7 | Cd3eap | 0.55 | 6.13E-29 | 1.27E-24 | 0.70 | 0.52 | 0.18 | 1.35 |
| 7 | Myl12b | 0.27 | 9.48E-29 | 1.97E-24 | 0.96 | 0.88 | 0.07 | 1.08 |
| 7 | Amt | 0.77 | 1.39E-28 | 2.89E-24 | 0.28 | 0.14 | 0.14 | 1.98 |
| 7 | Plxnb2 | 0.54 | 1.43E-28 | 2.97E-24 | 0.73 | 0.53 | 0.20 | 1.38 |
| 7 | Kbtbd8 | 1.45 | 1.45E-28 | 3.02E-24 | 0.11 | 0.04 | 0.07 | 3.09 |
| 7 | Erp29 | 0.32 | 1.65E-28 | 3.44E-24 | 0.97 | 0.88 | 0.09 | 1.10 |
| 7 | Mcm6 | 0.47 | 1.84E-28 | 3.82E-24 | 0.83 | 0.70 | 0.14 | 1.20 |
| 7 | Npepps | 0.46 | 2.49E-28 | 5.19E-24 | 0.79 | 0.64 | 0.15 | 1.24 |
| 7 | Cyb5r4 | 0.58 | 2.51E-28 | 5.23E-24 | 0.58 | 0.39 | 0.19 | 1.48 |
| 7 | Baz2b | 0.43 | 2.52E-28 | 5.24E-24 | 0.84 | 0.70 | 0.13 | 1.19 |
| 7 | Sema4d | 1.10 | 2.53E-28 | 5.26E-24 | 0.17 | 0.07 | 0.10 | 2.46 |
| 7 | Plpp2 | 0.68 | 2.56E-28 | 5.33E-24 | 0.43 | 0.26 | 0.18 | 1.68 |
| 7 | Meis3 | 0.65 | 2.64E-28 | 5.49E-24 | 0.36 | 0.19 | 0.16 | 1.84 |
| 7 | Thnsl1 | 1.15 | 3.61E-28 | 7.50E-24 | 0.17 | 0.07 | 0.10 | 2.43 |
| 7 | Ipo5 | 0.47 | 5.11E-28 | 1.06E-23 | 0.83 | 0.70 | 0.13 | 1.19 |
| 7 | Prxl2a | 0.54 | 5.60E-28 | 1.16E-23 | 0.57 | 0.37 | 0.20 | 1.53 |
| 7 | B9d1 | 0.48 | 7.45E-28 | 1.55E-23 | 0.74 | 0.56 | 0.18 | 1.33 |
| 7 | Spred3 | 0.79 | 7.72E-28 | 1.61E-23 | 0.20 | 0.09 | 0.11 | 2.27 |
| 7 | Ap1g2 | 1.62 | 8.29E-28 | 1.72E-23 | 0.11 | 0.04 | 0.08 | 2.92 |

**Supplementary Table 1. Marker genes for the original clusters of integrated clustering of E11.5 control and NotchGOF λ-junction cells.**

| | | | | | | | | |
|---|---|---|---|---|---|---|---|---|
| 7 | Mettl27 | 0.74 | 8.38E-28 | 1.74E-23 | 0.27 | 0.14 | 0.14 | 2.01 |
| 7 | Crip1 | 0.40 | 1.03E-27 | 2.15E-23 | 0.60 | 0.40 | 0.20 | 1.52 |
| 7 | Slc1a3 | 0.54 | 1.04E-27 | 2.17E-23 | 0.39 | 0.22 | 0.17 | 1.77 |
| 7 | Cdc34 | 0.37 | 1.06E-27 | 2.21E-23 | 0.90 | 0.77 | 0.13 | 1.17 |
| 7 | Elf1 | 0.54 | 1.48E-27 | 3.07E-23 | 0.43 | 0.26 | 0.18 | 1.68 |
| 7 | Arl6ip1 | 0.41 | 1.98E-27 | 4.11E-23 | 0.96 | 0.89 | 0.07 | 1.08 |
| 7 | Map2 | 0.61 | 2.14E-27 | 4.45E-23 | 0.38 | 0.21 | 0.17 | 1.78 |
| 7 | Fkbp4 | 0.25 | 2.73E-27 | 5.69E-23 | 0.99 | 0.97 | 0.02 | 1.02 |
| 7 | Ccpg1os | 0.72 | 3.85E-27 | 8.00E-23 | 0.30 | 0.15 | 0.14 | 1.94 |
| 7 | Lonrf1 | 0.93 | 4.43E-27 | 9.21E-23 | 0.20 | 0.09 | 0.11 | 2.23 |
| 7 | Stap2 | 1.06 | 4.63E-27 | 9.63E-23 | 0.18 | 0.08 | 0.10 | 2.31 |
| 7 | Sema3f | 0.69 | 8.04E-27 | 1.67E-22 | 0.46 | 0.29 | 0.18 | 1.61 |
| 7 | Erg28 | 0.45 | 8.46E-27 | 1.76E-22 | 0.80 | 0.68 | 0.13 | 1.19 |
| 7 | Sema3d | 0.36 | 8.75E-27 | 1.82E-22 | 0.24 | 0.11 | 0.13 | 2.12 |
| 7 | Tlcd1 | 0.97 | 9.46E-27 | 1.97E-22 | 0.19 | 0.09 | 0.11 | 2.26 |
| 7 | Kif13b | 0.79 | 9.83E-27 | 2.04E-22 | 0.30 | 0.16 | 0.14 | 1.89 |
| 7 | Baz2a | 0.56 | 1.13E-26 | 2.36E-22 | 0.57 | 0.38 | 0.19 | 1.49 |
| 7 | Larp4b | 0.50 | 1.17E-26 | 2.43E-22 | 0.74 | 0.61 | 0.14 | 1.22 |
| 7 | Dmrta1 | 1.22 | 1.27E-26 | 2.64E-22 | 0.11 | 0.04 | 0.07 | 2.97 |
| 7 | Podxl | 1.17 | 1.33E-26 | 2.76E-22 | 0.12 | 0.04 | 0.08 | 2.83 |
| 7 | Galnt7 | 0.60 | 1.33E-26 | 2.76E-22 | 0.53 | 0.34 | 0.19 | 1.56 |
| 7 | Mark2 | 0.52 | 1.36E-26 | 2.82E-22 | 0.64 | 0.46 | 0.18 | 1.40 |
| 7 | Thumpd3 | 0.55 | 1.47E-26 | 3.06E-22 | 0.56 | 0.38 | 0.19 | 1.50 |
| 7 | Prom1 | 0.56 | 1.63E-26 | 3.38E-22 | 0.59 | 0.40 | 0.19 | 1.48 |
| 7 | Grik5 | 0.71 | 1.65E-26 | 3.42E-22 | 0.29 | 0.15 | 0.14 | 1.93 |
| 7 | Pura | 0.51 | 1.66E-26 | 3.46E-22 | 0.65 | 0.47 | 0.19 | 1.40 |
| 7 | Rhpn1 | 1.44 | 1.86E-26 | 3.88E-22 | 0.11 | 0.04 | 0.07 | 3.00 |
| 7 | Arf3 | 0.57 | 1.92E-26 | 3.99E-22 | 0.49 | 0.31 | 0.18 | 1.60 |
| 7 | Cenpf | 0.48 | 2.18E-26 | 4.53E-22 | 0.83 | 0.70 | 0.13 | 1.19 |
| 7 | Pigp | 0.40 | 2.46E-26 | 5.12E-22 | 0.80 | 0.64 | 0.16 | 1.25 |
| 7 | Cask | 0.56 | 2.59E-26 | 5.39E-22 | 0.52 | 0.33 | 0.19 | 1.56 |
| 7 | Mettl1 | 0.40 | 2.81E-26 | 5.85E-22 | 0.77 | 0.58 | 0.19 | 1.33 |
| 7 | Prelid1 | 0.27 | 2.94E-26 | 6.11E-22 | 0.96 | 0.93 | 0.04 | 1.04 |
| 7 | H2afx | 0.33 | 2.94E-26 | 6.11E-22 | 0.97 | 0.93 | 0.04 | 1.04 |
| 7 | 1700025G04Rik | 0.54 | 3.38E-26 | 7.03E-22 | 0.52 | 0.33 | 0.19 | 1.57 |
| 7 | Ptprk | 0.52 | 3.53E-26 | 7.34E-22 | 0.63 | 0.43 | 0.20 | 1.46 |
| 7 | Dnajc2 | 0.32 | 3.61E-26 | 7.51E-22 | 0.95 | 0.86 | 0.09 | 1.10 |
| 7 | Smim4 | 0.53 | 3.74E-26 | 7.78E-22 | 0.61 | 0.42 | 0.19 | 1.45 |
| 7 | Aagab | 0.73 | 4.11E-26 | 8.55E-22 | 0.32 | 0.17 | 0.15 | 1.86 |
| 7 | Prtg | 0.71 | 4.27E-26 | 8.89E-22 | 0.31 | 0.17 | 0.15 | 1.87 |
| 7 | Zcchc2 | 1.02 | 4.44E-26 | 9.24E-22 | 0.20 | 0.09 | 0.11 | 2.19 |
| 7 | Itsn2 | 0.69 | 5.52E-26 | 1.15E-21 | 0.33 | 0.19 | 0.15 | 1.79 |
| 7 | Sertad2 | 0.56 | 5.72E-26 | 1.19E-21 | 0.58 | 0.40 | 0.18 | 1.46 |
| 7 | Klf3 | 0.52 | 6.41E-26 | 1.33E-21 | 0.67 | 0.49 | 0.18 | 1.36 |
| 7 | Ccdc12 | 0.38 | 6.51E-26 | 1.35E-21 | 0.83 | 0.70 | 0.13 | 1.19 |
| 7 | Ncor1 | 0.29 | 6.75E-26 | 1.40E-21 | 0.96 | 0.91 | 0.05 | 1.05 |
| 7 | Tom1 | 0.79 | 8.11E-26 | 1.69E-21 | 0.23 | 0.11 | 0.12 | 2.08 |
| 7 | 2310039H08Rik | 0.53 | 8.20E-26 | 1.70E-21 | 0.60 | 0.41 | 0.18 | 1.44 |
| 7 | Sinhcaf | 0.30 | 1.13E-25 | 2.36E-21 | 0.91 | 0.84 | 0.08 | 1.09 |
| 7 | Wdr76 | 0.64 | 1.39E-25 | 2.89E-21 | 0.50 | 0.32 | 0.18 | 1.55 |
| 7 | Ccdc112 | 0.44 | 1.48E-25 | 3.07E-21 | 0.70 | 0.51 | 0.19 | 1.37 |
| 7 | H1fx | 0.30 | 1.59E-25 | 3.30E-21 | 0.98 | 0.93 | 0.05 | 1.06 |
| 7 | Ruvbl2 | 0.43 | 1.60E-25 | 3.32E-21 | 0.80 | 0.65 | 0.15 | 1.23 |
| 7 | Cdon | 0.69 | 1.70E-25 | 3.53E-21 | 0.37 | 0.22 | 0.16 | 1.73 |
| 7 | Slc9a9 | 1.25 | 1.74E-25 | 3.61E-21 | 0.11 | 0.04 | 0.07 | 2.92 |
| 7 | Nop10 | 0.25 | 1.77E-25 | 3.69E-21 | 0.98 | 0.95 | 0.03 | 1.03 |

**Supplementary Table 1. Marker genes for the original clusters of integrated clustering of E11.5 control and NotchGOF λ-junction cells.**

| | | | | | | | | |
|---|---|---|---|---|---|---|---|---|
| 7 | Impdh2 | 0.29 | 1.80E-25 | 3.73E-21 | 0.95 | 0.89 | 0.06 | 1.06 |
| 7 | Polr2h | 0.37 | 1.89E-25 | 3.93E-21 | 0.90 | 0.78 | 0.12 | 1.15 |
| 7 | Bid | 0.73 | 2.18E-25 | 4.53E-21 | 0.24 | 0.12 | 0.12 | 2.03 |
| 7 | Ppan | 0.54 | 2.39E-25 | 4.97E-21 | 0.62 | 0.44 | 0.18 | 1.41 |
| 7 | Gmpr | 0.40 | 2.83E-25 | 5.88E-21 | 0.32 | 0.17 | 0.15 | 1.89 |
| 7 | 2700033N17Rik | 0.66 | 3.50E-25 | 7.29E-21 | 0.24 | 0.11 | 0.12 | 2.07 |
| 7 | Atp1b1 | 0.58 | 4.42E-25 | 9.19E-21 | 0.19 | 0.08 | 0.10 | 2.23 |
| 7 | Syt7 | 0.99 | 4.93E-25 | 1.02E-20 | 0.14 | 0.05 | 0.08 | 2.56 |
| 7 | Nsun4 | 0.80 | 5.00E-25 | 1.04E-20 | 0.29 | 0.15 | 0.13 | 1.88 |
| 7 | 2310002F09Rik | 1.31 | 5.57E-25 | 1.16E-20 | 0.10 | 0.04 | 0.07 | 2.94 |
| 7 | Sun1 | 0.68 | 5.75E-25 | 1.20E-20 | 0.44 | 0.27 | 0.17 | 1.61 |
| 7 | Hmgb2 | 0.28 | 5.79E-25 | 1.20E-20 | 0.99 | 0.98 | 0.01 | 1.01 |
| 7 | Tpp1 | 0.57 | 6.08E-25 | 1.26E-20 | 0.43 | 0.26 | 0.17 | 1.65 |
| 7 | Rnpepl1 | 0.60 | 6.49E-25 | 1.35E-20 | 0.42 | 0.25 | 0.17 | 1.69 |
| 7 | C130026L21Rik | 1.28 | 7.09E-25 | 1.47E-20 | 0.11 | 0.04 | 0.07 | 2.86 |
| 7 | Pde1c | 0.88 | 7.22E-25 | 1.50E-20 | 0.22 | 0.10 | 0.11 | 2.08 |
| 7 | Cep55 | 0.73 | 7.75E-25 | 1.61E-20 | 0.40 | 0.24 | 0.16 | 1.67 |
| 7 | Gipc1 | 0.40 | 8.47E-25 | 1.76E-20 | 0.83 | 0.68 | 0.15 | 1.21 |
| 7 | Tmed5 | 0.52 | 8.73E-25 | 1.82E-20 | 0.65 | 0.48 | 0.18 | 1.37 |
| 7 | Cd2ap | 0.32 | 1.01E-24 | 2.10E-20 | 0.77 | 0.62 | 0.16 | 1.25 |
| 7 | Cbx6 | 0.56 | 1.02E-24 | 2.12E-20 | 0.43 | 0.26 | 0.17 | 1.64 |
| 7 | Sel1l | 0.60 | 1.07E-24 | 2.22E-20 | 0.54 | 0.36 | 0.18 | 1.51 |
| 7 | Cep89 | 0.65 | 1.08E-24 | 2.26E-20 | 0.48 | 0.32 | 0.17 | 1.52 |
| 7 | Suclg2 | 0.47 | 1.34E-24 | 2.78E-20 | 0.67 | 0.49 | 0.18 | 1.37 |
| 7 | Rnf187 | 0.27 | 1.34E-24 | 2.79E-20 | 0.96 | 0.90 | 0.06 | 1.06 |
| 7 | Cul1 | 0.33 | 1.43E-24 | 2.97E-20 | 0.92 | 0.86 | 0.06 | 1.07 |
| 7 | H2-Ke6 | 0.65 | 1.48E-24 | 3.09E-20 | 0.39 | 0.23 | 0.16 | 1.69 |
| 7 | Sar1b | 0.38 | 1.52E-24 | 3.17E-20 | 0.83 | 0.69 | 0.13 | 1.19 |
| 7 | Barx2 | 0.76 | 1.53E-24 | 3.17E-20 | 0.11 | 0.04 | 0.07 | 2.79 |
| 7 | Prkci | 0.56 | 1.63E-24 | 3.39E-20 | 0.43 | 0.26 | 0.17 | 1.65 |
| 7 | Six4 | 0.46 | 1.64E-24 | 3.40E-20 | 0.65 | 0.45 | 0.19 | 1.43 |
| 7 | Ctf1 | 0.70 | 1.67E-24 | 3.48E-20 | 0.30 | 0.16 | 0.14 | 1.85 |
| 7 | Sema6a | 0.40 | 1.76E-24 | 3.65E-20 | 0.64 | 0.44 | 0.20 | 1.46 |
| 7 | Rnf165 | 0.61 | 1.84E-24 | 3.84E-20 | 0.40 | 0.24 | 0.16 | 1.68 |
| 7 | Nudt21 | 0.27 | 2.08E-24 | 4.33E-20 | 0.97 | 0.93 | 0.04 | 1.05 |
| 7 | Mgat4b | 0.50 | 3.17E-24 | 6.60E-20 | 0.57 | 0.39 | 0.18 | 1.46 |
| 7 | Ftsj3 | 0.50 | 3.82E-24 | 7.95E-20 | 0.72 | 0.57 | 0.16 | 1.27 |
| 7 | Fam13c | 1.07 | 4.31E-24 | 8.96E-20 | 0.16 | 0.07 | 0.09 | 2.35 |
| 7 | Cdv3 | 0.27 | 4.34E-24 | 9.03E-20 | 0.99 | 0.95 | 0.04 | 1.04 |
| 7 | Kif5c | 0.43 | 4.51E-24 | 9.38E-20 | 0.67 | 0.48 | 0.19 | 1.39 |
| 7 | Rundc3b | 1.30 | 4.57E-24 | 9.51E-20 | 0.11 | 0.04 | 0.07 | 2.77 |
| 7 | Klhl7 | 0.47 | 4.68E-24 | 9.74E-20 | 0.69 | 0.51 | 0.18 | 1.35 |
| 7 | Polr2l | 0.37 | 5.28E-24 | 1.10E-19 | 0.88 | 0.77 | 0.11 | 1.15 |
| 7 | Cfap298 | 0.52 | 6.22E-24 | 1.29E-19 | 0.54 | 0.36 | 0.18 | 1.50 |
| 7 | Chn2 | 1.00 | 6.59E-24 | 1.37E-19 | 0.16 | 0.07 | 0.09 | 2.32 |
| 7 | Bend4 | 0.84 | 8.68E-24 | 1.81E-19 | 0.23 | 0.11 | 0.11 | 2.01 |
| 7 | Zfp612 | 0.66 | 9.39E-24 | 1.95E-19 | 0.30 | 0.16 | 0.14 | 1.85 |
| 7 | Scamp2 | 0.57 | 9.39E-24 | 1.95E-19 | 0.48 | 0.31 | 0.17 | 1.55 |
| 7 | Pkia | 0.73 | 9.94E-24 | 2.07E-19 | 0.32 | 0.18 | 0.14 | 1.78 |
| 7 | Atic | 0.39 | 1.06E-23 | 2.21E-19 | 0.81 | 0.67 | 0.14 | 1.21 |
| 7 | Ank | 1.20 | 1.35E-23 | 2.81E-19 | 0.15 | 0.07 | 0.09 | 2.32 |
| 7 | Vamp8 | 0.37 | 1.38E-23 | 2.86E-19 | 0.78 | 0.61 | 0.17 | 1.29 |
| 7 | Letmd1 | 0.65 | 1.57E-23 | 3.28E-19 | 0.37 | 0.22 | 0.15 | 1.71 |
| 7 | Dalrd3 | 0.57 | 1.64E-23 | 3.42E-19 | 0.43 | 0.26 | 0.17 | 1.63 |
| 7 | 1700037H04Rik | 0.47 | 1.65E-23 | 3.44E-19 | 0.65 | 0.47 | 0.18 | 1.38 |
| 7 | Abcb8 | 0.63 | 2.07E-23 | 4.31E-19 | 0.32 | 0.18 | 0.14 | 1.81 |

**Supplementary Table 1. Marker genes for the original clusters of integrated clustering of E11.5 control and NotchGOF λ-junction cells.**

| | | | | | | | | |
|---|---|---|---|---|---|---|---|---|
| 7 | Otud4 | 0.54 | 2.28E-23 | 4.75E-19 | 0.47 | 0.30 | 0.17 | 1.55 |
| 7 | Poglut3 | 0.87 | 2.29E-23 | 4.77E-19 | 0.25 | 0.13 | 0.12 | 1.92 |
| 7 | Trib2 | 0.64 | 2.40E-23 | 5.00E-19 | 0.34 | 0.19 | 0.15 | 1.76 |
| 7 | Cd164 | 0.41 | 2.53E-23 | 5.26E-19 | 0.78 | 0.63 | 0.15 | 1.24 |
| 7 | Dipk2a | 0.84 | 3.96E-23 | 8.24E-19 | 0.38 | 0.23 | 0.15 | 1.62 |
| 7 | Lcor | 0.67 | 4.09E-23 | 8.51E-19 | 0.44 | 0.28 | 0.16 | 1.57 |
| 7 | Nfib | 0.27 | 4.22E-23 | 8.77E-19 | 0.89 | 0.74 | 0.15 | 1.20 |
| 7 | Ak3 | 0.53 | 4.34E-23 | 9.02E-19 | 0.55 | 0.38 | 0.18 | 1.47 |
| 7 | Camk2d | 0.56 | 4.75E-23 | 9.88E-19 | 0.34 | 0.19 | 0.15 | 1.77 |
| 7 | Ddx19b | 0.91 | 5.03E-23 | 1.05E-18 | 0.19 | 0.09 | 0.10 | 2.14 |
| 7 | Nrm | 0.62 | 5.47E-23 | 1.14E-18 | 0.52 | 0.35 | 0.17 | 1.48 |
| 7 | Pigt | 0.53 | 6.23E-23 | 1.30E-18 | 0.60 | 0.42 | 0.18 | 1.42 |
| 7 | Hat1 | 0.38 | 6.29E-23 | 1.31E-18 | 0.86 | 0.76 | 0.10 | 1.13 |
| 7 | Pgap1 | 0.73 | 6.51E-23 | 1.35E-18 | 0.27 | 0.15 | 0.13 | 1.86 |
| 7 | Ppil1 | 0.32 | 6.56E-23 | 1.36E-18 | 0.91 | 0.81 | 0.10 | 1.12 |
| 7 | C2cd3 | 0.55 | 6.66E-23 | 1.39E-18 | 0.47 | 0.30 | 0.17 | 1.56 |
| 7 | Atp2c1 | 0.50 | 7.11E-23 | 1.48E-18 | 0.58 | 0.39 | 0.19 | 1.47 |
| 7 | Prc1 | 0.54 | 7.66E-23 | 1.59E-18 | 0.67 | 0.52 | 0.15 | 1.29 |
| 7 | Camk2n2 | 0.97 | 8.47E-23 | 1.76E-18 | 0.20 | 0.09 | 0.10 | 2.10 |
| 7 | Dpy30 | 0.26 | 9.17E-23 | 1.91E-18 | 0.95 | 0.90 | 0.05 | 1.06 |
| 7 | Prkch | 0.81 | 9.71E-23 | 2.02E-18 | 0.18 | 0.08 | 0.10 | 2.15 |
| 7 | Nsun2 | 0.39 | 1.04E-22 | 2.16E-18 | 0.79 | 0.63 | 0.15 | 1.24 |
| 7 | Itgb8 | 0.62 | 1.22E-22 | 2.54E-18 | 0.25 | 0.13 | 0.12 | 1.95 |
| 7 | Tmem106c | 0.58 | 1.23E-22 | 2.55E-18 | 0.33 | 0.19 | 0.14 | 1.74 |
| 7 | Madd | 0.86 | 1.27E-22 | 2.65E-18 | 0.16 | 0.07 | 0.09 | 2.29 |
| 7 | Haus1 | 0.46 | 1.46E-22 | 3.04E-18 | 0.67 | 0.51 | 0.16 | 1.32 |
| 7 | Tagln2 | 0.30 | 1.61E-22 | 3.35E-18 | 0.44 | 0.29 | 0.15 | 1.51 |
| 7 | Plcb4 | 0.71 | 1.78E-22 | 3.70E-18 | 0.33 | 0.19 | 0.14 | 1.71 |
| 7 | Nipsnap1 | 0.40 | 1.93E-22 | 4.01E-18 | 0.73 | 0.55 | 0.18 | 1.32 |
| 7 | Tm9sf2 | 0.40 | 2.58E-22 | 5.36E-18 | 0.78 | 0.61 | 0.17 | 1.27 |
| 7 | Cetn3 | 0.28 | 2.70E-22 | 5.63E-18 | 0.93 | 0.88 | 0.05 | 1.06 |
| 7 | Rffl | 1.14 | 3.09E-22 | 6.43E-18 | 0.15 | 0.07 | 0.09 | 2.26 |
| 7 | Mogs | 0.56 | 3.23E-22 | 6.72E-18 | 0.51 | 0.34 | 0.17 | 1.49 |
| 7 | Rfxap | 0.44 | 3.32E-22 | 6.90E-18 | 0.68 | 0.53 | 0.16 | 1.30 |
| 7 | Dcps | 0.44 | 3.40E-22 | 7.08E-18 | 0.74 | 0.59 | 0.15 | 1.25 |
| 7 | Dpp3 | 0.40 | 3.64E-22 | 7.58E-18 | 0.74 | 0.57 | 0.17 | 1.29 |
| 7 | Dnph1 | 0.45 | 3.67E-22 | 7.64E-18 | 0.68 | 0.52 | 0.17 | 1.33 |
| 7 | Prr7 | 0.65 | 4.01E-22 | 8.34E-18 | 0.38 | 0.23 | 0.15 | 1.66 |
| 7 | Tmem121 | 0.60 | 4.33E-22 | 9.01E-18 | 0.38 | 0.23 | 0.15 | 1.66 |
| 7 | Zfp593 | 0.49 | 4.33E-22 | 9.01E-18 | 0.65 | 0.46 | 0.18 | 1.40 |
| 7 | Ssr4 | 0.28 | 4.53E-22 | 9.43E-18 | 0.91 | 0.84 | 0.08 | 1.09 |
| 7 | Lamb2 | 0.77 | 4.57E-22 | 9.51E-18 | 0.25 | 0.13 | 0.12 | 1.91 |
| 7 | Prag1 | 0.70 | 4.69E-22 | 9.76E-18 | 0.26 | 0.14 | 0.12 | 1.89 |
| 7 | Trip10 | 0.63 | 5.13E-22 | 1.07E-17 | 0.36 | 0.21 | 0.14 | 1.68 |
| 7 | Cd302 | 0.51 | 5.14E-22 | 1.07E-17 | 0.45 | 0.28 | 0.17 | 1.59 |
| 7 | Uaca | 0.63 | 5.66E-22 | 1.18E-17 | 0.49 | 0.33 | 0.16 | 1.49 |
| 7 | Jtb | 0.45 | 6.57E-22 | 1.37E-17 | 0.69 | 0.53 | 0.17 | 1.31 |
| 7 | Tmem205 | 0.54 | 7.01E-22 | 1.46E-17 | 0.43 | 0.27 | 0.16 | 1.59 |
| 7 | Nsdhl | 0.61 | 7.37E-22 | 1.53E-17 | 0.53 | 0.37 | 0.16 | 1.44 |
| 7 | Snx5 | 0.32 | 8.08E-22 | 1.68E-17 | 0.87 | 0.77 | 0.09 | 1.12 |
| 7 | Ect2 | 0.48 | 8.97E-22 | 1.87E-17 | 0.57 | 0.40 | 0.17 | 1.41 |
| 7 | Chka | 0.45 | 9.53E-22 | 1.98E-17 | 0.66 | 0.49 | 0.17 | 1.36 |
| 7 | Amer1 | 0.53 | 9.85E-22 | 2.05E-17 | 0.67 | 0.50 | 0.17 | 1.33 |
| 7 | Chst2 | 0.57 | 9.88E-22 | 2.06E-17 | 0.34 | 0.20 | 0.14 | 1.71 |
| 7 | Ptprd | 0.49 | 1.04E-21 | 2.16E-17 | 0.61 | 0.44 | 0.18 | 1.41 |
| 7 | Dnaaf5 | 0.83 | 1.09E-21 | 2.26E-17 | 0.19 | 0.09 | 0.10 | 2.11 |

**Supplementary Table 1. Marker genes for the original clusters of integrated clustering of E11.5 control and NotchGOF λ-junction cells.**

| | | | | | | | | |
|---|---|---|---|---|---|---|---|---|
| 7 | Chpf | 0.62 | 1.10E-21 | 2.29E-17 | 0.27 | 0.15 | 0.13 | 1.86 |
| 7 | Sdsl | 1.02 | 1.11E-21 | 2.31E-17 | 0.14 | 0.06 | 0.08 | 2.39 |
| 7 | Hspa5 | 0.27 | 1.18E-21 | 2.46E-17 | 1.00 | 0.97 | 0.03 | 1.03 |
| 7 | Nfs1 | 0.60 | 1.22E-21 | 2.54E-17 | 0.43 | 0.28 | 0.15 | 1.56 |
| 7 | Dcaf17 | 0.49 | 1.61E-21 | 3.36E-17 | 0.50 | 0.33 | 0.18 | 1.54 |
| 7 | Dcaf12l1 | 0.66 | 1.70E-21 | 3.54E-17 | 0.22 | 0.11 | 0.11 | 1.97 |
| 7 | Pak1ip1 | 0.40 | 1.71E-21 | 3.56E-17 | 0.78 | 0.65 | 0.13 | 1.20 |
| 7 | Racgap1 | 0.49 | 1.94E-21 | 4.03E-17 | 0.67 | 0.52 | 0.15 | 1.29 |
| 7 | Ybx2 | 0.90 | 2.03E-21 | 4.23E-17 | 0.16 | 0.07 | 0.09 | 2.24 |
| 7 | AU040320 | 0.96 | 2.32E-21 | 4.82E-17 | 0.21 | 0.11 | 0.10 | 1.97 |
| 7 | Jpt2 | 0.47 | 2.53E-21 | 5.27E-17 | 0.62 | 0.44 | 0.18 | 1.41 |
| 7 | Tmem63b | 0.45 | 2.65E-21 | 5.51E-17 | 0.51 | 0.33 | 0.18 | 1.54 |
| 7 | Bckdk | 0.47 | 2.82E-21 | 5.86E-17 | 0.64 | 0.49 | 0.16 | 1.33 |
| 7 | Higd1a | 0.34 | 2.84E-21 | 5.90E-17 | 0.91 | 0.87 | 0.04 | 1.04 |
| 7 | Pomgnt2 | 1.16 | 3.22E-21 | 6.69E-17 | 0.13 | 0.06 | 0.08 | 2.36 |
| 7 | Ppp1r9a | 0.52 | 3.48E-21 | 7.23E-17 | 0.56 | 0.39 | 0.17 | 1.43 |
| 7 | Akap12 | 0.64 | 3.74E-21 | 7.79E-17 | 0.32 | 0.19 | 0.13 | 1.68 |
| 7 | Nhlrc1 | 1.09 | 4.34E-21 | 9.02E-17 | 0.10 | 0.04 | 0.07 | 2.71 |
| 7 | Acadm | 0.52 | 4.40E-21 | 9.16E-17 | 0.49 | 0.33 | 0.16 | 1.50 |
| 7 | Rpn2 | 0.33 | 4.44E-21 | 9.24E-17 | 0.88 | 0.78 | 0.10 | 1.12 |
| 7 | Adipor2 | 0.52 | 4.72E-21 | 9.82E-17 | 0.50 | 0.34 | 0.16 | 1.47 |
| 7 | Nudt16 | 0.80 | 5.38E-21 | 1.12E-16 | 0.18 | 0.08 | 0.09 | 2.13 |
| 7 | Eya4 | 0.46 | 5.83E-21 | 1.21E-16 | 0.44 | 0.28 | 0.16 | 1.56 |
| 7 | Nop58 | 0.26 | 5.83E-21 | 1.21E-16 | 0.96 | 0.90 | 0.05 | 1.06 |
| 7 | Ier5 | 0.33 | 5.92E-21 | 1.23E-16 | 0.80 | 0.65 | 0.15 | 1.23 |
| 7 | Hist1h4i | 0.53 | 5.99E-21 | 1.25E-16 | 0.61 | 0.45 | 0.16 | 1.34 |
| 7 | Dph2 | 0.87 | 6.50E-21 | 1.35E-16 | 0.20 | 0.10 | 0.10 | 2.03 |
| 7 | Wnk2 | 0.98 | 7.77E-21 | 1.62E-16 | 0.12 | 0.05 | 0.07 | 2.52 |
| 7 | Cercam | 1.02 | 8.21E-21 | 1.71E-16 | 0.13 | 0.06 | 0.08 | 2.36 |
| 7 | 2610507B11Rik | 0.48 | 8.69E-21 | 1.81E-16 | 0.63 | 0.46 | 0.17 | 1.36 |
| 7 | Adgrl1 | 0.46 | 1.25E-20 | 2.60E-16 | 0.60 | 0.43 | 0.17 | 1.40 |
| 7 | Mccc2 | 0.55 | 1.26E-20 | 2.61E-16 | 0.43 | 0.27 | 0.16 | 1.58 |
| 7 | Mien1 | 0.36 | 1.42E-20 | 2.94E-16 | 0.73 | 0.56 | 0.17 | 1.31 |
| 7 | Spata24 | 0.54 | 1.50E-20 | 3.11E-16 | 0.43 | 0.28 | 0.15 | 1.55 |
| 7 | Pgf | 0.88 | 1.55E-20 | 3.22E-16 | 0.15 | 0.07 | 0.08 | 2.24 |
| 7 | Sft2d3 | 0.63 | 1.65E-20 | 3.43E-16 | 0.33 | 0.20 | 0.14 | 1.68 |
| 7 | Stim2 | 0.66 | 1.87E-20 | 3.88E-16 | 0.37 | 0.23 | 0.14 | 1.62 |
| 7 | Acly | 0.37 | 1.97E-20 | 4.09E-16 | 0.74 | 0.57 | 0.17 | 1.30 |
| 7 | Sephs2 | 0.48 | 1.99E-20 | 4.15E-16 | 0.58 | 0.42 | 0.17 | 1.40 |
| 7 | Wdr77 | 0.43 | 2.01E-20 | 4.19E-16 | 0.69 | 0.52 | 0.16 | 1.31 |
| 7 | Rangap1 | 0.37 | 2.13E-20 | 4.44E-16 | 0.83 | 0.72 | 0.12 | 1.16 |
| 7 | Mcoln3 | 0.92 | 2.26E-20 | 4.70E-16 | 0.13 | 0.05 | 0.07 | 2.40 |
| 7 | Cacnb2 | 1.05 | 2.32E-20 | 4.83E-16 | 0.16 | 0.07 | 0.09 | 2.18 |
| 7 | Os9 | 0.49 | 2.37E-20 | 4.92E-16 | 0.54 | 0.37 | 0.17 | 1.48 |
| 7 | Knl1 | 0.51 | 2.51E-20 | 5.23E-16 | 0.64 | 0.49 | 0.15 | 1.31 |
| 7 | Eml2 | 0.81 | 3.26E-20 | 6.78E-16 | 0.19 | 0.10 | 0.10 | 2.00 |
| 7 | Gnaz | 0.97 | 3.33E-20 | 6.92E-16 | 0.11 | 0.04 | 0.07 | 2.55 |
| 7 | Fbxl14 | 0.49 | 3.45E-20 | 7.17E-16 | 0.54 | 0.38 | 0.16 | 1.43 |
| 7 | Stamos | 0.97 | 3.48E-20 | 7.23E-16 | 0.11 | 0.05 | 0.07 | 2.51 |
| 7 | Zcchc18 | 0.43 | 4.24E-20 | 8.81E-16 | 0.50 | 0.33 | 0.17 | 1.51 |
| 7 | Ddx56 | 0.52 | 4.43E-20 | 9.21E-16 | 0.50 | 0.34 | 0.16 | 1.46 |
| 7 | Tfdp1 | 0.39 | 4.64E-20 | 9.64E-16 | 0.78 | 0.66 | 0.13 | 1.20 |
| 7 | Mpp2 | 0.68 | 4.65E-20 | 9.67E-16 | 0.29 | 0.17 | 0.12 | 1.74 |
| 7 | Sh3bp4 | 0.67 | 4.68E-20 | 9.74E-16 | 0.29 | 0.16 | 0.12 | 1.74 |
| 7 | Ldlr | 0.66 | 4.74E-20 | 9.85E-16 | 0.38 | 0.24 | 0.14 | 1.58 |
| 7 | Gphn | 0.62 | 4.98E-20 | 1.04E-15 | 0.31 | 0.18 | 0.13 | 1.71 |

**Supplementary Table 1. Marker genes for the original clusters of integrated clustering of E11.5 control and NotchGOF λ-junction cells.**

| | | | | | | | | |
|---|---|---|---|---|---|---|---|---|
| 7 | Rnpep | 0.44 | 5.26E-20 | 1.09E-15 | 0.61 | 0.43 | 0.18 | 1.41 |
| 7 | Bst2 | 0.86 | 5.35E-20 | 1.11E-15 | 0.13 | 0.06 | 0.08 | 2.36 |
| 7 | Znrd2 | 0.34 | 5.50E-20 | 1.14E-15 | 0.85 | 0.76 | 0.09 | 1.12 |
| 7 | Mrps36 | 0.31 | 5.70E-20 | 1.19E-15 | 0.88 | 0.77 | 0.11 | 1.14 |
| 7 | Usp1 | 0.30 | 6.20E-20 | 1.29E-15 | 0.89 | 0.81 | 0.08 | 1.10 |
| 7 | Nsd2 | 0.37 | 6.30E-20 | 1.31E-15 | 0.87 | 0.75 | 0.12 | 1.16 |
| 7 | Plekhb2 | 1.02 | 6.68E-20 | 1.39E-15 | 0.13 | 0.06 | 0.08 | 2.31 |
| 7 | Ggta1 | 0.98 | 6.75E-20 | 1.40E-15 | 0.11 | 0.04 | 0.07 | 2.55 |
| 7 | Yif1a | 0.42 | 7.46E-20 | 1.55E-15 | 0.64 | 0.47 | 0.16 | 1.34 |
| 7 | Wfs1 | 1.17 | 8.96E-20 | 1.86E-15 | 0.14 | 0.06 | 0.08 | 2.22 |
| 7 | Rhou | 0.50 | 9.63E-20 | 2.00E-15 | 0.57 | 0.41 | 0.16 | 1.39 |
| 7 | Hmga1b | 0.42 | 1.01E-19 | 2.11E-15 | 0.61 | 0.43 | 0.18 | 1.41 |
| 7 | Tmem42 | 0.66 | 1.03E-19 | 2.14E-15 | 0.27 | 0.15 | 0.12 | 1.82 |
| 7 | S100a11 | 0.41 | 1.05E-19 | 2.19E-15 | 0.84 | 0.76 | 0.08 | 1.11 |
| 7 | Wdfy1 | 0.43 | 1.09E-19 | 2.26E-15 | 0.51 | 0.35 | 0.17 | 1.48 |
| 7 | Rrp7a | 0.43 | 1.36E-19 | 2.83E-15 | 0.64 | 0.47 | 0.17 | 1.37 |
| 7 | Ociad2 | 0.60 | 1.43E-19 | 2.97E-15 | 0.31 | 0.18 | 0.13 | 1.72 |
| 7 | Tmem107 | 0.50 | 1.50E-19 | 3.12E-15 | 0.44 | 0.29 | 0.16 | 1.55 |
| 7 | Szrd1 | 0.42 | 1.69E-19 | 3.51E-15 | 0.66 | 0.50 | 0.16 | 1.31 |
| 7 | Tmem38b | 0.66 | 1.75E-19 | 3.63E-15 | 0.26 | 0.14 | 0.12 | 1.81 |
| 7 | Slco5a1 | 0.68 | 1.80E-19 | 3.74E-15 | 0.17 | 0.08 | 0.09 | 2.10 |
| 7 | Hspg2 | 0.38 | 1.90E-19 | 3.96E-15 | 0.37 | 0.22 | 0.14 | 1.65 |
| 7 | Lmo2 | 0.32 | 1.96E-19 | 4.08E-15 | 0.23 | 0.12 | 0.11 | 1.88 |
| 7 | Rfc5 | 0.38 | 1.99E-19 | 4.15E-15 | 0.73 | 0.58 | 0.14 | 1.25 |
| 7 | Cenpv | 0.35 | 2.08E-19 | 4.33E-15 | 0.66 | 0.50 | 0.17 | 1.34 |
| 7 | Sdf2l1 | 0.45 | 2.31E-19 | 4.80E-15 | 0.62 | 0.46 | 0.16 | 1.34 |
| 7 | Gpatch4 | 0.38 | 2.34E-19 | 4.86E-15 | 0.70 | 0.54 | 0.16 | 1.29 |
| 7 | Atp13a2 | 0.50 | 2.43E-19 | 5.05E-15 | 0.40 | 0.25 | 0.15 | 1.60 |
| 7 | Ppp1r26 | 0.99 | 2.45E-19 | 5.10E-15 | 0.11 | 0.04 | 0.06 | 2.52 |
| 7 | Klhl36 | 0.88 | 2.58E-19 | 5.36E-15 | 0.14 | 0.06 | 0.08 | 2.22 |
| 7 | Cdca8 | 0.33 | 2.77E-19 | 5.77E-15 | 0.87 | 0.75 | 0.12 | 1.16 |
| 7 | 1500011B03Rik | 0.33 | 3.08E-19 | 6.40E-15 | 0.81 | 0.66 | 0.14 | 1.22 |
| 7 | Shmt2 | 0.52 | 3.15E-19 | 6.55E-15 | 0.50 | 0.34 | 0.16 | 1.48 |
| 7 | Enpp5 | 1.10 | 3.21E-19 | 6.67E-15 | 0.11 | 0.04 | 0.06 | 2.50 |
| 7 | Rad23a | 0.46 | 3.21E-19 | 6.67E-15 | 0.61 | 0.45 | 0.16 | 1.35 |
| 7 | Snhg3 | 0.61 | 3.37E-19 | 7.00E-15 | 0.66 | 0.53 | 0.13 | 1.23 |
| 7 | Scap | 0.70 | 3.39E-19 | 7.05E-15 | 0.31 | 0.18 | 0.13 | 1.72 |
| 7 | Hook2 | 0.67 | 3.40E-19 | 7.07E-15 | 0.28 | 0.16 | 0.12 | 1.75 |
| 7 | Grhl3 | 0.63 | 3.65E-19 | 7.59E-15 | 0.13 | 0.05 | 0.07 | 2.38 |
| 7 | Pus1 | 0.58 | 4.36E-19 | 9.06E-15 | 0.44 | 0.29 | 0.15 | 1.53 |
| 7 | Zfp536 | 0.67 | 4.77E-19 | 9.92E-15 | 0.20 | 0.10 | 0.10 | 1.98 |
| 7 | Cldn11 | 0.52 | 5.12E-19 | 1.06E-14 | 0.24 | 0.13 | 0.11 | 1.87 |
| 7 | Rabggta | 0.77 | 5.21E-19 | 1.08E-14 | 0.23 | 0.13 | 0.11 | 1.85 |
| 7 | Galnt2 | 0.43 | 5.33E-19 | 1.11E-14 | 0.62 | 0.46 | 0.16 | 1.35 |
| 7 | Ubxn2a | 0.52 | 5.34E-19 | 1.11E-14 | 0.46 | 0.31 | 0.15 | 1.47 |
| 7 | Stox2 | 0.35 | 5.61E-19 | 1.17E-14 | 0.71 | 0.56 | 0.16 | 1.28 |
| 7 | Amfr | 0.51 | 5.97E-19 | 1.24E-14 | 0.51 | 0.36 | 0.16 | 1.44 |
| 7 | L2hgdh | 0.67 | 6.49E-19 | 1.35E-14 | 0.34 | 0.21 | 0.13 | 1.64 |
| 7 | Mcf2l | 0.86 | 7.11E-19 | 1.48E-14 | 0.15 | 0.07 | 0.08 | 2.20 |
| 7 | Prmt3 | 0.51 | 8.83E-19 | 1.84E-14 | 0.48 | 0.33 | 0.15 | 1.47 |
| 7 | Bola1 | 0.38 | 9.36E-19 | 1.95E-14 | 0.74 | 0.62 | 0.13 | 1.20 |
| 7 | Ppp2r1a | 0.28 | 1.00E-18 | 2.09E-14 | 0.89 | 0.78 | 0.11 | 1.14 |
| 7 | Kyat1 | 0.98 | 1.02E-18 | 2.12E-14 | 0.12 | 0.05 | 0.07 | 2.38 |
| 7 | Zc3h12c | 0.81 | 1.07E-18 | 2.22E-14 | 0.13 | 0.06 | 0.07 | 2.23 |
| 7 | Atp1a1 | 0.35 | 1.13E-18 | 2.34E-14 | 0.77 | 0.63 | 0.14 | 1.22 |
| 7 | Nmt2 | 0.51 | 1.20E-18 | 2.50E-14 | 0.49 | 0.34 | 0.15 | 1.44 |

**Supplementary Table 1. Marker genes for the original clusters of integrated clustering of E11.5 control and NotchGOF λ-junction cells.**

| | | | | | | | | |
|---|---|---|---|---|---|---|---|---|
| 7 | Crmp1 | 0.47 | 1.20E-18 | 2.50E-14 | 0.22 | 0.12 | 0.10 | 1.88 |
| 7 | Nudt11 | 0.47 | 1.23E-18 | 2.56E-14 | 0.30 | 0.17 | 0.13 | 1.73 |
| 7 | Lpin1 | 1.01 | 1.27E-18 | 2.64E-14 | 0.14 | 0.06 | 0.08 | 2.21 |
| 7 | Wdr46 | 0.43 | 1.37E-18 | 2.86E-14 | 0.62 | 0.44 | 0.18 | 1.40 |
| 7 | 2610306M01Rik | 0.58 | 1.45E-18 | 3.01E-14 | 0.28 | 0.16 | 0.12 | 1.75 |
| 7 | Khk | 0.80 | 1.49E-18 | 3.10E-14 | 0.14 | 0.06 | 0.08 | 2.22 |
| 7 | Tspan4 | 0.73 | 1.65E-18 | 3.44E-14 | 0.23 | 0.12 | 0.10 | 1.85 |
| 7 | Gnptab | 0.58 | 1.75E-18 | 3.64E-14 | 0.26 | 0.15 | 0.11 | 1.79 |
| 7 | Pex2 | 0.40 | 2.05E-18 | 4.27E-14 | 0.63 | 0.47 | 0.17 | 1.35 |
| 7 | Ring1 | 0.56 | 2.16E-18 | 4.48E-14 | 0.38 | 0.24 | 0.14 | 1.57 |
| 7 | Pop5 | 0.36 | 2.31E-18 | 4.80E-14 | 0.79 | 0.65 | 0.14 | 1.22 |
| 7 | Wdfy2 | 0.59 | 2.51E-18 | 5.23E-14 | 0.27 | 0.16 | 0.12 | 1.75 |
| 7 | Slc44a1 | 0.47 | 2.53E-18 | 5.25E-14 | 0.37 | 0.23 | 0.14 | 1.61 |
| 7 | Zfp800 | 0.50 | 2.78E-18 | 5.79E-14 | 0.40 | 0.26 | 0.14 | 1.55 |
| 7 | Celf4 | 0.51 | 2.84E-18 | 5.90E-14 | 0.36 | 0.22 | 0.14 | 1.62 |
| 7 | Mtss1 | 0.33 | 3.04E-18 | 6.32E-14 | 0.45 | 0.29 | 0.17 | 1.57 |
| 7 | Ift22 | 0.45 | 3.13E-18 | 6.52E-14 | 0.51 | 0.35 | 0.16 | 1.46 |
| 7 | Rbms1 | 0.28 | 3.27E-18 | 6.80E-14 | 0.88 | 0.78 | 0.10 | 1.13 |
| 7 | Srd5a1 | 0.57 | 3.38E-18 | 7.03E-14 | 0.30 | 0.18 | 0.12 | 1.69 |
| 7 | Dap | 0.41 | 3.49E-18 | 7.27E-14 | 0.71 | 0.56 | 0.15 | 1.27 |
| 7 | 2510002D24Rik | 0.40 | 3.51E-18 | 7.29E-14 | 0.55 | 0.39 | 0.16 | 1.42 |
| 7 | Polr3h | 0.45 | 4.27E-18 | 8.89E-14 | 0.53 | 0.38 | 0.16 | 1.42 |
| 7 | Abhd14a | 0.43 | 5.20E-18 | 1.08E-13 | 0.55 | 0.38 | 0.17 | 1.43 |
| 7 | Plpp6 | 0.73 | 5.47E-18 | 1.14E-13 | 0.14 | 0.07 | 0.08 | 2.17 |
| 7 | M6pr | 0.44 | 5.68E-18 | 1.18E-13 | 0.60 | 0.44 | 0.16 | 1.36 |
| 7 | Cyb5b | 0.43 | 5.74E-18 | 1.19E-13 | 0.65 | 0.50 | 0.15 | 1.30 |
| 7 | Slc22a17 | 0.48 | 6.01E-18 | 1.25E-13 | 0.50 | 0.34 | 0.16 | 1.47 |
| 7 | Rps6ka6 | 0.36 | 6.78E-18 | 1.41E-13 | 0.62 | 0.45 | 0.17 | 1.38 |
| 7 | Prkcd | 0.50 | 7.22E-18 | 1.50E-13 | 0.34 | 0.21 | 0.13 | 1.63 |
| 7 | Spc24 | 0.29 | 9.03E-18 | 1.88E-13 | 0.83 | 0.73 | 0.10 | 1.14 |
| 7 | Sh3glb2 | 0.50 | 9.04E-18 | 1.88E-13 | 0.47 | 0.32 | 0.15 | 1.49 |
| 7 | Mrpl44 | 0.41 | 9.17E-18 | 1.91E-13 | 0.57 | 0.41 | 0.16 | 1.39 |
| 7 | Zfp367 | 0.65 | 9.29E-18 | 1.93E-13 | 0.25 | 0.14 | 0.11 | 1.78 |
| 7 | Tmem179b | 0.54 | 9.30E-18 | 1.93E-13 | 0.43 | 0.29 | 0.14 | 1.49 |
| 7 | Diablo | 0.34 | 9.59E-18 | 2.00E-13 | 0.77 | 0.63 | 0.15 | 1.24 |
| 7 | Ggact | 1.09 | 1.04E-17 | 2.16E-13 | 0.11 | 0.05 | 0.06 | 2.36 |
| 7 | Mylk | 0.53 | 1.18E-17 | 2.46E-13 | 0.29 | 0.17 | 0.12 | 1.71 |
| 7 | Usp51 | 0.86 | 1.44E-17 | 2.99E-13 | 0.16 | 0.08 | 0.08 | 2.04 |
| 7 | Rnf121 | 0.45 | 1.69E-17 | 3.51E-13 | 0.52 | 0.36 | 0.16 | 1.43 |
| 7 | Mlec | 0.28 | 1.78E-17 | 3.69E-13 | 0.87 | 0.75 | 0.12 | 1.16 |
| 7 | Xylb | 0.97 | 1.79E-17 | 3.72E-13 | 0.13 | 0.06 | 0.07 | 2.21 |
| 7 | Clcc1 | 0.46 | 1.88E-17 | 3.91E-13 | 0.56 | 0.41 | 0.15 | 1.37 |
| 7 | Gid4 | 0.47 | 1.88E-17 | 3.91E-13 | 0.48 | 0.33 | 0.15 | 1.45 |
| 7 | Tctn2 | 0.79 | 1.96E-17 | 4.08E-13 | 0.18 | 0.09 | 0.09 | 1.97 |
| 7 | Ccdc18 | 0.51 | 2.09E-17 | 4.34E-13 | 0.44 | 0.30 | 0.14 | 1.48 |
| 7 | Ccpg1 | 0.69 | 2.10E-17 | 4.37E-13 | 0.27 | 0.16 | 0.11 | 1.71 |
| 7 | Yipf2 | 0.72 | 2.11E-17 | 4.39E-13 | 0.22 | 0.12 | 0.10 | 1.82 |
| 7 | Cryzl2 | 0.63 | 2.18E-17 | 4.54E-13 | 0.19 | 0.10 | 0.09 | 1.92 |
| 7 | Ebf4 | 1.01 | 2.26E-17 | 4.69E-13 | 0.13 | 0.06 | 0.07 | 2.23 |
| 7 | Maz | 0.27 | 2.30E-17 | 4.78E-13 | 0.89 | 0.81 | 0.08 | 1.10 |
| 7 | Hipk2 | 0.34 | 2.57E-17 | 5.35E-13 | 0.75 | 0.64 | 0.11 | 1.17 |
| 7 | Tmem237 | 0.44 | 2.69E-17 | 5.59E-13 | 0.54 | 0.39 | 0.15 | 1.38 |
| 7 | Cope | 0.27 | 2.97E-17 | 6.18E-13 | 0.86 | 0.77 | 0.10 | 1.12 |
| 7 | Gm45051 | 0.69 | 2.98E-17 | 6.21E-13 | 0.15 | 0.07 | 0.08 | 2.10 |
| 7 | Ccna2 | 0.37 | 3.23E-17 | 6.71E-13 | 0.76 | 0.63 | 0.13 | 1.21 |
| 7 | Ssc4d | 0.93 | 3.27E-17 | 6.81E-13 | 0.12 | 0.05 | 0.07 | 2.29 |

**Supplementary Table 1. Marker genes for the original clusters of integrated clustering of E11.5 control and NotchGOF λ-junction cells.**

| | | | | | | | | |
|---|---|---|---|---|---|---|---|---|
| 7 | Prrg3 | 0.73 | 3.44E-17 | 7.16E-13 | 0.20 | 0.11 | 0.09 | 1.87 |
| 7 | Setd6 | 0.80 | 3.48E-17 | 7.23E-13 | 0.22 | 0.12 | 0.10 | 1.80 |
| 7 | Iqsec1 | 0.77 | 3.58E-17 | 7.45E-13 | 0.24 | 0.13 | 0.10 | 1.76 |
| 7 | Slc20a2 | 0.50 | 3.79E-17 | 7.87E-13 | 0.37 | 0.23 | 0.14 | 1.58 |
| 7 | Smarcd3 | 0.46 | 3.83E-17 | 7.96E-13 | 0.42 | 0.27 | 0.14 | 1.52 |
| 7 | B4galt4 | 0.47 | 4.04E-17 | 8.40E-13 | 0.36 | 0.23 | 0.14 | 1.60 |
| 7 | Tspan7 | 0.27 | 4.11E-17 | 8.54E-13 | 0.43 | 0.28 | 0.15 | 1.56 |
| 7 | Rrs1 | 0.39 | 4.14E-17 | 8.61E-13 | 0.74 | 0.60 | 0.14 | 1.23 |
| 7 | Pisd | 0.57 | 4.54E-17 | 9.43E-13 | 0.33 | 0.21 | 0.13 | 1.61 |
| 7 | Rida | 0.40 | 4.62E-17 | 9.60E-13 | 0.57 | 0.41 | 0.16 | 1.38 |
| 7 | Intu | 0.69 | 4.64E-17 | 9.65E-13 | 0.20 | 0.10 | 0.09 | 1.90 |
| 7 | Kctd11 | 0.95 | 4.65E-17 | 9.68E-13 | 0.11 | 0.05 | 0.06 | 2.36 |
| 7 | Mrps34 | 0.31 | 5.06E-17 | 1.05E-12 | 0.75 | 0.62 | 0.14 | 1.22 |
| 7 | Fam98c | 0.76 | 5.18E-17 | 1.08E-12 | 0.21 | 0.11 | 0.10 | 1.84 |
| 7 | Ppp1r2 | 0.28 | 5.35E-17 | 1.11E-12 | 0.86 | 0.75 | 0.11 | 1.15 |
| 7 | Rps6ka1 | 0.45 | 5.78E-17 | 1.20E-12 | 0.38 | 0.24 | 0.14 | 1.58 |
| 7 | Mboat2 | 0.47 | 5.91E-17 | 1.23E-12 | 0.50 | 0.34 | 0.15 | 1.44 |
| 7 | Herc3 | 0.72 | 6.31E-17 | 1.31E-12 | 0.15 | 0.07 | 0.08 | 2.10 |
| 7 | Nr2c2ap | 0.48 | 6.68E-17 | 1.39E-12 | 0.36 | 0.23 | 0.13 | 1.57 |
| 7 | Galk1 | 0.30 | 6.83E-17 | 1.42E-12 | 0.83 | 0.71 | 0.12 | 1.18 |
| 7 | 1190007I07Rik | 0.47 | 7.09E-17 | 1.48E-12 | 0.42 | 0.27 | 0.14 | 1.52 |
| 7 | Cep57 | 0.35 | 7.11E-17 | 1.48E-12 | 0.73 | 0.59 | 0.14 | 1.23 |
| 7 | Larp7 | 0.33 | 7.23E-17 | 1.50E-12 | 0.86 | 0.77 | 0.10 | 1.12 |
| 7 | Wdr44 | 1.08 | 7.82E-17 | 1.63E-12 | 0.13 | 0.06 | 0.07 | 2.13 |
| 7 | Dhcr24 | 0.49 | 8.60E-17 | 1.79E-12 | 0.49 | 0.35 | 0.14 | 1.41 |
| 7 | Ankrd37 | 0.59 | 8.84E-17 | 1.84E-12 | 0.26 | 0.15 | 0.11 | 1.71 |
| 7 | 4933434E20Rik | 0.42 | 8.94E-17 | 1.86E-12 | 0.64 | 0.49 | 0.15 | 1.31 |
| 7 | Unc5b | 0.31 | 9.03E-17 | 1.88E-12 | 0.56 | 0.39 | 0.18 | 1.46 |
| 7 | Knstrn | 0.33 | 9.09E-17 | 1.89E-12 | 0.75 | 0.61 | 0.15 | 1.24 |
| 7 | Insr | 0.53 | 9.84E-17 | 2.05E-12 | 0.37 | 0.24 | 0.13 | 1.56 |
| 7 | Tma16 | 0.37 | 1.00E-16 | 2.08E-12 | 0.69 | 0.53 | 0.16 | 1.30 |
| 7 | Klf13 | 0.45 | 1.04E-16 | 2.17E-12 | 0.51 | 0.37 | 0.14 | 1.39 |
| 7 | Abr | 0.56 | 1.07E-16 | 2.23E-12 | 0.34 | 0.21 | 0.13 | 1.58 |
| 7 | Egln1 | 0.34 | 1.14E-16 | 2.36E-12 | 0.80 | 0.68 | 0.12 | 1.17 |
| 7 | Esco1 | 0.43 | 1.24E-16 | 2.58E-12 | 0.36 | 0.23 | 0.13 | 1.59 |
| 7 | Ext1 | 0.44 | 1.26E-16 | 2.62E-12 | 0.55 | 0.41 | 0.15 | 1.36 |
| 7 | Chmp2a | 0.32 | 1.52E-16 | 3.15E-12 | 0.78 | 0.66 | 0.13 | 1.19 |
| 7 | Ctnna2 | 0.96 | 1.53E-16 | 3.17E-12 | 0.11 | 0.05 | 0.06 | 2.32 |
| 7 | Pard3 | 0.39 | 1.86E-16 | 3.87E-12 | 0.71 | 0.57 | 0.13 | 1.23 |
| 7 | Cars | 0.46 | 1.87E-16 | 3.88E-12 | 0.44 | 0.30 | 0.14 | 1.48 |
| 7 | Atp11a | 0.56 | 1.90E-16 | 3.95E-12 | 0.28 | 0.17 | 0.11 | 1.68 |
| 7 | Top1mt | 1.02 | 2.06E-16 | 4.29E-12 | 0.12 | 0.05 | 0.07 | 2.20 |
| 7 | Micu3 | 0.59 | 2.12E-16 | 4.41E-12 | 0.28 | 0.17 | 0.11 | 1.65 |
| 7 | Golgb1 | 0.30 | 2.13E-16 | 4.44E-12 | 0.84 | 0.74 | 0.11 | 1.15 |
| 7 | Tamm41 | 0.40 | 2.17E-16 | 4.51E-12 | 0.49 | 0.34 | 0.15 | 1.45 |
| 7 | Pdrg1 | 0.31 | 2.44E-16 | 5.07E-12 | 0.68 | 0.53 | 0.15 | 1.29 |
| 7 | Gstz1 | 0.48 | 3.01E-16 | 6.25E-12 | 0.40 | 0.27 | 0.14 | 1.51 |
| 7 | Miga1 | 0.45 | 3.08E-16 | 6.41E-12 | 0.44 | 0.30 | 0.14 | 1.48 |
| 7 | Dkc1 | 0.30 | 3.26E-16 | 6.79E-12 | 0.87 | 0.79 | 0.08 | 1.11 |
| 7 | Erlin2 | 0.41 | 3.57E-16 | 7.42E-12 | 0.47 | 0.32 | 0.15 | 1.48 |
| 7 | Sp4 | 0.47 | 3.72E-16 | 7.75E-12 | 0.36 | 0.23 | 0.13 | 1.56 |
| 7 | Tsr1 | 0.46 | 3.80E-16 | 7.91E-12 | 0.50 | 0.36 | 0.14 | 1.40 |
| 7 | Ddb2 | 0.70 | 3.89E-16 | 8.09E-12 | 0.18 | 0.10 | 0.09 | 1.89 |
| 7 | Prmt5 | 0.33 | 3.92E-16 | 8.16E-12 | 0.74 | 0.60 | 0.14 | 1.24 |
| 7 | Rad51 | 0.34 | 4.18E-16 | 8.69E-12 | 0.69 | 0.54 | 0.15 | 1.27 |
| 7 | Clstn3 | 0.86 | 4.30E-16 | 8.94E-12 | 0.12 | 0.06 | 0.07 | 2.16 |

**Supplementary Table 1. Marker genes for the original clusters of integrated clustering of E11.5 control and NotchGOF λ-junction cells.**

| | | | | | | | | |
|---|---|---|---|---|---|---|---|---|
| 7 | Gstt2 | 0.79 | 4.32E-16 | 8.99E-12 | 0.15 | 0.07 | 0.08 | 2.04 |
| 7 | Rnf141 | 0.63 | 4.49E-16 | 9.34E-12 | 0.22 | 0.12 | 0.10 | 1.79 |
| 7 | Hif3a | 0.86 | 4.68E-16 | 9.72E-12 | 0.16 | 0.09 | 0.08 | 1.93 |
| 7 | Tfg | 0.27 | 4.80E-16 | 9.97E-12 | 0.85 | 0.73 | 0.13 | 1.17 |
| 7 | Apobec3 | 0.73 | 4.83E-16 | 1.00E-11 | 0.16 | 0.08 | 0.08 | 1.99 |
| 7 | Hmgcs1 | 0.31 | 5.09E-16 | 1.06E-11 | 0.87 | 0.77 | 0.11 | 1.14 |
| 7 | H2-T23 | 1.04 | 5.27E-16 | 1.10E-11 | 0.12 | 0.05 | 0.06 | 2.21 |
| 7 | Kcnc3 | 0.77 | 5.27E-16 | 1.10E-11 | 0.12 | 0.05 | 0.06 | 2.23 |
| 7 | Txndc16 | 0.79 | 5.46E-16 | 1.13E-11 | 0.24 | 0.14 | 0.10 | 1.71 |
| 7 | Dars2 | 0.57 | 5.63E-16 | 1.17E-11 | 0.32 | 0.20 | 0.12 | 1.60 |
| 7 | Cisd3 | 0.84 | 6.56E-16 | 1.36E-11 | 0.13 | 0.06 | 0.07 | 2.11 |
| 7 | Dazap2 | 0.32 | 6.69E-16 | 1.39E-11 | 0.71 | 0.57 | 0.14 | 1.25 |
| 7 | Ccdc173 | 0.51 | 6.95E-16 | 1.44E-11 | 0.21 | 0.12 | 0.10 | 1.83 |
| 7 | Setd7 | 0.44 | 7.09E-16 | 1.47E-11 | 0.33 | 0.20 | 0.13 | 1.63 |
| 7 | Map3k11 | 0.58 | 7.15E-16 | 1.49E-11 | 0.21 | 0.11 | 0.09 | 1.81 |
| 7 | Tmem120a | 0.44 | 7.60E-16 | 1.58E-11 | 0.43 | 0.30 | 0.14 | 1.46 |
| 7 | Plat | 0.67 | 7.64E-16 | 1.59E-11 | 0.44 | 0.31 | 0.13 | 1.40 |
| 7 | Pts | 0.35 | 7.77E-16 | 1.62E-11 | 0.56 | 0.40 | 0.16 | 1.39 |
| 7 | Ccdc51 | 0.69 | 8.06E-16 | 1.68E-11 | 0.21 | 0.12 | 0.09 | 1.79 |
| 7 | Ebna1bp2 | 0.27 | 8.18E-16 | 1.70E-11 | 0.88 | 0.79 | 0.09 | 1.11 |
| 7 | Aimp2 | 0.37 | 8.79E-16 | 1.83E-11 | 0.66 | 0.53 | 0.14 | 1.26 |
| 7 | C1galt1c1 | 0.53 | 9.05E-16 | 1.88E-11 | 0.26 | 0.16 | 0.11 | 1.69 |
| 7 | Slc39a3 | 0.78 | 9.53E-16 | 1.98E-11 | 0.22 | 0.12 | 0.10 | 1.78 |
| 7 | Pdia4 | 0.28 | 9.62E-16 | 2.00E-11 | 0.89 | 0.78 | 0.11 | 1.15 |
| 7 | Rbfa | 0.37 | 1.02E-15 | 2.13E-11 | 0.45 | 0.31 | 0.14 | 1.47 |
| 7 | Gm48226 | 0.57 | 1.05E-15 | 2.19E-11 | 0.29 | 0.17 | 0.11 | 1.65 |
| 7 | Pigl | 0.75 | 1.39E-15 | 2.89E-11 | 0.15 | 0.07 | 0.07 | 2.01 |
| 7 | Rtl6 | 0.56 | 1.43E-15 | 2.98E-11 | 0.20 | 0.11 | 0.09 | 1.83 |
| 7 | Slc25a13 | 0.36 | 1.44E-15 | 2.99E-11 | 0.60 | 0.44 | 0.16 | 1.37 |
| 7 | Fmn2 | 0.78 | 1.53E-15 | 3.18E-11 | 0.14 | 0.07 | 0.07 | 2.06 |
| 7 | Dok5 | 0.51 | 1.74E-15 | 3.63E-11 | 0.20 | 0.11 | 0.09 | 1.79 |
| 7 | C030006K11Rik | 0.72 | 1.78E-15 | 3.71E-11 | 0.17 | 0.09 | 0.08 | 1.92 |
| 7 | Thbs1 | 0.28 | 1.85E-15 | 3.85E-11 | 0.21 | 0.12 | 0.09 | 1.80 |
| 7 | Qtrt1 | 0.54 | 1.92E-15 | 4.00E-11 | 0.36 | 0.24 | 0.12 | 1.52 |
| 7 | Fam136a | 0.29 | 1.96E-15 | 4.08E-11 | 0.88 | 0.79 | 0.09 | 1.11 |
| 7 | Pdk3 | 0.41 | 2.10E-15 | 4.37E-11 | 0.55 | 0.40 | 0.15 | 1.36 |
| 7 | Jmjd1c | 0.36 | 2.10E-15 | 4.37E-11 | 0.63 | 0.48 | 0.16 | 1.32 |
| 7 | Kif15 | 0.31 | 2.12E-15 | 4.41E-11 | 0.76 | 0.63 | 0.13 | 1.21 |
| 7 | Stk4 | 0.36 | 2.12E-15 | 4.42E-11 | 0.51 | 0.36 | 0.15 | 1.43 |
| 7 | Stk11 | 0.40 | 2.23E-15 | 4.65E-11 | 0.61 | 0.48 | 0.14 | 1.29 |
| 7 | Ttc39b | 0.68 | 2.44E-15 | 5.08E-11 | 0.13 | 0.06 | 0.07 | 2.08 |
| 7 | Mrpl38 | 0.31 | 2.60E-15 | 5.40E-11 | 0.78 | 0.64 | 0.14 | 1.22 |
| 7 | Zfyve21 | 0.38 | 2.64E-15 | 5.50E-11 | 0.55 | 0.40 | 0.15 | 1.39 |
| 7 | Tomm40 | 0.27 | 2.68E-15 | 5.57E-11 | 0.88 | 0.81 | 0.08 | 1.10 |
| 7 | Oma1 | 0.53 | 2.75E-15 | 5.73E-11 | 0.29 | 0.18 | 0.11 | 1.63 |
| 7 | Pigs | 0.44 | 2.78E-15 | 5.79E-11 | 0.44 | 0.30 | 0.14 | 1.46 |
| 7 | Slc7a1 | 1.03 | 3.53E-15 | 7.34E-11 | 0.15 | 0.08 | 0.07 | 1.97 |
| 7 | Adsl | 0.35 | 3.75E-15 | 7.80E-11 | 0.62 | 0.46 | 0.16 | 1.34 |
| 7 | Cyb561d2 | 0.53 | 3.86E-15 | 8.03E-11 | 0.28 | 0.17 | 0.11 | 1.66 |
| 7 | Ddc | 0.87 | 4.11E-15 | 8.54E-11 | 0.11 | 0.05 | 0.06 | 2.19 |
| 7 | Tspyl5 | 0.73 | 4.13E-15 | 8.59E-11 | 0.13 | 0.07 | 0.07 | 2.06 |
| 7 | Hspbp1 | 0.38 | 4.15E-15 | 8.63E-11 | 0.54 | 0.40 | 0.15 | 1.37 |
| 7 | Tmem37 | 0.33 | 4.45E-15 | 9.25E-11 | 0.16 | 0.08 | 0.08 | 1.95 |
| 7 | Notch4 | 0.96 | 4.67E-15 | 9.72E-11 | 0.12 | 0.06 | 0.06 | 2.13 |
| 7 | Ammecr1 | 0.41 | 4.80E-15 | 9.99E-11 | 0.49 | 0.35 | 0.14 | 1.40 |
| 7 | Cit | 0.46 | 5.09E-15 | 1.06E-10 | 0.51 | 0.37 | 0.14 | 1.38 |

**Supplementary Table 1. Marker genes for the original clusters of integrated clustering of E11.5 control and NotchGOF λ-junction cells.**

| | | | | | | | | |
|---|---|---|---|---|---|---|---|---|
| 7 | Edem2 | 0.46 | 5.10E-15 | 1.06E-10 | 0.31 | 0.19 | 0.12 | 1.61 |
| 7 | 2010320M18Rik | 0.35 | 5.25E-15 | 1.09E-10 | 0.49 | 0.34 | 0.15 | 1.44 |
| 7 | Mid1 | 0.47 | 5.59E-15 | 1.16E-10 | 0.37 | 0.24 | 0.13 | 1.54 |
| 7 | Tmem176a | 0.36 | 5.68E-15 | 1.18E-10 | 0.55 | 0.40 | 0.15 | 1.37 |
| 7 | Acp6 | 0.46 | 6.17E-15 | 1.28E-10 | 0.34 | 0.22 | 0.12 | 1.55 |
| 7 | Gm3839 | 0.56 | 7.28E-15 | 1.51E-10 | 0.27 | 0.16 | 0.11 | 1.65 |
| 7 | Lrrc42 | 0.29 | 8.13E-15 | 1.69E-10 | 0.74 | 0.60 | 0.14 | 1.24 |
| 7 | Kxd1 | 0.34 | 8.71E-15 | 1.81E-10 | 0.64 | 0.50 | 0.15 | 1.30 |
| 7 | Tsc22d2 | 0.43 | 8.89E-15 | 1.85E-10 | 0.55 | 0.40 | 0.15 | 1.37 |
| 7 | Mef2d | 0.57 | 9.31E-15 | 1.94E-10 | 0.23 | 0.14 | 0.10 | 1.72 |
| 7 | Elovl2 | 0.71 | 9.39E-15 | 1.95E-10 | 0.19 | 0.11 | 0.09 | 1.79 |
| 7 | Gm9993 | 0.77 | 9.48E-15 | 1.97E-10 | 0.16 | 0.08 | 0.08 | 1.94 |
| 7 | Nek1 | 0.44 | 9.64E-15 | 2.01E-10 | 0.44 | 0.31 | 0.14 | 1.45 |
| 7 | Ltv1 | 0.38 | 1.02E-14 | 2.13E-10 | 0.60 | 0.46 | 0.14 | 1.31 |
| 7 | Ppie | 0.30 | 1.09E-14 | 2.27E-10 | 0.78 | 0.65 | 0.13 | 1.19 |
| 7 | Zadh2 | 0.73 | 1.15E-14 | 2.38E-10 | 0.19 | 0.10 | 0.08 | 1.80 |
| 7 | Ccdc43 | 0.42 | 1.16E-14 | 2.40E-10 | 0.55 | 0.40 | 0.15 | 1.36 |
| 7 | Kat2b | 1.00 | 1.16E-14 | 2.41E-10 | 0.10 | 0.05 | 0.06 | 2.24 |
| 7 | Agtrap | 0.61 | 1.25E-14 | 2.60E-10 | 0.24 | 0.14 | 0.10 | 1.69 |
| 7 | Ptpn4 | 0.50 | 1.29E-14 | 2.68E-10 | 0.26 | 0.16 | 0.10 | 1.67 |
| 7 | Pycard | 0.38 | 1.33E-14 | 2.77E-10 | 0.22 | 0.12 | 0.09 | 1.75 |
| 7 | Oscp1 | 0.50 | 1.36E-14 | 2.82E-10 | 0.31 | 0.19 | 0.11 | 1.59 |
| 7 | Mitd1 | 0.41 | 1.42E-14 | 2.95E-10 | 0.43 | 0.29 | 0.13 | 1.45 |
| 7 | Pigh | 0.37 | 1.52E-14 | 3.17E-10 | 0.41 | 0.27 | 0.14 | 1.50 |
| 7 | Wee1 | 0.42 | 1.54E-14 | 3.20E-10 | 0.59 | 0.45 | 0.14 | 1.30 |
| 7 | Trib1 | 0.62 | 1.54E-14 | 3.21E-10 | 0.28 | 0.18 | 0.11 | 1.60 |
| 7 | 2310033P09Rik | 0.29 | 1.57E-14 | 3.27E-10 | 0.75 | 0.62 | 0.14 | 1.22 |
| 7 | Vezt | 0.44 | 1.59E-14 | 3.31E-10 | 0.45 | 0.32 | 0.13 | 1.42 |
| 7 | Wdr4 | 0.50 | 1.60E-14 | 3.33E-10 | 0.39 | 0.26 | 0.13 | 1.49 |
| 7 | Zfp7 | 0.61 | 1.61E-14 | 3.36E-10 | 0.24 | 0.14 | 0.10 | 1.68 |
| 7 | Pot1a | 0.56 | 1.71E-14 | 3.56E-10 | 0.31 | 0.20 | 0.11 | 1.57 |
| 7 | Sfxn1 | 0.30 | 1.74E-14 | 3.62E-10 | 0.83 | 0.73 | 0.10 | 1.13 |
| 7 | Dcaf6 | 0.82 | 1.74E-14 | 3.62E-10 | 0.15 | 0.08 | 0.07 | 1.91 |
| 7 | Slc16a6 | 1.07 | 1.79E-14 | 3.72E-10 | 0.11 | 0.05 | 0.06 | 2.13 |
| 7 | Slc15a2 | 0.93 | 1.81E-14 | 3.76E-10 | 0.10 | 0.05 | 0.06 | 2.21 |
| 7 | Lrrc8d | 0.47 | 1.86E-14 | 3.87E-10 | 0.31 | 0.19 | 0.12 | 1.60 |
| 7 | B4galnt4 | 0.47 | 1.87E-14 | 3.88E-10 | 0.36 | 0.24 | 0.13 | 1.53 |
| 7 | Ap1s3 | 0.81 | 1.92E-14 | 3.99E-10 | 0.13 | 0.07 | 0.07 | 2.02 |
| 7 | Tshz2 | 0.50 | 2.03E-14 | 4.21E-10 | 0.46 | 0.34 | 0.12 | 1.37 |
| 7 | Ankrd13c | 0.51 | 2.05E-14 | 4.26E-10 | 0.36 | 0.24 | 0.12 | 1.51 |
| 7 | Mrps18b | 0.35 | 2.13E-14 | 4.42E-10 | 0.64 | 0.50 | 0.14 | 1.29 |
| 7 | Trappc13 | 0.38 | 2.14E-14 | 4.45E-10 | 0.57 | 0.42 | 0.15 | 1.36 |
| 7 | Thop1 | 0.38 | 2.17E-14 | 4.52E-10 | 0.65 | 0.51 | 0.14 | 1.28 |
| 7 | 2310015A10Rik | 0.66 | 2.36E-14 | 4.91E-10 | 0.14 | 0.07 | 0.07 | 1.97 |
| 7 | Phtf1 | 0.45 | 2.38E-14 | 4.95E-10 | 0.54 | 0.40 | 0.14 | 1.35 |
| 7 | Slc6a6 | 0.32 | 2.48E-14 | 5.15E-10 | 0.49 | 0.35 | 0.15 | 1.42 |
| 7 | Zfp704 | 0.31 | 2.61E-14 | 5.44E-10 | 0.62 | 0.46 | 0.15 | 1.33 |
| 7 | Chchd6 | 0.34 | 2.65E-14 | 5.52E-10 | 0.67 | 0.53 | 0.14 | 1.26 |
| 7 | Tfb1m | 0.64 | 2.73E-14 | 5.68E-10 | 0.21 | 0.12 | 0.09 | 1.75 |
| 7 | Acer3 | 0.59 | 2.77E-14 | 5.77E-10 | 0.26 | 0.16 | 0.10 | 1.64 |
| 7 | Sbno2 | 0.78 | 2.98E-14 | 6.19E-10 | 0.20 | 0.11 | 0.08 | 1.75 |
| 7 | Ubl3 | 0.34 | 3.22E-14 | 6.69E-10 | 0.51 | 0.37 | 0.14 | 1.39 |
| 7 | Snx1 | 0.37 | 3.40E-14 | 7.07E-10 | 0.58 | 0.44 | 0.14 | 1.31 |
| 7 | Slc37a3 | 0.40 | 3.43E-14 | 7.14E-10 | 0.39 | 0.26 | 0.13 | 1.50 |
| 7 | Hprt | 0.35 | 3.48E-14 | 7.24E-10 | 0.56 | 0.43 | 0.13 | 1.31 |
| 7 | Tram1 | 0.40 | 3.90E-14 | 8.12E-10 | 0.64 | 0.50 | 0.14 | 1.27 |

**Supplementary Table 1. Marker genes for the original clusters of integrated clustering of E11.5 control and NotchGOF λ-junction cells.**

| | | | | | | | | |
|---|---|---|---|---|---|---|---|---|
| 7 | Rcan3 | 0.67 | 3.96E-14 | 8.24E-10 | 0.15 | 0.08 | 0.07 | 1.95 |
| 7 | Kif13a | 0.48 | 4.05E-14 | 8.41E-10 | 0.32 | 0.21 | 0.12 | 1.56 |
| 7 | Gpr89 | 0.32 | 4.06E-14 | 8.44E-10 | 0.47 | 0.32 | 0.15 | 1.46 |
| 7 | Thtpa | 0.51 | 4.17E-14 | 8.68E-10 | 0.26 | 0.16 | 0.10 | 1.66 |
| 7 | Slc29a2 | 0.73 | 4.39E-14 | 9.13E-10 | 0.17 | 0.09 | 0.08 | 1.85 |
| 7 | Bmpr1b | 0.85 | 4.40E-14 | 9.15E-10 | 0.16 | 0.09 | 0.08 | 1.88 |
| 7 | Lockd | 0.34 | 4.50E-14 | 9.36E-10 | 0.50 | 0.36 | 0.14 | 1.40 |
| 7 | Pdia5 | 0.59 | 4.54E-14 | 9.44E-10 | 0.20 | 0.11 | 0.09 | 1.77 |
| 7 | Zdhhc13 | 0.47 | 4.73E-14 | 9.84E-10 | 0.23 | 0.14 | 0.10 | 1.71 |
| 7 | Rsph9 | 0.55 | 4.73E-14 | 9.84E-10 | 0.19 | 0.11 | 0.09 | 1.81 |
| 7 | Ninl | 0.51 | 5.15E-14 | 1.07E-09 | 0.35 | 0.24 | 0.12 | 1.49 |
| 7 | Sema4b | 0.69 | 5.27E-14 | 1.10E-09 | 0.16 | 0.09 | 0.08 | 1.90 |
| 7 | Bag4 | 0.35 | 5.28E-14 | 1.10E-09 | 0.54 | 0.39 | 0.15 | 1.38 |
| 7 | Lpcat3 | 0.40 | 5.47E-14 | 1.14E-09 | 0.43 | 0.30 | 0.13 | 1.42 |
| 7 | Pbx3 | 0.28 | 5.50E-14 | 1.14E-09 | 0.69 | 0.52 | 0.17 | 1.33 |
| 7 | Nacc1 | 0.38 | 5.50E-14 | 1.14E-09 | 0.53 | 0.39 | 0.14 | 1.36 |
| 7 | Fam185a | 0.82 | 5.66E-14 | 1.18E-09 | 0.12 | 0.06 | 0.06 | 2.11 |
| 7 | Iqcb1 | 0.28 | 5.82E-14 | 1.21E-09 | 0.67 | 0.52 | 0.16 | 1.31 |
| 7 | Fgfbp3 | 0.71 | 6.86E-14 | 1.43E-09 | 0.18 | 0.10 | 0.08 | 1.81 |
| 7 | Atpaf1 | 0.37 | 6.95E-14 | 1.44E-09 | 0.51 | 0.37 | 0.14 | 1.39 |
| 7 | Reep6 | 1.04 | 7.07E-14 | 1.47E-09 | 0.11 | 0.05 | 0.06 | 2.16 |
| 7 | Fry | 0.75 | 7.07E-14 | 1.47E-09 | 0.11 | 0.05 | 0.06 | 2.11 |
| 7 | Polr1e | 0.45 | 7.08E-14 | 1.47E-09 | 0.37 | 0.25 | 0.12 | 1.49 |
| 7 | Mindy2 | 0.61 | 7.13E-14 | 1.48E-09 | 0.27 | 0.17 | 0.10 | 1.59 |
| 7 | Wrnip1 | 0.48 | 7.26E-14 | 1.51E-09 | 0.31 | 0.19 | 0.11 | 1.58 |
| 7 | Nusap1 | 0.35 | 7.59E-14 | 1.58E-09 | 0.63 | 0.48 | 0.14 | 1.29 |
| 7 | Klf10 | 0.66 | 7.61E-14 | 1.58E-09 | 0.19 | 0.11 | 0.08 | 1.77 |
| 7 | Polr1c | 0.30 | 7.90E-14 | 1.64E-09 | 0.73 | 0.59 | 0.14 | 1.23 |
| 7 | Mfsd2a | 0.83 | 7.94E-14 | 1.65E-09 | 0.13 | 0.06 | 0.06 | 2.05 |
| 7 | Dolpp1 | 0.56 | 8.03E-14 | 1.67E-09 | 0.23 | 0.14 | 0.10 | 1.70 |
| 7 | Mark1 | 0.35 | 8.35E-14 | 1.74E-09 | 0.37 | 0.25 | 0.13 | 1.51 |
| 7 | Gm11361 | 0.38 | 8.92E-14 | 1.86E-09 | 0.54 | 0.41 | 0.14 | 1.34 |
| 7 | Noa1 | 0.40 | 8.96E-14 | 1.86E-09 | 0.44 | 0.31 | 0.13 | 1.43 |
| 7 | Agpat5 | 0.44 | 9.06E-14 | 1.89E-09 | 0.58 | 0.44 | 0.14 | 1.31 |
| 7 | Ccdc120 | 0.68 | 9.10E-14 | 1.89E-09 | 0.19 | 0.10 | 0.08 | 1.79 |
| 7 | Grwd1 | 0.36 | 9.40E-14 | 1.96E-09 | 0.57 | 0.42 | 0.14 | 1.34 |
| 7 | Serinc2 | 0.30 | 1.05E-13 | 2.18E-09 | 0.54 | 0.38 | 0.16 | 1.41 |
| 7 | Pak1 | 0.32 | 1.05E-13 | 2.18E-09 | 0.65 | 0.49 | 0.16 | 1.34 |
| 7 | Mblac2 | 0.98 | 1.06E-13 | 2.20E-09 | 0.11 | 0.05 | 0.06 | 2.16 |
| 7 | Slc35e3 | 0.59 | 1.09E-13 | 2.27E-09 | 0.22 | 0.13 | 0.09 | 1.71 |
| 7 | Ano10 | 0.49 | 1.15E-13 | 2.40E-09 | 0.22 | 0.13 | 0.09 | 1.73 |
| 7 | Sod2 | 0.28 | 1.16E-13 | 2.42E-09 | 0.76 | 0.62 | 0.14 | 1.23 |
| 7 | Sgsm1 | 0.71 | 1.17E-13 | 2.43E-09 | 0.15 | 0.08 | 0.07 | 1.90 |
| 7 | C1qtnf12 | 0.41 | 1.18E-13 | 2.46E-09 | 0.26 | 0.16 | 0.10 | 1.63 |
| 7 | E2f2 | 0.66 | 1.19E-13 | 2.47E-09 | 0.19 | 0.11 | 0.08 | 1.75 |
| 7 | Brca2 | 0.38 | 1.22E-13 | 2.54E-09 | 0.60 | 0.47 | 0.13 | 1.28 |
| 7 | Slc25a30 | 0.49 | 1.26E-13 | 2.62E-09 | 0.24 | 0.14 | 0.10 | 1.67 |
| 7 | Pgm1 | 0.37 | 1.28E-13 | 2.66E-09 | 0.45 | 0.31 | 0.14 | 1.45 |
| 7 | Hmga1 | 0.32 | 1.33E-13 | 2.77E-09 | 0.65 | 0.49 | 0.16 | 1.32 |
| 7 | Etfa | 0.28 | 1.45E-13 | 3.02E-09 | 0.84 | 0.73 | 0.11 | 1.14 |
| 7 | Coq4 | 0.38 | 1.48E-13 | 3.07E-09 | 0.46 | 0.33 | 0.13 | 1.40 |
| 7 | Tmem128 | 0.32 | 1.52E-13 | 3.16E-09 | 0.75 | 0.63 | 0.12 | 1.19 |
| 7 | Gchfr | 0.55 | 1.56E-13 | 3.24E-09 | 0.19 | 0.11 | 0.08 | 1.76 |
| 7 | Cib1 | 0.37 | 1.60E-13 | 3.33E-09 | 0.54 | 0.40 | 0.14 | 1.35 |
| 7 | Slc4a8 | 0.73 | 1.63E-13 | 3.38E-09 | 0.17 | 0.10 | 0.08 | 1.80 |
| 7 | Enpp4 | 0.79 | 1.67E-13 | 3.46E-09 | 0.13 | 0.06 | 0.06 | 2.03 |

**Supplementary Table 1. Marker genes for the original clusters of integrated clustering of E11.5 control and NotchGOF λ-junction cells.**

| | | | | | | | | |
|---|---|---|---|---|---|---|---|---|
| 7 | Pqlc3 | 0.86 | 1.74E-13 | 3.61E-09 | 0.11 | 0.05 | 0.06 | 2.09 |
| 7 | Hsdl1 | 0.50 | 1.85E-13 | 3.85E-09 | 0.25 | 0.15 | 0.10 | 1.64 |
| 7 | Fktn | 0.38 | 1.89E-13 | 3.92E-09 | 0.38 | 0.26 | 0.13 | 1.49 |
| 7 | Dbnl | 0.29 | 1.92E-13 | 3.98E-09 | 0.78 | 0.65 | 0.13 | 1.19 |
| 7 | Klhl2 | 0.64 | 2.02E-13 | 4.19E-09 | 0.20 | 0.12 | 0.09 | 1.71 |
| 7 | Exd2 | 0.52 | 2.05E-13 | 4.27E-09 | 0.25 | 0.15 | 0.10 | 1.65 |
| 7 | Ptrh2 | 0.30 | 2.09E-13 | 4.34E-09 | 0.79 | 0.66 | 0.13 | 1.19 |
| 7 | Wdr83os | 0.25 | 2.18E-13 | 4.53E-09 | 0.90 | 0.84 | 0.07 | 1.08 |
| 7 | Cdca3 | 0.31 | 2.59E-13 | 5.39E-09 | 0.76 | 0.65 | 0.11 | 1.17 |
| 7 | Marchf8 | 0.46 | 2.71E-13 | 5.63E-09 | 0.27 | 0.17 | 0.10 | 1.60 |
| 7 | Scrn2 | 0.42 | 2.78E-13 | 5.78E-09 | 0.51 | 0.37 | 0.14 | 1.38 |
| 7 | Lonp1 | 0.31 | 3.13E-13 | 6.51E-09 | 0.65 | 0.52 | 0.14 | 1.26 |
| 7 | Gm17018 | 0.30 | 3.14E-13 | 6.54E-09 | 0.66 | 0.52 | 0.14 | 1.27 |
| 7 | Agap1 | 0.34 | 3.15E-13 | 6.56E-09 | 0.56 | 0.41 | 0.14 | 1.35 |
| 7 | Eci1 | 0.35 | 3.17E-13 | 6.60E-09 | 0.66 | 0.52 | 0.14 | 1.27 |
| 7 | Tmem170b | 0.81 | 3.17E-13 | 6.60E-09 | 0.13 | 0.06 | 0.06 | 2.00 |
| 7 | Ankle1 | 0.81 | 3.34E-13 | 6.94E-09 | 0.16 | 0.09 | 0.07 | 1.79 |
| 7 | 1500004A13Rik | 0.61 | 4.10E-13 | 8.54E-09 | 0.14 | 0.07 | 0.07 | 1.94 |
| 7 | Mmgt2 | 0.37 | 4.24E-13 | 8.81E-09 | 0.57 | 0.43 | 0.14 | 1.32 |
| 7 | Enoph1 | 0.41 | 4.70E-13 | 9.77E-09 | 0.36 | 0.24 | 0.12 | 1.49 |
| 7 | Krcc1 | 0.31 | 5.12E-13 | 1.07E-08 | 0.71 | 0.57 | 0.14 | 1.24 |
| 7 | Slc45a4 | 0.51 | 5.23E-13 | 1.09E-08 | 0.22 | 0.13 | 0.09 | 1.69 |
| 7 | Fbxo16 | 0.57 | 5.68E-13 | 1.18E-08 | 0.19 | 0.11 | 0.08 | 1.76 |
| 7 | Exosc3 | 0.29 | 5.83E-13 | 1.21E-08 | 0.81 | 0.69 | 0.12 | 1.18 |
| 7 | Tmem97 | 0.37 | 6.03E-13 | 1.25E-08 | 0.60 | 0.46 | 0.14 | 1.29 |
| 7 | Pno1 | 0.32 | 6.05E-13 | 1.26E-08 | 0.72 | 0.59 | 0.12 | 1.21 |
| 7 | Ahdc1 | 0.43 | 6.06E-13 | 1.26E-08 | 0.25 | 0.15 | 0.10 | 1.63 |
| 7 | Snapin | 0.35 | 6.99E-13 | 1.45E-08 | 0.46 | 0.32 | 0.14 | 1.43 |
| 7 | Med9 | 0.30 | 7.54E-13 | 1.57E-08 | 0.69 | 0.55 | 0.14 | 1.26 |
| 7 | Arhgap21 | 0.48 | 7.71E-13 | 1.60E-08 | 0.33 | 0.22 | 0.11 | 1.51 |
| 7 | mt-Atp8 | 0.46 | 7.97E-13 | 1.66E-08 | 0.66 | 0.54 | 0.12 | 1.22 |
| 7 | Mif4gd | 0.46 | 8.17E-13 | 1.70E-08 | 0.26 | 0.17 | 0.10 | 1.59 |
| 7 | Ppt1 | 0.37 | 8.25E-13 | 1.72E-08 | 0.44 | 0.31 | 0.13 | 1.42 |
| 7 | Rnd2 | 0.64 | 8.66E-13 | 1.80E-08 | 0.16 | 0.09 | 0.07 | 1.85 |
| 7 | Tmub1 | 0.47 | 8.88E-13 | 1.85E-08 | 0.30 | 0.19 | 0.11 | 1.56 |
| 7 | Eif1a | 0.28 | 9.04E-13 | 1.88E-08 | 0.79 | 0.66 | 0.13 | 1.19 |
| 7 | Lss | 0.58 | 9.62E-13 | 2.00E-08 | 0.25 | 0.16 | 0.09 | 1.60 |
| 7 | Cad | 0.41 | 9.79E-13 | 2.04E-08 | 0.52 | 0.38 | 0.14 | 1.36 |
| 7 | Ero1l | 0.40 | 1.01E-12 | 2.09E-08 | 0.41 | 0.29 | 0.12 | 1.42 |
| 7 | Slfn9 | 0.41 | 1.02E-12 | 2.11E-08 | 0.37 | 0.25 | 0.12 | 1.48 |
| 7 | Ipo4 | 0.40 | 1.07E-12 | 2.23E-08 | 0.47 | 0.34 | 0.13 | 1.38 |
| 7 | 2310061I04Rik | 0.52 | 1.08E-12 | 2.24E-08 | 0.28 | 0.18 | 0.10 | 1.55 |
| 7 | Malt1 | 0.56 | 1.08E-12 | 2.25E-08 | 0.26 | 0.16 | 0.09 | 1.58 |
| 7 | Slc38a10 | 0.39 | 1.12E-12 | 2.34E-08 | 0.47 | 0.33 | 0.13 | 1.40 |
| 7 | Unc50 | 0.32 | 1.14E-12 | 2.37E-08 | 0.69 | 0.56 | 0.13 | 1.23 |
| 7 | Asl | 0.50 | 1.14E-12 | 2.38E-08 | 0.24 | 0.15 | 0.09 | 1.64 |
| 7 | Rrp15 | 0.26 | 1.27E-12 | 2.65E-08 | 0.79 | 0.67 | 0.12 | 1.19 |
| 7 | Psmb10 | 0.38 | 1.40E-12 | 2.92E-08 | 0.35 | 0.23 | 0.12 | 1.52 |
| 7 | Tmem170 | 0.58 | 1.43E-12 | 2.97E-08 | 0.17 | 0.09 | 0.07 | 1.80 |
| 7 | Trmt61a | 0.63 | 1.48E-12 | 3.07E-08 | 0.23 | 0.15 | 0.09 | 1.61 |
| 7 | Crls1 | 0.31 | 1.53E-12 | 3.19E-08 | 0.58 | 0.45 | 0.13 | 1.30 |
| 7 | Creld2 | 0.40 | 1.60E-12 | 3.32E-08 | 0.58 | 0.44 | 0.13 | 1.30 |
| 7 | Ext2 | 0.33 | 1.64E-12 | 3.41E-08 | 0.56 | 0.41 | 0.15 | 1.36 |
| 7 | Gaa | 0.44 | 1.64E-12 | 3.41E-08 | 0.28 | 0.18 | 0.10 | 1.56 |
| 7 | Iqgap3 | 0.56 | 1.65E-12 | 3.43E-08 | 0.32 | 0.22 | 0.10 | 1.47 |
| 7 | 9930014A18Rik | 0.59 | 1.71E-12 | 3.55E-08 | 0.10 | 0.05 | 0.06 | 2.15 |

**Supplementary Table 1. Marker genes for the original clusters of integrated clustering of E11.5 control and NotchGOF λ-junction cells.**

| | | | | | | | | |
|---|---|---|---|---|---|---|---|---|
| 7 | Hgh1 | 0.52 | 1.71E-12 | 3.55E-08 | 0.22 | 0.13 | 0.09 | 1.67 |
| 7 | Coq2 | 0.33 | 1.88E-12 | 3.91E-08 | 0.60 | 0.45 | 0.15 | 1.32 |
| 7 | Tnnt1 | 0.55 | 1.90E-12 | 3.96E-08 | 0.11 | 0.05 | 0.06 | 2.08 |
| 7 | Zfp651 | 0.56 | 1.94E-12 | 4.03E-08 | 0.25 | 0.15 | 0.09 | 1.62 |
| 7 | Cfap410 | 0.47 | 1.94E-12 | 4.04E-08 | 0.33 | 0.22 | 0.11 | 1.50 |
| 7 | Pter | 0.63 | 2.03E-12 | 4.23E-08 | 0.19 | 0.11 | 0.08 | 1.71 |
| 7 | Zc2hc1a | 0.38 | 2.16E-12 | 4.49E-08 | 0.56 | 0.41 | 0.14 | 1.34 |
| 7 | Txndc11 | 0.47 | 2.19E-12 | 4.55E-08 | 0.30 | 0.20 | 0.11 | 1.54 |
| 7 | Lrba | 0.33 | 2.22E-12 | 4.62E-08 | 0.34 | 0.22 | 0.11 | 1.51 |
| 7 | Alg8 | 0.59 | 2.23E-12 | 4.63E-08 | 0.24 | 0.15 | 0.09 | 1.62 |
| 7 | Lman2 | 0.28 | 2.23E-12 | 4.65E-08 | 0.76 | 0.62 | 0.15 | 1.24 |
| 7 | Vps13a | 0.31 | 2.42E-12 | 5.04E-08 | 0.47 | 0.34 | 0.13 | 1.39 |
| 7 | Lifr | 0.75 | 2.70E-12 | 5.62E-08 | 0.18 | 0.11 | 0.08 | 1.71 |
| 7 | Actr5 | 0.41 | 2.77E-12 | 5.75E-08 | 0.29 | 0.18 | 0.10 | 1.56 |
| 7 | Pex10 | 0.50 | 2.78E-12 | 5.78E-08 | 0.25 | 0.16 | 0.09 | 1.59 |
| 7 | Ostm1 | 0.55 | 3.03E-12 | 6.31E-08 | 0.13 | 0.07 | 0.06 | 1.91 |
| 7 | Efcab14 | 0.33 | 3.05E-12 | 6.35E-08 | 0.56 | 0.42 | 0.14 | 1.33 |
| 7 | Pex7 | 0.42 | 3.28E-12 | 6.82E-08 | 0.37 | 0.25 | 0.12 | 1.47 |
| 7 | Pltp | 0.34 | 3.40E-12 | 7.08E-08 | 0.25 | 0.16 | 0.10 | 1.61 |
| 7 | Tmem245 | 0.38 | 3.69E-12 | 7.67E-08 | 0.53 | 0.40 | 0.13 | 1.33 |
| 7 | Cdca5 | 0.34 | 3.77E-12 | 7.83E-08 | 0.46 | 0.33 | 0.13 | 1.38 |
| 7 | Tmc4 | 0.48 | 3.85E-12 | 8.01E-08 | 0.17 | 0.10 | 0.07 | 1.76 |
| 7 | Xpot | 0.37 | 3.92E-12 | 8.16E-08 | 0.54 | 0.41 | 0.13 | 1.33 |
| 7 | Bcap31 | 0.27 | 4.05E-12 | 8.41E-08 | 0.85 | 0.75 | 0.10 | 1.14 |
| 7 | Samhd1 | 0.31 | 4.10E-12 | 8.53E-08 | 0.52 | 0.38 | 0.14 | 1.37 |
| 7 | Mir99ahg | 0.25 | 4.17E-12 | 8.67E-08 | 0.63 | 0.48 | 0.15 | 1.30 |
| 7 | Kif23 | 0.30 | 4.34E-12 | 9.02E-08 | 0.57 | 0.44 | 0.13 | 1.29 |
| 7 | Atl2 | 0.43 | 4.39E-12 | 9.14E-08 | 0.36 | 0.25 | 0.12 | 1.46 |
| 7 | Lmbrd2 | 0.46 | 4.49E-12 | 9.34E-08 | 0.28 | 0.18 | 0.10 | 1.55 |
| 7 | Chchd4 | 0.34 | 4.60E-12 | 9.57E-08 | 0.61 | 0.48 | 0.13 | 1.27 |
| 7 | Ugcg | 0.44 | 4.68E-12 | 9.74E-08 | 0.49 | 0.37 | 0.13 | 1.35 |
| 7 | Pex26 | 0.66 | 4.80E-12 | 9.98E-08 | 0.15 | 0.08 | 0.07 | 1.84 |
| 7 | Lmtk2 | 0.61 | 4.85E-12 | 1.01E-07 | 0.17 | 0.10 | 0.07 | 1.76 |
| 7 | Scamp1 | 0.50 | 5.02E-12 | 1.04E-07 | 0.27 | 0.17 | 0.10 | 1.56 |
| 7 | Gns | 0.35 | 5.21E-12 | 1.08E-07 | 0.37 | 0.25 | 0.12 | 1.49 |
| 7 | Slc35b2 | 0.37 | 5.40E-12 | 1.12E-07 | 0.55 | 0.41 | 0.14 | 1.34 |
| 7 | 4930452B06Rik | 0.49 | 5.41E-12 | 1.13E-07 | 0.20 | 0.12 | 0.08 | 1.68 |
| 7 | Pla2g6 | 0.79 | 5.51E-12 | 1.15E-07 | 0.14 | 0.08 | 0.07 | 1.82 |
| 7 | Pigq | 0.47 | 6.11E-12 | 1.27E-07 | 0.34 | 0.24 | 0.11 | 1.46 |
| 7 | Nanp | 0.37 | 6.16E-12 | 1.28E-07 | 0.51 | 0.38 | 0.13 | 1.34 |
| 7 | Nav1 | 0.31 | 6.22E-12 | 1.29E-07 | 0.67 | 0.54 | 0.13 | 1.25 |
| 7 | Map9 | 0.38 | 6.42E-12 | 1.34E-07 | 0.28 | 0.18 | 0.10 | 1.57 |
| 7 | Eno1b | 0.37 | 6.57E-12 | 1.37E-07 | 0.47 | 0.34 | 0.13 | 1.37 |
| 7 | Lin37 | 0.42 | 6.61E-12 | 1.37E-07 | 0.34 | 0.23 | 0.11 | 1.46 |
| 7 | Hectd2 | 0.92 | 6.62E-12 | 1.38E-07 | 0.10 | 0.05 | 0.05 | 2.10 |
| 7 | Fbxo10 | 0.62 | 6.64E-12 | 1.38E-07 | 0.14 | 0.08 | 0.07 | 1.86 |
| 7 | Tsen15 | 0.29 | 6.85E-12 | 1.42E-07 | 0.76 | 0.63 | 0.13 | 1.20 |
| 7 | Qsox2 | 0.53 | 7.05E-12 | 1.47E-07 | 0.21 | 0.13 | 0.08 | 1.65 |
| 7 | Pepd | 0.37 | 7.24E-12 | 1.51E-07 | 0.55 | 0.41 | 0.14 | 1.33 |
| 7 | Rad51b | 0.61 | 7.39E-12 | 1.54E-07 | 0.19 | 0.11 | 0.08 | 1.69 |
| 7 | Stt3b | 0.36 | 7.60E-12 | 1.58E-07 | 0.62 | 0.51 | 0.12 | 1.23 |
| 7 | Msi2 | 0.26 | 7.71E-12 | 1.60E-07 | 0.78 | 0.68 | 0.10 | 1.15 |
| 7 | Atp9a | 0.46 | 7.85E-12 | 1.63E-07 | 0.27 | 0.17 | 0.10 | 1.55 |
| 7 | Erbb2 | 0.41 | 7.90E-12 | 1.64E-07 | 0.34 | 0.23 | 0.11 | 1.48 |
| 7 | Bclaf3 | 0.26 | 8.03E-12 | 1.67E-07 | 0.46 | 0.33 | 0.13 | 1.41 |
| 7 | Proser2 | 0.46 | 8.30E-12 | 1.73E-07 | 0.11 | 0.06 | 0.06 | 2.02 |

**Supplementary Table 1. Marker genes for the original clusters of integrated clustering of E11.5 control and NotchGOF λ-junction cells.**

| | | | | | | | | |
|---|---|---|---|---|---|---|---|---|
| 7 | Dgkz | 0.38 | 8.36E-12 | 1.74E-07 | 0.40 | 0.28 | 0.12 | 1.43 |
| 7 | Frat1 | 0.63 | 9.27E-12 | 1.93E-07 | 0.10 | 0.05 | 0.05 | 2.10 |
| 7 | Cdk7 | 0.28 | 9.33E-12 | 1.94E-07 | 0.62 | 0.48 | 0.14 | 1.28 |
| 7 | Mttp | 0.49 | 1.01E-11 | 2.10E-07 | 0.21 | 0.13 | 0.08 | 1.65 |
| 7 | Mad2l1 | 0.27 | 1.01E-11 | 2.11E-07 | 0.69 | 0.56 | 0.14 | 1.24 |
| 7 | Srgap1 | 0.37 | 1.05E-11 | 2.18E-07 | 0.46 | 0.34 | 0.13 | 1.37 |
| 7 | Kifc3 | 0.69 | 1.07E-11 | 2.23E-07 | 0.14 | 0.07 | 0.06 | 1.86 |
| 7 | Hltf | 0.31 | 1.10E-11 | 2.29E-07 | 0.44 | 0.32 | 0.13 | 1.40 |
| 7 | Zfp687 | 0.43 | 1.10E-11 | 2.30E-07 | 0.30 | 0.19 | 0.10 | 1.53 |
| 7 | Tmem141 | 0.34 | 1.12E-11 | 2.34E-07 | 0.30 | 0.19 | 0.11 | 1.55 |
| 7 | Atp13a3 | 0.37 | 1.13E-11 | 2.36E-07 | 0.53 | 0.40 | 0.13 | 1.31 |
| 7 | Znhit2 | 0.38 | 1.14E-11 | 2.38E-07 | 0.41 | 0.29 | 0.12 | 1.42 |
| 7 | Pim3 | 0.31 | 1.17E-11 | 2.44E-07 | 0.42 | 0.29 | 0.12 | 1.42 |
| 7 | Ctdspl | 0.32 | 1.18E-11 | 2.45E-07 | 0.47 | 0.34 | 0.13 | 1.38 |
| 7 | Xkr6 | 0.44 | 1.18E-11 | 2.46E-07 | 0.21 | 0.12 | 0.08 | 1.68 |
| 7 | Nfkbiz | 0.72 | 1.22E-11 | 2.54E-07 | 0.13 | 0.07 | 0.06 | 1.88 |
| 7 | Camkmt | 0.36 | 1.27E-11 | 2.65E-07 | 0.28 | 0.18 | 0.10 | 1.56 |
| 7 | D11Wsu47e | 0.58 | 1.28E-11 | 2.67E-07 | 0.16 | 0.09 | 0.07 | 1.76 |
| 7 | Socs3 | 0.53 | 1.32E-11 | 2.74E-07 | 0.27 | 0.17 | 0.10 | 1.56 |
| 7 | Uvrag | 0.49 | 1.39E-11 | 2.89E-07 | 0.33 | 0.22 | 0.11 | 1.47 |
| 7 | Il10rb | 0.68 | 1.40E-11 | 2.92E-07 | 0.15 | 0.08 | 0.07 | 1.79 |
| 7 | Srebf2 | 0.31 | 1.42E-11 | 2.95E-07 | 0.65 | 0.52 | 0.13 | 1.24 |
| 7 | Nudt12 | 0.82 | 1.46E-11 | 3.03E-07 | 0.10 | 0.05 | 0.05 | 2.02 |
| 7 | Arfgef1 | 0.29 | 1.55E-11 | 3.23E-07 | 0.61 | 0.48 | 0.13 | 1.27 |
| 7 | Rps6ka2 | 0.60 | 1.70E-11 | 3.53E-07 | 0.14 | 0.08 | 0.07 | 1.83 |
| 7 | Orai3 | 0.50 | 1.76E-11 | 3.65E-07 | 0.19 | 0.11 | 0.08 | 1.70 |
| 7 | Klc1 | 0.26 | 1.87E-11 | 3.88E-07 | 0.60 | 0.47 | 0.14 | 1.30 |
| 7 | Mgme1 | 0.40 | 1.87E-11 | 3.89E-07 | 0.34 | 0.23 | 0.11 | 1.47 |
| 7 | Acrbp | 0.65 | 1.92E-11 | 3.99E-07 | 0.18 | 0.10 | 0.07 | 1.72 |
| 7 | Traf3 | 0.38 | 1.94E-11 | 4.04E-07 | 0.29 | 0.19 | 0.10 | 1.53 |
| 7 | Dnajb2 | 0.35 | 2.00E-11 | 4.15E-07 | 0.39 | 0.27 | 0.12 | 1.45 |
| 7 | Slc37a4 | 0.75 | 2.08E-11 | 4.33E-07 | 0.11 | 0.06 | 0.06 | 1.95 |
| 7 | Zfyve19 | 0.36 | 2.09E-11 | 4.34E-07 | 0.35 | 0.23 | 0.11 | 1.48 |
| 7 | Stxbp5 | 0.55 | 2.18E-11 | 4.53E-07 | 0.14 | 0.08 | 0.07 | 1.84 |
| 7 | Ncoa2 | 0.32 | 2.25E-11 | 4.69E-07 | 0.41 | 0.29 | 0.12 | 1.42 |
| 7 | Lpcat1 | 0.33 | 2.42E-11 | 5.03E-07 | 0.59 | 0.47 | 0.13 | 1.27 |
| 7 | Arhgef26 | 0.57 | 2.48E-11 | 5.17E-07 | 0.20 | 0.12 | 0.08 | 1.65 |
| 7 | Rrp36 | 0.30 | 2.53E-11 | 5.27E-07 | 0.44 | 0.31 | 0.13 | 1.41 |
| 7 | Sil1 | 0.48 | 2.54E-11 | 5.27E-07 | 0.19 | 0.11 | 0.08 | 1.70 |
| 7 | Rbm14 | 0.38 | 2.55E-11 | 5.31E-07 | 0.56 | 0.43 | 0.13 | 1.30 |
| 7 | Dusp16 | 0.44 | 2.87E-11 | 5.96E-07 | 0.31 | 0.21 | 0.10 | 1.49 |
| 7 | Bag5 | 0.31 | 2.91E-11 | 6.06E-07 | 0.61 | 0.49 | 0.12 | 1.25 |
| 7 | Copg2 | 0.35 | 3.09E-11 | 6.42E-07 | 0.49 | 0.37 | 0.12 | 1.33 |
| 7 | Pan3 | 0.32 | 3.11E-11 | 6.47E-07 | 0.45 | 0.33 | 0.12 | 1.38 |
| 7 | Uxt | 0.38 | 3.14E-11 | 6.52E-07 | 0.50 | 0.38 | 0.12 | 1.33 |
| 7 | Ptpn21 | 0.54 | 3.18E-11 | 6.61E-07 | 0.19 | 0.11 | 0.08 | 1.67 |
| 7 | Fgd4 | 0.47 | 3.31E-11 | 6.88E-07 | 0.16 | 0.09 | 0.07 | 1.77 |
| 7 | Arhgef12 | 0.27 | 3.45E-11 | 7.19E-07 | 0.69 | 0.58 | 0.12 | 1.20 |
| 7 | Snx24 | 0.29 | 3.61E-11 | 7.52E-07 | 0.49 | 0.35 | 0.13 | 1.38 |
| 7 | Gm9856 | 0.37 | 3.70E-11 | 7.70E-07 | 0.24 | 0.15 | 0.09 | 1.60 |
| 7 | Ube2t | 0.39 | 3.75E-11 | 7.80E-07 | 0.44 | 0.32 | 0.12 | 1.37 |
| 7 | Pef1 | 0.36 | 3.76E-11 | 7.82E-07 | 0.48 | 0.35 | 0.13 | 1.35 |
| 7 | Pspc1 | 0.25 | 3.82E-11 | 7.94E-07 | 0.73 | 0.60 | 0.13 | 1.22 |
| 7 | Gspt2 | 0.46 | 3.83E-11 | 7.96E-07 | 0.24 | 0.15 | 0.09 | 1.57 |
| 7 | Mlycd | 0.47 | 3.86E-11 | 8.03E-07 | 0.22 | 0.14 | 0.08 | 1.60 |
| 7 | Wbp2 | 0.35 | 3.88E-11 | 8.07E-07 | 0.39 | 0.27 | 0.12 | 1.42 |

**Supplementary Table 1. Marker genes for the original clusters of integrated clustering of E11.5 control and NotchGOF λ-junction cells.**

| | | | | | | | | |
|---|---|---|---|---|---|---|---|---|
| 7 | mt-Nd4l | 0.25 | 3.91E-11 | 8.13E-07 | 0.97 | 0.90 | 0.07 | 1.08 |
| 7 | Rnf6 | 0.31 | 4.06E-11 | 8.45E-07 | 0.48 | 0.35 | 0.13 | 1.38 |
| 7 | Ift43 | 0.28 | 4.12E-11 | 8.57E-07 | 0.68 | 0.54 | 0.14 | 1.26 |
| 7 | Por | 0.39 | 4.20E-11 | 8.73E-07 | 0.59 | 0.47 | 0.13 | 1.27 |
| 7 | Usp7 | 0.26 | 4.39E-11 | 9.14E-07 | 0.79 | 0.67 | 0.12 | 1.17 |
| 7 | Insig1 | 0.44 | 4.41E-11 | 9.16E-07 | 0.42 | 0.30 | 0.12 | 1.38 |
| 7 | Sfmbt1 | 0.48 | 4.84E-11 | 1.01E-06 | 0.29 | 0.19 | 0.10 | 1.52 |
| 7 | AW146154 | 0.60 | 5.09E-11 | 1.06E-06 | 0.15 | 0.09 | 0.07 | 1.76 |
| 7 | Znrd1as | 0.44 | 5.44E-11 | 1.13E-06 | 0.20 | 0.13 | 0.08 | 1.63 |
| 7 | Apmap | 0.49 | 5.77E-11 | 1.20E-06 | 0.28 | 0.19 | 0.09 | 1.51 |
| 7 | Mvd | 0.36 | 6.07E-11 | 1.26E-06 | 0.41 | 0.30 | 0.12 | 1.40 |
| 7 | Slc44a2 | 0.40 | 6.17E-11 | 1.28E-06 | 0.39 | 0.27 | 0.11 | 1.41 |
| 7 | Oxnad1 | 0.40 | 6.24E-11 | 1.30E-06 | 0.28 | 0.19 | 0.10 | 1.52 |
| 7 | Paip2b | 0.35 | 6.32E-11 | 1.31E-06 | 0.41 | 0.29 | 0.11 | 1.39 |
| 7 | Fam214b | 0.61 | 6.42E-11 | 1.33E-06 | 0.16 | 0.10 | 0.07 | 1.72 |
| 7 | Uevld | 0.65 | 6.80E-11 | 1.41E-06 | 0.20 | 0.12 | 0.08 | 1.62 |
| 7 | Mettl5 | 0.34 | 6.82E-11 | 1.42E-06 | 0.43 | 0.31 | 0.12 | 1.37 |
| 7 | Oxsr1 | 0.38 | 6.91E-11 | 1.44E-06 | 0.43 | 0.32 | 0.12 | 1.38 |
| 7 | Ncstn | 0.32 | 7.08E-11 | 1.47E-06 | 0.40 | 0.29 | 0.12 | 1.41 |
| 7 | Trim3 | 0.64 | 7.46E-11 | 1.55E-06 | 0.16 | 0.09 | 0.07 | 1.72 |
| 7 | Vps54 | 0.30 | 7.54E-11 | 1.57E-06 | 0.43 | 0.31 | 0.12 | 1.40 |
| 7 | Ipo7 | 0.27 | 7.66E-11 | 1.59E-06 | 0.73 | 0.61 | 0.11 | 1.18 |
| 7 | Appbp2 | 0.25 | 8.15E-11 | 1.69E-06 | 0.77 | 0.65 | 0.12 | 1.18 |
| 7 | Zbtb8os | 0.30 | 8.17E-11 | 1.70E-06 | 0.62 | 0.50 | 0.12 | 1.25 |
| 7 | Ccdc86 | 0.27 | 8.19E-11 | 1.70E-06 | 0.69 | 0.55 | 0.14 | 1.25 |
| 7 | Mov10 | 0.46 | 8.29E-11 | 1.72E-06 | 0.29 | 0.19 | 0.09 | 1.49 |
| 7 | Plekha1 | 0.37 | 8.44E-11 | 1.76E-06 | 0.30 | 0.20 | 0.10 | 1.51 |
| 7 | Zfp316 | 0.46 | 8.49E-11 | 1.77E-06 | 0.23 | 0.14 | 0.08 | 1.60 |
| 7 | Rnaseh2b | 0.27 | 8.64E-11 | 1.80E-06 | 0.63 | 0.49 | 0.13 | 1.27 |
| 7 | Noc4l | 0.34 | 8.70E-11 | 1.81E-06 | 0.55 | 0.42 | 0.12 | 1.29 |
| 7 | Zswim1 | 0.47 | 8.74E-11 | 1.82E-06 | 0.19 | 0.12 | 0.08 | 1.65 |
| 7 | Fahd2a | 0.45 | 9.38E-11 | 1.95E-06 | 0.28 | 0.19 | 0.09 | 1.48 |
| 7 | F8a | 0.53 | 9.47E-11 | 1.97E-06 | 0.15 | 0.08 | 0.06 | 1.76 |
| 7 | Ncor2 | 0.39 | 9.66E-11 | 2.01E-06 | 0.47 | 0.35 | 0.12 | 1.35 |
| 7 | Tcirg1 | 0.67 | 9.72E-11 | 2.02E-06 | 0.15 | 0.09 | 0.07 | 1.75 |
| 7 | Ppip5k2 | 0.31 | 1.01E-10 | 2.09E-06 | 0.58 | 0.45 | 0.13 | 1.28 |
| 7 | Mecr | 0.31 | 1.01E-10 | 2.10E-06 | 0.43 | 0.31 | 0.12 | 1.40 |
| 7 | Pacs2 | 0.38 | 1.04E-10 | 2.16E-06 | 0.26 | 0.17 | 0.09 | 1.55 |
| 7 | Hint2 | 0.26 | 1.06E-10 | 2.21E-06 | 0.73 | 0.61 | 0.12 | 1.20 |
| 7 | Grsf1 | 0.27 | 1.08E-10 | 2.24E-06 | 0.71 | 0.60 | 0.10 | 1.17 |
| 7 | Pcbd2 | 0.30 | 1.12E-10 | 2.33E-06 | 0.56 | 0.43 | 0.13 | 1.29 |
| 7 | Wdr74 | 0.26 | 1.15E-10 | 2.40E-06 | 0.67 | 0.54 | 0.13 | 1.24 |
| 7 | Yars2 | 0.33 | 1.17E-10 | 2.43E-06 | 0.52 | 0.40 | 0.12 | 1.31 |
| 7 | Pih1d2 | 0.61 | 1.19E-10 | 2.47E-06 | 0.13 | 0.07 | 0.06 | 1.84 |
| 7 | Asah1 | 0.30 | 1.25E-10 | 2.60E-06 | 0.49 | 0.37 | 0.13 | 1.35 |
| 7 | Ttc13 | 0.41 | 1.26E-10 | 2.62E-06 | 0.33 | 0.23 | 0.10 | 1.45 |
| 7 | Vars | 0.30 | 1.30E-10 | 2.70E-06 | 0.72 | 0.60 | 0.12 | 1.19 |
| 7 | Cep152 | 0.55 | 1.32E-10 | 2.75E-06 | 0.20 | 0.12 | 0.08 | 1.61 |
| 7 | Xkr5 | 0.74 | 1.34E-10 | 2.79E-06 | 0.13 | 0.07 | 0.06 | 1.82 |
| 7 | Ccnq | 0.27 | 1.34E-10 | 2.79E-06 | 0.51 | 0.38 | 0.13 | 1.34 |
| 7 | Gpr27 | 0.42 | 1.35E-10 | 2.80E-06 | 0.22 | 0.14 | 0.08 | 1.60 |
| 7 | Flnb | 0.34 | 1.35E-10 | 2.80E-06 | 0.40 | 0.29 | 0.11 | 1.38 |
| 7 | Mettl14 | 0.42 | 1.36E-10 | 2.83E-06 | 0.38 | 0.27 | 0.11 | 1.43 |
| 7 | Dipk1b | 0.25 | 1.40E-10 | 2.90E-06 | 0.62 | 0.47 | 0.14 | 1.30 |
| 7 | Zdhhc6 | 0.26 | 1.46E-10 | 3.05E-06 | 0.58 | 0.45 | 0.13 | 1.30 |
| 7 | Adarb1 | 0.64 | 1.48E-10 | 3.08E-06 | 0.13 | 0.07 | 0.06 | 1.84 |

**Supplementary Table 1. Marker genes for the original clusters of integrated clustering of E11.5 control and NotchGOF λ-junction cells.**

| | | | | | | | | |
|---|---|---|---|---|---|---|---|---|
| 7 | Cep70 | 0.28 | 1.50E-10 | 3.11E-06 | 0.52 | 0.39 | 0.13 | 1.32 |
| 7 | Naa40 | 0.36 | 1.58E-10 | 3.29E-06 | 0.49 | 0.37 | 0.13 | 1.35 |
| 7 | Nup133 | 0.30 | 1.60E-10 | 3.32E-06 | 0.50 | 0.37 | 0.12 | 1.33 |
| 7 | Miip | 0.43 | 1.63E-10 | 3.39E-06 | 0.31 | 0.21 | 0.10 | 1.46 |
| 7 | Eed | 0.31 | 1.65E-10 | 3.43E-06 | 0.65 | 0.54 | 0.11 | 1.21 |
| 7 | Deaf1 | 0.37 | 1.65E-10 | 3.43E-06 | 0.32 | 0.22 | 0.10 | 1.47 |
| 7 | Rnf145 | 0.33 | 1.66E-10 | 3.46E-06 | 0.50 | 0.37 | 0.13 | 1.35 |
| 7 | Glis2 | 0.37 | 1.69E-10 | 3.51E-06 | 0.29 | 0.19 | 0.10 | 1.52 |
| 7 | Wdpcp | 0.60 | 1.85E-10 | 3.85E-06 | 0.14 | 0.08 | 0.06 | 1.78 |
| 7 | Nedd4l | 0.47 | 1.92E-10 | 3.99E-06 | 0.22 | 0.14 | 0.08 | 1.56 |
| 7 | Col4a5 | 0.37 | 1.98E-10 | 4.11E-06 | 0.21 | 0.13 | 0.08 | 1.63 |
| 7 | Tead3 | 0.35 | 1.98E-10 | 4.11E-06 | 0.35 | 0.24 | 0.11 | 1.44 |
| 7 | Btbd3 | 0.53 | 2.01E-10 | 4.19E-06 | 0.24 | 0.15 | 0.08 | 1.55 |
| 7 | Sac3d1 | 0.31 | 2.08E-10 | 4.33E-06 | 0.59 | 0.46 | 0.12 | 1.27 |
| 7 | Shmt1 | 0.37 | 2.22E-10 | 4.61E-06 | 0.48 | 0.35 | 0.12 | 1.35 |
| 7 | Cptp | 0.49 | 2.25E-10 | 4.69E-06 | 0.21 | 0.13 | 0.08 | 1.61 |
| 7 | Gm10282 | 0.38 | 2.28E-10 | 4.75E-06 | 0.24 | 0.15 | 0.09 | 1.58 |
| 7 | Pex16 | 0.48 | 2.37E-10 | 4.92E-06 | 0.27 | 0.18 | 0.09 | 1.49 |
| 7 | Zfp317 | 0.48 | 2.55E-10 | 5.31E-06 | 0.30 | 0.20 | 0.09 | 1.47 |
| 7 | Pknox1 | 0.32 | 2.59E-10 | 5.39E-06 | 0.34 | 0.23 | 0.11 | 1.47 |
| 7 | Klhl42 | 0.49 | 2.61E-10 | 5.42E-06 | 0.17 | 0.10 | 0.07 | 1.69 |
| 7 | Cabin1 | 0.31 | 2.67E-10 | 5.56E-06 | 0.58 | 0.45 | 0.13 | 1.29 |
| 7 | Homez | 0.48 | 2.71E-10 | 5.64E-06 | 0.16 | 0.09 | 0.07 | 1.71 |
| 7 | F2r | 0.25 | 2.76E-10 | 5.74E-06 | 0.71 | 0.56 | 0.14 | 1.25 |
| 7 | Cxxc5 | 0.28 | 2.91E-10 | 6.05E-06 | 0.58 | 0.45 | 0.13 | 1.29 |
| 7 | Rpp21 | 0.28 | 2.94E-10 | 6.12E-06 | 0.70 | 0.57 | 0.13 | 1.22 |
| 7 | Med25 | 0.26 | 3.17E-10 | 6.60E-06 | 0.61 | 0.49 | 0.12 | 1.25 |
| 7 | Tmem14a | 0.42 | 3.21E-10 | 6.67E-06 | 0.17 | 0.10 | 0.07 | 1.68 |
| 7 | Derl2 | 0.34 | 3.24E-10 | 6.73E-06 | 0.50 | 0.37 | 0.12 | 1.33 |
| 7 | Ubald1 | 0.31 | 3.35E-10 | 6.98E-06 | 0.37 | 0.26 | 0.11 | 1.41 |
| 7 | Odf2 | 0.37 | 3.38E-10 | 7.02E-06 | 0.57 | 0.45 | 0.11 | 1.25 |
| 7 | Prodh | 0.52 | 3.40E-10 | 7.07E-06 | 0.18 | 0.11 | 0.07 | 1.66 |
| 7 | Rab10os | 0.39 | 3.42E-10 | 7.11E-06 | 0.27 | 0.18 | 0.09 | 1.51 |
| 7 | Herpud2 | 0.30 | 3.61E-10 | 7.51E-06 | 0.39 | 0.28 | 0.12 | 1.41 |
| 7 | Cradd | 0.46 | 3.63E-10 | 7.56E-06 | 0.25 | 0.17 | 0.09 | 1.52 |
| 7 | Myo1e | 0.39 | 3.72E-10 | 7.75E-06 | 0.34 | 0.23 | 0.11 | 1.45 |
| 7 | Adam15 | 0.47 | 3.77E-10 | 7.84E-06 | 0.16 | 0.10 | 0.07 | 1.71 |
| 7 | Pdcd2 | 0.29 | 3.85E-10 | 8.00E-06 | 0.51 | 0.38 | 0.13 | 1.33 |
| 7 | Mob1b | 0.28 | 4.00E-10 | 8.32E-06 | 0.55 | 0.43 | 0.13 | 1.29 |
| 7 | Rel | 0.66 | 4.02E-10 | 8.37E-06 | 0.15 | 0.09 | 0.06 | 1.72 |
| 7 | Pcgf6 | 0.28 | 4.03E-10 | 8.38E-06 | 0.60 | 0.48 | 0.12 | 1.25 |
| 7 | Whamm | 0.62 | 4.28E-10 | 8.90E-06 | 0.16 | 0.10 | 0.07 | 1.67 |
| 7 | Tm2d3 | 0.30 | 4.33E-10 | 9.00E-06 | 0.63 | 0.50 | 0.13 | 1.26 |
| 7 | Insig2 | 0.33 | 4.45E-10 | 9.25E-06 | 0.38 | 0.27 | 0.11 | 1.40 |
| 7 | 1300002E11Rik | 0.29 | 4.51E-10 | 9.37E-06 | 0.44 | 0.32 | 0.12 | 1.38 |
| 7 | Kif1c | 0.48 | 4.92E-10 | 1.02E-05 | 0.24 | 0.16 | 0.08 | 1.54 |
| 7 | Ubac2 | 0.26 | 5.13E-10 | 1.07E-05 | 0.58 | 0.44 | 0.14 | 1.31 |
| 7 | Sec23ip | 0.36 | 5.21E-10 | 1.08E-05 | 0.41 | 0.30 | 0.11 | 1.36 |
| 7 | Inafm2 | 0.45 | 5.28E-10 | 1.10E-05 | 0.20 | 0.12 | 0.07 | 1.60 |
| 7 | Echdc2 | 0.32 | 5.42E-10 | 1.13E-05 | 0.36 | 0.25 | 0.11 | 1.43 |
| 7 | Mllt6 | 0.42 | 5.44E-10 | 1.13E-05 | 0.29 | 0.20 | 0.09 | 1.47 |
| 7 | Heatr1 | 0.31 | 5.55E-10 | 1.15E-05 | 0.42 | 0.30 | 0.12 | 1.39 |
| 7 | Dgcr2 | 0.31 | 5.56E-10 | 1.16E-05 | 0.59 | 0.46 | 0.12 | 1.27 |
| 7 | Med17 | 0.31 | 5.75E-10 | 1.20E-05 | 0.46 | 0.33 | 0.12 | 1.36 |
| 7 | Epb41l4b | 0.57 | 5.78E-10 | 1.20E-05 | 0.16 | 0.09 | 0.07 | 1.71 |
| 7 | Tor2a | 0.33 | 5.80E-10 | 1.21E-05 | 0.53 | 0.42 | 0.12 | 1.28 |

**Supplementary Table 1. Marker genes for the original clusters of integrated clustering of E11.5 control and NotchGOF λ-junction cells.**

| | | | | | | | | |
|---|---|---|---|---|---|---|---|---|
| 7 | Lpcat4 | 0.77 | 6.16E-10 | 1.28E-05 | 0.11 | 0.06 | 0.05 | 1.85 |
| 7 | Dnaaf2 | 0.28 | 6.40E-10 | 1.33E-05 | 0.64 | 0.51 | 0.13 | 1.25 |
| 7 | Suco | 0.27 | 6.44E-10 | 1.34E-05 | 0.61 | 0.49 | 0.12 | 1.24 |
| 7 | Gde1 | 0.27 | 6.45E-10 | 1.34E-05 | 0.64 | 0.52 | 0.13 | 1.25 |
| 7 | Nubp2 | 0.32 | 6.54E-10 | 1.36E-05 | 0.42 | 0.30 | 0.11 | 1.37 |
| 7 | Dclk1 | 0.28 | 6.58E-10 | 1.37E-05 | 0.36 | 0.25 | 0.11 | 1.45 |
| 7 | Epb41l3 | 0.26 | 6.73E-10 | 1.40E-05 | 0.58 | 0.44 | 0.14 | 1.32 |
| 7 | Npc1 | 0.26 | 6.81E-10 | 1.42E-05 | 0.37 | 0.26 | 0.11 | 1.44 |
| 7 | Cdk10 | 0.36 | 6.86E-10 | 1.43E-05 | 0.34 | 0.24 | 0.10 | 1.42 |
| 7 | Zxdb | 0.58 | 7.12E-10 | 1.48E-05 | 0.17 | 0.10 | 0.07 | 1.66 |
| 7 | Hacd2 | 0.28 | 7.39E-10 | 1.54E-05 | 0.49 | 0.37 | 0.12 | 1.34 |
| 7 | Dgat2 | 0.66 | 7.41E-10 | 1.54E-05 | 0.11 | 0.06 | 0.05 | 1.88 |
| 7 | Tmem219 | 0.36 | 7.78E-10 | 1.62E-05 | 0.24 | 0.15 | 0.08 | 1.55 |
| 7 | AV356131 | 0.45 | 7.83E-10 | 1.63E-05 | 0.14 | 0.08 | 0.06 | 1.79 |
| 7 | Mtr | 0.43 | 7.87E-10 | 1.64E-05 | 0.32 | 0.23 | 0.09 | 1.41 |
| 7 | Bag2 | 0.30 | 7.89E-10 | 1.64E-05 | 0.52 | 0.40 | 0.12 | 1.30 |
| 7 | Pik3cb | 0.50 | 7.91E-10 | 1.64E-05 | 0.11 | 0.06 | 0.05 | 1.87 |
| 7 | Aebp2 | 0.31 | 8.31E-10 | 1.73E-05 | 0.54 | 0.41 | 0.13 | 1.31 |
| 7 | Ankzf1 | 0.31 | 8.46E-10 | 1.76E-05 | 0.31 | 0.22 | 0.10 | 1.46 |
| 7 | Cldn12 | 0.49 | 8.48E-10 | 1.76E-05 | 0.18 | 0.11 | 0.07 | 1.64 |
| 7 | Enkd1 | 0.33 | 8.58E-10 | 1.78E-05 | 0.39 | 0.28 | 0.11 | 1.39 |
| 7 | Slc25a27 | 0.26 | 8.63E-10 | 1.79E-05 | 0.53 | 0.40 | 0.13 | 1.34 |
| 7 | Dtd2 | 0.35 | 8.88E-10 | 1.85E-05 | 0.44 | 0.32 | 0.12 | 1.36 |
| 7 | Tmem30a | 0.28 | 8.94E-10 | 1.86E-05 | 0.63 | 0.51 | 0.13 | 1.25 |
| 7 | Atmin | 0.42 | 9.26E-10 | 1.93E-05 | 0.21 | 0.14 | 0.08 | 1.57 |
| 7 | Fbxw7 | 0.35 | 9.27E-10 | 1.93E-05 | 0.49 | 0.38 | 0.11 | 1.30 |
| 7 | Wdr90 | 0.53 | 9.58E-10 | 1.99E-05 | 0.19 | 0.12 | 0.07 | 1.60 |
| 7 | Pmvk | 0.30 | 9.91E-10 | 2.06E-05 | 0.66 | 0.55 | 0.11 | 1.21 |
| 7 | Dnaaf4 | 0.73 | 9.98E-10 | 2.08E-05 | 0.11 | 0.06 | 0.05 | 1.88 |
| 7 | Uri1 | 0.25 | 1.04E-09 | 2.15E-05 | 0.72 | 0.61 | 0.10 | 1.17 |
| 7 | Mpv17l | 0.40 | 1.06E-09 | 2.19E-05 | 0.13 | 0.08 | 0.06 | 1.74 |
| 7 | Ralgapa1 | 0.35 | 1.07E-09 | 2.23E-05 | 0.45 | 0.34 | 0.11 | 1.32 |
| 7 | Nfxl1 | 0.34 | 1.08E-09 | 2.24E-05 | 0.45 | 0.34 | 0.11 | 1.34 |
| 7 | Ecpas | 0.30 | 1.10E-09 | 2.29E-05 | 0.54 | 0.42 | 0.12 | 1.29 |
| 7 | Zcchc7 | 0.28 | 1.27E-09 | 2.64E-05 | 0.64 | 0.52 | 0.12 | 1.23 |
| 7 | Ncoa6 | 0.33 | 1.28E-09 | 2.66E-05 | 0.47 | 0.35 | 0.12 | 1.34 |
| 7 | Rab28 | 0.28 | 1.29E-09 | 2.69E-05 | 0.62 | 0.50 | 0.13 | 1.26 |
| 7 | Pomk | 0.44 | 1.31E-09 | 2.73E-05 | 0.15 | 0.09 | 0.06 | 1.72 |
| 7 | Zfp703 | 0.39 | 1.34E-09 | 2.78E-05 | 0.35 | 0.25 | 0.09 | 1.37 |
| 7 | Slc30a1 | 0.75 | 1.37E-09 | 2.85E-05 | 0.14 | 0.08 | 0.06 | 1.71 |
| 7 | Mthfd1 | 0.31 | 1.42E-09 | 2.96E-05 | 0.50 | 0.38 | 0.12 | 1.31 |
| 7 | Mtmr4 | 0.38 | 1.43E-09 | 2.97E-05 | 0.39 | 0.29 | 0.11 | 1.37 |
| 7 | Wdr24 | 0.36 | 1.44E-09 | 2.99E-05 | 0.19 | 0.11 | 0.07 | 1.63 |
| 7 | Tbc1d1 | 0.28 | 1.44E-09 | 2.99E-05 | 0.49 | 0.37 | 0.12 | 1.33 |
| 7 | Serinc5 | 0.60 | 1.52E-09 | 3.16E-05 | 0.11 | 0.06 | 0.05 | 1.88 |
| 7 | Cdkl5 | 0.72 | 1.53E-09 | 3.18E-05 | 0.10 | 0.06 | 0.05 | 1.89 |
| 7 | Taf6l | 0.31 | 1.57E-09 | 3.26E-05 | 0.37 | 0.27 | 0.11 | 1.40 |
| 7 | Pex6 | 0.33 | 1.59E-09 | 3.32E-05 | 0.40 | 0.29 | 0.11 | 1.39 |
| 7 | Bpnt1 | 0.36 | 1.61E-09 | 3.34E-05 | 0.21 | 0.13 | 0.08 | 1.58 |
| 7 | Tctex1d2 | 0.27 | 1.79E-09 | 3.72E-05 | 0.56 | 0.43 | 0.12 | 1.29 |
| 7 | Gm3448 | 0.78 | 1.80E-09 | 3.74E-05 | 0.10 | 0.06 | 0.05 | 1.86 |
| 7 | Slc35a2 | 0.41 | 1.86E-09 | 3.88E-05 | 0.23 | 0.15 | 0.08 | 1.53 |
| 7 | 1600002K03Rik | 0.62 | 2.04E-09 | 4.24E-05 | 0.18 | 0.11 | 0.07 | 1.58 |
| 7 | Abcb7 | 0.36 | 2.05E-09 | 4.27E-05 | 0.29 | 0.20 | 0.09 | 1.47 |
| 7 | Trmt2b | 0.57 | 2.10E-09 | 4.36E-05 | 0.15 | 0.09 | 0.06 | 1.70 |
| 7 | G6pc3 | 0.36 | 2.15E-09 | 4.48E-05 | 0.37 | 0.27 | 0.10 | 1.38 |

**Supplementary Table 1. Marker genes for the original clusters of integrated clustering of E11.5 control and NotchGOF λ-junction cells.**

| | | | | | | | | |
|---|---|---|---|---|---|---|---|---|
| 7 | Coq10a | 0.49 | 2.16E-09 | 4.49E-05 | 0.14 | 0.08 | 0.06 | 1.73 |
| 7 | Snx16 | 0.40 | 2.25E-09 | 4.68E-05 | 0.28 | 0.19 | 0.09 | 1.47 |
| 7 | 2010204K13Rik | 0.30 | 2.39E-09 | 4.98E-05 | 0.36 | 0.25 | 0.11 | 1.42 |
| 7 | Cinp | 0.31 | 2.40E-09 | 5.00E-05 | 0.60 | 0.48 | 0.12 | 1.25 |
| 7 | Rcc1 | 0.26 | 2.45E-09 | 5.10E-05 | 0.63 | 0.51 | 0.12 | 1.23 |
| 7 | Ttll1 | 0.34 | 2.59E-09 | 5.39E-05 | 0.24 | 0.16 | 0.08 | 1.52 |
| 7 | Lipt1 | 0.65 | 2.62E-09 | 5.45E-05 | 0.11 | 0.06 | 0.05 | 1.88 |
| 7 | Uhrf1 | 0.27 | 2.70E-09 | 5.61E-05 | 0.78 | 0.68 | 0.10 | 1.15 |
| 7 | Scrib | 0.37 | 2.77E-09 | 5.76E-05 | 0.47 | 0.36 | 0.11 | 1.32 |
| 7 | Tapt1 | 0.47 | 2.86E-09 | 5.94E-05 | 0.23 | 0.15 | 0.08 | 1.52 |
| 7 | Zfp113 | 0.27 | 2.86E-09 | 5.94E-05 | 0.44 | 0.32 | 0.11 | 1.35 |
| 7 | Pex11a | 0.34 | 2.88E-09 | 5.99E-05 | 0.18 | 0.11 | 0.07 | 1.65 |
| 7 | Rfx3 | 0.37 | 2.88E-09 | 6.00E-05 | 0.31 | 0.21 | 0.10 | 1.47 |
| 7 | A430005L14Rik | 0.28 | 2.92E-09 | 6.06E-05 | 0.62 | 0.50 | 0.12 | 1.24 |
| 7 | 2310068J16Rik | 0.56 | 2.93E-09 | 6.10E-05 | 0.13 | 0.07 | 0.06 | 1.77 |
| 7 | Uck2 | 0.26 | 3.13E-09 | 6.51E-05 | 0.64 | 0.52 | 0.12 | 1.22 |
| 7 | Dennd6b | 0.46 | 3.20E-09 | 6.66E-05 | 0.15 | 0.09 | 0.06 | 1.68 |
| 7 | Alms1 | 0.41 | 3.24E-09 | 6.73E-05 | 0.35 | 0.26 | 0.10 | 1.38 |
| 7 | Pde5a | 0.37 | 3.24E-09 | 6.74E-05 | 0.27 | 0.18 | 0.09 | 1.48 |
| 7 | Piezo1 | 0.50 | 3.34E-09 | 6.96E-05 | 0.19 | 0.12 | 0.07 | 1.57 |
| 7 | Ptk7 | 0.32 | 3.41E-09 | 7.09E-05 | 0.48 | 0.36 | 0.12 | 1.33 |
| 7 | Urb2 | 0.53 | 3.43E-09 | 7.13E-05 | 0.14 | 0.08 | 0.06 | 1.70 |
| 7 | 0610010F05Rik | 0.26 | 3.48E-09 | 7.23E-05 | 0.35 | 0.24 | 0.10 | 1.43 |
| 7 | Gla | 0.51 | 3.48E-09 | 7.24E-05 | 0.17 | 0.10 | 0.07 | 1.64 |
| 7 | Tinf2 | 0.40 | 3.50E-09 | 7.28E-05 | 0.29 | 0.20 | 0.09 | 1.44 |
| 7 | Ccdc47 | 0.25 | 3.51E-09 | 7.29E-05 | 0.73 | 0.62 | 0.11 | 1.18 |
| 7 | Tor1b | 0.36 | 3.63E-09 | 7.55E-05 | 0.45 | 0.34 | 0.11 | 1.31 |
| 7 | Spryd7 | 0.39 | 3.80E-09 | 7.91E-05 | 0.23 | 0.15 | 0.08 | 1.52 |
| 7 | Acsl4 | 0.48 | 3.92E-09 | 8.15E-05 | 0.21 | 0.14 | 0.07 | 1.53 |
| 7 | Exoc4 | 0.27 | 4.02E-09 | 8.36E-05 | 0.53 | 0.40 | 0.12 | 1.30 |
| 7 | Rcan1 | 0.26 | 4.10E-09 | 8.53E-05 | 0.36 | 0.26 | 0.10 | 1.40 |
| 7 | Ppp1r13l | 0.43 | 4.21E-09 | 8.76E-05 | 0.13 | 0.08 | 0.06 | 1.73 |
| 7 | Rnf44 | 0.27 | 4.35E-09 | 9.05E-05 | 0.46 | 0.34 | 0.12 | 1.35 |
| 7 | Nr2c1 | 0.39 | 4.35E-09 | 9.05E-05 | 0.32 | 0.22 | 0.09 | 1.42 |
| 7 | Ggct | 0.31 | 4.40E-09 | 9.16E-05 | 0.22 | 0.14 | 0.08 | 1.56 |
| 7 | Fasn | 0.30 | 4.61E-09 | 9.59E-05 | 0.62 | 0.50 | 0.12 | 1.23 |
| 7 | Ganab | 0.25 | 4.62E-09 | 9.61E-05 | 0.66 | 0.53 | 0.13 | 1.25 |
| 7 | Smpdl3a | 0.48 | 4.62E-09 | 9.61E-05 | 0.20 | 0.13 | 0.07 | 1.55 |
| 7 | A830082K12Rik | 0.46 | 4.65E-09 | 9.68E-05 | 0.13 | 0.07 | 0.06 | 1.75 |
| 7 | Las1l | 0.28 | 4.69E-09 | 9.74E-05 | 0.64 | 0.51 | 0.12 | 1.24 |
| 7 | Alkbh5 | 0.27 | 4.82E-09 | 1.00E-04 | 0.65 | 0.54 | 0.11 | 1.20 |
| 7 | Taf5l | 0.31 | 4.83E-09 | 1.00E-04 | 0.38 | 0.27 | 0.11 | 1.39 |
| 7 | Flot2 | 0.38 | 4.95E-09 | 1.03E-04 | 0.30 | 0.21 | 0.09 | 1.42 |
| 7 | 2810408A11Rik | 0.32 | 4.99E-09 | 1.04E-04 | 0.23 | 0.15 | 0.08 | 1.54 |
| 7 | Aen | 0.28 | 5.28E-09 | 1.10E-04 | 0.51 | 0.39 | 0.12 | 1.30 |
| 7 | Pex12 | 0.47 | 5.56E-09 | 1.16E-04 | 0.15 | 0.09 | 0.06 | 1.66 |
| 7 | Atf6b | 0.29 | 5.64E-09 | 1.17E-04 | 0.52 | 0.40 | 0.12 | 1.30 |
| 7 | Trip6 | 0.26 | 5.66E-09 | 1.18E-04 | 0.59 | 0.46 | 0.13 | 1.28 |
| 7 | Senp8 | 0.27 | 5.73E-09 | 1.19E-04 | 0.30 | 0.21 | 0.09 | 1.45 |
| 7 | 1700037C18Rik | 0.50 | 5.82E-09 | 1.21E-04 | 0.12 | 0.06 | 0.05 | 1.80 |
| 7 | Zfp667 | 0.49 | 5.86E-09 | 1.22E-04 | 0.16 | 0.10 | 0.06 | 1.63 |
| 7 | Afmid | 0.72 | 5.93E-09 | 1.23E-04 | 0.11 | 0.06 | 0.05 | 1.78 |
| 7 | Eml1 | 0.32 | 6.04E-09 | 1.26E-04 | 0.25 | 0.17 | 0.08 | 1.50 |
| 7 | Bbc3 | 0.45 | 6.06E-09 | 1.26E-04 | 0.16 | 0.10 | 0.06 | 1.63 |
| 7 | Prrg2 | 0.38 | 6.30E-09 | 1.31E-04 | 0.33 | 0.24 | 0.09 | 1.40 |
| 7 | Slc39a9 | 0.40 | 6.36E-09 | 1.32E-04 | 0.23 | 0.15 | 0.08 | 1.52 |

**Supplementary Table 1. Marker genes for the original clusters of integrated clustering of E11.5 control and NotchGOF λ-junction cells.**

| | | | | | | | | |
|---|---|---|---|---|---|---|---|---|
| 7 | Gt(ROSA)26Sor | 0.39 | 6.64E-09 | 1.38E-04 | 0.33 | 0.24 | 0.09 | 1.39 |
| 7 | Eif2b2 | 0.27 | 6.89E-09 | 1.43E-04 | 0.60 | 0.49 | 0.12 | 1.24 |
| 7 | Minpp1 | 0.28 | 6.96E-09 | 1.45E-04 | 0.62 | 0.51 | 0.12 | 1.23 |
| 7 | Fam98a | 0.30 | 7.02E-09 | 1.46E-04 | 0.43 | 0.32 | 0.11 | 1.34 |
| 7 | AI413582 | 0.56 | 7.59E-09 | 1.58E-04 | 0.15 | 0.09 | 0.06 | 1.66 |
| 7 | Tulp1 | 0.32 | 7.84E-09 | 1.63E-04 | 0.14 | 0.09 | 0.06 | 1.69 |
| 7 | Gan | 0.41 | 8.13E-09 | 1.69E-04 | 0.33 | 0.23 | 0.09 | 1.39 |
| 7 | Reps1 | 0.26 | 8.40E-09 | 1.75E-04 | 0.46 | 0.35 | 0.11 | 1.32 |
| 7 | Nbn | 0.28 | 8.76E-09 | 1.82E-04 | 0.44 | 0.33 | 0.11 | 1.33 |
| 7 | Vmac | 0.37 | 8.88E-09 | 1.85E-04 | 0.15 | 0.09 | 0.06 | 1.66 |
| 7 | Fbxo44 | 0.36 | 8.89E-09 | 1.85E-04 | 0.15 | 0.09 | 0.06 | 1.66 |
| 7 | Fkbp14 | 0.34 | 8.91E-09 | 1.85E-04 | 0.21 | 0.13 | 0.08 | 1.56 |
| 7 | Dlg1 | 0.36 | 8.94E-09 | 1.86E-04 | 0.48 | 0.37 | 0.11 | 1.30 |
| 7 | Usp20 | 0.40 | 9.12E-09 | 1.90E-04 | 0.13 | 0.08 | 0.06 | 1.71 |
| 7 | Tmem123 | 0.27 | 9.19E-09 | 1.91E-04 | 0.44 | 0.32 | 0.12 | 1.37 |
| 7 | Tgoln1 | 0.25 | 9.83E-09 | 2.04E-04 | 0.39 | 0.29 | 0.11 | 1.37 |
| 7 | Cnpy3 | 0.29 | 9.98E-09 | 2.08E-04 | 0.46 | 0.35 | 0.11 | 1.33 |
| 7 | Scn8a | 0.64 | 1.01E-08 | 2.09E-04 | 0.12 | 0.07 | 0.05 | 1.76 |
| 7 | Tmem250-ps | 0.47 | 1.01E-08 | 2.10E-04 | 0.21 | 0.14 | 0.07 | 1.52 |
| 7 | Nfkbib | 0.26 | 1.01E-08 | 2.11E-04 | 0.34 | 0.24 | 0.10 | 1.40 |
| 7 | Zfp865 | 0.38 | 1.02E-08 | 2.12E-04 | 0.33 | 0.23 | 0.10 | 1.42 |
| 7 | Slf2 | 0.30 | 1.07E-08 | 2.22E-04 | 0.43 | 0.32 | 0.11 | 1.34 |
| 7 | Lrp10 | 0.36 | 1.09E-08 | 2.27E-04 | 0.25 | 0.17 | 0.08 | 1.48 |
| 7 | Zfp605 | 0.48 | 1.09E-08 | 2.27E-04 | 0.18 | 0.12 | 0.07 | 1.57 |
| 7 | Nek6 | 0.30 | 1.10E-08 | 2.29E-04 | 0.41 | 0.31 | 0.10 | 1.34 |
| 7 | Triobp | 0.28 | 1.14E-08 | 2.37E-04 | 0.42 | 0.31 | 0.11 | 1.35 |
| 7 | Amer2 | 0.33 | 1.14E-08 | 2.37E-04 | 0.17 | 0.11 | 0.06 | 1.60 |
| 7 | Gramd4 | 0.41 | 1.16E-08 | 2.42E-04 | 0.14 | 0.08 | 0.06 | 1.69 |
| 7 | Fam120a | 0.30 | 1.24E-08 | 2.59E-04 | 0.57 | 0.45 | 0.12 | 1.26 |
| 7 | Usp31 | 0.50 | 1.26E-08 | 2.63E-04 | 0.17 | 0.11 | 0.07 | 1.61 |
| 7 | Spcs3 | 0.32 | 1.29E-08 | 2.69E-04 | 0.49 | 0.38 | 0.11 | 1.30 |
| 7 | Rpe | 0.28 | 1.34E-08 | 2.78E-04 | 0.45 | 0.33 | 0.12 | 1.35 |
| 7 | Rars2 | 0.29 | 1.35E-08 | 2.81E-04 | 0.44 | 0.33 | 0.11 | 1.33 |
| 7 | BC004004 | 0.29 | 1.36E-08 | 2.83E-04 | 0.42 | 0.31 | 0.11 | 1.34 |
| 7 | Ascc1 | 0.25 | 1.40E-08 | 2.92E-04 | 0.51 | 0.39 | 0.12 | 1.31 |
| 7 | Tbc1d5 | 0.44 | 1.42E-08 | 2.95E-04 | 0.29 | 0.20 | 0.09 | 1.42 |
| 7 | Men1 | 0.26 | 1.47E-08 | 3.06E-04 | 0.50 | 0.38 | 0.12 | 1.31 |
| 7 | Slc1a5 | 0.46 | 1.52E-08 | 3.16E-04 | 0.28 | 0.20 | 0.08 | 1.41 |
| 7 | Pgap2 | 0.31 | 1.55E-08 | 3.21E-04 | 0.57 | 0.45 | 0.12 | 1.26 |
| 7 | Rab26os | 0.35 | 1.55E-08 | 3.22E-04 | 0.28 | 0.19 | 0.09 | 1.46 |
| 7 | Gclc | 0.27 | 1.55E-08 | 3.23E-04 | 0.40 | 0.29 | 0.11 | 1.36 |
| 7 | Slc25a10 | 0.27 | 1.57E-08 | 3.26E-04 | 0.33 | 0.23 | 0.10 | 1.41 |
| 7 | Aacs | 0.67 | 1.59E-08 | 3.30E-04 | 0.17 | 0.11 | 0.06 | 1.56 |
| 7 | Dph5 | 0.35 | 1.63E-08 | 3.39E-04 | 0.32 | 0.23 | 0.09 | 1.40 |
| 7 | Arhgap32 | 0.33 | 1.70E-08 | 3.54E-04 | 0.27 | 0.19 | 0.08 | 1.44 |
| 7 | Paxip1 | 0.28 | 1.78E-08 | 3.69E-04 | 0.44 | 0.33 | 0.10 | 1.31 |
| 7 | Ears2 | 0.33 | 1.79E-08 | 3.71E-04 | 0.23 | 0.16 | 0.08 | 1.50 |
| 7 | Immp2l | 0.51 | 1.84E-08 | 3.82E-04 | 0.14 | 0.08 | 0.06 | 1.68 |
| 7 | Pomt1 | 0.44 | 1.89E-08 | 3.93E-04 | 0.19 | 0.12 | 0.07 | 1.57 |
| 7 | Speg | 0.29 | 1.94E-08 | 4.04E-04 | 0.24 | 0.16 | 0.08 | 1.51 |
| 7 | Tmem87b | 0.36 | 1.97E-08 | 4.10E-04 | 0.30 | 0.21 | 0.09 | 1.41 |
| 7 | Man2c1 | 0.40 | 2.08E-08 | 4.33E-04 | 0.17 | 0.10 | 0.06 | 1.61 |
| 7 | Arrdc3 | 0.44 | 2.13E-08 | 4.43E-04 | 0.28 | 0.20 | 0.08 | 1.41 |
| 7 | Gm4890 | 0.52 | 2.14E-08 | 4.44E-04 | 0.12 | 0.07 | 0.05 | 1.75 |
| 7 | BC052040 | 0.32 | 2.14E-08 | 4.44E-04 | 0.26 | 0.17 | 0.08 | 1.47 |
| 7 | Oxld1 | 0.47 | 2.18E-08 | 4.54E-04 | 0.18 | 0.12 | 0.07 | 1.56 |

**Supplementary Table 1. Marker genes for the original clusters of integrated clustering of E11.5 control and NotchGOF λ-junction cells.**

| | | | | | | | | |
|---|---|---|---|---|---|---|---|---|
| 7 | Pigb | 0.68 | 2.19E-08 | 4.56E-04 | 0.13 | 0.07 | 0.05 | 1.71 |
| 7 | Atp6ap2 | 0.28 | 2.21E-08 | 4.59E-04 | 0.54 | 0.42 | 0.12 | 1.28 |
| 7 | Sdr39u1 | 0.40 | 2.24E-08 | 4.66E-04 | 0.23 | 0.16 | 0.08 | 1.49 |
| 7 | Sppl2b | 0.35 | 2.25E-08 | 4.67E-04 | 0.22 | 0.14 | 0.08 | 1.52 |
| 7 | Elk4 | 0.29 | 2.28E-08 | 4.74E-04 | 0.32 | 0.23 | 0.09 | 1.39 |
| 7 | Zfp217 | 0.32 | 2.32E-08 | 4.82E-04 | 0.28 | 0.20 | 0.09 | 1.44 |
| 7 | Wwox | 0.37 | 2.40E-08 | 4.99E-04 | 0.17 | 0.10 | 0.06 | 1.60 |
| 7 | Ppp1r16a | 0.25 | 2.43E-08 | 5.04E-04 | 0.27 | 0.18 | 0.09 | 1.47 |
| 7 | Fbxo36 | 0.54 | 2.45E-08 | 5.09E-04 | 0.15 | 0.09 | 0.06 | 1.64 |
| 7 | Ofd1 | 0.25 | 2.45E-08 | 5.10E-04 | 0.41 | 0.31 | 0.11 | 1.34 |
| 7 | Tmem158 | 0.34 | 2.51E-08 | 5.23E-04 | 0.20 | 0.13 | 0.07 | 1.53 |
| 7 | Skiv2l | 0.31 | 2.55E-08 | 5.31E-04 | 0.34 | 0.24 | 0.09 | 1.39 |
| 7 | Arhgap39 | 0.35 | 2.57E-08 | 5.34E-04 | 0.26 | 0.18 | 0.08 | 1.46 |
| 7 | Ccdc177 | 0.45 | 2.66E-08 | 5.54E-04 | 0.12 | 0.07 | 0.05 | 1.72 |
| 7 | 4932422M17Rik | 0.52 | 2.76E-08 | 5.74E-04 | 0.16 | 0.10 | 0.06 | 1.61 |
| 7 | Gpn2 | 0.29 | 2.77E-08 | 5.75E-04 | 0.29 | 0.20 | 0.09 | 1.44 |
| 7 | Sc5d | 0.30 | 2.80E-08 | 5.82E-04 | 0.48 | 0.38 | 0.11 | 1.28 |
| 7 | Nudt7 | 0.60 | 2.90E-08 | 6.04E-04 | 0.12 | 0.07 | 0.05 | 1.74 |
| 7 | Fig4 | 0.61 | 2.99E-08 | 6.23E-04 | 0.13 | 0.07 | 0.05 | 1.70 |
| 7 | LTO1 | 0.27 | 3.03E-08 | 6.31E-04 | 0.32 | 0.23 | 0.09 | 1.40 |
| 7 | Rabif | 0.35 | 3.09E-08 | 6.44E-04 | 0.34 | 0.25 | 0.09 | 1.37 |
| 7 | Nek2 | 0.30 | 3.15E-08 | 6.55E-04 | 0.33 | 0.24 | 0.09 | 1.38 |
| 7 | Dkk3 | 0.32 | 3.39E-08 | 7.05E-04 | 0.18 | 0.12 | 0.07 | 1.58 |
| 7 | Fam169a | 0.43 | 3.39E-08 | 7.06E-04 | 0.20 | 0.13 | 0.07 | 1.52 |
| 7 | Prepl | 0.50 | 3.42E-08 | 7.11E-04 | 0.19 | 0.12 | 0.07 | 1.54 |
| 7 | Ints6 | 0.32 | 3.54E-08 | 7.36E-04 | 0.46 | 0.36 | 0.10 | 1.28 |
| 7 | Selenoo | 0.35 | 3.66E-08 | 7.61E-04 | 0.22 | 0.14 | 0.07 | 1.50 |
| 7 | Hspbap1 | 0.28 | 3.67E-08 | 7.64E-04 | 0.29 | 0.20 | 0.09 | 1.44 |
| 7 | Gm16196 | 0.36 | 3.77E-08 | 7.84E-04 | 0.11 | 0.07 | 0.05 | 1.75 |
| 7 | Zbtb11os1 | 0.52 | 3.78E-08 | 7.87E-04 | 0.11 | 0.06 | 0.05 | 1.77 |
| 7 | Slc35c2 | 0.26 | 3.83E-08 | 7.98E-04 | 0.44 | 0.32 | 0.11 | 1.35 |
| 7 | Trmt1l | 0.32 | 3.86E-08 | 8.03E-04 | 0.36 | 0.26 | 0.10 | 1.37 |
| 7 | Nfix | 0.41 | 3.98E-08 | 8.27E-04 | 0.22 | 0.15 | 0.07 | 1.47 |
| 7 | Abhd6 | 0.35 | 4.10E-08 | 8.52E-04 | 0.24 | 0.17 | 0.08 | 1.47 |
| 7 | Cdk5rap1 | 0.62 | 4.28E-08 | 8.90E-04 | 0.14 | 0.09 | 0.05 | 1.61 |
| 7 | Ints1 | 0.29 | 4.30E-08 | 8.94E-04 | 0.39 | 0.29 | 0.10 | 1.35 |
| 7 | Taf4 | 0.45 | 4.31E-08 | 8.96E-04 | 0.21 | 0.14 | 0.07 | 1.52 |
| 7 | Epdr1 | 0.39 | 4.38E-08 | 9.11E-04 | 0.17 | 0.10 | 0.06 | 1.59 |
| 7 | Phkg2 | 0.29 | 4.78E-08 | 9.95E-04 | 0.33 | 0.24 | 0.09 | 1.39 |
| 7 | Lyrm9 | 0.33 | 5.08E-08 | 1.06E-03 | 0.25 | 0.17 | 0.08 | 1.45 |
| 7 | Aunip | 0.29 | 5.97E-08 | 1.24E-03 | 0.35 | 0.26 | 0.09 | 1.36 |
| 7 | Afg3l1 | 0.29 | 6.01E-08 | 1.25E-03 | 0.59 | 0.47 | 0.12 | 1.25 |
| 7 | Ankrd9 | 0.43 | 6.02E-08 | 1.25E-03 | 0.19 | 0.12 | 0.06 | 1.52 |
| 7 | Gm19426 | 0.50 | 6.14E-08 | 1.28E-03 | 0.19 | 0.12 | 0.07 | 1.53 |
| 7 | Nsun5 | 0.33 | 6.29E-08 | 1.31E-03 | 0.26 | 0.18 | 0.08 | 1.45 |
| 7 | Zfp949 | 0.31 | 6.37E-08 | 1.32E-03 | 0.18 | 0.12 | 0.07 | 1.55 |
| 7 | Nsun3 | 0.60 | 6.44E-08 | 1.34E-03 | 0.13 | 0.08 | 0.05 | 1.67 |
| 7 | Tut1 | 0.33 | 6.48E-08 | 1.35E-03 | 0.21 | 0.14 | 0.07 | 1.51 |
| 7 | Trappc2 | 0.37 | 6.78E-08 | 1.41E-03 | 0.24 | 0.16 | 0.07 | 1.45 |
| 7 | Ltbp1 | 0.26 | 7.01E-08 | 1.46E-03 | 0.31 | 0.22 | 0.09 | 1.41 |
| 7 | Zfp994 | 0.42 | 7.11E-08 | 1.48E-03 | 0.15 | 0.09 | 0.06 | 1.64 |
| 7 | Mpv17l2 | 0.28 | 7.54E-08 | 1.57E-03 | 0.31 | 0.22 | 0.09 | 1.38 |
| 7 | Rnf139 | 0.31 | 7.59E-08 | 1.58E-03 | 0.33 | 0.25 | 0.09 | 1.36 |
| 7 | Disp1 | 0.64 | 7.98E-08 | 1.66E-03 | 0.17 | 0.11 | 0.06 | 1.56 |
| 7 | Fntb | 0.26 | 8.22E-08 | 1.71E-03 | 0.31 | 0.22 | 0.09 | 1.41 |
| 7 | D3Ertd751e | 0.28 | 8.27E-08 | 1.72E-03 | 0.28 | 0.20 | 0.08 | 1.42 |

**Supplementary Table 1. Marker genes for the original clusters of integrated clustering of E11.5 control and NotchGOF λ-junction cells.**

| | | | | | | | | |
|---|---|---|---|---|---|---|---|---|
| 7 | Ppp2r3d | 0.27 | 8.41E-08 | 1.75E-03 | 0.31 | 0.22 | 0.09 | 1.40 |
| 7 | Fuom | 0.25 | 8.42E-08 | 1.75E-03 | 0.37 | 0.27 | 0.10 | 1.35 |
| 7 | Vav2 | 0.44 | 8.46E-08 | 1.76E-03 | 0.14 | 0.08 | 0.05 | 1.65 |
| 7 | Mboat7 | 0.29 | 9.06E-08 | 1.89E-03 | 0.43 | 0.33 | 0.10 | 1.31 |
| 7 | 2410022M11Rik | 0.42 | 9.16E-08 | 1.91E-03 | 0.17 | 0.11 | 0.06 | 1.55 |
| 7 | L3mbtl2 | 0.37 | 9.40E-08 | 1.95E-03 | 0.27 | 0.19 | 0.08 | 1.42 |
| 7 | Rpp38 | 0.33 | 1.00E-07 | 2.09E-03 | 0.25 | 0.17 | 0.08 | 1.45 |
| 7 | Hist1h1d | 0.27 | 1.04E-07 | 2.16E-03 | 0.45 | 0.35 | 0.10 | 1.28 |
| 7 | Zfp974 | 0.37 | 1.08E-07 | 2.24E-03 | 0.14 | 0.09 | 0.05 | 1.61 |
| 7 | Pdpr | 0.40 | 1.12E-07 | 2.32E-03 | 0.11 | 0.06 | 0.05 | 1.74 |
| 7 | Plekha5 | 0.28 | 1.12E-07 | 2.34E-03 | 0.56 | 0.46 | 0.10 | 1.21 |
| 7 | Rad54l | 0.28 | 1.14E-07 | 2.38E-03 | 0.40 | 0.31 | 0.10 | 1.32 |
| 7 | Mars | 0.29 | 1.17E-07 | 2.43E-03 | 0.46 | 0.36 | 0.10 | 1.28 |
| 7 | Eif4ebp3 | 0.29 | 1.20E-07 | 2.50E-03 | 0.27 | 0.19 | 0.08 | 1.43 |
| 7 | Whrn | 0.49 | 1.21E-07 | 2.52E-03 | 0.10 | 0.06 | 0.05 | 1.76 |
| 7 | Enox2 | 0.46 | 1.22E-07 | 2.53E-03 | 0.12 | 0.07 | 0.05 | 1.70 |
| 7 | Gm32856 | 0.55 | 1.23E-07 | 2.55E-03 | 0.12 | 0.07 | 0.05 | 1.68 |
| 7 | Pigf | 0.25 | 1.23E-07 | 2.56E-03 | 0.51 | 0.40 | 0.11 | 1.28 |
| 7 | Repin1 | 0.41 | 1.24E-07 | 2.57E-03 | 0.20 | 0.13 | 0.07 | 1.50 |
| 7 | Zc3h6 | 0.40 | 1.26E-07 | 2.61E-03 | 0.14 | 0.09 | 0.05 | 1.62 |
| 7 | C2cd5 | 0.38 | 1.35E-07 | 2.80E-03 | 0.33 | 0.24 | 0.09 | 1.36 |
| 7 | Siae | 0.47 | 1.43E-07 | 2.97E-03 | 0.12 | 0.07 | 0.05 | 1.69 |
| 7 | AU020206 | 0.33 | 1.44E-07 | 2.99E-03 | 0.31 | 0.22 | 0.09 | 1.40 |
| 7 | Bcdin3d | 0.31 | 1.55E-07 | 3.23E-03 | 0.34 | 0.25 | 0.09 | 1.35 |
| 7 | Tbc1d17 | 0.30 | 1.57E-07 | 3.26E-03 | 0.39 | 0.29 | 0.10 | 1.33 |
| 7 | Rccd1 | 0.43 | 1.61E-07 | 3.35E-03 | 0.22 | 0.15 | 0.07 | 1.45 |
| 7 | Dand5 | 0.72 | 1.64E-07 | 3.40E-03 | 0.10 | 0.06 | 0.04 | 1.73 |
| 7 | Tigar | 0.38 | 1.66E-07 | 3.46E-03 | 0.12 | 0.07 | 0.05 | 1.68 |
| 7 | Ift57 | 0.27 | 1.78E-07 | 3.70E-03 | 0.41 | 0.31 | 0.10 | 1.33 |
| 7 | Fads1 | 0.28 | 1.79E-07 | 3.73E-03 | 0.58 | 0.48 | 0.10 | 1.21 |
| 7 | Nek3 | 0.31 | 1.82E-07 | 3.79E-03 | 0.17 | 0.11 | 0.06 | 1.55 |
| 7 | Cds2 | 0.42 | 1.91E-07 | 3.97E-03 | 0.25 | 0.18 | 0.08 | 1.42 |
| 7 | Gbe1 | 0.46 | 1.92E-07 | 3.99E-03 | 0.16 | 0.10 | 0.06 | 1.56 |
| 7 | Golga1 | 0.27 | 1.94E-07 | 4.04E-03 | 0.38 | 0.28 | 0.10 | 1.35 |
| 7 | Ltbp3 | 0.38 | 2.05E-07 | 4.27E-03 | 0.13 | 0.08 | 0.05 | 1.63 |
| 7 | Msto1 | 0.32 | 2.05E-07 | 4.27E-03 | 0.30 | 0.22 | 0.08 | 1.38 |
| 7 | Pard6a | 0.31 | 2.06E-07 | 4.28E-03 | 0.24 | 0.16 | 0.07 | 1.45 |
| 7 | Atxn1l | 0.39 | 2.10E-07 | 4.37E-03 | 0.23 | 0.16 | 0.07 | 1.44 |
| 7 | Katnal2 | 0.41 | 2.22E-07 | 4.62E-03 | 0.11 | 0.06 | 0.05 | 1.73 |
| 7 | S1pr1 | 0.35 | 2.26E-07 | 4.70E-03 | 0.11 | 0.06 | 0.04 | 1.71 |
| 7 | 2510039O18Rik | 0.29 | 2.28E-07 | 4.75E-03 | 0.32 | 0.23 | 0.09 | 1.38 |
| 7 | Nr3c1 | 0.46 | 2.28E-07 | 4.75E-03 | 0.23 | 0.16 | 0.07 | 1.43 |
| 7 | Ttc27 | 0.30 | 2.33E-07 | 4.84E-03 | 0.27 | 0.19 | 0.08 | 1.41 |
| 7 | Lck | 0.50 | 2.43E-07 | 5.06E-03 | 0.11 | 0.07 | 0.05 | 1.68 |
| 7 | Srd5a3 | 0.27 | 2.47E-07 | 5.14E-03 | 0.29 | 0.21 | 0.08 | 1.40 |
| 7 | Rrp9 | 0.27 | 2.51E-07 | 5.23E-03 | 0.44 | 0.34 | 0.10 | 1.28 |
| 7 | Zfp937 | 0.54 | 2.65E-07 | 5.52E-03 | 0.16 | 0.10 | 0.06 | 1.56 |
| 7 | Hspe1-rs1 | 0.46 | 2.77E-07 | 5.76E-03 | 0.20 | 0.13 | 0.07 | 1.49 |
| 7 | Rad51d | 0.36 | 2.79E-07 | 5.80E-03 | 0.20 | 0.13 | 0.07 | 1.49 |
| 7 | 1810044D09Rik | 0.31 | 2.83E-07 | 5.88E-03 | 0.17 | 0.11 | 0.06 | 1.53 |
| 7 | Ogfod2 | 0.46 | 2.94E-07 | 6.11E-03 | 0.15 | 0.10 | 0.05 | 1.55 |
| 7 | Sec24a | 0.32 | 3.04E-07 | 6.32E-03 | 0.25 | 0.17 | 0.08 | 1.43 |
| 7 | Itfg1 | 0.25 | 3.06E-07 | 6.37E-03 | 0.45 | 0.35 | 0.10 | 1.30 |
| 7 | Smg9 | 0.31 | 3.08E-07 | 6.40E-03 | 0.20 | 0.14 | 0.07 | 1.48 |
| 7 | Lztfl1 | 0.26 | 3.15E-07 | 6.54E-03 | 0.59 | 0.49 | 0.11 | 1.22 |
| 7 | Nudcd1 | 0.25 | 3.20E-07 | 6.66E-03 | 0.51 | 0.41 | 0.11 | 1.26 |

**Supplementary Table 1. Marker genes for the original clusters of integrated clustering of E11.5 control and NotchGOF λ-junction cells.**

| | | | | | | | | |
|---|---|---|---|---|---|---|---|---|
| 7 | Skida1 | 0.34 | 3.35E-07 | 6.96E-03 | 0.22 | 0.15 | 0.07 | 1.46 |
| 7 | Errfi1 | 0.29 | 3.36E-07 | 6.99E-03 | 0.25 | 0.18 | 0.07 | 1.41 |
| 7 | Rmnd5b | 0.32 | 3.56E-07 | 7.41E-03 | 0.24 | 0.17 | 0.07 | 1.41 |
| 7 | Mfn2 | 0.27 | 3.63E-07 | 7.55E-03 | 0.23 | 0.16 | 0.07 | 1.45 |
| 7 | Chd1l | 0.27 | 3.69E-07 | 7.67E-03 | 0.28 | 0.20 | 0.08 | 1.40 |
| 7 | Ppip5k1 | 0.59 | 3.89E-07 | 8.10E-03 | 0.11 | 0.06 | 0.04 | 1.68 |
| 7 | Rce1 | 0.25 | 3.92E-07 | 8.15E-03 | 0.56 | 0.45 | 0.11 | 1.23 |
| 7 | Lbhd1 | 0.46 | 3.97E-07 | 8.25E-03 | 0.11 | 0.06 | 0.04 | 1.69 |
| 7 | Shld3 | 0.38 | 4.00E-07 | 8.33E-03 | 0.16 | 0.10 | 0.06 | 1.55 |
| 7 | 1700007L15Rik | 0.56 | 4.10E-07 | 8.53E-03 | 0.12 | 0.07 | 0.05 | 1.64 |
| 7 | Lyplal1 | 0.29 | 4.15E-07 | 8.64E-03 | 0.33 | 0.24 | 0.09 | 1.35 |
| 7 | Tti1 | 0.26 | 4.43E-07 | 9.22E-03 | 0.24 | 0.17 | 0.07 | 1.43 |
| 7 | Med16 | 0.30 | 4.95E-07 | 1.03E-02 | 0.39 | 0.29 | 0.10 | 1.33 |
| 7 | Slc19a1 | 0.28 | 5.19E-07 | 1.08E-02 | 0.22 | 0.15 | 0.07 | 1.46 |
| 7 | Src | 0.30 | 5.80E-07 | 1.21E-02 | 0.31 | 0.23 | 0.08 | 1.36 |
| 7 | Lpar2 | 0.36 | 5.89E-07 | 1.22E-02 | 0.12 | 0.08 | 0.05 | 1.64 |
| 7 | Mnd1 | 0.29 | 5.92E-07 | 1.23E-02 | 0.35 | 0.26 | 0.09 | 1.33 |
| 7 | Zmpste24 | 0.26 | 6.25E-07 | 1.30E-02 | 0.41 | 0.31 | 0.09 | 1.30 |
| 7 | Tmem265 | 0.36 | 6.52E-07 | 1.36E-02 | 0.16 | 0.10 | 0.06 | 1.53 |
| 7 | Mok | 0.46 | 6.70E-07 | 1.39E-02 | 0.10 | 0.06 | 0.04 | 1.70 |
| 7 | Spen | 0.25 | 6.74E-07 | 1.40E-02 | 0.56 | 0.46 | 0.10 | 1.22 |
| 7 | Plk1 | 0.26 | 7.12E-07 | 1.48E-02 | 0.55 | 0.46 | 0.09 | 1.19 |
| 7 | Tceal1 | 0.32 | 7.16E-07 | 1.49E-02 | 0.31 | 0.23 | 0.08 | 1.34 |
| 7 | Tmem251 | 0.33 | 7.16E-07 | 1.49E-02 | 0.34 | 0.26 | 0.08 | 1.32 |
| 7 | Polr3e | 0.28 | 7.33E-07 | 1.52E-02 | 0.29 | 0.21 | 0.08 | 1.37 |
| 7 | Pigw | 0.62 | 7.45E-07 | 1.55E-02 | 0.11 | 0.06 | 0.04 | 1.67 |
| 7 | Pkmyt1 | 0.36 | 7.76E-07 | 1.61E-02 | 0.24 | 0.17 | 0.07 | 1.40 |
| 7 | Btbd9 | 0.39 | 7.99E-07 | 1.66E-02 | 0.39 | 0.29 | 0.09 | 1.32 |
| 7 | 1600012H06Rik | 0.27 | 8.92E-07 | 1.85E-02 | 0.30 | 0.22 | 0.08 | 1.36 |
| 7 | Vangl1 | 0.34 | 9.07E-07 | 1.89E-02 | 0.22 | 0.15 | 0.07 | 1.44 |
| 7 | Dhx16 | 0.28 | 9.50E-07 | 1.98E-02 | 0.43 | 0.34 | 0.10 | 1.29 |
| 7 | Zfp446 | 0.56 | 1.08E-06 | 2.24E-02 | 0.12 | 0.08 | 0.05 | 1.58 |
| 7 | Arvcf | 0.27 | 1.15E-06 | 2.38E-02 | 0.33 | 0.24 | 0.09 | 1.36 |
| 7 | Mlh3 | 0.39 | 1.22E-06 | 2.53E-02 | 0.24 | 0.17 | 0.07 | 1.42 |
| 7 | Nin | 0.28 | 1.28E-06 | 2.66E-02 | 0.36 | 0.27 | 0.09 | 1.33 |
| 7 | Prss36 | 0.33 | 1.30E-06 | 2.70E-02 | 0.14 | 0.09 | 0.05 | 1.58 |
| 7 | Mcm9 | 0.36 | 1.38E-06 | 2.87E-02 | 0.23 | 0.16 | 0.07 | 1.43 |
| 7 | Gemin8 | 0.27 | 1.41E-06 | 2.92E-02 | 0.26 | 0.19 | 0.07 | 1.39 |
| 7 | Arhgef39 | 0.46 | 1.41E-06 | 2.92E-02 | 0.20 | 0.14 | 0.06 | 1.42 |
| 7 | Cstf2 | 0.25 | 1.41E-06 | 2.94E-02 | 0.44 | 0.34 | 0.10 | 1.28 |
| 7 | Per3 | 0.41 | 1.43E-06 | 2.98E-02 | 0.15 | 0.10 | 0.05 | 1.53 |
| 7 | Ralgapa2 | 0.27 | 1.49E-06 | 3.09E-02 | 0.20 | 0.14 | 0.06 | 1.46 |
| 7 | Mief1 | 0.30 | 1.55E-06 | 3.23E-02 | 0.23 | 0.16 | 0.07 | 1.41 |
| 7 | Donson | 0.33 | 1.62E-06 | 3.36E-02 | 0.32 | 0.24 | 0.08 | 1.32 |
| 7 | Hmces | 0.26 | 1.71E-06 | 3.55E-02 | 0.46 | 0.37 | 0.09 | 1.25 |
| 7 | Coq8b | 0.29 | 1.74E-06 | 3.62E-02 | 0.18 | 0.12 | 0.06 | 1.48 |
| 7 | Hira | 0.26 | 1.82E-06 | 3.78E-02 | 0.46 | 0.37 | 0.10 | 1.27 |
| 7 | Aste1 | 0.44 | 1.83E-06 | 3.80E-02 | 0.13 | 0.08 | 0.05 | 1.58 |
| 7 | Sptlc1 | 0.25 | 1.83E-06 | 3.80E-02 | 0.38 | 0.29 | 0.09 | 1.31 |
| 7 | Dolk | 0.34 | 1.98E-06 | 4.13E-02 | 0.22 | 0.15 | 0.06 | 1.41 |
| 7 | Tmem175 | 0.30 | 2.00E-06 | 4.16E-02 | 0.15 | 0.10 | 0.05 | 1.53 |
| 7 | Uhmk1 | 0.27 | 2.06E-06 | 4.29E-02 | 0.28 | 0.21 | 0.07 | 1.35 |
| 7 | Rassf1 | 0.29 | 2.14E-06 | 4.45E-02 | 0.48 | 0.40 | 0.09 | 1.21 |
| 7 | Zbtb9 | 0.34 | 2.14E-06 | 4.46E-02 | 0.16 | 0.11 | 0.05 | 1.50 |
| 7 | 2900060B14Rik | 0.39 | 2.35E-06 | 4.88E-02 | 0.12 | 0.08 | 0.05 | 1.58 |
| 7 | Tmem50b | 0.27 | 2.39E-06 | 4.97E-02 | 0.25 | 0.18 | 0.07 | 1.41 |

**Supplementary Table 1. Marker genes for the original clusters of integrated clustering of E11.5 control and NotchGOF λ-junction cells.**

| | | | | | | | | |
|---|---|---|---|---|---|---|---|---|
| 7 | Ctso | 0.37 | 2.40E-06 | 4.99E-02 | 0.11 | 0.06 | 0.04 | 1.66 |
| 8 | Foxf1 | 2.46 | 0 | 0 | 0.75 | 0.16 | 0.59 | 4.69 |
| 8 | Pax9 | 2.16 | 0 | 0 | 0.92 | 0.34 | 0.58 | 2.72 |
| 8 | Flrt2 | 2.18 | 0 | 0 | 0.97 | 0.43 | 0.53 | 2.23 |
| 8 | Isl2 | 2.53 | 0 | 0 | 0.56 | 0.11 | 0.44 | 4.93 |
| 8 | Fendrr | 2.70 | 0 | 0 | 0.48 | 0.08 | 0.40 | 5.94 |
| 8 | Frzb | 1.95 | 1.60E-285 | 3.33E-281 | 0.85 | 0.33 | 0.51 | 2.55 |
| 8 | Plagl1 | 1.81 | 1.26E-268 | 2.62E-264 | 0.90 | 0.41 | 0.49 | 2.18 |
| 8 | Hist1h2ap | 1.55 | 7.64E-267 | 1.59E-262 | 0.99 | 0.74 | 0.25 | 1.34 |
| 8 | Dkk1 | 2.44 | 2.88E-264 | 5.98E-260 | 0.74 | 0.26 | 0.48 | 2.84 |
| 8 | Unc5c | 1.74 | 1.23E-258 | 2.57E-254 | 0.84 | 0.32 | 0.52 | 2.62 |
| 8 | Sox6 | 2.36 | 2.81E-256 | 5.85E-252 | 0.50 | 0.11 | 0.39 | 4.54 |
| 8 | Hist1h1a | 1.61 | 1.11E-240 | 2.30E-236 | 0.89 | 0.40 | 0.49 | 2.21 |
| 8 | Epha3 | 1.59 | 9.40E-235 | 1.95E-230 | 0.84 | 0.34 | 0.50 | 2.47 |
| 8 | Hist1h1b | 1.39 | 3.85E-234 | 8.00E-230 | 0.99 | 0.63 | 0.36 | 1.58 |
| 8 | Hist1h1e | 1.33 | 3.60E-221 | 7.48E-217 | 0.98 | 0.63 | 0.35 | 1.55 |
| 8 | Hist1h2ae | 1.38 | 8.97E-221 | 1.87E-216 | 0.99 | 0.69 | 0.30 | 1.44 |
| 8 | Top2a | 1.18 | 1.03E-215 | 2.13E-211 | 1.00 | 0.78 | 0.22 | 1.28 |
| 8 | Fbln2 | 1.45 | 3.77E-204 | 7.85E-200 | 0.91 | 0.50 | 0.41 | 1.81 |
| 8 | Sall1 | 2.38 | 1.16E-191 | 2.42E-187 | 0.37 | 0.08 | 0.29 | 4.65 |
| 8 | Nrp2 | 1.62 | 3.57E-189 | 7.42E-185 | 0.79 | 0.35 | 0.43 | 2.22 |
| 8 | Col3a1 | 1.08 | 3.96E-189 | 8.23E-185 | 1.00 | 0.71 | 0.29 | 1.40 |
| 8 | Tenm3 | 1.27 | 6.40E-187 | 1.33E-182 | 0.93 | 0.63 | 0.30 | 1.48 |
| 8 | Fstl1 | 1.02 | 6.24E-181 | 1.30E-176 | 0.99 | 0.82 | 0.18 | 1.22 |
| 8 | Col9a1 | 1.55 | 8.25E-178 | 1.72E-173 | 0.69 | 0.27 | 0.42 | 2.52 |
| 8 | Sox9 | 1.68 | 3.42E-177 | 7.11E-173 | 0.89 | 0.58 | 0.31 | 1.54 |
| 8 | Piezo2 | 1.62 | 1.12E-168 | 2.32E-164 | 0.60 | 0.21 | 0.39 | 2.85 |
| 8 | Peg3 | 1.05 | 8.92E-168 | 1.86E-163 | 0.98 | 0.82 | 0.16 | 1.19 |
| 8 | Pclaf | 0.80 | 2.46E-162 | 5.12E-158 | 1.00 | 0.88 | 0.12 | 1.13 |
| 8 | Col9a3 | 2.14 | 1.52E-159 | 3.15E-155 | 0.45 | 0.13 | 0.32 | 3.42 |
| 8 | Hist1h4d | 1.30 | 2.52E-159 | 5.25E-155 | 0.85 | 0.44 | 0.41 | 1.92 |
| 8 | Hist1h3c | 1.20 | 1.05E-158 | 2.18E-154 | 0.93 | 0.58 | 0.35 | 1.59 |
| 8 | Hist1h2ab | 1.30 | 1.31E-156 | 2.73E-152 | 0.78 | 0.36 | 0.42 | 2.19 |
| 8 | Rspo3 | 1.71 | 2.30E-153 | 4.78E-149 | 0.71 | 0.33 | 0.38 | 2.17 |
| 8 | Rab32 | 2.44 | 3.90E-150 | 8.11E-146 | 0.34 | 0.08 | 0.26 | 4.11 |
| 8 | Hist1h1d | 1.40 | 2.96E-145 | 6.17E-141 | 0.73 | 0.34 | 0.39 | 2.15 |
| 8 | Neil3 | 1.86 | 2.25E-143 | 4.67E-139 | 0.55 | 0.22 | 0.33 | 2.54 |
| 8 | Itga8 | 1.72 | 9.87E-140 | 2.05E-135 | 0.51 | 0.18 | 0.33 | 2.88 |
| 8 | Alx3 | 1.30 | 2.43E-136 | 5.06E-132 | 0.67 | 0.28 | 0.39 | 2.39 |
| 8 | Ddit4l | 1.74 | 1.69E-134 | 3.52E-130 | 0.53 | 0.20 | 0.33 | 2.67 |
| 8 | Foxd1 | 2.33 | 2.38E-134 | 4.95E-130 | 0.36 | 0.10 | 0.26 | 3.63 |
| 8 | Hist1h3e | 1.31 | 1.35E-133 | 2.80E-129 | 0.77 | 0.41 | 0.36 | 1.87 |
| 8 | Cdh11 | 0.90 | 3.46E-133 | 7.19E-129 | 0.98 | 0.69 | 0.30 | 1.44 |
| 8 | Hist1h4h | 1.40 | 1.77E-131 | 3.69E-127 | 0.75 | 0.40 | 0.36 | 1.90 |
| 8 | Enpp2 | 1.80 | 9.51E-131 | 1.98E-126 | 0.37 | 0.10 | 0.27 | 3.59 |
| 8 | Phactr2 | 1.29 | 2.39E-130 | 4.96E-126 | 0.78 | 0.44 | 0.34 | 1.77 |
| 8 | Pdlim3 | 1.98 | 7.53E-130 | 1.57E-125 | 0.43 | 0.14 | 0.29 | 3.13 |
| 8 | Smc2 | 0.73 | 1.37E-127 | 2.85E-123 | 0.99 | 0.86 | 0.12 | 1.14 |
| 8 | Auts2 | 0.72 | 1.46E-125 | 3.04E-121 | 0.99 | 0.86 | 0.13 | 1.15 |
| 8 | Incenp | 1.01 | 8.88E-125 | 1.85E-120 | 0.91 | 0.58 | 0.33 | 1.56 |
| 8 | Gata2 | 1.21 | 7.74E-124 | 1.61E-119 | 0.79 | 0.43 | 0.36 | 1.84 |
| 8 | Ptch1 | 1.31 | 1.29E-123 | 2.69E-119 | 0.78 | 0.44 | 0.34 | 1.77 |
| 8 | Tox | 1.48 | 1.56E-122 | 3.25E-118 | 0.59 | 0.25 | 0.34 | 2.38 |
| 8 | Sobp | 1.34 | 2.06E-122 | 4.28E-118 | 0.68 | 0.33 | 0.35 | 2.08 |
| 8 | Fbxo5 | 0.99 | 3.41E-122 | 7.09E-118 | 0.91 | 0.58 | 0.33 | 1.57 |
| 8 | Thsd7a | 1.76 | 1.18E-121 | 2.46E-117 | 0.42 | 0.14 | 0.29 | 3.09 |

**Supplementary Table 1. Marker genes for the original clusters of integrated clustering of E11.5 control and NotchGOF λ-junction cells.**

| | | | | | | | | |
|---|---|---|---|---|---|---|---|---|
| 8 | Smc4 | 0.81 | 1.35E-121 | 2.80E-117 | 0.98 | 0.79 | 0.19 | 1.24 |
| 8 | Tril | 1.11 | 8.18E-120 | 1.70E-115 | 0.85 | 0.53 | 0.32 | 1.59 |
| 8 | Slitrk5 | 1.78 | 4.47E-115 | 9.30E-111 | 0.43 | 0.15 | 0.28 | 2.89 |
| 8 | Dek | 0.55 | 9.32E-115 | 1.94E-110 | 1.00 | 0.97 | 0.03 | 1.03 |
| 8 | Pcdh7 | 1.16 | 1.34E-114 | 2.79E-110 | 0.75 | 0.38 | 0.37 | 1.97 |
| 8 | Pex5l | 1.90 | 1.52E-114 | 3.16E-110 | 0.31 | 0.08 | 0.23 | 3.80 |
| 8 | Lgr5 | 1.77 | 3.95E-114 | 8.22E-110 | 0.49 | 0.20 | 0.30 | 2.51 |
| 8 | Kif26b | 1.17 | 1.71E-113 | 3.55E-109 | 0.75 | 0.39 | 0.36 | 1.91 |
| 8 | Hist1h1c | 1.19 | 1.05E-111 | 2.19E-107 | 0.74 | 0.41 | 0.33 | 1.82 |
| 8 | Hist2h2ac | 1.29 | 4.02E-109 | 8.36E-105 | 0.64 | 0.31 | 0.33 | 2.07 |
| 8 | Mecom | 1.28 | 1.00E-108 | 2.09E-104 | 0.72 | 0.39 | 0.33 | 1.83 |
| 8 | Pgm2l1 | 1.30 | 2.36E-108 | 4.92E-104 | 0.67 | 0.34 | 0.32 | 1.94 |
| 8 | Xist | 0.64 | 1.01E-106 | 2.11E-102 | 0.92 | 0.86 | 0.05 | 1.06 |
| 8 | Gata3 | 1.18 | 1.04E-106 | 2.17E-102 | 0.81 | 0.51 | 0.30 | 1.59 |
| 8 | Ppp3ca | 0.86 | 2.60E-105 | 5.41E-101 | 0.93 | 0.72 | 0.21 | 1.29 |
| 8 | Zbtb20 | 1.32 | 5.34E-105 | 1.11E-100 | 0.60 | 0.28 | 0.32 | 2.14 |
| 8 | Pcdh18 | 0.92 | 5.57E-105 | 1.16E-100 | 0.91 | 0.63 | 0.28 | 1.45 |
| 8 | Foxc1 | 1.25 | 1.02E-103 | 2.13E-99 | 0.60 | 0.28 | 0.32 | 2.17 |
| 8 | Arhgap28 | 1.03 | 5.55E-103 | 1.15E-98 | 0.81 | 0.52 | 0.29 | 1.56 |
| 8 | Hist1h2an | 1.29 | 1.31E-102 | 2.73E-98 | 0.61 | 0.29 | 0.32 | 2.08 |
| 8 | Col9a2 | 1.75 | 4.04E-102 | 8.40E-98 | 0.32 | 0.10 | 0.22 | 3.36 |
| 8 | Sned1 | 1.52 | 1.69E-101 | 3.51E-97 | 0.37 | 0.12 | 0.25 | 3.12 |
| 8 | Asap1 | 0.86 | 4.87E-101 | 1.01E-96 | 0.93 | 0.73 | 0.21 | 1.28 |
| 8 | Hist1h2be | 1.37 | 1.10E-99 | 2.29E-95 | 0.57 | 0.26 | 0.30 | 2.16 |
| 8 | Spc25 | 0.95 | 1.86E-99 | 3.86E-95 | 0.86 | 0.54 | 0.32 | 1.59 |
| 8 | Prrx1 | 0.80 | 1.87E-99 | 3.88E-95 | 0.94 | 0.64 | 0.30 | 1.46 |
| 8 | Rad51ap1 | 1.10 | 3.16E-99 | 6.56E-95 | 0.77 | 0.48 | 0.30 | 1.62 |
| 8 | Col1a2 | 0.73 | 1.07E-98 | 2.22E-94 | 0.99 | 0.73 | 0.26 | 1.35 |
| 8 | Oat | 0.90 | 3.29E-98 | 6.85E-94 | 0.90 | 0.67 | 0.23 | 1.34 |
| 8 | Arglu1 | 0.52 | 6.57E-98 | 1.37E-93 | 0.99 | 0.96 | 0.03 | 1.03 |
| 8 | Kif11 | 0.95 | 3.44E-97 | 7.16E-93 | 0.85 | 0.53 | 0.32 | 1.61 |
| 8 | Mki67 | 0.80 | 4.33E-97 | 9.00E-93 | 0.96 | 0.71 | 0.25 | 1.35 |
| 8 | Mfap4 | 0.63 | 6.03E-97 | 1.25E-92 | 0.99 | 0.84 | 0.15 | 1.17 |
| 8 | Tpr | 0.58 | 1.06E-96 | 2.20E-92 | 1.00 | 0.98 | 0.02 | 1.02 |
| 8 | Crym | 0.86 | 4.54E-96 | 9.44E-92 | 0.81 | 0.47 | 0.35 | 1.74 |
| 8 | Hist1h2bm | 1.29 | 2.96E-95 | 6.15E-91 | 0.60 | 0.30 | 0.30 | 2.02 |
| 8 | Col2a1 | 0.96 | 3.16E-95 | 6.57E-91 | 0.88 | 0.63 | 0.24 | 1.38 |
| 8 | Atad2 | 0.83 | 5.09E-95 | 1.06E-90 | 0.94 | 0.73 | 0.21 | 1.29 |
| 8 | Grin3a | 2.01 | 1.41E-94 | 2.92E-90 | 0.20 | 0.04 | 0.15 | 4.76 |
| 8 | Sfrp1 | 0.91 | 7.36E-94 | 1.53E-89 | 0.89 | 0.61 | 0.27 | 1.44 |
| 8 | Tyms | 0.70 | 7.22E-93 | 1.50E-88 | 0.97 | 0.83 | 0.14 | 1.17 |
| 8 | Epha5 | 2.11 | 1.17E-92 | 2.44E-88 | 0.24 | 0.06 | 0.18 | 3.83 |
| 8 | Esco2 | 1.08 | 2.41E-92 | 5.01E-88 | 0.74 | 0.44 | 0.30 | 1.69 |
| 8 | Sox5 | 1.17 | 1.44E-91 | 3.00E-87 | 0.69 | 0.38 | 0.30 | 1.79 |
| 8 | Zeb2 | 0.89 | 2.20E-91 | 4.57E-87 | 0.72 | 0.39 | 0.34 | 1.87 |
| 8 | Twist1 | 0.64 | 2.50E-91 | 5.20E-87 | 1.00 | 0.76 | 0.24 | 1.32 |
| 8 | Rrm1 | 0.69 | 2.53E-91 | 5.26E-87 | 0.97 | 0.81 | 0.16 | 1.20 |
| 8 | Tenm4 | 0.89 | 5.34E-91 | 1.11E-86 | 0.86 | 0.60 | 0.25 | 1.42 |
| 8 | Pdlim5 | 1.46 | 5.41E-91 | 1.12E-86 | 0.49 | 0.21 | 0.28 | 2.31 |
| 8 | Rhoj | 1.35 | 9.69E-91 | 2.02E-86 | 0.35 | 0.11 | 0.23 | 3.09 |
| 8 | Hist1h3b | 1.42 | 1.55E-90 | 3.22E-86 | 0.49 | 0.22 | 0.27 | 2.24 |
| 8 | Slit3 | 1.48 | 3.71E-90 | 7.72E-86 | 0.45 | 0.18 | 0.27 | 2.48 |
| 8 | Gpc3 | 0.66 | 6.02E-90 | 1.25E-85 | 0.99 | 0.83 | 0.16 | 1.20 |
| 8 | Hist2h2bb | 1.34 | 1.90E-89 | 3.96E-85 | 0.52 | 0.24 | 0.28 | 2.17 |
| 8 | Inhba | 1.44 | 3.65E-89 | 7.60E-85 | 0.41 | 0.16 | 0.26 | 2.61 |
| 8 | Itprid2 | 1.72 | 1.56E-88 | 3.24E-84 | 0.38 | 0.14 | 0.24 | 2.68 |

**Supplementary Table 1. Marker genes for the original clusters of integrated clustering of E11.5 control and NotchGOF λ-junction cells.**

| | | | | | | | | |
|---|---|---|---|---|---|---|---|---|
| 8 | Diaph3 | 1.26 | 3.25E-88 | 6.76E-84 | 0.64 | 0.36 | 0.28 | 1.77 |
| 8 | Luzp2 | 2.88 | 4.42E-88 | 9.19E-84 | 0.11 | 0.01 | 0.09 | 7.57 |
| 8 | Cped1 | 0.98 | 9.47E-88 | 1.97E-83 | 0.77 | 0.45 | 0.32 | 1.70 |
| 8 | Cdca2 | 1.02 | 1.60E-86 | 3.33E-82 | 0.72 | 0.42 | 0.31 | 1.74 |
| 8 | Hmgb2 | 0.50 | 1.44E-85 | 3.00E-81 | 1.00 | 0.98 | 0.02 | 1.02 |
| 8 | Tbx3 | 0.85 | 3.60E-85 | 7.49E-81 | 0.77 | 0.43 | 0.35 | 1.82 |
| 8 | Negr1 | 2.11 | 7.77E-85 | 1.62E-80 | 0.20 | 0.05 | 0.15 | 4.21 |
| 8 | Nid1 | 1.13 | 1.52E-84 | 3.15E-80 | 0.63 | 0.33 | 0.30 | 1.90 |
| 8 | Etv1 | 1.18 | 5.58E-84 | 1.16E-79 | 0.58 | 0.29 | 0.30 | 2.03 |
| 8 | Myo6 | 1.31 | 1.49E-83 | 3.10E-79 | 0.44 | 0.18 | 0.26 | 2.47 |
| 8 | Hjurp | 0.84 | 3.62E-83 | 7.53E-79 | 0.85 | 0.59 | 0.26 | 1.43 |
| 8 | Frem1 | 0.99 | 8.50E-83 | 1.77E-78 | 0.70 | 0.39 | 0.31 | 1.78 |
| 8 | Mis18bp1 | 0.87 | 1.16E-81 | 2.41E-77 | 0.80 | 0.50 | 0.30 | 1.59 |
| 8 | Rgs5 | 1.72 | 2.69E-80 | 5.59E-76 | 0.26 | 0.08 | 0.18 | 3.41 |
| 8 | Arid5b | 0.77 | 9.30E-80 | 1.93E-75 | 0.89 | 0.66 | 0.23 | 1.35 |
| 8 | Spc24 | 0.72 | 2.64E-79 | 5.49E-75 | 0.92 | 0.73 | 0.20 | 1.27 |
| 8 | Synpo | 1.30 | 2.89E-79 | 6.02E-75 | 0.50 | 0.23 | 0.27 | 2.17 |
| 8 | Plcl1 | 1.86 | 7.59E-79 | 1.58E-74 | 0.21 | 0.06 | 0.16 | 3.85 |
| 8 | Dnmt1 | 0.68 | 1.10E-78 | 2.28E-74 | 0.95 | 0.78 | 0.17 | 1.21 |
| 8 | Cacna1d | 1.26 | 2.88E-78 | 5.99E-74 | 0.49 | 0.23 | 0.27 | 2.19 |
| 8 | Prrx2 | 0.62 | 4.76E-78 | 9.89E-74 | 0.99 | 0.73 | 0.26 | 1.35 |
| 8 | Cenpq | 0.89 | 6.03E-78 | 1.25E-73 | 0.81 | 0.55 | 0.26 | 1.48 |
| 8 | Crispld1 | 1.16 | 1.44E-77 | 2.99E-73 | 0.60 | 0.32 | 0.29 | 1.90 |
| 8 | Gdf6 | 1.72 | 3.26E-77 | 6.78E-73 | 0.30 | 0.10 | 0.20 | 2.97 |
| 8 | Nusap1 | 0.79 | 4.10E-77 | 8.53E-73 | 0.80 | 0.48 | 0.32 | 1.67 |
| 8 | Aurkb | 0.85 | 6.20E-77 | 1.29E-72 | 0.79 | 0.48 | 0.31 | 1.64 |
| 8 | Kif15 | 0.79 | 4.66E-76 | 9.70E-72 | 0.87 | 0.62 | 0.25 | 1.40 |
| 8 | Daam2 | 1.14 | 5.15E-76 | 1.07E-71 | 0.56 | 0.29 | 0.28 | 1.96 |
| 8 | Fn1 | 0.77 | 6.18E-76 | 1.29E-71 | 0.93 | 0.78 | 0.15 | 1.19 |
| 8 | Tbx2 | 0.79 | 2.05E-75 | 4.26E-71 | 0.90 | 0.64 | 0.27 | 1.42 |
| 8 | Pla2g4a | 1.30 | 2.40E-75 | 5.00E-71 | 0.50 | 0.24 | 0.26 | 2.08 |
| 8 | Inmt | 2.19 | 1.31E-74 | 2.73E-70 | 0.13 | 0.02 | 0.10 | 5.48 |
| 8 | Dact3 | 0.87 | 2.34E-74 | 4.87E-70 | 0.76 | 0.48 | 0.28 | 1.58 |
| 8 | Tbx3os1 | 1.45 | 2.92E-74 | 6.07E-70 | 0.34 | 0.13 | 0.22 | 2.69 |
| 8 | Adamts5 | 1.54 | 7.07E-74 | 1.47E-69 | 0.30 | 0.10 | 0.20 | 2.91 |
| 8 | Limch1 | 0.81 | 8.35E-74 | 1.74E-69 | 0.89 | 0.67 | 0.23 | 1.34 |
| 8 | Foxf2 | 1.71 | 1.35E-73 | 2.81E-69 | 0.27 | 0.09 | 0.18 | 3.08 |
| 8 | Hist1h3i | 1.39 | 2.36E-73 | 4.91E-69 | 0.48 | 0.23 | 0.25 | 2.07 |
| 8 | Gabra1 | 2.00 | 3.00E-73 | 6.25E-69 | 0.19 | 0.05 | 0.14 | 3.92 |
| 8 | Clspn | 0.79 | 6.24E-73 | 1.30E-68 | 0.86 | 0.61 | 0.25 | 1.41 |
| 8 | Fbn2 | 0.67 | 9.05E-73 | 1.88E-68 | 0.94 | 0.73 | 0.21 | 1.28 |
| 8 | Ncapg | 0.92 | 1.55E-72 | 3.23E-68 | 0.73 | 0.47 | 0.26 | 1.55 |
| 8 | Lig1 | 0.74 | 8.56E-72 | 1.78E-67 | 0.93 | 0.78 | 0.15 | 1.19 |
| 8 | Cdk1 | 0.61 | 1.14E-71 | 2.37E-67 | 0.93 | 0.66 | 0.28 | 1.42 |
| 8 | Pid1 | 1.32 | 6.22E-71 | 1.29E-66 | 0.42 | 0.19 | 0.24 | 2.29 |
| 8 | Cbx5 | 0.44 | 8.40E-71 | 1.75E-66 | 0.99 | 0.95 | 0.04 | 1.04 |
| 8 | H1f0 | 0.72 | 8.65E-71 | 1.80E-66 | 0.96 | 0.85 | 0.12 | 1.14 |
| 8 | Tle4 | 0.80 | 8.67E-71 | 1.80E-66 | 0.81 | 0.58 | 0.23 | 1.40 |
| 8 | H2afz | 0.31 | 1.14E-70 | 2.37E-66 | 1.00 | 1.00 | 0.00 | 1.00 |
| 8 | Pde1c | 1.61 | 1.39E-70 | 2.90E-66 | 0.29 | 0.10 | 0.19 | 2.88 |
| 8 | Plod2 | 0.88 | 1.11E-69 | 2.30E-65 | 0.74 | 0.49 | 0.26 | 1.53 |
| 8 | Baz1b | 0.53 | 4.07E-69 | 8.46E-65 | 0.98 | 0.89 | 0.10 | 1.11 |
| 8 | Ncaph | 0.91 | 4.56E-69 | 9.48E-65 | 0.73 | 0.48 | 0.25 | 1.52 |
| 8 | Rcn2 | 0.49 | 7.09E-69 | 1.48E-64 | 0.99 | 0.91 | 0.08 | 1.09 |
| 8 | Vegfc | 1.17 | 7.09E-69 | 1.48E-64 | 0.51 | 0.26 | 0.26 | 2.01 |
| 8 | Pdgfrb | 0.87 | 1.96E-68 | 4.08E-64 | 0.72 | 0.44 | 0.28 | 1.62 |

**Supplementary Table 1. Marker genes for the original clusters of integrated clustering of E11.5 control and NotchGOF λ-junction cells.**

| | | | | | | | | |
|---|---|---|---|---|---|---|---|---|
| 8 | Angptl1 | 1.08 | 2.53E-68 | 5.26E-64 | 0.47 | 0.21 | 0.25 | 2.19 |
| 8 | Ube2e3 | 0.49 | 8.92E-68 | 1.85E-63 | 0.98 | 0.91 | 0.07 | 1.08 |
| 8 | Ttk | 1.05 | 7.79E-67 | 1.62E-62 | 0.61 | 0.35 | 0.26 | 1.72 |
| 8 | Rbms3 | 0.86 | 9.63E-67 | 2.00E-62 | 0.68 | 0.40 | 0.28 | 1.69 |
| 8 | Emilin1 | 0.79 | 1.37E-66 | 2.86E-62 | 0.79 | 0.52 | 0.27 | 1.52 |
| 8 | H3f3b | 0.27 | 1.50E-66 | 3.11E-62 | 1.00 | 1.00 | 0.00 | 1.00 |
| 8 | Prdm16 | 0.90 | 1.10E-65 | 2.29E-61 | 0.60 | 0.32 | 0.29 | 1.91 |
| 8 | Hist1h2bb | 1.36 | 1.50E-65 | 3.12E-61 | 0.42 | 0.20 | 0.22 | 2.13 |
| 8 | Zfp637 | 0.73 | 3.54E-65 | 7.37E-61 | 0.88 | 0.73 | 0.15 | 1.21 |
| 8 | App | 0.66 | 4.39E-65 | 9.14E-61 | 0.92 | 0.78 | 0.14 | 1.18 |
| 8 | Cdk5rap2 | 0.77 | 5.05E-65 | 1.05E-60 | 0.82 | 0.60 | 0.22 | 1.36 |
| 8 | Runx1t1 | 0.80 | 9.30E-65 | 1.93E-60 | 0.78 | 0.52 | 0.26 | 1.51 |
| 8 | Hist2h2aa1 | 1.41 | 3.45E-64 | 7.17E-60 | 0.38 | 0.17 | 0.21 | 2.27 |
| 8 | Pdlim4 | 0.73 | 3.48E-64 | 7.23E-60 | 0.84 | 0.62 | 0.22 | 1.36 |
| 8 | Cdk14 | 0.86 | 1.35E-63 | 2.80E-59 | 0.73 | 0.48 | 0.25 | 1.51 |
| 8 | Nuf2 | 0.86 | 7.46E-63 | 1.55E-58 | 0.72 | 0.47 | 0.25 | 1.54 |
| 8 | Chst15 | 1.28 | 8.92E-63 | 1.85E-58 | 0.38 | 0.16 | 0.21 | 2.32 |
| 8 | Hirip3 | 0.74 | 9.56E-63 | 1.99E-58 | 0.86 | 0.68 | 0.18 | 1.27 |
| 8 | Tmpo | 0.43 | 1.18E-62 | 2.46E-58 | 1.00 | 0.95 | 0.05 | 1.05 |
| 8 | Pdgfra | 0.71 | 1.45E-62 | 3.03E-58 | 0.80 | 0.53 | 0.27 | 1.51 |
| 8 | Ntrk2 | 2.20 | 4.09E-62 | 8.50E-58 | 0.15 | 0.03 | 0.11 | 4.26 |
| 8 | Prc1 | 0.67 | 4.88E-62 | 1.02E-57 | 0.81 | 0.52 | 0.29 | 1.56 |
| 8 | Igfbp7 | 1.03 | 3.31E-61 | 6.89E-57 | 0.50 | 0.25 | 0.25 | 1.99 |
| 8 | Rbm39 | 0.32 | 3.94E-61 | 8.19E-57 | 1.00 | 0.99 | 0.01 | 1.01 |
| 8 | Shcbp1 | 0.85 | 4.05E-61 | 8.43E-57 | 0.72 | 0.46 | 0.26 | 1.56 |
| 8 | Csrp2 | 0.42 | 5.08E-61 | 1.06E-56 | 1.00 | 0.92 | 0.07 | 1.08 |
| 8 | Efnb1 | 0.86 | 6.36E-61 | 1.32E-56 | 0.76 | 0.52 | 0.23 | 1.44 |
| 8 | Smc1a | 0.43 | 1.08E-60 | 2.25E-56 | 0.99 | 0.94 | 0.05 | 1.05 |
| 8 | Mxd3 | 1.31 | 3.87E-60 | 8.05E-56 | 0.45 | 0.23 | 0.22 | 1.93 |
| 8 | Fli1 | 0.88 | 5.61E-60 | 1.17E-55 | 0.64 | 0.36 | 0.28 | 1.76 |
| 8 | Ccdc34 | 0.45 | 1.36E-59 | 2.82E-55 | 0.98 | 0.92 | 0.06 | 1.07 |
| 8 | Foxd2 | 1.29 | 2.19E-59 | 4.56E-55 | 0.29 | 0.11 | 0.18 | 2.70 |
| 8 | Tmtc4 | 1.17 | 3.74E-59 | 7.77E-55 | 0.51 | 0.28 | 0.23 | 1.84 |
| 8 | Eml4 | 0.55 | 4.98E-59 | 1.03E-54 | 0.96 | 0.83 | 0.13 | 1.15 |
| 8 | Ndc80 | 0.85 | 1.94E-58 | 4.02E-54 | 0.66 | 0.41 | 0.25 | 1.61 |
| 8 | Cntln | 0.85 | 1.99E-58 | 4.14E-54 | 0.72 | 0.48 | 0.24 | 1.49 |
| 8 | Brca1 | 0.98 | 2.62E-58 | 5.46E-54 | 0.61 | 0.37 | 0.24 | 1.64 |
| 8 | Clmp | 0.61 | 4.08E-58 | 8.49E-54 | 0.90 | 0.63 | 0.27 | 1.43 |
| 8 | Kif4 | 0.81 | 5.87E-58 | 1.22E-53 | 0.72 | 0.46 | 0.25 | 1.55 |
| 8 | Basp1 | 0.35 | 6.21E-58 | 1.29E-53 | 1.00 | 0.95 | 0.05 | 1.05 |
| 8 | Pola1 | 0.84 | 9.13E-58 | 1.90E-53 | 0.72 | 0.48 | 0.23 | 1.48 |
| 8 | Hist1h2ak | 1.50 | 1.01E-57 | 2.10E-53 | 0.35 | 0.16 | 0.19 | 2.22 |
| 8 | Gmnn | 0.55 | 1.18E-57 | 2.45E-53 | 0.96 | 0.79 | 0.17 | 1.21 |
| 8 | Sulf1 | 0.61 | 1.36E-57 | 2.83E-53 | 0.80 | 0.53 | 0.27 | 1.52 |
| 8 | Hist1h3f | 1.10 | 1.53E-57 | 3.17E-53 | 0.49 | 0.26 | 0.23 | 1.87 |
| 8 | P3h3 | 1.01 | 2.94E-57 | 6.11E-53 | 0.54 | 0.30 | 0.24 | 1.82 |
| 8 | Mindy4 | 1.38 | 3.12E-57 | 6.49E-53 | 0.31 | 0.12 | 0.18 | 2.50 |
| 8 | Fign | 1.06 | 4.96E-57 | 1.03E-52 | 0.52 | 0.29 | 0.24 | 1.83 |
| 8 | Serpinf1 | 0.58 | 8.79E-57 | 1.83E-52 | 0.89 | 0.63 | 0.25 | 1.40 |
| 8 | 0610040J01Rik | 2.33 | 1.11E-56 | 2.30E-52 | 0.12 | 0.03 | 0.10 | 4.65 |
| 8 | Bgn | 1.06 | 2.36E-56 | 4.91E-52 | 0.49 | 0.25 | 0.23 | 1.92 |
| 8 | Malat1 | 0.26 | 4.08E-56 | 8.49E-52 | 1.00 | 1.00 | 0.01 | 1.01 |
| 8 | Crybg3 | 1.24 | 4.20E-56 | 8.74E-52 | 0.42 | 0.20 | 0.21 | 2.04 |
| 8 | Cenph | 0.69 | 8.28E-56 | 1.72E-51 | 0.83 | 0.63 | 0.21 | 1.33 |
| 8 | Ckap2l | 0.78 | 1.28E-55 | 2.66E-51 | 0.73 | 0.48 | 0.25 | 1.52 |
| 8 | Pth1r | 0.66 | 1.39E-55 | 2.89E-51 | 0.82 | 0.56 | 0.26 | 1.46 |

**Supplementary Table 1. Marker genes for the original clusters of integrated clustering of E11.5 control and NotchGOF λ-junction cells.**

| | | | | | | | | |
|---|---|---|---|---|---|---|---|---|
| 8 | Anln | 0.92 | 1.45E-55 | 3.01E-51 | 0.62 | 0.38 | 0.24 | 1.63 |
| 8 | Sgo1 | 0.80 | 1.63E-55 | 3.38E-51 | 0.68 | 0.44 | 0.24 | 1.55 |
| 8 | Ccdc88a | 0.56 | 1.74E-55 | 3.62E-51 | 0.94 | 0.81 | 0.14 | 1.17 |
| 8 | Tk1 | 0.75 | 2.38E-55 | 4.94E-51 | 0.78 | 0.54 | 0.24 | 1.44 |
| 8 | Reep5 | 0.59 | 2.46E-55 | 5.12E-51 | 0.92 | 0.78 | 0.14 | 1.18 |
| 8 | Kif20b | 0.69 | 2.86E-55 | 5.94E-51 | 0.81 | 0.55 | 0.26 | 1.47 |
| 8 | Oxct1 | 0.52 | 3.28E-55 | 6.82E-51 | 0.95 | 0.82 | 0.13 | 1.16 |
| 8 | Pbk | 0.72 | 3.65E-55 | 7.59E-51 | 0.76 | 0.53 | 0.23 | 1.43 |
| 8 | Slitrk4 | 1.94 | 4.53E-55 | 9.41E-51 | 0.17 | 0.05 | 0.12 | 3.46 |
| 8 | Cxxc4 | 1.00 | 4.69E-55 | 9.76E-51 | 0.53 | 0.30 | 0.23 | 1.78 |
| 8 | Bub3 | 0.51 | 9.30E-55 | 1.93E-50 | 0.96 | 0.82 | 0.13 | 1.16 |
| 8 | Smc6 | 0.50 | 1.72E-54 | 3.58E-50 | 0.96 | 0.87 | 0.10 | 1.11 |
| 8 | Arhgap11a | 0.86 | 3.46E-54 | 7.19E-50 | 0.63 | 0.40 | 0.23 | 1.59 |
| 8 | Pitpnc1 | 0.72 | 4.51E-54 | 9.38E-50 | 0.80 | 0.58 | 0.22 | 1.38 |
| 8 | Asxl3 | 0.82 | 5.12E-54 | 1.07E-49 | 0.70 | 0.45 | 0.24 | 1.53 |
| 8 | Plk4 | 0.66 | 7.02E-54 | 1.46E-49 | 0.76 | 0.54 | 0.22 | 1.41 |
| 8 | Kif22 | 0.72 | 7.49E-54 | 1.56E-49 | 0.73 | 0.49 | 0.24 | 1.50 |
| 8 | Mkx | 1.88 | 1.62E-53 | 3.36E-49 | 0.16 | 0.05 | 0.12 | 3.52 |
| 8 | Hs3st6 | 1.75 | 1.67E-53 | 3.48E-49 | 0.23 | 0.08 | 0.15 | 2.85 |
| 8 | Id2 | 0.53 | 5.93E-53 | 1.23E-48 | 0.97 | 0.86 | 0.11 | 1.13 |
| 8 | Raet1e | 1.08 | 7.88E-53 | 1.64E-48 | 0.44 | 0.22 | 0.22 | 1.98 |
| 8 | Mab21l2 | 0.40 | 7.95E-53 | 1.65E-48 | 0.82 | 0.48 | 0.34 | 1.70 |
| 8 | Bub1 | 0.68 | 9.03E-53 | 1.88E-48 | 0.80 | 0.56 | 0.24 | 1.43 |
| 8 | Sfrp2 | 0.38 | 9.83E-53 | 2.05E-48 | 0.82 | 0.60 | 0.21 | 1.35 |
| 8 | Cdca5 | 0.98 | 1.47E-52 | 3.05E-48 | 0.55 | 0.33 | 0.22 | 1.68 |
| 8 | Adgra2 | 0.99 | 4.41E-52 | 9.16E-48 | 0.45 | 0.23 | 0.22 | 1.96 |
| 8 | Pbx3 | 0.74 | 4.76E-52 | 9.91E-48 | 0.75 | 0.52 | 0.23 | 1.43 |
| 8 | Hist1h2bn | 1.41 | 3.00E-51 | 6.24E-47 | 0.32 | 0.14 | 0.18 | 2.23 |
| 8 | Cald1 | 0.40 | 5.82E-51 | 1.21E-46 | 1.00 | 0.89 | 0.11 | 1.12 |
| 8 | Scaf11 | 0.41 | 6.15E-51 | 1.28E-46 | 0.99 | 0.93 | 0.06 | 1.06 |
| 8 | Smc3 | 0.39 | 6.52E-51 | 1.36E-46 | 0.99 | 0.95 | 0.04 | 1.05 |
| 8 | Spdl1 | 0.79 | 6.96E-51 | 1.45E-46 | 0.66 | 0.43 | 0.23 | 1.55 |
| 8 | Hist1h4c | 1.35 | 7.85E-51 | 1.63E-46 | 0.32 | 0.14 | 0.18 | 2.25 |
| 8 | Ezh2 | 0.37 | 8.45E-51 | 1.76E-46 | 0.99 | 0.95 | 0.05 | 1.05 |
| 8 | Dnm3os | 0.61 | 9.33E-51 | 1.94E-46 | 0.77 | 0.51 | 0.25 | 1.49 |
| 8 | Prkar2b | 0.79 | 1.25E-50 | 2.60E-46 | 0.70 | 0.48 | 0.21 | 1.44 |
| 8 | Tubb5 | 0.26 | 1.60E-50 | 3.33E-46 | 1.00 | 1.00 | 0.00 | 1.00 |
| 8 | Hmmr | 0.65 | 2.00E-50 | 4.16E-46 | 0.79 | 0.52 | 0.26 | 1.50 |
| 8 | Etv5 | 0.99 | 2.07E-50 | 4.32E-46 | 0.53 | 0.31 | 0.22 | 1.71 |
| 8 | Wipi1 | 0.93 | 5.91E-50 | 1.23E-45 | 0.61 | 0.39 | 0.22 | 1.57 |
| 8 | Sgo2a | 0.73 | 5.99E-50 | 1.25E-45 | 0.71 | 0.47 | 0.24 | 1.51 |
| 8 | Exo1 | 0.79 | 6.40E-50 | 1.33E-45 | 0.68 | 0.44 | 0.24 | 1.53 |
| 8 | Six1 | 0.51 | 1.01E-49 | 2.09E-45 | 0.88 | 0.60 | 0.28 | 1.47 |
| 8 | Ddr2 | 0.69 | 1.21E-49 | 2.52E-45 | 0.74 | 0.48 | 0.26 | 1.55 |
| 8 | Mme | 0.76 | 1.31E-49 | 2.73E-45 | 0.70 | 0.46 | 0.24 | 1.52 |
| 8 | Igsf3 | 0.70 | 1.51E-49 | 3.14E-45 | 0.78 | 0.58 | 0.20 | 1.35 |
| 8 | H2afx | 0.45 | 1.78E-49 | 3.71E-45 | 0.99 | 0.93 | 0.07 | 1.07 |
| 8 | Pmepa1 | 0.81 | 6.65E-49 | 1.38E-44 | 0.69 | 0.47 | 0.22 | 1.47 |
| 8 | Lmo4 | 0.34 | 7.21E-49 | 1.50E-44 | 0.95 | 0.81 | 0.14 | 1.18 |
| 8 | Crlf1 | 0.55 | 8.22E-49 | 1.71E-44 | 0.87 | 0.63 | 0.24 | 1.38 |
| 8 | Hist1h2af | 1.44 | 8.93E-49 | 1.86E-44 | 0.31 | 0.14 | 0.17 | 2.22 |
| 8 | Smad7 | 0.60 | 1.25E-48 | 2.61E-44 | 0.86 | 0.69 | 0.17 | 1.25 |
| 8 | Mylip | 0.93 | 1.78E-48 | 3.70E-44 | 0.52 | 0.30 | 0.22 | 1.72 |
| 8 | Sncaip | 0.95 | 1.85E-48 | 3.85E-44 | 0.52 | 0.30 | 0.22 | 1.75 |
| 8 | Cthrc1 | 0.98 | 2.27E-48 | 4.72E-44 | 0.39 | 0.19 | 0.20 | 2.07 |
| 8 | Birc5 | 0.45 | 2.54E-48 | 5.29E-44 | 0.97 | 0.80 | 0.17 | 1.21 |

**Supplementary Table 1. Marker genes for the original clusters of integrated clustering of E11.5 control and NotchGOF λ-junction cells.**

| | | | | | | | | |
|---|---|---|---|---|---|---|---|---|
| 8 | Ptprd | 0.75 | 2.86E-48 | 5.95E-44 | 0.66 | 0.43 | 0.23 | 1.53 |
| 8 | Gjc1 | 0.53 | 5.76E-48 | 1.20E-43 | 0.94 | 0.80 | 0.14 | 1.17 |
| 8 | Hnrnpr | 0.34 | 6.13E-48 | 1.27E-43 | 0.99 | 0.97 | 0.03 | 1.03 |
| 8 | Wwtr1 | 0.79 | 6.16E-48 | 1.28E-43 | 0.64 | 0.43 | 0.22 | 1.51 |
| 8 | Melk | 0.81 | 9.25E-48 | 1.92E-43 | 0.69 | 0.46 | 0.22 | 1.48 |
| 8 | Cenpe | 0.53 | 9.32E-48 | 1.94E-43 | 0.88 | 0.65 | 0.24 | 1.37 |
| 8 | Tln2 | 0.68 | 1.72E-47 | 3.59E-43 | 0.78 | 0.58 | 0.20 | 1.34 |
| 8 | Tpd52l1 | 0.91 | 2.34E-47 | 4.87E-43 | 0.55 | 0.32 | 0.23 | 1.70 |
| 8 | Prss23 | 1.00 | 3.56E-47 | 7.40E-43 | 0.52 | 0.30 | 0.22 | 1.71 |
| 8 | Inka1 | 0.75 | 9.03E-47 | 1.88E-42 | 0.72 | 0.50 | 0.22 | 1.43 |
| 8 | Lima1 | 0.57 | 9.30E-47 | 1.93E-42 | 0.89 | 0.73 | 0.15 | 1.21 |
| 8 | Ednrb | 1.97 | 2.14E-46 | 4.46E-42 | 0.13 | 0.04 | 0.10 | 3.67 |
| 8 | Synpo2 | 1.31 | 2.42E-46 | 5.04E-42 | 0.20 | 0.07 | 0.13 | 2.90 |
| 8 | Septin11 | 0.39 | 3.70E-46 | 7.69E-42 | 0.98 | 0.87 | 0.11 | 1.12 |
| 8 | Foxd2os | 0.79 | 4.40E-46 | 9.14E-42 | 0.33 | 0.14 | 0.19 | 2.29 |
| 8 | Efnb2 | 0.83 | 8.20E-46 | 1.71E-41 | 0.66 | 0.44 | 0.22 | 1.51 |
| 8 | Septin7 | 0.32 | 1.25E-45 | 2.60E-41 | 0.99 | 0.97 | 0.02 | 1.02 |
| 8 | Stxbp6 | 1.32 | 1.27E-45 | 2.64E-41 | 0.29 | 0.13 | 0.16 | 2.25 |
| 8 | Rai14 | 0.59 | 2.12E-45 | 4.41E-41 | 0.84 | 0.64 | 0.20 | 1.31 |
| 8 | Hdac9 | 1.58 | 2.97E-45 | 6.17E-41 | 0.20 | 0.07 | 0.13 | 2.80 |
| 8 | Tut4 | 0.38 | 3.80E-45 | 7.91E-41 | 0.99 | 0.92 | 0.07 | 1.08 |
| 8 | Rps6ka3 | 0.74 | 4.43E-45 | 9.22E-41 | 0.67 | 0.45 | 0.22 | 1.49 |
| 8 | Kank4 | 0.99 | 5.54E-45 | 1.15E-40 | 0.39 | 0.19 | 0.19 | 2.00 |
| 8 | Haspin | 0.97 | 6.61E-45 | 1.37E-40 | 0.47 | 0.28 | 0.20 | 1.72 |
| 8 | Thrap3 | 0.40 | 7.84E-45 | 1.63E-40 | 0.99 | 0.93 | 0.06 | 1.07 |
| 8 | Zeb1 | 0.80 | 9.04E-45 | 1.88E-40 | 0.57 | 0.34 | 0.23 | 1.67 |
| 8 | Blm | 0.70 | 9.15E-45 | 1.90E-40 | 0.71 | 0.51 | 0.20 | 1.39 |
| 8 | E130114P18Rik | 0.95 | 9.76E-45 | 2.03E-40 | 0.43 | 0.22 | 0.21 | 1.94 |
| 8 | Acadl | 0.71 | 1.07E-44 | 2.23E-40 | 0.71 | 0.53 | 0.19 | 1.35 |
| 8 | Rfc4 | 0.50 | 1.11E-44 | 2.31E-40 | 0.92 | 0.79 | 0.13 | 1.17 |
| 8 | Fam110b | 0.96 | 2.28E-44 | 4.74E-40 | 0.51 | 0.30 | 0.21 | 1.69 |
| 8 | Fam111a | 0.92 | 2.83E-44 | 5.89E-40 | 0.53 | 0.32 | 0.21 | 1.65 |
| 8 | AI314278 | 1.68 | 3.88E-44 | 8.06E-40 | 0.15 | 0.05 | 0.11 | 3.33 |
| 8 | Nedd4 | 0.26 | 6.43E-44 | 1.34E-39 | 1.00 | 0.98 | 0.02 | 1.02 |
| 8 | Tsen34 | 0.45 | 1.44E-43 | 2.99E-39 | 0.96 | 0.84 | 0.12 | 1.15 |
| 8 | Ncapd3 | 0.70 | 1.73E-43 | 3.59E-39 | 0.72 | 0.53 | 0.19 | 1.36 |
| 8 | Kif2c | 0.68 | 2.00E-43 | 4.15E-39 | 0.69 | 0.46 | 0.23 | 1.50 |
| 8 | H2afv | 0.25 | 2.28E-43 | 4.74E-39 | 1.00 | 0.99 | 0.01 | 1.01 |
| 8 | Bora | 0.83 | 5.17E-43 | 1.08E-38 | 0.64 | 0.44 | 0.20 | 1.45 |
| 8 | Col13a1 | 1.30 | 5.20E-43 | 1.08E-38 | 0.25 | 0.10 | 0.15 | 2.45 |
| 8 | Vcan | 0.34 | 7.26E-43 | 1.51E-38 | 1.00 | 0.86 | 0.14 | 1.16 |
| 8 | Gpc6 | 0.56 | 7.85E-43 | 1.63E-38 | 0.79 | 0.58 | 0.21 | 1.36 |
| 8 | Tpx2 | 0.53 | 9.36E-43 | 1.95E-38 | 0.87 | 0.64 | 0.23 | 1.36 |
| 8 | Brip1 | 1.14 | 9.70E-43 | 2.02E-38 | 0.40 | 0.22 | 0.18 | 1.82 |
| 8 | Kank2 | 0.90 | 1.20E-42 | 2.50E-38 | 0.51 | 0.30 | 0.21 | 1.68 |
| 8 | Lor | 1.76 | 1.33E-42 | 2.77E-38 | 0.11 | 0.03 | 0.08 | 4.12 |
| 8 | Ttc28 | 0.63 | 1.33E-42 | 2.78E-38 | 0.79 | 0.59 | 0.20 | 1.34 |
| 8 | 2610035D17Rik | 1.75 | 1.74E-42 | 3.62E-38 | 0.16 | 0.05 | 0.11 | 3.10 |
| 8 | Lama2 | 1.05 | 1.86E-42 | 3.87E-38 | 0.31 | 0.14 | 0.17 | 2.21 |
| 8 | Cenpp | 0.87 | 2.07E-42 | 4.30E-38 | 0.59 | 0.40 | 0.19 | 1.49 |
| 8 | Cyp26b1 | 1.04 | 2.08E-42 | 4.33E-38 | 0.28 | 0.12 | 0.16 | 2.33 |
| 8 | Rfc1 | 0.45 | 2.23E-42 | 4.65E-38 | 0.95 | 0.86 | 0.09 | 1.11 |
| 8 | Epb41l2 | 0.54 | 5.28E-42 | 1.10E-37 | 0.85 | 0.66 | 0.18 | 1.28 |
| 8 | Mab21l1 | 0.97 | 5.91E-42 | 1.23E-37 | 0.34 | 0.16 | 0.18 | 2.12 |
| 8 | Frmd4a | 0.53 | 6.11E-42 | 1.27E-37 | 0.85 | 0.67 | 0.19 | 1.28 |
| 8 | Ldhb | 0.37 | 1.31E-41 | 2.72E-37 | 0.98 | 0.91 | 0.08 | 1.08 |

**Supplementary Table 1. Marker genes for the original clusters of integrated clustering of E11.5 control and NotchGOF λ-junction cells.**

| | | | | | | | | |
|---|---|---|---|---|---|---|---|---|
| 8 | Bambi | 0.47 | 1.63E-41 | 3.39E-37 | 0.82 | 0.63 | 0.19 | 1.30 |
| 8 | Snai1 | 0.68 | 1.99E-41 | 4.14E-37 | 0.64 | 0.41 | 0.23 | 1.58 |
| 8 | Fgfr3 | 1.97 | 2.61E-41 | 5.43E-37 | 0.12 | 0.03 | 0.09 | 3.62 |
| 8 | Cdca8 | 0.44 | 3.42E-41 | 7.12E-37 | 0.95 | 0.75 | 0.21 | 1.28 |
| 8 | Rbm46 | 1.51 | 4.99E-41 | 1.04E-36 | 0.21 | 0.08 | 0.13 | 2.62 |
| 8 | G2e3 | 0.74 | 5.57E-41 | 1.16E-36 | 0.67 | 0.46 | 0.20 | 1.44 |
| 8 | Dlx5 | 0.90 | 6.14E-41 | 1.28E-36 | 0.40 | 0.21 | 0.19 | 1.91 |
| 8 | Msn | 0.45 | 6.39E-41 | 1.33E-36 | 0.73 | 0.51 | 0.22 | 1.44 |
| 8 | Pla2g7 | 1.38 | 6.64E-41 | 1.38E-36 | 0.18 | 0.06 | 0.12 | 2.92 |
| 8 | 9130024F11Rik | 0.78 | 6.89E-41 | 1.43E-36 | 0.41 | 0.21 | 0.20 | 1.94 |
| 8 | Hist1h2br | 1.24 | 1.01E-40 | 2.10E-36 | 0.31 | 0.15 | 0.16 | 2.05 |
| 8 | Luc7l2 | 0.36 | 1.20E-40 | 2.50E-36 | 0.98 | 0.94 | 0.04 | 1.04 |
| 8 | Ptpdc1 | 1.13 | 1.21E-40 | 2.52E-36 | 0.33 | 0.16 | 0.17 | 2.06 |
| 8 | Scaper | 0.96 | 1.46E-40 | 3.04E-36 | 0.48 | 0.29 | 0.20 | 1.69 |
| 8 | Slc16a10 | 1.31 | 2.08E-40 | 4.33E-36 | 0.25 | 0.10 | 0.14 | 2.39 |
| 8 | Bub1b | 0.58 | 2.43E-40 | 5.06E-36 | 0.77 | 0.55 | 0.22 | 1.41 |
| 8 | Penk | 0.76 | 2.62E-40 | 5.45E-36 | 0.22 | 0.09 | 0.14 | 2.52 |
| 8 | Selenop | 0.59 | 2.89E-40 | 6.02E-36 | 0.37 | 0.18 | 0.18 | 2.01 |
| 8 | Eid1 | 0.33 | 3.17E-40 | 6.58E-36 | 0.99 | 0.97 | 0.03 | 1.03 |
| 8 | Kif18a | 0.78 | 3.63E-40 | 7.55E-36 | 0.54 | 0.34 | 0.21 | 1.61 |
| 8 | Scube1 | 0.78 | 3.66E-40 | 7.61E-36 | 0.59 | 0.37 | 0.22 | 1.60 |
| 8 | Brd8 | 0.53 | 4.58E-40 | 9.52E-36 | 0.88 | 0.75 | 0.13 | 1.18 |
| 8 | Rnf227 | 0.73 | 4.74E-40 | 9.86E-36 | 0.62 | 0.41 | 0.21 | 1.50 |
| 8 | Fxyd1 | 1.42 | 5.25E-40 | 1.09E-35 | 0.17 | 0.06 | 0.11 | 2.93 |
| 8 | Wnt5a | 0.53 | 9.71E-40 | 2.02E-35 | 0.84 | 0.63 | 0.21 | 1.33 |
| 8 | Lsm2 | 0.34 | 1.33E-39 | 2.76E-35 | 0.99 | 0.92 | 0.07 | 1.07 |
| 8 | Eya2 | 0.68 | 1.41E-39 | 2.94E-35 | 0.43 | 0.23 | 0.20 | 1.88 |
| 8 | Dock1 | 0.73 | 1.68E-39 | 3.49E-35 | 0.69 | 0.49 | 0.19 | 1.39 |
| 8 | Ghr | 1.14 | 1.70E-39 | 3.53E-35 | 0.29 | 0.13 | 0.16 | 2.16 |
| 8 | Supt16 | 0.33 | 1.91E-39 | 3.97E-35 | 0.99 | 0.96 | 0.03 | 1.03 |
| 8 | Knl1 | 0.68 | 3.39E-39 | 7.06E-35 | 0.69 | 0.48 | 0.21 | 1.43 |
| 8 | Prdx4 | 0.41 | 3.43E-39 | 7.14E-35 | 0.98 | 0.89 | 0.09 | 1.10 |
| 8 | Chaf1a | 0.56 | 3.99E-39 | 8.31E-35 | 0.83 | 0.64 | 0.18 | 1.29 |
| 8 | Nfatc4 | 0.64 | 4.06E-39 | 8.45E-35 | 0.73 | 0.55 | 0.19 | 1.34 |
| 8 | Bcl2 | 0.78 | 5.12E-39 | 1.07E-34 | 0.55 | 0.34 | 0.20 | 1.59 |
| 8 | Stag1 | 0.52 | 6.03E-39 | 1.26E-34 | 0.88 | 0.75 | 0.13 | 1.17 |
| 8 | Pcdh11x | 1.07 | 6.16E-39 | 1.28E-34 | 0.24 | 0.10 | 0.14 | 2.37 |
| 8 | Dennd5b | 0.72 | 1.08E-38 | 2.25E-34 | 0.58 | 0.38 | 0.20 | 1.53 |
| 8 | Ppp1r12a | 0.55 | 1.51E-38 | 3.13E-34 | 0.86 | 0.73 | 0.14 | 1.19 |
| 8 | Nsmce4a | 0.40 | 1.93E-38 | 4.01E-34 | 0.95 | 0.87 | 0.07 | 1.08 |
| 8 | Pex3 | 0.85 | 2.29E-38 | 4.75E-34 | 0.50 | 0.31 | 0.19 | 1.63 |
| 8 | Arhgef40 | 0.63 | 2.98E-38 | 6.21E-34 | 0.71 | 0.52 | 0.19 | 1.37 |
| 8 | Rad51 | 0.61 | 3.81E-38 | 7.93E-34 | 0.74 | 0.54 | 0.20 | 1.38 |
| 8 | Greb1 | 0.87 | 6.23E-38 | 1.30E-33 | 0.48 | 0.29 | 0.20 | 1.68 |
| 8 | Epc2 | 0.60 | 6.26E-38 | 1.30E-33 | 0.79 | 0.61 | 0.18 | 1.29 |
| 8 | Plk1 | 0.58 | 6.78E-38 | 1.41E-33 | 0.69 | 0.46 | 0.23 | 1.51 |
| 8 | Nts | 1.79 | 6.98E-38 | 1.45E-33 | 0.16 | 0.05 | 0.10 | 2.92 |
| 8 | Jph1 | 1.64 | 7.21E-38 | 1.50E-33 | 0.18 | 0.07 | 0.11 | 2.71 |
| 8 | Tra2a | 0.34 | 7.40E-38 | 1.54E-33 | 0.98 | 0.94 | 0.05 | 1.05 |
| 8 | Smarcc1 | 0.31 | 7.66E-38 | 1.59E-33 | 0.99 | 0.96 | 0.03 | 1.03 |
| 8 | Tnrc18 | 0.58 | 7.93E-38 | 1.65E-33 | 0.74 | 0.55 | 0.19 | 1.35 |
| 8 | Pakap.1 | 0.69 | 9.32E-38 | 1.94E-33 | 0.61 | 0.41 | 0.20 | 1.50 |
| 8 | Fxyd6 | 0.54 | 1.87E-37 | 3.89E-33 | 0.84 | 0.68 | 0.17 | 1.25 |
| 8 | Hmcn1 | 0.57 | 2.16E-37 | 4.49E-33 | 0.72 | 0.53 | 0.19 | 1.36 |
| 8 | Tceal6 | 1.86 | 3.20E-37 | 6.66E-33 | 0.12 | 0.03 | 0.08 | 3.52 |
| 8 | Depdc1a | 0.75 | 4.50E-37 | 9.35E-33 | 0.55 | 0.35 | 0.20 | 1.57 |

**Supplementary Table 1. Marker genes for the original clusters of integrated clustering of E11.5 control and NotchGOF λ-junction cells.**

| | | | | | | | | |
|---|---|---|---|---|---|---|---|---|
| 8 | Cks2 | 0.38 | 5.38E-37 | 1.12E-32 | 0.98 | 0.83 | 0.15 | 1.18 |
| 8 | Elf2 | 0.54 | 5.70E-37 | 1.18E-32 | 0.83 | 0.66 | 0.17 | 1.25 |
| 8 | Metrn | 0.73 | 7.33E-37 | 1.53E-32 | 0.69 | 0.51 | 0.18 | 1.35 |
| 8 | Golim4 | 0.62 | 7.67E-37 | 1.60E-32 | 0.77 | 0.61 | 0.17 | 1.27 |
| 8 | Ednra | 0.85 | 9.90E-37 | 2.06E-32 | 0.53 | 0.33 | 0.19 | 1.58 |
| 8 | Fndc3c1 | 0.66 | 1.14E-36 | 2.37E-32 | 0.65 | 0.46 | 0.19 | 1.42 |
| 8 | Nfic | 0.90 | 1.39E-36 | 2.90E-32 | 0.48 | 0.29 | 0.19 | 1.63 |
| 8 | Eva1b | 0.52 | 3.95E-36 | 8.22E-32 | 0.75 | 0.53 | 0.21 | 1.40 |
| 8 | Dnajc10 | 0.54 | 4.36E-36 | 9.07E-32 | 0.84 | 0.68 | 0.16 | 1.23 |
| 8 | Skp2 | 0.51 | 5.09E-36 | 1.06E-31 | 0.87 | 0.69 | 0.18 | 1.26 |
| 8 | Gja1 | 0.37 | 7.94E-36 | 1.65E-31 | 0.91 | 0.77 | 0.15 | 1.19 |
| 8 | Mis18a | 0.64 | 8.07E-36 | 1.68E-31 | 0.65 | 0.46 | 0.19 | 1.42 |
| 8 | Hhip | 1.73 | 8.47E-36 | 1.76E-31 | 0.14 | 0.05 | 0.09 | 3.04 |
| 8 | Gen1 | 0.88 | 1.06E-35 | 2.20E-31 | 0.45 | 0.27 | 0.18 | 1.66 |
| 8 | Dync2h1 | 0.83 | 1.11E-35 | 2.31E-31 | 0.56 | 0.37 | 0.19 | 1.51 |
| 8 | Nfat5 | 0.60 | 2.23E-35 | 4.64E-31 | 0.74 | 0.57 | 0.17 | 1.30 |
| 8 | Hist1h2bl | 1.35 | 2.25E-35 | 4.67E-31 | 0.23 | 0.10 | 0.13 | 2.28 |
| 8 | Ppfibp2 | 0.71 | 2.28E-35 | 4.74E-31 | 0.62 | 0.42 | 0.20 | 1.47 |
| 8 | Ctcf | 0.39 | 3.05E-35 | 6.34E-31 | 0.95 | 0.89 | 0.07 | 1.07 |
| 8 | Hist1h4a | 1.16 | 3.37E-35 | 7.00E-31 | 0.33 | 0.18 | 0.16 | 1.87 |
| 8 | Acvrl1 | 1.15 | 3.95E-35 | 8.21E-31 | 0.23 | 0.10 | 0.13 | 2.34 |
| 8 | Sass6 | 0.68 | 4.52E-35 | 9.39E-31 | 0.68 | 0.49 | 0.19 | 1.38 |
| 8 | Alx1 | 0.35 | 4.72E-35 | 9.81E-31 | 0.98 | 0.72 | 0.26 | 1.36 |
| 8 | Ube2e2 | 0.74 | 4.92E-35 | 1.02E-30 | 0.56 | 0.37 | 0.19 | 1.50 |
| 8 | Itga9 | 0.87 | 5.33E-35 | 1.11E-30 | 0.38 | 0.21 | 0.18 | 1.84 |
| 8 | Mdc1 | 0.80 | 6.63E-35 | 1.38E-30 | 0.56 | 0.38 | 0.18 | 1.47 |
| 8 | Dbf4 | 0.54 | 7.37E-35 | 1.53E-30 | 0.82 | 0.65 | 0.17 | 1.26 |
| 8 | Colec12 | 0.78 | 8.01E-35 | 1.67E-30 | 0.39 | 0.22 | 0.18 | 1.83 |
| 8 | Ccng2 | 0.70 | 9.01E-35 | 1.87E-30 | 0.61 | 0.43 | 0.18 | 1.43 |
| 8 | Zfp36l2 | 0.70 | 9.67E-35 | 2.01E-30 | 0.62 | 0.43 | 0.19 | 1.44 |
| 8 | Fgfr1 | 0.48 | 9.86E-35 | 2.05E-30 | 0.83 | 0.65 | 0.18 | 1.28 |
| 8 | Ahr | 1.10 | 9.97E-35 | 2.07E-30 | 0.21 | 0.08 | 0.12 | 2.45 |
| 8 | Prrc2c | 0.29 | 1.21E-34 | 2.53E-30 | 1.00 | 0.97 | 0.03 | 1.03 |
| 8 | Tmx4 | 0.66 | 1.43E-34 | 2.98E-30 | 0.64 | 0.46 | 0.18 | 1.39 |
| 8 | Slbp | 0.40 | 1.85E-34 | 3.84E-30 | 0.96 | 0.87 | 0.09 | 1.10 |
| 8 | Itgav | 0.79 | 1.98E-34 | 4.11E-30 | 0.45 | 0.27 | 0.19 | 1.69 |
| 8 | Map3k1 | 0.89 | 2.15E-34 | 4.46E-30 | 0.43 | 0.26 | 0.18 | 1.68 |
| 8 | Mcc | 1.10 | 2.62E-34 | 5.45E-30 | 0.27 | 0.13 | 0.14 | 2.14 |
| 8 | Ube2c | 0.29 | 2.65E-34 | 5.52E-30 | 0.92 | 0.73 | 0.19 | 1.26 |
| 8 | P4ha3 | 1.14 | 3.62E-34 | 7.52E-30 | 0.30 | 0.15 | 0.15 | 1.99 |
| 8 | Usp1 | 0.41 | 5.26E-34 | 1.09E-29 | 0.93 | 0.81 | 0.13 | 1.16 |
| 8 | Grb10 | 0.32 | 6.23E-34 | 1.30E-29 | 0.98 | 0.94 | 0.05 | 1.05 |
| 8 | Pde4a | 1.37 | 7.80E-34 | 1.62E-29 | 0.16 | 0.06 | 0.10 | 2.74 |
| 8 | Ercc6l | 0.81 | 7.90E-34 | 1.64E-29 | 0.53 | 0.35 | 0.17 | 1.50 |
| 8 | Ncapg2 | 0.70 | 8.61E-34 | 1.79E-29 | 0.59 | 0.40 | 0.19 | 1.48 |
| 8 | Nrp1 | 0.51 | 1.00E-33 | 2.09E-29 | 0.39 | 0.20 | 0.18 | 1.91 |
| 8 | Ing1 | 0.49 | 1.02E-33 | 2.13E-29 | 0.84 | 0.73 | 0.12 | 1.16 |
| 8 | Sfr1 | 0.29 | 1.65E-33 | 3.44E-29 | 0.99 | 0.98 | 0.01 | 1.01 |
| 8 | Zwilch | 0.88 | 1.67E-33 | 3.47E-29 | 0.49 | 0.31 | 0.18 | 1.57 |
| 8 | Reck | 0.87 | 1.81E-33 | 3.75E-29 | 0.42 | 0.24 | 0.18 | 1.75 |
| 8 | Mef2c | 0.54 | 1.95E-33 | 4.06E-29 | 0.34 | 0.18 | 0.16 | 1.92 |
| 8 | Slc35f2 | 1.51 | 2.22E-33 | 4.62E-29 | 0.16 | 0.06 | 0.10 | 2.75 |
| 8 | Ncapd2 | 0.56 | 2.43E-33 | 5.06E-29 | 0.73 | 0.55 | 0.18 | 1.34 |
| 8 | Camk2n1 | 0.73 | 2.69E-33 | 5.59E-29 | 0.35 | 0.18 | 0.17 | 1.90 |
| 8 | Cdca4 | 0.61 | 2.96E-33 | 6.15E-29 | 0.72 | 0.55 | 0.17 | 1.31 |
| 8 | Itga4 | 0.69 | 3.95E-33 | 8.22E-29 | 0.36 | 0.19 | 0.17 | 1.89 |

**Supplementary Table 1. Marker genes for the original clusters of integrated clustering of E11.5 control and NotchGOF λ-junction cells.**

| | | | | | | | | |
|---|---|---|---|---|---|---|---|---|
| 8 | Aebp1 | 1.04 | 6.73E-33 | 1.40E-28 | 0.36 | 0.20 | 0.16 | 1.80 |
| 8 | Phip | 0.37 | 7.45E-33 | 1.55E-28 | 0.96 | 0.88 | 0.08 | 1.09 |
| 8 | Cenpl | 0.86 | 1.04E-32 | 2.15E-28 | 0.44 | 0.27 | 0.17 | 1.63 |
| 8 | Hist1h3a | 1.03 | 1.07E-32 | 2.23E-28 | 0.34 | 0.18 | 0.15 | 1.83 |
| 8 | Prkg2 | 1.45 | 1.35E-32 | 2.80E-28 | 0.13 | 0.05 | 0.09 | 2.96 |
| 8 | Crtac1 | 1.34 | 1.58E-32 | 3.29E-28 | 0.15 | 0.05 | 0.10 | 2.81 |
| 8 | Bmpr2 | 0.59 | 1.70E-32 | 3.54E-28 | 0.72 | 0.55 | 0.17 | 1.31 |
| 8 | H1fx | 0.36 | 2.15E-32 | 4.48E-28 | 0.99 | 0.93 | 0.06 | 1.06 |
| 8 | Cenpf | 0.28 | 2.79E-32 | 5.81E-28 | 0.91 | 0.70 | 0.21 | 1.31 |
| 8 | Slco3a1 | 1.06 | 3.06E-32 | 6.36E-28 | 0.26 | 0.12 | 0.14 | 2.10 |
| 8 | Rad54b | 1.00 | 3.79E-32 | 7.89E-28 | 0.35 | 0.20 | 0.15 | 1.77 |
| 8 | Pwwp3a | 0.59 | 4.08E-32 | 8.50E-28 | 0.71 | 0.55 | 0.17 | 1.31 |
| 8 | Rell1 | 0.78 | 5.15E-32 | 1.07E-27 | 0.52 | 0.34 | 0.17 | 1.50 |
| 8 | Satb2 | 0.57 | 5.81E-32 | 1.21E-27 | 0.44 | 0.25 | 0.20 | 1.79 |
| 8 | Uhrf2 | 0.47 | 6.05E-32 | 1.26E-27 | 0.85 | 0.73 | 0.12 | 1.16 |
| 8 | Cep192 | 0.66 | 8.91E-32 | 1.85E-27 | 0.61 | 0.44 | 0.18 | 1.40 |
| 8 | Olfml3 | 0.71 | 1.01E-31 | 2.09E-27 | 0.47 | 0.28 | 0.18 | 1.65 |
| 8 | Tnrc6a | 0.47 | 1.05E-31 | 2.17E-27 | 0.90 | 0.77 | 0.12 | 1.16 |
| 8 | Olfm2 | 1.16 | 1.22E-31 | 2.53E-27 | 0.22 | 0.10 | 0.12 | 2.22 |
| 8 | Sp3 | 0.56 | 1.25E-31 | 2.61E-27 | 0.76 | 0.62 | 0.14 | 1.22 |
| 8 | Cenpm | 0.56 | 1.27E-31 | 2.64E-27 | 0.77 | 0.59 | 0.18 | 1.30 |
| 8 | Egflam | 1.21 | 1.40E-31 | 2.92E-27 | 0.18 | 0.07 | 0.11 | 2.52 |
| 8 | Pcsk5 | 0.62 | 1.75E-31 | 3.64E-27 | 0.56 | 0.36 | 0.20 | 1.54 |
| 8 | Mthfd2 | 0.67 | 2.10E-31 | 4.38E-27 | 0.64 | 0.47 | 0.17 | 1.37 |
| 8 | Ska1 | 0.66 | 2.49E-31 | 5.18E-27 | 0.59 | 0.41 | 0.18 | 1.43 |
| 8 | Mmp2 | 0.54 | 3.11E-31 | 6.46E-27 | 0.70 | 0.51 | 0.19 | 1.38 |
| 8 | Brd3 | 0.31 | 3.12E-31 | 6.49E-27 | 0.98 | 0.93 | 0.06 | 1.06 |
| 8 | Ncam1 | 0.59 | 3.93E-31 | 8.17E-27 | 0.54 | 0.36 | 0.19 | 1.53 |
| 8 | Plod1 | 0.97 | 5.40E-31 | 1.12E-26 | 0.35 | 0.20 | 0.16 | 1.79 |
| 8 | Apip | 0.52 | 6.01E-31 | 1.25E-26 | 0.77 | 0.60 | 0.17 | 1.28 |
| 8 | Alg13 | 0.93 | 8.13E-31 | 1.69E-26 | 0.42 | 0.26 | 0.16 | 1.59 |
| 8 | Apbb2 | 0.78 | 1.22E-30 | 2.54E-26 | 0.47 | 0.30 | 0.18 | 1.59 |
| 8 | Pds5b | 0.51 | 1.54E-30 | 3.21E-26 | 0.79 | 0.65 | 0.14 | 1.22 |
| 8 | Kcnj2 | 0.83 | 1.61E-30 | 3.35E-26 | 0.33 | 0.17 | 0.16 | 1.91 |
| 8 | Rest | 0.36 | 1.83E-30 | 3.80E-26 | 0.92 | 0.81 | 0.11 | 1.13 |
| 8 | Creb3l2 | 0.73 | 2.46E-30 | 5.12E-26 | 0.49 | 0.32 | 0.18 | 1.56 |
| 8 | Prim2 | 0.69 | 2.74E-30 | 5.69E-26 | 0.55 | 0.38 | 0.17 | 1.46 |
| 8 | Aurka | 0.56 | 2.83E-30 | 5.88E-26 | 0.64 | 0.45 | 0.20 | 1.44 |
| 8 | Plxdc2 | 0.77 | 3.08E-30 | 6.40E-26 | 0.41 | 0.24 | 0.17 | 1.69 |
| 8 | St3gal4 | 0.86 | 3.18E-30 | 6.60E-26 | 0.41 | 0.25 | 0.16 | 1.67 |
| 8 | Slc1a3 | 0.73 | 3.75E-30 | 7.80E-26 | 0.39 | 0.22 | 0.17 | 1.76 |
| 8 | Tbc1d15 | 0.71 | 4.48E-30 | 9.31E-26 | 0.66 | 0.51 | 0.16 | 1.31 |
| 8 | Tab2 | 0.52 | 5.35E-30 | 1.11E-25 | 0.81 | 0.67 | 0.14 | 1.21 |
| 8 | Mex3b | 0.46 | 5.99E-30 | 1.24E-25 | 0.83 | 0.69 | 0.14 | 1.20 |
| 8 | Cks1b | 0.27 | 7.64E-30 | 1.59E-25 | 0.99 | 0.96 | 0.04 | 1.04 |
| 8 | Kif23 | 0.55 | 8.78E-30 | 1.83E-25 | 0.63 | 0.44 | 0.19 | 1.44 |
| 8 | Prrg1 | 1.01 | 1.82E-29 | 3.79E-25 | 0.34 | 0.19 | 0.15 | 1.79 |
| 8 | H2-Q7 | 1.21 | 1.98E-29 | 4.13E-25 | 0.22 | 0.11 | 0.12 | 2.13 |
| 8 | Smyd3 | 0.90 | 2.20E-29 | 4.57E-25 | 0.39 | 0.23 | 0.16 | 1.68 |
| 8 | Sdc2 | 0.67 | 2.97E-29 | 6.18E-25 | 0.54 | 0.36 | 0.18 | 1.50 |
| 8 | 4930503L19Rik | 0.70 | 3.52E-29 | 7.31E-25 | 0.50 | 0.33 | 0.17 | 1.53 |
| 8 | Zcchc3 | 0.44 | 4.82E-29 | 1.00E-24 | 0.85 | 0.71 | 0.14 | 1.20 |
| 8 | Dach2 | 1.59 | 5.23E-29 | 1.09E-24 | 0.13 | 0.05 | 0.08 | 2.79 |
| 8 | Npnt | 0.73 | 7.09E-29 | 1.47E-24 | 0.40 | 0.23 | 0.17 | 1.73 |
| 8 | Ptprg | 0.70 | 7.53E-29 | 1.57E-24 | 0.53 | 0.36 | 0.17 | 1.45 |
| 8 | Cdc45 | 0.73 | 7.68E-29 | 1.60E-24 | 0.52 | 0.36 | 0.16 | 1.44 |

**Supplementary Table 1. Marker genes for the original clusters of integrated clustering of E11.5 control and NotchGOF λ-junction cells.**

| | | | | | | | | |
|---|---|---|---|---|---|---|---|---|
| 8 | Matr3 | 0.34 | 7.80E-29 | 1.62E-24 | 0.97 | 0.90 | 0.08 | 1.09 |
| 8 | Igf1r | 0.41 | 1.06E-28 | 2.21E-24 | 0.89 | 0.77 | 0.12 | 1.16 |
| 8 | Dst | 0.56 | 1.08E-28 | 2.24E-24 | 0.52 | 0.35 | 0.18 | 1.50 |
| 8 | Entpd1 | 1.31 | 1.12E-28 | 2.34E-24 | 0.15 | 0.06 | 0.09 | 2.59 |
| 8 | Ror2 | 0.69 | 1.19E-28 | 2.48E-24 | 0.47 | 0.29 | 0.17 | 1.59 |
| 8 | Map2k4 | 0.54 | 1.62E-28 | 3.37E-24 | 0.68 | 0.52 | 0.16 | 1.30 |
| 8 | Myh10 | 0.36 | 1.64E-28 | 3.42E-24 | 0.96 | 0.86 | 0.10 | 1.12 |
| 8 | Hapln1 | 1.01 | 1.76E-28 | 3.65E-24 | 0.12 | 0.04 | 0.08 | 2.93 |
| 8 | Limd2 | 0.31 | 1.87E-28 | 3.88E-24 | 0.99 | 0.90 | 0.09 | 1.10 |
| 8 | Fus | 0.26 | 1.87E-28 | 3.88E-24 | 0.99 | 0.97 | 0.03 | 1.03 |
| 8 | Bicc1 | 0.96 | 1.87E-28 | 3.89E-24 | 0.38 | 0.23 | 0.15 | 1.67 |
| 8 | Ccna2 | 0.41 | 2.07E-28 | 4.30E-24 | 0.83 | 0.62 | 0.21 | 1.33 |
| 8 | Osr2 | 0.80 | 2.09E-28 | 4.34E-24 | 0.20 | 0.09 | 0.11 | 2.32 |
| 8 | Actn1 | 0.39 | 2.40E-28 | 5.00E-24 | 0.86 | 0.72 | 0.14 | 1.19 |
| 8 | Rad21 | 0.37 | 2.72E-28 | 5.66E-24 | 0.91 | 0.80 | 0.10 | 1.13 |
| 8 | Kpna2 | 0.47 | 2.82E-28 | 5.87E-24 | 0.60 | 0.41 | 0.19 | 1.47 |
| 8 | Cpxm1 | 0.58 | 2.96E-28 | 6.15E-24 | 0.61 | 0.43 | 0.18 | 1.42 |
| 8 | Gulp1 | 0.82 | 3.28E-28 | 6.82E-24 | 0.40 | 0.24 | 0.16 | 1.66 |
| 8 | Ppig | 0.30 | 3.38E-28 | 7.02E-24 | 0.98 | 0.93 | 0.05 | 1.05 |
| 8 | Tiparp | 0.68 | 3.78E-28 | 7.86E-24 | 0.49 | 0.31 | 0.18 | 1.56 |
| 8 | Kmt2e | 0.34 | 4.04E-28 | 8.40E-24 | 0.93 | 0.84 | 0.09 | 1.10 |
| 8 | Macf1 | 0.36 | 4.45E-28 | 9.25E-24 | 0.93 | 0.84 | 0.09 | 1.11 |
| 8 | Trip13 | 0.50 | 5.68E-28 | 1.18E-23 | 0.71 | 0.53 | 0.18 | 1.34 |
| 8 | Dpep1 | 0.87 | 5.97E-28 | 1.24E-23 | 0.38 | 0.22 | 0.15 | 1.69 |
| 8 | Cmc2 | 0.48 | 7.00E-28 | 1.46E-23 | 0.75 | 0.60 | 0.15 | 1.25 |
| 8 | Cd248 | 0.64 | 7.46E-28 | 1.55E-23 | 0.42 | 0.25 | 0.17 | 1.70 |
| 8 | Vasn | 0.88 | 8.83E-28 | 1.84E-23 | 0.36 | 0.21 | 0.15 | 1.71 |
| 8 | Mtch1 | 0.27 | 1.31E-27 | 2.73E-23 | 0.99 | 0.96 | 0.04 | 1.04 |
| 8 | Srsf10 | 0.28 | 1.51E-27 | 3.15E-23 | 0.99 | 0.94 | 0.05 | 1.05 |
| 8 | Cdr1os | 1.28 | 1.57E-27 | 3.27E-23 | 0.15 | 0.06 | 0.09 | 2.58 |
| 8 | Plekha5 | 0.54 | 1.87E-27 | 3.90E-23 | 0.63 | 0.46 | 0.18 | 1.39 |
| 8 | Fen1 | 0.45 | 1.95E-27 | 4.06E-23 | 0.82 | 0.65 | 0.17 | 1.27 |
| 8 | Skil | 0.51 | 2.73E-27 | 5.69E-23 | 0.72 | 0.55 | 0.17 | 1.30 |
| 8 | Hs2st1 | 0.53 | 2.90E-27 | 6.03E-23 | 0.72 | 0.56 | 0.16 | 1.28 |
| 8 | Taf3 | 0.59 | 2.99E-27 | 6.21E-23 | 0.66 | 0.50 | 0.16 | 1.32 |
| 8 | Tcaf1 | 0.44 | 3.71E-27 | 7.72E-23 | 0.80 | 0.66 | 0.14 | 1.21 |
| 8 | Ddah1 | 1.05 | 4.32E-27 | 8.99E-23 | 0.26 | 0.14 | 0.13 | 1.94 |
| 8 | Rgma | 0.82 | 5.92E-27 | 1.23E-22 | 0.39 | 0.24 | 0.15 | 1.65 |
| 8 | Fzd4 | 0.76 | 7.58E-27 | 1.58E-22 | 0.40 | 0.25 | 0.16 | 1.64 |
| 8 | Rad18 | 0.68 | 8.08E-27 | 1.68E-22 | 0.53 | 0.36 | 0.17 | 1.45 |
| 8 | Stag2 | 0.48 | 8.18E-27 | 1.70E-22 | 0.79 | 0.65 | 0.14 | 1.22 |
| 8 | Rtf1 | 0.29 | 9.90E-27 | 2.06E-22 | 0.97 | 0.92 | 0.05 | 1.06 |
| 8 | Sox4 | 0.26 | 1.26E-26 | 2.62E-22 | 1.00 | 0.94 | 0.06 | 1.06 |
| 8 | Ankrd11 | 0.29 | 1.36E-26 | 2.83E-22 | 0.98 | 0.94 | 0.05 | 1.05 |
| 8 | Rif1 | 0.39 | 1.38E-26 | 2.88E-22 | 0.90 | 0.80 | 0.10 | 1.13 |
| 8 | 4930579G24Rik | 0.83 | 1.65E-26 | 3.43E-22 | 0.40 | 0.25 | 0.15 | 1.61 |
| 8 | Dnajc9 | 0.31 | 1.70E-26 | 3.53E-22 | 0.96 | 0.86 | 0.10 | 1.12 |
| 8 | Fam171b | 0.68 | 1.76E-26 | 3.66E-22 | 0.40 | 0.24 | 0.16 | 1.67 |
| 8 | Slc16a2 | 0.91 | 1.83E-26 | 3.80E-22 | 0.28 | 0.15 | 0.13 | 1.89 |
| 8 | Ticrr | 0.97 | 2.89E-26 | 6.02E-22 | 0.36 | 0.22 | 0.14 | 1.63 |
| 8 | Pcna | 0.38 | 2.92E-26 | 6.08E-22 | 0.94 | 0.83 | 0.11 | 1.13 |
| 8 | Dlc1 | 0.62 | 3.00E-26 | 6.23E-22 | 0.53 | 0.36 | 0.17 | 1.48 |
| 8 | Arhgef26 | 1.06 | 3.58E-26 | 7.45E-22 | 0.24 | 0.12 | 0.12 | 1.98 |
| 8 | Slf1 | 0.53 | 3.96E-26 | 8.24E-22 | 0.70 | 0.53 | 0.17 | 1.32 |
| 8 | Dcakd | 0.29 | 4.25E-26 | 8.84E-22 | 0.97 | 0.90 | 0.07 | 1.08 |
| 8 | Cspp1 | 0.47 | 4.32E-26 | 8.98E-22 | 0.81 | 0.68 | 0.12 | 1.18 |

**Supplementary Table 1. Marker genes for the original clusters of integrated clustering of E11.5 control and NotchGOF λ-junction cells.**

| | | | | | | | | |
|---|---|---|---|---|---|---|---|---|
| 8 | Sgpp1 | 1.16 | 4.60E-26 | 9.57E-22 | 0.21 | 0.10 | 0.11 | 2.10 |
| 8 | Rrm2 | 0.36 | 5.51E-26 | 1.15E-21 | 0.93 | 0.80 | 0.13 | 1.17 |
| 8 | Cul3 | 0.36 | 7.72E-26 | 1.61E-21 | 0.91 | 0.84 | 0.07 | 1.09 |
| 8 | Pdzrn3 | 0.89 | 8.19E-26 | 1.70E-21 | 0.30 | 0.17 | 0.14 | 1.81 |
| 8 | Nog | 1.31 | 9.70E-26 | 2.02E-21 | 0.20 | 0.09 | 0.10 | 2.14 |
| 8 | Serpine2 | 0.36 | 1.02E-25 | 2.12E-21 | 0.86 | 0.70 | 0.17 | 1.24 |
| 8 | Meaf6 | 0.41 | 1.03E-25 | 2.14E-21 | 0.82 | 0.66 | 0.16 | 1.23 |
| 8 | Rasl11b | 0.61 | 1.04E-25 | 2.16E-21 | 0.65 | 0.48 | 0.16 | 1.34 |
| 8 | Tnik | 0.73 | 1.08E-25 | 2.26E-21 | 0.39 | 0.24 | 0.16 | 1.65 |
| 8 | Traip | 0.79 | 1.28E-25 | 2.65E-21 | 0.41 | 0.27 | 0.14 | 1.53 |
| 8 | Ssbp2 | 0.41 | 1.53E-25 | 3.17E-21 | 0.84 | 0.70 | 0.14 | 1.19 |
| 8 | Tia1 | 0.31 | 2.22E-25 | 4.62E-21 | 0.95 | 0.87 | 0.08 | 1.09 |
| 8 | Edil3 | 0.96 | 3.01E-25 | 6.25E-21 | 0.30 | 0.17 | 0.13 | 1.79 |
| 8 | Slc38a2 | 0.29 | 4.07E-25 | 8.46E-21 | 0.97 | 0.90 | 0.06 | 1.07 |
| 8 | Lix1l | 0.52 | 4.21E-25 | 8.75E-21 | 0.56 | 0.37 | 0.19 | 1.51 |
| 8 | St6galnac5 | 1.46 | 4.59E-25 | 9.54E-21 | 0.13 | 0.05 | 0.08 | 2.66 |
| 8 | Tipin | 0.38 | 5.61E-25 | 1.17E-20 | 0.90 | 0.77 | 0.13 | 1.18 |
| 8 | Yaf2 | 0.38 | 6.93E-25 | 1.44E-20 | 0.82 | 0.70 | 0.13 | 1.18 |
| 8 | Zranb3 | 0.89 | 7.93E-25 | 1.65E-20 | 0.34 | 0.21 | 0.13 | 1.65 |
| 8 | Eme1 | 0.64 | 8.10E-25 | 1.68E-20 | 0.52 | 0.36 | 0.16 | 1.45 |
| 8 | Hist1h2bg | 1.27 | 8.34E-25 | 1.73E-20 | 0.20 | 0.10 | 0.10 | 2.08 |
| 8 | Ssbp4 | 0.43 | 9.80E-25 | 2.04E-20 | 0.80 | 0.65 | 0.16 | 1.24 |
| 8 | Tcf7l1 | 0.48 | 1.02E-24 | 2.13E-20 | 0.73 | 0.58 | 0.15 | 1.26 |
| 8 | Nipbl | 0.32 | 1.15E-24 | 2.38E-20 | 0.95 | 0.87 | 0.08 | 1.09 |
| 8 | Atad5 | 0.58 | 1.22E-24 | 2.53E-20 | 0.68 | 0.52 | 0.15 | 1.29 |
| 8 | Hist1h2bj | 0.84 | 1.30E-24 | 2.70E-20 | 0.38 | 0.24 | 0.14 | 1.58 |
| 8 | Alcam | 0.93 | 1.36E-24 | 2.82E-20 | 0.26 | 0.14 | 0.12 | 1.87 |
| 8 | Col25a1 | 0.69 | 1.63E-24 | 3.40E-20 | 0.34 | 0.19 | 0.15 | 1.76 |
| 8 | Slc40a1 | 1.37 | 1.99E-24 | 4.13E-20 | 0.14 | 0.06 | 0.08 | 2.44 |
| 8 | Osbpl1a | 0.62 | 2.03E-24 | 4.22E-20 | 0.54 | 0.38 | 0.16 | 1.41 |
| 8 | Kifc1 | 0.78 | 2.11E-24 | 4.39E-20 | 0.42 | 0.28 | 0.15 | 1.52 |
| 8 | E2f7 | 0.79 | 2.12E-24 | 4.40E-20 | 0.39 | 0.25 | 0.14 | 1.57 |
| 8 | Nsd3 | 0.46 | 2.31E-24 | 4.80E-20 | 0.76 | 0.63 | 0.13 | 1.21 |
| 8 | Kntc1 | 0.89 | 2.45E-24 | 5.09E-20 | 0.32 | 0.19 | 0.13 | 1.70 |
| 8 | Eif4g3 | 0.38 | 2.66E-24 | 5.54E-20 | 0.87 | 0.76 | 0.11 | 1.15 |
| 8 | Thsd4 | 1.46 | 2.91E-24 | 6.05E-20 | 0.11 | 0.04 | 0.07 | 2.79 |
| 8 | Pakap | 0.98 | 2.95E-24 | 6.13E-20 | 0.23 | 0.12 | 0.12 | 1.98 |
| 8 | Zwint | 0.36 | 3.17E-24 | 6.59E-20 | 0.93 | 0.87 | 0.06 | 1.07 |
| 8 | Ubap2l | 0.27 | 3.94E-24 | 8.20E-20 | 0.98 | 0.94 | 0.04 | 1.04 |
| 8 | Ect2 | 0.54 | 3.97E-24 | 8.25E-20 | 0.57 | 0.40 | 0.17 | 1.42 |
| 8 | Phactr3 | 1.16 | 3.98E-24 | 8.28E-20 | 0.17 | 0.07 | 0.09 | 2.26 |
| 8 | Sash1 | 0.64 | 4.85E-24 | 1.01E-19 | 0.55 | 0.40 | 0.15 | 1.39 |
| 8 | Cttnbp2 | 0.88 | 4.95E-24 | 1.03E-19 | 0.28 | 0.15 | 0.13 | 1.83 |
| 8 | Arhgdib | 0.47 | 5.01E-24 | 1.04E-19 | 0.66 | 0.49 | 0.17 | 1.34 |
| 8 | Kif20a | 0.46 | 5.49E-24 | 1.14E-19 | 0.64 | 0.46 | 0.18 | 1.39 |
| 8 | Cacna2d1 | 0.79 | 6.10E-24 | 1.27E-19 | 0.23 | 0.12 | 0.12 | 2.00 |
| 8 | Has2 | 0.79 | 6.22E-24 | 1.29E-19 | 0.33 | 0.19 | 0.14 | 1.72 |
| 8 | Unc5b | 0.62 | 6.27E-24 | 1.30E-19 | 0.55 | 0.39 | 0.16 | 1.41 |
| 8 | Pcnt | 0.54 | 6.87E-24 | 1.43E-19 | 0.62 | 0.47 | 0.16 | 1.34 |
| 8 | Sec24d | 1.17 | 8.63E-24 | 1.80E-19 | 0.18 | 0.08 | 0.10 | 2.22 |
| 8 | Ccp110 | 0.53 | 1.12E-23 | 2.33E-19 | 0.58 | 0.42 | 0.16 | 1.37 |
| 8 | Pde7b | 1.34 | 1.15E-23 | 2.38E-19 | 0.11 | 0.04 | 0.07 | 2.76 |
| 8 | Usp34 | 0.35 | 1.48E-23 | 3.09E-19 | 0.93 | 0.84 | 0.09 | 1.10 |
| 8 | Ubn1 | 0.39 | 1.55E-23 | 3.23E-19 | 0.84 | 0.76 | 0.08 | 1.10 |
| 8 | Elk3 | 0.45 | 1.57E-23 | 3.26E-19 | 0.59 | 0.41 | 0.17 | 1.42 |
| 8 | Spag5 | 0.71 | 1.72E-23 | 3.58E-19 | 0.44 | 0.30 | 0.14 | 1.48 |

**Supplementary Table 1. Marker genes for the original clusters of integrated clustering of E11.5 control and NotchGOF λ-junction cells.**

| | | | | | | | | |
|---|---|---|---|---|---|---|---|---|
| 8 | Rfc2 | 0.36 | 1.75E-23 | 3.64E-19 | 0.90 | 0.79 | 0.10 | 1.13 |
| 8 | Cdc25c | 0.62 | 1.83E-23 | 3.80E-19 | 0.50 | 0.34 | 0.15 | 1.44 |
| 8 | Bend4 | 1.14 | 2.06E-23 | 4.29E-19 | 0.22 | 0.11 | 0.11 | 1.95 |
| 8 | Dnm1 | 0.58 | 2.36E-23 | 4.90E-19 | 0.48 | 0.32 | 0.16 | 1.49 |
| 8 | Slc25a21 | 1.31 | 2.50E-23 | 5.19E-19 | 0.15 | 0.06 | 0.09 | 2.33 |
| 8 | Ptprs | 0.39 | 3.02E-23 | 6.27E-19 | 0.82 | 0.68 | 0.14 | 1.20 |
| 8 | Stil | 0.91 | 3.02E-23 | 6.28E-19 | 0.32 | 0.19 | 0.13 | 1.68 |
| 8 | E2f8 | 0.86 | 3.31E-23 | 6.88E-19 | 0.33 | 0.20 | 0.13 | 1.64 |
| 8 | Smarcc2 | 0.41 | 3.33E-23 | 6.93E-19 | 0.77 | 0.65 | 0.12 | 1.18 |
| 8 | Fam217b | 1.12 | 3.64E-23 | 7.57E-19 | 0.19 | 0.09 | 0.10 | 2.13 |
| 8 | Prim1 | 0.41 | 4.10E-23 | 8.53E-19 | 0.81 | 0.67 | 0.14 | 1.21 |
| 8 | Selenbp1 | 1.21 | 4.87E-23 | 1.01E-18 | 0.18 | 0.08 | 0.09 | 2.13 |
| 8 | Nhs | 0.66 | 5.06E-23 | 1.05E-18 | 0.49 | 0.33 | 0.15 | 1.46 |
| 8 | Ets2 | 0.71 | 5.56E-23 | 1.16E-18 | 0.39 | 0.25 | 0.14 | 1.57 |
| 8 | Stk3 | 0.57 | 5.86E-23 | 1.22E-18 | 0.57 | 0.42 | 0.15 | 1.35 |
| 8 | Adamts1 | 0.57 | 6.04E-23 | 1.26E-18 | 0.42 | 0.27 | 0.15 | 1.57 |
| 8 | Rfc5 | 0.48 | 6.16E-23 | 1.28E-18 | 0.72 | 0.58 | 0.14 | 1.24 |
| 8 | Mpzl1 | 0.39 | 7.52E-23 | 1.56E-18 | 0.77 | 0.63 | 0.14 | 1.22 |
| 8 | Ak1 | 1.09 | 8.07E-23 | 1.68E-18 | 0.18 | 0.08 | 0.09 | 2.16 |
| 8 | Ash1l | 0.38 | 8.60E-23 | 1.79E-18 | 0.84 | 0.73 | 0.11 | 1.16 |
| 8 | Sirt1 | 0.56 | 8.77E-23 | 1.82E-18 | 0.54 | 0.39 | 0.15 | 1.39 |
| 8 | Fuca2 | 0.69 | 8.83E-23 | 1.84E-18 | 0.38 | 0.23 | 0.15 | 1.63 |
| 8 | Cpq | 0.92 | 9.99E-23 | 2.08E-18 | 0.28 | 0.16 | 0.12 | 1.78 |
| 8 | Atrx | 0.26 | 1.03E-22 | 2.14E-18 | 0.99 | 0.96 | 0.03 | 1.03 |
| 8 | Ephb6 | 1.12 | 1.47E-22 | 3.06E-18 | 0.18 | 0.08 | 0.10 | 2.14 |
| 8 | Bcl9 | 0.40 | 1.52E-22 | 3.15E-18 | 0.80 | 0.68 | 0.12 | 1.17 |
| 8 | St6gal2 | 1.08 | 1.59E-22 | 3.30E-18 | 0.20 | 0.10 | 0.10 | 2.03 |
| 8 | Spidr | 0.68 | 1.61E-22 | 3.35E-18 | 0.45 | 0.30 | 0.14 | 1.47 |
| 8 | Asf1b | 0.62 | 2.02E-22 | 4.20E-18 | 0.57 | 0.43 | 0.14 | 1.32 |
| 8 | Eps8 | 0.86 | 2.04E-22 | 4.25E-18 | 0.28 | 0.16 | 0.12 | 1.77 |
| 8 | Fignl1 | 0.53 | 2.09E-22 | 4.34E-18 | 0.66 | 0.51 | 0.15 | 1.30 |
| 8 | Akt3 | 0.73 | 2.09E-22 | 4.35E-18 | 0.40 | 0.26 | 0.14 | 1.53 |
| 8 | Fancm | 0.62 | 2.15E-22 | 4.46E-18 | 0.51 | 0.36 | 0.15 | 1.43 |
| 8 | Adgrb3 | 1.34 | 2.27E-22 | 4.73E-18 | 0.14 | 0.06 | 0.08 | 2.32 |
| 8 | Cacng4 | 0.73 | 2.38E-22 | 4.94E-18 | 0.40 | 0.25 | 0.14 | 1.57 |
| 8 | Ypel2 | 0.99 | 2.39E-22 | 4.96E-18 | 0.24 | 0.13 | 0.11 | 1.90 |
| 8 | Smc5 | 0.45 | 2.50E-22 | 5.19E-18 | 0.75 | 0.64 | 0.11 | 1.18 |
| 8 | Anpep | 1.38 | 3.15E-22 | 6.56E-18 | 0.11 | 0.04 | 0.07 | 2.63 |
| 8 | Cenpk | 0.46 | 4.05E-22 | 8.43E-18 | 0.72 | 0.57 | 0.15 | 1.26 |
| 8 | Kctd12b | 0.84 | 4.25E-22 | 8.83E-18 | 0.30 | 0.18 | 0.13 | 1.72 |
| 8 | Gm3550 | 1.17 | 4.56E-22 | 9.49E-18 | 0.20 | 0.10 | 0.10 | 1.96 |
| 8 | Tns1 | 0.64 | 5.01E-22 | 1.04E-17 | 0.45 | 0.30 | 0.15 | 1.51 |
| 8 | Lin7a | 1.23 | 5.50E-22 | 1.14E-17 | 0.12 | 0.05 | 0.07 | 2.58 |
| 8 | Setd2 | 0.40 | 5.93E-22 | 1.23E-17 | 0.79 | 0.65 | 0.13 | 1.20 |
| 8 | Gli1 | 0.91 | 5.94E-22 | 1.24E-17 | 0.24 | 0.13 | 0.11 | 1.87 |
| 8 | Zfp644 | 0.30 | 6.30E-22 | 1.31E-17 | 0.95 | 0.86 | 0.09 | 1.10 |
| 8 | Foxm1 | 0.78 | 6.82E-22 | 1.42E-17 | 0.39 | 0.26 | 0.13 | 1.52 |
| 8 | Mmp16 | 0.65 | 6.86E-22 | 1.43E-17 | 0.37 | 0.23 | 0.14 | 1.60 |
| 8 | Fzd2 | 0.27 | 7.01E-22 | 1.46E-17 | 0.97 | 0.87 | 0.10 | 1.12 |
| 8 | Dab2ip | 0.43 | 7.02E-22 | 1.46E-17 | 0.74 | 0.60 | 0.14 | 1.23 |
| 8 | Spire1 | 0.66 | 7.98E-22 | 1.66E-17 | 0.49 | 0.34 | 0.14 | 1.42 |
| 8 | Mad2l2 | 0.58 | 8.00E-22 | 1.66E-17 | 0.56 | 0.41 | 0.15 | 1.37 |
| 8 | Tcf19 | 0.91 | 8.71E-22 | 1.81E-17 | 0.31 | 0.19 | 0.12 | 1.66 |
| 8 | Adamts10 | 0.64 | 1.04E-21 | 2.16E-17 | 0.47 | 0.32 | 0.15 | 1.46 |
| 8 | Erc1 | 0.48 | 1.11E-21 | 2.32E-17 | 0.70 | 0.55 | 0.15 | 1.28 |
| 8 | Sema3a | 0.40 | 1.15E-21 | 2.39E-17 | 0.57 | 0.40 | 0.17 | 1.41 |

**Supplementary Table 1. Marker genes for the original clusters of integrated clustering of E11.5 control and NotchGOF λ-junction cells.**

| | | | | | | | | |
|---|---|---|---|---|---|---|---|---|
| 8 | Dock11 | 0.76 | 1.34E-21 | 2.78E-17 | 0.35 | 0.22 | 0.13 | 1.58 |
| 8 | Dpy19l1 | 0.63 | 1.34E-21 | 2.79E-17 | 0.50 | 0.35 | 0.15 | 1.41 |
| 8 | Spsb4 | 0.69 | 1.38E-21 | 2.87E-17 | 0.34 | 0.20 | 0.13 | 1.66 |
| 8 | Pole | 0.71 | 1.50E-21 | 3.13E-17 | 0.43 | 0.29 | 0.13 | 1.46 |
| 8 | Smchd1 | 0.28 | 1.55E-21 | 3.22E-17 | 0.94 | 0.85 | 0.09 | 1.10 |
| 8 | 1810026B05Rik | 0.45 | 1.69E-21 | 3.52E-17 | 0.73 | 0.59 | 0.14 | 1.23 |
| 8 | Magi1 | 0.70 | 1.83E-21 | 3.81E-17 | 0.45 | 0.30 | 0.14 | 1.47 |
| 8 | Dock7 | 0.43 | 1.86E-21 | 3.87E-17 | 0.75 | 0.61 | 0.14 | 1.23 |
| 8 | Raet1d | 1.11 | 1.92E-21 | 3.99E-17 | 0.17 | 0.08 | 0.09 | 2.12 |
| 8 | Farp1 | 0.53 | 2.03E-21 | 4.23E-17 | 0.64 | 0.49 | 0.15 | 1.30 |
| 8 | Herc6 | 1.40 | 2.14E-21 | 4.46E-17 | 0.12 | 0.05 | 0.07 | 2.44 |
| 8 | Gatad2b | 0.42 | 2.35E-21 | 4.88E-17 | 0.76 | 0.64 | 0.13 | 1.20 |
| 8 | Mcm10 | 0.62 | 2.67E-21 | 5.55E-17 | 0.49 | 0.35 | 0.14 | 1.40 |
| 8 | Vkorc1 | 0.51 | 2.99E-21 | 6.22E-17 | 0.58 | 0.43 | 0.15 | 1.35 |
| 8 | Taf1 | 0.48 | 3.27E-21 | 6.80E-17 | 0.70 | 0.58 | 0.13 | 1.22 |
| 8 | Dhfr | 0.39 | 3.83E-21 | 7.96E-17 | 0.82 | 0.67 | 0.15 | 1.22 |
| 8 | Lrp4 | 1.07 | 4.27E-21 | 8.88E-17 | 0.19 | 0.10 | 0.10 | 2.01 |
| 8 | Brd4 | 0.35 | 4.43E-21 | 9.21E-17 | 0.88 | 0.80 | 0.08 | 1.10 |
| 8 | Hist1h3d | 0.89 | 4.65E-21 | 9.67E-17 | 0.30 | 0.18 | 0.12 | 1.66 |
| 8 | Fam193a | 0.43 | 5.03E-21 | 1.05E-16 | 0.75 | 0.61 | 0.14 | 1.22 |
| 8 | Oaf | 0.43 | 5.09E-21 | 1.06E-16 | 0.62 | 0.45 | 0.17 | 1.38 |
| 8 | Fancd2 | 1.09 | 5.53E-21 | 1.15E-16 | 0.23 | 0.12 | 0.10 | 1.84 |
| 8 | Anxa11 | 0.72 | 5.94E-21 | 1.24E-16 | 0.27 | 0.15 | 0.12 | 1.79 |
| 8 | Rbm5 | 0.33 | 6.15E-21 | 1.28E-16 | 0.91 | 0.82 | 0.09 | 1.11 |
| 8 | Vrk1 | 0.39 | 6.83E-21 | 1.42E-16 | 0.76 | 0.61 | 0.15 | 1.25 |
| 8 | Selenom | 0.28 | 7.77E-21 | 1.62E-16 | 0.67 | 0.52 | 0.15 | 1.30 |
| 8 | Pbrm1 | 0.26 | 8.56E-21 | 1.78E-16 | 0.97 | 0.94 | 0.04 | 1.04 |
| 8 | Rbl1 | 0.73 | 9.26E-21 | 1.93E-16 | 0.42 | 0.29 | 0.13 | 1.45 |
| 8 | Wnt5b | 0.69 | 9.59E-21 | 1.99E-16 | 0.39 | 0.25 | 0.14 | 1.56 |
| 8 | Cep295 | 0.48 | 1.00E-20 | 2.09E-16 | 0.63 | 0.48 | 0.15 | 1.32 |
| 8 | Dock4 | 0.58 | 1.04E-20 | 2.17E-16 | 0.46 | 0.31 | 0.15 | 1.49 |
| 8 | Rap1gds1 | 0.68 | 1.11E-20 | 2.31E-16 | 0.43 | 0.29 | 0.14 | 1.48 |
| 8 | Bmpr1b | 1.11 | 1.16E-20 | 2.41E-16 | 0.17 | 0.08 | 0.09 | 2.07 |
| 8 | Aard | 1.26 | 1.17E-20 | 2.43E-16 | 0.15 | 0.07 | 0.08 | 2.19 |
| 8 | Polr2a | 0.33 | 1.24E-20 | 2.57E-16 | 0.90 | 0.79 | 0.11 | 1.13 |
| 8 | Kifc5b | 0.96 | 1.27E-20 | 2.65E-16 | 0.26 | 0.15 | 0.11 | 1.72 |
| 8 | Usp29 | 0.76 | 1.33E-20 | 2.76E-16 | 0.33 | 0.20 | 0.13 | 1.63 |
| 8 | Arid1a | 0.33 | 1.41E-20 | 2.92E-16 | 0.91 | 0.81 | 0.09 | 1.12 |
| 8 | Pold3 | 0.46 | 1.44E-20 | 2.99E-16 | 0.70 | 0.57 | 0.13 | 1.23 |
| 8 | Med4 | 0.36 | 1.47E-20 | 3.06E-16 | 0.83 | 0.70 | 0.13 | 1.18 |
| 8 | Ntng1 | 0.88 | 1.71E-20 | 3.55E-16 | 0.23 | 0.12 | 0.11 | 1.90 |
| 8 | Cenpj | 0.65 | 1.78E-20 | 3.71E-16 | 0.51 | 0.37 | 0.15 | 1.39 |
| 8 | Zfpm1 | 0.58 | 2.38E-20 | 4.94E-16 | 0.45 | 0.31 | 0.15 | 1.47 |
| 8 | Fmnl3 | 0.72 | 2.57E-20 | 5.35E-16 | 0.33 | 0.20 | 0.13 | 1.63 |
| 8 | Orc6 | 0.30 | 2.58E-20 | 5.36E-16 | 0.90 | 0.79 | 0.11 | 1.14 |
| 8 | B2m | 0.35 | 2.58E-20 | 5.37E-16 | 0.68 | 0.53 | 0.15 | 1.28 |
| 8 | Gabra4 | 1.37 | 2.81E-20 | 5.85E-16 | 0.12 | 0.05 | 0.07 | 2.42 |
| 8 | Cdk2 | 0.50 | 2.93E-20 | 6.10E-16 | 0.56 | 0.41 | 0.15 | 1.38 |
| 8 | Dchs1 | 0.51 | 3.11E-20 | 6.46E-16 | 0.56 | 0.41 | 0.15 | 1.36 |
| 8 | Zfhx4 | 0.35 | 3.12E-20 | 6.49E-16 | 0.86 | 0.73 | 0.12 | 1.17 |
| 8 | Cenpn | 0.59 | 3.12E-20 | 6.50E-16 | 0.52 | 0.38 | 0.14 | 1.36 |
| 8 | Cacna1c | 1.08 | 3.19E-20 | 6.63E-16 | 0.18 | 0.09 | 0.09 | 2.03 |
| 8 | Senp6 | 0.29 | 3.25E-20 | 6.76E-16 | 0.94 | 0.88 | 0.07 | 1.08 |
| 8 | Pclo | 1.11 | 3.38E-20 | 7.04E-16 | 0.20 | 0.10 | 0.10 | 1.92 |
| 8 | Srek1 | 0.33 | 3.42E-20 | 7.12E-16 | 0.92 | 0.82 | 0.10 | 1.12 |
| 8 | Cpd | 0.53 | 3.48E-20 | 7.25E-16 | 0.48 | 0.33 | 0.15 | 1.45 |

**Supplementary Table 1. Marker genes for the original clusters of integrated clustering of E11.5 control and NotchGOF λ-junction cells.**

| | | | | | | | | |
|---|---|---|---|---|---|---|---|---|
| 8 | Septin6 | 0.47 | 3.58E-20 | 7.45E-16 | 0.71 | 0.56 | 0.14 | 1.25 |
| 8 | Prr11 | 0.87 | 4.64E-20 | 9.65E-16 | 0.31 | 0.19 | 0.12 | 1.62 |
| 8 | L1cam | 1.13 | 4.99E-20 | 1.04E-15 | 0.16 | 0.08 | 0.09 | 2.13 |
| 8 | Eif4e | 0.26 | 5.57E-20 | 1.16E-15 | 0.95 | 0.89 | 0.06 | 1.07 |
| 8 | Dscc1 | 0.63 | 5.64E-20 | 1.17E-15 | 0.43 | 0.30 | 0.13 | 1.45 |
| 8 | Chtf18 | 0.78 | 6.19E-20 | 1.29E-15 | 0.35 | 0.23 | 0.12 | 1.54 |
| 8 | Ncoa7 | 0.59 | 6.35E-20 | 1.32E-15 | 0.53 | 0.40 | 0.13 | 1.33 |
| 8 | Fam102b | 0.83 | 6.70E-20 | 1.39E-15 | 0.32 | 0.20 | 0.12 | 1.62 |
| 8 | Slc6a17 | 1.08 | 6.92E-20 | 1.44E-15 | 0.17 | 0.08 | 0.09 | 2.10 |
| 8 | Kif2a | 0.33 | 7.81E-20 | 1.63E-15 | 0.87 | 0.79 | 0.09 | 1.11 |
| 8 | Pip4k2b | 0.78 | 7.90E-20 | 1.64E-15 | 0.34 | 0.22 | 0.12 | 1.57 |
| 8 | Fmr1 | 0.43 | 9.08E-20 | 1.89E-15 | 0.76 | 0.66 | 0.10 | 1.16 |
| 8 | Smarcd1 | 0.31 | 1.00E-19 | 2.08E-15 | 0.90 | 0.84 | 0.07 | 1.08 |
| 8 | Gnb4 | 0.54 | 1.03E-19 | 2.14E-15 | 0.44 | 0.30 | 0.15 | 1.50 |
| 8 | Hspa13 | 0.45 | 1.04E-19 | 2.16E-15 | 0.64 | 0.49 | 0.15 | 1.30 |
| 8 | Wasf1 | 0.55 | 1.22E-19 | 2.54E-15 | 0.54 | 0.40 | 0.14 | 1.35 |
| 8 | Brd2 | 0.32 | 1.22E-19 | 2.55E-15 | 0.91 | 0.82 | 0.09 | 1.11 |
| 8 | Cnot6 | 0.32 | 1.27E-19 | 2.63E-15 | 0.87 | 0.77 | 0.10 | 1.14 |
| 8 | Iws1 | 0.39 | 1.33E-19 | 2.77E-15 | 0.81 | 0.70 | 0.11 | 1.15 |
| 8 | Nbea | 0.44 | 1.38E-19 | 2.86E-15 | 0.59 | 0.44 | 0.15 | 1.34 |
| 8 | Mybl2 | 0.76 | 1.39E-19 | 2.89E-15 | 0.36 | 0.24 | 0.12 | 1.50 |
| 8 | Tnks | 0.51 | 1.41E-19 | 2.94E-15 | 0.66 | 0.54 | 0.12 | 1.23 |
| 8 | Nfyb | 0.30 | 1.45E-19 | 3.02E-15 | 0.89 | 0.79 | 0.10 | 1.13 |
| 8 | Ilf3 | 0.30 | 1.56E-19 | 3.25E-15 | 0.91 | 0.79 | 0.12 | 1.14 |
| 8 | Inpp4b | 0.69 | 1.63E-19 | 3.38E-15 | 0.27 | 0.15 | 0.11 | 1.75 |
| 8 | Ift74 | 0.31 | 1.72E-19 | 3.58E-15 | 0.85 | 0.74 | 0.11 | 1.15 |
| 8 | Safb | 0.32 | 1.74E-19 | 3.62E-15 | 0.88 | 0.78 | 0.10 | 1.12 |
| 8 | Rspo2 | 0.31 | 1.88E-19 | 3.91E-15 | 0.54 | 0.36 | 0.18 | 1.48 |
| 8 | Sh3glb1 | 0.30 | 1.95E-19 | 4.06E-15 | 0.90 | 0.84 | 0.06 | 1.07 |
| 8 | Erbin | 0.40 | 1.96E-19 | 4.07E-15 | 0.75 | 0.63 | 0.12 | 1.20 |
| 8 | Frmd6 | 0.39 | 2.54E-19 | 5.29E-15 | 0.64 | 0.48 | 0.16 | 1.33 |
| 8 | Gulo | 0.75 | 2.65E-19 | 5.50E-15 | 0.19 | 0.09 | 0.09 | 2.01 |
| 8 | Rhobtb3 | 0.29 | 2.80E-19 | 5.81E-15 | 0.91 | 0.81 | 0.10 | 1.12 |
| 8 | Camk2d | 0.71 | 2.80E-19 | 5.82E-15 | 0.31 | 0.19 | 0.12 | 1.63 |
| 8 | Ttc39b | 1.22 | 3.11E-19 | 6.46E-15 | 0.14 | 0.06 | 0.08 | 2.19 |
| 8 | Gle1 | 0.54 | 3.14E-19 | 6.53E-15 | 0.60 | 0.48 | 0.12 | 1.26 |
| 8 | Rin2 | 0.64 | 3.28E-19 | 6.81E-15 | 0.32 | 0.20 | 0.13 | 1.63 |
| 8 | Casd1 | 0.47 | 3.28E-19 | 6.82E-15 | 0.64 | 0.49 | 0.15 | 1.30 |
| 8 | Ophn1 | 0.65 | 3.29E-19 | 6.83E-15 | 0.31 | 0.19 | 0.12 | 1.65 |
| 8 | Fat3 | 0.87 | 3.71E-19 | 7.71E-15 | 0.18 | 0.09 | 0.09 | 2.00 |
| 8 | Tmem26 | 0.56 | 3.77E-19 | 7.84E-15 | 0.39 | 0.25 | 0.14 | 1.55 |
| 8 | Cachd1 | 0.36 | 3.90E-19 | 8.12E-15 | 0.67 | 0.52 | 0.15 | 1.29 |
| 8 | Sae1 | 0.26 | 4.02E-19 | 8.36E-15 | 0.97 | 0.90 | 0.07 | 1.08 |
| 8 | Bmpr1a | 0.42 | 4.26E-19 | 8.85E-15 | 0.77 | 0.64 | 0.13 | 1.20 |
| 8 | Rftn2 | 0.62 | 4.67E-19 | 9.72E-15 | 0.36 | 0.22 | 0.14 | 1.61 |
| 8 | Ddx17 | 0.29 | 4.68E-19 | 9.73E-15 | 0.93 | 0.87 | 0.07 | 1.07 |
| 8 | Ckap5 | 0.31 | 5.15E-19 | 1.07E-14 | 0.88 | 0.77 | 0.11 | 1.14 |
| 8 | Nkain3 | 1.20 | 5.65E-19 | 1.18E-14 | 0.11 | 0.05 | 0.07 | 2.43 |
| 8 | Mns1 | 0.40 | 5.71E-19 | 1.19E-14 | 0.69 | 0.54 | 0.15 | 1.27 |
| 8 | Spry2 | 0.61 | 6.27E-19 | 1.30E-14 | 0.42 | 0.29 | 0.14 | 1.47 |
| 8 | Gpatch8 | 0.32 | 6.76E-19 | 1.41E-14 | 0.88 | 0.79 | 0.09 | 1.12 |
| 8 | Wwp2 | 0.70 | 7.01E-19 | 1.46E-14 | 0.43 | 0.30 | 0.13 | 1.43 |
| 8 | Pkn2 | 0.28 | 7.04E-19 | 1.46E-14 | 0.94 | 0.87 | 0.07 | 1.08 |
| 8 | Rad50 | 0.38 | 7.23E-19 | 1.50E-14 | 0.81 | 0.68 | 0.13 | 1.19 |
| 8 | Sclt1 | 0.53 | 8.11E-19 | 1.69E-14 | 0.56 | 0.42 | 0.14 | 1.33 |
| 8 | Sall4 | 0.67 | 8.19E-19 | 1.70E-14 | 0.31 | 0.19 | 0.12 | 1.66 |

**Supplementary Table 1. Marker genes for the original clusters of integrated clustering of E11.5 control and NotchGOF λ-junction cells.**

| | | | | | | | | |
|---|---|---|---|---|---|---|---|---|
| 8 | Mad2l1 | 0.41 | 1.00E-18 | 2.08E-14 | 0.70 | 0.56 | 0.14 | 1.25 |
| 8 | Heph | 0.64 | 1.11E-18 | 2.31E-14 | 0.38 | 0.25 | 0.13 | 1.51 |
| 8 | Nup85 | 0.61 | 1.13E-18 | 2.36E-14 | 0.49 | 0.36 | 0.13 | 1.35 |
| 8 | St6galnac6 | 0.74 | 1.20E-18 | 2.50E-14 | 0.27 | 0.16 | 0.11 | 1.72 |
| 8 | Lum | 0.76 | 1.20E-18 | 2.50E-14 | 0.27 | 0.15 | 0.11 | 1.72 |
| 8 | Prickle1 | 0.66 | 1.24E-18 | 2.58E-14 | 0.35 | 0.22 | 0.13 | 1.58 |
| 8 | Cnksr2 | 0.65 | 1.30E-18 | 2.70E-14 | 0.45 | 0.32 | 0.13 | 1.40 |
| 8 | Olfm1 | 0.74 | 1.33E-18 | 2.76E-14 | 0.35 | 0.22 | 0.12 | 1.54 |
| 8 | 0610010K14Rik | 0.30 | 1.34E-18 | 2.79E-14 | 0.90 | 0.81 | 0.09 | 1.11 |
| 8 | Hist1h2bp | 1.35 | 1.41E-18 | 2.94E-14 | 0.16 | 0.08 | 0.08 | 2.01 |
| 8 | Taf15 | 0.32 | 1.58E-18 | 3.28E-14 | 0.90 | 0.81 | 0.08 | 1.10 |
| 8 | Chpt1 | 0.51 | 1.67E-18 | 3.47E-14 | 0.52 | 0.39 | 0.14 | 1.36 |
| 8 | Suv39h1 | 0.48 | 1.73E-18 | 3.60E-14 | 0.60 | 0.46 | 0.14 | 1.30 |
| 8 | Eea1 | 0.50 | 1.76E-18 | 3.65E-14 | 0.57 | 0.42 | 0.15 | 1.34 |
| 8 | Cnot6l | 0.47 | 1.80E-18 | 3.74E-14 | 0.66 | 0.53 | 0.13 | 1.24 |
| 8 | Itpr1 | 0.65 | 1.81E-18 | 3.77E-14 | 0.33 | 0.21 | 0.12 | 1.59 |
| 8 | Klf4 | 0.83 | 1.84E-18 | 3.84E-14 | 0.32 | 0.21 | 0.12 | 1.58 |
| 8 | Ska3 | 0.68 | 2.51E-18 | 5.22E-14 | 0.40 | 0.28 | 0.12 | 1.44 |
| 8 | Ctc1 | 0.54 | 2.95E-18 | 6.14E-14 | 0.51 | 0.38 | 0.13 | 1.35 |
| 8 | Brca2 | 0.45 | 3.60E-18 | 7.48E-14 | 0.62 | 0.47 | 0.15 | 1.31 |
| 8 | Prrc2b | 0.32 | 3.67E-18 | 7.63E-14 | 0.85 | 0.74 | 0.11 | 1.15 |
| 8 | Nsl1 | 0.87 | 3.80E-18 | 7.89E-14 | 0.29 | 0.18 | 0.11 | 1.59 |
| 8 | Zfp24 | 0.41 | 3.89E-18 | 8.09E-14 | 0.71 | 0.58 | 0.13 | 1.23 |
| 8 | Pimreg | 0.28 | 4.19E-18 | 8.72E-14 | 0.70 | 0.51 | 0.19 | 1.37 |
| 8 | Pard3b | 0.92 | 4.81E-18 | 9.99E-14 | 0.21 | 0.12 | 0.10 | 1.81 |
| 8 | Zyx | 0.37 | 5.05E-18 | 1.05E-13 | 0.66 | 0.50 | 0.15 | 1.30 |
| 8 | Gfpt2 | 0.73 | 5.18E-18 | 1.08E-13 | 0.29 | 0.18 | 0.12 | 1.64 |
| 8 | Galnt1 | 0.34 | 6.35E-18 | 1.32E-13 | 0.78 | 0.66 | 0.12 | 1.18 |
| 8 | Ndn | 0.29 | 6.63E-18 | 1.38E-13 | 0.85 | 0.74 | 0.11 | 1.15 |
| 8 | Nsd2 | 0.31 | 6.90E-18 | 1.44E-13 | 0.88 | 0.75 | 0.13 | 1.17 |
| 8 | Trim24 | 0.34 | 7.34E-18 | 1.53E-13 | 0.82 | 0.71 | 0.11 | 1.16 |
| 8 | Ptgfrn | 0.54 | 7.46E-18 | 1.55E-13 | 0.47 | 0.33 | 0.14 | 1.42 |
| 8 | Iqgap3 | 0.72 | 8.73E-18 | 1.81E-13 | 0.33 | 0.22 | 0.11 | 1.52 |
| 8 | Gpr155 | 1.19 | 9.38E-18 | 1.95E-13 | 0.12 | 0.06 | 0.07 | 2.25 |
| 8 | Wee1 | 0.47 | 9.48E-18 | 1.97E-13 | 0.58 | 0.46 | 0.13 | 1.28 |
| 8 | Pkn1 | 0.55 | 9.63E-18 | 2.00E-13 | 0.48 | 0.35 | 0.13 | 1.38 |
| 8 | Topbp1 | 0.35 | 9.69E-18 | 2.01E-13 | 0.84 | 0.73 | 0.12 | 1.16 |
| 8 | Fam83d | 0.75 | 1.03E-17 | 2.15E-13 | 0.31 | 0.20 | 0.11 | 1.57 |
| 8 | Uba2 | 0.26 | 1.08E-17 | 2.24E-13 | 0.95 | 0.86 | 0.09 | 1.10 |
| 8 | Spred2 | 0.51 | 1.08E-17 | 2.24E-13 | 0.51 | 0.38 | 0.13 | 1.34 |
| 8 | Col5a2 | 0.36 | 1.24E-17 | 2.57E-13 | 0.68 | 0.52 | 0.16 | 1.30 |
| 8 | Smarca2 | 0.36 | 1.35E-17 | 2.81E-13 | 0.74 | 0.62 | 0.12 | 1.20 |
| 8 | Cyth3 | 0.53 | 1.42E-17 | 2.95E-13 | 0.52 | 0.39 | 0.14 | 1.35 |
| 8 | Acss3 | 1.10 | 1.74E-17 | 3.62E-13 | 0.13 | 0.06 | 0.07 | 2.20 |
| 8 | Pmaip1 | 0.50 | 1.79E-17 | 3.72E-13 | 0.37 | 0.24 | 0.13 | 1.55 |
| 8 | R3hdm1 | 0.31 | 1.87E-17 | 3.89E-13 | 0.83 | 0.73 | 0.10 | 1.14 |
| 8 | Pja2 | 0.42 | 1.89E-17 | 3.94E-13 | 0.67 | 0.55 | 0.13 | 1.23 |
| 8 | Lrr1 | 0.73 | 1.91E-17 | 3.96E-13 | 0.37 | 0.26 | 0.11 | 1.44 |
| 8 | Pum1 | 0.25 | 1.93E-17 | 4.01E-13 | 0.94 | 0.88 | 0.06 | 1.07 |
| 8 | Cit | 0.61 | 2.05E-17 | 4.25E-13 | 0.49 | 0.37 | 0.12 | 1.33 |
| 8 | Ypel3 | 0.37 | 2.21E-17 | 4.60E-13 | 0.69 | 0.57 | 0.13 | 1.22 |
| 8 | Tshz2 | 0.37 | 2.33E-17 | 4.84E-13 | 0.48 | 0.33 | 0.15 | 1.45 |
| 8 | Pcdh10 | 0.92 | 2.36E-17 | 4.91E-13 | 0.19 | 0.10 | 0.09 | 1.86 |
| 8 | Cttnbp2nl | 0.42 | 2.38E-17 | 4.96E-13 | 0.63 | 0.49 | 0.14 | 1.28 |
| 8 | Trim59 | 0.31 | 2.44E-17 | 5.08E-13 | 0.84 | 0.71 | 0.12 | 1.17 |
| 8 | Alx4 | 0.39 | 2.53E-17 | 5.27E-13 | 0.58 | 0.42 | 0.16 | 1.37 |

**Supplementary Table 1. Marker genes for the original clusters of integrated clustering of E11.5 control and NotchGOF λ-junction cells.**

| | | | | | | | | |
|---|---|---|---|---|---|---|---|---|
| 8 | Rprd1b | 0.49 | 2.79E-17 | 5.81E-13 | 0.65 | 0.53 | 0.12 | 1.23 |
| 8 | Prkce | 0.81 | 2.89E-17 | 6.02E-13 | 0.25 | 0.15 | 0.10 | 1.69 |
| 8 | Cdc42ep4 | 0.44 | 2.92E-17 | 6.07E-13 | 0.58 | 0.44 | 0.14 | 1.32 |
| 8 | Arhgap31 | 0.61 | 2.92E-17 | 6.08E-13 | 0.30 | 0.19 | 0.12 | 1.63 |
| 8 | Mllt3 | 0.35 | 2.95E-17 | 6.13E-13 | 0.78 | 0.66 | 0.13 | 1.20 |
| 8 | Gmeb1 | 0.53 | 2.99E-17 | 6.21E-13 | 0.53 | 0.41 | 0.13 | 1.31 |
| 8 | Srgap2 | 0.57 | 3.11E-17 | 6.47E-13 | 0.48 | 0.35 | 0.13 | 1.36 |
| 8 | Zfp101 | 0.57 | 3.44E-17 | 7.15E-13 | 0.48 | 0.36 | 0.13 | 1.35 |
| 8 | Siah1b | 0.64 | 3.65E-17 | 7.59E-13 | 0.41 | 0.29 | 0.12 | 1.42 |
| 8 | Aopep | 0.37 | 3.68E-17 | 7.66E-13 | 0.74 | 0.62 | 0.12 | 1.19 |
| 8 | Peak1 | 0.64 | 3.81E-17 | 7.93E-13 | 0.37 | 0.24 | 0.12 | 1.51 |
| 8 | Rab11b | 0.29 | 3.93E-17 | 8.17E-13 | 0.89 | 0.84 | 0.06 | 1.07 |
| 8 | Tdp2 | 0.41 | 4.31E-17 | 8.96E-13 | 0.69 | 0.56 | 0.12 | 1.21 |
| 8 | Fgfrl1 | 0.53 | 4.78E-17 | 9.94E-13 | 0.47 | 0.34 | 0.13 | 1.39 |
| 8 | Pias1 | 0.52 | 4.83E-17 | 1.00E-12 | 0.45 | 0.32 | 0.13 | 1.42 |
| 8 | Ankrd17 | 0.34 | 5.31E-17 | 1.10E-12 | 0.83 | 0.73 | 0.11 | 1.15 |
| 8 | Rbbp7 | 0.26 | 5.95E-17 | 1.24E-12 | 0.95 | 0.89 | 0.06 | 1.07 |
| 8 | Laptm4b | 0.43 | 5.96E-17 | 1.24E-12 | 0.68 | 0.55 | 0.13 | 1.23 |
| 8 | Pds5a | 0.38 | 5.97E-17 | 1.24E-12 | 0.78 | 0.67 | 0.12 | 1.17 |
| 8 | Gmds | 0.51 | 6.62E-17 | 1.38E-12 | 0.49 | 0.36 | 0.13 | 1.37 |
| 8 | Sp1 | 0.51 | 7.03E-17 | 1.46E-12 | 0.53 | 0.41 | 0.13 | 1.32 |
| 8 | Atxn1 | 0.88 | 7.66E-17 | 1.59E-12 | 0.17 | 0.09 | 0.08 | 1.93 |
| 8 | Phf3 | 0.31 | 7.79E-17 | 1.62E-12 | 0.88 | 0.81 | 0.07 | 1.08 |
| 8 | Iqgap2 | 0.52 | 7.86E-17 | 1.63E-12 | 0.49 | 0.36 | 0.13 | 1.37 |
| 8 | Ccnf | 0.59 | 7.97E-17 | 1.66E-12 | 0.46 | 0.33 | 0.12 | 1.37 |
| 8 | Cep128 | 0.90 | 8.41E-17 | 1.75E-12 | 0.27 | 0.17 | 0.10 | 1.58 |
| 8 | Gxylt1 | 0.44 | 8.74E-17 | 1.82E-12 | 0.59 | 0.46 | 0.13 | 1.28 |
| 8 | Mier1 | 0.31 | 8.79E-17 | 1.83E-12 | 0.87 | 0.79 | 0.08 | 1.10 |
| 8 | Numa1 | 0.33 | 9.02E-17 | 1.88E-12 | 0.81 | 0.72 | 0.09 | 1.13 |
| 8 | Pold1 | 0.56 | 9.54E-17 | 1.98E-12 | 0.46 | 0.34 | 0.12 | 1.36 |
| 8 | Slit2 | 0.32 | 9.75E-17 | 2.03E-12 | 0.51 | 0.36 | 0.14 | 1.40 |
| 8 | Taf5 | 0.50 | 1.01E-16 | 2.10E-12 | 0.55 | 0.43 | 0.13 | 1.29 |
| 8 | Mn1 | 0.64 | 1.09E-16 | 2.28E-12 | 0.40 | 0.28 | 0.12 | 1.44 |
| 8 | Car13 | 1.01 | 1.20E-16 | 2.49E-12 | 0.18 | 0.09 | 0.09 | 1.90 |
| 8 | Abcg2 | 0.46 | 1.22E-16 | 2.53E-12 | 0.48 | 0.34 | 0.14 | 1.40 |
| 8 | Arid1b | 0.45 | 1.30E-16 | 2.70E-12 | 0.62 | 0.48 | 0.14 | 1.28 |
| 8 | Rab21 | 0.67 | 1.48E-16 | 3.08E-12 | 0.57 | 0.46 | 0.11 | 1.24 |
| 8 | Haus3 | 0.46 | 1.66E-16 | 3.45E-12 | 0.62 | 0.49 | 0.12 | 1.25 |
| 8 | Fbxl7 | 0.43 | 1.66E-16 | 3.46E-12 | 0.60 | 0.46 | 0.14 | 1.30 |
| 8 | Trpm7 | 0.34 | 1.82E-16 | 3.79E-12 | 0.79 | 0.69 | 0.11 | 1.15 |
| 8 | Ehmt1 | 0.30 | 1.83E-16 | 3.82E-12 | 0.90 | 0.82 | 0.08 | 1.10 |
| 8 | Kcne1l | 0.79 | 1.84E-16 | 3.82E-12 | 0.22 | 0.12 | 0.10 | 1.77 |
| 8 | Cnrip1 | 0.75 | 2.76E-16 | 5.74E-12 | 0.24 | 0.14 | 0.10 | 1.72 |
| 8 | Usp51 | 1.06 | 2.83E-16 | 5.89E-12 | 0.16 | 0.08 | 0.08 | 1.97 |
| 8 | Akap11 | 0.41 | 2.85E-16 | 5.93E-12 | 0.61 | 0.48 | 0.13 | 1.28 |
| 8 | Loxl3 | 0.98 | 2.92E-16 | 6.07E-12 | 0.14 | 0.07 | 0.07 | 2.06 |
| 8 | Clec14a | 0.96 | 3.01E-16 | 6.25E-12 | 0.12 | 0.05 | 0.06 | 2.23 |
| 8 | E2f3 | 0.34 | 3.08E-16 | 6.40E-12 | 0.71 | 0.58 | 0.14 | 1.24 |
| 8 | Commd8 | 0.36 | 3.11E-16 | 6.47E-12 | 0.77 | 0.67 | 0.11 | 1.16 |
| 8 | Crebbp | 0.32 | 3.18E-16 | 6.61E-12 | 0.84 | 0.74 | 0.10 | 1.13 |
| 8 | Smarca1 | 0.44 | 3.19E-16 | 6.64E-12 | 0.59 | 0.46 | 0.13 | 1.29 |
| 8 | Nup35 | 0.47 | 3.32E-16 | 6.90E-12 | 0.59 | 0.47 | 0.12 | 1.25 |
| 8 | Timeless | 0.42 | 3.38E-16 | 7.04E-12 | 0.66 | 0.54 | 0.12 | 1.23 |
| 8 | Tacc3 | 0.31 | 3.65E-16 | 7.59E-12 | 0.75 | 0.60 | 0.15 | 1.26 |
| 8 | Rnf144a | 0.46 | 4.32E-16 | 8.99E-12 | 0.57 | 0.44 | 0.13 | 1.31 |
| 8 | Arhgap5 | 0.28 | 4.36E-16 | 9.06E-12 | 0.94 | 0.86 | 0.08 | 1.09 |

**Supplementary Table 1. Marker genes for the original clusters of integrated clustering of E11.5 control and NotchGOF λ-junction cells.**

| | | | | | | | | |
|---|---|---|---|---|---|---|---|---|
| 8 | Gtf2a1 | 0.45 | 4.66E-16 | 9.69E-12 | 0.61 | 0.49 | 0.12 | 1.25 |
| 8 | Slc44a1 | 0.64 | 5.22E-16 | 1.09E-11 | 0.35 | 0.23 | 0.12 | 1.50 |
| 8 | Arhgap19 | 0.63 | 5.35E-16 | 1.11E-11 | 0.40 | 0.29 | 0.11 | 1.39 |
| 8 | Mms22l | 0.50 | 5.58E-16 | 1.16E-11 | 0.48 | 0.35 | 0.13 | 1.36 |
| 8 | Tm9sf1 | 0.45 | 5.61E-16 | 1.17E-11 | 0.57 | 0.44 | 0.13 | 1.30 |
| 8 | Man1a2 | 0.31 | 5.78E-16 | 1.20E-11 | 0.79 | 0.69 | 0.11 | 1.16 |
| 8 | Ogfrl1 | 0.53 | 5.86E-16 | 1.22E-11 | 0.46 | 0.33 | 0.12 | 1.37 |
| 8 | Myof | 0.85 | 6.98E-16 | 1.45E-11 | 0.17 | 0.09 | 0.08 | 1.93 |
| 8 | Hist1h2ad | 1.03 | 7.00E-16 | 1.46E-11 | 0.18 | 0.10 | 0.08 | 1.84 |
| 8 | Pdzd8 | 0.46 | 7.00E-16 | 1.46E-11 | 0.57 | 0.45 | 0.13 | 1.29 |
| 8 | Rtkn | 0.64 | 7.13E-16 | 1.48E-11 | 0.32 | 0.21 | 0.11 | 1.54 |
| 8 | Vps13a | 0.63 | 7.65E-16 | 1.59E-11 | 0.45 | 0.34 | 0.12 | 1.34 |
| 8 | Eya4 | 0.41 | 7.70E-16 | 1.60E-11 | 0.42 | 0.28 | 0.14 | 1.49 |
| 8 | Mbd5 | 0.66 | 8.06E-16 | 1.68E-11 | 0.31 | 0.20 | 0.11 | 1.53 |
| 8 | Kctd1 | 0.35 | 8.11E-16 | 1.69E-11 | 0.60 | 0.46 | 0.14 | 1.29 |
| 8 | Prpf4 | 0.42 | 8.22E-16 | 1.71E-11 | 0.70 | 0.59 | 0.12 | 1.20 |
| 8 | Rad54l | 0.57 | 8.36E-16 | 1.74E-11 | 0.43 | 0.31 | 0.12 | 1.39 |
| 8 | Cdo1 | 0.29 | 8.44E-16 | 1.76E-11 | 0.64 | 0.50 | 0.14 | 1.28 |
| 8 | Mastl | 0.77 | 8.84E-16 | 1.84E-11 | 0.32 | 0.22 | 0.10 | 1.47 |
| 8 | Ss18 | 0.31 | 9.24E-16 | 1.92E-11 | 0.76 | 0.65 | 0.12 | 1.18 |
| 8 | N4bp2 | 0.34 | 1.04E-15 | 2.15E-11 | 0.77 | 0.67 | 0.10 | 1.15 |
| 8 | Atp1b3 | 0.32 | 1.05E-15 | 2.18E-11 | 0.81 | 0.72 | 0.10 | 1.13 |
| 8 | Ric1 | 0.55 | 1.11E-15 | 2.32E-11 | 0.45 | 0.33 | 0.12 | 1.38 |
| 8 | Rhno1 | 0.37 | 1.16E-15 | 2.40E-11 | 0.79 | 0.66 | 0.13 | 1.19 |
| 8 | Ints6l | 0.77 | 1.23E-15 | 2.56E-11 | 0.27 | 0.17 | 0.10 | 1.60 |
| 8 | Ctdspl2 | 0.38 | 1.43E-15 | 2.97E-11 | 0.70 | 0.57 | 0.12 | 1.21 |
| 8 | Nrm | 0.56 | 1.47E-15 | 3.05E-11 | 0.47 | 0.35 | 0.12 | 1.34 |
| 8 | Clec2l | 1.17 | 1.53E-15 | 3.19E-11 | 0.14 | 0.07 | 0.07 | 2.06 |
| 8 | R3hdm2 | 0.39 | 1.54E-15 | 3.20E-11 | 0.70 | 0.59 | 0.12 | 1.20 |
| 8 | Pten | 0.34 | 1.99E-15 | 4.13E-11 | 0.71 | 0.59 | 0.12 | 1.20 |
| 8 | Clcf1 | 1.12 | 1.99E-15 | 4.15E-11 | 0.12 | 0.06 | 0.07 | 2.16 |
| 8 | Ttc14 | 0.29 | 2.00E-15 | 4.16E-11 | 0.85 | 0.73 | 0.12 | 1.16 |
| 8 | Xiap | 0.49 | 2.08E-15 | 4.32E-11 | 0.60 | 0.49 | 0.11 | 1.23 |
| 8 | Dusp6 | 0.32 | 2.08E-15 | 4.34E-11 | 0.51 | 0.36 | 0.15 | 1.43 |
| 8 | Cenpu | 0.62 | 2.09E-15 | 4.34E-11 | 0.42 | 0.31 | 0.11 | 1.35 |
| 8 | Ttbk2 | 0.79 | 2.11E-15 | 4.38E-11 | 0.28 | 0.18 | 0.10 | 1.58 |
| 8 | A830018L16Rik | 0.85 | 2.17E-15 | 4.51E-11 | 0.21 | 0.12 | 0.09 | 1.75 |
| 8 | Pum2 | 0.31 | 2.18E-15 | 4.53E-11 | 0.85 | 0.75 | 0.11 | 1.14 |
| 8 | Dsel | 0.52 | 2.36E-15 | 4.91E-11 | 0.45 | 0.33 | 0.12 | 1.37 |
| 8 | Cand2 | 0.38 | 2.75E-15 | 5.72E-11 | 0.62 | 0.49 | 0.13 | 1.26 |
| 8 | Aspm | 0.31 | 2.84E-15 | 5.90E-11 | 0.56 | 0.41 | 0.15 | 1.37 |
| 8 | A330076H08Rik | 0.90 | 2.84E-15 | 5.91E-11 | 0.15 | 0.07 | 0.07 | 2.01 |
| 8 | Fgd1 | 0.56 | 2.87E-15 | 5.98E-11 | 0.44 | 0.32 | 0.12 | 1.37 |
| 8 | Plcb1 | 1.02 | 3.18E-15 | 6.61E-11 | 0.17 | 0.09 | 0.08 | 1.83 |
| 8 | Fbln1 | 0.26 | 3.18E-15 | 6.61E-11 | 0.92 | 0.79 | 0.13 | 1.17 |
| 8 | Ska2 | 0.40 | 3.21E-15 | 6.67E-11 | 0.65 | 0.54 | 0.11 | 1.21 |
| 8 | Adprh | 0.25 | 3.65E-15 | 7.58E-11 | 0.88 | 0.78 | 0.10 | 1.12 |
| 8 | Haus6 | 0.50 | 3.69E-15 | 7.67E-11 | 0.48 | 0.36 | 0.12 | 1.34 |
| 8 | Srcap | 0.44 | 3.92E-15 | 8.15E-11 | 0.61 | 0.50 | 0.12 | 1.23 |
| 8 | Peli2 | 0.66 | 4.45E-15 | 9.26E-11 | 0.29 | 0.18 | 0.11 | 1.58 |
| 8 | Usp48 | 0.42 | 4.60E-15 | 9.57E-11 | 0.64 | 0.52 | 0.13 | 1.24 |
| 8 | Col11a1 | 0.41 | 4.61E-15 | 9.59E-11 | 0.56 | 0.42 | 0.14 | 1.32 |
| 8 | Chd6 | 0.29 | 5.05E-15 | 1.05E-10 | 0.82 | 0.73 | 0.09 | 1.13 |
| 8 | Ctsf | 0.71 | 5.22E-15 | 1.09E-10 | 0.19 | 0.10 | 0.09 | 1.82 |
| 8 | Nck1 | 0.39 | 5.28E-15 | 1.10E-10 | 0.65 | 0.54 | 0.11 | 1.21 |
| 8 | Prex2 | 0.40 | 5.43E-15 | 1.13E-10 | 0.54 | 0.40 | 0.14 | 1.34 |

**Supplementary Table 1. Marker genes for the original clusters of integrated clustering of E11.5 control and NotchGOF λ-junction cells.**

| | | | | | | | | |
|---|---|---|---|---|---|---|---|---|
| 8 | Tbck | 0.54 | 5.54E-15 | 1.15E-10 | 0.40 | 0.28 | 0.12 | 1.43 |
| 8 | Zcwpw1 | 0.79 | 5.76E-15 | 1.20E-10 | 0.26 | 0.16 | 0.10 | 1.61 |
| 8 | Cep135 | 0.53 | 5.80E-15 | 1.21E-10 | 0.41 | 0.29 | 0.12 | 1.41 |
| 8 | Spred1 | 0.40 | 5.83E-15 | 1.21E-10 | 0.63 | 0.50 | 0.13 | 1.26 |
| 8 | Adnp | 0.26 | 5.99E-15 | 1.25E-10 | 0.91 | 0.84 | 0.07 | 1.08 |
| 8 | Slfn9 | 0.61 | 6.03E-15 | 1.25E-10 | 0.36 | 0.25 | 0.11 | 1.45 |
| 8 | Eif4a2 | 0.26 | 7.15E-15 | 1.49E-10 | 0.91 | 0.84 | 0.07 | 1.09 |
| 8 | Jam3 | 0.32 | 7.26E-15 | 1.51E-10 | 0.70 | 0.57 | 0.13 | 1.24 |
| 8 | Me1 | 0.61 | 7.45E-15 | 1.55E-10 | 0.36 | 0.25 | 0.11 | 1.45 |
| 8 | Gpbp1 | 0.27 | 7.48E-15 | 1.56E-10 | 0.88 | 0.78 | 0.10 | 1.12 |
| 8 | 5033430I15Rik | 1.17 | 7.63E-15 | 1.59E-10 | 0.11 | 0.05 | 0.06 | 2.17 |
| 8 | Washc2 | 0.37 | 8.68E-15 | 1.80E-10 | 0.65 | 0.52 | 0.13 | 1.25 |
| 8 | Yipf5 | 0.31 | 9.37E-15 | 1.95E-10 | 0.74 | 0.62 | 0.12 | 1.20 |
| 8 | Slc50a1 | 0.38 | 9.57E-15 | 1.99E-10 | 0.51 | 0.37 | 0.14 | 1.37 |
| 8 | Calu | 0.31 | 1.05E-14 | 2.18E-10 | 0.81 | 0.70 | 0.11 | 1.15 |
| 8 | Sdcbp | 0.33 | 1.06E-14 | 2.20E-10 | 0.77 | 0.69 | 0.08 | 1.12 |
| 8 | Sin3a | 0.35 | 1.07E-14 | 2.23E-10 | 0.65 | 0.52 | 0.13 | 1.25 |
| 8 | Cirbp | 0.31 | 1.19E-14 | 2.48E-10 | 0.81 | 0.70 | 0.10 | 1.15 |
| 8 | Six2 | 0.46 | 1.20E-14 | 2.49E-10 | 0.57 | 0.45 | 0.13 | 1.28 |
| 8 | Gps2 | 0.43 | 1.31E-14 | 2.72E-10 | 0.66 | 0.55 | 0.11 | 1.20 |
| 8 | Ttf2 | 0.56 | 1.33E-14 | 2.77E-10 | 0.42 | 0.32 | 0.11 | 1.35 |
| 8 | Uba7 | 0.84 | 1.42E-14 | 2.96E-10 | 0.19 | 0.11 | 0.08 | 1.78 |
| 8 | Trim44 | 0.26 | 1.45E-14 | 3.01E-10 | 0.82 | 0.71 | 0.12 | 1.16 |
| 8 | Lpin2 | 0.38 | 1.46E-14 | 3.04E-10 | 0.65 | 0.54 | 0.11 | 1.21 |
| 8 | Pkd2 | 0.47 | 1.46E-14 | 3.04E-10 | 0.50 | 0.38 | 0.12 | 1.31 |
| 8 | Gnaq | 0.41 | 1.54E-14 | 3.20E-10 | 0.64 | 0.52 | 0.12 | 1.22 |
| 8 | Gins4 | 0.35 | 1.55E-14 | 3.23E-10 | 0.76 | 0.64 | 0.12 | 1.19 |
| 8 | Pacs1 | 0.60 | 1.58E-14 | 3.29E-10 | 0.39 | 0.28 | 0.11 | 1.40 |
| 8 | Rarg | 0.36 | 1.63E-14 | 3.38E-10 | 0.65 | 0.52 | 0.13 | 1.25 |
| 8 | Trim35 | 0.29 | 1.63E-14 | 3.39E-10 | 0.81 | 0.70 | 0.11 | 1.16 |
| 8 | Bin1 | 0.36 | 1.65E-14 | 3.43E-10 | 0.73 | 0.62 | 0.11 | 1.18 |
| 8 | Nup155 | 0.48 | 1.78E-14 | 3.71E-10 | 0.52 | 0.40 | 0.12 | 1.30 |
| 8 | Mphosph8 | 0.28 | 1.94E-14 | 4.02E-10 | 0.86 | 0.76 | 0.10 | 1.13 |
| 8 | Chsy3 | 0.93 | 1.96E-14 | 4.08E-10 | 0.11 | 0.05 | 0.06 | 2.20 |
| 8 | Birc6 | 0.34 | 2.01E-14 | 4.19E-10 | 0.78 | 0.69 | 0.09 | 1.13 |
| 8 | 4932438A13Rik | 0.54 | 2.06E-14 | 4.28E-10 | 0.46 | 0.35 | 0.12 | 1.33 |
| 8 | C430049B03Rik | 0.44 | 2.15E-14 | 4.48E-10 | 0.47 | 0.35 | 0.12 | 1.34 |
| 8 | Chek1 | 0.37 | 2.27E-14 | 4.72E-10 | 0.67 | 0.55 | 0.13 | 1.23 |
| 8 | Emsy | 0.38 | 2.50E-14 | 5.20E-10 | 0.72 | 0.62 | 0.11 | 1.17 |
| 8 | Pcmtd1 | 0.45 | 2.54E-14 | 5.29E-10 | 0.58 | 0.46 | 0.11 | 1.24 |
| 8 | Cdc7 | 0.49 | 2.56E-14 | 5.31E-10 | 0.48 | 0.36 | 0.12 | 1.33 |
| 8 | Inpp1 | 0.71 | 2.59E-14 | 5.38E-10 | 0.26 | 0.16 | 0.10 | 1.59 |
| 8 | Lin54 | 0.70 | 2.59E-14 | 5.39E-10 | 0.33 | 0.23 | 0.10 | 1.43 |
| 8 | Hecw2 | 0.89 | 2.60E-14 | 5.42E-10 | 0.14 | 0.07 | 0.07 | 1.99 |
| 8 | Tox3 | 0.51 | 2.62E-14 | 5.45E-10 | 0.34 | 0.23 | 0.11 | 1.48 |
| 8 | Myo1b | 0.25 | 2.64E-14 | 5.49E-10 | 0.84 | 0.73 | 0.11 | 1.16 |
| 8 | Nsmce2 | 0.27 | 2.77E-14 | 5.76E-10 | 0.85 | 0.76 | 0.09 | 1.11 |
| 8 | Msantd2 | 0.43 | 2.79E-14 | 5.81E-10 | 0.56 | 0.44 | 0.12 | 1.27 |
| 8 | Ep400 | 0.30 | 2.87E-14 | 5.97E-10 | 0.77 | 0.67 | 0.10 | 1.15 |
| 8 | Zfp711 | 0.62 | 2.89E-14 | 6.01E-10 | 0.32 | 0.21 | 0.11 | 1.51 |
| 8 | Srpk2 | 0.29 | 3.19E-14 | 6.64E-10 | 0.81 | 0.71 | 0.10 | 1.14 |
| 8 | Etv4 | 0.38 | 3.20E-14 | 6.65E-10 | 0.46 | 0.33 | 0.14 | 1.41 |
| 8 | Lsamp | 1.17 | 3.20E-14 | 6.67E-10 | 0.10 | 0.05 | 0.06 | 2.24 |
| 8 | Ppp6r3 | 0.42 | 3.24E-14 | 6.75E-10 | 0.61 | 0.49 | 0.12 | 1.24 |
| 8 | Clcn5 | 0.44 | 3.33E-14 | 6.92E-10 | 0.51 | 0.39 | 0.12 | 1.32 |
| 8 | Pcmt1 | 0.34 | 3.42E-14 | 7.11E-10 | 0.84 | 0.74 | 0.10 | 1.13 |

**Supplementary Table 1. Marker genes for the original clusters of integrated clustering of E11.5 control and NotchGOF λ-junction cells.**

| | | | | | | | | |
|---|---|---|---|---|---|---|---|---|
| 8 | Prps1 | 0.41 | 3.45E-14 | 7.17E-10 | 0.56 | 0.44 | 0.12 | 1.27 |
| 8 | Sptbn1 | 0.26 | 3.47E-14 | 7.22E-10 | 0.86 | 0.76 | 0.11 | 1.14 |
| 8 | Ubl3 | 0.41 | 3.49E-14 | 7.25E-10 | 0.49 | 0.37 | 0.12 | 1.33 |
| 8 | Etfa | 0.28 | 3.50E-14 | 7.27E-10 | 0.81 | 0.73 | 0.08 | 1.11 |
| 8 | Pde4d | 0.58 | 3.64E-14 | 7.57E-10 | 0.38 | 0.26 | 0.12 | 1.46 |
| 8 | Pmf1 | 0.34 | 3.71E-14 | 7.72E-10 | 0.77 | 0.67 | 0.10 | 1.15 |
| 8 | Sik2 | 0.77 | 3.76E-14 | 7.82E-10 | 0.25 | 0.16 | 0.09 | 1.58 |
| 8 | Stard3nl | 0.33 | 4.02E-14 | 8.37E-10 | 0.70 | 0.59 | 0.12 | 1.20 |
| 8 | Mgp | 0.54 | 4.12E-14 | 8.58E-10 | 0.14 | 0.07 | 0.07 | 2.00 |
| 8 | Hist1h2ac | 0.98 | 4.24E-14 | 8.82E-10 | 0.18 | 0.10 | 0.08 | 1.76 |
| 8 | Gabpb2 | 0.45 | 4.27E-14 | 8.89E-10 | 0.53 | 0.42 | 0.12 | 1.28 |
| 8 | Znrf1 | 0.38 | 4.61E-14 | 9.58E-10 | 0.70 | 0.59 | 0.11 | 1.19 |
| 8 | Lpp | 0.31 | 4.69E-14 | 9.76E-10 | 0.68 | 0.55 | 0.13 | 1.23 |
| 8 | Zgrf1 | 0.68 | 4.86E-14 | 1.01E-09 | 0.30 | 0.20 | 0.10 | 1.49 |
| 8 | E2f6 | 0.41 | 5.17E-14 | 1.08E-09 | 0.64 | 0.53 | 0.11 | 1.22 |
| 8 | Tusc3 | 0.38 | 5.46E-14 | 1.14E-09 | 0.68 | 0.56 | 0.12 | 1.22 |
| 8 | Wwc2 | 0.35 | 5.70E-14 | 1.19E-09 | 0.70 | 0.58 | 0.12 | 1.21 |
| 8 | Fanci | 0.96 | 5.94E-14 | 1.24E-09 | 0.17 | 0.09 | 0.08 | 1.80 |
| 8 | Itih5 | 0.76 | 6.06E-14 | 1.26E-09 | 0.21 | 0.12 | 0.09 | 1.70 |
| 8 | Helz | 0.37 | 6.36E-14 | 1.32E-09 | 0.70 | 0.57 | 0.12 | 1.22 |
| 8 | Tmem45a | 0.87 | 6.37E-14 | 1.32E-09 | 0.15 | 0.08 | 0.07 | 1.90 |
| 8 | Dleu2 | 0.59 | 6.40E-14 | 1.33E-09 | 0.35 | 0.25 | 0.10 | 1.40 |
| 8 | Mis12 | 0.42 | 6.72E-14 | 1.40E-09 | 0.61 | 0.49 | 0.12 | 1.25 |
| 8 | Clasp2 | 0.67 | 7.06E-14 | 1.47E-09 | 0.34 | 0.24 | 0.10 | 1.42 |
| 8 | Glg1 | 0.29 | 7.74E-14 | 1.61E-09 | 0.79 | 0.71 | 0.09 | 1.12 |
| 8 | Nxt1 | 0.30 | 7.99E-14 | 1.66E-09 | 0.79 | 0.67 | 0.12 | 1.17 |
| 8 | Suz12 | 0.30 | 8.03E-14 | 1.67E-09 | 0.81 | 0.72 | 0.08 | 1.11 |
| 8 | Usp37 | 0.39 | 8.13E-14 | 1.69E-09 | 0.56 | 0.45 | 0.11 | 1.25 |
| 8 | Pkp4 | 0.45 | 9.66E-14 | 2.01E-09 | 0.50 | 0.38 | 0.13 | 1.33 |
| 8 | Arhgap24 | 0.77 | 9.70E-14 | 2.02E-09 | 0.20 | 0.12 | 0.08 | 1.71 |
| 8 | Osbpl6 | 0.71 | 1.01E-13 | 2.10E-09 | 0.24 | 0.15 | 0.09 | 1.62 |
| 8 | Fbln5 | 0.68 | 1.12E-13 | 2.34E-09 | 0.23 | 0.14 | 0.09 | 1.65 |
| 8 | Sox12 | 0.31 | 1.14E-13 | 2.36E-09 | 0.77 | 0.66 | 0.11 | 1.16 |
| 8 | Cdk12 | 0.29 | 1.16E-13 | 2.41E-09 | 0.81 | 0.73 | 0.09 | 1.12 |
| 8 | Slc43a3 | 0.53 | 1.22E-13 | 2.53E-09 | 0.39 | 0.28 | 0.11 | 1.39 |
| 8 | Dtx3 | 0.38 | 1.24E-13 | 2.59E-09 | 0.60 | 0.47 | 0.12 | 1.26 |
| 8 | Evi5 | 0.31 | 1.29E-13 | 2.69E-09 | 0.72 | 0.60 | 0.11 | 1.18 |
| 8 | Itsn1 | 0.45 | 1.30E-13 | 2.70E-09 | 0.54 | 0.43 | 0.11 | 1.25 |
| 8 | Wdr82 | 0.33 | 1.48E-13 | 3.07E-09 | 0.69 | 0.58 | 0.11 | 1.19 |
| 8 | Cenpw | 0.27 | 1.48E-13 | 3.09E-09 | 0.85 | 0.73 | 0.12 | 1.17 |
| 8 | Cir1 | 0.31 | 1.49E-13 | 3.11E-09 | 0.74 | 0.62 | 0.12 | 1.19 |
| 8 | Ppm1m | 0.60 | 1.52E-13 | 3.16E-09 | 0.35 | 0.24 | 0.11 | 1.43 |
| 8 | Itpripl2 | 0.76 | 1.53E-13 | 3.19E-09 | 0.22 | 0.13 | 0.09 | 1.67 |
| 8 | Prtg | 0.66 | 1.63E-13 | 3.38E-09 | 0.26 | 0.17 | 0.10 | 1.57 |
| 8 | Sgk3 | 0.68 | 1.64E-13 | 3.41E-09 | 0.24 | 0.15 | 0.09 | 1.61 |
| 8 | Prkacb | 0.34 | 1.65E-13 | 3.42E-09 | 0.65 | 0.52 | 0.12 | 1.23 |
| 8 | Gm16070 | 0.93 | 1.67E-13 | 3.47E-09 | 0.11 | 0.05 | 0.06 | 2.12 |
| 8 | Myo9a | 0.42 | 1.76E-13 | 3.66E-09 | 0.49 | 0.37 | 0.12 | 1.33 |
| 8 | Bcl2l11 | 0.36 | 1.77E-13 | 3.67E-09 | 0.47 | 0.35 | 0.12 | 1.35 |
| 8 | Tsix | 0.44 | 1.77E-13 | 3.68E-09 | 0.49 | 0.37 | 0.12 | 1.33 |
| 8 | Cep57 | 0.36 | 1.79E-13 | 3.72E-09 | 0.70 | 0.59 | 0.10 | 1.17 |
| 8 | Slc8a1 | 0.74 | 1.92E-13 | 4.00E-09 | 0.20 | 0.12 | 0.08 | 1.69 |
| 8 | B4galt6 | 0.66 | 1.97E-13 | 4.10E-09 | 0.33 | 0.23 | 0.10 | 1.45 |
| 8 | Hnrnpul1 | 0.31 | 2.00E-13 | 4.16E-09 | 0.77 | 0.68 | 0.09 | 1.13 |
| 8 | Ppp2r3a | 0.46 | 2.08E-13 | 4.33E-09 | 0.47 | 0.35 | 0.12 | 1.35 |
| 8 | Zranb1 | 0.34 | 2.11E-13 | 4.40E-09 | 0.71 | 0.61 | 0.10 | 1.17 |

**Supplementary Table 1. Marker genes for the original clusters of integrated clustering of E11.5 control and NotchGOF λ-junction cells.**

| | | | | | | | | |
|---|---|---|---|---|---|---|---|---|
| 8 | Astn1 | 0.77 | 2.27E-13 | 4.72E-09 | 0.20 | 0.12 | 0.08 | 1.70 |
| 8 | Lcorl | 0.37 | 2.28E-13 | 4.75E-09 | 0.60 | 0.48 | 0.12 | 1.25 |
| 8 | 1700025G04Rik | 0.44 | 2.39E-13 | 4.97E-09 | 0.45 | 0.34 | 0.12 | 1.34 |
| 8 | Pck2 | 0.58 | 2.43E-13 | 5.05E-09 | 0.30 | 0.20 | 0.10 | 1.49 |
| 8 | Man2a1 | 0.58 | 2.62E-13 | 5.45E-09 | 0.34 | 0.23 | 0.11 | 1.46 |
| 8 | Snx7 | 0.82 | 2.62E-13 | 5.46E-09 | 0.18 | 0.10 | 0.08 | 1.75 |
| 8 | Cbfb | 0.31 | 2.65E-13 | 5.50E-09 | 0.69 | 0.56 | 0.13 | 1.23 |
| 8 | Fnip1 | 0.62 | 2.70E-13 | 5.61E-09 | 0.34 | 0.24 | 0.10 | 1.44 |
| 8 | Rbm6 | 0.28 | 2.74E-13 | 5.69E-09 | 0.81 | 0.72 | 0.09 | 1.12 |
| 8 | Ipo9 | 0.39 | 2.74E-13 | 5.70E-09 | 0.64 | 0.54 | 0.11 | 1.20 |
| 8 | Ubr1 | 0.52 | 2.85E-13 | 5.94E-09 | 0.44 | 0.33 | 0.11 | 1.33 |
| 8 | Spry4 | 0.70 | 3.06E-13 | 6.35E-09 | 0.23 | 0.14 | 0.09 | 1.63 |
| 8 | Sh3gl1 | 0.30 | 3.08E-13 | 6.40E-09 | 0.73 | 0.62 | 0.11 | 1.18 |
| 8 | Cited2 | 0.36 | 3.16E-13 | 6.58E-09 | 0.73 | 0.64 | 0.09 | 1.15 |
| 8 | Dyrk2 | 0.48 | 3.39E-13 | 7.04E-09 | 0.49 | 0.38 | 0.12 | 1.31 |
| 8 | Ehmt2 | 0.35 | 3.39E-13 | 7.04E-09 | 0.67 | 0.56 | 0.11 | 1.19 |
| 8 | Terf1 | 0.38 | 3.64E-13 | 7.56E-09 | 0.64 | 0.52 | 0.12 | 1.22 |
| 8 | Zfp638 | 0.27 | 3.69E-13 | 7.68E-09 | 0.86 | 0.80 | 0.05 | 1.06 |
| 8 | Fxn | 0.37 | 3.69E-13 | 7.68E-09 | 0.65 | 0.55 | 0.10 | 1.19 |
| 8 | Pam | 0.66 | 3.78E-13 | 7.86E-09 | 0.24 | 0.15 | 0.09 | 1.61 |
| 8 | Med13l | 0.34 | 3.93E-13 | 8.17E-09 | 0.62 | 0.50 | 0.12 | 1.23 |
| 8 | Zcchc24 | 0.45 | 3.97E-13 | 8.27E-09 | 0.41 | 0.29 | 0.12 | 1.41 |
| 8 | Rnf122 | 0.34 | 3.99E-13 | 8.31E-09 | 0.65 | 0.53 | 0.12 | 1.23 |
| 8 | Tasor | 0.42 | 4.03E-13 | 8.37E-09 | 0.58 | 0.48 | 0.10 | 1.21 |
| 8 | Tmem131l | 0.36 | 4.18E-13 | 8.69E-09 | 0.66 | 0.54 | 0.11 | 1.21 |
| 8 | Nek1 | 0.57 | 4.19E-13 | 8.72E-09 | 0.41 | 0.31 | 0.10 | 1.33 |
| 8 | Suv39h2 | 0.41 | 4.29E-13 | 8.93E-09 | 0.53 | 0.41 | 0.12 | 1.30 |
| 8 | Ppt2 | 0.51 | 4.33E-13 | 9.00E-09 | 0.36 | 0.25 | 0.11 | 1.44 |
| 8 | Taf6 | 0.38 | 4.42E-13 | 9.19E-09 | 0.63 | 0.50 | 0.13 | 1.25 |
| 8 | Hist2h3b | 1.21 | 4.49E-13 | 9.33E-09 | 0.13 | 0.07 | 0.06 | 1.90 |
| 8 | Neo1 | 0.34 | 4.72E-13 | 9.81E-09 | 0.63 | 0.50 | 0.13 | 1.26 |
| 8 | Lrp6 | 0.34 | 4.83E-13 | 1.00E-08 | 0.68 | 0.57 | 0.10 | 1.18 |
| 8 | Abtb2 | 0.88 | 4.89E-13 | 1.02E-08 | 0.13 | 0.07 | 0.07 | 1.94 |
| 8 | Mbd4 | 0.54 | 4.90E-13 | 1.02E-08 | 0.43 | 0.33 | 0.11 | 1.33 |
| 8 | Aff3 | 0.52 | 4.97E-13 | 1.03E-08 | 0.32 | 0.22 | 0.11 | 1.49 |
| 8 | Dcaf8 | 0.27 | 5.01E-13 | 1.04E-08 | 0.84 | 0.74 | 0.10 | 1.13 |
| 8 | Rapgef6 | 0.48 | 5.16E-13 | 1.07E-08 | 0.53 | 0.44 | 0.10 | 1.22 |
| 8 | Cep350 | 0.38 | 5.17E-13 | 1.07E-08 | 0.56 | 0.45 | 0.11 | 1.24 |
| 8 | Hist1h2bk | 0.65 | 5.23E-13 | 1.09E-08 | 0.32 | 0.22 | 0.10 | 1.44 |
| 8 | Nid2 | 0.50 | 5.34E-13 | 1.11E-08 | 0.34 | 0.23 | 0.11 | 1.46 |
| 8 | Cdc14b | 0.44 | 5.95E-13 | 1.24E-08 | 0.51 | 0.40 | 0.11 | 1.28 |
| 8 | Foxp4 | 0.36 | 6.12E-13 | 1.27E-08 | 0.55 | 0.42 | 0.13 | 1.32 |
| 8 | Nfia | 0.31 | 6.19E-13 | 1.29E-08 | 0.58 | 0.47 | 0.12 | 1.25 |
| 8 | Gars | 0.30 | 6.78E-13 | 1.41E-08 | 0.78 | 0.67 | 0.11 | 1.17 |
| 8 | Rnf219 | 0.33 | 6.88E-13 | 1.43E-08 | 0.69 | 0.57 | 0.12 | 1.20 |
| 8 | Marveld1 | 0.47 | 7.31E-13 | 1.52E-08 | 0.35 | 0.24 | 0.11 | 1.47 |
| 8 | Pcolce | 0.29 | 7.60E-13 | 1.58E-08 | 0.60 | 0.46 | 0.15 | 1.33 |
| 8 | 4930452B06Rik | 0.85 | 7.75E-13 | 1.61E-08 | 0.20 | 0.12 | 0.08 | 1.64 |
| 8 | Gfpt1 | 0.43 | 8.38E-13 | 1.74E-08 | 0.49 | 0.38 | 0.11 | 1.30 |
| 8 | Pcgf2 | 0.41 | 8.93E-13 | 1.86E-08 | 0.49 | 0.38 | 0.11 | 1.29 |
| 8 | Eno3 | 0.28 | 9.29E-13 | 1.93E-08 | 0.50 | 0.37 | 0.13 | 1.34 |
| 8 | Pxdn | 0.30 | 9.34E-13 | 1.94E-08 | 0.64 | 0.51 | 0.13 | 1.25 |
| 8 | Xkr5 | 1.06 | 9.49E-13 | 1.97E-08 | 0.13 | 0.07 | 0.06 | 1.90 |
| 8 | Rab34 | 0.25 | 9.79E-13 | 2.04E-08 | 0.80 | 0.71 | 0.09 | 1.12 |
| 8 | Polr3gl | 0.51 | 9.98E-13 | 2.08E-08 | 0.42 | 0.31 | 0.11 | 1.36 |
| 8 | Mlf1 | 0.59 | 1.06E-12 | 2.21E-08 | 0.30 | 0.21 | 0.10 | 1.47 |

**Supplementary Table 1. Marker genes for the original clusters of integrated clustering of E11.5 control and NotchGOF λ-junction cells.**

| | | | | | | | | |
|---|---|---|---|---|---|---|---|---|
| 8 | Nckap1 | 0.36 | 1.08E-12 | 2.24E-08 | 0.67 | 0.56 | 0.10 | 1.18 |
| 8 | Sfswap | 0.34 | 1.09E-12 | 2.27E-08 | 0.69 | 0.57 | 0.12 | 1.22 |
| 8 | Lpar4 | 0.48 | 1.17E-12 | 2.42E-08 | 0.44 | 0.32 | 0.11 | 1.35 |
| 8 | Eloa | 0.35 | 1.18E-12 | 2.46E-08 | 0.58 | 0.46 | 0.12 | 1.26 |
| 8 | Spry1 | 0.45 | 1.19E-12 | 2.48E-08 | 0.39 | 0.29 | 0.11 | 1.38 |
| 8 | Nap1l3 | 0.84 | 1.20E-12 | 2.49E-08 | 0.14 | 0.07 | 0.07 | 1.89 |
| 8 | Cdc16 | 0.27 | 1.20E-12 | 2.49E-08 | 0.81 | 0.73 | 0.08 | 1.12 |
| 8 | Decr1 | 0.36 | 1.23E-12 | 2.55E-08 | 0.53 | 0.41 | 0.12 | 1.28 |
| 8 | Haus5 | 0.59 | 1.35E-12 | 2.81E-08 | 0.35 | 0.25 | 0.10 | 1.38 |
| 8 | Cep112 | 0.61 | 1.37E-12 | 2.84E-08 | 0.30 | 0.20 | 0.10 | 1.49 |
| 8 | Ppp2r2d | 0.27 | 1.37E-12 | 2.86E-08 | 0.79 | 0.67 | 0.11 | 1.17 |
| 8 | Hist1h2bf | 1.00 | 1.38E-12 | 2.86E-08 | 0.16 | 0.09 | 0.07 | 1.78 |
| 8 | Mpdz | 0.36 | 1.46E-12 | 3.03E-08 | 0.63 | 0.52 | 0.11 | 1.21 |
| 8 | Smco4 | 0.39 | 1.47E-12 | 3.05E-08 | 0.61 | 0.48 | 0.12 | 1.25 |
| 8 | Ahctf1 | 0.32 | 1.57E-12 | 3.27E-08 | 0.65 | 0.54 | 0.12 | 1.21 |
| 8 | Marchf6 | 0.41 | 1.60E-12 | 3.33E-08 | 0.50 | 0.39 | 0.11 | 1.29 |
| 8 | Prrc2a | 0.25 | 1.66E-12 | 3.46E-08 | 0.85 | 0.76 | 0.09 | 1.12 |
| 8 | Prkd1 | 0.78 | 1.68E-12 | 3.48E-08 | 0.21 | 0.13 | 0.08 | 1.61 |
| 8 | Rgs2 | 0.30 | 1.72E-12 | 3.59E-08 | 0.46 | 0.34 | 0.12 | 1.35 |
| 8 | Wdhd1 | 0.47 | 1.73E-12 | 3.59E-08 | 0.43 | 0.32 | 0.11 | 1.33 |
| 8 | Ylpm1 | 0.31 | 1.74E-12 | 3.61E-08 | 0.80 | 0.69 | 0.11 | 1.16 |
| 8 | Ubr5 | 0.28 | 1.76E-12 | 3.66E-08 | 0.76 | 0.66 | 0.10 | 1.14 |
| 8 | Trim8 | 0.45 | 1.80E-12 | 3.75E-08 | 0.41 | 0.30 | 0.11 | 1.38 |
| 8 | Dusp7 | 0.35 | 1.81E-12 | 3.76E-08 | 0.62 | 0.50 | 0.12 | 1.24 |
| 8 | Gins3 | 0.64 | 1.89E-12 | 3.93E-08 | 0.35 | 0.25 | 0.10 | 1.38 |
| 8 | Fibin | 0.59 | 1.89E-12 | 3.94E-08 | 0.20 | 0.12 | 0.08 | 1.67 |
| 8 | Sesn3 | 0.29 | 1.93E-12 | 4.02E-08 | 0.70 | 0.59 | 0.11 | 1.19 |
| 8 | Asap2 | 0.30 | 1.97E-12 | 4.10E-08 | 0.68 | 0.55 | 0.13 | 1.24 |
| 8 | Cacna1g | 0.40 | 2.07E-12 | 4.30E-08 | 0.45 | 0.33 | 0.12 | 1.37 |
| 8 | Cep170 | 0.33 | 2.13E-12 | 4.42E-08 | 0.62 | 0.51 | 0.11 | 1.22 |
| 8 | Rassf3 | 0.35 | 2.20E-12 | 4.57E-08 | 0.49 | 0.36 | 0.13 | 1.35 |
| 8 | Tbc1d31 | 0.62 | 2.21E-12 | 4.60E-08 | 0.31 | 0.22 | 0.09 | 1.43 |
| 8 | Zfp189 | 0.69 | 2.25E-12 | 4.67E-08 | 0.19 | 0.11 | 0.08 | 1.71 |
| 8 | Nlgn3 | 0.99 | 2.27E-12 | 4.71E-08 | 0.11 | 0.05 | 0.06 | 2.10 |
| 8 | Rasa2 | 0.47 | 2.27E-12 | 4.72E-08 | 0.46 | 0.35 | 0.11 | 1.33 |
| 8 | Irf2bp2 | 0.41 | 2.39E-12 | 4.97E-08 | 0.46 | 0.35 | 0.11 | 1.33 |
| 8 | Racgap1 | 0.29 | 2.40E-12 | 4.99E-08 | 0.65 | 0.52 | 0.13 | 1.25 |
| 8 | Stk24 | 0.40 | 2.50E-12 | 5.21E-08 | 0.57 | 0.46 | 0.11 | 1.24 |
| 8 | BC030867 | 0.79 | 2.54E-12 | 5.29E-08 | 0.22 | 0.14 | 0.08 | 1.58 |
| 8 | Capn6 | 0.29 | 2.61E-12 | 5.43E-08 | 0.67 | 0.54 | 0.13 | 1.23 |
| 8 | Srsf4 | 0.27 | 2.70E-12 | 5.62E-08 | 0.82 | 0.73 | 0.09 | 1.13 |
| 8 | Brwd3 | 0.40 | 2.71E-12 | 5.63E-08 | 0.57 | 0.46 | 0.11 | 1.23 |
| 8 | Tube1 | 0.80 | 2.74E-12 | 5.71E-08 | 0.26 | 0.17 | 0.08 | 1.48 |
| 8 | Sgcd | 0.52 | 2.80E-12 | 5.82E-08 | 0.32 | 0.22 | 0.10 | 1.47 |
| 8 | Hacd4 | 0.49 | 2.82E-12 | 5.87E-08 | 0.35 | 0.25 | 0.11 | 1.43 |
| 8 | Mmd | 0.46 | 2.84E-12 | 5.91E-08 | 0.46 | 0.35 | 0.11 | 1.31 |
| 8 | Gtse1 | 0.82 | 2.86E-12 | 5.96E-08 | 0.26 | 0.18 | 0.08 | 1.46 |
| 8 | Ugp2 | 0.36 | 3.03E-12 | 6.31E-08 | 0.60 | 0.49 | 0.11 | 1.21 |
| 8 | Eldr | 0.69 | 3.05E-12 | 6.33E-08 | 0.27 | 0.18 | 0.09 | 1.50 |
| 8 | Heca | 0.53 | 3.09E-12 | 6.42E-08 | 0.33 | 0.23 | 0.10 | 1.44 |
| 8 | Lmcd1 | 1.05 | 3.10E-12 | 6.46E-08 | 0.15 | 0.09 | 0.07 | 1.77 |
| 8 | Emp2 | 0.89 | 3.13E-12 | 6.51E-08 | 0.14 | 0.07 | 0.06 | 1.88 |
| 8 | Pxk | 0.66 | 3.28E-12 | 6.81E-08 | 0.25 | 0.16 | 0.09 | 1.54 |
| 8 | Hist1h2bh | 0.89 | 3.38E-12 | 7.03E-08 | 0.19 | 0.12 | 0.08 | 1.65 |
| 8 | Ctbp2 | 0.28 | 3.59E-12 | 7.46E-08 | 0.80 | 0.68 | 0.11 | 1.16 |
| 8 | Adam23 | 0.86 | 3.61E-12 | 7.52E-08 | 0.15 | 0.08 | 0.07 | 1.81 |

**Supplementary Table 1. Marker genes for the original clusters of integrated clustering of E11.5 control and NotchGOF λ-junction cells.**

| | | | | | | | | |
|---|---|---|---|---|---|---|---|---|
| 8 | Nbas | 0.58 | 3.65E-12 | 7.60E-08 | 0.31 | 0.22 | 0.10 | 1.44 |
| 8 | Pask | 0.72 | 3.84E-12 | 7.98E-08 | 0.24 | 0.15 | 0.08 | 1.55 |
| 8 | Dusp22 | 0.56 | 3.90E-12 | 8.10E-08 | 0.30 | 0.20 | 0.10 | 1.47 |
| 8 | Meis1 | 0.31 | 4.05E-12 | 8.42E-08 | 0.59 | 0.45 | 0.14 | 1.30 |
| 8 | Raph1 | 0.36 | 4.13E-12 | 8.59E-08 | 0.54 | 0.42 | 0.12 | 1.29 |
| 8 | Notch2 | 0.44 | 4.52E-12 | 9.39E-08 | 0.48 | 0.36 | 0.11 | 1.32 |
| 8 | Hist1h2ai | 1.21 | 5.04E-12 | 1.05E-07 | 0.11 | 0.06 | 0.05 | 1.98 |
| 8 | Trib2 | 0.66 | 5.60E-12 | 1.16E-07 | 0.29 | 0.19 | 0.09 | 1.48 |
| 8 | Vcam1 | 0.28 | 5.63E-12 | 1.17E-07 | 0.57 | 0.43 | 0.14 | 1.33 |
| 8 | Ythdc2 | 0.58 | 5.73E-12 | 1.19E-07 | 0.36 | 0.27 | 0.10 | 1.37 |
| 8 | Lnpk | 0.36 | 5.74E-12 | 1.19E-07 | 0.58 | 0.48 | 0.11 | 1.22 |
| 8 | Fer | 0.41 | 5.94E-12 | 1.23E-07 | 0.46 | 0.35 | 0.11 | 1.33 |
| 8 | Dlgap1 | 1.02 | 6.16E-12 | 1.28E-07 | 0.10 | 0.05 | 0.05 | 2.06 |
| 8 | Ddhd2 | 0.38 | 6.31E-12 | 1.31E-07 | 0.49 | 0.38 | 0.11 | 1.30 |
| 8 | Trim12c | 0.63 | 6.92E-12 | 1.44E-07 | 0.26 | 0.18 | 0.09 | 1.50 |
| 8 | Rtl8a | 0.37 | 7.01E-12 | 1.46E-07 | 0.50 | 0.39 | 0.11 | 1.29 |
| 8 | Clock | 0.50 | 7.09E-12 | 1.47E-07 | 0.42 | 0.31 | 0.11 | 1.35 |
| 8 | Slc7a2 | 0.92 | 7.16E-12 | 1.49E-07 | 0.13 | 0.07 | 0.06 | 1.89 |
| 8 | Colgalt1 | 0.38 | 7.54E-12 | 1.57E-07 | 0.58 | 0.47 | 0.11 | 1.24 |
| 8 | Magi2 | 0.57 | 7.71E-12 | 1.60E-07 | 0.28 | 0.18 | 0.09 | 1.51 |
| 8 | Cdk19 | 0.45 | 7.76E-12 | 1.61E-07 | 0.42 | 0.31 | 0.11 | 1.35 |
| 8 | Rgs12 | 0.38 | 7.79E-12 | 1.62E-07 | 0.56 | 0.45 | 0.11 | 1.25 |
| 8 | Anapc7 | 0.33 | 7.79E-12 | 1.62E-07 | 0.65 | 0.55 | 0.10 | 1.18 |
| 8 | Picalm | 0.28 | 7.81E-12 | 1.63E-07 | 0.72 | 0.61 | 0.11 | 1.17 |
| 8 | Prkg1 | 0.49 | 8.62E-12 | 1.79E-07 | 0.25 | 0.16 | 0.09 | 1.57 |
| 8 | Bbx | 0.29 | 8.77E-12 | 1.82E-07 | 0.70 | 0.58 | 0.11 | 1.19 |
| 8 | Nxpe3 | 0.81 | 9.02E-12 | 1.88E-07 | 0.17 | 0.10 | 0.07 | 1.70 |
| 8 | Ppp4r3a | 0.33 | 9.50E-12 | 1.98E-07 | 0.74 | 0.65 | 0.10 | 1.15 |
| 8 | Atf7ip | 0.31 | 9.66E-12 | 2.01E-07 | 0.72 | 0.62 | 0.10 | 1.15 |
| 8 | Zc3h14 | 0.27 | 9.76E-12 | 2.03E-07 | 0.84 | 0.77 | 0.08 | 1.10 |
| 8 | Appl2 | 0.61 | 9.77E-12 | 2.03E-07 | 0.23 | 0.15 | 0.08 | 1.56 |
| 8 | Lrp5 | 0.62 | 9.77E-12 | 2.03E-07 | 0.26 | 0.17 | 0.09 | 1.50 |
| 8 | Cast | 0.53 | 1.01E-11 | 2.10E-07 | 0.34 | 0.24 | 0.10 | 1.41 |
| 8 | Fbxo30 | 0.41 | 1.04E-11 | 2.16E-07 | 0.52 | 0.41 | 0.11 | 1.26 |
| 8 | Tmem119 | 0.30 | 1.06E-11 | 2.21E-07 | 0.59 | 0.45 | 0.13 | 1.30 |
| 8 | E130307A14Rik | 0.61 | 1.08E-11 | 2.24E-07 | 0.29 | 0.19 | 0.09 | 1.48 |
| 8 | Hook3 | 0.25 | 1.09E-11 | 2.27E-07 | 0.76 | 0.66 | 0.10 | 1.16 |
| 8 | Gnai1 | 0.62 | 1.10E-11 | 2.29E-07 | 0.24 | 0.15 | 0.09 | 1.57 |
| 8 | Gabbr1 | 0.47 | 1.15E-11 | 2.39E-07 | 0.34 | 0.24 | 0.10 | 1.42 |
| 8 | Tpst1 | 0.45 | 1.24E-11 | 2.58E-07 | 0.44 | 0.33 | 0.10 | 1.31 |
| 8 | Safb2 | 0.35 | 1.26E-11 | 2.62E-07 | 0.66 | 0.56 | 0.10 | 1.18 |
| 8 | Kat6a | 0.34 | 1.31E-11 | 2.72E-07 | 0.64 | 0.54 | 0.11 | 1.20 |
| 8 | Sh3kbp1 | 0.49 | 1.33E-11 | 2.78E-07 | 0.32 | 0.22 | 0.10 | 1.45 |
| 8 | Tmem263 | 0.35 | 1.41E-11 | 2.94E-07 | 0.58 | 0.47 | 0.11 | 1.23 |
| 8 | 9930021J03Rik | 0.36 | 1.41E-11 | 2.94E-07 | 0.64 | 0.55 | 0.09 | 1.16 |
| 8 | Samd1 | 0.40 | 1.47E-11 | 3.05E-07 | 0.51 | 0.40 | 0.11 | 1.26 |
| 8 | Fkbp9 | 0.33 | 1.47E-11 | 3.07E-07 | 0.56 | 0.44 | 0.12 | 1.28 |
| 8 | Olfml1 | 0.69 | 1.49E-11 | 3.10E-07 | 0.19 | 0.12 | 0.08 | 1.66 |
| 8 | Ift80 | 0.54 | 1.49E-11 | 3.10E-07 | 0.31 | 0.22 | 0.09 | 1.43 |
| 8 | Slc25a36 | 0.37 | 1.56E-11 | 3.24E-07 | 0.54 | 0.43 | 0.11 | 1.26 |
| 8 | Fut8 | 0.44 | 1.56E-11 | 3.24E-07 | 0.39 | 0.29 | 0.10 | 1.35 |
| 8 | Prdm16os | 0.50 | 1.57E-11 | 3.26E-07 | 0.20 | 0.12 | 0.08 | 1.68 |
| 8 | Cenpc1 | 0.33 | 1.58E-11 | 3.30E-07 | 0.57 | 0.45 | 0.12 | 1.25 |
| 8 | BC055324 | 0.57 | 1.61E-11 | 3.35E-07 | 0.32 | 0.23 | 0.09 | 1.39 |
| 8 | Pnpla8 | 0.34 | 1.62E-11 | 3.36E-07 | 0.59 | 0.50 | 0.10 | 1.19 |
| 8 | Nfatc2ip | 0.75 | 1.70E-11 | 3.54E-07 | 0.24 | 0.16 | 0.08 | 1.51 |

**Supplementary Table 1. Marker genes for the original clusters of integrated clustering of E11.5 control and NotchGOF λ-junction cells.**

| | | | | | | | | |
|---|---|---|---|---|---|---|---|---|
| 8 | Mre11a | 0.41 | 1.79E-11 | 3.71E-07 | 0.48 | 0.38 | 0.10 | 1.27 |
| 8 | Cfap97 | 0.37 | 1.80E-11 | 3.74E-07 | 0.55 | 0.45 | 0.11 | 1.23 |
| 8 | Pkmyt1 | 0.68 | 1.81E-11 | 3.76E-07 | 0.25 | 0.17 | 0.08 | 1.49 |
| 8 | Slc4a4 | 0.78 | 1.84E-11 | 3.83E-07 | 0.18 | 0.11 | 0.07 | 1.65 |
| 8 | Wdr86 | 0.33 | 1.92E-11 | 3.99E-07 | 0.64 | 0.51 | 0.12 | 1.24 |
| 8 | Ubr7 | 0.40 | 1.98E-11 | 4.12E-07 | 0.58 | 0.47 | 0.11 | 1.24 |
| 8 | Zfp318 | 0.33 | 2.05E-11 | 4.26E-07 | 0.58 | 0.47 | 0.11 | 1.24 |
| 8 | Mprip | 0.27 | 2.12E-11 | 4.41E-07 | 0.80 | 0.73 | 0.08 | 1.11 |
| 8 | Manea | 0.51 | 2.28E-11 | 4.73E-07 | 0.33 | 0.23 | 0.10 | 1.42 |
| 8 | Zfp982 | 0.72 | 2.28E-11 | 4.75E-07 | 0.17 | 0.10 | 0.07 | 1.72 |
| 8 | Plekho1 | 0.26 | 2.30E-11 | 4.78E-07 | 0.74 | 0.61 | 0.13 | 1.21 |
| 8 | Nynrin | 0.31 | 2.34E-11 | 4.86E-07 | 0.65 | 0.55 | 0.11 | 1.19 |
| 8 | Syce2 | 0.31 | 2.40E-11 | 4.98E-07 | 0.70 | 0.59 | 0.11 | 1.19 |
| 8 | Wsb1 | 0.31 | 2.46E-11 | 5.12E-07 | 0.77 | 0.69 | 0.08 | 1.11 |
| 8 | Mycbp2 | 0.25 | 2.49E-11 | 5.17E-07 | 0.77 | 0.70 | 0.08 | 1.11 |
| 8 | Timm50 | 0.26 | 2.52E-11 | 5.24E-07 | 0.84 | 0.73 | 0.10 | 1.14 |
| 8 | Fam168a | 0.33 | 2.56E-11 | 5.32E-07 | 0.65 | 0.54 | 0.11 | 1.20 |
| 8 | Nrk | 0.44 | 2.69E-11 | 5.60E-07 | 0.24 | 0.15 | 0.09 | 1.57 |
| 8 | Yeats2 | 0.30 | 2.71E-11 | 5.63E-07 | 0.69 | 0.57 | 0.12 | 1.21 |
| 8 | Lrrc4b | 0.60 | 2.72E-11 | 5.66E-07 | 0.21 | 0.13 | 0.08 | 1.63 |
| 8 | Ncoa2 | 0.45 | 2.74E-11 | 5.69E-07 | 0.39 | 0.29 | 0.10 | 1.34 |
| 8 | Ppp3cb | 0.31 | 2.74E-11 | 5.70E-07 | 0.72 | 0.62 | 0.11 | 1.17 |
| 8 | Chd9 | 0.31 | 2.76E-11 | 5.74E-07 | 0.63 | 0.53 | 0.10 | 1.20 |
| 8 | Ckap4 | 0.25 | 2.87E-11 | 5.97E-07 | 0.89 | 0.79 | 0.09 | 1.12 |
| 8 | Eri2 | 0.62 | 2.93E-11 | 6.09E-07 | 0.28 | 0.19 | 0.09 | 1.44 |
| 8 | Syde1 | 0.63 | 3.01E-11 | 6.25E-07 | 0.26 | 0.17 | 0.09 | 1.50 |
| 8 | Hunk | 0.34 | 3.03E-11 | 6.30E-07 | 0.60 | 0.50 | 0.11 | 1.21 |
| 8 | Rnaseh2b | 0.37 | 3.06E-11 | 6.36E-07 | 0.59 | 0.50 | 0.09 | 1.19 |
| 8 | Usp47 | 0.26 | 3.07E-11 | 6.37E-07 | 0.79 | 0.70 | 0.09 | 1.12 |
| 8 | Ifit2 | 0.75 | 3.46E-11 | 7.20E-07 | 0.16 | 0.09 | 0.07 | 1.73 |
| 8 | Six4 | 0.35 | 3.59E-11 | 7.46E-07 | 0.57 | 0.46 | 0.11 | 1.25 |
| 8 | Nup205 | 0.45 | 3.59E-11 | 7.48E-07 | 0.44 | 0.34 | 0.10 | 1.29 |
| 8 | Fam114a1 | 0.68 | 3.64E-11 | 7.57E-07 | 0.25 | 0.16 | 0.08 | 1.50 |
| 8 | Chd8 | 0.27 | 3.87E-11 | 8.04E-07 | 0.78 | 0.69 | 0.09 | 1.12 |
| 8 | Adat2 | 0.52 | 3.89E-11 | 8.08E-07 | 0.39 | 0.30 | 0.09 | 1.31 |
| 8 | Mark1 | 0.57 | 3.91E-11 | 8.13E-07 | 0.34 | 0.25 | 0.09 | 1.38 |
| 8 | Guf1 | 0.35 | 4.02E-11 | 8.35E-07 | 0.52 | 0.41 | 0.12 | 1.28 |
| 8 | Fanca | 0.95 | 4.15E-11 | 8.62E-07 | 0.18 | 0.11 | 0.07 | 1.61 |
| 8 | St6galnac4 | 0.55 | 4.19E-11 | 8.72E-07 | 0.29 | 0.20 | 0.09 | 1.47 |
| 8 | Arcn1 | 0.29 | 4.52E-11 | 9.41E-07 | 0.74 | 0.65 | 0.09 | 1.14 |
| 8 | Igf2r | 0.31 | 4.76E-11 | 9.90E-07 | 0.61 | 0.50 | 0.11 | 1.22 |
| 8 | Vopp1 | 0.51 | 4.80E-11 | 9.97E-07 | 0.37 | 0.27 | 0.10 | 1.35 |
| 8 | Zw10 | 0.53 | 4.88E-11 | 1.01E-06 | 0.41 | 0.32 | 0.09 | 1.29 |
| 8 | Tafa5 | 0.73 | 5.03E-11 | 1.05E-06 | 0.19 | 0.12 | 0.07 | 1.60 |
| 8 | Smndc1 | 0.27 | 5.27E-11 | 1.10E-06 | 0.76 | 0.68 | 0.08 | 1.11 |
| 8 | Prkca | 0.76 | 5.43E-11 | 1.13E-06 | 0.15 | 0.08 | 0.06 | 1.77 |
| 8 | Strn4 | 0.35 | 5.51E-11 | 1.15E-06 | 0.59 | 0.49 | 0.10 | 1.21 |
| 8 | Sspn | 0.81 | 5.56E-11 | 1.16E-06 | 0.13 | 0.07 | 0.06 | 1.85 |
| 8 | Lrp1 | 0.28 | 5.62E-11 | 1.17E-06 | 0.68 | 0.58 | 0.10 | 1.18 |
| 8 | Arid2 | 0.29 | 6.05E-11 | 1.26E-06 | 0.73 | 0.63 | 0.10 | 1.17 |
| 8 | Cenpi | 0.76 | 6.47E-11 | 1.35E-06 | 0.24 | 0.17 | 0.08 | 1.47 |
| 8 | Dhrs3 | 0.47 | 6.67E-11 | 1.39E-06 | 0.39 | 0.29 | 0.10 | 1.33 |
| 8 | Terf2 | 0.35 | 6.99E-11 | 1.45E-06 | 0.55 | 0.45 | 0.10 | 1.22 |
| 8 | Cenpo | 0.80 | 7.38E-11 | 1.53E-06 | 0.20 | 0.13 | 0.07 | 1.56 |
| 8 | Pik3ca | 0.39 | 7.48E-11 | 1.56E-06 | 0.54 | 0.43 | 0.11 | 1.25 |
| 8 | Bmp1 | 0.36 | 7.70E-11 | 1.60E-06 | 0.53 | 0.42 | 0.10 | 1.25 |

**Supplementary Table 1. Marker genes for the original clusters of integrated clustering of E11.5 control and NotchGOF λ-junction cells.**

| | | | | | | | | |
|---|---|---|---|---|---|---|---|---|
| 8 | Fam135a | 0.52 | 7.83E-11 | 1.63E-06 | 0.39 | 0.29 | 0.09 | 1.32 |
| 8 | Pwwp2a | 0.35 | 7.86E-11 | 1.63E-06 | 0.66 | 0.56 | 0.09 | 1.17 |
| 8 | Adgra3 | 0.32 | 7.96E-11 | 1.66E-06 | 0.66 | 0.56 | 0.11 | 1.19 |
| 8 | Plekhn1 | 0.59 | 7.97E-11 | 1.66E-06 | 0.29 | 0.20 | 0.09 | 1.44 |
| 8 | Mpp6 | 0.25 | 8.02E-11 | 1.67E-06 | 0.84 | 0.74 | 0.10 | 1.13 |
| 8 | Peg12 | 0.52 | 8.10E-11 | 1.68E-06 | 0.35 | 0.25 | 0.10 | 1.39 |
| 8 | Gng8 | 0.38 | 8.33E-11 | 1.73E-06 | 0.21 | 0.13 | 0.08 | 1.60 |
| 8 | Efemp2 | 0.34 | 8.33E-11 | 1.73E-06 | 0.58 | 0.47 | 0.10 | 1.22 |
| 8 | Atp7a | 0.45 | 8.36E-11 | 1.74E-06 | 0.33 | 0.23 | 0.10 | 1.41 |
| 8 | Dnaaf2 | 0.34 | 9.01E-11 | 1.87E-06 | 0.61 | 0.51 | 0.11 | 1.21 |
| 8 | Sec11c | 0.42 | 9.22E-11 | 1.92E-06 | 0.39 | 0.30 | 0.09 | 1.32 |
| 8 | Hyls1 | 0.71 | 9.27E-11 | 1.93E-06 | 0.25 | 0.17 | 0.08 | 1.46 |
| 8 | Syne1 | 0.58 | 9.48E-11 | 1.97E-06 | 0.21 | 0.13 | 0.08 | 1.60 |
| 8 | Cdan1 | 0.74 | 9.56E-11 | 1.99E-06 | 0.16 | 0.09 | 0.07 | 1.71 |
| 8 | Anapc1 | 0.31 | 9.65E-11 | 2.01E-06 | 0.68 | 0.58 | 0.10 | 1.18 |
| 8 | Cenpv | 0.26 | 9.80E-11 | 2.04E-06 | 0.62 | 0.50 | 0.13 | 1.25 |
| 8 | Fcho2 | 0.27 | 9.85E-11 | 2.05E-06 | 0.59 | 0.48 | 0.11 | 1.24 |
| 8 | Atf1 | 0.39 | 1.00E-10 | 2.09E-06 | 0.59 | 0.50 | 0.09 | 1.18 |
| 8 | Yap1 | 0.26 | 1.03E-10 | 2.13E-06 | 0.73 | 0.61 | 0.11 | 1.18 |
| 8 | Nup133 | 0.42 | 1.04E-10 | 2.16E-06 | 0.47 | 0.37 | 0.10 | 1.26 |
| 8 | Fbn1 | 0.39 | 1.06E-10 | 2.21E-06 | 0.43 | 0.33 | 0.10 | 1.31 |
| 8 | Wipf1 | 0.45 | 1.11E-10 | 2.30E-06 | 0.33 | 0.23 | 0.10 | 1.44 |
| 8 | Slc36a4 | 0.47 | 1.15E-10 | 2.39E-06 | 0.25 | 0.16 | 0.09 | 1.53 |
| 8 | B930095G15Rik | 0.55 | 1.18E-10 | 2.44E-06 | 0.26 | 0.18 | 0.08 | 1.48 |
| 8 | Atm | 0.50 | 1.20E-10 | 2.50E-06 | 0.41 | 0.32 | 0.09 | 1.30 |
| 8 | Ints12 | 0.51 | 1.22E-10 | 2.53E-06 | 0.39 | 0.30 | 0.09 | 1.31 |
| 8 | Rbm15 | 0.40 | 1.26E-10 | 2.62E-06 | 0.47 | 0.37 | 0.10 | 1.28 |
| 8 | Dxo | 0.36 | 1.27E-10 | 2.64E-06 | 0.50 | 0.40 | 0.10 | 1.25 |
| 8 | Polh | 0.97 | 1.30E-10 | 2.71E-06 | 0.15 | 0.09 | 0.06 | 1.70 |
| 8 | Sart3 | 0.29 | 1.32E-10 | 2.74E-06 | 0.66 | 0.56 | 0.10 | 1.18 |
| 8 | Mast2 | 0.44 | 1.37E-10 | 2.85E-06 | 0.38 | 0.29 | 0.10 | 1.33 |
| 8 | Gtf3c2 | 0.30 | 1.38E-10 | 2.87E-06 | 0.69 | 0.59 | 0.10 | 1.17 |
| 8 | Rttn | 0.66 | 1.41E-10 | 2.94E-06 | 0.24 | 0.16 | 0.08 | 1.49 |
| 8 | Nup62 | 0.33 | 1.44E-10 | 2.99E-06 | 0.64 | 0.55 | 0.09 | 1.15 |
| 8 | AI506816 | 0.38 | 1.49E-10 | 3.10E-06 | 0.64 | 0.53 | 0.10 | 1.19 |
| 8 | Cdc42bpa | 0.43 | 1.49E-10 | 3.10E-06 | 0.47 | 0.38 | 0.10 | 1.25 |
| 8 | Pcnx | 0.52 | 1.53E-10 | 3.18E-06 | 0.38 | 0.28 | 0.09 | 1.32 |
| 8 | Igsf10 | 0.40 | 1.61E-10 | 3.34E-06 | 0.33 | 0.23 | 0.10 | 1.41 |
| 8 | Ppp4r1 | 0.41 | 1.62E-10 | 3.38E-06 | 0.49 | 0.39 | 0.10 | 1.25 |
| 8 | Six5 | 0.56 | 1.68E-10 | 3.49E-06 | 0.28 | 0.20 | 0.09 | 1.45 |
| 8 | Ubr4 | 0.30 | 1.69E-10 | 3.51E-06 | 0.64 | 0.54 | 0.10 | 1.18 |
| 8 | Cnep1r1 | 0.36 | 1.72E-10 | 3.58E-06 | 0.64 | 0.54 | 0.10 | 1.19 |
| 8 | Aunip | 0.62 | 1.73E-10 | 3.59E-06 | 0.34 | 0.26 | 0.08 | 1.31 |
| 8 | Dgkh | 0.76 | 1.76E-10 | 3.66E-06 | 0.15 | 0.09 | 0.06 | 1.71 |
| 8 | Cdc27 | 0.33 | 1.76E-10 | 3.67E-06 | 0.62 | 0.52 | 0.10 | 1.19 |
| 8 | Ankmy2 | 0.45 | 1.80E-10 | 3.74E-06 | 0.38 | 0.28 | 0.09 | 1.33 |
| 8 | Cwf19l2 | 0.26 | 1.80E-10 | 3.74E-06 | 0.63 | 0.53 | 0.10 | 1.19 |
| 8 | Runx2 | 0.49 | 1.89E-10 | 3.93E-06 | 0.24 | 0.16 | 0.09 | 1.53 |
| 8 | Qdpr | 0.30 | 1.90E-10 | 3.95E-06 | 0.63 | 0.53 | 0.10 | 1.18 |
| 8 | Ppwd1 | 0.35 | 1.91E-10 | 3.96E-06 | 0.59 | 0.49 | 0.10 | 1.21 |
| 8 | Crot | 0.33 | 1.93E-10 | 4.02E-06 | 0.52 | 0.41 | 0.11 | 1.26 |
| 8 | Mcph1 | 0.58 | 1.95E-10 | 4.05E-06 | 0.39 | 0.30 | 0.09 | 1.30 |
| 8 | Dnajc30 | 0.36 | 1.99E-10 | 4.13E-06 | 0.54 | 0.44 | 0.10 | 1.22 |
| 8 | Mtrex | 0.30 | 2.07E-10 | 4.30E-06 | 0.72 | 0.62 | 0.10 | 1.17 |
| 8 | Pde3b | 0.52 | 2.12E-10 | 4.41E-06 | 0.33 | 0.24 | 0.09 | 1.38 |
| 8 | Rfwd3 | 0.41 | 2.20E-10 | 4.57E-06 | 0.47 | 0.37 | 0.10 | 1.26 |

**Supplementary Table 1. Marker genes for the original clusters of integrated clustering of E11.5 control and NotchGOF λ-junction cells.**

| | | | | | | | | |
|---|---|---|---|---|---|---|---|---|
| 8 | Adgrl1 | 0.39 | 2.24E-10 | 4.66E-06 | 0.53 | 0.44 | 0.10 | 1.22 |
| 8 | Tmem209 | 0.44 | 2.31E-10 | 4.80E-06 | 0.36 | 0.26 | 0.09 | 1.35 |
| 8 | Hmgxb4 | 0.30 | 2.51E-10 | 5.22E-06 | 0.71 | 0.62 | 0.09 | 1.15 |
| 8 | Setd1b | 0.44 | 2.68E-10 | 5.57E-06 | 0.52 | 0.43 | 0.09 | 1.21 |
| 8 | Exoc4 | 0.36 | 2.79E-10 | 5.81E-06 | 0.51 | 0.41 | 0.10 | 1.25 |
| 8 | Gse1 | 0.41 | 2.80E-10 | 5.82E-06 | 0.40 | 0.30 | 0.10 | 1.34 |
| 8 | Gpsm1 | 0.45 | 2.91E-10 | 6.05E-06 | 0.38 | 0.29 | 0.10 | 1.34 |
| 8 | Creb1 | 0.27 | 2.91E-10 | 6.06E-06 | 0.70 | 0.59 | 0.10 | 1.17 |
| 8 | Lhfpl2 | 0.36 | 2.97E-10 | 6.18E-06 | 0.44 | 0.33 | 0.11 | 1.33 |
| 8 | Kif18b | 0.78 | 3.04E-10 | 6.33E-06 | 0.23 | 0.16 | 0.07 | 1.46 |
| 8 | Nfx1 | 0.29 | 3.05E-10 | 6.34E-06 | 0.67 | 0.57 | 0.10 | 1.17 |
| 8 | Ddx11 | 0.54 | 3.08E-10 | 6.40E-06 | 0.33 | 0.25 | 0.09 | 1.35 |
| 8 | Rufy3 | 0.45 | 3.10E-10 | 6.45E-06 | 0.31 | 0.23 | 0.09 | 1.39 |
| 8 | Sestd1 | 0.32 | 3.15E-10 | 6.56E-06 | 0.55 | 0.44 | 0.11 | 1.25 |
| 8 | Tet2 | 0.40 | 3.19E-10 | 6.64E-06 | 0.35 | 0.25 | 0.10 | 1.41 |
| 8 | Hivep2 | 0.81 | 3.24E-10 | 6.73E-06 | 0.13 | 0.08 | 0.06 | 1.76 |
| 8 | Akap13 | 0.31 | 3.27E-10 | 6.80E-06 | 0.60 | 0.50 | 0.10 | 1.19 |
| 8 | Smad6 | 0.44 | 3.29E-10 | 6.84E-06 | 0.34 | 0.25 | 0.10 | 1.39 |
| 8 | Enox1 | 0.79 | 3.33E-10 | 6.92E-06 | 0.13 | 0.08 | 0.06 | 1.77 |
| 8 | Sirpa | 0.55 | 3.40E-10 | 7.07E-06 | 0.19 | 0.12 | 0.07 | 1.61 |
| 8 | Neto2 | 0.61 | 3.41E-10 | 7.10E-06 | 0.31 | 0.23 | 0.09 | 1.37 |
| 8 | Pdzd4 | 0.63 | 3.49E-10 | 7.25E-06 | 0.22 | 0.15 | 0.08 | 1.53 |
| 8 | Arhgef12 | 0.25 | 3.64E-10 | 7.57E-06 | 0.69 | 0.58 | 0.11 | 1.19 |
| 8 | Mpp5 | 0.45 | 3.67E-10 | 7.64E-06 | 0.47 | 0.37 | 0.09 | 1.25 |
| 8 | Rab12 | 0.32 | 3.68E-10 | 7.65E-06 | 0.58 | 0.48 | 0.10 | 1.21 |
| 8 | Ube2t | 0.46 | 3.70E-10 | 7.70E-06 | 0.42 | 0.33 | 0.09 | 1.28 |
| 8 | Lrrcc1 | 0.31 | 3.83E-10 | 7.96E-06 | 0.63 | 0.54 | 0.09 | 1.17 |
| 8 | Pabpc4l | 0.36 | 3.90E-10 | 8.12E-06 | 0.49 | 0.38 | 0.11 | 1.27 |
| 8 | Eed | 0.30 | 4.02E-10 | 8.37E-06 | 0.65 | 0.54 | 0.11 | 1.20 |
| 8 | Ggct | 0.68 | 4.06E-10 | 8.44E-06 | 0.22 | 0.15 | 0.07 | 1.51 |
| 8 | Rsbn1 | 0.31 | 4.11E-10 | 8.55E-06 | 0.64 | 0.55 | 0.09 | 1.16 |
| 8 | Arl15 | 0.56 | 4.38E-10 | 9.12E-06 | 0.25 | 0.17 | 0.08 | 1.48 |
| 8 | Cnih2 | 0.36 | 4.47E-10 | 9.30E-06 | 0.43 | 0.32 | 0.10 | 1.32 |
| 8 | Tiam1 | 0.34 | 4.74E-10 | 9.85E-06 | 0.41 | 0.31 | 0.10 | 1.32 |
| 8 | Ing3 | 0.45 | 4.86E-10 | 1.01E-05 | 0.45 | 0.36 | 0.10 | 1.27 |
| 8 | B3glct | 0.40 | 4.89E-10 | 1.02E-05 | 0.41 | 0.31 | 0.10 | 1.31 |
| 8 | Vcl | 0.40 | 5.05E-10 | 1.05E-05 | 0.49 | 0.39 | 0.10 | 1.24 |
| 8 | Tial1 | 0.31 | 5.12E-10 | 1.06E-05 | 0.74 | 0.67 | 0.07 | 1.11 |
| 8 | Zfp704 | 0.32 | 5.33E-10 | 1.11E-05 | 0.57 | 0.47 | 0.10 | 1.22 |
| 8 | Sh3d19 | 0.49 | 5.37E-10 | 1.12E-05 | 0.35 | 0.26 | 0.09 | 1.35 |
| 8 | Arhgef25 | 0.32 | 5.44E-10 | 1.13E-05 | 0.48 | 0.37 | 0.11 | 1.30 |
| 8 | Myo5a | 0.39 | 5.87E-10 | 1.22E-05 | 0.42 | 0.32 | 0.10 | 1.31 |
| 8 | Fam107b | 0.39 | 5.92E-10 | 1.23E-05 | 0.46 | 0.35 | 0.10 | 1.30 |
| 8 | Tmem243 | 0.55 | 5.94E-10 | 1.24E-05 | 0.30 | 0.21 | 0.09 | 1.41 |
| 8 | Lypla1 | 0.37 | 6.15E-10 | 1.28E-05 | 0.55 | 0.46 | 0.09 | 1.20 |
| 8 | Ski | 0.35 | 6.29E-10 | 1.31E-05 | 0.51 | 0.41 | 0.11 | 1.26 |
| 8 | Fam171a1 | 0.40 | 6.80E-10 | 1.42E-05 | 0.42 | 0.33 | 0.10 | 1.30 |
| 8 | Rab31 | 0.50 | 6.90E-10 | 1.44E-05 | 0.31 | 0.22 | 0.09 | 1.41 |
| 8 | Ctdspl | 0.34 | 6.91E-10 | 1.44E-05 | 0.45 | 0.34 | 0.11 | 1.31 |
| 8 | Specc1l | 0.32 | 6.99E-10 | 1.45E-05 | 0.53 | 0.42 | 0.11 | 1.25 |
| 8 | Kansl1l | 0.45 | 7.13E-10 | 1.48E-05 | 0.35 | 0.26 | 0.09 | 1.36 |
| 8 | Chm | 0.42 | 7.15E-10 | 1.49E-05 | 0.41 | 0.31 | 0.09 | 1.30 |
| 8 | Msl2 | 0.47 | 7.46E-10 | 1.55E-05 | 0.43 | 0.34 | 0.09 | 1.27 |
| 8 | Snx25 | 0.91 | 7.46E-10 | 1.55E-05 | 0.12 | 0.06 | 0.05 | 1.83 |
| 8 | Vps13d | 0.62 | 7.92E-10 | 1.65E-05 | 0.21 | 0.14 | 0.07 | 1.52 |
| 8 | Dusp4 | 0.63 | 8.03E-10 | 1.67E-05 | 0.14 | 0.08 | 0.06 | 1.73 |

**Supplementary Table 1. Marker genes for the original clusters of integrated clustering of E11.5 control and NotchGOF λ-junction cells.**

| | | | | | | | | |
|---|---|---|---|---|---|---|---|---|
| 8 | Rnf216 | 0.35 | 8.04E-10 | 1.67E-05 | 0.62 | 0.52 | 0.09 | 1.18 |
| 8 | Sap130 | 0.47 | 8.15E-10 | 1.69E-05 | 0.34 | 0.26 | 0.09 | 1.34 |
| 8 | Sgip1 | 0.59 | 8.27E-10 | 1.72E-05 | 0.18 | 0.11 | 0.07 | 1.63 |
| 8 | Ptpra | 0.30 | 8.33E-10 | 1.73E-05 | 0.70 | 0.60 | 0.10 | 1.16 |
| 8 | Rbmx2 | 0.34 | 8.36E-10 | 1.74E-05 | 0.52 | 0.43 | 0.09 | 1.20 |
| 8 | Slc30a5 | 0.39 | 8.72E-10 | 1.81E-05 | 0.40 | 0.31 | 0.09 | 1.30 |
| 8 | Diaph2 | 0.64 | 8.85E-10 | 1.84E-05 | 0.23 | 0.15 | 0.08 | 1.50 |
| 8 | Actr6 | 0.41 | 8.95E-10 | 1.86E-05 | 0.49 | 0.39 | 0.10 | 1.25 |
| 8 | Ntn3 | 0.60 | 9.04E-10 | 1.88E-05 | 0.23 | 0.16 | 0.08 | 1.48 |
| 8 | Ints14 | 0.46 | 9.42E-10 | 1.96E-05 | 0.48 | 0.39 | 0.09 | 1.22 |
| 8 | Fam184a | 0.71 | 9.71E-10 | 2.02E-05 | 0.18 | 0.11 | 0.07 | 1.60 |
| 8 | Espl1 | 0.54 | 1.02E-09 | 2.12E-05 | 0.30 | 0.22 | 0.08 | 1.36 |
| 8 | Col5a1 | 0.39 | 1.03E-09 | 2.14E-05 | 0.36 | 0.26 | 0.10 | 1.36 |
| 8 | Mafb | 0.28 | 1.05E-09 | 2.18E-05 | 0.31 | 0.22 | 0.09 | 1.42 |
| 8 | Gopc | 0.45 | 1.07E-09 | 2.23E-05 | 0.41 | 0.32 | 0.09 | 1.30 |
| 8 | Tmem246 | 0.71 | 1.09E-09 | 2.27E-05 | 0.20 | 0.13 | 0.07 | 1.54 |
| 8 | Cdkn2d | 0.32 | 1.10E-09 | 2.29E-05 | 0.48 | 0.38 | 0.10 | 1.26 |
| 8 | Zfp280d | 0.27 | 1.15E-09 | 2.39E-05 | 0.73 | 0.66 | 0.07 | 1.10 |
| 8 | Fzd1 | 0.33 | 1.18E-09 | 2.46E-05 | 0.44 | 0.34 | 0.11 | 1.32 |
| 8 | Dtnbp1 | 0.34 | 1.20E-09 | 2.49E-05 | 0.52 | 0.42 | 0.10 | 1.24 |
| 8 | Pms2 | 0.44 | 1.23E-09 | 2.55E-05 | 0.41 | 0.32 | 0.09 | 1.30 |
| 8 | Zcchc8 | 0.32 | 1.25E-09 | 2.59E-05 | 0.58 | 0.47 | 0.10 | 1.22 |
| 8 | Loxl2 | 0.28 | 1.27E-09 | 2.65E-05 | 0.54 | 0.44 | 0.11 | 1.25 |
| 8 | Tram2 | 0.67 | 1.35E-09 | 2.81E-05 | 0.16 | 0.10 | 0.06 | 1.62 |
| 8 | Psat1 | 0.30 | 1.36E-09 | 2.82E-05 | 0.55 | 0.45 | 0.10 | 1.23 |
| 8 | Tpm2 | 0.25 | 1.39E-09 | 2.89E-05 | 0.55 | 0.43 | 0.11 | 1.26 |
| 8 | Xrcc1 | 0.34 | 1.45E-09 | 3.02E-05 | 0.62 | 0.53 | 0.09 | 1.16 |
| 8 | St5 | 0.44 | 1.51E-09 | 3.13E-05 | 0.31 | 0.23 | 0.09 | 1.39 |
| 8 | Zfp871 | 0.29 | 1.51E-09 | 3.14E-05 | 0.67 | 0.58 | 0.09 | 1.15 |
| 8 | Vps16 | 0.45 | 1.57E-09 | 3.27E-05 | 0.38 | 0.29 | 0.09 | 1.31 |
| 8 | Rora | 0.58 | 1.69E-09 | 3.53E-05 | 0.26 | 0.18 | 0.08 | 1.46 |
| 8 | Col4a3bp | 0.44 | 1.72E-09 | 3.58E-05 | 0.41 | 0.32 | 0.09 | 1.29 |
| 8 | Spin4 | 0.75 | 1.74E-09 | 3.63E-05 | 0.19 | 0.12 | 0.07 | 1.56 |
| 8 | Adamts20 | 0.44 | 1.76E-09 | 3.66E-05 | 0.22 | 0.14 | 0.08 | 1.54 |
| 8 | Abhd10 | 0.47 | 1.77E-09 | 3.68E-05 | 0.41 | 0.32 | 0.09 | 1.27 |
| 8 | Lrch2 | 0.39 | 1.77E-09 | 3.68E-05 | 0.37 | 0.28 | 0.10 | 1.34 |
| 8 | Mcm8 | 0.49 | 1.83E-09 | 3.81E-05 | 0.34 | 0.26 | 0.08 | 1.32 |
| 8 | Atf2 | 0.29 | 1.85E-09 | 3.84E-05 | 0.63 | 0.54 | 0.09 | 1.18 |
| 8 | Gga2 | 0.25 | 1.87E-09 | 3.88E-05 | 0.73 | 0.64 | 0.09 | 1.14 |
| 8 | Zfp37 | 0.53 | 1.89E-09 | 3.93E-05 | 0.30 | 0.22 | 0.08 | 1.37 |
| 8 | Abi2 | 0.34 | 1.94E-09 | 4.04E-05 | 0.54 | 0.44 | 0.10 | 1.23 |
| 8 | Dclk2 | 0.58 | 2.07E-09 | 4.31E-05 | 0.18 | 0.11 | 0.07 | 1.60 |
| 8 | Zbtb10 | 0.57 | 2.08E-09 | 4.33E-05 | 0.27 | 0.20 | 0.08 | 1.40 |
| 8 | Brd9 | 0.30 | 2.18E-09 | 4.53E-05 | 0.65 | 0.55 | 0.10 | 1.17 |
| 8 | Arpin | 0.51 | 2.18E-09 | 4.54E-05 | 0.25 | 0.18 | 0.08 | 1.44 |
| 8 | Rexo5 | 0.78 | 2.20E-09 | 4.58E-05 | 0.18 | 0.12 | 0.06 | 1.55 |
| 8 | Lama1 | 0.70 | 2.21E-09 | 4.60E-05 | 0.17 | 0.10 | 0.06 | 1.61 |
| 8 | Vkorc1l1 | 0.38 | 2.26E-09 | 4.71E-05 | 0.49 | 0.40 | 0.09 | 1.23 |
| 8 | Maml3 | 0.35 | 2.27E-09 | 4.72E-05 | 0.49 | 0.40 | 0.09 | 1.24 |
| 8 | Nup160 | 0.44 | 2.37E-09 | 4.92E-05 | 0.39 | 0.31 | 0.09 | 1.28 |
| 8 | Tfdp1 | 0.25 | 2.38E-09 | 4.94E-05 | 0.75 | 0.66 | 0.10 | 1.15 |
| 8 | Enc1 | 0.46 | 2.40E-09 | 4.99E-05 | 0.30 | 0.22 | 0.09 | 1.39 |
| 8 | Prkd3 | 0.31 | 2.41E-09 | 5.01E-05 | 0.63 | 0.54 | 0.10 | 1.18 |
| 8 | Eri1 | 0.31 | 2.42E-09 | 5.03E-05 | 0.60 | 0.50 | 0.10 | 1.19 |
| 8 | Pogz | 0.39 | 2.46E-09 | 5.12E-05 | 0.46 | 0.38 | 0.09 | 1.23 |
| 8 | Ubr2 | 0.32 | 2.47E-09 | 5.13E-05 | 0.55 | 0.46 | 0.10 | 1.21 |

**Supplementary Table 1. Marker genes for the original clusters of integrated clustering of E11.5 control and NotchGOF λ-junction cells.**

| | | | | | | | | |
|---|---|---|---|---|---|---|---|---|
| 8 | Gm17092 | 0.46 | 2.50E-09 | 5.20E-05 | 0.33 | 0.24 | 0.08 | 1.34 |
| 8 | Fzd7 | 0.34 | 2.51E-09 | 5.21E-05 | 0.44 | 0.35 | 0.10 | 1.28 |
| 8 | Hcfc1 | 0.25 | 2.54E-09 | 5.29E-05 | 0.75 | 0.67 | 0.08 | 1.12 |
| 8 | Cand1 | 0.26 | 2.58E-09 | 5.36E-05 | 0.74 | 0.64 | 0.10 | 1.15 |
| 8 | Cdk13 | 0.32 | 2.59E-09 | 5.38E-05 | 0.63 | 0.54 | 0.08 | 1.15 |
| 8 | Mrc2 | 0.44 | 2.60E-09 | 5.41E-05 | 0.30 | 0.22 | 0.09 | 1.40 |
| 8 | Atad2b | 0.26 | 2.64E-09 | 5.50E-05 | 0.72 | 0.64 | 0.08 | 1.12 |
| 8 | Ilrun | 0.34 | 2.72E-09 | 5.65E-05 | 0.55 | 0.46 | 0.09 | 1.20 |
| 8 | Cpsf1 | 0.44 | 2.73E-09 | 5.68E-05 | 0.41 | 0.32 | 0.09 | 1.29 |
| 8 | Gclc | 0.49 | 2.74E-09 | 5.69E-05 | 0.38 | 0.29 | 0.08 | 1.29 |
| 8 | Yes1 | 0.41 | 2.95E-09 | 6.13E-05 | 0.44 | 0.35 | 0.10 | 1.27 |
| 8 | Dele1 | 0.39 | 2.98E-09 | 6.19E-05 | 0.39 | 0.30 | 0.09 | 1.31 |
| 8 | Fchsd2 | 0.38 | 3.03E-09 | 6.29E-05 | 0.42 | 0.33 | 0.09 | 1.28 |
| 8 | Fzr1 | 0.33 | 3.06E-09 | 6.37E-05 | 0.54 | 0.44 | 0.10 | 1.23 |
| 8 | Tubgcp6 | 0.59 | 3.08E-09 | 6.40E-05 | 0.21 | 0.14 | 0.07 | 1.52 |
| 8 | Copz2 | 0.35 | 3.14E-09 | 6.52E-05 | 0.35 | 0.26 | 0.09 | 1.36 |
| 8 | Uimc1 | 0.44 | 3.14E-09 | 6.54E-05 | 0.39 | 0.31 | 0.08 | 1.27 |
| 8 | Elovl5 | 0.41 | 3.14E-09 | 6.54E-05 | 0.39 | 0.30 | 0.09 | 1.30 |
| 8 | Tbc1d10b | 0.40 | 3.21E-09 | 6.68E-05 | 0.44 | 0.35 | 0.09 | 1.26 |
| 8 | Bach2 | 0.28 | 3.35E-09 | 6.97E-05 | 0.48 | 0.37 | 0.11 | 1.29 |
| 8 | Fndc4 | 0.32 | 3.36E-09 | 7.00E-05 | 0.51 | 0.41 | 0.10 | 1.25 |
| 8 | Fam76b | 0.29 | 3.49E-09 | 7.27E-05 | 0.65 | 0.55 | 0.10 | 1.18 |
| 8 | Fbxl17 | 0.44 | 3.51E-09 | 7.30E-05 | 0.28 | 0.20 | 0.08 | 1.42 |
| 8 | Lzts2 | 0.25 | 3.52E-09 | 7.33E-05 | 0.70 | 0.60 | 0.10 | 1.17 |
| 8 | Ndfip2 | 0.28 | 3.60E-09 | 7.48E-05 | 0.60 | 0.51 | 0.09 | 1.17 |
| 8 | Lrch1 | 0.73 | 3.67E-09 | 7.64E-05 | 0.13 | 0.08 | 0.06 | 1.71 |
| 8 | Nf1 | 0.36 | 3.81E-09 | 7.92E-05 | 0.49 | 0.40 | 0.09 | 1.23 |
| 8 | Zfp423 | 0.50 | 3.84E-09 | 7.98E-05 | 0.29 | 0.20 | 0.08 | 1.40 |
| 8 | P4ha2 | 0.36 | 4.01E-09 | 8.33E-05 | 0.36 | 0.27 | 0.09 | 1.34 |
| 8 | Ddias | 0.73 | 4.06E-09 | 8.43E-05 | 0.22 | 0.15 | 0.07 | 1.43 |
| 8 | Pom121 | 0.26 | 4.33E-09 | 9.01E-05 | 0.70 | 0.59 | 0.10 | 1.17 |
| 8 | Nup107 | 0.39 | 4.36E-09 | 9.07E-05 | 0.47 | 0.38 | 0.09 | 1.24 |
| 8 | Cnpy4 | 0.38 | 4.50E-09 | 9.36E-05 | 0.44 | 0.35 | 0.09 | 1.25 |
| 8 | Rpa1 | 0.41 | 4.54E-09 | 9.43E-05 | 0.50 | 0.42 | 0.08 | 1.19 |
| 8 | Platr25 | 0.44 | 4.78E-09 | 9.94E-05 | 0.31 | 0.22 | 0.08 | 1.38 |
| 8 | Minpp1 | 0.32 | 4.87E-09 | 1.01E-04 | 0.60 | 0.51 | 0.09 | 1.17 |
| 8 | Rad17 | 0.32 | 4.92E-09 | 1.02E-04 | 0.56 | 0.47 | 0.09 | 1.19 |
| 8 | Gucy1a2 | 0.89 | 4.99E-09 | 1.04E-04 | 0.12 | 0.07 | 0.05 | 1.74 |
| 8 | Fto | 0.31 | 5.09E-09 | 1.06E-04 | 0.57 | 0.47 | 0.10 | 1.22 |
| 8 | Ep300 | 0.31 | 5.15E-09 | 1.07E-04 | 0.63 | 0.54 | 0.09 | 1.16 |
| 8 | Itga5 | 0.43 | 5.20E-09 | 1.08E-04 | 0.30 | 0.21 | 0.08 | 1.39 |
| 8 | Zfand3 | 0.30 | 5.25E-09 | 1.09E-04 | 0.62 | 0.53 | 0.09 | 1.17 |
| 8 | Tor1aip1 | 0.32 | 5.32E-09 | 1.11E-04 | 0.53 | 0.44 | 0.10 | 1.22 |
| 8 | Pias4 | 0.35 | 5.40E-09 | 1.12E-04 | 0.53 | 0.44 | 0.09 | 1.20 |
| 8 | Clasp1 | 0.31 | 5.43E-09 | 1.13E-04 | 0.64 | 0.55 | 0.09 | 1.16 |
| 8 | Scarf2 | 0.37 | 5.49E-09 | 1.14E-04 | 0.35 | 0.26 | 0.09 | 1.35 |
| 8 | Cip2a | 0.35 | 5.68E-09 | 1.18E-04 | 0.47 | 0.39 | 0.09 | 1.23 |
| 8 | Mtbp | 0.56 | 5.94E-09 | 1.24E-04 | 0.30 | 0.22 | 0.08 | 1.35 |
| 8 | Wrn | 0.53 | 5.99E-09 | 1.25E-04 | 0.32 | 0.24 | 0.08 | 1.34 |
| 8 | Etaa1 | 0.28 | 6.12E-09 | 1.27E-04 | 0.62 | 0.53 | 0.09 | 1.17 |
| 8 | Fkbp7 | 0.33 | 6.13E-09 | 1.27E-04 | 0.38 | 0.28 | 0.10 | 1.35 |
| 8 | Sephs1 | 0.26 | 6.15E-09 | 1.28E-04 | 0.69 | 0.61 | 0.08 | 1.14 |
| 8 | 2700081O15Rik | 0.27 | 6.82E-09 | 1.42E-04 | 0.65 | 0.56 | 0.09 | 1.17 |
| 8 | Zfp148 | 0.26 | 6.88E-09 | 1.43E-04 | 0.77 | 0.67 | 0.10 | 1.14 |
| 8 | Gpm6b | 0.73 | 6.94E-09 | 1.44E-04 | 0.12 | 0.07 | 0.05 | 1.76 |
| 8 | Megf6 | 0.58 | 7.06E-09 | 1.47E-04 | 0.29 | 0.21 | 0.08 | 1.38 |

**Supplementary Table 1. Marker genes for the original clusters of integrated clustering of E11.5 control and NotchGOF λ-junction cells.**

| | | | | | | | | |
|---|---|---|---|---|---|---|---|---|
| 8 | Zmym5 | 0.25 | 7.20E-09 | 1.50E-04 | 0.69 | 0.61 | 0.08 | 1.13 |
| 8 | Spsb1 | 0.56 | 7.53E-09 | 1.57E-04 | 0.22 | 0.15 | 0.07 | 1.48 |
| 8 | Mosmo | 0.38 | 7.53E-09 | 1.57E-04 | 0.57 | 0.49 | 0.08 | 1.16 |
| 8 | Ugdh | 0.29 | 8.01E-09 | 1.67E-04 | 0.56 | 0.46 | 0.10 | 1.21 |
| 8 | Krit1 | 0.30 | 8.26E-09 | 1.72E-04 | 0.72 | 0.63 | 0.09 | 1.14 |
| 8 | Xpo4 | 0.35 | 8.43E-09 | 1.75E-04 | 0.53 | 0.43 | 0.09 | 1.21 |
| 8 | Tex30 | 0.33 | 8.70E-09 | 1.81E-04 | 0.56 | 0.47 | 0.09 | 1.19 |
| 8 | Chek2 | 0.57 | 9.12E-09 | 1.90E-04 | 0.31 | 0.23 | 0.07 | 1.31 |
| 8 | Rnf217 | 0.44 | 9.23E-09 | 1.92E-04 | 0.27 | 0.19 | 0.08 | 1.43 |
| 8 | Mapre2 | 0.30 | 9.38E-09 | 1.95E-04 | 0.50 | 0.40 | 0.10 | 1.25 |
| 8 | Ogdh | 0.30 | 9.46E-09 | 1.97E-04 | 0.61 | 0.53 | 0.08 | 1.15 |
| 8 | Uspl1 | 0.37 | 9.59E-09 | 1.99E-04 | 0.47 | 0.38 | 0.09 | 1.22 |
| 8 | Zfp281 | 0.31 | 9.60E-09 | 2.00E-04 | 0.55 | 0.46 | 0.10 | 1.21 |
| 8 | Dse | 0.36 | 9.64E-09 | 2.00E-04 | 0.36 | 0.27 | 0.09 | 1.34 |
| 8 | Shd | 0.39 | 9.64E-09 | 2.01E-04 | 0.25 | 0.17 | 0.08 | 1.46 |
| 8 | Tspan2 | 0.77 | 9.67E-09 | 2.01E-04 | 0.16 | 0.10 | 0.06 | 1.59 |
| 8 | Ubald2 | 0.27 | 9.85E-09 | 2.05E-04 | 0.77 | 0.69 | 0.08 | 1.12 |
| 8 | Lysmd2 | 0.50 | 9.91E-09 | 2.06E-04 | 0.30 | 0.22 | 0.08 | 1.37 |
| 8 | Chd1l | 0.55 | 9.92E-09 | 2.06E-04 | 0.27 | 0.20 | 0.08 | 1.38 |
| 8 | Mtss2 | 0.36 | 1.00E-08 | 2.09E-04 | 0.36 | 0.27 | 0.09 | 1.33 |
| 8 | Cables1 | 0.81 | 1.02E-08 | 2.12E-04 | 0.14 | 0.08 | 0.06 | 1.65 |
| 8 | Rpgrip1 | 0.42 | 1.02E-08 | 2.12E-04 | 0.35 | 0.27 | 0.09 | 1.33 |
| 8 | Tor1a | 0.30 | 1.02E-08 | 2.13E-04 | 0.61 | 0.52 | 0.09 | 1.18 |
| 8 | Myg1 | 0.26 | 1.04E-08 | 2.16E-04 | 0.72 | 0.62 | 0.10 | 1.16 |
| 8 | Zfp41 | 0.33 | 1.04E-08 | 2.16E-04 | 0.44 | 0.34 | 0.10 | 1.29 |
| 8 | Agap1 | 0.35 | 1.06E-08 | 2.21E-04 | 0.51 | 0.41 | 0.09 | 1.22 |
| 8 | Parp2 | 0.28 | 1.09E-08 | 2.27E-04 | 0.66 | 0.57 | 0.10 | 1.17 |
| 8 | Smg6 | 0.27 | 1.09E-08 | 2.28E-04 | 0.68 | 0.60 | 0.08 | 1.14 |
| 8 | Hsf2 | 0.27 | 1.10E-08 | 2.30E-04 | 0.58 | 0.48 | 0.10 | 1.21 |
| 8 | Nrf1 | 0.31 | 1.11E-08 | 2.31E-04 | 0.54 | 0.45 | 0.09 | 1.20 |
| 8 | Herc3 | 0.76 | 1.17E-08 | 2.43E-04 | 0.12 | 0.07 | 0.05 | 1.70 |
| 8 | Ehd3 | 0.74 | 1.20E-08 | 2.51E-04 | 0.17 | 0.11 | 0.06 | 1.55 |
| 8 | Dcaf7 | 0.29 | 1.21E-08 | 2.51E-04 | 0.59 | 0.49 | 0.10 | 1.21 |
| 8 | Chst3 | 0.79 | 1.22E-08 | 2.55E-04 | 0.11 | 0.06 | 0.05 | 1.77 |
| 8 | Arhgef2 | 0.29 | 1.25E-08 | 2.60E-04 | 0.53 | 0.42 | 0.11 | 1.25 |
| 8 | Lipt2 | 0.31 | 1.28E-08 | 2.67E-04 | 0.45 | 0.36 | 0.09 | 1.25 |
| 8 | Klhl13 | 0.44 | 1.29E-08 | 2.69E-04 | 0.35 | 0.27 | 0.09 | 1.32 |
| 8 | Tns3 | 0.27 | 1.29E-08 | 2.69E-04 | 0.59 | 0.49 | 0.10 | 1.21 |
| 8 | Meis2 | 0.30 | 1.31E-08 | 2.72E-04 | 0.75 | 0.67 | 0.08 | 1.12 |
| 8 | Lrch3 | 0.37 | 1.44E-08 | 2.99E-04 | 0.46 | 0.37 | 0.09 | 1.25 |
| 8 | Cep85l | 0.64 | 1.45E-08 | 3.01E-04 | 0.20 | 0.14 | 0.07 | 1.50 |
| 8 | Efna5 | 0.27 | 1.45E-08 | 3.01E-04 | 0.58 | 0.47 | 0.11 | 1.23 |
| 8 | Rab23 | 0.32 | 1.50E-08 | 3.12E-04 | 0.41 | 0.32 | 0.09 | 1.28 |
| 8 | AW554918 | 0.51 | 1.51E-08 | 3.13E-04 | 0.28 | 0.20 | 0.08 | 1.38 |
| 8 | Arl13b | 0.28 | 1.58E-08 | 3.29E-04 | 0.49 | 0.40 | 0.10 | 1.24 |
| 8 | Clcc1 | 0.27 | 1.62E-08 | 3.38E-04 | 0.51 | 0.41 | 0.10 | 1.24 |
| 8 | Ndc1 | 0.50 | 1.66E-08 | 3.45E-04 | 0.38 | 0.30 | 0.08 | 1.26 |
| 8 | Virma | 0.26 | 1.66E-08 | 3.46E-04 | 0.67 | 0.59 | 0.08 | 1.14 |
| 8 | Tdp1 | 0.40 | 1.69E-08 | 3.51E-04 | 0.38 | 0.30 | 0.08 | 1.27 |
| 8 | Ptpn9 | 0.27 | 1.71E-08 | 3.55E-04 | 0.59 | 0.49 | 0.10 | 1.21 |
| 8 | C230038L03Rik | 0.58 | 1.80E-08 | 3.74E-04 | 0.19 | 0.12 | 0.07 | 1.53 |
| 8 | Septin9 | 0.25 | 1.85E-08 | 3.85E-04 | 0.73 | 0.63 | 0.10 | 1.15 |
| 8 | Snx18 | 0.34 | 1.85E-08 | 3.85E-04 | 0.47 | 0.38 | 0.09 | 1.25 |
| 8 | Herc2 | 0.33 | 1.90E-08 | 3.94E-04 | 0.49 | 0.41 | 0.09 | 1.21 |
| 8 | Dzip3 | 0.27 | 1.92E-08 | 3.99E-04 | 0.69 | 0.61 | 0.08 | 1.14 |
| 8 | Tbc1d12 | 0.55 | 1.97E-08 | 4.09E-04 | 0.24 | 0.17 | 0.07 | 1.44 |

**Supplementary Table 1. Marker genes for the original clusters of integrated clustering of E11.5 control and NotchGOF λ-junction cells.**

| | | | | | | | | |
|---|---|---|---|---|---|---|---|---|
| 8 | Podxl2 | 0.26 | 2.03E-08 | 4.22E-04 | 0.52 | 0.42 | 0.10 | 1.24 |
| 8 | Herc1 | 0.27 | 2.07E-08 | 4.31E-04 | 0.64 | 0.56 | 0.08 | 1.14 |
| 8 | Slc35f5 | 0.51 | 2.09E-08 | 4.35E-04 | 0.20 | 0.13 | 0.07 | 1.51 |
| 8 | Tor1aip2 | 0.26 | 2.22E-08 | 4.61E-04 | 0.62 | 0.53 | 0.09 | 1.18 |
| 8 | Ccdc90b | 0.27 | 2.22E-08 | 4.62E-04 | 0.65 | 0.55 | 0.10 | 1.18 |
| 8 | 2610008E11Rik | 0.27 | 2.24E-08 | 4.66E-04 | 0.49 | 0.39 | 0.10 | 1.26 |
| 8 | Rgl1 | 0.54 | 2.28E-08 | 4.75E-04 | 0.22 | 0.15 | 0.07 | 1.46 |
| 8 | Adam9 | 0.51 | 2.33E-08 | 4.85E-04 | 0.21 | 0.14 | 0.07 | 1.49 |
| 8 | Gpaa1 | 0.34 | 2.41E-08 | 5.01E-04 | 0.50 | 0.42 | 0.09 | 1.21 |
| 8 | Scmh1 | 0.34 | 2.42E-08 | 5.03E-04 | 0.41 | 0.32 | 0.09 | 1.30 |
| 8 | Nck2 | 0.44 | 2.55E-08 | 5.31E-04 | 0.35 | 0.27 | 0.08 | 1.30 |
| 8 | Ahi1 | 0.28 | 2.62E-08 | 5.46E-04 | 0.52 | 0.42 | 0.10 | 1.24 |
| 8 | Sntb2 | 0.37 | 2.73E-08 | 5.68E-04 | 0.43 | 0.34 | 0.09 | 1.26 |
| 8 | Mrnip | 0.52 | 2.86E-08 | 5.95E-04 | 0.26 | 0.19 | 0.07 | 1.38 |
| 8 | Cep290 | 0.33 | 2.99E-08 | 6.23E-04 | 0.60 | 0.51 | 0.09 | 1.17 |
| 8 | Zmym1 | 0.42 | 3.02E-08 | 6.29E-04 | 0.41 | 0.33 | 0.08 | 1.23 |
| 8 | Angptl2 | 0.44 | 3.03E-08 | 6.31E-04 | 0.26 | 0.18 | 0.08 | 1.41 |
| 8 | Siah1a | 0.28 | 3.08E-08 | 6.40E-04 | 0.57 | 0.49 | 0.08 | 1.17 |
| 8 | Pdhx | 0.42 | 3.11E-08 | 6.46E-04 | 0.39 | 0.31 | 0.09 | 1.28 |
| 8 | Cep131 | 0.44 | 3.19E-08 | 6.64E-04 | 0.33 | 0.25 | 0.08 | 1.32 |
| 8 | Mettl15 | 0.94 | 3.25E-08 | 6.75E-04 | 0.12 | 0.07 | 0.05 | 1.70 |
| 8 | Tesk2 | 0.89 | 3.27E-08 | 6.81E-04 | 0.11 | 0.06 | 0.05 | 1.72 |
| 8 | Zzz3 | 0.25 | 3.28E-08 | 6.82E-04 | 0.65 | 0.54 | 0.10 | 1.19 |
| 8 | Zfc3h1 | 0.30 | 3.28E-08 | 6.82E-04 | 0.45 | 0.36 | 0.10 | 1.27 |
| 8 | Ctdsp2 | 0.26 | 3.39E-08 | 7.04E-04 | 0.65 | 0.56 | 0.09 | 1.16 |
| 8 | Kif14 | 0.63 | 3.41E-08 | 7.09E-04 | 0.25 | 0.19 | 0.07 | 1.36 |
| 8 | Gm36988 | 0.53 | 3.42E-08 | 7.12E-04 | 0.21 | 0.14 | 0.07 | 1.47 |
| 8 | Cdyl | 0.37 | 3.52E-08 | 7.32E-04 | 0.47 | 0.38 | 0.09 | 1.23 |
| 8 | Ttc21b | 0.56 | 3.55E-08 | 7.39E-04 | 0.30 | 0.23 | 0.08 | 1.33 |
| 8 | Cep55 | 0.43 | 3.61E-08 | 7.50E-04 | 0.32 | 0.24 | 0.08 | 1.33 |
| 8 | Eva1a | 0.40 | 3.88E-08 | 8.08E-04 | 0.33 | 0.24 | 0.08 | 1.34 |
| 8 | Abhd12 | 0.40 | 3.91E-08 | 8.12E-04 | 0.39 | 0.31 | 0.08 | 1.26 |
| 8 | Esyt2 | 0.53 | 3.91E-08 | 8.14E-04 | 0.29 | 0.21 | 0.07 | 1.35 |
| 8 | Hist1h4i | 0.33 | 4.17E-08 | 8.66E-04 | 0.53 | 0.46 | 0.08 | 1.16 |
| 8 | Smad5 | 0.28 | 4.29E-08 | 8.92E-04 | 0.57 | 0.49 | 0.08 | 1.16 |
| 8 | Dact1 | 0.27 | 4.40E-08 | 9.16E-04 | 0.40 | 0.31 | 0.10 | 1.31 |
| 8 | Lmnb2 | 0.34 | 4.42E-08 | 9.19E-04 | 0.46 | 0.37 | 0.09 | 1.24 |
| 8 | Emp3 | 0.39 | 4.47E-08 | 9.29E-04 | 0.20 | 0.14 | 0.07 | 1.50 |
| 8 | Cep44 | 0.35 | 4.55E-08 | 9.47E-04 | 0.47 | 0.38 | 0.09 | 1.22 |
| 8 | Pelp1 | 0.28 | 4.57E-08 | 9.51E-04 | 0.60 | 0.51 | 0.09 | 1.18 |
| 8 | Pi4k2b | 0.41 | 4.70E-08 | 9.78E-04 | 0.36 | 0.28 | 0.08 | 1.30 |
| 8 | Dennd1a | 0.47 | 4.71E-08 | 9.80E-04 | 0.28 | 0.21 | 0.08 | 1.36 |
| 8 | Baiap2l2 | 0.52 | 4.75E-08 | 9.89E-04 | 0.18 | 0.11 | 0.06 | 1.54 |
| 8 | Rassf8 | 0.42 | 4.88E-08 | 1.02E-03 | 0.26 | 0.18 | 0.08 | 1.41 |
| 8 | Slc18b1 | 0.46 | 5.29E-08 | 1.10E-03 | 0.19 | 0.12 | 0.07 | 1.53 |
| 8 | Slc20a2 | 0.39 | 5.44E-08 | 1.13E-03 | 0.31 | 0.24 | 0.08 | 1.34 |
| 8 | Hsd17b11 | 0.39 | 5.44E-08 | 1.13E-03 | 0.28 | 0.20 | 0.08 | 1.38 |
| 8 | Prss35 | 0.57 | 5.44E-08 | 1.13E-03 | 0.10 | 0.06 | 0.05 | 1.80 |
| 8 | Lin9 | 0.35 | 5.66E-08 | 1.18E-03 | 0.42 | 0.33 | 0.08 | 1.25 |
| 8 | Rnf157 | 0.54 | 5.69E-08 | 1.18E-03 | 0.22 | 0.15 | 0.07 | 1.45 |
| 8 | Tmem176a | 0.26 | 6.02E-08 | 1.25E-03 | 0.50 | 0.40 | 0.09 | 1.23 |
| 8 | Smpdl3a | 0.57 | 6.11E-08 | 1.27E-03 | 0.19 | 0.13 | 0.06 | 1.48 |
| 8 | Psmd5 | 0.27 | 6.14E-08 | 1.28E-03 | 0.60 | 0.52 | 0.09 | 1.17 |
| 8 | Gm11837 | 0.69 | 6.50E-08 | 1.35E-03 | 0.12 | 0.07 | 0.05 | 1.69 |
| 8 | Fkbp10 | 0.26 | 6.74E-08 | 1.40E-03 | 0.43 | 0.33 | 0.10 | 1.30 |
| 8 | Ankrd26 | 0.34 | 6.81E-08 | 1.42E-03 | 0.52 | 0.43 | 0.09 | 1.20 |

**Supplementary Table 1. Marker genes for the original clusters of integrated clustering of E11.5 control and NotchGOF λ-junction cells.**

| | | | | | | | | |
|---|---|---|---|---|---|---|---|---|
| 8 | Fancc | 0.65 | 6.88E-08 | 1.43E-03 | 0.16 | 0.10 | 0.06 | 1.55 |
| 8 | Prkar2a | 0.26 | 6.90E-08 | 1.44E-03 | 0.61 | 0.51 | 0.09 | 1.18 |
| 8 | Etnk1 | 0.29 | 6.97E-08 | 1.45E-03 | 0.54 | 0.46 | 0.08 | 1.18 |
| 8 | Adam22 | 0.73 | 7.03E-08 | 1.46E-03 | 0.12 | 0.07 | 0.05 | 1.66 |
| 8 | Map3k2 | 0.48 | 7.13E-08 | 1.48E-03 | 0.27 | 0.19 | 0.07 | 1.37 |
| 8 | Rrp1b | 0.29 | 7.19E-08 | 1.50E-03 | 0.62 | 0.53 | 0.09 | 1.18 |
| 8 | Pik3c3 | 0.50 | 7.21E-08 | 1.50E-03 | 0.24 | 0.17 | 0.07 | 1.39 |
| 8 | Acap2 | 0.31 | 7.22E-08 | 1.50E-03 | 0.55 | 0.46 | 0.09 | 1.20 |
| 8 | Apaf1 | 0.49 | 7.27E-08 | 1.51E-03 | 0.30 | 0.23 | 0.07 | 1.32 |
| 8 | Fam122a | 0.41 | 7.40E-08 | 1.54E-03 | 0.37 | 0.29 | 0.08 | 1.27 |
| 8 | Ptk2 | 0.28 | 7.52E-08 | 1.56E-03 | 0.62 | 0.53 | 0.08 | 1.16 |
| 8 | Pif1 | 0.61 | 7.67E-08 | 1.60E-03 | 0.20 | 0.14 | 0.06 | 1.46 |
| 8 | Azin1 | 0.28 | 7.76E-08 | 1.61E-03 | 0.69 | 0.62 | 0.07 | 1.11 |
| 8 | Gigyf2 | 0.26 | 7.83E-08 | 1.63E-03 | 0.67 | 0.59 | 0.08 | 1.14 |
| 8 | Lifr | 0.60 | 7.87E-08 | 1.64E-03 | 0.17 | 0.11 | 0.06 | 1.54 |
| 8 | S1pr2 | 0.40 | 8.15E-08 | 1.70E-03 | 0.40 | 0.32 | 0.09 | 1.27 |
| 8 | Ankrd44 | 0.27 | 8.25E-08 | 1.72E-03 | 0.37 | 0.28 | 0.09 | 1.32 |
| 8 | Top3a | 0.56 | 8.26E-08 | 1.72E-03 | 0.23 | 0.16 | 0.07 | 1.41 |
| 8 | St6gal1 | 0.44 | 8.60E-08 | 1.79E-03 | 0.26 | 0.19 | 0.07 | 1.39 |
| 8 | Iah1 | 0.27 | 8.71E-08 | 1.81E-03 | 0.60 | 0.52 | 0.08 | 1.15 |
| 8 | Twist2 | 0.33 | 8.74E-08 | 1.82E-03 | 0.31 | 0.23 | 0.08 | 1.36 |
| 8 | Usp13 | 0.87 | 8.77E-08 | 1.82E-03 | 0.12 | 0.07 | 0.05 | 1.64 |
| 8 | Ccny | 0.25 | 8.91E-08 | 1.85E-03 | 0.61 | 0.54 | 0.08 | 1.15 |
| 8 | Agps | 0.30 | 8.91E-08 | 1.85E-03 | 0.49 | 0.40 | 0.09 | 1.22 |
| 8 | Pwwp3b | 0.29 | 8.95E-08 | 1.86E-03 | 0.40 | 0.31 | 0.09 | 1.29 |
| 8 | Eml3 | 0.62 | 8.96E-08 | 1.86E-03 | 0.18 | 0.12 | 0.06 | 1.48 |
| 8 | Crtc3 | 0.35 | 9.05E-08 | 1.88E-03 | 0.33 | 0.24 | 0.08 | 1.34 |
| 8 | Stambpl1 | 0.67 | 9.08E-08 | 1.89E-03 | 0.11 | 0.07 | 0.05 | 1.71 |
| 8 | Sec23a | 0.33 | 9.12E-08 | 1.90E-03 | 0.42 | 0.34 | 0.08 | 1.24 |
| 8 | Tent4a | 0.50 | 9.27E-08 | 1.93E-03 | 0.33 | 0.26 | 0.07 | 1.27 |
| 8 | Atrip | 0.40 | 9.53E-08 | 1.98E-03 | 0.35 | 0.27 | 0.08 | 1.29 |
| 8 | Tmem109 | 0.29 | 9.57E-08 | 1.99E-03 | 0.45 | 0.36 | 0.09 | 1.24 |
| 8 | Fancb | 0.76 | 1.00E-07 | 2.08E-03 | 0.13 | 0.08 | 0.05 | 1.62 |
| 8 | Rapgef2 | 0.37 | 1.02E-07 | 2.12E-03 | 0.30 | 0.22 | 0.08 | 1.35 |
| 8 | Polq | 0.81 | 1.02E-07 | 2.12E-03 | 0.15 | 0.10 | 0.05 | 1.54 |
| 8 | Vrk2 | 0.56 | 1.03E-07 | 2.15E-03 | 0.30 | 0.23 | 0.07 | 1.31 |
| 8 | Phf13 | 0.40 | 1.06E-07 | 2.20E-03 | 0.35 | 0.27 | 0.08 | 1.30 |
| 8 | Rbp4 | 0.65 | 1.08E-07 | 2.24E-03 | 0.10 | 0.06 | 0.04 | 1.74 |
| 8 | Phkb | 0.45 | 1.09E-07 | 2.26E-03 | 0.34 | 0.27 | 0.08 | 1.29 |
| 8 | Map1a | 0.59 | 1.11E-07 | 2.32E-03 | 0.14 | 0.09 | 0.05 | 1.60 |
| 8 | Marf1 | 0.29 | 1.12E-07 | 2.32E-03 | 0.48 | 0.40 | 0.09 | 1.22 |
| 8 | Gm10184 | 0.68 | 1.13E-07 | 2.35E-03 | 0.20 | 0.14 | 0.06 | 1.42 |
| 8 | Casc3 | 0.28 | 1.16E-07 | 2.41E-03 | 0.61 | 0.53 | 0.08 | 1.14 |
| 8 | 2610020C07Rik | 0.47 | 1.17E-07 | 2.44E-03 | 0.25 | 0.18 | 0.07 | 1.38 |
| 8 | Thap2 | 0.28 | 1.18E-07 | 2.45E-03 | 0.60 | 0.51 | 0.09 | 1.17 |
| 8 | Arhgef9 | 0.43 | 1.28E-07 | 2.66E-03 | 0.26 | 0.18 | 0.07 | 1.39 |
| 8 | Lrrc40 | 0.33 | 1.28E-07 | 2.66E-03 | 0.41 | 0.33 | 0.08 | 1.25 |
| 8 | Stam | 0.37 | 1.31E-07 | 2.73E-03 | 0.37 | 0.29 | 0.08 | 1.27 |
| 8 | Rap2b | 0.25 | 1.36E-07 | 2.83E-03 | 0.57 | 0.48 | 0.09 | 1.18 |
| 8 | Rictor | 0.28 | 1.37E-07 | 2.86E-03 | 0.46 | 0.38 | 0.09 | 1.23 |
| 8 | Cntrob | 0.49 | 1.40E-07 | 2.92E-03 | 0.28 | 0.21 | 0.07 | 1.34 |
| 8 | S100pbp | 0.33 | 1.42E-07 | 2.95E-03 | 0.39 | 0.30 | 0.08 | 1.27 |
| 8 | Map1s | 0.29 | 1.48E-07 | 3.08E-03 | 0.46 | 0.37 | 0.09 | 1.25 |
| 8 | 1700066M21Rik | 0.78 | 1.48E-07 | 3.08E-03 | 0.13 | 0.08 | 0.05 | 1.60 |
| 8 | Mtmr2 | 0.29 | 1.49E-07 | 3.11E-03 | 0.46 | 0.38 | 0.09 | 1.23 |
| 8 | Lzic | 0.25 | 1.50E-07 | 3.13E-03 | 0.59 | 0.51 | 0.09 | 1.17 |

**Supplementary Table 1. Marker genes for the original clusters of integrated clustering of E11.5 control and NotchGOF λ-junction cells.**

| | | | | | | | | |
|---|---|---|---|---|---|---|---|---|
| 8 | Paxip1 | 0.35 | 1.51E-07 | 3.14E-03 | 0.42 | 0.34 | 0.08 | 1.25 |
| 8 | Antxr2 | 0.44 | 1.53E-07 | 3.18E-03 | 0.26 | 0.18 | 0.07 | 1.39 |
| 8 | 1700052K11Rik | 0.38 | 1.54E-07 | 3.21E-03 | 0.29 | 0.22 | 0.08 | 1.36 |
| 8 | Dvl2 | 0.38 | 1.57E-07 | 3.26E-03 | 0.46 | 0.38 | 0.08 | 1.20 |
| 8 | Dipk1a | 0.53 | 1.59E-07 | 3.31E-03 | 0.27 | 0.20 | 0.07 | 1.33 |
| 8 | Sacs | 0.48 | 1.65E-07 | 3.42E-03 | 0.24 | 0.17 | 0.07 | 1.40 |
| 8 | Dsn1 | 0.46 | 1.68E-07 | 3.49E-03 | 0.31 | 0.24 | 0.07 | 1.30 |
| 8 | 2810408I11Rik | 0.38 | 1.68E-07 | 3.50E-03 | 0.37 | 0.30 | 0.08 | 1.25 |
| 8 | Zfp407 | 0.39 | 1.69E-07 | 3.51E-03 | 0.39 | 0.31 | 0.08 | 1.26 |
| 8 | Wdr90 | 0.62 | 1.69E-07 | 3.51E-03 | 0.17 | 0.12 | 0.06 | 1.49 |
| 8 | Tubgcp5 | 0.30 | 1.71E-07 | 3.55E-03 | 0.43 | 0.34 | 0.09 | 1.25 |
| 8 | Pgm2 | 0.28 | 1.71E-07 | 3.55E-03 | 0.36 | 0.27 | 0.09 | 1.32 |
| 8 | Gm49797 | 0.49 | 1.73E-07 | 3.60E-03 | 0.31 | 0.24 | 0.07 | 1.30 |
| 8 | Hcfc2 | 0.52 | 1.75E-07 | 3.63E-03 | 0.21 | 0.15 | 0.06 | 1.43 |
| 8 | Ubxn7 | 0.30 | 1.75E-07 | 3.65E-03 | 0.55 | 0.47 | 0.08 | 1.18 |
| 8 | Zbed5 | 0.45 | 1.77E-07 | 3.68E-03 | 0.22 | 0.15 | 0.07 | 1.44 |
| 8 | Xndc1 | 0.42 | 1.82E-07 | 3.79E-03 | 0.32 | 0.24 | 0.08 | 1.32 |
| 8 | Mad1l1 | 0.32 | 1.83E-07 | 3.81E-03 | 0.44 | 0.35 | 0.09 | 1.25 |
| 8 | Ip6k1 | 0.33 | 1.84E-07 | 3.82E-03 | 0.45 | 0.37 | 0.08 | 1.22 |
| 8 | Piga | 0.56 | 1.84E-07 | 3.83E-03 | 0.22 | 0.16 | 0.06 | 1.39 |
| 8 | Map3k3 | 0.50 | 1.86E-07 | 3.86E-03 | 0.22 | 0.15 | 0.07 | 1.43 |
| 8 | Fem1c | 0.32 | 1.88E-07 | 3.91E-03 | 0.47 | 0.38 | 0.09 | 1.24 |
| 8 | Gbf1 | 0.30 | 1.89E-07 | 3.94E-03 | 0.52 | 0.44 | 0.09 | 1.20 |
| 8 | Gstm7 | 0.36 | 2.02E-07 | 4.19E-03 | 0.36 | 0.28 | 0.08 | 1.28 |
| 8 | Stard9 | 0.72 | 2.03E-07 | 4.22E-03 | 0.11 | 0.07 | 0.05 | 1.70 |
| 8 | Rprd2 | 0.30 | 2.03E-07 | 4.22E-03 | 0.52 | 0.44 | 0.09 | 1.19 |
| 8 | Tram1l1 | 0.46 | 2.07E-07 | 4.30E-03 | 0.24 | 0.18 | 0.07 | 1.39 |
| 8 | Dnajc5 | 0.28 | 2.14E-07 | 4.45E-03 | 0.60 | 0.52 | 0.08 | 1.15 |
| 8 | Kif24 | 0.69 | 2.16E-07 | 4.49E-03 | 0.15 | 0.10 | 0.05 | 1.54 |
| 8 | Mboat7 | 0.34 | 2.16E-07 | 4.50E-03 | 0.41 | 0.33 | 0.08 | 1.25 |
| 8 | Arhgap10 | 0.26 | 2.22E-07 | 4.61E-03 | 0.50 | 0.42 | 0.08 | 1.20 |
| 8 | Cbfa2t2 | 0.34 | 2.24E-07 | 4.65E-03 | 0.32 | 0.24 | 0.08 | 1.32 |
| 8 | Haus4 | 0.35 | 2.29E-07 | 4.77E-03 | 0.39 | 0.31 | 0.08 | 1.24 |
| 8 | Dzip1 | 0.25 | 2.42E-07 | 5.03E-03 | 0.42 | 0.33 | 0.09 | 1.27 |
| 8 | Siah2 | 0.64 | 2.48E-07 | 5.15E-03 | 0.12 | 0.07 | 0.05 | 1.65 |
| 8 | Uhrf1bp1l | 0.37 | 2.48E-07 | 5.15E-03 | 0.39 | 0.31 | 0.08 | 1.26 |
| 8 | Slc35b2 | 0.27 | 2.51E-07 | 5.22E-03 | 0.50 | 0.41 | 0.09 | 1.22 |
| 8 | B4galt1 | 0.41 | 2.58E-07 | 5.37E-03 | 0.30 | 0.23 | 0.07 | 1.33 |
| 8 | Nek7 | 0.31 | 2.59E-07 | 5.38E-03 | 0.37 | 0.29 | 0.08 | 1.27 |
| 8 | Usp42 | 0.31 | 2.66E-07 | 5.54E-03 | 0.44 | 0.36 | 0.08 | 1.22 |
| 8 | Pip5k1c | 0.40 | 2.72E-07 | 5.65E-03 | 0.30 | 0.23 | 0.07 | 1.32 |
| 8 | Cep57l1 | 0.34 | 2.78E-07 | 5.78E-03 | 0.35 | 0.28 | 0.08 | 1.27 |
| 8 | Zfp61 | 0.28 | 2.79E-07 | 5.81E-03 | 0.32 | 0.24 | 0.08 | 1.34 |
| 8 | Lmbrd1 | 0.34 | 2.84E-07 | 5.90E-03 | 0.30 | 0.22 | 0.08 | 1.35 |
| 8 | Ercc6l2 | 0.39 | 2.88E-07 | 6.00E-03 | 0.41 | 0.33 | 0.08 | 1.23 |
| 8 | C330007P06Rik | 0.30 | 2.94E-07 | 6.11E-03 | 0.51 | 0.43 | 0.08 | 1.18 |
| 8 | Zswim6 | 0.30 | 2.96E-07 | 6.15E-03 | 0.38 | 0.29 | 0.08 | 1.29 |
| 8 | Stx5a | 0.26 | 3.09E-07 | 6.44E-03 | 0.56 | 0.48 | 0.08 | 1.16 |
| 8 | Pcx | 0.57 | 3.15E-07 | 6.56E-03 | 0.19 | 0.13 | 0.06 | 1.46 |
| 8 | Dlx6 | 0.41 | 3.23E-07 | 6.71E-03 | 0.14 | 0.09 | 0.05 | 1.59 |
| 8 | Pias2 | 0.28 | 3.29E-07 | 6.84E-03 | 0.48 | 0.40 | 0.09 | 1.21 |
| 8 | Mfhas1 | 0.45 | 3.31E-07 | 6.89E-03 | 0.26 | 0.19 | 0.07 | 1.35 |
| 8 | Galnt7 | 0.30 | 3.32E-07 | 6.91E-03 | 0.42 | 0.34 | 0.08 | 1.23 |
| 8 | Klf12 | 0.46 | 3.38E-07 | 7.03E-03 | 0.22 | 0.16 | 0.07 | 1.42 |
| 8 | Chpf | 0.53 | 3.43E-07 | 7.13E-03 | 0.21 | 0.15 | 0.06 | 1.42 |
| 8 | Stn1 | 0.36 | 3.47E-07 | 7.21E-03 | 0.43 | 0.35 | 0.08 | 1.23 |

**Supplementary Table 1. Marker genes for the original clusters of integrated clustering of E11.5 control and NotchGOF λ-junction cells.**

| | | | | | | | | |
|---|---|---|---|---|---|---|---|---|
| 8 | Pcolce2 | 0.54 | 3.77E-07 | 7.84E-03 | 0.20 | 0.14 | 0.06 | 1.45 |
| 8 | Mon2 | 0.36 | 3.81E-07 | 7.93E-03 | 0.35 | 0.28 | 0.08 | 1.29 |
| 8 | Gm43728 | 0.67 | 3.90E-07 | 8.10E-03 | 0.10 | 0.06 | 0.04 | 1.72 |
| 8 | L3mbtl3 | 0.27 | 4.07E-07 | 8.46E-03 | 0.47 | 0.39 | 0.08 | 1.21 |
| 8 | Arnt | 0.45 | 4.07E-07 | 8.47E-03 | 0.32 | 0.25 | 0.08 | 1.30 |
| 8 | Dyrk1a | 0.33 | 4.10E-07 | 8.52E-03 | 0.46 | 0.38 | 0.08 | 1.21 |
| 8 | Macrod1 | 0.62 | 4.10E-07 | 8.54E-03 | 0.12 | 0.07 | 0.05 | 1.64 |
| 8 | Npat | 0.30 | 4.18E-07 | 8.69E-03 | 0.43 | 0.35 | 0.08 | 1.23 |
| 8 | Depdc5 | 0.49 | 4.21E-07 | 8.76E-03 | 0.27 | 0.20 | 0.07 | 1.32 |
| 8 | Pdpk1 | 0.31 | 4.35E-07 | 9.05E-03 | 0.48 | 0.40 | 0.08 | 1.19 |
| 8 | Dip2c | 0.38 | 4.36E-07 | 9.08E-03 | 0.31 | 0.24 | 0.07 | 1.31 |
| 8 | Plcg1 | 0.30 | 4.45E-07 | 9.25E-03 | 0.48 | 0.40 | 0.08 | 1.21 |
| 8 | Vps13b | 0.48 | 4.54E-07 | 9.43E-03 | 0.25 | 0.18 | 0.07 | 1.37 |
| 8 | Brip1os | 0.34 | 4.54E-07 | 9.44E-03 | 0.44 | 0.36 | 0.08 | 1.22 |
| 8 | Pgrmc2 | 0.29 | 4.55E-07 | 9.47E-03 | 0.65 | 0.58 | 0.07 | 1.12 |
| 8 | Plscr1 | 0.35 | 4.57E-07 | 9.50E-03 | 0.31 | 0.23 | 0.08 | 1.33 |
| 8 | Tbc1d5 | 0.44 | 4.64E-07 | 9.65E-03 | 0.27 | 0.20 | 0.07 | 1.33 |
| 8 | Vstm2b | 0.51 | 4.67E-07 | 9.71E-03 | 0.15 | 0.09 | 0.05 | 1.56 |
| 8 | Pros1 | 0.45 | 4.67E-07 | 9.72E-03 | 0.20 | 0.14 | 0.06 | 1.43 |
| 8 | Deaf1 | 0.36 | 4.71E-07 | 9.80E-03 | 0.29 | 0.22 | 0.07 | 1.32 |
| 8 | Cc2d2a | 0.43 | 4.76E-07 | 9.89E-03 | 0.28 | 0.21 | 0.07 | 1.34 |
| 8 | Homer1 | 0.40 | 4.87E-07 | 1.01E-02 | 0.36 | 0.28 | 0.07 | 1.26 |
| 8 | Eefsec | 0.36 | 4.87E-07 | 1.01E-02 | 0.46 | 0.39 | 0.07 | 1.19 |
| 8 | Plag1 | 0.45 | 4.97E-07 | 1.03E-02 | 0.26 | 0.19 | 0.07 | 1.36 |
| 8 | Kirrel | 0.37 | 5.05E-07 | 1.05E-02 | 0.29 | 0.22 | 0.07 | 1.33 |
| 8 | Nelfa | 0.27 | 5.10E-07 | 1.06E-02 | 0.61 | 0.54 | 0.08 | 1.14 |
| 8 | B3galt6 | 0.29 | 5.13E-07 | 1.07E-02 | 0.39 | 0.31 | 0.08 | 1.26 |
| 8 | Gng12 | 0.28 | 5.15E-07 | 1.07E-02 | 0.45 | 0.36 | 0.09 | 1.25 |
| 8 | Pola2 | 0.28 | 5.26E-07 | 1.09E-02 | 0.50 | 0.42 | 0.08 | 1.19 |
| 8 | Phf12 | 0.27 | 5.40E-07 | 1.12E-02 | 0.52 | 0.43 | 0.09 | 1.20 |
| 8 | Epb41l1 | 0.44 | 5.43E-07 | 1.13E-02 | 0.20 | 0.14 | 0.06 | 1.44 |
| 8 | Smtn | 0.43 | 5.51E-07 | 1.15E-02 | 0.20 | 0.14 | 0.06 | 1.42 |
| 8 | Tigd2 | 0.25 | 5.58E-07 | 1.16E-02 | 0.47 | 0.39 | 0.08 | 1.21 |
| 8 | 6030458C11Rik | 0.33 | 5.71E-07 | 1.19E-02 | 0.41 | 0.33 | 0.08 | 1.25 |
| 8 | Mllt1 | 0.41 | 5.72E-07 | 1.19E-02 | 0.37 | 0.30 | 0.07 | 1.25 |
| 8 | Msh3 | 0.28 | 5.78E-07 | 1.20E-02 | 0.49 | 0.41 | 0.08 | 1.19 |
| 8 | P3h4 | 0.30 | 6.03E-07 | 1.25E-02 | 0.43 | 0.35 | 0.08 | 1.23 |
| 8 | Ibtk | 0.28 | 6.04E-07 | 1.26E-02 | 0.46 | 0.38 | 0.08 | 1.22 |
| 8 | Trp53bp2 | 0.29 | 6.14E-07 | 1.28E-02 | 0.54 | 0.46 | 0.08 | 1.17 |
| 8 | Bard1 | 0.56 | 6.18E-07 | 1.29E-02 | 0.25 | 0.19 | 0.06 | 1.32 |
| 8 | Plcl2 | 0.58 | 6.20E-07 | 1.29E-02 | 0.17 | 0.11 | 0.05 | 1.49 |
| 8 | Snhg11 | 0.55 | 6.24E-07 | 1.30E-02 | 0.16 | 0.10 | 0.05 | 1.51 |
| 8 | Prkaca | 0.26 | 6.26E-07 | 1.30E-02 | 0.63 | 0.56 | 0.07 | 1.13 |
| 8 | Crip1 | 0.26 | 6.27E-07 | 1.30E-02 | 0.48 | 0.40 | 0.08 | 1.20 |
| 8 | Mtfr2 | 0.75 | 6.52E-07 | 1.36E-02 | 0.16 | 0.11 | 0.05 | 1.46 |
| 8 | Kdm6a | 0.29 | 6.63E-07 | 1.38E-02 | 0.50 | 0.42 | 0.07 | 1.17 |
| 8 | Sbf2 | 0.28 | 6.65E-07 | 1.38E-02 | 0.43 | 0.34 | 0.09 | 1.25 |
| 8 | A430005L14Rik | 0.28 | 6.81E-07 | 1.42E-02 | 0.58 | 0.50 | 0.08 | 1.16 |
| 8 | Nup153 | 0.31 | 6.83E-07 | 1.42E-02 | 0.51 | 0.43 | 0.07 | 1.17 |
| 8 | Setd1a | 0.39 | 6.87E-07 | 1.43E-02 | 0.37 | 0.30 | 0.07 | 1.23 |
| 8 | Cchcr1 | 0.59 | 6.88E-07 | 1.43E-02 | 0.20 | 0.15 | 0.06 | 1.39 |
| 8 | Tinf2 | 0.47 | 7.04E-07 | 1.46E-02 | 0.27 | 0.20 | 0.07 | 1.32 |
| 8 | Lpl | 0.41 | 7.08E-07 | 1.47E-02 | 0.22 | 0.16 | 0.06 | 1.38 |
| 8 | 2700049A03Rik | 0.31 | 7.16E-07 | 1.49E-02 | 0.39 | 0.31 | 0.08 | 1.25 |
| 8 | Pop4 | 0.26 | 7.18E-07 | 1.49E-02 | 0.53 | 0.46 | 0.07 | 1.16 |
| 8 | 2610037D02Rik | 0.31 | 7.24E-07 | 1.51E-02 | 0.41 | 0.33 | 0.08 | 1.24 |

**Supplementary Table 1. Marker genes for the original clusters of integrated clustering of E11.5 control and NotchGOF λ-junction cells.**

| | | | | | | | | |
|---|---|---|---|---|---|---|---|---|
| 8 | Fkbp5 | 0.29 | 7.33E-07 | 1.52E-02 | 0.50 | 0.42 | 0.08 | 1.20 |
| 8 | Lox | 0.58 | 7.38E-07 | 1.53E-02 | 0.11 | 0.07 | 0.04 | 1.64 |
| 8 | Cblb | 0.33 | 7.42E-07 | 1.54E-02 | 0.42 | 0.34 | 0.07 | 1.22 |
| 8 | Upp2 | 0.40 | 7.63E-07 | 1.59E-02 | 0.24 | 0.18 | 0.07 | 1.37 |
| 8 | Mphosph9 | 0.45 | 7.64E-07 | 1.59E-02 | 0.32 | 0.25 | 0.07 | 1.27 |
| 8 | Cdr2l | 0.44 | 7.79E-07 | 1.62E-02 | 0.26 | 0.19 | 0.07 | 1.35 |
| 8 | Hacd1 | 0.27 | 7.81E-07 | 1.63E-02 | 0.48 | 0.39 | 0.08 | 1.21 |
| 8 | Wwp1 | 0.29 | 7.88E-07 | 1.64E-02 | 0.49 | 0.41 | 0.08 | 1.19 |
| 8 | Arhgap42 | 0.33 | 8.00E-07 | 1.66E-02 | 0.45 | 0.37 | 0.08 | 1.22 |
| 8 | Stk39 | 0.42 | 8.01E-07 | 1.67E-02 | 0.33 | 0.26 | 0.07 | 1.27 |
| 8 | Troap | 0.52 | 8.10E-07 | 1.68E-02 | 0.24 | 0.18 | 0.06 | 1.33 |
| 8 | Impa2 | 0.41 | 8.20E-07 | 1.71E-02 | 0.36 | 0.29 | 0.07 | 1.23 |
| 8 | Rngtt | 0.31 | 8.21E-07 | 1.71E-02 | 0.48 | 0.40 | 0.08 | 1.21 |
| 8 | Pcca | 0.28 | 8.34E-07 | 1.73E-02 | 0.37 | 0.29 | 0.08 | 1.27 |
| 8 | Dna2 | 0.43 | 8.35E-07 | 1.74E-02 | 0.33 | 0.26 | 0.07 | 1.27 |
| 8 | Stau2 | 0.30 | 8.40E-07 | 1.75E-02 | 0.46 | 0.38 | 0.08 | 1.21 |
| 8 | Zfyve21 | 0.26 | 8.51E-07 | 1.77E-02 | 0.48 | 0.40 | 0.08 | 1.20 |
| 8 | Ltbp1 | 0.37 | 8.69E-07 | 1.81E-02 | 0.30 | 0.22 | 0.07 | 1.33 |
| 8 | Camk2n2 | 0.65 | 8.74E-07 | 1.82E-02 | 0.15 | 0.10 | 0.05 | 1.53 |
| 8 | Zcchc10 | 0.31 | 8.75E-07 | 1.82E-02 | 0.43 | 0.35 | 0.07 | 1.21 |
| 8 | Zdhhc17 | 0.37 | 8.80E-07 | 1.83E-02 | 0.33 | 0.25 | 0.07 | 1.29 |
| 8 | Crkl | 0.29 | 8.99E-07 | 1.87E-02 | 0.45 | 0.37 | 0.08 | 1.21 |
| 8 | Kdsr | 0.33 | 8.99E-07 | 1.87E-02 | 0.35 | 0.27 | 0.08 | 1.28 |
| 8 | Napg | 0.31 | 9.18E-07 | 1.91E-02 | 0.44 | 0.36 | 0.08 | 1.22 |
| 8 | Kbtbd2 | 0.36 | 9.43E-07 | 1.96E-02 | 0.41 | 0.33 | 0.08 | 1.23 |
| 8 | Acad9 | 0.36 | 9.79E-07 | 2.04E-02 | 0.32 | 0.25 | 0.07 | 1.29 |
| 8 | Tmem259 | 0.30 | 9.80E-07 | 2.04E-02 | 0.49 | 0.42 | 0.08 | 1.19 |
| 8 | Pou2f1 | 0.29 | 1.00E-06 | 2.09E-02 | 0.44 | 0.36 | 0.08 | 1.23 |
| 8 | Adprhl2 | 0.25 | 1.00E-06 | 2.09E-02 | 0.58 | 0.50 | 0.08 | 1.16 |
| 8 | Ret | 0.47 | 1.01E-06 | 2.10E-02 | 0.10 | 0.06 | 0.04 | 1.70 |
| 8 | Ing2 | 0.27 | 1.02E-06 | 2.12E-02 | 0.48 | 0.41 | 0.08 | 1.19 |
| 8 | B3galnt1 | 0.48 | 1.03E-06 | 2.14E-02 | 0.19 | 0.14 | 0.06 | 1.42 |
| 8 | Ice2 | 0.37 | 1.05E-06 | 2.17E-02 | 0.32 | 0.25 | 0.07 | 1.29 |
| 8 | Map7d3 | 0.72 | 1.05E-06 | 2.18E-02 | 0.11 | 0.06 | 0.04 | 1.66 |
| 8 | Cherp | 0.27 | 1.06E-06 | 2.20E-02 | 0.51 | 0.43 | 0.08 | 1.19 |
| 8 | Git2 | 0.37 | 1.09E-06 | 2.26E-02 | 0.41 | 0.34 | 0.07 | 1.21 |
| 8 | Pdss2 | 0.44 | 1.10E-06 | 2.28E-02 | 0.29 | 0.23 | 0.07 | 1.29 |
| 8 | Alg6 | 0.27 | 1.12E-06 | 2.33E-02 | 0.38 | 0.30 | 0.08 | 1.26 |
| 8 | Pxdc1 | 0.43 | 1.13E-06 | 2.35E-02 | 0.16 | 0.10 | 0.05 | 1.51 |
| 8 | Hus1 | 0.41 | 1.14E-06 | 2.37E-02 | 0.32 | 0.25 | 0.07 | 1.27 |
| 8 | Hdac5 | 0.40 | 1.15E-06 | 2.40E-02 | 0.24 | 0.17 | 0.07 | 1.38 |
| 8 | Trerf1 | 0.35 | 1.16E-06 | 2.40E-02 | 0.34 | 0.27 | 0.07 | 1.28 |
| 8 | Hmg20a | 0.26 | 1.18E-06 | 2.45E-02 | 0.51 | 0.42 | 0.09 | 1.20 |
| 8 | Crlf3 | 0.29 | 1.18E-06 | 2.46E-02 | 0.52 | 0.45 | 0.07 | 1.16 |
| 8 | Haus8 | 0.36 | 1.19E-06 | 2.48E-02 | 0.42 | 0.35 | 0.07 | 1.20 |
| 8 | Psd3 | 0.42 | 1.20E-06 | 2.49E-02 | 0.29 | 0.22 | 0.07 | 1.31 |
| 8 | Pank2 | 0.25 | 1.24E-06 | 2.57E-02 | 0.58 | 0.50 | 0.08 | 1.16 |
| 8 | Utp6 | 0.29 | 1.26E-06 | 2.61E-02 | 0.52 | 0.45 | 0.07 | 1.16 |
| 8 | Tsga10 | 0.60 | 1.30E-06 | 2.70E-02 | 0.15 | 0.10 | 0.05 | 1.51 |
| 8 | Map3k7 | 0.28 | 1.30E-06 | 2.70E-02 | 0.55 | 0.47 | 0.08 | 1.16 |
| 8 | Zfp827 | 0.33 | 1.30E-06 | 2.71E-02 | 0.38 | 0.31 | 0.08 | 1.25 |
| 8 | Rps6ka5 | 0.63 | 1.33E-06 | 2.76E-02 | 0.19 | 0.13 | 0.06 | 1.42 |
| 8 | Mepce | 0.36 | 1.33E-06 | 2.77E-02 | 0.41 | 0.34 | 0.08 | 1.22 |
| 8 | Insr | 0.40 | 1.37E-06 | 2.85E-02 | 0.31 | 0.24 | 0.07 | 1.29 |
| 8 | Hist1h2ag | 0.40 | 1.37E-06 | 2.85E-02 | 0.38 | 0.31 | 0.07 | 1.23 |
| 8 | Lancl1 | 0.31 | 1.37E-06 | 2.85E-02 | 0.44 | 0.35 | 0.08 | 1.23 |

**Supplementary Table 1. Marker genes for the original clusters of integrated clustering of E11.5 control and NotchGOF λ-junction cells.**

| | | | | | | | | |
|---|---|---|---|---|---|---|---|---|
| 8 | Tbl1xr1 | 0.28 | 1.38E-06 | 2.86E-02 | 0.49 | 0.42 | 0.08 | 1.18 |
| 8 | Kctd2 | 0.29 | 1.38E-06 | 2.87E-02 | 0.44 | 0.36 | 0.08 | 1.21 |
| 8 | Pxylp1 | 0.53 | 1.38E-06 | 2.88E-02 | 0.16 | 0.11 | 0.05 | 1.49 |
| 8 | Rnf26 | 0.32 | 1.40E-06 | 2.92E-02 | 0.43 | 0.36 | 0.07 | 1.20 |
| 8 | Zfp521 | 0.38 | 1.41E-06 | 2.94E-02 | 0.36 | 0.29 | 0.07 | 1.26 |
| 8 | Nr4a2 | 0.57 | 1.44E-06 | 2.99E-02 | 0.15 | 0.10 | 0.05 | 1.50 |
| 8 | Ifngr1 | 0.28 | 1.45E-06 | 3.01E-02 | 0.32 | 0.25 | 0.07 | 1.30 |
| 8 | Eln | 0.44 | 1.47E-06 | 3.06E-02 | 0.12 | 0.08 | 0.05 | 1.61 |
| 8 | Tgfbr3 | 0.51 | 1.52E-06 | 3.16E-02 | 0.13 | 0.08 | 0.05 | 1.58 |
| 8 | Tmem35a | 0.48 | 1.55E-06 | 3.22E-02 | 0.15 | 0.10 | 0.05 | 1.52 |
| 8 | Mapk8 | 0.27 | 1.62E-06 | 3.36E-02 | 0.51 | 0.42 | 0.08 | 1.20 |
| 8 | Cep162 | 0.36 | 1.62E-06 | 3.37E-02 | 0.33 | 0.26 | 0.07 | 1.28 |
| 8 | Zfp157 | 0.28 | 1.64E-06 | 3.40E-02 | 0.45 | 0.37 | 0.08 | 1.23 |
| 8 | Cep72 | 0.47 | 1.66E-06 | 3.45E-02 | 0.21 | 0.15 | 0.06 | 1.37 |
| 8 | Shf | 0.34 | 1.67E-06 | 3.47E-02 | 0.35 | 0.27 | 0.08 | 1.28 |
| 8 | Asxl1 | 0.27 | 1.69E-06 | 3.52E-02 | 0.57 | 0.50 | 0.08 | 1.16 |
| 8 | Cbx4 | 0.60 | 1.70E-06 | 3.53E-02 | 0.16 | 0.11 | 0.05 | 1.47 |
| 8 | Adcy4 | 0.67 | 1.72E-06 | 3.57E-02 | 0.10 | 0.06 | 0.04 | 1.65 |
| 8 | Btbd8 | 0.69 | 1.78E-06 | 3.71E-02 | 0.12 | 0.07 | 0.04 | 1.58 |
| 8 | Inpp5b | 0.70 | 1.81E-06 | 3.77E-02 | 0.17 | 0.12 | 0.05 | 1.43 |
| 8 | Zdhhc15 | 0.50 | 1.87E-06 | 3.90E-02 | 0.18 | 0.13 | 0.05 | 1.42 |
| 8 | Maoa | 0.29 | 1.91E-06 | 3.97E-02 | 0.34 | 0.27 | 0.07 | 1.28 |
| 8 | Cep250 | 0.33 | 1.94E-06 | 4.03E-02 | 0.33 | 0.26 | 0.07 | 1.27 |
| 8 | Gk | 0.76 | 1.96E-06 | 4.08E-02 | 0.12 | 0.08 | 0.04 | 1.55 |
| 8 | Gsn | 0.37 | 1.97E-06 | 4.10E-02 | 0.20 | 0.14 | 0.06 | 1.42 |
| 8 | Atr | 0.37 | 2.00E-06 | 4.16E-02 | 0.40 | 0.33 | 0.07 | 1.21 |
| 8 | Braf | 0.27 | 2.03E-06 | 4.21E-02 | 0.41 | 0.33 | 0.08 | 1.23 |
| 8 | Slc39a6 | 0.26 | 2.03E-06 | 4.22E-02 | 0.65 | 0.57 | 0.08 | 1.14 |
| 8 | Phrf1 | 0.27 | 2.04E-06 | 4.25E-02 | 0.60 | 0.53 | 0.07 | 1.13 |
| 8 | Ikzf5 | 0.30 | 2.05E-06 | 4.26E-02 | 0.41 | 0.33 | 0.08 | 1.23 |
| 8 | Phka2 | 0.55 | 2.06E-06 | 4.28E-02 | 0.18 | 0.13 | 0.05 | 1.43 |
| 8 | Ccser2 | 0.33 | 2.08E-06 | 4.32E-02 | 0.34 | 0.27 | 0.07 | 1.27 |
| 8 | Scrn3 | 0.57 | 2.12E-06 | 4.40E-02 | 0.21 | 0.15 | 0.06 | 1.37 |
| 8 | Zmat1 | 0.62 | 2.12E-06 | 4.41E-02 | 0.11 | 0.07 | 0.04 | 1.62 |
| 8 | Ppm1d | 0.32 | 2.16E-06 | 4.49E-02 | 0.46 | 0.38 | 0.08 | 1.20 |
| 8 | Cmtr1 | 0.32 | 2.19E-06 | 4.56E-02 | 0.41 | 0.34 | 0.07 | 1.22 |
| 8 | Arhgap17 | 0.29 | 2.22E-06 | 4.62E-02 | 0.44 | 0.37 | 0.07 | 1.20 |
| 8 | Ralgapa1 | 0.29 | 2.24E-06 | 4.65E-02 | 0.42 | 0.35 | 0.07 | 1.21 |
| 8 | 5330434G04Rik | 0.66 | 2.25E-06 | 4.67E-02 | 0.14 | 0.09 | 0.05 | 1.53 |
| 8 | Ift27 | 0.26 | 2.26E-06 | 4.71E-02 | 0.64 | 0.56 | 0.08 | 1.13 |
| 10 | Sfn | 2.63 | 0 | 0 | 0.97 | 0.18 | 0.79 | 5.42 |
| 10 | Perp | 3.13 | 0 | 0 | 0.90 | 0.11 | 0.78 | 7.99 |
| 10 | Dsp | 3.02 | 0 | 0 | 0.98 | 0.21 | 0.77 | 4.66 |
| 10 | Isl1 | 2.99 | 0 | 0 | 0.95 | 0.19 | 0.76 | 4.96 |
| 10 | Pitx1 | 2.42 | 0 | 0 | 0.91 | 0.16 | 0.75 | 5.84 |
| 10 | Foxe1 | 7.03 | 0 | 0 | 0.76 | 0.02 | 0.74 | 44.71 |
| 10 | Pitx2 | 4.13 | 0 | 0 | 0.86 | 0.12 | 0.74 | 7.43 |
| 10 | Bcam | 2.69 | 0 | 0 | 0.90 | 0.18 | 0.72 | 5.06 |
| 10 | Fermt1 | 2.96 | 0 | 0 | 0.83 | 0.12 | 0.72 | 7.23 |
| 10 | Sox2 | 2.48 | 0 | 0 | 0.88 | 0.17 | 0.71 | 5.28 |
| 10 | Pdgfa | 2.43 | 0 | 0 | 0.97 | 0.26 | 0.71 | 3.77 |
| 10 | Cldn6 | 2.19 | 0 | 0 | 0.90 | 0.21 | 0.70 | 4.39 |
| 10 | Spint2 | 2.17 | 0 | 0 | 0.97 | 0.29 | 0.69 | 3.39 |
| 10 | Krt18 | 2.62 | 0 | 0 | 0.96 | 0.28 | 0.68 | 3.48 |
| 10 | Wnt4 | 2.11 | 0 | 0 | 0.85 | 0.18 | 0.68 | 4.84 |
| 10 | Krt8 | 2.91 | 0 | 0 | 0.90 | 0.24 | 0.67 | 3.81 |

**Supplementary Table 1. Marker genes for the original clusters of integrated clustering of E11.5 control and NotchGOF λ-junction cells.**

| | | | | | | | | |
|---|---|---|---|---|---|---|---|---|
| 10 | Epcam | 2.12 | 0 | 0 | 0.99 | 0.32 | 0.66 | 3.05 |
| 10 | Cldn7 | 2.26 | 0 | 0 | 0.87 | 0.22 | 0.66 | 4.05 |
| 10 | Gm16136 | 2.70 | 0 | 0 | 0.80 | 0.15 | 0.65 | 5.32 |
| 10 | Fxyd3 | 3.89 | 0 | 0 | 0.70 | 0.07 | 0.63 | 10.31 |
| 10 | Ikzf2 | 2.11 | 0 | 0 | 0.89 | 0.26 | 0.63 | 3.41 |
| 10 | Trp63 | 2.59 | 0 | 0 | 0.74 | 0.11 | 0.63 | 6.60 |
| 10 | Ezr | 1.96 | 0 | 0 | 0.92 | 0.30 | 0.62 | 3.07 |
| 10 | Igfbp5 | 2.22 | 0 | 0 | 0.98 | 0.39 | 0.59 | 2.53 |
| 10 | Nectin4 | 2.90 | 0 | 0 | 0.70 | 0.11 | 0.59 | 6.41 |
| 10 | Klf5 | 3.24 | 0 | 0 | 0.69 | 0.10 | 0.59 | 6.75 |
| 10 | Mpzl2 | 2.15 | 0 | 0 | 0.77 | 0.19 | 0.59 | 4.15 |
| 10 | F11r | 2.51 | 0 | 0 | 0.74 | 0.16 | 0.58 | 4.65 |
| 10 | Anxa2 | 2.55 | 0 | 0 | 0.98 | 0.39 | 0.58 | 2.48 |
| 10 | Mycl | 2.41 | 0 | 0 | 0.85 | 0.28 | 0.57 | 3.02 |
| 10 | Sema3d | 3.03 | 0 | 0 | 0.66 | 0.10 | 0.56 | 6.80 |
| 10 | Frem2 | 2.40 | 0 | 0 | 0.74 | 0.17 | 0.56 | 4.25 |
| 10 | Esrp1 | 2.38 | 0 | 0 | 0.71 | 0.15 | 0.56 | 4.80 |
| 10 | Col18a1 | 1.91 | 0 | 0 | 0.91 | 0.34 | 0.56 | 2.64 |
| 10 | Cdh1 | 2.45 | 0 | 0 | 0.71 | 0.15 | 0.56 | 4.70 |
| 10 | Gm12446 | 7.40 | 0 | 0 | 0.55 | 0.01 | 0.55 | 92.33 |
| 10 | Hspb1 | 2.80 | 0 | 0 | 0.68 | 0.13 | 0.55 | 5.24 |
| 10 | Pltp | 2.57 | 0 | 0 | 0.68 | 0.14 | 0.54 | 4.88 |
| 10 | Dsg2 | 2.99 | 0 | 0 | 0.64 | 0.11 | 0.53 | 5.77 |
| 10 | Krt14 | 2.87 | 0 | 0 | 0.64 | 0.11 | 0.52 | 5.58 |
| 10 | Nrarp | 2.09 | 0 | 0 | 0.68 | 0.16 | 0.52 | 4.24 |
| 10 | Upk3bl | 4.16 | 0 | 0 | 0.57 | 0.05 | 0.52 | 10.57 |
| 10 | Runx1 | 2.09 | 0 | 0 | 0.69 | 0.18 | 0.52 | 3.95 |
| 10 | Hspb8 | 2.34 | 0 | 0 | 0.63 | 0.12 | 0.51 | 5.32 |
| 10 | Irf6 | 2.47 | 0 | 0 | 0.58 | 0.09 | 0.49 | 6.41 |
| 10 | Mir205hg | 2.45 | 0 | 0 | 0.59 | 0.10 | 0.49 | 5.73 |
| 10 | Fgf9 | 3.19 | 0 | 0 | 0.57 | 0.09 | 0.48 | 6.43 |
| 10 | Dapl1 | 3.47 | 0 | 0 | 0.52 | 0.05 | 0.47 | 9.77 |
| 10 | St14 | 2.70 | 0 | 0 | 0.57 | 0.10 | 0.46 | 5.50 |
| 10 | Tacstd2 | 3.30 | 0 | 0 | 0.50 | 0.05 | 0.45 | 9.45 |
| 10 | Cdkn1a | 1.89 | 0 | 0 | 0.54 | 0.10 | 0.44 | 5.65 |
| 10 | Adamts18 | 3.37 | 0 | 0 | 0.50 | 0.05 | 0.44 | 9.19 |
| 10 | S100a10 | 2.09 | 0 | 0 | 0.96 | 0.54 | 0.42 | 1.78 |
| 10 | Cryab | 5.08 | 0 | 0 | 0.45 | 0.03 | 0.42 | 13.67 |
| 10 | Bex1 | 1.87 | 0 | 0 | 0.99 | 0.58 | 0.41 | 1.72 |
| 10 | Krt19 | 4.26 | 0 | 0 | 0.47 | 0.06 | 0.41 | 7.92 |
| 10 | Krt7 | 3.85 | 0 | 0 | 0.43 | 0.04 | 0.39 | 11.26 |
| 10 | Trim29 | 3.87 | 0 | 0 | 0.42 | 0.03 | 0.39 | 12.79 |
| 10 | Barx2 | 3.93 | 0 | 0 | 0.42 | 0.03 | 0.39 | 15.96 |
| 10 | Hspb2 | 5.17 | 0 | 0 | 0.39 | 0.02 | 0.37 | 19.65 |
| 10 | Jph4 | 3.45 | 0 | 0 | 0.41 | 0.04 | 0.37 | 10.05 |
| 10 | Prss22 | 3.06 | 0 | 0 | 0.40 | 0.05 | 0.35 | 8.16 |
| 10 | Dmrt2 | 4.68 | 0 | 0 | 0.35 | 0.01 | 0.33 | 26.69 |
| 10 | Grhl3 | 3.26 | 0 | 0 | 0.38 | 0.04 | 0.33 | 8.52 |
| 10 | Lypd2 | 4.02 | 0 | 0 | 0.35 | 0.03 | 0.32 | 13.35 |
| 10 | Anxa8 | 4.96 | 0 | 0 | 0.32 | 0.01 | 0.31 | 29.36 |
| 10 | Lamc2 | 3.88 | 0 | 0 | 0.34 | 0.03 | 0.31 | 10.63 |
| 10 | Shh | 6.86 | 0 | 0 | 0.30 | 0.01 | 0.29 | 59.40 |
| 10 | S100a11 | 1.88 | 0 | 0 | 1.00 | 0.75 | 0.24 | 1.32 |
| 10 | Upk2 | 5.10 | 0 | 0 | 0.21 | 0.01 | 0.20 | 25.75 |
| 10 | Chl1 | 4.72 | 0 | 0 | 0.20 | 0.01 | 0.19 | 22.56 |
| 10 | H19 | 2.55 | 0 | 0 | 1.00 | 0.84 | 0.16 | 1.19 |

**Supplementary Table 1. Marker genes for the original clusters of integrated clustering of E11.5 control and NotchGOF λ-junction cells.**

| | | | | | | | | |
|---|---|---|---|---|---|---|---|---|
| 10 | Foxa1 | 5.39 | 0 | 0 | 0.13 | 0.00 | 0.13 | 33.25 |
| 10 | Siva1 | 1.73 | 0 | 0 | 1.00 | 0.95 | 0.05 | 1.05 |
| 10 | Atox1 | 1.66 | 0 | 0 | 1.00 | 0.95 | 0.05 | 1.05 |
| 10 | Fabp5 | 1.37 | 0 | 0 | 1.00 | 0.98 | 0.02 | 1.02 |
| 10 | Ly6g6e | 4.25 | 7.08E-295 | 1.47E-290 | 0.18 | 0.01 | 0.17 | 18.20 |
| 10 | Pdgfc | 1.99 | 1.29E-294 | 2.68E-290 | 0.81 | 0.26 | 0.55 | 3.09 |
| 10 | 2610016A17Rik | 4.79 | 5.04E-294 | 1.05E-289 | 0.14 | 0.01 | 0.14 | 28.20 |
| 10 | Wfdc2 | 2.28 | 7.76E-288 | 1.61E-283 | 0.91 | 0.44 | 0.47 | 2.06 |
| 10 | Amot | 2.59 | 3.87E-286 | 8.05E-282 | 0.88 | 0.47 | 0.42 | 1.89 |
| 10 | Ntf5 | 2.66 | 1.66E-284 | 3.46E-280 | 0.45 | 0.08 | 0.37 | 5.95 |
| 10 | Ap1m2 | 2.19 | 5.07E-282 | 1.06E-277 | 0.59 | 0.13 | 0.46 | 4.59 |
| 10 | Nav2 | 1.97 | 1.01E-280 | 2.10E-276 | 0.76 | 0.22 | 0.54 | 3.42 |
| 10 | 2310030G06Rik | 2.94 | 1.41E-278 | 2.93E-274 | 0.39 | 0.06 | 0.33 | 6.69 |
| 10 | Esrp2 | 2.88 | 2.70E-274 | 5.62E-270 | 0.37 | 0.05 | 0.32 | 6.89 |
| 10 | Stx3 | 2.51 | 7.19E-273 | 1.49E-268 | 0.53 | 0.11 | 0.42 | 4.73 |
| 10 | Lgi2 | 2.86 | 2.35E-272 | 4.89E-268 | 0.49 | 0.10 | 0.39 | 4.94 |
| 10 | Ptprf | 1.70 | 1.21E-270 | 2.51E-266 | 0.97 | 0.63 | 0.34 | 1.54 |
| 10 | Fgf8 | 2.40 | 5.05E-267 | 1.05E-262 | 0.41 | 0.07 | 0.34 | 6.09 |
| 10 | Ripk4 | 2.70 | 1.89E-266 | 3.93E-262 | 0.40 | 0.06 | 0.33 | 6.37 |
| 10 | Zfp385c | 3.31 | 3.39E-266 | 7.05E-262 | 0.25 | 0.03 | 0.23 | 10.12 |
| 10 | Map7 | 2.43 | 1.38E-263 | 2.87E-259 | 0.49 | 0.09 | 0.39 | 5.16 |
| 10 | Dsc2 | 3.18 | 2.05E-263 | 4.26E-259 | 0.34 | 0.05 | 0.29 | 7.19 |
| 10 | Cgnl1 | 3.18 | 2.29E-263 | 4.76E-259 | 0.50 | 0.11 | 0.39 | 4.58 |
| 10 | Slc9a3r1 | 1.73 | 1.23E-262 | 2.55E-258 | 0.76 | 0.22 | 0.54 | 3.42 |
| 10 | Capg | 2.46 | 6.61E-261 | 1.37E-256 | 0.57 | 0.14 | 0.43 | 4.21 |
| 10 | Rab27b | 2.45 | 1.02E-260 | 2.12E-256 | 0.45 | 0.08 | 0.37 | 5.41 |
| 10 | Tac2 | 4.32 | 2.86E-259 | 5.95E-255 | 0.13 | 0.01 | 0.12 | 25.60 |
| 10 | Sema3c | 2.60 | 5.68E-259 | 1.18E-254 | 0.44 | 0.08 | 0.36 | 5.53 |
| 10 | Gask1b | 3.74 | 1.76E-258 | 3.66E-254 | 0.21 | 0.02 | 0.19 | 12.41 |
| 10 | Hist3h2ba | 1.66 | 2.14E-256 | 4.44E-252 | 0.80 | 0.26 | 0.54 | 3.13 |
| 10 | Klc3 | 2.28 | 3.76E-256 | 7.83E-252 | 0.48 | 0.09 | 0.38 | 5.07 |
| 10 | Jag1 | 2.20 | 1.37E-255 | 2.85E-251 | 0.62 | 0.16 | 0.46 | 3.84 |
| 10 | Cers4 | 2.50 | 4.84E-254 | 1.01E-249 | 0.57 | 0.14 | 0.43 | 4.04 |
| 10 | Rab25 | 1.94 | 2.00E-253 | 4.16E-249 | 0.61 | 0.15 | 0.46 | 4.18 |
| 10 | Igf2 | 2.18 | 4.25E-253 | 8.84E-249 | 0.91 | 0.57 | 0.34 | 1.59 |
| 10 | Ctnnd2 | 2.37 | 5.50E-252 | 1.14E-247 | 0.50 | 0.11 | 0.40 | 4.75 |
| 10 | Bex4 | 1.54 | 3.07E-251 | 6.39E-247 | 0.97 | 0.64 | 0.33 | 1.51 |
| 10 | Stard10 | 1.95 | 4.17E-250 | 8.68E-246 | 0.79 | 0.28 | 0.50 | 2.78 |
| 10 | Cd9 | 1.91 | 2.59E-247 | 5.38E-243 | 0.78 | 0.28 | 0.50 | 2.77 |
| 10 | Gjb3 | 3.05 | 5.28E-246 | 1.10E-241 | 0.25 | 0.03 | 0.23 | 9.41 |
| 10 | Samd5 | 1.79 | 9.75E-246 | 2.03E-241 | 0.85 | 0.35 | 0.51 | 2.47 |
| 10 | Grhl2 | 2.24 | 1.05E-245 | 2.19E-241 | 0.52 | 0.12 | 0.41 | 4.51 |
| 10 | Cldn8 | 3.35 | 4.82E-241 | 1.00E-236 | 0.23 | 0.02 | 0.21 | 10.41 |
| 10 | Sdc4 | 1.94 | 4.07E-240 | 8.46E-236 | 0.81 | 0.33 | 0.48 | 2.46 |
| 10 | Pdlim1 | 1.68 | 5.01E-239 | 1.04E-234 | 0.77 | 0.27 | 0.50 | 2.82 |
| 10 | Six3 | 1.09 | 1.28E-236 | 2.66E-232 | 0.78 | 0.22 | 0.56 | 3.53 |
| 10 | Ccnd2 | 1.58 | 1.32E-233 | 2.75E-229 | 0.99 | 0.85 | 0.14 | 1.16 |
| 10 | Pof1b | 4.55 | 2.49E-233 | 5.17E-229 | 0.13 | 0.01 | 0.13 | 19.14 |
| 10 | Nr2f2 | 2.01 | 3.25E-232 | 6.77E-228 | 0.81 | 0.35 | 0.46 | 2.33 |
| 10 | Ablim1 | 2.11 | 4.30E-232 | 8.93E-228 | 0.67 | 0.21 | 0.46 | 3.20 |
| 10 | Ppl | 2.80 | 2.01E-231 | 4.18E-227 | 0.27 | 0.03 | 0.24 | 8.21 |
| 10 | Tpd52 | 1.51 | 5.20E-231 | 1.08E-226 | 0.77 | 0.24 | 0.52 | 3.14 |
| 10 | Kcnk1 | 2.12 | 3.00E-229 | 6.23E-225 | 0.61 | 0.17 | 0.44 | 3.57 |
| 10 | Rab15 | 2.18 | 3.19E-229 | 6.63E-225 | 0.51 | 0.12 | 0.40 | 4.32 |
| 10 | Fjx1 | 1.75 | 1.48E-227 | 3.07E-223 | 0.85 | 0.37 | 0.48 | 2.28 |
| 10 | Six3os1 | 1.10 | 1.38E-224 | 2.86E-220 | 0.74 | 0.21 | 0.53 | 3.48 |

**Supplementary Table 1. Marker genes for the original clusters of integrated clustering of E11.5 control and NotchGOF λ-junction cells.**

| | | | | | | | | |
|---|---|---|---|---|---|---|---|---|
| 10 | Lama5 | 2.28 | 1.06E-220 | 2.21E-216 | 0.47 | 0.10 | 0.37 | 4.56 |
| 10 | Npnt | 2.11 | 4.58E-219 | 9.53E-215 | 0.68 | 0.22 | 0.46 | 3.06 |
| 10 | Arhgef19 | 2.17 | 1.31E-218 | 2.72E-214 | 0.50 | 0.12 | 0.39 | 4.26 |
| 10 | Jup | 2.03 | 3.83E-217 | 7.96E-213 | 0.62 | 0.18 | 0.44 | 3.38 |
| 10 | Ston2 | 2.00 | 7.11E-217 | 1.48E-212 | 0.60 | 0.17 | 0.43 | 3.57 |
| 10 | Cldn4 | 2.30 | 1.30E-212 | 2.70E-208 | 0.44 | 0.09 | 0.34 | 4.74 |
| 10 | Ece1 | 2.21 | 4.52E-212 | 9.40E-208 | 0.64 | 0.22 | 0.43 | 2.98 |
| 10 | Igfbp2 | 1.28 | 1.42E-211 | 2.95E-207 | 0.98 | 0.75 | 0.23 | 1.31 |
| 10 | Il17re | 3.09 | 8.88E-211 | 1.85E-206 | 0.27 | 0.04 | 0.24 | 7.38 |
| 10 | Elf5 | 2.84 | 1.72E-210 | 3.58E-206 | 0.32 | 0.05 | 0.27 | 6.17 |
| 10 | Rxfp1 | 3.73 | 5.89E-208 | 1.23E-203 | 0.18 | 0.02 | 0.16 | 11.73 |
| 10 | Dsc3 | 4.50 | 1.32E-204 | 2.75E-200 | 0.16 | 0.01 | 0.14 | 12.92 |
| 10 | Dpysl5 | 2.20 | 4.03E-203 | 8.38E-199 | 0.44 | 0.10 | 0.34 | 4.55 |
| 10 | Emb | 2.00 | 1.34E-202 | 2.79E-198 | 0.81 | 0.39 | 0.42 | 2.07 |
| 10 | Klk8 | 1.84 | 2.47E-201 | 5.14E-197 | 0.64 | 0.20 | 0.44 | 3.23 |
| 10 | Alox12 | 2.77 | 3.80E-201 | 7.90E-197 | 0.19 | 0.02 | 0.17 | 10.94 |
| 10 | Lad1 | 2.29 | 9.89E-200 | 2.06E-195 | 0.39 | 0.08 | 0.31 | 4.97 |
| 10 | Cmtm8 | 1.95 | 1.23E-198 | 2.56E-194 | 0.57 | 0.16 | 0.41 | 3.49 |
| 10 | Ifitm3 | 2.48 | 4.16E-198 | 8.66E-194 | 0.70 | 0.28 | 0.41 | 2.46 |
| 10 | Macc1 | 2.76 | 1.85E-197 | 3.85E-193 | 0.30 | 0.05 | 0.25 | 6.34 |
| 10 | Plekha7 | 2.49 | 1.15E-196 | 2.40E-192 | 0.39 | 0.08 | 0.31 | 4.83 |
| 10 | Cldn23 | 3.14 | 7.75E-196 | 1.61E-191 | 0.24 | 0.03 | 0.21 | 7.71 |
| 10 | Srd5a1 | 2.27 | 1.86E-195 | 3.87E-191 | 0.56 | 0.17 | 0.39 | 3.30 |
| 10 | Fzd6 | 2.29 | 5.35E-195 | 1.11E-190 | 0.38 | 0.08 | 0.30 | 5.05 |
| 10 | Fam83f | 4.04 | 1.87E-194 | 3.90E-190 | 0.12 | 0.01 | 0.11 | 19.50 |
| 10 | Rps16 | 0.36 | 1.99E-194 | 4.15E-190 | 1.00 | 1.00 | 0.00 | 1.00 |
| 10 | Nrtn | 2.56 | 4.10E-193 | 8.52E-189 | 0.51 | 0.14 | 0.37 | 3.58 |
| 10 | Myh14 | 2.59 | 2.22E-192 | 4.61E-188 | 0.36 | 0.07 | 0.29 | 5.08 |
| 10 | Wnt6 | 1.23 | 1.65E-191 | 3.43E-187 | 0.42 | 0.09 | 0.34 | 4.76 |
| 10 | Etl4 | 1.80 | 3.61E-190 | 7.50E-186 | 0.73 | 0.30 | 0.43 | 2.47 |
| 10 | Jag2 | 2.24 | 5.51E-190 | 1.15E-185 | 0.37 | 0.07 | 0.29 | 5.08 |
| 10 | Tmem54 | 2.09 | 6.23E-190 | 1.30E-185 | 0.46 | 0.11 | 0.35 | 4.21 |
| 10 | Rps19 | 0.38 | 1.15E-188 | 2.40E-184 | 1.00 | 1.00 | 0.00 | 1.00 |
| 10 | Rps5 | 0.38 | 1.39E-187 | 2.90E-183 | 1.00 | 1.00 | 0.00 | 1.00 |
| 10 | Stk26 | 1.86 | 8.00E-187 | 1.66E-182 | 0.64 | 0.21 | 0.42 | 2.98 |
| 10 | Cgn | 2.37 | 5.89E-186 | 1.22E-181 | 0.42 | 0.10 | 0.32 | 4.39 |
| 10 | Itga6 | 1.45 | 7.32E-186 | 1.52E-181 | 0.78 | 0.30 | 0.48 | 2.63 |
| 10 | Rpl27a | 0.40 | 7.76E-186 | 1.61E-181 | 1.00 | 1.00 | 0.00 | 1.00 |
| 10 | Prom2 | 3.09 | 1.34E-185 | 2.80E-181 | 0.22 | 0.03 | 0.19 | 8.27 |
| 10 | Spint1 | 2.20 | 1.96E-185 | 4.08E-181 | 0.41 | 0.09 | 0.32 | 4.55 |
| 10 | Acaa2 | 1.64 | 2.43E-183 | 5.06E-179 | 0.85 | 0.47 | 0.38 | 1.82 |
| 10 | Foxq1 | 4.03 | 2.73E-182 | 5.68E-178 | 0.13 | 0.01 | 0.12 | 16.00 |
| 10 | Celf2 | 1.87 | 1.59E-181 | 3.32E-177 | 0.66 | 0.24 | 0.42 | 2.75 |
| 10 | Cdh3 | 1.71 | 5.94E-179 | 1.24E-174 | 0.61 | 0.20 | 0.41 | 3.11 |
| 10 | Pkp4 | 1.81 | 1.56E-178 | 3.24E-174 | 0.76 | 0.37 | 0.39 | 2.07 |
| 10 | Cadm1 | 1.51 | 5.33E-177 | 1.11E-172 | 0.80 | 0.37 | 0.43 | 2.15 |
| 10 | S100a6 | 2.84 | 7.06E-177 | 1.47E-172 | 0.51 | 0.15 | 0.35 | 3.28 |
| 10 | Acer2 | 1.34 | 7.27E-177 | 1.51E-172 | 0.84 | 0.38 | 0.46 | 2.21 |
| 10 | Cldn3 | 1.81 | 5.28E-175 | 1.10E-170 | 0.53 | 0.15 | 0.38 | 3.56 |
| 10 | Ano1 | 1.98 | 8.52E-175 | 1.77E-170 | 0.46 | 0.12 | 0.34 | 3.91 |
| 10 | Lmo7 | 2.20 | 1.30E-174 | 2.70E-170 | 0.51 | 0.15 | 0.36 | 3.42 |
| 10 | Arl4c | 2.14 | 3.10E-174 | 6.46E-170 | 0.71 | 0.32 | 0.39 | 2.25 |
| 10 | Plch2 | 3.19 | 3.22E-174 | 6.70E-170 | 0.20 | 0.02 | 0.17 | 8.52 |
| 10 | Tom1l1 | 1.97 | 1.23E-173 | 2.55E-169 | 0.51 | 0.15 | 0.36 | 3.45 |
| 10 | Notch1 | 1.55 | 1.32E-172 | 2.76E-168 | 0.65 | 0.23 | 0.42 | 2.82 |
| 10 | Mapk13 | 1.67 | 2.20E-172 | 4.57E-168 | 0.60 | 0.19 | 0.41 | 3.20 |

**Supplementary Table 1. Marker genes for the original clusters of integrated clustering of E11.5 control and NotchGOF λ-junction cells.**

| | | | | | | | | |
|---|---|---|---|---|---|---|---|---|
| 10 | Eps8l2 | 2.98 | 2.64E-172 | 5.49E-168 | 0.24 | 0.04 | 0.21 | 6.86 |
| 10 | Evpl | 3.12 | 1.45E-171 | 3.01E-167 | 0.23 | 0.03 | 0.20 | 7.29 |
| 10 | Erbb3 | 1.71 | 2.08E-171 | 4.33E-167 | 0.55 | 0.16 | 0.39 | 3.40 |
| 10 | Grb7 | 2.18 | 8.66E-170 | 1.80E-165 | 0.39 | 0.09 | 0.30 | 4.41 |
| 10 | Cd151 | 1.66 | 2.21E-169 | 4.59E-165 | 0.73 | 0.32 | 0.41 | 2.30 |
| 10 | Zfp750 | 2.48 | 9.76E-169 | 2.03E-164 | 0.26 | 0.04 | 0.22 | 6.26 |
| 10 | Exph5 | 3.08 | 3.79E-168 | 7.89E-164 | 0.22 | 0.03 | 0.19 | 7.48 |
| 10 | Socs2 | 1.42 | 5.44E-168 | 1.13E-163 | 0.77 | 0.33 | 0.44 | 2.33 |
| 10 | Platr21 | 3.35 | 1.00E-167 | 2.08E-163 | 0.17 | 0.02 | 0.16 | 9.16 |
| 10 | Anxa1 | 3.72 | 1.12E-167 | 2.34E-163 | 0.19 | 0.02 | 0.17 | 8.50 |
| 10 | Itga3 | 2.50 | 5.34E-167 | 1.11E-162 | 0.31 | 0.06 | 0.25 | 5.22 |
| 10 | Eppk1 | 3.40 | 4.77E-165 | 9.93E-161 | 0.16 | 0.02 | 0.14 | 10.00 |
| 10 | Arhgef16 | 2.64 | 1.82E-164 | 3.79E-160 | 0.28 | 0.05 | 0.23 | 5.57 |
| 10 | Rpl12 | 0.39 | 4.02E-164 | 8.35E-160 | 1.00 | 1.00 | 0.00 | 1.00 |
| 10 | Krt17 | 3.64 | 5.88E-164 | 1.22E-159 | 0.15 | 0.01 | 0.14 | 10.86 |
| 10 | Ppp2r2b | 2.09 | 1.02E-161 | 2.12E-157 | 0.38 | 0.09 | 0.29 | 4.36 |
| 10 | Rab38 | 2.54 | 8.38E-161 | 1.74E-156 | 0.31 | 0.06 | 0.25 | 5.12 |
| 10 | Pon2 | 2.11 | 1.92E-160 | 4.00E-156 | 0.52 | 0.17 | 0.35 | 3.10 |
| 10 | Sp6 | 2.94 | 5.15E-160 | 1.07E-155 | 0.19 | 0.02 | 0.17 | 8.08 |
| 10 | Dstn | 0.86 | 5.69E-160 | 1.18E-155 | 0.99 | 0.94 | 0.05 | 1.06 |
| 10 | Cers3 | 3.82 | 8.81E-160 | 1.83E-155 | 0.12 | 0.01 | 0.11 | 14.63 |
| 10 | Rps27 | 0.43 | 9.13E-160 | 1.90E-155 | 1.00 | 1.00 | 0.00 | 1.00 |
| 10 | Dok5 | 1.91 | 1.00E-159 | 2.09E-155 | 0.42 | 0.11 | 0.32 | 4.00 |
| 10 | Six6 | 2.19 | 3.29E-159 | 6.84E-155 | 0.31 | 0.06 | 0.25 | 5.13 |
| 10 | Krt15 | 2.54 | 5.55E-159 | 1.15E-154 | 0.23 | 0.03 | 0.19 | 6.68 |
| 10 | Krt5 | 2.01 | 5.55E-158 | 1.15E-153 | 0.36 | 0.08 | 0.28 | 4.40 |
| 10 | Rps14 | 0.34 | 2.34E-156 | 4.86E-152 | 1.00 | 1.00 | 0.00 | 1.00 |
| 10 | Clic6 | 2.50 | 7.50E-155 | 1.56E-150 | 0.27 | 0.05 | 0.22 | 5.66 |
| 10 | Hook1 | 1.65 | 2.83E-152 | 5.89E-148 | 0.57 | 0.19 | 0.38 | 3.01 |
| 10 | Cyb5r3 | 1.08 | 4.37E-152 | 9.09E-148 | 0.95 | 0.80 | 0.15 | 1.18 |
| 10 | Plp2 | 1.57 | 9.94E-152 | 2.07E-147 | 0.76 | 0.37 | 0.39 | 2.08 |
| 10 | Ccdc88c | 1.64 | 1.02E-151 | 2.11E-147 | 0.50 | 0.15 | 0.35 | 3.42 |
| 10 | Tbx1 | 2.80 | 1.93E-151 | 4.02E-147 | 0.18 | 0.02 | 0.16 | 7.96 |
| 10 | Rps26 | 0.38 | 1.71E-150 | 3.55E-146 | 1.00 | 1.00 | 0.00 | 1.00 |
| 10 | Khdrbs3 | 1.20 | 4.69E-150 | 9.74E-146 | 0.90 | 0.61 | 0.30 | 1.49 |
| 10 | Rps20 | 0.33 | 4.74E-150 | 9.85E-146 | 1.00 | 1.00 | 0.00 | 1.00 |
| 10 | Rps27a | 0.32 | 4.89E-150 | 1.02E-145 | 1.00 | 1.00 | 0.00 | 1.00 |
| 10 | Dlx2 | 2.23 | 1.33E-149 | 2.77E-145 | 0.43 | 0.12 | 0.31 | 3.61 |
| 10 | Smtnl2 | 2.52 | 1.86E-149 | 3.88E-145 | 0.31 | 0.07 | 0.25 | 4.74 |
| 10 | Rpl23 | 0.33 | 6.82E-149 | 1.42E-144 | 1.00 | 1.00 | 0.00 | 1.00 |
| 10 | Ocln | 2.37 | 1.20E-148 | 2.50E-144 | 0.32 | 0.07 | 0.25 | 4.62 |
| 10 | Kdf1 | 2.19 | 1.64E-148 | 3.41E-144 | 0.34 | 0.08 | 0.26 | 4.43 |
| 10 | Pkp3 | 2.29 | 3.64E-147 | 7.58E-143 | 0.29 | 0.06 | 0.24 | 4.98 |
| 10 | Rplp2 | 0.31 | 2.65E-146 | 5.51E-142 | 1.00 | 1.00 | 0.00 | 1.00 |
| 10 | Rpl26 | 0.33 | 4.11E-146 | 8.54E-142 | 1.00 | 1.00 | 0.00 | 1.00 |
| 10 | Col14a1 | 3.64 | 9.80E-146 | 2.04E-141 | 0.14 | 0.02 | 0.13 | 9.60 |
| 10 | Rpl38 | 0.39 | 1.01E-144 | 2.10E-140 | 1.00 | 1.00 | 0.00 | 1.00 |
| 10 | Unc5c | 1.46 | 1.62E-144 | 3.37E-140 | 0.74 | 0.33 | 0.41 | 2.26 |
| 10 | Lsr | 1.55 | 2.04E-144 | 4.25E-140 | 0.57 | 0.19 | 0.38 | 2.96 |
| 10 | Vwa2 | 2.95 | 5.96E-144 | 1.24E-139 | 0.20 | 0.03 | 0.17 | 6.77 |
| 10 | Lmo4 | 0.98 | 1.59E-143 | 3.30E-139 | 0.97 | 0.81 | 0.16 | 1.20 |
| 10 | Lratd2 | 1.91 | 2.52E-143 | 5.24E-139 | 0.39 | 0.10 | 0.29 | 3.86 |
| 10 | Hras | 0.97 | 4.60E-142 | 9.58E-138 | 0.96 | 0.80 | 0.16 | 1.20 |
| 10 | Tpt1 | 0.31 | 7.20E-142 | 1.50E-137 | 1.00 | 1.00 | 0.00 | 1.00 |
| 10 | Spats2l | 1.84 | 7.75E-142 | 1.61E-137 | 0.58 | 0.22 | 0.36 | 2.65 |
| 10 | Serpinb6b | 3.24 | 1.37E-141 | 2.85E-137 | 0.15 | 0.02 | 0.14 | 9.06 |

**Supplementary Table 1. Marker genes for the original clusters of integrated clustering of E11.5 control and NotchGOF λ-junction cells.**

| | | | | | | | | |
|---|---|---|---|---|---|---|---|---|
| 10 | Celsr2 | 2.18 | 4.87E-141 | 1.01E-136 | 0.35 | 0.08 | 0.26 | 4.18 |
| 10 | Shtn1 | 2.23 | 8.86E-140 | 1.84E-135 | 0.25 | 0.05 | 0.20 | 5.53 |
| 10 | Marchf9 | 1.77 | 1.46E-139 | 3.04E-135 | 0.46 | 0.14 | 0.32 | 3.34 |
| 10 | Efna1 | 1.72 | 4.28E-139 | 8.90E-135 | 0.45 | 0.13 | 0.32 | 3.47 |
| 10 | Rpl28 | 0.32 | 5.19E-139 | 1.08E-134 | 1.00 | 1.00 | 0.00 | 1.00 |
| 10 | Hip1r | 2.37 | 3.38E-136 | 7.02E-132 | 0.31 | 0.07 | 0.24 | 4.50 |
| 10 | Cystm1 | 2.03 | 4.01E-133 | 8.34E-129 | 0.34 | 0.08 | 0.26 | 4.15 |
| 10 | Afdn | 1.21 | 5.19E-133 | 1.08E-128 | 0.86 | 0.56 | 0.30 | 1.53 |
| 10 | Tcf7l2 | 1.43 | 1.06E-132 | 2.21E-128 | 0.86 | 0.60 | 0.26 | 1.43 |
| 10 | Epb41l5 | 1.67 | 1.07E-132 | 2.23E-128 | 0.60 | 0.24 | 0.36 | 2.49 |
| 10 | Wwc1 | 1.54 | 3.30E-131 | 6.86E-127 | 0.51 | 0.17 | 0.34 | 3.01 |
| 10 | Tpm1 | 1.11 | 3.52E-131 | 7.32E-127 | 0.98 | 0.91 | 0.07 | 1.08 |
| 10 | Rps13 | 0.31 | 4.31E-131 | 8.96E-127 | 1.00 | 1.00 | 0.00 | 1.00 |
| 10 | Slc12a2 | 1.54 | 1.10E-130 | 2.30E-126 | 0.77 | 0.42 | 0.35 | 1.82 |
| 10 | Rassf10 | 2.29 | 1.26E-130 | 2.62E-126 | 0.27 | 0.05 | 0.21 | 4.93 |
| 10 | Pqlc3 | 2.49 | 1.53E-130 | 3.17E-126 | 0.25 | 0.05 | 0.20 | 5.19 |
| 10 | Ptpru | 1.83 | 1.82E-130 | 3.79E-126 | 0.46 | 0.15 | 0.31 | 3.14 |
| 10 | Rpl32 | 0.31 | 7.80E-130 | 1.62E-125 | 1.00 | 1.00 | 0.00 | 1.00 |
| 10 | Nrxn3 | 2.93 | 8.68E-130 | 1.80E-125 | 0.17 | 0.02 | 0.15 | 7.35 |
| 10 | Myh9 | 1.39 | 1.74E-129 | 3.61E-125 | 0.81 | 0.50 | 0.31 | 1.63 |
| 10 | Arhgef5 | 1.67 | 2.26E-129 | 4.71E-125 | 0.53 | 0.19 | 0.34 | 2.79 |
| 10 | Ddr1 | 1.74 | 5.53E-129 | 1.15E-124 | 0.42 | 0.12 | 0.30 | 3.47 |
| 10 | Rpl14 | 0.29 | 7.37E-129 | 1.53E-124 | 1.00 | 1.00 | 0.00 | 1.00 |
| 10 | Pla2g12a | 1.51 | 1.91E-128 | 3.97E-124 | 0.68 | 0.32 | 0.36 | 2.14 |
| 10 | Rpl39 | 0.32 | 2.36E-128 | 4.92E-124 | 1.00 | 1.00 | 0.00 | 1.00 |
| 10 | Ptpn13 | 1.46 | 1.32E-127 | 2.75E-123 | 0.74 | 0.39 | 0.35 | 1.89 |
| 10 | Fau | 0.34 | 2.05E-126 | 4.26E-122 | 1.00 | 1.00 | 0.00 | 1.00 |
| 10 | Rps18 | 0.36 | 7.14E-126 | 1.48E-121 | 1.00 | 1.00 | 0.00 | 1.00 |
| 10 | Oxtr | 2.08 | 9.04E-126 | 1.88E-121 | 0.28 | 0.06 | 0.22 | 4.70 |
| 10 | Lrp2 | 2.46 | 1.16E-125 | 2.41E-121 | 0.24 | 0.05 | 0.20 | 5.08 |
| 10 | Crb3 | 1.83 | 1.87E-125 | 3.89E-121 | 0.38 | 0.10 | 0.27 | 3.71 |
| 10 | Rpl17 | 0.33 | 2.85E-125 | 5.93E-121 | 1.00 | 1.00 | 0.00 | 1.00 |
| 10 | Kif21a | 1.26 | 3.90E-125 | 8.11E-121 | 0.69 | 0.30 | 0.39 | 2.31 |
| 10 | Fgfr2 | 1.44 | 7.16E-125 | 1.49E-120 | 0.63 | 0.27 | 0.37 | 2.38 |
| 10 | Dmrta2 | 1.85 | 1.74E-124 | 3.61E-120 | 0.43 | 0.13 | 0.30 | 3.27 |
| 10 | Phldb3 | 3.12 | 2.40E-124 | 5.00E-120 | 0.14 | 0.02 | 0.13 | 8.47 |
| 10 | Rpl18a | 0.33 | 6.75E-123 | 1.40E-118 | 1.00 | 1.00 | 0.00 | 1.00 |
| 10 | Fras1 | 1.44 | 8.07E-123 | 1.68E-118 | 0.62 | 0.25 | 0.36 | 2.43 |
| 10 | Wnt7b | 1.78 | 2.24E-121 | 4.66E-117 | 0.21 | 0.04 | 0.18 | 5.73 |
| 10 | Ccdc68 | 2.40 | 2.34E-121 | 4.86E-117 | 0.24 | 0.05 | 0.19 | 5.15 |
| 10 | Arrdc1 | 1.89 | 6.44E-121 | 1.34E-116 | 0.37 | 0.10 | 0.27 | 3.60 |
| 10 | Krtcap3 | 1.90 | 1.44E-120 | 3.00E-116 | 0.37 | 0.10 | 0.27 | 3.56 |
| 10 | Itgb4 | 2.76 | 1.86E-119 | 3.87E-115 | 0.18 | 0.03 | 0.15 | 6.56 |
| 10 | Camsap3 | 2.19 | 2.56E-119 | 5.32E-115 | 0.30 | 0.07 | 0.23 | 4.11 |
| 10 | 2610528J11Rik | 2.16 | 5.08E-118 | 1.06E-113 | 0.26 | 0.06 | 0.20 | 4.71 |
| 10 | Frmd4b | 1.51 | 7.23E-118 | 1.50E-113 | 0.47 | 0.16 | 0.32 | 2.99 |
| 10 | Lypd3 | 2.71 | 2.10E-117 | 4.37E-113 | 0.16 | 0.02 | 0.13 | 7.38 |
| 10 | AI661453 | 2.63 | 4.97E-117 | 1.03E-112 | 0.19 | 0.03 | 0.16 | 6.20 |
| 10 | Plekhg3 | 2.07 | 1.25E-116 | 2.59E-112 | 0.30 | 0.07 | 0.23 | 4.09 |
| 10 | Rps28 | 0.33 | 1.63E-116 | 3.40E-112 | 1.00 | 1.00 | 0.00 | 1.00 |
| 10 | Rps21 | 0.30 | 1.80E-116 | 3.75E-112 | 1.00 | 1.00 | 0.00 | 1.00 |
| 10 | Csrp2 | 1.20 | 2.34E-116 | 4.87E-112 | 0.97 | 0.93 | 0.04 | 1.05 |
| 10 | Ktn1 | 0.86 | 2.59E-114 | 5.38E-110 | 0.94 | 0.82 | 0.12 | 1.15 |
| 10 | Ctnnb1 | 0.58 | 7.60E-114 | 1.58E-109 | 0.99 | 0.97 | 0.03 | 1.03 |
| 10 | Cacna1b | 2.17 | 1.05E-112 | 2.19E-108 | 0.30 | 0.07 | 0.22 | 4.04 |
| 10 | Pdpn | 1.76 | 1.09E-112 | 2.27E-108 | 0.48 | 0.18 | 0.31 | 2.75 |

**Supplementary Table 1. Marker genes for the original clusters of integrated clustering of E11.5 control and NotchGOF λ-junction cells.**

| | | | | | | | | |
|---|---|---|---|---|---|---|---|---|
| 10 | Cldn10 | 3.36 | 2.57E-112 | 5.35E-108 | 0.21 | 0.04 | 0.17 | 5.20 |
| 10 | Plec | 1.57 | 3.94E-112 | 8.19E-108 | 0.60 | 0.26 | 0.34 | 2.28 |
| 10 | Palmd | 3.20 | 4.07E-112 | 8.47E-108 | 0.14 | 0.02 | 0.12 | 7.72 |
| 10 | Aplp2 | 0.90 | 6.10E-112 | 1.27E-107 | 0.93 | 0.82 | 0.12 | 1.14 |
| 10 | Smim22 | 2.51 | 2.18E-111 | 4.54E-107 | 0.19 | 0.03 | 0.16 | 5.79 |
| 10 | Gca | 3.02 | 6.33E-111 | 1.32E-106 | 0.15 | 0.02 | 0.13 | 7.40 |
| 10 | Gm2694 | 3.09 | 8.79E-111 | 1.83E-106 | 0.13 | 0.02 | 0.11 | 8.60 |
| 10 | Hs6st1 | 1.66 | 1.53E-110 | 3.18E-106 | 0.46 | 0.16 | 0.30 | 2.83 |
| 10 | Tnk1 | 2.61 | 6.66E-110 | 1.38E-105 | 0.19 | 0.03 | 0.15 | 5.81 |
| 10 | Agrn | 1.28 | 1.18E-109 | 2.46E-105 | 0.72 | 0.38 | 0.34 | 1.89 |
| 10 | Egln3 | 2.18 | 1.94E-109 | 4.04E-105 | 0.28 | 0.07 | 0.21 | 4.06 |
| 10 | Prss8 | 1.77 | 6.19E-109 | 1.29E-104 | 0.33 | 0.09 | 0.24 | 3.72 |
| 10 | Hey1 | 1.04 | 8.13E-109 | 1.69E-104 | 0.65 | 0.29 | 0.37 | 2.26 |
| 10 | Slc2a3 | 1.77 | 1.65E-108 | 3.43E-104 | 0.40 | 0.13 | 0.27 | 3.07 |
| 10 | Ppp1r13l | 2.09 | 1.69E-108 | 3.52E-104 | 0.29 | 0.07 | 0.22 | 4.06 |
| 10 | Hes1 | 1.22 | 2.15E-108 | 4.48E-104 | 0.83 | 0.54 | 0.30 | 1.55 |
| 10 | Ndufa4l2 | 2.82 | 5.13E-108 | 1.07E-103 | 0.16 | 0.02 | 0.13 | 6.74 |
| 10 | Elf1 | 1.46 | 7.94E-108 | 1.65E-103 | 0.58 | 0.25 | 0.32 | 2.29 |
| 10 | Prkci | 1.47 | 2.01E-107 | 4.19E-103 | 0.58 | 0.25 | 0.33 | 2.30 |
| 10 | Cstb | 1.01 | 4.15E-107 | 8.62E-103 | 0.93 | 0.80 | 0.14 | 1.17 |
| 10 | Phlda3 | 1.62 | 1.20E-106 | 2.49E-102 | 0.71 | 0.40 | 0.31 | 1.77 |
| 10 | Pik3ip1 | 1.95 | 4.10E-105 | 8.53E-101 | 0.30 | 0.08 | 0.22 | 3.80 |
| 10 | Slc35g1 | 2.33 | 8.96E-105 | 1.86E-100 | 0.23 | 0.05 | 0.18 | 4.59 |
| 10 | Hsp90ab1 | 0.30 | 4.29E-104 | 8.91E-100 | 1.00 | 1.00 | 0.00 | 1.00 |
| 10 | Hpcal1 | 1.34 | 2.50E-103 | 5.20E-99 | 0.65 | 0.32 | 0.33 | 2.04 |
| 10 | Nr3c1 | 1.68 | 1.38E-102 | 2.87E-98 | 0.44 | 0.15 | 0.28 | 2.86 |
| 10 | Iqgap1 | 1.25 | 1.44E-102 | 3.00E-98 | 0.75 | 0.46 | 0.29 | 1.63 |
| 10 | Hs6st2 | 1.48 | 1.57E-102 | 3.26E-98 | 0.60 | 0.27 | 0.32 | 2.18 |
| 10 | 2900026A02Rik | 1.73 | 8.56E-102 | 1.78E-97 | 0.34 | 0.10 | 0.24 | 3.49 |
| 10 | Lmo2 | 1.70 | 9.27E-102 | 1.93E-97 | 0.37 | 0.12 | 0.25 | 3.15 |
| 10 | Cd59a | 1.84 | 1.31E-101 | 2.72E-97 | 0.51 | 0.21 | 0.30 | 2.45 |
| 10 | Plcd1 | 2.30 | 2.38E-101 | 4.94E-97 | 0.26 | 0.06 | 0.20 | 4.08 |
| 10 | Foxp1 | 0.78 | 2.88E-101 | 5.99E-97 | 0.94 | 0.59 | 0.35 | 1.60 |
| 10 | Nudt14 | 1.21 | 2.97E-101 | 6.17E-97 | 0.72 | 0.39 | 0.33 | 1.83 |
| 10 | Nebl | 1.63 | 1.17E-100 | 2.42E-96 | 0.34 | 0.10 | 0.24 | 3.40 |
| 10 | Palm3 | 2.10 | 4.25E-100 | 8.83E-96 | 0.40 | 0.14 | 0.26 | 2.84 |
| 10 | Pkp2 | 2.31 | 5.42E-99 | 1.13E-94 | 0.22 | 0.05 | 0.17 | 4.53 |
| 10 | Plce1 | 1.54 | 5.02E-98 | 1.04E-93 | 0.52 | 0.22 | 0.30 | 2.39 |
| 10 | Pdcd4 | 0.95 | 7.05E-98 | 1.47E-93 | 0.89 | 0.63 | 0.26 | 1.41 |
| 10 | F3 | 2.33 | 2.29E-97 | 4.76E-93 | 0.32 | 0.09 | 0.23 | 3.41 |
| 10 | Itm2b | 1.04 | 8.93E-97 | 1.86E-92 | 0.94 | 0.84 | 0.10 | 1.12 |
| 10 | Elovl7 | 2.75 | 1.06E-96 | 2.21E-92 | 0.15 | 0.02 | 0.12 | 6.35 |
| 10 | Tspan13 | 1.07 | 1.95E-96 | 4.06E-92 | 0.74 | 0.40 | 0.34 | 1.84 |
| 10 | Pik3r1 | 1.21 | 2.09E-96 | 4.35E-92 | 0.71 | 0.38 | 0.33 | 1.87 |
| 10 | Glrx | 1.92 | 5.18E-96 | 1.08E-91 | 0.51 | 0.23 | 0.28 | 2.23 |
| 10 | Ppp1r36 | 2.20 | 2.22E-95 | 4.61E-91 | 0.22 | 0.05 | 0.17 | 4.54 |
| 10 | Llgl2 | 2.25 | 6.07E-94 | 1.26E-89 | 0.21 | 0.04 | 0.16 | 4.68 |
| 10 | Marveld2 | 2.07 | 2.19E-93 | 4.55E-89 | 0.23 | 0.05 | 0.18 | 4.28 |
| 10 | Ovol2 | 2.64 | 4.91E-93 | 1.02E-88 | 0.15 | 0.03 | 0.13 | 6.00 |
| 10 | Mt2 | 0.52 | 3.45E-92 | 7.17E-88 | 0.38 | 0.12 | 0.26 | 3.12 |
| 10 | Zdhhc2 | 1.43 | 7.27E-92 | 1.51E-87 | 0.37 | 0.12 | 0.25 | 3.07 |
| 10 | Fam83b | 2.72 | 2.29E-91 | 4.77E-87 | 0.12 | 0.02 | 0.11 | 7.29 |
| 10 | Ldlrap1 | 1.56 | 3.16E-91 | 6.57E-87 | 0.45 | 0.17 | 0.27 | 2.57 |
| 10 | Ndrg2 | 1.68 | 4.42E-91 | 9.19E-87 | 0.38 | 0.13 | 0.25 | 2.96 |
| 10 | Celsr1 | 1.65 | 5.37E-91 | 1.12E-86 | 0.32 | 0.10 | 0.23 | 3.35 |
| 10 | Prr15l | 2.41 | 2.62E-90 | 5.45E-86 | 0.17 | 0.03 | 0.14 | 5.42 |

**Supplementary Table 1. Marker genes for the original clusters of integrated clustering of E11.5 control and NotchGOF λ-junction cells.**

| | | | | | | | | |
|---|---|---|---|---|---|---|---|---|
| 10 | Sinhcaf | 0.69 | 6.03E-90 | 1.25E-85 | 0.96 | 0.83 | 0.13 | 1.15 |
| 10 | Egfl6 | 1.89 | 1.05E-89 | 2.18E-85 | 0.32 | 0.10 | 0.22 | 3.31 |
| 10 | Rps10 | 0.25 | 1.38E-89 | 2.87E-85 | 1.00 | 1.00 | 0.00 | 1.00 |
| 10 | Cdo1 | 1.56 | 4.73E-89 | 9.84E-85 | 0.73 | 0.50 | 0.23 | 1.45 |
| 10 | Myo10 | 1.08 | 5.13E-89 | 1.07E-84 | 0.76 | 0.46 | 0.30 | 1.65 |
| 10 | Afap1l1 | 2.11 | 8.71E-89 | 1.81E-84 | 0.26 | 0.07 | 0.19 | 3.81 |
| 10 | Cdcp1 | 1.99 | 1.71E-88 | 3.56E-84 | 0.23 | 0.05 | 0.17 | 4.25 |
| 10 | Sptlc3 | 2.73 | 3.20E-87 | 6.65E-83 | 0.12 | 0.02 | 0.10 | 7.25 |
| 10 | Tmem238 | 1.50 | 3.77E-87 | 7.84E-83 | 0.44 | 0.17 | 0.27 | 2.58 |
| 10 | Gm9922 | 2.61 | 4.11E-87 | 8.54E-83 | 0.18 | 0.04 | 0.14 | 5.06 |
| 10 | Mdh1 | 0.56 | 1.40E-86 | 2.90E-82 | 0.98 | 0.95 | 0.04 | 1.04 |
| 10 | Hes5 | 0.95 | 2.53E-86 | 5.26E-82 | 0.35 | 0.11 | 0.24 | 3.17 |
| 10 | Cebpa | 2.54 | 2.73E-86 | 5.67E-82 | 0.14 | 0.02 | 0.12 | 6.13 |
| 10 | Arhgap29 | 1.42 | 9.48E-86 | 1.97E-81 | 0.63 | 0.36 | 0.28 | 1.77 |
| 10 | Arap2 | 2.27 | 1.25E-85 | 2.60E-81 | 0.21 | 0.05 | 0.16 | 4.31 |
| 10 | Synrg | 1.31 | 1.23E-84 | 2.56E-80 | 0.52 | 0.23 | 0.29 | 2.29 |
| 10 | A730020E08Rik | 2.49 | 1.34E-84 | 2.79E-80 | 0.18 | 0.04 | 0.14 | 4.89 |
| 10 | Cox4i2 | 1.39 | 1.52E-84 | 3.17E-80 | 0.54 | 0.26 | 0.29 | 2.13 |
| 10 | Zfp219 | 1.13 | 2.19E-84 | 4.56E-80 | 0.69 | 0.39 | 0.30 | 1.78 |
| 10 | Gkap1 | 1.07 | 3.86E-84 | 8.02E-80 | 0.73 | 0.42 | 0.31 | 1.72 |
| 10 | AW011738 | 1.75 | 4.25E-84 | 8.85E-80 | 0.29 | 0.08 | 0.20 | 3.43 |
| 10 | Rps25 | 0.27 | 4.52E-84 | 9.40E-80 | 1.00 | 1.00 | 0.00 | 1.00 |
| 10 | Uchl1 | 1.15 | 8.68E-84 | 1.81E-79 | 0.64 | 0.34 | 0.30 | 1.87 |
| 10 | Susd4 | 1.75 | 9.02E-84 | 1.88E-79 | 0.29 | 0.08 | 0.20 | 3.42 |
| 10 | Arhgap27os2 | 1.87 | 1.70E-83 | 3.54E-79 | 0.24 | 0.06 | 0.18 | 3.93 |
| 10 | Rps7 | 0.25 | 1.84E-83 | 3.82E-79 | 1.00 | 1.00 | 0.00 | 1.00 |
| 10 | Utrn | 1.08 | 1.92E-83 | 3.99E-79 | 0.83 | 0.61 | 0.22 | 1.36 |
| 10 | Dlk1 | 0.82 | 2.89E-83 | 6.00E-79 | 0.73 | 0.41 | 0.32 | 1.78 |
| 10 | Rpl30 | 0.26 | 3.81E-83 | 7.93E-79 | 1.00 | 1.00 | 0.00 | 1.00 |
| 10 | AI463170 | 2.78 | 4.44E-83 | 9.24E-79 | 0.12 | 0.02 | 0.11 | 6.53 |
| 10 | Rpl9 | 0.26 | 8.25E-83 | 1.71E-78 | 1.00 | 1.00 | 0.00 | 1.00 |
| 10 | Rpl15 | 0.25 | 1.37E-82 | 2.84E-78 | 1.00 | 1.00 | 0.00 | 1.00 |
| 10 | Rpl10a | 0.29 | 2.11E-82 | 4.39E-78 | 1.00 | 1.00 | 0.00 | 1.00 |
| 10 | Ntm | 2.16 | 3.01E-82 | 6.25E-78 | 0.31 | 0.10 | 0.21 | 3.08 |
| 10 | Ptprm | 2.33 | 5.93E-82 | 1.23E-77 | 0.20 | 0.05 | 0.15 | 4.36 |
| 10 | Pwwp2b | 1.94 | 9.46E-82 | 1.97E-77 | 0.25 | 0.07 | 0.18 | 3.72 |
| 10 | Fam83h | 2.55 | 1.06E-81 | 2.20E-77 | 0.15 | 0.03 | 0.13 | 5.46 |
| 10 | Tiam1 | 1.40 | 2.75E-81 | 5.72E-77 | 0.58 | 0.30 | 0.28 | 1.92 |
| 10 | Cd24a | 0.67 | 2.76E-81 | 5.74E-77 | 0.99 | 0.92 | 0.08 | 1.08 |
| 10 | Arg1 | 2.22 | 3.90E-81 | 8.12E-77 | 0.14 | 0.02 | 0.12 | 5.79 |
| 10 | Npm3 | 0.93 | 6.80E-81 | 1.41E-76 | 0.92 | 0.79 | 0.13 | 1.16 |
| 10 | Limk2 | 1.57 | 9.38E-81 | 1.95E-76 | 0.45 | 0.19 | 0.26 | 2.37 |
| 10 | Arhgap8 | 2.22 | 1.51E-80 | 3.13E-76 | 0.18 | 0.04 | 0.14 | 4.68 |
| 10 | Dag1 | 1.18 | 2.60E-80 | 5.41E-76 | 0.65 | 0.36 | 0.29 | 1.81 |
| 10 | Slain1 | 1.98 | 4.31E-80 | 8.96E-76 | 0.22 | 0.06 | 0.16 | 3.96 |
| 10 | Gpc4 | 1.76 | 4.37E-80 | 9.09E-76 | 0.36 | 0.13 | 0.23 | 2.73 |
| 10 | Foxg1 | 0.99 | 8.11E-80 | 1.69E-75 | 0.39 | 0.14 | 0.26 | 2.87 |
| 10 | Uba52 | 0.31 | 1.04E-79 | 2.16E-75 | 1.00 | 1.00 | 0.00 | 1.00 |
| 10 | Rpl36a | 0.28 | 1.42E-79 | 2.94E-75 | 1.00 | 1.00 | 0.00 | 1.00 |
| 10 | Rps15 | 0.26 | 1.46E-79 | 3.03E-75 | 1.00 | 1.00 | 0.00 | 1.00 |
| 10 | Epha1 | 2.06 | 1.53E-79 | 3.19E-75 | 0.22 | 0.05 | 0.16 | 4.00 |
| 10 | Hdgf | 0.47 | 1.86E-79 | 3.87E-75 | 1.00 | 0.97 | 0.02 | 1.02 |
| 10 | Nt5c | 0.98 | 2.41E-79 | 5.01E-75 | 0.78 | 0.53 | 0.26 | 1.48 |
| 10 | Myo5c | 2.37 | 3.61E-79 | 7.51E-75 | 0.17 | 0.03 | 0.13 | 4.91 |
| 10 | Dab1 | 1.28 | 6.83E-79 | 1.42E-74 | 0.40 | 0.15 | 0.25 | 2.71 |
| 10 | Gpc3 | 0.65 | 2.01E-77 | 4.18E-73 | 0.97 | 0.83 | 0.14 | 1.17 |

**Supplementary Table 1. Marker genes for the original clusters of integrated clustering of E11.5 control and NotchGOF λ-junction cells.**

| | | | | | | | | |
|---|---|---|---|---|---|---|---|---|
| 10 | Mpp7 | 1.29 | 2.67E-77 | 5.56E-73 | 0.63 | 0.36 | 0.26 | 1.72 |
| 10 | Pcdh7 | 0.95 | 2.76E-77 | 5.75E-73 | 0.70 | 0.38 | 0.31 | 1.82 |
| 10 | Gm32585 | 1.70 | 3.96E-77 | 8.24E-73 | 0.21 | 0.05 | 0.16 | 4.14 |
| 10 | Ctnnal1 | 1.28 | 1.60E-76 | 3.32E-72 | 0.53 | 0.25 | 0.28 | 2.08 |
| 10 | Bicd1 | 1.72 | 2.68E-76 | 5.58E-72 | 0.29 | 0.09 | 0.20 | 3.19 |
| 10 | Patj | 1.62 | 5.88E-76 | 1.22E-71 | 0.35 | 0.12 | 0.22 | 2.81 |
| 10 | Eef1b2 | 0.28 | 9.58E-76 | 1.99E-71 | 1.00 | 1.00 | 0.00 | 1.00 |
| 10 | Rnh1 | 0.95 | 1.32E-75 | 2.74E-71 | 0.79 | 0.55 | 0.24 | 1.45 |
| 10 | Dock9 | 1.88 | 1.74E-75 | 3.62E-71 | 0.24 | 0.07 | 0.17 | 3.54 |
| 10 | B130024G19Rik | 2.51 | 1.95E-75 | 4.05E-71 | 0.13 | 0.02 | 0.11 | 5.58 |
| 10 | Tjp3 | 2.16 | 2.05E-75 | 4.26E-71 | 0.19 | 0.04 | 0.14 | 4.27 |
| 10 | Chst5 | 2.99 | 3.54E-75 | 7.35E-71 | 0.10 | 0.02 | 0.09 | 6.93 |
| 10 | Slc15a2 | 2.19 | 5.21E-75 | 1.08E-70 | 0.19 | 0.04 | 0.14 | 4.25 |
| 10 | Ctsh | 1.93 | 5.61E-75 | 1.17E-70 | 0.22 | 0.06 | 0.16 | 3.81 |
| 10 | Usp43 | 2.97 | 6.08E-75 | 1.26E-70 | 0.10 | 0.01 | 0.09 | 7.21 |
| 10 | Usp25 | 1.11 | 7.84E-75 | 1.63E-70 | 0.69 | 0.43 | 0.26 | 1.61 |
| 10 | Dlx1 | 2.10 | 1.07E-74 | 2.22E-70 | 0.19 | 0.05 | 0.15 | 4.13 |
| 10 | Tpi1 | 0.52 | 1.19E-74 | 2.48E-70 | 0.99 | 0.96 | 0.04 | 1.04 |
| 10 | Fam187b | 2.21 | 2.78E-74 | 5.78E-70 | 0.16 | 0.03 | 0.13 | 4.79 |
| 10 | Ssx2ip | 1.04 | 5.85E-74 | 1.22E-69 | 0.64 | 0.35 | 0.29 | 1.83 |
| 10 | Dst | 1.41 | 7.05E-74 | 1.47E-69 | 0.61 | 0.35 | 0.26 | 1.75 |
| 10 | Pik3cb | 2.02 | 1.14E-73 | 2.38E-69 | 0.22 | 0.06 | 0.16 | 3.79 |
| 10 | Ctdspl | 1.10 | 1.50E-73 | 3.12E-69 | 0.63 | 0.34 | 0.29 | 1.85 |
| 10 | Ajuba | 1.26 | 6.35E-73 | 1.32E-68 | 0.54 | 0.27 | 0.27 | 1.99 |
| 10 | Tceal9 | 0.42 | 1.52E-72 | 3.16E-68 | 1.00 | 0.98 | 0.02 | 1.02 |
| 10 | Samd10 | 1.62 | 2.21E-72 | 4.59E-68 | 0.27 | 0.08 | 0.19 | 3.25 |
| 10 | Ly6h | 1.23 | 5.15E-72 | 1.07E-67 | 0.38 | 0.14 | 0.24 | 2.65 |
| 10 | Acot7 | 1.35 | 9.56E-72 | 1.99E-67 | 0.49 | 0.23 | 0.26 | 2.14 |
| 10 | Ephb1 | 1.61 | 1.05E-71 | 2.18E-67 | 0.29 | 0.09 | 0.20 | 3.11 |
| 10 | Cep170b | 1.49 | 2.49E-71 | 5.18E-67 | 0.34 | 0.12 | 0.22 | 2.79 |
| 10 | Tbc1d2 | 2.61 | 2.70E-71 | 5.62E-67 | 0.12 | 0.02 | 0.10 | 5.64 |
| 10 | Fry | 2.16 | 3.61E-71 | 7.51E-67 | 0.20 | 0.05 | 0.15 | 3.88 |
| 10 | Mfge8 | 1.12 | 5.19E-71 | 1.08E-66 | 0.75 | 0.52 | 0.24 | 1.46 |
| 10 | Frrs1 | 1.30 | 1.78E-70 | 3.71E-66 | 0.47 | 0.21 | 0.25 | 2.19 |
| 10 | Bend7 | 1.82 | 3.05E-70 | 6.34E-66 | 0.25 | 0.08 | 0.18 | 3.29 |
| 10 | 1700023F06Rik | 2.12 | 3.16E-70 | 6.58E-66 | 0.17 | 0.04 | 0.13 | 4.41 |
| 10 | Rpl36 | 0.25 | 5.45E-70 | 1.13E-65 | 1.00 | 1.00 | 0.00 | 1.00 |
| 10 | Fgf17 | 1.63 | 1.54E-69 | 3.21E-65 | 0.17 | 0.04 | 0.13 | 4.36 |
| 10 | B3gnt2 | 1.54 | 2.15E-69 | 4.47E-65 | 0.28 | 0.09 | 0.19 | 3.05 |
| 10 | Cyb561 | 2.10 | 4.14E-69 | 8.60E-65 | 0.20 | 0.05 | 0.15 | 3.86 |
| 10 | Prxl2a | 1.07 | 4.52E-69 | 9.41E-65 | 0.65 | 0.37 | 0.28 | 1.74 |
| 10 | Irf2 | 1.24 | 6.79E-69 | 1.41E-64 | 0.51 | 0.25 | 0.26 | 2.06 |
| 10 | Nrbp2 | 2.38 | 1.99E-68 | 4.14E-64 | 0.14 | 0.03 | 0.11 | 5.11 |
| 10 | Osbpl3 | 1.81 | 8.25E-68 | 1.71E-63 | 0.27 | 0.09 | 0.18 | 3.09 |
| 10 | Frmd5 | 1.96 | 2.22E-67 | 4.61E-63 | 0.19 | 0.05 | 0.14 | 4.04 |
| 10 | Cox7a2l | 0.50 | 2.59E-67 | 5.38E-63 | 0.99 | 0.96 | 0.03 | 1.03 |
| 10 | Dpysl3 | 1.09 | 4.47E-67 | 9.29E-63 | 0.69 | 0.44 | 0.24 | 1.55 |
| 10 | Cdca7l | 1.18 | 4.81E-67 | 1.00E-62 | 0.56 | 0.30 | 0.27 | 1.89 |
| 10 | Efcab1 | 2.08 | 6.42E-67 | 1.33E-62 | 0.17 | 0.04 | 0.13 | 4.14 |
| 10 | Cblc | 2.13 | 7.14E-67 | 1.48E-62 | 0.14 | 0.03 | 0.11 | 4.93 |
| 10 | Kif13a | 1.48 | 1.07E-66 | 2.23E-62 | 0.44 | 0.20 | 0.24 | 2.19 |
| 10 | Id1 | 0.63 | 1.09E-66 | 2.28E-62 | 0.97 | 0.86 | 0.12 | 1.14 |
| 10 | Tjp2 | 1.21 | 1.23E-66 | 2.57E-62 | 0.56 | 0.30 | 0.26 | 1.86 |
| 10 | Retreg1 | 1.71 | 1.92E-66 | 3.99E-62 | 0.27 | 0.09 | 0.18 | 3.08 |
| 10 | Fzd5 | 3.02 | 1.95E-66 | 4.06E-62 | 0.11 | 0.02 | 0.09 | 5.84 |
| 10 | Pabpc1 | 0.38 | 3.02E-66 | 6.28E-62 | 1.00 | 0.99 | 0.01 | 1.01 |

**Supplementary Table 1. Marker genes for the original clusters of integrated clustering of E11.5 control and NotchGOF λ-junction cells.**

| | | | | | | | | |
|---|---|---|---|---|---|---|---|---|
| 10 | Pkm | 0.37 | 4.31E-66 | 8.96E-62 | 1.00 | 0.98 | 0.02 | 1.02 |
| 10 | Dcaf4 | 1.41 | 4.46E-66 | 9.28E-62 | 0.48 | 0.23 | 0.24 | 2.05 |
| 10 | Sult5a1 | 1.94 | 5.73E-66 | 1.19E-61 | 0.21 | 0.06 | 0.15 | 3.68 |
| 10 | Cpne5 | 2.18 | 1.51E-65 | 3.15E-61 | 0.19 | 0.05 | 0.14 | 3.90 |
| 10 | Meg3 | 0.67 | 3.13E-65 | 6.52E-61 | 0.89 | 0.69 | 0.20 | 1.29 |
| 10 | Ier5 | 0.73 | 3.22E-65 | 6.70E-61 | 0.85 | 0.64 | 0.21 | 1.32 |
| 10 | Rnf208 | 1.79 | 4.09E-65 | 8.51E-61 | 0.22 | 0.06 | 0.16 | 3.49 |
| 10 | Foxo1 | 1.41 | 4.22E-65 | 8.78E-61 | 0.40 | 0.17 | 0.23 | 2.32 |
| 10 | Pawr | 1.39 | 4.28E-65 | 8.91E-61 | 0.33 | 0.12 | 0.21 | 2.70 |
| 10 | Pdk1 | 1.17 | 2.62E-64 | 5.46E-60 | 0.51 | 0.25 | 0.26 | 2.04 |
| 10 | Prkch | 1.70 | 2.96E-64 | 6.15E-60 | 0.26 | 0.08 | 0.17 | 3.11 |
| 10 | Fgd2 | 1.47 | 5.98E-64 | 1.24E-59 | 0.28 | 0.10 | 0.19 | 2.95 |
| 10 | Hes6 | 0.63 | 7.88E-64 | 1.64E-59 | 0.59 | 0.33 | 0.26 | 1.81 |
| 10 | Arid3b | 1.31 | 1.08E-63 | 2.25E-59 | 0.46 | 0.22 | 0.24 | 2.08 |
| 10 | 2310002F09Rik | 2.31 | 1.18E-63 | 2.46E-59 | 0.15 | 0.03 | 0.12 | 4.41 |
| 10 | Ralbp1 | 0.76 | 1.89E-63 | 3.93E-59 | 0.89 | 0.78 | 0.11 | 1.15 |
| 10 | Wipf3 | 1.96 | 2.17E-63 | 4.52E-59 | 0.20 | 0.06 | 0.15 | 3.59 |
| 10 | Cobl | 1.83 | 2.59E-63 | 5.38E-59 | 0.22 | 0.07 | 0.16 | 3.40 |
| 10 | Rflnb | 0.98 | 3.14E-63 | 6.53E-59 | 0.78 | 0.54 | 0.23 | 1.43 |
| 10 | Cecr2 | 1.24 | 4.00E-63 | 8.32E-59 | 0.46 | 0.22 | 0.24 | 2.13 |
| 10 | Cracr2b | 2.27 | 5.14E-63 | 1.07E-58 | 0.14 | 0.03 | 0.11 | 4.65 |
| 10 | Pgam1 | 0.41 | 9.24E-63 | 1.92E-58 | 1.00 | 0.97 | 0.03 | 1.03 |
| 10 | Kctd1 | 1.08 | 1.54E-62 | 3.19E-58 | 0.69 | 0.46 | 0.23 | 1.50 |
| 10 | Lrrc1 | 1.39 | 1.80E-62 | 3.73E-58 | 0.36 | 0.14 | 0.21 | 2.50 |
| 10 | Tspan12 | 1.17 | 2.70E-62 | 5.62E-58 | 0.45 | 0.20 | 0.25 | 2.21 |
| 10 | Kremen2 | 1.67 | 2.85E-62 | 5.93E-58 | 0.17 | 0.04 | 0.13 | 4.10 |
| 10 | Bspry | 1.71 | 2.93E-62 | 6.10E-58 | 0.21 | 0.06 | 0.15 | 3.48 |
| 10 | Mbp | 1.59 | 2.94E-62 | 6.11E-58 | 0.27 | 0.09 | 0.18 | 2.98 |
| 10 | Sptbn2 | 1.50 | 4.14E-62 | 8.60E-58 | 0.30 | 0.11 | 0.19 | 2.79 |
| 10 | Zfhx2 | 1.20 | 1.09E-61 | 2.27E-57 | 0.49 | 0.24 | 0.25 | 2.03 |
| 10 | Sorl1 | 1.39 | 1.17E-61 | 2.44E-57 | 0.36 | 0.14 | 0.21 | 2.47 |
| 10 | Trp53i11 | 1.06 | 1.61E-61 | 3.34E-57 | 0.46 | 0.22 | 0.25 | 2.12 |
| 10 | Lrrk1 | 1.52 | 9.20E-61 | 1.91E-56 | 0.35 | 0.14 | 0.21 | 2.44 |
| 10 | Tnfaip2 | 0.82 | 1.68E-60 | 3.50E-56 | 0.35 | 0.14 | 0.22 | 2.60 |
| 10 | Tmod3 | 0.93 | 3.30E-60 | 6.87E-56 | 0.80 | 0.63 | 0.17 | 1.27 |
| 10 | Nell2 | 2.05 | 3.74E-60 | 7.79E-56 | 0.18 | 0.05 | 0.13 | 3.83 |
| 10 | Eno1 | 0.40 | 4.09E-60 | 8.51E-56 | 1.00 | 0.97 | 0.03 | 1.03 |
| 10 | Sntb1 | 2.43 | 7.68E-60 | 1.60E-55 | 0.12 | 0.03 | 0.10 | 4.96 |
| 10 | H2afj | 0.58 | 7.95E-60 | 1.65E-55 | 0.94 | 0.81 | 0.13 | 1.16 |
| 10 | Lpar6 | 1.07 | 8.91E-60 | 1.85E-55 | 0.64 | 0.40 | 0.24 | 1.61 |
| 10 | Agl | 1.35 | 2.18E-59 | 4.53E-55 | 0.48 | 0.25 | 0.23 | 1.95 |
| 10 | Cd2ap | 0.81 | 3.10E-59 | 6.44E-55 | 0.81 | 0.62 | 0.20 | 1.32 |
| 10 | Igsf9 | 0.95 | 4.15E-59 | 8.63E-55 | 0.65 | 0.40 | 0.26 | 1.64 |
| 10 | Vwa1 | 2.01 | 6.77E-59 | 1.41E-54 | 0.19 | 0.05 | 0.14 | 3.56 |
| 10 | Apoc1 | 1.22 | 1.10E-58 | 2.29E-54 | 0.25 | 0.08 | 0.17 | 3.08 |
| 10 | Mtln | 1.05 | 1.70E-58 | 3.53E-54 | 0.57 | 0.32 | 0.25 | 1.77 |
| 10 | Ppp1r13b | 1.66 | 2.17E-58 | 4.51E-54 | 0.23 | 0.08 | 0.16 | 3.12 |
| 10 | Scube3 | 1.22 | 4.16E-58 | 8.65E-54 | 0.42 | 0.20 | 0.23 | 2.16 |
| 10 | Podxl | 1.88 | 6.01E-58 | 1.25E-53 | 0.16 | 0.04 | 0.12 | 3.98 |
| 10 | Dlx3 | 1.39 | 6.44E-58 | 1.34E-53 | 0.23 | 0.07 | 0.16 | 3.23 |
| 10 | Ccser1 | 2.28 | 7.12E-58 | 1.48E-53 | 0.13 | 0.03 | 0.10 | 4.59 |
| 10 | Barx1 | 0.89 | 3.73E-57 | 7.77E-53 | 0.46 | 0.21 | 0.25 | 2.18 |
| 10 | Hspg2 | 0.99 | 4.37E-57 | 9.09E-53 | 0.46 | 0.22 | 0.24 | 2.11 |
| 10 | Fhdc1 | 2.01 | 4.72E-57 | 9.82E-53 | 0.15 | 0.04 | 0.12 | 4.05 |
| 10 | Mgst1 | 0.94 | 7.62E-57 | 1.58E-52 | 0.62 | 0.37 | 0.26 | 1.69 |
| 10 | Krt1 | 1.88 | 9.26E-57 | 1.93E-52 | 0.17 | 0.05 | 0.13 | 3.70 |

**Supplementary Table 1. Marker genes for the original clusters of integrated clustering of E11.5 control and NotchGOF λ-junction cells.**

| | | | | | | | | |
|---|---|---|---|---|---|---|---|---|
| 10 | Ttc9 | 2.03 | 1.09E-56 | 2.26E-52 | 0.15 | 0.04 | 0.11 | 4.05 |
| 10 | Aig1 | 1.06 | 1.67E-56 | 3.48E-52 | 0.57 | 0.32 | 0.25 | 1.77 |
| 10 | Gldc | 2.15 | 3.19E-56 | 6.63E-52 | 0.15 | 0.04 | 0.12 | 4.03 |
| 10 | Nectin1 | 1.37 | 3.98E-56 | 8.28E-52 | 0.38 | 0.17 | 0.21 | 2.24 |
| 10 | Rab11fip1 | 1.55 | 4.12E-56 | 8.57E-52 | 0.29 | 0.11 | 0.18 | 2.62 |
| 10 | Rbms1 | 0.68 | 7.52E-56 | 1.56E-51 | 0.90 | 0.78 | 0.12 | 1.15 |
| 10 | Stxbp2 | 1.40 | 2.78E-55 | 5.79E-51 | 0.32 | 0.13 | 0.19 | 2.47 |
| 10 | S100a16 | 0.61 | 2.87E-55 | 5.98E-51 | 0.23 | 0.07 | 0.15 | 3.17 |
| 10 | Plcd3 | 2.17 | 3.42E-55 | 7.11E-51 | 0.13 | 0.03 | 0.10 | 4.63 |
| 10 | Cnksr1 | 2.47 | 5.45E-55 | 1.13E-50 | 0.10 | 0.02 | 0.08 | 5.37 |
| 10 | Mtcl1 | 1.43 | 6.04E-55 | 1.26E-50 | 0.30 | 0.12 | 0.18 | 2.58 |
| 10 | Prag1 | 1.40 | 1.04E-54 | 2.17E-50 | 0.33 | 0.13 | 0.19 | 2.45 |
| 10 | Errfi1 | 1.35 | 1.10E-54 | 2.30E-50 | 0.38 | 0.17 | 0.21 | 2.20 |
| 10 | Isoc1 | 0.83 | 1.69E-54 | 3.52E-50 | 0.83 | 0.72 | 0.11 | 1.15 |
| 10 | Usp53 | 1.77 | 2.22E-54 | 4.62E-50 | 0.22 | 0.07 | 0.15 | 3.11 |
| 10 | Kdm5a | 0.68 | 3.00E-54 | 6.24E-50 | 0.85 | 0.67 | 0.18 | 1.27 |
| 10 | Serpinb6a | 0.96 | 4.86E-54 | 1.01E-49 | 0.62 | 0.38 | 0.24 | 1.63 |
| 10 | Fosl2 | 2.29 | 5.10E-54 | 1.06E-49 | 0.12 | 0.03 | 0.10 | 4.59 |
| 10 | Igf2bp3 | 0.58 | 9.18E-54 | 1.91E-49 | 0.91 | 0.82 | 0.09 | 1.11 |
| 10 | Hpn | 2.20 | 1.30E-53 | 2.71E-49 | 0.14 | 0.04 | 0.11 | 4.09 |
| 10 | Large2 | 1.62 | 1.39E-53 | 2.89E-49 | 0.24 | 0.08 | 0.16 | 2.95 |
| 10 | Fhl1 | 1.51 | 1.40E-53 | 2.91E-49 | 0.33 | 0.14 | 0.19 | 2.40 |
| 10 | Acvr2b | 0.70 | 1.68E-53 | 3.49E-49 | 0.82 | 0.64 | 0.19 | 1.29 |
| 10 | Lgr4 | 0.99 | 2.02E-53 | 4.19E-49 | 0.45 | 0.22 | 0.23 | 2.03 |
| 10 | Dap | 0.82 | 2.27E-53 | 4.73E-49 | 0.76 | 0.56 | 0.20 | 1.35 |
| 10 | Slc2a1 | 0.89 | 4.68E-53 | 9.73E-49 | 0.66 | 0.42 | 0.24 | 1.57 |
| 10 | Atp6v1f | 0.48 | 8.40E-53 | 1.75E-48 | 0.96 | 0.90 | 0.06 | 1.06 |
| 10 | Fzd8 | 1.33 | 1.01E-52 | 2.11E-48 | 0.34 | 0.14 | 0.20 | 2.38 |
| 10 | Syt5 | 1.75 | 1.34E-52 | 2.79E-48 | 0.18 | 0.05 | 0.13 | 3.41 |
| 10 | Rnf43 | 1.86 | 1.51E-52 | 3.14E-48 | 0.17 | 0.05 | 0.12 | 3.63 |
| 10 | Slc44a2 | 1.12 | 1.80E-52 | 3.74E-48 | 0.49 | 0.27 | 0.22 | 1.83 |
| 10 | Pts | 0.91 | 2.84E-52 | 5.90E-48 | 0.64 | 0.40 | 0.24 | 1.59 |
| 10 | Spock2 | 1.50 | 3.07E-52 | 6.39E-48 | 0.14 | 0.04 | 0.11 | 4.00 |
| 10 | Ago1 | 0.99 | 5.95E-52 | 1.24E-47 | 0.59 | 0.36 | 0.24 | 1.66 |
| 10 | Map3k20 | 1.60 | 6.16E-52 | 1.28E-47 | 0.28 | 0.11 | 0.17 | 2.54 |
| 10 | Spred1 | 0.97 | 6.98E-52 | 1.45E-47 | 0.69 | 0.50 | 0.20 | 1.39 |
| 10 | Lamb1 | 0.86 | 7.09E-52 | 1.47E-47 | 0.63 | 0.38 | 0.25 | 1.67 |
| 10 | Marveld3 | 2.04 | 1.14E-51 | 2.36E-47 | 0.14 | 0.03 | 0.11 | 4.09 |
| 10 | Armc10 | 0.80 | 1.56E-51 | 3.24E-47 | 0.75 | 0.52 | 0.24 | 1.46 |
| 10 | Cyb5a | 0.46 | 3.54E-51 | 7.36E-47 | 0.97 | 0.92 | 0.05 | 1.05 |
| 10 | 2200002D01Rik | 2.25 | 3.64E-51 | 7.56E-47 | 0.14 | 0.03 | 0.10 | 4.12 |
| 10 | Svip | 1.50 | 5.45E-51 | 1.13E-46 | 0.24 | 0.09 | 0.16 | 2.78 |
| 10 | Ldha | 0.42 | 9.86E-51 | 2.05E-46 | 1.00 | 0.97 | 0.03 | 1.03 |
| 10 | Mtus1 | 1.47 | 1.35E-50 | 2.81E-46 | 0.27 | 0.10 | 0.17 | 2.66 |
| 10 | Ccn2 | 2.12 | 2.31E-50 | 4.81E-46 | 0.14 | 0.03 | 0.10 | 4.00 |
| 10 | Slc4a4 | 1.55 | 3.87E-50 | 8.05E-46 | 0.28 | 0.11 | 0.17 | 2.57 |
| 10 | Trpm4 | 1.43 | 8.10E-50 | 1.68E-45 | 0.25 | 0.09 | 0.16 | 2.73 |
| 10 | Hap1 | 1.78 | 1.21E-49 | 2.52E-45 | 0.17 | 0.05 | 0.12 | 3.48 |
| 10 | Mbnl2 | 0.69 | 2.02E-49 | 4.20E-45 | 0.85 | 0.67 | 0.18 | 1.26 |
| 10 | Ormdl2 | 0.89 | 2.09E-49 | 4.35E-45 | 0.58 | 0.34 | 0.24 | 1.70 |
| 10 | Ubald1 | 1.11 | 2.33E-49 | 4.85E-45 | 0.48 | 0.26 | 0.22 | 1.85 |
| 10 | Bmp7 | 0.92 | 2.89E-49 | 6.01E-45 | 0.61 | 0.38 | 0.23 | 1.62 |
| 10 | Crim1 | 1.23 | 3.32E-49 | 6.90E-45 | 0.40 | 0.19 | 0.21 | 2.07 |
| 10 | Tead1 | 0.78 | 3.84E-49 | 7.98E-45 | 0.78 | 0.60 | 0.18 | 1.30 |
| 10 | Cxadr | 0.78 | 1.16E-48 | 2.42E-44 | 0.70 | 0.46 | 0.24 | 1.53 |
| 10 | Endod1 | 1.65 | 1.25E-48 | 2.60E-44 | 0.24 | 0.09 | 0.15 | 2.73 |

**Supplementary Table 1. Marker genes for the original clusters of integrated clustering of E11.5 control and NotchGOF λ-junction cells.**

| | | | | | | | | |
|---|---|---|---|---|---|---|---|---|
| 10 | Gab1 | 1.21 | 3.53E-48 | 7.33E-44 | 0.39 | 0.18 | 0.20 | 2.09 |
| 10 | Lypd6b | 1.52 | 5.44E-48 | 1.13E-43 | 0.20 | 0.07 | 0.13 | 3.06 |
| 10 | Trmt9b | 1.56 | 5.82E-48 | 1.21E-43 | 0.20 | 0.07 | 0.13 | 3.06 |
| 10 | Ankrd6 | 1.59 | 8.61E-48 | 1.79E-43 | 0.26 | 0.10 | 0.16 | 2.59 |
| 10 | Gltp | 0.93 | 8.99E-48 | 1.87E-43 | 0.61 | 0.39 | 0.22 | 1.56 |
| 10 | Kif13b | 1.39 | 1.14E-47 | 2.36E-43 | 0.34 | 0.16 | 0.19 | 2.20 |
| 10 | Rab3ip | 0.98 | 1.65E-47 | 3.44E-43 | 0.52 | 0.30 | 0.22 | 1.75 |
| 10 | Cux1 | 0.62 | 2.46E-47 | 5.11E-43 | 0.86 | 0.69 | 0.17 | 1.24 |
| 10 | Isl2 | 1.01 | 2.99E-47 | 6.23E-43 | 0.31 | 0.13 | 0.19 | 2.47 |
| 10 | Syngr1 | 1.71 | 6.70E-47 | 1.39E-42 | 0.19 | 0.06 | 0.13 | 3.03 |
| 10 | Nup210 | 1.73 | 8.47E-47 | 1.76E-42 | 0.18 | 0.06 | 0.13 | 3.19 |
| 10 | Rhpn2 | 2.32 | 9.79E-47 | 2.04E-42 | 0.11 | 0.02 | 0.09 | 4.54 |
| 10 | Metrnl | 1.72 | 1.08E-46 | 2.24E-42 | 0.19 | 0.06 | 0.13 | 3.13 |
| 10 | Epn3 | 1.89 | 1.35E-46 | 2.80E-42 | 0.16 | 0.05 | 0.11 | 3.46 |
| 10 | Pax6 | 0.77 | 2.32E-46 | 4.82E-42 | 0.31 | 0.12 | 0.19 | 2.49 |
| 10 | Kdm5b | 0.50 | 2.85E-46 | 5.92E-42 | 0.95 | 0.86 | 0.09 | 1.10 |
| 10 | B230208H11Rik | 2.04 | 3.83E-46 | 7.96E-42 | 0.13 | 0.03 | 0.10 | 4.00 |
| 10 | Trappc6a | 0.77 | 4.09E-46 | 8.50E-42 | 0.72 | 0.52 | 0.20 | 1.39 |
| 10 | Marchf2 | 0.78 | 5.71E-46 | 1.19E-41 | 0.71 | 0.51 | 0.21 | 1.40 |
| 10 | Egfr | 1.23 | 6.32E-46 | 1.31E-41 | 0.26 | 0.10 | 0.16 | 2.66 |
| 10 | Pura | 0.82 | 8.19E-46 | 1.70E-41 | 0.67 | 0.47 | 0.21 | 1.45 |
| 10 | Met | 2.01 | 1.10E-45 | 2.29E-41 | 0.14 | 0.04 | 0.10 | 3.67 |
| 10 | Tmem79 | 2.26 | 1.11E-45 | 2.31E-41 | 0.12 | 0.03 | 0.09 | 4.26 |
| 10 | Syt7 | 1.73 | 1.80E-45 | 3.74E-41 | 0.17 | 0.05 | 0.12 | 3.26 |
| 10 | Chmp2b | 0.73 | 1.85E-45 | 3.85E-41 | 0.76 | 0.56 | 0.20 | 1.35 |
| 10 | Atp8b1 | 1.69 | 2.41E-45 | 5.01E-41 | 0.14 | 0.04 | 0.10 | 3.67 |
| 10 | Firre | 1.54 | 2.52E-45 | 5.24E-41 | 0.23 | 0.09 | 0.15 | 2.71 |
| 10 | Galnt3 | 1.93 | 2.62E-45 | 5.46E-41 | 0.12 | 0.03 | 0.09 | 4.07 |
| 10 | Ces1d | 1.87 | 2.81E-45 | 5.84E-41 | 0.15 | 0.04 | 0.11 | 3.63 |
| 10 | Ppm1l | 1.10 | 2.98E-45 | 6.20E-41 | 0.42 | 0.22 | 0.21 | 1.96 |
| 10 | Wls | 0.57 | 3.51E-45 | 7.30E-41 | 0.87 | 0.74 | 0.14 | 1.19 |
| 10 | Nkain3 | 1.82 | 4.22E-45 | 8.78E-41 | 0.15 | 0.04 | 0.11 | 3.50 |
| 10 | Ifi30 | 0.83 | 6.19E-45 | 1.29E-40 | 0.52 | 0.29 | 0.23 | 1.79 |
| 10 | Plekhg1 | 1.37 | 8.21E-45 | 1.71E-40 | 0.30 | 0.13 | 0.17 | 2.31 |
| 10 | Nr2f6 | 0.71 | 8.73E-45 | 1.81E-40 | 0.78 | 0.59 | 0.19 | 1.32 |
| 10 | Dnase2a | 1.04 | 1.09E-44 | 2.26E-40 | 0.42 | 0.21 | 0.21 | 1.98 |
| 10 | Pkp1 | 2.33 | 1.13E-44 | 2.34E-40 | 0.11 | 0.02 | 0.08 | 4.57 |
| 10 | Rbbp8 | 0.77 | 4.39E-44 | 9.13E-40 | 0.73 | 0.55 | 0.18 | 1.32 |
| 10 | Mfsd6 | 2.03 | 5.25E-44 | 1.09E-39 | 0.13 | 0.03 | 0.09 | 3.91 |
| 10 | Igdcc3 | 1.39 | 7.55E-44 | 1.57E-39 | 0.42 | 0.23 | 0.19 | 1.80 |
| 10 | Igfbp3 | 1.28 | 1.05E-43 | 2.17E-39 | 0.29 | 0.12 | 0.17 | 2.38 |
| 10 | Cdc42se1 | 0.85 | 1.16E-43 | 2.41E-39 | 0.63 | 0.42 | 0.21 | 1.49 |
| 10 | Pak2 | 0.48 | 1.19E-43 | 2.48E-39 | 0.93 | 0.83 | 0.10 | 1.11 |
| 10 | Kank1 | 1.19 | 1.37E-43 | 2.86E-39 | 0.34 | 0.15 | 0.18 | 2.19 |
| 10 | Cdk6 | 0.95 | 1.87E-43 | 3.89E-39 | 0.55 | 0.33 | 0.22 | 1.66 |
| 10 | Mest | 0.49 | 2.09E-43 | 4.35E-39 | 0.95 | 0.91 | 0.04 | 1.05 |
| 10 | Sparc | 0.47 | 2.18E-43 | 4.53E-39 | 0.92 | 0.74 | 0.18 | 1.24 |
| 10 | Plekhh1 | 1.70 | 2.85E-43 | 5.93E-39 | 0.19 | 0.06 | 0.12 | 3.00 |
| 10 | Usp46 | 0.64 | 3.46E-43 | 7.20E-39 | 0.81 | 0.62 | 0.19 | 1.31 |
| 10 | Tmem159 | 0.98 | 3.65E-43 | 7.60E-39 | 0.47 | 0.26 | 0.21 | 1.83 |
| 10 | Hist3h2a | 1.02 | 3.75E-43 | 7.79E-39 | 0.47 | 0.26 | 0.21 | 1.80 |
| 10 | Rhbdf1 | 1.43 | 7.54E-43 | 1.57E-38 | 0.25 | 0.10 | 0.15 | 2.55 |
| 10 | Alad | 0.30 | 1.00E-42 | 2.08E-38 | 0.81 | 0.63 | 0.19 | 1.29 |
| 10 | Gpr137c | 1.31 | 1.10E-42 | 2.29E-38 | 0.31 | 0.14 | 0.17 | 2.24 |
| 10 | Tuft1 | 2.12 | 1.24E-42 | 2.57E-38 | 0.13 | 0.04 | 0.10 | 3.71 |
| 10 | Ptch2 | 2.55 | 2.09E-42 | 4.35E-38 | 0.12 | 0.03 | 0.09 | 3.88 |

**Supplementary Table 1. Marker genes for the original clusters of integrated clustering of E11.5 control and NotchGOF λ-junction cells.**

| | | | | | | | | |
|---|---|---|---|---|---|---|---|---|
| 10 | Purb | 0.49 | 2.10E-42 | 4.37E-38 | 0.93 | 0.86 | 0.07 | 1.08 |
| 10 | Hspa2 | 1.70 | 2.53E-42 | 5.26E-38 | 0.17 | 0.05 | 0.11 | 3.15 |
| 10 | Pcbd1 | 1.26 | 2.83E-42 | 5.89E-38 | 0.28 | 0.12 | 0.16 | 2.38 |
| 10 | Sh3tc2 | 2.28 | 3.90E-42 | 8.12E-38 | 0.10 | 0.02 | 0.08 | 4.43 |
| 10 | Plxnb1 | 1.03 | 3.94E-42 | 8.20E-38 | 0.46 | 0.26 | 0.21 | 1.80 |
| 10 | Cntnap2 | 1.77 | 9.25E-42 | 1.92E-37 | 0.16 | 0.05 | 0.11 | 3.27 |
| 10 | Lrrfip1 | 1.02 | 1.09E-41 | 2.26E-37 | 0.53 | 0.33 | 0.20 | 1.60 |
| 10 | Churc1 | 0.73 | 1.14E-41 | 2.36E-37 | 0.77 | 0.58 | 0.19 | 1.33 |
| 10 | Mmp11 | 1.08 | 1.44E-41 | 2.99E-37 | 0.52 | 0.31 | 0.21 | 1.67 |
| 10 | Snx5 | 0.51 | 2.63E-41 | 5.48E-37 | 0.89 | 0.77 | 0.11 | 1.14 |
| 10 | Arf1 | 0.34 | 2.64E-41 | 5.49E-37 | 0.99 | 0.98 | 0.01 | 1.01 |
| 10 | Pycard | 1.19 | 3.62E-41 | 7.54E-37 | 0.28 | 0.12 | 0.16 | 2.32 |
| 10 | H2-Q4 | 1.34 | 1.18E-40 | 2.45E-36 | 0.25 | 0.10 | 0.15 | 2.46 |
| 10 | Nedd9 | 1.23 | 1.24E-40 | 2.58E-36 | 0.34 | 0.16 | 0.18 | 2.09 |
| 10 | Myl12b | 0.45 | 1.29E-40 | 2.68E-36 | 0.94 | 0.88 | 0.06 | 1.07 |
| 10 | Irx1 | 1.58 | 1.42E-40 | 2.96E-36 | 0.16 | 0.05 | 0.11 | 3.13 |
| 10 | Arhgdig | 0.50 | 1.49E-40 | 3.11E-36 | 0.21 | 0.08 | 0.13 | 2.70 |
| 10 | Nap1l1 | 0.27 | 1.85E-40 | 3.86E-36 | 1.00 | 1.00 | 0.00 | 1.00 |
| 10 | Gm17108 | 1.87 | 2.10E-40 | 4.36E-36 | 0.14 | 0.04 | 0.10 | 3.35 |
| 10 | Tinagl1 | 1.69 | 2.21E-40 | 4.60E-36 | 0.17 | 0.06 | 0.11 | 3.02 |
| 10 | Dazap2 | 0.67 | 2.25E-40 | 4.67E-36 | 0.76 | 0.57 | 0.19 | 1.34 |
| 10 | Cuedc1 | 0.85 | 2.45E-40 | 5.10E-36 | 0.59 | 0.38 | 0.21 | 1.55 |
| 10 | Id4 | 0.63 | 3.15E-40 | 6.55E-36 | 0.52 | 0.30 | 0.22 | 1.74 |
| 10 | 1810037I17Rik | 0.49 | 3.62E-40 | 7.53E-36 | 0.90 | 0.83 | 0.07 | 1.08 |
| 10 | Pcdh1 | 1.64 | 8.58E-40 | 1.78E-35 | 0.16 | 0.05 | 0.11 | 3.23 |
| 10 | Vamp8 | 0.57 | 1.51E-39 | 3.14E-35 | 0.79 | 0.61 | 0.19 | 1.31 |
| 10 | Spon2 | 1.93 | 1.58E-39 | 3.29E-35 | 0.11 | 0.03 | 0.09 | 3.93 |
| 10 | Rreb1 | 1.11 | 3.25E-39 | 6.77E-35 | 0.39 | 0.20 | 0.19 | 1.92 |
| 10 | Ptpn14 | 1.24 | 5.44E-39 | 1.13E-34 | 0.31 | 0.15 | 0.17 | 2.14 |
| 10 | Cpm | 0.43 | 6.20E-39 | 1.29E-34 | 0.36 | 0.17 | 0.19 | 2.11 |
| 10 | Fam83g | 1.97 | 1.05E-38 | 2.19E-34 | 0.12 | 0.03 | 0.09 | 3.67 |
| 10 | Slc39a10 | 0.62 | 1.08E-38 | 2.25E-34 | 0.81 | 0.68 | 0.14 | 1.20 |
| 10 | Rassf7 | 1.13 | 1.57E-38 | 3.26E-34 | 0.38 | 0.19 | 0.19 | 1.96 |
| 10 | Pard6b | 1.86 | 1.61E-38 | 3.34E-34 | 0.15 | 0.05 | 0.11 | 3.14 |
| 10 | Ell3 | 1.80 | 1.64E-38 | 3.42E-34 | 0.11 | 0.03 | 0.08 | 3.96 |
| 10 | Patz1 | 0.70 | 2.78E-38 | 5.78E-34 | 0.70 | 0.52 | 0.19 | 1.36 |
| 10 | Galnt18 | 1.68 | 2.78E-38 | 5.79E-34 | 0.12 | 0.03 | 0.09 | 3.74 |
| 10 | Pcgf5 | 1.49 | 2.85E-38 | 5.93E-34 | 0.21 | 0.08 | 0.13 | 2.57 |
| 10 | Golga4 | 0.61 | 3.38E-38 | 7.04E-34 | 0.79 | 0.64 | 0.15 | 1.24 |
| 10 | Plagl2 | 0.93 | 3.49E-38 | 7.27E-34 | 0.54 | 0.35 | 0.19 | 1.54 |
| 10 | Aktip | 0.90 | 3.94E-38 | 8.20E-34 | 0.49 | 0.28 | 0.20 | 1.72 |
| 10 | Junb | 1.29 | 4.81E-38 | 1.00E-33 | 0.26 | 0.11 | 0.15 | 2.32 |
| 10 | Col2a1 | 0.43 | 4.83E-38 | 1.00E-33 | 0.85 | 0.64 | 0.21 | 1.33 |
| 10 | Kazn | 1.08 | 5.61E-38 | 1.17E-33 | 0.36 | 0.18 | 0.18 | 2.01 |
| 10 | Pard3 | 0.65 | 6.35E-38 | 1.32E-33 | 0.75 | 0.57 | 0.17 | 1.30 |
| 10 | Fgfr3 | 1.73 | 6.50E-38 | 1.35E-33 | 0.13 | 0.04 | 0.09 | 3.57 |
| 10 | Fam3c | 0.90 | 6.78E-38 | 1.41E-33 | 0.55 | 0.36 | 0.20 | 1.54 |
| 10 | Amotl1 | 0.58 | 7.89E-38 | 1.64E-33 | 0.84 | 0.68 | 0.15 | 1.22 |
| 10 | Sik1 | 1.33 | 7.96E-38 | 1.66E-33 | 0.24 | 0.10 | 0.14 | 2.40 |
| 10 | Knop1 | 0.40 | 9.31E-38 | 1.94E-33 | 0.96 | 0.90 | 0.06 | 1.06 |
| 10 | Fzd3 | 0.64 | 2.35E-37 | 4.88E-33 | 0.76 | 0.58 | 0.18 | 1.31 |
| 10 | Actn4 | 0.64 | 2.81E-37 | 5.85E-33 | 0.82 | 0.70 | 0.12 | 1.17 |
| 10 | Kxd1 | 0.70 | 5.01E-37 | 1.04E-32 | 0.68 | 0.50 | 0.18 | 1.37 |
| 10 | Pgk1 | 0.37 | 5.71E-37 | 1.19E-32 | 0.99 | 0.95 | 0.03 | 1.03 |
| 10 | Trim2 | 0.80 | 6.14E-37 | 1.28E-32 | 0.62 | 0.41 | 0.21 | 1.51 |
| 10 | Slc22a23 | 1.17 | 6.88E-37 | 1.43E-32 | 0.23 | 0.09 | 0.14 | 2.49 |

**Supplementary Table 1. Marker genes for the original clusters of integrated clustering of E11.5 control and NotchGOF λ-junction cells.**

| | | | | | | | | |
|---|---|---|---|---|---|---|---|---|
| 10 | Lamc1 | 0.83 | 9.53E-37 | 1.98E-32 | 0.46 | 0.25 | 0.20 | 1.80 |
| 10 | Lurap1l | 1.07 | 1.65E-36 | 3.44E-32 | 0.20 | 0.07 | 0.13 | 2.71 |
| 10 | Arhgap18 | 0.85 | 2.11E-36 | 4.40E-32 | 0.27 | 0.12 | 0.15 | 2.26 |
| 10 | Zfp704 | 0.70 | 2.27E-36 | 4.71E-32 | 0.66 | 0.46 | 0.19 | 1.42 |
| 10 | Zbtb18 | 1.26 | 2.54E-36 | 5.29E-32 | 0.25 | 0.11 | 0.14 | 2.35 |
| 10 | Ankrd49 | 0.75 | 2.89E-36 | 6.02E-32 | 0.62 | 0.43 | 0.19 | 1.45 |
| 10 | Jade1 | 1.16 | 3.64E-36 | 7.58E-32 | 0.34 | 0.17 | 0.17 | 1.99 |
| 10 | Nol7 | 0.39 | 3.75E-36 | 7.80E-32 | 0.94 | 0.88 | 0.06 | 1.07 |
| 10 | Pomp | 0.30 | 5.09E-36 | 1.06E-31 | 0.99 | 0.98 | 0.01 | 1.01 |
| 10 | Nav3 | 0.79 | 5.48E-36 | 1.14E-31 | 0.28 | 0.12 | 0.16 | 2.27 |
| 10 | Gas7 | 1.49 | 5.62E-36 | 1.17E-31 | 0.15 | 0.05 | 0.10 | 3.10 |
| 10 | Tnks2 | 0.46 | 7.28E-36 | 1.51E-31 | 0.92 | 0.81 | 0.11 | 1.14 |
| 10 | Heyl | 1.31 | 8.15E-36 | 1.69E-31 | 0.16 | 0.05 | 0.11 | 3.00 |
| 10 | Otx1 | 0.95 | 8.19E-36 | 1.70E-31 | 0.26 | 0.11 | 0.15 | 2.37 |
| 10 | Lypd6 | 1.20 | 1.15E-35 | 2.40E-31 | 0.28 | 0.12 | 0.15 | 2.24 |
| 10 | Dmtn | 1.62 | 1.19E-35 | 2.47E-31 | 0.17 | 0.06 | 0.11 | 2.79 |
| 10 | Nckap5 | 1.24 | 1.64E-35 | 3.40E-31 | 0.22 | 0.09 | 0.13 | 2.49 |
| 10 | Ank3 | 0.76 | 1.64E-35 | 3.41E-31 | 0.67 | 0.48 | 0.19 | 1.38 |
| 10 | Cd164l2 | 1.88 | 1.66E-35 | 3.45E-31 | 0.13 | 0.04 | 0.09 | 3.37 |
| 10 | Scaper | 1.06 | 1.82E-35 | 3.79E-31 | 0.48 | 0.29 | 0.19 | 1.66 |
| 10 | Nedd4l | 1.23 | 2.06E-35 | 4.28E-31 | 0.30 | 0.14 | 0.16 | 2.11 |
| 10 | Tjp1 | 0.71 | 2.63E-35 | 5.46E-31 | 0.67 | 0.49 | 0.18 | 1.36 |
| 10 | Fgd4 | 1.18 | 3.61E-35 | 7.51E-31 | 0.22 | 0.09 | 0.13 | 2.51 |
| 10 | Ahr | 1.23 | 4.27E-35 | 8.89E-31 | 0.21 | 0.09 | 0.13 | 2.51 |
| 10 | Zfp568 | 1.06 | 5.20E-35 | 1.08E-30 | 0.44 | 0.27 | 0.18 | 1.67 |
| 10 | Fam89a | 1.31 | 5.41E-35 | 1.12E-30 | 0.19 | 0.07 | 0.12 | 2.69 |
| 10 | Mbnl3 | 1.21 | 6.12E-35 | 1.27E-30 | 0.27 | 0.12 | 0.15 | 2.21 |
| 10 | Plek2 | 1.59 | 6.33E-35 | 1.32E-30 | 0.17 | 0.06 | 0.11 | 2.84 |
| 10 | Ehbp1 | 0.90 | 1.55E-34 | 3.23E-30 | 0.42 | 0.23 | 0.19 | 1.79 |
| 10 | Dennd1c | 1.95 | 2.23E-34 | 4.63E-30 | 0.11 | 0.03 | 0.08 | 3.67 |
| 10 | Unc13b | 1.39 | 3.46E-34 | 7.19E-30 | 0.20 | 0.08 | 0.12 | 2.54 |
| 10 | Col17a1 | 1.55 | 3.71E-34 | 7.72E-30 | 0.12 | 0.03 | 0.08 | 3.44 |
| 10 | Ifitm1 | 1.49 | 3.76E-34 | 7.82E-30 | 0.34 | 0.18 | 0.16 | 1.85 |
| 10 | Emp1 | 0.98 | 4.07E-34 | 8.47E-30 | 0.28 | 0.13 | 0.15 | 2.20 |
| 10 | Ccdc112 | 0.67 | 4.95E-34 | 1.03E-29 | 0.69 | 0.51 | 0.18 | 1.36 |
| 10 | Cadps2 | 1.66 | 6.01E-34 | 1.25E-29 | 0.12 | 0.03 | 0.08 | 3.44 |
| 10 | Clip4 | 1.52 | 6.23E-34 | 1.30E-29 | 0.14 | 0.04 | 0.09 | 3.14 |
| 10 | Flot2 | 1.03 | 6.29E-34 | 1.31E-29 | 0.38 | 0.21 | 0.17 | 1.83 |
| 10 | Fam162a | 0.45 | 7.00E-34 | 1.46E-29 | 0.93 | 0.89 | 0.04 | 1.04 |
| 10 | Sh3d19 | 0.87 | 8.18E-34 | 1.70E-29 | 0.44 | 0.25 | 0.19 | 1.75 |
| 10 | Aprt | 0.38 | 9.34E-34 | 1.94E-29 | 0.98 | 0.92 | 0.05 | 1.06 |
| 10 | Imp3 | 0.49 | 1.09E-33 | 2.27E-29 | 0.88 | 0.79 | 0.09 | 1.12 |
| 10 | Ak4 | 1.30 | 1.39E-33 | 2.89E-29 | 0.23 | 0.10 | 0.13 | 2.32 |
| 10 | Cryl1 | 1.10 | 1.45E-33 | 3.03E-29 | 0.30 | 0.15 | 0.16 | 2.08 |
| 10 | Sft2d2 | 1.16 | 1.48E-33 | 3.08E-29 | 0.30 | 0.15 | 0.15 | 2.05 |
| 10 | Xpa | 0.72 | 2.34E-33 | 4.87E-29 | 0.63 | 0.43 | 0.20 | 1.46 |
| 10 | Ext1 | 0.84 | 2.96E-33 | 6.15E-29 | 0.59 | 0.41 | 0.18 | 1.45 |
| 10 | Adamts17 | 1.98 | 2.96E-33 | 6.16E-29 | 0.12 | 0.03 | 0.08 | 3.41 |
| 10 | Ppfibp1 | 0.60 | 4.05E-33 | 8.42E-29 | 0.63 | 0.42 | 0.21 | 1.49 |
| 10 | Tgfb1 | 0.85 | 4.16E-33 | 8.66E-29 | 0.24 | 0.10 | 0.14 | 2.31 |
| 10 | Ahnak | 0.84 | 4.37E-33 | 9.09E-29 | 0.63 | 0.45 | 0.17 | 1.38 |
| 10 | Capn6 | 0.71 | 5.39E-33 | 1.12E-28 | 0.70 | 0.54 | 0.16 | 1.30 |
| 10 | Krcc1 | 0.63 | 6.26E-33 | 1.30E-28 | 0.73 | 0.57 | 0.16 | 1.28 |
| 10 | Tnfrsf12a | 1.52 | 6.34E-33 | 1.32E-28 | 0.19 | 0.07 | 0.12 | 2.55 |
| 10 | Slc12a4 | 1.17 | 1.00E-32 | 2.08E-28 | 0.28 | 0.14 | 0.15 | 2.09 |
| 10 | Arfgef1 | 0.74 | 1.14E-32 | 2.37E-28 | 0.67 | 0.48 | 0.18 | 1.38 |

**Supplementary Table 1. Marker genes for the original clusters of integrated clustering of E11.5 control and NotchGOF λ-junction cells.**

| | | | | | | | | |
|---|---|---|---|---|---|---|---|---|
| 10 | Macrod2 | 0.96 | 1.27E-32 | 2.65E-28 | 0.39 | 0.22 | 0.17 | 1.80 |
| 10 | Exosc7 | 0.55 | 1.37E-32 | 2.85E-28 | 0.77 | 0.62 | 0.15 | 1.25 |
| 10 | Pfkl | 0.56 | 1.53E-32 | 3.19E-28 | 0.79 | 0.64 | 0.15 | 1.23 |
| 10 | Sostdc1 | 0.65 | 1.57E-32 | 3.27E-28 | 0.30 | 0.14 | 0.16 | 2.11 |
| 10 | Tpp1 | 0.92 | 1.81E-32 | 3.76E-28 | 0.44 | 0.26 | 0.18 | 1.68 |
| 10 | Clstn1 | 0.77 | 1.99E-32 | 4.14E-28 | 0.57 | 0.38 | 0.19 | 1.50 |
| 10 | Abcg1 | 1.10 | 2.42E-32 | 5.04E-28 | 0.20 | 0.08 | 0.12 | 2.49 |
| 10 | Hsdl2 | 1.08 | 2.52E-32 | 5.25E-28 | 0.43 | 0.26 | 0.17 | 1.64 |
| 10 | Itgb6 | 1.80 | 2.57E-32 | 5.35E-28 | 0.13 | 0.04 | 0.09 | 3.15 |
| 10 | Nectin3 | 0.67 | 2.82E-32 | 5.86E-28 | 0.70 | 0.54 | 0.16 | 1.29 |
| 10 | Ide | 0.87 | 3.40E-32 | 7.07E-28 | 0.45 | 0.26 | 0.18 | 1.69 |
| 10 | Fuom | 0.84 | 5.25E-32 | 1.09E-27 | 0.45 | 0.27 | 0.18 | 1.69 |
| 10 | Mark2 | 0.71 | 5.38E-32 | 1.12E-27 | 0.64 | 0.46 | 0.18 | 1.38 |
| 10 | Mettl27 | 1.05 | 9.55E-32 | 1.99E-27 | 0.29 | 0.14 | 0.15 | 2.08 |
| 10 | Pnck | 1.38 | 9.79E-32 | 2.04E-27 | 0.18 | 0.07 | 0.11 | 2.62 |
| 10 | Scp2 | 0.47 | 1.44E-31 | 3.00E-27 | 0.86 | 0.77 | 0.09 | 1.12 |
| 10 | Nudt18 | 1.46 | 1.68E-31 | 3.49E-27 | 0.16 | 0.06 | 0.10 | 2.76 |
| 10 | Fat1 | 0.67 | 1.71E-31 | 3.56E-27 | 0.68 | 0.51 | 0.18 | 1.35 |
| 10 | Plch1 | 1.01 | 1.98E-31 | 4.13E-27 | 0.23 | 0.10 | 0.13 | 2.34 |
| 10 | C330021F23Rik | 1.88 | 2.86E-31 | 5.94E-27 | 0.10 | 0.03 | 0.07 | 3.52 |
| 10 | Sfr1 | 0.28 | 3.11E-31 | 6.47E-27 | 0.99 | 0.98 | 0.01 | 1.01 |
| 10 | Prr5l | 0.93 | 3.39E-31 | 7.05E-27 | 0.32 | 0.16 | 0.16 | 1.98 |
| 10 | Ildr1 | 1.81 | 4.13E-31 | 8.60E-27 | 0.12 | 0.04 | 0.08 | 3.22 |
| 10 | Pnrc1 | 0.44 | 4.25E-31 | 8.84E-27 | 0.93 | 0.87 | 0.06 | 1.07 |
| 10 | Chchd10 | 0.93 | 5.19E-31 | 1.08E-26 | 0.31 | 0.16 | 0.16 | 2.01 |
| 10 | Plagl1 | 0.32 | 6.15E-31 | 1.28E-26 | 0.66 | 0.43 | 0.24 | 1.55 |
| 10 | Chst11 | 1.50 | 6.35E-31 | 1.32E-26 | 0.18 | 0.07 | 0.11 | 2.52 |
| 10 | Vmac | 1.20 | 6.42E-31 | 1.34E-26 | 0.21 | 0.09 | 0.12 | 2.34 |
| 10 | Tet3 | 0.60 | 9.01E-31 | 1.87E-26 | 0.72 | 0.57 | 0.16 | 1.28 |
| 10 | Erbb2 | 0.89 | 9.23E-31 | 1.92E-26 | 0.40 | 0.23 | 0.17 | 1.75 |
| 10 | Abhd14b | 1.49 | 9.63E-31 | 2.00E-26 | 0.16 | 0.06 | 0.10 | 2.75 |
| 10 | Ybx3 | 0.42 | 1.04E-30 | 2.16E-26 | 0.92 | 0.84 | 0.08 | 1.10 |
| 10 | Kctd11 | 1.61 | 1.25E-30 | 2.60E-26 | 0.13 | 0.04 | 0.09 | 3.02 |
| 10 | Mpst | 0.70 | 1.31E-30 | 2.72E-26 | 0.60 | 0.41 | 0.18 | 1.45 |
| 10 | Cemip2 | 0.74 | 1.53E-30 | 3.17E-26 | 0.61 | 0.45 | 0.17 | 1.37 |
| 10 | Pclo | 0.85 | 2.25E-30 | 4.69E-26 | 0.24 | 0.10 | 0.13 | 2.28 |
| 10 | Trim44 | 0.49 | 2.33E-30 | 4.85E-26 | 0.83 | 0.71 | 0.12 | 1.17 |
| 10 | Pls1 | 1.00 | 2.50E-30 | 5.20E-26 | 0.22 | 0.09 | 0.12 | 2.35 |
| 10 | Tmem123 | 0.76 | 2.64E-30 | 5.49E-26 | 0.50 | 0.32 | 0.18 | 1.57 |
| 10 | Plxnc1 | 1.63 | 2.86E-30 | 5.95E-26 | 0.13 | 0.04 | 0.09 | 3.13 |
| 10 | Col4a1 | 0.35 | 2.97E-30 | 6.17E-26 | 0.80 | 0.61 | 0.19 | 1.31 |
| 10 | Fam173a | 0.70 | 3.66E-30 | 7.62E-26 | 0.65 | 0.50 | 0.15 | 1.30 |
| 10 | Kremen1 | 1.01 | 3.92E-30 | 8.16E-26 | 0.29 | 0.14 | 0.15 | 2.04 |
| 10 | Micu2 | 0.64 | 4.87E-30 | 1.01E-25 | 0.67 | 0.50 | 0.17 | 1.34 |
| 10 | Psap | 0.51 | 5.20E-30 | 1.08E-25 | 0.77 | 0.61 | 0.16 | 1.25 |
| 10 | Acsl5 | 1.04 | 7.45E-30 | 1.55E-25 | 0.30 | 0.15 | 0.15 | 1.97 |
| 10 | Erich1 | 0.86 | 1.36E-29 | 2.83E-25 | 0.40 | 0.23 | 0.17 | 1.74 |
| 10 | Mcee | 0.67 | 1.45E-29 | 3.03E-25 | 0.62 | 0.43 | 0.19 | 1.43 |
| 10 | Prkaa2 | 1.72 | 1.78E-29 | 3.71E-25 | 0.10 | 0.03 | 0.07 | 3.48 |
| 10 | Prkcz | 1.18 | 2.79E-29 | 5.80E-25 | 0.23 | 0.10 | 0.13 | 2.23 |
| 10 | Slc38a4 | 1.08 | 2.90E-29 | 6.03E-25 | 0.43 | 0.27 | 0.16 | 1.59 |
| 10 | Pafah1b3 | 0.32 | 2.94E-29 | 6.11E-25 | 0.98 | 0.92 | 0.06 | 1.06 |
| 10 | Hspa4l | 1.15 | 2.99E-29 | 6.22E-25 | 0.23 | 0.11 | 0.13 | 2.17 |
| 10 | Rtn3 | 0.43 | 4.32E-29 | 8.98E-25 | 0.87 | 0.79 | 0.09 | 1.11 |
| 10 | Reep3 | 0.63 | 4.66E-29 | 9.69E-25 | 0.69 | 0.52 | 0.17 | 1.33 |
| 10 | Bnip3l | 0.32 | 4.73E-29 | 9.84E-25 | 0.97 | 0.93 | 0.03 | 1.04 |

**Supplementary Table 1. Marker genes for the original clusters of integrated clustering of E11.5 control and NotchGOF λ-junction cells.**

| | | | | | | | | |
|---|---|---|---|---|---|---|---|---|
| 10 | Sh3rf1 | 0.94 | 7.41E-29 | 1.54E-24 | 0.39 | 0.22 | 0.16 | 1.74 |
| 10 | Grip1 | 1.32 | 8.08E-29 | 1.68E-24 | 0.23 | 0.10 | 0.12 | 2.21 |
| 10 | Ebf4 | 1.41 | 8.81E-29 | 1.83E-24 | 0.15 | 0.06 | 0.10 | 2.73 |
| 10 | Maml2 | 1.18 | 1.16E-28 | 2.41E-24 | 0.21 | 0.09 | 0.12 | 2.33 |
| 10 | Rap2c | 0.60 | 1.26E-28 | 2.61E-24 | 0.74 | 0.59 | 0.15 | 1.25 |
| 10 | Prr36 | 1.26 | 1.50E-28 | 3.12E-24 | 0.20 | 0.09 | 0.11 | 2.33 |
| 10 | Dbp | 1.11 | 1.50E-28 | 3.12E-24 | 0.27 | 0.14 | 0.14 | 2.02 |
| 10 | Nhsl1 | 0.88 | 1.62E-28 | 3.37E-24 | 0.38 | 0.21 | 0.16 | 1.76 |
| 10 | Polr3g | 0.90 | 2.00E-28 | 4.17E-24 | 0.44 | 0.27 | 0.17 | 1.61 |
| 10 | Tbc1d8 | 1.82 | 2.24E-28 | 4.66E-24 | 0.10 | 0.03 | 0.07 | 3.37 |
| 10 | Rabgap1 | 0.61 | 3.62E-28 | 7.52E-24 | 0.72 | 0.56 | 0.15 | 1.27 |
| 10 | Bid | 1.09 | 4.21E-28 | 8.75E-24 | 0.25 | 0.12 | 0.13 | 2.11 |
| 10 | Cpz | 1.18 | 4.92E-28 | 1.02E-23 | 0.17 | 0.07 | 0.10 | 2.54 |
| 10 | Dcbld1 | 0.83 | 5.37E-28 | 1.12E-23 | 0.47 | 0.30 | 0.17 | 1.58 |
| 10 | Vwa5a | 1.30 | 6.98E-28 | 1.45E-23 | 0.20 | 0.08 | 0.11 | 2.35 |
| 10 | Bdh1 | 1.40 | 7.90E-28 | 1.64E-23 | 0.17 | 0.07 | 0.10 | 2.50 |
| 10 | Zdhhc12 | 0.82 | 1.12E-27 | 2.33E-23 | 0.43 | 0.26 | 0.17 | 1.67 |
| 10 | Tead3 | 0.81 | 1.51E-27 | 3.13E-23 | 0.41 | 0.24 | 0.17 | 1.69 |
| 10 | Vps36 | 0.48 | 1.72E-27 | 3.59E-23 | 0.88 | 0.79 | 0.09 | 1.11 |
| 10 | Pnpla2 | 1.26 | 1.82E-27 | 3.78E-23 | 0.20 | 0.09 | 0.11 | 2.29 |
| 10 | Lmo1 | 1.11 | 2.68E-27 | 5.57E-23 | 0.18 | 0.07 | 0.10 | 2.44 |
| 10 | Sipa1l3 | 1.18 | 3.15E-27 | 6.55E-23 | 0.22 | 0.10 | 0.12 | 2.18 |
| 10 | Aagab | 1.07 | 3.50E-27 | 7.28E-23 | 0.31 | 0.17 | 0.14 | 1.83 |
| 10 | Slc39a8 | 0.91 | 4.68E-27 | 9.74E-23 | 0.31 | 0.17 | 0.15 | 1.89 |
| 10 | Zfp36l1 | 0.38 | 5.08E-27 | 1.06E-22 | 0.94 | 0.85 | 0.09 | 1.11 |
| 10 | Hltf | 0.81 | 5.93E-27 | 1.23E-22 | 0.48 | 0.31 | 0.17 | 1.54 |
| 10 | Mid1ip1 | 1.07 | 6.82E-27 | 1.42E-22 | 0.37 | 0.21 | 0.15 | 1.71 |
| 10 | Lrrc8d | 0.99 | 8.63E-27 | 1.79E-22 | 0.34 | 0.19 | 0.15 | 1.78 |
| 10 | Prrg3 | 1.08 | 1.17E-26 | 2.43E-22 | 0.23 | 0.11 | 0.12 | 2.14 |
| 10 | Dmrt3 | 1.32 | 1.39E-26 | 2.89E-22 | 0.14 | 0.05 | 0.09 | 2.69 |
| 10 | Pard6g | 0.52 | 1.40E-26 | 2.91E-22 | 0.75 | 0.59 | 0.16 | 1.27 |
| 10 | Arid3a | 0.87 | 1.44E-26 | 3.00E-22 | 0.49 | 0.33 | 0.16 | 1.47 |
| 10 | Lonrf1 | 1.26 | 2.24E-26 | 4.66E-22 | 0.20 | 0.09 | 0.11 | 2.23 |
| 10 | Sh3gl2 | 1.52 | 2.46E-26 | 5.11E-22 | 0.11 | 0.04 | 0.08 | 3.03 |
| 10 | Zmiz2 | 0.74 | 2.69E-26 | 5.60E-22 | 0.47 | 0.31 | 0.17 | 1.55 |
| 10 | Ppp1r9a | 0.69 | 2.81E-26 | 5.85E-22 | 0.56 | 0.39 | 0.17 | 1.44 |
| 10 | Sox13 | 1.22 | 2.94E-26 | 6.11E-22 | 0.19 | 0.08 | 0.11 | 2.30 |
| 10 | Rcan1 | 0.83 | 2.99E-26 | 6.23E-22 | 0.42 | 0.26 | 0.16 | 1.63 |
| 10 | Cbx6 | 0.86 | 3.59E-26 | 7.47E-22 | 0.42 | 0.27 | 0.16 | 1.58 |
| 10 | Cdc42ep5 | 0.69 | 4.86E-26 | 1.01E-21 | 0.62 | 0.46 | 0.16 | 1.34 |
| 10 | Baz2a | 0.72 | 6.64E-26 | 1.38E-21 | 0.54 | 0.38 | 0.16 | 1.41 |
| 10 | Cmas | 0.64 | 7.99E-26 | 1.66E-21 | 0.71 | 0.59 | 0.12 | 1.20 |
| 10 | Sh3glb2 | 0.77 | 8.32E-26 | 1.73E-21 | 0.48 | 0.32 | 0.17 | 1.53 |
| 10 | Bnipl | 1.52 | 1.23E-25 | 2.56E-21 | 0.12 | 0.04 | 0.08 | 2.84 |
| 10 | Bcl11b | 0.54 | 1.25E-25 | 2.59E-21 | 0.74 | 0.59 | 0.16 | 1.27 |
| 10 | Mettl26 | 0.59 | 1.39E-25 | 2.89E-21 | 0.64 | 0.49 | 0.15 | 1.31 |
| 10 | Copg2 | 0.68 | 1.50E-25 | 3.12E-21 | 0.53 | 0.37 | 0.17 | 1.45 |
| 10 | Gtf3a | 0.50 | 2.05E-25 | 4.26E-21 | 0.76 | 0.64 | 0.12 | 1.19 |
| 10 | 5430403G16Rik | 1.41 | 2.70E-25 | 5.62E-21 | 0.16 | 0.07 | 0.10 | 2.41 |
| 10 | Txndc16 | 0.98 | 2.83E-25 | 5.89E-21 | 0.27 | 0.14 | 0.13 | 1.94 |
| 10 | Cdkl5 | 1.44 | 3.27E-25 | 6.80E-21 | 0.14 | 0.05 | 0.09 | 2.61 |
| 10 | Rian | 0.63 | 4.13E-25 | 8.59E-21 | 0.49 | 0.31 | 0.17 | 1.56 |
| 10 | Trim25 | 1.40 | 4.19E-25 | 8.71E-21 | 0.15 | 0.06 | 0.09 | 2.56 |
| 10 | Npepps | 0.49 | 5.80E-25 | 1.21E-20 | 0.76 | 0.64 | 0.12 | 1.18 |
| 10 | Tfap2a | 0.59 | 5.83E-25 | 1.21E-20 | 0.37 | 0.21 | 0.16 | 1.77 |
| 10 | Rassf9 | 0.95 | 6.22E-25 | 1.29E-20 | 0.30 | 0.16 | 0.14 | 1.86 |

**Supplementary Table 1. Marker genes for the original clusters of integrated clustering of E11.5 control and NotchGOF λ-junction cells.**

| | | | | | | | | |
|---|---|---|---|---|---|---|---|---|
| 10 | Elavl2 | 0.66 | 6.34E-25 | 1.32E-20 | 0.33 | 0.18 | 0.15 | 1.81 |
| 10 | Amt | 0.92 | 7.40E-25 | 1.54E-20 | 0.28 | 0.14 | 0.13 | 1.93 |
| 10 | Nab1 | 1.11 | 8.93E-25 | 1.86E-20 | 0.25 | 0.13 | 0.12 | 1.98 |
| 10 | Cbarp | 1.21 | 1.06E-24 | 2.21E-20 | 0.22 | 0.11 | 0.11 | 2.08 |
| 10 | Mtss1 | 0.73 | 1.26E-24 | 2.61E-20 | 0.46 | 0.29 | 0.17 | 1.58 |
| 10 | Alcam | 0.84 | 1.53E-24 | 3.17E-20 | 0.27 | 0.14 | 0.13 | 1.94 |
| 10 | E330009J07Rik | 1.09 | 1.82E-24 | 3.78E-20 | 0.23 | 0.12 | 0.12 | 2.03 |
| 10 | Ccdc120 | 1.07 | 2.48E-24 | 5.16E-20 | 0.22 | 0.10 | 0.11 | 2.11 |
| 10 | Cd3eap | 0.62 | 2.71E-24 | 5.64E-20 | 0.67 | 0.52 | 0.15 | 1.29 |
| 10 | Map3k5 | 1.71 | 3.13E-24 | 6.50E-20 | 0.10 | 0.03 | 0.07 | 3.06 |
| 10 | Pde5a | 0.95 | 3.25E-24 | 6.76E-20 | 0.32 | 0.18 | 0.14 | 1.78 |
| 10 | Rdm1 | 0.71 | 3.26E-24 | 6.78E-20 | 0.48 | 0.32 | 0.16 | 1.52 |
| 10 | Gm10260 | 0.38 | 3.89E-24 | 8.09E-20 | 0.92 | 0.85 | 0.07 | 1.08 |
| 10 | Skp1a | 0.28 | 3.92E-24 | 8.14E-20 | 0.98 | 0.96 | 0.03 | 1.03 |
| 10 | Plekhh3 | 1.74 | 4.88E-24 | 1.02E-19 | 0.10 | 0.03 | 0.07 | 3.00 |
| 10 | Syde2 | 1.31 | 5.32E-24 | 1.11E-19 | 0.17 | 0.07 | 0.10 | 2.35 |
| 10 | Stox2 | 0.52 | 5.33E-24 | 1.11E-19 | 0.72 | 0.56 | 0.16 | 1.29 |
| 10 | Glrb | 1.39 | 5.44E-24 | 1.13E-19 | 0.13 | 0.05 | 0.08 | 2.69 |
| 10 | Cpne3 | 0.57 | 6.14E-24 | 1.28E-19 | 0.66 | 0.52 | 0.15 | 1.28 |
| 10 | Arg2 | 1.26 | 6.14E-24 | 1.28E-19 | 0.13 | 0.05 | 0.08 | 2.63 |
| 10 | Cox14 | 0.35 | 6.28E-24 | 1.31E-19 | 0.89 | 0.80 | 0.09 | 1.11 |
| 10 | Kcnh8 | 1.55 | 6.62E-24 | 1.38E-19 | 0.10 | 0.03 | 0.07 | 3.13 |
| 10 | Ptpn4 | 0.95 | 8.17E-24 | 1.70E-19 | 0.29 | 0.16 | 0.13 | 1.85 |
| 10 | Fryl | 0.82 | 8.90E-24 | 1.85E-19 | 0.44 | 0.30 | 0.15 | 1.50 |
| 10 | Rims2 | 1.12 | 9.36E-24 | 1.95E-19 | 0.14 | 0.06 | 0.09 | 2.55 |
| 10 | Thbs1 | 0.54 | 1.15E-23 | 2.38E-19 | 0.24 | 0.12 | 0.12 | 2.05 |
| 10 | Arpc2 | 0.27 | 1.31E-23 | 2.72E-19 | 0.98 | 0.94 | 0.03 | 1.04 |
| 10 | Eef2kmt | 0.91 | 1.43E-23 | 2.97E-19 | 0.31 | 0.18 | 0.14 | 1.75 |
| 10 | Harbi1 | 0.88 | 1.81E-23 | 3.77E-19 | 0.32 | 0.18 | 0.14 | 1.76 |
| 10 | Baiap2 | 1.09 | 1.83E-23 | 3.80E-19 | 0.28 | 0.15 | 0.13 | 1.85 |
| 10 | Cacnb3 | 0.68 | 2.10E-23 | 4.37E-19 | 0.43 | 0.27 | 0.16 | 1.60 |
| 10 | Ptch1 | 0.50 | 2.20E-23 | 4.58E-19 | 0.63 | 0.45 | 0.18 | 1.41 |
| 10 | Rhov | 1.46 | 2.33E-23 | 4.84E-19 | 0.14 | 0.06 | 0.09 | 2.55 |
| 10 | Map7d2 | 1.11 | 2.36E-23 | 4.90E-19 | 0.18 | 0.08 | 0.10 | 2.25 |
| 10 | Tceal5 | 0.90 | 2.52E-23 | 5.24E-19 | 0.28 | 0.15 | 0.13 | 1.88 |
| 10 | Nsmaf | 0.75 | 2.82E-23 | 5.86E-19 | 0.41 | 0.26 | 0.15 | 1.56 |
| 10 | Npdc1 | 0.98 | 3.38E-23 | 7.02E-19 | 0.22 | 0.10 | 0.11 | 2.09 |
| 10 | Col4a2 | 0.45 | 3.48E-23 | 7.23E-19 | 0.60 | 0.42 | 0.18 | 1.42 |
| 10 | Tst | 1.08 | 3.79E-23 | 7.88E-19 | 0.24 | 0.12 | 0.12 | 1.94 |
| 10 | Xist | 0.31 | 4.11E-23 | 8.54E-19 | 0.96 | 0.86 | 0.10 | 1.11 |
| 10 | Agap1 | 0.67 | 4.28E-23 | 8.90E-19 | 0.57 | 0.41 | 0.15 | 1.37 |
| 10 | Esrra | 0.98 | 4.46E-23 | 9.27E-19 | 0.24 | 0.12 | 0.12 | 1.99 |
| 10 | Eci1 | 0.59 | 5.81E-23 | 1.21E-18 | 0.65 | 0.52 | 0.13 | 1.25 |
| 10 | Agap3 | 0.71 | 7.18E-23 | 1.49E-18 | 0.51 | 0.36 | 0.15 | 1.43 |
| 10 | Map3k1 | 0.70 | 7.37E-23 | 1.53E-18 | 0.41 | 0.26 | 0.15 | 1.60 |
| 10 | Ube2e1 | 0.36 | 8.00E-23 | 1.66E-18 | 0.91 | 0.84 | 0.07 | 1.09 |
| 10 | Pim3 | 0.69 | 8.06E-23 | 1.68E-18 | 0.44 | 0.29 | 0.15 | 1.52 |
| 10 | Ociad2 | 0.89 | 8.73E-23 | 1.82E-18 | 0.31 | 0.18 | 0.14 | 1.76 |
| 10 | Mkrn1 | 0.51 | 9.18E-23 | 1.91E-18 | 0.60 | 0.42 | 0.17 | 1.41 |
| 10 | Gid4 | 0.72 | 9.23E-23 | 1.92E-18 | 0.48 | 0.33 | 0.15 | 1.45 |
| 10 | Zfand1 | 0.73 | 9.78E-23 | 2.04E-18 | 0.50 | 0.34 | 0.15 | 1.45 |
| 10 | Fam214a | 1.01 | 1.11E-22 | 2.30E-18 | 0.21 | 0.10 | 0.11 | 2.11 |
| 10 | Igsf9b | 1.38 | 1.13E-22 | 2.34E-18 | 0.14 | 0.05 | 0.08 | 2.52 |
| 10 | Igf2os | 1.61 | 1.17E-22 | 2.44E-18 | 0.11 | 0.04 | 0.07 | 2.86 |
| 10 | H2-T23 | 1.53 | 1.39E-22 | 2.88E-18 | 0.13 | 0.05 | 0.08 | 2.52 |
| 10 | Acly | 0.52 | 1.80E-22 | 3.74E-18 | 0.73 | 0.57 | 0.15 | 1.26 |

**Supplementary Table 1. Marker genes for the original clusters of integrated clustering of E11.5 control and NotchGOF λ-junction cells.**

| | | | | | | | | |
|---|---|---|---|---|---|---|---|---|
| 10 | Exoc6 | 1.21 | 1.88E-22 | 3.91E-18 | 0.16 | 0.07 | 0.09 | 2.28 |
| 10 | Syngr2 | 0.97 | 2.26E-22 | 4.69E-18 | 0.26 | 0.14 | 0.12 | 1.89 |
| 10 | Aftph | 0.79 | 2.30E-22 | 4.78E-18 | 0.39 | 0.25 | 0.15 | 1.59 |
| 10 | Wdsub1 | 0.78 | 2.39E-22 | 4.97E-18 | 0.35 | 0.20 | 0.14 | 1.70 |
| 10 | Nfat5 | 0.50 | 2.43E-22 | 5.05E-18 | 0.72 | 0.57 | 0.14 | 1.25 |
| 10 | Rasef | 1.57 | 2.44E-22 | 5.07E-18 | 0.11 | 0.04 | 0.07 | 2.82 |
| 10 | Luzp1 | 0.40 | 2.55E-22 | 5.30E-18 | 0.86 | 0.77 | 0.10 | 1.13 |
| 10 | Zdhhc21 | 0.69 | 2.59E-22 | 5.39E-18 | 0.44 | 0.29 | 0.16 | 1.55 |
| 10 | Cyfip1 | 0.42 | 2.74E-22 | 5.69E-18 | 0.82 | 0.70 | 0.13 | 1.18 |
| 10 | Il10rb | 1.20 | 3.34E-22 | 6.95E-18 | 0.18 | 0.08 | 0.10 | 2.18 |
| 10 | Cndp2 | 0.67 | 4.16E-22 | 8.65E-18 | 0.52 | 0.37 | 0.15 | 1.40 |
| 10 | Hivep2 | 1.08 | 4.35E-22 | 9.05E-18 | 0.17 | 0.07 | 0.10 | 2.30 |
| 10 | Pak1 | 0.53 | 4.38E-22 | 9.11E-18 | 0.64 | 0.49 | 0.16 | 1.32 |
| 10 | Shroom2 | 1.30 | 4.51E-22 | 9.39E-18 | 0.17 | 0.08 | 0.09 | 2.22 |
| 10 | Gpi1 | 0.33 | 4.60E-22 | 9.57E-18 | 0.91 | 0.84 | 0.07 | 1.09 |
| 10 | Mttp | 1.01 | 4.86E-22 | 1.01E-17 | 0.25 | 0.13 | 0.12 | 1.90 |
| 10 | Bcl7a | 0.44 | 5.10E-22 | 1.06E-17 | 0.78 | 0.67 | 0.12 | 1.17 |
| 10 | Pex2 | 0.56 | 5.47E-22 | 1.14E-17 | 0.62 | 0.47 | 0.15 | 1.31 |
| 10 | Hmgn5 | 0.44 | 5.64E-22 | 1.17E-17 | 0.79 | 0.71 | 0.08 | 1.12 |
| 10 | Khk | 1.45 | 6.08E-22 | 1.27E-17 | 0.15 | 0.06 | 0.09 | 2.33 |
| 10 | Smco4 | 0.45 | 6.64E-22 | 1.38E-17 | 0.66 | 0.48 | 0.18 | 1.36 |
| 10 | Efna5 | 0.60 | 8.58E-22 | 1.79E-17 | 0.61 | 0.47 | 0.14 | 1.31 |
| 10 | Jph1 | 1.17 | 8.88E-22 | 1.85E-17 | 0.16 | 0.07 | 0.09 | 2.32 |
| 10 | Exosc5 | 0.47 | 1.06E-21 | 2.21E-17 | 0.72 | 0.58 | 0.14 | 1.24 |
| 10 | Tbrg1 | 0.38 | 1.25E-21 | 2.59E-17 | 0.90 | 0.83 | 0.08 | 1.09 |
| 10 | Usp31 | 1.11 | 1.32E-21 | 2.74E-17 | 0.21 | 0.11 | 0.11 | 2.01 |
| 10 | Herc3 | 1.12 | 1.51E-21 | 3.14E-17 | 0.16 | 0.07 | 0.09 | 2.30 |
| 10 | Adi1 | 0.73 | 1.62E-21 | 3.37E-17 | 0.46 | 0.31 | 0.14 | 1.46 |
| 10 | Mxi1 | 0.59 | 2.06E-21 | 4.29E-17 | 0.57 | 0.42 | 0.15 | 1.35 |
| 10 | Fgfr1op2 | 0.48 | 2.09E-21 | 4.35E-17 | 0.79 | 0.68 | 0.12 | 1.17 |
| 10 | Fam210b | 0.73 | 2.30E-21 | 4.79E-17 | 0.34 | 0.20 | 0.14 | 1.71 |
| 10 | Epb41l4a | 1.08 | 2.32E-21 | 4.82E-17 | 0.21 | 0.10 | 0.11 | 2.02 |
| 10 | Tmem205 | 0.73 | 2.38E-21 | 4.95E-17 | 0.43 | 0.27 | 0.15 | 1.55 |
| 10 | Snx10 | 0.84 | 2.54E-21 | 5.29E-17 | 0.28 | 0.15 | 0.13 | 1.81 |
| 10 | Tbc1d10a | 0.90 | 3.02E-21 | 6.28E-17 | 0.33 | 0.19 | 0.13 | 1.69 |
| 10 | Ralgps1 | 1.34 | 3.29E-21 | 6.85E-17 | 0.11 | 0.04 | 0.07 | 2.67 |
| 10 | Ctso | 1.26 | 3.91E-21 | 8.13E-17 | 0.15 | 0.06 | 0.09 | 2.35 |
| 10 | Hunk | 0.57 | 4.04E-21 | 8.40E-17 | 0.64 | 0.49 | 0.15 | 1.30 |
| 10 | Mtmr7 | 1.33 | 4.22E-21 | 8.79E-17 | 0.13 | 0.05 | 0.08 | 2.55 |
| 10 | Atp6v0e2 | 1.39 | 4.56E-21 | 9.49E-17 | 0.13 | 0.05 | 0.08 | 2.51 |
| 10 | Sos1 | 0.72 | 4.87E-21 | 1.01E-16 | 0.49 | 0.35 | 0.14 | 1.40 |
| 10 | Nrk | 0.91 | 6.24E-21 | 1.30E-16 | 0.27 | 0.15 | 0.12 | 1.82 |
| 10 | Frs2 | 0.82 | 8.90E-21 | 1.85E-16 | 0.41 | 0.27 | 0.14 | 1.52 |
| 10 | Smarcd2 | 0.62 | 9.81E-21 | 2.04E-16 | 0.44 | 0.29 | 0.15 | 1.53 |
| 10 | Plekhf2 | 0.72 | 1.02E-20 | 2.12E-16 | 0.37 | 0.23 | 0.14 | 1.62 |
| 10 | Ptpn3 | 1.44 | 1.21E-20 | 2.51E-16 | 0.10 | 0.04 | 0.07 | 2.76 |
| 10 | Epha4 | 0.49 | 1.26E-20 | 2.63E-16 | 0.78 | 0.65 | 0.14 | 1.21 |
| 10 | Hook2 | 0.80 | 1.31E-20 | 2.73E-16 | 0.29 | 0.16 | 0.13 | 1.78 |
| 10 | Gls | 0.75 | 1.41E-20 | 2.92E-16 | 0.54 | 0.41 | 0.13 | 1.33 |
| 10 | Igf1r | 0.37 | 1.51E-20 | 3.15E-16 | 0.87 | 0.77 | 0.10 | 1.13 |
| 10 | Smim10l2a | 1.22 | 1.78E-20 | 3.71E-16 | 0.16 | 0.07 | 0.09 | 2.25 |
| 10 | Mblac2 | 1.46 | 1.94E-20 | 4.03E-16 | 0.12 | 0.05 | 0.08 | 2.53 |
| 10 | Tnfrsf19 | 1.23 | 2.17E-20 | 4.52E-16 | 0.26 | 0.15 | 0.11 | 1.77 |
| 10 | Apc | 0.39 | 2.39E-20 | 4.96E-16 | 0.74 | 0.61 | 0.13 | 1.21 |
| 10 | Schip1 | 0.83 | 2.73E-20 | 5.67E-16 | 0.28 | 0.16 | 0.12 | 1.77 |
| 10 | Arhgap12 | 0.64 | 2.76E-20 | 5.74E-16 | 0.48 | 0.33 | 0.14 | 1.43 |

**Supplementary Table 1. Marker genes for the original clusters of integrated clustering of E11.5 control and NotchGOF λ-junction cells.**

| | | | | | | | | |
|---|---|---|---|---|---|---|---|---|
| 10 | Dnmbp | 0.89 | 2.92E-20 | 6.07E-16 | 0.24 | 0.13 | 0.11 | 1.90 |
| 10 | Taok1 | 0.47 | 3.21E-20 | 6.67E-16 | 0.72 | 0.60 | 0.12 | 1.20 |
| 10 | Akap9 | 0.35 | 3.39E-20 | 7.04E-16 | 0.88 | 0.78 | 0.10 | 1.12 |
| 10 | Dcaf17 | 0.68 | 3.93E-20 | 8.18E-16 | 0.47 | 0.33 | 0.14 | 1.42 |
| 10 | Armh4 | 1.14 | 4.78E-20 | 9.93E-16 | 0.26 | 0.15 | 0.11 | 1.78 |
| 10 | Elf3 | 1.08 | 5.83E-20 | 1.21E-15 | 0.16 | 0.07 | 0.09 | 2.21 |
| 10 | Ccdc18 | 0.71 | 6.90E-20 | 1.44E-15 | 0.44 | 0.30 | 0.14 | 1.47 |
| 10 | Brwd3 | 0.58 | 7.29E-20 | 1.52E-15 | 0.61 | 0.46 | 0.14 | 1.31 |
| 10 | Mtss2 | 0.72 | 8.02E-20 | 1.67E-15 | 0.41 | 0.27 | 0.15 | 1.54 |
| 10 | Ets2 | 0.63 | 9.16E-20 | 1.91E-15 | 0.39 | 0.25 | 0.14 | 1.57 |
| 10 | Notch3 | 0.66 | 9.59E-20 | 1.99E-15 | 0.54 | 0.40 | 0.14 | 1.36 |
| 10 | Gstz1 | 0.71 | 1.12E-19 | 2.32E-15 | 0.41 | 0.27 | 0.14 | 1.52 |
| 10 | Dbndd2 | 1.13 | 1.14E-19 | 2.36E-15 | 0.17 | 0.08 | 0.09 | 2.13 |
| 10 | 2010320M18Rik | 0.65 | 1.14E-19 | 2.37E-15 | 0.49 | 0.34 | 0.15 | 1.43 |
| 10 | Gipc2 | 0.99 | 1.14E-19 | 2.38E-15 | 0.14 | 0.06 | 0.08 | 2.34 |
| 10 | Ripply3 | 1.33 | 1.17E-19 | 2.44E-15 | 0.14 | 0.06 | 0.08 | 2.34 |
| 10 | Slc25a17 | 0.52 | 1.27E-19 | 2.64E-15 | 0.74 | 0.62 | 0.12 | 1.19 |
| 10 | Jdp2 | 1.36 | 1.30E-19 | 2.69E-15 | 0.10 | 0.04 | 0.06 | 2.78 |
| 10 | Tulp4 | 0.39 | 1.40E-19 | 2.91E-15 | 0.80 | 0.66 | 0.13 | 1.20 |
| 10 | 1700096K18Rik | 0.99 | 1.40E-19 | 2.92E-15 | 0.19 | 0.09 | 0.10 | 2.05 |
| 10 | Dtx3l | 1.53 | 1.67E-19 | 3.47E-15 | 0.11 | 0.04 | 0.07 | 2.69 |
| 10 | Rb1 | 0.82 | 1.95E-19 | 4.06E-15 | 0.28 | 0.16 | 0.12 | 1.76 |
| 10 | Inava | 0.70 | 1.96E-19 | 4.08E-15 | 0.36 | 0.23 | 0.14 | 1.60 |
| 10 | Per3 | 1.02 | 2.30E-19 | 4.79E-15 | 0.19 | 0.10 | 0.10 | 2.02 |
| 10 | Prr13 | 0.46 | 2.93E-19 | 6.10E-15 | 0.56 | 0.40 | 0.16 | 1.41 |
| 10 | Bmyc | 0.39 | 3.30E-19 | 6.86E-15 | 0.76 | 0.62 | 0.14 | 1.23 |
| 10 | Fam129b | 0.94 | 3.52E-19 | 7.31E-15 | 0.25 | 0.14 | 0.11 | 1.81 |
| 10 | Hprt | 0.52 | 4.17E-19 | 8.67E-15 | 0.56 | 0.43 | 0.13 | 1.30 |
| 10 | Ammecr1 | 0.64 | 4.50E-19 | 9.36E-15 | 0.49 | 0.35 | 0.14 | 1.41 |
| 10 | Gsr | 0.59 | 4.74E-19 | 9.85E-15 | 0.48 | 0.33 | 0.15 | 1.45 |
| 10 | Rhou | 0.60 | 4.82E-19 | 1.00E-14 | 0.55 | 0.41 | 0.14 | 1.35 |
| 10 | Hk2 | 0.70 | 5.78E-19 | 1.20E-14 | 0.39 | 0.25 | 0.14 | 1.55 |
| 10 | Aldh5a1 | 0.69 | 5.85E-19 | 1.22E-14 | 0.38 | 0.24 | 0.14 | 1.57 |
| 10 | Ddhd1 | 0.89 | 8.42E-19 | 1.75E-14 | 0.28 | 0.17 | 0.12 | 1.70 |
| 10 | Slc37a3 | 0.71 | 9.51E-19 | 1.98E-14 | 0.40 | 0.26 | 0.14 | 1.53 |
| 10 | Mgat4a | 1.03 | 1.04E-18 | 2.17E-14 | 0.18 | 0.09 | 0.09 | 2.02 |
| 10 | Fam120a | 0.54 | 1.06E-18 | 2.20E-14 | 0.58 | 0.45 | 0.13 | 1.28 |
| 10 | St7 | 0.83 | 1.08E-18 | 2.24E-14 | 0.35 | 0.22 | 0.13 | 1.57 |
| 10 | Jade3 | 1.01 | 1.14E-18 | 2.37E-14 | 0.23 | 0.12 | 0.10 | 1.84 |
| 10 | Vsig10 | 1.36 | 1.24E-18 | 2.57E-14 | 0.12 | 0.05 | 0.07 | 2.38 |
| 10 | Ptk7 | 0.60 | 1.39E-18 | 2.88E-14 | 0.51 | 0.36 | 0.15 | 1.40 |
| 10 | Anxa3 | 0.39 | 1.42E-18 | 2.95E-14 | 0.18 | 0.09 | 0.10 | 2.09 |
| 10 | Cd47 | 0.44 | 1.47E-18 | 3.05E-14 | 0.58 | 0.44 | 0.15 | 1.34 |
| 10 | Cc2d1b | 0.77 | 1.54E-18 | 3.20E-14 | 0.29 | 0.17 | 0.12 | 1.72 |
| 10 | Pak4 | 0.60 | 1.64E-18 | 3.40E-14 | 0.52 | 0.38 | 0.14 | 1.36 |
| 10 | Chd3 | 0.26 | 1.65E-18 | 3.44E-14 | 0.95 | 0.90 | 0.06 | 1.06 |
| 10 | Edaradd | 1.39 | 1.68E-18 | 3.50E-14 | 0.11 | 0.04 | 0.07 | 2.63 |
| 10 | Tmem256 | 0.25 | 1.71E-18 | 3.56E-14 | 0.93 | 0.89 | 0.04 | 1.05 |
| 10 | Vill | 1.12 | 1.75E-18 | 3.65E-14 | 0.17 | 0.08 | 0.09 | 2.09 |
| 10 | Rac3 | 0.61 | 2.04E-18 | 4.24E-14 | 0.47 | 0.32 | 0.14 | 1.44 |
| 10 | Ist1 | 0.51 | 2.18E-18 | 4.54E-14 | 0.60 | 0.46 | 0.15 | 1.32 |
| 10 | Epb41l3 | 0.51 | 2.19E-18 | 4.55E-14 | 0.60 | 0.44 | 0.15 | 1.35 |
| 10 | Tmem63b | 0.64 | 2.28E-18 | 4.73E-14 | 0.48 | 0.33 | 0.14 | 1.44 |
| 10 | Plpp2 | 0.74 | 2.48E-18 | 5.16E-14 | 0.39 | 0.26 | 0.14 | 1.52 |
| 10 | Chchd5 | 0.52 | 2.78E-18 | 5.79E-14 | 0.59 | 0.45 | 0.14 | 1.30 |
| 10 | Rnpepl1 | 0.68 | 3.01E-18 | 6.26E-14 | 0.39 | 0.25 | 0.13 | 1.53 |

**Supplementary Table 1. Marker genes for the original clusters of integrated clustering of E11.5 control and NotchGOF λ-junction cells.**

| | | | | | | | | |
|---|---|---|---|---|---|---|---|---|
| 10 | Lamtor2 | 0.27 | 3.29E-18 | 6.85E-14 | 0.95 | 0.89 | 0.06 | 1.06 |
| 10 | Carmil1 | 0.90 | 3.34E-18 | 6.95E-14 | 0.25 | 0.14 | 0.11 | 1.77 |
| 10 | Prickle3 | 1.27 | 4.16E-18 | 8.65E-14 | 0.14 | 0.06 | 0.08 | 2.20 |
| 10 | Ccdc115 | 0.46 | 4.65E-18 | 9.68E-14 | 0.64 | 0.49 | 0.15 | 1.31 |
| 10 | Nudt4 | 0.27 | 4.78E-18 | 9.94E-14 | 0.81 | 0.77 | 0.04 | 1.05 |
| 10 | Otud4 | 0.63 | 6.95E-18 | 1.45E-13 | 0.44 | 0.31 | 0.14 | 1.45 |
| 10 | Serinc5 | 1.32 | 7.35E-18 | 1.53E-13 | 0.13 | 0.06 | 0.07 | 2.33 |
| 10 | Shb | 0.75 | 8.58E-18 | 1.78E-13 | 0.32 | 0.19 | 0.12 | 1.64 |
| 10 | Prelid2 | 0.63 | 9.08E-18 | 1.89E-13 | 0.50 | 0.37 | 0.14 | 1.37 |
| 10 | Ccdc85c | 1.03 | 9.15E-18 | 1.90E-13 | 0.21 | 0.11 | 0.10 | 1.86 |
| 10 | Nipsnap1 | 0.42 | 1.21E-17 | 2.52E-13 | 0.69 | 0.56 | 0.13 | 1.24 |
| 10 | Chd7 | 0.37 | 1.25E-17 | 2.61E-13 | 0.79 | 0.65 | 0.13 | 1.21 |
| 10 | Klf10 | 1.00 | 1.28E-17 | 2.66E-13 | 0.20 | 0.11 | 0.10 | 1.91 |
| 10 | Nsun2 | 0.45 | 1.28E-17 | 2.67E-13 | 0.75 | 0.63 | 0.12 | 1.18 |
| 10 | Dynlt3 | 0.72 | 1.32E-17 | 2.74E-13 | 0.38 | 0.25 | 0.12 | 1.49 |
| 10 | Kit | 0.89 | 1.33E-17 | 2.78E-13 | 0.19 | 0.09 | 0.09 | 1.98 |
| 10 | Itgb8 | 0.82 | 1.61E-17 | 3.35E-13 | 0.24 | 0.13 | 0.11 | 1.82 |
| 10 | Lyrm9 | 0.92 | 1.75E-17 | 3.64E-13 | 0.28 | 0.17 | 0.11 | 1.66 |
| 10 | Ahdc1 | 0.78 | 2.06E-17 | 4.29E-13 | 0.27 | 0.15 | 0.11 | 1.73 |
| 10 | Sppl2a | 0.49 | 2.06E-17 | 4.29E-13 | 0.61 | 0.47 | 0.14 | 1.29 |
| 10 | Snhg14 | 0.89 | 2.13E-17 | 4.43E-13 | 0.29 | 0.18 | 0.11 | 1.64 |
| 10 | Cdca7 | 0.40 | 2.65E-17 | 5.52E-13 | 0.83 | 0.72 | 0.11 | 1.15 |
| 10 | Sft2d3 | 0.83 | 2.66E-17 | 5.53E-13 | 0.32 | 0.20 | 0.12 | 1.59 |
| 10 | Id3 | 0.26 | 2.67E-17 | 5.56E-13 | 0.99 | 0.96 | 0.03 | 1.03 |
| 10 | Abhd4 | 0.86 | 2.69E-17 | 5.59E-13 | 0.24 | 0.14 | 0.11 | 1.77 |
| 10 | Lmln | 1.35 | 2.89E-17 | 6.00E-13 | 0.12 | 0.05 | 0.07 | 2.30 |
| 10 | Abca3 | 1.22 | 3.03E-17 | 6.31E-13 | 0.14 | 0.06 | 0.08 | 2.19 |
| 10 | Tpbg | 0.60 | 3.40E-17 | 7.07E-13 | 0.44 | 0.30 | 0.14 | 1.47 |
| 10 | Kcnmb4 | 0.68 | 3.43E-17 | 7.13E-13 | 0.46 | 0.32 | 0.14 | 1.43 |
| 10 | Sesn2 | 1.10 | 3.58E-17 | 7.44E-13 | 0.15 | 0.07 | 0.08 | 2.15 |
| 10 | AC149090.1 | 0.66 | 3.85E-17 | 8.02E-13 | 0.41 | 0.28 | 0.13 | 1.47 |
| 10 | Jmy | 0.58 | 4.11E-17 | 8.56E-13 | 0.51 | 0.37 | 0.13 | 1.36 |
| 10 | Ppp1r26 | 1.39 | 4.14E-17 | 8.61E-13 | 0.11 | 0.04 | 0.06 | 2.50 |
| 10 | Itprid2 | 0.53 | 4.73E-17 | 9.84E-13 | 0.26 | 0.15 | 0.11 | 1.76 |
| 10 | Celf4 | 0.73 | 5.30E-17 | 1.10E-12 | 0.35 | 0.22 | 0.12 | 1.56 |
| 10 | Rps27rt | 0.38 | 5.56E-17 | 1.16E-12 | 0.82 | 0.73 | 0.09 | 1.13 |
| 10 | Pygl | 1.04 | 5.82E-17 | 1.21E-12 | 0.18 | 0.10 | 0.09 | 1.94 |
| 10 | Ssr4 | 0.30 | 6.04E-17 | 1.26E-12 | 0.90 | 0.84 | 0.06 | 1.07 |
| 10 | Ubl3 | 0.50 | 6.40E-17 | 1.33E-12 | 0.50 | 0.37 | 0.14 | 1.37 |
| 10 | Dynll2 | 0.27 | 6.50E-17 | 1.35E-12 | 0.90 | 0.84 | 0.06 | 1.07 |
| 10 | Larp4b | 0.44 | 7.42E-17 | 1.54E-12 | 0.72 | 0.61 | 0.11 | 1.18 |
| 10 | Arhgap23 | 1.03 | 1.03E-16 | 2.15E-12 | 0.18 | 0.09 | 0.09 | 1.95 |
| 10 | Pex7 | 0.67 | 1.50E-16 | 3.11E-12 | 0.38 | 0.25 | 0.12 | 1.49 |
| 10 | Npepl1 | 0.62 | 1.66E-16 | 3.44E-12 | 0.46 | 0.33 | 0.14 | 1.42 |
| 10 | 1810044D09Rik | 1.04 | 1.69E-16 | 3.51E-12 | 0.20 | 0.11 | 0.09 | 1.85 |
| 10 | Amer1 | 0.52 | 1.85E-16 | 3.84E-12 | 0.63 | 0.51 | 0.12 | 1.24 |
| 10 | Kdm3a | 0.56 | 1.89E-16 | 3.93E-12 | 0.50 | 0.37 | 0.14 | 1.37 |
| 10 | Ero1l | 0.67 | 1.90E-16 | 3.95E-12 | 0.42 | 0.29 | 0.13 | 1.43 |
| 10 | Naa38 | 0.35 | 2.01E-16 | 4.18E-12 | 0.82 | 0.74 | 0.08 | 1.11 |
| 10 | Tmc4 | 1.02 | 2.09E-16 | 4.34E-12 | 0.18 | 0.10 | 0.09 | 1.93 |
| 10 | Agpat3 | 0.59 | 2.63E-16 | 5.46E-12 | 0.39 | 0.26 | 0.13 | 1.51 |
| 10 | Nucb2 | 0.74 | 2.64E-16 | 5.50E-12 | 0.26 | 0.15 | 0.11 | 1.72 |
| 10 | Thumpd3 | 0.60 | 3.13E-16 | 6.50E-12 | 0.51 | 0.38 | 0.13 | 1.34 |
| 10 | Tvp23b | 0.53 | 3.45E-16 | 7.18E-12 | 0.54 | 0.41 | 0.13 | 1.33 |
| 10 | Nbeal1 | 0.50 | 4.06E-16 | 8.44E-12 | 0.58 | 0.45 | 0.13 | 1.29 |
| 10 | Gtf2i | 0.32 | 4.24E-16 | 8.82E-12 | 0.91 | 0.86 | 0.05 | 1.06 |

**Supplementary Table 1. Marker genes for the original clusters of integrated clustering of E11.5 control and NotchGOF λ-junction cells.**

| | | | | | | | | |
|---|---|---|---|---|---|---|---|---|
| 10 | Serinc2 | 0.48 | 4.66E-16 | 9.69E-12 | 0.53 | 0.39 | 0.14 | 1.37 |
| 10 | Anapc13 | 0.29 | 4.79E-16 | 9.97E-12 | 0.89 | 0.85 | 0.05 | 1.06 |
| 10 | Plcb3 | 0.67 | 5.33E-16 | 1.11E-11 | 0.41 | 0.29 | 0.13 | 1.45 |
| 10 | Med19 | 0.34 | 6.26E-16 | 1.30E-11 | 0.81 | 0.74 | 0.07 | 1.10 |
| 10 | Srgap3 | 0.89 | 6.47E-16 | 1.35E-11 | 0.23 | 0.13 | 0.10 | 1.76 |
| 10 | Akr1b10 | 0.88 | 6.64E-16 | 1.38E-11 | 0.21 | 0.11 | 0.09 | 1.82 |
| 10 | Pick1 | 0.62 | 7.09E-16 | 1.48E-11 | 0.38 | 0.25 | 0.13 | 1.50 |
| 10 | 1500009L16Rik | 0.55 | 1.05E-15 | 2.19E-11 | 0.67 | 0.58 | 0.10 | 1.17 |
| 10 | Bcl2l1 | 0.55 | 1.10E-15 | 2.28E-11 | 0.49 | 0.37 | 0.12 | 1.33 |
| 10 | Stxbp6 | 0.84 | 1.23E-15 | 2.55E-11 | 0.23 | 0.13 | 0.10 | 1.74 |
| 10 | Mvb12a | 0.48 | 1.30E-15 | 2.70E-11 | 0.57 | 0.45 | 0.12 | 1.26 |
| 10 | Zbtb12 | 0.33 | 1.48E-15 | 3.09E-11 | 0.80 | 0.72 | 0.08 | 1.11 |
| 10 | Kdsr | 0.67 | 1.56E-15 | 3.24E-11 | 0.39 | 0.27 | 0.12 | 1.45 |
| 10 | Smim4 | 0.51 | 2.13E-15 | 4.42E-11 | 0.55 | 0.42 | 0.13 | 1.30 |
| 10 | Sun1 | 0.61 | 2.41E-15 | 5.01E-11 | 0.40 | 0.27 | 0.13 | 1.46 |
| 10 | Otx2 | 0.58 | 2.53E-15 | 5.27E-11 | 0.19 | 0.10 | 0.09 | 1.91 |
| 10 | Elk4 | 0.72 | 2.79E-15 | 5.80E-11 | 0.34 | 0.23 | 0.12 | 1.50 |
| 10 | Avpi1 | 0.62 | 3.01E-15 | 6.27E-11 | 0.44 | 0.32 | 0.13 | 1.40 |
| 10 | Plekhb2 | 1.15 | 3.37E-15 | 7.01E-11 | 0.13 | 0.06 | 0.07 | 2.17 |
| 10 | Tmem53 | 0.94 | 3.59E-15 | 7.47E-11 | 0.17 | 0.09 | 0.08 | 1.96 |
| 10 | Efnb2 | 0.31 | 3.60E-15 | 7.48E-11 | 0.59 | 0.45 | 0.15 | 1.33 |
| 10 | Zbtb8os | 0.47 | 3.61E-15 | 7.52E-11 | 0.62 | 0.50 | 0.12 | 1.24 |
| 10 | Psme1 | 0.28 | 3.63E-15 | 7.54E-11 | 0.84 | 0.76 | 0.08 | 1.11 |
| 10 | Snrpn | 0.45 | 3.87E-15 | 8.05E-11 | 0.59 | 0.47 | 0.13 | 1.28 |
| 10 | Xpr1 | 0.61 | 3.91E-15 | 8.13E-11 | 0.39 | 0.27 | 0.12 | 1.44 |
| 10 | 4930523C07Rik | 1.20 | 4.05E-15 | 8.43E-11 | 0.15 | 0.07 | 0.07 | 2.00 |
| 10 | Cenpb | 0.40 | 4.41E-15 | 9.16E-11 | 0.75 | 0.66 | 0.09 | 1.13 |
| 10 | Ppp1r1a | 0.76 | 4.46E-15 | 9.28E-11 | 0.27 | 0.16 | 0.11 | 1.65 |
| 10 | Ddi2 | 0.52 | 4.61E-15 | 9.59E-11 | 0.54 | 0.41 | 0.13 | 1.32 |
| 10 | Plxdc2 | 0.65 | 4.73E-15 | 9.83E-11 | 0.37 | 0.25 | 0.13 | 1.51 |
| 10 | Ormdl1 | 0.46 | 4.81E-15 | 1.00E-10 | 0.57 | 0.44 | 0.13 | 1.29 |
| 10 | Wdpcp | 0.98 | 5.11E-15 | 1.06E-10 | 0.16 | 0.08 | 0.08 | 1.98 |
| 10 | Tmem141 | 0.64 | 5.16E-15 | 1.07E-10 | 0.31 | 0.19 | 0.11 | 1.59 |
| 10 | Acbd4 | 1.24 | 5.72E-15 | 1.19E-10 | 0.11 | 0.05 | 0.06 | 2.33 |
| 10 | F8a | 0.88 | 6.02E-15 | 1.25E-10 | 0.16 | 0.08 | 0.08 | 1.98 |
| 10 | Ctf1 | 0.82 | 6.15E-15 | 1.28E-10 | 0.27 | 0.16 | 0.10 | 1.64 |
| 10 | Kcnmb4os2 | 0.81 | 6.16E-15 | 1.28E-10 | 0.26 | 0.16 | 0.10 | 1.65 |
| 10 | Ago2 | 0.37 | 6.39E-15 | 1.33E-10 | 0.80 | 0.72 | 0.08 | 1.11 |
| 10 | Hebp2 | 0.75 | 6.49E-15 | 1.35E-10 | 0.35 | 0.24 | 0.11 | 1.46 |
| 10 | Rnf44 | 0.59 | 6.50E-15 | 1.35E-10 | 0.46 | 0.34 | 0.12 | 1.34 |
| 10 | Prrc2b | 0.34 | 6.71E-15 | 1.39E-10 | 0.83 | 0.74 | 0.09 | 1.12 |
| 10 | Pole3 | 0.35 | 6.78E-15 | 1.41E-10 | 0.85 | 0.76 | 0.09 | 1.12 |
| 10 | Tshz1 | 0.57 | 6.87E-15 | 1.43E-10 | 0.44 | 0.31 | 0.13 | 1.40 |
| 10 | Ldlrad3 | 0.57 | 7.12E-15 | 1.48E-10 | 0.43 | 0.31 | 0.13 | 1.41 |
| 10 | Stxbp3 | 0.63 | 7.76E-15 | 1.61E-10 | 0.35 | 0.23 | 0.12 | 1.50 |
| 10 | Os9 | 0.51 | 7.89E-15 | 1.64E-10 | 0.50 | 0.37 | 0.13 | 1.34 |
| 10 | Sdr39u1 | 0.75 | 8.21E-15 | 1.71E-10 | 0.26 | 0.16 | 0.10 | 1.66 |
| 10 | Nmt2 | 0.56 | 8.54E-15 | 1.78E-10 | 0.47 | 0.34 | 0.13 | 1.38 |
| 10 | Srcin1 | 1.20 | 8.70E-15 | 1.81E-10 | 0.11 | 0.05 | 0.06 | 2.30 |
| 10 | Tmem184a | 1.39 | 8.83E-15 | 1.84E-10 | 0.11 | 0.05 | 0.06 | 2.24 |
| 10 | Gpd2 | 0.65 | 9.56E-15 | 1.99E-10 | 0.34 | 0.23 | 0.12 | 1.51 |
| 10 | Lrrc42 | 0.37 | 9.58E-15 | 1.99E-10 | 0.72 | 0.60 | 0.12 | 1.20 |
| 10 | Erf | 0.60 | 1.01E-14 | 2.10E-10 | 0.42 | 0.30 | 0.12 | 1.41 |
| 10 | Zmynd8 | 0.38 | 1.09E-14 | 2.26E-10 | 0.71 | 0.60 | 0.11 | 1.19 |
| 10 | Prps2 | 0.63 | 1.09E-14 | 2.26E-10 | 0.27 | 0.17 | 0.11 | 1.63 |
| 10 | Smox | 0.71 | 1.16E-14 | 2.41E-10 | 0.35 | 0.24 | 0.11 | 1.47 |

**Supplementary Table 1. Marker genes for the original clusters of integrated clustering of E11.5 control and NotchGOF λ-junction cells.**

| | | | | | | | | |
|---|---|---|---|---|---|---|---|---|
| 10 | Hint2 | 0.42 | 1.19E-14 | 2.47E-10 | 0.70 | 0.61 | 0.09 | 1.14 |
| 10 | Sgsm2 | 1.15 | 1.22E-14 | 2.54E-10 | 0.12 | 0.06 | 0.07 | 2.18 |
| 10 | Hibadh | 0.48 | 1.25E-14 | 2.60E-10 | 0.66 | 0.55 | 0.11 | 1.20 |
| 10 | Vps4b | 0.38 | 1.36E-14 | 2.83E-10 | 0.73 | 0.62 | 0.10 | 1.17 |
| 10 | Pbx4 | 0.90 | 1.45E-14 | 3.01E-10 | 0.17 | 0.09 | 0.08 | 1.92 |
| 10 | Etfa | 0.32 | 1.69E-14 | 3.52E-10 | 0.80 | 0.73 | 0.07 | 1.09 |
| 10 | Jtb | 0.41 | 1.74E-14 | 3.62E-10 | 0.65 | 0.53 | 0.12 | 1.23 |
| 10 | Ube2e2 | 0.54 | 1.81E-14 | 3.77E-10 | 0.50 | 0.38 | 0.12 | 1.32 |
| 10 | Spop | 0.38 | 2.02E-14 | 4.19E-10 | 0.78 | 0.69 | 0.10 | 1.14 |
| 10 | Ppp1r12b | 1.02 | 2.09E-14 | 4.34E-10 | 0.20 | 0.11 | 0.09 | 1.77 |
| 10 | Prkag2 | 0.73 | 2.18E-14 | 4.54E-10 | 0.33 | 0.22 | 0.11 | 1.50 |
| 10 | Mif4gd | 0.77 | 2.55E-14 | 5.29E-10 | 0.27 | 0.17 | 0.10 | 1.60 |
| 10 | Tgif1 | 0.41 | 2.69E-14 | 5.60E-10 | 0.74 | 0.63 | 0.11 | 1.17 |
| 10 | Strn | 0.69 | 3.22E-14 | 6.70E-10 | 0.28 | 0.18 | 0.10 | 1.59 |
| 10 | Lpcat4 | 1.09 | 3.28E-14 | 6.82E-10 | 0.13 | 0.06 | 0.07 | 2.10 |
| 10 | Esco1 | 0.61 | 3.56E-14 | 7.41E-10 | 0.35 | 0.23 | 0.12 | 1.50 |
| 10 | Dipk2a | 0.67 | 3.58E-14 | 7.44E-10 | 0.35 | 0.24 | 0.11 | 1.47 |
| 10 | Gtf2ird1 | 0.48 | 3.58E-14 | 7.45E-10 | 0.58 | 0.47 | 0.12 | 1.25 |
| 10 | Ppip5k2 | 0.45 | 3.67E-14 | 7.63E-10 | 0.57 | 0.45 | 0.12 | 1.26 |
| 10 | Trim8 | 0.57 | 5.45E-14 | 1.13E-09 | 0.42 | 0.30 | 0.12 | 1.41 |
| 10 | Reep6 | 1.57 | 5.45E-14 | 1.13E-09 | 0.11 | 0.05 | 0.06 | 2.18 |
| 10 | Mbd2 | 0.42 | 5.65E-14 | 1.17E-09 | 0.66 | 0.56 | 0.10 | 1.19 |
| 10 | Itsn2 | 0.66 | 5.65E-14 | 1.18E-09 | 0.29 | 0.19 | 0.11 | 1.56 |
| 10 | Myo5b | 0.98 | 5.66E-14 | 1.18E-09 | 0.10 | 0.04 | 0.06 | 2.33 |
| 10 | Slk | 0.40 | 5.72E-14 | 1.19E-09 | 0.65 | 0.53 | 0.12 | 1.22 |
| 10 | Cyba | 0.30 | 6.21E-14 | 1.29E-09 | 0.58 | 0.45 | 0.13 | 1.28 |
| 10 | Saraf | 0.38 | 6.74E-14 | 1.40E-09 | 0.66 | 0.54 | 0.12 | 1.22 |
| 10 | Dtd1 | 0.46 | 7.29E-14 | 1.52E-09 | 0.54 | 0.41 | 0.12 | 1.30 |
| 10 | Ppil1 | 0.29 | 7.90E-14 | 1.64E-09 | 0.87 | 0.81 | 0.06 | 1.07 |
| 10 | Erlin2 | 0.53 | 8.03E-14 | 1.67E-09 | 0.44 | 0.32 | 0.12 | 1.37 |
| 10 | Strbp | 0.32 | 8.16E-14 | 1.70E-09 | 0.79 | 0.70 | 0.09 | 1.12 |
| 10 | Cep164 | 0.34 | 8.44E-14 | 1.76E-09 | 0.72 | 0.61 | 0.11 | 1.19 |
| 10 | Rpl9-ps6 | 0.31 | 9.75E-14 | 2.03E-09 | 0.86 | 0.78 | 0.08 | 1.10 |
| 10 | Pla2r1 | 1.01 | 1.02E-13 | 2.13E-09 | 0.15 | 0.07 | 0.07 | 1.96 |
| 10 | Naf1 | 0.48 | 1.02E-13 | 2.13E-09 | 0.52 | 0.39 | 0.12 | 1.31 |
| 10 | Eps15 | 0.59 | 1.03E-13 | 2.15E-09 | 0.42 | 0.30 | 0.12 | 1.39 |
| 10 | Tamm41 | 0.53 | 1.09E-13 | 2.26E-09 | 0.46 | 0.34 | 0.12 | 1.35 |
| 10 | Noc2l | 0.34 | 1.21E-13 | 2.51E-09 | 0.72 | 0.61 | 0.11 | 1.18 |
| 10 | Rnf6 | 0.47 | 1.23E-13 | 2.55E-09 | 0.47 | 0.35 | 0.12 | 1.35 |
| 10 | Vps54 | 0.56 | 1.29E-13 | 2.68E-09 | 0.43 | 0.31 | 0.12 | 1.38 |
| 10 | Col4a5 | 0.71 | 1.67E-13 | 3.47E-09 | 0.22 | 0.13 | 0.09 | 1.72 |
| 10 | Rtf2 | 0.34 | 1.72E-13 | 3.57E-09 | 0.76 | 0.66 | 0.11 | 1.16 |
| 10 | Tmem250-ps | 0.73 | 1.74E-13 | 3.62E-09 | 0.23 | 0.14 | 0.09 | 1.67 |
| 10 | Akap12 | 0.66 | 1.86E-13 | 3.88E-09 | 0.30 | 0.19 | 0.11 | 1.54 |
| 10 | Ccdc51 | 0.98 | 1.94E-13 | 4.03E-09 | 0.20 | 0.12 | 0.08 | 1.70 |
| 10 | Wtip | 0.41 | 2.03E-13 | 4.21E-09 | 0.67 | 0.56 | 0.11 | 1.19 |
| 10 | Wnk2 | 1.46 | 2.21E-13 | 4.59E-09 | 0.10 | 0.05 | 0.06 | 2.21 |
| 10 | Sec22a | 0.53 | 2.47E-13 | 5.14E-09 | 0.41 | 0.29 | 0.12 | 1.41 |
| 10 | 2610306M01Rik | 0.66 | 2.49E-13 | 5.19E-09 | 0.26 | 0.16 | 0.10 | 1.63 |
| 10 | Slc25a23 | 0.74 | 2.50E-13 | 5.20E-09 | 0.26 | 0.16 | 0.10 | 1.62 |
| 10 | Mgme1 | 0.70 | 2.60E-13 | 5.41E-09 | 0.34 | 0.23 | 0.11 | 1.45 |
| 10 | Gprc5c | 0.56 | 2.77E-13 | 5.76E-09 | 0.35 | 0.23 | 0.12 | 1.50 |
| 10 | Ift43 | 0.37 | 2.92E-13 | 6.07E-09 | 0.67 | 0.54 | 0.13 | 1.24 |
| 10 | Slc44a1 | 0.62 | 2.92E-13 | 6.08E-09 | 0.35 | 0.23 | 0.11 | 1.48 |
| 10 | Tspan7 | 0.59 | 3.21E-13 | 6.68E-09 | 0.39 | 0.28 | 0.11 | 1.41 |
| 10 | Prkd3 | 0.48 | 3.32E-13 | 6.91E-09 | 0.63 | 0.54 | 0.10 | 1.18 |

**Supplementary Table 1. Marker genes for the original clusters of integrated clustering of E11.5 control and NotchGOF λ-junction cells.**

| | | | | | | | | |
|---|---|---|---|---|---|---|---|---|
| 10 | Ankhd1 | 0.43 | 3.33E-13 | 6.92E-09 | 0.61 | 0.49 | 0.12 | 1.25 |
| 10 | Egln1 | 0.33 | 3.55E-13 | 7.38E-09 | 0.80 | 0.68 | 0.12 | 1.18 |
| 10 | Eif1a | 0.41 | 3.86E-13 | 8.03E-09 | 0.76 | 0.67 | 0.09 | 1.14 |
| 10 | Nlk | 0.53 | 4.27E-13 | 8.89E-09 | 0.39 | 0.27 | 0.12 | 1.44 |
| 10 | Parp1 | 0.27 | 4.77E-13 | 9.93E-09 | 0.91 | 0.88 | 0.03 | 1.04 |
| 10 | Abhd17c | 0.50 | 4.84E-13 | 1.01E-08 | 0.52 | 0.40 | 0.12 | 1.30 |
| 10 | Rnaseh2a | 0.42 | 5.08E-13 | 1.06E-08 | 0.67 | 0.57 | 0.11 | 1.19 |
| 10 | Acadm | 0.51 | 5.10E-13 | 1.06E-08 | 0.45 | 0.33 | 0.12 | 1.37 |
| 10 | Clcn3 | 0.41 | 5.21E-13 | 1.08E-08 | 0.50 | 0.38 | 0.12 | 1.32 |
| 10 | Plekha1 | 0.57 | 5.39E-13 | 1.12E-08 | 0.30 | 0.20 | 0.11 | 1.54 |
| 10 | Rpp21 | 0.36 | 5.70E-13 | 1.19E-08 | 0.66 | 0.57 | 0.09 | 1.16 |
| 10 | Arhgef28 | 0.87 | 5.78E-13 | 1.20E-08 | 0.18 | 0.10 | 0.08 | 1.79 |
| 10 | Yipf2 | 0.76 | 6.74E-13 | 1.40E-08 | 0.21 | 0.12 | 0.09 | 1.71 |
| 10 | Prxl2c | 0.59 | 7.25E-13 | 1.51E-08 | 0.40 | 0.29 | 0.11 | 1.40 |
| 10 | Thtpa | 0.74 | 7.52E-13 | 1.56E-08 | 0.25 | 0.16 | 0.09 | 1.60 |
| 10 | 1110032A03Rik | 0.46 | 8.51E-13 | 1.77E-08 | 0.44 | 0.32 | 0.12 | 1.38 |
| 10 | Prr5 | 0.85 | 8.77E-13 | 1.82E-08 | 0.20 | 0.12 | 0.08 | 1.70 |
| 10 | Inpp5f | 0.64 | 8.82E-13 | 1.83E-08 | 0.34 | 0.24 | 0.11 | 1.44 |
| 10 | Wrnip1 | 0.69 | 1.03E-12 | 2.14E-08 | 0.30 | 0.20 | 0.10 | 1.51 |
| 10 | Tmem106c | 0.65 | 1.04E-12 | 2.17E-08 | 0.29 | 0.19 | 0.10 | 1.52 |
| 10 | Dph6 | 0.42 | 1.05E-12 | 2.19E-08 | 0.53 | 0.41 | 0.12 | 1.29 |
| 10 | Tafa5 | 0.70 | 1.06E-12 | 2.21E-08 | 0.21 | 0.12 | 0.09 | 1.71 |
| 10 | Zbtb38 | 0.69 | 1.07E-12 | 2.22E-08 | 0.22 | 0.13 | 0.09 | 1.66 |
| 10 | Smad3 | 0.56 | 1.21E-12 | 2.52E-08 | 0.38 | 0.26 | 0.11 | 1.42 |
| 10 | Ccdc28a | 0.89 | 1.32E-12 | 2.74E-08 | 0.18 | 0.10 | 0.08 | 1.77 |
| 10 | Dennd1b | 0.65 | 1.35E-12 | 2.82E-08 | 0.25 | 0.16 | 0.09 | 1.60 |
| 10 | Pfn2 | 0.37 | 1.40E-12 | 2.91E-08 | 0.75 | 0.66 | 0.09 | 1.14 |
| 10 | Zfp467 | 0.49 | 1.49E-12 | 3.10E-08 | 0.40 | 0.29 | 0.12 | 1.40 |
| 10 | Ap1ar | 0.37 | 1.53E-12 | 3.18E-08 | 0.67 | 0.55 | 0.12 | 1.21 |
| 10 | Cnksr2 | 0.60 | 1.55E-12 | 3.23E-08 | 0.44 | 0.32 | 0.11 | 1.35 |
| 10 | Parva | 0.33 | 1.66E-12 | 3.45E-08 | 0.66 | 0.55 | 0.12 | 1.21 |
| 10 | Ddb1 | 0.26 | 1.72E-12 | 3.58E-08 | 0.91 | 0.88 | 0.03 | 1.03 |
| 10 | Arhgap10 | 0.46 | 1.83E-12 | 3.80E-08 | 0.55 | 0.42 | 0.13 | 1.31 |
| 10 | Mdm1 | 0.54 | 1.85E-12 | 3.84E-08 | 0.41 | 0.30 | 0.11 | 1.38 |
| 10 | Sdsl | 1.16 | 1.90E-12 | 3.95E-08 | 0.12 | 0.06 | 0.06 | 2.05 |
| 10 | Thnsl1 | 0.96 | 2.07E-12 | 4.31E-08 | 0.14 | 0.07 | 0.07 | 1.94 |
| 10 | Ralgapa2 | 0.74 | 2.09E-12 | 4.34E-08 | 0.22 | 0.14 | 0.09 | 1.65 |
| 10 | Txnip | 0.53 | 2.11E-12 | 4.39E-08 | 0.45 | 0.34 | 0.11 | 1.33 |
| 10 | Plpp6 | 0.93 | 2.16E-12 | 4.49E-08 | 0.13 | 0.07 | 0.06 | 1.97 |
| 10 | Arl6ip5 | 0.52 | 2.27E-12 | 4.72E-08 | 0.43 | 0.32 | 0.12 | 1.37 |
| 10 | Cyb5r1 | 0.61 | 2.30E-12 | 4.79E-08 | 0.36 | 0.25 | 0.11 | 1.42 |
| 10 | Ccdc191 | 0.61 | 2.32E-12 | 4.82E-08 | 0.32 | 0.22 | 0.10 | 1.47 |
| 10 | Pgap2 | 0.40 | 2.46E-12 | 5.11E-08 | 0.57 | 0.45 | 0.12 | 1.26 |
| 10 | Klhl8 | 0.97 | 2.47E-12 | 5.13E-08 | 0.16 | 0.09 | 0.07 | 1.79 |
| 10 | Map9 | 0.57 | 2.47E-12 | 5.15E-08 | 0.28 | 0.18 | 0.10 | 1.56 |
| 10 | Pigp | 0.28 | 2.58E-12 | 5.36E-08 | 0.75 | 0.64 | 0.11 | 1.17 |
| 10 | Eml5 | 0.88 | 2.71E-12 | 5.64E-08 | 0.16 | 0.08 | 0.07 | 1.85 |
| 10 | Bbx | 0.36 | 3.21E-12 | 6.67E-08 | 0.69 | 0.58 | 0.11 | 1.19 |
| 10 | Galnt7 | 0.53 | 3.57E-12 | 7.42E-08 | 0.45 | 0.34 | 0.11 | 1.32 |
| 10 | Zfp64 | 0.54 | 3.57E-12 | 7.42E-08 | 0.42 | 0.31 | 0.11 | 1.37 |
| 10 | Zbtb8a | 0.94 | 3.77E-12 | 7.83E-08 | 0.14 | 0.08 | 0.07 | 1.91 |
| 10 | Vps13a | 0.46 | 4.02E-12 | 8.36E-08 | 0.46 | 0.34 | 0.12 | 1.35 |
| 10 | Ythdf2 | 0.26 | 4.02E-12 | 8.36E-08 | 0.84 | 0.77 | 0.08 | 1.10 |
| 10 | Paip2b | 0.52 | 4.12E-12 | 8.57E-08 | 0.40 | 0.29 | 0.11 | 1.37 |
| 10 | Tspan5 | 0.48 | 4.19E-12 | 8.70E-08 | 0.38 | 0.26 | 0.12 | 1.44 |
| 10 | Malt1 | 0.63 | 4.27E-12 | 8.89E-08 | 0.26 | 0.16 | 0.10 | 1.58 |

**Supplementary Table 1. Marker genes for the original clusters of integrated clustering of E11.5 control and NotchGOF λ-junction cells.**

| | | | | | | | | |
|---|---|---|---|---|---|---|---|---|
| 10 | Atp9a | 0.63 | 4.39E-12 | 9.13E-08 | 0.27 | 0.17 | 0.10 | 1.55 |
| 10 | Zswim8 | 0.54 | 4.39E-12 | 9.14E-08 | 0.44 | 0.34 | 0.11 | 1.32 |
| 10 | Odc1 | 0.27 | 4.44E-12 | 9.23E-08 | 0.87 | 0.81 | 0.06 | 1.08 |
| 10 | Dcaf12l1 | 0.87 | 4.61E-12 | 9.58E-08 | 0.20 | 0.12 | 0.08 | 1.70 |
| 10 | Gadd45b | 0.99 | 4.69E-12 | 9.75E-08 | 0.22 | 0.14 | 0.08 | 1.58 |
| 10 | Prorp | 0.76 | 4.74E-12 | 9.85E-08 | 0.23 | 0.14 | 0.09 | 1.64 |
| 10 | Anxa4 | 0.89 | 4.96E-12 | 1.03E-07 | 0.19 | 0.11 | 0.08 | 1.72 |
| 10 | Poglut3 | 0.77 | 5.00E-12 | 1.04E-07 | 0.22 | 0.13 | 0.09 | 1.65 |
| 10 | Agpat2 | 0.62 | 5.15E-12 | 1.07E-07 | 0.26 | 0.16 | 0.09 | 1.57 |
| 10 | Golgb1 | 0.29 | 5.27E-12 | 1.10E-07 | 0.82 | 0.74 | 0.08 | 1.11 |
| 10 | Map4k3 | 0.57 | 5.77E-12 | 1.20E-07 | 0.39 | 0.28 | 0.11 | 1.39 |
| 10 | Ankra2 | 0.49 | 5.85E-12 | 1.22E-07 | 0.43 | 0.31 | 0.12 | 1.37 |
| 10 | Crebrf | 0.80 | 6.13E-12 | 1.27E-07 | 0.19 | 0.11 | 0.08 | 1.72 |
| 10 | Ralgps2 | 0.38 | 7.31E-12 | 1.52E-07 | 0.79 | 0.71 | 0.08 | 1.11 |
| 10 | Golim4 | 0.38 | 7.66E-12 | 1.59E-07 | 0.70 | 0.61 | 0.09 | 1.14 |
| 10 | Snapin | 0.49 | 7.71E-12 | 1.60E-07 | 0.43 | 0.32 | 0.11 | 1.34 |
| 10 | Mthfd1 | 0.48 | 7.87E-12 | 1.64E-07 | 0.49 | 0.38 | 0.11 | 1.28 |
| 10 | Mapkapk2 | 0.39 | 7.90E-12 | 1.64E-07 | 0.64 | 0.55 | 0.09 | 1.16 |
| 10 | Retreg2 | 0.52 | 8.76E-12 | 1.82E-07 | 0.39 | 0.29 | 0.11 | 1.38 |
| 10 | Prkcd | 0.61 | 8.82E-12 | 1.84E-07 | 0.31 | 0.21 | 0.10 | 1.47 |
| 10 | Ptprk | 0.34 | 9.21E-12 | 1.92E-07 | 0.57 | 0.44 | 0.13 | 1.29 |
| 10 | Tfap4 | 0.52 | 9.23E-12 | 1.92E-07 | 0.44 | 0.33 | 0.11 | 1.34 |
| 10 | mt-Nd6 | 0.49 | 9.35E-12 | 1.95E-07 | 0.47 | 0.35 | 0.11 | 1.32 |
| 10 | Minar2 | 1.07 | 9.75E-12 | 2.03E-07 | 0.17 | 0.09 | 0.07 | 1.78 |
| 10 | Oxsr1 | 0.51 | 1.00E-11 | 2.08E-07 | 0.42 | 0.32 | 0.11 | 1.34 |
| 10 | Tmem38a | 0.95 | 1.01E-11 | 2.10E-07 | 0.17 | 0.10 | 0.07 | 1.73 |
| 10 | Slc25a27 | 0.39 | 1.02E-11 | 2.13E-07 | 0.52 | 0.40 | 0.12 | 1.30 |
| 10 | Wdr31 | 0.90 | 1.03E-11 | 2.15E-07 | 0.16 | 0.09 | 0.07 | 1.80 |
| 10 | Nabp2 | 0.29 | 1.05E-11 | 2.17E-07 | 0.75 | 0.65 | 0.10 | 1.15 |
| 10 | Morc4 | 0.72 | 1.07E-11 | 2.22E-07 | 0.23 | 0.14 | 0.09 | 1.60 |
| 10 | Atic | 0.33 | 1.08E-11 | 2.25E-07 | 0.76 | 0.67 | 0.09 | 1.13 |
| 10 | Zfp612 | 0.62 | 1.10E-11 | 2.28E-07 | 0.26 | 0.17 | 0.09 | 1.56 |
| 10 | Msi2 | 0.32 | 1.14E-11 | 2.38E-07 | 0.76 | 0.68 | 0.08 | 1.12 |
| 10 | Fam114a2 | 0.30 | 1.30E-11 | 2.71E-07 | 0.78 | 0.67 | 0.11 | 1.16 |
| 10 | Miga1 | 0.54 | 1.39E-11 | 2.89E-07 | 0.41 | 0.30 | 0.11 | 1.37 |
| 10 | Nrg1 | 0.68 | 1.46E-11 | 3.04E-07 | 0.15 | 0.08 | 0.07 | 1.85 |
| 10 | Nr2c1 | 0.59 | 1.50E-11 | 3.13E-07 | 0.32 | 0.22 | 0.10 | 1.45 |
| 10 | Apobec3 | 0.70 | 1.52E-11 | 3.17E-07 | 0.15 | 0.08 | 0.07 | 1.85 |
| 10 | Cltb | 0.67 | 1.53E-11 | 3.18E-07 | 0.27 | 0.18 | 0.09 | 1.51 |
| 10 | Bcap31 | 0.27 | 1.56E-11 | 3.25E-07 | 0.81 | 0.75 | 0.06 | 1.09 |
| 10 | Rrs1 | 0.34 | 1.65E-11 | 3.43E-07 | 0.71 | 0.60 | 0.10 | 1.17 |
| 10 | Tgif2 | 0.42 | 1.71E-11 | 3.57E-07 | 0.56 | 0.45 | 0.11 | 1.24 |
| 10 | Comt | 0.40 | 1.76E-11 | 3.66E-07 | 0.63 | 0.53 | 0.10 | 1.18 |
| 10 | Kif1b | 0.35 | 1.83E-11 | 3.81E-07 | 0.74 | 0.66 | 0.08 | 1.13 |
| 10 | Lifr | 0.78 | 1.91E-11 | 3.98E-07 | 0.18 | 0.11 | 0.08 | 1.71 |
| 10 | Dusp28 | 0.68 | 2.03E-11 | 4.22E-07 | 0.19 | 0.11 | 0.08 | 1.70 |
| 10 | Ing4 | 0.25 | 2.05E-11 | 4.26E-07 | 0.84 | 0.76 | 0.08 | 1.11 |
| 10 | Mboat1 | 1.09 | 2.09E-11 | 4.34E-07 | 0.10 | 0.05 | 0.05 | 2.08 |
| 10 | Zfp651 | 0.65 | 2.32E-11 | 4.82E-07 | 0.24 | 0.15 | 0.09 | 1.57 |
| 10 | Edem2 | 0.64 | 2.35E-11 | 4.88E-07 | 0.29 | 0.19 | 0.09 | 1.49 |
| 10 | Kdm1b | 0.87 | 2.36E-11 | 4.92E-07 | 0.17 | 0.10 | 0.07 | 1.74 |
| 10 | Rnf114 | 0.40 | 2.44E-11 | 5.08E-07 | 0.58 | 0.48 | 0.11 | 1.22 |
| 10 | Bok | 0.58 | 2.52E-11 | 5.24E-07 | 0.29 | 0.20 | 0.09 | 1.47 |
| 10 | Sestd1 | 0.42 | 2.58E-11 | 5.37E-07 | 0.55 | 0.44 | 0.11 | 1.24 |
| 10 | Prdm10 | 0.75 | 2.70E-11 | 5.63E-07 | 0.19 | 0.11 | 0.08 | 1.70 |
| 10 | Dlg3 | 0.47 | 2.74E-11 | 5.70E-07 | 0.43 | 0.32 | 0.11 | 1.34 |

**Supplementary Table 1. Marker genes for the original clusters of integrated clustering of E11.5 control and NotchGOF λ-junction cells.**

| | | | | | | | | |
|---|---|---|---|---|---|---|---|---|
| 10 | Arhgap21 | 0.64 | 3.07E-11 | 6.39E-07 | 0.31 | 0.22 | 0.10 | 1.44 |
| 10 | Slc48a1 | 0.67 | 3.41E-11 | 7.09E-07 | 0.25 | 0.17 | 0.09 | 1.52 |
| 10 | Mplkip | 0.33 | 3.44E-11 | 7.16E-07 | 0.75 | 0.67 | 0.08 | 1.12 |
| 10 | Tug1 | 0.34 | 3.70E-11 | 7.70E-07 | 0.66 | 0.55 | 0.11 | 1.21 |
| 10 | Sbno2 | 0.75 | 4.01E-11 | 8.35E-07 | 0.19 | 0.11 | 0.08 | 1.69 |
| 10 | Cebpzos | 0.29 | 4.18E-11 | 8.69E-07 | 0.69 | 0.57 | 0.13 | 1.22 |
| 10 | Rif1 | 0.26 | 4.31E-11 | 8.97E-07 | 0.86 | 0.80 | 0.06 | 1.07 |
| 10 | Fkbp9 | 0.40 | 4.35E-11 | 9.05E-07 | 0.56 | 0.44 | 0.12 | 1.27 |
| 10 | Trpc1 | 0.65 | 4.42E-11 | 9.19E-07 | 0.19 | 0.11 | 0.08 | 1.69 |
| 10 | Ascc1 | 0.40 | 5.02E-11 | 1.04E-06 | 0.50 | 0.39 | 0.11 | 1.28 |
| 10 | Trit1 | 0.63 | 5.04E-11 | 1.05E-06 | 0.28 | 0.19 | 0.09 | 1.48 |
| 10 | 2510002D24Rik | 0.41 | 5.16E-11 | 1.07E-06 | 0.51 | 0.39 | 0.12 | 1.30 |
| 10 | Bcl2l11 | 0.41 | 5.29E-11 | 1.10E-06 | 0.46 | 0.35 | 0.11 | 1.33 |
| 10 | Zfp703 | 0.60 | 5.36E-11 | 1.12E-06 | 0.35 | 0.25 | 0.10 | 1.39 |
| 10 | Acvr2a | 0.36 | 5.57E-11 | 1.16E-06 | 0.38 | 0.27 | 0.11 | 1.42 |
| 10 | Dhrs4 | 0.35 | 5.97E-11 | 1.24E-06 | 0.71 | 0.64 | 0.08 | 1.12 |
| 10 | Cmip | 0.42 | 6.10E-11 | 1.27E-06 | 0.51 | 0.39 | 0.11 | 1.29 |
| 10 | Kif1a | 0.88 | 6.26E-11 | 1.30E-06 | 0.12 | 0.06 | 0.06 | 1.97 |
| 10 | Sort1 | 0.62 | 6.31E-11 | 1.31E-06 | 0.30 | 0.20 | 0.10 | 1.48 |
| 10 | Rnf38 | 0.48 | 6.53E-11 | 1.36E-06 | 0.37 | 0.27 | 0.10 | 1.37 |
| 10 | Sp8 | 0.28 | 6.63E-11 | 1.38E-06 | 0.23 | 0.14 | 0.09 | 1.64 |
| 10 | AW146154 | 0.85 | 6.64E-11 | 1.38E-06 | 0.15 | 0.09 | 0.07 | 1.79 |
| 10 | Atp2b4 | 0.58 | 6.66E-11 | 1.39E-06 | 0.30 | 0.20 | 0.10 | 1.49 |
| 10 | Arvcf | 0.48 | 6.70E-11 | 1.39E-06 | 0.35 | 0.24 | 0.10 | 1.43 |
| 10 | Rnf11 | 0.33 | 6.72E-11 | 1.40E-06 | 0.70 | 0.62 | 0.08 | 1.13 |
| 10 | Bckdha | 0.44 | 6.82E-11 | 1.42E-06 | 0.48 | 0.37 | 0.11 | 1.29 |
| 10 | Naxd | 0.28 | 6.84E-11 | 1.42E-06 | 0.75 | 0.64 | 0.11 | 1.17 |
| 10 | Zfp992 | 0.65 | 7.07E-11 | 1.47E-06 | 0.24 | 0.16 | 0.09 | 1.55 |
| 10 | Lasp1 | 0.42 | 7.30E-11 | 1.52E-06 | 0.53 | 0.43 | 0.11 | 1.25 |
| 10 | Fndc3b | 0.59 | 7.36E-11 | 1.53E-06 | 0.34 | 0.24 | 0.10 | 1.41 |
| 10 | Jrkl | 0.65 | 7.63E-11 | 1.59E-06 | 0.26 | 0.17 | 0.09 | 1.53 |
| 10 | Suclg2 | 0.38 | 7.70E-11 | 1.60E-06 | 0.60 | 0.49 | 0.11 | 1.22 |
| 10 | Rnf145 | 0.46 | 7.71E-11 | 1.60E-06 | 0.48 | 0.37 | 0.11 | 1.29 |
| 10 | Uaca | 0.57 | 7.75E-11 | 1.61E-06 | 0.43 | 0.34 | 0.09 | 1.28 |
| 10 | Tdrd3 | 0.47 | 7.79E-11 | 1.62E-06 | 0.45 | 0.34 | 0.11 | 1.32 |
| 10 | Atl2 | 0.69 | 8.65E-11 | 1.80E-06 | 0.34 | 0.25 | 0.09 | 1.36 |
| 10 | Abhd11 | 0.55 | 8.98E-11 | 1.87E-06 | 0.36 | 0.26 | 0.10 | 1.37 |
| 10 | Suco | 0.41 | 9.25E-11 | 1.92E-06 | 0.58 | 0.49 | 0.09 | 1.18 |
| 10 | Bmf | 0.68 | 9.60E-11 | 2.00E-06 | 0.18 | 0.11 | 0.07 | 1.70 |
| 10 | Fyco1 | 0.61 | 9.72E-11 | 2.02E-06 | 0.25 | 0.16 | 0.09 | 1.54 |
| 10 | Bag4 | 0.47 | 9.83E-11 | 2.04E-06 | 0.49 | 0.39 | 0.10 | 1.25 |
| 10 | Suz12 | 0.28 | 9.85E-11 | 2.05E-06 | 0.79 | 0.72 | 0.07 | 1.10 |
| 10 | Ostf1 | 0.32 | 9.93E-11 | 2.06E-06 | 0.38 | 0.27 | 0.11 | 1.38 |
| 10 | Cox6b2 | 0.34 | 1.05E-10 | 2.19E-06 | 0.58 | 0.47 | 0.12 | 1.25 |
| 10 | Bola1 | 0.33 | 1.09E-10 | 2.27E-06 | 0.71 | 0.62 | 0.09 | 1.14 |
| 10 | Lyplal1 | 0.53 | 1.10E-10 | 2.29E-06 | 0.35 | 0.24 | 0.10 | 1.42 |
| 10 | Ctnnd1 | 0.41 | 1.15E-10 | 2.40E-06 | 0.56 | 0.46 | 0.11 | 1.23 |
| 10 | Wdr43 | 0.27 | 1.22E-10 | 2.53E-06 | 0.77 | 0.69 | 0.08 | 1.12 |
| 10 | Gpbp1l1 | 0.48 | 1.24E-10 | 2.57E-06 | 0.49 | 0.38 | 0.11 | 1.28 |
| 10 | Dpm2 | 0.36 | 1.33E-10 | 2.76E-06 | 0.65 | 0.57 | 0.08 | 1.15 |
| 10 | 2610002M06Rik | 0.48 | 1.35E-10 | 2.81E-06 | 0.39 | 0.29 | 0.10 | 1.35 |
| 10 | Hacd2 | 0.55 | 1.38E-10 | 2.88E-06 | 0.47 | 0.37 | 0.09 | 1.25 |
| 10 | 1600014C10Rik | 0.81 | 1.40E-10 | 2.92E-06 | 0.19 | 0.11 | 0.07 | 1.65 |
| 10 | Pno1 | 0.34 | 1.41E-10 | 2.93E-06 | 0.68 | 0.60 | 0.08 | 1.13 |
| 10 | Fign | 0.43 | 1.55E-10 | 3.22E-06 | 0.40 | 0.29 | 0.10 | 1.36 |
| 10 | Dedd2 | 0.91 | 1.62E-10 | 3.37E-06 | 0.13 | 0.07 | 0.06 | 1.84 |

**Supplementary Table 1. Marker genes for the original clusters of integrated clustering of E11.5 control and NotchGOF λ-junction cells.**

| | | | | | | | | |
|---|---|---|---|---|---|---|---|---|
| 10 | Igf2r | 0.38 | 1.70E-10 | 3.54E-06 | 0.60 | 0.50 | 0.11 | 1.22 |
| 10 | Ddb2 | 0.79 | 1.76E-10 | 3.66E-06 | 0.17 | 0.10 | 0.07 | 1.70 |
| 10 | Zcchc7 | 0.33 | 1.81E-10 | 3.76E-06 | 0.63 | 0.52 | 0.11 | 1.22 |
| 10 | Lcor | 0.47 | 1.82E-10 | 3.79E-06 | 0.38 | 0.28 | 0.10 | 1.36 |
| 10 | Hdhd3 | 0.67 | 1.96E-10 | 4.08E-06 | 0.25 | 0.16 | 0.09 | 1.52 |
| 10 | Bphl | 0.41 | 1.97E-10 | 4.10E-06 | 0.48 | 0.38 | 0.11 | 1.28 |
| 10 | Mid1 | 0.55 | 2.03E-10 | 4.21E-06 | 0.34 | 0.24 | 0.10 | 1.39 |
| 10 | Rap2b | 0.36 | 2.12E-10 | 4.42E-06 | 0.59 | 0.48 | 0.11 | 1.23 |
| 10 | Myo1e | 0.56 | 2.38E-10 | 4.95E-06 | 0.33 | 0.23 | 0.10 | 1.41 |
| 10 | Sdf2 | 0.32 | 2.45E-10 | 5.09E-06 | 0.66 | 0.56 | 0.10 | 1.17 |
| 10 | Airn | 0.57 | 2.56E-10 | 5.33E-06 | 0.28 | 0.19 | 0.09 | 1.48 |
| 10 | 2010315B03Rik | 0.91 | 2.57E-10 | 5.34E-06 | 0.11 | 0.05 | 0.05 | 1.98 |
| 10 | Satb1 | 0.40 | 2.58E-10 | 5.37E-06 | 0.50 | 0.39 | 0.11 | 1.30 |
| 10 | Heca | 0.53 | 2.63E-10 | 5.47E-06 | 0.32 | 0.23 | 0.09 | 1.41 |
| 10 | Nme3 | 0.75 | 2.66E-10 | 5.53E-06 | 0.14 | 0.08 | 0.06 | 1.81 |
| 10 | Zfp7 | 0.61 | 2.81E-10 | 5.85E-06 | 0.22 | 0.14 | 0.08 | 1.57 |
| 10 | Trmo | 0.81 | 2.98E-10 | 6.21E-06 | 0.12 | 0.07 | 0.06 | 1.88 |
| 10 | AV356131 | 0.91 | 3.26E-10 | 6.79E-06 | 0.14 | 0.08 | 0.06 | 1.82 |
| 10 | Stk11 | 0.38 | 3.60E-10 | 7.48E-06 | 0.57 | 0.48 | 0.09 | 1.19 |
| 10 | Zfp809 | 0.89 | 3.84E-10 | 7.99E-06 | 0.13 | 0.07 | 0.06 | 1.82 |
| 10 | Ephb2 | 0.49 | 3.89E-10 | 8.10E-06 | 0.38 | 0.28 | 0.10 | 1.37 |
| 10 | Atp11a | 0.70 | 3.97E-10 | 8.26E-06 | 0.25 | 0.17 | 0.08 | 1.49 |
| 10 | Gm26917 | 0.38 | 4.14E-10 | 8.62E-06 | 0.64 | 0.56 | 0.08 | 1.14 |
| 10 | Hexim1 | 0.52 | 4.17E-10 | 8.67E-06 | 0.38 | 0.28 | 0.10 | 1.34 |
| 10 | Mycbp2 | 0.27 | 4.22E-10 | 8.77E-06 | 0.77 | 0.70 | 0.07 | 1.10 |
| 10 | 4931406P16Rik | 0.41 | 4.65E-10 | 9.68E-06 | 0.46 | 0.35 | 0.11 | 1.30 |
| 10 | Tmem35b | 0.96 | 4.98E-10 | 1.04E-05 | 0.11 | 0.05 | 0.05 | 1.96 |
| 10 | Tmem42 | 0.58 | 5.55E-10 | 1.15E-05 | 0.23 | 0.15 | 0.08 | 1.55 |
| 10 | Dcxr | 0.97 | 5.60E-10 | 1.17E-05 | 0.16 | 0.09 | 0.07 | 1.69 |
| 10 | Rida | 0.37 | 5.61E-10 | 1.17E-05 | 0.51 | 0.42 | 0.10 | 1.23 |
| 10 | Tsix | 0.41 | 5.66E-10 | 1.18E-05 | 0.48 | 0.37 | 0.11 | 1.29 |
| 10 | Hps1 | 0.66 | 5.94E-10 | 1.24E-05 | 0.20 | 0.13 | 0.08 | 1.59 |
| 10 | Usp8 | 0.37 | 6.17E-10 | 1.28E-05 | 0.55 | 0.46 | 0.09 | 1.20 |
| 10 | Zfp53 | 0.71 | 6.20E-10 | 1.29E-05 | 0.15 | 0.09 | 0.07 | 1.74 |
| 10 | Gnpda2 | 0.46 | 6.31E-10 | 1.31E-05 | 0.38 | 0.28 | 0.10 | 1.35 |
| 10 | Plcb4 | 0.52 | 6.51E-10 | 1.35E-05 | 0.28 | 0.20 | 0.09 | 1.45 |
| 10 | Msh2 | 0.30 | 6.60E-10 | 1.37E-05 | 0.74 | 0.67 | 0.07 | 1.10 |
| 10 | Fmc1 | 0.31 | 6.66E-10 | 1.39E-05 | 0.69 | 0.59 | 0.10 | 1.16 |
| 10 | Zc2hc1a | 0.40 | 6.88E-10 | 1.43E-05 | 0.52 | 0.42 | 0.11 | 1.25 |
| 10 | Asxl1 | 0.38 | 7.38E-10 | 1.53E-05 | 0.59 | 0.50 | 0.10 | 1.20 |
| 10 | Ski | 0.42 | 7.97E-10 | 1.66E-05 | 0.51 | 0.41 | 0.10 | 1.24 |
| 10 | Cep152 | 0.67 | 8.17E-10 | 1.70E-05 | 0.20 | 0.12 | 0.07 | 1.61 |
| 10 | Pik3c2a | 0.40 | 8.37E-10 | 1.74E-05 | 0.51 | 0.41 | 0.10 | 1.25 |
| 10 | Phactr4 | 0.40 | 9.00E-10 | 1.87E-05 | 0.59 | 0.49 | 0.10 | 1.20 |
| 10 | Faah | 0.86 | 9.27E-10 | 1.93E-05 | 0.11 | 0.06 | 0.05 | 1.93 |
| 10 | Myo1d | 0.88 | 1.00E-09 | 2.08E-05 | 0.10 | 0.05 | 0.05 | 1.96 |
| 10 | Fam117a | 0.60 | 1.02E-09 | 2.12E-05 | 0.32 | 0.23 | 0.09 | 1.38 |
| 10 | Ncaph2 | 0.31 | 1.06E-09 | 2.19E-05 | 0.68 | 0.59 | 0.09 | 1.15 |
| 10 | Scai | 0.41 | 1.07E-09 | 2.23E-05 | 0.45 | 0.35 | 0.10 | 1.29 |
| 10 | Echdc1 | 0.48 | 1.21E-09 | 2.51E-05 | 0.33 | 0.24 | 0.09 | 1.39 |
| 10 | Ddx58 | 1.14 | 1.26E-09 | 2.61E-05 | 0.10 | 0.05 | 0.05 | 1.93 |
| 10 | Fam241b | 0.58 | 1.27E-09 | 2.64E-05 | 0.25 | 0.17 | 0.08 | 1.50 |
| 10 | Baiap2l1 | 0.97 | 1.28E-09 | 2.67E-05 | 0.12 | 0.06 | 0.05 | 1.89 |
| 10 | Marchf8 | 0.63 | 1.31E-09 | 2.72E-05 | 0.25 | 0.17 | 0.08 | 1.49 |
| 10 | Zfp503 | 0.28 | 1.33E-09 | 2.76E-05 | 0.49 | 0.37 | 0.12 | 1.32 |
| 10 | Pisd | 0.50 | 1.39E-09 | 2.90E-05 | 0.30 | 0.21 | 0.09 | 1.44 |

**Supplementary Table 1. Marker genes for the original clusters of integrated clustering of E11.5 control and NotchGOF λ-junction cells.**

| | | | | | | | | |
|---|---|---|---|---|---|---|---|---|
| 10 | Tmc7 | 0.83 | 1.41E-09 | 2.92E-05 | 0.11 | 0.06 | 0.05 | 1.93 |
| 10 | Iqsec1 | 0.60 | 1.41E-09 | 2.94E-05 | 0.21 | 0.14 | 0.08 | 1.57 |
| 10 | Arhgap32 | 0.60 | 1.48E-09 | 3.08E-05 | 0.28 | 0.19 | 0.09 | 1.46 |
| 10 | Cybc1 | 0.58 | 1.51E-09 | 3.13E-05 | 0.31 | 0.22 | 0.09 | 1.40 |
| 10 | Alpl | 0.89 | 1.52E-09 | 3.16E-05 | 0.17 | 0.10 | 0.07 | 1.65 |
| 10 | Lrrc8b | 0.78 | 1.53E-09 | 3.19E-05 | 0.18 | 0.11 | 0.07 | 1.63 |
| 10 | Tradd | 0.78 | 1.74E-09 | 3.61E-05 | 0.18 | 0.11 | 0.07 | 1.60 |
| 10 | Mmgt2 | 0.40 | 1.77E-09 | 3.68E-05 | 0.53 | 0.43 | 0.10 | 1.23 |
| 10 | Ung | 0.57 | 1.80E-09 | 3.74E-05 | 0.43 | 0.34 | 0.09 | 1.27 |
| 10 | Slc12a6 | 0.69 | 1.84E-09 | 3.82E-05 | 0.21 | 0.13 | 0.08 | 1.57 |
| 10 | Zfp217 | 0.44 | 1.85E-09 | 3.84E-05 | 0.29 | 0.20 | 0.09 | 1.46 |
| 10 | Klf6 | 0.54 | 1.87E-09 | 3.88E-05 | 0.33 | 0.23 | 0.09 | 1.40 |
| 10 | Tedc1 | 0.63 | 1.99E-09 | 4.14E-05 | 0.20 | 0.13 | 0.07 | 1.58 |
| 10 | Oard1 | 0.27 | 2.11E-09 | 4.38E-05 | 0.75 | 0.67 | 0.08 | 1.12 |
| 10 | N4bp3 | 0.61 | 2.11E-09 | 4.39E-05 | 0.12 | 0.06 | 0.05 | 1.86 |
| 10 | Tfdp1 | 0.28 | 2.15E-09 | 4.48E-05 | 0.74 | 0.66 | 0.08 | 1.12 |
| 10 | Flywch2 | 0.32 | 2.15E-09 | 4.48E-05 | 0.56 | 0.45 | 0.11 | 1.24 |
| 10 | 1700025G04Rik | 0.47 | 2.28E-09 | 4.74E-05 | 0.43 | 0.34 | 0.10 | 1.29 |
| 10 | Zbtb7a | 0.45 | 2.29E-09 | 4.77E-05 | 0.43 | 0.33 | 0.10 | 1.29 |
| 10 | Jmjd1c | 0.32 | 2.30E-09 | 4.79E-05 | 0.59 | 0.48 | 0.11 | 1.22 |
| 10 | Rere | 0.29 | 2.56E-09 | 5.32E-05 | 0.68 | 0.59 | 0.09 | 1.16 |
| 10 | Poli | 0.85 | 2.70E-09 | 5.61E-05 | 0.11 | 0.06 | 0.05 | 1.87 |
| 10 | Cdc42ep3 | 0.38 | 2.80E-09 | 5.82E-05 | 0.25 | 0.16 | 0.08 | 1.51 |
| 10 | Emid1 | 0.82 | 2.87E-09 | 5.96E-05 | 0.15 | 0.09 | 0.06 | 1.69 |
| 10 | Mterf1a | 0.74 | 3.22E-09 | 6.70E-05 | 0.14 | 0.08 | 0.06 | 1.73 |
| 10 | C77080 | 0.90 | 3.25E-09 | 6.75E-05 | 0.11 | 0.06 | 0.05 | 1.91 |
| 10 | Rxrb | 0.34 | 3.58E-09 | 7.45E-05 | 0.52 | 0.41 | 0.11 | 1.27 |
| 10 | Nomo1 | 0.58 | 3.75E-09 | 7.81E-05 | 0.28 | 0.20 | 0.08 | 1.42 |
| 10 | Tfrc | 0.35 | 3.90E-09 | 8.11E-05 | 0.50 | 0.40 | 0.11 | 1.27 |
| 10 | Bclaf3 | 0.37 | 3.91E-09 | 8.12E-05 | 0.43 | 0.33 | 0.10 | 1.30 |
| 10 | Pik3c3 | 0.62 | 3.97E-09 | 8.25E-05 | 0.25 | 0.17 | 0.08 | 1.45 |
| 10 | 2310039H08Rik | 0.34 | 4.10E-09 | 8.52E-05 | 0.52 | 0.42 | 0.10 | 1.24 |
| 10 | Zdhhc13 | 0.72 | 4.21E-09 | 8.75E-05 | 0.21 | 0.14 | 0.07 | 1.53 |
| 10 | Asl | 0.60 | 4.38E-09 | 9.10E-05 | 0.22 | 0.15 | 0.08 | 1.51 |
| 10 | Ercc6l2 | 0.43 | 4.45E-09 | 9.26E-05 | 0.43 | 0.33 | 0.10 | 1.29 |
| 10 | Srebf1 | 0.54 | 4.47E-09 | 9.30E-05 | 0.28 | 0.19 | 0.09 | 1.45 |
| 10 | Acvr1b | 0.77 | 4.85E-09 | 1.01E-04 | 0.18 | 0.11 | 0.07 | 1.60 |
| 10 | Gm17251 | 0.64 | 4.90E-09 | 1.02E-04 | 0.23 | 0.15 | 0.08 | 1.50 |
| 10 | Ankzf1 | 0.50 | 5.15E-09 | 1.07E-04 | 0.31 | 0.22 | 0.09 | 1.42 |
| 10 | Rmnd5a | 0.37 | 5.51E-09 | 1.15E-04 | 0.49 | 0.40 | 0.09 | 1.23 |
| 10 | Sypl | 0.33 | 5.86E-09 | 1.22E-04 | 0.58 | 0.48 | 0.10 | 1.21 |
| 10 | Gcat | 0.37 | 5.93E-09 | 1.23E-04 | 0.57 | 0.48 | 0.09 | 1.19 |
| 10 | Zfp318 | 0.30 | 6.18E-09 | 1.29E-04 | 0.57 | 0.47 | 0.10 | 1.22 |
| 10 | Tlk1 | 0.30 | 6.28E-09 | 1.31E-04 | 0.71 | 0.65 | 0.07 | 1.10 |
| 10 | Pcolce2 | 0.91 | 6.62E-09 | 1.38E-04 | 0.20 | 0.14 | 0.07 | 1.50 |
| 10 | Ncor2 | 0.46 | 6.94E-09 | 1.44E-04 | 0.44 | 0.35 | 0.09 | 1.27 |
| 10 | Ift52 | 0.41 | 7.07E-09 | 1.47E-04 | 0.49 | 0.40 | 0.10 | 1.24 |
| 10 | BC005537 | 0.32 | 7.33E-09 | 1.52E-04 | 0.59 | 0.50 | 0.09 | 1.19 |
| 10 | Tmed5 | 0.42 | 7.79E-09 | 1.62E-04 | 0.57 | 0.48 | 0.09 | 1.19 |
| 10 | Mettl14 | 0.41 | 7.92E-09 | 1.65E-04 | 0.36 | 0.27 | 0.10 | 1.36 |
| 10 | Bmp2k | 0.46 | 8.41E-09 | 1.75E-04 | 0.23 | 0.15 | 0.08 | 1.50 |
| 10 | Stap2 | 0.70 | 8.65E-09 | 1.80E-04 | 0.14 | 0.08 | 0.06 | 1.73 |
| 10 | Hmga1 | 0.30 | 8.65E-09 | 1.80E-04 | 0.59 | 0.49 | 0.09 | 1.19 |
| 10 | Sp1 | 0.34 | 8.66E-09 | 1.80E-04 | 0.51 | 0.41 | 0.10 | 1.24 |
| 10 | Hdac8 | 0.60 | 9.10E-09 | 1.89E-04 | 0.31 | 0.23 | 0.09 | 1.38 |
| 10 | Wbp2 | 0.47 | 9.19E-09 | 1.91E-04 | 0.36 | 0.28 | 0.09 | 1.32 |

**Supplementary Table 1. Marker genes for the original clusters of integrated clustering of E11.5 control and NotchGOF λ-junction cells.**

| | | | | | | | | |
|---|---|---|---|---|---|---|---|---|
| 10 | Polr3h | 0.40 | 9.78E-09 | 2.03E-04 | 0.48 | 0.38 | 0.10 | 1.25 |
| 10 | Fbxl3 | 0.50 | 9.80E-09 | 2.04E-04 | 0.34 | 0.25 | 0.09 | 1.35 |
| 10 | Dip2b | 0.41 | 9.92E-09 | 2.06E-04 | 0.43 | 0.34 | 0.09 | 1.27 |
| 10 | Pcmtd1 | 0.29 | 1.04E-08 | 2.15E-04 | 0.56 | 0.46 | 0.10 | 1.21 |
| 10 | Acp6 | 0.48 | 1.06E-08 | 2.20E-04 | 0.31 | 0.22 | 0.09 | 1.38 |
| 10 | B230354K17Rik | 0.37 | 1.08E-08 | 2.25E-04 | 0.46 | 0.36 | 0.10 | 1.28 |
| 10 | Dgkz | 0.47 | 1.10E-08 | 2.30E-04 | 0.37 | 0.28 | 0.09 | 1.32 |
| 10 | Dapk1 | 0.46 | 1.20E-08 | 2.49E-04 | 0.37 | 0.28 | 0.09 | 1.33 |
| 10 | Iqcb1 | 0.31 | 1.22E-08 | 2.55E-04 | 0.62 | 0.52 | 0.10 | 1.19 |
| 10 | Tspan14 | 0.42 | 1.23E-08 | 2.57E-04 | 0.42 | 0.32 | 0.09 | 1.29 |
| 10 | Atp6v1d | 0.35 | 1.28E-08 | 2.66E-04 | 0.69 | 0.64 | 0.05 | 1.08 |
| 10 | Fbxw4 | 0.69 | 1.31E-08 | 2.72E-04 | 0.21 | 0.14 | 0.07 | 1.50 |
| 10 | Fam76b | 0.32 | 1.36E-08 | 2.82E-04 | 0.64 | 0.55 | 0.10 | 1.18 |
| 10 | Wwc2 | 0.42 | 1.37E-08 | 2.86E-04 | 0.64 | 0.58 | 0.06 | 1.10 |
| 10 | Zbtb11os1 | 0.73 | 1.40E-08 | 2.91E-04 | 0.12 | 0.07 | 0.05 | 1.78 |
| 10 | Creb3l2 | 0.39 | 1.40E-08 | 2.92E-04 | 0.41 | 0.32 | 0.10 | 1.30 |
| 10 | Fgd6 | 0.61 | 1.41E-08 | 2.92E-04 | 0.24 | 0.16 | 0.07 | 1.46 |
| 10 | 2810402E24Rik | 0.57 | 1.45E-08 | 3.01E-04 | 0.30 | 0.21 | 0.09 | 1.40 |
| 10 | Jpt2 | 0.37 | 1.47E-08 | 3.06E-04 | 0.53 | 0.45 | 0.09 | 1.19 |
| 10 | Zkscan7 | 0.80 | 1.53E-08 | 3.18E-04 | 0.13 | 0.08 | 0.06 | 1.72 |
| 10 | Ppfia3 | 0.45 | 1.55E-08 | 3.22E-04 | 0.20 | 0.13 | 0.07 | 1.55 |
| 10 | Zfp692 | 0.69 | 1.55E-08 | 3.23E-04 | 0.17 | 0.11 | 0.06 | 1.59 |
| 10 | Napa | 0.25 | 1.73E-08 | 3.59E-04 | 0.79 | 0.73 | 0.06 | 1.08 |
| 10 | 2310011J03Rik | 0.27 | 1.77E-08 | 3.69E-04 | 0.58 | 0.48 | 0.10 | 1.20 |
| 10 | Hat1 | 0.25 | 1.78E-08 | 3.69E-04 | 0.82 | 0.76 | 0.06 | 1.07 |
| 10 | Cmtm4 | 0.48 | 1.78E-08 | 3.71E-04 | 0.35 | 0.26 | 0.09 | 1.35 |
| 10 | Ltv1 | 0.35 | 1.80E-08 | 3.74E-04 | 0.54 | 0.46 | 0.08 | 1.18 |
| 10 | Proser2 | 0.88 | 1.87E-08 | 3.90E-04 | 0.10 | 0.06 | 0.05 | 1.86 |
| 10 | Nudt11 | 0.45 | 1.89E-08 | 3.92E-04 | 0.26 | 0.18 | 0.08 | 1.46 |
| 10 | Mindy2 | 0.62 | 1.90E-08 | 3.95E-04 | 0.25 | 0.17 | 0.07 | 1.42 |
| 10 | Nras | 0.25 | 1.95E-08 | 4.05E-04 | 0.77 | 0.69 | 0.08 | 1.12 |
| 10 | Plxna3 | 0.66 | 1.99E-08 | 4.14E-04 | 0.16 | 0.10 | 0.06 | 1.63 |
| 10 | Sorcs2 | 0.56 | 2.12E-08 | 4.40E-04 | 0.17 | 0.11 | 0.07 | 1.61 |
| 10 | Gm4890 | 0.71 | 2.19E-08 | 4.55E-04 | 0.12 | 0.07 | 0.05 | 1.78 |
| 10 | 3110039M20Rik | 0.40 | 2.26E-08 | 4.69E-04 | 0.13 | 0.08 | 0.06 | 1.72 |
| 10 | Atp1a1 | 0.32 | 2.30E-08 | 4.79E-04 | 0.71 | 0.63 | 0.07 | 1.12 |
| 10 | Znrf3 | 0.64 | 2.33E-08 | 4.84E-04 | 0.25 | 0.18 | 0.07 | 1.41 |
| 10 | Sirt7 | 0.51 | 2.34E-08 | 4.87E-04 | 0.25 | 0.17 | 0.08 | 1.45 |
| 10 | Klhl23 | 0.31 | 2.52E-08 | 5.24E-04 | 0.47 | 0.37 | 0.10 | 1.26 |
| 10 | Htatip2 | 0.87 | 2.55E-08 | 5.31E-04 | 0.13 | 0.07 | 0.05 | 1.71 |
| 10 | Cdk5rap3 | 0.37 | 2.56E-08 | 5.32E-04 | 0.53 | 0.43 | 0.10 | 1.23 |
| 10 | Arfip2 | 0.32 | 2.57E-08 | 5.34E-04 | 0.52 | 0.42 | 0.10 | 1.24 |
| 10 | 0610010F05Rik | 0.41 | 2.62E-08 | 5.44E-04 | 0.34 | 0.25 | 0.09 | 1.37 |
| 10 | Frat2 | 0.48 | 2.78E-08 | 5.78E-04 | 0.32 | 0.23 | 0.08 | 1.35 |
| 10 | Hsdl1 | 0.59 | 2.84E-08 | 5.91E-04 | 0.22 | 0.15 | 0.07 | 1.47 |
| 10 | Pdcd6ip | 0.36 | 2.85E-08 | 5.92E-04 | 0.58 | 0.48 | 0.10 | 1.20 |
| 10 | Axl | 0.48 | 2.85E-08 | 5.92E-04 | 0.23 | 0.15 | 0.08 | 1.50 |
| 10 | Adam22 | 0.71 | 2.86E-08 | 5.96E-04 | 0.13 | 0.07 | 0.05 | 1.73 |
| 10 | Gm17018 | 0.35 | 2.94E-08 | 6.12E-04 | 0.61 | 0.52 | 0.09 | 1.17 |
| 10 | Atp6v1b2 | 0.31 | 2.95E-08 | 6.15E-04 | 0.48 | 0.39 | 0.10 | 1.25 |
| 10 | Tmem189 | 0.43 | 3.12E-08 | 6.48E-04 | 0.39 | 0.30 | 0.09 | 1.30 |
| 10 | Chmp1b | 0.36 | 3.43E-08 | 7.13E-04 | 0.44 | 0.35 | 0.09 | 1.26 |
| 10 | Tmem19 | 0.55 | 3.55E-08 | 7.38E-04 | 0.27 | 0.19 | 0.08 | 1.41 |
| 10 | Osbpl6 | 0.54 | 3.56E-08 | 7.41E-04 | 0.22 | 0.15 | 0.07 | 1.50 |
| 10 | Stt3b | 0.32 | 3.72E-08 | 7.75E-04 | 0.60 | 0.51 | 0.09 | 1.18 |
| 10 | Obsl1 | 0.40 | 3.75E-08 | 7.79E-04 | 0.41 | 0.31 | 0.09 | 1.29 |

**Supplementary Table 1. Marker genes for the original clusters of integrated clustering of E11.5 control and NotchGOF λ-junction cells.**

| | | | | | | | | |
|---|---|---|---|---|---|---|---|---|
| 10 | Ccpg1os | 0.51 | 3.90E-08 | 8.10E-04 | 0.23 | 0.16 | 0.07 | 1.47 |
| 10 | Znrd1as | 0.51 | 4.12E-08 | 8.58E-04 | 0.19 | 0.13 | 0.07 | 1.54 |
| 10 | Pde7a | 0.30 | 4.22E-08 | 8.78E-04 | 0.62 | 0.55 | 0.08 | 1.14 |
| 10 | Ankrd13c | 0.43 | 4.32E-08 | 8.99E-04 | 0.32 | 0.24 | 0.09 | 1.36 |
| 10 | Krt10 | 0.35 | 4.69E-08 | 9.75E-04 | 0.52 | 0.43 | 0.09 | 1.21 |
| 10 | Pqlc1 | 0.61 | 4.73E-08 | 9.83E-04 | 0.20 | 0.14 | 0.07 | 1.49 |
| 10 | Zfp593 | 0.35 | 4.74E-08 | 9.85E-04 | 0.55 | 0.47 | 0.08 | 1.16 |
| 10 | 2700033N17Rik | 0.42 | 5.02E-08 | 1.04E-03 | 0.18 | 0.12 | 0.07 | 1.57 |
| 10 | Aco1 | 0.53 | 5.26E-08 | 1.09E-03 | 0.42 | 0.33 | 0.09 | 1.26 |
| 10 | Adamts6 | 0.62 | 5.28E-08 | 1.10E-03 | 0.12 | 0.07 | 0.05 | 1.76 |
| 10 | Rfxap | 0.36 | 5.46E-08 | 1.14E-03 | 0.60 | 0.53 | 0.07 | 1.13 |
| 10 | Pacsin3 | 0.48 | 5.57E-08 | 1.16E-03 | 0.30 | 0.22 | 0.08 | 1.37 |
| 10 | Gm45051 | 0.83 | 5.58E-08 | 1.16E-03 | 0.13 | 0.07 | 0.05 | 1.71 |
| 10 | Mob2 | 0.40 | 5.72E-08 | 1.19E-03 | 0.49 | 0.40 | 0.09 | 1.22 |
| 10 | Osbpl5 | 0.78 | 5.79E-08 | 1.20E-03 | 0.13 | 0.08 | 0.05 | 1.71 |
| 10 | Haghl | 0.41 | 5.81E-08 | 1.21E-03 | 0.43 | 0.35 | 0.09 | 1.25 |
| 10 | Ogfod3 | 0.61 | 5.81E-08 | 1.21E-03 | 0.21 | 0.14 | 0.07 | 1.50 |
| 10 | Lin7c | 0.28 | 5.98E-08 | 1.24E-03 | 0.59 | 0.50 | 0.09 | 1.19 |
| 10 | Tmem192 | 0.49 | 6.29E-08 | 1.31E-03 | 0.26 | 0.18 | 0.08 | 1.42 |
| 10 | Rimklb | 0.35 | 6.36E-08 | 1.32E-03 | 0.42 | 0.32 | 0.10 | 1.30 |
| 10 | Pgap1 | 0.64 | 6.56E-08 | 1.36E-03 | 0.22 | 0.15 | 0.07 | 1.45 |
| 10 | Magi3 | 0.39 | 6.67E-08 | 1.39E-03 | 0.27 | 0.19 | 0.08 | 1.43 |
| 10 | Lztfl1 | 0.27 | 6.72E-08 | 1.40E-03 | 0.59 | 0.49 | 0.10 | 1.20 |
| 10 | Rnpep | 0.27 | 6.97E-08 | 1.45E-03 | 0.54 | 0.43 | 0.11 | 1.25 |
| 10 | Arhgef12 | 0.32 | 6.97E-08 | 1.45E-03 | 0.66 | 0.58 | 0.08 | 1.13 |
| 10 | Meis3 | 0.49 | 7.30E-08 | 1.52E-03 | 0.28 | 0.20 | 0.08 | 1.41 |
| 10 | Scamp2 | 0.39 | 7.33E-08 | 1.53E-03 | 0.40 | 0.31 | 0.09 | 1.27 |
| 10 | Lrrc61 | 0.67 | 7.64E-08 | 1.59E-03 | 0.15 | 0.10 | 0.06 | 1.60 |
| 10 | Pou2f1 | 0.43 | 7.73E-08 | 1.61E-03 | 0.44 | 0.36 | 0.08 | 1.22 |
| 10 | Pigv | 0.71 | 7.78E-08 | 1.62E-03 | 0.15 | 0.09 | 0.06 | 1.60 |
| 10 | Zfyve27 | 0.65 | 7.82E-08 | 1.63E-03 | 0.19 | 0.13 | 0.07 | 1.52 |
| 10 | Slfn9 | 0.48 | 7.86E-08 | 1.63E-03 | 0.33 | 0.25 | 0.08 | 1.33 |
| 10 | Sfi1 | 0.90 | 7.96E-08 | 1.66E-03 | 0.12 | 0.07 | 0.05 | 1.72 |
| 10 | Ears2 | 0.54 | 8.27E-08 | 1.72E-03 | 0.23 | 0.16 | 0.07 | 1.46 |
| 10 | Cops7a | 0.25 | 8.72E-08 | 1.81E-03 | 0.68 | 0.59 | 0.08 | 1.14 |
| 10 | Aldh4a1 | 0.75 | 9.06E-08 | 1.88E-03 | 0.15 | 0.09 | 0.06 | 1.62 |
| 10 | 3110082I17Rik | 0.47 | 9.17E-08 | 1.91E-03 | 0.38 | 0.30 | 0.09 | 1.29 |
| 10 | Fam210a | 0.42 | 9.21E-08 | 1.92E-03 | 0.40 | 0.31 | 0.08 | 1.27 |
| 10 | Hbp1 | 0.28 | 9.38E-08 | 1.95E-03 | 0.65 | 0.56 | 0.09 | 1.16 |
| 10 | Rgs3 | 0.50 | 9.48E-08 | 1.97E-03 | 0.25 | 0.18 | 0.07 | 1.41 |
| 10 | Fuca2 | 0.37 | 9.62E-08 | 2.00E-03 | 0.32 | 0.23 | 0.09 | 1.37 |
| 10 | Parn | 0.47 | 1.00E-07 | 2.08E-03 | 0.34 | 0.26 | 0.08 | 1.30 |
| 10 | Sertad2 | 0.33 | 1.03E-07 | 2.14E-03 | 0.49 | 0.40 | 0.09 | 1.22 |
| 10 | Zfp800 | 0.39 | 1.03E-07 | 2.14E-03 | 0.35 | 0.27 | 0.08 | 1.31 |
| 10 | Brca2 | 0.39 | 1.15E-07 | 2.39E-03 | 0.54 | 0.47 | 0.07 | 1.15 |
| 10 | Zfp316 | 0.60 | 1.16E-07 | 2.42E-03 | 0.21 | 0.14 | 0.07 | 1.48 |
| 10 | Dvl1 | 0.60 | 1.31E-07 | 2.72E-03 | 0.20 | 0.13 | 0.07 | 1.50 |
| 10 | Ascc3 | 0.32 | 1.33E-07 | 2.77E-03 | 0.52 | 0.43 | 0.09 | 1.21 |
| 10 | Zfp113 | 0.38 | 1.37E-07 | 2.85E-03 | 0.41 | 0.32 | 0.09 | 1.27 |
| 10 | Upf1 | 0.35 | 1.38E-07 | 2.87E-03 | 0.55 | 0.47 | 0.08 | 1.16 |
| 10 | Ngrn | 0.29 | 1.47E-07 | 3.06E-03 | 0.55 | 0.46 | 0.08 | 1.17 |
| 10 | Eml2 | 0.57 | 1.48E-07 | 3.08E-03 | 0.16 | 0.10 | 0.06 | 1.58 |
| 10 | Gpalpp1 | 0.32 | 1.50E-07 | 3.12E-03 | 0.44 | 0.36 | 0.08 | 1.23 |
| 10 | Egr1 | 0.68 | 1.50E-07 | 3.12E-03 | 0.13 | 0.08 | 0.05 | 1.67 |
| 10 | Cep290 | 0.33 | 1.53E-07 | 3.19E-03 | 0.59 | 0.51 | 0.08 | 1.15 |
| 10 | Rcsd1 | 0.67 | 1.54E-07 | 3.21E-03 | 0.15 | 0.09 | 0.06 | 1.61 |

**Supplementary Table 1. Marker genes for the original clusters of integrated clustering of E11.5 control and NotchGOF λ-junction cells.**

| | | | | | | | | |
|---|---|---|---|---|---|---|---|---|
| 10 | Gucy1a2 | 0.72 | 1.57E-07 | 3.26E-03 | 0.12 | 0.07 | 0.05 | 1.72 |
| 10 | Renbp | 0.68 | 1.67E-07 | 3.48E-03 | 0.13 | 0.08 | 0.05 | 1.66 |
| 10 | Tab3 | 0.69 | 1.79E-07 | 3.72E-03 | 0.14 | 0.09 | 0.05 | 1.62 |
| 10 | Med18 | 0.61 | 1.86E-07 | 3.88E-03 | 0.18 | 0.12 | 0.06 | 1.51 |
| 10 | Bcl9l | 0.33 | 1.87E-07 | 3.89E-03 | 0.42 | 0.33 | 0.09 | 1.28 |
| 10 | Reps1 | 0.41 | 1.88E-07 | 3.92E-03 | 0.43 | 0.35 | 0.08 | 1.23 |
| 10 | Mettl1 | 0.26 | 1.90E-07 | 3.95E-03 | 0.67 | 0.58 | 0.09 | 1.15 |
| 10 | Brms1 | 0.36 | 1.91E-07 | 3.98E-03 | 0.43 | 0.35 | 0.08 | 1.24 |
| 10 | Shmt2 | 0.43 | 1.96E-07 | 4.08E-03 | 0.42 | 0.34 | 0.08 | 1.24 |
| 10 | Efs | 0.34 | 2.00E-07 | 4.16E-03 | 0.37 | 0.28 | 0.09 | 1.32 |
| 10 | Cacul1 | 0.41 | 2.03E-07 | 4.21E-03 | 0.37 | 0.28 | 0.08 | 1.29 |
| 10 | 1190007I07Rik | 0.41 | 2.14E-07 | 4.45E-03 | 0.36 | 0.28 | 0.09 | 1.31 |
| 10 | A430018G15Rik | 0.85 | 2.15E-07 | 4.48E-03 | 0.10 | 0.06 | 0.05 | 1.76 |
| 10 | Dck | 0.31 | 2.16E-07 | 4.50E-03 | 0.54 | 0.46 | 0.08 | 1.18 |
| 10 | Borcs8 | 0.26 | 2.19E-07 | 4.55E-03 | 0.64 | 0.55 | 0.09 | 1.16 |
| 10 | Rabggta | 0.62 | 2.26E-07 | 4.69E-03 | 0.19 | 0.13 | 0.06 | 1.49 |
| 10 | Pigt | 0.33 | 2.29E-07 | 4.77E-03 | 0.52 | 0.43 | 0.09 | 1.21 |
| 10 | Nudt8 | 0.45 | 2.33E-07 | 4.85E-03 | 0.30 | 0.22 | 0.08 | 1.35 |
| 10 | 2610507B11Rik | 0.31 | 2.40E-07 | 4.99E-03 | 0.54 | 0.46 | 0.08 | 1.17 |
| 10 | Mcf2l | 0.79 | 2.41E-07 | 5.02E-03 | 0.12 | 0.07 | 0.05 | 1.71 |
| 10 | Qars | 0.29 | 2.41E-07 | 5.02E-03 | 0.67 | 0.59 | 0.08 | 1.13 |
| 10 | Rabif | 0.37 | 2.53E-07 | 5.25E-03 | 0.33 | 0.25 | 0.08 | 1.32 |
| 10 | Zc3h11a | 0.92 | 2.54E-07 | 5.28E-03 | 0.11 | 0.06 | 0.05 | 1.73 |
| 10 | Haus2 | 0.42 | 2.56E-07 | 5.33E-03 | 0.35 | 0.27 | 0.08 | 1.31 |
| 10 | Pex16 | 0.50 | 2.60E-07 | 5.41E-03 | 0.25 | 0.18 | 0.07 | 1.39 |
| 10 | Nfkb1 | 0.36 | 2.66E-07 | 5.53E-03 | 0.39 | 0.30 | 0.09 | 1.31 |
| 10 | Ikbkg | 0.61 | 2.75E-07 | 5.72E-03 | 0.18 | 0.12 | 0.06 | 1.52 |
| 10 | Slc27a1 | 0.63 | 3.05E-07 | 6.35E-03 | 0.14 | 0.08 | 0.05 | 1.64 |
| 10 | Dtnb | 0.46 | 3.24E-07 | 6.74E-03 | 0.35 | 0.27 | 0.08 | 1.28 |
| 10 | Dtna | 0.39 | 3.42E-07 | 7.11E-03 | 0.21 | 0.14 | 0.07 | 1.48 |
| 10 | Pitpnm2 | 0.72 | 3.44E-07 | 7.15E-03 | 0.14 | 0.09 | 0.05 | 1.60 |
| 10 | Mrpl44 | 0.34 | 3.45E-07 | 7.17E-03 | 0.49 | 0.41 | 0.08 | 1.20 |
| 10 | Col4a3bp | 0.39 | 3.60E-07 | 7.48E-03 | 0.41 | 0.32 | 0.09 | 1.27 |
| 10 | Rasa3 | 0.67 | 3.68E-07 | 7.66E-03 | 0.13 | 0.08 | 0.05 | 1.64 |
| 10 | Mpp5 | 0.37 | 3.70E-07 | 7.70E-03 | 0.45 | 0.37 | 0.08 | 1.21 |
| 10 | Oxld1 | 0.58 | 3.71E-07 | 7.72E-03 | 0.18 | 0.12 | 0.06 | 1.51 |
| 10 | Kif16b | 0.55 | 3.78E-07 | 7.87E-03 | 0.22 | 0.15 | 0.07 | 1.43 |
| 10 | Tmem88 | 0.55 | 3.83E-07 | 7.96E-03 | 0.10 | 0.06 | 0.05 | 1.76 |
| 10 | Dcaf1 | 0.27 | 3.87E-07 | 8.05E-03 | 0.53 | 0.44 | 0.09 | 1.21 |
| 10 | Sirt5 | 0.46 | 3.95E-07 | 8.21E-03 | 0.35 | 0.27 | 0.08 | 1.31 |
| 10 | Kbtbd2 | 0.36 | 4.09E-07 | 8.51E-03 | 0.42 | 0.33 | 0.09 | 1.26 |
| 10 | Gm9856 | 0.50 | 4.20E-07 | 8.74E-03 | 0.22 | 0.15 | 0.07 | 1.43 |
| 10 | Ttf1 | 0.35 | 4.63E-07 | 9.63E-03 | 0.42 | 0.34 | 0.09 | 1.25 |
| 10 | Rrp7a | 0.27 | 4.80E-07 | 9.98E-03 | 0.56 | 0.48 | 0.09 | 1.19 |
| 10 | Zadh2 | 0.66 | 4.81E-07 | 1.00E-02 | 0.16 | 0.11 | 0.06 | 1.54 |
| 10 | Fntb | 0.40 | 5.01E-07 | 1.04E-02 | 0.30 | 0.22 | 0.08 | 1.34 |
| 10 | Rapgef5 | 0.43 | 5.02E-07 | 1.04E-02 | 0.18 | 0.12 | 0.06 | 1.52 |
| 10 | Nek7 | 0.36 | 5.05E-07 | 1.05E-02 | 0.38 | 0.29 | 0.08 | 1.28 |
| 10 | Sh3bp1 | 0.42 | 5.08E-07 | 1.06E-02 | 0.37 | 0.29 | 0.09 | 1.30 |
| 10 | Kidins220 | 0.41 | 5.15E-07 | 1.07E-02 | 0.36 | 0.28 | 0.08 | 1.28 |
| 10 | H2-K1 | 0.42 | 5.19E-07 | 1.08E-02 | 0.20 | 0.13 | 0.07 | 1.49 |
| 10 | Hnrnpul1 | 0.27 | 5.24E-07 | 1.09E-02 | 0.74 | 0.68 | 0.06 | 1.09 |
| 10 | Gmppa | 0.52 | 5.29E-07 | 1.10E-02 | 0.24 | 0.17 | 0.07 | 1.40 |
| 10 | Birc2 | 0.41 | 5.42E-07 | 1.13E-02 | 0.43 | 0.35 | 0.08 | 1.24 |
| 10 | Senp8 | 0.43 | 5.76E-07 | 1.20E-02 | 0.28 | 0.21 | 0.07 | 1.35 |
| 10 | Prmt3 | 0.34 | 5.86E-07 | 1.22E-02 | 0.41 | 0.33 | 0.08 | 1.25 |

**Supplementary Table 1. Marker genes for the original clusters of integrated clustering of E11.5 control and NotchGOF λ-junction cells.**

| | | | | | | | | |
|---|---|---|---|---|---|---|---|---|
| 10 | Prr14l | 0.41 | 5.86E-07 | 1.22E-02 | 0.32 | 0.24 | 0.08 | 1.32 |
| 10 | Zfp740 | 0.29 | 5.93E-07 | 1.23E-02 | 0.52 | 0.43 | 0.09 | 1.21 |
| 10 | Fbxl6 | 0.36 | 5.97E-07 | 1.24E-02 | 0.42 | 0.33 | 0.09 | 1.27 |
| 10 | Smim20 | 0.29 | 6.13E-07 | 1.28E-02 | 0.49 | 0.41 | 0.09 | 1.21 |
| 10 | Zfp458 | 0.69 | 6.15E-07 | 1.28E-02 | 0.11 | 0.06 | 0.04 | 1.71 |
| 10 | Dtd2 | 0.43 | 6.88E-07 | 1.43E-02 | 0.40 | 0.32 | 0.08 | 1.24 |
| 10 | Chd2 | 0.25 | 7.03E-07 | 1.46E-02 | 0.70 | 0.62 | 0.08 | 1.12 |
| 10 | Mnt | 0.50 | 7.70E-07 | 1.60E-02 | 0.24 | 0.17 | 0.07 | 1.39 |
| 10 | Pi4k2b | 0.47 | 7.70E-07 | 1.60E-02 | 0.36 | 0.28 | 0.08 | 1.27 |
| 10 | Rnf167 | 0.39 | 7.98E-07 | 1.66E-02 | 0.44 | 0.37 | 0.08 | 1.20 |
| 10 | Rnf24 | 0.42 | 7.98E-07 | 1.66E-02 | 0.37 | 0.30 | 0.08 | 1.25 |
| 10 | Tmem245 | 0.40 | 8.28E-07 | 1.72E-02 | 0.48 | 0.40 | 0.08 | 1.19 |
| 10 | Gan | 0.45 | 8.61E-07 | 1.79E-02 | 0.31 | 0.24 | 0.07 | 1.31 |
| 10 | Mgat5 | 0.42 | 8.67E-07 | 1.80E-02 | 0.34 | 0.26 | 0.08 | 1.30 |
| 10 | Cep70 | 0.35 | 8.78E-07 | 1.83E-02 | 0.47 | 0.40 | 0.07 | 1.18 |
| 10 | Gnb1l | 0.62 | 9.24E-07 | 1.92E-02 | 0.14 | 0.09 | 0.05 | 1.58 |
| 10 | Dpp3 | 0.25 | 9.36E-07 | 1.95E-02 | 0.64 | 0.58 | 0.06 | 1.11 |
| 10 | Prdm11 | 0.64 | 9.49E-07 | 1.97E-02 | 0.13 | 0.08 | 0.05 | 1.60 |
| 10 | 1700037H04Rik | 0.26 | 9.51E-07 | 1.98E-02 | 0.56 | 0.48 | 0.09 | 1.19 |
| 10 | Ptprz1 | 1.02 | 1.00E-06 | 2.09E-02 | 0.16 | 0.11 | 0.05 | 1.48 |
| 10 | C87436 | 0.44 | 1.06E-06 | 2.20E-02 | 0.37 | 0.29 | 0.08 | 1.26 |
| 10 | Stk4 | 0.34 | 1.06E-06 | 2.21E-02 | 0.44 | 0.36 | 0.08 | 1.22 |
| 10 | Pus1 | 0.36 | 1.09E-06 | 2.26E-02 | 0.37 | 0.29 | 0.08 | 1.27 |
| 10 | 1700007L15Rik | 0.72 | 1.12E-06 | 2.34E-02 | 0.12 | 0.07 | 0.05 | 1.65 |
| 10 | 2210016F16Rik | 0.37 | 1.12E-06 | 2.34E-02 | 0.43 | 0.35 | 0.08 | 1.22 |
| 10 | Rabgef1 | 0.59 | 1.13E-06 | 2.35E-02 | 0.23 | 0.16 | 0.06 | 1.38 |
| 10 | Gli1 | 0.50 | 1.13E-06 | 2.35E-02 | 0.19 | 0.13 | 0.06 | 1.46 |
| 10 | Dtx2 | 0.40 | 1.13E-06 | 2.35E-02 | 0.28 | 0.21 | 0.07 | 1.36 |
| 10 | Mga | 0.28 | 1.18E-06 | 2.46E-02 | 0.64 | 0.58 | 0.06 | 1.11 |
| 10 | Kif1c | 0.44 | 1.24E-06 | 2.59E-02 | 0.22 | 0.16 | 0.07 | 1.42 |
| 10 | Adipor1 | 0.27 | 1.24E-06 | 2.59E-02 | 0.63 | 0.56 | 0.07 | 1.13 |
| 10 | Sord | 0.49 | 1.27E-06 | 2.64E-02 | 0.21 | 0.15 | 0.06 | 1.41 |
| 10 | 1700012D14Rik | 0.52 | 1.29E-06 | 2.68E-02 | 0.19 | 0.13 | 0.06 | 1.47 |
| 10 | Aebp2 | 0.28 | 1.31E-06 | 2.73E-02 | 0.50 | 0.42 | 0.09 | 1.21 |
| 10 | Sgpl1 | 0.36 | 1.34E-06 | 2.78E-02 | 0.42 | 0.34 | 0.08 | 1.24 |
| 10 | 1600012H06Rik | 0.39 | 1.35E-06 | 2.81E-02 | 0.29 | 0.22 | 0.08 | 1.35 |
| 10 | Pea15a | 0.28 | 1.37E-06 | 2.84E-02 | 0.31 | 0.23 | 0.08 | 1.34 |
| 10 | Pank1 | 0.40 | 1.37E-06 | 2.85E-02 | 0.39 | 0.31 | 0.08 | 1.25 |
| 10 | Sephs2 | 0.34 | 1.38E-06 | 2.87E-02 | 0.49 | 0.42 | 0.07 | 1.18 |
| 10 | Wiz | 0.31 | 1.39E-06 | 2.90E-02 | 0.50 | 0.42 | 0.08 | 1.20 |
| 10 | Zfp367 | 0.60 | 1.40E-06 | 2.91E-02 | 0.20 | 0.14 | 0.06 | 1.42 |
| 10 | Dcaf6 | 0.62 | 1.43E-06 | 2.98E-02 | 0.13 | 0.08 | 0.05 | 1.60 |
| 10 | Sema4b | 0.70 | 1.44E-06 | 2.99E-02 | 0.14 | 0.09 | 0.05 | 1.57 |
| 10 | Fam172a | 0.34 | 1.50E-06 | 3.12E-02 | 0.43 | 0.35 | 0.08 | 1.23 |
| 10 | L3mbtl3 | 0.25 | 1.56E-06 | 3.24E-02 | 0.49 | 0.39 | 0.10 | 1.24 |
| 10 | Chd1l | 0.49 | 1.58E-06 | 3.29E-02 | 0.27 | 0.20 | 0.07 | 1.33 |
| 10 | Rad54b | 0.41 | 1.61E-06 | 3.35E-02 | 0.27 | 0.20 | 0.07 | 1.35 |
| 10 | Ccdc173 | 0.48 | 1.66E-06 | 3.45E-02 | 0.18 | 0.12 | 0.06 | 1.49 |
| 10 | Ube2t | 0.42 | 1.69E-06 | 3.52E-02 | 0.40 | 0.33 | 0.07 | 1.23 |
| 10 | Ankrd37 | 0.56 | 1.86E-06 | 3.86E-02 | 0.22 | 0.16 | 0.06 | 1.40 |
| 10 | Pmm1 | 0.36 | 1.86E-06 | 3.87E-02 | 0.35 | 0.27 | 0.08 | 1.29 |
| 10 | Etv3 | 0.46 | 1.89E-06 | 3.93E-02 | 0.29 | 0.22 | 0.07 | 1.32 |
| 10 | Rps6ka1 | 0.33 | 1.94E-06 | 4.04E-02 | 0.32 | 0.24 | 0.08 | 1.32 |
| 10 | C330018D20Rik | 0.48 | 2.02E-06 | 4.19E-02 | 0.25 | 0.18 | 0.07 | 1.35 |
| 10 | Maf | 0.37 | 2.13E-06 | 4.44E-02 | 0.12 | 0.07 | 0.04 | 1.60 |
| 10 | Slc39a11 | 0.63 | 2.14E-06 | 4.45E-02 | 0.11 | 0.07 | 0.04 | 1.65 |

**Supplementary Table 1. Marker genes for the original clusters of integrated clustering of E11.5 control and NotchGOF λ-junction cells.**

| | | | | | | | | |
|---|---|---|---|---|---|---|---|---|
| 10 | Ap3b1 | 0.30 | 2.18E-06 | 4.53E-02 | 0.54 | 0.47 | 0.07 | 1.14 |
| 10 | Slc25a36 | 0.31 | 2.18E-06 | 4.53E-02 | 0.51 | 0.43 | 0.08 | 1.17 |
| 10 | Mapk12 | 0.45 | 2.21E-06 | 4.59E-02 | 0.17 | 0.11 | 0.06 | 1.50 |
| 10 | Ppp1r21 | 0.39 | 2.28E-06 | 4.74E-02 | 0.28 | 0.21 | 0.07 | 1.34 |
| 11 | Sfn | 4.29 | 0 | 0 | 0.98 | 0.18 | 0.81 | 5.52 |
| 11 | Dsp | 3.15 | 0 | 0 | 0.97 | 0.21 | 0.76 | 4.59 |
| 11 | Nectin4 | 3.88 | 0 | 0 | 0.84 | 0.10 | 0.74 | 8.12 |
| 11 | Cldn6 | 2.81 | 0 | 0 | 0.94 | 0.20 | 0.74 | 4.62 |
| 11 | Hspb8 | 3.76 | 0 | 0 | 0.82 | 0.11 | 0.71 | 7.31 |
| 11 | Spint2 | 2.37 | 0 | 0 | 0.98 | 0.29 | 0.69 | 3.40 |
| 11 | Perp | 3.13 | 0 | 0 | 0.80 | 0.12 | 0.68 | 6.89 |
| 11 | Six3 | 2.31 | 0 | 0 | 0.90 | 0.22 | 0.68 | 4.18 |
| 11 | Cldn7 | 2.74 | 0 | 0 | 0.89 | 0.22 | 0.68 | 4.14 |
| 11 | Irf6 | 3.75 | 0 | 0 | 0.76 | 0.08 | 0.68 | 9.05 |
| 11 | Epcam | 2.40 | 0 | 0 | 1.00 | 0.32 | 0.67 | 3.08 |
| 11 | Gm16136 | 3.36 | 0 | 0 | 0.82 | 0.15 | 0.67 | 5.47 |
| 11 | Bcam | 2.59 | 0 | 0 | 0.84 | 0.18 | 0.66 | 4.68 |
| 11 | Six3os1 | 2.51 | 0 | 0 | 0.86 | 0.21 | 0.65 | 4.12 |
| 11 | Mpzl2 | 2.84 | 0 | 0 | 0.84 | 0.18 | 0.65 | 4.54 |
| 11 | Ikzf2 | 2.75 | 0 | 0 | 0.91 | 0.26 | 0.65 | 3.50 |
| 11 | Wnt4 | 2.59 | 0 | 0 | 0.82 | 0.18 | 0.65 | 4.63 |
| 11 | Fermt1 | 2.74 | 0 | 0 | 0.76 | 0.12 | 0.64 | 6.43 |
| 11 | Tpd52 | 2.24 | 0 | 0 | 0.88 | 0.24 | 0.64 | 3.67 |
| 11 | Cpm | 3.48 | 0 | 0 | 0.79 | 0.15 | 0.64 | 5.17 |
| 11 | Ezr | 2.26 | 0 | 0 | 0.93 | 0.30 | 0.63 | 3.13 |
| 11 | Cdh1 | 2.96 | 0 | 0 | 0.78 | 0.15 | 0.63 | 5.31 |
| 11 | Notch1 | 3.61 | 0 | 0 | 0.85 | 0.22 | 0.63 | 3.80 |
| 11 | Trp63 | 2.89 | 0 | 0 | 0.74 | 0.11 | 0.63 | 6.58 |
| 11 | Runx1 | 3.77 | 0 | 0 | 0.79 | 0.17 | 0.62 | 4.64 |
| 11 | Nav2 | 2.70 | 0 | 0 | 0.82 | 0.22 | 0.60 | 3.75 |
| 11 | Hist3h2ba | 1.91 | 0 | 0 | 0.86 | 0.25 | 0.60 | 3.38 |
| 11 | Isl1 | 2.32 | 0 | 0 | 0.80 | 0.20 | 0.60 | 4.01 |
| 11 | Cdkn1a | 4.41 | 0 | 0 | 0.69 | 0.09 | 0.60 | 7.70 |
| 11 | Hey1 | 2.41 | 0 | 0 | 0.88 | 0.28 | 0.60 | 3.12 |
| 11 | Igfbp5 | 2.85 | 0 | 0 | 0.98 | 0.39 | 0.59 | 2.53 |
| 11 | Nav3 | 3.57 | 0 | 0 | 0.70 | 0.11 | 0.59 | 6.58 |
| 11 | Krt5 | 4.52 | 0 | 0 | 0.65 | 0.07 | 0.58 | 9.33 |
| 11 | Wnt6 | 2.81 | 0 | 0 | 0.66 | 0.08 | 0.58 | 8.33 |
| 11 | Cldn3 | 2.66 | 0 | 0 | 0.72 | 0.14 | 0.58 | 5.03 |
| 11 | Frem2 | 2.61 | 0 | 0 | 0.75 | 0.17 | 0.57 | 4.31 |
| 11 | Slc9a3r1 | 2.12 | 0 | 0 | 0.79 | 0.22 | 0.57 | 3.59 |
| 11 | Esrp1 | 2.76 | 0 | 0 | 0.72 | 0.15 | 0.57 | 4.85 |
| 11 | Jag1 | 2.89 | 0 | 0 | 0.73 | 0.16 | 0.57 | 4.62 |
| 11 | Rab25 | 2.58 | 0 | 0 | 0.71 | 0.14 | 0.57 | 5.02 |
| 11 | Ly6h | 3.16 | 0 | 0 | 0.70 | 0.13 | 0.57 | 5.35 |
| 11 | Krt14 | 2.94 | 0 | 0 | 0.67 | 0.11 | 0.56 | 5.95 |
| 11 | Dab1 | 2.91 | 0 | 0 | 0.69 | 0.13 | 0.56 | 5.17 |
| 11 | F11r | 2.47 | 0 | 0 | 0.72 | 0.16 | 0.56 | 4.46 |
| 11 | Pdlim1 | 2.64 | 0 | 0 | 0.82 | 0.27 | 0.55 | 3.04 |
| 11 | Cd9 | 2.73 | 0 | 0 | 0.83 | 0.28 | 0.55 | 2.96 |
| 11 | Ablim1 | 2.49 | 0 | 0 | 0.75 | 0.21 | 0.54 | 3.62 |
| 11 | Jup | 2.65 | 0 | 0 | 0.72 | 0.18 | 0.54 | 4.01 |
| 11 | Anxa2 | 2.15 | 0 | 0 | 0.94 | 0.40 | 0.54 | 2.36 |
| 11 | Erbb3 | 2.44 | 0 | 0 | 0.69 | 0.16 | 0.54 | 4.41 |
| 11 | Nrarp | 2.36 | 0 | 0 | 0.70 | 0.16 | 0.53 | 4.32 |
| 11 | Ston2 | 2.45 | 0 | 0 | 0.70 | 0.16 | 0.53 | 4.28 |

**Supplementary Table 1. Marker genes for the original clusters of integrated clustering of E11.5 control and NotchGOF λ-junction cells.**

| | | | | | | | | |
|---|---|---|---|---|---|---|---|---|
| 11 | Dlx2 | 3.28 | 0 | 0 | 0.64 | 0.11 | 0.53 | 5.84 |
| 11 | Kcnk1 | 2.65 | 0 | 0 | 0.70 | 0.17 | 0.53 | 4.17 |
| 11 | Pik3r1 | 2.65 | 0 | 0 | 0.90 | 0.37 | 0.53 | 2.42 |
| 11 | Frmd4b | 2.64 | 0 | 0 | 0.67 | 0.15 | 0.52 | 4.46 |
| 11 | Wwc1 | 2.54 | 0 | 0 | 0.67 | 0.16 | 0.51 | 4.13 |
| 11 | Rab15 | 2.93 | 0 | 0 | 0.62 | 0.12 | 0.50 | 5.36 |
| 11 | Cacna1b | 3.58 | 0 | 0 | 0.56 | 0.06 | 0.49 | 8.84 |
| 11 | Efna1 | 2.57 | 0 | 0 | 0.61 | 0.12 | 0.49 | 4.98 |
| 11 | Zfp750 | 4.65 | 0 | 0 | 0.52 | 0.03 | 0.49 | 16.28 |
| 11 | Ppp2r2b | 3.27 | 0 | 0 | 0.57 | 0.08 | 0.49 | 7.19 |
| 11 | Ccdc88c | 2.53 | 0 | 0 | 0.63 | 0.14 | 0.49 | 4.49 |
| 11 | Dlx3 | 3.52 | 0 | 0 | 0.54 | 0.06 | 0.49 | 9.54 |
| 11 | Cldn4 | 3.33 | 0 | 0 | 0.57 | 0.09 | 0.48 | 6.55 |
| 11 | St14 | 2.85 | 0 | 0 | 0.58 | 0.10 | 0.48 | 5.73 |
| 11 | Ddr1 | 2.69 | 0 | 0 | 0.60 | 0.11 | 0.48 | 5.23 |
| 11 | Ap1m2 | 2.43 | 0 | 0 | 0.61 | 0.13 | 0.48 | 4.78 |
| 11 | Dok5 | 2.80 | 0 | 0 | 0.58 | 0.10 | 0.48 | 5.83 |
| 11 | Fgf8 | 4.81 | 0 | 0 | 0.54 | 0.06 | 0.48 | 8.69 |
| 11 | Dsg2 | 2.71 | 0 | 0 | 0.59 | 0.11 | 0.48 | 5.21 |
| 11 | Sp8 | 2.36 | 0 | 0 | 0.60 | 0.12 | 0.47 | 4.84 |
| 11 | Mir205hg | 2.58 | 0 | 0 | 0.57 | 0.10 | 0.47 | 5.50 |
| 11 | Arhgef19 | 2.62 | 0 | 0 | 0.58 | 0.12 | 0.47 | 5.04 |
| 11 | Grhl3 | 4.22 | 0 | 0 | 0.50 | 0.04 | 0.46 | 12.79 |
| 11 | Grhl2 | 2.57 | 0 | 0 | 0.57 | 0.11 | 0.46 | 5.01 |
| 11 | Lurap1l | 3.65 | 0 | 0 | 0.51 | 0.06 | 0.45 | 8.55 |
| 11 | Spint1 | 3.03 | 0 | 0 | 0.53 | 0.08 | 0.45 | 6.32 |
| 11 | Lad1 | 3.18 | 0 | 0 | 0.52 | 0.07 | 0.44 | 6.97 |
| 11 | Tacstd2 | 4.47 | 0 | 0 | 0.49 | 0.05 | 0.44 | 9.06 |
| 11 | Nebl | 3.47 | 0 | 0 | 0.53 | 0.09 | 0.43 | 5.67 |
| 11 | Klc3 | 2.52 | 0 | 0 | 0.52 | 0.09 | 0.42 | 5.55 |
| 11 | Fxyd3 | 2.88 | 0 | 0 | 0.48 | 0.08 | 0.41 | 6.27 |
| 11 | Ppl | 4.67 | 0 | 0 | 0.40 | 0.03 | 0.37 | 14.21 |
| 11 | Ptprf | 1.94 | 0 | 0 | 0.98 | 0.63 | 0.36 | 1.57 |
| 11 | Esrp2 | 3.32 | 0 | 0 | 0.41 | 0.05 | 0.36 | 7.72 |
| 11 | Cdkn2b | 5.59 | 0 | 0 | 0.37 | 0.01 | 0.36 | 30.58 |
| 11 | Rhov | 3.61 | 0 | 0 | 0.40 | 0.05 | 0.35 | 8.78 |
| 11 | Bspry | 3.12 | 0 | 0 | 0.40 | 0.05 | 0.35 | 7.55 |
| 11 | Tctex1d1 | 4.72 | 0 | 0 | 0.37 | 0.02 | 0.35 | 16.00 |
| 11 | Pkp1 | 5.09 | 0 | 0 | 0.36 | 0.01 | 0.34 | 27.31 |
| 11 | Proser2 | 3.57 | 0 | 0 | 0.38 | 0.05 | 0.33 | 8.38 |
| 11 | Cldn1 | 4.88 | 0 | 0 | 0.32 | 0.02 | 0.30 | 18.76 |
| 11 | D7Ertd443e | 4.50 | 0 | 0 | 0.29 | 0.01 | 0.28 | 20.64 |
| 11 | Gjb3 | 3.76 | 0 | 0 | 0.30 | 0.03 | 0.27 | 11.84 |
| 11 | Lratd1 | 4.17 | 0 | 0 | 0.29 | 0.03 | 0.27 | 11.60 |
| 11 | S100a11 | 2.52 | 0 | 0 | 1.00 | 0.75 | 0.25 | 1.33 |
| 11 | Lypd3 | 4.26 | 0 | 0 | 0.26 | 0.02 | 0.24 | 15.18 |
| 11 | Cldn8 | 3.81 | 0 | 0 | 0.26 | 0.02 | 0.24 | 12.43 |
| 11 | Ovol1 | 4.60 | 0 | 0 | 0.22 | 0.01 | 0.21 | 18.42 |
| 11 | Tgm1 | 5.06 | 0 | 0 | 0.19 | 0.01 | 0.18 | 23.50 |
| 11 | Rhcg | 4.64 | 0 | 0 | 0.15 | 0.01 | 0.14 | 29.60 |
| 11 | Gpr87 | 6.07 | 0 | 0 | 0.14 | 0.00 | 0.14 | 46.67 |
| 11 | Fabp5 | 1.69 | 1.27E-307 | 2.63E-303 | 1.00 | 0.98 | 0.02 | 1.02 |
| 11 | Col18a1 | 1.95 | 3.62E-302 | 7.53E-298 | 0.89 | 0.34 | 0.55 | 2.59 |
| 11 | Smtnl2 | 3.05 | 1.80E-301 | 3.75E-297 | 0.42 | 0.06 | 0.35 | 6.71 |
| 11 | Pdgfc | 2.10 | 2.01E-301 | 4.18E-297 | 0.82 | 0.26 | 0.55 | 3.11 |
| 11 | Diras2 | 4.23 | 3.62E-300 | 7.53E-296 | 0.22 | 0.02 | 0.20 | 13.75 |

**Supplementary Table 1. Marker genes for the original clusters of integrated clustering of E11.5 control and NotchGOF λ-junction cells.**

| | | | | | | | | |
|---|---|---|---|---|---|---|---|---|
| 11 | Itm2b | 2.65 | 2.18E-297 | 4.54E-293 | 0.99 | 0.84 | 0.15 | 1.18 |
| 11 | Klf5 | 2.63 | 1.52E-295 | 3.16E-291 | 0.54 | 0.11 | 0.43 | 5.02 |
| 11 | Sp6 | 3.64 | 3.02E-294 | 6.28E-290 | 0.25 | 0.02 | 0.23 | 11.55 |
| 11 | Lsr | 2.23 | 1.56E-292 | 3.24E-288 | 0.70 | 0.19 | 0.51 | 3.73 |
| 11 | Bex1 | 1.79 | 6.01E-292 | 1.25E-287 | 0.98 | 0.58 | 0.40 | 1.70 |
| 11 | Slc2a3 | 3.76 | 6.92E-292 | 1.44E-287 | 0.55 | 0.12 | 0.43 | 4.50 |
| 11 | Marveld2 | 3.17 | 1.49E-291 | 3.09E-287 | 0.36 | 0.05 | 0.32 | 7.58 |
| 11 | Igfbp2 | 1.92 | 7.24E-290 | 1.51E-285 | 1.00 | 0.75 | 0.25 | 1.34 |
| 11 | Tspan7 | 2.56 | 1.10E-288 | 2.30E-284 | 0.76 | 0.26 | 0.50 | 2.90 |
| 11 | Il17re | 3.16 | 1.61E-287 | 3.35E-283 | 0.31 | 0.04 | 0.28 | 8.72 |
| 11 | Gad2 | 3.24 | 5.00E-285 | 1.04E-280 | 0.41 | 0.06 | 0.34 | 6.53 |
| 11 | Socs2 | 2.47 | 5.24E-275 | 1.09E-270 | 0.82 | 0.33 | 0.50 | 2.52 |
| 11 | Itgb6 | 3.33 | 5.13E-272 | 1.07E-267 | 0.29 | 0.03 | 0.26 | 8.85 |
| 11 | Rab27b | 2.48 | 1.45E-269 | 3.02E-265 | 0.46 | 0.08 | 0.38 | 5.52 |
| 11 | Cdkn2a | 5.40 | 2.85E-269 | 5.93E-265 | 0.12 | 0.00 | 0.12 | 31.00 |
| 11 | Map7 | 2.61 | 8.45E-268 | 1.76E-263 | 0.49 | 0.10 | 0.39 | 5.11 |
| 11 | Apoc1 | 2.59 | 2.58E-266 | 5.37E-262 | 0.43 | 0.07 | 0.35 | 5.82 |
| 11 | Krt8 | 1.66 | 6.15E-266 | 1.28E-261 | 0.79 | 0.24 | 0.55 | 3.28 |
| 11 | Ripply3 | 3.13 | 8.41E-266 | 1.75E-261 | 0.35 | 0.05 | 0.30 | 7.08 |
| 11 | Stard10 | 2.00 | 3.36E-265 | 6.98E-261 | 0.80 | 0.28 | 0.51 | 2.82 |
| 11 | Krt18 | 1.52 | 6.22E-265 | 1.29E-260 | 0.85 | 0.28 | 0.57 | 3.05 |
| 11 | Ripk4 | 2.76 | 2.01E-264 | 4.18E-260 | 0.40 | 0.06 | 0.33 | 6.27 |
| 11 | Dstn | 1.28 | 1.33E-263 | 2.77E-259 | 0.99 | 0.94 | 0.06 | 1.06 |
| 11 | Fjx1 | 2.16 | 2.42E-263 | 5.03E-259 | 0.84 | 0.37 | 0.47 | 2.26 |
| 11 | Gad1 | 4.25 | 2.15E-261 | 4.46E-257 | 0.17 | 0.01 | 0.16 | 18.33 |
| 11 | Lmo7 | 2.94 | 1.18E-260 | 2.44E-256 | 0.58 | 0.15 | 0.43 | 3.92 |
| 11 | Crb3 | 2.42 | 4.18E-260 | 8.69E-256 | 0.49 | 0.10 | 0.39 | 5.04 |
| 11 | Hras | 1.49 | 1.58E-259 | 3.29E-255 | 0.96 | 0.80 | 0.17 | 1.21 |
| 11 | Lama5 | 2.54 | 4.87E-256 | 1.01E-251 | 0.49 | 0.10 | 0.39 | 4.84 |
| 11 | S100a10 | 2.20 | 6.74E-255 | 1.40E-250 | 0.92 | 0.54 | 0.38 | 1.70 |
| 11 | Fzd6 | 2.70 | 2.35E-254 | 4.90E-250 | 0.42 | 0.07 | 0.35 | 5.74 |
| 11 | Ppp1r13l | 3.01 | 1.13E-250 | 2.35E-246 | 0.39 | 0.07 | 0.33 | 5.87 |
| 11 | Cdh3 | 2.12 | 3.38E-249 | 7.04E-245 | 0.67 | 0.19 | 0.48 | 3.48 |
| 11 | Dapl1 | 2.68 | 1.47E-242 | 3.05E-238 | 0.37 | 0.06 | 0.31 | 6.27 |
| 11 | Ptprz1 | 3.01 | 3.75E-242 | 7.80E-238 | 0.45 | 0.09 | 0.36 | 4.87 |
| 11 | Tafa2 | 3.50 | 5.58E-242 | 1.16E-237 | 0.27 | 0.03 | 0.24 | 8.53 |
| 11 | Pdgfa | 1.41 | 1.50E-241 | 3.12E-237 | 0.83 | 0.26 | 0.57 | 3.16 |
| 11 | Duoxa1 | 5.52 | 1.82E-241 | 3.79E-237 | 0.10 | 0.00 | 0.10 | 34.33 |
| 11 | Tmem54 | 2.45 | 2.06E-240 | 4.28E-236 | 0.49 | 0.11 | 0.39 | 4.64 |
| 11 | Atox1 | 1.23 | 9.30E-240 | 1.93E-235 | 0.99 | 0.95 | 0.04 | 1.04 |
| 11 | Macc1 | 3.03 | 2.09E-238 | 4.35E-234 | 0.32 | 0.05 | 0.28 | 7.02 |
| 11 | Ocln | 2.72 | 2.46E-238 | 5.11E-234 | 0.39 | 0.07 | 0.32 | 5.86 |
| 11 | Ndufa4l2 | 3.76 | 2.21E-235 | 4.59E-231 | 0.22 | 0.02 | 0.20 | 10.85 |
| 11 | Snx31 | 4.59 | 9.04E-235 | 1.88E-230 | 0.13 | 0.01 | 0.13 | 22.17 |
| 11 | Mir9-3hg | 3.34 | 1.78E-234 | 3.70E-230 | 0.22 | 0.02 | 0.20 | 10.38 |
| 11 | Sema3d | 2.57 | 3.20E-231 | 6.65E-227 | 0.48 | 0.10 | 0.38 | 4.63 |
| 11 | Fgf9 | 2.63 | 1.96E-230 | 4.08E-226 | 0.46 | 0.09 | 0.36 | 4.85 |
| 11 | Cd2ap | 1.81 | 2.75E-230 | 5.72E-226 | 0.92 | 0.61 | 0.31 | 1.51 |
| 11 | Samd5 | 2.09 | 3.51E-229 | 7.31E-225 | 0.81 | 0.35 | 0.47 | 2.35 |
| 11 | Acer2 | 1.68 | 2.52E-228 | 5.24E-224 | 0.85 | 0.38 | 0.47 | 2.25 |
| 11 | Otx1 | 2.21 | 6.44E-228 | 1.34E-223 | 0.47 | 0.10 | 0.37 | 4.73 |
| 11 | Cox4i2 | 2.15 | 9.38E-228 | 1.95E-223 | 0.70 | 0.25 | 0.46 | 2.84 |
| 11 | Capg | 2.36 | 2.35E-226 | 4.88E-222 | 0.54 | 0.14 | 0.40 | 3.94 |
| 11 | Grb7 | 2.41 | 3.33E-226 | 6.94E-222 | 0.44 | 0.09 | 0.35 | 5.07 |
| 11 | Ier5 | 1.63 | 1.24E-224 | 2.58E-220 | 0.94 | 0.64 | 0.30 | 1.47 |
| 11 | Aplp2 | 1.45 | 2.91E-224 | 6.06E-220 | 0.97 | 0.81 | 0.16 | 1.20 |

**Supplementary Table 1. Marker genes for the original clusters of integrated clustering of E11.5 control and NotchGOF λ-junction cells.**

| | | | | | | | | |
|---|---|---|---|---|---|---|---|---|
| 11 | Nectin3 | 1.83 | 3.71E-223 | 7.71E-219 | 0.89 | 0.53 | 0.35 | 1.66 |
| 11 | Dlx1 | 3.02 | 4.24E-222 | 8.83E-218 | 0.30 | 0.04 | 0.26 | 6.98 |
| 11 | Plekhg3 | 2.86 | 1.63E-221 | 3.38E-217 | 0.38 | 0.07 | 0.31 | 5.41 |
| 11 | Pltp | 2.01 | 8.22E-221 | 1.71E-216 | 0.57 | 0.15 | 0.42 | 3.90 |
| 11 | Tgfb2 | 2.60 | 1.87E-218 | 3.88E-214 | 0.87 | 0.47 | 0.40 | 1.84 |
| 11 | Gjb2 | 3.61 | 4.44E-217 | 9.24E-213 | 0.28 | 0.04 | 0.24 | 7.29 |
| 11 | Hspb1 | 2.95 | 1.54E-215 | 3.20E-211 | 0.53 | 0.14 | 0.39 | 3.86 |
| 11 | Pkp3 | 2.63 | 1.41E-214 | 2.93E-210 | 0.34 | 0.06 | 0.29 | 6.00 |
| 11 | Ctnnd2 | 2.25 | 5.01E-209 | 1.04E-204 | 0.47 | 0.11 | 0.36 | 4.36 |
| 11 | Nfe2l3 | 3.23 | 7.49E-207 | 1.56E-202 | 0.23 | 0.03 | 0.21 | 8.67 |
| 11 | Gsta3 | 3.83 | 4.48E-204 | 9.32E-200 | 0.15 | 0.01 | 0.14 | 14.90 |
| 11 | AW011738 | 2.58 | 8.26E-203 | 1.72E-198 | 0.39 | 0.08 | 0.31 | 4.96 |
| 11 | Plp2 | 1.90 | 8.67E-201 | 1.80E-196 | 0.79 | 0.37 | 0.42 | 2.16 |
| 11 | Kcnj15 | 4.05 | 2.05E-199 | 4.26E-195 | 0.16 | 0.01 | 0.15 | 12.31 |
| 11 | Kif21a | 1.90 | 2.12E-199 | 4.42E-195 | 0.75 | 0.30 | 0.45 | 2.53 |
| 11 | Slc15a2 | 3.05 | 6.13E-199 | 1.27E-194 | 0.28 | 0.04 | 0.24 | 6.90 |
| 11 | Mycl | 1.89 | 8.72E-199 | 1.81E-194 | 0.72 | 0.29 | 0.44 | 2.51 |
| 11 | Ntf5 | 2.27 | 2.16E-197 | 4.48E-193 | 0.39 | 0.08 | 0.31 | 5.01 |
| 11 | Sptlc3 | 3.77 | 1.07E-196 | 2.23E-192 | 0.17 | 0.01 | 0.15 | 11.86 |
| 11 | Rab11fip1 | 2.43 | 1.20E-196 | 2.49E-192 | 0.45 | 0.11 | 0.34 | 4.22 |
| 11 | Eppk1 | 3.82 | 3.93E-196 | 8.17E-192 | 0.17 | 0.02 | 0.16 | 11.60 |
| 11 | Itga6 | 1.72 | 1.15E-194 | 2.39E-190 | 0.76 | 0.30 | 0.47 | 2.57 |
| 11 | Nxph3 | 3.50 | 3.14E-194 | 6.53E-190 | 0.17 | 0.02 | 0.16 | 11.33 |
| 11 | H2-T23 | 3.04 | 3.28E-193 | 6.83E-189 | 0.29 | 0.05 | 0.24 | 6.42 |
| 11 | Cblc | 3.09 | 2.34E-192 | 4.87E-188 | 0.22 | 0.03 | 0.20 | 8.50 |
| 11 | Gadd45g | 3.65 | 2.02E-191 | 4.21E-187 | 0.58 | 0.20 | 0.38 | 2.91 |
| 11 | Spock2 | 4.10 | 1.35E-190 | 2.80E-186 | 0.24 | 0.03 | 0.21 | 7.50 |
| 11 | Lmo2 | 2.53 | 1.94E-190 | 4.03E-186 | 0.46 | 0.12 | 0.34 | 3.98 |
| 11 | Fzd3 | 1.52 | 6.04E-190 | 1.26E-185 | 0.91 | 0.58 | 0.33 | 1.57 |
| 11 | Gucy1a1 | 2.98 | 1.42E-189 | 2.95E-185 | 0.35 | 0.07 | 0.28 | 5.23 |
| 11 | Palm3 | 2.43 | 2.39E-189 | 4.98E-185 | 0.50 | 0.14 | 0.36 | 3.67 |
| 11 | Dsc2 | 2.79 | 3.18E-189 | 6.61E-185 | 0.30 | 0.05 | 0.25 | 6.08 |
| 11 | Six6 | 2.25 | 2.03E-188 | 4.23E-184 | 0.34 | 0.06 | 0.28 | 5.51 |
| 11 | Afdn | 1.50 | 8.82E-188 | 1.83E-183 | 0.89 | 0.56 | 0.33 | 1.59 |
| 11 | Itgb4 | 3.23 | 2.42E-187 | 5.03E-183 | 0.22 | 0.03 | 0.19 | 8.60 |
| 11 | Lratd2 | 2.30 | 5.83E-187 | 1.21E-182 | 0.43 | 0.10 | 0.33 | 4.33 |
| 11 | Bex4 | 1.27 | 1.94E-186 | 4.04E-182 | 0.94 | 0.64 | 0.30 | 1.46 |
| 11 | Adamts9 | 3.08 | 5.55E-186 | 1.15E-181 | 0.49 | 0.14 | 0.35 | 3.56 |
| 11 | Cd151 | 1.77 | 1.87E-185 | 3.89E-181 | 0.75 | 0.31 | 0.43 | 2.37 |
| 11 | Mgst1 | 1.71 | 3.40E-185 | 7.06E-181 | 0.79 | 0.36 | 0.43 | 2.20 |
| 11 | Bicdl2 | 3.80 | 2.02E-183 | 4.20E-179 | 0.13 | 0.01 | 0.13 | 14.89 |
| 11 | Krt7 | 2.97 | 3.01E-183 | 6.26E-179 | 0.28 | 0.05 | 0.24 | 6.24 |
| 11 | Sh2d4a | 3.77 | 1.53E-182 | 3.18E-178 | 0.13 | 0.01 | 0.12 | 15.63 |
| 11 | Pdcd4 | 1.62 | 7.99E-182 | 1.66E-177 | 0.92 | 0.63 | 0.29 | 1.46 |
| 11 | Stx3 | 2.23 | 5.59E-181 | 1.16E-176 | 0.46 | 0.12 | 0.34 | 3.97 |
| 11 | 2900026A02Rik | 2.42 | 1.06E-180 | 2.21E-176 | 0.41 | 0.09 | 0.32 | 4.40 |
| 11 | Cndp1 | 4.24 | 2.00E-180 | 4.16E-176 | 0.11 | 0.01 | 0.11 | 18.50 |
| 11 | Sorl1 | 2.34 | 3.98E-180 | 8.28E-176 | 0.50 | 0.14 | 0.36 | 3.59 |
| 11 | Lgalsl | 2.52 | 8.51E-179 | 1.77E-174 | 0.70 | 0.30 | 0.40 | 2.31 |
| 11 | Elf5 | 2.48 | 1.06E-178 | 2.20E-174 | 0.30 | 0.05 | 0.25 | 5.72 |
| 11 | Arl4c | 1.96 | 1.14E-178 | 2.38E-174 | 0.72 | 0.32 | 0.41 | 2.29 |
| 11 | Thrb | 2.71 | 7.10E-178 | 1.48E-173 | 0.33 | 0.06 | 0.27 | 5.21 |
| 11 | Cpz | 2.64 | 1.24E-177 | 2.57E-173 | 0.33 | 0.06 | 0.27 | 5.34 |
| 11 | Aldh1a3 | 1.74 | 1.31E-177 | 2.72E-173 | 0.62 | 0.22 | 0.40 | 2.85 |
| 11 | Rnf43 | 2.87 | 6.31E-177 | 1.31E-172 | 0.27 | 0.04 | 0.23 | 6.36 |
| 11 | Gm32585 | 2.47 | 1.12E-176 | 2.33E-172 | 0.29 | 0.05 | 0.24 | 6.09 |

**Supplementary Table 1. Marker genes for the original clusters of integrated clustering of E11.5 control and NotchGOF λ-junction cells.**

| | | | | | | | | |
|---|---|---|---|---|---|---|---|---|
| 11 | Camsap3 | 2.46 | 2.67E-176 | 5.55E-172 | 0.35 | 0.07 | 0.28 | 4.92 |
| 11 | Cyb5a | 1.05 | 4.32E-176 | 8.99E-172 | 0.98 | 0.92 | 0.06 | 1.06 |
| 11 | Acaa2 | 1.86 | 1.22E-173 | 2.55E-169 | 0.81 | 0.47 | 0.34 | 1.74 |
| 11 | Wls | 1.26 | 1.51E-172 | 3.14E-168 | 0.95 | 0.73 | 0.21 | 1.29 |
| 11 | Elovl7 | 3.36 | 2.85E-172 | 5.93E-168 | 0.19 | 0.02 | 0.17 | 8.95 |
| 11 | Fam49a | 3.20 | 4.92E-172 | 1.02E-167 | 0.25 | 0.04 | 0.21 | 6.62 |
| 11 | Rhob | 1.92 | 1.54E-171 | 3.20E-167 | 0.72 | 0.33 | 0.39 | 2.18 |
| 11 | Phldb3 | 3.43 | 3.20E-171 | 6.65E-167 | 0.17 | 0.02 | 0.15 | 10.38 |
| 11 | Kdf1 | 2.39 | 1.99E-170 | 4.14E-166 | 0.36 | 0.08 | 0.28 | 4.74 |
| 11 | Etl4 | 1.78 | 6.29E-170 | 1.31E-165 | 0.71 | 0.30 | 0.41 | 2.38 |
| 11 | Myo10 | 1.72 | 2.60E-169 | 5.41E-165 | 0.82 | 0.46 | 0.36 | 1.79 |
| 11 | Cyb5r3 | 1.14 | 5.16E-169 | 1.07E-164 | 0.95 | 0.80 | 0.14 | 1.18 |
| 11 | Ahr | 2.56 | 6.00E-169 | 1.25E-164 | 0.36 | 0.08 | 0.28 | 4.59 |
| 11 | Trmt9b | 2.61 | 1.06E-168 | 2.21E-164 | 0.32 | 0.06 | 0.26 | 5.25 |
| 11 | Plce1 | 2.25 | 1.49E-168 | 3.11E-164 | 0.59 | 0.21 | 0.38 | 2.77 |
| 11 | Cebpb | 2.44 | 2.20E-168 | 4.58E-164 | 0.43 | 0.11 | 0.32 | 3.93 |
| 11 | Ppm1l | 2.09 | 2.28E-168 | 4.74E-164 | 0.59 | 0.21 | 0.38 | 2.81 |
| 11 | Tnfaip2 | 1.64 | 3.31E-167 | 6.88E-163 | 0.49 | 0.13 | 0.36 | 3.73 |
| 11 | Cgn | 2.22 | 6.04E-167 | 1.26E-162 | 0.40 | 0.10 | 0.31 | 4.21 |
| 11 | Evpl | 3.04 | 8.36E-167 | 1.74E-162 | 0.23 | 0.03 | 0.19 | 7.26 |
| 11 | Cux1 | 1.28 | 4.79E-166 | 9.96E-162 | 0.93 | 0.69 | 0.24 | 1.34 |
| 11 | Heyl | 2.60 | 1.27E-165 | 2.63E-161 | 0.28 | 0.05 | 0.23 | 5.76 |
| 11 | Zfp385c | 3.36 | 1.63E-165 | 3.40E-161 | 0.21 | 0.03 | 0.18 | 7.63 |
| 11 | Eaf2 | 3.23 | 7.96E-165 | 1.66E-160 | 0.19 | 0.02 | 0.17 | 8.39 |
| 11 | Myh9 | 1.61 | 1.32E-164 | 2.75E-160 | 0.84 | 0.49 | 0.35 | 1.71 |
| 11 | Eps8l2 | 2.71 | 1.85E-164 | 3.85E-160 | 0.24 | 0.04 | 0.20 | 6.80 |
| 11 | Myh14 | 2.24 | 4.75E-163 | 9.89E-159 | 0.34 | 0.07 | 0.27 | 4.78 |
| 11 | Klk8 | 1.73 | 1.75E-162 | 3.65E-158 | 0.60 | 0.20 | 0.40 | 2.97 |
| 11 | Tom1l1 | 1.95 | 2.72E-162 | 5.65E-158 | 0.50 | 0.15 | 0.35 | 3.37 |
| 11 | Ddx58 | 2.72 | 1.60E-160 | 3.33E-156 | 0.27 | 0.05 | 0.22 | 5.77 |
| 11 | Elf1 | 1.84 | 2.13E-160 | 4.44E-156 | 0.64 | 0.25 | 0.39 | 2.56 |
| 11 | Mapk13 | 1.75 | 2.19E-160 | 4.56E-156 | 0.57 | 0.19 | 0.38 | 3.04 |
| 11 | Hes1 | 1.74 | 7.37E-160 | 1.53E-155 | 0.84 | 0.54 | 0.30 | 1.57 |
| 11 | Kdm5b | 1.11 | 8.07E-160 | 1.68E-155 | 0.97 | 0.86 | 0.11 | 1.13 |
| 11 | Cstb | 1.44 | 4.08E-159 | 8.49E-155 | 0.93 | 0.80 | 0.14 | 1.17 |
| 11 | Cldn23 | 3.13 | 1.35E-158 | 2.81E-154 | 0.22 | 0.03 | 0.19 | 6.88 |
| 11 | Ptpn13 | 1.76 | 2.18E-158 | 4.53E-154 | 0.77 | 0.39 | 0.38 | 1.96 |
| 11 | Rpl7 | 0.41 | 3.33E-158 | 6.94E-154 | 1.00 | 1.00 | 0.00 | 1.00 |
| 11 | Clic6 | 2.70 | 4.58E-157 | 9.52E-153 | 0.27 | 0.05 | 0.22 | 5.70 |
| 11 | Rab38 | 2.68 | 3.20E-156 | 6.66E-152 | 0.31 | 0.06 | 0.25 | 5.08 |
| 11 | Bnipl | 2.82 | 3.26E-156 | 6.77E-152 | 0.24 | 0.04 | 0.20 | 6.34 |
| 11 | Platr21 | 3.24 | 4.56E-156 | 9.48E-152 | 0.17 | 0.02 | 0.15 | 8.95 |
| 11 | AI661453 | 3.09 | 7.34E-155 | 1.53E-150 | 0.21 | 0.03 | 0.18 | 7.21 |
| 11 | Kremen1 | 2.29 | 4.69E-154 | 9.74E-150 | 0.46 | 0.13 | 0.33 | 3.44 |
| 11 | Hspa2 | 2.67 | 8.52E-153 | 1.77E-148 | 0.27 | 0.05 | 0.22 | 5.60 |
| 11 | Prkch | 2.39 | 1.07E-152 | 2.22E-148 | 0.35 | 0.08 | 0.27 | 4.44 |
| 11 | Eml5 | 2.30 | 2.62E-152 | 5.46E-148 | 0.35 | 0.08 | 0.27 | 4.54 |
| 11 | 2310030G06Rik | 2.39 | 2.80E-151 | 5.83E-147 | 0.31 | 0.06 | 0.25 | 4.95 |
| 11 | Ppp1r36 | 2.57 | 2.88E-151 | 5.99E-147 | 0.26 | 0.05 | 0.22 | 5.72 |
| 11 | Npm3 | 1.27 | 3.02E-151 | 6.28E-147 | 0.94 | 0.79 | 0.15 | 1.19 |
| 11 | Marveld3 | 2.89 | 1.78E-150 | 3.70E-146 | 0.21 | 0.03 | 0.18 | 6.87 |
| 11 | Otx2 | 1.93 | 1.79E-150 | 3.72E-146 | 0.38 | 0.09 | 0.29 | 4.17 |
| 11 | Kif13b | 2.12 | 3.95E-150 | 8.21E-146 | 0.48 | 0.15 | 0.33 | 3.22 |
| 11 | Hs6st1 | 1.96 | 1.72E-149 | 3.57E-145 | 0.50 | 0.16 | 0.34 | 3.14 |
| 11 | Sox14 | 3.58 | 3.03E-149 | 6.30E-145 | 0.12 | 0.01 | 0.11 | 12.10 |
| 11 | Khdrbs3 | 1.50 | 8.77E-149 | 1.82E-144 | 0.88 | 0.61 | 0.27 | 1.45 |

**Supplementary Table 1. Marker genes for the original clusters of integrated clustering of E11.5 control and NotchGOF λ-junction cells.**

| | | | | | | | | |
|---|---|---|---|---|---|---|---|---|
| 11 | Nrip1 | 1.35 | 2.10E-148 | 4.36E-144 | 0.94 | 0.75 | 0.19 | 1.25 |
| 11 | Lmo4 | 1.18 | 3.81E-148 | 7.93E-144 | 0.95 | 0.81 | 0.14 | 1.17 |
| 11 | Ppfibp1 | 1.67 | 4.91E-147 | 1.02E-142 | 0.77 | 0.42 | 0.36 | 1.85 |
| 11 | Exph5 | 2.86 | 7.68E-147 | 1.60E-142 | 0.21 | 0.03 | 0.18 | 6.90 |
| 11 | Prag1 | 2.15 | 8.02E-147 | 1.67E-142 | 0.44 | 0.13 | 0.32 | 3.46 |
| 11 | Sdc4 | 1.76 | 2.59E-145 | 5.40E-141 | 0.70 | 0.33 | 0.36 | 2.09 |
| 11 | Pbx1 | 0.89 | 3.03E-144 | 6.30E-140 | 0.99 | 0.92 | 0.07 | 1.07 |
| 11 | Syt5 | 2.54 | 2.59E-143 | 5.39E-139 | 0.27 | 0.05 | 0.22 | 5.27 |
| 11 | Tfap2a | 1.81 | 2.75E-143 | 5.73E-139 | 0.56 | 0.20 | 0.36 | 2.78 |
| 11 | Retreg1 | 2.62 | 7.96E-143 | 1.65E-138 | 0.35 | 0.08 | 0.26 | 4.11 |
| 11 | Rps19 | 0.34 | 1.02E-142 | 2.11E-138 | 1.00 | 1.00 | 0.00 | 1.00 |
| 11 | Epn3 | 2.56 | 1.55E-142 | 3.23E-138 | 0.25 | 0.04 | 0.20 | 5.70 |
| 11 | Trim29 | 2.56 | 3.04E-142 | 6.32E-138 | 0.24 | 0.04 | 0.20 | 5.83 |
| 11 | Cd24a | 0.92 | 8.69E-142 | 1.81E-137 | 0.98 | 0.92 | 0.07 | 1.07 |
| 11 | Rnh1 | 1.34 | 1.24E-141 | 2.58E-137 | 0.84 | 0.54 | 0.29 | 1.54 |
| 11 | Ephb1 | 2.13 | 2.26E-141 | 4.71E-137 | 0.37 | 0.09 | 0.28 | 4.06 |
| 11 | Dcbld1 | 1.98 | 7.00E-141 | 1.46E-136 | 0.65 | 0.29 | 0.36 | 2.24 |
| 11 | Bcl11b | 1.36 | 4.38E-140 | 9.12E-136 | 0.87 | 0.58 | 0.29 | 1.49 |
| 11 | Tfap2b | 1.85 | 4.73E-140 | 9.84E-136 | 0.35 | 0.08 | 0.27 | 4.31 |
| 11 | Pdgfrl | 2.06 | 1.78E-139 | 3.70E-135 | 0.51 | 0.17 | 0.34 | 2.96 |
| 11 | Slc16a12 | 3.76 | 2.30E-139 | 4.77E-135 | 0.11 | 0.01 | 0.10 | 13.13 |
| 11 | Rbp1 | 1.57 | 2.44E-139 | 5.08E-135 | 0.96 | 0.91 | 0.05 | 1.06 |
| 11 | Plekhg1 | 2.20 | 3.53E-139 | 7.34E-135 | 0.43 | 0.13 | 0.30 | 3.40 |
| 11 | Tpm1 | 1.18 | 2.83E-138 | 5.89E-134 | 0.98 | 0.91 | 0.07 | 1.08 |
| 11 | Tspan13 | 1.40 | 7.34E-137 | 1.53E-132 | 0.77 | 0.40 | 0.37 | 1.92 |
| 11 | Pcdh17 | 2.06 | 1.42E-136 | 2.96E-132 | 0.54 | 0.19 | 0.35 | 2.78 |
| 11 | Cers3 | 3.58 | 3.58E-136 | 7.45E-132 | 0.11 | 0.01 | 0.10 | 12.22 |
| 11 | Fam89a | 2.48 | 1.27E-135 | 2.64E-131 | 0.30 | 0.07 | 0.23 | 4.55 |
| 11 | Fras1 | 1.62 | 1.56E-135 | 3.24E-131 | 0.63 | 0.25 | 0.37 | 2.46 |
| 11 | Hsp90ab1 | 0.37 | 3.43E-134 | 7.14E-130 | 1.00 | 1.00 | 0.00 | 1.00 |
| 11 | Fhdc1 | 2.65 | 6.92E-134 | 1.44E-129 | 0.22 | 0.04 | 0.18 | 6.17 |
| 11 | Pawr | 2.01 | 7.15E-134 | 1.49E-129 | 0.42 | 0.12 | 0.30 | 3.49 |
| 11 | Stk26 | 1.64 | 7.87E-134 | 1.64E-129 | 0.57 | 0.22 | 0.36 | 2.65 |
| 11 | Plekha7 | 2.05 | 1.27E-133 | 2.65E-129 | 0.34 | 0.08 | 0.26 | 4.14 |
| 11 | Arrdc1 | 1.93 | 3.27E-133 | 6.81E-129 | 0.38 | 0.10 | 0.28 | 3.75 |
| 11 | Gm35040 | 3.49 | 1.70E-132 | 3.53E-128 | 0.12 | 0.01 | 0.11 | 10.82 |
| 11 | Ermn | 2.48 | 1.78E-132 | 3.71E-128 | 0.22 | 0.04 | 0.18 | 5.84 |
| 11 | Slc2a1 | 1.43 | 3.21E-132 | 6.69E-128 | 0.77 | 0.42 | 0.36 | 1.86 |
| 11 | Rb1 | 2.38 | 7.97E-132 | 1.66E-127 | 0.45 | 0.15 | 0.30 | 3.04 |
| 11 | Prr15l | 2.77 | 1.94E-131 | 4.04E-127 | 0.20 | 0.03 | 0.17 | 6.57 |
| 11 | Tjp3 | 2.42 | 7.15E-131 | 1.49E-126 | 0.24 | 0.04 | 0.19 | 5.60 |
| 11 | Hook1 | 1.73 | 2.51E-130 | 5.22E-126 | 0.53 | 0.19 | 0.34 | 2.79 |
| 11 | Mtus1 | 2.11 | 7.57E-130 | 1.57E-125 | 0.36 | 0.10 | 0.27 | 3.79 |
| 11 | Tafa5 | 2.06 | 2.14E-128 | 4.46E-124 | 0.40 | 0.11 | 0.28 | 3.50 |
| 11 | Ptpru | 1.86 | 2.72E-128 | 5.67E-124 | 0.46 | 0.15 | 0.31 | 3.11 |
| 11 | Faah | 2.33 | 7.28E-128 | 1.51E-123 | 0.26 | 0.05 | 0.21 | 5.04 |
| 11 | Mbp | 2.03 | 8.22E-127 | 1.71E-122 | 0.34 | 0.09 | 0.26 | 3.97 |
| 11 | Mid1ip1 | 2.44 | 1.35E-126 | 2.80E-122 | 0.53 | 0.21 | 0.32 | 2.55 |
| 11 | Sinhcaf | 0.89 | 1.46E-126 | 3.04E-122 | 0.96 | 0.83 | 0.13 | 1.15 |
| 11 | Ly6g6e | 3.40 | 1.90E-126 | 3.95E-122 | 0.13 | 0.01 | 0.11 | 9.62 |
| 11 | Slc12a2 | 1.60 | 5.98E-126 | 1.24E-121 | 0.75 | 0.42 | 0.32 | 1.76 |
| 11 | Insyn2b | 3.16 | 2.28E-125 | 4.74E-121 | 0.15 | 0.02 | 0.13 | 8.59 |
| 11 | Ermp1 | 1.68 | 6.66E-125 | 1.38E-120 | 0.67 | 0.32 | 0.35 | 2.08 |
| 11 | Pcdh9 | 2.17 | 3.53E-124 | 7.34E-120 | 0.37 | 0.10 | 0.27 | 3.58 |
| 11 | Tmem178 | 2.78 | 9.67E-124 | 2.01E-119 | 0.18 | 0.03 | 0.16 | 6.78 |
| 11 | Prss8 | 2.01 | 1.01E-123 | 2.09E-119 | 0.35 | 0.09 | 0.26 | 3.88 |

**Supplementary Table 1. Marker genes for the original clusters of integrated clustering of E11.5 control and NotchGOF λ-junction cells.**

| | | | | | | | | |
|---|---|---|---|---|---|---|---|---|
| 11 | Sox2 | 0.88 | 3.48E-122 | 7.24E-118 | 0.55 | 0.18 | 0.37 | 3.04 |
| 11 | Nckap5 | 1.89 | 3.84E-121 | 7.98E-117 | 0.33 | 0.08 | 0.25 | 3.98 |
| 11 | Tmtc1 | 3.10 | 9.35E-121 | 1.94E-116 | 0.14 | 0.02 | 0.13 | 8.41 |
| 11 | Mtss2 | 1.69 | 2.37E-120 | 4.94E-116 | 0.60 | 0.26 | 0.34 | 2.32 |
| 11 | Celsr2 | 2.19 | 7.39E-120 | 1.54E-115 | 0.33 | 0.08 | 0.24 | 3.90 |
| 11 | Prxl2a | 1.47 | 2.39E-119 | 4.96E-115 | 0.71 | 0.37 | 0.34 | 1.92 |
| 11 | Abcg1 | 2.17 | 4.53E-118 | 9.42E-114 | 0.31 | 0.08 | 0.23 | 4.09 |
| 11 | 2610528J11Rik | 2.22 | 2.23E-117 | 4.63E-113 | 0.26 | 0.06 | 0.20 | 4.71 |
| 11 | Limk2 | 2.19 | 2.44E-117 | 5.07E-113 | 0.49 | 0.19 | 0.30 | 2.62 |
| 11 | Nab1 | 1.92 | 7.69E-117 | 1.60E-112 | 0.40 | 0.12 | 0.28 | 3.30 |
| 11 | Rpl17 | 0.30 | 2.04E-116 | 4.25E-112 | 1.00 | 1.00 | 0.00 | 1.00 |
| 11 | Ece1 | 1.65 | 2.34E-116 | 4.87E-112 | 0.55 | 0.22 | 0.33 | 2.50 |
| 11 | Prr5l | 1.86 | 2.86E-116 | 5.95E-112 | 0.45 | 0.16 | 0.30 | 2.92 |
| 11 | Meis2 | 0.86 | 2.90E-116 | 6.02E-112 | 0.98 | 0.66 | 0.31 | 1.47 |
| 11 | Glcci1 | 2.04 | 1.12E-114 | 2.32E-110 | 0.68 | 0.38 | 0.30 | 1.80 |
| 11 | Gja1 | 1.57 | 4.03E-114 | 8.39E-110 | 0.91 | 0.77 | 0.14 | 1.18 |
| 11 | Rdh10 | 2.41 | 4.86E-114 | 1.01E-109 | 0.43 | 0.15 | 0.28 | 2.94 |
| 11 | Ttc3 | 0.62 | 6.77E-114 | 1.41E-109 | 1.00 | 0.96 | 0.04 | 1.04 |
| 11 | Adamts18 | 2.38 | 1.17E-113 | 2.44E-109 | 0.28 | 0.06 | 0.21 | 4.31 |
| 11 | Arhgef5 | 1.73 | 1.29E-113 | 2.68E-109 | 0.51 | 0.19 | 0.32 | 2.68 |
| 11 | Plec | 1.63 | 1.66E-112 | 3.45E-108 | 0.59 | 0.26 | 0.33 | 2.26 |
| 11 | Smagp | 2.02 | 1.88E-112 | 3.91E-108 | 0.18 | 0.03 | 0.15 | 6.31 |
| 11 | Itga3 | 2.14 | 9.64E-112 | 2.00E-107 | 0.27 | 0.06 | 0.21 | 4.37 |
| 11 | Pygl | 2.50 | 4.65E-111 | 9.67E-107 | 0.32 | 0.09 | 0.23 | 3.60 |
| 11 | Ankrd22 | 3.48 | 9.07E-111 | 1.89E-106 | 0.12 | 0.01 | 0.10 | 9.58 |
| 11 | Satb1 | 1.42 | 9.69E-111 | 2.02E-106 | 0.70 | 0.38 | 0.32 | 1.86 |
| 11 | H19 | 1.31 | 1.69E-110 | 3.51E-106 | 0.94 | 0.84 | 0.10 | 1.11 |
| 11 | Rps16 | 0.25 | 1.75E-110 | 3.64E-106 | 1.00 | 1.00 | 0.00 | 1.00 |
| 11 | Mgll | 3.10 | 8.62E-110 | 1.79E-105 | 0.13 | 0.02 | 0.12 | 7.88 |
| 11 | Igfbp3 | 2.52 | 1.30E-109 | 2.70E-105 | 0.38 | 0.12 | 0.26 | 3.23 |
| 11 | Myl12b | 0.82 | 1.61E-109 | 3.34E-105 | 0.95 | 0.88 | 0.07 | 1.08 |
| 11 | Gadd45b | 2.15 | 3.03E-109 | 6.31E-105 | 0.41 | 0.13 | 0.27 | 3.04 |
| 11 | Scube3 | 1.71 | 4.03E-109 | 8.37E-105 | 0.50 | 0.19 | 0.31 | 2.61 |
| 11 | Rfk | 1.56 | 4.87E-108 | 1.01E-103 | 0.75 | 0.47 | 0.28 | 1.59 |
| 11 | Zfp503 | 1.44 | 1.46E-107 | 3.04E-103 | 0.71 | 0.36 | 0.34 | 1.95 |
| 11 | Sptbn2 | 1.95 | 1.60E-107 | 3.32E-103 | 0.36 | 0.11 | 0.25 | 3.39 |
| 11 | Gstt3 | 2.44 | 1.83E-107 | 3.81E-103 | 0.22 | 0.04 | 0.17 | 5.05 |
| 11 | Ano1 | 1.73 | 2.31E-107 | 4.80E-103 | 0.39 | 0.12 | 0.27 | 3.23 |
| 11 | Cmtm8 | 1.42 | 2.90E-107 | 6.03E-103 | 0.48 | 0.17 | 0.31 | 2.85 |
| 11 | Sema6a | 1.56 | 4.34E-107 | 9.03E-103 | 0.73 | 0.44 | 0.30 | 1.69 |
| 11 | Serinc5 | 2.56 | 5.40E-107 | 1.12E-102 | 0.24 | 0.05 | 0.18 | 4.61 |
| 11 | Syt1 | 2.60 | 5.64E-107 | 1.17E-102 | 0.19 | 0.03 | 0.15 | 5.78 |
| 11 | Id4 | 1.83 | 2.88E-106 | 5.99E-102 | 0.60 | 0.30 | 0.30 | 2.01 |
| 11 | Pax6 | 1.49 | 5.08E-106 | 1.06E-101 | 0.40 | 0.12 | 0.28 | 3.29 |
| 11 | Gldc | 2.63 | 2.02E-105 | 4.21E-101 | 0.20 | 0.04 | 0.16 | 5.42 |
| 11 | Fgd2 | 1.75 | 4.98E-105 | 1.04E-100 | 0.33 | 0.09 | 0.24 | 3.58 |
| 11 | Igsf9b | 2.35 | 1.31E-104 | 2.72E-100 | 0.23 | 0.05 | 0.18 | 4.68 |
| 11 | Synrg | 1.57 | 1.51E-104 | 3.13E-100 | 0.54 | 0.23 | 0.31 | 2.39 |
| 11 | Mtss1 | 1.62 | 1.94E-104 | 4.04E-100 | 0.60 | 0.29 | 0.31 | 2.10 |
| 11 | Klhl23 | 1.47 | 2.51E-104 | 5.22E-100 | 0.67 | 0.36 | 0.31 | 1.85 |
| 11 | Trpm3 | 2.35 | 1.68E-103 | 3.50E-99 | 0.21 | 0.04 | 0.17 | 5.10 |
| 11 | Anxa9 | 3.48 | 9.12E-103 | 1.90E-98 | 0.10 | 0.01 | 0.09 | 10.20 |
| 11 | Rpl28 | 0.27 | 1.29E-102 | 2.68E-98 | 1.00 | 1.00 | 0.00 | 1.00 |
| 11 | Cdcp1 | 2.22 | 2.54E-102 | 5.28E-98 | 0.24 | 0.05 | 0.19 | 4.49 |
| 11 | Smim5 | 3.04 | 6.23E-102 | 1.30E-97 | 0.15 | 0.02 | 0.13 | 6.73 |
| 11 | Zfhx3 | 0.72 | 8.24E-102 | 1.71E-97 | 0.98 | 0.81 | 0.17 | 1.22 |

**Supplementary Table 1. Marker genes for the original clusters of integrated clustering of E11.5 control and NotchGOF λ-junction cells.**

| | | | | | | | | |
|---|---|---|---|---|---|---|---|---|
| 11 | Dennd2c | 2.48 | 5.96E-101 | 1.24E-96 | 0.25 | 0.06 | 0.19 | 4.30 |
| 11 | Fau | 0.30 | 6.50E-101 | 1.35E-96 | 1.00 | 1.00 | 0.00 | 1.00 |
| 11 | Cystm1 | 1.65 | 9.14E-101 | 1.90E-96 | 0.31 | 0.08 | 0.23 | 3.71 |
| 11 | B3gnt2 | 1.84 | 1.95E-100 | 4.05E-96 | 0.32 | 0.09 | 0.23 | 3.51 |
| 11 | Tubb2b | 0.70 | 2.17E-100 | 4.51E-96 | 0.91 | 0.66 | 0.25 | 1.38 |
| 11 | Dnase2a | 1.73 | 2.24E-100 | 4.67E-96 | 0.51 | 0.21 | 0.30 | 2.43 |
| 11 | Mbnl2 | 1.08 | 8.09E-100 | 1.68E-95 | 0.89 | 0.67 | 0.22 | 1.33 |
| 11 | Susd4 | 1.89 | 2.97E-99 | 6.17E-95 | 0.31 | 0.08 | 0.22 | 3.63 |
| 11 | H3f3b | 0.41 | 4.16E-99 | 8.65E-95 | 1.00 | 1.00 | 0.00 | 1.00 |
| 11 | Tap1 | 2.71 | 6.88E-98 | 1.43E-93 | 0.16 | 0.03 | 0.13 | 6.24 |
| 11 | Arid3b | 1.67 | 8.01E-98 | 1.67E-93 | 0.52 | 0.22 | 0.30 | 2.36 |
| 11 | Eno1 | 0.54 | 9.41E-98 | 1.96E-93 | 1.00 | 0.97 | 0.02 | 1.02 |
| 11 | Tgfb1 | 1.78 | 1.28E-97 | 2.66E-93 | 0.33 | 0.10 | 0.23 | 3.33 |
| 11 | Ndrg2 | 1.79 | 1.72E-97 | 3.57E-93 | 0.38 | 0.13 | 0.26 | 3.02 |
| 11 | Slc7a11 | 2.32 | 3.83E-97 | 7.97E-93 | 0.28 | 0.07 | 0.20 | 3.79 |
| 11 | Dipk2a | 1.61 | 4.37E-97 | 9.10E-93 | 0.53 | 0.23 | 0.30 | 2.31 |
| 11 | Tmprss13 | 3.42 | 9.46E-97 | 1.97E-92 | 0.11 | 0.01 | 0.09 | 8.83 |
| 11 | Ktn1 | 0.83 | 1.01E-96 | 2.10E-92 | 0.95 | 0.82 | 0.13 | 1.16 |
| 11 | Hebp2 | 1.92 | 1.34E-96 | 2.79E-92 | 0.51 | 0.23 | 0.28 | 2.20 |
| 11 | Dmrta2 | 1.10 | 1.72E-96 | 3.58E-92 | 0.41 | 0.13 | 0.28 | 3.09 |
| 11 | Agrn | 1.28 | 1.46E-95 | 3.04E-91 | 0.69 | 0.38 | 0.31 | 1.81 |
| 11 | Metrnl | 2.51 | 1.51E-95 | 3.13E-91 | 0.25 | 0.06 | 0.19 | 4.15 |
| 11 | Siva1 | 0.72 | 1.80E-95 | 3.74E-91 | 0.98 | 0.95 | 0.03 | 1.03 |
| 11 | Id1 | 0.83 | 2.56E-95 | 5.31E-91 | 0.97 | 0.86 | 0.11 | 1.13 |
| 11 | Ostf1 | 1.33 | 2.91E-95 | 6.05E-91 | 0.57 | 0.27 | 0.31 | 2.16 |
| 11 | Id3 | 0.95 | 5.69E-95 | 1.18E-90 | 0.99 | 0.96 | 0.04 | 1.04 |
| 11 | Usp25 | 1.34 | 7.88E-95 | 1.64E-90 | 0.71 | 0.43 | 0.28 | 1.66 |
| 11 | Dgka | 2.22 | 1.17E-94 | 2.43E-90 | 0.24 | 0.06 | 0.18 | 4.27 |
| 11 | Tnfrsf12a | 2.15 | 1.53E-93 | 3.17E-89 | 0.27 | 0.07 | 0.20 | 3.84 |
| 11 | Tpi1 | 0.63 | 2.77E-93 | 5.77E-89 | 0.98 | 0.96 | 0.03 | 1.03 |
| 11 | Arf1 | 0.55 | 5.92E-93 | 1.23E-88 | 0.99 | 0.98 | 0.02 | 1.02 |
| 11 | Ifitm3 | 1.60 | 1.24E-92 | 2.58E-88 | 0.59 | 0.29 | 0.30 | 2.06 |
| 11 | Ctsh | 2.19 | 1.69E-92 | 3.52E-88 | 0.24 | 0.06 | 0.18 | 4.20 |
| 11 | Chmp4c | 3.17 | 3.54E-92 | 7.36E-88 | 0.11 | 0.01 | 0.10 | 7.93 |
| 11 | Churc1 | 1.23 | 4.09E-92 | 8.51E-88 | 0.81 | 0.58 | 0.23 | 1.40 |
| 11 | Rps14 | 0.25 | 9.12E-92 | 1.90E-87 | 1.00 | 1.00 | 0.00 | 1.00 |
| 11 | Prom2 | 2.56 | 1.00E-91 | 2.09E-87 | 0.16 | 0.03 | 0.13 | 5.75 |
| 11 | Unc5b | 1.28 | 1.76E-91 | 3.65E-87 | 0.69 | 0.38 | 0.31 | 1.81 |
| 11 | Znrf3 | 1.86 | 2.64E-91 | 5.49E-87 | 0.44 | 0.17 | 0.27 | 2.55 |
| 11 | Cyb561 | 2.24 | 3.21E-91 | 6.67E-87 | 0.22 | 0.05 | 0.17 | 4.40 |
| 11 | Gm2464 | 2.68 | 4.54E-91 | 9.45E-87 | 0.15 | 0.02 | 0.12 | 6.13 |
| 11 | Cebpa | 2.44 | 6.15E-91 | 1.28E-86 | 0.15 | 0.02 | 0.12 | 6.35 |
| 11 | Pclo | 1.62 | 1.35E-90 | 2.80E-86 | 0.33 | 0.10 | 0.23 | 3.33 |
| 11 | Nrtn | 1.87 | 2.78E-90 | 5.79E-86 | 0.40 | 0.15 | 0.25 | 2.71 |
| 11 | Ago1 | 1.34 | 2.97E-90 | 6.17E-86 | 0.66 | 0.35 | 0.31 | 1.87 |
| 11 | Rnf24 | 1.46 | 6.63E-90 | 1.38E-85 | 0.59 | 0.29 | 0.30 | 2.03 |
| 11 | Sult2b1 | 2.93 | 1.13E-89 | 2.35E-85 | 0.11 | 0.01 | 0.10 | 7.93 |
| 11 | Hspg2 | 1.45 | 1.70E-89 | 3.53E-85 | 0.51 | 0.22 | 0.29 | 2.34 |
| 11 | Prss22 | 1.99 | 2.12E-89 | 4.42E-85 | 0.23 | 0.06 | 0.18 | 4.14 |
| 11 | Epb41l5 | 1.48 | 4.83E-89 | 1.00E-84 | 0.53 | 0.24 | 0.29 | 2.19 |
| 11 | Nxn | 0.92 | 1.87E-88 | 3.89E-84 | 0.89 | 0.70 | 0.19 | 1.26 |
| 11 | Soga3 | 1.94 | 2.31E-88 | 4.80E-84 | 0.21 | 0.05 | 0.16 | 4.49 |
| 11 | Lrp2 | 2.13 | 3.05E-88 | 6.34E-84 | 0.22 | 0.05 | 0.17 | 4.39 |
| 11 | Rps5 | 0.25 | 5.35E-88 | 1.11E-83 | 1.00 | 1.00 | 0.00 | 1.00 |
| 11 | Nrcam | 2.12 | 6.21E-88 | 1.29E-83 | 0.21 | 0.05 | 0.16 | 4.52 |
| 11 | Ehbp1 | 1.48 | 2.07E-87 | 4.30E-83 | 0.52 | 0.23 | 0.29 | 2.24 |

**Supplementary Table 1. Marker genes for the original clusters of integrated clustering of E11.5 control and NotchGOF λ-junction cells.**

| | | | | | | | | |
|---|---|---|---|---|---|---|---|---|
| 11 | Arhgap27os2 | 1.99 | 3.01E-87 | 6.26E-83 | 0.24 | 0.06 | 0.18 | 4.02 |
| 11 | Ormdl2 | 1.28 | 1.74E-86 | 3.61E-82 | 0.63 | 0.34 | 0.29 | 1.84 |
| 11 | Acvr2a | 1.56 | 5.63E-86 | 1.17E-81 | 0.54 | 0.26 | 0.28 | 2.08 |
| 11 | Llgl2 | 2.22 | 7.80E-86 | 1.62E-81 | 0.20 | 0.04 | 0.16 | 4.57 |
| 11 | Ldha | 0.59 | 9.81E-86 | 2.04E-81 | 0.99 | 0.97 | 0.02 | 1.02 |
| 11 | Celf2 | 1.42 | 1.62E-85 | 3.37E-81 | 0.55 | 0.25 | 0.30 | 2.22 |
| 11 | Tead1 | 1.04 | 2.38E-85 | 4.95E-81 | 0.82 | 0.60 | 0.22 | 1.38 |
| 11 | Bmp4 | 1.35 | 3.61E-85 | 7.52E-81 | 0.78 | 0.51 | 0.27 | 1.54 |
| 11 | Atp6v1f | 0.65 | 3.98E-85 | 8.28E-81 | 0.95 | 0.90 | 0.05 | 1.06 |
| 11 | Pla2g12a | 1.27 | 4.61E-85 | 9.58E-81 | 0.61 | 0.32 | 0.29 | 1.92 |
| 11 | Mdh1 | 0.69 | 8.34E-85 | 1.73E-80 | 0.97 | 0.95 | 0.03 | 1.03 |
| 11 | Cxadr | 1.14 | 1.67E-84 | 3.47E-80 | 0.73 | 0.45 | 0.28 | 1.62 |
| 11 | Laptm4a | 0.68 | 1.91E-84 | 3.98E-80 | 0.98 | 0.94 | 0.04 | 1.04 |
| 11 | A730020E08Rik | 2.52 | 2.44E-84 | 5.07E-80 | 0.18 | 0.04 | 0.14 | 4.89 |
| 11 | Pla2r1 | 2.20 | 9.34E-84 | 1.94E-79 | 0.25 | 0.07 | 0.19 | 3.68 |
| 11 | Ell2 | 2.13 | 1.10E-83 | 2.28E-79 | 0.28 | 0.08 | 0.20 | 3.37 |
| 11 | Hip1r | 2.04 | 2.71E-83 | 5.64E-79 | 0.26 | 0.07 | 0.19 | 3.64 |
| 11 | Foxn3 | 1.23 | 3.28E-83 | 6.82E-79 | 0.85 | 0.68 | 0.17 | 1.25 |
| 11 | Trim2 | 1.34 | 3.57E-83 | 7.42E-79 | 0.66 | 0.41 | 0.26 | 1.63 |
| 11 | Fgfr2 | 1.16 | 4.88E-83 | 1.02E-78 | 0.58 | 0.27 | 0.31 | 2.15 |
| 11 | Gm28112 | 2.73 | 1.07E-82 | 2.22E-78 | 0.13 | 0.02 | 0.11 | 6.40 |
| 11 | Msx1 | 0.88 | 1.30E-82 | 2.71E-78 | 0.95 | 0.71 | 0.25 | 1.35 |
| 11 | Sox13 | 2.06 | 2.10E-82 | 4.36E-78 | 0.28 | 0.08 | 0.20 | 3.42 |
| 11 | Zfp704 | 1.18 | 3.30E-82 | 6.86E-78 | 0.72 | 0.46 | 0.26 | 1.58 |
| 11 | Kdm5a | 0.98 | 5.93E-82 | 1.23E-77 | 0.85 | 0.67 | 0.19 | 1.28 |
| 11 | Kcnh8 | 2.51 | 7.59E-82 | 1.58E-77 | 0.16 | 0.03 | 0.13 | 5.27 |
| 11 | Igf2 | 1.06 | 1.08E-81 | 2.25E-77 | 0.81 | 0.58 | 0.24 | 1.42 |
| 11 | Mpp7 | 1.30 | 3.06E-81 | 6.36E-77 | 0.64 | 0.36 | 0.28 | 1.77 |
| 11 | Lhfp | 1.53 | 3.52E-81 | 7.32E-77 | 0.50 | 0.22 | 0.27 | 2.22 |
| 11 | Dennd1c | 2.47 | 6.29E-81 | 1.31E-76 | 0.15 | 0.03 | 0.12 | 5.28 |
| 11 | Stxbp2 | 1.64 | 6.33E-81 | 1.32E-76 | 0.36 | 0.13 | 0.23 | 2.80 |
| 11 | Npdc1 | 1.62 | 6.64E-81 | 1.38E-76 | 0.31 | 0.10 | 0.22 | 3.17 |
| 11 | Irx2 | 3.06 | 6.77E-81 | 1.41E-76 | 0.11 | 0.02 | 0.10 | 7.33 |
| 11 | Tiam1 | 1.41 | 6.97E-81 | 1.45E-76 | 0.59 | 0.30 | 0.28 | 1.92 |
| 11 | Trappc6a | 1.05 | 7.91E-81 | 1.65E-76 | 0.77 | 0.52 | 0.26 | 1.50 |
| 11 | Foxp1 | 0.69 | 1.10E-80 | 2.29E-76 | 0.92 | 0.59 | 0.33 | 1.56 |
| 11 | Ptbp3 | 1.06 | 1.18E-80 | 2.45E-76 | 0.81 | 0.62 | 0.19 | 1.31 |
| 11 | Dlx5 | 0.99 | 1.34E-80 | 2.79E-76 | 0.50 | 0.21 | 0.30 | 2.43 |
| 11 | Notum | 3.01 | 1.37E-80 | 2.85E-76 | 0.17 | 0.03 | 0.13 | 4.88 |
| 11 | Krt17 | 2.67 | 1.76E-80 | 3.66E-76 | 0.11 | 0.02 | 0.10 | 7.00 |
| 11 | Marchf9 | 1.50 | 1.95E-80 | 4.06E-76 | 0.39 | 0.14 | 0.25 | 2.76 |
| 11 | Irf2 | 1.45 | 2.04E-80 | 4.25E-76 | 0.52 | 0.25 | 0.28 | 2.13 |
| 11 | Rps27 | 0.32 | 3.14E-80 | 6.52E-76 | 1.00 | 1.00 | 0.00 | 1.00 |
| 11 | Nectin1 | 1.62 | 3.21E-80 | 6.68E-76 | 0.42 | 0.17 | 0.25 | 2.52 |
| 11 | Tenm2 | 2.09 | 8.08E-80 | 1.68E-75 | 0.25 | 0.07 | 0.18 | 3.66 |
| 11 | Cep170b | 1.65 | 2.56E-79 | 5.33E-75 | 0.35 | 0.12 | 0.23 | 2.88 |
| 11 | S100a16 | 0.65 | 5.23E-79 | 1.09E-74 | 0.26 | 0.07 | 0.19 | 3.67 |
| 11 | Emp1 | 2.34 | 9.73E-79 | 2.02E-74 | 0.34 | 0.12 | 0.22 | 2.79 |
| 11 | Plcd1 | 2.25 | 2.22E-78 | 4.62E-74 | 0.24 | 0.07 | 0.17 | 3.68 |
| 11 | Rbms1 | 0.80 | 2.93E-78 | 6.10E-74 | 0.89 | 0.78 | 0.11 | 1.14 |
| 11 | Unc13b | 1.90 | 3.96E-78 | 8.24E-74 | 0.27 | 0.08 | 0.19 | 3.47 |
| 11 | Tbc1d9 | 1.97 | 1.90E-77 | 3.96E-73 | 0.25 | 0.07 | 0.18 | 3.57 |
| 11 | Epha1 | 2.19 | 3.18E-77 | 6.62E-73 | 0.22 | 0.05 | 0.16 | 3.98 |
| 11 | Cldn9 | 1.58 | 4.17E-77 | 8.68E-73 | 0.28 | 0.08 | 0.20 | 3.37 |
| 11 | Gab1 | 1.55 | 4.93E-77 | 1.03E-72 | 0.43 | 0.18 | 0.25 | 2.38 |
| 11 | Atp8b1 | 2.09 | 6.53E-77 | 1.36E-72 | 0.18 | 0.04 | 0.14 | 4.73 |

**Supplementary Table 1. Marker genes for the original clusters of integrated clustering of E11.5 control and NotchGOF λ-junction cells.**

| | | | | | | | | |
|---|---|---|---|---|---|---|---|---|
| 11 | Iqgap1 | 1.16 | 9.23E-77 | 1.92E-72 | 0.72 | 0.46 | 0.26 | 1.56 |
| 11 | Psap | 0.93 | 2.43E-76 | 5.04E-72 | 0.82 | 0.61 | 0.21 | 1.34 |
| 11 | Barx2 | 2.83 | 9.73E-76 | 2.02E-71 | 0.17 | 0.04 | 0.13 | 4.59 |
| 11 | Dcaf4 | 1.46 | 1.11E-75 | 2.31E-71 | 0.50 | 0.23 | 0.27 | 2.15 |
| 11 | Gtf2i | 0.70 | 1.65E-75 | 3.43E-71 | 0.95 | 0.85 | 0.09 | 1.11 |
| 11 | Hexb | 1.62 | 2.17E-75 | 4.50E-71 | 0.47 | 0.22 | 0.26 | 2.18 |
| 11 | Pgk1 | 0.54 | 1.64E-74 | 3.41E-70 | 0.99 | 0.95 | 0.03 | 1.03 |
| 11 | Arap2 | 2.15 | 2.04E-74 | 4.24E-70 | 0.20 | 0.05 | 0.15 | 4.04 |
| 11 | Junb | 1.49 | 3.71E-74 | 7.72E-70 | 0.32 | 0.11 | 0.21 | 2.95 |
| 11 | Tceal9 | 0.41 | 5.81E-74 | 1.21E-69 | 1.00 | 0.98 | 0.02 | 1.02 |
| 11 | Prrg4 | 2.66 | 7.42E-74 | 1.54E-69 | 0.12 | 0.02 | 0.10 | 6.00 |
| 11 | Large2 | 1.89 | 8.35E-74 | 1.74E-69 | 0.27 | 0.08 | 0.19 | 3.33 |
| 11 | Bdh1 | 2.09 | 1.32E-73 | 2.75E-69 | 0.24 | 0.07 | 0.17 | 3.56 |
| 11 | Foxg1 | 1.09 | 1.41E-73 | 2.94E-69 | 0.38 | 0.14 | 0.24 | 2.77 |
| 11 | 1700023F06Rik | 2.03 | 1.87E-73 | 3.89E-69 | 0.18 | 0.04 | 0.14 | 4.51 |
| 11 | Ldlrap1 | 1.55 | 5.58E-73 | 1.16E-68 | 0.42 | 0.18 | 0.24 | 2.39 |
| 11 | Oxtr | 1.70 | 6.65E-73 | 1.38E-68 | 0.23 | 0.06 | 0.17 | 3.67 |
| 11 | Casz1 | 2.61 | 8.63E-73 | 1.80E-68 | 0.14 | 0.03 | 0.12 | 5.30 |
| 11 | Tgif1 | 0.99 | 9.56E-73 | 1.99E-68 | 0.82 | 0.63 | 0.19 | 1.30 |
| 11 | Eya2 | 1.47 | 1.01E-72 | 2.10E-68 | 0.48 | 0.23 | 0.26 | 2.15 |
| 11 | Alox12 | 1.77 | 1.29E-72 | 2.68E-68 | 0.12 | 0.02 | 0.10 | 6.05 |
| 11 | Igsf9 | 1.13 | 4.55E-72 | 9.46E-68 | 0.67 | 0.40 | 0.28 | 1.69 |
| 11 | Zfhx2 | 1.27 | 5.45E-72 | 1.13E-67 | 0.51 | 0.24 | 0.27 | 2.11 |
| 11 | Ptk7 | 1.28 | 5.76E-72 | 1.20E-67 | 0.62 | 0.36 | 0.26 | 1.72 |
| 11 | Pgam1 | 0.48 | 6.03E-72 | 1.25E-67 | 0.99 | 0.97 | 0.03 | 1.03 |
| 11 | Vgll4 | 0.79 | 9.94E-72 | 2.07E-67 | 0.91 | 0.78 | 0.14 | 1.18 |
| 11 | Tmcc3 | 2.17 | 1.77E-71 | 3.68E-67 | 0.15 | 0.03 | 0.12 | 5.03 |
| 11 | Gas7 | 2.12 | 1.78E-71 | 3.71E-67 | 0.19 | 0.05 | 0.15 | 4.02 |
| 11 | Rcsd1 | 1.85 | 1.02E-70 | 2.11E-66 | 0.28 | 0.09 | 0.19 | 3.16 |
| 11 | Arhgef16 | 1.88 | 1.16E-70 | 2.42E-66 | 0.21 | 0.05 | 0.15 | 3.85 |
| 11 | Pdk1 | 1.36 | 1.97E-70 | 4.09E-66 | 0.51 | 0.25 | 0.26 | 2.04 |
| 11 | Wfdc2 | 0.78 | 2.02E-70 | 4.21E-66 | 0.72 | 0.45 | 0.27 | 1.60 |
| 11 | Krt1 | 1.92 | 2.05E-70 | 4.27E-66 | 0.19 | 0.05 | 0.14 | 4.13 |
| 11 | Alad | 0.65 | 3.34E-70 | 6.95E-66 | 0.81 | 0.63 | 0.18 | 1.29 |
| 11 | Foxo1 | 1.56 | 8.05E-70 | 1.68E-65 | 0.41 | 0.17 | 0.24 | 2.35 |
| 11 | Mtcl1 | 1.50 | 8.30E-70 | 1.73E-65 | 0.33 | 0.12 | 0.21 | 2.84 |
| 11 | Ass1 | 2.12 | 1.25E-69 | 2.59E-65 | 0.20 | 0.05 | 0.15 | 3.85 |
| 11 | Prr36 | 1.86 | 1.87E-69 | 3.88E-65 | 0.26 | 0.08 | 0.18 | 3.21 |
| 11 | Ctsc | 1.36 | 5.37E-69 | 1.12E-64 | 0.32 | 0.12 | 0.21 | 2.74 |
| 11 | Kdm6b | 0.97 | 1.80E-68 | 3.74E-64 | 0.83 | 0.65 | 0.18 | 1.27 |
| 11 | Ssx2ip | 1.07 | 2.53E-68 | 5.25E-64 | 0.62 | 0.35 | 0.27 | 1.77 |
| 11 | Mid2 | 2.62 | 3.04E-68 | 6.33E-64 | 0.13 | 0.02 | 0.11 | 5.38 |
| 11 | Dst | 1.41 | 5.29E-68 | 1.10E-63 | 0.60 | 0.35 | 0.26 | 1.74 |
| 11 | Slain1 | 1.92 | 6.10E-68 | 1.27E-63 | 0.21 | 0.06 | 0.15 | 3.76 |
| 11 | Gm13648 | 2.39 | 1.16E-67 | 2.41E-63 | 0.12 | 0.02 | 0.10 | 5.59 |
| 11 | Syt7 | 2.07 | 1.23E-67 | 2.56E-63 | 0.20 | 0.05 | 0.15 | 3.83 |
| 11 | Rnd3 | 0.84 | 1.33E-67 | 2.77E-63 | 0.85 | 0.64 | 0.21 | 1.34 |
| 11 | Zdhhc13 | 1.67 | 1.51E-67 | 3.15E-63 | 0.35 | 0.13 | 0.21 | 2.61 |
| 11 | Utrn | 0.91 | 1.83E-67 | 3.80E-63 | 0.81 | 0.61 | 0.20 | 1.33 |
| 11 | Erich1 | 1.35 | 3.23E-67 | 6.71E-63 | 0.48 | 0.23 | 0.25 | 2.12 |
| 11 | Eps8l1 | 2.06 | 4.66E-67 | 9.69E-63 | 0.16 | 0.04 | 0.13 | 4.57 |
| 11 | Nipal2 | 2.12 | 6.69E-67 | 1.39E-62 | 0.18 | 0.04 | 0.14 | 4.14 |
| 11 | 0610009E02Rik | 2.60 | 7.45E-67 | 1.55E-62 | 0.15 | 0.03 | 0.12 | 4.74 |
| 11 | Hspa1b | 2.80 | 1.25E-66 | 2.61E-62 | 0.10 | 0.02 | 0.09 | 6.44 |
| 11 | Calcoco1 | 1.46 | 8.19E-66 | 1.70E-61 | 0.45 | 0.21 | 0.24 | 2.17 |
| 11 | Cacng4 | 1.38 | 8.50E-66 | 1.77E-61 | 0.50 | 0.25 | 0.25 | 2.01 |

**Supplementary Table 1. Marker genes for the original clusters of integrated clustering of E11.5 control and NotchGOF λ-junction cells.**

| | | | | | | | | |
|---|---|---|---|---|---|---|---|---|
| 11 | Baiap2 | 1.95 | 1.54E-65 | 3.21E-61 | 0.36 | 0.15 | 0.21 | 2.42 |
| 11 | Sox9 | 0.79 | 2.37E-65 | 4.93E-61 | 0.83 | 0.58 | 0.26 | 1.44 |
| 11 | H2-Q4 | 1.64 | 3.03E-65 | 6.29E-61 | 0.29 | 0.10 | 0.19 | 2.89 |
| 11 | Kremen2 | 1.59 | 3.54E-65 | 7.37E-61 | 0.18 | 0.04 | 0.13 | 4.19 |
| 11 | Acpp | 2.73 | 3.72E-65 | 7.73E-61 | 0.10 | 0.02 | 0.09 | 6.67 |
| 11 | Slc25a27 | 1.12 | 1.26E-64 | 2.63E-60 | 0.64 | 0.39 | 0.24 | 1.62 |
| 11 | Tspan12 | 1.25 | 1.40E-64 | 2.92E-60 | 0.45 | 0.20 | 0.25 | 2.23 |
| 11 | Gnpda2 | 1.50 | 2.34E-64 | 4.87E-60 | 0.52 | 0.28 | 0.24 | 1.88 |
| 11 | Mettl27 | 1.54 | 3.35E-64 | 6.97E-60 | 0.35 | 0.13 | 0.21 | 2.58 |
| 11 | Arg1 | 2.43 | 3.92E-64 | 8.16E-60 | 0.13 | 0.02 | 0.10 | 5.25 |
| 11 | Tmem102 | 2.77 | 6.67E-64 | 1.39E-59 | 0.10 | 0.02 | 0.08 | 6.25 |
| 11 | Tpbg | 1.23 | 1.44E-63 | 3.00E-59 | 0.55 | 0.30 | 0.25 | 1.84 |
| 11 | Saraf | 1.03 | 1.55E-63 | 3.22E-59 | 0.75 | 0.54 | 0.21 | 1.39 |
| 11 | Kctd1 | 1.06 | 1.90E-63 | 3.95E-59 | 0.68 | 0.46 | 0.22 | 1.48 |
| 11 | Ralgps1 | 2.01 | 2.09E-63 | 4.34E-59 | 0.17 | 0.04 | 0.13 | 4.15 |
| 11 | Dazap2 | 0.89 | 2.54E-63 | 5.28E-59 | 0.77 | 0.57 | 0.20 | 1.35 |
| 11 | Tmem159 | 1.35 | 3.03E-63 | 6.29E-59 | 0.50 | 0.25 | 0.25 | 1.97 |
| 11 | Jag2 | 1.65 | 7.12E-63 | 1.48E-58 | 0.25 | 0.08 | 0.17 | 3.18 |
| 11 | Pard3 | 0.88 | 7.14E-63 | 1.49E-58 | 0.78 | 0.57 | 0.21 | 1.37 |
| 11 | Cdc42se1 | 1.11 | 8.48E-63 | 1.76E-58 | 0.65 | 0.42 | 0.23 | 1.55 |
| 11 | Aktip | 1.20 | 1.21E-62 | 2.52E-58 | 0.54 | 0.28 | 0.26 | 1.91 |
| 11 | 2310002F09Rik | 2.07 | 1.48E-62 | 3.08E-58 | 0.15 | 0.03 | 0.12 | 4.44 |
| 11 | Fosl2 | 2.60 | 1.90E-62 | 3.96E-58 | 0.13 | 0.03 | 0.11 | 4.89 |
| 11 | Zfp185 | 2.53 | 3.61E-62 | 7.51E-58 | 0.12 | 0.02 | 0.10 | 5.13 |
| 11 | Uchl1 | 0.94 | 4.06E-62 | 8.44E-58 | 0.60 | 0.34 | 0.26 | 1.76 |
| 11 | Tshz1 | 1.17 | 4.80E-62 | 9.97E-58 | 0.56 | 0.31 | 0.25 | 1.80 |
| 11 | Thsd7b | 2.75 | 5.11E-62 | 1.06E-57 | 0.11 | 0.02 | 0.09 | 5.60 |
| 11 | Ccser1 | 2.31 | 9.32E-62 | 1.94E-57 | 0.14 | 0.03 | 0.11 | 4.72 |
| 11 | Tbc1d2 | 2.56 | 1.01E-61 | 2.10E-57 | 0.12 | 0.02 | 0.10 | 5.32 |
| 11 | Pfkl | 0.85 | 1.08E-61 | 2.24E-57 | 0.82 | 0.64 | 0.18 | 1.28 |
| 11 | Ovol2 | 2.40 | 1.48E-61 | 3.08E-57 | 0.13 | 0.03 | 0.10 | 4.92 |
| 11 | Trpm4 | 1.74 | 1.53E-61 | 3.18E-57 | 0.27 | 0.09 | 0.18 | 2.95 |
| 11 | Cmip | 1.23 | 3.05E-61 | 6.35E-57 | 0.63 | 0.39 | 0.24 | 1.61 |
| 11 | Ctnnb1 | 0.50 | 4.38E-61 | 9.12E-57 | 0.99 | 0.97 | 0.02 | 1.02 |
| 11 | Bend7 | 1.67 | 4.71E-61 | 9.79E-57 | 0.24 | 0.08 | 0.17 | 3.17 |
| 11 | Ubald1 | 1.24 | 5.52E-61 | 1.15E-56 | 0.50 | 0.26 | 0.24 | 1.94 |
| 11 | Fat3 | 1.94 | 6.47E-61 | 1.35E-56 | 0.26 | 0.09 | 0.17 | 2.95 |
| 11 | Hs3st1 | 2.85 | 6.71E-61 | 1.39E-56 | 0.11 | 0.02 | 0.09 | 5.43 |
| 11 | Nhsl1 | 1.43 | 6.94E-61 | 1.44E-56 | 0.44 | 0.21 | 0.23 | 2.10 |
| 11 | Dpysl5 | 1.28 | 8.42E-61 | 1.75E-56 | 0.29 | 0.10 | 0.19 | 2.85 |
| 11 | Notch3 | 1.04 | 1.28E-60 | 2.65E-56 | 0.64 | 0.40 | 0.25 | 1.62 |
| 11 | Tjp1 | 0.90 | 1.30E-60 | 2.69E-56 | 0.71 | 0.49 | 0.23 | 1.47 |
| 11 | Pcdh1 | 1.86 | 1.65E-60 | 3.43E-56 | 0.18 | 0.05 | 0.13 | 3.85 |
| 11 | Adrb1 | 2.43 | 2.24E-59 | 4.66E-55 | 0.10 | 0.02 | 0.08 | 5.94 |
| 11 | Chmp2b | 0.86 | 2.41E-59 | 5.02E-55 | 0.77 | 0.56 | 0.21 | 1.38 |
| 11 | Dmbx1 | 1.79 | 5.53E-59 | 1.15E-54 | 0.16 | 0.04 | 0.12 | 4.08 |
| 11 | Trp53i11 | 1.07 | 9.63E-59 | 2.00E-54 | 0.46 | 0.22 | 0.24 | 2.11 |
| 11 | Cox7a2l | 0.48 | 1.33E-58 | 2.78E-54 | 0.98 | 0.96 | 0.03 | 1.03 |
| 11 | Cdc42ep5 | 1.02 | 1.49E-58 | 3.10E-54 | 0.67 | 0.46 | 0.21 | 1.46 |
| 11 | Trim16 | 2.79 | 2.82E-58 | 5.87E-54 | 0.11 | 0.02 | 0.09 | 5.45 |
| 11 | Kcnmb4os2 | 1.54 | 2.88E-58 | 5.99E-54 | 0.36 | 0.15 | 0.21 | 2.37 |
| 11 | Anxa1 | 2.54 | 3.63E-58 | 7.56E-54 | 0.12 | 0.02 | 0.10 | 5.04 |
| 11 | Pkp4 | 1.04 | 3.64E-58 | 7.57E-54 | 0.60 | 0.37 | 0.23 | 1.62 |
| 11 | Rasal2 | 1.29 | 3.93E-58 | 8.17E-54 | 0.48 | 0.24 | 0.24 | 1.98 |
| 11 | Lipg | 2.19 | 4.01E-58 | 8.34E-54 | 0.14 | 0.03 | 0.11 | 4.33 |
| 11 | Upk3bl | 1.26 | 4.94E-58 | 1.03E-53 | 0.23 | 0.07 | 0.16 | 3.31 |

**Supplementary Table 1. Marker genes for the original clusters of integrated clustering of E11.5 control and NotchGOF λ-junction cells.**

| | | | | | | | | |
|---|---|---|---|---|---|---|---|---|
| 11 | Lamb1 | 1.05 | 7.30E-58 | 1.52E-53 | 0.63 | 0.38 | 0.25 | 1.65 |
| 11 | Rap2c | 0.90 | 7.99E-58 | 1.66E-53 | 0.79 | 0.59 | 0.20 | 1.33 |
| 11 | Mkrn1 | 0.92 | 1.28E-57 | 2.66E-53 | 0.67 | 0.42 | 0.25 | 1.59 |
| 11 | Fam83h | 2.28 | 1.47E-57 | 3.06E-53 | 0.13 | 0.03 | 0.11 | 4.62 |
| 11 | Fam187b | 2.05 | 1.74E-57 | 3.63E-53 | 0.15 | 0.04 | 0.11 | 4.23 |
| 11 | Bnip3l | 0.49 | 2.52E-57 | 5.25E-53 | 0.97 | 0.93 | 0.04 | 1.04 |
| 11 | Ank3 | 0.94 | 2.54E-57 | 5.27E-53 | 0.70 | 0.48 | 0.22 | 1.46 |
| 11 | Minar2 | 1.78 | 2.55E-57 | 5.31E-53 | 0.26 | 0.09 | 0.17 | 2.88 |
| 11 | Mxi1 | 1.08 | 2.65E-57 | 5.52E-53 | 0.65 | 0.42 | 0.23 | 1.54 |
| 11 | Tmem79 | 2.30 | 4.18E-57 | 8.69E-53 | 0.13 | 0.03 | 0.10 | 4.67 |
| 11 | Celsr1 | 1.31 | 5.38E-57 | 1.12E-52 | 0.28 | 0.10 | 0.18 | 2.84 |
| 11 | Nedd9 | 1.61 | 5.47E-57 | 1.14E-52 | 0.37 | 0.16 | 0.21 | 2.27 |
| 11 | Hist3h2a | 1.22 | 5.55E-57 | 1.15E-52 | 0.49 | 0.26 | 0.24 | 1.91 |
| 11 | Lrrfip1 | 1.12 | 5.63E-57 | 1.17E-52 | 0.57 | 0.33 | 0.24 | 1.71 |
| 11 | C330021F23Rik | 2.12 | 1.26E-56 | 2.62E-52 | 0.13 | 0.03 | 0.10 | 4.61 |
| 11 | Vamp8 | 0.78 | 2.64E-56 | 5.48E-52 | 0.79 | 0.61 | 0.19 | 1.31 |
| 11 | Dock8 | 1.83 | 3.20E-56 | 6.66E-52 | 0.14 | 0.03 | 0.11 | 4.50 |
| 11 | Cab39 | 0.95 | 3.54E-56 | 7.36E-52 | 0.75 | 0.58 | 0.17 | 1.29 |
| 11 | Cxxc5 | 0.90 | 3.70E-56 | 7.69E-52 | 0.68 | 0.44 | 0.23 | 1.53 |
| 11 | Col17a1 | 1.94 | 3.98E-56 | 8.27E-52 | 0.14 | 0.03 | 0.11 | 4.30 |
| 11 | Lamc2 | 2.09 | 4.21E-56 | 8.77E-52 | 0.16 | 0.04 | 0.12 | 4.03 |
| 11 | Tjp2 | 1.21 | 5.00E-56 | 1.04E-51 | 0.53 | 0.30 | 0.23 | 1.74 |
| 11 | Gm48435 | 2.49 | 5.72E-56 | 1.19E-51 | 0.12 | 0.02 | 0.09 | 5.00 |
| 11 | Adi1 | 1.42 | 7.79E-56 | 1.62E-51 | 0.52 | 0.31 | 0.21 | 1.68 |
| 11 | Chchd10 | 1.14 | 1.07E-55 | 2.22E-51 | 0.37 | 0.15 | 0.22 | 2.40 |
| 11 | Dag1 | 1.09 | 1.50E-55 | 3.11E-51 | 0.60 | 0.36 | 0.23 | 1.64 |
| 11 | Bambi | 1.73 | 1.60E-55 | 3.34E-51 | 0.80 | 0.64 | 0.16 | 1.25 |
| 11 | Lonrf1 | 1.77 | 2.94E-55 | 6.11E-51 | 0.25 | 0.09 | 0.17 | 2.85 |
| 11 | Suco | 1.13 | 3.35E-55 | 6.96E-51 | 0.68 | 0.48 | 0.19 | 1.40 |
| 11 | Lrrk1 | 1.40 | 4.14E-55 | 8.62E-51 | 0.35 | 0.15 | 0.20 | 2.38 |
| 11 | Malat1 | 0.54 | 5.00E-55 | 1.04E-50 | 1.00 | 1.00 | 0.01 | 1.01 |
| 11 | Mtln | 1.10 | 7.29E-55 | 1.52E-50 | 0.56 | 0.32 | 0.24 | 1.74 |
| 11 | Zfp219 | 0.90 | 7.66E-55 | 1.59E-50 | 0.64 | 0.39 | 0.25 | 1.63 |
| 11 | Krt19 | 1.23 | 7.68E-55 | 1.60E-50 | 0.22 | 0.07 | 0.15 | 3.20 |
| 11 | Phlda3 | 1.24 | 7.80E-55 | 1.62E-50 | 0.63 | 0.40 | 0.23 | 1.56 |
| 11 | Aldoa | 0.48 | 1.40E-54 | 2.91E-50 | 0.98 | 0.95 | 0.03 | 1.03 |
| 11 | Grhl1 | 2.22 | 1.80E-54 | 3.75E-50 | 0.11 | 0.02 | 0.09 | 5.30 |
| 11 | Pqlc3 | 1.76 | 2.16E-54 | 4.48E-50 | 0.18 | 0.05 | 0.13 | 3.55 |
| 11 | Pura | 0.96 | 2.32E-54 | 4.82E-50 | 0.68 | 0.47 | 0.22 | 1.47 |
| 11 | AI463170 | 2.38 | 2.34E-54 | 4.87E-50 | 0.11 | 0.02 | 0.09 | 5.25 |
| 11 | Rnf149 | 1.60 | 2.85E-54 | 5.92E-50 | 0.43 | 0.21 | 0.21 | 1.99 |
| 11 | Cryab | 0.74 | 3.69E-54 | 7.68E-50 | 0.17 | 0.05 | 0.13 | 3.78 |
| 11 | Acbd4 | 2.01 | 3.72E-54 | 7.73E-50 | 0.16 | 0.04 | 0.12 | 3.77 |
| 11 | Nrg1 | 1.57 | 4.10E-54 | 8.53E-50 | 0.23 | 0.08 | 0.15 | 3.05 |
| 11 | Wnt7b | 1.21 | 5.34E-54 | 1.11E-49 | 0.16 | 0.04 | 0.12 | 4.00 |
| 11 | Ddhd1 | 1.52 | 8.99E-54 | 1.87E-49 | 0.36 | 0.16 | 0.20 | 2.23 |
| 11 | Pik3cb | 1.90 | 1.07E-53 | 2.22E-49 | 0.19 | 0.06 | 0.14 | 3.34 |
| 11 | Tmem205 | 1.13 | 1.12E-53 | 2.33E-49 | 0.51 | 0.27 | 0.24 | 1.87 |
| 11 | Hes5 | 0.56 | 1.28E-53 | 2.66E-49 | 0.30 | 0.11 | 0.19 | 2.70 |
| 11 | Meis1 | 0.82 | 1.57E-53 | 3.27E-49 | 0.71 | 0.45 | 0.26 | 1.58 |
| 11 | Pwwp2b | 1.74 | 2.46E-53 | 5.13E-49 | 0.22 | 0.07 | 0.15 | 3.13 |
| 11 | Klhl8 | 1.69 | 2.48E-53 | 5.15E-49 | 0.25 | 0.09 | 0.16 | 2.85 |
| 11 | Aspscr1 | 1.15 | 2.99E-53 | 6.21E-49 | 0.65 | 0.45 | 0.20 | 1.45 |
| 11 | Arhgap29 | 1.59 | 8.17E-53 | 1.70E-48 | 0.57 | 0.36 | 0.21 | 1.57 |
| 11 | Galnt18 | 1.92 | 1.11E-52 | 2.30E-48 | 0.13 | 0.03 | 0.10 | 4.40 |
| 11 | Cnksr1 | 2.66 | 1.48E-52 | 3.09E-48 | 0.10 | 0.02 | 0.08 | 5.32 |

**Supplementary Table 1. Marker genes for the original clusters of integrated clustering of E11.5 control and NotchGOF λ-junction cells.**

| | | | | | | | | |
|---|---|---|---|---|---|---|---|---|
| 11 | Nudt14 | 0.89 | 1.54E-52 | 3.20E-48 | 0.64 | 0.40 | 0.24 | 1.60 |
| 11 | Arg2 | 1.76 | 1.67E-52 | 3.47E-48 | 0.18 | 0.05 | 0.13 | 3.57 |
| 11 | Klf3 | 0.98 | 1.71E-52 | 3.57E-48 | 0.69 | 0.49 | 0.20 | 1.40 |
| 11 | Tbc1d30 | 2.07 | 5.32E-52 | 1.11E-47 | 0.13 | 0.03 | 0.10 | 4.46 |
| 11 | Golga4 | 0.79 | 5.63E-52 | 1.17E-47 | 0.80 | 0.64 | 0.16 | 1.26 |
| 11 | Pkm | 0.33 | 6.74E-52 | 1.40E-47 | 1.00 | 0.98 | 0.02 | 1.02 |
| 11 | 2700033N17Rik | 1.24 | 8.30E-52 | 1.73E-47 | 0.29 | 0.11 | 0.18 | 2.60 |
| 11 | Ildr1 | 2.08 | 1.55E-51 | 3.23E-47 | 0.14 | 0.04 | 0.11 | 4.00 |
| 11 | Trp53inp1 | 1.50 | 1.77E-51 | 3.67E-47 | 0.38 | 0.18 | 0.20 | 2.12 |
| 11 | Acvr2b | 0.71 | 2.08E-51 | 4.33E-47 | 0.82 | 0.64 | 0.18 | 1.28 |
| 11 | Slc44a2 | 1.19 | 2.23E-51 | 4.64E-47 | 0.49 | 0.27 | 0.23 | 1.84 |
| 11 | Mark2 | 0.92 | 2.35E-51 | 4.88E-47 | 0.67 | 0.46 | 0.21 | 1.46 |
| 11 | Mal2 | 2.44 | 2.53E-51 | 5.26E-47 | 0.10 | 0.02 | 0.08 | 5.15 |
| 11 | Ddi2 | 1.18 | 4.01E-51 | 8.33E-47 | 0.61 | 0.40 | 0.20 | 1.50 |
| 11 | 3110039M20Rik | 1.37 | 6.02E-51 | 1.25E-46 | 0.22 | 0.07 | 0.15 | 3.07 |
| 11 | Sparc | 0.63 | 6.46E-51 | 1.34E-46 | 0.91 | 0.74 | 0.17 | 1.23 |
| 11 | Rel | 1.82 | 8.31E-51 | 1.73E-46 | 0.24 | 0.09 | 0.15 | 2.80 |
| 11 | Rabgap1 | 0.81 | 8.71E-51 | 1.81E-46 | 0.75 | 0.56 | 0.19 | 1.33 |
| 11 | Fbxo32 | 2.70 | 1.47E-50 | 3.06E-46 | 0.12 | 0.03 | 0.09 | 4.44 |
| 11 | Errfi1 | 1.40 | 1.50E-50 | 3.12E-46 | 0.37 | 0.17 | 0.20 | 2.16 |
| 11 | Clcc1 | 1.08 | 1.76E-50 | 3.67E-46 | 0.61 | 0.41 | 0.21 | 1.51 |
| 11 | Krtcap3 | 1.54 | 2.08E-50 | 4.33E-46 | 0.28 | 0.11 | 0.17 | 2.56 |
| 11 | Ntn4 | 2.25 | 3.57E-50 | 7.41E-46 | 0.11 | 0.02 | 0.09 | 4.75 |
| 11 | Tmem30b | 1.74 | 4.92E-50 | 1.02E-45 | 0.33 | 0.14 | 0.18 | 2.27 |
| 11 | Etv3 | 1.31 | 5.73E-50 | 1.19E-45 | 0.42 | 0.22 | 0.21 | 1.95 |
| 11 | Ralbp1 | 0.71 | 6.58E-50 | 1.37E-45 | 0.86 | 0.78 | 0.08 | 1.10 |
| 11 | Mfsd6 | 2.03 | 7.12E-50 | 1.48E-45 | 0.13 | 0.03 | 0.10 | 4.26 |
| 11 | Fgf11 | 2.58 | 1.26E-49 | 2.61E-45 | 0.10 | 0.02 | 0.08 | 5.05 |
| 11 | Aopep | 1.00 | 2.50E-49 | 5.21E-45 | 0.79 | 0.62 | 0.17 | 1.28 |
| 11 | Cryl1 | 1.32 | 2.97E-49 | 6.18E-45 | 0.33 | 0.14 | 0.19 | 2.31 |
| 11 | Frrs1 | 1.27 | 3.20E-49 | 6.65E-45 | 0.42 | 0.22 | 0.21 | 1.97 |
| 11 | St8sia6 | 1.79 | 3.41E-49 | 7.09E-45 | 0.16 | 0.05 | 0.12 | 3.60 |
| 11 | Tor3a | 1.91 | 8.77E-49 | 1.83E-44 | 0.14 | 0.04 | 0.11 | 3.94 |
| 11 | Tagln | 3.23 | 1.02E-48 | 2.12E-44 | 0.13 | 0.03 | 0.10 | 4.03 |
| 11 | Patj | 1.41 | 1.04E-48 | 2.17E-44 | 0.30 | 0.13 | 0.18 | 2.43 |
| 11 | Frmd5 | 1.98 | 1.24E-48 | 2.59E-44 | 0.17 | 0.05 | 0.12 | 3.51 |
| 11 | Tmem123 | 1.18 | 1.97E-48 | 4.10E-44 | 0.53 | 0.32 | 0.21 | 1.68 |
| 11 | Dync1i1 | 1.83 | 2.31E-48 | 4.79E-44 | 0.19 | 0.06 | 0.13 | 3.16 |
| 11 | Galnt14 | 2.42 | 2.70E-48 | 5.61E-44 | 0.11 | 0.02 | 0.08 | 4.77 |
| 11 | Rhbdf1 | 1.57 | 2.79E-48 | 5.80E-44 | 0.25 | 0.10 | 0.16 | 2.65 |
| 11 | Armc10 | 0.80 | 2.92E-48 | 6.06E-44 | 0.72 | 0.52 | 0.20 | 1.39 |
| 11 | Lrrc8d | 1.30 | 4.12E-48 | 8.57E-44 | 0.39 | 0.19 | 0.20 | 2.06 |
| 11 | Gimap9 | 1.51 | 4.26E-48 | 8.85E-44 | 0.27 | 0.10 | 0.16 | 2.61 |
| 11 | Pigp | 0.72 | 4.31E-48 | 8.96E-44 | 0.80 | 0.64 | 0.16 | 1.25 |
| 11 | Plxdc2 | 1.08 | 4.49E-48 | 9.35E-44 | 0.46 | 0.24 | 0.22 | 1.91 |
| 11 | Slc16a7 | 2.63 | 6.63E-48 | 1.38E-43 | 0.10 | 0.02 | 0.08 | 4.81 |
| 11 | Gkap1 | 0.83 | 6.64E-48 | 1.38E-43 | 0.66 | 0.43 | 0.23 | 1.53 |
| 11 | Hap1 | 1.70 | 8.07E-48 | 1.68E-43 | 0.17 | 0.05 | 0.12 | 3.46 |
| 11 | Skil | 0.94 | 8.10E-48 | 1.69E-43 | 0.74 | 0.55 | 0.19 | 1.35 |
| 11 | Bcl7a | 0.68 | 8.73E-48 | 1.81E-43 | 0.82 | 0.67 | 0.15 | 1.23 |
| 11 | Cmas | 0.90 | 1.35E-47 | 2.80E-43 | 0.74 | 0.59 | 0.15 | 1.25 |
| 11 | Erc2 | 2.20 | 1.43E-47 | 2.97E-43 | 0.20 | 0.07 | 0.13 | 3.05 |
| 11 | Fbxw4 | 1.54 | 2.29E-47 | 4.75E-43 | 0.32 | 0.14 | 0.18 | 2.30 |
| 11 | Zdhhc21 | 1.12 | 2.57E-47 | 5.34E-43 | 0.50 | 0.28 | 0.22 | 1.78 |
| 11 | Slc44a1 | 1.15 | 2.57E-47 | 5.35E-43 | 0.44 | 0.23 | 0.21 | 1.93 |
| 11 | Sft2d2 | 1.45 | 2.73E-47 | 5.67E-43 | 0.33 | 0.14 | 0.18 | 2.26 |

**Supplementary Table 1. Marker genes for the original clusters of integrated clustering of E11.5 control and NotchGOF λ-junction cells.**

| | | | | | | | | |
|---|---|---|---|---|---|---|---|---|
| 11 | Chd3 | 0.58 | 2.74E-47 | 5.70E-43 | 0.96 | 0.89 | 0.06 | 1.07 |
| 11 | Pag1 | 2.06 | 3.11E-47 | 6.46E-43 | 0.15 | 0.04 | 0.11 | 3.62 |
| 11 | Smpdl3a | 1.33 | 3.15E-47 | 6.55E-43 | 0.31 | 0.13 | 0.18 | 2.42 |
| 11 | Creld1 | 1.41 | 6.84E-47 | 1.42E-42 | 0.30 | 0.12 | 0.18 | 2.41 |
| 11 | Tead3 | 1.07 | 9.03E-47 | 1.88E-42 | 0.46 | 0.24 | 0.22 | 1.91 |
| 11 | Hdgf | 0.36 | 1.44E-46 | 3.00E-42 | 0.99 | 0.97 | 0.01 | 1.01 |
| 11 | Ptprk | 0.90 | 1.74E-46 | 3.62E-42 | 0.66 | 0.43 | 0.22 | 1.51 |
| 11 | Tmem200a | 1.59 | 1.84E-46 | 3.82E-42 | 0.17 | 0.05 | 0.12 | 3.35 |
| 11 | Reep3 | 0.89 | 2.13E-46 | 4.42E-42 | 0.70 | 0.52 | 0.18 | 1.36 |
| 11 | Rgs2 | 1.63 | 2.45E-46 | 5.09E-42 | 0.53 | 0.34 | 0.19 | 1.56 |
| 11 | Tnks2 | 0.59 | 2.45E-46 | 5.09E-42 | 0.90 | 0.81 | 0.09 | 1.12 |
| 11 | Ptpn3 | 1.93 | 4.34E-46 | 9.02E-42 | 0.14 | 0.04 | 0.10 | 3.91 |
| 11 | Cacnb3 | 1.05 | 6.77E-46 | 1.41E-41 | 0.49 | 0.27 | 0.22 | 1.82 |
| 11 | Hpn | 2.04 | 6.78E-46 | 1.41E-41 | 0.14 | 0.04 | 0.10 | 3.86 |
| 11 | 1810037I17Rik | 0.55 | 7.40E-46 | 1.54E-41 | 0.91 | 0.83 | 0.08 | 1.09 |
| 11 | Phlpp1 | 1.31 | 7.54E-46 | 1.57E-41 | 0.30 | 0.13 | 0.18 | 2.39 |
| 11 | Dock9 | 1.60 | 1.41E-45 | 2.92E-41 | 0.20 | 0.07 | 0.14 | 2.96 |
| 11 | Emb | 0.83 | 1.44E-45 | 3.00E-41 | 0.63 | 0.40 | 0.23 | 1.57 |
| 11 | Sh3tc2 | 2.32 | 1.70E-45 | 3.54E-41 | 0.11 | 0.02 | 0.08 | 4.61 |
| 11 | Pomp | 0.35 | 1.84E-45 | 3.84E-41 | 0.98 | 0.98 | 0.01 | 1.01 |
| 11 | Nudt18 | 1.74 | 2.06E-45 | 4.29E-41 | 0.18 | 0.06 | 0.12 | 3.12 |
| 11 | Aftph | 1.16 | 2.49E-45 | 5.19E-41 | 0.45 | 0.25 | 0.21 | 1.84 |
| 11 | Svip | 1.36 | 2.95E-45 | 6.14E-41 | 0.24 | 0.09 | 0.15 | 2.71 |
| 11 | Pak3 | 1.29 | 6.67E-45 | 1.39E-40 | 0.31 | 0.13 | 0.18 | 2.33 |
| 11 | 2010300C02Rik | 2.03 | 8.48E-45 | 1.76E-40 | 0.10 | 0.02 | 0.08 | 4.68 |
| 11 | Gprc5c | 1.13 | 1.09E-44 | 2.26E-40 | 0.44 | 0.23 | 0.21 | 1.91 |
| 11 | Rcn1 | 0.56 | 1.15E-44 | 2.40E-40 | 0.91 | 0.82 | 0.09 | 1.10 |
| 11 | Crybg1 | 2.42 | 1.51E-44 | 3.14E-40 | 0.10 | 0.02 | 0.08 | 4.48 |
| 11 | Dusp2 | 1.93 | 2.01E-44 | 4.17E-40 | 0.12 | 0.03 | 0.09 | 4.10 |
| 11 | Wipf3 | 1.64 | 2.09E-44 | 4.35E-40 | 0.18 | 0.06 | 0.12 | 3.14 |
| 11 | Pak2 | 0.53 | 3.05E-44 | 6.35E-40 | 0.92 | 0.83 | 0.09 | 1.11 |
| 11 | Tet3 | 0.79 | 3.09E-44 | 6.43E-40 | 0.74 | 0.56 | 0.18 | 1.31 |
| 11 | Rims2 | 1.58 | 3.13E-44 | 6.50E-40 | 0.18 | 0.06 | 0.12 | 3.18 |
| 11 | Map1lc3b | 0.48 | 4.78E-44 | 9.94E-40 | 0.95 | 0.93 | 0.03 | 1.03 |
| 11 | Cobll1 | 2.03 | 1.30E-43 | 2.70E-39 | 0.12 | 0.03 | 0.09 | 4.00 |
| 11 | Pnrc1 | 0.58 | 1.36E-43 | 2.82E-39 | 0.94 | 0.87 | 0.07 | 1.08 |
| 11 | Kcnmb4 | 1.14 | 1.75E-43 | 3.64E-39 | 0.53 | 0.32 | 0.21 | 1.67 |
| 11 | Samd10 | 1.40 | 3.18E-43 | 6.61E-39 | 0.23 | 0.09 | 0.15 | 2.69 |
| 11 | Rreb1 | 1.14 | 3.77E-43 | 7.84E-39 | 0.40 | 0.20 | 0.20 | 1.97 |
| 11 | Sgpl1 | 1.01 | 4.29E-43 | 8.93E-39 | 0.55 | 0.33 | 0.22 | 1.65 |
| 11 | Fgd6 | 1.40 | 6.66E-43 | 1.39E-38 | 0.34 | 0.16 | 0.18 | 2.15 |
| 11 | Firre | 1.56 | 7.30E-43 | 1.52E-38 | 0.23 | 0.09 | 0.14 | 2.67 |
| 11 | Kxd1 | 0.77 | 7.37E-43 | 1.53E-38 | 0.69 | 0.50 | 0.19 | 1.39 |
| 11 | Jpt1 | 0.48 | 8.96E-43 | 1.86E-38 | 0.98 | 0.94 | 0.04 | 1.04 |
| 11 | Exoc3l4 | 1.38 | 9.07E-43 | 1.89E-38 | 0.20 | 0.07 | 0.13 | 2.94 |
| 11 | Arfgef1 | 0.88 | 1.11E-42 | 2.31E-38 | 0.66 | 0.48 | 0.18 | 1.37 |
| 11 | Ywhaz | 0.39 | 1.33E-42 | 2.77E-38 | 0.98 | 0.95 | 0.03 | 1.03 |
| 11 | Plch2 | 2.00 | 2.28E-42 | 4.75E-38 | 0.11 | 0.03 | 0.09 | 4.27 |
| 11 | Slc25a17 | 0.75 | 2.31E-42 | 4.81E-38 | 0.78 | 0.62 | 0.17 | 1.27 |
| 11 | Acsl5 | 1.25 | 2.59E-42 | 5.39E-38 | 0.33 | 0.15 | 0.18 | 2.16 |
| 11 | Erbb2 | 1.22 | 3.14E-42 | 6.53E-38 | 0.42 | 0.23 | 0.20 | 1.86 |
| 11 | Elf3 | 1.62 | 3.47E-42 | 7.21E-38 | 0.20 | 0.07 | 0.13 | 2.87 |
| 11 | Vgll3 | 2.13 | 4.34E-42 | 9.02E-38 | 0.11 | 0.03 | 0.08 | 4.19 |
| 11 | Echdc1 | 1.19 | 4.35E-42 | 9.06E-38 | 0.43 | 0.23 | 0.19 | 1.84 |
| 11 | Klhl24 | 0.78 | 4.40E-42 | 9.15E-38 | 0.77 | 0.60 | 0.17 | 1.29 |
| 11 | Fam162a | 0.60 | 4.45E-42 | 9.26E-38 | 0.94 | 0.89 | 0.05 | 1.05 |

**Supplementary Table 1. Marker genes for the original clusters of integrated clustering of E11.5 control and NotchGOF λ-junction cells.**

| | | | | | | | | |
|---|---|---|---|---|---|---|---|---|
| 11 | Prickle3 | 1.61 | 4.63E-42 | 9.62E-38 | 0.19 | 0.06 | 0.12 | 2.98 |
| 11 | Snn | 0.77 | 6.05E-42 | 1.26E-37 | 0.64 | 0.43 | 0.22 | 1.50 |
| 11 | Skp1a | 0.38 | 8.89E-42 | 1.85E-37 | 0.98 | 0.96 | 0.03 | 1.03 |
| 11 | Ddah2 | 0.38 | 9.89E-42 | 2.06E-37 | 0.99 | 0.95 | 0.04 | 1.05 |
| 11 | Cemip2 | 0.87 | 1.39E-41 | 2.90E-37 | 0.64 | 0.44 | 0.19 | 1.43 |
| 11 | Dach1 | 0.84 | 1.49E-41 | 3.10E-37 | 0.59 | 0.38 | 0.21 | 1.57 |
| 11 | Mcee | 0.84 | 2.35E-41 | 4.90E-37 | 0.64 | 0.43 | 0.20 | 1.47 |
| 11 | Efnb2 | 0.69 | 2.49E-41 | 5.18E-37 | 0.67 | 0.44 | 0.23 | 1.51 |
| 11 | Rnf44 | 0.97 | 3.42E-41 | 7.10E-37 | 0.54 | 0.34 | 0.20 | 1.59 |
| 11 | Ube2b | 0.43 | 5.21E-41 | 1.08E-36 | 0.96 | 0.93 | 0.04 | 1.04 |
| 11 | Sulf1 | 0.64 | 5.99E-41 | 1.24E-36 | 0.75 | 0.53 | 0.22 | 1.41 |
| 11 | Cited2 | 0.83 | 7.94E-41 | 1.65E-36 | 0.77 | 0.64 | 0.14 | 1.21 |
| 11 | Pkp2 | 1.72 | 1.16E-40 | 2.42E-36 | 0.16 | 0.05 | 0.11 | 3.18 |
| 11 | Dpysl3 | 0.75 | 1.52E-40 | 3.17E-36 | 0.65 | 0.45 | 0.21 | 1.46 |
| 11 | Nr2f2 | 0.58 | 1.54E-40 | 3.21E-36 | 0.59 | 0.36 | 0.23 | 1.65 |
| 11 | Dlgap4 | 0.67 | 1.98E-40 | 4.11E-36 | 0.80 | 0.66 | 0.14 | 1.21 |
| 11 | Ppfia3 | 1.19 | 2.92E-40 | 6.08E-36 | 0.29 | 0.13 | 0.16 | 2.31 |
| 11 | Ntm | 1.62 | 4.32E-40 | 8.98E-36 | 0.25 | 0.10 | 0.15 | 2.44 |
| 11 | Cuedc1 | 0.95 | 4.63E-40 | 9.63E-36 | 0.58 | 0.38 | 0.20 | 1.52 |
| 11 | Alpl | 1.66 | 4.87E-40 | 1.01E-35 | 0.24 | 0.10 | 0.14 | 2.46 |
| 11 | Stox2 | 0.71 | 5.12E-40 | 1.06E-35 | 0.72 | 0.56 | 0.17 | 1.30 |
| 11 | Tmem184a | 1.78 | 7.85E-40 | 1.63E-35 | 0.16 | 0.05 | 0.11 | 3.14 |
| 11 | Gipc1 | 0.62 | 8.81E-40 | 1.83E-35 | 0.82 | 0.68 | 0.14 | 1.20 |
| 11 | Map3k20 | 1.48 | 9.62E-40 | 2.00E-35 | 0.26 | 0.11 | 0.15 | 2.34 |
| 11 | Hs6st2 | 1.02 | 1.02E-39 | 2.12E-35 | 0.48 | 0.28 | 0.20 | 1.72 |
| 11 | Filip1 | 2.34 | 1.05E-39 | 2.19E-35 | 0.12 | 0.03 | 0.09 | 3.65 |
| 11 | Gabarapl2 | 0.51 | 1.27E-39 | 2.65E-35 | 0.90 | 0.82 | 0.08 | 1.10 |
| 11 | Mfsd1 | 1.12 | 1.40E-39 | 2.92E-35 | 0.55 | 0.36 | 0.19 | 1.54 |
| 11 | Ctdspl | 0.94 | 1.88E-39 | 3.90E-35 | 0.54 | 0.34 | 0.20 | 1.59 |
| 11 | Knop1 | 0.45 | 2.67E-39 | 5.56E-35 | 0.96 | 0.90 | 0.05 | 1.06 |
| 11 | Srd5a1 | 1.21 | 4.52E-39 | 9.40E-35 | 0.36 | 0.18 | 0.18 | 2.00 |
| 11 | Msx2 | 0.62 | 5.76E-39 | 1.20E-34 | 0.67 | 0.42 | 0.25 | 1.60 |
| 11 | Sppl2a | 0.89 | 6.93E-39 | 1.44E-34 | 0.65 | 0.47 | 0.18 | 1.38 |
| 11 | Mbd2 | 0.76 | 8.00E-39 | 1.66E-34 | 0.71 | 0.56 | 0.15 | 1.27 |
| 11 | Npepps | 0.60 | 1.31E-38 | 2.73E-34 | 0.79 | 0.64 | 0.15 | 1.23 |
| 11 | Pycard | 1.17 | 1.56E-38 | 3.25E-34 | 0.28 | 0.12 | 0.16 | 2.26 |
| 11 | Tmem63b | 1.01 | 1.63E-38 | 3.39E-34 | 0.53 | 0.33 | 0.20 | 1.60 |
| 11 | Nipal3 | 1.75 | 1.67E-38 | 3.46E-34 | 0.13 | 0.04 | 0.10 | 3.50 |
| 11 | Pak1 | 0.79 | 1.88E-38 | 3.90E-34 | 0.67 | 0.49 | 0.19 | 1.38 |
| 11 | Fat1 | 0.75 | 2.17E-38 | 4.51E-34 | 0.69 | 0.51 | 0.18 | 1.35 |
| 11 | Gm6169 | 2.15 | 2.34E-38 | 4.86E-34 | 0.12 | 0.03 | 0.09 | 3.62 |
| 11 | Ctnnal1 | 0.88 | 3.71E-38 | 7.71E-34 | 0.46 | 0.26 | 0.21 | 1.80 |
| 11 | Schip1 | 1.07 | 3.82E-38 | 7.96E-34 | 0.33 | 0.16 | 0.17 | 2.12 |
| 11 | Baz2b | 0.63 | 4.57E-38 | 9.51E-34 | 0.84 | 0.70 | 0.14 | 1.20 |
| 11 | Npnt | 0.89 | 5.87E-38 | 1.22E-33 | 0.43 | 0.23 | 0.20 | 1.86 |
| 11 | Pnpla2 | 1.38 | 6.88E-38 | 1.43E-33 | 0.22 | 0.09 | 0.13 | 2.56 |
| 11 | Vwa5a | 1.57 | 1.46E-37 | 3.05E-33 | 0.22 | 0.08 | 0.13 | 2.56 |
| 11 | Tspan5 | 1.07 | 2.92E-37 | 6.07E-33 | 0.45 | 0.26 | 0.19 | 1.72 |
| 11 | Dmrta1 | 2.03 | 3.01E-37 | 6.27E-33 | 0.13 | 0.04 | 0.09 | 3.47 |
| 11 | Sh3gl2 | 1.73 | 3.39E-37 | 7.04E-33 | 0.13 | 0.04 | 0.09 | 3.56 |
| 11 | Creb3l2 | 1.00 | 3.80E-37 | 7.91E-33 | 0.51 | 0.32 | 0.19 | 1.61 |
| 11 | Tulp4 | 0.64 | 4.25E-37 | 8.84E-33 | 0.79 | 0.66 | 0.13 | 1.19 |
| 11 | Ctsd | 0.52 | 4.30E-37 | 8.94E-33 | 0.75 | 0.60 | 0.15 | 1.25 |
| 11 | Parm1 | 1.47 | 4.40E-37 | 9.15E-33 | 0.32 | 0.16 | 0.16 | 2.03 |
| 11 | Tuft1 | 2.25 | 4.91E-37 | 1.02E-32 | 0.12 | 0.04 | 0.09 | 3.54 |
| 11 | Pacsin1 | 1.21 | 6.11E-37 | 1.27E-32 | 0.24 | 0.10 | 0.14 | 2.45 |

**Supplementary Table 1. Marker genes for the original clusters of integrated clustering of E11.5 control and NotchGOF λ-junction cells.**

| | | | | | | | | |
|---|---|---|---|---|---|---|---|---|
| 11 | Lypd6b | 1.36 | 7.68E-37 | 1.60E-32 | 0.18 | 0.07 | 0.12 | 2.83 |
| 11 | Ebf4 | 1.69 | 8.97E-37 | 1.86E-32 | 0.16 | 0.06 | 0.11 | 2.96 |
| 11 | Akap9 | 0.56 | 1.21E-36 | 2.52E-32 | 0.87 | 0.78 | 0.09 | 1.11 |
| 11 | Pik3ip1 | 1.55 | 1.24E-36 | 2.58E-32 | 0.21 | 0.08 | 0.13 | 2.52 |
| 11 | Vsig10 | 1.69 | 1.29E-36 | 2.67E-32 | 0.16 | 0.05 | 0.11 | 3.10 |
| 11 | Pnck | 1.43 | 1.57E-36 | 3.26E-32 | 0.19 | 0.07 | 0.12 | 2.76 |
| 11 | Lamp2 | 0.71 | 1.64E-36 | 3.42E-32 | 0.73 | 0.56 | 0.17 | 1.30 |
| 11 | Enc1 | 1.04 | 1.71E-36 | 3.57E-32 | 0.40 | 0.22 | 0.19 | 1.86 |
| 11 | Tpp1 | 0.97 | 1.90E-36 | 3.96E-32 | 0.45 | 0.26 | 0.19 | 1.73 |
| 11 | Zmiz2 | 0.93 | 2.61E-36 | 5.42E-32 | 0.50 | 0.30 | 0.19 | 1.63 |
| 11 | Adgrb3 | 1.53 | 2.62E-36 | 5.46E-32 | 0.18 | 0.06 | 0.11 | 2.87 |
| 11 | Rapgefl1 | 1.77 | 2.84E-36 | 5.90E-32 | 0.11 | 0.03 | 0.08 | 3.79 |
| 11 | Plekha1 | 1.15 | 3.74E-36 | 7.77E-32 | 0.37 | 0.20 | 0.18 | 1.91 |
| 11 | Syngr2 | 1.22 | 4.09E-36 | 8.50E-32 | 0.30 | 0.14 | 0.16 | 2.14 |
| 11 | Cox14 | 0.47 | 4.59E-36 | 9.54E-32 | 0.90 | 0.80 | 0.09 | 1.12 |
| 11 | Ank2 | 1.15 | 5.08E-36 | 1.06E-31 | 0.42 | 0.24 | 0.18 | 1.75 |
| 11 | Trps1 | 0.61 | 1.00E-35 | 2.09E-31 | 0.68 | 0.48 | 0.20 | 1.42 |
| 11 | C1qtnf12 | 1.51 | 1.02E-35 | 2.12E-31 | 0.32 | 0.16 | 0.16 | 1.99 |
| 11 | Lrrc1 | 1.18 | 1.22E-35 | 2.53E-31 | 0.31 | 0.14 | 0.16 | 2.12 |
| 11 | Ptpn4 | 1.18 | 1.48E-35 | 3.08E-31 | 0.32 | 0.15 | 0.16 | 2.06 |
| 11 | Podxl | 1.55 | 1.53E-35 | 3.17E-31 | 0.13 | 0.04 | 0.09 | 3.27 |
| 11 | Rasgef1b | 1.73 | 1.92E-35 | 3.99E-31 | 0.14 | 0.05 | 0.10 | 3.16 |
| 11 | Jdp2 | 1.70 | 2.05E-35 | 4.26E-31 | 0.12 | 0.04 | 0.09 | 3.51 |
| 11 | D230030E09Rik | 1.07 | 3.37E-35 | 7.02E-31 | 0.16 | 0.05 | 0.11 | 2.98 |
| 11 | Hint2 | 0.60 | 3.69E-35 | 7.68E-31 | 0.76 | 0.61 | 0.15 | 1.25 |
| 11 | Amotl1 | 0.75 | 4.79E-35 | 9.97E-31 | 0.79 | 0.69 | 0.11 | 1.15 |
| 11 | Hibadh | 0.81 | 6.03E-35 | 1.25E-30 | 0.69 | 0.55 | 0.14 | 1.25 |
| 11 | F3 | 1.28 | 6.32E-35 | 1.31E-30 | 0.23 | 0.10 | 0.14 | 2.39 |
| 11 | Xpr1 | 1.24 | 7.29E-35 | 1.52E-30 | 0.45 | 0.27 | 0.18 | 1.67 |
| 11 | Rrbp1 | 0.50 | 1.12E-34 | 2.32E-30 | 0.95 | 0.90 | 0.05 | 1.06 |
| 11 | Tdrp | 1.93 | 1.12E-34 | 2.32E-30 | 0.10 | 0.03 | 0.08 | 3.92 |
| 11 | Marchf7 | 0.56 | 1.56E-34 | 3.25E-30 | 0.83 | 0.72 | 0.10 | 1.14 |
| 11 | Smim22 | 1.62 | 2.66E-34 | 5.52E-30 | 0.12 | 0.04 | 0.09 | 3.42 |
| 11 | Crebrf | 1.29 | 2.68E-34 | 5.58E-30 | 0.25 | 0.11 | 0.14 | 2.32 |
| 11 | Mindy2 | 1.09 | 2.76E-34 | 5.74E-30 | 0.34 | 0.17 | 0.17 | 1.98 |
| 11 | Dclre1c | 2.23 | 2.86E-34 | 5.95E-30 | 0.15 | 0.05 | 0.10 | 2.96 |
| 11 | Epha2 | 1.94 | 3.16E-34 | 6.57E-30 | 0.11 | 0.03 | 0.08 | 3.58 |
| 11 | Fam32a | 0.51 | 3.23E-34 | 6.72E-30 | 0.86 | 0.77 | 0.09 | 1.12 |
| 11 | Sh3d19 | 1.00 | 3.30E-34 | 6.85E-30 | 0.43 | 0.25 | 0.18 | 1.72 |
| 11 | Eml2 | 1.40 | 3.49E-34 | 7.27E-30 | 0.23 | 0.10 | 0.13 | 2.38 |
| 11 | Renbp | 1.43 | 3.65E-34 | 7.59E-30 | 0.19 | 0.07 | 0.12 | 2.61 |
| 11 | Baz2a | 0.88 | 3.69E-34 | 7.67E-30 | 0.56 | 0.38 | 0.18 | 1.47 |
| 11 | Mcoln3 | 1.64 | 3.92E-34 | 8.15E-30 | 0.16 | 0.05 | 0.10 | 2.98 |
| 11 | Insr | 1.04 | 4.05E-34 | 8.43E-30 | 0.42 | 0.23 | 0.18 | 1.78 |
| 11 | Plxnb1 | 0.92 | 4.30E-34 | 8.94E-30 | 0.44 | 0.26 | 0.19 | 1.73 |
| 11 | Apc | 0.55 | 4.77E-34 | 9.92E-30 | 0.77 | 0.61 | 0.16 | 1.26 |
| 11 | Alcam | 1.07 | 5.07E-34 | 1.05E-29 | 0.30 | 0.14 | 0.16 | 2.14 |
| 11 | Kazn | 1.03 | 7.74E-34 | 1.61E-29 | 0.35 | 0.18 | 0.17 | 1.94 |
| 11 | Cobl | 1.28 | 9.85E-34 | 2.05E-29 | 0.18 | 0.07 | 0.11 | 2.70 |
| 11 | Ccdc68 | 1.57 | 1.05E-33 | 2.19E-29 | 0.15 | 0.05 | 0.10 | 2.98 |
| 11 | Gria3 | 1.20 | 1.49E-33 | 3.09E-29 | 0.32 | 0.16 | 0.16 | 2.01 |
| 11 | Vps4b | 0.62 | 2.13E-33 | 4.43E-29 | 0.77 | 0.62 | 0.15 | 1.23 |
| 11 | Nup210 | 1.56 | 3.05E-33 | 6.35E-29 | 0.16 | 0.06 | 0.11 | 2.81 |
| 11 | Fam117a | 1.03 | 3.62E-33 | 7.53E-29 | 0.41 | 0.23 | 0.18 | 1.76 |
| 11 | Atp2b1 | 0.61 | 4.08E-33 | 8.48E-29 | 0.88 | 0.84 | 0.04 | 1.05 |
| 11 | Pon2 | 0.93 | 4.93E-33 | 1.03E-28 | 0.34 | 0.18 | 0.17 | 1.95 |

**Supplementary Table 1. Marker genes for the original clusters of integrated clustering of E11.5 control and NotchGOF λ-junction cells.**

| | | | | | | | | |
|---|---|---|---|---|---|---|---|---|
| 11 | Sort1 | 1.04 | 5.89E-33 | 1.22E-28 | 0.37 | 0.20 | 0.17 | 1.86 |
| 11 | Patz1 | 0.69 | 6.57E-33 | 1.37E-28 | 0.68 | 0.52 | 0.16 | 1.31 |
| 11 | Egln3 | 1.28 | 6.94E-33 | 1.44E-28 | 0.19 | 0.07 | 0.12 | 2.63 |
| 11 | Dtx3l | 1.55 | 7.61E-33 | 1.58E-28 | 0.13 | 0.04 | 0.09 | 3.23 |
| 11 | Rsrp1 | 0.41 | 8.38E-33 | 1.74E-28 | 0.95 | 0.88 | 0.06 | 1.07 |
| 11 | Plxnb2 | 0.79 | 8.57E-33 | 1.78E-28 | 0.68 | 0.53 | 0.15 | 1.29 |
| 11 | Os9 | 0.85 | 9.16E-33 | 1.90E-28 | 0.55 | 0.37 | 0.18 | 1.50 |
| 11 | Rnf167 | 0.82 | 9.21E-33 | 1.92E-28 | 0.55 | 0.36 | 0.19 | 1.51 |
| 11 | Sypl | 0.71 | 1.00E-32 | 2.09E-28 | 0.65 | 0.48 | 0.17 | 1.36 |
| 11 | Gpi1 | 0.41 | 1.25E-32 | 2.59E-28 | 0.91 | 0.84 | 0.08 | 1.09 |
| 11 | Stk39 | 0.99 | 1.39E-32 | 2.89E-28 | 0.43 | 0.26 | 0.18 | 1.69 |
| 11 | Tbc1d10a | 1.14 | 1.40E-32 | 2.90E-28 | 0.36 | 0.19 | 0.17 | 1.85 |
| 11 | Hbp1 | 0.67 | 1.62E-32 | 3.37E-28 | 0.72 | 0.56 | 0.16 | 1.29 |
| 11 | Hdhd3 | 1.18 | 1.70E-32 | 3.53E-28 | 0.32 | 0.16 | 0.16 | 1.98 |
| 11 | Hunk | 0.72 | 2.03E-32 | 4.22E-28 | 0.67 | 0.49 | 0.18 | 1.36 |
| 11 | Gpr137c | 1.26 | 3.01E-32 | 6.27E-28 | 0.29 | 0.14 | 0.15 | 2.06 |
| 11 | Prelid2 | 0.87 | 4.07E-32 | 8.46E-28 | 0.54 | 0.36 | 0.18 | 1.49 |
| 11 | Sbno2 | 1.23 | 9.06E-32 | 1.88E-27 | 0.25 | 0.11 | 0.14 | 2.23 |
| 11 | Hsdl2 | 1.02 | 1.30E-31 | 2.70E-27 | 0.44 | 0.26 | 0.18 | 1.67 |
| 11 | Harbi1 | 1.11 | 1.41E-31 | 2.92E-27 | 0.34 | 0.18 | 0.16 | 1.88 |
| 11 | Rab6a | 0.60 | 1.47E-31 | 3.05E-27 | 0.81 | 0.71 | 0.09 | 1.13 |
| 11 | Arhgap8 | 1.41 | 1.60E-31 | 3.32E-27 | 0.13 | 0.04 | 0.09 | 3.20 |
| 11 | Pard6b | 1.48 | 1.69E-31 | 3.52E-27 | 0.15 | 0.05 | 0.10 | 2.92 |
| 11 | Plpp2 | 1.01 | 1.80E-31 | 3.74E-27 | 0.43 | 0.26 | 0.18 | 1.68 |
| 11 | Copg2 | 0.82 | 1.84E-31 | 3.82E-27 | 0.55 | 0.37 | 0.18 | 1.50 |
| 11 | Daam1 | 0.95 | 2.07E-31 | 4.31E-27 | 0.57 | 0.40 | 0.17 | 1.41 |
| 11 | Igf2r | 0.70 | 2.36E-31 | 4.91E-27 | 0.66 | 0.49 | 0.17 | 1.33 |
| 11 | Grip1 | 1.15 | 2.38E-31 | 4.94E-27 | 0.23 | 0.10 | 0.13 | 2.29 |
| 11 | Smim20 | 0.73 | 2.55E-31 | 5.30E-27 | 0.58 | 0.40 | 0.18 | 1.45 |
| 11 | Lnx1 | 1.89 | 2.91E-31 | 6.06E-27 | 0.11 | 0.03 | 0.08 | 3.53 |
| 11 | Nrbp2 | 1.98 | 3.50E-31 | 7.28E-27 | 0.10 | 0.03 | 0.07 | 3.52 |
| 11 | Syngr1 | 1.29 | 3.59E-31 | 7.47E-27 | 0.17 | 0.07 | 0.11 | 2.65 |
| 11 | Ccdc28a | 1.29 | 4.00E-31 | 8.32E-27 | 0.23 | 0.10 | 0.13 | 2.29 |
| 11 | Ak4 | 1.32 | 4.35E-31 | 9.05E-27 | 0.23 | 0.10 | 0.13 | 2.29 |
| 11 | Bckdha | 0.88 | 4.44E-31 | 9.24E-27 | 0.55 | 0.37 | 0.18 | 1.50 |
| 11 | Oga | 0.59 | 6.76E-31 | 1.41E-26 | 0.80 | 0.66 | 0.14 | 1.21 |
| 11 | Cd44 | 1.79 | 7.10E-31 | 1.48E-26 | 0.12 | 0.04 | 0.08 | 3.29 |
| 11 | Pde5a | 1.03 | 7.19E-31 | 1.49E-26 | 0.34 | 0.18 | 0.16 | 1.89 |
| 11 | Dap | 0.62 | 7.24E-31 | 1.51E-26 | 0.70 | 0.56 | 0.14 | 1.25 |
| 11 | Gipc2 | 1.56 | 7.43E-31 | 1.55E-26 | 0.16 | 0.06 | 0.10 | 2.72 |
| 11 | Cntnap2 | 1.43 | 7.83E-31 | 1.63E-26 | 0.14 | 0.05 | 0.09 | 2.92 |
| 11 | Mgat4a | 1.26 | 8.05E-31 | 1.68E-26 | 0.21 | 0.09 | 0.12 | 2.37 |
| 11 | Rgs9 | 1.30 | 8.47E-31 | 1.76E-26 | 0.20 | 0.08 | 0.12 | 2.45 |
| 11 | Ifi30 | 0.75 | 1.09E-30 | 2.26E-26 | 0.47 | 0.29 | 0.18 | 1.63 |
| 11 | Clip4 | 1.79 | 1.25E-30 | 2.60E-26 | 0.13 | 0.04 | 0.09 | 3.02 |
| 11 | Slc25a23 | 1.03 | 1.32E-30 | 2.75E-26 | 0.31 | 0.16 | 0.15 | 1.97 |
| 11 | Agpat3 | 0.96 | 1.50E-30 | 3.13E-26 | 0.43 | 0.26 | 0.17 | 1.65 |
| 11 | Ptpn14 | 1.18 | 1.88E-30 | 3.91E-26 | 0.30 | 0.15 | 0.15 | 2.01 |
| 11 | Skap2 | 1.63 | 1.96E-30 | 4.08E-26 | 0.15 | 0.05 | 0.10 | 2.83 |
| 11 | Cpne3 | 0.68 | 2.24E-30 | 4.65E-26 | 0.68 | 0.52 | 0.16 | 1.32 |
| 11 | Atxn1 | 2.17 | 2.29E-30 | 4.77E-26 | 0.20 | 0.09 | 0.11 | 2.30 |
| 11 | Cdk6 | 0.80 | 2.35E-30 | 4.89E-26 | 0.51 | 0.34 | 0.18 | 1.53 |
| 11 | Rgs12 | 0.94 | 2.58E-30 | 5.37E-26 | 0.60 | 0.45 | 0.15 | 1.34 |
| 11 | Plch1 | 1.12 | 3.25E-30 | 6.77E-26 | 0.23 | 0.10 | 0.13 | 2.33 |
| 11 | Taok1 | 0.62 | 4.26E-30 | 8.87E-26 | 0.73 | 0.60 | 0.13 | 1.22 |
| 11 | Scamp2 | 0.77 | 4.76E-30 | 9.91E-26 | 0.49 | 0.31 | 0.18 | 1.59 |

**Supplementary Table 1. Marker genes for the original clusters of integrated clustering of E11.5 control and NotchGOF λ-junction cells.**

| | | | | | | | | |
|---|---|---|---|---|---|---|---|---|
| 11 | Myo5b | 1.49 | 5.21E-30 | 1.08E-25 | 0.13 | 0.04 | 0.09 | 3.12 |
| 11 | Ccdc120 | 1.31 | 5.25E-30 | 1.09E-25 | 0.23 | 0.10 | 0.13 | 2.22 |
| 11 | Actn4 | 0.57 | 5.34E-30 | 1.11E-25 | 0.80 | 0.70 | 0.10 | 1.14 |
| 11 | Dnmbp | 1.09 | 7.78E-30 | 1.62E-25 | 0.26 | 0.12 | 0.14 | 2.12 |
| 11 | Mbtd1 | 0.51 | 1.05E-29 | 2.18E-25 | 0.84 | 0.73 | 0.11 | 1.16 |
| 11 | Anxa4 | 1.33 | 1.07E-29 | 2.23E-25 | 0.23 | 0.11 | 0.13 | 2.21 |
| 11 | Ptprd | 0.68 | 1.24E-29 | 2.58E-25 | 0.61 | 0.44 | 0.18 | 1.41 |
| 11 | Hspb2 | 1.09 | 1.43E-29 | 2.98E-25 | 0.11 | 0.03 | 0.08 | 3.45 |
| 11 | Chmp1b | 0.81 | 1.57E-29 | 3.27E-25 | 0.52 | 0.35 | 0.18 | 1.51 |
| 11 | Cd164l2 | 1.71 | 1.58E-29 | 3.28E-25 | 0.12 | 0.04 | 0.08 | 3.16 |
| 11 | Gltp | 0.74 | 1.63E-29 | 3.40E-25 | 0.56 | 0.39 | 0.17 | 1.43 |
| 11 | Trip12 | 0.58 | 1.87E-29 | 3.88E-25 | 0.77 | 0.66 | 0.11 | 1.17 |
| 11 | Nectin2 | 0.75 | 2.40E-29 | 4.99E-25 | 0.63 | 0.47 | 0.15 | 1.32 |
| 11 | Pls1 | 0.88 | 2.44E-29 | 5.07E-25 | 0.22 | 0.09 | 0.12 | 2.35 |
| 11 | Meis3 | 0.82 | 2.68E-29 | 5.58E-25 | 0.36 | 0.19 | 0.17 | 1.86 |
| 11 | Lef1 | 0.76 | 2.91E-29 | 6.06E-25 | 0.65 | 0.46 | 0.19 | 1.42 |
| 11 | Metap2 | 0.31 | 2.91E-29 | 6.06E-25 | 0.98 | 0.97 | 0.01 | 1.01 |
| 11 | St7 | 0.94 | 3.15E-29 | 6.55E-25 | 0.39 | 0.22 | 0.17 | 1.76 |
| 11 | Zcchc18 | 0.91 | 3.24E-29 | 6.74E-25 | 0.50 | 0.33 | 0.17 | 1.50 |
| 11 | Me1 | 1.07 | 3.70E-29 | 7.70E-25 | 0.41 | 0.24 | 0.17 | 1.68 |
| 11 | Kdm3a | 0.85 | 5.13E-29 | 1.07E-24 | 0.53 | 0.36 | 0.17 | 1.46 |
| 11 | Smim10l2a | 1.37 | 6.02E-29 | 1.25E-24 | 0.17 | 0.07 | 0.11 | 2.56 |
| 11 | Plxna2 | 0.91 | 6.16E-29 | 1.28E-24 | 0.39 | 0.23 | 0.17 | 1.73 |
| 11 | Zfp467 | 0.74 | 6.49E-29 | 1.35E-24 | 0.47 | 0.29 | 0.18 | 1.63 |
| 11 | Celf4 | 0.99 | 7.38E-29 | 1.54E-24 | 0.38 | 0.22 | 0.16 | 1.72 |
| 11 | Lamc1 | 0.94 | 7.66E-29 | 1.59E-24 | 0.42 | 0.26 | 0.17 | 1.66 |
| 11 | Rab21 | 0.62 | 1.07E-28 | 2.22E-24 | 0.63 | 0.46 | 0.17 | 1.36 |
| 11 | App | 0.39 | 1.42E-28 | 2.95E-24 | 0.89 | 0.78 | 0.11 | 1.14 |
| 11 | Jmy | 0.84 | 1.57E-28 | 3.26E-24 | 0.54 | 0.37 | 0.17 | 1.45 |
| 11 | B3galnt1 | 1.20 | 2.12E-28 | 4.40E-24 | 0.27 | 0.13 | 0.14 | 2.04 |
| 11 | mt-Co1 | 0.25 | 2.12E-28 | 4.42E-24 | 1.00 | 0.99 | 0.01 | 1.01 |
| 11 | Socs7 | 1.07 | 2.34E-28 | 4.87E-24 | 0.36 | 0.21 | 0.16 | 1.75 |
| 11 | Myo5c | 1.56 | 2.40E-28 | 4.98E-24 | 0.12 | 0.04 | 0.08 | 3.19 |
| 11 | Arhgap27 | 1.77 | 2.47E-28 | 5.13E-24 | 0.10 | 0.03 | 0.07 | 3.37 |
| 11 | Plxna1 | 1.14 | 2.85E-28 | 5.93E-24 | 0.35 | 0.20 | 0.15 | 1.75 |
| 11 | St6galnac5 | 1.52 | 3.45E-28 | 7.18E-24 | 0.14 | 0.05 | 0.09 | 2.87 |
| 11 | Sat1 | 0.84 | 3.67E-28 | 7.63E-24 | 0.46 | 0.30 | 0.16 | 1.52 |
| 11 | Sh3rf1 | 0.96 | 4.46E-28 | 9.27E-24 | 0.38 | 0.22 | 0.16 | 1.72 |
| 11 | Atp6v0e | 0.58 | 4.88E-28 | 1.02E-23 | 0.82 | 0.71 | 0.11 | 1.15 |
| 11 | Serinc2 | 0.71 | 5.93E-28 | 1.23E-23 | 0.57 | 0.38 | 0.18 | 1.48 |
| 11 | Tmem53 | 1.26 | 6.85E-28 | 1.42E-23 | 0.20 | 0.08 | 0.11 | 2.35 |
| 11 | Chst11 | 1.39 | 7.65E-28 | 1.59E-23 | 0.18 | 0.07 | 0.10 | 2.46 |
| 11 | Ankra2 | 0.76 | 7.89E-28 | 1.64E-23 | 0.48 | 0.31 | 0.17 | 1.55 |
| 11 | Ypel3 | 0.68 | 1.33E-27 | 2.76E-23 | 0.70 | 0.57 | 0.14 | 1.24 |
| 11 | Med19 | 0.50 | 1.45E-27 | 3.02E-23 | 0.82 | 0.74 | 0.08 | 1.11 |
| 11 | Atp11b | 0.74 | 1.51E-27 | 3.14E-23 | 0.58 | 0.42 | 0.17 | 1.40 |
| 11 | Kidins220 | 0.95 | 1.62E-27 | 3.37E-23 | 0.44 | 0.28 | 0.16 | 1.59 |
| 11 | Clta | 0.39 | 1.64E-27 | 3.42E-23 | 0.99 | 0.98 | 0.01 | 1.01 |
| 11 | Pcdh7 | 0.60 | 1.89E-27 | 3.92E-23 | 0.59 | 0.39 | 0.20 | 1.52 |
| 11 | Rassf7 | 0.97 | 1.90E-27 | 3.96E-23 | 0.35 | 0.20 | 0.15 | 1.78 |
| 11 | Arl6ip5 | 0.74 | 2.60E-27 | 5.41E-23 | 0.49 | 0.32 | 0.18 | 1.57 |
| 11 | Cds1 | 1.67 | 2.67E-27 | 5.55E-23 | 0.12 | 0.04 | 0.08 | 3.11 |
| 11 | Chmp2a | 0.49 | 2.67E-27 | 5.56E-23 | 0.78 | 0.66 | 0.12 | 1.19 |
| 11 | Itgb8 | 1.32 | 2.85E-27 | 5.93E-23 | 0.26 | 0.13 | 0.13 | 2.00 |
| 11 | Efs | 0.83 | 3.46E-27 | 7.19E-23 | 0.45 | 0.28 | 0.17 | 1.61 |
| 11 | Csnk1e | 0.43 | 3.55E-27 | 7.38E-23 | 0.92 | 0.84 | 0.08 | 1.09 |

**Supplementary Table 1. Marker genes for the original clusters of integrated clustering of E11.5 control and NotchGOF λ-junction cells.**

| | | | | | | | | |
|---|---|---|---|---|---|---|---|---|
| 11 | Vezt | 0.81 | 3.58E-27 | 7.45E-23 | 0.49 | 0.32 | 0.17 | 1.54 |
| 11 | Rnf11 | 0.56 | 3.79E-27 | 7.87E-23 | 0.75 | 0.62 | 0.13 | 1.21 |
| 11 | Abhd11 | 0.88 | 3.98E-27 | 8.28E-23 | 0.42 | 0.26 | 0.16 | 1.61 |
| 11 | Herc3 | 1.63 | 4.81E-27 | 9.99E-23 | 0.17 | 0.07 | 0.10 | 2.42 |
| 11 | Usp53 | 1.34 | 6.24E-27 | 1.30E-22 | 0.18 | 0.07 | 0.11 | 2.44 |
| 11 | Elk4 | 0.96 | 6.47E-27 | 1.35E-22 | 0.39 | 0.23 | 0.16 | 1.71 |
| 11 | Plekhf2 | 0.89 | 7.57E-27 | 1.57E-22 | 0.39 | 0.23 | 0.16 | 1.71 |
| 11 | Pnldc1 | 2.00 | 7.81E-27 | 1.62E-22 | 0.10 | 0.03 | 0.07 | 3.22 |
| 11 | Rps6ka6 | 0.66 | 8.98E-27 | 1.87E-22 | 0.62 | 0.45 | 0.17 | 1.37 |
| 11 | Miga1 | 0.84 | 9.21E-27 | 1.92E-22 | 0.46 | 0.30 | 0.16 | 1.55 |
| 11 | Rere | 0.73 | 9.38E-27 | 1.95E-22 | 0.72 | 0.59 | 0.13 | 1.21 |
| 11 | Tcn2 | 1.08 | 1.17E-26 | 2.43E-22 | 0.32 | 0.17 | 0.15 | 1.84 |
| 11 | 5430403G16Rik | 1.38 | 1.22E-26 | 2.55E-22 | 0.17 | 0.07 | 0.10 | 2.49 |
| 11 | 5033430I15Rik | 2.09 | 1.29E-26 | 2.68E-22 | 0.14 | 0.05 | 0.09 | 2.65 |
| 11 | Polr1d | 0.30 | 1.58E-26 | 3.29E-22 | 0.96 | 0.95 | 0.01 | 1.01 |
| 11 | Acot1 | 0.97 | 1.85E-26 | 3.84E-22 | 0.50 | 0.35 | 0.16 | 1.46 |
| 11 | Fam173a | 0.65 | 1.92E-26 | 4.00E-22 | 0.64 | 0.50 | 0.14 | 1.29 |
| 11 | Ces1d | 1.38 | 2.07E-26 | 4.31E-22 | 0.12 | 0.04 | 0.08 | 2.95 |
| 11 | Hk2 | 1.07 | 2.07E-26 | 4.31E-22 | 0.41 | 0.25 | 0.16 | 1.63 |
| 11 | Tvp23b | 0.71 | 2.34E-26 | 4.87E-22 | 0.56 | 0.40 | 0.16 | 1.39 |
| 11 | Dlx6 | 1.12 | 3.13E-26 | 6.51E-22 | 0.19 | 0.08 | 0.11 | 2.31 |
| 11 | Dynlt3 | 0.92 | 3.27E-26 | 6.79E-22 | 0.41 | 0.25 | 0.15 | 1.61 |
| 11 | Mlec | 0.46 | 3.47E-26 | 7.23E-22 | 0.84 | 0.75 | 0.09 | 1.13 |
| 11 | Inava | 0.81 | 3.51E-26 | 7.29E-22 | 0.39 | 0.23 | 0.16 | 1.72 |
| 11 | Galnt3 | 1.52 | 4.37E-26 | 9.10E-22 | 0.10 | 0.03 | 0.07 | 3.26 |
| 11 | Atp11a | 1.04 | 4.59E-26 | 9.54E-22 | 0.31 | 0.17 | 0.14 | 1.84 |
| 11 | Supt4a | 0.34 | 4.73E-26 | 9.83E-22 | 0.94 | 0.90 | 0.04 | 1.04 |
| 11 | Zdhhc12 | 0.88 | 4.95E-26 | 1.03E-21 | 0.42 | 0.26 | 0.16 | 1.63 |
| 11 | Fbxl20 | 0.84 | 5.41E-26 | 1.13E-21 | 0.37 | 0.21 | 0.16 | 1.74 |
| 11 | Mmp11 | 0.80 | 5.84E-26 | 1.21E-21 | 0.48 | 0.31 | 0.17 | 1.54 |
| 11 | Rbbp8 | 0.67 | 6.20E-26 | 1.29E-21 | 0.68 | 0.55 | 0.13 | 1.23 |
| 11 | Ccdc18 | 0.87 | 6.52E-26 | 1.36E-21 | 0.46 | 0.30 | 0.16 | 1.52 |
| 11 | Ide | 0.79 | 7.30E-26 | 1.52E-21 | 0.42 | 0.27 | 0.16 | 1.60 |
| 11 | Gsap | 1.69 | 7.56E-26 | 1.57E-21 | 0.11 | 0.04 | 0.07 | 3.11 |
| 11 | Maml2 | 1.35 | 7.63E-26 | 1.59E-21 | 0.20 | 0.09 | 0.11 | 2.22 |
| 11 | Enpp5 | 1.65 | 8.74E-26 | 1.82E-21 | 0.12 | 0.04 | 0.08 | 2.93 |
| 11 | Dtna | 0.86 | 1.05E-25 | 2.19E-21 | 0.28 | 0.14 | 0.14 | 1.99 |
| 11 | Med13 | 0.82 | 1.19E-25 | 2.47E-21 | 0.68 | 0.55 | 0.13 | 1.23 |
| 11 | Zcchc7 | 0.66 | 1.22E-25 | 2.53E-21 | 0.66 | 0.52 | 0.14 | 1.27 |
| 11 | Slc16a3 | 0.86 | 1.26E-25 | 2.63E-21 | 0.48 | 0.32 | 0.16 | 1.50 |
| 11 | Cpne5 | 1.40 | 1.34E-25 | 2.79E-21 | 0.14 | 0.05 | 0.09 | 2.74 |
| 11 | Abca3 | 1.49 | 1.38E-25 | 2.87E-21 | 0.16 | 0.06 | 0.09 | 2.52 |
| 11 | Airn | 0.91 | 1.48E-25 | 3.07E-21 | 0.33 | 0.19 | 0.15 | 1.79 |
| 11 | Sesn2 | 1.57 | 1.60E-25 | 3.34E-21 | 0.16 | 0.07 | 0.10 | 2.43 |
| 11 | Gtf3a | 0.51 | 1.63E-25 | 3.39E-21 | 0.76 | 0.64 | 0.12 | 1.19 |
| 11 | Vps36 | 0.47 | 1.98E-25 | 4.12E-21 | 0.84 | 0.79 | 0.05 | 1.07 |
| 11 | Magi3 | 1.12 | 2.32E-25 | 4.83E-21 | 0.33 | 0.19 | 0.14 | 1.76 |
| 11 | Cyb5r4 | 0.70 | 2.96E-25 | 6.15E-21 | 0.54 | 0.39 | 0.15 | 1.39 |
| 11 | Ube2h | 0.79 | 3.37E-25 | 7.01E-21 | 0.60 | 0.46 | 0.14 | 1.30 |
| 11 | Crim1 | 0.95 | 3.44E-25 | 7.16E-21 | 0.34 | 0.19 | 0.15 | 1.76 |
| 11 | Cd164 | 0.55 | 4.03E-25 | 8.37E-21 | 0.76 | 0.64 | 0.12 | 1.19 |
| 11 | Snx10 | 0.97 | 4.04E-25 | 8.40E-21 | 0.29 | 0.15 | 0.14 | 1.89 |
| 11 | Ankhd1 | 0.67 | 4.71E-25 | 9.79E-21 | 0.64 | 0.49 | 0.15 | 1.30 |
| 11 | Yipf2 | 1.03 | 4.78E-25 | 9.94E-21 | 0.24 | 0.12 | 0.13 | 2.06 |
| 11 | Sapcd1 | 1.70 | 4.87E-25 | 1.01E-20 | 0.11 | 0.04 | 0.07 | 3.03 |
| 11 | Ing4 | 0.42 | 5.27E-25 | 1.10E-20 | 0.85 | 0.76 | 0.09 | 1.12 |

**Supplementary Table 1. Marker genes for the original clusters of integrated clustering of E11.5 control and NotchGOF λ-junction cells.**

| | | | | | | | | |
|---|---|---|---|---|---|---|---|---|
| 11 | Rmnd5a | 0.69 | 5.33E-25 | 1.11E-20 | 0.56 | 0.40 | 0.16 | 1.40 |
| 11 | Acvr1b | 1.23 | 6.44E-25 | 1.34E-20 | 0.23 | 0.11 | 0.12 | 2.07 |
| 11 | Cnpy2 | 0.33 | 6.95E-25 | 1.45E-20 | 0.96 | 0.93 | 0.03 | 1.03 |
| 11 | Fam114a2 | 0.51 | 8.05E-25 | 1.67E-20 | 0.77 | 0.67 | 0.10 | 1.14 |
| 11 | Elavl2 | 0.64 | 8.65E-25 | 1.80E-20 | 0.33 | 0.18 | 0.15 | 1.83 |
| 11 | Map3k1 | 0.73 | 9.01E-25 | 1.87E-20 | 0.42 | 0.26 | 0.16 | 1.63 |
| 11 | Atp6ap2 | 0.73 | 9.59E-25 | 1.99E-20 | 0.57 | 0.42 | 0.15 | 1.36 |
| 11 | Lamtor2 | 0.33 | 1.04E-24 | 2.15E-20 | 0.93 | 0.89 | 0.03 | 1.04 |
| 11 | Shoc2 | 0.63 | 1.26E-24 | 2.62E-20 | 0.61 | 0.45 | 0.16 | 1.35 |
| 11 | Arhgap18 | 0.54 | 1.48E-24 | 3.07E-20 | 0.25 | 0.12 | 0.13 | 2.05 |
| 11 | Ddit4 | 1.03 | 1.48E-24 | 3.09E-20 | 0.52 | 0.37 | 0.15 | 1.39 |
| 11 | Hltf | 0.84 | 1.50E-24 | 3.11E-20 | 0.47 | 0.32 | 0.15 | 1.48 |
| 11 | H2-D1 | 0.93 | 1.52E-24 | 3.16E-20 | 0.39 | 0.24 | 0.15 | 1.63 |
| 11 | Gm15222 | 1.24 | 1.63E-24 | 3.40E-20 | 0.23 | 0.11 | 0.12 | 2.03 |
| 11 | Mien1 | 0.54 | 2.10E-24 | 4.38E-20 | 0.70 | 0.56 | 0.14 | 1.24 |
| 11 | Filip1l | 1.03 | 2.39E-24 | 4.97E-20 | 0.40 | 0.25 | 0.15 | 1.59 |
| 11 | Rhou | 0.74 | 2.63E-24 | 5.47E-20 | 0.56 | 0.41 | 0.15 | 1.36 |
| 11 | Rab3ip | 0.74 | 3.03E-24 | 6.31E-20 | 0.46 | 0.30 | 0.17 | 1.55 |
| 11 | 2010320M18Rik | 0.73 | 4.06E-24 | 8.45E-20 | 0.50 | 0.34 | 0.16 | 1.46 |
| 11 | Dusp16 | 0.90 | 4.42E-24 | 9.18E-20 | 0.36 | 0.21 | 0.15 | 1.71 |
| 11 | Zfp282 | 0.89 | 5.41E-24 | 1.12E-19 | 0.41 | 0.26 | 0.15 | 1.58 |
| 11 | Larp4b | 0.50 | 5.55E-24 | 1.15E-19 | 0.72 | 0.61 | 0.12 | 1.19 |
| 11 | Hook2 | 0.95 | 6.29E-24 | 1.31E-19 | 0.29 | 0.16 | 0.13 | 1.84 |
| 11 | Psme1 | 0.40 | 6.35E-24 | 1.32E-19 | 0.84 | 0.76 | 0.08 | 1.11 |
| 11 | Sash1 | 0.66 | 6.53E-24 | 1.36E-19 | 0.56 | 0.40 | 0.16 | 1.41 |
| 11 | Mtmr7 | 1.16 | 7.98E-24 | 1.66E-19 | 0.13 | 0.05 | 0.08 | 2.69 |
| 11 | Nudt16 | 1.21 | 8.72E-24 | 1.81E-19 | 0.19 | 0.08 | 0.10 | 2.21 |
| 11 | Rab11a | 0.44 | 8.90E-24 | 1.85E-19 | 0.76 | 0.64 | 0.12 | 1.19 |
| 11 | Prkci | 0.84 | 9.39E-24 | 1.95E-19 | 0.41 | 0.26 | 0.15 | 1.57 |
| 11 | Prr13 | 0.59 | 1.07E-23 | 2.23E-19 | 0.56 | 0.40 | 0.17 | 1.42 |
| 11 | Sik1 | 1.18 | 1.18E-23 | 2.46E-19 | 0.21 | 0.10 | 0.11 | 2.09 |
| 11 | Ubald2 | 0.50 | 1.34E-23 | 2.78E-19 | 0.79 | 0.69 | 0.10 | 1.15 |
| 11 | Borcs8 | 0.57 | 1.40E-23 | 2.92E-19 | 0.68 | 0.55 | 0.13 | 1.24 |
| 11 | Zfp36l1 | 0.46 | 1.43E-23 | 2.97E-19 | 0.90 | 0.85 | 0.05 | 1.06 |
| 11 | Acot7 | 0.78 | 1.47E-23 | 3.05E-19 | 0.38 | 0.23 | 0.15 | 1.66 |
| 11 | Slc35g1 | 1.57 | 1.50E-23 | 3.12E-19 | 0.14 | 0.06 | 0.08 | 2.53 |
| 11 | Cxcr4 | 1.31 | 1.51E-23 | 3.14E-19 | 0.13 | 0.05 | 0.08 | 2.72 |
| 11 | Rassf10 | 1.09 | 1.51E-23 | 3.15E-19 | 0.15 | 0.06 | 0.09 | 2.51 |
| 11 | Mgat5 | 0.82 | 1.51E-23 | 3.15E-19 | 0.41 | 0.26 | 0.15 | 1.60 |
| 11 | Nfat5 | 0.47 | 1.75E-23 | 3.64E-19 | 0.71 | 0.57 | 0.14 | 1.24 |
| 11 | Ephb2 | 0.85 | 2.12E-23 | 4.40E-19 | 0.43 | 0.28 | 0.15 | 1.54 |
| 11 | Prkcz | 1.09 | 2.20E-23 | 4.58E-19 | 0.22 | 0.10 | 0.11 | 2.10 |
| 11 | Prkg1 | 1.29 | 2.75E-23 | 5.72E-19 | 0.28 | 0.16 | 0.12 | 1.79 |
| 11 | Ccpg1 | 1.01 | 3.29E-23 | 6.84E-19 | 0.29 | 0.16 | 0.13 | 1.81 |
| 11 | Bsg | 0.26 | 4.01E-23 | 8.34E-19 | 1.00 | 0.98 | 0.02 | 1.02 |
| 11 | Ppp1r9a | 0.68 | 4.54E-23 | 9.44E-19 | 0.55 | 0.39 | 0.16 | 1.40 |
| 11 | Adcy1 | 1.45 | 4.75E-23 | 9.87E-19 | 0.11 | 0.04 | 0.07 | 2.89 |
| 11 | Scg5 | 1.02 | 4.95E-23 | 1.03E-18 | 0.13 | 0.05 | 0.08 | 2.63 |
| 11 | Mboat2 | 0.74 | 5.44E-23 | 1.13E-18 | 0.50 | 0.35 | 0.15 | 1.44 |
| 11 | Sema3f | 0.78 | 6.59E-23 | 1.37E-18 | 0.44 | 0.29 | 0.16 | 1.54 |
| 11 | mt-Nd6 | 0.82 | 6.83E-23 | 1.42E-18 | 0.50 | 0.35 | 0.16 | 1.44 |
| 11 | Lnx2 | 1.46 | 7.17E-23 | 1.49E-18 | 0.14 | 0.06 | 0.09 | 2.49 |
| 11 | Ccpg1os | 1.03 | 7.80E-23 | 1.62E-18 | 0.28 | 0.16 | 0.13 | 1.81 |
| 11 | Wdr26 | 0.46 | 8.44E-23 | 1.76E-18 | 0.82 | 0.71 | 0.11 | 1.15 |
| 11 | Tnfrsf1a | 0.89 | 8.46E-23 | 1.76E-18 | 0.44 | 0.30 | 0.15 | 1.49 |
| 11 | Suclg2 | 0.64 | 9.31E-23 | 1.94E-18 | 0.62 | 0.49 | 0.13 | 1.27 |

**Supplementary Table 1. Marker genes for the original clusters of integrated clustering of E11.5 control and NotchGOF λ-junction cells.**

| | | | | | | | | |
|---|---|---|---|---|---|---|---|---|
| 11 | Nr2f6 | 0.52 | 9.72E-23 | 2.02E-18 | 0.72 | 0.60 | 0.12 | 1.20 |
| 11 | Phyhipl | 1.65 | 9.75E-23 | 2.03E-18 | 0.11 | 0.04 | 0.07 | 2.86 |
| 11 | Wdsub1 | 0.76 | 9.83E-23 | 2.05E-18 | 0.35 | 0.20 | 0.15 | 1.73 |
| 11 | Wdpcp | 1.32 | 1.10E-22 | 2.29E-18 | 0.18 | 0.08 | 0.10 | 2.23 |
| 11 | Cd3eap | 0.66 | 1.40E-22 | 2.91E-18 | 0.65 | 0.52 | 0.13 | 1.24 |
| 11 | Nedd4l | 0.92 | 1.42E-22 | 2.96E-18 | 0.27 | 0.14 | 0.13 | 1.90 |
| 11 | Pim3 | 0.74 | 1.43E-22 | 2.97E-18 | 0.45 | 0.29 | 0.16 | 1.53 |
| 11 | Bmpr2 | 0.61 | 1.51E-22 | 3.14E-18 | 0.68 | 0.55 | 0.13 | 1.23 |
| 11 | Tnk1 | 1.50 | 1.59E-22 | 3.31E-18 | 0.10 | 0.04 | 0.07 | 2.94 |
| 11 | Ttc9 | 1.55 | 1.75E-22 | 3.64E-18 | 0.11 | 0.04 | 0.07 | 2.87 |
| 11 | Zfp385a | 0.91 | 1.88E-22 | 3.91E-18 | 0.39 | 0.25 | 0.14 | 1.56 |
| 11 | Txnip | 0.93 | 1.89E-22 | 3.93E-18 | 0.48 | 0.34 | 0.15 | 1.43 |
| 11 | Heca | 0.84 | 2.00E-22 | 4.15E-18 | 0.37 | 0.23 | 0.15 | 1.64 |
| 11 | Csnk1g1 | 0.75 | 2.29E-22 | 4.75E-18 | 0.51 | 0.36 | 0.15 | 1.42 |
| 11 | Cited4 | 0.32 | 2.47E-22 | 5.14E-18 | 0.14 | 0.06 | 0.09 | 2.57 |
| 11 | Ppp1r12b | 1.27 | 2.96E-22 | 6.16E-18 | 0.22 | 0.11 | 0.11 | 1.97 |
| 11 | Kdsr | 0.86 | 3.09E-22 | 6.42E-18 | 0.41 | 0.27 | 0.14 | 1.54 |
| 11 | Btbd7 | 0.50 | 3.09E-22 | 6.44E-18 | 0.77 | 0.67 | 0.10 | 1.14 |
| 11 | Ist1 | 0.69 | 3.25E-22 | 6.76E-18 | 0.59 | 0.46 | 0.13 | 1.29 |
| 11 | Pold4 | 0.98 | 3.55E-22 | 7.39E-18 | 0.32 | 0.18 | 0.13 | 1.72 |
| 11 | Acly | 0.56 | 3.65E-22 | 7.58E-18 | 0.69 | 0.58 | 0.12 | 1.20 |
| 11 | Arl6ip1 | 0.41 | 3.65E-22 | 7.60E-18 | 0.95 | 0.89 | 0.07 | 1.08 |
| 11 | Rtn4 | 0.30 | 5.03E-22 | 1.05E-17 | 0.97 | 0.94 | 0.04 | 1.04 |
| 11 | Rnf180 | 1.54 | 5.38E-22 | 1.12E-17 | 0.13 | 0.05 | 0.08 | 2.56 |
| 11 | Zkscan7 | 1.14 | 5.74E-22 | 1.19E-17 | 0.17 | 0.08 | 0.10 | 2.27 |
| 11 | Klf4 | 1.00 | 6.11E-22 | 1.27E-17 | 0.34 | 0.21 | 0.14 | 1.66 |
| 11 | Shtn1 | 0.98 | 6.74E-22 | 1.40E-17 | 0.13 | 0.05 | 0.08 | 2.58 |
| 11 | Mylk | 1.03 | 7.45E-22 | 1.55E-17 | 0.30 | 0.17 | 0.13 | 1.76 |
| 11 | Mapkapk3 | 1.56 | 7.71E-22 | 1.60E-17 | 0.10 | 0.04 | 0.07 | 2.91 |
| 11 | Igsf8 | 0.77 | 8.20E-22 | 1.71E-17 | 0.38 | 0.24 | 0.14 | 1.60 |
| 11 | Strbp | 0.42 | 8.66E-22 | 1.80E-17 | 0.81 | 0.70 | 0.10 | 1.15 |
| 11 | Rtn3 | 0.39 | 8.94E-22 | 1.86E-17 | 0.86 | 0.79 | 0.08 | 1.10 |
| 11 | 0610010F05Rik | 0.80 | 9.03E-22 | 1.88E-17 | 0.38 | 0.24 | 0.14 | 1.58 |
| 11 | Vwa2 | 1.63 | 1.01E-21 | 2.09E-17 | 0.10 | 0.03 | 0.07 | 2.94 |
| 11 | Neat1 | 1.51 | 1.03E-21 | 2.14E-17 | 0.13 | 0.05 | 0.08 | 2.49 |
| 11 | Rsbn1l | 0.49 | 1.16E-21 | 2.42E-17 | 0.73 | 0.63 | 0.09 | 1.15 |
| 11 | Fam129b | 0.97 | 1.24E-21 | 2.58E-17 | 0.26 | 0.14 | 0.12 | 1.88 |
| 11 | Cadm4 | 1.23 | 1.61E-21 | 3.35E-17 | 0.18 | 0.08 | 0.10 | 2.15 |
| 11 | H2afj | 0.35 | 1.62E-21 | 3.36E-17 | 0.88 | 0.81 | 0.07 | 1.08 |
| 11 | Usp46 | 0.59 | 1.68E-21 | 3.50E-17 | 0.73 | 0.63 | 0.11 | 1.17 |
| 11 | Atp13a2 | 0.86 | 1.71E-21 | 3.57E-17 | 0.40 | 0.25 | 0.14 | 1.56 |
| 11 | Ajuba | 0.87 | 1.84E-21 | 3.83E-17 | 0.42 | 0.28 | 0.14 | 1.52 |
| 11 | Hpcal1 | 0.78 | 1.87E-21 | 3.90E-17 | 0.47 | 0.33 | 0.14 | 1.44 |
| 11 | Prrg3 | 1.04 | 1.89E-21 | 3.93E-17 | 0.22 | 0.11 | 0.11 | 2.02 |
| 11 | Rap2b | 0.59 | 2.11E-21 | 4.39E-17 | 0.63 | 0.48 | 0.15 | 1.30 |
| 11 | Pcmtd2 | 0.70 | 4.28E-21 | 8.90E-17 | 0.52 | 0.37 | 0.14 | 1.39 |
| 11 | Imp3 | 0.38 | 4.38E-21 | 9.12E-17 | 0.85 | 0.79 | 0.06 | 1.08 |
| 11 | Adgrg1 | 1.35 | 4.40E-21 | 9.16E-17 | 0.11 | 0.04 | 0.07 | 2.75 |
| 11 | Palld | 0.69 | 5.08E-21 | 1.06E-16 | 0.72 | 0.62 | 0.10 | 1.15 |
| 11 | Lyplal1 | 0.81 | 5.51E-21 | 1.15E-16 | 0.39 | 0.24 | 0.14 | 1.59 |
| 11 | Arl8b | 0.62 | 5.57E-21 | 1.16E-16 | 0.61 | 0.47 | 0.14 | 1.30 |
| 11 | Fam192a | 0.47 | 6.62E-21 | 1.38E-16 | 0.76 | 0.65 | 0.11 | 1.17 |
| 11 | Slc48a1 | 0.88 | 6.68E-21 | 1.39E-16 | 0.29 | 0.16 | 0.13 | 1.77 |
| 11 | Slc22a23 | 0.97 | 6.92E-21 | 1.44E-16 | 0.19 | 0.09 | 0.10 | 2.09 |
| 11 | Nol7 | 0.31 | 6.93E-21 | 1.44E-16 | 0.92 | 0.88 | 0.04 | 1.05 |
| 11 | Arpc1a | 0.32 | 7.05E-21 | 1.47E-16 | 0.92 | 0.88 | 0.04 | 1.04 |

**Supplementary Table 1. Marker genes for the original clusters of integrated clustering of E11.5 control and NotchGOF λ-junction cells.**

| | | | | | | | | |
|---|---|---|---|---|---|---|---|---|
| 11 | Tmem238 | 0.75 | 7.20E-21 | 1.50E-16 | 0.31 | 0.18 | 0.13 | 1.76 |
| 11 | Nlk | 0.77 | 9.73E-21 | 2.02E-16 | 0.41 | 0.27 | 0.14 | 1.53 |
| 11 | Ccdc115 | 0.55 | 1.25E-20 | 2.60E-16 | 0.62 | 0.49 | 0.14 | 1.28 |
| 11 | Plk2 | 0.71 | 1.26E-20 | 2.63E-16 | 0.52 | 0.37 | 0.15 | 1.40 |
| 11 | Med12l | 1.38 | 1.28E-20 | 2.66E-16 | 0.11 | 0.04 | 0.07 | 2.65 |
| 11 | Tmem35b | 1.46 | 1.44E-20 | 3.00E-16 | 0.13 | 0.05 | 0.08 | 2.45 |
| 11 | Plekhh1 | 1.17 | 1.68E-20 | 3.50E-16 | 0.15 | 0.06 | 0.09 | 2.35 |
| 11 | Cers4 | 0.82 | 1.77E-20 | 3.68E-16 | 0.28 | 0.15 | 0.12 | 1.81 |
| 11 | Wdtc1 | 0.91 | 1.83E-20 | 3.81E-16 | 0.30 | 0.17 | 0.13 | 1.74 |
| 11 | Pick1 | 0.78 | 1.86E-20 | 3.86E-16 | 0.40 | 0.25 | 0.14 | 1.57 |
| 11 | Reps1 | 0.74 | 2.15E-20 | 4.47E-16 | 0.49 | 0.35 | 0.15 | 1.42 |
| 11 | Kank1 | 1.01 | 2.22E-20 | 4.61E-16 | 0.28 | 0.16 | 0.12 | 1.78 |
| 11 | Sp5 | 2.35 | 2.22E-20 | 4.62E-16 | 0.10 | 0.04 | 0.07 | 2.71 |
| 11 | Prrc2b | 0.41 | 2.35E-20 | 4.90E-16 | 0.82 | 0.74 | 0.08 | 1.11 |
| 11 | Mboat1 | 1.36 | 2.49E-20 | 5.18E-16 | 0.12 | 0.05 | 0.08 | 2.53 |
| 11 | Rdm1 | 0.69 | 2.50E-20 | 5.19E-16 | 0.46 | 0.32 | 0.14 | 1.44 |
| 11 | Slc12a4 | 1.12 | 2.77E-20 | 5.75E-16 | 0.25 | 0.14 | 0.11 | 1.82 |
| 11 | Gm46218 | 1.42 | 2.86E-20 | 5.95E-16 | 0.12 | 0.04 | 0.07 | 2.61 |
| 11 | Snapin | 0.64 | 3.40E-20 | 7.07E-16 | 0.47 | 0.32 | 0.15 | 1.47 |
| 11 | Cbarp | 1.00 | 3.89E-20 | 8.09E-16 | 0.21 | 0.11 | 0.11 | 2.00 |
| 11 | Egfr | 0.96 | 4.50E-20 | 9.36E-16 | 0.20 | 0.10 | 0.10 | 2.04 |
| 11 | Zfp113 | 0.71 | 4.97E-20 | 1.03E-15 | 0.47 | 0.32 | 0.14 | 1.45 |
| 11 | Marchf8 | 1.03 | 6.20E-20 | 1.29E-15 | 0.29 | 0.17 | 0.12 | 1.71 |
| 11 | Vwa1 | 1.63 | 6.54E-20 | 1.36E-15 | 0.13 | 0.06 | 0.08 | 2.35 |
| 11 | Axl | 0.91 | 7.13E-20 | 1.48E-15 | 0.27 | 0.15 | 0.12 | 1.79 |
| 11 | Tspan14 | 0.73 | 7.18E-20 | 1.49E-15 | 0.47 | 0.32 | 0.15 | 1.47 |
| 11 | Plpp6 | 1.11 | 7.37E-20 | 1.53E-15 | 0.15 | 0.07 | 0.08 | 2.29 |
| 11 | Zbtb7a | 0.76 | 8.30E-20 | 1.73E-15 | 0.46 | 0.33 | 0.14 | 1.41 |
| 11 | Sft2d3 | 0.83 | 8.75E-20 | 1.82E-15 | 0.33 | 0.20 | 0.13 | 1.66 |
| 11 | Cnot7 | 0.46 | 9.04E-20 | 1.88E-15 | 0.78 | 0.67 | 0.10 | 1.15 |
| 11 | Ormdl1 | 0.62 | 9.37E-20 | 1.95E-15 | 0.57 | 0.44 | 0.13 | 1.29 |
| 11 | Phc2 | 0.43 | 9.57E-20 | 1.99E-15 | 0.78 | 0.69 | 0.09 | 1.13 |
| 11 | Igf2os | 1.38 | 1.06E-19 | 2.19E-15 | 0.10 | 0.04 | 0.07 | 2.76 |
| 11 | Setd7 | 0.89 | 1.09E-19 | 2.26E-15 | 0.33 | 0.20 | 0.13 | 1.63 |
| 11 | Nsun4 | 0.93 | 1.17E-19 | 2.42E-15 | 0.27 | 0.15 | 0.12 | 1.77 |
| 11 | H1f0 | 0.39 | 1.63E-19 | 3.40E-15 | 0.92 | 0.85 | 0.07 | 1.09 |
| 11 | Nbr1 | 0.61 | 1.66E-19 | 3.45E-15 | 0.52 | 0.38 | 0.14 | 1.36 |
| 11 | Bcl2l11 | 1.04 | 1.70E-19 | 3.54E-15 | 0.47 | 0.35 | 0.13 | 1.37 |
| 11 | Tceal5 | 0.92 | 1.76E-19 | 3.66E-15 | 0.27 | 0.15 | 0.12 | 1.80 |
| 11 | Zfp395 | 0.80 | 1.79E-19 | 3.73E-15 | 0.37 | 0.23 | 0.13 | 1.58 |
| 11 | Abca1 | 1.23 | 1.88E-19 | 3.91E-15 | 0.15 | 0.07 | 0.08 | 2.24 |
| 11 | Txndc17 | 0.33 | 1.89E-19 | 3.92E-15 | 0.94 | 0.92 | 0.02 | 1.02 |
| 11 | Smarcd2 | 0.79 | 2.30E-19 | 4.78E-15 | 0.42 | 0.29 | 0.13 | 1.45 |
| 11 | Pak4 | 0.68 | 2.56E-19 | 5.32E-15 | 0.51 | 0.38 | 0.13 | 1.33 |
| 11 | Lgr4 | 0.54 | 2.62E-19 | 5.46E-15 | 0.36 | 0.23 | 0.14 | 1.61 |
| 11 | Aig1 | 0.55 | 2.76E-19 | 5.73E-15 | 0.48 | 0.33 | 0.16 | 1.47 |
| 11 | Dstyk | 0.87 | 2.94E-19 | 6.11E-15 | 0.36 | 0.23 | 0.13 | 1.58 |
| 11 | Ankrd49 | 0.59 | 3.27E-19 | 6.79E-15 | 0.56 | 0.43 | 0.13 | 1.31 |
| 11 | Abtb1 | 1.13 | 3.30E-19 | 6.86E-15 | 0.21 | 0.11 | 0.10 | 1.94 |
| 11 | Ccnd2 | 0.51 | 3.85E-19 | 8.01E-15 | 0.91 | 0.86 | 0.05 | 1.06 |
| 11 | Gucy1b1 | 1.42 | 4.16E-19 | 8.65E-15 | 0.10 | 0.04 | 0.06 | 2.68 |
| 11 | Nmt2 | 0.74 | 4.22E-19 | 8.77E-15 | 0.47 | 0.34 | 0.13 | 1.39 |
| 11 | Emid1 | 1.06 | 4.58E-19 | 9.53E-15 | 0.18 | 0.09 | 0.10 | 2.08 |
| 11 | Lrrc8b | 0.92 | 5.19E-19 | 1.08E-14 | 0.21 | 0.11 | 0.10 | 1.95 |
| 11 | Ctnna1 | 0.33 | 5.22E-19 | 1.08E-14 | 0.89 | 0.85 | 0.05 | 1.06 |
| 11 | Pqlc1 | 1.11 | 5.46E-19 | 1.13E-14 | 0.24 | 0.13 | 0.11 | 1.82 |

**Supplementary Table 1. Marker genes for the original clusters of integrated clustering of E11.5 control and NotchGOF λ-junction cells.**

| | | | | | | | | |
|---|---|---|---|---|---|---|---|---|
| 11 | Dynll2 | 0.34 | 6.95E-19 | 1.45E-14 | 0.91 | 0.84 | 0.07 | 1.08 |
| 11 | Agl | 0.82 | 7.64E-19 | 1.59E-14 | 0.38 | 0.25 | 0.13 | 1.54 |
| 11 | Atp6v0e2 | 1.30 | 7.70E-19 | 1.60E-14 | 0.12 | 0.05 | 0.07 | 2.43 |
| 11 | Maf1 | 0.53 | 7.74E-19 | 1.61E-14 | 0.65 | 0.53 | 0.12 | 1.23 |
| 11 | 1810058I24Rik | 0.35 | 7.83E-19 | 1.63E-14 | 0.96 | 0.94 | 0.02 | 1.02 |
| 11 | Pdcd6ip | 0.57 | 9.20E-19 | 1.91E-14 | 0.60 | 0.48 | 0.12 | 1.25 |
| 11 | Ralgps2 | 0.43 | 9.77E-19 | 2.03E-14 | 0.79 | 0.71 | 0.08 | 1.11 |
| 11 | Carmil1 | 0.87 | 1.05E-18 | 2.19E-14 | 0.26 | 0.14 | 0.11 | 1.80 |
| 11 | Gcat | 0.59 | 1.09E-18 | 2.27E-14 | 0.60 | 0.48 | 0.13 | 1.26 |
| 11 | Sqstm1 | 0.61 | 1.10E-18 | 2.28E-14 | 0.54 | 0.41 | 0.13 | 1.32 |
| 11 | Lbh | 1.03 | 1.17E-18 | 2.43E-14 | 0.32 | 0.20 | 0.12 | 1.61 |
| 11 | Ankrd50 | 0.79 | 1.17E-18 | 2.44E-14 | 0.36 | 0.23 | 0.13 | 1.56 |
| 11 | Iqcb1 | 0.46 | 1.23E-18 | 2.56E-14 | 0.66 | 0.52 | 0.14 | 1.26 |
| 11 | Plcb3 | 0.74 | 1.26E-18 | 2.62E-14 | 0.42 | 0.29 | 0.13 | 1.47 |
| 11 | Cdc14a | 0.86 | 1.30E-18 | 2.70E-14 | 0.39 | 0.25 | 0.13 | 1.53 |
| 11 | Mpst | 0.58 | 1.35E-18 | 2.81E-14 | 0.54 | 0.42 | 0.13 | 1.31 |
| 11 | 2410022M11Rik | 1.00 | 1.35E-18 | 2.81E-14 | 0.21 | 0.11 | 0.10 | 1.93 |
| 11 | Acp6 | 0.86 | 1.69E-18 | 3.53E-14 | 0.35 | 0.22 | 0.12 | 1.56 |
| 11 | Snca | 0.67 | 1.73E-18 | 3.61E-14 | 0.18 | 0.09 | 0.09 | 2.08 |
| 11 | Epb41l4a | 0.98 | 1.91E-18 | 3.97E-14 | 0.20 | 0.10 | 0.10 | 1.95 |
| 11 | Pcmtd1 | 0.65 | 1.95E-18 | 4.05E-14 | 0.58 | 0.46 | 0.12 | 1.25 |
| 11 | Pbxip1 | 0.83 | 2.11E-18 | 4.38E-14 | 0.32 | 0.20 | 0.13 | 1.63 |
| 11 | Sdf2 | 0.46 | 2.18E-18 | 4.53E-14 | 0.68 | 0.56 | 0.12 | 1.22 |
| 11 | Pgpep1 | 0.87 | 2.21E-18 | 4.60E-14 | 0.42 | 0.29 | 0.13 | 1.47 |
| 11 | Braf | 0.74 | 2.23E-18 | 4.65E-14 | 0.46 | 0.33 | 0.13 | 1.41 |
| 11 | Snx5 | 0.35 | 2.29E-18 | 4.76E-14 | 0.84 | 0.78 | 0.06 | 1.08 |
| 11 | Edem2 | 0.89 | 2.33E-18 | 4.86E-14 | 0.31 | 0.19 | 0.12 | 1.63 |
| 11 | Phtf2 | 0.88 | 2.34E-18 | 4.88E-14 | 0.32 | 0.20 | 0.12 | 1.63 |
| 11 | Nme3 | 1.27 | 2.35E-18 | 4.90E-14 | 0.16 | 0.07 | 0.08 | 2.12 |
| 11 | Ccdc191 | 0.78 | 2.37E-18 | 4.92E-14 | 0.34 | 0.22 | 0.13 | 1.59 |
| 11 | Cndp2 | 0.69 | 2.51E-18 | 5.23E-14 | 0.49 | 0.37 | 0.12 | 1.33 |
| 11 | Pcgf5 | 0.99 | 2.58E-18 | 5.36E-14 | 0.18 | 0.09 | 0.09 | 2.06 |
| 11 | Ube2e1 | 0.34 | 2.87E-18 | 5.97E-14 | 0.90 | 0.84 | 0.07 | 1.08 |
| 11 | Grb14 | 1.39 | 2.99E-18 | 6.23E-14 | 0.11 | 0.05 | 0.07 | 2.49 |
| 11 | Macrod2 | 1.12 | 4.03E-18 | 8.38E-14 | 0.34 | 0.22 | 0.12 | 1.56 |
| 11 | Aagab | 0.92 | 4.18E-18 | 8.69E-14 | 0.29 | 0.17 | 0.12 | 1.69 |
| 11 | Papss1 | 0.40 | 4.39E-18 | 9.13E-14 | 0.78 | 0.66 | 0.12 | 1.18 |
| 11 | Arpc5 | 0.32 | 4.77E-18 | 9.92E-14 | 0.84 | 0.76 | 0.08 | 1.10 |
| 11 | Naa80 | 1.11 | 4.86E-18 | 1.01E-13 | 0.26 | 0.16 | 0.11 | 1.70 |
| 11 | Luzp1 | 0.38 | 5.89E-18 | 1.22E-13 | 0.84 | 0.77 | 0.07 | 1.09 |
| 11 | Ppp2cb | 0.59 | 6.12E-18 | 1.27E-13 | 0.56 | 0.44 | 0.12 | 1.27 |
| 11 | Ppil2 | 0.44 | 6.39E-18 | 1.33E-13 | 0.71 | 0.60 | 0.11 | 1.19 |
| 11 | AC149090.1 | 0.65 | 6.60E-18 | 1.37E-13 | 0.42 | 0.28 | 0.14 | 1.49 |
| 11 | Micos13 | 0.26 | 7.45E-18 | 1.55E-13 | 0.95 | 0.94 | 0.01 | 1.01 |
| 11 | 2900097C17Rik | 0.37 | 7.67E-18 | 1.60E-13 | 0.84 | 0.74 | 0.10 | 1.14 |
| 11 | Gm42418 | 0.39 | 7.91E-18 | 1.65E-13 | 1.00 | 1.00 | 0.00 | 1.00 |
| 11 | Dapk1 | 0.68 | 8.46E-18 | 1.76E-13 | 0.41 | 0.27 | 0.14 | 1.50 |
| 11 | Tmem179b | 0.70 | 9.31E-18 | 1.94E-13 | 0.42 | 0.29 | 0.13 | 1.45 |
| 11 | Zdhhc2 | 0.43 | 9.65E-18 | 2.01E-13 | 0.24 | 0.13 | 0.11 | 1.88 |
| 11 | Rbpms | 0.60 | 9.73E-18 | 2.02E-13 | 0.64 | 0.54 | 0.10 | 1.18 |
| 11 | Kcnk2 | 0.72 | 1.43E-17 | 2.97E-13 | 0.43 | 0.30 | 0.14 | 1.46 |
| 11 | Espn | 0.57 | 1.63E-17 | 3.40E-13 | 0.21 | 0.11 | 0.10 | 1.95 |
| 11 | Wbp2 | 0.73 | 1.68E-17 | 3.50E-13 | 0.40 | 0.27 | 0.12 | 1.45 |
| 11 | Atp6v1g1 | 0.26 | 1.70E-17 | 3.54E-13 | 0.97 | 0.97 | 0.00 | 1.00 |
| 11 | Ncstn | 0.69 | 1.81E-17 | 3.76E-13 | 0.42 | 0.29 | 0.13 | 1.46 |
| 11 | Cyp39a1 | 0.89 | 1.95E-17 | 4.05E-13 | 0.25 | 0.15 | 0.11 | 1.74 |

**Supplementary Table 1. Marker genes for the original clusters of integrated clustering of E11.5 control and NotchGOF λ-junction cells.**

| | | | | | | | | |
|---|---|---|---|---|---|---|---|---|
| 11 | Ctsl | 0.43 | 2.02E-17 | 4.20E-13 | 0.90 | 0.82 | 0.08 | 1.09 |
| 11 | Pex2 | 0.56 | 2.21E-17 | 4.61E-13 | 0.60 | 0.47 | 0.12 | 1.26 |
| 11 | Smox | 0.70 | 2.45E-17 | 5.10E-13 | 0.37 | 0.24 | 0.13 | 1.55 |
| 11 | Chmp1a | 0.56 | 2.52E-17 | 5.24E-13 | 0.55 | 0.42 | 0.13 | 1.30 |
| 11 | Nmrk1 | 1.14 | 2.69E-17 | 5.59E-13 | 0.17 | 0.08 | 0.09 | 2.05 |
| 11 | Pard6g | 0.46 | 2.86E-17 | 5.94E-13 | 0.71 | 0.59 | 0.12 | 1.20 |
| 11 | Atg10 | 0.91 | 2.88E-17 | 5.99E-13 | 0.25 | 0.15 | 0.11 | 1.74 |
| 11 | Fryl | 0.71 | 3.04E-17 | 6.32E-13 | 0.43 | 0.30 | 0.13 | 1.44 |
| 11 | Chd7 | 0.38 | 3.27E-17 | 6.81E-13 | 0.76 | 0.65 | 0.11 | 1.17 |
| 11 | Fam214a | 0.88 | 3.30E-17 | 6.86E-13 | 0.19 | 0.10 | 0.10 | 1.98 |
| 11 | Fgf17 | 1.15 | 3.77E-17 | 7.85E-13 | 0.11 | 0.04 | 0.06 | 2.50 |
| 11 | Atp9a | 0.80 | 4.26E-17 | 8.87E-13 | 0.29 | 0.17 | 0.12 | 1.68 |
| 11 | Akirin2 | 0.37 | 4.33E-17 | 9.01E-13 | 0.87 | 0.81 | 0.06 | 1.08 |
| 11 | Dcaf17 | 0.63 | 4.59E-17 | 9.56E-13 | 0.46 | 0.33 | 0.13 | 1.39 |
| 11 | Ift43 | 0.42 | 4.73E-17 | 9.83E-13 | 0.67 | 0.54 | 0.13 | 1.24 |
| 11 | Retreg2 | 0.67 | 5.40E-17 | 1.12E-12 | 0.42 | 0.28 | 0.13 | 1.46 |
| 11 | Med13l | 0.56 | 5.87E-17 | 1.22E-12 | 0.60 | 0.50 | 0.10 | 1.20 |
| 11 | Cux2 | 0.97 | 6.76E-17 | 1.41E-12 | 0.19 | 0.10 | 0.09 | 1.96 |
| 11 | Wnt5a | 0.71 | 7.31E-17 | 1.52E-12 | 0.74 | 0.64 | 0.11 | 1.17 |
| 11 | Diablo | 0.41 | 7.91E-17 | 1.64E-12 | 0.73 | 0.63 | 0.10 | 1.17 |
| 11 | Spry1 | 0.85 | 8.29E-17 | 1.72E-12 | 0.41 | 0.29 | 0.13 | 1.45 |
| 11 | Map4k3 | 0.67 | 8.61E-17 | 1.79E-12 | 0.41 | 0.28 | 0.13 | 1.47 |
| 11 | Nanos3 | 1.21 | 8.77E-17 | 1.82E-12 | 0.12 | 0.05 | 0.07 | 2.32 |
| 11 | Nabp2 | 0.42 | 9.15E-17 | 1.90E-12 | 0.74 | 0.65 | 0.09 | 1.14 |
| 11 | Zbtb18 | 0.88 | 9.39E-17 | 1.95E-12 | 0.20 | 0.11 | 0.10 | 1.88 |
| 11 | Hs3st6 | 0.91 | 1.03E-16 | 2.15E-12 | 0.17 | 0.08 | 0.09 | 2.04 |
| 11 | Zfand1 | 0.58 | 1.08E-16 | 2.25E-12 | 0.48 | 0.34 | 0.14 | 1.39 |
| 11 | Lamtor5 | 0.35 | 1.19E-16 | 2.47E-12 | 0.85 | 0.79 | 0.07 | 1.09 |
| 11 | Myo1e | 0.75 | 1.21E-16 | 2.51E-12 | 0.36 | 0.23 | 0.12 | 1.54 |
| 11 | Cecr2 | 0.75 | 1.25E-16 | 2.60E-12 | 0.34 | 0.22 | 0.12 | 1.56 |
| 11 | Cltb | 0.79 | 1.26E-16 | 2.61E-12 | 0.29 | 0.18 | 0.11 | 1.63 |
| 11 | Pts | 0.57 | 1.28E-16 | 2.66E-12 | 0.54 | 0.40 | 0.13 | 1.33 |
| 11 | Slco5a1 | 1.05 | 1.31E-16 | 2.72E-12 | 0.16 | 0.08 | 0.08 | 2.01 |
| 11 | Zfp949 | 1.08 | 1.32E-16 | 2.75E-12 | 0.22 | 0.12 | 0.10 | 1.82 |
| 11 | Atp6v0b | 0.37 | 1.53E-16 | 3.19E-12 | 0.82 | 0.73 | 0.09 | 1.12 |
| 11 | Isl2 | 0.43 | 1.80E-16 | 3.75E-12 | 0.24 | 0.13 | 0.11 | 1.86 |
| 11 | Zmat4 | 1.16 | 1.86E-16 | 3.87E-12 | 0.12 | 0.05 | 0.07 | 2.28 |
| 11 | Arl5a | 0.48 | 2.27E-16 | 4.73E-12 | 0.69 | 0.60 | 0.09 | 1.15 |
| 11 | Pbx4 | 1.09 | 2.78E-16 | 5.78E-12 | 0.17 | 0.09 | 0.08 | 1.98 |
| 11 | Gm5617 | 0.59 | 2.79E-16 | 5.80E-12 | 0.51 | 0.39 | 0.12 | 1.32 |
| 11 | Nfkbia | 0.55 | 2.80E-16 | 5.83E-12 | 0.62 | 0.52 | 0.10 | 1.20 |
| 11 | Atp6v1d | 0.39 | 2.96E-16 | 6.15E-12 | 0.74 | 0.64 | 0.10 | 1.15 |
| 11 | Pink1 | 0.78 | 3.00E-16 | 6.24E-12 | 0.33 | 0.22 | 0.12 | 1.53 |
| 11 | Arfip1 | 0.82 | 3.04E-16 | 6.33E-12 | 0.30 | 0.18 | 0.11 | 1.63 |
| 11 | Slc20a2 | 0.80 | 3.14E-16 | 6.52E-12 | 0.36 | 0.23 | 0.13 | 1.53 |
| 11 | Ptar1 | 0.70 | 3.24E-16 | 6.75E-12 | 0.40 | 0.28 | 0.12 | 1.44 |
| 11 | Dcaf6 | 1.19 | 3.40E-16 | 7.07E-12 | 0.16 | 0.08 | 0.08 | 2.01 |
| 11 | Spop | 0.41 | 3.43E-16 | 7.13E-12 | 0.77 | 0.69 | 0.08 | 1.12 |
| 11 | Erf | 0.67 | 3.60E-16 | 7.49E-12 | 0.42 | 0.30 | 0.13 | 1.43 |
| 11 | Ctso | 1.16 | 3.71E-16 | 7.72E-12 | 0.14 | 0.06 | 0.07 | 2.17 |
| 11 | Zc3h6 | 0.88 | 4.21E-16 | 8.76E-12 | 0.17 | 0.09 | 0.09 | 2.00 |
| 11 | Rb1cc1 | 0.45 | 4.45E-16 | 9.25E-12 | 0.73 | 0.65 | 0.08 | 1.12 |
| 11 | Abracl | 0.32 | 4.89E-16 | 1.02E-11 | 0.96 | 0.93 | 0.03 | 1.04 |
| 11 | Gpbp1l1 | 0.61 | 5.50E-16 | 1.14E-11 | 0.51 | 0.38 | 0.13 | 1.34 |
| 11 | Scai | 0.61 | 5.78E-16 | 1.20E-11 | 0.48 | 0.35 | 0.13 | 1.36 |
| 11 | Gm26917 | 0.58 | 5.99E-16 | 1.25E-11 | 0.66 | 0.56 | 0.10 | 1.18 |

**Supplementary Table 1. Marker genes for the original clusters of integrated clustering of E11.5 control and NotchGOF λ-junction cells.**

| | | | | | | | | |
|---|---|---|---|---|---|---|---|---|
| 11 | Ier5l | 0.47 | 6.09E-16 | 1.27E-11 | 0.75 | 0.66 | 0.08 | 1.12 |
| 11 | Sox11 | 0.32 | 6.16E-16 | 1.28E-11 | 1.00 | 0.96 | 0.04 | 1.04 |
| 11 | Nagk | 0.78 | 6.26E-16 | 1.30E-11 | 0.31 | 0.20 | 0.11 | 1.58 |
| 11 | Prdm10 | 0.99 | 6.51E-16 | 1.35E-11 | 0.20 | 0.11 | 0.09 | 1.84 |
| 11 | Zswim8 | 0.69 | 6.58E-16 | 1.37E-11 | 0.46 | 0.34 | 0.12 | 1.36 |
| 11 | Abi1 | 0.53 | 6.70E-16 | 1.39E-11 | 0.51 | 0.38 | 0.13 | 1.33 |
| 11 | Sipa1l3 | 1.12 | 7.28E-16 | 1.51E-11 | 0.19 | 0.10 | 0.09 | 1.88 |
| 11 | Rnf208 | 0.97 | 7.35E-16 | 1.53E-11 | 0.14 | 0.07 | 0.08 | 2.15 |
| 11 | Tacc2 | 0.51 | 7.84E-16 | 1.63E-11 | 0.59 | 0.48 | 0.11 | 1.23 |
| 11 | Atp2b4 | 0.83 | 8.06E-16 | 1.68E-11 | 0.31 | 0.20 | 0.11 | 1.57 |
| 11 | Akap12 | 0.84 | 8.09E-16 | 1.68E-11 | 0.31 | 0.19 | 0.11 | 1.58 |
| 11 | Cir1 | 0.42 | 8.09E-16 | 1.68E-11 | 0.72 | 0.62 | 0.09 | 1.15 |
| 11 | Dok1 | 1.17 | 8.42E-16 | 1.75E-11 | 0.19 | 0.10 | 0.09 | 1.89 |
| 11 | Smad6 | 0.74 | 9.49E-16 | 1.97E-11 | 0.37 | 0.25 | 0.12 | 1.49 |
| 11 | Gdpd1 | 0.67 | 9.97E-16 | 2.07E-11 | 0.39 | 0.27 | 0.12 | 1.46 |
| 11 | Sppl3 | 0.51 | 1.01E-15 | 2.10E-11 | 0.71 | 0.62 | 0.09 | 1.15 |
| 11 | Bptf | 0.28 | 1.08E-15 | 2.25E-11 | 0.93 | 0.89 | 0.04 | 1.05 |
| 11 | Elmsan1 | 0.95 | 1.23E-15 | 2.55E-11 | 0.21 | 0.12 | 0.10 | 1.82 |
| 11 | Dpp3 | 0.44 | 1.23E-15 | 2.56E-11 | 0.67 | 0.58 | 0.10 | 1.17 |
| 11 | Pcm1 | 0.29 | 1.26E-15 | 2.61E-11 | 0.95 | 0.89 | 0.05 | 1.06 |
| 11 | 1700096K18Rik | 0.91 | 1.26E-15 | 2.63E-11 | 0.18 | 0.09 | 0.09 | 1.94 |
| 11 | Bcl2l1 | 0.55 | 1.38E-15 | 2.87E-11 | 0.49 | 0.37 | 0.13 | 1.34 |
| 11 | Cacna2d3 | 1.25 | 1.41E-15 | 2.94E-11 | 0.15 | 0.08 | 0.08 | 2.03 |
| 11 | Parva | 0.44 | 1.43E-15 | 2.97E-11 | 0.67 | 0.55 | 0.12 | 1.22 |
| 11 | Kifc3 | 0.95 | 1.43E-15 | 2.98E-11 | 0.15 | 0.07 | 0.08 | 2.07 |
| 11 | A130010J15Rik | 0.87 | 1.70E-15 | 3.53E-11 | 0.25 | 0.15 | 0.10 | 1.70 |
| 11 | Glrb | 1.36 | 1.82E-15 | 3.78E-11 | 0.11 | 0.05 | 0.06 | 2.29 |
| 11 | Casp8 | 0.70 | 1.94E-15 | 4.03E-11 | 0.30 | 0.18 | 0.12 | 1.64 |
| 11 | Dcxr | 0.98 | 1.94E-15 | 4.03E-11 | 0.18 | 0.09 | 0.09 | 1.92 |
| 11 | Ctnnd1 | 0.58 | 1.94E-15 | 4.03E-11 | 0.57 | 0.46 | 0.11 | 1.25 |
| 11 | Tmtc3 | 0.71 | 1.95E-15 | 4.05E-11 | 0.35 | 0.23 | 0.12 | 1.50 |
| 11 | Acyp1 | 0.52 | 2.05E-15 | 4.26E-11 | 0.55 | 0.42 | 0.12 | 1.29 |
| 11 | Cep70 | 0.57 | 2.07E-15 | 4.30E-11 | 0.52 | 0.39 | 0.12 | 1.31 |
| 11 | Ndufa6 | 0.29 | 2.20E-15 | 4.58E-11 | 0.96 | 0.93 | 0.02 | 1.03 |
| 11 | Gpalpp1 | 0.61 | 2.55E-15 | 5.29E-11 | 0.48 | 0.35 | 0.12 | 1.35 |
| 11 | Ddb1 | 0.28 | 2.63E-15 | 5.47E-11 | 0.91 | 0.88 | 0.03 | 1.04 |
| 11 | Kctd6 | 0.85 | 2.82E-15 | 5.86E-11 | 0.26 | 0.15 | 0.10 | 1.68 |
| 11 | Irs1 | 0.83 | 3.14E-15 | 6.54E-11 | 0.28 | 0.18 | 0.11 | 1.60 |
| 11 | Atxn3 | 0.84 | 3.27E-15 | 6.80E-11 | 0.29 | 0.19 | 0.11 | 1.57 |
| 11 | Ppm1b | 0.44 | 3.68E-15 | 7.65E-11 | 0.65 | 0.54 | 0.11 | 1.20 |
| 11 | Lpar6 | 0.54 | 3.91E-15 | 8.14E-11 | 0.53 | 0.40 | 0.13 | 1.32 |
| 11 | Rac3 | 0.57 | 4.40E-15 | 9.16E-11 | 0.45 | 0.33 | 0.13 | 1.39 |
| 11 | Irx1 | 0.89 | 4.53E-15 | 9.43E-11 | 0.12 | 0.05 | 0.07 | 2.26 |
| 11 | Bcl11a | 0.42 | 4.88E-15 | 1.02E-10 | 0.82 | 0.74 | 0.08 | 1.11 |
| 11 | Akr1b10 | 0.93 | 5.31E-15 | 1.10E-10 | 0.21 | 0.11 | 0.09 | 1.81 |
| 11 | A630072M18Rik | 0.90 | 5.38E-15 | 1.12E-10 | 0.19 | 0.10 | 0.09 | 1.84 |
| 11 | Sema3c | 0.72 | 5.38E-15 | 1.12E-10 | 0.18 | 0.09 | 0.09 | 1.93 |
| 11 | Stim2 | 0.80 | 5.80E-15 | 1.21E-10 | 0.34 | 0.23 | 0.11 | 1.47 |
| 11 | Arhgap32 | 0.73 | 6.49E-15 | 1.35E-10 | 0.30 | 0.19 | 0.11 | 1.59 |
| 11 | Tbrg1 | 0.31 | 6.52E-15 | 1.36E-10 | 0.88 | 0.83 | 0.05 | 1.06 |
| 11 | Zfp217 | 0.84 | 6.72E-15 | 1.40E-10 | 0.30 | 0.20 | 0.11 | 1.55 |
| 11 | Mcf2l | 1.02 | 7.34E-15 | 1.53E-10 | 0.14 | 0.07 | 0.07 | 2.09 |
| 11 | Ostm1 | 1.08 | 7.57E-15 | 1.57E-10 | 0.14 | 0.07 | 0.07 | 2.06 |
| 11 | Acrbp | 0.91 | 7.83E-15 | 1.63E-10 | 0.19 | 0.10 | 0.09 | 1.85 |
| 11 | Igf2bp3 | 0.30 | 8.12E-15 | 1.69E-10 | 0.87 | 0.82 | 0.05 | 1.06 |
| 11 | Ssr4 | 0.27 | 8.24E-15 | 1.71E-10 | 0.88 | 0.84 | 0.04 | 1.04 |

**Supplementary Table 1. Marker genes for the original clusters of integrated clustering of E11.5 control and NotchGOF λ-junction cells.**

| | | | | | | | | |
|---|---|---|---|---|---|---|---|---|
| 11 | Col4a3bp | 0.59 | 8.52E-15 | 1.77E-10 | 0.44 | 0.32 | 0.12 | 1.37 |
| 11 | Ankzf1 | 0.64 | 8.64E-15 | 1.80E-10 | 0.33 | 0.22 | 0.12 | 1.55 |
| 11 | Ddx6 | 0.29 | 8.95E-15 | 1.86E-10 | 0.89 | 0.84 | 0.05 | 1.06 |
| 11 | Syde2 | 0.94 | 1.02E-14 | 2.11E-10 | 0.15 | 0.07 | 0.08 | 2.07 |
| 11 | Cpne8 | 1.14 | 1.13E-14 | 2.35E-10 | 0.15 | 0.08 | 0.08 | 2.01 |
| 11 | Tmem141 | 0.75 | 1.14E-14 | 2.37E-10 | 0.30 | 0.19 | 0.11 | 1.58 |
| 11 | Sec22a | 0.62 | 1.16E-14 | 2.41E-10 | 0.41 | 0.29 | 0.12 | 1.41 |
| 11 | Slc3a2 | 0.53 | 1.16E-14 | 2.42E-10 | 0.69 | 0.62 | 0.06 | 1.10 |
| 11 | Sdsl | 1.08 | 1.21E-14 | 2.52E-10 | 0.13 | 0.06 | 0.07 | 2.16 |
| 11 | Ino80d | 0.62 | 1.23E-14 | 2.56E-10 | 0.46 | 0.34 | 0.12 | 1.34 |
| 11 | Plod2 | 0.52 | 1.34E-14 | 2.80E-10 | 0.62 | 0.49 | 0.13 | 1.25 |
| 11 | Reep6 | 1.39 | 1.39E-14 | 2.89E-10 | 0.11 | 0.05 | 0.06 | 2.27 |
| 11 | Arhgap12 | 0.50 | 1.40E-14 | 2.92E-10 | 0.46 | 0.33 | 0.13 | 1.38 |
| 11 | Clstn1 | 0.56 | 1.40E-14 | 2.92E-10 | 0.51 | 0.39 | 0.12 | 1.32 |
| 11 | Arvcf | 0.70 | 1.49E-14 | 3.09E-10 | 0.36 | 0.24 | 0.12 | 1.48 |
| 11 | 9930014A18Rik | 1.30 | 1.57E-14 | 3.26E-10 | 0.11 | 0.05 | 0.06 | 2.29 |
| 11 | Nr2c1 | 0.73 | 1.57E-14 | 3.27E-10 | 0.34 | 0.22 | 0.11 | 1.50 |
| 11 | Wtip | 0.47 | 1.58E-14 | 3.29E-10 | 0.65 | 0.56 | 0.10 | 1.17 |
| 11 | Pacsin3 | 0.68 | 1.67E-14 | 3.47E-10 | 0.33 | 0.22 | 0.12 | 1.53 |
| 11 | Aldh5a1 | 0.67 | 1.85E-14 | 3.85E-10 | 0.36 | 0.24 | 0.12 | 1.48 |
| 11 | Col11a1 | 0.65 | 1.91E-14 | 3.98E-10 | 0.54 | 0.42 | 0.11 | 1.26 |
| 11 | Stx4a | 0.49 | 1.93E-14 | 4.02E-10 | 0.44 | 0.31 | 0.12 | 1.39 |
| 11 | Ypel5 | 0.72 | 1.94E-14 | 4.04E-10 | 0.36 | 0.25 | 0.11 | 1.45 |
| 11 | Endod1 | 1.09 | 1.96E-14 | 4.07E-10 | 0.17 | 0.09 | 0.08 | 1.90 |
| 11 | Zdhhc5 | 0.71 | 1.99E-14 | 4.14E-10 | 0.43 | 0.32 | 0.11 | 1.34 |
| 11 | Pan3 | 0.63 | 1.99E-14 | 4.14E-10 | 0.45 | 0.33 | 0.12 | 1.38 |
| 11 | Ten1 | 0.42 | 2.09E-14 | 4.35E-10 | 0.67 | 0.56 | 0.10 | 1.18 |
| 11 | Stap2 | 0.92 | 2.19E-14 | 4.56E-10 | 0.16 | 0.08 | 0.08 | 1.96 |
| 11 | Ncoa3 | 0.77 | 2.20E-14 | 4.58E-10 | 0.33 | 0.22 | 0.11 | 1.49 |
| 11 | Hipk2 | 0.41 | 2.29E-14 | 4.76E-10 | 0.72 | 0.64 | 0.07 | 1.11 |
| 11 | Fam172a | 0.59 | 2.31E-14 | 4.81E-10 | 0.47 | 0.35 | 0.12 | 1.35 |
| 11 | Zkscan2 | 0.88 | 2.43E-14 | 5.05E-10 | 0.19 | 0.10 | 0.09 | 1.84 |
| 11 | Slc35b3 | 0.80 | 2.81E-14 | 5.84E-10 | 0.32 | 0.21 | 0.11 | 1.52 |
| 11 | Prdx5 | 0.54 | 2.86E-14 | 5.94E-10 | 0.84 | 0.80 | 0.04 | 1.05 |
| 11 | Rcan1 | 0.67 | 3.05E-14 | 6.34E-10 | 0.37 | 0.26 | 0.12 | 1.45 |
| 11 | Pcbd1 | 0.73 | 3.11E-14 | 6.47E-10 | 0.21 | 0.12 | 0.09 | 1.78 |
| 11 | Plagl2 | 0.63 | 3.18E-14 | 6.62E-10 | 0.47 | 0.36 | 0.12 | 1.33 |
| 11 | C130071C03Rik | 0.75 | 3.19E-14 | 6.64E-10 | 0.12 | 0.05 | 0.06 | 2.25 |
| 11 | Narf | 0.52 | 3.19E-14 | 6.64E-10 | 0.54 | 0.42 | 0.12 | 1.29 |
| 11 | Egfl6 | 0.82 | 3.32E-14 | 6.91E-10 | 0.19 | 0.10 | 0.09 | 1.85 |
| 11 | Zswim6 | 0.61 | 3.33E-14 | 6.93E-10 | 0.41 | 0.29 | 0.12 | 1.40 |
| 11 | Nucb2 | 0.93 | 3.34E-14 | 6.95E-10 | 0.25 | 0.15 | 0.10 | 1.65 |
| 11 | Cdk5rap1 | 1.02 | 3.51E-14 | 7.29E-10 | 0.17 | 0.09 | 0.08 | 1.91 |
| 11 | Eno1b | 0.57 | 3.61E-14 | 7.50E-10 | 0.47 | 0.34 | 0.13 | 1.37 |
| 11 | Gprc5b | 0.95 | 3.71E-14 | 7.71E-10 | 0.18 | 0.10 | 0.08 | 1.85 |
| 11 | Insig2 | 0.67 | 3.81E-14 | 7.92E-10 | 0.39 | 0.27 | 0.12 | 1.44 |
| 11 | Rtf2 | 0.37 | 3.90E-14 | 8.10E-10 | 0.74 | 0.66 | 0.08 | 1.12 |
| 11 | Pogk | 0.42 | 4.00E-14 | 8.32E-10 | 0.62 | 0.50 | 0.12 | 1.24 |
| 11 | Ociad2 | 0.73 | 4.01E-14 | 8.34E-10 | 0.29 | 0.18 | 0.11 | 1.59 |
| 11 | Hexim1 | 0.72 | 4.16E-14 | 8.65E-10 | 0.39 | 0.28 | 0.11 | 1.39 |
| 11 | Stxbp3 | 0.74 | 4.19E-14 | 8.72E-10 | 0.34 | 0.23 | 0.11 | 1.47 |
| 11 | H2-K1 | 0.87 | 4.20E-14 | 8.74E-10 | 0.23 | 0.13 | 0.09 | 1.71 |
| 11 | Ebpl | 0.46 | 4.34E-14 | 9.03E-10 | 0.65 | 0.54 | 0.11 | 1.19 |
| 11 | Olfm2 | 0.78 | 4.88E-14 | 1.02E-09 | 0.19 | 0.10 | 0.09 | 1.83 |
| 11 | Leprotl1 | 0.47 | 4.90E-14 | 1.02E-09 | 0.66 | 0.55 | 0.11 | 1.20 |
| 11 | Stx6 | 0.47 | 4.96E-14 | 1.03E-09 | 0.61 | 0.50 | 0.11 | 1.22 |

**Supplementary Table 1. Marker genes for the original clusters of integrated clustering of E11.5 control and NotchGOF λ-junction cells.**

| | | | | | | | | |
|---|---|---|---|---|---|---|---|---|
| 11 | Rnf38 | 0.70 | 5.15E-14 | 1.07E-09 | 0.39 | 0.27 | 0.12 | 1.43 |
| 11 | Nipsnap1 | 0.43 | 5.18E-14 | 1.08E-09 | 0.67 | 0.56 | 0.12 | 1.21 |
| 11 | Eef2kmt | 0.71 | 5.20E-14 | 1.08E-09 | 0.29 | 0.18 | 0.11 | 1.59 |
| 11 | Inhba | 0.95 | 5.27E-14 | 1.10E-09 | 0.26 | 0.17 | 0.10 | 1.59 |
| 11 | Tmbim1 | 1.08 | 5.44E-14 | 1.13E-09 | 0.18 | 0.10 | 0.08 | 1.84 |
| 11 | Dkk3 | 1.03 | 5.52E-14 | 1.15E-09 | 0.20 | 0.12 | 0.09 | 1.74 |
| 11 | Ccdc112 | 0.46 | 6.05E-14 | 1.26E-09 | 0.64 | 0.51 | 0.12 | 1.24 |
| 11 | Ammecr1 | 0.56 | 6.13E-14 | 1.27E-09 | 0.47 | 0.35 | 0.12 | 1.35 |
| 11 | Sirt7 | 0.78 | 7.22E-14 | 1.50E-09 | 0.27 | 0.17 | 0.10 | 1.59 |
| 11 | Pex7 | 0.70 | 7.94E-14 | 1.65E-09 | 0.36 | 0.25 | 0.11 | 1.44 |
| 11 | Lats1 | 0.67 | 8.10E-14 | 1.68E-09 | 0.37 | 0.26 | 0.11 | 1.43 |
| 11 | Ccdc173 | 0.86 | 8.29E-14 | 1.72E-09 | 0.20 | 0.12 | 0.09 | 1.76 |
| 11 | Unc5c | 0.34 | 8.39E-14 | 1.75E-09 | 0.48 | 0.34 | 0.14 | 1.42 |
| 11 | Furin | 0.70 | 8.44E-14 | 1.75E-09 | 0.31 | 0.20 | 0.11 | 1.55 |
| 11 | Sertad2 | 0.52 | 9.04E-14 | 1.88E-09 | 0.52 | 0.40 | 0.11 | 1.28 |
| 11 | Dbndd1 | 1.21 | 9.16E-14 | 1.91E-09 | 0.12 | 0.05 | 0.06 | 2.17 |
| 11 | Osbpl8 | 0.80 | 9.28E-14 | 1.93E-09 | 0.34 | 0.23 | 0.11 | 1.47 |
| 11 | Itgb5 | 0.46 | 9.72E-14 | 2.02E-09 | 0.68 | 0.58 | 0.10 | 1.17 |
| 11 | Gramd3 | 0.75 | 1.07E-13 | 2.22E-09 | 0.28 | 0.18 | 0.10 | 1.58 |
| 11 | Nras | 0.35 | 1.12E-13 | 2.32E-09 | 0.76 | 0.69 | 0.07 | 1.11 |
| 11 | Golgb1 | 0.36 | 1.30E-13 | 2.71E-09 | 0.81 | 0.74 | 0.07 | 1.10 |
| 11 | Bmp7 | 0.53 | 1.51E-13 | 3.15E-09 | 0.50 | 0.38 | 0.12 | 1.32 |
| 11 | Gm20186 | 1.04 | 1.54E-13 | 3.20E-09 | 0.13 | 0.06 | 0.07 | 2.06 |
| 11 | Kyat1 | 0.97 | 1.56E-13 | 3.23E-09 | 0.11 | 0.05 | 0.06 | 2.20 |
| 11 | Sdr39u1 | 0.75 | 1.74E-13 | 3.61E-09 | 0.25 | 0.16 | 0.10 | 1.63 |
| 11 | BC005537 | 0.43 | 1.80E-13 | 3.75E-09 | 0.61 | 0.50 | 0.12 | 1.24 |
| 11 | Vezf1 | 0.29 | 1.80E-13 | 3.75E-09 | 0.86 | 0.80 | 0.06 | 1.07 |
| 11 | Bclaf3 | 0.51 | 1.81E-13 | 3.76E-09 | 0.45 | 0.33 | 0.12 | 1.37 |
| 11 | Mllt11 | 0.46 | 1.86E-13 | 3.86E-09 | 0.37 | 0.26 | 0.11 | 1.44 |
| 11 | Malt1 | 0.73 | 1.87E-13 | 3.89E-09 | 0.26 | 0.16 | 0.10 | 1.61 |
| 11 | Slc25a11 | 0.36 | 1.90E-13 | 3.95E-09 | 0.76 | 0.69 | 0.07 | 1.10 |
| 11 | Fahd2a | 0.68 | 1.98E-13 | 4.12E-09 | 0.29 | 0.19 | 0.10 | 1.54 |
| 11 | Syf2 | 0.36 | 2.00E-13 | 4.17E-09 | 0.73 | 0.65 | 0.07 | 1.11 |
| 11 | Tmem59 | 0.29 | 2.05E-13 | 4.26E-09 | 0.93 | 0.88 | 0.04 | 1.05 |
| 11 | Tmf1 | 0.47 | 2.08E-13 | 4.33E-09 | 0.59 | 0.49 | 0.10 | 1.21 |
| 11 | Ccdc80 | 0.88 | 2.11E-13 | 4.39E-09 | 0.28 | 0.18 | 0.10 | 1.56 |
| 11 | A430018G15Rik | 1.02 | 2.13E-13 | 4.44E-09 | 0.12 | 0.06 | 0.07 | 2.12 |
| 11 | Ubl3 | 0.50 | 2.17E-13 | 4.52E-09 | 0.48 | 0.37 | 0.12 | 1.32 |
| 11 | Tmem106c | 0.72 | 2.20E-13 | 4.58E-09 | 0.30 | 0.19 | 0.10 | 1.53 |
| 11 | Peli1 | 0.38 | 2.24E-13 | 4.65E-09 | 0.76 | 0.68 | 0.09 | 1.13 |
| 11 | Krcc1 | 0.39 | 2.46E-13 | 5.11E-09 | 0.66 | 0.57 | 0.09 | 1.16 |
| 11 | Cd82 | 1.05 | 2.54E-13 | 5.28E-09 | 0.13 | 0.07 | 0.07 | 2.03 |
| 11 | Angel1 | 1.40 | 2.78E-13 | 5.77E-09 | 0.10 | 0.05 | 0.06 | 2.22 |
| 11 | Bnip3 | 0.80 | 2.79E-13 | 5.80E-09 | 0.51 | 0.41 | 0.10 | 1.24 |
| 11 | Mafg | 0.56 | 3.11E-13 | 6.46E-09 | 0.60 | 0.51 | 0.09 | 1.18 |
| 11 | Smad3 | 0.59 | 3.26E-13 | 6.77E-09 | 0.38 | 0.26 | 0.11 | 1.43 |
| 11 | 1700025G04Rik | 0.53 | 3.30E-13 | 6.87E-09 | 0.45 | 0.34 | 0.12 | 1.35 |
| 11 | Ap3b1 | 0.45 | 3.48E-13 | 7.23E-09 | 0.59 | 0.47 | 0.12 | 1.25 |
| 11 | Nell2 | 1.03 | 3.50E-13 | 7.27E-09 | 0.11 | 0.05 | 0.06 | 2.18 |
| 11 | Aamp | 0.31 | 3.89E-13 | 8.09E-09 | 0.85 | 0.80 | 0.05 | 1.07 |
| 11 | Prkcd | 0.68 | 4.02E-13 | 8.37E-09 | 0.32 | 0.21 | 0.11 | 1.51 |
| 11 | Aff4 | 0.42 | 4.05E-13 | 8.42E-09 | 0.71 | 0.63 | 0.08 | 1.13 |
| 11 | Zfp612 | 0.74 | 4.22E-13 | 8.78E-09 | 0.26 | 0.17 | 0.10 | 1.58 |
| 11 | Letmd1 | 0.67 | 4.31E-13 | 8.97E-09 | 0.32 | 0.22 | 0.11 | 1.48 |
| 11 | 1700012D14Rik | 0.88 | 4.48E-13 | 9.31E-09 | 0.21 | 0.13 | 0.09 | 1.68 |
| 11 | Csf2ra | 0.81 | 4.50E-13 | 9.36E-09 | 0.23 | 0.14 | 0.09 | 1.66 |

**Supplementary Table 1. Marker genes for the original clusters of integrated clustering of E11.5 control and NotchGOF λ-junction cells.**

| | | | | | | | | |
|---|---|---|---|---|---|---|---|---|
| 11 | Mllt10 | 0.37 | 4.63E-13 | 9.62E-09 | 0.72 | 0.63 | 0.09 | 1.14 |
| 11 | Zfand5 | 0.44 | 4.76E-13 | 9.89E-09 | 0.88 | 0.83 | 0.06 | 1.07 |
| 11 | Grn | 0.35 | 4.76E-13 | 9.90E-09 | 0.48 | 0.35 | 0.13 | 1.36 |
| 11 | 1600012H06Rik | 0.67 | 5.07E-13 | 1.06E-08 | 0.32 | 0.22 | 0.11 | 1.49 |
| 11 | Nr1h2 | 0.52 | 5.19E-13 | 1.08E-08 | 0.43 | 0.31 | 0.12 | 1.38 |
| 11 | Tmem245 | 0.55 | 5.38E-13 | 1.12E-08 | 0.51 | 0.40 | 0.11 | 1.28 |
| 11 | Fundc1 | 0.51 | 5.65E-13 | 1.18E-08 | 0.48 | 0.37 | 0.12 | 1.32 |
| 11 | Hyal1 | 0.94 | 6.50E-13 | 1.35E-08 | 0.17 | 0.09 | 0.08 | 1.85 |
| 11 | Nipbl | 0.25 | 6.72E-13 | 1.40E-08 | 0.92 | 0.87 | 0.05 | 1.06 |
| 11 | Nsmaf | 0.61 | 6.83E-13 | 1.42E-08 | 0.38 | 0.27 | 0.11 | 1.41 |
| 11 | Fnbp1l | 0.26 | 6.95E-13 | 1.45E-08 | 0.93 | 0.85 | 0.08 | 1.09 |
| 11 | Bcap31 | 0.34 | 6.98E-13 | 1.45E-08 | 0.80 | 0.75 | 0.05 | 1.07 |
| 11 | Mplkip | 0.34 | 7.00E-13 | 1.46E-08 | 0.75 | 0.67 | 0.08 | 1.11 |
| 11 | Slc35a5 | 1.00 | 7.36E-13 | 1.53E-08 | 0.17 | 0.10 | 0.08 | 1.80 |
| 11 | Rimklb | 0.47 | 7.50E-13 | 1.56E-08 | 0.44 | 0.32 | 0.12 | 1.37 |
| 11 | Fbxl3 | 0.64 | 7.80E-13 | 1.62E-08 | 0.36 | 0.25 | 0.11 | 1.43 |
| 11 | Zc2hc1a | 0.50 | 8.02E-13 | 1.67E-08 | 0.52 | 0.42 | 0.10 | 1.25 |
| 11 | Zfp292 | 0.27 | 8.09E-13 | 1.68E-08 | 0.90 | 0.86 | 0.04 | 1.05 |
| 11 | Scp2 | 0.30 | 8.56E-13 | 1.78E-08 | 0.84 | 0.77 | 0.06 | 1.08 |
| 11 | Smurf2 | 0.51 | 8.67E-13 | 1.80E-08 | 0.51 | 0.40 | 0.11 | 1.28 |
| 11 | Gpr107 | 0.64 | 8.75E-13 | 1.82E-08 | 0.42 | 0.31 | 0.11 | 1.34 |
| 11 | 2610002M06Rik | 0.57 | 8.86E-13 | 1.84E-08 | 0.40 | 0.29 | 0.11 | 1.38 |
| 11 | Ust | 0.97 | 9.75E-13 | 2.03E-08 | 0.12 | 0.06 | 0.06 | 2.07 |
| 11 | Rtn2 | 0.92 | 9.91E-13 | 2.06E-08 | 0.16 | 0.08 | 0.07 | 1.89 |
| 11 | Tmem87b | 0.70 | 1.01E-12 | 2.10E-08 | 0.32 | 0.21 | 0.11 | 1.50 |
| 11 | Thbs1 | 0.36 | 1.04E-12 | 2.17E-08 | 0.20 | 0.12 | 0.09 | 1.74 |
| 11 | Tmem19 | 0.65 | 1.06E-12 | 2.20E-08 | 0.29 | 0.19 | 0.10 | 1.54 |
| 11 | Arf3 | 0.58 | 1.09E-12 | 2.26E-08 | 0.42 | 0.31 | 0.11 | 1.36 |
| 11 | Pi4k2b | 0.60 | 1.10E-12 | 2.29E-08 | 0.39 | 0.28 | 0.11 | 1.40 |
| 11 | Bmyc | 0.35 | 1.20E-12 | 2.50E-08 | 0.72 | 0.62 | 0.10 | 1.16 |
| 11 | Rnf5 | 0.35 | 1.38E-12 | 2.86E-08 | 0.72 | 0.65 | 0.07 | 1.11 |
| 11 | Aass | 1.16 | 1.45E-12 | 3.02E-08 | 0.11 | 0.05 | 0.06 | 2.14 |
| 11 | Mex3a | 0.26 | 1.46E-12 | 3.04E-08 | 0.91 | 0.85 | 0.06 | 1.06 |
| 11 | Sema4d | 1.02 | 1.50E-12 | 3.12E-08 | 0.14 | 0.07 | 0.07 | 1.96 |
| 11 | Prkag2 | 0.82 | 1.55E-12 | 3.22E-08 | 0.32 | 0.22 | 0.10 | 1.45 |
| 11 | Maco1 | 0.38 | 1.55E-12 | 3.23E-08 | 0.77 | 0.69 | 0.08 | 1.11 |
| 11 | Ncoa2 | 0.50 | 1.65E-12 | 3.43E-08 | 0.41 | 0.29 | 0.12 | 1.41 |
| 11 | Nes | 0.52 | 1.70E-12 | 3.53E-08 | 0.52 | 0.42 | 0.10 | 1.23 |
| 11 | Pgap2 | 0.42 | 1.98E-12 | 4.12E-08 | 0.56 | 0.45 | 0.11 | 1.25 |
| 11 | Akap17b | 0.93 | 1.99E-12 | 4.13E-08 | 0.20 | 0.12 | 0.08 | 1.70 |
| 11 | Cdk17 | 0.65 | 2.04E-12 | 4.24E-08 | 0.31 | 0.21 | 0.10 | 1.51 |
| 11 | Map7d2 | 0.84 | 2.19E-12 | 4.56E-08 | 0.16 | 0.08 | 0.07 | 1.89 |
| 11 | Add1 | 0.37 | 2.20E-12 | 4.57E-08 | 0.73 | 0.65 | 0.08 | 1.12 |
| 11 | AU040320 | 0.90 | 2.36E-12 | 4.90E-08 | 0.19 | 0.11 | 0.08 | 1.73 |
| 11 | Slc4a2 | 0.72 | 2.41E-12 | 5.02E-08 | 0.32 | 0.22 | 0.10 | 1.47 |
| 11 | Purb | 0.25 | 2.52E-12 | 5.24E-08 | 0.90 | 0.87 | 0.03 | 1.04 |
| 11 | Gnptg | 0.67 | 2.61E-12 | 5.43E-08 | 0.30 | 0.20 | 0.10 | 1.50 |
| 11 | Spg21 | 0.34 | 2.68E-12 | 5.57E-08 | 0.72 | 0.64 | 0.08 | 1.12 |
| 11 | Ccndbp1 | 0.61 | 2.69E-12 | 5.59E-08 | 0.38 | 0.28 | 0.11 | 1.38 |
| 11 | Paip2b | 0.59 | 2.75E-12 | 5.72E-08 | 0.40 | 0.29 | 0.11 | 1.36 |
| 11 | Bcl9 | 0.35 | 2.75E-12 | 5.72E-08 | 0.75 | 0.68 | 0.07 | 1.10 |
| 11 | Tmtc2 | 0.61 | 2.85E-12 | 5.93E-08 | 0.30 | 0.20 | 0.10 | 1.53 |
| 11 | Cep152 | 0.79 | 2.87E-12 | 5.97E-08 | 0.21 | 0.12 | 0.08 | 1.69 |
| 11 | Chn2 | 1.01 | 2.90E-12 | 6.02E-08 | 0.14 | 0.07 | 0.07 | 1.93 |
| 11 | Strn | 0.72 | 3.28E-12 | 6.81E-08 | 0.27 | 0.18 | 0.10 | 1.55 |
| 11 | Agap3 | 0.55 | 3.32E-12 | 6.90E-08 | 0.46 | 0.36 | 0.10 | 1.29 |

**Supplementary Table 1. Marker genes for the original clusters of integrated clustering of E11.5 control and NotchGOF λ-junction cells.**

| | | | | | | | | |
|---|---|---|---|---|---|---|---|---|
| 11 | Eif4ebp3 | 0.72 | 3.48E-12 | 7.23E-08 | 0.28 | 0.19 | 0.10 | 1.51 |
| 11 | Shb | 0.67 | 3.61E-12 | 7.51E-08 | 0.29 | 0.19 | 0.10 | 1.50 |
| 11 | Uevld | 0.82 | 3.72E-12 | 7.74E-08 | 0.21 | 0.12 | 0.08 | 1.67 |
| 11 | Acsl3 | 0.36 | 3.79E-12 | 7.89E-08 | 0.79 | 0.74 | 0.05 | 1.07 |
| 11 | Usp22 | 0.34 | 3.82E-12 | 7.95E-08 | 0.72 | 0.62 | 0.10 | 1.16 |
| 11 | Vrk3 | 0.51 | 4.10E-12 | 8.53E-08 | 0.45 | 0.34 | 0.11 | 1.32 |
| 11 | 1810044D09Rik | 0.88 | 4.31E-12 | 8.97E-08 | 0.19 | 0.11 | 0.08 | 1.73 |
| 11 | Pdcd6 | 0.27 | 4.33E-12 | 9.00E-08 | 0.83 | 0.78 | 0.04 | 1.06 |
| 11 | Dcp1b | 0.86 | 4.44E-12 | 9.23E-08 | 0.16 | 0.09 | 0.07 | 1.84 |
| 11 | Micu2 | 0.39 | 4.58E-12 | 9.53E-08 | 0.62 | 0.51 | 0.11 | 1.22 |
| 11 | Tmed3 | 0.44 | 5.28E-12 | 1.10E-07 | 0.55 | 0.44 | 0.11 | 1.25 |
| 11 | Col9a3 | 0.60 | 5.34E-12 | 1.11E-07 | 0.23 | 0.14 | 0.09 | 1.64 |
| 11 | Hcfc1r1 | 0.41 | 5.37E-12 | 1.12E-07 | 0.66 | 0.58 | 0.08 | 1.15 |
| 11 | Ubn2 | 0.34 | 5.76E-12 | 1.20E-07 | 0.78 | 0.70 | 0.08 | 1.11 |
| 11 | Amn1 | 0.48 | 6.14E-12 | 1.28E-07 | 0.54 | 0.44 | 0.10 | 1.24 |
| 11 | Rlf | 0.40 | 6.44E-12 | 1.34E-07 | 0.67 | 0.57 | 0.10 | 1.17 |
| 11 | Fkbp1b | 0.67 | 7.39E-12 | 1.54E-07 | 0.23 | 0.14 | 0.09 | 1.63 |
| 11 | Slc38a4 | 0.72 | 7.39E-12 | 1.54E-07 | 0.38 | 0.27 | 0.10 | 1.38 |
| 11 | Fzd10 | 0.74 | 7.55E-12 | 1.57E-07 | 0.23 | 0.14 | 0.09 | 1.65 |
| 11 | Gdi1 | 0.50 | 8.00E-12 | 1.66E-07 | 0.53 | 0.42 | 0.11 | 1.26 |
| 11 | Mast4 | 0.47 | 8.21E-12 | 1.71E-07 | 0.55 | 0.45 | 0.11 | 1.24 |
| 11 | Cryzl2 | 0.85 | 8.23E-12 | 1.71E-07 | 0.18 | 0.10 | 0.08 | 1.76 |
| 11 | Cdca7l | 0.57 | 8.34E-12 | 1.73E-07 | 0.41 | 0.30 | 0.11 | 1.36 |
| 11 | F8a | 1.04 | 8.46E-12 | 1.76E-07 | 0.15 | 0.08 | 0.07 | 1.83 |
| 11 | Ypel1 | 0.42 | 9.02E-12 | 1.88E-07 | 0.60 | 0.50 | 0.10 | 1.21 |
| 11 | Sord | 0.73 | 9.72E-12 | 2.02E-07 | 0.23 | 0.14 | 0.09 | 1.60 |
| 11 | Sh3bp5 | 0.51 | 9.87E-12 | 2.05E-07 | 0.59 | 0.50 | 0.09 | 1.19 |
| 11 | Aptx | 0.54 | 1.03E-11 | 2.14E-07 | 0.36 | 0.26 | 0.10 | 1.40 |
| 11 | Tab3 | 0.90 | 1.03E-11 | 2.15E-07 | 0.16 | 0.09 | 0.07 | 1.80 |
| 11 | Hmga1 | 0.38 | 1.06E-11 | 2.20E-07 | 0.60 | 0.49 | 0.10 | 1.21 |
| 11 | Ubxn2a | 0.51 | 1.08E-11 | 2.25E-07 | 0.42 | 0.31 | 0.11 | 1.34 |
| 11 | Cpt1a | 0.89 | 1.14E-11 | 2.37E-07 | 0.17 | 0.10 | 0.08 | 1.76 |
| 11 | Zfp652 | 0.43 | 1.18E-11 | 2.46E-07 | 0.60 | 0.51 | 0.10 | 1.19 |
| 11 | Prmt2 | 0.52 | 1.25E-11 | 2.61E-07 | 0.42 | 0.31 | 0.11 | 1.36 |
| 11 | Zdbf2 | 0.46 | 1.32E-11 | 2.74E-07 | 0.52 | 0.41 | 0.11 | 1.27 |
| 11 | Epha4 | 0.28 | 1.54E-11 | 3.21E-07 | 0.77 | 0.65 | 0.12 | 1.19 |
| 11 | Tmc4 | 0.98 | 1.59E-11 | 3.30E-07 | 0.17 | 0.10 | 0.07 | 1.74 |
| 11 | Arhgef12 | 0.41 | 1.61E-11 | 3.34E-07 | 0.66 | 0.58 | 0.08 | 1.14 |
| 11 | Tmem14a | 0.83 | 1.71E-11 | 3.56E-07 | 0.18 | 0.10 | 0.08 | 1.74 |
| 11 | Ccdc85c | 0.70 | 1.88E-11 | 3.92E-07 | 0.19 | 0.11 | 0.08 | 1.71 |
| 11 | 3830406C13Rik | 0.58 | 2.24E-11 | 4.66E-07 | 0.34 | 0.24 | 0.10 | 1.42 |
| 11 | Chd2 | 0.35 | 2.33E-11 | 4.85E-07 | 0.71 | 0.62 | 0.09 | 1.14 |
| 11 | Heph | 0.53 | 2.38E-11 | 4.94E-07 | 0.36 | 0.25 | 0.11 | 1.42 |
| 11 | Tmed5 | 0.55 | 2.59E-11 | 5.39E-07 | 0.57 | 0.48 | 0.09 | 1.18 |
| 11 | Ncoa1 | 0.62 | 2.63E-11 | 5.46E-07 | 0.26 | 0.17 | 0.09 | 1.54 |
| 11 | Btg3 | 0.58 | 2.73E-11 | 5.67E-07 | 0.43 | 0.32 | 0.11 | 1.33 |
| 11 | Hectd1 | 0.25 | 2.77E-11 | 5.77E-07 | 0.84 | 0.81 | 0.04 | 1.05 |
| 11 | Jmjd1c | 0.39 | 2.82E-11 | 5.87E-07 | 0.59 | 0.48 | 0.11 | 1.23 |
| 11 | Dlg3 | 0.60 | 3.13E-11 | 6.50E-07 | 0.42 | 0.32 | 0.11 | 1.33 |
| 11 | Pdzrn3 | 0.73 | 3.13E-11 | 6.52E-07 | 0.26 | 0.17 | 0.09 | 1.54 |
| 11 | Tsc22d4 | 0.43 | 3.15E-11 | 6.54E-07 | 0.67 | 0.61 | 0.06 | 1.09 |
| 11 | Cnot6 | 0.26 | 3.21E-11 | 6.68E-07 | 0.83 | 0.77 | 0.05 | 1.07 |
| 11 | Mta3 | 0.52 | 3.32E-11 | 6.91E-07 | 0.48 | 0.38 | 0.10 | 1.26 |
| 11 | F2r | 0.36 | 3.34E-11 | 6.95E-07 | 0.67 | 0.57 | 0.11 | 1.19 |
| 11 | Thumpd3 | 0.50 | 3.41E-11 | 7.08E-07 | 0.48 | 0.38 | 0.10 | 1.26 |
| 11 | Ascc1 | 0.47 | 3.58E-11 | 7.45E-07 | 0.50 | 0.39 | 0.11 | 1.28 |

**Supplementary Table 1. Marker genes for the original clusters of integrated clustering of E11.5 control and NotchGOF λ-junction cells.**

| | | | | | | | | |
|---|---|---|---|---|---|---|---|---|
| 11 | Sidt2 | 0.78 | 3.65E-11 | 7.58E-07 | 0.27 | 0.18 | 0.09 | 1.49 |
| 11 | Rab10 | 0.31 | 3.68E-11 | 7.66E-07 | 0.82 | 0.79 | 0.03 | 1.04 |
| 11 | Chmp5 | 0.28 | 3.77E-11 | 7.85E-07 | 0.85 | 0.80 | 0.05 | 1.07 |
| 11 | Dcaf12 | 0.53 | 3.83E-11 | 7.96E-07 | 0.44 | 0.34 | 0.10 | 1.30 |
| 11 | Dedd2 | 1.09 | 3.83E-11 | 7.97E-07 | 0.14 | 0.07 | 0.06 | 1.85 |
| 11 | 2210016F16Rik | 0.53 | 3.91E-11 | 8.14E-07 | 0.45 | 0.35 | 0.10 | 1.28 |
| 11 | 2310039H08Rik | 0.45 | 4.01E-11 | 8.34E-07 | 0.52 | 0.42 | 0.10 | 1.24 |
| 11 | Rnf181 | 0.47 | 4.02E-11 | 8.35E-07 | 0.49 | 0.39 | 0.10 | 1.26 |
| 11 | Nos1ap | 1.06 | 4.21E-11 | 8.76E-07 | 0.10 | 0.05 | 0.05 | 2.08 |
| 11 | Arrb2 | 0.78 | 4.60E-11 | 9.56E-07 | 0.26 | 0.17 | 0.09 | 1.52 |
| 11 | Lin7c | 0.45 | 4.68E-11 | 9.74E-07 | 0.58 | 0.50 | 0.09 | 1.18 |
| 11 | Frs2 | 0.68 | 5.08E-11 | 1.06E-06 | 0.37 | 0.27 | 0.10 | 1.36 |
| 11 | Arl13b | 0.57 | 5.20E-11 | 1.08E-06 | 0.50 | 0.40 | 0.10 | 1.25 |
| 11 | B3galt6 | 0.48 | 5.28E-11 | 1.10E-06 | 0.42 | 0.31 | 0.11 | 1.37 |
| 11 | Chfr | 0.66 | 5.71E-11 | 1.19E-06 | 0.35 | 0.25 | 0.10 | 1.38 |
| 11 | Traf6 | 0.79 | 5.76E-11 | 1.20E-06 | 0.23 | 0.15 | 0.08 | 1.57 |
| 11 | Trit1 | 0.69 | 6.28E-11 | 1.31E-06 | 0.28 | 0.19 | 0.09 | 1.48 |
| 11 | Gid4 | 0.51 | 6.54E-11 | 1.36E-06 | 0.44 | 0.34 | 0.11 | 1.32 |
| 11 | Wiz | 0.47 | 6.60E-11 | 1.37E-06 | 0.51 | 0.42 | 0.10 | 1.23 |
| 11 | Gstt2 | 0.85 | 6.68E-11 | 1.39E-06 | 0.14 | 0.07 | 0.06 | 1.86 |
| 11 | Slmap | 0.55 | 7.06E-11 | 1.47E-06 | 0.43 | 0.33 | 0.10 | 1.31 |
| 11 | 1110032A03Rik | 0.48 | 7.61E-11 | 1.58E-06 | 0.42 | 0.32 | 0.11 | 1.33 |
| 11 | Nt5c | 0.41 | 7.78E-11 | 1.62E-06 | 0.63 | 0.53 | 0.09 | 1.17 |
| 11 | Cope | 0.30 | 8.08E-11 | 1.68E-06 | 0.80 | 0.77 | 0.03 | 1.04 |
| 11 | Fbxw7 | 0.44 | 8.29E-11 | 1.72E-06 | 0.48 | 0.38 | 0.11 | 1.28 |
| 11 | 2900060N12Rik | 0.89 | 8.29E-11 | 1.72E-06 | 0.13 | 0.07 | 0.06 | 1.91 |
| 11 | Dph6 | 0.41 | 8.43E-11 | 1.75E-06 | 0.52 | 0.41 | 0.11 | 1.26 |
| 11 | Zfp7 | 0.71 | 8.72E-11 | 1.81E-06 | 0.23 | 0.14 | 0.08 | 1.58 |
| 11 | Slc45a4 | 0.75 | 8.81E-11 | 1.83E-06 | 0.21 | 0.13 | 0.08 | 1.59 |
| 11 | Bod1l | 0.26 | 8.99E-11 | 1.87E-06 | 0.87 | 0.82 | 0.05 | 1.06 |
| 11 | Mdm2 | 0.43 | 9.85E-11 | 2.05E-06 | 0.53 | 0.43 | 0.10 | 1.23 |
| 11 | Ero1l | 0.51 | 1.00E-10 | 2.09E-06 | 0.40 | 0.29 | 0.11 | 1.37 |
| 11 | Wdr45 | 0.76 | 1.01E-10 | 2.11E-06 | 0.21 | 0.13 | 0.08 | 1.60 |
| 11 | Vps54 | 0.55 | 1.07E-10 | 2.23E-06 | 0.41 | 0.31 | 0.10 | 1.32 |
| 11 | Slc39a3 | 0.83 | 1.10E-10 | 2.29E-06 | 0.20 | 0.12 | 0.08 | 1.63 |
| 11 | Ube2e2 | 0.46 | 1.13E-10 | 2.35E-06 | 0.49 | 0.38 | 0.11 | 1.28 |
| 11 | Cxcl14 | 0.39 | 1.14E-10 | 2.37E-06 | 0.24 | 0.15 | 0.09 | 1.57 |
| 11 | Trim44 | 0.29 | 1.16E-10 | 2.42E-06 | 0.79 | 0.71 | 0.09 | 1.12 |
| 11 | Abhd17c | 0.49 | 1.17E-10 | 2.43E-06 | 0.50 | 0.40 | 0.10 | 1.24 |
| 11 | Cbx6 | 0.53 | 1.17E-10 | 2.44E-06 | 0.37 | 0.27 | 0.10 | 1.37 |
| 11 | Klf6 | 0.65 | 1.18E-10 | 2.46E-06 | 0.32 | 0.23 | 0.09 | 1.39 |
| 11 | Camk2d | 0.56 | 1.18E-10 | 2.46E-06 | 0.29 | 0.19 | 0.10 | 1.49 |
| 11 | Elovl1 | 0.54 | 1.37E-10 | 2.85E-06 | 0.36 | 0.26 | 0.10 | 1.39 |
| 11 | Vmac | 0.85 | 1.40E-10 | 2.91E-06 | 0.16 | 0.09 | 0.07 | 1.73 |
| 11 | Acadm | 0.50 | 1.41E-10 | 2.92E-06 | 0.43 | 0.33 | 0.10 | 1.31 |
| 11 | Arhgap23 | 0.87 | 1.44E-10 | 2.99E-06 | 0.16 | 0.09 | 0.07 | 1.73 |
| 11 | Nt5dc2 | 0.26 | 1.44E-10 | 3.00E-06 | 0.91 | 0.92 | -0.02 | 0.98 |
| 11 | Esco1 | 0.61 | 1.49E-10 | 3.09E-06 | 0.33 | 0.23 | 0.10 | 1.41 |
| 11 | Itpkc | 0.94 | 1.49E-10 | 3.11E-06 | 0.12 | 0.07 | 0.06 | 1.91 |
| 11 | Hps1 | 0.72 | 1.50E-10 | 3.12E-06 | 0.21 | 0.13 | 0.08 | 1.62 |
| 11 | Tnrc6b | 0.32 | 1.65E-10 | 3.44E-06 | 0.77 | 0.71 | 0.06 | 1.09 |
| 11 | Efcab14 | 0.47 | 1.74E-10 | 3.62E-06 | 0.52 | 0.42 | 0.10 | 1.23 |
| 11 | Klf7 | 0.46 | 1.77E-10 | 3.68E-06 | 0.32 | 0.23 | 0.10 | 1.43 |
| 11 | Gm4890 | 0.92 | 1.99E-10 | 4.15E-06 | 0.13 | 0.07 | 0.06 | 1.88 |
| 11 | Aldh4a1 | 0.93 | 2.00E-10 | 4.16E-06 | 0.16 | 0.09 | 0.07 | 1.72 |
| 11 | Lars2 | 0.48 | 2.02E-10 | 4.20E-06 | 0.55 | 0.47 | 0.08 | 1.17 |

**Supplementary Table 1. Marker genes for the original clusters of integrated clustering of E11.5 control and NotchGOF λ-junction cells.**

| | | | | | | | | |
|---|---|---|---|---|---|---|---|---|
| 11 | Acad11 | 0.86 | 2.05E-10 | 4.27E-06 | 0.15 | 0.09 | 0.07 | 1.76 |
| 11 | 2510039O18Rik | 0.56 | 2.06E-10 | 4.28E-06 | 0.33 | 0.23 | 0.10 | 1.41 |
| 11 | Zfp458 | 0.88 | 2.14E-10 | 4.45E-06 | 0.12 | 0.06 | 0.06 | 1.95 |
| 11 | C1galt1c1 | 0.67 | 2.19E-10 | 4.56E-06 | 0.24 | 0.16 | 0.09 | 1.54 |
| 11 | Htatip2 | 1.34 | 2.21E-10 | 4.59E-06 | 0.13 | 0.07 | 0.06 | 1.81 |
| 11 | Prtg | 0.79 | 2.23E-10 | 4.63E-06 | 0.25 | 0.17 | 0.09 | 1.50 |
| 11 | Tmem189 | 0.49 | 2.29E-10 | 4.76E-06 | 0.40 | 0.30 | 0.10 | 1.33 |
| 11 | Kcnj2 | 0.87 | 2.36E-10 | 4.92E-06 | 0.26 | 0.17 | 0.09 | 1.50 |
| 11 | Pank1 | 0.49 | 2.39E-10 | 4.96E-06 | 0.42 | 0.31 | 0.11 | 1.34 |
| 11 | Ttc5 | 0.36 | 2.40E-10 | 5.00E-06 | 0.64 | 0.54 | 0.09 | 1.17 |
| 11 | Ptgr2 | 0.81 | 2.46E-10 | 5.12E-06 | 0.23 | 0.15 | 0.08 | 1.53 |
| 11 | Rabgef1 | 0.69 | 2.50E-10 | 5.21E-06 | 0.25 | 0.16 | 0.08 | 1.51 |
| 11 | Tulp1 | 0.75 | 2.59E-10 | 5.38E-06 | 0.15 | 0.09 | 0.07 | 1.78 |
| 11 | Arl5b | 0.51 | 2.59E-10 | 5.39E-06 | 0.55 | 0.45 | 0.10 | 1.23 |
| 11 | Plscr3 | 0.34 | 2.68E-10 | 5.57E-06 | 0.66 | 0.56 | 0.10 | 1.18 |
| 11 | Tmem170 | 0.83 | 2.68E-10 | 5.57E-06 | 0.16 | 0.09 | 0.07 | 1.73 |
| 11 | Ammecr1l | 0.53 | 2.72E-10 | 5.66E-06 | 0.40 | 0.29 | 0.10 | 1.34 |
| 11 | Fam98c | 0.76 | 2.74E-10 | 5.70E-06 | 0.19 | 0.11 | 0.07 | 1.65 |
| 11 | Lrch1 | 0.95 | 2.82E-10 | 5.86E-06 | 0.14 | 0.08 | 0.06 | 1.79 |
| 11 | Cradd | 0.70 | 2.91E-10 | 6.05E-06 | 0.25 | 0.17 | 0.09 | 1.51 |
| 11 | Vps13a | 0.50 | 2.92E-10 | 6.06E-06 | 0.44 | 0.34 | 0.10 | 1.29 |
| 11 | Selenot | 0.38 | 2.94E-10 | 6.11E-06 | 0.63 | 0.54 | 0.09 | 1.17 |
| 11 | Stim1 | 0.81 | 3.01E-10 | 6.26E-06 | 0.22 | 0.14 | 0.08 | 1.54 |
| 11 | Sptlc1 | 0.52 | 3.04E-10 | 6.32E-06 | 0.39 | 0.29 | 0.10 | 1.35 |
| 11 | BC031181 | 0.27 | 3.06E-10 | 6.36E-06 | 0.79 | 0.73 | 0.06 | 1.08 |
| 11 | Dbndd2 | 0.71 | 3.13E-10 | 6.52E-06 | 0.15 | 0.08 | 0.07 | 1.80 |
| 11 | Nsun3 | 0.79 | 3.16E-10 | 6.57E-06 | 0.14 | 0.07 | 0.06 | 1.85 |
| 11 | Mterf4 | 0.48 | 3.32E-10 | 6.90E-06 | 0.44 | 0.33 | 0.10 | 1.31 |
| 11 | Lcor | 0.45 | 3.35E-10 | 6.96E-06 | 0.38 | 0.28 | 0.10 | 1.36 |
| 11 | Zfp994 | 0.81 | 3.41E-10 | 7.10E-06 | 0.16 | 0.09 | 0.07 | 1.74 |
| 11 | 2610307P16Rik | 0.55 | 3.48E-10 | 7.25E-06 | 0.26 | 0.17 | 0.09 | 1.51 |
| 11 | Fyco1 | 0.67 | 3.58E-10 | 7.45E-06 | 0.25 | 0.16 | 0.08 | 1.52 |
| 11 | Kmt5b | 0.40 | 3.78E-10 | 7.87E-06 | 0.51 | 0.41 | 0.10 | 1.24 |
| 11 | Stambp | 0.57 | 3.80E-10 | 7.90E-06 | 0.32 | 0.22 | 0.10 | 1.43 |
| 11 | Disp1 | 0.79 | 3.90E-10 | 8.12E-06 | 0.18 | 0.11 | 0.07 | 1.68 |
| 11 | Dhrs3 | 0.72 | 3.91E-10 | 8.14E-06 | 0.38 | 0.29 | 0.09 | 1.31 |
| 11 | Sowahc | 0.93 | 4.00E-10 | 8.33E-06 | 0.21 | 0.14 | 0.08 | 1.55 |
| 11 | Zbtb34 | 0.99 | 4.49E-10 | 9.35E-06 | 0.15 | 0.09 | 0.06 | 1.70 |
| 11 | 4931406P16Rik | 0.47 | 4.87E-10 | 1.01E-05 | 0.46 | 0.35 | 0.10 | 1.29 |
| 11 | Zfp729a | 0.64 | 4.99E-10 | 1.04E-05 | 0.22 | 0.14 | 0.08 | 1.56 |
| 11 | Tmem51 | 0.63 | 5.08E-10 | 1.06E-05 | 0.17 | 0.10 | 0.07 | 1.70 |
| 11 | Dlg1 | 0.48 | 5.40E-10 | 1.12E-05 | 0.46 | 0.37 | 0.09 | 1.25 |
| 11 | Sh3bp4 | 0.66 | 5.45E-10 | 1.13E-05 | 0.25 | 0.17 | 0.08 | 1.49 |
| 11 | Dgkz | 0.53 | 5.49E-10 | 1.14E-05 | 0.38 | 0.28 | 0.09 | 1.33 |
| 11 | Wsb2 | 0.55 | 5.72E-10 | 1.19E-05 | 0.41 | 0.31 | 0.10 | 1.32 |
| 11 | Rnf114 | 0.46 | 5.86E-10 | 1.22E-05 | 0.56 | 0.48 | 0.08 | 1.17 |
| 11 | Spty2d1 | 0.66 | 5.99E-10 | 1.25E-05 | 0.34 | 0.25 | 0.09 | 1.36 |
| 11 | Mtpn | 0.26 | 6.01E-10 | 1.25E-05 | 0.85 | 0.81 | 0.04 | 1.05 |
| 11 | Sc5d | 0.48 | 6.41E-10 | 1.33E-05 | 0.48 | 0.38 | 0.10 | 1.26 |
| 11 | Ehd4 | 0.49 | 6.43E-10 | 1.34E-05 | 0.26 | 0.17 | 0.08 | 1.48 |
| 11 | Zadh2 | 0.80 | 6.81E-10 | 1.42E-05 | 0.17 | 0.11 | 0.07 | 1.66 |
| 11 | Kdm7a | 0.65 | 6.87E-10 | 1.43E-05 | 0.44 | 0.35 | 0.09 | 1.25 |
| 11 | Zfp937 | 0.73 | 7.03E-10 | 1.46E-05 | 0.17 | 0.10 | 0.07 | 1.69 |
| 11 | Podxl2 | 0.44 | 7.22E-10 | 1.50E-05 | 0.52 | 0.42 | 0.10 | 1.23 |
| 11 | Arhgef17 | 0.44 | 7.58E-10 | 1.58E-05 | 0.46 | 0.36 | 0.10 | 1.27 |
| 11 | Ascc3 | 0.45 | 7.74E-10 | 1.61E-05 | 0.51 | 0.43 | 0.08 | 1.19 |

**Supplementary Table 1. Marker genes for the original clusters of integrated clustering of E11.5 control and NotchGOF λ-junction cells.**

| | | | | | | | | |
|---|---|---|---|---|---|---|---|---|
| 11 | Gsr | 0.43 | 7.99E-10 | 1.66E-05 | 0.44 | 0.33 | 0.10 | 1.31 |
| 11 | Axin2 | 0.38 | 8.03E-10 | 1.67E-05 | 0.45 | 0.34 | 0.11 | 1.33 |
| 11 | Jrkl | 0.65 | 8.15E-10 | 1.69E-05 | 0.25 | 0.17 | 0.09 | 1.51 |
| 11 | Stxbp5 | 0.92 | 8.25E-10 | 1.72E-05 | 0.14 | 0.08 | 0.06 | 1.77 |
| 11 | Bmf | 0.83 | 8.27E-10 | 1.72E-05 | 0.18 | 0.11 | 0.07 | 1.65 |
| 11 | Pacs2 | 0.61 | 8.31E-10 | 1.73E-05 | 0.25 | 0.17 | 0.09 | 1.51 |
| 11 | Pef1 | 0.46 | 8.48E-10 | 1.76E-05 | 0.45 | 0.36 | 0.09 | 1.26 |
| 11 | Clcn3 | 0.45 | 8.50E-10 | 1.77E-05 | 0.47 | 0.38 | 0.09 | 1.25 |
| 11 | Qars | 0.35 | 8.95E-10 | 1.86E-05 | 0.66 | 0.59 | 0.07 | 1.11 |
| 11 | Rgs3 | 0.77 | 9.12E-10 | 1.90E-05 | 0.26 | 0.18 | 0.08 | 1.46 |
| 11 | Rnpep | 0.46 | 9.22E-10 | 1.92E-05 | 0.52 | 0.43 | 0.09 | 1.21 |
| 11 | Ncaph2 | 0.34 | 9.34E-10 | 1.94E-05 | 0.66 | 0.59 | 0.07 | 1.11 |
| 11 | Otud5 | 0.44 | 9.82E-10 | 2.04E-05 | 0.46 | 0.37 | 0.09 | 1.24 |
| 11 | Reep4 | 0.58 | 9.83E-10 | 2.04E-05 | 0.41 | 0.33 | 0.09 | 1.26 |
| 11 | Cdk2ap2 | 0.36 | 1.00E-09 | 2.08E-05 | 0.55 | 0.46 | 0.09 | 1.20 |
| 11 | Penk | 0.47 | 1.00E-09 | 2.09E-05 | 0.16 | 0.09 | 0.07 | 1.71 |
| 11 | Ccdc181 | 0.96 | 1.02E-09 | 2.11E-05 | 0.12 | 0.06 | 0.06 | 1.86 |
| 11 | Mlycd | 0.76 | 1.04E-09 | 2.17E-05 | 0.22 | 0.14 | 0.08 | 1.54 |
| 11 | Abhd4 | 0.78 | 1.11E-09 | 2.31E-05 | 0.21 | 0.14 | 0.07 | 1.53 |
| 11 | Pld3 | 0.75 | 1.17E-09 | 2.43E-05 | 0.29 | 0.21 | 0.09 | 1.41 |
| 11 | Npc1 | 0.56 | 1.17E-09 | 2.44E-05 | 0.35 | 0.26 | 0.09 | 1.37 |
| 11 | Zmym2 | 0.36 | 1.19E-09 | 2.47E-05 | 0.62 | 0.53 | 0.09 | 1.17 |
| 11 | Hscb | 0.48 | 1.47E-09 | 3.05E-05 | 0.42 | 0.32 | 0.09 | 1.29 |
| 11 | Fgfr1op2 | 0.27 | 1.51E-09 | 3.13E-05 | 0.75 | 0.68 | 0.07 | 1.11 |
| 11 | Thnsl1 | 0.87 | 1.54E-09 | 3.19E-05 | 0.13 | 0.07 | 0.06 | 1.79 |
| 11 | Tinagl1 | 0.97 | 1.71E-09 | 3.56E-05 | 0.11 | 0.06 | 0.05 | 1.90 |
| 11 | Amot | 0.38 | 1.84E-09 | 3.83E-05 | 0.57 | 0.48 | 0.09 | 1.19 |
| 11 | Dph5 | 0.61 | 1.87E-09 | 3.89E-05 | 0.31 | 0.23 | 0.09 | 1.39 |
| 11 | Slc29a4 | 0.75 | 1.93E-09 | 4.01E-05 | 0.17 | 0.10 | 0.07 | 1.65 |
| 11 | Slc38a10 | 0.50 | 1.93E-09 | 4.02E-05 | 0.43 | 0.33 | 0.09 | 1.28 |
| 11 | Itpripl1 | 0.71 | 2.04E-09 | 4.24E-05 | 0.16 | 0.09 | 0.07 | 1.71 |
| 11 | Smim4 | 0.44 | 2.04E-09 | 4.25E-05 | 0.51 | 0.42 | 0.09 | 1.21 |
| 11 | Fzd10os | 0.48 | 2.22E-09 | 4.61E-05 | 0.29 | 0.20 | 0.09 | 1.46 |
| 11 | Dusp11 | 0.28 | 2.30E-09 | 4.78E-05 | 0.72 | 0.66 | 0.06 | 1.09 |
| 11 | Tgfbr1 | 0.37 | 2.39E-09 | 4.96E-05 | 0.64 | 0.55 | 0.08 | 1.15 |
| 11 | Ccnjl | 0.45 | 2.39E-09 | 4.97E-05 | 0.50 | 0.41 | 0.09 | 1.22 |
| 11 | Rida | 0.41 | 2.39E-09 | 4.98E-05 | 0.51 | 0.42 | 0.09 | 1.22 |
| 11 | Col4a2 | 0.36 | 2.42E-09 | 5.04E-05 | 0.52 | 0.42 | 0.10 | 1.23 |
| 11 | Spaca6 | 0.51 | 2.48E-09 | 5.16E-05 | 0.42 | 0.33 | 0.10 | 1.30 |
| 11 | Rfx3 | 0.61 | 2.48E-09 | 5.16E-05 | 0.30 | 0.21 | 0.09 | 1.41 |
| 11 | Pigt | 0.42 | 2.50E-09 | 5.21E-05 | 0.52 | 0.43 | 0.09 | 1.22 |
| 11 | Tmsb15b1 | 0.56 | 2.97E-09 | 6.18E-05 | 0.30 | 0.21 | 0.09 | 1.41 |
| 11 | Znrf2 | 0.59 | 3.03E-09 | 6.31E-05 | 0.27 | 0.18 | 0.08 | 1.45 |
| 11 | Ro60 | 0.32 | 3.20E-09 | 6.66E-05 | 0.65 | 0.57 | 0.08 | 1.14 |
| 11 | Tm9sf2 | 0.28 | 3.29E-09 | 6.84E-05 | 0.69 | 0.62 | 0.08 | 1.13 |
| 11 | Irf2bp2 | 0.43 | 3.30E-09 | 6.87E-05 | 0.45 | 0.35 | 0.11 | 1.31 |
| 11 | Pole3 | 0.27 | 3.39E-09 | 7.05E-05 | 0.78 | 0.76 | 0.02 | 1.03 |
| 11 | AW146154 | 0.85 | 3.44E-09 | 7.15E-05 | 0.15 | 0.09 | 0.06 | 1.72 |
| 11 | Sptan1 | 0.35 | 3.60E-09 | 7.49E-05 | 0.66 | 0.56 | 0.10 | 1.17 |
| 11 | Adam10 | 0.35 | 3.65E-09 | 7.59E-05 | 0.61 | 0.53 | 0.07 | 1.14 |
| 11 | Mterf1a | 0.82 | 3.65E-09 | 7.59E-05 | 0.14 | 0.08 | 0.06 | 1.73 |
| 11 | AW549877 | 0.57 | 3.86E-09 | 8.03E-05 | 0.30 | 0.21 | 0.09 | 1.42 |
| 11 | Chd6 | 0.28 | 3.92E-09 | 8.15E-05 | 0.78 | 0.73 | 0.05 | 1.07 |
| 11 | Tmod3 | 0.31 | 4.00E-09 | 8.31E-05 | 0.71 | 0.64 | 0.07 | 1.12 |
| 11 | Pigv | 0.85 | 4.25E-09 | 8.84E-05 | 0.16 | 0.09 | 0.06 | 1.68 |
| 11 | Nsg1 | 0.32 | 4.38E-09 | 9.10E-05 | 0.75 | 0.67 | 0.08 | 1.12 |

**Supplementary Table 1. Marker genes for the original clusters of integrated clustering of E11.5 control and NotchGOF λ-junction cells.**

| | | | | | | | | |
|---|---|---|---|---|---|---|---|---|
| 11 | Mmp15 | 0.71 | 4.71E-09 | 9.79E-05 | 0.15 | 0.09 | 0.06 | 1.71 |
| 11 | Atg4d | 0.64 | 4.72E-09 | 9.81E-05 | 0.29 | 0.20 | 0.08 | 1.40 |
| 11 | Fam241b | 0.55 | 4.82E-09 | 1.00E-04 | 0.25 | 0.17 | 0.08 | 1.49 |
| 11 | Stk11 | 0.36 | 5.16E-09 | 1.07E-04 | 0.57 | 0.48 | 0.09 | 1.18 |
| 11 | Itsn2 | 0.52 | 5.20E-09 | 1.08E-04 | 0.27 | 0.19 | 0.08 | 1.44 |
| 11 | Rcan3 | 0.76 | 5.26E-09 | 1.09E-04 | 0.13 | 0.08 | 0.06 | 1.76 |
| 11 | Glmp | 0.37 | 5.75E-09 | 1.20E-04 | 0.57 | 0.47 | 0.10 | 1.21 |
| 11 | Cd47 | 0.34 | 5.78E-09 | 1.20E-04 | 0.54 | 0.44 | 0.11 | 1.24 |
| 11 | Rps6ka2 | 0.87 | 5.94E-09 | 1.24E-04 | 0.14 | 0.08 | 0.06 | 1.73 |
| 11 | Galc | 0.64 | 6.14E-09 | 1.28E-04 | 0.25 | 0.17 | 0.08 | 1.44 |
| 11 | Fam8a1 | 0.50 | 6.35E-09 | 1.32E-04 | 0.36 | 0.27 | 0.09 | 1.34 |
| 11 | Dbt | 0.50 | 6.50E-09 | 1.35E-04 | 0.38 | 0.29 | 0.09 | 1.31 |
| 11 | Cdk19 | 0.51 | 6.78E-09 | 1.41E-04 | 0.40 | 0.31 | 0.09 | 1.29 |
| 11 | Hyal2 | 0.35 | 6.92E-09 | 1.44E-04 | 0.61 | 0.53 | 0.08 | 1.16 |
| 11 | Fry | 0.81 | 6.99E-09 | 1.45E-04 | 0.10 | 0.05 | 0.05 | 1.91 |
| 11 | Slc30a4 | 0.84 | 7.42E-09 | 1.54E-04 | 0.20 | 0.13 | 0.07 | 1.55 |
| 11 | Tob2 | 0.47 | 7.46E-09 | 1.55E-04 | 0.33 | 0.24 | 0.09 | 1.38 |
| 11 | Trim25 | 0.94 | 7.49E-09 | 1.56E-04 | 0.11 | 0.06 | 0.05 | 1.85 |
| 11 | Osbpl5 | 0.90 | 7.69E-09 | 1.60E-04 | 0.13 | 0.08 | 0.06 | 1.73 |
| 11 | Gorasp2 | 0.38 | 7.70E-09 | 1.60E-04 | 0.60 | 0.53 | 0.07 | 1.13 |
| 11 | Fbxo10 | 0.76 | 7.72E-09 | 1.61E-04 | 0.13 | 0.08 | 0.06 | 1.74 |
| 11 | Fndc3b | 0.57 | 7.74E-09 | 1.61E-04 | 0.33 | 0.24 | 0.08 | 1.34 |
| 11 | Ahdc1 | 0.66 | 7.81E-09 | 1.62E-04 | 0.23 | 0.16 | 0.08 | 1.49 |
| 11 | 4732440D04Rik | 0.84 | 8.27E-09 | 1.72E-04 | 0.16 | 0.10 | 0.06 | 1.64 |
| 11 | Pex11g | 0.78 | 8.40E-09 | 1.75E-04 | 0.15 | 0.09 | 0.06 | 1.68 |
| 11 | Fgd4 | 0.66 | 8.47E-09 | 1.76E-04 | 0.15 | 0.09 | 0.06 | 1.67 |
| 11 | Dctn3 | 0.26 | 9.04E-09 | 1.88E-04 | 0.82 | 0.78 | 0.04 | 1.05 |
| 11 | Phyh | 0.50 | 9.23E-09 | 1.92E-04 | 0.38 | 0.29 | 0.09 | 1.32 |
| 11 | Atg13 | 0.62 | 9.57E-09 | 1.99E-04 | 0.22 | 0.15 | 0.08 | 1.51 |
| 11 | Gabarapl1 | 0.44 | 9.67E-09 | 2.01E-04 | 0.42 | 0.33 | 0.09 | 1.28 |
| 11 | Ssc4d | 1.09 | 9.78E-09 | 2.03E-04 | 0.10 | 0.05 | 0.05 | 1.87 |
| 11 | Map4k5 | 0.50 | 9.96E-09 | 2.07E-04 | 0.39 | 0.31 | 0.09 | 1.28 |
| 11 | Fbxo25 | 0.64 | 1.01E-08 | 2.11E-04 | 0.23 | 0.15 | 0.08 | 1.50 |
| 11 | Ndfip2 | 0.37 | 1.14E-08 | 2.37E-04 | 0.59 | 0.51 | 0.08 | 1.16 |
| 11 | Nxt2 | 0.77 | 1.14E-08 | 2.38E-04 | 0.13 | 0.07 | 0.06 | 1.74 |
| 11 | Atxn7 | 0.54 | 1.15E-08 | 2.39E-04 | 0.30 | 0.21 | 0.08 | 1.40 |
| 11 | Pip4k2c | 0.61 | 1.22E-08 | 2.53E-04 | 0.26 | 0.18 | 0.08 | 1.43 |
| 11 | Ppp1r13b | 0.62 | 1.22E-08 | 2.53E-04 | 0.14 | 0.08 | 0.06 | 1.73 |
| 11 | Rnf6 | 0.47 | 1.31E-08 | 2.72E-04 | 0.44 | 0.35 | 0.09 | 1.25 |
| 11 | Mif4gd | 0.57 | 1.39E-08 | 2.90E-04 | 0.24 | 0.17 | 0.08 | 1.46 |
| 11 | Asl | 0.61 | 1.39E-08 | 2.90E-04 | 0.22 | 0.15 | 0.07 | 1.50 |
| 11 | Brwd3 | 0.43 | 1.48E-08 | 3.08E-04 | 0.54 | 0.46 | 0.08 | 1.17 |
| 11 | Zfp593 | 0.35 | 1.53E-08 | 3.18E-04 | 0.55 | 0.47 | 0.08 | 1.17 |
| 11 | Etfa | 0.27 | 1.61E-08 | 3.34E-04 | 0.77 | 0.74 | 0.03 | 1.05 |
| 11 | Poli | 0.92 | 1.66E-08 | 3.45E-04 | 0.11 | 0.06 | 0.05 | 1.83 |
| 11 | Kantr | 0.64 | 1.66E-08 | 3.46E-04 | 0.25 | 0.17 | 0.08 | 1.47 |
| 11 | Gm26930 | 0.84 | 1.69E-08 | 3.51E-04 | 0.15 | 0.09 | 0.06 | 1.67 |
| 11 | Decr2 | 0.67 | 1.70E-08 | 3.53E-04 | 0.22 | 0.15 | 0.07 | 1.49 |
| 11 | Hif1a | 0.30 | 1.73E-08 | 3.59E-04 | 0.81 | 0.75 | 0.06 | 1.08 |
| 11 | Atf6b | 0.34 | 1.73E-08 | 3.60E-04 | 0.50 | 0.40 | 0.10 | 1.25 |
| 11 | Cnksr2 | 0.55 | 1.77E-08 | 3.69E-04 | 0.41 | 0.32 | 0.08 | 1.26 |
| 11 | Fubp3 | 0.38 | 1.82E-08 | 3.79E-04 | 0.54 | 0.45 | 0.09 | 1.20 |
| 11 | Mettl23 | 0.29 | 1.84E-08 | 3.84E-04 | 0.68 | 0.62 | 0.06 | 1.10 |
| 11 | Slk | 0.36 | 1.87E-08 | 3.88E-04 | 0.61 | 0.53 | 0.08 | 1.15 |
| 11 | Apcdd1 | 0.37 | 1.87E-08 | 3.88E-04 | 0.30 | 0.22 | 0.09 | 1.41 |
| 11 | Apba2 | 0.85 | 1.88E-08 | 3.90E-04 | 0.11 | 0.06 | 0.05 | 1.83 |

**Supplementary Table 1. Marker genes for the original clusters of integrated clustering of E11.5 control and NotchGOF λ-junction cells.**

| | | | | | | | | |
|---|---|---|---|---|---|---|---|---|
| 11 | Naxd | 0.31 | 1.94E-08 | 4.03E-04 | 0.70 | 0.64 | 0.06 | 1.10 |
| 11 | Oxsr1 | 0.51 | 1.96E-08 | 4.08E-04 | 0.41 | 0.32 | 0.09 | 1.28 |
| 11 | Tmub2 | 0.63 | 1.98E-08 | 4.11E-04 | 0.31 | 0.23 | 0.08 | 1.33 |
| 11 | Btbd9 | 0.45 | 2.01E-08 | 4.18E-04 | 0.38 | 0.29 | 0.09 | 1.30 |
| 11 | Nr1d2 | 0.96 | 2.02E-08 | 4.20E-04 | 0.11 | 0.06 | 0.05 | 1.83 |
| 11 | Ccdc51 | 0.73 | 2.11E-08 | 4.38E-04 | 0.18 | 0.12 | 0.07 | 1.56 |
| 11 | Gm43445 | 0.82 | 2.12E-08 | 4.40E-04 | 0.15 | 0.09 | 0.06 | 1.67 |
| 11 | Med14 | 0.56 | 2.19E-08 | 4.54E-04 | 0.34 | 0.26 | 0.08 | 1.31 |
| 11 | Tmem38a | 0.74 | 2.19E-08 | 4.55E-04 | 0.16 | 0.10 | 0.06 | 1.62 |
| 11 | Phactr4 | 0.36 | 2.20E-08 | 4.58E-04 | 0.57 | 0.49 | 0.08 | 1.17 |
| 11 | Brms1 | 0.46 | 2.21E-08 | 4.61E-04 | 0.43 | 0.35 | 0.09 | 1.26 |
| 11 | 0610009L18Rik | 0.82 | 2.27E-08 | 4.72E-04 | 0.13 | 0.07 | 0.05 | 1.74 |
| 11 | Nudt11 | 0.56 | 2.27E-08 | 4.72E-04 | 0.26 | 0.18 | 0.08 | 1.44 |
| 11 | Dck | 0.36 | 2.27E-08 | 4.73E-04 | 0.54 | 0.46 | 0.08 | 1.18 |
| 11 | Ulk1 | 0.57 | 2.45E-08 | 5.09E-04 | 0.35 | 0.27 | 0.08 | 1.31 |
| 11 | Klhl7 | 0.30 | 2.46E-08 | 5.12E-04 | 0.60 | 0.52 | 0.08 | 1.16 |
| 11 | Ext1 | 0.37 | 2.55E-08 | 5.31E-04 | 0.50 | 0.41 | 0.09 | 1.22 |
| 11 | Atp6v0d1 | 0.38 | 2.66E-08 | 5.54E-04 | 0.60 | 0.54 | 0.07 | 1.13 |
| 11 | Araf | 0.42 | 2.67E-08 | 5.55E-04 | 0.49 | 0.40 | 0.09 | 1.21 |
| 11 | Kdm6a | 0.42 | 2.76E-08 | 5.74E-04 | 0.51 | 0.42 | 0.08 | 1.20 |
| 11 | Frmd8 | 0.54 | 2.97E-08 | 6.18E-04 | 0.28 | 0.20 | 0.08 | 1.40 |
| 11 | Rbbp9 | 0.36 | 3.09E-08 | 6.42E-04 | 0.48 | 0.39 | 0.09 | 1.23 |
| 11 | Klhl36 | 0.83 | 3.10E-08 | 6.45E-04 | 0.12 | 0.07 | 0.05 | 1.76 |
| 11 | E330009J07Rik | 0.66 | 3.20E-08 | 6.66E-04 | 0.18 | 0.12 | 0.07 | 1.56 |
| 11 | Hivep2 | 0.74 | 3.33E-08 | 6.92E-04 | 0.13 | 0.08 | 0.05 | 1.72 |
| 11 | Mapk12 | 0.34 | 3.35E-08 | 6.96E-04 | 0.18 | 0.11 | 0.07 | 1.60 |
| 11 | Fmn2 | 0.95 | 3.48E-08 | 7.24E-04 | 0.12 | 0.07 | 0.05 | 1.74 |
| 11 | Smim8 | 0.41 | 3.52E-08 | 7.33E-04 | 0.52 | 0.42 | 0.09 | 1.22 |
| 11 | 1700001O22Rik | 0.82 | 3.56E-08 | 7.41E-04 | 0.12 | 0.07 | 0.05 | 1.76 |
| 11 | Tmed4 | 0.29 | 3.65E-08 | 7.58E-04 | 0.65 | 0.56 | 0.09 | 1.16 |
| 11 | Slc25a13 | 0.39 | 3.69E-08 | 7.68E-04 | 0.53 | 0.44 | 0.08 | 1.19 |
| 11 | Dennd4c | 0.68 | 3.70E-08 | 7.70E-04 | 0.24 | 0.17 | 0.07 | 1.44 |
| 11 | Siae | 0.88 | 3.71E-08 | 7.72E-04 | 0.12 | 0.07 | 0.05 | 1.75 |
| 11 | Cyfip2 | 0.59 | 3.81E-08 | 7.92E-04 | 0.15 | 0.09 | 0.06 | 1.66 |
| 11 | Zbtb8os | 0.34 | 3.86E-08 | 8.04E-04 | 0.58 | 0.50 | 0.08 | 1.16 |
| 11 | Tgoln1 | 0.52 | 3.87E-08 | 8.05E-04 | 0.38 | 0.29 | 0.09 | 1.30 |
| 11 | Tnrc18 | 0.30 | 4.10E-08 | 8.53E-04 | 0.64 | 0.56 | 0.08 | 1.15 |
| 11 | Etfbkmt | 0.83 | 4.12E-08 | 8.57E-04 | 0.12 | 0.07 | 0.05 | 1.73 |
| 11 | Tnfrsf19 | 0.53 | 4.13E-08 | 8.59E-04 | 0.22 | 0.15 | 0.07 | 1.48 |
| 11 | Nit1 | 0.36 | 4.21E-08 | 8.75E-04 | 0.44 | 0.35 | 0.09 | 1.27 |
| 11 | 4933434E20Rik | 0.34 | 4.24E-08 | 8.82E-04 | 0.58 | 0.49 | 0.09 | 1.18 |
| 11 | Esrra | 0.66 | 4.25E-08 | 8.84E-04 | 0.19 | 0.12 | 0.07 | 1.54 |
| 11 | Fam126b | 0.53 | 4.41E-08 | 9.16E-04 | 0.27 | 0.19 | 0.08 | 1.42 |
| 11 | Baiap2l1 | 0.76 | 4.42E-08 | 9.19E-04 | 0.11 | 0.06 | 0.05 | 1.79 |
| 11 | Olfm1 | 0.39 | 4.49E-08 | 9.35E-04 | 0.31 | 0.23 | 0.09 | 1.38 |
| 11 | Rxrb | 0.36 | 4.54E-08 | 9.43E-04 | 0.50 | 0.41 | 0.09 | 1.21 |
| 11 | Ski | 0.40 | 4.59E-08 | 9.56E-04 | 0.50 | 0.41 | 0.09 | 1.23 |
| 11 | Prss36 | 0.61 | 4.60E-08 | 9.57E-04 | 0.15 | 0.09 | 0.06 | 1.67 |
| 11 | Pfn2 | 0.38 | 4.66E-08 | 9.68E-04 | 0.73 | 0.66 | 0.07 | 1.10 |
| 11 | Vps37d | 0.49 | 4.77E-08 | 9.91E-04 | 0.37 | 0.29 | 0.08 | 1.28 |
| 11 | Dyrk1b | 0.65 | 4.77E-08 | 9.92E-04 | 0.21 | 0.14 | 0.07 | 1.49 |
| 11 | Tox4 | 0.27 | 4.99E-08 | 1.04E-03 | 0.61 | 0.52 | 0.09 | 1.17 |
| 11 | Prkx | 0.64 | 5.07E-08 | 1.05E-03 | 0.22 | 0.15 | 0.07 | 1.46 |
| 11 | Jkamp | 0.32 | 5.13E-08 | 1.07E-03 | 0.62 | 0.53 | 0.09 | 1.17 |
| 11 | Amfr | 0.38 | 5.21E-08 | 1.08E-03 | 0.45 | 0.36 | 0.09 | 1.25 |
| 11 | Fmnl2 | 0.56 | 5.29E-08 | 1.10E-03 | 0.52 | 0.44 | 0.08 | 1.17 |

**Supplementary Table 1. Marker genes for the original clusters of integrated clustering of E11.5 control and NotchGOF λ-junction cells.**

| | | | | | | | | |
|---|---|---|---|---|---|---|---|---|
| 11 | Slc39a10 | 0.29 | 5.35E-08 | 1.11E-03 | 0.74 | 0.68 | 0.06 | 1.08 |
| 11 | Bid | 0.62 | 5.38E-08 | 1.12E-03 | 0.19 | 0.12 | 0.07 | 1.54 |
| 11 | Zmym5 | 0.34 | 5.41E-08 | 1.12E-03 | 0.67 | 0.61 | 0.05 | 1.08 |
| 11 | B9d2 | 0.62 | 5.61E-08 | 1.17E-03 | 0.35 | 0.27 | 0.08 | 1.30 |
| 11 | Gnl3l | 0.46 | 5.68E-08 | 1.18E-03 | 0.40 | 0.32 | 0.08 | 1.26 |
| 11 | Unc50 | 0.33 | 5.74E-08 | 1.19E-03 | 0.64 | 0.56 | 0.08 | 1.14 |
| 11 | Sil1 | 0.65 | 5.90E-08 | 1.23E-03 | 0.18 | 0.11 | 0.06 | 1.58 |
| 11 | Zfyve1 | 0.71 | 6.24E-08 | 1.30E-03 | 0.22 | 0.15 | 0.07 | 1.46 |
| 11 | Rab5a | 0.28 | 6.34E-08 | 1.32E-03 | 0.70 | 0.63 | 0.06 | 1.10 |
| 11 | Rrp36 | 0.46 | 6.36E-08 | 1.32E-03 | 0.40 | 0.32 | 0.09 | 1.27 |
| 11 | Zfp608 | 0.28 | 6.46E-08 | 1.34E-03 | 0.63 | 0.53 | 0.09 | 1.17 |
| 11 | Pip4p1 | 0.36 | 6.53E-08 | 1.36E-03 | 0.45 | 0.36 | 0.09 | 1.26 |
| 11 | Map3k11 | 0.60 | 6.62E-08 | 1.38E-03 | 0.18 | 0.12 | 0.06 | 1.56 |
| 11 | E2f8 | 0.69 | 6.71E-08 | 1.40E-03 | 0.28 | 0.20 | 0.07 | 1.36 |
| 11 | Lrrc56 | 0.66 | 6.96E-08 | 1.45E-03 | 0.16 | 0.10 | 0.06 | 1.63 |
| 11 | Zdhhc20 | 0.31 | 6.99E-08 | 1.45E-03 | 0.66 | 0.58 | 0.08 | 1.13 |
| 11 | Golga1 | 0.48 | 7.02E-08 | 1.46E-03 | 0.37 | 0.28 | 0.09 | 1.30 |
| 11 | Pnpla6 | 0.91 | 7.15E-08 | 1.49E-03 | 0.14 | 0.09 | 0.06 | 1.63 |
| 11 | Fbxo44 | 0.72 | 7.61E-08 | 1.58E-03 | 0.15 | 0.09 | 0.06 | 1.62 |
| 11 | Fggy | 0.82 | 7.80E-08 | 1.62E-03 | 0.13 | 0.08 | 0.05 | 1.67 |
| 11 | Msi2 | 0.29 | 7.86E-08 | 1.64E-03 | 0.74 | 0.68 | 0.06 | 1.09 |
| 11 | Klhl13 | 0.32 | 7.95E-08 | 1.65E-03 | 0.36 | 0.27 | 0.09 | 1.35 |
| 11 | Asah1 | 0.35 | 7.97E-08 | 1.66E-03 | 0.46 | 0.37 | 0.09 | 1.24 |
| 11 | Trappc12 | 0.47 | 8.07E-08 | 1.68E-03 | 0.29 | 0.21 | 0.08 | 1.39 |
| 11 | Twsg1 | 0.40 | 8.25E-08 | 1.71E-03 | 0.57 | 0.49 | 0.08 | 1.17 |
| 11 | Parn | 0.48 | 8.86E-08 | 1.84E-03 | 0.34 | 0.26 | 0.08 | 1.31 |
| 11 | Dusp5 | 0.69 | 8.93E-08 | 1.86E-03 | 0.18 | 0.12 | 0.06 | 1.54 |
| 11 | Slu7 | 0.37 | 9.33E-08 | 1.94E-03 | 0.58 | 0.49 | 0.08 | 1.17 |
| 11 | Mier2 | 0.44 | 9.59E-08 | 1.99E-03 | 0.28 | 0.20 | 0.08 | 1.41 |
| 11 | Bin3 | 0.51 | 9.61E-08 | 2.00E-03 | 0.37 | 0.29 | 0.09 | 1.30 |
| 11 | Tasor | 0.36 | 9.69E-08 | 2.01E-03 | 0.56 | 0.48 | 0.08 | 1.17 |
| 11 | Rela | 0.47 | 1.03E-07 | 2.13E-03 | 0.33 | 0.24 | 0.08 | 1.34 |
| 11 | Pttg1ip | 0.36 | 1.10E-07 | 2.29E-03 | 0.48 | 0.39 | 0.09 | 1.23 |
| 11 | Sipa1l1 | 0.30 | 1.13E-07 | 2.35E-03 | 0.59 | 0.51 | 0.08 | 1.15 |
| 11 | Adipor1 | 0.29 | 1.13E-07 | 2.36E-03 | 0.64 | 0.56 | 0.08 | 1.13 |
| 11 | Zfp316 | 0.58 | 1.20E-07 | 2.50E-03 | 0.21 | 0.14 | 0.07 | 1.49 |
| 11 | Shroom2 | 0.65 | 1.21E-07 | 2.51E-03 | 0.13 | 0.08 | 0.05 | 1.69 |
| 11 | Stk4 | 0.42 | 1.27E-07 | 2.63E-03 | 0.44 | 0.36 | 0.09 | 1.24 |
| 11 | Abcd3 | 0.38 | 1.28E-07 | 2.65E-03 | 0.50 | 0.42 | 0.08 | 1.20 |
| 11 | Rnf103 | 0.61 | 1.30E-07 | 2.71E-03 | 0.25 | 0.18 | 0.07 | 1.41 |
| 11 | Zc3h12c | 0.68 | 1.33E-07 | 2.76E-03 | 0.11 | 0.06 | 0.05 | 1.79 |
| 11 | Znrd1as | 0.59 | 1.39E-07 | 2.88E-03 | 0.19 | 0.13 | 0.06 | 1.50 |
| 11 | Hspa4l | 0.55 | 1.41E-07 | 2.94E-03 | 0.17 | 0.11 | 0.06 | 1.55 |
| 11 | Rabac1 | 0.30 | 1.41E-07 | 2.94E-03 | 0.63 | 0.55 | 0.08 | 1.14 |
| 11 | Zfp536 | 0.58 | 1.42E-07 | 2.94E-03 | 0.16 | 0.10 | 0.06 | 1.59 |
| 11 | Pxmp4 | 0.47 | 1.43E-07 | 2.98E-03 | 0.25 | 0.18 | 0.07 | 1.41 |
| 11 | Sh3bp1 | 0.42 | 1.43E-07 | 2.98E-03 | 0.37 | 0.29 | 0.09 | 1.30 |
| 11 | Fut8 | 0.48 | 1.45E-07 | 3.02E-03 | 0.37 | 0.29 | 0.08 | 1.28 |
| 11 | Slc37a3 | 0.49 | 1.47E-07 | 3.06E-03 | 0.34 | 0.26 | 0.08 | 1.30 |
| 11 | Myo9b | 0.54 | 1.52E-07 | 3.15E-03 | 0.30 | 0.23 | 0.07 | 1.32 |
| 11 | Jade1 | 0.58 | 1.57E-07 | 3.26E-03 | 0.24 | 0.17 | 0.07 | 1.41 |
| 11 | Bok | 0.52 | 1.60E-07 | 3.32E-03 | 0.28 | 0.20 | 0.08 | 1.39 |
| 11 | Clcn4 | 0.43 | 1.60E-07 | 3.33E-03 | 0.38 | 0.30 | 0.08 | 1.26 |
| 11 | Samd4b | 0.29 | 1.69E-07 | 3.52E-03 | 0.59 | 0.50 | 0.09 | 1.18 |
| 11 | Atp2c1 | 0.39 | 1.76E-07 | 3.67E-03 | 0.48 | 0.40 | 0.08 | 1.21 |
| 11 | Glod4 | 0.25 | 1.77E-07 | 3.67E-03 | 0.74 | 0.69 | 0.05 | 1.07 |

**Supplementary Table 1. Marker genes for the original clusters of integrated clustering of E11.5 control and NotchGOF λ-junction cells.**

| | | | | | | | | |
|---|---|---|---|---|---|---|---|---|
| 11 | D030056L22Rik | 0.40 | 1.78E-07 | 3.70E-03 | 0.55 | 0.48 | 0.07 | 1.14 |
| 11 | Sbk1 | 0.30 | 1.78E-07 | 3.71E-03 | 0.60 | 0.52 | 0.08 | 1.15 |
| 11 | Etfrf1 | 0.51 | 1.80E-07 | 3.75E-03 | 0.29 | 0.22 | 0.07 | 1.34 |
| 11 | Exosc5 | 0.30 | 1.82E-07 | 3.77E-03 | 0.64 | 0.58 | 0.06 | 1.10 |
| 11 | Pign | 0.67 | 1.82E-07 | 3.79E-03 | 0.21 | 0.14 | 0.07 | 1.45 |
| 11 | Marchf2 | 0.37 | 1.83E-07 | 3.80E-03 | 0.58 | 0.51 | 0.07 | 1.13 |
| 11 | Tprgl | 0.26 | 1.83E-07 | 3.81E-03 | 0.67 | 0.60 | 0.06 | 1.11 |
| 11 | Abhd14b | 0.77 | 1.86E-07 | 3.87E-03 | 0.11 | 0.06 | 0.05 | 1.78 |
| 11 | Sesn3 | 0.29 | 1.92E-07 | 3.99E-03 | 0.66 | 0.59 | 0.07 | 1.12 |
| 11 | Slc12a6 | 0.66 | 1.92E-07 | 3.99E-03 | 0.20 | 0.13 | 0.07 | 1.49 |
| 11 | Mvb12a | 0.30 | 1.93E-07 | 4.02E-03 | 0.54 | 0.46 | 0.09 | 1.19 |
| 11 | Ak3 | 0.37 | 1.93E-07 | 4.02E-03 | 0.47 | 0.38 | 0.09 | 1.23 |
| 11 | Kif16b | 0.52 | 1.94E-07 | 4.03E-03 | 0.22 | 0.15 | 0.07 | 1.45 |
| 11 | Pisd | 0.45 | 1.98E-07 | 4.12E-03 | 0.29 | 0.21 | 0.08 | 1.38 |
| 11 | Asxl2 | 0.40 | 1.99E-07 | 4.14E-03 | 0.49 | 0.41 | 0.08 | 1.19 |
| 11 | Yif1a | 0.31 | 2.05E-07 | 4.27E-03 | 0.56 | 0.48 | 0.08 | 1.17 |
| 11 | Numb | 0.53 | 2.06E-07 | 4.29E-03 | 0.40 | 0.32 | 0.08 | 1.25 |
| 11 | Ogfod2 | 0.70 | 2.14E-07 | 4.45E-03 | 0.16 | 0.10 | 0.06 | 1.58 |
| 11 | Lrrc61 | 0.71 | 2.27E-07 | 4.73E-03 | 0.15 | 0.10 | 0.06 | 1.58 |
| 11 | Numbl | 0.49 | 2.27E-07 | 4.73E-03 | 0.20 | 0.13 | 0.07 | 1.51 |
| 11 | B230354K17Rik | 0.41 | 2.31E-07 | 4.80E-03 | 0.43 | 0.36 | 0.08 | 1.22 |
| 11 | Trak1 | 0.66 | 2.34E-07 | 4.87E-03 | 0.18 | 0.12 | 0.06 | 1.52 |
| 11 | Pard3b | 0.59 | 2.39E-07 | 4.97E-03 | 0.18 | 0.12 | 0.06 | 1.53 |
| 11 | CAAA01118383.1 | 0.49 | 2.41E-07 | 5.01E-03 | 0.31 | 0.23 | 0.08 | 1.33 |
| 11 | Flad1 | 0.71 | 2.41E-07 | 5.02E-03 | 0.16 | 0.10 | 0.06 | 1.56 |
| 11 | Taz | 0.45 | 2.44E-07 | 5.08E-03 | 0.48 | 0.40 | 0.08 | 1.20 |
| 11 | Rdh14 | 0.41 | 2.47E-07 | 5.14E-03 | 0.36 | 0.28 | 0.08 | 1.30 |
| 11 | Nfe2l1 | 0.36 | 2.50E-07 | 5.19E-03 | 0.60 | 0.53 | 0.07 | 1.13 |
| 11 | Xylt1 | 0.78 | 2.55E-07 | 5.31E-03 | 0.11 | 0.06 | 0.05 | 1.75 |
| 11 | Cybc1 | 0.49 | 2.57E-07 | 5.35E-03 | 0.30 | 0.22 | 0.08 | 1.35 |
| 11 | 1190007I07Rik | 0.46 | 2.62E-07 | 5.44E-03 | 0.36 | 0.28 | 0.08 | 1.29 |
| 11 | Rab5b | 0.45 | 2.65E-07 | 5.50E-03 | 0.35 | 0.27 | 0.08 | 1.29 |
| 11 | A430105J06Rik | 0.92 | 2.67E-07 | 5.56E-03 | 0.10 | 0.06 | 0.04 | 1.79 |
| 11 | Usp6nl | 0.43 | 2.76E-07 | 5.74E-03 | 0.30 | 0.22 | 0.08 | 1.35 |
| 11 | Ppp6r1 | 0.41 | 2.92E-07 | 6.08E-03 | 0.33 | 0.25 | 0.08 | 1.32 |
| 11 | Sft2d1 | 0.51 | 3.00E-07 | 6.23E-03 | 0.30 | 0.22 | 0.08 | 1.34 |
| 11 | Sfi1 | 0.77 | 3.03E-07 | 6.31E-03 | 0.12 | 0.07 | 0.05 | 1.71 |
| 11 | Chst3 | 1.01 | 3.04E-07 | 6.33E-03 | 0.11 | 0.06 | 0.05 | 1.73 |
| 11 | Tst | 0.47 | 3.05E-07 | 6.35E-03 | 0.19 | 0.13 | 0.06 | 1.51 |
| 11 | Top3b | 0.39 | 3.36E-07 | 6.98E-03 | 0.42 | 0.34 | 0.08 | 1.24 |
| 11 | Tdrd3 | 0.39 | 3.38E-07 | 7.03E-03 | 0.43 | 0.34 | 0.09 | 1.26 |
| 11 | Rad9b | 0.69 | 3.45E-07 | 7.17E-03 | 0.13 | 0.08 | 0.05 | 1.63 |
| 11 | Nudt7 | 0.61 | 3.47E-07 | 7.21E-03 | 0.12 | 0.07 | 0.05 | 1.70 |
| 11 | Pdcl | 0.34 | 3.49E-07 | 7.26E-03 | 0.52 | 0.43 | 0.09 | 1.20 |
| 11 | Necap2 | 0.41 | 3.49E-07 | 7.26E-03 | 0.47 | 0.39 | 0.08 | 1.20 |
| 11 | Zmynd8 | 0.31 | 3.53E-07 | 7.35E-03 | 0.66 | 0.60 | 0.06 | 1.09 |
| 11 | Atl2 | 0.47 | 3.56E-07 | 7.41E-03 | 0.33 | 0.25 | 0.08 | 1.31 |
| 11 | Wdr31 | 0.53 | 3.57E-07 | 7.43E-03 | 0.14 | 0.09 | 0.06 | 1.63 |
| 11 | Sun1 | 0.42 | 3.58E-07 | 7.44E-03 | 0.36 | 0.27 | 0.09 | 1.31 |
| 11 | Pygb | 0.43 | 3.73E-07 | 7.76E-03 | 0.31 | 0.23 | 0.08 | 1.33 |
| 11 | Mbd6 | 0.45 | 3.92E-07 | 8.14E-03 | 0.31 | 0.23 | 0.08 | 1.34 |
| 11 | Man2c1 | 0.59 | 3.97E-07 | 8.25E-03 | 0.16 | 0.11 | 0.06 | 1.54 |
| 11 | Gm4924 | 0.62 | 3.98E-07 | 8.28E-03 | 0.20 | 0.14 | 0.06 | 1.47 |
| 11 | Letm2 | 0.50 | 4.02E-07 | 8.36E-03 | 0.23 | 0.16 | 0.07 | 1.44 |
| 11 | Rabggta | 0.70 | 4.06E-07 | 8.45E-03 | 0.19 | 0.13 | 0.06 | 1.47 |
| 11 | 2810408A11Rik | 0.53 | 4.09E-07 | 8.50E-03 | 0.21 | 0.15 | 0.07 | 1.44 |

**Supplementary Table 1. Marker genes for the original clusters of integrated clustering of E11.5 control and NotchGOF λ-junction cells.**

| | | | | | | | | |
|---|---|---|---|---|---|---|---|---|
| 11 | Svil | 0.37 | 4.14E-07 | 8.62E-03 | 0.46 | 0.38 | 0.08 | 1.20 |
| 11 | Efhd2 | 0.34 | 4.35E-07 | 9.04E-03 | 0.27 | 0.20 | 0.07 | 1.36 |
| 11 | Commd4 | 0.29 | 4.35E-07 | 9.05E-03 | 0.61 | 0.53 | 0.08 | 1.16 |
| 11 | 0610009B22Rik | 0.28 | 4.42E-07 | 9.19E-03 | 0.60 | 0.52 | 0.08 | 1.14 |
| 11 | Megf9 | 0.43 | 4.46E-07 | 9.28E-03 | 0.26 | 0.19 | 0.08 | 1.40 |
| 11 | Mycbp2 | 0.28 | 4.63E-07 | 9.63E-03 | 0.76 | 0.70 | 0.06 | 1.08 |
| 11 | Pgap1 | 0.48 | 4.65E-07 | 9.67E-03 | 0.22 | 0.15 | 0.07 | 1.45 |
| 11 | Ikbkg | 0.58 | 4.79E-07 | 9.97E-03 | 0.18 | 0.12 | 0.06 | 1.52 |
| 11 | Spry2 | 0.41 | 5.10E-07 | 1.06E-02 | 0.37 | 0.29 | 0.08 | 1.28 |
| 11 | Kif13a | 0.50 | 5.20E-07 | 1.08E-02 | 0.28 | 0.21 | 0.08 | 1.36 |
| 11 | Txlng | 0.49 | 5.20E-07 | 1.08E-02 | 0.29 | 0.22 | 0.07 | 1.33 |
| 11 | Mob1a | 0.32 | 5.34E-07 | 1.11E-02 | 0.53 | 0.45 | 0.07 | 1.16 |
| 11 | Amt | 0.56 | 5.40E-07 | 1.12E-02 | 0.21 | 0.15 | 0.07 | 1.44 |
| 11 | Zfp800 | 0.46 | 5.45E-07 | 1.13E-02 | 0.34 | 0.27 | 0.08 | 1.28 |
| 11 | Osbpl2 | 0.52 | 5.47E-07 | 1.14E-02 | 0.22 | 0.15 | 0.07 | 1.44 |
| 11 | Ofd1 | 0.47 | 5.55E-07 | 1.15E-02 | 0.38 | 0.31 | 0.07 | 1.23 |
| 11 | Tug1 | 0.33 | 5.56E-07 | 1.16E-02 | 0.62 | 0.55 | 0.07 | 1.13 |
| 11 | Mpp5 | 0.41 | 5.61E-07 | 1.17E-02 | 0.45 | 0.37 | 0.07 | 1.19 |
| 11 | Commd9 | 0.29 | 5.85E-07 | 1.22E-02 | 0.57 | 0.49 | 0.08 | 1.16 |
| 11 | Cflar | 0.40 | 5.86E-07 | 1.22E-02 | 0.39 | 0.31 | 0.08 | 1.25 |
| 11 | Cyb561d2 | 0.64 | 6.11E-07 | 1.27E-02 | 0.23 | 0.17 | 0.07 | 1.38 |
| 11 | Tnpo1 | 0.25 | 6.29E-07 | 1.31E-02 | 0.74 | 0.68 | 0.06 | 1.08 |
| 11 | Ift52 | 0.41 | 6.70E-07 | 1.39E-02 | 0.47 | 0.40 | 0.08 | 1.19 |
| 11 | Usp32 | 0.63 | 6.83E-07 | 1.42E-02 | 0.22 | 0.15 | 0.06 | 1.41 |
| 11 | Jtb | 0.34 | 6.94E-07 | 1.44E-02 | 0.60 | 0.53 | 0.07 | 1.13 |
| 11 | Map9 | 0.51 | 6.97E-07 | 1.45E-02 | 0.25 | 0.18 | 0.07 | 1.39 |
| 11 | Zfp974 | 0.65 | 7.23E-07 | 1.50E-02 | 0.14 | 0.09 | 0.05 | 1.59 |
| 11 | Ntmt1 | 0.27 | 7.33E-07 | 1.52E-02 | 0.69 | 0.62 | 0.06 | 1.10 |
| 11 | Ddx19b | 0.75 | 7.35E-07 | 1.53E-02 | 0.14 | 0.09 | 0.05 | 1.57 |
| 11 | BC052040 | 0.53 | 7.72E-07 | 1.61E-02 | 0.24 | 0.18 | 0.07 | 1.39 |
| 11 | Exd2 | 0.53 | 7.82E-07 | 1.63E-02 | 0.22 | 0.15 | 0.07 | 1.43 |
| 11 | Cdkn1b | 0.40 | 7.85E-07 | 1.63E-02 | 0.46 | 0.39 | 0.08 | 1.19 |
| 11 | Fam120a | 0.32 | 8.01E-07 | 1.66E-02 | 0.53 | 0.46 | 0.08 | 1.17 |
| 11 | Rps6ka1 | 0.44 | 8.09E-07 | 1.68E-02 | 0.31 | 0.24 | 0.07 | 1.30 |
| 11 | Nub1 | 0.35 | 8.12E-07 | 1.69E-02 | 0.45 | 0.37 | 0.08 | 1.21 |
| 11 | 2610306M01Rik | 0.61 | 8.94E-07 | 1.86E-02 | 0.23 | 0.16 | 0.06 | 1.40 |
| 11 | Tefm | 0.62 | 9.18E-07 | 1.91E-02 | 0.25 | 0.19 | 0.07 | 1.35 |
| 11 | Ccdc47 | 0.27 | 9.26E-07 | 1.93E-02 | 0.68 | 0.62 | 0.06 | 1.09 |
| 11 | Micall1 | 0.55 | 1.01E-06 | 2.10E-02 | 0.18 | 0.12 | 0.06 | 1.49 |
| 11 | Nomo1 | 0.49 | 1.11E-06 | 2.32E-02 | 0.27 | 0.20 | 0.07 | 1.36 |
| 11 | 2810402E24Rik | 0.40 | 1.24E-06 | 2.59E-02 | 0.29 | 0.21 | 0.08 | 1.35 |
| 11 | Zfp667 | 0.64 | 1.25E-06 | 2.60E-02 | 0.15 | 0.10 | 0.05 | 1.54 |
| 11 | Oard1 | 0.28 | 1.26E-06 | 2.62E-02 | 0.70 | 0.67 | 0.03 | 1.04 |
| 11 | Coa4 | 0.37 | 1.26E-06 | 2.62E-02 | 0.34 | 0.27 | 0.08 | 1.29 |
| 11 | Kmt2d | 0.56 | 1.27E-06 | 2.65E-02 | 0.32 | 0.25 | 0.07 | 1.28 |
| 11 | Slc29a3 | 0.77 | 1.28E-06 | 2.66E-02 | 0.15 | 0.10 | 0.05 | 1.53 |
| 11 | Fzd7 | 0.42 | 1.31E-06 | 2.72E-02 | 0.42 | 0.35 | 0.08 | 1.22 |
| 11 | Btbd1 | 0.27 | 1.31E-06 | 2.72E-02 | 0.67 | 0.63 | 0.04 | 1.06 |
| 11 | Cd59a | 0.56 | 1.32E-06 | 2.74E-02 | 0.29 | 0.22 | 0.07 | 1.31 |
| 11 | Atf7ip | 0.29 | 1.33E-06 | 2.77E-02 | 0.68 | 0.62 | 0.06 | 1.09 |
| 11 | Cep95 | 0.56 | 1.35E-06 | 2.80E-02 | 0.23 | 0.17 | 0.07 | 1.39 |
| 11 | Dhrs4 | 0.26 | 1.39E-06 | 2.88E-02 | 0.69 | 0.64 | 0.05 | 1.08 |
| 11 | Usp20 | 0.72 | 1.41E-06 | 2.93E-02 | 0.13 | 0.08 | 0.05 | 1.62 |
| 11 | Khsrp | 0.28 | 1.44E-06 | 2.99E-02 | 0.61 | 0.56 | 0.05 | 1.10 |
| 11 | Tmem127 | 0.43 | 1.44E-06 | 2.99E-02 | 0.26 | 0.19 | 0.07 | 1.37 |
| 11 | Ccdc92 | 0.63 | 1.48E-06 | 3.08E-02 | 0.13 | 0.08 | 0.05 | 1.59 |

**Supplementary Table 1. Marker genes for the original clusters of integrated clustering of E11.5 control and NotchGOF λ-junction cells.**

| | | | | | | | | |
|---|---|---|---|---|---|---|---|---|
| 11 | Exoc4 | 0.31 | 1.53E-06 | 3.17E-02 | 0.49 | 0.41 | 0.08 | 1.20 |
| 11 | Map2k1 | 0.41 | 1.60E-06 | 3.32E-02 | 0.43 | 0.34 | 0.08 | 1.24 |
| 11 | Tor1b | 0.40 | 1.60E-06 | 3.33E-02 | 0.42 | 0.35 | 0.07 | 1.21 |
| 11 | A930005H10Rik | 0.52 | 1.61E-06 | 3.34E-02 | 0.30 | 0.23 | 0.07 | 1.31 |
| 11 | Zfp236 | 0.49 | 1.66E-06 | 3.45E-02 | 0.26 | 0.19 | 0.07 | 1.36 |
| 11 | Poglut3 | 0.47 | 1.67E-06 | 3.46E-02 | 0.20 | 0.13 | 0.06 | 1.47 |
| 11 | Gchfr | 0.41 | 1.70E-06 | 3.53E-02 | 0.16 | 0.11 | 0.06 | 1.52 |
| 11 | Chchd5 | 0.33 | 1.70E-06 | 3.53E-02 | 0.53 | 0.46 | 0.07 | 1.15 |
| 11 | Kmt2b | 0.31 | 1.73E-06 | 3.60E-02 | 0.56 | 0.49 | 0.07 | 1.15 |
| 11 | Dennd1b | 0.44 | 1.74E-06 | 3.61E-02 | 0.22 | 0.16 | 0.07 | 1.41 |
| 11 | Zfp740 | 0.32 | 1.77E-06 | 3.68E-02 | 0.51 | 0.43 | 0.08 | 1.19 |
| 11 | Mitd1 | 0.38 | 1.78E-06 | 3.70E-02 | 0.37 | 0.30 | 0.08 | 1.26 |
| 11 | Dnajb9 | 0.48 | 1.78E-06 | 3.70E-02 | 0.34 | 0.27 | 0.07 | 1.27 |
| 11 | Nr2c2ap | 0.37 | 1.80E-06 | 3.74E-02 | 0.31 | 0.23 | 0.08 | 1.33 |
| 11 | Wfs1 | 0.67 | 1.81E-06 | 3.77E-02 | 0.11 | 0.07 | 0.04 | 1.67 |
| 11 | Apmap | 0.50 | 1.82E-06 | 3.78E-02 | 0.26 | 0.19 | 0.07 | 1.36 |
| 11 | Zc3h11a | 0.81 | 1.82E-06 | 3.79E-02 | 0.11 | 0.06 | 0.04 | 1.68 |
| 11 | Xrn1 | 0.29 | 1.86E-06 | 3.88E-02 | 0.59 | 0.52 | 0.07 | 1.14 |
| 11 | Dtd2 | 0.37 | 1.94E-06 | 4.04E-02 | 0.40 | 0.32 | 0.08 | 1.24 |
| 11 | Mon1b | 0.79 | 2.05E-06 | 4.27E-02 | 0.10 | 0.06 | 0.04 | 1.69 |
| 11 | Camta1 | 0.29 | 2.23E-06 | 4.64E-02 | 0.67 | 0.62 | 0.05 | 1.08 |
| 11 | Efna5 | 0.37 | 2.24E-06 | 4.65E-02 | 0.54 | 0.47 | 0.07 | 1.15 |
| 11 | Grhpr | 0.37 | 2.24E-06 | 4.65E-02 | 0.52 | 0.44 | 0.08 | 1.19 |
| 11 | Avl9 | 0.39 | 2.27E-06 | 4.72E-02 | 0.42 | 0.35 | 0.08 | 1.22 |
| 11 | Mettl26 | 0.30 | 2.31E-06 | 4.81E-02 | 0.57 | 0.49 | 0.08 | 1.15 |
| 11 | Abhd13 | 0.46 | 2.32E-06 | 4.82E-02 | 0.26 | 0.19 | 0.07 | 1.37 |
| 11 | Zfp809 | 0.71 | 2.32E-06 | 4.83E-02 | 0.12 | 0.07 | 0.05 | 1.63 |
| 11 | Tspan15 | 0.52 | 2.37E-06 | 4.93E-02 | 0.26 | 0.19 | 0.07 | 1.36 |
| 12 | Gypa | 7.34 | 0 | 0 | 0.99 | 0.02 | 0.97 | 54.78 |
| 12 | Slc4a1 | 6.91 | 0 | 0 | 1.00 | 0.03 | 0.97 | 32.19 |
| 12 | Alas2 | 7.31 | 0 | 0 | 0.99 | 0.03 | 0.97 | 38.15 |
| 12 | Rhd | 7.19 | 0 | 0 | 0.98 | 0.02 | 0.96 | 54.56 |
| 12 | Hemgn | 7.15 | 0 | 0 | 0.98 | 0.02 | 0.95 | 40.67 |
| 12 | Trim10 | 7.50 | 0 | 0 | 0.96 | 0.01 | 0.95 | 87.45 |
| 12 | Ermap | 7.14 | 0 | 0 | 0.96 | 0.01 | 0.95 | 73.92 |
| 12 | Cited4 | 6.28 | 0 | 0 | 0.97 | 0.03 | 0.94 | 33.28 |
| 12 | Klf1 | 6.33 | 0 | 0 | 0.96 | 0.03 | 0.93 | 36.77 |
| 12 | Acp5 | 5.89 | 0 | 0 | 0.98 | 0.06 | 0.92 | 16.37 |
| 12 | Pdzk1ip1 | 7.14 | 0 | 0 | 0.93 | 0.02 | 0.92 | 58.25 |
| 12 | Car2 | 6.70 | 0 | 0 | 1.00 | 0.09 | 0.91 | 11.34 |
| 12 | Ncf4 | 6.73 | 0 | 0 | 0.92 | 0.02 | 0.91 | 54.29 |
| 12 | Icam4 | 7.07 | 0 | 0 | 0.92 | 0.01 | 0.91 | 70.62 |
| 12 | Rhag | 7.15 | 0 | 0 | 0.92 | 0.01 | 0.91 | 91.50 |
| 12 | Tspan33 | 5.65 | 0 | 0 | 0.96 | 0.06 | 0.90 | 17.42 |
| 12 | Spta1 | 6.90 | 0 | 0 | 0.90 | 0.01 | 0.89 | 75.25 |
| 12 | Snca | 4.62 | 0 | 0 | 0.94 | 0.06 | 0.88 | 15.48 |
| 12 | Mt2 | 5.65 | 0 | 0 | 0.98 | 0.10 | 0.88 | 9.58 |
| 12 | Tal1 | 6.09 | 0 | 0 | 0.89 | 0.02 | 0.87 | 44.50 |
| 12 | Sptb | 6.25 | 0 | 0 | 0.89 | 0.02 | 0.86 | 42.14 |
| 12 | Ube2l6 | 4.41 | 0 | 0 | 0.97 | 0.11 | 0.86 | 9.05 |
| 12 | Kel | 6.82 | 0 | 0 | 0.87 | 0.01 | 0.86 | 67.00 |
| 12 | Pnpo | 4.44 | 0 | 0 | 0.97 | 0.15 | 0.82 | 6.68 |
| 12 | Add2 | 4.44 | 0 | 0 | 0.88 | 0.06 | 0.82 | 14.03 |
| 12 | Smim1 | 4.56 | 0 | 0 | 0.96 | 0.14 | 0.81 | 6.73 |
| 12 | Epb42 | 7.06 | 0 | 0 | 0.82 | 0.01 | 0.81 | 82.10 |
| 12 | Hsd3b6 | 7.39 | 0 | 0 | 0.81 | 0.01 | 0.80 | 81.40 |

**Supplementary Table 1. Marker genes for the original clusters of integrated clustering of E11.5 control and NotchGOF λ-junction cells.**

| | | | | | | | | |
|---|---|---|---|---|---|---|---|---|
| 12 | Slc39a8 | 3.82 | 0 | 0 | 0.95 | 0.15 | 0.80 | 6.54 |
| 12 | Tuba4a | 3.86 | 0 | 0 | 0.85 | 0.04 | 0.80 | 19.67 |
| 12 | Fam210b | 3.97 | 0 | 0 | 0.98 | 0.18 | 0.80 | 5.56 |
| 12 | Slc38a5 | 6.95 | 0 | 0 | 0.80 | 0.01 | 0.79 | 80.30 |
| 12 | Cldn13 | 7.12 | 0 | 0 | 0.80 | 0.01 | 0.79 | 88.44 |
| 12 | Hbb-bs | 7.33 | 0 | 0 | 0.99 | 0.21 | 0.78 | 4.64 |
| 12 | Slc25a37 | 5.33 | 0 | 0 | 0.99 | 0.22 | 0.78 | 4.60 |
| 12 | Marchf2 | 3.39 | 0 | 0 | 0.96 | 0.19 | 0.78 | 5.10 |
| 12 | Tspan32 | 5.51 | 0 | 0 | 0.78 | 0.01 | 0.77 | 60.31 |
| 12 | Gmpr | 3.71 | 0 | 0 | 0.92 | 0.15 | 0.77 | 6.04 |
| 12 | Chchd10 | 3.11 | 0 | 0 | 0.90 | 0.14 | 0.76 | 6.57 |
| 12 | Cd59a | 3.14 | 0 | 0 | 0.96 | 0.20 | 0.76 | 4.87 |
| 12 | Mt1 | 5.75 | 0 | 0 | 0.99 | 0.24 | 0.75 | 4.07 |
| 12 | Hbq1b | 7.23 | 0 | 0 | 0.76 | 0.01 | 0.75 | 108.00 |
| 12 | Ptp4a3 | 3.18 | 0 | 0 | 0.97 | 0.23 | 0.74 | 4.23 |
| 12 | Fech | 3.99 | 0 | 0 | 0.99 | 0.25 | 0.74 | 3.94 |
| 12 | Epor | 4.19 | 0 | 0 | 0.79 | 0.05 | 0.73 | 14.81 |
| 12 | Hebp1 | 3.99 | 0 | 0 | 0.97 | 0.25 | 0.73 | 3.97 |
| 12 | Atp8a1 | 3.75 | 0 | 0 | 0.79 | 0.06 | 0.73 | 12.36 |
| 12 | 1300017J02Rik | 6.83 | 0 | 0 | 0.73 | 0.01 | 0.73 | 81.56 |
| 12 | Hbb-bt | 7.28 | 0 | 0 | 1.00 | 0.28 | 0.72 | 3.58 |
| 12 | Slc30a10 | 5.47 | 0 | 0 | 0.74 | 0.02 | 0.72 | 41.00 |
| 12 | Plek2 | 4.64 | 0 | 0 | 0.76 | 0.04 | 0.72 | 19.44 |
| 12 | Gm16867 | 5.27 | 0 | 0 | 0.74 | 0.03 | 0.72 | 29.72 |
| 12 | Tent5c | 5.75 | 0 | 0 | 0.73 | 0.02 | 0.72 | 40.78 |
| 12 | Ell2 | 3.91 | 0 | 0 | 0.78 | 0.07 | 0.71 | 11.66 |
| 12 | Abcb10 | 3.11 | 0 | 0 | 0.92 | 0.21 | 0.71 | 4.46 |
| 12 | Rbm38 | 2.79 | 0 | 0 | 0.92 | 0.21 | 0.71 | 4.30 |
| 12 | Gata1 | 6.20 | 0 | 0 | 0.72 | 0.01 | 0.71 | 71.60 |
| 12 | Cfap57 | 7.15 | 0 | 0 | 0.71 | 0.01 | 0.70 | 88.50 |
| 12 | Ank1 | 6.13 | 0 | 0 | 0.70 | 0.01 | 0.69 | 58.58 |
| 12 | Kcnn4 | 6.53 | 0 | 0 | 0.70 | 0.01 | 0.69 | 70.00 |
| 12 | Cpox | 4.23 | 0 | 0 | 0.99 | 0.30 | 0.69 | 3.33 |
| 12 | Nfe2 | 5.54 | 0 | 0 | 0.70 | 0.01 | 0.69 | 53.85 |
| 12 | Lmx1a | 7.02 | 0 | 0 | 0.69 | 0.01 | 0.68 | 85.63 |
| 12 | Asb17os | 7.45 | 0 | 0 | 0.68 | 0.01 | 0.67 | 135.20 |
| 12 | Uros | 2.79 | 0 | 0 | 0.82 | 0.15 | 0.67 | 5.47 |
| 12 | Ctse | 6.65 | 0 | 0 | 0.68 | 0.01 | 0.67 | 84.38 |
| 12 | Blvrb | 4.90 | 0 | 0 | 0.99 | 0.32 | 0.67 | 3.06 |
| 12 | Grina | 2.34 | 0 | 0 | 0.89 | 0.23 | 0.66 | 3.87 |
| 12 | Creg1 | 3.39 | 0 | 0 | 0.98 | 0.33 | 0.65 | 2.95 |
| 12 | Ucp2 | 3.06 | 0 | 0 | 0.99 | 0.34 | 0.65 | 2.93 |
| 12 | Tmod1 | 4.18 | 0 | 0 | 0.69 | 0.04 | 0.64 | 16.31 |
| 12 | Rfesd | 2.81 | 0 | 0 | 0.93 | 0.29 | 0.64 | 3.22 |
| 12 | Dmtn | 3.94 | 0 | 0 | 0.68 | 0.05 | 0.63 | 14.67 |
| 12 | Serinc3 | 2.71 | 0 | 0 | 0.97 | 0.34 | 0.63 | 2.83 |
| 12 | Chac2 | 2.99 | 0 | 0 | 0.74 | 0.11 | 0.62 | 6.46 |
| 12 | Ncoa4 | 2.32 | 0 | 0 | 0.92 | 0.30 | 0.62 | 3.09 |
| 12 | Aqp3 | 5.17 | 0 | 0 | 0.64 | 0.02 | 0.62 | 39.88 |
| 12 | Gclm | 3.56 | 0 | 0 | 0.98 | 0.37 | 0.61 | 2.66 |
| 12 | St3gal5 | 2.21 | 0 | 0 | 0.84 | 0.22 | 0.61 | 3.76 |
| 12 | Grap2 | 6.68 | 0 | 0 | 0.62 | 0.01 | 0.61 | 77.13 |
| 12 | Cpeb4 | 3.42 | 0 | 0 | 0.68 | 0.08 | 0.61 | 8.97 |
| 12 | Tspo | 2.28 | 0 | 0 | 0.94 | 0.33 | 0.60 | 2.81 |
| 12 | Slc25a21 | 3.54 | 0 | 0 | 0.65 | 0.05 | 0.60 | 13.50 |
| 12 | Hbb-bh1 | 7.55 | 0 | 0 | 1.00 | 0.40 | 0.60 | 2.49 |

**Supplementary Table 1. Marker genes for the original clusters of integrated clustering of E11.5 control and NotchGOF λ-junction cells.**

| | | | | | | | | |
|---|---|---|---|---|---|---|---|---|
| 12 | Fam241a | 2.83 | 0 | 0 | 0.69 | 0.10 | 0.59 | 6.86 |
| 12 | Vangl1 | 2.43 | 0 | 0 | 0.71 | 0.14 | 0.58 | 5.20 |
| 12 | Josd2 | 3.18 | 0 | 0 | 0.98 | 0.40 | 0.58 | 2.43 |
| 12 | Lyl1 | 4.70 | 0 | 0 | 0.58 | 0.02 | 0.56 | 32.28 |
| 12 | Xk | 5.19 | 0 | 0 | 0.58 | 0.02 | 0.56 | 34.00 |
| 12 | Slc43a1 | 2.37 | 0 | 0 | 0.70 | 0.14 | 0.56 | 4.94 |
| 12 | Dhrs11 | 4.26 | 0 | 0 | 0.58 | 0.03 | 0.55 | 20.79 |
| 12 | Bmp2k | 2.36 | 0 | 0 | 0.68 | 0.13 | 0.55 | 5.09 |
| 12 | Gm2061 | 7.12 | 0 | 0 | 0.55 | 0.01 | 0.55 | 92.00 |
| 12 | Tinagl1 | 3.57 | 0 | 0 | 0.59 | 0.04 | 0.54 | 13.93 |
| 12 | Bpgm | 3.12 | 0 | 0 | 0.98 | 0.44 | 0.54 | 2.22 |
| 12 | Arrb1 | 2.75 | 0 | 0 | 0.63 | 0.10 | 0.54 | 6.47 |
| 12 | Gm867 | 6.98 | 0 | 0 | 0.53 | 0.01 | 0.53 | 89.00 |
| 12 | Reln | 4.48 | 0 | 0 | 0.53 | 0.02 | 0.51 | 24.23 |
| 12 | Rec114 | 5.64 | 0 | 0 | 0.51 | 0.01 | 0.50 | 46.18 |
| 12 | Fbxo6 | 3.13 | 0 | 0 | 0.54 | 0.05 | 0.50 | 11.31 |
| 12 | Mgst3 | 4.41 | 0 | 0 | 0.99 | 0.49 | 0.50 | 2.01 |
| 12 | Pkhd1l1 | 7.03 | 0 | 0 | 0.50 | 0.01 | 0.49 | 82.50 |
| 12 | Tmem184a | 3.64 | 0 | 0 | 0.52 | 0.04 | 0.49 | 13.76 |
| 12 | Gpr146 | 4.78 | 0 | 0 | 0.50 | 0.02 | 0.48 | 33.07 |
| 12 | Svip | 2.19 | 0 | 0 | 0.55 | 0.08 | 0.47 | 7.16 |
| 12 | Pklr | 6.78 | 0 | 0 | 0.47 | 0.01 | 0.47 | 94.40 |
| 12 | Dapk2 | 3.65 | 0 | 0 | 0.50 | 0.03 | 0.47 | 15.12 |
| 12 | Steap3 | 5.57 | 0 | 0 | 0.46 | 0.01 | 0.45 | 51.22 |
| 12 | Htatip2 | 2.79 | 0 | 0 | 0.51 | 0.06 | 0.45 | 8.47 |
| 12 | Gm32051 | 7.16 | 0 | 0 | 0.45 | 0.00 | 0.44 | 111.25 |
| 12 | Urod | 2.68 | 0 | 0 | 0.99 | 0.55 | 0.44 | 1.79 |
| 12 | Ikzf1 | 4.62 | 0 | 0 | 0.45 | 0.01 | 0.43 | 31.86 |
| 12 | Aqp8 | 7.31 | 0 | 0 | 0.43 | 0.01 | 0.43 | 86.80 |
| 12 | Slc6a9 | 3.61 | 0 | 0 | 0.46 | 0.03 | 0.42 | 14.22 |
| 12 | Gfi1b | 6.04 | 0 | 0 | 0.43 | 0.01 | 0.42 | 85.00 |
| 12 | Mylk3 | 4.90 | 0 | 0 | 0.43 | 0.01 | 0.42 | 33.31 |
| 12 | Hmbs | 3.41 | 0 | 0 | 0.99 | 0.57 | 0.42 | 1.72 |
| 12 | Nmnat3 | 3.38 | 0 | 0 | 0.45 | 0.03 | 0.42 | 13.58 |
| 12 | 6030468B19Rik | 7.59 | 0 | 0 | 0.40 | 0.00 | 0.40 | 134.67 |
| 12 | Gm14275 | 7.27 | 0 | 0 | 0.40 | 0.00 | 0.40 | 101.00 |
| 12 | Slc22a4 | 6.39 | 0 | 0 | 0.41 | 0.01 | 0.40 | 67.50 |
| 12 | Cmpk2 | 3.87 | 0 | 0 | 0.42 | 0.03 | 0.40 | 16.84 |
| 12 | Pheta2 | 2.76 | 0 | 0 | 0.44 | 0.05 | 0.39 | 8.90 |
| 12 | Redrum | 5.01 | 0 | 0 | 0.40 | 0.01 | 0.39 | 39.50 |
| 12 | Slfn14 | 7.08 | 0 | 0 | 0.39 | 0.00 | 0.38 | 128.67 |
| 12 | Ccdc141 | 5.20 | 0 | 0 | 0.39 | 0.01 | 0.38 | 30.15 |
| 12 | Cd59b | 3.20 | 0 | 0 | 0.41 | 0.03 | 0.38 | 12.84 |
| 12 | Gm38619.1 | 7.52 | 0 | 0 | 0.38 | 0.00 | 0.37 | 125.67 |
| 12 | Alad | 3.38 | 0 | 0 | 1.00 | 0.62 | 0.37 | 1.59 |
| 12 | Asb17 | 6.58 | 0 | 0 | 0.37 | 0.00 | 0.37 | 93.00 |
| 12 | Hba-a2 | 7.23 | 0 | 0 | 1.00 | 0.65 | 0.35 | 1.54 |
| 12 | Ifi35 | 3.88 | 0 | 0 | 0.37 | 0.02 | 0.35 | 19.53 |
| 12 | Cdkn2c | 3.40 | 0 | 0 | 0.38 | 0.03 | 0.35 | 13.10 |
| 12 | Rnf212 | 7.26 | 0 | 0 | 0.35 | 0.00 | 0.35 | 118.00 |
| 12 | Zc3hav1 | 4.25 | 0 | 0 | 0.36 | 0.02 | 0.35 | 21.35 |
| 12 | Spns2 | 3.94 | 0 | 0 | 0.36 | 0.02 | 0.35 | 20.17 |
| 12 | Mrap | 7.07 | 0 | 0 | 0.33 | 0.00 | 0.33 | 111.00 |
| 12 | Ccdc92b | 6.22 | 0 | 0 | 0.33 | 0.01 | 0.33 | 66.80 |
| 12 | Gm43154 | 6.82 | 0 | 0 | 0.33 | 0.00 | 0.33 | 110.33 |
| 12 | Adra2b | 6.87 | 0 | 0 | 0.33 | 0.00 | 0.32 | 81.75 |

**Supplementary Table 1. Marker genes for the original clusters of integrated clustering of E11.5 control and NotchGOF λ-junction cells.**

| | | | | | | | | |
|---|---|---|---|---|---|---|---|---|
| 12 | Ache | 5.01 | 0 | 0 | 0.33 | 0.01 | 0.32 | 41.38 |
| 12 | Hba-a1 | 7.55 | 0 | 0 | 1.00 | 0.69 | 0.32 | 1.46 |
| 12 | 5430401H09Rik | 6.87 | 0 | 0 | 0.32 | 0.00 | 0.31 | 105.00 |
| 12 | Tmem86b | 3.76 | 0 | 0 | 0.33 | 0.02 | 0.31 | 15.76 |
| 12 | Mfsd2b | 4.94 | 0 | 0 | 0.31 | 0.01 | 0.30 | 44.14 |
| 12 | Wnk4 | 3.06 | 0 | 0 | 0.32 | 0.03 | 0.30 | 12.38 |
| 12 | Acss1 | 4.04 | 0 | 0 | 0.31 | 0.02 | 0.29 | 20.60 |
| 12 | Gm20161 | 7.15 | 0 | 0 | 0.30 | 0.00 | 0.29 | 98.33 |
| 12 | Treml2 | 4.60 | 0 | 0 | 0.28 | 0.01 | 0.27 | 39.29 |
| 12 | Art4 | 3.27 | 0 | 0 | 0.28 | 0.02 | 0.26 | 13.52 |
| 12 | Trib3 | 4.11 | 0 | 0 | 0.27 | 0.01 | 0.26 | 19.43 |
| 12 | Igtp | 3.74 | 0 | 0 | 0.27 | 0.01 | 0.25 | 22.17 |
| 12 | Gm9895 | 5.38 | 0 | 0 | 0.25 | 0.01 | 0.25 | 50.80 |
| 12 | Tmem56 | 3.91 | 0 | 0 | 0.26 | 0.02 | 0.24 | 17.27 |
| 12 | Popdc2 | 5.11 | 0 | 0 | 0.25 | 0.01 | 0.24 | 49.40 |
| 12 | Flt3l | 5.14 | 0 | 0 | 0.24 | 0.01 | 0.23 | 47.60 |
| 12 | Gm10390 | 6.27 | 0 | 0 | 0.23 | 0.00 | 0.23 | 77.67 |
| 12 | Nudt4 | 3.15 | 0 | 0 | 1.00 | 0.77 | 0.23 | 1.30 |
| 12 | Ccrl2 | 4.82 | 0 | 0 | 0.22 | 0.01 | 0.22 | 44.40 |
| 12 | Gm38619 | 7.49 | 0 | 0 | 0.21 | 0.00 | 0.21 | 207.00 |
| 12 | Arsg | 3.87 | 0 | 0 | 0.22 | 0.01 | 0.21 | 21.50 |
| 12 | Ackr1 | 4.05 | 0 | 0 | 0.21 | 0.01 | 0.20 | 20.90 |
| 12 | Gm10603 | 5.29 | 0 | 0 | 0.20 | 0.01 | 0.20 | 40.20 |
| 12 | Lrrc39 | 5.59 | 0 | 0 | 0.19 | 0.00 | 0.18 | 61.67 |
| 12 | Csf2rb | 4.43 | 0 | 0 | 0.18 | 0.01 | 0.17 | 29.83 |
| 12 | 1810053B23Rik | 7.31 | 0 | 0 | 0.17 | 0.00 | 0.16 | 165.00 |
| 12 | Lefty1 | 4.27 | 0 | 0 | 0.16 | 0.01 | 0.15 | 31.20 |
| 12 | Nxpe2 | 5.40 | 0 | 0 | 0.15 | 0.00 | 0.15 | 51.00 |
| 12 | H2-T24 | 5.06 | 0 | 0 | 0.15 | 0.00 | 0.15 | 49.33 |
| 12 | Apol8 | 5.18 | 0 | 0 | 0.15 | 0.00 | 0.14 | 49.00 |
| 12 | Il1bos | 7.34 | 0 | 0 | 0.13 | 0.00 | 0.13 | 132.00 |
| 12 | Gpr150 | 5.71 | 0 | 0 | 0.13 | 0.00 | 0.13 | 65.00 |
| 12 | Cyp4b1 | 7.09 | 0 | 0 | 0.11 | 0.00 | 0.11 | 113.00 |
| 12 | Hba-x | 7.49 | 0 | 0 | 1.00 | 0.89 | 0.11 | 1.12 |
| 12 | Glrx5 | 1.97 | 0 | 0 | 0.99 | 0.88 | 0.11 | 1.12 |
| 12 | Pdia2 | 6.29 | 0 | 0 | 0.11 | 0.00 | 0.11 | 107.00 |
| 12 | Acmsd | 6.26 | 0 | 0 | 0.10 | 0.00 | 0.10 | 103.00 |
| 12 | Hbq1a | 6.27 | 0 | 0 | 0.10 | 0.00 | 0.10 | 100.00 |
| 12 | Tmem14c | 1.98 | 0 | 0 | 0.99 | 0.92 | 0.07 | 1.07 |
| 12 | Gpx1 | 3.45 | 0 | 0 | 1.00 | 0.95 | 0.05 | 1.05 |
| 12 | Hbb-y | 7.42 | 0 | 0 | 1.00 | 0.98 | 0.02 | 1.02 |
| 12 | Ftl1 | 0.78 | 0 | 0 | 1.00 | 1.00 | 0.00 | 1.00 |
| 12 | Prdx2 | 1.08 | 0 | 0 | 1.00 | 1.00 | 0.00 | 1.00 |
| 12 | Fth1 | 3.27 | 0 | 0 | 1.00 | 1.00 | 0.00 | 1.00 |
| 12 | Ypel4 | 3.68 | 6.90E-305 | 1.43E-300 | 0.27 | 0.02 | 0.25 | 12.90 |
| 12 | Hopx | 3.48 | 1.32E-302 | 2.75E-298 | 0.19 | 0.01 | 0.18 | 21.33 |
| 12 | Ttc39a | 2.53 | 9.88E-293 | 2.06E-288 | 0.40 | 0.05 | 0.35 | 8.20 |
| 12 | Zfpm1 | 2.07 | 9.99E-289 | 2.08E-284 | 0.89 | 0.30 | 0.60 | 3.03 |
| 12 | Mst1 | 5.63 | 1.01E-286 | 2.11E-282 | 0.10 | 0.00 | 0.10 | 50.50 |
| 12 | Tgm2 | 3.54 | 3.76E-286 | 7.83E-282 | 0.19 | 0.01 | 0.18 | 20.67 |
| 12 | Cox17 | 1.62 | 4.41E-286 | 9.18E-282 | 0.98 | 0.76 | 0.22 | 1.29 |
| 12 | Smim5 | 3.13 | 1.58E-281 | 3.29E-277 | 0.25 | 0.02 | 0.23 | 13.21 |
| 12 | Prr13 | 2.13 | 2.74E-279 | 5.69E-275 | 0.93 | 0.38 | 0.55 | 2.42 |
| 12 | 4930455G09Rik | 3.73 | 1.62E-278 | 3.38E-274 | 0.18 | 0.01 | 0.17 | 19.44 |
| 12 | Ifrd2 | 2.09 | 2.36E-278 | 4.90E-274 | 0.89 | 0.31 | 0.57 | 2.83 |
| 12 | Lyrm9 | 2.13 | 4.37E-277 | 9.09E-273 | 0.70 | 0.16 | 0.54 | 4.49 |

**Supplementary Table 1. Marker genes for the original clusters of integrated clustering of E11.5 control and NotchGOF λ-junction cells.**

| | | | | | | | | |
|---|---|---|---|---|---|---|---|---|
| 12 | Abcg4 | 3.47 | 4.81E-277 | 1.00E-272 | 0.21 | 0.01 | 0.19 | 15.85 |
| 12 | Larp1b | 2.40 | 1.15E-275 | 2.39E-271 | 0.45 | 0.07 | 0.39 | 6.94 |
| 12 | Cox6b2 | 2.16 | 1.53E-273 | 3.19E-269 | 0.94 | 0.45 | 0.49 | 2.08 |
| 12 | Tmem238 | 1.96 | 6.22E-268 | 1.29E-263 | 0.71 | 0.16 | 0.55 | 4.36 |
| 12 | St3gal6 | 2.21 | 7.80E-257 | 1.62E-252 | 0.54 | 0.10 | 0.45 | 5.54 |
| 12 | Atp1b2 | 3.48 | 2.38E-251 | 4.95E-247 | 0.19 | 0.01 | 0.18 | 16.17 |
| 12 | Ubac1 | 1.78 | 2.02E-249 | 4.19E-245 | 0.95 | 0.56 | 0.39 | 1.69 |
| 12 | Samd11 | 4.39 | 8.42E-248 | 1.75E-243 | 0.14 | 0.01 | 0.13 | 27.20 |
| 12 | Lmo2 | 0.99 | 5.86E-244 | 1.22E-239 | 0.59 | 0.11 | 0.48 | 5.29 |
| 12 | Adrb2 | 3.62 | 2.09E-243 | 4.34E-239 | 0.16 | 0.01 | 0.15 | 18.00 |
| 12 | Ankrd9 | 2.02 | 8.04E-239 | 1.67E-234 | 0.56 | 0.11 | 0.45 | 5.09 |
| 12 | Rgs10 | 1.55 | 2.34E-237 | 4.86E-233 | 0.46 | 0.08 | 0.38 | 5.99 |
| 12 | Acyp2 | 2.17 | 3.58E-237 | 7.44E-233 | 0.51 | 0.09 | 0.41 | 5.45 |
| 12 | Ccnd3 | 1.28 | 1.02E-234 | 2.13E-230 | 0.99 | 0.85 | 0.14 | 1.16 |
| 12 | Mkrn1 | 1.75 | 2.24E-228 | 4.66E-224 | 0.93 | 0.41 | 0.52 | 2.26 |
| 12 | Fam117a | 1.97 | 1.89E-227 | 3.94E-223 | 0.76 | 0.22 | 0.54 | 3.48 |
| 12 | Marchf3 | 4.46 | 4.88E-227 | 1.02E-222 | 0.11 | 0.00 | 0.10 | 35.33 |
| 12 | Btg2 | 1.52 | 9.54E-226 | 1.98E-221 | 0.82 | 0.26 | 0.57 | 3.21 |
| 12 | Txnrd2 | 1.88 | 1.15E-224 | 2.39E-220 | 0.73 | 0.20 | 0.53 | 3.71 |
| 12 | Nt5c3 | 1.86 | 1.71E-222 | 3.57E-218 | 0.91 | 0.38 | 0.52 | 2.36 |
| 12 | 4930523C07Rik | 2.61 | 6.39E-221 | 1.33E-216 | 0.40 | 0.07 | 0.34 | 6.22 |
| 12 | Gm42418 | 0.92 | 2.34E-220 | 4.86E-216 | 1.00 | 1.00 | 0.00 | 1.00 |
| 12 | Isca1 | 1.80 | 9.24E-219 | 1.92E-214 | 0.85 | 0.30 | 0.56 | 2.88 |
| 12 | E2f2 | 2.18 | 9.45E-219 | 1.97E-214 | 0.51 | 0.10 | 0.41 | 5.15 |
| 12 | Pigq | 1.94 | 1.98E-217 | 4.11E-213 | 0.77 | 0.22 | 0.55 | 3.48 |
| 12 | Tfrc | 2.01 | 2.62E-215 | 5.44E-211 | 0.89 | 0.38 | 0.51 | 2.33 |
| 12 | Cdc42ep3 | 1.39 | 1.99E-212 | 4.14E-208 | 0.65 | 0.15 | 0.50 | 4.32 |
| 12 | Pim1 | 1.78 | 1.51E-206 | 3.14E-202 | 0.93 | 0.46 | 0.48 | 2.04 |
| 12 | Nrtn | 1.54 | 8.66E-204 | 1.80E-199 | 0.61 | 0.14 | 0.47 | 4.33 |
| 12 | Ubash3a | 4.27 | 1.20E-203 | 2.49E-199 | 0.10 | 0.00 | 0.10 | 33.67 |
| 12 | Dnaja4 | 2.38 | 6.32E-203 | 1.31E-198 | 0.32 | 0.04 | 0.28 | 7.30 |
| 12 | Spaca9 | 2.58 | 7.88E-203 | 1.64E-198 | 0.25 | 0.03 | 0.22 | 9.19 |
| 12 | Fam78a | 3.65 | 9.65E-203 | 2.01E-198 | 0.15 | 0.01 | 0.14 | 16.44 |
| 12 | Tmcc2 | 2.56 | 1.54E-202 | 3.20E-198 | 0.42 | 0.07 | 0.34 | 5.68 |
| 12 | Ppox | 1.77 | 4.27E-202 | 8.87E-198 | 0.74 | 0.23 | 0.52 | 3.29 |
| 12 | Gcnt1 | 3.45 | 9.54E-202 | 1.98E-197 | 0.24 | 0.03 | 0.21 | 9.52 |
| 12 | Tfr2 | 3.14 | 2.50E-201 | 5.20E-197 | 0.16 | 0.01 | 0.15 | 15.60 |
| 12 | Slc40a1 | 1.86 | 1.21E-199 | 2.52E-195 | 0.34 | 0.05 | 0.29 | 6.86 |
| 12 | Prdx3 | 1.54 | 1.54E-196 | 3.19E-192 | 0.94 | 0.52 | 0.42 | 1.80 |
| 12 | Stom | 2.69 | 1.74E-194 | 3.63E-190 | 0.29 | 0.04 | 0.25 | 7.61 |
| 12 | Fads3 | 2.17 | 7.66E-194 | 1.59E-189 | 0.33 | 0.05 | 0.28 | 6.83 |
| 12 | Pip5k1b | 2.50 | 2.09E-193 | 4.35E-189 | 0.25 | 0.03 | 0.22 | 8.66 |
| 12 | C1qtnf4 | 1.58 | 1.65E-190 | 3.44E-186 | 0.81 | 0.27 | 0.54 | 2.99 |
| 12 | Slc29a1 | 1.60 | 3.02E-188 | 6.29E-184 | 0.51 | 0.11 | 0.40 | 4.66 |
| 12 | Exoc6 | 1.97 | 1.95E-187 | 4.05E-183 | 0.38 | 0.07 | 0.32 | 5.86 |
| 12 | Pfkfb1 | 2.37 | 1.25E-180 | 2.59E-176 | 0.27 | 0.04 | 0.24 | 7.53 |
| 12 | Abcg2 | 1.60 | 6.10E-180 | 1.27E-175 | 0.84 | 0.33 | 0.51 | 2.52 |
| 12 | D030028A08Rik | 2.14 | 2.43E-174 | 5.05E-170 | 0.33 | 0.05 | 0.28 | 6.29 |
| 12 | Asb1 | 1.95 | 8.87E-174 | 1.84E-169 | 0.35 | 0.06 | 0.29 | 5.87 |
| 12 | Prokr1 | 3.23 | 5.43E-171 | 1.13E-166 | 0.14 | 0.01 | 0.13 | 14.10 |
| 12 | Cltb | 1.62 | 4.49E-166 | 9.34E-162 | 0.62 | 0.17 | 0.45 | 3.62 |
| 12 | 5430431A17Rik | 3.64 | 2.03E-165 | 4.22E-161 | 0.11 | 0.01 | 0.10 | 21.40 |
| 12 | Arhgap23 | 1.80 | 1.89E-163 | 3.93E-159 | 0.42 | 0.09 | 0.33 | 4.92 |
| 12 | Dnajb3 | 1.59 | 2.81E-162 | 5.85E-158 | 0.55 | 0.14 | 0.41 | 4.00 |
| 12 | Lockd | 1.42 | 3.46E-162 | 7.19E-158 | 0.86 | 0.35 | 0.51 | 2.45 |
| 12 | Slc20a1 | 1.79 | 1.40E-158 | 2.90E-154 | 0.81 | 0.31 | 0.50 | 2.64 |

**Supplementary Table 1. Marker genes for the original clusters of integrated clustering of E11.5 control and NotchGOF λ-junction cells.**

| | | | | | | | | |
|---|---|---|---|---|---|---|---|---|
| 12 | Bola3 | 1.08 | 2.02E-158 | 4.20E-154 | 0.97 | 0.74 | 0.23 | 1.31 |
| 12 | Slc16a6 | 2.38 | 2.40E-154 | 4.98E-150 | 0.29 | 0.05 | 0.24 | 6.17 |
| 12 | Ppp1r15a | 1.48 | 7.20E-154 | 1.50E-149 | 0.73 | 0.24 | 0.49 | 3.09 |
| 12 | Tango2 | 1.73 | 1.56E-153 | 3.24E-149 | 0.55 | 0.14 | 0.40 | 3.81 |
| 12 | Ssx2ip | 1.27 | 2.86E-153 | 5.94E-149 | 0.84 | 0.35 | 0.50 | 2.44 |
| 12 | Tuft1 | 2.26 | 3.08E-153 | 6.41E-149 | 0.24 | 0.03 | 0.21 | 7.44 |
| 12 | Endod1 | 1.69 | 7.04E-152 | 1.46E-147 | 0.40 | 0.08 | 0.32 | 4.89 |
| 12 | Fbxo7 | 1.69 | 2.16E-150 | 4.49E-146 | 0.51 | 0.12 | 0.38 | 4.07 |
| 12 | Cdr2 | 1.60 | 2.12E-148 | 4.40E-144 | 0.66 | 0.21 | 0.45 | 3.19 |
| 12 | 5033406O09Rik | 3.22 | 5.19E-148 | 1.08E-143 | 0.12 | 0.01 | 0.11 | 13.67 |
| 12 | H2afj | 0.92 | 3.39E-147 | 7.05E-143 | 0.98 | 0.81 | 0.18 | 1.22 |
| 12 | Slc43a3 | 1.54 | 2.15E-145 | 4.47E-141 | 0.75 | 0.27 | 0.48 | 2.77 |
| 12 | Ocel1 | 1.96 | 2.87E-145 | 5.96E-141 | 0.28 | 0.05 | 0.24 | 6.04 |
| 12 | Nacad | 3.38 | 4.93E-145 | 1.03E-140 | 0.10 | 0.01 | 0.09 | 16.67 |
| 12 | Ehd1 | 1.25 | 7.63E-143 | 1.59E-138 | 0.93 | 0.51 | 0.41 | 1.81 |
| 12 | Lmna | 1.49 | 2.14E-138 | 4.44E-134 | 0.71 | 0.26 | 0.45 | 2.72 |
| 12 | Cdc25b | 1.70 | 1.50E-136 | 3.11E-132 | 0.60 | 0.19 | 0.41 | 3.11 |
| 12 | Mindy2 | 1.39 | 2.63E-134 | 5.47E-130 | 0.57 | 0.16 | 0.40 | 3.47 |
| 12 | Hsd17b14 | 3.13 | 8.49E-134 | 1.77E-129 | 0.13 | 0.01 | 0.12 | 12.00 |
| 12 | Hesx1 | 3.11 | 2.47E-133 | 5.13E-129 | 0.11 | 0.01 | 0.10 | 15.29 |
| 12 | Rexo2 | 1.14 | 8.08E-132 | 1.68E-127 | 0.94 | 0.58 | 0.35 | 1.60 |
| 12 | Pde10a | 2.00 | 4.54E-131 | 9.45E-127 | 0.24 | 0.04 | 0.20 | 6.37 |
| 12 | Sgk3 | 1.45 | 2.00E-128 | 4.16E-124 | 0.51 | 0.14 | 0.37 | 3.65 |
| 12 | Slc25a39 | 1.01 | 3.36E-127 | 6.99E-123 | 0.97 | 0.70 | 0.27 | 1.38 |
| 12 | Apeh | 1.42 | 4.88E-126 | 1.01E-121 | 0.70 | 0.26 | 0.44 | 2.71 |
| 12 | Ehbp1l1 | 1.86 | 1.51E-125 | 3.14E-121 | 0.30 | 0.06 | 0.24 | 5.23 |
| 12 | Cdkn3 | 1.41 | 4.32E-123 | 8.98E-119 | 0.71 | 0.28 | 0.43 | 2.52 |
| 12 | Slc48a1 | 1.46 | 3.41E-122 | 7.09E-118 | 0.53 | 0.16 | 0.38 | 3.42 |
| 12 | Gadd45a | 1.16 | 5.87E-122 | 1.22E-117 | 0.67 | 0.23 | 0.44 | 2.88 |
| 12 | Trak2 | 1.66 | 2.20E-118 | 4.58E-114 | 0.46 | 0.13 | 0.33 | 3.63 |
| 12 | Unc13d | 2.31 | 4.07E-118 | 8.47E-114 | 0.18 | 0.02 | 0.16 | 7.58 |
| 12 | Cd55 | 2.19 | 3.21E-117 | 6.68E-113 | 0.16 | 0.02 | 0.14 | 8.15 |
| 12 | Slc41a3 | 2.31 | 3.66E-116 | 7.61E-112 | 0.16 | 0.02 | 0.14 | 8.15 |
| 12 | Taldo1 | 0.76 | 3.80E-116 | 7.90E-112 | 0.98 | 0.89 | 0.09 | 1.10 |
| 12 | Slc16a10 | 1.39 | 4.59E-116 | 9.54E-112 | 0.40 | 0.10 | 0.30 | 4.06 |
| 12 | St3gal1 | 1.60 | 1.14E-114 | 2.36E-110 | 0.38 | 0.09 | 0.29 | 4.13 |
| 12 | Dyrk3 | 1.52 | 2.47E-111 | 5.15E-107 | 0.37 | 0.09 | 0.28 | 4.09 |
| 12 | Fzd5 | 2.46 | 2.67E-111 | 5.56E-107 | 0.15 | 0.02 | 0.13 | 8.39 |
| 12 | Pon2 | 1.05 | 7.11E-111 | 1.48E-106 | 0.56 | 0.17 | 0.39 | 3.28 |
| 12 | Brpf3 | 1.68 | 1.77E-105 | 3.69E-101 | 0.42 | 0.12 | 0.31 | 3.66 |
| 12 | Cercam | 1.51 | 2.14E-105 | 4.44E-101 | 0.27 | 0.05 | 0.21 | 5.00 |
| 12 | Sh3yl1 | 1.38 | 2.70E-102 | 5.61E-98 | 0.56 | 0.18 | 0.38 | 3.05 |
| 12 | Tubb4b | 0.91 | 4.09E-99 | 8.51E-95 | 0.97 | 0.85 | 0.12 | 1.14 |
| 12 | Mylpf | 1.50 | 5.64E-99 | 1.17E-94 | 0.30 | 0.07 | 0.23 | 4.43 |
| 12 | Dusp8 | 1.81 | 9.09E-99 | 1.89E-94 | 0.13 | 0.02 | 0.12 | 8.80 |
| 12 | Fermt3 | 0.63 | 1.22E-96 | 2.54E-92 | 0.23 | 0.04 | 0.18 | 5.28 |
| 12 | Uba7 | 1.46 | 5.39E-94 | 1.12E-89 | 0.37 | 0.10 | 0.27 | 3.72 |
| 12 | S100a13 | 0.87 | 6.44E-92 | 1.34E-87 | 0.37 | 0.10 | 0.27 | 3.75 |
| 12 | Vamp8 | 0.89 | 2.37E-91 | 4.93E-87 | 0.91 | 0.60 | 0.31 | 1.51 |
| 12 | Arl4a | 1.01 | 1.20E-90 | 2.50E-86 | 0.56 | 0.19 | 0.37 | 2.89 |
| 12 | Atpif1 | 0.39 | 6.18E-90 | 1.28E-85 | 1.00 | 1.00 | 0.00 | 1.00 |
| 12 | Arid3a | 1.09 | 4.45E-89 | 9.26E-85 | 0.75 | 0.32 | 0.43 | 2.35 |
| 12 | Katnb1 | 1.13 | 1.16E-88 | 2.42E-84 | 0.58 | 0.21 | 0.37 | 2.76 |
| 12 | Fam110c | 2.71 | 3.41E-88 | 7.09E-84 | 0.10 | 0.01 | 0.09 | 10.00 |
| 12 | Ppp1r9b | 1.19 | 2.26E-87 | 4.70E-83 | 0.64 | 0.24 | 0.39 | 2.60 |
| 12 | Slc30a1 | 1.54 | 6.85E-87 | 1.42E-82 | 0.30 | 0.08 | 0.23 | 3.95 |

**Supplementary Table 1. Marker genes for the original clusters of integrated clustering of E11.5 control and NotchGOF λ-junction cells.**

| | | | | | | | | |
|---|---|---|---|---|---|---|---|---|
| 12 | Hscb | 0.98 | 6.26E-86 | 1.30E-81 | 0.74 | 0.31 | 0.43 | 2.37 |
| 12 | Hagh | 0.99 | 1.77E-83 | 3.68E-79 | 0.62 | 0.23 | 0.38 | 2.64 |
| 12 | G430095P16Rik | 1.22 | 6.32E-83 | 1.31E-78 | 0.35 | 0.10 | 0.25 | 3.62 |
| 12 | Capn1 | 1.61 | 4.68E-82 | 9.73E-78 | 0.22 | 0.05 | 0.17 | 4.80 |
| 12 | Reep6 | 1.58 | 8.11E-82 | 1.69E-77 | 0.22 | 0.05 | 0.17 | 4.76 |
| 12 | Mbnl1 | 1.05 | 2.43E-81 | 5.05E-77 | 0.86 | 0.46 | 0.40 | 1.86 |
| 12 | Itpka | 2.24 | 6.13E-81 | 1.27E-76 | 0.13 | 0.02 | 0.11 | 7.17 |
| 12 | Fos | 0.67 | 1.93E-80 | 4.01E-76 | 0.41 | 0.13 | 0.28 | 3.15 |
| 12 | Gypc | 1.08 | 3.15E-78 | 6.55E-74 | 0.57 | 0.21 | 0.36 | 2.73 |
| 12 | Rbks | 1.62 | 7.87E-77 | 1.64E-72 | 0.17 | 0.03 | 0.14 | 5.53 |
| 12 | Mki67 | 0.93 | 8.82E-77 | 1.84E-72 | 0.95 | 0.71 | 0.24 | 1.33 |
| 12 | 2600006K01Rik | 1.80 | 1.37E-76 | 2.84E-72 | 0.13 | 0.02 | 0.11 | 6.65 |
| 12 | Rnf123 | 1.22 | 1.04E-75 | 2.17E-71 | 0.38 | 0.12 | 0.26 | 3.26 |
| 12 | Tmc6 | 1.36 | 2.78E-75 | 5.78E-71 | 0.27 | 0.07 | 0.20 | 3.91 |
| 12 | Nudt19 | 1.03 | 5.46E-75 | 1.14E-70 | 0.68 | 0.31 | 0.37 | 2.18 |
| 12 | 1700037H04Rik | 0.86 | 2.71E-73 | 5.63E-69 | 0.86 | 0.47 | 0.40 | 1.85 |
| 12 | Rffl | 1.35 | 4.86E-73 | 1.01E-68 | 0.26 | 0.07 | 0.19 | 3.94 |
| 12 | Dnase1l1 | 1.41 | 5.91E-69 | 1.23E-64 | 0.20 | 0.04 | 0.15 | 4.53 |
| 12 | Ankle1 | 1.35 | 1.25E-68 | 2.60E-64 | 0.30 | 0.09 | 0.21 | 3.46 |
| 12 | Ybx3 | 0.61 | 4.55E-68 | 9.47E-64 | 0.97 | 0.83 | 0.14 | 1.17 |
| 12 | S100a1 | 1.20 | 5.69E-68 | 1.18E-63 | 0.24 | 0.06 | 0.18 | 3.98 |
| 12 | Tom1l1 | 0.72 | 8.06E-68 | 1.68E-63 | 0.45 | 0.15 | 0.29 | 2.91 |
| 12 | Skap2 | 0.99 | 9.48E-68 | 1.97E-63 | 0.22 | 0.05 | 0.17 | 4.30 |
| 12 | Ifi30 | 0.76 | 1.04E-67 | 2.16E-63 | 0.65 | 0.28 | 0.37 | 2.30 |
| 12 | Arf5 | 0.36 | 3.14E-64 | 6.54E-60 | 0.99 | 0.98 | 0.01 | 1.01 |
| 12 | Ampd3 | 2.19 | 4.83E-64 | 1.01E-59 | 0.10 | 0.02 | 0.09 | 6.87 |
| 12 | Ift140 | 1.03 | 8.85E-64 | 1.84E-59 | 0.43 | 0.15 | 0.28 | 2.80 |
| 12 | Nipa1 | 0.86 | 7.89E-63 | 1.64E-58 | 0.40 | 0.14 | 0.26 | 2.93 |
| 12 | Mpst | 0.80 | 2.22E-62 | 4.63E-58 | 0.79 | 0.41 | 0.38 | 1.93 |
| 12 | Pop5 | 0.68 | 4.59E-61 | 9.55E-57 | 0.92 | 0.64 | 0.27 | 1.42 |
| 12 | Dennd2c | 1.16 | 1.79E-60 | 3.72E-56 | 0.23 | 0.06 | 0.17 | 3.85 |
| 12 | Slc25a38 | 0.98 | 1.97E-59 | 4.09E-55 | 0.60 | 0.26 | 0.33 | 2.26 |
| 12 | Pycr2 | 0.72 | 2.42E-59 | 5.04E-55 | 0.91 | 0.60 | 0.30 | 1.50 |
| 12 | Ftl1-ps1 | 0.74 | 3.90E-59 | 8.11E-55 | 0.78 | 0.40 | 0.39 | 1.98 |
| 12 | Stx2 | 0.87 | 1.53E-58 | 3.18E-54 | 0.52 | 0.21 | 0.31 | 2.45 |
| 12 | Nars | 0.61 | 7.28E-58 | 1.51E-53 | 0.96 | 0.80 | 0.16 | 1.20 |
| 12 | Tuba1c | 0.96 | 3.99E-57 | 8.30E-53 | 0.74 | 0.43 | 0.31 | 1.71 |
| 12 | Rad23a | 0.83 | 6.40E-57 | 1.33E-52 | 0.83 | 0.45 | 0.39 | 1.86 |
| 12 | H2-T23 | 1.12 | 1.15E-56 | 2.39E-52 | 0.20 | 0.05 | 0.15 | 3.96 |
| 12 | Nadk2 | 0.82 | 1.91E-56 | 3.97E-52 | 0.70 | 0.34 | 0.35 | 2.03 |
| 12 | Gch1 | 0.92 | 2.93E-56 | 6.10E-52 | 0.25 | 0.07 | 0.18 | 3.49 |
| 12 | Hspa4l | 0.82 | 4.19E-56 | 8.72E-52 | 0.32 | 0.11 | 0.22 | 3.06 |
| 12 | Pabpc4 | 0.77 | 1.60E-55 | 3.32E-51 | 0.92 | 0.64 | 0.28 | 1.43 |
| 12 | Zfp367 | 1.02 | 3.90E-53 | 8.10E-49 | 0.37 | 0.14 | 0.23 | 2.70 |
| 12 | Mafk | 1.50 | 1.28E-52 | 2.67E-48 | 0.16 | 0.04 | 0.12 | 4.30 |
| 12 | Tmem179b | 0.72 | 3.45E-52 | 7.17E-48 | 0.62 | 0.29 | 0.34 | 2.18 |
| 12 | Cep76 | 0.87 | 2.73E-51 | 5.67E-47 | 0.41 | 0.16 | 0.25 | 2.61 |
| 12 | Hist1h4i | 0.79 | 7.47E-50 | 1.55E-45 | 0.79 | 0.45 | 0.34 | 1.76 |
| 12 | Cdkn2d | 1.00 | 8.84E-49 | 1.84E-44 | 0.65 | 0.38 | 0.28 | 1.74 |
| 12 | Smox | 0.72 | 4.21E-48 | 8.75E-44 | 0.53 | 0.23 | 0.30 | 2.28 |
| 12 | Rab3il1 | 0.61 | 1.25E-46 | 2.59E-42 | 0.62 | 0.29 | 0.34 | 2.17 |
| 12 | Adcy7 | 1.85 | 1.37E-46 | 2.86E-42 | 0.11 | 0.02 | 0.09 | 5.19 |
| 12 | Pcgf5 | 0.96 | 2.28E-46 | 4.74E-42 | 0.25 | 0.08 | 0.17 | 3.06 |
| 12 | Nqo1 | 0.91 | 3.68E-46 | 7.65E-42 | 0.28 | 0.09 | 0.18 | 2.96 |
| 12 | Optn | 1.46 | 3.93E-46 | 8.18E-42 | 0.12 | 0.03 | 0.10 | 4.73 |
| 12 | Smim4 | 0.62 | 6.34E-46 | 1.32E-41 | 0.77 | 0.42 | 0.35 | 1.84 |

**Supplementary Table 1. Marker genes for the original clusters of integrated clustering of E11.5 control and NotchGOF λ-junction cells.**

| | | | | | | | | |
|---|---|---|---|---|---|---|---|---|
| 12 | Aff1 | 1.32 | 9.22E-46 | 1.92E-41 | 0.22 | 0.07 | 0.15 | 3.27 |
| 12 | Nup210 | 1.58 | 2.42E-45 | 5.04E-41 | 0.20 | 0.06 | 0.14 | 3.41 |
| 12 | Fbxo9 | 0.67 | 5.91E-44 | 1.23E-39 | 0.60 | 0.28 | 0.31 | 2.11 |
| 12 | Spsb2 | 0.80 | 8.16E-44 | 1.70E-39 | 0.23 | 0.07 | 0.16 | 3.19 |
| 12 | Stradb | 0.83 | 2.11E-43 | 4.40E-39 | 0.44 | 0.19 | 0.25 | 2.36 |
| 12 | Asns | 0.70 | 4.95E-43 | 1.03E-38 | 0.76 | 0.43 | 0.32 | 1.75 |
| 12 | Iba57 | 1.37 | 9.14E-43 | 1.90E-38 | 0.14 | 0.03 | 0.11 | 4.18 |
| 12 | Rap1a | 0.48 | 4.99E-42 | 1.04E-37 | 0.93 | 0.71 | 0.22 | 1.30 |
| 12 | Frrs1 | 0.74 | 1.31E-41 | 2.73E-37 | 0.47 | 0.22 | 0.26 | 2.20 |
| 12 | Yod1 | 0.78 | 7.31E-41 | 1.52E-36 | 0.35 | 0.14 | 0.21 | 2.53 |
| 12 | Capn5 | 1.66 | 1.12E-40 | 2.33E-36 | 0.11 | 0.03 | 0.09 | 4.52 |
| 12 | Skiv2l | 0.89 | 1.63E-40 | 3.40E-36 | 0.52 | 0.24 | 0.28 | 2.18 |
| 12 | Glrx | 0.48 | 1.75E-40 | 3.63E-36 | 0.50 | 0.23 | 0.27 | 2.20 |
| 12 | Rmc1 | 0.88 | 1.90E-40 | 3.95E-36 | 0.23 | 0.08 | 0.16 | 3.03 |
| 12 | Hspa12b | 0.66 | 2.00E-39 | 4.15E-35 | 0.33 | 0.13 | 0.20 | 2.58 |
| 12 | Tmem120b | 0.92 | 1.25E-38 | 2.59E-34 | 0.19 | 0.06 | 0.13 | 3.30 |
| 12 | Akip1 | 0.58 | 1.27E-38 | 2.65E-34 | 0.44 | 0.19 | 0.25 | 2.30 |
| 12 | Abcb6 | 0.95 | 2.69E-38 | 5.59E-34 | 0.30 | 0.11 | 0.18 | 2.61 |
| 12 | Tarsl2 | 0.77 | 3.74E-38 | 7.77E-34 | 0.26 | 0.09 | 0.17 | 2.79 |
| 12 | Pir | 0.79 | 4.07E-38 | 8.47E-34 | 0.31 | 0.12 | 0.19 | 2.54 |
| 12 | Psme3 | 0.61 | 4.72E-38 | 9.83E-34 | 0.82 | 0.49 | 0.33 | 1.68 |
| 12 | Tatdn3 | 0.86 | 7.99E-38 | 1.66E-33 | 0.27 | 0.10 | 0.17 | 2.69 |
| 12 | Narf | 0.55 | 6.89E-37 | 1.43E-32 | 0.72 | 0.41 | 0.31 | 1.75 |
| 12 | Ubl4a | 0.52 | 6.97E-37 | 1.45E-32 | 0.87 | 0.57 | 0.30 | 1.53 |
| 12 | Bcor | 0.68 | 1.24E-36 | 2.57E-32 | 0.71 | 0.41 | 0.31 | 1.76 |
| 12 | Slc11a2 | 0.68 | 1.41E-36 | 2.93E-32 | 0.39 | 0.17 | 0.23 | 2.35 |
| 12 | Pdk2 | 0.86 | 4.54E-36 | 9.44E-32 | 0.23 | 0.08 | 0.15 | 2.88 |
| 12 | Dleu2 | 0.72 | 9.66E-36 | 2.01E-31 | 0.51 | 0.25 | 0.26 | 2.06 |
| 12 | Zfand2b | 0.53 | 1.24E-35 | 2.59E-31 | 0.52 | 0.25 | 0.27 | 2.10 |
| 12 | Ank | 1.00 | 1.84E-35 | 3.83E-31 | 0.20 | 0.07 | 0.13 | 3.00 |
| 12 | Eif2ak1 | 0.58 | 4.46E-35 | 9.28E-31 | 0.73 | 0.41 | 0.32 | 1.79 |
| 12 | Ctsb | 0.30 | 7.22E-35 | 1.50E-30 | 0.95 | 0.75 | 0.21 | 1.28 |
| 12 | Sord | 0.52 | 8.16E-35 | 1.70E-30 | 0.34 | 0.14 | 0.20 | 2.41 |
| 12 | Foxo3 | 1.00 | 2.59E-34 | 5.39E-30 | 0.30 | 0.12 | 0.18 | 2.47 |
| 12 | Abhd5 | 0.48 | 1.84E-33 | 3.83E-29 | 0.43 | 0.19 | 0.24 | 2.22 |
| 12 | Irak2 | 0.78 | 3.02E-33 | 6.27E-29 | 0.16 | 0.05 | 0.11 | 3.26 |
| 12 | Stxbp2 | 0.42 | 4.09E-33 | 8.51E-29 | 0.32 | 0.13 | 0.19 | 2.43 |
| 12 | Fam174b | 0.65 | 8.83E-33 | 1.84E-28 | 0.27 | 0.11 | 0.17 | 2.51 |
| 12 | Gm35082 | 1.41 | 9.44E-33 | 1.96E-28 | 0.11 | 0.03 | 0.08 | 4.08 |
| 12 | Mthfd2 | 0.52 | 1.52E-32 | 3.16E-28 | 0.78 | 0.46 | 0.31 | 1.67 |
| 12 | Ogfrl1 | 0.56 | 2.11E-32 | 4.38E-28 | 0.62 | 0.33 | 0.29 | 1.89 |
| 12 | Fryl | 0.60 | 6.21E-32 | 1.29E-27 | 0.57 | 0.29 | 0.28 | 1.95 |
| 12 | Clk3 | 0.60 | 1.32E-31 | 2.75E-27 | 0.67 | 0.37 | 0.30 | 1.80 |
| 12 | Tnfrsf21 | 0.70 | 2.07E-31 | 4.30E-27 | 0.32 | 0.14 | 0.18 | 2.34 |
| 12 | Ier3 | 0.99 | 1.92E-30 | 3.98E-26 | 0.20 | 0.07 | 0.13 | 2.78 |
| 12 | Dedd2 | 0.82 | 2.26E-30 | 4.70E-26 | 0.20 | 0.07 | 0.13 | 2.82 |
| 12 | Pabpc1 | 0.31 | 3.91E-30 | 8.14E-26 | 1.00 | 0.99 | 0.01 | 1.01 |
| 12 | Khk | 0.55 | 6.45E-30 | 1.34E-25 | 0.18 | 0.06 | 0.12 | 2.90 |
| 12 | Podxl | 0.71 | 2.26E-28 | 4.70E-24 | 0.14 | 0.04 | 0.09 | 3.29 |
| 12 | Ap1g2 | 0.93 | 5.07E-28 | 1.05E-23 | 0.13 | 0.04 | 0.09 | 3.25 |
| 12 | Mbd2 | 0.47 | 1.03E-27 | 2.15E-23 | 0.84 | 0.55 | 0.29 | 1.53 |
| 12 | Taok3 | 0.47 | 2.23E-27 | 4.65E-23 | 0.25 | 0.10 | 0.15 | 2.45 |
| 12 | Mvb12b | 0.45 | 2.77E-27 | 5.77E-23 | 0.45 | 0.22 | 0.23 | 2.02 |
| 12 | Stat5a | 1.30 | 3.48E-27 | 7.23E-23 | 0.10 | 0.03 | 0.07 | 3.70 |
| 12 | Il10rb | 0.51 | 3.74E-27 | 7.78E-23 | 0.21 | 0.08 | 0.13 | 2.60 |
| 12 | Atg4d | 0.50 | 5.35E-27 | 1.11E-22 | 0.42 | 0.20 | 0.22 | 2.08 |

**Supplementary Table 1. Marker genes for the original clusters of integrated clustering of E11.5 control and NotchGOF λ-junction cells.**

| | | | | | | | | |
|---|---|---|---|---|---|---|---|---|
| 12 | Retreg2 | 0.44 | 7.15E-27 | 1.49E-22 | 0.54 | 0.28 | 0.26 | 1.93 |
| 12 | Snx15 | 0.42 | 1.41E-26 | 2.93E-22 | 0.56 | 0.30 | 0.26 | 1.87 |
| 12 | E2f8 | 0.48 | 2.29E-26 | 4.76E-22 | 0.41 | 0.20 | 0.21 | 2.04 |
| 12 | Ccndbp1 | 0.28 | 3.43E-26 | 7.13E-22 | 0.54 | 0.27 | 0.27 | 1.97 |
| 12 | 1110008P14Rik | 0.39 | 9.63E-26 | 2.00E-21 | 0.77 | 0.46 | 0.31 | 1.68 |
| 12 | Zfp524 | 0.53 | 1.39E-25 | 2.90E-21 | 0.24 | 0.10 | 0.14 | 2.40 |
| 12 | Micall2 | 0.72 | 5.20E-25 | 1.08E-20 | 0.17 | 0.06 | 0.11 | 2.72 |
| 12 | Ccdc82 | 0.45 | 5.99E-25 | 1.25E-20 | 0.27 | 0.11 | 0.15 | 2.32 |
| 12 | Rgcc | 0.55 | 6.02E-25 | 1.25E-20 | 0.22 | 0.09 | 0.13 | 2.43 |
| 12 | Plin2 | 0.47 | 4.13E-24 | 8.60E-20 | 0.79 | 0.52 | 0.26 | 1.50 |
| 12 | Neat1 | 0.78 | 5.81E-24 | 1.21E-19 | 0.15 | 0.05 | 0.10 | 2.79 |
| 12 | Cacna2d3 | 0.33 | 8.17E-24 | 1.70E-19 | 0.19 | 0.07 | 0.12 | 2.58 |
| 12 | Ptgr2 | 0.35 | 1.09E-23 | 2.27E-19 | 0.32 | 0.15 | 0.17 | 2.17 |
| 12 | Synj1 | 0.90 | 1.81E-23 | 3.77E-19 | 0.24 | 0.10 | 0.14 | 2.32 |
| 12 | Atp2b4 | 0.49 | 2.71E-23 | 5.64E-19 | 0.40 | 0.20 | 0.20 | 2.01 |
| 12 | Chst10 | 0.73 | 2.91E-23 | 6.06E-19 | 0.20 | 0.08 | 0.12 | 2.46 |
| 12 | Fbxo34 | 0.44 | 3.05E-23 | 6.34E-19 | 0.53 | 0.29 | 0.24 | 1.83 |
| 12 | Mad2l1bp | 0.33 | 7.94E-23 | 1.65E-18 | 0.45 | 0.23 | 0.22 | 1.95 |
| 12 | Selenbp1 | 0.44 | 9.27E-23 | 1.93E-18 | 0.20 | 0.08 | 0.12 | 2.45 |
| 12 | Eipr1 | 0.40 | 1.37E-22 | 2.84E-18 | 0.45 | 0.23 | 0.22 | 1.93 |
| 12 | Tlcd1 | 0.83 | 1.38E-22 | 2.88E-18 | 0.21 | 0.09 | 0.12 | 2.41 |
| 12 | Afg3l2 | 0.35 | 1.79E-22 | 3.73E-18 | 0.60 | 0.34 | 0.27 | 1.80 |
| 12 | Fam104a | 0.29 | 3.44E-22 | 7.14E-18 | 0.97 | 0.88 | 0.09 | 1.10 |
| 12 | Cmas | 0.40 | 4.05E-22 | 8.42E-18 | 0.84 | 0.59 | 0.25 | 1.43 |
| 12 | Tapbp | 0.59 | 4.49E-22 | 9.35E-18 | 0.19 | 0.08 | 0.12 | 2.46 |
| 12 | Agfg2 | 0.38 | 8.21E-22 | 1.71E-17 | 0.41 | 0.21 | 0.20 | 1.97 |
| 12 | D1Ertd622e | 0.54 | 2.33E-21 | 4.84E-17 | 0.27 | 0.13 | 0.14 | 2.13 |
| 12 | Wbp2 | 0.30 | 6.35E-21 | 1.32E-16 | 0.51 | 0.27 | 0.24 | 1.87 |
| 12 | Slc23a2 | 0.73 | 1.53E-20 | 3.17E-16 | 0.18 | 0.07 | 0.11 | 2.43 |
| 12 | Dusp1 | 0.49 | 1.69E-20 | 3.52E-16 | 0.51 | 0.29 | 0.23 | 1.79 |
| 12 | Hist1h2bj | 0.47 | 2.27E-20 | 4.73E-16 | 0.45 | 0.24 | 0.21 | 1.86 |
| 12 | Adipor1 | 0.34 | 8.87E-20 | 1.84E-15 | 0.84 | 0.56 | 0.28 | 1.51 |
| 12 | Zbtb7a | 0.38 | 1.50E-19 | 3.12E-15 | 0.56 | 0.33 | 0.24 | 1.73 |
| 12 | Pip4k2a | 0.55 | 1.88E-19 | 3.92E-15 | 0.24 | 0.11 | 0.13 | 2.16 |
| 12 | Adgrg1 | 0.76 | 2.15E-19 | 4.48E-15 | 0.12 | 0.04 | 0.08 | 2.88 |
| 12 | Pik3cb | 0.55 | 2.49E-19 | 5.19E-15 | 0.15 | 0.06 | 0.09 | 2.51 |
| 12 | 1600002K03Rik | 0.39 | 8.06E-19 | 1.68E-14 | 0.24 | 0.11 | 0.13 | 2.13 |
| 12 | Sod2 | 0.30 | 1.00E-18 | 2.08E-14 | 0.87 | 0.62 | 0.25 | 1.41 |
| 12 | Tacc1 | 0.31 | 1.43E-18 | 2.98E-14 | 0.55 | 0.31 | 0.24 | 1.76 |
| 12 | Gm4013 | 0.62 | 5.04E-18 | 1.05E-13 | 0.13 | 0.05 | 0.08 | 2.68 |
| 12 | Noct | 0.38 | 5.64E-18 | 1.17E-13 | 0.35 | 0.19 | 0.17 | 1.90 |
| 12 | Hif3a | 0.52 | 1.15E-17 | 2.38E-13 | 0.19 | 0.09 | 0.11 | 2.25 |
| 12 | Clcn3 | 0.58 | 1.28E-17 | 2.67E-13 | 0.64 | 0.37 | 0.27 | 1.72 |
| 12 | Napepld | 0.48 | 3.46E-17 | 7.20E-13 | 0.14 | 0.06 | 0.09 | 2.48 |
| 12 | Emc9 | 0.57 | 3.92E-17 | 8.16E-13 | 0.12 | 0.05 | 0.08 | 2.63 |
| 12 | Bcat2 | 0.26 | 4.77E-17 | 9.93E-13 | 0.50 | 0.29 | 0.22 | 1.77 |
| 12 | Hipk1 | 0.36 | 7.22E-17 | 1.50E-12 | 0.61 | 0.36 | 0.25 | 1.70 |
| 12 | Traf5 | 0.68 | 9.15E-17 | 1.90E-12 | 0.11 | 0.04 | 0.07 | 2.74 |
| 12 | Cbfa2t3 | 0.91 | 1.67E-16 | 3.47E-12 | 0.13 | 0.05 | 0.08 | 2.49 |
| 12 | Wdr81 | 0.74 | 1.94E-16 | 4.03E-12 | 0.12 | 0.05 | 0.08 | 2.56 |
| 12 | Cln8 | 0.26 | 2.39E-16 | 4.97E-12 | 0.37 | 0.20 | 0.17 | 1.86 |
| 12 | Snrnp25 | 0.38 | 6.51E-16 | 1.35E-11 | 0.82 | 0.61 | 0.21 | 1.35 |
| 12 | Myc | 0.35 | 7.33E-16 | 1.52E-11 | 0.30 | 0.16 | 0.14 | 1.88 |
| 12 | Pet100 | 0.34 | 1.04E-15 | 2.16E-11 | 0.81 | 0.55 | 0.26 | 1.48 |
| 12 | Polh | 0.46 | 1.35E-15 | 2.82E-11 | 0.19 | 0.09 | 0.10 | 2.16 |
| 12 | Piezo1 | 0.46 | 1.87E-15 | 3.89E-11 | 0.24 | 0.12 | 0.12 | 1.99 |

**Supplementary Table 1. Marker genes for the original clusters of integrated clustering of E11.5 control and NotchGOF λ-junction cells.**

| | | | | | | | | |
|---|---|---|---|---|---|---|---|---|
| 12 | Rab8b | 0.34 | 5.70E-15 | 1.19E-10 | 0.42 | 0.24 | 0.18 | 1.76 |
| 12 | Ogfod2 | 0.33 | 1.40E-14 | 2.92E-10 | 0.20 | 0.10 | 0.10 | 2.06 |
| 12 | Slc7a5 | 0.56 | 1.44E-14 | 2.99E-10 | 0.12 | 0.05 | 0.07 | 2.50 |
| 12 | Otub2 | 0.56 | 3.09E-14 | 6.43E-10 | 0.12 | 0.05 | 0.07 | 2.42 |
| 12 | Arrdc2 | 0.90 | 3.37E-14 | 7.01E-10 | 0.11 | 0.04 | 0.07 | 2.48 |
| 12 | Man1a | 0.89 | 1.06E-13 | 2.20E-09 | 0.14 | 0.06 | 0.08 | 2.27 |
| 12 | Flvcr1 | 0.57 | 1.81E-13 | 3.77E-09 | 0.19 | 0.10 | 0.10 | 2.01 |
| 12 | Ddit3 | 0.38 | 1.98E-13 | 4.13E-09 | 0.35 | 0.20 | 0.15 | 1.74 |
| 12 | Eif2b3 | 0.30 | 6.68E-13 | 1.39E-08 | 0.70 | 0.48 | 0.23 | 1.48 |
| 12 | Grtp1 | 0.27 | 1.49E-12 | 3.10E-08 | 0.26 | 0.14 | 0.12 | 1.88 |
| 12 | Sppl2b | 0.34 | 1.57E-12 | 3.26E-08 | 0.27 | 0.14 | 0.12 | 1.84 |
| 12 | Jak2 | 0.29 | 1.91E-12 | 3.98E-08 | 0.28 | 0.15 | 0.13 | 1.82 |
| 12 | Marchf8 | 0.39 | 6.05E-12 | 1.26E-07 | 0.29 | 0.17 | 0.13 | 1.75 |
| 12 | Ninl | 0.42 | 1.05E-11 | 2.18E-07 | 0.40 | 0.24 | 0.16 | 1.67 |
| 12 | Ddc | 0.55 | 1.15E-11 | 2.39E-07 | 0.12 | 0.05 | 0.06 | 2.19 |
| 12 | Kat2b | 0.63 | 1.54E-10 | 3.20E-06 | 0.10 | 0.05 | 0.06 | 2.24 |
| 12 | Hipk2 | 0.29 | 1.93E-10 | 4.01E-06 | 0.85 | 0.64 | 0.21 | 1.33 |
| 12 | Slc7a1 | 0.32 | 2.46E-10 | 5.12E-06 | 0.15 | 0.08 | 0.07 | 1.96 |
| 12 | Rhbdd1 | 0.35 | 4.54E-10 | 9.43E-06 | 0.11 | 0.05 | 0.06 | 2.18 |
| 12 | Cish | 0.56 | 4.75E-10 | 9.87E-06 | 0.13 | 0.06 | 0.06 | 2.03 |
| 12 | Znfx1 | 0.34 | 3.85E-09 | 8.01E-05 | 0.12 | 0.06 | 0.06 | 2.03 |
| 12 | Zfp874b | 0.26 | 7.10E-08 | 1.48E-03 | 0.10 | 0.05 | 0.05 | 1.98 |
| 12 | Fam234b | 0.35 | 1.34E-07 | 2.79E-03 | 0.11 | 0.06 | 0.05 | 1.95 |
| 12 | Spata13 | 0.76 | 1.62E-06 | 3.36E-02 | 0.13 | 0.08 | 0.05 | 1.71 |
| 13 | Aldh1a3 | 4.21 | 0 | 0 | 0.97 | 0.21 | 0.75 | 4.57 |
| 13 | Six3 | 2.77 | 0 | 0 | 0.97 | 0.22 | 0.75 | 4.43 |
| 13 | Pdgfa | 2.34 | 0 | 0 | 0.99 | 0.26 | 0.73 | 3.77 |
| 13 | Cldn6 | 2.27 | 0 | 0 | 0.94 | 0.21 | 0.73 | 4.44 |
| 13 | Six3os1 | 2.96 | 0 | 0 | 0.93 | 0.21 | 0.72 | 4.40 |
| 13 | Isl1 | 2.54 | 0 | 0 | 0.89 | 0.20 | 0.69 | 4.43 |
| 13 | Krt8 | 1.76 | 0 | 0 | 0.93 | 0.24 | 0.69 | 3.84 |
| 13 | Spint2 | 2.01 | 0 | 0 | 0.98 | 0.29 | 0.69 | 3.34 |
| 13 | Cldn7 | 2.24 | 0 | 0 | 0.90 | 0.22 | 0.67 | 4.04 |
| 13 | Pitx1 | 2.96 | 0 | 0 | 0.84 | 0.16 | 0.67 | 5.10 |
| 13 | Krt18 | 2.11 | 0 | 0 | 0.95 | 0.28 | 0.66 | 3.35 |
| 13 | Epcam | 2.33 | 0 | 0 | 0.99 | 0.33 | 0.66 | 3.01 |
| 13 | Sox2 | 2.16 | 0 | 0 | 0.84 | 0.18 | 0.66 | 4.77 |
| 13 | Foxg1 | 2.80 | 0 | 0 | 0.78 | 0.13 | 0.66 | 6.17 |
| 13 | Ezr | 2.19 | 0 | 0 | 0.96 | 0.30 | 0.65 | 3.16 |
| 13 | Hist3h2ba | 2.16 | 0 | 0 | 0.90 | 0.26 | 0.64 | 3.49 |
| 13 | Fgf9 | 3.25 | 0 | 0 | 0.71 | 0.09 | 0.62 | 8.01 |
| 13 | Dmrta2 | 2.81 | 0 | 0 | 0.74 | 0.13 | 0.61 | 5.89 |
| 13 | Bcam | 2.49 | 0 | 0 | 0.79 | 0.19 | 0.60 | 4.21 |
| 13 | Sp8 | 2.73 | 0 | 0 | 0.72 | 0.12 | 0.60 | 5.88 |
| 13 | Pax6 | 2.57 | 0 | 0 | 0.66 | 0.11 | 0.55 | 5.81 |
| 13 | Cldn9 | 3.08 | 0 | 0 | 0.59 | 0.08 | 0.51 | 7.81 |
| 13 | Nrg1 | 3.29 | 0 | 0 | 0.55 | 0.07 | 0.49 | 8.39 |
| 13 | D230030E09Rik | 4.34 | 0 | 0 | 0.53 | 0.04 | 0.48 | 12.26 |
| 13 | Dmrt3 | 3.47 | 0 | 0 | 0.44 | 0.04 | 0.40 | 10.28 |
| 13 | Fgf12 | 3.56 | 0 | 0 | 0.41 | 0.04 | 0.37 | 10.49 |
| 13 | Unc13c | 4.94 | 0 | 0 | 0.28 | 0.01 | 0.27 | 25.45 |
| 13 | Fam107a | 3.35 | 2.34E-307 | 4.88E-303 | 0.36 | 0.04 | 0.32 | 9.92 |
| 13 | Otx1 | 2.57 | 5.35E-307 | 1.11E-302 | 0.59 | 0.10 | 0.49 | 5.90 |
| 13 | 3110039M20Rik | 2.68 | 6.41E-303 | 1.33E-298 | 0.48 | 0.07 | 0.41 | 7.24 |
| 13 | Ikzf2 | 2.15 | 7.12E-299 | 1.48E-294 | 0.90 | 0.27 | 0.64 | 3.39 |
| 13 | Rab25 | 2.18 | 6.32E-298 | 1.31E-293 | 0.72 | 0.15 | 0.57 | 4.91 |

**Supplementary Table 1. Marker genes for the original clusters of integrated clustering of E11.5 control and NotchGOF λ-junction cells.**

| | | | | | | | | |
|---|---|---|---|---|---|---|---|---|
| 13 | Cadm1 | 2.46 | 1.08E-293 | 2.24E-289 | 0.92 | 0.37 | 0.55 | 2.47 |
| 13 | Nrxn3 | 4.14 | 1.16E-289 | 2.41E-285 | 0.27 | 0.02 | 0.25 | 12.86 |
| 13 | Espn | 2.91 | 3.58E-285 | 7.45E-281 | 0.55 | 0.10 | 0.46 | 5.69 |
| 13 | Pdgfc | 2.12 | 4.76E-284 | 9.91E-280 | 0.88 | 0.27 | 0.61 | 3.31 |
| 13 | Cdh1 | 2.26 | 1.07E-282 | 2.23E-278 | 0.72 | 0.16 | 0.56 | 4.63 |
| 13 | Tmeff2 | 4.69 | 3.84E-282 | 7.99E-278 | 0.19 | 0.01 | 0.18 | 19.30 |
| 13 | Ap1m2 | 2.31 | 1.75E-281 | 3.63E-277 | 0.65 | 0.13 | 0.52 | 5.02 |
| 13 | Prss22 | 3.18 | 1.10E-277 | 2.29E-273 | 0.41 | 0.05 | 0.35 | 7.79 |
| 13 | Wnt4 | 2.20 | 3.46E-274 | 7.19E-270 | 0.76 | 0.19 | 0.57 | 4.08 |
| 13 | Cldn3 | 2.04 | 1.10E-271 | 2.30E-267 | 0.70 | 0.15 | 0.55 | 4.68 |
| 13 | Fgf17 | 3.43 | 1.65E-267 | 3.42E-263 | 0.33 | 0.04 | 0.29 | 9.40 |
| 13 | Tpd52 | 1.86 | 2.63E-267 | 5.48E-263 | 0.86 | 0.25 | 0.61 | 3.49 |
| 13 | Igfbp2 | 1.92 | 6.28E-264 | 1.31E-259 | 1.00 | 0.75 | 0.25 | 1.33 |
| 13 | Pcp4 | 4.36 | 3.39E-259 | 7.04E-255 | 0.20 | 0.01 | 0.18 | 16.33 |
| 13 | Igfbp5 | 2.34 | 4.27E-256 | 8.89E-252 | 0.98 | 0.39 | 0.58 | 2.48 |
| 13 | Ccdc88c | 2.28 | 7.12E-252 | 1.48E-247 | 0.65 | 0.14 | 0.51 | 4.52 |
| 13 | Crb3 | 2.37 | 2.32E-250 | 4.83E-246 | 0.54 | 0.10 | 0.44 | 5.45 |
| 13 | Scn9a | 4.26 | 2.85E-246 | 5.92E-242 | 0.16 | 0.01 | 0.15 | 22.14 |
| 13 | Pla2r1 | 2.60 | 6.19E-246 | 1.29E-241 | 0.43 | 0.07 | 0.37 | 6.66 |
| 13 | Ly6h | 2.70 | 1.89E-242 | 3.93E-238 | 0.61 | 0.14 | 0.48 | 4.44 |
| 13 | Ano1 | 2.30 | 2.94E-242 | 6.12E-238 | 0.58 | 0.12 | 0.46 | 4.98 |
| 13 | Prss8 | 2.52 | 3.04E-238 | 6.33E-234 | 0.49 | 0.09 | 0.41 | 5.67 |
| 13 | Slc9a3r1 | 1.86 | 9.34E-234 | 1.94E-229 | 0.79 | 0.23 | 0.57 | 3.50 |
| 13 | Bex1 | 1.78 | 2.07E-233 | 4.30E-229 | 0.98 | 0.58 | 0.40 | 1.69 |
| 13 | Cldn4 | 2.34 | 5.86E-233 | 1.22E-228 | 0.51 | 0.09 | 0.41 | 5.45 |
| 13 | Gad2 | 2.84 | 2.11E-232 | 4.40E-228 | 0.42 | 0.07 | 0.36 | 6.46 |
| 13 | Ctxn3 | 3.81 | 1.57E-231 | 3.26E-227 | 0.26 | 0.02 | 0.23 | 10.63 |
| 13 | Dsp | 1.51 | 1.86E-230 | 3.86E-226 | 0.82 | 0.22 | 0.60 | 3.71 |
| 13 | Lypd6b | 2.98 | 2.29E-229 | 4.77E-225 | 0.40 | 0.06 | 0.34 | 6.58 |
| 13 | Esrp1 | 2.04 | 1.30E-226 | 2.71E-222 | 0.66 | 0.16 | 0.50 | 4.21 |
| 13 | Upk3bl | 2.54 | 5.56E-226 | 1.16E-221 | 0.41 | 0.06 | 0.35 | 6.44 |
| 13 | Mapk13 | 2.04 | 2.50E-224 | 5.20E-220 | 0.71 | 0.19 | 0.52 | 3.79 |
| 13 | Acer2 | 1.81 | 1.16E-219 | 2.42E-215 | 0.91 | 0.38 | 0.53 | 2.38 |
| 13 | Lad1 | 2.54 | 6.63E-215 | 1.38E-210 | 0.45 | 0.08 | 0.37 | 5.64 |
| 13 | Grhl2 | 2.14 | 2.24E-214 | 4.66E-210 | 0.56 | 0.12 | 0.44 | 4.69 |
| 13 | Lama5 | 2.33 | 3.76E-208 | 7.83E-204 | 0.51 | 0.11 | 0.41 | 4.90 |
| 13 | Mpzl2 | 1.69 | 7.40E-208 | 1.54E-203 | 0.73 | 0.19 | 0.53 | 3.74 |
| 13 | Kif21a | 1.82 | 1.00E-207 | 2.09E-203 | 0.83 | 0.30 | 0.53 | 2.79 |
| 13 | Ptchd1 | 3.36 | 3.29E-202 | 6.84E-198 | 0.21 | 0.02 | 0.20 | 11.26 |
| 13 | Frem2 | 2.10 | 3.57E-202 | 7.43E-198 | 0.67 | 0.18 | 0.49 | 3.69 |
| 13 | Prr15 | 3.15 | 2.93E-200 | 6.09E-196 | 0.26 | 0.03 | 0.23 | 9.11 |
| 13 | Ccdc171 | 2.66 | 5.33E-197 | 1.11E-192 | 0.45 | 0.09 | 0.36 | 5.17 |
| 13 | Eya2 | 2.14 | 6.54E-197 | 1.36E-192 | 0.72 | 0.22 | 0.50 | 3.27 |
| 13 | Ctnnd2 | 2.21 | 3.44E-195 | 7.16E-191 | 0.51 | 0.11 | 0.40 | 4.65 |
| 13 | Kdf1 | 2.39 | 3.68E-195 | 7.66E-191 | 0.43 | 0.08 | 0.35 | 5.52 |
| 13 | Gm16136 | 1.49 | 9.83E-195 | 2.04E-190 | 0.65 | 0.16 | 0.49 | 4.03 |
| 13 | Fgf16 | 3.88 | 6.59E-194 | 1.37E-189 | 0.18 | 0.01 | 0.17 | 12.93 |
| 13 | Lsr | 1.91 | 3.99E-193 | 8.29E-189 | 0.68 | 0.19 | 0.49 | 3.52 |
| 13 | Tubb2b | 1.52 | 5.25E-193 | 1.09E-188 | 0.97 | 0.66 | 0.31 | 1.47 |
| 13 | Arhgef19 | 2.05 | 6.30E-190 | 1.31E-185 | 0.54 | 0.12 | 0.41 | 4.42 |
| 13 | Ces1d | 3.08 | 7.48E-189 | 1.56E-184 | 0.29 | 0.04 | 0.25 | 7.70 |
| 13 | Rragd | 2.07 | 4.17E-188 | 8.68E-184 | 0.68 | 0.21 | 0.47 | 3.22 |
| 13 | Jag2 | 2.66 | 7.13E-188 | 1.48E-183 | 0.40 | 0.07 | 0.33 | 5.51 |
| 13 | Bex4 | 1.45 | 2.01E-187 | 4.19E-183 | 0.97 | 0.64 | 0.33 | 1.52 |
| 13 | Gm32585 | 2.84 | 2.06E-184 | 4.29E-180 | 0.32 | 0.05 | 0.28 | 6.73 |
| 13 | Abcg1 | 2.49 | 5.38E-183 | 1.12E-178 | 0.40 | 0.07 | 0.33 | 5.46 |

**Supplementary Table 1. Marker genes for the original clusters of integrated clustering of E11.5 control and NotchGOF λ-junction cells.**

| | | | | | | | | |
|---|---|---|---|---|---|---|---|---|
| 13 | Emb | 1.63 | 5.55E-183 | 1.15E-178 | 0.89 | 0.39 | 0.49 | 2.25 |
| 13 | Ptprf | 1.48 | 9.07E-182 | 1.89E-177 | 0.97 | 0.63 | 0.34 | 1.54 |
| 13 | Uchl1 | 1.73 | 1.56E-181 | 3.25E-177 | 0.82 | 0.34 | 0.48 | 2.42 |
| 13 | Sema3c | 2.67 | 5.21E-180 | 1.08E-175 | 0.42 | 0.08 | 0.34 | 5.04 |
| 13 | Celsr1 | 2.38 | 2.83E-179 | 5.88E-175 | 0.45 | 0.09 | 0.36 | 4.80 |
| 13 | Otx2 | 2.22 | 4.78E-179 | 9.95E-175 | 0.45 | 0.09 | 0.36 | 4.85 |
| 13 | Grb7 | 2.23 | 6.46E-179 | 1.34E-174 | 0.44 | 0.09 | 0.35 | 4.98 |
| 13 | Bnipl | 2.94 | 6.28E-178 | 1.31E-173 | 0.29 | 0.04 | 0.25 | 7.31 |
| 13 | Krtcap3 | 2.32 | 1.01E-173 | 2.11E-169 | 0.47 | 0.10 | 0.37 | 4.57 |
| 13 | 2610528J11Rik | 2.55 | 1.35E-171 | 2.81E-167 | 0.34 | 0.06 | 0.28 | 6.11 |
| 13 | Wfdc2 | 1.49 | 7.65E-168 | 1.59E-163 | 0.89 | 0.45 | 0.44 | 1.97 |
| 13 | Tmem54 | 1.91 | 1.79E-166 | 3.73E-162 | 0.49 | 0.11 | 0.38 | 4.44 |
| 13 | Dpysl5 | 2.12 | 2.13E-166 | 4.43E-162 | 0.46 | 0.10 | 0.36 | 4.64 |
| 13 | Erbb3 | 1.74 | 3.77E-165 | 7.85E-161 | 0.61 | 0.16 | 0.44 | 3.71 |
| 13 | Cystm1 | 1.94 | 5.03E-165 | 1.05E-160 | 0.41 | 0.08 | 0.33 | 5.02 |
| 13 | Lhfp | 2.12 | 5.02E-164 | 1.04E-159 | 0.66 | 0.22 | 0.44 | 2.98 |
| 13 | Hook1 | 1.90 | 2.30E-161 | 4.78E-157 | 0.63 | 0.19 | 0.44 | 3.31 |
| 13 | Fgd2 | 2.04 | 2.53E-159 | 5.27E-155 | 0.43 | 0.09 | 0.34 | 4.67 |
| 13 | Tjp3 | 2.58 | 1.46E-158 | 3.04E-154 | 0.28 | 0.04 | 0.24 | 6.74 |
| 13 | Samd5 | 1.86 | 5.54E-156 | 1.15E-151 | 0.80 | 0.35 | 0.45 | 2.28 |
| 13 | Ntf5 | 2.18 | 3.92E-155 | 8.15E-151 | 0.40 | 0.08 | 0.32 | 4.94 |
| 13 | Zdhhc2 | 2.05 | 5.08E-154 | 1.06E-149 | 0.49 | 0.12 | 0.37 | 4.07 |
| 13 | Adarb2 | 4.06 | 5.69E-153 | 1.18E-148 | 0.15 | 0.01 | 0.14 | 12.25 |
| 13 | Wipf3 | 2.53 | 5.79E-150 | 1.21E-145 | 0.31 | 0.05 | 0.26 | 5.76 |
| 13 | Susd4 | 2.18 | 1.51E-146 | 3.14E-142 | 0.39 | 0.08 | 0.31 | 4.70 |
| 13 | Six2 | 1.42 | 3.30E-146 | 6.86E-142 | 0.88 | 0.44 | 0.44 | 1.99 |
| 13 | Stard10 | 1.57 | 5.09E-145 | 1.06E-140 | 0.75 | 0.29 | 0.46 | 2.60 |
| 13 | Wwc1 | 1.71 | 8.18E-144 | 1.70E-139 | 0.58 | 0.17 | 0.41 | 3.39 |
| 13 | 2010300C02Rik | 3.35 | 1.04E-143 | 2.17E-139 | 0.19 | 0.02 | 0.17 | 8.86 |
| 13 | Meis2 | 1.20 | 1.11E-143 | 2.30E-139 | 0.99 | 0.66 | 0.33 | 1.50 |
| 13 | Spint1 | 1.99 | 1.22E-142 | 2.53E-138 | 0.41 | 0.09 | 0.32 | 4.48 |
| 13 | Dlx3 | 2.40 | 1.42E-142 | 2.95E-138 | 0.35 | 0.07 | 0.28 | 5.09 |
| 13 | Trp63 | 1.71 | 2.32E-142 | 4.83E-138 | 0.49 | 0.13 | 0.37 | 3.93 |
| 13 | Mtus1 | 2.08 | 4.99E-142 | 1.04E-137 | 0.42 | 0.10 | 0.32 | 4.33 |
| 13 | Hs6st2 | 1.87 | 7.23E-141 | 1.50E-136 | 0.70 | 0.27 | 0.43 | 2.55 |
| 13 | Cpz | 2.54 | 2.91E-140 | 6.05E-136 | 0.33 | 0.06 | 0.27 | 5.22 |
| 13 | Pltp | 2.10 | 1.37E-139 | 2.84E-135 | 0.52 | 0.15 | 0.37 | 3.46 |
| 13 | Fras1 | 1.72 | 3.72E-138 | 7.73E-134 | 0.69 | 0.26 | 0.43 | 2.70 |
| 13 | Runx1 | 1.74 | 1.41E-137 | 2.94E-133 | 0.58 | 0.18 | 0.40 | 3.19 |
| 13 | Nr2e1 | 2.64 | 4.95E-137 | 1.03E-132 | 0.25 | 0.04 | 0.21 | 6.55 |
| 13 | Hdgf | 0.72 | 2.45E-136 | 5.09E-132 | 0.99 | 0.97 | 0.02 | 1.02 |
| 13 | Cxcr4 | 2.62 | 7.24E-135 | 1.51E-130 | 0.27 | 0.04 | 0.22 | 6.07 |
| 13 | Ephb1 | 2.07 | 2.45E-134 | 5.09E-130 | 0.40 | 0.09 | 0.31 | 4.35 |
| 13 | Krt19 | 2.32 | 6.80E-133 | 1.41E-128 | 0.33 | 0.07 | 0.27 | 4.97 |
| 13 | Atox1 | 0.82 | 2.76E-132 | 5.74E-128 | 0.99 | 0.95 | 0.04 | 1.04 |
| 13 | Plch1 | 2.01 | 6.60E-132 | 1.37E-127 | 0.40 | 0.09 | 0.31 | 4.30 |
| 13 | Krt14 | 1.84 | 1.89E-131 | 3.94E-127 | 0.47 | 0.13 | 0.34 | 3.73 |
| 13 | Mbp | 2.20 | 4.52E-131 | 9.40E-127 | 0.38 | 0.09 | 0.30 | 4.41 |
| 13 | Ntn1 | 2.86 | 5.78E-131 | 1.20E-126 | 0.20 | 0.03 | 0.17 | 7.84 |
| 13 | Arrdc1 | 2.08 | 8.02E-131 | 1.67E-126 | 0.42 | 0.10 | 0.32 | 4.06 |
| 13 | Marchf9 | 1.79 | 1.19E-128 | 2.47E-124 | 0.50 | 0.14 | 0.36 | 3.58 |
| 13 | Kcnk1 | 1.57 | 2.10E-128 | 4.36E-124 | 0.58 | 0.18 | 0.40 | 3.28 |
| 13 | Bspry | 2.16 | 3.10E-128 | 6.45E-124 | 0.31 | 0.06 | 0.25 | 5.25 |
| 13 | Col18a1 | 1.11 | 5.55E-127 | 1.16E-122 | 0.83 | 0.35 | 0.47 | 2.35 |
| 13 | Ston2 | 1.63 | 7.42E-127 | 1.54E-122 | 0.56 | 0.17 | 0.39 | 3.24 |
| 13 | Cdh2 | 1.42 | 2.88E-126 | 5.98E-122 | 0.83 | 0.41 | 0.41 | 2.00 |

**Supplementary Table 1. Marker genes for the original clusters of integrated clustering of E11.5 control and NotchGOF λ-junction cells.**

| | | | | | | | | |
|---|---|---|---|---|---|---|---|---|
| 13 | Cgn | 1.96 | 2.27E-125 | 4.72E-121 | 0.41 | 0.10 | 0.31 | 4.09 |
| 13 | Fzd6 | 2.15 | 7.05E-125 | 1.47E-120 | 0.36 | 0.08 | 0.28 | 4.56 |
| 13 | Khdrbs3 | 1.18 | 1.95E-124 | 4.06E-120 | 0.93 | 0.61 | 0.32 | 1.53 |
| 13 | Dtna | 1.92 | 1.15E-123 | 2.38E-119 | 0.48 | 0.13 | 0.34 | 3.55 |
| 13 | Rdh10 | 2.38 | 1.36E-123 | 2.82E-119 | 0.48 | 0.15 | 0.33 | 3.27 |
| 13 | Map7 | 1.87 | 1.94E-123 | 4.04E-119 | 0.41 | 0.10 | 0.31 | 4.06 |
| 13 | Dsg2 | 1.74 | 1.83E-122 | 3.81E-118 | 0.46 | 0.12 | 0.34 | 3.79 |
| 13 | Atp8b1 | 2.60 | 2.59E-121 | 5.39E-117 | 0.23 | 0.04 | 0.20 | 6.27 |
| 13 | Ocln | 1.95 | 2.79E-120 | 5.80E-116 | 0.33 | 0.07 | 0.26 | 4.70 |
| 13 | Vgll3 | 3.07 | 1.06E-119 | 2.21E-115 | 0.19 | 0.02 | 0.16 | 7.71 |
| 13 | Cmtm8 | 1.47 | 1.33E-119 | 2.77E-115 | 0.55 | 0.17 | 0.38 | 3.25 |
| 13 | Pcdh19 | 2.11 | 1.93E-119 | 4.02E-115 | 0.50 | 0.16 | 0.34 | 3.14 |
| 13 | Kitl | 2.27 | 5.64E-119 | 1.17E-114 | 0.32 | 0.07 | 0.25 | 4.65 |
| 13 | Nt5dc2 | 0.88 | 3.61E-118 | 7.50E-114 | 0.98 | 0.92 | 0.06 | 1.07 |
| 13 | Ssx2ip | 1.49 | 1.99E-117 | 4.13E-113 | 0.75 | 0.35 | 0.40 | 2.14 |
| 13 | Tom1l1 | 1.73 | 2.08E-117 | 4.33E-113 | 0.50 | 0.15 | 0.35 | 3.30 |
| 13 | Apoc1 | 1.88 | 1.27E-116 | 2.65E-112 | 0.35 | 0.08 | 0.27 | 4.44 |
| 13 | Nav2 | 1.27 | 3.30E-116 | 6.87E-112 | 0.64 | 0.23 | 0.41 | 2.80 |
| 13 | Rnf43 | 2.60 | 5.04E-116 | 1.05E-111 | 0.25 | 0.04 | 0.21 | 5.68 |
| 13 | Pkp2 | 2.35 | 7.46E-116 | 1.55E-111 | 0.27 | 0.05 | 0.22 | 5.41 |
| 13 | Mir205hg | 1.61 | 8.06E-116 | 1.68E-111 | 0.43 | 0.11 | 0.31 | 3.78 |
| 13 | Fermt1 | 1.51 | 9.08E-115 | 1.89E-110 | 0.47 | 0.13 | 0.34 | 3.54 |
| 13 | Acvr2a | 1.81 | 1.35E-114 | 2.80E-110 | 0.63 | 0.26 | 0.37 | 2.43 |
| 13 | Cd151 | 1.51 | 1.58E-114 | 3.29E-110 | 0.72 | 0.32 | 0.40 | 2.27 |
| 13 | Ildr1 | 2.66 | 1.96E-114 | 4.07E-110 | 0.21 | 0.03 | 0.18 | 6.48 |
| 13 | Hsp90ab1 | 0.36 | 2.23E-113 | 4.64E-109 | 1.00 | 1.00 | 0.00 | 1.00 |
| 13 | St14 | 1.69 | 3.38E-113 | 7.03E-109 | 0.42 | 0.11 | 0.31 | 3.78 |
| 13 | Adamts18 | 2.23 | 2.93E-112 | 6.09E-108 | 0.31 | 0.07 | 0.24 | 4.69 |
| 13 | Dmrta2os | 2.95 | 5.99E-112 | 1.25E-107 | 0.15 | 0.02 | 0.14 | 8.50 |
| 13 | Hap1 | 2.39 | 1.89E-111 | 3.93E-107 | 0.25 | 0.05 | 0.21 | 5.48 |
| 13 | Slain1 | 2.13 | 4.55E-111 | 9.47E-107 | 0.28 | 0.05 | 0.22 | 5.13 |
| 13 | Cd24a | 0.84 | 1.08E-110 | 2.25E-106 | 0.99 | 0.92 | 0.07 | 1.08 |
| 13 | Arg2 | 2.29 | 1.63E-110 | 3.39E-106 | 0.26 | 0.05 | 0.21 | 5.35 |
| 13 | Rprm | 2.01 | 2.75E-110 | 5.72E-106 | 0.48 | 0.15 | 0.33 | 3.14 |
| 13 | Slc44a3 | 3.59 | 6.71E-110 | 1.40E-105 | 0.11 | 0.01 | 0.10 | 12.22 |
| 13 | Shtn1 | 1.92 | 3.59E-109 | 7.48E-105 | 0.25 | 0.05 | 0.21 | 5.40 |
| 13 | Lmod1 | 2.95 | 3.79E-109 | 7.88E-105 | 0.14 | 0.02 | 0.13 | 9.33 |
| 13 | F11r | 1.44 | 5.85E-109 | 1.22E-104 | 0.54 | 0.17 | 0.37 | 3.13 |
| 13 | Igfbp3 | 1.63 | 2.49E-107 | 5.18E-103 | 0.42 | 0.12 | 0.31 | 3.62 |
| 13 | Schip1 | 1.75 | 6.80E-107 | 1.41E-102 | 0.48 | 0.15 | 0.33 | 3.14 |
| 13 | Chchd10 | 1.57 | 6.82E-107 | 1.42E-102 | 0.49 | 0.15 | 0.34 | 3.24 |
| 13 | Lmo4 | 1.20 | 1.44E-106 | 2.99E-102 | 0.97 | 0.81 | 0.16 | 1.20 |
| 13 | Zfhx3 | 0.98 | 6.28E-106 | 1.31E-101 | 0.98 | 0.81 | 0.17 | 1.21 |
| 13 | Plekha7 | 1.97 | 1.23E-105 | 2.56E-101 | 0.35 | 0.09 | 0.26 | 4.06 |
| 13 | Gsta4 | 1.09 | 1.35E-105 | 2.81E-101 | 0.93 | 0.75 | 0.18 | 1.24 |
| 13 | Podn | 3.45 | 1.37E-105 | 2.84E-101 | 0.13 | 0.01 | 0.12 | 9.29 |
| 13 | Cdkn1a | 1.53 | 3.48E-105 | 7.23E-101 | 0.39 | 0.10 | 0.29 | 3.75 |
| 13 | Pla2g12a | 1.41 | 3.61E-105 | 7.51E-101 | 0.71 | 0.32 | 0.39 | 2.23 |
| 13 | Ppfia3 | 1.88 | 3.33E-104 | 6.92E-100 | 0.42 | 0.12 | 0.30 | 3.44 |
| 13 | Hspb8 | 1.37 | 3.78E-104 | 7.86E-100 | 0.45 | 0.13 | 0.32 | 3.45 |
| 13 | Meis1 | 1.29 | 6.27E-104 | 1.30E-99 | 0.81 | 0.45 | 0.36 | 1.80 |
| 13 | Rpl7 | 0.40 | 1.29E-103 | 2.67E-99 | 1.00 | 1.00 | 0.00 | 1.00 |
| 13 | Bicd1 | 1.98 | 2.12E-103 | 4.40E-99 | 0.36 | 0.09 | 0.27 | 3.95 |
| 13 | Ppp2r2b | 1.78 | 2.82E-103 | 5.87E-99 | 0.36 | 0.09 | 0.27 | 3.99 |
| 13 | Itga3 | 2.26 | 3.79E-103 | 7.89E-99 | 0.29 | 0.06 | 0.23 | 4.60 |
| 13 | Celf2 | 1.56 | 2.88E-102 | 6.00E-98 | 0.61 | 0.25 | 0.37 | 2.49 |

**Supplementary Table 1. Marker genes for the original clusters of integrated clustering of E11.5 control and NotchGOF λ-junction cells.**

| | | | | | | | | |
|---|---|---|---|---|---|---|---|---|
| 13 | Arhgap8 | 2.31 | 5.41E-102 | 1.13E-97 | 0.22 | 0.04 | 0.18 | 5.76 |
| 13 | Klf5 | 1.80 | 7.84E-102 | 1.63E-97 | 0.41 | 0.12 | 0.30 | 3.52 |
| 13 | Bend7 | 2.01 | 8.78E-101 | 1.83E-96 | 0.32 | 0.08 | 0.25 | 4.22 |
| 13 | Met | 2.65 | 8.41E-100 | 1.75E-95 | 0.21 | 0.04 | 0.18 | 5.63 |
| 13 | Smim22 | 2.49 | 1.23E-99 | 2.55E-95 | 0.20 | 0.03 | 0.17 | 6.00 |
| 13 | Sfn | 1.16 | 4.77E-98 | 9.91E-94 | 0.56 | 0.20 | 0.36 | 2.79 |
| 13 | Pcbp4 | 0.79 | 7.91E-97 | 1.65E-92 | 0.98 | 0.86 | 0.12 | 1.13 |
| 13 | Galm | 1.73 | 1.71E-96 | 3.55E-92 | 0.53 | 0.20 | 0.33 | 2.63 |
| 13 | Gapdh | 0.44 | 3.56E-96 | 7.41E-92 | 1.00 | 1.00 | 0.00 | 1.00 |
| 13 | Hs6st1 | 1.70 | 4.16E-96 | 8.65E-92 | 0.49 | 0.16 | 0.32 | 2.98 |
| 13 | Ddr1 | 1.68 | 5.57E-96 | 1.16E-91 | 0.42 | 0.12 | 0.29 | 3.37 |
| 13 | Gm48435 | 2.83 | 6.16E-96 | 1.28E-91 | 0.16 | 0.02 | 0.14 | 7.27 |
| 13 | Psip1 | 0.94 | 1.31E-95 | 2.73E-91 | 1.00 | 0.98 | 0.02 | 1.02 |
| 13 | Epb41l5 | 1.53 | 2.14E-95 | 4.45E-91 | 0.60 | 0.24 | 0.35 | 2.45 |
| 13 | Tnnt1 | 2.38 | 3.37E-95 | 7.00E-91 | 0.24 | 0.05 | 0.19 | 4.94 |
| 13 | Has3 | 3.60 | 9.96E-95 | 2.07E-90 | 0.10 | 0.01 | 0.09 | 11.33 |
| 13 | Cdh3 | 1.45 | 1.19E-94 | 2.47E-90 | 0.55 | 0.20 | 0.35 | 2.75 |
| 13 | Klk8 | 1.36 | 2.72E-94 | 5.66E-90 | 0.56 | 0.21 | 0.36 | 2.73 |
| 13 | Clu | 2.31 | 3.18E-94 | 6.61E-90 | 0.52 | 0.20 | 0.32 | 2.59 |
| 13 | Kcnd2 | 3.14 | 1.35E-93 | 2.80E-89 | 0.14 | 0.02 | 0.12 | 8.44 |
| 13 | Cd9 | 0.95 | 3.41E-93 | 7.08E-89 | 0.69 | 0.29 | 0.40 | 2.37 |
| 13 | Elf5 | 2.15 | 5.72E-93 | 1.19E-88 | 0.26 | 0.06 | 0.20 | 4.64 |
| 13 | Mdh1 | 0.71 | 2.16E-92 | 4.50E-88 | 0.99 | 0.95 | 0.04 | 1.05 |
| 13 | Pak3 | 1.90 | 2.66E-92 | 5.54E-88 | 0.42 | 0.13 | 0.29 | 3.19 |
| 13 | Nrip1 | 1.12 | 1.23E-91 | 2.55E-87 | 0.93 | 0.75 | 0.18 | 1.24 |
| 13 | Dmrt2 | 3.11 | 1.49E-91 | 3.09E-87 | 0.16 | 0.02 | 0.14 | 7.14 |
| 13 | Afdn | 1.05 | 1.82E-91 | 3.79E-87 | 0.89 | 0.56 | 0.32 | 1.57 |
| 13 | Slco5a1 | 2.02 | 5.10E-91 | 1.06E-86 | 0.31 | 0.08 | 0.23 | 4.00 |
| 13 | Cdcp1 | 2.17 | 1.08E-90 | 2.25E-86 | 0.25 | 0.05 | 0.20 | 4.67 |
| 13 | Rps19 | 0.26 | 1.09E-90 | 2.27E-86 | 1.00 | 1.00 | 0.00 | 1.00 |
| 13 | Rasef | 2.46 | 1.38E-90 | 2.87E-86 | 0.20 | 0.04 | 0.17 | 5.51 |
| 13 | Myh14 | 1.81 | 1.72E-90 | 3.57E-86 | 0.31 | 0.08 | 0.23 | 4.11 |
| 13 | S100a11 | 1.29 | 1.89E-90 | 3.93E-86 | 0.91 | 0.76 | 0.15 | 1.20 |
| 13 | Stxbp2 | 1.69 | 2.03E-90 | 4.22E-86 | 0.42 | 0.13 | 0.29 | 3.19 |
| 13 | Trim2 | 1.29 | 2.79E-89 | 5.80E-85 | 0.76 | 0.41 | 0.35 | 1.87 |
| 13 | Gadd45g | 1.61 | 7.58E-89 | 1.58E-84 | 0.52 | 0.20 | 0.32 | 2.55 |
| 13 | Rassf10 | 2.01 | 8.03E-89 | 1.67E-84 | 0.26 | 0.06 | 0.20 | 4.59 |
| 13 | Stx3 | 1.62 | 2.71E-88 | 5.63E-84 | 0.40 | 0.12 | 0.28 | 3.33 |
| 13 | Cacna1b | 1.69 | 4.22E-88 | 8.77E-84 | 0.31 | 0.08 | 0.23 | 4.07 |
| 13 | Ormdl2 | 1.31 | 5.10E-88 | 1.06E-83 | 0.69 | 0.34 | 0.35 | 2.03 |
| 13 | Svip | 2.18 | 7.30E-88 | 1.52E-83 | 0.32 | 0.09 | 0.23 | 3.72 |
| 13 | Stra6 | 1.46 | 2.58E-87 | 5.37E-83 | 0.68 | 0.34 | 0.35 | 2.04 |
| 13 | F3 | 2.27 | 4.99E-87 | 1.04E-82 | 0.34 | 0.10 | 0.24 | 3.53 |
| 13 | Pcdh7 | 1.27 | 2.69E-86 | 5.61E-82 | 0.75 | 0.38 | 0.37 | 1.96 |
| 13 | Gpm6a | 2.27 | 4.91E-86 | 1.02E-81 | 0.22 | 0.04 | 0.18 | 5.02 |
| 13 | Lmo7 | 1.46 | 5.34E-86 | 1.11E-81 | 0.45 | 0.15 | 0.30 | 2.94 |
| 13 | Pik3ap1 | 2.70 | 1.14E-85 | 2.37E-81 | 0.13 | 0.02 | 0.11 | 8.33 |
| 13 | Ago1 | 1.29 | 3.85E-85 | 8.01E-81 | 0.70 | 0.36 | 0.35 | 1.98 |
| 13 | Ktn1 | 0.79 | 1.75E-84 | 3.63E-80 | 0.96 | 0.82 | 0.14 | 1.17 |
| 13 | Jade1 | 1.77 | 1.89E-84 | 3.92E-80 | 0.46 | 0.17 | 0.30 | 2.79 |
| 13 | Mal2 | 2.72 | 3.00E-84 | 6.25E-80 | 0.14 | 0.02 | 0.12 | 7.10 |
| 13 | Dusp5 | 1.87 | 3.01E-84 | 6.27E-80 | 0.37 | 0.11 | 0.26 | 3.34 |
| 13 | Siva1 | 0.71 | 3.03E-84 | 6.31E-80 | 0.99 | 0.95 | 0.04 | 1.05 |
| 13 | Chmp2b | 1.12 | 3.35E-84 | 6.97E-80 | 0.85 | 0.56 | 0.30 | 1.53 |
| 13 | Prr36 | 1.75 | 8.15E-84 | 1.69E-79 | 0.31 | 0.08 | 0.23 | 3.82 |
| 13 | Rpl10a | 0.30 | 1.34E-83 | 2.79E-79 | 1.00 | 1.00 | 0.00 | 1.00 |

**Supplementary Table 1. Marker genes for the original clusters of integrated clustering of E11.5 control and NotchGOF λ-junction cells.**

| | | | | | | | | |
|---|---|---|---|---|---|---|---|---|
| 13 | Marveld2 | 1.98 | 2.58E-83 | 5.36E-79 | 0.25 | 0.06 | 0.19 | 4.49 |
| 13 | Plekhg3 | 2.03 | 5.98E-83 | 1.24E-78 | 0.30 | 0.08 | 0.22 | 3.86 |
| 13 | Pkm | 0.47 | 3.35E-82 | 6.96E-78 | 1.00 | 0.98 | 0.02 | 1.02 |
| 13 | Olfm2 | 1.71 | 3.13E-81 | 6.50E-77 | 0.34 | 0.10 | 0.25 | 3.47 |
| 13 | Lurap1l | 1.82 | 3.47E-81 | 7.23E-77 | 0.29 | 0.07 | 0.21 | 3.96 |
| 13 | Sema3d | 1.56 | 3.32E-80 | 6.91E-76 | 0.37 | 0.11 | 0.26 | 3.32 |
| 13 | Hey1 | 0.84 | 3.76E-80 | 7.83E-76 | 0.67 | 0.29 | 0.38 | 2.31 |
| 13 | Prr5l | 1.65 | 2.05E-79 | 4.27E-75 | 0.44 | 0.16 | 0.29 | 2.80 |
| 13 | Rnh1 | 1.06 | 1.01E-78 | 2.10E-74 | 0.85 | 0.55 | 0.30 | 1.55 |
| 13 | Clic5 | 3.01 | 1.04E-78 | 2.16E-74 | 0.12 | 0.02 | 0.10 | 7.80 |
| 13 | Sparc | 1.07 | 2.27E-78 | 4.72E-74 | 0.94 | 0.74 | 0.20 | 1.28 |
| 13 | Igsf9 | 1.28 | 3.87E-78 | 8.04E-74 | 0.73 | 0.40 | 0.33 | 1.82 |
| 13 | Mycl | 1.18 | 6.95E-78 | 1.44E-73 | 0.65 | 0.29 | 0.35 | 2.20 |
| 13 | Trpm3 | 2.26 | 7.02E-78 | 1.46E-73 | 0.21 | 0.04 | 0.17 | 4.84 |
| 13 | Tmem30b | 1.70 | 7.75E-78 | 1.61E-73 | 0.42 | 0.14 | 0.28 | 2.94 |
| 13 | Chd3 | 0.75 | 2.97E-77 | 6.18E-73 | 0.99 | 0.89 | 0.09 | 1.11 |
| 13 | Syt5 | 2.13 | 3.56E-77 | 7.40E-73 | 0.24 | 0.05 | 0.18 | 4.37 |
| 13 | Pbx1 | 0.74 | 3.60E-77 | 7.48E-73 | 0.99 | 0.92 | 0.07 | 1.08 |
| 13 | Slc12a2 | 1.22 | 3.78E-77 | 7.86E-73 | 0.75 | 0.43 | 0.32 | 1.76 |
| 13 | Tspan13 | 0.98 | 5.16E-77 | 1.07E-72 | 0.76 | 0.40 | 0.36 | 1.89 |
| 13 | Fam20c | 2.61 | 5.65E-77 | 1.17E-72 | 0.14 | 0.02 | 0.12 | 6.75 |
| 13 | Hes6 | 1.16 | 6.44E-77 | 1.34E-72 | 0.64 | 0.33 | 0.32 | 1.98 |
| 13 | Cyb5r3 | 0.77 | 1.96E-76 | 4.07E-72 | 0.95 | 0.81 | 0.15 | 1.19 |
| 13 | Npdc1 | 1.48 | 3.86E-76 | 8.03E-72 | 0.34 | 0.10 | 0.24 | 3.41 |
| 13 | Zfp467 | 1.31 | 1.32E-75 | 2.75E-71 | 0.61 | 0.28 | 0.33 | 2.17 |
| 13 | Klc3 | 1.29 | 2.08E-75 | 4.32E-71 | 0.34 | 0.10 | 0.24 | 3.37 |
| 13 | Ascl1 | 3.20 | 2.88E-75 | 5.99E-71 | 0.10 | 0.01 | 0.09 | 9.09 |
| 13 | Synrg | 1.40 | 5.53E-75 | 1.15E-70 | 0.54 | 0.23 | 0.31 | 2.38 |
| 13 | Thbs1 | 1.36 | 1.27E-74 | 2.63E-70 | 0.36 | 0.11 | 0.24 | 3.16 |
| 13 | Ablim1 | 1.29 | 2.35E-74 | 4.88E-70 | 0.54 | 0.22 | 0.32 | 2.46 |
| 13 | Syndig1 | 2.46 | 2.93E-74 | 6.09E-70 | 0.13 | 0.02 | 0.11 | 7.17 |
| 13 | Rnf208 | 1.87 | 9.26E-74 | 1.93E-69 | 0.26 | 0.06 | 0.19 | 3.98 |
| 13 | Nudt14 | 1.14 | 1.04E-73 | 2.17E-69 | 0.72 | 0.40 | 0.33 | 1.83 |
| 13 | Rpl38 | 0.28 | 3.40E-73 | 7.08E-69 | 1.00 | 1.00 | 0.00 | 1.00 |
| 13 | Tead1 | 1.02 | 7.29E-73 | 1.52E-68 | 0.87 | 0.60 | 0.27 | 1.45 |
| 13 | Skp1a | 0.62 | 1.08E-72 | 2.24E-68 | 0.99 | 0.96 | 0.04 | 1.04 |
| 13 | Rps27 | 0.28 | 1.15E-72 | 2.40E-68 | 1.00 | 1.00 | 0.00 | 1.00 |
| 13 | Tmem200c | 2.66 | 3.00E-72 | 6.23E-68 | 0.14 | 0.02 | 0.12 | 6.22 |
| 13 | Eno1 | 0.50 | 3.48E-71 | 7.23E-67 | 0.98 | 0.97 | 0.01 | 1.01 |
| 13 | Epha1 | 1.99 | 3.78E-71 | 7.86E-67 | 0.23 | 0.06 | 0.18 | 4.20 |
| 13 | Gm2694 | 2.32 | 4.00E-71 | 8.32E-67 | 0.12 | 0.02 | 0.10 | 7.06 |
| 13 | Ece1 | 1.33 | 4.52E-71 | 9.41E-67 | 0.53 | 0.22 | 0.31 | 2.39 |
| 13 | Crim1 | 1.62 | 4.85E-71 | 1.01E-66 | 0.47 | 0.19 | 0.28 | 2.48 |
| 13 | Itga6 | 0.58 | 8.17E-71 | 1.70E-66 | 0.69 | 0.30 | 0.38 | 2.27 |
| 13 | Esrp2 | 1.83 | 2.41E-70 | 5.00E-66 | 0.25 | 0.06 | 0.18 | 4.02 |
| 13 | Mtmr7 | 1.93 | 2.70E-70 | 5.61E-66 | 0.21 | 0.05 | 0.16 | 4.49 |
| 13 | Sox21 | 2.11 | 4.67E-70 | 9.72E-66 | 0.16 | 0.03 | 0.13 | 5.37 |
| 13 | Prr15l | 2.20 | 5.09E-70 | 1.06E-65 | 0.17 | 0.03 | 0.14 | 5.31 |
| 13 | Trim29 | 2.30 | 1.66E-69 | 3.46E-65 | 0.20 | 0.04 | 0.16 | 4.52 |
| 13 | Ldha | 0.54 | 4.00E-69 | 8.32E-65 | 0.99 | 0.97 | 0.02 | 1.02 |
| 13 | Zfp219 | 1.11 | 8.60E-69 | 1.79E-64 | 0.71 | 0.39 | 0.32 | 1.82 |
| 13 | Cers4 | 1.48 | 9.53E-69 | 1.98E-64 | 0.42 | 0.15 | 0.27 | 2.76 |
| 13 | Tnfrsf12a | 1.81 | 1.97E-68 | 4.10E-64 | 0.27 | 0.07 | 0.20 | 3.71 |
| 13 | Gpi1 | 0.70 | 8.39E-68 | 1.74E-63 | 0.96 | 0.84 | 0.13 | 1.15 |
| 13 | Cep170b | 1.60 | 8.40E-68 | 1.75E-63 | 0.36 | 0.12 | 0.24 | 3.01 |
| 13 | Metrnl | 1.91 | 1.35E-67 | 2.81E-63 | 0.24 | 0.06 | 0.18 | 3.95 |

**Supplementary Table 1. Marker genes for the original clusters of integrated clustering of E11.5 control and NotchGOF λ-junction cells.**

| | | | | | | | | |
|---|---|---|---|---|---|---|---|---|
| 13 | Cd2ap | 0.89 | 2.48E-67 | 5.16E-63 | 0.86 | 0.62 | 0.24 | 1.39 |
| 13 | Rps18 | 0.26 | 2.52E-67 | 5.25E-63 | 1.00 | 1.00 | 0.00 | 1.00 |
| 13 | Dstn | 0.55 | 2.88E-67 | 5.98E-63 | 0.99 | 0.94 | 0.05 | 1.06 |
| 13 | Frmd5 | 2.04 | 4.66E-67 | 9.68E-63 | 0.21 | 0.05 | 0.16 | 4.38 |
| 13 | Patj | 1.57 | 5.34E-67 | 1.11E-62 | 0.37 | 0.13 | 0.24 | 2.94 |
| 13 | Fzd8 | 1.65 | 6.85E-67 | 1.42E-62 | 0.39 | 0.14 | 0.25 | 2.75 |
| 13 | Maob | 2.23 | 8.25E-67 | 1.72E-62 | 0.19 | 0.04 | 0.15 | 4.61 |
| 13 | Dsc2 | 1.95 | 1.10E-66 | 2.28E-62 | 0.22 | 0.05 | 0.17 | 4.17 |
| 13 | Tmem238 | 1.51 | 1.19E-66 | 2.47E-62 | 0.44 | 0.17 | 0.27 | 2.55 |
| 13 | Sorl1 | 1.51 | 2.30E-66 | 4.77E-62 | 0.40 | 0.15 | 0.26 | 2.76 |
| 13 | Plp2 | 1.02 | 3.17E-66 | 6.60E-62 | 0.71 | 0.37 | 0.34 | 1.91 |
| 13 | Camsap3 | 1.69 | 3.34E-66 | 6.94E-62 | 0.28 | 0.08 | 0.20 | 3.57 |
| 13 | Sptbn2 | 1.61 | 5.83E-66 | 1.21E-61 | 0.34 | 0.11 | 0.23 | 3.11 |
| 13 | Efna1 | 1.37 | 1.17E-65 | 2.43E-61 | 0.39 | 0.13 | 0.25 | 2.88 |
| 13 | Syngr1 | 1.71 | 1.37E-65 | 2.85E-61 | 0.24 | 0.06 | 0.18 | 3.87 |
| 13 | Hist3h2a | 1.30 | 2.56E-65 | 5.32E-61 | 0.56 | 0.26 | 0.30 | 2.16 |
| 13 | Dok5 | 1.42 | 3.50E-65 | 7.28E-61 | 0.34 | 0.11 | 0.23 | 3.07 |
| 13 | Rbm47 | 2.29 | 3.76E-65 | 7.82E-61 | 0.16 | 0.03 | 0.13 | 5.19 |
| 13 | Fat3 | 2.09 | 4.13E-65 | 8.60E-61 | 0.29 | 0.09 | 0.20 | 3.27 |
| 13 | 2700033N17Rik | 1.61 | 5.05E-65 | 1.05E-60 | 0.34 | 0.11 | 0.23 | 3.04 |
| 13 | Epn3 | 1.86 | 7.35E-65 | 1.53E-60 | 0.20 | 0.05 | 0.16 | 4.37 |
| 13 | Pkp3 | 1.79 | 8.60E-65 | 1.79E-60 | 0.24 | 0.06 | 0.18 | 3.83 |
| 13 | Frat2 | 1.28 | 1.09E-64 | 2.27E-60 | 0.52 | 0.23 | 0.29 | 2.29 |
| 13 | Slc22a23 | 1.66 | 1.44E-64 | 3.00E-60 | 0.30 | 0.09 | 0.21 | 3.31 |
| 13 | Kit | 1.80 | 1.48E-64 | 3.08E-60 | 0.30 | 0.09 | 0.21 | 3.27 |
| 13 | Perp | 0.68 | 3.61E-64 | 7.52E-60 | 0.40 | 0.14 | 0.27 | 2.96 |
| 13 | Egr1 | 2.60 | 3.68E-64 | 7.66E-60 | 0.25 | 0.07 | 0.18 | 3.52 |
| 13 | Scube3 | 1.29 | 4.81E-64 | 1.00E-59 | 0.48 | 0.20 | 0.28 | 2.44 |
| 13 | Zfhx2 | 1.28 | 5.26E-64 | 1.09E-59 | 0.54 | 0.24 | 0.29 | 2.20 |
| 13 | Zfp608 | 1.08 | 5.43E-64 | 1.13E-59 | 0.79 | 0.53 | 0.27 | 1.50 |
| 13 | Npnt | 1.24 | 1.03E-63 | 2.14E-59 | 0.53 | 0.23 | 0.30 | 2.28 |
| 13 | Col11a1 | 1.29 | 1.10E-63 | 2.30E-59 | 0.71 | 0.42 | 0.29 | 1.69 |
| 13 | 4930426D05Rik | 2.43 | 3.47E-63 | 7.22E-59 | 0.15 | 0.03 | 0.12 | 5.28 |
| 13 | Gkap1 | 0.95 | 1.46E-62 | 3.03E-58 | 0.74 | 0.43 | 0.31 | 1.74 |
| 13 | Pgam1 | 0.45 | 5.11E-62 | 1.06E-57 | 1.00 | 0.97 | 0.03 | 1.03 |
| 13 | Six1 | 0.79 | 7.59E-62 | 1.58E-57 | 0.90 | 0.61 | 0.30 | 1.49 |
| 13 | Crip2 | 0.54 | 9.88E-62 | 2.05E-57 | 0.44 | 0.17 | 0.27 | 2.63 |
| 13 | Cryl1 | 1.55 | 2.25E-61 | 4.68E-57 | 0.38 | 0.14 | 0.24 | 2.67 |
| 13 | Eya1 | 1.01 | 4.67E-61 | 9.71E-57 | 0.79 | 0.49 | 0.30 | 1.60 |
| 13 | Nuak2 | 2.20 | 5.73E-61 | 1.19E-56 | 0.17 | 0.03 | 0.13 | 4.85 |
| 13 | Pdlim1 | 0.84 | 2.23E-60 | 4.64E-56 | 0.59 | 0.28 | 0.30 | 2.06 |
| 13 | Tmem108 | 1.66 | 2.32E-60 | 4.83E-56 | 0.48 | 0.21 | 0.26 | 2.25 |
| 13 | Nfib | 0.77 | 2.73E-60 | 5.68E-56 | 0.92 | 0.74 | 0.18 | 1.24 |
| 13 | Aldoa | 0.52 | 4.28E-60 | 8.91E-56 | 0.99 | 0.95 | 0.04 | 1.04 |
| 13 | Cyb5a | 0.50 | 5.40E-60 | 1.12E-55 | 0.97 | 0.92 | 0.05 | 1.06 |
| 13 | Smim4 | 1.06 | 1.08E-59 | 2.24E-55 | 0.72 | 0.42 | 0.30 | 1.72 |
| 13 | Il17re | 2.06 | 1.27E-59 | 2.63E-55 | 0.19 | 0.04 | 0.14 | 4.40 |
| 13 | Acsl5 | 1.45 | 2.53E-59 | 5.27E-55 | 0.40 | 0.15 | 0.24 | 2.60 |
| 13 | Ociad2 | 1.42 | 3.19E-59 | 6.64E-55 | 0.44 | 0.18 | 0.26 | 2.48 |
| 13 | Grk5 | 1.61 | 5.57E-59 | 1.16E-54 | 0.33 | 0.12 | 0.22 | 2.90 |
| 13 | Myh9 | 1.08 | 6.27E-59 | 1.30E-54 | 0.77 | 0.50 | 0.27 | 1.54 |
| 13 | Ell3 | 2.15 | 1.46E-58 | 3.05E-54 | 0.15 | 0.03 | 0.12 | 5.18 |
| 13 | Arhgef16 | 1.85 | 1.50E-58 | 3.12E-54 | 0.21 | 0.06 | 0.16 | 3.89 |
| 13 | Errfi1 | 1.48 | 1.98E-58 | 4.12E-54 | 0.42 | 0.17 | 0.25 | 2.45 |
| 13 | Rab3ip | 1.17 | 3.45E-58 | 7.17E-54 | 0.59 | 0.30 | 0.29 | 1.99 |
| 13 | Arhgef5 | 1.16 | 3.50E-58 | 7.28E-54 | 0.47 | 0.19 | 0.28 | 2.44 |

**Supplementary Table 1. Marker genes for the original clusters of integrated clustering of E11.5 control and NotchGOF λ-junction cells.**

| | | | | | | | | |
|---|---|---|---|---|---|---|---|---|
| 13 | Galnt7 | 1.19 | 4.72E-58 | 9.82E-54 | 0.62 | 0.34 | 0.29 | 1.85 |
| 13 | Asprv1 | 2.67 | 5.47E-58 | 1.14E-53 | 0.11 | 0.02 | 0.09 | 6.56 |
| 13 | Dnase2a | 1.16 | 5.63E-58 | 1.17E-53 | 0.49 | 0.21 | 0.28 | 2.31 |
| 13 | Ckmt1 | 1.23 | 5.97E-58 | 1.24E-53 | 0.18 | 0.04 | 0.14 | 4.40 |
| 13 | Samd10 | 1.50 | 7.52E-58 | 1.56E-53 | 0.28 | 0.09 | 0.19 | 3.26 |
| 13 | Llgl2 | 1.88 | 7.71E-58 | 1.60E-53 | 0.19 | 0.05 | 0.15 | 4.15 |
| 13 | Mkrn1 | 0.97 | 7.92E-58 | 1.65E-53 | 0.72 | 0.42 | 0.30 | 1.71 |
| 13 | Fam187b | 2.00 | 9.68E-58 | 2.01E-53 | 0.17 | 0.04 | 0.13 | 4.71 |
| 13 | Pgk1 | 0.51 | 1.06E-57 | 2.21E-53 | 0.99 | 0.95 | 0.03 | 1.04 |
| 13 | Pdpn | 1.50 | 1.26E-57 | 2.63E-53 | 0.44 | 0.18 | 0.25 | 2.40 |
| 13 | Cpm | 0.95 | 1.40E-57 | 2.91E-53 | 0.43 | 0.17 | 0.26 | 2.53 |
| 13 | Ddc | 2.17 | 1.44E-57 | 3.00E-53 | 0.20 | 0.05 | 0.15 | 3.98 |
| 13 | Zfp536 | 1.54 | 1.58E-57 | 3.28E-53 | 0.30 | 0.10 | 0.20 | 3.07 |
| 13 | Gldc | 2.14 | 1.87E-57 | 3.88E-53 | 0.17 | 0.04 | 0.13 | 4.50 |
| 13 | Fgfr2 | 1.34 | 2.19E-57 | 4.56E-53 | 0.54 | 0.27 | 0.27 | 1.99 |
| 13 | Mgat4a | 1.88 | 2.95E-57 | 6.14E-53 | 0.28 | 0.09 | 0.19 | 3.12 |
| 13 | Diras2 | 2.18 | 6.08E-57 | 1.26E-52 | 0.12 | 0.02 | 0.10 | 5.71 |
| 13 | Creld1 | 1.48 | 1.01E-56 | 2.09E-52 | 0.35 | 0.12 | 0.22 | 2.81 |
| 13 | Junb | 1.77 | 2.35E-56 | 4.89E-52 | 0.32 | 0.11 | 0.21 | 2.85 |
| 13 | Scd2 | 0.64 | 3.64E-56 | 7.58E-52 | 0.99 | 0.91 | 0.08 | 1.08 |
| 13 | Frrs1 | 1.23 | 3.84E-56 | 7.99E-52 | 0.49 | 0.22 | 0.27 | 2.26 |
| 13 | Chn2 | 1.86 | 4.29E-56 | 8.92E-52 | 0.24 | 0.07 | 0.17 | 3.52 |
| 13 | Jup | 1.03 | 9.81E-56 | 2.04E-51 | 0.47 | 0.19 | 0.27 | 2.41 |
| 13 | Ralgapa2 | 1.63 | 1.05E-55 | 2.18E-51 | 0.35 | 0.13 | 0.22 | 2.67 |
| 13 | Rims2 | 1.74 | 2.36E-55 | 4.90E-51 | 0.21 | 0.06 | 0.15 | 3.80 |
| 13 | Eml5 | 1.61 | 2.59E-55 | 5.39E-51 | 0.27 | 0.08 | 0.18 | 3.27 |
| 13 | Atp6v0e2 | 1.65 | 3.22E-55 | 6.69E-51 | 0.20 | 0.05 | 0.15 | 3.96 |
| 13 | Eps8l1 | 2.29 | 3.93E-55 | 8.18E-51 | 0.16 | 0.04 | 0.13 | 4.53 |
| 13 | Gm266 | 1.02 | 3.97E-55 | 8.25E-51 | 0.57 | 0.30 | 0.27 | 1.88 |
| 13 | Rab15 | 1.35 | 5.16E-55 | 1.07E-50 | 0.35 | 0.13 | 0.23 | 2.76 |
| 13 | Ripply3 | 1.94 | 5.51E-55 | 1.15E-50 | 0.21 | 0.06 | 0.16 | 3.74 |
| 13 | Cd47 | 1.11 | 5.73E-55 | 1.19E-50 | 0.71 | 0.43 | 0.28 | 1.64 |
| 13 | Mgp | 2.46 | 6.51E-55 | 1.35E-50 | 0.23 | 0.07 | 0.16 | 3.41 |
| 13 | D430041D05Rik | 2.76 | 1.02E-54 | 2.13E-50 | 0.11 | 0.02 | 0.09 | 6.11 |
| 13 | Gm13648 | 2.18 | 1.35E-54 | 2.80E-50 | 0.13 | 0.02 | 0.10 | 5.43 |
| 13 | Tenm2 | 1.69 | 1.84E-54 | 3.83E-50 | 0.24 | 0.07 | 0.17 | 3.43 |
| 13 | Dipk2a | 1.61 | 1.93E-54 | 4.01E-50 | 0.49 | 0.23 | 0.26 | 2.09 |
| 13 | Pard6b | 1.84 | 2.14E-54 | 4.45E-50 | 0.19 | 0.05 | 0.15 | 3.96 |
| 13 | Clic6 | 1.75 | 2.16E-54 | 4.48E-50 | 0.20 | 0.05 | 0.15 | 3.90 |
| 13 | Cblc | 1.97 | 2.36E-54 | 4.90E-50 | 0.15 | 0.03 | 0.12 | 4.90 |
| 13 | Igf2 | 0.72 | 3.30E-54 | 6.87E-50 | 0.85 | 0.58 | 0.27 | 1.47 |
| 13 | Pls1 | 1.39 | 3.44E-54 | 7.15E-50 | 0.28 | 0.09 | 0.19 | 3.11 |
| 13 | Pqlc3 | 1.97 | 7.56E-54 | 1.57E-49 | 0.20 | 0.05 | 0.15 | 3.88 |
| 13 | Adamts17 | 2.08 | 9.90E-54 | 2.06E-49 | 0.16 | 0.03 | 0.12 | 4.56 |
| 13 | Atp2b1 | 0.73 | 1.01E-53 | 2.11E-49 | 0.95 | 0.84 | 0.11 | 1.13 |
| 13 | Unc13b | 1.58 | 3.61E-53 | 7.52E-49 | 0.26 | 0.08 | 0.18 | 3.28 |
| 13 | Fjx1 | 0.97 | 4.00E-53 | 8.32E-49 | 0.67 | 0.38 | 0.29 | 1.75 |
| 13 | Tpi1 | 0.48 | 9.49E-53 | 1.97E-48 | 0.99 | 0.96 | 0.03 | 1.04 |
| 13 | Tbc1d1 | 1.14 | 1.16E-52 | 2.40E-48 | 0.63 | 0.37 | 0.27 | 1.74 |
| 13 | Celsr2 | 1.43 | 1.54E-52 | 3.21E-48 | 0.28 | 0.09 | 0.19 | 3.13 |
| 13 | Srcin1 | 1.91 | 1.88E-52 | 3.90E-48 | 0.18 | 0.04 | 0.14 | 4.09 |
| 13 | 2900026A02Rik | 1.44 | 2.43E-52 | 5.05E-48 | 0.30 | 0.10 | 0.20 | 2.95 |
| 13 | Ripk4 | 1.48 | 2.56E-52 | 5.33E-48 | 0.24 | 0.07 | 0.17 | 3.41 |
| 13 | Bcl11a | 0.74 | 2.88E-52 | 6.00E-48 | 0.91 | 0.74 | 0.17 | 1.24 |
| 13 | Wscd1 | 2.43 | 3.88E-52 | 8.08E-48 | 0.12 | 0.02 | 0.10 | 5.39 |
| 13 | Zfp385c | 2.11 | 6.90E-52 | 1.43E-47 | 0.14 | 0.03 | 0.11 | 4.77 |

**Supplementary Table 1. Marker genes for the original clusters of integrated clustering of E11.5 control and NotchGOF λ-junction cells.**

| | | | | | | | | |
|---|---|---|---|---|---|---|---|---|
| 13 | Cyb561 | 1.68 | 7.49E-52 | 1.56E-47 | 0.20 | 0.05 | 0.15 | 3.81 |
| 13 | Ntm | 1.47 | 9.05E-52 | 1.88E-47 | 0.30 | 0.10 | 0.20 | 2.91 |
| 13 | Rbbp8 | 0.99 | 1.65E-51 | 3.42E-47 | 0.79 | 0.55 | 0.24 | 1.43 |
| 13 | Trpm4 | 1.59 | 2.52E-51 | 5.23E-47 | 0.28 | 0.09 | 0.19 | 2.98 |
| 13 | Cacnb3 | 1.03 | 7.41E-51 | 1.54E-46 | 0.55 | 0.27 | 0.28 | 2.06 |
| 13 | Serinc2 | 0.97 | 8.16E-51 | 1.70E-46 | 0.68 | 0.38 | 0.30 | 1.78 |
| 13 | Dlx6 | 1.45 | 9.52E-51 | 1.98E-46 | 0.26 | 0.08 | 0.18 | 3.13 |
| 13 | Itm2b | 1.17 | 1.08E-50 | 2.24E-46 | 0.96 | 0.84 | 0.12 | 1.14 |
| 13 | Atp6v1f | 0.54 | 1.35E-50 | 2.80E-46 | 0.96 | 0.90 | 0.06 | 1.07 |
| 13 | Etl4 | 1.05 | 3.79E-50 | 7.88E-46 | 0.59 | 0.30 | 0.29 | 1.94 |
| 13 | Tbc1d30 | 2.09 | 4.83E-50 | 1.00E-45 | 0.14 | 0.03 | 0.11 | 4.72 |
| 13 | Bphl | 1.03 | 7.37E-50 | 1.53E-45 | 0.65 | 0.37 | 0.28 | 1.74 |
| 13 | Tceal9 | 0.38 | 9.69E-50 | 2.01E-45 | 1.00 | 0.98 | 0.02 | 1.02 |
| 13 | Arhgap27os2 | 1.60 | 1.01E-49 | 2.10E-45 | 0.22 | 0.06 | 0.15 | 3.44 |
| 13 | Nfasc | 2.51 | 1.34E-49 | 2.79E-45 | 0.12 | 0.02 | 0.10 | 5.08 |
| 13 | Ttc3 | 0.51 | 1.39E-49 | 2.90E-45 | 1.00 | 0.96 | 0.04 | 1.04 |
| 13 | Dhx32 | 1.32 | 1.49E-49 | 3.11E-45 | 0.46 | 0.21 | 0.24 | 2.15 |
| 13 | Ovol2 | 2.33 | 1.86E-49 | 3.87E-45 | 0.13 | 0.03 | 0.10 | 4.81 |
| 13 | Dcx | 1.52 | 2.37E-49 | 4.93E-45 | 0.13 | 0.03 | 0.10 | 5.08 |
| 13 | Acvr2b | 0.73 | 2.73E-49 | 5.67E-45 | 0.86 | 0.64 | 0.22 | 1.34 |
| 13 | Slc25a23 | 1.29 | 5.90E-49 | 1.23E-44 | 0.38 | 0.16 | 0.23 | 2.46 |
| 13 | Lrp2 | 1.64 | 1.37E-48 | 2.85E-44 | 0.19 | 0.05 | 0.14 | 3.75 |
| 13 | Agrn | 1.04 | 2.95E-48 | 6.14E-44 | 0.66 | 0.38 | 0.27 | 1.71 |
| 13 | Lgi2 | 1.77 | 4.42E-48 | 9.19E-44 | 0.30 | 0.11 | 0.19 | 2.71 |
| 13 | Sft2d2 | 1.30 | 5.10E-48 | 1.06E-43 | 0.36 | 0.15 | 0.22 | 2.50 |
| 13 | Cxadr | 0.89 | 6.20E-48 | 1.29E-43 | 0.73 | 0.46 | 0.27 | 1.60 |
| 13 | Marveld3 | 1.88 | 8.14E-48 | 1.69E-43 | 0.15 | 0.04 | 0.12 | 4.29 |
| 13 | Vwa2 | 2.03 | 9.96E-48 | 2.07E-43 | 0.15 | 0.03 | 0.11 | 4.45 |
| 13 | Elf3 | 1.75 | 1.16E-47 | 2.41E-43 | 0.23 | 0.07 | 0.16 | 3.24 |
| 13 | Pawr | 1.29 | 1.30E-47 | 2.71E-43 | 0.33 | 0.13 | 0.21 | 2.65 |
| 13 | Knop1 | 0.49 | 1.34E-47 | 2.79E-43 | 0.97 | 0.90 | 0.06 | 1.07 |
| 13 | Anxa2 | 0.96 | 1.52E-47 | 3.17E-43 | 0.68 | 0.41 | 0.27 | 1.65 |
| 13 | Stk26 | 1.04 | 2.14E-47 | 4.45E-43 | 0.48 | 0.22 | 0.26 | 2.16 |
| 13 | AI661453 | 1.95 | 2.20E-47 | 4.57E-43 | 0.15 | 0.03 | 0.11 | 4.39 |
| 13 | Nedd9 | 1.40 | 1.09E-46 | 2.28E-42 | 0.38 | 0.16 | 0.22 | 2.36 |
| 13 | Dennd1c | 2.04 | 2.06E-46 | 4.29E-42 | 0.14 | 0.03 | 0.11 | 4.57 |
| 13 | Scg5 | 1.45 | 3.15E-46 | 6.55E-42 | 0.19 | 0.05 | 0.14 | 3.72 |
| 13 | Pcdh17 | 1.19 | 4.43E-46 | 9.22E-42 | 0.44 | 0.20 | 0.24 | 2.19 |
| 13 | Irs1 | 1.28 | 5.44E-46 | 1.13E-41 | 0.40 | 0.17 | 0.23 | 2.30 |
| 13 | Ltbp1 | 1.21 | 5.64E-46 | 1.17E-41 | 0.46 | 0.22 | 0.24 | 2.11 |
| 13 | Ppm1l | 1.25 | 5.71E-46 | 1.19E-41 | 0.46 | 0.22 | 0.24 | 2.12 |
| 13 | Ptpru | 1.13 | 5.93E-46 | 1.23E-41 | 0.37 | 0.15 | 0.22 | 2.44 |
| 13 | Cntnap2 | 1.77 | 6.03E-46 | 1.25E-41 | 0.18 | 0.05 | 0.13 | 3.75 |
| 13 | Fgf8 | 0.92 | 1.24E-45 | 2.58E-41 | 0.24 | 0.08 | 0.16 | 3.12 |
| 13 | Calm2 | 0.42 | 1.33E-45 | 2.77E-41 | 1.00 | 0.98 | 0.02 | 1.02 |
| 13 | Elf1 | 1.05 | 2.13E-45 | 4.43E-41 | 0.51 | 0.26 | 0.26 | 2.00 |
| 13 | Cemip2 | 1.01 | 2.31E-45 | 4.81E-41 | 0.69 | 0.44 | 0.24 | 1.55 |
| 13 | Ttc9 | 1.96 | 2.79E-45 | 5.81E-41 | 0.15 | 0.04 | 0.12 | 4.03 |
| 13 | Rgs9 | 1.67 | 4.45E-45 | 9.25E-41 | 0.25 | 0.08 | 0.16 | 3.02 |
| 13 | Chd7 | 0.73 | 6.55E-45 | 1.36E-40 | 0.85 | 0.65 | 0.19 | 1.30 |
| 13 | Ifi30 | 0.93 | 6.93E-45 | 1.44E-40 | 0.55 | 0.29 | 0.26 | 1.91 |
| 13 | Fam162a | 0.59 | 8.22E-45 | 1.71E-40 | 0.95 | 0.89 | 0.06 | 1.07 |
| 13 | Asb7 | 1.55 | 1.03E-44 | 2.15E-40 | 0.43 | 0.20 | 0.22 | 2.09 |
| 13 | Tedc1 | 1.44 | 1.98E-44 | 4.12E-40 | 0.32 | 0.13 | 0.19 | 2.54 |
| 13 | Krt15 | 0.77 | 2.04E-44 | 4.25E-40 | 0.16 | 0.04 | 0.12 | 4.08 |
| 13 | Il17rd | 0.88 | 2.22E-44 | 4.63E-40 | 0.74 | 0.50 | 0.24 | 1.49 |

**Supplementary Table 1. Marker genes for the original clusters of integrated clustering of E11.5 control and NotchGOF λ-junction cells.**

| | | | | | | | | |
|---|---|---|---|---|---|---|---|---|
| 13 | Bcl11b | 1.02 | 2.98E-44 | 6.20E-40 | 0.78 | 0.59 | 0.20 | 1.34 |
| 13 | Crabp2 | 0.46 | 3.11E-44 | 6.48E-40 | 0.99 | 0.91 | 0.08 | 1.09 |
| 13 | Dapl1 | 1.39 | 3.65E-44 | 7.60E-40 | 0.22 | 0.07 | 0.15 | 3.22 |
| 13 | Capsl | 2.26 | 4.12E-44 | 8.56E-40 | 0.12 | 0.02 | 0.09 | 4.88 |
| 13 | Rasgef1b | 2.02 | 5.57E-44 | 1.16E-39 | 0.17 | 0.05 | 0.12 | 3.73 |
| 13 | Tshz2 | 1.16 | 1.13E-43 | 2.34E-39 | 0.59 | 0.33 | 0.25 | 1.76 |
| 13 | Nol7 | 0.51 | 1.29E-43 | 2.67E-39 | 0.95 | 0.88 | 0.08 | 1.09 |
| 13 | Ppargc1a | 2.20 | 1.34E-43 | 2.79E-39 | 0.12 | 0.03 | 0.10 | 4.69 |
| 13 | Ostf1 | 0.98 | 2.28E-43 | 4.74E-39 | 0.52 | 0.27 | 0.25 | 1.92 |
| 13 | Pcgf5 | 1.48 | 2.35E-43 | 4.90E-39 | 0.25 | 0.08 | 0.16 | 2.95 |
| 13 | AI463170 | 2.49 | 3.54E-43 | 7.37E-39 | 0.11 | 0.02 | 0.09 | 5.25 |
| 13 | Gsr | 1.04 | 4.90E-43 | 1.02E-38 | 0.58 | 0.33 | 0.25 | 1.76 |
| 13 | Tacstd2 | 0.85 | 1.09E-42 | 2.26E-38 | 0.22 | 0.07 | 0.15 | 3.22 |
| 13 | Foxp1 | 0.50 | 2.45E-42 | 5.09E-38 | 0.90 | 0.59 | 0.30 | 1.51 |
| 13 | Akap9 | 0.70 | 2.76E-42 | 5.75E-38 | 0.92 | 0.78 | 0.15 | 1.19 |
| 13 | Hip1r | 1.57 | 3.12E-42 | 6.48E-38 | 0.22 | 0.07 | 0.15 | 3.07 |
| 13 | Pard3b | 1.28 | 4.19E-42 | 8.71E-38 | 0.30 | 0.12 | 0.19 | 2.62 |
| 13 | Pdcd4 | 0.80 | 5.21E-42 | 1.08E-37 | 0.83 | 0.64 | 0.20 | 1.31 |
| 13 | Gdpd1 | 1.00 | 5.26E-42 | 1.09E-37 | 0.51 | 0.26 | 0.25 | 1.94 |
| 13 | Jag1 | 0.79 | 6.50E-42 | 1.35E-37 | 0.41 | 0.17 | 0.23 | 2.35 |
| 13 | Tnk1 | 1.92 | 7.42E-42 | 1.54E-37 | 0.14 | 0.04 | 0.11 | 4.06 |
| 13 | Parva | 0.80 | 9.10E-42 | 1.89E-37 | 0.76 | 0.54 | 0.22 | 1.40 |
| 13 | Lratd2 | 1.19 | 1.04E-41 | 2.16E-37 | 0.29 | 0.11 | 0.18 | 2.68 |
| 13 | Arhgef28 | 1.42 | 1.06E-41 | 2.20E-37 | 0.27 | 0.10 | 0.17 | 2.76 |
| 13 | Gchfr | 1.22 | 1.17E-41 | 2.44E-37 | 0.28 | 0.10 | 0.18 | 2.71 |
| 13 | Ermn | 1.76 | 2.46E-41 | 5.13E-37 | 0.16 | 0.04 | 0.12 | 3.83 |
| 13 | Tmem47 | 1.01 | 6.34E-41 | 1.32E-36 | 0.61 | 0.36 | 0.25 | 1.68 |
| 13 | Uqcr11 | 0.33 | 6.79E-41 | 1.41E-36 | 0.99 | 0.97 | 0.01 | 1.01 |
| 13 | H2-Q4 | 1.39 | 7.61E-41 | 1.58E-36 | 0.28 | 0.10 | 0.17 | 2.69 |
| 13 | Vamp8 | 0.69 | 8.70E-41 | 1.81E-36 | 0.83 | 0.61 | 0.22 | 1.36 |
| 13 | H2afj | 0.54 | 1.11E-40 | 2.31E-36 | 0.93 | 0.81 | 0.12 | 1.15 |
| 13 | 1700023F06Rik | 1.65 | 1.14E-40 | 2.38E-36 | 0.16 | 0.04 | 0.12 | 3.83 |
| 13 | Rnf24 | 1.11 | 1.29E-40 | 2.69E-36 | 0.54 | 0.29 | 0.24 | 1.83 |
| 13 | Armc10 | 0.75 | 1.97E-40 | 4.09E-36 | 0.76 | 0.52 | 0.24 | 1.46 |
| 13 | Shroom3 | 1.27 | 2.01E-40 | 4.18E-36 | 0.35 | 0.15 | 0.20 | 2.34 |
| 13 | Tjp2 | 1.03 | 2.22E-40 | 4.62E-36 | 0.55 | 0.31 | 0.24 | 1.80 |
| 13 | Phgdh | 0.57 | 3.00E-40 | 6.24E-36 | 0.93 | 0.82 | 0.11 | 1.13 |
| 13 | Ctnna2 | 1.78 | 5.16E-40 | 1.07E-35 | 0.17 | 0.05 | 0.12 | 3.61 |
| 13 | Slc2a1 | 0.86 | 5.44E-40 | 1.13E-35 | 0.67 | 0.42 | 0.24 | 1.58 |
| 13 | Id4 | 0.88 | 5.46E-40 | 1.14E-35 | 0.55 | 0.30 | 0.25 | 1.82 |
| 13 | Grb10 | 0.53 | 6.10E-40 | 1.27E-35 | 0.98 | 0.94 | 0.04 | 1.05 |
| 13 | Mxi1 | 0.95 | 1.38E-39 | 2.88E-35 | 0.66 | 0.42 | 0.23 | 1.55 |
| 13 | Fzd3 | 0.66 | 1.39E-39 | 2.89E-35 | 0.81 | 0.58 | 0.23 | 1.39 |
| 13 | Cyfip2 | 1.24 | 2.32E-39 | 4.83E-35 | 0.25 | 0.09 | 0.16 | 2.82 |
| 13 | Wnt6 | 0.52 | 2.73E-39 | 5.68E-35 | 0.27 | 0.10 | 0.17 | 2.77 |
| 13 | Malat1 | 0.54 | 2.82E-39 | 5.87E-35 | 1.00 | 1.00 | 0.00 | 1.00 |
| 13 | Vps36 | 0.57 | 2.94E-39 | 6.11E-35 | 0.92 | 0.79 | 0.13 | 1.16 |
| 13 | Zfp503 | 0.73 | 3.06E-39 | 6.36E-35 | 0.63 | 0.37 | 0.26 | 1.72 |
| 13 | Mtln | 0.92 | 3.78E-39 | 7.86E-35 | 0.57 | 0.32 | 0.24 | 1.75 |
| 13 | Tpbg | 0.92 | 4.11E-39 | 8.55E-35 | 0.55 | 0.30 | 0.25 | 1.84 |
| 13 | Tmem176b | 0.82 | 6.39E-39 | 1.33E-34 | 0.77 | 0.56 | 0.21 | 1.37 |
| 13 | Macc1 | 1.48 | 8.69E-39 | 1.81E-34 | 0.18 | 0.05 | 0.13 | 3.40 |
| 13 | Tspan12 | 1.03 | 9.00E-39 | 1.87E-34 | 0.43 | 0.21 | 0.23 | 2.10 |
| 13 | 2310002F09Rik | 1.72 | 1.83E-38 | 3.81E-34 | 0.14 | 0.04 | 0.10 | 3.94 |
| 13 | Plxnb1 | 1.04 | 2.35E-38 | 4.88E-34 | 0.49 | 0.26 | 0.24 | 1.93 |
| 13 | Itgb4 | 2.06 | 3.32E-38 | 6.90E-34 | 0.13 | 0.03 | 0.10 | 4.17 |

**Supplementary Table 1. Marker genes for the original clusters of integrated clustering of E11.5 control and NotchGOF λ-junction cells.**

| | | | | | | | | |
|---|---|---|---|---|---|---|---|---|
| 13 | Abca3 | 1.58 | 3.47E-38 | 7.21E-34 | 0.19 | 0.06 | 0.13 | 3.13 |
| 13 | Tst | 1.23 | 5.59E-38 | 1.16E-33 | 0.30 | 0.12 | 0.18 | 2.46 |
| 13 | Slc35g1 | 1.54 | 8.97E-38 | 1.87E-33 | 0.18 | 0.05 | 0.13 | 3.33 |
| 13 | Pcm1 | 0.47 | 1.09E-37 | 2.28E-33 | 0.96 | 0.89 | 0.07 | 1.08 |
| 13 | Syde2 | 1.47 | 1.58E-37 | 3.28E-33 | 0.21 | 0.07 | 0.14 | 3.01 |
| 13 | Mcoln3 | 1.70 | 2.27E-37 | 4.73E-33 | 0.18 | 0.05 | 0.12 | 3.30 |
| 13 | Prkcz | 1.23 | 2.30E-37 | 4.79E-33 | 0.27 | 0.10 | 0.17 | 2.64 |
| 13 | Psma7 | 0.31 | 2.37E-37 | 4.93E-33 | 1.00 | 0.99 | 0.01 | 1.01 |
| 13 | Dab1 | 0.83 | 2.37E-37 | 4.93E-33 | 0.35 | 0.15 | 0.20 | 2.35 |
| 13 | Tafa2 | 1.70 | 2.41E-37 | 5.02E-33 | 0.15 | 0.04 | 0.11 | 3.72 |
| 13 | 2310030G06Rik | 1.27 | 2.45E-37 | 5.10E-33 | 0.21 | 0.07 | 0.14 | 3.07 |
| 13 | Sema6a | 1.09 | 3.18E-37 | 6.60E-33 | 0.66 | 0.44 | 0.22 | 1.49 |
| 13 | 1700096K18Rik | 1.32 | 3.53E-37 | 7.35E-33 | 0.25 | 0.09 | 0.16 | 2.74 |
| 13 | Slc35f1 | 1.87 | 3.86E-37 | 8.03E-33 | 0.13 | 0.03 | 0.10 | 4.10 |
| 13 | Lrig3 | 0.99 | 4.19E-37 | 8.71E-33 | 0.45 | 0.22 | 0.23 | 2.02 |
| 13 | Macrod2 | 1.07 | 4.76E-37 | 9.91E-33 | 0.43 | 0.22 | 0.22 | 1.99 |
| 13 | Zdhhc12 | 0.96 | 8.84E-37 | 1.84E-32 | 0.49 | 0.26 | 0.23 | 1.91 |
| 13 | Gm28112 | 2.05 | 9.19E-37 | 1.91E-32 | 0.10 | 0.02 | 0.08 | 4.64 |
| 13 | Rab27b | 1.15 | 1.03E-36 | 2.14E-32 | 0.25 | 0.09 | 0.16 | 2.68 |
| 13 | Sinhcaf | 0.49 | 1.14E-36 | 2.37E-32 | 0.96 | 0.84 | 0.12 | 1.14 |
| 13 | Large2 | 1.31 | 1.51E-36 | 3.13E-32 | 0.23 | 0.08 | 0.15 | 2.80 |
| 13 | Efnb2 | 0.81 | 1.76E-36 | 3.66E-32 | 0.69 | 0.44 | 0.25 | 1.56 |
| 13 | Ptprk | 0.82 | 1.81E-36 | 3.76E-32 | 0.68 | 0.44 | 0.24 | 1.56 |
| 13 | Npm3 | 0.54 | 2.16E-36 | 4.50E-32 | 0.91 | 0.79 | 0.11 | 1.14 |
| 13 | Baiap2l1 | 1.58 | 2.53E-36 | 5.26E-32 | 0.19 | 0.06 | 0.13 | 3.13 |
| 13 | Hpn | 1.87 | 3.15E-36 | 6.55E-32 | 0.14 | 0.04 | 0.10 | 3.81 |
| 13 | Mpst | 0.80 | 3.27E-36 | 6.79E-32 | 0.65 | 0.41 | 0.24 | 1.58 |
| 13 | Zmat4 | 1.50 | 3.55E-36 | 7.38E-32 | 0.17 | 0.05 | 0.12 | 3.26 |
| 13 | Ankrd6 | 1.61 | 3.60E-36 | 7.48E-32 | 0.26 | 0.10 | 0.16 | 2.52 |
| 13 | Ralgps1 | 1.95 | 3.80E-36 | 7.91E-32 | 0.15 | 0.04 | 0.11 | 3.61 |
| 13 | Ide | 0.98 | 4.06E-36 | 8.44E-32 | 0.49 | 0.26 | 0.23 | 1.86 |
| 13 | Npepps | 0.63 | 5.12E-36 | 1.06E-31 | 0.84 | 0.64 | 0.19 | 1.30 |
| 13 | Enpp5 | 1.70 | 5.53E-36 | 1.15E-31 | 0.15 | 0.04 | 0.11 | 3.61 |
| 13 | Daam1 | 1.01 | 5.58E-36 | 1.16E-31 | 0.62 | 0.40 | 0.22 | 1.55 |
| 13 | Tubb4a | 1.50 | 6.42E-36 | 1.34E-31 | 0.14 | 0.04 | 0.10 | 3.86 |
| 13 | Plekhf2 | 1.05 | 7.74E-36 | 1.61E-31 | 0.44 | 0.23 | 0.21 | 1.95 |
| 13 | Sord | 1.09 | 8.36E-36 | 1.74E-31 | 0.33 | 0.14 | 0.19 | 2.31 |
| 13 | Dag1 | 0.81 | 9.10E-36 | 1.89E-31 | 0.61 | 0.37 | 0.25 | 1.67 |
| 13 | Mgst1 | 0.67 | 1.15E-35 | 2.40E-31 | 0.64 | 0.37 | 0.27 | 1.73 |
| 13 | Ubash3b | 1.06 | 2.54E-35 | 5.28E-31 | 0.44 | 0.23 | 0.21 | 1.91 |
| 13 | Cux1 | 0.59 | 2.54E-35 | 5.28E-31 | 0.87 | 0.69 | 0.17 | 1.25 |
| 13 | Kif13b | 1.13 | 3.55E-35 | 7.39E-31 | 0.35 | 0.16 | 0.19 | 2.21 |
| 13 | Fbln7 | 2.10 | 4.30E-35 | 8.94E-31 | 0.10 | 0.02 | 0.08 | 4.64 |
| 13 | Ssc4d | 1.56 | 4.43E-35 | 9.20E-31 | 0.17 | 0.05 | 0.12 | 3.27 |
| 13 | Apba2 | 1.39 | 4.57E-35 | 9.50E-31 | 0.18 | 0.06 | 0.13 | 3.16 |
| 13 | Lypd2 | 1.62 | 5.89E-35 | 1.22E-30 | 0.14 | 0.04 | 0.10 | 3.75 |
| 13 | Tmem205 | 0.97 | 6.32E-35 | 1.31E-30 | 0.50 | 0.27 | 0.23 | 1.85 |
| 13 | Tent5a | 1.71 | 6.97E-35 | 1.45E-30 | 0.14 | 0.04 | 0.10 | 3.70 |
| 13 | Dmbx1 | 1.61 | 7.32E-35 | 1.52E-30 | 0.15 | 0.04 | 0.11 | 3.52 |
| 13 | Sox3 | 1.44 | 8.05E-35 | 1.67E-30 | 0.14 | 0.04 | 0.10 | 3.68 |
| 13 | Cmtm4 | 0.94 | 9.43E-35 | 1.96E-30 | 0.47 | 0.25 | 0.22 | 1.87 |
| 13 | Sult4a1 | 1.88 | 1.21E-34 | 2.52E-30 | 0.13 | 0.04 | 0.10 | 3.69 |
| 13 | Mapk12 | 1.20 | 1.35E-34 | 2.80E-30 | 0.28 | 0.11 | 0.16 | 2.48 |
| 13 | Tsc22d2 | 0.92 | 1.64E-34 | 3.41E-30 | 0.63 | 0.40 | 0.23 | 1.56 |
| 13 | Dock9 | 1.49 | 1.78E-34 | 3.71E-30 | 0.20 | 0.07 | 0.13 | 2.91 |
| 13 | Lgr4 | 0.82 | 1.82E-34 | 3.80E-30 | 0.45 | 0.22 | 0.22 | 2.00 |

**Supplementary Table 1. Marker genes for the original clusters of integrated clustering of E11.5 control and NotchGOF λ-junction cells.**

| | | | | | | | | |
|---|---|---|---|---|---|---|---|---|
| 13 | Kcnh8 | 1.89 | 1.89E-34 | 3.93E-30 | 0.13 | 0.03 | 0.09 | 3.91 |
| 13 | Ralyl | 1.80 | 3.08E-34 | 6.40E-30 | 0.11 | 0.03 | 0.08 | 4.28 |
| 13 | Dcxr | 1.42 | 3.54E-34 | 7.36E-30 | 0.24 | 0.09 | 0.15 | 2.62 |
| 13 | Aig1 | 0.87 | 4.27E-34 | 8.89E-30 | 0.57 | 0.33 | 0.24 | 1.74 |
| 13 | Dlx5 | 0.82 | 4.28E-34 | 8.91E-30 | 0.43 | 0.21 | 0.22 | 2.04 |
| 13 | Nedd4l | 1.19 | 4.68E-34 | 9.73E-30 | 0.32 | 0.14 | 0.18 | 2.26 |
| 13 | Wls | 0.78 | 5.12E-34 | 1.06E-29 | 0.85 | 0.74 | 0.11 | 1.15 |
| 13 | Ccdc92 | 1.39 | 6.31E-34 | 1.31E-29 | 0.22 | 0.08 | 0.14 | 2.78 |
| 13 | Cacnb2 | 1.49 | 7.90E-34 | 1.64E-29 | 0.21 | 0.07 | 0.13 | 2.86 |
| 13 | Clstn1 | 0.81 | 8.05E-34 | 1.67E-29 | 0.62 | 0.38 | 0.24 | 1.62 |
| 13 | Matk | 2.07 | 1.02E-33 | 2.12E-29 | 0.11 | 0.03 | 0.08 | 4.28 |
| 13 | Aftph | 1.09 | 1.02E-33 | 2.13E-29 | 0.46 | 0.25 | 0.21 | 1.86 |
| 13 | Itgb6 | 1.68 | 1.23E-33 | 2.55E-29 | 0.14 | 0.04 | 0.10 | 3.55 |
| 13 | Cdca7l | 0.95 | 2.00E-33 | 4.15E-29 | 0.52 | 0.30 | 0.22 | 1.73 |
| 13 | Tox3 | 1.03 | 2.07E-33 | 4.30E-29 | 0.44 | 0.23 | 0.21 | 1.92 |
| 13 | Arid3b | 0.98 | 2.34E-33 | 4.86E-29 | 0.44 | 0.22 | 0.21 | 1.95 |
| 13 | Socs2 | 0.56 | 2.50E-33 | 5.19E-29 | 0.58 | 0.34 | 0.24 | 1.71 |
| 13 | Sorbs2 | 0.73 | 2.64E-33 | 5.50E-29 | 0.44 | 0.22 | 0.22 | 2.03 |
| 13 | Ptpn4 | 1.20 | 2.99E-33 | 6.21E-29 | 0.34 | 0.16 | 0.18 | 2.17 |
| 13 | Ctnnal1 | 0.76 | 3.20E-33 | 6.65E-29 | 0.49 | 0.26 | 0.23 | 1.89 |
| 13 | Lamc2 | 1.49 | 3.94E-33 | 8.20E-29 | 0.14 | 0.04 | 0.10 | 3.49 |
| 13 | Fam189a2 | 2.02 | 5.26E-33 | 1.09E-28 | 0.10 | 0.02 | 0.08 | 4.35 |
| 13 | Acot7 | 0.94 | 5.74E-33 | 1.19E-28 | 0.45 | 0.23 | 0.21 | 1.93 |
| 13 | Nudt16 | 1.33 | 7.85E-33 | 1.63E-28 | 0.22 | 0.08 | 0.14 | 2.70 |
| 13 | Sulf1 | 0.77 | 8.24E-33 | 1.71E-28 | 0.76 | 0.53 | 0.23 | 1.42 |
| 13 | Alad | 0.39 | 1.01E-32 | 2.10E-28 | 0.83 | 0.63 | 0.20 | 1.32 |
| 13 | Frat1 | 1.50 | 1.04E-32 | 2.17E-28 | 0.16 | 0.05 | 0.11 | 3.34 |
| 13 | Nipsnap1 | 0.62 | 1.13E-32 | 2.35E-28 | 0.78 | 0.56 | 0.22 | 1.40 |
| 13 | Map7d2 | 1.39 | 4.33E-32 | 9.02E-28 | 0.22 | 0.08 | 0.14 | 2.68 |
| 13 | Card19 | 1.08 | 4.85E-32 | 1.01E-27 | 0.45 | 0.25 | 0.20 | 1.82 |
| 13 | Pdgfrl | 1.06 | 5.70E-32 | 1.19E-27 | 0.37 | 0.18 | 0.19 | 2.07 |
| 13 | Rasal2 | 1.04 | 6.08E-32 | 1.26E-27 | 0.45 | 0.24 | 0.21 | 1.85 |
| 13 | Fxyd3 | 1.11 | 6.33E-32 | 1.32E-27 | 0.23 | 0.09 | 0.14 | 2.61 |
| 13 | Ptpn14 | 1.33 | 7.45E-32 | 1.55E-27 | 0.32 | 0.15 | 0.17 | 2.15 |
| 13 | Rimklb | 0.83 | 1.10E-31 | 2.28E-27 | 0.55 | 0.32 | 0.23 | 1.71 |
| 13 | Cltb | 1.01 | 1.71E-31 | 3.56E-27 | 0.37 | 0.18 | 0.19 | 2.04 |
| 13 | Rrp12 | 1.08 | 1.76E-31 | 3.67E-27 | 0.38 | 0.19 | 0.19 | 1.97 |
| 13 | Dennd1b | 1.10 | 2.00E-31 | 4.15E-27 | 0.34 | 0.16 | 0.18 | 2.17 |
| 13 | Churc1 | 0.70 | 2.11E-31 | 4.38E-27 | 0.79 | 0.58 | 0.20 | 1.35 |
| 13 | Irf6 | 0.61 | 2.45E-31 | 5.10E-27 | 0.27 | 0.11 | 0.16 | 2.50 |
| 13 | Tdrd7 | 2.12 | 3.67E-31 | 7.63E-27 | 0.11 | 0.03 | 0.08 | 4.04 |
| 13 | Nup210 | 1.48 | 4.04E-31 | 8.41E-27 | 0.18 | 0.06 | 0.12 | 2.97 |
| 13 | Myc | 1.31 | 4.62E-31 | 9.60E-27 | 0.33 | 0.16 | 0.17 | 2.08 |
| 13 | Lamb1 | 0.85 | 6.53E-31 | 1.36E-26 | 0.60 | 0.38 | 0.22 | 1.57 |
| 13 | Nectin2 | 0.75 | 7.90E-31 | 1.64E-26 | 0.69 | 0.47 | 0.22 | 1.46 |
| 13 | Hif3a | 1.22 | 8.65E-31 | 1.80E-26 | 0.22 | 0.08 | 0.14 | 2.63 |
| 13 | Palm3 | 0.98 | 9.53E-31 | 1.98E-26 | 0.32 | 0.14 | 0.18 | 2.22 |
| 13 | Cracr2b | 1.66 | 1.04E-30 | 2.16E-26 | 0.12 | 0.03 | 0.09 | 3.70 |
| 13 | Mien1 | 0.66 | 1.84E-30 | 3.82E-26 | 0.75 | 0.56 | 0.19 | 1.35 |
| 13 | Nipal3 | 1.82 | 2.46E-30 | 5.11E-26 | 0.13 | 0.04 | 0.09 | 3.41 |
| 13 | Baz2b | 0.58 | 2.72E-30 | 5.66E-26 | 0.86 | 0.70 | 0.16 | 1.23 |
| 13 | Myo5c | 1.76 | 2.92E-30 | 6.08E-26 | 0.13 | 0.04 | 0.09 | 3.49 |
| 13 | Tdrkh | 1.58 | 3.05E-30 | 6.35E-26 | 0.17 | 0.06 | 0.11 | 2.95 |
| 13 | B3gnt2 | 0.99 | 3.11E-30 | 6.48E-26 | 0.24 | 0.10 | 0.15 | 2.54 |
| 13 | Atp7b | 1.83 | 3.38E-30 | 7.02E-26 | 0.12 | 0.03 | 0.08 | 3.71 |
| 13 | Rassf7 | 0.90 | 3.75E-30 | 7.81E-26 | 0.39 | 0.19 | 0.20 | 2.03 |

**Supplementary Table 1. Marker genes for the original clusters of integrated clustering of E11.5 control and NotchGOF λ-junction cells.**

| | | | | | | | | |
|---|---|---|---|---|---|---|---|---|
| 13 | Strbp | 0.53 | 3.77E-30 | 7.85E-26 | 0.84 | 0.70 | 0.14 | 1.20 |
| 13 | Rel | 1.40 | 4.24E-30 | 8.82E-26 | 0.22 | 0.09 | 0.14 | 2.55 |
| 13 | Ajuba | 0.99 | 6.35E-30 | 1.32E-25 | 0.48 | 0.28 | 0.20 | 1.74 |
| 13 | Tspan7 | 0.81 | 6.86E-30 | 1.43E-25 | 0.50 | 0.28 | 0.22 | 1.80 |
| 13 | Prxl2a | 0.76 | 7.55E-30 | 1.57E-25 | 0.60 | 0.38 | 0.23 | 1.60 |
| 13 | Elavl2 | 0.80 | 7.96E-30 | 1.65E-25 | 0.37 | 0.18 | 0.19 | 2.04 |
| 13 | Rab11fip1 | 1.04 | 7.96E-30 | 1.66E-25 | 0.27 | 0.11 | 0.16 | 2.37 |
| 13 | Rtn3 | 0.49 | 8.29E-30 | 1.72E-25 | 0.91 | 0.79 | 0.13 | 1.16 |
| 13 | Frmd4b | 0.85 | 8.71E-30 | 1.81E-25 | 0.35 | 0.17 | 0.18 | 2.12 |
| 13 | Pdk1 | 0.80 | 9.53E-30 | 1.98E-25 | 0.47 | 0.25 | 0.21 | 1.85 |
| 13 | Cdon | 1.10 | 1.33E-29 | 2.77E-25 | 0.41 | 0.22 | 0.19 | 1.87 |
| 13 | Rbms1 | 0.59 | 1.52E-29 | 3.17E-25 | 0.89 | 0.78 | 0.11 | 1.14 |
| 13 | Myo10 | 0.74 | 1.90E-29 | 3.95E-25 | 0.68 | 0.47 | 0.21 | 1.45 |
| 13 | Myl12b | 0.37 | 2.30E-29 | 4.78E-25 | 0.95 | 0.88 | 0.07 | 1.08 |
| 13 | Klhl23 | 0.77 | 2.32E-29 | 4.83E-25 | 0.59 | 0.37 | 0.23 | 1.61 |
| 13 | Dedd2 | 1.29 | 2.50E-29 | 5.20E-25 | 0.20 | 0.07 | 0.12 | 2.72 |
| 13 | Gpc4 | 1.15 | 2.59E-29 | 5.39E-25 | 0.30 | 0.14 | 0.16 | 2.21 |
| 13 | Prps2 | 1.05 | 2.84E-29 | 5.91E-25 | 0.34 | 0.17 | 0.17 | 2.05 |
| 13 | Ehbp1 | 0.82 | 3.07E-29 | 6.39E-25 | 0.44 | 0.24 | 0.21 | 1.87 |
| 13 | Fbxo27 | 1.28 | 3.44E-29 | 7.15E-25 | 0.18 | 0.06 | 0.12 | 2.86 |
| 13 | Lamc1 | 0.81 | 3.60E-29 | 7.49E-25 | 0.46 | 0.26 | 0.21 | 1.81 |
| 13 | Fgd4 | 1.19 | 5.61E-29 | 1.17E-24 | 0.22 | 0.09 | 0.14 | 2.55 |
| 13 | Erc2 | 1.39 | 5.64E-29 | 1.17E-24 | 0.19 | 0.07 | 0.12 | 2.78 |
| 13 | Clip4 | 1.31 | 5.72E-29 | 1.19E-24 | 0.14 | 0.04 | 0.10 | 3.30 |
| 13 | Gtf3a | 0.57 | 5.85E-29 | 1.22E-24 | 0.79 | 0.64 | 0.15 | 1.23 |
| 13 | Sh3gl2 | 1.53 | 6.24E-29 | 1.30E-24 | 0.13 | 0.04 | 0.09 | 3.49 |
| 13 | Zswim6 | 0.89 | 1.02E-28 | 2.13E-24 | 0.50 | 0.29 | 0.21 | 1.72 |
| 13 | Gltp | 0.80 | 1.04E-28 | 2.17E-24 | 0.60 | 0.39 | 0.20 | 1.52 |
| 13 | St8sia6 | 1.18 | 1.08E-28 | 2.24E-24 | 0.15 | 0.05 | 0.10 | 3.15 |
| 13 | Pfkl | 0.53 | 1.50E-28 | 3.11E-24 | 0.83 | 0.64 | 0.19 | 1.29 |
| 13 | Scamp2 | 0.80 | 1.50E-28 | 3.12E-24 | 0.53 | 0.31 | 0.22 | 1.70 |
| 13 | Myo5b | 1.74 | 1.79E-28 | 3.73E-24 | 0.14 | 0.04 | 0.10 | 3.26 |
| 13 | Snn | 0.71 | 1.83E-28 | 3.82E-24 | 0.65 | 0.43 | 0.22 | 1.52 |
| 13 | Stox2 | 0.67 | 1.89E-28 | 3.94E-24 | 0.74 | 0.56 | 0.18 | 1.33 |
| 13 | Trp53i11 | 0.61 | 2.03E-28 | 4.23E-24 | 0.43 | 0.22 | 0.21 | 1.92 |
| 13 | Smim10l2a | 1.34 | 2.06E-28 | 4.29E-24 | 0.19 | 0.07 | 0.12 | 2.76 |
| 13 | Egln3 | 1.24 | 2.21E-28 | 4.61E-24 | 0.20 | 0.07 | 0.12 | 2.68 |
| 13 | Dsel | 0.86 | 2.48E-28 | 5.16E-24 | 0.53 | 0.33 | 0.21 | 1.63 |
| 13 | Plcb4 | 0.98 | 2.50E-28 | 5.20E-24 | 0.38 | 0.19 | 0.18 | 1.94 |
| 13 | Med19 | 0.49 | 2.73E-28 | 5.67E-24 | 0.88 | 0.74 | 0.14 | 1.18 |
| 13 | Galnt3 | 1.69 | 3.34E-28 | 6.95E-24 | 0.11 | 0.03 | 0.08 | 3.68 |
| 13 | Usp25 | 0.67 | 3.50E-28 | 7.27E-24 | 0.65 | 0.43 | 0.22 | 1.51 |
| 13 | AC149090.1 | 0.92 | 3.56E-28 | 7.40E-24 | 0.48 | 0.28 | 0.20 | 1.72 |
| 13 | Nucb2 | 1.10 | 3.62E-28 | 7.54E-24 | 0.32 | 0.15 | 0.17 | 2.11 |
| 13 | Arhgdig | 0.29 | 4.19E-28 | 8.71E-24 | 0.20 | 0.08 | 0.13 | 2.58 |
| 13 | Zdhhc21 | 0.80 | 4.31E-28 | 8.96E-24 | 0.50 | 0.29 | 0.21 | 1.75 |
| 13 | Grip1 | 1.21 | 6.50E-28 | 1.35E-23 | 0.24 | 0.10 | 0.14 | 2.37 |
| 13 | Os9 | 0.76 | 6.64E-28 | 1.38E-23 | 0.59 | 0.37 | 0.22 | 1.60 |
| 13 | Dclre1c | 1.80 | 8.92E-28 | 1.85E-23 | 0.15 | 0.05 | 0.10 | 2.98 |
| 13 | Dmtn | 1.54 | 1.00E-27 | 2.08E-23 | 0.18 | 0.06 | 0.11 | 2.78 |
| 13 | mt-Nd6 | 0.85 | 1.11E-27 | 2.31E-23 | 0.56 | 0.35 | 0.21 | 1.61 |
| 13 | Dazap2 | 0.60 | 1.33E-27 | 2.77E-23 | 0.77 | 0.57 | 0.20 | 1.36 |
| 13 | P4ha3 | 0.97 | 1.35E-27 | 2.80E-23 | 0.32 | 0.15 | 0.17 | 2.10 |
| 13 | Krt7 | 1.23 | 1.39E-27 | 2.89E-23 | 0.15 | 0.05 | 0.10 | 3.00 |
| 13 | Nabp2 | 0.52 | 1.48E-27 | 3.07E-23 | 0.83 | 0.65 | 0.18 | 1.27 |
| 13 | Pick1 | 0.80 | 1.95E-27 | 4.05E-23 | 0.46 | 0.25 | 0.20 | 1.81 |

**Supplementary Table 1. Marker genes for the original clusters of integrated clustering of E11.5 control and NotchGOF λ-junction cells.**

| | | | | | | | | |
|---|---|---|---|---|---|---|---|---|
| 13 | Lrrc1 | 1.10 | 2.18E-27 | 4.53E-23 | 0.31 | 0.15 | 0.16 | 2.10 |
| 13 | Tle5 | 0.34 | 2.28E-27 | 4.74E-23 | 0.98 | 0.94 | 0.05 | 1.05 |
| 13 | Ier5 | 0.75 | 2.34E-27 | 4.86E-23 | 0.80 | 0.65 | 0.15 | 1.23 |
| 13 | Nme3 | 1.29 | 2.36E-27 | 4.90E-23 | 0.19 | 0.07 | 0.12 | 2.62 |
| 13 | Pura | 0.67 | 2.76E-27 | 5.74E-23 | 0.68 | 0.47 | 0.21 | 1.45 |
| 13 | Slc48a1 | 0.96 | 2.80E-27 | 5.82E-23 | 0.33 | 0.16 | 0.17 | 2.05 |
| 13 | Syngr2 | 0.95 | 3.33E-27 | 6.92E-23 | 0.30 | 0.14 | 0.16 | 2.16 |
| 13 | Lipg | 1.73 | 4.82E-27 | 1.00E-22 | 0.12 | 0.03 | 0.09 | 3.50 |
| 13 | Smtnl2 | 1.14 | 5.24E-27 | 1.09E-22 | 0.19 | 0.07 | 0.12 | 2.64 |
| 13 | Bnip3 | 0.84 | 5.36E-27 | 1.12E-22 | 0.61 | 0.41 | 0.20 | 1.48 |
| 13 | Ndrg2 | 1.06 | 5.89E-27 | 1.22E-22 | 0.29 | 0.13 | 0.16 | 2.18 |
| 13 | Prkci | 0.85 | 6.30E-27 | 1.31E-22 | 0.46 | 0.26 | 0.20 | 1.75 |
| 13 | Erich1 | 0.98 | 6.47E-27 | 1.35E-22 | 0.41 | 0.23 | 0.18 | 1.79 |
| 13 | Zdbf2 | 0.69 | 6.83E-27 | 1.42E-22 | 0.62 | 0.40 | 0.22 | 1.55 |
| 13 | Pts | 0.69 | 7.05E-27 | 1.47E-22 | 0.63 | 0.40 | 0.23 | 1.56 |
| 13 | Nrtn | 1.02 | 7.64E-27 | 1.59E-22 | 0.31 | 0.15 | 0.16 | 2.06 |
| 13 | Trp53inp1 | 0.89 | 7.95E-27 | 1.65E-22 | 0.36 | 0.18 | 0.18 | 1.99 |
| 13 | Zswim5 | 1.70 | 9.40E-27 | 1.96E-22 | 0.11 | 0.03 | 0.08 | 3.55 |
| 13 | Tdrp | 1.68 | 9.70E-27 | 2.02E-22 | 0.10 | 0.03 | 0.08 | 3.78 |
| 13 | Gucy1a1 | 1.74 | 9.96E-27 | 2.07E-22 | 0.19 | 0.07 | 0.12 | 2.55 |
| 13 | Aktip | 0.75 | 1.02E-26 | 2.11E-22 | 0.49 | 0.28 | 0.21 | 1.74 |
| 13 | Agpat3 | 0.76 | 1.11E-26 | 2.32E-22 | 0.46 | 0.26 | 0.20 | 1.78 |
| 13 | Smim20 | 0.71 | 1.20E-26 | 2.49E-22 | 0.61 | 0.40 | 0.21 | 1.52 |
| 13 | Cmip | 0.74 | 1.78E-26 | 3.70E-22 | 0.60 | 0.39 | 0.20 | 1.52 |
| 13 | Kcnk2 | 1.21 | 1.80E-26 | 3.73E-22 | 0.48 | 0.30 | 0.18 | 1.61 |
| 13 | Edem2 | 1.04 | 2.07E-26 | 4.31E-22 | 0.36 | 0.19 | 0.17 | 1.92 |
| 13 | Pard3 | 0.58 | 2.10E-26 | 4.37E-22 | 0.76 | 0.57 | 0.19 | 1.33 |
| 13 | Kdm5a | 0.63 | 2.26E-26 | 4.69E-22 | 0.81 | 0.67 | 0.14 | 1.21 |
| 13 | Zfp703 | 0.87 | 2.28E-26 | 4.74E-22 | 0.44 | 0.25 | 0.19 | 1.77 |
| 13 | Retreg1 | 1.22 | 2.42E-26 | 5.04E-22 | 0.22 | 0.09 | 0.13 | 2.43 |
| 13 | Necab2 | 1.34 | 2.57E-26 | 5.34E-22 | 0.36 | 0.20 | 0.16 | 1.79 |
| 13 | Id1 | 0.51 | 2.80E-26 | 5.83E-22 | 0.95 | 0.86 | 0.09 | 1.11 |
| 13 | Hspg2 | 0.70 | 3.12E-26 | 6.50E-22 | 0.42 | 0.22 | 0.19 | 1.87 |
| 13 | Tmem53 | 1.17 | 4.23E-26 | 8.79E-22 | 0.21 | 0.08 | 0.13 | 2.51 |
| 13 | Gprc5c | 0.82 | 4.86E-26 | 1.01E-21 | 0.43 | 0.23 | 0.19 | 1.84 |
| 13 | Gcat | 0.65 | 5.38E-26 | 1.12E-21 | 0.68 | 0.48 | 0.20 | 1.43 |
| 13 | Jpt1 | 0.36 | 7.44E-26 | 1.55E-21 | 0.98 | 0.94 | 0.04 | 1.04 |
| 13 | Spats2l | 1.16 | 9.25E-26 | 1.92E-21 | 0.40 | 0.23 | 0.17 | 1.76 |
| 13 | Homer2 | 0.95 | 9.53E-26 | 1.98E-21 | 0.38 | 0.21 | 0.18 | 1.86 |
| 13 | Lrrn3 | 1.48 | 1.27E-25 | 2.64E-21 | 0.18 | 0.07 | 0.11 | 2.65 |
| 13 | Dynll2 | 0.43 | 1.37E-25 | 2.84E-21 | 0.93 | 0.84 | 0.09 | 1.10 |
| 13 | Cdc42se1 | 0.62 | 1.40E-25 | 2.91E-21 | 0.65 | 0.42 | 0.23 | 1.54 |
| 13 | Tead3 | 0.85 | 1.44E-25 | 3.00E-21 | 0.43 | 0.24 | 0.19 | 1.79 |
| 13 | Arfgef1 | 0.68 | 1.48E-25 | 3.07E-21 | 0.68 | 0.48 | 0.20 | 1.41 |
| 13 | Eps8l2 | 1.46 | 1.60E-25 | 3.32E-21 | 0.13 | 0.04 | 0.09 | 3.23 |
| 13 | Hist2h2be | 1.72 | 1.61E-25 | 3.36E-21 | 0.12 | 0.04 | 0.08 | 3.34 |
| 13 | Mtcl1 | 1.00 | 1.68E-25 | 3.50E-21 | 0.27 | 0.12 | 0.15 | 2.23 |
| 13 | Cd164 | 0.56 | 2.10E-25 | 4.38E-21 | 0.80 | 0.64 | 0.17 | 1.27 |
| 13 | Thrb | 1.37 | 2.26E-25 | 4.70E-21 | 0.18 | 0.07 | 0.11 | 2.59 |
| 13 | mt-Nd5 | 0.42 | 2.76E-25 | 5.74E-21 | 0.98 | 0.96 | 0.02 | 1.02 |
| 13 | Nmrk1 | 1.25 | 2.97E-25 | 6.18E-21 | 0.20 | 0.08 | 0.12 | 2.48 |
| 13 | Cadps2 | 1.32 | 3.19E-25 | 6.63E-21 | 0.12 | 0.04 | 0.08 | 3.34 |
| 13 | Cep152 | 1.04 | 3.58E-25 | 7.44E-21 | 0.27 | 0.12 | 0.14 | 2.19 |
| 13 | Setd7 | 1.00 | 4.47E-25 | 9.30E-21 | 0.37 | 0.20 | 0.17 | 1.86 |
| 13 | Hspa8 | 0.27 | 4.66E-25 | 9.69E-21 | 1.00 | 1.00 | 0.00 | 1.00 |
| 13 | Nr3c1 | 1.12 | 5.32E-25 | 1.11E-20 | 0.32 | 0.16 | 0.16 | 1.99 |

**Supplementary Table 1. Marker genes for the original clusters of integrated clustering of E11.5 control and NotchGOF λ-junction cells.**

| | | | | | | | | |
|---|---|---|---|---|---|---|---|---|
| 13 | Arl6ip5 | 0.72 | 5.55E-25 | 1.15E-20 | 0.52 | 0.32 | 0.21 | 1.65 |
| 13 | Nectin1 | 0.92 | 5.68E-25 | 1.18E-20 | 0.34 | 0.17 | 0.17 | 1.98 |
| 13 | Hltf | 0.81 | 7.78E-25 | 1.62E-20 | 0.51 | 0.32 | 0.19 | 1.62 |
| 13 | Unc5b | 0.75 | 8.83E-25 | 1.84E-20 | 0.60 | 0.39 | 0.21 | 1.54 |
| 13 | Pbx4 | 1.20 | 9.35E-25 | 1.94E-20 | 0.21 | 0.09 | 0.12 | 2.45 |
| 13 | Zfp612 | 0.97 | 1.16E-24 | 2.40E-20 | 0.33 | 0.17 | 0.16 | 1.98 |
| 13 | Lgalsl | 0.62 | 1.17E-24 | 2.44E-20 | 0.51 | 0.31 | 0.20 | 1.64 |
| 13 | Dgka | 1.37 | 1.20E-24 | 2.50E-20 | 0.16 | 0.06 | 0.10 | 2.73 |
| 13 | mt-Co1 | 0.28 | 1.22E-24 | 2.53E-20 | 0.99 | 0.99 | 0.00 | 1.00 |
| 13 | Gsap | 1.56 | 1.53E-24 | 3.19E-20 | 0.12 | 0.04 | 0.08 | 3.25 |
| 13 | Abhd11 | 0.77 | 1.93E-24 | 4.01E-20 | 0.45 | 0.26 | 0.19 | 1.72 |
| 13 | Grhl3 | 0.82 | 2.07E-24 | 4.30E-20 | 0.15 | 0.05 | 0.10 | 2.83 |
| 13 | Tmem189 | 0.84 | 2.17E-24 | 4.51E-20 | 0.49 | 0.30 | 0.20 | 1.65 |
| 13 | Slc16a1 | 0.81 | 2.70E-24 | 5.62E-20 | 0.49 | 0.30 | 0.19 | 1.66 |
| 13 | Nog | 1.18 | 2.81E-24 | 5.84E-20 | 0.22 | 0.09 | 0.12 | 2.35 |
| 13 | Ucp2 | 0.37 | 3.41E-24 | 7.09E-20 | 0.56 | 0.35 | 0.21 | 1.60 |
| 13 | Lmcd1 | 1.32 | 3.96E-24 | 8.24E-20 | 0.21 | 0.09 | 0.12 | 2.40 |
| 13 | Cpne8 | 1.58 | 4.17E-24 | 8.67E-20 | 0.19 | 0.08 | 0.11 | 2.47 |
| 13 | Cacna2d3 | 1.43 | 4.22E-24 | 8.78E-20 | 0.19 | 0.08 | 0.11 | 2.48 |
| 13 | Abhd14b | 1.42 | 4.37E-24 | 9.09E-20 | 0.16 | 0.06 | 0.10 | 2.72 |
| 13 | Grtp1 | 0.94 | 5.37E-24 | 1.12E-19 | 0.29 | 0.14 | 0.15 | 2.10 |
| 13 | Adam22 | 1.56 | 5.38E-24 | 1.12E-19 | 0.18 | 0.07 | 0.11 | 2.51 |
| 13 | Faah | 1.01 | 5.49E-24 | 1.14E-19 | 0.16 | 0.06 | 0.10 | 2.80 |
| 13 | Rnf121 | 0.80 | 5.81E-24 | 1.21E-19 | 0.55 | 0.36 | 0.19 | 1.53 |
| 13 | Tm7sf2 | 1.04 | 6.80E-24 | 1.41E-19 | 0.29 | 0.14 | 0.15 | 2.04 |
| 13 | Rb1 | 1.05 | 7.30E-24 | 1.52E-19 | 0.31 | 0.16 | 0.15 | 1.97 |
| 13 | Rhob | 0.77 | 7.80E-24 | 1.62E-19 | 0.53 | 0.34 | 0.19 | 1.56 |
| 13 | Bst2 | 1.21 | 1.04E-23 | 2.17E-19 | 0.16 | 0.06 | 0.10 | 2.77 |
| 13 | Cecr2 | 0.75 | 1.07E-23 | 2.22E-19 | 0.40 | 0.22 | 0.18 | 1.82 |
| 13 | Nt5c | 0.54 | 1.33E-23 | 2.76E-19 | 0.74 | 0.53 | 0.20 | 1.38 |
| 13 | Harbi1 | 0.97 | 1.60E-23 | 3.32E-19 | 0.35 | 0.18 | 0.16 | 1.89 |
| 13 | Ggact | 1.36 | 1.62E-23 | 3.36E-19 | 0.14 | 0.05 | 0.09 | 2.94 |
| 13 | Snx10 | 0.95 | 1.66E-23 | 3.46E-19 | 0.31 | 0.16 | 0.16 | 2.00 |
| 13 | H2-T23 | 1.53 | 2.33E-23 | 4.85E-19 | 0.15 | 0.05 | 0.09 | 2.79 |
| 13 | Gpd2 | 0.90 | 2.67E-23 | 5.55E-19 | 0.40 | 0.23 | 0.17 | 1.76 |
| 13 | Slc36a1 | 1.41 | 2.79E-23 | 5.81E-19 | 0.15 | 0.05 | 0.10 | 2.83 |
| 13 | Pgap1 | 1.05 | 2.92E-23 | 6.08E-19 | 0.30 | 0.15 | 0.15 | 2.01 |
| 13 | Limk2 | 0.77 | 3.00E-23 | 6.23E-19 | 0.36 | 0.19 | 0.17 | 1.88 |
| 13 | Meg3 | 0.56 | 3.24E-23 | 6.74E-19 | 0.84 | 0.69 | 0.15 | 1.21 |
| 13 | Dlx6os1 | 1.15 | 3.47E-23 | 7.22E-19 | 0.12 | 0.04 | 0.08 | 3.13 |
| 13 | Psap | 0.53 | 4.37E-23 | 9.09E-19 | 0.79 | 0.61 | 0.18 | 1.29 |
| 13 | Igf1r | 0.52 | 4.79E-23 | 9.97E-19 | 0.89 | 0.77 | 0.12 | 1.16 |
| 13 | Imp3 | 0.42 | 5.28E-23 | 1.10E-18 | 0.90 | 0.79 | 0.11 | 1.15 |
| 13 | Mfsd6 | 1.58 | 5.53E-23 | 1.15E-18 | 0.11 | 0.03 | 0.08 | 3.30 |
| 13 | Suclg2 | 0.57 | 5.77E-23 | 1.20E-18 | 0.68 | 0.49 | 0.19 | 1.38 |
| 13 | Adamts9 | 1.28 | 6.32E-23 | 1.31E-18 | 0.29 | 0.15 | 0.15 | 1.99 |
| 13 | Tet3 | 0.59 | 7.25E-23 | 1.51E-18 | 0.74 | 0.57 | 0.18 | 1.31 |
| 13 | Mid1ip1 | 0.91 | 7.59E-23 | 1.58E-18 | 0.38 | 0.22 | 0.17 | 1.77 |
| 13 | Ell2 | 1.04 | 8.91E-23 | 1.85E-18 | 0.21 | 0.09 | 0.12 | 2.37 |
| 13 | Trim25 | 1.34 | 9.00E-23 | 1.87E-18 | 0.16 | 0.06 | 0.10 | 2.67 |
| 13 | Sdsl | 1.39 | 1.14E-22 | 2.37E-18 | 0.16 | 0.06 | 0.10 | 2.67 |
| 13 | Gclm | 0.33 | 1.28E-22 | 2.67E-18 | 0.60 | 0.38 | 0.21 | 1.56 |
| 13 | H19 | 0.45 | 1.51E-22 | 3.15E-18 | 0.89 | 0.84 | 0.04 | 1.05 |
| 13 | mt-Atp6 | 0.26 | 1.59E-22 | 3.31E-18 | 1.00 | 0.99 | 0.00 | 1.00 |
| 13 | Mettl27 | 1.06 | 1.60E-22 | 3.33E-18 | 0.28 | 0.14 | 0.14 | 2.04 |
| 13 | Tmem184a | 1.44 | 1.64E-22 | 3.42E-18 | 0.14 | 0.05 | 0.09 | 2.78 |

**Supplementary Table 1. Marker genes for the original clusters of integrated clustering of E11.5 control and NotchGOF λ-junction cells.**

| | | | | | | | | |
|---|---|---|---|---|---|---|---|---|
| 13 | Atp2c1 | 0.70 | 1.67E-22 | 3.47E-18 | 0.59 | 0.40 | 0.19 | 1.48 |
| 13 | Tpp1 | 0.71 | 1.69E-22 | 3.52E-18 | 0.45 | 0.26 | 0.19 | 1.73 |
| 13 | Slit2 | 0.42 | 2.36E-22 | 4.90E-18 | 0.57 | 0.36 | 0.21 | 1.58 |
| 13 | Syne4 | 0.87 | 3.52E-22 | 7.32E-18 | 0.36 | 0.19 | 0.16 | 1.85 |
| 13 | Clcn3 | 0.70 | 3.72E-22 | 7.73E-18 | 0.57 | 0.38 | 0.20 | 1.52 |
| 13 | Ak4 | 1.18 | 4.32E-22 | 8.99E-18 | 0.23 | 0.10 | 0.13 | 2.24 |
| 13 | Pnpla2 | 1.16 | 4.97E-22 | 1.03E-17 | 0.20 | 0.09 | 0.11 | 2.31 |
| 13 | Jmy | 0.66 | 4.98E-22 | 1.04E-17 | 0.58 | 0.37 | 0.20 | 1.54 |
| 13 | Nek7 | 0.85 | 5.05E-22 | 1.05E-17 | 0.48 | 0.29 | 0.19 | 1.64 |
| 13 | Apc | 0.54 | 6.09E-22 | 1.27E-17 | 0.77 | 0.61 | 0.16 | 1.26 |
| 13 | Nxn | 0.56 | 6.41E-22 | 1.33E-17 | 0.82 | 0.71 | 0.11 | 1.16 |
| 13 | Mettl26 | 0.53 | 7.76E-22 | 1.61E-17 | 0.67 | 0.49 | 0.18 | 1.37 |
| 13 | Map3k5 | 1.58 | 8.46E-22 | 1.76E-17 | 0.11 | 0.03 | 0.07 | 3.24 |
| 13 | Lrrk1 | 0.90 | 8.62E-22 | 1.79E-17 | 0.29 | 0.15 | 0.15 | 1.98 |
| 13 | Fam173a | 0.57 | 9.38E-22 | 1.95E-17 | 0.69 | 0.50 | 0.19 | 1.38 |
| 13 | Klhl13 | 0.74 | 9.39E-22 | 1.95E-17 | 0.45 | 0.26 | 0.18 | 1.69 |
| 13 | Hspa4l | 0.87 | 1.42E-21 | 2.95E-17 | 0.24 | 0.11 | 0.13 | 2.17 |
| 13 | Acsl3 | 0.47 | 1.49E-21 | 3.10E-17 | 0.87 | 0.73 | 0.13 | 1.18 |
| 13 | Lrrfip1 | 0.63 | 1.54E-21 | 3.20E-17 | 0.53 | 0.34 | 0.20 | 1.59 |
| 13 | Ccn2 | 1.56 | 1.54E-21 | 3.21E-17 | 0.11 | 0.04 | 0.08 | 3.14 |
| 13 | Fbxl20 | 0.76 | 1.58E-21 | 3.29E-17 | 0.39 | 0.21 | 0.17 | 1.80 |
| 13 | Pclo | 0.68 | 1.60E-21 | 3.32E-17 | 0.23 | 0.11 | 0.13 | 2.21 |
| 13 | Gm16141 | 1.26 | 1.85E-21 | 3.84E-17 | 0.11 | 0.03 | 0.08 | 3.21 |
| 13 | Kcnmb4 | 0.89 | 2.54E-21 | 5.29E-17 | 0.49 | 0.32 | 0.18 | 1.55 |
| 13 | Ppp1r1a | 0.82 | 3.04E-21 | 6.32E-17 | 0.31 | 0.16 | 0.15 | 1.94 |
| 13 | Aldh5a1 | 0.80 | 3.91E-21 | 8.12E-17 | 0.41 | 0.24 | 0.17 | 1.71 |
| 13 | Igsf9b | 1.19 | 4.37E-21 | 9.08E-17 | 0.15 | 0.06 | 0.09 | 2.67 |
| 13 | Rnf167 | 0.74 | 5.17E-21 | 1.08E-16 | 0.55 | 0.37 | 0.18 | 1.49 |
| 13 | Nfe2l3 | 1.51 | 5.20E-21 | 1.08E-16 | 0.11 | 0.03 | 0.07 | 3.18 |
| 13 | Fam214a | 0.96 | 5.75E-21 | 1.20E-16 | 0.22 | 0.10 | 0.12 | 2.24 |
| 13 | Tshz1 | 0.60 | 6.90E-21 | 1.44E-16 | 0.50 | 0.31 | 0.19 | 1.61 |
| 13 | Wdsub1 | 0.89 | 7.32E-21 | 1.52E-16 | 0.36 | 0.20 | 0.16 | 1.77 |
| 13 | Plec | 0.79 | 7.46E-21 | 1.55E-16 | 0.44 | 0.27 | 0.17 | 1.63 |
| 13 | Nr2f6 | 0.53 | 7.46E-21 | 1.55E-16 | 0.76 | 0.60 | 0.16 | 1.27 |
| 13 | Tnks2 | 0.41 | 9.79E-21 | 2.04E-16 | 0.92 | 0.81 | 0.12 | 1.14 |
| 13 | Cldn23 | 1.24 | 9.86E-21 | 2.05E-16 | 0.11 | 0.04 | 0.08 | 3.03 |
| 13 | Lcor | 0.73 | 1.02E-20 | 2.12E-16 | 0.46 | 0.28 | 0.18 | 1.63 |
| 13 | Bnip3l | 0.31 | 1.26E-20 | 2.63E-16 | 0.98 | 0.93 | 0.05 | 1.05 |
| 13 | Patz1 | 0.52 | 1.34E-20 | 2.79E-16 | 0.70 | 0.52 | 0.18 | 1.35 |
| 13 | Wnk2 | 1.41 | 1.41E-20 | 2.93E-16 | 0.13 | 0.05 | 0.08 | 2.83 |
| 13 | Baz2a | 0.65 | 1.44E-20 | 3.00E-16 | 0.57 | 0.38 | 0.18 | 1.48 |
| 13 | Mov10 | 0.81 | 1.60E-20 | 3.34E-16 | 0.35 | 0.19 | 0.16 | 1.82 |
| 13 | Exosc7 | 0.47 | 1.96E-20 | 4.08E-16 | 0.76 | 0.62 | 0.14 | 1.22 |
| 13 | Npepl1 | 0.65 | 2.14E-20 | 4.44E-16 | 0.51 | 0.33 | 0.18 | 1.56 |
| 13 | Prkag2 | 0.79 | 2.38E-20 | 4.96E-16 | 0.38 | 0.22 | 0.16 | 1.74 |
| 13 | Tspan5 | 0.72 | 2.68E-20 | 5.57E-16 | 0.44 | 0.26 | 0.17 | 1.66 |
| 13 | mt-Nd4 | 0.26 | 2.74E-20 | 5.69E-16 | 0.99 | 0.99 | 0.01 | 1.01 |
| 13 | Eno1b | 0.64 | 3.75E-20 | 7.81E-16 | 0.52 | 0.34 | 0.18 | 1.54 |
| 13 | Rabgap1 | 0.50 | 4.09E-20 | 8.51E-16 | 0.75 | 0.56 | 0.18 | 1.33 |
| 13 | Dnajc1 | 0.55 | 4.43E-20 | 9.21E-16 | 0.71 | 0.56 | 0.16 | 1.28 |
| 13 | Ppp1r36 | 1.29 | 4.46E-20 | 9.27E-16 | 0.14 | 0.05 | 0.09 | 2.65 |
| 13 | Aprt | 0.30 | 4.59E-20 | 9.55E-16 | 0.96 | 0.92 | 0.04 | 1.05 |
| 13 | Ppp1r9a | 0.62 | 5.21E-20 | 1.08E-15 | 0.59 | 0.39 | 0.20 | 1.50 |
| 13 | Zfp36l1 | 0.44 | 5.79E-20 | 1.20E-15 | 0.93 | 0.85 | 0.08 | 1.09 |
| 13 | Phlda3 | 0.59 | 8.28E-20 | 1.72E-15 | 0.58 | 0.41 | 0.17 | 1.42 |
| 13 | Atp13a2 | 0.70 | 8.31E-20 | 1.73E-15 | 0.43 | 0.25 | 0.17 | 1.68 |

**Supplementary Table 1. Marker genes for the original clusters of integrated clustering of E11.5 control and NotchGOF λ-junction cells.**

| | | | | | | | | |
|---|---|---|---|---|---|---|---|---|
| 13 | Zcchc18 | 0.71 | 8.52E-20 | 1.77E-15 | 0.51 | 0.34 | 0.17 | 1.52 |
| 13 | Asl | 0.85 | 9.27E-20 | 1.93E-15 | 0.29 | 0.15 | 0.14 | 1.95 |
| 13 | Nrcam | 1.11 | 1.17E-19 | 2.43E-15 | 0.14 | 0.05 | 0.09 | 2.70 |
| 13 | Foxn3 | 0.58 | 1.21E-19 | 2.51E-15 | 0.81 | 0.68 | 0.13 | 1.19 |
| 13 | Sp4 | 0.81 | 1.48E-19 | 3.07E-15 | 0.39 | 0.23 | 0.16 | 1.69 |
| 13 | Saraf | 0.69 | 1.53E-19 | 3.18E-15 | 0.70 | 0.54 | 0.16 | 1.30 |
| 13 | 2310039H08Rik | 0.61 | 1.59E-19 | 3.31E-15 | 0.60 | 0.42 | 0.18 | 1.44 |
| 13 | Slc35d1 | 0.99 | 1.71E-19 | 3.56E-15 | 0.29 | 0.15 | 0.14 | 1.90 |
| 13 | Zfp7 | 0.92 | 1.98E-19 | 4.12E-15 | 0.28 | 0.14 | 0.13 | 1.95 |
| 13 | Col2a1 | 0.25 | 2.06E-19 | 4.29E-15 | 0.85 | 0.64 | 0.21 | 1.33 |
| 13 | Mllt6 | 0.87 | 2.35E-19 | 4.88E-15 | 0.34 | 0.20 | 0.15 | 1.76 |
| 13 | Prr13 | 0.48 | 2.42E-19 | 5.03E-15 | 0.60 | 0.40 | 0.20 | 1.51 |
| 13 | Sdc4 | 0.52 | 2.44E-19 | 5.08E-15 | 0.53 | 0.34 | 0.19 | 1.56 |
| 13 | Dmd | 1.05 | 2.48E-19 | 5.16E-15 | 0.20 | 0.09 | 0.11 | 2.22 |
| 13 | Kank1 | 0.86 | 2.49E-19 | 5.18E-15 | 0.30 | 0.16 | 0.14 | 1.90 |
| 13 | Plagl2 | 0.66 | 2.67E-19 | 5.54E-15 | 0.54 | 0.36 | 0.18 | 1.52 |
| 13 | Ctso | 1.14 | 3.27E-19 | 6.79E-15 | 0.16 | 0.06 | 0.09 | 2.49 |
| 13 | Tnfrsf19 | 0.60 | 3.54E-19 | 7.36E-15 | 0.29 | 0.15 | 0.14 | 1.95 |
| 13 | Sesn3 | 0.52 | 3.70E-19 | 7.70E-15 | 0.75 | 0.59 | 0.16 | 1.27 |
| 13 | Prag1 | 0.87 | 3.88E-19 | 8.07E-15 | 0.27 | 0.14 | 0.13 | 1.95 |
| 13 | Thtpa | 0.83 | 3.99E-19 | 8.30E-15 | 0.30 | 0.16 | 0.14 | 1.90 |
| 13 | Vmac | 1.04 | 4.00E-19 | 8.33E-15 | 0.20 | 0.09 | 0.11 | 2.21 |
| 13 | Avl9 | 0.71 | 4.25E-19 | 8.84E-15 | 0.52 | 0.34 | 0.17 | 1.50 |
| 13 | Srebf1 | 0.93 | 4.27E-19 | 8.87E-15 | 0.34 | 0.19 | 0.15 | 1.79 |
| 13 | Nr2c1 | 0.80 | 4.48E-19 | 9.31E-15 | 0.38 | 0.22 | 0.16 | 1.71 |
| 13 | Sgsm2 | 1.06 | 4.84E-19 | 1.01E-14 | 0.15 | 0.06 | 0.09 | 2.59 |
| 13 | Raver2 | 0.86 | 5.13E-19 | 1.07E-14 | 0.30 | 0.16 | 0.14 | 1.86 |
| 13 | Lrrn1 | 1.44 | 5.55E-19 | 1.15E-14 | 0.12 | 0.04 | 0.08 | 2.77 |
| 13 | Smco4 | 0.45 | 6.85E-19 | 1.42E-14 | 0.68 | 0.48 | 0.20 | 1.40 |
| 13 | Klf3 | 0.59 | 7.62E-19 | 1.59E-14 | 0.66 | 0.50 | 0.16 | 1.32 |
| 13 | Ddx58 | 1.03 | 8.05E-19 | 1.67E-14 | 0.14 | 0.05 | 0.09 | 2.60 |
| 13 | Fbxo44 | 0.95 | 8.55E-19 | 1.78E-14 | 0.20 | 0.09 | 0.11 | 2.21 |
| 13 | 1810037I17Rik | 0.35 | 1.19E-18 | 2.48E-14 | 0.92 | 0.83 | 0.09 | 1.10 |
| 13 | Dpysl3 | 0.85 | 1.23E-18 | 2.56E-14 | 0.61 | 0.45 | 0.16 | 1.36 |
| 13 | Kdm7a | 0.68 | 1.32E-18 | 2.74E-14 | 0.52 | 0.35 | 0.17 | 1.49 |
| 13 | Slc9a9 | 1.51 | 1.35E-18 | 2.81E-14 | 0.11 | 0.04 | 0.07 | 2.89 |
| 13 | Nmt2 | 0.67 | 1.57E-18 | 3.26E-14 | 0.51 | 0.34 | 0.17 | 1.51 |
| 13 | Eml2 | 0.91 | 1.60E-18 | 3.33E-14 | 0.21 | 0.10 | 0.11 | 2.15 |
| 13 | Ptpn6 | 0.87 | 1.62E-18 | 3.38E-14 | 0.11 | 0.04 | 0.07 | 2.92 |
| 13 | Slc37a3 | 0.61 | 1.73E-18 | 3.60E-14 | 0.44 | 0.26 | 0.18 | 1.68 |
| 13 | Klf9 | 1.34 | 2.04E-18 | 4.24E-14 | 0.13 | 0.05 | 0.08 | 2.69 |
| 13 | Cd82 | 1.07 | 2.08E-18 | 4.32E-14 | 0.16 | 0.06 | 0.09 | 2.45 |
| 13 | Ppa1 | 0.43 | 2.20E-18 | 4.58E-14 | 0.91 | 0.90 | 0.02 | 1.02 |
| 13 | Kdsr | 0.78 | 2.23E-18 | 4.63E-14 | 0.43 | 0.27 | 0.16 | 1.61 |
| 13 | Taok1 | 0.48 | 2.36E-18 | 4.92E-14 | 0.75 | 0.60 | 0.15 | 1.25 |
| 13 | Mettl23 | 0.46 | 2.81E-18 | 5.84E-14 | 0.77 | 0.62 | 0.16 | 1.26 |
| 13 | Gm6169 | 1.41 | 2.81E-18 | 5.85E-14 | 0.10 | 0.04 | 0.07 | 2.97 |
| 13 | Trappc6a | 0.59 | 2.86E-18 | 5.94E-14 | 0.68 | 0.52 | 0.16 | 1.30 |
| 13 | Tspan14 | 0.67 | 3.25E-18 | 6.75E-14 | 0.50 | 0.32 | 0.18 | 1.55 |
| 13 | Glrb | 1.38 | 3.35E-18 | 6.96E-14 | 0.13 | 0.05 | 0.08 | 2.63 |
| 13 | Dnmbp | 0.80 | 3.38E-18 | 7.03E-14 | 0.25 | 0.13 | 0.13 | 2.02 |
| 13 | Cadm4 | 1.00 | 3.45E-18 | 7.17E-14 | 0.19 | 0.08 | 0.10 | 2.24 |
| 13 | Zfp217 | 0.78 | 3.88E-18 | 8.06E-14 | 0.34 | 0.20 | 0.15 | 1.76 |
| 13 | Mcf2l | 1.20 | 4.30E-18 | 8.95E-14 | 0.16 | 0.07 | 0.09 | 2.39 |
| 13 | Foxe1 | 1.01 | 4.42E-18 | 9.19E-14 | 0.12 | 0.04 | 0.08 | 2.73 |
| 13 | Rundc3b | 1.30 | 4.53E-18 | 9.43E-14 | 0.11 | 0.04 | 0.07 | 2.80 |

**Supplementary Table 1. Marker genes for the original clusters of integrated clustering of E11.5 control and NotchGOF λ-junction cells.**

| | | | | | | | | |
|---|---|---|---|---|---|---|---|---|
| 13 | Calcoco1 | 0.75 | 4.54E-18 | 9.44E-14 | 0.36 | 0.21 | 0.15 | 1.70 |
| 13 | Pigp | 0.49 | 4.77E-18 | 9.92E-14 | 0.78 | 0.64 | 0.14 | 1.21 |
| 13 | Ccdc112 | 0.54 | 4.79E-18 | 9.95E-14 | 0.69 | 0.51 | 0.17 | 1.34 |
| 13 | Ephb2 | 0.68 | 4.87E-18 | 1.01E-13 | 0.44 | 0.28 | 0.16 | 1.59 |
| 13 | Golga4 | 0.47 | 5.11E-18 | 1.06E-13 | 0.77 | 0.64 | 0.13 | 1.20 |
| 13 | Ptpn3 | 1.21 | 6.84E-18 | 1.42E-13 | 0.11 | 0.04 | 0.07 | 2.89 |
| 13 | Nexn | 1.40 | 7.85E-18 | 1.63E-13 | 0.12 | 0.04 | 0.07 | 2.74 |
| 13 | Ostm1 | 1.22 | 7.88E-18 | 1.64E-13 | 0.16 | 0.07 | 0.09 | 2.30 |
| 13 | Pafah1b3 | 0.28 | 8.18E-18 | 1.70E-13 | 0.97 | 0.92 | 0.05 | 1.05 |
| 13 | C130071C03Rik | 0.80 | 8.28E-18 | 1.72E-13 | 0.13 | 0.05 | 0.08 | 2.61 |
| 13 | Cdc42ep5 | 0.56 | 8.58E-18 | 1.78E-13 | 0.64 | 0.47 | 0.18 | 1.38 |
| 13 | Btg2 | 1.19 | 8.82E-18 | 1.83E-13 | 0.41 | 0.27 | 0.14 | 1.52 |
| 13 | Zmiz2 | 0.69 | 9.18E-18 | 1.91E-13 | 0.48 | 0.31 | 0.17 | 1.56 |
| 13 | Arf3 | 0.58 | 9.76E-18 | 2.03E-13 | 0.48 | 0.31 | 0.18 | 1.57 |
| 13 | Nudt18 | 1.14 | 1.02E-17 | 2.12E-13 | 0.15 | 0.06 | 0.09 | 2.45 |
| 13 | Znrd1as | 0.90 | 1.03E-17 | 2.14E-13 | 0.24 | 0.13 | 0.12 | 1.95 |
| 13 | Ust | 1.20 | 1.11E-17 | 2.31E-13 | 0.15 | 0.06 | 0.09 | 2.45 |
| 13 | Frs2 | 0.94 | 1.16E-17 | 2.41E-13 | 0.42 | 0.27 | 0.15 | 1.56 |
| 13 | Gm26917 | 0.68 | 1.30E-17 | 2.71E-13 | 0.70 | 0.56 | 0.14 | 1.26 |
| 13 | 0610010F05Rik | 0.65 | 1.31E-17 | 2.73E-13 | 0.41 | 0.24 | 0.16 | 1.66 |
| 13 | Rnf145 | 0.63 | 1.32E-17 | 2.74E-13 | 0.55 | 0.37 | 0.18 | 1.48 |
| 13 | Pnck | 1.03 | 1.48E-17 | 3.08E-13 | 0.16 | 0.07 | 0.09 | 2.33 |
| 13 | Amer1 | 0.66 | 1.81E-17 | 3.76E-13 | 0.67 | 0.51 | 0.16 | 1.31 |
| 13 | Nectin3 | 0.47 | 2.29E-17 | 4.76E-13 | 0.69 | 0.54 | 0.15 | 1.28 |
| 13 | Rps27rt | 0.37 | 2.46E-17 | 5.11E-13 | 0.85 | 0.73 | 0.12 | 1.16 |
| 13 | Slc6a6 | 0.60 | 2.52E-17 | 5.24E-13 | 0.53 | 0.35 | 0.18 | 1.52 |
| 13 | Satb1 | 0.53 | 2.80E-17 | 5.82E-13 | 0.57 | 0.38 | 0.19 | 1.49 |
| 13 | Zfp704 | 0.54 | 3.01E-17 | 6.25E-13 | 0.65 | 0.46 | 0.19 | 1.40 |
| 13 | Exph5 | 1.53 | 3.21E-17 | 6.68E-13 | 0.10 | 0.04 | 0.07 | 2.86 |
| 13 | Mthfd1 | 0.66 | 3.22E-17 | 6.69E-13 | 0.55 | 0.38 | 0.17 | 1.45 |
| 13 | Adgrg1 | 1.09 | 3.30E-17 | 6.87E-13 | 0.11 | 0.04 | 0.07 | 2.73 |
| 13 | Igf2bp3 | 0.33 | 3.33E-17 | 6.92E-13 | 0.91 | 0.82 | 0.09 | 1.11 |
| 13 | Map3k11 | 0.81 | 3.35E-17 | 6.98E-13 | 0.23 | 0.11 | 0.12 | 2.01 |
| 13 | Mmgt2 | 0.49 | 3.54E-17 | 7.37E-13 | 0.62 | 0.43 | 0.19 | 1.44 |
| 13 | Kcnmb4os2 | 0.86 | 3.56E-17 | 7.41E-13 | 0.29 | 0.16 | 0.13 | 1.84 |
| 13 | Tex15 | 0.66 | 3.63E-17 | 7.55E-13 | 0.22 | 0.11 | 0.11 | 2.05 |
| 13 | Jund | 0.29 | 3.80E-17 | 7.91E-13 | 0.97 | 0.94 | 0.03 | 1.03 |
| 13 | Thumpd3 | 0.56 | 3.91E-17 | 8.13E-13 | 0.55 | 0.38 | 0.17 | 1.46 |
| 13 | Ccdc28a | 1.02 | 3.91E-17 | 8.14E-13 | 0.21 | 0.10 | 0.11 | 2.05 |
| 13 | Hook2 | 0.73 | 4.34E-17 | 9.03E-13 | 0.30 | 0.16 | 0.13 | 1.83 |
| 13 | Mtch1 | 0.25 | 4.55E-17 | 9.47E-13 | 0.98 | 0.96 | 0.03 | 1.03 |
| 13 | Vsig10 | 1.28 | 4.81E-17 | 1.00E-12 | 0.13 | 0.05 | 0.08 | 2.54 |
| 13 | Firre | 1.12 | 4.93E-17 | 1.03E-12 | 0.19 | 0.09 | 0.10 | 2.14 |
| 13 | Bmf | 0.91 | 6.31E-17 | 1.31E-12 | 0.22 | 0.11 | 0.11 | 2.06 |
| 13 | Dpp3 | 0.48 | 6.98E-17 | 1.45E-12 | 0.73 | 0.57 | 0.16 | 1.27 |
| 13 | Epha5 | 0.59 | 7.17E-17 | 1.49E-12 | 0.16 | 0.07 | 0.09 | 2.35 |
| 13 | Abr | 0.76 | 7.52E-17 | 1.56E-12 | 0.36 | 0.22 | 0.15 | 1.69 |
| 13 | Zbtb18 | 0.86 | 7.95E-17 | 1.65E-12 | 0.22 | 0.11 | 0.11 | 2.03 |
| 13 | Ccng1 | 0.77 | 9.12E-17 | 1.90E-12 | 0.34 | 0.20 | 0.14 | 1.71 |
| 13 | Sft2d3 | 0.76 | 9.60E-17 | 2.00E-12 | 0.34 | 0.20 | 0.14 | 1.72 |
| 13 | Ndufa6 | 0.27 | 1.00E-16 | 2.09E-12 | 0.96 | 0.93 | 0.03 | 1.03 |
| 13 | Snx5 | 0.34 | 1.06E-16 | 2.21E-12 | 0.88 | 0.77 | 0.10 | 1.13 |
| 13 | Rapgef5 | 0.78 | 1.09E-16 | 2.27E-12 | 0.23 | 0.12 | 0.12 | 2.00 |
| 13 | Cndp2 | 0.55 | 1.10E-16 | 2.29E-12 | 0.53 | 0.37 | 0.17 | 1.45 |
| 13 | Abhd17c | 0.49 | 1.14E-16 | 2.37E-12 | 0.58 | 0.40 | 0.18 | 1.45 |
| 13 | Ypel1 | 0.50 | 1.16E-16 | 2.42E-12 | 0.66 | 0.50 | 0.17 | 1.34 |

**Supplementary Table 1. Marker genes for the original clusters of integrated clustering of E11.5 control and NotchGOF λ-junction cells.**

| | | | | | | | | |
|---|---|---|---|---|---|---|---|---|
| 13 | Fam32a | 0.34 | 1.37E-16 | 2.86E-12 | 0.86 | 0.77 | 0.10 | 1.13 |
| 13 | Hras | 0.29 | 1.39E-16 | 2.89E-12 | 0.88 | 0.80 | 0.08 | 1.10 |
| 13 | Ppip5k2 | 0.49 | 1.60E-16 | 3.32E-12 | 0.62 | 0.45 | 0.17 | 1.37 |
| 13 | Pik3cb | 1.04 | 1.84E-16 | 3.83E-12 | 0.15 | 0.06 | 0.08 | 2.38 |
| 13 | Tmem141 | 0.69 | 2.00E-16 | 4.17E-12 | 0.33 | 0.19 | 0.14 | 1.73 |
| 13 | Ctf1 | 0.77 | 2.11E-16 | 4.39E-12 | 0.29 | 0.16 | 0.13 | 1.81 |
| 13 | Khk | 1.10 | 2.20E-16 | 4.58E-12 | 0.15 | 0.06 | 0.09 | 2.34 |
| 13 | Tgif2 | 0.61 | 2.44E-16 | 5.07E-12 | 0.61 | 0.45 | 0.15 | 1.34 |
| 13 | Sort1 | 0.75 | 2.83E-16 | 5.88E-12 | 0.34 | 0.20 | 0.14 | 1.70 |
| 13 | Flrt3 | 0.52 | 3.34E-16 | 6.95E-12 | 0.61 | 0.44 | 0.17 | 1.38 |
| 13 | D030056L22Rik | 0.50 | 3.57E-16 | 7.42E-12 | 0.62 | 0.48 | 0.15 | 1.31 |
| 13 | Fam98c | 0.82 | 3.59E-16 | 7.46E-12 | 0.23 | 0.11 | 0.11 | 1.98 |
| 13 | Capg | 0.60 | 3.90E-16 | 8.11E-12 | 0.28 | 0.15 | 0.13 | 1.85 |
| 13 | Sertad2 | 0.51 | 3.92E-16 | 8.15E-12 | 0.58 | 0.40 | 0.18 | 1.44 |
| 13 | Stim2 | 0.58 | 4.34E-16 | 9.02E-12 | 0.39 | 0.23 | 0.16 | 1.67 |
| 13 | Plekhg1 | 0.64 | 4.45E-16 | 9.25E-12 | 0.26 | 0.14 | 0.12 | 1.90 |
| 13 | Axl | 0.84 | 4.61E-16 | 9.59E-12 | 0.27 | 0.15 | 0.12 | 1.81 |
| 13 | Mpp7 | 0.51 | 4.72E-16 | 9.81E-12 | 0.54 | 0.37 | 0.17 | 1.47 |
| 13 | Kif5c | 0.51 | 5.80E-16 | 1.21E-11 | 0.63 | 0.48 | 0.15 | 1.31 |
| 13 | Amt | 0.75 | 5.95E-16 | 1.24E-11 | 0.27 | 0.15 | 0.12 | 1.86 |
| 13 | Dlg1 | 0.63 | 6.17E-16 | 1.28E-11 | 0.53 | 0.37 | 0.16 | 1.43 |
| 13 | Cyb5r4 | 0.54 | 6.51E-16 | 1.35E-11 | 0.56 | 0.39 | 0.16 | 1.41 |
| 13 | Atp1b1 | 1.24 | 6.91E-16 | 1.44E-11 | 0.18 | 0.09 | 0.09 | 2.09 |
| 13 | Tjp1 | 0.48 | 7.44E-16 | 1.55E-11 | 0.66 | 0.49 | 0.18 | 1.36 |
| 13 | Pnrc1 | 0.34 | 7.53E-16 | 1.57E-11 | 0.93 | 0.87 | 0.06 | 1.07 |
| 13 | Psme1 | 0.36 | 7.65E-16 | 1.59E-11 | 0.85 | 0.76 | 0.09 | 1.11 |
| 13 | Letmd1 | 0.71 | 7.86E-16 | 1.64E-11 | 0.36 | 0.22 | 0.14 | 1.65 |
| 13 | Bclaf3 | 0.59 | 8.15E-16 | 1.70E-11 | 0.50 | 0.33 | 0.17 | 1.51 |
| 13 | Irf2 | 0.74 | 9.38E-16 | 1.95E-11 | 0.40 | 0.25 | 0.15 | 1.60 |
| 13 | Aplp2 | 0.33 | 1.10E-15 | 2.28E-11 | 0.89 | 0.82 | 0.07 | 1.09 |
| 13 | Herc3 | 1.14 | 1.21E-15 | 2.53E-11 | 0.16 | 0.07 | 0.09 | 2.22 |
| 13 | Cpe | 0.68 | 1.31E-15 | 2.72E-11 | 0.58 | 0.41 | 0.16 | 1.39 |
| 13 | Usp53 | 1.23 | 1.59E-15 | 3.30E-11 | 0.16 | 0.08 | 0.09 | 2.17 |
| 13 | Glcci1 | 0.50 | 1.64E-15 | 3.41E-11 | 0.56 | 0.39 | 0.17 | 1.45 |
| 13 | Rffl | 1.16 | 1.66E-15 | 3.46E-11 | 0.16 | 0.07 | 0.09 | 2.21 |
| 13 | 2410022M11Rik | 0.82 | 1.67E-15 | 3.47E-11 | 0.22 | 0.11 | 0.11 | 1.99 |
| 13 | Rsrp1 | 0.27 | 1.67E-15 | 3.47E-11 | 0.93 | 0.89 | 0.04 | 1.05 |
| 13 | Meis3 | 0.65 | 1.68E-15 | 3.49E-11 | 0.34 | 0.20 | 0.14 | 1.73 |
| 13 | Ctsz | 0.37 | 1.72E-15 | 3.58E-11 | 0.74 | 0.57 | 0.16 | 1.28 |
| 13 | Nuak1 | 0.79 | 1.74E-15 | 3.61E-11 | 0.34 | 0.20 | 0.14 | 1.67 |
| 13 | Mfsd3 | 1.32 | 1.88E-15 | 3.90E-11 | 0.11 | 0.04 | 0.07 | 2.66 |
| 13 | Larp4b | 0.46 | 1.88E-15 | 3.91E-11 | 0.73 | 0.61 | 0.12 | 1.20 |
| 13 | Usp46 | 0.42 | 2.23E-15 | 4.64E-11 | 0.78 | 0.62 | 0.16 | 1.25 |
| 13 | Cib1 | 0.53 | 2.39E-15 | 4.97E-11 | 0.56 | 0.40 | 0.16 | 1.41 |
| 13 | Gstm1 | 0.47 | 2.44E-15 | 5.07E-11 | 0.39 | 0.24 | 0.15 | 1.60 |
| 13 | Sapcd1 | 1.27 | 2.48E-15 | 5.15E-11 | 0.10 | 0.04 | 0.06 | 2.70 |
| 13 | Plekha1 | 0.67 | 2.54E-15 | 5.28E-11 | 0.33 | 0.20 | 0.14 | 1.68 |
| 13 | Tubb2a | 0.40 | 2.65E-15 | 5.50E-11 | 0.46 | 0.29 | 0.16 | 1.56 |
| 13 | Slc44a2 | 0.58 | 3.28E-15 | 6.82E-11 | 0.43 | 0.27 | 0.15 | 1.56 |
| 13 | Cryzl2 | 0.83 | 3.67E-15 | 7.63E-11 | 0.20 | 0.10 | 0.10 | 2.01 |
| 13 | Mcee | 0.60 | 3.70E-15 | 7.70E-11 | 0.59 | 0.44 | 0.15 | 1.35 |
| 13 | Lrba | 0.61 | 3.80E-15 | 7.90E-11 | 0.37 | 0.22 | 0.14 | 1.65 |
| 13 | Tmem245 | 0.57 | 4.24E-15 | 8.81E-11 | 0.56 | 0.40 | 0.16 | 1.40 |
| 13 | Vwa5a | 0.90 | 4.26E-15 | 8.86E-11 | 0.18 | 0.09 | 0.09 | 2.09 |
| 13 | Ppm1b | 0.53 | 4.29E-15 | 8.91E-11 | 0.68 | 0.54 | 0.14 | 1.25 |
| 13 | Arap2 | 1.08 | 4.59E-15 | 9.55E-11 | 0.13 | 0.05 | 0.08 | 2.43 |

**Supplementary Table 1. Marker genes for the original clusters of integrated clustering of E11.5 control and NotchGOF λ-junction cells.**

| | | | | | | | | |
|---|---|---|---|---|---|---|---|---|
| 13 | Arhgef12 | 0.44 | 4.61E-15 | 9.59E-11 | 0.73 | 0.58 | 0.16 | 1.27 |
| 13 | Mtss1 | 0.54 | 4.78E-15 | 9.94E-11 | 0.45 | 0.29 | 0.16 | 1.54 |
| 13 | Zkscan7 | 0.93 | 5.37E-15 | 1.12E-10 | 0.17 | 0.08 | 0.09 | 2.17 |
| 13 | Pogk | 0.52 | 5.81E-15 | 1.21E-10 | 0.66 | 0.50 | 0.17 | 1.33 |
| 13 | Rreb1 | 0.69 | 5.97E-15 | 1.24E-10 | 0.34 | 0.21 | 0.14 | 1.67 |
| 13 | Klhl8 | 1.15 | 6.58E-15 | 1.37E-10 | 0.19 | 0.09 | 0.09 | 2.03 |
| 13 | Kif26a | 0.81 | 6.65E-15 | 1.38E-10 | 0.22 | 0.11 | 0.11 | 1.94 |
| 13 | Micu2 | 0.45 | 6.88E-15 | 1.43E-10 | 0.67 | 0.51 | 0.16 | 1.32 |
| 13 | Fmc1 | 0.45 | 6.89E-15 | 1.43E-10 | 0.73 | 0.59 | 0.14 | 1.23 |
| 13 | Pik3ip1 | 0.79 | 7.07E-15 | 1.47E-10 | 0.18 | 0.09 | 0.09 | 2.09 |
| 13 | Nomo1 | 0.66 | 7.21E-15 | 1.50E-10 | 0.33 | 0.20 | 0.14 | 1.69 |
| 13 | Ldlrap1 | 0.77 | 7.72E-15 | 1.61E-10 | 0.31 | 0.18 | 0.13 | 1.71 |
| 13 | Sgpl1 | 0.57 | 8.00E-15 | 1.66E-10 | 0.49 | 0.34 | 0.16 | 1.47 |
| 13 | Psmg4 | 0.37 | 1.01E-14 | 2.11E-10 | 0.83 | 0.74 | 0.09 | 1.12 |
| 13 | Copg2 | 0.57 | 1.05E-14 | 2.18E-10 | 0.53 | 0.37 | 0.16 | 1.43 |
| 13 | Miga1 | 0.53 | 1.16E-14 | 2.42E-10 | 0.46 | 0.30 | 0.16 | 1.54 |
| 13 | Jtb | 0.43 | 1.17E-14 | 2.44E-10 | 0.68 | 0.53 | 0.15 | 1.28 |
| 13 | Tulp4 | 0.47 | 1.18E-14 | 2.46E-10 | 0.79 | 0.67 | 0.12 | 1.18 |
| 13 | Retreg2 | 0.70 | 1.19E-14 | 2.47E-10 | 0.43 | 0.29 | 0.14 | 1.50 |
| 13 | Cdv3 | 0.31 | 1.21E-14 | 2.51E-10 | 0.97 | 0.95 | 0.02 | 1.02 |
| 13 | Pisd | 0.71 | 1.36E-14 | 2.84E-10 | 0.34 | 0.21 | 0.13 | 1.64 |
| 13 | Srsf5 | 0.25 | 1.39E-14 | 2.88E-10 | 0.99 | 0.94 | 0.04 | 1.05 |
| 13 | Lgr5 | 0.32 | 1.45E-14 | 3.02E-10 | 0.34 | 0.21 | 0.13 | 1.65 |
| 13 | Trpc1 | 0.89 | 1.51E-14 | 3.14E-10 | 0.21 | 0.11 | 0.10 | 1.93 |
| 13 | Slc25a27 | 0.56 | 1.64E-14 | 3.41E-10 | 0.56 | 0.40 | 0.17 | 1.42 |
| 13 | Ubxn2a | 0.53 | 1.66E-14 | 3.46E-10 | 0.47 | 0.31 | 0.16 | 1.50 |
| 13 | Sun1 | 0.56 | 1.66E-14 | 3.46E-10 | 0.42 | 0.27 | 0.15 | 1.55 |
| 13 | Bmyc | 0.39 | 1.74E-14 | 3.63E-10 | 0.78 | 0.62 | 0.16 | 1.26 |
| 13 | Gse1 | 0.56 | 1.93E-14 | 4.01E-10 | 0.46 | 0.30 | 0.15 | 1.51 |
| 13 | Kxd1 | 0.44 | 2.02E-14 | 4.21E-10 | 0.66 | 0.50 | 0.17 | 1.33 |
| 13 | Maoa | 0.63 | 2.14E-14 | 4.45E-10 | 0.41 | 0.27 | 0.14 | 1.53 |
| 13 | Man2a1 | 0.76 | 2.30E-14 | 4.77E-10 | 0.36 | 0.23 | 0.13 | 1.58 |
| 13 | Cdk6 | 0.48 | 2.33E-14 | 4.85E-10 | 0.50 | 0.34 | 0.16 | 1.47 |
| 13 | Zfp809 | 1.18 | 2.41E-14 | 5.02E-10 | 0.16 | 0.07 | 0.08 | 2.15 |
| 13 | Efcab14 | 0.58 | 2.55E-14 | 5.31E-10 | 0.57 | 0.42 | 0.15 | 1.37 |
| 13 | Slc4a7 | 0.49 | 2.64E-14 | 5.49E-10 | 0.68 | 0.52 | 0.17 | 1.32 |
| 13 | Hk2 | 0.51 | 2.91E-14 | 6.05E-10 | 0.40 | 0.25 | 0.15 | 1.59 |
| 13 | Gipc2 | 0.94 | 3.02E-14 | 6.29E-10 | 0.14 | 0.06 | 0.08 | 2.26 |
| 13 | Slc38a1 | 0.30 | 3.06E-14 | 6.36E-10 | 0.88 | 0.77 | 0.12 | 1.15 |
| 13 | Agl | 0.65 | 3.06E-14 | 6.36E-10 | 0.39 | 0.25 | 0.14 | 1.57 |
| 13 | Isyna1 | 0.41 | 3.46E-14 | 7.19E-10 | 0.77 | 0.68 | 0.09 | 1.14 |
| 13 | Cask | 0.57 | 3.47E-14 | 7.22E-10 | 0.49 | 0.34 | 0.16 | 1.47 |
| 13 | Gm4890 | 1.20 | 3.50E-14 | 7.27E-10 | 0.15 | 0.07 | 0.08 | 2.18 |
| 13 | Fmn2 | 1.03 | 3.53E-14 | 7.34E-10 | 0.15 | 0.07 | 0.08 | 2.17 |
| 13 | Alcam | 1.01 | 3.54E-14 | 7.37E-10 | 0.25 | 0.14 | 0.11 | 1.76 |
| 13 | Tmub1 | 0.65 | 3.70E-14 | 7.70E-10 | 0.33 | 0.20 | 0.13 | 1.67 |
| 13 | Zfp318 | 0.39 | 3.70E-14 | 7.70E-10 | 0.64 | 0.47 | 0.17 | 1.36 |
| 13 | Lmo1 | 0.36 | 3.77E-14 | 7.83E-10 | 0.16 | 0.07 | 0.09 | 2.16 |
| 13 | Irf2bpl | 0.52 | 4.45E-14 | 9.25E-10 | 0.53 | 0.38 | 0.15 | 1.38 |
| 13 | Marchf7 | 0.37 | 4.64E-14 | 9.66E-10 | 0.82 | 0.73 | 0.10 | 1.13 |
| 13 | Pradc1 | 0.72 | 4.87E-14 | 1.01E-09 | 0.35 | 0.22 | 0.13 | 1.62 |
| 13 | Rab27a | 1.40 | 4.90E-14 | 1.02E-09 | 0.11 | 0.05 | 0.07 | 2.43 |
| 13 | Pex16 | 0.70 | 5.03E-14 | 1.05E-09 | 0.31 | 0.18 | 0.13 | 1.69 |
| 13 | Scnm1 | 0.53 | 5.06E-14 | 1.05E-09 | 0.58 | 0.41 | 0.16 | 1.39 |
| 13 | Lrrc8d | 0.67 | 5.21E-14 | 1.08E-09 | 0.32 | 0.19 | 0.13 | 1.66 |
| 13 | Mlec | 0.33 | 5.44E-14 | 1.13E-09 | 0.85 | 0.75 | 0.10 | 1.14 |

**Supplementary Table 1. Marker genes for the original clusters of integrated clustering of E11.5 control and NotchGOF λ-junction cells.**

| | | | | | | | | |
|---|---|---|---|---|---|---|---|---|
| 13 | Gstz1 | 0.66 | 5.65E-14 | 1.18E-09 | 0.41 | 0.27 | 0.14 | 1.53 |
| 13 | Rhbdf1 | 0.84 | 5.86E-14 | 1.22E-09 | 0.19 | 0.10 | 0.10 | 1.96 |
| 13 | Ogfod3 | 0.75 | 6.02E-14 | 1.25E-09 | 0.25 | 0.14 | 0.11 | 1.80 |
| 13 | Scai | 0.49 | 6.06E-14 | 1.26E-09 | 0.51 | 0.35 | 0.16 | 1.45 |
| 13 | Erf | 0.53 | 6.33E-14 | 1.32E-09 | 0.46 | 0.30 | 0.16 | 1.53 |
| 13 | Pak1 | 0.42 | 6.40E-14 | 1.33E-09 | 0.66 | 0.49 | 0.17 | 1.35 |
| 13 | Arfip2 | 0.49 | 6.53E-14 | 1.36E-09 | 0.58 | 0.42 | 0.16 | 1.39 |
| 13 | Reep3 | 0.47 | 6.63E-14 | 1.38E-09 | 0.67 | 0.52 | 0.16 | 1.30 |
| 13 | Tmem191c | 1.31 | 6.77E-14 | 1.41E-09 | 0.10 | 0.04 | 0.06 | 2.55 |
| 13 | Mrps36 | 0.34 | 7.14E-14 | 1.48E-09 | 0.86 | 0.77 | 0.08 | 1.10 |
| 13 | Ankrd49 | 0.46 | 7.38E-14 | 1.54E-09 | 0.59 | 0.43 | 0.16 | 1.37 |
| 13 | Atp11a | 0.64 | 7.51E-14 | 1.56E-09 | 0.29 | 0.17 | 0.12 | 1.74 |
| 13 | Rnf180 | 1.14 | 8.15E-14 | 1.69E-09 | 0.12 | 0.05 | 0.07 | 2.35 |
| 13 | Msrb3 | 0.87 | 8.63E-14 | 1.80E-09 | 0.21 | 0.11 | 0.10 | 1.88 |
| 13 | Hexb | 0.67 | 9.33E-14 | 1.94E-09 | 0.35 | 0.22 | 0.13 | 1.57 |
| 13 | Trim3 | 1.02 | 9.40E-14 | 1.95E-09 | 0.18 | 0.09 | 0.09 | 1.99 |
| 13 | Inava | 0.66 | 9.84E-14 | 2.05E-09 | 0.36 | 0.23 | 0.13 | 1.58 |
| 13 | Cpd | 0.67 | 1.03E-13 | 2.14E-09 | 0.49 | 0.34 | 0.15 | 1.46 |
| 13 | Dap | 0.40 | 1.16E-13 | 2.41E-09 | 0.72 | 0.56 | 0.16 | 1.28 |
| 13 | Ifitm3 | 0.99 | 1.20E-13 | 2.50E-09 | 0.41 | 0.30 | 0.11 | 1.39 |
| 13 | Bid | 0.78 | 1.27E-13 | 2.64E-09 | 0.22 | 0.12 | 0.10 | 1.87 |
| 13 | Camkmt | 0.70 | 1.31E-13 | 2.72E-09 | 0.30 | 0.18 | 0.12 | 1.67 |
| 13 | Ybx2 | 1.04 | 1.32E-13 | 2.74E-09 | 0.15 | 0.07 | 0.08 | 2.13 |
| 13 | Pcyox1l | 0.67 | 1.35E-13 | 2.80E-09 | 0.33 | 0.20 | 0.13 | 1.65 |
| 13 | Kif16b | 0.82 | 1.36E-13 | 2.83E-09 | 0.26 | 0.15 | 0.11 | 1.75 |
| 13 | Rida | 0.49 | 1.58E-13 | 3.29E-09 | 0.57 | 0.42 | 0.15 | 1.36 |
| 13 | Polr2h | 0.32 | 1.72E-13 | 3.58E-09 | 0.88 | 0.79 | 0.09 | 1.12 |
| 13 | Phlpp1 | 0.67 | 1.80E-13 | 3.74E-09 | 0.24 | 0.13 | 0.11 | 1.82 |
| 13 | Gnpda2 | 0.59 | 1.90E-13 | 3.95E-09 | 0.42 | 0.28 | 0.14 | 1.51 |
| 13 | Llph | 0.27 | 1.93E-13 | 4.02E-09 | 0.91 | 0.86 | 0.05 | 1.06 |
| 13 | Rnf5 | 0.39 | 2.00E-13 | 4.16E-09 | 0.77 | 0.65 | 0.13 | 1.20 |
| 13 | Sema4d | 0.95 | 2.21E-13 | 4.59E-09 | 0.15 | 0.07 | 0.08 | 2.14 |
| 13 | Dcaf6 | 0.89 | 2.24E-13 | 4.66E-09 | 0.17 | 0.08 | 0.09 | 2.05 |
| 13 | Zdhhc13 | 0.94 | 2.30E-13 | 4.79E-09 | 0.24 | 0.14 | 0.11 | 1.77 |
| 13 | Exosc5 | 0.39 | 2.43E-13 | 5.05E-09 | 0.73 | 0.58 | 0.15 | 1.25 |
| 13 | Immp2l | 0.91 | 2.45E-13 | 5.11E-09 | 0.17 | 0.08 | 0.09 | 2.05 |
| 13 | Dapk1 | 0.55 | 2.98E-13 | 6.20E-09 | 0.42 | 0.27 | 0.14 | 1.53 |
| 13 | Grb14 | 1.09 | 3.14E-13 | 6.53E-09 | 0.11 | 0.05 | 0.06 | 2.39 |
| 13 | Tmem175 | 0.88 | 3.31E-13 | 6.88E-09 | 0.19 | 0.10 | 0.09 | 1.93 |
| 13 | Zfp395 | 0.60 | 3.44E-13 | 7.16E-09 | 0.37 | 0.23 | 0.13 | 1.57 |
| 13 | Mapk8ip1 | 0.52 | 4.01E-13 | 8.34E-09 | 0.40 | 0.26 | 0.15 | 1.57 |
| 13 | Grik5 | 0.71 | 4.08E-13 | 8.49E-09 | 0.27 | 0.16 | 0.12 | 1.74 |
| 13 | Wdr31 | 0.87 | 4.23E-13 | 8.81E-09 | 0.18 | 0.09 | 0.09 | 2.00 |
| 13 | Cep70 | 0.55 | 4.30E-13 | 8.93E-09 | 0.53 | 0.40 | 0.14 | 1.34 |
| 13 | Ybx3 | 0.30 | 4.30E-13 | 8.95E-09 | 0.91 | 0.84 | 0.07 | 1.08 |
| 13 | Fhl1 | 0.68 | 4.34E-13 | 9.02E-09 | 0.25 | 0.14 | 0.11 | 1.77 |
| 13 | Nell2 | 1.03 | 4.72E-13 | 9.81E-09 | 0.12 | 0.05 | 0.07 | 2.35 |
| 13 | Fkbp8 | 0.27 | 4.76E-13 | 9.89E-09 | 0.89 | 0.82 | 0.07 | 1.08 |
| 13 | Snapin | 0.53 | 4.85E-13 | 1.01E-08 | 0.47 | 0.32 | 0.15 | 1.46 |
| 13 | Sh3bp4 | 0.73 | 5.00E-13 | 1.04E-08 | 0.28 | 0.17 | 0.12 | 1.70 |
| 13 | Ebf1 | 0.26 | 5.03E-13 | 1.05E-08 | 0.44 | 0.29 | 0.14 | 1.49 |
| 13 | Ormdl1 | 0.50 | 5.33E-13 | 1.11E-08 | 0.60 | 0.44 | 0.16 | 1.35 |
| 13 | Atg4d | 0.66 | 5.46E-13 | 1.14E-08 | 0.33 | 0.20 | 0.13 | 1.62 |
| 13 | Yipf2 | 0.72 | 5.86E-13 | 1.22E-08 | 0.22 | 0.12 | 0.10 | 1.84 |
| 13 | Sypl | 0.42 | 6.25E-13 | 1.30E-08 | 0.63 | 0.48 | 0.16 | 1.32 |
| 13 | M6pr | 0.47 | 6.29E-13 | 1.31E-08 | 0.60 | 0.44 | 0.16 | 1.36 |

**Supplementary Table 1. Marker genes for the original clusters of integrated clustering of E11.5 control and NotchGOF λ-junction cells.**

| | | | | | | | | |
|---|---|---|---|---|---|---|---|---|
| 13 | Ccdc181 | 0.95 | 6.38E-13 | 1.33E-08 | 0.14 | 0.06 | 0.07 | 2.16 |
| 13 | Map3k20 | 0.85 | 6.39E-13 | 1.33E-08 | 0.21 | 0.12 | 0.10 | 1.82 |
| 13 | Tmc4 | 0.98 | 6.80E-13 | 1.41E-08 | 0.19 | 0.10 | 0.09 | 1.93 |
| 13 | Fam185a | 1.24 | 7.08E-13 | 1.47E-08 | 0.13 | 0.06 | 0.07 | 2.23 |
| 13 | Abcd3 | 0.52 | 7.13E-13 | 1.48E-08 | 0.57 | 0.42 | 0.15 | 1.37 |
| 13 | S100a10 | 0.39 | 8.05E-13 | 1.67E-08 | 0.69 | 0.55 | 0.14 | 1.25 |
| 13 | Disp1 | 0.87 | 8.26E-13 | 1.72E-08 | 0.20 | 0.11 | 0.09 | 1.88 |
| 13 | Ccndbp1 | 0.62 | 8.57E-13 | 1.78E-08 | 0.41 | 0.28 | 0.13 | 1.47 |
| 13 | Lonrf1 | 0.94 | 9.51E-13 | 1.98E-08 | 0.18 | 0.09 | 0.09 | 1.94 |
| 13 | Commd6 | 0.39 | 1.01E-12 | 2.11E-08 | 0.77 | 0.65 | 0.12 | 1.18 |
| 13 | Ift43 | 0.37 | 1.06E-12 | 2.21E-08 | 0.70 | 0.54 | 0.16 | 1.29 |
| 13 | Ptprd | 0.45 | 1.08E-12 | 2.25E-08 | 0.60 | 0.44 | 0.16 | 1.36 |
| 13 | Sec22a | 0.55 | 1.09E-12 | 2.26E-08 | 0.43 | 0.29 | 0.14 | 1.49 |
| 13 | Pak4 | 0.47 | 1.09E-12 | 2.28E-08 | 0.54 | 0.38 | 0.15 | 1.40 |
| 13 | Six4 | 0.44 | 1.12E-12 | 2.32E-08 | 0.62 | 0.46 | 0.17 | 1.36 |
| 13 | Cuedc2 | 0.32 | 1.17E-12 | 2.44E-08 | 0.87 | 0.77 | 0.10 | 1.13 |
| 13 | Cpne5 | 1.24 | 1.19E-12 | 2.47E-08 | 0.12 | 0.05 | 0.07 | 2.29 |
| 13 | Slc12a4 | 0.97 | 1.21E-12 | 2.52E-08 | 0.24 | 0.14 | 0.10 | 1.73 |
| 13 | Ly75 | 0.69 | 1.22E-12 | 2.54E-08 | 0.26 | 0.15 | 0.11 | 1.72 |
| 13 | Bag4 | 0.52 | 1.24E-12 | 2.58E-08 | 0.54 | 0.39 | 0.15 | 1.38 |
| 13 | Vps37d | 0.57 | 1.25E-12 | 2.60E-08 | 0.43 | 0.29 | 0.14 | 1.49 |
| 13 | Dyrk3 | 0.79 | 1.29E-12 | 2.68E-08 | 0.19 | 0.10 | 0.09 | 1.92 |
| 13 | Msl3 | 0.46 | 1.35E-12 | 2.81E-08 | 0.57 | 0.42 | 0.15 | 1.35 |
| 13 | Rtkn2 | 0.75 | 1.37E-12 | 2.84E-08 | 0.16 | 0.08 | 0.08 | 2.09 |
| 13 | Fam192a | 0.32 | 1.39E-12 | 2.88E-08 | 0.79 | 0.65 | 0.14 | 1.22 |
| 13 | Ears2 | 0.65 | 1.39E-12 | 2.89E-08 | 0.27 | 0.16 | 0.11 | 1.74 |
| 13 | Jmjd1c | 0.44 | 1.40E-12 | 2.91E-08 | 0.64 | 0.48 | 0.16 | 1.33 |
| 13 | Eif2s3y | 1.04 | 1.48E-12 | 3.07E-08 | 0.16 | 0.08 | 0.08 | 2.00 |
| 13 | Aagab | 0.67 | 1.57E-12 | 3.26E-08 | 0.29 | 0.17 | 0.11 | 1.66 |
| 13 | Klf10 | 0.77 | 1.64E-12 | 3.41E-08 | 0.20 | 0.11 | 0.09 | 1.86 |
| 13 | Cul1 | 0.32 | 1.73E-12 | 3.59E-08 | 0.90 | 0.87 | 0.04 | 1.04 |
| 13 | Plekhh1 | 0.81 | 1.89E-12 | 3.94E-08 | 0.14 | 0.06 | 0.07 | 2.16 |
| 13 | C1qtnf4 | 0.58 | 1.93E-12 | 4.01E-08 | 0.43 | 0.28 | 0.14 | 1.51 |
| 13 | Prepl | 0.66 | 1.94E-12 | 4.03E-08 | 0.22 | 0.12 | 0.10 | 1.82 |
| 13 | Rps6ka1 | 0.58 | 1.96E-12 | 4.08E-08 | 0.37 | 0.24 | 0.13 | 1.55 |
| 13 | Acly | 0.40 | 1.97E-12 | 4.10E-08 | 0.70 | 0.58 | 0.13 | 1.22 |
| 13 | Nhs | 0.46 | 1.99E-12 | 4.13E-08 | 0.48 | 0.34 | 0.15 | 1.44 |
| 13 | Cdk2ap2 | 0.43 | 2.02E-12 | 4.20E-08 | 0.61 | 0.46 | 0.15 | 1.34 |
| 13 | Sbk1 | 0.38 | 2.03E-12 | 4.22E-08 | 0.67 | 0.52 | 0.15 | 1.29 |
| 13 | Cybrd1 | 1.09 | 2.03E-12 | 4.23E-08 | 0.11 | 0.05 | 0.07 | 2.33 |
| 13 | Tradd | 0.91 | 2.10E-12 | 4.38E-08 | 0.21 | 0.11 | 0.09 | 1.81 |
| 13 | Ckb | 0.52 | 2.20E-12 | 4.57E-08 | 0.46 | 0.33 | 0.13 | 1.40 |
| 13 | Pigt | 0.45 | 2.48E-12 | 5.16E-08 | 0.58 | 0.43 | 0.15 | 1.35 |
| 13 | Marchf8 | 0.69 | 2.49E-12 | 5.18E-08 | 0.28 | 0.17 | 0.11 | 1.65 |
| 13 | Lrrc42 | 0.38 | 2.70E-12 | 5.62E-08 | 0.77 | 0.60 | 0.17 | 1.28 |
| 13 | Olfm1 | 0.52 | 2.77E-12 | 5.77E-08 | 0.36 | 0.23 | 0.13 | 1.58 |
| 13 | Mvb12a | 0.42 | 2.88E-12 | 5.99E-08 | 0.60 | 0.45 | 0.15 | 1.33 |
| 13 | Shoc2 | 0.46 | 3.06E-12 | 6.37E-08 | 0.59 | 0.46 | 0.14 | 1.30 |
| 13 | Tbl1x | 0.28 | 3.10E-12 | 6.46E-08 | 0.88 | 0.80 | 0.09 | 1.11 |
| 13 | Pcbd1 | 0.68 | 3.14E-12 | 6.52E-08 | 0.22 | 0.12 | 0.10 | 1.81 |
| 13 | Tmem176a | 0.42 | 3.47E-12 | 7.23E-08 | 0.56 | 0.40 | 0.15 | 1.37 |
| 13 | Dcaf17 | 0.51 | 3.61E-12 | 7.52E-08 | 0.48 | 0.33 | 0.15 | 1.45 |
| 13 | Spata13 | 0.93 | 3.66E-12 | 7.62E-08 | 0.15 | 0.08 | 0.08 | 2.04 |
| 13 | Dusp14 | 0.92 | 3.96E-12 | 8.23E-08 | 0.12 | 0.05 | 0.07 | 2.30 |
| 13 | Tmem63b | 0.52 | 4.13E-12 | 8.58E-08 | 0.47 | 0.33 | 0.14 | 1.42 |
| 13 | Ica1 | 0.76 | 4.23E-12 | 8.80E-08 | 0.22 | 0.12 | 0.10 | 1.84 |

**Supplementary Table 1. Marker genes for the original clusters of integrated clustering of E11.5 control and NotchGOF λ-junction cells.**

| | | | | | | | | |
|---|---|---|---|---|---|---|---|---|
| 13 | Snx14 | 0.63 | 4.45E-12 | 9.25E-08 | 0.27 | 0.16 | 0.11 | 1.70 |
| 13 | Kdm6b | 0.46 | 4.65E-12 | 9.67E-08 | 0.77 | 0.65 | 0.12 | 1.19 |
| 13 | Amer2 | 0.76 | 4.76E-12 | 9.91E-08 | 0.20 | 0.11 | 0.09 | 1.87 |
| 13 | Cthrc1 | 0.61 | 4.77E-12 | 9.92E-08 | 0.31 | 0.19 | 0.12 | 1.61 |
| 13 | Cpne3 | 0.37 | 4.80E-12 | 9.98E-08 | 0.68 | 0.52 | 0.16 | 1.31 |
| 13 | Rab11fip4 | 0.72 | 5.54E-12 | 1.15E-07 | 0.24 | 0.13 | 0.10 | 1.76 |
| 13 | Pan3 | 0.57 | 6.31E-12 | 1.31E-07 | 0.46 | 0.33 | 0.14 | 1.42 |
| 13 | Nhsl1 | 0.58 | 6.62E-12 | 1.38E-07 | 0.34 | 0.22 | 0.13 | 1.59 |
| 13 | Mitd1 | 0.52 | 6.74E-12 | 1.40E-07 | 0.44 | 0.29 | 0.14 | 1.48 |
| 13 | Ppp1r26 | 1.04 | 6.82E-12 | 1.42E-07 | 0.10 | 0.04 | 0.06 | 2.37 |
| 13 | Cdk7 | 0.45 | 6.93E-12 | 1.44E-07 | 0.63 | 0.48 | 0.15 | 1.30 |
| 13 | Pacsin1 | 0.96 | 7.56E-12 | 1.57E-07 | 0.19 | 0.10 | 0.09 | 1.85 |
| 13 | Zfp672 | 0.60 | 7.71E-12 | 1.60E-07 | 0.28 | 0.17 | 0.11 | 1.67 |
| 13 | Hdac1 | 0.33 | 8.27E-12 | 1.72E-07 | 0.86 | 0.80 | 0.07 | 1.08 |
| 13 | Foxo1 | 0.69 | 9.01E-12 | 1.87E-07 | 0.29 | 0.18 | 0.11 | 1.63 |
| 13 | Mllt10 | 0.36 | 9.35E-12 | 1.94E-07 | 0.75 | 0.63 | 0.12 | 1.18 |
| 13 | Pkp4 | 0.29 | 9.36E-12 | 1.95E-07 | 0.54 | 0.38 | 0.16 | 1.43 |
| 13 | Dkk3 | 0.77 | 9.73E-12 | 2.02E-07 | 0.21 | 0.12 | 0.10 | 1.82 |
| 13 | Cdc14a | 0.53 | 9.89E-12 | 2.06E-07 | 0.38 | 0.25 | 0.13 | 1.50 |
| 13 | Rian | 0.42 | 1.06E-11 | 2.21E-07 | 0.46 | 0.32 | 0.14 | 1.46 |
| 13 | Pycard | 0.49 | 1.11E-11 | 2.31E-07 | 0.22 | 0.13 | 0.10 | 1.76 |
| 13 | Nceh1 | 1.06 | 1.17E-11 | 2.43E-07 | 0.11 | 0.05 | 0.06 | 2.28 |
| 13 | Dbp | 0.64 | 1.19E-11 | 2.48E-07 | 0.24 | 0.14 | 0.10 | 1.74 |
| 13 | Pbxip1 | 0.59 | 1.20E-11 | 2.49E-07 | 0.32 | 0.20 | 0.12 | 1.58 |
| 13 | Rnpep | 0.39 | 1.23E-11 | 2.56E-07 | 0.58 | 0.43 | 0.15 | 1.34 |
| 13 | Tafa5 | 0.63 | 1.25E-11 | 2.60E-07 | 0.22 | 0.12 | 0.10 | 1.80 |
| 13 | Nop53 | 0.32 | 1.37E-11 | 2.85E-07 | 0.84 | 0.74 | 0.10 | 1.14 |
| 13 | Ermp1 | 0.68 | 1.38E-11 | 2.87E-07 | 0.46 | 0.33 | 0.13 | 1.38 |
| 13 | Uchl3 | 0.30 | 1.40E-11 | 2.92E-07 | 0.87 | 0.80 | 0.07 | 1.08 |
| 13 | 4931406P16Rik | 0.47 | 1.43E-11 | 2.98E-07 | 0.50 | 0.35 | 0.15 | 1.42 |
| 13 | Ccdc8 | 0.46 | 1.47E-11 | 3.06E-07 | 0.47 | 0.33 | 0.15 | 1.45 |
| 13 | Nr1h2 | 0.50 | 1.56E-11 | 3.25E-07 | 0.45 | 0.31 | 0.14 | 1.44 |
| 13 | Hmga1 | 0.35 | 1.61E-11 | 3.34E-07 | 0.65 | 0.49 | 0.16 | 1.33 |
| 13 | Rabgef1 | 0.70 | 1.87E-11 | 3.89E-07 | 0.27 | 0.16 | 0.11 | 1.67 |
| 13 | mt-Nd4l | 0.31 | 1.92E-11 | 4.00E-07 | 0.94 | 0.90 | 0.04 | 1.04 |
| 13 | Gid4 | 0.48 | 1.93E-11 | 4.01E-07 | 0.48 | 0.34 | 0.14 | 1.42 |
| 13 | Zfp800 | 0.61 | 1.93E-11 | 4.01E-07 | 0.39 | 0.26 | 0.12 | 1.46 |
| 13 | Zfp458 | 0.95 | 2.05E-11 | 4.27E-07 | 0.13 | 0.06 | 0.07 | 2.13 |
| 13 | Fry | 1.01 | 2.15E-11 | 4.47E-07 | 0.12 | 0.05 | 0.07 | 2.20 |
| 13 | Fuom | 0.51 | 2.15E-11 | 4.48E-07 | 0.40 | 0.27 | 0.13 | 1.49 |
| 13 | mt-Atp8 | 0.50 | 2.22E-11 | 4.63E-07 | 0.67 | 0.54 | 0.13 | 1.24 |
| 13 | Sh3d19 | 0.54 | 2.38E-11 | 4.95E-07 | 0.38 | 0.26 | 0.13 | 1.50 |
| 13 | Bckdha | 0.45 | 2.39E-11 | 4.97E-07 | 0.52 | 0.37 | 0.15 | 1.41 |
| 13 | Mycbp2 | 0.36 | 2.40E-11 | 4.99E-07 | 0.81 | 0.70 | 0.11 | 1.16 |
| 13 | Echdc2 | 0.52 | 2.48E-11 | 5.16E-07 | 0.38 | 0.25 | 0.13 | 1.51 |
| 13 | Dcp1b | 0.86 | 2.55E-11 | 5.31E-07 | 0.17 | 0.09 | 0.08 | 1.91 |
| 13 | Ddx19b | 0.84 | 2.58E-11 | 5.37E-07 | 0.17 | 0.09 | 0.08 | 1.91 |
| 13 | Tiam1 | 0.42 | 2.59E-11 | 5.38E-07 | 0.45 | 0.31 | 0.14 | 1.45 |
| 13 | Grn | 0.38 | 2.64E-11 | 5.50E-07 | 0.48 | 0.35 | 0.14 | 1.39 |
| 13 | Fos | 1.62 | 2.77E-11 | 5.76E-07 | 0.22 | 0.14 | 0.08 | 1.61 |
| 13 | Trak1 | 0.59 | 2.79E-11 | 5.80E-07 | 0.21 | 0.12 | 0.10 | 1.81 |
| 13 | Nfkbiz | 1.03 | 2.85E-11 | 5.93E-07 | 0.14 | 0.07 | 0.07 | 2.03 |
| 13 | Rragb | 0.93 | 2.86E-11 | 5.94E-07 | 0.16 | 0.09 | 0.08 | 1.92 |
| 13 | Apobec3 | 0.68 | 3.03E-11 | 6.30E-07 | 0.16 | 0.08 | 0.08 | 1.96 |
| 13 | Plxna3 | 0.88 | 3.29E-11 | 6.84E-07 | 0.18 | 0.10 | 0.08 | 1.86 |
| 13 | Marchf2 | 0.40 | 3.32E-11 | 6.90E-07 | 0.65 | 0.51 | 0.13 | 1.26 |

**Supplementary Table 1. Marker genes for the original clusters of integrated clustering of E11.5 control and NotchGOF λ-junction cells.**

| | | | | | | | | |
|---|---|---|---|---|---|---|---|---|
| 13 | Dcaf12l1 | 0.76 | 3.46E-11 | 7.19E-07 | 0.21 | 0.12 | 0.09 | 1.81 |
| 13 | Gpr137c | 0.58 | 3.74E-11 | 7.78E-07 | 0.25 | 0.14 | 0.10 | 1.70 |
| 13 | Ppp2r3d | 0.58 | 3.93E-11 | 8.18E-07 | 0.35 | 0.22 | 0.12 | 1.55 |
| 13 | Noct | 0.54 | 3.98E-11 | 8.29E-07 | 0.30 | 0.19 | 0.11 | 1.61 |
| 13 | Palld | 0.32 | 3.99E-11 | 8.29E-07 | 0.75 | 0.62 | 0.13 | 1.21 |
| 13 | Zcchc7 | 0.45 | 4.18E-11 | 8.69E-07 | 0.63 | 0.52 | 0.12 | 1.22 |
| 13 | Ank | 0.90 | 4.35E-11 | 9.05E-07 | 0.14 | 0.07 | 0.07 | 2.03 |
| 13 | Sppl2a | 0.44 | 4.37E-11 | 9.10E-07 | 0.62 | 0.47 | 0.15 | 1.32 |
| 13 | Shroom2 | 0.87 | 4.58E-11 | 9.53E-07 | 0.15 | 0.08 | 0.08 | 1.97 |
| 13 | Plpp2 | 0.50 | 4.75E-11 | 9.87E-07 | 0.39 | 0.26 | 0.13 | 1.50 |
| 13 | Cic | 0.48 | 4.95E-11 | 1.03E-06 | 0.51 | 0.37 | 0.14 | 1.37 |
| 13 | Gm43445 | 0.92 | 5.25E-11 | 1.09E-06 | 0.17 | 0.09 | 0.08 | 1.89 |
| 13 | Tanc2 | 0.80 | 5.83E-11 | 1.21E-06 | 0.34 | 0.23 | 0.11 | 1.48 |
| 13 | Utp14b | 0.47 | 5.88E-11 | 1.22E-06 | 0.52 | 0.38 | 0.14 | 1.37 |
| 13 | Akap1 | 0.57 | 5.88E-11 | 1.22E-06 | 0.30 | 0.19 | 0.11 | 1.59 |
| 13 | Erlin2 | 0.47 | 6.14E-11 | 1.28E-06 | 0.45 | 0.32 | 0.13 | 1.41 |
| 13 | Trmt1l | 0.45 | 6.18E-11 | 1.29E-06 | 0.40 | 0.26 | 0.13 | 1.50 |
| 13 | Txndc16 | 0.74 | 6.21E-11 | 1.29E-06 | 0.24 | 0.14 | 0.10 | 1.70 |
| 13 | 3110082I17Rik | 0.50 | 6.43E-11 | 1.34E-06 | 0.43 | 0.29 | 0.13 | 1.45 |
| 13 | E330009J07Rik | 0.86 | 6.54E-11 | 1.36E-06 | 0.21 | 0.12 | 0.09 | 1.76 |
| 13 | Ddb2 | 0.62 | 6.62E-11 | 1.38E-06 | 0.18 | 0.10 | 0.08 | 1.87 |
| 13 | Etaa1os | 0.88 | 6.71E-11 | 1.39E-06 | 0.13 | 0.06 | 0.07 | 2.08 |
| 13 | Rpp21 | 0.34 | 7.40E-11 | 1.54E-06 | 0.71 | 0.57 | 0.14 | 1.24 |
| 13 | Ptrh2 | 0.35 | 7.45E-11 | 1.55E-06 | 0.78 | 0.66 | 0.12 | 1.18 |
| 13 | Zc3h6 | 0.88 | 7.56E-11 | 1.57E-06 | 0.17 | 0.09 | 0.08 | 1.90 |
| 13 | Crebrf | 0.79 | 7.58E-11 | 1.58E-06 | 0.19 | 0.11 | 0.09 | 1.78 |
| 13 | Cdk5rap3 | 0.43 | 7.91E-11 | 1.65E-06 | 0.59 | 0.43 | 0.16 | 1.37 |
| 13 | Kdm5d | 1.33 | 8.00E-11 | 1.66E-06 | 0.10 | 0.05 | 0.06 | 2.26 |
| 13 | Xpa | 0.50 | 8.11E-11 | 1.69E-06 | 0.57 | 0.44 | 0.13 | 1.29 |
| 13 | Trim37 | 0.49 | 8.35E-11 | 1.74E-06 | 0.52 | 0.39 | 0.13 | 1.33 |
| 13 | Ttc5 | 0.35 | 8.37E-11 | 1.74E-06 | 0.69 | 0.54 | 0.15 | 1.28 |
| 13 | Acvr1b | 0.74 | 8.49E-11 | 1.77E-06 | 0.20 | 0.11 | 0.09 | 1.79 |
| 13 | Kat2b | 1.21 | 8.93E-11 | 1.86E-06 | 0.10 | 0.05 | 0.06 | 2.27 |
| 13 | Uaca | 0.47 | 9.28E-11 | 1.93E-06 | 0.47 | 0.33 | 0.14 | 1.42 |
| 13 | Smpdl3a | 0.78 | 9.95E-11 | 2.07E-06 | 0.22 | 0.13 | 0.09 | 1.69 |
| 13 | Shb | 0.73 | 1.03E-10 | 2.14E-06 | 0.30 | 0.20 | 0.11 | 1.54 |
| 13 | Rbbp9 | 0.48 | 1.03E-10 | 2.15E-06 | 0.52 | 0.39 | 0.13 | 1.34 |
| 13 | Myo6 | 0.59 | 1.06E-10 | 2.21E-06 | 0.30 | 0.19 | 0.11 | 1.61 |
| 13 | Utrn | 0.49 | 1.09E-10 | 2.27E-06 | 0.72 | 0.61 | 0.11 | 1.17 |
| 13 | Coasy | 0.51 | 1.11E-10 | 2.32E-06 | 0.45 | 0.31 | 0.13 | 1.42 |
| 13 | Chfr | 0.57 | 1.12E-10 | 2.33E-06 | 0.37 | 0.25 | 0.12 | 1.46 |
| 13 | Syne2 | 0.32 | 1.20E-10 | 2.50E-06 | 0.81 | 0.72 | 0.09 | 1.12 |
| 13 | Nlk | 0.38 | 1.23E-10 | 2.56E-06 | 0.40 | 0.27 | 0.13 | 1.48 |
| 13 | Mir99ahg | 0.36 | 1.33E-10 | 2.78E-06 | 0.63 | 0.48 | 0.15 | 1.31 |
| 13 | Gtf2ird1 | 0.47 | 1.41E-10 | 2.93E-06 | 0.60 | 0.47 | 0.13 | 1.29 |
| 13 | Magi3 | 0.52 | 1.61E-10 | 3.34E-06 | 0.30 | 0.19 | 0.11 | 1.60 |
| 13 | Naa80 | 0.61 | 1.72E-10 | 3.57E-06 | 0.26 | 0.16 | 0.10 | 1.63 |
| 13 | Rb1cc1 | 0.34 | 1.86E-10 | 3.86E-06 | 0.78 | 0.65 | 0.13 | 1.19 |
| 13 | Micu3 | 0.51 | 1.94E-10 | 4.03E-06 | 0.28 | 0.17 | 0.11 | 1.62 |
| 13 | Pim3 | 0.50 | 2.00E-10 | 4.15E-06 | 0.42 | 0.30 | 0.13 | 1.42 |
| 13 | Lpin1 | 0.79 | 2.02E-10 | 4.20E-06 | 0.13 | 0.07 | 0.07 | 2.03 |
| 13 | Impad1 | 0.31 | 2.10E-10 | 4.37E-06 | 0.84 | 0.72 | 0.12 | 1.17 |
| 13 | Mtmr4 | 0.42 | 2.14E-10 | 4.45E-06 | 0.42 | 0.29 | 0.14 | 1.48 |
| 13 | Ddx3y | 0.90 | 2.20E-10 | 4.57E-06 | 0.15 | 0.07 | 0.07 | 1.96 |
| 13 | Lbh | 0.65 | 2.23E-10 | 4.63E-06 | 0.31 | 0.20 | 0.11 | 1.53 |
| 13 | St7 | 0.68 | 2.26E-10 | 4.70E-06 | 0.33 | 0.22 | 0.11 | 1.49 |

**Supplementary Table 1. Marker genes for the original clusters of integrated clustering of E11.5 control and NotchGOF λ-junction cells.**

| | | | | | | | | |
|---|---|---|---|---|---|---|---|---|
| 13 | Noc2l | 0.39 | 2.30E-10 | 4.79E-06 | 0.72 | 0.61 | 0.11 | 1.17 |
| 13 | Tlcd1 | 0.74 | 2.44E-10 | 5.07E-06 | 0.17 | 0.09 | 0.08 | 1.88 |
| 13 | 2010320M18Rik | 0.44 | 2.46E-10 | 5.12E-06 | 0.47 | 0.34 | 0.13 | 1.38 |
| 13 | Aspscr1 | 0.42 | 2.54E-10 | 5.29E-06 | 0.59 | 0.45 | 0.14 | 1.30 |
| 13 | Hsdl1 | 0.62 | 2.74E-10 | 5.70E-06 | 0.25 | 0.15 | 0.10 | 1.64 |
| 13 | Eef2kmt | 0.53 | 2.80E-10 | 5.82E-06 | 0.29 | 0.18 | 0.11 | 1.59 |
| 13 | Ptprz1 | 1.00 | 2.80E-10 | 5.83E-06 | 0.19 | 0.11 | 0.08 | 1.76 |
| 13 | Atp13a3 | 0.48 | 2.86E-10 | 5.95E-06 | 0.54 | 0.41 | 0.13 | 1.32 |
| 13 | Hscb | 0.44 | 3.06E-10 | 6.35E-06 | 0.46 | 0.32 | 0.13 | 1.42 |
| 13 | Sec62 | 0.33 | 3.14E-10 | 6.54E-06 | 0.96 | 0.93 | 0.03 | 1.03 |
| 13 | Xkr6 | 0.47 | 3.33E-10 | 6.92E-06 | 0.22 | 0.13 | 0.09 | 1.75 |
| 13 | Gm45051 | 0.83 | 3.54E-10 | 7.36E-06 | 0.14 | 0.07 | 0.07 | 1.96 |
| 13 | Tbl1xr1 | 0.46 | 3.57E-10 | 7.43E-06 | 0.54 | 0.42 | 0.13 | 1.30 |
| 13 | Sox13 | 0.60 | 3.66E-10 | 7.62E-06 | 0.16 | 0.09 | 0.08 | 1.90 |
| 13 | Tmem38a | 0.62 | 3.85E-10 | 8.01E-06 | 0.18 | 0.10 | 0.08 | 1.83 |
| 13 | Vill | 0.68 | 3.93E-10 | 8.16E-06 | 0.16 | 0.08 | 0.07 | 1.91 |
| 13 | Ankrd13c | 0.49 | 3.96E-10 | 8.24E-06 | 0.36 | 0.24 | 0.12 | 1.49 |
| 13 | Stk39 | 0.49 | 3.98E-10 | 8.29E-06 | 0.38 | 0.26 | 0.13 | 1.48 |
| 13 | Lman2 | 0.32 | 4.04E-10 | 8.41E-06 | 0.75 | 0.62 | 0.14 | 1.22 |
| 13 | Hspa2 | 1.03 | 4.07E-10 | 8.46E-06 | 0.12 | 0.06 | 0.06 | 2.09 |
| 13 | Acbd5 | 0.55 | 4.14E-10 | 8.61E-06 | 0.48 | 0.35 | 0.12 | 1.35 |
| 13 | Krba1 | 0.61 | 4.28E-10 | 8.90E-06 | 0.28 | 0.18 | 0.11 | 1.59 |
| 13 | Slc25a17 | 0.36 | 4.31E-10 | 8.96E-06 | 0.73 | 0.62 | 0.11 | 1.18 |
| 13 | Asah1 | 0.35 | 4.32E-10 | 8.99E-06 | 0.51 | 0.37 | 0.14 | 1.38 |
| 13 | Prickle3 | 0.84 | 4.36E-10 | 9.07E-06 | 0.13 | 0.07 | 0.07 | 2.00 |
| 13 | Chka | 0.42 | 4.41E-10 | 9.18E-06 | 0.61 | 0.49 | 0.12 | 1.25 |
| 13 | Kremen1 | 0.61 | 4.49E-10 | 9.35E-06 | 0.24 | 0.14 | 0.10 | 1.67 |
| 13 | Ubald1 | 0.52 | 4.64E-10 | 9.65E-06 | 0.38 | 0.26 | 0.12 | 1.45 |
| 13 | Ccpg1 | 0.65 | 4.66E-10 | 9.69E-06 | 0.26 | 0.16 | 0.10 | 1.60 |
| 13 | Rrp36 | 0.45 | 4.79E-10 | 9.96E-06 | 0.45 | 0.32 | 0.13 | 1.41 |
| 13 | Rps6ka6 | 0.35 | 4.83E-10 | 1.00E-05 | 0.59 | 0.45 | 0.14 | 1.31 |
| 13 | Ring1 | 0.53 | 4.93E-10 | 1.03E-05 | 0.36 | 0.24 | 0.12 | 1.49 |
| 13 | Mccc2 | 0.51 | 5.00E-10 | 1.04E-05 | 0.40 | 0.28 | 0.13 | 1.46 |
| 13 | Mbnl2 | 0.30 | 5.04E-10 | 1.05E-05 | 0.78 | 0.68 | 0.10 | 1.15 |
| 13 | Tcf7l2 | 0.41 | 5.41E-10 | 1.13E-05 | 0.72 | 0.61 | 0.11 | 1.19 |
| 13 | Iqgap1 | 0.32 | 5.49E-10 | 1.14E-05 | 0.62 | 0.47 | 0.15 | 1.31 |
| 13 | Malt1 | 0.59 | 5.64E-10 | 1.17E-05 | 0.26 | 0.16 | 0.10 | 1.61 |
| 13 | Tmem179b | 0.46 | 5.67E-10 | 1.18E-05 | 0.42 | 0.29 | 0.13 | 1.45 |
| 13 | 2210016F16Rik | 0.46 | 5.71E-10 | 1.19E-05 | 0.48 | 0.35 | 0.13 | 1.36 |
| 13 | Fam114a2 | 0.33 | 5.72E-10 | 1.19E-05 | 0.77 | 0.68 | 0.10 | 1.14 |
| 13 | Mterf1a | 0.84 | 6.48E-10 | 1.35E-05 | 0.15 | 0.08 | 0.07 | 1.89 |
| 13 | C1galt1c1 | 0.62 | 6.51E-10 | 1.35E-05 | 0.26 | 0.16 | 0.10 | 1.64 |
| 13 | Asxl1 | 0.42 | 6.77E-10 | 1.41E-05 | 0.62 | 0.50 | 0.12 | 1.24 |
| 13 | Lmbr1l | 0.56 | 6.82E-10 | 1.42E-05 | 0.28 | 0.18 | 0.10 | 1.59 |
| 13 | Zfp53 | 0.67 | 7.50E-10 | 1.56E-05 | 0.16 | 0.09 | 0.08 | 1.85 |
| 13 | Krt1 | 0.89 | 7.63E-10 | 1.59E-05 | 0.11 | 0.05 | 0.06 | 2.14 |
| 13 | Leprotl1 | 0.35 | 7.77E-10 | 1.62E-05 | 0.68 | 0.55 | 0.14 | 1.26 |
| 13 | Dnajc21 | 0.34 | 7.83E-10 | 1.63E-05 | 0.72 | 0.59 | 0.13 | 1.22 |
| 13 | Zfp516 | 0.43 | 7.91E-10 | 1.64E-05 | 0.62 | 0.49 | 0.13 | 1.26 |
| 13 | Hbp1 | 0.34 | 8.03E-10 | 1.67E-05 | 0.70 | 0.56 | 0.13 | 1.24 |
| 13 | Ncor2 | 0.47 | 8.08E-10 | 1.68E-05 | 0.48 | 0.35 | 0.13 | 1.37 |
| 13 | Wdr26 | 0.29 | 8.12E-10 | 1.69E-05 | 0.81 | 0.71 | 0.10 | 1.14 |
| 13 | Esco1 | 0.54 | 8.22E-10 | 1.71E-05 | 0.34 | 0.23 | 0.11 | 1.49 |
| 13 | Fcf1 | 0.32 | 8.92E-10 | 1.86E-05 | 0.78 | 0.67 | 0.11 | 1.16 |
| 13 | Efs | 0.45 | 9.45E-10 | 1.96E-05 | 0.41 | 0.28 | 0.13 | 1.46 |
| 13 | Ccser2 | 0.52 | 9.56E-10 | 1.99E-05 | 0.39 | 0.27 | 0.12 | 1.45 |

**Supplementary Table 1. Marker genes for the original clusters of integrated clustering of E11.5 control and NotchGOF λ-junction cells.**

| | | | | | | | | |
|---|---|---|---|---|---|---|---|---|
| 13 | Slc20a2 | 0.50 | 9.58E-10 | 1.99E-05 | 0.35 | 0.24 | 0.12 | 1.49 |
| 13 | Gnptg | 0.57 | 1.01E-09 | 2.11E-05 | 0.31 | 0.20 | 0.11 | 1.54 |
| 13 | Kctd6 | 0.58 | 1.04E-09 | 2.17E-05 | 0.25 | 0.16 | 0.10 | 1.62 |
| 13 | Atp6ap2 | 0.40 | 1.04E-09 | 2.17E-05 | 0.56 | 0.42 | 0.13 | 1.31 |
| 13 | Mpv17l2 | 0.53 | 1.05E-09 | 2.18E-05 | 0.33 | 0.23 | 0.11 | 1.48 |
| 13 | Ppil1 | 0.25 | 1.06E-09 | 2.21E-05 | 0.87 | 0.81 | 0.06 | 1.07 |
| 13 | Ctdsp2 | 0.34 | 1.06E-09 | 2.21E-05 | 0.70 | 0.56 | 0.13 | 1.24 |
| 13 | Magi2 | 0.57 | 1.09E-09 | 2.27E-05 | 0.29 | 0.19 | 0.11 | 1.57 |
| 13 | Fbrsl1 | 0.41 | 1.23E-09 | 2.55E-05 | 0.50 | 0.37 | 0.13 | 1.36 |
| 13 | Fam3c | 0.40 | 1.25E-09 | 2.59E-05 | 0.50 | 0.36 | 0.14 | 1.38 |
| 13 | Plekhb2 | 0.69 | 1.28E-09 | 2.66E-05 | 0.12 | 0.06 | 0.06 | 2.03 |
| 13 | Dynlt3 | 0.52 | 1.30E-09 | 2.70E-05 | 0.37 | 0.26 | 0.11 | 1.44 |
| 13 | H2-K1 | 0.57 | 1.30E-09 | 2.70E-05 | 0.22 | 0.13 | 0.09 | 1.66 |
| 13 | 4933434E20Rik | 0.36 | 1.32E-09 | 2.75E-05 | 0.63 | 0.49 | 0.14 | 1.28 |
| 13 | Bckdk | 0.38 | 1.32E-09 | 2.75E-05 | 0.62 | 0.49 | 0.13 | 1.27 |
| 13 | Cyb561d2 | 0.62 | 1.38E-09 | 2.87E-05 | 0.27 | 0.17 | 0.10 | 1.58 |
| 13 | Etaa1 | 0.39 | 1.43E-09 | 2.98E-05 | 0.65 | 0.53 | 0.11 | 1.21 |
| 13 | Psph | 0.41 | 1.46E-09 | 3.05E-05 | 0.40 | 0.28 | 0.13 | 1.45 |
| 13 | Pex2 | 0.36 | 1.49E-09 | 3.09E-05 | 0.61 | 0.47 | 0.14 | 1.29 |
| 13 | Mnt | 0.73 | 1.49E-09 | 3.11E-05 | 0.27 | 0.17 | 0.09 | 1.54 |
| 13 | Pank1 | 0.43 | 1.52E-09 | 3.16E-05 | 0.44 | 0.31 | 0.12 | 1.39 |
| 13 | Txnip | 0.49 | 1.53E-09 | 3.18E-05 | 0.47 | 0.34 | 0.13 | 1.37 |
| 13 | Spop | 0.30 | 1.54E-09 | 3.20E-05 | 0.78 | 0.69 | 0.09 | 1.13 |
| 13 | Mtss2 | 0.42 | 1.56E-09 | 3.25E-05 | 0.39 | 0.27 | 0.12 | 1.44 |
| 13 | Lpar6 | 0.52 | 1.60E-09 | 3.33E-05 | 0.52 | 0.40 | 0.12 | 1.29 |
| 13 | Hint2 | 0.35 | 1.68E-09 | 3.49E-05 | 0.72 | 0.61 | 0.11 | 1.18 |
| 13 | Slk | 0.39 | 1.70E-09 | 3.54E-05 | 0.66 | 0.53 | 0.13 | 1.24 |
| 13 | Mark2 | 0.36 | 1.71E-09 | 3.56E-05 | 0.60 | 0.46 | 0.14 | 1.30 |
| 13 | Tmed4 | 0.33 | 1.74E-09 | 3.61E-05 | 0.69 | 0.56 | 0.13 | 1.23 |
| 13 | Sirt7 | 0.65 | 1.74E-09 | 3.62E-05 | 0.27 | 0.17 | 0.10 | 1.57 |
| 13 | Fbxo46 | 0.65 | 1.77E-09 | 3.68E-05 | 0.21 | 0.13 | 0.09 | 1.69 |
| 13 | Sipa1l3 | 0.72 | 1.86E-09 | 3.88E-05 | 0.18 | 0.10 | 0.08 | 1.76 |
| 13 | Stk4 | 0.41 | 1.87E-09 | 3.88E-05 | 0.49 | 0.36 | 0.13 | 1.36 |
| 13 | Amotl2 | 0.55 | 1.89E-09 | 3.93E-05 | 0.45 | 0.34 | 0.11 | 1.33 |
| 13 | Fuca2 | 0.47 | 1.99E-09 | 4.13E-05 | 0.35 | 0.23 | 0.12 | 1.51 |
| 13 | Rfx3 | 0.51 | 2.01E-09 | 4.17E-05 | 0.32 | 0.21 | 0.11 | 1.51 |
| 13 | Fmnl2 | 0.47 | 2.02E-09 | 4.21E-05 | 0.56 | 0.44 | 0.13 | 1.29 |
| 13 | Atg101 | 0.33 | 2.05E-09 | 4.26E-05 | 0.63 | 0.49 | 0.14 | 1.30 |
| 13 | Ncstn | 0.46 | 2.07E-09 | 4.30E-05 | 0.41 | 0.29 | 0.12 | 1.43 |
| 13 | Arvcf | 0.47 | 2.07E-09 | 4.31E-05 | 0.36 | 0.24 | 0.12 | 1.48 |
| 13 | Ccdc173 | 0.63 | 2.20E-09 | 4.57E-05 | 0.20 | 0.12 | 0.08 | 1.72 |
| 13 | Haghl | 0.40 | 2.27E-09 | 4.71E-05 | 0.49 | 0.35 | 0.14 | 1.40 |
| 13 | Pex7 | 0.43 | 2.27E-09 | 4.71E-05 | 0.37 | 0.25 | 0.12 | 1.47 |
| 13 | Lpcat3 | 0.42 | 2.31E-09 | 4.81E-05 | 0.43 | 0.30 | 0.13 | 1.42 |
| 13 | Rgcc | 0.54 | 2.33E-09 | 4.84E-05 | 0.17 | 0.09 | 0.08 | 1.82 |
| 13 | Wdfy1 | 0.43 | 2.37E-09 | 4.93E-05 | 0.48 | 0.35 | 0.13 | 1.37 |
| 13 | Lrrc8b | 0.67 | 2.38E-09 | 4.95E-05 | 0.19 | 0.11 | 0.08 | 1.72 |
| 13 | Tmx2 | 0.38 | 2.41E-09 | 5.01E-05 | 0.61 | 0.49 | 0.12 | 1.25 |
| 13 | Hmga1b | 0.30 | 2.42E-09 | 5.03E-05 | 0.58 | 0.43 | 0.15 | 1.35 |
| 13 | St3gal5 | 0.44 | 2.45E-09 | 5.09E-05 | 0.35 | 0.24 | 0.11 | 1.48 |
| 13 | Rnf44 | 0.42 | 2.63E-09 | 5.47E-05 | 0.47 | 0.34 | 0.13 | 1.37 |
| 13 | Gart | 0.39 | 2.77E-09 | 5.77E-05 | 0.67 | 0.56 | 0.11 | 1.19 |
| 13 | Zfp827 | 0.45 | 2.87E-09 | 5.96E-05 | 0.43 | 0.31 | 0.12 | 1.40 |
| 13 | Ccdc85c | 0.69 | 2.88E-09 | 5.98E-05 | 0.19 | 0.11 | 0.08 | 1.72 |
| 13 | Cuedc1 | 0.44 | 3.14E-09 | 6.53E-05 | 0.51 | 0.38 | 0.12 | 1.32 |
| 13 | Wfs1 | 0.69 | 3.19E-09 | 6.64E-05 | 0.13 | 0.07 | 0.06 | 1.98 |

**Supplementary Table 1. Marker genes for the original clusters of integrated clustering of E11.5 control and NotchGOF λ-junction cells.**

| | | | | | | | | |
|---|---|---|---|---|---|---|---|---|
| 13 | C2cd3 | 0.43 | 3.20E-09 | 6.66E-05 | 0.43 | 0.30 | 0.12 | 1.40 |
| 13 | Mrps34 | 0.35 | 3.22E-09 | 6.70E-05 | 0.72 | 0.62 | 0.10 | 1.16 |
| 13 | Egfr | 0.69 | 3.28E-09 | 6.83E-05 | 0.18 | 0.10 | 0.08 | 1.76 |
| 13 | Col4a3bp | 0.59 | 3.33E-09 | 6.93E-05 | 0.44 | 0.32 | 0.12 | 1.37 |
| 13 | Fgf13 | 0.64 | 3.46E-09 | 7.19E-05 | 0.19 | 0.11 | 0.08 | 1.74 |
| 13 | Col4a5 | 0.70 | 3.63E-09 | 7.55E-05 | 0.21 | 0.13 | 0.09 | 1.66 |
| 13 | Tgoln1 | 0.48 | 4.00E-09 | 8.32E-05 | 0.41 | 0.29 | 0.12 | 1.41 |
| 13 | Whamm | 0.66 | 4.15E-09 | 8.64E-05 | 0.18 | 0.10 | 0.08 | 1.77 |
| 13 | Gabra4 | 0.84 | 4.16E-09 | 8.65E-05 | 0.11 | 0.05 | 0.06 | 2.10 |
| 13 | Fgd6 | 0.66 | 4.21E-09 | 8.76E-05 | 0.25 | 0.16 | 0.09 | 1.57 |
| 13 | Rnd2 | 0.66 | 4.22E-09 | 8.77E-05 | 0.16 | 0.09 | 0.07 | 1.82 |
| 13 | Bcl7a | 0.26 | 4.29E-09 | 8.92E-05 | 0.80 | 0.67 | 0.13 | 1.19 |
| 13 | Rfx7 | 0.40 | 4.46E-09 | 9.27E-05 | 0.59 | 0.46 | 0.13 | 1.28 |
| 13 | Zfp667 | 0.69 | 4.66E-09 | 9.69E-05 | 0.17 | 0.10 | 0.08 | 1.77 |
| 13 | Fam49b | 0.29 | 4.68E-09 | 9.73E-05 | 0.76 | 0.65 | 0.11 | 1.16 |
| 13 | Atraid | 0.33 | 4.74E-09 | 9.85E-05 | 0.64 | 0.50 | 0.13 | 1.26 |
| 13 | Larp1b | 0.78 | 4.93E-09 | 1.02E-04 | 0.14 | 0.08 | 0.07 | 1.87 |
| 13 | Fam172a | 0.42 | 4.93E-09 | 1.03E-04 | 0.48 | 0.35 | 0.13 | 1.38 |
| 13 | Ppa2 | 0.33 | 4.95E-09 | 1.03E-04 | 0.70 | 0.55 | 0.14 | 1.26 |
| 13 | Agap3 | 0.44 | 5.19E-09 | 1.08E-04 | 0.48 | 0.36 | 0.13 | 1.36 |
| 13 | Dhcr24 | 0.54 | 5.34E-09 | 1.11E-04 | 0.47 | 0.35 | 0.12 | 1.35 |
| 13 | Zmpste24 | 0.41 | 5.43E-09 | 1.13E-04 | 0.44 | 0.31 | 0.12 | 1.40 |
| 13 | Tm2d3 | 0.32 | 5.59E-09 | 1.16E-04 | 0.64 | 0.50 | 0.14 | 1.28 |
| 13 | Pink1 | 0.49 | 5.62E-09 | 1.17E-04 | 0.33 | 0.22 | 0.11 | 1.49 |
| 13 | Wdpcp | 0.79 | 5.68E-09 | 1.18E-04 | 0.15 | 0.08 | 0.07 | 1.83 |
| 13 | Echdc1 | 0.48 | 5.76E-09 | 1.20E-04 | 0.34 | 0.24 | 0.11 | 1.46 |
| 13 | Actn4 | 0.29 | 5.88E-09 | 1.22E-04 | 0.82 | 0.70 | 0.13 | 1.18 |
| 13 | Zfp444 | 0.30 | 5.90E-09 | 1.23E-04 | 0.56 | 0.42 | 0.14 | 1.34 |
| 13 | Pla2g6 | 0.64 | 5.96E-09 | 1.24E-04 | 0.15 | 0.08 | 0.07 | 1.85 |
| 13 | Acrbp | 0.58 | 6.05E-09 | 1.26E-04 | 0.18 | 0.10 | 0.08 | 1.75 |
| 13 | Ptar1 | 0.46 | 6.36E-09 | 1.32E-04 | 0.39 | 0.28 | 0.11 | 1.41 |
| 13 | Stap2 | 0.72 | 6.62E-09 | 1.38E-04 | 0.15 | 0.08 | 0.07 | 1.83 |
| 13 | Ppt1 | 0.39 | 6.65E-09 | 1.38E-04 | 0.44 | 0.31 | 0.13 | 1.40 |
| 13 | Mpv17l | 0.77 | 6.83E-09 | 1.42E-04 | 0.14 | 0.08 | 0.07 | 1.86 |
| 13 | Rrp7a | 0.34 | 6.94E-09 | 1.44E-04 | 0.61 | 0.47 | 0.13 | 1.28 |
| 13 | Maml2 | 0.63 | 7.06E-09 | 1.47E-04 | 0.17 | 0.09 | 0.07 | 1.78 |
| 13 | Exoc6 | 0.88 | 7.13E-09 | 1.48E-04 | 0.14 | 0.07 | 0.06 | 1.85 |
| 13 | Herpud2 | 0.41 | 7.31E-09 | 1.52E-04 | 0.40 | 0.28 | 0.12 | 1.42 |
| 13 | Ccdc80 | 0.56 | 7.45E-09 | 1.55E-04 | 0.28 | 0.18 | 0.10 | 1.53 |
| 13 | Pacsin3 | 0.45 | 7.46E-09 | 1.55E-04 | 0.33 | 0.22 | 0.11 | 1.50 |
| 13 | Map9 | 0.50 | 7.55E-09 | 1.57E-04 | 0.28 | 0.18 | 0.10 | 1.54 |
| 13 | Sirt5 | 0.38 | 7.67E-09 | 1.60E-04 | 0.38 | 0.26 | 0.12 | 1.45 |
| 13 | Tm9sf2 | 0.31 | 7.81E-09 | 1.63E-04 | 0.72 | 0.62 | 0.11 | 1.17 |
| 13 | Mterf4 | 0.40 | 7.82E-09 | 1.63E-04 | 0.47 | 0.33 | 0.13 | 1.39 |
| 13 | Cnot7 | 0.29 | 7.83E-09 | 1.63E-04 | 0.78 | 0.68 | 0.10 | 1.15 |
| 13 | Chst11 | 0.77 | 8.13E-09 | 1.69E-04 | 0.14 | 0.07 | 0.06 | 1.88 |
| 13 | Pole3 | 0.26 | 8.26E-09 | 1.72E-04 | 0.84 | 0.76 | 0.08 | 1.10 |
| 13 | Gm3839 | 0.53 | 8.40E-09 | 1.75E-04 | 0.26 | 0.16 | 0.09 | 1.57 |
| 13 | Zfp646 | 0.37 | 8.41E-09 | 1.75E-04 | 0.54 | 0.41 | 0.13 | 1.32 |
| 13 | Diaph1 | 0.40 | 9.84E-09 | 2.05E-04 | 0.55 | 0.42 | 0.13 | 1.30 |
| 13 | Nat2 | 0.95 | 1.02E-08 | 2.13E-04 | 0.11 | 0.06 | 0.06 | 2.00 |
| 13 | Wdr89 | 0.26 | 1.05E-08 | 2.19E-04 | 0.85 | 0.77 | 0.09 | 1.11 |
| 13 | Cope | 0.26 | 1.07E-08 | 2.23E-04 | 0.85 | 0.77 | 0.08 | 1.11 |
| 13 | Ebpl | 0.44 | 1.10E-08 | 2.30E-04 | 0.64 | 0.55 | 0.10 | 1.18 |
| 13 | Deaf1 | 0.49 | 1.13E-08 | 2.35E-04 | 0.33 | 0.22 | 0.11 | 1.48 |
| 13 | Lmln | 0.91 | 1.14E-08 | 2.38E-04 | 0.11 | 0.06 | 0.06 | 2.00 |

**Supplementary Table 1. Marker genes for the original clusters of integrated clustering of E11.5 control and NotchGOF λ-junction cells.**

| | | | | | | | | |
|---|---|---|---|---|---|---|---|---|
| 13 | Tmem170b | 0.86 | 1.15E-08 | 2.38E-04 | 0.12 | 0.06 | 0.06 | 1.94 |
| 13 | Ank3 | 0.29 | 1.15E-08 | 2.39E-04 | 0.63 | 0.49 | 0.14 | 1.29 |
| 13 | Prom1 | 0.43 | 1.18E-08 | 2.45E-04 | 0.54 | 0.41 | 0.13 | 1.33 |
| 13 | Mtmr1 | 0.86 | 1.19E-08 | 2.48E-04 | 0.13 | 0.07 | 0.06 | 1.89 |
| 13 | Wdr45 | 0.49 | 1.20E-08 | 2.49E-04 | 0.21 | 0.13 | 0.09 | 1.66 |
| 13 | Lypd6 | 0.61 | 1.21E-08 | 2.51E-04 | 0.21 | 0.13 | 0.08 | 1.64 |
| 13 | Nanos3 | 0.91 | 1.25E-08 | 2.59E-04 | 0.11 | 0.05 | 0.06 | 2.02 |
| 13 | Cenpb | 0.25 | 1.26E-08 | 2.62E-04 | 0.77 | 0.66 | 0.11 | 1.16 |
| 13 | Rabggta | 0.60 | 1.27E-08 | 2.65E-04 | 0.21 | 0.13 | 0.08 | 1.65 |
| 13 | Cc2d1b | 0.55 | 1.28E-08 | 2.66E-04 | 0.27 | 0.17 | 0.10 | 1.56 |
| 13 | Chd1l | 0.52 | 1.29E-08 | 2.68E-04 | 0.30 | 0.20 | 0.10 | 1.52 |
| 13 | Gns | 0.50 | 1.30E-08 | 2.71E-04 | 0.36 | 0.25 | 0.11 | 1.44 |
| 13 | Zc2hc1a | 0.40 | 1.36E-08 | 2.82E-04 | 0.54 | 0.42 | 0.12 | 1.29 |
| 13 | Flot2 | 0.48 | 1.44E-08 | 2.99E-04 | 0.32 | 0.21 | 0.10 | 1.49 |
| 13 | Cobl | 0.57 | 1.52E-08 | 3.16E-04 | 0.13 | 0.07 | 0.06 | 1.91 |
| 13 | Hdhd3 | 0.57 | 1.53E-08 | 3.18E-04 | 0.25 | 0.16 | 0.09 | 1.55 |
| 13 | Dvl1 | 0.63 | 1.58E-08 | 3.28E-04 | 0.21 | 0.13 | 0.08 | 1.63 |
| 13 | Chmp2a | 0.26 | 1.58E-08 | 3.30E-04 | 0.76 | 0.66 | 0.10 | 1.15 |
| 13 | Hpcal1 | 0.37 | 1.60E-08 | 3.32E-04 | 0.45 | 0.33 | 0.12 | 1.36 |
| 13 | Etv3 | 0.53 | 1.66E-08 | 3.45E-04 | 0.33 | 0.22 | 0.11 | 1.48 |
| 13 | Ttc32 | 0.39 | 1.66E-08 | 3.45E-04 | 0.37 | 0.25 | 0.11 | 1.44 |
| 13 | Pomgnt2 | 0.78 | 1.69E-08 | 3.51E-04 | 0.11 | 0.06 | 0.06 | 1.97 |
| 13 | Qars | 0.31 | 1.71E-08 | 3.56E-04 | 0.70 | 0.59 | 0.11 | 1.19 |
| 13 | Myef2 | 0.26 | 1.73E-08 | 3.59E-04 | 0.78 | 0.69 | 0.10 | 1.14 |
| 13 | Klhl24 | 0.39 | 1.78E-08 | 3.70E-04 | 0.70 | 0.60 | 0.10 | 1.17 |
| 13 | Dipk1b | 0.29 | 1.87E-08 | 3.88E-04 | 0.61 | 0.48 | 0.14 | 1.29 |
| 13 | Eipr1 | 0.48 | 1.88E-08 | 3.91E-04 | 0.35 | 0.24 | 0.11 | 1.45 |
| 13 | Atp11b | 0.34 | 2.00E-08 | 4.16E-04 | 0.54 | 0.42 | 0.12 | 1.29 |
| 13 | Bcar3 | 0.85 | 2.00E-08 | 4.16E-04 | 0.10 | 0.05 | 0.05 | 2.04 |
| 13 | Enc1 | 0.28 | 2.03E-08 | 4.22E-04 | 0.33 | 0.22 | 0.11 | 1.48 |
| 13 | Ydjc | 0.61 | 2.06E-08 | 4.28E-04 | 0.15 | 0.08 | 0.07 | 1.81 |
| 13 | Sdf2 | 0.35 | 2.06E-08 | 4.29E-04 | 0.69 | 0.56 | 0.13 | 1.23 |
| 13 | Ino80d | 0.37 | 2.09E-08 | 4.34E-04 | 0.47 | 0.34 | 0.13 | 1.38 |
| 13 | Socs7 | 0.42 | 2.09E-08 | 4.35E-04 | 0.32 | 0.21 | 0.11 | 1.50 |
| 13 | Esrra | 0.66 | 2.10E-08 | 4.36E-04 | 0.20 | 0.12 | 0.08 | 1.64 |
| 13 | Ccpg1os | 0.47 | 2.14E-08 | 4.45E-04 | 0.25 | 0.16 | 0.09 | 1.57 |
| 13 | Traf6 | 0.67 | 2.16E-08 | 4.49E-04 | 0.23 | 0.15 | 0.09 | 1.59 |
| 13 | Vps13b | 0.50 | 2.20E-08 | 4.57E-04 | 0.28 | 0.18 | 0.10 | 1.53 |
| 13 | Med13 | 0.36 | 2.24E-08 | 4.65E-04 | 0.67 | 0.55 | 0.12 | 1.21 |
| 13 | Tjap1 | 0.44 | 2.28E-08 | 4.73E-04 | 0.46 | 0.34 | 0.12 | 1.35 |
| 13 | 2510002D24Rik | 0.41 | 2.36E-08 | 4.91E-04 | 0.51 | 0.39 | 0.12 | 1.30 |
| 13 | Ric3 | 0.93 | 2.39E-08 | 4.98E-04 | 0.10 | 0.05 | 0.05 | 2.04 |
| 13 | Slc45a4 | 0.45 | 2.41E-08 | 5.01E-04 | 0.22 | 0.13 | 0.09 | 1.65 |
| 13 | Sall2 | 0.37 | 2.43E-08 | 5.06E-04 | 0.52 | 0.39 | 0.13 | 1.33 |
| 13 | Plxdc2 | 0.47 | 2.50E-08 | 5.20E-04 | 0.36 | 0.25 | 0.11 | 1.44 |
| 13 | Smox | 0.53 | 2.60E-08 | 5.40E-04 | 0.34 | 0.24 | 0.10 | 1.44 |
| 13 | Mrpl44 | 0.39 | 2.69E-08 | 5.59E-04 | 0.53 | 0.41 | 0.11 | 1.28 |
| 13 | Prss36 | 0.53 | 2.70E-08 | 5.62E-04 | 0.16 | 0.09 | 0.07 | 1.78 |
| 13 | Rnf38 | 0.46 | 2.71E-08 | 5.63E-04 | 0.38 | 0.27 | 0.11 | 1.41 |
| 13 | Rnf6 | 0.34 | 2.72E-08 | 5.65E-04 | 0.48 | 0.35 | 0.13 | 1.36 |
| 13 | Msi1 | 0.36 | 2.72E-08 | 5.66E-04 | 0.51 | 0.37 | 0.14 | 1.37 |
| 13 | Kifc3 | 0.86 | 2.84E-08 | 5.92E-04 | 0.14 | 0.07 | 0.06 | 1.82 |
| 13 | Kdm3a | 0.37 | 2.88E-08 | 6.00E-04 | 0.49 | 0.37 | 0.12 | 1.33 |
| 13 | Atmin | 0.57 | 2.93E-08 | 6.10E-04 | 0.22 | 0.14 | 0.08 | 1.60 |
| 13 | Reep4 | 0.45 | 3.00E-08 | 6.24E-04 | 0.44 | 0.33 | 0.11 | 1.35 |
| 13 | Vps13a | 0.36 | 3.03E-08 | 6.29E-04 | 0.47 | 0.34 | 0.13 | 1.37 |

**Supplementary Table 1. Marker genes for the original clusters of integrated clustering of E11.5 control and NotchGOF λ-junction cells.**

| | | | | | | | | |
|---|---|---|---|---|---|---|---|---|
| 13 | Ino80b | 0.38 | 3.09E-08 | 6.42E-04 | 0.57 | 0.45 | 0.11 | 1.25 |
| 13 | Bdh1 | 0.61 | 3.11E-08 | 6.46E-04 | 0.13 | 0.07 | 0.06 | 1.86 |
| 13 | Zfand1 | 0.40 | 3.14E-08 | 6.52E-04 | 0.47 | 0.35 | 0.12 | 1.35 |
| 13 | Vezt | 0.33 | 3.17E-08 | 6.59E-04 | 0.44 | 0.32 | 0.12 | 1.38 |
| 13 | Usp20 | 0.88 | 3.21E-08 | 6.69E-04 | 0.14 | 0.08 | 0.06 | 1.82 |
| 13 | Nr2c2ap | 0.43 | 3.23E-08 | 6.72E-04 | 0.33 | 0.23 | 0.10 | 1.45 |
| 13 | Itsn2 | 0.43 | 3.26E-08 | 6.78E-04 | 0.29 | 0.19 | 0.10 | 1.52 |
| 13 | Tvp23b | 0.36 | 3.34E-08 | 6.95E-04 | 0.54 | 0.41 | 0.13 | 1.33 |
| 13 | Stk11 | 0.34 | 3.39E-08 | 7.05E-04 | 0.60 | 0.48 | 0.12 | 1.26 |
| 13 | AW146154 | 0.81 | 3.45E-08 | 7.17E-04 | 0.15 | 0.09 | 0.06 | 1.74 |
| 13 | Aasdh | 0.74 | 3.48E-08 | 7.23E-04 | 0.16 | 0.09 | 0.07 | 1.74 |
| 13 | Dcaf13 | 0.30 | 3.54E-08 | 7.36E-04 | 0.59 | 0.46 | 0.13 | 1.28 |
| 13 | Gm26930 | 0.86 | 3.61E-08 | 7.51E-04 | 0.16 | 0.09 | 0.07 | 1.74 |
| 13 | Cbarp | 0.74 | 3.62E-08 | 7.53E-04 | 0.18 | 0.11 | 0.07 | 1.67 |
| 13 | Parp8 | 0.64 | 3.68E-08 | 7.65E-04 | 0.22 | 0.14 | 0.08 | 1.60 |
| 13 | Ebf4 | 0.79 | 3.72E-08 | 7.74E-04 | 0.11 | 0.06 | 0.06 | 1.96 |
| 13 | Abcb7 | 0.54 | 3.74E-08 | 7.78E-04 | 0.30 | 0.20 | 0.10 | 1.49 |
| 13 | BC052040 | 0.46 | 3.82E-08 | 7.94E-04 | 0.27 | 0.18 | 0.09 | 1.54 |
| 13 | Plek2 | 0.64 | 3.93E-08 | 8.17E-04 | 0.12 | 0.06 | 0.06 | 1.92 |
| 13 | Rnaseh2a | 0.31 | 4.02E-08 | 8.37E-04 | 0.67 | 0.57 | 0.10 | 1.18 |
| 13 | Rmnd5b | 0.49 | 4.08E-08 | 8.49E-04 | 0.26 | 0.17 | 0.09 | 1.54 |
| 13 | 1300002E11Rik | 0.40 | 4.11E-08 | 8.55E-04 | 0.42 | 0.32 | 0.10 | 1.32 |
| 13 | Sh3glb2 | 0.41 | 4.11E-08 | 8.56E-04 | 0.43 | 0.32 | 0.12 | 1.36 |
| 13 | Use1 | 0.28 | 4.19E-08 | 8.71E-04 | 0.75 | 0.65 | 0.10 | 1.16 |
| 13 | Wbp2 | 0.40 | 4.32E-08 | 8.99E-04 | 0.38 | 0.28 | 0.11 | 1.39 |
| 13 | Ccdc12 | 0.25 | 4.36E-08 | 9.08E-04 | 0.79 | 0.70 | 0.09 | 1.13 |
| 13 | Lats1 | 0.44 | 4.38E-08 | 9.11E-04 | 0.36 | 0.26 | 0.10 | 1.40 |
| 13 | Mmp11 | 0.40 | 4.46E-08 | 9.27E-04 | 0.43 | 0.32 | 0.11 | 1.36 |
| 13 | Wrnip1 | 0.55 | 4.47E-08 | 9.30E-04 | 0.29 | 0.20 | 0.10 | 1.48 |
| 13 | Adamts6 | 0.92 | 4.60E-08 | 9.56E-04 | 0.12 | 0.07 | 0.06 | 1.88 |
| 13 | Mid1 | 0.45 | 4.79E-08 | 9.96E-04 | 0.35 | 0.24 | 0.11 | 1.44 |
| 13 | Hipk2 | 0.28 | 4.85E-08 | 1.01E-03 | 0.75 | 0.64 | 0.10 | 1.16 |
| 13 | Cux2 | 0.54 | 4.88E-08 | 1.02E-03 | 0.17 | 0.10 | 0.07 | 1.73 |
| 13 | Mrpl36 | 0.29 | 4.95E-08 | 1.03E-03 | 0.78 | 0.72 | 0.07 | 1.10 |
| 13 | Cpt1a | 0.70 | 4.99E-08 | 1.04E-03 | 0.17 | 0.10 | 0.07 | 1.70 |
| 13 | Hsdl2 | 0.49 | 5.03E-08 | 1.05E-03 | 0.37 | 0.27 | 0.10 | 1.39 |
| 13 | Madd | 0.82 | 5.05E-08 | 1.05E-03 | 0.13 | 0.07 | 0.06 | 1.83 |
| 13 | 4930513N10Rik | 0.96 | 5.18E-08 | 1.08E-03 | 0.10 | 0.05 | 0.05 | 2.00 |
| 13 | Rcan3 | 0.69 | 5.29E-08 | 1.10E-03 | 0.14 | 0.08 | 0.06 | 1.79 |
| 13 | AU040320 | 0.62 | 5.38E-08 | 1.12E-03 | 0.18 | 0.11 | 0.07 | 1.66 |
| 13 | Lin37 | 0.37 | 5.68E-08 | 1.18E-03 | 0.34 | 0.24 | 0.10 | 1.44 |
| 13 | Otud4 | 0.52 | 5.81E-08 | 1.21E-03 | 0.42 | 0.31 | 0.11 | 1.36 |
| 13 | Imp4 | 0.29 | 5.85E-08 | 1.22E-03 | 0.67 | 0.53 | 0.14 | 1.26 |
| 13 | Mboat2 | 0.35 | 5.98E-08 | 1.24E-03 | 0.47 | 0.35 | 0.13 | 1.37 |
| 13 | Mga | 0.28 | 6.09E-08 | 1.27E-03 | 0.70 | 0.58 | 0.13 | 1.22 |
| 13 | Aldh7a1 | 0.31 | 6.16E-08 | 1.28E-03 | 0.68 | 0.56 | 0.12 | 1.22 |
| 13 | Nes | 0.29 | 6.28E-08 | 1.31E-03 | 0.55 | 0.42 | 0.13 | 1.30 |
| 13 | Fahd2a | 0.51 | 6.32E-08 | 1.31E-03 | 0.28 | 0.19 | 0.09 | 1.47 |
| 13 | Fam169a | 0.46 | 6.38E-08 | 1.33E-03 | 0.22 | 0.13 | 0.08 | 1.62 |
| 13 | Strn3 | 0.27 | 6.44E-08 | 1.34E-03 | 0.84 | 0.75 | 0.10 | 1.13 |
| 13 | Pigs | 0.38 | 6.61E-08 | 1.37E-03 | 0.42 | 0.30 | 0.12 | 1.40 |
| 13 | Nudt11 | 0.39 | 6.70E-08 | 1.39E-03 | 0.27 | 0.18 | 0.09 | 1.53 |
| 13 | Zfp626 | 0.57 | 6.88E-08 | 1.43E-03 | 0.21 | 0.13 | 0.08 | 1.60 |
| 13 | Phkg2 | 0.46 | 6.97E-08 | 1.45E-03 | 0.34 | 0.24 | 0.10 | 1.43 |
| 13 | Gpbp1l1 | 0.39 | 7.06E-08 | 1.47E-03 | 0.50 | 0.38 | 0.12 | 1.31 |
| 13 | Rbfa | 0.37 | 7.07E-08 | 1.47E-03 | 0.42 | 0.31 | 0.12 | 1.37 |

**Supplementary Table 1. Marker genes for the original clusters of integrated clustering of E11.5 control and NotchGOF λ-junction cells.**

| | | | | | | | | |
|---|---|---|---|---|---|---|---|---|
| 13 | Lin28b | 0.43 | 7.09E-08 | 1.47E-03 | 0.31 | 0.21 | 0.10 | 1.49 |
| 13 | Rdh14 | 0.37 | 7.23E-08 | 1.50E-03 | 0.39 | 0.28 | 0.12 | 1.42 |
| 13 | Scarb1 | 0.61 | 7.25E-08 | 1.51E-03 | 0.24 | 0.16 | 0.09 | 1.54 |
| 13 | Rab21 | 0.33 | 7.33E-08 | 1.53E-03 | 0.60 | 0.46 | 0.13 | 1.28 |
| 13 | Tmod1 | 0.95 | 7.66E-08 | 1.59E-03 | 0.12 | 0.06 | 0.05 | 1.85 |
| 13 | Utp4 | 0.38 | 7.68E-08 | 1.60E-03 | 0.41 | 0.31 | 0.11 | 1.36 |
| 13 | Nudt7 | 0.72 | 7.71E-08 | 1.60E-03 | 0.13 | 0.07 | 0.06 | 1.84 |
| 13 | Chd2 | 0.32 | 7.75E-08 | 1.61E-03 | 0.72 | 0.62 | 0.10 | 1.16 |
| 13 | Hcfc1r1 | 0.33 | 7.77E-08 | 1.62E-03 | 0.69 | 0.58 | 0.11 | 1.19 |
| 13 | Jmjd8 | 0.30 | 7.96E-08 | 1.65E-03 | 0.40 | 0.28 | 0.12 | 1.42 |
| 13 | Zadh2 | 0.60 | 8.42E-08 | 1.75E-03 | 0.18 | 0.11 | 0.07 | 1.66 |
| 13 | Spry1 | 0.32 | 8.84E-08 | 1.84E-03 | 0.40 | 0.29 | 0.12 | 1.41 |
| 13 | Snx1 | 0.34 | 9.18E-08 | 1.91E-03 | 0.56 | 0.44 | 0.12 | 1.26 |
| 13 | Macrod1 | 0.67 | 9.25E-08 | 1.92E-03 | 0.13 | 0.07 | 0.06 | 1.82 |
| 13 | Tada3 | 0.39 | 9.60E-08 | 2.00E-03 | 0.47 | 0.36 | 0.12 | 1.32 |
| 13 | Paf1 | 0.39 | 9.73E-08 | 2.02E-03 | 0.54 | 0.43 | 0.11 | 1.26 |
| 13 | B4galt4 | 0.43 | 9.74E-08 | 2.03E-03 | 0.33 | 0.23 | 0.10 | 1.45 |
| 13 | Zfp652 | 0.29 | 1.01E-07 | 2.11E-03 | 0.62 | 0.51 | 0.11 | 1.22 |
| 13 | Zfp113 | 0.35 | 1.04E-07 | 2.16E-03 | 0.44 | 0.32 | 0.12 | 1.36 |
| 13 | Pcdh11x | 0.33 | 1.04E-07 | 2.17E-03 | 0.18 | 0.11 | 0.07 | 1.68 |
| 13 | Cdkl5 | 0.85 | 1.07E-07 | 2.22E-03 | 0.11 | 0.06 | 0.05 | 1.91 |
| 13 | Pde7a | 0.29 | 1.07E-07 | 2.23E-03 | 0.69 | 0.54 | 0.14 | 1.26 |
| 13 | Ccdc43 | 0.33 | 1.11E-07 | 2.31E-03 | 0.54 | 0.41 | 0.13 | 1.32 |
| 13 | Ankra2 | 0.36 | 1.16E-07 | 2.40E-03 | 0.43 | 0.32 | 0.12 | 1.37 |
| 13 | Prkcd | 0.48 | 1.22E-07 | 2.53E-03 | 0.31 | 0.21 | 0.09 | 1.44 |
| 13 | Aptx | 0.48 | 1.27E-07 | 2.64E-03 | 0.36 | 0.26 | 0.10 | 1.40 |
| 13 | Kyat1 | 0.78 | 1.27E-07 | 2.65E-03 | 0.10 | 0.05 | 0.05 | 1.96 |
| 13 | Dhdds | 0.34 | 1.29E-07 | 2.68E-03 | 0.40 | 0.28 | 0.11 | 1.40 |
| 13 | AW549877 | 0.50 | 1.34E-07 | 2.78E-03 | 0.31 | 0.21 | 0.10 | 1.46 |
| 13 | Tmem14a | 0.67 | 1.45E-07 | 3.02E-03 | 0.17 | 0.10 | 0.07 | 1.67 |
| 13 | Zfyve1 | 0.47 | 1.47E-07 | 3.05E-03 | 0.23 | 0.15 | 0.08 | 1.55 |
| 13 | Atf6b | 0.38 | 1.48E-07 | 3.08E-03 | 0.53 | 0.40 | 0.13 | 1.32 |
| 13 | Cdk17 | 0.46 | 1.58E-07 | 3.30E-03 | 0.30 | 0.21 | 0.09 | 1.45 |
| 13 | Cbx6 | 0.37 | 1.62E-07 | 3.37E-03 | 0.38 | 0.27 | 0.11 | 1.42 |
| 13 | Ncoa2 | 0.42 | 1.63E-07 | 3.39E-03 | 0.40 | 0.29 | 0.11 | 1.39 |
| 13 | Brms1 | 0.33 | 1.69E-07 | 3.51E-03 | 0.46 | 0.35 | 0.11 | 1.33 |
| 13 | Idnk | 0.33 | 1.73E-07 | 3.59E-03 | 0.52 | 0.41 | 0.12 | 1.28 |
| 13 | Il10rb | 0.88 | 1.76E-07 | 3.66E-03 | 0.15 | 0.09 | 0.06 | 1.73 |
| 13 | Trmt2b | 0.52 | 1.79E-07 | 3.72E-03 | 0.15 | 0.09 | 0.07 | 1.74 |
| 13 | Nras | 0.28 | 1.79E-07 | 3.73E-03 | 0.79 | 0.69 | 0.11 | 1.15 |
| 13 | Polr3h | 0.31 | 1.81E-07 | 3.76E-03 | 0.50 | 0.38 | 0.12 | 1.31 |
| 13 | Sbno2 | 0.49 | 1.90E-07 | 3.96E-03 | 0.19 | 0.11 | 0.07 | 1.65 |
| 13 | Cyp4f16 | 0.67 | 1.91E-07 | 3.98E-03 | 0.17 | 0.10 | 0.07 | 1.68 |
| 13 | Mindy2 | 0.55 | 1.91E-07 | 3.98E-03 | 0.26 | 0.17 | 0.09 | 1.49 |
| 13 | Zfp692 | 0.65 | 1.92E-07 | 4.00E-03 | 0.18 | 0.11 | 0.07 | 1.65 |
| 13 | Tmem170 | 0.57 | 1.94E-07 | 4.04E-03 | 0.16 | 0.09 | 0.07 | 1.70 |
| 13 | Atg13 | 0.57 | 1.95E-07 | 4.06E-03 | 0.23 | 0.15 | 0.08 | 1.53 |
| 13 | Fbxo10 | 0.59 | 1.96E-07 | 4.07E-03 | 0.14 | 0.08 | 0.06 | 1.78 |
| 13 | Gcdh | 0.37 | 1.96E-07 | 4.07E-03 | 0.41 | 0.30 | 0.11 | 1.37 |
| 13 | Rpe | 0.41 | 1.97E-07 | 4.11E-03 | 0.44 | 0.34 | 0.11 | 1.31 |
| 13 | Rab38 | 0.39 | 1.99E-07 | 4.13E-03 | 0.13 | 0.07 | 0.06 | 1.84 |
| 13 | Flad1 | 0.56 | 1.99E-07 | 4.14E-03 | 0.17 | 0.10 | 0.07 | 1.66 |
| 13 | Ap1s3 | 0.60 | 2.01E-07 | 4.17E-03 | 0.12 | 0.07 | 0.06 | 1.85 |
| 13 | Ap2b1 | 0.29 | 2.02E-07 | 4.19E-03 | 0.60 | 0.49 | 0.11 | 1.22 |
| 13 | Mettl17 | 0.42 | 2.09E-07 | 4.34E-03 | 0.27 | 0.18 | 0.09 | 1.51 |
| 13 | Dbndd1 | 0.79 | 2.13E-07 | 4.42E-03 | 0.10 | 0.05 | 0.05 | 1.93 |

**Supplementary Table 1. Marker genes for the original clusters of integrated clustering of E11.5 control and NotchGOF λ-junction cells.**

| | | | | | | | | |
|---|---|---|---|---|---|---|---|---|
| 13 | Cs | 0.25 | 2.15E-07 | 4.46E-03 | 0.81 | 0.72 | 0.09 | 1.13 |
| 13 | Ppan | 0.33 | 2.16E-07 | 4.50E-03 | 0.56 | 0.45 | 0.11 | 1.25 |
| 13 | 1700025G04Rik | 0.36 | 2.17E-07 | 4.52E-03 | 0.45 | 0.34 | 0.12 | 1.34 |
| 13 | Dstyk | 0.43 | 2.17E-07 | 4.52E-03 | 0.33 | 0.23 | 0.10 | 1.44 |
| 13 | Slc44a1 | 0.43 | 2.22E-07 | 4.61E-03 | 0.33 | 0.24 | 0.10 | 1.42 |
| 13 | Atxn1 | 0.62 | 2.22E-07 | 4.61E-03 | 0.16 | 0.09 | 0.06 | 1.70 |
| 13 | Itprip | 0.75 | 2.31E-07 | 4.80E-03 | 0.14 | 0.08 | 0.06 | 1.75 |
| 13 | Nfia | 0.28 | 2.31E-07 | 4.81E-03 | 0.58 | 0.47 | 0.11 | 1.24 |
| 13 | Bok | 0.33 | 2.32E-07 | 4.82E-03 | 0.29 | 0.20 | 0.09 | 1.46 |
| 13 | Cdc73 | 0.30 | 2.32E-07 | 4.82E-03 | 0.59 | 0.48 | 0.12 | 1.25 |
| 13 | Socs3 | 0.46 | 2.32E-07 | 4.83E-03 | 0.26 | 0.17 | 0.09 | 1.50 |
| 13 | Zfp651 | 0.34 | 2.37E-07 | 4.92E-03 | 0.24 | 0.15 | 0.09 | 1.55 |
| 13 | Aldh4a1 | 0.73 | 2.38E-07 | 4.94E-03 | 0.16 | 0.09 | 0.06 | 1.69 |
| 13 | Ttc13 | 0.40 | 2.43E-07 | 5.05E-03 | 0.33 | 0.23 | 0.10 | 1.44 |
| 13 | Ttc39b | 0.91 | 2.46E-07 | 5.11E-03 | 0.12 | 0.07 | 0.06 | 1.85 |
| 13 | Prorp | 0.62 | 2.49E-07 | 5.18E-03 | 0.22 | 0.14 | 0.08 | 1.55 |
| 13 | Poll | 0.42 | 2.52E-07 | 5.24E-03 | 0.26 | 0.17 | 0.09 | 1.51 |
| 13 | Nfat5 | 0.34 | 2.54E-07 | 5.28E-03 | 0.68 | 0.57 | 0.11 | 1.18 |
| 13 | Proser2 | 0.62 | 2.61E-07 | 5.44E-03 | 0.11 | 0.06 | 0.05 | 1.91 |
| 13 | Gprc5b | 0.63 | 2.65E-07 | 5.52E-03 | 0.17 | 0.10 | 0.07 | 1.68 |
| 13 | Zfp995 | 0.68 | 2.67E-07 | 5.56E-03 | 0.14 | 0.08 | 0.06 | 1.75 |
| 13 | Nsmaf | 0.43 | 2.70E-07 | 5.62E-03 | 0.37 | 0.27 | 0.10 | 1.39 |
| 13 | Fam8a1 | 0.44 | 2.76E-07 | 5.73E-03 | 0.37 | 0.27 | 0.10 | 1.38 |
| 13 | Exd2 | 0.60 | 2.89E-07 | 6.02E-03 | 0.23 | 0.15 | 0.08 | 1.52 |
| 13 | Zfp622 | 0.31 | 2.94E-07 | 6.11E-03 | 0.53 | 0.42 | 0.11 | 1.27 |
| 13 | Hivep2 | 0.64 | 3.00E-07 | 6.24E-03 | 0.13 | 0.08 | 0.06 | 1.77 |
| 13 | Bin3 | 0.40 | 3.01E-07 | 6.26E-03 | 0.39 | 0.29 | 0.10 | 1.35 |
| 13 | Sema3f | 0.41 | 3.01E-07 | 6.26E-03 | 0.39 | 0.29 | 0.10 | 1.35 |
| 13 | Nnat | 0.50 | 3.05E-07 | 6.35E-03 | 0.99 | 0.94 | 0.05 | 1.05 |
| 13 | Slc7a6 | 0.48 | 3.10E-07 | 6.45E-03 | 0.38 | 0.28 | 0.10 | 1.36 |
| 13 | Map2 | 0.38 | 3.12E-07 | 6.49E-03 | 0.31 | 0.22 | 0.09 | 1.43 |
| 13 | Reps1 | 0.37 | 3.13E-07 | 6.51E-03 | 0.46 | 0.35 | 0.11 | 1.32 |
| 13 | Ankhd1 | 0.30 | 3.16E-07 | 6.57E-03 | 0.60 | 0.49 | 0.11 | 1.22 |
| 13 | Slc38a10 | 0.36 | 3.20E-07 | 6.66E-03 | 0.45 | 0.34 | 0.11 | 1.34 |
| 13 | Bmt2 | 0.37 | 3.29E-07 | 6.84E-03 | 0.43 | 0.32 | 0.11 | 1.35 |
| 13 | Ccl27a | 0.36 | 3.29E-07 | 6.85E-03 | 0.40 | 0.30 | 0.11 | 1.35 |
| 13 | Klhl2 | 0.60 | 3.31E-07 | 6.88E-03 | 0.19 | 0.12 | 0.07 | 1.60 |
| 13 | Tapt1 | 0.68 | 3.39E-07 | 7.05E-03 | 0.23 | 0.15 | 0.08 | 1.51 |
| 13 | Fuz | 0.63 | 3.51E-07 | 7.31E-03 | 0.17 | 0.11 | 0.07 | 1.65 |
| 13 | Thoc6 | 0.31 | 3.52E-07 | 7.32E-03 | 0.59 | 0.48 | 0.11 | 1.23 |
| 13 | Pex12 | 0.72 | 3.54E-07 | 7.35E-03 | 0.16 | 0.09 | 0.06 | 1.67 |
| 13 | Minar2 | 0.56 | 3.59E-07 | 7.47E-03 | 0.16 | 0.10 | 0.07 | 1.68 |
| 13 | Ero1l | 0.44 | 3.71E-07 | 7.71E-03 | 0.40 | 0.29 | 0.11 | 1.36 |
| 13 | Gba | 0.74 | 3.82E-07 | 7.95E-03 | 0.16 | 0.10 | 0.06 | 1.66 |
| 13 | Fam120b | 0.37 | 3.82E-07 | 7.95E-03 | 0.36 | 0.26 | 0.10 | 1.38 |
| 13 | Snx16 | 0.40 | 3.85E-07 | 8.01E-03 | 0.28 | 0.19 | 0.09 | 1.47 |
| 13 | Zmym2 | 0.30 | 3.89E-07 | 8.10E-03 | 0.65 | 0.53 | 0.12 | 1.23 |
| 13 | Plpp6 | 0.72 | 4.06E-07 | 8.43E-03 | 0.12 | 0.07 | 0.05 | 1.82 |
| 13 | Pou2f1 | 0.37 | 4.14E-07 | 8.61E-03 | 0.46 | 0.36 | 0.11 | 1.29 |
| 13 | Wdr44 | 0.81 | 4.28E-07 | 8.89E-03 | 0.11 | 0.06 | 0.05 | 1.84 |
| 13 | Bbc3 | 0.69 | 4.30E-07 | 8.94E-03 | 0.17 | 0.10 | 0.06 | 1.63 |
| 13 | Aebp2 | 0.29 | 4.33E-07 | 9.01E-03 | 0.54 | 0.42 | 0.12 | 1.29 |
| 13 | Erp44 | 0.30 | 4.45E-07 | 9.25E-03 | 0.63 | 0.51 | 0.13 | 1.25 |
| 13 | Zfp654 | 0.53 | 4.47E-07 | 9.29E-03 | 0.22 | 0.15 | 0.08 | 1.51 |
| 13 | Zc3h12c | 0.95 | 4.62E-07 | 9.61E-03 | 0.11 | 0.06 | 0.05 | 1.84 |
| 13 | Alpl | 0.47 | 4.72E-07 | 9.82E-03 | 0.17 | 0.10 | 0.07 | 1.67 |

**Supplementary Table 1. Marker genes for the original clusters of integrated clustering of E11.5 control and NotchGOF λ-junction cells.**

| | | | | | | | | |
|---|---|---|---|---|---|---|---|---|
| 13 | Nsun2 | 0.28 | 4.79E-07 | 9.97E-03 | 0.74 | 0.64 | 0.10 | 1.16 |
| 13 | Usp49 | 0.38 | 4.80E-07 | 9.99E-03 | 0.19 | 0.11 | 0.07 | 1.64 |
| 13 | Slc29a3 | 0.54 | 4.87E-07 | 1.01E-02 | 0.17 | 0.10 | 0.07 | 1.64 |
| 13 | Mybl1 | 0.90 | 4.91E-07 | 1.02E-02 | 0.11 | 0.06 | 0.05 | 1.85 |
| 13 | Btbd9 | 0.29 | 4.94E-07 | 1.03E-02 | 0.41 | 0.29 | 0.11 | 1.38 |
| 13 | Glb1 | 0.59 | 5.12E-07 | 1.07E-02 | 0.24 | 0.16 | 0.08 | 1.49 |
| 13 | Rab11fip2 | 0.34 | 5.14E-07 | 1.07E-02 | 0.48 | 0.37 | 0.12 | 1.32 |
| 13 | Ankzf1 | 0.45 | 5.48E-07 | 1.14E-02 | 0.31 | 0.22 | 0.09 | 1.42 |
| 13 | Commd9 | 0.25 | 5.64E-07 | 1.17E-02 | 0.62 | 0.49 | 0.13 | 1.26 |
| 13 | Senp8 | 0.43 | 5.95E-07 | 1.24E-02 | 0.30 | 0.21 | 0.09 | 1.44 |
| 13 | Zfp568 | 0.39 | 6.01E-07 | 1.25E-02 | 0.37 | 0.27 | 0.10 | 1.37 |
| 13 | Tmem192 | 0.58 | 6.07E-07 | 1.26E-02 | 0.27 | 0.18 | 0.09 | 1.47 |
| 13 | Nbeal1 | 0.37 | 6.09E-07 | 1.27E-02 | 0.56 | 0.45 | 0.11 | 1.24 |
| 13 | Ascc1 | 0.33 | 6.10E-07 | 1.27E-02 | 0.51 | 0.39 | 0.11 | 1.29 |
| 13 | Chmp1b | 0.33 | 6.11E-07 | 1.27E-02 | 0.46 | 0.35 | 0.11 | 1.32 |
| 13 | 2310061I04Rik | 0.49 | 6.11E-07 | 1.27E-02 | 0.27 | 0.19 | 0.09 | 1.46 |
| 13 | Ccdc18 | 0.39 | 6.14E-07 | 1.28E-02 | 0.40 | 0.30 | 0.10 | 1.33 |
| 13 | Fam184a | 0.53 | 6.19E-07 | 1.29E-02 | 0.19 | 0.12 | 0.07 | 1.61 |
| 13 | Ddhd1 | 0.59 | 6.23E-07 | 1.29E-02 | 0.25 | 0.17 | 0.08 | 1.47 |
| 13 | Atp9a | 0.50 | 6.26E-07 | 1.30E-02 | 0.26 | 0.18 | 0.08 | 1.46 |
| 13 | 4930452B06Rik | 0.53 | 6.29E-07 | 1.31E-02 | 0.19 | 0.12 | 0.07 | 1.58 |
| 13 | Dph6 | 0.37 | 6.31E-07 | 1.31E-02 | 0.52 | 0.41 | 0.11 | 1.27 |
| 13 | Kdm4c | 0.59 | 6.33E-07 | 1.32E-02 | 0.18 | 0.11 | 0.07 | 1.59 |
| 13 | Cyb5b | 0.38 | 6.33E-07 | 1.32E-02 | 0.61 | 0.50 | 0.11 | 1.21 |
| 13 | Ank2 | 0.37 | 6.42E-07 | 1.34E-02 | 0.33 | 0.24 | 0.09 | 1.37 |
| 13 | Tab3 | 0.67 | 6.50E-07 | 1.35E-02 | 0.15 | 0.09 | 0.06 | 1.68 |
| 13 | Pea15a | 0.32 | 6.64E-07 | 1.38E-02 | 0.33 | 0.23 | 0.10 | 1.41 |
| 13 | Mlh3 | 0.46 | 6.81E-07 | 1.42E-02 | 0.25 | 0.17 | 0.08 | 1.48 |
| 13 | Paip2b | 0.39 | 7.09E-07 | 1.48E-02 | 0.40 | 0.29 | 0.10 | 1.35 |
| 13 | 2610306M01Rik | 0.47 | 7.10E-07 | 1.48E-02 | 0.24 | 0.16 | 0.08 | 1.49 |
| 13 | Alkbh6 | 0.26 | 7.12E-07 | 1.48E-02 | 0.56 | 0.44 | 0.13 | 1.29 |
| 13 | Fbxl6 | 0.38 | 7.35E-07 | 1.53E-02 | 0.43 | 0.33 | 0.10 | 1.31 |
| 13 | Kantr | 0.47 | 7.42E-07 | 1.54E-02 | 0.25 | 0.17 | 0.08 | 1.48 |
| 13 | Tigar | 0.77 | 7.44E-07 | 1.55E-02 | 0.13 | 0.07 | 0.05 | 1.76 |
| 13 | Mdm2 | 0.29 | 7.55E-07 | 1.57E-02 | 0.57 | 0.43 | 0.13 | 1.31 |
| 13 | Tgif1 | 0.30 | 7.73E-07 | 1.61E-02 | 0.73 | 0.63 | 0.10 | 1.16 |
| 13 | Pdcd2 | 0.34 | 7.91E-07 | 1.64E-02 | 0.49 | 0.39 | 0.11 | 1.28 |
| 13 | Gm2a | 0.53 | 8.32E-07 | 1.73E-02 | 0.13 | 0.07 | 0.06 | 1.76 |
| 13 | Rnf141 | 0.59 | 8.34E-07 | 1.73E-02 | 0.19 | 0.12 | 0.07 | 1.57 |
| 13 | Mmp15 | 0.55 | 8.53E-07 | 1.77E-02 | 0.15 | 0.09 | 0.06 | 1.68 |
| 13 | Prkx | 0.54 | 8.63E-07 | 1.79E-02 | 0.23 | 0.15 | 0.08 | 1.50 |
| 13 | Txnrd3 | 0.45 | 8.67E-07 | 1.80E-02 | 0.19 | 0.12 | 0.07 | 1.59 |
| 13 | Clstn3 | 0.77 | 8.68E-07 | 1.80E-02 | 0.11 | 0.06 | 0.05 | 1.84 |
| 13 | Airn | 0.32 | 9.00E-07 | 1.87E-02 | 0.28 | 0.19 | 0.09 | 1.48 |
| 13 | Fryl | 0.38 | 9.08E-07 | 1.89E-02 | 0.40 | 0.30 | 0.10 | 1.33 |
| 13 | Slc38a9 | 0.39 | 9.09E-07 | 1.89E-02 | 0.27 | 0.19 | 0.09 | 1.46 |
| 13 | Dtnb | 0.41 | 9.15E-07 | 1.90E-02 | 0.37 | 0.27 | 0.10 | 1.35 |
| 13 | Sptlc1 | 0.32 | 9.61E-07 | 2.00E-02 | 0.40 | 0.29 | 0.11 | 1.38 |
| 13 | Tert | 0.82 | 9.80E-07 | 2.04E-02 | 0.17 | 0.10 | 0.06 | 1.63 |
| 13 | Cxxc5 | 0.30 | 9.84E-07 | 2.05E-02 | 0.56 | 0.45 | 0.11 | 1.25 |
| 13 | Tdrd3 | 0.32 | 1.00E-06 | 2.08E-02 | 0.45 | 0.34 | 0.11 | 1.33 |
| 13 | Fktn | 0.27 | 1.00E-06 | 2.08E-02 | 0.36 | 0.26 | 0.11 | 1.41 |
| 13 | Maf1 | 0.28 | 1.03E-06 | 2.14E-02 | 0.63 | 0.53 | 0.10 | 1.20 |
| 13 | Abca1 | 0.30 | 1.04E-06 | 2.16E-02 | 0.12 | 0.07 | 0.05 | 1.77 |
| 13 | Slc22a17 | 0.28 | 1.08E-06 | 2.24E-02 | 0.46 | 0.34 | 0.11 | 1.33 |
| 13 | Srgap1 | 0.30 | 1.08E-06 | 2.25E-02 | 0.45 | 0.34 | 0.11 | 1.32 |

**Supplementary Table 1. Marker genes for the original clusters of integrated clustering of E11.5 control and NotchGOF λ-junction cells.**

| | | | | | | | | |
|---|---|---|---|---|---|---|---|---|
| 13 | Zfp367 | 0.58 | 1.13E-06 | 2.36E-02 | 0.22 | 0.14 | 0.07 | 1.51 |
| 13 | Cep164 | 0.33 | 1.14E-06 | 2.36E-02 | 0.71 | 0.61 | 0.10 | 1.16 |
| 13 | Abi1 | 0.34 | 1.14E-06 | 2.37E-02 | 0.50 | 0.39 | 0.11 | 1.30 |
| 13 | Cnot1 | 0.26 | 1.14E-06 | 2.38E-02 | 0.76 | 0.66 | 0.10 | 1.16 |
| 13 | Plxnb2 | 0.30 | 1.15E-06 | 2.38E-02 | 0.65 | 0.53 | 0.12 | 1.22 |
| 13 | Dcaf4 | 0.42 | 1.15E-06 | 2.38E-02 | 0.33 | 0.24 | 0.09 | 1.38 |
| 13 | Ano10 | 0.62 | 1.15E-06 | 2.39E-02 | 0.20 | 0.13 | 0.07 | 1.56 |
| 13 | Ddah1 | 0.46 | 1.15E-06 | 2.40E-02 | 0.21 | 0.14 | 0.08 | 1.54 |
| 13 | Gm3448 | 0.80 | 1.16E-06 | 2.41E-02 | 0.10 | 0.06 | 0.05 | 1.86 |
| 13 | Abhd4 | 0.50 | 1.17E-06 | 2.42E-02 | 0.21 | 0.14 | 0.07 | 1.53 |
| 13 | Atpsckmt | 0.32 | 1.19E-06 | 2.48E-02 | 0.38 | 0.28 | 0.10 | 1.38 |
| 13 | Pnpla6 | 0.66 | 1.21E-06 | 2.51E-02 | 0.15 | 0.09 | 0.06 | 1.67 |
| 13 | Pik3c2a | 0.40 | 1.26E-06 | 2.61E-02 | 0.52 | 0.41 | 0.11 | 1.26 |
| 13 | Tmem203 | 0.44 | 1.29E-06 | 2.69E-02 | 0.31 | 0.22 | 0.09 | 1.39 |
| 13 | Lamb2 | 0.42 | 1.31E-06 | 2.72E-02 | 0.20 | 0.13 | 0.07 | 1.56 |
| 13 | Mief1 | 0.45 | 1.32E-06 | 2.74E-02 | 0.24 | 0.16 | 0.08 | 1.48 |
| 13 | Lmbrd1 | 0.45 | 1.33E-06 | 2.77E-02 | 0.31 | 0.22 | 0.09 | 1.39 |
| 13 | Uhmk1 | 0.48 | 1.34E-06 | 2.79E-02 | 0.29 | 0.21 | 0.09 | 1.41 |
| 13 | 5033430I15Rik | 0.51 | 1.34E-06 | 2.79E-02 | 0.10 | 0.05 | 0.05 | 1.85 |
| 13 | Atp6ap1 | 0.38 | 1.34E-06 | 2.80E-02 | 0.46 | 0.35 | 0.10 | 1.29 |
| 13 | Dtd2 | 0.35 | 1.34E-06 | 2.80E-02 | 0.42 | 0.32 | 0.10 | 1.31 |
| 13 | Etfrf1 | 0.37 | 1.37E-06 | 2.84E-02 | 0.31 | 0.22 | 0.09 | 1.41 |
| 13 | Grwd1 | 0.28 | 1.37E-06 | 2.85E-02 | 0.55 | 0.43 | 0.12 | 1.28 |
| 13 | Pgpep1 | 0.34 | 1.40E-06 | 2.92E-02 | 0.39 | 0.29 | 0.10 | 1.34 |
| 13 | Mlycd | 0.34 | 1.48E-06 | 3.07E-02 | 0.22 | 0.14 | 0.08 | 1.55 |
| 13 | Tmem159 | 0.26 | 1.48E-06 | 3.08E-02 | 0.37 | 0.26 | 0.11 | 1.41 |
| 13 | Dtd1 | 0.28 | 1.48E-06 | 3.08E-02 | 0.53 | 0.41 | 0.12 | 1.29 |
| 13 | Smad3 | 0.41 | 1.49E-06 | 3.09E-02 | 0.36 | 0.27 | 0.10 | 1.37 |
| 13 | Tmem38b | 0.40 | 1.49E-06 | 3.10E-02 | 0.23 | 0.15 | 0.08 | 1.54 |
| 13 | Zfp511 | 0.35 | 1.55E-06 | 3.22E-02 | 0.48 | 0.38 | 0.11 | 1.28 |
| 13 | Trmt1 | 0.30 | 1.55E-06 | 3.23E-02 | 0.54 | 0.43 | 0.12 | 1.27 |
| 13 | Tmem39b | 0.31 | 1.55E-06 | 3.23E-02 | 0.46 | 0.36 | 0.11 | 1.30 |
| 13 | Zscan21 | 0.27 | 1.61E-06 | 3.34E-02 | 0.52 | 0.40 | 0.12 | 1.29 |
| 13 | Dusp28 | 0.53 | 1.64E-06 | 3.40E-02 | 0.18 | 0.11 | 0.07 | 1.59 |
| 13 | Gm10130 | 0.51 | 1.64E-06 | 3.42E-02 | 0.15 | 0.09 | 0.06 | 1.65 |
| 13 | Gas2l3 | 0.52 | 1.65E-06 | 3.43E-02 | 0.26 | 0.18 | 0.08 | 1.42 |
| 13 | Skil | 0.36 | 1.65E-06 | 3.43E-02 | 0.65 | 0.55 | 0.10 | 1.18 |
| 13 | Aplf | 0.49 | 1.66E-06 | 3.45E-02 | 0.17 | 0.11 | 0.07 | 1.60 |
| 13 | Tbc1d23 | 0.40 | 1.67E-06 | 3.48E-02 | 0.35 | 0.26 | 0.09 | 1.36 |
| 13 | Otud7b | 0.34 | 1.81E-06 | 3.75E-02 | 0.49 | 0.38 | 0.10 | 1.27 |
| 13 | Zfp992 | 0.51 | 1.84E-06 | 3.82E-02 | 0.23 | 0.16 | 0.08 | 1.48 |
| 13 | Nol12 | 0.25 | 1.85E-06 | 3.84E-02 | 0.65 | 0.55 | 0.10 | 1.18 |
| 13 | BC004004 | 0.27 | 1.85E-06 | 3.84E-02 | 0.42 | 0.32 | 0.11 | 1.34 |
| 13 | Jmjd4 | 0.74 | 1.86E-06 | 3.86E-02 | 0.18 | 0.12 | 0.07 | 1.56 |
| 13 | Hyal2 | 0.25 | 1.92E-06 | 3.99E-02 | 0.64 | 0.53 | 0.11 | 1.21 |
| 13 | Surf2 | 0.30 | 1.94E-06 | 4.03E-02 | 0.50 | 0.38 | 0.11 | 1.30 |
| 13 | Galnt2 | 0.28 | 1.98E-06 | 4.11E-02 | 0.57 | 0.46 | 0.11 | 1.24 |
| 13 | Smpd1 | 0.37 | 1.98E-06 | 4.12E-02 | 0.28 | 0.19 | 0.09 | 1.46 |
| 13 | Gpr89 | 0.32 | 2.03E-06 | 4.23E-02 | 0.43 | 0.33 | 0.11 | 1.32 |
| 13 | Plin3 | 0.39 | 2.06E-06 | 4.29E-02 | 0.62 | 0.55 | 0.08 | 1.14 |
| 13 | Eif4ebp3 | 0.33 | 2.08E-06 | 4.32E-02 | 0.28 | 0.19 | 0.09 | 1.46 |
| 13 | Fbxo16 | 0.41 | 2.13E-06 | 4.43E-02 | 0.17 | 0.11 | 0.07 | 1.60 |
| 13 | Enox2 | 0.72 | 2.16E-06 | 4.49E-02 | 0.12 | 0.07 | 0.05 | 1.71 |
| 13 | Ikbkb | 0.38 | 2.24E-06 | 4.67E-02 | 0.41 | 0.32 | 0.10 | 1.31 |
| 13 | Trip11 | 0.31 | 2.27E-06 | 4.72E-02 | 0.61 | 0.49 | 0.12 | 1.23 |
| 13 | Adsl | 0.35 | 2.31E-06 | 4.79E-02 | 0.56 | 0.47 | 0.09 | 1.20 |

**Supplementary Table 1. Marker genes for the original clusters of integrated clustering of E11.5 control and NotchGOF λ-junction cells.**

| | | | | | | | | |
|---|---|---|---|---|---|---|---|---|
| 13 | Nbr1 | 0.40 | 2.32E-06 | 4.82E-02 | 0.49 | 0.39 | 0.10 | 1.26 |
| 13 | Tmem123 | 0.26 | 2.34E-06 | 4.86E-02 | 0.42 | 0.32 | 0.10 | 1.32 |
| 13 | Jkamp | 0.30 | 2.35E-06 | 4.88E-02 | 0.64 | 0.53 | 0.11 | 1.21 |
| 13 | Acadm | 0.33 | 2.36E-06 | 4.91E-02 | 0.44 | 0.33 | 0.10 | 1.31 |
| 14 | Wnt6 | 5.34 | 0 | 0 | 0.94 | 0.08 | 0.86 | 12.37 |
| 14 | Cxcl14 | 5.46 | 0 | 0 | 0.96 | 0.13 | 0.83 | 7.52 |
| 14 | Tfap2b | 5.56 | 0 | 0 | 0.90 | 0.07 | 0.83 | 13.61 |
| 14 | Trp63 | 3.88 | 0 | 0 | 0.88 | 0.11 | 0.76 | 7.68 |
| 14 | Perp | 3.77 | 0 | 0 | 0.88 | 0.12 | 0.76 | 7.34 |
| 14 | Wnt3 | 7.82 | 0 | 0 | 0.74 | 0.01 | 0.73 | 92.75 |
| 14 | Sostdc1 | 4.88 | 0 | 0 | 0.84 | 0.13 | 0.72 | 6.67 |
| 14 | Fermt1 | 3.55 | 0 | 0 | 0.84 | 0.12 | 0.71 | 6.85 |
| 14 | Pdgfa | 3.02 | 0 | 0 | 0.97 | 0.26 | 0.70 | 3.66 |
| 14 | Krt14 | 3.60 | 0 | 0 | 0.80 | 0.11 | 0.69 | 7.05 |
| 14 | Wnt4 | 3.46 | 0 | 0 | 0.86 | 0.18 | 0.68 | 4.69 |
| 14 | Pdlim1 | 3.38 | 0 | 0 | 0.93 | 0.27 | 0.66 | 3.41 |
| 14 | Mir205hg | 3.95 | 0 | 0 | 0.74 | 0.10 | 0.64 | 7.19 |
| 14 | Bcam | 2.67 | 0 | 0 | 0.81 | 0.19 | 0.62 | 4.34 |
| 14 | Kremen2 | 5.71 | 0 | 0 | 0.65 | 0.03 | 0.62 | 22.45 |
| 14 | Nav2 | 3.00 | 0 | 0 | 0.84 | 0.22 | 0.62 | 3.75 |
| 14 | Wnt9b | 6.89 | 0 | 0 | 0.63 | 0.01 | 0.62 | 48.46 |
| 14 | Tfap2a | 3.43 | 0 | 0 | 0.81 | 0.20 | 0.61 | 4.09 |
| 14 | Wnt7b | 5.95 | 0 | 0 | 0.62 | 0.03 | 0.59 | 23.85 |
| 14 | Hspb1 | 3.30 | 0 | 0 | 0.69 | 0.13 | 0.56 | 5.18 |
| 14 | Dlx2 | 3.35 | 0 | 0 | 0.67 | 0.11 | 0.55 | 5.84 |
| 14 | Tnfrsf19 | 3.57 | 0 | 0 | 0.67 | 0.14 | 0.53 | 4.90 |
| 14 | Dok5 | 3.35 | 0 | 0 | 0.63 | 0.10 | 0.53 | 6.18 |
| 14 | Jag2 | 3.86 | 0 | 0 | 0.59 | 0.07 | 0.52 | 8.66 |
| 14 | Krt15 | 5.60 | 0 | 0 | 0.52 | 0.03 | 0.49 | 20.00 |
| 14 | Pcp4l1 | 6.76 | 0 | 0 | 0.50 | 0.01 | 0.48 | 45.00 |
| 14 | Col17a1 | 5.38 | 0 | 0 | 0.50 | 0.02 | 0.47 | 21.52 |
| 14 | Ntf5 | 3.36 | 0 | 0 | 0.53 | 0.08 | 0.46 | 7.00 |
| 14 | Dapl1 | 3.55 | 0 | 0 | 0.49 | 0.06 | 0.43 | 8.45 |
| 14 | Gm32585 | 3.89 | 0 | 0 | 0.45 | 0.04 | 0.41 | 10.30 |
| 14 | Gria2 | 5.76 | 0 | 0 | 0.37 | 0.02 | 0.35 | 20.44 |
| 14 | Rprml | 7.97 | 0 | 0 | 0.34 | 0.00 | 0.34 | 86.00 |
| 14 | Cobll1 | 4.83 | 0 | 0 | 0.36 | 0.02 | 0.34 | 15.70 |
| 14 | Jdp2 | 4.38 | 0 | 0 | 0.33 | 0.03 | 0.31 | 11.52 |
| 14 | Insyn2b | 4.70 | 0 | 0 | 0.27 | 0.02 | 0.25 | 17.93 |
| 14 | Lgals7 | 5.25 | 0 | 0 | 0.25 | 0.01 | 0.24 | 21.08 |
| 14 | Npr3 | 5.32 | 0 | 0 | 0.25 | 0.01 | 0.24 | 22.64 |
| 14 | Wnt10a | 7.00 | 0 | 0 | 0.18 | 0.00 | 0.18 | 90.50 |
| 14 | C130021I20Rik | 6.85 | 0 | 0 | 0.17 | 0.00 | 0.17 | 57.33 |
| 14 | Plbd1 | 5.66 | 0 | 0 | 0.17 | 0.01 | 0.16 | 33.20 |
| 14 | Irx4 | 8.01 | 0 | 0 | 0.15 | 0.00 | 0.14 | 145.00 |
| 14 | Lmx1b | 6.37 | 0 | 0 | 0.15 | 0.00 | 0.14 | 48.33 |
| 14 | Plxna2 | 3.33 | 3.63E-303 | 7.56E-299 | 0.76 | 0.22 | 0.55 | 3.52 |
| 14 | Otx1 | 3.35 | 8.98E-303 | 1.87E-298 | 0.57 | 0.10 | 0.46 | 5.59 |
| 14 | Dock8 | 4.01 | 1.87E-300 | 3.90E-296 | 0.30 | 0.03 | 0.28 | 11.65 |
| 14 | Sfn | 1.71 | 1.64E-299 | 3.40E-295 | 0.82 | 0.19 | 0.63 | 4.29 |
| 14 | Pdgfc | 2.75 | 9.10E-293 | 1.89E-288 | 0.83 | 0.27 | 0.56 | 3.08 |
| 14 | Igfbp5 | 2.44 | 4.93E-287 | 1.02E-282 | 0.96 | 0.40 | 0.56 | 2.42 |
| 14 | Trp73 | 5.79 | 1.69E-282 | 3.52E-278 | 0.14 | 0.00 | 0.13 | 34.25 |
| 14 | Fgfr2 | 2.97 | 5.53E-282 | 1.15E-277 | 0.80 | 0.26 | 0.53 | 3.03 |
| 14 | Pltp | 2.93 | 7.47E-279 | 1.55E-274 | 0.65 | 0.15 | 0.51 | 4.46 |
| 14 | Lmo1 | 3.76 | 1.30E-278 | 2.70E-274 | 0.44 | 0.07 | 0.38 | 6.82 |

**Supplementary Table 1. Marker genes for the original clusters of integrated clustering of E11.5 control and NotchGOF λ-junction cells.**

| | | | | | | | | |
|---|---|---|---|---|---|---|---|---|
| 14 | Bnc1 | 4.86 | 4.19E-276 | 8.71E-272 | 0.16 | 0.01 | 0.16 | 26.83 |
| 14 | Tmem178 | 4.19 | 8.67E-275 | 1.80E-270 | 0.29 | 0.03 | 0.26 | 11.08 |
| 14 | Irf6 | 3.15 | 3.32E-269 | 6.91E-265 | 0.54 | 0.10 | 0.44 | 5.49 |
| 14 | Pla2r1 | 3.55 | 4.91E-267 | 1.02E-262 | 0.44 | 0.07 | 0.38 | 6.77 |
| 14 | Col18a1 | 2.62 | 3.32E-264 | 6.91E-260 | 0.87 | 0.35 | 0.52 | 2.47 |
| 14 | Hras | 1.94 | 7.49E-263 | 1.56E-258 | 0.97 | 0.80 | 0.17 | 1.22 |
| 14 | Krt5 | 3.14 | 1.06E-261 | 2.20E-257 | 0.49 | 0.08 | 0.40 | 5.99 |
| 14 | Sox14 | 4.86 | 2.84E-261 | 5.90E-257 | 0.18 | 0.01 | 0.17 | 19.67 |
| 14 | Nckap5 | 3.51 | 2.51E-258 | 5.22E-254 | 0.47 | 0.08 | 0.39 | 5.84 |
| 14 | Cdh3 | 2.91 | 4.28E-258 | 8.89E-254 | 0.70 | 0.20 | 0.51 | 3.59 |
| 14 | Igfbp2 | 2.38 | 5.33E-258 | 1.11E-253 | 0.97 | 0.75 | 0.22 | 1.30 |
| 14 | Edaradd | 3.73 | 6.76E-247 | 1.41E-242 | 0.31 | 0.04 | 0.28 | 8.94 |
| 14 | Frmd4b | 2.87 | 1.26E-245 | 2.63E-241 | 0.63 | 0.16 | 0.48 | 4.06 |
| 14 | Ccnd2 | 1.99 | 1.75E-243 | 3.65E-239 | 1.00 | 0.85 | 0.14 | 1.17 |
| 14 | Slc7a11 | 3.78 | 9.76E-236 | 2.03E-231 | 0.43 | 0.07 | 0.36 | 6.07 |
| 14 | Lratd2 | 3.25 | 2.80E-234 | 5.82E-230 | 0.50 | 0.10 | 0.40 | 5.03 |
| 14 | Wnt10b | 5.99 | 1.70E-233 | 3.54E-229 | 0.11 | 0.00 | 0.10 | 35.00 |
| 14 | Anxa2 | 2.06 | 2.07E-233 | 4.31E-229 | 0.91 | 0.40 | 0.50 | 2.25 |
| 14 | S100a11 | 1.71 | 4.46E-226 | 9.27E-222 | 0.98 | 0.75 | 0.23 | 1.30 |
| 14 | Pthlh | 4.41 | 7.22E-226 | 1.50E-221 | 0.22 | 0.02 | 0.20 | 12.71 |
| 14 | Brinp1 | 4.40 | 2.66E-221 | 5.53E-217 | 0.15 | 0.01 | 0.15 | 19.25 |
| 14 | Spint2 | 1.74 | 1.62E-219 | 3.37E-215 | 0.88 | 0.30 | 0.58 | 2.97 |
| 14 | Dlx3 | 3.13 | 3.08E-218 | 6.40E-214 | 0.41 | 0.07 | 0.34 | 6.14 |
| 14 | Dsp | 1.93 | 1.15E-216 | 2.39E-212 | 0.77 | 0.22 | 0.55 | 3.46 |
| 14 | Ikzf2 | 2.21 | 1.92E-215 | 3.99E-211 | 0.79 | 0.27 | 0.52 | 2.92 |
| 14 | Stk26 | 2.61 | 4.74E-214 | 9.85E-210 | 0.69 | 0.22 | 0.47 | 3.18 |
| 14 | Galnt18 | 3.98 | 4.71E-211 | 9.79E-207 | 0.26 | 0.03 | 0.23 | 9.56 |
| 14 | Chst11 | 3.36 | 8.87E-210 | 1.84E-205 | 0.39 | 0.07 | 0.33 | 6.00 |
| 14 | Dlx1 | 3.52 | 1.04E-207 | 2.17E-203 | 0.32 | 0.04 | 0.28 | 7.36 |
| 14 | Ddr1 | 2.57 | 1.61E-200 | 3.35E-196 | 0.53 | 0.12 | 0.41 | 4.36 |
| 14 | Ehbp1 | 2.62 | 3.43E-200 | 7.14E-196 | 0.68 | 0.23 | 0.45 | 2.97 |
| 14 | Slc2a3 | 2.78 | 4.61E-198 | 9.59E-194 | 0.53 | 0.13 | 0.40 | 4.14 |
| 14 | Irx5 | 2.14 | 4.32E-196 | 8.99E-192 | 0.76 | 0.27 | 0.49 | 2.82 |
| 14 | Ston2 | 2.33 | 3.91E-195 | 8.13E-191 | 0.62 | 0.17 | 0.45 | 3.63 |
| 14 | Pkp3 | 3.25 | 1.35E-193 | 2.80E-189 | 0.36 | 0.06 | 0.30 | 6.12 |
| 14 | Piezo2 | 2.61 | 1.18E-192 | 2.45E-188 | 0.66 | 0.22 | 0.45 | 3.07 |
| 14 | Epcam | 1.44 | 9.79E-192 | 2.04E-187 | 0.92 | 0.33 | 0.58 | 2.76 |
| 14 | H19 | 1.78 | 1.03E-190 | 2.14E-186 | 0.98 | 0.84 | 0.13 | 1.16 |
| 14 | Txndc17 | 1.55 | 5.75E-185 | 1.20E-180 | 0.96 | 0.92 | 0.03 | 1.04 |
| 14 | Frem2 | 2.27 | 1.90E-182 | 3.94E-178 | 0.62 | 0.18 | 0.44 | 3.43 |
| 14 | Adamts18 | 2.91 | 1.90E-180 | 3.96E-176 | 0.37 | 0.06 | 0.30 | 5.79 |
| 14 | Sema3d | 2.53 | 1.50E-179 | 3.12E-175 | 0.48 | 0.11 | 0.37 | 4.43 |
| 14 | Ccdc68 | 3.36 | 6.81E-176 | 1.42E-171 | 0.31 | 0.05 | 0.26 | 6.65 |
| 14 | Gm16136 | 1.95 | 1.44E-174 | 3.00E-170 | 0.60 | 0.16 | 0.44 | 3.69 |
| 14 | Susd4 | 3.03 | 6.34E-174 | 1.32E-169 | 0.41 | 0.08 | 0.32 | 4.88 |
| 14 | Cd24a | 1.18 | 1.22E-173 | 2.53E-169 | 0.98 | 0.92 | 0.07 | 1.07 |
| 14 | Tpd52 | 1.91 | 1.74E-170 | 3.62E-166 | 0.72 | 0.25 | 0.47 | 2.86 |
| 14 | Rab38 | 3.07 | 8.17E-170 | 1.70E-165 | 0.35 | 0.06 | 0.29 | 5.70 |
| 14 | Wls | 1.71 | 9.41E-168 | 1.96E-163 | 0.91 | 0.74 | 0.18 | 1.24 |
| 14 | Htra1 | 3.70 | 4.02E-165 | 8.36E-161 | 0.22 | 0.03 | 0.19 | 8.68 |
| 14 | Gadd45g | 2.49 | 3.46E-161 | 7.20E-157 | 0.60 | 0.20 | 0.40 | 2.99 |
| 14 | Ripk4 | 2.91 | 2.37E-159 | 4.92E-155 | 0.36 | 0.07 | 0.29 | 5.30 |
| 14 | Dkk4 | 4.47 | 4.32E-158 | 8.98E-154 | 0.10 | 0.01 | 0.10 | 20.80 |
| 14 | Esrp1 | 2.02 | 9.15E-156 | 1.90E-151 | 0.56 | 0.16 | 0.40 | 3.52 |
| 14 | Pgam1 | 0.91 | 3.03E-155 | 6.29E-151 | 0.99 | 0.97 | 0.03 | 1.03 |
| 14 | Mpzl2 | 1.94 | 8.44E-152 | 1.76E-147 | 0.62 | 0.20 | 0.42 | 3.15 |

**Supplementary Table 1. Marker genes for the original clusters of integrated clustering of E11.5 control and NotchGOF λ-junction cells.**

| | | | | | | | | |
|---|---|---|---|---|---|---|---|---|
| 14 | Ctsl | 1.58 | 1.57E-150 | 3.26E-146 | 0.91 | 0.82 | 0.09 | 1.11 |
| 14 | F11r | 2.07 | 2.42E-150 | 5.03E-146 | 0.57 | 0.17 | 0.40 | 3.32 |
| 14 | Fth1 | 0.35 | 1.01E-149 | 2.10E-145 | 1.00 | 1.00 | 0.00 | 1.00 |
| 14 | Hist3h2ba | 1.84 | 2.72E-149 | 5.65E-145 | 0.71 | 0.26 | 0.45 | 2.70 |
| 14 | Acer2 | 1.75 | 3.83E-149 | 7.96E-145 | 0.81 | 0.39 | 0.42 | 2.09 |
| 14 | Npnt | 2.25 | 1.85E-148 | 3.85E-144 | 0.62 | 0.23 | 0.40 | 2.74 |
| 14 | Fjx1 | 2.02 | 2.41E-147 | 5.01E-143 | 0.78 | 0.38 | 0.40 | 2.05 |
| 14 | Hspb8 | 2.24 | 6.85E-145 | 1.43E-140 | 0.49 | 0.13 | 0.36 | 3.78 |
| 14 | Igf2 | 1.67 | 1.33E-143 | 2.77E-139 | 0.88 | 0.58 | 0.30 | 1.52 |
| 14 | Nebl | 2.74 | 3.84E-142 | 7.99E-138 | 0.42 | 0.10 | 0.32 | 4.12 |
| 14 | Fgd2 | 2.52 | 1.04E-139 | 2.16E-135 | 0.40 | 0.09 | 0.31 | 4.28 |
| 14 | Arhgap22 | 4.21 | 3.37E-139 | 7.01E-135 | 0.13 | 0.01 | 0.12 | 14.11 |
| 14 | Cbr2 | 4.25 | 1.23E-136 | 2.55E-132 | 0.11 | 0.01 | 0.11 | 14.25 |
| 14 | Grhl2 | 2.40 | 1.51E-135 | 3.14E-131 | 0.45 | 0.12 | 0.33 | 3.71 |
| 14 | Itga6 | 1.54 | 6.53E-135 | 1.36E-130 | 0.74 | 0.30 | 0.44 | 2.45 |
| 14 | Calml3 | 4.41 | 4.49E-133 | 9.35E-129 | 0.12 | 0.01 | 0.12 | 13.78 |
| 14 | Ier5 | 1.51 | 2.81E-132 | 5.84E-128 | 0.90 | 0.65 | 0.25 | 1.39 |
| 14 | Ap1m2 | 2.17 | 1.42E-131 | 2.95E-127 | 0.48 | 0.14 | 0.34 | 3.53 |
| 14 | Art4 | 3.71 | 1.06E-130 | 2.20E-126 | 0.19 | 0.02 | 0.17 | 8.00 |
| 14 | Irx3 | 1.98 | 7.98E-130 | 1.66E-125 | 0.65 | 0.26 | 0.39 | 2.51 |
| 14 | Ptprf | 1.39 | 4.81E-129 | 1.00E-124 | 0.89 | 0.63 | 0.26 | 1.41 |
| 14 | Barx2 | 3.21 | 1.31E-128 | 2.73E-124 | 0.23 | 0.04 | 0.20 | 6.50 |
| 14 | Fam162a | 1.21 | 1.39E-128 | 2.89E-124 | 0.95 | 0.89 | 0.06 | 1.06 |
| 14 | Klk8 | 2.00 | 1.57E-127 | 3.27E-123 | 0.58 | 0.21 | 0.38 | 2.84 |
| 14 | Artn | 4.30 | 1.64E-123 | 3.41E-119 | 0.11 | 0.01 | 0.10 | 13.75 |
| 14 | Plxna4 | 3.71 | 3.01E-123 | 6.26E-119 | 0.17 | 0.02 | 0.15 | 8.79 |
| 14 | Sema3a | 2.34 | 1.27E-122 | 2.64E-118 | 0.71 | 0.40 | 0.32 | 1.79 |
| 14 | Fzd3 | 1.67 | 2.77E-122 | 5.76E-118 | 0.84 | 0.58 | 0.26 | 1.45 |
| 14 | Fabp5 | 0.78 | 9.43E-119 | 1.96E-114 | 0.99 | 0.98 | 0.01 | 1.01 |
| 14 | Ctnnal1 | 2.32 | 4.12E-117 | 8.56E-113 | 0.60 | 0.26 | 0.34 | 2.33 |
| 14 | Rnase4 | 2.44 | 3.94E-116 | 8.19E-112 | 0.36 | 0.09 | 0.27 | 4.00 |
| 14 | Ajuba | 2.03 | 4.51E-114 | 9.38E-110 | 0.61 | 0.27 | 0.34 | 2.26 |
| 14 | Fzd6 | 2.74 | 1.10E-112 | 2.29E-108 | 0.33 | 0.08 | 0.25 | 4.22 |
| 14 | Sema3c | 2.91 | 2.85E-109 | 5.92E-105 | 0.34 | 0.09 | 0.26 | 3.95 |
| 14 | Bex1 | 1.12 | 5.62E-108 | 1.17E-103 | 0.90 | 0.58 | 0.31 | 1.54 |
| 14 | Tmem54 | 2.13 | 2.80E-107 | 5.83E-103 | 0.40 | 0.11 | 0.29 | 3.55 |
| 14 | Siva1 | 0.74 | 3.32E-107 | 6.91E-103 | 0.98 | 0.95 | 0.03 | 1.03 |
| 14 | S100a10 | 1.24 | 3.63E-107 | 7.56E-103 | 0.84 | 0.55 | 0.29 | 1.54 |
| 14 | Cldn1 | 2.94 | 1.18E-106 | 2.44E-102 | 0.18 | 0.02 | 0.15 | 7.33 |
| 14 | Ezr | 1.10 | 7.51E-106 | 1.56E-101 | 0.75 | 0.31 | 0.44 | 2.41 |
| 14 | Rxfp1 | 3.70 | 1.86E-105 | 3.88E-101 | 0.15 | 0.02 | 0.13 | 8.65 |
| 14 | Gm42418 | 1.08 | 3.04E-105 | 6.32E-101 | 1.00 | 1.00 | 0.00 | 1.00 |
| 14 | Fry | 2.86 | 3.56E-105 | 7.40E-101 | 0.25 | 0.05 | 0.20 | 5.08 |
| 14 | Lama5 | 2.20 | 3.74E-105 | 7.78E-101 | 0.39 | 0.11 | 0.28 | 3.58 |
| 14 | Rab27b | 2.29 | 1.60E-103 | 3.33E-99 | 0.35 | 0.09 | 0.26 | 3.87 |
| 14 | Itga3 | 2.74 | 9.28E-103 | 1.93E-98 | 0.28 | 0.06 | 0.22 | 4.51 |
| 14 | Nrep | 1.07 | 9.56E-103 | 1.99E-98 | 0.92 | 0.82 | 0.10 | 1.13 |
| 14 | Meis2 | 0.96 | 1.14E-102 | 2.37E-98 | 0.97 | 0.67 | 0.30 | 1.45 |
| 14 | Socs2 | 1.41 | 2.80E-102 | 5.83E-98 | 0.71 | 0.34 | 0.38 | 2.13 |
| 14 | Adamts17 | 3.27 | 3.01E-102 | 6.26E-98 | 0.20 | 0.03 | 0.17 | 6.22 |
| 14 | Pdpn | 2.12 | 6.48E-102 | 1.35E-97 | 0.49 | 0.18 | 0.31 | 2.75 |
| 14 | Serpinb6b | 3.60 | 9.32E-102 | 1.94E-97 | 0.15 | 0.02 | 0.13 | 7.95 |
| 14 | Prps2 | 2.49 | 1.34E-101 | 2.79E-97 | 0.46 | 0.16 | 0.30 | 2.84 |
| 14 | 2610204G07Rik | 3.98 | 1.87E-100 | 3.89E-96 | 0.11 | 0.01 | 0.10 | 10.90 |
| 14 | Ngf | 3.63 | 3.76E-100 | 7.83E-96 | 0.15 | 0.02 | 0.13 | 7.60 |
| 14 | Tdrp | 3.26 | 4.68E-100 | 9.74E-96 | 0.17 | 0.03 | 0.15 | 6.88 |

**Supplementary Table 1. Marker genes for the original clusters of integrated clustering of E11.5 control and NotchGOF λ-junction cells.**

| | | | | | | | | |
|---|---|---|---|---|---|---|---|---|
| 14 | Itm2b | 1.43 | 5.32E-99 | 1.11E-94 | 0.90 | 0.84 | 0.06 | 1.07 |
| 14 | Tpt1 | 0.33 | 7.65E-99 | 1.59E-94 | 1.00 | 1.00 | 0.00 | 1.00 |
| 14 | Dst | 1.95 | 1.84E-98 | 3.82E-94 | 0.66 | 0.35 | 0.32 | 1.91 |
| 14 | Stx3 | 2.08 | 1.92E-97 | 3.99E-93 | 0.40 | 0.12 | 0.28 | 3.32 |
| 14 | Robo2 | 2.15 | 2.75E-97 | 5.72E-93 | 0.49 | 0.18 | 0.31 | 2.67 |
| 14 | Malat1 | 0.93 | 3.84E-97 | 7.98E-93 | 0.98 | 1.00 | -0.01 | 0.99 |
| 14 | Mbnl2 | 1.34 | 4.01E-97 | 8.34E-93 | 0.86 | 0.67 | 0.18 | 1.27 |
| 14 | Meis1 | 1.51 | 1.41E-96 | 2.94E-92 | 0.76 | 0.45 | 0.32 | 1.70 |
| 14 | Arhgef19 | 2.01 | 2.26E-96 | 4.70E-92 | 0.41 | 0.13 | 0.29 | 3.28 |
| 14 | Gjb3 | 2.86 | 2.24E-94 | 4.67E-90 | 0.19 | 0.03 | 0.16 | 6.10 |
| 14 | Ablim1 | 1.89 | 6.14E-94 | 1.28E-89 | 0.54 | 0.22 | 0.32 | 2.46 |
| 14 | Celsr2 | 2.53 | 6.86E-94 | 1.43E-89 | 0.32 | 0.09 | 0.24 | 3.72 |
| 14 | Rps27 | 0.44 | 7.07E-94 | 1.47E-89 | 1.00 | 1.00 | 0.00 | 1.00 |
| 14 | Rnd3 | 1.28 | 3.91E-93 | 8.13E-89 | 0.84 | 0.64 | 0.21 | 1.32 |
| 14 | Dag1 | 1.85 | 7.05E-93 | 1.47E-88 | 0.65 | 0.36 | 0.29 | 1.80 |
| 14 | Calm2 | 0.66 | 7.39E-92 | 1.54E-87 | 0.99 | 0.98 | 0.01 | 1.01 |
| 14 | Cd59a | 2.03 | 1.53E-91 | 3.19E-87 | 0.53 | 0.21 | 0.31 | 2.47 |
| 14 | Acaa2 | 1.61 | 2.20E-91 | 4.59E-87 | 0.75 | 0.47 | 0.27 | 1.58 |
| 14 | Lratd1 | 2.95 | 6.97E-91 | 1.45E-86 | 0.18 | 0.03 | 0.15 | 5.94 |
| 14 | Efna1 | 2.09 | 1.02E-89 | 2.11E-85 | 0.41 | 0.13 | 0.28 | 3.06 |
| 14 | Foxp1 | 1.08 | 6.46E-89 | 1.34E-84 | 0.89 | 0.59 | 0.30 | 1.50 |
| 14 | Atxn10 | 0.99 | 3.65E-86 | 7.59E-82 | 0.92 | 0.89 | 0.03 | 1.04 |
| 14 | Anxa8 | 3.23 | 3.17E-85 | 6.59E-81 | 0.14 | 0.02 | 0.12 | 7.10 |
| 14 | Itgb5 | 1.48 | 3.80E-85 | 7.91E-81 | 0.79 | 0.58 | 0.21 | 1.36 |
| 14 | Kdf1 | 2.30 | 3.18E-84 | 6.61E-80 | 0.30 | 0.08 | 0.22 | 3.72 |
| 14 | Gjb6 | 3.87 | 5.02E-83 | 1.04E-78 | 0.12 | 0.01 | 0.11 | 8.50 |
| 14 | Znrf3 | 2.23 | 2.44E-82 | 5.08E-78 | 0.44 | 0.17 | 0.27 | 2.55 |
| 14 | Dab1 | 2.02 | 8.64E-82 | 1.80E-77 | 0.42 | 0.15 | 0.27 | 2.84 |
| 14 | Gjb2 | 2.93 | 1.27E-81 | 2.64E-77 | 0.21 | 0.04 | 0.17 | 4.93 |
| 14 | Atp2c2 | 3.58 | 1.96E-81 | 4.07E-77 | 0.11 | 0.01 | 0.10 | 8.46 |
| 14 | Rpl23 | 0.30 | 3.03E-80 | 6.30E-76 | 1.00 | 1.00 | 0.00 | 1.00 |
| 14 | Kif21a | 1.55 | 4.44E-80 | 9.24E-76 | 0.62 | 0.30 | 0.32 | 2.04 |
| 14 | Afdn | 1.26 | 1.34E-78 | 2.80E-74 | 0.79 | 0.57 | 0.23 | 1.40 |
| 14 | Slc15a2 | 2.85 | 9.64E-78 | 2.00E-73 | 0.21 | 0.04 | 0.17 | 4.75 |
| 14 | Rnf43 | 2.59 | 1.25E-77 | 2.60E-73 | 0.21 | 0.05 | 0.17 | 4.71 |
| 14 | Rb1 | 2.13 | 5.53E-77 | 1.15E-72 | 0.41 | 0.15 | 0.26 | 2.69 |
| 14 | Cd151 | 1.56 | 1.28E-76 | 2.67E-72 | 0.61 | 0.32 | 0.29 | 1.90 |
| 14 | Cntnap2 | 2.84 | 1.62E-75 | 3.37E-71 | 0.21 | 0.05 | 0.17 | 4.51 |
| 14 | Fam89a | 2.46 | 6.60E-75 | 1.37E-70 | 0.26 | 0.07 | 0.19 | 3.81 |
| 14 | Sdc4 | 1.55 | 1.69E-74 | 3.52E-70 | 0.62 | 0.34 | 0.29 | 1.84 |
| 14 | Cpz | 2.25 | 1.77E-74 | 3.68E-70 | 0.26 | 0.07 | 0.19 | 3.91 |
| 14 | Efcab1 | 2.59 | 1.55E-73 | 3.23E-69 | 0.20 | 0.04 | 0.16 | 4.74 |
| 14 | Cdh1 | 1.29 | 4.09E-72 | 8.51E-68 | 0.45 | 0.16 | 0.29 | 2.76 |
| 14 | Rps27a | 0.28 | 7.07E-72 | 1.47E-67 | 1.00 | 1.00 | 0.00 | 1.00 |
| 14 | Mtcl1 | 2.14 | 1.29E-71 | 2.68E-67 | 0.34 | 0.12 | 0.23 | 2.97 |
| 14 | Rps19 | 0.30 | 2.67E-71 | 5.54E-67 | 1.00 | 1.00 | 0.00 | 1.00 |
| 14 | Pdcd4 | 1.18 | 2.74E-71 | 5.69E-67 | 0.82 | 0.64 | 0.19 | 1.29 |
| 14 | Rps20 | 0.28 | 8.02E-71 | 1.67E-66 | 1.00 | 1.00 | 0.00 | 1.00 |
| 14 | Large2 | 2.42 | 2.09E-70 | 4.34E-66 | 0.28 | 0.08 | 0.20 | 3.43 |
| 14 | Rpl7 | 0.31 | 3.06E-70 | 6.37E-66 | 1.00 | 1.00 | 0.00 | 1.00 |
| 14 | Dnase2a | 1.86 | 4.86E-70 | 1.01E-65 | 0.48 | 0.21 | 0.26 | 2.24 |
| 14 | Wwc1 | 1.58 | 9.28E-70 | 1.93E-65 | 0.45 | 0.18 | 0.27 | 2.54 |
| 14 | Klhl13 | 1.70 | 9.38E-70 | 1.95E-65 | 0.53 | 0.26 | 0.27 | 2.03 |
| 14 | Rapgefl1 | 2.84 | 9.66E-70 | 2.01E-65 | 0.16 | 0.03 | 0.13 | 5.57 |
| 14 | Cldn6 | 0.76 | 1.07E-69 | 2.22E-65 | 0.58 | 0.22 | 0.35 | 2.59 |
| 14 | Elf1 | 1.70 | 1.14E-69 | 2.37E-65 | 0.53 | 0.26 | 0.28 | 2.07 |

**Supplementary Table 1. Marker genes for the original clusters of integrated clustering of E11.5 control and NotchGOF λ-junction cells.**

| | | | | | | | | |
|---|---|---|---|---|---|---|---|---|
| 14 | Rbbp8 | 1.35 | 4.92E-69 | 1.02E-64 | 0.74 | 0.55 | 0.19 | 1.33 |
| 14 | Krt17 | 3.22 | 1.92E-68 | 3.99E-64 | 0.12 | 0.02 | 0.10 | 6.88 |
| 14 | Adk | 1.24 | 2.06E-68 | 4.29E-64 | 0.76 | 0.61 | 0.16 | 1.26 |
| 14 | Cox7a2l | 0.71 | 9.40E-68 | 1.96E-63 | 0.96 | 0.96 | 0.00 | 1.00 |
| 14 | Knop1 | 0.69 | 1.12E-67 | 2.34E-63 | 0.95 | 0.91 | 0.04 | 1.05 |
| 14 | Rps7 | 0.29 | 3.76E-67 | 7.81E-63 | 1.00 | 1.00 | 0.00 | 1.00 |
| 14 | Slc2a1 | 1.52 | 4.31E-67 | 8.95E-63 | 0.67 | 0.42 | 0.24 | 1.58 |
| 14 | Fau | 0.32 | 4.76E-67 | 9.90E-63 | 1.00 | 1.00 | 0.00 | 1.00 |
| 14 | Col4a2 | 1.35 | 8.94E-67 | 1.86E-62 | 0.66 | 0.42 | 0.24 | 1.58 |
| 14 | 2310002F09Rik | 2.83 | 2.04E-66 | 4.24E-62 | 0.17 | 0.03 | 0.14 | 4.97 |
| 14 | Runx1 | 1.52 | 1.26E-65 | 2.62E-61 | 0.45 | 0.19 | 0.26 | 2.39 |
| 14 | Col12a1 | 2.50 | 2.88E-65 | 5.99E-61 | 0.23 | 0.06 | 0.17 | 3.90 |
| 14 | Epha1 | 2.46 | 5.18E-65 | 1.08E-60 | 0.22 | 0.06 | 0.16 | 3.98 |
| 14 | Cd2ap | 1.11 | 1.54E-64 | 3.21E-60 | 0.79 | 0.62 | 0.17 | 1.28 |
| 14 | Mycl | 1.41 | 6.01E-64 | 1.25E-59 | 0.57 | 0.30 | 0.27 | 1.92 |
| 14 | Slc35b3 | 2.01 | 6.41E-64 | 1.33E-59 | 0.45 | 0.21 | 0.25 | 2.18 |
| 14 | Plk2 | 1.49 | 8.45E-64 | 1.76E-59 | 0.63 | 0.37 | 0.26 | 1.71 |
| 14 | Nav3 | 2.41 | 2.03E-63 | 4.22E-59 | 0.34 | 0.12 | 0.22 | 2.76 |
| 14 | D7Ertd443e | 3.42 | 2.85E-63 | 5.93E-59 | 0.13 | 0.02 | 0.11 | 5.86 |
| 14 | Sinhcaf | 0.79 | 3.20E-63 | 6.65E-59 | 0.92 | 0.84 | 0.09 | 1.10 |
| 14 | Castor1 | 3.06 | 7.99E-63 | 1.66E-58 | 0.14 | 0.03 | 0.12 | 5.46 |
| 14 | Tspan18 | 2.10 | 2.44E-62 | 5.07E-58 | 0.18 | 0.04 | 0.14 | 4.63 |
| 14 | Zfp503 | 1.32 | 2.42E-61 | 5.03E-57 | 0.64 | 0.37 | 0.27 | 1.73 |
| 14 | Irx2 | 3.37 | 6.40E-61 | 1.33E-56 | 0.11 | 0.02 | 0.09 | 6.81 |
| 14 | Luzp1 | 0.87 | 3.52E-60 | 7.32E-56 | 0.86 | 0.77 | 0.09 | 1.12 |
| 14 | Map7 | 1.83 | 5.62E-60 | 1.17E-55 | 0.31 | 0.10 | 0.21 | 2.97 |
| 14 | Pcdh7 | 1.27 | 1.54E-59 | 3.20E-55 | 0.65 | 0.39 | 0.26 | 1.67 |
| 14 | Afap1l1 | 2.27 | 1.59E-59 | 3.31E-55 | 0.24 | 0.07 | 0.17 | 3.49 |
| 14 | Rps14 | 0.28 | 3.59E-59 | 7.47E-55 | 1.00 | 1.00 | 0.00 | 1.00 |
| 14 | Wnk2 | 2.54 | 5.21E-59 | 1.08E-54 | 0.19 | 0.05 | 0.14 | 4.16 |
| 14 | Lgi2 | 1.96 | 4.51E-58 | 9.39E-54 | 0.31 | 0.11 | 0.20 | 2.85 |
| 14 | Sgk3 | 2.06 | 6.74E-58 | 1.40E-53 | 0.36 | 0.15 | 0.22 | 2.49 |
| 14 | Myh9 | 1.16 | 1.40E-57 | 2.91E-53 | 0.71 | 0.50 | 0.21 | 1.41 |
| 14 | Rpl17 | 0.30 | 2.36E-57 | 4.91E-53 | 1.00 | 1.00 | 0.00 | 1.00 |
| 14 | Kitl | 2.23 | 3.39E-57 | 7.04E-53 | 0.24 | 0.07 | 0.17 | 3.41 |
| 14 | Msx2 | 1.09 | 4.08E-57 | 8.48E-53 | 0.71 | 0.42 | 0.29 | 1.68 |
| 14 | Pnrc1 | 0.76 | 1.02E-56 | 2.11E-52 | 0.93 | 0.87 | 0.06 | 1.07 |
| 14 | Palmd | 2.92 | 1.50E-56 | 3.12E-52 | 0.12 | 0.02 | 0.10 | 5.85 |
| 14 | Rpl27a | 0.26 | 4.64E-56 | 9.64E-52 | 1.00 | 1.00 | 0.00 | 1.00 |
| 14 | Nudt11 | 1.81 | 1.63E-54 | 3.39E-50 | 0.39 | 0.17 | 0.22 | 2.26 |
| 14 | Rap2c | 1.17 | 2.10E-54 | 4.37E-50 | 0.74 | 0.60 | 0.15 | 1.25 |
| 14 | Nell2 | 2.31 | 2.86E-54 | 5.94E-50 | 0.19 | 0.05 | 0.14 | 3.96 |
| 14 | Arhgef5 | 1.65 | 1.22E-53 | 2.53E-49 | 0.42 | 0.20 | 0.23 | 2.17 |
| 14 | Agrn | 1.38 | 1.31E-53 | 2.72E-49 | 0.61 | 0.39 | 0.23 | 1.59 |
| 14 | Rpl38 | 0.30 | 1.97E-53 | 4.10E-49 | 1.00 | 1.00 | 0.00 | 1.00 |
| 14 | Cox4i2 | 2.02 | 1.99E-53 | 4.15E-49 | 0.49 | 0.26 | 0.23 | 1.87 |
| 14 | Fam83b | 3.00 | 2.81E-53 | 5.85E-49 | 0.11 | 0.02 | 0.09 | 5.79 |
| 14 | Pqlc3 | 2.42 | 4.15E-53 | 8.62E-49 | 0.19 | 0.05 | 0.14 | 3.73 |
| 14 | Parm1 | 1.77 | 8.82E-53 | 1.83E-48 | 0.38 | 0.16 | 0.22 | 2.39 |
| 14 | Lsr | 1.51 | 1.13E-52 | 2.35E-48 | 0.44 | 0.20 | 0.24 | 2.18 |
| 14 | Stard10 | 1.13 | 2.65E-52 | 5.52E-48 | 0.56 | 0.29 | 0.27 | 1.90 |
| 14 | Erich1 | 1.74 | 9.30E-52 | 1.94E-47 | 0.46 | 0.23 | 0.23 | 1.98 |
| 14 | Diras2 | 2.60 | 1.95E-51 | 4.06E-47 | 0.12 | 0.02 | 0.09 | 5.48 |
| 14 | Card10 | 3.25 | 2.17E-51 | 4.52E-47 | 0.11 | 0.02 | 0.09 | 5.60 |
| 14 | Dnajc1 | 1.44 | 3.71E-51 | 7.71E-47 | 0.69 | 0.56 | 0.14 | 1.25 |
| 14 | Tom1l1 | 1.66 | 4.43E-51 | 9.22E-47 | 0.37 | 0.16 | 0.21 | 2.36 |

**Supplementary Table 1. Marker genes for the original clusters of integrated clustering of E11.5 control and NotchGOF λ-junction cells.**

| | | | | | | | | |
|---|---|---|---|---|---|---|---|---|
| 14 | Ptpn13 | 1.26 | 4.85E-51 | 1.01E-46 | 0.62 | 0.40 | 0.22 | 1.56 |
| 14 | Tmem123 | 1.48 | 1.11E-50 | 2.32E-46 | 0.54 | 0.32 | 0.22 | 1.69 |
| 14 | Plekha7 | 1.79 | 1.71E-50 | 3.56E-46 | 0.26 | 0.09 | 0.18 | 3.00 |
| 14 | Hook1 | 1.45 | 5.85E-50 | 1.22E-45 | 0.43 | 0.20 | 0.23 | 2.18 |
| 14 | Trp53i11 | 1.42 | 9.44E-50 | 1.96E-45 | 0.45 | 0.22 | 0.23 | 2.04 |
| 14 | Sp6 | 2.77 | 1.43E-49 | 2.97E-45 | 0.13 | 0.03 | 0.11 | 4.79 |
| 14 | 1700023F06Rik | 2.44 | 2.90E-49 | 6.03E-45 | 0.17 | 0.04 | 0.13 | 4.05 |
| 14 | Ctnnb1 | 0.55 | 5.30E-49 | 1.10E-44 | 0.97 | 0.97 | 0.01 | 1.01 |
| 14 | Pkm | 0.44 | 9.04E-49 | 1.88E-44 | 0.99 | 0.98 | 0.01 | 1.01 |
| 14 | Arf5 | 0.43 | 1.16E-48 | 2.41E-44 | 0.97 | 0.98 | -0.02 | 0.98 |
| 14 | Krt7 | 1.89 | 3.33E-48 | 6.92E-44 | 0.19 | 0.05 | 0.14 | 3.72 |
| 14 | Cpne5 | 2.65 | 4.14E-48 | 8.62E-44 | 0.18 | 0.05 | 0.13 | 3.64 |
| 14 | Rps26 | 0.29 | 5.84E-48 | 1.21E-43 | 1.00 | 1.00 | 0.00 | 1.00 |
| 14 | Tpi1 | 0.56 | 1.45E-47 | 3.02E-43 | 0.97 | 0.96 | 0.01 | 1.01 |
| 14 | Hs6st1 | 1.65 | 2.12E-47 | 4.42E-43 | 0.38 | 0.17 | 0.21 | 2.26 |
| 14 | Ipmk | 1.84 | 2.18E-47 | 4.53E-43 | 0.39 | 0.18 | 0.20 | 2.13 |
| 14 | Lef1 | 0.96 | 2.40E-47 | 4.99E-43 | 0.70 | 0.46 | 0.25 | 1.55 |
| 14 | Vwa1 | 2.17 | 4.04E-47 | 8.41E-43 | 0.19 | 0.06 | 0.14 | 3.53 |
| 14 | Btg1 | 1.07 | 8.06E-47 | 1.68E-42 | 0.83 | 0.79 | 0.05 | 1.06 |
| 14 | Fzd10 | 2.04 | 2.04E-46 | 4.25E-42 | 0.32 | 0.13 | 0.19 | 2.40 |
| 14 | Tafa2 | 2.34 | 2.67E-46 | 5.56E-42 | 0.16 | 0.04 | 0.12 | 4.11 |
| 14 | Rps4x | 0.27 | 4.91E-46 | 1.02E-41 | 1.00 | 1.00 | 0.00 | 1.00 |
| 14 | Tmcc3 | 2.60 | 5.24E-46 | 1.09E-41 | 0.14 | 0.03 | 0.11 | 4.34 |
| 14 | Crnde | 1.68 | 6.85E-46 | 1.42E-41 | 0.34 | 0.15 | 0.20 | 2.36 |
| 14 | Rps5 | 0.26 | 1.13E-45 | 2.36E-41 | 1.00 | 1.00 | 0.00 | 1.00 |
| 14 | Filip1l | 1.49 | 1.45E-45 | 3.02E-41 | 0.47 | 0.25 | 0.22 | 1.86 |
| 14 | Vwa5a | 2.00 | 1.74E-45 | 3.61E-41 | 0.24 | 0.08 | 0.16 | 2.90 |
| 14 | Nxn | 0.83 | 2.33E-45 | 4.84E-41 | 0.81 | 0.71 | 0.10 | 1.14 |
| 14 | Pcdh9 | 2.00 | 9.42E-45 | 1.96E-40 | 0.28 | 0.11 | 0.18 | 2.61 |
| 14 | Fras1 | 1.43 | 1.23E-44 | 2.55E-40 | 0.49 | 0.26 | 0.22 | 1.85 |
| 14 | Ctdspl | 1.33 | 1.46E-44 | 3.04E-40 | 0.55 | 0.34 | 0.21 | 1.61 |
| 14 | Nudt10 | 1.93 | 2.31E-44 | 4.81E-40 | 0.28 | 0.11 | 0.17 | 2.60 |
| 14 | Cdcp1 | 2.37 | 2.71E-44 | 5.64E-40 | 0.19 | 0.06 | 0.13 | 3.38 |
| 14 | Eif3f | 0.41 | 3.09E-44 | 6.44E-40 | 0.99 | 0.99 | -0.01 | 0.99 |
| 14 | Abca1 | 1.98 | 3.48E-44 | 7.24E-40 | 0.21 | 0.07 | 0.15 | 3.16 |
| 14 | Rpl37 | 0.27 | 3.51E-44 | 7.30E-40 | 1.00 | 1.00 | 0.00 | 1.00 |
| 14 | Nt5c | 1.10 | 4.06E-44 | 8.45E-40 | 0.69 | 0.53 | 0.16 | 1.29 |
| 14 | Sox11 | 0.56 | 5.30E-44 | 1.10E-39 | 0.99 | 0.97 | 0.03 | 1.03 |
| 14 | Cacnb3 | 1.36 | 5.65E-44 | 1.17E-39 | 0.49 | 0.27 | 0.22 | 1.80 |
| 14 | Celf2 | 1.43 | 6.51E-44 | 1.35E-39 | 0.47 | 0.25 | 0.22 | 1.88 |
| 14 | Dstn | 0.45 | 1.66E-43 | 3.46E-39 | 0.96 | 0.94 | 0.02 | 1.02 |
| 14 | Pik3r1 | 1.62 | 2.84E-43 | 5.91E-39 | 0.58 | 0.39 | 0.19 | 1.50 |
| 14 | Llgl2 | 2.37 | 3.44E-43 | 7.16E-39 | 0.17 | 0.05 | 0.12 | 3.60 |
| 14 | Itgb6 | 2.24 | 4.97E-43 | 1.03E-38 | 0.15 | 0.04 | 0.11 | 3.85 |
| 14 | mt-Co1 | 0.38 | 8.00E-43 | 1.66E-38 | 0.95 | 0.99 | -0.04 | 0.96 |
| 14 | Ppp1r36 | 2.26 | 9.79E-43 | 2.04E-38 | 0.18 | 0.05 | 0.13 | 3.47 |
| 14 | Osbpl3 | 2.20 | 1.50E-42 | 3.12E-38 | 0.24 | 0.09 | 0.16 | 2.74 |
| 14 | St14 | 1.48 | 1.77E-42 | 3.68E-38 | 0.30 | 0.12 | 0.18 | 2.55 |
| 14 | Rhpn2 | 2.88 | 2.84E-42 | 5.91E-38 | 0.12 | 0.03 | 0.09 | 4.60 |
| 14 | Etaa1os | 2.26 | 3.93E-42 | 8.17E-38 | 0.19 | 0.06 | 0.13 | 3.23 |
| 14 | Atox1 | 0.38 | 4.08E-42 | 8.49E-38 | 0.96 | 0.95 | 0.01 | 1.01 |
| 14 | Bcl11b | 0.94 | 4.63E-42 | 9.62E-38 | 0.75 | 0.59 | 0.17 | 1.28 |
| 14 | Pard6g | 0.88 | 1.30E-41 | 2.70E-37 | 0.73 | 0.59 | 0.14 | 1.24 |
| 14 | Ube2e3 | 0.62 | 2.60E-41 | 5.40E-37 | 0.93 | 0.92 | 0.01 | 1.01 |
| 14 | Kcnk1 | 1.20 | 3.11E-41 | 6.47E-37 | 0.40 | 0.18 | 0.21 | 2.17 |
| 14 | Mcee | 1.12 | 5.51E-41 | 1.15E-36 | 0.62 | 0.44 | 0.18 | 1.41 |

**Supplementary Table 1. Marker genes for the original clusters of integrated clustering of E11.5 control and NotchGOF λ-junction cells.**

| | | | | | | | | |
|---|---|---|---|---|---|---|---|---|
| 14 | Arl4c | 1.11 | 6.18E-41 | 1.29E-36 | 0.54 | 0.33 | 0.22 | 1.67 |
| 14 | Ucp2 | 1.01 | 1.01E-40 | 2.10E-36 | 0.57 | 0.35 | 0.22 | 1.63 |
| 14 | Prag1 | 1.72 | 1.41E-40 | 2.93E-36 | 0.31 | 0.14 | 0.18 | 2.32 |
| 14 | Mdh1 | 0.51 | 3.63E-40 | 7.54E-36 | 0.94 | 0.95 | -0.01 | 0.99 |
| 14 | Apoc1 | 1.56 | 3.97E-40 | 8.26E-36 | 0.23 | 0.08 | 0.15 | 2.85 |
| 14 | Cdkn1a | 1.30 | 7.17E-40 | 1.49E-35 | 0.28 | 0.11 | 0.17 | 2.57 |
| 14 | H2afj | 0.62 | 8.06E-40 | 1.68E-35 | 0.87 | 0.81 | 0.06 | 1.07 |
| 14 | Dclre1c | 2.36 | 1.12E-39 | 2.32E-35 | 0.17 | 0.05 | 0.12 | 3.42 |
| 14 | Atxn7l3b | 0.37 | 1.22E-39 | 2.55E-35 | 0.99 | 0.98 | 0.01 | 1.01 |
| 14 | Eno1 | 0.44 | 1.24E-39 | 2.58E-35 | 0.98 | 0.97 | 0.01 | 1.01 |
| 14 | Lmo4 | 0.67 | 1.51E-39 | 3.13E-35 | 0.88 | 0.81 | 0.07 | 1.08 |
| 14 | Itgb4 | 2.43 | 8.84E-39 | 1.84E-34 | 0.13 | 0.03 | 0.10 | 4.17 |
| 14 | Emp1 | 1.61 | 9.12E-39 | 1.90E-34 | 0.30 | 0.13 | 0.17 | 2.37 |
| 14 | Arhgap27os2 | 2.24 | 1.15E-38 | 2.38E-34 | 0.19 | 0.06 | 0.13 | 3.08 |
| 14 | Mbd2 | 0.93 | 1.29E-38 | 2.69E-34 | 0.69 | 0.56 | 0.13 | 1.23 |
| 14 | Mid2 | 2.56 | 1.46E-38 | 3.04E-34 | 0.11 | 0.03 | 0.09 | 4.38 |
| 14 | Phlda3 | 1.14 | 1.93E-38 | 4.01E-34 | 0.58 | 0.41 | 0.18 | 1.44 |
| 14 | Gask1b | 2.75 | 3.12E-38 | 6.49E-34 | 0.11 | 0.02 | 0.08 | 4.77 |
| 14 | Prkci | 1.48 | 5.97E-38 | 1.24E-33 | 0.45 | 0.26 | 0.19 | 1.73 |
| 14 | Bdh1 | 2.10 | 6.35E-38 | 1.32E-33 | 0.20 | 0.07 | 0.13 | 2.97 |
| 14 | Pgk1 | 0.55 | 7.51E-38 | 1.56E-33 | 0.96 | 0.96 | 0.01 | 1.01 |
| 14 | Grb7 | 1.72 | 1.35E-37 | 2.82E-33 | 0.25 | 0.10 | 0.15 | 2.62 |
| 14 | Lgalsl | 1.23 | 2.12E-37 | 4.41E-33 | 0.51 | 0.31 | 0.20 | 1.64 |
| 14 | Kdm5a | 0.83 | 2.38E-37 | 4.94E-33 | 0.78 | 0.67 | 0.11 | 1.17 |
| 14 | Plekhg1 | 1.83 | 2.49E-37 | 5.18E-33 | 0.30 | 0.13 | 0.17 | 2.26 |
| 14 | Gstk1 | 1.77 | 2.99E-37 | 6.21E-33 | 0.36 | 0.17 | 0.18 | 2.05 |
| 14 | Cacng4 | 1.31 | 3.76E-37 | 7.82E-33 | 0.45 | 0.25 | 0.20 | 1.78 |
| 14 | Erf | 1.34 | 4.12E-37 | 8.56E-33 | 0.49 | 0.30 | 0.19 | 1.65 |
| 14 | Serpinb6a | 1.20 | 4.29E-37 | 8.92E-33 | 0.56 | 0.39 | 0.17 | 1.45 |
| 14 | Arpc2 | 0.44 | 4.98E-37 | 1.04E-32 | 0.95 | 0.94 | 0.01 | 1.01 |
| 14 | Chmp2b | 0.93 | 2.73E-36 | 5.68E-32 | 0.70 | 0.56 | 0.14 | 1.25 |
| 14 | Tenm2 | 1.86 | 3.62E-36 | 7.53E-32 | 0.20 | 0.07 | 0.13 | 2.89 |
| 14 | Aldoa | 0.51 | 1.09E-35 | 2.26E-31 | 0.95 | 0.95 | 0.00 | 1.00 |
| 14 | Sparc | 0.63 | 1.41E-35 | 2.93E-31 | 0.84 | 0.74 | 0.11 | 1.14 |
| 14 | Ass1 | 2.22 | 1.93E-35 | 4.01E-31 | 0.17 | 0.06 | 0.12 | 3.11 |
| 14 | Ccdc88c | 1.33 | 2.18E-35 | 4.54E-31 | 0.34 | 0.15 | 0.18 | 2.18 |
| 14 | Ktn1 | 0.69 | 2.21E-35 | 4.61E-31 | 0.85 | 0.83 | 0.03 | 1.03 |
| 14 | Ctnnd2 | 1.49 | 2.27E-35 | 4.73E-31 | 0.28 | 0.12 | 0.16 | 2.40 |
| 14 | Ank3 | 0.93 | 6.31E-35 | 1.31E-30 | 0.66 | 0.49 | 0.18 | 1.36 |
| 14 | Pygl | 2.00 | 8.47E-35 | 1.76E-30 | 0.24 | 0.09 | 0.14 | 2.52 |
| 14 | Spats2l | 1.28 | 8.84E-35 | 1.84E-30 | 0.42 | 0.23 | 0.19 | 1.85 |
| 14 | Dennd2c | 2.12 | 1.13E-34 | 2.35E-30 | 0.18 | 0.06 | 0.12 | 2.98 |
| 14 | Vamp8 | 0.83 | 1.39E-34 | 2.89E-30 | 0.74 | 0.61 | 0.13 | 1.21 |
| 14 | Ifitm3 | 1.32 | 1.65E-34 | 3.43E-30 | 0.48 | 0.29 | 0.19 | 1.64 |
| 14 | Krt1 | 2.06 | 2.35E-34 | 4.89E-30 | 0.16 | 0.05 | 0.11 | 3.24 |
| 14 | Kit | 1.90 | 2.61E-34 | 5.43E-30 | 0.24 | 0.09 | 0.14 | 2.54 |
| 14 | Lbh | 1.48 | 3.36E-34 | 6.99E-30 | 0.38 | 0.20 | 0.18 | 1.89 |
| 14 | Rtn4ip1 | 2.10 | 3.70E-34 | 7.70E-30 | 0.19 | 0.07 | 0.12 | 2.91 |
| 14 | B3gnt2 | 1.79 | 4.56E-34 | 9.48E-30 | 0.24 | 0.10 | 0.15 | 2.51 |
| 14 | Tspan13 | 0.93 | 9.63E-34 | 2.00E-29 | 0.60 | 0.41 | 0.19 | 1.46 |
| 14 | Hist3h2a | 1.40 | 1.17E-33 | 2.43E-29 | 0.44 | 0.26 | 0.18 | 1.69 |
| 14 | Marchf9 | 1.44 | 1.35E-33 | 2.80E-29 | 0.31 | 0.15 | 0.17 | 2.17 |
| 14 | Itgav | 1.39 | 1.56E-33 | 3.24E-29 | 0.45 | 0.27 | 0.18 | 1.66 |
| 14 | Cdkn1c | 0.61 | 1.57E-33 | 3.26E-29 | 0.96 | 0.93 | 0.03 | 1.03 |
| 14 | Fosl2 | 2.59 | 2.31E-33 | 4.81E-29 | 0.11 | 0.03 | 0.09 | 4.07 |
| 14 | Pdk1 | 1.31 | 2.59E-33 | 5.39E-29 | 0.44 | 0.25 | 0.18 | 1.73 |

**Supplementary Table 1. Marker genes for the original clusters of integrated clustering of E11.5 control and NotchGOF λ-junction cells.**

| | | | | | | | | |
|---|---|---|---|---|---|---|---|---|
| 14 | Arid3b | 1.36 | 3.93E-33 | 8.17E-29 | 0.41 | 0.23 | 0.18 | 1.80 |
| 14 | Tulp4 | 0.79 | 4.33E-33 | 9.01E-29 | 0.77 | 0.67 | 0.11 | 1.16 |
| 14 | Pak2 | 0.57 | 4.74E-33 | 9.86E-29 | 0.88 | 0.84 | 0.04 | 1.05 |
| 14 | Gm9908 | 2.19 | 1.26E-32 | 2.63E-28 | 0.11 | 0.03 | 0.09 | 3.93 |
| 14 | Kank1 | 1.69 | 3.04E-32 | 6.33E-28 | 0.32 | 0.16 | 0.16 | 2.04 |
| 14 | Dsg2 | 1.47 | 3.26E-32 | 6.79E-28 | 0.29 | 0.13 | 0.16 | 2.25 |
| 14 | Sord | 1.70 | 3.40E-32 | 7.07E-28 | 0.30 | 0.14 | 0.16 | 2.11 |
| 14 | Stox2 | 0.95 | 5.36E-32 | 1.11E-27 | 0.69 | 0.56 | 0.13 | 1.23 |
| 14 | Hdgf | 0.36 | 7.07E-32 | 1.47E-27 | 0.96 | 0.98 | -0.01 | 0.99 |
| 14 | Trappc6a | 0.96 | 9.42E-32 | 1.96E-27 | 0.66 | 0.52 | 0.14 | 1.26 |
| 14 | Trp53inp1 | 1.61 | 1.02E-31 | 2.12E-27 | 0.35 | 0.18 | 0.17 | 1.92 |
| 14 | Cacna1b | 1.75 | 1.34E-31 | 2.79E-27 | 0.21 | 0.08 | 0.13 | 2.65 |
| 14 | Mgst1 | 0.88 | 1.79E-31 | 3.71E-27 | 0.58 | 0.37 | 0.21 | 1.56 |
| 14 | Gabarapl1 | 1.28 | 2.60E-31 | 5.40E-27 | 0.49 | 0.33 | 0.17 | 1.51 |
| 14 | Synrg | 1.33 | 2.61E-31 | 5.43E-27 | 0.41 | 0.23 | 0.18 | 1.77 |
| 14 | Pwwp2b | 1.97 | 3.04E-31 | 6.32E-27 | 0.19 | 0.07 | 0.12 | 2.70 |
| 14 | Ctso | 2.00 | 3.61E-31 | 7.51E-27 | 0.18 | 0.06 | 0.12 | 2.84 |
| 14 | Cux1 | 0.73 | 4.54E-31 | 9.45E-27 | 0.77 | 0.70 | 0.08 | 1.11 |
| 14 | AY036118 | 1.02 | 4.89E-31 | 1.02E-26 | 0.72 | 0.61 | 0.11 | 1.18 |
| 14 | Ttc7 | 2.29 | 7.27E-31 | 1.51E-26 | 0.12 | 0.03 | 0.09 | 3.72 |
| 14 | Phlda2 | 2.65 | 1.05E-30 | 2.19E-26 | 0.13 | 0.04 | 0.09 | 3.42 |
| 14 | Prxl2a | 1.10 | 1.12E-30 | 2.34E-26 | 0.55 | 0.38 | 0.17 | 1.45 |
| 14 | Clic6 | 2.15 | 1.13E-30 | 2.35E-26 | 0.16 | 0.05 | 0.11 | 3.06 |
| 14 | Rimklb | 1.21 | 1.94E-30 | 4.03E-26 | 0.49 | 0.32 | 0.17 | 1.52 |
| 14 | Rbms1 | 0.64 | 3.64E-30 | 7.58E-26 | 0.82 | 0.78 | 0.04 | 1.05 |
| 14 | Arhgef16 | 1.75 | 3.67E-30 | 7.63E-26 | 0.17 | 0.06 | 0.11 | 2.93 |
| 14 | Atp6v1f | 0.52 | 3.82E-30 | 7.94E-26 | 0.91 | 0.90 | 0.00 | 1.00 |
| 14 | 2900026A02Rik | 1.56 | 4.54E-30 | 9.45E-26 | 0.24 | 0.10 | 0.14 | 2.39 |
| 14 | Cldn7 | 0.26 | 5.65E-30 | 1.17E-25 | 0.47 | 0.24 | 0.24 | 2.00 |
| 14 | Laptm4a | 0.51 | 7.41E-30 | 1.54E-25 | 0.92 | 0.95 | -0.02 | 0.98 |
| 14 | Purb | 0.57 | 9.53E-30 | 1.98E-25 | 0.87 | 0.87 | 0.00 | 1.00 |
| 14 | Eps8l2 | 2.33 | 1.02E-29 | 2.13E-25 | 0.13 | 0.04 | 0.09 | 3.35 |
| 14 | AI661453 | 2.21 | 1.24E-29 | 2.59E-25 | 0.12 | 0.03 | 0.09 | 3.53 |
| 14 | Eif3h | 0.32 | 1.52E-29 | 3.17E-25 | 0.98 | 0.99 | -0.01 | 0.99 |
| 14 | Slc39a10 | 0.83 | 2.00E-29 | 4.16E-25 | 0.75 | 0.68 | 0.07 | 1.11 |
| 14 | Acvr2a | 1.38 | 2.93E-29 | 6.10E-25 | 0.43 | 0.27 | 0.17 | 1.62 |
| 14 | Cox6b2 | 0.93 | 3.50E-29 | 7.29E-25 | 0.62 | 0.47 | 0.15 | 1.32 |
| 14 | Tgif1 | 0.80 | 4.76E-29 | 9.89E-25 | 0.73 | 0.63 | 0.10 | 1.16 |
| 14 | Scube3 | 1.29 | 4.78E-29 | 9.94E-25 | 0.37 | 0.20 | 0.17 | 1.83 |
| 14 | Bri3 | 0.41 | 5.45E-29 | 1.13E-24 | 0.96 | 0.96 | 0.00 | 1.00 |
| 14 | Ldha | 0.44 | 1.14E-28 | 2.37E-24 | 0.98 | 0.97 | 0.01 | 1.01 |
| 14 | Hspg2 | 1.16 | 3.16E-28 | 6.57E-24 | 0.39 | 0.22 | 0.17 | 1.75 |
| 14 | Sik1 | 1.80 | 3.69E-28 | 7.67E-24 | 0.24 | 0.10 | 0.13 | 2.31 |
| 14 | Pik3ip1 | 1.80 | 4.02E-28 | 8.36E-24 | 0.21 | 0.09 | 0.13 | 2.48 |
| 14 | Bambi | 1.04 | 4.17E-28 | 8.67E-24 | 0.71 | 0.64 | 0.07 | 1.11 |
| 14 | Ddit4 | 1.34 | 7.13E-28 | 1.48E-23 | 0.53 | 0.37 | 0.15 | 1.40 |
| 14 | Tafa5 | 1.63 | 9.62E-28 | 2.00E-23 | 0.26 | 0.12 | 0.14 | 2.16 |
| 14 | Igsf9 | 1.05 | 1.01E-27 | 2.10E-23 | 0.56 | 0.40 | 0.15 | 1.38 |
| 14 | Ednrb | 1.77 | 2.76E-27 | 5.74E-23 | 0.13 | 0.04 | 0.09 | 3.38 |
| 14 | Crim1 | 1.37 | 4.39E-27 | 9.13E-23 | 0.35 | 0.20 | 0.15 | 1.79 |
| 14 | Fat1 | 0.89 | 6.97E-27 | 1.45E-22 | 0.64 | 0.51 | 0.13 | 1.25 |
| 14 | Slc3a2 | 0.89 | 8.14E-27 | 1.69E-22 | 0.71 | 0.62 | 0.08 | 1.13 |
| 14 | Id1 | 0.51 | 9.69E-27 | 2.01E-22 | 0.93 | 0.86 | 0.08 | 1.09 |
| 14 | Gprc5b | 1.70 | 1.01E-26 | 2.10E-22 | 0.22 | 0.10 | 0.13 | 2.31 |
| 14 | Btbd11 | 2.24 | 1.05E-26 | 2.18E-22 | 0.12 | 0.03 | 0.08 | 3.38 |
| 14 | Ptpn14 | 1.43 | 1.14E-26 | 2.36E-22 | 0.29 | 0.15 | 0.15 | 1.97 |

**Supplementary Table 1. Marker genes for the original clusters of integrated clustering of E11.5 control and NotchGOF λ-junction cells.**

| | | | | | | | | |
|---|---|---|---|---|---|---|---|---|
| 14 | Ovol2 | 2.14 | 1.68E-26 | 3.50E-22 | 0.10 | 0.03 | 0.08 | 3.78 |
| 14 | Efnb2 | 0.83 | 2.10E-26 | 4.36E-22 | 0.61 | 0.45 | 0.16 | 1.36 |
| 14 | Klc3 | 1.18 | 3.43E-26 | 7.13E-22 | 0.24 | 0.11 | 0.13 | 2.28 |
| 14 | Cpe | 1.00 | 4.44E-26 | 9.24E-22 | 0.56 | 0.42 | 0.14 | 1.34 |
| 14 | Nudt14 | 0.95 | 4.53E-26 | 9.42E-22 | 0.55 | 0.40 | 0.14 | 1.36 |
| 14 | Selenow | 0.28 | 4.61E-26 | 9.59E-22 | 0.99 | 0.99 | -0.01 | 0.99 |
| 14 | Imp3 | 0.57 | 5.25E-26 | 1.09E-21 | 0.83 | 0.79 | 0.04 | 1.05 |
| 14 | Kctd1 | 0.94 | 8.00E-26 | 1.66E-21 | 0.60 | 0.47 | 0.14 | 1.29 |
| 14 | Mllt10 | 0.70 | 1.19E-25 | 2.48E-21 | 0.73 | 0.63 | 0.09 | 1.15 |
| 14 | Cldn8 | 2.42 | 1.26E-25 | 2.61E-21 | 0.10 | 0.03 | 0.07 | 3.64 |
| 14 | Tead1 | 0.74 | 1.28E-25 | 2.67E-21 | 0.70 | 0.60 | 0.10 | 1.16 |
| 14 | Fam49a | 2.00 | 1.46E-25 | 3.03E-21 | 0.13 | 0.04 | 0.09 | 3.10 |
| 14 | Bnip3 | 1.08 | 1.55E-25 | 3.23E-21 | 0.55 | 0.41 | 0.14 | 1.34 |
| 14 | Tpbg | 1.11 | 1.58E-25 | 3.28E-21 | 0.46 | 0.30 | 0.16 | 1.52 |
| 14 | Tpm1 | 0.50 | 2.09E-25 | 4.35E-21 | 0.91 | 0.91 | 0.00 | 1.00 |
| 14 | P4ha1 | 1.02 | 2.12E-25 | 4.40E-21 | 0.62 | 0.52 | 0.11 | 1.20 |
| 14 | Sfr1 | 0.34 | 2.14E-25 | 4.45E-21 | 0.98 | 0.98 | -0.01 | 0.99 |
| 14 | Vgll4 | 0.63 | 2.37E-25 | 4.93E-21 | 0.82 | 0.78 | 0.04 | 1.05 |
| 14 | Tfap2c | 1.05 | 2.80E-25 | 5.82E-21 | 0.44 | 0.27 | 0.17 | 1.61 |
| 14 | Ahcyl2 | 1.10 | 3.40E-25 | 7.08E-21 | 0.44 | 0.29 | 0.15 | 1.52 |
| 14 | Clip4 | 1.97 | 4.39E-25 | 9.12E-21 | 0.13 | 0.04 | 0.09 | 3.00 |
| 14 | Spint1 | 1.36 | 4.63E-25 | 9.62E-21 | 0.22 | 0.10 | 0.13 | 2.29 |
| 14 | Trim29 | 1.70 | 4.85E-25 | 1.01E-20 | 0.14 | 0.05 | 0.09 | 2.98 |
| 14 | Ackr3 | 1.11 | 7.18E-25 | 1.49E-20 | 0.54 | 0.42 | 0.12 | 1.30 |
| 14 | Tspan2 | 1.64 | 8.69E-25 | 1.81E-20 | 0.22 | 0.10 | 0.12 | 2.26 |
| 14 | Ndrg2 | 1.40 | 9.14E-25 | 1.90E-20 | 0.27 | 0.13 | 0.14 | 2.02 |
| 14 | Klhl8 | 1.60 | 1.04E-24 | 2.15E-20 | 0.21 | 0.09 | 0.12 | 2.32 |
| 14 | Bex4 | 0.45 | 4.93E-24 | 1.02E-19 | 0.78 | 0.65 | 0.14 | 1.21 |
| 14 | Cebpb | 1.74 | 5.18E-24 | 1.08E-19 | 0.24 | 0.12 | 0.13 | 2.07 |
| 14 | Stxbp2 | 1.56 | 5.92E-24 | 1.23E-19 | 0.27 | 0.13 | 0.13 | 2.00 |
| 14 | Samd5 | 0.48 | 7.68E-24 | 1.60E-19 | 0.57 | 0.36 | 0.21 | 1.58 |
| 14 | Lmo7 | 1.64 | 8.37E-24 | 1.74E-19 | 0.30 | 0.16 | 0.14 | 1.86 |
| 14 | Plec | 1.23 | 9.09E-24 | 1.89E-19 | 0.42 | 0.27 | 0.15 | 1.56 |
| 14 | Plxdc2 | 1.19 | 9.96E-24 | 2.07E-19 | 0.40 | 0.25 | 0.15 | 1.62 |
| 14 | Cep164 | 0.80 | 1.08E-23 | 2.24E-19 | 0.69 | 0.61 | 0.08 | 1.12 |
| 14 | Sorcs2 | 1.65 | 2.45E-23 | 5.10E-19 | 0.23 | 0.11 | 0.12 | 2.13 |
| 14 | Gm16141 | 2.07 | 2.51E-23 | 5.21E-19 | 0.11 | 0.03 | 0.08 | 3.24 |
| 14 | Fxyd3 | 1.06 | 2.54E-23 | 5.29E-19 | 0.21 | 0.09 | 0.12 | 2.34 |
| 14 | Acyp1 | 0.91 | 2.98E-23 | 6.20E-19 | 0.56 | 0.43 | 0.14 | 1.32 |
| 14 | Grb14 | 2.17 | 3.03E-23 | 6.29E-19 | 0.13 | 0.05 | 0.09 | 2.89 |
| 14 | Minar2 | 1.56 | 3.12E-23 | 6.48E-19 | 0.21 | 0.09 | 0.12 | 2.24 |
| 14 | Mbp | 1.55 | 3.13E-23 | 6.51E-19 | 0.21 | 0.09 | 0.11 | 2.23 |
| 14 | Osbpl8 | 1.19 | 3.76E-23 | 7.82E-19 | 0.38 | 0.23 | 0.15 | 1.63 |
| 14 | Hunk | 0.83 | 4.02E-23 | 8.36E-19 | 0.62 | 0.50 | 0.12 | 1.24 |
| 14 | Rhob | 1.26 | 4.92E-23 | 1.02E-18 | 0.48 | 0.34 | 0.13 | 1.39 |
| 14 | Nr2f2 | 0.64 | 5.00E-23 | 1.04E-18 | 0.54 | 0.36 | 0.17 | 1.48 |
| 14 | Socs7 | 1.31 | 7.18E-23 | 1.49E-18 | 0.35 | 0.21 | 0.14 | 1.66 |
| 14 | Cited2 | 0.75 | 9.03E-23 | 1.88E-18 | 0.73 | 0.64 | 0.10 | 1.15 |
| 14 | Eya2 | 1.00 | 9.06E-23 | 1.89E-18 | 0.39 | 0.23 | 0.16 | 1.70 |
| 14 | Tceal9 | 0.31 | 1.18E-22 | 2.46E-18 | 0.98 | 0.98 | 0.00 | 1.00 |
| 14 | Etl4 | 1.02 | 1.80E-22 | 3.74E-18 | 0.46 | 0.31 | 0.16 | 1.50 |
| 14 | Map4k5 | 1.18 | 2.00E-22 | 4.15E-18 | 0.45 | 0.31 | 0.14 | 1.46 |
| 14 | Rap2b | 0.87 | 2.11E-22 | 4.38E-18 | 0.60 | 0.48 | 0.11 | 1.23 |
| 14 | Ly6h | 0.87 | 2.19E-22 | 4.55E-18 | 0.29 | 0.15 | 0.14 | 1.97 |
| 14 | Cxcr4 | 1.75 | 2.36E-22 | 4.91E-18 | 0.14 | 0.05 | 0.09 | 2.81 |
| 14 | Tspan7 | 1.03 | 2.61E-22 | 5.43E-18 | 0.42 | 0.28 | 0.15 | 1.52 |

**Supplementary Table 1. Marker genes for the original clusters of integrated clustering of E11.5 control and NotchGOF λ-junction cells.**

| | | | | | | | | |
|---|---|---|---|---|---|---|---|---|
| 14 | Notch1 | 1.28 | 3.22E-22 | 6.71E-18 | 0.39 | 0.25 | 0.15 | 1.59 |
| 14 | Trps1 | 0.74 | 4.25E-22 | 8.84E-18 | 0.61 | 0.48 | 0.13 | 1.26 |
| 14 | Mycn | 1.19 | 5.00E-22 | 1.04E-17 | 0.50 | 0.38 | 0.13 | 1.33 |
| 14 | Syt1 | 1.61 | 5.04E-22 | 1.05E-17 | 0.11 | 0.04 | 0.08 | 3.11 |
| 14 | Adgrg1 | 1.88 | 5.76E-22 | 1.20E-17 | 0.12 | 0.04 | 0.08 | 3.00 |
| 14 | Crppa | 1.85 | 7.03E-22 | 1.46E-17 | 0.14 | 0.05 | 0.09 | 2.74 |
| 14 | Slc16a13 | 1.58 | 7.42E-22 | 1.54E-17 | 0.20 | 0.09 | 0.11 | 2.22 |
| 14 | Lrp2 | 1.67 | 7.63E-22 | 1.59E-17 | 0.14 | 0.05 | 0.09 | 2.68 |
| 14 | Slc25a27 | 0.95 | 8.74E-22 | 1.82E-17 | 0.54 | 0.40 | 0.14 | 1.35 |
| 14 | Avpi1 | 1.16 | 8.77E-22 | 1.82E-17 | 0.44 | 0.32 | 0.13 | 1.40 |
| 14 | Rabgap1 | 0.71 | 9.70E-22 | 2.02E-17 | 0.66 | 0.57 | 0.10 | 1.17 |
| 14 | Jup | 1.03 | 2.05E-21 | 4.27E-17 | 0.35 | 0.20 | 0.15 | 1.76 |
| 14 | Fam192a | 0.62 | 2.19E-21 | 4.55E-17 | 0.71 | 0.65 | 0.06 | 1.09 |
| 14 | Lars2 | 0.83 | 2.21E-21 | 4.60E-17 | 0.60 | 0.47 | 0.13 | 1.28 |
| 14 | Celsr1 | 1.40 | 2.50E-21 | 5.20E-17 | 0.22 | 0.10 | 0.12 | 2.17 |
| 14 | Ptpru | 1.45 | 2.85E-21 | 5.93E-17 | 0.29 | 0.16 | 0.13 | 1.83 |
| 14 | Gimap9 | 1.39 | 3.00E-21 | 6.24E-17 | 0.22 | 0.11 | 0.12 | 2.10 |
| 14 | Zfp704 | 0.82 | 3.21E-21 | 6.68E-17 | 0.59 | 0.47 | 0.13 | 1.27 |
| 14 | Acvr2b | 0.60 | 3.57E-21 | 7.43E-17 | 0.73 | 0.64 | 0.08 | 1.13 |
| 14 | Bnip3l | 0.40 | 4.03E-21 | 8.39E-17 | 0.92 | 0.93 | -0.01 | 0.99 |
| 14 | Slc25a17 | 0.70 | 5.94E-21 | 1.24E-16 | 0.70 | 0.62 | 0.08 | 1.13 |
| 14 | Isl1 | 0.36 | 9.72E-21 | 2.02E-16 | 0.39 | 0.22 | 0.18 | 1.81 |
| 14 | Mtmr7 | 1.81 | 9.99E-21 | 2.08E-16 | 0.13 | 0.05 | 0.08 | 2.68 |
| 14 | Clta | 0.44 | 1.10E-20 | 2.29E-16 | 0.96 | 0.98 | -0.02 | 0.98 |
| 14 | Cxadr | 0.83 | 1.12E-20 | 2.32E-16 | 0.57 | 0.46 | 0.11 | 1.24 |
| 14 | Prkch | 1.68 | 1.43E-20 | 2.98E-16 | 0.19 | 0.09 | 0.10 | 2.22 |
| 14 | Adgrl2 | 0.71 | 1.70E-20 | 3.53E-16 | 0.74 | 0.69 | 0.06 | 1.08 |
| 14 | Chst15 | 1.19 | 1.77E-20 | 3.67E-16 | 0.30 | 0.17 | 0.13 | 1.78 |
| 14 | Mfge8 | 0.83 | 2.15E-20 | 4.47E-16 | 0.62 | 0.52 | 0.10 | 1.18 |
| 14 | Zfp462 | 0.55 | 2.78E-20 | 5.78E-16 | 0.79 | 0.73 | 0.06 | 1.09 |
| 14 | Nectin4 | 1.11 | 3.23E-20 | 6.72E-16 | 0.26 | 0.13 | 0.13 | 1.98 |
| 14 | Slc9a3r1 | 0.69 | 3.44E-20 | 7.16E-16 | 0.41 | 0.24 | 0.17 | 1.72 |
| 14 | Ahnak | 0.90 | 3.53E-20 | 7.34E-16 | 0.58 | 0.46 | 0.13 | 1.28 |
| 14 | Nectin1 | 1.34 | 4.44E-20 | 9.23E-16 | 0.30 | 0.17 | 0.13 | 1.73 |
| 14 | Lamb1 | 0.94 | 4.56E-20 | 9.48E-16 | 0.51 | 0.39 | 0.13 | 1.32 |
| 14 | 2610528J11Rik | 1.63 | 6.10E-20 | 1.27E-15 | 0.15 | 0.06 | 0.09 | 2.48 |
| 14 | Edf1 | 0.38 | 6.19E-20 | 1.29E-15 | 0.94 | 0.96 | -0.02 | 0.98 |
| 14 | Rere | 0.74 | 6.74E-20 | 1.40E-15 | 0.68 | 0.59 | 0.09 | 1.15 |
| 14 | Phc2 | 0.62 | 8.83E-20 | 1.84E-15 | 0.74 | 0.69 | 0.05 | 1.07 |
| 14 | Nabp2 | 0.58 | 1.47E-19 | 3.06E-15 | 0.73 | 0.65 | 0.08 | 1.12 |
| 14 | 2310030G06Rik | 1.55 | 1.62E-19 | 3.36E-15 | 0.16 | 0.07 | 0.09 | 2.35 |
| 14 | Serinc2 | 0.78 | 2.02E-19 | 4.19E-15 | 0.52 | 0.39 | 0.14 | 1.35 |
| 14 | Retreg1 | 1.49 | 3.22E-19 | 6.69E-15 | 0.19 | 0.09 | 0.10 | 2.13 |
| 14 | Ptprd | 0.82 | 3.23E-19 | 6.71E-15 | 0.57 | 0.44 | 0.13 | 1.29 |
| 14 | Syt5 | 1.71 | 4.06E-19 | 8.44E-15 | 0.14 | 0.06 | 0.09 | 2.49 |
| 14 | Zfp219 | 0.79 | 4.18E-19 | 8.69E-15 | 0.54 | 0.40 | 0.14 | 1.35 |
| 14 | Pde10a | 2.13 | 5.07E-19 | 1.06E-14 | 0.12 | 0.04 | 0.07 | 2.72 |
| 14 | Gm9922 | 1.98 | 7.74E-19 | 1.61E-14 | 0.11 | 0.04 | 0.07 | 2.82 |
| 14 | Cmip | 0.82 | 1.00E-18 | 2.08E-14 | 0.52 | 0.39 | 0.13 | 1.32 |
| 14 | Mkrn1 | 0.73 | 1.28E-18 | 2.65E-14 | 0.56 | 0.43 | 0.13 | 1.32 |
| 14 | Trmt9b | 1.70 | 2.02E-18 | 4.21E-14 | 0.16 | 0.07 | 0.09 | 2.34 |
| 14 | Usp25 | 0.76 | 2.16E-18 | 4.50E-14 | 0.55 | 0.43 | 0.11 | 1.26 |
| 14 | Gpc4 | 1.32 | 3.01E-18 | 6.26E-14 | 0.25 | 0.14 | 0.12 | 1.85 |
| 14 | Tbc1d10a | 1.15 | 3.40E-18 | 7.08E-14 | 0.32 | 0.20 | 0.12 | 1.62 |
| 14 | Glrb | 1.66 | 3.90E-18 | 8.12E-14 | 0.13 | 0.05 | 0.08 | 2.59 |
| 14 | Gm20186 | 1.77 | 3.94E-18 | 8.20E-14 | 0.15 | 0.06 | 0.09 | 2.36 |

**Supplementary Table 1. Marker genes for the original clusters of integrated clustering of E11.5 control and NotchGOF λ-junction cells.**

| | | | | | | | | |
|---|---|---|---|---|---|---|---|---|
| 14 | Kdm5b | 0.46 | 4.21E-18 | 8.75E-14 | 0.88 | 0.86 | 0.02 | 1.02 |
| 14 | Armc10 | 0.73 | 4.36E-18 | 9.08E-14 | 0.62 | 0.52 | 0.09 | 1.18 |
| 14 | Zfp385a | 1.15 | 4.38E-18 | 9.12E-14 | 0.38 | 0.25 | 0.13 | 1.51 |
| 14 | Igf2bp3 | 0.43 | 5.08E-18 | 1.06E-13 | 0.85 | 0.82 | 0.03 | 1.03 |
| 14 | Scp2 | 0.52 | 5.99E-18 | 1.25E-13 | 0.81 | 0.77 | 0.04 | 1.05 |
| 14 | Rnf180 | 1.72 | 7.55E-18 | 1.57E-13 | 0.13 | 0.05 | 0.08 | 2.58 |
| 14 | Polr3g | 1.09 | 7.78E-18 | 1.62E-13 | 0.40 | 0.27 | 0.12 | 1.44 |
| 14 | Rac3 | 1.01 | 9.49E-18 | 1.97E-13 | 0.45 | 0.33 | 0.12 | 1.37 |
| 14 | Pts | 0.88 | 1.03E-17 | 2.15E-13 | 0.52 | 0.41 | 0.12 | 1.29 |
| 14 | A930004D18Rik | 1.96 | 1.31E-17 | 2.73E-13 | 0.12 | 0.05 | 0.08 | 2.58 |
| 14 | Iqgap1 | 0.80 | 1.48E-17 | 3.07E-13 | 0.58 | 0.47 | 0.10 | 1.22 |
| 14 | Rassf7 | 1.17 | 1.48E-17 | 3.08E-13 | 0.32 | 0.20 | 0.12 | 1.62 |
| 14 | Fryl | 1.06 | 1.62E-17 | 3.37E-13 | 0.42 | 0.30 | 0.12 | 1.39 |
| 14 | Myo10 | 0.82 | 2.00E-17 | 4.16E-13 | 0.57 | 0.47 | 0.10 | 1.21 |
| 14 | Dipk2a | 1.13 | 2.23E-17 | 4.63E-13 | 0.36 | 0.24 | 0.12 | 1.51 |
| 14 | Cib2 | 0.99 | 2.64E-17 | 5.49E-13 | 0.39 | 0.25 | 0.13 | 1.52 |
| 14 | Pik3cb | 1.68 | 2.69E-17 | 5.59E-13 | 0.14 | 0.06 | 0.08 | 2.36 |
| 14 | Gkap1 | 0.79 | 2.81E-17 | 5.85E-13 | 0.56 | 0.43 | 0.12 | 1.28 |
| 14 | Txnip | 1.01 | 3.08E-17 | 6.40E-13 | 0.46 | 0.34 | 0.12 | 1.35 |
| 14 | Fhdc1 | 2.07 | 4.31E-17 | 8.97E-13 | 0.11 | 0.04 | 0.07 | 2.73 |
| 14 | Ak4 | 1.49 | 5.27E-17 | 1.10E-12 | 0.20 | 0.10 | 0.10 | 1.98 |
| 14 | Hk2 | 1.12 | 5.29E-17 | 1.10E-12 | 0.38 | 0.25 | 0.12 | 1.49 |
| 14 | Espn | 1.02 | 5.53E-17 | 1.15E-12 | 0.22 | 0.11 | 0.11 | 2.00 |
| 14 | Kxd1 | 0.77 | 6.20E-17 | 1.29E-12 | 0.61 | 0.50 | 0.11 | 1.21 |
| 14 | Baiap2 | 1.50 | 6.73E-17 | 1.40E-12 | 0.26 | 0.15 | 0.11 | 1.72 |
| 14 | Slc25a33 | 1.18 | 7.13E-17 | 1.48E-12 | 0.28 | 0.17 | 0.11 | 1.69 |
| 14 | Satb1 | 0.84 | 8.65E-17 | 1.80E-12 | 0.50 | 0.39 | 0.12 | 1.30 |
| 14 | Gpi1 | 0.41 | 9.79E-17 | 2.04E-12 | 0.87 | 0.84 | 0.03 | 1.03 |
| 14 | Ppp1r3b | 1.91 | 1.15E-16 | 2.40E-12 | 0.11 | 0.04 | 0.07 | 2.66 |
| 14 | Ppp1r12b | 1.80 | 1.59E-16 | 3.30E-12 | 0.21 | 0.11 | 0.10 | 1.88 |
| 14 | Schip1 | 1.14 | 1.60E-16 | 3.34E-12 | 0.28 | 0.16 | 0.12 | 1.74 |
| 14 | Snai2 | 0.76 | 1.86E-16 | 3.86E-12 | 0.58 | 0.46 | 0.12 | 1.26 |
| 14 | Mapk13 | 0.74 | 2.15E-16 | 4.47E-12 | 0.33 | 0.20 | 0.14 | 1.68 |
| 14 | Plekhg3 | 1.30 | 2.20E-16 | 4.57E-12 | 0.17 | 0.08 | 0.09 | 2.12 |
| 14 | Pxn | 1.27 | 2.70E-16 | 5.62E-12 | 0.30 | 0.18 | 0.11 | 1.62 |
| 14 | Nab1 | 1.32 | 2.90E-16 | 6.03E-12 | 0.23 | 0.13 | 0.10 | 1.81 |
| 14 | Antxr1 | 1.06 | 2.91E-16 | 6.05E-12 | 0.42 | 0.31 | 0.11 | 1.35 |
| 14 | Dcxr | 1.50 | 3.12E-16 | 6.48E-12 | 0.19 | 0.09 | 0.09 | 1.99 |
| 14 | Myl12b | 0.33 | 4.13E-16 | 8.58E-12 | 0.89 | 0.89 | 0.00 | 1.00 |
| 14 | Zbtb34 | 1.45 | 4.33E-16 | 9.02E-12 | 0.18 | 0.09 | 0.09 | 2.04 |
| 14 | Cuedc1 | 0.83 | 4.46E-16 | 9.28E-12 | 0.50 | 0.38 | 0.11 | 1.29 |
| 14 | Dazap2 | 0.66 | 4.59E-16 | 9.54E-12 | 0.65 | 0.57 | 0.07 | 1.13 |
| 14 | Sowahc | 1.39 | 4.98E-16 | 1.04E-11 | 0.24 | 0.14 | 0.11 | 1.79 |
| 14 | Glrx3 | 0.31 | 5.71E-16 | 1.19E-11 | 0.95 | 0.96 | -0.01 | 0.99 |
| 14 | Nsmaf | 1.17 | 6.83E-16 | 1.42E-11 | 0.38 | 0.27 | 0.11 | 1.42 |
| 14 | Cstb | 0.43 | 7.46E-16 | 1.55E-11 | 0.80 | 0.80 | 0.00 | 1.00 |
| 14 | Ifi30 | 0.90 | 7.52E-16 | 1.56E-11 | 0.41 | 0.29 | 0.12 | 1.40 |
| 14 | Cth | 1.30 | 8.77E-16 | 1.82E-11 | 0.31 | 0.21 | 0.11 | 1.52 |
| 14 | Ttc3 | 0.29 | 8.85E-16 | 1.84E-11 | 0.96 | 0.96 | 0.01 | 1.01 |
| 14 | Cd9 | 0.43 | 1.43E-15 | 2.97E-11 | 0.44 | 0.30 | 0.14 | 1.47 |
| 14 | Spock2 | 1.53 | 1.45E-15 | 3.01E-11 | 0.10 | 0.04 | 0.06 | 2.68 |
| 14 | Pura | 0.78 | 1.48E-15 | 3.08E-11 | 0.56 | 0.47 | 0.09 | 1.19 |
| 14 | Rftn1 | 2.36 | 1.82E-15 | 3.78E-11 | 0.10 | 0.04 | 0.06 | 2.63 |
| 14 | Arrdc1 | 1.14 | 1.85E-15 | 3.85E-11 | 0.21 | 0.11 | 0.10 | 1.90 |
| 14 | Frrs1 | 1.08 | 1.85E-15 | 3.85E-11 | 0.34 | 0.22 | 0.12 | 1.53 |
| 14 | Mast4 | 0.80 | 1.91E-15 | 3.97E-11 | 0.54 | 0.45 | 0.09 | 1.20 |

**Supplementary Table 1. Marker genes for the original clusters of integrated clustering of E11.5 control and NotchGOF λ-junction cells.**

| | | | | | | | | |
|---|---|---|---|---|---|---|---|---|
| 14 | Rcan1 | 1.16 | 1.98E-15 | 4.12E-11 | 0.37 | 0.26 | 0.11 | 1.41 |
| 14 | Myo5b | 1.48 | 2.17E-15 | 4.50E-11 | 0.11 | 0.04 | 0.07 | 2.56 |
| 14 | Rsrp1 | 0.40 | 2.54E-15 | 5.28E-11 | 0.87 | 0.89 | -0.02 | 0.98 |
| 14 | Utrn | 0.64 | 2.65E-15 | 5.52E-11 | 0.67 | 0.61 | 0.06 | 1.10 |
| 14 | Col4a1 | 0.43 | 2.68E-15 | 5.58E-11 | 0.69 | 0.62 | 0.07 | 1.12 |
| 14 | Cep170b | 1.26 | 2.79E-15 | 5.81E-11 | 0.23 | 0.13 | 0.10 | 1.83 |
| 14 | Ist1 | 0.72 | 3.46E-15 | 7.19E-11 | 0.55 | 0.46 | 0.09 | 1.20 |
| 14 | Cdc42se1 | 0.73 | 4.04E-15 | 8.41E-11 | 0.52 | 0.42 | 0.09 | 1.22 |
| 14 | Skap2 | 1.73 | 6.87E-15 | 1.43E-10 | 0.13 | 0.05 | 0.07 | 2.31 |
| 14 | Cd164l2 | 1.89 | 1.04E-14 | 2.15E-10 | 0.10 | 0.04 | 0.06 | 2.55 |
| 14 | Spop | 0.62 | 1.15E-14 | 2.38E-10 | 0.71 | 0.69 | 0.02 | 1.02 |
| 14 | Plekha1 | 1.16 | 1.15E-14 | 2.39E-10 | 0.31 | 0.20 | 0.11 | 1.56 |
| 14 | Gas5 | 0.31 | 1.18E-14 | 2.44E-10 | 0.98 | 0.99 | -0.01 | 0.99 |
| 14 | Ptpn4 | 1.17 | 1.21E-14 | 2.51E-10 | 0.26 | 0.16 | 0.11 | 1.68 |
| 14 | Commd3 | 0.44 | 1.34E-14 | 2.79E-10 | 0.80 | 0.81 | -0.01 | 0.99 |
| 14 | B3gnt7 | 1.92 | 1.43E-14 | 2.97E-10 | 0.11 | 0.04 | 0.06 | 2.50 |
| 14 | Fhit | 1.68 | 2.15E-14 | 4.48E-10 | 0.13 | 0.06 | 0.07 | 2.23 |
| 14 | Rnf187 | 0.32 | 2.56E-14 | 5.32E-10 | 0.90 | 0.91 | 0.00 | 1.00 |
| 14 | Lypd6b | 1.46 | 2.68E-14 | 5.58E-10 | 0.15 | 0.07 | 0.08 | 2.13 |
| 14 | Hapln1 | 1.69 | 2.79E-14 | 5.80E-10 | 0.11 | 0.04 | 0.07 | 2.48 |
| 14 | Cnot7 | 0.51 | 3.68E-14 | 7.66E-10 | 0.71 | 0.68 | 0.03 | 1.05 |
| 14 | Eef1e1 | 0.53 | 3.70E-14 | 7.69E-10 | 0.68 | 0.65 | 0.03 | 1.05 |
| 14 | Fgf8 | 1.41 | 4.26E-14 | 8.87E-10 | 0.16 | 0.08 | 0.08 | 2.06 |
| 14 | Foxo1 | 1.27 | 5.53E-14 | 1.15E-09 | 0.28 | 0.18 | 0.10 | 1.58 |
| 14 | Med19 | 0.46 | 6.23E-14 | 1.30E-09 | 0.77 | 0.74 | 0.02 | 1.03 |
| 14 | Trim32 | 0.95 | 6.52E-14 | 1.36E-09 | 0.44 | 0.34 | 0.10 | 1.29 |
| 14 | Hltf | 0.88 | 7.01E-14 | 1.46E-09 | 0.42 | 0.32 | 0.10 | 1.32 |
| 14 | Esrra | 1.36 | 7.45E-14 | 1.55E-09 | 0.22 | 0.12 | 0.09 | 1.76 |
| 14 | Cisd1 | 0.41 | 7.80E-14 | 1.62E-09 | 0.76 | 0.75 | 0.01 | 1.02 |
| 14 | Kif13b | 1.07 | 8.01E-14 | 1.67E-09 | 0.26 | 0.16 | 0.10 | 1.65 |
| 14 | Acvr1b | 1.45 | 1.10E-13 | 2.29E-09 | 0.20 | 0.11 | 0.09 | 1.81 |
| 14 | Cnr1 | 1.26 | 1.55E-13 | 3.22E-09 | 0.17 | 0.09 | 0.09 | 1.98 |
| 14 | Etaa1 | 0.81 | 1.67E-13 | 3.48E-09 | 0.58 | 0.54 | 0.05 | 1.09 |
| 14 | Dynll2 | 0.42 | 2.28E-13 | 4.74E-09 | 0.82 | 0.84 | -0.03 | 0.97 |
| 14 | Calcoco1 | 1.07 | 2.35E-13 | 4.88E-09 | 0.32 | 0.21 | 0.10 | 1.49 |
| 14 | Crebrf | 1.36 | 2.40E-13 | 4.99E-09 | 0.20 | 0.11 | 0.09 | 1.83 |
| 14 | Thbs1 | 0.72 | 2.90E-13 | 6.04E-09 | 0.21 | 0.12 | 0.09 | 1.80 |
| 14 | Rab25 | 0.52 | 3.01E-13 | 6.25E-09 | 0.28 | 0.16 | 0.12 | 1.74 |
| 14 | Ermp1 | 1.00 | 3.13E-13 | 6.50E-09 | 0.43 | 0.33 | 0.10 | 1.29 |
| 14 | Ddx6 | 0.39 | 3.66E-13 | 7.61E-09 | 0.84 | 0.85 | -0.01 | 0.99 |
| 14 | Prss22 | 1.49 | 3.80E-13 | 7.91E-09 | 0.13 | 0.06 | 0.07 | 2.20 |
| 14 | Akap9 | 0.54 | 3.89E-13 | 8.10E-09 | 0.77 | 0.78 | -0.01 | 0.99 |
| 14 | 2700033N17Rik | 1.05 | 3.96E-13 | 8.24E-09 | 0.21 | 0.12 | 0.10 | 1.81 |
| 14 | Smim20 | 0.76 | 4.13E-13 | 8.59E-09 | 0.50 | 0.41 | 0.09 | 1.22 |
| 14 | Krt19 | 0.78 | 4.23E-13 | 8.80E-09 | 0.15 | 0.07 | 0.08 | 2.08 |
| 14 | Slc12a4 | 1.29 | 4.79E-13 | 9.96E-09 | 0.23 | 0.14 | 0.09 | 1.68 |
| 14 | Pawr | 0.96 | 4.87E-13 | 1.01E-08 | 0.22 | 0.13 | 0.10 | 1.74 |
| 14 | Errfi1 | 1.08 | 4.99E-13 | 1.04E-08 | 0.28 | 0.18 | 0.10 | 1.56 |
| 14 | Patj | 1.19 | 5.13E-13 | 1.07E-08 | 0.22 | 0.13 | 0.10 | 1.74 |
| 14 | Tinagl1 | 1.39 | 6.08E-13 | 1.27E-08 | 0.13 | 0.06 | 0.07 | 2.19 |
| 14 | Gtf3a | 0.48 | 6.41E-13 | 1.33E-08 | 0.69 | 0.64 | 0.04 | 1.07 |
| 14 | Cyb5r3 | 0.34 | 8.02E-13 | 1.67E-08 | 0.82 | 0.81 | 0.01 | 1.01 |
| 14 | Mbnl1 | 0.81 | 9.41E-13 | 1.96E-08 | 0.54 | 0.47 | 0.07 | 1.14 |
| 14 | Kcnq1ot1 | 0.43 | 1.08E-12 | 2.25E-08 | 0.86 | 0.92 | -0.06 | 0.93 |
| 14 | Amotl1 | 0.50 | 1.12E-12 | 2.33E-08 | 0.74 | 0.69 | 0.05 | 1.08 |
| 14 | Sh3d19 | 0.91 | 1.21E-12 | 2.51E-08 | 0.36 | 0.26 | 0.10 | 1.41 |

**Supplementary Table 1. Marker genes for the original clusters of integrated clustering of E11.5 control and NotchGOF λ-junction cells.**

| | | | | | | | | |
|---|---|---|---|---|---|---|---|---|
| 14 | Zdhhc13 | 1.31 | 1.28E-12 | 2.66E-08 | 0.23 | 0.14 | 0.09 | 1.66 |
| 14 | Mcoln3 | 1.52 | 1.34E-12 | 2.78E-08 | 0.12 | 0.05 | 0.07 | 2.22 |
| 14 | Prickle3 | 1.52 | 1.62E-12 | 3.36E-08 | 0.14 | 0.07 | 0.07 | 2.08 |
| 14 | Sppl2a | 0.73 | 1.65E-12 | 3.43E-08 | 0.53 | 0.47 | 0.06 | 1.12 |
| 14 | Crb3 | 0.95 | 1.76E-12 | 3.66E-08 | 0.20 | 0.11 | 0.09 | 1.81 |
| 14 | Lrrc1 | 1.21 | 1.96E-12 | 4.08E-08 | 0.24 | 0.15 | 0.09 | 1.63 |
| 14 | Proser2 | 1.51 | 2.21E-12 | 4.60E-08 | 0.12 | 0.06 | 0.07 | 2.18 |
| 14 | Tceal5 | 1.17 | 2.62E-12 | 5.45E-08 | 0.25 | 0.15 | 0.10 | 1.64 |
| 14 | Rps27rt | 0.46 | 2.82E-12 | 5.87E-08 | 0.76 | 0.73 | 0.03 | 1.04 |
| 14 | Pex2 | 0.61 | 2.83E-12 | 5.88E-08 | 0.55 | 0.47 | 0.07 | 1.16 |
| 14 | Comtd1 | 1.20 | 3.13E-12 | 6.51E-08 | 0.23 | 0.14 | 0.09 | 1.65 |
| 14 | Wtip | 0.55 | 3.18E-12 | 6.60E-08 | 0.62 | 0.56 | 0.06 | 1.11 |
| 14 | Pkp4 | 0.71 | 3.94E-12 | 8.20E-08 | 0.47 | 0.38 | 0.09 | 1.24 |
| 14 | Cryl1 | 1.22 | 4.60E-12 | 9.57E-08 | 0.24 | 0.15 | 0.09 | 1.60 |
| 14 | Helb | 1.26 | 4.71E-12 | 9.80E-08 | 0.22 | 0.13 | 0.09 | 1.68 |
| 14 | Tmem132a | 0.59 | 5.49E-12 | 1.14E-07 | 0.67 | 0.64 | 0.03 | 1.05 |
| 14 | Baz1a | 0.58 | 5.68E-12 | 1.18E-07 | 0.60 | 0.56 | 0.04 | 1.08 |
| 14 | Plppr1 | 1.86 | 7.10E-12 | 1.48E-07 | 0.10 | 0.04 | 0.06 | 2.32 |
| 14 | Slc4a2 | 1.02 | 7.65E-12 | 1.59E-07 | 0.31 | 0.22 | 0.10 | 1.45 |
| 14 | Pcyt1b | 1.47 | 8.13E-12 | 1.69E-07 | 0.20 | 0.12 | 0.09 | 1.71 |
| 14 | Smarcad1 | 0.44 | 8.32E-12 | 1.73E-07 | 0.76 | 0.76 | 0.00 | 1.01 |
| 14 | Gprc5c | 1.00 | 1.06E-11 | 2.21E-07 | 0.33 | 0.23 | 0.10 | 1.41 |
| 14 | Esrp2 | 1.24 | 1.56E-11 | 3.25E-07 | 0.13 | 0.07 | 0.07 | 2.03 |
| 14 | Tnks2 | 0.38 | 1.96E-11 | 4.08E-07 | 0.81 | 0.81 | 0.00 | 1.00 |
| 14 | Atg10 | 1.00 | 2.17E-11 | 4.52E-07 | 0.24 | 0.15 | 0.09 | 1.60 |
| 14 | Hmga2 | 0.31 | 2.22E-11 | 4.63E-07 | 0.95 | 0.93 | 0.01 | 1.02 |
| 14 | 2010320M18Rik | 0.75 | 2.26E-11 | 4.70E-07 | 0.43 | 0.34 | 0.09 | 1.26 |
| 14 | Glcci1 | 1.09 | 2.58E-11 | 5.36E-07 | 0.46 | 0.39 | 0.07 | 1.18 |
| 14 | Ppp2r2b | 1.29 | 3.49E-11 | 7.26E-07 | 0.17 | 0.10 | 0.08 | 1.81 |
| 14 | 2610307P16Rik | 1.02 | 3.81E-11 | 7.93E-07 | 0.27 | 0.18 | 0.09 | 1.53 |
| 14 | Mlec | 0.57 | 4.09E-11 | 8.50E-07 | 0.73 | 0.75 | -0.02 | 0.97 |
| 14 | Sdr39u1 | 0.98 | 4.41E-11 | 9.17E-07 | 0.25 | 0.16 | 0.09 | 1.58 |
| 14 | Ldb1 | 0.49 | 4.67E-11 | 9.70E-07 | 0.67 | 0.64 | 0.03 | 1.05 |
| 14 | Smim10l2a | 1.32 | 4.93E-11 | 1.02E-06 | 0.14 | 0.07 | 0.07 | 1.96 |
| 14 | Bcl7a | 0.44 | 4.99E-11 | 1.04E-06 | 0.72 | 0.67 | 0.05 | 1.07 |
| 14 | Slc7a1 | 1.70 | 5.62E-11 | 1.17E-06 | 0.15 | 0.08 | 0.07 | 1.88 |
| 14 | Cd109 | 1.53 | 5.86E-11 | 1.22E-06 | 0.11 | 0.05 | 0.06 | 2.15 |
| 14 | Aass | 1.61 | 6.39E-11 | 1.33E-06 | 0.11 | 0.05 | 0.06 | 2.18 |
| 14 | Zfp992 | 1.24 | 6.49E-11 | 1.35E-06 | 0.24 | 0.16 | 0.09 | 1.55 |
| 14 | Cyb561 | 1.41 | 7.62E-11 | 1.59E-06 | 0.12 | 0.06 | 0.06 | 2.09 |
| 14 | Sox6 | 0.76 | 8.31E-11 | 1.73E-06 | 0.21 | 0.12 | 0.09 | 1.71 |
| 14 | Trpm3 | 1.71 | 8.73E-11 | 1.82E-06 | 0.10 | 0.05 | 0.06 | 2.22 |
| 14 | Rreb1 | 0.99 | 9.26E-11 | 1.93E-06 | 0.30 | 0.21 | 0.09 | 1.44 |
| 14 | Vps4b | 0.51 | 9.36E-11 | 1.95E-06 | 0.66 | 0.63 | 0.04 | 1.06 |
| 14 | Yap1 | 0.49 | 9.82E-11 | 2.04E-06 | 0.67 | 0.62 | 0.05 | 1.09 |
| 14 | Tmem53 | 1.29 | 1.03E-10 | 2.13E-06 | 0.16 | 0.09 | 0.07 | 1.85 |
| 14 | Rasa3 | 1.26 | 1.03E-10 | 2.14E-06 | 0.14 | 0.08 | 0.07 | 1.89 |
| 14 | Grip1 | 1.26 | 1.26E-10 | 2.62E-06 | 0.18 | 0.11 | 0.08 | 1.73 |
| 14 | Zfp664 | 0.39 | 1.27E-10 | 2.65E-06 | 0.77 | 0.75 | 0.02 | 1.03 |
| 14 | Limk2 | 0.86 | 1.28E-10 | 2.66E-06 | 0.29 | 0.20 | 0.09 | 1.48 |
| 14 | Kdm3a | 0.76 | 1.35E-10 | 2.80E-06 | 0.45 | 0.37 | 0.08 | 1.21 |
| 14 | Nox4 | 1.50 | 1.38E-10 | 2.87E-06 | 0.15 | 0.08 | 0.07 | 1.89 |
| 14 | Snap47 | 0.49 | 1.44E-10 | 2.99E-06 | 0.69 | 0.65 | 0.04 | 1.05 |
| 14 | Fam172a | 0.81 | 1.52E-10 | 3.16E-06 | 0.43 | 0.35 | 0.08 | 1.24 |
| 14 | Baz2b | 0.43 | 1.59E-10 | 3.31E-06 | 0.73 | 0.71 | 0.02 | 1.03 |
| 14 | Tead3 | 0.98 | 1.63E-10 | 3.39E-06 | 0.34 | 0.25 | 0.09 | 1.38 |

**Supplementary Table 1. Marker genes for the original clusters of integrated clustering of E11.5 control and NotchGOF λ-junction cells.**

| | | | | | | | | |
|---|---|---|---|---|---|---|---|---|
| 14 | Scd2 | 0.33 | 1.68E-10 | 3.50E-06 | 0.91 | 0.92 | -0.01 | 0.99 |
| 14 | Acp6 | 1.06 | 1.96E-10 | 4.07E-06 | 0.31 | 0.22 | 0.09 | 1.38 |
| 14 | Flywch2 | 0.62 | 1.98E-10 | 4.12E-06 | 0.54 | 0.46 | 0.08 | 1.18 |
| 14 | Arl2 | 0.46 | 2.08E-10 | 4.32E-06 | 0.67 | 0.65 | 0.02 | 1.03 |
| 14 | Ogt | 0.46 | 2.15E-10 | 4.47E-06 | 0.77 | 0.78 | -0.01 | 0.99 |
| 14 | Tiam1 | 0.81 | 2.29E-10 | 4.76E-06 | 0.40 | 0.31 | 0.09 | 1.28 |
| 14 | Khk | 1.50 | 2.30E-10 | 4.79E-06 | 0.13 | 0.07 | 0.06 | 1.95 |
| 14 | Nudt18 | 1.56 | 2.43E-10 | 5.06E-06 | 0.12 | 0.06 | 0.06 | 2.00 |
| 14 | Atp6v1a | 0.62 | 2.84E-10 | 5.90E-06 | 0.57 | 0.53 | 0.04 | 1.08 |
| 14 | Kif13a | 0.90 | 2.95E-10 | 6.14E-06 | 0.30 | 0.21 | 0.09 | 1.44 |
| 14 | Col4a5 | 1.20 | 3.14E-10 | 6.54E-06 | 0.21 | 0.13 | 0.08 | 1.63 |
| 14 | Abhd4 | 1.02 | 3.34E-10 | 6.94E-06 | 0.22 | 0.14 | 0.08 | 1.60 |
| 14 | Zfp612 | 0.98 | 3.63E-10 | 7.56E-06 | 0.25 | 0.17 | 0.09 | 1.51 |
| 14 | Hip1r | 1.24 | 3.75E-10 | 7.80E-06 | 0.14 | 0.08 | 0.07 | 1.89 |
| 14 | Pard3 | 0.51 | 3.79E-10 | 7.89E-06 | 0.63 | 0.58 | 0.05 | 1.09 |
| 14 | Fhl1 | 1.19 | 4.08E-10 | 8.48E-06 | 0.22 | 0.14 | 0.08 | 1.56 |
| 14 | Prdx5 | 0.41 | 4.10E-10 | 8.53E-06 | 0.79 | 0.80 | -0.01 | 0.99 |
| 14 | Tgfb1 | 1.04 | 4.12E-10 | 8.57E-06 | 0.18 | 0.11 | 0.08 | 1.72 |
| 14 | Ift43 | 0.47 | 4.55E-10 | 9.47E-06 | 0.62 | 0.55 | 0.08 | 1.14 |
| 14 | 1700025G04Rik | 0.91 | 4.61E-10 | 9.58E-06 | 0.41 | 0.34 | 0.08 | 1.23 |
| 14 | Copg2 | 0.73 | 4.78E-10 | 9.94E-06 | 0.45 | 0.37 | 0.08 | 1.22 |
| 14 | Llph | 0.34 | 4.82E-10 | 1.00E-05 | 0.84 | 0.86 | -0.02 | 0.97 |
| 14 | Ppfibp1 | 0.74 | 4.90E-10 | 1.02E-05 | 0.51 | 0.43 | 0.08 | 1.18 |
| 14 | Cecr2 | 0.87 | 5.24E-10 | 1.09E-05 | 0.31 | 0.22 | 0.09 | 1.39 |
| 14 | Fam32a | 0.40 | 5.74E-10 | 1.19E-05 | 0.76 | 0.77 | -0.01 | 0.99 |
| 14 | Kif5c | 0.61 | 5.83E-10 | 1.21E-05 | 0.54 | 0.49 | 0.06 | 1.11 |
| 14 | Bnc2 | 0.94 | 6.35E-10 | 1.32E-05 | 0.26 | 0.18 | 0.09 | 1.48 |
| 14 | Msi2 | 0.41 | 6.50E-10 | 1.35E-05 | 0.71 | 0.68 | 0.03 | 1.04 |
| 14 | Zbtb43 | 0.97 | 7.13E-10 | 1.48E-05 | 0.31 | 0.23 | 0.09 | 1.38 |
| 14 | Sema4g | 1.10 | 7.36E-10 | 1.53E-05 | 0.22 | 0.14 | 0.08 | 1.58 |
| 14 | Actn4 | 0.43 | 7.48E-10 | 1.56E-05 | 0.71 | 0.70 | 0.01 | 1.01 |
| 14 | Sertad2 | 0.68 | 7.68E-10 | 1.60E-05 | 0.47 | 0.41 | 0.07 | 1.17 |
| 14 | Ccdc126 | 1.15 | 8.28E-10 | 1.72E-05 | 0.16 | 0.09 | 0.07 | 1.79 |
| 14 | Arhgap12 | 0.73 | 8.93E-10 | 1.86E-05 | 0.42 | 0.34 | 0.08 | 1.24 |
| 14 | Ldlrap1 | 0.93 | 1.13E-09 | 2.36E-05 | 0.27 | 0.18 | 0.09 | 1.47 |
| 14 | Zfhx2 | 0.81 | 1.15E-09 | 2.39E-05 | 0.34 | 0.25 | 0.09 | 1.36 |
| 14 | Cyp4f17 | 1.22 | 1.21E-09 | 2.52E-05 | 0.13 | 0.07 | 0.06 | 1.95 |
| 14 | Ttc5 | 0.47 | 1.30E-09 | 2.70E-05 | 0.60 | 0.54 | 0.05 | 1.10 |
| 14 | Tamm41 | 0.81 | 1.41E-09 | 2.93E-05 | 0.42 | 0.34 | 0.08 | 1.22 |
| 14 | Gcat | 0.61 | 1.46E-09 | 3.05E-05 | 0.54 | 0.48 | 0.06 | 1.11 |
| 14 | Dcbld1 | 1.02 | 1.53E-09 | 3.18E-05 | 0.38 | 0.30 | 0.08 | 1.25 |
| 14 | 6330403K07Rik | 0.33 | 1.54E-09 | 3.19E-05 | 0.86 | 0.85 | 0.00 | 1.00 |
| 14 | Igsf9b | 1.50 | 1.57E-09 | 3.26E-05 | 0.11 | 0.06 | 0.06 | 2.00 |
| 14 | Exoc3l4 | 1.25 | 1.63E-09 | 3.38E-05 | 0.13 | 0.07 | 0.06 | 1.89 |
| 14 | Mettl27 | 1.04 | 1.68E-09 | 3.49E-05 | 0.22 | 0.14 | 0.08 | 1.58 |
| 14 | Chmp1b | 0.73 | 1.73E-09 | 3.60E-05 | 0.43 | 0.35 | 0.08 | 1.21 |
| 14 | Cdc42ep3 | 0.72 | 1.81E-09 | 3.76E-05 | 0.25 | 0.16 | 0.09 | 1.53 |
| 14 | 5430403G16Rik | 1.54 | 1.84E-09 | 3.82E-05 | 0.13 | 0.07 | 0.06 | 1.89 |
| 14 | Syde2 | 1.29 | 1.91E-09 | 3.97E-05 | 0.14 | 0.07 | 0.06 | 1.85 |
| 14 | Ncoa3 | 1.01 | 2.00E-09 | 4.16E-05 | 0.31 | 0.23 | 0.08 | 1.36 |
| 14 | 1700012D14Rik | 1.19 | 2.01E-09 | 4.18E-05 | 0.20 | 0.13 | 0.08 | 1.59 |
| 14 | Ilrun | 0.65 | 2.21E-09 | 4.60E-05 | 0.52 | 0.46 | 0.06 | 1.13 |
| 14 | Lgr4 | 0.62 | 2.30E-09 | 4.78E-05 | 0.32 | 0.23 | 0.09 | 1.40 |
| 14 | Klhl24 | 0.58 | 2.53E-09 | 5.26E-05 | 0.64 | 0.60 | 0.04 | 1.07 |
| 14 | Cgn | 1.02 | 2.59E-09 | 5.39E-05 | 0.18 | 0.11 | 0.07 | 1.69 |
| 14 | Timp3 | 1.02 | 2.62E-09 | 5.44E-05 | 0.25 | 0.17 | 0.08 | 1.48 |

**Supplementary Table 1. Marker genes for the original clusters of integrated clustering of E11.5 control and NotchGOF λ-junction cells.**

| | | | | | | | | |
|---|---|---|---|---|---|---|---|---|
| 14 | Mblac2 | 1.48 | 2.66E-09 | 5.53E-05 | 0.10 | 0.05 | 0.05 | 2.08 |
| 14 | Abracl | 0.25 | 2.74E-09 | 5.69E-05 | 0.92 | 0.93 | 0.00 | 1.00 |
| 14 | Supt4a | 0.29 | 2.75E-09 | 5.73E-05 | 0.88 | 0.90 | -0.03 | 0.97 |
| 14 | Etv3 | 0.97 | 3.01E-09 | 6.26E-05 | 0.30 | 0.22 | 0.08 | 1.36 |
| 14 | Map4k3 | 0.87 | 3.12E-09 | 6.50E-05 | 0.36 | 0.28 | 0.08 | 1.29 |
| 14 | Naxd | 0.44 | 3.22E-09 | 6.69E-05 | 0.66 | 0.64 | 0.02 | 1.03 |
| 14 | Gltp | 0.64 | 3.48E-09 | 7.24E-05 | 0.47 | 0.40 | 0.08 | 1.19 |
| 14 | 1810058I24Rik | 0.26 | 3.58E-09 | 7.44E-05 | 0.92 | 0.94 | -0.02 | 0.98 |
| 14 | 9930014A18Rik | 1.32 | 3.66E-09 | 7.60E-05 | 0.10 | 0.05 | 0.05 | 2.08 |
| 14 | Fam120a | 0.66 | 3.70E-09 | 7.69E-05 | 0.51 | 0.46 | 0.05 | 1.12 |
| 14 | Pak1 | 0.65 | 3.83E-09 | 7.96E-05 | 0.55 | 0.49 | 0.06 | 1.13 |
| 14 | Arpc3 | 0.25 | 3.85E-09 | 8.01E-05 | 0.89 | 0.90 | -0.02 | 0.98 |
| 14 | Bod1l | 0.33 | 4.33E-09 | 9.00E-05 | 0.80 | 0.82 | -0.02 | 0.97 |
| 14 | Pank1 | 0.80 | 4.60E-09 | 9.56E-05 | 0.39 | 0.31 | 0.08 | 1.25 |
| 14 | Ing4 | 0.40 | 4.93E-09 | 1.03E-04 | 0.76 | 0.76 | 0.00 | 1.00 |
| 14 | Zfp458 | 1.45 | 5.04E-09 | 1.05E-04 | 0.12 | 0.06 | 0.06 | 1.92 |
| 14 | Ammecr1 | 0.76 | 5.46E-09 | 1.14E-04 | 0.42 | 0.35 | 0.07 | 1.21 |
| 14 | Tmem205 | 0.84 | 5.79E-09 | 1.20E-04 | 0.36 | 0.28 | 0.09 | 1.31 |
| 14 | Abcg1 | 1.05 | 6.56E-09 | 1.36E-04 | 0.15 | 0.08 | 0.07 | 1.79 |
| 14 | Slc26a2 | 0.96 | 6.69E-09 | 1.39E-04 | 0.23 | 0.16 | 0.08 | 1.50 |
| 14 | Capg | 0.95 | 6.83E-09 | 1.42E-04 | 0.23 | 0.15 | 0.08 | 1.53 |
| 14 | Lpar6 | 0.63 | 7.68E-09 | 1.60E-04 | 0.47 | 0.40 | 0.07 | 1.16 |
| 14 | Gm17251 | 1.03 | 7.97E-09 | 1.66E-04 | 0.23 | 0.15 | 0.08 | 1.51 |
| 14 | Chd6 | 0.40 | 8.04E-09 | 1.67E-04 | 0.75 | 0.73 | 0.02 | 1.02 |
| 14 | Micall1 | 0.99 | 8.37E-09 | 1.74E-04 | 0.19 | 0.12 | 0.07 | 1.62 |
| 14 | Ago1 | 0.66 | 8.63E-09 | 1.80E-04 | 0.44 | 0.36 | 0.08 | 1.21 |
| 14 | Lad1 | 1.12 | 9.42E-09 | 1.96E-04 | 0.15 | 0.09 | 0.07 | 1.73 |
| 14 | Ptar1 | 0.84 | 9.75E-09 | 2.03E-04 | 0.35 | 0.28 | 0.07 | 1.27 |
| 14 | Fundc2 | 0.40 | 1.13E-08 | 2.36E-04 | 0.75 | 0.78 | -0.03 | 0.97 |
| 14 | Cep70 | 0.64 | 1.14E-08 | 2.37E-04 | 0.46 | 0.40 | 0.07 | 1.17 |
| 14 | Sidt2 | 1.05 | 1.16E-08 | 2.42E-04 | 0.26 | 0.18 | 0.08 | 1.43 |
| 14 | Airn | 0.85 | 1.21E-08 | 2.51E-04 | 0.27 | 0.19 | 0.08 | 1.42 |
| 14 | Zkscan7 | 1.29 | 1.22E-08 | 2.54E-04 | 0.14 | 0.08 | 0.06 | 1.78 |
| 14 | Kidins220 | 0.86 | 1.26E-08 | 2.62E-04 | 0.36 | 0.28 | 0.08 | 1.28 |
| 14 | Grn | 0.52 | 1.28E-08 | 2.67E-04 | 0.43 | 0.35 | 0.08 | 1.23 |
| 14 | Nlk | 0.76 | 1.29E-08 | 2.68E-04 | 0.35 | 0.27 | 0.08 | 1.29 |
| 14 | Dcaf17 | 0.86 | 1.31E-08 | 2.73E-04 | 0.40 | 0.33 | 0.06 | 1.19 |
| 14 | Rfx7 | 0.59 | 1.44E-08 | 2.99E-04 | 0.52 | 0.46 | 0.06 | 1.12 |
| 14 | Cldn4 | 0.79 | 1.52E-08 | 3.15E-04 | 0.17 | 0.10 | 0.07 | 1.67 |
| 14 | Ppip5k2 | 0.60 | 1.63E-08 | 3.39E-04 | 0.51 | 0.46 | 0.05 | 1.12 |
| 14 | Lamc1 | 0.78 | 1.64E-08 | 3.41E-04 | 0.34 | 0.26 | 0.08 | 1.31 |
| 14 | Mtln | 0.63 | 1.70E-08 | 3.53E-04 | 0.41 | 0.33 | 0.08 | 1.23 |
| 14 | Cab39 | 0.58 | 1.70E-08 | 3.54E-04 | 0.61 | 0.58 | 0.03 | 1.05 |
| 14 | Hbp1 | 0.56 | 1.70E-08 | 3.55E-04 | 0.60 | 0.57 | 0.04 | 1.07 |
| 14 | Gnpda2 | 0.79 | 1.91E-08 | 3.96E-04 | 0.36 | 0.28 | 0.08 | 1.28 |
| 14 | Inava | 0.70 | 2.09E-08 | 4.35E-04 | 0.31 | 0.23 | 0.08 | 1.35 |
| 14 | Armcx1 | 0.74 | 2.13E-08 | 4.42E-04 | 0.41 | 0.34 | 0.07 | 1.20 |
| 14 | Nipsnap1 | 0.46 | 2.14E-08 | 4.45E-04 | 0.61 | 0.56 | 0.04 | 1.08 |
| 14 | Snhg11 | 1.14 | 2.19E-08 | 4.55E-04 | 0.17 | 0.10 | 0.07 | 1.66 |
| 14 | Ap2b1 | 0.52 | 2.23E-08 | 4.64E-04 | 0.54 | 0.49 | 0.05 | 1.11 |
| 14 | Arhgap29 | 0.70 | 2.27E-08 | 4.72E-04 | 0.43 | 0.37 | 0.06 | 1.17 |
| 14 | Map1lc3b | 0.29 | 2.28E-08 | 4.73E-04 | 0.90 | 0.93 | -0.03 | 0.97 |
| 14 | Ccdc112 | 0.57 | 2.49E-08 | 5.18E-04 | 0.56 | 0.52 | 0.05 | 1.09 |
| 14 | Ctsc | 0.72 | 2.57E-08 | 5.35E-04 | 0.19 | 0.12 | 0.07 | 1.56 |
| 14 | Bbx | 0.47 | 2.76E-08 | 5.74E-04 | 0.63 | 0.59 | 0.04 | 1.07 |
| 14 | Xpr1 | 0.87 | 2.94E-08 | 6.12E-04 | 0.35 | 0.27 | 0.08 | 1.29 |

**Supplementary Table 1. Marker genes for the original clusters of integrated clustering of E11.5 control and NotchGOF λ-junction cells.**

| | | | | | | | | |
|---|---|---|---|---|---|---|---|---|
| 14 | Cript | 0.32 | 2.99E-08 | 6.22E-04 | 0.78 | 0.80 | -0.02 | 0.97 |
| 14 | Amotl2 | 0.78 | 3.05E-08 | 6.34E-04 | 0.41 | 0.34 | 0.07 | 1.20 |
| 14 | Zfp36l2 | 0.90 | 3.22E-08 | 6.69E-04 | 0.48 | 0.43 | 0.04 | 1.09 |
| 14 | Mpst | 0.62 | 3.29E-08 | 6.84E-04 | 0.48 | 0.42 | 0.06 | 1.15 |
| 14 | Apbb2 | 0.79 | 3.67E-08 | 7.63E-04 | 0.37 | 0.30 | 0.07 | 1.22 |
| 14 | Megf6 | 0.91 | 4.20E-08 | 8.74E-04 | 0.29 | 0.21 | 0.08 | 1.37 |
| 14 | Gnai1 | 1.01 | 4.28E-08 | 8.90E-04 | 0.22 | 0.15 | 0.07 | 1.47 |
| 14 | Iqsec1 | 1.06 | 4.40E-08 | 9.15E-04 | 0.21 | 0.14 | 0.07 | 1.50 |
| 14 | Wipf2 | 0.79 | 4.87E-08 | 1.01E-03 | 0.33 | 0.25 | 0.07 | 1.29 |
| 14 | Rab12 | 0.58 | 5.09E-08 | 1.06E-03 | 0.54 | 0.49 | 0.05 | 1.10 |
| 14 | Apba2 | 1.59 | 5.14E-08 | 1.07E-03 | 0.11 | 0.06 | 0.05 | 1.84 |
| 14 | Foxn3 | 0.47 | 5.17E-08 | 1.07E-03 | 0.69 | 0.69 | 0.00 | 1.00 |
| 14 | Egfr | 1.17 | 5.28E-08 | 1.10E-03 | 0.17 | 0.10 | 0.07 | 1.64 |
| 14 | Nrarp | 0.45 | 5.57E-08 | 1.16E-03 | 0.27 | 0.18 | 0.09 | 1.50 |
| 14 | Cox14 | 0.33 | 5.74E-08 | 1.19E-03 | 0.79 | 0.81 | -0.01 | 0.98 |
| 14 | Ankrd49 | 0.66 | 6.26E-08 | 1.30E-03 | 0.49 | 0.44 | 0.05 | 1.12 |
| 14 | Ece1 | 0.67 | 6.40E-08 | 1.33E-03 | 0.31 | 0.23 | 0.08 | 1.35 |
| 14 | Tmem63b | 0.67 | 6.68E-08 | 1.39E-03 | 0.41 | 0.34 | 0.07 | 1.21 |
| 14 | Galc | 0.93 | 6.77E-08 | 1.41E-03 | 0.25 | 0.17 | 0.07 | 1.42 |
| 14 | Acrbp | 1.26 | 6.95E-08 | 1.45E-03 | 0.17 | 0.10 | 0.06 | 1.62 |
| 14 | Dbp | 1.00 | 7.01E-08 | 1.46E-03 | 0.21 | 0.14 | 0.07 | 1.51 |
| 14 | Ap4s1 | 0.73 | 7.03E-08 | 1.46E-03 | 0.46 | 0.42 | 0.04 | 1.11 |
| 14 | Reps1 | 0.71 | 7.03E-08 | 1.46E-03 | 0.42 | 0.35 | 0.06 | 1.18 |
| 14 | Arhgap42 | 0.66 | 7.67E-08 | 1.60E-03 | 0.44 | 0.37 | 0.06 | 1.17 |
| 14 | Hpcal1 | 0.71 | 8.02E-08 | 1.67E-03 | 0.40 | 0.33 | 0.07 | 1.20 |
| 14 | Pfkl | 0.39 | 8.15E-08 | 1.69E-03 | 0.66 | 0.65 | 0.01 | 1.02 |
| 14 | Mvb12a | 0.58 | 8.68E-08 | 1.81E-03 | 0.51 | 0.46 | 0.05 | 1.12 |
| 14 | Prr13 | 0.51 | 8.68E-08 | 1.81E-03 | 0.46 | 0.40 | 0.06 | 1.15 |
| 14 | Jmy | 0.68 | 8.80E-08 | 1.83E-03 | 0.44 | 0.38 | 0.06 | 1.16 |
| 14 | Usp46 | 0.41 | 8.94E-08 | 1.86E-03 | 0.65 | 0.63 | 0.02 | 1.03 |
| 14 | 1110012L19Rik | 0.80 | 8.97E-08 | 1.87E-03 | 0.35 | 0.28 | 0.07 | 1.26 |
| 14 | Hspa2 | 1.23 | 9.56E-08 | 1.99E-03 | 0.11 | 0.06 | 0.05 | 1.88 |
| 14 | Irs1 | 0.79 | 9.83E-08 | 2.04E-03 | 0.25 | 0.18 | 0.08 | 1.42 |
| 14 | Fbxl20 | 0.73 | 1.02E-07 | 2.11E-03 | 0.30 | 0.22 | 0.08 | 1.36 |
| 14 | H2-Q4 | 1.05 | 1.02E-07 | 2.12E-03 | 0.17 | 0.11 | 0.07 | 1.61 |
| 14 | Rdm1 | 0.74 | 1.02E-07 | 2.12E-03 | 0.39 | 0.32 | 0.07 | 1.22 |
| 14 | Dlgap4 | 0.46 | 1.14E-07 | 2.37E-03 | 0.67 | 0.67 | 0.00 | 1.00 |
| 14 | Nectin2 | 0.60 | 1.16E-07 | 2.40E-03 | 0.52 | 0.48 | 0.04 | 1.09 |
| 14 | Csnk1g1 | 0.62 | 1.19E-07 | 2.47E-03 | 0.43 | 0.36 | 0.06 | 1.17 |
| 14 | Npdc1 | 0.90 | 1.27E-07 | 2.65E-03 | 0.17 | 0.11 | 0.06 | 1.61 |
| 14 | Fam114a2 | 0.40 | 1.32E-07 | 2.74E-03 | 0.69 | 0.68 | 0.01 | 1.02 |
| 14 | Grasp | 0.87 | 1.33E-07 | 2.76E-03 | 0.26 | 0.19 | 0.07 | 1.36 |
| 14 | Sdhaf4 | 0.42 | 1.47E-07 | 3.07E-03 | 0.65 | 0.65 | 0.00 | 1.00 |
| 14 | Man1a | 1.14 | 1.48E-07 | 3.07E-03 | 0.11 | 0.06 | 0.05 | 1.84 |
| 14 | Ppp1r13l | 1.26 | 1.50E-07 | 3.13E-03 | 0.13 | 0.08 | 0.06 | 1.72 |
| 14 | Egln1 | 0.39 | 1.57E-07 | 3.26E-03 | 0.69 | 0.69 | 0.01 | 1.01 |
| 14 | Mrpl36 | 0.39 | 1.69E-07 | 3.52E-03 | 0.71 | 0.72 | -0.01 | 0.98 |
| 14 | Amn1 | 0.62 | 1.70E-07 | 3.53E-03 | 0.49 | 0.44 | 0.05 | 1.11 |
| 14 | Parp11 | 0.90 | 1.74E-07 | 3.61E-03 | 0.23 | 0.16 | 0.07 | 1.45 |
| 14 | Cebpzos | 0.44 | 1.75E-07 | 3.64E-03 | 0.60 | 0.57 | 0.03 | 1.05 |
| 14 | Polr3k | 0.32 | 1.75E-07 | 3.65E-03 | 0.82 | 0.85 | -0.03 | 0.97 |
| 14 | Zfp182 | 1.23 | 1.79E-07 | 3.72E-03 | 0.11 | 0.06 | 0.05 | 1.83 |
| 14 | Rab15 | 0.92 | 1.84E-07 | 3.82E-03 | 0.20 | 0.13 | 0.07 | 1.52 |
| 14 | Gpc2 | 0.62 | 1.89E-07 | 3.93E-03 | 0.46 | 0.40 | 0.07 | 1.17 |
| 14 | Ctr9 | 0.42 | 2.05E-07 | 4.26E-03 | 0.69 | 0.68 | 0.01 | 1.01 |
| 14 | Harbi1 | 0.95 | 2.24E-07 | 4.65E-03 | 0.25 | 0.19 | 0.07 | 1.37 |

**Supplementary Table 1. Marker genes for the original clusters of integrated clustering of E11.5 control and NotchGOF λ-junction cells.**

| | | | | | | | | |
|---|---|---|---|---|---|---|---|---|
| 14 | Ephb2 | 0.73 | 2.32E-07 | 4.82E-03 | 0.35 | 0.28 | 0.07 | 1.25 |
| 14 | Pbxip1 | 0.81 | 2.40E-07 | 4.99E-03 | 0.28 | 0.20 | 0.07 | 1.37 |
| 14 | Myef2 | 0.40 | 2.42E-07 | 5.04E-03 | 0.70 | 0.69 | 0.01 | 1.01 |
| 14 | Pcgf5 | 1.47 | 2.46E-07 | 5.13E-03 | 0.14 | 0.09 | 0.06 | 1.63 |
| 14 | Dlg3 | 0.60 | 2.48E-07 | 5.17E-03 | 0.40 | 0.32 | 0.08 | 1.24 |
| 14 | Bcl11a | 0.29 | 2.60E-07 | 5.41E-03 | 0.76 | 0.74 | 0.02 | 1.03 |
| 14 | Mterf1a | 1.10 | 2.62E-07 | 5.45E-03 | 0.14 | 0.08 | 0.06 | 1.69 |
| 14 | Prkx | 0.86 | 2.71E-07 | 5.64E-03 | 0.22 | 0.15 | 0.07 | 1.45 |
| 14 | Ppm1l | 0.79 | 2.95E-07 | 6.14E-03 | 0.30 | 0.22 | 0.07 | 1.33 |
| 14 | Blvra | 0.79 | 2.99E-07 | 6.21E-03 | 0.27 | 0.20 | 0.07 | 1.34 |
| 14 | Bmp7 | 0.65 | 3.13E-07 | 6.51E-03 | 0.45 | 0.38 | 0.06 | 1.16 |
| 14 | Tshz1 | 0.74 | 3.26E-07 | 6.79E-03 | 0.39 | 0.32 | 0.07 | 1.21 |
| 14 | Scamp2 | 0.56 | 3.32E-07 | 6.90E-03 | 0.39 | 0.31 | 0.07 | 1.23 |
| 14 | Gm16286 | 0.30 | 3.35E-07 | 6.98E-03 | 0.79 | 0.81 | -0.02 | 0.98 |
| 14 | Mterf4 | 0.71 | 3.41E-07 | 7.09E-03 | 0.40 | 0.34 | 0.06 | 1.17 |
| 14 | Pik3c2a | 0.61 | 3.45E-07 | 7.18E-03 | 0.46 | 0.41 | 0.05 | 1.13 |
| 14 | Slain1 | 1.21 | 3.45E-07 | 7.18E-03 | 0.11 | 0.06 | 0.05 | 1.82 |
| 14 | Emb | 0.30 | 3.51E-07 | 7.30E-03 | 0.49 | 0.41 | 0.08 | 1.19 |
| 14 | Gab1 | 0.90 | 3.67E-07 | 7.64E-03 | 0.26 | 0.19 | 0.07 | 1.38 |
| 14 | Axl | 0.87 | 3.89E-07 | 8.09E-03 | 0.22 | 0.15 | 0.07 | 1.44 |
| 14 | AC149090.1 | 0.69 | 4.63E-07 | 9.64E-03 | 0.35 | 0.28 | 0.07 | 1.24 |
| 14 | Ypel3 | 0.46 | 4.76E-07 | 9.91E-03 | 0.60 | 0.57 | 0.03 | 1.05 |
| 14 | 1810044D09Rik | 1.13 | 4.81E-07 | 1.00E-02 | 0.17 | 0.11 | 0.06 | 1.53 |
| 14 | Inafm2 | 0.98 | 5.24E-07 | 1.09E-02 | 0.19 | 0.13 | 0.06 | 1.50 |
| 14 | Firre | 1.25 | 5.34E-07 | 1.11E-02 | 0.14 | 0.09 | 0.06 | 1.62 |
| 14 | Sdsl | 1.18 | 5.89E-07 | 1.23E-02 | 0.11 | 0.06 | 0.05 | 1.81 |
| 14 | Pbx3 | 0.42 | 5.91E-07 | 1.23E-02 | 0.57 | 0.53 | 0.04 | 1.08 |
| 14 | Rb1cc1 | 0.43 | 6.61E-07 | 1.37E-02 | 0.66 | 0.66 | 0.00 | 1.00 |
| 14 | Ypel1 | 0.53 | 6.61E-07 | 1.37E-02 | 0.54 | 0.50 | 0.04 | 1.08 |
| 14 | Esco1 | 0.76 | 6.65E-07 | 1.38E-02 | 0.30 | 0.23 | 0.07 | 1.29 |
| 14 | Tgfbr1 | 0.49 | 6.66E-07 | 1.39E-02 | 0.57 | 0.56 | 0.02 | 1.03 |
| 14 | Nras | 0.37 | 6.69E-07 | 1.39E-02 | 0.70 | 0.69 | 0.01 | 1.01 |
| 14 | 4732440D04Rik | 1.12 | 6.80E-07 | 1.41E-02 | 0.16 | 0.10 | 0.06 | 1.59 |
| 14 | Slc4a4 | 1.23 | 7.02E-07 | 1.46E-02 | 0.17 | 0.11 | 0.06 | 1.53 |
| 14 | Pcyox1l | 0.88 | 7.09E-07 | 1.47E-02 | 0.27 | 0.20 | 0.07 | 1.33 |
| 14 | Dpm2 | 0.45 | 7.17E-07 | 1.49E-02 | 0.61 | 0.57 | 0.03 | 1.06 |
| 14 | Heph | 0.84 | 7.38E-07 | 1.53E-02 | 0.32 | 0.25 | 0.07 | 1.26 |
| 14 | Edem2 | 0.80 | 7.60E-07 | 1.58E-02 | 0.26 | 0.19 | 0.07 | 1.35 |
| 14 | Kifc3 | 1.28 | 7.86E-07 | 1.63E-02 | 0.13 | 0.07 | 0.05 | 1.69 |
| 14 | Os9 | 0.76 | 7.89E-07 | 1.64E-02 | 0.42 | 0.37 | 0.05 | 1.14 |
| 14 | Ankrd13c | 0.89 | 7.90E-07 | 1.64E-02 | 0.30 | 0.24 | 0.06 | 1.26 |
| 14 | Med13 | 0.49 | 7.96E-07 | 1.66E-02 | 0.58 | 0.55 | 0.02 | 1.04 |
| 14 | Grina | 0.65 | 8.08E-07 | 1.68E-02 | 0.32 | 0.25 | 0.07 | 1.28 |
| 14 | Nop53 | 0.41 | 8.46E-07 | 1.76E-02 | 0.71 | 0.74 | -0.03 | 0.96 |
| 14 | Ociad2 | 0.84 | 9.41E-07 | 1.96E-02 | 0.25 | 0.18 | 0.07 | 1.37 |
| 14 | Sgpl1 | 0.74 | 9.54E-07 | 1.98E-02 | 0.40 | 0.34 | 0.06 | 1.17 |
| 14 | Ide | 0.71 | 9.94E-07 | 2.07E-02 | 0.34 | 0.27 | 0.07 | 1.26 |
| 14 | Tprgl | 0.49 | 1.01E-06 | 2.10E-02 | 0.62 | 0.60 | 0.01 | 1.02 |
| 14 | Setd5 | 0.33 | 1.03E-06 | 2.15E-02 | 0.73 | 0.71 | 0.02 | 1.02 |
| 14 | Irx1 | 0.95 | 1.04E-06 | 2.16E-02 | 0.10 | 0.06 | 0.05 | 1.82 |
| 14 | Zcchc7 | 0.54 | 1.05E-06 | 2.18E-02 | 0.55 | 0.52 | 0.03 | 1.05 |
| 14 | Lrrc42 | 0.41 | 1.11E-06 | 2.31E-02 | 0.64 | 0.61 | 0.03 | 1.05 |
| 14 | Ptk7 | 0.72 | 1.11E-06 | 2.31E-02 | 0.42 | 0.37 | 0.05 | 1.14 |
| 14 | Bmyc | 0.48 | 1.14E-06 | 2.38E-02 | 0.65 | 0.62 | 0.02 | 1.04 |
| 14 | Suco | 0.60 | 1.23E-06 | 2.55E-02 | 0.52 | 0.49 | 0.03 | 1.06 |
| 14 | Atp6v0e | 0.34 | 1.25E-06 | 2.60E-02 | 0.72 | 0.72 | 0.00 | 1.00 |

**Supplementary Table 1. Marker genes for the original clusters of integrated clustering of E11.5 control and NotchGOF λ-junction cells.**

| | | | | | | | | |
|---|---|---|---|---|---|---|---|---|
| 14 | Macc1 | 0.90 | 1.25E-06 | 2.60E-02 | 0.10 | 0.06 | 0.05 | 1.82 |
| 14 | Ufl1 | 0.53 | 1.28E-06 | 2.66E-02 | 0.47 | 0.42 | 0.05 | 1.12 |
| 14 | Tradd | 0.88 | 1.30E-06 | 2.71E-02 | 0.17 | 0.12 | 0.06 | 1.51 |
| 14 | Tmem19 | 0.88 | 1.31E-06 | 2.72E-02 | 0.26 | 0.19 | 0.07 | 1.34 |
| 14 | Neto2 | 0.70 | 1.32E-06 | 2.74E-02 | 0.30 | 0.23 | 0.07 | 1.31 |
| 14 | Scai | 0.59 | 1.38E-06 | 2.87E-02 | 0.42 | 0.35 | 0.06 | 1.18 |
| 14 | Slc25a11 | 0.34 | 1.46E-06 | 3.03E-02 | 0.68 | 0.69 | -0.01 | 0.99 |
| 14 | Ndufaf4 | 0.62 | 1.51E-06 | 3.15E-02 | 0.59 | 0.60 | -0.02 | 0.97 |
| 14 | B230307C23Rik | 1.08 | 1.57E-06 | 3.27E-02 | 0.23 | 0.16 | 0.06 | 1.38 |
| 14 | Kantr | 0.76 | 1.62E-06 | 3.36E-02 | 0.23 | 0.17 | 0.07 | 1.39 |
| 14 | Uri1 | 0.37 | 1.74E-06 | 3.61E-02 | 0.63 | 0.62 | 0.01 | 1.01 |
| 14 | Fam173a | 0.50 | 1.76E-06 | 3.65E-02 | 0.53 | 0.51 | 0.02 | 1.04 |
| 14 | Rnf114 | 0.59 | 1.79E-06 | 3.72E-02 | 0.51 | 0.48 | 0.03 | 1.07 |
| 14 | C1qtnf4 | 0.62 | 1.85E-06 | 3.85E-02 | 0.36 | 0.29 | 0.07 | 1.24 |
| 14 | Scmh1 | 0.62 | 1.88E-06 | 3.92E-02 | 0.38 | 0.32 | 0.06 | 1.19 |
| 14 | Nsg1 | 0.42 | 1.93E-06 | 4.01E-02 | 0.69 | 0.67 | 0.02 | 1.03 |
| 14 | Snhg20 | 0.53 | 1.97E-06 | 4.11E-02 | 0.43 | 0.38 | 0.06 | 1.15 |
| 14 | 1500011B03Rik | 0.34 | 2.09E-06 | 4.34E-02 | 0.68 | 0.67 | 0.01 | 1.01 |
| 14 | Hes1 | 0.56 | 2.22E-06 | 4.63E-02 | 0.58 | 0.55 | 0.03 | 1.06 |
| 14 | Pick1 | 0.63 | 2.29E-06 | 4.76E-02 | 0.32 | 0.26 | 0.07 | 1.25 |
| 15 | Pdgfa | 0.93 | 1.34E-149 | 2.79E-145 | 0.93 | 0.27 | 0.66 | 3.42 |
| 15 | Six3os1 | 1.22 | 3.75E-148 | 7.80E-144 | 0.79 | 0.22 | 0.57 | 3.57 |
| 15 | Six3 | 1.09 | 1.60E-147 | 3.33E-143 | 0.82 | 0.23 | 0.59 | 3.59 |
| 15 | Dsp | 0.84 | 6.07E-147 | 1.26E-142 | 0.85 | 0.23 | 0.62 | 3.72 |
| 15 | Epcam | 0.98 | 3.52E-143 | 7.31E-139 | 0.98 | 0.34 | 0.65 | 2.93 |
| 15 | Spint2 | 0.96 | 3.60E-131 | 7.50E-127 | 0.94 | 0.30 | 0.64 | 3.14 |
| 15 | Igfbp5 | 0.94 | 7.27E-129 | 1.51E-124 | 0.98 | 0.40 | 0.58 | 2.44 |
| 15 | Sox2 | 1.03 | 3.82E-128 | 7.94E-124 | 0.69 | 0.18 | 0.51 | 3.77 |
| 15 | Cldn6 | 0.90 | 1.62E-127 | 3.36E-123 | 0.80 | 0.22 | 0.58 | 3.63 |
| 15 | Cldn7 | 0.90 | 1.11E-123 | 2.30E-119 | 0.80 | 0.23 | 0.57 | 3.46 |
| 15 | Pitx1 | 1.06 | 1.46E-118 | 3.04E-114 | 0.65 | 0.18 | 0.48 | 3.73 |
| 15 | Cdh1 | 1.04 | 4.43E-117 | 9.22E-113 | 0.63 | 0.16 | 0.47 | 3.91 |
| 15 | Krt8 | 0.77 | 1.96E-116 | 4.07E-112 | 0.81 | 0.25 | 0.56 | 3.23 |
| 15 | Ezr | 0.94 | 8.01E-116 | 1.67E-111 | 0.90 | 0.31 | 0.59 | 2.90 |
| 15 | Mpzl2 | 0.99 | 3.68E-112 | 7.66E-108 | 0.71 | 0.20 | 0.51 | 3.57 |
| 15 | Ikzf2 | 0.89 | 5.28E-112 | 1.10E-107 | 0.85 | 0.27 | 0.57 | 3.08 |
| 15 | Krt18 | 0.82 | 1.09E-111 | 2.26E-107 | 0.85 | 0.29 | 0.56 | 2.91 |
| 15 | Wnt4 | 0.93 | 6.77E-110 | 1.41E-105 | 0.67 | 0.19 | 0.48 | 3.47 |
| 15 | Bcam | 0.98 | 9.75E-106 | 2.03E-101 | 0.68 | 0.20 | 0.49 | 3.49 |
| 15 | Isl1 | 0.96 | 6.01E-104 | 1.25E-99 | 0.71 | 0.21 | 0.49 | 3.33 |
| 15 | Sp8 | 1.13 | 1.28E-100 | 2.67E-96 | 0.53 | 0.13 | 0.40 | 3.97 |
| 15 | Pax6 | 1.27 | 1.79E-100 | 3.73E-96 | 0.50 | 0.12 | 0.38 | 4.05 |
| 15 | Nav2 | 0.98 | 9.76E-99 | 2.03E-94 | 0.73 | 0.23 | 0.50 | 3.14 |
| 15 | Foxg1 | 1.12 | 8.95E-98 | 1.86E-93 | 0.53 | 0.14 | 0.39 | 3.83 |
| 15 | Cpm | 1.18 | 5.66E-97 | 1.18E-92 | 0.58 | 0.17 | 0.41 | 3.41 |
| 15 | Trp63 | 1.01 | 1.11E-95 | 2.31E-91 | 0.50 | 0.13 | 0.37 | 3.92 |
| 15 | Slc9a3r1 | 0.95 | 3.99E-94 | 8.31E-90 | 0.72 | 0.23 | 0.49 | 3.10 |
| 15 | Itga6 | 0.91 | 1.12E-93 | 2.32E-89 | 0.82 | 0.30 | 0.52 | 2.72 |
| 15 | Ano1 | 1.30 | 3.08E-93 | 6.41E-89 | 0.48 | 0.12 | 0.36 | 3.93 |
| 15 | Hist3h2ba | 0.71 | 1.39E-91 | 2.89E-87 | 0.79 | 0.27 | 0.53 | 2.99 |
| 15 | Ephb1 | 1.41 | 1.89E-91 | 3.93E-87 | 0.41 | 0.09 | 0.32 | 4.36 |
| 15 | Dmrta2 | 1.26 | 3.98E-91 | 8.28E-87 | 0.51 | 0.14 | 0.37 | 3.71 |
| 15 | Socs2 | 0.97 | 1.07E-88 | 2.23E-84 | 0.82 | 0.34 | 0.48 | 2.44 |
| 15 | Sfn | 0.45 | 9.30E-88 | 1.94E-83 | 0.64 | 0.20 | 0.44 | 3.18 |
| 15 | Cldn3 | 0.70 | 2.25E-87 | 4.68E-83 | 0.56 | 0.16 | 0.41 | 3.58 |
| 15 | Hey1 | 0.93 | 1.09E-86 | 2.27E-82 | 0.77 | 0.29 | 0.48 | 2.62 |

**Supplementary Table 1. Marker genes for the original clusters of integrated clustering of E11.5 control and NotchGOF λ-junction cells.**

| | | | | | | | | |
|---|---|---|---|---|---|---|---|---|
| 15 | Fermt1 | 0.91 | 2.25E-86 | 4.68E-82 | 0.51 | 0.14 | 0.37 | 3.76 |
| 15 | Krt14 | 0.98 | 5.51E-86 | 1.15E-81 | 0.47 | 0.13 | 0.34 | 3.66 |
| 15 | Tpd52 | 0.69 | 9.39E-86 | 1.95E-81 | 0.76 | 0.25 | 0.51 | 2.99 |
| 15 | Lama5 | 1.16 | 2.66E-84 | 5.53E-80 | 0.44 | 0.11 | 0.33 | 3.98 |
| 15 | St14 | 1.17 | 3.92E-83 | 8.16E-79 | 0.45 | 0.11 | 0.33 | 3.92 |
| 15 | Aldh1a3 | 1.22 | 1.50E-82 | 3.11E-78 | 0.62 | 0.23 | 0.39 | 2.74 |
| 15 | Frem2 | 0.91 | 1.10E-81 | 2.28E-77 | 0.61 | 0.19 | 0.42 | 3.27 |
| 15 | Esrp1 | 0.89 | 1.48E-81 | 3.07E-77 | 0.56 | 0.16 | 0.40 | 3.47 |
| 15 | Igfbp2 | 0.70 | 4.08E-80 | 8.49E-76 | 1.00 | 0.75 | 0.25 | 1.33 |
| 15 | Cldn9 | 1.34 | 4.21E-80 | 8.76E-76 | 0.36 | 0.09 | 0.28 | 4.27 |
| 15 | Kcnk1 | 1.03 | 1.16E-79 | 2.41E-75 | 0.58 | 0.18 | 0.40 | 3.26 |
| 15 | Rab25 | 0.81 | 4.68E-79 | 9.74E-75 | 0.54 | 0.16 | 0.39 | 3.50 |
| 15 | Otx2 | 1.31 | 5.00E-79 | 1.04E-74 | 0.39 | 0.10 | 0.29 | 4.03 |
| 15 | Bex1 | 0.75 | 1.01E-78 | 2.11E-74 | 0.98 | 0.58 | 0.40 | 1.69 |
| 15 | Gm16136 | 0.53 | 1.04E-78 | 2.16E-74 | 0.57 | 0.17 | 0.40 | 3.41 |
| 15 | Nrarp | 0.94 | 1.61E-77 | 3.35E-73 | 0.56 | 0.17 | 0.39 | 3.25 |
| 15 | Tenm2 | 1.85 | 2.38E-75 | 4.95E-71 | 0.31 | 0.07 | 0.24 | 4.46 |
| 15 | Pdgfc | 0.79 | 1.10E-74 | 2.29E-70 | 0.74 | 0.27 | 0.47 | 2.70 |
| 15 | Ccdc88c | 0.91 | 1.43E-74 | 2.97E-70 | 0.52 | 0.15 | 0.37 | 3.42 |
| 15 | Runx1 | 1.01 | 2.92E-74 | 6.07E-70 | 0.56 | 0.19 | 0.37 | 2.98 |
| 15 | Ptprf | 0.76 | 6.95E-74 | 1.45E-69 | 0.98 | 0.63 | 0.35 | 1.55 |
| 15 | Espn | 1.25 | 3.42E-73 | 7.11E-69 | 0.40 | 0.11 | 0.29 | 3.77 |
| 15 | Fgf9 | 1.21 | 5.51E-73 | 1.15E-68 | 0.39 | 0.10 | 0.29 | 3.81 |
| 15 | Jag1 | 1.09 | 8.67E-73 | 1.80E-68 | 0.54 | 0.17 | 0.37 | 3.15 |
| 15 | Ly6h | 0.95 | 2.87E-72 | 5.96E-68 | 0.48 | 0.15 | 0.34 | 3.32 |
| 15 | Col18a1 | 0.62 | 6.59E-72 | 1.37E-67 | 0.85 | 0.35 | 0.50 | 2.41 |
| 15 | Grhl2 | 1.12 | 2.69E-71 | 5.60E-67 | 0.44 | 0.13 | 0.32 | 3.55 |
| 15 | Fjx1 | 0.86 | 6.44E-71 | 1.34E-66 | 0.84 | 0.38 | 0.46 | 2.20 |
| 15 | Erbb3 | 0.82 | 3.75E-70 | 7.79E-66 | 0.55 | 0.17 | 0.38 | 3.23 |
| 15 | Otx1 | 1.06 | 4.10E-69 | 8.52E-65 | 0.40 | 0.11 | 0.29 | 3.70 |
| 15 | Sema3d | 1.17 | 1.01E-68 | 2.11E-64 | 0.41 | 0.11 | 0.29 | 3.60 |
| 15 | Igfbp3 | 1.10 | 1.16E-68 | 2.41E-64 | 0.42 | 0.12 | 0.30 | 3.48 |
| 15 | Kif21a | 0.82 | 1.49E-68 | 3.09E-64 | 0.76 | 0.30 | 0.46 | 2.51 |
| 15 | Wwc1 | 0.92 | 2.02E-68 | 4.21E-64 | 0.55 | 0.18 | 0.37 | 3.14 |
| 15 | Ston2 | 0.81 | 5.40E-68 | 1.12E-63 | 0.55 | 0.18 | 0.38 | 3.15 |
| 15 | Dsg2 | 1.10 | 4.73E-67 | 9.83E-63 | 0.43 | 0.13 | 0.31 | 3.46 |
| 15 | Hes5 | 1.25 | 8.77E-67 | 1.82E-62 | 0.39 | 0.11 | 0.28 | 3.49 |
| 15 | Cadm1 | 0.87 | 1.09E-66 | 2.27E-62 | 0.82 | 0.38 | 0.44 | 2.16 |
| 15 | Ap1m2 | 0.84 | 2.88E-65 | 5.99E-61 | 0.47 | 0.14 | 0.33 | 3.38 |
| 15 | Wfdc2 | 0.74 | 6.31E-65 | 1.31E-60 | 0.87 | 0.45 | 0.42 | 1.92 |
| 15 | C130071C03Rik | 1.65 | 9.36E-65 | 1.95E-60 | 0.24 | 0.05 | 0.19 | 4.82 |
| 15 | Lsr | 0.91 | 8.44E-63 | 1.76E-58 | 0.58 | 0.20 | 0.38 | 2.88 |
| 15 | Nectin4 | 0.62 | 8.64E-63 | 1.80E-58 | 0.43 | 0.13 | 0.31 | 3.44 |
| 15 | Tnfaip2 | 1.19 | 9.57E-63 | 1.99E-58 | 0.44 | 0.14 | 0.30 | 3.15 |
| 15 | Hspb8 | 0.87 | 1.00E-61 | 2.08E-57 | 0.44 | 0.13 | 0.31 | 3.29 |
| 15 | Perp | 0.51 | 1.51E-61 | 3.14E-57 | 0.45 | 0.14 | 0.31 | 3.30 |
| 15 | Stx3 | 1.07 | 2.61E-60 | 5.43E-56 | 0.41 | 0.12 | 0.29 | 3.38 |
| 15 | Arhgef19 | 0.96 | 5.98E-60 | 1.24E-55 | 0.42 | 0.13 | 0.30 | 3.34 |
| 15 | Frmd4b | 0.85 | 2.00E-58 | 4.15E-54 | 0.49 | 0.16 | 0.33 | 3.01 |
| 15 | Dlx2 | 1.05 | 5.06E-58 | 1.05E-53 | 0.40 | 0.13 | 0.28 | 3.20 |
| 15 | Wnt6 | 0.84 | 5.33E-58 | 1.11E-53 | 0.34 | 0.10 | 0.25 | 3.54 |
| 15 | Dab1 | 0.74 | 5.45E-58 | 1.13E-53 | 0.47 | 0.15 | 0.32 | 3.12 |
| 15 | Crb3 | 1.04 | 6.99E-58 | 1.45E-53 | 0.37 | 0.11 | 0.27 | 3.52 |
| 15 | Cdkn1a | 0.85 | 3.35E-57 | 6.96E-53 | 0.37 | 0.11 | 0.26 | 3.42 |
| 15 | Plch1 | 1.32 | 6.11E-57 | 1.27E-52 | 0.34 | 0.10 | 0.25 | 3.54 |
| 15 | Pltp | 0.82 | 7.61E-56 | 1.58E-51 | 0.47 | 0.15 | 0.32 | 3.05 |

**Supplementary Table 1. Marker genes for the original clusters of integrated clustering of E11.5 control and NotchGOF λ-junction cells.**

| | | | | | | | | |
|---|---|---|---|---|---|---|---|---|
| 15 | Fgd2 | 1.03 | 1.25E-55 | 2.59E-51 | 0.35 | 0.10 | 0.25 | 3.58 |
| 15 | Cmtm8 | 0.70 | 5.14E-55 | 1.07E-50 | 0.51 | 0.17 | 0.34 | 2.96 |
| 15 | Ddr1 | 0.88 | 5.52E-55 | 1.15E-50 | 0.41 | 0.13 | 0.29 | 3.24 |
| 15 | Lad1 | 1.13 | 1.96E-54 | 4.07E-50 | 0.32 | 0.09 | 0.23 | 3.69 |
| 15 | Id4 | 0.78 | 3.68E-54 | 7.66E-50 | 0.69 | 0.30 | 0.39 | 2.31 |
| 15 | Samd5 | 0.82 | 6.02E-54 | 1.25E-49 | 0.76 | 0.36 | 0.40 | 2.12 |
| 15 | Pawr | 1.10 | 8.80E-54 | 1.83E-49 | 0.40 | 0.13 | 0.27 | 3.17 |
| 15 | Krt5 | 0.30 | 1.57E-53 | 3.26E-49 | 0.32 | 0.09 | 0.23 | 3.60 |
| 15 | Tmem54 | 0.75 | 1.62E-53 | 3.37E-49 | 0.39 | 0.12 | 0.27 | 3.34 |
| 15 | Notch1 | 0.71 | 3.84E-53 | 7.98E-49 | 0.60 | 0.24 | 0.36 | 2.51 |
| 15 | Ctnnd2 | 0.92 | 6.47E-53 | 1.35E-48 | 0.38 | 0.12 | 0.27 | 3.31 |
| 15 | Sox3 | 1.66 | 1.84E-52 | 3.83E-48 | 0.19 | 0.04 | 0.15 | 4.95 |
| 15 | Mapk13 | 0.79 | 8.61E-52 | 1.79E-47 | 0.54 | 0.20 | 0.34 | 2.75 |
| 15 | Gad2 | 1.02 | 1.84E-51 | 3.82E-47 | 0.27 | 0.07 | 0.20 | 3.85 |
| 15 | Acer2 | 0.58 | 3.03E-51 | 6.29E-47 | 0.82 | 0.39 | 0.43 | 2.12 |
| 15 | Map7 | 0.95 | 4.12E-51 | 8.57E-47 | 0.35 | 0.11 | 0.25 | 3.35 |
| 15 | Dok5 | 0.95 | 6.29E-51 | 1.31E-46 | 0.37 | 0.11 | 0.25 | 3.27 |
| 15 | Lratd2 | 1.01 | 7.83E-51 | 1.63E-46 | 0.35 | 0.11 | 0.25 | 3.31 |
| 15 | Emb | 0.73 | 8.05E-51 | 1.67E-46 | 0.80 | 0.40 | 0.40 | 2.00 |
| 15 | Spint1 | 0.91 | 2.26E-50 | 4.71E-46 | 0.33 | 0.10 | 0.24 | 3.47 |
| 15 | Myh14 | 1.04 | 2.50E-50 | 5.20E-46 | 0.29 | 0.08 | 0.21 | 3.71 |
| 15 | Bex4 | 0.63 | 5.95E-50 | 1.24E-45 | 0.95 | 0.65 | 0.31 | 1.47 |
| 15 | Pla2r1 | 1.09 | 1.15E-49 | 2.39E-45 | 0.27 | 0.07 | 0.20 | 3.79 |
| 15 | Nebl | 0.80 | 1.83E-49 | 3.81E-45 | 0.35 | 0.11 | 0.24 | 3.29 |
| 15 | 3110039M20Rik | 1.09 | 1.98E-49 | 4.12E-45 | 0.28 | 0.07 | 0.20 | 3.72 |
| 15 | D230030E09Rik | 1.22 | 1.12E-48 | 2.33E-44 | 0.23 | 0.05 | 0.17 | 4.20 |
| 15 | Sorbs2 | 0.88 | 2.03E-48 | 4.21E-44 | 0.55 | 0.21 | 0.34 | 2.57 |
| 15 | 2900026A02Rik | 1.00 | 2.65E-48 | 5.52E-44 | 0.34 | 0.10 | 0.24 | 3.36 |
| 15 | Dpysl5 | 0.93 | 3.74E-48 | 7.77E-44 | 0.34 | 0.10 | 0.24 | 3.30 |
| 15 | Hook1 | 0.78 | 5.17E-48 | 1.07E-43 | 0.53 | 0.20 | 0.33 | 2.66 |
| 15 | Dmbx1 | 1.51 | 9.91E-48 | 2.06E-43 | 0.19 | 0.04 | 0.15 | 4.57 |
| 15 | Plekha7 | 1.04 | 1.56E-47 | 3.25E-43 | 0.31 | 0.09 | 0.22 | 3.46 |
| 15 | Irf6 | 0.83 | 2.96E-47 | 6.16E-43 | 0.34 | 0.11 | 0.24 | 3.24 |
| 15 | Oxtr | 1.19 | 3.29E-47 | 6.83E-43 | 0.25 | 0.07 | 0.19 | 3.85 |
| 15 | Pdlim1 | 0.54 | 3.70E-47 | 7.70E-43 | 0.65 | 0.29 | 0.37 | 2.29 |
| 15 | Grb7 | 0.94 | 4.11E-47 | 8.55E-43 | 0.32 | 0.10 | 0.23 | 3.39 |
| 15 | Ablim1 | 0.78 | 4.35E-47 | 9.04E-43 | 0.55 | 0.22 | 0.33 | 2.50 |
| 15 | Cdcp1 | 1.21 | 6.44E-47 | 1.34E-42 | 0.23 | 0.06 | 0.17 | 4.09 |
| 15 | Slc22a23 | 1.19 | 1.04E-46 | 2.17E-42 | 0.31 | 0.09 | 0.22 | 3.35 |
| 15 | Kdf1 | 1.04 | 1.14E-46 | 2.37E-42 | 0.29 | 0.08 | 0.21 | 3.54 |
| 15 | Lgr4 | 0.83 | 1.52E-46 | 3.15E-42 | 0.56 | 0.22 | 0.34 | 2.50 |
| 15 | Gm266 | 1.00 | 1.61E-46 | 3.35E-42 | 0.64 | 0.31 | 0.33 | 2.09 |
| 15 | Celf2 | 0.76 | 2.70E-46 | 5.61E-42 | 0.60 | 0.25 | 0.35 | 2.41 |
| 15 | Mir205hg | 0.62 | 4.63E-46 | 9.63E-42 | 0.37 | 0.12 | 0.25 | 3.13 |
| 15 | Klk8 | 0.65 | 6.75E-46 | 1.40E-41 | 0.55 | 0.21 | 0.34 | 2.61 |
| 15 | Fgf8 | 1.13 | 1.73E-45 | 3.61E-41 | 0.27 | 0.08 | 0.19 | 3.50 |
| 15 | Nr2e1 | 1.34 | 5.14E-45 | 1.07E-40 | 0.19 | 0.04 | 0.14 | 4.51 |
| 15 | Rab15 | 0.75 | 5.83E-45 | 1.21E-40 | 0.39 | 0.13 | 0.26 | 3.03 |
| 15 | Kremen2 | 1.10 | 5.98E-45 | 1.24E-40 | 0.19 | 0.04 | 0.15 | 4.41 |
| 15 | Celsr2 | 1.01 | 2.59E-44 | 5.39E-40 | 0.30 | 0.09 | 0.21 | 3.38 |
| 15 | Scube3 | 0.77 | 3.47E-44 | 7.21E-40 | 0.52 | 0.20 | 0.32 | 2.62 |
| 15 | Prss22 | 1.27 | 3.77E-44 | 7.85E-40 | 0.23 | 0.06 | 0.17 | 3.88 |
| 15 | Faah | 1.12 | 5.52E-44 | 1.15E-39 | 0.22 | 0.06 | 0.17 | 3.96 |
| 15 | Celsr1 | 0.88 | 7.79E-44 | 1.62E-39 | 0.32 | 0.10 | 0.22 | 3.24 |
| 15 | Stard10 | 0.52 | 8.59E-44 | 1.79E-39 | 0.69 | 0.29 | 0.40 | 2.37 |
| 15 | Cldn4 | 0.76 | 1.62E-43 | 3.36E-39 | 0.33 | 0.10 | 0.23 | 3.23 |

**Supplementary Table 1. Marker genes for the original clusters of integrated clustering of E11.5 control and NotchGOF λ-junction cells.**

| | | | | | | | | |
|---|---|---|---|---|---|---|---|---|
| 15 | Cdh3 | 0.70 | 1.70E-43 | 3.54E-39 | 0.52 | 0.20 | 0.32 | 2.56 |
| 15 | Lmo7 | 0.87 | 3.55E-43 | 7.39E-39 | 0.43 | 0.16 | 0.27 | 2.70 |
| 15 | Dlk1 | 0.75 | 4.95E-43 | 1.03E-38 | 0.76 | 0.42 | 0.35 | 1.84 |
| 15 | Dlx3 | 0.96 | 5.05E-43 | 1.05E-38 | 0.26 | 0.07 | 0.19 | 3.60 |
| 15 | Cgn | 0.94 | 5.70E-43 | 1.19E-38 | 0.33 | 0.10 | 0.22 | 3.17 |
| 15 | Khdrbs3 | 0.64 | 9.03E-43 | 1.88E-38 | 0.93 | 0.61 | 0.32 | 1.52 |
| 15 | Fras1 | 0.73 | 1.08E-42 | 2.24E-38 | 0.61 | 0.26 | 0.35 | 2.34 |
| 15 | Dtna | 1.00 | 2.34E-42 | 4.88E-38 | 0.39 | 0.14 | 0.26 | 2.83 |
| 15 | AW011738 | 1.13 | 3.53E-42 | 7.34E-38 | 0.29 | 0.09 | 0.20 | 3.30 |
| 15 | Tubb2b | 0.49 | 1.32E-41 | 2.74E-37 | 0.95 | 0.66 | 0.29 | 1.43 |
| 15 | Abcg1 | 1.12 | 1.56E-41 | 3.25E-37 | 0.27 | 0.08 | 0.19 | 3.39 |
| 15 | Itgb6 | 1.42 | 3.97E-41 | 8.25E-37 | 0.18 | 0.04 | 0.14 | 4.40 |
| 15 | Nrcam | 1.52 | 4.02E-41 | 8.36E-37 | 0.20 | 0.05 | 0.15 | 3.98 |
| 15 | Fgf17 | 1.22 | 4.35E-41 | 9.05E-37 | 0.18 | 0.04 | 0.14 | 4.34 |
| 15 | Prss8 | 0.95 | 1.44E-40 | 3.00E-36 | 0.30 | 0.10 | 0.21 | 3.19 |
| 15 | Scg5 | 1.12 | 2.10E-40 | 4.37E-36 | 0.20 | 0.05 | 0.15 | 4.00 |
| 15 | Rprm | 0.82 | 2.88E-40 | 5.98E-36 | 0.41 | 0.16 | 0.25 | 2.59 |
| 15 | Cdh2 | 0.65 | 4.46E-40 | 9.27E-36 | 0.78 | 0.42 | 0.36 | 1.86 |
| 15 | Schip1 | 0.72 | 7.87E-40 | 1.64E-35 | 0.43 | 0.16 | 0.27 | 2.71 |
| 15 | Cpz | 1.20 | 1.02E-39 | 2.13E-35 | 0.24 | 0.07 | 0.17 | 3.54 |
| 15 | Jup | 0.66 | 2.05E-39 | 4.27E-35 | 0.49 | 0.20 | 0.30 | 2.52 |
| 15 | Zdhhc2 | 1.09 | 2.47E-39 | 5.14E-35 | 0.35 | 0.13 | 0.23 | 2.79 |
| 15 | Rragd | 0.69 | 4.02E-39 | 8.36E-35 | 0.53 | 0.22 | 0.31 | 2.40 |
| 15 | Slc12a2 | 0.62 | 4.78E-39 | 9.94E-35 | 0.81 | 0.43 | 0.38 | 1.88 |
| 15 | F11r | 0.42 | 6.74E-39 | 1.40E-34 | 0.47 | 0.18 | 0.29 | 2.67 |
| 15 | Trpm3 | 1.73 | 6.93E-39 | 1.44E-34 | 0.18 | 0.05 | 0.14 | 4.07 |
| 15 | Wipf3 | 1.27 | 1.40E-38 | 2.91E-34 | 0.22 | 0.06 | 0.16 | 3.71 |
| 15 | Pbx1 | 0.47 | 1.94E-38 | 4.04E-34 | 1.00 | 0.92 | 0.08 | 1.09 |
| 15 | Hs6st2 | 0.70 | 2.50E-38 | 5.19E-34 | 0.61 | 0.28 | 0.33 | 2.18 |
| 15 | Tex15 | 0.80 | 4.89E-38 | 1.02E-33 | 0.32 | 0.11 | 0.21 | 2.96 |
| 15 | Tom1l1 | 0.69 | 5.68E-38 | 1.18E-33 | 0.42 | 0.16 | 0.26 | 2.66 |
| 15 | Tspan13 | 0.54 | 7.43E-38 | 1.55E-33 | 0.81 | 0.41 | 0.41 | 2.00 |
| 15 | Galm | 0.78 | 1.11E-37 | 2.30E-33 | 0.50 | 0.21 | 0.29 | 2.39 |
| 15 | Klf5 | 0.68 | 1.20E-37 | 2.49E-33 | 0.35 | 0.12 | 0.23 | 2.87 |
| 15 | Sptbn2 | 0.91 | 1.50E-37 | 3.13E-33 | 0.33 | 0.11 | 0.22 | 2.96 |
| 15 | Zfhx2 | 0.72 | 1.82E-37 | 3.79E-33 | 0.57 | 0.25 | 0.32 | 2.31 |
| 15 | 2310030G06Rik | 1.04 | 2.26E-37 | 4.70E-33 | 0.24 | 0.07 | 0.17 | 3.47 |
| 15 | Hap1 | 1.32 | 3.40E-37 | 7.08E-33 | 0.19 | 0.05 | 0.14 | 3.92 |
| 15 | Meg3 | 0.57 | 3.87E-37 | 8.05E-33 | 0.95 | 0.69 | 0.26 | 1.38 |
| 15 | Nav3 | 0.68 | 3.90E-37 | 8.12E-33 | 0.35 | 0.12 | 0.22 | 2.78 |
| 15 | Esrp2 | 1.15 | 5.28E-37 | 1.10E-32 | 0.23 | 0.06 | 0.16 | 3.57 |
| 15 | Nckap5 | 0.78 | 9.51E-37 | 1.98E-32 | 0.28 | 0.09 | 0.19 | 3.16 |
| 15 | Ntf5 | 0.90 | 1.36E-36 | 2.83E-32 | 0.27 | 0.09 | 0.19 | 3.17 |
| 15 | Ermn | 1.34 | 1.90E-36 | 3.96E-32 | 0.17 | 0.04 | 0.13 | 4.07 |
| 15 | Tfap2b | 0.51 | 2.39E-36 | 4.98E-32 | 0.27 | 0.09 | 0.18 | 3.11 |
| 15 | Ago1 | 0.68 | 5.88E-36 | 1.22E-31 | 0.72 | 0.36 | 0.36 | 2.01 |
| 15 | Ocln | 1.10 | 6.94E-36 | 1.44E-31 | 0.25 | 0.08 | 0.17 | 3.31 |
| 15 | Adamts18 | 0.90 | 8.72E-36 | 1.81E-31 | 0.23 | 0.07 | 0.17 | 3.39 |
| 15 | Jag2 | 1.02 | 1.54E-35 | 3.20E-31 | 0.26 | 0.08 | 0.18 | 3.19 |
| 15 | Acvr2a | 0.71 | 1.87E-35 | 3.89E-31 | 0.58 | 0.27 | 0.32 | 2.20 |
| 15 | Bicd1 | 0.89 | 2.08E-35 | 4.33E-31 | 0.29 | 0.10 | 0.19 | 3.04 |
| 15 | Tgfb2 | 0.78 | 2.31E-35 | 4.80E-31 | 0.80 | 0.48 | 0.32 | 1.66 |
| 15 | Pkp2 | 1.30 | 2.48E-35 | 5.15E-31 | 0.19 | 0.05 | 0.14 | 3.73 |
| 15 | Epb41l5 | 0.61 | 2.92E-35 | 6.08E-31 | 0.57 | 0.25 | 0.32 | 2.30 |
| 15 | Pkp3 | 0.86 | 3.87E-35 | 8.05E-31 | 0.23 | 0.07 | 0.16 | 3.46 |
| 15 | Pcdh7 | 0.49 | 4.03E-35 | 8.39E-31 | 0.78 | 0.39 | 0.39 | 2.00 |

**Supplementary Table 1. Marker genes for the original clusters of integrated clustering of E11.5 control and NotchGOF λ-junction cells.**

| | | | | | | | | |
|---|---|---|---|---|---|---|---|---|
| 15 | Fzd3 | 0.55 | 6.14E-35 | 1.28E-30 | 0.91 | 0.58 | 0.33 | 1.57 |
| 15 | Itga3 | 1.08 | 6.89E-35 | 1.43E-30 | 0.23 | 0.07 | 0.16 | 3.39 |
| 15 | Susd4 | 0.88 | 7.57E-35 | 1.57E-30 | 0.27 | 0.09 | 0.19 | 3.10 |
| 15 | Marveld2 | 1.15 | 9.50E-35 | 1.98E-30 | 0.21 | 0.06 | 0.15 | 3.55 |
| 15 | Cep170b | 0.72 | 9.50E-35 | 1.98E-30 | 0.35 | 0.12 | 0.22 | 2.80 |
| 15 | Cbln2 | 1.55 | 3.18E-34 | 6.60E-30 | 0.16 | 0.04 | 0.12 | 4.10 |
| 15 | Etl4 | 0.59 | 9.77E-34 | 2.03E-29 | 0.65 | 0.31 | 0.34 | 2.12 |
| 15 | Hs6st1 | 0.70 | 1.32E-33 | 2.74E-29 | 0.42 | 0.17 | 0.25 | 2.51 |
| 15 | Pcdh17 | 0.54 | 1.58E-33 | 3.28E-29 | 0.48 | 0.20 | 0.28 | 2.36 |
| 15 | Efna1 | 0.59 | 2.12E-33 | 4.41E-29 | 0.37 | 0.14 | 0.23 | 2.69 |
| 15 | Nuak2 | 1.40 | 2.36E-33 | 4.92E-29 | 0.15 | 0.04 | 0.11 | 4.17 |
| 15 | Adamts9 | 0.79 | 2.53E-33 | 5.27E-29 | 0.38 | 0.15 | 0.23 | 2.58 |
| 15 | Syt1 | 1.09 | 2.89E-33 | 6.01E-29 | 0.15 | 0.04 | 0.11 | 4.17 |
| 15 | Ppm1l | 0.77 | 3.05E-33 | 6.33E-29 | 0.50 | 0.22 | 0.28 | 2.29 |
| 15 | Heyl | 1.32 | 3.48E-33 | 7.24E-29 | 0.20 | 0.06 | 0.14 | 3.58 |
| 15 | Ripk4 | 0.77 | 3.67E-33 | 7.63E-29 | 0.24 | 0.07 | 0.16 | 3.28 |
| 15 | Lurap1l | 0.91 | 4.36E-33 | 9.07E-29 | 0.24 | 0.08 | 0.16 | 3.17 |
| 15 | Synrg | 0.56 | 4.78E-33 | 9.95E-29 | 0.53 | 0.23 | 0.30 | 2.30 |
| 15 | Rims2 | 1.06 | 8.78E-33 | 1.83E-28 | 0.20 | 0.06 | 0.14 | 3.55 |
| 15 | Rdh10 | 0.92 | 9.48E-33 | 1.97E-28 | 0.38 | 0.15 | 0.23 | 2.51 |
| 15 | Nectin1 | 0.75 | 1.27E-32 | 2.64E-28 | 0.42 | 0.17 | 0.25 | 2.48 |
| 15 | Eml5 | 0.98 | 1.34E-32 | 2.78E-28 | 0.26 | 0.08 | 0.17 | 3.10 |
| 15 | Elavl2 | 0.56 | 1.61E-32 | 3.36E-28 | 0.44 | 0.18 | 0.26 | 2.42 |
| 15 | Cdk6 | 0.66 | 1.62E-32 | 3.37E-28 | 0.67 | 0.34 | 0.33 | 1.99 |
| 15 | Dapl1 | 0.87 | 2.95E-32 | 6.14E-28 | 0.22 | 0.07 | 0.15 | 3.26 |
| 15 | Pdk1 | 0.61 | 3.00E-32 | 6.24E-28 | 0.56 | 0.25 | 0.31 | 2.24 |
| 15 | Prr15 | 1.56 | 3.71E-32 | 7.72E-28 | 0.14 | 0.03 | 0.11 | 4.34 |
| 15 | Six6 | 0.80 | 6.01E-32 | 1.25E-27 | 0.22 | 0.07 | 0.15 | 3.26 |
| 15 | Slc25a23 | 0.79 | 6.71E-32 | 1.39E-27 | 0.40 | 0.16 | 0.24 | 2.51 |
| 15 | Prr36 | 0.99 | 7.27E-32 | 1.51E-27 | 0.26 | 0.09 | 0.17 | 3.05 |
| 15 | Arrdc1 | 0.74 | 7.50E-32 | 1.56E-27 | 0.31 | 0.11 | 0.20 | 2.83 |
| 15 | Col2a1 | 0.44 | 8.55E-32 | 1.78E-27 | 0.94 | 0.64 | 0.30 | 1.47 |
| 15 | Jph4 | 1.05 | 1.08E-31 | 2.24E-27 | 0.19 | 0.05 | 0.14 | 3.55 |
| 15 | Nrip1 | 0.50 | 1.58E-31 | 3.28E-27 | 0.96 | 0.75 | 0.21 | 1.28 |
| 15 | 4930426D05Rik | 1.50 | 1.60E-31 | 3.32E-27 | 0.13 | 0.03 | 0.10 | 4.32 |
| 15 | Dmrt3 | 0.91 | 1.88E-31 | 3.92E-27 | 0.19 | 0.05 | 0.14 | 3.62 |
| 15 | Gm19938 | 1.85 | 2.12E-31 | 4.42E-27 | 0.11 | 0.02 | 0.09 | 4.83 |
| 15 | Clu | 0.63 | 2.70E-31 | 5.62E-27 | 0.47 | 0.20 | 0.26 | 2.28 |
| 15 | Nrg1 | 0.81 | 3.65E-31 | 7.58E-27 | 0.24 | 0.08 | 0.16 | 3.09 |
| 15 | Unc13b | 0.83 | 4.00E-31 | 8.32E-27 | 0.25 | 0.08 | 0.17 | 3.09 |
| 15 | Arid3b | 0.74 | 5.00E-31 | 1.04E-26 | 0.50 | 0.22 | 0.27 | 2.22 |
| 15 | Pla2g12a | 0.62 | 5.95E-31 | 1.24E-26 | 0.64 | 0.32 | 0.32 | 1.99 |
| 15 | Stk26 | 0.54 | 6.35E-31 | 1.32E-26 | 0.51 | 0.22 | 0.28 | 2.26 |
| 15 | Uchl1 | 0.40 | 1.14E-30 | 2.38E-26 | 0.68 | 0.35 | 0.33 | 1.96 |
| 15 | Mbp | 0.69 | 1.15E-30 | 2.39E-26 | 0.27 | 0.09 | 0.18 | 2.95 |
| 15 | Mapk12 | 1.03 | 1.16E-30 | 2.42E-26 | 0.30 | 0.11 | 0.19 | 2.71 |
| 15 | Large2 | 0.93 | 1.35E-30 | 2.80E-26 | 0.25 | 0.08 | 0.17 | 3.00 |
| 15 | Pkp4 | 0.55 | 1.67E-30 | 3.46E-26 | 0.73 | 0.37 | 0.36 | 1.97 |
| 15 | Nectin3 | 0.57 | 2.26E-30 | 4.71E-26 | 0.85 | 0.54 | 0.31 | 1.57 |
| 15 | Gkap1 | 0.49 | 2.59E-30 | 5.39E-26 | 0.80 | 0.43 | 0.37 | 1.86 |
| 15 | Mtcl1 | 0.83 | 2.74E-30 | 5.70E-26 | 0.32 | 0.12 | 0.20 | 2.68 |
| 15 | S100a11 | 0.32 | 3.26E-30 | 6.77E-26 | 0.97 | 0.76 | 0.21 | 1.28 |
| 15 | 2610528J11Rik | 0.99 | 4.44E-30 | 9.24E-26 | 0.20 | 0.06 | 0.14 | 3.35 |
| 15 | Dipk2a | 0.63 | 4.48E-30 | 9.31E-26 | 0.51 | 0.24 | 0.28 | 2.18 |
| 15 | Dmrta2os | 1.69 | 4.64E-30 | 9.65E-26 | 0.10 | 0.02 | 0.08 | 5.10 |
| 15 | Stxbp2 | 0.70 | 6.11E-30 | 1.27E-25 | 0.35 | 0.13 | 0.21 | 2.57 |

**Supplementary Table 1. Marker genes for the original clusters of integrated clustering of E11.5 control and NotchGOF λ-junction cells.**

| | | | | | | | | |
|---|---|---|---|---|---|---|---|---|
| 15 | Gjb2 | 1.06 | 8.40E-30 | 1.75E-25 | 0.16 | 0.05 | 0.12 | 3.64 |
| 15 | Tspan12 | 0.59 | 1.07E-29 | 2.23E-25 | 0.48 | 0.21 | 0.27 | 2.31 |
| 15 | Cd151 | 0.46 | 1.18E-29 | 2.45E-25 | 0.67 | 0.32 | 0.35 | 2.06 |
| 15 | Ptpru | 0.68 | 1.21E-29 | 2.51E-25 | 0.38 | 0.16 | 0.23 | 2.46 |
| 15 | Satb1 | 0.58 | 1.65E-29 | 3.44E-25 | 0.73 | 0.38 | 0.35 | 1.91 |
| 15 | Slain1 | 1.12 | 1.69E-29 | 3.50E-25 | 0.19 | 0.06 | 0.14 | 3.34 |
| 15 | Ehbp1 | 0.54 | 1.79E-29 | 3.72E-25 | 0.52 | 0.24 | 0.28 | 2.20 |
| 15 | Mgst1 | 0.34 | 2.03E-29 | 4.22E-25 | 0.73 | 0.37 | 0.36 | 1.98 |
| 15 | Tnfrsf19 | 0.73 | 4.70E-29 | 9.78E-25 | 0.36 | 0.15 | 0.21 | 2.46 |
| 15 | Zfp219 | 0.55 | 4.71E-29 | 9.80E-25 | 0.74 | 0.39 | 0.35 | 1.89 |
| 15 | Apoc1 | 0.56 | 7.00E-29 | 1.46E-24 | 0.25 | 0.08 | 0.16 | 2.95 |
| 15 | Cbarp | 0.80 | 8.36E-29 | 1.74E-24 | 0.29 | 0.11 | 0.19 | 2.75 |
| 15 | Vwa2 | 1.47 | 8.78E-29 | 1.83E-24 | 0.14 | 0.04 | 0.10 | 3.97 |
| 15 | Dsc2 | 0.93 | 8.88E-29 | 1.85E-24 | 0.19 | 0.06 | 0.13 | 3.36 |
| 15 | Lamc1 | 0.56 | 1.14E-28 | 2.37E-24 | 0.55 | 0.26 | 0.30 | 2.16 |
| 15 | Rassf10 | 0.94 | 2.52E-28 | 5.24E-24 | 0.19 | 0.06 | 0.14 | 3.29 |
| 15 | Lratd1 | 1.19 | 2.63E-28 | 5.47E-24 | 0.13 | 0.03 | 0.10 | 4.06 |
| 15 | Rnf43 | 1.21 | 2.74E-28 | 5.71E-24 | 0.17 | 0.05 | 0.12 | 3.52 |
| 15 | Igsf9 | 0.55 | 2.87E-28 | 5.97E-24 | 0.74 | 0.40 | 0.34 | 1.85 |
| 15 | Dhx32 | 0.62 | 3.33E-28 | 6.92E-24 | 0.48 | 0.21 | 0.26 | 2.23 |
| 15 | Gpd2 | 0.66 | 3.98E-28 | 8.28E-24 | 0.49 | 0.22 | 0.27 | 2.20 |
| 15 | Meis2 | 0.37 | 4.02E-28 | 8.35E-24 | 0.98 | 0.67 | 0.32 | 1.48 |
| 15 | Bspry | 0.96 | 4.18E-28 | 8.70E-24 | 0.20 | 0.06 | 0.14 | 3.19 |
| 15 | Sdc4 | 0.49 | 4.55E-28 | 9.45E-24 | 0.67 | 0.34 | 0.33 | 1.98 |
| 15 | Apc | 0.57 | 5.17E-28 | 1.08E-23 | 0.89 | 0.61 | 0.28 | 1.46 |
| 15 | Ece1 | 0.59 | 5.42E-28 | 1.13E-23 | 0.50 | 0.23 | 0.27 | 2.19 |
| 15 | Agrn | 0.54 | 8.35E-28 | 1.74E-23 | 0.74 | 0.38 | 0.35 | 1.92 |
| 15 | Gucy1a1 | 0.83 | 9.73E-28 | 2.02E-23 | 0.22 | 0.07 | 0.15 | 3.00 |
| 15 | Lmo2 | 0.70 | 1.00E-27 | 2.08E-23 | 0.31 | 0.12 | 0.19 | 2.52 |
| 15 | Gm48435 | 1.55 | 1.00E-27 | 2.09E-23 | 0.11 | 0.03 | 0.09 | 4.44 |
| 15 | Ppfia3 | 0.68 | 1.14E-27 | 2.37E-23 | 0.33 | 0.13 | 0.20 | 2.57 |
| 15 | Plekhg3 | 0.89 | 1.23E-27 | 2.56E-23 | 0.24 | 0.08 | 0.16 | 2.95 |
| 15 | Grip1 | 0.87 | 1.27E-27 | 2.63E-23 | 0.28 | 0.10 | 0.18 | 2.72 |
| 15 | Lypd6b | 0.98 | 1.50E-27 | 3.12E-23 | 0.21 | 0.07 | 0.14 | 3.10 |
| 15 | St8sia6 | 1.32 | 1.91E-27 | 3.98E-23 | 0.16 | 0.05 | 0.12 | 3.49 |
| 15 | Cers4 | 0.64 | 2.33E-27 | 4.84E-23 | 0.37 | 0.16 | 0.22 | 2.41 |
| 15 | Mgat4a | 0.96 | 2.65E-27 | 5.50E-23 | 0.26 | 0.09 | 0.16 | 2.80 |
| 15 | Clic6 | 1.05 | 2.71E-27 | 5.63E-23 | 0.18 | 0.05 | 0.13 | 3.36 |
| 15 | Cxcr4 | 1.08 | 3.31E-27 | 6.89E-23 | 0.17 | 0.05 | 0.12 | 3.48 |
| 15 | Wnt9b | 1.11 | 3.39E-27 | 7.05E-23 | 0.12 | 0.03 | 0.09 | 4.24 |
| 15 | Prr5l | 0.62 | 3.59E-27 | 7.46E-23 | 0.39 | 0.16 | 0.23 | 2.39 |
| 15 | Tmem200c | 1.61 | 4.11E-27 | 8.54E-23 | 0.11 | 0.03 | 0.09 | 4.44 |
| 15 | Slc2a3 | 0.45 | 5.57E-27 | 1.16E-22 | 0.33 | 0.14 | 0.20 | 2.47 |
| 15 | Pls1 | 0.92 | 7.75E-27 | 1.61E-22 | 0.26 | 0.09 | 0.16 | 2.76 |
| 15 | Mllt6 | 0.66 | 9.34E-27 | 1.94E-22 | 0.44 | 0.19 | 0.25 | 2.27 |
| 15 | Ces1d | 0.91 | 1.01E-26 | 2.09E-22 | 0.15 | 0.04 | 0.11 | 3.64 |
| 15 | Fzd6 | 0.89 | 1.01E-26 | 2.11E-22 | 0.24 | 0.08 | 0.15 | 2.83 |
| 15 | Chd7 | 0.48 | 1.02E-26 | 2.12E-22 | 0.92 | 0.65 | 0.27 | 1.41 |
| 15 | Fgfr2 | 0.51 | 1.38E-26 | 2.88E-22 | 0.55 | 0.27 | 0.28 | 2.03 |
| 15 | Ntm | 0.97 | 1.63E-26 | 3.40E-22 | 0.28 | 0.10 | 0.17 | 2.64 |
| 15 | Tmem108 | 0.59 | 2.05E-26 | 4.26E-22 | 0.47 | 0.22 | 0.25 | 2.18 |
| 15 | Ide | 0.60 | 2.07E-26 | 4.31E-22 | 0.54 | 0.27 | 0.28 | 2.05 |
| 15 | Myh9 | 0.42 | 2.18E-26 | 4.54E-22 | 0.84 | 0.50 | 0.34 | 1.68 |
| 15 | Ssx2ip | 0.40 | 2.75E-26 | 5.72E-22 | 0.68 | 0.36 | 0.32 | 1.90 |
| 15 | Klhl13 | 0.61 | 4.17E-26 | 8.66E-22 | 0.54 | 0.26 | 0.28 | 2.05 |
| 15 | Dlx6os1 | 1.18 | 5.87E-26 | 1.22E-21 | 0.14 | 0.04 | 0.10 | 3.60 |

**Supplementary Table 1. Marker genes for the original clusters of integrated clustering of E11.5 control and NotchGOF λ-junction cells.**

| | | | | | | | | |
|---|---|---|---|---|---|---|---|---|
| 15 | Camsap3 | 0.73 | 7.46E-26 | 1.55E-21 | 0.23 | 0.08 | 0.15 | 2.89 |
| 15 | Rian | 0.57 | 7.51E-26 | 1.56E-21 | 0.60 | 0.31 | 0.28 | 1.89 |
| 15 | Ndrg2 | 0.66 | 7.65E-26 | 1.59E-21 | 0.33 | 0.13 | 0.20 | 2.47 |
| 15 | Frrs1 | 0.60 | 7.69E-26 | 1.60E-21 | 0.47 | 0.22 | 0.25 | 2.16 |
| 15 | 2310002F09Rik | 1.20 | 7.91E-26 | 1.64E-21 | 0.14 | 0.04 | 0.10 | 3.81 |
| 15 | Fgf12 | 1.13 | 8.14E-26 | 1.69E-21 | 0.16 | 0.05 | 0.12 | 3.45 |
| 15 | Nr2f2 | 0.42 | 9.71E-26 | 2.02E-21 | 0.69 | 0.36 | 0.33 | 1.92 |
| 15 | Npnt | 0.38 | 9.95E-26 | 2.07E-21 | 0.50 | 0.23 | 0.27 | 2.14 |
| 15 | Znrf3 | 0.68 | 1.01E-25 | 2.11E-21 | 0.40 | 0.18 | 0.22 | 2.26 |
| 15 | Cacna1b | 0.64 | 1.03E-25 | 2.14E-21 | 0.23 | 0.08 | 0.15 | 2.90 |
| 15 | Ctnnal1 | 0.51 | 1.11E-25 | 2.31E-21 | 0.54 | 0.26 | 0.28 | 2.07 |
| 15 | Mycl | 0.43 | 1.12E-25 | 2.34E-21 | 0.60 | 0.30 | 0.30 | 1.99 |
| 15 | Patj | 0.70 | 1.17E-25 | 2.43E-21 | 0.32 | 0.13 | 0.19 | 2.49 |
| 15 | Tafa5 | 0.59 | 1.37E-25 | 2.85E-21 | 0.31 | 0.12 | 0.19 | 2.58 |
| 15 | Marchf9 | 0.58 | 1.68E-25 | 3.49E-21 | 0.35 | 0.15 | 0.21 | 2.43 |
| 15 | Trpc1 | 0.83 | 1.84E-25 | 3.83E-21 | 0.29 | 0.11 | 0.18 | 2.59 |
| 15 | Cxadr | 0.52 | 2.20E-25 | 4.57E-21 | 0.80 | 0.46 | 0.34 | 1.75 |
| 15 | Socs7 | 0.55 | 2.61E-25 | 5.44E-21 | 0.46 | 0.21 | 0.25 | 2.20 |
| 15 | Dusp5 | 0.83 | 2.81E-25 | 5.84E-21 | 0.29 | 0.11 | 0.18 | 2.56 |
| 15 | Hspg2 | 0.52 | 4.17E-25 | 8.67E-21 | 0.48 | 0.22 | 0.26 | 2.15 |
| 15 | Macc1 | 0.90 | 5.14E-25 | 1.07E-20 | 0.18 | 0.06 | 0.12 | 3.20 |
| 15 | Dsel | 0.50 | 5.76E-25 | 1.20E-20 | 0.64 | 0.33 | 0.32 | 1.97 |
| 15 | Ttc9 | 1.08 | 8.01E-25 | 1.67E-20 | 0.14 | 0.04 | 0.10 | 3.62 |
| 15 | Syt5 | 0.78 | 9.05E-25 | 1.88E-20 | 0.18 | 0.06 | 0.12 | 3.18 |
| 15 | Lrig3 | 0.48 | 1.36E-24 | 2.83E-20 | 0.48 | 0.22 | 0.26 | 2.15 |
| 15 | Ppp2r2b | 0.77 | 1.39E-24 | 2.89E-20 | 0.25 | 0.10 | 0.16 | 2.65 |
| 15 | Clstn1 | 0.51 | 1.39E-24 | 2.90E-20 | 0.72 | 0.38 | 0.33 | 1.86 |
| 15 | Nrxn3 | 1.41 | 1.46E-24 | 3.03E-20 | 0.11 | 0.03 | 0.08 | 4.11 |
| 15 | Sox21 | 1.13 | 1.59E-24 | 3.32E-20 | 0.12 | 0.03 | 0.09 | 3.84 |
| 15 | Mid1ip1 | 0.56 | 1.83E-24 | 3.80E-20 | 0.46 | 0.22 | 0.24 | 2.12 |
| 15 | Msi1 | 0.59 | 1.84E-24 | 3.83E-20 | 0.68 | 0.37 | 0.31 | 1.84 |
| 15 | Lrrk1 | 0.71 | 1.92E-24 | 4.00E-20 | 0.35 | 0.15 | 0.20 | 2.36 |
| 15 | Npas3 | 0.98 | 2.02E-24 | 4.19E-20 | 0.18 | 0.06 | 0.12 | 3.10 |
| 15 | Cryab | 1.50 | 2.02E-24 | 4.19E-20 | 0.16 | 0.05 | 0.11 | 3.34 |
| 15 | Npdc1 | 0.65 | 2.23E-24 | 4.63E-20 | 0.27 | 0.10 | 0.17 | 2.63 |
| 15 | Firre | 0.74 | 2.41E-24 | 5.01E-20 | 0.24 | 0.09 | 0.15 | 2.77 |
| 15 | Zdbf2 | 0.53 | 2.79E-24 | 5.79E-20 | 0.72 | 0.40 | 0.32 | 1.79 |
| 15 | Itm2b | 0.43 | 3.04E-24 | 6.32E-20 | 0.98 | 0.84 | 0.14 | 1.17 |
| 15 | Lamb1 | 0.50 | 3.30E-24 | 6.86E-20 | 0.69 | 0.38 | 0.31 | 1.81 |
| 15 | Sema6a | 0.53 | 3.48E-24 | 7.25E-20 | 0.76 | 0.44 | 0.32 | 1.74 |
| 15 | Lgi2 | 0.63 | 3.80E-24 | 7.91E-20 | 0.29 | 0.11 | 0.17 | 2.54 |
| 15 | Mid1 | 0.57 | 4.12E-24 | 8.58E-20 | 0.50 | 0.24 | 0.26 | 2.09 |
| 15 | Thbs1 | 0.43 | 5.23E-24 | 1.09E-19 | 0.29 | 0.12 | 0.17 | 2.49 |
| 15 | Ctxn3 | 1.33 | 5.30E-24 | 1.10E-19 | 0.12 | 0.03 | 0.09 | 4.00 |
| 15 | Lmo1 | 0.59 | 5.67E-24 | 1.18E-19 | 0.21 | 0.07 | 0.14 | 2.89 |
| 15 | Sparc | 0.32 | 7.30E-24 | 1.52E-19 | 0.98 | 0.74 | 0.24 | 1.33 |
| 15 | Ppp1r1a | 0.52 | 7.86E-24 | 1.64E-19 | 0.38 | 0.16 | 0.21 | 2.33 |
| 15 | Kif26a | 0.64 | 8.73E-24 | 1.82E-19 | 0.29 | 0.11 | 0.17 | 2.54 |
| 15 | Gpc4 | 0.66 | 8.75E-24 | 1.82E-19 | 0.33 | 0.14 | 0.19 | 2.38 |
| 15 | Rasef | 1.06 | 1.01E-23 | 2.10E-19 | 0.14 | 0.04 | 0.10 | 3.53 |
| 15 | Tspan7 | 0.60 | 1.01E-23 | 2.11E-19 | 0.54 | 0.28 | 0.27 | 1.97 |
| 15 | Hspa4l | 0.83 | 1.40E-23 | 2.91E-19 | 0.27 | 0.11 | 0.16 | 2.50 |
| 15 | Cobll1 | 0.97 | 1.75E-23 | 3.64E-19 | 0.12 | 0.03 | 0.09 | 3.87 |
| 15 | Dmd | 0.82 | 1.79E-23 | 3.71E-19 | 0.24 | 0.09 | 0.15 | 2.67 |
| 15 | Wnk2 | 1.13 | 2.29E-23 | 4.77E-19 | 0.16 | 0.05 | 0.11 | 3.30 |
| 15 | Adamts17 | 1.18 | 2.62E-23 | 5.44E-19 | 0.13 | 0.04 | 0.10 | 3.71 |

**Supplementary Table 1. Marker genes for the original clusters of integrated clustering of E11.5 control and NotchGOF λ-junction cells.**

| | | | | | | | | |
|---|---|---|---|---|---|---|---|---|
| 15 | Lmo4 | 0.36 | 2.84E-23 | 5.91E-19 | 0.98 | 0.81 | 0.17 | 1.21 |
| 15 | Ktn1 | 0.37 | 2.89E-23 | 6.01E-19 | 0.98 | 0.82 | 0.16 | 1.19 |
| 15 | Cdca7l | 0.53 | 3.04E-23 | 6.33E-19 | 0.59 | 0.30 | 0.28 | 1.94 |
| 15 | Hip1r | 0.91 | 3.36E-23 | 7.00E-19 | 0.21 | 0.07 | 0.14 | 2.85 |
| 15 | Zfp385c | 1.30 | 3.51E-23 | 7.29E-19 | 0.12 | 0.03 | 0.09 | 3.75 |
| 15 | Acvr2b | 0.44 | 3.57E-23 | 7.43E-19 | 0.92 | 0.64 | 0.28 | 1.43 |
| 15 | Pdpn | 0.56 | 3.82E-23 | 7.95E-19 | 0.40 | 0.18 | 0.22 | 2.19 |
| 15 | Cystm1 | 0.52 | 4.15E-23 | 8.62E-19 | 0.24 | 0.09 | 0.15 | 2.70 |
| 15 | Arhgef5 | 0.44 | 4.15E-23 | 8.63E-19 | 0.43 | 0.20 | 0.23 | 2.18 |
| 15 | Prps2 | 0.56 | 4.60E-23 | 9.56E-19 | 0.37 | 0.17 | 0.21 | 2.25 |
| 15 | Syngr1 | 0.82 | 4.84E-23 | 1.01E-18 | 0.19 | 0.07 | 0.13 | 2.94 |
| 15 | Snx10 | 0.59 | 4.98E-23 | 1.04E-18 | 0.36 | 0.16 | 0.20 | 2.30 |
| 15 | Palld | 0.42 | 6.08E-23 | 1.26E-18 | 0.92 | 0.62 | 0.31 | 1.49 |
| 15 | Tmem238 | 0.57 | 6.32E-23 | 1.31E-18 | 0.39 | 0.18 | 0.21 | 2.20 |
| 15 | Syt11 | 0.46 | 6.35E-23 | 1.32E-18 | 0.58 | 0.30 | 0.28 | 1.93 |
| 15 | Sft2d2 | 0.68 | 6.75E-23 | 1.40E-18 | 0.34 | 0.15 | 0.19 | 2.31 |
| 15 | Elf5 | 0.86 | 7.27E-23 | 1.51E-18 | 0.18 | 0.06 | 0.12 | 3.02 |
| 15 | Gm20342 | 0.63 | 1.28E-22 | 2.67E-18 | 0.33 | 0.14 | 0.19 | 2.33 |
| 15 | Slc2a1 | 0.44 | 1.44E-22 | 2.99E-18 | 0.73 | 0.42 | 0.30 | 1.72 |
| 15 | Ak4 | 0.74 | 1.47E-22 | 3.06E-18 | 0.26 | 0.10 | 0.16 | 2.57 |
| 15 | Cox4i2 | 0.40 | 1.76E-22 | 3.65E-18 | 0.52 | 0.26 | 0.26 | 2.00 |
| 15 | Eya2 | 0.33 | 1.85E-22 | 3.86E-18 | 0.48 | 0.23 | 0.25 | 2.07 |
| 15 | Hras | 0.31 | 2.16E-22 | 4.49E-18 | 0.97 | 0.80 | 0.17 | 1.22 |
| 15 | Mtus1 | 0.56 | 2.29E-22 | 4.76E-18 | 0.26 | 0.10 | 0.16 | 2.56 |
| 15 | Ptpn14 | 0.71 | 2.49E-22 | 5.18E-18 | 0.34 | 0.15 | 0.19 | 2.28 |
| 15 | Epb41l4a | 0.59 | 2.60E-22 | 5.40E-18 | 0.27 | 0.10 | 0.16 | 2.56 |
| 15 | Gm32585 | 0.45 | 2.69E-22 | 5.60E-18 | 0.17 | 0.05 | 0.12 | 3.13 |
| 15 | Pkp1 | 1.34 | 2.82E-22 | 5.86E-18 | 0.10 | 0.03 | 0.08 | 4.08 |
| 15 | Vwa1 | 0.99 | 2.96E-22 | 6.16E-18 | 0.17 | 0.06 | 0.12 | 3.05 |
| 15 | Smim22 | 0.91 | 3.01E-22 | 6.25E-18 | 0.13 | 0.04 | 0.10 | 3.57 |
| 15 | Myo10 | 0.47 | 3.03E-22 | 6.31E-18 | 0.78 | 0.47 | 0.31 | 1.66 |
| 15 | Zmat4 | 0.90 | 3.37E-22 | 7.01E-18 | 0.17 | 0.05 | 0.11 | 3.09 |
| 15 | Myo5c | 1.13 | 3.38E-22 | 7.02E-18 | 0.13 | 0.04 | 0.10 | 3.57 |
| 15 | Tafa2 | 1.12 | 3.47E-22 | 7.22E-18 | 0.14 | 0.04 | 0.10 | 3.43 |
| 15 | Tmem63b | 0.55 | 3.65E-22 | 7.59E-18 | 0.62 | 0.33 | 0.29 | 1.88 |
| 15 | Crb2 | 1.43 | 3.90E-22 | 8.12E-18 | 0.10 | 0.03 | 0.08 | 4.08 |
| 15 | Wnt3 | 0.59 | 4.04E-22 | 8.40E-18 | 0.11 | 0.03 | 0.08 | 3.83 |
| 15 | Sema3c | 0.64 | 4.27E-22 | 8.88E-18 | 0.24 | 0.09 | 0.15 | 2.62 |
| 15 | Crim1 | 0.57 | 5.87E-22 | 1.22E-17 | 0.41 | 0.20 | 0.22 | 2.11 |
| 15 | Usp25 | 0.49 | 5.94E-22 | 1.24E-17 | 0.73 | 0.43 | 0.30 | 1.70 |
| 15 | Serinc2 | 0.43 | 7.26E-22 | 1.51E-17 | 0.70 | 0.38 | 0.32 | 1.82 |
| 15 | Gadd45g | 0.83 | 8.38E-22 | 1.74E-17 | 0.41 | 0.21 | 0.20 | 1.97 |
| 15 | Zdhhc21 | 0.56 | 8.61E-22 | 1.79E-17 | 0.55 | 0.29 | 0.27 | 1.93 |
| 15 | Cd2ap | 0.42 | 8.73E-22 | 1.82E-17 | 0.90 | 0.62 | 0.29 | 1.46 |
| 15 | Krtcap3 | 0.57 | 8.93E-22 | 1.86E-17 | 0.28 | 0.11 | 0.16 | 2.48 |
| 15 | Zfp750 | 0.69 | 9.95E-22 | 2.07E-17 | 0.16 | 0.05 | 0.11 | 3.23 |
| 15 | Zfp536 | 0.70 | 1.13E-21 | 2.35E-17 | 0.25 | 0.10 | 0.15 | 2.50 |
| 15 | Rassf9 | 0.53 | 1.15E-21 | 2.38E-17 | 0.36 | 0.16 | 0.20 | 2.23 |
| 15 | Chst11 | 0.81 | 1.28E-21 | 2.66E-17 | 0.20 | 0.07 | 0.13 | 2.79 |
| 15 | Nup210 | 0.99 | 1.55E-21 | 3.22E-17 | 0.18 | 0.06 | 0.12 | 2.93 |
| 15 | Kremen1 | 0.53 | 1.56E-21 | 3.25E-17 | 0.33 | 0.14 | 0.19 | 2.32 |
| 15 | Fat1 | 0.46 | 1.56E-21 | 3.25E-17 | 0.81 | 0.51 | 0.30 | 1.59 |
| 15 | Errfi1 | 0.53 | 1.64E-21 | 3.40E-17 | 0.38 | 0.18 | 0.21 | 2.18 |
| 15 | Plekhg1 | 0.55 | 1.74E-21 | 3.61E-17 | 0.32 | 0.13 | 0.18 | 2.35 |
| 15 | Rasal2 | 0.46 | 1.87E-21 | 3.88E-17 | 0.50 | 0.25 | 0.26 | 2.04 |
| 15 | Ccdc80 | 0.49 | 1.90E-21 | 3.96E-17 | 0.40 | 0.18 | 0.22 | 2.20 |

**Supplementary Table 1. Marker genes for the original clusters of integrated clustering of E11.5 control and NotchGOF λ-junction cells.**

| | | | | | | | | |
|---|---|---|---|---|---|---|---|---|
| 15 | Bcl11b | 0.39 | 2.03E-21 | 4.23E-17 | 0.86 | 0.59 | 0.27 | 1.46 |
| 15 | Trpm4 | 0.72 | 2.08E-21 | 4.32E-17 | 0.25 | 0.10 | 0.15 | 2.55 |
| 15 | Pdgfrl | 0.56 | 2.18E-21 | 4.54E-17 | 0.39 | 0.18 | 0.21 | 2.16 |
| 15 | Hist3h2a | 0.26 | 2.28E-21 | 4.75E-17 | 0.53 | 0.26 | 0.27 | 2.02 |
| 15 | Arhgef16 | 0.63 | 2.37E-21 | 4.93E-17 | 0.17 | 0.06 | 0.12 | 3.00 |
| 15 | Cyb5r3 | 0.31 | 2.73E-21 | 5.68E-17 | 0.97 | 0.81 | 0.16 | 1.20 |
| 15 | Aftph | 0.46 | 2.83E-21 | 5.88E-17 | 0.50 | 0.25 | 0.25 | 2.02 |
| 15 | Egfr | 0.74 | 2.88E-21 | 6.00E-17 | 0.25 | 0.10 | 0.15 | 2.53 |
| 15 | Pnck | 0.78 | 2.90E-21 | 6.03E-17 | 0.20 | 0.07 | 0.13 | 2.81 |
| 15 | Metrnl | 0.76 | 2.93E-21 | 6.10E-17 | 0.19 | 0.06 | 0.12 | 2.89 |
| 15 | Nuak1 | 0.54 | 3.11E-21 | 6.47E-17 | 0.42 | 0.20 | 0.22 | 2.08 |
| 15 | 1700023F06Rik | 0.81 | 4.04E-21 | 8.40E-17 | 0.14 | 0.04 | 0.10 | 3.28 |
| 15 | Dlx1 | 0.76 | 4.20E-21 | 8.73E-17 | 0.16 | 0.05 | 0.11 | 3.14 |
| 15 | Ttc3 | 0.27 | 4.70E-21 | 9.79E-17 | 1.00 | 0.96 | 0.04 | 1.04 |
| 15 | Furin | 0.49 | 4.86E-21 | 1.01E-16 | 0.42 | 0.20 | 0.22 | 2.13 |
| 15 | Fut9 | 1.33 | 4.89E-21 | 1.02E-16 | 0.11 | 0.03 | 0.08 | 3.79 |
| 15 | Airn | 0.58 | 5.64E-21 | 1.17E-16 | 0.40 | 0.19 | 0.21 | 2.12 |
| 15 | Stox2 | 0.49 | 5.77E-21 | 1.20E-16 | 0.83 | 0.56 | 0.28 | 1.50 |
| 15 | Sostdc1 | 0.37 | 6.18E-21 | 1.29E-16 | 0.32 | 0.14 | 0.17 | 2.19 |
| 15 | Tyro3 | 0.61 | 6.51E-21 | 1.35E-16 | 0.37 | 0.17 | 0.20 | 2.20 |
| 15 | Mest | 0.35 | 7.60E-21 | 1.58E-16 | 0.99 | 0.91 | 0.08 | 1.09 |
| 15 | Dag1 | 0.42 | 8.25E-21 | 1.72E-16 | 0.66 | 0.37 | 0.29 | 1.79 |
| 15 | Zfhx3 | 0.27 | 8.55E-21 | 1.78E-16 | 0.99 | 0.81 | 0.18 | 1.22 |
| 15 | Upk3bl | 0.44 | 8.69E-21 | 1.81E-16 | 0.20 | 0.07 | 0.13 | 2.76 |
| 15 | Eps8l2 | 0.93 | 9.03E-21 | 1.88E-16 | 0.14 | 0.04 | 0.10 | 3.34 |
| 15 | Dnase2a | 0.39 | 1.05E-20 | 2.19E-16 | 0.44 | 0.22 | 0.23 | 2.06 |
| 15 | Lgalsl | 0.50 | 1.16E-20 | 2.42E-16 | 0.58 | 0.31 | 0.27 | 1.87 |
| 15 | Foxp1 | 0.31 | 1.19E-20 | 2.47E-16 | 0.91 | 0.60 | 0.32 | 1.53 |
| 15 | Trim2 | 0.43 | 1.22E-20 | 2.55E-16 | 0.71 | 0.41 | 0.30 | 1.73 |
| 15 | Tshz2 | 0.42 | 1.38E-20 | 2.87E-16 | 0.61 | 0.34 | 0.28 | 1.83 |
| 15 | Phlpp1 | 0.54 | 1.42E-20 | 2.95E-16 | 0.31 | 0.13 | 0.18 | 2.35 |
| 15 | Sorl1 | 0.52 | 1.47E-20 | 3.07E-16 | 0.34 | 0.15 | 0.19 | 2.27 |
| 15 | Rhov | 0.84 | 1.52E-20 | 3.17E-16 | 0.17 | 0.06 | 0.11 | 2.98 |
| 15 | Gnao1 | 0.74 | 1.52E-20 | 3.17E-16 | 0.22 | 0.08 | 0.13 | 2.62 |
| 15 | Galnt7 | 0.44 | 1.67E-20 | 3.47E-16 | 0.62 | 0.34 | 0.28 | 1.83 |
| 15 | Ppfibp1 | 0.45 | 1.72E-20 | 3.57E-16 | 0.74 | 0.43 | 0.31 | 1.74 |
| 15 | Fzd8 | 0.42 | 1.73E-20 | 3.61E-16 | 0.33 | 0.15 | 0.19 | 2.28 |
| 15 | Kcnh8 | 1.07 | 2.05E-20 | 4.27E-16 | 0.12 | 0.03 | 0.09 | 3.58 |
| 15 | Lrp2 | 0.98 | 2.20E-20 | 4.58E-16 | 0.16 | 0.05 | 0.11 | 3.02 |
| 15 | Shroom3 | 0.65 | 2.24E-20 | 4.66E-16 | 0.34 | 0.15 | 0.19 | 2.23 |
| 15 | Ptprd | 0.48 | 2.29E-20 | 4.75E-16 | 0.73 | 0.44 | 0.30 | 1.68 |
| 15 | Retreg1 | 0.74 | 2.29E-20 | 4.77E-16 | 0.23 | 0.09 | 0.14 | 2.54 |
| 15 | Prag1 | 0.49 | 2.33E-20 | 4.86E-16 | 0.32 | 0.14 | 0.18 | 2.30 |
| 15 | Frat2 | 0.58 | 2.47E-20 | 5.13E-16 | 0.46 | 0.23 | 0.22 | 1.97 |
| 15 | Ntn1 | 1.10 | 2.56E-20 | 5.33E-16 | 0.11 | 0.03 | 0.08 | 3.79 |
| 15 | Unc5b | 0.42 | 3.09E-20 | 6.43E-16 | 0.68 | 0.39 | 0.30 | 1.76 |
| 15 | Hes1 | 0.34 | 3.09E-20 | 6.43E-16 | 0.84 | 0.54 | 0.30 | 1.55 |
| 15 | Trim29 | 0.84 | 3.11E-20 | 6.47E-16 | 0.15 | 0.05 | 0.10 | 3.17 |
| 15 | Adgrb3 | 0.97 | 3.31E-20 | 6.88E-16 | 0.18 | 0.06 | 0.12 | 2.83 |
| 15 | Rhbdf1 | 0.71 | 3.34E-20 | 6.94E-16 | 0.25 | 0.10 | 0.15 | 2.47 |
| 15 | Setd7 | 0.56 | 3.34E-20 | 6.95E-16 | 0.42 | 0.20 | 0.22 | 2.07 |
| 15 | Dync1i1 | 0.75 | 3.34E-20 | 6.95E-16 | 0.18 | 0.06 | 0.12 | 2.83 |
| 15 | Cadm4 | 0.57 | 3.58E-20 | 7.44E-16 | 0.22 | 0.09 | 0.14 | 2.61 |
| 15 | H2-Q4 | 0.57 | 3.64E-20 | 7.57E-16 | 0.26 | 0.11 | 0.15 | 2.47 |
| 15 | H2-K1 | 0.52 | 4.01E-20 | 8.33E-16 | 0.30 | 0.13 | 0.17 | 2.30 |
| 15 | Afdn | 0.31 | 4.09E-20 | 8.51E-16 | 0.88 | 0.57 | 0.31 | 1.55 |

**Supplementary Table 1. Marker genes for the original clusters of integrated clustering of E11.5 control and NotchGOF λ-junction cells.**

| | | | | | | | | |
|---|---|---|---|---|---|---|---|---|
| 15 | Ephb2 | 0.48 | 4.81E-20 | 1.00E-15 | 0.54 | 0.28 | 0.26 | 1.94 |
| 15 | Jade1 | 0.54 | 5.22E-20 | 1.09E-15 | 0.36 | 0.17 | 0.19 | 2.14 |
| 15 | Arhgap32 | 0.54 | 5.47E-20 | 1.14E-15 | 0.40 | 0.19 | 0.21 | 2.11 |
| 15 | Acrbp | 0.59 | 5.54E-20 | 1.15E-15 | 0.25 | 0.10 | 0.15 | 2.47 |
| 15 | Wdsub1 | 0.54 | 5.58E-20 | 1.16E-15 | 0.42 | 0.20 | 0.22 | 2.06 |
| 15 | Vezt | 0.47 | 6.59E-20 | 1.37E-15 | 0.60 | 0.32 | 0.28 | 1.87 |
| 15 | Gdpd1 | 0.40 | 8.29E-20 | 1.72E-15 | 0.52 | 0.27 | 0.26 | 1.97 |
| 15 | Gm9856 | 0.39 | 8.62E-20 | 1.79E-15 | 0.34 | 0.15 | 0.19 | 2.27 |
| 15 | Cemip2 | 0.41 | 9.61E-20 | 2.00E-15 | 0.74 | 0.45 | 0.29 | 1.66 |
| 15 | Pclo | 0.34 | 9.98E-20 | 2.08E-15 | 0.26 | 0.11 | 0.15 | 2.45 |
| 15 | Gstm1 | 0.56 | 1.00E-19 | 2.09E-15 | 0.48 | 0.24 | 0.23 | 1.95 |
| 15 | Rab11fip1 | 0.62 | 1.10E-19 | 2.29E-15 | 0.27 | 0.12 | 0.16 | 2.35 |
| 15 | Tpp1 | 0.40 | 1.18E-19 | 2.46E-15 | 0.51 | 0.26 | 0.25 | 1.95 |
| 15 | Bend7 | 0.74 | 1.26E-19 | 2.62E-15 | 0.21 | 0.08 | 0.13 | 2.60 |
| 15 | Gldc | 1.01 | 1.27E-19 | 2.65E-15 | 0.13 | 0.04 | 0.09 | 3.30 |
| 15 | Rhob | 0.71 | 1.32E-19 | 2.75E-15 | 0.61 | 0.34 | 0.26 | 1.77 |
| 15 | Mtss1 | 0.38 | 1.34E-19 | 2.80E-15 | 0.55 | 0.29 | 0.26 | 1.90 |
| 15 | Ormdl2 | 0.43 | 1.40E-19 | 2.92E-15 | 0.61 | 0.35 | 0.27 | 1.77 |
| 15 | Adi1 | 0.35 | 1.44E-19 | 2.99E-15 | 0.59 | 0.31 | 0.28 | 1.88 |
| 15 | Ybx2 | 0.76 | 1.89E-19 | 3.93E-15 | 0.19 | 0.07 | 0.12 | 2.69 |
| 15 | Arl4c | 0.31 | 1.90E-19 | 3.95E-15 | 0.60 | 0.33 | 0.28 | 1.85 |
| 15 | Tubb4a | 0.91 | 1.91E-19 | 3.96E-15 | 0.12 | 0.04 | 0.09 | 3.42 |
| 15 | Cacnb3 | 0.37 | 2.20E-19 | 4.58E-15 | 0.53 | 0.27 | 0.26 | 1.96 |
| 15 | Col4a5 | 0.53 | 2.29E-19 | 4.76E-15 | 0.29 | 0.13 | 0.17 | 2.31 |
| 15 | Slc22a17 | 0.32 | 2.49E-19 | 5.18E-15 | 0.65 | 0.34 | 0.31 | 1.91 |
| 15 | Zfp612 | 0.48 | 2.57E-19 | 5.35E-15 | 0.36 | 0.17 | 0.20 | 2.17 |
| 15 | Hdgf | 0.25 | 2.68E-19 | 5.58E-15 | 1.00 | 0.98 | 0.02 | 1.02 |
| 15 | Arg2 | 0.78 | 2.91E-19 | 6.06E-15 | 0.16 | 0.05 | 0.10 | 2.98 |
| 15 | AC149090.1 | 0.38 | 3.15E-19 | 6.55E-15 | 0.54 | 0.28 | 0.26 | 1.92 |
| 15 | Tpi1 | 0.25 | 3.17E-19 | 6.60E-15 | 1.00 | 0.96 | 0.04 | 1.04 |
| 15 | Plp2 | 0.31 | 3.45E-19 | 7.17E-15 | 0.67 | 0.38 | 0.29 | 1.78 |
| 15 | Nhs | 0.47 | 3.90E-19 | 8.11E-15 | 0.60 | 0.33 | 0.27 | 1.80 |
| 15 | Nfe2l3 | 1.15 | 4.16E-19 | 8.65E-15 | 0.12 | 0.03 | 0.08 | 3.41 |
| 15 | AI661453 | 0.94 | 4.43E-19 | 9.21E-15 | 0.12 | 0.03 | 0.08 | 3.47 |
| 15 | Srd5a1 | 0.40 | 4.68E-19 | 9.73E-15 | 0.38 | 0.18 | 0.20 | 2.12 |
| 15 | Ermp1 | 0.39 | 4.76E-19 | 9.89E-15 | 0.61 | 0.33 | 0.28 | 1.85 |
| 15 | Ldha | 0.25 | 4.87E-19 | 1.01E-14 | 1.00 | 0.97 | 0.03 | 1.03 |
| 15 | Fxyd3 | 0.33 | 4.87E-19 | 1.01E-14 | 0.23 | 0.09 | 0.14 | 2.52 |
| 15 | Ifitm3 | 0.40 | 5.07E-19 | 1.05E-14 | 0.53 | 0.29 | 0.24 | 1.81 |
| 15 | Rreb1 | 0.45 | 5.09E-19 | 1.06E-14 | 0.42 | 0.21 | 0.22 | 2.04 |
| 15 | Pik3r1 | 0.48 | 5.39E-19 | 1.12E-14 | 0.64 | 0.39 | 0.25 | 1.65 |
| 15 | Nr1h5 | 0.57 | 5.49E-19 | 1.14E-14 | 0.30 | 0.13 | 0.17 | 2.28 |
| 15 | Eya1 | 0.40 | 5.52E-19 | 1.15E-14 | 0.79 | 0.49 | 0.29 | 1.59 |
| 15 | Tiam1 | 0.43 | 5.98E-19 | 1.24E-14 | 0.58 | 0.31 | 0.27 | 1.86 |
| 15 | Nomo1 | 0.53 | 6.13E-19 | 1.27E-14 | 0.40 | 0.20 | 0.21 | 2.07 |
| 15 | Wipf2 | 0.44 | 6.29E-19 | 1.31E-14 | 0.49 | 0.25 | 0.24 | 1.96 |
| 15 | Tpbg | 0.55 | 6.47E-19 | 1.35E-14 | 0.58 | 0.30 | 0.28 | 1.92 |
| 15 | Pcdh19 | 0.60 | 6.65E-19 | 1.38E-14 | 0.35 | 0.17 | 0.18 | 2.09 |
| 15 | Ptprz1 | 0.63 | 7.48E-19 | 1.56E-14 | 0.25 | 0.10 | 0.15 | 2.40 |
| 15 | Fam126a | 0.52 | 7.49E-19 | 1.56E-14 | 0.48 | 0.25 | 0.23 | 1.92 |
| 15 | Zfp113 | 0.43 | 7.55E-19 | 1.57E-14 | 0.60 | 0.32 | 0.27 | 1.85 |
| 15 | Lin28b | 0.54 | 7.78E-19 | 1.62E-14 | 0.41 | 0.20 | 0.21 | 2.02 |
| 15 | Cux1 | 0.36 | 8.13E-19 | 1.69E-14 | 0.93 | 0.69 | 0.23 | 1.34 |
| 15 | Mpp7 | 0.37 | 9.18E-19 | 1.91E-14 | 0.66 | 0.37 | 0.29 | 1.80 |
| 15 | Notch3 | 0.46 | 9.77E-19 | 2.03E-14 | 0.69 | 0.40 | 0.29 | 1.72 |
| 15 | Gpc1 | 0.47 | 9.79E-19 | 2.04E-14 | 0.67 | 0.40 | 0.28 | 1.70 |

**Supplementary Table 1. Marker genes for the original clusters of integrated clustering of E11.5 control and NotchGOF λ-junction cells.**

| | | | | | | | | |
|---|---|---|---|---|---|---|---|---|
| 15 | Trim32 | 0.44 | 1.27E-18 | 2.64E-14 | 0.61 | 0.34 | 0.27 | 1.81 |
| 15 | Tfap2a | 0.34 | 1.27E-18 | 2.64E-14 | 0.42 | 0.21 | 0.21 | 1.98 |
| 15 | Otud4 | 0.45 | 1.38E-18 | 2.87E-14 | 0.57 | 0.31 | 0.26 | 1.86 |
| 15 | Tmem178 | 0.81 | 1.39E-18 | 2.90E-14 | 0.11 | 0.03 | 0.08 | 3.47 |
| 15 | Nab1 | 0.56 | 1.42E-18 | 2.95E-14 | 0.29 | 0.13 | 0.16 | 2.24 |
| 15 | Shtn1 | 0.59 | 1.58E-18 | 3.29E-14 | 0.15 | 0.05 | 0.10 | 2.94 |
| 15 | Nes | 0.40 | 1.75E-18 | 3.64E-14 | 0.70 | 0.42 | 0.28 | 1.66 |
| 15 | Col11a1 | 0.37 | 1.78E-18 | 3.69E-14 | 0.72 | 0.42 | 0.29 | 1.69 |
| 15 | Prkci | 0.40 | 1.94E-18 | 4.04E-14 | 0.50 | 0.26 | 0.24 | 1.92 |
| 15 | Rab27b | 0.43 | 2.14E-18 | 4.45E-14 | 0.23 | 0.10 | 0.14 | 2.43 |
| 15 | Gga3 | 0.54 | 2.29E-18 | 4.77E-14 | 0.31 | 0.14 | 0.17 | 2.23 |
| 15 | Ctf1 | 0.50 | 2.50E-18 | 5.21E-14 | 0.35 | 0.16 | 0.19 | 2.14 |
| 15 | Creld1 | 0.52 | 2.56E-18 | 5.32E-14 | 0.29 | 0.13 | 0.16 | 2.26 |
| 15 | Aig1 | 0.41 | 2.58E-18 | 5.37E-14 | 0.60 | 0.33 | 0.27 | 1.84 |
| 15 | Arfgef1 | 0.39 | 2.62E-18 | 5.45E-14 | 0.79 | 0.48 | 0.31 | 1.63 |
| 15 | Pcgf5 | 0.73 | 2.78E-18 | 5.79E-14 | 0.21 | 0.09 | 0.13 | 2.48 |
| 15 | Samd10 | 0.69 | 3.14E-18 | 6.52E-14 | 0.22 | 0.09 | 0.13 | 2.47 |
| 15 | Map7d2 | 0.64 | 3.25E-18 | 6.75E-14 | 0.21 | 0.08 | 0.13 | 2.54 |
| 15 | Plec | 0.39 | 3.27E-18 | 6.79E-14 | 0.51 | 0.27 | 0.24 | 1.90 |
| 15 | Pak3 | 0.46 | 3.64E-18 | 7.57E-14 | 0.30 | 0.14 | 0.17 | 2.23 |
| 15 | Zswim6 | 0.44 | 3.67E-18 | 7.63E-14 | 0.54 | 0.29 | 0.24 | 1.84 |
| 15 | Ccpg1 | 0.51 | 3.71E-18 | 7.72E-14 | 0.34 | 0.16 | 0.18 | 2.13 |
| 15 | Fam107a | 1.05 | 3.76E-18 | 7.81E-14 | 0.13 | 0.04 | 0.09 | 3.12 |
| 15 | Lrba | 0.38 | 3.81E-18 | 7.93E-14 | 0.44 | 0.22 | 0.22 | 2.00 |
| 15 | D930016D06Rik | 0.44 | 3.83E-18 | 7.97E-14 | 0.46 | 0.24 | 0.23 | 1.97 |
| 15 | Srcin1 | 1.06 | 4.23E-18 | 8.80E-14 | 0.14 | 0.05 | 0.09 | 3.02 |
| 15 | Ccdc120 | 0.83 | 4.25E-18 | 8.85E-14 | 0.25 | 0.11 | 0.14 | 2.36 |
| 15 | Tnpo2 | 0.43 | 4.99E-18 | 1.04E-13 | 0.46 | 0.23 | 0.23 | 1.97 |
| 15 | Plxnb1 | 0.44 | 5.21E-18 | 1.08E-13 | 0.50 | 0.26 | 0.24 | 1.93 |
| 15 | Fam214a | 0.57 | 5.34E-18 | 1.11E-13 | 0.24 | 0.10 | 0.14 | 2.41 |
| 15 | Cxxc5 | 0.37 | 5.65E-18 | 1.17E-13 | 0.73 | 0.45 | 0.28 | 1.62 |
| 15 | Gnpda2 | 0.36 | 6.19E-18 | 1.29E-13 | 0.53 | 0.28 | 0.25 | 1.90 |
| 15 | Rnf208 | 0.63 | 6.29E-18 | 1.31E-13 | 0.18 | 0.07 | 0.11 | 2.70 |
| 15 | Spen | 0.43 | 6.35E-18 | 1.32E-13 | 0.75 | 0.46 | 0.29 | 1.63 |
| 15 | Npepps | 0.41 | 6.43E-18 | 1.34E-13 | 0.90 | 0.64 | 0.26 | 1.40 |
| 15 | St8sia2 | 0.43 | 7.04E-18 | 1.46E-13 | 0.39 | 0.19 | 0.20 | 2.07 |
| 15 | Klhl23 | 0.41 | 7.21E-18 | 1.50E-13 | 0.64 | 0.37 | 0.28 | 1.75 |
| 15 | Scrn1 | 0.83 | 7.39E-18 | 1.54E-13 | 0.13 | 0.04 | 0.09 | 3.07 |
| 15 | Rif1 | 0.34 | 7.97E-18 | 1.66E-13 | 0.95 | 0.80 | 0.15 | 1.19 |
| 15 | Zfp704 | 0.39 | 8.00E-18 | 1.66E-13 | 0.77 | 0.46 | 0.31 | 1.66 |
| 15 | Cdkl1 | 1.16 | 8.31E-18 | 1.73E-13 | 0.11 | 0.03 | 0.08 | 3.32 |
| 15 | Palm3 | 0.38 | 8.69E-18 | 1.81E-13 | 0.32 | 0.15 | 0.17 | 2.17 |
| 15 | Acsl5 | 0.48 | 8.92E-18 | 1.86E-13 | 0.33 | 0.16 | 0.18 | 2.12 |
| 15 | Wnt7b | 0.57 | 9.25E-18 | 1.92E-13 | 0.13 | 0.04 | 0.09 | 3.10 |
| 15 | Gnb5 | 0.35 | 9.66E-18 | 2.01E-13 | 0.49 | 0.25 | 0.24 | 1.99 |
| 15 | Glcci1 | 0.40 | 9.85E-18 | 2.05E-13 | 0.66 | 0.38 | 0.28 | 1.72 |
| 15 | Ly75 | 0.62 | 1.00E-17 | 2.08E-13 | 0.32 | 0.15 | 0.17 | 2.15 |
| 15 | Ext1 | 0.40 | 1.02E-17 | 2.12E-13 | 0.70 | 0.41 | 0.30 | 1.73 |
| 15 | Usp40 | 0.37 | 1.04E-17 | 2.16E-13 | 0.62 | 0.33 | 0.28 | 1.85 |
| 15 | Atp6v0e2 | 0.77 | 1.09E-17 | 2.26E-13 | 0.15 | 0.05 | 0.10 | 2.88 |
| 15 | Prkcd | 0.33 | 1.13E-17 | 2.34E-13 | 0.43 | 0.21 | 0.22 | 2.04 |
| 15 | Atp2b1 | 0.32 | 1.13E-17 | 2.34E-13 | 0.97 | 0.84 | 0.13 | 1.15 |
| 15 | Hmga2 | 0.30 | 1.14E-17 | 2.36E-13 | 1.00 | 0.93 | 0.06 | 1.07 |
| 15 | Prkcz | 0.50 | 1.14E-17 | 2.37E-13 | 0.25 | 0.10 | 0.14 | 2.38 |
| 15 | Igdcc3 | 0.45 | 1.22E-17 | 2.53E-13 | 0.45 | 0.24 | 0.22 | 1.92 |
| 15 | B4galnt4 | 0.45 | 1.22E-17 | 2.54E-13 | 0.46 | 0.24 | 0.23 | 1.95 |

**Supplementary Table 1. Marker genes for the original clusters of integrated clustering of E11.5 control and NotchGOF λ-junction cells.**

| | | | | | | | | |
|---|---|---|---|---|---|---|---|---|
| 15 | Kif13b | 0.55 | 1.25E-17 | 2.60E-13 | 0.34 | 0.16 | 0.18 | 2.10 |
| 15 | Atp11a | 0.51 | 1.29E-17 | 2.67E-13 | 0.35 | 0.17 | 0.18 | 2.10 |
| 15 | Tdrp | 1.09 | 1.40E-17 | 2.91E-13 | 0.10 | 0.03 | 0.07 | 3.57 |
| 15 | Mpp2 | 0.54 | 1.41E-17 | 2.92E-13 | 0.35 | 0.17 | 0.18 | 2.08 |
| 15 | Elf1 | 0.25 | 1.46E-17 | 3.04E-13 | 0.50 | 0.26 | 0.24 | 1.93 |
| 15 | Ripply3 | 0.69 | 1.49E-17 | 3.10E-13 | 0.16 | 0.06 | 0.10 | 2.73 |
| 15 | Med13l | 0.45 | 1.51E-17 | 3.15E-13 | 0.78 | 0.50 | 0.28 | 1.56 |
| 15 | Cuedc1 | 0.39 | 1.52E-17 | 3.17E-13 | 0.67 | 0.38 | 0.29 | 1.76 |
| 15 | Tshz1 | 0.40 | 1.61E-17 | 3.34E-13 | 0.57 | 0.31 | 0.26 | 1.81 |
| 15 | Cadps2 | 1.12 | 1.66E-17 | 3.44E-13 | 0.12 | 0.04 | 0.08 | 3.22 |
| 15 | Acvr1b | 0.68 | 1.74E-17 | 3.61E-13 | 0.26 | 0.11 | 0.14 | 2.30 |
| 15 | Cecr2 | 0.32 | 1.94E-17 | 4.04E-13 | 0.44 | 0.22 | 0.22 | 2.00 |
| 15 | Npat | 0.36 | 1.97E-17 | 4.11E-13 | 0.63 | 0.34 | 0.28 | 1.83 |
| 15 | Gpm6a | 0.74 | 2.04E-17 | 4.23E-13 | 0.14 | 0.05 | 0.09 | 2.94 |
| 15 | Mnt | 0.47 | 2.15E-17 | 4.48E-13 | 0.36 | 0.17 | 0.18 | 2.07 |
| 15 | Kif13a | 0.54 | 2.22E-17 | 4.61E-13 | 0.41 | 0.21 | 0.20 | 1.97 |
| 15 | Kank1 | 0.47 | 2.22E-17 | 4.63E-13 | 0.33 | 0.16 | 0.18 | 2.12 |
| 15 | Pank1 | 0.41 | 2.25E-17 | 4.68E-13 | 0.57 | 0.31 | 0.26 | 1.85 |
| 15 | Slit2 | 0.26 | 2.40E-17 | 4.99E-13 | 0.62 | 0.36 | 0.26 | 1.72 |
| 15 | Psen2 | 0.59 | 2.40E-17 | 5.00E-13 | 0.19 | 0.08 | 0.12 | 2.55 |
| 15 | Irs1 | 0.41 | 2.46E-17 | 5.11E-13 | 0.37 | 0.18 | 0.19 | 2.07 |
| 15 | Dcbld1 | 0.47 | 2.47E-17 | 5.13E-13 | 0.54 | 0.30 | 0.24 | 1.80 |
| 15 | Ier5 | 0.33 | 2.52E-17 | 5.23E-13 | 0.88 | 0.65 | 0.24 | 1.37 |
| 15 | Mttp | 0.45 | 3.28E-17 | 6.82E-13 | 0.29 | 0.13 | 0.16 | 2.22 |
| 15 | Pqlc3 | 0.67 | 3.60E-17 | 7.50E-13 | 0.15 | 0.05 | 0.10 | 2.83 |
| 15 | Dst | 0.41 | 3.62E-17 | 7.53E-13 | 0.59 | 0.35 | 0.24 | 1.68 |
| 15 | Nfib | 0.33 | 3.64E-17 | 7.57E-13 | 0.96 | 0.74 | 0.22 | 1.29 |
| 15 | Npm3 | 0.33 | 3.78E-17 | 7.87E-13 | 0.97 | 0.79 | 0.18 | 1.22 |
| 15 | Ipo4 | 0.46 | 4.16E-17 | 8.66E-13 | 0.60 | 0.34 | 0.26 | 1.75 |
| 15 | Lrrc8b | 0.47 | 4.26E-17 | 8.87E-13 | 0.26 | 0.11 | 0.15 | 2.32 |
| 15 | Gigyf1 | 0.46 | 4.34E-17 | 9.03E-13 | 0.61 | 0.35 | 0.26 | 1.73 |
| 15 | Mtln | 0.34 | 4.79E-17 | 9.97E-13 | 0.58 | 0.33 | 0.26 | 1.79 |
| 15 | Slc35e2 | 0.45 | 4.86E-17 | 1.01E-12 | 0.41 | 0.20 | 0.20 | 2.00 |
| 15 | Gm6710 | 0.67 | 4.94E-17 | 1.03E-12 | 0.23 | 0.10 | 0.13 | 2.38 |
| 15 | Lrrc8d | 0.31 | 4.99E-17 | 1.04E-12 | 0.40 | 0.19 | 0.20 | 2.05 |
| 15 | Fsd1 | 0.32 | 5.11E-17 | 1.06E-12 | 0.47 | 0.24 | 0.23 | 1.97 |
| 15 | Zswim5 | 1.31 | 5.17E-17 | 1.07E-12 | 0.11 | 0.03 | 0.07 | 3.31 |
| 15 | Pard3 | 0.37 | 5.43E-17 | 1.13E-12 | 0.87 | 0.57 | 0.30 | 1.51 |
| 15 | Atxn7 | 0.41 | 5.64E-17 | 1.17E-12 | 0.42 | 0.21 | 0.21 | 1.99 |
| 15 | Nedd4l | 0.39 | 5.91E-17 | 1.23E-12 | 0.31 | 0.14 | 0.17 | 2.17 |
| 15 | Gab1 | 0.53 | 5.94E-17 | 1.24E-12 | 0.38 | 0.19 | 0.19 | 2.01 |
| 15 | Pik3ip1 | 0.56 | 6.70E-17 | 1.39E-12 | 0.21 | 0.09 | 0.13 | 2.45 |
| 15 | Selenoo | 0.45 | 6.93E-17 | 1.44E-12 | 0.31 | 0.14 | 0.17 | 2.17 |
| 15 | Edem2 | 0.44 | 7.40E-17 | 1.54E-12 | 0.39 | 0.19 | 0.20 | 2.04 |
| 15 | Snn | 0.32 | 7.52E-17 | 1.56E-12 | 0.72 | 0.43 | 0.29 | 1.67 |
| 15 | Pard3b | 0.56 | 7.52E-17 | 1.56E-12 | 0.26 | 0.12 | 0.15 | 2.24 |
| 15 | Lhfp | 0.40 | 7.87E-17 | 1.64E-12 | 0.44 | 0.23 | 0.21 | 1.91 |
| 15 | Nudt14 | 0.31 | 7.94E-17 | 1.65E-12 | 0.70 | 0.40 | 0.31 | 1.77 |
| 15 | Agl | 0.39 | 8.59E-17 | 1.79E-12 | 0.48 | 0.25 | 0.23 | 1.90 |
| 15 | Tcf7l2 | 0.35 | 8.70E-17 | 1.81E-12 | 0.88 | 0.61 | 0.27 | 1.45 |
| 15 | Msi2 | 0.37 | 8.84E-17 | 1.84E-12 | 0.91 | 0.68 | 0.23 | 1.34 |
| 15 | Srgap3 | 0.58 | 8.85E-17 | 1.84E-12 | 0.29 | 0.13 | 0.16 | 2.17 |
| 15 | Gcat | 0.39 | 8.86E-17 | 1.84E-12 | 0.77 | 0.48 | 0.29 | 1.61 |
| 15 | Col17a1 | 0.83 | 9.55E-17 | 1.99E-12 | 0.11 | 0.04 | 0.08 | 3.23 |
| 15 | Rimklb | 0.35 | 9.65E-17 | 2.01E-12 | 0.59 | 0.32 | 0.27 | 1.83 |
| 15 | Mtmr7 | 0.83 | 9.82E-17 | 2.04E-12 | 0.14 | 0.05 | 0.09 | 2.88 |

**Supplementary Table 1. Marker genes for the original clusters of integrated clustering of E11.5 control and NotchGOF λ-junction cells.**

| | | | | | | | | |
|---|---|---|---|---|---|---|---|---|
| 15 | Plce1 | 0.38 | 1.02E-16 | 2.13E-12 | 0.43 | 0.22 | 0.20 | 1.91 |
| 15 | Usp38 | 0.44 | 1.03E-16 | 2.15E-12 | 0.41 | 0.21 | 0.20 | 1.98 |
| 15 | Llgl2 | 0.55 | 1.05E-16 | 2.18E-12 | 0.14 | 0.05 | 0.09 | 2.94 |
| 15 | Gsap | 1.04 | 1.05E-16 | 2.19E-12 | 0.12 | 0.04 | 0.08 | 3.22 |
| 15 | Prkg1 | 0.49 | 1.07E-16 | 2.23E-12 | 0.32 | 0.16 | 0.17 | 2.08 |
| 15 | Lmtk2 | 0.67 | 1.09E-16 | 2.27E-12 | 0.23 | 0.10 | 0.13 | 2.37 |
| 15 | Gm43696 | 0.54 | 1.11E-16 | 2.32E-12 | 0.26 | 0.11 | 0.14 | 2.27 |
| 15 | Gsr | 0.42 | 1.22E-16 | 2.54E-12 | 0.58 | 0.33 | 0.25 | 1.75 |
| 15 | Tedc1 | 0.54 | 1.26E-16 | 2.63E-12 | 0.28 | 0.13 | 0.15 | 2.20 |
| 15 | Lypd6 | 0.37 | 1.28E-16 | 2.66E-12 | 0.28 | 0.13 | 0.15 | 2.22 |
| 15 | Lrrc1 | 0.43 | 1.29E-16 | 2.68E-12 | 0.32 | 0.15 | 0.17 | 2.14 |
| 15 | Cmtm4 | 0.38 | 1.39E-16 | 2.90E-12 | 0.49 | 0.25 | 0.23 | 1.92 |
| 15 | Myo9b | 0.44 | 1.42E-16 | 2.95E-12 | 0.44 | 0.23 | 0.21 | 1.94 |
| 15 | Ccdc85c | 0.64 | 1.44E-16 | 3.00E-12 | 0.25 | 0.11 | 0.14 | 2.25 |
| 15 | Dkk3 | 0.59 | 1.47E-16 | 3.06E-12 | 0.26 | 0.12 | 0.15 | 2.26 |
| 15 | Arhgap27os2 | 0.66 | 1.50E-16 | 3.11E-12 | 0.17 | 0.07 | 0.11 | 2.63 |
| 15 | Sema3a | 0.35 | 1.56E-16 | 3.24E-12 | 0.67 | 0.40 | 0.27 | 1.68 |
| 15 | Arnt2 | 0.42 | 1.59E-16 | 3.30E-12 | 0.38 | 0.18 | 0.19 | 2.05 |
| 15 | Tmem30b | 0.55 | 1.59E-16 | 3.31E-12 | 0.31 | 0.15 | 0.16 | 2.11 |
| 15 | Miga1 | 0.31 | 1.61E-16 | 3.34E-12 | 0.56 | 0.30 | 0.26 | 1.87 |
| 15 | Prpf40b | 0.40 | 1.69E-16 | 3.51E-12 | 0.45 | 0.24 | 0.21 | 1.91 |
| 15 | Ajuba | 0.34 | 1.82E-16 | 3.79E-12 | 0.52 | 0.28 | 0.24 | 1.87 |
| 15 | Zfp949 | 0.45 | 1.82E-16 | 3.79E-12 | 0.27 | 0.12 | 0.15 | 2.25 |
| 15 | Repin1 | 0.47 | 1.91E-16 | 3.98E-12 | 0.29 | 0.13 | 0.16 | 2.19 |
| 15 | Pygb | 0.36 | 1.98E-16 | 4.11E-12 | 0.45 | 0.23 | 0.22 | 1.97 |
| 15 | Usp24 | 0.40 | 2.06E-16 | 4.29E-12 | 0.55 | 0.30 | 0.25 | 1.82 |
| 15 | Fbrsl1 | 0.35 | 2.08E-16 | 4.34E-12 | 0.65 | 0.37 | 0.28 | 1.76 |
| 15 | Plcb4 | 0.38 | 2.29E-16 | 4.75E-12 | 0.39 | 0.20 | 0.19 | 1.99 |
| 15 | Kantr | 0.41 | 2.54E-16 | 5.29E-12 | 0.35 | 0.17 | 0.18 | 2.07 |
| 15 | Pkia | 0.44 | 2.67E-16 | 5.54E-12 | 0.37 | 0.18 | 0.19 | 2.02 |
| 15 | Iqsec1 | 0.44 | 2.73E-16 | 5.67E-12 | 0.29 | 0.14 | 0.16 | 2.16 |
| 15 | Golga4 | 0.37 | 2.75E-16 | 5.71E-12 | 0.90 | 0.64 | 0.26 | 1.41 |
| 15 | Slc35f1 | 1.04 | 2.79E-16 | 5.80E-12 | 0.11 | 0.03 | 0.07 | 3.31 |
| 15 | Sbf2 | 0.45 | 2.81E-16 | 5.85E-12 | 0.60 | 0.34 | 0.26 | 1.78 |
| 15 | Tnfrsf12a | 0.63 | 2.83E-16 | 5.88E-12 | 0.19 | 0.08 | 0.11 | 2.50 |
| 15 | Rab11fip4 | 0.53 | 2.91E-16 | 6.05E-12 | 0.29 | 0.13 | 0.16 | 2.17 |
| 15 | Tenm3 | 0.28 | 3.11E-16 | 6.47E-12 | 0.90 | 0.63 | 0.26 | 1.42 |
| 15 | Nedd9 | 0.47 | 3.13E-16 | 6.52E-12 | 0.34 | 0.17 | 0.17 | 2.05 |
| 15 | Pex6 | 0.41 | 3.30E-16 | 6.86E-12 | 0.53 | 0.29 | 0.24 | 1.83 |
| 15 | Mark2 | 0.38 | 3.45E-16 | 7.19E-12 | 0.76 | 0.46 | 0.30 | 1.65 |
| 15 | Hgh1 | 0.58 | 3.58E-16 | 7.44E-12 | 0.29 | 0.13 | 0.15 | 2.16 |
| 15 | B3gnt7 | 1.14 | 3.60E-16 | 7.48E-12 | 0.13 | 0.04 | 0.08 | 2.98 |
| 15 | Arfip2 | 0.39 | 3.64E-16 | 7.56E-12 | 0.69 | 0.42 | 0.28 | 1.66 |
| 15 | Ldlr | 0.41 | 3.69E-16 | 7.67E-12 | 0.45 | 0.24 | 0.21 | 1.88 |
| 15 | Atp9a | 0.38 | 3.83E-16 | 7.96E-12 | 0.36 | 0.17 | 0.18 | 2.05 |
| 15 | Acot7 | 0.36 | 3.90E-16 | 8.10E-12 | 0.45 | 0.23 | 0.22 | 1.94 |
| 15 | Setd1a | 0.40 | 3.92E-16 | 8.16E-12 | 0.54 | 0.30 | 0.25 | 1.83 |
| 15 | Smtnl2 | 0.38 | 3.96E-16 | 8.23E-12 | 0.19 | 0.07 | 0.11 | 2.54 |
| 15 | Pura | 0.36 | 3.96E-16 | 8.23E-12 | 0.75 | 0.47 | 0.28 | 1.61 |
| 15 | Gsta4 | 0.30 | 4.06E-16 | 8.45E-12 | 0.96 | 0.75 | 0.20 | 1.27 |
| 15 | Srgap1 | 0.28 | 4.11E-16 | 8.55E-12 | 0.61 | 0.34 | 0.27 | 1.81 |
| 15 | Ulk1 | 0.42 | 4.34E-16 | 9.03E-12 | 0.50 | 0.26 | 0.23 | 1.88 |
| 15 | Ncam1 | 0.28 | 4.44E-16 | 9.23E-12 | 0.62 | 0.36 | 0.26 | 1.73 |
| 15 | Rnf44 | 0.40 | 4.79E-16 | 9.95E-12 | 0.60 | 0.34 | 0.26 | 1.77 |
| 15 | Kdm5a | 0.35 | 4.83E-16 | 1.00E-11 | 0.91 | 0.67 | 0.25 | 1.37 |
| 15 | Rb1 | 0.47 | 4.84E-16 | 1.01E-11 | 0.32 | 0.16 | 0.17 | 2.05 |

**Supplementary Table 1. Marker genes for the original clusters of integrated clustering of E11.5 control and NotchGOF λ-junction cells.**

| | | | | | | | | |
|---|---|---|---|---|---|---|---|---|
| 15 | Il17rd | 0.36 | 5.12E-16 | 1.06E-11 | 0.78 | 0.50 | 0.28 | 1.57 |
| 15 | Tead1 | 0.35 | 5.49E-16 | 1.14E-11 | 0.86 | 0.60 | 0.26 | 1.44 |
| 15 | 2700033N17Rik | 0.41 | 5.58E-16 | 1.16E-11 | 0.26 | 0.12 | 0.15 | 2.24 |
| 15 | Fam89a | 0.54 | 5.64E-16 | 1.17E-11 | 0.18 | 0.07 | 0.11 | 2.54 |
| 15 | Mfsd13a | 1.05 | 5.78E-16 | 1.20E-11 | 0.12 | 0.04 | 0.08 | 3.14 |
| 15 | Stra6 | 0.33 | 5.95E-16 | 1.24E-11 | 0.60 | 0.34 | 0.26 | 1.76 |
| 15 | Egr1 | 0.65 | 6.05E-16 | 1.26E-11 | 0.18 | 0.07 | 0.11 | 2.47 |
| 15 | Larp4b | 0.38 | 6.23E-16 | 1.30E-11 | 0.87 | 0.61 | 0.26 | 1.43 |
| 15 | Proser2 | 1.02 | 6.28E-16 | 1.31E-11 | 0.15 | 0.06 | 0.09 | 2.68 |
| 15 | Nek1 | 0.33 | 6.31E-16 | 1.31E-11 | 0.56 | 0.31 | 0.25 | 1.81 |
| 15 | Dennd1c | 1.11 | 6.50E-16 | 1.35E-11 | 0.10 | 0.03 | 0.07 | 3.25 |
| 15 | Foxj3 | 0.32 | 6.55E-16 | 1.36E-11 | 0.53 | 0.28 | 0.25 | 1.88 |
| 15 | Rnf24 | 0.42 | 6.77E-16 | 1.41E-11 | 0.53 | 0.30 | 0.24 | 1.80 |
| 15 | Paqr4 | 1.04 | 6.98E-16 | 1.45E-11 | 0.11 | 0.03 | 0.08 | 3.21 |
| 15 | Nr2f1 | 0.40 | 7.44E-16 | 1.55E-11 | 0.57 | 0.33 | 0.24 | 1.72 |
| 15 | Phtf2 | 0.42 | 7.52E-16 | 1.56E-11 | 0.39 | 0.20 | 0.19 | 1.97 |
| 15 | Rbp1 | 0.31 | 7.70E-16 | 1.60E-11 | 1.00 | 0.91 | 0.09 | 1.09 |
| 15 | Cryl1 | 0.44 | 7.71E-16 | 1.60E-11 | 0.31 | 0.15 | 0.16 | 2.09 |
| 15 | Bnipl | 0.82 | 7.82E-16 | 1.63E-11 | 0.13 | 0.04 | 0.09 | 2.95 |
| 15 | Rbms1 | 0.31 | 8.43E-16 | 1.75E-11 | 0.96 | 0.78 | 0.18 | 1.23 |
| 15 | E330009J07Rik | 0.56 | 9.62E-16 | 2.00E-11 | 0.26 | 0.12 | 0.14 | 2.21 |
| 15 | Tmem201 | 0.38 | 1.04E-15 | 2.17E-11 | 0.47 | 0.25 | 0.22 | 1.89 |
| 15 | Macrod2 | 0.35 | 1.08E-15 | 2.24E-11 | 0.43 | 0.22 | 0.21 | 1.94 |
| 15 | Kdm7a | 0.40 | 1.13E-15 | 2.35E-11 | 0.59 | 0.35 | 0.24 | 1.71 |
| 15 | Rab38 | 0.42 | 1.14E-15 | 2.36E-11 | 0.17 | 0.07 | 0.11 | 2.56 |
| 15 | Zfp651 | 0.43 | 1.15E-15 | 2.40E-11 | 0.32 | 0.15 | 0.17 | 2.08 |
| 15 | Magi3 | 0.38 | 1.18E-15 | 2.46E-11 | 0.38 | 0.19 | 0.19 | 1.99 |
| 15 | Scrib | 0.33 | 1.26E-15 | 2.61E-11 | 0.63 | 0.36 | 0.28 | 1.78 |
| 15 | Zmiz2 | 0.37 | 1.30E-15 | 2.71E-11 | 0.55 | 0.31 | 0.24 | 1.78 |
| 15 | Kcnmb4os2 | 0.41 | 1.32E-15 | 2.75E-11 | 0.32 | 0.16 | 0.17 | 2.08 |
| 15 | Brf1 | 0.43 | 1.38E-15 | 2.87E-11 | 0.47 | 0.25 | 0.22 | 1.88 |
| 15 | Tjp2 | 0.32 | 1.45E-15 | 3.02E-11 | 0.56 | 0.31 | 0.25 | 1.82 |
| 15 | Dedd2 | 0.93 | 1.48E-15 | 3.08E-11 | 0.18 | 0.07 | 0.11 | 2.48 |
| 15 | Ncstn | 0.34 | 1.53E-15 | 3.19E-11 | 0.53 | 0.29 | 0.24 | 1.84 |
| 15 | Bcar3 | 0.95 | 1.54E-15 | 3.21E-11 | 0.14 | 0.05 | 0.09 | 2.78 |
| 15 | Gprc5c | 0.36 | 1.58E-15 | 3.28E-11 | 0.44 | 0.23 | 0.21 | 1.91 |
| 15 | Slc29a4 | 0.55 | 1.60E-15 | 3.32E-11 | 0.23 | 0.10 | 0.13 | 2.31 |
| 15 | Zbtb5 | 0.35 | 1.67E-15 | 3.48E-11 | 0.54 | 0.29 | 0.25 | 1.84 |
| 15 | Neu1 | 0.33 | 1.70E-15 | 3.53E-11 | 0.28 | 0.13 | 0.15 | 2.19 |
| 15 | Btbd9 | 0.40 | 1.75E-15 | 3.64E-11 | 0.53 | 0.29 | 0.23 | 1.80 |
| 15 | Mau2 | 0.39 | 1.84E-15 | 3.82E-11 | 0.69 | 0.40 | 0.29 | 1.72 |
| 15 | Wls | 0.29 | 1.88E-15 | 3.91E-11 | 0.93 | 0.74 | 0.20 | 1.27 |
| 15 | Pea15a | 0.35 | 1.95E-15 | 4.05E-11 | 0.43 | 0.23 | 0.20 | 1.88 |
| 15 | Psph | 0.40 | 2.00E-15 | 4.17E-11 | 0.51 | 0.28 | 0.23 | 1.83 |
| 15 | Thbs3 | 0.50 | 2.06E-15 | 4.28E-11 | 0.23 | 0.10 | 0.13 | 2.32 |
| 15 | Fbxl18 | 0.36 | 2.07E-15 | 4.30E-11 | 0.35 | 0.17 | 0.18 | 2.06 |
| 15 | Fam160a2 | 0.32 | 2.16E-15 | 4.49E-11 | 0.36 | 0.18 | 0.18 | 2.04 |
| 15 | Slc6a6 | 0.31 | 2.24E-15 | 4.66E-11 | 0.60 | 0.35 | 0.25 | 1.72 |
| 15 | Rassf7 | 0.35 | 2.24E-15 | 4.67E-11 | 0.39 | 0.20 | 0.19 | 1.98 |
| 15 | Syde2 | 0.65 | 2.29E-15 | 4.76E-11 | 0.18 | 0.07 | 0.11 | 2.51 |
| 15 | Sik1 | 0.47 | 2.35E-15 | 4.88E-11 | 0.24 | 0.10 | 0.13 | 2.29 |
| 15 | Ptprk | 0.36 | 2.46E-15 | 5.12E-11 | 0.71 | 0.44 | 0.27 | 1.63 |
| 15 | Gm26930 | 0.61 | 2.48E-15 | 5.17E-11 | 0.21 | 0.09 | 0.12 | 2.36 |
| 15 | Tbc1d31 | 0.38 | 2.53E-15 | 5.27E-11 | 0.42 | 0.22 | 0.20 | 1.94 |
| 15 | Qpct | 0.98 | 2.65E-15 | 5.50E-11 | 0.11 | 0.04 | 0.08 | 3.05 |
| 15 | Dach1 | 0.32 | 2.72E-15 | 5.67E-11 | 0.65 | 0.38 | 0.27 | 1.70 |

**Supplementary Table 1. Marker genes for the original clusters of integrated clustering of E11.5 control and NotchGOF λ-junction cells.**

| | | | | | | | | |
|---|---|---|---|---|---|---|---|---|
| 15 | Inava | 0.45 | 2.76E-15 | 5.73E-11 | 0.44 | 0.23 | 0.21 | 1.92 |
| 15 | Irs2 | 0.38 | 2.77E-15 | 5.76E-11 | 0.69 | 0.43 | 0.26 | 1.59 |
| 15 | Dnmt3b | 0.46 | 2.88E-15 | 5.99E-11 | 0.40 | 0.21 | 0.19 | 1.89 |
| 15 | Notum | 0.93 | 2.95E-15 | 6.14E-11 | 0.11 | 0.04 | 0.08 | 3.05 |
| 15 | Per3 | 0.54 | 2.99E-15 | 6.23E-11 | 0.23 | 0.10 | 0.13 | 2.30 |
| 15 | Zfp697 | 0.45 | 3.04E-15 | 6.33E-11 | 0.33 | 0.16 | 0.17 | 2.05 |
| 15 | Ghdc | 0.84 | 3.04E-15 | 6.33E-11 | 0.18 | 0.07 | 0.11 | 2.47 |
| 15 | Kazn | 0.39 | 3.28E-15 | 6.83E-11 | 0.36 | 0.18 | 0.18 | 1.99 |
| 15 | Rai1 | 0.38 | 3.32E-15 | 6.91E-11 | 0.67 | 0.40 | 0.27 | 1.68 |
| 15 | Plpp2 | 0.37 | 3.41E-15 | 7.09E-11 | 0.48 | 0.26 | 0.22 | 1.85 |
| 15 | Myo5b | 0.85 | 3.66E-15 | 7.62E-11 | 0.13 | 0.04 | 0.08 | 2.91 |
| 15 | Axl | 0.38 | 4.03E-15 | 8.39E-11 | 0.32 | 0.15 | 0.17 | 2.10 |
| 15 | Nr2f6 | 0.33 | 4.11E-15 | 8.55E-11 | 0.87 | 0.59 | 0.27 | 1.46 |
| 15 | Sh3gl2 | 1.02 | 4.15E-15 | 8.62E-11 | 0.12 | 0.04 | 0.08 | 3.05 |
| 15 | Hspb2 | 0.77 | 4.17E-15 | 8.67E-11 | 0.10 | 0.03 | 0.07 | 3.15 |
| 15 | Necab2 | 0.42 | 4.50E-15 | 9.37E-11 | 0.39 | 0.20 | 0.19 | 1.91 |
| 15 | Igf2r | 0.39 | 4.51E-15 | 9.38E-11 | 0.77 | 0.49 | 0.28 | 1.56 |
| 15 | Mex3a | 0.31 | 4.56E-15 | 9.49E-11 | 0.99 | 0.85 | 0.14 | 1.16 |
| 15 | Zfp503 | 0.34 | 4.60E-15 | 9.56E-11 | 0.62 | 0.37 | 0.25 | 1.67 |
| 15 | Limk2 | 0.41 | 4.68E-15 | 9.74E-11 | 0.38 | 0.19 | 0.18 | 1.93 |
| 15 | Stxbp5 | 0.56 | 4.69E-15 | 9.76E-11 | 0.19 | 0.08 | 0.11 | 2.44 |
| 15 | Sox13 | 0.75 | 5.01E-15 | 1.04E-10 | 0.20 | 0.09 | 0.12 | 2.34 |
| 15 | Fhl1 | 0.52 | 5.19E-15 | 1.08E-10 | 0.29 | 0.14 | 0.15 | 2.07 |
| 15 | Epha1 | 0.79 | 5.30E-15 | 1.10E-10 | 0.15 | 0.06 | 0.10 | 2.64 |
| 15 | Exph5 | 0.89 | 5.32E-15 | 1.11E-10 | 0.11 | 0.04 | 0.07 | 3.11 |
| 15 | Vps11 | 0.50 | 5.34E-15 | 1.11E-10 | 0.32 | 0.15 | 0.16 | 2.05 |
| 15 | Echdc1 | 0.34 | 5.38E-15 | 1.12E-10 | 0.44 | 0.23 | 0.21 | 1.88 |
| 15 | Ssh2 | 0.35 | 5.65E-15 | 1.18E-10 | 0.43 | 0.23 | 0.21 | 1.91 |
| 15 | Haus2 | 0.34 | 5.77E-15 | 1.20E-10 | 0.50 | 0.27 | 0.23 | 1.87 |
| 15 | Plxna2 | 0.26 | 6.24E-15 | 1.30E-10 | 0.42 | 0.23 | 0.19 | 1.85 |
| 15 | Ltbp1 | 0.37 | 6.59E-15 | 1.37E-10 | 0.42 | 0.22 | 0.20 | 1.91 |
| 15 | Cdon | 0.38 | 6.60E-15 | 1.37E-10 | 0.42 | 0.22 | 0.20 | 1.91 |
| 15 | Xpr1 | 0.36 | 6.65E-15 | 1.38E-10 | 0.49 | 0.27 | 0.22 | 1.81 |
| 15 | Cdh24 | 0.54 | 6.69E-15 | 1.39E-10 | 0.28 | 0.13 | 0.15 | 2.12 |
| 15 | Exoc3l4 | 0.59 | 6.79E-15 | 1.41E-10 | 0.17 | 0.07 | 0.11 | 2.52 |
| 15 | Cdh4 | 0.92 | 6.79E-15 | 1.41E-10 | 0.14 | 0.05 | 0.09 | 2.74 |
| 15 | Rnpepl1 | 0.42 | 7.02E-15 | 1.46E-10 | 0.46 | 0.25 | 0.21 | 1.84 |
| 15 | Agap1 | 0.37 | 7.11E-15 | 1.48E-10 | 0.69 | 0.41 | 0.27 | 1.66 |
| 15 | Hk2 | 0.27 | 7.41E-15 | 1.54E-10 | 0.47 | 0.25 | 0.22 | 1.86 |
| 15 | Nup62 | 0.39 | 7.91E-15 | 1.64E-10 | 0.84 | 0.55 | 0.29 | 1.52 |
| 15 | Grtp1 | 0.46 | 7.95E-15 | 1.65E-10 | 0.29 | 0.14 | 0.15 | 2.08 |
| 15 | Ubash3b | 0.33 | 8.24E-15 | 1.71E-10 | 0.44 | 0.23 | 0.20 | 1.86 |
| 15 | Zfp687 | 0.37 | 8.29E-15 | 1.72E-10 | 0.38 | 0.19 | 0.19 | 1.97 |
| 15 | Slc25a33 | 0.40 | 8.33E-15 | 1.73E-10 | 0.33 | 0.16 | 0.17 | 2.03 |
| 15 | Prxl2a | 0.27 | 8.36E-15 | 1.74E-10 | 0.64 | 0.38 | 0.27 | 1.71 |
| 15 | Trak1 | 0.52 | 8.49E-15 | 1.77E-10 | 0.26 | 0.12 | 0.14 | 2.18 |
| 15 | Cybrd1 | 0.66 | 8.80E-15 | 1.83E-10 | 0.14 | 0.05 | 0.09 | 2.80 |
| 15 | Chn2 | 0.69 | 9.25E-15 | 1.92E-10 | 0.17 | 0.07 | 0.10 | 2.49 |
| 15 | Ptpn13 | 0.31 | 9.47E-15 | 1.97E-10 | 0.67 | 0.40 | 0.27 | 1.69 |
| 15 | Arhgef39 | 0.57 | 9.69E-15 | 2.02E-10 | 0.29 | 0.14 | 0.15 | 2.03 |
| 15 | Pde5a | 0.25 | 9.94E-15 | 2.07E-10 | 0.36 | 0.18 | 0.18 | 2.01 |
| 15 | Zfp467 | 0.25 | 1.00E-14 | 2.08E-10 | 0.53 | 0.29 | 0.24 | 1.84 |
| 15 | Kdm3a | 0.33 | 1.01E-14 | 2.10E-10 | 0.63 | 0.37 | 0.26 | 1.72 |
| 15 | Ric3 | 0.92 | 1.06E-14 | 2.21E-10 | 0.14 | 0.05 | 0.09 | 2.74 |
| 15 | Slc45a4 | 0.40 | 1.07E-14 | 2.23E-10 | 0.28 | 0.13 | 0.15 | 2.12 |
| 15 | Tmem191c | 0.80 | 1.08E-14 | 2.25E-10 | 0.12 | 0.04 | 0.08 | 2.95 |

**Supplementary Table 1. Marker genes for the original clusters of integrated clustering of E11.5 control and NotchGOF λ-junction cells.**

| | | | | | | | | |
|---|---|---|---|---|---|---|---|---|
| 15 | Alpl | 0.65 | 1.17E-14 | 2.43E-10 | 0.23 | 0.10 | 0.12 | 2.23 |
| 15 | Pitx2 | 0.43 | 1.17E-14 | 2.44E-10 | 0.29 | 0.14 | 0.14 | 2.02 |
| 15 | Spata13 | 0.71 | 1.22E-14 | 2.53E-10 | 0.18 | 0.08 | 0.11 | 2.41 |
| 15 | Tmem8b | 0.66 | 1.23E-14 | 2.55E-10 | 0.20 | 0.09 | 0.11 | 2.34 |
| 15 | Osbpl3 | 0.48 | 1.25E-14 | 2.61E-10 | 0.21 | 0.09 | 0.12 | 2.32 |
| 15 | Tmcc3 | 0.92 | 1.28E-14 | 2.65E-10 | 0.10 | 0.03 | 0.07 | 3.06 |
| 15 | Mtmr3 | 0.38 | 1.29E-14 | 2.68E-10 | 0.48 | 0.26 | 0.22 | 1.83 |
| 15 | Cux2 | 0.52 | 1.31E-14 | 2.72E-10 | 0.22 | 0.10 | 0.12 | 2.27 |
| 15 | Zfp518a | 0.36 | 1.37E-14 | 2.85E-10 | 0.52 | 0.29 | 0.23 | 1.79 |
| 15 | Slc25a27 | 0.28 | 1.48E-14 | 3.09E-10 | 0.68 | 0.40 | 0.29 | 1.72 |
| 15 | Ptk7 | 0.34 | 1.49E-14 | 3.09E-10 | 0.63 | 0.36 | 0.26 | 1.72 |
| 15 | Ust | 0.89 | 1.54E-14 | 3.21E-10 | 0.16 | 0.06 | 0.09 | 2.54 |
| 15 | Arhgap35 | 0.37 | 1.55E-14 | 3.22E-10 | 0.73 | 0.44 | 0.28 | 1.64 |
| 15 | Gramd4 | 0.73 | 1.57E-14 | 3.27E-10 | 0.19 | 0.08 | 0.11 | 2.34 |
| 15 | Zxdb | 0.53 | 1.65E-14 | 3.42E-10 | 0.23 | 0.10 | 0.13 | 2.25 |
| 15 | Mmab | 0.40 | 1.66E-14 | 3.45E-10 | 0.39 | 0.20 | 0.19 | 1.94 |
| 15 | Dnmbp | 0.50 | 1.67E-14 | 3.46E-10 | 0.27 | 0.13 | 0.14 | 2.11 |
| 15 | Armc8 | 0.41 | 1.74E-14 | 3.61E-10 | 0.41 | 0.22 | 0.20 | 1.89 |
| 15 | Bmf | 0.35 | 1.77E-14 | 3.68E-10 | 0.24 | 0.11 | 0.13 | 2.25 |
| 15 | Pdp2 | 0.52 | 1.78E-14 | 3.71E-10 | 0.34 | 0.17 | 0.17 | 1.95 |
| 15 | Daam1 | 0.33 | 1.80E-14 | 3.75E-10 | 0.67 | 0.40 | 0.27 | 1.66 |
| 15 | Zkscan2 | 0.46 | 1.82E-14 | 3.79E-10 | 0.23 | 0.10 | 0.13 | 2.25 |
| 15 | Micu3 | 0.39 | 1.83E-14 | 3.81E-10 | 0.34 | 0.17 | 0.17 | 1.99 |
| 15 | Slc7a11 | 0.67 | 1.89E-14 | 3.94E-10 | 0.19 | 0.08 | 0.11 | 2.37 |
| 15 | Phlda3 | 0.34 | 1.91E-14 | 3.98E-10 | 0.65 | 0.41 | 0.24 | 1.60 |
| 15 | Efs | 0.36 | 1.93E-14 | 4.01E-10 | 0.51 | 0.28 | 0.23 | 1.80 |
| 15 | Bptf | 0.28 | 1.95E-14 | 4.05E-10 | 0.99 | 0.88 | 0.11 | 1.12 |
| 15 | Tshz3 | 0.58 | 1.96E-14 | 4.08E-10 | 0.23 | 0.10 | 0.13 | 2.20 |
| 15 | Zfp335 | 0.58 | 1.99E-14 | 4.13E-10 | 0.21 | 0.09 | 0.12 | 2.29 |
| 15 | Fbxo10 | 0.76 | 2.00E-14 | 4.17E-10 | 0.18 | 0.08 | 0.11 | 2.38 |
| 15 | Rbbp8 | 0.29 | 2.17E-14 | 4.52E-10 | 0.83 | 0.55 | 0.28 | 1.50 |
| 15 | Itgb8 | 0.51 | 2.26E-14 | 4.71E-10 | 0.28 | 0.13 | 0.14 | 2.10 |
| 15 | Mta1 | 0.34 | 2.40E-14 | 5.00E-10 | 0.93 | 0.69 | 0.24 | 1.34 |
| 15 | Fbrs | 0.35 | 2.42E-14 | 5.04E-10 | 0.39 | 0.20 | 0.19 | 1.94 |
| 15 | Ccdc28a | 0.41 | 2.45E-14 | 5.10E-10 | 0.23 | 0.10 | 0.13 | 2.25 |
| 15 | Syne4 | 0.35 | 2.52E-14 | 5.24E-10 | 0.38 | 0.19 | 0.18 | 1.94 |
| 15 | Rnf38 | 0.31 | 2.63E-14 | 5.46E-10 | 0.50 | 0.27 | 0.23 | 1.83 |
| 15 | Uvssa | 0.36 | 2.73E-14 | 5.68E-10 | 0.43 | 0.23 | 0.20 | 1.85 |
| 15 | Usp54 | 0.44 | 2.76E-14 | 5.74E-10 | 0.27 | 0.13 | 0.14 | 2.12 |
| 15 | Ammecr1 | 0.46 | 2.79E-14 | 5.80E-10 | 0.58 | 0.35 | 0.23 | 1.66 |
| 15 | Mgat5 | 0.36 | 2.82E-14 | 5.87E-10 | 0.47 | 0.26 | 0.21 | 1.82 |
| 15 | Abhd13 | 0.35 | 2.85E-14 | 5.92E-10 | 0.37 | 0.19 | 0.18 | 1.96 |
| 15 | Grik5 | 0.32 | 3.01E-14 | 6.27E-10 | 0.32 | 0.16 | 0.16 | 2.05 |
| 15 | Cds1 | 1.02 | 3.04E-14 | 6.32E-10 | 0.11 | 0.04 | 0.07 | 2.90 |
| 15 | Marchf8 | 0.38 | 3.16E-14 | 6.58E-10 | 0.33 | 0.17 | 0.17 | 1.98 |
| 15 | Cpne5 | 0.68 | 3.19E-14 | 6.64E-10 | 0.14 | 0.05 | 0.09 | 2.67 |
| 15 | Kitl | 0.51 | 3.20E-14 | 6.65E-10 | 0.18 | 0.07 | 0.10 | 2.41 |
| 15 | Mbd6 | 0.32 | 3.31E-14 | 6.87E-10 | 0.44 | 0.23 | 0.20 | 1.88 |
| 15 | Igsf9b | 0.64 | 3.31E-14 | 6.88E-10 | 0.15 | 0.06 | 0.09 | 2.61 |
| 15 | Rps6ka6 | 0.33 | 3.55E-14 | 7.39E-10 | 0.74 | 0.45 | 0.29 | 1.64 |
| 15 | Vps13c | 0.36 | 3.58E-14 | 7.44E-10 | 0.37 | 0.19 | 0.18 | 1.92 |
| 15 | mt-Nd6 | 0.32 | 3.69E-14 | 7.68E-10 | 0.60 | 0.35 | 0.25 | 1.72 |
| 15 | Lamc2 | 0.71 | 3.69E-14 | 7.68E-10 | 0.12 | 0.04 | 0.08 | 2.86 |
| 15 | Plagl2 | 0.38 | 3.85E-14 | 8.00E-10 | 0.60 | 0.36 | 0.25 | 1.70 |
| 15 | Marveld3 | 0.94 | 3.89E-14 | 8.09E-10 | 0.11 | 0.04 | 0.07 | 2.95 |
| 15 | Kcnmb4 | 0.35 | 3.95E-14 | 8.22E-10 | 0.55 | 0.32 | 0.23 | 1.73 |

**Supplementary Table 1. Marker genes for the original clusters of integrated clustering of E11.5 control and NotchGOF λ-junction cells.**

| | | | | | | | | |
|---|---|---|---|---|---|---|---|---|
| 15 | Sh3d19 | 0.29 | 3.99E-14 | 8.30E-10 | 0.47 | 0.25 | 0.22 | 1.85 |
| 15 | Patz1 | 0.30 | 4.05E-14 | 8.42E-10 | 0.81 | 0.52 | 0.29 | 1.57 |
| 15 | Vgll4 | 0.26 | 4.14E-14 | 8.61E-10 | 0.95 | 0.78 | 0.18 | 1.23 |
| 15 | Ppip5k2 | 0.34 | 4.19E-14 | 8.72E-10 | 0.72 | 0.45 | 0.27 | 1.59 |
| 15 | Hivep2 | 0.81 | 4.39E-14 | 9.14E-10 | 0.18 | 0.08 | 0.10 | 2.37 |
| 15 | Slc12a7 | 0.34 | 4.42E-14 | 9.20E-10 | 0.43 | 0.23 | 0.20 | 1.89 |
| 15 | Gcnt2 | 0.74 | 4.45E-14 | 9.25E-10 | 0.13 | 0.05 | 0.08 | 2.76 |
| 15 | Suco | 0.37 | 4.56E-14 | 9.49E-10 | 0.75 | 0.49 | 0.26 | 1.53 |
| 15 | F3 | 0.49 | 4.58E-14 | 9.54E-10 | 0.22 | 0.10 | 0.12 | 2.20 |
| 15 | Wdr13 | 0.41 | 4.66E-14 | 9.70E-10 | 0.23 | 0.11 | 0.13 | 2.21 |
| 15 | Inpp5f | 0.42 | 4.68E-14 | 9.74E-10 | 0.43 | 0.24 | 0.19 | 1.81 |
| 15 | Srebf1 | 0.35 | 4.79E-14 | 9.97E-10 | 0.37 | 0.19 | 0.18 | 1.94 |
| 15 | Zfp654 | 0.48 | 4.84E-14 | 1.01E-09 | 0.30 | 0.15 | 0.15 | 2.01 |
| 15 | Fktn | 0.32 | 4.84E-14 | 1.01E-09 | 0.47 | 0.26 | 0.21 | 1.83 |
| 15 | Dot1l | 0.36 | 4.95E-14 | 1.03E-09 | 0.70 | 0.43 | 0.27 | 1.62 |
| 15 | Ipo5 | 0.32 | 4.98E-14 | 1.04E-09 | 0.92 | 0.70 | 0.23 | 1.33 |
| 15 | Armc10 | 0.28 | 5.14E-14 | 1.07E-09 | 0.83 | 0.52 | 0.31 | 1.60 |
| 15 | Wdr78 | 0.45 | 5.14E-14 | 1.07E-09 | 0.24 | 0.11 | 0.13 | 2.20 |
| 15 | Marchf7 | 0.30 | 5.15E-14 | 1.07E-09 | 0.93 | 0.72 | 0.21 | 1.29 |
| 15 | Mms19 | 0.29 | 5.28E-14 | 1.10E-09 | 0.48 | 0.26 | 0.22 | 1.85 |
| 15 | Sema3f | 0.30 | 5.31E-14 | 1.10E-09 | 0.52 | 0.29 | 0.23 | 1.79 |
| 15 | Tbc1d1 | 0.28 | 5.33E-14 | 1.11E-09 | 0.63 | 0.37 | 0.27 | 1.73 |
| 15 | Echdc2 | 0.29 | 5.34E-14 | 1.11E-09 | 0.47 | 0.25 | 0.22 | 1.86 |
| 15 | Wee1 | 0.34 | 5.42E-14 | 1.13E-09 | 0.71 | 0.46 | 0.26 | 1.57 |
| 15 | Phyhipl | 0.89 | 5.50E-14 | 1.14E-09 | 0.11 | 0.04 | 0.07 | 2.92 |
| 15 | Kif5c | 0.27 | 5.59E-14 | 1.16E-09 | 0.78 | 0.48 | 0.29 | 1.61 |
| 15 | Fam187b | 0.83 | 5.61E-14 | 1.17E-09 | 0.11 | 0.04 | 0.07 | 2.92 |
| 15 | Aplp2 | 0.25 | 5.63E-14 | 1.17E-09 | 0.98 | 0.82 | 0.16 | 1.19 |
| 15 | Frmd5 | 0.79 | 5.66E-14 | 1.18E-09 | 0.13 | 0.05 | 0.08 | 2.68 |
| 15 | Ddah1 | 0.28 | 5.67E-14 | 1.18E-09 | 0.29 | 0.14 | 0.15 | 2.07 |
| 15 | Tsr1 | 0.34 | 5.94E-14 | 1.24E-09 | 0.61 | 0.36 | 0.25 | 1.71 |
| 15 | Asb7 | 0.35 | 5.97E-14 | 1.24E-09 | 0.39 | 0.21 | 0.18 | 1.89 |
| 15 | Ubox5 | 0.45 | 6.15E-14 | 1.28E-09 | 0.21 | 0.09 | 0.12 | 2.27 |
| 15 | Foxm1 | 0.33 | 6.33E-14 | 1.32E-09 | 0.47 | 0.26 | 0.21 | 1.82 |
| 15 | Tmem127 | 0.36 | 6.67E-14 | 1.39E-09 | 0.36 | 0.19 | 0.17 | 1.93 |
| 15 | Slc16a3 | 0.31 | 6.67E-14 | 1.39E-09 | 0.56 | 0.32 | 0.23 | 1.72 |
| 15 | Zfp609 | 0.37 | 6.69E-14 | 1.39E-09 | 0.69 | 0.42 | 0.27 | 1.65 |
| 15 | Pdzd8 | 0.37 | 6.74E-14 | 1.40E-09 | 0.73 | 0.45 | 0.28 | 1.63 |
| 15 | Col9a2 | 0.38 | 6.74E-14 | 1.40E-09 | 0.23 | 0.10 | 0.12 | 2.21 |
| 15 | Ddx58 | 0.67 | 6.90E-14 | 1.43E-09 | 0.14 | 0.05 | 0.09 | 2.61 |
| 15 | Akap12 | 0.35 | 6.91E-14 | 1.44E-09 | 0.37 | 0.19 | 0.18 | 1.91 |
| 15 | Mtss2 | 0.38 | 7.00E-14 | 1.45E-09 | 0.48 | 0.27 | 0.21 | 1.77 |
| 15 | Agpat3 | 0.29 | 7.08E-14 | 1.47E-09 | 0.48 | 0.26 | 0.22 | 1.83 |
| 15 | Pgap1 | 0.40 | 7.12E-14 | 1.48E-09 | 0.30 | 0.15 | 0.15 | 2.02 |
| 15 | Zbed4 | 0.38 | 7.22E-14 | 1.50E-09 | 0.51 | 0.29 | 0.22 | 1.77 |
| 15 | Prkdc | 0.33 | 7.22E-14 | 1.50E-09 | 0.40 | 0.21 | 0.19 | 1.90 |
| 15 | Zfp7 | 0.45 | 7.29E-14 | 1.52E-09 | 0.29 | 0.14 | 0.15 | 2.04 |
| 15 | Pcbd1 | 0.39 | 7.30E-14 | 1.52E-09 | 0.26 | 0.12 | 0.14 | 2.12 |
| 15 | Med26 | 0.46 | 7.35E-14 | 1.53E-09 | 0.27 | 0.13 | 0.14 | 2.10 |
| 15 | Stk11 | 0.36 | 7.50E-14 | 1.56E-09 | 0.75 | 0.48 | 0.27 | 1.57 |
| 15 | Traf6 | 0.45 | 7.70E-14 | 1.60E-09 | 0.29 | 0.15 | 0.15 | 2.03 |
| 15 | Leng8 | 0.33 | 8.01E-14 | 1.67E-09 | 0.59 | 0.34 | 0.25 | 1.73 |
| 15 | Tmem189 | 0.36 | 8.05E-14 | 1.67E-09 | 0.53 | 0.30 | 0.23 | 1.76 |
| 15 | Baz2b | 0.31 | 8.22E-14 | 1.71E-09 | 0.94 | 0.70 | 0.24 | 1.34 |
| 15 | Psap | 0.30 | 8.46E-14 | 1.76E-09 | 0.87 | 0.61 | 0.25 | 1.41 |
| 15 | Apba2 | 0.56 | 8.51E-14 | 1.77E-09 | 0.15 | 0.06 | 0.09 | 2.55 |

**Supplementary Table 1. Marker genes for the original clusters of integrated clustering of E11.5 control and NotchGOF λ-junction cells.**

| | | | | | | | | |
|---|---|---|---|---|---|---|---|---|
| 15 | Rhbdd2 | 0.33 | 8.63E-14 | 1.80E-09 | 0.47 | 0.26 | 0.21 | 1.82 |
| 15 | Esrra | 0.46 | 8.74E-14 | 1.82E-09 | 0.26 | 0.12 | 0.14 | 2.11 |
| 15 | Abhd14b | 0.62 | 9.25E-14 | 1.92E-09 | 0.15 | 0.06 | 0.09 | 2.54 |
| 15 | Mtmr4 | 0.36 | 9.44E-14 | 1.96E-09 | 0.51 | 0.29 | 0.22 | 1.78 |
| 15 | Slc4a4 | 0.54 | 9.55E-14 | 1.99E-09 | 0.24 | 0.11 | 0.13 | 2.14 |
| 15 | Atg13 | 0.33 | 9.71E-14 | 2.02E-09 | 0.30 | 0.15 | 0.15 | 2.05 |
| 15 | Slc38a10 | 0.27 | 1.00E-13 | 2.08E-09 | 0.59 | 0.33 | 0.26 | 1.77 |
| 15 | Zfp180 | 0.39 | 1.01E-13 | 2.09E-09 | 0.38 | 0.20 | 0.18 | 1.88 |
| 15 | 1700017B05Rik | 0.39 | 1.03E-13 | 2.14E-09 | 0.35 | 0.18 | 0.17 | 1.94 |
| 15 | Igf1r | 0.29 | 1.03E-13 | 2.14E-09 | 0.97 | 0.77 | 0.20 | 1.26 |
| 15 | Lonrf1 | 0.58 | 1.06E-13 | 2.20E-09 | 0.21 | 0.09 | 0.12 | 2.24 |
| 15 | Dnajc1 | 0.29 | 1.06E-13 | 2.20E-09 | 0.81 | 0.55 | 0.25 | 1.46 |
| 15 | Ppp1r36 | 0.64 | 1.06E-13 | 2.21E-09 | 0.14 | 0.05 | 0.09 | 2.62 |
| 15 | Sgsm2 | 0.86 | 1.08E-13 | 2.25E-09 | 0.15 | 0.06 | 0.09 | 2.56 |
| 15 | Igf2bp2 | 0.30 | 1.08E-13 | 2.25E-09 | 0.95 | 0.79 | 0.17 | 1.21 |
| 15 | Men1 | 0.38 | 1.09E-13 | 2.27E-09 | 0.64 | 0.38 | 0.26 | 1.67 |
| 15 | Champ1 | 0.36 | 1.19E-13 | 2.46E-09 | 0.60 | 0.35 | 0.25 | 1.73 |
| 15 | Jmjd4 | 0.50 | 1.20E-13 | 2.51E-09 | 0.25 | 0.12 | 0.13 | 2.14 |
| 15 | Malt1 | 0.40 | 1.21E-13 | 2.51E-09 | 0.32 | 0.16 | 0.16 | 1.98 |
| 15 | Wbp2 | 0.27 | 1.22E-13 | 2.53E-09 | 0.50 | 0.27 | 0.22 | 1.82 |
| 15 | Calcoco1 | 0.27 | 1.24E-13 | 2.57E-09 | 0.40 | 0.21 | 0.19 | 1.89 |
| 15 | Cbx6 | 0.29 | 1.24E-13 | 2.59E-09 | 0.48 | 0.27 | 0.22 | 1.82 |
| 15 | Gm9922 | 0.86 | 1.26E-13 | 2.62E-09 | 0.11 | 0.04 | 0.07 | 2.90 |
| 15 | Ubald1 | 0.26 | 1.26E-13 | 2.62E-09 | 0.48 | 0.26 | 0.22 | 1.83 |
| 15 | Mrtfa | 0.44 | 1.28E-13 | 2.66E-09 | 0.26 | 0.12 | 0.14 | 2.12 |
| 15 | Tet3 | 0.38 | 1.30E-13 | 2.69E-09 | 0.81 | 0.57 | 0.25 | 1.43 |
| 15 | Trp53inp1 | 0.26 | 1.34E-13 | 2.78E-09 | 0.36 | 0.18 | 0.18 | 1.95 |
| 15 | Dcaf17 | 0.29 | 1.35E-13 | 2.80E-09 | 0.58 | 0.33 | 0.25 | 1.76 |
| 15 | Scap | 0.46 | 1.35E-13 | 2.82E-09 | 0.35 | 0.18 | 0.17 | 1.92 |
| 15 | Agap3 | 0.28 | 1.36E-13 | 2.83E-09 | 0.62 | 0.36 | 0.26 | 1.73 |
| 15 | Hist2h2be | 0.89 | 1.40E-13 | 2.91E-09 | 0.11 | 0.04 | 0.07 | 2.94 |
| 15 | Bcl11a | 0.27 | 1.46E-13 | 3.04E-09 | 0.94 | 0.74 | 0.20 | 1.27 |
| 15 | Mapk8ip1 | 0.28 | 1.48E-13 | 3.07E-09 | 0.47 | 0.26 | 0.22 | 1.84 |
| 15 | Cdc14a | 0.39 | 1.50E-13 | 3.12E-09 | 0.45 | 0.25 | 0.20 | 1.77 |
| 15 | Zfp719 | 0.47 | 1.51E-13 | 3.14E-09 | 0.27 | 0.13 | 0.14 | 2.07 |
| 15 | Raver2 | 0.38 | 1.61E-13 | 3.36E-09 | 0.32 | 0.16 | 0.16 | 1.97 |
| 15 | Zfp36l1 | 0.27 | 1.64E-13 | 3.41E-09 | 0.98 | 0.85 | 0.14 | 1.16 |
| 15 | Snhg14 | 0.39 | 1.68E-13 | 3.50E-09 | 0.34 | 0.18 | 0.17 | 1.95 |
| 15 | Zfp703 | 0.37 | 1.69E-13 | 3.51E-09 | 0.45 | 0.25 | 0.20 | 1.78 |
| 15 | Snhg16 | 0.38 | 1.80E-13 | 3.75E-09 | 0.41 | 0.22 | 0.19 | 1.85 |
| 15 | Letmd1 | 0.32 | 1.85E-13 | 3.86E-09 | 0.41 | 0.22 | 0.19 | 1.88 |
| 15 | Hunk | 0.29 | 1.88E-13 | 3.92E-09 | 0.77 | 0.49 | 0.28 | 1.56 |
| 15 | Irf2bp2 | 0.27 | 1.89E-13 | 3.93E-09 | 0.60 | 0.35 | 0.26 | 1.74 |
| 15 | E130308A19Rik | 0.29 | 1.90E-13 | 3.96E-09 | 0.59 | 0.34 | 0.25 | 1.72 |
| 15 | Snhg20 | 0.29 | 1.95E-13 | 4.05E-09 | 0.62 | 0.37 | 0.25 | 1.68 |
| 15 | Rtl6 | 0.28 | 1.95E-13 | 4.05E-09 | 0.24 | 0.11 | 0.13 | 2.18 |
| 15 | Sp4 | 0.29 | 2.04E-13 | 4.25E-09 | 0.44 | 0.23 | 0.20 | 1.86 |
| 15 | Arhgap33 | 0.50 | 2.13E-13 | 4.44E-09 | 0.27 | 0.13 | 0.14 | 2.05 |
| 15 | Pold1 | 0.30 | 2.16E-13 | 4.49E-09 | 0.58 | 0.34 | 0.24 | 1.72 |
| 15 | H2-Ke6 | 0.37 | 2.17E-13 | 4.52E-09 | 0.42 | 0.23 | 0.19 | 1.82 |
| 15 | Lamb2 | 0.38 | 2.18E-13 | 4.54E-09 | 0.27 | 0.13 | 0.14 | 2.08 |
| 15 | Gm42595 | 0.74 | 2.19E-13 | 4.56E-09 | 0.12 | 0.04 | 0.08 | 2.81 |
| 15 | 2810002D19Rik | 0.35 | 2.24E-13 | 4.65E-09 | 0.30 | 0.15 | 0.15 | 2.05 |
| 15 | Pacsin1 | 0.49 | 2.27E-13 | 4.71E-09 | 0.22 | 0.10 | 0.12 | 2.18 |
| 15 | Ppp4r1 | 0.33 | 2.27E-13 | 4.71E-09 | 0.66 | 0.39 | 0.26 | 1.67 |
| 15 | Nat10 | 0.37 | 2.31E-13 | 4.81E-09 | 0.37 | 0.19 | 0.17 | 1.91 |

**Supplementary Table 1. Marker genes for the original clusters of integrated clustering of E11.5 control and NotchGOF λ-junction cells.**

| | | | | | | | | |
|---|---|---|---|---|---|---|---|---|
| 15 | Cmip | 0.29 | 2.34E-13 | 4.86E-09 | 0.65 | 0.39 | 0.26 | 1.66 |
| 15 | Tst | 0.50 | 2.37E-13 | 4.93E-09 | 0.26 | 0.13 | 0.13 | 2.06 |
| 15 | Zcchc18 | 0.32 | 2.57E-13 | 5.34E-09 | 0.57 | 0.34 | 0.24 | 1.71 |
| 15 | Ppl | 0.39 | 2.60E-13 | 5.40E-09 | 0.12 | 0.04 | 0.08 | 2.83 |
| 15 | Dbt | 0.34 | 2.62E-13 | 5.45E-09 | 0.51 | 0.29 | 0.22 | 1.75 |
| 15 | Ncor2 | 0.30 | 2.63E-13 | 5.48E-09 | 0.60 | 0.35 | 0.25 | 1.72 |
| 15 | Arid1a | 0.30 | 2.68E-13 | 5.58E-09 | 0.96 | 0.81 | 0.15 | 1.19 |
| 15 | Vac14 | 0.29 | 2.68E-13 | 5.58E-09 | 0.41 | 0.22 | 0.19 | 1.88 |
| 15 | Ubl3 | 0.32 | 2.70E-13 | 5.61E-09 | 0.63 | 0.37 | 0.27 | 1.72 |
| 15 | Irf2bpl | 0.30 | 2.70E-13 | 5.62E-09 | 0.63 | 0.38 | 0.25 | 1.65 |
| 15 | Slc38a1 | 0.26 | 2.74E-13 | 5.70E-09 | 0.96 | 0.76 | 0.19 | 1.25 |
| 15 | Pard6g | 0.31 | 2.76E-13 | 5.74E-09 | 0.86 | 0.59 | 0.27 | 1.46 |
| 15 | Cdip1 | 0.35 | 2.80E-13 | 5.83E-09 | 0.73 | 0.47 | 0.27 | 1.57 |
| 15 | Arvcf | 0.39 | 2.83E-13 | 5.89E-09 | 0.43 | 0.24 | 0.19 | 1.80 |
| 15 | Nectin2 | 0.31 | 2.91E-13 | 6.05E-09 | 0.74 | 0.47 | 0.27 | 1.56 |
| 15 | Ahcyl2 | 0.34 | 2.93E-13 | 6.09E-09 | 0.51 | 0.29 | 0.22 | 1.75 |
| 15 | Large1 | 0.31 | 2.97E-13 | 6.17E-09 | 0.37 | 0.19 | 0.18 | 1.92 |
| 15 | Tmem69 | 0.61 | 3.01E-13 | 6.26E-09 | 0.19 | 0.08 | 0.11 | 2.29 |
| 15 | Actn4 | 0.31 | 3.05E-13 | 6.34E-09 | 0.92 | 0.70 | 0.22 | 1.31 |
| 15 | Mapk8ip2 | 0.97 | 3.11E-13 | 6.46E-09 | 0.11 | 0.04 | 0.07 | 2.87 |
| 15 | Dtnb | 0.35 | 3.23E-13 | 6.71E-09 | 0.49 | 0.27 | 0.21 | 1.79 |
| 15 | Tjp1 | 0.26 | 3.23E-13 | 6.71E-09 | 0.77 | 0.49 | 0.28 | 1.57 |
| 15 | Epn3 | 0.69 | 3.24E-13 | 6.73E-09 | 0.13 | 0.05 | 0.08 | 2.65 |
| 15 | Khnyn | 0.27 | 3.32E-13 | 6.91E-09 | 0.43 | 0.23 | 0.21 | 1.90 |
| 15 | Slc12a4 | 0.45 | 3.34E-13 | 6.94E-09 | 0.28 | 0.14 | 0.14 | 2.01 |
| 15 | 2010320M18Rik | 0.30 | 3.36E-13 | 6.99E-09 | 0.57 | 0.34 | 0.23 | 1.69 |
| 15 | Exd2 | 0.35 | 3.37E-13 | 7.00E-09 | 0.30 | 0.15 | 0.15 | 2.01 |
| 15 | Tctex1d1 | 1.12 | 3.46E-13 | 7.19E-09 | 0.10 | 0.04 | 0.07 | 2.91 |
| 15 | Cipc | 0.41 | 3.50E-13 | 7.27E-09 | 0.40 | 0.22 | 0.18 | 1.83 |
| 15 | Ticrr | 0.39 | 3.53E-13 | 7.34E-09 | 0.41 | 0.23 | 0.19 | 1.83 |
| 15 | Clcn5 | 0.33 | 3.78E-13 | 7.86E-09 | 0.63 | 0.39 | 0.25 | 1.64 |
| 15 | Rnf166 | 0.27 | 3.97E-13 | 8.26E-09 | 0.47 | 0.26 | 0.21 | 1.83 |
| 15 | Rrp12 | 0.28 | 4.06E-13 | 8.44E-09 | 0.37 | 0.20 | 0.18 | 1.90 |
| 15 | Insr | 0.37 | 4.09E-13 | 8.51E-09 | 0.43 | 0.24 | 0.20 | 1.83 |
| 15 | Rnf167 | 0.29 | 4.09E-13 | 8.52E-09 | 0.62 | 0.37 | 0.26 | 1.71 |
| 15 | Gfm2 | 0.38 | 4.22E-13 | 8.79E-09 | 0.44 | 0.25 | 0.20 | 1.80 |
| 15 | Rcan3 | 0.52 | 4.33E-13 | 9.00E-09 | 0.18 | 0.08 | 0.10 | 2.34 |
| 15 | Neto2 | 0.30 | 4.41E-13 | 9.16E-09 | 0.42 | 0.23 | 0.20 | 1.85 |
| 15 | Tmem161b | 0.40 | 4.56E-13 | 9.48E-09 | 0.30 | 0.15 | 0.15 | 1.97 |
| 15 | Pou2f1 | 0.32 | 4.71E-13 | 9.80E-09 | 0.60 | 0.36 | 0.24 | 1.69 |
| 15 | Nr3c1 | 0.37 | 4.76E-13 | 9.91E-09 | 0.31 | 0.16 | 0.15 | 1.94 |
| 15 | Rnpep | 0.26 | 4.80E-13 | 9.98E-09 | 0.72 | 0.43 | 0.29 | 1.66 |
| 15 | Cnnm3 | 0.42 | 5.02E-13 | 1.04E-08 | 0.34 | 0.18 | 0.16 | 1.90 |
| 15 | Mthfr | 0.63 | 5.05E-13 | 1.05E-08 | 0.21 | 0.10 | 0.11 | 2.19 |
| 15 | Trim37 | 0.30 | 5.05E-13 | 1.05E-08 | 0.65 | 0.39 | 0.26 | 1.66 |
| 15 | Adam22 | 0.61 | 5.07E-13 | 1.06E-08 | 0.17 | 0.07 | 0.10 | 2.35 |
| 15 | Sort1 | 0.33 | 5.12E-13 | 1.06E-08 | 0.38 | 0.20 | 0.18 | 1.88 |
| 15 | Tesk1 | 0.26 | 5.13E-13 | 1.07E-08 | 0.62 | 0.37 | 0.26 | 1.70 |
| 15 | Fam227a | 0.72 | 5.15E-13 | 1.07E-08 | 0.13 | 0.05 | 0.08 | 2.64 |
| 15 | Nsmaf | 0.28 | 5.18E-13 | 1.08E-08 | 0.48 | 0.27 | 0.21 | 1.80 |
| 15 | Kdm5b | 0.25 | 5.22E-13 | 1.08E-08 | 0.98 | 0.86 | 0.12 | 1.13 |
| 15 | Amt | 0.36 | 5.27E-13 | 1.10E-08 | 0.29 | 0.15 | 0.15 | 2.01 |
| 15 | Pcdh9 | 0.50 | 5.36E-13 | 1.12E-08 | 0.23 | 0.11 | 0.12 | 2.09 |
| 15 | Pqlc2 | 0.43 | 5.47E-13 | 1.14E-08 | 0.22 | 0.10 | 0.12 | 2.16 |
| 15 | Thrb | 0.40 | 5.48E-13 | 1.14E-08 | 0.17 | 0.07 | 0.10 | 2.38 |
| 15 | Nucb2 | 0.38 | 5.48E-13 | 1.14E-08 | 0.30 | 0.15 | 0.15 | 1.98 |

**Supplementary Table 1. Marker genes for the original clusters of integrated clustering of E11.5 control and NotchGOF λ-junction cells.**

| | | | | | | | | |
|---|---|---|---|---|---|---|---|---|
| 15 | Rgs9 | 0.36 | 5.55E-13 | 1.15E-08 | 0.19 | 0.08 | 0.11 | 2.29 |
| 15 | Ncoa2 | 0.35 | 5.60E-13 | 1.16E-08 | 0.50 | 0.29 | 0.21 | 1.74 |
| 15 | Ubr4 | 0.30 | 5.65E-13 | 1.17E-08 | 0.82 | 0.54 | 0.28 | 1.53 |
| 15 | Zbtb34 | 0.53 | 5.65E-13 | 1.18E-08 | 0.20 | 0.09 | 0.11 | 2.23 |
| 15 | Trub2 | 0.31 | 5.80E-13 | 1.21E-08 | 0.53 | 0.30 | 0.23 | 1.75 |
| 15 | Usp36 | 0.37 | 5.92E-13 | 1.23E-08 | 0.41 | 0.22 | 0.19 | 1.85 |
| 15 | Gns | 0.33 | 5.98E-13 | 1.24E-08 | 0.45 | 0.25 | 0.20 | 1.80 |
| 15 | Ccdc136 | 0.30 | 6.07E-13 | 1.26E-08 | 0.49 | 0.28 | 0.21 | 1.77 |
| 15 | Ormdl1 | 0.28 | 6.08E-13 | 1.27E-08 | 0.71 | 0.44 | 0.27 | 1.62 |
| 15 | Slc16a1 | 0.34 | 6.17E-13 | 1.28E-08 | 0.52 | 0.30 | 0.22 | 1.74 |
| 15 | Ankrd26 | 0.26 | 6.26E-13 | 1.30E-08 | 0.69 | 0.43 | 0.27 | 1.62 |
| 15 | Gtf3c1 | 0.34 | 6.35E-13 | 1.32E-08 | 0.83 | 0.57 | 0.26 | 1.45 |
| 15 | Klf3 | 0.28 | 6.37E-13 | 1.32E-08 | 0.77 | 0.50 | 0.27 | 1.54 |
| 15 | Snx8 | 0.36 | 6.41E-13 | 1.33E-08 | 0.27 | 0.13 | 0.14 | 2.05 |
| 15 | G630016G05Rik | 0.74 | 6.51E-13 | 1.35E-08 | 0.10 | 0.03 | 0.07 | 3.00 |
| 15 | Zfp217 | 0.41 | 6.56E-13 | 1.36E-08 | 0.37 | 0.20 | 0.17 | 1.87 |
| 15 | Ankrd50 | 0.28 | 6.67E-13 | 1.39E-08 | 0.42 | 0.23 | 0.19 | 1.85 |
| 15 | Spock2 | 0.57 | 6.88E-13 | 1.43E-08 | 0.11 | 0.04 | 0.07 | 2.79 |
| 15 | Cacng4 | 0.27 | 6.91E-13 | 1.44E-08 | 0.46 | 0.26 | 0.20 | 1.79 |
| 15 | Parp8 | 0.33 | 6.93E-13 | 1.44E-08 | 0.28 | 0.14 | 0.14 | 2.04 |
| 15 | Phlpp2 | 0.51 | 6.99E-13 | 1.45E-08 | 0.25 | 0.12 | 0.13 | 2.07 |
| 15 | Pbx4 | 0.33 | 7.02E-13 | 1.46E-08 | 0.20 | 0.09 | 0.11 | 2.29 |
| 15 | Mmp14 | 0.28 | 7.10E-13 | 1.48E-08 | 0.88 | 0.63 | 0.26 | 1.41 |
| 15 | Magi1 | 0.26 | 7.11E-13 | 1.48E-08 | 0.54 | 0.30 | 0.23 | 1.76 |
| 15 | Utrn | 0.30 | 7.23E-13 | 1.50E-08 | 0.86 | 0.61 | 0.25 | 1.41 |
| 15 | Vill | 0.50 | 7.28E-13 | 1.51E-08 | 0.19 | 0.08 | 0.10 | 2.28 |
| 15 | Abca3 | 0.57 | 7.30E-13 | 1.52E-08 | 0.16 | 0.06 | 0.09 | 2.42 |
| 15 | Lrrn3 | 0.83 | 7.34E-13 | 1.53E-08 | 0.16 | 0.07 | 0.09 | 2.31 |
| 15 | Efna5 | 0.34 | 7.35E-13 | 1.53E-08 | 0.72 | 0.47 | 0.25 | 1.54 |
| 15 | Gm29170 | 0.61 | 7.38E-13 | 1.53E-08 | 0.18 | 0.08 | 0.10 | 2.31 |
| 15 | Avl9 | 0.30 | 7.41E-13 | 1.54E-08 | 0.59 | 0.34 | 0.24 | 1.71 |
| 15 | D430042O09Rik | 0.32 | 7.43E-13 | 1.54E-08 | 0.48 | 0.27 | 0.21 | 1.77 |
| 15 | Dennd1b | 0.35 | 7.45E-13 | 1.55E-08 | 0.31 | 0.16 | 0.15 | 1.96 |
| 15 | Vps52 | 0.33 | 7.46E-13 | 1.55E-08 | 0.45 | 0.25 | 0.20 | 1.79 |
| 15 | Cask | 0.28 | 7.51E-13 | 1.56E-08 | 0.57 | 0.34 | 0.23 | 1.69 |
| 15 | BC051226 | 0.44 | 7.61E-13 | 1.58E-08 | 0.26 | 0.13 | 0.13 | 2.05 |
| 15 | Poll | 0.34 | 7.66E-13 | 1.59E-08 | 0.33 | 0.17 | 0.16 | 1.95 |
| 15 | Zfp292 | 0.27 | 7.74E-13 | 1.61E-08 | 0.98 | 0.86 | 0.13 | 1.15 |
| 15 | Impact | 0.26 | 7.78E-13 | 1.62E-08 | 0.57 | 0.32 | 0.24 | 1.75 |
| 15 | Slc30a4 | 0.45 | 7.81E-13 | 1.62E-08 | 0.26 | 0.13 | 0.13 | 2.06 |
| 15 | Mphosph9 | 0.35 | 7.91E-13 | 1.64E-08 | 0.44 | 0.25 | 0.19 | 1.78 |
| 15 | Unc119b | 0.29 | 8.10E-13 | 1.68E-08 | 0.59 | 0.34 | 0.25 | 1.74 |
| 15 | Peli2 | 0.37 | 8.12E-13 | 1.69E-08 | 0.35 | 0.18 | 0.16 | 1.90 |
| 15 | Plxnb2 | 0.30 | 8.13E-13 | 1.69E-08 | 0.80 | 0.53 | 0.27 | 1.51 |
| 15 | Ildr1 | 0.78 | 8.31E-13 | 1.73E-08 | 0.11 | 0.04 | 0.07 | 2.86 |
| 15 | Egln3 | 0.57 | 8.32E-13 | 1.73E-08 | 0.17 | 0.08 | 0.10 | 2.32 |
| 15 | Ep400 | 0.35 | 8.37E-13 | 1.74E-08 | 0.89 | 0.67 | 0.22 | 1.33 |
| 15 | Polr1b | 0.34 | 8.38E-13 | 1.74E-08 | 0.33 | 0.17 | 0.16 | 1.95 |
| 15 | Dvl1 | 0.44 | 8.39E-13 | 1.75E-08 | 0.27 | 0.13 | 0.14 | 2.03 |
| 15 | Sun1 | 0.28 | 8.42E-13 | 1.75E-08 | 0.49 | 0.27 | 0.21 | 1.78 |
| 15 | Baiap2 | 0.35 | 8.48E-13 | 1.76E-08 | 0.30 | 0.15 | 0.15 | 1.98 |
| 15 | Tmem184b | 0.36 | 8.52E-13 | 1.77E-08 | 0.35 | 0.19 | 0.17 | 1.90 |
| 15 | B230354K17Rik | 0.35 | 8.58E-13 | 1.79E-08 | 0.59 | 0.35 | 0.24 | 1.68 |
| 15 | Dcaf4 | 0.27 | 8.70E-13 | 1.81E-08 | 0.44 | 0.24 | 0.20 | 1.83 |
| 15 | Naa80 | 0.28 | 8.74E-13 | 1.82E-08 | 0.31 | 0.16 | 0.15 | 1.97 |
| 15 | Cyb561 | 0.74 | 8.75E-13 | 1.82E-08 | 0.14 | 0.06 | 0.08 | 2.53 |

**Supplementary Table 1. Marker genes for the original clusters of integrated clustering of E11.5 control and NotchGOF λ-junction cells.**

| | | | | | | | | |
|---|---|---|---|---|---|---|---|---|
| 15 | Tm7sf2 | 0.33 | 9.13E-13 | 1.90E-08 | 0.29 | 0.14 | 0.15 | 2.01 |
| 15 | Flnb | 0.26 | 9.38E-13 | 1.95E-08 | 0.52 | 0.29 | 0.23 | 1.78 |
| 15 | Nudt11 | 0.30 | 9.44E-13 | 1.96E-08 | 0.34 | 0.18 | 0.17 | 1.94 |
| 15 | Mapk14 | 0.28 | 9.57E-13 | 1.99E-08 | 0.57 | 0.32 | 0.24 | 1.75 |
| 15 | Gaa | 0.33 | 9.62E-13 | 2.00E-08 | 0.35 | 0.18 | 0.17 | 1.92 |
| 15 | Retreg2 | 0.32 | 9.71E-13 | 2.02E-08 | 0.50 | 0.29 | 0.22 | 1.75 |
| 15 | Clcn4 | 0.29 | 9.76E-13 | 2.03E-08 | 0.52 | 0.30 | 0.22 | 1.74 |
| 15 | Gtf3c2 | 0.35 | 1.00E-12 | 2.08E-08 | 0.84 | 0.59 | 0.25 | 1.43 |
| 15 | Fgd4 | 0.30 | 1.01E-12 | 2.09E-08 | 0.20 | 0.09 | 0.11 | 2.24 |
| 15 | Dhx40 | 0.30 | 1.03E-12 | 2.14E-08 | 0.70 | 0.43 | 0.28 | 1.66 |
| 15 | Ank | 0.60 | 1.06E-12 | 2.21E-08 | 0.16 | 0.07 | 0.09 | 2.38 |
| 15 | Gm49700 | 0.33 | 1.07E-12 | 2.22E-08 | 0.31 | 0.16 | 0.15 | 1.97 |
| 15 | Klhl8 | 0.46 | 1.07E-12 | 2.23E-08 | 0.20 | 0.09 | 0.11 | 2.21 |
| 15 | Gstt3 | 0.63 | 1.10E-12 | 2.29E-08 | 0.13 | 0.05 | 0.08 | 2.65 |
| 15 | Gria3 | 0.46 | 1.11E-12 | 2.30E-08 | 0.31 | 0.16 | 0.15 | 1.93 |
| 15 | 2310068J16Rik | 0.43 | 1.11E-12 | 2.31E-08 | 0.17 | 0.07 | 0.10 | 2.35 |
| 15 | Pogk | 0.29 | 1.12E-12 | 2.32E-08 | 0.78 | 0.50 | 0.28 | 1.57 |
| 15 | Fmn2 | 0.57 | 1.13E-12 | 2.35E-08 | 0.16 | 0.07 | 0.10 | 2.38 |
| 15 | Sec16a | 0.36 | 1.13E-12 | 2.36E-08 | 0.50 | 0.29 | 0.21 | 1.74 |
| 15 | Cd44 | 0.94 | 1.14E-12 | 2.36E-08 | 0.10 | 0.04 | 0.07 | 2.89 |
| 15 | Parva | 0.26 | 1.15E-12 | 2.39E-08 | 0.83 | 0.54 | 0.29 | 1.53 |
| 15 | Gm16536 | 0.37 | 1.24E-12 | 2.57E-08 | 0.30 | 0.15 | 0.15 | 1.98 |
| 15 | Zfp629 | 0.26 | 1.28E-12 | 2.66E-08 | 0.48 | 0.27 | 0.21 | 1.79 |
| 15 | Irx1 | 0.50 | 1.31E-12 | 2.73E-08 | 0.14 | 0.05 | 0.08 | 2.54 |
| 15 | Mier2 | 0.30 | 1.32E-12 | 2.74E-08 | 0.37 | 0.20 | 0.17 | 1.89 |
| 15 | Nipsnap1 | 0.30 | 1.33E-12 | 2.76E-08 | 0.83 | 0.56 | 0.27 | 1.49 |
| 15 | Tkfc | 0.33 | 1.36E-12 | 2.83E-08 | 0.38 | 0.20 | 0.17 | 1.87 |
| 15 | Rgl2 | 0.34 | 1.37E-12 | 2.85E-08 | 0.37 | 0.20 | 0.17 | 1.87 |
| 15 | Med16 | 0.34 | 1.40E-12 | 2.91E-08 | 0.50 | 0.29 | 0.21 | 1.73 |
| 15 | Prr14 | 0.45 | 1.40E-12 | 2.92E-08 | 0.37 | 0.20 | 0.17 | 1.86 |
| 15 | Zscan12 | 0.30 | 1.42E-12 | 2.95E-08 | 0.44 | 0.24 | 0.20 | 1.84 |
| 15 | Tmem245 | 0.28 | 1.44E-12 | 2.99E-08 | 0.66 | 0.40 | 0.26 | 1.65 |
| 15 | Il17re | 0.68 | 1.44E-12 | 2.99E-08 | 0.12 | 0.05 | 0.08 | 2.67 |
| 15 | Tnk1 | 0.90 | 1.47E-12 | 3.05E-08 | 0.10 | 0.04 | 0.07 | 2.81 |
| 15 | Fam120a | 0.31 | 1.47E-12 | 3.06E-08 | 0.73 | 0.45 | 0.28 | 1.61 |
| 15 | Kcnj2 | 0.26 | 1.47E-12 | 3.06E-08 | 0.33 | 0.17 | 0.16 | 1.90 |
| 15 | Dock9 | 0.53 | 1.49E-12 | 3.09E-08 | 0.17 | 0.07 | 0.10 | 2.35 |
| 15 | Rabgap1 | 0.29 | 1.50E-12 | 3.12E-08 | 0.81 | 0.56 | 0.25 | 1.44 |
| 15 | Nol6 | 0.30 | 1.51E-12 | 3.13E-08 | 0.39 | 0.21 | 0.18 | 1.87 |
| 15 | Aldh5a1 | 0.36 | 1.54E-12 | 3.20E-08 | 0.44 | 0.24 | 0.20 | 1.81 |
| 15 | Fblim1 | 0.27 | 1.55E-12 | 3.23E-08 | 0.72 | 0.46 | 0.26 | 1.56 |
| 15 | Mybbp1a | 0.29 | 1.56E-12 | 3.23E-08 | 0.96 | 0.77 | 0.18 | 1.24 |
| 15 | Clcn6 | 0.49 | 1.60E-12 | 3.32E-08 | 0.19 | 0.09 | 0.11 | 2.23 |
| 15 | Slc6a8 | 0.37 | 1.66E-12 | 3.46E-08 | 0.35 | 0.19 | 0.17 | 1.88 |
| 15 | Strbp | 0.29 | 1.67E-12 | 3.48E-08 | 0.91 | 0.70 | 0.21 | 1.30 |
| 15 | 2610301B20Rik | 0.25 | 1.68E-12 | 3.50E-08 | 0.50 | 0.28 | 0.22 | 1.79 |
| 15 | Snx14 | 0.34 | 1.72E-12 | 3.58E-08 | 0.32 | 0.16 | 0.15 | 1.94 |
| 15 | Iqgap2 | 0.34 | 1.73E-12 | 3.59E-08 | 0.58 | 0.36 | 0.22 | 1.61 |
| 15 | Slc35a3 | 0.40 | 1.73E-12 | 3.61E-08 | 0.25 | 0.12 | 0.13 | 2.05 |
| 15 | 2610306M01Rik | 0.29 | 1.74E-12 | 3.61E-08 | 0.32 | 0.16 | 0.15 | 1.96 |
| 15 | Rtn3 | 0.27 | 1.74E-12 | 3.63E-08 | 0.95 | 0.79 | 0.17 | 1.21 |
| 15 | Rhou | 0.26 | 1.82E-12 | 3.79E-08 | 0.69 | 0.41 | 0.28 | 1.69 |
| 15 | Glis2 | 0.38 | 1.82E-12 | 3.79E-08 | 0.36 | 0.19 | 0.17 | 1.87 |
| 15 | Spry1 | 0.32 | 1.83E-12 | 3.81E-08 | 0.48 | 0.29 | 0.20 | 1.69 |
| 15 | Tfrc | 0.26 | 1.84E-12 | 3.83E-08 | 0.66 | 0.39 | 0.26 | 1.66 |
| 15 | Dpp9 | 0.31 | 1.88E-12 | 3.90E-08 | 0.51 | 0.29 | 0.22 | 1.75 |

**Supplementary Table 1. Marker genes for the original clusters of integrated clustering of E11.5 control and NotchGOF λ-junction cells.**

| | | | | | | | | |
|---|---|---|---|---|---|---|---|---|
| 15 | 2500004C02Rik | 0.32 | 1.91E-12 | 3.97E-08 | 0.28 | 0.14 | 0.14 | 2.01 |
| 15 | Vegfa | 0.32 | 1.92E-12 | 3.99E-08 | 0.48 | 0.27 | 0.21 | 1.77 |
| 15 | Smad3 | 0.28 | 1.93E-12 | 4.02E-08 | 0.47 | 0.26 | 0.21 | 1.79 |
| 15 | Habp4 | 0.27 | 1.95E-12 | 4.05E-08 | 0.34 | 0.18 | 0.16 | 1.92 |
| 15 | Syvn1 | 0.37 | 2.01E-12 | 4.19E-08 | 0.30 | 0.16 | 0.15 | 1.94 |
| 15 | Atr | 0.36 | 2.02E-12 | 4.19E-08 | 0.56 | 0.33 | 0.23 | 1.70 |
| 15 | Ube3b | 0.33 | 2.18E-12 | 4.54E-08 | 0.42 | 0.23 | 0.19 | 1.81 |
| 15 | Parm1 | 0.33 | 2.18E-12 | 4.54E-08 | 0.31 | 0.16 | 0.15 | 1.93 |
| 15 | Pacsin3 | 0.28 | 2.20E-12 | 4.57E-08 | 0.40 | 0.22 | 0.18 | 1.84 |
| 15 | Mlycd | 0.32 | 2.20E-12 | 4.58E-08 | 0.28 | 0.14 | 0.14 | 2.00 |
| 15 | Zc2hc1a | 0.28 | 2.23E-12 | 4.65E-08 | 0.69 | 0.41 | 0.28 | 1.67 |
| 15 | Gemin5 | 0.28 | 2.24E-12 | 4.65E-08 | 0.62 | 0.36 | 0.26 | 1.71 |
| 15 | Gpr137c | 0.35 | 2.24E-12 | 4.66E-08 | 0.29 | 0.14 | 0.14 | 1.98 |
| 15 | Chmp2b | 0.26 | 2.24E-12 | 4.67E-08 | 0.84 | 0.56 | 0.28 | 1.49 |
| 15 | Togaram1 | 0.26 | 2.25E-12 | 4.68E-08 | 0.47 | 0.26 | 0.21 | 1.79 |
| 15 | Bicc1 | 0.35 | 2.27E-12 | 4.72E-08 | 0.41 | 0.23 | 0.18 | 1.79 |
| 15 | Zfp516 | 0.32 | 2.28E-12 | 4.73E-08 | 0.76 | 0.49 | 0.27 | 1.54 |
| 15 | Mif4gd | 0.34 | 2.29E-12 | 4.76E-08 | 0.32 | 0.17 | 0.15 | 1.91 |
| 15 | Gnaq | 0.33 | 2.31E-12 | 4.81E-08 | 0.77 | 0.52 | 0.25 | 1.48 |
| 15 | E430018J23Rik | 0.51 | 2.31E-12 | 4.81E-08 | 0.16 | 0.06 | 0.09 | 2.42 |
| 15 | Pigu | 0.31 | 2.34E-12 | 4.86E-08 | 0.37 | 0.20 | 0.17 | 1.87 |
| 15 | Pcnx3 | 0.33 | 2.39E-12 | 4.96E-08 | 0.54 | 0.32 | 0.22 | 1.69 |
| 15 | Tspan5 | 0.28 | 2.40E-12 | 4.98E-08 | 0.47 | 0.26 | 0.21 | 1.79 |
| 15 | Uvrag | 0.32 | 2.45E-12 | 5.09E-08 | 0.41 | 0.23 | 0.19 | 1.83 |
| 15 | 2900076A07Rik | 0.28 | 2.48E-12 | 5.15E-08 | 0.53 | 0.30 | 0.23 | 1.75 |
| 15 | Patl1 | 0.29 | 2.50E-12 | 5.21E-08 | 0.44 | 0.24 | 0.19 | 1.80 |
| 15 | Fbxo27 | 0.59 | 2.52E-12 | 5.23E-08 | 0.16 | 0.07 | 0.09 | 2.38 |
| 15 | Dgcr2 | 0.29 | 2.55E-12 | 5.31E-08 | 0.74 | 0.46 | 0.28 | 1.60 |
| 15 | Thumpd3 | 0.32 | 2.57E-12 | 5.35E-08 | 0.61 | 0.38 | 0.24 | 1.62 |
| 15 | Ipo13 | 0.37 | 2.63E-12 | 5.48E-08 | 0.30 | 0.15 | 0.15 | 1.97 |
| 15 | Smurf1 | 0.35 | 2.69E-12 | 5.59E-08 | 0.28 | 0.14 | 0.14 | 2.00 |
| 15 | Nup133 | 0.30 | 2.72E-12 | 5.66E-08 | 0.61 | 0.37 | 0.24 | 1.64 |
| 15 | Lrfn4 | 0.27 | 2.73E-12 | 5.68E-08 | 0.46 | 0.26 | 0.20 | 1.79 |
| 15 | Rpgrip1 | 0.47 | 2.75E-12 | 5.73E-08 | 0.46 | 0.27 | 0.19 | 1.72 |
| 15 | Mok | 0.69 | 2.79E-12 | 5.81E-08 | 0.15 | 0.06 | 0.09 | 2.43 |
| 15 | Cobl | 0.59 | 2.82E-12 | 5.87E-08 | 0.16 | 0.07 | 0.09 | 2.35 |
| 15 | Zdhhc13 | 0.37 | 2.85E-12 | 5.92E-08 | 0.27 | 0.14 | 0.14 | 1.99 |
| 15 | Rnf139 | 0.29 | 2.87E-12 | 5.97E-08 | 0.44 | 0.25 | 0.20 | 1.80 |
| 15 | Enpp5 | 0.67 | 2.95E-12 | 6.13E-08 | 0.12 | 0.04 | 0.07 | 2.70 |
| 15 | Zfp667 | 0.43 | 2.96E-12 | 6.16E-08 | 0.21 | 0.10 | 0.11 | 2.15 |
| 15 | Adgrl1 | 0.31 | 2.99E-12 | 6.22E-08 | 0.71 | 0.43 | 0.27 | 1.63 |
| 15 | Klhl28 | 0.32 | 2.99E-12 | 6.22E-08 | 0.37 | 0.20 | 0.17 | 1.85 |
| 15 | Tmem50b | 0.28 | 3.01E-12 | 6.27E-08 | 0.34 | 0.18 | 0.16 | 1.92 |
| 15 | Ago2 | 0.30 | 3.03E-12 | 6.29E-08 | 0.90 | 0.72 | 0.18 | 1.25 |
| 15 | Ccnd1 | 0.28 | 3.10E-12 | 6.45E-08 | 0.96 | 0.80 | 0.16 | 1.20 |
| 15 | Fads1 | 0.28 | 3.11E-12 | 6.47E-08 | 0.75 | 0.48 | 0.27 | 1.56 |
| 15 | 2410002F23Rik | 0.32 | 3.12E-12 | 6.48E-08 | 0.82 | 0.57 | 0.25 | 1.45 |
| 15 | Gprc5b | 0.41 | 3.13E-12 | 6.51E-08 | 0.21 | 0.10 | 0.11 | 2.15 |
| 15 | Src | 0.29 | 3.18E-12 | 6.61E-08 | 0.41 | 0.23 | 0.19 | 1.82 |
| 15 | Trip10 | 0.27 | 3.26E-12 | 6.78E-08 | 0.40 | 0.21 | 0.18 | 1.86 |
| 15 | Ccnt1 | 0.33 | 3.29E-12 | 6.85E-08 | 0.83 | 0.58 | 0.26 | 1.44 |
| 15 | Ptpn4 | 0.39 | 3.36E-12 | 6.98E-08 | 0.30 | 0.16 | 0.14 | 1.92 |
| 15 | Acsl3 | 0.26 | 3.50E-12 | 7.27E-08 | 0.95 | 0.73 | 0.21 | 1.29 |
| 15 | Foxn3 | 0.31 | 3.54E-12 | 7.35E-08 | 0.89 | 0.68 | 0.21 | 1.31 |
| 15 | Nek7 | 0.27 | 3.78E-12 | 7.85E-08 | 0.51 | 0.29 | 0.22 | 1.75 |
| 15 | Pik3r4 | 0.33 | 3.88E-12 | 8.07E-08 | 0.38 | 0.21 | 0.17 | 1.83 |

**Supplementary Table 1. Marker genes for the original clusters of integrated clustering of E11.5 control and NotchGOF λ-junction cells.**

| | | | | | | | | |
|---|---|---|---|---|---|---|---|---|
| 15 | Myo5a | 0.27 | 3.93E-12 | 8.17E-08 | 0.54 | 0.32 | 0.23 | 1.71 |
| 15 | Sh3bp4 | 0.29 | 3.94E-12 | 8.19E-08 | 0.32 | 0.17 | 0.16 | 1.94 |
| 15 | Fgd6 | 0.31 | 3.94E-12 | 8.19E-08 | 0.31 | 0.16 | 0.15 | 1.94 |
| 15 | Trim3 | 0.52 | 3.99E-12 | 8.30E-08 | 0.20 | 0.09 | 0.11 | 2.18 |
| 15 | Tle3 | 0.25 | 4.11E-12 | 8.55E-08 | 0.70 | 0.43 | 0.27 | 1.63 |
| 15 | Tefm | 0.29 | 4.11E-12 | 8.56E-08 | 0.35 | 0.19 | 0.17 | 1.89 |
| 15 | Plxna3 | 0.45 | 4.14E-12 | 8.61E-08 | 0.21 | 0.10 | 0.11 | 2.14 |
| 15 | Pgap2 | 0.29 | 4.30E-12 | 8.95E-08 | 0.71 | 0.45 | 0.26 | 1.58 |
| 15 | Cpd | 0.28 | 4.41E-12 | 9.17E-08 | 0.57 | 0.34 | 0.23 | 1.69 |
| 15 | Ctns | 0.48 | 4.44E-12 | 9.24E-08 | 0.21 | 0.10 | 0.11 | 2.15 |
| 15 | Chml | 0.44 | 4.45E-12 | 9.25E-08 | 0.29 | 0.15 | 0.14 | 1.95 |
| 15 | Acadm | 0.27 | 4.60E-12 | 9.58E-08 | 0.56 | 0.33 | 0.23 | 1.70 |
| 15 | Fryl | 0.32 | 4.84E-12 | 1.01E-07 | 0.51 | 0.30 | 0.22 | 1.73 |
| 15 | Bcl2l2 | 0.35 | 4.93E-12 | 1.03E-07 | 0.20 | 0.09 | 0.11 | 2.21 |
| 15 | Swsap1 | 0.46 | 5.18E-12 | 1.08E-07 | 0.21 | 0.10 | 0.11 | 2.14 |
| 15 | Nfrkb | 0.27 | 5.23E-12 | 1.09E-07 | 0.56 | 0.33 | 0.24 | 1.72 |
| 15 | Efna3 | 0.31 | 5.29E-12 | 1.10E-07 | 0.15 | 0.06 | 0.09 | 2.43 |
| 15 | Gas2l3 | 0.38 | 5.33E-12 | 1.11E-07 | 0.33 | 0.18 | 0.15 | 1.83 |
| 15 | Crtc2 | 0.35 | 5.55E-12 | 1.15E-07 | 0.24 | 0.12 | 0.12 | 2.07 |
| 15 | Asxl2 | 0.28 | 5.55E-12 | 1.15E-07 | 0.67 | 0.41 | 0.26 | 1.64 |
| 15 | Zbtb21 | 0.45 | 5.57E-12 | 1.16E-07 | 0.27 | 0.13 | 0.13 | 1.99 |
| 15 | Usp6nl | 0.27 | 5.57E-12 | 1.16E-07 | 0.41 | 0.22 | 0.19 | 1.84 |
| 15 | Utp25 | 0.32 | 5.81E-12 | 1.21E-07 | 0.47 | 0.26 | 0.21 | 1.78 |
| 15 | Gdf11 | 0.25 | 5.97E-12 | 1.24E-07 | 0.54 | 0.32 | 0.23 | 1.72 |
| 15 | Tab3 | 0.45 | 5.99E-12 | 1.25E-07 | 0.19 | 0.09 | 0.11 | 2.21 |
| 15 | Plxna1 | 0.28 | 6.10E-12 | 1.27E-07 | 0.38 | 0.20 | 0.17 | 1.85 |
| 15 | Map3k20 | 0.51 | 6.26E-12 | 1.30E-07 | 0.24 | 0.12 | 0.12 | 2.03 |
| 15 | Map2k7 | 0.31 | 6.36E-12 | 1.32E-07 | 0.45 | 0.26 | 0.19 | 1.74 |
| 15 | Gen1 | 0.33 | 6.38E-12 | 1.33E-07 | 0.47 | 0.27 | 0.20 | 1.73 |
| 15 | Myadm | 0.33 | 6.47E-12 | 1.35E-07 | 0.25 | 0.12 | 0.13 | 2.03 |
| 15 | Slc39a14 | 0.39 | 6.61E-12 | 1.37E-07 | 0.22 | 0.10 | 0.12 | 2.12 |
| 15 | Vldlr | 0.36 | 6.85E-12 | 1.42E-07 | 0.24 | 0.12 | 0.12 | 2.05 |
| 15 | Proser3 | 0.31 | 6.93E-12 | 1.44E-07 | 0.33 | 0.17 | 0.16 | 1.92 |
| 15 | Gtf3c4 | 0.31 | 6.95E-12 | 1.45E-07 | 0.59 | 0.36 | 0.24 | 1.67 |
| 15 | Adipor2 | 0.28 | 6.96E-12 | 1.45E-07 | 0.58 | 0.35 | 0.24 | 1.69 |
| 15 | Cldn23 | 0.63 | 7.05E-12 | 1.47E-07 | 0.10 | 0.04 | 0.07 | 2.81 |
| 15 | Mink1 | 0.30 | 7.15E-12 | 1.49E-07 | 0.32 | 0.17 | 0.15 | 1.91 |
| 15 | Scamp1 | 0.30 | 7.24E-12 | 1.50E-07 | 0.33 | 0.17 | 0.16 | 1.90 |
| 15 | Nr2c2ap | 0.26 | 7.24E-12 | 1.51E-07 | 0.41 | 0.23 | 0.19 | 1.81 |
| 15 | Ppp1r9a | 0.25 | 7.40E-12 | 1.54E-07 | 0.65 | 0.39 | 0.26 | 1.66 |
| 15 | Map3k3 | 0.41 | 7.49E-12 | 1.56E-07 | 0.29 | 0.15 | 0.14 | 1.92 |
| 15 | Svip | 0.44 | 7.53E-12 | 1.57E-07 | 0.19 | 0.09 | 0.10 | 2.13 |
| 15 | Zfp58 | 0.28 | 7.61E-12 | 1.58E-07 | 0.26 | 0.13 | 0.13 | 2.03 |
| 15 | Ubxn2a | 0.28 | 7.62E-12 | 1.58E-07 | 0.53 | 0.31 | 0.22 | 1.69 |
| 15 | Tbce | 0.26 | 7.64E-12 | 1.59E-07 | 0.51 | 0.29 | 0.22 | 1.74 |
| 15 | Rnf165 | 0.30 | 7.65E-12 | 1.59E-07 | 0.43 | 0.25 | 0.19 | 1.77 |
| 15 | Fam53c | 0.28 | 7.70E-12 | 1.60E-07 | 0.37 | 0.20 | 0.17 | 1.86 |
| 15 | Prkd3 | 0.29 | 7.88E-12 | 1.64E-07 | 0.80 | 0.53 | 0.27 | 1.50 |
| 15 | Trim56 | 0.51 | 7.99E-12 | 1.66E-07 | 0.27 | 0.14 | 0.13 | 1.95 |
| 15 | Sos2 | 0.37 | 8.26E-12 | 1.72E-07 | 0.36 | 0.20 | 0.16 | 1.84 |
| 15 | Gba2 | 0.40 | 8.28E-12 | 1.72E-07 | 0.25 | 0.12 | 0.12 | 2.02 |
| 15 | Carmil1 | 0.36 | 8.41E-12 | 1.75E-07 | 0.28 | 0.14 | 0.14 | 1.96 |
| 15 | Sp1 | 0.27 | 8.48E-12 | 1.76E-07 | 0.67 | 0.41 | 0.26 | 1.65 |
| 15 | Cnksr2 | 0.27 | 8.50E-12 | 1.77E-07 | 0.55 | 0.32 | 0.22 | 1.70 |
| 15 | Slc20a2 | 0.33 | 8.58E-12 | 1.79E-07 | 0.42 | 0.23 | 0.19 | 1.79 |
| 15 | Six5 | 0.33 | 8.61E-12 | 1.79E-07 | 0.36 | 0.20 | 0.17 | 1.85 |

**Supplementary Table 1. Marker genes for the original clusters of integrated clustering of E11.5 control and NotchGOF λ-junction cells.**

| | | | | | | | | |
|---|---|---|---|---|---|---|---|---|
| 15 | Mdn1 | 0.33 | 8.69E-12 | 1.81E-07 | 0.64 | 0.41 | 0.23 | 1.57 |
| 15 | Uso1 | 0.27 | 8.74E-12 | 1.82E-07 | 0.63 | 0.38 | 0.25 | 1.64 |
| 15 | Sord | 0.28 | 8.76E-12 | 1.82E-07 | 0.28 | 0.14 | 0.14 | 1.94 |
| 15 | Taok2 | 0.30 | 8.82E-12 | 1.84E-07 | 0.53 | 0.31 | 0.21 | 1.69 |
| 15 | Wnk1 | 0.27 | 8.87E-12 | 1.84E-07 | 0.96 | 0.78 | 0.18 | 1.24 |
| 15 | Dtx3l | 0.93 | 8.96E-12 | 1.86E-07 | 0.11 | 0.04 | 0.07 | 2.66 |
| 15 | Zxdc | 0.50 | 9.09E-12 | 1.89E-07 | 0.22 | 0.10 | 0.11 | 2.09 |
| 15 | Akap9 | 0.26 | 9.32E-12 | 1.94E-07 | 0.96 | 0.78 | 0.18 | 1.24 |
| 15 | Stim1 | 0.37 | 9.43E-12 | 1.96E-07 | 0.28 | 0.14 | 0.14 | 1.96 |
| 15 | Osbpl8 | 0.30 | 9.52E-12 | 1.98E-07 | 0.41 | 0.23 | 0.18 | 1.77 |
| 15 | Gm43445 | 0.63 | 9.52E-12 | 1.98E-07 | 0.19 | 0.09 | 0.10 | 2.16 |
| 15 | Bcl2l13 | 0.34 | 9.52E-12 | 1.98E-07 | 0.60 | 0.37 | 0.23 | 1.62 |
| 15 | Rps6ka1 | 0.29 | 9.77E-12 | 2.03E-07 | 0.42 | 0.24 | 0.18 | 1.77 |
| 15 | Nfia | 0.25 | 9.79E-12 | 2.04E-07 | 0.72 | 0.47 | 0.26 | 1.55 |
| 15 | Smim10l2a | 0.49 | 9.93E-12 | 2.07E-07 | 0.16 | 0.07 | 0.09 | 2.31 |
| 15 | Ap4e1 | 0.58 | 9.94E-12 | 2.07E-07 | 0.22 | 0.11 | 0.11 | 2.06 |
| 15 | Abhd11 | 0.33 | 1.02E-11 | 2.13E-07 | 0.46 | 0.26 | 0.19 | 1.73 |
| 15 | Mboat1 | 0.65 | 1.03E-11 | 2.14E-07 | 0.13 | 0.05 | 0.08 | 2.54 |
| 15 | Zkscan7 | 0.60 | 1.03E-11 | 2.15E-07 | 0.17 | 0.08 | 0.10 | 2.25 |
| 15 | Carf | 0.44 | 1.03E-11 | 2.15E-07 | 0.22 | 0.11 | 0.11 | 2.08 |
| 15 | Arid2 | 0.28 | 1.05E-11 | 2.17E-07 | 0.86 | 0.62 | 0.24 | 1.38 |
| 15 | Gm43560 | 0.79 | 1.06E-11 | 2.21E-07 | 0.12 | 0.05 | 0.07 | 2.62 |
| 15 | Soga3 | 0.36 | 1.08E-11 | 2.25E-07 | 0.13 | 0.05 | 0.08 | 2.50 |
| 15 | Ctdspl | 0.28 | 1.10E-11 | 2.28E-07 | 0.58 | 0.34 | 0.24 | 1.69 |
| 15 | Dgkd | 0.37 | 1.11E-11 | 2.31E-07 | 0.41 | 0.23 | 0.18 | 1.78 |
| 15 | Plpp5 | 0.30 | 1.15E-11 | 2.40E-07 | 0.35 | 0.19 | 0.16 | 1.85 |
| 15 | Gm17108 | 0.68 | 1.16E-11 | 2.41E-07 | 0.12 | 0.05 | 0.07 | 2.62 |
| 15 | Kcnc3 | 0.58 | 1.16E-11 | 2.41E-07 | 0.13 | 0.05 | 0.08 | 2.49 |
| 15 | Ppip5k1 | 0.76 | 1.16E-11 | 2.42E-07 | 0.15 | 0.06 | 0.09 | 2.35 |
| 15 | Pptc7 | 0.30 | 1.16E-11 | 2.42E-07 | 0.35 | 0.19 | 0.16 | 1.87 |
| 15 | Ext2 | 0.34 | 1.17E-11 | 2.43E-07 | 0.66 | 0.42 | 0.24 | 1.58 |
| 15 | Klf13 | 0.28 | 1.17E-11 | 2.43E-07 | 0.59 | 0.37 | 0.22 | 1.60 |
| 15 | B230118H07Rik | 0.33 | 1.20E-11 | 2.50E-07 | 0.74 | 0.50 | 0.24 | 1.48 |
| 15 | Smarcd3 | 0.27 | 1.23E-11 | 2.56E-07 | 0.48 | 0.28 | 0.20 | 1.73 |
| 15 | Mirg | 0.89 | 1.24E-11 | 2.58E-07 | 0.12 | 0.05 | 0.07 | 2.55 |
| 15 | Ccdc92 | 0.51 | 1.24E-11 | 2.59E-07 | 0.18 | 0.08 | 0.10 | 2.23 |
| 15 | Lrrc61 | 0.46 | 1.29E-11 | 2.69E-07 | 0.20 | 0.10 | 0.11 | 2.13 |
| 15 | Tmem43 | 0.27 | 1.30E-11 | 2.70E-07 | 0.38 | 0.21 | 0.18 | 1.85 |
| 15 | Atraid | 0.26 | 1.35E-11 | 2.80E-07 | 0.78 | 0.50 | 0.27 | 1.55 |
| 15 | Lbh | 0.32 | 1.36E-11 | 2.82E-07 | 0.36 | 0.20 | 0.16 | 1.82 |
| 15 | Mical3 | 0.30 | 1.39E-11 | 2.89E-07 | 0.49 | 0.28 | 0.21 | 1.74 |
| 15 | Med13 | 0.29 | 1.41E-11 | 2.94E-07 | 0.82 | 0.55 | 0.27 | 1.49 |
| 15 | AU040320 | 0.32 | 1.42E-11 | 2.96E-07 | 0.23 | 0.11 | 0.12 | 2.08 |
| 15 | Klf10 | 0.34 | 1.43E-11 | 2.98E-07 | 0.22 | 0.11 | 0.12 | 2.07 |
| 15 | Ccdc171 | 0.35 | 1.46E-11 | 3.03E-07 | 0.20 | 0.09 | 0.11 | 2.14 |
| 15 | Rictor | 0.29 | 1.46E-11 | 3.03E-07 | 0.61 | 0.38 | 0.23 | 1.62 |
| 15 | Urb1 | 0.32 | 1.47E-11 | 3.06E-07 | 0.26 | 0.13 | 0.13 | 2.00 |
| 15 | Vps54 | 0.26 | 1.49E-11 | 3.09E-07 | 0.54 | 0.31 | 0.23 | 1.72 |
| 15 | Zcchc7 | 0.27 | 1.49E-11 | 3.09E-07 | 0.78 | 0.52 | 0.27 | 1.52 |
| 15 | Uhrf1bp1l | 0.27 | 1.50E-11 | 3.11E-07 | 0.53 | 0.31 | 0.22 | 1.71 |
| 15 | Vangl2 | 0.27 | 1.51E-11 | 3.15E-07 | 0.84 | 0.57 | 0.27 | 1.48 |
| 15 | Zfp141 | 0.45 | 1.52E-11 | 3.17E-07 | 0.26 | 0.14 | 0.13 | 1.96 |
| 15 | Pik3cb | 0.65 | 1.54E-11 | 3.20E-07 | 0.15 | 0.06 | 0.08 | 2.35 |
| 15 | Pard6b | 0.52 | 1.55E-11 | 3.22E-07 | 0.13 | 0.05 | 0.08 | 2.50 |
| 15 | Odf2l | 0.30 | 1.55E-11 | 3.23E-07 | 0.58 | 0.35 | 0.23 | 1.64 |
| 15 | Nhsl1 | 0.29 | 1.58E-11 | 3.28E-07 | 0.39 | 0.22 | 0.17 | 1.80 |

**Supplementary Table 1. Marker genes for the original clusters of integrated clustering of E11.5 control and NotchGOF λ-junction cells.**

| | | | | | | | | |
|---|---|---|---|---|---|---|---|---|
| 15 | Sh3bgrl2 | 0.55 | 1.62E-11 | 3.37E-07 | 0.18 | 0.08 | 0.10 | 2.23 |
| 15 | Trim12c | 0.30 | 1.63E-11 | 3.38E-07 | 0.33 | 0.18 | 0.15 | 1.86 |
| 15 | Zfp692 | 0.42 | 1.63E-11 | 3.39E-07 | 0.22 | 0.11 | 0.11 | 2.08 |
| 15 | Prkx | 0.34 | 1.64E-11 | 3.41E-07 | 0.29 | 0.15 | 0.14 | 1.93 |
| 15 | Zcchc14 | 0.27 | 1.65E-11 | 3.42E-07 | 0.54 | 0.32 | 0.22 | 1.69 |
| 15 | Chkb | 0.26 | 1.67E-11 | 3.47E-07 | 0.41 | 0.23 | 0.18 | 1.77 |
| 15 | Rhebl1 | 0.28 | 1.70E-11 | 3.53E-07 | 0.35 | 0.19 | 0.16 | 1.86 |
| 15 | Rnf11 | 0.28 | 1.70E-11 | 3.54E-07 | 0.85 | 0.62 | 0.23 | 1.37 |
| 15 | Tpra1 | 0.39 | 1.81E-11 | 3.77E-07 | 0.25 | 0.12 | 0.12 | 2.00 |
| 15 | Celf1 | 0.26 | 1.84E-11 | 3.82E-07 | 0.95 | 0.81 | 0.14 | 1.17 |
| 15 | Syne2 | 0.27 | 1.87E-11 | 3.89E-07 | 0.95 | 0.72 | 0.23 | 1.32 |
| 15 | Sipa1l3 | 0.45 | 1.92E-11 | 4.00E-07 | 0.21 | 0.10 | 0.11 | 2.09 |
| 15 | Gm20045 | 0.27 | 1.93E-11 | 4.02E-07 | 0.28 | 0.14 | 0.14 | 1.94 |
| 15 | Armh4 | 0.48 | 1.94E-11 | 4.04E-07 | 0.28 | 0.15 | 0.13 | 1.91 |
| 15 | Mettl27 | 0.34 | 1.97E-11 | 4.10E-07 | 0.27 | 0.14 | 0.13 | 1.95 |
| 15 | Zfp618 | 0.27 | 2.00E-11 | 4.15E-07 | 0.80 | 0.52 | 0.28 | 1.54 |
| 15 | Slc35d1 | 0.41 | 2.01E-11 | 4.17E-07 | 0.29 | 0.15 | 0.14 | 1.90 |
| 15 | Ephb4 | 0.26 | 2.01E-11 | 4.19E-07 | 0.64 | 0.39 | 0.25 | 1.64 |
| 15 | Kif16b | 0.34 | 2.04E-11 | 4.25E-07 | 0.29 | 0.15 | 0.14 | 1.91 |
| 15 | Six4 | 0.30 | 2.08E-11 | 4.33E-07 | 0.70 | 0.46 | 0.25 | 1.54 |
| 15 | Txlng | 0.38 | 2.09E-11 | 4.34E-07 | 0.38 | 0.22 | 0.17 | 1.77 |
| 15 | Ebf4 | 0.47 | 2.17E-11 | 4.52E-07 | 0.14 | 0.06 | 0.08 | 2.44 |
| 15 | Ubxn7 | 0.26 | 2.18E-11 | 4.53E-07 | 0.73 | 0.47 | 0.26 | 1.56 |
| 15 | Cep104 | 0.27 | 2.19E-11 | 4.56E-07 | 0.38 | 0.21 | 0.17 | 1.83 |
| 15 | Csf2ra | 0.25 | 2.29E-11 | 4.75E-07 | 0.28 | 0.14 | 0.14 | 1.97 |
| 15 | Grb14 | 0.72 | 2.32E-11 | 4.82E-07 | 0.12 | 0.05 | 0.07 | 2.57 |
| 15 | Tom1 | 0.34 | 2.32E-11 | 4.82E-07 | 0.23 | 0.11 | 0.12 | 2.06 |
| 15 | Polr3b | 0.31 | 2.32E-11 | 4.82E-07 | 0.35 | 0.19 | 0.16 | 1.83 |
| 15 | Klhl21 | 0.64 | 2.38E-11 | 4.94E-07 | 0.12 | 0.05 | 0.08 | 2.56 |
| 15 | Eif4g1 | 0.25 | 2.40E-11 | 5.00E-07 | 1.00 | 0.91 | 0.08 | 1.09 |
| 15 | Slc25a16 | 0.35 | 2.41E-11 | 5.02E-07 | 0.30 | 0.16 | 0.14 | 1.89 |
| 15 | Klhl36 | 0.48 | 2.46E-11 | 5.12E-07 | 0.15 | 0.07 | 0.09 | 2.32 |
| 15 | Pygo2 | 0.25 | 2.49E-11 | 5.18E-07 | 0.39 | 0.21 | 0.18 | 1.83 |
| 15 | Atg2a | 0.52 | 2.54E-11 | 5.29E-07 | 0.22 | 0.11 | 0.11 | 2.04 |
| 15 | Cnnm4 | 0.36 | 2.54E-11 | 5.29E-07 | 0.19 | 0.09 | 0.10 | 2.19 |
| 15 | Phf21a | 0.25 | 2.60E-11 | 5.40E-07 | 0.74 | 0.47 | 0.27 | 1.57 |
| 15 | Cit | 0.33 | 2.62E-11 | 5.45E-07 | 0.59 | 0.37 | 0.22 | 1.59 |
| 15 | Prickle3 | 0.61 | 2.65E-11 | 5.52E-07 | 0.15 | 0.07 | 0.09 | 2.31 |
| 15 | Lpar6 | 0.27 | 2.67E-11 | 5.55E-07 | 0.65 | 0.40 | 0.25 | 1.62 |
| 15 | Olfm2 | 0.33 | 2.69E-11 | 5.59E-07 | 0.22 | 0.10 | 0.11 | 2.07 |
| 15 | Aagab | 0.25 | 2.70E-11 | 5.61E-07 | 0.33 | 0.17 | 0.15 | 1.87 |
| 15 | Zbtb43 | 0.26 | 2.71E-11 | 5.65E-07 | 0.41 | 0.23 | 0.18 | 1.78 |
| 15 | Zcchc2 | 0.41 | 2.77E-11 | 5.76E-07 | 0.20 | 0.09 | 0.11 | 2.14 |
| 15 | Noc2l | 0.26 | 2.78E-11 | 5.78E-07 | 0.85 | 0.61 | 0.25 | 1.41 |
| 15 | H2-T22 | 0.35 | 2.78E-11 | 5.79E-07 | 0.21 | 0.10 | 0.11 | 2.12 |
| 15 | Lmcd1 | 0.54 | 2.84E-11 | 5.90E-07 | 0.19 | 0.09 | 0.10 | 2.14 |
| 15 | Pcdh11x | 0.63 | 2.88E-11 | 5.99E-07 | 0.22 | 0.11 | 0.11 | 2.03 |
| 15 | Tnk2 | 0.32 | 2.96E-11 | 6.16E-07 | 0.28 | 0.14 | 0.13 | 1.94 |
| 15 | Pitpnm2 | 0.53 | 3.09E-11 | 6.42E-07 | 0.18 | 0.08 | 0.10 | 2.18 |
| 15 | Smpdl3a | 0.26 | 3.13E-11 | 6.52E-07 | 0.26 | 0.13 | 0.13 | 1.98 |
| 15 | Mast2 | 0.30 | 3.14E-11 | 6.53E-07 | 0.50 | 0.29 | 0.21 | 1.71 |
| 15 | Zscan29 | 0.33 | 3.15E-11 | 6.55E-07 | 0.23 | 0.12 | 0.12 | 2.03 |
| 15 | Tecpr1 | 0.40 | 3.16E-11 | 6.57E-07 | 0.27 | 0.14 | 0.13 | 1.94 |
| 15 | Tulp1 | 0.39 | 3.17E-11 | 6.59E-07 | 0.19 | 0.09 | 0.10 | 2.18 |
| 15 | Fam160b2 | 0.34 | 3.19E-11 | 6.63E-07 | 0.27 | 0.14 | 0.13 | 1.96 |
| 15 | Kat2a | 0.27 | 3.23E-11 | 6.72E-07 | 0.58 | 0.35 | 0.23 | 1.67 |

**Supplementary Table 1. Marker genes for the original clusters of integrated clustering of E11.5 control and NotchGOF λ-junction cells.**

| | | | | | | | | |
|---|---|---|---|---|---|---|---|---|
| 15 | Tox3 | 0.25 | 3.31E-11 | 6.89E-07 | 0.40 | 0.23 | 0.17 | 1.75 |
| 15 | Scamp5 | 0.76 | 3.35E-11 | 6.98E-07 | 0.11 | 0.04 | 0.07 | 2.58 |
| 15 | Ccnd2 | 0.30 | 3.37E-11 | 7.01E-07 | 0.98 | 0.86 | 0.12 | 1.15 |
| 15 | Pacs2 | 0.31 | 3.43E-11 | 7.14E-07 | 0.32 | 0.17 | 0.15 | 1.89 |
| 15 | Ccdc191 | 0.25 | 3.48E-11 | 7.23E-07 | 0.39 | 0.22 | 0.17 | 1.80 |
| 15 | Pomgnt2 | 0.46 | 3.48E-11 | 7.24E-07 | 0.14 | 0.06 | 0.08 | 2.40 |
| 15 | Wdr4 | 0.26 | 3.48E-11 | 7.24E-07 | 0.46 | 0.26 | 0.20 | 1.76 |
| 15 | Suclg2 | 0.25 | 3.49E-11 | 7.26E-07 | 0.77 | 0.49 | 0.28 | 1.57 |
| 15 | Fbxl14 | 0.32 | 3.50E-11 | 7.28E-07 | 0.61 | 0.38 | 0.23 | 1.61 |
| 15 | Rmdn1 | 0.25 | 3.50E-11 | 7.28E-07 | 0.32 | 0.17 | 0.15 | 1.87 |
| 15 | Arhgef10 | 0.47 | 3.59E-11 | 7.46E-07 | 0.26 | 0.13 | 0.13 | 1.96 |
| 15 | Cd3eap | 0.26 | 3.59E-11 | 7.47E-07 | 0.78 | 0.52 | 0.26 | 1.50 |
| 15 | Tbl1x | 0.26 | 3.60E-11 | 7.48E-07 | 0.96 | 0.79 | 0.16 | 1.21 |
| 15 | Numbl | 0.26 | 3.62E-11 | 7.52E-07 | 0.26 | 0.13 | 0.13 | 1.98 |
| 15 | Smox | 0.30 | 3.64E-11 | 7.57E-07 | 0.42 | 0.24 | 0.18 | 1.75 |
| 15 | Hmgxb3 | 0.31 | 3.69E-11 | 7.67E-07 | 0.37 | 0.20 | 0.16 | 1.80 |
| 15 | Fam53b | 0.35 | 3.72E-11 | 7.74E-07 | 0.32 | 0.18 | 0.15 | 1.84 |
| 15 | Tmub1 | 0.37 | 3.75E-11 | 7.80E-07 | 0.35 | 0.20 | 0.16 | 1.80 |
| 15 | Amfr | 0.26 | 3.78E-11 | 7.86E-07 | 0.60 | 0.36 | 0.24 | 1.67 |
| 15 | Cldn10 | 0.64 | 3.83E-11 | 7.97E-07 | 0.12 | 0.05 | 0.07 | 2.51 |
| 15 | Gabre | 0.58 | 3.87E-11 | 8.04E-07 | 0.11 | 0.04 | 0.07 | 2.63 |
| 15 | Antxr1 | 0.29 | 3.90E-11 | 8.10E-07 | 0.52 | 0.31 | 0.21 | 1.68 |
| 15 | C030006K11Rik | 0.46 | 3.91E-11 | 8.14E-07 | 0.19 | 0.09 | 0.10 | 2.16 |
| 15 | Fbxo30 | 0.25 | 3.93E-11 | 8.17E-07 | 0.66 | 0.41 | 0.25 | 1.62 |
| 15 | Dlg4 | 0.29 | 4.08E-11 | 8.49E-07 | 0.52 | 0.31 | 0.21 | 1.68 |
| 15 | Gcn1 | 0.26 | 4.09E-11 | 8.51E-07 | 0.78 | 0.51 | 0.27 | 1.54 |
| 15 | Nsun4 | 0.25 | 4.10E-11 | 8.52E-07 | 0.30 | 0.16 | 0.14 | 1.91 |
| 15 | Sgpl1 | 0.28 | 4.14E-11 | 8.62E-07 | 0.56 | 0.34 | 0.22 | 1.66 |
| 15 | Etv3 | 0.26 | 4.25E-11 | 8.83E-07 | 0.39 | 0.22 | 0.17 | 1.77 |
| 15 | B930095G15Rik | 0.26 | 4.25E-11 | 8.84E-07 | 0.33 | 0.18 | 0.16 | 1.88 |
| 15 | Slc25a13 | 0.26 | 4.31E-11 | 8.97E-07 | 0.70 | 0.44 | 0.26 | 1.58 |
| 15 | Dbndd2 | 0.57 | 4.42E-11 | 9.19E-07 | 0.18 | 0.08 | 0.10 | 2.17 |
| 15 | Ubqln4 | 0.27 | 4.49E-11 | 9.35E-07 | 0.60 | 0.37 | 0.24 | 1.64 |
| 15 | Rel | 0.39 | 4.59E-11 | 9.54E-07 | 0.19 | 0.09 | 0.10 | 2.13 |
| 15 | Tlcd1 | 0.45 | 4.59E-11 | 9.55E-07 | 0.19 | 0.09 | 0.10 | 2.14 |
| 15 | Sin3a | 0.27 | 4.63E-11 | 9.63E-07 | 0.77 | 0.52 | 0.25 | 1.49 |
| 15 | Ect2 | 0.29 | 4.75E-11 | 9.87E-07 | 0.63 | 0.41 | 0.22 | 1.54 |
| 15 | Pan3 | 0.27 | 4.77E-11 | 9.93E-07 | 0.54 | 0.33 | 0.22 | 1.66 |
| 15 | Syngr2 | 0.30 | 4.95E-11 | 1.03E-06 | 0.27 | 0.14 | 0.13 | 1.92 |
| 15 | Trmt1l | 0.28 | 4.99E-11 | 1.04E-06 | 0.45 | 0.26 | 0.19 | 1.72 |
| 15 | Ass1 | 0.65 | 5.03E-11 | 1.05E-06 | 0.14 | 0.06 | 0.08 | 2.36 |
| 15 | Trmt61a | 0.45 | 5.04E-11 | 1.05E-06 | 0.28 | 0.15 | 0.13 | 1.88 |
| 15 | Reps1 | 0.33 | 5.07E-11 | 1.05E-06 | 0.57 | 0.35 | 0.22 | 1.64 |
| 15 | Itsn2 | 0.31 | 5.08E-11 | 1.06E-06 | 0.34 | 0.19 | 0.15 | 1.81 |
| 15 | Mlh1 | 0.32 | 5.18E-11 | 1.08E-06 | 0.37 | 0.20 | 0.16 | 1.81 |
| 15 | Ncoa1 | 0.27 | 5.33E-11 | 1.11E-06 | 0.32 | 0.17 | 0.15 | 1.85 |
| 15 | Shmt2 | 0.33 | 5.41E-11 | 1.13E-06 | 0.56 | 0.34 | 0.22 | 1.65 |
| 15 | Gramd3 | 0.30 | 5.43E-11 | 1.13E-06 | 0.33 | 0.18 | 0.15 | 1.85 |
| 15 | Nhlrc3 | 0.44 | 5.45E-11 | 1.13E-06 | 0.24 | 0.12 | 0.12 | 1.98 |
| 15 | Smyd5 | 0.27 | 5.55E-11 | 1.15E-06 | 0.54 | 0.32 | 0.22 | 1.66 |
| 15 | Ralgps1 | 0.68 | 5.58E-11 | 1.16E-06 | 0.11 | 0.04 | 0.07 | 2.58 |
| 15 | Lpar2 | 0.57 | 5.78E-11 | 1.20E-06 | 0.17 | 0.08 | 0.09 | 2.23 |
| 15 | Pcnx | 0.29 | 5.78E-11 | 1.20E-06 | 0.48 | 0.28 | 0.20 | 1.71 |
| 15 | Tnrc18 | 0.27 | 5.89E-11 | 1.23E-06 | 0.80 | 0.55 | 0.24 | 1.44 |
| 15 | Ralgapa2 | 0.35 | 5.94E-11 | 1.24E-06 | 0.26 | 0.14 | 0.13 | 1.94 |
| 15 | Fam3c | 0.32 | 6.00E-11 | 1.25E-06 | 0.58 | 0.36 | 0.22 | 1.61 |

**Supplementary Table 1. Marker genes for the original clusters of integrated clustering of E11.5 control and NotchGOF λ-junction cells.**

| | | | | | | | | |
|---|---|---|---|---|---|---|---|---|
| 15 | Jmjd7 | 0.27 | 6.11E-11 | 1.27E-06 | 0.32 | 0.17 | 0.15 | 1.86 |
| 15 | Ldlrap1 | 0.34 | 6.17E-11 | 1.28E-06 | 0.33 | 0.18 | 0.15 | 1.81 |
| 15 | Fasn | 0.28 | 6.30E-11 | 1.31E-06 | 0.76 | 0.50 | 0.26 | 1.52 |
| 15 | Clec16a | 0.47 | 6.45E-11 | 1.34E-06 | 0.19 | 0.09 | 0.10 | 2.11 |
| 15 | Adck1 | 0.32 | 6.46E-11 | 1.34E-06 | 0.30 | 0.16 | 0.14 | 1.88 |
| 15 | Mcoln3 | 0.53 | 6.47E-11 | 1.35E-06 | 0.13 | 0.06 | 0.08 | 2.40 |
| 15 | Rnf13 | 0.28 | 6.52E-11 | 1.36E-06 | 0.42 | 0.24 | 0.18 | 1.75 |
| 15 | Inpp5e | 0.32 | 6.54E-11 | 1.36E-06 | 0.28 | 0.14 | 0.13 | 1.93 |
| 15 | Rfx7 | 0.28 | 6.66E-11 | 1.39E-06 | 0.71 | 0.46 | 0.25 | 1.54 |
| 15 | Mief1 | 0.38 | 6.68E-11 | 1.39E-06 | 0.30 | 0.16 | 0.14 | 1.86 |
| 15 | Nabp2 | 0.28 | 6.68E-11 | 1.39E-06 | 0.88 | 0.65 | 0.22 | 1.34 |
| 15 | Tmem38a | 0.35 | 6.88E-11 | 1.43E-06 | 0.21 | 0.10 | 0.11 | 2.08 |
| 15 | Rbbp9 | 0.28 | 7.17E-11 | 1.49E-06 | 0.63 | 0.39 | 0.24 | 1.63 |
| 15 | Rasgef1b | 0.55 | 7.20E-11 | 1.50E-06 | 0.12 | 0.05 | 0.07 | 2.51 |
| 15 | Gm28791 | 0.30 | 7.34E-11 | 1.53E-06 | 0.26 | 0.13 | 0.13 | 1.98 |
| 15 | Mga | 0.26 | 7.37E-11 | 1.53E-06 | 0.84 | 0.57 | 0.26 | 1.46 |
| 15 | Cc2d1a | 0.43 | 7.43E-11 | 1.55E-06 | 0.20 | 0.09 | 0.10 | 2.10 |
| 15 | Zfp940 | 0.32 | 7.45E-11 | 1.55E-06 | 0.24 | 0.12 | 0.12 | 2.01 |
| 15 | Noct | 0.25 | 7.45E-11 | 1.55E-06 | 0.34 | 0.19 | 0.16 | 1.83 |
| 15 | Fndc3b | 0.30 | 7.61E-11 | 1.58E-06 | 0.42 | 0.24 | 0.18 | 1.74 |
| 15 | Hic2 | 0.28 | 7.69E-11 | 1.60E-06 | 0.50 | 0.29 | 0.21 | 1.71 |
| 15 | Gm1976 | 0.30 | 7.77E-11 | 1.62E-06 | 0.23 | 0.11 | 0.12 | 2.03 |
| 15 | Tob2 | 0.25 | 7.82E-11 | 1.63E-06 | 0.42 | 0.24 | 0.18 | 1.77 |
| 15 | Klhl42 | 0.40 | 7.91E-11 | 1.64E-06 | 0.21 | 0.10 | 0.11 | 2.08 |
| 15 | Galnt2 | 0.27 | 8.02E-11 | 1.67E-06 | 0.72 | 0.46 | 0.26 | 1.57 |
| 15 | Phc3 | 0.28 | 8.08E-11 | 1.68E-06 | 0.36 | 0.20 | 0.16 | 1.79 |
| 15 | Scaf1 | 0.28 | 8.26E-11 | 1.72E-06 | 0.65 | 0.41 | 0.25 | 1.60 |
| 15 | Bicral | 0.40 | 8.35E-11 | 1.74E-06 | 0.21 | 0.10 | 0.11 | 2.06 |
| 15 | Epdr1 | 0.28 | 8.41E-11 | 1.75E-06 | 0.22 | 0.10 | 0.11 | 2.07 |
| 15 | Esyt2 | 0.27 | 8.59E-11 | 1.79E-06 | 0.38 | 0.21 | 0.17 | 1.79 |
| 15 | Iffo2 | 0.35 | 8.60E-11 | 1.79E-06 | 0.26 | 0.13 | 0.13 | 1.95 |
| 15 | Zdhhc20 | 0.26 | 8.64E-11 | 1.80E-06 | 0.85 | 0.58 | 0.27 | 1.47 |
| 15 | Zc3h10 | 0.40 | 8.69E-11 | 1.81E-06 | 0.26 | 0.14 | 0.13 | 1.93 |
| 15 | Zfp617 | 0.27 | 8.94E-11 | 1.86E-06 | 0.51 | 0.30 | 0.21 | 1.69 |
| 15 | Lin54 | 0.28 | 8.94E-11 | 1.86E-06 | 0.41 | 0.23 | 0.18 | 1.76 |
| 15 | Phf21b | 0.84 | 9.18E-11 | 1.91E-06 | 0.11 | 0.04 | 0.07 | 2.59 |
| 15 | Slc48a1 | 0.31 | 9.30E-11 | 1.94E-06 | 0.31 | 0.17 | 0.14 | 1.84 |
| 15 | Stk39 | 0.30 | 9.32E-11 | 1.94E-06 | 0.44 | 0.26 | 0.19 | 1.71 |
| 15 | Pnpla2 | 0.45 | 9.36E-11 | 1.95E-06 | 0.19 | 0.09 | 0.10 | 2.10 |
| 15 | Arhgef12 | 0.27 | 9.46E-11 | 1.97E-06 | 0.81 | 0.58 | 0.23 | 1.40 |
| 15 | Slc35e3 | 0.32 | 9.78E-11 | 2.03E-06 | 0.26 | 0.13 | 0.13 | 1.96 |
| 15 | Clip4 | 0.64 | 1.01E-10 | 2.09E-06 | 0.11 | 0.05 | 0.07 | 2.51 |
| 15 | Foxo1 | 0.28 | 1.01E-10 | 2.10E-06 | 0.33 | 0.18 | 0.15 | 1.83 |
| 15 | Sh3rf1 | 0.27 | 1.01E-10 | 2.11E-06 | 0.40 | 0.23 | 0.17 | 1.77 |
| 15 | Prep | 0.26 | 1.01E-10 | 2.11E-06 | 0.63 | 0.39 | 0.24 | 1.62 |
| 15 | Kmt2c | 0.25 | 1.02E-10 | 2.12E-06 | 0.74 | 0.49 | 0.25 | 1.51 |
| 15 | Adgrl2 | 0.27 | 1.04E-10 | 2.16E-06 | 0.91 | 0.69 | 0.23 | 1.33 |
| 15 | Qtrt2 | 0.32 | 1.07E-10 | 2.23E-06 | 0.43 | 0.25 | 0.18 | 1.73 |
| 15 | Cdkl5 | 0.71 | 1.07E-10 | 2.23E-06 | 0.13 | 0.06 | 0.08 | 2.36 |
| 15 | Fam185a | 0.51 | 1.09E-10 | 2.27E-06 | 0.13 | 0.06 | 0.08 | 2.39 |
| 15 | Smyd4 | 0.31 | 1.10E-10 | 2.30E-06 | 0.32 | 0.17 | 0.15 | 1.85 |
| 15 | Zfp426 | 0.25 | 1.11E-10 | 2.31E-06 | 0.38 | 0.21 | 0.17 | 1.79 |
| 15 | Emc1 | 0.26 | 1.12E-10 | 2.33E-06 | 0.32 | 0.17 | 0.15 | 1.85 |
| 15 | Zfp26 | 0.27 | 1.12E-10 | 2.33E-06 | 0.58 | 0.36 | 0.23 | 1.64 |
| 15 | 2810021J22Rik | 0.60 | 1.14E-10 | 2.38E-06 | 0.17 | 0.08 | 0.09 | 2.18 |
| 15 | Cited2 | 0.25 | 1.15E-10 | 2.38E-06 | 0.88 | 0.64 | 0.25 | 1.39 |

**Supplementary Table 1. Marker genes for the original clusters of integrated clustering of E11.5 control and NotchGOF λ-junction cells.**

| | | | | | | | | |
|---|---|---|---|---|---|---|---|---|
| 15 | Cad | 0.32 | 1.16E-10 | 2.42E-06 | 0.61 | 0.38 | 0.23 | 1.60 |
| 15 | Zbtb39 | 0.27 | 1.18E-10 | 2.45E-06 | 0.26 | 0.13 | 0.13 | 1.96 |
| 15 | Sema4d | 0.64 | 1.18E-10 | 2.46E-06 | 0.16 | 0.07 | 0.09 | 2.21 |
| 15 | Trmt12 | 0.28 | 1.19E-10 | 2.48E-06 | 0.35 | 0.19 | 0.15 | 1.81 |
| 15 | Sdk1 | 0.44 | 1.21E-10 | 2.51E-06 | 0.18 | 0.08 | 0.09 | 2.15 |
| 15 | Eif4ebp2 | 0.26 | 1.23E-10 | 2.55E-06 | 0.90 | 0.70 | 0.21 | 1.30 |
| 15 | Tent5a | 0.84 | 1.26E-10 | 2.61E-06 | 0.10 | 0.04 | 0.06 | 2.62 |
| 15 | Sipa1l1 | 0.28 | 1.26E-10 | 2.61E-06 | 0.77 | 0.51 | 0.26 | 1.51 |
| 15 | Zfp101 | 0.27 | 1.27E-10 | 2.64E-06 | 0.59 | 0.36 | 0.23 | 1.64 |
| 15 | Ldb1 | 0.29 | 1.33E-10 | 2.77E-06 | 0.86 | 0.64 | 0.22 | 1.35 |
| 15 | Slc15a2 | 0.60 | 1.34E-10 | 2.79E-06 | 0.12 | 0.05 | 0.07 | 2.46 |
| 15 | Smad6 | 0.29 | 1.34E-10 | 2.79E-06 | 0.43 | 0.25 | 0.18 | 1.73 |
| 15 | Nanos3 | 0.56 | 1.35E-10 | 2.81E-06 | 0.13 | 0.05 | 0.08 | 2.41 |
| 15 | Hsf1 | 0.25 | 1.36E-10 | 2.83E-06 | 0.57 | 0.34 | 0.23 | 1.67 |
| 15 | Ubn2 | 0.26 | 1.38E-10 | 2.88E-06 | 0.93 | 0.70 | 0.23 | 1.34 |
| 15 | Tcf7l1 | 0.28 | 1.40E-10 | 2.91E-06 | 0.83 | 0.58 | 0.25 | 1.43 |
| 15 | Atp9b | 0.28 | 1.40E-10 | 2.92E-06 | 0.34 | 0.19 | 0.15 | 1.82 |
| 15 | Zmat3 | 0.27 | 1.41E-10 | 2.92E-06 | 0.35 | 0.19 | 0.16 | 1.82 |
| 15 | Herc3 | 0.39 | 1.42E-10 | 2.94E-06 | 0.16 | 0.07 | 0.09 | 2.22 |
| 15 | Senp3 | 0.29 | 1.46E-10 | 3.03E-06 | 0.71 | 0.47 | 0.24 | 1.51 |
| 15 | Zmynd8 | 0.27 | 1.46E-10 | 3.04E-06 | 0.85 | 0.60 | 0.25 | 1.42 |
| 15 | Ppp6r2 | 0.30 | 1.48E-10 | 3.08E-06 | 0.29 | 0.15 | 0.13 | 1.88 |
| 15 | Tagap1 | 0.30 | 1.48E-10 | 3.08E-06 | 0.27 | 0.14 | 0.13 | 1.91 |
| 15 | 2810402E24Rik | 0.27 | 1.51E-10 | 3.14E-06 | 0.38 | 0.21 | 0.16 | 1.76 |
| 15 | Rgs12 | 0.29 | 1.52E-10 | 3.16E-06 | 0.69 | 0.45 | 0.25 | 1.56 |
| 15 | Ddx10 | 0.31 | 1.58E-10 | 3.28E-06 | 0.73 | 0.48 | 0.25 | 1.52 |
| 15 | Slc35f6 | 0.41 | 1.58E-10 | 3.28E-06 | 0.22 | 0.11 | 0.11 | 2.05 |
| 15 | Nudt16 | 0.46 | 1.59E-10 | 3.31E-06 | 0.18 | 0.09 | 0.10 | 2.13 |
| 15 | Gm43813 | 0.55 | 1.63E-10 | 3.38E-06 | 0.18 | 0.09 | 0.10 | 2.10 |
| 15 | Lmbrd2 | 0.34 | 1.64E-10 | 3.41E-06 | 0.33 | 0.18 | 0.15 | 1.82 |
| 15 | Zfp994 | 0.37 | 1.65E-10 | 3.43E-06 | 0.19 | 0.09 | 0.10 | 2.09 |
| 15 | Azin2 | 0.37 | 1.65E-10 | 3.43E-06 | 0.22 | 0.11 | 0.11 | 2.02 |
| 15 | Pwp2 | 0.35 | 1.65E-10 | 3.43E-06 | 0.29 | 0.15 | 0.14 | 1.90 |
| 15 | Setd6 | 0.39 | 1.66E-10 | 3.46E-06 | 0.24 | 0.13 | 0.12 | 1.94 |
| 15 | Hectd2 | 0.74 | 1.67E-10 | 3.47E-06 | 0.12 | 0.05 | 0.07 | 2.45 |
| 15 | Rasa1 | 0.29 | 1.68E-10 | 3.49E-06 | 0.85 | 0.61 | 0.24 | 1.40 |
| 15 | Cdc25c | 0.34 | 1.68E-10 | 3.50E-06 | 0.55 | 0.35 | 0.20 | 1.58 |
| 15 | Hlcs | 0.40 | 1.69E-10 | 3.52E-06 | 0.25 | 0.13 | 0.12 | 1.94 |
| 15 | Grk3 | 0.47 | 1.70E-10 | 3.53E-06 | 0.23 | 0.11 | 0.11 | 1.99 |
| 15 | Eps8l1 | 0.76 | 1.71E-10 | 3.55E-06 | 0.10 | 0.04 | 0.06 | 2.62 |
| 15 | Megf8 | 0.38 | 1.71E-10 | 3.55E-06 | 0.30 | 0.16 | 0.14 | 1.84 |
| 15 | Sec31a | 0.26 | 1.72E-10 | 3.57E-06 | 0.81 | 0.55 | 0.26 | 1.48 |
| 15 | Zgrf1 | 0.29 | 1.75E-10 | 3.64E-06 | 0.36 | 0.20 | 0.16 | 1.80 |
| 15 | Zfp605 | 0.38 | 1.76E-10 | 3.67E-06 | 0.23 | 0.12 | 0.11 | 1.99 |
| 15 | Trmt9b | 0.48 | 1.79E-10 | 3.71E-06 | 0.15 | 0.07 | 0.09 | 2.25 |
| 15 | Atf7ip | 0.25 | 1.79E-10 | 3.73E-06 | 0.87 | 0.62 | 0.25 | 1.40 |
| 15 | Atf7 | 0.29 | 1.84E-10 | 3.82E-06 | 0.35 | 0.20 | 0.16 | 1.80 |
| 15 | Fbxw7 | 0.27 | 1.86E-10 | 3.87E-06 | 0.61 | 0.38 | 0.23 | 1.61 |
| 15 | Adam11 | 0.42 | 1.87E-10 | 3.88E-06 | 0.18 | 0.08 | 0.10 | 2.14 |
| 15 | Ttc7b | 0.27 | 1.92E-10 | 4.00E-06 | 0.30 | 0.16 | 0.14 | 1.89 |
| 15 | Pcdhga12 | 0.31 | 1.93E-10 | 4.01E-06 | 0.24 | 0.12 | 0.12 | 1.96 |
| 15 | Inpp5k | 0.39 | 1.94E-10 | 4.03E-06 | 0.22 | 0.11 | 0.11 | 1.98 |
| 15 | Ankrd52 | 0.40 | 1.94E-10 | 4.03E-06 | 0.22 | 0.11 | 0.11 | 2.00 |
| 15 | Rad9b | 0.43 | 1.97E-10 | 4.09E-06 | 0.18 | 0.08 | 0.09 | 2.15 |
| 15 | Skida1 | 0.31 | 1.97E-10 | 4.10E-06 | 0.29 | 0.15 | 0.13 | 1.88 |
| 15 | Rnf103 | 0.33 | 2.00E-10 | 4.16E-06 | 0.32 | 0.18 | 0.14 | 1.81 |

**Supplementary Table 1. Marker genes for the original clusters of integrated clustering of E11.5 control and NotchGOF λ-junction cells.**

| | | | | | | | | |
|---|---|---|---|---|---|---|---|---|
| 15 | Mon1b | 0.54 | 2.00E-10 | 4.16E-06 | 0.14 | 0.06 | 0.08 | 2.31 |
| 15 | Trmt2b | 0.39 | 2.03E-10 | 4.22E-06 | 0.19 | 0.09 | 0.10 | 2.10 |
| 15 | Megf9 | 0.30 | 2.06E-10 | 4.29E-06 | 0.34 | 0.19 | 0.15 | 1.81 |
| 15 | Sema4g | 0.27 | 2.09E-10 | 4.35E-06 | 0.27 | 0.14 | 0.13 | 1.91 |
| 15 | Sfxn3 | 0.58 | 2.10E-10 | 4.37E-06 | 0.14 | 0.06 | 0.08 | 2.29 |
| 15 | Wrb | 0.26 | 2.12E-10 | 4.42E-06 | 0.41 | 0.24 | 0.17 | 1.73 |
| 15 | Lrrc56 | 0.41 | 2.12E-10 | 4.42E-06 | 0.20 | 0.10 | 0.10 | 2.05 |
| 15 | Fbxo38 | 0.25 | 2.16E-10 | 4.49E-06 | 0.41 | 0.24 | 0.18 | 1.74 |
| 15 | Wdfy3 | 0.27 | 2.17E-10 | 4.51E-06 | 0.58 | 0.36 | 0.22 | 1.62 |
| 15 | Fgfbp3 | 0.33 | 2.20E-10 | 4.57E-06 | 0.20 | 0.10 | 0.11 | 2.06 |
| 15 | Faap24 | 0.32 | 2.22E-10 | 4.62E-06 | 0.30 | 0.16 | 0.14 | 1.86 |
| 15 | Sbsn | 0.29 | 2.24E-10 | 4.65E-06 | 0.26 | 0.14 | 0.13 | 1.91 |
| 15 | Deaf1 | 0.26 | 2.25E-10 | 4.68E-06 | 0.39 | 0.22 | 0.17 | 1.78 |
| 15 | Dcaf7 | 0.28 | 2.27E-10 | 4.72E-06 | 0.73 | 0.48 | 0.25 | 1.51 |
| 15 | Mettl26 | 0.26 | 2.38E-10 | 4.95E-06 | 0.73 | 0.49 | 0.24 | 1.49 |
| 15 | Mtmr10 | 0.38 | 2.39E-10 | 4.96E-06 | 0.16 | 0.08 | 0.09 | 2.19 |
| 15 | Tmem47 | 0.29 | 2.45E-10 | 5.09E-06 | 0.57 | 0.36 | 0.21 | 1.58 |
| 15 | Kdm4c | 0.52 | 2.47E-10 | 5.14E-06 | 0.23 | 0.11 | 0.11 | 1.99 |
| 15 | Fst | 0.54 | 2.51E-10 | 5.22E-06 | 0.23 | 0.12 | 0.11 | 1.91 |
| 15 | Abcc1 | 0.34 | 2.57E-10 | 5.35E-06 | 0.26 | 0.14 | 0.12 | 1.90 |
| 15 | Zfp384 | 0.30 | 2.59E-10 | 5.38E-06 | 0.29 | 0.16 | 0.14 | 1.88 |
| 15 | Eif4ebp3 | 0.31 | 2.59E-10 | 5.39E-06 | 0.34 | 0.19 | 0.15 | 1.80 |
| 15 | Ttc13 | 0.27 | 2.61E-10 | 5.42E-06 | 0.40 | 0.23 | 0.17 | 1.75 |
| 15 | Iqgap3 | 0.32 | 2.64E-10 | 5.49E-06 | 0.38 | 0.22 | 0.16 | 1.75 |
| 15 | Arhgap39 | 0.33 | 2.66E-10 | 5.52E-06 | 0.32 | 0.18 | 0.14 | 1.81 |
| 15 | Fam98c | 0.34 | 2.74E-10 | 5.71E-06 | 0.23 | 0.12 | 0.11 | 1.97 |
| 15 | Fbxo44 | 0.29 | 2.74E-10 | 5.71E-06 | 0.19 | 0.09 | 0.10 | 2.09 |
| 15 | Zfp868 | 0.44 | 2.76E-10 | 5.74E-06 | 0.27 | 0.14 | 0.13 | 1.88 |
| 15 | Trim8 | 0.27 | 2.79E-10 | 5.79E-06 | 0.49 | 0.30 | 0.19 | 1.65 |
| 15 | Pygl | 0.39 | 2.83E-10 | 5.89E-06 | 0.20 | 0.10 | 0.10 | 2.03 |
| 15 | Gm47320 | 0.58 | 2.86E-10 | 5.94E-06 | 0.13 | 0.06 | 0.08 | 2.36 |
| 15 | Tbc1d9 | 0.71 | 2.86E-10 | 5.95E-06 | 0.16 | 0.08 | 0.09 | 2.16 |
| 15 | Mrm1 | 0.72 | 2.86E-10 | 5.96E-06 | 0.11 | 0.04 | 0.06 | 2.52 |
| 15 | Gmeb2 | 0.27 | 2.88E-10 | 5.99E-06 | 0.27 | 0.14 | 0.13 | 1.90 |
| 15 | Fam126b | 0.29 | 2.98E-10 | 6.19E-06 | 0.34 | 0.19 | 0.15 | 1.80 |
| 15 | Cd82 | 0.33 | 3.04E-10 | 6.33E-06 | 0.15 | 0.07 | 0.08 | 2.28 |
| 15 | Tctn2 | 0.41 | 3.15E-10 | 6.56E-06 | 0.19 | 0.09 | 0.10 | 2.09 |
| 15 | Dnajc28 | 0.51 | 3.29E-10 | 6.84E-06 | 0.12 | 0.05 | 0.07 | 2.47 |
| 15 | Tmem51 | 0.26 | 3.32E-10 | 6.90E-06 | 0.20 | 0.10 | 0.10 | 2.05 |
| 15 | Osbpl2 | 0.31 | 3.32E-10 | 6.90E-06 | 0.29 | 0.15 | 0.13 | 1.86 |
| 15 | Ncoa3 | 0.30 | 3.39E-10 | 7.05E-06 | 0.39 | 0.23 | 0.16 | 1.73 |
| 15 | Kdm2a | 0.28 | 3.51E-10 | 7.29E-06 | 0.87 | 0.62 | 0.25 | 1.40 |
| 15 | Jmjd1c | 0.27 | 3.54E-10 | 7.36E-06 | 0.74 | 0.48 | 0.26 | 1.54 |
| 15 | Arhgef17 | 0.28 | 3.59E-10 | 7.47E-06 | 0.59 | 0.36 | 0.23 | 1.64 |
| 15 | Rad51d | 0.30 | 3.63E-10 | 7.54E-06 | 0.26 | 0.13 | 0.12 | 1.93 |
| 15 | Zfp319 | 0.36 | 3.63E-10 | 7.56E-06 | 0.29 | 0.16 | 0.13 | 1.84 |
| 15 | Mrpl37 | 0.25 | 3.66E-10 | 7.61E-06 | 0.63 | 0.39 | 0.23 | 1.59 |
| 15 | Fam168a | 0.28 | 3.70E-10 | 7.70E-06 | 0.80 | 0.54 | 0.26 | 1.48 |
| 15 | Kbtbd7 | 0.34 | 3.74E-10 | 7.78E-06 | 0.21 | 0.10 | 0.11 | 2.02 |
| 15 | Fut10 | 0.33 | 3.81E-10 | 7.93E-06 | 0.25 | 0.13 | 0.12 | 1.92 |
| 15 | Lpin1 | 0.60 | 3.85E-10 | 8.00E-06 | 0.15 | 0.07 | 0.08 | 2.25 |
| 15 | Gm10138 | 0.26 | 3.91E-10 | 8.14E-06 | 0.36 | 0.20 | 0.16 | 1.77 |
| 15 | Tceal5 | 0.31 | 3.93E-10 | 8.17E-06 | 0.28 | 0.15 | 0.13 | 1.87 |
| 15 | Spry4 | 0.35 | 3.98E-10 | 8.28E-06 | 0.27 | 0.14 | 0.13 | 1.87 |
| 15 | Krt15 | 0.60 | 4.01E-10 | 8.35E-06 | 0.10 | 0.04 | 0.06 | 2.55 |
| 15 | Aldh6a1 | 0.49 | 4.02E-10 | 8.37E-06 | 0.22 | 0.11 | 0.11 | 1.97 |

**Supplementary Table 1. Marker genes for the original clusters of integrated clustering of E11.5 control and NotchGOF λ-junction cells.**

| | | | | | | | | |
|---|---|---|---|---|---|---|---|---|
| 15 | Kcnb1 | 0.45 | 4.03E-10 | 8.39E-06 | 0.14 | 0.06 | 0.08 | 2.28 |
| 15 | Tekt1 | 0.34 | 4.17E-10 | 8.67E-06 | 0.19 | 0.09 | 0.10 | 2.09 |
| 15 | Gas7 | 0.64 | 4.43E-10 | 9.22E-06 | 0.12 | 0.05 | 0.07 | 2.37 |
| 15 | Rnf19b | 0.29 | 4.44E-10 | 9.24E-06 | 0.26 | 0.13 | 0.12 | 1.92 |
| 15 | Thtpa | 0.39 | 4.52E-10 | 9.40E-06 | 0.29 | 0.16 | 0.13 | 1.83 |
| 15 | Bmp2 | 0.52 | 4.62E-10 | 9.61E-06 | 0.15 | 0.07 | 0.08 | 2.25 |
| 15 | Hps1 | 0.37 | 4.71E-10 | 9.80E-06 | 0.24 | 0.13 | 0.12 | 1.91 |
| 15 | Clip1 | 0.27 | 4.73E-10 | 9.85E-06 | 0.73 | 0.49 | 0.23 | 1.47 |
| 15 | Rfx1 | 0.39 | 4.77E-10 | 9.91E-06 | 0.23 | 0.12 | 0.11 | 1.93 |
| 15 | Cables2 | 0.27 | 4.86E-10 | 1.01E-05 | 0.33 | 0.18 | 0.15 | 1.81 |
| 15 | Tbl2 | 0.34 | 4.89E-10 | 1.02E-05 | 0.23 | 0.12 | 0.11 | 1.97 |
| 15 | Tmem181a | 0.38 | 4.91E-10 | 1.02E-05 | 0.20 | 0.10 | 0.10 | 2.02 |
| 15 | Zkscan17 | 0.25 | 4.91E-10 | 1.02E-05 | 0.43 | 0.25 | 0.18 | 1.71 |
| 15 | Mrs2 | 0.29 | 5.04E-10 | 1.05E-05 | 0.37 | 0.21 | 0.16 | 1.75 |
| 15 | Prdm10 | 0.31 | 5.05E-10 | 1.05E-05 | 0.22 | 0.11 | 0.11 | 1.98 |
| 15 | BC052040 | 0.26 | 5.14E-10 | 1.07E-05 | 0.32 | 0.17 | 0.14 | 1.82 |
| 15 | Rtel1 | 0.39 | 5.17E-10 | 1.07E-05 | 0.32 | 0.18 | 0.14 | 1.77 |
| 15 | Acad11 | 0.33 | 5.26E-10 | 1.09E-05 | 0.18 | 0.09 | 0.10 | 2.10 |
| 15 | 2900060N12Rik | 0.44 | 5.35E-10 | 1.11E-05 | 0.15 | 0.07 | 0.08 | 2.24 |
| 15 | Man2c1 | 0.50 | 5.52E-10 | 1.15E-05 | 0.21 | 0.11 | 0.10 | 1.98 |
| 15 | Ccdc181 | 0.52 | 5.62E-10 | 1.17E-05 | 0.14 | 0.06 | 0.08 | 2.25 |
| 15 | Zscan18 | 0.30 | 5.78E-10 | 1.20E-05 | 0.23 | 0.11 | 0.11 | 1.97 |
| 15 | Ganab | 0.27 | 5.83E-10 | 1.21E-05 | 0.79 | 0.53 | 0.26 | 1.48 |
| 15 | Arhgap23 | 0.49 | 5.86E-10 | 1.22E-05 | 0.19 | 0.09 | 0.10 | 2.02 |
| 15 | Ydjc | 0.47 | 5.88E-10 | 1.22E-05 | 0.17 | 0.08 | 0.09 | 2.10 |
| 15 | Dennd4c | 0.33 | 6.02E-10 | 1.25E-05 | 0.30 | 0.17 | 0.13 | 1.81 |
| 15 | Pnpla6 | 0.42 | 6.08E-10 | 1.26E-05 | 0.18 | 0.09 | 0.09 | 2.08 |
| 15 | Zfp142 | 0.29 | 6.18E-10 | 1.29E-05 | 0.29 | 0.16 | 0.14 | 1.85 |
| 15 | Sgpp1 | 0.26 | 6.21E-10 | 1.29E-05 | 0.21 | 0.10 | 0.10 | 2.02 |
| 15 | 1700096K18Rik | 0.50 | 6.36E-10 | 1.32E-05 | 0.19 | 0.09 | 0.10 | 2.02 |
| 15 | Wdr59 | 0.39 | 6.40E-10 | 1.33E-05 | 0.19 | 0.09 | 0.10 | 2.08 |
| 15 | Dyrk3 | 0.57 | 6.42E-10 | 1.34E-05 | 0.20 | 0.10 | 0.10 | 1.99 |
| 15 | Zfp598 | 0.25 | 6.48E-10 | 1.35E-05 | 0.49 | 0.30 | 0.20 | 1.65 |
| 15 | Xpo6 | 0.26 | 6.49E-10 | 1.35E-05 | 0.59 | 0.37 | 0.22 | 1.61 |
| 15 | Tab2 | 0.27 | 6.81E-10 | 1.42E-05 | 0.88 | 0.67 | 0.20 | 1.30 |
| 15 | Slco5a1 | 0.47 | 7.08E-10 | 1.47E-05 | 0.17 | 0.08 | 0.09 | 2.09 |
| 15 | Yipf2 | 0.28 | 7.18E-10 | 1.49E-05 | 0.23 | 0.12 | 0.11 | 1.93 |
| 15 | Zfp592 | 0.28 | 7.45E-10 | 1.55E-05 | 0.28 | 0.15 | 0.13 | 1.85 |
| 15 | Rragb | 0.35 | 7.51E-10 | 1.56E-05 | 0.18 | 0.09 | 0.09 | 2.09 |
| 15 | Etfbkmt | 0.48 | 7.53E-10 | 1.57E-05 | 0.15 | 0.07 | 0.08 | 2.19 |
| 15 | Prss12 | 0.73 | 7.81E-10 | 1.62E-05 | 0.12 | 0.05 | 0.07 | 2.35 |
| 15 | Tanc2 | 0.26 | 7.97E-10 | 1.66E-05 | 0.39 | 0.23 | 0.16 | 1.71 |
| 15 | Tspyl2 | 0.26 | 8.35E-10 | 1.74E-05 | 0.32 | 0.18 | 0.14 | 1.80 |
| 15 | Adamts7 | 0.29 | 8.47E-10 | 1.76E-05 | 0.19 | 0.10 | 0.10 | 2.04 |
| 15 | Slc4a7 | 0.25 | 8.72E-10 | 1.81E-05 | 0.77 | 0.52 | 0.25 | 1.49 |
| 15 | Mtr | 0.25 | 8.86E-10 | 1.84E-05 | 0.40 | 0.23 | 0.17 | 1.73 |
| 15 | Egln1 | 0.26 | 9.06E-10 | 1.88E-05 | 0.91 | 0.68 | 0.23 | 1.33 |
| 15 | Prepl | 0.37 | 9.08E-10 | 1.89E-05 | 0.24 | 0.12 | 0.11 | 1.92 |
| 15 | 1700012D14Rik | 0.33 | 9.27E-10 | 1.93E-05 | 0.24 | 0.13 | 0.12 | 1.90 |
| 15 | Stat1 | 0.40 | 9.46E-10 | 1.97E-05 | 0.19 | 0.10 | 0.10 | 2.00 |
| 15 | Kri1 | 0.35 | 9.91E-10 | 2.06E-05 | 0.32 | 0.18 | 0.14 | 1.78 |
| 15 | C1qtnf12 | 0.29 | 1.00E-09 | 2.08E-05 | 0.30 | 0.16 | 0.13 | 1.80 |
| 15 | Fam214b | 0.40 | 1.02E-09 | 2.13E-05 | 0.19 | 0.10 | 0.10 | 2.00 |
| 15 | Sema4b | 0.47 | 1.03E-09 | 2.13E-05 | 0.18 | 0.09 | 0.09 | 2.06 |
| 15 | Mbnl3 | 0.31 | 1.04E-09 | 2.16E-05 | 0.24 | 0.13 | 0.12 | 1.93 |
| 15 | Stxbp6 | 0.26 | 1.06E-09 | 2.20E-05 | 0.25 | 0.13 | 0.12 | 1.88 |

**Supplementary Table 1. Marker genes for the original clusters of integrated clustering of E11.5 control and NotchGOF λ-junction cells.**

| | | | | | | | | |
|---|---|---|---|---|---|---|---|---|
| 15 | Tsc22d2 | 0.26 | 1.10E-09 | 2.28E-05 | 0.62 | 0.40 | 0.22 | 1.54 |
| 15 | Fto | 0.27 | 1.10E-09 | 2.29E-05 | 0.72 | 0.47 | 0.25 | 1.53 |
| 15 | Baiap2l1 | 0.52 | 1.11E-09 | 2.30E-05 | 0.14 | 0.06 | 0.08 | 2.24 |
| 15 | Gm4890 | 0.41 | 1.13E-09 | 2.35E-05 | 0.15 | 0.07 | 0.08 | 2.17 |
| 15 | Zfp626 | 0.44 | 1.13E-09 | 2.35E-05 | 0.25 | 0.13 | 0.12 | 1.88 |
| 15 | Nhsl2 | 0.26 | 1.15E-09 | 2.38E-05 | 0.29 | 0.16 | 0.13 | 1.83 |
| 15 | Aasdh | 0.36 | 1.18E-09 | 2.45E-05 | 0.19 | 0.09 | 0.10 | 2.04 |
| 15 | Rps6ka2 | 0.52 | 1.18E-09 | 2.45E-05 | 0.17 | 0.08 | 0.09 | 2.11 |
| 15 | Lpcat3 | 0.29 | 1.21E-09 | 2.52E-05 | 0.49 | 0.30 | 0.19 | 1.62 |
| 15 | Jmy | 0.25 | 1.25E-09 | 2.59E-05 | 0.60 | 0.37 | 0.22 | 1.59 |
| 15 | Akap1 | 0.27 | 1.25E-09 | 2.61E-05 | 0.33 | 0.19 | 0.15 | 1.77 |
| 15 | Ccdc68 | 0.44 | 1.26E-09 | 2.62E-05 | 0.12 | 0.05 | 0.07 | 2.37 |
| 15 | Dclre1a | 0.28 | 1.30E-09 | 2.70E-05 | 0.28 | 0.16 | 0.13 | 1.82 |
| 15 | Uckl1 | 0.30 | 1.32E-09 | 2.75E-05 | 0.23 | 0.12 | 0.11 | 1.94 |
| 15 | Zhx2 | 0.60 | 1.32E-09 | 2.75E-05 | 0.10 | 0.04 | 0.06 | 2.55 |
| 15 | Xylt1 | 0.59 | 1.34E-09 | 2.78E-05 | 0.13 | 0.06 | 0.07 | 2.23 |
| 15 | Trappc8 | 0.27 | 1.34E-09 | 2.79E-05 | 0.41 | 0.24 | 0.17 | 1.71 |
| 15 | Emid1 | 0.44 | 1.35E-09 | 2.81E-05 | 0.18 | 0.09 | 0.09 | 2.03 |
| 15 | 1810044D09Rik | 0.37 | 1.38E-09 | 2.87E-05 | 0.22 | 0.11 | 0.10 | 1.94 |
| 15 | Zc3h11a | 0.49 | 1.40E-09 | 2.92E-05 | 0.14 | 0.06 | 0.08 | 2.24 |
| 15 | Tarbp1 | 0.80 | 1.45E-09 | 3.02E-05 | 0.12 | 0.05 | 0.07 | 2.36 |
| 15 | Serinc5 | 0.78 | 1.46E-09 | 3.04E-05 | 0.13 | 0.06 | 0.07 | 2.27 |
| 15 | Ddx51 | 0.31 | 1.56E-09 | 3.24E-05 | 0.30 | 0.17 | 0.13 | 1.78 |
| 15 | Arap2 | 0.45 | 1.56E-09 | 3.25E-05 | 0.13 | 0.05 | 0.07 | 2.31 |
| 15 | Tbc1d23 | 0.28 | 1.58E-09 | 3.29E-05 | 0.43 | 0.26 | 0.17 | 1.66 |
| 15 | Nacc1 | 0.26 | 1.64E-09 | 3.41E-05 | 0.61 | 0.39 | 0.22 | 1.56 |
| 15 | Abcb8 | 0.25 | 1.66E-09 | 3.45E-05 | 0.32 | 0.18 | 0.14 | 1.78 |
| 15 | 2410131K14Rik | 0.28 | 1.67E-09 | 3.47E-05 | 0.28 | 0.15 | 0.13 | 1.83 |
| 15 | Tmem177 | 0.29 | 1.69E-09 | 3.52E-05 | 0.22 | 0.11 | 0.11 | 1.95 |
| 15 | Snx16 | 0.30 | 1.70E-09 | 3.55E-05 | 0.33 | 0.19 | 0.14 | 1.75 |
| 15 | Bahcc1 | 0.25 | 1.71E-09 | 3.55E-05 | 0.46 | 0.27 | 0.18 | 1.68 |
| 15 | Zc3h12c | 0.60 | 1.71E-09 | 3.55E-05 | 0.14 | 0.06 | 0.08 | 2.25 |
| 15 | Glrb | 0.46 | 1.76E-09 | 3.66E-05 | 0.12 | 0.05 | 0.07 | 2.36 |
| 15 | Elmsan1 | 0.40 | 1.77E-09 | 3.69E-05 | 0.23 | 0.12 | 0.11 | 1.91 |
| 15 | Zbtb2 | 0.35 | 1.78E-09 | 3.70E-05 | 0.23 | 0.12 | 0.11 | 1.92 |
| 15 | Plekhm2 | 0.26 | 1.79E-09 | 3.72E-05 | 0.35 | 0.20 | 0.16 | 1.78 |
| 15 | Pkmyt1 | 0.29 | 1.81E-09 | 3.77E-05 | 0.31 | 0.17 | 0.14 | 1.80 |
| 15 | Jph1 | 0.43 | 1.87E-09 | 3.88E-05 | 0.15 | 0.07 | 0.08 | 2.14 |
| 15 | Zfp81 | 0.25 | 1.87E-09 | 3.90E-05 | 0.42 | 0.25 | 0.17 | 1.67 |
| 15 | Srebf2 | 0.25 | 1.89E-09 | 3.93E-05 | 0.78 | 0.52 | 0.25 | 1.48 |
| 15 | Klhl2 | 0.26 | 1.94E-09 | 4.04E-05 | 0.23 | 0.12 | 0.11 | 1.91 |
| 15 | Dmac2l | 0.31 | 1.96E-09 | 4.09E-05 | 0.26 | 0.14 | 0.12 | 1.85 |
| 15 | Bicdl1 | 0.63 | 2.05E-09 | 4.27E-05 | 0.10 | 0.04 | 0.06 | 2.50 |
| 15 | Entpd5 | 0.31 | 2.11E-09 | 4.39E-05 | 0.16 | 0.08 | 0.09 | 2.10 |
| 15 | Ddi2 | 0.30 | 2.14E-09 | 4.45E-05 | 0.63 | 0.41 | 0.23 | 1.56 |
| 15 | Sbno2 | 0.41 | 2.16E-09 | 4.50E-05 | 0.22 | 0.11 | 0.11 | 1.93 |
| 15 | Slc25a10 | 0.25 | 2.17E-09 | 4.51E-05 | 0.40 | 0.23 | 0.16 | 1.69 |
| 15 | Cnot1 | 0.25 | 2.19E-09 | 4.56E-05 | 0.88 | 0.66 | 0.22 | 1.34 |
| 15 | 1190007I07Rik | 0.27 | 2.20E-09 | 4.58E-05 | 0.45 | 0.28 | 0.18 | 1.64 |
| 15 | 2900052L18Rik | 0.30 | 2.27E-09 | 4.73E-05 | 0.28 | 0.15 | 0.13 | 1.82 |
| 15 | Stc1 | 0.34 | 2.37E-09 | 4.93E-05 | 0.15 | 0.07 | 0.08 | 2.18 |
| 15 | Cep350 | 0.28 | 2.38E-09 | 4.96E-05 | 0.69 | 0.45 | 0.24 | 1.53 |
| 15 | Tnnt1 | 0.36 | 2.44E-09 | 5.08E-05 | 0.12 | 0.05 | 0.07 | 2.32 |
| 15 | Ylpm1 | 0.26 | 2.58E-09 | 5.37E-05 | 0.90 | 0.69 | 0.21 | 1.31 |
| 15 | Mtg2 | 0.27 | 2.59E-09 | 5.38E-05 | 0.44 | 0.26 | 0.18 | 1.67 |
| 15 | Snhg4 | 0.25 | 2.61E-09 | 5.42E-05 | 0.68 | 0.46 | 0.21 | 1.46 |

**Supplementary Table 1. Marker genes for the original clusters of integrated clustering of E11.5 control and NotchGOF λ-junction cells.**

| | | | | | | | | |
|---|---|---|---|---|---|---|---|---|
| 15 | Tbc1d22b | 0.47 | 2.65E-09 | 5.51E-05 | 0.20 | 0.10 | 0.10 | 1.95 |
| 15 | Rasa3 | 0.38 | 2.65E-09 | 5.52E-05 | 0.16 | 0.08 | 0.08 | 2.11 |
| 15 | Vps13b | 0.35 | 2.68E-09 | 5.58E-05 | 0.32 | 0.18 | 0.14 | 1.75 |
| 15 | Map10 | 0.46 | 2.72E-09 | 5.65E-05 | 0.13 | 0.05 | 0.07 | 2.31 |
| 15 | Piga | 0.29 | 2.77E-09 | 5.76E-05 | 0.29 | 0.16 | 0.13 | 1.81 |
| 15 | Limd1 | 0.31 | 2.81E-09 | 5.84E-05 | 0.73 | 0.51 | 0.23 | 1.44 |
| 15 | Txnrd3 | 0.35 | 2.86E-09 | 5.94E-05 | 0.22 | 0.12 | 0.11 | 1.90 |
| 15 | A930004D18Rik | 0.48 | 2.94E-09 | 6.11E-05 | 0.12 | 0.05 | 0.07 | 2.37 |
| 15 | Ipo7 | 0.27 | 2.95E-09 | 6.14E-05 | 0.82 | 0.61 | 0.21 | 1.34 |
| 15 | Gspt2 | 0.32 | 3.00E-09 | 6.25E-05 | 0.28 | 0.15 | 0.13 | 1.82 |
| 15 | Tyk2 | 0.39 | 3.02E-09 | 6.28E-05 | 0.13 | 0.06 | 0.07 | 2.28 |
| 15 | Sowahc | 0.37 | 3.04E-09 | 6.32E-05 | 0.25 | 0.14 | 0.11 | 1.84 |
| 15 | Snx33 | 0.39 | 3.05E-09 | 6.34E-05 | 0.20 | 0.10 | 0.10 | 1.95 |
| 15 | Sdad1 | 0.29 | 3.10E-09 | 6.45E-05 | 0.70 | 0.47 | 0.23 | 1.48 |
| 15 | Adarb1 | 0.40 | 3.12E-09 | 6.50E-05 | 0.15 | 0.07 | 0.08 | 2.14 |
| 15 | Kdm6a | 0.26 | 3.15E-09 | 6.54E-05 | 0.64 | 0.42 | 0.22 | 1.52 |
| 15 | Tnfsf9 | 0.32 | 3.34E-09 | 6.94E-05 | 0.16 | 0.08 | 0.08 | 2.11 |
| 15 | Zfp729a | 0.33 | 3.34E-09 | 6.95E-05 | 0.26 | 0.14 | 0.12 | 1.83 |
| 15 | Pter | 0.37 | 3.40E-09 | 7.07E-05 | 0.22 | 0.11 | 0.10 | 1.90 |
| 15 | Otud3 | 0.27 | 3.40E-09 | 7.07E-05 | 0.22 | 0.11 | 0.11 | 1.95 |
| 15 | Cul9 | 0.40 | 3.46E-09 | 7.20E-05 | 0.11 | 0.04 | 0.06 | 2.41 |
| 15 | Spred3 | 0.27 | 3.49E-09 | 7.26E-05 | 0.19 | 0.09 | 0.10 | 2.02 |
| 15 | Klhl15 | 0.51 | 3.51E-09 | 7.29E-05 | 0.14 | 0.06 | 0.08 | 2.21 |
| 15 | Nudt7 | 0.44 | 3.52E-09 | 7.32E-05 | 0.15 | 0.07 | 0.08 | 2.14 |
| 15 | Akap10 | 0.32 | 3.56E-09 | 7.41E-05 | 0.25 | 0.13 | 0.11 | 1.87 |
| 15 | Grk5 | 0.31 | 3.58E-09 | 7.44E-05 | 0.23 | 0.12 | 0.11 | 1.91 |
| 15 | Tsacc | 0.31 | 3.69E-09 | 7.68E-05 | 0.15 | 0.07 | 0.08 | 2.15 |
| 15 | Dusp28 | 0.34 | 3.73E-09 | 7.75E-05 | 0.22 | 0.11 | 0.11 | 1.93 |
| 15 | Atg4d | 0.26 | 3.74E-09 | 7.79E-05 | 0.35 | 0.20 | 0.15 | 1.73 |
| 15 | Elf3 | 0.36 | 3.76E-09 | 7.81E-05 | 0.16 | 0.07 | 0.08 | 2.12 |
| 15 | Hspa2 | 0.60 | 3.87E-09 | 8.04E-05 | 0.13 | 0.06 | 0.07 | 2.23 |
| 15 | Usp51 | 0.41 | 3.87E-09 | 8.05E-05 | 0.17 | 0.08 | 0.09 | 2.06 |
| 15 | Aaas | 0.27 | 3.88E-09 | 8.08E-05 | 0.52 | 0.33 | 0.19 | 1.59 |
| 15 | Prkch | 0.30 | 3.92E-09 | 8.14E-05 | 0.18 | 0.09 | 0.09 | 2.02 |
| 15 | Ids | 0.42 | 4.12E-09 | 8.56E-05 | 0.17 | 0.08 | 0.09 | 2.06 |
| 15 | Nin | 0.30 | 4.17E-09 | 8.68E-05 | 0.44 | 0.27 | 0.17 | 1.64 |
| 15 | Bahd1 | 0.65 | 4.37E-09 | 9.08E-05 | 0.10 | 0.04 | 0.06 | 2.43 |
| 15 | Ctdp1 | 0.25 | 4.42E-09 | 9.19E-05 | 0.35 | 0.20 | 0.15 | 1.72 |
| 15 | Poglut3 | 0.40 | 4.47E-09 | 9.29E-05 | 0.25 | 0.13 | 0.11 | 1.84 |
| 15 | Mllt10 | 0.25 | 4.48E-09 | 9.31E-05 | 0.85 | 0.63 | 0.22 | 1.34 |
| 15 | Pla2g6 | 0.46 | 4.50E-09 | 9.37E-05 | 0.16 | 0.08 | 0.08 | 2.05 |
| 15 | Mtmr9 | 0.38 | 4.66E-09 | 9.69E-05 | 0.26 | 0.15 | 0.12 | 1.82 |
| 15 | Aldh4a1 | 0.31 | 4.70E-09 | 9.77E-05 | 0.19 | 0.09 | 0.09 | 1.99 |
| 15 | Cpt1a | 0.29 | 4.77E-09 | 9.93E-05 | 0.20 | 0.10 | 0.10 | 1.97 |
| 15 | Gipc2 | 0.40 | 4.78E-09 | 9.95E-05 | 0.14 | 0.06 | 0.07 | 2.17 |
| 15 | Bbs2 | 0.27 | 4.84E-09 | 1.01E-04 | 0.19 | 0.09 | 0.09 | 2.01 |
| 15 | Maml2 | 0.39 | 4.95E-09 | 1.03E-04 | 0.19 | 0.09 | 0.09 | 1.99 |
| 15 | AI987944 | 0.28 | 4.97E-09 | 1.03E-04 | 0.24 | 0.13 | 0.11 | 1.87 |
| 15 | Nudt18 | 0.35 | 5.15E-09 | 1.07E-04 | 0.13 | 0.06 | 0.07 | 2.20 |
| 15 | Gadd45b | 0.35 | 5.16E-09 | 1.07E-04 | 0.26 | 0.14 | 0.11 | 1.80 |
| 15 | Xylt2 | 0.37 | 5.30E-09 | 1.10E-04 | 0.22 | 0.11 | 0.10 | 1.91 |
| 15 | Alkal2 | 0.27 | 5.32E-09 | 1.11E-04 | 0.17 | 0.08 | 0.09 | 2.04 |
| 15 | Dip2a | 0.43 | 5.34E-09 | 1.11E-04 | 0.19 | 0.10 | 0.09 | 1.96 |
| 15 | Prodh | 0.32 | 5.38E-09 | 1.12E-04 | 0.21 | 0.11 | 0.10 | 1.94 |
| 15 | Arsk | 0.27 | 5.46E-09 | 1.14E-04 | 0.24 | 0.13 | 0.11 | 1.89 |
| 15 | Endod1 | 0.45 | 5.54E-09 | 1.15E-04 | 0.18 | 0.09 | 0.09 | 1.98 |

**Supplementary Table 1. Marker genes for the original clusters of integrated clustering of E11.5 control and NotchGOF λ-junction cells.**

| | | | | | | | | |
|---|---|---|---|---|---|---|---|---|
| 15 | Gnaz | 0.51 | 5.59E-09 | 1.16E-04 | 0.10 | 0.04 | 0.06 | 2.42 |
| 15 | Ggnbp1 | 0.51 | 5.63E-09 | 1.17E-04 | 0.13 | 0.06 | 0.07 | 2.24 |
| 15 | Cpne8 | 0.43 | 5.78E-09 | 1.20E-04 | 0.16 | 0.08 | 0.08 | 2.05 |
| 15 | Arhgap19 | 0.26 | 6.05E-09 | 1.26E-04 | 0.46 | 0.29 | 0.17 | 1.60 |
| 15 | Pex11a | 0.27 | 6.35E-09 | 1.32E-04 | 0.21 | 0.11 | 0.10 | 1.94 |
| 15 | Palb2 | 0.28 | 6.53E-09 | 1.36E-04 | 0.25 | 0.13 | 0.11 | 1.86 |
| 15 | Slc25a44 | 0.28 | 6.55E-09 | 1.36E-04 | 0.28 | 0.15 | 0.12 | 1.81 |
| 15 | Lins1 | 0.27 | 6.57E-09 | 1.37E-04 | 0.19 | 0.10 | 0.10 | 1.96 |
| 15 | Rundc3b | 0.60 | 6.72E-09 | 1.40E-04 | 0.10 | 0.04 | 0.06 | 2.44 |
| 15 | Wdr19 | 0.31 | 6.76E-09 | 1.41E-04 | 0.20 | 0.10 | 0.10 | 1.97 |
| 15 | Slc4a2 | 0.25 | 6.84E-09 | 1.42E-04 | 0.36 | 0.22 | 0.15 | 1.68 |
| 15 | Mcf2l | 0.40 | 6.95E-09 | 1.44E-04 | 0.15 | 0.07 | 0.08 | 2.15 |
| 15 | Upf1 | 0.26 | 7.01E-09 | 1.46E-04 | 0.70 | 0.47 | 0.23 | 1.49 |
| 15 | Sil1 | 0.32 | 7.13E-09 | 1.48E-04 | 0.21 | 0.11 | 0.10 | 1.90 |
| 15 | Whrn | 0.42 | 7.17E-09 | 1.49E-04 | 0.13 | 0.06 | 0.07 | 2.20 |
| 15 | Zfp27 | 0.31 | 7.57E-09 | 1.57E-04 | 0.17 | 0.08 | 0.09 | 2.06 |
| 15 | Cers6 | 0.28 | 7.61E-09 | 1.58E-04 | 0.40 | 0.24 | 0.16 | 1.66 |
| 15 | Ern1 | 0.77 | 7.73E-09 | 1.61E-04 | 0.10 | 0.04 | 0.06 | 2.44 |
| 15 | Inafm2 | 0.28 | 7.76E-09 | 1.61E-04 | 0.23 | 0.12 | 0.11 | 1.86 |
| 15 | Gm20186 | 0.30 | 7.82E-09 | 1.63E-04 | 0.14 | 0.07 | 0.08 | 2.17 |
| 15 | Zfp959 | 0.33 | 7.86E-09 | 1.63E-04 | 0.22 | 0.12 | 0.10 | 1.88 |
| 15 | Dpp7 | 0.44 | 8.03E-09 | 1.67E-04 | 0.15 | 0.07 | 0.08 | 2.08 |
| 15 | Usp30 | 0.43 | 8.07E-09 | 1.68E-04 | 0.21 | 0.11 | 0.10 | 1.88 |
| 15 | Zmym6 | 0.44 | 8.23E-09 | 1.71E-04 | 0.16 | 0.08 | 0.08 | 2.07 |
| 15 | 2610507I01Rik | 0.41 | 8.28E-09 | 1.72E-04 | 0.14 | 0.06 | 0.07 | 2.17 |
| 15 | Klf9 | 0.55 | 8.45E-09 | 1.76E-04 | 0.11 | 0.05 | 0.06 | 2.31 |
| 15 | Sfi1 | 0.41 | 8.51E-09 | 1.77E-04 | 0.15 | 0.07 | 0.08 | 2.15 |
| 15 | Tango6 | 0.37 | 8.51E-09 | 1.77E-04 | 0.22 | 0.11 | 0.10 | 1.89 |
| 15 | Cdk5rap1 | 0.38 | 8.79E-09 | 1.83E-04 | 0.18 | 0.09 | 0.09 | 2.00 |
| 15 | Bdh1 | 0.49 | 8.90E-09 | 1.85E-04 | 0.15 | 0.07 | 0.08 | 2.08 |
| 15 | Dgke | 0.48 | 8.93E-09 | 1.86E-04 | 0.15 | 0.07 | 0.08 | 2.11 |
| 15 | Cplane1 | 0.28 | 9.00E-09 | 1.87E-04 | 0.54 | 0.35 | 0.20 | 1.56 |
| 15 | Ggct | 0.26 | 9.10E-09 | 1.89E-04 | 0.26 | 0.15 | 0.12 | 1.82 |
| 15 | Sall1 | 0.29 | 9.87E-09 | 2.05E-04 | 0.18 | 0.09 | 0.09 | 1.99 |
| 15 | Zfr2 | 0.45 | 1.01E-08 | 2.10E-04 | 0.10 | 0.04 | 0.06 | 2.43 |
| 15 | Adgrb2 | 0.46 | 1.01E-08 | 2.10E-04 | 0.10 | 0.04 | 0.06 | 2.36 |
| 15 | Mlst8 | 0.30 | 1.02E-08 | 2.12E-04 | 0.24 | 0.13 | 0.11 | 1.84 |
| 15 | Caskin2 | 0.34 | 1.03E-08 | 2.14E-04 | 0.21 | 0.11 | 0.10 | 1.90 |
| 15 | Hdac4 | 0.29 | 1.04E-08 | 2.16E-04 | 0.40 | 0.24 | 0.16 | 1.64 |
| 15 | Itpripl1 | 0.36 | 1.06E-08 | 2.20E-04 | 0.18 | 0.09 | 0.09 | 1.97 |
| 15 | Slc35g1 | 0.55 | 1.06E-08 | 2.21E-04 | 0.13 | 0.06 | 0.07 | 2.19 |
| 15 | Sv2a | 0.36 | 1.08E-08 | 2.25E-04 | 0.16 | 0.08 | 0.08 | 2.05 |
| 15 | Frk | 0.31 | 1.12E-08 | 2.33E-04 | 0.26 | 0.15 | 0.11 | 1.79 |
| 15 | Dand5 | 0.53 | 1.13E-08 | 2.35E-04 | 0.13 | 0.06 | 0.07 | 2.16 |
| 15 | 9930014A18Rik | 0.51 | 1.14E-08 | 2.37E-04 | 0.11 | 0.05 | 0.06 | 2.31 |
| 15 | Ago4 | 0.27 | 1.14E-08 | 2.38E-04 | 0.15 | 0.07 | 0.08 | 2.11 |
| 15 | Dclre1c | 0.36 | 1.17E-08 | 2.42E-04 | 0.12 | 0.05 | 0.07 | 2.27 |
| 15 | Nsun3 | 0.41 | 1.17E-08 | 2.44E-04 | 0.16 | 0.08 | 0.08 | 2.07 |
| 15 | Trrap | 0.26 | 1.20E-08 | 2.50E-04 | 0.83 | 0.58 | 0.25 | 1.43 |
| 15 | 5430403G16Rik | 0.34 | 1.21E-08 | 2.51E-04 | 0.15 | 0.07 | 0.08 | 2.11 |
| 15 | Tjp3 | 0.52 | 1.21E-08 | 2.52E-04 | 0.11 | 0.05 | 0.06 | 2.31 |
| 15 | Poli | 0.46 | 1.21E-08 | 2.53E-04 | 0.13 | 0.06 | 0.07 | 2.16 |
| 15 | Dpy19l3 | 0.28 | 1.22E-08 | 2.54E-04 | 0.27 | 0.15 | 0.12 | 1.79 |
| 15 | Aph1b | 0.29 | 1.24E-08 | 2.57E-04 | 0.18 | 0.09 | 0.09 | 1.98 |
| 15 | Ubiad1 | 0.39 | 1.26E-08 | 2.63E-04 | 0.15 | 0.07 | 0.08 | 2.12 |
| 15 | Tmem200a | 0.48 | 1.38E-08 | 2.88E-04 | 0.12 | 0.06 | 0.07 | 2.24 |

**Supplementary Table 1. Marker genes for the original clusters of integrated clustering of E11.5 control and NotchGOF λ-junction cells.**

| | | | | | | | | |
|---|---|---|---|---|---|---|---|---|
| 15 | Tmem94 | 0.25 | 1.43E-08 | 2.96E-04 | 0.31 | 0.18 | 0.13 | 1.73 |
| 15 | Gm46218 | 0.49 | 1.43E-08 | 2.98E-04 | 0.11 | 0.05 | 0.06 | 2.36 |
| 15 | Kdm5c | 0.25 | 1.44E-08 | 3.00E-04 | 0.75 | 0.52 | 0.23 | 1.44 |
| 15 | Fam217b | 0.27 | 1.45E-08 | 3.02E-04 | 0.18 | 0.09 | 0.09 | 1.99 |
| 15 | Mfsd2a | 0.57 | 1.45E-08 | 3.02E-04 | 0.13 | 0.06 | 0.07 | 2.16 |
| 15 | Madd | 0.31 | 1.48E-08 | 3.08E-04 | 0.15 | 0.07 | 0.08 | 2.08 |
| 15 | Dcaf6 | 0.38 | 1.48E-08 | 3.08E-04 | 0.16 | 0.08 | 0.08 | 2.00 |
| 15 | Mast3 | 0.28 | 1.50E-08 | 3.12E-04 | 0.21 | 0.11 | 0.10 | 1.91 |
| 15 | Mon1a | 0.50 | 1.51E-08 | 3.15E-04 | 0.14 | 0.07 | 0.07 | 2.11 |
| 15 | Ccdc51 | 0.31 | 1.53E-08 | 3.19E-04 | 0.22 | 0.12 | 0.10 | 1.86 |
| 15 | Usp53 | 0.41 | 1.56E-08 | 3.24E-04 | 0.16 | 0.08 | 0.08 | 2.04 |
| 15 | Slc33a1 | 0.26 | 1.62E-08 | 3.37E-04 | 0.21 | 0.11 | 0.10 | 1.90 |
| 15 | Tmem231 | 0.32 | 1.63E-08 | 3.39E-04 | 0.22 | 0.12 | 0.10 | 1.86 |
| 15 | Col12a1 | 0.34 | 1.64E-08 | 3.42E-04 | 0.13 | 0.06 | 0.07 | 2.16 |
| 15 | Wdr62 | 0.60 | 1.66E-08 | 3.45E-04 | 0.11 | 0.05 | 0.06 | 2.27 |
| 15 | Wdpcp | 0.34 | 1.67E-08 | 3.48E-04 | 0.16 | 0.08 | 0.08 | 2.02 |
| 15 | Coq10a | 0.45 | 1.78E-08 | 3.69E-04 | 0.16 | 0.08 | 0.08 | 2.00 |
| 15 | Chtf8 | 0.28 | 1.83E-08 | 3.80E-04 | 0.23 | 0.13 | 0.11 | 1.85 |
| 15 | Kdm6b | 0.27 | 1.83E-08 | 3.81E-04 | 0.87 | 0.65 | 0.22 | 1.33 |
| 15 | Gm12940 | 0.38 | 1.85E-08 | 3.85E-04 | 0.20 | 0.11 | 0.10 | 1.90 |
| 15 | Zbtb9 | 0.25 | 1.88E-08 | 3.91E-04 | 0.21 | 0.11 | 0.10 | 1.91 |
| 15 | Zfp764 | 0.53 | 1.89E-08 | 3.93E-04 | 0.13 | 0.06 | 0.07 | 2.16 |
| 15 | Zfp128 | 0.59 | 1.90E-08 | 3.94E-04 | 0.11 | 0.05 | 0.06 | 2.27 |
| 15 | C130036L24Rik | 0.26 | 2.00E-08 | 4.17E-04 | 0.17 | 0.09 | 0.09 | 2.01 |
| 15 | Ankrd16 | 0.27 | 2.08E-08 | 4.32E-04 | 0.37 | 0.22 | 0.15 | 1.67 |
| 15 | Gm43462 | 0.30 | 2.13E-08 | 4.42E-04 | 0.20 | 0.10 | 0.10 | 1.91 |
| 15 | C2cd2 | 0.30 | 2.13E-08 | 4.43E-04 | 0.19 | 0.10 | 0.09 | 1.92 |
| 15 | Rbl2 | 0.37 | 2.16E-08 | 4.50E-04 | 0.17 | 0.09 | 0.09 | 1.98 |
| 15 | Rfng | 0.27 | 2.24E-08 | 4.66E-04 | 0.31 | 0.18 | 0.13 | 1.73 |
| 15 | Wdr45 | 0.27 | 2.28E-08 | 4.74E-04 | 0.24 | 0.13 | 0.11 | 1.82 |
| 15 | Hdx | 0.26 | 2.32E-08 | 4.82E-04 | 0.19 | 0.10 | 0.09 | 1.94 |
| 15 | BC025920 | 0.45 | 2.38E-08 | 4.94E-04 | 0.12 | 0.06 | 0.07 | 2.20 |
| 15 | Fam220a.1 | 0.30 | 2.38E-08 | 4.95E-04 | 0.19 | 0.10 | 0.09 | 1.92 |
| 15 | Zbtb8a | 0.36 | 2.39E-08 | 4.97E-04 | 0.16 | 0.08 | 0.08 | 2.04 |
| 15 | Immp2l | 0.25 | 2.60E-08 | 5.40E-04 | 0.17 | 0.08 | 0.08 | 2.01 |
| 15 | Coro1b | 0.34 | 2.63E-08 | 5.46E-04 | 0.14 | 0.07 | 0.08 | 2.09 |
| 15 | Asl | 0.28 | 2.64E-08 | 5.50E-04 | 0.26 | 0.15 | 0.12 | 1.80 |
| 15 | Trim45 | 0.26 | 2.66E-08 | 5.52E-04 | 0.21 | 0.11 | 0.10 | 1.89 |
| 15 | Fam184a | 0.29 | 2.67E-08 | 5.55E-04 | 0.22 | 0.12 | 0.10 | 1.87 |
| 15 | Dolpp1 | 0.29 | 2.67E-08 | 5.55E-04 | 0.25 | 0.14 | 0.11 | 1.80 |
| 15 | 4933408B17Rik | 0.65 | 2.75E-08 | 5.72E-04 | 0.11 | 0.05 | 0.06 | 2.27 |
| 15 | Itpkc | 0.37 | 2.76E-08 | 5.74E-04 | 0.14 | 0.07 | 0.07 | 2.11 |
| 15 | Tsc1 | 0.30 | 2.81E-08 | 5.85E-04 | 0.36 | 0.22 | 0.14 | 1.66 |
| 15 | 5033430I15Rik | 0.56 | 2.83E-08 | 5.88E-04 | 0.12 | 0.05 | 0.06 | 2.19 |
| 15 | Afmid | 0.30 | 2.84E-08 | 5.90E-04 | 0.14 | 0.06 | 0.07 | 2.14 |
| 15 | Cntnap2 | 0.60 | 2.85E-08 | 5.92E-04 | 0.11 | 0.05 | 0.06 | 2.22 |
| 15 | Nfkbiz | 0.35 | 3.04E-08 | 6.31E-04 | 0.15 | 0.07 | 0.08 | 2.09 |
| 15 | Timmdc1 | 0.28 | 3.07E-08 | 6.38E-04 | 0.38 | 0.23 | 0.15 | 1.64 |
| 15 | Napb | 0.56 | 3.18E-08 | 6.62E-04 | 0.12 | 0.05 | 0.06 | 2.19 |
| 15 | Ptprm | 0.54 | 3.19E-08 | 6.63E-04 | 0.11 | 0.05 | 0.06 | 2.27 |
| 15 | Elac1 | 0.52 | 3.19E-08 | 6.64E-04 | 0.13 | 0.06 | 0.07 | 2.15 |
| 15 | Garem1 | 0.42 | 3.22E-08 | 6.71E-04 | 0.17 | 0.09 | 0.08 | 1.96 |
| 15 | Evi5l | 0.27 | 3.32E-08 | 6.91E-04 | 0.21 | 0.11 | 0.10 | 1.87 |
| 15 | Zfp46 | 0.27 | 3.38E-08 | 7.04E-04 | 0.24 | 0.14 | 0.11 | 1.80 |
| 15 | Syt7 | 0.50 | 3.44E-08 | 7.16E-04 | 0.12 | 0.06 | 0.07 | 2.16 |
| 15 | Coro2b | 0.73 | 3.46E-08 | 7.19E-04 | 0.11 | 0.05 | 0.06 | 2.26 |

**Supplementary Table 1. Marker genes for the original clusters of integrated clustering of E11.5 control and NotchGOF λ-junction cells.**

| | | | | | | | | |
|---|---|---|---|---|---|---|---|---|
| 15 | Plekhh1 | 0.39 | 3.47E-08 | 7.22E-04 | 0.14 | 0.07 | 0.07 | 2.11 |
| 15 | Smg9 | 0.34 | 3.56E-08 | 7.40E-04 | 0.25 | 0.14 | 0.11 | 1.78 |
| 15 | Mblac2 | 0.50 | 3.61E-08 | 7.50E-04 | 0.11 | 0.05 | 0.06 | 2.22 |
| 15 | Lck | 0.40 | 3.70E-08 | 7.69E-04 | 0.14 | 0.07 | 0.07 | 2.11 |
| 15 | Urb2 | 0.27 | 3.70E-08 | 7.70E-04 | 0.17 | 0.09 | 0.08 | 1.99 |
| 15 | 2010315B03Rik | 0.45 | 3.70E-08 | 7.70E-04 | 0.12 | 0.06 | 0.07 | 2.18 |
| 15 | Trp53inp2 | 0.26 | 3.71E-08 | 7.71E-04 | 0.19 | 0.10 | 0.09 | 1.93 |
| 15 | Erc2 | 0.60 | 3.87E-08 | 8.05E-04 | 0.14 | 0.07 | 0.07 | 2.04 |
| 15 | Ddhd1 | 0.30 | 3.88E-08 | 8.08E-04 | 0.29 | 0.17 | 0.12 | 1.71 |
| 15 | Gk | 0.49 | 4.11E-08 | 8.56E-04 | 0.15 | 0.08 | 0.08 | 2.01 |
| 15 | Btbd8 | 0.42 | 4.21E-08 | 8.75E-04 | 0.15 | 0.07 | 0.08 | 2.03 |
| 15 | Atxn1 | 0.31 | 4.24E-08 | 8.82E-04 | 0.18 | 0.09 | 0.09 | 1.93 |
| 15 | Ltbp3 | 0.31 | 4.32E-08 | 8.99E-04 | 0.16 | 0.08 | 0.08 | 1.98 |
| 15 | Zfp563 | 0.35 | 4.34E-08 | 9.03E-04 | 0.16 | 0.08 | 0.08 | 2.00 |
| 15 | Pigb | 0.26 | 4.51E-08 | 9.38E-04 | 0.15 | 0.07 | 0.08 | 2.05 |
| 15 | Prrg3 | 0.27 | 4.86E-08 | 1.01E-03 | 0.21 | 0.11 | 0.10 | 1.87 |
| 15 | Kat2b | 0.65 | 4.95E-08 | 1.03E-03 | 0.10 | 0.05 | 0.06 | 2.26 |
| 15 | Pign | 0.27 | 5.00E-08 | 1.04E-03 | 0.26 | 0.14 | 0.11 | 1.77 |
| 15 | Arfgef2 | 0.34 | 5.34E-08 | 1.11E-03 | 0.18 | 0.09 | 0.09 | 1.93 |
| 15 | Tmem38b | 0.26 | 5.37E-08 | 1.12E-03 | 0.26 | 0.15 | 0.11 | 1.76 |
| 15 | Lysmd4 | 0.34 | 5.48E-08 | 1.14E-03 | 0.16 | 0.08 | 0.08 | 1.99 |
| 15 | Rubcn | 0.45 | 5.55E-08 | 1.15E-03 | 0.16 | 0.08 | 0.08 | 1.98 |
| 15 | Rfx5 | 0.25 | 5.61E-08 | 1.17E-03 | 0.20 | 0.11 | 0.10 | 1.87 |
| 15 | Rnf180 | 0.56 | 5.61E-08 | 1.17E-03 | 0.11 | 0.05 | 0.06 | 2.22 |
| 15 | Cfp | 0.60 | 5.63E-08 | 1.17E-03 | 0.10 | 0.04 | 0.06 | 2.32 |
| 15 | Pde1c | 0.27 | 5.75E-08 | 1.20E-03 | 0.20 | 0.11 | 0.09 | 1.86 |
| 15 | B830012L14Rik | 0.38 | 5.87E-08 | 1.22E-03 | 0.12 | 0.05 | 0.06 | 2.19 |
| 15 | Gm5577 | 0.29 | 6.00E-08 | 1.25E-03 | 0.15 | 0.07 | 0.08 | 2.03 |
| 15 | Tmc7 | 0.30 | 6.05E-08 | 1.26E-03 | 0.12 | 0.06 | 0.07 | 2.18 |
| 15 | Gm17491 | 0.31 | 6.07E-08 | 1.26E-03 | 0.18 | 0.09 | 0.09 | 1.93 |
| 15 | Nudt10 | 0.28 | 6.21E-08 | 1.29E-03 | 0.21 | 0.11 | 0.10 | 1.84 |
| 15 | 9330104G04Rik | 0.32 | 6.24E-08 | 1.30E-03 | 0.15 | 0.07 | 0.08 | 2.06 |
| 15 | A830082K12Rik | 0.64 | 6.72E-08 | 1.40E-03 | 0.15 | 0.07 | 0.07 | 2.00 |
| 15 | Cdan1 | 0.39 | 6.95E-08 | 1.45E-03 | 0.18 | 0.09 | 0.08 | 1.91 |
| 15 | Gm45051 | 0.40 | 7.29E-08 | 1.52E-03 | 0.15 | 0.07 | 0.07 | 2.00 |
| 15 | Izumo4 | 0.26 | 7.50E-08 | 1.56E-03 | 0.13 | 0.06 | 0.07 | 2.16 |
| 15 | Rph3al | 0.38 | 7.51E-08 | 1.56E-03 | 0.11 | 0.05 | 0.06 | 2.22 |
| 15 | Slc37a4 | 0.36 | 7.51E-08 | 1.56E-03 | 0.13 | 0.06 | 0.07 | 2.12 |
| 15 | Tdrkh | 0.30 | 7.59E-08 | 1.58E-03 | 0.13 | 0.06 | 0.07 | 2.12 |
| 15 | Cdk20 | 0.44 | 7.73E-08 | 1.61E-03 | 0.12 | 0.05 | 0.06 | 2.19 |
| 15 | Fanci | 0.27 | 8.28E-08 | 1.72E-03 | 0.18 | 0.10 | 0.09 | 1.93 |
| 15 | Zfp597 | 0.42 | 8.36E-08 | 1.74E-03 | 0.15 | 0.08 | 0.08 | 2.00 |
| 15 | 4930563E22Rik | 0.42 | 8.54E-08 | 1.78E-03 | 0.13 | 0.06 | 0.07 | 2.12 |
| 15 | Myo19 | 0.33 | 9.42E-08 | 1.96E-03 | 0.16 | 0.08 | 0.08 | 1.95 |
| 15 | Tapbp | 0.30 | 9.84E-08 | 2.05E-03 | 0.16 | 0.08 | 0.08 | 1.98 |
| 15 | Alg1 | 0.29 | 9.92E-08 | 2.06E-03 | 0.15 | 0.08 | 0.08 | 1.99 |
| 15 | Eif2ak3 | 0.46 | 9.98E-08 | 2.07E-03 | 0.14 | 0.07 | 0.07 | 2.01 |
| 15 | Gm14418 | 0.49 | 1.01E-07 | 2.10E-03 | 0.12 | 0.06 | 0.06 | 2.15 |
| 15 | Gm14296 | 0.32 | 1.02E-07 | 2.12E-03 | 0.14 | 0.07 | 0.07 | 2.08 |
| 15 | Slc9a6 | 0.28 | 1.04E-07 | 2.17E-03 | 0.22 | 0.12 | 0.10 | 1.82 |
| 15 | Kctd11 | 0.32 | 1.07E-07 | 2.22E-03 | 0.10 | 0.05 | 0.06 | 2.26 |
| 15 | Apobec3 | 0.27 | 1.07E-07 | 2.22E-03 | 0.16 | 0.08 | 0.08 | 1.95 |
| 15 | Akr1b10 | 0.32 | 1.07E-07 | 2.22E-03 | 0.21 | 0.12 | 0.10 | 1.82 |
| 15 | Mutyh | 0.29 | 1.09E-07 | 2.26E-03 | 0.21 | 0.11 | 0.10 | 1.84 |
| 15 | Nyap1 | 0.32 | 1.09E-07 | 2.26E-03 | 0.17 | 0.09 | 0.08 | 1.94 |
| 15 | Lpcat4 | 0.50 | 1.09E-07 | 2.27E-03 | 0.13 | 0.06 | 0.07 | 2.08 |

**Supplementary Table 1. Marker genes for the original clusters of integrated clustering of E11.5 control and NotchGOF λ-junction cells.**

| | | | | | | | | |
|---|---|---|---|---|---|---|---|---|
| 15 | Tmem184a | 0.42 | 1.10E-07 | 2.28E-03 | 0.11 | 0.05 | 0.06 | 2.13 |
| 15 | Casp9 | 0.32 | 1.10E-07 | 2.29E-03 | 0.18 | 0.10 | 0.09 | 1.89 |
| 15 | Gm5141 | 0.29 | 1.12E-07 | 2.34E-03 | 0.19 | 0.10 | 0.09 | 1.89 |
| 15 | Top1mt | 0.54 | 1.13E-07 | 2.36E-03 | 0.12 | 0.06 | 0.06 | 2.15 |
| 15 | Lama1 | 0.40 | 1.17E-07 | 2.42E-03 | 0.19 | 0.10 | 0.09 | 1.85 |
| 15 | Pcdhgc4 | 0.31 | 1.19E-07 | 2.47E-03 | 0.15 | 0.07 | 0.07 | 2.00 |
| 15 | Pwwp2b | 0.32 | 1.21E-07 | 2.53E-03 | 0.15 | 0.07 | 0.07 | 2.00 |
| 15 | Gpm6b | 0.29 | 1.22E-07 | 2.54E-03 | 0.14 | 0.07 | 0.07 | 2.04 |
| 15 | Ddx19b | 0.33 | 1.22E-07 | 2.54E-03 | 0.17 | 0.09 | 0.08 | 1.90 |
| 15 | Tepsin | 0.46 | 1.24E-07 | 2.57E-03 | 0.14 | 0.07 | 0.07 | 2.04 |
| 15 | A630089N07Rik | 0.31 | 1.24E-07 | 2.59E-03 | 0.24 | 0.14 | 0.11 | 1.77 |
| 15 | Trmo | 0.61 | 1.25E-07 | 2.60E-03 | 0.13 | 0.07 | 0.07 | 2.03 |
| 15 | D030045P18Rik | 0.45 | 1.34E-07 | 2.78E-03 | 0.10 | 0.05 | 0.06 | 2.27 |
| 15 | Gucy1a2 | 0.46 | 1.34E-07 | 2.79E-03 | 0.14 | 0.07 | 0.07 | 2.01 |
| 15 | Loxl1 | 0.35 | 1.40E-07 | 2.90E-03 | 0.15 | 0.07 | 0.07 | 2.00 |
| 15 | D430020J02Rik | 0.26 | 1.44E-07 | 2.99E-03 | 0.16 | 0.08 | 0.08 | 1.96 |
| 15 | Fitm2 | 0.29 | 1.46E-07 | 3.04E-03 | 0.17 | 0.09 | 0.08 | 1.92 |
| 15 | Cish | 0.31 | 1.46E-07 | 3.04E-03 | 0.13 | 0.06 | 0.07 | 2.10 |
| 15 | Slc12a9 | 0.44 | 1.46E-07 | 3.04E-03 | 0.16 | 0.08 | 0.08 | 1.94 |
| 15 | Nt5dc1 | 0.30 | 1.50E-07 | 3.12E-03 | 0.20 | 0.11 | 0.09 | 1.84 |
| 15 | Pgap3 | 0.47 | 1.61E-07 | 3.35E-03 | 0.10 | 0.05 | 0.06 | 2.27 |
| 15 | Nell2 | 0.35 | 1.64E-07 | 3.42E-03 | 0.11 | 0.05 | 0.06 | 2.18 |
| 15 | Nxt2 | 0.26 | 1.68E-07 | 3.48E-03 | 0.15 | 0.07 | 0.07 | 2.00 |
| 15 | N4bp2os | 0.51 | 1.73E-07 | 3.60E-03 | 0.10 | 0.04 | 0.06 | 2.27 |
| 15 | D11Wsu47e | 0.30 | 1.78E-07 | 3.70E-03 | 0.18 | 0.10 | 0.08 | 1.87 |
| 15 | Kif5a | 0.27 | 1.81E-07 | 3.76E-03 | 0.10 | 0.05 | 0.06 | 2.21 |
| 15 | Cyp4f17 | 0.38 | 1.88E-07 | 3.91E-03 | 0.13 | 0.07 | 0.07 | 2.03 |
| 15 | Spef1 | 0.32 | 1.89E-07 | 3.93E-03 | 0.13 | 0.06 | 0.07 | 2.08 |
| 15 | Tmem250-ps | 0.34 | 1.91E-07 | 3.97E-03 | 0.25 | 0.14 | 0.10 | 1.73 |
| 15 | Ap1s3 | 0.36 | 1.96E-07 | 4.08E-03 | 0.14 | 0.07 | 0.07 | 2.04 |
| 15 | Slc38a7 | 0.38 | 2.02E-07 | 4.20E-03 | 0.16 | 0.08 | 0.08 | 1.93 |
| 15 | Cyb5rl | 0.42 | 2.09E-07 | 4.35E-03 | 0.13 | 0.06 | 0.07 | 2.05 |
| 15 | Lipe | 0.32 | 2.13E-07 | 4.44E-03 | 0.16 | 0.08 | 0.08 | 1.95 |
| 15 | Ap5z1 | 0.35 | 2.25E-07 | 4.69E-03 | 0.11 | 0.05 | 0.06 | 2.18 |
| 15 | Wfs1 | 0.30 | 2.26E-07 | 4.71E-03 | 0.13 | 0.07 | 0.07 | 2.03 |
| 15 | Zbtb4 | 0.37 | 2.28E-07 | 4.75E-03 | 0.13 | 0.07 | 0.07 | 2.03 |
| 15 | Rbak | 0.33 | 2.33E-07 | 4.85E-03 | 0.23 | 0.13 | 0.10 | 1.75 |
| 15 | Khk | 0.47 | 2.34E-07 | 4.86E-03 | 0.13 | 0.07 | 0.07 | 2.00 |
| 15 | Polq | 0.32 | 2.36E-07 | 4.91E-03 | 0.18 | 0.10 | 0.09 | 1.87 |
| 15 | Rcsd1 | 0.28 | 2.39E-07 | 4.96E-03 | 0.17 | 0.09 | 0.08 | 1.87 |
| 15 | Bbs1 | 0.36 | 2.40E-07 | 4.99E-03 | 0.12 | 0.06 | 0.06 | 2.11 |
| 15 | Acvr1 | 0.26 | 2.44E-07 | 5.07E-03 | 0.32 | 0.19 | 0.13 | 1.66 |
| 15 | Dph2 | 0.33 | 2.46E-07 | 5.12E-03 | 0.18 | 0.10 | 0.08 | 1.85 |
| 15 | Ttc30b | 0.32 | 2.58E-07 | 5.36E-03 | 0.16 | 0.08 | 0.08 | 1.94 |
| 15 | Krt1 | 0.48 | 2.62E-07 | 5.44E-03 | 0.11 | 0.05 | 0.06 | 2.14 |
| 15 | 4930452B06Rik | 0.35 | 2.69E-07 | 5.60E-03 | 0.22 | 0.12 | 0.10 | 1.77 |
| 15 | Gbe1 | 0.41 | 2.73E-07 | 5.68E-03 | 0.19 | 0.10 | 0.08 | 1.83 |
| 15 | Scn8a | 0.38 | 2.73E-07 | 5.69E-03 | 0.13 | 0.07 | 0.07 | 2.03 |
| 15 | Bicra | 0.27 | 2.73E-07 | 5.69E-03 | 0.25 | 0.15 | 0.11 | 1.71 |
| 15 | Snx29 | 0.30 | 2.76E-07 | 5.74E-03 | 0.14 | 0.07 | 0.07 | 2.01 |
| 15 | Vps39 | 0.26 | 2.80E-07 | 5.82E-03 | 0.26 | 0.15 | 0.11 | 1.72 |
| 15 | Pcdhgc5 | 0.35 | 2.80E-07 | 5.82E-03 | 0.10 | 0.05 | 0.06 | 2.21 |
| 15 | Fry | 0.35 | 2.92E-07 | 6.07E-03 | 0.12 | 0.06 | 0.06 | 2.11 |
| 15 | Ccdc15 | 0.26 | 2.94E-07 | 6.12E-03 | 0.21 | 0.11 | 0.09 | 1.81 |
| 15 | Dnaaf5 | 0.29 | 2.98E-07 | 6.20E-03 | 0.17 | 0.09 | 0.08 | 1.88 |
| 15 | Dnal1 | 0.36 | 3.02E-07 | 6.28E-03 | 0.16 | 0.09 | 0.08 | 1.91 |

**Supplementary Table 1. Marker genes for the original clusters of integrated clustering of E11.5 control and NotchGOF λ-junction cells.**

| | | | | | | | | |
|---|---|---|---|---|---|---|---|---|
| 15 | F8a | 0.29 | 3.18E-07 | 6.61E-03 | 0.16 | 0.08 | 0.08 | 1.90 |
| 15 | Tctn3 | 0.34 | 3.24E-07 | 6.75E-03 | 0.12 | 0.06 | 0.06 | 2.11 |
| 15 | Wdr24 | 0.27 | 3.30E-07 | 6.87E-03 | 0.21 | 0.12 | 0.09 | 1.79 |
| 15 | Tep1 | 0.37 | 3.35E-07 | 6.97E-03 | 0.12 | 0.06 | 0.06 | 2.07 |
| 15 | Scfd2 | 0.39 | 3.37E-07 | 7.02E-03 | 0.13 | 0.06 | 0.07 | 2.03 |
| 15 | Lrp8 | 0.52 | 3.38E-07 | 7.04E-03 | 0.18 | 0.10 | 0.08 | 1.79 |
| 15 | Rundc1 | 0.30 | 3.45E-07 | 7.18E-03 | 0.15 | 0.08 | 0.07 | 1.94 |
| 15 | Dcxr | 0.25 | 3.63E-07 | 7.54E-03 | 0.18 | 0.10 | 0.08 | 1.85 |
| 15 | Bbs10 | 0.27 | 3.66E-07 | 7.61E-03 | 0.13 | 0.07 | 0.07 | 2.03 |
| 15 | 9030025P20Rik | 0.43 | 3.71E-07 | 7.71E-03 | 0.11 | 0.05 | 0.06 | 2.14 |
| 15 | Aldh3a2 | 0.28 | 4.01E-07 | 8.35E-03 | 0.16 | 0.08 | 0.08 | 1.94 |
| 15 | B4galt5 | 0.33 | 4.04E-07 | 8.40E-03 | 0.19 | 0.11 | 0.09 | 1.81 |
| 15 | Itprip | 0.30 | 4.06E-07 | 8.45E-03 | 0.15 | 0.08 | 0.07 | 1.94 |
| 15 | Zbtb40 | 0.36 | 4.08E-07 | 8.49E-03 | 0.15 | 0.08 | 0.07 | 1.95 |
| 15 | Tle2 | 0.28 | 4.19E-07 | 8.71E-03 | 0.21 | 0.12 | 0.09 | 1.77 |
| 15 | Snx19 | 0.27 | 4.33E-07 | 9.02E-03 | 0.27 | 0.16 | 0.11 | 1.69 |
| 15 | Ehd3 | 0.30 | 4.77E-07 | 9.93E-03 | 0.19 | 0.11 | 0.09 | 1.78 |
| 15 | Lifr | 0.28 | 5.07E-07 | 1.05E-02 | 0.19 | 0.11 | 0.09 | 1.80 |
| 15 | Pcdhgc3 | 0.29 | 5.37E-07 | 1.12E-02 | 0.15 | 0.08 | 0.07 | 1.95 |
| 15 | Dop1b | 0.42 | 5.48E-07 | 1.14E-02 | 0.12 | 0.06 | 0.06 | 2.03 |
| 15 | Per1 | 0.31 | 5.67E-07 | 1.18E-02 | 0.13 | 0.06 | 0.07 | 2.03 |
| 15 | Zfp870 | 0.26 | 5.83E-07 | 1.21E-02 | 0.17 | 0.09 | 0.08 | 1.86 |
| 15 | Pex26 | 0.25 | 5.97E-07 | 1.24E-02 | 0.16 | 0.08 | 0.07 | 1.89 |
| 15 | 4632404H12Rik | 0.44 | 6.08E-07 | 1.26E-02 | 0.10 | 0.05 | 0.05 | 2.17 |
| 15 | Frs3 | 0.27 | 6.10E-07 | 1.27E-02 | 0.10 | 0.05 | 0.06 | 2.17 |
| 15 | Mapk11 | 0.43 | 6.24E-07 | 1.30E-02 | 0.14 | 0.07 | 0.07 | 1.96 |
| 15 | Spata2 | 0.27 | 6.61E-07 | 1.37E-02 | 0.21 | 0.12 | 0.09 | 1.76 |
| 15 | Thnsl1 | 0.32 | 6.63E-07 | 1.38E-02 | 0.14 | 0.07 | 0.07 | 1.96 |
| 15 | Gm17435 | 0.33 | 6.75E-07 | 1.40E-02 | 0.14 | 0.07 | 0.07 | 1.96 |
| 15 | Gtse1 | 0.30 | 6.95E-07 | 1.45E-02 | 0.29 | 0.18 | 0.11 | 1.62 |
| 15 | Gm38115 | 0.43 | 7.03E-07 | 1.46E-02 | 0.12 | 0.06 | 0.06 | 2.03 |
| 15 | Ankrd45 | 0.34 | 7.45E-07 | 1.55E-02 | 0.12 | 0.06 | 0.06 | 2.05 |
| 15 | Fbxl22 | 0.52 | 7.45E-07 | 1.55E-02 | 0.10 | 0.05 | 0.05 | 2.13 |
| 15 | H2-T23 | 0.34 | 7.85E-07 | 1.63E-02 | 0.11 | 0.05 | 0.06 | 2.06 |
| 15 | Ogdhl | 0.36 | 7.97E-07 | 1.66E-02 | 0.14 | 0.07 | 0.07 | 1.96 |
| 15 | Plpp6 | 0.26 | 8.07E-07 | 1.68E-02 | 0.13 | 0.07 | 0.07 | 1.97 |
| 15 | Nudt12 | 0.38 | 8.11E-07 | 1.69E-02 | 0.11 | 0.05 | 0.06 | 2.10 |
| 15 | Dgat2 | 0.40 | 8.15E-07 | 1.69E-02 | 0.12 | 0.06 | 0.06 | 2.03 |
| 15 | Gdpd5 | 0.61 | 8.15E-07 | 1.69E-02 | 0.10 | 0.05 | 0.05 | 2.17 |
| 15 | Enpp4 | 0.31 | 8.17E-07 | 1.70E-02 | 0.13 | 0.06 | 0.06 | 1.98 |
| 15 | Cercam | 0.50 | 8.25E-07 | 1.72E-02 | 0.12 | 0.06 | 0.06 | 2.03 |
| 15 | Sdsl | 0.29 | 8.39E-07 | 1.74E-02 | 0.12 | 0.06 | 0.06 | 2.03 |
| 15 | Ptpn21 | 0.27 | 8.42E-07 | 1.75E-02 | 0.20 | 0.11 | 0.09 | 1.76 |
| 15 | Zscan22 | 0.29 | 9.14E-07 | 1.90E-02 | 0.18 | 0.10 | 0.08 | 1.81 |
| 15 | C9orf72 | 0.29 | 9.40E-07 | 1.96E-02 | 0.20 | 0.11 | 0.09 | 1.77 |
| 15 | Slc29a2 | 0.28 | 9.60E-07 | 2.00E-02 | 0.17 | 0.10 | 0.08 | 1.83 |
| 15 | Pcyt1b | 0.27 | 1.18E-06 | 2.46E-02 | 0.21 | 0.12 | 0.09 | 1.73 |
| 15 | Tcea2 | 0.37 | 1.24E-06 | 2.58E-02 | 0.10 | 0.05 | 0.05 | 2.13 |
| 15 | Slc25a19 | 0.25 | 1.25E-06 | 2.61E-02 | 0.20 | 0.12 | 0.09 | 1.76 |
| 15 | Efcab7 | 0.26 | 1.29E-06 | 2.68E-02 | 0.17 | 0.09 | 0.08 | 1.84 |
| 15 | Stk38l | 0.31 | 1.31E-06 | 2.72E-02 | 0.20 | 0.12 | 0.09 | 1.74 |
| 15 | Xylb | 0.33 | 1.33E-06 | 2.76E-02 | 0.12 | 0.06 | 0.06 | 2.00 |
| 15 | Fam122b | 0.39 | 1.39E-06 | 2.90E-02 | 0.15 | 0.08 | 0.07 | 1.87 |
| 15 | Plekhm3 | 0.28 | 1.41E-06 | 2.93E-02 | 0.14 | 0.07 | 0.07 | 1.93 |
| 15 | Polg2 | 0.29 | 1.50E-06 | 3.12E-02 | 0.17 | 0.10 | 0.08 | 1.81 |
| 15 | Slc7a5 | 0.57 | 1.56E-06 | 3.24E-02 | 0.10 | 0.05 | 0.05 | 2.13 |

**Supplementary Table 1. Marker genes for the original clusters of integrated clustering of E11.5 control and NotchGOF λ-junction cells.**

| | | | | | | | | |
|---|---|---|---|---|---|---|---|---|
| 15 | Zfp202 | 0.49 | 1.71E-06 | 3.55E-02 | 0.10 | 0.05 | 0.05 | 2.08 |
| 15 | Trim68 | 0.29 | 1.71E-06 | 3.56E-02 | 0.17 | 0.09 | 0.08 | 1.84 |
| 15 | Tom1l2 | 0.40 | 1.77E-06 | 3.68E-02 | 0.11 | 0.05 | 0.05 | 2.04 |
| 15 | Mybl1 | 0.32 | 1.81E-06 | 3.77E-02 | 0.12 | 0.06 | 0.06 | 2.00 |
| 15 | Gm16196 | 0.45 | 1.85E-06 | 3.86E-02 | 0.13 | 0.07 | 0.06 | 1.92 |
| 15 | Ctu1 | 0.28 | 2.00E-06 | 4.16E-02 | 0.19 | 0.11 | 0.08 | 1.75 |
| 15 | Lipt1 | 0.31 | 2.10E-06 | 4.37E-02 | 0.12 | 0.06 | 0.06 | 2.04 |
| 15 | Gm2237 | 0.38 | 2.22E-06 | 4.62E-02 | 0.10 | 0.05 | 0.05 | 2.13 |
| 15 | Zfp94 | 0.42 | 2.23E-06 | 4.63E-02 | 0.12 | 0.06 | 0.06 | 1.97 |
| 15 | Slc25a53 | 0.31 | 2.28E-06 | 4.75E-02 | 0.19 | 0.11 | 0.08 | 1.76 |
| 15 | Fancf | 0.25 | 2.30E-06 | 4.78E-02 | 0.23 | 0.14 | 0.09 | 1.68 |
| 16 | Sox2 | 3.80 | 0 | 0 | 0.98 | 0.18 | 0.80 | 5.46 |
| 16 | Six3os1 | 3.32 | 0 | 0 | 0.98 | 0.22 | 0.76 | 4.44 |
| 16 | Six3 | 3.21 | 0 | 0 | 0.98 | 0.23 | 0.76 | 4.32 |
| 16 | Dlx5 | 3.41 | 0 | 0 | 0.91 | 0.21 | 0.70 | 4.41 |
| 16 | Dlx6os1 | 5.79 | 0 | 0 | 0.72 | 0.03 | 0.69 | 24.76 |
| 16 | Sp8 | 3.19 | 0 | 0 | 0.81 | 0.13 | 0.68 | 6.29 |
| 16 | Otx2 | 3.50 | 0 | 0 | 0.69 | 0.09 | 0.60 | 7.51 |
| 16 | Gad2 | 3.57 | 0 | 0 | 0.64 | 0.07 | 0.58 | 9.86 |
| 16 | Six6 | 3.61 | 0 | 0 | 0.59 | 0.06 | 0.53 | 9.69 |
| 16 | Ermn | 4.37 | 0 | 0 | 0.53 | 0.04 | 0.50 | 15.23 |
| 16 | Dync1i1 | 4.19 | 0 | 0 | 0.53 | 0.06 | 0.47 | 9.10 |
| 16 | Syt1 | 4.49 | 0 | 0 | 0.48 | 0.03 | 0.45 | 16.00 |
| 16 | Fezf1 | 4.06 | 0 | 0 | 0.43 | 0.02 | 0.41 | 25.29 |
| 16 | Mir9-3hg | 5.01 | 0 | 0 | 0.42 | 0.02 | 0.40 | 19.86 |
| 16 | Fezf2 | 6.79 | 0 | 0 | 0.23 | 0.00 | 0.23 | 76.67 |
| 16 | C230057A21Rik | 5.41 | 0 | 0 | 0.21 | 0.01 | 0.20 | 30.14 |
| 16 | Slitrk1 | 5.58 | 0 | 0 | 0.19 | 0.01 | 0.18 | 37.40 |
| 16 | Helt | 7.62 | 0 | 0 | 0.11 | 0.00 | 0.11 | 108.00 |
| 16 | Lgr4 | 3.63 | 6.48E-299 | 1.35E-294 | 0.88 | 0.22 | 0.67 | 4.07 |
| 16 | Foxg1 | 2.99 | 3.22E-292 | 6.70E-288 | 0.78 | 0.14 | 0.64 | 5.76 |
| 16 | Krt8 | 2.63 | 3.65E-288 | 7.60E-284 | 0.98 | 0.25 | 0.73 | 3.93 |
| 16 | Hes5 | 3.36 | 6.33E-277 | 1.32E-272 | 0.68 | 0.11 | 0.57 | 6.28 |
| 16 | Dlk1 | 4.08 | 7.66E-275 | 1.59E-270 | 0.99 | 0.41 | 0.58 | 2.41 |
| 16 | Cldn7 | 2.48 | 4.39E-267 | 9.14E-263 | 0.94 | 0.23 | 0.71 | 4.11 |
| 16 | Cldn3 | 2.64 | 1.39E-266 | 2.90E-262 | 0.81 | 0.15 | 0.65 | 5.26 |
| 16 | Hey1 | 3.00 | 2.11E-263 | 4.39E-259 | 0.95 | 0.29 | 0.66 | 3.26 |
| 16 | Cldn6 | 2.46 | 1.49E-259 | 3.10E-255 | 0.93 | 0.22 | 0.71 | 4.23 |
| 16 | Rprm | 3.66 | 6.93E-259 | 1.44E-254 | 0.75 | 0.15 | 0.60 | 4.96 |
| 16 | Mir670hg | 4.45 | 9.43E-259 | 1.96E-254 | 0.25 | 0.01 | 0.24 | 17.93 |
| 16 | Gm35040 | 4.70 | 1.17E-254 | 2.43E-250 | 0.22 | 0.01 | 0.21 | 20.36 |
| 16 | Epcam | 2.53 | 4.54E-245 | 9.44E-241 | 1.00 | 0.34 | 0.66 | 2.97 |
| 16 | Nr2e1 | 3.73 | 4.80E-244 | 9.99E-240 | 0.39 | 0.04 | 0.35 | 10.49 |
| 16 | Krt18 | 2.43 | 5.52E-240 | 1.15E-235 | 0.97 | 0.29 | 0.68 | 3.35 |
| 16 | Spint2 | 2.38 | 4.12E-239 | 8.56E-235 | 0.98 | 0.30 | 0.68 | 3.24 |
| 16 | Sox3 | 3.88 | 3.29E-234 | 6.84E-230 | 0.37 | 0.04 | 0.33 | 10.49 |
| 16 | C130071C03Rik | 3.69 | 2.05E-228 | 4.27E-224 | 0.42 | 0.05 | 0.37 | 8.87 |
| 16 | Gpm6a | 3.78 | 3.66E-228 | 7.61E-224 | 0.40 | 0.04 | 0.36 | 9.33 |
| 16 | Pdgfa | 2.14 | 2.01E-227 | 4.19E-223 | 0.98 | 0.27 | 0.71 | 3.60 |
| 16 | Nckap5 | 3.29 | 1.77E-216 | 3.69E-212 | 0.54 | 0.08 | 0.46 | 6.44 |
| 16 | Socs2 | 2.44 | 1.57E-209 | 3.27E-205 | 0.94 | 0.34 | 0.60 | 2.80 |
| 16 | Espn | 3.01 | 3.41E-208 | 7.10E-204 | 0.59 | 0.10 | 0.48 | 5.75 |
| 16 | Dlx6 | 3.21 | 8.20E-207 | 1.71E-202 | 0.52 | 0.08 | 0.44 | 6.46 |
| 16 | Ezr | 2.16 | 1.55E-197 | 3.22E-193 | 0.94 | 0.31 | 0.63 | 3.03 |
| 16 | Pax6 | 2.56 | 1.65E-196 | 3.43E-192 | 0.63 | 0.12 | 0.51 | 5.23 |
| 16 | Plch1 | 3.13 | 1.01E-195 | 2.11E-191 | 0.54 | 0.09 | 0.45 | 5.79 |

**Supplementary Table 1. Marker genes for the original clusters of integrated clustering of E11.5 control and NotchGOF λ-junction cells.**

| | | | | | | | | |
|---|---|---|---|---|---|---|---|---|
| 16 | Mapk12 | 3.07 | 8.88E-194 | 1.85E-189 | 0.57 | 0.11 | 0.47 | 5.36 |
| 16 | Cldn9 | 3.00 | 3.69E-191 | 7.67E-187 | 0.51 | 0.08 | 0.43 | 6.24 |
| 16 | Wfdc2 | 2.51 | 8.77E-190 | 1.82E-185 | 0.96 | 0.45 | 0.51 | 2.12 |
| 16 | Pitx1 | 2.40 | 9.19E-187 | 1.91E-182 | 0.74 | 0.17 | 0.57 | 4.28 |
| 16 | Fzd8 | 3.15 | 1.17E-182 | 2.43E-178 | 0.63 | 0.14 | 0.49 | 4.52 |
| 16 | Rab25 | 2.37 | 4.10E-182 | 8.52E-178 | 0.69 | 0.15 | 0.53 | 4.48 |
| 16 | Gna14 | 4.62 | 4.64E-181 | 9.65E-177 | 0.14 | 0.01 | 0.13 | 23.33 |
| 16 | Nebl | 2.59 | 1.86E-180 | 3.86E-176 | 0.55 | 0.10 | 0.45 | 5.49 |
| 16 | Nrarp | 2.51 | 3.99E-179 | 8.30E-175 | 0.71 | 0.17 | 0.54 | 4.15 |
| 16 | Slc9a3r1 | 2.34 | 1.19E-178 | 2.47E-174 | 0.81 | 0.23 | 0.58 | 3.50 |
| 16 | Igfbp2 | 2.23 | 1.28E-177 | 2.66E-173 | 1.00 | 0.75 | 0.25 | 1.33 |
| 16 | Scg5 | 3.34 | 2.06E-177 | 4.28E-173 | 0.38 | 0.05 | 0.33 | 7.85 |
| 16 | Sox2ot | 4.54 | 1.69E-173 | 3.51E-169 | 0.21 | 0.01 | 0.19 | 14.71 |
| 16 | Stmn3 | 2.84 | 3.44E-172 | 7.16E-168 | 0.30 | 0.03 | 0.27 | 9.71 |
| 16 | Kif21a | 2.26 | 1.73E-160 | 3.59E-156 | 0.83 | 0.30 | 0.53 | 2.75 |
| 16 | Nav3 | 2.74 | 4.19E-159 | 8.72E-155 | 0.57 | 0.12 | 0.45 | 4.69 |
| 16 | Mest | 1.92 | 4.87E-159 | 1.01E-154 | 1.00 | 0.91 | 0.09 | 1.10 |
| 16 | Mpzl2 | 2.34 | 1.91E-158 | 3.98E-154 | 0.73 | 0.20 | 0.53 | 3.66 |
| 16 | Car2 | 0.51 | 7.25E-158 | 1.51E-153 | 0.56 | 0.11 | 0.45 | 5.13 |
| 16 | Sox21 | 3.61 | 1.34E-157 | 2.79E-153 | 0.28 | 0.03 | 0.25 | 9.55 |
| 16 | Abcg1 | 3.19 | 3.51E-157 | 7.29E-153 | 0.45 | 0.08 | 0.37 | 5.83 |
| 16 | Id4 | 2.42 | 2.46E-156 | 5.11E-152 | 0.82 | 0.30 | 0.52 | 2.74 |
| 16 | Tgfb2 | 1.97 | 6.97E-155 | 1.45E-150 | 0.95 | 0.48 | 0.47 | 1.98 |
| 16 | Fgf17 | 3.87 | 7.84E-155 | 1.63E-150 | 0.32 | 0.04 | 0.28 | 8.13 |
| 16 | Lrrn3 | 3.46 | 3.00E-153 | 6.24E-149 | 0.41 | 0.07 | 0.34 | 6.20 |
| 16 | Hist3h2ba | 1.97 | 1.57E-152 | 3.26E-148 | 0.83 | 0.27 | 0.57 | 3.12 |
| 16 | Nrcam | 3.13 | 1.01E-149 | 2.10E-145 | 0.35 | 0.05 | 0.30 | 7.36 |
| 16 | Meg3 | 2.14 | 9.08E-149 | 1.89E-144 | 0.96 | 0.69 | 0.27 | 1.39 |
| 16 | Pls1 | 2.76 | 1.34E-148 | 2.79E-144 | 0.48 | 0.09 | 0.39 | 5.34 |
| 16 | Tox2 | 3.45 | 1.98E-148 | 4.12E-144 | 0.29 | 0.03 | 0.26 | 8.53 |
| 16 | Igfbp5 | 1.53 | 7.92E-148 | 1.65E-143 | 0.98 | 0.40 | 0.58 | 2.45 |
| 16 | Cystm1 | 2.66 | 1.04E-146 | 2.16E-142 | 0.46 | 0.08 | 0.38 | 5.50 |
| 16 | Cdh2 | 2.26 | 8.55E-146 | 1.78E-141 | 0.88 | 0.42 | 0.46 | 2.11 |
| 16 | Stmn2 | 2.17 | 2.02E-145 | 4.19E-141 | 0.48 | 0.09 | 0.39 | 5.14 |
| 16 | Isl1 | 1.65 | 4.56E-145 | 9.49E-141 | 0.78 | 0.21 | 0.57 | 3.67 |
| 16 | Npdc1 | 2.73 | 4.03E-142 | 8.38E-138 | 0.49 | 0.10 | 0.39 | 4.93 |
| 16 | Nuak2 | 3.22 | 7.57E-140 | 1.57E-135 | 0.28 | 0.03 | 0.25 | 8.29 |
| 16 | Cd9 | 1.96 | 1.21E-137 | 2.51E-133 | 0.81 | 0.29 | 0.52 | 2.77 |
| 16 | Ssx2ip | 2.12 | 6.97E-134 | 1.45E-129 | 0.83 | 0.35 | 0.47 | 2.33 |
| 16 | Shtn1 | 3.17 | 2.10E-131 | 4.38E-127 | 0.33 | 0.05 | 0.28 | 6.81 |
| 16 | Prss8 | 2.66 | 3.98E-131 | 8.28E-127 | 0.46 | 0.09 | 0.37 | 4.99 |
| 16 | Tmem54 | 2.37 | 4.56E-131 | 9.48E-127 | 0.52 | 0.11 | 0.41 | 4.56 |
| 16 | Tmem47 | 2.57 | 6.62E-130 | 1.38E-125 | 0.80 | 0.36 | 0.44 | 2.21 |
| 16 | Khdrbs3 | 1.76 | 1.38E-129 | 2.86E-125 | 0.94 | 0.61 | 0.33 | 1.53 |
| 16 | Galm | 2.51 | 5.32E-127 | 1.11E-122 | 0.65 | 0.20 | 0.45 | 3.21 |
| 16 | Myh14 | 2.88 | 1.25E-124 | 2.61E-120 | 0.41 | 0.08 | 0.33 | 5.34 |
| 16 | Ap1m2 | 2.23 | 3.64E-124 | 7.57E-120 | 0.56 | 0.14 | 0.42 | 4.05 |
| 16 | Crb3 | 2.56 | 7.67E-124 | 1.60E-119 | 0.48 | 0.11 | 0.38 | 4.60 |
| 16 | Bex1 | 1.57 | 3.05E-123 | 6.35E-119 | 0.96 | 0.59 | 0.37 | 1.64 |
| 16 | 3110039M20Rik | 2.48 | 6.52E-122 | 1.36E-117 | 0.40 | 0.07 | 0.32 | 5.50 |
| 16 | Lmo1 | 2.65 | 6.74E-121 | 1.40E-116 | 0.39 | 0.07 | 0.32 | 5.50 |
| 16 | Dpysl5 | 2.41 | 8.11E-117 | 1.69E-112 | 0.46 | 0.10 | 0.36 | 4.55 |
| 16 | Rian | 2.24 | 9.10E-117 | 1.89E-112 | 0.76 | 0.31 | 0.44 | 2.42 |
| 16 | Ccdc88c | 2.21 | 9.59E-116 | 1.99E-111 | 0.57 | 0.15 | 0.42 | 3.79 |
| 16 | Ptprf | 1.52 | 1.22E-115 | 2.54E-111 | 0.95 | 0.64 | 0.32 | 1.50 |
| 16 | Gpc4 | 2.60 | 1.24E-115 | 2.57E-111 | 0.52 | 0.13 | 0.39 | 3.91 |

**Supplementary Table 1. Marker genes for the original clusters of integrated clustering of E11.5 control and NotchGOF λ-junction cells.**

| | | | | | | | | |
|---|---|---|---|---|---|---|---|---|
| 16 | Wwc1 | 2.12 | 2.81E-115 | 5.84E-111 | 0.61 | 0.18 | 0.43 | 3.48 |
| 16 | Lmo4 | 1.51 | 7.09E-110 | 1.48E-105 | 0.98 | 0.81 | 0.17 | 1.21 |
| 16 | Esrp1 | 2.01 | 1.60E-109 | 3.32E-105 | 0.59 | 0.16 | 0.43 | 3.61 |
| 16 | Mecom | 1.86 | 2.05E-109 | 4.27E-105 | 0.86 | 0.40 | 0.46 | 2.14 |
| 16 | Pawr | 2.47 | 2.38E-109 | 4.94E-105 | 0.50 | 0.12 | 0.37 | 4.00 |
| 16 | Mapk13 | 1.94 | 1.02E-108 | 2.12E-104 | 0.64 | 0.19 | 0.44 | 3.28 |
| 16 | Pdgfc | 1.77 | 1.37E-108 | 2.84E-104 | 0.76 | 0.28 | 0.48 | 2.75 |
| 16 | Dsp | 1.20 | 3.61E-108 | 7.51E-104 | 0.75 | 0.23 | 0.52 | 3.24 |
| 16 | Rasef | 3.36 | 1.33E-107 | 2.78E-103 | 0.26 | 0.04 | 0.23 | 6.95 |
| 16 | Cmtm8 | 2.09 | 7.53E-107 | 1.57E-102 | 0.59 | 0.17 | 0.42 | 3.42 |
| 16 | Rragd | 2.26 | 1.48E-106 | 3.08E-102 | 0.64 | 0.22 | 0.42 | 2.93 |
| 16 | Syt5 | 2.91 | 2.33E-106 | 4.85E-102 | 0.32 | 0.06 | 0.26 | 5.80 |
| 16 | Crip2 | 1.48 | 2.49E-106 | 5.19E-102 | 0.58 | 0.17 | 0.42 | 3.49 |
| 16 | Bex4 | 1.42 | 1.23E-105 | 2.55E-101 | 0.93 | 0.65 | 0.28 | 1.43 |
| 16 | Cadps2 | 3.27 | 1.48E-105 | 3.09E-101 | 0.25 | 0.03 | 0.21 | 7.42 |
| 16 | Rhcg | 4.62 | 1.73E-105 | 3.59E-101 | 0.12 | 0.01 | 0.12 | 15.50 |
| 16 | Itga6 | 2.03 | 3.63E-104 | 7.56E-100 | 0.75 | 0.31 | 0.45 | 2.46 |
| 16 | Tubb2b | 1.39 | 5.88E-103 | 1.22E-98 | 0.95 | 0.66 | 0.28 | 1.42 |
| 16 | Emb | 2.09 | 1.95E-102 | 4.06E-98 | 0.81 | 0.40 | 0.41 | 2.01 |
| 16 | Gnai2 | 0.85 | 1.33E-99 | 2.77E-95 | 0.99 | 0.97 | 0.02 | 1.02 |
| 16 | Ikzf2 | 1.62 | 4.71E-99 | 9.79E-95 | 0.75 | 0.28 | 0.47 | 2.70 |
| 16 | Rims2 | 2.81 | 4.86E-99 | 1.01E-94 | 0.31 | 0.06 | 0.25 | 5.62 |
| 16 | Lrp2 | 3.26 | 1.00E-98 | 2.09E-94 | 0.30 | 0.05 | 0.25 | 5.80 |
| 16 | Gad1 | 3.66 | 2.22E-98 | 4.62E-94 | 0.15 | 0.01 | 0.14 | 11.38 |
| 16 | Tpd52 | 1.59 | 4.31E-98 | 8.96E-94 | 0.72 | 0.26 | 0.46 | 2.80 |
| 16 | Stx3 | 2.24 | 1.30E-96 | 2.71E-92 | 0.47 | 0.12 | 0.35 | 3.87 |
| 16 | Frmd3 | 3.62 | 1.10E-95 | 2.29E-91 | 0.17 | 0.02 | 0.15 | 9.56 |
| 16 | Macc1 | 2.89 | 3.17E-95 | 6.60E-91 | 0.30 | 0.05 | 0.25 | 5.62 |
| 16 | Psip1 | 0.92 | 5.47E-95 | 1.14E-90 | 0.99 | 0.98 | 0.01 | 1.01 |
| 16 | Kcnd2 | 3.77 | 2.44E-94 | 5.07E-90 | 0.17 | 0.02 | 0.15 | 9.76 |
| 16 | Lsr | 1.87 | 4.84E-94 | 1.01E-89 | 0.62 | 0.20 | 0.42 | 3.08 |
| 16 | Mgst1 | 1.74 | 1.53E-93 | 3.18E-89 | 0.78 | 0.37 | 0.41 | 2.12 |
| 16 | Ephb1 | 2.35 | 1.82E-93 | 3.79E-89 | 0.41 | 0.10 | 0.31 | 4.31 |
| 16 | Adrb1 | 3.92 | 5.07E-93 | 1.06E-88 | 0.17 | 0.02 | 0.15 | 9.22 |
| 16 | Dmbx1 | 2.99 | 8.03E-93 | 1.67E-88 | 0.26 | 0.04 | 0.22 | 6.32 |
| 16 | Fam107a | 2.98 | 4.17E-92 | 8.67E-88 | 0.26 | 0.04 | 0.22 | 6.32 |
| 16 | Col2a1 | 1.35 | 4.68E-92 | 9.74E-88 | 0.94 | 0.64 | 0.30 | 1.47 |
| 16 | Erbb3 | 2.01 | 2.38E-91 | 4.96E-87 | 0.56 | 0.17 | 0.39 | 3.28 |
| 16 | Nav2 | 1.55 | 1.11E-90 | 2.30E-86 | 0.67 | 0.23 | 0.43 | 2.85 |
| 16 | Ocln | 2.52 | 3.88E-90 | 8.08E-86 | 0.35 | 0.07 | 0.28 | 4.74 |
| 16 | Fgf3 | 4.35 | 6.18E-90 | 1.29E-85 | 0.10 | 0.01 | 0.10 | 14.71 |
| 16 | St14 | 2.24 | 1.44E-89 | 2.99E-85 | 0.45 | 0.12 | 0.33 | 3.88 |
| 16 | Acer2 | 1.58 | 2.50E-89 | 5.20E-85 | 0.81 | 0.39 | 0.42 | 2.07 |
| 16 | Frmd4b | 2.09 | 3.19E-89 | 6.64E-85 | 0.53 | 0.16 | 0.37 | 3.25 |
| 16 | Pcdh11x | 2.24 | 6.66E-89 | 1.39E-84 | 0.42 | 0.10 | 0.32 | 4.08 |
| 16 | Dab1 | 1.97 | 1.04E-88 | 2.17E-84 | 0.52 | 0.15 | 0.37 | 3.46 |
| 16 | Slc22a23 | 2.51 | 2.05E-87 | 4.27E-83 | 0.39 | 0.09 | 0.29 | 4.23 |
| 16 | Dtna | 2.25 | 7.17E-87 | 1.49E-82 | 0.48 | 0.14 | 0.34 | 3.46 |
| 16 | Elavl2 | 2.17 | 1.01E-86 | 2.09E-82 | 0.55 | 0.18 | 0.37 | 3.02 |
| 16 | Grhl2 | 2.02 | 2.16E-86 | 4.50E-82 | 0.46 | 0.13 | 0.34 | 3.68 |
| 16 | Gm16136 | 1.49 | 2.38E-85 | 4.94E-81 | 0.56 | 0.17 | 0.39 | 3.31 |
| 16 | Kcnk1 | 1.98 | 3.80E-85 | 7.91E-81 | 0.56 | 0.18 | 0.38 | 3.09 |
| 16 | Col18a1 | 1.38 | 4.34E-85 | 9.02E-81 | 0.80 | 0.36 | 0.44 | 2.23 |
| 16 | Rtn3 | 1.18 | 1.01E-84 | 2.10E-80 | 0.94 | 0.79 | 0.16 | 1.20 |
| 16 | Phlpp1 | 2.37 | 1.55E-84 | 3.22E-80 | 0.45 | 0.13 | 0.32 | 3.52 |
| 16 | Ano1 | 2.02 | 1.77E-84 | 3.69E-80 | 0.46 | 0.12 | 0.34 | 3.70 |

**Supplementary Table 1. Marker genes for the original clusters of integrated clustering of E11.5 control and NotchGOF λ-junction cells.**

| | | | | | | | | |
|---|---|---|---|---|---|---|---|---|
| 16 | Gm266 | 1.54 | 3.44E-83 | 7.16E-79 | 0.70 | 0.30 | 0.40 | 2.32 |
| 16 | Arhgef19 | 2.03 | 1.15E-82 | 2.39E-78 | 0.46 | 0.13 | 0.33 | 3.61 |
| 16 | Cdh1 | 1.72 | 2.85E-81 | 5.92E-77 | 0.54 | 0.17 | 0.37 | 3.25 |
| 16 | Notch1 | 1.41 | 2.83E-79 | 5.89E-75 | 0.64 | 0.24 | 0.40 | 2.65 |
| 16 | Myo5b | 2.89 | 7.65E-79 | 1.59E-74 | 0.24 | 0.04 | 0.20 | 5.85 |
| 16 | Otx1 | 2.01 | 2.37E-78 | 4.94E-74 | 0.41 | 0.11 | 0.31 | 3.80 |
| 16 | Pbx1 | 0.91 | 6.65E-78 | 1.38E-73 | 0.98 | 0.92 | 0.06 | 1.07 |
| 16 | Stard10 | 1.59 | 3.08E-77 | 6.41E-73 | 0.68 | 0.29 | 0.39 | 2.32 |
| 16 | Slit2 | 1.69 | 4.67E-77 | 9.71E-73 | 0.75 | 0.36 | 0.39 | 2.08 |
| 16 | Pkp2 | 2.80 | 4.64E-76 | 9.65E-72 | 0.27 | 0.05 | 0.22 | 5.22 |
| 16 | Cobl | 2.43 | 1.49E-75 | 3.11E-71 | 0.31 | 0.07 | 0.24 | 4.61 |
| 16 | Pcdh17 | 2.45 | 1.34E-74 | 2.80E-70 | 0.55 | 0.20 | 0.35 | 2.72 |
| 16 | Slc6a15 | 2.95 | 1.16E-73 | 2.42E-69 | 0.20 | 0.03 | 0.17 | 6.60 |
| 16 | Marchf9 | 1.99 | 1.40E-73 | 2.92E-69 | 0.46 | 0.14 | 0.32 | 3.22 |
| 16 | Celf2 | 1.72 | 5.20E-73 | 1.08E-68 | 0.62 | 0.25 | 0.37 | 2.47 |
| 16 | Cyb5a | 0.86 | 5.89E-73 | 1.22E-68 | 0.96 | 0.92 | 0.04 | 1.04 |
| 16 | 2900026A02Rik | 2.29 | 1.53E-72 | 3.19E-68 | 0.38 | 0.10 | 0.28 | 3.77 |
| 16 | Tmem238 | 2.03 | 7.39E-72 | 1.54E-67 | 0.51 | 0.18 | 0.33 | 2.90 |
| 16 | Syt11 | 2.22 | 6.56E-71 | 1.36E-66 | 0.63 | 0.30 | 0.33 | 2.10 |
| 16 | Gipc2 | 2.78 | 1.55E-70 | 3.22E-66 | 0.28 | 0.06 | 0.22 | 4.67 |
| 16 | Adgrg6 | 3.94 | 4.77E-70 | 9.91E-66 | 0.10 | 0.01 | 0.09 | 12.50 |
| 16 | Gucy1a1 | 2.41 | 1.83E-69 | 3.80E-65 | 0.31 | 0.07 | 0.24 | 4.26 |
| 16 | Bcam | 1.48 | 2.86E-69 | 5.95E-65 | 0.57 | 0.20 | 0.37 | 2.86 |
| 16 | Fgd2 | 2.08 | 3.73E-69 | 7.75E-65 | 0.37 | 0.10 | 0.27 | 3.78 |
| 16 | Sorbs2 | 1.75 | 5.66E-69 | 1.18E-64 | 0.57 | 0.22 | 0.35 | 2.63 |
| 16 | Syngr1 | 2.55 | 8.63E-69 | 1.79E-64 | 0.29 | 0.07 | 0.23 | 4.46 |
| 16 | Dpysl3 | 1.92 | 2.15E-68 | 4.48E-64 | 0.75 | 0.45 | 0.30 | 1.67 |
| 16 | Id3 | 0.99 | 5.74E-68 | 1.19E-63 | 0.98 | 0.96 | 0.03 | 1.03 |
| 16 | Clic5 | 3.36 | 6.28E-68 | 1.31E-63 | 0.14 | 0.02 | 0.12 | 8.44 |
| 16 | Grb7 | 2.10 | 1.39E-67 | 2.89E-63 | 0.36 | 0.10 | 0.26 | 3.78 |
| 16 | 2310030G06Rik | 2.35 | 3.54E-66 | 7.36E-62 | 0.29 | 0.07 | 0.23 | 4.37 |
| 16 | Aplp2 | 1.16 | 4.77E-66 | 9.92E-62 | 0.94 | 0.82 | 0.12 | 1.15 |
| 16 | F11r | 1.75 | 5.58E-66 | 1.16E-61 | 0.51 | 0.18 | 0.34 | 2.91 |
| 16 | Fjx1 | 1.53 | 2.50E-65 | 5.20E-61 | 0.73 | 0.39 | 0.34 | 1.89 |
| 16 | Jag1 | 1.68 | 1.23E-64 | 2.55E-60 | 0.50 | 0.17 | 0.33 | 2.88 |
| 16 | Pclo | 1.93 | 6.08E-64 | 1.26E-59 | 0.37 | 0.10 | 0.27 | 3.61 |
| 16 | Ndrg2 | 2.17 | 7.46E-64 | 1.55E-59 | 0.42 | 0.13 | 0.29 | 3.16 |
| 16 | Ralyl | 2.99 | 1.01E-63 | 2.10E-59 | 0.17 | 0.03 | 0.14 | 6.64 |
| 16 | Plp2 | 1.44 | 7.19E-63 | 1.49E-58 | 0.71 | 0.38 | 0.34 | 1.89 |
| 16 | Cgn | 2.25 | 1.44E-62 | 2.99E-58 | 0.36 | 0.10 | 0.26 | 3.53 |
| 16 | Spint1 | 2.06 | 1.84E-62 | 3.83E-58 | 0.35 | 0.10 | 0.26 | 3.65 |
| 16 | Tspan13 | 1.46 | 2.42E-62 | 5.04E-58 | 0.74 | 0.41 | 0.33 | 1.82 |
| 16 | Rassf10 | 2.41 | 2.83E-62 | 5.88E-58 | 0.26 | 0.06 | 0.21 | 4.55 |
| 16 | Sdc4 | 1.54 | 5.59E-62 | 1.16E-57 | 0.68 | 0.34 | 0.34 | 1.99 |
| 16 | D030055H07Rik | 3.17 | 9.60E-62 | 2.00E-57 | 0.14 | 0.02 | 0.12 | 7.94 |
| 16 | Rnf208 | 2.43 | 2.11E-61 | 4.39E-57 | 0.28 | 0.07 | 0.22 | 4.31 |
| 16 | Cbln2 | 2.65 | 2.13E-61 | 4.43E-57 | 0.21 | 0.04 | 0.17 | 5.47 |
| 16 | Aig1 | 1.66 | 2.78E-61 | 5.78E-57 | 0.66 | 0.33 | 0.33 | 2.01 |
| 16 | Cldn4 | 1.89 | 5.60E-61 | 1.16E-56 | 0.36 | 0.10 | 0.26 | 3.57 |
| 16 | Faah | 2.41 | 3.42E-60 | 7.10E-56 | 0.25 | 0.06 | 0.20 | 4.52 |
| 16 | Pea15a | 1.80 | 5.00E-60 | 1.04E-55 | 0.54 | 0.23 | 0.32 | 2.39 |
| 16 | Cdcp1 | 2.43 | 1.15E-59 | 2.40E-55 | 0.25 | 0.06 | 0.20 | 4.52 |
| 16 | Fam214a | 2.38 | 2.06E-59 | 4.29E-55 | 0.34 | 0.10 | 0.24 | 3.54 |
| 16 | Ckmt1 | 1.91 | 6.21E-59 | 1.29E-54 | 0.21 | 0.04 | 0.17 | 5.15 |
| 16 | Ktn1 | 0.92 | 6.60E-59 | 1.37E-54 | 0.95 | 0.82 | 0.12 | 1.15 |
| 16 | Dmrta1 | 2.80 | 1.53E-57 | 3.19E-53 | 0.20 | 0.04 | 0.16 | 5.42 |

**Supplementary Table 1. Marker genes for the original clusters of integrated clustering of E11.5 control and NotchGOF λ-junction cells.**

| | | | | | | | | |
|---|---|---|---|---|---|---|---|---|
| 16 | Hook1 | 1.65 | 1.70E-57 | 3.54E-53 | 0.52 | 0.20 | 0.32 | 2.60 |
| 16 | Ddr1 | 1.89 | 2.76E-57 | 5.74E-53 | 0.40 | 0.13 | 0.27 | 3.13 |
| 16 | Pla2g12a | 1.52 | 6.64E-57 | 1.38E-52 | 0.65 | 0.32 | 0.32 | 1.99 |
| 16 | Itgb6 | 2.66 | 1.11E-56 | 2.30E-52 | 0.21 | 0.04 | 0.17 | 5.15 |
| 16 | Cyb561 | 2.42 | 1.16E-56 | 2.40E-52 | 0.24 | 0.05 | 0.19 | 4.53 |
| 16 | Nectin3 | 1.29 | 3.11E-56 | 6.47E-52 | 0.81 | 0.54 | 0.27 | 1.49 |
| 16 | Sall2 | 1.56 | 3.82E-56 | 7.94E-52 | 0.68 | 0.39 | 0.29 | 1.75 |
| 16 | Nes | 1.42 | 4.99E-56 | 1.04E-51 | 0.73 | 0.42 | 0.31 | 1.72 |
| 16 | Marveld3 | 2.80 | 6.28E-56 | 1.31E-51 | 0.19 | 0.04 | 0.16 | 5.43 |
| 16 | Tcf7l2 | 1.26 | 1.65E-55 | 3.44E-51 | 0.83 | 0.61 | 0.22 | 1.36 |
| 16 | Id1 | 0.93 | 2.12E-55 | 4.42E-51 | 0.97 | 0.86 | 0.11 | 1.13 |
| 16 | Lurap1l | 2.37 | 2.46E-55 | 5.11E-51 | 0.29 | 0.07 | 0.21 | 3.89 |
| 16 | Ehbp1 | 1.64 | 6.55E-55 | 1.36E-50 | 0.54 | 0.24 | 0.31 | 2.31 |
| 16 | Mapk11 | 2.31 | 8.75E-55 | 1.82E-50 | 0.27 | 0.07 | 0.21 | 3.97 |
| 16 | Klk8 | 1.56 | 3.32E-54 | 6.90E-50 | 0.53 | 0.21 | 0.31 | 2.49 |
| 16 | Hist3h2a | 1.79 | 1.45E-53 | 3.01E-49 | 0.56 | 0.26 | 0.30 | 2.13 |
| 16 | Fam187b | 2.62 | 2.16E-53 | 4.50E-49 | 0.19 | 0.04 | 0.15 | 5.28 |
| 16 | Six1 | 1.02 | 3.34E-53 | 6.94E-49 | 0.90 | 0.61 | 0.29 | 1.47 |
| 16 | Uchl1 | 1.30 | 7.52E-53 | 1.56E-48 | 0.67 | 0.35 | 0.32 | 1.93 |
| 16 | Kcnh8 | 2.68 | 1.70E-52 | 3.54E-48 | 0.18 | 0.03 | 0.15 | 5.53 |
| 16 | Dnajc1 | 1.36 | 1.86E-52 | 3.88E-48 | 0.78 | 0.56 | 0.22 | 1.40 |
| 16 | Slc25a13 | 1.30 | 3.63E-52 | 7.54E-48 | 0.72 | 0.44 | 0.28 | 1.63 |
| 16 | Klc3 | 1.78 | 6.98E-52 | 1.45E-47 | 0.35 | 0.11 | 0.24 | 3.30 |
| 16 | Unc13b | 2.27 | 8.33E-52 | 1.73E-47 | 0.29 | 0.08 | 0.21 | 3.62 |
| 16 | AW011738 | 2.07 | 1.28E-51 | 2.67E-47 | 0.31 | 0.09 | 0.22 | 3.52 |
| 16 | Myo5c | 2.91 | 1.37E-51 | 2.85E-47 | 0.19 | 0.04 | 0.15 | 5.05 |
| 16 | Tspan12 | 1.80 | 2.22E-51 | 4.62E-47 | 0.50 | 0.21 | 0.29 | 2.40 |
| 16 | Lratd1 | 2.25 | 2.85E-51 | 5.93E-47 | 0.18 | 0.03 | 0.14 | 5.36 |
| 16 | Sema3d | 2.01 | 8.63E-51 | 1.79E-46 | 0.36 | 0.11 | 0.25 | 3.15 |
| 16 | Ppfia3 | 1.92 | 1.64E-50 | 3.42E-46 | 0.38 | 0.13 | 0.25 | 2.98 |
| 16 | Nectin4 | 1.35 | 1.99E-50 | 4.15E-46 | 0.40 | 0.13 | 0.27 | 3.09 |
| 16 | Ston2 | 1.48 | 2.52E-50 | 5.24E-46 | 0.47 | 0.18 | 0.29 | 2.64 |
| 16 | Ppp1r1a | 2.03 | 2.58E-50 | 5.36E-46 | 0.42 | 0.16 | 0.26 | 2.62 |
| 16 | Map7 | 1.92 | 2.91E-50 | 6.05E-46 | 0.34 | 0.11 | 0.24 | 3.24 |
| 16 | Ormdl2 | 1.45 | 3.35E-50 | 6.97E-46 | 0.64 | 0.35 | 0.30 | 1.86 |
| 16 | Plekha7 | 2.07 | 3.49E-50 | 7.26E-46 | 0.31 | 0.09 | 0.22 | 3.47 |
| 16 | Hsp90ab1 | 0.30 | 1.86E-49 | 3.87E-45 | 1.00 | 1.00 | 0.00 | 1.00 |
| 16 | Kdf1 | 1.98 | 2.38E-49 | 4.95E-45 | 0.30 | 0.08 | 0.21 | 3.57 |
| 16 | Ctnnd2 | 1.89 | 4.55E-49 | 9.46E-45 | 0.36 | 0.12 | 0.24 | 3.06 |
| 16 | Fzd3 | 1.10 | 5.42E-49 | 1.13E-44 | 0.83 | 0.59 | 0.24 | 1.42 |
| 16 | Klf5 | 1.56 | 6.24E-49 | 1.30E-44 | 0.38 | 0.12 | 0.25 | 3.10 |
| 16 | Scrn1 | 2.32 | 1.13E-48 | 2.35E-44 | 0.20 | 0.04 | 0.16 | 4.71 |
| 16 | Ttyh1 | 3.09 | 2.04E-48 | 4.24E-44 | 0.14 | 0.02 | 0.12 | 6.09 |
| 16 | Samd10 | 1.99 | 3.76E-48 | 7.81E-44 | 0.30 | 0.09 | 0.21 | 3.42 |
| 16 | Sfrp1 | 1.28 | 4.18E-48 | 8.69E-44 | 0.84 | 0.62 | 0.22 | 1.35 |
| 16 | Rnh1 | 1.05 | 4.70E-48 | 9.77E-44 | 0.79 | 0.55 | 0.23 | 1.42 |
| 16 | Ildr1 | 2.61 | 2.32E-47 | 4.83E-43 | 0.18 | 0.04 | 0.14 | 4.97 |
| 16 | Wipf3 | 2.32 | 2.46E-47 | 5.11E-43 | 0.24 | 0.06 | 0.18 | 4.05 |
| 16 | Cpm | 1.03 | 1.29E-46 | 2.69E-42 | 0.46 | 0.17 | 0.29 | 2.65 |
| 16 | Lmo2 | 2.21 | 1.69E-46 | 3.51E-42 | 0.36 | 0.12 | 0.24 | 2.91 |
| 16 | Rtn2 | 2.15 | 1.99E-46 | 4.15E-42 | 0.28 | 0.08 | 0.20 | 3.48 |
| 16 | Syndig1 | 3.44 | 2.52E-46 | 5.25E-42 | 0.13 | 0.02 | 0.11 | 6.35 |
| 16 | Atox1 | 0.67 | 5.60E-46 | 1.17E-41 | 0.97 | 0.95 | 0.03 | 1.03 |
| 16 | Cyb5r3 | 0.87 | 7.62E-46 | 1.59E-41 | 0.90 | 0.81 | 0.09 | 1.11 |
| 16 | Crip1 | 1.23 | 9.61E-46 | 2.00E-41 | 0.68 | 0.40 | 0.28 | 1.71 |
| 16 | Chchd10 | 1.77 | 1.00E-45 | 2.08E-41 | 0.42 | 0.16 | 0.26 | 2.68 |

**Supplementary Table 1. Marker genes for the original clusters of integrated clustering of E11.5 control and NotchGOF λ-junction cells.**

| | | | | | | | | |
|---|---|---|---|---|---|---|---|---|
| 16 | Tex15 | 1.54 | 1.36E-45 | 2.83E-41 | 0.34 | 0.11 | 0.23 | 3.13 |
| 16 | Dstn | 0.58 | 2.71E-45 | 5.64E-41 | 0.98 | 0.94 | 0.04 | 1.04 |
| 16 | Trpc1 | 1.97 | 4.43E-45 | 9.21E-41 | 0.33 | 0.11 | 0.22 | 3.02 |
| 16 | Kif5c | 1.28 | 1.55E-44 | 3.23E-40 | 0.72 | 0.48 | 0.24 | 1.49 |
| 16 | Lypd6b | 2.10 | 2.05E-44 | 4.26E-40 | 0.25 | 0.07 | 0.18 | 3.76 |
| 16 | Dock9 | 2.17 | 5.25E-44 | 1.09E-39 | 0.26 | 0.07 | 0.19 | 3.61 |
| 16 | Slain1 | 2.31 | 6.49E-44 | 1.35E-39 | 0.23 | 0.06 | 0.17 | 3.91 |
| 16 | Wdfy1 | 1.42 | 8.96E-44 | 1.86E-39 | 0.63 | 0.35 | 0.28 | 1.80 |
| 16 | Sem1 | 0.47 | 1.08E-43 | 2.26E-39 | 1.00 | 1.00 | 0.01 | 1.01 |
| 16 | Crabp2 | 0.61 | 1.32E-43 | 2.74E-39 | 0.99 | 0.91 | 0.08 | 1.09 |
| 16 | Gnao1 | 2.18 | 1.51E-43 | 3.14E-39 | 0.27 | 0.08 | 0.19 | 3.38 |
| 16 | Epn3 | 2.38 | 1.57E-43 | 3.26E-39 | 0.20 | 0.05 | 0.16 | 4.23 |
| 16 | Dsg2 | 1.59 | 2.58E-43 | 5.36E-39 | 0.36 | 0.13 | 0.24 | 2.87 |
| 16 | Dipk2a | 1.73 | 2.67E-43 | 5.55E-39 | 0.50 | 0.24 | 0.27 | 2.12 |
| 16 | Rundc3a | 2.35 | 2.97E-43 | 6.18E-39 | 0.16 | 0.03 | 0.13 | 5.10 |
| 16 | Smim22 | 2.68 | 3.41E-43 | 7.09E-39 | 0.17 | 0.04 | 0.14 | 4.70 |
| 16 | Pdgfrl | 1.60 | 4.45E-43 | 9.26E-39 | 0.44 | 0.18 | 0.26 | 2.46 |
| 16 | Fam3c | 1.51 | 6.30E-43 | 1.31E-38 | 0.63 | 0.36 | 0.27 | 1.74 |
| 16 | Hdgf | 0.51 | 1.51E-42 | 3.15E-38 | 0.98 | 0.98 | 0.00 | 1.00 |
| 16 | 1500004A13Rik | 2.39 | 1.96E-42 | 4.07E-38 | 0.25 | 0.07 | 0.18 | 3.54 |
| 16 | Rab11fip1 | 1.82 | 3.90E-42 | 8.11E-38 | 0.34 | 0.12 | 0.22 | 2.91 |
| 16 | Dmd | 2.11 | 5.58E-42 | 1.16E-37 | 0.29 | 0.09 | 0.20 | 3.24 |
| 16 | Arrdc1 | 1.93 | 7.25E-42 | 1.51E-37 | 0.32 | 0.11 | 0.21 | 2.95 |
| 16 | Adgrv1 | 2.93 | 8.75E-42 | 1.82E-37 | 0.13 | 0.02 | 0.11 | 5.86 |
| 16 | Xpr1 | 1.72 | 1.90E-41 | 3.95E-37 | 0.53 | 0.27 | 0.26 | 1.96 |
| 16 | Krtcap3 | 2.04 | 1.92E-41 | 4.00E-37 | 0.32 | 0.11 | 0.21 | 2.90 |
| 16 | Cadm1 | 1.09 | 2.67E-41 | 5.54E-37 | 0.68 | 0.38 | 0.29 | 1.77 |
| 16 | Mettl26 | 1.16 | 4.86E-41 | 1.01E-36 | 0.73 | 0.49 | 0.24 | 1.49 |
| 16 | Atp6v0e2 | 2.20 | 1.13E-40 | 2.35E-36 | 0.21 | 0.05 | 0.16 | 4.04 |
| 16 | Nfasc | 2.86 | 1.44E-40 | 2.99E-36 | 0.14 | 0.02 | 0.11 | 5.63 |
| 16 | Serinc2 | 1.21 | 1.45E-40 | 3.02E-36 | 0.67 | 0.39 | 0.28 | 1.74 |
| 16 | Ttc9 | 2.39 | 2.03E-40 | 4.22E-36 | 0.17 | 0.04 | 0.14 | 4.58 |
| 16 | Mip | 2.78 | 2.89E-40 | 6.02E-36 | 0.13 | 0.02 | 0.11 | 5.50 |
| 16 | Sptbn2 | 1.86 | 2.97E-40 | 6.18E-36 | 0.32 | 0.11 | 0.21 | 2.90 |
| 16 | Fam126a | 1.72 | 3.06E-40 | 6.37E-36 | 0.50 | 0.25 | 0.25 | 2.01 |
| 16 | Nrbp2 | 2.64 | 5.99E-40 | 1.25E-35 | 0.15 | 0.03 | 0.12 | 5.10 |
| 16 | Atp2b1 | 0.75 | 6.07E-40 | 1.26E-35 | 0.94 | 0.84 | 0.10 | 1.12 |
| 16 | Mfsd6 | 2.64 | 6.83E-40 | 1.42E-35 | 0.16 | 0.03 | 0.13 | 4.79 |
| 16 | Scube3 | 1.65 | 1.52E-39 | 3.17E-35 | 0.45 | 0.20 | 0.25 | 2.26 |
| 16 | Mapk8ip2 | 2.46 | 1.55E-39 | 3.23E-35 | 0.17 | 0.04 | 0.13 | 4.57 |
| 16 | Bicd1 | 2.15 | 3.35E-39 | 6.97E-35 | 0.29 | 0.10 | 0.19 | 3.02 |
| 16 | Eya1 | 1.11 | 3.48E-39 | 7.23E-35 | 0.74 | 0.50 | 0.24 | 1.49 |
| 16 | Celsr1 | 1.93 | 6.50E-39 | 1.35E-34 | 0.30 | 0.10 | 0.20 | 3.00 |
| 16 | Heyl | 1.97 | 9.24E-39 | 1.92E-34 | 0.21 | 0.06 | 0.16 | 3.89 |
| 16 | Srcin1 | 2.47 | 1.10E-38 | 2.28E-34 | 0.19 | 0.05 | 0.14 | 4.16 |
| 16 | Prr36 | 1.99 | 1.10E-38 | 2.29E-34 | 0.27 | 0.09 | 0.19 | 3.16 |
| 16 | Srgap3 | 1.79 | 1.54E-38 | 3.21E-34 | 0.35 | 0.13 | 0.22 | 2.66 |
| 16 | Zfp219 | 1.20 | 1.85E-38 | 3.85E-34 | 0.65 | 0.40 | 0.25 | 1.63 |
| 16 | Phyhipl | 2.42 | 3.05E-38 | 6.34E-34 | 0.17 | 0.04 | 0.13 | 4.49 |
| 16 | Baiap2l1 | 2.18 | 3.27E-38 | 6.81E-34 | 0.22 | 0.06 | 0.16 | 3.64 |
| 16 | Cep170b | 1.83 | 5.50E-38 | 1.14E-33 | 0.34 | 0.12 | 0.22 | 2.74 |
| 16 | Ago1 | 1.21 | 6.58E-38 | 1.37E-33 | 0.63 | 0.36 | 0.26 | 1.73 |
| 16 | Notch3 | 1.27 | 7.65E-38 | 1.59E-33 | 0.65 | 0.40 | 0.25 | 1.61 |
| 16 | Bnipl | 2.16 | 2.10E-37 | 4.36E-33 | 0.18 | 0.04 | 0.14 | 4.14 |
| 16 | Cxxc5 | 1.19 | 2.45E-37 | 5.10E-33 | 0.69 | 0.45 | 0.24 | 1.54 |
| 16 | A330048O09Rik | 2.77 | 2.61E-37 | 5.44E-33 | 0.10 | 0.02 | 0.09 | 6.44 |

**Supplementary Table 1. Marker genes for the original clusters of integrated clustering of E11.5 control and NotchGOF λ-junction cells.**

| | | | | | | | | |
|---|---|---|---|---|---|---|---|---|
| 16 | Cd151 | 1.28 | 3.06E-37 | 6.36E-33 | 0.59 | 0.33 | 0.27 | 1.81 |
| 16 | Exph5 | 2.31 | 4.62E-37 | 9.61E-33 | 0.16 | 0.03 | 0.12 | 4.65 |
| 16 | Nup210 | 2.42 | 6.08E-37 | 1.27E-32 | 0.21 | 0.06 | 0.16 | 3.63 |
| 16 | Acvr2a | 1.50 | 6.61E-37 | 1.37E-32 | 0.52 | 0.27 | 0.25 | 1.94 |
| 16 | Epb41l5 | 1.26 | 1.53E-36 | 3.19E-32 | 0.51 | 0.25 | 0.26 | 2.04 |
| 16 | Acpp | 3.15 | 2.09E-36 | 4.34E-32 | 0.11 | 0.02 | 0.09 | 6.24 |
| 16 | Hap1 | 2.31 | 2.14E-36 | 4.45E-32 | 0.19 | 0.05 | 0.14 | 3.86 |
| 16 | Misp | 2.92 | 2.15E-36 | 4.47E-32 | 0.11 | 0.02 | 0.09 | 6.24 |
| 16 | Mir99ahg | 1.25 | 3.05E-36 | 6.34E-32 | 0.72 | 0.48 | 0.24 | 1.49 |
| 16 | Apc | 1.02 | 3.37E-36 | 7.01E-32 | 0.79 | 0.61 | 0.18 | 1.29 |
| 16 | Tnfrsf12a | 2.08 | 4.48E-36 | 9.32E-32 | 0.25 | 0.08 | 0.17 | 3.27 |
| 16 | Tnfaip2 | 1.20 | 5.44E-36 | 1.13E-31 | 0.37 | 0.14 | 0.23 | 2.62 |
| 16 | Gsta4 | 0.77 | 5.93E-36 | 1.23E-31 | 0.89 | 0.75 | 0.14 | 1.18 |
| 16 | Map7d2 | 2.00 | 6.42E-36 | 1.33E-31 | 0.26 | 0.08 | 0.18 | 3.16 |
| 16 | Tnk1 | 2.50 | 1.05E-35 | 2.18E-31 | 0.16 | 0.04 | 0.12 | 4.39 |
| 16 | Camsap3 | 1.76 | 1.52E-35 | 3.17E-31 | 0.26 | 0.08 | 0.18 | 3.20 |
| 16 | Plppr1 | 2.44 | 1.73E-35 | 3.61E-31 | 0.18 | 0.04 | 0.13 | 4.12 |
| 16 | Saraf | 1.25 | 1.92E-35 | 3.99E-31 | 0.73 | 0.54 | 0.19 | 1.35 |
| 16 | Hspa4l | 1.86 | 2.41E-35 | 5.01E-31 | 0.30 | 0.11 | 0.19 | 2.78 |
| 16 | Afdn | 0.90 | 3.41E-35 | 7.09E-31 | 0.81 | 0.57 | 0.24 | 1.41 |
| 16 | Sox13 | 2.05 | 4.75E-35 | 9.87E-31 | 0.26 | 0.09 | 0.18 | 3.07 |
| 16 | Nabp2 | 0.90 | 5.29E-35 | 1.10E-30 | 0.80 | 0.65 | 0.15 | 1.23 |
| 16 | Lama5 | 1.83 | 5.49E-35 | 1.14E-30 | 0.32 | 0.11 | 0.20 | 2.78 |
| 16 | Nog | 1.90 | 7.71E-35 | 1.60E-30 | 0.27 | 0.09 | 0.18 | 2.98 |
| 16 | Bend7 | 1.93 | 7.85E-35 | 1.63E-30 | 0.25 | 0.08 | 0.17 | 3.12 |
| 16 | Atp6v1f | 0.58 | 8.82E-35 | 1.83E-30 | 0.95 | 0.90 | 0.04 | 1.05 |
| 16 | Tspan3 | 0.61 | 1.11E-34 | 2.31E-30 | 0.96 | 0.92 | 0.05 | 1.05 |
| 16 | Syt7 | 2.31 | 3.14E-34 | 6.53E-30 | 0.20 | 0.06 | 0.15 | 3.65 |
| 16 | Gm13648 | 2.48 | 4.75E-34 | 9.88E-30 | 0.12 | 0.02 | 0.10 | 5.17 |
| 16 | Fras1 | 1.22 | 5.29E-34 | 1.10E-29 | 0.52 | 0.26 | 0.26 | 1.97 |
| 16 | Frat2 | 1.54 | 6.68E-34 | 1.39E-29 | 0.47 | 0.23 | 0.24 | 2.03 |
| 16 | Sord | 2.23 | 8.51E-34 | 1.77E-29 | 0.34 | 0.14 | 0.20 | 2.40 |
| 16 | Rassf9 | 1.54 | 1.54E-33 | 3.21E-29 | 0.39 | 0.16 | 0.22 | 2.39 |
| 16 | Dag1 | 1.15 | 1.55E-33 | 3.22E-29 | 0.62 | 0.37 | 0.25 | 1.68 |
| 16 | Flrt2 | 0.67 | 1.56E-33 | 3.25E-29 | 0.76 | 0.45 | 0.30 | 1.67 |
| 16 | Pcbp4 | 0.62 | 2.48E-33 | 5.16E-29 | 0.94 | 0.87 | 0.07 | 1.08 |
| 16 | Ldha | 0.52 | 2.57E-33 | 5.35E-29 | 1.00 | 0.97 | 0.03 | 1.03 |
| 16 | Acsl5 | 1.64 | 3.70E-33 | 7.69E-29 | 0.37 | 0.16 | 0.21 | 2.37 |
| 16 | Acvr2b | 0.88 | 5.43E-33 | 1.13E-28 | 0.79 | 0.64 | 0.15 | 1.23 |
| 16 | H2-K1 | 1.68 | 6.80E-33 | 1.41E-28 | 0.34 | 0.13 | 0.20 | 2.54 |
| 16 | Ptpru | 1.80 | 9.38E-33 | 1.95E-28 | 0.37 | 0.16 | 0.21 | 2.37 |
| 16 | Schip1 | 1.44 | 2.35E-32 | 4.89E-28 | 0.38 | 0.16 | 0.22 | 2.39 |
| 16 | Tom1l1 | 1.51 | 6.06E-32 | 1.26E-27 | 0.37 | 0.16 | 0.21 | 2.35 |
| 16 | Frem2 | 1.07 | 6.43E-32 | 1.34E-27 | 0.44 | 0.19 | 0.25 | 2.29 |
| 16 | Klhl7 | 1.04 | 7.75E-32 | 1.61E-27 | 0.73 | 0.52 | 0.21 | 1.41 |
| 16 | Eml5 | 1.72 | 8.55E-32 | 1.78E-27 | 0.25 | 0.08 | 0.17 | 3.01 |
| 16 | Srd5a1 | 1.59 | 8.57E-32 | 1.78E-27 | 0.39 | 0.18 | 0.21 | 2.17 |
| 16 | Gm19938 | 2.65 | 1.34E-31 | 2.80E-27 | 0.12 | 0.02 | 0.09 | 5.04 |
| 16 | Marveld2 | 2.09 | 2.10E-31 | 4.37E-27 | 0.20 | 0.06 | 0.14 | 3.47 |
| 16 | Mirg | 2.28 | 2.36E-31 | 4.91E-27 | 0.17 | 0.05 | 0.13 | 3.78 |
| 16 | Cers4 | 1.67 | 2.37E-31 | 4.92E-27 | 0.36 | 0.16 | 0.21 | 2.33 |
| 16 | Nr3c1 | 1.57 | 2.47E-31 | 5.13E-27 | 0.37 | 0.16 | 0.21 | 2.33 |
| 16 | Tnnt1 | 2.04 | 2.83E-31 | 5.88E-27 | 0.19 | 0.05 | 0.14 | 3.60 |
| 16 | Hes6 | 1.43 | 2.84E-31 | 5.90E-27 | 0.57 | 0.33 | 0.23 | 1.71 |
| 16 | Sypl | 1.11 | 3.63E-31 | 7.56E-27 | 0.70 | 0.48 | 0.22 | 1.45 |
| 16 | Pkm | 0.38 | 3.76E-31 | 7.82E-27 | 1.00 | 0.98 | 0.02 | 1.02 |

**Supplementary Table 1. Marker genes for the original clusters of integrated clustering of E11.5 control and NotchGOF λ-junction cells.**

| | | | | | | | | |
|---|---|---|---|---|---|---|---|---|
| 16 | Wnt4 | 0.95 | 4.14E-31 | 8.61E-27 | 0.45 | 0.20 | 0.25 | 2.26 |
| 16 | Oxtr | 1.98 | 5.09E-31 | 1.06E-26 | 0.22 | 0.07 | 0.15 | 3.27 |
| 16 | Galnt2 | 1.17 | 5.71E-31 | 1.19E-26 | 0.67 | 0.46 | 0.21 | 1.46 |
| 16 | Tctex1d1 | 2.31 | 5.87E-31 | 1.22E-26 | 0.15 | 0.03 | 0.11 | 4.26 |
| 16 | Wdr66 | 1.94 | 6.94E-31 | 1.44E-26 | 0.10 | 0.02 | 0.08 | 5.56 |
| 16 | Nrxn1 | 2.01 | 7.41E-31 | 1.54E-26 | 0.12 | 0.03 | 0.10 | 4.84 |
| 16 | Cadm4 | 2.07 | 7.45E-31 | 1.55E-26 | 0.25 | 0.08 | 0.16 | 2.95 |
| 16 | A930004D18Rik | 2.25 | 9.62E-31 | 2.00E-26 | 0.18 | 0.05 | 0.13 | 3.69 |
| 16 | Lmo3 | 2.36 | 1.14E-30 | 2.37E-26 | 0.16 | 0.04 | 0.12 | 4.00 |
| 16 | Mpped1 | 2.27 | 1.44E-30 | 3.00E-26 | 0.16 | 0.04 | 0.12 | 4.00 |
| 16 | Kazald1 | 2.48 | 1.46E-30 | 3.03E-26 | 0.13 | 0.03 | 0.10 | 4.70 |
| 16 | Zswim5 | 2.49 | 1.61E-30 | 3.35E-26 | 0.14 | 0.03 | 0.11 | 4.42 |
| 16 | Eef2 | 0.33 | 2.08E-30 | 4.32E-26 | 1.00 | 1.00 | 0.00 | 1.00 |
| 16 | Tpi1 | 0.47 | 3.60E-30 | 7.48E-26 | 0.98 | 0.96 | 0.02 | 1.02 |
| 16 | Tmem184a | 2.18 | 4.59E-30 | 9.55E-26 | 0.18 | 0.05 | 0.13 | 3.57 |
| 16 | Tubb4a | 2.28 | 6.75E-30 | 1.40E-25 | 0.15 | 0.04 | 0.11 | 4.11 |
| 16 | Nhs | 1.13 | 9.70E-30 | 2.02E-25 | 0.56 | 0.34 | 0.22 | 1.66 |
| 16 | Arhgef5 | 1.26 | 1.03E-29 | 2.14E-25 | 0.42 | 0.20 | 0.23 | 2.14 |
| 16 | Gm2464 | 2.60 | 1.08E-29 | 2.24E-25 | 0.12 | 0.03 | 0.10 | 4.59 |
| 16 | Hes1 | 0.96 | 1.17E-29 | 2.42E-25 | 0.75 | 0.54 | 0.20 | 1.37 |
| 16 | Smtnl2 | 1.69 | 3.23E-29 | 6.71E-25 | 0.23 | 0.07 | 0.15 | 3.11 |
| 16 | Mycl | 1.09 | 4.25E-29 | 8.83E-25 | 0.54 | 0.30 | 0.24 | 1.80 |
| 16 | Bex2 | 0.63 | 7.17E-29 | 1.49E-24 | 0.92 | 0.85 | 0.07 | 1.08 |
| 16 | Dhx32 | 1.59 | 7.97E-29 | 1.66E-24 | 0.42 | 0.22 | 0.20 | 1.93 |
| 16 | Dmrt3 | 2.01 | 1.17E-28 | 2.43E-24 | 0.18 | 0.05 | 0.13 | 3.43 |
| 16 | Ctnnb1 | 0.44 | 1.29E-28 | 2.69E-24 | 0.99 | 0.97 | 0.02 | 1.02 |
| 16 | Igf2 | 0.77 | 1.35E-28 | 2.81E-24 | 0.81 | 0.58 | 0.22 | 1.39 |
| 16 | Gkap1 | 1.02 | 4.06E-28 | 8.43E-24 | 0.66 | 0.43 | 0.23 | 1.53 |
| 16 | Auts2 | 0.63 | 5.00E-28 | 1.04E-23 | 0.95 | 0.87 | 0.08 | 1.10 |
| 16 | Synrg | 1.32 | 5.53E-28 | 1.15E-23 | 0.45 | 0.23 | 0.22 | 1.95 |
| 16 | Ugcg | 1.17 | 6.13E-28 | 1.27E-23 | 0.58 | 0.37 | 0.21 | 1.58 |
| 16 | Mtch1 | 0.41 | 6.70E-28 | 1.39E-23 | 0.98 | 0.96 | 0.02 | 1.02 |
| 16 | Zfhx2 | 1.29 | 7.92E-28 | 1.65E-23 | 0.47 | 0.25 | 0.22 | 1.89 |
| 16 | Palm3 | 1.44 | 7.99E-28 | 1.66E-23 | 0.34 | 0.15 | 0.19 | 2.33 |
| 16 | Retreg1 | 1.67 | 1.14E-27 | 2.37E-23 | 0.25 | 0.09 | 0.16 | 2.78 |
| 16 | Mfng | 1.36 | 1.20E-27 | 2.49E-23 | 0.16 | 0.04 | 0.12 | 3.73 |
| 16 | Sparc | 0.72 | 1.23E-27 | 2.56E-23 | 0.88 | 0.74 | 0.14 | 1.19 |
| 16 | AI463170 | 2.68 | 1.34E-27 | 2.79E-23 | 0.11 | 0.02 | 0.09 | 5.05 |
| 16 | Pla2r1 | 1.76 | 1.43E-27 | 2.97E-23 | 0.22 | 0.07 | 0.15 | 3.00 |
| 16 | Sinhcaf | 0.56 | 1.47E-27 | 3.06E-23 | 0.94 | 0.84 | 0.10 | 1.12 |
| 16 | Vwa2 | 2.22 | 1.60E-27 | 3.32E-23 | 0.14 | 0.04 | 0.11 | 4.00 |
| 16 | 2610528J11Rik | 1.95 | 1.70E-27 | 3.53E-23 | 0.20 | 0.06 | 0.13 | 3.20 |
| 16 | Ctnnal1 | 1.28 | 2.78E-27 | 5.77E-23 | 0.48 | 0.26 | 0.22 | 1.84 |
| 16 | Hs6st1 | 1.54 | 3.84E-27 | 7.99E-23 | 0.36 | 0.17 | 0.20 | 2.15 |
| 16 | Slc35f1 | 2.56 | 5.33E-27 | 1.11E-22 | 0.13 | 0.03 | 0.10 | 4.13 |
| 16 | Pcdh7 | 0.90 | 5.92E-27 | 1.23E-22 | 0.64 | 0.39 | 0.25 | 1.64 |
| 16 | Wasl | 0.94 | 6.36E-27 | 1.32E-22 | 0.72 | 0.56 | 0.17 | 1.30 |
| 16 | Mdh1 | 0.48 | 6.60E-27 | 1.37E-22 | 0.96 | 0.95 | 0.02 | 1.02 |
| 16 | Clic6 | 1.95 | 9.24E-27 | 1.92E-22 | 0.18 | 0.05 | 0.13 | 3.38 |
| 16 | Pcgf5 | 1.69 | 9.30E-27 | 1.93E-22 | 0.24 | 0.09 | 0.15 | 2.79 |
| 16 | Dach1 | 1.00 | 1.02E-26 | 2.12E-22 | 0.60 | 0.38 | 0.21 | 1.56 |
| 16 | Myl12b | 0.51 | 1.05E-26 | 2.18E-22 | 0.93 | 0.89 | 0.05 | 1.05 |
| 16 | Apoc1 | 1.43 | 1.06E-26 | 2.20E-22 | 0.24 | 0.08 | 0.15 | 2.82 |
| 16 | Vill | 1.85 | 1.19E-26 | 2.48E-22 | 0.23 | 0.08 | 0.15 | 2.88 |
| 16 | Igfbp3 | 0.77 | 1.25E-26 | 2.60E-22 | 0.31 | 0.12 | 0.19 | 2.51 |
| 16 | Mtus1 | 1.58 | 1.26E-26 | 2.62E-22 | 0.27 | 0.10 | 0.17 | 2.61 |

**Supplementary Table 1. Marker genes for the original clusters of integrated clustering of E11.5 control and NotchGOF λ-junction cells.**

| | | | | | | | | |
|---|---|---|---|---|---|---|---|---|
| 16 | Satb1 | 1.09 | 1.52E-26 | 3.17E-22 | 0.59 | 0.39 | 0.21 | 1.54 |
| 16 | Upk3bl | 1.49 | 1.78E-26 | 3.69E-22 | 0.22 | 0.07 | 0.14 | 3.00 |
| 16 | Msi1 | 1.11 | 2.20E-26 | 4.57E-22 | 0.58 | 0.37 | 0.21 | 1.57 |
| 16 | Sostdc1 | 0.74 | 2.35E-26 | 4.89E-22 | 0.34 | 0.14 | 0.20 | 2.36 |
| 16 | S100a6 | 1.35 | 2.37E-26 | 4.93E-22 | 0.36 | 0.16 | 0.20 | 2.20 |
| 16 | Adgrg1 | 2.15 | 3.47E-26 | 7.21E-22 | 0.15 | 0.04 | 0.11 | 3.66 |
| 16 | Ccdc171 | 1.88 | 8.06E-26 | 1.68E-21 | 0.25 | 0.09 | 0.15 | 2.64 |
| 16 | Mkrn1 | 0.95 | 8.33E-26 | 1.73E-21 | 0.64 | 0.43 | 0.21 | 1.50 |
| 16 | Galnt3 | 2.25 | 9.20E-26 | 1.91E-21 | 0.13 | 0.03 | 0.10 | 4.03 |
| 16 | Ripk4 | 1.47 | 1.00E-25 | 2.09E-21 | 0.22 | 0.07 | 0.14 | 2.96 |
| 16 | Ier5 | 0.76 | 1.10E-25 | 2.28E-21 | 0.79 | 0.65 | 0.14 | 1.22 |
| 16 | Rasal2 | 1.31 | 1.11E-25 | 2.31E-21 | 0.45 | 0.25 | 0.20 | 1.83 |
| 16 | Agrn | 1.10 | 2.16E-25 | 4.50E-21 | 0.59 | 0.39 | 0.20 | 1.52 |
| 16 | Tox3 | 1.22 | 2.20E-25 | 4.57E-21 | 0.44 | 0.23 | 0.21 | 1.93 |
| 16 | Zdhhc2 | 1.41 | 3.59E-25 | 7.46E-21 | 0.30 | 0.13 | 0.18 | 2.37 |
| 16 | Negr1 | 1.81 | 4.21E-25 | 8.76E-21 | 0.17 | 0.05 | 0.12 | 3.37 |
| 16 | Cxadr | 0.87 | 4.48E-25 | 9.32E-21 | 0.66 | 0.46 | 0.20 | 1.44 |
| 16 | Nol7 | 0.56 | 6.74E-25 | 1.40E-20 | 0.89 | 0.88 | 0.02 | 1.02 |
| 16 | Arhgef16 | 1.90 | 7.86E-25 | 1.64E-20 | 0.18 | 0.06 | 0.12 | 3.14 |
| 16 | Mtmr7 | 2.10 | 8.29E-25 | 1.72E-20 | 0.17 | 0.05 | 0.12 | 3.32 |
| 16 | Cpz | 2.06 | 9.18E-25 | 1.91E-20 | 0.20 | 0.07 | 0.13 | 2.94 |
| 16 | Dsel | 1.28 | 1.10E-24 | 2.28E-20 | 0.53 | 0.33 | 0.20 | 1.61 |
| 16 | Hmgn2 | 0.37 | 1.73E-24 | 3.61E-20 | 0.99 | 0.98 | 0.01 | 1.01 |
| 16 | Ptchd1 | 2.50 | 1.95E-24 | 4.05E-20 | 0.11 | 0.02 | 0.08 | 4.61 |
| 16 | mt-Cytb | 0.35 | 2.14E-24 | 4.44E-20 | 1.00 | 0.99 | 0.01 | 1.01 |
| 16 | Ndfip1 | 0.58 | 3.01E-24 | 6.27E-20 | 0.89 | 0.83 | 0.06 | 1.07 |
| 16 | Mbp | 1.66 | 3.50E-24 | 7.28E-20 | 0.24 | 0.09 | 0.15 | 2.61 |
| 16 | Gas5 | 0.38 | 3.81E-24 | 7.92E-20 | 1.00 | 0.99 | 0.01 | 1.01 |
| 16 | Malat1 | 0.51 | 4.92E-24 | 1.02E-19 | 1.00 | 1.00 | 0.00 | 1.00 |
| 16 | Olfm2 | 1.66 | 5.75E-24 | 1.20E-19 | 0.26 | 0.10 | 0.16 | 2.51 |
| 16 | Fzd6 | 1.40 | 9.59E-24 | 2.00E-19 | 0.23 | 0.08 | 0.15 | 2.74 |
| 16 | Nr2f6 | 0.80 | 1.11E-23 | 2.31E-19 | 0.72 | 0.60 | 0.13 | 1.21 |
| 16 | Cybrd1 | 2.26 | 1.18E-23 | 2.46E-19 | 0.16 | 0.05 | 0.11 | 3.29 |
| 16 | Gatm | 1.50 | 1.22E-23 | 2.53E-19 | 0.14 | 0.04 | 0.10 | 3.64 |
| 16 | Lypd2 | 1.93 | 1.24E-23 | 2.59E-19 | 0.14 | 0.04 | 0.10 | 3.70 |
| 16 | Cdh3 | 1.01 | 1.33E-23 | 2.77E-19 | 0.42 | 0.21 | 0.21 | 2.01 |
| 16 | Plxnb1 | 1.22 | 1.39E-23 | 2.89E-19 | 0.46 | 0.26 | 0.20 | 1.78 |
| 16 | Rcn1 | 0.65 | 1.76E-23 | 3.65E-19 | 0.91 | 0.82 | 0.08 | 1.10 |
| 16 | Egln3 | 1.84 | 2.16E-23 | 4.50E-19 | 0.21 | 0.07 | 0.13 | 2.81 |
| 16 | Clu | 1.42 | 2.37E-23 | 4.94E-19 | 0.39 | 0.21 | 0.19 | 1.90 |
| 16 | Zc3h12c | 1.74 | 2.65E-23 | 5.51E-19 | 0.19 | 0.06 | 0.12 | 3.03 |
| 16 | Pcbd1 | 1.61 | 3.00E-23 | 6.25E-19 | 0.28 | 0.12 | 0.16 | 2.33 |
| 16 | Fat1 | 1.03 | 3.99E-23 | 8.29E-19 | 0.68 | 0.51 | 0.17 | 1.33 |
| 16 | Tjp3 | 1.83 | 4.56E-23 | 9.47E-19 | 0.16 | 0.05 | 0.11 | 3.29 |
| 16 | Tmx2 | 0.92 | 5.04E-23 | 1.05E-18 | 0.66 | 0.49 | 0.17 | 1.35 |
| 16 | Arap2 | 2.02 | 7.09E-23 | 1.47E-18 | 0.17 | 0.05 | 0.12 | 3.13 |
| 16 | Ppp1r9a | 1.15 | 1.06E-22 | 2.20E-18 | 0.59 | 0.40 | 0.20 | 1.50 |
| 16 | Crb2 | 2.60 | 1.75E-22 | 3.64E-18 | 0.11 | 0.03 | 0.08 | 4.24 |
| 16 | Meis2 | 0.36 | 1.79E-22 | 3.72E-18 | 0.96 | 0.67 | 0.30 | 1.44 |
| 16 | Sema6a | 0.97 | 1.90E-22 | 3.95E-18 | 0.64 | 0.44 | 0.19 | 1.44 |
| 16 | H2afj | 0.55 | 2.00E-22 | 4.17E-18 | 0.90 | 0.81 | 0.09 | 1.11 |
| 16 | Stc1 | 1.86 | 2.37E-22 | 4.92E-18 | 0.19 | 0.07 | 0.13 | 2.84 |
| 16 | Zdhhc12 | 1.15 | 2.43E-22 | 5.05E-18 | 0.46 | 0.26 | 0.20 | 1.77 |
| 16 | Pdpn | 1.23 | 2.77E-22 | 5.75E-18 | 0.37 | 0.19 | 0.19 | 2.01 |
| 16 | Igsf9 | 0.92 | 3.66E-22 | 7.62E-18 | 0.60 | 0.41 | 0.20 | 1.49 |
| 16 | Irf2bpl | 1.11 | 3.81E-22 | 7.93E-18 | 0.56 | 0.38 | 0.18 | 1.47 |

**Supplementary Table 1. Marker genes for the original clusters of integrated clustering of E11.5 control and NotchGOF λ-junction cells.**

| | | | | | | | | |
|---|---|---|---|---|---|---|---|---|
| 16 | Gpm6b | 1.77 | 4.01E-22 | 8.35E-18 | 0.19 | 0.07 | 0.12 | 2.88 |
| 16 | H2-Q4 | 1.62 | 4.30E-22 | 8.94E-18 | 0.26 | 0.11 | 0.15 | 2.42 |
| 16 | Gsr | 0.99 | 9.20E-22 | 1.91E-17 | 0.53 | 0.33 | 0.20 | 1.59 |
| 16 | Ust | 1.75 | 1.16E-21 | 2.41E-17 | 0.18 | 0.06 | 0.12 | 2.93 |
| 16 | Nudt4 | 0.28 | 1.37E-21 | 2.84E-17 | 0.86 | 0.77 | 0.09 | 1.11 |
| 16 | C330021F23Rik | 2.18 | 1.40E-21 | 2.92E-17 | 0.12 | 0.03 | 0.09 | 3.87 |
| 16 | Gm15417 | 2.27 | 1.50E-21 | 3.13E-17 | 0.10 | 0.03 | 0.08 | 4.12 |
| 16 | Etl4 | 1.08 | 1.87E-21 | 3.90E-17 | 0.51 | 0.31 | 0.20 | 1.65 |
| 16 | Gpd2 | 1.33 | 1.95E-21 | 4.06E-17 | 0.40 | 0.23 | 0.18 | 1.78 |
| 16 | Nap1l1 | 0.27 | 2.18E-21 | 4.54E-17 | 1.00 | 1.00 | 0.00 | 1.00 |
| 16 | Creld1 | 1.63 | 2.19E-21 | 4.56E-17 | 0.29 | 0.13 | 0.16 | 2.23 |
| 16 | Plch2 | 2.33 | 2.76E-21 | 5.75E-17 | 0.11 | 0.03 | 0.08 | 3.96 |
| 16 | Tceal3 | 1.36 | 3.21E-21 | 6.67E-17 | 0.27 | 0.12 | 0.16 | 2.32 |
| 16 | Ovol2 | 2.12 | 3.55E-21 | 7.38E-17 | 0.11 | 0.03 | 0.08 | 3.96 |
| 16 | B830012L14Rik | 2.05 | 3.56E-21 | 7.40E-17 | 0.16 | 0.05 | 0.11 | 3.09 |
| 16 | Abhd14b | 1.83 | 3.70E-21 | 7.71E-17 | 0.17 | 0.06 | 0.12 | 2.95 |
| 16 | Arl4c | 0.93 | 3.81E-21 | 7.93E-17 | 0.53 | 0.33 | 0.20 | 1.60 |
| 16 | Dnase2a | 1.22 | 4.31E-21 | 8.96E-17 | 0.40 | 0.22 | 0.18 | 1.84 |
| 16 | Mapk8ip1 | 1.16 | 4.47E-21 | 9.30E-17 | 0.44 | 0.26 | 0.19 | 1.72 |
| 16 | Mettl27 | 1.40 | 4.58E-21 | 9.53E-17 | 0.30 | 0.14 | 0.16 | 2.15 |
| 16 | Abca3 | 1.87 | 5.32E-21 | 1.11E-16 | 0.18 | 0.06 | 0.12 | 2.89 |
| 16 | Armc10 | 0.81 | 8.39E-21 | 1.74E-16 | 0.68 | 0.52 | 0.16 | 1.30 |
| 16 | Fgf15 | 2.06 | 8.59E-21 | 1.79E-16 | 0.10 | 0.03 | 0.08 | 4.12 |
| 16 | Tafa2 | 1.71 | 8.85E-21 | 1.84E-16 | 0.14 | 0.04 | 0.10 | 3.43 |
| 16 | Fkbp4 | 0.34 | 8.86E-21 | 1.84E-16 | 0.98 | 0.97 | 0.01 | 1.01 |
| 16 | Btg2 | 1.55 | 9.08E-21 | 1.89E-16 | 0.45 | 0.27 | 0.18 | 1.66 |
| 16 | B230118H07Rik | 0.90 | 9.58E-21 | 1.99E-16 | 0.65 | 0.50 | 0.15 | 1.30 |
| 16 | Acot7 | 1.14 | 1.05E-20 | 2.18E-16 | 0.41 | 0.23 | 0.18 | 1.77 |
| 16 | Ctf1 | 1.32 | 1.29E-20 | 2.68E-16 | 0.33 | 0.16 | 0.17 | 2.02 |
| 16 | Bcar3 | 2.01 | 1.31E-20 | 2.73E-16 | 0.16 | 0.05 | 0.11 | 3.12 |
| 16 | Ccdc160 | 2.31 | 1.46E-20 | 3.04E-16 | 0.11 | 0.03 | 0.08 | 3.86 |
| 16 | Rgs17 | 2.09 | 1.63E-20 | 3.39E-16 | 0.12 | 0.04 | 0.09 | 3.54 |
| 16 | Pnpla2 | 1.58 | 1.69E-20 | 3.51E-16 | 0.22 | 0.09 | 0.13 | 2.52 |
| 16 | Clstn1 | 1.04 | 1.87E-20 | 3.89E-16 | 0.57 | 0.39 | 0.18 | 1.47 |
| 16 | Bst2 | 1.60 | 2.66E-20 | 5.54E-16 | 0.17 | 0.06 | 0.11 | 2.96 |
| 16 | Ntn1 | 2.60 | 3.12E-20 | 6.48E-16 | 0.11 | 0.03 | 0.08 | 3.86 |
| 16 | Itgb8 | 1.52 | 3.32E-20 | 6.90E-16 | 0.29 | 0.13 | 0.15 | 2.18 |
| 16 | Cltb | 1.35 | 3.50E-20 | 7.28E-16 | 0.35 | 0.18 | 0.17 | 1.92 |
| 16 | Col11a1 | 0.91 | 4.02E-20 | 8.37E-16 | 0.61 | 0.43 | 0.18 | 1.43 |
| 16 | Mtcl1 | 1.42 | 4.69E-20 | 9.76E-16 | 0.27 | 0.12 | 0.15 | 2.27 |
| 16 | Epb41l4a | 1.52 | 5.13E-20 | 1.07E-15 | 0.25 | 0.11 | 0.14 | 2.36 |
| 16 | Furin | 1.44 | 6.89E-20 | 1.43E-15 | 0.36 | 0.20 | 0.17 | 1.83 |
| 16 | Elf5 | 1.56 | 7.52E-20 | 1.56E-15 | 0.17 | 0.06 | 0.11 | 2.90 |
| 16 | Pard3b | 1.61 | 8.13E-20 | 1.69E-15 | 0.27 | 0.12 | 0.15 | 2.25 |
| 16 | Gadd45b | 1.66 | 8.91E-20 | 1.85E-15 | 0.30 | 0.14 | 0.16 | 2.11 |
| 16 | Hmgcll1 | 2.33 | 1.07E-19 | 2.22E-15 | 0.12 | 0.03 | 0.09 | 3.61 |
| 16 | Mtln | 1.08 | 1.15E-19 | 2.40E-15 | 0.51 | 0.33 | 0.18 | 1.56 |
| 16 | Lrp11 | 2.10 | 1.54E-19 | 3.20E-15 | 0.12 | 0.03 | 0.09 | 3.56 |
| 16 | Fgd4 | 1.72 | 1.54E-19 | 3.21E-15 | 0.22 | 0.09 | 0.13 | 2.46 |
| 16 | Pgrmc1 | 0.47 | 1.73E-19 | 3.60E-15 | 0.95 | 0.92 | 0.04 | 1.04 |
| 16 | Frrs1 | 1.23 | 1.78E-19 | 3.70E-15 | 0.39 | 0.22 | 0.17 | 1.79 |
| 16 | Cblc | 1.92 | 2.30E-19 | 4.79E-15 | 0.12 | 0.03 | 0.08 | 3.63 |
| 16 | Fermt1 | 0.90 | 2.61E-19 | 5.44E-15 | 0.31 | 0.14 | 0.17 | 2.21 |
| 16 | Epha5 | 1.49 | 3.15E-19 | 6.54E-15 | 0.19 | 0.07 | 0.12 | 2.71 |
| 16 | Cmtm4 | 1.17 | 3.58E-19 | 7.44E-15 | 0.43 | 0.26 | 0.17 | 1.67 |
| 16 | Pgk1 | 0.39 | 4.28E-19 | 8.91E-15 | 0.97 | 0.96 | 0.01 | 1.01 |

**Supplementary Table 1. Marker genes for the original clusters of integrated clustering of E11.5 control and NotchGOF λ-junction cells.**

| | | | | | | | | |
|---|---|---|---|---|---|---|---|---|
| 16 | 4930426D05Rik | 2.34 | 4.30E-19 | 8.94E-15 | 0.11 | 0.03 | 0.08 | 3.53 |
| 16 | Ric3 | 2.06 | 5.82E-19 | 1.21E-14 | 0.15 | 0.05 | 0.10 | 3.00 |
| 16 | Cdc42ep5 | 0.86 | 5.88E-19 | 1.22E-14 | 0.63 | 0.47 | 0.16 | 1.34 |
| 16 | Cd47 | 0.84 | 5.90E-19 | 1.23E-14 | 0.60 | 0.44 | 0.16 | 1.37 |
| 16 | Aktip | 1.08 | 6.11E-19 | 1.27E-14 | 0.46 | 0.29 | 0.18 | 1.61 |
| 16 | Pacsin3 | 1.13 | 6.16E-19 | 1.28E-14 | 0.39 | 0.22 | 0.18 | 1.80 |
| 16 | Slc12a2 | 0.88 | 6.60E-19 | 1.37E-14 | 0.61 | 0.43 | 0.17 | 1.40 |
| 16 | Cbarp | 1.50 | 7.85E-19 | 1.63E-14 | 0.25 | 0.11 | 0.14 | 2.32 |
| 16 | B3gnt7 | 2.12 | 8.30E-19 | 1.73E-14 | 0.14 | 0.04 | 0.09 | 3.21 |
| 16 | H2-T23 | 1.81 | 8.97E-19 | 1.87E-14 | 0.16 | 0.05 | 0.10 | 2.94 |
| 16 | Prr13 | 0.81 | 9.80E-19 | 2.04E-14 | 0.58 | 0.40 | 0.18 | 1.45 |
| 16 | Arhgap8 | 1.85 | 1.02E-18 | 2.12E-14 | 0.14 | 0.04 | 0.09 | 3.21 |
| 16 | Adgrl2 | 0.69 | 1.07E-18 | 2.22E-14 | 0.81 | 0.69 | 0.12 | 1.17 |
| 16 | Celsr2 | 1.36 | 1.11E-18 | 2.31E-14 | 0.22 | 0.09 | 0.13 | 2.46 |
| 16 | Jup | 1.04 | 1.15E-18 | 2.39E-14 | 0.38 | 0.20 | 0.18 | 1.89 |
| 16 | Cpt1a | 1.64 | 1.42E-18 | 2.96E-14 | 0.23 | 0.10 | 0.13 | 2.32 |
| 16 | Eps8l1 | 2.18 | 1.54E-18 | 3.20E-14 | 0.13 | 0.04 | 0.09 | 3.26 |
| 16 | Dennd1c | 1.95 | 1.77E-18 | 3.68E-14 | 0.11 | 0.03 | 0.08 | 3.53 |
| 16 | Bmp4 | 0.71 | 1.98E-18 | 4.12E-14 | 0.69 | 0.51 | 0.18 | 1.35 |
| 16 | Bsg | 0.33 | 2.50E-18 | 5.21E-14 | 0.98 | 0.98 | 0.00 | 1.00 |
| 16 | B3gnt2 | 1.34 | 3.40E-18 | 7.06E-14 | 0.23 | 0.10 | 0.13 | 2.37 |
| 16 | Txnip | 0.96 | 4.17E-18 | 8.68E-14 | 0.53 | 0.34 | 0.19 | 1.54 |
| 16 | Baz2b | 0.66 | 4.67E-18 | 9.72E-14 | 0.81 | 0.71 | 0.10 | 1.14 |
| 16 | Prr15l | 1.95 | 5.56E-18 | 1.16E-13 | 0.12 | 0.04 | 0.08 | 3.40 |
| 16 | Zdbf2 | 0.93 | 5.75E-18 | 1.20E-13 | 0.58 | 0.41 | 0.17 | 1.41 |
| 16 | Pnck | 1.65 | 6.82E-18 | 1.42E-13 | 0.19 | 0.07 | 0.12 | 2.64 |
| 16 | Arl6ip5 | 1.01 | 8.52E-18 | 1.77E-13 | 0.50 | 0.32 | 0.18 | 1.55 |
| 16 | Pard6b | 1.87 | 9.34E-18 | 1.94E-13 | 0.15 | 0.05 | 0.10 | 2.88 |
| 16 | Fut9 | 2.34 | 9.38E-18 | 1.95E-13 | 0.10 | 0.03 | 0.08 | 3.68 |
| 16 | Snx10 | 1.23 | 9.68E-18 | 2.01E-13 | 0.31 | 0.16 | 0.15 | 1.97 |
| 16 | Mrps28 | 0.62 | 1.11E-17 | 2.32E-13 | 0.72 | 0.62 | 0.10 | 1.16 |
| 16 | Adgrb3 | 1.71 | 1.33E-17 | 2.76E-13 | 0.17 | 0.06 | 0.11 | 2.73 |
| 16 | 4930523C07Rik | 1.54 | 1.46E-17 | 3.04E-13 | 0.19 | 0.07 | 0.12 | 2.57 |
| 16 | Scd2 | 0.49 | 1.54E-17 | 3.20E-13 | 0.98 | 0.91 | 0.07 | 1.07 |
| 16 | Ces1d | 2.05 | 1.63E-17 | 3.39E-13 | 0.13 | 0.04 | 0.09 | 3.07 |
| 16 | Arhgap27os2 | 1.64 | 1.66E-17 | 3.46E-13 | 0.17 | 0.07 | 0.11 | 2.68 |
| 16 | Pgam1 | 0.30 | 1.72E-17 | 3.58E-13 | 0.98 | 0.97 | 0.01 | 1.01 |
| 16 | Klf3 | 0.85 | 1.81E-17 | 3.77E-13 | 0.65 | 0.50 | 0.15 | 1.29 |
| 16 | Slc25a23 | 1.26 | 2.26E-17 | 4.69E-13 | 0.31 | 0.16 | 0.15 | 1.96 |
| 16 | Bag2 | 0.85 | 2.90E-17 | 6.03E-13 | 0.57 | 0.40 | 0.17 | 1.42 |
| 16 | 2010320M18Rik | 0.99 | 2.99E-17 | 6.23E-13 | 0.51 | 0.34 | 0.17 | 1.48 |
| 16 | Frmd5 | 1.91 | 3.35E-17 | 6.98E-13 | 0.15 | 0.05 | 0.10 | 2.90 |
| 16 | Hey2 | 1.51 | 4.66E-17 | 9.68E-13 | 0.18 | 0.07 | 0.11 | 2.59 |
| 16 | 2310002F09Rik | 2.04 | 5.83E-17 | 1.21E-12 | 0.12 | 0.04 | 0.08 | 3.22 |
| 16 | Chd7 | 0.65 | 5.90E-17 | 1.23E-12 | 0.77 | 0.66 | 0.11 | 1.17 |
| 16 | Fgfbp3 | 1.58 | 9.08E-17 | 1.89E-12 | 0.22 | 0.10 | 0.13 | 2.26 |
| 16 | Pon2 | 1.23 | 1.07E-16 | 2.22E-12 | 0.34 | 0.18 | 0.16 | 1.87 |
| 16 | Erp29 | 0.40 | 1.15E-16 | 2.39E-12 | 0.91 | 0.88 | 0.02 | 1.03 |
| 16 | Dusp5 | 1.45 | 1.20E-16 | 2.50E-12 | 0.25 | 0.12 | 0.13 | 2.14 |
| 16 | 1500011B03Rik | 0.61 | 1.49E-16 | 3.10E-12 | 0.77 | 0.67 | 0.10 | 1.16 |
| 16 | Ssr4 | 0.47 | 1.61E-16 | 3.34E-12 | 0.89 | 0.84 | 0.05 | 1.06 |
| 16 | Irf2 | 1.11 | 1.80E-16 | 3.73E-12 | 0.42 | 0.25 | 0.16 | 1.64 |
| 16 | S100a11 | 0.39 | 1.80E-16 | 3.75E-12 | 0.84 | 0.76 | 0.08 | 1.11 |
| 16 | Zswim6 | 1.05 | 1.88E-16 | 3.91E-12 | 0.46 | 0.29 | 0.17 | 1.58 |
| 16 | Fgf13 | 1.45 | 2.04E-16 | 4.24E-12 | 0.24 | 0.11 | 0.13 | 2.18 |
| 16 | Svip | 1.30 | 2.06E-16 | 4.28E-12 | 0.21 | 0.09 | 0.12 | 2.32 |

**Supplementary Table 1. Marker genes for the original clusters of integrated clustering of E11.5 control and NotchGOF λ-junction cells.**

| | | | | | | | | |
|---|---|---|---|---|---|---|---|---|
| 16 | Arhgdig | 0.52 | 2.29E-16 | 4.75E-12 | 0.20 | 0.08 | 0.12 | 2.44 |
| 16 | Cd164 | 0.72 | 2.38E-16 | 4.95E-12 | 0.73 | 0.64 | 0.09 | 1.14 |
| 16 | Nudt14 | 0.81 | 2.66E-16 | 5.53E-12 | 0.57 | 0.40 | 0.17 | 1.41 |
| 16 | Ccdc85c | 1.44 | 2.69E-16 | 5.59E-12 | 0.24 | 0.11 | 0.13 | 2.15 |
| 16 | Cox7a2l | 0.32 | 3.22E-16 | 6.71E-12 | 0.96 | 0.96 | 0.01 | 1.01 |
| 16 | Hpn | 1.85 | 3.62E-16 | 7.52E-12 | 0.12 | 0.04 | 0.08 | 3.13 |
| 16 | Igsf9b | 1.66 | 3.72E-16 | 7.74E-12 | 0.15 | 0.06 | 0.10 | 2.73 |
| 16 | Sgce | 1.01 | 4.00E-16 | 8.31E-12 | 0.49 | 0.32 | 0.16 | 1.50 |
| 16 | Myo10 | 0.75 | 4.06E-16 | 8.45E-12 | 0.63 | 0.47 | 0.16 | 1.33 |
| 16 | Ahcyl2 | 1.21 | 4.20E-16 | 8.73E-12 | 0.44 | 0.29 | 0.15 | 1.53 |
| 16 | H2-D1 | 1.00 | 4.84E-16 | 1.01E-11 | 0.40 | 0.24 | 0.16 | 1.66 |
| 16 | Chd3 | 0.46 | 5.29E-16 | 1.10E-11 | 0.94 | 0.90 | 0.05 | 1.05 |
| 16 | Gpc1 | 0.85 | 5.76E-16 | 1.20E-11 | 0.57 | 0.40 | 0.17 | 1.42 |
| 16 | Aftph | 1.07 | 5.81E-16 | 1.21E-11 | 0.40 | 0.25 | 0.15 | 1.62 |
| 16 | Dynll2 | 0.47 | 6.37E-16 | 1.32E-11 | 0.89 | 0.84 | 0.05 | 1.05 |
| 16 | Stxbp2 | 1.24 | 6.66E-16 | 1.39E-11 | 0.27 | 0.14 | 0.14 | 2.01 |
| 16 | Gabarapl2 | 0.43 | 7.11E-16 | 1.48E-11 | 0.86 | 0.82 | 0.04 | 1.04 |
| 16 | Nfe2l3 | 1.83 | 7.30E-16 | 1.52E-11 | 0.11 | 0.03 | 0.08 | 3.26 |
| 16 | Pura | 0.77 | 7.50E-16 | 1.56E-11 | 0.62 | 0.47 | 0.15 | 1.31 |
| 16 | Prrg2 | 1.32 | 8.20E-16 | 1.70E-11 | 0.39 | 0.24 | 0.15 | 1.64 |
| 16 | Plekhf2 | 1.08 | 9.47E-16 | 1.97E-11 | 0.38 | 0.23 | 0.15 | 1.65 |
| 16 | Pdlim1 | 0.42 | 9.72E-16 | 2.02E-11 | 0.48 | 0.29 | 0.19 | 1.65 |
| 16 | Nnat | 0.44 | 1.01E-15 | 2.11E-11 | 0.99 | 0.94 | 0.05 | 1.05 |
| 16 | Pbx3 | 0.68 | 1.01E-15 | 2.11E-11 | 0.67 | 0.53 | 0.14 | 1.27 |
| 16 | C130026L21Rik | 2.08 | 1.05E-15 | 2.19E-11 | 0.12 | 0.04 | 0.08 | 3.05 |
| 16 | Tle5 | 0.38 | 1.12E-15 | 2.34E-11 | 0.98 | 0.94 | 0.04 | 1.05 |
| 16 | Ccdc92 | 1.42 | 1.34E-15 | 2.78E-11 | 0.19 | 0.08 | 0.11 | 2.41 |
| 16 | Dnajc21 | 0.63 | 1.57E-15 | 3.26E-11 | 0.71 | 0.60 | 0.11 | 1.19 |
| 16 | Ralgps1 | 1.99 | 2.26E-15 | 4.71E-11 | 0.13 | 0.04 | 0.08 | 2.95 |
| 16 | Cdca7l | 0.96 | 2.36E-15 | 4.91E-11 | 0.46 | 0.31 | 0.15 | 1.50 |
| 16 | Gpi1 | 0.44 | 2.37E-15 | 4.94E-11 | 0.91 | 0.84 | 0.07 | 1.08 |
| 16 | Pbx4 | 1.38 | 2.76E-15 | 5.75E-11 | 0.20 | 0.09 | 0.11 | 2.31 |
| 16 | Susd4 | 1.57 | 2.93E-15 | 6.09E-11 | 0.21 | 0.09 | 0.12 | 2.29 |
| 16 | Patj | 1.33 | 2.99E-15 | 6.21E-11 | 0.26 | 0.13 | 0.13 | 2.03 |
| 16 | Dock10 | 1.58 | 3.18E-15 | 6.62E-11 | 0.20 | 0.08 | 0.11 | 2.32 |
| 16 | Thbs3 | 1.50 | 3.76E-15 | 7.83E-11 | 0.22 | 0.10 | 0.12 | 2.18 |
| 16 | Klf13 | 0.94 | 4.13E-15 | 8.58E-11 | 0.52 | 0.37 | 0.15 | 1.40 |
| 16 | Grtp1 | 1.33 | 4.20E-15 | 8.73E-11 | 0.27 | 0.14 | 0.13 | 1.95 |
| 16 | Pqlc3 | 1.83 | 7.09E-15 | 1.47E-10 | 0.15 | 0.05 | 0.09 | 2.69 |
| 16 | Bpgm | 0.74 | 7.25E-15 | 1.51E-10 | 0.60 | 0.46 | 0.15 | 1.32 |
| 16 | Tmc4 | 1.74 | 1.03E-14 | 2.13E-10 | 0.21 | 0.10 | 0.11 | 2.18 |
| 16 | Llgl2 | 1.71 | 1.23E-14 | 2.56E-10 | 0.14 | 0.05 | 0.09 | 2.76 |
| 16 | Iqgap1 | 0.71 | 1.24E-14 | 2.58E-10 | 0.62 | 0.47 | 0.15 | 1.32 |
| 16 | Pak1 | 0.70 | 1.30E-14 | 2.70E-10 | 0.62 | 0.49 | 0.13 | 1.26 |
| 16 | Gnptg | 1.16 | 1.31E-14 | 2.72E-10 | 0.35 | 0.20 | 0.15 | 1.72 |
| 16 | Nectin1 | 1.13 | 1.33E-14 | 2.78E-10 | 0.32 | 0.17 | 0.14 | 1.82 |
| 16 | Ppp1r36 | 1.58 | 1.60E-14 | 3.33E-10 | 0.14 | 0.05 | 0.09 | 2.68 |
| 16 | Lgi2 | 1.24 | 1.74E-14 | 3.61E-10 | 0.24 | 0.11 | 0.12 | 2.10 |
| 16 | Prom2 | 1.96 | 1.75E-14 | 3.64E-10 | 0.10 | 0.03 | 0.07 | 3.22 |
| 16 | Wwc2 | 0.65 | 2.22E-14 | 4.61E-10 | 0.70 | 0.58 | 0.12 | 1.21 |
| 16 | Cyfip2 | 1.38 | 2.28E-14 | 4.74E-10 | 0.20 | 0.09 | 0.11 | 2.23 |
| 16 | Kazn | 1.18 | 2.37E-14 | 4.92E-10 | 0.33 | 0.18 | 0.14 | 1.78 |
| 16 | Ncor1 | 0.38 | 2.70E-14 | 5.61E-10 | 0.93 | 0.91 | 0.02 | 1.02 |
| 16 | Clip4 | 1.74 | 2.87E-14 | 5.97E-10 | 0.13 | 0.05 | 0.08 | 2.82 |
| 16 | Slc1a4 | 1.73 | 2.93E-14 | 6.09E-10 | 0.17 | 0.07 | 0.10 | 2.41 |
| 16 | C530008M17Rik | 0.97 | 3.09E-14 | 6.43E-10 | 0.56 | 0.42 | 0.14 | 1.34 |

**Supplementary Table 1. Marker genes for the original clusters of integrated clustering of E11.5 control and NotchGOF λ-junction cells.**

| | | | | | | | | |
|---|---|---|---|---|---|---|---|---|
| 16 | Arid3b | 1.13 | 3.11E-14 | 6.46E-10 | 0.37 | 0.23 | 0.14 | 1.62 |
| 16 | Cacna1b | 1.27 | 3.15E-14 | 6.56E-10 | 0.19 | 0.08 | 0.11 | 2.34 |
| 16 | Ass1 | 1.78 | 3.23E-14 | 6.72E-10 | 0.15 | 0.06 | 0.09 | 2.55 |
| 16 | Cryab | 0.63 | 3.28E-14 | 6.81E-10 | 0.14 | 0.05 | 0.09 | 2.81 |
| 16 | Elf3 | 1.46 | 3.54E-14 | 7.37E-10 | 0.17 | 0.07 | 0.10 | 2.38 |
| 16 | Trim37 | 0.80 | 3.68E-14 | 7.65E-10 | 0.54 | 0.39 | 0.15 | 1.38 |
| 16 | Lrrk1 | 1.18 | 4.33E-14 | 9.00E-10 | 0.29 | 0.15 | 0.14 | 1.90 |
| 16 | Xylt1 | 1.92 | 4.37E-14 | 9.10E-10 | 0.15 | 0.06 | 0.09 | 2.50 |
| 16 | Tedc1 | 1.41 | 4.47E-14 | 9.30E-10 | 0.25 | 0.13 | 0.13 | 1.98 |
| 16 | Gltp | 0.73 | 4.85E-14 | 1.01E-09 | 0.55 | 0.39 | 0.16 | 1.39 |
| 16 | Amt | 1.10 | 5.62E-14 | 1.17E-09 | 0.28 | 0.15 | 0.13 | 1.90 |
| 16 | Adam10 | 0.82 | 5.93E-14 | 1.23E-09 | 0.65 | 0.53 | 0.12 | 1.22 |
| 16 | Ptprk | 0.86 | 6.61E-14 | 1.37E-09 | 0.58 | 0.44 | 0.14 | 1.31 |
| 16 | Jund | 0.38 | 6.63E-14 | 1.38E-09 | 0.97 | 0.94 | 0.03 | 1.03 |
| 16 | Rab15 | 0.97 | 7.26E-14 | 1.51E-09 | 0.26 | 0.13 | 0.13 | 2.00 |
| 16 | Elf1 | 0.93 | 8.25E-14 | 1.72E-09 | 0.42 | 0.26 | 0.16 | 1.60 |
| 16 | Tpp1 | 1.09 | 8.41E-14 | 1.75E-09 | 0.41 | 0.27 | 0.14 | 1.53 |
| 16 | Pard3 | 0.64 | 9.97E-14 | 2.07E-09 | 0.68 | 0.58 | 0.10 | 1.18 |
| 16 | Prkcz | 1.22 | 1.00E-13 | 2.09E-09 | 0.22 | 0.11 | 0.12 | 2.11 |
| 16 | Zfp467 | 0.97 | 1.46E-13 | 3.04E-09 | 0.44 | 0.29 | 0.15 | 1.51 |
| 16 | 1810037I17Rik | 0.45 | 1.59E-13 | 3.32E-09 | 0.86 | 0.84 | 0.03 | 1.03 |
| 16 | Sh3gl2 | 1.57 | 1.60E-13 | 3.32E-09 | 0.11 | 0.04 | 0.07 | 2.90 |
| 16 | Med19 | 0.51 | 1.62E-13 | 3.37E-09 | 0.82 | 0.74 | 0.07 | 1.10 |
| 16 | Pde1b | 1.81 | 1.76E-13 | 3.66E-09 | 0.11 | 0.04 | 0.07 | 3.00 |
| 16 | Ctsd | 0.51 | 1.77E-13 | 3.69E-09 | 0.70 | 0.61 | 0.09 | 1.15 |
| 16 | Gstm1 | 1.04 | 1.81E-13 | 3.76E-09 | 0.39 | 0.25 | 0.15 | 1.60 |
| 16 | Itga3 | 1.49 | 1.93E-13 | 4.02E-09 | 0.16 | 0.07 | 0.10 | 2.41 |
| 16 | Zbtb18 | 1.41 | 1.95E-13 | 4.05E-09 | 0.22 | 0.11 | 0.12 | 2.06 |
| 16 | Ggt7 | 1.83 | 2.02E-13 | 4.20E-09 | 0.10 | 0.03 | 0.07 | 3.13 |
| 16 | Gtf3a | 0.55 | 2.46E-13 | 5.11E-09 | 0.74 | 0.64 | 0.10 | 1.15 |
| 16 | Ppa1 | 0.35 | 2.56E-13 | 5.33E-09 | 0.94 | 0.90 | 0.04 | 1.05 |
| 16 | Cyba | 0.60 | 2.59E-13 | 5.40E-09 | 0.59 | 0.45 | 0.14 | 1.30 |
| 16 | Ptpn3 | 1.71 | 2.91E-13 | 6.05E-09 | 0.11 | 0.04 | 0.07 | 2.92 |
| 16 | Lrrc8d | 1.16 | 3.22E-13 | 6.69E-09 | 0.33 | 0.20 | 0.14 | 1.69 |
| 16 | Gstt3 | 1.72 | 3.35E-13 | 6.96E-09 | 0.13 | 0.05 | 0.08 | 2.69 |
| 16 | Rnf182 | 0.90 | 3.42E-13 | 7.12E-09 | 0.36 | 0.21 | 0.15 | 1.69 |
| 16 | Tead1 | 0.60 | 3.45E-13 | 7.17E-09 | 0.72 | 0.60 | 0.11 | 1.18 |
| 16 | Gm42418 | 0.61 | 3.58E-13 | 7.44E-09 | 1.00 | 1.00 | 0.00 | 1.00 |
| 16 | Glo1 | 0.38 | 3.71E-13 | 7.71E-09 | 0.89 | 0.85 | 0.04 | 1.05 |
| 16 | Phc2 | 0.51 | 3.92E-13 | 8.14E-09 | 0.78 | 0.69 | 0.09 | 1.13 |
| 16 | Tmem178 | 1.38 | 4.56E-13 | 9.48E-09 | 0.10 | 0.03 | 0.07 | 3.13 |
| 16 | Dedd2 | 1.39 | 4.68E-13 | 9.74E-09 | 0.17 | 0.07 | 0.10 | 2.32 |
| 16 | Klf10 | 1.38 | 4.78E-13 | 9.94E-09 | 0.22 | 0.11 | 0.11 | 2.03 |
| 16 | Galnt7 | 0.85 | 4.97E-13 | 1.03E-08 | 0.49 | 0.34 | 0.14 | 1.41 |
| 16 | Npepl1 | 0.86 | 5.10E-13 | 1.06E-08 | 0.46 | 0.33 | 0.13 | 1.41 |
| 16 | Fmc1 | 0.63 | 5.17E-13 | 1.08E-08 | 0.69 | 0.59 | 0.10 | 1.16 |
| 16 | Rfxap | 0.64 | 6.19E-13 | 1.29E-08 | 0.64 | 0.53 | 0.11 | 1.21 |
| 16 | Shroom3 | 1.24 | 7.06E-13 | 1.47E-08 | 0.28 | 0.15 | 0.12 | 1.81 |
| 16 | Aldoa | 0.35 | 7.11E-13 | 1.48E-08 | 0.97 | 0.95 | 0.02 | 1.02 |
| 16 | Rhov | 1.18 | 7.18E-13 | 1.49E-08 | 0.15 | 0.06 | 0.09 | 2.54 |
| 16 | Cux1 | 0.51 | 7.46E-13 | 1.55E-08 | 0.78 | 0.70 | 0.08 | 1.12 |
| 16 | Kif13b | 1.26 | 7.48E-13 | 1.56E-08 | 0.29 | 0.16 | 0.13 | 1.79 |
| 16 | Tmem205 | 0.87 | 7.83E-13 | 1.63E-08 | 0.43 | 0.28 | 0.15 | 1.53 |
| 16 | Ncam1 | 1.08 | 7.96E-13 | 1.66E-08 | 0.48 | 0.36 | 0.12 | 1.34 |
| 16 | Lrrc8b | 1.50 | 8.05E-13 | 1.67E-08 | 0.22 | 0.11 | 0.11 | 2.00 |
| 16 | Sulf1 | 0.44 | 8.47E-13 | 1.76E-08 | 0.70 | 0.54 | 0.16 | 1.30 |

**Supplementary Table 1. Marker genes for the original clusters of integrated clustering of E11.5 control and NotchGOF λ-junction cells.**

| | | | | | | | | |
|---|---|---|---|---|---|---|---|---|
| 16 | Soga3 | 1.33 | 1.05E-12 | 2.17E-08 | 0.14 | 0.05 | 0.08 | 2.60 |
| 16 | Rgs9 | 1.49 | 1.14E-12 | 2.37E-08 | 0.19 | 0.08 | 0.10 | 2.20 |
| 16 | Dgat2 | 1.87 | 1.22E-12 | 2.54E-08 | 0.14 | 0.06 | 0.09 | 2.49 |
| 16 | Eya4 | 0.83 | 1.67E-12 | 3.47E-08 | 0.44 | 0.29 | 0.15 | 1.52 |
| 16 | Palld | 0.56 | 1.73E-12 | 3.60E-08 | 0.73 | 0.62 | 0.10 | 1.16 |
| 16 | Skp1a | 0.29 | 1.93E-12 | 4.01E-08 | 0.97 | 0.96 | 0.01 | 1.01 |
| 16 | Akap9 | 0.51 | 2.01E-12 | 4.18E-08 | 0.86 | 0.78 | 0.07 | 1.09 |
| 16 | Tst | 1.34 | 2.12E-12 | 4.40E-08 | 0.24 | 0.13 | 0.11 | 1.89 |
| 16 | Tspan33 | 0.56 | 2.14E-12 | 4.46E-08 | 0.19 | 0.08 | 0.10 | 2.25 |
| 16 | Crmp1 | 1.02 | 2.69E-12 | 5.60E-08 | 0.24 | 0.12 | 0.11 | 1.94 |
| 16 | Unc5b | 0.65 | 3.12E-12 | 6.48E-08 | 0.55 | 0.39 | 0.16 | 1.40 |
| 16 | Hspa2 | 1.75 | 3.13E-12 | 6.50E-08 | 0.14 | 0.06 | 0.08 | 2.45 |
| 16 | Shisa2 | 1.11 | 3.17E-12 | 6.60E-08 | 0.22 | 0.11 | 0.11 | 2.04 |
| 16 | Dgka | 1.52 | 3.31E-12 | 6.88E-08 | 0.15 | 0.06 | 0.09 | 2.42 |
| 16 | Plcb4 | 1.03 | 3.44E-12 | 7.16E-08 | 0.33 | 0.20 | 0.13 | 1.66 |
| 16 | Etv5 | 1.05 | 3.55E-12 | 7.38E-08 | 0.44 | 0.32 | 0.13 | 1.41 |
| 16 | Scp2 | 0.48 | 4.25E-12 | 8.83E-08 | 0.82 | 0.77 | 0.04 | 1.05 |
| 16 | Esrp2 | 1.34 | 4.69E-12 | 9.76E-08 | 0.15 | 0.07 | 0.09 | 2.35 |
| 16 | Tmem106c | 1.04 | 4.83E-12 | 1.00E-07 | 0.33 | 0.20 | 0.13 | 1.68 |
| 16 | Tm7sf2 | 1.18 | 4.99E-12 | 1.04E-07 | 0.26 | 0.15 | 0.12 | 1.82 |
| 16 | Cdc42se1 | 0.74 | 5.34E-12 | 1.11E-07 | 0.56 | 0.42 | 0.14 | 1.32 |
| 16 | Ptms | 0.27 | 5.59E-12 | 1.16E-07 | 0.98 | 0.95 | 0.03 | 1.03 |
| 16 | Reln | 2.11 | 5.64E-12 | 1.17E-07 | 0.11 | 0.04 | 0.07 | 2.79 |
| 16 | Strbp | 0.48 | 5.80E-12 | 1.21E-07 | 0.79 | 0.71 | 0.09 | 1.12 |
| 16 | Paqr4 | 1.81 | 5.84E-12 | 1.21E-07 | 0.10 | 0.04 | 0.07 | 2.86 |
| 16 | Gm45051 | 1.39 | 6.60E-12 | 1.37E-07 | 0.17 | 0.07 | 0.09 | 2.24 |
| 16 | Phlda1 | 0.84 | 6.93E-12 | 1.44E-07 | 0.55 | 0.42 | 0.14 | 1.33 |
| 16 | Nipal3 | 1.74 | 7.02E-12 | 1.46E-07 | 0.11 | 0.04 | 0.07 | 2.71 |
| 16 | Grip1 | 1.25 | 7.60E-12 | 1.58E-07 | 0.21 | 0.11 | 0.11 | 2.01 |
| 16 | Eya2 | 0.80 | 7.82E-12 | 1.63E-07 | 0.37 | 0.23 | 0.14 | 1.60 |
| 16 | Ifi30 | 0.75 | 7.83E-12 | 1.63E-07 | 0.43 | 0.29 | 0.14 | 1.46 |
| 16 | Camkmt | 1.03 | 7.85E-12 | 1.63E-07 | 0.31 | 0.18 | 0.13 | 1.69 |
| 16 | AC149090.1 | 1.07 | 7.99E-12 | 1.66E-07 | 0.42 | 0.28 | 0.14 | 1.50 |
| 16 | Dnmbp | 1.31 | 8.30E-12 | 1.73E-07 | 0.24 | 0.13 | 0.11 | 1.89 |
| 16 | Cebpb | 0.73 | 8.63E-12 | 1.80E-07 | 0.24 | 0.12 | 0.12 | 1.97 |
| 16 | Olfm1 | 1.10 | 8.96E-12 | 1.86E-07 | 0.36 | 0.23 | 0.13 | 1.57 |
| 16 | Bspry | 1.38 | 9.19E-12 | 1.91E-07 | 0.15 | 0.07 | 0.09 | 2.35 |
| 16 | Pkp3 | 1.21 | 9.24E-12 | 1.92E-07 | 0.16 | 0.07 | 0.09 | 2.36 |
| 16 | Gchfr | 1.23 | 9.42E-12 | 1.96E-07 | 0.21 | 0.11 | 0.11 | 2.00 |
| 16 | Sec22a | 0.93 | 9.51E-12 | 1.98E-07 | 0.43 | 0.29 | 0.14 | 1.47 |
| 16 | Eno1 | 0.29 | 9.59E-12 | 2.00E-07 | 0.98 | 0.97 | 0.01 | 1.01 |
| 16 | Hs6st2 | 0.76 | 9.99E-12 | 2.08E-07 | 0.44 | 0.28 | 0.15 | 1.54 |
| 16 | Arpc5 | 0.33 | 1.03E-11 | 2.14E-07 | 0.82 | 0.76 | 0.05 | 1.07 |
| 16 | Rab28 | 0.65 | 1.03E-11 | 2.15E-07 | 0.61 | 0.50 | 0.11 | 1.21 |
| 16 | Gldc | 1.73 | 1.05E-11 | 2.18E-07 | 0.11 | 0.04 | 0.07 | 2.71 |
| 16 | Tenm3 | 0.47 | 1.12E-11 | 2.33E-07 | 0.73 | 0.64 | 0.10 | 1.15 |
| 16 | Pde5a | 1.24 | 1.18E-11 | 2.46E-07 | 0.30 | 0.18 | 0.12 | 1.66 |
| 16 | Os9 | 0.82 | 1.24E-11 | 2.58E-07 | 0.51 | 0.37 | 0.14 | 1.37 |
| 16 | Card19 | 1.08 | 1.25E-11 | 2.60E-07 | 0.38 | 0.25 | 0.13 | 1.51 |
| 16 | Kxd1 | 0.67 | 1.27E-11 | 2.64E-07 | 0.60 | 0.50 | 0.10 | 1.19 |
| 16 | Kctd6 | 1.14 | 1.40E-11 | 2.90E-07 | 0.28 | 0.16 | 0.12 | 1.78 |
| 16 | Golga4 | 0.54 | 1.41E-11 | 2.94E-07 | 0.75 | 0.64 | 0.10 | 1.16 |
| 16 | Prtg | 1.12 | 1.42E-11 | 2.95E-07 | 0.29 | 0.17 | 0.12 | 1.72 |
| 16 | Rragb | 1.47 | 1.58E-11 | 3.28E-07 | 0.18 | 0.09 | 0.09 | 2.11 |
| 16 | Usp53 | 1.39 | 1.90E-11 | 3.95E-07 | 0.17 | 0.08 | 0.09 | 2.18 |
| 16 | Arf5 | 0.26 | 2.14E-11 | 4.44E-07 | 1.00 | 0.98 | 0.01 | 1.01 |

**Supplementary Table 1. Marker genes for the original clusters of integrated clustering of E11.5 control and NotchGOF λ-junction cells.**

| | | | | | | | | |
|---|---|---|---|---|---|---|---|---|
| 16 | Tmem200a | 1.69 | 2.22E-11 | 4.62E-07 | 0.14 | 0.06 | 0.08 | 2.45 |
| 16 | Cask | 0.81 | 2.25E-11 | 4.67E-07 | 0.47 | 0.34 | 0.13 | 1.39 |
| 16 | Chmp2b | 0.58 | 2.28E-11 | 4.73E-07 | 0.66 | 0.56 | 0.10 | 1.17 |
| 16 | Exosc7 | 0.57 | 2.32E-11 | 4.83E-07 | 0.70 | 0.62 | 0.08 | 1.13 |
| 16 | Dusp14 | 1.54 | 2.50E-11 | 5.21E-07 | 0.13 | 0.05 | 0.08 | 2.49 |
| 16 | Ssh2 | 1.07 | 2.59E-11 | 5.38E-07 | 0.36 | 0.23 | 0.13 | 1.57 |
| 16 | Maf | 1.09 | 2.77E-11 | 5.75E-07 | 0.16 | 0.07 | 0.09 | 2.21 |
| 16 | Arl6ip1 | 0.49 | 3.23E-11 | 6.71E-07 | 0.94 | 0.89 | 0.05 | 1.05 |
| 16 | Cox4i2 | 0.78 | 3.28E-11 | 6.82E-07 | 0.40 | 0.26 | 0.14 | 1.52 |
| 16 | Acp6 | 0.97 | 3.36E-11 | 6.98E-07 | 0.35 | 0.22 | 0.13 | 1.57 |
| 16 | Epha1 | 1.32 | 3.68E-11 | 7.66E-07 | 0.14 | 0.06 | 0.08 | 2.37 |
| 16 | Acsl3 | 0.46 | 4.18E-11 | 8.70E-07 | 0.81 | 0.74 | 0.07 | 1.09 |
| 16 | Tpbg | 0.79 | 4.33E-11 | 9.01E-07 | 0.45 | 0.31 | 0.14 | 1.47 |
| 16 | Rnf24 | 0.90 | 4.45E-11 | 9.25E-07 | 0.43 | 0.30 | 0.14 | 1.46 |
| 16 | Chmp2a | 0.46 | 4.54E-11 | 9.45E-07 | 0.75 | 0.66 | 0.09 | 1.14 |
| 16 | Ccdc115 | 0.62 | 4.61E-11 | 9.58E-07 | 0.61 | 0.49 | 0.12 | 1.24 |
| 16 | Trpm4 | 1.37 | 4.71E-11 | 9.80E-07 | 0.20 | 0.10 | 0.10 | 2.01 |
| 16 | Dtx3l | 1.65 | 5.29E-11 | 1.10E-06 | 0.11 | 0.04 | 0.07 | 2.63 |
| 16 | B4galnt4 | 0.82 | 5.54E-11 | 1.15E-06 | 0.38 | 0.24 | 0.13 | 1.56 |
| 16 | Pgf | 1.18 | 5.54E-11 | 1.15E-06 | 0.16 | 0.07 | 0.09 | 2.26 |
| 16 | Map9 | 1.09 | 5.86E-11 | 1.22E-06 | 0.30 | 0.18 | 0.12 | 1.66 |
| 16 | Syngr2 | 1.01 | 6.07E-11 | 1.26E-06 | 0.26 | 0.14 | 0.12 | 1.82 |
| 16 | Pnp | 0.52 | 6.63E-11 | 1.38E-06 | 0.73 | 0.66 | 0.07 | 1.11 |
| 16 | Spata13 | 1.34 | 6.63E-11 | 1.38E-06 | 0.16 | 0.08 | 0.09 | 2.16 |
| 16 | Srebf1 | 1.04 | 6.63E-11 | 1.38E-06 | 0.31 | 0.19 | 0.12 | 1.64 |
| 16 | Emc10 | 0.48 | 7.21E-11 | 1.50E-06 | 0.77 | 0.75 | 0.02 | 1.03 |
| 16 | Fgf9 | 0.48 | 7.40E-11 | 1.54E-06 | 0.22 | 0.11 | 0.11 | 2.04 |
| 16 | Peg3 | 0.52 | 8.03E-11 | 1.67E-06 | 0.90 | 0.83 | 0.08 | 1.09 |
| 16 | Smim1 | 0.25 | 8.09E-11 | 1.68E-06 | 0.30 | 0.17 | 0.13 | 1.80 |
| 16 | Scamp2 | 0.88 | 8.48E-11 | 1.76E-06 | 0.44 | 0.31 | 0.13 | 1.41 |
| 16 | Pick1 | 0.83 | 8.89E-11 | 1.85E-06 | 0.39 | 0.26 | 0.14 | 1.53 |
| 16 | Ube2b | 0.28 | 9.51E-11 | 1.98E-06 | 0.94 | 0.93 | 0.01 | 1.01 |
| 16 | Marchf2 | 0.76 | 9.65E-11 | 2.01E-06 | 0.62 | 0.51 | 0.11 | 1.21 |
| 16 | Lrig3 | 0.93 | 9.71E-11 | 2.02E-06 | 0.35 | 0.23 | 0.13 | 1.55 |
| 16 | Cd2ap | 0.52 | 9.96E-11 | 2.07E-06 | 0.72 | 0.62 | 0.10 | 1.16 |
| 16 | Dtx4 | 1.78 | 1.21E-10 | 2.52E-06 | 0.11 | 0.04 | 0.07 | 2.65 |
| 16 | Fbxo27 | 1.50 | 1.30E-10 | 2.70E-06 | 0.15 | 0.07 | 0.08 | 2.24 |
| 16 | Slc48a1 | 0.96 | 1.33E-10 | 2.77E-06 | 0.28 | 0.17 | 0.12 | 1.70 |
| 16 | Phtf2 | 1.06 | 1.44E-10 | 2.99E-06 | 0.31 | 0.20 | 0.11 | 1.57 |
| 16 | Eps8l2 | 1.71 | 1.51E-10 | 3.14E-06 | 0.11 | 0.04 | 0.07 | 2.57 |
| 16 | Mboat2 | 0.91 | 1.59E-10 | 3.32E-06 | 0.46 | 0.35 | 0.12 | 1.33 |
| 16 | Dsc2 | 1.16 | 1.67E-10 | 3.48E-06 | 0.14 | 0.06 | 0.08 | 2.37 |
| 16 | Wdsub1 | 0.99 | 1.73E-10 | 3.61E-06 | 0.33 | 0.21 | 0.12 | 1.58 |
| 16 | Vamp8 | 0.47 | 1.94E-10 | 4.04E-06 | 0.70 | 0.61 | 0.09 | 1.15 |
| 16 | Nudt18 | 1.42 | 2.02E-10 | 4.20E-06 | 0.14 | 0.06 | 0.08 | 2.30 |
| 16 | Miga1 | 0.90 | 2.16E-10 | 4.50E-06 | 0.43 | 0.30 | 0.12 | 1.41 |
| 16 | Vsig10 | 1.54 | 2.29E-10 | 4.77E-06 | 0.13 | 0.05 | 0.07 | 2.40 |
| 16 | Mir205hg | 0.80 | 2.37E-10 | 4.93E-06 | 0.23 | 0.12 | 0.11 | 1.89 |
| 16 | Rundc3b | 1.67 | 2.39E-10 | 4.98E-06 | 0.11 | 0.04 | 0.07 | 2.59 |
| 16 | Atp6ap2 | 0.63 | 2.52E-10 | 5.24E-06 | 0.54 | 0.43 | 0.12 | 1.28 |
| 16 | 1700023F06Rik | 1.56 | 2.69E-10 | 5.60E-06 | 0.11 | 0.04 | 0.07 | 2.58 |
| 16 | Cradd | 0.99 | 2.96E-10 | 6.16E-06 | 0.29 | 0.17 | 0.12 | 1.70 |
| 16 | Ext1 | 0.68 | 3.28E-10 | 6.83E-06 | 0.53 | 0.41 | 0.12 | 1.30 |
| 16 | H1fx | 0.34 | 3.41E-10 | 7.09E-06 | 0.97 | 0.93 | 0.04 | 1.04 |
| 16 | Psap | 0.47 | 3.65E-10 | 7.60E-06 | 0.72 | 0.62 | 0.10 | 1.16 |
| 16 | Gdpd1 | 0.86 | 3.81E-10 | 7.92E-06 | 0.39 | 0.27 | 0.12 | 1.46 |

**Supplementary Table 1. Marker genes for the original clusters of integrated clustering of E11.5 control and NotchGOF λ-junction cells.**

| | | | | | | | | |
|---|---|---|---|---|---|---|---|---|
| 16 | Fbxo44 | 1.17 | 3.82E-10 | 7.94E-06 | 0.19 | 0.09 | 0.09 | 1.99 |
| 16 | Rtn1 | 0.53 | 4.01E-10 | 8.33E-06 | 0.38 | 0.26 | 0.12 | 1.46 |
| 16 | Mllt10 | 0.50 | 4.06E-10 | 8.45E-06 | 0.72 | 0.64 | 0.08 | 1.13 |
| 16 | Slc2a1 | 0.63 | 4.08E-10 | 8.49E-06 | 0.55 | 0.43 | 0.13 | 1.30 |
| 16 | Atraid | 0.59 | 4.13E-10 | 8.60E-06 | 0.62 | 0.51 | 0.12 | 1.23 |
| 16 | Apba2 | 1.39 | 4.34E-10 | 9.02E-06 | 0.14 | 0.06 | 0.08 | 2.25 |
| 16 | Tmem128 | 0.49 | 4.52E-10 | 9.40E-06 | 0.72 | 0.63 | 0.09 | 1.13 |
| 16 | Ccdc112 | 0.58 | 4.74E-10 | 9.87E-06 | 0.62 | 0.52 | 0.11 | 1.21 |
| 16 | Myh9 | 0.56 | 5.22E-10 | 1.09E-05 | 0.61 | 0.51 | 0.11 | 1.21 |
| 16 | 1700096K18Rik | 1.26 | 5.30E-10 | 1.10E-05 | 0.19 | 0.10 | 0.09 | 1.97 |
| 16 | Mtss1 | 0.84 | 5.49E-10 | 1.14E-05 | 0.42 | 0.29 | 0.13 | 1.44 |
| 16 | Ide | 0.88 | 5.56E-10 | 1.16E-05 | 0.39 | 0.27 | 0.12 | 1.44 |
| 16 | Ly75 | 1.05 | 5.56E-10 | 1.16E-05 | 0.26 | 0.15 | 0.11 | 1.72 |
| 16 | Stbd1 | 1.31 | 5.91E-10 | 1.23E-05 | 0.20 | 0.11 | 0.10 | 1.91 |
| 16 | Xkr6 | 1.04 | 6.18E-10 | 1.29E-05 | 0.23 | 0.13 | 0.10 | 1.83 |
| 16 | Snrpn | 0.69 | 6.44E-10 | 1.34E-05 | 0.56 | 0.47 | 0.09 | 1.20 |
| 16 | Tshz2 | 0.93 | 6.87E-10 | 1.43E-05 | 0.45 | 0.34 | 0.12 | 1.34 |
| 16 | Cxcr4 | 1.39 | 6.87E-10 | 1.43E-05 | 0.12 | 0.05 | 0.07 | 2.43 |
| 16 | Gm9922 | 1.58 | 7.06E-10 | 1.47E-05 | 0.10 | 0.04 | 0.06 | 2.58 |
| 16 | mt-Nd3 | 0.27 | 7.06E-10 | 1.47E-05 | 0.98 | 0.97 | 0.01 | 1.01 |
| 16 | Thnsl1 | 1.47 | 7.33E-10 | 1.53E-05 | 0.15 | 0.07 | 0.08 | 2.13 |
| 16 | Sema4d | 1.43 | 7.78E-10 | 1.62E-05 | 0.15 | 0.07 | 0.08 | 2.14 |
| 16 | C1qtnf4 | 0.94 | 8.20E-10 | 1.71E-05 | 0.41 | 0.29 | 0.13 | 1.45 |
| 16 | Glrb | 1.50 | 8.40E-10 | 1.75E-05 | 0.12 | 0.05 | 0.07 | 2.38 |
| 16 | Lmo7 | 0.64 | 8.61E-10 | 1.79E-05 | 0.28 | 0.16 | 0.12 | 1.74 |
| 16 | Zfp800 | 0.84 | 8.75E-10 | 1.82E-05 | 0.39 | 0.27 | 0.12 | 1.45 |
| 16 | Klhl5 | 1.07 | 8.82E-10 | 1.83E-05 | 0.28 | 0.17 | 0.11 | 1.65 |
| 16 | Baz2a | 0.70 | 8.89E-10 | 1.85E-05 | 0.50 | 0.39 | 0.12 | 1.31 |
| 16 | Rnf187 | 0.28 | 1.02E-09 | 2.13E-05 | 0.92 | 0.91 | 0.01 | 1.02 |
| 16 | Tmem161b | 1.20 | 1.04E-09 | 2.17E-05 | 0.26 | 0.16 | 0.11 | 1.68 |
| 16 | Slc22a17 | 0.85 | 1.09E-09 | 2.27E-05 | 0.47 | 0.35 | 0.12 | 1.35 |
| 16 | Atp8b1 | 1.32 | 1.14E-09 | 2.36E-05 | 0.11 | 0.04 | 0.06 | 2.52 |
| 16 | Dlx3 | 1.04 | 1.14E-09 | 2.37E-05 | 0.16 | 0.08 | 0.08 | 2.11 |
| 16 | Smim20 | 0.63 | 1.19E-09 | 2.47E-05 | 0.53 | 0.41 | 0.12 | 1.31 |
| 16 | Spred3 | 1.16 | 1.21E-09 | 2.51E-05 | 0.19 | 0.09 | 0.09 | 1.97 |
| 16 | Efcab14 | 0.65 | 1.31E-09 | 2.73E-05 | 0.54 | 0.42 | 0.11 | 1.27 |
| 16 | Cirbp | 0.42 | 1.31E-09 | 2.73E-05 | 0.78 | 0.70 | 0.07 | 1.10 |
| 16 | Pcm1 | 0.36 | 1.32E-09 | 2.75E-05 | 0.92 | 0.90 | 0.03 | 1.03 |
| 16 | Sft2d2 | 1.01 | 1.40E-09 | 2.91E-05 | 0.26 | 0.15 | 0.11 | 1.72 |
| 16 | Ckb | 1.47 | 1.41E-09 | 2.93E-05 | 0.44 | 0.33 | 0.10 | 1.31 |
| 16 | Cacnb3 | 0.69 | 1.43E-09 | 2.98E-05 | 0.40 | 0.27 | 0.13 | 1.47 |
| 16 | Zdhhc21 | 0.85 | 1.44E-09 | 2.99E-05 | 0.40 | 0.29 | 0.11 | 1.39 |
| 16 | Psen2 | 1.33 | 1.65E-09 | 3.44E-05 | 0.16 | 0.08 | 0.08 | 2.05 |
| 16 | Ppm1l | 0.81 | 1.75E-09 | 3.64E-05 | 0.34 | 0.22 | 0.12 | 1.52 |
| 16 | Cenpb | 0.51 | 1.91E-09 | 3.98E-05 | 0.73 | 0.66 | 0.07 | 1.10 |
| 16 | Bend4 | 1.21 | 2.19E-09 | 4.55E-05 | 0.21 | 0.12 | 0.10 | 1.83 |
| 16 | Kcnc3 | 1.68 | 2.25E-09 | 4.69E-05 | 0.12 | 0.05 | 0.07 | 2.28 |
| 16 | Jade1 | 1.05 | 2.32E-09 | 4.83E-05 | 0.28 | 0.17 | 0.11 | 1.63 |
| 16 | Gpr137c | 1.23 | 2.38E-09 | 4.95E-05 | 0.25 | 0.15 | 0.10 | 1.69 |
| 16 | Lamb2 | 1.24 | 2.45E-09 | 5.09E-05 | 0.23 | 0.13 | 0.10 | 1.76 |
| 16 | F2r | 0.56 | 2.55E-09 | 5.31E-05 | 0.67 | 0.57 | 0.10 | 1.17 |
| 16 | Prkg1 | 1.20 | 2.59E-09 | 5.40E-05 | 0.26 | 0.16 | 0.10 | 1.64 |
| 16 | Bcap31 | 0.37 | 2.93E-09 | 6.10E-05 | 0.81 | 0.75 | 0.06 | 1.07 |
| 16 | Slc44a2 | 0.92 | 3.40E-09 | 7.08E-05 | 0.39 | 0.28 | 0.12 | 1.42 |
| 16 | Cpne5 | 1.20 | 3.41E-09 | 7.09E-05 | 0.12 | 0.05 | 0.07 | 2.28 |
| 16 | Eef2kmt | 1.08 | 3.41E-09 | 7.09E-05 | 0.29 | 0.18 | 0.11 | 1.59 |

**Supplementary Table 1. Marker genes for the original clusters of integrated clustering of E11.5 control and NotchGOF λ-junction cells.**

| | | | | | | | | |
|---|---|---|---|---|---|---|---|---|
| 16 | Lrrc42 | 0.45 | 3.44E-09 | 7.15E-05 | 0.70 | 0.61 | 0.09 | 1.15 |
| 16 | Ly6h | 0.49 | 3.48E-09 | 7.24E-05 | 0.26 | 0.15 | 0.11 | 1.75 |
| 16 | Dvl1 | 1.04 | 3.50E-09 | 7.29E-05 | 0.23 | 0.13 | 0.10 | 1.76 |
| 16 | Ssbp2 | 0.49 | 3.68E-09 | 7.66E-05 | 0.78 | 0.71 | 0.07 | 1.10 |
| 16 | Gclm | 0.27 | 3.80E-09 | 7.91E-05 | 0.52 | 0.39 | 0.13 | 1.33 |
| 16 | Klf9 | 1.51 | 3.84E-09 | 7.99E-05 | 0.12 | 0.05 | 0.07 | 2.37 |
| 16 | Fam171a2 | 0.71 | 3.95E-09 | 8.23E-05 | 0.55 | 0.45 | 0.09 | 1.21 |
| 16 | Patz1 | 0.56 | 4.32E-09 | 8.98E-05 | 0.62 | 0.52 | 0.09 | 1.18 |
| 16 | Upp2 | 1.14 | 4.47E-09 | 9.30E-05 | 0.28 | 0.18 | 0.11 | 1.59 |
| 16 | Map2 | 0.77 | 4.50E-09 | 9.36E-05 | 0.33 | 0.22 | 0.11 | 1.53 |
| 16 | Jpt1 | 0.34 | 4.75E-09 | 9.88E-05 | 0.95 | 0.94 | 0.00 | 1.00 |
| 16 | D230030E09Rik | 0.66 | 5.10E-09 | 1.06E-04 | 0.13 | 0.06 | 0.07 | 2.26 |
| 16 | Bbc3 | 1.35 | 5.81E-09 | 1.21E-04 | 0.19 | 0.10 | 0.09 | 1.85 |
| 16 | Apobec3 | 1.01 | 6.06E-09 | 1.26E-04 | 0.17 | 0.08 | 0.08 | 2.00 |
| 16 | Pip4p2 | 0.54 | 6.68E-09 | 1.39E-04 | 0.65 | 0.57 | 0.08 | 1.15 |
| 16 | Igf1r | 0.49 | 6.82E-09 | 1.42E-04 | 0.80 | 0.77 | 0.03 | 1.04 |
| 16 | Usp25 | 0.55 | 6.92E-09 | 1.44E-04 | 0.55 | 0.44 | 0.12 | 1.27 |
| 16 | Pgap1 | 1.13 | 7.35E-09 | 1.53E-04 | 0.25 | 0.15 | 0.10 | 1.66 |
| 16 | Pik3ip1 | 1.39 | 7.39E-09 | 1.54E-04 | 0.17 | 0.09 | 0.08 | 1.94 |
| 16 | Pde9a | 1.54 | 8.68E-09 | 1.80E-04 | 0.11 | 0.05 | 0.06 | 2.31 |
| 16 | Dazap2 | 0.45 | 8.76E-09 | 1.82E-04 | 0.65 | 0.57 | 0.08 | 1.13 |
| 16 | Trp63 | 0.81 | 9.25E-09 | 1.92E-04 | 0.24 | 0.14 | 0.10 | 1.74 |
| 16 | Snhg14 | 0.93 | 9.35E-09 | 1.94E-04 | 0.28 | 0.18 | 0.10 | 1.58 |
| 16 | Sort1 | 0.94 | 9.83E-09 | 2.04E-04 | 0.31 | 0.20 | 0.11 | 1.55 |
| 16 | Rpia | 0.84 | 1.05E-08 | 2.19E-04 | 0.40 | 0.30 | 0.11 | 1.37 |
| 16 | Npepps | 0.44 | 1.08E-08 | 2.25E-04 | 0.72 | 0.65 | 0.07 | 1.11 |
| 16 | Ptprm | 1.45 | 1.08E-08 | 2.25E-04 | 0.11 | 0.05 | 0.06 | 2.31 |
| 16 | Ntm | 1.03 | 1.10E-08 | 2.29E-04 | 0.20 | 0.11 | 0.09 | 1.82 |
| 16 | Daam1 | 0.80 | 1.17E-08 | 2.44E-04 | 0.49 | 0.41 | 0.09 | 1.21 |
| 16 | Tmx4 | 0.73 | 1.19E-08 | 2.47E-04 | 0.57 | 0.47 | 0.11 | 1.23 |
| 16 | Xist | 0.36 | 1.19E-08 | 2.48E-04 | 0.91 | 0.86 | 0.05 | 1.06 |
| 16 | Mpst | 0.69 | 1.19E-08 | 2.48E-04 | 0.52 | 0.42 | 0.10 | 1.23 |
| 16 | Sec11a | 0.31 | 1.27E-08 | 2.64E-04 | 0.93 | 0.90 | 0.04 | 1.04 |
| 16 | Tuba1c | 0.80 | 1.28E-08 | 2.66E-04 | 0.53 | 0.44 | 0.10 | 1.22 |
| 16 | Leprotl1 | 0.50 | 1.29E-08 | 2.67E-04 | 0.64 | 0.55 | 0.10 | 1.18 |
| 16 | Lrrc1 | 1.07 | 1.29E-08 | 2.68E-04 | 0.25 | 0.15 | 0.10 | 1.66 |
| 16 | Smim10l2a | 1.21 | 1.31E-08 | 2.72E-04 | 0.15 | 0.07 | 0.07 | 2.04 |
| 16 | Shroom2 | 1.31 | 1.32E-08 | 2.74E-04 | 0.16 | 0.08 | 0.08 | 2.00 |
| 16 | Cox14 | 0.34 | 1.34E-08 | 2.79E-04 | 0.83 | 0.81 | 0.03 | 1.03 |
| 16 | Pkn2 | 0.32 | 1.46E-08 | 3.03E-04 | 0.88 | 0.87 | 0.01 | 1.01 |
| 16 | Ifitm1 | 0.81 | 1.57E-08 | 3.27E-04 | 0.29 | 0.19 | 0.10 | 1.54 |
| 16 | Snapin | 0.71 | 1.67E-08 | 3.47E-04 | 0.44 | 0.33 | 0.12 | 1.36 |
| 16 | Tle1 | 0.55 | 1.70E-08 | 3.53E-04 | 0.62 | 0.53 | 0.09 | 1.17 |
| 16 | Celf4 | 0.85 | 1.97E-08 | 4.09E-04 | 0.33 | 0.23 | 0.11 | 1.48 |
| 16 | Siva1 | 0.32 | 2.07E-08 | 4.31E-04 | 0.96 | 0.95 | 0.01 | 1.01 |
| 16 | Ccdc12 | 0.37 | 2.21E-08 | 4.60E-04 | 0.75 | 0.71 | 0.04 | 1.06 |
| 16 | Sun1 | 0.85 | 2.35E-08 | 4.89E-04 | 0.38 | 0.28 | 0.11 | 1.39 |
| 16 | Agpat2 | 0.98 | 2.40E-08 | 4.99E-04 | 0.27 | 0.17 | 0.10 | 1.60 |
| 16 | Plekhg3 | 1.25 | 2.53E-08 | 5.25E-04 | 0.16 | 0.08 | 0.08 | 1.93 |
| 16 | Kcnb1 | 1.51 | 2.53E-08 | 5.27E-04 | 0.13 | 0.06 | 0.07 | 2.11 |
| 16 | Pigp | 0.43 | 2.64E-08 | 5.49E-04 | 0.72 | 0.65 | 0.08 | 1.12 |
| 16 | Cdh4 | 1.60 | 2.74E-08 | 5.71E-04 | 0.11 | 0.05 | 0.06 | 2.22 |
| 16 | Acvr1 | 1.03 | 2.91E-08 | 6.06E-04 | 0.30 | 0.19 | 0.10 | 1.53 |
| 16 | Sdsl | 1.35 | 3.11E-08 | 6.48E-04 | 0.13 | 0.06 | 0.07 | 2.15 |
| 16 | Arfip2 | 0.57 | 3.17E-08 | 6.59E-04 | 0.52 | 0.42 | 0.10 | 1.24 |
| 16 | Tmem42 | 1.02 | 3.30E-08 | 6.85E-04 | 0.25 | 0.15 | 0.10 | 1.63 |

**Supplementary Table 1. Marker genes for the original clusters of integrated clustering of E11.5 control and NotchGOF λ-junction cells.**

| | | | | | | | | |
|---|---|---|---|---|---|---|---|---|
| 16 | Ak4 | 1.23 | 3.50E-08 | 7.29E-04 | 0.19 | 0.10 | 0.08 | 1.80 |
| 16 | Sft2d3 | 0.91 | 3.53E-08 | 7.34E-04 | 0.31 | 0.20 | 0.10 | 1.51 |
| 16 | Sp4 | 0.87 | 3.61E-08 | 7.52E-04 | 0.34 | 0.24 | 0.10 | 1.44 |
| 16 | Lyplal1 | 0.75 | 3.72E-08 | 7.74E-04 | 0.35 | 0.25 | 0.11 | 1.44 |
| 16 | Canx | 0.29 | 3.76E-08 | 7.81E-04 | 0.97 | 0.93 | 0.04 | 1.04 |
| 16 | Zcchc7 | 0.55 | 3.76E-08 | 7.82E-04 | 0.61 | 0.52 | 0.09 | 1.17 |
| 16 | Tmem30b | 0.79 | 3.92E-08 | 8.16E-04 | 0.25 | 0.15 | 0.10 | 1.66 |
| 16 | 1700001O22Rik | 1.10 | 3.94E-08 | 8.20E-04 | 0.15 | 0.07 | 0.07 | 2.04 |
| 16 | Fndc3b | 0.76 | 3.99E-08 | 8.30E-04 | 0.35 | 0.25 | 0.11 | 1.43 |
| 16 | Pradc1 | 0.95 | 4.01E-08 | 8.35E-04 | 0.32 | 0.22 | 0.10 | 1.47 |
| 16 | Ist1 | 0.63 | 4.33E-08 | 9.01E-04 | 0.55 | 0.46 | 0.09 | 1.20 |
| 16 | Plin3 | 0.58 | 4.58E-08 | 9.52E-04 | 0.62 | 0.55 | 0.07 | 1.13 |
| 16 | mt-Nd5 | 0.32 | 4.67E-08 | 9.70E-04 | 0.96 | 0.96 | 0.01 | 1.01 |
| 16 | Zfp612 | 0.97 | 4.67E-08 | 9.72E-04 | 0.27 | 0.17 | 0.10 | 1.57 |
| 16 | Pak2 | 0.31 | 4.69E-08 | 9.76E-04 | 0.86 | 0.84 | 0.02 | 1.03 |
| 16 | Fbxl14 | 0.55 | 4.99E-08 | 1.04E-03 | 0.50 | 0.38 | 0.12 | 1.31 |
| 16 | Pgpep1 | 0.78 | 5.18E-08 | 1.08E-03 | 0.39 | 0.29 | 0.10 | 1.34 |
| 16 | Prr7 | 0.86 | 5.30E-08 | 1.10E-03 | 0.33 | 0.23 | 0.10 | 1.44 |
| 16 | Igdcc3 | 0.85 | 5.43E-08 | 1.13E-03 | 0.34 | 0.24 | 0.10 | 1.44 |
| 16 | Stox2 | 0.55 | 5.48E-08 | 1.14E-03 | 0.65 | 0.56 | 0.09 | 1.16 |
| 16 | Tmed9 | 0.28 | 5.79E-08 | 1.20E-03 | 0.96 | 0.94 | 0.03 | 1.03 |
| 16 | Arg2 | 1.22 | 5.93E-08 | 1.23E-03 | 0.12 | 0.05 | 0.06 | 2.19 |
| 16 | Madd | 1.12 | 6.04E-08 | 1.26E-03 | 0.15 | 0.07 | 0.07 | 1.99 |
| 16 | Sorl1 | 1.05 | 6.32E-08 | 1.31E-03 | 0.24 | 0.15 | 0.09 | 1.61 |
| 16 | Ing4 | 0.36 | 6.64E-08 | 1.38E-03 | 0.79 | 0.76 | 0.03 | 1.04 |
| 16 | Large2 | 1.10 | 6.70E-08 | 1.39E-03 | 0.16 | 0.09 | 0.08 | 1.91 |
| 16 | Cd82 | 1.25 | 6.99E-08 | 1.45E-03 | 0.14 | 0.07 | 0.07 | 2.05 |
| 16 | Sms | 0.29 | 7.28E-08 | 1.51E-03 | 0.88 | 0.86 | 0.02 | 1.02 |
| 16 | Cyb5r4 | 0.60 | 7.33E-08 | 1.52E-03 | 0.50 | 0.40 | 0.10 | 1.26 |
| 16 | Tmem107 | 0.67 | 8.04E-08 | 1.67E-03 | 0.40 | 0.29 | 0.10 | 1.36 |
| 16 | Slc38a1 | 0.35 | 8.66E-08 | 1.80E-03 | 0.83 | 0.77 | 0.06 | 1.08 |
| 16 | Hltf | 0.66 | 9.16E-08 | 1.91E-03 | 0.43 | 0.32 | 0.11 | 1.33 |
| 16 | Atp1a1 | 0.46 | 9.37E-08 | 1.95E-03 | 0.70 | 0.64 | 0.06 | 1.10 |
| 16 | Vmac | 1.23 | 9.58E-08 | 1.99E-03 | 0.17 | 0.09 | 0.08 | 1.83 |
| 16 | Pdk1 | 0.76 | 9.84E-08 | 2.05E-03 | 0.36 | 0.26 | 0.11 | 1.41 |
| 16 | Pbxip1 | 1.12 | 1.02E-07 | 2.12E-03 | 0.30 | 0.20 | 0.10 | 1.48 |
| 16 | Larp4b | 0.44 | 1.06E-07 | 2.21E-03 | 0.68 | 0.61 | 0.07 | 1.11 |
| 16 | Ino80b | 0.58 | 1.11E-07 | 2.30E-03 | 0.54 | 0.46 | 0.08 | 1.18 |
| 16 | Rab27b | 0.51 | 1.15E-07 | 2.40E-03 | 0.18 | 0.10 | 0.08 | 1.86 |
| 16 | Bid | 1.15 | 1.18E-07 | 2.46E-03 | 0.21 | 0.12 | 0.09 | 1.70 |
| 16 | Fam136a | 0.30 | 1.19E-07 | 2.47E-03 | 0.84 | 0.79 | 0.04 | 1.06 |
| 16 | Abhd11 | 0.84 | 1.27E-07 | 2.64E-03 | 0.36 | 0.27 | 0.10 | 1.37 |
| 16 | Bphl | 0.58 | 1.29E-07 | 2.69E-03 | 0.48 | 0.38 | 0.10 | 1.27 |
| 16 | Igsf8 | 0.80 | 1.33E-07 | 2.76E-03 | 0.35 | 0.24 | 0.11 | 1.43 |
| 16 | Ccpg1 | 1.18 | 1.35E-07 | 2.81E-03 | 0.25 | 0.16 | 0.09 | 1.56 |
| 16 | Taok1 | 0.42 | 1.45E-07 | 3.01E-03 | 0.67 | 0.60 | 0.07 | 1.12 |
| 16 | Irs1 | 0.87 | 1.51E-07 | 3.13E-03 | 0.28 | 0.18 | 0.10 | 1.55 |
| 16 | Tulp1 | 1.12 | 1.54E-07 | 3.20E-03 | 0.16 | 0.09 | 0.08 | 1.87 |
| 16 | Fam173a | 0.53 | 1.56E-07 | 3.24E-03 | 0.59 | 0.51 | 0.08 | 1.16 |
| 16 | Slc35g1 | 1.43 | 1.61E-07 | 3.34E-03 | 0.12 | 0.06 | 0.06 | 2.09 |
| 16 | Nipsnap1 | 0.47 | 1.61E-07 | 3.35E-03 | 0.64 | 0.56 | 0.08 | 1.15 |
| 16 | Aplp1 | 0.85 | 1.73E-07 | 3.60E-03 | 0.32 | 0.21 | 0.11 | 1.50 |
| 16 | Cecr2 | 0.82 | 1.78E-07 | 3.70E-03 | 0.32 | 0.22 | 0.10 | 1.44 |
| 16 | Tmem141 | 0.84 | 1.88E-07 | 3.91E-03 | 0.29 | 0.19 | 0.10 | 1.49 |
| 16 | Plxnb2 | 0.55 | 1.88E-07 | 3.91E-03 | 0.62 | 0.53 | 0.09 | 1.16 |
| 16 | Dyrk3 | 1.22 | 1.92E-07 | 4.00E-03 | 0.18 | 0.10 | 0.08 | 1.79 |

**Supplementary Table 1. Marker genes for the original clusters of integrated clustering of E11.5 control and NotchGOF λ-junction cells.**

| | | | | | | | | |
|---|---|---|---|---|---|---|---|---|
| 16 | Rsrp1 | 0.27 | 2.00E-07 | 4.16E-03 | 0.91 | 0.89 | 0.02 | 1.02 |
| 16 | 1700037H04Rik | 0.58 | 2.00E-07 | 4.17E-03 | 0.55 | 0.48 | 0.07 | 1.15 |
| 16 | Cryzl2 | 1.11 | 2.03E-07 | 4.22E-03 | 0.18 | 0.10 | 0.08 | 1.77 |
| 16 | Sod1 | 0.31 | 2.10E-07 | 4.36E-03 | 0.83 | 0.80 | 0.03 | 1.04 |
| 16 | Plekha1 | 0.83 | 2.16E-07 | 4.50E-03 | 0.30 | 0.20 | 0.10 | 1.49 |
| 16 | Uvrag | 0.86 | 2.26E-07 | 4.71E-03 | 0.33 | 0.23 | 0.10 | 1.43 |
| 16 | Plekhb2 | 1.31 | 2.34E-07 | 4.86E-03 | 0.12 | 0.06 | 0.06 | 2.03 |
| 16 | Grb14 | 1.33 | 2.37E-07 | 4.92E-03 | 0.10 | 0.05 | 0.06 | 2.19 |
| 16 | Tm9sf3 | 0.29 | 2.39E-07 | 4.97E-03 | 0.89 | 0.86 | 0.03 | 1.03 |
| 16 | Proser2 | 1.24 | 2.51E-07 | 5.22E-03 | 0.12 | 0.06 | 0.06 | 2.09 |
| 16 | Pex2 | 0.56 | 2.52E-07 | 5.24E-03 | 0.56 | 0.48 | 0.09 | 1.18 |
| 16 | Carmil3 | 1.17 | 2.62E-07 | 5.44E-03 | 0.12 | 0.06 | 0.06 | 2.11 |
| 16 | Dmrta2 | 0.45 | 2.83E-07 | 5.89E-03 | 0.24 | 0.14 | 0.10 | 1.69 |
| 16 | Fubp3 | 0.53 | 2.83E-07 | 5.89E-03 | 0.54 | 0.45 | 0.09 | 1.20 |
| 16 | Slc35d1 | 1.05 | 2.86E-07 | 5.95E-03 | 0.24 | 0.15 | 0.09 | 1.57 |
| 16 | Clcn3 | 0.63 | 2.87E-07 | 5.97E-03 | 0.47 | 0.38 | 0.09 | 1.25 |
| 16 | Ablim1 | 0.60 | 2.91E-07 | 6.06E-03 | 0.33 | 0.23 | 0.11 | 1.47 |
| 16 | Mgat4b | 0.54 | 3.00E-07 | 6.23E-03 | 0.49 | 0.40 | 0.09 | 1.24 |
| 16 | Tmem9 | 0.46 | 3.04E-07 | 6.33E-03 | 0.69 | 0.65 | 0.05 | 1.08 |
| 16 | Notch4 | 1.23 | 3.06E-07 | 6.36E-03 | 0.12 | 0.06 | 0.06 | 2.09 |
| 16 | Gipc1 | 0.38 | 3.20E-07 | 6.65E-03 | 0.74 | 0.69 | 0.06 | 1.08 |
| 16 | Fam13c | 1.27 | 3.37E-07 | 7.01E-03 | 0.14 | 0.07 | 0.07 | 1.96 |
| 16 | Tjp1 | 0.46 | 3.38E-07 | 7.02E-03 | 0.59 | 0.49 | 0.10 | 1.19 |
| 16 | Fuz | 1.02 | 3.47E-07 | 7.21E-03 | 0.19 | 0.11 | 0.08 | 1.75 |
| 16 | Yif1a | 0.47 | 3.60E-07 | 7.49E-03 | 0.58 | 0.48 | 0.10 | 1.20 |
| 16 | Plpp2 | 0.75 | 3.68E-07 | 7.66E-03 | 0.36 | 0.26 | 0.10 | 1.37 |
| 16 | Rimklb | 0.59 | 3.68E-07 | 7.66E-03 | 0.43 | 0.33 | 0.10 | 1.31 |
| 16 | Agpat3 | 0.78 | 3.69E-07 | 7.68E-03 | 0.36 | 0.26 | 0.10 | 1.36 |
| 16 | Ubxn4 | 0.32 | 4.12E-07 | 8.57E-03 | 0.87 | 0.84 | 0.03 | 1.03 |
| 16 | Plekhh1 | 1.18 | 4.22E-07 | 8.77E-03 | 0.13 | 0.07 | 0.06 | 1.98 |
| 16 | Cetn3 | 0.26 | 4.48E-07 | 9.31E-03 | 0.89 | 0.89 | 0.01 | 1.01 |
| 16 | Il17rd | 0.47 | 4.48E-07 | 9.32E-03 | 0.60 | 0.50 | 0.10 | 1.21 |
| 16 | Atp6v0e | 0.39 | 4.63E-07 | 9.63E-03 | 0.76 | 0.72 | 0.04 | 1.06 |
| 16 | Ctnna2 | 1.14 | 4.88E-07 | 1.01E-02 | 0.11 | 0.05 | 0.06 | 2.16 |
| 16 | Nectin2 | 0.50 | 4.98E-07 | 1.04E-02 | 0.57 | 0.48 | 0.09 | 1.18 |
| 16 | Ormdl1 | 0.56 | 5.06E-07 | 1.05E-02 | 0.53 | 0.44 | 0.09 | 1.19 |
| 16 | Fut8 | 0.75 | 5.09E-07 | 1.06E-02 | 0.39 | 0.29 | 0.10 | 1.33 |
| 16 | Zfp7 | 0.92 | 5.12E-07 | 1.06E-02 | 0.23 | 0.14 | 0.09 | 1.60 |
| 16 | Morn2 | 0.57 | 5.34E-07 | 1.11E-02 | 0.44 | 0.33 | 0.11 | 1.32 |
| 16 | Egfr | 1.03 | 5.40E-07 | 1.12E-02 | 0.18 | 0.10 | 0.08 | 1.77 |
| 16 | Gpr27 | 0.92 | 5.58E-07 | 1.16E-02 | 0.23 | 0.14 | 0.09 | 1.62 |
| 16 | Tmem87b | 0.78 | 5.90E-07 | 1.23E-02 | 0.31 | 0.21 | 0.10 | 1.45 |
| 16 | Mpdu1 | 0.54 | 6.30E-07 | 1.31E-02 | 0.59 | 0.50 | 0.09 | 1.17 |
| 16 | Rtkn2 | 1.20 | 6.78E-07 | 1.41E-02 | 0.14 | 0.08 | 0.07 | 1.87 |
| 16 | Mcf2l | 1.40 | 6.98E-07 | 1.45E-02 | 0.13 | 0.07 | 0.06 | 1.91 |
| 16 | Tmem170 | 1.36 | 6.98E-07 | 1.45E-02 | 0.17 | 0.10 | 0.07 | 1.75 |
| 16 | Tm2d3 | 0.57 | 7.01E-07 | 1.46E-02 | 0.57 | 0.50 | 0.07 | 1.14 |
| 16 | Dbndd2 | 1.29 | 7.13E-07 | 1.48E-02 | 0.15 | 0.08 | 0.07 | 1.83 |
| 16 | Dmtn | 1.42 | 7.47E-07 | 1.55E-02 | 0.13 | 0.07 | 0.06 | 1.92 |
| 16 | mt-Nd6 | 0.72 | 7.65E-07 | 1.59E-02 | 0.45 | 0.35 | 0.10 | 1.27 |
| 16 | Ctsz | 0.33 | 7.66E-07 | 1.59E-02 | 0.64 | 0.58 | 0.06 | 1.11 |
| 16 | Efna1 | 0.63 | 7.88E-07 | 1.64E-02 | 0.23 | 0.14 | 0.09 | 1.64 |
| 16 | Chd2 | 0.44 | 7.93E-07 | 1.65E-02 | 0.69 | 0.62 | 0.06 | 1.10 |
| 16 | Lgalsl | 0.57 | 8.13E-07 | 1.69E-02 | 0.41 | 0.32 | 0.10 | 1.31 |
| 16 | Prkcd | 0.86 | 8.14E-07 | 1.69E-02 | 0.31 | 0.21 | 0.10 | 1.45 |
| 16 | Oard1 | 0.38 | 8.26E-07 | 1.72E-02 | 0.72 | 0.67 | 0.04 | 1.07 |

**Supplementary Table 1. Marker genes for the original clusters of integrated clustering of E11.5 control and NotchGOF λ-junction cells.**

| | | | | | | | | |
|---|---|---|---|---|---|---|---|---|
| 16 | Plcl1 | 1.27 | 8.30E-07 | 1.73E-02 | 0.12 | 0.06 | 0.06 | 1.98 |
| 16 | Hlcs | 0.91 | 8.46E-07 | 1.76E-02 | 0.21 | 0.13 | 0.08 | 1.63 |
| 16 | Acadm | 0.71 | 8.50E-07 | 1.77E-02 | 0.42 | 0.33 | 0.09 | 1.26 |
| 16 | Rhbdf1 | 1.30 | 8.51E-07 | 1.77E-02 | 0.17 | 0.10 | 0.07 | 1.74 |
| 16 | Zfp667 | 1.14 | 8.62E-07 | 1.79E-02 | 0.17 | 0.10 | 0.07 | 1.74 |
| 16 | Ift43 | 0.41 | 8.73E-07 | 1.82E-02 | 0.63 | 0.55 | 0.09 | 1.16 |
| 16 | Atp11b | 0.63 | 8.73E-07 | 1.82E-02 | 0.50 | 0.42 | 0.08 | 1.19 |
| 16 | Commd6 | 0.37 | 9.02E-07 | 1.88E-02 | 0.71 | 0.66 | 0.06 | 1.09 |
| 16 | Jtb | 0.47 | 9.80E-07 | 2.04E-02 | 0.59 | 0.53 | 0.05 | 1.10 |
| 16 | Nfib | 0.31 | 9.95E-07 | 2.07E-02 | 0.83 | 0.74 | 0.09 | 1.12 |
| 16 | Dcaf17 | 0.68 | 1.06E-06 | 2.20E-02 | 0.42 | 0.33 | 0.09 | 1.27 |
| 16 | Vangl2 | 0.47 | 1.13E-06 | 2.35E-02 | 0.64 | 0.57 | 0.07 | 1.13 |
| 16 | Cdkn1a | 0.62 | 1.13E-06 | 2.36E-02 | 0.19 | 0.11 | 0.08 | 1.71 |
| 16 | Cdkl5 | 1.31 | 1.19E-06 | 2.46E-02 | 0.11 | 0.06 | 0.06 | 2.02 |
| 16 | 2310015A10Rik | 1.14 | 1.20E-06 | 2.49E-02 | 0.14 | 0.08 | 0.07 | 1.87 |
| 16 | 1110020A21Rik | 1.29 | 1.21E-06 | 2.51E-02 | 0.13 | 0.07 | 0.06 | 1.92 |
| 16 | Rassf7 | 0.82 | 1.21E-06 | 2.52E-02 | 0.29 | 0.20 | 0.09 | 1.46 |
| 16 | Rida | 0.65 | 1.26E-06 | 2.62E-02 | 0.50 | 0.42 | 0.08 | 1.18 |
| 16 | Ssc4d | 1.38 | 1.38E-06 | 2.87E-02 | 0.11 | 0.05 | 0.06 | 2.06 |
| 16 | F8a | 0.99 | 1.39E-06 | 2.89E-02 | 0.15 | 0.08 | 0.07 | 1.82 |
| 16 | Dnajb2 | 0.77 | 1.49E-06 | 3.10E-02 | 0.36 | 0.27 | 0.09 | 1.34 |
| 16 | Msi2 | 0.40 | 1.50E-06 | 3.12E-02 | 0.72 | 0.68 | 0.04 | 1.06 |
| 16 | Rsbn1l | 0.43 | 1.50E-06 | 3.12E-02 | 0.70 | 0.64 | 0.06 | 1.10 |
| 16 | Dusp6 | 0.46 | 1.52E-06 | 3.17E-02 | 0.47 | 0.36 | 0.11 | 1.31 |
| 16 | Amer2 | 1.01 | 1.53E-06 | 3.17E-02 | 0.18 | 0.11 | 0.08 | 1.70 |
| 16 | Rab3ip | 0.57 | 1.67E-06 | 3.48E-02 | 0.40 | 0.30 | 0.10 | 1.32 |
| 16 | Nme3 | 1.26 | 1.68E-06 | 3.49E-02 | 0.14 | 0.08 | 0.06 | 1.84 |
| 16 | Tfrc | 0.54 | 1.70E-06 | 3.54E-02 | 0.48 | 0.40 | 0.08 | 1.21 |
| 16 | Prss22 | 1.14 | 1.71E-06 | 3.56E-02 | 0.12 | 0.06 | 0.06 | 1.98 |
| 16 | Tmem189 | 0.73 | 1.77E-06 | 3.68E-02 | 0.39 | 0.30 | 0.09 | 1.30 |
| 16 | Syne4 | 0.82 | 1.78E-06 | 3.69E-02 | 0.29 | 0.20 | 0.09 | 1.46 |
| 16 | Selenot | 0.48 | 1.88E-06 | 3.90E-02 | 0.60 | 0.54 | 0.05 | 1.10 |
| 16 | Bcl2l1 | 0.64 | 1.88E-06 | 3.92E-02 | 0.46 | 0.37 | 0.09 | 1.23 |
| 16 | Gm4258 | 0.62 | 1.99E-06 | 4.14E-02 | 0.57 | 0.50 | 0.08 | 1.15 |
| 16 | Echdc1 | 0.63 | 2.00E-06 | 4.16E-02 | 0.33 | 0.24 | 0.09 | 1.39 |
| 16 | Trim25 | 1.34 | 2.01E-06 | 4.19E-02 | 0.12 | 0.06 | 0.06 | 1.97 |
| 16 | Whrn | 1.22 | 2.01E-06 | 4.19E-02 | 0.12 | 0.06 | 0.06 | 1.98 |
| 16 | Entpd5 | 1.27 | 2.07E-06 | 4.30E-02 | 0.14 | 0.08 | 0.06 | 1.82 |
| 16 | Gm43445 | 1.36 | 2.12E-06 | 4.41E-02 | 0.16 | 0.09 | 0.07 | 1.75 |
| 16 | Trim44 | 0.38 | 2.15E-06 | 4.47E-02 | 0.77 | 0.71 | 0.06 | 1.08 |
| 17 | Krt18 | 2.47 | 2.36E-138 | 4.90E-134 | 0.82 | 0.29 | 0.52 | 2.78 |
| 17 | Rps26 | 0.78 | 1.20E-134 | 2.50E-130 | 1.00 | 1.00 | 0.00 | 1.00 |
| 17 | Six3 | 2.43 | 1.04E-128 | 2.16E-124 | 0.74 | 0.23 | 0.50 | 3.16 |
| 17 | Aldh1a3 | 3.01 | 1.02E-119 | 2.12E-115 | 0.67 | 0.23 | 0.45 | 2.97 |
| 17 | Krt8 | 2.12 | 1.78E-105 | 3.70E-101 | 0.72 | 0.26 | 0.47 | 2.83 |
| 17 | Rps8 | 0.44 | 2.71E-103 | 5.63E-99 | 1.00 | 1.00 | 0.00 | 1.00 |
| 17 | Fau | 0.55 | 7.79E-102 | 1.62E-97 | 0.99 | 1.00 | -0.01 | 0.99 |
| 17 | Rpl27a | 0.54 | 1.90E-101 | 3.95E-97 | 1.00 | 1.00 | 0.00 | 1.00 |
| 17 | Bex1 | 2.07 | 1.32E-98 | 2.74E-94 | 0.86 | 0.59 | 0.27 | 1.46 |
| 17 | Gapdh | 0.76 | 1.21E-97 | 2.51E-93 | 1.00 | 1.00 | 0.00 | 1.00 |
| 17 | Rps27a | 0.47 | 1.74E-94 | 3.62E-90 | 1.00 | 1.00 | 0.00 | 1.00 |
| 17 | Pdgfa | 1.92 | 5.73E-93 | 1.19E-88 | 0.75 | 0.28 | 0.47 | 2.70 |
| 17 | Rps4x | 0.53 | 2.50E-92 | 5.20E-88 | 1.00 | 1.00 | 0.00 | 1.00 |
| 17 | Rpl28 | 0.51 | 3.77E-92 | 7.84E-88 | 0.99 | 1.00 | -0.01 | 0.99 |
| 17 | Rps19 | 0.53 | 2.67E-91 | 5.55E-87 | 1.00 | 1.00 | 0.00 | 1.00 |
| 17 | Rps12 | 0.55 | 4.65E-90 | 9.67E-86 | 1.00 | 1.00 | 0.00 | 1.00 |

**Supplementary Table 1. Marker genes for the original clusters of integrated clustering of E11.5 control and NotchGOF λ-junction cells.**

| | | | | | | | | |
|---|---|---|---|---|---|---|---|---|
| 17 | Six3os1 | 2.11 | 6.81E-89 | 1.42E-84 | 0.65 | 0.23 | 0.42 | 2.84 |
| 17 | Rps16 | 0.44 | 1.88E-88 | 3.92E-84 | 1.00 | 1.00 | 0.00 | 1.00 |
| 17 | Rpl32 | 0.48 | 2.60E-88 | 5.42E-84 | 1.00 | 1.00 | 0.00 | 1.00 |
| 17 | Rps11 | 0.44 | 2.62E-88 | 5.45E-84 | 1.00 | 1.00 | 0.00 | 1.00 |
| 17 | Rpl38 | 0.62 | 3.09E-88 | 6.43E-84 | 0.99 | 1.00 | -0.01 | 0.99 |
| 17 | Rpl11 | 0.47 | 1.00E-86 | 2.08E-82 | 1.00 | 1.00 | 0.00 | 1.00 |
| 17 | Rps21 | 0.51 | 2.26E-86 | 4.69E-82 | 0.98 | 1.00 | -0.02 | 0.98 |
| 17 | Rpl10a | 0.59 | 1.03E-84 | 2.15E-80 | 0.99 | 1.00 | -0.01 | 0.99 |
| 17 | Rplp0 | 0.48 | 5.63E-83 | 1.17E-78 | 0.99 | 1.00 | -0.01 | 0.99 |
| 17 | Rps3a1 | 0.44 | 1.07E-82 | 2.23E-78 | 1.00 | 1.00 | 0.00 | 1.00 |
| 17 | Rpl13 | 0.38 | 1.11E-82 | 2.30E-78 | 1.00 | 1.00 | 0.00 | 1.00 |
| 17 | Rpl21 | 0.43 | 3.13E-82 | 6.50E-78 | 1.00 | 1.00 | 0.00 | 1.00 |
| 17 | Rps10 | 0.45 | 1.09E-81 | 2.27E-77 | 1.00 | 1.00 | 0.00 | 1.00 |
| 17 | Sox2 | 2.41 | 3.39E-81 | 7.04E-77 | 0.57 | 0.19 | 0.38 | 2.99 |
| 17 | Rps7 | 0.46 | 6.24E-81 | 1.30E-76 | 1.00 | 1.00 | 0.00 | 1.00 |
| 17 | Rpl7 | 0.55 | 9.76E-78 | 2.03E-73 | 1.00 | 1.00 | 0.00 | 1.00 |
| 17 | Rps20 | 0.43 | 2.27E-77 | 4.73E-73 | 1.00 | 1.00 | 0.00 | 1.00 |
| 17 | Rpl6 | 0.48 | 3.10E-77 | 6.46E-73 | 1.00 | 1.00 | 0.00 | 1.00 |
| 17 | Rps28 | 0.55 | 1.53E-76 | 3.18E-72 | 0.99 | 1.00 | -0.01 | 0.99 |
| 17 | Rpl18a | 0.48 | 6.44E-76 | 1.34E-71 | 1.00 | 1.00 | 0.00 | 1.00 |
| 17 | Hsp90ab1 | 0.48 | 3.96E-75 | 8.23E-71 | 1.00 | 1.00 | 0.00 | 1.00 |
| 17 | Rpl17 | 0.48 | 2.03E-74 | 4.21E-70 | 1.00 | 1.00 | 0.00 | 1.00 |
| 17 | Rps13 | 0.45 | 5.64E-74 | 1.17E-69 | 0.99 | 1.00 | -0.01 | 0.99 |
| 17 | Tubb2b | 1.70 | 1.19E-72 | 2.48E-68 | 0.86 | 0.67 | 0.19 | 1.29 |
| 17 | Rps5 | 0.44 | 1.39E-72 | 2.90E-68 | 0.99 | 1.00 | -0.01 | 0.99 |
| 17 | Rpl26 | 0.44 | 1.41E-72 | 2.94E-68 | 1.00 | 1.00 | 0.00 | 1.00 |
| 17 | Rpl37 | 0.51 | 1.76E-71 | 3.66E-67 | 0.99 | 1.00 | -0.01 | 0.99 |
| 17 | Rpl22 | 0.54 | 1.09E-70 | 2.26E-66 | 0.99 | 1.00 | -0.01 | 0.99 |
| 17 | Rpl23 | 0.44 | 2.15E-70 | 4.47E-66 | 1.00 | 1.00 | 0.00 | 1.00 |
| 17 | Atox1 | 1.32 | 2.95E-70 | 6.13E-66 | 0.89 | 0.95 | -0.06 | 0.93 |
| 17 | Rpl36 | 0.47 | 1.61E-69 | 3.35E-65 | 0.99 | 1.00 | -0.01 | 0.99 |
| 17 | Rps9 | 0.44 | 6.56E-68 | 1.36E-63 | 1.00 | 1.00 | 0.00 | 1.00 |
| 17 | Pitx1 | 2.28 | 4.10E-67 | 8.52E-63 | 0.51 | 0.18 | 0.33 | 2.85 |
| 17 | Rpl24 | 0.43 | 1.04E-63 | 2.17E-59 | 0.99 | 1.00 | -0.01 | 0.99 |
| 17 | Rps27 | 0.53 | 2.06E-63 | 4.29E-59 | 0.98 | 1.00 | -0.02 | 0.98 |
| 17 | Rps18 | 0.45 | 1.05E-62 | 2.19E-58 | 0.98 | 1.00 | -0.02 | 0.98 |
| 17 | Rpl35a | 0.44 | 2.48E-62 | 5.16E-58 | 0.99 | 1.00 | -0.01 | 0.99 |
| 17 | Rplp2 | 0.38 | 2.75E-62 | 5.71E-58 | 1.00 | 1.00 | 0.00 | 1.00 |
| 17 | Rpl15 | 0.42 | 1.22E-60 | 2.54E-56 | 0.99 | 1.00 | -0.01 | 0.99 |
| 17 | Rpl36a | 0.53 | 2.60E-60 | 5.40E-56 | 0.98 | 1.00 | -0.02 | 0.98 |
| 17 | Eef1a1 | 0.43 | 2.98E-60 | 6.19E-56 | 1.00 | 1.00 | 0.00 | 1.00 |
| 17 | Bex4 | 1.73 | 3.83E-60 | 7.96E-56 | 0.79 | 0.65 | 0.14 | 1.21 |
| 17 | Rpl14 | 0.41 | 1.09E-59 | 2.27E-55 | 1.00 | 1.00 | 0.00 | 1.00 |
| 17 | Rpl39 | 0.42 | 1.84E-59 | 3.83E-55 | 0.99 | 1.00 | -0.01 | 0.99 |
| 17 | Rpl41 | 0.37 | 2.41E-59 | 5.00E-55 | 1.00 | 1.00 | 0.00 | 1.00 |
| 17 | Naca | 0.45 | 7.55E-59 | 1.57E-54 | 0.99 | 1.00 | -0.01 | 0.99 |
| 17 | Rps29 | 0.38 | 7.72E-59 | 1.61E-54 | 0.99 | 1.00 | -0.01 | 0.99 |
| 17 | Rpl37a | 0.40 | 6.65E-57 | 1.38E-52 | 0.99 | 1.00 | -0.01 | 0.99 |
| 17 | Epcam | 1.06 | 1.73E-56 | 3.61E-52 | 0.75 | 0.34 | 0.41 | 2.20 |
| 17 | Rpl9 | 0.38 | 2.73E-56 | 5.68E-52 | 1.00 | 1.00 | 0.00 | 1.00 |
| 17 | Ezr | 1.96 | 7.27E-56 | 1.51E-51 | 0.63 | 0.32 | 0.32 | 1.99 |
| 17 | Rps14 | 0.36 | 5.56E-55 | 1.16E-50 | 0.99 | 1.00 | -0.01 | 0.99 |
| 17 | Isl1 | 1.86 | 5.32E-54 | 1.11E-49 | 0.53 | 0.22 | 0.32 | 2.46 |
| 17 | Rpsa | 0.33 | 9.28E-54 | 1.93E-49 | 1.00 | 1.00 | 0.00 | 1.00 |
| 17 | Rpl3 | 0.39 | 8.39E-52 | 1.74E-47 | 0.99 | 1.00 | -0.01 | 0.99 |
| 17 | Acer2 | 2.13 | 9.77E-51 | 2.03E-46 | 0.63 | 0.39 | 0.23 | 1.59 |

**Supplementary Table 1. Marker genes for the original clusters of integrated clustering of E11.5 control and NotchGOF λ-junction cells.**

| | | | | | | | | |
|---|---|---|---|---|---|---|---|---|
| 17 | Pkm | 0.65 | 9.49E-50 | 1.97E-45 | 0.96 | 0.98 | -0.02 | 0.98 |
| 17 | Tpt1 | 0.36 | 1.40E-48 | 2.91E-44 | 0.99 | 1.00 | -0.01 | 0.99 |
| 17 | Rplp1 | 0.39 | 5.12E-48 | 1.07E-43 | 1.00 | 1.00 | 0.00 | 1.00 |
| 17 | Hist3h2ba | 2.10 | 2.80E-47 | 5.82E-43 | 0.55 | 0.27 | 0.28 | 2.01 |
| 17 | Rpl12 | 0.43 | 4.41E-47 | 9.18E-43 | 1.00 | 1.00 | 0.00 | 1.00 |
| 17 | Rps23 | 0.32 | 4.49E-47 | 9.35E-43 | 0.99 | 1.00 | -0.01 | 0.99 |
| 17 | Eno1 | 0.81 | 7.92E-47 | 1.65E-42 | 0.93 | 0.97 | -0.05 | 0.95 |
| 17 | Rps2 | 0.41 | 9.45E-47 | 1.97E-42 | 1.00 | 1.00 | 0.00 | 1.00 |
| 17 | Rpl34 | 0.33 | 1.74E-46 | 3.63E-42 | 0.99 | 1.00 | -0.01 | 0.99 |
| 17 | Rpl5 | 0.46 | 1.89E-46 | 3.94E-42 | 0.99 | 1.00 | -0.01 | 0.99 |
| 17 | Tpi1 | 0.88 | 2.66E-46 | 5.52E-42 | 0.90 | 0.96 | -0.06 | 0.94 |
| 17 | Rpl18 | 0.31 | 7.29E-46 | 1.52E-41 | 1.00 | 1.00 | 0.00 | 1.00 |
| 17 | Rps24 | 0.27 | 9.79E-46 | 2.04E-41 | 0.99 | 1.00 | -0.01 | 0.99 |
| 17 | Rps15a | 0.31 | 1.22E-45 | 2.54E-41 | 0.99 | 1.00 | -0.01 | 0.99 |
| 17 | Ldha | 0.82 | 1.70E-45 | 3.54E-41 | 0.94 | 0.97 | -0.04 | 0.96 |
| 17 | Khdrbs3 | 1.73 | 4.09E-45 | 8.51E-41 | 0.73 | 0.62 | 0.11 | 1.19 |
| 17 | Rack1 | 0.40 | 8.53E-45 | 1.77E-40 | 0.99 | 1.00 | -0.01 | 0.99 |
| 17 | Rpl10 | 0.42 | 9.67E-45 | 2.01E-40 | 0.99 | 1.00 | -0.01 | 0.99 |
| 17 | Psip1 | 1.01 | 3.54E-44 | 7.37E-40 | 0.97 | 0.98 | -0.02 | 0.98 |
| 17 | Rpl7a | 0.36 | 4.59E-44 | 9.54E-40 | 0.99 | 1.00 | -0.02 | 0.99 |
| 17 | Rpl8 | 0.31 | 4.69E-44 | 9.76E-40 | 0.99 | 1.00 | -0.01 | 0.99 |
| 17 | Hey1 | 2.19 | 7.76E-44 | 1.61E-39 | 0.56 | 0.30 | 0.26 | 1.86 |
| 17 | Rpl35 | 0.38 | 1.27E-43 | 2.63E-39 | 0.98 | 1.00 | -0.02 | 0.98 |
| 17 | Rps25 | 0.38 | 3.99E-43 | 8.29E-39 | 0.98 | 1.00 | -0.02 | 0.98 |
| 17 | Rpl30 | 0.35 | 1.17E-42 | 2.44E-38 | 0.99 | 1.00 | -0.01 | 0.99 |
| 17 | Tpd52 | 2.02 | 1.21E-42 | 2.51E-38 | 0.52 | 0.26 | 0.26 | 2.00 |
| 17 | Rps3 | 0.30 | 7.23E-42 | 1.50E-37 | 0.99 | 1.00 | -0.01 | 0.99 |
| 17 | Rpl27 | 0.33 | 8.71E-42 | 1.81E-37 | 0.99 | 1.00 | -0.02 | 0.99 |
| 17 | Mdh1 | 0.95 | 1.37E-40 | 2.85E-36 | 0.85 | 0.95 | -0.10 | 0.89 |
| 17 | Rpl36al | 0.52 | 1.92E-40 | 3.99E-36 | 0.94 | 1.00 | -0.06 | 0.94 |
| 17 | Dstn | 1.04 | 4.29E-40 | 8.92E-36 | 0.87 | 0.94 | -0.08 | 0.92 |
| 17 | Rps6 | 0.41 | 5.52E-40 | 1.15E-35 | 0.99 | 1.00 | -0.01 | 0.99 |
| 17 | Sfn | 1.88 | 1.53E-38 | 3.18E-34 | 0.46 | 0.21 | 0.25 | 2.23 |
| 17 | Rpl19 | 0.28 | 8.49E-38 | 1.76E-33 | 1.00 | 1.00 | 0.00 | 1.00 |
| 17 | Pgam1 | 0.66 | 3.30E-37 | 6.86E-33 | 0.89 | 0.97 | -0.08 | 0.92 |
| 17 | Sp8 | 2.63 | 5.92E-37 | 1.23E-32 | 0.35 | 0.14 | 0.21 | 2.53 |
| 17 | Rpl29 | 0.41 | 6.03E-37 | 1.25E-32 | 0.97 | 1.00 | -0.03 | 0.97 |
| 17 | Rab25 | 2.55 | 1.24E-36 | 2.59E-32 | 0.38 | 0.16 | 0.22 | 2.34 |
| 17 | Foxg1 | 2.32 | 6.01E-35 | 1.25E-30 | 0.36 | 0.14 | 0.21 | 2.47 |
| 17 | S100a11 | 1.61 | 9.20E-35 | 1.91E-30 | 0.75 | 0.76 | -0.01 | 0.99 |
| 17 | Uba52 | 0.42 | 1.64E-34 | 3.42E-30 | 0.96 | 1.00 | -0.04 | 0.96 |
| 17 | Ppia | 0.31 | 1.73E-34 | 3.59E-30 | 1.00 | 1.00 | 0.00 | 1.00 |
| 17 | Pgk1 | 0.81 | 7.06E-34 | 1.47E-29 | 0.88 | 0.96 | -0.08 | 0.92 |
| 17 | Tceal9 | 0.60 | 7.59E-34 | 1.58E-29 | 0.93 | 0.98 | -0.06 | 0.94 |
| 17 | Gm10076 | 0.33 | 1.58E-30 | 3.28E-26 | 0.99 | 1.00 | -0.01 | 0.99 |
| 17 | Gm16136 | 2.05 | 2.06E-30 | 4.28E-26 | 0.38 | 0.17 | 0.21 | 2.22 |
| 17 | Spint2 | 1.43 | 2.95E-30 | 6.14E-26 | 0.56 | 0.31 | 0.25 | 1.82 |
| 17 | Cyb5a | 0.99 | 3.58E-30 | 7.45E-26 | 0.82 | 0.92 | -0.10 | 0.89 |
| 17 | Rps17 | 0.32 | 1.06E-29 | 2.20E-25 | 0.99 | 1.00 | -0.01 | 0.99 |
| 17 | Pomp | 0.62 | 4.87E-28 | 1.01E-23 | 0.88 | 0.98 | -0.10 | 0.90 |
| 17 | Cldn7 | 1.55 | 1.31E-27 | 2.73E-23 | 0.47 | 0.24 | 0.23 | 1.96 |
| 17 | Mif | 0.44 | 3.05E-27 | 6.35E-23 | 0.99 | 0.99 | -0.01 | 0.99 |
| 17 | Atp5d | 0.59 | 1.05E-26 | 2.18E-22 | 0.90 | 0.99 | -0.09 | 0.91 |
| 17 | Rpl31 | 0.37 | 1.44E-25 | 2.99E-21 | 0.96 | 1.00 | -0.04 | 0.96 |
| 17 | Rpl13a | 0.26 | 4.36E-25 | 9.06E-21 | 0.99 | 1.00 | -0.01 | 0.99 |
| 17 | Ap1m2 | 2.29 | 1.43E-24 | 2.96E-20 | 0.31 | 0.14 | 0.17 | 2.17 |

**Supplementary Table 1. Marker genes for the original clusters of integrated clustering of E11.5 control and NotchGOF λ-junction cells.**

| | | | | | | | | |
|---|---|---|---|---|---|---|---|---|
| 17 | Aldoa | 0.69 | 2.76E-24 | 5.74E-20 | 0.88 | 0.96 | -0.08 | 0.92 |
| 17 | Siva1 | 0.88 | 1.04E-23 | 2.16E-19 | 0.83 | 0.95 | -0.12 | 0.88 |
| 17 | Atp6v1f | 0.94 | 1.07E-23 | 2.23E-19 | 0.79 | 0.91 | -0.12 | 0.87 |
| 17 | Rprm | 2.47 | 1.15E-23 | 2.40E-19 | 0.33 | 0.16 | 0.17 | 2.07 |
| 17 | Slc9a3r1 | 1.75 | 2.15E-23 | 4.46E-19 | 0.42 | 0.24 | 0.18 | 1.76 |
| 17 | Eef1b2 | 0.27 | 3.74E-23 | 7.78E-19 | 0.98 | 1.00 | -0.02 | 0.98 |
| 17 | Eif3f | 0.43 | 6.19E-23 | 1.29E-18 | 0.93 | 0.99 | -0.06 | 0.94 |
| 17 | Prdx1 | 0.48 | 1.67E-22 | 3.48E-18 | 0.94 | 1.00 | -0.06 | 0.94 |
| 17 | Mapk13 | 2.13 | 3.24E-22 | 6.74E-18 | 0.36 | 0.20 | 0.16 | 1.82 |
| 17 | Atp5g2 | 0.41 | 1.67E-21 | 3.48E-17 | 0.95 | 1.00 | -0.05 | 0.95 |
| 17 | Calm2 | 0.61 | 2.57E-21 | 5.34E-17 | 0.91 | 0.98 | -0.07 | 0.93 |
| 17 | Dlx6os1 | 2.94 | 1.79E-20 | 3.73E-16 | 0.14 | 0.04 | 0.10 | 3.45 |
| 17 | Atp5e | 0.34 | 2.55E-20 | 5.30E-16 | 0.93 | 1.00 | -0.07 | 0.93 |
| 17 | Psma7 | 0.48 | 2.61E-20 | 5.43E-16 | 0.91 | 0.99 | -0.09 | 0.91 |
| 17 | Rbp1 | 0.97 | 2.67E-20 | 5.56E-16 | 0.87 | 0.91 | -0.04 | 0.95 |
| 17 | Gng5 | 0.30 | 2.80E-20 | 5.83E-16 | 0.96 | 1.00 | -0.04 | 0.96 |
| 17 | Sub1 | 0.42 | 4.45E-20 | 9.25E-16 | 0.90 | 0.99 | -0.09 | 0.91 |
| 17 | Rpl23a | 0.27 | 7.84E-20 | 1.63E-15 | 0.99 | 1.00 | -0.01 | 0.99 |
| 17 | Skp1a | 0.66 | 1.40E-19 | 2.91E-15 | 0.83 | 0.96 | -0.13 | 0.87 |
| 17 | Hnrnpa1 | 0.34 | 1.52E-19 | 3.16E-15 | 0.98 | 0.99 | -0.01 | 0.99 |
| 17 | Stard10 | 2.01 | 4.97E-19 | 1.03E-14 | 0.43 | 0.30 | 0.13 | 1.45 |
| 17 | D230030E09Rik | 2.70 | 3.58E-18 | 7.44E-14 | 0.16 | 0.06 | 0.11 | 2.88 |
| 17 | Btf3 | 0.27 | 7.34E-18 | 1.53E-13 | 0.97 | 1.00 | -0.03 | 0.97 |
| 17 | Tle5 | 0.61 | 9.93E-18 | 2.07E-13 | 0.85 | 0.94 | -0.09 | 0.90 |
| 17 | 3110039M20Rik | 2.94 | 1.28E-17 | 2.66E-13 | 0.19 | 0.08 | 0.12 | 2.51 |
| 17 | Fabp5 | 0.63 | 8.88E-17 | 1.85E-12 | 0.92 | 0.98 | -0.06 | 0.94 |
| 17 | H3f3b | 0.31 | 9.52E-17 | 1.98E-12 | 0.99 | 1.00 | -0.01 | 0.99 |
| 17 | Ermn | 3.00 | 2.94E-16 | 6.11E-12 | 0.13 | 0.04 | 0.09 | 3.07 |
| 17 | Slc25a3 | 0.40 | 3.02E-16 | 6.27E-12 | 0.94 | 0.99 | -0.05 | 0.95 |
| 17 | Rpl22l1 | 0.29 | 5.96E-16 | 1.24E-11 | 0.96 | 1.00 | -0.04 | 0.96 |
| 17 | Rbmx2 | 0.32 | 7.13E-16 | 1.48E-11 | 0.16 | 0.44 | -0.28 | 0.37 |
| 17 | Wnt4 | 1.15 | 1.65E-15 | 3.44E-11 | 0.36 | 0.20 | 0.16 | 1.80 |
| 17 | Eif3k | 0.39 | 1.81E-15 | 3.76E-11 | 0.88 | 0.99 | -0.11 | 0.89 |
| 17 | Pebp1 | 0.36 | 1.82E-15 | 3.79E-11 | 0.94 | 0.99 | -0.05 | 0.95 |
| 17 | Id1 | 0.82 | 3.14E-15 | 6.52E-11 | 0.82 | 0.86 | -0.04 | 0.95 |
| 17 | Id3 | 0.77 | 3.29E-15 | 6.85E-11 | 0.91 | 0.96 | -0.04 | 0.95 |
| 17 | Crabp2 | 0.56 | 5.86E-15 | 1.22E-10 | 0.88 | 0.92 | -0.04 | 0.96 |
| 17 | Jpt1 | 0.74 | 6.14E-15 | 1.28E-10 | 0.81 | 0.95 | -0.14 | 0.86 |
| 17 | Cldn3 | 1.63 | 9.70E-15 | 2.02E-10 | 0.30 | 0.16 | 0.14 | 1.85 |
| 17 | Gpi1 | 0.88 | 1.43E-14 | 2.97E-10 | 0.73 | 0.84 | -0.11 | 0.87 |
| 17 | Hmgn2 | 0.49 | 1.73E-14 | 3.60E-10 | 0.89 | 0.98 | -0.09 | 0.91 |
| 17 | Serf2 | 0.28 | 2.42E-14 | 5.03E-10 | 0.97 | 1.00 | -0.03 | 0.97 |
| 17 | Dsp | 1.05 | 4.76E-14 | 9.90E-10 | 0.40 | 0.24 | 0.16 | 1.68 |
| 17 | Rbm10 | 0.34 | 1.03E-13 | 2.14E-09 | 0.27 | 0.60 | -0.33 | 0.46 |
| 17 | Mob2 | 0.35 | 1.07E-13 | 2.22E-09 | 0.15 | 0.41 | -0.26 | 0.37 |
| 17 | Sem1 | 0.33 | 1.07E-13 | 2.22E-09 | 0.94 | 1.00 | -0.06 | 0.94 |
| 17 | Stk25 | 0.40 | 1.19E-13 | 2.46E-09 | 0.19 | 0.46 | -0.27 | 0.41 |
| 17 | Eps15 | 0.35 | 2.09E-13 | 4.35E-09 | 0.10 | 0.31 | -0.21 | 0.32 |
| 17 | Myl12b | 0.82 | 2.45E-13 | 5.10E-09 | 0.73 | 0.89 | -0.16 | 0.82 |
| 17 | Dok5 | 2.10 | 2.50E-13 | 5.19E-09 | 0.23 | 0.12 | 0.11 | 1.97 |
| 17 | Hdgf | 0.51 | 3.51E-13 | 7.30E-09 | 0.85 | 0.98 | -0.13 | 0.86 |
| 17 | Ndufa6 | 0.69 | 3.98E-13 | 8.27E-09 | 0.76 | 0.94 | -0.18 | 0.81 |
| 17 | Dlx5 | 2.18 | 4.26E-13 | 8.85E-09 | 0.33 | 0.22 | 0.12 | 1.54 |
| 17 | Uqcr11 | 0.48 | 5.38E-13 | 1.12E-08 | 0.84 | 0.98 | -0.14 | 0.86 |
| 17 | Nup37 | 0.31 | 5.66E-13 | 1.18E-08 | 0.24 | 0.54 | -0.30 | 0.45 |
| 17 | Msantd3 | 0.31 | 6.50E-13 | 1.35E-08 | 0.23 | 0.51 | -0.29 | 0.44 |

**Supplementary Table 1. Marker genes for the original clusters of integrated clustering of E11.5 control and NotchGOF λ-junction cells.**

| | | | | | | | | |
|---|---|---|---|---|---|---|---|---|
| 17 | Ubl5 | 0.45 | 7.16E-13 | 1.49E-08 | 0.86 | 0.99 | -0.13 | 0.87 |
| 17 | Myl6 | 0.39 | 7.18E-13 | 1.49E-08 | 0.89 | 0.99 | -0.11 | 0.89 |
| 17 | Hspb8 | 1.91 | 9.95E-13 | 2.07E-08 | 0.25 | 0.14 | 0.11 | 1.83 |
| 17 | Hspa8 | 0.36 | 1.12E-12 | 2.32E-08 | 0.99 | 1.00 | -0.01 | 0.99 |
| 17 | Dmrta2 | 1.72 | 1.36E-12 | 2.83E-08 | 0.26 | 0.14 | 0.12 | 1.84 |
| 17 | Gm266 | 1.90 | 1.78E-12 | 3.70E-08 | 0.41 | 0.31 | 0.10 | 1.31 |
| 17 | Pax6 | 2.19 | 1.78E-12 | 3.70E-08 | 0.24 | 0.13 | 0.11 | 1.86 |
| 17 | Espn | 2.39 | 2.08E-12 | 4.32E-08 | 0.21 | 0.11 | 0.11 | 1.96 |
| 17 | Socs2 | 1.35 | 2.24E-12 | 4.67E-08 | 0.45 | 0.35 | 0.11 | 1.31 |
| 17 | Trim37 | 0.27 | 2.50E-12 | 5.19E-08 | 0.16 | 0.40 | -0.24 | 0.40 |
| 17 | Eef2 | 0.32 | 2.61E-12 | 5.43E-08 | 0.93 | 1.00 | -0.07 | 0.93 |
| 17 | Ttc37 | 0.36 | 3.67E-12 | 7.62E-08 | 0.12 | 0.33 | -0.21 | 0.36 |
| 17 | Ppfia1 | 0.31 | 4.95E-12 | 1.03E-07 | 0.27 | 0.57 | -0.30 | 0.47 |
| 17 | Ercc3 | 0.49 | 8.21E-12 | 1.71E-07 | 0.07 | 0.25 | -0.18 | 0.29 |
| 17 | Cox6b1 | 0.29 | 1.06E-11 | 2.21E-07 | 0.92 | 1.00 | -0.07 | 0.93 |
| 17 | Gsta4 | 1.09 | 1.17E-11 | 2.44E-07 | 0.66 | 0.76 | -0.10 | 0.87 |
| 17 | Nudt16l1 | 0.32 | 1.33E-11 | 2.77E-07 | 0.23 | 0.52 | -0.29 | 0.44 |
| 17 | Lurap1l | 2.55 | 1.51E-11 | 3.13E-07 | 0.17 | 0.08 | 0.09 | 2.17 |
| 17 | Tmsb4x | 0.28 | 1.81E-11 | 3.77E-07 | 0.98 | 0.99 | -0.01 | 0.99 |
| 17 | Uqcrh | 0.29 | 1.97E-11 | 4.09E-07 | 0.92 | 1.00 | -0.08 | 0.92 |
| 17 | Dimt1 | 0.41 | 2.10E-11 | 4.36E-07 | 0.17 | 0.42 | -0.24 | 0.42 |
| 17 | Fgf17 | 3.01 | 2.25E-11 | 4.68E-07 | 0.11 | 0.04 | 0.07 | 2.65 |
| 17 | Crb3 | 2.25 | 2.45E-11 | 5.10E-07 | 0.21 | 0.11 | 0.10 | 1.87 |
| 17 | Edf1 | 0.56 | 2.69E-11 | 5.60E-07 | 0.77 | 0.96 | -0.19 | 0.80 |
| 17 | Atp6v1b2 | 0.27 | 2.71E-11 | 5.63E-07 | 0.16 | 0.39 | -0.24 | 0.40 |
| 17 | Ado | 0.59 | 3.22E-11 | 6.70E-07 | 0.16 | 0.40 | -0.23 | 0.41 |
| 17 | Crnkl1 | 0.26 | 3.60E-11 | 7.48E-07 | 0.26 | 0.56 | -0.30 | 0.46 |
| 17 | Cdk4 | 0.27 | 3.76E-11 | 7.82E-07 | 0.93 | 0.99 | -0.06 | 0.94 |
| 17 | Cenpb | 0.26 | 3.97E-11 | 8.25E-07 | 0.33 | 0.67 | -0.34 | 0.49 |
| 17 | Nsa2 | 0.49 | 4.00E-11 | 8.31E-07 | 0.84 | 0.97 | -0.13 | 0.86 |
| 17 | Tmem47 | 0.26 | 4.39E-11 | 9.12E-07 | 0.15 | 0.37 | -0.22 | 0.40 |
| 17 | Rpe | 0.51 | 4.47E-11 | 9.29E-07 | 0.13 | 0.34 | -0.21 | 0.38 |
| 17 | Gad2 | 2.49 | 4.97E-11 | 1.03E-06 | 0.16 | 0.07 | 0.09 | 2.18 |
| 17 | Nipsnap2 | 0.34 | 4.97E-11 | 1.03E-06 | 0.21 | 0.46 | -0.25 | 0.46 |
| 17 | Chmp6 | 0.33 | 4.97E-11 | 1.03E-06 | 0.23 | 0.51 | -0.28 | 0.45 |
| 17 | Commd7 | 0.26 | 5.19E-11 | 1.08E-06 | 0.28 | 0.58 | -0.30 | 0.48 |
| 17 | Pno1 | 0.30 | 6.16E-11 | 1.28E-06 | 0.29 | 0.60 | -0.31 | 0.49 |
| 17 | Cep120 | 0.26 | 6.86E-11 | 1.43E-06 | 0.07 | 0.24 | -0.17 | 0.30 |
| 17 | Ppp6r1 | 0.37 | 7.17E-11 | 1.49E-06 | 0.08 | 0.26 | -0.17 | 0.32 |
| 17 | Pdgfc | 1.41 | 8.82E-11 | 1.83E-06 | 0.39 | 0.28 | 0.10 | 1.37 |
| 17 | Champ1 | 0.37 | 9.00E-11 | 1.87E-06 | 0.14 | 0.36 | -0.22 | 0.40 |
| 17 | Fam107a | 2.58 | 9.13E-11 | 1.90E-06 | 0.11 | 0.04 | 0.07 | 2.59 |
| 17 | Tbca | 0.38 | 9.17E-11 | 1.91E-06 | 0.86 | 0.99 | -0.13 | 0.87 |
| 17 | Tatdn1 | 0.28 | 1.00E-10 | 2.09E-06 | 0.16 | 0.38 | -0.22 | 0.42 |
| 17 | Ndufaf3 | 0.55 | 1.24E-10 | 2.58E-06 | 0.18 | 0.42 | -0.24 | 0.43 |
| 17 | Eif2d | 0.28 | 1.25E-10 | 2.60E-06 | 0.11 | 0.30 | -0.19 | 0.37 |
| 17 | Uap1 | 0.31 | 1.52E-10 | 3.17E-06 | 0.12 | 0.32 | -0.20 | 0.38 |
| 17 | Kctd17 | 0.34 | 1.55E-10 | 3.23E-06 | 0.10 | 0.28 | -0.18 | 0.36 |
| 17 | Cdkn1a | 1.46 | 1.61E-10 | 3.34E-06 | 0.21 | 0.11 | 0.10 | 1.90 |
| 17 | Nup85 | 0.29 | 1.67E-10 | 3.48E-06 | 0.15 | 0.37 | -0.22 | 0.41 |
| 17 | Pinx1 | 0.36 | 1.72E-10 | 3.59E-06 | 0.27 | 0.55 | -0.29 | 0.48 |
| 17 | Oxsr1 | 0.29 | 1.90E-10 | 3.94E-06 | 0.12 | 0.32 | -0.20 | 0.38 |
| 17 | Gnl2 | 0.25 | 2.13E-10 | 4.42E-06 | 0.25 | 0.53 | -0.28 | 0.47 |
| 17 | Szrd1 | 0.26 | 2.19E-10 | 4.56E-06 | 0.24 | 0.52 | -0.27 | 0.47 |
| 17 | Aip | 0.27 | 2.26E-10 | 4.71E-06 | 0.18 | 0.42 | -0.24 | 0.44 |
| 17 | Lmo4 | 1.03 | 2.33E-10 | 4.84E-06 | 0.68 | 0.82 | -0.13 | 0.84 |

**Supplementary Table 1. Marker genes for the original clusters of integrated clustering of E11.5 control and NotchGOF λ-junction cells.**

| | | | | | | | | |
|---|---|---|---|---|---|---|---|---|
| 17 | Nol7 | 0.78 | 2.36E-10 | 4.91E-06 | 0.71 | 0.88 | -0.17 | 0.81 |
| 17 | Slc39a1 | 0.42 | 2.40E-10 | 4.99E-06 | 0.09 | 0.26 | -0.17 | 0.35 |
| 17 | Kifc1 | 0.34 | 2.43E-10 | 5.06E-06 | 0.11 | 0.29 | -0.18 | 0.37 |
| 17 | Cdk2 | 0.28 | 2.51E-10 | 5.21E-06 | 0.18 | 0.42 | -0.24 | 0.43 |
| 17 | Fgd2 | 2.19 | 2.71E-10 | 5.63E-06 | 0.19 | 0.10 | 0.09 | 1.89 |
| 17 | Coq2 | 0.27 | 3.29E-10 | 6.84E-06 | 0.21 | 0.46 | -0.25 | 0.45 |
| 17 | Elf5 | 2.91 | 3.62E-10 | 7.52E-06 | 0.14 | 0.06 | 0.08 | 2.26 |
| 17 | Fam117b | 0.40 | 3.98E-10 | 8.28E-06 | 0.12 | 0.31 | -0.19 | 0.38 |
| 17 | 2700062C07Rik | 0.37 | 4.01E-10 | 8.34E-06 | 0.17 | 0.39 | -0.22 | 0.44 |
| 17 | Ppat | 0.35 | 4.02E-10 | 8.36E-06 | 0.25 | 0.51 | -0.26 | 0.49 |
| 17 | Wfdc2 | 1.13 | 4.06E-10 | 8.44E-06 | 0.52 | 0.46 | 0.05 | 1.12 |
| 17 | Rrs1 | 0.26 | 4.30E-10 | 8.94E-06 | 0.31 | 0.61 | -0.31 | 0.50 |
| 17 | Cox5b | 0.38 | 4.32E-10 | 8.98E-06 | 0.86 | 0.99 | -0.13 | 0.87 |
| 17 | Wdr74 | 0.29 | 4.70E-10 | 9.76E-06 | 0.26 | 0.55 | -0.29 | 0.48 |
| 17 | Rpf1 | 0.38 | 4.92E-10 | 1.02E-05 | 0.26 | 0.54 | -0.28 | 0.49 |
| 17 | Ech1 | 0.34 | 4.99E-10 | 1.04E-05 | 0.24 | 0.52 | -0.28 | 0.47 |
| 17 | Cluap1 | 0.39 | 5.06E-10 | 1.05E-05 | 0.17 | 0.39 | -0.22 | 0.44 |
| 17 | Tmem54 | 2.12 | 5.27E-10 | 1.10E-05 | 0.21 | 0.12 | 0.09 | 1.78 |
| 17 | Usp8 | 0.45 | 5.70E-10 | 1.18E-05 | 0.21 | 0.47 | -0.26 | 0.45 |
| 17 | Tjp3 | 2.97 | 5.83E-10 | 1.21E-05 | 0.12 | 0.05 | 0.07 | 2.39 |
| 17 | Cystm1 | 2.16 | 6.06E-10 | 1.26E-05 | 0.18 | 0.09 | 0.09 | 1.96 |
| 17 | Dtnbp1 | 0.45 | 6.26E-10 | 1.30E-05 | 0.19 | 0.43 | -0.23 | 0.45 |
| 17 | Cmtm4 | 0.34 | 7.36E-10 | 1.53E-05 | 0.09 | 0.26 | -0.17 | 0.36 |
| 17 | Fam207a | 0.48 | 7.50E-10 | 1.56E-05 | 0.14 | 0.35 | -0.20 | 0.41 |
| 17 | Rexo4 | 0.48 | 7.58E-10 | 1.58E-05 | 0.21 | 0.46 | -0.25 | 0.46 |
| 17 | Nmt2 | 0.30 | 7.93E-10 | 1.65E-05 | 0.15 | 0.35 | -0.20 | 0.42 |
| 17 | Ndc80 | 0.29 | 8.14E-10 | 1.69E-05 | 0.20 | 0.43 | -0.22 | 0.47 |
| 17 | Fermt1 | 1.49 | 8.79E-10 | 1.83E-05 | 0.24 | 0.14 | 0.10 | 1.72 |
| 17 | Necap2 | 0.49 | 8.93E-10 | 1.86E-05 | 0.18 | 0.40 | -0.22 | 0.44 |
| 17 | Cstf2 | 0.25 | 9.71E-10 | 2.02E-05 | 0.15 | 0.35 | -0.20 | 0.42 |
| 17 | Bcl7b | 0.30 | 1.01E-09 | 2.11E-05 | 0.18 | 0.41 | -0.22 | 0.45 |
| 17 | Arl4a | 0.39 | 1.07E-09 | 2.23E-05 | 0.06 | 0.21 | -0.15 | 0.30 |
| 17 | Spata24 | 0.40 | 1.10E-09 | 2.28E-05 | 0.11 | 0.29 | -0.18 | 0.38 |
| 17 | Lasp1 | 0.37 | 1.10E-09 | 2.28E-05 | 0.20 | 0.44 | -0.23 | 0.46 |
| 17 | Dpm2 | 0.28 | 1.11E-09 | 2.32E-05 | 0.29 | 0.58 | -0.29 | 0.50 |
| 17 | Lamtor3 | 0.32 | 1.15E-09 | 2.39E-05 | 0.22 | 0.48 | -0.25 | 0.47 |
| 17 | Fam3c | 0.27 | 1.18E-09 | 2.45E-05 | 0.16 | 0.37 | -0.21 | 0.44 |
| 17 | Mtch1 | 0.44 | 1.22E-09 | 2.53E-05 | 0.81 | 0.96 | -0.15 | 0.85 |
| 17 | Zfp560 | 0.28 | 1.24E-09 | 2.57E-05 | 0.16 | 0.38 | -0.21 | 0.44 |
| 17 | Dlx6 | 2.47 | 1.28E-09 | 2.66E-05 | 0.17 | 0.09 | 0.08 | 1.95 |
| 17 | Tprgl | 0.29 | 1.32E-09 | 2.75E-05 | 0.31 | 0.61 | -0.30 | 0.50 |
| 17 | Ubl3 | 0.28 | 1.33E-09 | 2.76E-05 | 0.17 | 0.38 | -0.21 | 0.44 |
| 17 | Fastk | 0.39 | 1.33E-09 | 2.77E-05 | 0.20 | 0.43 | -0.24 | 0.45 |
| 17 | Mfap1b | 0.31 | 1.43E-09 | 2.98E-05 | 0.15 | 0.36 | -0.20 | 0.43 |
| 17 | Ube2g1 | 0.28 | 1.45E-09 | 3.01E-05 | 0.31 | 0.61 | -0.31 | 0.50 |
| 17 | Spindoc | 0.34 | 1.52E-09 | 3.16E-05 | 0.12 | 0.31 | -0.19 | 0.40 |
| 17 | Znhit1 | 0.26 | 1.66E-09 | 3.46E-05 | 0.33 | 0.66 | -0.33 | 0.50 |
| 17 | Tusc2 | 0.36 | 1.67E-09 | 3.47E-05 | 0.18 | 0.40 | -0.22 | 0.45 |
| 17 | Hibadh | 0.27 | 1.81E-09 | 3.77E-05 | 0.27 | 0.56 | -0.29 | 0.49 |
| 17 | Mark1 | 0.44 | 1.94E-09 | 4.03E-05 | 0.09 | 0.25 | -0.16 | 0.36 |
| 17 | Fbxl17 | 0.28 | 1.94E-09 | 4.04E-05 | 0.06 | 0.20 | -0.14 | 0.29 |
| 17 | Gtpbp10 | 0.38 | 1.97E-09 | 4.09E-05 | 0.13 | 0.32 | -0.19 | 0.41 |
| 17 | Tmem120a | 0.28 | 2.00E-09 | 4.15E-05 | 0.12 | 0.30 | -0.18 | 0.40 |
| 17 | Pigyl | 0.26 | 2.00E-09 | 4.17E-05 | 0.24 | 0.49 | -0.26 | 0.48 |
| 17 | Mrpl10 | 0.35 | 2.06E-09 | 4.28E-05 | 0.23 | 0.49 | -0.26 | 0.47 |
| 17 | Rad18 | 0.27 | 2.11E-09 | 4.38E-05 | 0.17 | 0.37 | -0.21 | 0.45 |

**Supplementary Table 1. Marker genes for the original clusters of integrated clustering of E11.5 control and NotchGOF λ-junction cells.**

| | | | | | | | | |
|---|---|---|---|---|---|---|---|---|
| 17 | Nif3l1 | 0.26 | 2.15E-09 | 4.47E-05 | 0.16 | 0.36 | -0.21 | 0.43 |
| 17 | Nbdy | 0.25 | 2.33E-09 | 4.85E-05 | 0.32 | 0.63 | -0.31 | 0.51 |
| 17 | Bag5 | 0.27 | 2.34E-09 | 4.86E-05 | 0.24 | 0.50 | -0.26 | 0.47 |
| 17 | Usp39 | 0.44 | 2.36E-09 | 4.90E-05 | 0.30 | 0.59 | -0.30 | 0.50 |
| 17 | Ate1 | 0.28 | 2.46E-09 | 5.12E-05 | 0.21 | 0.45 | -0.24 | 0.47 |
| 17 | Rnf138 | 0.30 | 2.48E-09 | 5.16E-05 | 0.26 | 0.53 | -0.27 | 0.50 |
| 17 | Tomm7 | 0.39 | 2.52E-09 | 5.24E-05 | 0.86 | 0.99 | -0.13 | 0.87 |
| 17 | Cox8a | 0.27 | 2.53E-09 | 5.25E-05 | 0.93 | 1.00 | -0.07 | 0.93 |
| 17 | Tfip11 | 0.30 | 2.70E-09 | 5.62E-05 | 0.09 | 0.25 | -0.16 | 0.36 |
| 17 | Xrcc6 | 0.28 | 2.85E-09 | 5.93E-05 | 0.20 | 0.44 | -0.24 | 0.46 |
| 17 | Herc4 | 0.30 | 2.91E-09 | 6.05E-05 | 0.12 | 0.30 | -0.18 | 0.40 |
| 17 | Ap1ar | 0.25 | 3.16E-09 | 6.58E-05 | 0.28 | 0.56 | -0.28 | 0.51 |
| 17 | Golga5 | 0.39 | 3.18E-09 | 6.61E-05 | 0.10 | 0.26 | -0.16 | 0.37 |
| 17 | Sall2 | 0.29 | 3.20E-09 | 6.66E-05 | 0.18 | 0.40 | -0.22 | 0.46 |
| 17 | Arid4a | 0.37 | 3.31E-09 | 6.88E-05 | 0.23 | 0.48 | -0.25 | 0.48 |
| 17 | Riok2 | 0.32 | 3.44E-09 | 7.15E-05 | 0.20 | 0.42 | -0.22 | 0.47 |
| 17 | 2700049A03Rik | 0.42 | 3.51E-09 | 7.30E-05 | 0.14 | 0.32 | -0.18 | 0.43 |
| 17 | Klhdc4 | 0.40 | 3.61E-09 | 7.51E-05 | 0.15 | 0.35 | -0.20 | 0.43 |
| 17 | Six1 | 0.81 | 3.71E-09 | 7.72E-05 | 0.62 | 0.61 | 0.01 | 1.01 |
| 17 | Sacm1l | 0.30 | 3.97E-09 | 8.25E-05 | 0.14 | 0.33 | -0.19 | 0.43 |
| 17 | Gprasp2 | 0.27 | 3.97E-09 | 8.26E-05 | 0.13 | 0.30 | -0.18 | 0.41 |
| 17 | Otud6b | 0.35 | 4.08E-09 | 8.49E-05 | 0.19 | 0.41 | -0.22 | 0.46 |
| 17 | Eif2b4 | 0.31 | 4.08E-09 | 8.49E-05 | 0.16 | 0.37 | -0.21 | 0.44 |
| 17 | BC005624 | 0.31 | 4.12E-09 | 8.57E-05 | 0.30 | 0.61 | -0.31 | 0.49 |
| 17 | Rpl4 | 0.27 | 4.44E-09 | 9.23E-05 | 0.97 | 1.00 | -0.02 | 0.98 |
| 17 | Arfrp1 | 0.30 | 4.68E-09 | 9.74E-05 | 0.28 | 0.56 | -0.28 | 0.50 |
| 17 | Prxl2c | 0.54 | 4.78E-09 | 9.93E-05 | 0.12 | 0.30 | -0.18 | 0.41 |
| 17 | Ppfia3 | 2.31 | 5.22E-09 | 1.09E-04 | 0.21 | 0.13 | 0.08 | 1.65 |
| 17 | Sars | 0.45 | 5.42E-09 | 1.13E-04 | 0.37 | 0.72 | -0.35 | 0.52 |
| 17 | Fam174a | 0.29 | 5.80E-09 | 1.21E-04 | 0.14 | 0.31 | -0.18 | 0.43 |
| 17 | Ube2h | 0.30 | 5.83E-09 | 1.21E-04 | 0.23 | 0.47 | -0.25 | 0.48 |
| 17 | Nrbp1 | 0.25 | 6.02E-09 | 1.25E-04 | 0.28 | 0.57 | -0.28 | 0.50 |
| 17 | Nob1 | 0.27 | 6.47E-09 | 1.35E-04 | 0.26 | 0.52 | -0.26 | 0.50 |
| 17 | Pithd1 | 0.48 | 6.47E-09 | 1.35E-04 | 0.22 | 0.46 | -0.24 | 0.48 |
| 17 | Akt2 | 0.39 | 6.56E-09 | 1.36E-04 | 0.16 | 0.36 | -0.20 | 0.44 |
| 17 | Rnf6 | 0.32 | 6.71E-09 | 1.40E-04 | 0.16 | 0.36 | -0.20 | 0.44 |
| 17 | Gtpbp6 | 0.50 | 6.81E-09 | 1.42E-04 | 0.11 | 0.27 | -0.16 | 0.39 |
| 17 | Clp1 | 0.32 | 7.88E-09 | 1.64E-04 | 0.15 | 0.35 | -0.19 | 0.44 |
| 17 | Dexi | 0.30 | 8.23E-09 | 1.71E-04 | 0.18 | 0.39 | -0.21 | 0.46 |
| 17 | Asb8 | 0.30 | 8.26E-09 | 1.72E-04 | 0.13 | 0.30 | -0.18 | 0.41 |
| 17 | Swi5 | 0.33 | 8.29E-09 | 1.72E-04 | 0.89 | 0.99 | -0.10 | 0.90 |
| 17 | Kri1 | 0.35 | 8.59E-09 | 1.79E-04 | 0.05 | 0.18 | -0.13 | 0.29 |
| 17 | Eif2b2 | 0.42 | 8.88E-09 | 1.85E-04 | 0.25 | 0.50 | -0.25 | 0.49 |
| 17 | 2310011J03Rik | 0.47 | 8.94E-09 | 1.86E-04 | 0.24 | 0.49 | -0.25 | 0.49 |
| 17 | Mrpl52 | 0.47 | 9.25E-09 | 1.92E-04 | 0.80 | 0.96 | -0.17 | 0.83 |
| 17 | Lzic | 0.44 | 9.57E-09 | 1.99E-04 | 0.26 | 0.52 | -0.26 | 0.49 |
| 17 | Yju2 | 0.35 | 9.62E-09 | 2.00E-04 | 0.09 | 0.25 | -0.16 | 0.37 |
| 17 | Nt5dc2 | 0.76 | 9.65E-09 | 2.01E-04 | 0.75 | 0.92 | -0.18 | 0.81 |
| 17 | Psmd5 | 0.29 | 9.99E-09 | 2.08E-04 | 0.26 | 0.52 | -0.27 | 0.49 |
| 17 | Ndufaf7 | 0.42 | 1.01E-08 | 2.10E-04 | 0.14 | 0.32 | -0.18 | 0.43 |
| 17 | Gnl1 | 0.53 | 1.02E-08 | 2.12E-04 | 0.23 | 0.48 | -0.25 | 0.48 |
| 17 | Upf3b | 0.26 | 1.02E-08 | 2.12E-04 | 0.37 | 0.71 | -0.33 | 0.53 |
| 17 | Atp6v0e | 0.26 | 1.07E-08 | 2.22E-04 | 0.38 | 0.72 | -0.34 | 0.53 |
| 17 | Abitram | 0.45 | 1.10E-08 | 2.28E-04 | 0.09 | 0.23 | -0.15 | 0.36 |
| 17 | Nudcd1 | 0.27 | 1.12E-08 | 2.33E-04 | 0.19 | 0.41 | -0.22 | 0.47 |
| 17 | Hexim1 | 0.34 | 1.13E-08 | 2.35E-04 | 0.12 | 0.29 | -0.17 | 0.41 |

**Supplementary Table 1. Marker genes for the original clusters of integrated clustering of E11.5 control and NotchGOF λ-junction cells.**

| | | | | | | | | |
|---|---|---|---|---|---|---|---|---|
| 17 | Zmynd19 | 0.36 | 1.13E-08 | 2.35E-04 | 0.24 | 0.49 | -0.25 | 0.49 |
| 17 | Dph6 | 0.32 | 1.17E-08 | 2.43E-04 | 0.19 | 0.42 | -0.22 | 0.46 |
| 17 | Tysnd1 | 0.43 | 1.18E-08 | 2.45E-04 | 0.15 | 0.34 | -0.19 | 0.45 |
| 17 | Hist1h4i | 0.26 | 1.19E-08 | 2.48E-04 | 0.23 | 0.46 | -0.23 | 0.50 |
| 17 | Nme6 | 0.26 | 1.29E-08 | 2.68E-04 | 0.21 | 0.44 | -0.23 | 0.48 |
| 17 | Tmem179b | 0.35 | 1.32E-08 | 2.74E-04 | 0.13 | 0.30 | -0.17 | 0.42 |
| 17 | Cdkn2d | 0.28 | 1.32E-08 | 2.75E-04 | 0.18 | 0.39 | -0.21 | 0.46 |
| 17 | Prelid2 | 0.29 | 1.33E-08 | 2.77E-04 | 0.17 | 0.37 | -0.20 | 0.46 |
| 17 | Cyb5b | 0.31 | 1.36E-08 | 2.82E-04 | 0.25 | 0.51 | -0.26 | 0.49 |
| 17 | Nectin1 | 0.44 | 1.44E-08 | 3.00E-04 | 0.05 | 0.18 | -0.13 | 0.30 |
| 17 | Krt14 | 1.59 | 1.48E-08 | 3.08E-04 | 0.23 | 0.13 | 0.10 | 1.71 |
| 17 | Cnot10 | 0.34 | 1.49E-08 | 3.10E-04 | 0.23 | 0.47 | -0.24 | 0.49 |
| 17 | Vps51 | 0.46 | 1.51E-08 | 3.15E-04 | 0.07 | 0.21 | -0.14 | 0.34 |
| 17 | Magoh | 0.47 | 1.53E-08 | 3.18E-04 | 0.78 | 0.94 | -0.16 | 0.83 |
| 17 | Ppdpf | 0.45 | 1.57E-08 | 3.26E-04 | 0.82 | 0.97 | -0.14 | 0.85 |
| 17 | Cep57l1 | 0.46 | 1.60E-08 | 3.33E-04 | 0.12 | 0.28 | -0.17 | 0.41 |
| 17 | Ccdc102a | 0.29 | 1.65E-08 | 3.42E-04 | 0.11 | 0.28 | -0.17 | 0.41 |
| 17 | Cfap36 | 0.27 | 1.65E-08 | 3.43E-04 | 0.28 | 0.56 | -0.27 | 0.51 |
| 17 | Alas1 | 0.27 | 1.77E-08 | 3.68E-04 | 0.16 | 0.34 | -0.19 | 0.45 |
| 17 | Anxa2 | 1.08 | 1.80E-08 | 3.75E-04 | 0.48 | 0.42 | 0.06 | 1.15 |
| 17 | Kti12 | 0.35 | 1.96E-08 | 4.08E-04 | 0.21 | 0.45 | -0.24 | 0.47 |
| 17 | Ppp1r35 | 0.32 | 2.00E-08 | 4.16E-04 | 0.29 | 0.56 | -0.27 | 0.52 |
| 17 | Ppm1b | 0.33 | 2.06E-08 | 4.29E-04 | 0.28 | 0.55 | -0.27 | 0.51 |
| 17 | Gas5 | 0.34 | 2.14E-08 | 4.45E-04 | 0.89 | 0.99 | -0.10 | 0.90 |
| 17 | Ubl7 | 0.51 | 2.19E-08 | 4.55E-04 | 0.18 | 0.39 | -0.21 | 0.47 |
| 17 | Scamp2 | 0.32 | 2.19E-08 | 4.56E-04 | 0.14 | 0.32 | -0.18 | 0.44 |
| 17 | Thumpd3 | 0.38 | 2.23E-08 | 4.64E-04 | 0.18 | 0.39 | -0.21 | 0.47 |
| 17 | Snrpd2 | 0.25 | 2.27E-08 | 4.71E-04 | 0.91 | 0.99 | -0.09 | 0.91 |
| 17 | 1810037I17Rik | 0.84 | 2.27E-08 | 4.72E-04 | 0.65 | 0.84 | -0.19 | 0.78 |
| 17 | Ccdc71 | 0.43 | 2.35E-08 | 4.89E-04 | 0.14 | 0.32 | -0.18 | 0.44 |
| 17 | Tpgs1 | 0.45 | 2.36E-08 | 4.92E-04 | 0.21 | 0.44 | -0.23 | 0.48 |
| 17 | Zbtb33 | 0.31 | 2.52E-08 | 5.24E-04 | 0.11 | 0.27 | -0.16 | 0.41 |
| 17 | Coa6 | 0.28 | 2.53E-08 | 5.25E-04 | 0.21 | 0.44 | -0.22 | 0.49 |
| 17 | Krcc1 | 0.26 | 2.64E-08 | 5.49E-04 | 0.30 | 0.58 | -0.29 | 0.51 |
| 17 | Aldh16a1 | 0.35 | 2.67E-08 | 5.55E-04 | 0.08 | 0.23 | -0.14 | 0.36 |
| 17 | 2510002D24Rik | 0.34 | 2.72E-08 | 5.67E-04 | 0.19 | 0.40 | -0.21 | 0.48 |
| 17 | Fbxw7 | 0.36 | 2.85E-08 | 5.93E-04 | 0.18 | 0.39 | -0.21 | 0.46 |
| 17 | Park7 | 0.36 | 2.93E-08 | 6.10E-04 | 0.85 | 0.99 | -0.13 | 0.87 |
| 17 | Trappc13 | 0.35 | 2.94E-08 | 6.11E-04 | 0.21 | 0.43 | -0.23 | 0.48 |
| 17 | Sap30bp | 0.48 | 2.95E-08 | 6.14E-04 | 0.26 | 0.52 | -0.26 | 0.50 |
| 17 | Tbcc | 0.34 | 2.97E-08 | 6.17E-04 | 0.23 | 0.47 | -0.24 | 0.49 |
| 17 | Rragc | 0.26 | 2.99E-08 | 6.22E-04 | 0.23 | 0.46 | -0.23 | 0.50 |
| 17 | Taf10 | 0.54 | 3.03E-08 | 6.30E-04 | 0.75 | 0.93 | -0.18 | 0.80 |
| 17 | Rangrf | 0.29 | 3.12E-08 | 6.49E-04 | 0.28 | 0.54 | -0.26 | 0.52 |
| 17 | Dgkz | 0.31 | 3.12E-08 | 6.49E-04 | 0.12 | 0.29 | -0.17 | 0.42 |
| 17 | Nosip | 0.50 | 3.30E-08 | 6.86E-04 | 0.15 | 0.34 | -0.19 | 0.45 |
| 17 | Rab5b | 0.43 | 3.45E-08 | 7.17E-04 | 0.11 | 0.28 | -0.16 | 0.41 |
| 17 | Rad54b | 0.40 | 3.46E-08 | 7.21E-04 | 0.07 | 0.21 | -0.13 | 0.35 |
| 17 | Gnl3l | 0.35 | 3.48E-08 | 7.24E-04 | 0.15 | 0.33 | -0.18 | 0.45 |
| 17 | Zfp277 | 0.33 | 3.52E-08 | 7.31E-04 | 0.22 | 0.43 | -0.22 | 0.50 |
| 17 | Camk1 | 0.32 | 3.80E-08 | 7.90E-04 | 0.21 | 0.43 | -0.22 | 0.49 |
| 17 | Dram2 | 0.40 | 4.07E-08 | 8.46E-04 | 0.07 | 0.21 | -0.14 | 0.35 |
| 17 | Mycbp | 0.29 | 4.07E-08 | 8.47E-04 | 0.26 | 0.52 | -0.26 | 0.51 |
| 17 | Xab2 | 0.60 | 4.15E-08 | 8.63E-04 | 0.13 | 0.30 | -0.17 | 0.43 |
| 17 | Sugp1 | 0.58 | 4.18E-08 | 8.68E-04 | 0.12 | 0.28 | -0.16 | 0.42 |
| 17 | Oxr1 | 0.30 | 4.20E-08 | 8.73E-04 | 0.16 | 0.33 | -0.18 | 0.46 |

**Supplementary Table 1. Marker genes for the original clusters of integrated clustering of E11.5 control and NotchGOF λ-junction cells.**

| | | | | | | | | |
|---|---|---|---|---|---|---|---|---|
| 17 | Smim19 | 0.36 | 4.20E-08 | 8.73E-04 | 0.21 | 0.42 | -0.22 | 0.49 |
| 17 | Sec24a | 0.46 | 4.27E-08 | 8.89E-04 | 0.06 | 0.18 | -0.12 | 0.31 |
| 17 | Eya1 | 0.28 | 4.31E-08 | 8.96E-04 | 0.26 | 0.50 | -0.24 | 0.52 |
| 17 | Pop5 | 0.28 | 4.39E-08 | 9.13E-04 | 0.35 | 0.66 | -0.31 | 0.53 |
| 17 | Mgat4b | 0.29 | 4.67E-08 | 9.72E-04 | 0.19 | 0.40 | -0.21 | 0.48 |
| 17 | Dlg3 | 0.43 | 4.69E-08 | 9.76E-04 | 0.15 | 0.33 | -0.18 | 0.45 |
| 17 | Slc25a13 | 0.31 | 4.95E-08 | 1.03E-03 | 0.22 | 0.45 | -0.23 | 0.49 |
| 17 | Anks1 | 0.27 | 5.08E-08 | 1.06E-03 | 0.05 | 0.16 | -0.12 | 0.29 |
| 17 | Fdx1 | 0.35 | 5.24E-08 | 1.09E-03 | 0.28 | 0.54 | -0.26 | 0.51 |
| 17 | Ubxn2b | 0.29 | 5.33E-08 | 1.11E-03 | 0.10 | 0.25 | -0.15 | 0.40 |
| 17 | Arhgap8 | 2.97 | 5.38E-08 | 1.12E-03 | 0.10 | 0.04 | 0.06 | 2.33 |
| 17 | Rrp9 | 0.29 | 5.45E-08 | 1.13E-03 | 0.16 | 0.35 | -0.19 | 0.47 |
| 17 | Lztfl1 | 0.40 | 5.63E-08 | 1.17E-03 | 0.26 | 0.50 | -0.24 | 0.51 |
| 17 | Eri3 | 0.30 | 5.68E-08 | 1.18E-03 | 0.31 | 0.60 | -0.29 | 0.52 |
| 17 | Nle1 | 0.28 | 5.72E-08 | 1.19E-03 | 0.09 | 0.24 | -0.14 | 0.39 |
| 17 | Prpsap2 | 0.52 | 5.90E-08 | 1.23E-03 | 0.11 | 0.27 | -0.16 | 0.41 |
| 17 | Zfp800 | 0.38 | 6.13E-08 | 1.28E-03 | 0.11 | 0.27 | -0.16 | 0.42 |
| 17 | Zfand1 | 0.27 | 6.26E-08 | 1.30E-03 | 0.16 | 0.35 | -0.19 | 0.46 |
| 17 | Crcp | 0.58 | 6.37E-08 | 1.32E-03 | 0.18 | 0.38 | -0.20 | 0.47 |
| 17 | Eif3h | 0.35 | 6.69E-08 | 1.39E-03 | 0.88 | 0.99 | -0.11 | 0.89 |
| 17 | Cldn6 | 0.95 | 6.87E-08 | 1.43E-03 | 0.34 | 0.23 | 0.11 | 1.48 |
| 17 | Pcbd2 | 0.32 | 6.99E-08 | 1.45E-03 | 0.21 | 0.44 | -0.23 | 0.49 |
| 17 | Prim1 | 0.40 | 7.17E-08 | 1.49E-03 | 0.36 | 0.68 | -0.32 | 0.54 |
| 17 | Coq9 | 0.30 | 7.18E-08 | 1.49E-03 | 0.11 | 0.27 | -0.16 | 0.42 |
| 17 | Aatf | 0.44 | 7.30E-08 | 1.52E-03 | 0.18 | 0.38 | -0.20 | 0.47 |
| 17 | Ubc | 0.46 | 7.56E-08 | 1.57E-03 | 0.87 | 0.98 | -0.11 | 0.89 |
| 17 | Thap7 | 0.28 | 8.40E-08 | 1.75E-03 | 0.16 | 0.34 | -0.18 | 0.46 |
| 17 | Thoc5 | 0.32 | 8.64E-08 | 1.80E-03 | 0.17 | 0.36 | -0.19 | 0.47 |
| 17 | Cdk5rap3 | 0.37 | 8.66E-08 | 1.80E-03 | 0.22 | 0.44 | -0.22 | 0.50 |
| 17 | Arpin | 0.29 | 8.86E-08 | 1.84E-03 | 0.06 | 0.18 | -0.12 | 0.33 |
| 17 | Btbd10 | 0.69 | 9.39E-08 | 1.95E-03 | 0.10 | 0.25 | -0.15 | 0.41 |
| 17 | Wtip | 0.28 | 9.71E-08 | 2.02E-03 | 0.29 | 0.57 | -0.27 | 0.52 |
| 17 | Vps28 | 0.25 | 9.84E-08 | 2.05E-03 | 0.33 | 0.63 | -0.29 | 0.53 |
| 17 | Flot1 | 0.27 | 1.00E-07 | 2.08E-03 | 0.14 | 0.31 | -0.17 | 0.46 |
| 17 | Sh3yl1 | 0.32 | 1.01E-07 | 2.10E-03 | 0.07 | 0.20 | -0.13 | 0.35 |
| 17 | Pip4p1 | 0.50 | 1.04E-07 | 2.16E-03 | 0.17 | 0.37 | -0.19 | 0.47 |
| 17 | Cpsf3 | 0.58 | 1.06E-07 | 2.20E-03 | 0.24 | 0.47 | -0.24 | 0.50 |
| 17 | Cep85 | 0.28 | 1.07E-07 | 2.23E-03 | 0.05 | 0.17 | -0.12 | 0.32 |
| 17 | Kctd3 | 0.61 | 1.08E-07 | 2.24E-03 | 0.13 | 0.29 | -0.17 | 0.44 |
| 17 | Ubash3b | 0.44 | 1.09E-07 | 2.27E-03 | 0.10 | 0.24 | -0.14 | 0.40 |
| 17 | Pcbp4 | 0.66 | 1.10E-07 | 2.29E-03 | 0.72 | 0.87 | -0.15 | 0.83 |
| 17 | Hdac8 | 0.28 | 1.11E-07 | 2.31E-03 | 0.09 | 0.23 | -0.14 | 0.39 |
| 17 | Rnaseh2b | 0.39 | 1.11E-07 | 2.32E-03 | 0.26 | 0.50 | -0.25 | 0.51 |
| 17 | Ddx51 | 0.57 | 1.13E-07 | 2.34E-03 | 0.06 | 0.18 | -0.12 | 0.32 |
| 17 | Pdcd2 | 0.30 | 1.14E-07 | 2.37E-03 | 0.19 | 0.39 | -0.20 | 0.49 |
| 17 | Apoc1 | 1.93 | 1.14E-07 | 2.37E-03 | 0.16 | 0.09 | 0.07 | 1.86 |
| 17 | Gid8 | 0.32 | 1.14E-07 | 2.37E-03 | 0.28 | 0.55 | -0.27 | 0.51 |
| 17 | Yrdc | 0.41 | 1.18E-07 | 2.46E-03 | 0.21 | 0.42 | -0.22 | 0.49 |
| 17 | Hes5 | 1.82 | 1.21E-07 | 2.52E-03 | 0.20 | 0.12 | 0.08 | 1.70 |
| 17 | Med17 | 0.27 | 1.24E-07 | 2.59E-03 | 0.16 | 0.34 | -0.18 | 0.47 |
| 17 | Aggf1 | 0.30 | 1.26E-07 | 2.61E-03 | 0.31 | 0.60 | -0.28 | 0.53 |
| 17 | Cstf1 | 0.52 | 1.32E-07 | 2.74E-03 | 0.12 | 0.27 | -0.16 | 0.43 |
| 17 | Tomm6 | 0.38 | 1.36E-07 | 2.82E-03 | 0.80 | 0.97 | -0.17 | 0.82 |
| 17 | Cog7 | 0.25 | 1.39E-07 | 2.89E-03 | 0.04 | 0.14 | -0.10 | 0.25 |
| 17 | Bnip3l | 0.53 | 1.41E-07 | 2.93E-03 | 0.74 | 0.94 | -0.20 | 0.79 |
| 17 | Ccdc117 | 0.42 | 1.43E-07 | 2.98E-03 | 0.08 | 0.21 | -0.13 | 0.37 |

**Supplementary Table 1. Marker genes for the original clusters of integrated clustering of E11.5 control and NotchGOF λ-junction cells.**

| | | | | | | | | |
|---|---|---|---|---|---|---|---|---|
| 17 | Vasp | 0.37 | 1.52E-07 | 3.15E-03 | 0.25 | 0.49 | -0.24 | 0.51 |
| 17 | Dcaf17 | 0.31 | 1.54E-07 | 3.20E-03 | 0.16 | 0.34 | -0.18 | 0.47 |
| 17 | Toe1 | 0.39 | 1.66E-07 | 3.45E-03 | 0.16 | 0.33 | -0.18 | 0.47 |
| 17 | Erf | 0.28 | 1.66E-07 | 3.46E-03 | 0.14 | 0.31 | -0.17 | 0.45 |
| 17 | Pus10 | 0.47 | 1.67E-07 | 3.48E-03 | 0.15 | 0.32 | -0.17 | 0.46 |
| 17 | Taf6l | 0.55 | 1.68E-07 | 3.49E-03 | 0.12 | 0.27 | -0.15 | 0.44 |
| 17 | Mrpl9 | 0.31 | 1.70E-07 | 3.53E-03 | 0.31 | 0.58 | -0.28 | 0.53 |
| 17 | Zfp358 | 0.34 | 1.75E-07 | 3.65E-03 | 0.22 | 0.44 | -0.22 | 0.50 |
| 17 | Rnh1 | 1.36 | 1.77E-07 | 3.68E-03 | 0.50 | 0.56 | -0.06 | 0.90 |
| 17 | Fgf9 | 1.44 | 1.83E-07 | 3.80E-03 | 0.19 | 0.11 | 0.08 | 1.76 |
| 17 | Med11 | 0.36 | 1.85E-07 | 3.85E-03 | 0.13 | 0.30 | -0.16 | 0.45 |
| 17 | Cd24a | 0.57 | 1.88E-07 | 3.91E-03 | 0.78 | 0.92 | -0.14 | 0.85 |
| 17 | Esrp1 | 1.44 | 1.89E-07 | 3.93E-03 | 0.26 | 0.17 | 0.09 | 1.51 |
| 17 | Gins1 | 0.56 | 1.92E-07 | 3.99E-03 | 0.19 | 0.40 | -0.21 | 0.48 |
| 17 | Stk19 | 0.28 | 1.99E-07 | 4.13E-03 | 0.11 | 0.26 | -0.15 | 0.43 |
| 17 | Septin10 | 0.29 | 1.99E-07 | 4.14E-03 | 0.18 | 0.38 | -0.19 | 0.49 |
| 17 | Ifi30 | 0.42 | 2.00E-07 | 4.15E-03 | 0.14 | 0.30 | -0.16 | 0.46 |
| 17 | Rgs3 | 0.29 | 2.00E-07 | 4.17E-03 | 0.07 | 0.19 | -0.12 | 0.35 |
| 17 | Rpia | 0.38 | 2.05E-07 | 4.26E-03 | 0.14 | 0.30 | -0.17 | 0.45 |
| 17 | Pih1d1 | 0.40 | 2.13E-07 | 4.43E-03 | 0.29 | 0.56 | -0.27 | 0.52 |
| 17 | Cat | 0.42 | 2.13E-07 | 4.43E-03 | 0.22 | 0.44 | -0.22 | 0.50 |
| 17 | Nacc1 | 0.54 | 2.18E-07 | 4.53E-03 | 0.20 | 0.40 | -0.20 | 0.49 |
| 17 | Airn | 0.27 | 2.20E-07 | 4.57E-03 | 0.07 | 0.19 | -0.12 | 0.36 |
| 17 | Stk11 | 0.35 | 2.26E-07 | 4.69E-03 | 0.25 | 0.49 | -0.23 | 0.52 |
| 17 | Cpne3 | 0.27 | 2.28E-07 | 4.75E-03 | 0.28 | 0.53 | -0.25 | 0.53 |
| 17 | Aif1l | 0.50 | 2.29E-07 | 4.76E-03 | 0.19 | 0.39 | -0.20 | 0.49 |
| 17 | Lsm11 | 0.32 | 2.36E-07 | 4.92E-03 | 0.07 | 0.20 | -0.13 | 0.37 |
| 17 | Abcf3 | 0.40 | 2.40E-07 | 4.98E-03 | 0.13 | 0.30 | -0.16 | 0.45 |
| 17 | Dtd2 | 0.26 | 2.42E-07 | 5.04E-03 | 0.16 | 0.33 | -0.17 | 0.47 |
| 17 | Ska3 | 0.57 | 2.45E-07 | 5.09E-03 | 0.13 | 0.29 | -0.16 | 0.45 |
| 17 | 1110051M20Rik | 0.35 | 2.68E-07 | 5.57E-03 | 0.17 | 0.35 | -0.18 | 0.48 |
| 17 | Sorbs3 | 0.32 | 2.77E-07 | 5.75E-03 | 0.04 | 0.14 | -0.10 | 0.27 |
| 17 | Khsrp | 0.34 | 2.84E-07 | 5.91E-03 | 0.30 | 0.57 | -0.27 | 0.53 |
| 17 | Nelfb | 0.56 | 3.22E-07 | 6.69E-03 | 0.29 | 0.56 | -0.27 | 0.52 |
| 17 | Tbc1d7 | 0.42 | 3.23E-07 | 6.71E-03 | 0.20 | 0.40 | -0.20 | 0.49 |
| 17 | Ddx28 | 0.28 | 3.30E-07 | 6.87E-03 | 0.09 | 0.22 | -0.13 | 0.40 |
| 17 | Pccb | 0.46 | 3.47E-07 | 7.22E-03 | 0.08 | 0.21 | -0.13 | 0.39 |
| 17 | Crls1 | 0.34 | 3.57E-07 | 7.43E-03 | 0.24 | 0.46 | -0.22 | 0.51 |
| 17 | Gatd3a | 0.36 | 3.58E-07 | 7.45E-03 | 0.20 | 0.41 | -0.21 | 0.50 |
| 17 | Ngrn | 0.30 | 3.60E-07 | 7.48E-03 | 0.24 | 0.47 | -0.23 | 0.52 |
| 17 | Bckdk | 0.27 | 3.61E-07 | 7.50E-03 | 0.26 | 0.50 | -0.24 | 0.52 |
| 17 | Dus1l | 0.70 | 3.65E-07 | 7.58E-03 | 0.20 | 0.41 | -0.21 | 0.49 |
| 17 | Vps41 | 0.32 | 3.67E-07 | 7.62E-03 | 0.23 | 0.45 | -0.22 | 0.51 |
| 17 | Mta3 | 0.39 | 3.74E-07 | 7.77E-03 | 0.19 | 0.38 | -0.19 | 0.50 |
| 17 | Rbms2 | 0.44 | 3.80E-07 | 7.90E-03 | 0.24 | 0.47 | -0.23 | 0.51 |
| 17 | Amz2 | 0.45 | 3.82E-07 | 7.95E-03 | 0.18 | 0.37 | -0.19 | 0.48 |
| 17 | Pole2 | 0.33 | 3.90E-07 | 8.10E-03 | 0.13 | 0.29 | -0.16 | 0.45 |
| 17 | Proser3 | 0.25 | 4.00E-07 | 8.32E-03 | 0.06 | 0.18 | -0.11 | 0.35 |
| 17 | Zbtb25 | 0.28 | 4.03E-07 | 8.37E-03 | 0.03 | 0.13 | -0.10 | 0.25 |
| 17 | Znhit3 | 0.40 | 4.04E-07 | 8.40E-03 | 0.30 | 0.58 | -0.28 | 0.52 |
| 17 | Dcaf13 | 0.52 | 4.04E-07 | 8.41E-03 | 0.24 | 0.47 | -0.23 | 0.51 |
| 17 | Ywhae | 0.26 | 4.05E-07 | 8.43E-03 | 0.97 | 1.00 | -0.03 | 0.97 |
| 17 | L2hgdh | 0.33 | 4.18E-07 | 8.69E-03 | 0.09 | 0.22 | -0.13 | 0.40 |
| 17 | Mvd | 0.42 | 4.23E-07 | 8.79E-03 | 0.14 | 0.31 | -0.16 | 0.47 |
| 17 | Smtn | 0.40 | 4.33E-07 | 9.00E-03 | 0.04 | 0.15 | -0.10 | 0.30 |
| 17 | Gfod2 | 0.28 | 4.36E-07 | 9.07E-03 | 0.14 | 0.31 | -0.17 | 0.47 |

**Supplementary Table 1. Marker genes for the original clusters of integrated clustering of E11.5 control and NotchGOF λ-junction cells.**

| | | | | | | | | |
|---|---|---|---|---|---|---|---|---|
| 17 | Tinf2 | 0.29 | 4.38E-07 | 9.12E-03 | 0.08 | 0.21 | -0.13 | 0.39 |
| 17 | Riox2 | 0.39 | 4.49E-07 | 9.34E-03 | 0.12 | 0.27 | -0.15 | 0.44 |
| 17 | Txndc16 | 0.37 | 4.65E-07 | 9.66E-03 | 0.04 | 0.15 | -0.10 | 0.30 |
| 17 | Umps | 0.34 | 4.67E-07 | 9.71E-03 | 0.33 | 0.61 | -0.28 | 0.54 |
| 17 | Mettl5 | 0.47 | 4.69E-07 | 9.75E-03 | 0.15 | 0.32 | -0.17 | 0.47 |
| 17 | Tars2 | 0.50 | 4.80E-07 | 9.99E-03 | 0.11 | 0.25 | -0.14 | 0.44 |
| 17 | Rfk | 0.48 | 4.83E-07 | 1.00E-02 | 0.25 | 0.49 | -0.24 | 0.51 |
| 17 | Ttc33 | 0.31 | 4.83E-07 | 1.00E-02 | 0.16 | 0.34 | -0.18 | 0.47 |
| 17 | Atg12 | 0.33 | 4.90E-07 | 1.02E-02 | 0.28 | 0.52 | -0.24 | 0.53 |
| 17 | Bcar1 | 0.32 | 4.93E-07 | 1.03E-02 | 0.11 | 0.25 | -0.14 | 0.43 |
| 17 | Rnf34 | 0.39 | 4.96E-07 | 1.03E-02 | 0.25 | 0.48 | -0.23 | 0.52 |
| 17 | Mphosph6 | 0.45 | 5.08E-07 | 1.06E-02 | 0.26 | 0.50 | -0.25 | 0.51 |
| 17 | Zfp703 | 0.44 | 5.09E-07 | 1.06E-02 | 0.12 | 0.26 | -0.14 | 0.45 |
| 17 | Ghitm | 0.26 | 5.13E-07 | 1.07E-02 | 0.36 | 0.68 | -0.31 | 0.54 |
| 17 | Ccdc9 | 0.25 | 5.16E-07 | 1.07E-02 | 0.14 | 0.31 | -0.16 | 0.47 |
| 17 | Mov10 | 0.54 | 5.31E-07 | 1.10E-02 | 0.08 | 0.20 | -0.12 | 0.38 |
| 17 | Nphp1 | 0.60 | 5.45E-07 | 1.13E-02 | 0.14 | 0.31 | -0.16 | 0.47 |
| 17 | Uchl1 | 1.38 | 5.49E-07 | 1.14E-02 | 0.39 | 0.35 | 0.04 | 1.11 |
| 17 | Cfap298 | 0.60 | 5.75E-07 | 1.20E-02 | 0.19 | 0.37 | -0.19 | 0.50 |
| 17 | Tspan7 | 0.36 | 5.88E-07 | 1.22E-02 | 0.14 | 0.29 | -0.15 | 0.47 |
| 17 | 2010204K13Rik | 0.25 | 5.96E-07 | 1.24E-02 | 0.11 | 0.26 | -0.14 | 0.44 |
| 17 | Isca2 | 0.29 | 5.98E-07 | 1.24E-02 | 0.31 | 0.59 | -0.27 | 0.53 |
| 17 | Serinc2 | 0.30 | 6.11E-07 | 1.27E-02 | 0.21 | 0.39 | -0.19 | 0.52 |
| 17 | Tsen2 | 0.37 | 6.13E-07 | 1.28E-02 | 0.12 | 0.27 | -0.15 | 0.45 |
| 17 | Cox7a2l | 0.50 | 6.40E-07 | 1.33E-02 | 0.78 | 0.96 | -0.18 | 0.81 |
| 17 | Tacc3 | 0.38 | 6.43E-07 | 1.34E-02 | 0.35 | 0.61 | -0.26 | 0.58 |
| 17 | Zbtb43 | 0.55 | 6.47E-07 | 1.35E-02 | 0.10 | 0.23 | -0.13 | 0.43 |
| 17 | Scyl2 | 0.52 | 6.47E-07 | 1.35E-02 | 0.13 | 0.28 | -0.15 | 0.45 |
| 17 | Pde5a | 0.43 | 6.52E-07 | 1.36E-02 | 0.07 | 0.19 | -0.12 | 0.38 |
| 17 | Dxo | 0.30 | 6.54E-07 | 1.36E-02 | 0.21 | 0.41 | -0.20 | 0.51 |
| 17 | Bloc1s5 | 0.45 | 6.67E-07 | 1.39E-02 | 0.16 | 0.33 | -0.17 | 0.48 |
| 17 | Ahcy | 0.36 | 6.93E-07 | 1.44E-02 | 0.13 | 0.28 | -0.15 | 0.46 |
| 17 | Ly6h | 1.51 | 7.03E-07 | 1.46E-02 | 0.23 | 0.15 | 0.08 | 1.54 |
| 17 | Ccdc18 | 0.38 | 7.17E-07 | 1.49E-02 | 0.15 | 0.31 | -0.16 | 0.48 |
| 17 | 2610002M06Rik | 0.28 | 7.19E-07 | 1.50E-02 | 0.14 | 0.30 | -0.15 | 0.48 |
| 17 | Rwdd4a | 0.28 | 7.51E-07 | 1.56E-02 | 0.32 | 0.59 | -0.27 | 0.54 |
| 17 | Prcc | 0.53 | 7.62E-07 | 1.58E-02 | 0.21 | 0.41 | -0.20 | 0.50 |
| 17 | Ap3m1 | 0.31 | 7.64E-07 | 1.59E-02 | 0.20 | 0.40 | -0.20 | 0.51 |
| 17 | Chkb | 0.35 | 7.65E-07 | 1.59E-02 | 0.10 | 0.24 | -0.14 | 0.43 |
| 17 | Carmil1 | 0.44 | 7.83E-07 | 1.63E-02 | 0.05 | 0.15 | -0.10 | 0.32 |
| 17 | Hspbap1 | 0.37 | 7.89E-07 | 1.64E-02 | 0.09 | 0.21 | -0.13 | 0.40 |
| 17 | Qars | 0.44 | 7.92E-07 | 1.65E-02 | 0.33 | 0.60 | -0.27 | 0.55 |
| 17 | Cep70 | 0.40 | 8.07E-07 | 1.68E-02 | 0.21 | 0.40 | -0.20 | 0.51 |
| 17 | Arf3 | 0.36 | 8.13E-07 | 1.69E-02 | 0.16 | 0.32 | -0.16 | 0.49 |
| 17 | Aldh7a1 | 0.36 | 8.40E-07 | 1.75E-02 | 0.30 | 0.56 | -0.26 | 0.54 |
| 17 | Ctps2 | 0.28 | 8.48E-07 | 1.76E-02 | 0.14 | 0.29 | -0.15 | 0.47 |
| 17 | Eci1 | 0.28 | 8.62E-07 | 1.79E-02 | 0.28 | 0.53 | -0.25 | 0.53 |
| 17 | Smim8 | 0.43 | 8.73E-07 | 1.82E-02 | 0.22 | 0.43 | -0.21 | 0.52 |
| 17 | Haus2 | 0.55 | 9.00E-07 | 1.87E-02 | 0.13 | 0.28 | -0.15 | 0.46 |
| 17 | E2f8 | 0.29 | 9.01E-07 | 1.87E-02 | 0.09 | 0.21 | -0.12 | 0.41 |
| 17 | Rnf219 | 0.25 | 9.08E-07 | 1.89E-02 | 0.31 | 0.58 | -0.27 | 0.54 |
| 17 | Rfc5 | 0.30 | 9.12E-07 | 1.90E-02 | 0.32 | 0.59 | -0.27 | 0.54 |
| 17 | Trnt1 | 0.32 | 9.50E-07 | 1.98E-02 | 0.27 | 0.51 | -0.24 | 0.53 |
| 17 | Idh3g | 0.42 | 9.53E-07 | 1.98E-02 | 0.35 | 0.67 | -0.31 | 0.53 |
| 17 | Rgs19 | 0.44 | 9.56E-07 | 1.99E-02 | 0.24 | 0.46 | -0.22 | 0.52 |
| 17 | H2afj | 0.81 | 9.60E-07 | 2.00E-02 | 0.64 | 0.82 | -0.17 | 0.79 |

**Supplementary Table 1. Marker genes for the original clusters of integrated clustering of E11.5 control and NotchGOF λ-junction cells.**

| | | | | | | | | |
|---|---|---|---|---|---|---|---|---|
| 17 | Lgr4 | 0.51 | 9.69E-07 | 2.02E-02 | 0.10 | 0.23 | -0.13 | 0.44 |
| 17 | Sars2 | 0.37 | 9.84E-07 | 2.05E-02 | 0.07 | 0.18 | -0.11 | 0.37 |
| 17 | Sertad3 | 0.29 | 9.85E-07 | 2.05E-02 | 0.11 | 0.25 | -0.14 | 0.45 |
| 17 | Vps4b | 0.32 | 1.02E-06 | 2.12E-02 | 0.35 | 0.63 | -0.29 | 0.55 |
| 17 | Taf1a | 0.31 | 1.05E-06 | 2.17E-02 | 0.03 | 0.12 | -0.09 | 0.25 |
| 17 | Kctd1 | 0.40 | 1.08E-06 | 2.25E-02 | 0.25 | 0.47 | -0.22 | 0.53 |
| 17 | Zdhhc12 | 0.31 | 1.09E-06 | 2.26E-02 | 0.12 | 0.26 | -0.14 | 0.45 |
| 17 | Inava | 0.53 | 1.09E-06 | 2.27E-02 | 0.10 | 0.24 | -0.13 | 0.44 |
| 17 | Knstrn | 0.26 | 1.11E-06 | 2.31E-02 | 0.36 | 0.62 | -0.26 | 0.58 |
| 17 | Cab39 | 0.35 | 1.12E-06 | 2.32E-02 | 0.32 | 0.59 | -0.27 | 0.55 |
| 17 | Cdca5 | 0.53 | 1.12E-06 | 2.34E-02 | 0.17 | 0.34 | -0.17 | 0.49 |
| 17 | Glrx3 | 0.41 | 1.16E-06 | 2.42E-02 | 0.80 | 0.96 | -0.17 | 0.83 |
| 17 | Cetn2 | 0.34 | 1.23E-06 | 2.55E-02 | 0.36 | 0.65 | -0.29 | 0.56 |
| 17 | Map2k3 | 0.60 | 1.24E-06 | 2.59E-02 | 0.25 | 0.49 | -0.24 | 0.52 |
| 17 | Usp46 | 0.28 | 1.25E-06 | 2.60E-02 | 0.36 | 0.63 | -0.28 | 0.56 |
| 17 | Tada1 | 0.46 | 1.26E-06 | 2.62E-02 | 0.20 | 0.38 | -0.19 | 0.51 |
| 17 | Pkia | 0.29 | 1.29E-06 | 2.69E-02 | 0.07 | 0.19 | -0.12 | 0.39 |
| 17 | 2210016L21Rik | 0.32 | 1.31E-06 | 2.73E-02 | 0.36 | 0.68 | -0.31 | 0.54 |
| 17 | Dnaja3 | 0.26 | 1.34E-06 | 2.79E-02 | 0.17 | 0.34 | -0.17 | 0.50 |
| 17 | Slc25a46 | 0.26 | 1.34E-06 | 2.80E-02 | 0.14 | 0.29 | -0.15 | 0.47 |
| 17 | Gpatch4 | 0.37 | 1.38E-06 | 2.87E-02 | 0.30 | 0.56 | -0.26 | 0.54 |
| 17 | Agap3 | 0.26 | 1.39E-06 | 2.89E-02 | 0.19 | 0.36 | -0.18 | 0.51 |
| 17 | Cln3 | 0.31 | 1.42E-06 | 2.94E-02 | 0.05 | 0.15 | -0.10 | 0.32 |
| 17 | Pfdn5 | 0.26 | 1.43E-06 | 2.97E-02 | 0.89 | 0.99 | -0.10 | 0.90 |
| 17 | Gga3 | 0.43 | 1.49E-06 | 3.10E-02 | 0.05 | 0.14 | -0.10 | 0.33 |
| 17 | Uck2 | 0.28 | 1.50E-06 | 3.13E-02 | 0.29 | 0.53 | -0.24 | 0.54 |
| 17 | Fam219b | 0.33 | 1.51E-06 | 3.14E-02 | 0.05 | 0.15 | -0.10 | 0.34 |
| 17 | Patz1 | 0.42 | 1.51E-06 | 3.14E-02 | 0.28 | 0.53 | -0.25 | 0.53 |
| 17 | Epb41l4aos | 0.37 | 1.52E-06 | 3.17E-02 | 0.30 | 0.56 | -0.26 | 0.54 |
| 17 | Mlh1 | 0.60 | 1.56E-06 | 3.25E-02 | 0.09 | 0.21 | -0.12 | 0.41 |
| 17 | Mat2b | 0.36 | 1.57E-06 | 3.27E-02 | 0.33 | 0.60 | -0.27 | 0.55 |
| 17 | Snx1 | 0.41 | 1.65E-06 | 3.44E-02 | 0.24 | 0.45 | -0.22 | 0.52 |
| 17 | Asb7 | 0.50 | 1.67E-06 | 3.48E-02 | 0.09 | 0.21 | -0.12 | 0.43 |
| 17 | Aen | 0.44 | 1.67E-06 | 3.48E-02 | 0.21 | 0.40 | -0.20 | 0.51 |
| 17 | Id4 | 1.58 | 1.69E-06 | 3.51E-02 | 0.36 | 0.31 | 0.05 | 1.16 |
| 17 | Fkbp3 | 0.28 | 1.69E-06 | 3.52E-02 | 0.91 | 0.99 | -0.07 | 0.93 |
| 17 | Cdk6 | 0.36 | 1.76E-06 | 3.66E-02 | 0.18 | 0.35 | -0.17 | 0.52 |
| 17 | Ciao2b | 0.29 | 1.79E-06 | 3.73E-02 | 0.41 | 0.74 | -0.33 | 0.56 |
| 17 | Armh3 | 0.39 | 1.80E-06 | 3.75E-02 | 0.02 | 0.11 | -0.08 | 0.22 |
| 17 | Cldn9 | 2.11 | 1.80E-06 | 3.75E-02 | 0.16 | 0.09 | 0.07 | 1.72 |
| 17 | Fam216a | 0.27 | 1.81E-06 | 3.76E-02 | 0.28 | 0.53 | -0.25 | 0.53 |
| 17 | Hinfp | 0.44 | 1.81E-06 | 3.77E-02 | 0.06 | 0.16 | -0.10 | 0.35 |
| 17 | Mrpl35 | 0.29 | 1.84E-06 | 3.82E-02 | 0.41 | 0.74 | -0.33 | 0.56 |
| 17 | Zfp317 | 0.59 | 1.87E-06 | 3.90E-02 | 0.09 | 0.21 | -0.12 | 0.43 |
| 17 | Adi1 | 0.52 | 1.88E-06 | 3.91E-02 | 0.16 | 0.32 | -0.17 | 0.49 |
| 17 | Ppa1 | 0.63 | 1.90E-06 | 3.96E-02 | 0.72 | 0.90 | -0.18 | 0.80 |
| 17 | Fam45a | 0.35 | 1.94E-06 | 4.04E-02 | 0.09 | 0.21 | -0.12 | 0.43 |
| 17 | Grb2 | 0.36 | 1.95E-06 | 4.06E-02 | 0.36 | 0.64 | -0.29 | 0.56 |
| 17 | Anp32a | 0.30 | 1.96E-06 | 4.07E-02 | 0.88 | 0.99 | -0.11 | 0.89 |
| 17 | Ier5 | 1.18 | 1.97E-06 | 4.11E-02 | 0.55 | 0.65 | -0.10 | 0.84 |
| 17 | Gmeb2 | 0.65 | 1.99E-06 | 4.15E-02 | 0.05 | 0.14 | -0.10 | 0.33 |
| 17 | Apeh | 0.42 | 2.00E-06 | 4.15E-02 | 0.13 | 0.28 | -0.15 | 0.47 |
| 17 | Trim32 | 0.46 | 2.01E-06 | 4.18E-02 | 0.17 | 0.35 | -0.17 | 0.50 |
| 17 | Gpatch1 | 0.49 | 2.02E-06 | 4.21E-02 | 0.11 | 0.24 | -0.13 | 0.45 |
| 17 | Ing2 | 0.49 | 2.07E-06 | 4.31E-02 | 0.21 | 0.41 | -0.20 | 0.52 |
| 17 | Chmp1b | 0.32 | 2.14E-06 | 4.45E-02 | 0.18 | 0.36 | -0.18 | 0.50 |

**Supplementary Table 1. Marker genes for the original clusters of integrated clustering of E11.5 control and NotchGOF λ-junction cells.**

| | | | | | | | | |
|---|---|---|---|---|---|---|---|---|
| 17 | Mrps23 | 0.31 | 2.15E-06 | 4.46E-02 | 0.34 | 0.61 | -0.27 | 0.55 |
| 17 | Gm32585 | 2.20 | 2.18E-06 | 4.54E-02 | 0.11 | 0.06 | 0.06 | 1.98 |
| 17 | Mkrn2 | 0.53 | 2.19E-06 | 4.54E-02 | 0.15 | 0.30 | -0.16 | 0.48 |
| 17 | Aftph | 0.67 | 2.24E-06 | 4.66E-02 | 0.12 | 0.26 | -0.14 | 0.46 |
| 17 | Gphn | 0.48 | 2.30E-06 | 4.78E-02 | 0.08 | 0.19 | -0.11 | 0.40 |
| 17 | Bhlhb9 | 0.40 | 2.34E-06 | 4.87E-02 | 0.27 | 0.51 | -0.24 | 0.53 |
| 17 | Fundc1 | 0.39 | 2.35E-06 | 4.89E-02 | 0.19 | 0.37 | -0.18 | 0.52 |
| 17 | Dsn1 | 0.49 | 2.36E-06 | 4.91E-02 | 0.11 | 0.24 | -0.13 | 0.45 |
| 17 | Rras2 | 0.25 | 2.39E-06 | 4.98E-02 | 0.13 | 0.28 | -0.15 | 0.47 |
| 18 | Cldn5 | 9.58 | 0 | 0 | 1.00 | 0.01 | 0.99 | 142.86 |
| 18 | Cdh5 | 9.52 | 0 | 0 | 1.00 | 0.01 | 0.99 | 111.11 |
| 18 | Emcn | 9.05 | 0 | 0 | 1.00 | 0.01 | 0.99 | 76.92 |
| 18 | Kdr | 9.04 | 0 | 0 | 1.00 | 0.01 | 0.99 | 99.70 |
| 18 | Ecscr | 9.08 | 0 | 0 | 1.00 | 0.01 | 0.99 | 99.70 |
| 18 | Cd93 | 9.15 | 0 | 0 | 0.99 | 0.00 | 0.98 | 246.50 |
| 18 | Pecam1 | 8.45 | 0 | 0 | 1.00 | 0.02 | 0.98 | 66.47 |
| 18 | Icam2 | 9.05 | 0 | 0 | 0.99 | 0.01 | 0.98 | 164.33 |
| 18 | Flt1 | 9.59 | 0 | 0 | 0.98 | 0.01 | 0.97 | 163.33 |
| 18 | Esam | 8.14 | 0 | 0 | 0.99 | 0.01 | 0.97 | 75.85 |
| 18 | Plvap | 8.28 | 0 | 0 | 1.00 | 0.03 | 0.97 | 35.71 |
| 18 | Aplnr | 9.17 | 0 | 0 | 0.97 | 0.01 | 0.97 | 162.17 |
| 18 | Rasip1 | 6.56 | 0 | 0 | 0.99 | 0.03 | 0.96 | 31.94 |
| 18 | Col15a1 | 8.38 | 0 | 0 | 0.97 | 0.01 | 0.96 | 107.33 |
| 18 | Gimap6 | 7.34 | 0 | 0 | 0.96 | 0.01 | 0.95 | 68.50 |
| 18 | Plxnd1 | 6.97 | 0 | 0 | 0.99 | 0.05 | 0.95 | 22.00 |
| 18 | Tmem255a | 7.45 | 0 | 0 | 0.96 | 0.02 | 0.94 | 63.93 |
| 18 | Vamp5 | 6.25 | 0 | 0 | 0.99 | 0.05 | 0.94 | 19.86 |
| 18 | Tspan18 | 6.44 | 0 | 0 | 0.97 | 0.03 | 0.94 | 34.50 |
| 18 | Sox18 | 7.50 | 0 | 0 | 0.94 | 0.01 | 0.94 | 134.57 |
| 18 | Eng | 6.37 | 0 | 0 | 0.96 | 0.03 | 0.93 | 33.07 |
| 18 | S100a16 | 6.15 | 0 | 0 | 0.98 | 0.06 | 0.92 | 15.60 |
| 18 | Gngt2 | 6.08 | 0 | 0 | 0.95 | 0.03 | 0.92 | 36.38 |
| 18 | Clec1b | 6.25 | 0 | 0 | 0.92 | 0.01 | 0.92 | 184.40 |
| 18 | Ctla2a | 5.93 | 0 | 0 | 0.92 | 0.01 | 0.92 | 153.67 |
| 18 | Fxyd5 | 6.11 | 0 | 0 | 0.93 | 0.02 | 0.91 | 58.06 |
| 18 | Pde8a | 7.12 | 0 | 0 | 0.92 | 0.01 | 0.90 | 83.18 |
| 18 | Lcp1 | 4.89 | 0 | 0 | 0.92 | 0.03 | 0.89 | 36.72 |
| 18 | Adgrf5 | 8.92 | 0 | 0 | 0.90 | 0.00 | 0.89 | 298.33 |
| 18 | Anxa3 | 5.19 | 0 | 0 | 0.97 | 0.08 | 0.89 | 12.42 |
| 18 | Mfng | 4.41 | 0 | 0 | 0.92 | 0.03 | 0.88 | 26.91 |
| 18 | Cd34 | 5.92 | 0 | 0 | 1.00 | 0.12 | 0.88 | 8.31 |
| 18 | Tie1 | 9.11 | 0 | 0 | 0.88 | 0.00 | 0.88 | 220.25 |
| 18 | Gimap4 | 7.78 | 0 | 0 | 0.87 | 0.00 | 0.87 | 218.50 |
| 18 | Tgfb1 | 4.18 | 0 | 0 | 0.97 | 0.10 | 0.87 | 10.06 |
| 18 | Flt4 | 9.34 | 0 | 0 | 0.87 | 0.00 | 0.86 | 289.00 |
| 18 | Gmfg | 4.98 | 0 | 0 | 0.87 | 0.02 | 0.86 | 58.27 |
| 18 | Tfpi | 4.24 | 0 | 0 | 0.96 | 0.10 | 0.86 | 9.47 |
| 18 | S100a13 | 4.22 | 0 | 0 | 0.95 | 0.10 | 0.85 | 9.89 |
| 18 | Ehd2 | 5.33 | 0 | 0 | 0.89 | 0.04 | 0.85 | 22.77 |
| 18 | Arhgap18 | 4.64 | 0 | 0 | 0.96 | 0.11 | 0.85 | 8.45 |
| 18 | Dok4 | 5.03 | 0 | 0 | 0.92 | 0.07 | 0.85 | 13.11 |
| 18 | She | 6.57 | 0 | 0 | 0.86 | 0.01 | 0.85 | 77.91 |
| 18 | Rasgrp3 | 7.99 | 0 | 0 | 0.85 | 0.01 | 0.85 | 170.00 |
| 18 | Klhl4 | 8.49 | 0 | 0 | 0.85 | 0.00 | 0.84 | 282.33 |
| 18 | Sema6d | 5.13 | 0 | 0 | 0.90 | 0.05 | 0.84 | 17.21 |
| 18 | Smagp | 5.70 | 0 | 0 | 0.86 | 0.02 | 0.84 | 37.57 |

**Supplementary Table 1. Marker genes for the original clusters of integrated clustering of E11.5 control and NotchGOF λ-junction cells.**

| | | | | | | | | |
|---|---|---|---|---|---|---|---|---|
| 18 | Lmo2 | 3.73 | 0 | 0 | 0.96 | 0.12 | 0.84 | 8.24 |
| 18 | Tmem88 | 5.23 | 0 | 0 | 0.89 | 0.05 | 0.84 | 18.50 |
| 18 | N4bp3 | 5.10 | 0 | 0 | 0.89 | 0.05 | 0.84 | 16.81 |
| 18 | Mmrn2 | 5.56 | 0 | 0 | 0.88 | 0.04 | 0.84 | 21.95 |
| 18 | Cavin3 | 4.96 | 0 | 0 | 0.95 | 0.11 | 0.84 | 8.47 |
| 18 | Slc29a1 | 4.62 | 0 | 0 | 0.95 | 0.11 | 0.84 | 8.52 |
| 18 | Stab1 | 5.27 | 0 | 0 | 0.84 | 0.01 | 0.83 | 76.73 |
| 18 | Gimap1 | 7.34 | 0 | 0 | 0.84 | 0.01 | 0.83 | 167.40 |
| 18 | Crip2 | 5.14 | 0 | 0 | 0.99 | 0.16 | 0.83 | 6.09 |
| 18 | Ramp2 | 5.74 | 0 | 0 | 1.00 | 0.17 | 0.82 | 5.73 |
| 18 | Ets1 | 4.74 | 0 | 0 | 0.99 | 0.17 | 0.82 | 5.89 |
| 18 | Mef2c | 4.01 | 0 | 0 | 0.99 | 0.17 | 0.82 | 5.69 |
| 18 | Rhoj | 3.95 | 0 | 0 | 0.92 | 0.11 | 0.81 | 8.38 |
| 18 | Rasgrp2 | 4.71 | 0 | 0 | 0.85 | 0.04 | 0.81 | 21.79 |
| 18 | Myzap | 5.70 | 0 | 0 | 0.83 | 0.02 | 0.81 | 39.52 |
| 18 | Dysf | 5.45 | 0 | 0 | 0.83 | 0.03 | 0.81 | 32.04 |
| 18 | Myct1 | 7.60 | 0 | 0 | 0.81 | 0.00 | 0.81 | 202.50 |
| 18 | Arhgef15 | 5.78 | 0 | 0 | 0.82 | 0.02 | 0.81 | 48.41 |
| 18 | Tubb6 | 4.37 | 0 | 0 | 0.91 | 0.10 | 0.81 | 8.82 |
| 18 | Afap1l1 | 4.27 | 0 | 0 | 0.87 | 0.06 | 0.80 | 13.76 |
| 18 | Arap3 | 5.32 | 0 | 0 | 0.83 | 0.03 | 0.80 | 25.24 |
| 18 | Pon2 | 3.57 | 0 | 0 | 0.97 | 0.17 | 0.80 | 5.67 |
| 18 | Timp3 | 4.63 | 0 | 0 | 0.96 | 0.16 | 0.80 | 5.94 |
| 18 | Thsd1 | 7.32 | 0 | 0 | 0.80 | 0.01 | 0.79 | 114.14 |
| 18 | Gpr183 | 6.84 | 0 | 0 | 0.80 | 0.01 | 0.79 | 113.71 |
| 18 | Egfl7 | 5.52 | 0 | 0 | 1.00 | 0.21 | 0.79 | 4.77 |
| 18 | Mcam | 5.34 | 0 | 0 | 0.82 | 0.03 | 0.79 | 25.63 |
| 18 | Sparcl1 | 7.14 | 0 | 0 | 0.80 | 0.02 | 0.79 | 47.24 |
| 18 | Lama4 | 3.84 | 0 | 0 | 0.91 | 0.12 | 0.79 | 7.38 |
| 18 | Pde2a | 8.37 | 0 | 0 | 0.78 | 0.00 | 0.78 | 260.67 |
| 18 | Hapln1 | 5.62 | 0 | 0 | 0.81 | 0.03 | 0.78 | 23.91 |
| 18 | Adgrl4 | 8.41 | 0 | 0 | 0.78 | 0.00 | 0.78 | 259.67 |
| 18 | Tek | 7.15 | 0 | 0 | 0.78 | 0.01 | 0.77 | 97.75 |
| 18 | Kitl | 4.44 | 0 | 0 | 0.84 | 0.07 | 0.77 | 12.88 |
| 18 | Kank3 | 4.77 | 0 | 0 | 0.83 | 0.06 | 0.77 | 15.04 |
| 18 | Emp1 | 3.80 | 0 | 0 | 0.89 | 0.12 | 0.77 | 7.40 |
| 18 | Kcne3 | 6.31 | 0 | 0 | 0.79 | 0.03 | 0.77 | 31.72 |
| 18 | Tbx1 | 5.81 | 0 | 0 | 0.79 | 0.02 | 0.77 | 43.67 |
| 18 | Mmrn1 | 7.09 | 0 | 0 | 0.77 | 0.00 | 0.77 | 256.33 |
| 18 | D030007L05Rik | 8.25 | 0 | 0 | 0.76 | 0.00 | 0.76 | 762.00 |
| 18 | Efna1 | 3.43 | 0 | 0 | 0.89 | 0.13 | 0.76 | 6.78 |
| 18 | Sipa1 | 3.60 | 0 | 0 | 0.92 | 0.17 | 0.76 | 5.55 |
| 18 | Ptpn18 | 3.60 | 0 | 0 | 0.82 | 0.07 | 0.75 | 12.24 |
| 18 | Tm6sf1 | 4.52 | 0 | 0 | 0.80 | 0.05 | 0.75 | 17.69 |
| 18 | Lyn | 4.86 | 0 | 0 | 0.77 | 0.02 | 0.75 | 35.09 |
| 18 | Gm16104 | 4.84 | 0 | 0 | 0.78 | 0.03 | 0.75 | 24.44 |
| 18 | Scarf1 | 8.20 | 0 | 0 | 0.75 | 0.00 | 0.74 | 248.33 |
| 18 | S1pr1 | 4.49 | 0 | 0 | 0.79 | 0.05 | 0.74 | 14.89 |
| 18 | Cavin1 | 6.53 | 0 | 0 | 0.75 | 0.01 | 0.74 | 82.78 |
| 18 | Adam15 | 3.81 | 0 | 0 | 0.82 | 0.09 | 0.73 | 9.43 |
| 18 | Tmem204 | 7.81 | 0 | 0 | 0.74 | 0.00 | 0.73 | 245.00 |
| 18 | Dusp2 | 5.21 | 0 | 0 | 0.75 | 0.02 | 0.73 | 32.70 |
| 18 | Klf2 | 6.43 | 0 | 0 | 0.76 | 0.03 | 0.73 | 26.96 |
| 18 | Fam167b | 7.15 | 0 | 0 | 0.73 | 0.00 | 0.73 | 364.00 |
| 18 | Grrp1 | 4.51 | 0 | 0 | 0.77 | 0.05 | 0.72 | 15.76 |
| 18 | Fgd5 | 5.98 | 0 | 0 | 0.73 | 0.01 | 0.72 | 52.21 |

**Supplementary Table 1. Marker genes for the original clusters of integrated clustering of E11.5 control and NotchGOF λ-junction cells.**

| | | | | | | | | |
|---|---|---|---|---|---|---|---|---|
| 18 | Klhl6 | 5.82 | 0 | 0 | 0.72 | 0.01 | 0.72 | 80.44 |
| 18 | Crmp1 | 3.18 | 0 | 0 | 0.82 | 0.11 | 0.71 | 7.35 |
| 18 | Cyyr1 | 4.70 | 0 | 0 | 0.75 | 0.04 | 0.71 | 17.49 |
| 18 | Pcdh12 | 8.72 | 0 | 0 | 0.71 | 0.00 | 0.71 | 707.00 |
| 18 | Dusp3 | 4.55 | 0 | 0 | 0.75 | 0.05 | 0.71 | 16.00 |
| 18 | Gchfr | 3.45 | 0 | 0 | 0.80 | 0.10 | 0.70 | 8.11 |
| 18 | Ptprb | 8.19 | 0 | 0 | 0.71 | 0.00 | 0.70 | 235.67 |
| 18 | Robo4 | 9.43 | 0 | 0 | 0.70 | 0.00 | 0.70 | 704.00 |
| 18 | Apold1 | 6.95 | 0 | 0 | 0.71 | 0.01 | 0.70 | 79.00 |
| 18 | Sox17 | 6.96 | 0 | 0 | 0.70 | 0.00 | 0.70 | 176.00 |
| 18 | Grap | 6.98 | 0 | 0 | 0.70 | 0.01 | 0.70 | 117.33 |
| 18 | Junb | 3.42 | 0 | 0 | 0.80 | 0.11 | 0.69 | 7.47 |
| 18 | 2610203C22Rik | 7.50 | 0 | 0 | 0.69 | 0.00 | 0.69 | 230.00 |
| 18 | Podxl | 5.23 | 0 | 0 | 0.72 | 0.03 | 0.69 | 21.21 |
| 18 | Erg | 5.21 | 0 | 0 | 0.69 | 0.02 | 0.68 | 38.56 |
| 18 | Pdgfb | 6.79 | 0 | 0 | 0.68 | 0.01 | 0.67 | 112.83 |
| 18 | Clec14a | 4.43 | 0 | 0 | 0.71 | 0.05 | 0.67 | 15.87 |
| 18 | Pcdh1 | 4.33 | 0 | 0 | 0.71 | 0.04 | 0.67 | 16.53 |
| 18 | Epas1 | 7.45 | 0 | 0 | 0.67 | 0.00 | 0.67 | 167.50 |
| 18 | Tgfbr2 | 3.68 | 0 | 0 | 0.74 | 0.08 | 0.67 | 9.75 |
| 18 | Cd38 | 4.35 | 0 | 0 | 0.70 | 0.04 | 0.66 | 16.69 |
| 18 | Sh2d3c | 3.37 | 0 | 0 | 0.75 | 0.09 | 0.66 | 8.13 |
| 18 | Exoc3l4 | 3.77 | 0 | 0 | 0.72 | 0.06 | 0.66 | 11.58 |
| 18 | Npr1 | 6.58 | 0 | 0 | 0.65 | 0.01 | 0.65 | 93.29 |
| 18 | Bcl6b | 6.26 | 0 | 0 | 0.66 | 0.01 | 0.65 | 65.60 |
| 18 | Ppp1r13b | 3.61 | 0 | 0 | 0.72 | 0.07 | 0.65 | 9.97 |
| 18 | Abi3 | 6.26 | 0 | 0 | 0.65 | 0.01 | 0.65 | 81.63 |
| 18 | Bst2 | 3.90 | 0 | 0 | 0.69 | 0.05 | 0.65 | 14.16 |
| 18 | BC028528 | 3.71 | 0 | 0 | 0.71 | 0.07 | 0.64 | 10.46 |
| 18 | Nos3 | 6.89 | 0 | 0 | 0.64 | 0.01 | 0.64 | 107.17 |
| 18 | Prkch | 3.30 | 0 | 0 | 0.71 | 0.08 | 0.64 | 9.04 |
| 18 | Ushbp1 | 7.73 | 0 | 0 | 0.64 | 0.00 | 0.63 | 318.00 |
| 18 | Mmp15 | 3.39 | 0 | 0 | 0.71 | 0.08 | 0.63 | 8.73 |
| 18 | Nrros | 4.12 | 0 | 0 | 0.65 | 0.03 | 0.62 | 21.77 |
| 18 | Cd40 | 5.18 | 0 | 0 | 0.64 | 0.02 | 0.62 | 37.41 |
| 18 | Pald1 | 3.49 | 0 | 0 | 0.69 | 0.08 | 0.62 | 9.25 |
| 18 | Ptprm | 4.42 | 0 | 0 | 0.66 | 0.04 | 0.62 | 16.10 |
| 18 | Upp1 | 7.00 | 0 | 0 | 0.62 | 0.00 | 0.62 | 155.50 |
| 18 | Icam1 | 6.50 | 0 | 0 | 0.63 | 0.01 | 0.62 | 69.56 |
| 18 | Itga2 | 6.96 | 0 | 0 | 0.62 | 0.01 | 0.62 | 124.40 |
| 18 | Sncg | 7.29 | 0 | 0 | 0.62 | 0.00 | 0.62 | 309.50 |
| 18 | Cd109 | 4.04 | 0 | 0 | 0.66 | 0.05 | 0.61 | 14.58 |
| 18 | Arhgap27 | 5.02 | 0 | 0 | 0.62 | 0.02 | 0.60 | 25.92 |
| 18 | Elmo1 | 4.12 | 0 | 0 | 0.64 | 0.04 | 0.60 | 16.74 |
| 18 | Adora2a | 7.05 | 0 | 0 | 0.59 | 0.00 | 0.59 | 148.00 |
| 18 | Calcrl | 4.45 | 0 | 0 | 0.61 | 0.03 | 0.59 | 22.67 |
| 18 | Cdc42ep1 | 3.62 | 0 | 0 | 0.64 | 0.05 | 0.59 | 11.83 |
| 18 | Fam43a | 3.73 | 0 | 0 | 0.63 | 0.05 | 0.58 | 13.98 |
| 18 | Cdc42ep2 | 4.25 | 0 | 0 | 0.62 | 0.04 | 0.58 | 17.19 |
| 18 | Samsn1 | 5.99 | 0 | 0 | 0.59 | 0.01 | 0.58 | 98.00 |
| 18 | Exoc3l | 7.65 | 0 | 0 | 0.58 | 0.00 | 0.58 | 291.00 |
| 18 | Ripply3 | 3.27 | 0 | 0 | 0.63 | 0.05 | 0.58 | 11.72 |
| 18 | Vav3 | 3.51 | 0 | 0 | 0.61 | 0.05 | 0.56 | 12.00 |
| 18 | Cav2 | 5.52 | 0 | 0 | 0.57 | 0.01 | 0.56 | 43.92 |
| 18 | Itga1 | 4.56 | 0 | 0 | 0.58 | 0.03 | 0.55 | 19.27 |
| 18 | Cda | 4.75 | 0 | 0 | 0.57 | 0.02 | 0.55 | 27.05 |

**Supplementary Table 1. Marker genes for the original clusters of integrated clustering of E11.5 control and NotchGOF λ-junction cells.**

| | | | | | | | | |
|---|---|---|---|---|---|---|---|---|
| 18 | Notch4 | 3.64 | 0 | 0 | 0.60 | 0.05 | 0.55 | 11.90 |
| 18 | Plac1 | 5.00 | 0 | 0 | 0.55 | 0.02 | 0.54 | 34.63 |
| 18 | Hhex | 5.68 | 0 | 0 | 0.54 | 0.01 | 0.54 | 60.44 |
| 18 | Gdpd5 | 3.94 | 0 | 0 | 0.57 | 0.04 | 0.53 | 14.64 |
| 18 | Ccm2l | 7.97 | 0 | 0 | 0.53 | 0.00 | 0.53 | 527.00 |
| 18 | Sox7 | 7.57 | 0 | 0 | 0.52 | 0.00 | 0.52 | 524.00 |
| 18 | Bik | 5.27 | 0 | 0 | 0.54 | 0.02 | 0.52 | 30.06 |
| 18 | Rapgef3 | 6.30 | 0 | 0 | 0.53 | 0.01 | 0.52 | 105.40 |
| 18 | Cav1 | 7.03 | 0 | 0 | 0.52 | 0.00 | 0.51 | 129.25 |
| 18 | Nova2 | 4.70 | 0 | 0 | 0.53 | 0.02 | 0.51 | 27.74 |
| 18 | Enpp3 | 4.54 | 0 | 0 | 0.53 | 0.03 | 0.51 | 20.54 |
| 18 | C130074G19Rik | 7.68 | 0 | 0 | 0.51 | 0.00 | 0.51 | 253.50 |
| 18 | Exoc3l2 | 6.92 | 0 | 0 | 0.50 | 0.00 | 0.50 | 167.67 |
| 18 | Rin3 | 4.63 | 0 | 0 | 0.51 | 0.02 | 0.50 | 27.05 |
| 18 | Ggta1 | 3.74 | 0 | 0 | 0.53 | 0.04 | 0.49 | 13.97 |
| 18 | Spns2 | 4.73 | 0 | 0 | 0.51 | 0.02 | 0.49 | 23.05 |
| 18 | Lbp | 4.05 | 0 | 0 | 0.51 | 0.02 | 0.48 | 21.13 |
| 18 | Hlx | 6.22 | 0 | 0 | 0.48 | 0.00 | 0.48 | 120.75 |
| 18 | Ppp1r16b | 6.91 | 0 | 0 | 0.47 | 0.00 | 0.47 | 236.50 |
| 18 | Cracr2b | 3.91 | 0 | 0 | 0.50 | 0.03 | 0.47 | 17.24 |
| 18 | Adgrg3 | 7.85 | 0 | 0 | 0.47 | 0.00 | 0.47 | 234.50 |
| 18 | Atf3 | 3.84 | 0 | 0 | 0.48 | 0.02 | 0.46 | 24.15 |
| 18 | Adam12 | 5.59 | 0 | 0 | 0.46 | 0.01 | 0.46 | 57.88 |
| 18 | Zfpm2 | 6.82 | 0 | 0 | 0.46 | 0.01 | 0.45 | 91.80 |
| 18 | Eogt | 3.63 | 0 | 0 | 0.49 | 0.04 | 0.45 | 13.24 |
| 18 | Gm13889 | 5.71 | 0 | 0 | 0.46 | 0.01 | 0.45 | 57.00 |
| 18 | Tm4sf1 | 5.57 | 0 | 0 | 0.46 | 0.01 | 0.45 | 41.45 |
| 18 | Card10 | 4.34 | 0 | 0 | 0.46 | 0.02 | 0.44 | 28.69 |
| 18 | Shank3 | 4.72 | 0 | 0 | 0.46 | 0.01 | 0.44 | 32.57 |
| 18 | Pon3 | 3.88 | 0 | 0 | 0.47 | 0.03 | 0.44 | 17.92 |
| 18 | Lat | 4.73 | 0 | 0 | 0.45 | 0.01 | 0.44 | 44.90 |
| 18 | Cd200 | 4.09 | 0 | 0 | 0.46 | 0.02 | 0.44 | 21.86 |
| 18 | Abcc9 | 7.14 | 0 | 0 | 0.44 | 0.00 | 0.44 | 146.33 |
| 18 | Jak3 | 3.99 | 0 | 0 | 0.46 | 0.02 | 0.44 | 19.13 |
| 18 | Filip1 | 3.58 | 0 | 0 | 0.46 | 0.03 | 0.43 | 14.94 |
| 18 | Man1c1 | 3.64 | 0 | 0 | 0.45 | 0.03 | 0.42 | 16.14 |
| 18 | Ctla2b | 4.34 | 0 | 0 | 0.43 | 0.01 | 0.42 | 85.80 |
| 18 | Sqor | 4.71 | 0 | 0 | 0.43 | 0.01 | 0.42 | 30.64 |
| 18 | Epha2 | 4.23 | 0 | 0 | 0.44 | 0.03 | 0.41 | 15.54 |
| 18 | Inpp5d | 4.49 | 0 | 0 | 0.42 | 0.01 | 0.41 | 46.11 |
| 18 | Apln | 4.77 | 0 | 0 | 0.43 | 0.02 | 0.41 | 21.25 |
| 18 | Dll4 | 8.32 | 0 | 0 | 0.41 | 0.00 | 0.40 | 405.00 |
| 18 | Pear1 | 3.50 | 0 | 0 | 0.43 | 0.03 | 0.40 | 17.00 |
| 18 | Rgs7bp | 4.43 | 0 | 0 | 0.42 | 0.02 | 0.40 | 23.22 |
| 18 | Thbd | 5.82 | 0 | 0 | 0.41 | 0.01 | 0.40 | 67.50 |
| 18 | Tbxa2r | 5.35 | 0 | 0 | 0.41 | 0.01 | 0.40 | 40.80 |
| 18 | Car8 | 4.45 | 0 | 0 | 0.41 | 0.02 | 0.40 | 27.47 |
| 18 | Elf4 | 4.21 | 0 | 0 | 0.41 | 0.02 | 0.39 | 21.68 |
| 18 | Prl3b1 | 10.79 | 0 | 0 | 0.37 | 0.00 | 0.37 | #NA |
| 18 | Gm32688 | 4.68 | 0 | 0 | 0.37 | 0.01 | 0.36 | 37.40 |
| 18 | Laptm5 | 2.42 | 0 | 0 | 0.37 | 0.02 | 0.36 | 24.73 |
| 18 | Rapgef4 | 5.48 | 0 | 0 | 0.36 | 0.01 | 0.35 | 45.13 |
| 18 | Csgalnact1 | 6.13 | 0 | 0 | 0.35 | 0.00 | 0.35 | 88.50 |
| 18 | Gm9917 | 4.86 | 0 | 0 | 0.35 | 0.01 | 0.35 | 39.33 |
| 18 | Gimap5 | 7.30 | 0 | 0 | 0.34 | 0.00 | 0.34 | 344.00 |
| 18 | Trim47 | 4.40 | 0 | 0 | 0.34 | 0.01 | 0.33 | 24.29 |

**Supplementary Table 1. Marker genes for the original clusters of integrated clustering of E11.5 control and NotchGOF λ-junction cells.**

| | | | | | | | | |
|---|---|---|---|---|---|---|---|---|
| 18 | Clec1a | 8.61 | 0 | 0 | 0.32 | 0.00 | 0.32 | 323.00 |
| 18 | F2rl3 | 6.06 | 0 | 0 | 0.32 | 0.00 | 0.32 | 106.67 |
| 18 | Zfp366 | 7.65 | 0 | 0 | 0.32 | 0.00 | 0.32 | 316.00 |
| 18 | Procr | 4.38 | 0 | 0 | 0.32 | 0.01 | 0.31 | 24.62 |
| 18 | Sh3tc1 | 6.45 | 0 | 0 | 0.31 | 0.00 | 0.31 | 103.33 |
| 18 | Prnd | 8.74 | 0 | 0 | 0.29 | 0.00 | 0.29 | 293.00 |
| 18 | Trpc6 | 5.42 | 0 | 0 | 0.29 | 0.00 | 0.29 | 146.50 |
| 18 | Gpihbp1 | 8.55 | 0 | 0 | 0.29 | 0.00 | 0.29 | 289.00 |
| 18 | Prdm1 | 4.90 | 0 | 0 | 0.29 | 0.01 | 0.29 | 41.86 |
| 18 | Depp1 | 7.75 | 0 | 0 | 0.27 | 0.00 | 0.27 | #NA |
| 18 | Pik3r6 | 6.02 | 0 | 0 | 0.26 | 0.00 | 0.26 | 86.33 |
| 18 | Hcls1 | 3.93 | 0 | 0 | 0.25 | 0.01 | 0.24 | 35.43 |
| 18 | Bace2 | 4.92 | 0 | 0 | 0.25 | 0.01 | 0.24 | 35.43 |
| 18 | Arhgap25 | 4.19 | 0 | 0 | 0.25 | 0.01 | 0.24 | 31.00 |
| 18 | 9230102K24Rik | 7.98 | 0 | 0 | 0.24 | 0.00 | 0.24 | #NA |
| 18 | Madcam1 | 8.79 | 0 | 0 | 0.21 | 0.00 | 0.21 | #NA |
| 18 | Ppm1j | 4.94 | 0 | 0 | 0.22 | 0.01 | 0.21 | 43.60 |
| 18 | Ptpn7 | 4.14 | 0 | 0 | 0.22 | 0.01 | 0.21 | 36.33 |
| 18 | Rsad2 | 6.50 | 0 | 0 | 0.21 | 0.00 | 0.21 | 105.50 |
| 18 | Dusp27 | 4.91 | 0 | 0 | 0.21 | 0.01 | 0.21 | 42.20 |
| 18 | Gimap8 | 6.52 | 0 | 0 | 0.21 | 0.00 | 0.21 | 207.00 |
| 18 | Adamtsl2 | 7.37 | 0 | 0 | 0.20 | 0.00 | 0.20 | 201.00 |
| 18 | Colgalt2 | 5.00 | 0 | 0 | 0.19 | 0.00 | 0.19 | 64.67 |
| 18 | Tfpi2 | 7.64 | 0 | 0 | 0.19 | 0.00 | 0.19 | #NA |
| 18 | Unc45b | 5.25 | 0 | 0 | 0.18 | 0.00 | 0.18 | 88.50 |
| 18 | AU021092 | 7.91 | 0 | 0 | 0.17 | 0.00 | 0.17 | 170.00 |
| 18 | Gpr182 | 9.67 | 0 | 0 | 0.17 | 0.00 | 0.17 | #NA |
| 18 | Gm13056 | 7.58 | 0 | 0 | 0.16 | 0.00 | 0.16 | #NA |
| 18 | Nos2 | 6.01 | 0 | 0 | 0.16 | 0.00 | 0.16 | #NA |
| 18 | Kcnq1 | 7.18 | 0 | 0 | 0.16 | 0.00 | 0.16 | 156.00 |
| 18 | Mmp25 | 5.87 | 0 | 0 | 0.16 | 0.00 | 0.15 | 78.00 |
| 18 | Esm1 | 7.43 | 0 | 0 | 0.15 | 0.00 | 0.15 | #NA |
| 18 | Sema3g | 6.51 | 0 | 0 | 0.15 | 0.00 | 0.15 | 146.00 |
| 18 | Gm38414 | 5.93 | 0 | 0 | 0.14 | 0.00 | 0.14 | 143.00 |
| 18 | D330037F02Rik | 8.79 | 0 | 0 | 0.14 | 0.00 | 0.14 | #NA |
| 18 | Gm33370 | 6.46 | 0 | 0 | 0.13 | 0.00 | 0.13 | 133.00 |
| 18 | Gm38832 | 6.91 | 0 | 0 | 0.13 | 0.00 | 0.13 | 129.00 |
| 18 | 4930542D17Rik | 9.71 | 0 | 0 | 0.12 | 0.00 | 0.12 | #NA |
| 18 | Gm17745 | 6.19 | 0 | 0 | 0.12 | 0.00 | 0.12 | 122.00 |
| 18 | Tmem252 | 7.70 | 0 | 0 | 0.12 | 0.00 | 0.12 | #NA |
| 18 | Endou | 7.02 | 0 | 0 | 0.12 | 0.00 | 0.12 | #NA |
| 18 | 4930555A03Rik | 6.70 | 0 | 0 | 0.12 | 0.00 | 0.12 | 116.00 |
| 18 | S1pr4 | 6.98 | 0 | 0 | 0.12 | 0.00 | 0.12 | 116.00 |
| 18 | Dipk2b | 9.91 | 0 | 0 | 0.11 | 0.00 | 0.11 | #NA |
| 18 | Fam89a | 3.39 | 1.60E-304 | 3.32E-300 | 0.64 | 0.07 | 0.58 | 9.74 |
| 18 | Tnfaip8l1 | 3.57 | 3.12E-304 | 6.50E-300 | 0.65 | 0.07 | 0.58 | 9.33 |
| 18 | Hecw2 | 3.31 | 1.71E-303 | 3.56E-299 | 0.64 | 0.07 | 0.57 | 9.78 |
| 18 | Jcad | 3.77 | 2.57E-302 | 5.34E-298 | 0.56 | 0.05 | 0.51 | 11.16 |
| 18 | Fam78a | 4.67 | 3.88E-302 | 8.08E-298 | 0.26 | 0.01 | 0.25 | 26.20 |
| 18 | Tnnt2 | 5.33 | 1.81E-301 | 3.77E-297 | 0.14 | 0.00 | 0.14 | 71.50 |
| 18 | BC049762 | 5.26 | 7.54E-301 | 1.57E-296 | 0.15 | 0.00 | 0.15 | 51.00 |
| 18 | Rcsd1 | 3.22 | 2.70E-300 | 5.62E-296 | 0.72 | 0.09 | 0.63 | 8.45 |
| 18 | Scn1b | 3.50 | 1.52E-297 | 3.17E-293 | 0.36 | 0.02 | 0.34 | 18.05 |
| 18 | Rapgef5 | 3.38 | 3.64E-296 | 7.57E-292 | 0.79 | 0.11 | 0.68 | 7.21 |
| 18 | Hspg2 | 3.69 | 1.35E-295 | 2.82E-291 | 0.97 | 0.22 | 0.75 | 4.47 |
| 18 | Slc31a2 | 3.43 | 1.87E-295 | 3.89E-291 | 0.63 | 0.07 | 0.56 | 9.68 |

**Supplementary Table 1. Marker genes for the original clusters of integrated clustering of E11.5 control and NotchGOF λ-junction cells.**

| | | | | | | | | |
|---|---|---|---|---|---|---|---|---|
| 18 | Plau | 4.45 | 7.60E-295 | 1.58E-290 | 0.22 | 0.01 | 0.22 | 32.00 |
| 18 | Dock6 | 3.38 | 2.08E-294 | 4.33E-290 | 0.75 | 0.10 | 0.65 | 7.63 |
| 18 | B3gnt2 | 3.43 | 9.23E-292 | 1.92E-287 | 0.72 | 0.09 | 0.63 | 7.96 |
| 18 | Igfbp3 | 4.16 | 1.63E-291 | 3.39E-287 | 0.79 | 0.12 | 0.68 | 6.84 |
| 18 | Acvrl1 | 3.17 | 3.54E-290 | 7.37E-286 | 0.74 | 0.09 | 0.65 | 7.97 |
| 18 | Cgnl1 | 3.19 | 1.27E-283 | 2.65E-279 | 0.79 | 0.11 | 0.68 | 6.92 |
| 18 | Gja4 | 6.70 | 1.75E-283 | 3.65E-279 | 0.19 | 0.01 | 0.19 | 38.80 |
| 18 | Ecm1 | 3.90 | 3.74E-283 | 7.79E-279 | 0.35 | 0.02 | 0.33 | 18.26 |
| 18 | Rassf2 | 3.93 | 9.42E-283 | 1.96E-278 | 0.34 | 0.02 | 0.32 | 18.72 |
| 18 | Cmtm8 | 3.11 | 1.86E-282 | 3.87E-278 | 0.92 | 0.17 | 0.75 | 5.43 |
| 18 | Lyl1 | 3.73 | 3.44E-282 | 7.16E-278 | 0.43 | 0.03 | 0.40 | 13.84 |
| 18 | Stk10 | 3.80 | 2.92E-279 | 6.08E-275 | 0.38 | 0.02 | 0.36 | 15.88 |
| 18 | Ramp3 | 3.77 | 1.49E-278 | 3.10E-274 | 0.40 | 0.03 | 0.37 | 15.19 |
| 18 | Tagln2 | 4.32 | 1.56E-273 | 3.24E-269 | 0.99 | 0.29 | 0.70 | 3.43 |
| 18 | Gng11 | 5.39 | 4.86E-273 | 1.01E-268 | 0.99 | 0.31 | 0.69 | 3.25 |
| 18 | Sema6b | 4.69 | 8.16E-273 | 1.70E-268 | 0.20 | 0.01 | 0.20 | 33.50 |
| 18 | Dock9 | 3.33 | 3.38E-272 | 7.04E-268 | 0.61 | 0.07 | 0.54 | 9.23 |
| 18 | Dnah2os | 5.77 | 3.13E-266 | 6.51E-262 | 0.13 | 0.00 | 0.12 | 63.00 |
| 18 | Trpc3 | 3.63 | 3.02E-264 | 6.27E-260 | 0.41 | 0.03 | 0.38 | 13.97 |
| 18 | Shroom2 | 3.27 | 5.36E-264 | 1.12E-259 | 0.62 | 0.07 | 0.55 | 8.76 |
| 18 | Anxa6 | 3.28 | 1.31E-263 | 2.72E-259 | 0.88 | 0.16 | 0.71 | 5.35 |
| 18 | Samd9l | 5.59 | 1.83E-262 | 3.81E-258 | 0.13 | 0.00 | 0.13 | 66.50 |
| 18 | Tlr3 | 4.77 | 2.47E-261 | 5.13E-257 | 0.19 | 0.01 | 0.18 | 37.40 |
| 18 | Ulbp1 | 4.00 | 4.86E-261 | 1.01E-256 | 0.34 | 0.02 | 0.32 | 16.85 |
| 18 | Ednrb | 3.08 | 1.97E-259 | 4.09E-255 | 0.44 | 0.03 | 0.40 | 12.79 |
| 18 | Nostrin | 5.91 | 4.31E-257 | 8.96E-253 | 0.12 | 0.00 | 0.11 | 58.00 |
| 18 | Bok | 3.28 | 2.34E-256 | 4.87E-252 | 0.90 | 0.19 | 0.71 | 4.72 |
| 18 | 6430548M08Rik | 3.91 | 6.74E-256 | 1.40E-251 | 0.26 | 0.01 | 0.25 | 21.58 |
| 18 | Ptgs1 | 3.03 | 1.12E-255 | 2.32E-251 | 0.44 | 0.04 | 0.41 | 12.28 |
| 18 | Lrrc8c | 3.15 | 1.53E-255 | 3.19E-251 | 0.77 | 0.12 | 0.65 | 6.52 |
| 18 | Tinagl1 | 3.73 | 8.70E-254 | 1.81E-249 | 0.52 | 0.05 | 0.47 | 9.89 |
| 18 | Arhgap31 | 3.19 | 9.79E-252 | 2.04E-247 | 0.89 | 0.18 | 0.71 | 4.95 |
| 18 | Fam124b | 4.96 | 1.61E-251 | 3.35E-247 | 0.13 | 0.00 | 0.13 | 64.50 |
| 18 | Rasa3 | 3.01 | 1.56E-250 | 3.25E-246 | 0.61 | 0.07 | 0.54 | 8.64 |
| 18 | Coro2b | 3.38 | 2.11E-248 | 4.38E-244 | 0.47 | 0.04 | 0.43 | 11.17 |
| 18 | Cbfa2t3 | 3.45 | 4.71E-248 | 9.79E-244 | 0.50 | 0.05 | 0.45 | 10.27 |
| 18 | Bvht | 3.02 | 1.33E-245 | 2.78E-241 | 0.52 | 0.05 | 0.47 | 10.14 |
| 18 | Il16 | 4.47 | 1.06E-243 | 2.20E-239 | 0.18 | 0.01 | 0.17 | 35.40 |
| 18 | F11r | 2.68 | 1.36E-243 | 2.83E-239 | 0.90 | 0.17 | 0.73 | 5.24 |
| 18 | Prcp | 3.10 | 1.34E-240 | 2.80E-236 | 0.96 | 0.25 | 0.71 | 3.87 |
| 18 | Klf7 | 2.98 | 1.06E-239 | 2.20E-235 | 0.93 | 0.22 | 0.71 | 4.24 |
| 18 | Akap12 | 3.08 | 2.12E-238 | 4.40E-234 | 0.89 | 0.19 | 0.70 | 4.76 |
| 18 | Pcdh17 | 3.04 | 1.91E-233 | 3.97E-229 | 0.90 | 0.20 | 0.70 | 4.54 |
| 18 | Gm36028 | 4.98 | 2.26E-233 | 4.71E-229 | 0.13 | 0.00 | 0.13 | 43.00 |
| 18 | H2-D1 | 3.11 | 3.07E-233 | 6.38E-229 | 0.94 | 0.23 | 0.70 | 4.01 |
| 18 | Tead4 | 4.02 | 1.18E-230 | 2.45E-226 | 0.25 | 0.01 | 0.24 | 20.67 |
| 18 | Tbkbp1 | 3.03 | 4.10E-230 | 8.53E-226 | 0.53 | 0.06 | 0.47 | 9.32 |
| 18 | Asb4 | 2.82 | 1.83E-226 | 3.80E-222 | 0.99 | 0.29 | 0.70 | 3.41 |
| 18 | Slfn5 | 4.28 | 1.05E-225 | 2.19E-221 | 0.19 | 0.01 | 0.19 | 27.71 |
| 18 | Nrp1 | 2.53 | 4.05E-225 | 8.43E-221 | 0.93 | 0.20 | 0.73 | 4.66 |
| 18 | Tmem173 | 3.27 | 1.16E-224 | 2.42E-220 | 0.49 | 0.05 | 0.44 | 9.61 |
| 18 | P2ry10b | 5.60 | 7.76E-224 | 1.61E-219 | 0.11 | 0.00 | 0.10 | 52.50 |
| 18 | Arpc1b | 3.05 | 3.14E-223 | 6.52E-219 | 0.98 | 0.34 | 0.64 | 2.89 |
| 18 | Orai1 | 2.80 | 9.45E-223 | 1.97E-218 | 0.83 | 0.16 | 0.67 | 5.24 |
| 18 | Nfatc1 | 3.38 | 1.32E-222 | 2.75E-218 | 0.37 | 0.03 | 0.34 | 13.11 |
| 18 | Ostf1 | 2.98 | 2.04E-221 | 4.24E-217 | 0.94 | 0.27 | 0.68 | 3.53 |

**Supplementary Table 1. Marker genes for the original clusters of integrated clustering of E11.5 control and NotchGOF λ-junction cells.**

| | | | | | | | | |
|---|---|---|---|---|---|---|---|---|
| 18 | Gm10687 | 5.27 | 8.42E-220 | 1.75E-215 | 0.12 | 0.00 | 0.12 | 61.00 |
| 18 | Rnf125 | 3.77 | 1.12E-219 | 2.33E-215 | 0.25 | 0.01 | 0.23 | 20.42 |
| 18 | Sardh | 2.76 | 7.40E-219 | 1.54E-214 | 0.14 | 0.00 | 0.14 | 35.75 |
| 18 | Arhgef3 | 3.51 | 8.53E-218 | 1.77E-213 | 0.27 | 0.02 | 0.25 | 17.93 |
| 18 | Cmbl | 3.76 | 4.37E-216 | 9.09E-212 | 0.26 | 0.01 | 0.24 | 18.21 |
| 18 | Ipo11 | 2.98 | 2.28E-215 | 4.74E-211 | 0.97 | 0.33 | 0.65 | 2.98 |
| 18 | Ets2 | 3.04 | 8.87E-215 | 1.85E-210 | 0.92 | 0.25 | 0.68 | 3.75 |
| 18 | Cdkl1 | 2.84 | 1.50E-214 | 3.12E-210 | 0.38 | 0.03 | 0.35 | 12.19 |
| 18 | Fes | 3.10 | 3.13E-214 | 6.52E-210 | 0.42 | 0.04 | 0.38 | 10.92 |
| 18 | Col4a2 | 3.52 | 1.89E-213 | 3.94E-209 | 0.98 | 0.42 | 0.56 | 2.35 |
| 18 | Hapln3 | 4.01 | 2.38E-213 | 4.94E-209 | 0.25 | 0.01 | 0.24 | 19.38 |
| 18 | Map3k11 | 2.87 | 4.13E-213 | 8.58E-209 | 0.69 | 0.11 | 0.58 | 6.39 |
| 18 | 5033406O09Rik | 4.30 | 2.80E-210 | 5.83E-206 | 0.21 | 0.01 | 0.20 | 23.44 |
| 18 | Ripor1 | 2.89 | 1.51E-208 | 3.15E-204 | 0.61 | 0.08 | 0.52 | 7.38 |
| 18 | Tspan4 | 2.81 | 7.98E-208 | 1.66E-203 | 0.72 | 0.12 | 0.60 | 6.03 |
| 18 | Slc7a7 | 3.53 | 5.48E-207 | 1.14E-202 | 0.26 | 0.02 | 0.24 | 17.27 |
| 18 | Piezo1 | 2.90 | 8.00E-206 | 1.66E-201 | 0.70 | 0.12 | 0.58 | 5.99 |
| 18 | Slc9a3r2 | 3.26 | 1.24E-205 | 2.58E-201 | 0.92 | 0.26 | 0.67 | 3.59 |
| 18 | Piezo2 | 2.49 | 1.29E-204 | 2.69E-200 | 0.93 | 0.22 | 0.71 | 4.22 |
| 18 | Cotl1 | 3.03 | 5.54E-204 | 1.15E-199 | 0.99 | 0.44 | 0.56 | 2.27 |
| 18 | Ptp4a3 | 3.23 | 7.15E-204 | 1.49E-199 | 0.90 | 0.25 | 0.65 | 3.67 |
| 18 | Elk3 | 2.93 | 5.03E-203 | 1.05E-198 | 0.99 | 0.41 | 0.58 | 2.40 |
| 18 | Gas6 | 3.23 | 2.10E-200 | 4.36E-196 | 0.32 | 0.02 | 0.30 | 13.33 |
| 18 | Col4a1 | 3.63 | 3.44E-199 | 7.14E-195 | 1.00 | 0.61 | 0.39 | 1.63 |
| 18 | Slc25a45 | 3.98 | 3.59E-199 | 7.46E-195 | 0.20 | 0.01 | 0.19 | 22.33 |
| 18 | Msn | 3.22 | 1.94E-198 | 4.04E-194 | 1.00 | 0.51 | 0.49 | 1.96 |
| 18 | Plaur | 4.67 | 7.67E-196 | 1.60E-191 | 0.17 | 0.01 | 0.17 | 24.71 |
| 18 | Slc43a2 | 3.05 | 1.36E-195 | 2.82E-191 | 0.39 | 0.04 | 0.36 | 10.86 |
| 18 | Card6 | 3.62 | 2.46E-195 | 5.11E-191 | 0.22 | 0.01 | 0.21 | 18.67 |
| 18 | Gm47664 | 3.96 | 2.98E-195 | 6.20E-191 | 0.21 | 0.01 | 0.20 | 21.10 |
| 18 | Pea15a | 2.66 | 1.07E-194 | 2.23E-190 | 0.89 | 0.22 | 0.66 | 3.96 |
| 18 | Col18a1 | 2.72 | 1.72E-193 | 3.58E-189 | 0.99 | 0.36 | 0.64 | 2.79 |
| 18 | Abcg2 | 2.72 | 9.35E-191 | 1.94E-186 | 0.97 | 0.34 | 0.63 | 2.84 |
| 18 | Jup | 2.50 | 2.61E-190 | 5.43E-186 | 0.85 | 0.19 | 0.66 | 4.43 |
| 18 | Zfp69 | 3.33 | 4.09E-190 | 8.50E-186 | 0.38 | 0.04 | 0.34 | 10.80 |
| 18 | Map4k2 | 2.96 | 9.97E-190 | 2.07E-185 | 0.51 | 0.06 | 0.45 | 7.97 |
| 18 | Hid1 | 3.94 | 3.51E-189 | 7.31E-185 | 0.21 | 0.01 | 0.20 | 20.70 |
| 18 | Hbegf | 2.99 | 4.78E-189 | 9.94E-185 | 0.55 | 0.08 | 0.48 | 7.19 |
| 18 | Ccdc85b | 2.91 | 1.44E-188 | 3.00E-184 | 0.99 | 0.58 | 0.41 | 1.71 |
| 18 | Tlr4 | 5.30 | 5.84E-188 | 1.21E-183 | 0.11 | 0.00 | 0.10 | 52.50 |
| 18 | Rftn1 | 2.97 | 2.25E-187 | 4.68E-183 | 0.37 | 0.04 | 0.34 | 10.69 |
| 18 | 1110032F04Rik | 3.81 | 9.20E-187 | 1.91E-182 | 0.28 | 0.02 | 0.26 | 14.53 |
| 18 | Vim | 3.24 | 3.78E-186 | 7.87E-182 | 1.00 | 0.84 | 0.16 | 1.19 |
| 18 | Myo6 | 2.56 | 1.14E-185 | 2.37E-181 | 0.81 | 0.18 | 0.63 | 4.53 |
| 18 | Ica1 | 2.62 | 5.09E-184 | 1.06E-179 | 0.67 | 0.11 | 0.56 | 5.88 |
| 18 | Gnai2 | 1.89 | 3.59E-183 | 7.47E-179 | 1.00 | 0.97 | 0.03 | 1.03 |
| 18 | Sacs | 2.73 | 3.94E-183 | 8.19E-179 | 0.79 | 0.17 | 0.62 | 4.71 |
| 18 | Cfap45 | 4.38 | 4.38E-182 | 9.11E-178 | 0.17 | 0.01 | 0.16 | 24.29 |
| 18 | Fkbp1a | 2.02 | 6.62E-182 | 1.38E-177 | 1.00 | 0.94 | 0.06 | 1.06 |
| 18 | Zc3hav1 | 3.49 | 8.79E-182 | 1.83E-177 | 0.31 | 0.02 | 0.28 | 12.75 |
| 18 | Npdc1 | 2.49 | 1.19E-181 | 2.47E-177 | 0.63 | 0.10 | 0.53 | 6.35 |
| 18 | Cmklr1 | 4.30 | 3.80E-181 | 7.90E-177 | 0.13 | 0.00 | 0.13 | 33.25 |
| 18 | St8sia1 | 3.33 | 1.41E-180 | 2.93E-176 | 0.32 | 0.03 | 0.29 | 12.31 |
| 18 | Itga6 | 2.29 | 2.73E-178 | 5.67E-174 | 0.98 | 0.30 | 0.68 | 3.22 |
| 18 | Cd81 | 2.19 | 5.79E-178 | 1.20E-173 | 1.00 | 0.89 | 0.11 | 1.12 |
| 18 | St8sia4 | 2.84 | 2.62E-177 | 5.45E-173 | 0.58 | 0.09 | 0.49 | 6.61 |

**Supplementary Table 1. Marker genes for the original clusters of integrated clustering of E11.5 control and NotchGOF λ-junction cells.**

| | | | | | | | | |
|---|---|---|---|---|---|---|---|---|
| 18 | Tmsb4x | 2.54 | 5.39E-177 | 1.12E-172 | 1.00 | 0.99 | 0.01 | 1.01 |
| 18 | Fam217a | 4.54 | 7.01E-177 | 1.46E-172 | 0.12 | 0.00 | 0.11 | 38.67 |
| 18 | Gprc5a | 3.95 | 7.73E-177 | 1.61E-172 | 0.22 | 0.01 | 0.21 | 17.00 |
| 18 | Sh2b3 | 2.69 | 6.47E-176 | 1.35E-171 | 0.52 | 0.07 | 0.45 | 7.39 |
| 18 | Ralb | 2.61 | 1.96E-175 | 4.08E-171 | 0.98 | 0.43 | 0.55 | 2.30 |
| 18 | Myo18a | 2.64 | 2.07E-175 | 4.31E-171 | 0.86 | 0.23 | 0.63 | 3.79 |
| 18 | Sparc | 2.53 | 2.24E-175 | 4.65E-171 | 1.00 | 0.74 | 0.26 | 1.36 |
| 18 | Mest | 2.14 | 1.07E-174 | 2.24E-170 | 1.00 | 0.91 | 0.09 | 1.10 |
| 18 | Gimap9 | 2.57 | 1.34E-174 | 2.79E-170 | 0.62 | 0.10 | 0.52 | 6.16 |
| 18 | Ttc9 | 3.04 | 6.25E-174 | 1.30E-169 | 0.37 | 0.04 | 0.33 | 10.19 |
| 18 | Kcnk6 | 4.15 | 1.05E-173 | 2.19E-169 | 0.17 | 0.01 | 0.16 | 24.29 |
| 18 | Arhgap29 | 2.65 | 2.27E-173 | 4.73E-169 | 0.95 | 0.36 | 0.59 | 2.64 |
| 18 | Notch1 | 2.01 | 4.55E-173 | 9.47E-169 | 0.93 | 0.24 | 0.69 | 3.89 |
| 18 | Trim25 | 3.04 | 9.11E-173 | 1.90E-168 | 0.45 | 0.06 | 0.39 | 8.16 |
| 18 | Pfkp | 3.05 | 1.06E-172 | 2.19E-168 | 0.42 | 0.05 | 0.37 | 8.89 |
| 18 | Rras | 2.47 | 3.93E-172 | 8.18E-168 | 0.84 | 0.21 | 0.63 | 4.02 |
| 18 | Plpp1 | 2.65 | 5.31E-172 | 1.11E-167 | 0.71 | 0.14 | 0.57 | 5.09 |
| 18 | Casq1 | 4.87 | 4.22E-171 | 8.78E-167 | 0.12 | 0.00 | 0.11 | 38.67 |
| 18 | Nes | 2.65 | 4.59E-168 | 9.54E-164 | 0.96 | 0.42 | 0.55 | 2.30 |
| 18 | Lxn | 2.64 | 6.99E-168 | 1.45E-163 | 0.95 | 0.36 | 0.59 | 2.65 |
| 18 | Clic1 | 1.85 | 1.45E-167 | 3.02E-163 | 1.00 | 0.93 | 0.07 | 1.08 |
| 18 | Pde4b | 3.41 | 8.59E-167 | 1.79E-162 | 0.37 | 0.04 | 0.33 | 9.51 |
| 18 | Anxa2 | 2.36 | 1.04E-166 | 2.16E-162 | 0.99 | 0.41 | 0.58 | 2.42 |
| 18 | Tgm2 | 4.23 | 1.32E-166 | 2.75E-162 | 0.21 | 0.01 | 0.20 | 17.58 |
| 18 | Cmtm3 | 2.44 | 1.02E-165 | 2.12E-161 | 0.98 | 0.41 | 0.56 | 2.36 |
| 18 | Tubb2a | 2.69 | 1.23E-165 | 2.56E-161 | 0.91 | 0.29 | 0.62 | 3.14 |
| 18 | Calm1 | 1.36 | 8.52E-165 | 1.77E-160 | 1.00 | 0.99 | 0.01 | 1.01 |
| 18 | Zbtb46 | 3.00 | 1.23E-164 | 2.56E-160 | 0.36 | 0.04 | 0.32 | 9.76 |
| 18 | Pkn3 | 2.92 | 1.66E-164 | 3.45E-160 | 0.40 | 0.04 | 0.35 | 8.98 |
| 18 | Ppic | 2.18 | 2.50E-163 | 5.20E-159 | 1.00 | 0.71 | 0.29 | 1.42 |
| 18 | Klk8 | 2.31 | 1.75E-161 | 3.64E-157 | 0.84 | 0.21 | 0.63 | 4.06 |
| 18 | Il2rg | 4.65 | 8.17E-160 | 1.70E-155 | 0.13 | 0.00 | 0.13 | 32.25 |
| 18 | Slco3a1 | 2.82 | 5.20E-159 | 1.08E-154 | 0.65 | 0.12 | 0.52 | 5.30 |
| 18 | Sh3bgrl3 | 1.69 | 7.45E-159 | 1.55E-154 | 1.00 | 0.91 | 0.09 | 1.10 |
| 18 | Arpc3 | 1.76 | 8.12E-159 | 1.69E-154 | 1.00 | 0.90 | 0.10 | 1.11 |
| 18 | Disc1 | 3.26 | 3.43E-158 | 7.14E-154 | 0.23 | 0.02 | 0.21 | 15.20 |
| 18 | Adcy4 | 2.77 | 8.41E-158 | 1.75E-153 | 0.45 | 0.06 | 0.39 | 7.66 |
| 18 | Sptbn1 | 2.17 | 1.29E-157 | 2.68E-153 | 1.00 | 0.76 | 0.24 | 1.32 |
| 18 | Plk2 | 2.42 | 2.19E-157 | 4.56E-153 | 0.95 | 0.37 | 0.58 | 2.58 |
| 18 | Rgl1 | 2.47 | 4.02E-156 | 8.37E-152 | 0.70 | 0.15 | 0.56 | 4.82 |
| 18 | Dgkh | 2.68 | 6.07E-156 | 1.26E-151 | 0.54 | 0.08 | 0.45 | 6.47 |
| 18 | Nid1 | 2.38 | 1.30E-155 | 2.70E-151 | 0.95 | 0.34 | 0.61 | 2.81 |
| 18 | Tspan9 | 2.42 | 1.54E-155 | 3.21E-151 | 0.79 | 0.19 | 0.60 | 4.20 |
| 18 | Gnas | 1.18 | 2.34E-154 | 4.87E-150 | 1.00 | 1.00 | 0.00 | 1.00 |
| 18 | Nlrc3 | 3.57 | 7.10E-154 | 1.48E-149 | 0.24 | 0.02 | 0.22 | 13.82 |
| 18 | Rhob | 2.45 | 1.52E-153 | 3.15E-149 | 0.93 | 0.34 | 0.59 | 2.75 |
| 18 | Cx3cl1 | 3.79 | 6.47E-153 | 1.34E-148 | 0.17 | 0.01 | 0.16 | 18.89 |
| 18 | Tns2 | 2.82 | 1.34E-152 | 2.78E-148 | 0.41 | 0.05 | 0.36 | 8.16 |
| 18 | Parvb | 2.42 | 2.18E-152 | 4.54E-148 | 0.65 | 0.12 | 0.52 | 5.30 |
| 18 | Ece1 | 2.38 | 2.89E-152 | 6.00E-148 | 0.83 | 0.22 | 0.61 | 3.71 |
| 18 | Cebpd | 2.59 | 3.56E-152 | 7.39E-148 | 0.56 | 0.09 | 0.47 | 6.03 |
| 18 | Ppp1r2 | 1.89 | 7.19E-151 | 1.49E-146 | 0.99 | 0.75 | 0.25 | 1.33 |
| 18 | Abcg1 | 2.32 | 7.55E-151 | 1.57E-146 | 0.52 | 0.08 | 0.44 | 6.63 |
| 18 | Smtn | 2.54 | 3.24E-150 | 6.74E-146 | 0.68 | 0.14 | 0.54 | 4.94 |
| 18 | Actb | 0.95 | 1.12E-149 | 2.32E-145 | 1.00 | 1.00 | 0.00 | 1.00 |
| 18 | Hebp1 | 1.89 | 3.68E-148 | 7.66E-144 | 0.89 | 0.26 | 0.63 | 3.42 |

**Supplementary Table 1. Marker genes for the original clusters of integrated clustering of E11.5 control and NotchGOF λ-junction cells.**

| | | | | | | | | |
|---|---|---|---|---|---|---|---|---|
| 18 | Tpst2 | 2.36 | 4.50E-147 | 9.35E-143 | 0.83 | 0.23 | 0.60 | 3.63 |
| 18 | Plagl1 | 2.10 | 2.04E-146 | 4.24E-142 | 0.98 | 0.43 | 0.55 | 2.28 |
| 18 | Mgat4b | 2.18 | 3.04E-145 | 6.32E-141 | 0.96 | 0.39 | 0.57 | 2.46 |
| 18 | Cyth4 | 2.50 | 4.01E-145 | 8.34E-141 | 0.17 | 0.01 | 0.16 | 19.22 |
| 18 | Tspan17 | 3.08 | 4.04E-145 | 8.41E-141 | 0.31 | 0.03 | 0.28 | 10.20 |
| 18 | Sgk1 | 2.45 | 1.04E-144 | 2.17E-140 | 0.79 | 0.20 | 0.59 | 3.91 |
| 18 | Ppp1r14a | 1.74 | 1.46E-144 | 3.04E-140 | 0.47 | 0.07 | 0.40 | 7.00 |
| 18 | Vsir | 3.89 | 3.58E-144 | 7.45E-140 | 0.20 | 0.01 | 0.19 | 15.69 |
| 18 | Prex1 | 2.28 | 6.08E-144 | 1.26E-139 | 0.77 | 0.19 | 0.58 | 4.09 |
| 18 | Skap2 | 2.41 | 8.86E-144 | 1.84E-139 | 0.40 | 0.05 | 0.35 | 7.86 |
| 18 | Tjp1 | 2.31 | 1.24E-143 | 2.59E-139 | 0.96 | 0.49 | 0.47 | 1.96 |
| 18 | Fyn | 2.15 | 5.76E-143 | 1.20E-138 | 0.95 | 0.44 | 0.51 | 2.16 |
| 18 | Rbpms | 2.05 | 8.09E-143 | 1.68E-138 | 0.98 | 0.54 | 0.45 | 1.83 |
| 18 | Lamc1 | 2.30 | 2.03E-142 | 4.22E-138 | 0.86 | 0.25 | 0.61 | 3.43 |
| 18 | Myl6 | 1.10 | 2.72E-142 | 5.66E-138 | 1.00 | 0.99 | 0.01 | 1.01 |
| 18 | Trf | 2.44 | 2.36E-140 | 4.90E-136 | 0.35 | 0.04 | 0.31 | 8.90 |
| 18 | Tal1 | 2.11 | 2.39E-140 | 4.97E-136 | 0.37 | 0.04 | 0.33 | 8.50 |
| 18 | Grasp | 1.98 | 3.04E-140 | 6.32E-136 | 0.78 | 0.19 | 0.59 | 4.17 |
| 18 | Myo1c | 2.56 | 5.11E-140 | 1.06E-135 | 0.53 | 0.09 | 0.44 | 5.80 |
| 18 | Fam13c | 2.70 | 3.96E-139 | 8.24E-135 | 0.45 | 0.06 | 0.38 | 6.97 |
| 18 | Kctd12b | 2.45 | 1.14E-138 | 2.38E-134 | 0.72 | 0.17 | 0.55 | 4.16 |
| 18 | Mef2a | 2.28 | 1.56E-138 | 3.25E-134 | 0.92 | 0.39 | 0.53 | 2.38 |
| 18 | Itsn2 | 2.33 | 2.77E-138 | 5.75E-134 | 0.75 | 0.18 | 0.57 | 4.09 |
| 18 | Csrp1 | 2.16 | 3.43E-138 | 7.14E-134 | 0.92 | 0.33 | 0.58 | 2.74 |
| 18 | Pfn1 | 0.95 | 7.87E-138 | 1.64E-133 | 1.00 | 1.00 | 0.00 | 1.00 |
| 18 | Selenop | 1.95 | 1.27E-137 | 2.64E-133 | 0.75 | 0.18 | 0.57 | 4.11 |
| 18 | Slc2a1 | 2.32 | 6.80E-137 | 1.41E-132 | 0.96 | 0.42 | 0.54 | 2.28 |
| 18 | Srgn | 1.39 | 1.39E-136 | 2.89E-132 | 0.17 | 0.01 | 0.16 | 17.00 |
| 18 | Il4ra | 3.13 | 3.01E-136 | 6.26E-132 | 0.21 | 0.02 | 0.20 | 14.07 |
| 18 | Abhd17a | 1.96 | 1.65E-135 | 3.44E-131 | 0.98 | 0.70 | 0.28 | 1.40 |
| 18 | Rcan3 | 2.46 | 2.00E-135 | 4.17E-131 | 0.47 | 0.07 | 0.40 | 6.48 |
| 18 | 2900026A02Rik | 2.19 | 2.08E-135 | 4.32E-131 | 0.56 | 0.10 | 0.46 | 5.67 |
| 18 | Tgfb1i1 | 2.31 | 9.26E-135 | 1.93E-130 | 0.64 | 0.13 | 0.51 | 4.89 |
| 18 | Slco2b1 | 3.53 | 9.65E-135 | 2.01E-130 | 0.12 | 0.00 | 0.11 | 29.00 |
| 18 | Sigirr | 3.11 | 1.50E-134 | 3.13E-130 | 0.26 | 0.02 | 0.24 | 10.92 |
| 18 | Syk | 2.64 | 3.00E-134 | 6.24E-130 | 0.27 | 0.03 | 0.25 | 10.88 |
| 18 | Arpc5 | 1.64 | 3.53E-134 | 7.34E-130 | 0.99 | 0.76 | 0.23 | 1.31 |
| 18 | Lfng | 3.64 | 8.85E-134 | 1.84E-129 | 0.21 | 0.02 | 0.20 | 13.38 |
| 18 | Ndrg1 | 3.21 | 7.00E-131 | 1.46E-126 | 0.22 | 0.02 | 0.21 | 12.44 |
| 18 | Rab11a | 1.92 | 9.94E-131 | 2.07E-126 | 0.98 | 0.64 | 0.34 | 1.53 |
| 18 | Degs1 | 2.19 | 2.87E-130 | 5.96E-126 | 0.94 | 0.39 | 0.54 | 2.39 |
| 18 | Zfp979 | 2.84 | 3.45E-130 | 7.17E-126 | 0.30 | 0.03 | 0.27 | 9.55 |
| 18 | Peg13 | 2.67 | 5.69E-130 | 1.18E-125 | 0.36 | 0.05 | 0.32 | 7.91 |
| 18 | Hsd3b7 | 2.90 | 6.73E-130 | 1.40E-125 | 0.30 | 0.03 | 0.27 | 9.55 |
| 18 | Trim16 | 2.94 | 1.03E-129 | 2.13E-125 | 0.24 | 0.02 | 0.22 | 11.90 |
| 18 | Vasp | 2.05 | 2.54E-129 | 5.28E-125 | 0.95 | 0.48 | 0.47 | 1.99 |
| 18 | Lrrc32 | 4.74 | 4.32E-129 | 8.99E-125 | 0.12 | 0.01 | 0.11 | 23.80 |
| 18 | Mtss1 | 2.09 | 5.36E-129 | 1.11E-124 | 0.86 | 0.29 | 0.58 | 3.00 |
| 18 | Atg9b | 4.45 | 1.25E-127 | 2.60E-123 | 0.10 | 0.00 | 0.10 | 34.00 |
| 18 | Tmsb10 | 0.81 | 6.87E-127 | 1.43E-122 | 1.00 | 1.00 | 0.00 | 1.00 |
| 18 | Cd9 | 1.80 | 1.20E-125 | 2.49E-121 | 0.90 | 0.29 | 0.60 | 3.07 |
| 18 | Abca1 | 2.09 | 3.06E-125 | 6.35E-121 | 0.44 | 0.07 | 0.37 | 6.59 |
| 18 | Gm13431 | 3.73 | 3.85E-125 | 8.00E-121 | 0.15 | 0.01 | 0.14 | 18.75 |
| 18 | Acot9 | 2.19 | 1.31E-124 | 2.73E-120 | 0.67 | 0.15 | 0.52 | 4.39 |
| 18 | Tpm4 | 1.20 | 5.57E-124 | 1.16E-119 | 1.00 | 0.93 | 0.06 | 1.07 |
| 18 | Tmem37 | 2.35 | 2.15E-123 | 4.48E-119 | 0.48 | 0.08 | 0.40 | 5.95 |

**Supplementary Table 1. Marker genes for the original clusters of integrated clustering of E11.5 control and NotchGOF λ-junction cells.**

| | | | | | | | | |
|---|---|---|---|---|---|---|---|---|
| 18 | Ehd4 | 1.91 | 2.41E-123 | 5.02E-119 | 0.70 | 0.17 | 0.53 | 4.12 |
| 18 | Itm2b | 1.39 | 6.48E-123 | 1.35E-118 | 1.00 | 0.84 | 0.16 | 1.18 |
| 18 | Slc25a5 | 1.27 | 2.10E-122 | 4.36E-118 | 1.00 | 0.93 | 0.06 | 1.07 |
| 18 | Ripk1 | 2.25 | 1.19E-120 | 2.48E-116 | 0.58 | 0.12 | 0.46 | 4.89 |
| 18 | Adam19 | 2.18 | 1.64E-120 | 3.42E-116 | 0.66 | 0.15 | 0.51 | 4.37 |
| 18 | Fzd4 | 1.93 | 1.36E-119 | 2.83E-115 | 0.83 | 0.25 | 0.59 | 3.40 |
| 18 | Itpkb | 2.17 | 1.61E-119 | 3.35E-115 | 0.72 | 0.19 | 0.53 | 3.86 |
| 18 | Zeb1 | 2.04 | 1.95E-119 | 4.05E-115 | 0.89 | 0.34 | 0.55 | 2.60 |
| 18 | Rgs3 | 2.21 | 1.41E-118 | 2.93E-114 | 0.70 | 0.18 | 0.52 | 3.98 |
| 18 | Serpinh1 | 1.06 | 1.89E-118 | 3.92E-114 | 1.00 | 0.94 | 0.06 | 1.06 |
| 18 | Galnt18 | 3.04 | 3.73E-118 | 7.76E-114 | 0.28 | 0.03 | 0.25 | 9.30 |
| 18 | Snx3 | 1.24 | 7.44E-118 | 1.55E-113 | 0.99 | 0.89 | 0.10 | 1.11 |
| 18 | Snx9 | 1.97 | 1.92E-117 | 3.99E-113 | 0.87 | 0.31 | 0.56 | 2.83 |
| 18 | Pnp | 1.71 | 2.31E-117 | 4.81E-113 | 0.97 | 0.66 | 0.31 | 1.47 |
| 18 | Mid2 | 3.03 | 2.32E-117 | 4.82E-113 | 0.25 | 0.03 | 0.23 | 10.08 |
| 18 | Gng12 | 2.00 | 2.48E-117 | 5.15E-113 | 0.87 | 0.36 | 0.51 | 2.44 |
| 18 | Frmd4b | 1.81 | 3.22E-117 | 6.70E-113 | 0.68 | 0.16 | 0.52 | 4.20 |
| 18 | Stard8 | 2.22 | 2.03E-116 | 4.23E-112 | 0.48 | 0.08 | 0.39 | 5.73 |
| 18 | Svil | 2.03 | 2.42E-116 | 5.04E-112 | 0.92 | 0.38 | 0.54 | 2.41 |
| 18 | Galnt12 | 2.91 | 1.13E-115 | 2.35E-111 | 0.23 | 0.02 | 0.21 | 11.00 |
| 18 | Cnn2 | 1.61 | 1.96E-115 | 4.07E-111 | 0.99 | 0.63 | 0.36 | 1.58 |
| 18 | Ppp1r18 | 1.96 | 2.32E-115 | 4.83E-111 | 0.81 | 0.25 | 0.57 | 3.31 |
| 18 | Ldb2 | 1.98 | 1.39E-114 | 2.90E-110 | 0.82 | 0.27 | 0.55 | 3.08 |
| 18 | Anxa5 | 1.75 | 1.73E-114 | 3.60E-110 | 0.97 | 0.60 | 0.36 | 1.60 |
| 18 | Serf2 | 0.66 | 3.07E-114 | 6.39E-110 | 1.00 | 1.00 | 0.00 | 1.00 |
| 18 | Rrbp1 | 1.30 | 8.42E-114 | 1.75E-109 | 0.99 | 0.90 | 0.10 | 1.11 |
| 18 | Cdc42 | 0.89 | 9.00E-114 | 1.87E-109 | 0.99 | 0.98 | 0.01 | 1.01 |
| 18 | Ppm1f | 2.28 | 3.22E-113 | 6.69E-109 | 0.52 | 0.10 | 0.42 | 5.14 |
| 18 | Gmpr | 1.80 | 5.21E-113 | 1.08E-108 | 0.69 | 0.17 | 0.52 | 4.06 |
| 18 | Cfl1 | 0.67 | 3.27E-112 | 6.79E-108 | 1.00 | 1.00 | 0.00 | 1.00 |
| 18 | Itga5 | 2.05 | 3.63E-112 | 7.55E-108 | 0.76 | 0.21 | 0.55 | 3.63 |
| 18 | Chst1 | 2.66 | 1.18E-111 | 2.46E-107 | 0.33 | 0.04 | 0.28 | 7.60 |
| 18 | Vat1 | 2.27 | 1.31E-111 | 2.73E-107 | 0.59 | 0.13 | 0.46 | 4.49 |
| 18 | Igf2 | 1.98 | 4.18E-111 | 8.69E-107 | 0.97 | 0.58 | 0.39 | 1.68 |
| 18 | Kif26a | 2.08 | 1.30E-110 | 2.71E-106 | 0.55 | 0.11 | 0.44 | 4.94 |
| 18 | B2m | 1.61 | 2.97E-110 | 6.18E-106 | 0.98 | 0.53 | 0.45 | 1.84 |
| 18 | Caskin2 | 2.30 | 4.18E-110 | 8.70E-106 | 0.53 | 0.11 | 0.42 | 4.92 |
| 18 | Lifr | 2.31 | 5.22E-110 | 1.09E-105 | 0.52 | 0.10 | 0.42 | 5.00 |
| 18 | Arl4a | 1.95 | 1.83E-109 | 3.80E-105 | 0.73 | 0.20 | 0.53 | 3.69 |
| 18 | Tcim | 2.27 | 3.39E-109 | 7.04E-105 | 0.88 | 0.36 | 0.52 | 2.43 |
| 18 | Dapk2 | 2.90 | 3.77E-109 | 7.84E-105 | 0.33 | 0.05 | 0.28 | 7.27 |
| 18 | Dlc1 | 1.82 | 6.12E-109 | 1.27E-104 | 0.89 | 0.35 | 0.53 | 2.51 |
| 18 | Maff | 2.52 | 6.27E-109 | 1.30E-104 | 0.32 | 0.04 | 0.28 | 7.62 |
| 18 | C430049B03Rik | 1.97 | 7.51E-109 | 1.56E-104 | 0.88 | 0.35 | 0.54 | 2.55 |
| 18 | Sh3bp5 | 2.00 | 1.14E-108 | 2.36E-104 | 0.93 | 0.50 | 0.43 | 1.85 |
| 18 | Ubtd1 | 2.01 | 2.09E-108 | 4.34E-104 | 0.62 | 0.14 | 0.48 | 4.32 |
| 18 | Sh3glb1 | 1.30 | 4.16E-108 | 8.65E-104 | 0.99 | 0.84 | 0.15 | 1.18 |
| 18 | Pitpna | 1.48 | 4.28E-108 | 8.91E-104 | 0.98 | 0.70 | 0.28 | 1.40 |
| 18 | Myo7a | 3.32 | 9.02E-108 | 1.88E-103 | 0.16 | 0.01 | 0.15 | 13.58 |
| 18 | Heg1 | 2.27 | 9.52E-108 | 1.98E-103 | 0.62 | 0.15 | 0.47 | 4.22 |
| 18 | Bgn | 1.90 | 1.05E-107 | 2.18E-103 | 0.80 | 0.26 | 0.54 | 3.12 |
| 18 | Nab1 | 1.87 | 1.06E-107 | 2.20E-103 | 0.58 | 0.12 | 0.45 | 4.66 |
| 18 | Kcna5 | 3.51 | 1.73E-107 | 3.60E-103 | 0.20 | 0.02 | 0.18 | 11.59 |
| 18 | Elovl1 | 2.08 | 1.89E-107 | 3.94E-103 | 0.79 | 0.25 | 0.53 | 3.12 |
| 18 | Tacc1 | 1.84 | 2.66E-107 | 5.54E-103 | 0.86 | 0.31 | 0.55 | 2.78 |
| 18 | Fam129b | 1.97 | 2.74E-107 | 5.70E-103 | 0.61 | 0.14 | 0.47 | 4.42 |

**Supplementary Table 1. Marker genes for the original clusters of integrated clustering of E11.5 control and NotchGOF λ-junction cells.**

| | | | | | | | | |
|---|---|---|---|---|---|---|---|---|
| 18 | Als2cl | 3.54 | 4.70E-107 | 9.78E-103 | 0.14 | 0.01 | 0.13 | 15.89 |
| 18 | Magi3 | 2.05 | 1.24E-106 | 2.58E-102 | 0.69 | 0.18 | 0.50 | 3.73 |
| 18 | Dgkz | 1.87 | 1.59E-106 | 3.31E-102 | 0.83 | 0.28 | 0.55 | 2.97 |
| 18 | Gm26862 | 3.34 | 1.72E-106 | 3.58E-102 | 0.14 | 0.01 | 0.13 | 15.89 |
| 18 | Lrp10 | 2.22 | 5.31E-106 | 1.10E-101 | 0.65 | 0.17 | 0.48 | 3.92 |
| 18 | Tcf15 | 1.88 | 6.59E-106 | 1.37E-101 | 0.83 | 0.28 | 0.55 | 3.00 |
| 18 | Itgb1 | 1.28 | 6.89E-106 | 1.43E-101 | 1.00 | 0.85 | 0.15 | 1.17 |
| 18 | Ly6h | 1.59 | 6.95E-106 | 1.44E-101 | 0.63 | 0.15 | 0.48 | 4.29 |
| 18 | Ndufa8 | 1.21 | 9.91E-106 | 2.06E-101 | 0.98 | 0.90 | 0.08 | 1.09 |
| 18 | Marchf2 | 1.82 | 9.92E-106 | 2.06E-101 | 0.97 | 0.51 | 0.46 | 1.90 |
| 18 | Pam | 1.90 | 9.98E-106 | 2.08E-101 | 0.62 | 0.15 | 0.48 | 4.29 |
| 18 | Nid2 | 2.08 | 1.26E-105 | 2.62E-101 | 0.77 | 0.23 | 0.54 | 3.33 |
| 18 | Ddah1 | 2.11 | 2.48E-105 | 5.16E-101 | 0.59 | 0.13 | 0.45 | 4.37 |
| 18 | Actg1 | 0.69 | 6.28E-105 | 1.31E-100 | 1.00 | 1.00 | 0.00 | 1.00 |
| 18 | S100a10 | 1.57 | 1.26E-104 | 2.62E-100 | 0.97 | 0.55 | 0.42 | 1.76 |
| 18 | Paqr4 | 2.59 | 4.32E-104 | 8.99E-100 | 0.27 | 0.03 | 0.24 | 8.50 |
| 18 | Uaca | 1.92 | 4.49E-104 | 9.34E-100 | 0.85 | 0.33 | 0.52 | 2.57 |
| 18 | Fmnl3 | 1.96 | 4.77E-104 | 9.92E-100 | 0.71 | 0.20 | 0.51 | 3.55 |
| 18 | Pgm2 | 1.91 | 6.65E-104 | 1.38E-99 | 0.81 | 0.27 | 0.54 | 3.02 |
| 18 | Gpx3 | 1.83 | 9.03E-104 | 1.88E-99 | 0.75 | 0.21 | 0.53 | 3.51 |
| 18 | Maged2 | 1.60 | 1.74E-103 | 3.62E-99 | 0.99 | 0.82 | 0.17 | 1.21 |
| 18 | 4931406P16Rik | 1.93 | 2.09E-103 | 4.36E-99 | 0.87 | 0.35 | 0.52 | 2.49 |
| 18 | Lpar4 | 2.01 | 6.87E-103 | 1.43E-98 | 0.85 | 0.32 | 0.53 | 2.66 |
| 18 | Arpc4 | 1.27 | 1.30E-102 | 2.70E-98 | 0.98 | 0.82 | 0.16 | 1.19 |
| 18 | Rhoa | 0.92 | 2.13E-102 | 4.44E-98 | 1.00 | 0.97 | 0.03 | 1.04 |
| 18 | Gnb2 | 0.93 | 2.49E-102 | 5.18E-98 | 1.00 | 0.96 | 0.04 | 1.04 |
| 18 | Ppfibp1 | 1.67 | 3.19E-102 | 6.64E-98 | 0.93 | 0.42 | 0.50 | 2.18 |
| 18 | Rasal2 | 1.86 | 4.66E-102 | 9.68E-98 | 0.77 | 0.24 | 0.53 | 3.18 |
| 18 | Cabp1 | 3.48 | 8.02E-102 | 1.67E-97 | 0.16 | 0.01 | 0.15 | 13.33 |
| 18 | Peg10 | 1.73 | 4.24E-101 | 8.81E-97 | 0.92 | 0.43 | 0.49 | 2.13 |
| 18 | Dram2 | 1.86 | 7.40E-101 | 1.54E-96 | 0.72 | 0.20 | 0.52 | 3.61 |
| 18 | Fhod1 | 2.18 | 1.32E-100 | 2.74E-96 | 0.45 | 0.08 | 0.37 | 5.41 |
| 18 | Plekha1 | 2.02 | 1.38E-100 | 2.88E-96 | 0.69 | 0.19 | 0.50 | 3.58 |
| 18 | Schip1 | 1.80 | 1.79E-100 | 3.72E-96 | 0.64 | 0.16 | 0.48 | 4.10 |
| 18 | 9930012K11Rik | 2.33 | 2.38E-100 | 4.94E-96 | 0.37 | 0.06 | 0.31 | 6.33 |
| 18 | Rasd1 | 2.66 | 4.91E-100 | 1.02E-95 | 0.30 | 0.04 | 0.26 | 7.59 |
| 18 | Arhgef5 | 1.87 | 5.18E-100 | 1.08E-95 | 0.69 | 0.19 | 0.50 | 3.56 |
| 18 | H2-K1 | 2.15 | 7.49E-99 | 1.56E-94 | 0.56 | 0.13 | 0.43 | 4.33 |
| 18 | Rap1a | 1.39 | 1.71E-98 | 3.56E-94 | 0.98 | 0.71 | 0.27 | 1.37 |
| 18 | Slc43a3 | 1.85 | 2.13E-98 | 4.42E-94 | 0.80 | 0.28 | 0.52 | 2.86 |
| 18 | Mpp1 | 2.00 | 5.43E-98 | 1.13E-93 | 0.74 | 0.23 | 0.51 | 3.18 |
| 18 | Crip1 | 2.01 | 9.89E-98 | 2.06E-93 | 0.88 | 0.40 | 0.49 | 2.22 |
| 18 | Klhl36 | 2.23 | 1.05E-97 | 2.18E-93 | 0.38 | 0.06 | 0.32 | 6.05 |
| 18 | Bmp2k | 1.82 | 1.26E-97 | 2.62E-93 | 0.60 | 0.15 | 0.46 | 4.12 |
| 18 | Chst7 | 3.34 | 4.03E-97 | 8.38E-93 | 0.17 | 0.01 | 0.16 | 12.36 |
| 18 | Aprt | 1.13 | 4.93E-97 | 1.03E-92 | 1.00 | 0.92 | 0.08 | 1.08 |
| 18 | Slc7a1 | 2.09 | 5.41E-97 | 1.12E-92 | 0.42 | 0.07 | 0.34 | 5.61 |
| 18 | Lamb1 | 1.71 | 5.82E-97 | 1.21E-92 | 0.90 | 0.38 | 0.51 | 2.35 |
| 18 | Pitpnm2 | 2.09 | 6.31E-96 | 1.31E-91 | 0.44 | 0.08 | 0.35 | 5.37 |
| 18 | Hdac7 | 1.85 | 6.94E-96 | 1.44E-91 | 0.81 | 0.28 | 0.53 | 2.90 |
| 18 | Tecpr1 | 2.09 | 1.13E-95 | 2.36E-91 | 0.57 | 0.13 | 0.43 | 4.22 |
| 18 | H19 | 1.33 | 1.91E-95 | 3.97E-91 | 0.99 | 0.84 | 0.15 | 1.18 |
| 18 | Kcnj15 | 2.93 | 7.66E-95 | 1.59E-90 | 0.18 | 0.02 | 0.16 | 11.25 |
| 18 | Fam102b | 1.91 | 8.55E-95 | 1.78E-90 | 0.68 | 0.19 | 0.49 | 3.53 |
| 18 | Hprt | 1.62 | 1.33E-94 | 2.76E-90 | 0.92 | 0.43 | 0.49 | 2.13 |
| 18 | Slc6a6 | 1.99 | 1.05E-93 | 2.18E-89 | 0.85 | 0.35 | 0.50 | 2.45 |

**Supplementary Table 1. Marker genes for the original clusters of integrated clustering of E11.5 control and NotchGOF λ-junction cells.**

| | | | | | | | | |
|---|---|---|---|---|---|---|---|---|
| 18 | Coro1c | 1.82 | 1.10E-93 | 2.28E-89 | 0.84 | 0.34 | 0.50 | 2.45 |
| 18 | Inf2 | 2.49 | 3.41E-93 | 7.08E-89 | 0.31 | 0.05 | 0.27 | 6.80 |
| 18 | Tax1bp3 | 1.45 | 3.92E-93 | 8.15E-89 | 0.97 | 0.64 | 0.33 | 1.51 |
| 18 | Tnfrsf1a | 1.76 | 4.22E-93 | 8.77E-89 | 0.82 | 0.30 | 0.52 | 2.76 |
| 18 | Actr3 | 1.16 | 4.71E-93 | 9.79E-89 | 0.99 | 0.82 | 0.17 | 1.21 |
| 18 | Map4k5 | 1.96 | 7.37E-93 | 1.53E-88 | 0.80 | 0.30 | 0.49 | 2.63 |
| 18 | Adm | 3.32 | 1.15E-92 | 2.40E-88 | 0.11 | 0.01 | 0.10 | 21.00 |
| 18 | Klf6 | 1.99 | 1.69E-92 | 3.52E-88 | 0.71 | 0.23 | 0.48 | 3.10 |
| 18 | Parp4 | 2.97 | 3.83E-92 | 7.97E-88 | 0.18 | 0.02 | 0.17 | 10.82 |
| 18 | Pdlim5 | 1.75 | 5.63E-92 | 1.17E-87 | 0.72 | 0.22 | 0.50 | 3.29 |
| 18 | Ehbp1l1 | 2.27 | 1.28E-91 | 2.67E-87 | 0.36 | 0.06 | 0.30 | 5.92 |
| 18 | Tpm3 | 0.92 | 3.27E-91 | 6.81E-87 | 0.99 | 0.96 | 0.04 | 1.04 |
| 18 | Arpc2 | 0.93 | 3.31E-91 | 6.88E-87 | 1.00 | 0.94 | 0.06 | 1.06 |
| 18 | Prkd2 | 2.13 | 4.59E-91 | 9.54E-87 | 0.48 | 0.10 | 0.38 | 4.71 |
| 18 | Kctd17 | 1.69 | 5.58E-91 | 1.16E-86 | 0.79 | 0.27 | 0.52 | 2.96 |
| 18 | Mapk12 | 1.82 | 1.34E-90 | 2.79E-86 | 0.50 | 0.11 | 0.39 | 4.57 |
| 18 | Fli1 | 1.67 | 1.45E-90 | 3.01E-86 | 0.89 | 0.37 | 0.52 | 2.43 |
| 18 | Rsu1 | 1.52 | 2.27E-90 | 4.72E-86 | 0.94 | 0.48 | 0.46 | 1.96 |
| 18 | Myh9 | 1.73 | 3.29E-90 | 6.84E-86 | 0.92 | 0.50 | 0.42 | 1.84 |
| 18 | Tspan15 | 1.86 | 4.13E-90 | 8.58E-86 | 0.66 | 0.19 | 0.48 | 3.57 |
| 18 | Rps2 | 0.50 | 5.30E-90 | 1.10E-85 | 1.00 | 1.00 | 0.00 | 1.00 |
| 18 | Pxn | 1.79 | 5.67E-90 | 1.18E-85 | 0.65 | 0.18 | 0.47 | 3.63 |
| 18 | Spata13 | 2.20 | 1.02E-89 | 2.12E-85 | 0.40 | 0.07 | 0.32 | 5.41 |
| 18 | Cpne8 | 1.90 | 1.37E-89 | 2.86E-85 | 0.41 | 0.08 | 0.33 | 5.40 |
| 18 | Nexmif | 2.39 | 1.43E-89 | 2.97E-85 | 0.25 | 0.03 | 0.22 | 7.88 |
| 18 | Abcc4 | 3.03 | 6.74E-89 | 1.40E-84 | 0.19 | 0.02 | 0.17 | 10.00 |
| 18 | Gng5 | 0.55 | 8.39E-89 | 1.74E-84 | 1.00 | 1.00 | 0.00 | 1.00 |
| 18 | Ccdc82 | 2.00 | 1.00E-88 | 2.09E-84 | 0.50 | 0.11 | 0.39 | 4.45 |
| 18 | Klf10 | 1.98 | 1.91E-88 | 3.96E-84 | 0.48 | 0.10 | 0.38 | 4.62 |
| 18 | Frmd8 | 1.82 | 1.95E-88 | 4.06E-84 | 0.68 | 0.20 | 0.48 | 3.44 |
| 18 | Gng2 | 1.43 | 2.59E-88 | 5.38E-84 | 0.96 | 0.57 | 0.39 | 1.69 |
| 18 | St6galnac4 | 1.74 | 1.34E-87 | 2.78E-83 | 0.67 | 0.19 | 0.48 | 3.47 |
| 18 | Mxra7 | 1.70 | 1.79E-87 | 3.73E-83 | 0.73 | 0.23 | 0.50 | 3.21 |
| 18 | Tnfaip2 | 0.83 | 3.15E-87 | 6.56E-83 | 0.59 | 0.14 | 0.45 | 4.29 |
| 18 | Tdrd7 | 2.54 | 5.07E-87 | 1.06E-82 | 0.22 | 0.03 | 0.20 | 8.50 |
| 18 | Dock11 | 1.87 | 5.87E-87 | 1.22E-82 | 0.70 | 0.22 | 0.47 | 3.13 |
| 18 | Ppp2r5a | 1.82 | 9.57E-87 | 1.99E-82 | 0.80 | 0.31 | 0.48 | 2.54 |
| 18 | Prr5l | 1.84 | 3.46E-86 | 7.19E-82 | 0.60 | 0.16 | 0.44 | 3.76 |
| 18 | Sptan1 | 1.62 | 4.30E-86 | 8.94E-82 | 0.95 | 0.56 | 0.39 | 1.69 |
| 18 | Tln1 | 1.17 | 6.72E-86 | 1.40E-81 | 0.99 | 0.74 | 0.25 | 1.33 |
| 18 | Cd151 | 1.52 | 1.13E-85 | 2.35E-81 | 0.86 | 0.32 | 0.54 | 2.67 |
| 18 | Pik3c2b | 2.29 | 1.59E-85 | 3.30E-81 | 0.37 | 0.07 | 0.30 | 5.48 |
| 18 | Prkce | 1.80 | 2.78E-85 | 5.78E-81 | 0.58 | 0.15 | 0.44 | 3.96 |
| 18 | Mpzl1 | 1.46 | 7.00E-85 | 1.46E-80 | 0.96 | 0.63 | 0.33 | 1.53 |
| 18 | Dock4 | 1.66 | 1.16E-84 | 2.41E-80 | 0.81 | 0.31 | 0.50 | 2.64 |
| 18 | Slco2a1 | 2.85 | 1.44E-84 | 2.99E-80 | 0.20 | 0.02 | 0.18 | 8.87 |
| 18 | Acsl5 | 1.85 | 1.65E-84 | 3.44E-80 | 0.58 | 0.15 | 0.43 | 3.80 |
| 18 | Pold4 | 1.71 | 1.69E-84 | 3.52E-80 | 0.65 | 0.18 | 0.47 | 3.57 |
| 18 | Ctbs | 2.15 | 2.11E-84 | 4.38E-80 | 0.43 | 0.09 | 0.34 | 4.80 |
| 18 | Plec | 1.71 | 2.30E-84 | 4.78E-80 | 0.77 | 0.27 | 0.50 | 2.88 |
| 18 | Snrk | 1.71 | 3.64E-84 | 7.56E-80 | 0.76 | 0.26 | 0.50 | 2.96 |
| 18 | Ywhab | 0.75 | 3.80E-84 | 7.91E-80 | 1.00 | 0.97 | 0.03 | 1.03 |
| 18 | Lpar6 | 1.59 | 4.40E-84 | 9.16E-80 | 0.89 | 0.40 | 0.49 | 2.23 |
| 18 | Prickle3 | 2.11 | 1.33E-83 | 2.77E-79 | 0.35 | 0.06 | 0.29 | 5.71 |
| 18 | Baiap2 | 1.63 | 1.66E-83 | 3.45E-79 | 0.58 | 0.15 | 0.43 | 3.91 |
| 18 | Cpne2 | 2.27 | 5.19E-83 | 1.08E-78 | 0.31 | 0.05 | 0.26 | 6.38 |

**Supplementary Table 1. Marker genes for the original clusters of integrated clustering of E11.5 control and NotchGOF λ-junction cells.**

| | | | | | | | | |
|---|---|---|---|---|---|---|---|---|
| 18 | Hfe | 2.89 | 5.73E-83 | 1.19E-78 | 0.15 | 0.01 | 0.14 | 11.77 |
| 18 | D5Ertd605e | 3.45 | 1.62E-82 | 3.37E-78 | 0.11 | 0.01 | 0.11 | 16.00 |
| 18 | Lipt2 | 2.13 | 3.42E-82 | 7.12E-78 | 0.80 | 0.35 | 0.44 | 2.25 |
| 18 | Srgap1 | 1.61 | 7.29E-82 | 1.52E-77 | 0.83 | 0.34 | 0.49 | 2.45 |
| 18 | Eif1 | 0.46 | 1.92E-81 | 3.98E-77 | 1.00 | 1.00 | 0.00 | 1.00 |
| 18 | Pdia5 | 2.07 | 2.17E-81 | 4.52E-77 | 0.48 | 0.11 | 0.37 | 4.29 |
| 18 | Mfge8 | 1.44 | 3.28E-81 | 6.82E-77 | 0.96 | 0.52 | 0.44 | 1.85 |
| 18 | Reep1 | 2.41 | 3.32E-81 | 6.90E-77 | 0.29 | 0.05 | 0.24 | 6.42 |
| 18 | Actn4 | 1.29 | 5.54E-81 | 1.15E-76 | 0.97 | 0.70 | 0.27 | 1.39 |
| 18 | Ly6c1 | 4.01 | 7.13E-81 | 1.48E-76 | 0.10 | 0.01 | 0.10 | 17.00 |
| 18 | Ywhah | 0.91 | 1.43E-80 | 2.98E-76 | 1.00 | 0.91 | 0.09 | 1.10 |
| 18 | Mall | 3.40 | 1.45E-80 | 3.02E-76 | 0.10 | 0.01 | 0.10 | 17.00 |
| 18 | Afdn | 1.30 | 2.19E-80 | 4.55E-76 | 0.96 | 0.57 | 0.40 | 1.70 |
| 18 | Nrarp | 1.08 | 5.20E-80 | 1.08E-75 | 0.65 | 0.18 | 0.48 | 3.71 |
| 18 | Casp7 | 1.87 | 7.29E-80 | 1.52E-75 | 0.59 | 0.17 | 0.42 | 3.56 |
| 18 | Arhgef28 | 1.92 | 7.34E-80 | 1.53E-75 | 0.45 | 0.10 | 0.35 | 4.58 |
| 18 | Fam78b | 2.44 | 7.43E-80 | 1.54E-75 | 0.22 | 0.03 | 0.19 | 7.89 |
| 18 | Ripk3 | 1.65 | 1.14E-79 | 2.37E-75 | 0.60 | 0.16 | 0.44 | 3.72 |
| 18 | Yes1 | 1.63 | 5.60E-79 | 1.16E-74 | 0.83 | 0.34 | 0.49 | 2.42 |
| 18 | Rgl2 | 1.73 | 7.39E-79 | 1.54E-74 | 0.66 | 0.20 | 0.46 | 3.33 |
| 18 | Gnb4 | 1.58 | 1.21E-78 | 2.51E-74 | 0.79 | 0.29 | 0.50 | 2.70 |
| 18 | Rinl | 2.94 | 1.84E-78 | 3.82E-74 | 0.16 | 0.02 | 0.15 | 10.19 |
| 18 | Arl6ip5 | 1.54 | 1.92E-78 | 3.99E-74 | 0.83 | 0.31 | 0.51 | 2.63 |
| 18 | Fermt2 | 1.15 | 2.75E-78 | 5.72E-74 | 0.98 | 0.72 | 0.26 | 1.36 |
| 18 | Wasf2 | 1.20 | 3.38E-78 | 7.03E-74 | 0.98 | 0.71 | 0.28 | 1.39 |
| 18 | Nhp2 | 1.03 | 3.72E-78 | 7.73E-74 | 0.99 | 0.88 | 0.11 | 1.12 |
| 18 | Ssh2 | 1.64 | 7.84E-78 | 1.63E-73 | 0.70 | 0.23 | 0.47 | 3.10 |
| 18 | Kifc3 | 2.06 | 8.07E-78 | 1.68E-73 | 0.37 | 0.07 | 0.30 | 5.23 |
| 18 | Git1 | 1.62 | 1.19E-77 | 2.48E-73 | 0.74 | 0.26 | 0.48 | 2.89 |
| 18 | Casp8 | 1.79 | 1.44E-77 | 2.99E-73 | 0.61 | 0.18 | 0.43 | 3.40 |
| 18 | Arhgef1 | 1.67 | 1.50E-77 | 3.11E-73 | 0.73 | 0.26 | 0.47 | 2.79 |
| 18 | Itpr2 | 2.16 | 2.45E-77 | 5.09E-73 | 0.28 | 0.04 | 0.24 | 6.41 |
| 18 | Pomp | 0.75 | 2.81E-77 | 5.85E-73 | 1.00 | 0.98 | 0.02 | 1.02 |
| 18 | Cd79b | 2.67 | 3.26E-77 | 6.78E-73 | 0.14 | 0.01 | 0.13 | 11.58 |
| 18 | Smpd5 | 3.01 | 3.48E-77 | 7.24E-73 | 0.13 | 0.01 | 0.12 | 12.90 |
| 18 | Mapk3 | 1.52 | 4.53E-77 | 9.41E-73 | 0.90 | 0.44 | 0.45 | 2.02 |
| 18 | Ebf3 | 1.39 | 5.30E-77 | 1.10E-72 | 0.61 | 0.17 | 0.44 | 3.60 |
| 18 | Myl12a | 0.82 | 6.88E-77 | 1.43E-72 | 1.00 | 0.95 | 0.05 | 1.05 |
| 18 | Rab13 | 1.56 | 7.93E-77 | 1.65E-72 | 0.89 | 0.42 | 0.47 | 2.12 |
| 18 | Palm | 1.51 | 1.20E-76 | 2.49E-72 | 0.89 | 0.46 | 0.43 | 1.95 |
| 18 | Aplp2 | 1.08 | 1.61E-76 | 3.35E-72 | 0.99 | 0.82 | 0.17 | 1.21 |
| 18 | Adgre5 | 1.77 | 1.16E-75 | 2.41E-71 | 0.50 | 0.12 | 0.38 | 4.07 |
| 18 | Cdipt | 1.66 | 1.64E-75 | 3.41E-71 | 0.84 | 0.35 | 0.49 | 2.39 |
| 18 | Rap1b | 1.09 | 1.79E-75 | 3.72E-71 | 0.97 | 0.76 | 0.21 | 1.28 |
| 18 | Peak1 | 1.66 | 2.47E-75 | 5.14E-71 | 0.71 | 0.24 | 0.47 | 2.95 |
| 18 | Sptlc2 | 1.60 | 3.41E-75 | 7.10E-71 | 0.81 | 0.32 | 0.49 | 2.51 |
| 18 | Sp140 | 3.03 | 6.68E-75 | 1.39E-70 | 0.13 | 0.01 | 0.12 | 12.60 |
| 18 | Stard4 | 1.55 | 7.45E-75 | 1.55E-70 | 0.76 | 0.27 | 0.49 | 2.78 |
| 18 | Rhoc | 1.16 | 1.15E-74 | 2.38E-70 | 0.98 | 0.79 | 0.19 | 1.24 |
| 18 | Mef2d | 1.75 | 2.59E-74 | 5.39E-70 | 0.52 | 0.13 | 0.39 | 3.88 |
| 18 | Pxdc1 | 1.63 | 3.01E-74 | 6.26E-70 | 0.45 | 0.10 | 0.35 | 4.42 |
| 18 | Ap1s2 | 1.91 | 3.31E-74 | 6.89E-70 | 0.50 | 0.13 | 0.37 | 3.87 |
| 18 | Myadm | 1.92 | 1.57E-73 | 3.26E-69 | 0.49 | 0.12 | 0.37 | 4.02 |
| 18 | Scarb1 | 1.84 | 1.84E-73 | 3.82E-69 | 0.55 | 0.15 | 0.39 | 3.56 |
| 18 | Nr4a1 | 2.64 | 2.24E-73 | 4.66E-69 | 0.26 | 0.04 | 0.22 | 6.48 |
| 18 | Sorbs1 | 1.91 | 2.26E-73 | 4.69E-69 | 0.39 | 0.08 | 0.31 | 4.85 |

**Supplementary Table 1. Marker genes for the original clusters of integrated clustering of E11.5 control and NotchGOF λ-junction cells.**

| | | | | | | | | |
|---|---|---|---|---|---|---|---|---|
| 18 | Nrp2 | 1.18 | 3.00E-73 | 6.25E-69 | 0.88 | 0.37 | 0.52 | 2.42 |
| 18 | Cmtm6 | 1.82 | 3.56E-73 | 7.41E-69 | 0.50 | 0.13 | 0.37 | 3.97 |
| 18 | Wdr1 | 1.31 | 3.58E-73 | 7.44E-69 | 0.92 | 0.54 | 0.38 | 1.70 |
| 18 | Tspan6 | 1.22 | 5.59E-73 | 1.16E-68 | 0.97 | 0.74 | 0.23 | 1.31 |
| 18 | Lrrfip1 | 1.51 | 7.00E-73 | 1.46E-68 | 0.82 | 0.34 | 0.49 | 2.46 |
| 18 | Rbfox3 | 3.36 | 8.19E-73 | 1.70E-68 | 0.11 | 0.01 | 0.10 | 14.00 |
| 18 | Stim1 | 1.72 | 8.66E-73 | 1.80E-68 | 0.53 | 0.14 | 0.39 | 3.79 |
| 18 | Nckap5l | 1.73 | 8.87E-73 | 1.85E-68 | 0.53 | 0.14 | 0.39 | 3.82 |
| 18 | Nr2f2 | 1.38 | 1.47E-72 | 3.07E-68 | 0.84 | 0.36 | 0.48 | 2.33 |
| 18 | Ltbr | 1.60 | 1.54E-72 | 3.19E-68 | 0.69 | 0.23 | 0.47 | 3.08 |
| 18 | Bnip2 | 1.17 | 1.67E-72 | 3.46E-68 | 0.96 | 0.68 | 0.28 | 1.40 |
| 18 | Gale | 1.83 | 5.51E-72 | 1.15E-67 | 0.48 | 0.12 | 0.36 | 3.99 |
| 18 | Ccser2 | 1.52 | 6.28E-72 | 1.31E-67 | 0.74 | 0.27 | 0.47 | 2.77 |
| 18 | Wscd1 | 2.35 | 1.70E-71 | 3.53E-67 | 0.19 | 0.02 | 0.17 | 8.08 |
| 18 | Sntb1 | 2.37 | 5.45E-71 | 1.13E-66 | 0.20 | 0.03 | 0.18 | 7.85 |
| 18 | Foxo1 | 1.56 | 6.92E-71 | 1.44E-66 | 0.60 | 0.18 | 0.42 | 3.36 |
| 18 | Ifi35 | 2.75 | 9.36E-71 | 1.95E-66 | 0.21 | 0.03 | 0.18 | 7.54 |
| 18 | Ctsz | 1.33 | 1.36E-70 | 2.82E-66 | 0.94 | 0.57 | 0.37 | 1.64 |
| 18 | Sypl | 1.37 | 1.88E-70 | 3.90E-66 | 0.91 | 0.48 | 0.43 | 1.90 |
| 18 | Usp43 | 2.78 | 2.71E-70 | 5.64E-66 | 0.15 | 0.02 | 0.14 | 10.20 |
| 18 | Arhgdia | 0.85 | 1.81E-69 | 3.76E-65 | 0.99 | 0.88 | 0.11 | 1.12 |
| 18 | Peli2 | 1.44 | 1.85E-69 | 3.84E-65 | 0.62 | 0.18 | 0.44 | 3.44 |
| 18 | Stx2 | 1.58 | 3.81E-69 | 7.93E-65 | 0.66 | 0.22 | 0.44 | 3.04 |
| 18 | Fam102a | 1.94 | 1.82E-68 | 3.79E-64 | 0.52 | 0.15 | 0.37 | 3.54 |
| 18 | Pros1 | 1.72 | 3.61E-68 | 7.51E-64 | 0.51 | 0.14 | 0.37 | 3.67 |
| 18 | Hspa12b | 1.72 | 4.30E-68 | 8.95E-64 | 0.49 | 0.13 | 0.36 | 3.80 |
| 18 | Nav3 | 1.15 | 4.99E-68 | 1.04E-63 | 0.49 | 0.12 | 0.37 | 3.95 |
| 18 | Eva1b | 1.29 | 6.53E-68 | 1.36E-63 | 0.93 | 0.54 | 0.39 | 1.73 |
| 18 | Actr2 | 1.16 | 8.59E-68 | 1.79E-63 | 0.95 | 0.63 | 0.33 | 1.52 |
| 18 | Krtcap2 | 0.77 | 1.43E-67 | 2.97E-63 | 0.99 | 0.94 | 0.05 | 1.05 |
| 18 | Nectin2 | 1.34 | 1.83E-67 | 3.80E-63 | 0.92 | 0.47 | 0.44 | 1.94 |
| 18 | Wsb2 | 1.52 | 1.88E-67 | 3.92E-63 | 0.78 | 0.31 | 0.47 | 2.54 |
| 18 | Btg2 | 1.31 | 5.32E-67 | 1.11E-62 | 0.73 | 0.27 | 0.46 | 2.73 |
| 18 | Prr5 | 1.70 | 5.66E-67 | 1.18E-62 | 0.46 | 0.12 | 0.35 | 4.03 |
| 18 | Flnb | 1.57 | 5.74E-67 | 1.19E-62 | 0.75 | 0.29 | 0.46 | 2.57 |
| 18 | Ccnyl1 | 1.60 | 6.21E-67 | 1.29E-62 | 0.59 | 0.18 | 0.41 | 3.29 |
| 18 | Stim2 | 1.57 | 7.67E-67 | 1.60E-62 | 0.68 | 0.23 | 0.45 | 2.96 |
| 18 | Fhl2 | 2.07 | 7.91E-67 | 1.64E-62 | 0.38 | 0.08 | 0.29 | 4.50 |
| 18 | Iqgap1 | 1.34 | 1.10E-66 | 2.28E-62 | 0.89 | 0.47 | 0.42 | 1.90 |
| 18 | Spata6 | 1.57 | 1.54E-66 | 3.20E-62 | 0.62 | 0.20 | 0.42 | 3.16 |
| 18 | Arid3a | 1.47 | 2.58E-66 | 5.37E-62 | 0.78 | 0.33 | 0.45 | 2.37 |
| 18 | Twf2 | 1.28 | 3.64E-66 | 7.56E-62 | 0.88 | 0.42 | 0.46 | 2.10 |
| 18 | Ptpn1 | 1.30 | 4.68E-66 | 9.74E-62 | 0.92 | 0.52 | 0.40 | 1.77 |
| 18 | Rac1 | 0.80 | 8.39E-66 | 1.74E-61 | 0.99 | 0.90 | 0.09 | 1.10 |
| 18 | Nmi | 1.80 | 1.31E-65 | 2.73E-61 | 0.39 | 0.09 | 0.30 | 4.41 |
| 18 | Prnp | 1.56 | 1.32E-65 | 2.74E-61 | 0.57 | 0.17 | 0.40 | 3.34 |
| 18 | Nol4l | 1.60 | 1.67E-65 | 3.48E-61 | 0.56 | 0.16 | 0.40 | 3.44 |
| 18 | Bag3 | 1.66 | 2.27E-65 | 4.72E-61 | 0.44 | 0.11 | 0.33 | 4.10 |
| 18 | Plekhg5 | 1.78 | 2.55E-65 | 5.31E-61 | 0.41 | 0.10 | 0.31 | 4.29 |
| 18 | Cdc42bpb | 1.39 | 5.13E-65 | 1.07E-60 | 0.81 | 0.35 | 0.46 | 2.32 |
| 18 | Pim3 | 1.50 | 5.86E-65 | 1.22E-60 | 0.74 | 0.29 | 0.44 | 2.52 |
| 18 | Asah2 | 2.08 | 1.14E-64 | 2.38E-60 | 0.26 | 0.04 | 0.22 | 5.95 |
| 18 | Lrrc58 | 1.14 | 2.30E-64 | 4.77E-60 | 0.95 | 0.63 | 0.33 | 1.52 |
| 18 | Tspan13 | 1.41 | 3.17E-64 | 6.58E-60 | 0.86 | 0.41 | 0.45 | 2.11 |
| 18 | Ifngr1 | 1.50 | 4.60E-64 | 9.57E-60 | 0.68 | 0.24 | 0.44 | 2.81 |
| 18 | Gm43260 | 2.29 | 5.92E-64 | 1.23E-59 | 0.22 | 0.03 | 0.19 | 6.70 |

**Supplementary Table 1. Marker genes for the original clusters of integrated clustering of E11.5 control and NotchGOF λ-junction cells.**

| | | | | | | | | |
|---|---|---|---|---|---|---|---|---|
| 18 | Trib1 | 1.60 | 6.01E-64 | 1.25E-59 | 0.58 | 0.18 | 0.40 | 3.27 |
| 18 | Nfic | 1.28 | 1.02E-63 | 2.12E-59 | 0.78 | 0.29 | 0.48 | 2.64 |
| 18 | Tmed5 | 1.33 | 1.09E-63 | 2.27E-59 | 0.90 | 0.48 | 0.42 | 1.87 |
| 18 | Cap1 | 1.23 | 2.24E-63 | 4.67E-59 | 0.90 | 0.54 | 0.36 | 1.67 |
| 18 | Spns1 | 1.49 | 2.27E-63 | 4.72E-59 | 0.84 | 0.41 | 0.43 | 2.07 |
| 18 | Arsa | 1.98 | 2.53E-63 | 5.25E-59 | 0.32 | 0.06 | 0.26 | 5.00 |
| 18 | Slc43a1 | 1.54 | 2.66E-63 | 5.54E-59 | 0.54 | 0.16 | 0.38 | 3.46 |
| 18 | Cyp20a1 | 1.44 | 4.47E-63 | 9.29E-59 | 0.71 | 0.25 | 0.46 | 2.84 |
| 18 | Dcbld1 | 1.31 | 4.94E-63 | 1.03E-58 | 0.77 | 0.30 | 0.48 | 2.62 |
| 18 | Hivep3 | 2.45 | 1.28E-62 | 2.67E-58 | 0.16 | 0.02 | 0.14 | 8.58 |
| 18 | Btbd3 | 1.71 | 2.51E-62 | 5.22E-58 | 0.52 | 0.15 | 0.37 | 3.40 |
| 18 | Glud1 | 1.19 | 2.54E-62 | 5.28E-58 | 0.94 | 0.65 | 0.29 | 1.45 |
| 18 | Sri | 1.01 | 6.17E-62 | 1.28E-57 | 0.96 | 0.77 | 0.19 | 1.24 |
| 18 | Triobp | 1.48 | 6.60E-62 | 1.37E-57 | 0.76 | 0.31 | 0.45 | 2.45 |
| 18 | Aim2 | 2.17 | 7.35E-62 | 1.53E-57 | 0.30 | 0.06 | 0.24 | 5.05 |
| 18 | Prrg3 | 1.82 | 1.00E-61 | 2.09E-57 | 0.43 | 0.11 | 0.32 | 3.94 |
| 18 | Lrp11 | 2.23 | 1.43E-61 | 2.98E-57 | 0.22 | 0.03 | 0.19 | 6.61 |
| 18 | Chst2 | 1.45 | 2.85E-61 | 5.92E-57 | 0.61 | 0.20 | 0.41 | 3.03 |
| 18 | Rpl28 | 0.29 | 4.58E-61 | 9.53E-57 | 1.00 | 1.00 | 0.00 | 1.00 |
| 18 | D1Ertd622e | 1.72 | 8.27E-61 | 1.72E-56 | 0.47 | 0.13 | 0.34 | 3.67 |
| 18 | Ppia | 0.35 | 8.41E-61 | 1.75E-56 | 1.00 | 1.00 | 0.00 | 1.00 |
| 18 | Vwa1 | 2.17 | 1.31E-60 | 2.72E-56 | 0.29 | 0.06 | 0.23 | 5.16 |
| 18 | Tmem120b | 1.93 | 1.40E-60 | 2.90E-56 | 0.30 | 0.06 | 0.24 | 5.10 |
| 18 | Jak1 | 1.23 | 1.53E-60 | 3.17E-56 | 0.93 | 0.60 | 0.33 | 1.54 |
| 18 | Itga9 | 1.50 | 1.69E-60 | 3.52E-56 | 0.63 | 0.21 | 0.42 | 3.03 |
| 18 | Stxbp1 | 1.72 | 2.15E-60 | 4.47E-56 | 0.39 | 0.09 | 0.30 | 4.26 |
| 18 | Rgs2 | 1.47 | 3.19E-60 | 6.64E-56 | 0.77 | 0.34 | 0.43 | 2.27 |
| 18 | Cdc42ep3 | 1.47 | 4.58E-60 | 9.52E-56 | 0.53 | 0.16 | 0.37 | 3.30 |
| 18 | Fam111a | 1.46 | 4.73E-60 | 9.84E-56 | 0.76 | 0.32 | 0.44 | 2.36 |
| 18 | Rap2a | 1.45 | 6.08E-60 | 1.26E-55 | 0.76 | 0.32 | 0.44 | 2.40 |
| 18 | Phactr2 | 1.23 | 6.41E-60 | 1.33E-55 | 0.88 | 0.45 | 0.44 | 1.97 |
| 18 | Dusp6 | 1.14 | 6.86E-60 | 1.43E-55 | 0.85 | 0.36 | 0.49 | 2.39 |
| 18 | Crem | 1.58 | 1.98E-59 | 4.13E-55 | 0.56 | 0.18 | 0.38 | 3.19 |
| 18 | Plp2 | 1.30 | 2.32E-59 | 4.83E-55 | 0.83 | 0.38 | 0.46 | 2.21 |
| 18 | Sec11c | 1.31 | 2.60E-59 | 5.42E-55 | 0.74 | 0.30 | 0.44 | 2.48 |
| 18 | Cyba | 1.28 | 4.05E-59 | 8.42E-55 | 0.86 | 0.45 | 0.41 | 1.91 |
| 18 | Lnx2 | 1.87 | 5.04E-59 | 1.05E-54 | 0.29 | 0.06 | 0.23 | 5.07 |
| 18 | Pls3 | 1.44 | 7.26E-59 | 1.51E-54 | 0.73 | 0.29 | 0.44 | 2.48 |
| 18 | Rtl8a | 1.31 | 7.39E-59 | 1.54E-54 | 0.83 | 0.38 | 0.45 | 2.17 |
| 18 | Wipf1 | 1.33 | 7.73E-59 | 1.61E-54 | 0.66 | 0.23 | 0.43 | 2.92 |
| 18 | Tmc6 | 2.08 | 8.77E-59 | 1.82E-54 | 0.33 | 0.07 | 0.26 | 4.52 |
| 18 | Ocel1 | 1.95 | 1.11E-58 | 2.31E-54 | 0.27 | 0.05 | 0.22 | 5.23 |
| 18 | Irgm1 | 2.82 | 1.15E-58 | 2.39E-54 | 0.12 | 0.01 | 0.11 | 10.17 |
| 18 | Cdk17 | 1.52 | 1.98E-58 | 4.11E-54 | 0.60 | 0.20 | 0.40 | 2.95 |
| 18 | Rab12 | 1.26 | 3.26E-58 | 6.77E-54 | 0.88 | 0.48 | 0.40 | 1.84 |
| 18 | Rock2 | 1.11 | 5.78E-58 | 1.20E-53 | 0.95 | 0.66 | 0.29 | 1.43 |
| 18 | Sox13 | 1.59 | 9.29E-58 | 1.93E-53 | 0.36 | 0.09 | 0.28 | 4.28 |
| 18 | Rpsa | 0.29 | 1.32E-57 | 2.75E-53 | 1.00 | 1.00 | 0.00 | 1.00 |
| 18 | Npr3 | 2.49 | 1.33E-57 | 2.76E-53 | 0.14 | 0.02 | 0.13 | 8.94 |
| 18 | Uvrag | 1.54 | 1.89E-57 | 3.93E-53 | 0.62 | 0.22 | 0.40 | 2.79 |
| 18 | Prag1 | 1.48 | 2.64E-57 | 5.49E-53 | 0.48 | 0.14 | 0.34 | 3.53 |
| 18 | Nt5c | 1.16 | 2.69E-57 | 5.59E-53 | 0.90 | 0.53 | 0.37 | 1.69 |
| 18 | Strn | 1.59 | 2.90E-57 | 6.04E-53 | 0.55 | 0.17 | 0.38 | 3.18 |
| 18 | Gap43 | 1.38 | 3.24E-57 | 6.73E-53 | 0.83 | 0.41 | 0.41 | 2.00 |
| 18 | Chst15 | 1.30 | 4.20E-57 | 8.74E-53 | 0.55 | 0.17 | 0.38 | 3.30 |
| 18 | Arhgef7 | 1.41 | 4.71E-57 | 9.80E-53 | 0.74 | 0.31 | 0.43 | 2.37 |

**Supplementary Table 1. Marker genes for the original clusters of integrated clustering of E11.5 control and NotchGOF λ-junction cells.**

| | | | | | | | | |
|---|---|---|---|---|---|---|---|---|
| 18 | Man1a | 1.67 | 4.85E-57 | 1.01E-52 | 0.29 | 0.06 | 0.23 | 4.97 |
| 18 | Fez2 | 1.30 | 5.08E-57 | 1.06E-52 | 0.75 | 0.31 | 0.45 | 2.46 |
| 18 | Rps6ka1 | 1.45 | 5.14E-57 | 1.07E-52 | 0.66 | 0.24 | 0.43 | 2.79 |
| 18 | Nqo2 | 1.61 | 5.82E-57 | 1.21E-52 | 0.46 | 0.13 | 0.33 | 3.62 |
| 18 | Mydgf | 0.94 | 1.65E-56 | 3.43E-52 | 0.97 | 0.73 | 0.24 | 1.33 |
| 18 | St3gal5 | 1.35 | 2.40E-56 | 5.00E-52 | 0.65 | 0.24 | 0.42 | 2.77 |
| 18 | Rgcc | 1.79 | 3.80E-56 | 7.91E-52 | 0.37 | 0.09 | 0.28 | 4.12 |
| 18 | Marchf2 | 1.17 | 4.41E-56 | 9.18E-52 | 0.62 | 0.21 | 0.41 | 2.96 |
| 18 | Casd1 | 1.18 | 4.45E-56 | 9.26E-52 | 0.92 | 0.50 | 0.42 | 1.85 |
| 18 | Git2 | 1.36 | 4.90E-56 | 1.02E-51 | 0.78 | 0.33 | 0.44 | 2.32 |
| 18 | Med10 | 0.96 | 5.50E-56 | 1.14E-51 | 0.97 | 0.72 | 0.25 | 1.35 |
| 18 | Gpsm3 | 1.60 | 6.55E-56 | 1.36E-51 | 0.36 | 0.08 | 0.27 | 4.25 |
| 18 | Irf2 | 1.34 | 7.97E-56 | 1.66E-51 | 0.67 | 0.25 | 0.42 | 2.68 |
| 18 | Map3k20 | 1.56 | 8.64E-56 | 1.80E-51 | 0.43 | 0.11 | 0.31 | 3.73 |
| 18 | Tmbim1 | 1.81 | 1.04E-55 | 2.16E-51 | 0.38 | 0.09 | 0.28 | 4.02 |
| 18 | Cnrip1 | 1.29 | 1.28E-55 | 2.67E-51 | 0.49 | 0.14 | 0.35 | 3.57 |
| 18 | Stk32c | 2.82 | 1.34E-55 | 2.78E-51 | 0.12 | 0.01 | 0.11 | 9.38 |
| 18 | Odc1 | 1.49 | 3.04E-55 | 6.32E-51 | 0.94 | 0.81 | 0.13 | 1.15 |
| 18 | Cdc42se1 | 1.23 | 9.31E-55 | 1.94E-50 | 0.84 | 0.42 | 0.42 | 2.00 |
| 18 | Cpd | 1.31 | 1.32E-54 | 2.74E-50 | 0.77 | 0.33 | 0.44 | 2.32 |
| 18 | A730020E08Rik | 1.94 | 1.76E-54 | 3.66E-50 | 0.22 | 0.04 | 0.19 | 5.74 |
| 18 | Plod1 | 1.46 | 2.43E-54 | 5.05E-50 | 0.58 | 0.20 | 0.39 | 2.97 |
| 18 | Syngr2 | 1.43 | 2.47E-54 | 5.15E-50 | 0.48 | 0.14 | 0.34 | 3.40 |
| 18 | Kit | 1.83 | 2.49E-54 | 5.19E-50 | 0.37 | 0.09 | 0.28 | 3.95 |
| 18 | Sytl2 | 2.56 | 2.63E-54 | 5.46E-50 | 0.10 | 0.01 | 0.09 | 11.33 |
| 18 | Pelo | 1.38 | 3.95E-54 | 8.22E-50 | 0.79 | 0.39 | 0.40 | 2.03 |
| 18 | Cast | 1.31 | 5.97E-54 | 1.24E-49 | 0.65 | 0.24 | 0.41 | 2.72 |
| 18 | Tspan14 | 1.27 | 8.36E-54 | 1.74E-49 | 0.75 | 0.32 | 0.43 | 2.35 |
| 18 | Rdx | 0.91 | 1.05E-53 | 2.17E-49 | 1.00 | 0.96 | 0.04 | 1.04 |
| 18 | Gamt | 1.57 | 1.64E-53 | 3.42E-49 | 0.36 | 0.09 | 0.28 | 4.09 |
| 18 | Ptbp3 | 1.10 | 1.68E-53 | 3.50E-49 | 0.93 | 0.62 | 0.31 | 1.51 |
| 18 | Swap70 | 1.37 | 1.98E-53 | 4.12E-49 | 0.75 | 0.32 | 0.42 | 2.31 |
| 18 | Tcf4 | 0.81 | 2.32E-53 | 4.83E-49 | 0.99 | 0.90 | 0.09 | 1.10 |
| 18 | Mtus1 | 1.48 | 2.37E-53 | 4.94E-49 | 0.40 | 0.10 | 0.29 | 3.87 |
| 18 | Usp12 | 1.36 | 3.14E-53 | 6.53E-49 | 0.59 | 0.20 | 0.39 | 2.93 |
| 18 | Esyt1 | 1.73 | 4.19E-53 | 8.72E-49 | 0.32 | 0.07 | 0.25 | 4.45 |
| 18 | Pakap.1 | 1.24 | 4.23E-53 | 8.81E-49 | 0.84 | 0.41 | 0.43 | 2.05 |
| 18 | Rbms1 | 0.87 | 6.18E-53 | 1.29E-48 | 0.97 | 0.78 | 0.19 | 1.24 |
| 18 | Adamts9 | 1.17 | 7.23E-53 | 1.50E-48 | 0.49 | 0.15 | 0.34 | 3.36 |
| 18 | Rab8b | 1.35 | 8.99E-53 | 1.87E-48 | 0.64 | 0.24 | 0.41 | 2.70 |
| 18 | Raph1 | 1.33 | 1.20E-52 | 2.50E-48 | 0.82 | 0.42 | 0.40 | 1.95 |
| 18 | Ybx3 | 0.85 | 2.11E-52 | 4.38E-48 | 0.98 | 0.84 | 0.14 | 1.17 |
| 18 | Lgals8 | 1.76 | 2.80E-52 | 5.82E-48 | 0.35 | 0.08 | 0.26 | 4.13 |
| 18 | F2r | 1.17 | 3.20E-52 | 6.66E-48 | 0.93 | 0.57 | 0.36 | 1.64 |
| 18 | Chmp2a | 1.00 | 3.65E-52 | 7.58E-48 | 0.95 | 0.66 | 0.29 | 1.44 |
| 18 | Stat3 | 1.50 | 4.10E-52 | 8.52E-48 | 0.45 | 0.13 | 0.32 | 3.48 |
| 18 | Sdcbp | 0.97 | 4.12E-52 | 8.58E-48 | 0.93 | 0.69 | 0.24 | 1.35 |
| 18 | Il10rb | 1.78 | 4.72E-52 | 9.81E-48 | 0.34 | 0.08 | 0.26 | 4.10 |
| 18 | Entpd1 | 1.96 | 4.86E-52 | 1.01E-47 | 0.28 | 0.06 | 0.22 | 4.73 |
| 18 | Vps29 | 0.85 | 7.47E-52 | 1.55E-47 | 0.97 | 0.78 | 0.19 | 1.25 |
| 18 | Ndel1 | 1.27 | 7.96E-52 | 1.66E-47 | 0.74 | 0.31 | 0.43 | 2.39 |
| 18 | Slc38a10 | 1.35 | 1.17E-51 | 2.43E-47 | 0.76 | 0.33 | 0.43 | 2.30 |
| 18 | Mgat2 | 1.17 | 1.18E-51 | 2.46E-47 | 0.85 | 0.46 | 0.40 | 1.86 |
| 18 | Wwtr1 | 1.19 | 1.67E-51 | 3.46E-47 | 0.82 | 0.43 | 0.39 | 1.92 |
| 18 | Emilin1 | 1.12 | 1.89E-51 | 3.94E-47 | 0.92 | 0.53 | 0.39 | 1.75 |
| 18 | Ap2s1 | 0.71 | 1.99E-51 | 4.13E-47 | 0.99 | 0.88 | 0.10 | 1.12 |

**Supplementary Table 1. Marker genes for the original clusters of integrated clustering of E11.5 control and NotchGOF λ-junction cells.**

| | | | | | | | | |
|---|---|---|---|---|---|---|---|---|
| 18 | Gnai1 | 1.40 | 2.11E-51 | 4.38E-47 | 0.49 | 0.15 | 0.34 | 3.29 |
| 18 | Neu1 | 1.55 | 3.18E-51 | 6.62E-47 | 0.44 | 0.13 | 0.31 | 3.45 |
| 18 | Lhx6 | 1.63 | 3.26E-51 | 6.78E-47 | 0.28 | 0.06 | 0.22 | 4.70 |
| 18 | Eif4ebp1 | 1.03 | 3.63E-51 | 7.55E-47 | 0.93 | 0.69 | 0.24 | 1.34 |
| 18 | Pqlc1 | 1.54 | 3.91E-51 | 8.13E-47 | 0.45 | 0.13 | 0.32 | 3.37 |
| 18 | Nin | 1.37 | 3.96E-51 | 8.23E-47 | 0.67 | 0.27 | 0.40 | 2.52 |
| 18 | Acy3 | 2.47 | 5.90E-51 | 1.23E-46 | 0.12 | 0.01 | 0.11 | 9.15 |
| 18 | Xbp1 | 1.18 | 9.02E-51 | 1.88E-46 | 0.86 | 0.47 | 0.39 | 1.83 |
| 18 | Sorbs3 | 1.34 | 9.14E-51 | 1.90E-46 | 0.46 | 0.13 | 0.33 | 3.45 |
| 18 | Ppargc1b | 2.24 | 1.13E-50 | 2.36E-46 | 0.18 | 0.03 | 0.15 | 6.56 |
| 18 | Itpripl2 | 1.36 | 1.29E-50 | 2.69E-46 | 0.45 | 0.13 | 0.32 | 3.51 |
| 18 | Slc23a2 | 1.88 | 1.33E-50 | 2.76E-46 | 0.32 | 0.07 | 0.24 | 4.27 |
| 18 | Atp8b2 | 1.54 | 1.62E-50 | 3.38E-46 | 0.34 | 0.08 | 0.26 | 4.14 |
| 18 | B4galt5 | 1.72 | 1.65E-50 | 3.43E-46 | 0.38 | 0.10 | 0.28 | 3.73 |
| 18 | Klhl3 | 2.19 | 1.76E-50 | 3.67E-46 | 0.17 | 0.02 | 0.14 | 6.96 |
| 18 | Cdr2l | 1.38 | 3.25E-50 | 6.75E-46 | 0.56 | 0.19 | 0.37 | 2.95 |
| 18 | Rhog | 1.32 | 3.45E-50 | 7.17E-46 | 0.76 | 0.35 | 0.41 | 2.18 |
| 18 | Clta | 0.49 | 3.63E-50 | 7.55E-46 | 1.00 | 0.98 | 0.02 | 1.02 |
| 18 | Cr1l | 1.24 | 3.73E-50 | 7.75E-46 | 0.74 | 0.32 | 0.42 | 2.33 |
| 18 | Tlnrd1 | 1.15 | 4.17E-50 | 8.68E-46 | 0.90 | 0.50 | 0.40 | 1.81 |
| 18 | Ctnnb1 | 0.55 | 5.42E-50 | 1.13E-45 | 1.00 | 0.97 | 0.03 | 1.03 |
| 18 | Commd7 | 0.99 | 9.04E-50 | 1.88E-45 | 0.91 | 0.57 | 0.34 | 1.60 |
| 18 | Ptk2 | 1.22 | 9.24E-50 | 1.92E-45 | 0.88 | 0.53 | 0.35 | 1.67 |
| 18 | Manba | 2.68 | 1.92E-49 | 4.00E-45 | 0.13 | 0.02 | 0.11 | 8.06 |
| 18 | Ost4 | 0.61 | 2.32E-49 | 4.83E-45 | 1.00 | 0.95 | 0.04 | 1.05 |
| 18 | Ptpn12 | 1.29 | 3.64E-49 | 7.56E-45 | 0.75 | 0.35 | 0.40 | 2.14 |
| 18 | Prelid1 | 0.68 | 3.68E-49 | 7.66E-45 | 0.99 | 0.93 | 0.06 | 1.07 |
| 18 | Rps3 | 0.30 | 4.28E-49 | 8.90E-45 | 1.00 | 1.00 | 0.00 | 1.00 |
| 18 | Zfp36l2 | 1.15 | 4.30E-49 | 8.93E-45 | 0.84 | 0.43 | 0.42 | 1.97 |
| 18 | Acsf2 | 1.50 | 4.51E-49 | 9.39E-45 | 0.46 | 0.14 | 0.32 | 3.33 |
| 18 | Klf13 | 1.17 | 5.52E-49 | 1.15E-44 | 0.79 | 0.37 | 0.42 | 2.14 |
| 18 | Kctd10 | 1.21 | 5.54E-49 | 1.15E-44 | 0.71 | 0.29 | 0.42 | 2.43 |
| 18 | Creg1 | 0.94 | 6.68E-49 | 1.39E-44 | 0.79 | 0.35 | 0.44 | 2.27 |
| 18 | Sft2d1 | 1.33 | 7.53E-49 | 1.57E-44 | 0.60 | 0.22 | 0.38 | 2.75 |
| 18 | Dnajc3 | 1.13 | 7.76E-49 | 1.61E-44 | 0.90 | 0.52 | 0.38 | 1.73 |
| 18 | Hmgn5 | 1.10 | 9.36E-49 | 1.95E-44 | 0.95 | 0.71 | 0.24 | 1.34 |
| 18 | Iffo2 | 1.31 | 9.57E-49 | 1.99E-44 | 0.45 | 0.13 | 0.32 | 3.43 |
| 18 | Cdyl2 | 1.67 | 1.04E-48 | 2.17E-44 | 0.25 | 0.05 | 0.20 | 4.96 |
| 18 | Fkbp10 | 1.17 | 1.17E-48 | 2.43E-44 | 0.76 | 0.33 | 0.43 | 2.29 |
| 18 | Ldlrap1 | 1.29 | 1.62E-48 | 3.37E-44 | 0.54 | 0.18 | 0.36 | 2.98 |
| 18 | Shc1 | 1.28 | 1.70E-48 | 3.53E-44 | 0.63 | 0.23 | 0.39 | 2.68 |
| 18 | Tmbim6 | 0.78 | 1.89E-48 | 3.94E-44 | 0.99 | 0.84 | 0.15 | 1.18 |
| 18 | Mgat1 | 1.47 | 2.41E-48 | 5.02E-44 | 0.57 | 0.20 | 0.36 | 2.77 |
| 18 | Gm42702 | 2.12 | 2.96E-48 | 6.15E-44 | 0.20 | 0.04 | 0.17 | 5.67 |
| 18 | Mindy2 | 1.47 | 3.47E-48 | 7.23E-44 | 0.51 | 0.17 | 0.34 | 2.97 |
| 18 | Arrb1 | 1.57 | 3.64E-48 | 7.57E-44 | 0.40 | 0.11 | 0.29 | 3.55 |
| 18 | Il3ra | 1.83 | 4.44E-48 | 9.23E-44 | 0.25 | 0.05 | 0.20 | 5.00 |
| 18 | Trp53i11 | 1.21 | 5.17E-48 | 1.08E-43 | 0.60 | 0.22 | 0.38 | 2.70 |
| 18 | Tnfaip1 | 1.01 | 6.31E-48 | 1.31E-43 | 0.92 | 0.62 | 0.30 | 1.49 |
| 18 | Nme2 | 0.41 | 6.98E-48 | 1.45E-43 | 1.00 | 0.98 | 0.01 | 1.01 |
| 18 | Vangl1 | 1.57 | 7.38E-48 | 1.53E-43 | 0.47 | 0.15 | 0.32 | 3.09 |
| 18 | Endod1 | 1.72 | 7.45E-48 | 1.55E-43 | 0.34 | 0.09 | 0.25 | 3.86 |
| 18 | Mogat2 | 1.46 | 1.48E-47 | 3.07E-43 | 0.41 | 0.12 | 0.29 | 3.52 |
| 18 | Acyp2 | 1.55 | 2.55E-47 | 5.30E-43 | 0.38 | 0.10 | 0.28 | 3.67 |
| 18 | Ubald2 | 0.95 | 2.89E-47 | 6.02E-43 | 0.95 | 0.69 | 0.26 | 1.38 |
| 18 | Slc12a2 | 0.95 | 2.95E-47 | 6.14E-43 | 0.85 | 0.43 | 0.42 | 1.99 |

**Supplementary Table 1. Marker genes for the original clusters of integrated clustering of E11.5 control and NotchGOF λ-junction cells.**

| | | | | | | | | |
|---|---|---|---|---|---|---|---|---|
| 18 | Gm43573 | 2.25 | 3.12E-47 | 6.48E-43 | 0.18 | 0.03 | 0.15 | 6.10 |
| 18 | Hmox1 | 0.95 | 3.34E-47 | 6.96E-43 | 0.55 | 0.19 | 0.37 | 2.99 |
| 18 | Prkar1a | 0.69 | 3.71E-47 | 7.72E-43 | 1.00 | 0.88 | 0.12 | 1.13 |
| 18 | Lrrk2 | 1.88 | 3.88E-47 | 8.08E-43 | 0.18 | 0.03 | 0.15 | 6.13 |
| 18 | Fam171a2 | 1.15 | 4.42E-47 | 9.20E-43 | 0.84 | 0.45 | 0.39 | 1.88 |
| 18 | Rell1 | 1.33 | 4.48E-47 | 9.31E-43 | 0.77 | 0.35 | 0.42 | 2.22 |
| 18 | Rabac1 | 1.04 | 4.60E-47 | 9.57E-43 | 0.91 | 0.55 | 0.36 | 1.65 |
| 18 | Homer3 | 1.06 | 7.11E-47 | 1.48E-42 | 0.85 | 0.48 | 0.38 | 1.79 |
| 18 | Cd302 | 1.15 | 1.21E-46 | 2.51E-42 | 0.70 | 0.28 | 0.42 | 2.48 |
| 18 | Necap2 | 1.15 | 1.41E-46 | 2.93E-42 | 0.80 | 0.39 | 0.41 | 2.05 |
| 18 | Apobec3 | 1.45 | 1.79E-46 | 3.73E-42 | 0.33 | 0.08 | 0.25 | 4.04 |
| 18 | Prdx1 | 0.44 | 1.93E-46 | 4.01E-42 | 1.00 | 0.99 | 0.01 | 1.01 |
| 18 | Smap2 | 1.20 | 2.12E-46 | 4.41E-42 | 0.77 | 0.37 | 0.40 | 2.09 |
| 18 | Relb | 2.07 | 2.60E-46 | 5.40E-42 | 0.19 | 0.03 | 0.16 | 5.76 |
| 18 | Itpr3 | 1.98 | 3.05E-46 | 6.34E-42 | 0.24 | 0.05 | 0.19 | 4.86 |
| 18 | Fzd10 | 1.52 | 3.61E-46 | 7.52E-42 | 0.44 | 0.14 | 0.30 | 3.22 |
| 18 | Zfp710 | 1.30 | 3.71E-46 | 7.72E-42 | 0.52 | 0.18 | 0.34 | 2.94 |
| 18 | Iqcg | 1.82 | 4.26E-46 | 8.85E-42 | 0.20 | 0.04 | 0.16 | 5.63 |
| 18 | Npepl1 | 1.10 | 4.29E-46 | 8.92E-42 | 0.76 | 0.32 | 0.43 | 2.33 |
| 18 | Acadl | 0.96 | 6.07E-46 | 1.26E-41 | 0.91 | 0.53 | 0.38 | 1.71 |
| 18 | Rilpl1 | 1.29 | 7.06E-46 | 1.47E-41 | 0.69 | 0.29 | 0.40 | 2.41 |
| 18 | Fscn1 | 0.69 | 8.09E-46 | 1.68E-41 | 1.00 | 0.91 | 0.09 | 1.10 |
| 18 | Gm17018 | 1.07 | 8.22E-46 | 1.71E-41 | 0.87 | 0.52 | 0.35 | 1.67 |
| 18 | Arhgap23 | 1.89 | 9.57E-46 | 1.99E-41 | 0.34 | 0.09 | 0.25 | 3.74 |
| 18 | Ccnd1 | 0.81 | 1.73E-45 | 3.59E-41 | 0.97 | 0.80 | 0.18 | 1.22 |
| 18 | Hras | 0.71 | 1.92E-45 | 4.00E-41 | 0.98 | 0.80 | 0.18 | 1.22 |
| 18 | Adam10 | 1.12 | 2.29E-45 | 4.77E-41 | 0.89 | 0.53 | 0.36 | 1.67 |
| 18 | Plekho1 | 0.96 | 4.20E-45 | 8.73E-41 | 0.93 | 0.61 | 0.32 | 1.52 |
| 18 | Ktn1 | 0.76 | 5.14E-45 | 1.07E-40 | 0.98 | 0.82 | 0.16 | 1.19 |
| 18 | Qk | 0.66 | 8.43E-45 | 1.75E-40 | 0.99 | 0.90 | 0.09 | 1.10 |
| 18 | St5 | 1.28 | 8.59E-45 | 1.79E-40 | 0.60 | 0.22 | 0.38 | 2.70 |
| 18 | Ccser1 | 2.24 | 8.93E-45 | 1.86E-40 | 0.18 | 0.03 | 0.15 | 5.81 |
| 18 | Numb | 1.14 | 1.11E-44 | 2.31E-40 | 0.72 | 0.31 | 0.41 | 2.30 |
| 18 | Nfkb1 | 1.25 | 1.66E-44 | 3.45E-40 | 0.68 | 0.30 | 0.38 | 2.30 |
| 18 | Egln3 | 1.64 | 2.02E-44 | 4.21E-40 | 0.30 | 0.07 | 0.23 | 4.04 |
| 18 | Vamp8 | 0.97 | 2.19E-44 | 4.56E-40 | 0.93 | 0.61 | 0.32 | 1.53 |
| 18 | Prex2 | 1.19 | 2.71E-44 | 5.64E-40 | 0.81 | 0.40 | 0.41 | 2.03 |
| 18 | Slk | 1.08 | 4.31E-44 | 8.95E-40 | 0.88 | 0.53 | 0.35 | 1.66 |
| 18 | Dennd5b | 1.13 | 4.39E-44 | 9.12E-40 | 0.79 | 0.38 | 0.41 | 2.07 |
| 18 | Myo10 | 1.15 | 4.55E-44 | 9.46E-40 | 0.83 | 0.47 | 0.36 | 1.78 |
| 18 | Abhd5 | 1.28 | 4.85E-44 | 1.01E-39 | 0.54 | 0.20 | 0.35 | 2.78 |
| 18 | Dhrs7 | 1.34 | 5.68E-44 | 1.18E-39 | 0.42 | 0.12 | 0.29 | 3.35 |
| 18 | Gm26632 | 2.09 | 8.01E-44 | 1.67E-39 | 0.17 | 0.03 | 0.14 | 6.19 |
| 18 | Mgmt | 1.21 | 9.50E-44 | 1.98E-39 | 0.56 | 0.20 | 0.36 | 2.75 |
| 18 | Gpr146 | 2.34 | 1.07E-43 | 2.24E-39 | 0.17 | 0.03 | 0.14 | 5.86 |
| 18 | Carhsp1 | 0.99 | 1.42E-43 | 2.96E-39 | 0.93 | 0.59 | 0.34 | 1.57 |
| 18 | Plekha3 | 1.23 | 1.61E-43 | 3.34E-39 | 0.63 | 0.25 | 0.38 | 2.53 |
| 18 | Mrpl17 | 0.83 | 1.61E-43 | 3.35E-39 | 0.95 | 0.79 | 0.15 | 1.19 |
| 18 | Gm42670 | 2.39 | 1.95E-43 | 4.06E-39 | 0.13 | 0.02 | 0.11 | 7.41 |
| 18 | Gdi2 | 0.55 | 2.29E-43 | 4.77E-39 | 0.99 | 0.95 | 0.05 | 1.05 |
| 18 | Tbc1d4 | 1.49 | 2.76E-43 | 5.74E-39 | 0.38 | 0.11 | 0.27 | 3.43 |
| 18 | Renbp | 1.40 | 3.30E-43 | 6.86E-39 | 0.30 | 0.08 | 0.23 | 4.04 |
| 18 | Hexa | 0.98 | 6.69E-43 | 1.39E-38 | 0.72 | 0.31 | 0.41 | 2.31 |
| 18 | Jag2 | 1.05 | 7.38E-43 | 1.53E-38 | 0.32 | 0.08 | 0.24 | 3.95 |
| 18 | Mast4 | 1.24 | 1.04E-42 | 2.16E-38 | 0.82 | 0.44 | 0.38 | 1.85 |
| 18 | Anxa4 | 1.36 | 1.55E-42 | 3.23E-38 | 0.37 | 0.11 | 0.27 | 3.50 |

**Supplementary Table 1. Marker genes for the original clusters of integrated clustering of E11.5 control and NotchGOF λ-junction cells.**

| | | | | | | | | |
|---|---|---|---|---|---|---|---|---|
| 18 | Ephb4 | 1.23 | 1.73E-42 | 3.60E-38 | 0.79 | 0.39 | 0.40 | 2.01 |
| 18 | Ier2 | 0.80 | 1.93E-42 | 4.00E-38 | 0.96 | 0.79 | 0.18 | 1.23 |
| 18 | Igfbp4 | 0.64 | 2.89E-42 | 6.02E-38 | 1.00 | 0.90 | 0.10 | 1.11 |
| 18 | Rhbdf1 | 1.39 | 3.63E-42 | 7.54E-38 | 0.35 | 0.10 | 0.26 | 3.61 |
| 18 | Gatm | 1.50 | 4.08E-42 | 8.48E-38 | 0.20 | 0.04 | 0.16 | 5.15 |
| 18 | Acot7 | 1.07 | 1.08E-41 | 2.25E-37 | 0.61 | 0.23 | 0.38 | 2.64 |
| 18 | Tstd3 | 1.21 | 1.35E-41 | 2.81E-37 | 0.55 | 0.21 | 0.35 | 2.69 |
| 18 | Bahcc1 | 1.32 | 1.70E-41 | 3.54E-37 | 0.63 | 0.27 | 0.36 | 2.34 |
| 18 | Siah2 | 1.52 | 1.75E-41 | 3.64E-37 | 0.29 | 0.07 | 0.22 | 4.01 |
| 18 | Nek6 | 1.06 | 2.07E-41 | 4.31E-37 | 0.70 | 0.31 | 0.39 | 2.28 |
| 18 | Zfp948 | 1.42 | 2.14E-41 | 4.45E-37 | 0.33 | 0.09 | 0.24 | 3.66 |
| 18 | Dad1 | 0.59 | 2.93E-41 | 6.09E-37 | 1.00 | 0.94 | 0.06 | 1.06 |
| 18 | Stard13 | 2.06 | 3.07E-41 | 6.38E-37 | 0.15 | 0.02 | 0.13 | 6.25 |
| 18 | Tmem63a | 2.37 | 3.74E-41 | 7.77E-37 | 0.15 | 0.03 | 0.13 | 6.12 |
| 18 | Atl1 | 1.73 | 8.94E-41 | 1.86E-36 | 0.22 | 0.05 | 0.17 | 4.70 |
| 18 | Tprgl | 0.91 | 1.17E-40 | 2.44E-36 | 0.89 | 0.60 | 0.29 | 1.48 |
| 18 | Amotl1 | 0.81 | 1.31E-40 | 2.73E-36 | 0.93 | 0.69 | 0.25 | 1.36 |
| 18 | Gne | 1.47 | 1.52E-40 | 3.16E-36 | 0.30 | 0.08 | 0.22 | 3.88 |
| 18 | Bambi | 0.74 | 1.53E-40 | 3.19E-36 | 0.94 | 0.64 | 0.30 | 1.47 |
| 18 | Hspa8 | 0.49 | 1.57E-40 | 3.27E-36 | 1.00 | 1.00 | 0.00 | 1.00 |
| 18 | Exoc6 | 1.78 | 1.93E-40 | 4.02E-36 | 0.28 | 0.07 | 0.21 | 3.86 |
| 18 | Pde10a | 1.83 | 2.21E-40 | 4.59E-36 | 0.21 | 0.04 | 0.17 | 4.93 |
| 18 | Taok2 | 1.14 | 2.26E-40 | 4.69E-36 | 0.70 | 0.31 | 0.39 | 2.25 |
| 18 | Bola2 | 0.63 | 3.36E-40 | 6.99E-36 | 0.98 | 0.88 | 0.09 | 1.10 |
| 18 | Mthfd1l | 1.43 | 3.50E-40 | 7.29E-36 | 0.46 | 0.16 | 0.30 | 2.96 |
| 18 | Gmds | 1.02 | 4.56E-40 | 9.49E-36 | 0.75 | 0.36 | 0.39 | 2.11 |
| 18 | C2cd2 | 1.67 | 4.63E-40 | 9.63E-36 | 0.35 | 0.10 | 0.25 | 3.51 |
| 18 | Nipsnap3b | 0.99 | 4.65E-40 | 9.67E-36 | 0.82 | 0.43 | 0.39 | 1.89 |
| 18 | Cers4 | 1.28 | 4.99E-40 | 1.04E-35 | 0.46 | 0.16 | 0.30 | 2.96 |
| 18 | Mark4 | 1.23 | 5.15E-40 | 1.07E-35 | 0.55 | 0.21 | 0.34 | 2.65 |
| 18 | Pdzd2 | 1.93 | 6.32E-40 | 1.31E-35 | 0.18 | 0.03 | 0.14 | 5.36 |
| 18 | Ssu72 | 0.72 | 7.04E-40 | 1.46E-35 | 0.98 | 0.83 | 0.15 | 1.18 |
| 18 | Tmem184b | 1.35 | 7.34E-40 | 1.53E-35 | 0.51 | 0.18 | 0.33 | 2.77 |
| 18 | Capns1 | 0.71 | 1.10E-39 | 2.29E-35 | 0.96 | 0.80 | 0.16 | 1.20 |
| 18 | Chmp3 | 0.96 | 1.66E-39 | 3.44E-35 | 0.83 | 0.45 | 0.38 | 1.83 |
| 18 | Enc1 | 0.99 | 2.38E-39 | 4.95E-35 | 0.57 | 0.22 | 0.35 | 2.62 |
| 18 | Ldlrad4 | 2.00 | 2.96E-39 | 6.17E-35 | 0.17 | 0.03 | 0.14 | 5.48 |
| 18 | Ctnnd1 | 1.01 | 3.65E-39 | 7.59E-35 | 0.82 | 0.46 | 0.36 | 1.79 |
| 18 | Kdelr2 | 0.68 | 3.86E-39 | 8.03E-35 | 0.98 | 0.83 | 0.14 | 1.17 |
| 18 | Vash1 | 2.30 | 3.86E-39 | 8.03E-35 | 0.13 | 0.02 | 0.11 | 7.00 |
| 18 | Bcar1 | 1.23 | 6.60E-39 | 1.37E-34 | 0.60 | 0.24 | 0.36 | 2.49 |
| 18 | Myc | 1.29 | 7.49E-39 | 1.56E-34 | 0.46 | 0.16 | 0.30 | 2.87 |
| 18 | Fam219a | 2.00 | 8.39E-39 | 1.75E-34 | 0.17 | 0.03 | 0.14 | 5.41 |
| 18 | Tuba1a | 0.44 | 8.59E-39 | 1.79E-34 | 1.00 | 0.98 | 0.02 | 1.02 |
| 18 | Ddx41 | 0.95 | 9.44E-39 | 1.96E-34 | 0.88 | 0.56 | 0.32 | 1.58 |
| 18 | Stap2 | 1.41 | 1.19E-38 | 2.48E-34 | 0.30 | 0.08 | 0.22 | 3.78 |
| 18 | Msrb3 | 1.48 | 1.20E-38 | 2.50E-34 | 0.37 | 0.11 | 0.26 | 3.31 |
| 18 | Ccdc125 | 1.54 | 1.23E-38 | 2.55E-34 | 0.25 | 0.06 | 0.19 | 4.20 |
| 18 | Tspan2 | 1.33 | 1.36E-38 | 2.82E-34 | 0.34 | 0.10 | 0.24 | 3.51 |
| 18 | Kti12 | 1.02 | 1.44E-38 | 3.00E-34 | 0.80 | 0.44 | 0.35 | 1.80 |
| 18 | Colgalt1 | 0.99 | 2.02E-38 | 4.20E-34 | 0.85 | 0.47 | 0.38 | 1.80 |
| 18 | Acsl1 | 1.28 | 2.79E-38 | 5.81E-34 | 0.35 | 0.10 | 0.25 | 3.43 |
| 18 | Ltv1 | 1.08 | 3.32E-38 | 6.90E-34 | 0.81 | 0.46 | 0.35 | 1.75 |
| 18 | Rps6ka2 | 1.63 | 3.98E-38 | 8.27E-34 | 0.29 | 0.08 | 0.22 | 3.76 |
| 18 | Tmed3 | 0.94 | 4.24E-38 | 8.83E-34 | 0.83 | 0.44 | 0.39 | 1.88 |
| 18 | Capza2 | 0.61 | 5.20E-38 | 1.08E-33 | 0.99 | 0.86 | 0.13 | 1.15 |

**Supplementary Table 1. Marker genes for the original clusters of integrated clustering of E11.5 control and NotchGOF λ-junction cells.**

| | | | | | | | | |
|---|---|---|---|---|---|---|---|---|
| 18 | Zfp715 | 1.50 | 5.39E-38 | 1.12E-33 | 0.30 | 0.08 | 0.22 | 3.70 |
| 18 | Dusp1 | 1.23 | 5.43E-38 | 1.13E-33 | 0.64 | 0.29 | 0.35 | 2.20 |
| 18 | Gm10638 | 1.62 | 6.09E-38 | 1.27E-33 | 0.22 | 0.05 | 0.17 | 4.48 |
| 18 | Nhsl2 | 1.22 | 7.08E-38 | 1.47E-33 | 0.46 | 0.16 | 0.30 | 2.89 |
| 18 | Ahr | 1.07 | 7.61E-38 | 1.58E-33 | 0.32 | 0.09 | 0.23 | 3.67 |
| 18 | Plcg1 | 1.04 | 1.08E-37 | 2.25E-33 | 0.77 | 0.40 | 0.37 | 1.95 |
| 18 | Ctnna1 | 0.67 | 1.33E-37 | 2.76E-33 | 0.98 | 0.85 | 0.13 | 1.16 |
| 18 | Kbtbd11 | 2.00 | 1.51E-37 | 3.14E-33 | 0.16 | 0.03 | 0.13 | 5.57 |
| 18 | Fam3c | 1.01 | 1.58E-37 | 3.29E-33 | 0.75 | 0.36 | 0.39 | 2.09 |
| 18 | Plcb4 | 1.11 | 1.83E-37 | 3.81E-33 | 0.52 | 0.19 | 0.33 | 2.70 |
| 18 | Mmd | 1.04 | 2.27E-37 | 4.71E-33 | 0.72 | 0.35 | 0.38 | 2.09 |
| 18 | Hivep2 | 1.60 | 2.61E-37 | 5.43E-33 | 0.28 | 0.07 | 0.21 | 3.81 |
| 18 | Txnip | 0.95 | 3.27E-37 | 6.81E-33 | 0.74 | 0.34 | 0.40 | 2.18 |
| 18 | Npl | 1.08 | 4.54E-37 | 9.44E-33 | 0.33 | 0.10 | 0.24 | 3.47 |
| 18 | Gabarapl1 | 1.09 | 5.07E-37 | 1.05E-32 | 0.69 | 0.33 | 0.37 | 2.13 |
| 18 | Naa20 | 0.83 | 5.42E-37 | 1.13E-32 | 0.95 | 0.72 | 0.23 | 1.32 |
| 18 | Sh3tc2 | 1.96 | 5.56E-37 | 1.16E-32 | 0.15 | 0.03 | 0.12 | 5.84 |
| 18 | Xpnpep1 | 0.94 | 5.74E-37 | 1.19E-32 | 0.86 | 0.52 | 0.34 | 1.65 |
| 18 | Rtl8b | 1.04 | 6.73E-37 | 1.40E-32 | 0.70 | 0.33 | 0.37 | 2.13 |
| 18 | Gab2 | 1.70 | 7.17E-37 | 1.49E-32 | 0.20 | 0.04 | 0.16 | 4.64 |
| 18 | Gas7 | 1.27 | 9.20E-37 | 1.91E-32 | 0.22 | 0.05 | 0.17 | 4.39 |
| 18 | Lck | 1.59 | 1.31E-36 | 2.72E-32 | 0.26 | 0.07 | 0.19 | 3.98 |
| 18 | Mylk | 0.99 | 1.35E-36 | 2.80E-32 | 0.48 | 0.17 | 0.31 | 2.84 |
| 18 | Reep3 | 0.91 | 1.55E-36 | 3.22E-32 | 0.89 | 0.52 | 0.37 | 1.72 |
| 18 | Desi2 | 0.97 | 2.19E-36 | 4.54E-32 | 0.80 | 0.44 | 0.35 | 1.80 |
| 18 | Map3k3 | 1.23 | 2.50E-36 | 5.19E-32 | 0.44 | 0.15 | 0.29 | 2.89 |
| 18 | Sertad1 | 1.22 | 3.35E-36 | 6.97E-32 | 0.55 | 0.22 | 0.33 | 2.50 |
| 18 | Eef1akmt4 | 1.08 | 3.93E-36 | 8.18E-32 | 0.57 | 0.23 | 0.34 | 2.48 |
| 18 | Dnm2 | 1.01 | 4.08E-36 | 8.48E-32 | 0.77 | 0.38 | 0.38 | 2.00 |
| 18 | Rexo2 | 0.87 | 4.21E-36 | 8.75E-32 | 0.88 | 0.59 | 0.29 | 1.49 |
| 18 | Lta4h | 1.03 | 4.34E-36 | 9.03E-32 | 0.80 | 0.46 | 0.34 | 1.74 |
| 18 | Pdcd6ip | 0.94 | 4.39E-36 | 9.13E-32 | 0.82 | 0.48 | 0.33 | 1.69 |
| 18 | Dmtn | 1.57 | 7.73E-36 | 1.61E-31 | 0.25 | 0.06 | 0.19 | 3.94 |
| 18 | Zfp36 | 1.42 | 7.88E-36 | 1.64E-31 | 0.27 | 0.07 | 0.20 | 3.79 |
| 18 | Gch1 | 1.54 | 8.97E-36 | 1.87E-31 | 0.28 | 0.08 | 0.20 | 3.72 |
| 18 | Nedd9 | 1.14 | 1.40E-35 | 2.91E-31 | 0.46 | 0.17 | 0.29 | 2.77 |
| 18 | Pak2 | 0.61 | 1.44E-35 | 2.99E-31 | 0.98 | 0.84 | 0.14 | 1.17 |
| 18 | Stat1 | 1.44 | 1.50E-35 | 3.12E-31 | 0.32 | 0.10 | 0.23 | 3.36 |
| 18 | Lrrc8b | 1.36 | 2.01E-35 | 4.19E-31 | 0.35 | 0.11 | 0.25 | 3.25 |
| 18 | Nckipsd | 1.46 | 2.09E-35 | 4.35E-31 | 0.26 | 0.07 | 0.19 | 3.92 |
| 18 | Map4 | 0.95 | 2.51E-35 | 5.22E-31 | 0.78 | 0.39 | 0.39 | 1.99 |
| 18 | Pip4k2a | 1.30 | 3.20E-35 | 6.65E-31 | 0.36 | 0.11 | 0.25 | 3.17 |
| 18 | Ksr1 | 2.11 | 3.53E-35 | 7.35E-31 | 0.17 | 0.03 | 0.13 | 5.06 |
| 18 | Crlf2 | 1.50 | 3.95E-35 | 8.22E-31 | 0.26 | 0.07 | 0.19 | 3.92 |
| 18 | Rnpep | 0.98 | 4.09E-35 | 8.50E-31 | 0.79 | 0.43 | 0.36 | 1.83 |
| 18 | Fkbp7 | 0.97 | 4.29E-35 | 8.92E-31 | 0.65 | 0.28 | 0.37 | 2.32 |
| 18 | Pdia6 | 0.53 | 4.79E-35 | 9.96E-31 | 1.00 | 0.96 | 0.05 | 1.05 |
| 18 | Ccdc126 | 1.32 | 5.55E-35 | 1.15E-30 | 0.30 | 0.09 | 0.22 | 3.52 |
| 18 | Rmc1 | 1.44 | 6.45E-35 | 1.34E-30 | 0.29 | 0.08 | 0.21 | 3.62 |
| 18 | Fam57a | 1.45 | 1.08E-34 | 2.25E-30 | 0.40 | 0.14 | 0.26 | 2.91 |
| 18 | Mospd1 | 1.06 | 1.26E-34 | 2.61E-30 | 0.59 | 0.24 | 0.35 | 2.42 |
| 18 | Iqsec1 | 1.32 | 1.64E-34 | 3.41E-30 | 0.40 | 0.14 | 0.27 | 2.97 |
| 18 | Cdc42bpa | 0.98 | 2.17E-34 | 4.52E-30 | 0.74 | 0.37 | 0.37 | 1.98 |
| 18 | Rreb1 | 1.06 | 2.25E-34 | 4.68E-30 | 0.53 | 0.21 | 0.32 | 2.56 |
| 18 | Cers2 | 0.87 | 2.69E-34 | 5.60E-30 | 0.88 | 0.54 | 0.35 | 1.65 |
| 18 | Cd47 | 0.83 | 4.89E-34 | 1.02E-29 | 0.82 | 0.44 | 0.39 | 1.89 |

**Supplementary Table 1. Marker genes for the original clusters of integrated clustering of E11.5 control and NotchGOF λ-junction cells.**

| | | | | | | | | |
|---|---|---|---|---|---|---|---|---|
| 18 | Litaf | 0.90 | 4.95E-34 | 1.03E-29 | 0.84 | 0.50 | 0.34 | 1.69 |
| 18 | Gm43259 | 2.15 | 5.23E-34 | 1.09E-29 | 0.14 | 0.02 | 0.11 | 5.91 |
| 18 | C1qtnf6 | 1.22 | 5.38E-34 | 1.12E-29 | 0.44 | 0.16 | 0.28 | 2.80 |
| 18 | Naa38 | 0.69 | 5.61E-34 | 1.17E-29 | 0.95 | 0.74 | 0.21 | 1.28 |
| 18 | Prodh | 1.24 | 6.21E-34 | 1.29E-29 | 0.34 | 0.11 | 0.24 | 3.25 |
| 18 | Abca3 | 1.54 | 6.37E-34 | 1.32E-29 | 0.25 | 0.06 | 0.18 | 3.89 |
| 18 | Srgap2 | 0.96 | 6.39E-34 | 1.33E-29 | 0.73 | 0.35 | 0.37 | 2.06 |
| 18 | A330009N23Rik | 1.78 | 8.05E-34 | 1.67E-29 | 0.14 | 0.02 | 0.11 | 5.91 |
| 18 | Septin4 | 2.19 | 9.54E-34 | 1.98E-29 | 0.13 | 0.02 | 0.11 | 6.14 |
| 18 | Gypc | 1.07 | 1.04E-33 | 2.17E-29 | 0.53 | 0.22 | 0.32 | 2.48 |
| 18 | Mvb12b | 1.06 | 1.05E-33 | 2.18E-29 | 0.55 | 0.23 | 0.33 | 2.45 |
| 18 | Zyx | 0.87 | 1.06E-33 | 2.20E-29 | 0.85 | 0.51 | 0.34 | 1.67 |
| 18 | Enox2 | 1.43 | 1.20E-33 | 2.49E-29 | 0.26 | 0.07 | 0.19 | 3.81 |
| 18 | Rit1 | 1.04 | 1.23E-33 | 2.56E-29 | 0.58 | 0.25 | 0.33 | 2.33 |
| 18 | Ssbp4 | 0.76 | 1.23E-33 | 2.57E-29 | 0.93 | 0.65 | 0.28 | 1.44 |
| 18 | Dpagt1 | 1.09 | 1.24E-33 | 2.57E-29 | 0.64 | 0.28 | 0.35 | 2.25 |
| 18 | Ccnjl | 0.93 | 1.38E-33 | 2.88E-29 | 0.78 | 0.41 | 0.37 | 1.90 |
| 18 | Ppib | 0.41 | 1.42E-33 | 2.95E-29 | 1.00 | 0.98 | 0.02 | 1.02 |
| 18 | Hipk3 | 1.09 | 1.65E-33 | 3.43E-29 | 0.43 | 0.15 | 0.28 | 2.88 |
| 18 | Limch1 | 0.74 | 2.70E-33 | 5.61E-29 | 0.96 | 0.67 | 0.28 | 1.42 |
| 18 | Def6 | 1.67 | 4.53E-33 | 9.41E-29 | 0.18 | 0.04 | 0.14 | 4.74 |
| 18 | Lats2 | 1.07 | 4.60E-33 | 9.57E-29 | 0.49 | 0.18 | 0.30 | 2.66 |
| 18 | Slc44a2 | 0.98 | 4.87E-33 | 1.01E-28 | 0.63 | 0.27 | 0.36 | 2.30 |
| 18 | Fam53b | 1.07 | 6.45E-33 | 1.34E-28 | 0.47 | 0.18 | 0.29 | 2.68 |
| 18 | Man2a1 | 1.00 | 6.77E-33 | 1.41E-28 | 0.56 | 0.23 | 0.33 | 2.45 |
| 18 | Ube2a | 0.78 | 8.35E-33 | 1.74E-28 | 0.92 | 0.71 | 0.22 | 1.31 |
| 18 | Pkn1 | 0.93 | 1.10E-32 | 2.29E-28 | 0.72 | 0.35 | 0.37 | 2.06 |
| 18 | Aph1a | 0.90 | 1.21E-32 | 2.52E-28 | 0.86 | 0.50 | 0.37 | 1.73 |
| 18 | Ifnar2 | 0.95 | 1.92E-32 | 3.99E-28 | 0.68 | 0.33 | 0.35 | 2.08 |
| 18 | Sgms1 | 0.98 | 2.30E-32 | 4.78E-28 | 0.46 | 0.17 | 0.29 | 2.75 |
| 18 | Tmem123 | 0.86 | 2.44E-32 | 5.07E-28 | 0.68 | 0.32 | 0.37 | 2.15 |
| 18 | Ccdc102a | 1.00 | 2.64E-32 | 5.48E-28 | 0.63 | 0.27 | 0.35 | 2.29 |
| 18 | Slc41a3 | 2.22 | 2.69E-32 | 5.58E-28 | 0.13 | 0.02 | 0.11 | 5.78 |
| 18 | Gpi1 | 0.59 | 2.87E-32 | 5.96E-28 | 0.99 | 0.84 | 0.15 | 1.18 |
| 18 | Rab5c | 0.80 | 3.18E-32 | 6.62E-28 | 0.88 | 0.65 | 0.24 | 1.37 |
| 18 | Tmem241 | 1.86 | 3.35E-32 | 6.96E-28 | 0.18 | 0.04 | 0.14 | 4.66 |
| 18 | Utp11 | 0.65 | 4.87E-32 | 1.01E-27 | 0.96 | 0.77 | 0.19 | 1.25 |
| 18 | Chmp6 | 0.92 | 5.22E-32 | 1.09E-27 | 0.83 | 0.51 | 0.33 | 1.64 |
| 18 | Sash1 | 0.97 | 5.67E-32 | 1.18E-27 | 0.76 | 0.40 | 0.36 | 1.92 |
| 18 | Cd59a | 0.55 | 5.71E-32 | 1.19E-27 | 0.55 | 0.22 | 0.34 | 2.55 |
| 18 | Rnf19a | 1.03 | 6.79E-32 | 1.41E-27 | 0.53 | 0.22 | 0.31 | 2.45 |
| 18 | Rap1gds1 | 0.92 | 7.38E-32 | 1.53E-27 | 0.65 | 0.29 | 0.36 | 2.26 |
| 18 | Rnasek | 0.69 | 8.10E-32 | 1.68E-27 | 0.95 | 0.72 | 0.23 | 1.33 |
| 18 | Tbrg1 | 0.60 | 1.09E-31 | 2.28E-27 | 0.95 | 0.83 | 0.12 | 1.15 |
| 18 | Ssr4 | 0.58 | 1.42E-31 | 2.95E-27 | 0.95 | 0.84 | 0.11 | 1.13 |
| 18 | Polr1d | 0.49 | 1.42E-31 | 2.96E-27 | 0.99 | 0.95 | 0.04 | 1.05 |
| 18 | Ptrhd1 | 0.99 | 1.43E-31 | 2.98E-27 | 0.71 | 0.36 | 0.35 | 1.99 |
| 18 | Dock8 | 1.36 | 1.91E-31 | 3.97E-27 | 0.16 | 0.03 | 0.13 | 5.00 |
| 18 | Slc8b1 | 1.93 | 2.46E-31 | 5.12E-27 | 0.13 | 0.02 | 0.11 | 5.54 |
| 18 | Tfec | 2.04 | 2.73E-31 | 5.68E-27 | 0.10 | 0.02 | 0.09 | 6.80 |
| 18 | Txn1 | 0.36 | 2.99E-31 | 6.22E-27 | 1.00 | 0.99 | 0.01 | 1.01 |
| 18 | Shisa5 | 0.87 | 3.38E-31 | 7.02E-27 | 0.57 | 0.24 | 0.33 | 2.41 |
| 18 | 2200002D01Rik | 1.91 | 3.66E-31 | 7.62E-27 | 0.17 | 0.04 | 0.13 | 4.64 |
| 18 | Apbb2 | 1.04 | 3.76E-31 | 7.82E-27 | 0.64 | 0.30 | 0.34 | 2.14 |
| 18 | B4galt7 | 0.95 | 3.77E-31 | 7.84E-27 | 0.50 | 0.19 | 0.31 | 2.60 |
| 18 | Pcbp3 | 1.14 | 3.94E-31 | 8.20E-27 | 0.45 | 0.17 | 0.28 | 2.64 |

**Supplementary Table 1. Marker genes for the original clusters of integrated clustering of E11.5 control and NotchGOF λ-junction cells.**

| | | | | | | | | |
|---|---|---|---|---|---|---|---|---|
| 18 | Cd2ap | 0.71 | 3.98E-31 | 8.27E-27 | 0.91 | 0.62 | 0.29 | 1.47 |
| 18 | Tmem109 | 1.02 | 4.05E-31 | 8.43E-27 | 0.71 | 0.36 | 0.35 | 1.98 |
| 18 | Cers5 | 0.87 | 4.26E-31 | 8.87E-27 | 0.72 | 0.34 | 0.38 | 2.12 |
| 18 | Eps15 | 0.90 | 4.57E-31 | 9.50E-27 | 0.65 | 0.30 | 0.35 | 2.16 |
| 18 | Gmppb | 1.00 | 5.21E-31 | 1.08E-26 | 0.48 | 0.19 | 0.30 | 2.60 |
| 18 | Ninj1 | 1.03 | 5.22E-31 | 1.09E-26 | 0.48 | 0.19 | 0.29 | 2.56 |
| 18 | Cerk | 1.01 | 1.17E-30 | 2.42E-26 | 0.41 | 0.15 | 0.27 | 2.82 |
| 18 | Tram1 | 0.90 | 1.23E-30 | 2.56E-26 | 0.82 | 0.50 | 0.32 | 1.63 |
| 18 | Cltb | 1.00 | 1.37E-30 | 2.85E-26 | 0.47 | 0.18 | 0.29 | 2.61 |
| 18 | Idh2 | 0.52 | 1.57E-30 | 3.26E-26 | 0.99 | 0.90 | 0.10 | 1.11 |
| 18 | Comt | 0.79 | 1.75E-30 | 3.64E-26 | 0.86 | 0.53 | 0.33 | 1.62 |
| 18 | Capn2 | 0.80 | 1.78E-30 | 3.71E-26 | 0.81 | 0.45 | 0.36 | 1.79 |
| 18 | Top1mt | 1.61 | 1.81E-30 | 3.75E-26 | 0.21 | 0.05 | 0.16 | 3.96 |
| 18 | Spag9 | 0.78 | 1.87E-30 | 3.89E-26 | 0.88 | 0.62 | 0.26 | 1.42 |
| 18 | Hmg20b | 0.62 | 2.00E-30 | 4.16E-26 | 0.98 | 0.80 | 0.18 | 1.22 |
| 18 | Tgds | 1.00 | 2.40E-30 | 4.99E-26 | 0.49 | 0.20 | 0.30 | 2.52 |
| 18 | Tmod3 | 0.79 | 2.43E-30 | 5.05E-26 | 0.91 | 0.64 | 0.27 | 1.43 |
| 18 | Ass1 | 1.56 | 2.44E-30 | 5.07E-26 | 0.22 | 0.06 | 0.16 | 3.88 |
| 18 | Gmip | 1.66 | 2.48E-30 | 5.16E-26 | 0.21 | 0.05 | 0.16 | 3.96 |
| 18 | Rab2a | 0.50 | 2.52E-30 | 5.23E-26 | 0.99 | 0.91 | 0.07 | 1.08 |
| 18 | Calm2 | 0.43 | 2.73E-30 | 5.67E-26 | 1.00 | 0.98 | 0.02 | 1.02 |
| 18 | Gna12 | 0.98 | 2.83E-30 | 5.89E-26 | 0.64 | 0.30 | 0.34 | 2.12 |
| 18 | Lman2 | 0.75 | 3.27E-30 | 6.81E-26 | 0.89 | 0.62 | 0.27 | 1.44 |
| 18 | Acsl4 | 1.16 | 3.30E-30 | 6.85E-26 | 0.39 | 0.14 | 0.25 | 2.81 |
| 18 | Snx7 | 1.18 | 3.56E-30 | 7.40E-26 | 0.32 | 0.10 | 0.22 | 3.17 |
| 18 | Pacc1 | 1.15 | 3.76E-30 | 7.83E-26 | 0.30 | 0.09 | 0.21 | 3.26 |
| 18 | Uchl5 | 0.84 | 4.62E-30 | 9.62E-26 | 0.84 | 0.54 | 0.31 | 1.57 |
| 18 | Cdr2 | 1.13 | 5.46E-30 | 1.13E-25 | 0.51 | 0.22 | 0.30 | 2.38 |
| 18 | Atox1 | 0.32 | 6.00E-30 | 1.25E-25 | 1.00 | 0.95 | 0.05 | 1.05 |
| 18 | Timp1 | 0.99 | 6.72E-30 | 1.40E-25 | 0.45 | 0.17 | 0.28 | 2.69 |
| 18 | Myd88 | 1.46 | 7.16E-30 | 1.49E-25 | 0.21 | 0.05 | 0.16 | 3.98 |
| 18 | Manf | 0.56 | 8.28E-30 | 1.72E-25 | 0.99 | 0.90 | 0.09 | 1.10 |
| 18 | Wdfy1 | 0.79 | 8.45E-30 | 1.76E-25 | 0.72 | 0.35 | 0.37 | 2.07 |
| 18 | Anxa11 | 0.80 | 8.51E-30 | 1.77E-25 | 0.42 | 0.15 | 0.27 | 2.79 |
| 18 | Nagk | 1.11 | 8.68E-30 | 1.81E-25 | 0.48 | 0.20 | 0.29 | 2.48 |
| 18 | 4933415A04Rik | 1.41 | 1.08E-29 | 2.25E-25 | 0.25 | 0.07 | 0.18 | 3.55 |
| 18 | Bcl2 | 1.02 | 1.18E-29 | 2.45E-25 | 0.68 | 0.35 | 0.33 | 1.95 |
| 18 | Igf2r | 0.81 | 1.33E-29 | 2.76E-25 | 0.85 | 0.49 | 0.36 | 1.72 |
| 18 | Lpp | 0.78 | 1.42E-29 | 2.96E-25 | 0.86 | 0.56 | 0.30 | 1.54 |
| 18 | Rps6ka3 | 0.75 | 1.76E-29 | 3.66E-25 | 0.82 | 0.45 | 0.37 | 1.81 |
| 18 | Gm42701 | 2.60 | 1.95E-29 | 4.06E-25 | 0.10 | 0.02 | 0.09 | 6.38 |
| 18 | Emc7 | 0.85 | 2.05E-29 | 4.27E-25 | 0.82 | 0.49 | 0.33 | 1.67 |
| 18 | Pdia3 | 0.46 | 2.35E-29 | 4.90E-25 | 1.00 | 0.96 | 0.04 | 1.04 |
| 18 | Map4k4 | 0.52 | 3.04E-29 | 6.32E-25 | 0.98 | 0.90 | 0.08 | 1.09 |
| 18 | Cables1 | 1.25 | 3.06E-29 | 6.37E-25 | 0.28 | 0.08 | 0.20 | 3.32 |
| 18 | Ampd2 | 0.88 | 3.53E-29 | 7.34E-25 | 0.43 | 0.16 | 0.27 | 2.75 |
| 18 | Arf1 | 0.39 | 3.93E-29 | 8.17E-25 | 0.99 | 0.98 | 0.02 | 1.02 |
| 18 | Nras | 0.72 | 4.16E-29 | 8.65E-25 | 0.95 | 0.69 | 0.26 | 1.38 |
| 18 | Alad | 0.42 | 4.27E-29 | 8.88E-25 | 0.91 | 0.63 | 0.27 | 1.43 |
| 18 | Mbd2 | 0.77 | 4.45E-29 | 9.26E-25 | 0.88 | 0.56 | 0.32 | 1.58 |
| 18 | Slc39a8 | 0.71 | 5.01E-29 | 1.04E-24 | 0.45 | 0.17 | 0.28 | 2.67 |
| 18 | Capza1 | 0.75 | 5.89E-29 | 1.22E-24 | 0.90 | 0.64 | 0.26 | 1.40 |
| 18 | Rassf1 | 0.92 | 5.93E-29 | 1.23E-24 | 0.74 | 0.40 | 0.34 | 1.86 |
| 18 | Riox1 | 0.98 | 6.48E-29 | 1.35E-24 | 0.60 | 0.28 | 0.32 | 2.17 |
| 18 | Lrp5 | 0.94 | 7.34E-29 | 1.53E-24 | 0.45 | 0.17 | 0.28 | 2.60 |
| 18 | Nagpa | 1.11 | 8.56E-29 | 1.78E-24 | 0.31 | 0.10 | 0.21 | 3.16 |

**Supplementary Table 1. Marker genes for the original clusters of integrated clustering of E11.5 control and NotchGOF λ-junction cells.**

| | | | | | | | | |
|---|---|---|---|---|---|---|---|---|
| 18 | Irak4 | 1.61 | 9.15E-29 | 1.90E-24 | 0.19 | 0.05 | 0.14 | 4.13 |
| 18 | Dnaja1 | 0.37 | 9.32E-29 | 1.94E-24 | 1.00 | 0.99 | 0.01 | 1.01 |
| 18 | Ophn1 | 0.91 | 1.01E-28 | 2.09E-24 | 0.48 | 0.19 | 0.29 | 2.56 |
| 18 | Shroom4 | 1.30 | 1.07E-28 | 2.23E-24 | 0.28 | 0.09 | 0.20 | 3.28 |
| 18 | Lims1 | 0.79 | 1.11E-28 | 2.31E-24 | 0.86 | 0.57 | 0.29 | 1.52 |
| 18 | Btbd10 | 0.83 | 1.24E-28 | 2.58E-24 | 0.57 | 0.25 | 0.32 | 2.32 |
| 18 | Fry | 1.21 | 1.27E-28 | 2.64E-24 | 0.21 | 0.05 | 0.16 | 3.91 |
| 18 | Slc35g2 | 2.15 | 1.42E-28 | 2.94E-24 | 0.12 | 0.02 | 0.10 | 5.67 |
| 18 | Myo1b | 0.72 | 1.44E-28 | 2.99E-24 | 0.93 | 0.73 | 0.20 | 1.28 |
| 18 | Ccdc12 | 0.70 | 1.46E-28 | 3.04E-24 | 0.93 | 0.70 | 0.23 | 1.33 |
| 18 | Ifitm3 | 0.49 | 1.56E-28 | 3.24E-24 | 0.65 | 0.29 | 0.36 | 2.21 |
| 18 | Dnajb11 | 0.71 | 1.72E-28 | 3.57E-24 | 0.93 | 0.68 | 0.25 | 1.37 |
| 18 | Ctso | 1.35 | 1.78E-28 | 3.70E-24 | 0.23 | 0.06 | 0.17 | 3.61 |
| 18 | Rmnd5b | 0.92 | 1.92E-28 | 4.00E-24 | 0.44 | 0.17 | 0.27 | 2.61 |
| 18 | Nrbp1 | 0.76 | 1.93E-28 | 4.01E-24 | 0.87 | 0.56 | 0.31 | 1.56 |
| 18 | Grpel1 | 0.61 | 2.08E-28 | 4.32E-24 | 0.95 | 0.83 | 0.12 | 1.15 |
| 18 | Rab1a | 0.66 | 2.33E-28 | 4.84E-24 | 0.95 | 0.71 | 0.24 | 1.34 |
| 18 | Trak2 | 1.10 | 2.33E-28 | 4.85E-24 | 0.37 | 0.13 | 0.24 | 2.79 |
| 18 | Lamp1 | 0.55 | 3.15E-28 | 6.55E-24 | 0.97 | 0.86 | 0.12 | 1.13 |
| 18 | Stc1 | 1.40 | 3.28E-28 | 6.82E-24 | 0.24 | 0.07 | 0.17 | 3.55 |
| 18 | Cgrrf1 | 0.89 | 3.45E-28 | 7.17E-24 | 0.58 | 0.25 | 0.32 | 2.27 |
| 18 | Adarb1 | 1.51 | 3.69E-28 | 7.67E-24 | 0.24 | 0.07 | 0.17 | 3.49 |
| 18 | Saraf | 0.70 | 3.93E-28 | 8.17E-24 | 0.87 | 0.54 | 0.33 | 1.61 |
| 18 | Fam122b | 1.43 | 4.14E-28 | 8.60E-24 | 0.27 | 0.08 | 0.18 | 3.27 |
| 18 | Plekhg1 | 1.10 | 4.48E-28 | 9.31E-24 | 0.37 | 0.14 | 0.24 | 2.77 |
| 18 | Rabep2 | 0.98 | 5.05E-28 | 1.05E-23 | 0.47 | 0.19 | 0.28 | 2.49 |
| 18 | Nudt9 | 0.80 | 5.87E-28 | 1.22E-23 | 0.81 | 0.48 | 0.33 | 1.70 |
| 18 | Gls | 0.95 | 5.91E-28 | 1.23E-23 | 0.73 | 0.41 | 0.33 | 1.80 |
| 18 | Rbm43 | 0.96 | 5.96E-28 | 1.24E-23 | 0.33 | 0.11 | 0.22 | 3.06 |
| 18 | Slc30a1 | 1.29 | 7.49E-28 | 1.56E-23 | 0.27 | 0.08 | 0.19 | 3.28 |
| 18 | Pla2g6 | 1.32 | 8.16E-28 | 1.70E-23 | 0.26 | 0.08 | 0.18 | 3.32 |
| 18 | Zdhhc12 | 0.97 | 8.96E-28 | 1.86E-23 | 0.57 | 0.26 | 0.31 | 2.22 |
| 18 | Zbtb38 | 1.00 | 9.70E-28 | 2.02E-23 | 0.37 | 0.13 | 0.24 | 2.79 |
| 18 | Stx4a | 0.88 | 1.33E-27 | 2.76E-23 | 0.65 | 0.31 | 0.34 | 2.07 |
| 18 | Capg | 0.83 | 1.58E-27 | 3.28E-23 | 0.40 | 0.15 | 0.25 | 2.69 |
| 18 | Snx4 | 0.79 | 1.58E-27 | 3.29E-23 | 0.78 | 0.42 | 0.36 | 1.86 |
| 18 | Dennd2c | 1.34 | 2.30E-27 | 4.78E-23 | 0.22 | 0.06 | 0.16 | 3.61 |
| 18 | Cltc | 0.69 | 2.55E-27 | 5.30E-23 | 0.95 | 0.75 | 0.20 | 1.27 |
| 18 | Orai3 | 1.10 | 2.70E-27 | 5.62E-23 | 0.33 | 0.11 | 0.22 | 2.92 |
| 18 | Mgll | 2.47 | 3.70E-27 | 7.69E-23 | 0.11 | 0.02 | 0.09 | 5.60 |
| 18 | Ufm1 | 0.65 | 4.14E-27 | 8.62E-23 | 0.92 | 0.69 | 0.23 | 1.34 |
| 18 | Rab35 | 0.82 | 4.38E-27 | 9.10E-23 | 0.70 | 0.36 | 0.34 | 1.93 |
| 18 | Anxa1 | 1.66 | 5.44E-27 | 1.13E-22 | 0.13 | 0.03 | 0.11 | 4.93 |
| 18 | Gab1 | 1.04 | 5.70E-27 | 1.18E-22 | 0.46 | 0.19 | 0.27 | 2.44 |
| 18 | Impdh1 | 1.08 | 6.27E-27 | 1.30E-22 | 0.51 | 0.22 | 0.30 | 2.37 |
| 18 | Map3k1 | 0.86 | 9.27E-27 | 1.93E-22 | 0.57 | 0.26 | 0.31 | 2.20 |
| 18 | Efnb2 | 0.80 | 9.97E-27 | 2.07E-22 | 0.79 | 0.45 | 0.35 | 1.78 |
| 18 | Cmtm7 | 0.70 | 1.07E-26 | 2.23E-22 | 0.83 | 0.48 | 0.35 | 1.72 |
| 18 | Dnajc13 | 0.87 | 1.13E-26 | 2.34E-22 | 0.71 | 0.37 | 0.34 | 1.90 |
| 18 | Rwdd2a | 1.79 | 1.17E-26 | 2.44E-22 | 0.12 | 0.02 | 0.10 | 5.52 |
| 18 | Eef2kmt | 0.92 | 1.28E-26 | 2.66E-22 | 0.45 | 0.18 | 0.27 | 2.51 |
| 18 | Tgfbr3 | 1.26 | 1.29E-26 | 2.69E-22 | 0.26 | 0.08 | 0.18 | 3.28 |
| 18 | Arhgap1 | 0.84 | 1.62E-26 | 3.37E-22 | 0.57 | 0.26 | 0.31 | 2.21 |
| 18 | Haus4 | 0.80 | 1.79E-26 | 3.72E-22 | 0.64 | 0.31 | 0.33 | 2.06 |
| 18 | Septin7 | 0.38 | 1.83E-26 | 3.81E-22 | 1.00 | 0.97 | 0.03 | 1.03 |
| 18 | Inka1 | 0.70 | 1.84E-26 | 3.83E-22 | 0.85 | 0.51 | 0.35 | 1.68 |

**Supplementary Table 1. Marker genes for the original clusters of integrated clustering of E11.5 control and NotchGOF λ-junction cells.**

| | | | | | | | | |
|---|---|---|---|---|---|---|---|---|
| 18 | Plcb1 | 1.17 | 1.95E-26 | 4.06E-22 | 0.29 | 0.09 | 0.20 | 3.07 |
| 18 | Vps26c | 0.83 | 2.23E-26 | 4.64E-22 | 0.53 | 0.22 | 0.30 | 2.35 |
| 18 | Mob3c | 1.43 | 2.39E-26 | 4.96E-22 | 0.20 | 0.05 | 0.15 | 3.79 |
| 18 | Pygb | 0.85 | 2.46E-26 | 5.11E-22 | 0.53 | 0.23 | 0.30 | 2.32 |
| 18 | Fignl2 | 1.10 | 2.61E-26 | 5.43E-22 | 0.34 | 0.12 | 0.22 | 2.80 |
| 18 | Hps3 | 1.20 | 2.63E-26 | 5.48E-22 | 0.26 | 0.08 | 0.18 | 3.23 |
| 18 | Tbc1d1 | 0.89 | 2.70E-26 | 5.63E-22 | 0.70 | 0.37 | 0.33 | 1.89 |
| 18 | Vdac2 | 0.41 | 2.74E-26 | 5.70E-22 | 0.99 | 0.96 | 0.03 | 1.03 |
| 18 | Trappc2 | 1.12 | 2.98E-26 | 6.20E-22 | 0.41 | 0.16 | 0.25 | 2.56 |
| 18 | Mars2 | 1.44 | 3.24E-26 | 6.73E-22 | 0.24 | 0.07 | 0.17 | 3.41 |
| 18 | Ydjc | 1.38 | 3.32E-26 | 6.90E-22 | 0.26 | 0.08 | 0.18 | 3.20 |
| 18 | Fam107b | 0.65 | 3.33E-26 | 6.92E-22 | 0.72 | 0.35 | 0.37 | 2.05 |
| 18 | Vcl | 0.79 | 4.28E-26 | 8.90E-22 | 0.74 | 0.39 | 0.35 | 1.88 |
| 18 | Dusp23 | 0.89 | 4.37E-26 | 9.09E-22 | 0.43 | 0.17 | 0.26 | 2.54 |
| 18 | Zdhhc20 | 0.75 | 4.40E-26 | 9.16E-22 | 0.90 | 0.58 | 0.32 | 1.55 |
| 18 | Lrrc59 | 0.60 | 4.61E-26 | 9.58E-22 | 0.95 | 0.79 | 0.16 | 1.20 |
| 18 | Armcx4 | 0.88 | 4.80E-26 | 9.99E-22 | 0.70 | 0.36 | 0.34 | 1.94 |
| 18 | Bicd2 | 0.87 | 5.21E-26 | 1.08E-21 | 0.62 | 0.29 | 0.34 | 2.18 |
| 18 | Leprot | 0.78 | 5.57E-26 | 1.16E-21 | 0.82 | 0.52 | 0.30 | 1.58 |
| 18 | St3gal1 | 1.19 | 7.87E-26 | 1.64E-21 | 0.30 | 0.10 | 0.20 | 2.96 |
| 18 | Ppp1r14b | 0.35 | 9.97E-26 | 2.07E-21 | 1.00 | 0.99 | 0.01 | 1.01 |
| 18 | Airn | 0.99 | 1.10E-25 | 2.29E-21 | 0.46 | 0.19 | 0.28 | 2.48 |
| 18 | Adgra2 | 1.10 | 1.12E-25 | 2.33E-21 | 0.53 | 0.23 | 0.30 | 2.29 |
| 18 | Sat1 | 0.65 | 1.68E-25 | 3.49E-21 | 0.63 | 0.30 | 0.33 | 2.07 |
| 18 | Plpp2 | 0.89 | 3.37E-25 | 7.01E-21 | 0.56 | 0.26 | 0.30 | 2.15 |
| 18 | Hk1 | 0.83 | 3.41E-25 | 7.09E-21 | 0.55 | 0.25 | 0.30 | 2.23 |
| 18 | Prelid2 | 0.77 | 3.58E-25 | 7.44E-21 | 0.69 | 0.37 | 0.33 | 1.90 |
| 18 | Fnbp1 | 0.86 | 3.90E-25 | 8.11E-21 | 0.53 | 0.23 | 0.30 | 2.27 |
| 18 | Cst3 | 0.41 | 4.13E-25 | 8.59E-21 | 0.99 | 0.92 | 0.07 | 1.08 |
| 18 | Babam1 | 0.68 | 4.90E-25 | 1.02E-20 | 0.87 | 0.58 | 0.29 | 1.51 |
| 18 | Ntan1 | 0.61 | 5.35E-25 | 1.11E-20 | 0.91 | 0.66 | 0.25 | 1.37 |
| 18 | Tsc22d4 | 0.64 | 5.44E-25 | 1.13E-20 | 0.89 | 0.61 | 0.28 | 1.46 |
| 18 | Atp2b4 | 1.16 | 6.28E-25 | 1.31E-20 | 0.47 | 0.20 | 0.27 | 2.34 |
| 18 | Map3k14 | 1.66 | 7.21E-25 | 1.50E-20 | 0.15 | 0.04 | 0.12 | 4.25 |
| 18 | Erp29 | 0.46 | 7.27E-25 | 1.51E-20 | 0.98 | 0.88 | 0.10 | 1.11 |
| 18 | Klhl2 | 1.09 | 7.65E-25 | 1.59E-20 | 0.33 | 0.12 | 0.21 | 2.78 |
| 18 | Mrps18b | 0.76 | 7.67E-25 | 1.60E-20 | 0.82 | 0.50 | 0.32 | 1.64 |
| 18 | Ucp2 | 0.33 | 1.10E-24 | 2.28E-20 | 0.71 | 0.35 | 0.36 | 2.02 |
| 18 | Stx6 | 0.79 | 1.25E-24 | 2.59E-20 | 0.82 | 0.50 | 0.32 | 1.64 |
| 18 | Ddrgk1 | 0.71 | 1.42E-24 | 2.95E-20 | 0.84 | 0.55 | 0.29 | 1.53 |
| 18 | Arf5 | 0.34 | 1.46E-24 | 3.03E-20 | 1.00 | 0.98 | 0.01 | 1.01 |
| 18 | Mcl1 | 0.53 | 1.60E-24 | 3.32E-20 | 0.96 | 0.82 | 0.15 | 1.18 |
| 18 | 0610012G03Rik | 0.50 | 1.66E-24 | 3.44E-20 | 0.98 | 0.86 | 0.12 | 1.13 |
| 18 | Hadhb | 0.68 | 1.69E-24 | 3.52E-20 | 0.88 | 0.62 | 0.27 | 1.43 |
| 18 | Exoc5 | 0.69 | 1.78E-24 | 3.71E-20 | 0.86 | 0.57 | 0.29 | 1.51 |
| 18 | Twf1 | 0.70 | 1.86E-24 | 3.88E-20 | 0.82 | 0.49 | 0.33 | 1.68 |
| 18 | Inpp5a | 0.96 | 2.03E-24 | 4.23E-20 | 0.39 | 0.15 | 0.24 | 2.60 |
| 18 | Atp6v0e | 0.59 | 2.31E-24 | 4.80E-20 | 0.92 | 0.72 | 0.20 | 1.28 |
| 18 | Ctsd | 0.53 | 2.31E-24 | 4.81E-20 | 0.87 | 0.61 | 0.27 | 1.44 |
| 18 | Dync1li1 | 0.69 | 2.50E-24 | 5.20E-20 | 0.81 | 0.52 | 0.29 | 1.57 |
| 18 | Arpc5l | 0.56 | 2.86E-24 | 5.96E-20 | 0.97 | 0.81 | 0.16 | 1.20 |
| 18 | St3gal6 | 1.15 | 2.89E-24 | 6.01E-20 | 0.31 | 0.11 | 0.20 | 2.82 |
| 18 | Stk25 | 0.76 | 2.92E-24 | 6.08E-20 | 0.79 | 0.45 | 0.34 | 1.75 |
| 18 | Akt1 | 0.65 | 3.34E-24 | 6.96E-20 | 0.93 | 0.72 | 0.21 | 1.30 |
| 18 | Tpcn1 | 1.20 | 3.49E-24 | 7.26E-20 | 0.27 | 0.09 | 0.18 | 3.02 |
| 18 | Ly96 | 1.23 | 3.68E-24 | 7.65E-20 | 0.20 | 0.06 | 0.15 | 3.58 |

**Supplementary Table 1. Marker genes for the original clusters of integrated clustering of E11.5 control and NotchGOF λ-junction cells.**

| | | | | | | | | |
|---|---|---|---|---|---|---|---|---|
| 18 | Tcn2 | 1.05 | 3.99E-24 | 8.30E-20 | 0.43 | 0.18 | 0.25 | 2.43 |
| 18 | Rgs10 | 0.50 | 4.02E-24 | 8.37E-20 | 0.27 | 0.09 | 0.18 | 3.05 |
| 18 | B4galt1 | 0.89 | 4.07E-24 | 8.46E-20 | 0.51 | 0.23 | 0.28 | 2.24 |
| 18 | Add2 | 0.93 | 4.29E-24 | 8.92E-20 | 0.27 | 0.09 | 0.18 | 3.06 |
| 18 | Chst12 | 1.11 | 4.61E-24 | 9.60E-20 | 0.26 | 0.08 | 0.18 | 3.12 |
| 18 | Lpcat1 | 0.79 | 5.36E-24 | 1.12E-19 | 0.77 | 0.47 | 0.30 | 1.64 |
| 18 | Rel | 1.04 | 6.13E-24 | 1.28E-19 | 0.27 | 0.09 | 0.18 | 3.06 |
| 18 | Spryd3 | 1.38 | 6.32E-24 | 1.32E-19 | 0.23 | 0.07 | 0.16 | 3.25 |
| 18 | Dnmbp | 0.88 | 6.60E-24 | 1.37E-19 | 0.34 | 0.13 | 0.22 | 2.73 |
| 18 | Hip1 | 0.78 | 6.82E-24 | 1.42E-19 | 0.77 | 0.44 | 0.32 | 1.72 |
| 18 | Mri1 | 0.75 | 7.38E-24 | 1.53E-19 | 0.58 | 0.27 | 0.31 | 2.15 |
| 18 | Diaph2 | 0.93 | 8.03E-24 | 1.67E-19 | 0.39 | 0.15 | 0.24 | 2.55 |
| 18 | Eif5a | 0.31 | 8.90E-24 | 1.85E-19 | 1.00 | 1.00 | 0.00 | 1.00 |
| 18 | Trim3 | 1.11 | 9.21E-24 | 1.91E-19 | 0.28 | 0.09 | 0.18 | 3.00 |
| 18 | Sbf2 | 0.80 | 1.09E-23 | 2.27E-19 | 0.65 | 0.34 | 0.31 | 1.91 |
| 18 | Sik3 | 0.76 | 1.09E-23 | 2.27E-19 | 0.61 | 0.28 | 0.32 | 2.15 |
| 18 | Stx7 | 0.78 | 1.13E-23 | 2.35E-19 | 0.62 | 0.31 | 0.31 | 2.01 |
| 18 | Egr1 | 0.75 | 1.26E-23 | 2.62E-19 | 0.24 | 0.07 | 0.16 | 3.22 |
| 18 | Uprt | 2.13 | 1.87E-23 | 3.89E-19 | 0.10 | 0.02 | 0.08 | 5.37 |
| 18 | Lrrc8a | 0.85 | 2.05E-23 | 4.26E-19 | 0.57 | 0.27 | 0.30 | 2.11 |
| 18 | Cd55 | 1.57 | 2.09E-23 | 4.34E-19 | 0.12 | 0.02 | 0.09 | 5.04 |
| 18 | Acat1 | 0.73 | 2.10E-23 | 4.37E-19 | 0.77 | 0.46 | 0.31 | 1.67 |
| 18 | Acap2 | 0.71 | 2.14E-23 | 4.44E-19 | 0.79 | 0.46 | 0.33 | 1.71 |
| 18 | D830030K20Rik | 1.75 | 2.20E-23 | 4.58E-19 | 0.13 | 0.03 | 0.10 | 4.67 |
| 18 | Zswim6 | 0.78 | 2.30E-23 | 4.79E-19 | 0.61 | 0.29 | 0.32 | 2.09 |
| 18 | Coq10b | 0.82 | 2.41E-23 | 5.01E-19 | 0.55 | 0.25 | 0.30 | 2.17 |
| 18 | Jmjd1c | 0.70 | 2.85E-23 | 5.93E-19 | 0.81 | 0.48 | 0.33 | 1.69 |
| 18 | Yrdc | 0.80 | 3.02E-23 | 6.28E-19 | 0.74 | 0.41 | 0.32 | 1.78 |
| 18 | Tmem14c | 0.32 | 3.07E-23 | 6.38E-19 | 0.98 | 0.92 | 0.06 | 1.07 |
| 18 | Serpinb6a | 0.75 | 3.09E-23 | 6.43E-19 | 0.71 | 0.39 | 0.32 | 1.84 |
| 18 | Gask1b | 1.74 | 3.11E-23 | 6.47E-19 | 0.12 | 0.02 | 0.09 | 4.83 |
| 18 | Nav1 | 0.67 | 3.12E-23 | 6.50E-19 | 0.84 | 0.54 | 0.30 | 1.56 |
| 18 | Pdcd10 | 0.61 | 3.30E-23 | 6.86E-19 | 0.91 | 0.72 | 0.19 | 1.26 |
| 18 | Zfp36l1 | 0.51 | 3.65E-23 | 7.59E-19 | 0.97 | 0.85 | 0.12 | 1.14 |
| 18 | Brk1 | 0.42 | 3.91E-23 | 8.12E-19 | 0.99 | 0.92 | 0.07 | 1.07 |
| 18 | Alg1 | 1.07 | 3.99E-23 | 8.30E-19 | 0.24 | 0.08 | 0.17 | 3.17 |
| 18 | Ttc7b | 1.06 | 4.21E-23 | 8.76E-19 | 0.39 | 0.16 | 0.23 | 2.47 |
| 18 | Cyfip1 | 0.58 | 4.79E-23 | 9.96E-19 | 0.90 | 0.70 | 0.20 | 1.29 |
| 18 | Ppp1ca | 0.33 | 5.05E-23 | 1.05E-18 | 0.99 | 0.98 | 0.01 | 1.01 |
| 18 | Tbc1d2b | 1.44 | 5.08E-23 | 1.06E-18 | 0.25 | 0.08 | 0.17 | 3.07 |
| 18 | Ctsl | 0.46 | 5.18E-23 | 1.08E-18 | 0.97 | 0.82 | 0.15 | 1.18 |
| 18 | Mrpl36 | 0.55 | 5.35E-23 | 1.11E-18 | 0.93 | 0.71 | 0.22 | 1.31 |
| 18 | Irak2 | 1.39 | 5.81E-23 | 1.21E-18 | 0.19 | 0.05 | 0.14 | 3.60 |
| 18 | Smpdl3b | 1.36 | 6.46E-23 | 1.34E-18 | 0.22 | 0.07 | 0.15 | 3.30 |
| 18 | Poglut3 | 0.99 | 6.68E-23 | 1.39E-18 | 0.35 | 0.13 | 0.21 | 2.61 |
| 18 | Ndfip2 | 0.75 | 9.07E-23 | 1.89E-18 | 0.82 | 0.51 | 0.31 | 1.60 |
| 18 | Cfap20 | 0.60 | 9.18E-23 | 1.91E-18 | 0.92 | 0.75 | 0.17 | 1.23 |
| 18 | Aacs | 0.93 | 1.01E-22 | 2.10E-18 | 0.31 | 0.11 | 0.20 | 2.79 |
| 18 | Fam241a | 0.97 | 1.12E-22 | 2.33E-18 | 0.32 | 0.12 | 0.20 | 2.71 |
| 18 | Fkbp15 | 0.98 | 1.20E-22 | 2.50E-18 | 0.34 | 0.13 | 0.21 | 2.63 |
| 18 | Ccnd3 | 0.49 | 1.45E-22 | 3.02E-18 | 0.97 | 0.85 | 0.12 | 1.14 |
| 18 | Ap1s1 | 0.54 | 1.47E-22 | 3.07E-18 | 0.95 | 0.76 | 0.19 | 1.24 |
| 18 | Hsd17b11 | 0.83 | 1.74E-22 | 3.62E-18 | 0.48 | 0.20 | 0.27 | 2.34 |
| 18 | Iqsec2 | 1.79 | 1.77E-22 | 3.69E-18 | 0.11 | 0.02 | 0.09 | 4.87 |
| 18 | Stat6 | 1.23 | 1.78E-22 | 3.70E-18 | 0.19 | 0.05 | 0.14 | 3.52 |
| 18 | Fam49b | 0.59 | 1.82E-22 | 3.78E-18 | 0.90 | 0.65 | 0.25 | 1.38 |

**Supplementary Table 1. Marker genes for the original clusters of integrated clustering of E11.5 control and NotchGOF λ-junction cells.**

| | | | | | | | | |
|---|---|---|---|---|---|---|---|---|
| 18 | Pde4d | 0.90 | 1.83E-22 | 3.81E-18 | 0.54 | 0.26 | 0.28 | 2.08 |
| 18 | Camk4 | 1.62 | 2.13E-22 | 4.42E-18 | 0.11 | 0.02 | 0.08 | 5.00 |
| 18 | Tram2 | 0.93 | 2.18E-22 | 4.52E-18 | 0.29 | 0.10 | 0.19 | 2.86 |
| 18 | Mcc | 1.06 | 2.60E-22 | 5.41E-18 | 0.33 | 0.13 | 0.21 | 2.60 |
| 18 | Filip1l | 0.76 | 2.66E-22 | 5.53E-18 | 0.54 | 0.25 | 0.28 | 2.11 |
| 18 | Lmf2 | 0.90 | 2.80E-22 | 5.82E-18 | 0.44 | 0.19 | 0.25 | 2.36 |
| 18 | Nr2c2ap | 0.80 | 2.92E-22 | 6.07E-18 | 0.51 | 0.23 | 0.28 | 2.23 |
| 18 | Ubl3 | 0.70 | 3.23E-22 | 6.71E-18 | 0.69 | 0.37 | 0.33 | 1.89 |
| 18 | Rrm2b | 0.91 | 3.23E-22 | 6.72E-18 | 0.33 | 0.12 | 0.21 | 2.68 |
| 18 | Clic4 | 0.60 | 3.56E-22 | 7.41E-18 | 0.92 | 0.68 | 0.25 | 1.37 |
| 18 | Rpl36al | 0.26 | 3.82E-22 | 7.95E-18 | 1.00 | 1.00 | 0.00 | 1.00 |
| 18 | Cds2 | 0.88 | 4.05E-22 | 8.42E-18 | 0.43 | 0.18 | 0.25 | 2.40 |
| 18 | Tnip1 | 1.03 | 4.42E-22 | 9.19E-18 | 0.29 | 0.10 | 0.19 | 2.83 |
| 18 | Crtap | 0.82 | 4.56E-22 | 9.49E-18 | 0.56 | 0.27 | 0.29 | 2.05 |
| 18 | Golm1 | 0.73 | 4.57E-22 | 9.51E-18 | 0.82 | 0.55 | 0.27 | 1.49 |
| 18 | Atp6v0a2 | 0.86 | 4.92E-22 | 1.02E-17 | 0.55 | 0.26 | 0.29 | 2.12 |
| 18 | Tro | 0.81 | 4.99E-22 | 1.04E-17 | 0.74 | 0.42 | 0.32 | 1.77 |
| 18 | Blvra | 0.87 | 5.14E-22 | 1.07E-17 | 0.46 | 0.20 | 0.25 | 2.26 |
| 18 | Gmpr2 | 0.74 | 5.23E-22 | 1.09E-17 | 0.72 | 0.41 | 0.30 | 1.73 |
| 18 | Bcat2 | 0.74 | 5.55E-22 | 1.15E-17 | 0.59 | 0.29 | 0.30 | 2.06 |
| 18 | Fosl2 | 1.49 | 5.66E-22 | 1.18E-17 | 0.13 | 0.03 | 0.10 | 4.45 |
| 18 | Cdc37 | 0.35 | 5.73E-22 | 1.19E-17 | 0.99 | 0.96 | 0.03 | 1.03 |
| 18 | Rgs19 | 0.69 | 5.93E-22 | 1.23E-17 | 0.76 | 0.45 | 0.31 | 1.69 |
| 18 | Gm43328 | 1.84 | 6.05E-22 | 1.26E-17 | 0.12 | 0.03 | 0.10 | 4.52 |
| 18 | Rpp21 | 0.66 | 6.27E-22 | 1.30E-17 | 0.85 | 0.57 | 0.28 | 1.48 |
| 18 | Septin10 | 0.73 | 7.59E-22 | 1.58E-17 | 0.68 | 0.37 | 0.31 | 1.85 |
| 18 | Ggh | 0.84 | 7.86E-22 | 1.64E-17 | 0.53 | 0.25 | 0.28 | 2.11 |
| 18 | Mrrf | 0.72 | 7.99E-22 | 1.66E-17 | 0.67 | 0.36 | 0.31 | 1.87 |
| 18 | Rab5if | 0.42 | 8.24E-22 | 1.71E-17 | 0.97 | 0.91 | 0.06 | 1.07 |
| 18 | Psat1 | 0.69 | 9.71E-22 | 2.02E-17 | 0.77 | 0.45 | 0.32 | 1.71 |
| 18 | Fkbp9 | 0.66 | 1.00E-21 | 2.08E-17 | 0.78 | 0.44 | 0.34 | 1.78 |
| 18 | Tmem181a | 0.97 | 1.20E-21 | 2.49E-17 | 0.29 | 0.10 | 0.19 | 2.83 |
| 18 | Irf2bp2 | 0.74 | 1.23E-21 | 2.55E-17 | 0.67 | 0.35 | 0.33 | 1.95 |
| 18 | Zbtb1 | 0.77 | 1.26E-21 | 2.61E-17 | 0.67 | 0.35 | 0.32 | 1.92 |
| 18 | Slc20a1 | 0.71 | 1.40E-21 | 2.92E-17 | 0.63 | 0.32 | 0.31 | 1.98 |
| 18 | Arf6 | 0.80 | 1.45E-21 | 3.02E-17 | 0.70 | 0.39 | 0.30 | 1.77 |
| 18 | Arhgef2 | 0.74 | 1.60E-21 | 3.32E-17 | 0.73 | 0.42 | 0.31 | 1.73 |
| 18 | Tgoln1 | 0.88 | 2.10E-21 | 4.37E-17 | 0.58 | 0.29 | 0.29 | 2.01 |
| 18 | Cdk1 | 0.75 | 2.27E-21 | 4.72E-17 | 0.90 | 0.67 | 0.24 | 1.35 |
| 18 | 0610010K14Rik | 0.48 | 2.27E-21 | 4.72E-17 | 0.96 | 0.81 | 0.14 | 1.18 |
| 18 | Rer1 | 0.47 | 2.30E-21 | 4.78E-17 | 0.96 | 0.83 | 0.13 | 1.16 |
| 18 | Slc39a10 | 0.63 | 2.69E-21 | 5.60E-17 | 0.92 | 0.68 | 0.24 | 1.36 |
| 18 | Alg3 | 0.78 | 2.78E-21 | 5.78E-17 | 0.57 | 0.28 | 0.30 | 2.07 |
| 18 | Tsen15 | 0.57 | 3.03E-21 | 6.30E-17 | 0.88 | 0.63 | 0.25 | 1.39 |
| 18 | Bcl10 | 0.67 | 3.05E-21 | 6.35E-17 | 0.80 | 0.49 | 0.31 | 1.62 |
| 18 | Map7d1 | 0.63 | 3.29E-21 | 6.84E-17 | 0.87 | 0.62 | 0.26 | 1.42 |
| 18 | Cemip2 | 0.62 | 3.30E-21 | 6.86E-17 | 0.76 | 0.45 | 0.31 | 1.70 |
| 18 | Layn | 1.67 | 4.14E-21 | 8.62E-17 | 0.12 | 0.03 | 0.09 | 4.36 |
| 18 | Itpr1 | 0.83 | 4.33E-21 | 9.01E-17 | 0.47 | 0.21 | 0.25 | 2.20 |
| 18 | Cflar | 0.72 | 4.88E-21 | 1.01E-16 | 0.61 | 0.31 | 0.30 | 1.96 |
| 18 | Surf4 | 0.60 | 5.00E-21 | 1.04E-16 | 0.90 | 0.59 | 0.31 | 1.54 |
| 18 | Nfkbie | 1.68 | 6.08E-21 | 1.27E-16 | 0.15 | 0.04 | 0.11 | 3.85 |
| 18 | Selenof | 0.36 | 6.15E-21 | 1.28E-16 | 0.99 | 0.96 | 0.03 | 1.04 |
| 18 | Szrd1 | 0.67 | 6.55E-21 | 1.36E-16 | 0.81 | 0.51 | 0.31 | 1.61 |
| 18 | Gemin7 | 0.57 | 6.88E-21 | 1.43E-16 | 0.92 | 0.70 | 0.21 | 1.30 |
| 18 | Mrtfb | 0.83 | 7.20E-21 | 1.50E-16 | 0.52 | 0.24 | 0.28 | 2.14 |

**Supplementary Table 1. Marker genes for the original clusters of integrated clustering of E11.5 control and NotchGOF λ-junction cells.**

| | | | | | | | | |
|---|---|---|---|---|---|---|---|---|
| 18 | Nfe2l2 | 0.62 | 7.29E-21 | 1.52E-16 | 0.79 | 0.44 | 0.34 | 1.77 |
| 18 | Hexb | 0.52 | 7.49E-21 | 1.56E-16 | 0.50 | 0.22 | 0.27 | 2.23 |
| 18 | Nfkbil1 | 0.87 | 7.76E-21 | 1.61E-16 | 0.56 | 0.28 | 0.28 | 2.02 |
| 18 | Ddx39 | 0.56 | 8.62E-21 | 1.79E-16 | 0.95 | 0.79 | 0.16 | 1.20 |
| 18 | Slc35d2 | 1.57 | 1.05E-20 | 2.19E-16 | 0.12 | 0.03 | 0.09 | 4.46 |
| 18 | Slc25a30 | 1.02 | 1.06E-20 | 2.20E-16 | 0.35 | 0.15 | 0.21 | 2.44 |
| 18 | Zdhhc9 | 1.04 | 1.06E-20 | 2.21E-16 | 0.26 | 0.09 | 0.17 | 2.90 |
| 18 | Pisd | 0.91 | 1.25E-20 | 2.60E-16 | 0.45 | 0.21 | 0.25 | 2.18 |
| 18 | Arxes1 | 0.65 | 1.32E-20 | 2.74E-16 | 0.68 | 0.37 | 0.31 | 1.84 |
| 18 | Cptp | 0.94 | 1.42E-20 | 2.96E-16 | 0.33 | 0.13 | 0.20 | 2.52 |
| 18 | Atp1b3 | 0.54 | 1.43E-20 | 2.98E-16 | 0.95 | 0.72 | 0.23 | 1.33 |
| 18 | Srp9 | 0.42 | 1.45E-20 | 3.01E-16 | 0.98 | 0.92 | 0.06 | 1.07 |
| 18 | Zdhhc7 | 0.92 | 1.45E-20 | 3.01E-16 | 0.34 | 0.14 | 0.21 | 2.52 |
| 18 | Tifa | 1.72 | 1.48E-20 | 3.07E-16 | 0.12 | 0.03 | 0.09 | 4.36 |
| 18 | Chn2 | 1.12 | 1.65E-20 | 3.43E-16 | 0.22 | 0.07 | 0.15 | 3.11 |
| 18 | Btbd19 | 1.64 | 1.67E-20 | 3.47E-16 | 0.12 | 0.03 | 0.09 | 4.41 |
| 18 | Gem | 1.27 | 1.80E-20 | 3.75E-16 | 0.15 | 0.04 | 0.11 | 3.85 |
| 18 | Fundc2 | 0.49 | 1.89E-20 | 3.93E-16 | 0.93 | 0.77 | 0.16 | 1.21 |
| 18 | Prdx6 | 0.34 | 1.91E-20 | 3.97E-16 | 1.00 | 0.97 | 0.04 | 1.04 |
| 18 | 2810006K23Rik | 0.70 | 1.97E-20 | 4.09E-16 | 0.66 | 0.35 | 0.31 | 1.89 |
| 18 | Pabpc1l | 1.66 | 2.34E-20 | 4.86E-16 | 0.12 | 0.03 | 0.09 | 4.41 |
| 18 | Nomo1 | 0.82 | 2.43E-20 | 5.06E-16 | 0.44 | 0.20 | 0.25 | 2.26 |
| 18 | Yaf2 | 0.52 | 2.50E-20 | 5.21E-16 | 0.92 | 0.70 | 0.22 | 1.31 |
| 18 | Vamp3 | 0.61 | 2.69E-20 | 5.58E-16 | 0.86 | 0.57 | 0.30 | 1.52 |
| 18 | Gpd1l | 1.26 | 3.42E-20 | 7.12E-16 | 0.22 | 0.07 | 0.15 | 3.07 |
| 18 | Ankrd37 | 1.17 | 3.72E-20 | 7.74E-16 | 0.36 | 0.16 | 0.21 | 2.33 |
| 18 | Prkaa1 | 0.95 | 3.74E-20 | 7.78E-16 | 0.37 | 0.15 | 0.22 | 2.41 |
| 18 | Mapk11 | 1.08 | 3.79E-20 | 7.87E-16 | 0.22 | 0.07 | 0.15 | 3.07 |
| 18 | Pigt | 0.70 | 3.97E-20 | 8.25E-16 | 0.72 | 0.43 | 0.29 | 1.69 |
| 18 | Dynlt3 | 0.77 | 4.87E-20 | 1.01E-15 | 0.52 | 0.26 | 0.27 | 2.04 |
| 18 | Trdmt1 | 1.29 | 5.17E-20 | 1.07E-15 | 0.18 | 0.05 | 0.13 | 3.40 |
| 18 | Cox18 | 0.70 | 5.49E-20 | 1.14E-15 | 0.60 | 0.30 | 0.29 | 1.97 |
| 18 | Tbc1d9b | 0.83 | 5.57E-20 | 1.16E-15 | 0.47 | 0.21 | 0.26 | 2.20 |
| 18 | Rpn2 | 0.48 | 6.63E-20 | 1.38E-15 | 0.96 | 0.79 | 0.17 | 1.22 |
| 18 | Akap13 | 0.64 | 7.15E-20 | 1.49E-15 | 0.81 | 0.50 | 0.31 | 1.62 |
| 18 | Mitd1 | 0.66 | 7.70E-20 | 1.60E-15 | 0.58 | 0.29 | 0.29 | 1.98 |
| 18 | Nme1 | 0.32 | 7.86E-20 | 1.64E-15 | 1.00 | 0.99 | 0.01 | 1.01 |
| 18 | Wdr45b | 0.60 | 8.02E-20 | 1.67E-15 | 0.86 | 0.58 | 0.28 | 1.48 |
| 18 | Tjp2 | 0.66 | 8.04E-20 | 1.67E-15 | 0.61 | 0.31 | 0.30 | 1.97 |
| 18 | Pgs1 | 0.71 | 8.17E-20 | 1.70E-15 | 0.59 | 0.30 | 0.29 | 1.98 |
| 18 | Epn2 | 0.73 | 8.23E-20 | 1.71E-15 | 0.55 | 0.27 | 0.28 | 2.02 |
| 18 | Mapre1 | 0.44 | 8.47E-20 | 1.76E-15 | 0.99 | 0.91 | 0.08 | 1.09 |
| 18 | Man2b1 | 0.63 | 8.50E-20 | 1.77E-15 | 0.49 | 0.23 | 0.27 | 2.16 |
| 18 | Spr | 0.61 | 9.27E-20 | 1.93E-15 | 0.80 | 0.50 | 0.30 | 1.61 |
| 18 | Whamm | 1.06 | 9.34E-20 | 1.94E-15 | 0.27 | 0.10 | 0.17 | 2.72 |
| 18 | Gramd4 | 1.65 | 1.02E-19 | 2.13E-15 | 0.24 | 0.08 | 0.16 | 2.87 |
| 18 | Imp4 | 0.63 | 1.11E-19 | 2.32E-15 | 0.82 | 0.53 | 0.29 | 1.54 |
| 18 | Parp9 | 1.53 | 1.25E-19 | 2.61E-15 | 0.13 | 0.03 | 0.10 | 4.06 |
| 18 | Ankrd50 | 0.77 | 1.27E-19 | 2.63E-15 | 0.48 | 0.23 | 0.26 | 2.12 |
| 18 | Crybg3 | 0.75 | 1.27E-19 | 2.64E-15 | 0.46 | 0.21 | 0.25 | 2.19 |
| 18 | Rrp7a | 0.67 | 1.36E-19 | 2.83E-15 | 0.76 | 0.47 | 0.29 | 1.60 |
| 18 | Slc9a9 | 1.39 | 1.52E-19 | 3.17E-15 | 0.14 | 0.04 | 0.11 | 3.86 |
| 18 | Arhgef12 | 0.58 | 1.67E-19 | 3.48E-15 | 0.86 | 0.58 | 0.29 | 1.50 |
| 18 | Sdf2l1 | 0.70 | 1.82E-19 | 3.79E-15 | 0.77 | 0.47 | 0.30 | 1.64 |
| 18 | Mfsd13a | 1.32 | 1.92E-19 | 3.98E-15 | 0.14 | 0.04 | 0.11 | 3.76 |
| 18 | Pgp | 0.54 | 2.11E-19 | 4.39E-15 | 0.91 | 0.66 | 0.26 | 1.39 |

**Supplementary Table 1. Marker genes for the original clusters of integrated clustering of E11.5 control and NotchGOF λ-junction cells.**

| | | | | | | | | |
|---|---|---|---|---|---|---|---|---|
| 18 | Hmox2 | 0.59 | 2.19E-19 | 4.56E-15 | 0.87 | 0.64 | 0.23 | 1.35 |
| 18 | Herc6 | 1.25 | 2.23E-19 | 4.64E-15 | 0.17 | 0.05 | 0.12 | 3.39 |
| 18 | Plekhh3 | 1.55 | 2.37E-19 | 4.92E-15 | 0.14 | 0.04 | 0.10 | 3.89 |
| 18 | Cln5 | 0.73 | 2.41E-19 | 5.01E-15 | 0.46 | 0.21 | 0.25 | 2.19 |
| 18 | Tomm34 | 0.60 | 2.51E-19 | 5.22E-15 | 0.74 | 0.41 | 0.32 | 1.78 |
| 18 | Lipa | 0.75 | 2.69E-19 | 5.58E-15 | 0.47 | 0.22 | 0.25 | 2.18 |
| 18 | Slc41a1 | 0.81 | 2.87E-19 | 5.97E-15 | 0.35 | 0.14 | 0.21 | 2.45 |
| 18 | Ap2m1 | 0.44 | 2.94E-19 | 6.12E-15 | 0.97 | 0.86 | 0.11 | 1.13 |
| 18 | Edf1 | 0.32 | 3.21E-19 | 6.67E-15 | 0.98 | 0.96 | 0.03 | 1.03 |
| 18 | Mpdu1 | 0.69 | 3.45E-19 | 7.18E-15 | 0.78 | 0.50 | 0.28 | 1.56 |
| 18 | Nck1 | 0.63 | 3.66E-19 | 7.62E-15 | 0.80 | 0.54 | 0.27 | 1.50 |
| 18 | Txndc17 | 0.33 | 3.67E-19 | 7.64E-15 | 0.98 | 0.92 | 0.06 | 1.06 |
| 18 | Hspa4l | 0.90 | 3.69E-19 | 7.67E-15 | 0.29 | 0.11 | 0.18 | 2.60 |
| 18 | Pxk | 0.87 | 4.46E-19 | 9.28E-15 | 0.38 | 0.17 | 0.22 | 2.31 |
| 18 | Pwwp2b | 0.92 | 5.71E-19 | 1.19E-14 | 0.22 | 0.07 | 0.15 | 3.03 |
| 18 | Atp5g1 | 0.31 | 6.26E-19 | 1.30E-14 | 1.00 | 0.99 | 0.01 | 1.01 |
| 18 | Slc22a15 | 0.92 | 6.31E-19 | 1.31E-14 | 0.26 | 0.09 | 0.17 | 2.76 |
| 18 | Mfsd5 | 0.81 | 6.39E-19 | 1.33E-14 | 0.49 | 0.24 | 0.25 | 2.06 |
| 18 | Serhl | 0.68 | 7.29E-19 | 1.52E-14 | 0.39 | 0.17 | 0.22 | 2.34 |
| 18 | Scamp2 | 0.55 | 8.24E-19 | 1.71E-14 | 0.63 | 0.31 | 0.31 | 2.01 |
| 18 | Trmt10a | 0.63 | 9.57E-19 | 1.99E-14 | 0.64 | 0.34 | 0.30 | 1.89 |
| 18 | 2010204K13Rik | 0.84 | 9.62E-19 | 2.00E-14 | 0.50 | 0.25 | 0.25 | 1.99 |
| 18 | Ccny | 0.65 | 1.12E-18 | 2.33E-14 | 0.80 | 0.53 | 0.27 | 1.50 |
| 18 | Nans | 0.57 | 1.32E-18 | 2.74E-14 | 0.82 | 0.60 | 0.22 | 1.36 |
| 18 | Hspa5 | 0.37 | 1.38E-18 | 2.87E-14 | 1.00 | 0.97 | 0.03 | 1.03 |
| 18 | Esyt2 | 0.83 | 1.39E-18 | 2.90E-14 | 0.45 | 0.21 | 0.24 | 2.11 |
| 18 | Cox5a | 0.27 | 1.42E-18 | 2.95E-14 | 1.00 | 0.99 | 0.01 | 1.01 |
| 18 | Cish | 1.11 | 1.44E-18 | 3.01E-14 | 0.19 | 0.06 | 0.13 | 3.13 |
| 18 | Atf7 | 0.66 | 1.54E-18 | 3.21E-14 | 0.44 | 0.20 | 0.24 | 2.23 |
| 18 | Kbtbd2 | 0.62 | 1.59E-18 | 3.30E-14 | 0.63 | 0.33 | 0.29 | 1.89 |
| 18 | Limd1 | 0.59 | 1.76E-18 | 3.66E-14 | 0.81 | 0.51 | 0.30 | 1.58 |
| 18 | Engase | 1.11 | 1.77E-18 | 3.69E-14 | 0.15 | 0.04 | 0.11 | 3.56 |
| 18 | Xiap | 0.63 | 1.80E-18 | 3.74E-14 | 0.77 | 0.49 | 0.28 | 1.58 |
| 18 | Pus1 | 0.71 | 1.95E-18 | 4.05E-14 | 0.57 | 0.29 | 0.27 | 1.94 |
| 18 | Slc16a13 | 1.03 | 2.37E-18 | 4.94E-14 | 0.25 | 0.09 | 0.16 | 2.70 |
| 18 | Gars | 0.53 | 2.60E-18 | 5.40E-14 | 0.91 | 0.67 | 0.24 | 1.35 |
| 18 | Wwc2 | 0.55 | 2.66E-18 | 5.53E-14 | 0.86 | 0.58 | 0.28 | 1.49 |
| 18 | Tnfrsf21 | 0.79 | 2.86E-18 | 5.95E-14 | 0.34 | 0.14 | 0.20 | 2.41 |
| 18 | Inpp4b | 0.51 | 3.09E-18 | 6.42E-14 | 0.37 | 0.15 | 0.22 | 2.41 |
| 18 | Cttnbp2 | 0.80 | 3.11E-18 | 6.46E-14 | 0.36 | 0.15 | 0.21 | 2.34 |
| 18 | Pcbp1 | 0.29 | 3.15E-18 | 6.55E-14 | 0.99 | 0.98 | 0.01 | 1.01 |
| 18 | 4921524J17Rik | 0.65 | 3.20E-18 | 6.67E-14 | 0.75 | 0.49 | 0.26 | 1.54 |
| 18 | Hyal2 | 0.60 | 3.53E-18 | 7.33E-14 | 0.84 | 0.53 | 0.32 | 1.60 |
| 18 | Elf2 | 0.57 | 3.53E-18 | 7.34E-14 | 0.87 | 0.66 | 0.21 | 1.31 |
| 18 | Cep55 | 0.92 | 3.63E-18 | 7.56E-14 | 0.47 | 0.24 | 0.23 | 1.97 |
| 18 | C2cd2l | 1.39 | 3.64E-18 | 7.57E-14 | 0.17 | 0.05 | 0.12 | 3.27 |
| 18 | Dubr | 1.32 | 3.82E-18 | 7.95E-14 | 0.14 | 0.04 | 0.10 | 3.78 |
| 18 | Tdrp | 1.48 | 4.10E-18 | 8.52E-14 | 0.12 | 0.03 | 0.09 | 4.14 |
| 18 | Ran | 0.27 | 4.37E-18 | 9.08E-14 | 1.00 | 1.00 | 0.00 | 1.00 |
| 18 | Vdac1 | 0.33 | 4.39E-18 | 9.14E-14 | 0.98 | 0.95 | 0.03 | 1.04 |
| 18 | Ptpre | 1.10 | 4.48E-18 | 9.32E-14 | 0.14 | 0.04 | 0.10 | 3.78 |
| 18 | Sh3pxd2b | 0.69 | 4.76E-18 | 9.89E-14 | 0.57 | 0.29 | 0.28 | 1.98 |
| 18 | Mtln | 0.60 | 4.79E-18 | 9.96E-14 | 0.62 | 0.33 | 0.29 | 1.89 |
| 18 | Gorasp2 | 0.63 | 5.09E-18 | 1.06E-13 | 0.81 | 0.53 | 0.28 | 1.53 |
| 18 | Creld2 | 0.60 | 5.11E-18 | 1.06E-13 | 0.75 | 0.44 | 0.31 | 1.69 |
| 18 | Creb3 | 0.77 | 5.21E-18 | 1.08E-13 | 0.41 | 0.19 | 0.23 | 2.20 |

**Supplementary Table 1. Marker genes for the original clusters of integrated clustering of E11.5 control and NotchGOF λ-junction cells.**

| | | | | | | | | |
|---|---|---|---|---|---|---|---|---|
| 18 | B130024G19Rik | 1.47 | 5.26E-18 | 1.09E-13 | 0.11 | 0.03 | 0.09 | 4.15 |
| 18 | Stx12 | 0.66 | 5.97E-18 | 1.24E-13 | 0.80 | 0.52 | 0.28 | 1.54 |
| 18 | Tubb4b | 0.45 | 6.08E-18 | 1.26E-13 | 0.97 | 0.85 | 0.12 | 1.14 |
| 18 | Septin2 | 0.76 | 6.10E-18 | 1.27E-13 | 0.40 | 0.17 | 0.22 | 2.27 |
| 18 | Tm2d3 | 0.60 | 6.34E-18 | 1.32E-13 | 0.80 | 0.50 | 0.30 | 1.61 |
| 18 | 2610507B11Rik | 0.57 | 6.51E-18 | 1.35E-13 | 0.76 | 0.46 | 0.30 | 1.65 |
| 18 | Faap100 | 0.81 | 6.54E-18 | 1.36E-13 | 0.49 | 0.24 | 0.25 | 2.05 |
| 18 | Flna | 0.49 | 6.81E-18 | 1.42E-13 | 0.95 | 0.79 | 0.16 | 1.20 |
| 18 | Picalm | 0.55 | 7.16E-18 | 1.49E-13 | 0.86 | 0.62 | 0.24 | 1.39 |
| 18 | Pxdn | 0.64 | 7.50E-18 | 1.56E-13 | 0.81 | 0.51 | 0.30 | 1.58 |
| 18 | Tex261 | 0.60 | 8.26E-18 | 1.72E-13 | 0.86 | 0.60 | 0.26 | 1.44 |
| 18 | Cox5b | 0.26 | 8.27E-18 | 1.72E-13 | 0.99 | 0.99 | 0.01 | 1.01 |
| 18 | Txnrd2 | 0.69 | 8.56E-18 | 1.78E-13 | 0.45 | 0.21 | 0.24 | 2.14 |
| 18 | Gusb | 0.54 | 8.77E-18 | 1.82E-13 | 0.60 | 0.31 | 0.29 | 1.95 |
| 18 | Slc1a5 | 0.81 | 8.86E-18 | 1.84E-13 | 0.43 | 0.20 | 0.22 | 2.11 |
| 18 | Dph5 | 0.77 | 8.89E-18 | 1.85E-13 | 0.46 | 0.23 | 0.24 | 2.06 |
| 18 | Xaf1 | 0.96 | 9.69E-18 | 2.02E-13 | 0.18 | 0.06 | 0.12 | 3.10 |
| 18 | Ppp1r15b | 0.64 | 9.72E-18 | 2.02E-13 | 0.59 | 0.30 | 0.28 | 1.92 |
| 18 | Gpr180 | 0.66 | 1.14E-17 | 2.38E-13 | 0.58 | 0.30 | 0.29 | 1.97 |
| 18 | Srm | 0.61 | 1.14E-17 | 2.38E-13 | 0.89 | 0.68 | 0.21 | 1.31 |
| 18 | Stox1 | 1.40 | 1.46E-17 | 3.03E-13 | 0.11 | 0.03 | 0.08 | 4.19 |
| 18 | BC029722 | 0.94 | 1.68E-17 | 3.49E-13 | 0.29 | 0.11 | 0.17 | 2.53 |
| 18 | Hmbox1 | 0.76 | 1.88E-17 | 3.91E-13 | 0.60 | 0.33 | 0.28 | 1.85 |
| 18 | Cnn3 | 0.35 | 1.94E-17 | 4.03E-13 | 0.99 | 0.94 | 0.05 | 1.06 |
| 18 | Psen2 | 0.89 | 1.97E-17 | 4.10E-13 | 0.22 | 0.08 | 0.14 | 2.87 |
| 18 | Yif1b | 0.56 | 2.00E-17 | 4.16E-13 | 0.89 | 0.67 | 0.22 | 1.33 |
| 18 | Abl2 | 0.82 | 2.04E-17 | 4.25E-13 | 0.41 | 0.19 | 0.22 | 2.18 |
| 18 | Cyth1 | 0.76 | 2.18E-17 | 4.53E-13 | 0.34 | 0.14 | 0.20 | 2.36 |
| 18 | Micall2 | 1.03 | 2.31E-17 | 4.80E-13 | 0.20 | 0.07 | 0.13 | 2.98 |
| 18 | Fam81a | 1.78 | 2.35E-17 | 4.89E-13 | 0.11 | 0.03 | 0.08 | 4.20 |
| 18 | Gramd1a | 0.58 | 2.35E-17 | 4.89E-13 | 0.50 | 0.24 | 0.26 | 2.09 |
| 18 | Fosb | 1.63 | 2.47E-17 | 5.13E-13 | 0.12 | 0.03 | 0.09 | 3.81 |
| 18 | Pfkfb3 | 0.93 | 2.52E-17 | 5.24E-13 | 0.27 | 0.10 | 0.17 | 2.59 |
| 18 | Zfp961 | 0.81 | 2.55E-17 | 5.31E-13 | 0.38 | 0.17 | 0.21 | 2.24 |
| 18 | Ptpn4 | 0.95 | 2.72E-17 | 5.65E-13 | 0.35 | 0.16 | 0.20 | 2.25 |
| 18 | Prkab1 | 0.70 | 2.78E-17 | 5.78E-13 | 0.60 | 0.33 | 0.27 | 1.84 |
| 18 | Grk5 | 0.83 | 2.79E-17 | 5.81E-13 | 0.30 | 0.12 | 0.18 | 2.49 |
| 18 | Bop1 | 0.63 | 2.81E-17 | 5.85E-13 | 0.67 | 0.37 | 0.30 | 1.82 |
| 18 | Sema6a | 0.55 | 2.90E-17 | 6.04E-13 | 0.77 | 0.44 | 0.33 | 1.74 |
| 18 | Lix1l | 0.56 | 3.03E-17 | 6.29E-13 | 0.68 | 0.38 | 0.31 | 1.81 |
| 18 | Klhl8 | 0.84 | 3.06E-17 | 6.36E-13 | 0.25 | 0.09 | 0.15 | 2.69 |
| 18 | Ginm1 | 0.69 | 3.07E-17 | 6.38E-13 | 0.56 | 0.29 | 0.27 | 1.95 |
| 18 | Tnk2 | 0.85 | 3.27E-17 | 6.79E-13 | 0.33 | 0.14 | 0.19 | 2.35 |
| 18 | Stard3 | 0.66 | 3.37E-17 | 7.01E-13 | 0.48 | 0.23 | 0.25 | 2.06 |
| 18 | Polg | 0.78 | 4.04E-17 | 8.41E-13 | 0.47 | 0.22 | 0.24 | 2.08 |
| 18 | Rab1b | 0.55 | 4.06E-17 | 8.44E-13 | 0.83 | 0.53 | 0.30 | 1.57 |
| 18 | Mtmr6 | 0.76 | 4.47E-17 | 9.31E-13 | 0.32 | 0.13 | 0.19 | 2.41 |
| 18 | L3hypdh | 0.62 | 4.61E-17 | 9.59E-13 | 0.41 | 0.19 | 0.23 | 2.23 |
| 18 | Uqcrb | 0.26 | 4.71E-17 | 9.81E-13 | 0.99 | 0.98 | 0.01 | 1.01 |
| 18 | Efhd2 | 0.51 | 4.81E-17 | 1.00E-12 | 0.43 | 0.20 | 0.23 | 2.13 |
| 18 | Ski | 0.59 | 4.85E-17 | 1.01E-12 | 0.71 | 0.41 | 0.30 | 1.74 |
| 18 | Mdga2 | 0.79 | 5.06E-17 | 1.05E-12 | 0.20 | 0.07 | 0.14 | 2.96 |
| 18 | Zfpm1 | 0.56 | 5.49E-17 | 1.14E-12 | 0.59 | 0.31 | 0.27 | 1.88 |
| 18 | Mtcl1 | 0.80 | 5.82E-17 | 1.21E-12 | 0.30 | 0.12 | 0.18 | 2.45 |
| 18 | Cfap298 | 0.56 | 6.02E-17 | 1.25E-12 | 0.65 | 0.36 | 0.29 | 1.79 |
| 18 | Tspan12 | 0.55 | 6.08E-17 | 1.26E-12 | 0.45 | 0.21 | 0.24 | 2.14 |

**Supplementary Table 1. Marker genes for the original clusters of integrated clustering of E11.5 control and NotchGOF λ-junction cells.**

| | | | | | | | | |
|---|---|---|---|---|---|---|---|---|
| 18 | Spindoc | 0.68 | 6.20E-17 | 1.29E-12 | 0.57 | 0.30 | 0.27 | 1.88 |
| 18 | Zfp560 | 0.61 | 6.23E-17 | 1.30E-12 | 0.67 | 0.37 | 0.30 | 1.81 |
| 18 | Sav1 | 0.58 | 6.48E-17 | 1.35E-12 | 0.75 | 0.45 | 0.30 | 1.68 |
| 18 | Tiam1 | 0.57 | 6.57E-17 | 1.37E-12 | 0.59 | 0.31 | 0.28 | 1.90 |
| 18 | Urm1 | 0.60 | 6.75E-17 | 1.40E-12 | 0.66 | 0.37 | 0.29 | 1.77 |
| 18 | Katnbl1 | 0.69 | 7.25E-17 | 1.51E-12 | 0.63 | 0.35 | 0.28 | 1.79 |
| 18 | Sco2 | 0.75 | 8.16E-17 | 1.70E-12 | 0.40 | 0.19 | 0.21 | 2.14 |
| 18 | Rheb | 0.41 | 8.49E-17 | 1.77E-12 | 0.99 | 0.87 | 0.12 | 1.14 |
| 18 | Dock1 | 0.63 | 8.50E-17 | 1.77E-12 | 0.78 | 0.50 | 0.28 | 1.57 |
| 18 | Mrpl57 | 0.47 | 8.80E-17 | 1.83E-12 | 0.94 | 0.78 | 0.17 | 1.22 |
| 18 | Baz1a | 0.50 | 9.02E-17 | 1.88E-12 | 0.83 | 0.56 | 0.28 | 1.50 |
| 18 | Mcur1 | 0.78 | 9.54E-17 | 1.98E-12 | 0.44 | 0.22 | 0.22 | 2.02 |
| 18 | Cpeb4 | 0.90 | 9.69E-17 | 2.02E-12 | 0.25 | 0.09 | 0.15 | 2.61 |
| 18 | Mtmr11 | 0.98 | 9.69E-17 | 2.02E-12 | 0.26 | 0.10 | 0.16 | 2.56 |
| 18 | Svbp | 0.42 | 1.01E-16 | 2.11E-12 | 0.97 | 0.87 | 0.11 | 1.12 |
| 18 | Fam120a | 0.70 | 1.02E-16 | 2.12E-12 | 0.76 | 0.46 | 0.30 | 1.66 |
| 18 | Pip5k1c | 0.62 | 1.03E-16 | 2.15E-12 | 0.47 | 0.23 | 0.25 | 2.10 |
| 18 | Nus1 | 0.60 | 1.03E-16 | 2.15E-12 | 0.76 | 0.48 | 0.27 | 1.56 |
| 18 | Lysmd2 | 0.61 | 1.09E-16 | 2.27E-12 | 0.46 | 0.22 | 0.24 | 2.12 |
| 18 | Macf1 | 0.48 | 1.26E-16 | 2.62E-12 | 0.96 | 0.85 | 0.12 | 1.14 |
| 18 | Crim1 | 0.60 | 1.31E-16 | 2.72E-12 | 0.42 | 0.20 | 0.23 | 2.15 |
| 18 | Tmbim4 | 0.57 | 1.50E-16 | 3.12E-12 | 0.78 | 0.49 | 0.28 | 1.57 |
| 18 | Sipa1l2 | 0.75 | 1.67E-16 | 3.48E-12 | 0.32 | 0.13 | 0.18 | 2.39 |
| 18 | Jph1 | 0.89 | 1.77E-16 | 3.68E-12 | 0.20 | 0.07 | 0.13 | 2.91 |
| 18 | Gnb1 | 0.29 | 1.79E-16 | 3.73E-12 | 0.98 | 0.96 | 0.02 | 1.02 |
| 18 | Inpp1 | 0.66 | 1.79E-16 | 3.73E-12 | 0.37 | 0.16 | 0.21 | 2.26 |
| 18 | Ankrd39 | 0.71 | 1.81E-16 | 3.77E-12 | 0.35 | 0.16 | 0.20 | 2.28 |
| 18 | Rnf213 | 1.71 | 2.03E-16 | 4.23E-12 | 0.11 | 0.03 | 0.08 | 4.04 |
| 18 | Nxpe4 | 1.31 | 2.14E-16 | 4.45E-12 | 0.14 | 0.04 | 0.10 | 3.40 |
| 18 | Luzp1 | 0.47 | 2.15E-16 | 4.47E-12 | 0.94 | 0.77 | 0.17 | 1.23 |
| 18 | Gm12166 | 0.85 | 2.16E-16 | 4.49E-12 | 0.31 | 0.13 | 0.18 | 2.37 |
| 18 | Edem2 | 0.78 | 2.21E-16 | 4.59E-12 | 0.41 | 0.19 | 0.22 | 2.14 |
| 18 | Edem1 | 0.63 | 2.29E-16 | 4.76E-12 | 0.48 | 0.24 | 0.24 | 2.01 |
| 18 | Ccdc32 | 0.60 | 2.30E-16 | 4.79E-12 | 0.58 | 0.31 | 0.27 | 1.86 |
| 18 | F8 | 1.27 | 2.35E-16 | 4.88E-12 | 0.14 | 0.04 | 0.10 | 3.49 |
| 18 | B4galt4 | 0.57 | 2.47E-16 | 5.13E-12 | 0.47 | 0.23 | 0.25 | 2.07 |
| 18 | Ap3m2 | 1.33 | 2.53E-16 | 5.25E-12 | 0.13 | 0.04 | 0.10 | 3.50 |
| 18 | Xylt1 | 1.05 | 2.77E-16 | 5.76E-12 | 0.18 | 0.06 | 0.12 | 3.00 |
| 18 | Faap24 | 0.87 | 2.89E-16 | 6.00E-12 | 0.36 | 0.16 | 0.20 | 2.20 |
| 18 | Pofut2 | 0.68 | 2.95E-16 | 6.14E-12 | 0.76 | 0.48 | 0.28 | 1.59 |
| 18 | Ubxn8 | 0.79 | 3.28E-16 | 6.81E-12 | 0.41 | 0.20 | 0.21 | 2.08 |
| 18 | 2510039O18Rik | 0.68 | 3.32E-16 | 6.90E-12 | 0.47 | 0.23 | 0.24 | 2.04 |
| 18 | Tradd | 0.90 | 3.40E-16 | 7.06E-12 | 0.28 | 0.11 | 0.17 | 2.45 |
| 18 | 2810408I11Rik | 0.61 | 3.43E-16 | 7.12E-12 | 0.57 | 0.30 | 0.27 | 1.90 |
| 18 | Elk4 | 0.65 | 3.59E-16 | 7.47E-12 | 0.47 | 0.23 | 0.24 | 2.03 |
| 18 | Mtbp | 0.61 | 3.69E-16 | 7.68E-12 | 0.45 | 0.22 | 0.23 | 2.06 |
| 18 | Crtc2 | 0.86 | 3.81E-16 | 7.92E-12 | 0.28 | 0.12 | 0.17 | 2.45 |
| 18 | Pttg1ip | 0.63 | 3.87E-16 | 8.05E-12 | 0.69 | 0.39 | 0.30 | 1.77 |
| 18 | Ptpra | 0.54 | 3.92E-16 | 8.15E-12 | 0.86 | 0.60 | 0.26 | 1.44 |
| 18 | Amdhd2 | 0.56 | 3.97E-16 | 8.26E-12 | 0.50 | 0.24 | 0.25 | 2.04 |
| 18 | Ebna1bp2 | 0.44 | 4.16E-16 | 8.65E-12 | 0.94 | 0.79 | 0.15 | 1.18 |
| 18 | Dennd5a | 0.62 | 4.64E-16 | 9.64E-12 | 0.63 | 0.34 | 0.29 | 1.83 |
| 18 | Crlf3 | 0.64 | 4.73E-16 | 9.84E-12 | 0.73 | 0.45 | 0.28 | 1.62 |
| 18 | Bbc3 | 0.95 | 4.94E-16 | 1.03E-11 | 0.26 | 0.10 | 0.15 | 2.52 |
| 18 | Xrcc4 | 0.79 | 5.23E-16 | 1.09E-11 | 0.39 | 0.18 | 0.21 | 2.13 |
| 18 | Tpm3-rs7 | 0.93 | 5.65E-16 | 1.17E-11 | 0.27 | 0.11 | 0.16 | 2.49 |

**Supplementary Table 1. Marker genes for the original clusters of integrated clustering of E11.5 control and NotchGOF λ-junction cells.**

| | | | | | | | | |
|---|---|---|---|---|---|---|---|---|
| 18 | Czib | 0.62 | 6.28E-16 | 1.31E-11 | 0.79 | 0.53 | 0.26 | 1.49 |
| 18 | Brms1 | 0.57 | 6.62E-16 | 1.38E-11 | 0.62 | 0.34 | 0.28 | 1.81 |
| 18 | Ssna1 | 0.39 | 6.81E-16 | 1.42E-11 | 0.96 | 0.84 | 0.11 | 1.13 |
| 18 | Ppp2cb | 0.57 | 7.63E-16 | 1.59E-11 | 0.72 | 0.44 | 0.28 | 1.64 |
| 18 | Sec14l2 | 1.70 | 7.66E-16 | 1.59E-11 | 0.10 | 0.03 | 0.08 | 4.08 |
| 18 | Fbxl12 | 0.60 | 7.76E-16 | 1.61E-11 | 0.42 | 0.20 | 0.22 | 2.12 |
| 18 | Lsm11 | 0.75 | 7.93E-16 | 1.65E-11 | 0.41 | 0.19 | 0.21 | 2.10 |
| 18 | Tmcc1 | 0.57 | 8.07E-16 | 1.68E-11 | 0.74 | 0.44 | 0.30 | 1.69 |
| 18 | Akirin1 | 0.53 | 9.00E-16 | 1.87E-11 | 0.84 | 0.57 | 0.27 | 1.47 |
| 18 | Hdac10 | 1.18 | 9.10E-16 | 1.89E-11 | 0.13 | 0.04 | 0.10 | 3.50 |
| 18 | Plaat3 | 0.53 | 9.18E-16 | 1.91E-11 | 0.46 | 0.22 | 0.24 | 2.09 |
| 18 | Ric8a | 0.56 | 9.31E-16 | 1.94E-11 | 0.50 | 0.25 | 0.25 | 1.98 |
| 18 | Ube2d3 | 0.30 | 9.73E-16 | 2.02E-11 | 1.00 | 0.97 | 0.03 | 1.03 |
| 18 | Fhl3 | 0.89 | 1.00E-15 | 2.08E-11 | 0.26 | 0.11 | 0.16 | 2.47 |
| 18 | Sec24d | 0.73 | 1.02E-15 | 2.11E-11 | 0.22 | 0.08 | 0.14 | 2.73 |
| 18 | Smdt1 | 0.30 | 1.03E-15 | 2.13E-11 | 0.98 | 0.94 | 0.04 | 1.04 |
| 18 | Gosr1 | 0.70 | 1.07E-15 | 2.22E-11 | 0.37 | 0.17 | 0.20 | 2.20 |
| 18 | Cryzl2 | 0.70 | 1.08E-15 | 2.24E-11 | 0.26 | 0.10 | 0.16 | 2.54 |
| 18 | Mmp2 | 0.54 | 1.09E-15 | 2.26E-11 | 0.82 | 0.52 | 0.30 | 1.58 |
| 18 | Ranbp2 | 0.59 | 1.09E-15 | 2.27E-11 | 0.90 | 0.66 | 0.24 | 1.37 |
| 18 | Cd276 | 0.57 | 1.13E-15 | 2.35E-11 | 0.60 | 0.33 | 0.28 | 1.85 |
| 18 | Nlk | 0.54 | 1.14E-15 | 2.37E-11 | 0.53 | 0.27 | 0.26 | 1.94 |
| 18 | Icmt | 0.84 | 1.16E-15 | 2.41E-11 | 0.30 | 0.13 | 0.17 | 2.35 |
| 18 | Mcm3 | 0.50 | 1.21E-15 | 2.51E-11 | 0.95 | 0.79 | 0.16 | 1.21 |
| 18 | Sbds | 0.52 | 1.25E-15 | 2.59E-11 | 0.79 | 0.52 | 0.28 | 1.53 |
| 18 | Kif1c | 0.76 | 1.25E-15 | 2.60E-11 | 0.35 | 0.16 | 0.19 | 2.22 |
| 18 | Sec11a | 0.37 | 1.28E-15 | 2.67E-11 | 0.98 | 0.90 | 0.08 | 1.09 |
| 18 | Cog8 | 0.60 | 1.32E-15 | 2.75E-11 | 0.71 | 0.42 | 0.30 | 1.72 |
| 18 | Nfkbib | 0.65 | 1.36E-15 | 2.82E-11 | 0.48 | 0.24 | 0.24 | 1.98 |
| 18 | Galns | 1.38 | 1.39E-15 | 2.89E-11 | 0.12 | 0.03 | 0.09 | 3.72 |
| 18 | Lrch1 | 0.88 | 1.40E-15 | 2.91E-11 | 0.21 | 0.08 | 0.14 | 2.71 |
| 18 | Stt3b | 0.56 | 1.44E-15 | 2.99E-11 | 0.80 | 0.51 | 0.29 | 1.57 |
| 18 | Cyb5a | 0.28 | 1.45E-15 | 3.02E-11 | 0.98 | 0.92 | 0.06 | 1.06 |
| 18 | Vsig10 | 1.06 | 1.52E-15 | 3.16E-11 | 0.16 | 0.05 | 0.11 | 3.08 |
| 18 | Cycs | 0.32 | 1.58E-15 | 3.28E-11 | 0.99 | 0.96 | 0.03 | 1.03 |
| 18 | Mapre2 | 0.56 | 1.68E-15 | 3.49E-11 | 0.68 | 0.40 | 0.28 | 1.70 |
| 18 | Arhgap17 | 0.54 | 1.70E-15 | 3.53E-11 | 0.65 | 0.37 | 0.29 | 1.77 |
| 18 | Itga3 | 0.83 | 1.75E-15 | 3.64E-11 | 0.19 | 0.07 | 0.13 | 2.85 |
| 18 | Utrn | 0.46 | 1.77E-15 | 3.67E-11 | 0.87 | 0.61 | 0.26 | 1.43 |
| 18 | Atg3 | 0.48 | 1.95E-15 | 4.05E-11 | 0.74 | 0.44 | 0.30 | 1.67 |
| 18 | Uqcc2 | 0.31 | 2.03E-15 | 4.21E-11 | 0.99 | 0.95 | 0.03 | 1.03 |
| 18 | Adipor1 | 0.59 | 2.16E-15 | 4.49E-11 | 0.81 | 0.56 | 0.25 | 1.44 |
| 18 | Sub1 | 0.26 | 2.17E-15 | 4.52E-11 | 1.00 | 0.99 | 0.01 | 1.01 |
| 18 | Znrf2 | 0.64 | 2.24E-15 | 4.66E-11 | 0.39 | 0.18 | 0.21 | 2.14 |
| 18 | Myl7 | 1.49 | 2.36E-15 | 4.91E-11 | 0.11 | 0.03 | 0.08 | 3.89 |
| 18 | Il13ra1 | 1.10 | 2.61E-15 | 5.42E-11 | 0.15 | 0.05 | 0.10 | 3.24 |
| 18 | Nts | 0.90 | 2.67E-15 | 5.55E-11 | 0.17 | 0.06 | 0.11 | 2.98 |
| 18 | Pcbd1 | 0.68 | 2.72E-15 | 5.66E-11 | 0.29 | 0.12 | 0.17 | 2.39 |
| 18 | Fbxo6 | 1.07 | 2.73E-15 | 5.67E-11 | 0.18 | 0.06 | 0.12 | 2.86 |
| 18 | Pvr | 1.05 | 2.75E-15 | 5.72E-11 | 0.19 | 0.07 | 0.12 | 2.84 |
| 18 | Cenpx | 0.40 | 2.83E-15 | 5.88E-11 | 0.93 | 0.79 | 0.14 | 1.18 |
| 18 | Pla2g15 | 1.21 | 2.90E-15 | 6.03E-11 | 0.19 | 0.07 | 0.12 | 2.84 |
| 18 | Sharpin | 0.67 | 2.92E-15 | 6.06E-11 | 0.37 | 0.17 | 0.20 | 2.20 |
| 18 | Ints6l | 0.72 | 2.94E-15 | 6.11E-11 | 0.37 | 0.17 | 0.20 | 2.17 |
| 18 | Fbxw11 | 0.57 | 2.94E-15 | 6.12E-11 | 0.60 | 0.33 | 0.27 | 1.83 |
| 18 | Lbr | 0.64 | 2.96E-15 | 6.15E-11 | 0.78 | 0.50 | 0.28 | 1.55 |

**Supplementary Table 1. Marker genes for the original clusters of integrated clustering of E11.5 control and NotchGOF λ-junction cells.**

| | | | | | | | | |
|---|---|---|---|---|---|---|---|---|
| 18 | Isca1 | 0.58 | 3.09E-15 | 6.43E-11 | 0.58 | 0.31 | 0.27 | 1.85 |
| 18 | Fam160b1 | 0.65 | 3.19E-15 | 6.64E-11 | 0.53 | 0.28 | 0.25 | 1.88 |
| 18 | Fkbp14 | 0.68 | 3.27E-15 | 6.80E-11 | 0.31 | 0.14 | 0.18 | 2.32 |
| 18 | 2510009E07Rik | 0.91 | 3.31E-15 | 6.88E-11 | 0.18 | 0.06 | 0.12 | 2.90 |
| 18 | Trip10 | 0.80 | 3.43E-15 | 7.14E-11 | 0.43 | 0.22 | 0.21 | 2.00 |
| 18 | Socs5 | 0.71 | 3.47E-15 | 7.21E-11 | 0.32 | 0.14 | 0.18 | 2.30 |
| 18 | Glb1 | 0.68 | 3.49E-15 | 7.26E-11 | 0.35 | 0.16 | 0.19 | 2.19 |
| 18 | Trim71 | 0.66 | 3.53E-15 | 7.34E-11 | 0.43 | 0.21 | 0.22 | 2.06 |
| 18 | Mfhas1 | 0.62 | 3.63E-15 | 7.54E-11 | 0.41 | 0.19 | 0.22 | 2.15 |
| 18 | Btbd1 | 0.47 | 3.69E-15 | 7.68E-11 | 0.88 | 0.63 | 0.25 | 1.39 |
| 18 | Grk2 | 0.60 | 3.97E-15 | 8.25E-11 | 0.60 | 0.33 | 0.27 | 1.83 |
| 18 | Ralgps2 | 0.52 | 4.21E-15 | 8.76E-11 | 0.91 | 0.71 | 0.20 | 1.28 |
| 18 | Ccn2 | 1.40 | 4.44E-15 | 9.23E-11 | 0.13 | 0.04 | 0.09 | 3.50 |
| 18 | Rhobtb1 | 0.66 | 4.85E-15 | 1.01E-10 | 0.33 | 0.15 | 0.19 | 2.28 |
| 18 | Pigk | 0.62 | 4.96E-15 | 1.03E-10 | 0.55 | 0.29 | 0.25 | 1.86 |
| 18 | Fuca1 | 0.54 | 5.00E-15 | 1.04E-10 | 0.64 | 0.35 | 0.28 | 1.80 |
| 18 | Pacsin2 | 0.53 | 5.19E-15 | 1.08E-10 | 0.62 | 0.35 | 0.27 | 1.79 |
| 18 | Slc9a1 | 0.70 | 5.32E-15 | 1.11E-10 | 0.27 | 0.11 | 0.16 | 2.45 |
| 18 | Uck2 | 0.56 | 5.62E-15 | 1.17E-10 | 0.80 | 0.52 | 0.27 | 1.52 |
| 18 | Tmem65 | 0.53 | 5.74E-15 | 1.19E-10 | 0.58 | 0.31 | 0.27 | 1.87 |
| 18 | Mindy3 | 0.67 | 5.74E-15 | 1.19E-10 | 0.52 | 0.28 | 0.25 | 1.89 |
| 18 | Zfp287 | 0.88 | 5.90E-15 | 1.23E-10 | 0.20 | 0.07 | 0.13 | 2.79 |
| 18 | Maml2 | 0.75 | 5.97E-15 | 1.24E-10 | 0.24 | 0.09 | 0.15 | 2.56 |
| 18 | Adamts1 | 0.50 | 6.72E-15 | 1.40E-10 | 0.52 | 0.27 | 0.25 | 1.90 |
| 18 | Card19 | 0.55 | 6.77E-15 | 1.41E-10 | 0.49 | 0.25 | 0.24 | 1.96 |
| 18 | Sec14l1 | 0.62 | 7.50E-15 | 1.56E-10 | 0.59 | 0.32 | 0.26 | 1.81 |
| 18 | Sft2d2 | 0.91 | 7.81E-15 | 1.62E-10 | 0.32 | 0.15 | 0.17 | 2.17 |
| 18 | Arap2 | 0.90 | 7.90E-15 | 1.64E-10 | 0.16 | 0.05 | 0.11 | 3.02 |
| 18 | Ctnnbip1 | 0.42 | 8.05E-15 | 1.67E-10 | 0.96 | 0.81 | 0.15 | 1.18 |
| 18 | Slc31a1 | 0.64 | 8.30E-15 | 1.73E-10 | 0.61 | 0.34 | 0.26 | 1.76 |
| 18 | Spcs3 | 0.57 | 9.11E-15 | 1.89E-10 | 0.66 | 0.38 | 0.28 | 1.75 |
| 18 | Cul7 | 0.54 | 9.24E-15 | 1.92E-10 | 0.65 | 0.36 | 0.29 | 1.80 |
| 18 | Sil1 | 0.77 | 9.50E-15 | 1.98E-10 | 0.27 | 0.11 | 0.16 | 2.42 |
| 18 | Cbl | 0.54 | 9.72E-15 | 2.02E-10 | 0.75 | 0.47 | 0.28 | 1.59 |
| 18 | Plpp3 | 0.29 | 9.81E-15 | 2.04E-10 | 0.94 | 0.66 | 0.28 | 1.42 |
| 18 | Frg1 | 0.40 | 1.04E-14 | 2.17E-10 | 0.91 | 0.74 | 0.17 | 1.23 |
| 18 | Rassf4 | 1.07 | 1.06E-14 | 2.21E-10 | 0.11 | 0.03 | 0.08 | 3.76 |
| 18 | Tmem150a | 1.15 | 1.06E-14 | 2.21E-10 | 0.18 | 0.06 | 0.12 | 2.85 |
| 18 | Snap23 | 0.51 | 1.10E-14 | 2.29E-10 | 0.83 | 0.56 | 0.27 | 1.48 |
| 18 | Zmpste24 | 0.58 | 1.11E-14 | 2.31E-10 | 0.58 | 0.31 | 0.26 | 1.84 |
| 18 | Mtpn | 0.36 | 1.18E-14 | 2.46E-10 | 0.94 | 0.81 | 0.13 | 1.16 |
| 18 | A930006K02Rik | 1.28 | 1.18E-14 | 2.46E-10 | 0.14 | 0.04 | 0.10 | 3.23 |
| 18 | Pigq | 0.57 | 1.24E-14 | 2.58E-10 | 0.47 | 0.24 | 0.23 | 1.97 |
| 18 | Slc25a25 | 0.66 | 1.45E-14 | 3.01E-10 | 0.38 | 0.18 | 0.20 | 2.11 |
| 18 | Plod3 | 0.55 | 1.51E-14 | 3.15E-10 | 0.60 | 0.32 | 0.27 | 1.85 |
| 18 | Armc6 | 0.63 | 1.56E-14 | 3.25E-10 | 0.32 | 0.14 | 0.18 | 2.29 |
| 18 | Nckap1 | 0.47 | 1.73E-14 | 3.59E-10 | 0.84 | 0.56 | 0.27 | 1.49 |
| 18 | Psma6 | 0.31 | 1.73E-14 | 3.60E-10 | 0.99 | 0.94 | 0.05 | 1.05 |
| 18 | Tspo | 0.32 | 1.76E-14 | 3.66E-10 | 0.62 | 0.35 | 0.27 | 1.78 |
| 18 | Tbc1d12 | 0.70 | 1.82E-14 | 3.78E-10 | 0.35 | 0.17 | 0.19 | 2.13 |
| 18 | Smarcd2 | 0.49 | 1.82E-14 | 3.80E-10 | 0.54 | 0.29 | 0.26 | 1.89 |
| 18 | Adamts4 | 0.87 | 1.84E-14 | 3.83E-10 | 0.17 | 0.06 | 0.11 | 2.93 |
| 18 | Nucb2 | 0.73 | 1.89E-14 | 3.93E-10 | 0.34 | 0.15 | 0.19 | 2.21 |
| 18 | Zfp994 | 0.88 | 1.91E-14 | 3.97E-10 | 0.23 | 0.09 | 0.14 | 2.51 |
| 18 | Ammecr1 | 0.68 | 1.94E-14 | 4.04E-10 | 0.62 | 0.35 | 0.27 | 1.77 |
| 18 | Ano10 | 0.75 | 1.95E-14 | 4.06E-10 | 0.29 | 0.13 | 0.17 | 2.31 |

**Supplementary Table 1. Marker genes for the original clusters of integrated clustering of E11.5 control and NotchGOF λ-junction cells.**

| | | | | | | | | |
|---|---|---|---|---|---|---|---|---|
| 18 | Mrps16 | 0.37 | 1.99E-14 | 4.13E-10 | 0.95 | 0.84 | 0.12 | 1.14 |
| 18 | Rab31 | 0.62 | 2.16E-14 | 4.50E-10 | 0.45 | 0.22 | 0.23 | 2.02 |
| 18 | Morc4 | 0.60 | 2.32E-14 | 4.82E-10 | 0.33 | 0.15 | 0.18 | 2.26 |
| 18 | Nudt16 | 0.75 | 2.39E-14 | 4.96E-10 | 0.22 | 0.09 | 0.14 | 2.57 |
| 18 | Lin28b | 0.71 | 2.39E-14 | 4.96E-10 | 0.42 | 0.21 | 0.21 | 2.01 |
| 18 | Shkbp1 | 0.63 | 2.44E-14 | 5.08E-10 | 0.59 | 0.34 | 0.25 | 1.75 |
| 18 | Grb2 | 0.42 | 2.47E-14 | 5.15E-10 | 0.89 | 0.64 | 0.26 | 1.40 |
| 18 | Tusc2 | 0.53 | 2.49E-14 | 5.18E-10 | 0.66 | 0.39 | 0.27 | 1.70 |
| 18 | Amfr | 0.63 | 2.54E-14 | 5.29E-10 | 0.64 | 0.36 | 0.28 | 1.78 |
| 18 | Tmem131 | 0.54 | 2.63E-14 | 5.47E-10 | 0.67 | 0.40 | 0.27 | 1.67 |
| 18 | Traf4 | 0.43 | 2.86E-14 | 5.95E-10 | 0.86 | 0.63 | 0.24 | 1.37 |
| 18 | Sap18 | 0.34 | 2.87E-14 | 5.97E-10 | 0.98 | 0.85 | 0.13 | 1.15 |
| 18 | Rybp | 0.49 | 3.02E-14 | 6.28E-10 | 0.69 | 0.42 | 0.28 | 1.66 |
| 18 | Sirt7 | 0.65 | 3.22E-14 | 6.69E-10 | 0.36 | 0.17 | 0.19 | 2.12 |
| 18 | Otud4 | 0.54 | 3.23E-14 | 6.71E-10 | 0.57 | 0.31 | 0.26 | 1.85 |
| 18 | Tmem30a | 0.56 | 3.26E-14 | 6.78E-10 | 0.78 | 0.51 | 0.27 | 1.53 |
| 18 | Spry4 | 0.66 | 3.61E-14 | 7.50E-10 | 0.32 | 0.14 | 0.18 | 2.22 |
| 18 | Naa60 | 0.63 | 3.69E-14 | 7.68E-10 | 0.49 | 0.25 | 0.24 | 1.97 |
| 18 | Bcap31 | 0.44 | 3.90E-14 | 8.11E-10 | 0.93 | 0.75 | 0.18 | 1.25 |
| 18 | Septin11 | 0.64 | 3.98E-14 | 8.27E-10 | 0.44 | 0.22 | 0.22 | 1.97 |
| 18 | Hdlbp | 0.35 | 4.00E-14 | 8.32E-10 | 0.96 | 0.86 | 0.10 | 1.12 |
| 18 | Sh3bp4 | 0.74 | 4.01E-14 | 8.35E-10 | 0.35 | 0.17 | 0.18 | 2.10 |
| 18 | Dmwd | 0.54 | 4.31E-14 | 8.96E-10 | 0.68 | 0.44 | 0.24 | 1.54 |
| 18 | Arrdc1 | 0.47 | 4.55E-14 | 9.47E-10 | 0.27 | 0.11 | 0.16 | 2.41 |
| 18 | Akip1 | 0.61 | 4.85E-14 | 1.01E-09 | 0.40 | 0.20 | 0.20 | 2.04 |
| 18 | Ankrd13a | 0.56 | 4.89E-14 | 1.02E-09 | 0.53 | 0.29 | 0.25 | 1.85 |
| 18 | Hpn | 1.26 | 4.91E-14 | 1.02E-09 | 0.13 | 0.04 | 0.09 | 3.32 |
| 18 | Wbp2 | 0.52 | 5.06E-14 | 1.05E-09 | 0.52 | 0.28 | 0.24 | 1.88 |
| 18 | Macrod2 | 0.45 | 5.17E-14 | 1.07E-09 | 0.45 | 0.22 | 0.23 | 2.02 |
| 18 | Cttnbp2nl | 0.48 | 5.24E-14 | 1.09E-09 | 0.76 | 0.50 | 0.27 | 1.54 |
| 18 | Nln | 0.56 | 5.29E-14 | 1.10E-09 | 0.53 | 0.29 | 0.25 | 1.86 |
| 18 | Dip2b | 0.58 | 5.32E-14 | 1.11E-09 | 0.60 | 0.34 | 0.26 | 1.77 |
| 18 | Mmp14 | 0.47 | 5.56E-14 | 1.16E-09 | 0.86 | 0.63 | 0.23 | 1.36 |
| 18 | Tmc4 | 0.56 | 5.78E-14 | 1.20E-09 | 0.24 | 0.10 | 0.14 | 2.48 |
| 18 | Agfg1 | 0.57 | 5.80E-14 | 1.21E-09 | 0.83 | 0.59 | 0.24 | 1.41 |
| 18 | Vapb | 0.47 | 6.04E-14 | 1.26E-09 | 0.84 | 0.61 | 0.24 | 1.39 |
| 18 | Spty2d1 | 0.65 | 6.16E-14 | 1.28E-09 | 0.48 | 0.25 | 0.23 | 1.92 |
| 18 | Ddi2 | 0.63 | 6.33E-14 | 1.32E-09 | 0.66 | 0.41 | 0.25 | 1.61 |
| 18 | Epn1 | 0.48 | 6.57E-14 | 1.37E-09 | 0.86 | 0.62 | 0.24 | 1.39 |
| 18 | Tnnt1 | 0.97 | 7.18E-14 | 1.49E-09 | 0.16 | 0.05 | 0.10 | 2.94 |
| 18 | Map2k1 | 0.49 | 7.28E-14 | 1.51E-09 | 0.61 | 0.34 | 0.27 | 1.77 |
| 18 | Fut4 | 1.04 | 7.32E-14 | 1.52E-09 | 0.17 | 0.06 | 0.11 | 2.83 |
| 18 | Acvr1 | 0.64 | 7.45E-14 | 1.55E-09 | 0.40 | 0.19 | 0.20 | 2.06 |
| 18 | Tnip2 | 1.47 | 7.56E-14 | 1.57E-09 | 0.10 | 0.03 | 0.08 | 3.78 |
| 18 | Fnip2 | 0.92 | 7.62E-14 | 1.59E-09 | 0.24 | 0.10 | 0.14 | 2.41 |
| 18 | Dgke | 0.93 | 7.64E-14 | 1.59E-09 | 0.19 | 0.07 | 0.12 | 2.71 |
| 18 | Tmem50a | 0.39 | 7.71E-14 | 1.60E-09 | 0.95 | 0.81 | 0.14 | 1.17 |
| 18 | Pdcl3 | 0.44 | 7.80E-14 | 1.62E-09 | 0.91 | 0.70 | 0.20 | 1.29 |
| 18 | Retreg1 | 0.74 | 8.44E-14 | 1.76E-09 | 0.23 | 0.09 | 0.14 | 2.48 |
| 18 | Uckl1 | 0.89 | 8.50E-14 | 1.77E-09 | 0.27 | 0.12 | 0.15 | 2.30 |
| 18 | Tomm5 | 0.33 | 8.50E-14 | 1.77E-09 | 0.99 | 0.90 | 0.09 | 1.09 |
| 18 | Sgsm3 | 0.95 | 8.67E-14 | 1.80E-09 | 0.24 | 0.10 | 0.14 | 2.39 |
| 18 | Ctsa | 0.55 | 9.03E-14 | 1.88E-09 | 0.71 | 0.43 | 0.28 | 1.64 |
| 18 | Gsr | 0.54 | 9.37E-14 | 1.95E-09 | 0.59 | 0.33 | 0.25 | 1.76 |
| 18 | Dnmt3b | 0.57 | 9.52E-14 | 1.98E-09 | 0.42 | 0.21 | 0.21 | 1.98 |
| 18 | Sgpp1 | 0.72 | 9.57E-14 | 1.99E-09 | 0.25 | 0.10 | 0.15 | 2.43 |

**Supplementary Table 1. Marker genes for the original clusters of integrated clustering of E11.5 control and NotchGOF λ-junction cells.**

| | | | | | | | | |
|---|---|---|---|---|---|---|---|---|
| 18 | Add1 | 0.47 | 9.65E-14 | 2.01E-09 | 0.87 | 0.65 | 0.22 | 1.33 |
| 18 | Gm26532 | 1.11 | 1.00E-13 | 2.08E-09 | 0.13 | 0.04 | 0.09 | 3.32 |
| 18 | Antxr2 | 0.57 | 1.06E-13 | 2.20E-09 | 0.38 | 0.18 | 0.20 | 2.09 |
| 18 | Slc35e1 | 0.69 | 1.06E-13 | 2.21E-09 | 0.35 | 0.17 | 0.19 | 2.09 |
| 18 | Prkab2 | 0.54 | 1.12E-13 | 2.32E-09 | 0.29 | 0.13 | 0.16 | 2.31 |
| 18 | Pcgf5 | 0.82 | 1.14E-13 | 2.37E-09 | 0.22 | 0.09 | 0.13 | 2.53 |
| 18 | Zdhhc17 | 0.54 | 1.18E-13 | 2.46E-09 | 0.49 | 0.25 | 0.23 | 1.92 |
| 18 | Tyw5 | 0.57 | 1.18E-13 | 2.46E-09 | 0.44 | 0.22 | 0.22 | 1.99 |
| 18 | Cstf2 | 0.54 | 1.20E-13 | 2.49E-09 | 0.61 | 0.34 | 0.27 | 1.79 |
| 18 | Nmb | 0.89 | 1.22E-13 | 2.53E-09 | 0.20 | 0.08 | 0.12 | 2.61 |
| 18 | Flii | 0.59 | 1.23E-13 | 2.55E-09 | 0.62 | 0.37 | 0.26 | 1.70 |
| 18 | Cxxc5 | 0.42 | 1.30E-13 | 2.70E-09 | 0.74 | 0.45 | 0.29 | 1.64 |
| 18 | Tomm40l | 0.72 | 1.32E-13 | 2.74E-09 | 0.34 | 0.16 | 0.18 | 2.12 |
| 18 | Fam149a | 0.57 | 1.32E-13 | 2.75E-09 | 0.32 | 0.15 | 0.18 | 2.23 |
| 18 | Jmjd6 | 0.47 | 1.39E-13 | 2.89E-09 | 0.78 | 0.52 | 0.26 | 1.51 |
| 18 | Plpp5 | 0.56 | 1.42E-13 | 2.96E-09 | 0.39 | 0.19 | 0.20 | 2.07 |
| 18 | Nudt14 | 0.51 | 1.53E-13 | 3.18E-09 | 0.68 | 0.40 | 0.28 | 1.70 |
| 18 | Klhl5 | 0.57 | 1.53E-13 | 3.18E-09 | 0.36 | 0.17 | 0.19 | 2.12 |
| 18 | Actr1a | 0.58 | 1.56E-13 | 3.25E-09 | 0.72 | 0.46 | 0.26 | 1.56 |
| 18 | Rnf14 | 0.50 | 1.57E-13 | 3.27E-09 | 0.62 | 0.35 | 0.27 | 1.77 |
| 18 | Gpatch4 | 0.55 | 1.61E-13 | 3.35E-09 | 0.78 | 0.55 | 0.23 | 1.42 |
| 18 | Atp6v0b | 0.39 | 1.67E-13 | 3.48E-09 | 0.92 | 0.73 | 0.18 | 1.25 |
| 18 | Mtmr12 | 0.68 | 1.71E-13 | 3.56E-09 | 0.27 | 0.11 | 0.15 | 2.35 |
| 18 | Lamp2 | 0.47 | 1.80E-13 | 3.75E-09 | 0.83 | 0.56 | 0.26 | 1.47 |
| 18 | Aldoa | 0.31 | 1.82E-13 | 3.79E-09 | 0.99 | 0.95 | 0.04 | 1.04 |
| 18 | Pdk1 | 0.38 | 1.86E-13 | 3.88E-09 | 0.49 | 0.25 | 0.24 | 1.94 |
| 18 | Glrx3 | 0.28 | 1.89E-13 | 3.93E-09 | 0.99 | 0.96 | 0.03 | 1.04 |
| 18 | Atraid | 0.50 | 1.91E-13 | 3.97E-09 | 0.78 | 0.50 | 0.27 | 1.54 |
| 18 | Nhs | 0.53 | 1.94E-13 | 4.03E-09 | 0.59 | 0.34 | 0.25 | 1.75 |
| 18 | Slc38a4 | 0.50 | 1.96E-13 | 4.07E-09 | 0.52 | 0.28 | 0.24 | 1.88 |
| 18 | Pmepa1 | 0.57 | 2.06E-13 | 4.29E-09 | 0.74 | 0.48 | 0.26 | 1.55 |
| 18 | Slc25a24 | 0.58 | 2.23E-13 | 4.63E-09 | 0.61 | 0.35 | 0.26 | 1.75 |
| 18 | Ppt2 | 0.56 | 2.33E-13 | 4.85E-09 | 0.48 | 0.25 | 0.23 | 1.91 |
| 18 | Sh3pxd2a | 0.53 | 2.46E-13 | 5.12E-09 | 0.61 | 0.36 | 0.25 | 1.68 |
| 18 | Ero1l | 0.50 | 2.46E-13 | 5.13E-09 | 0.54 | 0.29 | 0.24 | 1.83 |
| 18 | Ttpal | 0.81 | 2.48E-13 | 5.15E-09 | 0.26 | 0.11 | 0.15 | 2.30 |
| 18 | Pcbd2 | 0.55 | 2.56E-13 | 5.32E-09 | 0.70 | 0.43 | 0.26 | 1.61 |
| 18 | Acap3 | 0.71 | 2.62E-13 | 5.44E-09 | 0.19 | 0.07 | 0.12 | 2.66 |
| 18 | Asl | 0.57 | 2.68E-13 | 5.58E-09 | 0.32 | 0.15 | 0.18 | 2.20 |
| 18 | Bysl | 0.54 | 2.80E-13 | 5.83E-09 | 0.59 | 0.34 | 0.26 | 1.76 |
| 18 | Stard9 | 0.92 | 2.86E-13 | 5.94E-09 | 0.18 | 0.07 | 0.11 | 2.73 |
| 18 | Zdhhc18 | 0.55 | 2.91E-13 | 6.05E-09 | 0.32 | 0.15 | 0.18 | 2.21 |
| 18 | Fndc3b | 0.46 | 2.93E-13 | 6.09E-09 | 0.48 | 0.24 | 0.23 | 1.95 |
| 18 | Ppp1r11 | 0.35 | 2.98E-13 | 6.19E-09 | 0.97 | 0.81 | 0.16 | 1.20 |
| 18 | Gipc2 | 0.68 | 3.42E-13 | 7.11E-09 | 0.17 | 0.06 | 0.11 | 2.75 |
| 18 | Bak1 | 0.54 | 3.54E-13 | 7.36E-09 | 0.67 | 0.41 | 0.27 | 1.65 |
| 18 | Atl3 | 0.58 | 3.56E-13 | 7.40E-09 | 0.43 | 0.22 | 0.21 | 1.96 |
| 18 | Plcb3 | 0.50 | 3.58E-13 | 7.45E-09 | 0.53 | 0.29 | 0.24 | 1.84 |
| 18 | Dip2a | 0.96 | 3.61E-13 | 7.52E-09 | 0.24 | 0.10 | 0.14 | 2.40 |
| 18 | Atp9a | 0.52 | 3.63E-13 | 7.55E-09 | 0.37 | 0.18 | 0.19 | 2.10 |
| 18 | Gm6169 | 1.16 | 3.89E-13 | 8.10E-09 | 0.12 | 0.04 | 0.08 | 3.31 |
| 18 | Cmpk2 | 1.37 | 3.98E-13 | 8.27E-09 | 0.12 | 0.04 | 0.08 | 3.22 |
| 18 | Eif3b | 0.38 | 4.01E-13 | 8.34E-09 | 0.94 | 0.81 | 0.13 | 1.15 |
| 18 | Acadm | 0.49 | 4.12E-13 | 8.57E-09 | 0.59 | 0.33 | 0.26 | 1.78 |
| 18 | Ifngr2 | 0.47 | 4.26E-13 | 8.87E-09 | 0.64 | 0.37 | 0.27 | 1.74 |
| 18 | Zyg11b | 0.51 | 4.41E-13 | 9.18E-09 | 0.60 | 0.34 | 0.25 | 1.74 |

**Supplementary Table 1. Marker genes for the original clusters of integrated clustering of E11.5 control and NotchGOF λ-junction cells.**

| | | | | | | | | |
|---|---|---|---|---|---|---|---|---|
| 18 | Ppp1r12c | 0.54 | 4.45E-13 | 9.25E-09 | 0.56 | 0.32 | 0.24 | 1.75 |
| 18 | Ttc32 | 0.46 | 4.88E-13 | 1.02E-08 | 0.49 | 0.25 | 0.23 | 1.91 |
| 18 | Ppp6c | 0.50 | 5.03E-13 | 1.05E-08 | 0.73 | 0.47 | 0.26 | 1.57 |
| 18 | Uggt1 | 0.60 | 5.08E-13 | 1.06E-08 | 0.62 | 0.35 | 0.27 | 1.76 |
| 18 | Magt1 | 0.46 | 5.16E-13 | 1.07E-08 | 0.94 | 0.75 | 0.19 | 1.25 |
| 18 | Tmem97 | 0.56 | 5.19E-13 | 1.08E-08 | 0.71 | 0.46 | 0.25 | 1.55 |
| 18 | Snap29 | 0.57 | 5.24E-13 | 1.09E-08 | 0.39 | 0.19 | 0.20 | 2.01 |
| 18 | Marcks | 0.26 | 5.25E-13 | 1.09E-08 | 1.00 | 0.97 | 0.03 | 1.03 |
| 18 | Scyl1 | 0.56 | 5.34E-13 | 1.11E-08 | 0.51 | 0.28 | 0.23 | 1.83 |
| 18 | Rnf141 | 0.72 | 5.41E-13 | 1.13E-08 | 0.28 | 0.12 | 0.15 | 2.26 |
| 18 | Rnf13 | 0.51 | 5.62E-13 | 1.17E-08 | 0.47 | 0.24 | 0.23 | 1.94 |
| 18 | Ppp1r15a | 0.61 | 5.64E-13 | 1.17E-08 | 0.46 | 0.25 | 0.22 | 1.87 |
| 18 | Cox7a2l | 0.27 | 5.96E-13 | 1.24E-08 | 0.99 | 0.96 | 0.04 | 1.04 |
| 18 | Synj1 | 0.81 | 6.26E-13 | 1.30E-08 | 0.25 | 0.11 | 0.14 | 2.32 |
| 18 | Mrps6 | 0.45 | 6.41E-13 | 1.33E-08 | 0.86 | 0.63 | 0.23 | 1.37 |
| 18 | Cnih4 | 0.38 | 6.45E-13 | 1.34E-08 | 0.92 | 0.79 | 0.13 | 1.16 |
| 18 | Vps28 | 0.44 | 6.48E-13 | 1.35E-08 | 0.85 | 0.62 | 0.23 | 1.37 |
| 18 | Ralgapa2 | 0.69 | 6.64E-13 | 1.38E-08 | 0.30 | 0.14 | 0.16 | 2.18 |
| 18 | Mvp | 0.87 | 6.86E-13 | 1.43E-08 | 0.17 | 0.06 | 0.11 | 2.78 |
| 18 | Sh3glb2 | 0.55 | 6.91E-13 | 1.44E-08 | 0.58 | 0.32 | 0.26 | 1.82 |
| 18 | Srp19 | 0.32 | 6.93E-13 | 1.44E-08 | 0.97 | 0.88 | 0.09 | 1.11 |
| 18 | Marchf2 | 0.54 | 7.62E-13 | 1.59E-08 | 0.35 | 0.17 | 0.19 | 2.09 |
| 18 | Gm11772 | 0.94 | 7.74E-13 | 1.61E-08 | 0.14 | 0.05 | 0.09 | 3.02 |
| 18 | Arl8a | 0.40 | 7.83E-13 | 1.63E-08 | 0.71 | 0.43 | 0.29 | 1.68 |
| 18 | Kpna4 | 0.41 | 8.75E-13 | 1.82E-08 | 0.84 | 0.61 | 0.24 | 1.39 |
| 18 | Ilk | 0.39 | 8.76E-13 | 1.82E-08 | 0.86 | 0.61 | 0.26 | 1.42 |
| 18 | Pcyox1l | 0.55 | 8.96E-13 | 1.86E-08 | 0.41 | 0.20 | 0.20 | 2.01 |
| 18 | Ciapin1 | 0.42 | 9.12E-13 | 1.90E-08 | 0.91 | 0.73 | 0.17 | 1.23 |
| 18 | Rbfa | 0.44 | 9.36E-13 | 1.95E-08 | 0.56 | 0.31 | 0.25 | 1.81 |
| 18 | Plcl1 | 0.82 | 9.58E-13 | 1.99E-08 | 0.17 | 0.06 | 0.11 | 2.78 |
| 18 | Btg3 | 0.54 | 9.59E-13 | 1.99E-08 | 0.56 | 0.32 | 0.24 | 1.75 |
| 18 | Septin11 | 0.30 | 9.70E-13 | 2.02E-08 | 0.98 | 0.87 | 0.11 | 1.13 |
| 18 | Sgce | 0.66 | 1.04E-12 | 2.16E-08 | 0.55 | 0.32 | 0.23 | 1.72 |
| 18 | Lypla2 | 0.45 | 1.05E-12 | 2.18E-08 | 0.62 | 0.36 | 0.27 | 1.75 |
| 18 | Elovl5 | 0.60 | 1.09E-12 | 2.26E-08 | 0.54 | 0.30 | 0.24 | 1.81 |
| 18 | Arpp19 | 0.26 | 1.15E-12 | 2.38E-08 | 0.99 | 0.95 | 0.04 | 1.05 |
| 18 | Alg14 | 0.59 | 1.16E-12 | 2.42E-08 | 0.55 | 0.31 | 0.24 | 1.77 |
| 18 | Ccm2 | 0.49 | 1.17E-12 | 2.43E-08 | 0.67 | 0.40 | 0.28 | 1.70 |
| 18 | Tmem144 | 1.05 | 1.19E-12 | 2.47E-08 | 0.14 | 0.05 | 0.09 | 2.96 |
| 18 | Atp6ap1 | 0.44 | 1.21E-12 | 2.52E-08 | 0.62 | 0.35 | 0.26 | 1.75 |
| 18 | Trmt61a | 0.69 | 1.23E-12 | 2.56E-08 | 0.31 | 0.15 | 0.17 | 2.13 |
| 18 | Camkk2 | 0.74 | 1.33E-12 | 2.77E-08 | 0.45 | 0.23 | 0.21 | 1.92 |
| 18 | Clcn4 | 0.50 | 1.38E-12 | 2.87E-08 | 0.54 | 0.30 | 0.24 | 1.80 |
| 18 | Dazap2 | 0.43 | 1.43E-12 | 2.97E-08 | 0.83 | 0.57 | 0.26 | 1.45 |
| 18 | Gyg | 0.46 | 1.46E-12 | 3.03E-08 | 0.69 | 0.42 | 0.27 | 1.65 |
| 18 | Slc39a11 | 0.94 | 1.54E-12 | 3.19E-08 | 0.18 | 0.07 | 0.11 | 2.65 |
| 18 | Ptprg | 0.49 | 1.55E-12 | 3.23E-08 | 0.63 | 0.37 | 0.26 | 1.71 |
| 18 | Pgm3 | 0.58 | 1.78E-12 | 3.70E-08 | 0.52 | 0.29 | 0.23 | 1.81 |
| 18 | Selenom | 0.31 | 1.84E-12 | 3.83E-08 | 0.78 | 0.52 | 0.26 | 1.49 |
| 18 | Bmpr2 | 0.41 | 1.87E-12 | 3.89E-08 | 0.81 | 0.55 | 0.26 | 1.47 |
| 18 | Emc3 | 0.48 | 1.89E-12 | 3.93E-08 | 0.83 | 0.55 | 0.28 | 1.51 |
| 18 | Mlx | 0.50 | 1.89E-12 | 3.93E-08 | 0.54 | 0.30 | 0.24 | 1.79 |
| 18 | Fars2 | 0.51 | 2.06E-12 | 4.29E-08 | 0.44 | 0.23 | 0.21 | 1.91 |
| 18 | Smad1 | 0.50 | 2.08E-12 | 4.32E-08 | 0.74 | 0.47 | 0.26 | 1.56 |
| 18 | Nmd3 | 0.52 | 2.16E-12 | 4.49E-08 | 0.70 | 0.45 | 0.25 | 1.55 |
| 18 | Rnf31 | 0.85 | 2.33E-12 | 4.84E-08 | 0.18 | 0.07 | 0.11 | 2.64 |

**Supplementary Table 1. Marker genes for the original clusters of integrated clustering of E11.5 control and NotchGOF λ-junction cells.**

| | | | | | | | | |
|---|---|---|---|---|---|---|---|---|
| 18 | Mcts1 | 0.41 | 2.36E-12 | 4.92E-08 | 0.75 | 0.47 | 0.28 | 1.60 |
| 18 | Neurl2 | 0.75 | 2.36E-12 | 4.92E-08 | 0.18 | 0.07 | 0.11 | 2.64 |
| 18 | Tcirg1 | 0.73 | 2.44E-12 | 5.08E-08 | 0.21 | 0.09 | 0.13 | 2.43 |
| 18 | Slc12a6 | 0.71 | 2.50E-12 | 5.20E-08 | 0.29 | 0.13 | 0.15 | 2.15 |
| 18 | Dapk3 | 0.52 | 2.50E-12 | 5.20E-08 | 0.71 | 0.47 | 0.24 | 1.51 |
| 18 | Tigar | 0.91 | 2.60E-12 | 5.40E-08 | 0.18 | 0.07 | 0.11 | 2.59 |
| 18 | Snx2 | 0.40 | 2.69E-12 | 5.60E-08 | 0.82 | 0.58 | 0.25 | 1.43 |
| 18 | Tatdn3 | 0.77 | 2.73E-12 | 5.68E-08 | 0.24 | 0.11 | 0.14 | 2.30 |
| 18 | Eea1 | 0.43 | 2.99E-12 | 6.21E-08 | 0.68 | 0.43 | 0.26 | 1.61 |
| 18 | Zfp524 | 0.77 | 3.03E-12 | 6.31E-08 | 0.24 | 0.10 | 0.13 | 2.29 |
| 18 | Trabd | 0.60 | 3.07E-12 | 6.39E-08 | 0.65 | 0.42 | 0.23 | 1.55 |
| 18 | Sgf29 | 0.50 | 3.16E-12 | 6.57E-08 | 0.57 | 0.32 | 0.24 | 1.74 |
| 18 | Slc25a32 | 0.75 | 3.20E-12 | 6.65E-08 | 0.21 | 0.09 | 0.13 | 2.45 |
| 18 | Baz2a | 0.46 | 3.21E-12 | 6.67E-08 | 0.65 | 0.38 | 0.26 | 1.68 |
| 18 | Rasa1 | 0.44 | 3.26E-12 | 6.78E-08 | 0.86 | 0.61 | 0.25 | 1.41 |
| 18 | Dera | 0.40 | 3.36E-12 | 6.99E-08 | 0.86 | 0.66 | 0.19 | 1.29 |
| 18 | Fam98a | 0.49 | 3.37E-12 | 7.00E-08 | 0.58 | 0.33 | 0.25 | 1.77 |
| 18 | Prelid3b | 0.43 | 3.39E-12 | 7.04E-08 | 0.86 | 0.65 | 0.21 | 1.33 |
| 18 | Lurap1l | 0.37 | 3.40E-12 | 7.07E-08 | 0.19 | 0.08 | 0.12 | 2.52 |
| 18 | Cacul1 | 0.44 | 3.46E-12 | 7.19E-08 | 0.52 | 0.28 | 0.24 | 1.84 |
| 18 | Shmt2 | 0.50 | 3.56E-12 | 7.41E-08 | 0.58 | 0.34 | 0.24 | 1.72 |
| 18 | Phldb1 | 0.53 | 3.74E-12 | 7.78E-08 | 0.40 | 0.20 | 0.20 | 1.98 |
| 18 | Sema4c | 0.62 | 3.88E-12 | 8.07E-08 | 0.48 | 0.27 | 0.21 | 1.80 |
| 18 | Rcan1 | 0.45 | 3.94E-12 | 8.20E-08 | 0.49 | 0.26 | 0.23 | 1.88 |
| 18 | Septin8 | 0.52 | 3.98E-12 | 8.29E-08 | 0.42 | 0.22 | 0.21 | 1.94 |
| 18 | Armcx3 | 0.49 | 3.99E-12 | 8.31E-08 | 0.67 | 0.43 | 0.25 | 1.57 |
| 18 | Asah1 | 0.48 | 4.16E-12 | 8.66E-08 | 0.63 | 0.37 | 0.27 | 1.72 |
| 18 | Glmp | 0.56 | 4.19E-12 | 8.72E-08 | 0.72 | 0.47 | 0.26 | 1.55 |
| 18 | Ralgds | 0.48 | 4.22E-12 | 8.77E-08 | 0.43 | 0.22 | 0.21 | 1.93 |
| 18 | Rcan2 | 1.37 | 4.26E-12 | 8.85E-08 | 0.10 | 0.03 | 0.07 | 3.40 |
| 18 | Map1lc3b | 0.27 | 4.48E-12 | 9.32E-08 | 0.97 | 0.93 | 0.05 | 1.05 |
| 18 | Med27 | 0.56 | 4.54E-12 | 9.44E-08 | 0.47 | 0.25 | 0.22 | 1.87 |
| 18 | Ncln | 0.50 | 4.56E-12 | 9.48E-08 | 0.63 | 0.37 | 0.26 | 1.70 |
| 18 | Entr1 | 0.45 | 4.57E-12 | 9.50E-08 | 0.71 | 0.43 | 0.28 | 1.64 |
| 18 | Tcta | 0.89 | 4.70E-12 | 9.78E-08 | 0.16 | 0.06 | 0.10 | 2.67 |
| 18 | Mvb12a | 0.44 | 4.71E-12 | 9.80E-08 | 0.72 | 0.45 | 0.27 | 1.59 |
| 18 | Chd7 | 0.37 | 4.83E-12 | 1.00E-07 | 0.89 | 0.65 | 0.24 | 1.36 |
| 18 | Fos | 0.45 | 5.04E-12 | 1.05E-07 | 0.29 | 0.14 | 0.15 | 2.09 |
| 18 | Dnajc18 | 0.48 | 5.06E-12 | 1.05E-07 | 0.36 | 0.18 | 0.18 | 2.04 |
| 18 | Reep5 | 0.26 | 5.20E-12 | 1.08E-07 | 0.96 | 0.78 | 0.17 | 1.22 |
| 18 | Traf5 | 1.32 | 5.29E-12 | 1.10E-07 | 0.12 | 0.04 | 0.08 | 3.05 |
| 18 | Yars | 0.51 | 5.45E-12 | 1.13E-07 | 0.61 | 0.37 | 0.24 | 1.65 |
| 18 | Pter | 0.69 | 5.64E-12 | 1.17E-07 | 0.26 | 0.11 | 0.14 | 2.26 |
| 18 | Serinc5 | 1.21 | 5.73E-12 | 1.19E-07 | 0.15 | 0.06 | 0.10 | 2.73 |
| 18 | Tmem170 | 0.60 | 5.77E-12 | 1.20E-07 | 0.22 | 0.09 | 0.13 | 2.38 |
| 18 | Gpatch3 | 0.84 | 5.94E-12 | 1.23E-07 | 0.18 | 0.07 | 0.11 | 2.60 |
| 18 | Acot8 | 0.55 | 6.20E-12 | 1.29E-07 | 0.31 | 0.15 | 0.17 | 2.11 |
| 18 | Ppid | 0.35 | 6.30E-12 | 1.31E-07 | 0.96 | 0.88 | 0.08 | 1.09 |
| 18 | Lcorl | 0.40 | 6.31E-12 | 1.31E-07 | 0.75 | 0.48 | 0.27 | 1.56 |
| 18 | Ccdc85c | 0.93 | 6.38E-12 | 1.33E-07 | 0.25 | 0.11 | 0.14 | 2.23 |
| 18 | Gas2l3 | 0.70 | 6.64E-12 | 1.38E-07 | 0.35 | 0.18 | 0.17 | 1.96 |
| 18 | Btg1 | 0.37 | 6.67E-12 | 1.39E-07 | 0.94 | 0.79 | 0.15 | 1.19 |
| 18 | Mcu | 0.76 | 6.85E-12 | 1.42E-07 | 0.32 | 0.15 | 0.16 | 2.08 |
| 18 | Zfp423 | 0.53 | 7.31E-12 | 1.52E-07 | 0.40 | 0.21 | 0.19 | 1.94 |
| 18 | Rufy3 | 0.41 | 7.33E-12 | 1.52E-07 | 0.44 | 0.23 | 0.21 | 1.92 |
| 18 | Ifrd2 | 0.52 | 7.77E-12 | 1.61E-07 | 0.57 | 0.33 | 0.24 | 1.73 |

**Supplementary Table 1. Marker genes for the original clusters of integrated clustering of E11.5 control and NotchGOF λ-junction cells.**

| | | | | | | | | |
|---|---|---|---|---|---|---|---|---|
| 18 | Gatb | 0.43 | 8.00E-12 | 1.66E-07 | 0.35 | 0.17 | 0.18 | 2.08 |
| 18 | Dcaf12 | 0.42 | 8.24E-12 | 1.71E-07 | 0.59 | 0.34 | 0.26 | 1.76 |
| 18 | Irf1 | 1.00 | 8.38E-12 | 1.74E-07 | 0.12 | 0.04 | 0.08 | 3.13 |
| 18 | Anln | 0.55 | 8.44E-12 | 1.76E-07 | 0.61 | 0.39 | 0.22 | 1.56 |
| 18 | Sgcb | 0.49 | 8.65E-12 | 1.80E-07 | 0.41 | 0.21 | 0.20 | 1.97 |
| 18 | Bach1 | 0.46 | 8.73E-12 | 1.81E-07 | 0.74 | 0.47 | 0.26 | 1.55 |
| 18 | Cpq | 0.48 | 8.77E-12 | 1.82E-07 | 0.33 | 0.16 | 0.17 | 2.06 |
| 18 | Dnajc2 | 0.32 | 9.03E-12 | 1.88E-07 | 0.97 | 0.86 | 0.11 | 1.13 |
| 18 | Upp2 | 0.43 | 9.19E-12 | 1.91E-07 | 0.36 | 0.18 | 0.18 | 2.02 |
| 18 | Riok3 | 0.44 | 9.50E-12 | 1.98E-07 | 0.75 | 0.50 | 0.25 | 1.50 |
| 18 | Rassf3 | 0.35 | 9.55E-12 | 1.99E-07 | 0.64 | 0.37 | 0.27 | 1.75 |
| 18 | Armcx1 | 0.43 | 9.69E-12 | 2.01E-07 | 0.59 | 0.34 | 0.25 | 1.75 |
| 18 | Setx | 0.43 | 9.99E-12 | 2.08E-07 | 0.71 | 0.45 | 0.27 | 1.60 |
| 18 | 1600002K03Rik | 0.64 | 1.00E-11 | 2.09E-07 | 0.26 | 0.11 | 0.14 | 2.24 |
| 18 | Mfsd1 | 0.46 | 1.12E-11 | 2.33E-07 | 0.63 | 0.36 | 0.27 | 1.74 |
| 18 | Golt1b | 0.46 | 1.15E-11 | 2.40E-07 | 0.73 | 0.45 | 0.28 | 1.61 |
| 18 | Ubc | 0.26 | 1.18E-11 | 2.45E-07 | 0.99 | 0.98 | 0.02 | 1.02 |
| 18 | Psap | 0.36 | 1.19E-11 | 2.48E-07 | 0.86 | 0.62 | 0.24 | 1.39 |
| 18 | Coa3 | 0.31 | 1.21E-11 | 2.51E-07 | 0.97 | 0.90 | 0.07 | 1.08 |
| 18 | B230307C23Rik | 0.58 | 1.21E-11 | 2.51E-07 | 0.33 | 0.16 | 0.17 | 2.04 |
| 18 | Suclg2 | 0.50 | 1.23E-11 | 2.55E-07 | 0.72 | 0.49 | 0.23 | 1.47 |
| 18 | Ogfod2 | 0.73 | 1.30E-11 | 2.70E-07 | 0.23 | 0.10 | 0.13 | 2.33 |
| 18 | Cep68 | 0.42 | 1.32E-11 | 2.74E-07 | 0.52 | 0.29 | 0.24 | 1.83 |
| 18 | Ttc13 | 0.51 | 1.35E-11 | 2.81E-07 | 0.43 | 0.23 | 0.20 | 1.88 |
| 18 | Stam2 | 0.43 | 1.36E-11 | 2.82E-07 | 0.55 | 0.31 | 0.24 | 1.79 |
| 18 | Galnt4 | 1.21 | 1.36E-11 | 2.84E-07 | 0.12 | 0.04 | 0.08 | 2.98 |
| 18 | Pnpt1 | 0.57 | 1.37E-11 | 2.85E-07 | 0.50 | 0.29 | 0.21 | 1.73 |
| 18 | Usp10 | 0.45 | 1.59E-11 | 3.31E-07 | 0.85 | 0.69 | 0.17 | 1.24 |
| 18 | Tor1aip1 | 0.53 | 1.61E-11 | 3.36E-07 | 0.70 | 0.44 | 0.26 | 1.59 |
| 18 | Atf6 | 0.56 | 1.66E-11 | 3.46E-07 | 0.30 | 0.14 | 0.16 | 2.13 |
| 18 | Larp4 | 0.47 | 1.71E-11 | 3.55E-07 | 0.83 | 0.60 | 0.23 | 1.39 |
| 18 | Fpgs | 0.59 | 1.73E-11 | 3.61E-07 | 0.36 | 0.19 | 0.18 | 1.97 |
| 18 | Ptdss1 | 0.54 | 1.74E-11 | 3.63E-07 | 0.39 | 0.20 | 0.19 | 1.93 |
| 18 | Pan3 | 0.50 | 1.77E-11 | 3.68E-07 | 0.58 | 0.33 | 0.25 | 1.76 |
| 18 | Wdr11 | 0.47 | 1.79E-11 | 3.71E-07 | 0.45 | 0.24 | 0.21 | 1.89 |
| 18 | CAAA01118383.1 | 0.50 | 1.84E-11 | 3.83E-07 | 0.43 | 0.23 | 0.20 | 1.87 |
| 18 | Nox4 | 0.66 | 1.89E-11 | 3.92E-07 | 0.20 | 0.08 | 0.12 | 2.43 |
| 18 | Zfp800 | 0.48 | 1.92E-11 | 3.99E-07 | 0.49 | 0.27 | 0.23 | 1.85 |
| 18 | Ablim1 | 0.46 | 1.99E-11 | 4.14E-07 | 0.42 | 0.23 | 0.19 | 1.86 |
| 18 | Rab21 | 0.31 | 2.02E-11 | 4.19E-07 | 0.71 | 0.46 | 0.25 | 1.53 |
| 18 | Prkx | 0.55 | 2.02E-11 | 4.20E-07 | 0.32 | 0.15 | 0.16 | 2.07 |
| 18 | Them4 | 0.69 | 2.02E-11 | 4.21E-07 | 0.27 | 0.13 | 0.14 | 2.13 |
| 18 | Rnf135 | 1.25 | 2.09E-11 | 4.35E-07 | 0.11 | 0.04 | 0.08 | 3.11 |
| 18 | Gna11 | 0.38 | 2.11E-11 | 4.39E-07 | 0.91 | 0.67 | 0.24 | 1.35 |
| 18 | Snn | 0.41 | 2.13E-11 | 4.42E-07 | 0.69 | 0.43 | 0.26 | 1.59 |
| 18 | Apmap | 0.59 | 2.18E-11 | 4.53E-07 | 0.37 | 0.19 | 0.18 | 1.96 |
| 18 | S100a1 | 0.73 | 2.31E-11 | 4.80E-07 | 0.17 | 0.06 | 0.10 | 2.61 |
| 18 | Tmem164 | 0.52 | 2.34E-11 | 4.87E-07 | 0.54 | 0.31 | 0.23 | 1.73 |
| 18 | Ssr2 | 0.26 | 2.36E-11 | 4.92E-07 | 0.99 | 0.95 | 0.04 | 1.04 |
| 18 | Bag1 | 0.27 | 2.37E-11 | 4.92E-07 | 0.98 | 0.92 | 0.06 | 1.07 |
| 18 | Adss | 0.41 | 2.43E-11 | 5.06E-07 | 0.88 | 0.72 | 0.16 | 1.22 |
| 18 | Zfp335 | 0.61 | 2.48E-11 | 5.15E-07 | 0.22 | 0.09 | 0.13 | 2.34 |
| 18 | Mtmr9 | 0.45 | 2.65E-11 | 5.51E-07 | 0.31 | 0.15 | 0.16 | 2.11 |
| 18 | Cit | 0.40 | 2.70E-11 | 5.61E-07 | 0.63 | 0.37 | 0.26 | 1.69 |
| 18 | Tmem248 | 0.49 | 2.71E-11 | 5.64E-07 | 0.59 | 0.35 | 0.24 | 1.68 |
| 18 | Sugt1 | 0.29 | 2.85E-11 | 5.93E-07 | 0.98 | 0.87 | 0.11 | 1.12 |

**Supplementary Table 1. Marker genes for the original clusters of integrated clustering of E11.5 control and NotchGOF λ-junction cells.**

| | | | | | | | | |
|---|---|---|---|---|---|---|---|---|
| 18 | Ebp | 0.45 | 2.88E-11 | 5.98E-07 | 0.65 | 0.40 | 0.25 | 1.62 |
| 18 | Paox | 0.41 | 3.07E-11 | 6.38E-07 | 0.33 | 0.16 | 0.17 | 2.08 |
| 18 | Abhd16a | 0.43 | 3.14E-11 | 6.54E-07 | 0.50 | 0.27 | 0.23 | 1.84 |
| 18 | Cables2 | 0.56 | 3.21E-11 | 6.68E-07 | 0.36 | 0.18 | 0.18 | 1.96 |
| 18 | Herc2 | 0.51 | 3.29E-11 | 6.85E-07 | 0.65 | 0.41 | 0.24 | 1.60 |
| 18 | Ube2g2 | 0.46 | 3.36E-11 | 6.99E-07 | 0.57 | 0.33 | 0.24 | 1.73 |
| 18 | Zbtb11 | 0.49 | 3.36E-11 | 6.99E-07 | 0.43 | 0.23 | 0.20 | 1.87 |
| 18 | Slc17a5 | 0.54 | 3.41E-11 | 7.09E-07 | 0.36 | 0.18 | 0.18 | 2.00 |
| 18 | Arhgap26 | 1.19 | 3.49E-11 | 7.26E-07 | 0.11 | 0.04 | 0.08 | 3.11 |
| 18 | Afg1l | 0.92 | 3.50E-11 | 7.27E-07 | 0.13 | 0.05 | 0.08 | 2.87 |
| 18 | Entpd6 | 0.66 | 3.51E-11 | 7.30E-07 | 0.30 | 0.15 | 0.15 | 2.05 |
| 18 | Plekhm1 | 1.07 | 3.51E-11 | 7.31E-07 | 0.16 | 0.06 | 0.10 | 2.60 |
| 18 | Ppat | 0.42 | 3.53E-11 | 7.34E-07 | 0.75 | 0.50 | 0.25 | 1.49 |
| 18 | Ptpn14 | 0.34 | 3.69E-11 | 7.68E-07 | 0.32 | 0.15 | 0.17 | 2.09 |
| 18 | Ppp2r2a | 0.33 | 3.72E-11 | 7.74E-07 | 0.94 | 0.79 | 0.14 | 1.18 |
| 18 | Fbxl17 | 0.41 | 4.13E-11 | 8.58E-07 | 0.38 | 0.20 | 0.19 | 1.97 |
| 18 | Fh1 | 0.44 | 4.13E-11 | 8.59E-07 | 0.91 | 0.72 | 0.19 | 1.26 |
| 18 | Creld1 | 0.53 | 4.33E-11 | 9.01E-07 | 0.28 | 0.13 | 0.15 | 2.14 |
| 18 | Foxn2 | 0.75 | 4.48E-11 | 9.33E-07 | 0.32 | 0.16 | 0.15 | 1.95 |
| 18 | Fam114a1 | 0.47 | 4.76E-11 | 9.91E-07 | 0.33 | 0.17 | 0.17 | 2.02 |
| 18 | Yipf3 | 0.44 | 4.87E-11 | 1.01E-06 | 0.68 | 0.41 | 0.26 | 1.64 |
| 18 | Klhl21 | 0.94 | 5.01E-11 | 1.04E-06 | 0.14 | 0.05 | 0.09 | 2.78 |
| 18 | Agpat4 | 0.42 | 5.05E-11 | 1.05E-06 | 0.68 | 0.44 | 0.25 | 1.57 |
| 18 | Malt1 | 0.50 | 5.15E-11 | 1.07E-06 | 0.33 | 0.16 | 0.17 | 2.01 |
| 18 | Nat10 | 0.60 | 5.17E-11 | 1.08E-06 | 0.37 | 0.19 | 0.17 | 1.90 |
| 18 | Rad51b | 0.77 | 5.71E-11 | 1.19E-06 | 0.25 | 0.12 | 0.13 | 2.16 |
| 18 | Cyc1 | 0.29 | 5.86E-11 | 1.22E-06 | 0.97 | 0.90 | 0.07 | 1.08 |
| 18 | Eif4h | 0.26 | 5.93E-11 | 1.23E-06 | 0.98 | 0.93 | 0.06 | 1.06 |
| 18 | Dnajc1 | 0.27 | 6.15E-11 | 1.28E-06 | 0.81 | 0.56 | 0.25 | 1.46 |
| 18 | Galk1 | 0.41 | 6.22E-11 | 1.29E-06 | 0.89 | 0.71 | 0.18 | 1.26 |
| 18 | Eva1a | 0.57 | 6.22E-11 | 1.29E-06 | 0.44 | 0.24 | 0.20 | 1.81 |
| 18 | Mcrip2 | 1.48 | 6.29E-11 | 1.31E-06 | 0.17 | 0.07 | 0.10 | 2.46 |
| 18 | Rep15 | 0.76 | 6.32E-11 | 1.31E-06 | 0.11 | 0.04 | 0.08 | 3.03 |
| 18 | Faim | 0.43 | 6.36E-11 | 1.32E-06 | 0.70 | 0.46 | 0.24 | 1.52 |
| 18 | Fdx2 | 0.40 | 6.47E-11 | 1.35E-06 | 0.83 | 0.59 | 0.24 | 1.41 |
| 18 | Ubash3b | 0.30 | 6.74E-11 | 1.40E-06 | 0.44 | 0.24 | 0.21 | 1.87 |
| 18 | Gipc1 | 0.37 | 6.77E-11 | 1.41E-06 | 0.88 | 0.69 | 0.20 | 1.29 |
| 18 | Fbxl7 | 0.34 | 7.31E-11 | 1.52E-06 | 0.72 | 0.46 | 0.26 | 1.56 |
| 18 | Erich1 | 0.37 | 7.32E-11 | 1.52E-06 | 0.44 | 0.23 | 0.20 | 1.87 |
| 18 | Etv6 | 0.57 | 7.53E-11 | 1.57E-06 | 0.48 | 0.28 | 0.20 | 1.73 |
| 18 | Klhdc10 | 0.47 | 7.70E-11 | 1.60E-06 | 0.68 | 0.44 | 0.24 | 1.55 |
| 18 | Nfya | 0.42 | 7.71E-11 | 1.60E-06 | 0.55 | 0.32 | 0.23 | 1.73 |
| 18 | Cln3 | 0.46 | 7.77E-11 | 1.62E-06 | 0.30 | 0.14 | 0.16 | 2.10 |
| 18 | Gabarapl2 | 0.28 | 7.88E-11 | 1.64E-06 | 0.95 | 0.82 | 0.13 | 1.15 |
| 18 | Tmed9 | 0.28 | 7.96E-11 | 1.66E-06 | 1.00 | 0.94 | 0.06 | 1.07 |
| 18 | Sh3kbp1 | 0.48 | 8.02E-11 | 1.67E-06 | 0.42 | 0.22 | 0.20 | 1.88 |
| 18 | Ndufa5 | 0.26 | 8.10E-11 | 1.68E-06 | 0.99 | 0.92 | 0.07 | 1.07 |
| 18 | Heatr5a | 0.57 | 8.12E-11 | 1.69E-06 | 0.36 | 0.19 | 0.17 | 1.92 |
| 18 | Oaz2 | 0.43 | 8.44E-11 | 1.76E-06 | 0.78 | 0.55 | 0.23 | 1.43 |
| 18 | Mad2l1bp | 0.44 | 8.69E-11 | 1.81E-06 | 0.43 | 0.23 | 0.20 | 1.85 |
| 18 | Smurf1 | 0.68 | 8.85E-11 | 1.84E-06 | 0.29 | 0.14 | 0.15 | 2.09 |
| 18 | Tmem106b | 0.45 | 8.87E-11 | 1.85E-06 | 0.48 | 0.26 | 0.21 | 1.80 |
| 18 | Emc2 | 0.38 | 8.98E-11 | 1.87E-06 | 0.83 | 0.59 | 0.24 | 1.41 |
| 18 | Plekho2 | 0.67 | 9.03E-11 | 1.88E-06 | 0.17 | 0.07 | 0.10 | 2.51 |
| 18 | Tpgs1 | 0.44 | 9.16E-11 | 1.90E-06 | 0.67 | 0.43 | 0.24 | 1.55 |
| 18 | Dennd4a | 0.68 | 9.36E-11 | 1.95E-06 | 0.19 | 0.08 | 0.11 | 2.40 |

**Supplementary Table 1. Marker genes for the original clusters of integrated clustering of E11.5 control and NotchGOF λ-junction cells.**

| | | | | | | | | |
|---|---|---|---|---|---|---|---|---|
| 18 | Tpra1 | 0.57 | 9.38E-11 | 1.95E-06 | 0.27 | 0.13 | 0.14 | 2.12 |
| 18 | Polk | 0.76 | 9.53E-11 | 1.98E-06 | 0.26 | 0.12 | 0.14 | 2.18 |
| 18 | A630072M18Rik | 0.71 | 9.63E-11 | 2.00E-06 | 0.23 | 0.10 | 0.13 | 2.22 |
| 18 | Tnfsf12 | 0.74 | 9.65E-11 | 2.01E-06 | 0.17 | 0.07 | 0.10 | 2.47 |
| 18 | Mafb | 0.43 | 9.74E-11 | 2.03E-06 | 0.41 | 0.22 | 0.19 | 1.84 |
| 18 | Alg5 | 0.38 | 9.81E-11 | 2.04E-06 | 0.62 | 0.36 | 0.25 | 1.69 |
| 18 | Phf7 | 0.57 | 9.85E-11 | 2.05E-06 | 0.30 | 0.15 | 0.15 | 2.04 |
| 18 | Aida | 0.45 | 9.92E-11 | 2.06E-06 | 0.73 | 0.50 | 0.23 | 1.47 |
| 18 | Snhg15 | 0.45 | 9.96E-11 | 2.07E-06 | 0.51 | 0.29 | 0.22 | 1.77 |
| 18 | Micall1 | 0.76 | 1.04E-10 | 2.15E-06 | 0.25 | 0.12 | 0.13 | 2.14 |
| 18 | Prps2 | 0.45 | 1.05E-10 | 2.19E-06 | 0.33 | 0.17 | 0.16 | 1.96 |
| 18 | Gopc | 0.43 | 1.07E-10 | 2.22E-06 | 0.55 | 0.32 | 0.23 | 1.74 |
| 18 | Ufl1 | 0.45 | 1.09E-10 | 2.26E-06 | 0.67 | 0.42 | 0.26 | 1.62 |
| 18 | Agap3 | 0.51 | 1.15E-10 | 2.38E-06 | 0.59 | 0.36 | 0.23 | 1.63 |
| 18 | Crtc3 | 0.50 | 1.16E-10 | 2.41E-06 | 0.45 | 0.24 | 0.21 | 1.85 |
| 18 | Gbe1 | 0.56 | 1.17E-10 | 2.44E-06 | 0.23 | 0.10 | 0.13 | 2.26 |
| 18 | Esco1 | 0.42 | 1.19E-10 | 2.48E-06 | 0.43 | 0.23 | 0.20 | 1.85 |
| 18 | Bpnt1 | 0.49 | 1.22E-10 | 2.54E-06 | 0.28 | 0.14 | 0.15 | 2.09 |
| 18 | Mrps18a | 0.35 | 1.23E-10 | 2.55E-06 | 0.91 | 0.73 | 0.18 | 1.25 |
| 18 | Bend7 | 0.69 | 1.23E-10 | 2.56E-06 | 0.19 | 0.08 | 0.11 | 2.37 |
| 18 | Pigm | 0.50 | 1.26E-10 | 2.62E-06 | 0.45 | 0.25 | 0.20 | 1.81 |
| 18 | Sh3gl1 | 0.41 | 1.26E-10 | 2.62E-06 | 0.88 | 0.62 | 0.26 | 1.42 |
| 18 | Chfr | 0.37 | 1.27E-10 | 2.65E-06 | 0.47 | 0.26 | 0.21 | 1.83 |
| 18 | Tmem86a | 0.62 | 1.29E-10 | 2.67E-06 | 0.23 | 0.10 | 0.13 | 2.22 |
| 18 | Nop2 | 0.40 | 1.30E-10 | 2.70E-06 | 0.56 | 0.33 | 0.23 | 1.71 |
| 18 | Retreg2 | 0.37 | 1.33E-10 | 2.76E-06 | 0.51 | 0.29 | 0.23 | 1.80 |
| 18 | Tmem219 | 0.45 | 1.34E-10 | 2.78E-06 | 0.31 | 0.15 | 0.16 | 2.03 |
| 18 | Rwdd2b | 0.74 | 1.38E-10 | 2.86E-06 | 0.25 | 0.12 | 0.13 | 2.16 |
| 18 | Timm10b | 0.35 | 1.38E-10 | 2.87E-06 | 0.94 | 0.85 | 0.09 | 1.11 |
| 18 | Col5a2 | 0.29 | 1.38E-10 | 2.87E-06 | 0.81 | 0.53 | 0.28 | 1.53 |
| 18 | Gm49066 | 0.96 | 1.40E-10 | 2.92E-06 | 0.13 | 0.04 | 0.08 | 2.86 |
| 18 | Rapgef2 | 0.57 | 1.46E-10 | 3.03E-06 | 0.41 | 0.22 | 0.18 | 1.83 |
| 18 | Uba1 | 0.32 | 1.46E-10 | 3.04E-06 | 0.94 | 0.76 | 0.17 | 1.23 |
| 18 | Cars | 0.59 | 1.59E-10 | 3.30E-06 | 0.51 | 0.30 | 0.22 | 1.72 |
| 18 | Nsmce1 | 0.38 | 1.59E-10 | 3.30E-06 | 0.90 | 0.70 | 0.20 | 1.28 |
| 18 | Tbrg4 | 0.64 | 1.63E-10 | 3.39E-06 | 0.35 | 0.19 | 0.17 | 1.89 |
| 18 | Tyk2 | 0.82 | 1.64E-10 | 3.42E-06 | 0.15 | 0.06 | 0.09 | 2.59 |
| 18 | Tcea2 | 0.87 | 1.90E-10 | 3.96E-06 | 0.13 | 0.05 | 0.08 | 2.74 |
| 18 | Ndufa9 | 0.40 | 1.93E-10 | 4.02E-06 | 0.88 | 0.71 | 0.17 | 1.23 |
| 18 | Aldh3a2 | 0.65 | 1.98E-10 | 4.11E-06 | 0.19 | 0.08 | 0.11 | 2.38 |
| 18 | Zfp28 | 0.61 | 2.09E-10 | 4.35E-06 | 0.19 | 0.08 | 0.11 | 2.38 |
| 18 | Dtd2 | 0.34 | 2.10E-10 | 4.37E-06 | 0.57 | 0.32 | 0.24 | 1.75 |
| 18 | Fancg | 0.97 | 2.15E-10 | 4.48E-06 | 0.14 | 0.05 | 0.09 | 2.67 |
| 18 | Prdx4 | 0.30 | 2.16E-10 | 4.48E-06 | 0.99 | 0.89 | 0.09 | 1.11 |
| 18 | Prr13 | 0.28 | 2.20E-10 | 4.58E-06 | 0.66 | 0.40 | 0.26 | 1.65 |
| 18 | Ppp5c | 0.41 | 2.23E-10 | 4.64E-06 | 0.76 | 0.54 | 0.22 | 1.42 |
| 18 | Ddx54 | 0.37 | 2.32E-10 | 4.82E-06 | 0.79 | 0.56 | 0.24 | 1.43 |
| 18 | Tpk1 | 0.68 | 2.32E-10 | 4.84E-06 | 0.15 | 0.06 | 0.09 | 2.63 |
| 18 | Zfp568 | 0.48 | 2.42E-10 | 5.03E-06 | 0.48 | 0.27 | 0.21 | 1.79 |
| 18 | Itgb1bp1 | 0.40 | 2.42E-10 | 5.04E-06 | 0.64 | 0.39 | 0.25 | 1.63 |
| 18 | Pop5 | 0.41 | 2.48E-10 | 5.16E-06 | 0.85 | 0.65 | 0.20 | 1.30 |
| 18 | Trp53inp2 | 0.70 | 2.52E-10 | 5.24E-06 | 0.21 | 0.10 | 0.12 | 2.23 |
| 18 | Emc6 | 0.34 | 2.58E-10 | 5.36E-06 | 0.96 | 0.83 | 0.13 | 1.16 |
| 18 | Acox1 | 0.47 | 2.60E-10 | 5.40E-06 | 0.29 | 0.14 | 0.15 | 2.05 |
| 18 | Nop16 | 0.39 | 2.62E-10 | 5.44E-06 | 0.84 | 0.61 | 0.24 | 1.40 |
| 18 | Agpat3 | 0.48 | 2.65E-10 | 5.52E-06 | 0.47 | 0.26 | 0.21 | 1.78 |

**Supplementary Table 1. Marker genes for the original clusters of integrated clustering of E11.5 control and NotchGOF λ-junction cells.**

| | | | | | | | | |
|---|---|---|---|---|---|---|---|---|
| 18 | Sys1 | 0.38 | 2.73E-10 | 5.67E-06 | 0.88 | 0.70 | 0.18 | 1.26 |
| 18 | Tysnd1 | 0.45 | 2.75E-10 | 5.71E-06 | 0.56 | 0.33 | 0.23 | 1.70 |
| 18 | Zdhhc15 | 0.45 | 2.77E-10 | 5.76E-06 | 0.27 | 0.13 | 0.14 | 2.11 |
| 18 | Smco4 | 0.54 | 2.79E-10 | 5.80E-06 | 0.70 | 0.49 | 0.22 | 1.45 |
| 18 | Snx25 | 0.80 | 2.83E-10 | 5.88E-06 | 0.16 | 0.06 | 0.10 | 2.50 |
| 18 | Gstk1 | 0.54 | 2.92E-10 | 6.07E-06 | 0.34 | 0.18 | 0.16 | 1.93 |
| 18 | Brox | 0.38 | 3.05E-10 | 6.35E-06 | 0.42 | 0.22 | 0.20 | 1.87 |
| 18 | Samm50 | 0.34 | 3.15E-10 | 6.55E-06 | 0.90 | 0.72 | 0.18 | 1.25 |
| 18 | Ropn1l | 0.72 | 3.24E-10 | 6.73E-06 | 0.16 | 0.06 | 0.10 | 2.54 |
| 18 | Mak16 | 0.41 | 3.28E-10 | 6.82E-06 | 0.86 | 0.66 | 0.20 | 1.30 |
| 18 | Yipf6 | 0.55 | 3.30E-10 | 6.86E-06 | 0.37 | 0.19 | 0.18 | 1.93 |
| 18 | Rftn2 | 0.37 | 3.42E-10 | 7.10E-06 | 0.42 | 0.23 | 0.20 | 1.86 |
| 18 | Phlda3 | 0.37 | 3.43E-10 | 7.13E-06 | 0.65 | 0.41 | 0.25 | 1.60 |
| 18 | Pigl | 0.75 | 3.48E-10 | 7.23E-06 | 0.18 | 0.07 | 0.10 | 2.39 |
| 18 | Pik3cb | 0.75 | 3.51E-10 | 7.31E-06 | 0.16 | 0.06 | 0.09 | 2.52 |
| 18 | Gmfb | 0.42 | 3.60E-10 | 7.50E-06 | 0.79 | 0.55 | 0.24 | 1.43 |
| 18 | Txndc11 | 0.48 | 3.62E-10 | 7.53E-06 | 0.38 | 0.20 | 0.18 | 1.88 |
| 18 | 1700017B05Rik | 0.50 | 3.69E-10 | 7.68E-06 | 0.34 | 0.18 | 0.17 | 1.92 |
| 18 | Pnpla2 | 0.67 | 3.74E-10 | 7.78E-06 | 0.20 | 0.09 | 0.11 | 2.28 |
| 18 | Zfp787 | 0.47 | 3.75E-10 | 7.79E-06 | 0.54 | 0.33 | 0.22 | 1.66 |
| 18 | Aen | 0.40 | 3.78E-10 | 7.86E-06 | 0.64 | 0.39 | 0.25 | 1.63 |
| 18 | Ttyh3 | 0.47 | 3.79E-10 | 7.89E-06 | 0.78 | 0.53 | 0.25 | 1.46 |
| 18 | Mprip | 0.37 | 3.84E-10 | 7.99E-06 | 0.91 | 0.73 | 0.18 | 1.25 |
| 18 | Echdc1 | 0.38 | 3.86E-10 | 8.03E-06 | 0.43 | 0.24 | 0.20 | 1.83 |
| 18 | Atg4a | 0.65 | 3.95E-10 | 8.21E-06 | 0.20 | 0.09 | 0.11 | 2.32 |
| 18 | Slc16a6 | 1.10 | 4.01E-10 | 8.34E-06 | 0.14 | 0.05 | 0.09 | 2.57 |
| 18 | Hif3a | 0.67 | 4.09E-10 | 8.51E-06 | 0.20 | 0.09 | 0.11 | 2.26 |
| 18 | Mknk2 | 0.33 | 4.10E-10 | 8.53E-06 | 0.67 | 0.41 | 0.26 | 1.63 |
| 18 | Nup93 | 0.42 | 4.34E-10 | 9.03E-06 | 0.70 | 0.47 | 0.24 | 1.50 |
| 18 | Pcsk7 | 0.49 | 4.36E-10 | 9.06E-06 | 0.29 | 0.14 | 0.15 | 2.04 |
| 18 | Bloc1s4 | 0.36 | 4.38E-10 | 9.11E-06 | 0.57 | 0.32 | 0.25 | 1.77 |
| 18 | Eif2s1 | 0.27 | 4.51E-10 | 9.37E-06 | 0.95 | 0.90 | 0.05 | 1.05 |
| 18 | Pmp22 | 0.52 | 4.75E-10 | 9.89E-06 | 0.35 | 0.18 | 0.17 | 1.91 |
| 18 | Klf9 | 0.92 | 4.76E-10 | 9.89E-06 | 0.13 | 0.05 | 0.08 | 2.71 |
| 18 | St6galnac6 | 0.44 | 4.92E-10 | 1.02E-05 | 0.31 | 0.16 | 0.16 | 1.98 |
| 18 | Acox3 | 0.89 | 5.10E-10 | 1.06E-05 | 0.17 | 0.07 | 0.10 | 2.39 |
| 18 | Nmt1 | 0.40 | 5.15E-10 | 1.07E-05 | 0.83 | 0.61 | 0.22 | 1.36 |
| 18 | Ssr3 | 0.29 | 5.34E-10 | 1.11E-05 | 0.99 | 0.88 | 0.11 | 1.13 |
| 18 | Pop7 | 0.40 | 5.62E-10 | 1.17E-05 | 0.75 | 0.52 | 0.23 | 1.44 |
| 18 | Cep85l | 0.43 | 5.62E-10 | 1.17E-05 | 0.28 | 0.14 | 0.15 | 2.07 |
| 18 | Gcnt2 | 1.10 | 5.69E-10 | 1.18E-05 | 0.13 | 0.05 | 0.08 | 2.68 |
| 18 | Bud31 | 0.28 | 5.92E-10 | 1.23E-05 | 0.96 | 0.83 | 0.13 | 1.16 |
| 18 | Wbp1l | 0.43 | 6.07E-10 | 1.26E-05 | 0.49 | 0.28 | 0.21 | 1.75 |
| 18 | Krcc1 | 0.36 | 6.22E-10 | 1.29E-05 | 0.81 | 0.57 | 0.24 | 1.42 |
| 18 | Mafk | 1.24 | 6.62E-10 | 1.38E-05 | 0.11 | 0.04 | 0.07 | 2.80 |
| 18 | Lbh | 0.37 | 6.75E-10 | 1.40E-05 | 0.38 | 0.20 | 0.18 | 1.88 |
| 18 | Gtpbp6 | 0.27 | 6.78E-10 | 1.41E-05 | 0.48 | 0.26 | 0.22 | 1.83 |
| 18 | Fzd10os | 0.43 | 6.98E-10 | 1.45E-05 | 0.37 | 0.20 | 0.17 | 1.86 |
| 18 | Taf9b | 0.96 | 7.42E-10 | 1.54E-05 | 0.11 | 0.04 | 0.07 | 2.87 |
| 18 | Lamtor5 | 0.29 | 7.61E-10 | 1.58E-05 | 0.95 | 0.79 | 0.16 | 1.20 |
| 18 | Rab7 | 0.33 | 8.11E-10 | 1.69E-05 | 0.87 | 0.69 | 0.19 | 1.27 |
| 18 | Fam174a | 0.35 | 8.30E-10 | 1.73E-05 | 0.53 | 0.31 | 0.23 | 1.75 |
| 18 | Os9 | 0.31 | 8.44E-10 | 1.75E-05 | 0.61 | 0.37 | 0.24 | 1.65 |
| 18 | Sars | 0.35 | 9.25E-10 | 1.92E-05 | 0.86 | 0.71 | 0.15 | 1.21 |
| 18 | Ufc1 | 0.29 | 9.34E-10 | 1.94E-05 | 0.95 | 0.83 | 0.12 | 1.14 |
| 18 | Prdx3 | 0.35 | 9.76E-10 | 2.03E-05 | 0.78 | 0.53 | 0.24 | 1.46 |

**Supplementary Table 1. Marker genes for the original clusters of integrated clustering of E11.5 control and NotchGOF λ-junction cells.**

| | | | | | | | | |
|---|---|---|---|---|---|---|---|---|
| 18 | Slc26a2 | 0.36 | 9.92E-10 | 2.06E-05 | 0.31 | 0.16 | 0.16 | 2.00 |
| 18 | 1600012H06Rik | 0.38 | 9.95E-10 | 2.07E-05 | 0.41 | 0.22 | 0.19 | 1.85 |
| 18 | Kif1b | 0.36 | 1.01E-09 | 2.10E-05 | 0.87 | 0.66 | 0.22 | 1.33 |
| 18 | Rtkn | 0.39 | 1.03E-09 | 2.15E-05 | 0.39 | 0.21 | 0.18 | 1.84 |
| 18 | Tpp2 | 0.38 | 1.04E-09 | 2.16E-05 | 0.83 | 0.60 | 0.23 | 1.39 |
| 18 | Eif6 | 0.29 | 1.04E-09 | 2.17E-05 | 0.94 | 0.87 | 0.07 | 1.08 |
| 18 | Dusp22 | 0.36 | 1.05E-09 | 2.18E-05 | 0.38 | 0.20 | 0.18 | 1.87 |
| 18 | Uqcc3 | 0.37 | 1.06E-09 | 2.20E-05 | 0.85 | 0.67 | 0.19 | 1.28 |
| 18 | Atp6v0d1 | 0.35 | 1.08E-09 | 2.24E-05 | 0.77 | 0.53 | 0.24 | 1.44 |
| 18 | Trappc8 | 0.39 | 1.10E-09 | 2.29E-05 | 0.44 | 0.24 | 0.20 | 1.81 |
| 18 | Rwdd1 | 0.26 | 1.11E-09 | 2.31E-05 | 0.98 | 0.93 | 0.05 | 1.05 |
| 18 | Tsen54 | 0.43 | 1.12E-09 | 2.32E-05 | 0.32 | 0.17 | 0.16 | 1.96 |
| 18 | Sin3b | 0.28 | 1.13E-09 | 2.34E-05 | 0.95 | 0.80 | 0.15 | 1.18 |
| 18 | Gpr108 | 0.49 | 1.16E-09 | 2.42E-05 | 0.31 | 0.16 | 0.15 | 1.96 |
| 18 | Ano6 | 0.42 | 1.18E-09 | 2.45E-05 | 0.44 | 0.25 | 0.20 | 1.80 |
| 18 | Kidins220 | 0.34 | 1.18E-09 | 2.45E-05 | 0.49 | 0.28 | 0.21 | 1.76 |
| 18 | Babam2 | 0.40 | 1.19E-09 | 2.47E-05 | 0.57 | 0.35 | 0.22 | 1.63 |
| 18 | Qsox1 | 0.51 | 1.20E-09 | 2.51E-05 | 0.25 | 0.12 | 0.13 | 2.11 |
| 18 | Dgkd | 0.44 | 1.26E-09 | 2.62E-05 | 0.42 | 0.23 | 0.19 | 1.80 |
| 18 | Rabl3 | 0.30 | 1.26E-09 | 2.62E-05 | 0.36 | 0.19 | 0.17 | 1.93 |
| 18 | Clptm1 | 0.45 | 1.37E-09 | 2.84E-05 | 0.70 | 0.44 | 0.26 | 1.58 |
| 18 | Srd5a3 | 0.48 | 1.48E-09 | 3.08E-05 | 0.38 | 0.21 | 0.17 | 1.82 |
| 18 | Gnpda2 | 0.33 | 1.49E-09 | 3.09E-05 | 0.49 | 0.28 | 0.21 | 1.75 |
| 18 | Trappc6a | 0.34 | 1.61E-09 | 3.34E-05 | 0.77 | 0.52 | 0.25 | 1.48 |
| 18 | Mrps34 | 0.32 | 1.69E-09 | 3.51E-05 | 0.85 | 0.62 | 0.23 | 1.37 |
| 18 | Cdc34 | 0.30 | 1.80E-09 | 3.75E-05 | 0.93 | 0.77 | 0.16 | 1.20 |
| 18 | Btbd9 | 0.36 | 1.82E-09 | 3.78E-05 | 0.50 | 0.29 | 0.21 | 1.72 |
| 18 | Hook3 | 0.34 | 1.83E-09 | 3.81E-05 | 0.87 | 0.66 | 0.21 | 1.32 |
| 18 | Desi1 | 0.40 | 1.84E-09 | 3.83E-05 | 0.61 | 0.38 | 0.23 | 1.61 |
| 18 | Klf3 | 0.50 | 1.85E-09 | 3.84E-05 | 0.74 | 0.50 | 0.24 | 1.47 |
| 18 | Hadha | 0.30 | 1.87E-09 | 3.88E-05 | 0.95 | 0.80 | 0.14 | 1.18 |
| 18 | 4732440D04Rik | 0.47 | 1.97E-09 | 4.11E-05 | 0.22 | 0.10 | 0.12 | 2.18 |
| 18 | Ech1 | 0.35 | 2.06E-09 | 4.28E-05 | 0.75 | 0.51 | 0.24 | 1.46 |
| 18 | Zfp277 | 0.39 | 2.06E-09 | 4.28E-05 | 0.67 | 0.42 | 0.24 | 1.57 |
| 18 | Galnt1 | 0.43 | 2.08E-09 | 4.33E-05 | 0.86 | 0.66 | 0.21 | 1.31 |
| 18 | Slc25a26 | 0.73 | 2.10E-09 | 4.37E-05 | 0.20 | 0.09 | 0.11 | 2.21 |
| 18 | Dda1 | 0.39 | 2.12E-09 | 4.40E-05 | 0.80 | 0.57 | 0.23 | 1.41 |
| 18 | P4ha1 | 0.36 | 2.17E-09 | 4.52E-05 | 0.77 | 0.51 | 0.25 | 1.49 |
| 18 | Rpe | 0.28 | 2.22E-09 | 4.61E-05 | 0.59 | 0.33 | 0.25 | 1.75 |
| 18 | Rabggtb | 0.35 | 2.23E-09 | 4.64E-05 | 0.81 | 0.58 | 0.23 | 1.40 |
| 18 | Gatad2a | 0.35 | 2.24E-09 | 4.65E-05 | 0.86 | 0.67 | 0.19 | 1.28 |
| 18 | Pemt | 0.88 | 2.30E-09 | 4.78E-05 | 0.12 | 0.04 | 0.08 | 2.70 |
| 18 | Esrra | 0.43 | 2.30E-09 | 4.78E-05 | 0.26 | 0.12 | 0.13 | 2.06 |
| 18 | Fcho2 | 0.38 | 2.46E-09 | 5.12E-05 | 0.72 | 0.48 | 0.24 | 1.51 |
| 18 | Pkp4 | 0.28 | 2.48E-09 | 5.15E-05 | 0.62 | 0.38 | 0.24 | 1.63 |
| 18 | Immp2l | 0.75 | 2.50E-09 | 5.19E-05 | 0.19 | 0.08 | 0.10 | 2.25 |
| 18 | Ggct | 0.57 | 2.54E-09 | 5.28E-05 | 0.29 | 0.15 | 0.14 | 1.96 |
| 18 | Tom1l2 | 0.95 | 2.59E-09 | 5.39E-05 | 0.13 | 0.05 | 0.08 | 2.56 |
| 18 | Eif1a | 0.34 | 2.63E-09 | 5.47E-05 | 0.86 | 0.67 | 0.19 | 1.29 |
| 18 | Cd1d1 | 0.43 | 2.70E-09 | 5.61E-05 | 0.32 | 0.16 | 0.16 | 1.95 |
| 18 | Tomm22 | 0.25 | 2.75E-09 | 5.72E-05 | 0.97 | 0.90 | 0.06 | 1.07 |
| 18 | Dcaf6 | 0.70 | 2.87E-09 | 5.96E-05 | 0.18 | 0.08 | 0.10 | 2.24 |
| 18 | Atg101 | 0.38 | 2.87E-09 | 5.97E-05 | 0.74 | 0.49 | 0.25 | 1.51 |
| 18 | Fam98c | 0.59 | 2.92E-09 | 6.06E-05 | 0.24 | 0.12 | 0.12 | 2.05 |
| 18 | Utp4 | 0.55 | 2.92E-09 | 6.07E-05 | 0.52 | 0.31 | 0.22 | 1.72 |
| 18 | Gnl3 | 0.37 | 3.00E-09 | 6.24E-05 | 0.89 | 0.75 | 0.14 | 1.19 |

**Supplementary Table 1. Marker genes for the original clusters of integrated clustering of E11.5 control and NotchGOF λ-junction cells.**

| | | | | | | | | |
|---|---|---|---|---|---|---|---|---|
| 18 | Tbc1d19 | 0.38 | 3.08E-09 | 6.41E-05 | 0.39 | 0.21 | 0.18 | 1.85 |
| 18 | Smim12 | 0.38 | 3.09E-09 | 6.42E-05 | 0.76 | 0.52 | 0.24 | 1.47 |
| 18 | Rpn1 | 0.36 | 3.14E-09 | 6.53E-05 | 0.93 | 0.76 | 0.16 | 1.21 |
| 18 | Marveld2 | 0.72 | 3.16E-09 | 6.58E-05 | 0.15 | 0.06 | 0.09 | 2.43 |
| 18 | Mtg1 | 0.39 | 3.34E-09 | 6.94E-05 | 0.29 | 0.15 | 0.15 | 1.98 |
| 18 | Ogfod3 | 0.52 | 3.49E-09 | 7.25E-05 | 0.28 | 0.14 | 0.14 | 1.99 |
| 18 | Btd | 0.47 | 3.65E-09 | 7.59E-05 | 0.25 | 0.12 | 0.13 | 2.07 |
| 18 | Ypel2 | 0.49 | 3.71E-09 | 7.73E-05 | 0.26 | 0.13 | 0.13 | 2.02 |
| 18 | Cbx4 | 0.54 | 3.77E-09 | 7.83E-05 | 0.22 | 0.11 | 0.12 | 2.11 |
| 18 | Irf3 | 0.32 | 3.78E-09 | 7.86E-05 | 0.56 | 0.33 | 0.23 | 1.70 |
| 18 | AI504432 | 0.75 | 3.79E-09 | 7.87E-05 | 0.12 | 0.04 | 0.07 | 2.76 |
| 18 | Eif4e2 | 0.25 | 3.83E-09 | 7.96E-05 | 0.95 | 0.86 | 0.09 | 1.10 |
| 18 | Agtrap | 0.50 | 4.06E-09 | 8.45E-05 | 0.29 | 0.15 | 0.14 | 1.95 |
| 18 | Inpp5k | 0.55 | 4.29E-09 | 8.91E-05 | 0.23 | 0.11 | 0.12 | 2.08 |
| 18 | Cib1 | 0.34 | 4.39E-09 | 9.14E-05 | 0.65 | 0.40 | 0.25 | 1.62 |
| 18 | Vps33a | 0.55 | 4.40E-09 | 9.14E-05 | 0.27 | 0.14 | 0.13 | 1.96 |
| 18 | Dohh | 0.42 | 4.50E-09 | 9.35E-05 | 0.74 | 0.52 | 0.22 | 1.43 |
| 18 | Amacr | 0.42 | 4.56E-09 | 9.49E-05 | 0.28 | 0.14 | 0.14 | 1.97 |
| 18 | Phlpp1 | 0.40 | 4.70E-09 | 9.78E-05 | 0.27 | 0.13 | 0.13 | 2.01 |
| 18 | Lepr | 0.95 | 4.75E-09 | 9.89E-05 | 0.12 | 0.05 | 0.08 | 2.60 |
| 18 | Zbtb7a | 0.38 | 4.85E-09 | 1.01E-04 | 0.54 | 0.33 | 0.21 | 1.65 |
| 18 | Trmt1 | 0.43 | 4.93E-09 | 1.02E-04 | 0.68 | 0.43 | 0.25 | 1.58 |
| 18 | Dynlt1a | 0.38 | 4.94E-09 | 1.03E-04 | 0.51 | 0.30 | 0.21 | 1.68 |
| 18 | Arl4d | 0.78 | 4.96E-09 | 1.03E-04 | 0.11 | 0.04 | 0.07 | 2.73 |
| 18 | Actr1b | 0.35 | 5.08E-09 | 1.06E-04 | 0.77 | 0.52 | 0.25 | 1.47 |
| 18 | Tmem243 | 0.31 | 5.16E-09 | 1.07E-04 | 0.39 | 0.21 | 0.18 | 1.85 |
| 18 | Rnf185 | 0.43 | 5.22E-09 | 1.08E-04 | 0.34 | 0.18 | 0.16 | 1.89 |
| 18 | Dph2 | 0.60 | 5.24E-09 | 1.09E-04 | 0.21 | 0.10 | 0.11 | 2.13 |
| 18 | Ankrd13c | 0.35 | 5.36E-09 | 1.11E-04 | 0.43 | 0.24 | 0.19 | 1.77 |
| 18 | Ranbp9 | 0.36 | 5.38E-09 | 1.12E-04 | 0.57 | 0.34 | 0.23 | 1.67 |
| 18 | Xylt2 | 0.52 | 5.44E-09 | 1.13E-04 | 0.24 | 0.11 | 0.12 | 2.09 |
| 18 | Prrg2 | 0.35 | 5.44E-09 | 1.13E-04 | 0.42 | 0.24 | 0.19 | 1.78 |
| 18 | Zfp984 | 0.51 | 5.60E-09 | 1.17E-04 | 0.17 | 0.07 | 0.10 | 2.33 |
| 18 | Eif2b2 | 0.38 | 5.60E-09 | 1.17E-04 | 0.72 | 0.49 | 0.23 | 1.47 |
| 18 | Stn1 | 0.32 | 5.62E-09 | 1.17E-04 | 0.59 | 0.35 | 0.24 | 1.67 |
| 18 | BC037039 | 0.53 | 5.77E-09 | 1.20E-04 | 0.17 | 0.07 | 0.10 | 2.35 |
| 18 | Prtg | 0.54 | 5.88E-09 | 1.22E-04 | 0.32 | 0.17 | 0.15 | 1.88 |
| 18 | Atp6v1a | 0.35 | 6.15E-09 | 1.28E-04 | 0.77 | 0.53 | 0.24 | 1.46 |
| 18 | Snx8 | 0.51 | 6.17E-09 | 1.28E-04 | 0.27 | 0.13 | 0.13 | 1.99 |
| 18 | Ddhd1 | 0.41 | 6.25E-09 | 1.30E-04 | 0.32 | 0.17 | 0.15 | 1.89 |
| 18 | Dhrs13 | 0.44 | 6.28E-09 | 1.31E-04 | 0.35 | 0.19 | 0.16 | 1.85 |
| 18 | Emp3 | 0.35 | 6.39E-09 | 1.33E-04 | 0.27 | 0.14 | 0.13 | 1.98 |
| 18 | Pcyt1a | 0.31 | 6.56E-09 | 1.36E-04 | 0.53 | 0.32 | 0.22 | 1.70 |
| 18 | Klf16 | 0.57 | 6.70E-09 | 1.39E-04 | 0.18 | 0.08 | 0.10 | 2.27 |
| 18 | Naa25 | 0.45 | 6.77E-09 | 1.41E-04 | 0.55 | 0.36 | 0.20 | 1.55 |
| 18 | Ankrd40 | 0.35 | 6.82E-09 | 1.42E-04 | 0.74 | 0.47 | 0.27 | 1.57 |
| 18 | Tmf1 | 0.35 | 6.88E-09 | 1.43E-04 | 0.73 | 0.49 | 0.24 | 1.49 |
| 18 | Mink1 | 0.29 | 6.90E-09 | 1.44E-04 | 0.32 | 0.17 | 0.16 | 1.93 |
| 18 | Laptm4b | 0.32 | 7.00E-09 | 1.46E-04 | 0.79 | 0.55 | 0.24 | 1.43 |
| 18 | Psmc4 | 0.25 | 7.32E-09 | 1.52E-04 | 0.99 | 0.91 | 0.07 | 1.08 |
| 18 | Zpr1 | 0.39 | 7.40E-09 | 1.54E-04 | 0.65 | 0.43 | 0.22 | 1.52 |
| 18 | Pprc1 | 0.29 | 7.41E-09 | 1.54E-04 | 0.45 | 0.25 | 0.20 | 1.79 |
| 18 | Abi1 | 0.42 | 7.56E-09 | 1.57E-04 | 0.61 | 0.39 | 0.22 | 1.58 |
| 18 | Crk | 0.37 | 7.58E-09 | 1.58E-04 | 0.80 | 0.57 | 0.23 | 1.41 |
| 18 | Rnaseh1 | 0.39 | 7.63E-09 | 1.59E-04 | 0.47 | 0.28 | 0.20 | 1.71 |
| 18 | Traf7 | 0.37 | 7.88E-09 | 1.64E-04 | 0.77 | 0.54 | 0.23 | 1.43 |

**Supplementary Table 1. Marker genes for the original clusters of integrated clustering of E11.5 control and NotchGOF λ-junction cells.**

| | | | | | | | | |
|---|---|---|---|---|---|---|---|---|
| 18 | Atg16l1 | 0.42 | 7.90E-09 | 1.64E-04 | 0.37 | 0.20 | 0.17 | 1.84 |
| 18 | Sec24a | 0.35 | 8.04E-09 | 1.67E-04 | 0.33 | 0.17 | 0.16 | 1.90 |
| 18 | Cyb561d2 | 0.46 | 8.17E-09 | 1.70E-04 | 0.32 | 0.17 | 0.15 | 1.87 |
| 18 | Leprotl1 | 0.35 | 8.25E-09 | 1.72E-04 | 0.77 | 0.55 | 0.23 | 1.41 |
| 18 | Prxl2c | 0.39 | 8.59E-09 | 1.79E-04 | 0.48 | 0.29 | 0.19 | 1.67 |
| 18 | Mob3a | 0.50 | 8.61E-09 | 1.79E-04 | 0.30 | 0.16 | 0.15 | 1.92 |
| 18 | Tubd1 | 0.64 | 8.62E-09 | 1.79E-04 | 0.20 | 0.10 | 0.11 | 2.13 |
| 18 | Gbf1 | 0.33 | 8.70E-09 | 1.81E-04 | 0.68 | 0.44 | 0.25 | 1.56 |
| 18 | Mrps11 | 0.30 | 8.73E-09 | 1.81E-04 | 0.90 | 0.70 | 0.20 | 1.28 |
| 18 | Nfkb2 | 0.82 | 9.44E-09 | 1.96E-04 | 0.16 | 0.07 | 0.09 | 2.33 |
| 18 | Vac14 | 0.44 | 9.45E-09 | 1.97E-04 | 0.38 | 0.22 | 0.17 | 1.76 |
| 18 | Usp14 | 0.28 | 9.70E-09 | 2.02E-04 | 0.95 | 0.79 | 0.15 | 1.19 |
| 18 | Pde12 | 0.39 | 9.71E-09 | 2.02E-04 | 0.47 | 0.28 | 0.20 | 1.71 |
| 18 | Shq1 | 0.41 | 9.75E-09 | 2.03E-04 | 0.35 | 0.19 | 0.16 | 1.83 |
| 18 | Aimp2 | 0.41 | 9.80E-09 | 2.04E-04 | 0.74 | 0.53 | 0.21 | 1.39 |
| 18 | Fbxo4 | 0.73 | 9.97E-09 | 2.07E-04 | 0.12 | 0.05 | 0.08 | 2.60 |
| 18 | Dpy19l1 | 0.33 | 1.03E-08 | 2.15E-04 | 0.59 | 0.36 | 0.23 | 1.64 |
| 18 | Trak1 | 0.50 | 1.08E-08 | 2.24E-04 | 0.24 | 0.12 | 0.12 | 2.03 |
| 18 | Nr2f6 | 0.33 | 1.09E-08 | 2.26E-04 | 0.83 | 0.60 | 0.23 | 1.39 |
| 18 | Cenpt | 0.27 | 1.09E-08 | 2.26E-04 | 0.44 | 0.25 | 0.19 | 1.77 |
| 18 | Elmsan1 | 0.42 | 1.09E-08 | 2.26E-04 | 0.24 | 0.12 | 0.12 | 2.04 |
| 18 | Msra | 0.59 | 1.10E-08 | 2.29E-04 | 0.31 | 0.16 | 0.14 | 1.89 |
| 18 | 1110059E24Rik | 0.33 | 1.13E-08 | 2.34E-04 | 0.79 | 0.56 | 0.23 | 1.42 |
| 18 | Rnf215 | 0.33 | 1.17E-08 | 2.43E-04 | 0.44 | 0.25 | 0.19 | 1.74 |
| 18 | Fkbp5 | 0.43 | 1.18E-08 | 2.45E-04 | 0.64 | 0.42 | 0.22 | 1.53 |
| 18 | Idh3a | 0.34 | 1.19E-08 | 2.48E-04 | 0.79 | 0.56 | 0.23 | 1.40 |
| 18 | Abhd17b | 0.41 | 1.21E-08 | 2.51E-04 | 0.59 | 0.37 | 0.22 | 1.61 |
| 18 | Rab43 | 1.08 | 1.21E-08 | 2.52E-04 | 0.15 | 0.07 | 0.09 | 2.32 |
| 18 | Coro1b | 0.62 | 1.24E-08 | 2.58E-04 | 0.16 | 0.07 | 0.09 | 2.32 |
| 18 | Col4a3bp | 0.41 | 1.28E-08 | 2.67E-04 | 0.52 | 0.32 | 0.20 | 1.64 |
| 18 | Tmem165 | 0.30 | 1.30E-08 | 2.70E-04 | 0.95 | 0.80 | 0.14 | 1.18 |
| 18 | Znrf1 | 0.31 | 1.35E-08 | 2.81E-04 | 0.82 | 0.60 | 0.22 | 1.38 |
| 18 | C1d | 0.27 | 1.37E-08 | 2.85E-04 | 0.93 | 0.84 | 0.09 | 1.11 |
| 18 | Ap2a2 | 0.34 | 1.39E-08 | 2.89E-04 | 0.77 | 0.54 | 0.23 | 1.42 |
| 18 | Zcchc2 | 0.56 | 1.41E-08 | 2.93E-04 | 0.20 | 0.09 | 0.11 | 2.16 |
| 18 | Dhrs7b | 0.39 | 1.42E-08 | 2.96E-04 | 0.52 | 0.32 | 0.20 | 1.61 |
| 18 | Llph | 0.28 | 1.45E-08 | 3.01E-04 | 0.97 | 0.86 | 0.11 | 1.13 |
| 18 | Wdr37 | 0.49 | 1.46E-08 | 3.03E-04 | 0.37 | 0.21 | 0.17 | 1.82 |
| 18 | Mcf2l | 0.47 | 1.59E-08 | 3.30E-04 | 0.16 | 0.07 | 0.09 | 2.32 |
| 18 | Lmtk2 | 0.55 | 1.62E-08 | 3.36E-04 | 0.20 | 0.10 | 0.11 | 2.13 |
| 18 | Nt5dc1 | 0.56 | 1.62E-08 | 3.37E-04 | 0.22 | 0.11 | 0.11 | 2.07 |
| 18 | Rab4b | 0.33 | 1.63E-08 | 3.40E-04 | 0.62 | 0.39 | 0.23 | 1.59 |
| 18 | Fam221a | 0.92 | 1.63E-08 | 3.40E-04 | 0.16 | 0.07 | 0.09 | 2.26 |
| 18 | Uqcrc1 | 0.29 | 1.64E-08 | 3.41E-04 | 0.96 | 0.87 | 0.09 | 1.10 |
| 18 | Lactb2 | 0.33 | 1.64E-08 | 3.41E-04 | 0.33 | 0.18 | 0.16 | 1.89 |
| 18 | Alkbh3 | 0.39 | 1.64E-08 | 3.42E-04 | 0.36 | 0.20 | 0.16 | 1.82 |
| 18 | Ppm1m | 0.30 | 1.68E-08 | 3.49E-04 | 0.44 | 0.25 | 0.19 | 1.77 |
| 18 | Ankrd54 | 0.29 | 1.71E-08 | 3.55E-04 | 0.55 | 0.33 | 0.22 | 1.66 |
| 18 | Cactin | 0.42 | 1.71E-08 | 3.55E-04 | 0.51 | 0.32 | 0.20 | 1.63 |
| 18 | Man2c1 | 0.59 | 1.71E-08 | 3.56E-04 | 0.22 | 0.11 | 0.11 | 2.08 |
| 18 | Pum3 | 0.32 | 1.72E-08 | 3.57E-04 | 0.81 | 0.61 | 0.20 | 1.33 |
| 18 | AI413582 | 0.34 | 1.72E-08 | 3.57E-04 | 0.20 | 0.09 | 0.11 | 2.16 |
| 18 | Mtx1 | 0.36 | 1.75E-08 | 3.64E-04 | 0.77 | 0.55 | 0.22 | 1.39 |
| 18 | Ddx31 | 0.42 | 1.76E-08 | 3.65E-04 | 0.25 | 0.13 | 0.13 | 2.00 |
| 18 | Limk2 | 0.31 | 1.77E-08 | 3.69E-04 | 0.35 | 0.20 | 0.16 | 1.81 |
| 18 | Polr3c | 0.32 | 1.78E-08 | 3.70E-04 | 0.52 | 0.31 | 0.21 | 1.69 |

**Supplementary Table 1. Marker genes for the original clusters of integrated clustering of E11.5 control and NotchGOF λ-junction cells.**

| | | | | | | | | |
|---|---|---|---|---|---|---|---|---|
| 18 | Dtx3l | 1.05 | 1.78E-08 | 3.70E-04 | 0.11 | 0.04 | 0.07 | 2.66 |
| 18 | 2410006H16Rik | 0.26 | 1.78E-08 | 3.71E-04 | 0.96 | 0.86 | 0.10 | 1.12 |
| 18 | Mtmr2 | 0.36 | 1.79E-08 | 3.72E-04 | 0.60 | 0.38 | 0.22 | 1.59 |
| 18 | Commd1 | 0.26 | 2.10E-08 | 4.38E-04 | 0.96 | 0.88 | 0.08 | 1.10 |
| 18 | Ifnar1 | 0.36 | 2.21E-08 | 4.60E-04 | 0.56 | 0.35 | 0.22 | 1.62 |
| 18 | Alg8 | 0.44 | 2.24E-08 | 4.65E-04 | 0.28 | 0.15 | 0.13 | 1.91 |
| 18 | R3hcc1l | 0.44 | 2.24E-08 | 4.66E-04 | 0.35 | 0.19 | 0.16 | 1.82 |
| 18 | Trappc5 | 0.34 | 2.26E-08 | 4.71E-04 | 0.86 | 0.66 | 0.20 | 1.30 |
| 18 | 4930503L19Rik | 0.30 | 2.28E-08 | 4.74E-04 | 0.55 | 0.33 | 0.22 | 1.66 |
| 18 | Vrk3 | 0.26 | 2.30E-08 | 4.78E-04 | 0.57 | 0.34 | 0.23 | 1.68 |
| 18 | Slc12a9 | 0.39 | 2.32E-08 | 4.82E-04 | 0.18 | 0.08 | 0.10 | 2.22 |
| 18 | Snx1 | 0.35 | 2.39E-08 | 4.96E-04 | 0.69 | 0.44 | 0.25 | 1.55 |
| 18 | Sdccag8 | 0.47 | 2.40E-08 | 5.00E-04 | 0.25 | 0.13 | 0.12 | 1.98 |
| 18 | Rxylt1 | 0.35 | 2.52E-08 | 5.25E-04 | 0.73 | 0.47 | 0.26 | 1.55 |
| 18 | Pomt2 | 0.63 | 2.58E-08 | 5.37E-04 | 0.19 | 0.09 | 0.10 | 2.11 |
| 18 | Sec23ip | 0.41 | 2.71E-08 | 5.64E-04 | 0.50 | 0.31 | 0.20 | 1.64 |
| 18 | Hikeshi | 0.28 | 2.75E-08 | 5.72E-04 | 0.90 | 0.77 | 0.13 | 1.17 |
| 18 | 2500002B13Rik | 0.96 | 2.76E-08 | 5.73E-04 | 0.12 | 0.05 | 0.07 | 2.53 |
| 18 | Zfp704 | 0.31 | 2.80E-08 | 5.82E-04 | 0.70 | 0.47 | 0.24 | 1.51 |
| 18 | Tusc3 | 0.30 | 2.82E-08 | 5.86E-04 | 0.80 | 0.56 | 0.24 | 1.42 |
| 18 | Trim56 | 0.40 | 2.83E-08 | 5.89E-04 | 0.27 | 0.14 | 0.13 | 1.96 |
| 18 | Zfp931 | 0.31 | 2.84E-08 | 5.90E-04 | 0.35 | 0.19 | 0.16 | 1.83 |
| 18 | Atg7 | 0.41 | 2.86E-08 | 5.95E-04 | 0.25 | 0.13 | 0.12 | 1.98 |
| 18 | Cat | 0.36 | 2.88E-08 | 6.00E-04 | 0.67 | 0.43 | 0.24 | 1.55 |
| 18 | Zc3h7a | 0.33 | 2.92E-08 | 6.06E-04 | 0.54 | 0.34 | 0.21 | 1.60 |
| 18 | Pgm1 | 0.27 | 2.95E-08 | 6.14E-04 | 0.53 | 0.31 | 0.22 | 1.70 |
| 18 | Bcar3 | 0.77 | 2.96E-08 | 6.15E-04 | 0.13 | 0.05 | 0.08 | 2.47 |
| 18 | Rnf165 | 0.47 | 3.02E-08 | 6.28E-04 | 0.42 | 0.25 | 0.17 | 1.70 |
| 18 | Stx1a | 0.32 | 3.03E-08 | 6.30E-04 | 0.20 | 0.09 | 0.11 | 2.14 |
| 18 | Ahnak | 0.46 | 3.17E-08 | 6.58E-04 | 0.65 | 0.46 | 0.19 | 1.41 |
| 18 | Lap3 | 0.32 | 3.17E-08 | 6.60E-04 | 0.85 | 0.67 | 0.18 | 1.27 |
| 18 | Shc2 | 0.53 | 3.30E-08 | 6.86E-04 | 0.18 | 0.08 | 0.10 | 2.17 |
| 18 | Tollip | 0.44 | 3.33E-08 | 6.92E-04 | 0.30 | 0.16 | 0.14 | 1.86 |
| 18 | Map4k3 | 0.37 | 3.38E-08 | 7.04E-04 | 0.46 | 0.28 | 0.19 | 1.67 |
| 18 | Rtn3 | 0.28 | 3.53E-08 | 7.33E-04 | 0.94 | 0.79 | 0.16 | 1.20 |
| 18 | Prkag1 | 0.37 | 3.58E-08 | 7.45E-04 | 0.68 | 0.45 | 0.23 | 1.51 |
| 18 | Dhx33 | 0.32 | 3.59E-08 | 7.46E-04 | 0.28 | 0.14 | 0.14 | 1.94 |
| 18 | Trip11 | 0.29 | 3.63E-08 | 7.55E-04 | 0.74 | 0.49 | 0.25 | 1.50 |
| 18 | Lrch4 | 0.27 | 3.65E-08 | 7.58E-04 | 0.31 | 0.16 | 0.15 | 1.90 |
| 18 | Mrpl38 | 0.33 | 3.70E-08 | 7.69E-04 | 0.85 | 0.64 | 0.21 | 1.32 |
| 18 | Obsl1 | 0.33 | 3.75E-08 | 7.80E-04 | 0.51 | 0.31 | 0.20 | 1.62 |
| 18 | Trip12 | 0.34 | 3.87E-08 | 8.04E-04 | 0.89 | 0.66 | 0.23 | 1.34 |
| 18 | 4930402H24Rik | 0.26 | 4.01E-08 | 8.35E-04 | 0.44 | 0.25 | 0.19 | 1.74 |
| 18 | Nosip | 0.38 | 4.02E-08 | 8.36E-04 | 0.54 | 0.33 | 0.21 | 1.64 |
| 18 | mt-Nd4l | 0.35 | 4.03E-08 | 8.39E-04 | 0.99 | 0.90 | 0.09 | 1.10 |
| 18 | Zfp597 | 1.02 | 4.07E-08 | 8.47E-04 | 0.17 | 0.08 | 0.09 | 2.20 |
| 18 | Tmem128 | 0.33 | 4.17E-08 | 8.67E-04 | 0.85 | 0.63 | 0.22 | 1.34 |
| 18 | Erf | 0.37 | 4.23E-08 | 8.79E-04 | 0.50 | 0.30 | 0.20 | 1.66 |
| 18 | Slc35c1 | 1.18 | 4.25E-08 | 8.85E-04 | 0.11 | 0.04 | 0.07 | 2.63 |
| 18 | Spryd7 | 0.45 | 4.28E-08 | 8.91E-04 | 0.29 | 0.15 | 0.13 | 1.87 |
| 18 | Chek2 | 0.44 | 4.31E-08 | 8.96E-04 | 0.40 | 0.23 | 0.16 | 1.70 |
| 18 | Ptgr2 | 0.37 | 4.43E-08 | 9.22E-04 | 0.28 | 0.15 | 0.13 | 1.89 |
| 18 | Adipor2 | 0.37 | 4.43E-08 | 9.22E-04 | 0.56 | 0.35 | 0.21 | 1.62 |
| 18 | Mybl2 | 0.41 | 4.47E-08 | 9.30E-04 | 0.42 | 0.24 | 0.17 | 1.71 |
| 18 | Mphosph6 | 0.36 | 4.57E-08 | 9.50E-04 | 0.71 | 0.50 | 0.22 | 1.43 |
| 18 | Zfp622 | 0.29 | 4.60E-08 | 9.56E-04 | 0.67 | 0.42 | 0.25 | 1.61 |

**Supplementary Table 1. Marker genes for the original clusters of integrated clustering of E11.5 control and NotchGOF λ-junction cells.**

| | | | | | | | | |
|---|---|---|---|---|---|---|---|---|
| 18 | Pxmp2 | 0.63 | 4.69E-08 | 9.74E-04 | 0.16 | 0.07 | 0.09 | 2.23 |
| 18 | Ugp2 | 0.29 | 4.83E-08 | 1.00E-03 | 0.72 | 0.50 | 0.23 | 1.45 |
| 18 | Slc35a3 | 0.46 | 4.84E-08 | 1.01E-03 | 0.24 | 0.12 | 0.12 | 1.96 |
| 18 | Tmem177 | 0.59 | 4.90E-08 | 1.02E-03 | 0.22 | 0.11 | 0.11 | 2.01 |
| 18 | Med9 | 0.28 | 5.04E-08 | 1.05E-03 | 0.77 | 0.55 | 0.22 | 1.40 |
| 18 | Fam171a1 | 0.28 | 5.17E-08 | 1.08E-03 | 0.54 | 0.33 | 0.22 | 1.66 |
| 18 | Borcs6 | 0.41 | 5.19E-08 | 1.08E-03 | 0.28 | 0.15 | 0.13 | 1.90 |
| 18 | Paxx | 0.46 | 5.28E-08 | 1.10E-03 | 0.23 | 0.11 | 0.11 | 2.00 |
| 18 | Arhgef10 | 0.49 | 5.29E-08 | 1.10E-03 | 0.26 | 0.13 | 0.12 | 1.93 |
| 18 | Mapk1 | 0.33 | 5.31E-08 | 1.10E-03 | 0.84 | 0.65 | 0.19 | 1.30 |
| 18 | Ggact | 0.58 | 5.41E-08 | 1.13E-03 | 0.12 | 0.05 | 0.07 | 2.49 |
| 18 | Atp6v0a1 | 0.43 | 5.43E-08 | 1.13E-03 | 0.35 | 0.20 | 0.15 | 1.78 |
| 18 | Dab2ip | 0.32 | 5.85E-08 | 1.22E-03 | 0.84 | 0.60 | 0.25 | 1.41 |
| 18 | Agpat5 | 0.40 | 6.10E-08 | 1.27E-03 | 0.68 | 0.45 | 0.23 | 1.52 |
| 18 | Pnkd | 0.29 | 6.19E-08 | 1.29E-03 | 0.91 | 0.74 | 0.17 | 1.23 |
| 18 | Ube2f | 0.35 | 6.22E-08 | 1.29E-03 | 0.78 | 0.56 | 0.21 | 1.38 |
| 18 | Cog1 | 0.28 | 6.24E-08 | 1.30E-03 | 0.31 | 0.17 | 0.15 | 1.87 |
| 18 | Hint3 | 0.38 | 6.37E-08 | 1.32E-03 | 0.41 | 0.24 | 0.17 | 1.73 |
| 18 | Ncoa3 | 0.43 | 6.62E-08 | 1.38E-03 | 0.38 | 0.23 | 0.16 | 1.70 |
| 18 | Vps35l | 0.37 | 6.70E-08 | 1.39E-03 | 0.44 | 0.26 | 0.18 | 1.67 |
| 18 | Nrbf2 | 0.32 | 6.70E-08 | 1.39E-03 | 0.67 | 0.43 | 0.23 | 1.54 |
| 18 | Dpp8 | 0.31 | 6.72E-08 | 1.40E-03 | 0.62 | 0.40 | 0.21 | 1.53 |
| 18 | Chml | 0.44 | 6.75E-08 | 1.40E-03 | 0.28 | 0.15 | 0.13 | 1.89 |
| 18 | Drg1 | 0.35 | 6.75E-08 | 1.40E-03 | 0.81 | 0.61 | 0.20 | 1.32 |
| 18 | Arfgef2 | 0.61 | 6.98E-08 | 1.45E-03 | 0.19 | 0.09 | 0.10 | 2.07 |
| 18 | Fads3 | 0.77 | 7.10E-08 | 1.48E-03 | 0.13 | 0.06 | 0.08 | 2.38 |
| 18 | Me2 | 0.35 | 7.29E-08 | 1.52E-03 | 0.68 | 0.45 | 0.23 | 1.51 |
| 18 | Zfp820 | 0.38 | 7.29E-08 | 1.52E-03 | 0.36 | 0.20 | 0.15 | 1.76 |
| 18 | Zzef1 | 0.32 | 7.49E-08 | 1.56E-03 | 0.31 | 0.17 | 0.14 | 1.84 |
| 18 | 1810037I17Rik | 0.28 | 7.71E-08 | 1.60E-03 | 0.94 | 0.83 | 0.11 | 1.13 |
| 18 | Pa2g4 | 0.25 | 7.72E-08 | 1.61E-03 | 0.96 | 0.91 | 0.05 | 1.06 |
| 18 | Nle1 | 0.38 | 7.76E-08 | 1.61E-03 | 0.40 | 0.23 | 0.17 | 1.71 |
| 18 | Pdss1 | 0.48 | 8.01E-08 | 1.67E-03 | 0.30 | 0.16 | 0.14 | 1.84 |
| 18 | Sidt2 | 0.33 | 8.05E-08 | 1.67E-03 | 0.33 | 0.18 | 0.15 | 1.82 |
| 18 | Ncdn | 0.33 | 8.18E-08 | 1.70E-03 | 0.26 | 0.13 | 0.13 | 1.95 |
| 18 | Ogdh | 0.37 | 8.32E-08 | 1.73E-03 | 0.75 | 0.53 | 0.22 | 1.42 |
| 18 | Hgsnat | 0.32 | 8.33E-08 | 1.73E-03 | 0.35 | 0.20 | 0.16 | 1.80 |
| 18 | Fam83d | 0.42 | 8.44E-08 | 1.75E-03 | 0.34 | 0.20 | 0.14 | 1.73 |
| 18 | Tm9sf2 | 0.35 | 8.60E-08 | 1.79E-03 | 0.81 | 0.62 | 0.20 | 1.32 |
| 18 | Dna2 | 0.45 | 8.71E-08 | 1.81E-03 | 0.43 | 0.26 | 0.17 | 1.67 |
| 18 | Pggt1b | 0.29 | 8.74E-08 | 1.82E-03 | 0.77 | 0.49 | 0.27 | 1.56 |
| 18 | Sar1b | 0.29 | 8.80E-08 | 1.83E-03 | 0.91 | 0.70 | 0.21 | 1.30 |
| 18 | Pmm2 | 0.30 | 8.83E-08 | 1.84E-03 | 0.52 | 0.32 | 0.21 | 1.66 |
| 18 | Mbnl2 | 0.26 | 8.86E-08 | 1.84E-03 | 0.88 | 0.68 | 0.21 | 1.31 |
| 18 | Noa1 | 0.32 | 8.88E-08 | 1.85E-03 | 0.51 | 0.31 | 0.20 | 1.64 |
| 18 | Mtch2 | 0.26 | 8.93E-08 | 1.86E-03 | 0.94 | 0.78 | 0.15 | 1.20 |
| 18 | Pnpla6 | 0.86 | 8.96E-08 | 1.86E-03 | 0.18 | 0.09 | 0.10 | 2.09 |
| 18 | Vps8 | 0.62 | 9.07E-08 | 1.89E-03 | 0.13 | 0.06 | 0.08 | 2.38 |
| 18 | Yipf1 | 0.29 | 9.51E-08 | 1.98E-03 | 0.57 | 0.35 | 0.21 | 1.60 |
| 18 | As3mt | 0.32 | 9.58E-08 | 1.99E-03 | 0.59 | 0.37 | 0.22 | 1.60 |
| 18 | Pctp | 0.34 | 9.81E-08 | 2.04E-03 | 0.23 | 0.12 | 0.11 | 1.97 |
| 18 | Sntb2 | 0.34 | 9.96E-08 | 2.07E-03 | 0.55 | 0.34 | 0.21 | 1.62 |
| 18 | Slc35f5 | 0.30 | 1.00E-07 | 2.09E-03 | 0.26 | 0.14 | 0.13 | 1.94 |
| 18 | Fbxw8 | 0.34 | 1.02E-07 | 2.13E-03 | 0.46 | 0.28 | 0.19 | 1.67 |
| 18 | Slc12a4 | 0.33 | 1.02E-07 | 2.13E-03 | 0.27 | 0.14 | 0.13 | 1.91 |
| 18 | Utp20 | 0.41 | 1.03E-07 | 2.15E-03 | 0.53 | 0.33 | 0.20 | 1.60 |

**Supplementary Table 1. Marker genes for the original clusters of integrated clustering of E11.5 control and NotchGOF λ-junction cells.**

| | | | | | | | | |
|---|---|---|---|---|---|---|---|---|
| 18 | Gm9844 | 0.73 | 1.05E-07 | 2.18E-03 | 0.13 | 0.05 | 0.07 | 2.42 |
| 18 | Gosr2 | 0.27 | 1.07E-07 | 2.22E-03 | 0.58 | 0.35 | 0.23 | 1.64 |
| 18 | Cluh | 0.25 | 1.07E-07 | 2.23E-03 | 0.40 | 0.23 | 0.17 | 1.76 |
| 18 | Arhgef9 | 0.26 | 1.08E-07 | 2.24E-03 | 0.34 | 0.19 | 0.15 | 1.82 |
| 18 | Vps37b | 0.26 | 1.12E-07 | 2.34E-03 | 0.49 | 0.29 | 0.20 | 1.67 |
| 18 | Ints11 | 0.33 | 1.15E-07 | 2.38E-03 | 0.71 | 0.49 | 0.23 | 1.46 |
| 18 | Mrpl48 | 0.37 | 1.17E-07 | 2.43E-03 | 0.75 | 0.53 | 0.22 | 1.41 |
| 18 | Eaf1 | 0.39 | 1.17E-07 | 2.43E-03 | 0.34 | 0.19 | 0.15 | 1.78 |
| 18 | Klhdc4 | 0.27 | 1.17E-07 | 2.44E-03 | 0.57 | 0.35 | 0.22 | 1.63 |
| 18 | Tspyl3 | 0.76 | 1.20E-07 | 2.50E-03 | 0.11 | 0.05 | 0.07 | 2.49 |
| 18 | Tmem159 | 0.29 | 1.20E-07 | 2.50E-03 | 0.45 | 0.26 | 0.19 | 1.72 |
| 18 | Mon2 | 0.48 | 1.22E-07 | 2.54E-03 | 0.45 | 0.28 | 0.18 | 1.64 |
| 18 | Eif2b1 | 0.30 | 1.23E-07 | 2.55E-03 | 0.70 | 0.49 | 0.21 | 1.42 |
| 18 | Ube2w | 0.34 | 1.26E-07 | 2.62E-03 | 0.56 | 0.35 | 0.21 | 1.59 |
| 18 | L3mbtl3 | 0.29 | 1.27E-07 | 2.64E-03 | 0.61 | 0.39 | 0.22 | 1.57 |
| 18 | Derl2 | 0.34 | 1.27E-07 | 2.64E-03 | 0.60 | 0.38 | 0.22 | 1.59 |
| 18 | Tatdn2 | 0.35 | 1.29E-07 | 2.68E-03 | 0.29 | 0.15 | 0.13 | 1.88 |
| 18 | Smim26 | 0.29 | 1.37E-07 | 2.84E-03 | 0.65 | 0.43 | 0.22 | 1.51 |
| 18 | Zfp46 | 0.49 | 1.38E-07 | 2.86E-03 | 0.26 | 0.14 | 0.12 | 1.89 |
| 18 | Copg1 | 0.31 | 1.39E-07 | 2.89E-03 | 0.75 | 0.53 | 0.22 | 1.41 |
| 18 | Ccdc22 | 0.27 | 1.40E-07 | 2.92E-03 | 0.50 | 0.30 | 0.20 | 1.68 |
| 18 | Becn1 | 0.29 | 1.42E-07 | 2.95E-03 | 0.69 | 0.45 | 0.23 | 1.51 |
| 18 | Fmc1 | 0.26 | 1.42E-07 | 2.96E-03 | 0.81 | 0.59 | 0.22 | 1.37 |
| 18 | Zhx3 | 0.40 | 1.43E-07 | 2.96E-03 | 0.15 | 0.07 | 0.09 | 2.25 |
| 18 | Uimc1 | 0.32 | 1.44E-07 | 3.00E-03 | 0.50 | 0.31 | 0.19 | 1.61 |
| 18 | Fitm2 | 0.53 | 1.49E-07 | 3.10E-03 | 0.18 | 0.09 | 0.10 | 2.09 |
| 18 | Exosc1 | 0.39 | 1.50E-07 | 3.13E-03 | 0.74 | 0.55 | 0.20 | 1.36 |
| 18 | Trappc13 | 0.31 | 1.53E-07 | 3.18E-03 | 0.66 | 0.42 | 0.23 | 1.55 |
| 18 | Nexn | 0.75 | 1.55E-07 | 3.22E-03 | 0.11 | 0.04 | 0.07 | 2.53 |
| 18 | Zdhhc1 | 0.59 | 1.57E-07 | 3.26E-03 | 0.19 | 0.09 | 0.10 | 2.05 |
| 18 | Pmvk | 0.31 | 1.58E-07 | 3.28E-03 | 0.79 | 0.55 | 0.24 | 1.43 |
| 18 | Donson | 0.47 | 1.67E-07 | 3.47E-03 | 0.41 | 0.24 | 0.17 | 1.68 |
| 18 | Ext2 | 0.38 | 1.67E-07 | 3.48E-03 | 0.62 | 0.42 | 0.20 | 1.48 |
| 18 | Asap1 | 0.26 | 1.72E-07 | 3.58E-03 | 0.92 | 0.73 | 0.18 | 1.25 |
| 18 | Rbm19 | 0.36 | 1.74E-07 | 3.62E-03 | 0.45 | 0.27 | 0.18 | 1.67 |
| 18 | Stx8 | 0.31 | 1.75E-07 | 3.65E-03 | 0.65 | 0.44 | 0.20 | 1.46 |
| 18 | Nub1 | 0.27 | 1.85E-07 | 3.85E-03 | 0.59 | 0.37 | 0.22 | 1.59 |
| 18 | Klc1 | 0.32 | 1.86E-07 | 3.86E-03 | 0.68 | 0.47 | 0.22 | 1.46 |
| 18 | Lin54 | 0.38 | 1.94E-07 | 4.03E-03 | 0.40 | 0.23 | 0.16 | 1.70 |
| 18 | Stat5b | 0.35 | 1.98E-07 | 4.11E-03 | 0.25 | 0.13 | 0.12 | 1.93 |
| 18 | Rab6a | 0.27 | 1.99E-07 | 4.14E-03 | 0.89 | 0.71 | 0.18 | 1.25 |
| 18 | Slc37a4 | 0.48 | 2.01E-07 | 4.18E-03 | 0.14 | 0.06 | 0.08 | 2.31 |
| 18 | Nsf | 0.40 | 2.03E-07 | 4.21E-03 | 0.53 | 0.34 | 0.20 | 1.58 |
| 18 | Armc5 | 0.39 | 2.04E-07 | 4.24E-03 | 0.27 | 0.14 | 0.12 | 1.88 |
| 18 | Eri1 | 0.30 | 2.06E-07 | 4.28E-03 | 0.70 | 0.50 | 0.20 | 1.40 |
| 18 | Eif3j1 | 0.31 | 2.07E-07 | 4.31E-03 | 0.90 | 0.73 | 0.17 | 1.23 |
| 18 | Cdk9 | 0.32 | 2.08E-07 | 4.33E-03 | 0.85 | 0.63 | 0.23 | 1.36 |
| 18 | Dbndd2 | 0.45 | 2.09E-07 | 4.35E-03 | 0.17 | 0.08 | 0.09 | 2.11 |
| 18 | Susd6 | 0.47 | 2.12E-07 | 4.41E-03 | 0.34 | 0.20 | 0.14 | 1.72 |
| 18 | Utp18 | 0.32 | 2.12E-07 | 4.42E-03 | 0.78 | 0.58 | 0.20 | 1.34 |
| 18 | Tspan5 | 0.32 | 2.13E-07 | 4.42E-03 | 0.44 | 0.27 | 0.17 | 1.66 |
| 18 | Dync1h1 | 0.27 | 2.18E-07 | 4.54E-03 | 0.87 | 0.71 | 0.17 | 1.24 |
| 18 | Xkr8 | 0.59 | 2.22E-07 | 4.62E-03 | 0.12 | 0.05 | 0.07 | 2.47 |
| 18 | Mrpl2 | 0.32 | 2.25E-07 | 4.68E-03 | 0.86 | 0.68 | 0.18 | 1.27 |
| 18 | Eif2ak3 | 0.62 | 2.29E-07 | 4.76E-03 | 0.15 | 0.07 | 0.08 | 2.19 |
| 18 | Mphosph9 | 0.35 | 2.29E-07 | 4.77E-03 | 0.42 | 0.25 | 0.17 | 1.68 |

**Supplementary Table 1. Marker genes for the original clusters of integrated clustering of E11.5 control and NotchGOF λ-junction cells.**

| | | | | | | | | |
|---|---|---|---|---|---|---|---|---|
| 18 | Tmx1 | 0.30 | 2.31E-07 | 4.80E-03 | 0.81 | 0.59 | 0.22 | 1.38 |
| 18 | Dpm2 | 0.26 | 2.37E-07 | 4.93E-03 | 0.80 | 0.57 | 0.23 | 1.40 |
| 18 | Myo1e | 0.34 | 2.40E-07 | 4.98E-03 | 0.40 | 0.23 | 0.17 | 1.71 |
| 18 | Hyou1 | 0.29 | 2.48E-07 | 5.15E-03 | 0.48 | 0.29 | 0.19 | 1.67 |
| 18 | Arhgap21 | 0.33 | 2.72E-07 | 5.67E-03 | 0.37 | 0.22 | 0.16 | 1.72 |
| 18 | Paqr8 | 0.51 | 2.89E-07 | 6.02E-03 | 0.14 | 0.06 | 0.08 | 2.28 |
| 18 | Cyb5r1 | 0.25 | 2.94E-07 | 6.11E-03 | 0.43 | 0.25 | 0.18 | 1.70 |
| 18 | Thop1 | 0.32 | 2.99E-07 | 6.21E-03 | 0.75 | 0.51 | 0.23 | 1.46 |
| 18 | Yif1a | 0.29 | 3.02E-07 | 6.27E-03 | 0.70 | 0.48 | 0.23 | 1.47 |
| 18 | Fig4 | 0.63 | 3.03E-07 | 6.30E-03 | 0.16 | 0.08 | 0.09 | 2.13 |
| 18 | Aip | 0.30 | 3.05E-07 | 6.35E-03 | 0.61 | 0.41 | 0.20 | 1.49 |
| 18 | Erp44 | 0.31 | 3.06E-07 | 6.36E-03 | 0.72 | 0.51 | 0.21 | 1.42 |
| 18 | Taf4b | 0.60 | 3.07E-07 | 6.38E-03 | 0.12 | 0.05 | 0.07 | 2.35 |
| 18 | Nbas | 0.36 | 3.12E-07 | 6.50E-03 | 0.38 | 0.22 | 0.16 | 1.73 |
| 18 | Ogfod1 | 0.60 | 3.19E-07 | 6.63E-03 | 0.21 | 0.11 | 0.10 | 1.91 |
| 18 | Cpt1a | 0.65 | 3.20E-07 | 6.65E-03 | 0.20 | 0.10 | 0.10 | 1.99 |
| 18 | Myh10 | 0.27 | 3.28E-07 | 6.83E-03 | 0.97 | 0.87 | 0.11 | 1.12 |
| 18 | Atp13a1 | 0.48 | 3.36E-07 | 7.00E-03 | 0.39 | 0.23 | 0.16 | 1.68 |
| 18 | Erlin1 | 0.43 | 3.38E-07 | 7.03E-03 | 0.28 | 0.15 | 0.12 | 1.82 |
| 18 | Cnpy3 | 0.27 | 3.38E-07 | 7.03E-03 | 0.57 | 0.35 | 0.22 | 1.62 |
| 18 | Rapgef1 | 0.48 | 3.40E-07 | 7.08E-03 | 0.43 | 0.27 | 0.16 | 1.59 |
| 18 | Scml2 | 0.64 | 3.42E-07 | 7.10E-03 | 0.12 | 0.05 | 0.07 | 2.35 |
| 18 | Kat2a | 0.34 | 3.50E-07 | 7.28E-03 | 0.55 | 0.35 | 0.20 | 1.57 |
| 18 | Washc4 | 0.33 | 3.59E-07 | 7.46E-03 | 0.51 | 0.31 | 0.19 | 1.61 |
| 18 | Klhdc3 | 0.30 | 3.69E-07 | 7.66E-03 | 0.67 | 0.43 | 0.24 | 1.56 |
| 18 | Snd1 | 0.31 | 3.71E-07 | 7.72E-03 | 0.73 | 0.51 | 0.21 | 1.42 |
| 18 | Slc39a1 | 0.44 | 3.76E-07 | 7.82E-03 | 0.42 | 0.26 | 0.17 | 1.65 |
| 18 | Agtpbp1 | 0.39 | 3.83E-07 | 7.97E-03 | 0.31 | 0.18 | 0.14 | 1.77 |
| 18 | Pkmyt1 | 0.35 | 3.85E-07 | 8.00E-03 | 0.31 | 0.17 | 0.14 | 1.79 |
| 18 | 4932438A13Rik | 0.27 | 3.92E-07 | 8.16E-03 | 0.56 | 0.35 | 0.21 | 1.59 |
| 18 | Dusp4 | 0.30 | 3.94E-07 | 8.20E-03 | 0.18 | 0.09 | 0.09 | 2.08 |
| 18 | Mtor | 0.38 | 3.95E-07 | 8.21E-03 | 0.40 | 0.23 | 0.16 | 1.69 |
| 18 | Tada2b | 0.32 | 4.03E-07 | 8.39E-03 | 0.26 | 0.14 | 0.12 | 1.88 |
| 18 | Selenot | 0.26 | 4.21E-07 | 8.76E-03 | 0.76 | 0.54 | 0.22 | 1.41 |
| 18 | Rnf41 | 0.30 | 4.32E-07 | 8.98E-03 | 0.44 | 0.26 | 0.17 | 1.65 |
| 18 | Chpf2 | 0.51 | 4.45E-07 | 9.25E-03 | 0.20 | 0.10 | 0.10 | 1.97 |
| 18 | Gga3 | 0.42 | 4.52E-07 | 9.40E-03 | 0.26 | 0.14 | 0.12 | 1.84 |
| 18 | Tomm70a | 0.29 | 4.58E-07 | 9.52E-03 | 0.87 | 0.71 | 0.16 | 1.23 |
| 18 | Rab10os | 0.30 | 4.71E-07 | 9.79E-03 | 0.32 | 0.18 | 0.14 | 1.76 |
| 18 | Spopl | 0.29 | 4.82E-07 | 1.00E-02 | 0.44 | 0.26 | 0.18 | 1.69 |
| 18 | Tmem127 | 0.47 | 4.82E-07 | 1.00E-02 | 0.33 | 0.19 | 0.14 | 1.75 |
| 18 | Mtrr | 0.49 | 4.84E-07 | 1.01E-02 | 0.22 | 0.12 | 0.11 | 1.91 |
| 18 | Pradc1 | 0.33 | 4.89E-07 | 1.02E-02 | 0.37 | 0.22 | 0.15 | 1.71 |
| 18 | Exosc9 | 0.29 | 4.92E-07 | 1.02E-02 | 0.75 | 0.51 | 0.24 | 1.47 |
| 18 | Klf4 | 0.67 | 5.02E-07 | 1.05E-02 | 0.34 | 0.21 | 0.13 | 1.63 |
| 18 | Rgmb | 0.28 | 5.26E-07 | 1.09E-02 | 0.24 | 0.13 | 0.11 | 1.90 |
| 18 | Cyp4f16 | 0.58 | 5.32E-07 | 1.11E-02 | 0.20 | 0.10 | 0.10 | 1.95 |
| 18 | Gm20721 | 0.87 | 5.33E-07 | 1.11E-02 | 0.12 | 0.05 | 0.07 | 2.33 |
| 18 | Ice1 | 0.30 | 5.44E-07 | 1.13E-02 | 0.77 | 0.54 | 0.23 | 1.43 |
| 18 | Gtdc1 | 0.44 | 5.45E-07 | 1.13E-02 | 0.24 | 0.13 | 0.11 | 1.88 |
| 18 | Pigw | 0.57 | 5.49E-07 | 1.14E-02 | 0.14 | 0.07 | 0.08 | 2.20 |
| 18 | Pgd | 0.29 | 5.54E-07 | 1.15E-02 | 0.83 | 0.58 | 0.25 | 1.43 |
| 18 | Rab3gap1 | 0.26 | 5.64E-07 | 1.17E-02 | 0.54 | 0.33 | 0.21 | 1.62 |
| 18 | Slc39a9 | 0.28 | 5.65E-07 | 1.17E-02 | 0.28 | 0.15 | 0.13 | 1.84 |
| 18 | Fer | 0.29 | 5.67E-07 | 1.18E-02 | 0.54 | 0.35 | 0.19 | 1.55 |
| 18 | Poll | 0.33 | 5.74E-07 | 1.19E-02 | 0.31 | 0.17 | 0.13 | 1.78 |

**Supplementary Table 1. Marker genes for the original clusters of integrated clustering of E11.5 control and NotchGOF λ-junction cells.**

| | | | | | | | | |
|---|---|---|---|---|---|---|---|---|
| 18 | Abcb7 | 0.47 | 5.77E-07 | 1.20E-02 | 0.34 | 0.20 | 0.14 | 1.70 |
| 18 | Dlst | 0.27 | 5.91E-07 | 1.23E-02 | 0.79 | 0.57 | 0.23 | 1.40 |
| 18 | Dok1 | 0.56 | 5.95E-07 | 1.24E-02 | 0.20 | 0.10 | 0.10 | 1.97 |
| 18 | Gprc5b | 0.30 | 5.96E-07 | 1.24E-02 | 0.20 | 0.10 | 0.10 | 1.99 |
| 18 | Tmem44 | 0.31 | 6.00E-07 | 1.25E-02 | 0.37 | 0.21 | 0.15 | 1.71 |
| 18 | Snf8 | 0.25 | 6.26E-07 | 1.30E-02 | 0.92 | 0.75 | 0.17 | 1.23 |
| 18 | Sbno2 | 0.54 | 6.30E-07 | 1.31E-02 | 0.22 | 0.11 | 0.10 | 1.91 |
| 18 | Tmem43 | 0.35 | 6.44E-07 | 1.34E-02 | 0.36 | 0.21 | 0.15 | 1.71 |
| 18 | Abhd4 | 0.28 | 6.47E-07 | 1.35E-02 | 0.26 | 0.14 | 0.12 | 1.85 |
| 18 | 8030462N17Rik | 0.42 | 6.74E-07 | 1.40E-02 | 0.31 | 0.18 | 0.13 | 1.76 |
| 18 | Spata2 | 0.44 | 6.91E-07 | 1.44E-02 | 0.22 | 0.12 | 0.11 | 1.90 |
| 18 | Med21 | 0.27 | 7.13E-07 | 1.48E-02 | 0.91 | 0.74 | 0.17 | 1.23 |
| 18 | Clpb | 0.32 | 7.29E-07 | 1.52E-02 | 0.54 | 0.35 | 0.19 | 1.55 |
| 18 | Lztr1 | 0.32 | 7.47E-07 | 1.55E-02 | 0.50 | 0.32 | 0.18 | 1.58 |
| 18 | Slc15a4 | 0.56 | 7.87E-07 | 1.64E-02 | 0.27 | 0.15 | 0.12 | 1.80 |
| 18 | Eif2ak2 | 0.30 | 8.56E-07 | 1.78E-02 | 0.45 | 0.28 | 0.17 | 1.60 |
| 18 | Hus1 | 0.42 | 8.76E-07 | 1.82E-02 | 0.41 | 0.25 | 0.16 | 1.63 |
| 18 | Primpol | 0.27 | 8.79E-07 | 1.83E-02 | 0.21 | 0.11 | 0.10 | 1.95 |
| 18 | Rapgef6 | 0.34 | 8.94E-07 | 1.86E-02 | 0.63 | 0.44 | 0.20 | 1.45 |
| 18 | Tmem214 | 0.26 | 9.05E-07 | 1.88E-02 | 0.47 | 0.29 | 0.18 | 1.61 |
| 18 | Uhmk1 | 0.33 | 9.19E-07 | 1.91E-02 | 0.35 | 0.21 | 0.14 | 1.70 |
| 18 | Alg12 | 0.43 | 9.21E-07 | 1.91E-02 | 0.19 | 0.09 | 0.09 | 1.99 |
| 18 | Exosc10 | 0.27 | 9.43E-07 | 1.96E-02 | 0.65 | 0.43 | 0.21 | 1.49 |
| 18 | Bahd1 | 0.88 | 9.52E-07 | 1.98E-02 | 0.10 | 0.04 | 0.06 | 2.43 |
| 18 | Akt1s1 | 0.28 | 9.57E-07 | 1.99E-02 | 0.82 | 0.62 | 0.20 | 1.32 |
| 18 | Aven | 0.26 | 9.68E-07 | 2.01E-02 | 0.61 | 0.39 | 0.21 | 1.54 |
| 18 | Ap2b1 | 0.27 | 1.01E-06 | 2.09E-02 | 0.70 | 0.49 | 0.22 | 1.44 |
| 18 | Ykt6 | 0.29 | 1.07E-06 | 2.23E-02 | 0.63 | 0.41 | 0.23 | 1.56 |
| 18 | Ckap2 | 0.37 | 1.10E-06 | 2.28E-02 | 0.58 | 0.40 | 0.18 | 1.46 |
| 18 | Mtfr2 | 0.41 | 1.10E-06 | 2.30E-02 | 0.21 | 0.11 | 0.10 | 1.93 |
| 18 | Mpv17l2 | 0.28 | 1.13E-06 | 2.34E-02 | 0.38 | 0.23 | 0.16 | 1.70 |
| 18 | Dnajc5 | 0.31 | 1.14E-06 | 2.36E-02 | 0.72 | 0.52 | 0.20 | 1.38 |
| 18 | Il2 | 0.51 | 1.17E-06 | 2.43E-02 | 0.11 | 0.04 | 0.06 | 2.39 |
| 18 | C1galt1c1 | 0.26 | 1.19E-06 | 2.48E-02 | 0.29 | 0.16 | 0.13 | 1.80 |
| 18 | Trmo | 0.45 | 1.20E-06 | 2.49E-02 | 0.14 | 0.07 | 0.08 | 2.17 |
| 18 | Psmb10 | 0.27 | 1.24E-06 | 2.58E-02 | 0.39 | 0.23 | 0.16 | 1.69 |
| 18 | Lemd2 | 0.37 | 1.24E-06 | 2.58E-02 | 0.50 | 0.31 | 0.18 | 1.59 |
| 18 | Plekhb2 | 0.64 | 1.26E-06 | 2.63E-02 | 0.13 | 0.06 | 0.07 | 2.18 |
| 18 | Get4 | 0.37 | 1.26E-06 | 2.63E-02 | 0.74 | 0.55 | 0.19 | 1.35 |
| 18 | Oma1 | 0.35 | 1.29E-06 | 2.68E-02 | 0.31 | 0.18 | 0.13 | 1.72 |
| 18 | Dcps | 0.25 | 1.30E-06 | 2.70E-02 | 0.82 | 0.60 | 0.22 | 1.37 |
| 18 | Vma21 | 0.30 | 1.31E-06 | 2.71E-02 | 0.72 | 0.52 | 0.20 | 1.38 |
| 18 | Etf1 | 0.25 | 1.35E-06 | 2.80E-02 | 0.93 | 0.80 | 0.13 | 1.16 |
| 18 | Sec31a | 0.28 | 1.36E-06 | 2.83E-02 | 0.77 | 0.55 | 0.22 | 1.39 |
| 18 | Tdrkh | 0.48 | 1.38E-06 | 2.87E-02 | 0.13 | 0.06 | 0.07 | 2.22 |
| 18 | Ncstn | 0.41 | 1.39E-06 | 2.88E-02 | 0.46 | 0.29 | 0.17 | 1.57 |
| 18 | Dnajc16 | 0.57 | 1.41E-06 | 2.93E-02 | 0.13 | 0.06 | 0.07 | 2.25 |
| 18 | Wwox | 0.46 | 1.44E-06 | 3.00E-02 | 0.20 | 0.10 | 0.10 | 1.93 |
| 18 | Cyb5b | 0.36 | 1.45E-06 | 3.01E-02 | 0.70 | 0.50 | 0.20 | 1.40 |
| 18 | Kras | 0.27 | 1.50E-06 | 3.13E-02 | 0.93 | 0.85 | 0.09 | 1.10 |
| 18 | Gak | 0.33 | 1.52E-06 | 3.16E-02 | 0.37 | 0.23 | 0.14 | 1.63 |
| 18 | Map11 | 0.35 | 1.57E-06 | 3.27E-02 | 0.20 | 0.11 | 0.10 | 1.94 |
| 18 | Plxnc1 | 0.90 | 1.62E-06 | 3.36E-02 | 0.10 | 0.04 | 0.06 | 2.37 |
| 18 | Apex2 | 0.28 | 1.67E-06 | 3.47E-02 | 0.39 | 0.23 | 0.16 | 1.68 |
| 18 | Fbxl3 | 0.36 | 1.70E-06 | 3.53E-02 | 0.40 | 0.25 | 0.15 | 1.60 |
| 18 | Atad3a | 0.26 | 1.76E-06 | 3.67E-02 | 0.79 | 0.57 | 0.22 | 1.38 |

**Supplementary Table 1. Marker genes for the original clusters of integrated clustering of E11.5 control and NotchGOF λ-junction cells.**

| | | | | | | | | |
|---|---|---|---|---|---|---|---|---|
| 18 | Cnot8 | 0.26 | 1.78E-06 | 3.70E-02 | 0.70 | 0.48 | 0.22 | 1.46 |
| 18 | Kif3b | 0.30 | 1.81E-06 | 3.76E-02 | 0.32 | 0.19 | 0.13 | 1.71 |
| 18 | Wrnip1 | 0.29 | 1.82E-06 | 3.78E-02 | 0.34 | 0.20 | 0.14 | 1.70 |
| 18 | Exoc8 | 0.43 | 1.90E-06 | 3.96E-02 | 0.19 | 0.10 | 0.09 | 1.95 |
| 18 | Dctn1 | 0.28 | 1.93E-06 | 4.02E-02 | 0.73 | 0.52 | 0.21 | 1.41 |
| 18 | Sec22b | 0.31 | 1.94E-06 | 4.04E-02 | 0.82 | 0.60 | 0.22 | 1.37 |
| 18 | Pip5k1a | 0.26 | 1.95E-06 | 4.05E-02 | 0.68 | 0.46 | 0.21 | 1.46 |
| 18 | Ldlr | 0.34 | 1.96E-06 | 4.08E-02 | 0.40 | 0.24 | 0.15 | 1.63 |
| 18 | Rab29 | 0.77 | 2.00E-06 | 4.16E-02 | 0.12 | 0.05 | 0.07 | 2.27 |
| 18 | Secisbp2 | 0.33 | 2.03E-06 | 4.22E-02 | 0.46 | 0.29 | 0.17 | 1.58 |
| 18 | Ugcg | 0.32 | 2.04E-06 | 4.24E-02 | 0.57 | 0.37 | 0.20 | 1.54 |
| 18 | Gm9774 | 0.35 | 2.20E-06 | 4.59E-02 | 0.16 | 0.08 | 0.08 | 2.08 |
| 18 | Gnl3l | 0.31 | 2.22E-06 | 4.61E-02 | 0.49 | 0.32 | 0.17 | 1.53 |
| 18 | Rlim | 0.34 | 2.22E-06 | 4.61E-02 | 0.76 | 0.54 | 0.22 | 1.41 |
| 18 | Sdhaf2 | 0.25 | 2.30E-06 | 4.79E-02 | 0.67 | 0.44 | 0.22 | 1.50 |
| 18 | Rnpepl1 | 0.27 | 2.36E-06 | 4.91E-02 | 0.41 | 0.25 | 0.16 | 1.62 |
| 18 | Cep135 | 0.34 | 2.39E-06 | 4.98E-02 | 0.47 | 0.29 | 0.17 | 1.59 |
| 19 | Lhx2 | 10.28 | 0 | 0 | 0.88 | 0.02 | 0.86 | 55.00 |
| 19 | Insm1 | 9.09 | 0 | 0 | 0.86 | 0.01 | 0.85 | 85.50 |
| 19 | Elavl4 | 8.32 | 0 | 0 | 0.87 | 0.06 | 0.81 | 14.95 |
| 19 | Nhlh2 | 10.84 | 0 | 0 | 0.81 | 0.00 | 0.80 | 269.00 |
| 19 | Neurod1 | 11.75 | 0 | 0 | 0.80 | 0.00 | 0.80 | 267.00 |
| 19 | Igfbpl1 | 10.67 | 0 | 0 | 0.80 | 0.01 | 0.80 | 133.50 |
| 19 | Elavl3 | 10.25 | 0 | 0 | 0.80 | 0.01 | 0.79 | 99.38 |
| 19 | Ebf2 | 6.67 | 0 | 0 | 0.81 | 0.04 | 0.78 | 23.23 |
| 19 | Ppp1r14a | 6.64 | 0 | 0 | 0.84 | 0.07 | 0.77 | 12.49 |
| 19 | Srrm4 | 10.32 | 0 | 0 | 0.77 | 0.00 | 0.76 | 255.00 |
| 19 | Mfng | 6.75 | 0 | 0 | 0.80 | 0.04 | 0.76 | 20.03 |
| 19 | Neurog1 | 11.14 | 0 | 0 | 0.75 | 0.00 | 0.75 | 373.50 |
| 19 | Wdr66 | 7.77 | 0 | 0 | 0.70 | 0.01 | 0.69 | 49.93 |
| 19 | Nhlh1 | 11.43 | 0 | 0 | 0.66 | 0.00 | 0.66 | 657.00 |
| 19 | Soga3 | 6.00 | 0 | 0 | 0.70 | 0.05 | 0.65 | 14.56 |
| 19 | Fezf1 | 6.92 | 0 | 0 | 0.63 | 0.02 | 0.61 | 33.32 |
| 19 | Btbd17 | 9.11 | 0 | 0 | 0.62 | 0.01 | 0.61 | 103.33 |
| 19 | Dcx | 6.64 | 0 | 0 | 0.63 | 0.02 | 0.61 | 27.52 |
| 19 | Miat | 7.31 | 0 | 0 | 0.60 | 0.01 | 0.59 | 60.20 |
| 19 | Ckmt1 | 6.70 | 0 | 0 | 0.63 | 0.04 | 0.59 | 16.08 |
| 19 | Sowaha | 10.59 | 0 | 0 | 0.59 | 0.00 | 0.59 | 295.00 |
| 19 | Stmn3 | 7.43 | 0 | 0 | 0.61 | 0.03 | 0.58 | 19.00 |
| 19 | Syp | 7.04 | 0 | 0 | 0.58 | 0.01 | 0.57 | 41.71 |
| 19 | Kif19a | 7.67 | 0 | 0 | 0.56 | 0.01 | 0.55 | 70.00 |
| 19 | Cdhr1 | 9.99 | 0 | 0 | 0.54 | 0.00 | 0.54 | 180.67 |
| 19 | St18 | 11.39 | 0 | 0 | 0.52 | 0.00 | 0.52 | 518.00 |
| 19 | Rbm24 | 7.24 | 0 | 0 | 0.52 | 0.01 | 0.51 | 43.67 |
| 19 | Syt13 | 7.04 | 0 | 0 | 0.51 | 0.01 | 0.50 | 51.20 |
| 19 | Chga | 8.62 | 0 | 0 | 0.51 | 0.01 | 0.50 | 56.22 |
| 19 | Ina | 10.90 | 0 | 0 | 0.50 | 0.01 | 0.50 | 100.00 |
| 19 | Coro2a | 6.10 | 0 | 0 | 0.51 | 0.02 | 0.49 | 29.76 |
| 19 | Thsd7b | 6.00 | 0 | 0 | 0.49 | 0.02 | 0.47 | 24.70 |
| 19 | Uncx | 6.80 | 0 | 0 | 0.49 | 0.02 | 0.47 | 23.52 |
| 19 | Nrxn1 | 6.27 | 0 | 0 | 0.49 | 0.02 | 0.47 | 21.48 |
| 19 | Slc17a6 | 11.56 | 0 | 0 | 0.46 | 0.00 | 0.46 | #NA |
| 19 | Dcc | 7.62 | 0 | 0 | 0.47 | 0.01 | 0.46 | 58.75 |
| 19 | Gdap1l1 | 6.99 | 0 | 0 | 0.47 | 0.01 | 0.46 | 33.57 |
| 19 | Olig2 | 11.82 | 0 | 0 | 0.44 | 0.00 | 0.44 | #NA |
| 19 | Celf3 | 8.96 | 0 | 0 | 0.43 | 0.00 | 0.43 | 217.00 |

**Supplementary Table 1. Marker genes for the original clusters of integrated clustering of E11.5 control and NotchGOF λ-junction cells.**

| | | | | | | | | |
|---|---|---|---|---|---|---|---|---|
| 19 | Foxo6 | 7.05 | 0 | 0 | 0.44 | 0.01 | 0.43 | 40.00 |
| 19 | Brsk2 | 6.38 | 0 | 0 | 0.42 | 0.01 | 0.41 | 42.20 |
| 19 | Myt1l | 10.99 | 0 | 0 | 0.40 | 0.00 | 0.40 | 398.00 |
| 19 | Atp8b4 | 9.83 | 0 | 0 | 0.40 | 0.00 | 0.40 | 199.00 |
| 19 | Scg3 | 8.16 | 0 | 0 | 0.40 | 0.01 | 0.39 | 79.60 |
| 19 | Ttc9b | 8.31 | 0 | 0 | 0.38 | 0.00 | 0.38 | 95.00 |
| 19 | Syt4 | 6.80 | 0 | 0 | 0.37 | 0.01 | 0.36 | 61.17 |
| 19 | Inka2 | 6.64 | 0 | 0 | 0.36 | 0.01 | 0.35 | 45.13 |
| 19 | Scn9a | 6.49 | 0 | 0 | 0.36 | 0.01 | 0.35 | 40.11 |
| 19 | Shisal1 | 7.06 | 0 | 0 | 0.36 | 0.01 | 0.35 | 59.17 |
| 19 | Fstl5 | 8.25 | 0 | 0 | 0.34 | 0.01 | 0.34 | 42.88 |
| 19 | Dscaml1 | 9.46 | 0 | 0 | 0.32 | 0.00 | 0.32 | 319.00 |
| 19 | Nxph4 | 8.54 | 0 | 0 | 0.32 | 0.00 | 0.32 | 159.50 |
| 19 | Pcdh8 | 7.16 | 0 | 0 | 0.32 | 0.01 | 0.31 | 53.17 |
| 19 | Akap6 | 8.15 | 0 | 0 | 0.30 | 0.01 | 0.30 | 60.20 |
| 19 | Olig1 | 9.83 | 0 | 0 | 0.29 | 0.00 | 0.29 | 289.00 |
| 19 | Rgs20 | 7.71 | 0 | 0 | 0.27 | 0.00 | 0.27 | 90.33 |
| 19 | Clvs1 | 8.15 | 0 | 0 | 0.27 | 0.00 | 0.27 | 90.33 |
| 19 | Igsf21 | 9.22 | 0 | 0 | 0.27 | 0.00 | 0.26 | 265.00 |
| 19 | Stmn4 | 8.17 | 0 | 0 | 0.27 | 0.00 | 0.26 | 66.25 |
| 19 | Dner | 6.49 | 0 | 0 | 0.26 | 0.00 | 0.26 | 64.75 |
| 19 | Mast1 | 6.52 | 0 | 0 | 0.26 | 0.00 | 0.26 | 64.75 |
| 19 | Nol4 | 6.87 | 0 | 0 | 0.26 | 0.01 | 0.25 | 51.80 |
| 19 | Myt1 | 9.91 | 0 | 0 | 0.24 | 0.00 | 0.24 | #NA |
| 19 | Chgb | 8.24 | 0 | 0 | 0.23 | 0.00 | 0.23 | 76.33 |
| 19 | Slc17a7 | 7.80 | 0 | 0 | 0.22 | 0.00 | 0.22 | 223.00 |
| 19 | Smim18 | 10.63 | 0 | 0 | 0.22 | 0.00 | 0.22 | #NA |
| 19 | Pcsk2 | 8.89 | 0 | 0 | 0.22 | 0.00 | 0.22 | 217.00 |
| 19 | Actl6b | 11.39 | 0 | 0 | 0.21 | 0.00 | 0.21 | #NA |
| 19 | 5730403I07Rik | 9.53 | 0 | 0 | 0.21 | 0.00 | 0.21 | 211.00 |
| 19 | Pirt | 11.07 | 0 | 0 | 0.19 | 0.00 | 0.19 | #NA |
| 19 | A230051N06Rik | 10.58 | 0 | 0 | 0.18 | 0.00 | 0.18 | #NA |
| 19 | Chrna3 | 8.51 | 0 | 0 | 0.18 | 0.00 | 0.17 | 175.00 |
| 19 | Gfy | 10.30 | 0 | 0 | 0.17 | 0.00 | 0.17 | #NA |
| 19 | Ppp1r17 | 10.39 | 0 | 0 | 0.16 | 0.00 | 0.16 | 157.00 |
| 19 | Gm3234 | 7.97 | 0 | 0 | 0.16 | 0.00 | 0.16 | 157.00 |
| 19 | Tmprss6 | 8.87 | 0 | 0 | 0.16 | 0.00 | 0.16 | 157.00 |
| 19 | Gda | 8.64 | 0 | 0 | 0.16 | 0.00 | 0.16 | 157.00 |
| 19 | Rd3 | 9.94 | 0 | 0 | 0.15 | 0.00 | 0.15 | #NA |
| 19 | Gm27197 | 10.50 | 0 | 0 | 0.13 | 0.00 | 0.13 | #NA |
| 19 | Gm31819 | 9.65 | 0 | 0 | 0.11 | 0.00 | 0.11 | #NA |
| 19 | Eef1a2 | 8.96 | 0 | 0 | 0.10 | 0.00 | 0.10 | #NA |
| 19 | Kcnk10 | 8.89 | 0 | 0 | 0.10 | 0.00 | 0.10 | #NA |
| 19 | Islr2 | 7.80 | 1.83E-295 | 3.80E-291 | 0.16 | 0.00 | 0.16 | 81.50 |
| 19 | Srrm3 | 6.19 | 6.35E-292 | 1.32E-287 | 0.28 | 0.01 | 0.27 | 39.57 |
| 19 | Rnf112 | 6.49 | 1.99E-288 | 4.14E-284 | 0.19 | 0.00 | 0.19 | 64.33 |
| 19 | Syt2 | 6.89 | 5.08E-285 | 1.06E-280 | 0.22 | 0.00 | 0.21 | 54.25 |
| 19 | Mreg | 6.29 | 1.01E-284 | 2.10E-280 | 0.34 | 0.01 | 0.33 | 30.64 |
| 19 | Gm17322 | 8.40 | 5.90E-283 | 1.23E-278 | 0.16 | 0.00 | 0.16 | 78.50 |
| 19 | Svop | 8.11 | 4.12E-278 | 8.58E-274 | 0.11 | 0.00 | 0.11 | 108.00 |
| 19 | Tesc | 8.40 | 1.34E-277 | 2.79E-273 | 0.26 | 0.01 | 0.25 | 43.17 |
| 19 | Map6 | 6.60 | 2.41E-277 | 5.01E-273 | 0.34 | 0.01 | 0.33 | 28.08 |
| 19 | Gm13589 | 7.14 | 4.54E-277 | 9.44E-273 | 0.19 | 0.00 | 0.18 | 62.33 |
| 19 | Ccdc187 | 8.00 | 8.96E-266 | 1.86E-261 | 0.10 | 0.00 | 0.10 | 102.00 |
| 19 | Nsg2 | 6.49 | 4.21E-265 | 8.75E-261 | 0.33 | 0.01 | 0.32 | 27.58 |
| 19 | Spock1 | 7.09 | 2.52E-259 | 5.25E-255 | 0.19 | 0.00 | 0.19 | 64.33 |

**Supplementary Table 1. Marker genes for the original clusters of integrated clustering of E11.5 control and NotchGOF λ-junction cells.**

| | | | | | | | | |
|---|---|---|---|---|---|---|---|---|
| 19 | Sez6l2 | 7.12 | 2.95E-257 | 6.13E-253 | 0.18 | 0.00 | 0.17 | 58.33 |
| 19 | Rundc3a | 5.62 | 4.59E-257 | 9.54E-253 | 0.50 | 0.03 | 0.47 | 17.24 |
| 19 | Ntrk1 | 8.40 | 2.10E-252 | 4.36E-248 | 0.11 | 0.00 | 0.11 | 108.00 |
| 19 | Scrt1 | 6.45 | 3.89E-252 | 8.09E-248 | 0.23 | 0.01 | 0.22 | 45.80 |
| 19 | Lingo2 | 6.94 | 1.50E-247 | 3.13E-243 | 0.22 | 0.01 | 0.22 | 44.60 |
| 19 | Ndrg4 | 5.44 | 3.53E-242 | 7.35E-238 | 0.49 | 0.03 | 0.46 | 15.94 |
| 19 | Cpeb1 | 8.69 | 5.27E-231 | 1.10E-226 | 0.11 | 0.00 | 0.11 | 108.00 |
| 19 | Cntn2 | 6.16 | 3.52E-229 | 7.33E-225 | 0.34 | 0.02 | 0.33 | 22.87 |
| 19 | Tubb3 | 7.61 | 1.66E-226 | 3.44E-222 | 0.98 | 0.17 | 0.81 | 5.64 |
| 19 | Nrn1 | 7.47 | 1.21E-222 | 2.52E-218 | 0.22 | 0.01 | 0.22 | 37.17 |
| 19 | Prmt8 | 8.25 | 4.62E-220 | 9.62E-216 | 0.11 | 0.00 | 0.11 | 114.00 |
| 19 | Asic1 | 6.28 | 8.79E-219 | 1.83E-214 | 0.18 | 0.00 | 0.17 | 58.33 |
| 19 | Brinp2 | 6.13 | 3.44E-216 | 7.16E-212 | 0.19 | 0.00 | 0.19 | 48.25 |
| 19 | Scg5 | 4.74 | 3.91E-212 | 8.14E-208 | 0.58 | 0.05 | 0.53 | 11.68 |
| 19 | D930020B18Rik | 5.86 | 4.56E-212 | 9.48E-208 | 0.36 | 0.02 | 0.34 | 19.72 |
| 19 | Fam57b | 6.41 | 6.21E-212 | 1.29E-207 | 0.20 | 0.01 | 0.19 | 39.80 |
| 19 | Shisa7 | 6.00 | 8.99E-210 | 1.87E-205 | 0.21 | 0.01 | 0.21 | 35.17 |
| 19 | Vwa5b2 | 5.76 | 1.77E-205 | 3.68E-201 | 0.17 | 0.00 | 0.17 | 56.33 |
| 19 | Gdap1 | 5.29 | 8.28E-205 | 1.72E-200 | 0.36 | 0.02 | 0.34 | 19.00 |
| 19 | Sct | 7.64 | 1.55E-200 | 3.22E-196 | 0.42 | 0.03 | 0.39 | 16.00 |
| 19 | Stmn2 | 6.83 | 1.48E-199 | 3.09E-195 | 0.75 | 0.10 | 0.65 | 7.78 |
| 19 | Cntn1 | 6.22 | 4.98E-187 | 1.04E-182 | 0.21 | 0.01 | 0.20 | 34.17 |
| 19 | Ap3b2 | 5.43 | 1.62E-185 | 3.37E-181 | 0.34 | 0.02 | 0.32 | 18.72 |
| 19 | Dll1 | 6.66 | 3.42E-185 | 7.10E-181 | 0.16 | 0.00 | 0.15 | 52.33 |
| 19 | Myo7a | 5.83 | 2.41E-181 | 5.00E-177 | 0.27 | 0.01 | 0.26 | 22.58 |
| 19 | Gm2694 | 5.52 | 3.67E-179 | 7.63E-175 | 0.33 | 0.02 | 0.31 | 19.12 |
| 19 | Tuba4a | 4.49 | 2.06E-178 | 4.28E-174 | 0.61 | 0.07 | 0.54 | 9.35 |
| 19 | Snap25 | 6.09 | 2.86E-178 | 5.94E-174 | 0.23 | 0.01 | 0.22 | 28.63 |
| 19 | Arhgdig | 7.11 | 6.09E-174 | 1.27E-169 | 0.64 | 0.08 | 0.56 | 8.19 |
| 19 | 4930578M01Rik | 5.24 | 5.09E-173 | 1.06E-168 | 0.37 | 0.02 | 0.35 | 15.54 |
| 19 | Gm16364 | 7.74 | 1.21E-171 | 2.51E-167 | 0.10 | 0.00 | 0.10 | 102.00 |
| 19 | Lrrc10b | 6.00 | 2.90E-170 | 6.04E-166 | 0.17 | 0.00 | 0.17 | 42.25 |
| 19 | Dll3 | 5.20 | 8.65E-157 | 1.80E-152 | 0.29 | 0.02 | 0.27 | 18.06 |
| 19 | Pnma2 | 5.48 | 2.43E-152 | 5.06E-148 | 0.24 | 0.01 | 0.23 | 23.50 |
| 19 | Lamp5 | 5.96 | 9.41E-152 | 1.96E-147 | 0.18 | 0.01 | 0.18 | 30.17 |
| 19 | Mapk10 | 6.04 | 4.10E-149 | 8.53E-145 | 0.18 | 0.01 | 0.18 | 30.17 |
| 19 | Atp1a3 | 6.56 | 2.20E-142 | 4.59E-138 | 0.13 | 0.00 | 0.12 | 42.33 |
| 19 | Gpx2 | 6.90 | 3.46E-141 | 7.19E-137 | 0.10 | 0.00 | 0.10 | 51.00 |
| 19 | Emx2 | 5.07 | 6.42E-140 | 1.34E-135 | 0.48 | 0.05 | 0.43 | 9.52 |
| 19 | Dpysl5 | 4.06 | 1.63E-137 | 3.40E-133 | 0.67 | 0.11 | 0.56 | 6.37 |
| 19 | Apba1 | 6.84 | 1.13E-135 | 2.35E-131 | 0.10 | 0.00 | 0.10 | 51.00 |
| 19 | Rgs11 | 5.17 | 1.88E-135 | 3.90E-131 | 0.23 | 0.01 | 0.22 | 20.82 |
| 19 | Mapk8ip2 | 4.78 | 7.20E-135 | 1.50E-130 | 0.40 | 0.04 | 0.37 | 10.92 |
| 19 | Rap1gap2 | 5.22 | 6.75E-133 | 1.40E-128 | 0.25 | 0.01 | 0.24 | 18.07 |
| 19 | Celf6 | 6.77 | 3.93E-132 | 8.16E-128 | 0.11 | 0.00 | 0.11 | 54.00 |
| 19 | Kcnmb2 | 5.59 | 2.04E-130 | 4.24E-126 | 0.30 | 0.02 | 0.28 | 14.33 |
| 19 | Rtn1 | 5.66 | 5.78E-130 | 1.20E-125 | 0.89 | 0.25 | 0.64 | 3.51 |
| 19 | Klc3 | 4.57 | 3.88E-129 | 8.07E-125 | 0.65 | 0.11 | 0.55 | 6.20 |
| 19 | Celsr3 | 5.83 | 1.98E-128 | 4.13E-124 | 0.13 | 0.00 | 0.12 | 42.33 |
| 19 | Nrg1 | 4.06 | 1.84E-127 | 3.84E-123 | 0.57 | 0.08 | 0.49 | 7.35 |
| 19 | Dkkl1 | 5.32 | 2.92E-127 | 6.08E-123 | 0.19 | 0.01 | 0.18 | 23.38 |
| 19 | Gng3 | 5.62 | 1.24E-126 | 2.58E-122 | 0.52 | 0.07 | 0.46 | 7.59 |
| 19 | Galnt17 | 5.09 | 3.91E-126 | 8.14E-122 | 0.19 | 0.01 | 0.19 | 24.13 |
| 19 | Kcnh8 | 4.47 | 2.69E-124 | 5.59E-120 | 0.37 | 0.03 | 0.34 | 11.47 |
| 19 | Cxcr4 | 4.32 | 4.04E-123 | 8.41E-119 | 0.44 | 0.05 | 0.39 | 9.36 |
| 19 | Dusp8 | 5.63 | 3.88E-121 | 8.08E-117 | 0.27 | 0.02 | 0.25 | 15.59 |

**Supplementary Table 1. Marker genes for the original clusters of integrated clustering of E11.5 control and NotchGOF λ-junction cells.**

| | | | | | | | | |
|---|---|---|---|---|---|---|---|---|
| 19 | Ell3 | 4.75 | 7.41E-121 | 1.54E-116 | 0.34 | 0.03 | 0.31 | 11.83 |
| 19 | Gm13648 | 4.91 | 2.02E-119 | 4.20E-115 | 0.31 | 0.02 | 0.28 | 12.79 |
| 19 | Shtn1 | 4.86 | 6.97E-119 | 1.45E-114 | 0.44 | 0.05 | 0.39 | 8.80 |
| 19 | Adgrg1 | 4.04 | 2.27E-118 | 4.72E-114 | 0.40 | 0.04 | 0.36 | 9.95 |
| 19 | Trp53i11 | 4.07 | 1.24E-116 | 2.57E-112 | 0.83 | 0.22 | 0.61 | 3.73 |
| 19 | Cplx2 | 4.95 | 1.04E-115 | 2.15E-111 | 0.74 | 0.17 | 0.57 | 4.44 |
| 19 | Crmp1 | 4.62 | 3.47E-114 | 7.22E-110 | 0.65 | 0.12 | 0.53 | 5.47 |
| 19 | Ccdc184 | 6.88 | 1.22E-111 | 2.54E-107 | 0.11 | 0.00 | 0.11 | 38.00 |
| 19 | Elavl2 | 3.89 | 1.54E-109 | 3.20E-105 | 0.76 | 0.18 | 0.58 | 4.15 |
| 19 | Tpd52 | 3.27 | 2.00E-108 | 4.15E-104 | 0.88 | 0.26 | 0.62 | 3.38 |
| 19 | Rem2 | 4.70 | 1.47E-107 | 3.06E-103 | 0.47 | 0.06 | 0.41 | 7.58 |
| 19 | Cadps | 5.08 | 6.84E-107 | 1.42E-102 | 0.28 | 0.02 | 0.26 | 13.19 |
| 19 | Hes6 | 5.35 | 1.64E-106 | 3.40E-102 | 0.89 | 0.33 | 0.56 | 2.68 |
| 19 | Fgd4 | 3.84 | 2.21E-106 | 4.59E-102 | 0.55 | 0.09 | 0.47 | 6.37 |
| 19 | Tmcc3 | 4.24 | 1.46E-103 | 3.04E-99 | 0.34 | 0.03 | 0.30 | 10.21 |
| 19 | Cotl1 | 4.30 | 2.45E-103 | 5.10E-99 | 0.95 | 0.44 | 0.50 | 2.14 |
| 19 | P2rx3 | 5.60 | 4.07E-103 | 8.46E-99 | 0.13 | 0.00 | 0.12 | 31.75 |
| 19 | Cystm1 | 4.47 | 7.74E-103 | 1.61E-98 | 0.54 | 0.09 | 0.45 | 6.16 |
| 19 | Gm14226 | 5.45 | 2.29E-102 | 4.77E-98 | 0.12 | 0.00 | 0.12 | 30.00 |
| 19 | Sult4a1 | 4.46 | 1.09E-100 | 2.27E-96 | 0.35 | 0.04 | 0.31 | 9.69 |
| 19 | Art5 | 4.66 | 1.46E-100 | 3.03E-96 | 0.20 | 0.01 | 0.19 | 18.09 |
| 19 | Cadm3 | 4.77 | 2.09E-100 | 4.35E-96 | 0.25 | 0.02 | 0.23 | 13.32 |
| 19 | Tmsb4x | 2.62 | 1.61E-99 | 3.36E-95 | 1.00 | 0.99 | 0.01 | 1.01 |
| 19 | Spock2 | 3.98 | 2.41E-99 | 5.02E-95 | 0.36 | 0.04 | 0.32 | 9.59 |
| 19 | Fam110a | 4.13 | 4.31E-98 | 8.95E-94 | 0.82 | 0.26 | 0.56 | 3.17 |
| 19 | Pclo | 3.98 | 3.36E-97 | 6.99E-93 | 0.57 | 0.11 | 0.47 | 5.45 |
| 19 | Atp1b1 | 4.73 | 8.95E-96 | 1.86E-91 | 0.51 | 0.09 | 0.43 | 6.02 |
| 19 | Gfra1 | 4.26 | 1.08E-94 | 2.25E-90 | 0.37 | 0.04 | 0.33 | 8.48 |
| 19 | Dusp26 | 5.50 | 6.55E-94 | 1.36E-89 | 0.15 | 0.01 | 0.14 | 24.17 |
| 19 | Gng8 | 4.69 | 1.16E-93 | 2.42E-89 | 0.62 | 0.13 | 0.49 | 4.66 |
| 19 | Uchl1 | 4.15 | 7.09E-92 | 1.47E-87 | 0.89 | 0.35 | 0.54 | 2.54 |
| 19 | Hspa4l | 3.69 | 1.39E-91 | 2.89E-87 | 0.57 | 0.11 | 0.46 | 5.25 |
| 19 | Necab3 | 6.65 | 2.06E-90 | 4.28E-86 | 0.15 | 0.01 | 0.14 | 20.71 |
| 19 | A330009N23Rik | 4.78 | 4.56E-90 | 9.48E-86 | 0.27 | 0.02 | 0.24 | 11.52 |
| 19 | Gnrh1 | 9.90 | 1.40E-89 | 2.91E-85 | 0.13 | 0.01 | 0.13 | 22.17 |
| 19 | Gm26871 | 4.97 | 1.43E-89 | 2.97E-85 | 0.18 | 0.01 | 0.17 | 17.50 |
| 19 | Foxg1 | 3.28 | 2.56E-89 | 5.32E-85 | 0.65 | 0.14 | 0.50 | 4.51 |
| 19 | Camk2n1 | 4.16 | 3.30E-89 | 6.86E-85 | 0.69 | 0.19 | 0.51 | 3.71 |
| 19 | Nfasc | 4.53 | 2.28E-88 | 4.75E-84 | 0.27 | 0.03 | 0.25 | 10.84 |
| 19 | Nacad | 5.29 | 2.90E-88 | 6.02E-84 | 0.15 | 0.01 | 0.14 | 21.57 |
| 19 | Eya2 | 3.40 | 3.18E-88 | 6.61E-84 | 0.78 | 0.23 | 0.55 | 3.40 |
| 19 | Cbfa2t3 | 3.74 | 2.42E-87 | 5.02E-83 | 0.40 | 0.05 | 0.35 | 7.51 |
| 19 | Sgpp2 | 3.93 | 6.00E-87 | 1.25E-82 | 0.30 | 0.03 | 0.27 | 10.03 |
| 19 | 9330159F19Rik | 4.08 | 7.14E-87 | 1.48E-82 | 0.27 | 0.02 | 0.24 | 11.04 |
| 19 | Unc13a | 4.63 | 2.17E-86 | 4.50E-82 | 0.18 | 0.01 | 0.17 | 16.45 |
| 19 | Pcsk1n | 6.66 | 8.96E-86 | 1.86E-81 | 0.19 | 0.01 | 0.18 | 15.58 |
| 19 | Serping1 | 4.02 | 2.13E-85 | 4.42E-81 | 0.40 | 0.06 | 0.35 | 7.21 |
| 19 | Tubb2b | 3.34 | 2.29E-85 | 4.76E-81 | 0.97 | 0.67 | 0.30 | 1.45 |
| 19 | Trim36 | 4.25 | 6.96E-84 | 1.45E-79 | 0.28 | 0.03 | 0.26 | 10.11 |
| 19 | Rab6b | 5.15 | 1.19E-82 | 2.48E-78 | 0.33 | 0.04 | 0.29 | 8.28 |
| 19 | Pnmal1 | 5.43 | 4.15E-82 | 8.64E-78 | 0.20 | 0.01 | 0.19 | 14.21 |
| 19 | Cep152 | 3.58 | 6.23E-82 | 1.30E-77 | 0.57 | 0.12 | 0.45 | 4.69 |
| 19 | Psip1 | 1.70 | 1.17E-81 | 2.43E-77 | 0.98 | 0.98 | 0.00 | 1.00 |
| 19 | Grasp | 4.30 | 1.32E-81 | 2.74E-77 | 0.68 | 0.19 | 0.48 | 3.52 |
| 19 | Hist3h2ba | 3.37 | 2.45E-81 | 5.09E-77 | 0.83 | 0.27 | 0.56 | 3.06 |
| 19 | Gnas | 1.60 | 4.99E-81 | 1.04E-76 | 1.00 | 1.00 | 0.00 | 1.00 |

**Supplementary Table 1. Marker genes for the original clusters of integrated clustering of E11.5 control and NotchGOF λ-junction cells.**

| | | | | | | | | |
|---|---|---|---|---|---|---|---|---|
| 19 | Snca | 4.11 | 1.42E-80 | 2.95E-76 | 0.49 | 0.09 | 0.41 | 5.68 |
| 19 | Cyfip2 | 4.12 | 5.83E-80 | 1.21E-75 | 0.48 | 0.09 | 0.39 | 5.48 |
| 19 | Gadd45a | 4.36 | 6.72E-80 | 1.40E-75 | 0.74 | 0.24 | 0.49 | 3.02 |
| 19 | 3110039M20Rik | 3.88 | 2.30E-79 | 4.79E-75 | 0.45 | 0.08 | 0.38 | 6.03 |
| 19 | Gprin1 | 4.19 | 2.76E-79 | 5.75E-75 | 0.25 | 0.02 | 0.22 | 10.74 |
| 19 | Kif5a | 4.35 | 3.21E-79 | 6.67E-75 | 0.35 | 0.05 | 0.30 | 7.76 |
| 19 | Plk3 | 4.82 | 1.72E-78 | 3.57E-74 | 0.24 | 0.02 | 0.22 | 10.95 |
| 19 | Map1b | 3.90 | 1.93E-78 | 4.01E-74 | 0.95 | 0.63 | 0.32 | 1.51 |
| 19 | Nova2 | 4.33 | 2.18E-78 | 4.53E-74 | 0.26 | 0.03 | 0.23 | 10.36 |
| 19 | Btg2 | 3.96 | 4.21E-78 | 8.76E-74 | 0.77 | 0.27 | 0.50 | 2.85 |
| 19 | Ssc5d | 4.87 | 1.33E-77 | 2.76E-73 | 0.15 | 0.01 | 0.14 | 16.78 |
| 19 | Dpysl3 | 3.44 | 5.89E-77 | 1.23E-72 | 0.89 | 0.45 | 0.44 | 1.98 |
| 19 | Fam169b | 5.14 | 1.05E-76 | 2.19E-72 | 0.10 | 0.00 | 0.10 | 25.50 |
| 19 | Prnp | 3.62 | 1.16E-75 | 2.42E-71 | 0.65 | 0.17 | 0.48 | 3.76 |
| 19 | Apc2 | 4.58 | 1.24E-74 | 2.58E-70 | 0.24 | 0.02 | 0.21 | 10.68 |
| 19 | Tuba1a | 3.06 | 1.42E-74 | 2.95E-70 | 0.99 | 0.98 | 0.00 | 1.00 |
| 19 | Stmn1 | 1.45 | 9.19E-74 | 1.91E-69 | 0.99 | 1.00 | 0.00 | 1.00 |
| 19 | Flrt1 | 4.27 | 1.12E-73 | 2.34E-69 | 0.24 | 0.02 | 0.21 | 10.68 |
| 19 | Dusp4 | 3.70 | 7.05E-73 | 1.47E-68 | 0.46 | 0.08 | 0.38 | 5.52 |
| 19 | Scg2 | 4.08 | 9.15E-73 | 1.90E-68 | 0.15 | 0.01 | 0.14 | 16.78 |
| 19 | Cadps2 | 4.34 | 1.14E-72 | 2.38E-68 | 0.30 | 0.04 | 0.26 | 8.43 |
| 19 | Map2 | 3.63 | 1.23E-72 | 2.57E-68 | 0.69 | 0.21 | 0.47 | 3.21 |
| 19 | Ebf3 | 4.04 | 1.87E-72 | 3.90E-68 | 0.63 | 0.17 | 0.46 | 3.70 |
| 19 | Riiad1 | 4.91 | 1.32E-71 | 2.74E-67 | 0.15 | 0.01 | 0.14 | 16.11 |
| 19 | Tmem163 | 4.99 | 8.21E-71 | 1.71E-66 | 0.30 | 0.04 | 0.26 | 7.92 |
| 19 | Mapt | 5.08 | 1.71E-70 | 3.55E-66 | 0.31 | 0.04 | 0.27 | 7.63 |
| 19 | Rab3c | 5.02 | 1.68E-69 | 3.49E-65 | 0.16 | 0.01 | 0.15 | 15.70 |
| 19 | Syt1 | 4.12 | 2.80E-69 | 5.82E-65 | 0.30 | 0.04 | 0.26 | 8.19 |
| 19 | Kif1a | 4.10 | 4.39E-69 | 9.12E-65 | 0.37 | 0.06 | 0.32 | 6.43 |
| 19 | Ckb | 3.75 | 8.91E-68 | 1.85E-63 | 0.78 | 0.33 | 0.45 | 2.36 |
| 19 | Rims2 | 3.86 | 1.11E-67 | 2.30E-63 | 0.37 | 0.06 | 0.31 | 6.44 |
| 19 | Gadd45g | 3.38 | 2.33E-67 | 4.84E-63 | 0.68 | 0.21 | 0.47 | 3.24 |
| 19 | Mllt11 | 4.27 | 2.86E-67 | 5.95E-63 | 0.72 | 0.26 | 0.47 | 2.80 |
| 19 | Cldn3 | 2.89 | 4.58E-67 | 9.52E-63 | 0.63 | 0.16 | 0.47 | 3.91 |
| 19 | Ripor2 | 5.10 | 5.23E-66 | 1.09E-61 | 0.13 | 0.01 | 0.12 | 18.14 |
| 19 | Arfgef3 | 5.05 | 8.07E-66 | 1.68E-61 | 0.19 | 0.02 | 0.17 | 11.69 |
| 19 | Phactr1 | 4.22 | 1.40E-65 | 2.92E-61 | 0.24 | 0.03 | 0.21 | 9.40 |
| 19 | Ebf1 | 3.33 | 2.75E-65 | 5.73E-61 | 0.75 | 0.30 | 0.46 | 2.55 |
| 19 | Rassf4 | 4.26 | 3.70E-65 | 7.70E-61 | 0.25 | 0.03 | 0.23 | 9.04 |
| 19 | Cerkl | 4.31 | 1.14E-64 | 2.37E-60 | 0.14 | 0.01 | 0.13 | 15.44 |
| 19 | Tagln3 | 4.16 | 1.97E-64 | 4.11E-60 | 0.72 | 0.26 | 0.46 | 2.75 |
| 19 | Cdyl2 | 4.55 | 8.88E-64 | 1.85E-59 | 0.33 | 0.05 | 0.28 | 6.49 |
| 19 | Cadm1 | 2.93 | 3.16E-63 | 6.57E-59 | 0.80 | 0.39 | 0.42 | 2.08 |
| 19 | Lmo1 | 3.74 | 5.56E-63 | 1.16E-58 | 0.41 | 0.07 | 0.34 | 5.62 |
| 19 | Trim9 | 5.07 | 5.60E-63 | 1.16E-58 | 0.10 | 0.01 | 0.10 | 20.40 |
| 19 | Klf7 | 3.78 | 6.49E-62 | 1.35E-57 | 0.67 | 0.23 | 0.44 | 2.95 |
| 19 | Tox3 | 3.19 | 7.89E-62 | 1.64E-57 | 0.68 | 0.23 | 0.45 | 2.96 |
| 19 | Atf5 | 4.45 | 7.89E-62 | 1.64E-57 | 0.62 | 0.19 | 0.43 | 3.21 |
| 19 | Nkain4 | 4.62 | 2.32E-61 | 4.83E-57 | 0.22 | 0.02 | 0.20 | 9.29 |
| 19 | 1500004A13Rik | 4.19 | 5.51E-61 | 1.15E-56 | 0.39 | 0.07 | 0.32 | 5.51 |
| 19 | Cacna2d1 | 3.97 | 7.55E-61 | 1.57E-56 | 0.49 | 0.12 | 0.38 | 4.22 |
| 19 | Hecw1 | 3.72 | 1.00E-60 | 2.08E-56 | 0.24 | 0.03 | 0.21 | 9.04 |
| 19 | Ucp2 | 3.16 | 1.65E-60 | 3.43E-56 | 0.80 | 0.36 | 0.45 | 2.26 |
| 19 | Nrsn1 | 5.26 | 2.13E-60 | 4.43E-56 | 0.15 | 0.01 | 0.14 | 14.50 |
| 19 | Hid1 | 4.60 | 3.19E-60 | 6.63E-56 | 0.16 | 0.01 | 0.15 | 13.08 |
| 19 | Fam20c | 3.95 | 5.83E-60 | 1.21E-55 | 0.21 | 0.02 | 0.19 | 9.59 |

**Supplementary Table 1. Marker genes for the original clusters of integrated clustering of E11.5 control and NotchGOF λ-junction cells.**

| | | | | | | | | |
|---|---|---|---|---|---|---|---|---|
| 19 | Basp1 | 1.93 | 6.02E-60 | 1.25E-55 | 0.96 | 0.96 | 0.01 | 1.01 |
| 19 | Efna3 | 4.74 | 7.59E-60 | 1.58E-55 | 0.35 | 0.06 | 0.29 | 5.92 |
| 19 | Numbl | 3.29 | 2.46E-59 | 5.13E-55 | 0.52 | 0.13 | 0.39 | 4.03 |
| 19 | Zc3h12c | 3.36 | 8.37E-59 | 1.74E-54 | 0.36 | 0.06 | 0.29 | 5.82 |
| 19 | Npdc1 | 3.23 | 1.07E-58 | 2.22E-54 | 0.47 | 0.10 | 0.37 | 4.52 |
| 19 | Ksr2 | 4.71 | 1.10E-58 | 2.29E-54 | 0.15 | 0.01 | 0.13 | 13.18 |
| 19 | Lrp11 | 3.60 | 1.80E-58 | 3.75E-54 | 0.27 | 0.03 | 0.23 | 7.79 |
| 19 | Serpini1 | 3.44 | 3.92E-58 | 8.15E-54 | 0.36 | 0.06 | 0.29 | 5.82 |
| 19 | Ndufa4l2 | 4.25 | 3.93E-57 | 8.18E-53 | 0.23 | 0.03 | 0.20 | 8.48 |
| 19 | Mical2 | 3.64 | 1.15E-56 | 2.40E-52 | 0.22 | 0.03 | 0.20 | 8.92 |
| 19 | Nrxn2 | 3.81 | 1.64E-56 | 3.41E-52 | 0.30 | 0.05 | 0.26 | 6.54 |
| 19 | AC109619.1 | 4.71 | 5.93E-56 | 1.23E-51 | 0.10 | 0.01 | 0.10 | 20.40 |
| 19 | Smpd3 | 4.01 | 6.03E-56 | 1.25E-51 | 0.15 | 0.01 | 0.14 | 12.58 |
| 19 | Cemip2 | 3.00 | 2.51E-55 | 5.23E-51 | 0.79 | 0.45 | 0.34 | 1.76 |
| 19 | Emx2os | 3.87 | 3.90E-55 | 8.10E-51 | 0.18 | 0.02 | 0.16 | 10.94 |
| 19 | Map1a | 3.21 | 4.53E-55 | 9.42E-51 | 0.42 | 0.09 | 0.33 | 4.69 |
| 19 | Aplp1 | 3.55 | 6.79E-55 | 1.41E-50 | 0.62 | 0.21 | 0.41 | 2.95 |
| 19 | Fam124a | 4.29 | 6.90E-55 | 1.43E-50 | 0.14 | 0.01 | 0.13 | 13.90 |
| 19 | Gjb5 | 5.24 | 2.43E-54 | 5.06E-50 | 0.11 | 0.01 | 0.11 | 16.29 |
| 19 | Crip2 | 2.61 | 1.03E-53 | 2.15E-49 | 0.60 | 0.17 | 0.43 | 3.49 |
| 19 | Ptms | 1.56 | 1.42E-53 | 2.95E-49 | 0.95 | 0.95 | -0.01 | 0.99 |
| 19 | Art1 | 5.47 | 5.46E-53 | 1.14E-48 | 0.12 | 0.01 | 0.11 | 15.00 |
| 19 | Spint2 | 1.96 | 7.88E-53 | 1.64E-48 | 0.81 | 0.31 | 0.50 | 2.60 |
| 19 | Rgs16 | 3.99 | 2.20E-52 | 4.57E-48 | 0.29 | 0.05 | 0.24 | 6.42 |
| 19 | Slc27a2 | 4.16 | 7.94E-52 | 1.65E-47 | 0.16 | 0.01 | 0.14 | 11.21 |
| 19 | Sel1l3 | 4.06 | 9.97E-52 | 2.07E-47 | 0.18 | 0.02 | 0.16 | 10.29 |
| 19 | Kdm7a | 2.58 | 1.52E-51 | 3.17E-47 | 0.75 | 0.35 | 0.41 | 2.16 |
| 19 | Atl1 | 3.29 | 2.03E-51 | 4.23E-47 | 0.30 | 0.05 | 0.25 | 6.28 |
| 19 | Hpca | 5.58 | 2.43E-51 | 5.04E-47 | 0.11 | 0.01 | 0.10 | 15.43 |
| 19 | Chrnb2 | 4.47 | 6.46E-51 | 1.34E-46 | 0.11 | 0.01 | 0.10 | 15.43 |
| 19 | Snap91 | 3.48 | 1.36E-50 | 2.83E-46 | 0.22 | 0.03 | 0.19 | 8.35 |
| 19 | Nnat | 2.69 | 2.40E-50 | 5.00E-46 | 0.97 | 0.94 | 0.03 | 1.03 |
| 19 | Ada | 3.52 | 1.00E-49 | 2.08E-45 | 0.38 | 0.08 | 0.30 | 4.81 |
| 19 | Mtcl1 | 2.51 | 1.13E-48 | 2.34E-44 | 0.48 | 0.12 | 0.36 | 3.97 |
| 19 | Apba2 | 3.56 | 1.57E-48 | 3.27E-44 | 0.33 | 0.06 | 0.27 | 5.42 |
| 19 | Gnao1 | 3.56 | 5.70E-48 | 1.19E-43 | 0.38 | 0.08 | 0.30 | 4.63 |
| 19 | Col9a2 | 4.16 | 8.36E-48 | 1.74E-43 | 0.42 | 0.10 | 0.32 | 4.14 |
| 19 | Robo2 | 2.92 | 8.97E-48 | 1.86E-43 | 0.58 | 0.19 | 0.39 | 3.04 |
| 19 | Mxi1 | 2.51 | 2.13E-47 | 4.43E-43 | 0.79 | 0.43 | 0.36 | 1.85 |
| 19 | Phyhipl | 3.46 | 2.49E-47 | 5.17E-43 | 0.25 | 0.04 | 0.22 | 6.66 |
| 19 | Pcbp4 | 1.58 | 2.70E-47 | 5.62E-43 | 0.92 | 0.87 | 0.05 | 1.06 |
| 19 | Myo5b | 3.30 | 5.04E-47 | 1.05E-42 | 0.27 | 0.04 | 0.23 | 6.30 |
| 19 | Bcl2l1 | 2.43 | 2.31E-46 | 4.80E-42 | 0.75 | 0.37 | 0.38 | 2.02 |
| 19 | Fam78b | 4.06 | 3.84E-46 | 7.99E-42 | 0.22 | 0.03 | 0.19 | 7.48 |
| 19 | Tenm1 | 4.07 | 7.63E-46 | 1.59E-41 | 0.14 | 0.01 | 0.13 | 11.58 |
| 19 | Map9 | 2.82 | 8.16E-46 | 1.70E-41 | 0.55 | 0.18 | 0.37 | 3.08 |
| 19 | Tmem198 | 4.94 | 1.30E-45 | 2.70E-41 | 0.10 | 0.01 | 0.10 | 14.57 |
| 19 | B630019K06Rik | 3.33 | 1.81E-45 | 3.77E-41 | 0.40 | 0.10 | 0.31 | 4.25 |
| 19 | Kif5c | 2.80 | 6.47E-45 | 1.35E-40 | 0.80 | 0.49 | 0.32 | 1.65 |
| 19 | Ddr1 | 2.51 | 9.36E-45 | 1.95E-40 | 0.48 | 0.13 | 0.35 | 3.71 |
| 19 | Svip | 3.23 | 1.19E-44 | 2.47E-40 | 0.39 | 0.09 | 0.30 | 4.31 |
| 19 | Parp6 | 3.10 | 1.40E-44 | 2.91E-40 | 0.43 | 0.11 | 0.32 | 3.88 |
| 19 | Selenom | 3.85 | 4.01E-44 | 8.33E-40 | 0.78 | 0.52 | 0.26 | 1.50 |
| 19 | Plekha6 | 4.23 | 5.08E-44 | 1.06E-39 | 0.16 | 0.02 | 0.14 | 9.81 |
| 19 | Nlk | 2.86 | 5.49E-44 | 1.14E-39 | 0.64 | 0.27 | 0.37 | 2.35 |
| 19 | Nuak2 | 3.54 | 6.26E-44 | 1.30E-39 | 0.24 | 0.04 | 0.20 | 6.51 |

**Supplementary Table 1. Marker genes for the original clusters of integrated clustering of E11.5 control and NotchGOF λ-junction cells.**

| | | | | | | | | |
|---|---|---|---|---|---|---|---|---|
| 19 | Calm2 | 1.14 | 8.03E-44 | 1.67E-39 | 0.96 | 0.98 | -0.02 | 0.98 |
| 19 | Sptbn2 | 2.94 | 2.14E-43 | 4.44E-39 | 0.43 | 0.11 | 0.32 | 3.88 |
| 19 | Rnf182 | 3.73 | 5.06E-43 | 1.05E-38 | 0.57 | 0.21 | 0.35 | 2.67 |
| 19 | Ptma | 0.76 | 1.02E-42 | 2.13E-38 | 1.00 | 1.00 | 0.00 | 1.00 |
| 19 | Plk4 | 2.75 | 6.37E-42 | 1.33E-37 | 0.79 | 0.55 | 0.24 | 1.43 |
| 19 | Stk33 | 4.92 | 9.06E-42 | 1.88E-37 | 0.11 | 0.01 | 0.11 | 12.67 |
| 19 | Abcd2 | 3.05 | 1.54E-41 | 3.20E-37 | 0.34 | 0.07 | 0.27 | 4.68 |
| 19 | Zbtb18 | 3.00 | 1.54E-41 | 3.21E-37 | 0.42 | 0.11 | 0.31 | 3.82 |
| 19 | Mturn | 3.13 | 1.65E-41 | 3.42E-37 | 0.30 | 0.06 | 0.24 | 5.10 |
| 19 | Jpt1 | 1.35 | 2.01E-41 | 4.18E-37 | 0.94 | 0.94 | 0.00 | 1.00 |
| 19 | Amotl2 | 2.51 | 3.65E-41 | 7.60E-37 | 0.69 | 0.34 | 0.35 | 2.04 |
| 19 | Rgmb | 3.19 | 7.09E-41 | 1.47E-36 | 0.45 | 0.12 | 0.32 | 3.60 |
| 19 | Pak3 | 2.86 | 8.04E-41 | 1.67E-36 | 0.46 | 0.14 | 0.33 | 3.41 |
| 19 | Rgs4 | 4.38 | 1.29E-40 | 2.68E-36 | 0.23 | 0.04 | 0.19 | 6.36 |
| 19 | Amer2 | 2.69 | 2.28E-40 | 4.74E-36 | 0.42 | 0.11 | 0.31 | 3.92 |
| 19 | Mbp | 3.03 | 2.63E-40 | 5.47E-36 | 0.39 | 0.09 | 0.29 | 4.11 |
| 19 | Gal | 3.16 | 3.77E-40 | 7.84E-36 | 0.13 | 0.01 | 0.12 | 10.23 |
| 19 | Plcxd3 | 4.80 | 7.70E-40 | 1.60E-35 | 0.11 | 0.01 | 0.10 | 12.00 |
| 19 | Col16a1 | 4.80 | 1.12E-39 | 2.33E-35 | 0.27 | 0.05 | 0.22 | 5.52 |
| 19 | Ndufaf3 | 2.61 | 1.61E-39 | 3.34E-35 | 0.72 | 0.42 | 0.31 | 1.74 |
| 19 | Rbfox3 | 4.18 | 2.98E-39 | 6.20E-35 | 0.11 | 0.01 | 0.10 | 12.00 |
| 19 | Eif4e3 | 2.47 | 3.33E-39 | 6.93E-35 | 0.64 | 0.28 | 0.36 | 2.30 |
| 19 | Ctnna2 | 3.23 | 3.88E-39 | 8.07E-35 | 0.27 | 0.05 | 0.22 | 5.52 |
| 19 | Ptpn6 | 3.14 | 5.46E-39 | 1.13E-34 | 0.23 | 0.04 | 0.19 | 6.36 |
| 19 | Sult2b1 | 3.89 | 6.74E-39 | 1.40E-34 | 0.15 | 0.02 | 0.13 | 8.88 |
| 19 | Fam107a | 2.71 | 1.21E-38 | 2.51E-34 | 0.25 | 0.04 | 0.21 | 5.75 |
| 19 | Ssx2ip | 2.29 | 1.87E-38 | 3.89E-34 | 0.72 | 0.36 | 0.36 | 1.99 |
| 19 | Olfm1 | 2.80 | 4.31E-38 | 8.96E-34 | 0.57 | 0.23 | 0.35 | 2.53 |
| 19 | Bcl7a | 1.86 | 6.98E-38 | 1.45E-33 | 0.83 | 0.67 | 0.16 | 1.24 |
| 19 | Fam219a | 3.37 | 1.13E-37 | 2.35E-33 | 0.21 | 0.03 | 0.18 | 6.39 |
| 19 | Sox11 | 0.90 | 1.28E-37 | 2.66E-33 | 0.95 | 0.97 | -0.02 | 0.98 |
| 19 | Kcnn2 | 3.94 | 1.36E-37 | 2.82E-33 | 0.12 | 0.01 | 0.11 | 10.91 |
| 19 | Cyth2 | 1.87 | 1.47E-37 | 3.06E-33 | 0.83 | 0.67 | 0.16 | 1.23 |
| 19 | Cecr2 | 2.31 | 1.50E-37 | 3.12E-33 | 0.58 | 0.22 | 0.36 | 2.60 |
| 19 | Cldn9 | 2.44 | 1.75E-37 | 3.63E-33 | 0.37 | 0.09 | 0.28 | 4.17 |
| 19 | Slc43a2 | 3.17 | 1.93E-37 | 4.02E-33 | 0.24 | 0.04 | 0.20 | 5.88 |
| 19 | Fam131a | 3.71 | 2.06E-37 | 4.28E-33 | 0.16 | 0.02 | 0.14 | 8.15 |
| 19 | C530008M17Rik | 2.36 | 2.15E-37 | 4.48E-33 | 0.73 | 0.42 | 0.31 | 1.74 |
| 19 | Nat14 | 3.20 | 2.76E-37 | 5.74E-33 | 0.24 | 0.04 | 0.20 | 5.74 |
| 19 | Baz1a | 2.81 | 3.33E-37 | 6.92E-33 | 0.80 | 0.56 | 0.24 | 1.43 |
| 19 | Wdr47 | 2.66 | 7.03E-37 | 1.46E-32 | 0.37 | 0.09 | 0.28 | 3.97 |
| 19 | Gabarapl2 | 1.49 | 1.26E-36 | 2.63E-32 | 0.88 | 0.82 | 0.06 | 1.07 |
| 19 | Wscd1 | 4.03 | 1.55E-36 | 3.23E-32 | 0.18 | 0.03 | 0.16 | 7.24 |
| 19 | Smarcd2 | 2.72 | 3.93E-36 | 8.18E-32 | 0.63 | 0.29 | 0.34 | 2.17 |
| 19 | Jag2 | 2.35 | 1.20E-35 | 2.50E-31 | 0.34 | 0.08 | 0.26 | 4.23 |
| 19 | Rhbdl3 | 3.90 | 1.88E-35 | 3.91E-31 | 0.11 | 0.01 | 0.10 | 10.80 |
| 19 | Ascl1 | 4.41 | 2.46E-35 | 5.11E-31 | 0.13 | 0.01 | 0.11 | 9.77 |
| 19 | Ank2 | 3.64 | 1.20E-34 | 2.50E-30 | 0.56 | 0.25 | 0.32 | 2.29 |
| 19 | Syt11 | 3.03 | 2.15E-34 | 4.48E-30 | 0.63 | 0.31 | 0.32 | 2.06 |
| 19 | Dlx6 | 3.38 | 2.98E-34 | 6.20E-30 | 0.34 | 0.09 | 0.26 | 3.99 |
| 19 | Ccser2 | 2.35 | 3.68E-34 | 7.64E-30 | 0.60 | 0.27 | 0.33 | 2.24 |
| 19 | Nol4l | 2.55 | 4.29E-34 | 8.91E-30 | 0.48 | 0.17 | 0.32 | 2.92 |
| 19 | Trnp1 | 4.11 | 5.48E-34 | 1.14E-29 | 0.18 | 0.03 | 0.15 | 7.00 |
| 19 | Rtn2 | 2.85 | 5.73E-34 | 1.19E-29 | 0.34 | 0.08 | 0.25 | 4.06 |
| 19 | Otx2 | 2.80 | 7.86E-34 | 1.63E-29 | 0.38 | 0.10 | 0.28 | 3.76 |
| 19 | Lipg | 3.30 | 9.91E-34 | 2.06E-29 | 0.21 | 0.04 | 0.18 | 5.86 |

**Supplementary Table 1. Marker genes for the original clusters of integrated clustering of E11.5 control and NotchGOF λ-junction cells.**

| | | | | | | | | |
|---|---|---|---|---|---|---|---|---|
| 19 | Tekt2 | 4.15 | 1.21E-33 | 2.51E-29 | 0.12 | 0.01 | 0.11 | 9.23 |
| 19 | Krt8 | 1.59 | 1.33E-33 | 2.77E-29 | 0.68 | 0.26 | 0.42 | 2.62 |
| 19 | Mkrn1 | 1.99 | 2.35E-33 | 4.89E-29 | 0.74 | 0.43 | 0.32 | 1.74 |
| 19 | Jakmip2 | 2.95 | 2.81E-33 | 5.84E-29 | 0.43 | 0.14 | 0.29 | 3.01 |
| 19 | Mapk8ip1 | 2.65 | 2.91E-33 | 6.05E-29 | 0.58 | 0.26 | 0.33 | 2.27 |
| 19 | Grik2 | 3.62 | 4.14E-33 | 8.60E-29 | 0.16 | 0.02 | 0.14 | 7.41 |
| 19 | Actb | 0.61 | 5.11E-33 | 1.06E-28 | 1.00 | 1.00 | 0.00 | 1.00 |
| 19 | Map1lc3b | 1.31 | 9.99E-33 | 2.08E-28 | 0.90 | 0.93 | -0.02 | 0.98 |
| 19 | Syt5 | 3.10 | 1.62E-32 | 3.36E-28 | 0.27 | 0.06 | 0.21 | 4.67 |
| 19 | Mbd2 | 2.04 | 1.65E-32 | 3.44E-28 | 0.79 | 0.56 | 0.23 | 1.41 |
| 19 | Trim62 | 2.91 | 1.87E-32 | 3.89E-28 | 0.36 | 0.10 | 0.26 | 3.61 |
| 19 | Pcdh20 | 3.84 | 3.39E-32 | 7.05E-28 | 0.13 | 0.01 | 0.11 | 9.07 |
| 19 | Rab3b | 3.93 | 3.90E-32 | 8.10E-28 | 0.27 | 0.06 | 0.21 | 4.44 |
| 19 | Cdkn1a | 2.14 | 4.38E-32 | 9.11E-28 | 0.40 | 0.11 | 0.29 | 3.62 |
| 19 | Mdm1 | 2.81 | 5.52E-32 | 1.15E-27 | 0.60 | 0.30 | 0.30 | 2.02 |
| 19 | Hpcal1 | 2.31 | 5.65E-32 | 1.17E-27 | 0.66 | 0.33 | 0.33 | 1.98 |
| 19 | Trp53inp2 | 2.66 | 6.54E-32 | 1.36E-27 | 0.36 | 0.10 | 0.26 | 3.70 |
| 19 | Onecut2 | 4.50 | 1.10E-31 | 2.29E-27 | 0.15 | 0.02 | 0.13 | 7.55 |
| 19 | Ildr1 | 3.09 | 1.31E-31 | 2.72E-27 | 0.21 | 0.04 | 0.17 | 5.70 |
| 19 | Prr36 | 2.45 | 1.94E-31 | 4.04E-27 | 0.34 | 0.09 | 0.25 | 3.87 |
| 19 | Abcg1 | 2.77 | 2.72E-31 | 5.66E-27 | 0.33 | 0.08 | 0.24 | 3.96 |
| 19 | A330084C13Rik | 3.72 | 2.89E-31 | 6.02E-27 | 0.10 | 0.01 | 0.09 | 10.20 |
| 19 | Rab3a | 3.15 | 4.09E-31 | 8.50E-27 | 0.42 | 0.14 | 0.28 | 3.08 |
| 19 | Scrn1 | 3.79 | 5.69E-31 | 1.18E-26 | 0.22 | 0.04 | 0.18 | 5.19 |
| 19 | Dbndd1 | 2.99 | 6.20E-31 | 1.29E-26 | 0.25 | 0.05 | 0.20 | 4.69 |
| 19 | Samd10 | 2.63 | 9.55E-31 | 1.99E-26 | 0.34 | 0.09 | 0.25 | 3.81 |
| 19 | Eid1 | 0.97 | 1.87E-30 | 3.90E-26 | 0.94 | 0.97 | -0.03 | 0.97 |
| 19 | Ccdc88c | 2.01 | 2.49E-30 | 5.19E-26 | 0.46 | 0.16 | 0.31 | 2.96 |
| 19 | Shb | 2.55 | 6.37E-30 | 1.33E-25 | 0.50 | 0.20 | 0.30 | 2.55 |
| 19 | Epcam | 0.95 | 7.20E-30 | 1.50E-25 | 0.78 | 0.35 | 0.44 | 2.26 |
| 19 | Gdi1 | 2.26 | 9.08E-30 | 1.89E-25 | 0.70 | 0.42 | 0.28 | 1.67 |
| 19 | Tomm34 | 2.03 | 9.56E-30 | 1.99E-25 | 0.69 | 0.42 | 0.28 | 1.67 |
| 19 | Ube2b | 1.11 | 1.28E-29 | 2.67E-25 | 0.91 | 0.93 | -0.02 | 0.98 |
| 19 | Tfeb | 3.36 | 2.15E-29 | 4.48E-25 | 0.16 | 0.02 | 0.13 | 6.83 |
| 19 | Sp8 | 1.82 | 2.26E-29 | 4.70E-25 | 0.44 | 0.14 | 0.30 | 3.17 |
| 19 | Slc26a10 | 3.28 | 2.32E-29 | 4.82E-25 | 0.13 | 0.02 | 0.11 | 8.47 |
| 19 | Cspg5 | 3.88 | 2.96E-29 | 6.16E-25 | 0.14 | 0.02 | 0.12 | 7.32 |
| 19 | Gdpd1 | 2.49 | 3.53E-29 | 7.35E-25 | 0.58 | 0.27 | 0.31 | 2.15 |
| 19 | Zmat4 | 3.67 | 4.69E-29 | 9.76E-25 | 0.25 | 0.06 | 0.19 | 4.49 |
| 19 | Ccdc96 | 3.28 | 5.59E-29 | 1.16E-24 | 0.15 | 0.02 | 0.13 | 6.86 |
| 19 | Celf5 | 3.59 | 9.91E-29 | 2.06E-24 | 0.15 | 0.02 | 0.13 | 6.86 |
| 19 | Tdrkh | 2.91 | 9.96E-29 | 2.07E-24 | 0.26 | 0.06 | 0.20 | 4.32 |
| 19 | Tmcc2 | 2.42 | 1.07E-28 | 2.23E-24 | 0.31 | 0.08 | 0.23 | 3.82 |
| 19 | Hspa12a | 3.14 | 1.29E-28 | 2.68E-24 | 0.18 | 0.03 | 0.15 | 6.03 |
| 19 | Dlx6os1 | 3.16 | 1.81E-28 | 3.75E-24 | 0.21 | 0.04 | 0.17 | 5.28 |
| 19 | Rcor2 | 1.83 | 1.82E-28 | 3.78E-24 | 0.72 | 0.49 | 0.23 | 1.48 |
| 19 | Prex1 | 2.33 | 2.09E-28 | 4.34E-24 | 0.49 | 0.19 | 0.30 | 2.53 |
| 19 | Mcf2l | 2.87 | 3.31E-28 | 6.89E-24 | 0.28 | 0.07 | 0.21 | 4.07 |
| 19 | Pitx1 | 1.69 | 3.91E-28 | 8.13E-24 | 0.51 | 0.18 | 0.33 | 2.81 |
| 19 | Gse1 | 2.12 | 5.44E-28 | 1.13E-23 | 0.60 | 0.30 | 0.29 | 1.97 |
| 19 | Zfhx2 | 1.91 | 6.07E-28 | 1.26E-23 | 0.57 | 0.25 | 0.32 | 2.25 |
| 19 | Stox1 | 4.03 | 7.41E-28 | 1.54E-23 | 0.16 | 0.03 | 0.14 | 6.27 |
| 19 | Hap1 | 3.31 | 7.82E-28 | 1.63E-23 | 0.24 | 0.05 | 0.18 | 4.61 |
| 19 | Pnmal2 | 3.03 | 9.53E-28 | 1.98E-23 | 0.27 | 0.06 | 0.20 | 4.14 |
| 19 | Nefm | 4.38 | 9.73E-28 | 2.02E-23 | 0.16 | 0.03 | 0.14 | 6.27 |
| 19 | Lrrn3 | 2.71 | 1.53E-27 | 3.18E-23 | 0.28 | 0.07 | 0.21 | 3.99 |

**Supplementary Table 1. Marker genes for the original clusters of integrated clustering of E11.5 control and NotchGOF λ-junction cells.**

| | | | | | | | | |
|---|---|---|---|---|---|---|---|---|
| 19 | Fyn | 1.76 | 7.96E-27 | 1.66E-22 | 0.72 | 0.44 | 0.28 | 1.64 |
| 19 | Ddit4 | 2.21 | 8.25E-27 | 1.72E-22 | 0.68 | 0.38 | 0.30 | 1.80 |
| 19 | Stxbp1 | 2.62 | 8.57E-27 | 1.78E-22 | 0.33 | 0.09 | 0.23 | 3.46 |
| 19 | Kif21a | 1.99 | 8.71E-27 | 1.81E-22 | 0.63 | 0.31 | 0.32 | 2.02 |
| 19 | Ralgds | 2.18 | 8.87E-27 | 1.85E-22 | 0.52 | 0.22 | 0.30 | 2.35 |
| 19 | Srgap1 | 2.04 | 1.23E-26 | 2.56E-22 | 0.63 | 0.34 | 0.29 | 1.86 |
| 19 | Gm19938 | 3.31 | 1.25E-26 | 2.61E-22 | 0.15 | 0.02 | 0.13 | 6.57 |
| 19 | Gck | 3.37 | 1.35E-26 | 2.81E-22 | 0.16 | 0.03 | 0.13 | 6.28 |
| 19 | Rnf5 | 1.74 | 1.72E-26 | 3.57E-22 | 0.81 | 0.65 | 0.16 | 1.25 |
| 19 | Calm1 | 1.17 | 1.95E-26 | 4.06E-22 | 0.95 | 0.99 | -0.05 | 0.95 |
| 19 | Lzts1 | 2.85 | 3.28E-26 | 6.82E-22 | 0.20 | 0.04 | 0.16 | 5.10 |
| 19 | Mycl | 2.06 | 4.02E-26 | 8.36E-22 | 0.60 | 0.30 | 0.29 | 1.97 |
| 19 | Eif1 | 0.49 | 4.93E-26 | 1.03E-21 | 0.98 | 1.00 | -0.02 | 0.98 |
| 19 | Marchf9 | 2.24 | 6.52E-26 | 1.36E-21 | 0.42 | 0.15 | 0.27 | 2.85 |
| 19 | Adgrb2 | 2.76 | 6.58E-26 | 1.37E-21 | 0.21 | 0.04 | 0.17 | 4.91 |
| 19 | Mybl1 | 3.48 | 7.08E-26 | 1.47E-21 | 0.25 | 0.06 | 0.19 | 4.19 |
| 19 | Lpcat2 | 2.91 | 9.36E-26 | 1.95E-21 | 0.19 | 0.04 | 0.16 | 5.08 |
| 19 | Fgf12 | 2.95 | 1.02E-25 | 2.12E-21 | 0.22 | 0.05 | 0.18 | 4.65 |
| 19 | Ddx5 | 0.53 | 1.24E-25 | 2.58E-21 | 0.99 | 1.00 | -0.01 | 0.99 |
| 19 | Acer2 | 1.61 | 1.55E-25 | 3.22E-21 | 0.71 | 0.40 | 0.32 | 1.80 |
| 19 | Ppp1r14b | 0.63 | 3.01E-25 | 6.27E-21 | 0.95 | 0.99 | -0.04 | 0.96 |
| 19 | Neurl1a | 2.46 | 3.12E-25 | 6.49E-21 | 0.26 | 0.06 | 0.20 | 4.05 |
| 19 | Elmo1 | 3.53 | 3.78E-25 | 7.85E-21 | 0.21 | 0.05 | 0.17 | 4.69 |
| 19 | Rasgef1b | 3.08 | 3.78E-25 | 7.85E-21 | 0.22 | 0.05 | 0.17 | 4.62 |
| 19 | Gabrb3 | 3.45 | 3.90E-25 | 8.12E-21 | 0.10 | 0.01 | 0.09 | 8.50 |
| 19 | Kdm4b | 2.22 | 5.45E-25 | 1.13E-20 | 0.44 | 0.17 | 0.27 | 2.65 |
| 19 | Stil | 2.53 | 5.66E-25 | 1.18E-20 | 0.45 | 0.19 | 0.26 | 2.34 |
| 19 | Kdm1a | 0.96 | 7.00E-25 | 1.46E-20 | 0.88 | 0.92 | -0.04 | 0.95 |
| 19 | Mfsd6 | 2.82 | 7.94E-25 | 1.65E-20 | 0.18 | 0.03 | 0.15 | 5.32 |
| 19 | Casp3 | 1.67 | 1.02E-24 | 2.12E-20 | 0.82 | 0.72 | 0.10 | 1.13 |
| 19 | Slco5a1 | 2.55 | 1.11E-24 | 2.30E-20 | 0.30 | 0.08 | 0.21 | 3.60 |
| 19 | Rb1 | 2.06 | 1.28E-24 | 2.65E-20 | 0.43 | 0.16 | 0.28 | 2.73 |
| 19 | Arpc5 | 1.40 | 1.55E-24 | 3.22E-20 | 0.84 | 0.76 | 0.08 | 1.11 |
| 19 | Chd7 | 1.62 | 1.67E-24 | 3.47E-20 | 0.77 | 0.66 | 0.11 | 1.17 |
| 19 | Rufy3 | 3.31 | 1.72E-24 | 3.58E-20 | 0.49 | 0.23 | 0.26 | 2.15 |
| 19 | Actg1 | 0.64 | 1.78E-24 | 3.71E-20 | 0.99 | 1.00 | -0.01 | 0.99 |
| 19 | Prdx2 | 0.56 | 2.23E-24 | 4.63E-20 | 1.00 | 1.00 | 0.00 | 1.00 |
| 19 | Tle1 | 1.79 | 2.43E-24 | 5.05E-20 | 0.77 | 0.53 | 0.23 | 1.44 |
| 19 | Cdh2 | 1.35 | 2.84E-24 | 5.91E-20 | 0.71 | 0.42 | 0.29 | 1.68 |
| 19 | Synrg | 1.96 | 3.39E-24 | 7.04E-20 | 0.52 | 0.24 | 0.29 | 2.23 |
| 19 | Olfm2 | 2.47 | 3.50E-24 | 7.27E-20 | 0.34 | 0.10 | 0.23 | 3.24 |
| 19 | Tshz1 | 2.41 | 4.25E-24 | 8.85E-20 | 0.58 | 0.32 | 0.27 | 1.84 |
| 19 | Tsg101 | 1.21 | 4.82E-24 | 1.00E-19 | 0.87 | 0.85 | 0.02 | 1.03 |
| 19 | Atp6v1g1 | 0.92 | 5.37E-24 | 1.12E-19 | 0.90 | 0.97 | -0.06 | 0.94 |
| 19 | Nab1 | 2.09 | 5.73E-24 | 1.19E-19 | 0.38 | 0.13 | 0.25 | 2.95 |
| 19 | Napb | 2.71 | 6.15E-24 | 1.28E-19 | 0.23 | 0.05 | 0.18 | 4.24 |
| 19 | Sertad2 | 1.87 | 1.15E-23 | 2.39E-19 | 0.66 | 0.41 | 0.26 | 1.63 |
| 19 | Nedd4l | 2.31 | 1.24E-23 | 2.59E-19 | 0.39 | 0.14 | 0.25 | 2.72 |
| 19 | Rapgef5 | 2.52 | 1.71E-23 | 3.55E-19 | 0.36 | 0.12 | 0.24 | 3.03 |
| 19 | Spsb4 | 2.17 | 2.03E-23 | 4.23E-19 | 0.48 | 0.21 | 0.28 | 2.34 |
| 19 | Dctn3 | 1.37 | 2.17E-23 | 4.52E-19 | 0.81 | 0.79 | 0.03 | 1.04 |
| 19 | 1700088E04Rik | 2.86 | 2.41E-23 | 5.02E-19 | 0.22 | 0.05 | 0.17 | 4.21 |
| 19 | Klf13 | 2.02 | 8.50E-23 | 1.77E-18 | 0.61 | 0.37 | 0.24 | 1.65 |
| 19 | Pdrg1 | 2.09 | 9.05E-23 | 1.88E-18 | 0.74 | 0.54 | 0.20 | 1.37 |
| 19 | Saraf | 2.07 | 1.87E-22 | 3.88E-18 | 0.71 | 0.54 | 0.17 | 1.31 |
| 19 | Bex2 | 1.05 | 2.18E-22 | 4.53E-18 | 0.91 | 0.85 | 0.06 | 1.07 |

**Supplementary Table 1. Marker genes for the original clusters of integrated clustering of E11.5 control and NotchGOF λ-junction cells.**

| | | | | | | | | |
|---|---|---|---|---|---|---|---|---|
| 19 | Fdps | 1.26 | 2.18E-22 | 4.53E-18 | 0.86 | 0.82 | 0.04 | 1.05 |
| 19 | Prox1 | 3.37 | 2.49E-22 | 5.19E-18 | 0.13 | 0.02 | 0.11 | 6.05 |
| 19 | Sh3bgrl | 1.16 | 5.30E-22 | 1.10E-17 | 0.87 | 0.86 | 0.02 | 1.02 |
| 19 | Ube2s | 1.12 | 5.68E-22 | 1.18E-17 | 0.94 | 0.97 | -0.03 | 0.97 |
| 19 | Fry | 2.67 | 6.90E-22 | 1.43E-17 | 0.22 | 0.06 | 0.17 | 4.05 |
| 19 | Kcnma1 | 4.18 | 7.51E-22 | 1.56E-17 | 0.11 | 0.02 | 0.09 | 7.20 |
| 19 | Rnf165 | 2.00 | 8.43E-22 | 1.75E-17 | 0.51 | 0.25 | 0.27 | 2.08 |
| 19 | D230030E09Rik | 2.74 | 1.01E-21 | 2.11E-17 | 0.23 | 0.06 | 0.17 | 4.02 |
| 19 | Kif5b | 0.81 | 1.71E-21 | 3.55E-17 | 0.91 | 0.94 | -0.03 | 0.97 |
| 19 | Atp6v0b | 1.40 | 1.97E-21 | 4.09E-17 | 0.80 | 0.74 | 0.06 | 1.09 |
| 19 | Gm5468 | 3.43 | 2.13E-21 | 4.43E-17 | 0.10 | 0.01 | 0.09 | 7.29 |
| 19 | Add2 | 2.39 | 2.74E-21 | 5.70E-17 | 0.29 | 0.09 | 0.20 | 3.25 |
| 19 | 2310040G24Rik | 3.76 | 3.45E-21 | 7.18E-17 | 0.16 | 0.03 | 0.13 | 4.94 |
| 19 | Atp8a1 | 2.68 | 5.12E-21 | 1.07E-16 | 0.28 | 0.09 | 0.20 | 3.25 |
| 19 | Espn | 2.27 | 6.63E-21 | 1.38E-16 | 0.33 | 0.11 | 0.22 | 3.04 |
| 19 | Ttll7 | 2.83 | 7.75E-21 | 1.61E-16 | 0.25 | 0.07 | 0.18 | 3.47 |
| 19 | Slain1 | 2.78 | 8.23E-21 | 1.71E-16 | 0.23 | 0.06 | 0.17 | 3.82 |
| 19 | Rab39b | 2.19 | 8.37E-21 | 1.74E-16 | 0.40 | 0.16 | 0.25 | 2.56 |
| 19 | Ubb | 0.47 | 1.07E-20 | 2.22E-16 | 0.99 | 1.00 | -0.01 | 0.99 |
| 19 | Ccdc92 | 2.54 | 1.27E-20 | 2.65E-16 | 0.27 | 0.08 | 0.19 | 3.35 |
| 19 | Ppp2r5b | 2.25 | 1.32E-20 | 2.75E-16 | 0.29 | 0.09 | 0.20 | 3.25 |
| 19 | Mien1 | 1.51 | 1.43E-20 | 2.98E-16 | 0.74 | 0.56 | 0.17 | 1.30 |
| 19 | Ncam1 | 3.45 | 1.72E-20 | 3.57E-16 | 0.58 | 0.36 | 0.22 | 1.61 |
| 19 | Ap3m2 | 2.75 | 1.98E-20 | 4.11E-16 | 0.18 | 0.04 | 0.14 | 4.61 |
| 19 | Enc1 | 2.61 | 2.08E-20 | 4.32E-16 | 0.46 | 0.22 | 0.24 | 2.10 |
| 19 | Mfap4 | 1.32 | 2.23E-20 | 4.63E-16 | 0.87 | 0.85 | 0.02 | 1.02 |
| 19 | Tmem200a | 2.89 | 2.35E-20 | 4.90E-16 | 0.22 | 0.06 | 0.16 | 3.88 |
| 19 | Cend1 | 2.96 | 2.42E-20 | 5.04E-16 | 0.22 | 0.06 | 0.16 | 3.78 |
| 19 | Rprm | 1.55 | 3.23E-20 | 6.71E-16 | 0.42 | 0.16 | 0.26 | 2.58 |
| 19 | C130071C03Rik | 2.51 | 3.25E-20 | 6.76E-16 | 0.21 | 0.05 | 0.16 | 4.06 |
| 19 | Bicd1 | 2.54 | 3.85E-20 | 8.00E-16 | 0.30 | 0.10 | 0.20 | 3.07 |
| 19 | Tcta | 2.63 | 4.60E-20 | 9.56E-16 | 0.23 | 0.06 | 0.17 | 3.75 |
| 19 | Eif4g2 | 0.49 | 5.06E-20 | 1.05E-15 | 0.93 | 0.99 | -0.05 | 0.94 |
| 19 | Cmpk1 | 1.14 | 5.20E-20 | 1.08E-15 | 0.83 | 0.79 | 0.04 | 1.05 |
| 19 | Lsr | 1.59 | 5.79E-20 | 1.21E-15 | 0.48 | 0.21 | 0.27 | 2.31 |
| 19 | Nabp1 | 2.87 | 5.94E-20 | 1.24E-15 | 0.15 | 0.03 | 0.12 | 5.03 |
| 19 | Tspan13 | 1.65 | 6.09E-20 | 1.27E-15 | 0.66 | 0.41 | 0.25 | 1.61 |
| 19 | Phf21b | 3.20 | 1.00E-19 | 2.09E-15 | 0.18 | 0.04 | 0.14 | 4.41 |
| 19 | Dubr | 2.76 | 1.31E-19 | 2.73E-15 | 0.17 | 0.04 | 0.13 | 4.57 |
| 19 | Ccsap | 2.58 | 1.42E-19 | 2.96E-15 | 0.32 | 0.11 | 0.21 | 2.87 |
| 19 | Gpm6b | 3.00 | 1.78E-19 | 3.70E-15 | 0.24 | 0.07 | 0.17 | 3.51 |
| 19 | 0610010F05Rik | 1.82 | 1.79E-19 | 3.72E-15 | 0.50 | 0.25 | 0.25 | 2.02 |
| 19 | Stx1a | 2.29 | 3.12E-19 | 6.49E-15 | 0.29 | 0.09 | 0.20 | 3.07 |
| 19 | Ralyl | 4.17 | 4.56E-19 | 9.48E-15 | 0.14 | 0.03 | 0.11 | 5.35 |
| 19 | Arid3b | 1.86 | 4.62E-19 | 9.61E-15 | 0.48 | 0.23 | 0.25 | 2.11 |
| 19 | Afap1 | 2.20 | 4.73E-19 | 9.85E-15 | 0.49 | 0.26 | 0.24 | 1.94 |
| 19 | H3f3b | 0.41 | 7.75E-19 | 1.61E-14 | 1.00 | 1.00 | 0.00 | 1.00 |
| 19 | Rab3d | 3.25 | 8.63E-19 | 1.80E-14 | 0.12 | 0.02 | 0.10 | 5.71 |
| 19 | Apbb1 | 2.10 | 9.23E-19 | 1.92E-14 | 0.55 | 0.32 | 0.23 | 1.71 |
| 19 | Ccp110 | 2.02 | 1.04E-18 | 2.16E-14 | 0.63 | 0.43 | 0.20 | 1.48 |
| 19 | Hist3h2a | 2.28 | 1.05E-18 | 2.18E-14 | 0.51 | 0.27 | 0.25 | 1.93 |
| 19 | Dpysl2 | 1.40 | 1.29E-18 | 2.69E-14 | 0.78 | 0.70 | 0.09 | 1.12 |
| 19 | BC031181 | 1.30 | 1.33E-18 | 2.78E-14 | 0.80 | 0.74 | 0.06 | 1.08 |
| 19 | Ccpg1 | 1.96 | 1.34E-18 | 2.78E-14 | 0.40 | 0.16 | 0.24 | 2.46 |
| 19 | Ackr1 | 3.57 | 1.59E-18 | 3.31E-14 | 0.10 | 0.02 | 0.09 | 6.38 |
| 19 | Dbn1 | 0.99 | 1.69E-18 | 3.51E-14 | 0.83 | 0.88 | -0.06 | 0.94 |

**Supplementary Table 1. Marker genes for the original clusters of integrated clustering of E11.5 control and NotchGOF λ-junction cells.**

| | | | | | | | | |
|---|---|---|---|---|---|---|---|---|
| 19 | Pou2f2 | 3.33 | 2.13E-18 | 4.43E-14 | 0.14 | 0.03 | 0.11 | 5.15 |
| 19 | Mtmr7 | 2.57 | 2.53E-18 | 5.27E-14 | 0.20 | 0.05 | 0.15 | 3.90 |
| 19 | Dpysl4 | 2.07 | 2.70E-18 | 5.62E-14 | 0.41 | 0.18 | 0.23 | 2.25 |
| 19 | Baz2b | 1.36 | 2.76E-18 | 5.75E-14 | 0.79 | 0.71 | 0.08 | 1.12 |
| 19 | Dstn | 0.73 | 3.43E-18 | 7.13E-14 | 0.91 | 0.94 | -0.03 | 0.97 |
| 19 | Mospd1 | 2.11 | 4.01E-18 | 8.33E-14 | 0.49 | 0.25 | 0.24 | 1.98 |
| 19 | Ralgps1 | 2.48 | 4.18E-18 | 8.69E-14 | 0.18 | 0.04 | 0.14 | 4.21 |
| 19 | Ddah2 | 0.89 | 4.84E-18 | 1.01E-13 | 0.93 | 0.95 | -0.02 | 0.98 |
| 19 | Scn3a | 3.32 | 5.01E-18 | 1.04E-13 | 0.21 | 0.06 | 0.15 | 3.66 |
| 19 | Tmem176b | 1.49 | 5.41E-18 | 1.13E-13 | 0.72 | 0.57 | 0.15 | 1.27 |
| 19 | Frrs1l | 2.85 | 6.72E-18 | 1.40E-13 | 0.15 | 0.03 | 0.12 | 4.83 |
| 19 | Tubb5 | 0.50 | 7.02E-18 | 1.46E-13 | 1.00 | 1.00 | 0.00 | 1.00 |
| 19 | Tecpr1 | 1.88 | 7.70E-18 | 1.60E-13 | 0.36 | 0.14 | 0.22 | 2.57 |
| 19 | Actr3b | 2.92 | 9.92E-18 | 2.06E-13 | 0.14 | 0.03 | 0.11 | 4.96 |
| 19 | Ttll1 | 2.29 | 9.97E-18 | 2.07E-13 | 0.39 | 0.16 | 0.23 | 2.43 |
| 19 | Akap17b | 2.68 | 1.27E-17 | 2.64E-13 | 0.32 | 0.12 | 0.20 | 2.64 |
| 19 | Riok3 | 1.51 | 1.33E-17 | 2.77E-13 | 0.68 | 0.50 | 0.18 | 1.35 |
| 19 | Pdgfa | 1.08 | 1.74E-17 | 3.63E-13 | 0.59 | 0.28 | 0.31 | 2.08 |
| 19 | Aldh1b1 | 3.08 | 1.76E-17 | 3.67E-13 | 0.11 | 0.02 | 0.09 | 6.00 |
| 19 | Smarca5 | 0.61 | 2.12E-17 | 4.41E-13 | 0.93 | 0.98 | -0.05 | 0.95 |
| 19 | Zfp386 | 1.81 | 3.28E-17 | 6.83E-13 | 0.65 | 0.41 | 0.23 | 1.56 |
| 19 | Pcp4l1 | 2.91 | 3.42E-17 | 7.10E-13 | 0.13 | 0.02 | 0.10 | 5.29 |
| 19 | Malat1 | 1.13 | 6.58E-17 | 1.37E-12 | 0.92 | 1.00 | -0.07 | 0.93 |
| 19 | Ypel3 | 1.71 | 7.11E-17 | 1.48E-12 | 0.71 | 0.57 | 0.14 | 1.25 |
| 19 | Fabp5 | 0.51 | 7.13E-17 | 1.48E-12 | 0.95 | 0.98 | -0.03 | 0.97 |
| 19 | Gm10544 | 3.04 | 7.69E-17 | 1.60E-12 | 0.15 | 0.03 | 0.11 | 4.68 |
| 19 | Cldn7 | 0.75 | 1.05E-16 | 2.19E-12 | 0.55 | 0.24 | 0.31 | 2.28 |
| 19 | Cox6a1 | 0.68 | 1.11E-16 | 2.31E-12 | 0.93 | 0.96 | -0.03 | 0.97 |
| 19 | Carhsp1 | 1.52 | 1.20E-16 | 2.50E-12 | 0.72 | 0.59 | 0.12 | 1.21 |
| 19 | Rsph9 | 2.29 | 1.22E-16 | 2.54E-12 | 0.30 | 0.11 | 0.19 | 2.76 |
| 19 | Tppp3 | 4.41 | 1.58E-16 | 3.28E-12 | 0.16 | 0.04 | 0.12 | 4.08 |
| 19 | Ywhag | 1.52 | 1.98E-16 | 4.11E-12 | 0.73 | 0.63 | 0.10 | 1.15 |
| 19 | Racgap1 | 1.49 | 2.05E-16 | 4.25E-12 | 0.71 | 0.53 | 0.19 | 1.35 |
| 19 | Amigo1 | 2.82 | 2.09E-16 | 4.36E-12 | 0.11 | 0.02 | 0.09 | 5.43 |
| 19 | Cxxc5 | 1.74 | 2.39E-16 | 4.96E-12 | 0.65 | 0.45 | 0.20 | 1.44 |
| 19 | Ago1 | 1.56 | 2.45E-16 | 5.09E-12 | 0.58 | 0.36 | 0.22 | 1.60 |
| 19 | Atp8a2 | 3.09 | 2.79E-16 | 5.81E-12 | 0.12 | 0.02 | 0.10 | 5.22 |
| 19 | Gm49066 | 2.40 | 2.80E-16 | 5.82E-12 | 0.18 | 0.04 | 0.13 | 3.98 |
| 19 | Trim24 | 1.34 | 3.30E-16 | 6.86E-12 | 0.77 | 0.71 | 0.05 | 1.07 |
| 19 | Sdk1 | 2.86 | 3.99E-16 | 8.30E-12 | 0.25 | 0.08 | 0.17 | 3.01 |
| 19 | Pim3 | 2.54 | 4.02E-16 | 8.36E-12 | 0.51 | 0.30 | 0.22 | 1.72 |
| 19 | Pacsin1 | 1.94 | 4.13E-16 | 8.58E-12 | 0.29 | 0.10 | 0.19 | 2.81 |
| 19 | Unc13b | 2.04 | 4.57E-16 | 9.50E-12 | 0.25 | 0.08 | 0.17 | 3.01 |
| 19 | Cep120 | 1.72 | 5.06E-16 | 1.05E-11 | 0.46 | 0.24 | 0.23 | 1.96 |
| 19 | Gm30015 | 4.20 | 5.50E-16 | 1.14E-11 | 0.10 | 0.02 | 0.08 | 5.67 |
| 19 | Dnajb6 | 0.96 | 5.99E-16 | 1.25E-11 | 0.88 | 0.89 | -0.01 | 0.99 |
| 19 | Cep97 | 2.34 | 6.13E-16 | 1.28E-11 | 0.41 | 0.21 | 0.21 | 2.00 |
| 19 | Selenok | 0.99 | 6.15E-16 | 1.28E-11 | 0.86 | 0.92 | -0.06 | 0.93 |
| 19 | Pick1 | 1.87 | 6.48E-16 | 1.35E-11 | 0.49 | 0.26 | 0.23 | 1.91 |
| 19 | Gm4651 | 2.50 | 7.87E-16 | 1.64E-11 | 0.13 | 0.03 | 0.10 | 4.88 |
| 19 | Pou3f1 | 3.08 | 7.95E-16 | 1.65E-11 | 0.13 | 0.03 | 0.10 | 4.88 |
| 19 | Cd55 | 2.56 | 8.92E-16 | 1.86E-11 | 0.12 | 0.02 | 0.10 | 5.00 |
| 19 | Rmst | 2.27 | 9.73E-16 | 2.02E-11 | 0.24 | 0.08 | 0.16 | 3.13 |
| 19 | Rnf32 | 2.81 | 9.77E-16 | 2.03E-11 | 0.12 | 0.02 | 0.10 | 5.00 |
| 19 | Msi2 | 1.11 | 1.22E-15 | 2.53E-11 | 0.78 | 0.68 | 0.10 | 1.14 |
| 19 | Dtna | 1.97 | 1.44E-15 | 3.00E-11 | 0.34 | 0.14 | 0.20 | 2.40 |

**Supplementary Table 1. Marker genes for the original clusters of integrated clustering of E11.5 control and NotchGOF λ-junction cells.**

| | | | | | | | | |
|---|---|---|---|---|---|---|---|---|
| 19 | Ache | 3.98 | 1.48E-15 | 3.07E-11 | 0.10 | 0.02 | 0.08 | 5.67 |
| 19 | Tmem108 | 1.84 | 1.64E-15 | 3.42E-11 | 0.44 | 0.22 | 0.22 | 2.02 |
| 19 | Sh2d3c | 2.06 | 1.65E-15 | 3.44E-11 | 0.28 | 0.10 | 0.18 | 2.77 |
| 19 | Kalrn | 1.99 | 1.84E-15 | 3.83E-11 | 0.40 | 0.20 | 0.21 | 2.05 |
| 19 | Fmn2 | 2.35 | 1.88E-15 | 3.92E-11 | 0.22 | 0.07 | 0.15 | 3.19 |
| 19 | Iqgap1 | 1.28 | 2.01E-15 | 4.18E-11 | 0.70 | 0.47 | 0.23 | 1.48 |
| 19 | Srp14 | 0.76 | 2.10E-15 | 4.36E-11 | 0.90 | 0.95 | -0.05 | 0.95 |
| 19 | Selenow | 0.65 | 2.94E-15 | 6.12E-11 | 0.95 | 0.99 | -0.04 | 0.96 |
| 19 | Kdm5b | 0.84 | 2.96E-15 | 6.15E-11 | 0.86 | 0.86 | 0.00 | 1.00 |
| 19 | Atp6v0e2 | 3.18 | 3.27E-15 | 6.80E-11 | 0.19 | 0.05 | 0.13 | 3.53 |
| 19 | Prkar1a | 1.05 | 3.45E-15 | 7.17E-11 | 0.88 | 0.88 | 0.00 | 1.00 |
| 19 | Jund | 0.87 | 3.58E-15 | 7.45E-11 | 0.89 | 0.94 | -0.05 | 0.94 |
| 19 | Dlx5 | 2.16 | 3.90E-15 | 8.11E-11 | 0.44 | 0.22 | 0.22 | 2.03 |
| 19 | Rtn3 | 0.93 | 4.00E-15 | 8.32E-11 | 0.81 | 0.79 | 0.02 | 1.03 |
| 19 | Tubb4a | 3.36 | 4.02E-15 | 8.37E-11 | 0.15 | 0.04 | 0.11 | 4.08 |
| 19 | Coro1a | 1.95 | 4.70E-15 | 9.78E-11 | 0.35 | 0.15 | 0.20 | 2.36 |
| 19 | Ubald2 | 1.16 | 5.47E-15 | 1.14E-10 | 0.78 | 0.69 | 0.09 | 1.13 |
| 19 | Septin3 | 1.55 | 6.48E-15 | 1.35E-10 | 0.61 | 0.44 | 0.17 | 1.39 |
| 19 | Cep170 | 1.34 | 7.54E-15 | 1.57E-10 | 0.68 | 0.51 | 0.17 | 1.33 |
| 19 | Gm15417 | 2.82 | 7.93E-15 | 1.65E-10 | 0.12 | 0.03 | 0.10 | 4.80 |
| 19 | Chfr | 1.62 | 8.88E-15 | 1.85E-10 | 0.48 | 0.26 | 0.22 | 1.86 |
| 19 | Gpi1 | 1.06 | 9.07E-15 | 1.89E-10 | 0.86 | 0.84 | 0.01 | 1.02 |
| 19 | Syt7 | 3.04 | 1.02E-14 | 2.13E-10 | 0.19 | 0.06 | 0.14 | 3.39 |
| 19 | Nfe2l3 | 2.43 | 1.15E-14 | 2.40E-10 | 0.15 | 0.04 | 0.11 | 4.14 |
| 19 | Ank3 | 2.01 | 1.23E-14 | 2.56E-10 | 0.65 | 0.49 | 0.16 | 1.33 |
| 19 | Maml3 | 1.32 | 1.35E-14 | 2.80E-10 | 0.61 | 0.40 | 0.21 | 1.53 |
| 19 | Tgfbr1 | 1.83 | 1.50E-14 | 3.13E-10 | 0.68 | 0.55 | 0.12 | 1.22 |
| 19 | Pkig | 1.06 | 1.62E-14 | 3.37E-10 | 0.86 | 0.79 | 0.07 | 1.09 |
| 19 | Rgs17 | 2.65 | 1.84E-14 | 3.83E-10 | 0.15 | 0.04 | 0.11 | 4.14 |
| 19 | Map7d2 | 2.08 | 2.01E-14 | 4.17E-10 | 0.24 | 0.08 | 0.16 | 2.87 |
| 19 | Gm43260 | 2.48 | 2.06E-14 | 4.29E-10 | 0.15 | 0.04 | 0.11 | 4.14 |
| 19 | Hsp90aa1 | 0.40 | 2.14E-14 | 4.45E-10 | 0.97 | 1.00 | -0.03 | 0.97 |
| 19 | Ctxn1 | 1.26 | 2.38E-14 | 4.95E-10 | 0.80 | 0.78 | 0.01 | 1.02 |
| 19 | Fbxl16 | 3.30 | 2.46E-14 | 5.12E-10 | 0.13 | 0.03 | 0.10 | 4.54 |
| 19 | Fgf15 | 3.67 | 2.58E-14 | 5.37E-10 | 0.12 | 0.03 | 0.09 | 4.62 |
| 19 | Marveld2 | 1.97 | 2.74E-14 | 5.71E-10 | 0.20 | 0.06 | 0.14 | 3.32 |
| 19 | Ccdc171 | 2.08 | 2.77E-14 | 5.75E-10 | 0.27 | 0.10 | 0.17 | 2.79 |
| 19 | Marchf8 | 1.88 | 2.81E-14 | 5.85E-10 | 0.37 | 0.17 | 0.20 | 2.16 |
| 19 | B3gnt4 | 2.56 | 3.11E-14 | 6.46E-10 | 0.16 | 0.04 | 0.12 | 3.93 |
| 19 | Mtch1 | 0.68 | 3.40E-14 | 7.07E-10 | 0.93 | 0.96 | -0.02 | 0.98 |
| 19 | Nefl | 4.36 | 4.22E-14 | 8.77E-10 | 0.13 | 0.03 | 0.10 | 4.29 |
| 19 | Igf1r | 1.19 | 5.16E-14 | 1.07E-09 | 0.79 | 0.77 | 0.02 | 1.02 |
| 19 | Hook1 | 1.45 | 5.55E-14 | 1.15E-09 | 0.43 | 0.20 | 0.23 | 2.12 |
| 19 | Snx32 | 2.82 | 6.08E-14 | 1.26E-09 | 0.11 | 0.02 | 0.09 | 4.75 |
| 19 | Zcchc18 | 1.66 | 6.59E-14 | 1.37E-09 | 0.54 | 0.34 | 0.20 | 1.60 |
| 19 | Gldc | 2.59 | 7.12E-14 | 1.48E-09 | 0.16 | 0.04 | 0.12 | 3.83 |
| 19 | Gap43 | 3.34 | 7.99E-14 | 1.66E-09 | 0.58 | 0.42 | 0.16 | 1.38 |
| 19 | Lmo2 | 2.35 | 8.36E-14 | 1.74E-09 | 0.31 | 0.13 | 0.19 | 2.46 |
| 19 | Baz2a | 1.23 | 8.83E-14 | 1.84E-09 | 0.61 | 0.39 | 0.22 | 1.58 |
| 19 | Klc1 | 1.97 | 9.12E-14 | 1.90E-09 | 0.63 | 0.47 | 0.16 | 1.33 |
| 19 | Six3 | 0.87 | 1.04E-13 | 2.16E-09 | 0.51 | 0.24 | 0.27 | 2.11 |
| 19 | Msi1 | 1.45 | 1.11E-13 | 2.32E-09 | 0.58 | 0.37 | 0.21 | 1.55 |
| 19 | Ift43 | 1.66 | 1.33E-13 | 2.76E-09 | 0.71 | 0.55 | 0.16 | 1.29 |
| 19 | Prr15 | 2.18 | 1.50E-13 | 3.11E-09 | 0.14 | 0.03 | 0.11 | 4.09 |
| 19 | Ssh2 | 1.54 | 1.50E-13 | 3.11E-09 | 0.43 | 0.23 | 0.20 | 1.89 |
| 19 | Rtn4 | 0.81 | 1.83E-13 | 3.80E-09 | 0.83 | 0.94 | -0.11 | 0.88 |

**Supplementary Table 1. Marker genes for the original clusters of integrated clustering of E11.5 control and NotchGOF λ-junction cells.**

| | | | | | | | | |
|---|---|---|---|---|---|---|---|---|
| 19 | Gpr137c | 1.53 | 1.84E-13 | 3.83E-09 | 0.34 | 0.15 | 0.19 | 2.32 |
| 19 | Gfod1 | 2.99 | 1.85E-13 | 3.85E-09 | 0.10 | 0.02 | 0.08 | 4.86 |
| 19 | Tubb2a | 1.89 | 2.22E-13 | 4.61E-09 | 0.50 | 0.30 | 0.20 | 1.68 |
| 19 | Dusp14 | 3.12 | 2.22E-13 | 4.61E-09 | 0.18 | 0.05 | 0.12 | 3.43 |
| 19 | Lcorl | 1.58 | 2.73E-13 | 5.69E-09 | 0.64 | 0.48 | 0.16 | 1.33 |
| 19 | Cpe | 1.50 | 2.88E-13 | 5.98E-09 | 0.60 | 0.42 | 0.18 | 1.43 |
| 19 | Tulp4 | 1.11 | 3.41E-13 | 7.09E-09 | 0.77 | 0.67 | 0.10 | 1.15 |
| 19 | Nrcam | 2.56 | 3.82E-13 | 7.95E-09 | 0.18 | 0.05 | 0.12 | 3.37 |
| 19 | C9orf72 | 2.48 | 4.05E-13 | 8.42E-09 | 0.28 | 0.11 | 0.17 | 2.47 |
| 19 | Csnk1e | 0.94 | 4.19E-13 | 8.72E-09 | 0.84 | 0.84 | -0.01 | 0.99 |
| 19 | Rac3 | 1.95 | 4.27E-13 | 8.88E-09 | 0.51 | 0.33 | 0.18 | 1.54 |
| 19 | Dmxl2 | 2.19 | 4.41E-13 | 9.18E-09 | 0.22 | 0.07 | 0.14 | 2.93 |
| 19 | Rala | 0.82 | 4.66E-13 | 9.69E-09 | 0.84 | 0.85 | -0.02 | 0.98 |
| 19 | Arhgap33 | 1.74 | 4.79E-13 | 9.97E-09 | 0.31 | 0.13 | 0.18 | 2.34 |
| 19 | Dynll1 | 0.51 | 4.85E-13 | 1.01E-08 | 0.99 | 1.00 | -0.01 | 0.99 |
| 19 | Acap3 | 2.01 | 5.13E-13 | 1.07E-08 | 0.22 | 0.07 | 0.14 | 2.93 |
| 19 | C2cd2l | 2.49 | 5.28E-13 | 1.10E-08 | 0.18 | 0.05 | 0.12 | 3.30 |
| 19 | Tanc2 | 1.60 | 5.50E-13 | 1.14E-08 | 0.43 | 0.23 | 0.20 | 1.87 |
| 19 | Pabpc1l | 2.76 | 5.81E-13 | 1.21E-08 | 0.12 | 0.03 | 0.09 | 4.29 |
| 19 | Ankrd37 | 1.62 | 5.93E-13 | 1.23E-08 | 0.35 | 0.16 | 0.19 | 2.21 |
| 19 | Ptprf | 0.76 | 6.28E-13 | 1.31E-08 | 0.80 | 0.64 | 0.16 | 1.24 |
| 19 | Them6 | 2.93 | 6.36E-13 | 1.32E-08 | 0.10 | 0.02 | 0.08 | 4.86 |
| 19 | Cep63 | 1.43 | 7.06E-13 | 1.47E-08 | 0.52 | 0.32 | 0.19 | 1.60 |
| 19 | Six1 | 1.11 | 7.82E-13 | 1.63E-08 | 0.75 | 0.61 | 0.14 | 1.23 |
| 19 | Zfp667 | 2.12 | 7.91E-13 | 1.64E-08 | 0.26 | 0.10 | 0.16 | 2.62 |
| 19 | Tnfrsf12a | 1.79 | 9.94E-13 | 2.07E-08 | 0.22 | 0.08 | 0.15 | 2.90 |
| 19 | Ccpg1os | 1.79 | 1.13E-12 | 2.34E-08 | 0.34 | 0.16 | 0.19 | 2.17 |
| 19 | Mapk11 | 2.12 | 1.18E-12 | 2.46E-08 | 0.21 | 0.07 | 0.14 | 2.93 |
| 19 | Nectin1 | 1.79 | 1.22E-12 | 2.54E-08 | 0.36 | 0.18 | 0.19 | 2.06 |
| 19 | Meg3 | 1.89 | 1.23E-12 | 2.57E-08 | 0.74 | 0.69 | 0.04 | 1.06 |
| 19 | Ttc5 | 1.17 | 1.31E-12 | 2.73E-08 | 0.69 | 0.54 | 0.14 | 1.26 |
| 19 | Kctd13 | 1.70 | 1.44E-12 | 2.99E-08 | 0.33 | 0.15 | 0.18 | 2.16 |
| 19 | Crabp2 | 0.65 | 1.48E-12 | 3.08E-08 | 0.95 | 0.92 | 0.03 | 1.03 |
| 19 | Cpeb4 | 2.09 | 1.59E-12 | 3.31E-08 | 0.25 | 0.10 | 0.15 | 2.60 |
| 19 | Ncoa1 | 2.16 | 1.65E-12 | 3.42E-08 | 0.35 | 0.17 | 0.18 | 2.03 |
| 19 | Bcl11b | 1.89 | 1.82E-12 | 3.79E-08 | 0.63 | 0.59 | 0.04 | 1.06 |
| 19 | Parm1 | 2.26 | 1.97E-12 | 4.09E-08 | 0.34 | 0.16 | 0.18 | 2.12 |
| 19 | Rian | 1.89 | 2.06E-12 | 4.29E-08 | 0.51 | 0.32 | 0.19 | 1.61 |
| 19 | Eml1 | 1.78 | 2.25E-12 | 4.68E-08 | 0.35 | 0.17 | 0.18 | 2.09 |
| 19 | Chchd10 | 1.64 | 2.28E-12 | 4.75E-08 | 0.35 | 0.16 | 0.19 | 2.17 |
| 19 | Fam187b | 1.89 | 2.32E-12 | 4.84E-08 | 0.15 | 0.04 | 0.11 | 3.72 |
| 19 | Card10 | 2.90 | 2.80E-12 | 5.81E-08 | 0.10 | 0.02 | 0.08 | 4.64 |
| 19 | Rab33a | 3.06 | 3.40E-12 | 7.07E-08 | 0.13 | 0.03 | 0.10 | 4.10 |
| 19 | Myo1b | 0.91 | 3.45E-12 | 7.18E-08 | 0.80 | 0.73 | 0.06 | 1.09 |
| 19 | Ric3 | 2.17 | 3.47E-12 | 7.23E-08 | 0.17 | 0.05 | 0.12 | 3.31 |
| 19 | Cox7a1 | 3.70 | 3.59E-12 | 7.47E-08 | 0.10 | 0.02 | 0.08 | 4.64 |
| 19 | Mex3a | 0.83 | 3.66E-12 | 7.62E-08 | 0.83 | 0.85 | -0.03 | 0.97 |
| 19 | Nav1 | 1.21 | 3.78E-12 | 7.86E-08 | 0.66 | 0.54 | 0.12 | 1.22 |
| 19 | Wfs1 | 1.81 | 3.85E-12 | 8.02E-08 | 0.20 | 0.07 | 0.13 | 3.02 |
| 19 | Rnasek | 1.00 | 4.07E-12 | 8.47E-08 | 0.76 | 0.72 | 0.04 | 1.05 |
| 19 | Phyh | 1.61 | 4.99E-12 | 1.04E-07 | 0.48 | 0.29 | 0.19 | 1.65 |
| 19 | Plch1 | 1.51 | 6.16E-12 | 1.28E-07 | 0.26 | 0.10 | 0.16 | 2.56 |
| 19 | Psma7 | 0.55 | 6.32E-12 | 1.31E-07 | 0.96 | 0.99 | -0.03 | 0.97 |
| 19 | Gabarap | 0.61 | 6.78E-12 | 1.41E-07 | 0.91 | 0.98 | -0.07 | 0.93 |
| 19 | Prmt2 | 1.70 | 7.43E-12 | 1.54E-07 | 0.48 | 0.31 | 0.17 | 1.55 |
| 19 | Ralgapa2 | 1.79 | 7.51E-12 | 1.56E-07 | 0.30 | 0.14 | 0.16 | 2.18 |

**Supplementary Table 1. Marker genes for the original clusters of integrated clustering of E11.5 control and NotchGOF λ-junction cells.**

| | | | | | | | | |
|---|---|---|---|---|---|---|---|---|
| 19 | Gm5860 | 3.19 | 9.22E-12 | 1.92E-07 | 0.13 | 0.03 | 0.09 | 3.85 |
| 19 | Tex15 | 4.24 | 9.54E-12 | 1.98E-07 | 0.26 | 0.11 | 0.15 | 2.35 |
| 19 | Usp29 | 1.52 | 9.81E-12 | 2.04E-07 | 0.40 | 0.21 | 0.19 | 1.93 |
| 19 | Mapre3 | 2.43 | 1.06E-11 | 2.20E-07 | 0.27 | 0.11 | 0.16 | 2.38 |
| 19 | Gm42702 | 2.25 | 1.06E-11 | 2.20E-07 | 0.14 | 0.04 | 0.10 | 3.66 |
| 19 | Tmem178 | 2.89 | 1.13E-11 | 2.34E-07 | 0.13 | 0.03 | 0.09 | 3.85 |
| 19 | Snrpn | 1.78 | 1.14E-11 | 2.36E-07 | 0.63 | 0.47 | 0.16 | 1.34 |
| 19 | Dync1i2 | 0.73 | 1.37E-11 | 2.84E-07 | 0.84 | 0.89 | -0.05 | 0.94 |
| 19 | Lrrc42 | 1.51 | 1.47E-11 | 3.05E-07 | 0.68 | 0.61 | 0.07 | 1.11 |
| 19 | Lman2l | 1.52 | 1.62E-11 | 3.37E-07 | 0.53 | 0.35 | 0.18 | 1.53 |
| 19 | Atp6v1e1 | 1.09 | 1.89E-11 | 3.93E-07 | 0.72 | 0.68 | 0.04 | 1.07 |
| 19 | Unc45a | 1.80 | 2.13E-11 | 4.43E-07 | 0.45 | 0.27 | 0.18 | 1.67 |
| 19 | Jade1 | 1.98 | 2.57E-11 | 5.35E-07 | 0.34 | 0.17 | 0.17 | 1.98 |
| 19 | Tmeff1 | 1.11 | 2.67E-11 | 5.56E-07 | 0.74 | 0.69 | 0.05 | 1.07 |
| 19 | Arpc3 | 0.70 | 2.74E-11 | 5.70E-07 | 0.87 | 0.90 | -0.03 | 0.97 |
| 19 | Kit | 2.58 | 2.82E-11 | 5.86E-07 | 0.24 | 0.10 | 0.15 | 2.51 |
| 19 | Prkx | 1.69 | 2.84E-11 | 5.90E-07 | 0.32 | 0.15 | 0.17 | 2.07 |
| 19 | Fkbp1a | 0.56 | 3.06E-11 | 6.37E-07 | 0.89 | 0.94 | -0.05 | 0.94 |
| 19 | Atp6v0d1 | 1.31 | 3.19E-11 | 6.64E-07 | 0.69 | 0.54 | 0.15 | 1.28 |
| 19 | Akna | 1.96 | 3.22E-11 | 6.69E-07 | 0.26 | 0.11 | 0.15 | 2.44 |
| 19 | Frmd5 | 2.03 | 3.40E-11 | 7.07E-07 | 0.16 | 0.05 | 0.11 | 3.20 |
| 19 | Dnaja1 | 0.38 | 3.42E-11 | 7.11E-07 | 0.93 | 0.99 | -0.06 | 0.94 |
| 19 | Gamt | 2.85 | 3.45E-11 | 7.18E-07 | 0.24 | 0.09 | 0.14 | 2.55 |
| 19 | 4930452B06Rik | 2.46 | 3.67E-11 | 7.63E-07 | 0.28 | 0.12 | 0.16 | 2.28 |
| 19 | Mgst3 | 1.45 | 3.74E-11 | 7.78E-07 | 0.63 | 0.51 | 0.13 | 1.25 |
| 19 | Tnik | 1.77 | 3.75E-11 | 7.81E-07 | 0.42 | 0.25 | 0.17 | 1.70 |
| 19 | Dda1 | 1.32 | 4.15E-11 | 8.63E-07 | 0.69 | 0.57 | 0.12 | 1.21 |
| 19 | 2810455O05Rik | 2.74 | 4.71E-11 | 9.79E-07 | 0.10 | 0.02 | 0.08 | 4.25 |
| 19 | Rnd2 | 2.26 | 4.77E-11 | 9.92E-07 | 0.23 | 0.09 | 0.14 | 2.60 |
| 19 | Snta1 | 2.23 | 5.13E-11 | 1.07E-06 | 0.16 | 0.05 | 0.11 | 3.20 |
| 19 | Hist2h2be | 2.57 | 5.21E-11 | 1.08E-06 | 0.13 | 0.04 | 0.10 | 3.59 |
| 19 | Tnfrsf21 | 1.78 | 5.66E-11 | 1.18E-06 | 0.31 | 0.14 | 0.17 | 2.16 |
| 19 | Strbp | 1.06 | 5.67E-11 | 1.18E-06 | 0.75 | 0.71 | 0.04 | 1.06 |
| 19 | Jup | 1.17 | 5.71E-11 | 1.19E-06 | 0.40 | 0.20 | 0.20 | 2.00 |
| 19 | 2900093K20Rik | 1.99 | 5.91E-11 | 1.23E-06 | 0.22 | 0.09 | 0.14 | 2.59 |
| 19 | Tmcc1 | 1.24 | 6.07E-11 | 1.26E-06 | 0.60 | 0.44 | 0.16 | 1.37 |
| 19 | Cyb5r4 | 1.40 | 6.23E-11 | 1.29E-06 | 0.57 | 0.40 | 0.17 | 1.43 |
| 19 | Kras | 0.87 | 7.34E-11 | 1.53E-06 | 0.82 | 0.85 | -0.03 | 0.97 |
| 19 | Fam122b | 1.94 | 7.57E-11 | 1.57E-06 | 0.22 | 0.08 | 0.13 | 2.61 |
| 19 | Ezr | 0.71 | 8.04E-11 | 1.67E-06 | 0.57 | 0.32 | 0.25 | 1.76 |
| 19 | Daam1 | 1.33 | 8.57E-11 | 1.78E-06 | 0.58 | 0.41 | 0.18 | 1.43 |
| 19 | Mob3b | 1.80 | 9.02E-11 | 1.88E-06 | 0.23 | 0.09 | 0.14 | 2.54 |
| 19 | Cyb561 | 2.30 | 1.00E-10 | 2.08E-06 | 0.17 | 0.06 | 0.11 | 3.02 |
| 19 | Dram2 | 1.43 | 1.02E-10 | 2.12E-06 | 0.39 | 0.21 | 0.19 | 1.91 |
| 19 | Lurap1l | 1.71 | 1.21E-10 | 2.51E-06 | 0.21 | 0.08 | 0.13 | 2.74 |
| 19 | Myef2 | 0.88 | 1.21E-10 | 2.53E-06 | 0.75 | 0.69 | 0.06 | 1.09 |
| 19 | Cdkn2d | 1.71 | 1.29E-10 | 2.69E-06 | 0.53 | 0.38 | 0.15 | 1.38 |
| 19 | Kcnn1 | 2.18 | 1.33E-10 | 2.77E-06 | 0.15 | 0.04 | 0.10 | 3.37 |
| 19 | Susd6 | 1.68 | 1.35E-10 | 2.80E-06 | 0.37 | 0.20 | 0.17 | 1.86 |
| 19 | Lbh | 1.97 | 1.49E-10 | 3.09E-06 | 0.37 | 0.20 | 0.17 | 1.85 |
| 19 | 1700023F06Rik | 2.63 | 1.52E-10 | 3.17E-06 | 0.15 | 0.04 | 0.10 | 3.30 |
| 19 | Arpc1a | 0.73 | 1.54E-10 | 3.20E-06 | 0.85 | 0.88 | -0.04 | 0.96 |
| 19 | Gdf11 | 1.44 | 1.55E-10 | 3.22E-06 | 0.49 | 0.32 | 0.17 | 1.53 |
| 19 | Twf1 | 1.92 | 1.58E-10 | 3.29E-06 | 0.61 | 0.49 | 0.12 | 1.24 |
| 19 | Ubl5 | 0.53 | 1.59E-10 | 3.30E-06 | 0.93 | 0.98 | -0.05 | 0.95 |
| 19 | Cdc42ep3 | 1.57 | 1.66E-10 | 3.44E-06 | 0.34 | 0.17 | 0.17 | 2.04 |

**Supplementary Table 1. Marker genes for the original clusters of integrated clustering of E11.5 control and NotchGOF λ-junction cells.**

| | | | | | | | | |
|---|---|---|---|---|---|---|---|---|
| 19 | Rundc3b | 2.40 | 1.75E-10 | 3.64E-06 | 0.14 | 0.04 | 0.10 | 3.39 |
| 19 | Kif3c | 1.82 | 1.89E-10 | 3.93E-06 | 0.27 | 0.12 | 0.15 | 2.25 |
| 19 | Gm42372 | 2.14 | 1.90E-10 | 3.94E-06 | 0.16 | 0.05 | 0.11 | 3.14 |
| 19 | Dctn1 | 1.46 | 2.30E-10 | 4.78E-06 | 0.62 | 0.52 | 0.10 | 1.19 |
| 19 | Dnajc6 | 2.54 | 2.38E-10 | 4.94E-06 | 0.10 | 0.03 | 0.08 | 4.08 |
| 19 | Camsap3 | 2.08 | 2.49E-10 | 5.17E-06 | 0.21 | 0.08 | 0.13 | 2.57 |
| 19 | Srcin1 | 2.03 | 2.56E-10 | 5.33E-06 | 0.15 | 0.05 | 0.10 | 3.21 |
| 19 | Trp53inp1 | 1.42 | 2.64E-10 | 5.48E-06 | 0.37 | 0.19 | 0.19 | 2.01 |
| 19 | Fam222a | 2.97 | 2.81E-10 | 5.85E-06 | 0.11 | 0.03 | 0.08 | 3.68 |
| 19 | Vamp2 | 1.80 | 3.29E-10 | 6.85E-06 | 0.45 | 0.30 | 0.15 | 1.52 |
| 19 | Zfp664 | 0.85 | 3.30E-10 | 6.86E-06 | 0.78 | 0.75 | 0.03 | 1.04 |
| 19 | Sbk1 | 1.19 | 3.34E-10 | 6.95E-06 | 0.62 | 0.52 | 0.10 | 1.18 |
| 19 | Id4 | 2.64 | 3.71E-10 | 7.71E-06 | 0.47 | 0.31 | 0.16 | 1.53 |
| 19 | Ppp1r11 | 0.84 | 3.78E-10 | 7.87E-06 | 0.83 | 0.81 | 0.02 | 1.03 |
| 19 | Gm2464 | 3.29 | 3.84E-10 | 7.98E-06 | 0.11 | 0.03 | 0.08 | 3.86 |
| 19 | Celsr2 | 1.70 | 3.90E-10 | 8.11E-06 | 0.23 | 0.09 | 0.14 | 2.46 |
| 19 | Cxxc4 | 1.79 | 4.14E-10 | 8.62E-06 | 0.46 | 0.31 | 0.15 | 1.50 |
| 19 | Cdk5r1 | 2.17 | 4.17E-10 | 8.68E-06 | 0.25 | 0.11 | 0.14 | 2.29 |
| 19 | Map2k1 | 1.64 | 4.43E-10 | 9.20E-06 | 0.50 | 0.35 | 0.15 | 1.45 |
| 19 | Dynll2 | 0.86 | 4.69E-10 | 9.76E-06 | 0.81 | 0.84 | -0.04 | 0.96 |
| 19 | Kcnh2 | 1.95 | 5.12E-10 | 1.06E-05 | 0.15 | 0.05 | 0.10 | 3.22 |
| 19 | Sdccag8 | 2.01 | 5.41E-10 | 1.13E-05 | 0.28 | 0.13 | 0.15 | 2.20 |
| 19 | Ciao2b | 0.96 | 5.53E-10 | 1.15E-05 | 0.75 | 0.73 | 0.02 | 1.02 |
| 19 | Map3k13 | 3.11 | 5.73E-10 | 1.19E-05 | 0.12 | 0.03 | 0.09 | 3.53 |
| 19 | Rbfox2 | 0.88 | 5.93E-10 | 1.23E-05 | 0.78 | 0.77 | 0.02 | 1.02 |
| 19 | Gng2 | 1.20 | 6.52E-10 | 1.36E-05 | 0.67 | 0.57 | 0.10 | 1.17 |
| 19 | Armh4 | 2.47 | 7.75E-10 | 1.61E-05 | 0.30 | 0.15 | 0.15 | 1.97 |
| 19 | Abca3 | 1.87 | 8.09E-10 | 1.68E-05 | 0.18 | 0.07 | 0.12 | 2.78 |
| 19 | Nbea | 1.20 | 8.54E-10 | 1.78E-05 | 0.59 | 0.44 | 0.15 | 1.33 |
| 19 | Ift22 | 1.27 | 8.74E-10 | 1.82E-05 | 0.51 | 0.35 | 0.15 | 1.43 |
| 19 | Cib1 | 1.36 | 9.60E-10 | 2.00E-05 | 0.55 | 0.40 | 0.15 | 1.38 |
| 19 | Dync1li1 | 1.21 | 1.00E-09 | 2.08E-05 | 0.64 | 0.52 | 0.12 | 1.23 |
| 19 | Pbx3 | 1.23 | 1.14E-09 | 2.37E-05 | 0.65 | 0.53 | 0.12 | 1.23 |
| 19 | Plxnc1 | 2.02 | 1.23E-09 | 2.57E-05 | 0.14 | 0.04 | 0.10 | 3.23 |
| 19 | 1700037H04Rik | 1.59 | 1.27E-09 | 2.64E-05 | 0.62 | 0.48 | 0.14 | 1.30 |
| 19 | Plppr2 | 2.11 | 1.36E-09 | 2.83E-05 | 0.18 | 0.06 | 0.11 | 2.78 |
| 19 | Rcan1 | 2.08 | 1.40E-09 | 2.90E-05 | 0.42 | 0.26 | 0.16 | 1.62 |
| 19 | Peg13 | 2.11 | 1.58E-09 | 3.29E-05 | 0.15 | 0.05 | 0.10 | 3.02 |
| 19 | Gnaz | 2.32 | 1.60E-09 | 3.32E-05 | 0.14 | 0.04 | 0.10 | 3.16 |
| 19 | Arl8a | 1.69 | 1.67E-09 | 3.47E-05 | 0.54 | 0.43 | 0.11 | 1.26 |
| 19 | Zc2hc1a | 1.26 | 1.67E-09 | 3.48E-05 | 0.57 | 0.42 | 0.15 | 1.37 |
| 19 | Zbtb20 | 2.04 | 1.71E-09 | 3.56E-05 | 0.44 | 0.29 | 0.15 | 1.50 |
| 19 | Hspa2 | 2.07 | 1.75E-09 | 3.64E-05 | 0.16 | 0.06 | 0.11 | 2.91 |
| 19 | Akr1b8 | 1.78 | 1.75E-09 | 3.64E-05 | 0.20 | 0.08 | 0.12 | 2.58 |
| 19 | Mindy2 | 1.73 | 1.80E-09 | 3.74E-05 | 0.33 | 0.18 | 0.15 | 1.86 |
| 19 | Crtc3 | 1.75 | 1.81E-09 | 3.77E-05 | 0.40 | 0.25 | 0.15 | 1.62 |
| 19 | H13 | 0.92 | 1.84E-09 | 3.82E-05 | 0.72 | 0.63 | 0.08 | 1.13 |
| 19 | Rab6a | 1.14 | 1.95E-09 | 4.06E-05 | 0.74 | 0.72 | 0.03 | 1.03 |
| 19 | Clta | 0.55 | 2.03E-09 | 4.22E-05 | 0.94 | 0.98 | -0.04 | 0.96 |
| 19 | Gnptg | 1.65 | 2.26E-09 | 4.69E-05 | 0.37 | 0.20 | 0.17 | 1.82 |
| 19 | Cfap36 | 1.32 | 2.32E-09 | 4.82E-05 | 0.69 | 0.55 | 0.14 | 1.25 |
| 19 | Arg2 | 2.00 | 2.41E-09 | 5.01E-05 | 0.16 | 0.05 | 0.10 | 2.96 |
| 19 | Cfl1 | 0.31 | 2.44E-09 | 5.08E-05 | 0.99 | 1.00 | -0.01 | 0.99 |
| 19 | Ttc39b | 2.00 | 2.45E-09 | 5.09E-05 | 0.18 | 0.07 | 0.12 | 2.74 |
| 19 | Ccdc28b | 1.35 | 2.53E-09 | 5.25E-05 | 0.53 | 0.38 | 0.16 | 1.41 |
| 19 | Eml5 | 1.89 | 2.61E-09 | 5.44E-05 | 0.21 | 0.09 | 0.13 | 2.45 |

**Supplementary Table 1. Marker genes for the original clusters of integrated clustering of E11.5 control and NotchGOF λ-junction cells.**

| | | | | | | | | |
|---|---|---|---|---|---|---|---|---|
| 19 | Mapk8 | 1.05 | 2.78E-09 | 5.78E-05 | 0.59 | 0.43 | 0.17 | 1.39 |
| 19 | Atp6v1h | 1.04 | 2.88E-09 | 5.99E-05 | 0.54 | 0.36 | 0.17 | 1.48 |
| 19 | Rasal2 | 1.81 | 2.92E-09 | 6.08E-05 | 0.40 | 0.25 | 0.16 | 1.62 |
| 19 | Dedd2 | 1.90 | 3.00E-09 | 6.23E-05 | 0.19 | 0.08 | 0.12 | 2.57 |
| 19 | Stx7 | 2.22 | 3.07E-09 | 6.38E-05 | 0.45 | 0.31 | 0.14 | 1.45 |
| 19 | B830012L14Rik | 2.28 | 3.08E-09 | 6.40E-05 | 0.16 | 0.06 | 0.10 | 2.85 |
| 19 | Mterf2 | 1.51 | 3.27E-09 | 6.81E-05 | 0.37 | 0.21 | 0.16 | 1.77 |
| 19 | Smim22 | 3.07 | 3.43E-09 | 7.14E-05 | 0.13 | 0.04 | 0.09 | 3.26 |
| 19 | Palm | 1.46 | 3.52E-09 | 7.33E-05 | 0.58 | 0.46 | 0.12 | 1.25 |
| 19 | Clcn2 | 2.06 | 4.03E-09 | 8.38E-05 | 0.20 | 0.08 | 0.12 | 2.55 |
| 19 | Akirin1 | 1.31 | 4.14E-09 | 8.61E-05 | 0.65 | 0.58 | 0.07 | 1.12 |
| 19 | Slc24a5 | 1.08 | 4.54E-09 | 9.43E-05 | 0.57 | 0.40 | 0.16 | 1.40 |
| 19 | Gde1 | 1.17 | 4.67E-09 | 9.70E-05 | 0.63 | 0.52 | 0.11 | 1.20 |
| 19 | Bdh1 | 2.23 | 4.69E-09 | 9.76E-05 | 0.19 | 0.07 | 0.12 | 2.60 |
| 19 | Ndufb1-ps | 0.63 | 4.81E-09 | 9.99E-05 | 0.89 | 0.94 | -0.05 | 0.95 |
| 19 | Lcor | 1.26 | 4.83E-09 | 1.00E-04 | 0.45 | 0.28 | 0.16 | 1.57 |
| 19 | Garnl3 | 2.31 | 5.09E-09 | 1.06E-04 | 0.11 | 0.03 | 0.08 | 3.60 |
| 19 | Dmrta2 | 1.58 | 5.18E-09 | 1.08E-04 | 0.29 | 0.14 | 0.15 | 2.04 |
| 19 | Hspa8 | 0.32 | 5.24E-09 | 1.09E-04 | 0.98 | 1.00 | -0.02 | 0.98 |
| 19 | Cep250 | 1.40 | 5.27E-09 | 1.10E-04 | 0.42 | 0.26 | 0.16 | 1.60 |
| 19 | Mgat5b | 2.04 | 5.29E-09 | 1.10E-04 | 0.13 | 0.04 | 0.09 | 3.26 |
| 19 | Mapk3 | 1.32 | 5.57E-09 | 1.16E-04 | 0.58 | 0.45 | 0.13 | 1.28 |
| 19 | Ciapin1 | 0.92 | 6.30E-09 | 1.31E-04 | 0.78 | 0.74 | 0.05 | 1.06 |
| 19 | Pfdn4 | 0.99 | 6.30E-09 | 1.31E-04 | 0.81 | 0.75 | 0.05 | 1.07 |
| 19 | Wbp2 | 1.55 | 6.55E-09 | 1.36E-04 | 0.43 | 0.28 | 0.16 | 1.57 |
| 19 | Ppp2cb | 1.79 | 7.03E-09 | 1.46E-04 | 0.57 | 0.44 | 0.13 | 1.30 |
| 19 | Mycbp2 | 0.92 | 7.22E-09 | 1.50E-04 | 0.73 | 0.70 | 0.03 | 1.04 |
| 19 | Lpin1 | 1.78 | 7.31E-09 | 1.52E-04 | 0.18 | 0.07 | 0.11 | 2.65 |
| 19 | Enox1 | 2.27 | 7.35E-09 | 1.53E-04 | 0.19 | 0.08 | 0.12 | 2.51 |
| 19 | Cnih2 | 2.01 | 7.36E-09 | 1.53E-04 | 0.46 | 0.33 | 0.14 | 1.42 |
| 19 | Esrp1 | 0.75 | 7.37E-09 | 1.53E-04 | 0.35 | 0.17 | 0.18 | 2.07 |
| 19 | Wnk2 | 1.81 | 8.05E-09 | 1.68E-04 | 0.15 | 0.05 | 0.10 | 3.02 |
| 19 | Akap9 | 0.80 | 8.23E-09 | 1.71E-04 | 0.77 | 0.78 | -0.01 | 0.98 |
| 19 | Gm36839 | 2.24 | 8.33E-09 | 1.73E-04 | 0.11 | 0.03 | 0.08 | 3.35 |
| 19 | Micu3 | 2.30 | 8.37E-09 | 1.74E-04 | 0.33 | 0.17 | 0.15 | 1.87 |
| 19 | Vps28 | 1.14 | 8.47E-09 | 1.76E-04 | 0.71 | 0.62 | 0.09 | 1.14 |
| 19 | Cacnb3 | 1.04 | 8.63E-09 | 1.79E-04 | 0.45 | 0.27 | 0.18 | 1.65 |
| 19 | Vav2 | 1.76 | 8.95E-09 | 1.86E-04 | 0.21 | 0.09 | 0.12 | 2.41 |
| 19 | Zdhhc12 | 1.65 | 1.06E-08 | 2.20E-04 | 0.42 | 0.26 | 0.16 | 1.62 |
| 19 | Dynlt1f | 0.91 | 1.08E-08 | 2.24E-04 | 0.80 | 0.77 | 0.02 | 1.03 |
| 19 | Rsrp1 | 0.73 | 1.15E-08 | 2.39E-04 | 0.83 | 0.89 | -0.06 | 0.94 |
| 19 | Mdm2 | 0.94 | 1.20E-08 | 2.50E-04 | 0.59 | 0.44 | 0.15 | 1.35 |
| 19 | Hmgcll1 | 2.14 | 1.33E-08 | 2.76E-04 | 0.11 | 0.03 | 0.08 | 3.35 |
| 19 | Trappc2l | 1.21 | 1.40E-08 | 2.90E-04 | 0.70 | 0.62 | 0.08 | 1.13 |
| 19 | Ywhah | 0.79 | 1.40E-08 | 2.91E-04 | 0.88 | 0.91 | -0.03 | 0.97 |
| 19 | Med12l | 3.27 | 1.40E-08 | 2.91E-04 | 0.13 | 0.05 | 0.09 | 2.96 |
| 19 | Tspyl4 | 1.82 | 1.41E-08 | 2.92E-04 | 0.38 | 0.24 | 0.14 | 1.59 |
| 19 | Fgd2 | 2.02 | 1.44E-08 | 2.99E-04 | 0.23 | 0.10 | 0.13 | 2.27 |
| 19 | Btbd10 | 1.57 | 1.45E-08 | 3.01E-04 | 0.40 | 0.25 | 0.16 | 1.63 |
| 19 | Zdhhc2 | 1.27 | 1.47E-08 | 3.06E-04 | 0.28 | 0.13 | 0.15 | 2.13 |
| 19 | Hdac2 | 0.66 | 1.57E-08 | 3.27E-04 | 0.86 | 0.90 | -0.05 | 0.95 |
| 19 | Celf4 | 1.72 | 1.59E-08 | 3.30E-04 | 0.37 | 0.23 | 0.15 | 1.65 |
| 19 | Myl6 | 0.41 | 1.63E-08 | 3.40E-04 | 0.92 | 0.99 | -0.08 | 0.92 |
| 19 | Ccndbp1 | 1.76 | 1.64E-08 | 3.42E-04 | 0.45 | 0.28 | 0.17 | 1.59 |
| 19 | Grcc10 | 0.65 | 1.76E-08 | 3.65E-04 | 0.93 | 0.98 | -0.04 | 0.96 |
| 19 | Nuak1 | 1.61 | 1.79E-08 | 3.72E-04 | 0.36 | 0.20 | 0.15 | 1.74 |

**Supplementary Table 1. Marker genes for the original clusters of integrated clustering of E11.5 control and NotchGOF λ-junction cells.**

| | | | | | | | | |
|---|---|---|---|---|---|---|---|---|
| 19 | AI413582 | 1.96 | 1.80E-08 | 3.74E-04 | 0.21 | 0.09 | 0.12 | 2.29 |
| 19 | Enho | 1.19 | 1.85E-08 | 3.84E-04 | 0.58 | 0.45 | 0.13 | 1.28 |
| 19 | Ank | 2.14 | 1.92E-08 | 3.99E-04 | 0.18 | 0.07 | 0.11 | 2.54 |
| 19 | Atp6v1b2 | 1.39 | 1.94E-08 | 4.04E-04 | 0.55 | 0.39 | 0.16 | 1.42 |
| 19 | Pak7 | 2.24 | 1.96E-08 | 4.09E-04 | 0.14 | 0.05 | 0.09 | 2.90 |
| 19 | Cebpg | 1.21 | 2.10E-08 | 4.36E-04 | 0.65 | 0.55 | 0.10 | 1.19 |
| 19 | Katnal2 | 1.89 | 2.21E-08 | 4.60E-04 | 0.17 | 0.06 | 0.11 | 2.68 |
| 19 | Fgf9 | 0.66 | 2.24E-08 | 4.65E-04 | 0.25 | 0.11 | 0.14 | 2.31 |
| 19 | Rnd3 | 1.08 | 2.24E-08 | 4.66E-04 | 0.70 | 0.64 | 0.06 | 1.09 |
| 19 | Trak1 | 1.49 | 2.29E-08 | 4.77E-04 | 0.25 | 0.12 | 0.13 | 2.13 |
| 19 | Igfbp2 | 0.50 | 2.30E-08 | 4.78E-04 | 0.78 | 0.76 | 0.03 | 1.04 |
| 19 | Chmp1b | 1.34 | 2.35E-08 | 4.89E-04 | 0.51 | 0.35 | 0.15 | 1.44 |
| 19 | Carmil3 | 2.25 | 2.47E-08 | 5.14E-04 | 0.15 | 0.06 | 0.10 | 2.75 |
| 19 | Ankrd13b | 1.49 | 2.48E-08 | 5.16E-04 | 0.34 | 0.19 | 0.15 | 1.80 |
| 19 | Snx11 | 1.66 | 2.51E-08 | 5.23E-04 | 0.30 | 0.16 | 0.14 | 1.91 |
| 19 | Dennd1a | 1.22 | 2.53E-08 | 5.26E-04 | 0.36 | 0.21 | 0.15 | 1.73 |
| 19 | Pax6 | 1.33 | 2.66E-08 | 5.53E-04 | 0.27 | 0.13 | 0.14 | 2.08 |
| 19 | Wsb2 | 1.35 | 3.13E-08 | 6.50E-04 | 0.46 | 0.31 | 0.15 | 1.47 |
| 19 | Smim20 | 1.37 | 3.18E-08 | 6.61E-04 | 0.55 | 0.41 | 0.15 | 1.36 |
| 19 | Acrbp | 1.83 | 3.19E-08 | 6.64E-04 | 0.23 | 0.10 | 0.13 | 2.20 |
| 19 | Rc3h2 | 1.15 | 3.34E-08 | 6.94E-04 | 0.63 | 0.53 | 0.11 | 1.20 |
| 19 | Pls1 | 2.17 | 3.35E-08 | 6.96E-04 | 0.22 | 0.10 | 0.12 | 2.26 |
| 19 | Mdh2 | 0.64 | 3.42E-08 | 7.11E-04 | 0.89 | 0.95 | -0.07 | 0.93 |
| 19 | Gm42418 | 0.29 | 3.56E-08 | 7.40E-04 | 0.99 | 1.00 | 0.00 | 1.00 |
| 19 | B3gnt2 | 2.13 | 3.58E-08 | 7.44E-04 | 0.22 | 0.10 | 0.12 | 2.17 |
| 19 | Ap2m1 | 0.66 | 3.72E-08 | 7.74E-04 | 0.82 | 0.86 | -0.04 | 0.95 |
| 19 | Ptprz1 | 2.07 | 3.84E-08 | 8.00E-04 | 0.23 | 0.11 | 0.12 | 2.14 |
| 19 | Psmc5 | 0.53 | 4.89E-08 | 1.02E-03 | 0.85 | 0.92 | -0.07 | 0.93 |
| 19 | Snn | 0.93 | 5.16E-08 | 1.07E-03 | 0.58 | 0.44 | 0.15 | 1.34 |
| 19 | Zfp428 | 0.85 | 5.25E-08 | 1.09E-03 | 0.74 | 0.69 | 0.04 | 1.06 |
| 19 | Ttc3 | 0.44 | 5.95E-08 | 1.24E-03 | 0.86 | 0.96 | -0.10 | 0.90 |
| 19 | Mtus2 | 2.40 | 6.00E-08 | 1.25E-03 | 0.12 | 0.04 | 0.08 | 3.00 |
| 19 | Sirt7 | 1.73 | 6.01E-08 | 1.25E-03 | 0.31 | 0.17 | 0.14 | 1.82 |
| 19 | Fbxo32 | 2.01 | 6.17E-08 | 1.28E-03 | 0.10 | 0.03 | 0.07 | 3.40 |
| 19 | Dennd1b | 1.54 | 6.18E-08 | 1.28E-03 | 0.30 | 0.16 | 0.14 | 1.88 |
| 19 | Khdrbs2 | 1.90 | 6.42E-08 | 1.33E-03 | 0.11 | 0.04 | 0.08 | 3.17 |
| 19 | Slc25a33 | 1.65 | 6.61E-08 | 1.37E-03 | 0.30 | 0.17 | 0.13 | 1.80 |
| 19 | Psmc2 | 0.65 | 6.75E-08 | 1.40E-03 | 0.82 | 0.88 | -0.06 | 0.93 |
| 19 | Smchd1 | 1.59 | 6.81E-08 | 1.42E-03 | 0.79 | 0.86 | -0.07 | 0.92 |
| 19 | Fzd3 | 0.85 | 6.84E-08 | 1.42E-03 | 0.68 | 0.59 | 0.09 | 1.15 |
| 19 | Chmp5 | 0.67 | 6.87E-08 | 1.43E-03 | 0.80 | 0.80 | 0.00 | 1.00 |
| 19 | Gsk3b | 0.76 | 6.87E-08 | 1.43E-03 | 0.82 | 0.89 | -0.07 | 0.92 |
| 19 | Hsdl1 | 1.72 | 7.35E-08 | 1.53E-03 | 0.29 | 0.15 | 0.14 | 1.88 |
| 19 | Esyt1 | 1.84 | 7.48E-08 | 1.56E-03 | 0.18 | 0.07 | 0.11 | 2.45 |
| 19 | Ntm | 1.87 | 7.58E-08 | 1.58E-03 | 0.23 | 0.11 | 0.12 | 2.14 |
| 19 | Cog1 | 1.61 | 7.89E-08 | 1.64E-03 | 0.31 | 0.17 | 0.14 | 1.83 |
| 19 | Hnrnpa2b1 | 0.33 | 8.02E-08 | 1.67E-03 | 0.98 | 1.00 | -0.02 | 0.98 |
| 19 | Rb1cc1 | 0.80 | 8.20E-08 | 1.70E-03 | 0.72 | 0.66 | 0.07 | 1.10 |
| 19 | Amn1 | 1.04 | 9.05E-08 | 1.88E-03 | 0.59 | 0.44 | 0.15 | 1.34 |
| 19 | Rab3ip | 1.44 | 9.28E-08 | 1.93E-03 | 0.45 | 0.31 | 0.14 | 1.46 |
| 19 | 1700096K18Rik | 1.72 | 9.63E-08 | 2.00E-03 | 0.21 | 0.10 | 0.12 | 2.22 |
| 19 | Chchd6 | 1.17 | 9.73E-08 | 2.02E-03 | 0.64 | 0.54 | 0.10 | 1.19 |
| 19 | Katnal1 | 1.56 | 9.93E-08 | 2.06E-03 | 0.32 | 0.17 | 0.15 | 1.83 |
| 19 | Chn2 | 1.69 | 1.01E-07 | 2.09E-03 | 0.18 | 0.07 | 0.10 | 2.46 |
| 19 | Isl1 | 1.31 | 1.10E-07 | 2.28E-03 | 0.40 | 0.22 | 0.18 | 1.80 |
| 19 | Six6 | 3.34 | 1.10E-07 | 2.29E-03 | 0.17 | 0.07 | 0.10 | 2.38 |

**Supplementary Table 1. Marker genes for the original clusters of integrated clustering of E11.5 control and NotchGOF λ-junction cells.**

| | | | | | | | | |
|---|---|---|---|---|---|---|---|---|
| 19 | Ypel5 | 1.13 | 1.16E-07 | 2.42E-03 | 0.42 | 0.25 | 0.17 | 1.66 |
| 19 | Capn10 | 1.12 | 1.17E-07 | 2.44E-03 | 0.37 | 0.22 | 0.15 | 1.69 |
| 19 | L1cam | 2.12 | 1.17E-07 | 2.44E-03 | 0.19 | 0.08 | 0.11 | 2.34 |
| 19 | Cmpk2 | 2.11 | 1.19E-07 | 2.47E-03 | 0.11 | 0.04 | 0.08 | 3.08 |
| 19 | Rtl8a | 1.72 | 1.22E-07 | 2.55E-03 | 0.51 | 0.39 | 0.12 | 1.31 |
| 19 | Jarid2 | 0.97 | 1.24E-07 | 2.58E-03 | 0.66 | 0.61 | 0.05 | 1.09 |
| 19 | Dynlrb1 | 0.65 | 1.30E-07 | 2.70E-03 | 0.91 | 0.95 | -0.03 | 0.96 |
| 19 | Wwc1 | 0.99 | 1.43E-07 | 2.98E-03 | 0.34 | 0.18 | 0.16 | 1.85 |
| 19 | Nebl | 1.01 | 1.60E-07 | 3.32E-03 | 0.24 | 0.11 | 0.13 | 2.16 |
| 19 | Klhl29 | 1.80 | 1.62E-07 | 3.37E-03 | 0.13 | 0.05 | 0.09 | 2.83 |
| 19 | Leprotl1 | 0.79 | 1.63E-07 | 3.39E-03 | 0.65 | 0.55 | 0.10 | 1.17 |
| 19 | Dclk1 | 1.76 | 1.66E-07 | 3.45E-03 | 0.39 | 0.26 | 0.13 | 1.51 |
| 19 | Dst | 1.35 | 1.71E-07 | 3.55E-03 | 0.48 | 0.36 | 0.13 | 1.35 |
| 19 | Evl | 0.94 | 1.86E-07 | 3.88E-03 | 0.78 | 0.76 | 0.01 | 1.02 |
| 19 | Acvr2a | 1.01 | 1.90E-07 | 3.94E-03 | 0.43 | 0.27 | 0.16 | 1.59 |
| 19 | Septin5 | 2.01 | 1.93E-07 | 4.02E-03 | 0.16 | 0.07 | 0.10 | 2.43 |
| 19 | Sort1 | 1.67 | 1.97E-07 | 4.10E-03 | 0.34 | 0.20 | 0.14 | 1.68 |
| 19 | Epb41 | 0.94 | 1.98E-07 | 4.11E-03 | 0.74 | 0.76 | -0.02 | 0.98 |
| 19 | Dyrk1b | 1.22 | 2.14E-07 | 4.46E-03 | 0.28 | 0.14 | 0.13 | 1.94 |
| 19 | Cdh4 | 2.29 | 2.25E-07 | 4.67E-03 | 0.14 | 0.05 | 0.09 | 2.73 |
| 19 | Chmp2a | 0.92 | 2.26E-07 | 4.71E-03 | 0.72 | 0.66 | 0.06 | 1.09 |
| 19 | Csnk1g3 | 0.91 | 2.39E-07 | 4.97E-03 | 0.75 | 0.68 | 0.06 | 1.09 |
| 19 | Krtcap3 | 1.38 | 2.39E-07 | 4.97E-03 | 0.24 | 0.11 | 0.12 | 2.06 |
| 19 | Fam204a | 1.01 | 2.41E-07 | 5.01E-03 | 0.73 | 0.65 | 0.08 | 1.12 |
| 19 | Akap5 | 2.03 | 2.46E-07 | 5.11E-03 | 0.12 | 0.04 | 0.08 | 2.93 |
| 19 | Kbtbd8 | 2.01 | 2.51E-07 | 5.22E-03 | 0.11 | 0.04 | 0.08 | 3.00 |
| 19 | Afdn | 0.74 | 2.70E-07 | 5.62E-03 | 0.71 | 0.57 | 0.13 | 1.23 |
| 19 | Vasp | 1.20 | 2.75E-07 | 5.73E-03 | 0.58 | 0.49 | 0.10 | 1.20 |
| 19 | Fam227a | 1.87 | 2.84E-07 | 5.91E-03 | 0.14 | 0.05 | 0.09 | 2.73 |
| 19 | Dmrt3 | 1.45 | 3.24E-07 | 6.75E-03 | 0.15 | 0.05 | 0.09 | 2.69 |
| 19 | Dach1 | 1.19 | 3.34E-07 | 6.95E-03 | 0.53 | 0.39 | 0.15 | 1.38 |
| 19 | Zswim5 | 2.22 | 3.42E-07 | 7.11E-03 | 0.10 | 0.03 | 0.07 | 3.09 |
| 19 | Bex1 | 0.35 | 3.46E-07 | 7.19E-03 | 0.77 | 0.59 | 0.17 | 1.29 |
| 19 | Dctn2 | 0.90 | 4.03E-07 | 8.37E-03 | 0.80 | 0.83 | -0.03 | 0.96 |
| 19 | Pik3ip1 | 1.90 | 4.14E-07 | 8.62E-03 | 0.19 | 0.09 | 0.11 | 2.19 |
| 19 | Chn1 | 1.42 | 4.39E-07 | 9.12E-03 | 0.52 | 0.40 | 0.12 | 1.29 |
| 19 | Ufm1 | 0.93 | 4.58E-07 | 9.52E-03 | 0.72 | 0.69 | 0.03 | 1.04 |
| 19 | Ggt7 | 2.04 | 4.60E-07 | 9.57E-03 | 0.10 | 0.03 | 0.07 | 3.09 |
| 19 | Nin | 1.29 | 4.77E-07 | 9.92E-03 | 0.41 | 0.27 | 0.14 | 1.51 |
| 19 | Peli1 | 0.83 | 4.86E-07 | 1.01E-02 | 0.71 | 0.68 | 0.03 | 1.05 |
| 19 | Cep290 | 1.03 | 4.94E-07 | 1.03E-02 | 0.61 | 0.51 | 0.09 | 1.18 |
| 19 | Ly6h | 1.98 | 5.06E-07 | 1.05E-02 | 0.28 | 0.15 | 0.13 | 1.82 |
| 19 | Nell2 | 2.81 | 5.13E-07 | 1.07E-02 | 0.13 | 0.05 | 0.08 | 2.66 |
| 19 | Gipc2 | 1.67 | 5.14E-07 | 1.07E-02 | 0.16 | 0.06 | 0.09 | 2.45 |
| 19 | 1110006O24Rik | 2.83 | 5.16E-07 | 1.07E-02 | 0.11 | 0.04 | 0.08 | 2.92 |
| 19 | Ssbp3 | 0.47 | 5.40E-07 | 1.12E-02 | 0.86 | 0.94 | -0.08 | 0.92 |
| 19 | Gls | 1.03 | 5.48E-07 | 1.14E-02 | 0.52 | 0.41 | 0.12 | 1.28 |
| 19 | Znrf1 | 0.98 | 5.80E-07 | 1.21E-02 | 0.68 | 0.60 | 0.08 | 1.13 |
| 19 | Madd | 1.84 | 5.95E-07 | 1.24E-02 | 0.17 | 0.07 | 0.10 | 2.32 |
| 19 | 2700097O09Rik | 1.43 | 5.99E-07 | 1.25E-02 | 0.46 | 0.35 | 0.11 | 1.33 |
| 19 | Pcdh17 | 1.54 | 6.10E-07 | 1.27E-02 | 0.34 | 0.21 | 0.14 | 1.66 |
| 19 | Kcne1l | 2.23 | 6.65E-07 | 1.38E-02 | 0.24 | 0.13 | 0.12 | 1.91 |
| 19 | Rbm18 | 0.97 | 6.92E-07 | 1.44E-02 | 0.64 | 0.55 | 0.09 | 1.16 |
| 19 | Minar2 | 1.86 | 6.92E-07 | 1.44E-02 | 0.21 | 0.10 | 0.11 | 2.14 |
| 19 | BC005537 | 0.91 | 7.08E-07 | 1.47E-02 | 0.59 | 0.50 | 0.09 | 1.18 |
| 19 | Lrrn1 | 1.69 | 7.31E-07 | 1.52E-02 | 0.13 | 0.05 | 0.08 | 2.76 |

**Supplementary Table 1. Marker genes for the original clusters of integrated clustering of E11.5 control and NotchGOF λ-junction cells.**

| | | | | | | | | |
|---|---|---|---|---|---|---|---|---|
| 19 | Rnf121 | 1.16 | 7.44E-07 | 1.55E-02 | 0.49 | 0.37 | 0.12 | 1.33 |
| 19 | Slc43a1 | 1.36 | 7.53E-07 | 1.57E-02 | 0.29 | 0.16 | 0.13 | 1.81 |
| 19 | Psen2 | 2.41 | 8.00E-07 | 1.66E-02 | 0.18 | 0.08 | 0.10 | 2.27 |
| 19 | Stx2 | 1.38 | 8.11E-07 | 1.69E-02 | 0.34 | 0.22 | 0.12 | 1.55 |
| 19 | Ncstn | 1.30 | 8.51E-07 | 1.77E-02 | 0.42 | 0.29 | 0.13 | 1.45 |
| 19 | Prr13 | 1.06 | 8.58E-07 | 1.79E-02 | 0.54 | 0.40 | 0.14 | 1.35 |
| 19 | Hmgn5 | 1.05 | 8.59E-07 | 1.79E-02 | 0.71 | 0.71 | -0.01 | 0.99 |
| 19 | Synj1 | 2.00 | 8.77E-07 | 1.82E-02 | 0.22 | 0.11 | 0.11 | 2.01 |
| 19 | Slc9a3r1 | 0.62 | 8.98E-07 | 1.87E-02 | 0.42 | 0.24 | 0.17 | 1.72 |
| 19 | Brsk1 | 1.72 | 9.13E-07 | 1.90E-02 | 0.18 | 0.08 | 0.10 | 2.23 |
| 19 | Gkap1 | 1.20 | 9.21E-07 | 1.92E-02 | 0.57 | 0.44 | 0.14 | 1.31 |
| 19 | Meak7 | 2.17 | 9.29E-07 | 1.93E-02 | 0.15 | 0.06 | 0.09 | 2.54 |
| 19 | Tnnt1 | 1.86 | 9.55E-07 | 1.99E-02 | 0.14 | 0.05 | 0.09 | 2.57 |
| 19 | Cep70 | 0.91 | 9.61E-07 | 2.00E-02 | 0.54 | 0.40 | 0.14 | 1.35 |
| 19 | Rnf38 | 1.17 | 1.09E-06 | 2.26E-02 | 0.40 | 0.27 | 0.13 | 1.47 |
| 19 | Ell2 | 1.62 | 1.10E-06 | 2.29E-02 | 0.19 | 0.09 | 0.10 | 2.17 |
| 19 | Atp6v0e | 0.78 | 1.14E-06 | 2.38E-02 | 0.70 | 0.72 | -0.02 | 0.97 |
| 19 | Trappc6b | 0.89 | 1.16E-06 | 2.41E-02 | 0.63 | 0.54 | 0.09 | 1.17 |
| 19 | Pard6a | 1.76 | 1.28E-06 | 2.65E-02 | 0.29 | 0.17 | 0.12 | 1.73 |
| 19 | Thnsl1 | 1.62 | 1.29E-06 | 2.68E-02 | 0.17 | 0.07 | 0.10 | 2.32 |
| 19 | Smarcc2 | 0.89 | 1.37E-06 | 2.84E-02 | 0.71 | 0.65 | 0.06 | 1.09 |
| 19 | Hs6st1 | 1.27 | 1.59E-06 | 3.31E-02 | 0.30 | 0.17 | 0.13 | 1.75 |
| 19 | Calm3 | 0.70 | 1.60E-06 | 3.32E-02 | 0.87 | 0.91 | -0.05 | 0.95 |
| 19 | Meis3 | 2.06 | 1.61E-06 | 3.35E-02 | 0.33 | 0.20 | 0.13 | 1.63 |
| 19 | Fam8a1 | 1.38 | 1.65E-06 | 3.43E-02 | 0.39 | 0.27 | 0.12 | 1.46 |
| 19 | Ooep | 2.44 | 1.65E-06 | 3.44E-02 | 0.11 | 0.04 | 0.07 | 2.84 |
| 19 | Reep1 | 1.73 | 1.67E-06 | 3.47E-02 | 0.13 | 0.05 | 0.08 | 2.65 |
| 19 | Ndfip2 | 0.99 | 1.74E-06 | 3.62E-02 | 0.63 | 0.51 | 0.12 | 1.22 |
| 19 | Dysf | 1.59 | 1.75E-06 | 3.63E-02 | 0.11 | 0.04 | 0.07 | 2.84 |
| 19 | Kctd10 | 1.30 | 1.75E-06 | 3.63E-02 | 0.43 | 0.30 | 0.14 | 1.46 |
| 19 | Spa17 | 1.58 | 1.76E-06 | 3.65E-02 | 0.28 | 0.15 | 0.12 | 1.81 |
| 19 | Mpc1 | 1.08 | 1.78E-06 | 3.70E-02 | 0.68 | 0.63 | 0.06 | 1.09 |
| 19 | Atp6v1f | 0.55 | 1.81E-06 | 3.77E-02 | 0.90 | 0.90 | -0.01 | 0.99 |
| 19 | Morn4 | 1.37 | 1.88E-06 | 3.92E-02 | 0.29 | 0.16 | 0.13 | 1.76 |
| 19 | Map3k12 | 1.14 | 1.95E-06 | 4.05E-02 | 0.34 | 0.21 | 0.13 | 1.61 |
| 19 | Rab15 | 1.27 | 1.96E-06 | 4.09E-02 | 0.25 | 0.13 | 0.12 | 1.89 |
| 19 | Atp6v1d | 0.85 | 2.08E-06 | 4.33E-02 | 0.68 | 0.64 | 0.04 | 1.06 |
| 19 | Arl6 | 1.37 | 2.15E-06 | 4.48E-02 | 0.54 | 0.42 | 0.12 | 1.28 |
| 19 | 1110008P14Rik | 1.19 | 2.24E-06 | 4.65E-02 | 0.56 | 0.47 | 0.09 | 1.20 |
| 19 | Prepl | 1.17 | 2.28E-06 | 4.73E-02 | 0.24 | 0.13 | 0.12 | 1.93 |
| 19 | Whamm | 1.45 | 2.38E-06 | 4.95E-02 | 0.21 | 0.10 | 0.11 | 2.05 |
| 20 | Rhd | 4.17 | 0 | 0 | 0.99 | 0.04 | 0.95 | 23.09 |
| 20 | Alas2 | 4.10 | 0 | 0 | 1.00 | 0.05 | 0.95 | 19.23 |
| 20 | Hemgn | 4.16 | 0 | 0 | 0.99 | 0.05 | 0.94 | 20.27 |
| 20 | Gypa | 4.14 | 0 | 0 | 0.99 | 0.04 | 0.94 | 22.93 |
| 20 | Slc4a1 | 3.95 | 0 | 0 | 1.00 | 0.06 | 0.94 | 17.54 |
| 20 | Ermap | 4.11 | 0 | 0 | 0.97 | 0.04 | 0.94 | 25.61 |
| 20 | Cited4 | 3.77 | 0 | 0 | 0.97 | 0.05 | 0.92 | 18.36 |
| 20 | Klf1 | 3.92 | 0 | 0 | 0.97 | 0.05 | 0.92 | 19.32 |
| 20 | Acp5 | 3.56 | 0 | 0 | 0.99 | 0.08 | 0.91 | 11.96 |
| 20 | Icam4 | 4.13 | 0 | 0 | 0.95 | 0.04 | 0.91 | 25.57 |
| 20 | Tspan33 | 3.52 | 0 | 0 | 0.97 | 0.08 | 0.90 | 12.47 |
| 20 | Sptb | 4.04 | 0 | 0 | 0.93 | 0.04 | 0.89 | 21.67 |
| 20 | Trim10 | 3.90 | 0 | 0 | 0.93 | 0.04 | 0.89 | 25.69 |
| 20 | Spta1 | 4.37 | 0 | 0 | 0.92 | 0.04 | 0.88 | 26.23 |
| 20 | Pdzk1ip1 | 3.99 | 0 | 0 | 0.91 | 0.04 | 0.87 | 22.80 |

**Supplementary Table 1. Marker genes for the original clusters of integrated clustering of E11.5 control and NotchGOF λ-junction cells.**

| | | | | | | | | |
|---|---|---|---|---|---|---|---|---|
| 20 | Ncf4 | 3.93 | 0 | 0 | 0.91 | 0.04 | 0.86 | 22.07 |
| 20 | Rhag | 4.28 | 0 | 0 | 0.89 | 0.03 | 0.86 | 26.21 |
| 20 | Tal1 | 3.93 | 0 | 0 | 0.90 | 0.04 | 0.86 | 20.88 |
| 20 | Cldn13 | 4.25 | 0 | 0 | 0.88 | 0.03 | 0.85 | 30.28 |
| 20 | Kel | 3.88 | 0 | 0 | 0.84 | 0.04 | 0.80 | 23.25 |
| 20 | Epb42 | 4.08 | 0 | 0 | 0.80 | 0.03 | 0.77 | 25.90 |
| 20 | Slc38a5 | 4.17 | 0 | 0 | 0.80 | 0.03 | 0.77 | 26.53 |
| 20 | Hsd3b6 | 4.02 | 0 | 0 | 0.78 | 0.03 | 0.75 | 25.03 |
| 20 | Kcnn4 | 4.16 | 0 | 0 | 0.74 | 0.03 | 0.71 | 26.46 |
| 20 | Cfap57 | 4.22 | 0 | 0 | 0.73 | 0.03 | 0.70 | 28.00 |
| 20 | Tspan32 | 3.61 | 0 | 0 | 0.73 | 0.03 | 0.69 | 21.41 |
| 20 | Gata1 | 3.94 | 0 | 0 | 0.72 | 0.03 | 0.69 | 25.75 |
| 20 | Hbq1b | 4.14 | 0 | 0 | 0.71 | 0.03 | 0.68 | 26.19 |
| 20 | Tent5c | 3.80 | 0 | 0 | 0.71 | 0.04 | 0.68 | 19.30 |
| 20 | Ctse | 4.29 | 0 | 0 | 0.68 | 0.03 | 0.66 | 27.20 |
| 20 | Slc30a10 | 3.75 | 0 | 0 | 0.69 | 0.04 | 0.65 | 18.57 |
| 20 | Grap2 | 3.94 | 0 | 0 | 0.67 | 0.02 | 0.65 | 29.26 |
| 20 | Ank1 | 4.06 | 0 | 0 | 0.67 | 0.03 | 0.64 | 22.43 |
| 20 | Nfe2 | 3.72 | 0 | 0 | 0.67 | 0.03 | 0.64 | 21.71 |
| 20 | 1300017J02Rik | 3.78 | 0 | 0 | 0.67 | 0.03 | 0.64 | 23.00 |
| 20 | Lmx1a | 3.92 | 0 | 0 | 0.66 | 0.03 | 0.63 | 25.38 |
| 20 | Gm867 | 4.22 | 0 | 0 | 0.55 | 0.02 | 0.53 | 29.00 |
| 20 | Pkhd1l1 | 4.09 | 0 | 0 | 0.53 | 0.02 | 0.51 | 29.50 |
| 20 | Aqp8 | 4.09 | 0 | 0 | 0.46 | 0.02 | 0.44 | 28.50 |
| 20 | Gm2061 | 3.88 | 3.27E-304 | 6.81E-300 | 0.50 | 0.02 | 0.48 | 24.85 |
| 20 | Asb17os | 3.89 | 2.93E-290 | 6.09E-286 | 0.52 | 0.02 | 0.50 | 21.83 |
| 20 | Aqp3 | 3.43 | 1.69E-284 | 3.52E-280 | 0.61 | 0.03 | 0.57 | 18.33 |
| 20 | Ikzf1 | 3.59 | 2.46E-276 | 5.12E-272 | 0.52 | 0.02 | 0.49 | 21.54 |
| 20 | Car2 | 4.01 | 4.00E-270 | 8.31E-266 | 1.00 | 0.11 | 0.89 | 8.93 |
| 20 | Gm20161 | 4.47 | 1.09E-266 | 2.26E-262 | 0.33 | 0.01 | 0.32 | 33.30 |
| 20 | Snca | 3.22 | 9.69E-266 | 2.02E-261 | 0.91 | 0.09 | 0.83 | 10.73 |
| 20 | Tuba4a | 2.52 | 1.50E-264 | 3.13E-260 | 0.82 | 0.06 | 0.75 | 12.75 |
| 20 | Gm16867 | 3.43 | 2.97E-258 | 6.17E-254 | 0.67 | 0.04 | 0.62 | 15.16 |
| 20 | Pklr | 4.27 | 2.76E-252 | 5.74E-248 | 0.42 | 0.02 | 0.40 | 24.41 |
| 20 | Gm14275 | 4.21 | 9.14E-248 | 1.90E-243 | 0.37 | 0.01 | 0.36 | 26.71 |
| 20 | Mt2 | 3.59 | 1.96E-243 | 4.08E-239 | 1.00 | 0.13 | 0.88 | 8.00 |
| 20 | Epor | 3.06 | 9.19E-241 | 1.91E-236 | 0.83 | 0.07 | 0.76 | 11.53 |
| 20 | Xk | 3.48 | 1.59E-240 | 3.31E-236 | 0.54 | 0.03 | 0.51 | 17.00 |
| 20 | Gm32051 | 4.18 | 3.53E-240 | 7.34E-236 | 0.39 | 0.02 | 0.37 | 24.25 |
| 20 | Mylk3 | 3.52 | 2.81E-239 | 5.83E-235 | 0.47 | 0.02 | 0.45 | 20.39 |
| 20 | Redrum | 4.06 | 1.34E-232 | 2.80E-228 | 0.43 | 0.02 | 0.41 | 21.45 |
| 20 | Adra2b | 4.08 | 5.53E-231 | 1.15E-226 | 0.33 | 0.01 | 0.32 | 27.75 |
| 20 | Gpr146 | 3.32 | 1.67E-228 | 3.48E-224 | 0.50 | 0.03 | 0.47 | 18.41 |
| 20 | Plek2 | 3.22 | 1.01E-227 | 2.10E-223 | 0.71 | 0.06 | 0.66 | 12.31 |
| 20 | Add2 | 3.11 | 7.85E-223 | 1.63E-218 | 0.85 | 0.09 | 0.77 | 10.00 |
| 20 | Steap3 | 3.89 | 6.33E-221 | 1.32E-216 | 0.43 | 0.02 | 0.41 | 20.43 |
| 20 | Slfn14 | 4.15 | 3.52E-217 | 7.32E-213 | 0.34 | 0.01 | 0.33 | 26.15 |
| 20 | Dhrs11 | 3.26 | 9.11E-215 | 1.89E-210 | 0.60 | 0.04 | 0.56 | 14.26 |
| 20 | Dmtn | 3.03 | 4.59E-213 | 9.54E-209 | 0.72 | 0.06 | 0.66 | 11.63 |
| 20 | 6030468B19Rik | 3.88 | 3.76E-206 | 7.83E-202 | 0.34 | 0.01 | 0.33 | 24.29 |
| 20 | Gfi1b | 4.08 | 4.43E-205 | 9.21E-201 | 0.37 | 0.02 | 0.35 | 21.59 |
| 20 | Gm10390 | 4.66 | 1.21E-203 | 2.51E-199 | 0.27 | 0.01 | 0.26 | 29.44 |
| 20 | Ube2l6 | 3.10 | 1.69E-202 | 3.52E-198 | 0.97 | 0.13 | 0.84 | 7.49 |
| 20 | Asb17 | 4.07 | 3.59E-199 | 7.46E-195 | 0.33 | 0.01 | 0.32 | 23.79 |
| 20 | Tmod1 | 2.98 | 9.81E-195 | 2.04E-190 | 0.67 | 0.06 | 0.61 | 11.41 |
| 20 | Ache | 3.39 | 3.30E-190 | 6.87E-186 | 0.35 | 0.02 | 0.34 | 20.82 |

**Supplementary Table 1. Marker genes for the original clusters of integrated clustering of E11.5 control and NotchGOF λ-junction cells.**

| | | | | | | | | |
|---|---|---|---|---|---|---|---|---|
| 20 | Ell2 | 2.89 | 7.31E-186 | 1.52E-181 | 0.79 | 0.09 | 0.70 | 9.28 |
| 20 | Slc22a4 | 3.52 | 1.42E-183 | 2.95E-179 | 0.35 | 0.02 | 0.33 | 20.41 |
| 20 | Lyl1 | 3.10 | 1.69E-181 | 3.51E-177 | 0.49 | 0.03 | 0.46 | 14.41 |
| 20 | Reln | 3.18 | 7.80E-176 | 1.62E-171 | 0.50 | 0.04 | 0.46 | 13.81 |
| 20 | Rec114 | 3.44 | 1.45E-175 | 3.01E-171 | 0.41 | 0.02 | 0.38 | 17.00 |
| 20 | Pnpo | 3.12 | 2.96E-171 | 6.15E-167 | 0.98 | 0.17 | 0.81 | 5.90 |
| 20 | Mrap | 3.92 | 3.40E-170 | 7.07E-166 | 0.29 | 0.01 | 0.27 | 23.83 |
| 20 | Dapk2 | 2.93 | 1.56E-168 | 3.24E-164 | 0.55 | 0.05 | 0.51 | 12.24 |
| 20 | Smim1 | 3.15 | 1.55E-164 | 3.23E-160 | 0.95 | 0.16 | 0.78 | 5.80 |
| 20 | Ccdc92b | 3.64 | 1.80E-163 | 3.74E-159 | 0.30 | 0.01 | 0.29 | 21.36 |
| 20 | Atp8a1 | 2.59 | 1.86E-158 | 3.87E-154 | 0.74 | 0.08 | 0.65 | 8.75 |
| 20 | Rnf212 | 3.97 | 3.26E-157 | 6.77E-153 | 0.28 | 0.01 | 0.27 | 21.46 |
| 20 | Cmpk2 | 2.98 | 4.05E-155 | 8.43E-151 | 0.46 | 0.04 | 0.43 | 13.23 |
| 20 | Hbb-bs | 4.11 | 2.02E-154 | 4.20E-150 | 0.99 | 0.23 | 0.76 | 4.24 |
| 20 | Slc39a8 | 2.76 | 1.05E-153 | 2.19E-149 | 0.96 | 0.17 | 0.79 | 5.78 |
| 20 | Slc25a37 | 3.55 | 2.95E-153 | 6.13E-149 | 1.00 | 0.24 | 0.76 | 4.24 |
| 20 | Fam210b | 2.88 | 9.79E-149 | 2.04E-144 | 0.98 | 0.20 | 0.78 | 4.97 |
| 20 | Treml2 | 3.28 | 1.08E-144 | 2.24E-140 | 0.29 | 0.01 | 0.27 | 20.43 |
| 20 | Zc3hav1 | 3.46 | 2.03E-141 | 4.22E-137 | 0.38 | 0.03 | 0.36 | 14.65 |
| 20 | Mt1 | 3.62 | 3.54E-138 | 7.36E-134 | 0.99 | 0.26 | 0.73 | 3.76 |
| 20 | Gmpr | 2.70 | 1.01E-137 | 2.10E-133 | 0.94 | 0.17 | 0.77 | 5.46 |
| 20 | Gm38619.1 | 3.44 | 1.77E-134 | 3.67E-130 | 0.27 | 0.01 | 0.25 | 20.38 |
| 20 | Spns2 | 3.08 | 4.21E-134 | 8.75E-130 | 0.37 | 0.03 | 0.35 | 13.85 |
| 20 | Hsd3b1 | 5.22 | 5.27E-131 | 1.10E-126 | 0.13 | 0.00 | 0.13 | 43.00 |
| 20 | Hbb-bt | 4.14 | 4.15E-130 | 8.63E-126 | 0.99 | 0.30 | 0.70 | 3.33 |
| 20 | 5430401H09Rik | 3.71 | 1.35E-129 | 2.81E-125 | 0.25 | 0.01 | 0.23 | 20.42 |
| 20 | Fech | 2.91 | 1.77E-129 | 3.68E-125 | 1.00 | 0.27 | 0.73 | 3.70 |
| 20 | Mfsd2b | 3.03 | 3.03E-128 | 6.31E-124 | 0.28 | 0.02 | 0.26 | 18.60 |
| 20 | Hebp1 | 2.84 | 7.01E-128 | 1.46E-123 | 0.99 | 0.26 | 0.73 | 3.78 |
| 20 | Ifi35 | 3.02 | 3.62E-123 | 7.54E-119 | 0.37 | 0.03 | 0.35 | 12.90 |
| 20 | Tinagl1 | 2.40 | 8.07E-123 | 1.68E-118 | 0.54 | 0.06 | 0.48 | 9.42 |
| 20 | Slc25a21 | 2.36 | 4.23E-121 | 8.79E-117 | 0.57 | 0.06 | 0.51 | 8.92 |
| 20 | Cpox | 3.05 | 1.42E-120 | 2.96E-116 | 0.99 | 0.31 | 0.68 | 3.16 |
| 20 | Gm43154 | 4.02 | 2.47E-119 | 5.13E-115 | 0.24 | 0.01 | 0.23 | 19.83 |
| 20 | Ccdc141 | 3.00 | 2.40E-118 | 4.99E-114 | 0.33 | 0.02 | 0.30 | 14.22 |
| 20 | Chchd10 | 2.24 | 1.06E-117 | 2.20E-113 | 0.88 | 0.16 | 0.73 | 5.63 |
| 20 | Marchf2 | 2.41 | 8.62E-117 | 1.79E-112 | 0.95 | 0.21 | 0.74 | 4.50 |
| 20 | Blvrb | 3.36 | 1.55E-116 | 3.23E-112 | 0.99 | 0.34 | 0.65 | 2.91 |
| 20 | Fam241a | 2.36 | 3.80E-116 | 7.91E-112 | 0.76 | 0.12 | 0.65 | 6.57 |
| 20 | Cd59a | 2.23 | 1.49E-111 | 3.11E-107 | 0.96 | 0.22 | 0.74 | 4.42 |
| 20 | Lrrc39 | 4.30 | 4.75E-109 | 9.87E-105 | 0.18 | 0.01 | 0.18 | 23.00 |
| 20 | Trib3 | 2.98 | 7.78E-108 | 1.62E-103 | 0.30 | 0.02 | 0.28 | 14.24 |
| 20 | Hbb-bh1 | 3.94 | 3.61E-106 | 7.50E-102 | 1.00 | 0.42 | 0.58 | 2.40 |
| 20 | Slc6a9 | 2.60 | 4.08E-106 | 8.49E-102 | 0.44 | 0.04 | 0.39 | 10.12 |
| 20 | Creg1 | 2.53 | 4.76E-106 | 9.90E-102 | 1.00 | 0.35 | 0.65 | 2.86 |
| 20 | Cpeb4 | 2.19 | 7.77E-104 | 1.62E-99 | 0.65 | 0.09 | 0.55 | 7.02 |
| 20 | Gclm | 2.59 | 5.11E-103 | 1.06E-98 | 0.99 | 0.39 | 0.61 | 2.57 |
| 20 | Abcb10 | 2.25 | 5.51E-102 | 1.15E-97 | 0.95 | 0.22 | 0.72 | 4.22 |
| 20 | Chac2 | 2.29 | 1.69E-100 | 3.51E-96 | 0.75 | 0.13 | 0.62 | 5.75 |
| 20 | Fbxo6 | 2.29 | 4.32E-100 | 8.98E-96 | 0.51 | 0.06 | 0.45 | 8.36 |
| 20 | Tmem184a | 2.39 | 2.89E-98 | 6.01E-94 | 0.46 | 0.05 | 0.41 | 8.94 |
| 20 | Ptp4a3 | 2.17 | 2.17E-97 | 4.51E-93 | 0.95 | 0.25 | 0.70 | 3.81 |
| 20 | Mgst3 | 3.04 | 1.63E-94 | 3.39E-90 | 0.99 | 0.50 | 0.49 | 1.97 |
| 20 | Uros | 2.09 | 2.05E-94 | 4.27E-90 | 0.85 | 0.17 | 0.68 | 5.09 |
| 20 | Cdkn2c | 2.61 | 6.63E-93 | 1.38E-88 | 0.38 | 0.04 | 0.34 | 10.03 |
| 20 | Arrb1 | 2.05 | 1.15E-92 | 2.40E-88 | 0.69 | 0.11 | 0.58 | 6.13 |

**Supplementary Table 1. Marker genes for the original clusters of integrated clustering of E11.5 control and NotchGOF λ-junction cells.**

| | | | | | | | | |
|---|---|---|---|---|---|---|---|---|
| 20 | Ucp2 | 2.16 | 3.06E-91 | 6.37E-87 | 0.99 | 0.35 | 0.63 | 2.79 |
| 20 | Hmbs | 2.51 | 3.78E-90 | 7.86E-86 | 0.99 | 0.58 | 0.41 | 1.70 |
| 20 | Apol8 | 3.94 | 5.60E-90 | 1.16E-85 | 0.16 | 0.01 | 0.15 | 22.29 |
| 20 | Ccrl2 | 3.47 | 6.39E-90 | 1.33E-85 | 0.20 | 0.01 | 0.19 | 17.91 |
| 20 | Hba-a2 | 3.88 | 1.36E-88 | 2.83E-84 | 1.00 | 0.66 | 0.34 | 1.52 |
| 20 | Tmem86b | 2.70 | 1.68E-88 | 3.50E-84 | 0.32 | 0.03 | 0.29 | 11.03 |
| 20 | Josd2 | 2.26 | 5.58E-88 | 1.16E-83 | 0.99 | 0.42 | 0.57 | 2.37 |
| 20 | Hba-a1 | 4.00 | 1.18E-87 | 2.46E-83 | 1.00 | 0.69 | 0.31 | 1.44 |
| 20 | Adrb2 | 3.15 | 2.88E-87 | 5.99E-83 | 0.20 | 0.01 | 0.19 | 17.00 |
| 20 | Nudt4 | 2.35 | 6.50E-87 | 1.35E-82 | 1.00 | 0.77 | 0.23 | 1.30 |
| 20 | Nxpe2 | 4.10 | 8.83E-87 | 1.84E-82 | 0.16 | 0.01 | 0.15 | 22.29 |
| 20 | Gpx1 | 2.49 | 1.79E-85 | 3.72E-81 | 1.00 | 0.95 | 0.05 | 1.05 |
| 20 | Fth1 | 2.38 | 9.98E-85 | 2.08E-80 | 1.00 | 1.00 | 0.00 | 1.00 |
| 20 | Htatip2 | 2.11 | 2.74E-84 | 5.69E-80 | 0.52 | 0.07 | 0.45 | 7.18 |
| 20 | Alad | 2.32 | 3.25E-84 | 6.75E-80 | 0.99 | 0.63 | 0.36 | 1.57 |
| 20 | Hba-x | 4.00 | 3.55E-84 | 7.38E-80 | 1.00 | 0.89 | 0.11 | 1.12 |
| 20 | Hbb-y | 4.07 | 2.16E-83 | 4.48E-79 | 1.00 | 0.98 | 0.02 | 1.02 |
| 20 | Popdc2 | 3.52 | 9.80E-83 | 2.04E-78 | 0.19 | 0.01 | 0.18 | 17.27 |
| 20 | Gm38619 | 4.15 | 4.41E-81 | 9.17E-77 | 0.15 | 0.01 | 0.14 | 21.43 |
| 20 | Art4 | 2.34 | 9.71E-81 | 2.02E-76 | 0.30 | 0.03 | 0.27 | 11.07 |
| 20 | Tmem56 | 2.85 | 2.02E-80 | 4.20E-76 | 0.26 | 0.02 | 0.24 | 12.33 |
| 20 | E2f2 | 1.93 | 2.14E-80 | 4.45E-76 | 0.64 | 0.11 | 0.53 | 5.86 |
| 20 | Gm9895 | 3.38 | 3.20E-79 | 6.65E-75 | 0.19 | 0.01 | 0.18 | 15.83 |
| 20 | Urod | 1.95 | 9.67E-79 | 2.01E-74 | 0.99 | 0.56 | 0.43 | 1.77 |
| 20 | 4930455G09Rik | 3.32 | 1.16E-78 | 2.41E-74 | 0.20 | 0.01 | 0.18 | 15.15 |
| 20 | Pheta2 | 2.02 | 3.16E-78 | 6.57E-74 | 0.45 | 0.06 | 0.39 | 7.61 |
| 20 | Bpgm | 2.20 | 1.05E-75 | 2.19E-71 | 0.99 | 0.46 | 0.53 | 2.16 |
| 20 | Arsg | 2.84 | 1.53E-75 | 3.19E-71 | 0.21 | 0.02 | 0.20 | 14.07 |
| 20 | 2810030D12Rik | 3.99 | 1.88E-75 | 3.90E-71 | 0.12 | 0.01 | 0.12 | 24.40 |
| 20 | Ackr1 | 3.11 | 5.59E-75 | 1.16E-70 | 0.21 | 0.02 | 0.20 | 14.07 |
| 20 | Rbm38 | 1.80 | 1.43E-74 | 2.98E-70 | 0.90 | 0.23 | 0.67 | 3.85 |
| 20 | St3gal5 | 1.91 | 3.27E-74 | 6.80E-70 | 0.89 | 0.24 | 0.65 | 3.76 |
| 20 | Pdia2 | 4.47 | 3.51E-74 | 7.31E-70 | 0.11 | 0.00 | 0.11 | 27.25 |
| 20 | Prdx2 | 0.76 | 7.17E-74 | 1.49E-69 | 1.00 | 1.00 | 0.00 | 1.00 |
| 20 | Glrx5 | 1.41 | 1.19E-73 | 2.47E-69 | 1.00 | 0.89 | 0.12 | 1.13 |
| 20 | Grina | 1.73 | 1.81E-73 | 3.77E-69 | 0.93 | 0.25 | 0.69 | 3.77 |
| 20 | Mst1 | 4.23 | 3.01E-73 | 6.26E-69 | 0.11 | 0.00 | 0.11 | 27.25 |
| 20 | Rfesd | 2.01 | 9.68E-73 | 2.01E-68 | 0.94 | 0.31 | 0.63 | 3.08 |
| 20 | 1810053B23Rik | 3.71 | 1.60E-72 | 3.33E-68 | 0.13 | 0.01 | 0.12 | 21.50 |
| 20 | Wnk4 | 2.02 | 1.86E-72 | 3.87E-68 | 0.32 | 0.03 | 0.29 | 9.41 |
| 20 | Tspo | 1.77 | 3.89E-72 | 8.09E-68 | 0.98 | 0.35 | 0.63 | 2.82 |
| 20 | Serinc3 | 1.93 | 1.40E-70 | 2.91E-66 | 0.97 | 0.36 | 0.61 | 2.70 |
| 20 | Slc43a1 | 1.80 | 1.76E-70 | 3.67E-66 | 0.74 | 0.16 | 0.58 | 4.72 |
| 20 | Flt3l | 3.31 | 4.23E-68 | 8.79E-64 | 0.18 | 0.01 | 0.17 | 14.75 |
| 20 | Acss1 | 2.57 | 3.61E-67 | 7.50E-63 | 0.25 | 0.02 | 0.23 | 10.96 |
| 20 | Cyp4b1 | 4.42 | 3.85E-67 | 8.00E-63 | 0.10 | 0.00 | 0.10 | 25.50 |
| 20 | Igtp | 2.56 | 5.02E-67 | 1.04E-62 | 0.22 | 0.02 | 0.21 | 11.79 |
| 20 | Tmem14c | 1.41 | 1.41E-66 | 2.92E-62 | 1.00 | 0.92 | 0.08 | 1.08 |
| 20 | 4930523C07Rik | 2.01 | 1.71E-66 | 3.56E-62 | 0.47 | 0.07 | 0.40 | 6.34 |
| 20 | Smim5 | 2.22 | 6.90E-65 | 1.43E-60 | 0.26 | 0.03 | 0.23 | 10.36 |
| 20 | Abcg4 | 2.89 | 4.23E-64 | 8.80E-60 | 0.21 | 0.02 | 0.19 | 11.72 |
| 20 | Nmnat3 | 2.17 | 3.34E-63 | 6.95E-59 | 0.35 | 0.05 | 0.30 | 7.71 |
| 20 | Slc16a6 | 1.99 | 1.13E-62 | 2.36E-58 | 0.38 | 0.05 | 0.33 | 7.19 |
| 20 | Gm10603 | 3.17 | 4.62E-62 | 9.60E-58 | 0.16 | 0.01 | 0.15 | 15.60 |
| 20 | Gpr150 | 4.03 | 1.57E-61 | 3.27E-57 | 0.12 | 0.01 | 0.11 | 19.33 |
| 20 | Tmcc2 | 1.93 | 5.79E-61 | 1.21E-56 | 0.48 | 0.08 | 0.40 | 5.88 |

**Supplementary Table 1. Marker genes for the original clusters of integrated clustering of E11.5 control and NotchGOF λ-junction cells.**

| | | | | | | | | |
|---|---|---|---|---|---|---|---|---|
| 20 | Ubash3a | 3.91 | 1.57E-59 | 3.27E-55 | 0.12 | 0.01 | 0.11 | 19.33 |
| 20 | Ncoa4 | 1.59 | 4.24E-59 | 8.82E-55 | 0.95 | 0.31 | 0.63 | 3.01 |
| 20 | Bmp2k | 1.61 | 9.81E-59 | 2.04E-54 | 0.67 | 0.15 | 0.52 | 4.48 |
| 20 | Vangl1 | 1.80 | 1.58E-57 | 3.28E-53 | 0.66 | 0.15 | 0.51 | 4.34 |
| 20 | Slc40a1 | 1.46 | 2.93E-57 | 6.09E-53 | 0.39 | 0.06 | 0.33 | 6.69 |
| 20 | Rgs10 | 0.93 | 7.29E-57 | 1.52E-52 | 0.49 | 0.09 | 0.40 | 5.70 |
| 20 | Lmo2 | 0.52 | 1.46E-56 | 3.04E-52 | 0.62 | 0.13 | 0.49 | 4.95 |
| 20 | Gcnt1 | 2.06 | 3.01E-56 | 6.27E-52 | 0.27 | 0.03 | 0.24 | 8.83 |
| 20 | Dpf3 | 3.43 | 4.23E-56 | 8.80E-52 | 0.11 | 0.01 | 0.10 | 21.80 |
| 20 | Lyrm9 | 1.46 | 1.71E-55 | 3.55E-51 | 0.72 | 0.17 | 0.55 | 4.24 |
| 20 | 5430431A17Rik | 3.34 | 1.38E-54 | 2.86E-50 | 0.13 | 0.01 | 0.12 | 16.13 |
| 20 | Ppox | 1.56 | 1.11E-53 | 2.31E-49 | 0.84 | 0.24 | 0.60 | 3.50 |
| 20 | Pip5k1b | 2.06 | 1.34E-53 | 2.78E-49 | 0.28 | 0.03 | 0.25 | 8.21 |
| 20 | Zfpm1 | 1.49 | 3.59E-53 | 7.46E-49 | 0.91 | 0.31 | 0.60 | 2.93 |
| 20 | Ftl1 | 0.49 | 1.22E-52 | 2.53E-48 | 1.00 | 1.00 | 0.00 | 1.00 |
| 20 | Larp1b | 1.57 | 7.12E-52 | 1.48E-47 | 0.44 | 0.08 | 0.36 | 5.72 |
| 20 | Tmem238 | 1.28 | 3.34E-51 | 6.95E-47 | 0.72 | 0.18 | 0.54 | 4.07 |
| 20 | Ttc39a | 1.67 | 5.83E-50 | 1.21E-45 | 0.37 | 0.06 | 0.31 | 6.33 |
| 20 | Ifrd2 | 1.54 | 1.18E-49 | 2.45E-45 | 0.90 | 0.33 | 0.57 | 2.74 |
| 20 | Ypel4 | 1.91 | 1.75E-49 | 3.65E-45 | 0.24 | 0.03 | 0.21 | 8.81 |
| 20 | Samd11 | 2.78 | 3.67E-49 | 7.63E-45 | 0.13 | 0.01 | 0.12 | 14.33 |
| 20 | Ankrd9 | 1.37 | 2.73E-48 | 5.68E-44 | 0.57 | 0.12 | 0.44 | 4.67 |
| 20 | Fzd5 | 2.14 | 3.61E-48 | 7.51E-44 | 0.20 | 0.02 | 0.18 | 9.71 |
| 20 | Dnaja4 | 1.67 | 8.37E-48 | 1.74E-43 | 0.33 | 0.05 | 0.28 | 6.53 |
| 20 | Fads3 | 1.82 | 8.87E-48 | 1.84E-43 | 0.35 | 0.06 | 0.29 | 6.31 |
| 20 | Prr13 | 1.32 | 1.05E-47 | 2.19E-43 | 0.97 | 0.40 | 0.58 | 2.44 |
| 20 | Acyp2 | 1.55 | 4.42E-47 | 9.19E-43 | 0.50 | 0.10 | 0.40 | 4.84 |
| 20 | Stom | 1.96 | 2.54E-46 | 5.28E-42 | 0.30 | 0.04 | 0.26 | 6.80 |
| 20 | Pigq | 1.42 | 3.61E-46 | 7.52E-42 | 0.79 | 0.24 | 0.55 | 3.34 |
| 20 | Svip | 1.34 | 1.84E-45 | 3.82E-41 | 0.46 | 0.09 | 0.37 | 5.07 |
| 20 | H2-T24 | 3.52 | 2.96E-45 | 6.16E-41 | 0.11 | 0.01 | 0.10 | 15.57 |
| 20 | Fam117a | 1.27 | 2.87E-44 | 5.97E-40 | 0.80 | 0.23 | 0.56 | 3.42 |
| 20 | Cdc25b | 1.47 | 2.91E-44 | 6.06E-40 | 0.71 | 0.20 | 0.51 | 3.50 |
| 20 | Capn1 | 1.66 | 3.69E-44 | 7.68E-40 | 0.31 | 0.05 | 0.26 | 6.39 |
| 20 | Trak2 | 1.42 | 4.34E-44 | 9.03E-40 | 0.57 | 0.13 | 0.44 | 4.26 |
| 20 | 1700034P13Rik | 3.10 | 1.39E-43 | 2.89E-39 | 0.11 | 0.01 | 0.10 | 15.57 |
| 20 | Ehbp1l1 | 1.61 | 1.74E-43 | 3.61E-39 | 0.36 | 0.06 | 0.30 | 5.73 |
| 20 | Txnrd2 | 1.29 | 2.59E-43 | 5.38E-39 | 0.75 | 0.21 | 0.54 | 3.56 |
| 20 | Mkrn1 | 1.27 | 3.46E-43 | 7.20E-39 | 0.95 | 0.43 | 0.53 | 2.24 |
| 20 | Cox6b2 | 1.43 | 5.67E-43 | 1.18E-38 | 0.97 | 0.47 | 0.50 | 2.07 |
| 20 | Btg2 | 1.02 | 6.19E-43 | 1.29E-38 | 0.84 | 0.27 | 0.57 | 3.11 |
| 20 | Csf2rb | 2.84 | 8.91E-43 | 1.85E-38 | 0.14 | 0.01 | 0.13 | 12.36 |
| 20 | Tango2 | 1.27 | 1.65E-42 | 3.43E-38 | 0.62 | 0.15 | 0.47 | 4.02 |
| 20 | Mindy2 | 1.30 | 7.43E-42 | 1.55E-37 | 0.65 | 0.17 | 0.48 | 3.77 |
| 20 | 5033406O09Rik | 2.71 | 2.24E-41 | 4.66E-37 | 0.14 | 0.01 | 0.12 | 11.33 |
| 20 | St3gal6 | 1.39 | 2.97E-41 | 6.18E-37 | 0.50 | 0.11 | 0.39 | 4.52 |
| 20 | St3gal1 | 1.28 | 3.49E-41 | 7.27E-37 | 0.47 | 0.10 | 0.37 | 4.69 |
| 20 | Tfr2 | 2.68 | 4.42E-41 | 9.18E-37 | 0.15 | 0.01 | 0.14 | 10.71 |
| 20 | Pfkfb1 | 1.83 | 9.50E-40 | 1.98E-35 | 0.27 | 0.04 | 0.23 | 6.48 |
| 20 | Cltb | 1.30 | 1.41E-39 | 2.94E-35 | 0.65 | 0.18 | 0.47 | 3.61 |
| 20 | Dnase1l1 | 1.55 | 9.35E-39 | 1.94E-34 | 0.29 | 0.05 | 0.24 | 6.22 |
| 20 | Fbxo7 | 1.19 | 1.14E-38 | 2.37E-34 | 0.55 | 0.13 | 0.42 | 4.11 |
| 20 | Lmna | 1.29 | 1.55E-38 | 3.22E-34 | 0.79 | 0.27 | 0.52 | 2.92 |
| 20 | Cd59b | 1.91 | 4.23E-38 | 8.79E-34 | 0.27 | 0.04 | 0.23 | 6.33 |
| 20 | D030028A08Rik | 1.86 | 4.46E-38 | 9.27E-34 | 0.33 | 0.06 | 0.27 | 5.54 |
| 20 | Uba7 | 1.20 | 5.80E-38 | 1.21E-33 | 0.47 | 0.11 | 0.36 | 4.42 |

**Supplementary Table 1. Marker genes for the original clusters of integrated clustering of E11.5 control and NotchGOF λ-junction cells.**

| | | | | | | | | |
|---|---|---|---|---|---|---|---|---|
| 20 | Ppp1r15a | 1.13 | 1.56E-36 | 3.24E-32 | 0.78 | 0.25 | 0.53 | 3.15 |
| 20 | Brpf3 | 1.37 | 4.33E-36 | 9.00E-32 | 0.50 | 0.12 | 0.38 | 4.09 |
| 20 | Ccnd3 | 0.84 | 4.39E-36 | 9.12E-32 | 1.00 | 0.85 | 0.15 | 1.17 |
| 20 | Pim1 | 1.27 | 4.97E-36 | 1.03E-31 | 0.94 | 0.47 | 0.47 | 2.00 |
| 20 | Ocel1 | 1.73 | 7.51E-36 | 1.56E-31 | 0.30 | 0.05 | 0.25 | 5.64 |
| 20 | Hsd17b14 | 2.52 | 8.48E-36 | 1.76E-31 | 0.14 | 0.01 | 0.13 | 10.21 |
| 20 | Slc43a3 | 1.05 | 1.02E-35 | 2.12E-31 | 0.84 | 0.28 | 0.56 | 2.98 |
| 20 | Exoc6 | 1.65 | 1.38E-35 | 2.86E-31 | 0.36 | 0.07 | 0.29 | 4.88 |
| 20 | Ubac1 | 1.15 | 4.12E-35 | 8.57E-31 | 0.97 | 0.57 | 0.39 | 1.69 |
| 20 | Atp1b2 | 2.20 | 7.35E-35 | 1.53E-30 | 0.16 | 0.02 | 0.14 | 9.18 |
| 20 | Prdx3 | 1.11 | 4.33E-34 | 9.00E-30 | 0.97 | 0.53 | 0.44 | 1.82 |
| 20 | Cdkn3 | 1.09 | 9.51E-34 | 1.98E-29 | 0.82 | 0.29 | 0.53 | 2.83 |
| 20 | Fam78a | 2.76 | 9.77E-34 | 2.03E-29 | 0.13 | 0.01 | 0.12 | 9.92 |
| 20 | Cox17 | 0.91 | 1.13E-33 | 2.36E-29 | 0.98 | 0.77 | 0.21 | 1.28 |
| 20 | Endod1 | 1.24 | 2.50E-32 | 5.20E-28 | 0.40 | 0.09 | 0.31 | 4.44 |
| 20 | Hopx | 2.59 | 4.65E-32 | 9.67E-28 | 0.14 | 0.02 | 0.12 | 9.07 |
| 20 | Sgk3 | 1.20 | 5.33E-32 | 1.11E-27 | 0.54 | 0.15 | 0.39 | 3.63 |
| 20 | Lefty1 | 2.75 | 5.36E-32 | 1.12E-27 | 0.11 | 0.01 | 0.10 | 10.90 |
| 20 | Dnajb3 | 1.01 | 7.44E-32 | 1.55E-27 | 0.55 | 0.15 | 0.40 | 3.72 |
| 20 | Prokr1 | 2.39 | 9.83E-32 | 2.04E-27 | 0.13 | 0.01 | 0.12 | 9.92 |
| 20 | Slc16a10 | 0.92 | 2.19E-31 | 4.57E-27 | 0.44 | 0.11 | 0.34 | 4.13 |
| 20 | C1qtnf4 | 1.01 | 2.97E-31 | 6.18E-27 | 0.80 | 0.29 | 0.52 | 2.82 |
| 20 | Dyrk3 | 1.12 | 6.48E-31 | 1.35E-26 | 0.42 | 0.10 | 0.32 | 4.23 |
| 20 | Reep6 | 1.43 | 1.12E-30 | 2.34E-26 | 0.27 | 0.05 | 0.22 | 5.44 |
| 20 | Isca1 | 1.06 | 1.86E-30 | 3.86E-26 | 0.84 | 0.31 | 0.53 | 2.70 |
| 20 | Gadd45a | 0.71 | 2.82E-30 | 5.88E-26 | 0.76 | 0.24 | 0.51 | 3.09 |
| 20 | Lockd | 0.97 | 2.95E-30 | 6.13E-26 | 0.88 | 0.36 | 0.52 | 2.43 |
| 20 | Cdr2 | 1.11 | 7.08E-30 | 1.47E-25 | 0.67 | 0.22 | 0.45 | 3.07 |
| 20 | Ssx2ip | 0.82 | 2.27E-29 | 4.72E-25 | 0.89 | 0.36 | 0.53 | 2.48 |
| 20 | Rnf123 | 0.93 | 2.97E-29 | 6.17E-25 | 0.47 | 0.12 | 0.35 | 3.88 |
| 20 | Slc46a3 | 2.38 | 4.55E-29 | 9.46E-25 | 0.12 | 0.01 | 0.10 | 9.67 |
| 20 | Nrtn | 0.84 | 5.85E-29 | 1.22E-24 | 0.54 | 0.15 | 0.38 | 3.49 |
| 20 | Abcg2 | 1.06 | 6.19E-29 | 1.29E-24 | 0.84 | 0.35 | 0.50 | 2.43 |
| 20 | Cdc42ep3 | 0.53 | 6.74E-29 | 1.40E-24 | 0.57 | 0.16 | 0.41 | 3.50 |
| 20 | Tgm2 | 2.47 | 6.74E-29 | 1.40E-24 | 0.13 | 0.02 | 0.11 | 8.60 |
| 20 | Nt5c3 | 1.12 | 8.67E-29 | 1.80E-24 | 0.87 | 0.40 | 0.47 | 2.19 |
| 20 | Slc20a1 | 0.95 | 8.73E-29 | 1.82E-24 | 0.86 | 0.32 | 0.54 | 2.69 |
| 20 | Fam174b | 1.06 | 1.49E-28 | 3.09E-24 | 0.44 | 0.11 | 0.32 | 3.88 |
| 20 | Ppp1r9b | 0.91 | 2.21E-28 | 4.60E-24 | 0.76 | 0.25 | 0.50 | 2.98 |
| 20 | Spaca9 | 1.58 | 2.55E-28 | 5.31E-24 | 0.20 | 0.03 | 0.17 | 6.18 |
| 20 | Asb1 | 1.45 | 2.56E-28 | 5.33E-24 | 0.31 | 0.07 | 0.25 | 4.60 |
| 20 | Slc29a1 | 0.87 | 3.58E-28 | 7.44E-24 | 0.46 | 0.12 | 0.34 | 3.77 |
| 20 | Unc13d | 1.76 | 2.63E-27 | 5.47E-23 | 0.18 | 0.03 | 0.16 | 6.57 |
| 20 | Apeh | 0.92 | 3.76E-27 | 7.81E-23 | 0.76 | 0.27 | 0.49 | 2.82 |
| 20 | Tatdn3 | 0.89 | 4.34E-27 | 9.02E-23 | 0.42 | 0.11 | 0.31 | 3.95 |
| 20 | G430095P16Rik | 1.06 | 2.88E-26 | 5.99E-22 | 0.40 | 0.10 | 0.30 | 3.86 |
| 20 | Tmc6 | 1.01 | 2.94E-26 | 6.11E-22 | 0.33 | 0.08 | 0.25 | 4.36 |
| 20 | Mki67 | 0.89 | 3.42E-26 | 7.12E-22 | 1.00 | 0.72 | 0.28 | 1.39 |
| 20 | Tfrc | 1.01 | 6.79E-26 | 1.41E-21 | 0.87 | 0.40 | 0.47 | 2.19 |
| 20 | Tom1l1 | 0.65 | 1.04E-25 | 2.17E-21 | 0.54 | 0.16 | 0.38 | 3.36 |
| 20 | Arhgap23 | 1.24 | 1.54E-25 | 3.21E-21 | 0.37 | 0.09 | 0.27 | 3.90 |
| 20 | Sh3yl1 | 0.71 | 4.44E-25 | 9.22E-21 | 0.61 | 0.19 | 0.42 | 3.17 |
| 20 | Ift140 | 0.76 | 4.74E-25 | 9.86E-21 | 0.54 | 0.16 | 0.38 | 3.36 |
| 20 | Adcy7 | 1.76 | 7.08E-25 | 1.47E-20 | 0.16 | 0.02 | 0.13 | 6.78 |
| 20 | Slc48a1 | 0.76 | 8.41E-25 | 1.75E-20 | 0.54 | 0.17 | 0.38 | 3.28 |
| 20 | Tuba1c | 0.89 | 3.15E-24 | 6.54E-20 | 0.88 | 0.44 | 0.44 | 2.01 |

**Supplementary Table 1. Marker genes for the original clusters of integrated clustering of E11.5 control and NotchGOF λ-junction cells.**

| | | | | | | | | |
|---|---|---|---|---|---|---|---|---|
| 20 | Ehd1 | 0.86 | 3.35E-24 | 6.98E-20 | 0.97 | 0.52 | 0.45 | 1.86 |
| 20 | Pdk2 | 0.85 | 3.43E-24 | 7.13E-20 | 0.34 | 0.08 | 0.26 | 4.10 |
| 20 | Mafk | 1.37 | 5.43E-24 | 1.13E-19 | 0.21 | 0.04 | 0.17 | 5.41 |
| 20 | Pon2 | 0.58 | 5.74E-24 | 1.19E-19 | 0.57 | 0.18 | 0.39 | 3.14 |
| 20 | Tubb4b | 0.73 | 6.61E-24 | 1.38E-19 | 0.99 | 0.85 | 0.14 | 1.17 |
| 20 | Slc41a3 | 1.97 | 7.53E-24 | 1.57E-19 | 0.16 | 0.02 | 0.13 | 6.50 |
| 20 | Pir | 0.93 | 8.51E-24 | 1.77E-19 | 0.44 | 0.13 | 0.32 | 3.51 |
| 20 | Khk | 1.15 | 1.20E-23 | 2.50E-19 | 0.29 | 0.07 | 0.22 | 4.40 |
| 20 | Optn | 1.62 | 2.59E-23 | 5.39E-19 | 0.17 | 0.03 | 0.14 | 6.07 |
| 20 | Hagh | 0.81 | 7.71E-23 | 1.60E-18 | 0.69 | 0.24 | 0.44 | 2.83 |
| 20 | Foxo3 | 0.72 | 1.47E-22 | 3.06E-18 | 0.44 | 0.13 | 0.31 | 3.45 |
| 20 | Rexo2 | 0.81 | 1.51E-22 | 3.13E-18 | 0.97 | 0.59 | 0.37 | 1.63 |
| 20 | Pde10a | 1.19 | 2.85E-22 | 5.93E-18 | 0.22 | 0.04 | 0.17 | 4.95 |
| 20 | Slc30a1 | 0.88 | 4.48E-22 | 9.31E-18 | 0.33 | 0.08 | 0.24 | 3.94 |
| 20 | Iba57 | 1.36 | 8.75E-22 | 1.82E-17 | 0.19 | 0.04 | 0.15 | 5.28 |
| 20 | Gm35082 | 1.54 | 1.07E-21 | 2.22E-17 | 0.16 | 0.03 | 0.14 | 5.82 |
| 20 | Bola3 | 0.70 | 1.99E-21 | 4.14E-17 | 1.00 | 0.75 | 0.25 | 1.34 |
| 20 | Fam110c | 2.06 | 3.62E-21 | 7.53E-17 | 0.10 | 0.01 | 0.09 | 8.50 |
| 20 | Mbnl1 | 0.85 | 4.27E-21 | 8.88E-17 | 0.93 | 0.47 | 0.46 | 1.97 |
| 20 | Fos | 0.40 | 4.48E-21 | 9.31E-17 | 0.44 | 0.14 | 0.30 | 3.15 |
| 20 | Aff1 | 1.16 | 5.45E-21 | 1.13E-16 | 0.29 | 0.07 | 0.22 | 4.09 |
| 20 | Gypc | 0.65 | 6.15E-21 | 1.28E-16 | 0.63 | 0.22 | 0.41 | 2.90 |
| 20 | Nudt19 | 0.75 | 9.84E-21 | 2.05E-16 | 0.80 | 0.32 | 0.48 | 2.50 |
| 20 | Piezo1 | 0.87 | 1.17E-20 | 2.42E-16 | 0.42 | 0.12 | 0.29 | 3.37 |
| 20 | Dennd2c | 1.00 | 1.71E-20 | 3.56E-16 | 0.27 | 0.06 | 0.20 | 4.21 |
| 20 | Frrs1 | 0.80 | 3.43E-20 | 7.12E-16 | 0.61 | 0.22 | 0.39 | 2.78 |
| 20 | Taldo1 | 0.56 | 3.97E-20 | 8.25E-16 | 1.00 | 0.89 | 0.11 | 1.12 |
| 20 | Skap2 | 0.75 | 7.76E-20 | 1.61E-15 | 0.24 | 0.06 | 0.18 | 4.33 |
| 20 | Neat1 | 0.46 | 8.54E-20 | 1.78E-15 | 0.24 | 0.06 | 0.18 | 4.33 |
| 20 | Ankle1 | 0.99 | 8.89E-20 | 1.85E-15 | 0.33 | 0.09 | 0.24 | 3.62 |
| 20 | Rab3il1 | 0.59 | 1.53E-19 | 3.17E-15 | 0.76 | 0.30 | 0.47 | 2.58 |
| 20 | Mylpf | 1.11 | 2.88E-19 | 5.99E-15 | 0.29 | 0.08 | 0.21 | 3.81 |
| 20 | Ank | 0.92 | 3.59E-19 | 7.47E-15 | 0.27 | 0.07 | 0.20 | 3.94 |
| 20 | Ogfrl1 | 0.70 | 3.93E-19 | 8.17E-15 | 0.80 | 0.34 | 0.47 | 2.40 |
| 20 | S100a13 | 0.28 | 4.34E-19 | 9.02E-15 | 0.37 | 0.11 | 0.26 | 3.43 |
| 20 | Nipa1 | 0.75 | 4.46E-19 | 9.27E-15 | 0.44 | 0.14 | 0.30 | 3.13 |
| 20 | Arl4a | 0.75 | 6.12E-19 | 1.27E-14 | 0.56 | 0.20 | 0.36 | 2.75 |
| 20 | Tarsl2 | 0.61 | 7.23E-19 | 1.50E-14 | 0.34 | 0.10 | 0.24 | 3.54 |
| 20 | Slc11a2 | 0.66 | 9.77E-19 | 2.03E-14 | 0.51 | 0.17 | 0.34 | 2.97 |
| 20 | Itpka | 1.73 | 1.08E-18 | 2.24E-14 | 0.13 | 0.02 | 0.11 | 6.14 |
| 20 | Slc25a39 | 0.63 | 1.24E-18 | 2.57E-14 | 0.99 | 0.71 | 0.28 | 1.39 |
| 20 | Yod1 | 0.86 | 1.62E-18 | 3.36E-14 | 0.44 | 0.14 | 0.30 | 3.07 |
| 20 | Ston2 | 0.44 | 5.41E-18 | 1.12E-13 | 0.52 | 0.18 | 0.34 | 2.86 |
| 20 | Junb | 0.36 | 8.62E-18 | 1.79E-13 | 0.37 | 0.12 | 0.26 | 3.22 |
| 20 | Otub2 | 1.44 | 1.01E-17 | 2.11E-13 | 0.21 | 0.05 | 0.16 | 4.31 |
| 20 | Rmc1 | 0.84 | 1.58E-17 | 3.28E-13 | 0.29 | 0.08 | 0.21 | 3.62 |
| 20 | Zfp524 | 0.71 | 1.90E-17 | 3.94E-13 | 0.35 | 0.10 | 0.24 | 3.34 |
| 20 | Gch1 | 0.92 | 2.56E-17 | 5.32E-13 | 0.28 | 0.08 | 0.20 | 3.67 |
| 20 | Spsb2 | 0.79 | 3.47E-17 | 7.22E-13 | 0.28 | 0.08 | 0.20 | 3.67 |
| 20 | Tuft1 | 1.40 | 3.56E-17 | 7.40E-13 | 0.18 | 0.04 | 0.14 | 4.66 |
| 20 | Chst10 | 0.88 | 3.64E-17 | 7.57E-13 | 0.29 | 0.08 | 0.21 | 3.57 |
| 20 | Fam214b | 0.64 | 5.32E-17 | 1.11E-12 | 0.33 | 0.10 | 0.23 | 3.41 |
| 20 | Hspa4l | 0.75 | 5.53E-17 | 1.15E-12 | 0.35 | 0.11 | 0.24 | 3.19 |
| 20 | Rad23a | 0.71 | 7.92E-17 | 1.65E-12 | 0.91 | 0.46 | 0.45 | 1.98 |
| 20 | Akip1 | 0.60 | 1.02E-16 | 2.11E-12 | 0.54 | 0.20 | 0.35 | 2.76 |
| 20 | Capn5 | 1.57 | 1.23E-16 | 2.56E-12 | 0.14 | 0.03 | 0.12 | 5.30 |

**Supplementary Table 1. Marker genes for the original clusters of integrated clustering of E11.5 control and NotchGOF λ-junction cells.**

| | | | | | | | | |
|---|---|---|---|---|---|---|---|---|
| 20 | Smox | 0.46 | 1.63E-16 | 3.40E-12 | 0.63 | 0.24 | 0.39 | 2.65 |
| 20 | 2600006K01Rik | 1.45 | 1.66E-16 | 3.46E-12 | 0.13 | 0.02 | 0.11 | 5.61 |
| 20 | Katnb1 | 0.69 | 2.01E-16 | 4.18E-12 | 0.57 | 0.22 | 0.35 | 2.62 |
| 20 | Fbxo9 | 0.50 | 2.09E-16 | 4.34E-12 | 0.73 | 0.29 | 0.44 | 2.52 |
| 20 | Atp2b4 | 0.55 | 2.13E-16 | 4.43E-12 | 0.54 | 0.20 | 0.34 | 2.71 |
| 20 | Rgcc | 0.86 | 3.92E-16 | 8.15E-12 | 0.31 | 0.09 | 0.21 | 3.33 |
| 20 | Stradb | 0.47 | 4.09E-16 | 8.51E-12 | 0.53 | 0.19 | 0.34 | 2.78 |
| 20 | Cep76 | 0.55 | 4.79E-16 | 9.95E-12 | 0.47 | 0.16 | 0.31 | 2.88 |
| 20 | Dleu2 | 0.62 | 5.84E-16 | 1.21E-11 | 0.63 | 0.25 | 0.37 | 2.48 |
| 20 | Sord | 0.54 | 6.20E-16 | 1.29E-11 | 0.42 | 0.15 | 0.28 | 2.91 |
| 20 | S100a1 | 0.47 | 6.52E-16 | 1.36E-11 | 0.25 | 0.07 | 0.18 | 3.77 |
| 20 | H2afj | 0.49 | 1.00E-15 | 2.09E-11 | 0.99 | 0.81 | 0.18 | 1.22 |
| 20 | Taok3 | 0.56 | 1.13E-15 | 2.34E-11 | 0.33 | 0.10 | 0.23 | 3.20 |
| 20 | Tlcd1 | 0.67 | 1.14E-15 | 2.38E-11 | 0.30 | 0.09 | 0.21 | 3.36 |
| 20 | Ifi30 | 0.39 | 1.26E-15 | 2.62E-11 | 0.71 | 0.29 | 0.41 | 2.41 |
| 20 | Hscb | 0.58 | 1.30E-15 | 2.69E-11 | 0.75 | 0.32 | 0.43 | 2.32 |
| 20 | Tmem179b | 0.50 | 1.52E-15 | 3.16E-11 | 0.71 | 0.29 | 0.42 | 2.44 |
| 20 | Nup210 | 0.81 | 1.54E-15 | 3.21E-11 | 0.23 | 0.06 | 0.17 | 3.79 |
| 20 | Cdkn2d | 0.79 | 1.98E-15 | 4.11E-11 | 0.78 | 0.38 | 0.40 | 2.04 |
| 20 | Rbks | 1.37 | 2.26E-15 | 4.69E-11 | 0.16 | 0.03 | 0.12 | 4.73 |
| 20 | Mvb12b | 0.51 | 2.34E-15 | 4.87E-11 | 0.59 | 0.23 | 0.36 | 2.58 |
| 20 | D1Ertd622e | 0.58 | 2.96E-15 | 6.16E-11 | 0.39 | 0.13 | 0.26 | 2.98 |
| 20 | Arid3a | 0.56 | 3.96E-15 | 8.24E-11 | 0.74 | 0.33 | 0.40 | 2.21 |
| 20 | Mpst | 0.65 | 4.16E-15 | 8.64E-11 | 0.83 | 0.42 | 0.41 | 1.99 |
| 20 | Nadk2 | 0.59 | 5.65E-15 | 1.18E-10 | 0.78 | 0.35 | 0.43 | 2.23 |
| 20 | Rffl | 0.96 | 6.29E-15 | 1.31E-10 | 0.25 | 0.07 | 0.18 | 3.55 |
| 20 | Dclre1b | 0.51 | 6.41E-15 | 1.33E-10 | 0.36 | 0.12 | 0.24 | 3.09 |
| 20 | E2f8 | 0.42 | 6.45E-15 | 1.34E-10 | 0.54 | 0.20 | 0.33 | 2.65 |
| 20 | Lrrc8c | 0.27 | 7.26E-15 | 1.51E-10 | 0.38 | 0.13 | 0.26 | 3.02 |
| 20 | Dusp8 | 1.20 | 7.82E-15 | 1.63E-10 | 0.11 | 0.02 | 0.09 | 5.74 |
| 20 | Napepld | 0.88 | 9.63E-15 | 2.00E-10 | 0.22 | 0.06 | 0.16 | 3.73 |
| 20 | Tnfrsf21 | 0.64 | 1.09E-14 | 2.28E-10 | 0.40 | 0.14 | 0.26 | 2.84 |
| 20 | Zbtb7a | 0.49 | 1.24E-14 | 2.59E-10 | 0.76 | 0.33 | 0.43 | 2.31 |
| 20 | Pcgf5 | 0.65 | 1.30E-14 | 2.71E-10 | 0.29 | 0.09 | 0.20 | 3.29 |
| 20 | Hspa12b | 0.56 | 1.46E-14 | 3.04E-10 | 0.39 | 0.13 | 0.26 | 2.94 |
| 20 | Mthfd2 | 0.65 | 1.50E-14 | 3.12E-10 | 0.89 | 0.47 | 0.42 | 1.90 |
| 20 | Depdc1b | 0.68 | 1.60E-14 | 3.32E-10 | 0.35 | 0.12 | 0.24 | 3.03 |
| 20 | Abhd5 | 0.57 | 1.75E-14 | 3.64E-10 | 0.52 | 0.20 | 0.32 | 2.61 |
| 20 | Glrx | 0.43 | 2.26E-14 | 4.70E-10 | 0.59 | 0.24 | 0.36 | 2.52 |
| 20 | Slc7a1 | 0.58 | 2.77E-14 | 5.76E-10 | 0.27 | 0.08 | 0.19 | 3.40 |
| 20 | Asns | 0.63 | 3.34E-14 | 6.94E-10 | 0.84 | 0.44 | 0.41 | 1.92 |
| 20 | Eif2ak1 | 0.57 | 3.40E-14 | 7.08E-10 | 0.86 | 0.42 | 0.44 | 2.05 |
| 20 | Nqo1 | 0.63 | 4.79E-14 | 9.97E-10 | 0.31 | 0.10 | 0.21 | 3.15 |
| 20 | Vamp8 | 0.55 | 7.47E-14 | 1.55E-09 | 0.97 | 0.61 | 0.36 | 1.58 |
| 20 | Cd55 | 1.27 | 8.29E-14 | 1.72E-09 | 0.12 | 0.02 | 0.10 | 5.08 |
| 20 | Pop5 | 0.56 | 8.91E-14 | 1.85E-09 | 0.97 | 0.65 | 0.32 | 1.50 |
| 20 | Homer2 | 0.35 | 9.16E-14 | 1.91E-09 | 0.54 | 0.21 | 0.33 | 2.58 |
| 20 | Dusp1 | 0.67 | 9.38E-14 | 1.95E-09 | 0.65 | 0.29 | 0.36 | 2.24 |
| 20 | Slc25a38 | 0.58 | 1.16E-13 | 2.41E-09 | 0.63 | 0.27 | 0.36 | 2.33 |
| 20 | Tacc1 | 0.47 | 1.31E-13 | 2.73E-09 | 0.71 | 0.32 | 0.40 | 2.26 |
| 20 | Noct | 0.39 | 1.40E-13 | 2.90E-09 | 0.50 | 0.19 | 0.31 | 2.64 |
| 20 | Itga2b | 0.44 | 1.60E-13 | 3.33E-09 | 0.10 | 0.02 | 0.08 | 5.67 |
| 20 | Rab8b | 0.39 | 2.19E-13 | 4.55E-09 | 0.60 | 0.24 | 0.36 | 2.49 |
| 20 | Ap1g2 | 0.89 | 2.27E-13 | 4.72E-09 | 0.17 | 0.04 | 0.13 | 4.05 |
| 20 | Wbp2 | 0.37 | 2.59E-13 | 5.38E-09 | 0.67 | 0.28 | 0.39 | 2.42 |
| 20 | Aftph | 0.35 | 3.18E-13 | 6.61E-09 | 0.61 | 0.25 | 0.36 | 2.45 |

**Supplementary Table 1. Marker genes for the original clusters of integrated clustering of E11.5 control and NotchGOF λ-junction cells.**

| | | | | | | | | |
|---|---|---|---|---|---|---|---|---|
| 20 | Ptgr2 | 0.43 | 3.33E-13 | 6.92E-09 | 0.41 | 0.15 | 0.26 | 2.74 |
| 20 | Stxbp2 | 0.38 | 4.28E-13 | 8.89E-09 | 0.38 | 0.14 | 0.24 | 2.78 |
| 20 | Hipk2 | 0.56 | 4.61E-13 | 9.60E-09 | 0.96 | 0.64 | 0.32 | 1.49 |
| 20 | Fryl | 0.53 | 4.67E-13 | 9.71E-09 | 0.67 | 0.30 | 0.37 | 2.23 |
| 20 | Il10rb | 0.47 | 5.60E-13 | 1.17E-08 | 0.27 | 0.09 | 0.19 | 3.20 |
| 20 | Skiv2l | 0.42 | 5.82E-13 | 1.21E-08 | 0.59 | 0.24 | 0.35 | 2.44 |
| 20 | Trim56 | 0.37 | 6.03E-13 | 1.25E-08 | 0.39 | 0.14 | 0.25 | 2.77 |
| 20 | Chek2 | 0.43 | 7.43E-13 | 1.55E-08 | 0.57 | 0.23 | 0.34 | 2.44 |
| 20 | Mad2l1bp | 0.37 | 8.19E-13 | 1.70E-08 | 0.58 | 0.23 | 0.34 | 2.47 |
| 20 | Cdkl1 | 0.72 | 9.11E-13 | 1.90E-08 | 0.15 | 0.04 | 0.12 | 4.29 |
| 20 | Ninl | 0.47 | 1.07E-12 | 2.23E-08 | 0.58 | 0.24 | 0.34 | 2.43 |
| 20 | Jak2 | 0.36 | 1.11E-12 | 2.30E-08 | 0.42 | 0.15 | 0.26 | 2.69 |
| 20 | Pycr2 | 0.54 | 1.20E-12 | 2.49E-08 | 0.94 | 0.61 | 0.33 | 1.54 |
| 20 | Srxn1 | 0.47 | 1.36E-12 | 2.82E-08 | 0.33 | 0.11 | 0.22 | 2.92 |
| 20 | Tmem9b | 0.38 | 1.48E-12 | 3.07E-08 | 0.38 | 0.14 | 0.24 | 2.76 |
| 20 | Ogfod2 | 0.40 | 1.55E-12 | 3.23E-08 | 0.30 | 0.10 | 0.20 | 3.02 |
| 20 | Pabpc1 | 0.28 | 1.65E-12 | 3.44E-08 | 1.00 | 0.99 | 0.01 | 1.01 |
| 20 | Stat5a | 1.12 | 1.85E-12 | 3.84E-08 | 0.13 | 0.03 | 0.10 | 4.61 |
| 20 | Grtp1 | 0.34 | 2.37E-12 | 4.94E-08 | 0.38 | 0.14 | 0.24 | 2.74 |
| 20 | Tapbp | 0.55 | 2.67E-12 | 5.56E-08 | 0.26 | 0.08 | 0.18 | 3.16 |
| 20 | Selenbp1 | 0.40 | 2.81E-12 | 5.84E-08 | 0.27 | 0.09 | 0.18 | 3.12 |
| 20 | Plin2 | 0.45 | 3.01E-12 | 6.26E-08 | 0.91 | 0.53 | 0.38 | 1.72 |
| 20 | Agfg2 | 0.35 | 3.09E-12 | 6.43E-08 | 0.53 | 0.21 | 0.32 | 2.50 |
| 20 | Prox1 | 1.58 | 3.60E-12 | 7.49E-08 | 0.11 | 0.02 | 0.09 | 4.95 |
| 20 | Hipk1 | 0.42 | 3.75E-12 | 7.79E-08 | 0.80 | 0.37 | 0.43 | 2.17 |
| 20 | 4930481A15Rik | 1.17 | 3.87E-12 | 8.05E-08 | 0.12 | 0.02 | 0.09 | 4.83 |
| 20 | Zfp367 | 0.33 | 4.25E-12 | 8.83E-08 | 0.39 | 0.14 | 0.24 | 2.69 |
| 20 | Nars | 0.46 | 4.34E-12 | 9.02E-08 | 0.99 | 0.80 | 0.19 | 1.24 |
| 20 | Cercam | 0.61 | 4.56E-12 | 9.48E-08 | 0.20 | 0.06 | 0.15 | 3.46 |
| 20 | Slc7a5 | 0.92 | 4.94E-12 | 1.03E-07 | 0.18 | 0.05 | 0.13 | 3.69 |
| 20 | Adck2 | 0.45 | 7.00E-12 | 1.46E-07 | 0.31 | 0.11 | 0.20 | 2.89 |
| 20 | Pip4k2a | 0.44 | 7.40E-12 | 1.54E-07 | 0.33 | 0.12 | 0.21 | 2.82 |
| 20 | Ccdc82 | 0.58 | 7.71E-12 | 1.60E-07 | 0.33 | 0.12 | 0.21 | 2.77 |
| 20 | Itsn2 | 0.29 | 8.89E-12 | 1.85E-07 | 0.48 | 0.19 | 0.29 | 2.51 |
| 20 | Scrn3 | 0.28 | 1.00E-11 | 2.08E-07 | 0.40 | 0.15 | 0.25 | 2.66 |
| 20 | Ampd2 | 0.57 | 1.00E-11 | 2.08E-07 | 0.41 | 0.16 | 0.25 | 2.58 |
| 20 | Marchf8 | 0.31 | 1.08E-11 | 2.26E-07 | 0.44 | 0.17 | 0.27 | 2.57 |
| 20 | Atg4d | 0.44 | 1.39E-11 | 2.89E-07 | 0.50 | 0.21 | 0.29 | 2.42 |
| 20 | Zfand2b | 0.37 | 1.42E-11 | 2.96E-07 | 0.59 | 0.26 | 0.34 | 2.31 |
| 20 | Hist1h2bj | 0.44 | 1.46E-11 | 3.03E-07 | 0.57 | 0.25 | 0.32 | 2.31 |
| 20 | Fbxo34 | 0.35 | 1.47E-11 | 3.05E-07 | 0.67 | 0.30 | 0.38 | 2.28 |
| 20 | Sppl2b | 0.31 | 1.58E-11 | 3.28E-07 | 0.39 | 0.15 | 0.24 | 2.66 |
| 20 | Podxl | 0.75 | 1.80E-11 | 3.74E-07 | 0.16 | 0.04 | 0.12 | 3.79 |
| 20 | Kat2b | 0.81 | 2.17E-11 | 4.52E-07 | 0.17 | 0.05 | 0.12 | 3.70 |
| 20 | Abcb6 | 0.44 | 2.95E-11 | 6.14E-07 | 0.33 | 0.12 | 0.21 | 2.75 |
| 20 | Wdr81 | 0.78 | 3.17E-11 | 6.59E-07 | 0.18 | 0.05 | 0.13 | 3.54 |
| 20 | Nfix | 0.26 | 3.40E-11 | 7.07E-07 | 0.40 | 0.15 | 0.24 | 2.62 |
| 20 | Stk17b | 0.51 | 3.71E-11 | 7.72E-07 | 0.35 | 0.14 | 0.22 | 2.62 |
| 20 | Ybx3 | 0.40 | 4.02E-11 | 8.36E-07 | 0.99 | 0.84 | 0.16 | 1.19 |
| 20 | Retreg2 | 0.35 | 4.26E-11 | 8.86E-07 | 0.65 | 0.29 | 0.36 | 2.25 |
| 20 | Afg3l2 | 0.39 | 4.36E-11 | 9.08E-07 | 0.76 | 0.34 | 0.41 | 2.21 |
| 20 | Inpp5k | 0.27 | 5.35E-11 | 1.11E-06 | 0.31 | 0.11 | 0.20 | 2.79 |
| 20 | Synj1 | 0.43 | 7.95E-11 | 1.65E-06 | 0.30 | 0.11 | 0.19 | 2.77 |
| 20 | Pip4k2c | 0.25 | 7.98E-11 | 1.66E-06 | 0.46 | 0.18 | 0.27 | 2.48 |
| 20 | Psme3 | 0.51 | 8.70E-11 | 1.81E-06 | 0.90 | 0.50 | 0.40 | 1.81 |
| 20 | Gm43378 | 1.39 | 9.87E-11 | 2.05E-06 | 0.10 | 0.02 | 0.08 | 4.64 |

**Supplementary Table 1. Marker genes for the original clusters of integrated clustering of E11.5 control and NotchGOF λ-junction cells.**

| | | | | | | | | |
|---|---|---|---|---|---|---|---|---|
| 20 | Snx15 | 0.36 | 1.03E-10 | 2.14E-06 | 0.67 | 0.30 | 0.36 | 2.19 |
| 20 | Bcat2 | 0.44 | 1.15E-10 | 2.39E-06 | 0.64 | 0.29 | 0.35 | 2.20 |
| 20 | Narf | 0.37 | 1.28E-10 | 2.65E-06 | 0.84 | 0.42 | 0.42 | 2.00 |
| 20 | Mon1a | 0.56 | 1.46E-10 | 3.03E-06 | 0.21 | 0.07 | 0.15 | 3.20 |
| 20 | Hist1h4i | 0.49 | 1.53E-10 | 3.18E-06 | 0.88 | 0.46 | 0.43 | 1.93 |
| 20 | Il17ra | 0.97 | 1.70E-10 | 3.53E-06 | 0.12 | 0.03 | 0.09 | 4.30 |
| 20 | Adipor1 | 0.46 | 2.67E-10 | 5.56E-06 | 0.95 | 0.56 | 0.39 | 1.69 |
| 20 | Ehd3 | 0.26 | 3.61E-10 | 7.51E-06 | 0.30 | 0.11 | 0.19 | 2.74 |
| 20 | Clk3 | 0.42 | 3.67E-10 | 7.64E-06 | 0.78 | 0.38 | 0.40 | 2.04 |
| 20 | Hspa2 | 0.60 | 4.24E-10 | 8.81E-06 | 0.18 | 0.06 | 0.13 | 3.29 |
| 20 | Dclk2 | 0.41 | 4.29E-10 | 8.92E-06 | 0.31 | 0.11 | 0.19 | 2.68 |
| 20 | Smim4 | 0.36 | 4.68E-10 | 9.74E-06 | 0.84 | 0.43 | 0.41 | 1.97 |
| 20 | Hint3 | 0.25 | 4.85E-10 | 1.01E-05 | 0.54 | 0.24 | 0.31 | 2.31 |
| 20 | Clcn3 | 0.30 | 5.94E-10 | 1.24E-05 | 0.78 | 0.38 | 0.40 | 2.05 |
| 20 | Ftl1-ps1 | 0.40 | 6.50E-10 | 1.35E-05 | 0.81 | 0.41 | 0.40 | 2.00 |
| 20 | Faap24 | 0.46 | 8.51E-10 | 1.77E-05 | 0.40 | 0.16 | 0.23 | 2.41 |
| 20 | Pnpla7 | 0.80 | 9.47E-10 | 1.97E-05 | 0.14 | 0.04 | 0.10 | 3.78 |
| 20 | Ranbp10 | 0.32 | 9.83E-10 | 2.04E-05 | 0.64 | 0.30 | 0.34 | 2.14 |
| 20 | Zfp809 | 0.55 | 1.45E-09 | 3.01E-05 | 0.22 | 0.07 | 0.14 | 2.95 |
| 20 | Hk1 | 0.27 | 1.69E-09 | 3.51E-05 | 0.56 | 0.25 | 0.31 | 2.24 |
| 20 | Cmas | 0.39 | 1.85E-09 | 3.85E-05 | 0.97 | 0.60 | 0.38 | 1.64 |
| 20 | Rfx2 | 0.88 | 2.46E-09 | 5.12E-05 | 0.14 | 0.04 | 0.10 | 3.58 |
| 20 | Znfx1 | 0.77 | 3.12E-09 | 6.49E-05 | 0.18 | 0.06 | 0.13 | 3.12 |
| 20 | Pabpc4 | 0.46 | 3.36E-09 | 6.99E-05 | 0.95 | 0.65 | 0.30 | 1.46 |
| 20 | Ddhd1 | 0.41 | 3.60E-09 | 7.48E-05 | 0.40 | 0.17 | 0.23 | 2.39 |
| 20 | Btrc | 0.27 | 4.99E-09 | 1.04E-04 | 0.71 | 0.34 | 0.37 | 2.09 |
| 20 | Slc1a5 | 0.27 | 6.07E-09 | 1.26E-04 | 0.46 | 0.20 | 0.26 | 2.29 |
| 20 | Cacna2d3 | 0.42 | 6.22E-09 | 1.29E-04 | 0.22 | 0.08 | 0.14 | 2.83 |
| 20 | Msh5 | 0.80 | 9.10E-09 | 1.89E-04 | 0.13 | 0.04 | 0.09 | 3.58 |
| 20 | Hist1h3f | 0.34 | 9.58E-09 | 1.99E-04 | 0.55 | 0.27 | 0.28 | 2.03 |
| 20 | 1700037H04Rik | 0.42 | 1.09E-08 | 2.28E-04 | 0.82 | 0.48 | 0.35 | 1.73 |
| 20 | Rhbdd1 | 0.76 | 1.10E-08 | 2.29E-04 | 0.16 | 0.05 | 0.11 | 3.13 |
| 20 | Ghitm | 0.43 | 1.11E-08 | 2.30E-04 | 0.96 | 0.67 | 0.29 | 1.43 |
| 20 | Mrm1 | 0.89 | 1.21E-08 | 2.52E-04 | 0.14 | 0.04 | 0.10 | 3.40 |
| 20 | Dedd2 | 0.48 | 1.37E-08 | 2.84E-04 | 0.21 | 0.08 | 0.14 | 2.81 |
| 20 | Garem1 | 0.43 | 1.81E-08 | 3.76E-04 | 0.23 | 0.09 | 0.15 | 2.72 |
| 20 | Wdr62 | 0.43 | 2.65E-08 | 5.51E-04 | 0.16 | 0.05 | 0.11 | 3.18 |
| 20 | Herc1 | 0.36 | 2.76E-08 | 5.74E-04 | 0.95 | 0.56 | 0.38 | 1.68 |
| 20 | Ccna2 | 0.36 | 4.36E-08 | 9.07E-04 | 0.92 | 0.63 | 0.29 | 1.46 |
| 20 | Proser2 | 0.59 | 4.78E-08 | 9.95E-04 | 0.17 | 0.06 | 0.11 | 2.98 |
| 20 | H2-T23 | 0.39 | 5.74E-08 | 1.19E-03 | 0.16 | 0.05 | 0.11 | 3.02 |
| 20 | Fam234b | 0.49 | 6.09E-08 | 1.27E-03 | 0.17 | 0.06 | 0.11 | 2.98 |
| 20 | Polh | 0.64 | 6.17E-08 | 1.28E-03 | 0.23 | 0.09 | 0.14 | 2.63 |
| 20 | Tmem120b | 0.49 | 7.42E-08 | 1.54E-03 | 0.18 | 0.06 | 0.12 | 2.90 |
| 20 | Kif14 | 0.29 | 7.55E-08 | 1.57E-03 | 0.41 | 0.19 | 0.22 | 2.17 |
| 20 | Mtmr1 | 0.43 | 7.91E-08 | 1.65E-03 | 0.20 | 0.07 | 0.13 | 2.77 |
| 20 | Helb | 0.27 | 1.14E-07 | 2.38E-03 | 0.31 | 0.13 | 0.18 | 2.37 |
| 20 | Neil3 | 0.26 | 1.32E-07 | 2.75E-03 | 0.48 | 0.23 | 0.25 | 2.07 |
| 20 | 1110008P14Rik | 0.34 | 1.33E-07 | 2.76E-03 | 0.83 | 0.47 | 0.36 | 1.78 |
| 20 | Tprgl | 0.35 | 1.41E-07 | 2.94E-03 | 0.94 | 0.60 | 0.34 | 1.56 |
| 20 | Tmed8 | 0.50 | 1.71E-07 | 3.56E-03 | 0.16 | 0.05 | 0.10 | 2.94 |
| 20 | Atg9a | 0.26 | 1.75E-07 | 3.64E-03 | 0.26 | 0.11 | 0.15 | 2.47 |
| 20 | Bcor | 0.44 | 2.17E-07 | 4.52E-03 | 0.74 | 0.41 | 0.32 | 1.78 |
| 20 | Mapkapk3 | 0.82 | 2.18E-07 | 4.54E-03 | 0.12 | 0.04 | 0.09 | 3.30 |
| 20 | Naga | 0.26 | 2.21E-07 | 4.60E-03 | 0.28 | 0.12 | 0.16 | 2.38 |
| 20 | Relt | 0.81 | 2.31E-07 | 4.80E-03 | 0.11 | 0.03 | 0.08 | 3.52 |

**Supplementary Table 1. Marker genes for the original clusters of integrated clustering of E11.5 control and NotchGOF λ-junction cells.**

| | | | | | | | | |
|---|---|---|---|---|---|---|---|---|
| 20 | Wdr44 | 0.35 | 2.34E-07 | 4.86E-03 | 0.18 | 0.06 | 0.11 | 2.81 |
| 20 | Cish | 0.36 | 2.56E-07 | 5.33E-03 | 0.18 | 0.06 | 0.11 | 2.81 |
| 20 | Hist4h4 | 0.25 | 3.38E-07 | 7.03E-03 | 0.25 | 0.10 | 0.15 | 2.47 |
| 20 | Ubl4a | 0.35 | 3.68E-07 | 7.65E-03 | 0.97 | 0.57 | 0.39 | 1.68 |
| 20 | Rnf141 | 0.32 | 4.67E-07 | 9.71E-03 | 0.29 | 0.12 | 0.16 | 2.33 |
| 20 | Spata13 | 0.50 | 5.40E-07 | 1.12E-02 | 0.20 | 0.08 | 0.12 | 2.59 |
| 20 | Gm8251 | 0.36 | 5.45E-07 | 1.13E-02 | 0.19 | 0.07 | 0.12 | 2.60 |
| 20 | Fhdc1 | 0.57 | 6.30E-07 | 1.31E-02 | 0.13 | 0.04 | 0.09 | 3.07 |
| 20 | Ier3 | 0.27 | 7.09E-07 | 1.48E-02 | 0.20 | 0.08 | 0.12 | 2.56 |
| 20 | Asf1b | 0.27 | 7.23E-07 | 1.50E-02 | 0.78 | 0.44 | 0.34 | 1.78 |
| 20 | Hist2h4 | 0.75 | 7.41E-07 | 1.54E-02 | 0.10 | 0.03 | 0.07 | 3.52 |
| 20 | Adgrg1 | 0.50 | 7.51E-07 | 1.56E-02 | 0.13 | 0.04 | 0.09 | 3.07 |
| 20 | Eif2b3 | 0.33 | 8.36E-07 | 1.74E-02 | 0.82 | 0.48 | 0.34 | 1.71 |
| 20 | 1700007L15Rik | 0.32 | 1.12E-06 | 2.33E-02 | 0.19 | 0.07 | 0.12 | 2.60 |
| 20 | Commd5 | 0.35 | 1.33E-06 | 2.77E-02 | 0.18 | 0.07 | 0.11 | 2.63 |
| 20 | Cdan1 | 0.63 | 1.96E-06 | 4.08E-02 | 0.22 | 0.09 | 0.13 | 2.41 |
| 20 | Klhl15 | 0.41 | 1.98E-06 | 4.11E-02 | 0.17 | 0.06 | 0.11 | 2.66 |
| 21 | Tyrobp | 12.38 | 0 | 0 | 1.00 | 0.01 | 1.00 | 200.00 |
| 21 | Rac2 | 10.45 | 0 | 0 | 1.00 | 0.01 | 0.99 | 166.67 |
| 21 | Fcer1g | 9.28 | 0 | 0 | 0.99 | 0.02 | 0.98 | 66.20 |
| 21 | Laptm5 | 9.00 | 0 | 0 | 0.99 | 0.01 | 0.97 | 75.85 |
| 21 | Spi1 | 11.82 | 0 | 0 | 0.96 | 0.00 | 0.96 | 479.00 |
| 21 | Cyth4 | 9.28 | 0 | 0 | 0.96 | 0.01 | 0.95 | 191.60 |
| 21 | Lcp1 | 7.11 | 0 | 0 | 0.98 | 0.03 | 0.95 | 30.59 |
| 21 | Cd52 | 12.36 | 0 | 0 | 0.94 | 0.00 | 0.94 | 944.00 |
| 21 | Gmfg | 6.53 | 0 | 0 | 0.96 | 0.02 | 0.94 | 45.62 |
| 21 | Lst1 | 9.49 | 0 | 0 | 0.94 | 0.01 | 0.94 | 134.86 |
| 21 | Ncf2 | 8.22 | 0 | 0 | 0.94 | 0.01 | 0.93 | 78.67 |
| 21 | Ptprc | 11.48 | 0 | 0 | 0.92 | 0.00 | 0.92 | 923.00 |
| 21 | Ptpn6 | 6.50 | 0 | 0 | 0.95 | 0.03 | 0.92 | 30.68 |
| 21 | Cd53 | 11.63 | 0 | 0 | 0.92 | 0.00 | 0.92 | 916.00 |
| 21 | Ptpn18 | 6.32 | 0 | 0 | 0.99 | 0.07 | 0.91 | 13.69 |
| 21 | Fermt3 | 5.60 | 0 | 0 | 0.95 | 0.04 | 0.91 | 22.64 |
| 21 | Bin2 | 6.86 | 0 | 0 | 0.89 | 0.01 | 0.88 | 177.60 |
| 21 | Fyb | 6.87 | 0 | 0 | 0.92 | 0.04 | 0.88 | 25.44 |
| 21 | Nrros | 6.02 | 0 | 0 | 0.90 | 0.03 | 0.87 | 27.33 |
| 21 | Apbb1ip | 8.25 | 0 | 0 | 0.87 | 0.01 | 0.87 | 109.25 |
| 21 | Tm6sf1 | 5.32 | 0 | 0 | 0.92 | 0.05 | 0.87 | 18.32 |
| 21 | Fcgr3 | 11.84 | 0 | 0 | 0.87 | 0.00 | 0.87 | 433.50 |
| 21 | Evi2a | 9.92 | 0 | 0 | 0.87 | 0.00 | 0.87 | 433.50 |
| 21 | Cx3cr1 | 11.25 | 0 | 0 | 0.87 | 0.00 | 0.86 | 216.75 |
| 21 | Lyn | 6.15 | 0 | 0 | 0.89 | 0.03 | 0.86 | 32.89 |
| 21 | Pou2f2 | 6.69 | 0 | 0 | 0.88 | 0.02 | 0.86 | 40.05 |
| 21 | Arhgap30 | 9.86 | 0 | 0 | 0.86 | 0.00 | 0.86 | 430.00 |
| 21 | Csf1r | 11.74 | 0 | 0 | 0.86 | 0.00 | 0.86 | 430.00 |
| 21 | Tnfaip8l2 | 10.95 | 0 | 0 | 0.85 | 0.00 | 0.85 | 853.00 |
| 21 | Ly86 | 11.90 | 0 | 0 | 0.85 | 0.00 | 0.85 | 426.50 |
| 21 | Rgs10 | 5.42 | 0 | 0 | 0.93 | 0.08 | 0.85 | 11.20 |
| 21 | Psmb8 | 11.24 | 0 | 0 | 0.85 | 0.00 | 0.85 | 846.00 |
| 21 | Inpp5d | 7.60 | 0 | 0 | 0.85 | 0.01 | 0.84 | 94.00 |
| 21 | Fxyd5 | 5.75 | 0 | 0 | 0.86 | 0.02 | 0.84 | 35.83 |
| 21 | Nckap1l | 8.63 | 0 | 0 | 0.84 | 0.00 | 0.84 | 209.75 |
| 21 | Alox5ap | 6.05 | 0 | 0 | 0.87 | 0.03 | 0.83 | 25.50 |
| 21 | Pld4 | 7.44 | 0 | 0 | 0.85 | 0.02 | 0.83 | 56.40 |
| 21 | Unc93b1 | 6.28 | 0 | 0 | 0.87 | 0.04 | 0.83 | 20.33 |
| 21 | Ctss | 12.45 | 0 | 0 | 0.83 | 0.00 | 0.82 | 825.00 |

**Supplementary Table 1. Marker genes for the original clusters of integrated clustering of E11.5 control and NotchGOF λ-junction cells.**

| | | | | | | | | |
|---|---|---|---|---|---|---|---|---|
| 21 | Lyz2 | 10.19 | 0 | 0 | 0.83 | 0.00 | 0.82 | 206.25 |
| 21 | Srgn | 7.73 | 0 | 0 | 0.83 | 0.01 | 0.82 | 137.50 |
| 21 | Plek | 5.91 | 0 | 0 | 0.83 | 0.01 | 0.82 | 117.86 |
| 21 | Ighm | 12.37 | 0 | 0 | 0.82 | 0.00 | 0.82 | 818.00 |
| 21 | Zfp36 | 5.82 | 0 | 0 | 0.88 | 0.07 | 0.81 | 13.15 |
| 21 | Irf8 | 7.32 | 0 | 0 | 0.83 | 0.02 | 0.81 | 48.53 |
| 21 | Hcls1 | 8.34 | 0 | 0 | 0.81 | 0.01 | 0.81 | 162.20 |
| 21 | Ncf1 | 8.69 | 0 | 0 | 0.80 | 0.01 | 0.80 | 160.80 |
| 21 | Ms4a6c | 10.93 | 0 | 0 | 0.80 | 0.00 | 0.79 | 199.25 |
| 21 | Skap2 | 5.08 | 0 | 0 | 0.84 | 0.05 | 0.79 | 16.78 |
| 21 | Dock8 | 5.45 | 0 | 0 | 0.82 | 0.03 | 0.79 | 28.21 |
| 21 | Was | 8.48 | 0 | 0 | 0.79 | 0.00 | 0.79 | 395.00 |
| 21 | Lgals9 | 9.30 | 0 | 0 | 0.78 | 0.00 | 0.78 | 391.50 |
| 21 | Otulinl | 4.72 | 0 | 0 | 0.83 | 0.05 | 0.78 | 15.70 |
| 21 | Ptgs1 | 5.44 | 0 | 0 | 0.81 | 0.04 | 0.78 | 22.53 |
| 21 | Lcp2 | 8.14 | 0 | 0 | 0.78 | 0.00 | 0.77 | 388.00 |
| 21 | Hpgds | 6.79 | 0 | 0 | 0.80 | 0.03 | 0.77 | 24.36 |
| 21 | Aif1 | 11.26 | 0 | 0 | 0.77 | 0.00 | 0.77 | 384.50 |
| 21 | Vav1 | 8.87 | 0 | 0 | 0.77 | 0.00 | 0.77 | 384.50 |
| 21 | Rassf4 | 5.82 | 0 | 0 | 0.79 | 0.03 | 0.77 | 31.60 |
| 21 | Plcg2 | 6.53 | 0 | 0 | 0.78 | 0.01 | 0.76 | 59.69 |
| 21 | Myo1f | 11.70 | 0 | 0 | 0.76 | 0.00 | 0.76 | #NA |
| 21 | Mpeg1 | 11.67 | 0 | 0 | 0.76 | 0.00 | 0.75 | 755.00 |
| 21 | Cd68 | 6.07 | 0 | 0 | 0.78 | 0.04 | 0.75 | 21.75 |
| 21 | Trem2 | 11.91 | 0 | 0 | 0.75 | 0.00 | 0.75 | 748.00 |
| 21 | Ccl3 | 11.00 | 0 | 0 | 0.75 | 0.00 | 0.75 | 249.33 |
| 21 | Bst2 | 4.82 | 0 | 0 | 0.80 | 0.05 | 0.74 | 14.76 |
| 21 | Fcgr1 | 12.89 | 0 | 0 | 0.73 | 0.00 | 0.73 | #NA |
| 21 | Klhl6 | 5.80 | 0 | 0 | 0.75 | 0.02 | 0.73 | 49.87 |
| 21 | Ms4a6b | 11.74 | 0 | 0 | 0.73 | 0.00 | 0.73 | 734.00 |
| 21 | C3ar1 | 11.25 | 0 | 0 | 0.73 | 0.00 | 0.73 | 367.00 |
| 21 | Dock2 | 9.16 | 0 | 0 | 0.73 | 0.00 | 0.73 | 367.00 |
| 21 | Fes | 4.84 | 0 | 0 | 0.77 | 0.04 | 0.73 | 20.24 |
| 21 | C1qb | 12.65 | 0 | 0 | 0.73 | 0.01 | 0.73 | 146.80 |
| 21 | Cybb | 12.11 | 0 | 0 | 0.73 | 0.00 | 0.73 | 727.00 |
| 21 | Cd300a | 7.73 | 0 | 0 | 0.73 | 0.01 | 0.73 | 81.56 |
| 21 | Neat1 | 5.70 | 0 | 0 | 0.78 | 0.05 | 0.73 | 15.22 |
| 21 | Ncf4 | 3.66 | 0 | 0 | 0.76 | 0.04 | 0.72 | 18.14 |
| 21 | Cd79b | 7.28 | 0 | 0 | 0.73 | 0.01 | 0.72 | 90.88 |
| 21 | Ikzf1 | 6.06 | 0 | 0 | 0.74 | 0.02 | 0.72 | 32.22 |
| 21 | C1qc | 12.45 | 0 | 0 | 0.72 | 0.00 | 0.72 | 180.00 |
| 21 | Milr1 | 7.41 | 0 | 0 | 0.72 | 0.01 | 0.71 | 102.86 |
| 21 | Gatm | 6.58 | 0 | 0 | 0.75 | 0.04 | 0.71 | 20.78 |
| 21 | C1qa | 12.93 | 0 | 0 | 0.71 | 0.00 | 0.71 | 237.67 |
| 21 | Hck | 8.75 | 0 | 0 | 0.71 | 0.00 | 0.70 | 235.33 |
| 21 | Slfn2 | 7.19 | 0 | 0 | 0.71 | 0.01 | 0.70 | 59.42 |
| 21 | Fcrls | 11.82 | 0 | 0 | 0.70 | 0.00 | 0.70 | 349.50 |
| 21 | BC035044 | 10.85 | 0 | 0 | 0.69 | 0.00 | 0.69 | 692.00 |
| 21 | Cfh | 9.83 | 0 | 0 | 0.69 | 0.00 | 0.69 | 346.00 |
| 21 | Gngt2 | 4.36 | 0 | 0 | 0.72 | 0.03 | 0.69 | 21.18 |
| 21 | Gpr34 | 12.10 | 0 | 0 | 0.69 | 0.00 | 0.69 | #NA |
| 21 | Adgre1 | 10.87 | 0 | 0 | 0.69 | 0.00 | 0.68 | 685.00 |
| 21 | Ms4a6d | 11.48 | 0 | 0 | 0.69 | 0.00 | 0.68 | 685.00 |
| 21 | P2ry6 | 9.79 | 0 | 0 | 0.69 | 0.00 | 0.68 | 342.50 |
| 21 | Traf3ip3 | 10.41 | 0 | 0 | 0.68 | 0.00 | 0.68 | 678.00 |
| 21 | Btk | 7.74 | 0 | 0 | 0.68 | 0.00 | 0.68 | 339.00 |

**Supplementary Table 1. Marker genes for the original clusters of integrated clustering of E11.5 control and NotchGOF λ-junction cells.**

| | | | | | | | | |
|---|---|---|---|---|---|---|---|---|
| 21 | Hhex | 5.82 | 0 | 0 | 0.68 | 0.01 | 0.67 | 56.50 |
| 21 | Psmb9 | 6.66 | 0 | 0 | 0.67 | 0.01 | 0.66 | 74.56 |
| 21 | Nfam1 | 7.76 | 0 | 0 | 0.66 | 0.01 | 0.66 | 110.67 |
| 21 | Pik3ap1 | 6.33 | 0 | 0 | 0.67 | 0.01 | 0.66 | 47.93 |
| 21 | Stab1 | 7.40 | 0 | 0 | 0.67 | 0.02 | 0.65 | 37.28 |
| 21 | Irf5 | 8.14 | 0 | 0 | 0.65 | 0.00 | 0.65 | 216.67 |
| 21 | Vsir | 6.50 | 0 | 0 | 0.66 | 0.01 | 0.65 | 54.75 |
| 21 | Lpcat2 | 4.98 | 0 | 0 | 0.68 | 0.03 | 0.64 | 19.94 |
| 21 | Il6ra | 10.18 | 0 | 0 | 0.64 | 0.00 | 0.64 | 643.00 |
| 21 | Ramp1 | 7.86 | 0 | 0 | 0.64 | 0.01 | 0.64 | 128.60 |
| 21 | Mrc1 | 11.33 | 0 | 0 | 0.64 | 0.00 | 0.63 | 318.00 |
| 21 | Lrmp | 6.97 | 0 | 0 | 0.64 | 0.01 | 0.63 | 106.00 |
| 21 | Adcy7 | 5.46 | 0 | 0 | 0.65 | 0.02 | 0.63 | 32.50 |
| 21 | Sp140 | 6.93 | 0 | 0 | 0.64 | 0.01 | 0.63 | 90.86 |
| 21 | Fmnl1 | 8.78 | 0 | 0 | 0.63 | 0.00 | 0.63 | 314.50 |
| 21 | Ccr1 | 11.79 | 0 | 0 | 0.62 | 0.00 | 0.62 | #NA |
| 21 | Arhgef6 | 6.09 | 0 | 0 | 0.64 | 0.01 | 0.62 | 45.43 |
| 21 | Cd300c2 | 10.48 | 0 | 0 | 0.62 | 0.00 | 0.62 | 622.00 |
| 21 | Glipr1 | 9.94 | 0 | 0 | 0.62 | 0.00 | 0.62 | 311.00 |
| 21 | Tnfrsf13b | 7.24 | 0 | 0 | 0.62 | 0.01 | 0.62 | 124.40 |
| 21 | Selplg | 9.96 | 0 | 0 | 0.61 | 0.00 | 0.61 | 608.00 |
| 21 | Trim30a | 10.31 | 0 | 0 | 0.60 | 0.00 | 0.60 | 601.00 |
| 21 | Slc7a8 | 8.00 | 0 | 0 | 0.60 | 0.00 | 0.60 | 200.33 |
| 21 | Uap1l1 | 5.03 | 0 | 0 | 0.63 | 0.03 | 0.60 | 19.66 |
| 21 | Il4ra | 6.45 | 0 | 0 | 0.61 | 0.01 | 0.59 | 43.43 |
| 21 | Sash3 | 11.23 | 0 | 0 | 0.59 | 0.00 | 0.59 | #NA |
| 21 | Itgam | 12.17 | 0 | 0 | 0.59 | 0.00 | 0.59 | #NA |
| 21 | Cd86 | 11.35 | 0 | 0 | 0.59 | 0.00 | 0.59 | #NA |
| 21 | Apobec1 | 8.09 | 0 | 0 | 0.59 | 0.01 | 0.58 | 97.83 |
| 21 | Cd33 | 11.70 | 0 | 0 | 0.58 | 0.00 | 0.58 | #NA |
| 21 | Lrrc25 | 11.58 | 0 | 0 | 0.58 | 0.00 | 0.58 | #NA |
| 21 | Gpr183 | 5.73 | 0 | 0 | 0.59 | 0.01 | 0.58 | 42.43 |
| 21 | Igsf6 | 8.01 | 0 | 0 | 0.58 | 0.00 | 0.58 | 193.33 |
| 21 | Syk | 4.90 | 0 | 0 | 0.60 | 0.03 | 0.58 | 24.04 |
| 21 | Abi3 | 5.56 | 0 | 0 | 0.59 | 0.01 | 0.57 | 45.15 |
| 21 | Tifab | 11.28 | 0 | 0 | 0.57 | 0.00 | 0.57 | #NA |
| 21 | Ifi204 | 10.41 | 0 | 0 | 0.57 | 0.00 | 0.57 | 573.00 |
| 21 | Snx20 | 5.89 | 0 | 0 | 0.59 | 0.02 | 0.57 | 39.13 |
| 21 | Il17ra | 5.41 | 0 | 0 | 0.59 | 0.02 | 0.57 | 25.83 |
| 21 | Samsn1 | 6.36 | 0 | 0 | 0.58 | 0.01 | 0.57 | 52.73 |
| 21 | Csf2rb | 6.91 | 0 | 0 | 0.57 | 0.01 | 0.57 | 71.63 |
| 21 | Parvg | 8.88 | 0 | 0 | 0.57 | 0.00 | 0.57 | 566.00 |
| 21 | Sp110 | 8.97 | 0 | 0 | 0.57 | 0.00 | 0.56 | 283.00 |
| 21 | Ptpn7 | 7.35 | 0 | 0 | 0.57 | 0.01 | 0.56 | 113.20 |
| 21 | Ifi202b | 7.27 | 0 | 0 | 0.57 | 0.01 | 0.56 | 94.33 |
| 21 | Ccl9 | 11.39 | 0 | 0 | 0.56 | 0.00 | 0.56 | #NA |
| 21 | Rhoh | 9.13 | 0 | 0 | 0.56 | 0.00 | 0.56 | 559.00 |
| 21 | Sla | 7.45 | 0 | 0 | 0.56 | 0.00 | 0.56 | 279.50 |
| 21 | Zc3hav1 | 5.06 | 0 | 0 | 0.58 | 0.03 | 0.56 | 23.20 |
| 21 | P2rx4 | 4.78 | 0 | 0 | 0.58 | 0.03 | 0.55 | 22.31 |
| 21 | Gpr65 | 11.23 | 0 | 0 | 0.55 | 0.00 | 0.55 | #NA |
| 21 | Slc15a3 | 6.37 | 0 | 0 | 0.57 | 0.01 | 0.55 | 40.43 |
| 21 | Csf3r | 10.39 | 0 | 0 | 0.55 | 0.00 | 0.55 | 552.00 |
| 21 | Pik3cd | 5.57 | 0 | 0 | 0.57 | 0.02 | 0.55 | 37.73 |
| 21 | Tnfrsf11a | 7.86 | 0 | 0 | 0.55 | 0.00 | 0.55 | 138.00 |
| 21 | Cd14 | 5.82 | 0 | 0 | 0.56 | 0.02 | 0.54 | 27.95 |

**Supplementary Table 1. Marker genes for the original clusters of integrated clustering of E11.5 control and NotchGOF λ-junction cells.**

| | | | | | | | | |
|---|---|---|---|---|---|---|---|---|
| 21 | Myo1g | 10.80 | 0 | 0 | 0.53 | 0.00 | 0.53 | #NA |
| 21 | Slc25a45 | 6.05 | 0 | 0 | 0.53 | 0.01 | 0.52 | 66.38 |
| 21 | Ppfia4 | 5.08 | 0 | 0 | 0.55 | 0.02 | 0.52 | 22.71 |
| 21 | Tfec | 5.83 | 0 | 0 | 0.53 | 0.01 | 0.52 | 44.25 |
| 21 | Gbp7 | 5.78 | 0 | 0 | 0.53 | 0.02 | 0.51 | 31.24 |
| 21 | Icam1 | 5.03 | 0 | 0 | 0.52 | 0.01 | 0.51 | 37.43 |
| 21 | 9930111J21Rik2 | 11.07 | 0 | 0 | 0.51 | 0.00 | 0.51 | #NA |
| 21 | Ccr5 | 9.60 | 0 | 0 | 0.51 | 0.00 | 0.51 | 510.00 |
| 21 | Tnfrsf1b | 8.03 | 0 | 0 | 0.51 | 0.00 | 0.51 | 170.00 |
| 21 | Ifi207 | 11.83 | 0 | 0 | 0.50 | 0.00 | 0.50 | #NA |
| 21 | Adap2 | 11.16 | 0 | 0 | 0.50 | 0.00 | 0.50 | #NA |
| 21 | Rgs1 | 10.14 | 0 | 0 | 0.50 | 0.00 | 0.50 | 503.00 |
| 21 | Slco2b1 | 9.08 | 0 | 0 | 0.50 | 0.00 | 0.50 | 251.50 |
| 21 | Dok2 | 5.22 | 0 | 0 | 0.52 | 0.02 | 0.50 | 32.31 |
| 21 | Cela1 | 7.83 | 0 | 0 | 0.50 | 0.00 | 0.50 | 251.50 |
| 21 | Camk1d | 5.87 | 0 | 0 | 0.51 | 0.01 | 0.50 | 51.00 |
| 21 | Tnni2 | 8.46 | 0 | 0 | 0.50 | 0.00 | 0.50 | 167.67 |
| 21 | Pik3cg | 7.31 | 0 | 0 | 0.50 | 0.00 | 0.50 | 167.67 |
| 21 | Clec4a2 | 11.76 | 0 | 0 | 0.50 | 0.00 | 0.50 | #NA |
| 21 | Itgb2 | 8.31 | 0 | 0 | 0.50 | 0.00 | 0.50 | 497.00 |
| 21 | Ccl4 | 11.25 | 0 | 0 | 0.50 | 0.00 | 0.50 | 248.50 |
| 21 | Slc11a1 | 11.75 | 0 | 0 | 0.49 | 0.00 | 0.49 | #NA |
| 21 | Cd84 | 9.18 | 0 | 0 | 0.49 | 0.00 | 0.49 | 490.00 |
| 21 | Casp1 | 10.48 | 0 | 0 | 0.48 | 0.00 | 0.48 | #NA |
| 21 | Bcl2a1b | 12.24 | 0 | 0 | 0.48 | 0.00 | 0.48 | #NA |
| 21 | Cd180 | 10.83 | 0 | 0 | 0.48 | 0.00 | 0.48 | #NA |
| 21 | Tmem273 | 13.08 | 0 | 0 | 0.48 | 0.00 | 0.48 | #NA |
| 21 | P2ry12 | 9.66 | 0 | 0 | 0.48 | 0.00 | 0.48 | 483.00 |
| 21 | Wdfy4 | 10.10 | 0 | 0 | 0.48 | 0.00 | 0.48 | #NA |
| 21 | Arhgap25 | 6.13 | 0 | 0 | 0.48 | 0.01 | 0.48 | 60.38 |
| 21 | Lair1 | 10.56 | 0 | 0 | 0.48 | 0.00 | 0.48 | 476.00 |
| 21 | F13a1 | 9.85 | 0 | 0 | 0.48 | 0.00 | 0.48 | 476.00 |
| 21 | Arid5a | 5.13 | 0 | 0 | 0.48 | 0.01 | 0.46 | 36.62 |
| 21 | Arl11 | 9.79 | 0 | 0 | 0.45 | 0.00 | 0.45 | 448.00 |
| 21 | Tmem8 | 5.53 | 0 | 0 | 0.46 | 0.02 | 0.45 | 28.88 |
| 21 | Arhgap4 | 5.65 | 0 | 0 | 0.46 | 0.01 | 0.44 | 41.36 |
| 21 | Hfe | 5.88 | 0 | 0 | 0.46 | 0.01 | 0.44 | 37.92 |
| 21 | Cytip | 11.90 | 0 | 0 | 0.44 | 0.00 | 0.44 | #NA |
| 21 | Cryba4 | 10.31 | 0 | 0 | 0.44 | 0.00 | 0.44 | 441.00 |
| 21 | Il10ra | 9.10 | 0 | 0 | 0.43 | 0.00 | 0.43 | 434.00 |
| 21 | Ccl6 | 10.99 | 0 | 0 | 0.43 | 0.00 | 0.43 | 434.00 |
| 21 | Clec12a | 8.44 | 0 | 0 | 0.43 | 0.00 | 0.43 | 144.67 |
| 21 | A630001G21Rik | 5.32 | 0 | 0 | 0.44 | 0.01 | 0.43 | 40.09 |
| 21 | P2ry13 | 10.98 | 0 | 0 | 0.43 | 0.00 | 0.43 | #NA |
| 21 | Parp14 | 7.20 | 0 | 0 | 0.43 | 0.00 | 0.42 | 106.75 |
| 21 | Tlr7 | 10.51 | 0 | 0 | 0.42 | 0.00 | 0.42 | #NA |
| 21 | F630028O10Rik | 8.87 | 0 | 0 | 0.42 | 0.00 | 0.42 | 420.00 |
| 21 | Rassf5 | 8.48 | 0 | 0 | 0.41 | 0.00 | 0.41 | 413.00 |
| 21 | Hlx | 5.61 | 0 | 0 | 0.42 | 0.01 | 0.41 | 52.50 |
| 21 | Lilrb4a | 10.92 | 0 | 0 | 0.41 | 0.00 | 0.41 | 413.00 |
| 21 | Rapsn | 7.76 | 0 | 0 | 0.41 | 0.00 | 0.41 | 206.50 |
| 21 | Gna15 | 6.45 | 0 | 0 | 0.41 | 0.01 | 0.41 | 82.60 |
| 21 | Ang | 5.15 | 0 | 0 | 0.42 | 0.01 | 0.41 | 32.31 |
| 21 | Themis2 | 8.74 | 0 | 0 | 0.41 | 0.00 | 0.41 | 406.00 |
| 21 | Il21r | 8.52 | 0 | 0 | 0.41 | 0.00 | 0.41 | 406.00 |
| 21 | I830077J02Rik | 6.73 | 0 | 0 | 0.41 | 0.00 | 0.40 | 203.00 |

**Supplementary Table 1. Marker genes for the original clusters of integrated clustering of E11.5 control and NotchGOF λ-junction cells.**

| | | | | | | | | |
|---|---|---|---|---|---|---|---|---|
| 21 | Psd4 | 6.82 | 0 | 0 | 0.41 | 0.00 | 0.40 | 101.50 |
| 21 | Arhgap15 | 10.57 | 0 | 0 | 0.40 | 0.00 | 0.40 | #NA |
| 21 | Ccl24 | 12.20 | 0 | 0 | 0.40 | 0.00 | 0.40 | #NA |
| 21 | Abca9 | 7.12 | 0 | 0 | 0.40 | 0.00 | 0.40 | 99.75 |
| 21 | Susd3 | 5.14 | 0 | 0 | 0.41 | 0.01 | 0.39 | 33.83 |
| 21 | Pilra | 10.95 | 0 | 0 | 0.39 | 0.00 | 0.39 | #NA |
| 21 | Nupr1 | 7.90 | 0 | 0 | 0.39 | 0.00 | 0.39 | 196.00 |
| 21 | Il16 | 6.94 | 0 | 0 | 0.39 | 0.01 | 0.39 | 78.40 |
| 21 | Adap1 | 5.96 | 0 | 0 | 0.39 | 0.01 | 0.39 | 65.33 |
| 21 | AB124611 | 8.85 | 0 | 0 | 0.39 | 0.00 | 0.38 | 385.00 |
| 21 | Trpv2 | 6.19 | 0 | 0 | 0.38 | 0.01 | 0.37 | 75.60 |
| 21 | Ripor2 | 5.66 | 0 | 0 | 0.38 | 0.01 | 0.37 | 75.60 |
| 21 | Ptafr | 10.85 | 0 | 0 | 0.37 | 0.00 | 0.37 | #NA |
| 21 | Rgs18 | 5.30 | 0 | 0 | 0.37 | 0.00 | 0.37 | 185.50 |
| 21 | Irgm1 | 5.32 | 0 | 0 | 0.38 | 0.01 | 0.37 | 34.36 |
| 21 | Siglech | 12.50 | 0 | 0 | 0.36 | 0.00 | 0.36 | #NA |
| 21 | Ms4a7 | 11.81 | 0 | 0 | 0.36 | 0.00 | 0.36 | #NA |
| 21 | Rasal3 | 8.71 | 0 | 0 | 0.36 | 0.00 | 0.36 | 364.00 |
| 21 | Cysltr1 | 5.57 | 0 | 0 | 0.37 | 0.01 | 0.36 | 46.38 |
| 21 | Ptk2b | 6.25 | 0 | 0 | 0.36 | 0.00 | 0.36 | 91.00 |
| 21 | Cd37 | 6.97 | 0 | 0 | 0.36 | 0.01 | 0.36 | 72.80 |
| 21 | Clec5a | 10.80 | 0 | 0 | 0.36 | 0.00 | 0.36 | #NA |
| 21 | C5ar1 | 11.18 | 0 | 0 | 0.36 | 0.00 | 0.36 | #NA |
| 21 | Pkib | 9.42 | 0 | 0 | 0.36 | 0.00 | 0.36 | 357.00 |
| 21 | Ctla2b | 4.75 | 0 | 0 | 0.36 | 0.01 | 0.36 | 40.44 |
| 21 | Ccdc88b | 6.43 | 0 | 0 | 0.35 | 0.00 | 0.35 | 87.50 |
| 21 | Cd48 | 11.18 | 0 | 0 | 0.34 | 0.00 | 0.34 | #NA |
| 21 | Epsti1 | 11.21 | 0 | 0 | 0.34 | 0.00 | 0.34 | #NA |
| 21 | Cd74 | 6.46 | 0 | 0 | 0.34 | 0.00 | 0.34 | 85.75 |
| 21 | 2310001H17Rik | 7.76 | 0 | 0 | 0.34 | 0.00 | 0.33 | 168.00 |
| 21 | Neurl3 | 6.96 | 0 | 0 | 0.34 | 0.00 | 0.33 | 112.00 |
| 21 | Hk3 | 10.78 | 0 | 0 | 0.32 | 0.00 | 0.32 | #NA |
| 21 | Evi2 | 6.93 | 0 | 0 | 0.32 | 0.00 | 0.32 | 161.00 |
| 21 | Asb2 | 10.20 | 0 | 0 | 0.32 | 0.00 | 0.32 | #NA |
| 21 | Clec4a1 | 9.55 | 0 | 0 | 0.32 | 0.00 | 0.31 | 315.00 |
| 21 | Il18 | 6.26 | 0 | 0 | 0.32 | 0.01 | 0.31 | 63.00 |
| 21 | Il2rg | 6.43 | 0 | 0 | 0.31 | 0.00 | 0.30 | 77.00 |
| 21 | Pstpip1 | 5.85 | 0 | 0 | 0.31 | 0.01 | 0.30 | 51.33 |
| 21 | Tlr13 | 11.04 | 0 | 0 | 0.30 | 0.00 | 0.30 | #NA |
| 21 | Hcst | 7.13 | 0 | 0 | 0.30 | 0.00 | 0.30 | 301.00 |
| 21 | Ptpro | 6.85 | 0 | 0 | 0.30 | 0.00 | 0.30 | 150.50 |
| 21 | Prkcb | 6.82 | 0 | 0 | 0.30 | 0.00 | 0.30 | 100.33 |
| 21 | Rnase6 | 7.69 | 0 | 0 | 0.30 | 0.00 | 0.30 | 100.33 |
| 21 | Lgals3 | 8.20 | 0 | 0 | 0.30 | 0.00 | 0.30 | 75.25 |
| 21 | Nlrp3 | 11.74 | 0 | 0 | 0.29 | 0.00 | 0.29 | #NA |
| 21 | Serpinb1a | 9.36 | 0 | 0 | 0.29 | 0.00 | 0.29 | 294.00 |
| 21 | Plcb2 | 6.85 | 0 | 0 | 0.29 | 0.00 | 0.29 | 147.00 |
| 21 | Tap2 | 5.58 | 0 | 0 | 0.29 | 0.01 | 0.29 | 49.00 |
| 21 | Ifi211 | 11.00 | 0 | 0 | 0.29 | 0.00 | 0.29 | #NA |
| 21 | 5430427O19Rik | 9.33 | 0 | 0 | 0.29 | 0.00 | 0.29 | #NA |
| 21 | Marchf1 | 8.18 | 0 | 0 | 0.29 | 0.00 | 0.29 | 287.00 |
| 21 | Tbxas1 | 6.34 | 0 | 0 | 0.29 | 0.00 | 0.28 | 95.67 |
| 21 | Fcgr4 | 10.47 | 0 | 0 | 0.28 | 0.00 | 0.28 | #NA |
| 21 | Mndal | 12.19 | 0 | 0 | 0.28 | 0.00 | 0.28 | #NA |
| 21 | Tlr1 | 9.95 | 0 | 0 | 0.28 | 0.00 | 0.28 | #NA |
| 21 | Napsa | 8.04 | 0 | 0 | 0.28 | 0.00 | 0.28 | 140.00 |

**Supplementary Table 1. Marker genes for the original clusters of integrated clustering of E11.5 control and NotchGOF λ-junction cells.**

| | | | | | | | | |
|---|---|---|---|---|---|---|---|---|
| 21 | Sp100 | 9.90 | 0 | 0 | 0.27 | 0.00 | 0.27 | #NA |
| 21 | Cd300lf | 8.22 | 0 | 0 | 0.27 | 0.00 | 0.27 | 273.00 |
| 21 | Ticam2 | 6.33 | 0 | 0 | 0.27 | 0.00 | 0.27 | 91.00 |
| 21 | Stap1 | 6.63 | 0 | 0 | 0.27 | 0.00 | 0.27 | 68.25 |
| 21 | Apoc2 | 11.55 | 0 | 0 | 0.27 | 0.00 | 0.27 | #NA |
| 21 | Cd200r1 | 7.87 | 0 | 0 | 0.27 | 0.00 | 0.27 | 266.00 |
| 21 | Itgal | 6.28 | 0 | 0 | 0.27 | 0.00 | 0.26 | 66.50 |
| 21 | Gm44751 | 5.82 | 0 | 0 | 0.27 | 0.01 | 0.26 | 53.20 |
| 21 | Aoah | 9.84 | 0 | 0 | 0.26 | 0.00 | 0.26 | #NA |
| 21 | Ltc4s | 7.90 | 0 | 0 | 0.26 | 0.00 | 0.26 | 129.50 |
| 21 | AI467606 | 6.86 | 0 | 0 | 0.26 | 0.00 | 0.26 | 86.33 |
| 21 | AI662270 | 6.81 | 0 | 0 | 0.26 | 0.00 | 0.26 | 86.33 |
| 21 | Bank1 | 9.86 | 0 | 0 | 0.25 | 0.00 | 0.25 | #NA |
| 21 | Clec4a3 | 10.65 | 0 | 0 | 0.25 | 0.00 | 0.25 | #NA |
| 21 | Phf11b | 12.21 | 0 | 0 | 0.25 | 0.00 | 0.25 | #NA |
| 21 | Gbp3 | 6.80 | 0 | 0 | 0.25 | 0.00 | 0.25 | 126.00 |
| 21 | P2rx7 | 9.33 | 0 | 0 | 0.25 | 0.00 | 0.25 | #NA |
| 21 | Hp | 10.55 | 0 | 0 | 0.25 | 0.00 | 0.25 | #NA |
| 21 | Wfdc17 | 12.12 | 0 | 0 | 0.25 | 0.00 | 0.25 | #NA |
| 21 | Naip2 | 8.17 | 0 | 0 | 0.25 | 0.00 | 0.24 | 245.00 |
| 21 | Fgl2 | 7.04 | 0 | 0 | 0.25 | 0.00 | 0.24 | 122.50 |
| 21 | P2rx1 | 5.45 | 0 | 0 | 0.25 | 0.00 | 0.24 | 122.50 |
| 21 | Alox5 | 7.82 | 0 | 0 | 0.24 | 0.00 | 0.24 | 238.00 |
| 21 | Ptprcap | 8.87 | 0 | 0 | 0.24 | 0.00 | 0.24 | 119.00 |
| 21 | Lpxn | 6.89 | 0 | 0 | 0.24 | 0.00 | 0.24 | 119.00 |
| 21 | Gpr171 | 8.29 | 0 | 0 | 0.23 | 0.00 | 0.23 | 231.00 |
| 21 | Clec10a | 9.41 | 0 | 0 | 0.23 | 0.00 | 0.23 | 231.00 |
| 21 | Ccl12 | 10.51 | 0 | 0 | 0.23 | 0.00 | 0.23 | 231.00 |
| 21 | Mpo | 10.54 | 0 | 0 | 0.23 | 0.00 | 0.23 | 77.00 |
| 21 | Cd69 | 10.58 | 0 | 0 | 0.22 | 0.00 | 0.22 | #NA |
| 21 | Bex6 | 11.63 | 0 | 0 | 0.22 | 0.00 | 0.22 | #NA |
| 21 | Rab7b | 7.24 | 0 | 0 | 0.22 | 0.00 | 0.22 | 112.00 |
| 21 | Atp2a3 | 6.03 | 0 | 0 | 0.22 | 0.00 | 0.22 | 112.00 |
| 21 | Ltb | 7.24 | 0 | 0 | 0.22 | 0.00 | 0.22 | 74.67 |
| 21 | Clec4n | 11.70 | 0 | 0 | 0.22 | 0.00 | 0.22 | #NA |
| 21 | Tagap | 8.77 | 0 | 0 | 0.22 | 0.00 | 0.22 | #NA |
| 21 | Gm26740 | 7.02 | 0 | 0 | 0.22 | 0.00 | 0.22 | 217.00 |
| 21 | Tlr4 | 6.86 | 0 | 0 | 0.22 | 0.00 | 0.22 | 108.50 |
| 21 | Ccr2 | 11.94 | 0 | 0 | 0.21 | 0.00 | 0.21 | #NA |
| 21 | Tespa1 | 10.69 | 0 | 0 | 0.21 | 0.00 | 0.21 | #NA |
| 21 | Card9 | 8.14 | 0 | 0 | 0.21 | 0.00 | 0.21 | 210.00 |
| 21 | Cmah | 5.85 | 0 | 0 | 0.21 | 0.00 | 0.21 | 105.00 |
| 21 | Abcc3 | 7.18 | 0 | 0 | 0.21 | 0.00 | 0.21 | 70.00 |
| 21 | Cd28 | 10.80 | 0 | 0 | 0.20 | 0.00 | 0.20 | #NA |
| 21 | Tmem156 | 9.93 | 0 | 0 | 0.20 | 0.00 | 0.20 | #NA |
| 21 | Gpr141 | 10.50 | 0 | 0 | 0.20 | 0.00 | 0.20 | #NA |
| 21 | Mcoln2 | 8.05 | 0 | 0 | 0.20 | 0.00 | 0.20 | 203.00 |
| 21 | Mir142hg | 7.59 | 0 | 0 | 0.20 | 0.00 | 0.20 | 203.00 |
| 21 | Ifi209 | 11.88 | 0 | 0 | 0.20 | 0.00 | 0.20 | #NA |
| 21 | Msr1 | 9.32 | 0 | 0 | 0.20 | 0.00 | 0.20 | #NA |
| 21 | Cfb | 9.75 | 0 | 0 | 0.20 | 0.00 | 0.20 | #NA |
| 21 | Tnf | 8.83 | 0 | 0 | 0.20 | 0.00 | 0.20 | #NA |
| 21 | Ifi203 | 10.12 | 0 | 0 | 0.19 | 0.00 | 0.19 | #NA |
| 21 | Ms4a4c | 11.32 | 0 | 0 | 0.19 | 0.00 | 0.19 | #NA |
| 21 | Crybb1 | 9.51 | 0 | 0 | 0.18 | 0.00 | 0.18 | #NA |
| 21 | Ccl2 | 9.91 | 0 | 0 | 0.18 | 0.00 | 0.18 | #NA |

**Supplementary Table 1. Marker genes for the original clusters of integrated clustering of E11.5 control and NotchGOF λ-junction cells.**

| | | | | | | | | |
|---|---|---|---|---|---|---|---|---|
| 21 | Serpina3g | 8.84 | 0 | 0 | 0.18 | 0.00 | 0.18 | 182.00 |
| 21 | Ly9 | 10.79 | 0 | 0 | 0.18 | 0.00 | 0.18 | #NA |
| 21 | Siglece | 7.98 | 0 | 0 | 0.18 | 0.00 | 0.18 | #NA |
| 21 | Tpbgl | 9.64 | 0 | 0 | 0.18 | 0.00 | 0.18 | #NA |
| 21 | Il1b | 8.75 | 0 | 0 | 0.18 | 0.00 | 0.17 | 175.00 |
| 21 | Kcnk12 | 7.37 | 0 | 0 | 0.17 | 0.00 | 0.17 | 168.00 |
| 21 | Fcgr2b | 9.26 | 0 | 0 | 0.16 | 0.00 | 0.16 | #NA |
| 21 | Trim30d | 9.38 | 0 | 0 | 0.16 | 0.00 | 0.16 | #NA |
| 21 | Slamf9 | 8.32 | 0 | 0 | 0.16 | 0.00 | 0.16 | 161.00 |
| 21 | Gpr84 | 8.00 | 0 | 0 | 0.16 | 0.00 | 0.16 | 161.00 |
| 21 | Spn | 9.36 | 0 | 0 | 0.15 | 0.00 | 0.15 | #NA |
| 21 | Cyp4f18 | 8.16 | 0 | 0 | 0.15 | 0.00 | 0.15 | #NA |
| 21 | Lilr4b | 10.64 | 0 | 0 | 0.15 | 0.00 | 0.15 | #NA |
| 21 | Oas1a | 7.49 | 0 | 0 | 0.15 | 0.00 | 0.15 | 154.00 |
| 21 | Trbc1 | 9.25 | 0 | 0 | 0.15 | 0.00 | 0.15 | 154.00 |
| 21 | Havcr2 | 8.21 | 0 | 0 | 0.15 | 0.00 | 0.15 | 154.00 |
| 21 | Slfn8 | 7.03 | 0 | 0 | 0.15 | 0.00 | 0.15 | 154.00 |
| 21 | Gm38843 | 9.39 | 0 | 0 | 0.15 | 0.00 | 0.15 | #NA |
| 21 | Abcg3 | 8.17 | 0 | 0 | 0.15 | 0.00 | 0.15 | 147.00 |
| 21 | Cst7 | 9.33 | 0 | 0 | 0.14 | 0.00 | 0.14 | #NA |
| 21 | Gsg1 | 7.33 | 0 | 0 | 0.14 | 0.00 | 0.14 | 140.00 |
| 21 | 1600010M07Rik | 9.76 | 0 | 0 | 0.13 | 0.00 | 0.13 | #NA |
| 21 | Il7r | 10.54 | 0 | 0 | 0.13 | 0.00 | 0.13 | #NA |
| 21 | Cd300ld | 8.33 | 0 | 0 | 0.13 | 0.00 | 0.13 | 133.00 |
| 21 | Gm5547 | 8.53 | 0 | 0 | 0.13 | 0.00 | 0.13 | #NA |
| 21 | Gdf3 | 10.12 | 0 | 0 | 0.13 | 0.00 | 0.13 | #NA |
| 21 | Bcl2a1d | 10.38 | 0 | 0 | 0.13 | 0.00 | 0.13 | #NA |
| 21 | Gm5086 | 9.68 | 0 | 0 | 0.13 | 0.00 | 0.13 | #NA |
| 21 | Pglyrp2 | 10.66 | 0 | 0 | 0.13 | 0.00 | 0.13 | #NA |
| 21 | C130050O18Rik | 7.91 | 0 | 0 | 0.13 | 0.00 | 0.13 | 126.00 |
| 21 | Scimp | 10.72 | 0 | 0 | 0.12 | 0.00 | 0.12 | #NA |
| 21 | Adap2os | 9.12 | 0 | 0 | 0.12 | 0.00 | 0.12 | #NA |
| 21 | Flt3 | 8.41 | 0 | 0 | 0.11 | 0.00 | 0.11 | #NA |
| 21 | 2010013B24Rik | 8.91 | 0 | 0 | 0.11 | 0.00 | 0.11 | #NA |
| 21 | Gapt | 10.25 | 0 | 0 | 0.11 | 0.00 | 0.11 | #NA |
| 21 | Ebi3 | 4.95 | 4.72E-308 | 9.83E-304 | 0.43 | 0.01 | 0.41 | 30.50 |
| 21 | Elmo1 | 4.25 | 1.91E-306 | 3.97E-302 | 0.70 | 0.04 | 0.66 | 16.64 |
| 21 | Tlr2 | 4.87 | 1.63E-300 | 3.39E-296 | 0.51 | 0.02 | 0.49 | 23.18 |
| 21 | Ptpn22 | 6.51 | 4.33E-298 | 9.01E-294 | 0.15 | 0.00 | 0.15 | 147.00 |
| 21 | Smagp | 4.44 | 1.07E-295 | 2.23E-291 | 0.60 | 0.03 | 0.57 | 19.39 |
| 21 | Parp10 | 6.39 | 3.82E-295 | 7.94E-291 | 0.19 | 0.00 | 0.19 | 94.50 |
| 21 | Slc7a7 | 5.25 | 2.69E-293 | 5.59E-289 | 0.43 | 0.02 | 0.41 | 28.47 |
| 21 | Gas6 | 6.32 | 4.41E-290 | 9.16E-286 | 0.52 | 0.03 | 0.50 | 20.96 |
| 21 | Ptpre | 4.85 | 5.69E-290 | 1.18E-285 | 0.61 | 0.03 | 0.57 | 17.88 |
| 21 | Klf2 | 4.53 | 3.63E-289 | 7.54E-285 | 0.62 | 0.03 | 0.58 | 18.09 |
| 21 | Hdc | 9.64 | 4.44E-288 | 9.24E-284 | 0.12 | 0.00 | 0.12 | 119.00 |
| 21 | Gm44067 | 7.66 | 4.69E-288 | 9.76E-284 | 0.12 | 0.00 | 0.12 | 119.00 |
| 21 | Cd44 | 4.60 | 2.30E-287 | 4.79E-283 | 0.61 | 0.03 | 0.57 | 17.88 |
| 21 | Tifa | 4.92 | 2.71E-286 | 5.64E-282 | 0.54 | 0.03 | 0.51 | 20.69 |
| 21 | Tmem106a | 5.61 | 1.96E-277 | 4.07E-273 | 0.33 | 0.01 | 0.32 | 36.56 |
| 21 | Gpr160 | 5.08 | 3.18E-273 | 6.62E-269 | 0.33 | 0.01 | 0.32 | 36.56 |
| 21 | Spns3 | 7.54 | 1.52E-269 | 3.16E-265 | 0.11 | 0.00 | 0.10 | 105.00 |
| 21 | Dapp1 | 4.47 | 1.59E-268 | 3.31E-264 | 0.53 | 0.03 | 0.50 | 19.67 |
| 21 | Coro2a | 4.82 | 9.18E-267 | 1.91E-262 | 0.43 | 0.02 | 0.42 | 24.11 |
| 21 | Paqr7 | 4.96 | 3.06E-263 | 6.36E-259 | 0.32 | 0.01 | 0.31 | 35.00 |
| 21 | Plac8 | 7.80 | 1.29E-262 | 2.69E-258 | 0.43 | 0.02 | 0.41 | 23.72 |

**Supplementary Table 1. Marker genes for the original clusters of integrated clustering of E11.5 control and NotchGOF λ-junction cells.**

| | | | | | | | | |
|---|---|---|---|---|---|---|---|---|
| 21 | Csf2rb2 | 7.55 | 3.57E-262 | 7.42E-258 | 0.13 | 0.00 | 0.13 | 133.00 |
| 21 | Myo7a | 4.43 | 1.19E-260 | 2.49E-256 | 0.35 | 0.01 | 0.34 | 31.82 |
| 21 | Slc46a3 | 5.44 | 1.94E-257 | 4.04E-253 | 0.33 | 0.01 | 0.32 | 32.90 |
| 21 | Ctsc | 6.22 | 9.57E-256 | 1.99E-251 | 0.97 | 0.12 | 0.85 | 8.10 |
| 21 | Egr1 | 4.65 | 1.58E-255 | 3.29E-251 | 0.80 | 0.07 | 0.73 | 11.32 |
| 21 | Spp1 | 10.79 | 2.23E-253 | 4.64E-249 | 0.14 | 0.00 | 0.14 | 140.00 |
| 21 | Maf | 6.52 | 7.37E-253 | 1.53E-248 | 0.77 | 0.07 | 0.70 | 11.14 |
| 21 | Pycard | 5.27 | 1.51E-252 | 3.13E-248 | 0.97 | 0.12 | 0.85 | 7.90 |
| 21 | Grap | 4.34 | 2.91E-249 | 6.06E-245 | 0.37 | 0.01 | 0.36 | 28.54 |
| 21 | Tgfb1 | 4.07 | 3.18E-249 | 6.62E-245 | 0.94 | 0.10 | 0.83 | 9.10 |
| 21 | Atf3 | 5.71 | 2.12E-246 | 4.40E-242 | 0.48 | 0.02 | 0.45 | 19.83 |
| 21 | Arhgap9 | 4.33 | 1.33E-244 | 2.77E-240 | 0.57 | 0.04 | 0.54 | 16.37 |
| 21 | Lgals3bp | 5.87 | 3.17E-244 | 6.60E-240 | 0.26 | 0.01 | 0.25 | 43.17 |
| 21 | Gpsm3 | 4.28 | 8.70E-244 | 1.81E-239 | 0.85 | 0.08 | 0.76 | 10.32 |
| 21 | Abcb1b | 5.39 | 1.33E-243 | 2.76E-239 | 0.24 | 0.01 | 0.23 | 47.60 |
| 21 | Nfkbid | 5.12 | 4.37E-243 | 9.08E-239 | 0.28 | 0.01 | 0.27 | 35.00 |
| 21 | Coro1a | 6.28 | 1.19E-241 | 2.48E-237 | 1.00 | 0.14 | 0.86 | 6.99 |
| 21 | Tec | 4.81 | 4.94E-240 | 1.03E-235 | 0.33 | 0.01 | 0.32 | 29.91 |
| 21 | Plxna4os1 | 6.76 | 9.83E-233 | 2.04E-228 | 0.14 | 0.00 | 0.14 | 70.00 |
| 21 | Ehbp1l1 | 4.23 | 3.33E-232 | 6.92E-228 | 0.72 | 0.06 | 0.66 | 11.80 |
| 21 | Ifi35 | 4.55 | 3.73E-231 | 7.75E-227 | 0.50 | 0.03 | 0.47 | 17.75 |
| 21 | Ifi27 | 5.48 | 6.39E-230 | 1.33E-225 | 0.29 | 0.01 | 0.28 | 31.89 |
| 21 | Gas7 | 4.82 | 1.78E-228 | 3.71E-224 | 0.64 | 0.05 | 0.59 | 13.12 |
| 21 | Pirb | 6.05 | 3.78E-228 | 7.87E-224 | 0.18 | 0.00 | 0.18 | 60.67 |
| 21 | Arhgap45 | 3.85 | 4.18E-228 | 8.68E-224 | 0.72 | 0.06 | 0.66 | 12.00 |
| 21 | Lyl1 | 4.48 | 1.74E-227 | 3.63E-223 | 0.54 | 0.03 | 0.51 | 16.30 |
| 21 | Trf | 5.73 | 1.22E-226 | 2.53E-222 | 0.58 | 0.04 | 0.54 | 14.50 |
| 21 | Anxa3 | 4.27 | 1.24E-226 | 2.57E-222 | 0.83 | 0.09 | 0.74 | 9.59 |
| 21 | S100a13 | 3.99 | 1.34E-224 | 2.79E-220 | 0.90 | 0.10 | 0.79 | 8.69 |
| 21 | Abhd15 | 6.54 | 2.50E-220 | 5.19E-216 | 0.12 | 0.00 | 0.12 | 119.00 |
| 21 | Rasgrp3 | 3.97 | 2.62E-220 | 5.45E-216 | 0.36 | 0.02 | 0.35 | 24.27 |
| 21 | Irf7 | 7.05 | 5.17E-220 | 1.07E-215 | 0.15 | 0.00 | 0.15 | 77.00 |
| 21 | Serpinb8 | 5.14 | 1.24E-218 | 2.58E-214 | 0.20 | 0.00 | 0.20 | 50.75 |
| 21 | Slc22a3 | 8.05 | 3.48E-218 | 7.23E-214 | 0.11 | 0.00 | 0.11 | 112.00 |
| 21 | Ptprj | 4.43 | 7.38E-218 | 1.53E-213 | 0.54 | 0.04 | 0.50 | 15.37 |
| 21 | Npl | 5.21 | 2.43E-217 | 5.05E-213 | 0.84 | 0.09 | 0.75 | 8.93 |
| 21 | Atp1a3 | 5.43 | 9.12E-217 | 1.90E-212 | 0.17 | 0.00 | 0.17 | 56.00 |
| 21 | Haao | 6.48 | 1.89E-216 | 3.93E-212 | 0.11 | 0.00 | 0.10 | 105.00 |
| 21 | Slc43a2 | 4.48 | 8.63E-215 | 1.79E-210 | 0.56 | 0.04 | 0.52 | 14.71 |
| 21 | Lat2 | 4.90 | 1.60E-214 | 3.33E-210 | 0.34 | 0.01 | 0.33 | 24.50 |
| 21 | Parp9 | 4.11 | 2.32E-213 | 4.83E-209 | 0.49 | 0.03 | 0.46 | 16.90 |
| 21 | Dok3 | 4.63 | 4.76E-213 | 9.91E-209 | 0.39 | 0.02 | 0.37 | 22.65 |
| 21 | Treml2 | 4.82 | 6.52E-211 | 1.36E-206 | 0.34 | 0.01 | 0.33 | 24.50 |
| 21 | Elf4 | 4.39 | 7.08E-211 | 1.47E-206 | 0.43 | 0.02 | 0.41 | 19.41 |
| 21 | Cebpb | 4.89 | 1.53E-208 | 3.17E-204 | 0.89 | 0.12 | 0.77 | 7.66 |
| 21 | Fam129a | 6.12 | 3.12E-206 | 6.48E-202 | 0.20 | 0.00 | 0.19 | 49.00 |
| 21 | Pik3r6 | 5.57 | 3.64E-206 | 7.58E-202 | 0.21 | 0.01 | 0.21 | 42.00 |
| 21 | Cd83 | 5.22 | 4.78E-206 | 9.95E-202 | 0.40 | 0.02 | 0.38 | 19.95 |
| 21 | D5Ertd605e | 4.92 | 8.30E-205 | 1.73E-200 | 0.25 | 0.01 | 0.24 | 35.00 |
| 21 | Tlr6 | 6.29 | 3.32E-204 | 6.90E-200 | 0.12 | 0.00 | 0.12 | 119.00 |
| 21 | Sall3 | 6.32 | 8.93E-204 | 1.86E-199 | 0.14 | 0.00 | 0.14 | 70.00 |
| 21 | Kynu | 6.23 | 1.05E-203 | 2.18E-199 | 0.14 | 0.00 | 0.14 | 70.00 |
| 21 | Zeb2os | 4.33 | 1.38E-201 | 2.87E-197 | 0.58 | 0.04 | 0.54 | 13.18 |
| 21 | Rnase4 | 5.38 | 1.86E-198 | 3.88E-194 | 0.79 | 0.09 | 0.70 | 8.49 |
| 21 | Sirpa | 4.48 | 2.79E-198 | 5.79E-194 | 0.88 | 0.11 | 0.77 | 7.73 |
| 21 | Trim14 | 6.50 | 5.25E-197 | 1.09E-192 | 0.11 | 0.00 | 0.10 | 105.00 |

**Supplementary Table 1. Marker genes for the original clusters of integrated clustering of E11.5 control and NotchGOF λ-junction cells.**

| | | | | | | | | |
|---|---|---|---|---|---|---|---|---|
| 21 | Junb | 4.13 | 4.25E-195 | 8.83E-191 | 0.87 | 0.11 | 0.76 | 7.74 |
| 21 | Ctsh | 4.87 | 5.68E-195 | 1.18E-190 | 0.64 | 0.06 | 0.58 | 10.90 |
| 21 | Stk10 | 4.23 | 1.73E-194 | 3.59E-190 | 0.45 | 0.03 | 0.42 | 17.23 |
| 21 | Pparg | 6.11 | 6.71E-194 | 1.40E-189 | 0.13 | 0.00 | 0.13 | 66.50 |
| 21 | Aim2 | 3.59 | 1.09E-193 | 2.26E-189 | 0.67 | 0.06 | 0.61 | 11.18 |
| 21 | Smim40 | 6.17 | 8.88E-192 | 1.85E-187 | 0.16 | 0.00 | 0.16 | 53.67 |
| 21 | Kcnk6 | 4.99 | 1.65E-191 | 3.43E-187 | 0.25 | 0.01 | 0.24 | 31.50 |
| 21 | Emp3 | 4.30 | 1.56E-190 | 3.24E-186 | 0.90 | 0.13 | 0.77 | 6.83 |
| 21 | Fos | 5.54 | 3.46E-189 | 7.19E-185 | 0.90 | 0.14 | 0.76 | 6.63 |
| 21 | Gm2a | 3.73 | 7.27E-189 | 1.51E-184 | 0.71 | 0.07 | 0.64 | 10.23 |
| 21 | Pf4 | 1.24 | 3.83E-187 | 7.96E-183 | 0.43 | 0.03 | 0.41 | 16.69 |
| 21 | Ifi27l2a | 6.85 | 2.19E-185 | 4.56E-181 | 0.21 | 0.01 | 0.20 | 35.00 |
| 21 | Taok3 | 3.85 | 2.48E-185 | 5.16E-181 | 0.82 | 0.10 | 0.72 | 8.18 |
| 21 | Cgas | 4.03 | 2.68E-184 | 5.56E-180 | 0.40 | 0.02 | 0.38 | 18.14 |
| 21 | Cmklr1 | 5.62 | 3.46E-183 | 7.19E-179 | 0.19 | 0.01 | 0.18 | 37.80 |
| 21 | Dusp2 | 3.82 | 1.14E-182 | 2.36E-178 | 0.47 | 0.03 | 0.44 | 15.13 |
| 21 | Jak3 | 3.90 | 2.74E-180 | 5.69E-176 | 0.44 | 0.03 | 0.41 | 16.33 |
| 21 | Man1c1 | 4.16 | 5.98E-179 | 1.24E-174 | 0.47 | 0.03 | 0.44 | 15.13 |
| 21 | Ttc7 | 3.95 | 1.95E-178 | 4.06E-174 | 0.47 | 0.03 | 0.44 | 14.66 |
| 21 | H2-K1 | 3.88 | 1.49E-176 | 3.11E-172 | 0.90 | 0.13 | 0.77 | 6.88 |
| 21 | Plxnc1 | 3.95 | 5.18E-176 | 1.08E-171 | 0.52 | 0.04 | 0.48 | 13.10 |
| 21 | Slc16a6 | 3.97 | 9.17E-174 | 1.91E-169 | 0.58 | 0.05 | 0.53 | 11.37 |
| 21 | Erap1 | 3.84 | 7.07E-173 | 1.47E-168 | 0.48 | 0.04 | 0.45 | 13.80 |
| 21 | Arsb | 4.16 | 6.08E-170 | 1.26E-165 | 0.63 | 0.06 | 0.57 | 9.98 |
| 21 | Rgs14 | 4.93 | 2.61E-169 | 5.42E-165 | 0.16 | 0.00 | 0.16 | 53.67 |
| 21 | B4galnt1 | 4.13 | 1.14E-167 | 2.38E-163 | 0.52 | 0.04 | 0.48 | 12.19 |
| 21 | Batf | 3.70 | 3.25E-167 | 6.76E-163 | 0.46 | 0.03 | 0.43 | 14.44 |
| 21 | Rinl | 4.30 | 5.39E-165 | 1.12E-160 | 0.32 | 0.02 | 0.31 | 20.13 |
| 21 | Rasa4 | 4.33 | 7.08E-165 | 1.47E-160 | 0.31 | 0.01 | 0.29 | 22.00 |
| 21 | Mef2c | 4.61 | 2.58E-164 | 5.36E-160 | 0.94 | 0.18 | 0.76 | 5.18 |
| 21 | Sh3tc1 | 5.23 | 7.63E-164 | 1.59E-159 | 0.20 | 0.01 | 0.20 | 33.83 |
| 21 | Tent5a | 4.49 | 1.49E-163 | 3.10E-159 | 0.48 | 0.04 | 0.45 | 13.05 |
| 21 | Pip4k2a | 3.64 | 6.17E-163 | 1.28E-158 | 0.81 | 0.11 | 0.70 | 7.24 |
| 21 | E230029C05Rik | 4.73 | 7.77E-162 | 1.62E-157 | 0.39 | 0.02 | 0.37 | 16.33 |
| 21 | Abca1 | 5.26 | 1.69E-160 | 3.51E-156 | 0.63 | 0.07 | 0.56 | 9.25 |
| 21 | Apobec3 | 3.64 | 1.15E-159 | 2.40E-155 | 0.69 | 0.08 | 0.61 | 8.65 |
| 21 | Def6 | 3.73 | 4.96E-159 | 1.03E-154 | 0.48 | 0.04 | 0.45 | 13.05 |
| 21 | Ier3 | 4.45 | 9.48E-159 | 1.97E-154 | 0.65 | 0.07 | 0.58 | 8.90 |
| 21 | Cbfa2t3 | 4.15 | 1.11E-158 | 2.31E-154 | 0.56 | 0.05 | 0.51 | 10.75 |
| 21 | Anxa1 | 3.77 | 1.66E-157 | 3.46E-153 | 0.40 | 0.03 | 0.37 | 15.35 |
| 21 | Tbc1d10c | 5.49 | 5.50E-157 | 1.14E-152 | 0.19 | 0.01 | 0.18 | 37.80 |
| 21 | Rnf213 | 4.43 | 1.32E-156 | 2.74E-152 | 0.40 | 0.03 | 0.37 | 15.35 |
| 21 | Ehd4 | 4.71 | 5.98E-154 | 1.24E-149 | 0.91 | 0.17 | 0.74 | 5.28 |
| 21 | Xlr | 6.76 | 6.39E-154 | 1.33E-149 | 0.13 | 0.00 | 0.12 | 63.00 |
| 21 | Apoc1 | 5.15 | 1.04E-153 | 2.17E-149 | 0.68 | 0.08 | 0.60 | 8.27 |
| 21 | Renbp | 3.55 | 2.40E-153 | 4.99E-149 | 0.66 | 0.07 | 0.59 | 8.97 |
| 21 | Clec2d | 4.12 | 3.54E-152 | 7.36E-148 | 0.36 | 0.02 | 0.34 | 17.00 |
| 21 | Parp12 | 5.19 | 1.01E-151 | 2.09E-147 | 0.21 | 0.01 | 0.20 | 30.00 |
| 21 | Dock10 | 3.49 | 7.37E-150 | 1.53E-145 | 0.69 | 0.08 | 0.60 | 8.35 |
| 21 | Efhd2 | 4.51 | 1.63E-149 | 3.39E-145 | 0.94 | 0.20 | 0.74 | 4.71 |
| 21 | Slc37a2 | 4.82 | 6.73E-147 | 1.40E-142 | 0.32 | 0.02 | 0.30 | 17.89 |
| 21 | Lsp1 | 4.78 | 3.47E-146 | 7.22E-142 | 0.85 | 0.16 | 0.69 | 5.46 |
| 21 | Tmem140 | 4.63 | 2.72E-143 | 5.65E-139 | 0.24 | 0.01 | 0.23 | 23.80 |
| 21 | Samd9l | 6.01 | 2.12E-142 | 4.41E-138 | 0.14 | 0.00 | 0.14 | 46.67 |
| 21 | St8sia4 | 3.70 | 3.39E-142 | 7.06E-138 | 0.69 | 0.09 | 0.60 | 7.69 |
| 21 | Acap1 | 6.03 | 3.20E-141 | 6.65E-137 | 0.15 | 0.00 | 0.14 | 36.75 |

**Supplementary Table 1. Marker genes for the original clusters of integrated clustering of E11.5 control and NotchGOF λ-junction cells.**

| | | | | | | | | |
|---|---|---|---|---|---|---|---|---|
| 21 | Tgfbr2 | 3.57 | 1.29E-140 | 2.69E-136 | 0.66 | 0.08 | 0.58 | 8.20 |
| 21 | Trim47 | 4.60 | 1.33E-140 | 2.77E-136 | 0.31 | 0.02 | 0.29 | 18.12 |
| 21 | Il18rap | 4.42 | 1.16E-139 | 2.42E-135 | 0.27 | 0.01 | 0.25 | 20.46 |
| 21 | P2ry10b | 6.17 | 2.13E-139 | 4.43E-135 | 0.12 | 0.00 | 0.12 | 59.50 |
| 21 | Phf11d | 6.37 | 1.79E-138 | 3.72E-134 | 0.11 | 0.00 | 0.10 | 52.50 |
| 21 | Pstpip2 | 4.44 | 4.61E-138 | 9.60E-134 | 0.25 | 0.01 | 0.23 | 22.27 |
| 21 | Man2b1 | 3.69 | 6.19E-138 | 1.29E-133 | 0.96 | 0.23 | 0.73 | 4.24 |
| 21 | Trim25 | 3.44 | 9.22E-137 | 1.92E-132 | 0.55 | 0.06 | 0.50 | 9.68 |
| 21 | Tmem173 | 3.57 | 1.37E-136 | 2.86E-132 | 0.54 | 0.05 | 0.48 | 9.96 |
| 21 | Ccdc152 | 5.75 | 1.55E-135 | 3.22E-131 | 0.16 | 0.01 | 0.16 | 32.20 |
| 21 | Rin3 | 3.69 | 1.98E-135 | 4.12E-131 | 0.36 | 0.02 | 0.33 | 14.88 |
| 21 | Selenop | 5.37 | 3.83E-135 | 7.97E-131 | 0.90 | 0.19 | 0.72 | 4.88 |
| 21 | Arap1 | 3.52 | 6.80E-135 | 1.41E-130 | 0.59 | 0.07 | 0.53 | 8.74 |
| 21 | Plxnd1 | 2.24 | 1.27E-134 | 2.64E-130 | 0.55 | 0.06 | 0.50 | 10.04 |
| 21 | Man1a | 3.73 | 6.33E-133 | 1.32E-128 | 0.55 | 0.06 | 0.49 | 9.36 |
| 21 | Ostf1 | 3.72 | 6.56E-133 | 1.36E-128 | 0.98 | 0.27 | 0.71 | 3.60 |
| 21 | Gpr146 | 4.12 | 5.68E-131 | 1.18E-126 | 0.38 | 0.03 | 0.35 | 13.50 |
| 21 | Sema4d | 3.55 | 9.36E-131 | 1.95E-126 | 0.59 | 0.07 | 0.52 | 8.63 |
| 21 | Manba | 4.45 | 3.62E-130 | 7.52E-126 | 0.28 | 0.02 | 0.27 | 18.67 |
| 21 | Ifi47 | 5.44 | 1.34E-129 | 2.79E-125 | 0.11 | 0.00 | 0.11 | 56.00 |
| 21 | Ocel1 | 3.83 | 1.39E-129 | 2.88E-125 | 0.51 | 0.05 | 0.46 | 9.81 |
| 21 | Blnk | 4.40 | 2.34E-127 | 4.87E-123 | 0.25 | 0.01 | 0.24 | 19.38 |
| 21 | Apobr | 4.43 | 9.46E-127 | 1.97E-122 | 0.25 | 0.01 | 0.23 | 20.42 |
| 21 | Arpc1b | 4.78 | 1.58E-126 | 3.29E-122 | 0.99 | 0.34 | 0.65 | 2.89 |
| 21 | Syngr2 | 3.42 | 2.86E-126 | 5.94E-122 | 0.79 | 0.14 | 0.65 | 5.64 |
| 21 | Cebpa | 4.07 | 2.88E-126 | 5.99E-122 | 0.36 | 0.03 | 0.33 | 13.73 |
| 21 | Pnpla7 | 3.65 | 2.96E-126 | 6.15E-122 | 0.41 | 0.03 | 0.38 | 12.15 |
| 21 | Myd88 | 3.59 | 4.84E-126 | 1.01E-121 | 0.50 | 0.05 | 0.45 | 9.86 |
| 21 | Plekha2 | 3.62 | 6.48E-124 | 1.35E-119 | 0.47 | 0.05 | 0.42 | 10.42 |
| 21 | Bmp2k | 3.34 | 1.81E-123 | 3.76E-119 | 0.81 | 0.15 | 0.66 | 5.48 |
| 21 | Pygm | 5.41 | 3.79E-122 | 7.88E-118 | 0.13 | 0.00 | 0.13 | 44.33 |
| 21 | Tor3a | 3.56 | 8.56E-122 | 1.78E-117 | 0.43 | 0.04 | 0.39 | 11.24 |
| 21 | Capn3 | 4.60 | 7.28E-119 | 1.51E-114 | 0.11 | 0.00 | 0.11 | 56.00 |
| 21 | Amz1 | 4.24 | 2.61E-118 | 5.42E-114 | 0.21 | 0.01 | 0.20 | 23.33 |
| 21 | Runx1 | 2.56 | 4.03E-118 | 8.38E-114 | 0.92 | 0.19 | 0.73 | 4.82 |
| 21 | Il10rb | 3.70 | 5.79E-118 | 1.21E-113 | 0.61 | 0.08 | 0.53 | 7.33 |
| 21 | Ifih1 | 4.46 | 8.64E-118 | 1.80E-113 | 0.22 | 0.01 | 0.21 | 21.70 |
| 21 | Fosb | 3.83 | 3.09E-117 | 6.43E-113 | 0.38 | 0.03 | 0.35 | 12.19 |
| 21 | Morrbid | 4.86 | 9.10E-117 | 1.89E-112 | 0.14 | 0.00 | 0.14 | 35.00 |
| 21 | Map3k8 | 4.03 | 1.59E-116 | 3.30E-112 | 0.30 | 0.02 | 0.28 | 15.05 |
| 21 | Rrad | 4.42 | 2.47E-116 | 5.14E-112 | 0.25 | 0.01 | 0.23 | 18.85 |
| 21 | Rassf2 | 4.05 | 1.29E-114 | 2.68E-110 | 0.31 | 0.02 | 0.29 | 14.67 |
| 21 | Ggta1 | 3.60 | 2.71E-113 | 5.64E-109 | 0.43 | 0.04 | 0.39 | 10.33 |
| 21 | Lmo2 | 2.44 | 3.67E-113 | 7.64E-109 | 0.76 | 0.12 | 0.63 | 6.09 |
| 21 | Serinc3 | 3.80 | 5.72E-112 | 1.19E-107 | 0.98 | 0.36 | 0.62 | 2.73 |
| 21 | Hivep3 | 4.10 | 2.40E-111 | 4.98E-107 | 0.29 | 0.02 | 0.27 | 14.70 |
| 21 | Slc16a7 | 4.33 | 4.27E-111 | 8.89E-107 | 0.31 | 0.02 | 0.29 | 14.00 |
| 21 | Rnasel | 3.51 | 1.69E-110 | 3.52E-106 | 0.60 | 0.08 | 0.52 | 7.24 |
| 21 | Hexb | 4.03 | 6.02E-110 | 1.25E-105 | 0.89 | 0.22 | 0.67 | 4.00 |
| 21 | Ifngr1 | 3.32 | 2.42E-109 | 5.04E-105 | 0.92 | 0.24 | 0.68 | 3.80 |
| 21 | Stxbp2 | 2.86 | 1.01E-108 | 2.09E-104 | 0.76 | 0.13 | 0.62 | 5.63 |
| 21 | Pik3r5 | 4.62 | 1.06E-108 | 2.21E-104 | 0.18 | 0.01 | 0.17 | 25.00 |
| 21 | Sat1 | 4.34 | 1.22E-108 | 2.53E-104 | 0.96 | 0.30 | 0.65 | 3.15 |
| 21 | Gmip | 3.29 | 3.90E-108 | 8.11E-104 | 0.48 | 0.05 | 0.43 | 9.11 |
| 21 | Stat1 | 3.17 | 5.25E-108 | 1.09E-103 | 0.64 | 0.10 | 0.54 | 6.69 |
| 21 | Gsdmd | 3.84 | 1.98E-107 | 4.12E-103 | 0.32 | 0.02 | 0.30 | 13.42 |

**Supplementary Table 1. Marker genes for the original clusters of integrated clustering of E11.5 control and NotchGOF λ-junction cells.**

| | | | | | | | | |
|---|---|---|---|---|---|---|---|---|
| 21 | Dclre1c | 3.13 | 2.44E-107 | 5.08E-103 | 0.47 | 0.05 | 0.42 | 9.20 |
| 21 | H2-D1 | 3.05 | 3.37E-107 | 7.02E-103 | 0.92 | 0.24 | 0.68 | 3.83 |
| 21 | Rab32 | 3.07 | 6.88E-107 | 1.43E-102 | 0.62 | 0.09 | 0.53 | 6.91 |
| 21 | Rbm47 | 3.81 | 9.65E-107 | 2.01E-102 | 0.37 | 0.03 | 0.34 | 11.24 |
| 21 | Rcsd1 | 2.86 | 1.22E-106 | 2.53E-102 | 0.63 | 0.09 | 0.54 | 6.91 |
| 21 | AI413582 | 3.10 | 1.56E-106 | 3.25E-102 | 0.62 | 0.09 | 0.53 | 6.91 |
| 21 | Gsn | 3.42 | 4.06E-106 | 8.45E-102 | 0.75 | 0.14 | 0.61 | 5.30 |
| 21 | Cyba | 3.97 | 7.29E-106 | 1.52E-101 | 0.99 | 0.45 | 0.54 | 2.21 |
| 21 | Adrb2 | 5.24 | 8.96E-106 | 1.86E-101 | 0.22 | 0.01 | 0.21 | 18.67 |
| 21 | Psmb10 | 3.03 | 9.32E-106 | 1.94E-101 | 0.90 | 0.23 | 0.67 | 3.93 |
| 21 | Reps2 | 3.79 | 2.26E-105 | 4.71E-101 | 0.34 | 0.03 | 0.32 | 12.25 |
| 21 | Tspo | 3.91 | 2.88E-105 | 5.99E-101 | 0.97 | 0.35 | 0.62 | 2.79 |
| 21 | Dram1 | 4.83 | 5.29E-105 | 1.10E-100 | 0.13 | 0.00 | 0.13 | 33.25 |
| 21 | Tpd52 | 3.08 | 5.90E-105 | 1.23E-100 | 0.93 | 0.26 | 0.67 | 3.58 |
| 21 | Celf2 | 2.75 | 4.12E-104 | 8.58E-100 | 0.93 | 0.25 | 0.68 | 3.68 |
| 21 | Rtp4 | 4.66 | 8.61E-104 | 1.79E-99 | 0.20 | 0.01 | 0.19 | 21.78 |
| 21 | Nfatc1 | 3.68 | 9.96E-104 | 2.07E-99 | 0.36 | 0.03 | 0.33 | 11.52 |
| 21 | Pla2g15 | 3.22 | 2.72E-103 | 5.66E-99 | 0.52 | 0.07 | 0.46 | 7.94 |
| 21 | Nostrin | 5.75 | 6.83E-103 | 1.42E-98 | 0.11 | 0.00 | 0.10 | 52.50 |
| 21 | Tmem37 | 3.59 | 9.90E-103 | 2.06E-98 | 0.58 | 0.08 | 0.50 | 6.99 |
| 21 | Plekho2 | 3.34 | 1.45E-102 | 3.03E-98 | 0.52 | 0.07 | 0.46 | 7.71 |
| 21 | Phyhd1 | 3.95 | 2.34E-102 | 4.87E-98 | 0.29 | 0.02 | 0.27 | 13.36 |
| 21 | Nagpa | 3.02 | 4.18E-102 | 8.69E-98 | 0.63 | 0.10 | 0.53 | 6.48 |
| 21 | Stat6 | 3.06 | 1.99E-101 | 4.14E-97 | 0.47 | 0.05 | 0.42 | 8.85 |
| 21 | 2610203C22Rik | 3.93 | 2.24E-101 | 4.66E-97 | 0.22 | 0.01 | 0.21 | 18.08 |
| 21 | Guca1a | 3.45 | 2.79E-101 | 5.80E-97 | 0.36 | 0.03 | 0.33 | 11.03 |
| 21 | Sec11c | 3.06 | 3.90E-101 | 8.10E-97 | 0.92 | 0.30 | 0.63 | 3.10 |
| 21 | Cotl1 | 3.22 | 6.56E-100 | 1.36E-95 | 1.00 | 0.44 | 0.56 | 2.26 |
| 21 | Gm36738 | 3.82 | 2.36E-99 | 4.90E-95 | 0.22 | 0.01 | 0.21 | 18.08 |
| 21 | Zbtb7b | 3.97 | 2.54E-99 | 5.28E-95 | 0.27 | 0.02 | 0.25 | 14.37 |
| 21 | B2m | 3.75 | 2.11E-98 | 4.40E-94 | 1.00 | 0.54 | 0.46 | 1.86 |
| 21 | Sh2b3 | 3.11 | 3.45E-98 | 7.17E-94 | 0.54 | 0.07 | 0.47 | 7.37 |
| 21 | Gusb | 3.21 | 3.45E-98 | 7.18E-94 | 0.92 | 0.30 | 0.62 | 3.04 |
| 21 | Nr4a1 | 3.74 | 4.47E-98 | 9.30E-94 | 0.40 | 0.04 | 0.36 | 9.73 |
| 21 | Khk | 3.14 | 9.71E-98 | 2.02E-93 | 0.50 | 0.06 | 0.44 | 7.86 |
| 21 | Anxa6 | 2.78 | 1.27E-97 | 2.65E-93 | 0.80 | 0.17 | 0.63 | 4.69 |
| 21 | Arhgdib | 3.27 | 2.46E-97 | 5.12E-93 | 0.99 | 0.50 | 0.50 | 1.99 |
| 21 | Tmem229b | 3.51 | 8.30E-97 | 1.73E-92 | 0.30 | 0.02 | 0.28 | 13.09 |
| 21 | Pon3 | 3.50 | 1.32E-96 | 2.75E-92 | 0.34 | 0.03 | 0.31 | 11.06 |
| 21 | Adam15 | 3.02 | 2.29E-96 | 4.76E-92 | 0.61 | 0.10 | 0.51 | 6.40 |
| 21 | Ckb | 4.27 | 2.79E-96 | 5.81E-92 | 0.94 | 0.33 | 0.61 | 2.86 |
| 21 | Fgd2 | 3.01 | 4.98E-96 | 1.04E-91 | 0.62 | 0.10 | 0.52 | 6.28 |
| 21 | Nabp1 | 3.81 | 5.28E-96 | 1.10E-91 | 0.33 | 0.03 | 0.30 | 11.34 |
| 21 | Ubash3a | 4.80 | 1.96E-95 | 4.07E-91 | 0.15 | 0.01 | 0.14 | 24.50 |
| 21 | Hexa | 3.76 | 6.13E-95 | 1.27E-90 | 0.92 | 0.31 | 0.60 | 2.93 |
| 21 | Mvp | 3.26 | 7.87E-95 | 1.64E-90 | 0.48 | 0.06 | 0.42 | 8.07 |
| 21 | Ucp2 | 3.04 | 2.60E-94 | 5.41E-90 | 0.97 | 0.35 | 0.61 | 2.73 |
| 21 | Sipa1 | 2.60 | 8.35E-94 | 1.74E-89 | 0.80 | 0.17 | 0.63 | 4.65 |
| 21 | Hebp1 | 2.74 | 5.08E-93 | 1.06E-88 | 0.93 | 0.26 | 0.67 | 3.52 |
| 21 | Stard8 | 3.64 | 8.30E-93 | 1.73E-88 | 0.56 | 0.09 | 0.47 | 6.50 |
| 21 | Daglb | 3.05 | 2.56E-92 | 5.33E-88 | 0.63 | 0.11 | 0.52 | 5.93 |
| 21 | Arrb2 | 2.81 | 2.87E-92 | 5.96E-88 | 0.78 | 0.17 | 0.61 | 4.62 |
| 21 | Klf6 | 3.44 | 5.04E-92 | 1.05E-87 | 0.85 | 0.23 | 0.62 | 3.68 |
| 21 | Ninj1 | 3.12 | 6.29E-92 | 1.31E-87 | 0.80 | 0.19 | 0.62 | 4.25 |
| 21 | Abcd1 | 3.06 | 7.98E-92 | 1.66E-87 | 0.48 | 0.06 | 0.42 | 7.79 |
| 21 | Lpl | 4.65 | 1.30E-91 | 2.71E-87 | 0.71 | 0.16 | 0.55 | 4.50 |

**Supplementary Table 1. Marker genes for the original clusters of integrated clustering of E11.5 control and NotchGOF λ-junction cells.**

| | | | | | | | | |
|---|---|---|---|---|---|---|---|---|
| 21 | Ap1g2 | 3.14 | 2.12E-91 | 4.41E-87 | 0.39 | 0.04 | 0.35 | 9.63 |
| 21 | Gimap1 | 3.50 | 5.81E-91 | 1.21E-86 | 0.24 | 0.02 | 0.22 | 14.88 |
| 21 | Ifnar2 | 2.90 | 8.06E-91 | 1.68E-86 | 0.94 | 0.33 | 0.62 | 2.88 |
| 21 | Casp8 | 2.68 | 9.26E-91 | 1.93E-86 | 0.80 | 0.18 | 0.62 | 4.40 |
| 21 | Enpp1 | 4.91 | 6.88E-90 | 1.43E-85 | 0.14 | 0.01 | 0.14 | 28.00 |
| 21 | Tmsb4x | 3.78 | 1.28E-89 | 2.66E-85 | 1.00 | 0.99 | 0.01 | 1.01 |
| 21 | Stat3 | 2.73 | 6.59E-88 | 1.37E-83 | 0.69 | 0.13 | 0.56 | 5.31 |
| 21 | Tfeb | 3.43 | 6.95E-88 | 1.45E-83 | 0.28 | 0.02 | 0.26 | 12.73 |
| 21 | Mt1 | 2.98 | 9.78E-88 | 2.03E-83 | 0.90 | 0.27 | 0.64 | 3.40 |
| 21 | E030030I06Rik | 4.97 | 1.14E-87 | 2.36E-83 | 0.13 | 0.00 | 0.12 | 31.50 |
| 21 | Rasa3 | 3.19 | 1.77E-87 | 3.68E-83 | 0.52 | 0.08 | 0.44 | 6.89 |
| 21 | Padi2 | 5.96 | 4.03E-87 | 8.39E-83 | 0.11 | 0.00 | 0.10 | 35.00 |
| 21 | Creg1 | 2.88 | 1.71E-86 | 3.55E-82 | 0.94 | 0.35 | 0.59 | 2.68 |
| 21 | Sh3bgrl3 | 2.21 | 1.76E-86 | 3.67E-82 | 1.00 | 0.91 | 0.09 | 1.10 |
| 21 | Fam78a | 4.34 | 2.62E-86 | 5.44E-82 | 0.20 | 0.01 | 0.19 | 16.92 |
| 21 | Tagln2 | 3.20 | 1.45E-85 | 3.02E-81 | 0.88 | 0.30 | 0.59 | 2.98 |
| 21 | Lrmda | 4.06 | 1.49E-85 | 3.09E-81 | 0.18 | 0.01 | 0.17 | 19.44 |
| 21 | Rasgrp2 | 2.66 | 1.85E-85 | 3.84E-81 | 0.41 | 0.05 | 0.36 | 8.43 |
| 21 | Nceh1 | 3.44 | 3.84E-85 | 7.98E-81 | 0.41 | 0.05 | 0.36 | 8.26 |
| 21 | Naaa | 3.73 | 5.40E-85 | 1.12E-80 | 0.27 | 0.02 | 0.25 | 12.67 |
| 21 | Ifi30 | 3.82 | 1.53E-84 | 3.19E-80 | 0.87 | 0.29 | 0.58 | 2.99 |
| 21 | Serpinb6b | 3.57 | 2.33E-84 | 4.84E-80 | 0.27 | 0.02 | 0.25 | 12.67 |
| 21 | Acss1 | 3.52 | 2.50E-84 | 5.19E-80 | 0.28 | 0.02 | 0.26 | 12.17 |
| 21 | Egr3 | 5.08 | 1.33E-83 | 2.77E-79 | 0.11 | 0.00 | 0.10 | 35.00 |
| 21 | Ppp1r18 | 2.57 | 1.45E-83 | 3.02E-79 | 0.88 | 0.25 | 0.63 | 3.55 |
| 21 | Mtus1 | 3.12 | 2.66E-83 | 5.52E-79 | 0.59 | 0.10 | 0.48 | 5.70 |
| 21 | Arpc2 | 1.89 | 3.75E-83 | 7.81E-79 | 1.00 | 0.94 | 0.06 | 1.06 |
| 21 | Lacc1 | 3.21 | 6.48E-83 | 1.35E-78 | 0.54 | 0.09 | 0.45 | 6.18 |
| 21 | Apoe | 7.13 | 3.75E-82 | 7.79E-78 | 0.94 | 0.43 | 0.51 | 2.18 |
| 21 | Sqor | 4.02 | 5.28E-82 | 1.10E-77 | 0.25 | 0.02 | 0.23 | 12.89 |
| 21 | Galns | 3.33 | 8.46E-81 | 1.76E-76 | 0.32 | 0.03 | 0.29 | 10.06 |
| 21 | Ctsz | 3.38 | 1.02E-80 | 2.12E-76 | 0.99 | 0.58 | 0.41 | 1.71 |
| 21 | Tmem9b | 2.58 | 1.63E-80 | 3.39E-76 | 0.69 | 0.14 | 0.55 | 5.04 |
| 21 | Rgs7bp | 3.48 | 1.87E-80 | 3.88E-76 | 0.27 | 0.02 | 0.24 | 12.09 |
| 21 | Gbp9 | 4.72 | 2.39E-80 | 4.97E-76 | 0.11 | 0.00 | 0.10 | 35.00 |
| 21 | Tbc1d8 | 3.17 | 5.77E-80 | 1.20E-75 | 0.32 | 0.03 | 0.28 | 10.16 |
| 21 | Rnf144b | 4.53 | 4.48E-79 | 9.32E-75 | 0.15 | 0.01 | 0.14 | 21.00 |
| 21 | Arpc5 | 2.37 | 5.51E-79 | 1.15E-74 | 0.99 | 0.76 | 0.23 | 1.30 |
| 21 | Mctp1 | 4.25 | 8.34E-79 | 1.74E-74 | 0.15 | 0.01 | 0.14 | 21.00 |
| 21 | Mbnl1 | 2.65 | 1.34E-78 | 2.78E-74 | 0.97 | 0.47 | 0.50 | 2.07 |
| 21 | Cst3 | 3.22 | 1.78E-78 | 3.70E-74 | 1.00 | 0.92 | 0.08 | 1.09 |
| 21 | Tmcc3 | 3.35 | 3.38E-78 | 7.02E-74 | 0.32 | 0.03 | 0.29 | 9.76 |
| 21 | Prkcd | 2.76 | 6.09E-78 | 1.27E-73 | 0.80 | 0.21 | 0.59 | 3.80 |
| 21 | Ly6e | 3.25 | 9.30E-78 | 1.93E-73 | 0.94 | 0.45 | 0.49 | 2.09 |
| 21 | Tapbp | 2.84 | 1.02E-77 | 2.12E-73 | 0.51 | 0.08 | 0.43 | 6.38 |
| 21 | Arid3a | 2.64 | 2.37E-77 | 4.93E-73 | 0.90 | 0.33 | 0.57 | 2.73 |
| 21 | S100a1 | 2.99 | 5.01E-77 | 1.04E-72 | 0.45 | 0.06 | 0.39 | 7.11 |
| 21 | Cebpd | 2.95 | 5.18E-77 | 1.08E-72 | 0.56 | 0.10 | 0.46 | 5.76 |
| 21 | Ulbp1 | 3.43 | 8.33E-77 | 1.73E-72 | 0.27 | 0.02 | 0.24 | 11.57 |
| 21 | Nbeal2 | 4.10 | 9.76E-77 | 2.03E-72 | 0.15 | 0.01 | 0.15 | 19.25 |
| 21 | Btg2 | 2.98 | 1.30E-76 | 2.70E-72 | 0.85 | 0.27 | 0.58 | 3.12 |
| 21 | Hprt | 2.54 | 1.32E-76 | 2.74E-72 | 0.94 | 0.43 | 0.50 | 2.16 |
| 21 | Cmtm7 | 2.85 | 1.41E-76 | 2.94E-72 | 0.97 | 0.48 | 0.48 | 2.00 |
| 21 | Capg | 2.96 | 1.54E-76 | 3.21E-72 | 0.69 | 0.15 | 0.54 | 4.60 |
| 21 | Gpx1 | 1.77 | 1.69E-76 | 3.51E-72 | 1.00 | 0.95 | 0.05 | 1.05 |
| 21 | Fcgrt | 3.73 | 3.23E-76 | 6.72E-72 | 0.65 | 0.14 | 0.51 | 4.61 |

**Supplementary Table 1. Marker genes for the original clusters of integrated clustering of E11.5 control and NotchGOF λ-junction cells.**

| | | | | | | | | |
|---|---|---|---|---|---|---|---|---|
| 21 | Gcnt1 | 3.62 | 3.23E-76 | 6.72E-72 | 0.30 | 0.03 | 0.27 | 10.03 |
| 21 | Tacc1 | 2.42 | 3.41E-76 | 7.09E-72 | 0.90 | 0.32 | 0.58 | 2.84 |
| 21 | Ctsd | 4.50 | 4.31E-76 | 8.95E-72 | 0.99 | 0.61 | 0.39 | 1.64 |
| 21 | Fam111a | 2.71 | 4.94E-76 | 1.03E-71 | 0.90 | 0.32 | 0.57 | 2.77 |
| 21 | Tnip2 | 3.11 | 5.09E-76 | 1.06E-71 | 0.29 | 0.03 | 0.26 | 10.63 |
| 21 | Actr3 | 1.98 | 5.60E-76 | 1.16E-71 | 0.99 | 0.82 | 0.18 | 1.22 |
| 21 | Atp13a2 | 2.52 | 7.95E-76 | 1.65E-71 | 0.86 | 0.26 | 0.61 | 3.37 |
| 21 | Pld1 | 3.62 | 1.07E-75 | 2.22E-71 | 0.24 | 0.02 | 0.22 | 12.53 |
| 21 | Ccrl2 | 4.83 | 1.79E-75 | 3.73E-71 | 0.18 | 0.01 | 0.17 | 16.55 |
| 21 | Erp29 | 2.07 | 1.93E-75 | 4.02E-71 | 0.99 | 0.88 | 0.11 | 1.13 |
| 21 | Rhog | 2.56 | 2.27E-75 | 4.72E-71 | 0.93 | 0.35 | 0.58 | 2.66 |
| 21 | Arpc4 | 1.82 | 4.05E-75 | 8.42E-71 | 1.00 | 0.83 | 0.18 | 1.21 |
| 21 | Mknk1 | 2.67 | 9.58E-75 | 1.99E-70 | 0.54 | 0.09 | 0.45 | 5.98 |
| 21 | Lgmn | 5.05 | 1.08E-74 | 2.25E-70 | 0.80 | 0.29 | 0.52 | 2.82 |
| 21 | Actb | 1.50 | 1.28E-74 | 2.67E-70 | 1.00 | 1.00 | 0.00 | 1.00 |
| 21 | Tubb2a | 2.90 | 1.44E-74 | 2.99E-70 | 0.90 | 0.30 | 0.60 | 3.03 |
| 21 | Rps6ka1 | 2.58 | 3.89E-74 | 8.10E-70 | 0.82 | 0.24 | 0.58 | 3.41 |
| 21 | Clic1 | 1.71 | 6.42E-74 | 1.33E-69 | 1.00 | 0.93 | 0.07 | 1.08 |
| 21 | Kctd12 | 2.62 | 8.30E-74 | 1.73E-69 | 0.89 | 0.30 | 0.59 | 3.01 |
| 21 | Hmox1 | 4.94 | 1.37E-73 | 2.86E-69 | 0.72 | 0.19 | 0.53 | 3.85 |
| 21 | Tnnt1 | 2.72 | 6.42E-73 | 1.33E-68 | 0.40 | 0.05 | 0.35 | 7.67 |
| 21 | Rftn1 | 3.03 | 7.78E-73 | 1.62E-68 | 0.34 | 0.04 | 0.30 | 8.84 |
| 21 | Pdgfb | 3.64 | 7.85E-73 | 1.63E-68 | 0.20 | 0.01 | 0.19 | 14.50 |
| 21 | Clta | 1.71 | 1.83E-72 | 3.81E-68 | 1.00 | 0.98 | 0.02 | 1.02 |
| 21 | Exoc6 | 2.82 | 2.36E-72 | 4.90E-68 | 0.47 | 0.07 | 0.40 | 6.42 |
| 21 | Fth1 | 2.31 | 5.36E-72 | 1.11E-67 | 1.00 | 1.00 | 0.00 | 1.00 |
| 21 | Pold4 | 2.45 | 8.05E-72 | 1.67E-67 | 0.75 | 0.18 | 0.56 | 4.07 |
| 21 | Tmem104 | 3.46 | 1.77E-71 | 3.69E-67 | 0.27 | 0.03 | 0.24 | 10.64 |
| 21 | Itpr2 | 3.11 | 2.88E-71 | 6.00E-67 | 0.36 | 0.05 | 0.32 | 8.09 |
| 21 | Mpp1 | 2.87 | 4.73E-71 | 9.83E-67 | 0.80 | 0.24 | 0.56 | 3.36 |
| 21 | Ednrb | 3.65 | 5.91E-71 | 1.23E-66 | 0.33 | 0.04 | 0.29 | 8.66 |
| 21 | Irak4 | 2.72 | 1.69E-70 | 3.51E-66 | 0.37 | 0.05 | 0.33 | 8.07 |
| 21 | Arap3 | 2.80 | 1.75E-70 | 3.63E-66 | 0.36 | 0.04 | 0.31 | 8.30 |
| 21 | Pfn1 | 1.17 | 3.34E-70 | 6.95E-66 | 0.99 | 1.00 | 0.00 | 1.00 |
| 21 | Gnai2 | 1.48 | 3.97E-70 | 8.26E-66 | 1.00 | 0.97 | 0.03 | 1.03 |
| 21 | Itm2b | 2.83 | 4.84E-70 | 1.01E-65 | 1.00 | 0.84 | 0.16 | 1.19 |
| 21 | Ctsa | 2.83 | 5.89E-70 | 1.23E-65 | 0.94 | 0.44 | 0.51 | 2.17 |
| 21 | Iqgap1 | 2.25 | 1.54E-69 | 3.21E-65 | 0.97 | 0.47 | 0.49 | 2.05 |
| 21 | Slc9a9 | 3.40 | 1.81E-69 | 3.75E-65 | 0.32 | 0.04 | 0.29 | 8.70 |
| 21 | Acsl5 | 2.59 | 3.85E-69 | 8.00E-65 | 0.68 | 0.16 | 0.52 | 4.35 |
| 21 | Atp8a1 | 2.69 | 5.48E-69 | 1.14E-64 | 0.50 | 0.09 | 0.42 | 5.92 |
| 21 | Vamp8 | 2.14 | 5.65E-69 | 1.18E-64 | 0.97 | 0.61 | 0.36 | 1.59 |
| 21 | Scamp2 | 2.43 | 5.71E-69 | 1.19E-64 | 0.89 | 0.31 | 0.58 | 2.85 |
| 21 | Slfn5 | 5.18 | 1.24E-68 | 2.59E-64 | 0.15 | 0.01 | 0.15 | 17.11 |
| 21 | Aldh3b1 | 4.58 | 1.27E-68 | 2.65E-64 | 0.11 | 0.00 | 0.10 | 26.25 |
| 21 | Orai1 | 2.35 | 3.43E-68 | 7.13E-64 | 0.71 | 0.17 | 0.54 | 4.28 |
| 21 | Arpc3 | 1.57 | 6.03E-68 | 1.25E-63 | 0.99 | 0.90 | 0.09 | 1.10 |
| 21 | Gm16096 | 4.68 | 6.27E-68 | 1.30E-63 | 0.11 | 0.00 | 0.10 | 26.25 |
| 21 | Icos | 4.10 | 2.87E-67 | 5.97E-63 | 0.15 | 0.01 | 0.14 | 18.38 |
| 21 | Car9 | 3.97 | 3.96E-67 | 8.23E-63 | 0.15 | 0.01 | 0.14 | 18.38 |
| 21 | Gpr157 | 3.98 | 8.23E-67 | 1.71E-62 | 0.18 | 0.01 | 0.16 | 14.58 |
| 21 | Msn | 2.15 | 1.08E-66 | 2.24E-62 | 0.98 | 0.51 | 0.47 | 1.91 |
| 21 | Entpd1 | 3.27 | 2.50E-66 | 5.20E-62 | 0.41 | 0.06 | 0.35 | 6.88 |
| 21 | Ptgs2 | 3.75 | 2.66E-66 | 5.53E-62 | 0.13 | 0.01 | 0.12 | 21.00 |
| 21 | Dusp3 | 2.65 | 5.12E-66 | 1.07E-61 | 0.39 | 0.06 | 0.34 | 7.13 |
| 21 | Scamp5 | 3.29 | 6.20E-66 | 1.29E-61 | 0.34 | 0.04 | 0.29 | 8.00 |

**Supplementary Table 1. Marker genes for the original clusters of integrated clustering of E11.5 control and NotchGOF λ-junction cells.**

| | | | | | | | | |
|---|---|---|---|---|---|---|---|---|
| 21 | Capza2 | 1.72 | 8.42E-66 | 1.75E-61 | 1.00 | 0.86 | 0.14 | 1.16 |
| 21 | Rel | 2.45 | 1.23E-65 | 2.55E-61 | 0.50 | 0.09 | 0.42 | 5.72 |
| 21 | Bhlhe41 | 4.07 | 1.39E-65 | 2.90E-61 | 0.18 | 0.01 | 0.16 | 14.58 |
| 21 | Atp6v0b | 1.95 | 1.71E-65 | 3.55E-61 | 0.98 | 0.74 | 0.24 | 1.33 |
| 21 | Susd1 | 3.83 | 1.97E-65 | 4.09E-61 | 0.19 | 0.01 | 0.18 | 13.50 |
| 21 | Vwa5a | 2.71 | 4.37E-65 | 9.09E-61 | 0.49 | 0.09 | 0.40 | 5.70 |
| 21 | Rgs2 | 2.55 | 5.62E-65 | 1.17E-60 | 0.89 | 0.34 | 0.55 | 2.60 |
| 21 | Tln1 | 1.73 | 1.22E-64 | 2.54E-60 | 0.99 | 0.74 | 0.25 | 1.33 |
| 21 | Trim12a | 3.08 | 1.48E-64 | 3.08E-60 | 0.24 | 0.02 | 0.22 | 10.82 |
| 21 | Pon2 | 2.03 | 1.03E-63 | 2.14E-59 | 0.72 | 0.18 | 0.54 | 4.02 |
| 21 | Psap | 2.97 | 1.11E-63 | 2.31E-59 | 0.97 | 0.62 | 0.35 | 1.56 |
| 21 | Nfatc2 | 3.91 | 1.39E-63 | 2.89E-59 | 0.19 | 0.01 | 0.18 | 13.50 |
| 21 | Camk2d | 2.60 | 2.36E-63 | 4.92E-59 | 0.71 | 0.19 | 0.52 | 3.68 |
| 21 | Itsn2 | 2.29 | 7.59E-63 | 1.58E-58 | 0.71 | 0.19 | 0.52 | 3.77 |
| 21 | Grn | 3.86 | 1.34E-62 | 2.80E-58 | 0.84 | 0.35 | 0.49 | 2.40 |
| 21 | Zc3h12a | 4.04 | 2.23E-62 | 4.63E-58 | 0.15 | 0.01 | 0.14 | 15.40 |
| 21 | P2ry14 | 3.57 | 2.69E-62 | 5.59E-58 | 0.24 | 0.02 | 0.22 | 10.35 |
| 21 | Elf1 | 2.21 | 2.96E-62 | 6.16E-58 | 0.82 | 0.26 | 0.56 | 3.13 |
| 21 | Nup210 | 2.76 | 3.00E-62 | 6.23E-58 | 0.40 | 0.06 | 0.34 | 6.65 |
| 21 | Fkbp15 | 2.57 | 4.34E-62 | 9.02E-58 | 0.60 | 0.13 | 0.47 | 4.59 |
| 21 | N4bp2l1 | 2.87 | 4.60E-62 | 9.57E-58 | 0.29 | 0.03 | 0.26 | 8.91 |
| 21 | H2afj | 1.62 | 9.13E-62 | 1.90E-57 | 0.99 | 0.81 | 0.18 | 1.22 |
| 21 | Plaur | 4.17 | 4.45E-61 | 9.25E-57 | 0.14 | 0.01 | 0.13 | 17.50 |
| 21 | Psme2b | 4.01 | 4.89E-61 | 1.02E-56 | 0.13 | 0.01 | 0.13 | 19.00 |
| 21 | Tpp1 | 2.71 | 5.89E-61 | 1.23E-56 | 0.80 | 0.26 | 0.53 | 3.03 |
| 21 | Arl6ip5 | 2.15 | 6.67E-61 | 1.39E-56 | 0.87 | 0.32 | 0.55 | 2.73 |
| 21 | Disc1 | 3.42 | 6.70E-61 | 1.39E-56 | 0.20 | 0.02 | 0.19 | 11.94 |
| 21 | Edem1 | 2.33 | 6.97E-61 | 1.45E-56 | 0.79 | 0.24 | 0.55 | 3.29 |
| 21 | Cpeb2 | 3.51 | 8.01E-61 | 1.66E-56 | 0.20 | 0.02 | 0.18 | 12.25 |
| 21 | Slc48a1 | 2.58 | 8.34E-61 | 1.74E-56 | 0.66 | 0.17 | 0.49 | 3.98 |
| 21 | Abcd2 | 2.74 | 9.59E-61 | 1.99E-56 | 0.43 | 0.07 | 0.36 | 6.03 |
| 21 | Gm13212 | 3.05 | 1.25E-60 | 2.60E-56 | 0.26 | 0.03 | 0.23 | 9.59 |
| 21 | Blvra | 2.32 | 1.25E-60 | 2.61E-56 | 0.73 | 0.20 | 0.53 | 3.60 |
| 21 | Myo5a | 2.45 | 2.22E-60 | 4.62E-56 | 0.87 | 0.32 | 0.56 | 2.74 |
| 21 | Ptger3 | 3.35 | 3.24E-60 | 6.73E-56 | 0.17 | 0.01 | 0.16 | 14.00 |
| 21 | Itga6 | 1.97 | 3.89E-60 | 8.09E-56 | 0.91 | 0.31 | 0.60 | 2.93 |
| 21 | Sh3kbp1 | 2.25 | 3.91E-60 | 8.14E-56 | 0.77 | 0.22 | 0.55 | 3.48 |
| 21 | Fam49b | 1.74 | 4.72E-60 | 9.83E-56 | 0.99 | 0.65 | 0.33 | 1.51 |
| 21 | Klk8 | 2.34 | 5.38E-60 | 1.12E-55 | 0.74 | 0.21 | 0.53 | 3.48 |
| 21 | Rogdi | 2.51 | 5.51E-60 | 1.15E-55 | 0.76 | 0.25 | 0.52 | 3.10 |
| 21 | Madd | 2.50 | 5.92E-60 | 1.23E-55 | 0.43 | 0.07 | 0.36 | 6.11 |
| 21 | Glrx | 2.32 | 1.14E-59 | 2.38E-55 | 0.76 | 0.23 | 0.53 | 3.27 |
| 21 | Stk17b | 2.75 | 1.84E-59 | 3.82E-55 | 0.58 | 0.13 | 0.45 | 4.36 |
| 21 | Ssr4 | 1.46 | 2.16E-59 | 4.49E-55 | 0.99 | 0.84 | 0.15 | 1.18 |
| 21 | Rab27a | 2.94 | 2.84E-59 | 5.91E-55 | 0.34 | 0.05 | 0.30 | 7.30 |
| 21 | Dtx3l | 3.15 | 4.71E-59 | 9.80E-55 | 0.32 | 0.04 | 0.28 | 7.88 |
| 21 | Tcirg1 | 2.63 | 5.02E-59 | 1.04E-54 | 0.48 | 0.09 | 0.39 | 5.47 |
| 21 | Rap1b | 1.75 | 5.94E-59 | 1.24E-54 | 0.99 | 0.76 | 0.23 | 1.30 |
| 21 | Tep1 | 2.75 | 7.25E-59 | 1.51E-54 | 0.38 | 0.06 | 0.32 | 6.63 |
| 21 | Gba | 2.54 | 8.17E-59 | 1.70E-54 | 0.50 | 0.09 | 0.40 | 5.29 |
| 21 | Zeb2 | 2.26 | 9.62E-59 | 2.00E-54 | 0.92 | 0.40 | 0.52 | 2.30 |
| 21 | Pygl | 2.21 | 1.51E-58 | 3.14E-54 | 0.51 | 0.10 | 0.41 | 5.31 |
| 21 | Slc8b1 | 3.05 | 1.55E-58 | 3.23E-54 | 0.24 | 0.02 | 0.21 | 9.92 |
| 21 | Csf2ra | 2.46 | 1.76E-58 | 3.66E-54 | 0.60 | 0.14 | 0.46 | 4.26 |
| 21 | Rap1a | 1.65 | 2.55E-58 | 5.31E-54 | 0.99 | 0.72 | 0.27 | 1.38 |
| 21 | 1600014C10Rik | 3.05 | 6.84E-58 | 1.42E-53 | 0.54 | 0.11 | 0.43 | 4.76 |

**Supplementary Table 1. Marker genes for the original clusters of integrated clustering of E11.5 control and NotchGOF λ-junction cells.**

| | | | | | | | | |
|---|---|---|---|---|---|---|---|---|
| 21 | Lamp1 | 2.03 | 1.16E-57 | 2.42E-53 | 0.98 | 0.86 | 0.12 | 1.14 |
| 21 | Tmc8 | 5.55 | 2.46E-57 | 5.12E-53 | 0.12 | 0.01 | 0.11 | 19.83 |
| 21 | Runx3 | 4.27 | 2.71E-57 | 5.63E-53 | 0.20 | 0.02 | 0.18 | 11.53 |
| 21 | Lrch4 | 2.51 | 5.00E-57 | 1.04E-52 | 0.64 | 0.16 | 0.48 | 3.98 |
| 21 | Snx5 | 1.78 | 5.11E-57 | 1.06E-52 | 0.98 | 0.78 | 0.20 | 1.26 |
| 21 | Prps2 | 2.11 | 5.60E-57 | 1.16E-52 | 0.66 | 0.17 | 0.50 | 3.98 |
| 21 | Plin2 | 3.23 | 5.87E-57 | 1.22E-52 | 0.94 | 0.53 | 0.41 | 1.77 |
| 21 | Rnf130 | 1.76 | 6.05E-57 | 1.26E-52 | 0.97 | 0.56 | 0.41 | 1.73 |
| 21 | Slc8a1 | 2.45 | 9.91E-57 | 2.06E-52 | 0.56 | 0.12 | 0.44 | 4.62 |
| 21 | Twf2 | 1.97 | 1.91E-56 | 3.97E-52 | 0.90 | 0.42 | 0.47 | 2.12 |
| 21 | A330040F15Rik | 3.68 | 2.61E-56 | 5.43E-52 | 0.15 | 0.01 | 0.14 | 14.70 |
| 21 | Dusp27 | 4.16 | 2.85E-56 | 5.93E-52 | 0.13 | 0.01 | 0.12 | 18.00 |
| 21 | BC028528 | 2.42 | 4.03E-56 | 8.37E-52 | 0.43 | 0.08 | 0.36 | 5.79 |
| 21 | Spaca9 | 3.19 | 8.17E-56 | 1.70E-51 | 0.27 | 0.03 | 0.24 | 8.53 |
| 21 | Stx7 | 2.10 | 8.31E-56 | 1.73E-51 | 0.86 | 0.31 | 0.55 | 2.78 |
| 21 | Grb2 | 1.72 | 1.19E-55 | 2.48E-51 | 0.99 | 0.64 | 0.35 | 1.55 |
| 21 | Rnpep | 2.11 | 1.29E-55 | 2.69E-51 | 0.92 | 0.43 | 0.48 | 2.11 |
| 21 | Hacd4 | 2.65 | 1.36E-55 | 2.84E-51 | 0.75 | 0.25 | 0.50 | 3.03 |
| 21 | Flt3l | 4.26 | 2.53E-55 | 5.25E-51 | 0.16 | 0.01 | 0.15 | 13.42 |
| 21 | Atp6v0e | 1.51 | 2.75E-55 | 5.71E-51 | 0.98 | 0.72 | 0.26 | 1.37 |
| 21 | Rab8b | 2.16 | 8.22E-55 | 1.71E-50 | 0.77 | 0.24 | 0.53 | 3.20 |
| 21 | Cd47 | 2.36 | 8.97E-55 | 1.87E-50 | 0.92 | 0.44 | 0.48 | 2.09 |
| 21 | H2-Oa | 4.24 | 2.06E-54 | 4.27E-50 | 0.12 | 0.01 | 0.11 | 17.00 |
| 21 | Tpcn1 | 2.36 | 2.41E-54 | 5.02E-50 | 0.47 | 0.09 | 0.38 | 5.33 |
| 21 | Fgd4 | 2.50 | 3.13E-54 | 6.51E-50 | 0.46 | 0.09 | 0.37 | 5.25 |
| 21 | Igtp | 3.73 | 3.45E-54 | 7.18E-50 | 0.20 | 0.02 | 0.18 | 10.68 |
| 21 | Tap1 | 3.06 | 4.12E-54 | 8.57E-50 | 0.25 | 0.03 | 0.22 | 8.69 |
| 21 | Grk2 | 1.93 | 9.26E-54 | 1.93E-49 | 0.85 | 0.33 | 0.52 | 2.60 |
| 21 | Actr2 | 1.69 | 1.08E-53 | 2.24E-49 | 0.97 | 0.63 | 0.34 | 1.55 |
| 21 | Ptbp3 | 1.59 | 1.20E-53 | 2.49E-49 | 0.97 | 0.62 | 0.35 | 1.57 |
| 21 | S100a4 | 4.03 | 1.30E-53 | 2.71E-49 | 0.17 | 0.01 | 0.16 | 12.92 |
| 21 | Lipa | 2.57 | 1.95E-53 | 4.05E-49 | 0.70 | 0.22 | 0.48 | 3.24 |
| 21 | Sigirr | 2.88 | 2.07E-53 | 4.31E-49 | 0.24 | 0.03 | 0.21 | 9.15 |
| 21 | Ripk1 | 2.38 | 2.89E-53 | 6.01E-49 | 0.55 | 0.12 | 0.42 | 4.43 |
| 21 | Rab3il1 | 2.65 | 3.64E-53 | 7.58E-49 | 0.78 | 0.30 | 0.48 | 2.63 |
| 21 | Gamt | 2.41 | 3.82E-53 | 7.94E-49 | 0.47 | 0.09 | 0.38 | 5.21 |
| 21 | Pecam1 | 1.49 | 4.21E-53 | 8.75E-49 | 0.25 | 0.03 | 0.22 | 9.00 |
| 21 | Cd36 | 4.04 | 7.53E-53 | 1.57E-48 | 0.15 | 0.01 | 0.14 | 14.00 |
| 21 | Sh3bp1 | 1.94 | 1.23E-52 | 2.55E-48 | 0.80 | 0.29 | 0.52 | 2.81 |
| 21 | Asah1 | 2.40 | 1.39E-52 | 2.88E-48 | 0.86 | 0.37 | 0.49 | 2.34 |
| 21 | Parp4 | 3.30 | 1.41E-52 | 2.93E-48 | 0.20 | 0.02 | 0.18 | 10.89 |
| 21 | Samhd1 | 2.11 | 2.47E-52 | 5.13E-48 | 0.88 | 0.39 | 0.50 | 2.28 |
| 21 | Dram2 | 2.29 | 3.00E-52 | 6.23E-48 | 0.69 | 0.20 | 0.49 | 3.41 |
| 21 | Ssh2 | 2.29 | 3.59E-52 | 7.46E-48 | 0.73 | 0.23 | 0.51 | 3.22 |
| 21 | B3gnt8 | 3.53 | 3.76E-52 | 7.81E-48 | 0.18 | 0.02 | 0.16 | 11.67 |
| 21 | Tfpi | 1.94 | 4.26E-52 | 8.87E-48 | 0.52 | 0.11 | 0.41 | 4.76 |
| 21 | Abca3 | 2.64 | 4.44E-52 | 9.23E-48 | 0.38 | 0.06 | 0.32 | 6.00 |
| 21 | Tspan4 | 2.23 | 6.93E-52 | 1.44E-47 | 0.55 | 0.13 | 0.43 | 4.42 |
| 21 | Cap1 | 1.94 | 6.94E-52 | 1.44E-47 | 0.93 | 0.54 | 0.39 | 1.72 |
| 21 | Psme1 | 1.50 | 1.12E-51 | 2.33E-47 | 0.98 | 0.76 | 0.22 | 1.29 |
| 21 | Neu1 | 2.25 | 1.13E-51 | 2.34E-47 | 0.56 | 0.13 | 0.43 | 4.33 |
| 21 | Tpst2 | 2.12 | 1.87E-51 | 3.90E-47 | 0.74 | 0.23 | 0.51 | 3.17 |
| 21 | Abhd12 | 2.71 | 3.69E-51 | 7.67E-47 | 0.79 | 0.31 | 0.48 | 2.55 |
| 21 | Anxa4 | 2.64 | 4.14E-51 | 8.61E-47 | 0.50 | 0.11 | 0.39 | 4.60 |
| 21 | Slc25a5 | 1.26 | 8.51E-51 | 1.77E-46 | 1.00 | 0.93 | 0.07 | 1.07 |
| 21 | Snx2 | 2.34 | 1.24E-50 | 2.57E-46 | 0.93 | 0.58 | 0.35 | 1.61 |

**Supplementary Table 1. Marker genes for the original clusters of integrated clustering of E11.5 control and NotchGOF λ-junction cells.**

| | | | | | | | | |
|---|---|---|---|---|---|---|---|---|
| 21 | Icam2 | 1.07 | 1.63E-50 | 3.38E-46 | 0.20 | 0.02 | 0.18 | 10.15 |
| 21 | Mfng | 1.45 | 4.43E-50 | 9.21E-46 | 0.32 | 0.05 | 0.27 | 7.00 |
| 21 | Ntpcr | 2.36 | 1.14E-49 | 2.37E-45 | 0.71 | 0.23 | 0.48 | 3.14 |
| 21 | Cyp4f13 | 3.74 | 1.17E-49 | 2.44E-45 | 0.15 | 0.01 | 0.14 | 13.36 |
| 21 | Rplp1 | 0.58 | 1.64E-49 | 3.41E-45 | 1.00 | 1.00 | 0.00 | 1.00 |
| 21 | Coro7 | 2.15 | 3.89E-49 | 8.10E-45 | 0.49 | 0.10 | 0.39 | 4.71 |
| 21 | Arhgap27 | 2.66 | 3.91E-49 | 8.12E-45 | 0.25 | 0.03 | 0.22 | 8.13 |
| 21 | Hps3 | 2.55 | 4.17E-49 | 8.68E-45 | 0.42 | 0.08 | 0.34 | 5.19 |
| 21 | Ccdc69 | 3.64 | 4.99E-49 | 1.04E-44 | 0.11 | 0.01 | 0.10 | 17.50 |
| 21 | Pros1 | 2.14 | 7.98E-49 | 1.66E-44 | 0.58 | 0.14 | 0.44 | 4.06 |
| 21 | Tmem63a | 3.27 | 9.35E-49 | 1.94E-44 | 0.22 | 0.03 | 0.20 | 8.96 |
| 21 | Tdrd7 | 3.02 | 4.03E-48 | 8.39E-44 | 0.23 | 0.03 | 0.20 | 8.56 |
| 21 | Lgals8 | 2.32 | 4.92E-48 | 1.02E-43 | 0.43 | 0.09 | 0.35 | 5.11 |
| 21 | Irf9 | 2.49 | 7.21E-48 | 1.50E-43 | 0.37 | 0.06 | 0.31 | 5.80 |
| 21 | Rtn4rl1 | 4.49 | 7.76E-48 | 1.61E-43 | 0.11 | 0.01 | 0.11 | 16.00 |
| 21 | Tal1 | 2.29 | 8.40E-48 | 1.75E-43 | 0.32 | 0.05 | 0.27 | 6.70 |
| 21 | Gpx3 | 2.50 | 8.89E-48 | 1.85E-43 | 0.69 | 0.22 | 0.47 | 3.17 |
| 21 | Ncoa3 | 1.94 | 1.85E-47 | 3.86E-43 | 0.71 | 0.23 | 0.49 | 3.17 |
| 21 | Prex1 | 1.93 | 2.10E-47 | 4.36E-43 | 0.67 | 0.19 | 0.48 | 3.49 |
| 21 | Cib1 | 1.79 | 2.21E-47 | 4.59E-43 | 0.87 | 0.40 | 0.47 | 2.17 |
| 21 | Gab3 | 3.72 | 2.55E-47 | 5.30E-43 | 0.13 | 0.01 | 0.12 | 14.00 |
| 21 | Ctsb | 4.03 | 3.03E-47 | 6.30E-43 | 0.95 | 0.75 | 0.20 | 1.27 |
| 21 | Man2b2 | 3.52 | 4.63E-47 | 9.63E-43 | 0.17 | 0.02 | 0.15 | 11.20 |
| 21 | Capn1 | 2.84 | 5.50E-47 | 1.14E-42 | 0.32 | 0.05 | 0.27 | 6.43 |
| 21 | Arhgap17 | 1.97 | 5.82E-47 | 1.21E-42 | 0.86 | 0.37 | 0.49 | 2.33 |
| 21 | Ptp4a3 | 1.99 | 5.98E-47 | 1.24E-42 | 0.75 | 0.25 | 0.50 | 2.98 |
| 21 | Mir22hg | 3.94 | 1.06E-46 | 2.20E-42 | 0.15 | 0.01 | 0.14 | 12.25 |
| 21 | Nfkbiz | 2.77 | 1.19E-46 | 2.48E-42 | 0.38 | 0.07 | 0.31 | 5.40 |
| 21 | Itgb7 | 4.72 | 1.55E-46 | 3.23E-42 | 0.12 | 0.01 | 0.11 | 14.88 |
| 21 | Mapkapk3 | 2.82 | 2.46E-46 | 5.11E-42 | 0.27 | 0.04 | 0.23 | 7.39 |
| 21 | Mfsd12 | 2.98 | 3.51E-46 | 7.30E-42 | 0.34 | 0.06 | 0.28 | 5.89 |
| 21 | Zfp36l2 | 1.97 | 3.57E-46 | 7.42E-42 | 0.87 | 0.43 | 0.44 | 2.02 |
| 21 | Sgk3 | 2.01 | 3.58E-46 | 7.44E-42 | 0.58 | 0.15 | 0.43 | 3.92 |
| 21 | Kat2b | 2.60 | 3.65E-46 | 7.59E-42 | 0.30 | 0.05 | 0.26 | 6.69 |
| 21 | Vav3 | 2.48 | 4.23E-46 | 8.80E-42 | 0.34 | 0.06 | 0.29 | 6.02 |
| 21 | Slc6a6 | 1.95 | 4.30E-46 | 8.94E-42 | 0.86 | 0.35 | 0.51 | 2.45 |
| 21 | Pdxk | 2.11 | 6.65E-46 | 1.38E-41 | 0.50 | 0.11 | 0.39 | 4.41 |
| 21 | Scarb2 | 2.42 | 7.25E-46 | 1.51E-41 | 0.62 | 0.18 | 0.44 | 3.42 |
| 21 | Elovl1 | 1.95 | 1.07E-45 | 2.23E-41 | 0.73 | 0.26 | 0.48 | 2.86 |
| 21 | Eng | 1.61 | 1.12E-45 | 2.33E-41 | 0.29 | 0.04 | 0.25 | 7.00 |
| 21 | Serp1 | 1.88 | 1.30E-45 | 2.70E-41 | 0.97 | 0.76 | 0.22 | 1.29 |
| 21 | Fau | 0.53 | 1.35E-45 | 2.80E-41 | 1.00 | 1.00 | 0.00 | 1.00 |
| 21 | Shisa5 | 2.01 | 1.69E-45 | 3.52E-41 | 0.73 | 0.24 | 0.50 | 3.07 |
| 21 | Ttyh2 | 3.22 | 1.94E-45 | 4.03E-41 | 0.22 | 0.03 | 0.20 | 8.30 |
| 21 | Tmed5 | 1.63 | 3.75E-45 | 7.79E-41 | 0.93 | 0.48 | 0.45 | 1.92 |
| 21 | Aldoa | 1.09 | 4.77E-45 | 9.92E-41 | 0.99 | 0.95 | 0.04 | 1.04 |
| 21 | Ap1s2 | 2.28 | 5.46E-45 | 1.14E-40 | 0.53 | 0.13 | 0.40 | 3.99 |
| 21 | Slc38a7 | 2.64 | 6.06E-45 | 1.26E-40 | 0.41 | 0.08 | 0.33 | 5.04 |
| 21 | Syngr1 | 3.22 | 6.54E-45 | 1.36E-40 | 0.36 | 0.07 | 0.30 | 5.43 |
| 21 | Gm47664 | 3.55 | 7.40E-45 | 1.54E-40 | 0.15 | 0.01 | 0.14 | 12.25 |
| 21 | Picalm | 2.04 | 7.45E-45 | 1.55E-40 | 0.96 | 0.62 | 0.34 | 1.56 |
| 21 | Cxcl16 | 5.13 | 7.68E-45 | 1.60E-40 | 0.11 | 0.01 | 0.11 | 16.00 |
| 21 | Calm1 | 0.82 | 1.28E-44 | 2.67E-40 | 1.00 | 0.99 | 0.01 | 1.01 |
| 21 | Crlf2 | 2.23 | 1.75E-44 | 3.63E-40 | 0.37 | 0.07 | 0.30 | 5.54 |
| 21 | Ly96 | 2.54 | 1.81E-44 | 3.77E-40 | 0.34 | 0.06 | 0.28 | 5.89 |
| 21 | Dpp7 | 2.94 | 2.88E-44 | 5.98E-40 | 0.38 | 0.07 | 0.31 | 5.32 |

**Supplementary Table 1. Marker genes for the original clusters of integrated clustering of E11.5 control and NotchGOF λ-junction cells.**

| | | | | | | | | |
|---|---|---|---|---|---|---|---|---|
| 21 | Tubb6 | 2.07 | 3.48E-44 | 7.24E-40 | 0.49 | 0.11 | 0.38 | 4.34 |
| 21 | Tsc22d4 | 1.52 | 3.75E-44 | 7.80E-40 | 0.97 | 0.61 | 0.35 | 1.58 |
| 21 | Apold1 | 2.69 | 6.18E-44 | 1.29E-39 | 0.18 | 0.02 | 0.16 | 10.11 |
| 21 | Gna12 | 2.06 | 6.41E-44 | 1.33E-39 | 0.78 | 0.31 | 0.48 | 2.56 |
| 21 | Prr5 | 2.42 | 7.97E-44 | 1.66E-39 | 0.49 | 0.12 | 0.37 | 4.15 |
| 21 | Myl6 | 0.92 | 9.50E-44 | 1.98E-39 | 0.99 | 0.99 | 0.00 | 1.00 |
| 21 | Aup1 | 1.60 | 9.95E-44 | 2.07E-39 | 0.91 | 0.44 | 0.47 | 2.05 |
| 21 | Rab5if | 1.07 | 1.28E-43 | 2.66E-39 | 0.99 | 0.91 | 0.08 | 1.08 |
| 21 | Xaf1 | 1.53 | 1.48E-43 | 3.08E-39 | 0.34 | 0.06 | 0.28 | 5.89 |
| 21 | Hdac9 | 2.49 | 2.18E-43 | 4.53E-39 | 0.39 | 0.07 | 0.31 | 5.20 |
| 21 | Atp6v1b2 | 2.03 | 2.68E-43 | 5.58E-39 | 0.85 | 0.39 | 0.47 | 2.22 |
| 21 | Akna | 2.07 | 2.84E-43 | 5.90E-39 | 0.47 | 0.11 | 0.36 | 4.47 |
| 21 | 0610040J01Rik | 2.59 | 3.00E-43 | 6.24E-39 | 0.23 | 0.03 | 0.20 | 7.97 |
| 21 | Arhgap18 | 1.62 | 3.02E-43 | 6.28E-39 | 0.52 | 0.12 | 0.39 | 4.17 |
| 21 | S1pr1 | 2.88 | 3.84E-43 | 7.99E-39 | 0.34 | 0.06 | 0.28 | 5.53 |
| 21 | Snta1 | 2.61 | 3.86E-43 | 8.02E-39 | 0.31 | 0.05 | 0.26 | 6.16 |
| 21 | Naglu | 2.95 | 3.87E-43 | 8.06E-39 | 0.23 | 0.03 | 0.20 | 7.70 |
| 21 | Mgat4a | 2.21 | 4.06E-43 | 8.45E-39 | 0.43 | 0.09 | 0.34 | 4.72 |
| 21 | Cfl1 | 0.67 | 4.36E-43 | 9.07E-39 | 1.00 | 1.00 | 0.00 | 1.00 |
| 21 | Gm15417 | 2.82 | 4.58E-43 | 9.53E-39 | 0.21 | 0.03 | 0.19 | 8.40 |
| 21 | Akap13 | 1.65 | 5.23E-43 | 1.09E-38 | 0.92 | 0.50 | 0.42 | 1.83 |
| 21 | Isg15 | 4.57 | 6.90E-43 | 1.44E-38 | 0.13 | 0.01 | 0.12 | 12.09 |
| 21 | Gpi1 | 1.20 | 9.49E-43 | 1.97E-38 | 0.98 | 0.84 | 0.14 | 1.17 |
| 21 | Itga4 | 1.78 | 1.45E-42 | 3.01E-38 | 0.66 | 0.19 | 0.46 | 3.40 |
| 21 | Relt | 3.11 | 1.48E-42 | 3.07E-38 | 0.23 | 0.03 | 0.20 | 7.70 |
| 21 | Wdr1 | 1.73 | 1.61E-42 | 3.35E-38 | 0.90 | 0.55 | 0.36 | 1.66 |
| 21 | Ptpn1 | 1.72 | 1.82E-42 | 3.79E-38 | 0.92 | 0.52 | 0.40 | 1.76 |
| 21 | Prr13 | 1.68 | 2.32E-42 | 4.82E-38 | 0.90 | 0.40 | 0.50 | 2.24 |
| 21 | Irf2bp2 | 1.75 | 2.49E-42 | 5.18E-38 | 0.83 | 0.35 | 0.49 | 2.40 |
| 21 | Myo1c | 2.17 | 3.16E-42 | 6.58E-38 | 0.44 | 0.10 | 0.35 | 4.59 |
| 21 | Pnp | 1.55 | 3.46E-42 | 7.20E-38 | 0.97 | 0.66 | 0.31 | 1.47 |
| 21 | Rhob | 2.28 | 3.72E-42 | 7.73E-38 | 0.80 | 0.35 | 0.45 | 2.31 |
| 21 | Tspan32 | 2.68 | 4.96E-42 | 1.03E-37 | 0.26 | 0.04 | 0.22 | 7.00 |
| 21 | Sdf2l1 | 1.89 | 6.83E-42 | 1.42E-37 | 0.87 | 0.47 | 0.41 | 1.86 |
| 21 | Acp5 | 1.53 | 1.22E-41 | 2.54E-37 | 0.43 | 0.09 | 0.34 | 4.85 |
| 21 | Cd93 | 1.21 | 1.47E-41 | 3.06E-37 | 0.18 | 0.02 | 0.16 | 9.72 |
| 21 | Gltp | 1.79 | 1.55E-41 | 3.23E-37 | 0.87 | 0.39 | 0.47 | 2.20 |
| 21 | Emb | 2.06 | 1.80E-41 | 3.74E-37 | 0.84 | 0.41 | 0.43 | 2.07 |
| 21 | Htatip2 | 2.48 | 2.31E-41 | 4.80E-37 | 0.37 | 0.07 | 0.30 | 5.08 |
| 21 | Dcxr | 2.13 | 2.72E-41 | 5.66E-37 | 0.43 | 0.09 | 0.34 | 4.62 |
| 21 | Atf6 | 2.11 | 3.21E-41 | 6.68E-37 | 0.53 | 0.14 | 0.39 | 3.82 |
| 21 | Map3k5 | 2.72 | 3.56E-41 | 7.40E-37 | 0.25 | 0.03 | 0.21 | 7.21 |
| 21 | Cryzl2 | 2.16 | 4.13E-41 | 8.60E-37 | 0.45 | 0.10 | 0.35 | 4.39 |
| 21 | Cdc42 | 0.82 | 4.25E-41 | 8.84E-37 | 0.99 | 0.98 | 0.01 | 1.01 |
| 21 | Cd9 | 1.42 | 4.41E-41 | 9.18E-37 | 0.81 | 0.30 | 0.51 | 2.73 |
| 21 | Zfp710 | 2.19 | 5.27E-41 | 1.10E-36 | 0.60 | 0.18 | 0.42 | 3.36 |
| 21 | Map3k11 | 2.13 | 6.36E-41 | 1.32E-36 | 0.48 | 0.11 | 0.36 | 4.18 |
| 21 | Mfsd1 | 1.81 | 7.59E-41 | 1.58E-36 | 0.83 | 0.36 | 0.47 | 2.31 |
| 21 | Plekhb2 | 2.56 | 8.26E-41 | 1.72E-36 | 0.33 | 0.06 | 0.27 | 5.48 |
| 21 | Slc29a1 | 1.77 | 8.86E-41 | 1.84E-36 | 0.50 | 0.12 | 0.38 | 4.11 |
| 21 | Znrf2 | 2.02 | 9.51E-41 | 1.98E-36 | 0.61 | 0.18 | 0.43 | 3.32 |
| 21 | Ncoa4 | 1.99 | 1.79E-40 | 3.72E-36 | 0.77 | 0.32 | 0.45 | 2.44 |
| 21 | Ppm1h | 2.33 | 3.20E-40 | 6.65E-36 | 0.39 | 0.08 | 0.31 | 4.78 |
| 21 | Abca7 | 3.12 | 3.31E-40 | 6.88E-36 | 0.19 | 0.02 | 0.17 | 8.59 |
| 21 | Vamp5 | 1.90 | 3.43E-40 | 7.13E-36 | 0.34 | 0.06 | 0.27 | 5.42 |
| 21 | Irf1 | 2.33 | 3.86E-40 | 8.03E-36 | 0.26 | 0.04 | 0.22 | 6.82 |

**Supplementary Table 1. Marker genes for the original clusters of integrated clustering of E11.5 control and NotchGOF λ-junction cells.**

| | | | | | | | | |
|---|---|---|---|---|---|---|---|---|
| 21 | Il3ra | 2.41 | 4.83E-40 | 1.01E-35 | 0.30 | 0.05 | 0.25 | 6.02 |
| 21 | Mcur1 | 1.75 | 7.47E-40 | 1.55E-35 | 0.68 | 0.22 | 0.46 | 3.15 |
| 21 | Dab2 | 3.68 | 8.81E-40 | 1.83E-35 | 0.71 | 0.35 | 0.36 | 2.02 |
| 21 | Cltc | 1.73 | 1.29E-39 | 2.69E-35 | 0.94 | 0.75 | 0.20 | 1.26 |
| 21 | Esyt1 | 2.23 | 1.49E-39 | 3.10E-35 | 0.37 | 0.07 | 0.30 | 5.08 |
| 21 | Dtx4 | 2.82 | 5.33E-39 | 1.11E-34 | 0.26 | 0.04 | 0.22 | 6.48 |
| 21 | Rps29 | 0.40 | 6.97E-39 | 1.45E-34 | 1.00 | 1.00 | 0.00 | 1.00 |
| 21 | Rab29 | 2.71 | 9.16E-39 | 1.91E-34 | 0.29 | 0.05 | 0.24 | 5.88 |
| 21 | Slc16a10 | 2.16 | 1.14E-38 | 2.38E-34 | 0.45 | 0.11 | 0.34 | 4.19 |
| 21 | Fnbp1 | 1.84 | 1.22E-38 | 2.53E-34 | 0.68 | 0.24 | 0.44 | 2.89 |
| 21 | Prdx5 | 1.40 | 1.58E-38 | 3.29E-34 | 0.99 | 0.80 | 0.19 | 1.24 |
| 21 | Gns | 2.33 | 2.36E-38 | 4.90E-34 | 0.69 | 0.25 | 0.43 | 2.72 |
| 21 | Prdm1 | 3.79 | 3.09E-38 | 6.43E-34 | 0.13 | 0.01 | 0.12 | 11.45 |
| 21 | Ap5b1 | 3.65 | 3.98E-38 | 8.27E-34 | 0.11 | 0.01 | 0.10 | 14.00 |
| 21 | Slc9a3r1 | 1.24 | 4.01E-38 | 8.35E-34 | 0.75 | 0.24 | 0.51 | 3.12 |
| 21 | 2310040G24Rik | 2.33 | 4.21E-38 | 8.76E-34 | 0.23 | 0.03 | 0.20 | 7.22 |
| 21 | Rreb1 | 1.83 | 4.42E-38 | 9.19E-34 | 0.64 | 0.21 | 0.44 | 3.11 |
| 21 | Sdcbp | 1.81 | 5.67E-38 | 1.18E-33 | 0.94 | 0.69 | 0.25 | 1.36 |
| 21 | Cdc42se1 | 1.53 | 6.08E-38 | 1.27E-33 | 0.91 | 0.42 | 0.49 | 2.14 |
| 21 | Lamp2 | 1.74 | 6.85E-38 | 1.42E-33 | 0.92 | 0.56 | 0.35 | 1.62 |
| 21 | Cln6 | 2.04 | 7.19E-38 | 1.50E-33 | 0.43 | 0.10 | 0.33 | 4.40 |
| 21 | Tmem179b | 1.54 | 8.82E-38 | 1.83E-33 | 0.78 | 0.29 | 0.48 | 2.65 |
| 21 | Rbfa | 1.86 | 1.12E-37 | 2.34E-33 | 0.74 | 0.31 | 0.43 | 2.41 |
| 21 | Txnip | 1.83 | 1.52E-37 | 3.16E-33 | 0.80 | 0.34 | 0.46 | 2.34 |
| 21 | Sirt7 | 1.87 | 1.82E-37 | 3.79E-33 | 0.57 | 0.17 | 0.40 | 3.37 |
| 21 | Snx30 | 2.03 | 3.17E-37 | 6.58E-33 | 0.53 | 0.15 | 0.38 | 3.56 |
| 21 | Mindy2 | 1.96 | 3.61E-37 | 7.50E-33 | 0.58 | 0.17 | 0.41 | 3.33 |
| 21 | D1Ertd622e | 2.06 | 4.42E-37 | 9.19E-33 | 0.49 | 0.13 | 0.36 | 3.80 |
| 21 | Asl | 1.79 | 4.67E-37 | 9.71E-33 | 0.54 | 0.15 | 0.39 | 3.66 |
| 21 | Tmem176a | 2.23 | 4.87E-37 | 1.01E-32 | 0.83 | 0.41 | 0.42 | 2.04 |
| 21 | Exoc3l2 | 3.29 | 5.58E-37 | 1.16E-32 | 0.12 | 0.01 | 0.11 | 11.90 |
| 21 | Cox7a1 | 2.66 | 1.02E-36 | 2.11E-32 | 0.18 | 0.02 | 0.16 | 8.27 |
| 21 | Itpripl2 | 1.93 | 1.52E-36 | 3.16E-32 | 0.50 | 0.13 | 0.37 | 3.82 |
| 21 | Nfkb1 | 1.80 | 1.75E-36 | 3.64E-32 | 0.76 | 0.30 | 0.46 | 2.55 |
| 21 | Slc40a1 | 4.23 | 2.49E-36 | 5.18E-32 | 0.30 | 0.06 | 0.24 | 5.19 |
| 21 | Ppp2r5a | 1.52 | 3.48E-36 | 7.24E-32 | 0.79 | 0.32 | 0.47 | 2.48 |
| 21 | Irf2 | 1.72 | 4.74E-36 | 9.86E-32 | 0.69 | 0.25 | 0.44 | 2.72 |
| 21 | Acer3 | 1.97 | 5.24E-36 | 1.09E-31 | 0.55 | 0.16 | 0.38 | 3.39 |
| 21 | Rnaset2a | 1.49 | 5.28E-36 | 1.10E-31 | 0.92 | 0.52 | 0.40 | 1.77 |
| 21 | Agtrap | 2.01 | 7.49E-36 | 1.56E-31 | 0.52 | 0.15 | 0.37 | 3.54 |
| 21 | Atp6v0c | 1.26 | 7.89E-36 | 1.64E-31 | 1.00 | 0.96 | 0.04 | 1.04 |
| 21 | Tmem219 | 1.94 | 8.69E-36 | 1.81E-31 | 0.53 | 0.15 | 0.38 | 3.45 |
| 21 | Chst12 | 1.90 | 1.04E-35 | 2.16E-31 | 0.39 | 0.08 | 0.30 | 4.64 |
| 21 | Myh9 | 1.31 | 1.54E-35 | 3.20E-31 | 0.93 | 0.51 | 0.43 | 1.84 |
| 21 | Serf2 | 0.59 | 1.55E-35 | 3.23E-31 | 1.00 | 1.00 | 0.00 | 1.00 |
| 21 | Wdr26 | 1.37 | 1.59E-35 | 3.30E-31 | 0.94 | 0.71 | 0.23 | 1.33 |
| 21 | Frmd4b | 1.76 | 1.66E-35 | 3.45E-31 | 0.57 | 0.17 | 0.40 | 3.37 |
| 21 | Dgkz | 1.81 | 1.68E-35 | 3.49E-31 | 0.73 | 0.28 | 0.45 | 2.58 |
| 21 | Retreg1 | 2.17 | 2.35E-35 | 4.89E-31 | 0.40 | 0.09 | 0.31 | 4.34 |
| 21 | Bscl2 | 1.49 | 2.38E-35 | 4.95E-31 | 0.65 | 0.21 | 0.44 | 3.11 |
| 21 | Blvrb | 1.60 | 2.59E-35 | 5.38E-31 | 0.80 | 0.34 | 0.46 | 2.34 |
| 21 | Pxn | 1.70 | 2.90E-35 | 6.03E-31 | 0.60 | 0.18 | 0.42 | 3.27 |
| 21 | Engase | 2.50 | 3.11E-35 | 6.48E-31 | 0.26 | 0.04 | 0.22 | 6.02 |
| 21 | Fgd3 | 3.19 | 3.14E-35 | 6.53E-31 | 0.13 | 0.01 | 0.12 | 10.23 |
| 21 | Nrgn | 3.88 | 3.39E-35 | 7.04E-31 | 0.27 | 0.05 | 0.22 | 5.66 |
| 21 | Rnasek | 1.25 | 3.57E-35 | 7.43E-31 | 0.96 | 0.72 | 0.24 | 1.33 |

**Supplementary Table 1. Marker genes for the original clusters of integrated clustering of E11.5 control and NotchGOF λ-junction cells.**

| | | | | | | | | |
|---|---|---|---|---|---|---|---|---|
| 21 | Mroh1 | 2.23 | 3.63E-35 | 7.55E-31 | 0.36 | 0.08 | 0.28 | 4.70 |
| 21 | Cndp2 | 2.12 | 4.41E-35 | 9.18E-31 | 0.79 | 0.37 | 0.42 | 2.14 |
| 21 | Slc3a2 | 1.42 | 4.85E-35 | 1.01E-30 | 0.94 | 0.62 | 0.32 | 1.52 |
| 21 | Lfng | 2.58 | 5.29E-35 | 1.10E-30 | 0.16 | 0.02 | 0.14 | 8.94 |
| 21 | Ftl1 | 1.90 | 8.69E-35 | 1.81E-30 | 1.00 | 1.00 | 0.00 | 1.00 |
| 21 | Pkn1 | 1.52 | 1.05E-34 | 2.18E-30 | 0.83 | 0.35 | 0.48 | 2.35 |
| 21 | Snx3 | 1.03 | 1.06E-34 | 2.21E-30 | 0.99 | 0.89 | 0.10 | 1.11 |
| 21 | G6pdx | 1.79 | 1.14E-34 | 2.38E-30 | 0.66 | 0.24 | 0.42 | 2.74 |
| 21 | Git2 | 1.54 | 1.17E-34 | 2.43E-30 | 0.80 | 0.34 | 0.46 | 2.36 |
| 21 | Rgl1 | 2.16 | 1.18E-34 | 2.45E-30 | 0.51 | 0.15 | 0.36 | 3.38 |
| 21 | Zfp992 | 1.81 | 1.31E-34 | 2.72E-30 | 0.54 | 0.16 | 0.38 | 3.45 |
| 21 | Cyb561a3 | 2.42 | 1.73E-34 | 3.61E-30 | 0.30 | 0.06 | 0.24 | 5.19 |
| 21 | Nmi | 2.14 | 1.97E-34 | 4.10E-30 | 0.39 | 0.09 | 0.30 | 4.36 |
| 21 | Adam17 | 1.80 | 2.29E-34 | 4.77E-30 | 0.79 | 0.33 | 0.46 | 2.37 |
| 21 | Tmem86a | 3.01 | 2.63E-34 | 5.46E-30 | 0.41 | 0.10 | 0.30 | 3.90 |
| 21 | Synj1 | 2.44 | 2.64E-34 | 5.50E-30 | 0.43 | 0.11 | 0.32 | 3.99 |
| 21 | Hsd3b7 | 2.53 | 3.08E-34 | 6.40E-30 | 0.22 | 0.03 | 0.19 | 6.59 |
| 21 | Gabarap | 1.14 | 3.10E-34 | 6.45E-30 | 1.00 | 0.98 | 0.02 | 1.02 |
| 21 | Slc12a9 | 2.27 | 3.36E-34 | 6.98E-30 | 0.36 | 0.08 | 0.28 | 4.55 |
| 21 | Tmem144 | 2.72 | 4.62E-34 | 9.61E-30 | 0.27 | 0.05 | 0.22 | 5.66 |
| 21 | Rab43 | 2.28 | 5.48E-34 | 1.14E-29 | 0.32 | 0.07 | 0.26 | 4.95 |
| 21 | Adssl1 | 2.50 | 8.79E-34 | 1.83E-29 | 0.23 | 0.04 | 0.20 | 6.42 |
| 21 | Frrs1 | 1.79 | 9.19E-34 | 1.91E-29 | 0.63 | 0.22 | 0.41 | 2.86 |
| 21 | Slc6a13 | 3.69 | 1.10E-33 | 2.28E-29 | 0.15 | 0.02 | 0.13 | 9.19 |
| 21 | Mitf | 2.96 | 1.11E-33 | 2.31E-29 | 0.17 | 0.02 | 0.15 | 8.40 |
| 21 | Lrp10 | 1.68 | 1.13E-33 | 2.35E-29 | 0.57 | 0.17 | 0.40 | 3.33 |
| 21 | Necap2 | 1.61 | 1.16E-33 | 2.41E-29 | 0.82 | 0.39 | 0.43 | 2.09 |
| 21 | Cyfip1 | 1.71 | 2.43E-33 | 5.06E-29 | 0.94 | 0.70 | 0.24 | 1.35 |
| 21 | Tradd | 1.83 | 3.07E-33 | 6.38E-29 | 0.45 | 0.11 | 0.33 | 3.93 |
| 21 | Mob3c | 2.27 | 3.42E-33 | 7.12E-29 | 0.29 | 0.05 | 0.23 | 5.31 |
| 21 | Slc36a1 | 2.19 | 3.75E-33 | 7.80E-29 | 0.29 | 0.05 | 0.23 | 5.42 |
| 21 | Map4k1 | 2.15 | 4.58E-33 | 9.53E-29 | 0.26 | 0.04 | 0.22 | 5.89 |
| 21 | Rps12 | 0.44 | 4.96E-33 | 1.03E-28 | 1.00 | 1.00 | 0.00 | 1.00 |
| 21 | Ubl3 | 1.51 | 5.04E-33 | 1.05E-28 | 0.80 | 0.37 | 0.43 | 2.17 |
| 21 | Chmp2a | 1.19 | 7.83E-33 | 1.63E-28 | 0.95 | 0.66 | 0.29 | 1.44 |
| 21 | Ifngr2 | 1.70 | 8.75E-33 | 1.82E-28 | 0.78 | 0.37 | 0.41 | 2.11 |
| 21 | Pitpna | 1.21 | 8.93E-33 | 1.86E-28 | 0.94 | 0.70 | 0.23 | 1.33 |
| 21 | Plekho1 | 1.42 | 9.64E-33 | 2.00E-28 | 0.94 | 0.62 | 0.33 | 1.53 |
| 21 | Pag1 | 2.10 | 1.15E-32 | 2.39E-28 | 0.26 | 0.05 | 0.21 | 5.76 |
| 21 | Pla2g4a | 1.51 | 1.28E-32 | 2.66E-28 | 0.68 | 0.25 | 0.43 | 2.72 |
| 21 | Lrrfip1 | 1.54 | 2.13E-32 | 4.44E-28 | 0.76 | 0.34 | 0.42 | 2.25 |
| 21 | Dleu2 | 1.95 | 2.48E-32 | 5.17E-28 | 0.65 | 0.25 | 0.40 | 2.58 |
| 21 | Fosl2 | 2.53 | 2.59E-32 | 5.38E-28 | 0.20 | 0.03 | 0.17 | 6.77 |
| 21 | Rac1 | 0.92 | 2.70E-32 | 5.62E-28 | 0.99 | 0.90 | 0.08 | 1.09 |
| 21 | Arhgdia | 0.96 | 3.32E-32 | 6.90E-28 | 0.99 | 0.88 | 0.11 | 1.12 |
| 21 | Wdr81 | 2.48 | 4.05E-32 | 8.43E-28 | 0.27 | 0.05 | 0.22 | 5.43 |
| 21 | Pqlc2 | 2.47 | 5.27E-32 | 1.10E-27 | 0.41 | 0.10 | 0.30 | 3.94 |
| 21 | Pnpla2 | 1.92 | 5.50E-32 | 1.14E-27 | 0.38 | 0.09 | 0.29 | 4.30 |
| 21 | Ndufv3 | 0.99 | 7.07E-32 | 1.47E-27 | 0.98 | 0.84 | 0.14 | 1.17 |
| 21 | Vps13c | 1.84 | 7.28E-32 | 1.51E-27 | 0.58 | 0.20 | 0.39 | 2.97 |
| 21 | H2-T23 | 2.44 | 7.36E-32 | 1.53E-27 | 0.28 | 0.05 | 0.23 | 5.28 |
| 21 | Adgre5 | 2.09 | 1.04E-31 | 2.16E-27 | 0.46 | 0.13 | 0.33 | 3.64 |
| 21 | Slc7a1 | 2.22 | 1.27E-31 | 2.64E-27 | 0.34 | 0.08 | 0.27 | 4.40 |
| 21 | Ap1b1 | 1.80 | 1.49E-31 | 3.10E-27 | 0.64 | 0.24 | 0.40 | 2.64 |
| 21 | Lamtor4 | 1.30 | 1.62E-31 | 3.38E-27 | 0.93 | 0.66 | 0.27 | 1.41 |
| 21 | Rap1gap2 | 2.37 | 1.71E-31 | 3.56E-27 | 0.14 | 0.02 | 0.13 | 9.33 |

**Supplementary Table 1. Marker genes for the original clusters of integrated clustering of E11.5 control and NotchGOF λ-junction cells.**

| | | | | | | | | |
|---|---|---|---|---|---|---|---|---|
| 21 | M6pr | 1.50 | 1.74E-31 | 3.61E-27 | 0.85 | 0.44 | 0.41 | 1.92 |
| 21 | Dennd4a | 1.94 | 1.95E-31 | 4.06E-27 | 0.36 | 0.08 | 0.28 | 4.41 |
| 21 | Nhlrc3 | 1.87 | 2.14E-31 | 4.45E-27 | 0.46 | 0.12 | 0.33 | 3.73 |
| 21 | Dazap2 | 1.30 | 2.45E-31 | 5.09E-27 | 0.91 | 0.57 | 0.34 | 1.59 |
| 21 | Stxbp5 | 1.90 | 2.45E-31 | 5.10E-27 | 0.35 | 0.08 | 0.27 | 4.49 |
| 21 | Vasp | 1.34 | 2.77E-31 | 5.77E-27 | 0.88 | 0.48 | 0.40 | 1.83 |
| 21 | Tmbim1 | 2.04 | 3.09E-31 | 6.42E-27 | 0.39 | 0.10 | 0.30 | 4.08 |
| 21 | Cryl1 | 1.85 | 3.11E-31 | 6.46E-27 | 0.50 | 0.15 | 0.35 | 3.34 |
| 21 | Acox3 | 2.07 | 3.16E-31 | 6.58E-27 | 0.33 | 0.07 | 0.26 | 4.63 |
| 21 | Tmem170b | 2.18 | 3.22E-31 | 6.70E-27 | 0.31 | 0.06 | 0.24 | 4.81 |
| 21 | Cd34 | 1.80 | 3.74E-31 | 7.78E-27 | 0.46 | 0.13 | 0.32 | 3.47 |
| 21 | Cdkn2c | 2.79 | 3.99E-31 | 8.31E-27 | 0.23 | 0.04 | 0.19 | 5.92 |
| 21 | Cln3 | 2.11 | 5.03E-31 | 1.05E-26 | 0.48 | 0.14 | 0.34 | 3.43 |
| 21 | Nrp1 | 1.23 | 7.12E-31 | 1.48E-26 | 0.62 | 0.21 | 0.41 | 2.99 |
| 21 | Rhbdf2 | 2.63 | 9.38E-31 | 1.95E-26 | 0.20 | 0.03 | 0.17 | 6.55 |
| 21 | Sh3glb1 | 1.09 | 1.47E-30 | 3.06E-26 | 0.97 | 0.84 | 0.13 | 1.16 |
| 21 | Gdi2 | 0.80 | 1.49E-30 | 3.09E-26 | 0.99 | 0.95 | 0.04 | 1.04 |
| 21 | Glb1 | 1.56 | 1.68E-30 | 3.50E-26 | 0.54 | 0.16 | 0.38 | 3.32 |
| 21 | Atp6v0d1 | 1.34 | 2.04E-30 | 4.25E-26 | 0.90 | 0.54 | 0.37 | 1.69 |
| 21 | Rnf141 | 2.16 | 2.12E-30 | 4.41E-26 | 0.44 | 0.12 | 0.32 | 3.61 |
| 21 | Edem2 | 1.62 | 2.62E-30 | 5.44E-26 | 0.58 | 0.19 | 0.39 | 3.02 |
| 21 | Sesn1 | 1.89 | 3.22E-30 | 6.70E-26 | 0.61 | 0.23 | 0.38 | 2.70 |
| 21 | Ftl1-ps1 | 1.92 | 3.46E-30 | 7.20E-26 | 0.78 | 0.41 | 0.37 | 1.91 |
| 21 | Slc12a6 | 1.91 | 3.79E-30 | 7.88E-26 | 0.46 | 0.13 | 0.33 | 3.47 |
| 21 | Ctla2a | 1.30 | 4.55E-30 | 9.46E-26 | 0.15 | 0.02 | 0.14 | 8.11 |
| 21 | Pdcd6ip | 1.30 | 5.96E-30 | 1.24E-25 | 0.87 | 0.48 | 0.38 | 1.79 |
| 21 | Capzb | 0.86 | 6.07E-30 | 1.26E-25 | 0.99 | 0.92 | 0.07 | 1.08 |
| 21 | Rnf13 | 1.68 | 6.54E-30 | 1.36E-25 | 0.64 | 0.24 | 0.40 | 2.67 |
| 21 | Pcgf5 | 1.75 | 7.38E-30 | 1.53E-25 | 0.36 | 0.09 | 0.28 | 4.23 |
| 21 | Rsu1 | 1.38 | 9.10E-30 | 1.89E-25 | 0.89 | 0.48 | 0.41 | 1.84 |
| 21 | Wasf2 | 1.22 | 9.62E-30 | 2.00E-25 | 0.96 | 0.71 | 0.25 | 1.35 |
| 21 | Znfx1 | 2.22 | 9.66E-30 | 2.01E-25 | 0.29 | 0.06 | 0.23 | 4.86 |
| 21 | Capza1 | 1.14 | 1.20E-29 | 2.49E-25 | 0.92 | 0.65 | 0.28 | 1.43 |
| 21 | Heatr5a | 1.65 | 1.30E-29 | 2.69E-25 | 0.56 | 0.19 | 0.37 | 3.01 |
| 21 | Slc29a3 | 1.89 | 1.50E-29 | 3.12E-25 | 0.39 | 0.10 | 0.29 | 3.92 |
| 21 | Mapk3 | 1.38 | 1.94E-29 | 4.04E-25 | 0.83 | 0.45 | 0.38 | 1.86 |
| 21 | Diaph2 | 1.72 | 2.47E-29 | 5.14E-25 | 0.50 | 0.15 | 0.35 | 3.29 |
| 21 | H6pd | 2.47 | 3.19E-29 | 6.63E-25 | 0.20 | 0.03 | 0.17 | 6.53 |
| 21 | Sptlc2 | 1.57 | 3.60E-29 | 7.49E-25 | 0.76 | 0.33 | 0.44 | 2.33 |
| 21 | Emc9 | 2.35 | 3.67E-29 | 7.64E-25 | 0.25 | 0.05 | 0.21 | 5.36 |
| 21 | Glmp | 1.25 | 3.70E-29 | 7.70E-25 | 0.91 | 0.47 | 0.44 | 1.95 |
| 21 | Arrdc1 | 1.52 | 4.02E-29 | 8.36E-25 | 0.42 | 0.11 | 0.31 | 3.82 |
| 21 | Tfe3 | 1.80 | 4.72E-29 | 9.82E-25 | 0.54 | 0.18 | 0.36 | 2.99 |
| 21 | Nudt16 | 1.90 | 8.54E-29 | 1.78E-24 | 0.36 | 0.09 | 0.27 | 4.15 |
| 21 | Mob1a | 1.35 | 1.11E-28 | 2.31E-24 | 0.85 | 0.45 | 0.39 | 1.87 |
| 21 | Tmem192 | 1.70 | 1.13E-28 | 2.34E-24 | 0.55 | 0.18 | 0.36 | 2.98 |
| 21 | Nfe2l2 | 1.58 | 1.13E-28 | 2.36E-24 | 0.84 | 0.45 | 0.39 | 1.88 |
| 21 | Vps18 | 1.71 | 1.23E-28 | 2.56E-24 | 0.46 | 0.14 | 0.33 | 3.40 |
| 21 | Ccdc82 | 1.74 | 1.45E-28 | 3.02E-24 | 0.43 | 0.12 | 0.31 | 3.65 |
| 21 | Stk4 | 1.51 | 1.97E-28 | 4.10E-24 | 0.76 | 0.36 | 0.40 | 2.10 |
| 21 | Ostm1 | 2.14 | 2.10E-28 | 4.36E-24 | 0.32 | 0.07 | 0.24 | 4.38 |
| 21 | Npc2 | 0.95 | 3.15E-28 | 6.55E-24 | 0.99 | 0.93 | 0.06 | 1.07 |
| 21 | Vps8 | 2.13 | 3.38E-28 | 7.03E-24 | 0.27 | 0.06 | 0.22 | 4.96 |
| 21 | Calhm6 | 2.86 | 4.61E-28 | 9.59E-24 | 0.17 | 0.02 | 0.14 | 7.00 |
| 21 | Cenpx | 1.00 | 4.89E-28 | 1.02E-23 | 0.95 | 0.79 | 0.17 | 1.21 |
| 21 | Scarb1 | 1.75 | 4.94E-28 | 1.03E-23 | 0.49 | 0.16 | 0.33 | 3.12 |

**Supplementary Table 1. Marker genes for the original clusters of integrated clustering of E11.5 control and NotchGOF λ-junction cells.**

| | | | | | | | | |
|---|---|---|---|---|---|---|---|---|
| 21 | Tuba1c | 1.25 | 5.49E-28 | 1.14E-23 | 0.84 | 0.44 | 0.40 | 1.92 |
| 21 | Camk1 | 1.57 | 6.75E-28 | 1.40E-23 | 0.82 | 0.42 | 0.40 | 1.94 |
| 21 | Dhx58 | 3.20 | 9.26E-28 | 1.93E-23 | 0.11 | 0.01 | 0.10 | 10.50 |
| 21 | Ost4 | 0.65 | 1.32E-27 | 2.74E-23 | 0.98 | 0.95 | 0.03 | 1.03 |
| 21 | Aff1 | 2.06 | 1.33E-27 | 2.77E-23 | 0.31 | 0.07 | 0.24 | 4.40 |
| 21 | Nlrc3 | 2.78 | 1.47E-27 | 3.06E-23 | 0.15 | 0.02 | 0.13 | 7.74 |
| 21 | Cd302 | 1.59 | 1.50E-27 | 3.12E-23 | 0.67 | 0.29 | 0.39 | 2.35 |
| 21 | Gnb2 | 0.65 | 1.60E-27 | 3.32E-23 | 0.99 | 0.96 | 0.02 | 1.02 |
| 21 | Gab2 | 2.35 | 1.72E-27 | 3.58E-23 | 0.24 | 0.05 | 0.19 | 5.29 |
| 21 | Coa5 | 1.40 | 1.79E-27 | 3.72E-23 | 0.68 | 0.27 | 0.41 | 2.55 |
| 21 | Fli1 | 1.21 | 1.80E-27 | 3.73E-23 | 0.82 | 0.37 | 0.45 | 2.21 |
| 21 | Dnase2a | 1.57 | 1.82E-27 | 3.78E-23 | 0.60 | 0.22 | 0.38 | 2.77 |
| 21 | Stat5a | 2.56 | 2.05E-27 | 4.27E-23 | 0.18 | 0.03 | 0.15 | 6.50 |
| 21 | Eps15 | 1.50 | 2.10E-27 | 4.37E-23 | 0.70 | 0.31 | 0.39 | 2.29 |
| 21 | Atp6v1f | 0.79 | 2.22E-27 | 4.63E-23 | 1.00 | 0.90 | 0.10 | 1.11 |
| 21 | Rnf135 | 2.24 | 2.49E-27 | 5.18E-23 | 0.21 | 0.04 | 0.17 | 5.83 |
| 21 | Calhm2 | 2.01 | 2.62E-27 | 5.46E-23 | 0.21 | 0.04 | 0.18 | 6.00 |
| 21 | Stim1 | 1.49 | 2.65E-27 | 5.52E-23 | 0.48 | 0.14 | 0.33 | 3.35 |
| 21 | Lamtor1 | 0.98 | 4.02E-27 | 8.37E-23 | 0.97 | 0.86 | 0.10 | 1.12 |
| 21 | Gpd1l | 1.74 | 5.18E-27 | 1.08E-22 | 0.32 | 0.07 | 0.24 | 4.38 |
| 21 | Atp6ap2 | 1.33 | 5.40E-27 | 1.12E-22 | 0.84 | 0.43 | 0.41 | 1.97 |
| 21 | 5033406O09Rik | 3.01 | 5.68E-27 | 1.18E-22 | 0.11 | 0.01 | 0.10 | 9.33 |
| 21 | Tubgcp5 | 1.71 | 5.83E-27 | 1.21E-22 | 0.75 | 0.34 | 0.40 | 2.17 |
| 21 | Fam174a | 1.44 | 5.87E-27 | 1.22E-22 | 0.72 | 0.31 | 0.41 | 2.35 |
| 21 | St6galnac4 | 1.56 | 5.94E-27 | 1.23E-22 | 0.56 | 0.20 | 0.36 | 2.85 |
| 21 | Zfp120 | 1.85 | 5.96E-27 | 1.24E-22 | 0.45 | 0.13 | 0.32 | 3.37 |
| 21 | Pgd | 1.26 | 6.26E-27 | 1.30E-22 | 0.89 | 0.59 | 0.30 | 1.52 |
| 21 | Rp2 | 1.93 | 7.08E-27 | 1.47E-22 | 0.50 | 0.17 | 0.33 | 2.91 |
| 21 | Tcn2 | 1.94 | 7.22E-27 | 1.50E-22 | 0.51 | 0.18 | 0.33 | 2.90 |
| 21 | Aagab | 1.56 | 8.50E-27 | 1.77E-22 | 0.53 | 0.17 | 0.36 | 3.05 |
| 21 | Ptgr2 | 1.75 | 1.04E-26 | 2.16E-22 | 0.47 | 0.15 | 0.32 | 3.17 |
| 21 | Matk | 2.45 | 1.21E-26 | 2.52E-22 | 0.18 | 0.03 | 0.15 | 6.48 |
| 21 | Glud1 | 1.03 | 1.44E-26 | 3.00E-22 | 0.92 | 0.65 | 0.27 | 1.41 |
| 21 | Bid | 1.75 | 1.64E-26 | 3.41E-22 | 0.42 | 0.12 | 0.30 | 3.50 |
| 21 | Cmtm6 | 1.66 | 1.65E-26 | 3.44E-22 | 0.44 | 0.13 | 0.31 | 3.39 |
| 21 | Nek7 | 1.50 | 1.72E-26 | 3.58E-22 | 0.67 | 0.29 | 0.38 | 2.28 |
| 21 | Il15 | 2.57 | 1.89E-26 | 3.93E-22 | 0.16 | 0.02 | 0.14 | 7.00 |
| 21 | Rab5c | 1.23 | 1.96E-26 | 4.07E-22 | 0.94 | 0.65 | 0.30 | 1.46 |
| 21 | Pld2 | 2.69 | 2.14E-26 | 4.46E-22 | 0.15 | 0.02 | 0.13 | 7.33 |
| 21 | Pea15a | 1.67 | 2.40E-26 | 4.99E-22 | 0.60 | 0.23 | 0.37 | 2.60 |
| 21 | Ppp1r3b | 2.23 | 2.42E-26 | 5.04E-22 | 0.22 | 0.04 | 0.18 | 5.46 |
| 21 | Cfp | 2.48 | 2.46E-26 | 5.11E-22 | 0.23 | 0.04 | 0.19 | 5.25 |
| 21 | Akr1b10 | 1.66 | 2.93E-26 | 6.09E-22 | 0.41 | 0.12 | 0.30 | 3.56 |
| 21 | Ada | 1.82 | 2.98E-26 | 6.20E-22 | 0.33 | 0.08 | 0.25 | 4.11 |
| 21 | Arhgef1 | 1.42 | 3.25E-26 | 6.76E-22 | 0.66 | 0.27 | 0.39 | 2.47 |
| 21 | Comtd1 | 1.65 | 3.49E-26 | 7.26E-22 | 0.46 | 0.14 | 0.32 | 3.25 |
| 21 | Mthfs | 1.88 | 3.57E-26 | 7.43E-22 | 0.46 | 0.14 | 0.32 | 3.30 |
| 21 | Acsl1 | 1.70 | 3.68E-26 | 7.64E-22 | 0.39 | 0.10 | 0.28 | 3.74 |
| 21 | Rpl10a | 0.37 | 3.96E-26 | 8.23E-22 | 0.99 | 1.00 | -0.01 | 0.99 |
| 21 | Gnpda1 | 2.00 | 4.87E-26 | 1.01E-21 | 0.52 | 0.18 | 0.33 | 2.83 |
| 21 | Tor1aip1 | 1.35 | 5.09E-26 | 1.06E-21 | 0.85 | 0.44 | 0.41 | 1.94 |
| 21 | Psme2 | 1.01 | 5.20E-26 | 1.08E-21 | 0.96 | 0.76 | 0.20 | 1.26 |
| 21 | Washc4 | 1.47 | 6.13E-26 | 1.28E-21 | 0.72 | 0.31 | 0.41 | 2.29 |
| 21 | Atp6ap1 | 1.28 | 7.03E-26 | 1.46E-21 | 0.76 | 0.35 | 0.41 | 2.15 |
| 21 | Cox17 | 0.88 | 7.38E-26 | 1.53E-21 | 0.98 | 0.77 | 0.21 | 1.28 |
| 21 | Gla | 2.23 | 8.53E-26 | 1.77E-21 | 0.38 | 0.10 | 0.28 | 3.71 |

**Supplementary Table 1. Marker genes for the original clusters of integrated clustering of E11.5 control and NotchGOF λ-junction cells.**

| | | | | | | | | |
|---|---|---|---|---|---|---|---|---|
| 21 | Eif2ak2 | 1.41 | 8.87E-26 | 1.85E-21 | 0.67 | 0.28 | 0.39 | 2.42 |
| 21 | Cnp | 1.61 | 1.08E-25 | 2.25E-21 | 0.48 | 0.16 | 0.33 | 3.12 |
| 21 | Pfkp | 2.04 | 1.13E-25 | 2.35E-21 | 0.25 | 0.05 | 0.20 | 4.94 |
| 21 | Il13ra1 | 2.31 | 1.21E-25 | 2.52E-21 | 0.23 | 0.05 | 0.19 | 5.13 |
| 21 | Slc35c1 | 2.20 | 1.24E-25 | 2.58E-21 | 0.22 | 0.04 | 0.18 | 5.43 |
| 21 | Rpl35a | 0.31 | 1.40E-25 | 2.92E-21 | 0.99 | 1.00 | -0.01 | 0.99 |
| 21 | Smap2 | 1.31 | 1.50E-25 | 3.12E-21 | 0.77 | 0.37 | 0.40 | 2.07 |
| 21 | Csk | 1.15 | 1.51E-25 | 3.15E-21 | 0.87 | 0.54 | 0.33 | 1.61 |
| 21 | Mcl1 | 0.96 | 1.65E-25 | 3.44E-21 | 0.99 | 0.82 | 0.17 | 1.21 |
| 21 | Dhrs3 | 1.66 | 1.69E-25 | 3.52E-21 | 0.66 | 0.29 | 0.36 | 2.23 |
| 21 | Tpm2 | 1.99 | 2.32E-25 | 4.82E-21 | 0.78 | 0.44 | 0.35 | 1.80 |
| 21 | Swap70 | 1.34 | 2.33E-25 | 4.84E-21 | 0.72 | 0.33 | 0.39 | 2.20 |
| 21 | Marchf2 | 1.25 | 2.41E-25 | 5.01E-21 | 0.59 | 0.21 | 0.37 | 2.76 |
| 21 | Colgalt1 | 1.22 | 2.55E-25 | 5.31E-21 | 0.87 | 0.47 | 0.39 | 1.83 |
| 21 | Aph1c | 2.40 | 2.57E-25 | 5.35E-21 | 0.21 | 0.04 | 0.17 | 5.53 |
| 21 | Zfp979 | 2.24 | 2.61E-25 | 5.44E-21 | 0.20 | 0.03 | 0.16 | 5.76 |
| 21 | Nr3c1 | 1.59 | 2.74E-25 | 5.69E-21 | 0.49 | 0.16 | 0.33 | 3.02 |
| 21 | Ormdl2 | 1.22 | 3.04E-25 | 6.32E-21 | 0.75 | 0.35 | 0.40 | 2.14 |
| 21 | Vps33a | 1.64 | 3.37E-25 | 7.01E-21 | 0.45 | 0.14 | 0.31 | 3.25 |
| 21 | Ints6l | 2.00 | 3.43E-25 | 7.13E-21 | 0.48 | 0.17 | 0.31 | 2.84 |
| 21 | Mef2a | 1.46 | 3.58E-25 | 7.44E-21 | 0.78 | 0.39 | 0.39 | 2.01 |
| 21 | Cpt1a | 1.93 | 3.79E-25 | 7.89E-21 | 0.36 | 0.10 | 0.26 | 3.64 |
| 21 | Acat1 | 1.20 | 4.45E-25 | 9.26E-21 | 0.85 | 0.46 | 0.38 | 1.82 |
| 21 | Ubtd1 | 1.56 | 4.63E-25 | 9.63E-21 | 0.47 | 0.15 | 0.32 | 3.15 |
| 21 | Gdpd3 | 3.98 | 5.07E-25 | 1.05E-20 | 0.12 | 0.01 | 0.11 | 8.50 |
| 21 | Taldo1 | 0.84 | 5.18E-25 | 1.08E-20 | 0.96 | 0.89 | 0.07 | 1.08 |
| 21 | Dnase1l1 | 2.37 | 6.82E-25 | 1.42E-20 | 0.23 | 0.05 | 0.19 | 5.02 |
| 21 | Prcp | 1.49 | 6.86E-25 | 1.43E-20 | 0.64 | 0.26 | 0.38 | 2.48 |
| 21 | Dennd4b | 2.33 | 7.32E-25 | 1.52E-20 | 0.24 | 0.05 | 0.19 | 4.86 |
| 21 | Cdkn1a | 0.82 | 8.76E-25 | 1.82E-20 | 0.40 | 0.11 | 0.29 | 3.59 |
| 21 | BC005537 | 1.38 | 9.10E-25 | 1.89E-20 | 0.82 | 0.50 | 0.32 | 1.65 |
| 21 | Fuca1 | 1.40 | 9.12E-25 | 1.90E-20 | 0.75 | 0.35 | 0.39 | 2.11 |
| 21 | Slc2a5 | 2.81 | 9.66E-25 | 2.01E-20 | 0.13 | 0.02 | 0.11 | 7.88 |
| 21 | Hpse | 2.78 | 9.91E-25 | 2.06E-20 | 0.13 | 0.02 | 0.12 | 7.82 |
| 21 | Litaf | 1.67 | 1.08E-24 | 2.24E-20 | 0.85 | 0.50 | 0.34 | 1.68 |
| 21 | Cltb | 1.52 | 1.24E-24 | 2.58E-20 | 0.52 | 0.18 | 0.34 | 2.88 |
| 21 | B4galt1 | 1.45 | 1.26E-24 | 2.61E-20 | 0.60 | 0.23 | 0.37 | 2.65 |
| 21 | H2-DMa | 2.20 | 1.34E-24 | 2.78E-20 | 0.25 | 0.05 | 0.20 | 4.67 |
| 21 | Rhoa | 0.75 | 1.66E-24 | 3.45E-20 | 1.00 | 0.97 | 0.03 | 1.04 |
| 21 | Unc13d | 2.33 | 1.90E-24 | 3.96E-20 | 0.18 | 0.03 | 0.15 | 6.03 |
| 21 | Fbxo6 | 2.31 | 1.94E-24 | 4.03E-20 | 0.27 | 0.06 | 0.21 | 4.33 |
| 21 | Dnm2 | 1.27 | 2.15E-24 | 4.47E-20 | 0.80 | 0.39 | 0.41 | 2.06 |
| 21 | Kif21b | 1.54 | 2.34E-24 | 4.87E-20 | 0.49 | 0.16 | 0.33 | 3.02 |
| 21 | Lpcat3 | 1.31 | 2.87E-24 | 5.97E-20 | 0.71 | 0.31 | 0.40 | 2.31 |
| 21 | Jmjd1c | 1.32 | 3.08E-24 | 6.40E-20 | 0.85 | 0.48 | 0.36 | 1.76 |
| 21 | Osbpl11 | 1.54 | 3.40E-24 | 7.07E-20 | 0.64 | 0.28 | 0.36 | 2.30 |
| 21 | Nlrx1 | 2.63 | 3.59E-24 | 7.47E-20 | 0.12 | 0.01 | 0.11 | 8.50 |
| 21 | Rnaset2b | 1.42 | 3.80E-24 | 7.90E-20 | 0.57 | 0.21 | 0.36 | 2.68 |
| 21 | Ptrhd1 | 1.28 | 4.20E-24 | 8.73E-20 | 0.76 | 0.36 | 0.40 | 2.10 |
| 21 | Ccdc12 | 0.90 | 4.27E-24 | 8.87E-20 | 0.94 | 0.71 | 0.23 | 1.33 |
| 21 | Arf3 | 1.34 | 5.07E-24 | 1.06E-19 | 0.71 | 0.31 | 0.40 | 2.29 |
| 21 | Mcrip2 | 1.93 | 5.22E-24 | 1.09E-19 | 0.29 | 0.07 | 0.22 | 4.16 |
| 21 | Etfb | 0.79 | 5.26E-24 | 1.09E-19 | 0.98 | 0.89 | 0.09 | 1.10 |
| 21 | Shfl | 2.73 | 5.66E-24 | 1.18E-19 | 0.12 | 0.02 | 0.10 | 7.93 |
| 21 | Slc45a4 | 1.74 | 6.02E-24 | 1.25E-19 | 0.43 | 0.13 | 0.29 | 3.19 |
| 21 | Fcho1 | 3.07 | 6.11E-24 | 1.27E-19 | 0.13 | 0.02 | 0.12 | 7.39 |

**Supplementary Table 1. Marker genes for the original clusters of integrated clustering of E11.5 control and NotchGOF λ-junction cells.**

| | | | | | | | | |
|---|---|---|---|---|---|---|---|---|
| 21 | Fuca2 | 1.42 | 6.56E-24 | 1.36E-19 | 0.59 | 0.24 | 0.36 | 2.53 |
| 21 | Tmbim4 | 1.18 | 6.91E-24 | 1.44E-19 | 0.83 | 0.49 | 0.34 | 1.68 |
| 21 | Iqgap2 | 1.33 | 7.05E-24 | 1.47E-19 | 0.73 | 0.36 | 0.37 | 2.02 |
| 21 | Ppp6r1 | 1.37 | 7.90E-24 | 1.64E-19 | 0.62 | 0.25 | 0.37 | 2.46 |
| 21 | Jdp2 | 2.00 | 8.00E-24 | 1.66E-19 | 0.20 | 0.04 | 0.17 | 5.49 |
| 21 | Wbp2 | 1.32 | 1.02E-23 | 2.12E-19 | 0.65 | 0.28 | 0.37 | 2.36 |
| 21 | Piezo1 | 1.66 | 1.25E-23 | 2.59E-19 | 0.41 | 0.12 | 0.28 | 3.30 |
| 21 | Tpm3 | 0.59 | 1.35E-23 | 2.80E-19 | 0.99 | 0.96 | 0.04 | 1.04 |
| 21 | Dgkd | 1.59 | 1.37E-23 | 2.85E-19 | 0.57 | 0.24 | 0.34 | 2.44 |
| 21 | Rpl36al | 0.44 | 1.47E-23 | 3.05E-19 | 0.99 | 1.00 | 0.00 | 1.00 |
| 21 | AU020206 | 1.39 | 1.49E-23 | 3.10E-19 | 0.59 | 0.22 | 0.37 | 2.66 |
| 21 | 5031439G07Rik | 1.53 | 1.51E-23 | 3.13E-19 | 0.59 | 0.25 | 0.34 | 2.38 |
| 21 | Pde4b | 2.16 | 1.58E-23 | 3.28E-19 | 0.22 | 0.04 | 0.17 | 5.05 |
| 21 | Brk1 | 0.72 | 1.59E-23 | 3.31E-19 | 0.99 | 0.92 | 0.07 | 1.08 |
| 21 | Rps11 | 0.26 | 1.64E-23 | 3.42E-19 | 1.00 | 1.00 | 0.00 | 1.00 |
| 21 | Tbc1d22a | 1.78 | 1.88E-23 | 3.92E-19 | 0.30 | 0.07 | 0.23 | 4.07 |
| 21 | Gng2 | 0.90 | 1.92E-23 | 3.99E-19 | 0.93 | 0.57 | 0.36 | 1.63 |
| 21 | Tmem50a | 0.93 | 2.00E-23 | 4.17E-19 | 0.99 | 0.81 | 0.19 | 1.23 |
| 21 | Dok1 | 1.77 | 2.18E-23 | 4.53E-19 | 0.36 | 0.10 | 0.26 | 3.57 |
| 21 | Fam98c | 1.64 | 2.24E-23 | 4.66E-19 | 0.39 | 0.12 | 0.28 | 3.41 |
| 21 | Psen1 | 1.51 | 2.58E-23 | 5.37E-19 | 0.62 | 0.25 | 0.37 | 2.49 |
| 21 | Mef2d | 1.65 | 2.60E-23 | 5.41E-19 | 0.43 | 0.14 | 0.30 | 3.14 |
| 21 | Sord | 1.46 | 2.76E-23 | 5.74E-19 | 0.45 | 0.15 | 0.30 | 3.09 |
| 21 | Ankrd13a | 1.37 | 2.98E-23 | 6.19E-19 | 0.66 | 0.29 | 0.38 | 2.30 |
| 21 | Fam20c | 3.92 | 3.02E-23 | 6.28E-19 | 0.15 | 0.02 | 0.13 | 6.68 |
| 21 | Ciao2a | 0.84 | 3.07E-23 | 6.38E-19 | 0.95 | 0.85 | 0.10 | 1.12 |
| 21 | Gng12 | 1.56 | 3.44E-23 | 7.16E-19 | 0.72 | 0.36 | 0.36 | 2.00 |
| 21 | Plcl2 | 1.60 | 3.48E-23 | 7.24E-19 | 0.39 | 0.11 | 0.27 | 3.44 |
| 21 | 1700017B05Rik | 1.65 | 3.54E-23 | 7.37E-19 | 0.51 | 0.18 | 0.33 | 2.85 |
| 21 | Rilpl2 | 1.33 | 4.05E-23 | 8.41E-19 | 0.67 | 0.28 | 0.40 | 2.43 |
| 21 | Nptn | 1.32 | 4.41E-23 | 9.17E-19 | 0.80 | 0.45 | 0.34 | 1.76 |
| 21 | Orai2 | 2.22 | 5.03E-23 | 1.05E-18 | 0.22 | 0.04 | 0.17 | 4.93 |
| 21 | Cyb561d2 | 1.52 | 5.21E-23 | 1.08E-18 | 0.49 | 0.17 | 0.32 | 2.90 |
| 21 | Rpl39 | 0.30 | 5.36E-23 | 1.12E-18 | 1.00 | 1.00 | 0.00 | 1.00 |
| 21 | Frmd8 | 1.39 | 5.51E-23 | 1.15E-18 | 0.55 | 0.20 | 0.34 | 2.70 |
| 21 | Mgat2 | 1.35 | 5.59E-23 | 1.16E-18 | 0.80 | 0.46 | 0.34 | 1.74 |
| 21 | Rab39 | 3.00 | 5.89E-23 | 1.22E-18 | 0.11 | 0.01 | 0.09 | 8.75 |
| 21 | Sbno2 | 1.62 | 6.91E-23 | 1.44E-18 | 0.39 | 0.11 | 0.27 | 3.38 |
| 21 | Psen2 | 1.87 | 8.71E-23 | 1.81E-18 | 0.30 | 0.08 | 0.22 | 3.91 |
| 21 | Lgals1 | 1.41 | 8.83E-23 | 1.84E-18 | 0.98 | 0.83 | 0.15 | 1.18 |
| 21 | Ogfrl1 | 1.55 | 1.35E-22 | 2.80E-18 | 0.67 | 0.34 | 0.34 | 2.00 |
| 21 | Zdhhc14 | 2.91 | 1.71E-22 | 3.56E-18 | 0.16 | 0.03 | 0.13 | 5.96 |
| 21 | Stom | 2.22 | 1.95E-22 | 4.05E-18 | 0.22 | 0.05 | 0.17 | 4.82 |
| 21 | Pik3cb | 1.77 | 1.99E-22 | 4.13E-18 | 0.27 | 0.06 | 0.20 | 4.29 |
| 21 | Fnip2 | 2.14 | 2.00E-22 | 4.16E-18 | 0.34 | 0.10 | 0.24 | 3.43 |
| 21 | Msrb1 | 1.46 | 2.03E-22 | 4.22E-18 | 0.83 | 0.49 | 0.34 | 1.69 |
| 21 | Zyx | 1.15 | 2.08E-22 | 4.33E-18 | 0.84 | 0.51 | 0.33 | 1.65 |
| 21 | Ralb | 1.20 | 2.52E-22 | 5.24E-18 | 0.80 | 0.43 | 0.37 | 1.85 |
| 21 | Plk3 | 2.68 | 3.24E-22 | 6.75E-18 | 0.15 | 0.02 | 0.12 | 6.39 |
| 21 | Rnh1 | 0.89 | 3.31E-22 | 6.89E-18 | 0.90 | 0.55 | 0.34 | 1.62 |
| 21 | Tnfrsf1a | 1.32 | 3.39E-22 | 7.05E-18 | 0.69 | 0.30 | 0.39 | 2.30 |
| 21 | Herc6 | 1.84 | 3.41E-22 | 7.09E-18 | 0.24 | 0.05 | 0.19 | 4.67 |
| 21 | Ripk3 | 1.43 | 3.66E-22 | 7.61E-18 | 0.48 | 0.16 | 0.32 | 2.95 |
| 21 | Bcl3 | 2.19 | 3.69E-22 | 7.67E-18 | 0.27 | 0.07 | 0.21 | 4.14 |
| 21 | Hspa4l | 1.31 | 3.85E-22 | 8.02E-18 | 0.38 | 0.11 | 0.27 | 3.41 |
| 21 | Slc15a4 | 1.60 | 5.07E-22 | 1.05E-17 | 0.44 | 0.15 | 0.30 | 3.02 |

**Supplementary Table 1. Marker genes for the original clusters of integrated clustering of E11.5 control and NotchGOF λ-junction cells.**

| | | | | | | | | |
|---|---|---|---|---|---|---|---|---|
| 21 | Arfgef2 | 1.56 | 6.45E-22 | 1.34E-17 | 0.34 | 0.09 | 0.25 | 3.69 |
| 21 | Ap2s1 | 0.90 | 6.47E-22 | 1.35E-17 | 0.99 | 0.89 | 0.11 | 1.12 |
| 21 | Cln5 | 1.38 | 6.52E-22 | 1.36E-17 | 0.55 | 0.21 | 0.34 | 2.59 |
| 21 | Sft2d1 | 1.43 | 6.64E-22 | 1.38E-17 | 0.55 | 0.22 | 0.33 | 2.49 |
| 21 | Glul | 1.69 | 7.80E-22 | 1.62E-17 | 0.87 | 0.62 | 0.25 | 1.41 |
| 21 | Atp6v1a | 1.24 | 9.28E-22 | 1.93E-17 | 0.88 | 0.53 | 0.36 | 1.67 |
| 21 | Npepl1 | 1.13 | 9.28E-22 | 1.93E-17 | 0.71 | 0.33 | 0.39 | 2.18 |
| 21 | H2-M3 | 2.86 | 9.36E-22 | 1.95E-17 | 0.13 | 0.02 | 0.11 | 7.00 |
| 21 | Cpeb4 | 1.65 | 1.06E-21 | 2.21E-17 | 0.34 | 0.09 | 0.24 | 3.57 |
| 21 | Tmem14c | 0.56 | 1.14E-21 | 2.38E-17 | 0.99 | 0.92 | 0.06 | 1.07 |
| 21 | Rin2 | 1.79 | 1.19E-21 | 2.47E-17 | 0.51 | 0.20 | 0.31 | 2.54 |
| 21 | Fmn1 | 2.83 | 1.25E-21 | 2.59E-17 | 0.13 | 0.02 | 0.11 | 7.00 |
| 21 | Psat1 | 1.23 | 1.38E-21 | 2.87E-17 | 0.78 | 0.45 | 0.33 | 1.72 |
| 21 | Ophn1 | 1.84 | 1.49E-21 | 3.10E-17 | 0.50 | 0.19 | 0.31 | 2.60 |
| 21 | Tram1 | 1.15 | 1.58E-21 | 3.28E-17 | 0.83 | 0.51 | 0.32 | 1.63 |
| 21 | Orai3 | 1.58 | 1.65E-21 | 3.42E-17 | 0.38 | 0.11 | 0.26 | 3.32 |
| 21 | Derl2 | 1.22 | 1.79E-21 | 3.72E-17 | 0.74 | 0.38 | 0.37 | 1.97 |
| 21 | Mapkapk2 | 1.11 | 1.98E-21 | 4.12E-17 | 0.85 | 0.55 | 0.30 | 1.55 |
| 21 | mt-Co1 | 0.44 | 2.07E-21 | 4.31E-17 | 1.00 | 0.99 | 0.01 | 1.01 |
| 21 | Nmt1 | 1.07 | 2.13E-21 | 4.42E-17 | 0.90 | 0.61 | 0.29 | 1.47 |
| 21 | Amdhd2 | 1.45 | 2.41E-21 | 5.02E-17 | 0.59 | 0.25 | 0.35 | 2.42 |
| 21 | Eef2kmt | 1.37 | 2.68E-21 | 5.58E-17 | 0.50 | 0.18 | 0.32 | 2.76 |
| 21 | Ppp1r21 | 1.40 | 2.72E-21 | 5.66E-17 | 0.54 | 0.21 | 0.33 | 2.55 |
| 21 | Rab7 | 1.00 | 2.87E-21 | 5.97E-17 | 0.96 | 0.69 | 0.27 | 1.39 |
| 21 | Ctbs | 1.84 | 2.97E-21 | 6.18E-17 | 0.33 | 0.09 | 0.24 | 3.50 |
| 21 | Cox4i1 | 0.38 | 3.14E-21 | 6.52E-17 | 1.00 | 1.00 | 0.00 | 1.00 |
| 21 | Tmem176b | 1.68 | 3.22E-21 | 6.69E-17 | 0.81 | 0.57 | 0.24 | 1.43 |
| 21 | Agpat5 | 1.25 | 3.51E-21 | 7.31E-17 | 0.78 | 0.45 | 0.34 | 1.76 |
| 21 | Morc3 | 1.16 | 3.54E-21 | 7.37E-17 | 0.79 | 0.42 | 0.37 | 1.87 |
| 21 | Pip4p1 | 1.26 | 3.82E-21 | 7.95E-17 | 0.72 | 0.36 | 0.36 | 2.00 |
| 21 | Zzef1 | 1.55 | 3.94E-21 | 8.18E-17 | 0.48 | 0.17 | 0.31 | 2.80 |
| 21 | Lactb | 1.33 | 4.01E-21 | 8.34E-17 | 0.69 | 0.34 | 0.35 | 2.04 |
| 21 | Ccdc22 | 1.16 | 4.05E-21 | 8.43E-17 | 0.69 | 0.30 | 0.39 | 2.32 |
| 21 | D6Wsu163e | 1.72 | 4.15E-21 | 8.63E-17 | 0.34 | 0.10 | 0.24 | 3.54 |
| 21 | Wipf1 | 1.35 | 4.33E-21 | 9.00E-17 | 0.57 | 0.23 | 0.34 | 2.49 |
| 21 | Jak1 | 0.98 | 4.61E-21 | 9.60E-17 | 0.93 | 0.61 | 0.32 | 1.53 |
| 21 | Ldlrad4 | 2.09 | 4.62E-21 | 9.61E-17 | 0.18 | 0.03 | 0.14 | 5.47 |
| 21 | Paqr4 | 2.46 | 4.66E-21 | 9.69E-17 | 0.18 | 0.04 | 0.15 | 5.20 |
| 21 | B3gnt2 | 1.50 | 4.72E-21 | 9.81E-17 | 0.34 | 0.10 | 0.24 | 3.46 |
| 21 | Cstb | 1.30 | 5.71E-21 | 1.19E-16 | 0.93 | 0.80 | 0.13 | 1.16 |
| 21 | Rbm43 | 1.56 | 6.73E-21 | 1.40E-16 | 0.36 | 0.11 | 0.26 | 3.37 |
| 21 | Pomp | 0.54 | 6.79E-21 | 1.41E-16 | 0.99 | 0.98 | 0.02 | 1.02 |
| 21 | Rgs19 | 1.11 | 7.50E-21 | 1.56E-16 | 0.80 | 0.45 | 0.35 | 1.78 |
| 21 | Abr | 1.19 | 8.48E-21 | 1.76E-16 | 0.57 | 0.22 | 0.36 | 2.64 |
| 21 | Nubp2 | 1.21 | 8.81E-21 | 1.83E-16 | 0.67 | 0.31 | 0.37 | 2.19 |
| 21 | Qsox1 | 1.51 | 9.74E-21 | 2.03E-16 | 0.38 | 0.12 | 0.26 | 3.26 |
| 21 | Chmp3 | 1.05 | 1.08E-20 | 2.24E-16 | 0.83 | 0.46 | 0.37 | 1.81 |
| 21 | Spcs3 | 1.12 | 1.14E-20 | 2.36E-16 | 0.76 | 0.38 | 0.38 | 2.02 |
| 21 | Myo9b | 1.26 | 1.16E-20 | 2.41E-16 | 0.58 | 0.23 | 0.35 | 2.52 |
| 21 | Tbc1d14 | 1.45 | 1.23E-20 | 2.55E-16 | 0.51 | 0.19 | 0.32 | 2.64 |
| 21 | Sfxn5 | 2.28 | 1.23E-20 | 2.56E-16 | 0.20 | 0.04 | 0.16 | 4.90 |
| 21 | Tbc1d9 | 1.98 | 1.38E-20 | 2.87E-16 | 0.29 | 0.08 | 0.21 | 3.78 |
| 21 | Arf6 | 1.14 | 1.49E-20 | 3.10E-16 | 0.75 | 0.40 | 0.35 | 1.89 |
| 21 | Dse | 1.46 | 1.63E-20 | 3.40E-16 | 0.59 | 0.27 | 0.33 | 2.22 |
| 21 | Cln8 | 1.69 | 1.69E-20 | 3.51E-16 | 0.52 | 0.21 | 0.32 | 2.56 |
| 21 | Stx4a | 1.30 | 1.78E-20 | 3.71E-16 | 0.69 | 0.32 | 0.37 | 2.17 |

**Supplementary Table 1. Marker genes for the original clusters of integrated clustering of E11.5 control and NotchGOF λ-junction cells.**

| | | | | | | | | |
|---|---|---|---|---|---|---|---|---|
| 21 | Akr1a1 | 0.77 | 2.07E-20 | 4.30E-16 | 1.00 | 0.96 | 0.04 | 1.04 |
| 21 | Myadm | 1.54 | 2.12E-20 | 4.41E-16 | 0.39 | 0.13 | 0.27 | 3.14 |
| 21 | Vps29 | 0.81 | 2.47E-20 | 5.14E-16 | 0.94 | 0.78 | 0.17 | 1.21 |
| 21 | 2510039O18Rik | 1.30 | 2.81E-20 | 5.84E-16 | 0.57 | 0.23 | 0.33 | 2.43 |
| 21 | Bola2 | 0.72 | 3.10E-20 | 6.46E-16 | 0.97 | 0.88 | 0.08 | 1.09 |
| 21 | Snapc5 | 0.97 | 3.59E-20 | 7.47E-16 | 0.86 | 0.55 | 0.31 | 1.56 |
| 21 | B4galt5 | 1.67 | 3.62E-20 | 7.53E-16 | 0.35 | 0.11 | 0.25 | 3.33 |
| 21 | Lman2 | 0.99 | 4.03E-20 | 8.39E-16 | 0.86 | 0.62 | 0.24 | 1.39 |
| 21 | Ech1 | 1.00 | 4.26E-20 | 8.87E-16 | 0.86 | 0.51 | 0.35 | 1.68 |
| 21 | Pde3b | 1.32 | 4.73E-20 | 9.85E-16 | 0.59 | 0.24 | 0.35 | 2.44 |
| 21 | Atxn3 | 1.23 | 5.93E-20 | 1.23E-15 | 0.50 | 0.19 | 0.31 | 2.66 |
| 21 | Rpl10 | 0.32 | 6.26E-20 | 1.30E-15 | 1.00 | 1.00 | 0.00 | 1.00 |
| 21 | Cebpzos | 1.36 | 6.43E-20 | 1.34E-15 | 0.87 | 0.57 | 0.30 | 1.53 |
| 21 | Bsn | 2.10 | 6.65E-20 | 1.38E-15 | 0.18 | 0.03 | 0.14 | 5.30 |
| 21 | Herpud1 | 1.30 | 6.73E-20 | 1.40E-15 | 0.60 | 0.25 | 0.35 | 2.41 |
| 21 | Actg1 | 0.44 | 8.61E-20 | 1.79E-15 | 1.00 | 1.00 | 0.00 | 1.00 |
| 21 | Pign | 1.26 | 8.66E-20 | 1.80E-15 | 0.43 | 0.14 | 0.29 | 3.01 |
| 21 | Mertk | 1.90 | 8.67E-20 | 1.80E-15 | 0.43 | 0.16 | 0.27 | 2.70 |
| 21 | Mob3a | 1.41 | 8.95E-20 | 1.86E-15 | 0.46 | 0.16 | 0.30 | 2.88 |
| 21 | Emilin2 | 1.55 | 9.21E-20 | 1.92E-15 | 0.58 | 0.26 | 0.33 | 2.27 |
| 21 | Acadm | 1.28 | 1.05E-19 | 2.19E-15 | 0.67 | 0.33 | 0.34 | 2.02 |
| 21 | Dynlt3 | 1.19 | 1.08E-19 | 2.25E-15 | 0.61 | 0.26 | 0.35 | 2.38 |
| 21 | Rab3d | 2.33 | 1.08E-19 | 2.25E-15 | 0.13 | 0.02 | 0.11 | 6.33 |
| 21 | Abi1 | 1.18 | 1.25E-19 | 2.60E-15 | 0.76 | 0.39 | 0.37 | 1.96 |
| 21 | Ppp1r13b | 1.58 | 1.36E-19 | 2.83E-15 | 0.29 | 0.08 | 0.21 | 3.68 |
| 21 | Derl1 | 1.09 | 1.42E-19 | 2.96E-15 | 0.80 | 0.49 | 0.32 | 1.65 |
| 21 | Hgf | 2.67 | 1.52E-19 | 3.16E-15 | 0.13 | 0.02 | 0.11 | 6.63 |
| 21 | Pts | 1.13 | 1.66E-19 | 3.44E-15 | 0.78 | 0.41 | 0.38 | 1.93 |
| 21 | Fig4 | 1.76 | 2.07E-19 | 4.30E-15 | 0.28 | 0.08 | 0.21 | 3.73 |
| 21 | Atg3 | 1.16 | 2.19E-19 | 4.55E-15 | 0.78 | 0.45 | 0.33 | 1.74 |
| 21 | 0610012G03Rik | 0.84 | 2.54E-19 | 5.28E-15 | 0.99 | 0.86 | 0.12 | 1.14 |
| 21 | Tnfrsf21 | 1.51 | 2.58E-19 | 5.36E-15 | 0.41 | 0.14 | 0.27 | 2.93 |
| 21 | Cdk6 | 1.43 | 2.74E-19 | 5.69E-15 | 0.68 | 0.34 | 0.34 | 1.99 |
| 21 | Trappc6a | 0.97 | 2.77E-19 | 5.77E-15 | 0.89 | 0.52 | 0.37 | 1.70 |
| 21 | Gimap6 | 1.57 | 3.04E-19 | 6.31E-15 | 0.15 | 0.03 | 0.13 | 5.70 |
| 21 | Ap5z1 | 1.74 | 3.17E-19 | 6.60E-15 | 0.22 | 0.05 | 0.17 | 4.31 |
| 21 | Malt1 | 1.46 | 3.53E-19 | 7.34E-15 | 0.45 | 0.17 | 0.28 | 2.72 |
| 21 | Dnajb14 | 1.35 | 4.09E-19 | 8.52E-15 | 0.67 | 0.32 | 0.35 | 2.09 |
| 21 | Card19 | 1.07 | 4.20E-19 | 8.74E-15 | 0.60 | 0.25 | 0.35 | 2.39 |
| 21 | Plgrkt | 1.18 | 4.26E-19 | 8.87E-15 | 0.81 | 0.48 | 0.33 | 1.70 |
| 21 | Hps1 | 1.38 | 5.99E-19 | 1.25E-14 | 0.39 | 0.13 | 0.26 | 3.06 |
| 21 | Osbpl8 | 1.24 | 6.16E-19 | 1.28E-14 | 0.56 | 0.23 | 0.33 | 2.41 |
| 21 | Commd4 | 0.96 | 6.46E-19 | 1.34E-14 | 0.87 | 0.53 | 0.34 | 1.64 |
| 21 | Cds1 | 1.86 | 7.20E-19 | 1.50E-14 | 0.19 | 0.04 | 0.15 | 4.85 |
| 21 | Sys1 | 0.82 | 7.21E-19 | 1.50E-14 | 0.94 | 0.70 | 0.23 | 1.33 |
| 21 | Plod1 | 1.51 | 8.11E-19 | 1.69E-14 | 0.50 | 0.20 | 0.30 | 2.49 |
| 21 | Tapbpl | 2.64 | 8.32E-19 | 1.73E-14 | 0.11 | 0.02 | 0.10 | 7.00 |
| 21 | Nfe2 | 2.12 | 9.11E-19 | 1.89E-14 | 0.18 | 0.04 | 0.14 | 5.00 |
| 21 | Ebp | 1.22 | 9.55E-19 | 1.99E-14 | 0.73 | 0.40 | 0.32 | 1.80 |
| 21 | 2610507B11Rik | 1.06 | 9.61E-19 | 2.00E-14 | 0.81 | 0.46 | 0.35 | 1.75 |
| 21 | Slc41a3 | 2.73 | 9.73E-19 | 2.02E-14 | 0.14 | 0.02 | 0.12 | 5.83 |
| 21 | Wwp2 | 1.37 | 9.80E-19 | 2.04E-14 | 0.64 | 0.31 | 0.33 | 2.09 |
| 21 | Fam173a | 1.03 | 9.82E-19 | 2.04E-14 | 0.84 | 0.50 | 0.34 | 1.66 |
| 21 | Uba7 | 1.50 | 9.87E-19 | 2.05E-14 | 0.34 | 0.11 | 0.24 | 3.21 |
| 21 | Ppt1 | 1.29 | 1.14E-18 | 2.37E-14 | 0.65 | 0.31 | 0.34 | 2.07 |
| 21 | Dennd1c | 1.90 | 1.20E-18 | 2.49E-14 | 0.17 | 0.03 | 0.14 | 5.09 |

**Supplementary Table 1. Marker genes for the original clusters of integrated clustering of E11.5 control and NotchGOF λ-junction cells.**

| | | | | | | | | |
|---|---|---|---|---|---|---|---|---|
| 21 | Timm10b | 0.69 | 1.33E-18 | 2.77E-14 | 0.96 | 0.85 | 0.11 | 1.13 |
| 21 | Tie1 | 1.53 | 1.39E-18 | 2.90E-14 | 0.11 | 0.02 | 0.10 | 7.00 |
| 21 | Mgat1 | 1.36 | 1.54E-18 | 3.21E-14 | 0.51 | 0.21 | 0.30 | 2.46 |
| 21 | Vac14 | 1.54 | 1.60E-18 | 3.32E-14 | 0.52 | 0.22 | 0.31 | 2.40 |
| 21 | Uchl5 | 1.13 | 1.62E-18 | 3.37E-14 | 0.79 | 0.54 | 0.25 | 1.46 |
| 21 | Os9 | 0.97 | 1.66E-18 | 3.45E-14 | 0.75 | 0.37 | 0.38 | 2.02 |
| 21 | Ctse | 3.17 | 1.70E-18 | 3.54E-14 | 0.15 | 0.03 | 0.13 | 5.31 |
| 21 | Agps | 1.15 | 1.75E-18 | 3.65E-14 | 0.76 | 0.41 | 0.36 | 1.88 |
| 21 | Use1 | 0.94 | 2.02E-18 | 4.21E-14 | 0.91 | 0.65 | 0.26 | 1.40 |
| 21 | Acot9 | 1.34 | 2.41E-18 | 5.01E-14 | 0.43 | 0.16 | 0.28 | 2.76 |
| 21 | Ppard | 1.62 | 2.45E-18 | 5.09E-14 | 0.33 | 0.10 | 0.23 | 3.26 |
| 21 | Slc31a2 | 1.95 | 2.62E-18 | 5.46E-14 | 0.27 | 0.07 | 0.19 | 3.69 |
| 21 | Rpsa | 0.33 | 2.66E-18 | 5.53E-14 | 1.00 | 1.00 | 0.00 | 1.00 |
| 21 | Ptpra | 1.13 | 2.66E-18 | 5.53E-14 | 0.87 | 0.60 | 0.26 | 1.44 |
| 21 | Serpinb6a | 1.98 | 2.78E-18 | 5.79E-14 | 0.71 | 0.39 | 0.32 | 1.81 |
| 21 | Slc35c2 | 1.15 | 2.93E-18 | 6.10E-14 | 0.68 | 0.33 | 0.35 | 2.07 |
| 21 | Tuba4a | 1.85 | 3.31E-18 | 6.87E-14 | 0.25 | 0.07 | 0.18 | 3.65 |
| 21 | Tango2 | 1.33 | 3.54E-18 | 7.37E-14 | 0.43 | 0.16 | 0.27 | 2.74 |
| 21 | Gdap2 | 1.40 | 3.61E-18 | 7.51E-14 | 0.36 | 0.11 | 0.24 | 3.13 |
| 21 | Dhrs7 | 1.37 | 3.66E-18 | 7.60E-14 | 0.39 | 0.13 | 0.26 | 3.03 |
| 21 | Sharpin | 1.23 | 3.82E-18 | 7.95E-14 | 0.46 | 0.17 | 0.29 | 2.71 |
| 21 | Flot1 | 1.10 | 4.96E-18 | 1.03E-13 | 0.64 | 0.30 | 0.34 | 2.12 |
| 21 | Trim12c | 1.47 | 5.05E-18 | 1.05E-13 | 0.46 | 0.18 | 0.29 | 2.61 |
| 21 | 4931406C07Rik | 1.34 | 5.08E-18 | 1.06E-13 | 0.58 | 0.26 | 0.32 | 2.24 |
| 21 | Enox2 | 1.80 | 5.40E-18 | 1.12E-13 | 0.26 | 0.07 | 0.19 | 3.70 |
| 21 | Pnpo | 0.79 | 5.98E-18 | 1.24E-13 | 0.47 | 0.17 | 0.30 | 2.76 |
| 21 | Sorl1 | 1.48 | 8.72E-18 | 1.81E-13 | 0.41 | 0.15 | 0.26 | 2.74 |
| 21 | Tmbim6 | 0.71 | 9.24E-18 | 1.92E-13 | 0.97 | 0.84 | 0.13 | 1.15 |
| 21 | Relch | 1.25 | 1.07E-17 | 2.22E-13 | 0.55 | 0.23 | 0.32 | 2.37 |
| 21 | Ptpmt1 | 0.93 | 1.13E-17 | 2.34E-13 | 0.85 | 0.56 | 0.29 | 1.52 |
| 21 | Snap29 | 1.43 | 1.21E-17 | 2.53E-13 | 0.48 | 0.19 | 0.29 | 2.49 |
| 21 | Nagk | 1.53 | 1.23E-17 | 2.56E-13 | 0.48 | 0.20 | 0.29 | 2.45 |
| 21 | Atp6v1e1 | 0.86 | 1.24E-17 | 2.59E-13 | 0.90 | 0.68 | 0.23 | 1.33 |
| 21 | Snx14 | 1.22 | 1.26E-17 | 2.63E-13 | 0.45 | 0.16 | 0.29 | 2.75 |
| 21 | Capns1 | 0.79 | 1.29E-17 | 2.68E-13 | 0.97 | 0.80 | 0.16 | 1.20 |
| 21 | Atp6v0a1 | 1.51 | 1.41E-17 | 2.93E-13 | 0.49 | 0.20 | 0.29 | 2.49 |
| 21 | Tmco1 | 0.84 | 1.47E-17 | 3.06E-13 | 0.92 | 0.70 | 0.23 | 1.32 |
| 21 | Gmpr | 1.01 | 1.48E-17 | 3.07E-13 | 0.47 | 0.18 | 0.29 | 2.68 |
| 21 | Phka2 | 1.55 | 1.50E-17 | 3.13E-13 | 0.37 | 0.13 | 0.24 | 2.92 |
| 21 | Alg1 | 1.73 | 1.57E-17 | 3.26E-13 | 0.27 | 0.08 | 0.20 | 3.55 |
| 21 | Mpv17l2 | 1.23 | 1.57E-17 | 3.27E-13 | 0.54 | 0.23 | 0.31 | 2.38 |
| 21 | Lrpap1 | 0.93 | 1.93E-17 | 4.02E-13 | 0.85 | 0.49 | 0.36 | 1.73 |
| 21 | Ap2m1 | 0.78 | 1.96E-17 | 4.07E-13 | 0.97 | 0.86 | 0.11 | 1.13 |
| 21 | Mien1 | 0.85 | 2.24E-17 | 4.66E-13 | 0.88 | 0.56 | 0.32 | 1.56 |
| 21 | 2510009E07Rik | 1.85 | 2.54E-17 | 5.29E-13 | 0.24 | 0.06 | 0.18 | 3.84 |
| 21 | Trappc2l | 0.88 | 2.57E-17 | 5.35E-13 | 0.90 | 0.62 | 0.28 | 1.45 |
| 21 | Lyplal1 | 1.19 | 2.75E-17 | 5.72E-13 | 0.57 | 0.25 | 0.32 | 2.30 |
| 21 | Mafk | 1.86 | 2.83E-17 | 5.89E-13 | 0.18 | 0.04 | 0.14 | 4.55 |
| 21 | Mif4gd | 1.34 | 2.85E-17 | 5.94E-13 | 0.45 | 0.17 | 0.28 | 2.67 |
| 21 | Trim21 | 2.29 | 2.90E-17 | 6.04E-13 | 0.16 | 0.03 | 0.13 | 4.88 |
| 21 | Galk2 | 1.06 | 3.07E-17 | 6.38E-13 | 0.61 | 0.27 | 0.34 | 2.23 |
| 21 | Atp6v1c1 | 1.23 | 3.19E-17 | 6.63E-13 | 0.69 | 0.36 | 0.33 | 1.92 |
| 21 | Gls | 1.21 | 3.27E-17 | 6.80E-13 | 0.73 | 0.41 | 0.32 | 1.78 |
| 21 | Flii | 1.16 | 3.34E-17 | 6.94E-13 | 0.73 | 0.37 | 0.36 | 1.99 |
| 21 | Efr3a | 0.95 | 3.42E-17 | 7.12E-13 | 0.73 | 0.36 | 0.37 | 2.04 |
| 21 | Mid1ip1 | 1.39 | 3.64E-17 | 7.57E-13 | 0.51 | 0.22 | 0.29 | 2.34 |

**Supplementary Table 1. Marker genes for the original clusters of integrated clustering of E11.5 control and NotchGOF λ-junction cells.**

| | | | | | | | | |
|---|---|---|---|---|---|---|---|---|
| 21 | Ifnar1 | 1.12 | 3.74E-17 | 7.77E-13 | 0.69 | 0.35 | 0.35 | 1.99 |
| 21 | Arl4c | 1.36 | 3.94E-17 | 8.20E-13 | 0.64 | 0.33 | 0.31 | 1.95 |
| 21 | Tbkbp1 | 1.33 | 3.95E-17 | 8.21E-13 | 0.25 | 0.06 | 0.18 | 3.89 |
| 21 | Vav2 | 1.58 | 4.21E-17 | 8.76E-13 | 0.29 | 0.08 | 0.20 | 3.42 |
| 21 | St3gal6 | 2.01 | 4.29E-17 | 8.92E-13 | 0.33 | 0.11 | 0.22 | 2.94 |
| 21 | Fam53b | 1.34 | 4.49E-17 | 9.35E-13 | 0.46 | 0.18 | 0.28 | 2.57 |
| 21 | Npr1 | 2.09 | 4.94E-17 | 1.03E-12 | 0.11 | 0.02 | 0.09 | 7.00 |
| 21 | Grina | 1.13 | 5.09E-17 | 1.06E-12 | 0.57 | 0.25 | 0.32 | 2.30 |
| 21 | Tyk2 | 2.00 | 5.56E-17 | 1.16E-12 | 0.22 | 0.06 | 0.17 | 3.86 |
| 21 | Sri | 0.82 | 6.16E-17 | 1.28E-12 | 0.97 | 0.77 | 0.20 | 1.26 |
| 21 | Twf1 | 1.16 | 7.26E-17 | 1.51E-12 | 0.80 | 0.49 | 0.31 | 1.63 |
| 21 | Ap2a2 | 1.23 | 7.36E-17 | 1.53E-12 | 0.83 | 0.54 | 0.29 | 1.54 |
| 21 | Cast | 1.19 | 8.37E-17 | 1.74E-12 | 0.56 | 0.24 | 0.32 | 2.30 |
| 21 | Ddx58 | 1.84 | 8.97E-17 | 1.86E-12 | 0.22 | 0.05 | 0.16 | 4.02 |
| 21 | Mt2 | 1.23 | 1.01E-16 | 2.11E-12 | 0.36 | 0.13 | 0.24 | 2.82 |
| 21 | Layn | 2.05 | 1.07E-16 | 2.23E-12 | 0.15 | 0.03 | 0.12 | 5.25 |
| 21 | Pak2 | 0.63 | 1.14E-16 | 2.38E-12 | 0.99 | 0.84 | 0.15 | 1.18 |
| 21 | Mthfsl | 1.14 | 1.15E-16 | 2.38E-12 | 0.55 | 0.23 | 0.32 | 2.39 |
| 21 | Acsl4 | 1.28 | 1.23E-16 | 2.55E-12 | 0.39 | 0.14 | 0.25 | 2.80 |
| 21 | Hsd17b11 | 1.12 | 1.29E-16 | 2.67E-12 | 0.51 | 0.21 | 0.31 | 2.49 |
| 21 | Otulin | 0.96 | 1.37E-16 | 2.84E-12 | 0.73 | 0.41 | 0.32 | 1.79 |
| 21 | Fabp3 | 2.59 | 1.39E-16 | 2.89E-12 | 0.14 | 0.03 | 0.11 | 5.19 |
| 21 | Wdr41 | 1.51 | 1.46E-16 | 3.03E-12 | 0.32 | 0.10 | 0.22 | 3.16 |
| 21 | Mfsd11 | 1.52 | 1.49E-16 | 3.10E-12 | 0.41 | 0.15 | 0.25 | 2.64 |
| 21 | Akr1b3 | 0.77 | 1.51E-16 | 3.14E-12 | 0.89 | 0.68 | 0.21 | 1.31 |
| 21 | Washc2 | 0.90 | 1.83E-16 | 3.81E-12 | 0.83 | 0.52 | 0.31 | 1.60 |
| 21 | Flt1 | 0.72 | 1.85E-16 | 3.85E-12 | 0.12 | 0.02 | 0.10 | 5.95 |
| 21 | Abhd6 | 1.36 | 1.98E-16 | 4.11E-12 | 0.44 | 0.17 | 0.27 | 2.63 |
| 21 | Ppcdc | 1.37 | 1.99E-16 | 4.13E-12 | 0.38 | 0.13 | 0.24 | 2.82 |
| 21 | Hk2 | 1.18 | 1.99E-16 | 4.14E-12 | 0.57 | 0.25 | 0.31 | 2.23 |
| 21 | Coro1c | 1.11 | 2.04E-16 | 4.24E-12 | 0.68 | 0.35 | 0.33 | 1.95 |
| 21 | Cerk | 1.62 | 2.13E-16 | 4.42E-12 | 0.40 | 0.15 | 0.25 | 2.68 |
| 21 | Fchsd2 | 1.22 | 2.17E-16 | 4.52E-12 | 0.66 | 0.33 | 0.33 | 1.99 |
| 21 | Fam49a | 1.90 | 2.19E-16 | 4.55E-12 | 0.19 | 0.04 | 0.15 | 4.30 |
| 21 | Rab1a | 0.86 | 2.24E-16 | 4.67E-12 | 0.94 | 0.71 | 0.23 | 1.33 |
| 21 | Tnfaip3 | 2.84 | 2.26E-16 | 4.69E-12 | 0.11 | 0.02 | 0.09 | 6.56 |
| 21 | Ssbp4 | 0.79 | 2.27E-16 | 4.73E-12 | 0.92 | 0.65 | 0.27 | 1.41 |
| 21 | Haus4 | 1.00 | 2.44E-16 | 5.08E-12 | 0.66 | 0.32 | 0.34 | 2.09 |
| 21 | Birc3 | 1.94 | 2.49E-16 | 5.17E-12 | 0.19 | 0.04 | 0.15 | 4.30 |
| 21 | Emid1 | 1.82 | 2.50E-16 | 5.21E-12 | 0.29 | 0.09 | 0.20 | 3.23 |
| 21 | Arhgef3 | 2.46 | 2.54E-16 | 5.28E-12 | 0.11 | 0.02 | 0.09 | 6.22 |
| 21 | Crlf3 | 0.98 | 2.55E-16 | 5.30E-12 | 0.79 | 0.45 | 0.34 | 1.75 |
| 21 | Nfic | 1.17 | 2.64E-16 | 5.48E-12 | 0.62 | 0.30 | 0.32 | 2.08 |
| 21 | Arl5c | 2.84 | 2.95E-16 | 6.13E-12 | 0.11 | 0.02 | 0.09 | 6.56 |
| 21 | Emc7 | 0.90 | 3.05E-16 | 6.34E-12 | 0.80 | 0.50 | 0.30 | 1.61 |
| 21 | Sfmbt2 | 2.10 | 3.07E-16 | 6.38E-12 | 0.14 | 0.03 | 0.11 | 5.19 |
| 21 | Rps20 | 0.27 | 3.27E-16 | 6.81E-12 | 1.00 | 1.00 | 0.00 | 1.00 |
| 21 | Ndufb1-ps | 0.57 | 3.47E-16 | 7.22E-12 | 0.98 | 0.94 | 0.04 | 1.04 |
| 21 | Calm3 | 0.57 | 3.50E-16 | 7.27E-12 | 0.99 | 0.91 | 0.08 | 1.09 |
| 21 | Synrg | 0.93 | 3.54E-16 | 7.37E-12 | 0.55 | 0.24 | 0.32 | 2.35 |
| 21 | Arhgap31 | 1.29 | 3.92E-16 | 8.14E-12 | 0.47 | 0.19 | 0.28 | 2.49 |
| 21 | Rpl12 | 0.26 | 3.92E-16 | 8.15E-12 | 1.00 | 1.00 | 0.00 | 1.00 |
| 21 | Uvrag | 1.06 | 4.03E-16 | 8.39E-12 | 0.54 | 0.23 | 0.31 | 2.37 |
| 21 | Baz1a | 0.92 | 4.95E-16 | 1.03E-11 | 0.82 | 0.56 | 0.26 | 1.47 |
| 21 | Fez2 | 1.24 | 4.96E-16 | 1.03E-11 | 0.65 | 0.31 | 0.34 | 2.10 |
| 21 | Pnkp | 1.18 | 5.36E-16 | 1.11E-11 | 0.52 | 0.22 | 0.30 | 2.33 |

**Supplementary Table 1. Marker genes for the original clusters of integrated clustering of E11.5 control and NotchGOF λ-junction cells.**

| | | | | | | | | |
|---|---|---|---|---|---|---|---|---|
| 21 | Helz2 | 2.51 | 5.56E-16 | 1.16E-11 | 0.11 | 0.02 | 0.09 | 6.56 |
| 21 | Fbxw4 | 1.13 | 6.19E-16 | 1.29E-11 | 0.40 | 0.14 | 0.26 | 2.79 |
| 21 | AC149090.1 | 1.13 | 6.46E-16 | 1.34E-11 | 0.61 | 0.28 | 0.33 | 2.16 |
| 21 | Mast3 | 1.37 | 6.76E-16 | 1.41E-11 | 0.33 | 0.11 | 0.22 | 3.02 |
| 21 | Uqcrq | 0.46 | 7.76E-16 | 1.61E-11 | 1.00 | 0.97 | 0.03 | 1.03 |
| 21 | Cracr2b | 2.13 | 8.35E-16 | 1.74E-11 | 0.16 | 0.04 | 0.13 | 4.60 |
| 21 | Fam102b | 1.31 | 8.62E-16 | 1.79E-11 | 0.48 | 0.20 | 0.28 | 2.38 |
| 21 | Pacc1 | 1.87 | 8.94E-16 | 1.86E-11 | 0.30 | 0.09 | 0.21 | 3.20 |
| 21 | Rela | 1.14 | 9.22E-16 | 1.92E-11 | 0.54 | 0.25 | 0.29 | 2.20 |
| 21 | Gak | 1.22 | 1.02E-15 | 2.11E-11 | 0.52 | 0.23 | 0.30 | 2.31 |
| 21 | Atp1b3 | 0.79 | 1.04E-15 | 2.17E-11 | 0.93 | 0.72 | 0.21 | 1.30 |
| 21 | Foxo3 | 1.16 | 1.12E-15 | 2.34E-11 | 0.36 | 0.13 | 0.24 | 2.89 |
| 21 | Rbl2 | 1.39 | 1.24E-15 | 2.58E-11 | 0.29 | 0.09 | 0.20 | 3.26 |
| 21 | Ampd3 | 2.65 | 1.30E-15 | 2.71E-11 | 0.11 | 0.02 | 0.09 | 6.18 |
| 21 | Mtr | 1.39 | 1.34E-15 | 2.79E-11 | 0.50 | 0.23 | 0.27 | 2.18 |
| 21 | Clptm1 | 0.99 | 1.50E-15 | 3.11E-11 | 0.76 | 0.45 | 0.32 | 1.71 |
| 21 | Foxj2 | 1.65 | 1.53E-15 | 3.19E-11 | 0.25 | 0.07 | 0.18 | 3.55 |
| 21 | Rhof | 2.26 | 1.56E-15 | 3.25E-11 | 0.11 | 0.02 | 0.09 | 6.18 |
| 21 | Klhl5 | 1.14 | 1.64E-15 | 3.41E-11 | 0.44 | 0.17 | 0.27 | 2.58 |
| 21 | Pald1 | 1.54 | 1.77E-15 | 3.69E-11 | 0.27 | 0.08 | 0.19 | 3.29 |
| 21 | Mtss1 | 1.10 | 1.96E-15 | 4.08E-11 | 0.62 | 0.30 | 0.32 | 2.08 |
| 21 | Nqo2 | 1.29 | 2.07E-15 | 4.30E-11 | 0.36 | 0.13 | 0.24 | 2.82 |
| 21 | Ydjc | 1.68 | 2.21E-15 | 4.60E-11 | 0.27 | 0.08 | 0.19 | 3.29 |
| 21 | 1600002K03Rik | 1.34 | 2.21E-15 | 4.61E-11 | 0.34 | 0.12 | 0.22 | 2.92 |
| 21 | Cflar | 1.00 | 2.42E-15 | 5.03E-11 | 0.63 | 0.31 | 0.32 | 2.03 |
| 21 | Slc2a3 | 0.56 | 2.43E-15 | 5.05E-11 | 0.39 | 0.14 | 0.25 | 2.79 |
| 21 | Nudt18 | 1.65 | 2.58E-15 | 5.38E-11 | 0.22 | 0.06 | 0.16 | 3.67 |
| 21 | Ccpg1 | 1.20 | 2.72E-15 | 5.65E-11 | 0.42 | 0.16 | 0.26 | 2.59 |
| 21 | Slc38a6 | 1.58 | 2.91E-15 | 6.05E-11 | 0.39 | 0.15 | 0.24 | 2.57 |
| 21 | Iqsec2 | 2.22 | 3.23E-15 | 6.71E-11 | 0.13 | 0.02 | 0.10 | 5.48 |
| 21 | Ranbp2 | 0.77 | 3.24E-15 | 6.75E-11 | 0.90 | 0.66 | 0.25 | 1.38 |
| 21 | Trappc8 | 1.12 | 3.96E-15 | 8.25E-11 | 0.54 | 0.24 | 0.30 | 2.21 |
| 21 | Ptger4 | 2.11 | 4.11E-15 | 8.54E-11 | 0.13 | 0.03 | 0.11 | 5.12 |
| 21 | Inpp5b | 1.34 | 4.43E-15 | 9.21E-11 | 0.34 | 0.12 | 0.22 | 2.90 |
| 21 | Ankrd44 | 1.09 | 4.55E-15 | 9.46E-11 | 0.61 | 0.28 | 0.33 | 2.18 |
| 21 | Adam9 | 1.67 | 4.62E-15 | 9.61E-11 | 0.36 | 0.14 | 0.22 | 2.60 |
| 21 | Rab10os | 1.20 | 4.62E-15 | 9.61E-11 | 0.45 | 0.18 | 0.26 | 2.43 |
| 21 | Pcbd2 | 0.84 | 5.02E-15 | 1.04E-10 | 0.78 | 0.43 | 0.35 | 1.80 |
| 21 | Snx18 | 1.12 | 5.05E-15 | 1.05E-10 | 0.69 | 0.38 | 0.32 | 1.84 |
| 21 | Eml3 | 1.29 | 5.17E-15 | 1.08E-10 | 0.35 | 0.12 | 0.23 | 2.82 |
| 21 | Heatr1 | 1.16 | 5.22E-15 | 1.09E-10 | 0.62 | 0.31 | 0.31 | 2.01 |
| 21 | Inpp5k | 1.51 | 5.32E-15 | 1.11E-10 | 0.32 | 0.11 | 0.21 | 2.88 |
| 21 | Nab1 | 1.19 | 5.35E-15 | 1.11E-10 | 0.36 | 0.13 | 0.24 | 2.82 |
| 21 | Aldh2 | 0.79 | 5.38E-15 | 1.12E-10 | 0.92 | 0.78 | 0.15 | 1.19 |
| 21 | Atp6v1h | 1.06 | 5.68E-15 | 1.18E-10 | 0.68 | 0.36 | 0.32 | 1.88 |
| 21 | Ubc | 0.56 | 5.77E-15 | 1.20E-10 | 1.00 | 0.98 | 0.03 | 1.03 |
| 21 | Lyrm9 | 1.05 | 5.79E-15 | 1.20E-10 | 0.44 | 0.17 | 0.27 | 2.56 |
| 21 | Smarcd2 | 0.92 | 5.89E-15 | 1.23E-10 | 0.62 | 0.29 | 0.33 | 2.12 |
| 21 | Bcap29 | 0.92 | 5.90E-15 | 1.23E-10 | 0.69 | 0.34 | 0.34 | 2.00 |
| 21 | Arhgap19 | 1.05 | 5.94E-15 | 1.23E-10 | 0.59 | 0.29 | 0.31 | 2.07 |
| 21 | Map3k1 | 1.03 | 6.35E-15 | 1.32E-10 | 0.57 | 0.26 | 0.31 | 2.17 |
| 21 | Tmem141 | 1.28 | 7.13E-15 | 1.48E-10 | 0.46 | 0.19 | 0.27 | 2.38 |
| 21 | Cbl | 0.83 | 7.15E-15 | 1.49E-10 | 0.81 | 0.47 | 0.34 | 1.73 |
| 21 | Man2c1 | 1.41 | 7.74E-15 | 1.61E-10 | 0.32 | 0.11 | 0.21 | 2.97 |
| 21 | Rbks | 1.83 | 7.91E-15 | 1.65E-10 | 0.15 | 0.03 | 0.12 | 4.67 |
| 21 | Erlin1 | 1.19 | 8.29E-15 | 1.72E-10 | 0.40 | 0.15 | 0.25 | 2.63 |

**Supplementary Table 1. Marker genes for the original clusters of integrated clustering of E11.5 control and NotchGOF λ-junction cells.**

| | | | | | | | | |
|---|---|---|---|---|---|---|---|---|
| 21 | Itpkb | 1.17 | 8.31E-15 | 1.73E-10 | 0.47 | 0.19 | 0.28 | 2.43 |
| 21 | Gm42670 | 2.36 | 8.59E-15 | 1.79E-10 | 0.11 | 0.02 | 0.09 | 6.18 |
| 21 | Slc38a10 | 1.06 | 8.76E-15 | 1.82E-10 | 0.66 | 0.34 | 0.33 | 1.98 |
| 21 | Prtn3 | 4.40 | 9.29E-15 | 1.93E-10 | 0.34 | 0.14 | 0.20 | 2.45 |
| 21 | Rab11a | 0.81 | 9.45E-15 | 1.97E-10 | 0.90 | 0.64 | 0.26 | 1.41 |
| 21 | Dnajc3 | 0.90 | 9.69E-15 | 2.02E-10 | 0.83 | 0.52 | 0.31 | 1.59 |
| 21 | Slc35d2 | 1.79 | 1.02E-14 | 2.12E-10 | 0.13 | 0.03 | 0.11 | 5.12 |
| 21 | Emc2 | 0.72 | 1.05E-14 | 2.18E-10 | 0.88 | 0.59 | 0.29 | 1.49 |
| 21 | Map3k14 | 1.97 | 1.14E-14 | 2.36E-10 | 0.16 | 0.04 | 0.12 | 4.35 |
| 21 | Gaa | 1.31 | 1.15E-14 | 2.39E-10 | 0.45 | 0.18 | 0.26 | 2.43 |
| 21 | Isoc2b | 2.18 | 1.25E-14 | 2.60E-10 | 0.13 | 0.02 | 0.10 | 5.25 |
| 21 | Stard3 | 1.12 | 1.30E-14 | 2.69E-10 | 0.52 | 0.24 | 0.29 | 2.23 |
| 21 | Fbxl8 | 2.05 | 1.30E-14 | 2.71E-10 | 0.12 | 0.02 | 0.10 | 5.41 |
| 21 | Map3k3 | 1.21 | 1.33E-14 | 2.77E-10 | 0.40 | 0.15 | 0.25 | 2.59 |
| 21 | Tet2 | 1.09 | 1.46E-14 | 3.03E-10 | 0.55 | 0.25 | 0.30 | 2.19 |
| 21 | Fxyd2 | 2.89 | 1.51E-14 | 3.15E-10 | 0.11 | 0.02 | 0.09 | 5.60 |
| 21 | Foxn2 | 1.23 | 1.61E-14 | 3.34E-10 | 0.41 | 0.16 | 0.25 | 2.55 |
| 21 | Mpc2 | 0.75 | 1.61E-14 | 3.36E-10 | 0.93 | 0.79 | 0.14 | 1.18 |
| 21 | Trim65 | 1.85 | 1.71E-14 | 3.55E-10 | 0.16 | 0.04 | 0.12 | 4.35 |
| 21 | Selenbp1 | 1.31 | 1.73E-14 | 3.59E-10 | 0.27 | 0.09 | 0.19 | 3.21 |
| 21 | Dtnbp1 | 0.91 | 1.85E-14 | 3.85E-10 | 0.75 | 0.42 | 0.33 | 1.79 |
| 21 | Ssu72 | 0.58 | 1.88E-14 | 3.92E-10 | 0.96 | 0.84 | 0.12 | 1.15 |
| 21 | Idua | 2.09 | 1.89E-14 | 3.93E-10 | 0.15 | 0.03 | 0.12 | 4.53 |
| 21 | Ap3b1 | 0.90 | 1.99E-14 | 4.14E-10 | 0.79 | 0.47 | 0.32 | 1.68 |
| 21 | Arl6ip1 | 0.41 | 2.13E-14 | 4.44E-10 | 0.99 | 0.89 | 0.10 | 1.12 |
| 21 | Mthfd1l | 1.20 | 2.16E-14 | 4.50E-10 | 0.41 | 0.16 | 0.25 | 2.57 |
| 21 | Gyg | 1.08 | 2.29E-14 | 4.76E-10 | 0.73 | 0.42 | 0.31 | 1.74 |
| 21 | Dnajc13 | 1.10 | 2.34E-14 | 4.88E-10 | 0.67 | 0.38 | 0.30 | 1.78 |
| 21 | Klf13 | 1.04 | 2.36E-14 | 4.91E-10 | 0.71 | 0.37 | 0.34 | 1.92 |
| 21 | Cdk2ap2 | 0.84 | 2.36E-14 | 4.92E-10 | 0.78 | 0.46 | 0.32 | 1.71 |
| 21 | Slc17a5 | 1.19 | 2.39E-14 | 4.97E-10 | 0.44 | 0.18 | 0.26 | 2.41 |
| 21 | Sft2d2 | 1.35 | 2.44E-14 | 5.07E-10 | 0.39 | 0.15 | 0.24 | 2.61 |
| 21 | Ten1 | 0.78 | 2.55E-14 | 5.30E-10 | 0.86 | 0.57 | 0.30 | 1.52 |
| 21 | Parp8 | 1.26 | 2.76E-14 | 5.74E-10 | 0.37 | 0.14 | 0.23 | 2.71 |
| 21 | Sppl2a | 0.87 | 2.79E-14 | 5.80E-10 | 0.83 | 0.47 | 0.36 | 1.76 |
| 21 | Aftph | 1.08 | 2.90E-14 | 6.03E-10 | 0.54 | 0.25 | 0.29 | 2.14 |
| 21 | Taf10 | 0.45 | 3.14E-14 | 6.53E-10 | 0.99 | 0.93 | 0.06 | 1.06 |
| 21 | Tmem268 | 1.56 | 3.21E-14 | 6.68E-10 | 0.27 | 0.08 | 0.18 | 3.17 |
| 21 | Mafb | 1.46 | 3.30E-14 | 6.86E-10 | 0.49 | 0.22 | 0.27 | 2.22 |
| 21 | Shmt2 | 1.14 | 3.76E-14 | 7.82E-10 | 0.62 | 0.34 | 0.28 | 1.83 |
| 21 | Siah2 | 1.29 | 3.77E-14 | 7.83E-10 | 0.25 | 0.08 | 0.18 | 3.36 |
| 21 | Rb1 | 0.96 | 3.77E-14 | 7.84E-10 | 0.41 | 0.16 | 0.25 | 2.55 |
| 21 | Zfp385a | 0.99 | 3.92E-14 | 8.16E-10 | 0.54 | 0.25 | 0.28 | 2.12 |
| 21 | Aldh9a1 | 1.12 | 4.05E-14 | 8.43E-10 | 0.62 | 0.32 | 0.31 | 1.97 |
| 21 | Mtmr1 | 1.63 | 4.17E-14 | 8.66E-10 | 0.24 | 0.07 | 0.17 | 3.35 |
| 21 | Fhod1 | 1.50 | 4.35E-14 | 9.04E-10 | 0.27 | 0.09 | 0.19 | 3.14 |
| 21 | Zbtb7a | 1.01 | 4.43E-14 | 9.21E-10 | 0.63 | 0.33 | 0.30 | 1.90 |
| 21 | Bnip2 | 0.74 | 5.00E-14 | 1.04E-09 | 0.92 | 0.69 | 0.23 | 1.34 |
| 21 | Sh2d3c | 1.21 | 5.15E-14 | 1.07E-09 | 0.30 | 0.10 | 0.20 | 3.01 |
| 21 | Edf1 | 0.39 | 5.18E-14 | 1.08E-09 | 0.99 | 0.96 | 0.03 | 1.03 |
| 21 | Pepd | 0.91 | 5.68E-14 | 1.18E-09 | 0.76 | 0.42 | 0.34 | 1.81 |
| 21 | Gcnt2 | 1.99 | 5.70E-14 | 1.19E-09 | 0.18 | 0.05 | 0.14 | 3.87 |
| 21 | Galc | 1.60 | 5.96E-14 | 1.24E-09 | 0.42 | 0.18 | 0.25 | 2.40 |
| 21 | Spty2d1 | 0.99 | 6.00E-14 | 1.25E-09 | 0.54 | 0.25 | 0.29 | 2.14 |
| 21 | Ctso | 1.82 | 6.73E-14 | 1.40E-09 | 0.22 | 0.07 | 0.16 | 3.45 |
| 21 | Plxnb2 | 1.01 | 6.84E-14 | 1.42E-09 | 0.74 | 0.53 | 0.21 | 1.39 |

**Supplementary Table 1. Marker genes for the original clusters of integrated clustering of E11.5 control and NotchGOF λ-junction cells.**

| | | | | | | | | |
|---|---|---|---|---|---|---|---|---|
| 21 | Ankfy1 | 1.01 | 7.29E-14 | 1.52E-09 | 0.59 | 0.28 | 0.31 | 2.08 |
| 21 | Creb3 | 1.10 | 9.15E-14 | 1.90E-09 | 0.45 | 0.19 | 0.26 | 2.37 |
| 21 | Vmp1 | 0.83 | 9.45E-14 | 1.97E-09 | 0.87 | 0.59 | 0.28 | 1.48 |
| 21 | Tmem175 | 1.19 | 1.03E-13 | 2.13E-09 | 0.30 | 0.10 | 0.20 | 2.98 |
| 21 | Vat1 | 1.47 | 1.08E-13 | 2.25E-09 | 0.36 | 0.14 | 0.22 | 2.63 |
| 21 | Rnf31 | 1.33 | 1.09E-13 | 2.28E-09 | 0.23 | 0.07 | 0.16 | 3.45 |
| 21 | Comt | 0.88 | 1.12E-13 | 2.34E-09 | 0.83 | 0.54 | 0.29 | 1.54 |
| 21 | Calr | 0.65 | 1.15E-13 | 2.38E-09 | 0.99 | 0.97 | 0.02 | 1.02 |
| 21 | Agpat2 | 1.00 | 1.16E-13 | 2.41E-09 | 0.42 | 0.17 | 0.25 | 2.53 |
| 21 | Hist1h2bc | 1.57 | 1.21E-13 | 2.53E-09 | 0.52 | 0.26 | 0.26 | 1.99 |
| 21 | Ghitm | 0.72 | 1.22E-13 | 2.53E-09 | 0.90 | 0.67 | 0.23 | 1.35 |
| 21 | Oplah | 2.44 | 1.31E-13 | 2.71E-09 | 0.11 | 0.02 | 0.09 | 5.33 |
| 21 | Gpr108 | 1.17 | 1.35E-13 | 2.81E-09 | 0.40 | 0.16 | 0.24 | 2.49 |
| 21 | Cd82 | 1.98 | 1.39E-13 | 2.89E-09 | 0.22 | 0.07 | 0.16 | 3.39 |
| 21 | Ilk | 0.86 | 1.39E-13 | 2.90E-09 | 0.89 | 0.61 | 0.28 | 1.46 |
| 21 | Tdrkh | 1.72 | 1.41E-13 | 2.93E-09 | 0.21 | 0.06 | 0.15 | 3.50 |
| 21 | Atg7 | 1.23 | 1.44E-13 | 3.00E-09 | 0.34 | 0.13 | 0.22 | 2.74 |
| 21 | Acaa1a | 1.01 | 1.51E-13 | 3.13E-09 | 0.75 | 0.46 | 0.29 | 1.63 |
| 21 | Arhgap1 | 0.90 | 1.53E-13 | 3.19E-09 | 0.56 | 0.26 | 0.30 | 2.16 |
| 21 | Ufsp2 | 0.80 | 1.57E-13 | 3.26E-09 | 0.80 | 0.49 | 0.31 | 1.62 |
| 21 | Nsf | 0.99 | 1.59E-13 | 3.30E-09 | 0.64 | 0.34 | 0.31 | 1.91 |
| 21 | Ldlrap1 | 1.33 | 1.62E-13 | 3.37E-09 | 0.43 | 0.18 | 0.25 | 2.37 |
| 21 | Gm26532 | 1.88 | 1.91E-13 | 3.97E-09 | 0.16 | 0.04 | 0.12 | 4.13 |
| 21 | Mov10 | 0.87 | 1.92E-13 | 4.00E-09 | 0.47 | 0.19 | 0.28 | 2.42 |
| 21 | Myc | 1.51 | 2.09E-13 | 4.35E-09 | 0.39 | 0.16 | 0.23 | 2.40 |
| 21 | Iscu | 0.78 | 2.15E-13 | 4.46E-09 | 0.90 | 0.64 | 0.25 | 1.39 |
| 21 | Tmem256 | 0.63 | 2.16E-13 | 4.50E-09 | 0.99 | 0.89 | 0.10 | 1.11 |
| 21 | Hpgd | 0.73 | 2.25E-13 | 4.67E-09 | 0.67 | 0.35 | 0.32 | 1.93 |
| 21 | Paox | 1.33 | 2.26E-13 | 4.69E-09 | 0.39 | 0.16 | 0.23 | 2.48 |
| 21 | Egfl7 | 0.75 | 2.28E-13 | 4.74E-09 | 0.48 | 0.22 | 0.26 | 2.17 |
| 21 | Slc23a2 | 1.63 | 2.49E-13 | 5.18E-09 | 0.25 | 0.08 | 0.17 | 3.18 |
| 21 | Atox1 | 0.29 | 2.50E-13 | 5.19E-09 | 1.00 | 0.95 | 0.05 | 1.05 |
| 21 | Rnf17 | 1.48 | 2.60E-13 | 5.40E-09 | 0.15 | 0.03 | 0.11 | 4.45 |
| 21 | Pacsin2 | 0.96 | 2.63E-13 | 5.47E-09 | 0.65 | 0.35 | 0.30 | 1.87 |
| 21 | Fam107b | 0.85 | 2.68E-13 | 5.57E-09 | 0.65 | 0.36 | 0.30 | 1.83 |
| 21 | Ppp1r9b | 1.14 | 2.68E-13 | 5.57E-09 | 0.53 | 0.26 | 0.28 | 2.08 |
| 21 | Mcts1 | 0.86 | 2.71E-13 | 5.64E-09 | 0.75 | 0.47 | 0.28 | 1.58 |
| 21 | Limd2 | 0.55 | 2.75E-13 | 5.72E-09 | 0.99 | 0.90 | 0.09 | 1.10 |
| 21 | Prelid1 | 0.49 | 2.89E-13 | 6.00E-09 | 0.99 | 0.93 | 0.07 | 1.07 |
| 21 | Tbc1d5 | 1.07 | 3.05E-13 | 6.34E-09 | 0.47 | 0.21 | 0.26 | 2.29 |
| 21 | Arid4a | 0.82 | 3.05E-13 | 6.34E-09 | 0.78 | 0.47 | 0.31 | 1.65 |
| 21 | Arl8b | 0.88 | 3.19E-13 | 6.64E-09 | 0.76 | 0.47 | 0.29 | 1.61 |
| 21 | Zfp984 | 1.65 | 3.28E-13 | 6.82E-09 | 0.24 | 0.07 | 0.17 | 3.26 |
| 21 | Myb | 2.49 | 3.37E-13 | 7.00E-09 | 0.25 | 0.08 | 0.17 | 3.00 |
| 21 | Ncstn | 1.03 | 3.60E-13 | 7.48E-09 | 0.58 | 0.29 | 0.29 | 1.99 |
| 21 | Vrk2 | 0.88 | 3.64E-13 | 7.57E-09 | 0.51 | 0.23 | 0.28 | 2.26 |
| 21 | Apmap | 1.38 | 3.68E-13 | 7.66E-09 | 0.44 | 0.19 | 0.25 | 2.35 |
| 21 | Myom1 | 1.71 | 3.71E-13 | 7.71E-09 | 0.14 | 0.03 | 0.11 | 4.52 |
| 21 | Epg5 | 1.50 | 3.92E-13 | 8.15E-09 | 0.17 | 0.04 | 0.13 | 4.00 |
| 21 | Susd6 | 1.16 | 3.98E-13 | 8.27E-09 | 0.46 | 0.20 | 0.26 | 2.31 |
| 21 | Arsa | 1.47 | 4.14E-13 | 8.60E-09 | 0.22 | 0.07 | 0.16 | 3.34 |
| 21 | Fkbp2 | 0.73 | 4.34E-13 | 9.02E-09 | 0.91 | 0.65 | 0.26 | 1.39 |
| 21 | Epn1 | 0.87 | 4.65E-13 | 9.67E-09 | 0.87 | 0.62 | 0.25 | 1.40 |
| 21 | Ptges3l | 1.49 | 4.93E-13 | 1.03E-08 | 0.21 | 0.06 | 0.15 | 3.50 |
| 21 | Ripor1 | 1.55 | 4.98E-13 | 1.04E-08 | 0.27 | 0.09 | 0.18 | 2.99 |
| 21 | Rffl | 1.55 | 5.03E-13 | 1.05E-08 | 0.23 | 0.07 | 0.16 | 3.25 |

**Supplementary Table 1. Marker genes for the original clusters of integrated clustering of E11.5 control and NotchGOF λ-junction cells.**

| | | | | | | | | |
|---|---|---|---|---|---|---|---|---|
| 21 | Mavs | 1.53 | 5.11E-13 | 1.06E-08 | 0.27 | 0.09 | 0.18 | 2.99 |
| 21 | Ctsl | 1.24 | 5.28E-13 | 1.10E-08 | 0.93 | 0.82 | 0.11 | 1.13 |
| 21 | Shtn1 | 1.60 | 5.37E-13 | 1.12E-08 | 0.19 | 0.05 | 0.14 | 3.63 |
| 21 | Wdfy2 | 1.15 | 5.40E-13 | 1.12E-08 | 0.40 | 0.16 | 0.24 | 2.48 |
| 21 | Smg9 | 1.09 | 5.74E-13 | 1.19E-08 | 0.36 | 0.14 | 0.23 | 2.62 |
| 21 | Hltf | 0.83 | 5.93E-13 | 1.23E-08 | 0.64 | 0.32 | 0.32 | 2.00 |
| 21 | Depdc7 | 1.84 | 6.02E-13 | 1.25E-08 | 0.14 | 0.03 | 0.11 | 4.52 |
| 21 | Jun | 0.84 | 6.39E-13 | 1.33E-08 | 0.90 | 0.67 | 0.23 | 1.34 |
| 21 | Pbxip1 | 1.04 | 6.84E-13 | 1.42E-08 | 0.47 | 0.20 | 0.27 | 2.32 |
| 21 | Bach1 | 0.97 | 6.93E-13 | 1.44E-08 | 0.79 | 0.48 | 0.31 | 1.66 |
| 21 | Pgap1 | 1.31 | 7.02E-13 | 1.46E-08 | 0.37 | 0.15 | 0.22 | 2.46 |
| 21 | Smcr8 | 1.72 | 7.29E-13 | 1.52E-08 | 0.25 | 0.08 | 0.17 | 3.07 |
| 21 | Ppp1ca | 0.36 | 7.46E-13 | 1.55E-08 | 1.00 | 0.98 | 0.02 | 1.02 |
| 21 | Ecpas | 0.85 | 7.69E-13 | 1.60E-08 | 0.75 | 0.42 | 0.33 | 1.77 |
| 21 | Adrb1 | 2.18 | 8.32E-13 | 1.73E-08 | 0.11 | 0.02 | 0.09 | 5.25 |
| 21 | Nfkb2 | 1.73 | 8.41E-13 | 1.75E-08 | 0.22 | 0.07 | 0.16 | 3.34 |
| 21 | Mapk14 | 0.98 | 8.56E-13 | 1.78E-08 | 0.62 | 0.33 | 0.29 | 1.88 |
| 21 | Commd7 | 0.71 | 9.11E-13 | 1.89E-08 | 0.84 | 0.57 | 0.27 | 1.47 |
| 21 | Dusp22 | 0.99 | 9.29E-13 | 1.93E-08 | 0.46 | 0.20 | 0.26 | 2.28 |
| 21 | Eea1 | 0.79 | 9.46E-13 | 1.97E-08 | 0.78 | 0.43 | 0.35 | 1.82 |
| 21 | Irak1 | 0.84 | 9.64E-13 | 2.00E-08 | 0.75 | 0.44 | 0.31 | 1.70 |
| 21 | Casp7 | 1.09 | 9.88E-13 | 2.05E-08 | 0.41 | 0.17 | 0.24 | 2.44 |
| 21 | Acot8 | 1.07 | 1.04E-12 | 2.16E-08 | 0.38 | 0.15 | 0.23 | 2.54 |
| 21 | Crat | 1.09 | 1.06E-12 | 2.21E-08 | 0.46 | 0.20 | 0.25 | 2.23 |
| 21 | Tbc1d4 | 1.62 | 1.08E-12 | 2.24E-08 | 0.31 | 0.11 | 0.19 | 2.70 |
| 21 | Wnk1 | 0.73 | 1.10E-12 | 2.28E-08 | 0.96 | 0.78 | 0.18 | 1.23 |
| 21 | Vps26a | 0.96 | 1.14E-12 | 2.37E-08 | 0.73 | 0.43 | 0.30 | 1.70 |
| 21 | Cyld | 1.15 | 1.15E-12 | 2.38E-08 | 0.48 | 0.22 | 0.26 | 2.16 |
| 21 | Ciao3 | 1.13 | 1.15E-12 | 2.40E-08 | 0.40 | 0.17 | 0.23 | 2.40 |
| 21 | Rmc1 | 1.26 | 1.17E-12 | 2.44E-08 | 0.25 | 0.08 | 0.17 | 3.11 |
| 21 | Rras | 1.14 | 1.19E-12 | 2.48E-08 | 0.48 | 0.22 | 0.26 | 2.18 |
| 21 | Vps35l | 1.08 | 1.25E-12 | 2.61E-08 | 0.54 | 0.26 | 0.28 | 2.07 |
| 21 | Trpc4ap | 0.85 | 1.30E-12 | 2.70E-08 | 0.82 | 0.53 | 0.29 | 1.54 |
| 21 | Tmem189 | 0.89 | 1.32E-12 | 2.74E-08 | 0.62 | 0.30 | 0.31 | 2.04 |
| 21 | Adgrg1 | 2.89 | 1.43E-12 | 2.98E-08 | 0.16 | 0.04 | 0.12 | 3.83 |
| 21 | Marchf2 | 0.73 | 1.44E-12 | 3.00E-08 | 0.82 | 0.51 | 0.30 | 1.59 |
| 21 | Snx10 | 1.16 | 1.75E-12 | 3.63E-08 | 0.39 | 0.16 | 0.23 | 2.44 |
| 21 | Slc30a5 | 0.94 | 1.79E-12 | 3.72E-08 | 0.61 | 0.31 | 0.30 | 1.95 |
| 21 | Lamtor2 | 0.55 | 1.98E-12 | 4.11E-08 | 0.97 | 0.89 | 0.07 | 1.08 |
| 21 | Rnft1 | 0.94 | 2.11E-12 | 4.38E-08 | 0.50 | 0.23 | 0.27 | 2.21 |
| 21 | Map11 | 1.31 | 2.12E-12 | 4.41E-08 | 0.29 | 0.11 | 0.19 | 2.80 |
| 21 | Atp6v0a2 | 0.97 | 2.24E-12 | 4.67E-08 | 0.55 | 0.26 | 0.29 | 2.11 |
| 21 | Rab11fip5 | 2.02 | 2.44E-12 | 5.08E-08 | 0.14 | 0.03 | 0.11 | 4.24 |
| 21 | Ppp1r15a | 1.09 | 2.57E-12 | 5.35E-08 | 0.51 | 0.25 | 0.26 | 2.04 |
| 21 | Adipor1 | 0.76 | 2.62E-12 | 5.45E-08 | 0.87 | 0.56 | 0.31 | 1.55 |
| 21 | Etv3 | 1.07 | 2.71E-12 | 5.65E-08 | 0.49 | 0.22 | 0.27 | 2.20 |
| 21 | Raph1 | 1.01 | 2.94E-12 | 6.10E-08 | 0.73 | 0.42 | 0.31 | 1.72 |
| 21 | Bcap31 | 0.56 | 2.95E-12 | 6.14E-08 | 0.95 | 0.75 | 0.20 | 1.27 |
| 21 | Usp12 | 0.97 | 3.14E-12 | 6.52E-08 | 0.46 | 0.21 | 0.26 | 2.25 |
| 21 | Agfg2 | 0.86 | 3.20E-12 | 6.66E-08 | 0.48 | 0.21 | 0.26 | 2.25 |
| 21 | Cars2 | 1.06 | 3.37E-12 | 7.01E-08 | 0.42 | 0.18 | 0.24 | 2.37 |
| 21 | Rap2a | 1.00 | 3.49E-12 | 7.27E-08 | 0.61 | 0.32 | 0.29 | 1.88 |
| 21 | Ifitm3 | 2.04 | 3.76E-12 | 7.83E-08 | 0.54 | 0.30 | 0.24 | 1.81 |
| 21 | Vim | 0.71 | 4.14E-12 | 8.61E-08 | 0.99 | 0.84 | 0.15 | 1.18 |
| 21 | Ggh | 0.95 | 4.26E-12 | 8.85E-08 | 0.54 | 0.25 | 0.28 | 2.12 |
| 21 | Exosc5 | 0.86 | 4.31E-12 | 8.97E-08 | 0.80 | 0.58 | 0.22 | 1.38 |

**Supplementary Table 1. Marker genes for the original clusters of integrated clustering of E11.5 control and NotchGOF λ-junction cells.**

| | | | | | | | | |
|---|---|---|---|---|---|---|---|---|
| 21 | Adar | 1.19 | 4.46E-12 | 9.28E-08 | 0.37 | 0.15 | 0.22 | 2.44 |
| 21 | Chmp4b | 0.57 | 4.98E-12 | 1.04E-07 | 0.97 | 0.85 | 0.12 | 1.14 |
| 21 | Cept1 | 0.84 | 5.05E-12 | 1.05E-07 | 0.62 | 0.33 | 0.29 | 1.87 |
| 21 | Tox4 | 0.76 | 5.34E-12 | 1.11E-07 | 0.81 | 0.52 | 0.29 | 1.56 |
| 21 | Dip2b | 0.91 | 5.41E-12 | 1.13E-07 | 0.64 | 0.34 | 0.29 | 1.85 |
| 21 | Hes6 | 0.51 | 5.93E-12 | 1.23E-07 | 0.62 | 0.33 | 0.29 | 1.87 |
| 21 | Steap3 | 2.54 | 6.20E-12 | 1.29E-07 | 0.11 | 0.02 | 0.09 | 4.67 |
| 21 | Sprtn | 1.34 | 6.46E-12 | 1.34E-07 | 0.26 | 0.09 | 0.17 | 2.94 |
| 21 | Lnpep | 0.79 | 6.63E-12 | 1.38E-07 | 0.59 | 0.29 | 0.30 | 2.04 |
| 21 | Cpd | 1.39 | 6.68E-12 | 1.39E-07 | 0.59 | 0.34 | 0.26 | 1.76 |
| 21 | Chid1 | 0.92 | 6.71E-12 | 1.40E-07 | 0.53 | 0.25 | 0.28 | 2.09 |
| 21 | Irf3 | 0.81 | 7.00E-12 | 1.46E-07 | 0.63 | 0.33 | 0.30 | 1.91 |
| 21 | Fbrs | 0.85 | 7.07E-12 | 1.47E-07 | 0.46 | 0.20 | 0.26 | 2.30 |
| 21 | Flcn | 1.35 | 7.21E-12 | 1.50E-07 | 0.30 | 0.11 | 0.19 | 2.66 |
| 21 | Bahd1 | 1.64 | 7.36E-12 | 1.53E-07 | 0.16 | 0.04 | 0.12 | 3.83 |
| 21 | Ogfod2 | 1.17 | 8.36E-12 | 1.74E-07 | 0.28 | 0.10 | 0.18 | 2.83 |
| 21 | Nrm | 0.95 | 8.38E-12 | 1.74E-07 | 0.64 | 0.35 | 0.28 | 1.80 |
| 21 | Acox1 | 1.43 | 8.48E-12 | 1.76E-07 | 0.35 | 0.14 | 0.21 | 2.43 |
| 21 | Zbtb38 | 1.13 | 8.55E-12 | 1.78E-07 | 0.34 | 0.14 | 0.21 | 2.52 |
| 21 | Xbp1 | 0.84 | 9.20E-12 | 1.91E-07 | 0.77 | 0.48 | 0.29 | 1.61 |
| 21 | Klf7 | 1.00 | 9.46E-12 | 1.97E-07 | 0.48 | 0.23 | 0.25 | 2.11 |
| 21 | Ecm1 | 2.41 | 9.54E-12 | 1.98E-07 | 0.11 | 0.02 | 0.09 | 4.67 |
| 21 | Usp8 | 0.80 | 1.11E-11 | 2.30E-07 | 0.78 | 0.46 | 0.32 | 1.68 |
| 21 | Tmem87b | 1.03 | 1.11E-11 | 2.31E-07 | 0.47 | 0.21 | 0.26 | 2.20 |
| 21 | Pkig | 0.56 | 1.13E-11 | 2.35E-07 | 0.94 | 0.79 | 0.16 | 1.20 |
| 21 | Tbc1d2b | 1.46 | 1.21E-11 | 2.51E-07 | 0.25 | 0.08 | 0.16 | 2.92 |
| 21 | Krcc1 | 0.70 | 1.22E-11 | 2.53E-07 | 0.85 | 0.58 | 0.28 | 1.48 |
| 21 | Bad | 0.72 | 1.28E-11 | 2.66E-07 | 0.81 | 0.53 | 0.28 | 1.54 |
| 21 | Galnt12 | 2.24 | 1.30E-11 | 2.70E-07 | 0.11 | 0.02 | 0.09 | 4.67 |
| 21 | Soat1 | 1.24 | 1.30E-11 | 2.71E-07 | 0.69 | 0.44 | 0.24 | 1.55 |
| 21 | Auh | 0.90 | 1.36E-11 | 2.82E-07 | 0.55 | 0.28 | 0.27 | 1.95 |
| 21 | Cyfip2 | 1.28 | 1.37E-11 | 2.85E-07 | 0.26 | 0.09 | 0.17 | 2.88 |
| 21 | Hras | 0.50 | 1.43E-11 | 2.97E-07 | 0.94 | 0.80 | 0.14 | 1.18 |
| 21 | Tmed3 | 0.79 | 1.56E-11 | 3.25E-07 | 0.72 | 0.45 | 0.27 | 1.61 |
| 21 | BC004004 | 0.94 | 1.64E-11 | 3.41E-07 | 0.59 | 0.32 | 0.27 | 1.85 |
| 21 | Dennd1b | 1.05 | 1.74E-11 | 3.61E-07 | 0.39 | 0.16 | 0.23 | 2.42 |
| 21 | Zdhhc9 | 1.30 | 1.77E-11 | 3.68E-07 | 0.26 | 0.09 | 0.17 | 2.91 |
| 21 | Gm26782 | 1.63 | 1.82E-11 | 3.78E-07 | 0.21 | 0.07 | 0.14 | 3.18 |
| 21 | Slk | 0.69 | 1.82E-11 | 3.80E-07 | 0.80 | 0.53 | 0.27 | 1.51 |
| 21 | Retsat | 1.56 | 1.83E-11 | 3.81E-07 | 0.18 | 0.05 | 0.13 | 3.43 |
| 21 | Hspa8 | 0.38 | 1.87E-11 | 3.89E-07 | 1.00 | 1.00 | 0.00 | 1.00 |
| 21 | Nin | 0.88 | 1.89E-11 | 3.93E-07 | 0.54 | 0.27 | 0.27 | 1.99 |
| 21 | Hivep2 | 1.42 | 1.90E-11 | 3.95E-07 | 0.23 | 0.08 | 0.16 | 3.04 |
| 21 | Uckl1 | 0.88 | 2.03E-11 | 4.22E-07 | 0.32 | 0.12 | 0.20 | 2.69 |
| 21 | Nuak2 | 1.56 | 2.03E-11 | 4.23E-07 | 0.15 | 0.04 | 0.11 | 3.97 |
| 21 | 1810037I17Rik | 0.55 | 2.14E-11 | 4.45E-07 | 0.94 | 0.84 | 0.10 | 1.12 |
| 21 | Rexo2 | 0.74 | 2.17E-11 | 4.52E-07 | 0.81 | 0.59 | 0.22 | 1.37 |
| 21 | Traf5 | 1.80 | 2.19E-11 | 4.56E-07 | 0.15 | 0.04 | 0.11 | 3.76 |
| 21 | Reep5 | 0.48 | 2.43E-11 | 5.05E-07 | 0.97 | 0.78 | 0.19 | 1.24 |
| 21 | Vps13d | 0.97 | 2.48E-11 | 5.15E-07 | 0.35 | 0.14 | 0.21 | 2.52 |
| 21 | Cr1l | 0.77 | 2.63E-11 | 5.46E-07 | 0.60 | 0.32 | 0.28 | 1.88 |
| 21 | Npc1 | 1.03 | 2.71E-11 | 5.64E-07 | 0.53 | 0.26 | 0.27 | 2.05 |
| 21 | Cox5b | 0.34 | 2.72E-11 | 5.66E-07 | 0.99 | 0.99 | 0.01 | 1.01 |
| 21 | Cybc1 | 0.95 | 2.77E-11 | 5.76E-07 | 0.48 | 0.22 | 0.25 | 2.14 |
| 21 | Nr2c2ap | 0.98 | 2.79E-11 | 5.80E-07 | 0.48 | 0.23 | 0.25 | 2.06 |
| 21 | Dusp1 | 1.69 | 2.81E-11 | 5.85E-07 | 0.52 | 0.29 | 0.22 | 1.76 |

**Supplementary Table 1. Marker genes for the original clusters of integrated clustering of E11.5 control and NotchGOF λ-junction cells.**

| | | | | | | | | |
|---|---|---|---|---|---|---|---|---|
| 21 | Rmnd5b | 1.37 | 2.92E-11 | 6.08E-07 | 0.39 | 0.17 | 0.22 | 2.31 |
| 21 | Mbd2 | 0.64 | 3.06E-11 | 6.36E-07 | 0.83 | 0.56 | 0.27 | 1.49 |
| 21 | Map2k1 | 0.97 | 3.07E-11 | 6.38E-07 | 0.60 | 0.35 | 0.26 | 1.74 |
| 21 | Ier2 | 0.64 | 3.15E-11 | 6.55E-07 | 0.94 | 0.79 | 0.15 | 1.19 |
| 21 | BC029722 | 1.02 | 3.25E-11 | 6.76E-07 | 0.31 | 0.12 | 0.19 | 2.68 |
| 21 | Jtb | 0.78 | 3.33E-11 | 6.92E-07 | 0.80 | 0.53 | 0.26 | 1.50 |
| 21 | Tmem181a | 1.15 | 3.55E-11 | 7.38E-07 | 0.28 | 0.10 | 0.18 | 2.75 |
| 21 | Bcl10 | 0.85 | 3.62E-11 | 7.52E-07 | 0.76 | 0.50 | 0.27 | 1.53 |
| 21 | Oma1 | 0.99 | 3.64E-11 | 7.57E-07 | 0.41 | 0.18 | 0.23 | 2.27 |
| 21 | Prorsd1 | 0.80 | 4.07E-11 | 8.46E-07 | 0.67 | 0.38 | 0.29 | 1.76 |
| 21 | Qprt | 1.47 | 4.07E-11 | 8.46E-07 | 0.16 | 0.04 | 0.12 | 3.66 |
| 21 | Naprt | 2.02 | 4.20E-11 | 8.74E-07 | 0.11 | 0.03 | 0.09 | 4.48 |
| 21 | Rrbp1 | 0.50 | 4.22E-11 | 8.77E-07 | 0.99 | 0.90 | 0.10 | 1.11 |
| 21 | Nipsnap3b | 0.86 | 4.26E-11 | 8.86E-07 | 0.70 | 0.44 | 0.26 | 1.61 |
| 21 | Tnip1 | 1.20 | 4.38E-11 | 9.10E-07 | 0.28 | 0.10 | 0.18 | 2.72 |
| 21 | Pigt | 0.82 | 4.42E-11 | 9.20E-07 | 0.69 | 0.43 | 0.26 | 1.62 |
| 21 | Zfp800 | 0.91 | 4.68E-11 | 9.73E-07 | 0.53 | 0.27 | 0.27 | 2.00 |
| 21 | Tob2 | 0.98 | 4.79E-11 | 9.96E-07 | 0.50 | 0.24 | 0.26 | 2.07 |
| 21 | Itpripl1 | 1.37 | 4.84E-11 | 1.01E-06 | 0.26 | 0.09 | 0.17 | 2.76 |
| 21 | Rnf166 | 0.85 | 4.88E-11 | 1.01E-06 | 0.53 | 0.26 | 0.27 | 2.04 |
| 21 | Mospd3 | 1.16 | 5.13E-11 | 1.07E-06 | 0.39 | 0.17 | 0.22 | 2.28 |
| 21 | Gpt | 1.91 | 5.55E-11 | 1.15E-06 | 0.15 | 0.04 | 0.11 | 3.67 |
| 21 | Tbk1 | 1.06 | 5.66E-11 | 1.18E-06 | 0.54 | 0.28 | 0.26 | 1.91 |
| 21 | Mrpl36 | 0.69 | 6.06E-11 | 1.26E-06 | 0.92 | 0.72 | 0.20 | 1.28 |
| 21 | Csrnp1 | 2.00 | 6.69E-11 | 1.39E-06 | 0.13 | 0.03 | 0.10 | 4.03 |
| 21 | Slc35b2 | 0.73 | 6.74E-11 | 1.40E-06 | 0.72 | 0.41 | 0.31 | 1.74 |
| 21 | Spns2 | 2.92 | 6.98E-11 | 1.45E-06 | 0.12 | 0.03 | 0.09 | 4.25 |
| 21 | Rab31 | 0.96 | 7.11E-11 | 1.48E-06 | 0.47 | 0.22 | 0.25 | 2.09 |
| 21 | Ltbr | 0.90 | 7.89E-11 | 1.64E-06 | 0.48 | 0.23 | 0.25 | 2.10 |
| 21 | Elmsan1 | 1.12 | 8.03E-11 | 1.67E-06 | 0.31 | 0.12 | 0.19 | 2.59 |
| 21 | Rit1 | 0.93 | 8.25E-11 | 1.72E-06 | 0.52 | 0.25 | 0.26 | 2.04 |
| 21 | Snx9 | 0.87 | 8.37E-11 | 1.74E-06 | 0.59 | 0.31 | 0.27 | 1.87 |
| 21 | Tbc1d13 | 0.97 | 8.42E-11 | 1.75E-06 | 0.34 | 0.14 | 0.20 | 2.47 |
| 21 | Sqstm1 | 0.80 | 8.54E-11 | 1.78E-06 | 0.71 | 0.41 | 0.30 | 1.73 |
| 21 | Anxa5 | 1.00 | 8.66E-11 | 1.80E-06 | 0.83 | 0.61 | 0.23 | 1.38 |
| 21 | Usp24 | 0.91 | 8.86E-11 | 1.84E-06 | 0.57 | 0.31 | 0.27 | 1.87 |
| 21 | Iqsec1 | 1.15 | 8.91E-11 | 1.85E-06 | 0.34 | 0.14 | 0.20 | 2.43 |
| 21 | Coq8b | 0.99 | 9.00E-11 | 1.87E-06 | 0.32 | 0.12 | 0.19 | 2.58 |
| 21 | Rap1gds1 | 0.96 | 9.22E-11 | 1.92E-06 | 0.56 | 0.29 | 0.27 | 1.91 |
| 21 | Max | 0.80 | 9.39E-11 | 1.95E-06 | 0.69 | 0.40 | 0.28 | 1.70 |
| 21 | Itgav | 1.07 | 9.90E-11 | 2.06E-06 | 0.53 | 0.27 | 0.26 | 1.94 |
| 21 | Gclm | 0.75 | 1.03E-10 | 2.14E-06 | 0.69 | 0.39 | 0.30 | 1.78 |
| 21 | Akap10 | 1.20 | 1.08E-10 | 2.25E-06 | 0.32 | 0.13 | 0.19 | 2.44 |
| 21 | Slc39a1 | 0.72 | 1.09E-10 | 2.26E-06 | 0.53 | 0.26 | 0.28 | 2.07 |
| 21 | Pim2 | 2.11 | 1.10E-10 | 2.29E-06 | 0.11 | 0.03 | 0.09 | 4.48 |
| 21 | Myl12b | 0.52 | 1.28E-10 | 2.65E-06 | 0.98 | 0.89 | 0.09 | 1.10 |
| 21 | Nans | 0.65 | 1.32E-10 | 2.74E-06 | 0.86 | 0.60 | 0.26 | 1.42 |
| 21 | Atp2b1 | 0.70 | 1.35E-10 | 2.81E-06 | 0.96 | 0.85 | 0.11 | 1.13 |
| 21 | Lin28b | 0.93 | 1.44E-10 | 3.01E-06 | 0.43 | 0.21 | 0.23 | 2.10 |
| 21 | Atp13a3 | 1.05 | 1.45E-10 | 3.01E-06 | 0.68 | 0.41 | 0.27 | 1.66 |
| 21 | Dna2 | 0.83 | 1.47E-10 | 3.06E-06 | 0.52 | 0.26 | 0.26 | 2.00 |
| 21 | Alkbh3 | 0.86 | 1.48E-10 | 3.08E-06 | 0.43 | 0.20 | 0.24 | 2.18 |
| 21 | Ubl4a | 0.76 | 1.51E-10 | 3.14E-06 | 0.80 | 0.58 | 0.22 | 1.39 |
| 21 | Commd8 | 0.62 | 1.52E-10 | 3.15E-06 | 0.88 | 0.67 | 0.21 | 1.32 |
| 21 | Pfkfb3 | 1.38 | 1.52E-10 | 3.16E-06 | 0.27 | 0.11 | 0.17 | 2.60 |
| 21 | Sgpp1 | 1.05 | 1.61E-10 | 3.34E-06 | 0.28 | 0.10 | 0.18 | 2.72 |

**Supplementary Table 1. Marker genes for the original clusters of integrated clustering of E11.5 control and NotchGOF λ-junction cells.**

| | | | | | | | | |
|---|---|---|---|---|---|---|---|---|
| 21 | Mfsd10 | 0.86 | 1.61E-10 | 3.34E-06 | 0.60 | 0.35 | 0.25 | 1.72 |
| 21 | Idh3g | 0.60 | 1.62E-10 | 3.36E-06 | 0.86 | 0.66 | 0.20 | 1.31 |
| 21 | Spryd7 | 0.90 | 1.63E-10 | 3.39E-06 | 0.36 | 0.15 | 0.21 | 2.38 |
| 21 | Pxk | 0.80 | 1.63E-10 | 3.40E-06 | 0.39 | 0.17 | 0.23 | 2.36 |
| 21 | Mydgf | 0.56 | 1.76E-10 | 3.66E-06 | 0.94 | 0.73 | 0.20 | 1.28 |
| 21 | Eef1akmt4 | 0.88 | 1.81E-10 | 3.76E-06 | 0.48 | 0.23 | 0.25 | 2.07 |
| 21 | Mlec | 0.70 | 1.82E-10 | 3.79E-06 | 0.94 | 0.75 | 0.19 | 1.26 |
| 21 | Slc25a26 | 1.06 | 1.83E-10 | 3.80E-06 | 0.25 | 0.09 | 0.16 | 2.83 |
| 21 | Tm9sf2 | 0.61 | 1.90E-10 | 3.95E-06 | 0.89 | 0.62 | 0.27 | 1.44 |
| 21 | Anxa11 | 0.69 | 1.93E-10 | 4.02E-06 | 0.37 | 0.15 | 0.22 | 2.41 |
| 21 | Rufy1 | 0.93 | 1.98E-10 | 4.11E-06 | 0.50 | 0.25 | 0.25 | 2.02 |
| 21 | Tmem35b | 1.27 | 2.15E-10 | 4.48E-06 | 0.18 | 0.06 | 0.13 | 3.31 |
| 21 | Irak2 | 1.68 | 2.17E-10 | 4.52E-06 | 0.18 | 0.05 | 0.12 | 3.30 |
| 21 | Tmem123 | 0.70 | 2.18E-10 | 4.53E-06 | 0.62 | 0.32 | 0.29 | 1.92 |
| 21 | Hacd1 | 0.85 | 2.21E-10 | 4.60E-06 | 0.65 | 0.39 | 0.26 | 1.65 |
| 21 | Coro1b | 1.31 | 2.25E-10 | 4.69E-06 | 0.21 | 0.07 | 0.14 | 3.04 |
| 21 | Colca2 | 1.49 | 2.32E-10 | 4.82E-06 | 0.14 | 0.04 | 0.10 | 3.78 |
| 21 | Plbd2 | 0.99 | 2.36E-10 | 4.90E-06 | 0.48 | 0.24 | 0.24 | 1.98 |
| 21 | Ttll3 | 2.42 | 2.41E-10 | 5.02E-06 | 0.12 | 0.03 | 0.09 | 4.10 |
| 21 | Vps52 | 0.92 | 2.44E-10 | 5.07E-06 | 0.50 | 0.25 | 0.25 | 1.99 |
| 21 | Zfp991 | 1.28 | 2.48E-10 | 5.15E-06 | 0.25 | 0.09 | 0.16 | 2.78 |
| 21 | Selenot | 0.64 | 2.48E-10 | 5.17E-06 | 0.84 | 0.54 | 0.30 | 1.55 |
| 21 | Impa2 | 0.87 | 2.54E-10 | 5.29E-06 | 0.55 | 0.30 | 0.25 | 1.85 |
| 21 | 4833438C02Rik | 1.64 | 2.59E-10 | 5.38E-06 | 0.13 | 0.03 | 0.10 | 4.06 |
| 21 | Ccdc115 | 0.70 | 2.59E-10 | 5.38E-06 | 0.73 | 0.49 | 0.25 | 1.50 |
| 21 | Sap25 | 1.72 | 2.69E-10 | 5.59E-06 | 0.13 | 0.03 | 0.10 | 3.91 |
| 21 | Tmem250-ps | 0.98 | 2.70E-10 | 5.62E-06 | 0.34 | 0.14 | 0.20 | 2.40 |
| 21 | Fam120a | 0.81 | 2.78E-10 | 5.78E-06 | 0.71 | 0.46 | 0.26 | 1.56 |
| 21 | Dnajc5 | 0.71 | 3.21E-10 | 6.68E-06 | 0.83 | 0.52 | 0.30 | 1.58 |
| 21 | Atp5l | 0.29 | 3.21E-10 | 6.68E-06 | 0.99 | 0.99 | 0.00 | 1.00 |
| 21 | Snf8 | 0.58 | 3.42E-10 | 7.11E-06 | 0.89 | 0.75 | 0.14 | 1.18 |
| 21 | Gna13 | 0.83 | 3.44E-10 | 7.16E-06 | 0.69 | 0.43 | 0.26 | 1.61 |
| 21 | Gm12166 | 1.02 | 3.55E-10 | 7.38E-06 | 0.32 | 0.13 | 0.19 | 2.44 |
| 21 | Dipk1a | 0.87 | 3.87E-10 | 8.04E-06 | 0.44 | 0.20 | 0.24 | 2.16 |
| 21 | Ccng1 | 0.79 | 3.97E-10 | 8.25E-06 | 0.43 | 0.20 | 0.23 | 2.16 |
| 21 | 2900026A02Rik | 1.18 | 4.12E-10 | 8.57E-06 | 0.27 | 0.10 | 0.17 | 2.63 |
| 21 | C2cd2l | 1.32 | 4.13E-10 | 8.58E-06 | 0.18 | 0.05 | 0.12 | 3.30 |
| 21 | Cox8a | 0.25 | 4.19E-10 | 8.72E-06 | 0.99 | 1.00 | 0.00 | 1.00 |
| 21 | Man2a1 | 0.93 | 4.23E-10 | 8.79E-06 | 0.47 | 0.23 | 0.24 | 2.02 |
| 21 | Zmpste24 | 0.82 | 4.23E-10 | 8.79E-06 | 0.59 | 0.32 | 0.27 | 1.86 |
| 21 | Brms1 | 0.83 | 4.25E-10 | 8.84E-06 | 0.62 | 0.35 | 0.27 | 1.78 |
| 21 | Tmem135 | 0.93 | 4.90E-10 | 1.02E-05 | 0.32 | 0.13 | 0.19 | 2.46 |
| 21 | Calcrl | 1.60 | 5.00E-10 | 1.04E-05 | 0.13 | 0.04 | 0.10 | 3.80 |
| 21 | 3830406C13Rik | 0.81 | 5.14E-10 | 1.07E-05 | 0.50 | 0.24 | 0.26 | 2.05 |
| 21 | Lifr | 1.08 | 5.40E-10 | 1.12E-05 | 0.28 | 0.11 | 0.17 | 2.57 |
| 21 | Necap1 | 0.97 | 5.82E-10 | 1.21E-05 | 0.48 | 0.24 | 0.24 | 2.02 |
| 21 | Ykt6 | 0.75 | 5.89E-10 | 1.22E-05 | 0.69 | 0.41 | 0.28 | 1.70 |
| 21 | Klf10 | 1.06 | 6.00E-10 | 1.25E-05 | 0.28 | 0.11 | 0.17 | 2.57 |
| 21 | Helb | 0.93 | 6.02E-10 | 1.25E-05 | 0.32 | 0.13 | 0.19 | 2.44 |
| 21 | Ctsf | 1.30 | 6.12E-10 | 1.27E-05 | 0.27 | 0.11 | 0.17 | 2.58 |
| 21 | Rragc | 0.89 | 6.25E-10 | 1.30E-05 | 0.74 | 0.45 | 0.29 | 1.63 |
| 21 | Aph1a | 0.69 | 6.28E-10 | 1.31E-05 | 0.79 | 0.50 | 0.29 | 1.57 |
| 21 | Scn1b | 2.24 | 6.52E-10 | 1.36E-05 | 0.11 | 0.02 | 0.08 | 4.38 |
| 21 | Hmox2 | 0.66 | 6.75E-10 | 1.40E-05 | 0.88 | 0.64 | 0.24 | 1.37 |
| 21 | Slc35a3 | 0.94 | 6.84E-10 | 1.42E-05 | 0.31 | 0.12 | 0.19 | 2.50 |
| 21 | Apaf1 | 0.83 | 6.96E-10 | 1.45E-05 | 0.47 | 0.23 | 0.24 | 2.08 |

**Supplementary Table 1. Marker genes for the original clusters of integrated clustering of E11.5 control and NotchGOF λ-junction cells.**

| | | | | | | | | |
|---|---|---|---|---|---|---|---|---|
| 21 | Ywhag | 0.76 | 7.02E-10 | 1.46E-05 | 0.82 | 0.63 | 0.19 | 1.29 |
| 21 | Acap2 | 0.70 | 7.30E-10 | 1.52E-05 | 0.75 | 0.46 | 0.29 | 1.62 |
| 21 | Sh3bp2 | 1.35 | 7.54E-10 | 1.57E-05 | 0.22 | 0.08 | 0.15 | 2.84 |
| 21 | Map3k20 | 1.41 | 7.55E-10 | 1.57E-05 | 0.29 | 0.12 | 0.18 | 2.51 |
| 21 | Anxa7 | 0.72 | 7.60E-10 | 1.58E-05 | 0.63 | 0.36 | 0.27 | 1.74 |
| 21 | Abracl | 0.36 | 8.07E-10 | 1.68E-05 | 0.99 | 0.93 | 0.06 | 1.06 |
| 21 | Speg | 1.43 | 8.39E-10 | 1.75E-05 | 0.36 | 0.16 | 0.19 | 2.18 |
| 21 | Nmrk1 | 1.29 | 8.48E-10 | 1.76E-05 | 0.23 | 0.08 | 0.15 | 2.78 |
| 21 | Esrra | 1.03 | 9.33E-10 | 1.94E-05 | 0.31 | 0.13 | 0.18 | 2.46 |
| 21 | Cdk18 | 2.01 | 9.44E-10 | 1.96E-05 | 0.11 | 0.02 | 0.08 | 4.38 |
| 21 | Srd5a3 | 0.87 | 9.69E-10 | 2.01E-05 | 0.44 | 0.21 | 0.23 | 2.08 |
| 21 | Dock11 | 0.83 | 1.05E-09 | 2.19E-05 | 0.47 | 0.23 | 0.24 | 2.06 |
| 21 | Vps35 | 0.66 | 1.06E-09 | 2.21E-05 | 0.91 | 0.76 | 0.15 | 1.19 |
| 21 | Vti1b | 0.57 | 1.10E-09 | 2.29E-05 | 0.83 | 0.52 | 0.31 | 1.59 |
| 21 | Kit | 1.37 | 1.11E-09 | 2.31E-05 | 0.25 | 0.10 | 0.16 | 2.60 |
| 21 | Otud1 | 1.91 | 1.11E-09 | 2.31E-05 | 0.12 | 0.03 | 0.09 | 3.97 |
| 21 | Cnn2 | 0.61 | 1.22E-09 | 2.53E-05 | 0.87 | 0.63 | 0.25 | 1.39 |
| 21 | Prr14l | 0.69 | 1.24E-09 | 2.58E-05 | 0.49 | 0.24 | 0.25 | 2.03 |
| 21 | Pqlc3 | 1.40 | 1.25E-09 | 2.61E-05 | 0.18 | 0.06 | 0.12 | 3.18 |
| 21 | Igsf8 | 0.78 | 1.28E-09 | 2.66E-05 | 0.49 | 0.24 | 0.25 | 2.01 |
| 21 | Rock2 | 0.73 | 1.32E-09 | 2.75E-05 | 0.87 | 0.66 | 0.20 | 1.31 |
| 21 | Uqcr10 | 0.32 | 1.37E-09 | 2.84E-05 | 0.99 | 0.98 | 0.01 | 1.01 |
| 21 | Vps28 | 0.59 | 1.37E-09 | 2.85E-05 | 0.89 | 0.62 | 0.27 | 1.43 |
| 21 | Tbc1d9b | 0.88 | 1.38E-09 | 2.87E-05 | 0.44 | 0.22 | 0.23 | 2.05 |
| 21 | Usp2 | 1.17 | 1.47E-09 | 3.06E-05 | 0.33 | 0.14 | 0.19 | 2.33 |
| 21 | Pigk | 0.77 | 1.58E-09 | 3.29E-05 | 0.55 | 0.30 | 0.26 | 1.86 |
| 21 | Zfp524 | 1.13 | 1.62E-09 | 3.38E-05 | 0.27 | 0.10 | 0.16 | 2.56 |
| 21 | Sun2 | 1.09 | 1.70E-09 | 3.53E-05 | 0.37 | 0.17 | 0.20 | 2.13 |
| 21 | Snx1 | 0.69 | 1.75E-09 | 3.64E-05 | 0.72 | 0.45 | 0.27 | 1.61 |
| 21 | Sgsh | 2.05 | 1.78E-09 | 3.71E-05 | 0.11 | 0.03 | 0.08 | 4.20 |
| 21 | Klhl18 | 1.19 | 1.80E-09 | 3.75E-05 | 0.23 | 0.08 | 0.15 | 2.75 |
| 21 | Gm20707 | 1.58 | 1.80E-09 | 3.75E-05 | 0.13 | 0.04 | 0.10 | 3.69 |
| 21 | Stxbp3 | 0.85 | 1.86E-09 | 3.87E-05 | 0.48 | 0.24 | 0.24 | 2.02 |
| 21 | Naga | 1.07 | 1.88E-09 | 3.92E-05 | 0.29 | 0.12 | 0.17 | 2.45 |
| 21 | Atg10 | 0.78 | 1.92E-09 | 3.99E-05 | 0.35 | 0.15 | 0.20 | 2.35 |
| 21 | Nipa2 | 0.70 | 2.01E-09 | 4.19E-05 | 0.79 | 0.54 | 0.25 | 1.46 |
| 21 | Slc26a2 | 1.09 | 2.06E-09 | 4.28E-05 | 0.35 | 0.16 | 0.19 | 2.24 |
| 21 | Abcg1 | 1.49 | 2.08E-09 | 4.32E-05 | 0.22 | 0.08 | 0.14 | 2.70 |
| 21 | Gm43328 | 2.02 | 2.16E-09 | 4.50E-05 | 0.11 | 0.03 | 0.08 | 4.00 |
| 21 | Htt | 0.75 | 2.20E-09 | 4.57E-05 | 0.57 | 0.30 | 0.27 | 1.89 |
| 21 | Slc25a12 | 1.05 | 2.25E-09 | 4.67E-05 | 0.38 | 0.18 | 0.20 | 2.14 |
| 21 | Ube2f | 0.57 | 2.42E-09 | 5.04E-05 | 0.81 | 0.56 | 0.25 | 1.44 |
| 21 | Neurl2 | 1.53 | 2.46E-09 | 5.13E-05 | 0.20 | 0.07 | 0.13 | 2.93 |
| 21 | Fkbp8 | 0.56 | 2.53E-09 | 5.25E-05 | 0.98 | 0.82 | 0.16 | 1.19 |
| 21 | Pex7 | 0.83 | 2.55E-09 | 5.30E-05 | 0.49 | 0.25 | 0.24 | 1.93 |
| 21 | Mbp | 0.80 | 2.73E-09 | 5.68E-05 | 0.25 | 0.10 | 0.16 | 2.65 |
| 21 | Pgk1 | 0.43 | 2.76E-09 | 5.75E-05 | 0.98 | 0.96 | 0.02 | 1.03 |
| 21 | Ttc32 | 0.81 | 2.84E-09 | 5.90E-05 | 0.50 | 0.26 | 0.24 | 1.94 |
| 21 | AU040320 | 1.10 | 2.86E-09 | 5.94E-05 | 0.27 | 0.11 | 0.16 | 2.46 |
| 21 | Atp1a1 | 0.63 | 2.88E-09 | 5.98E-05 | 0.87 | 0.63 | 0.23 | 1.37 |
| 21 | Cdkn3 | 1.06 | 2.94E-09 | 6.12E-05 | 0.52 | 0.29 | 0.22 | 1.76 |
| 21 | Fancg | 1.36 | 2.98E-09 | 6.19E-05 | 0.17 | 0.05 | 0.12 | 3.17 |
| 21 | Churc1 | 0.59 | 2.98E-09 | 6.20E-05 | 0.85 | 0.59 | 0.27 | 1.45 |
| 21 | Ppp1r12c | 0.72 | 2.99E-09 | 6.21E-05 | 0.58 | 0.32 | 0.26 | 1.82 |
| 21 | Zmiz2 | 0.77 | 3.15E-09 | 6.54E-05 | 0.59 | 0.31 | 0.28 | 1.89 |
| 21 | Alkbh2 | 0.91 | 3.16E-09 | 6.58E-05 | 0.37 | 0.17 | 0.21 | 2.23 |

**Supplementary Table 1. Marker genes for the original clusters of integrated clustering of E11.5 control and NotchGOF λ-junction cells.**

| | | | | | | | | |
|---|---|---|---|---|---|---|---|---|
| 21 | Tgfbrap1 | 1.22 | 3.36E-09 | 6.98E-05 | 0.29 | 0.12 | 0.17 | 2.37 |
| 21 | Sel1l | 0.70 | 3.47E-09 | 7.21E-05 | 0.64 | 0.37 | 0.27 | 1.74 |
| 21 | Sgpl1 | 1.04 | 3.49E-09 | 7.25E-05 | 0.55 | 0.34 | 0.21 | 1.60 |
| 21 | Slc30a1 | 1.44 | 3.49E-09 | 7.26E-05 | 0.22 | 0.08 | 0.14 | 2.67 |
| 21 | Ccdc86 | 0.84 | 3.61E-09 | 7.50E-05 | 0.80 | 0.55 | 0.25 | 1.45 |
| 21 | Arhgef2 | 0.73 | 3.90E-09 | 8.11E-05 | 0.68 | 0.43 | 0.25 | 1.60 |
| 21 | Ide | 0.77 | 3.90E-09 | 8.12E-05 | 0.50 | 0.27 | 0.23 | 1.87 |
| 21 | Ptgr1 | 0.85 | 4.00E-09 | 8.32E-05 | 0.51 | 0.26 | 0.25 | 1.93 |
| 21 | Ncln | 0.68 | 4.08E-09 | 8.50E-05 | 0.64 | 0.37 | 0.26 | 1.71 |
| 21 | Plekhf2 | 0.67 | 4.09E-09 | 8.50E-05 | 0.47 | 0.23 | 0.24 | 2.03 |
| 21 | B230307C23Rik | 0.83 | 4.37E-09 | 9.09E-05 | 0.36 | 0.16 | 0.20 | 2.22 |
| 21 | Rsrp1 | 0.49 | 4.54E-09 | 9.45E-05 | 0.97 | 0.89 | 0.08 | 1.09 |
| 21 | Zbtb18 | 1.08 | 4.56E-09 | 9.48E-05 | 0.27 | 0.11 | 0.16 | 2.48 |
| 21 | Trafd1 | 0.74 | 4.87E-09 | 1.01E-04 | 0.55 | 0.30 | 0.25 | 1.85 |
| 21 | Gmppa | 0.89 | 4.87E-09 | 1.01E-04 | 0.37 | 0.17 | 0.20 | 2.17 |
| 21 | Axl | 1.66 | 4.94E-09 | 1.03E-04 | 0.32 | 0.15 | 0.17 | 2.12 |
| 21 | Mrps26 | 0.68 | 5.37E-09 | 1.12E-04 | 0.78 | 0.65 | 0.13 | 1.20 |
| 21 | Wwp1 | 0.79 | 5.45E-09 | 1.13E-04 | 0.69 | 0.41 | 0.27 | 1.66 |
| 21 | Mknk2 | 0.72 | 5.49E-09 | 1.14E-04 | 0.69 | 0.41 | 0.27 | 1.67 |
| 21 | Specc1 | 0.66 | 5.51E-09 | 1.15E-04 | 0.70 | 0.42 | 0.28 | 1.65 |
| 21 | Hipk3 | 0.91 | 5.54E-09 | 1.15E-04 | 0.34 | 0.15 | 0.19 | 2.26 |
| 21 | Hsdl1 | 1.00 | 5.58E-09 | 1.16E-04 | 0.34 | 0.15 | 0.19 | 2.23 |
| 21 | Ccdc50 | 0.62 | 5.99E-09 | 1.25E-04 | 0.90 | 0.76 | 0.14 | 1.19 |
| 21 | Rabac1 | 0.64 | 5.99E-09 | 1.25E-04 | 0.79 | 0.55 | 0.24 | 1.43 |
| 21 | Pcyox1 | 0.85 | 6.05E-09 | 1.26E-04 | 0.54 | 0.30 | 0.24 | 1.82 |
| 21 | Brox | 0.87 | 6.09E-09 | 1.27E-04 | 0.45 | 0.22 | 0.22 | 2.00 |
| 21 | Ppargc1b | 1.69 | 6.17E-09 | 1.28E-04 | 0.11 | 0.03 | 0.08 | 3.86 |
| 21 | B3galt4 | 1.59 | 6.27E-09 | 1.30E-04 | 0.14 | 0.04 | 0.10 | 3.41 |
| 21 | Neurl1a | 1.23 | 6.40E-09 | 1.33E-04 | 0.19 | 0.07 | 0.12 | 2.91 |
| 21 | Smim4 | 0.75 | 6.41E-09 | 1.33E-04 | 0.68 | 0.43 | 0.25 | 1.59 |
| 21 | Camta2 | 1.39 | 6.57E-09 | 1.37E-04 | 0.17 | 0.06 | 0.11 | 3.05 |
| 21 | Usp15 | 0.63 | 6.72E-09 | 1.40E-04 | 0.80 | 0.57 | 0.23 | 1.40 |
| 21 | Vps26c | 0.86 | 6.87E-09 | 1.43E-04 | 0.45 | 0.23 | 0.22 | 1.97 |
| 21 | Stx2 | 0.89 | 7.73E-09 | 1.61E-04 | 0.44 | 0.22 | 0.22 | 2.00 |
| 21 | Hps5 | 0.84 | 7.76E-09 | 1.61E-04 | 0.33 | 0.14 | 0.19 | 2.32 |
| 21 | Cyp4v3 | 1.78 | 8.01E-09 | 1.67E-04 | 0.13 | 0.04 | 0.09 | 3.60 |
| 21 | Ap3m2 | 1.81 | 8.18E-09 | 1.70E-04 | 0.13 | 0.04 | 0.10 | 3.50 |
| 21 | Trmt2b | 1.34 | 8.29E-09 | 1.72E-04 | 0.23 | 0.09 | 0.14 | 2.60 |
| 21 | Atxn7l1 | 0.79 | 8.42E-09 | 1.75E-04 | 0.36 | 0.17 | 0.20 | 2.21 |
| 21 | Mogs | 0.75 | 8.43E-09 | 1.75E-04 | 0.59 | 0.35 | 0.25 | 1.71 |
| 21 | Scly | 0.96 | 8.47E-09 | 1.76E-04 | 0.27 | 0.11 | 0.16 | 2.48 |
| 21 | Pop5 | 0.64 | 8.50E-09 | 1.77E-04 | 0.83 | 0.65 | 0.18 | 1.28 |
| 21 | Mxd4 | 0.65 | 8.54E-09 | 1.78E-04 | 0.73 | 0.47 | 0.26 | 1.55 |
| 21 | Spsb2 | 1.31 | 8.72E-09 | 1.81E-04 | 0.21 | 0.08 | 0.13 | 2.76 |
| 21 | Hdac5 | 0.78 | 8.86E-09 | 1.84E-04 | 0.38 | 0.17 | 0.20 | 2.17 |
| 21 | Nat9 | 0.79 | 9.14E-09 | 1.90E-04 | 0.42 | 0.20 | 0.22 | 2.06 |
| 21 | Adam10 | 0.65 | 9.16E-09 | 1.91E-04 | 0.80 | 0.54 | 0.27 | 1.50 |
| 21 | Hpn | 1.67 | 9.33E-09 | 1.94E-04 | 0.13 | 0.04 | 0.10 | 3.50 |
| 21 | Zfp592 | 0.95 | 9.50E-09 | 1.98E-04 | 0.34 | 0.15 | 0.19 | 2.23 |
| 21 | Rabgap1l | 1.00 | 9.56E-09 | 1.99E-04 | 0.31 | 0.14 | 0.17 | 2.23 |
| 21 | Crebrf | 1.17 | 9.66E-09 | 2.01E-04 | 0.27 | 0.11 | 0.16 | 2.40 |
| 21 | Pomt1 | 0.97 | 1.04E-08 | 2.16E-04 | 0.29 | 0.12 | 0.17 | 2.39 |
| 21 | Jup | 0.54 | 1.05E-08 | 2.18E-04 | 0.42 | 0.20 | 0.22 | 2.11 |
| 21 | Atl3 | 0.79 | 1.08E-08 | 2.26E-04 | 0.44 | 0.22 | 0.22 | 1.99 |
| 21 | Wdr91 | 0.98 | 1.12E-08 | 2.32E-04 | 0.26 | 0.10 | 0.16 | 2.51 |
| 21 | Degs1 | 0.74 | 1.12E-08 | 2.34E-04 | 0.66 | 0.40 | 0.27 | 1.67 |

**Supplementary Table 1. Marker genes for the original clusters of integrated clustering of E11.5 control and NotchGOF λ-junction cells.**

| | | | | | | | | |
|---|---|---|---|---|---|---|---|---|
| 21 | Ccdc93 | 0.76 | 1.18E-08 | 2.45E-04 | 0.37 | 0.17 | 0.20 | 2.20 |
| 21 | Acvr1b | 0.97 | 1.21E-08 | 2.52E-04 | 0.27 | 0.11 | 0.16 | 2.42 |
| 21 | Stard9 | 1.52 | 1.36E-08 | 2.82E-04 | 0.19 | 0.07 | 0.12 | 2.82 |
| 21 | Chkb | 0.70 | 1.36E-08 | 2.83E-04 | 0.47 | 0.24 | 0.23 | 2.00 |
| 21 | Anapc15 | 0.55 | 1.36E-08 | 2.83E-04 | 0.84 | 0.60 | 0.24 | 1.39 |
| 21 | Clptm1l | 0.67 | 1.44E-08 | 3.00E-04 | 0.72 | 0.48 | 0.24 | 1.49 |
| 21 | Kidins220 | 0.71 | 1.50E-08 | 3.11E-04 | 0.52 | 0.28 | 0.24 | 1.87 |
| 21 | Alg5 | 0.64 | 1.51E-08 | 3.14E-04 | 0.64 | 0.37 | 0.27 | 1.74 |
| 21 | Vill | 1.23 | 1.51E-08 | 3.15E-04 | 0.22 | 0.08 | 0.14 | 2.65 |
| 21 | Dusp5 | 1.35 | 1.55E-08 | 3.21E-04 | 0.27 | 0.12 | 0.16 | 2.33 |
| 21 | Rnpepl1 | 0.75 | 1.59E-08 | 3.31E-04 | 0.48 | 0.26 | 0.23 | 1.89 |
| 21 | Sh2b2 | 0.97 | 1.62E-08 | 3.36E-04 | 0.34 | 0.16 | 0.18 | 2.13 |
| 21 | Tmem97 | 0.76 | 1.76E-08 | 3.66E-04 | 0.69 | 0.46 | 0.23 | 1.49 |
| 21 | Pim1 | 0.68 | 1.79E-08 | 3.71E-04 | 0.69 | 0.47 | 0.22 | 1.47 |
| 21 | Dpm3 | 0.45 | 1.92E-08 | 4.00E-04 | 0.92 | 0.75 | 0.17 | 1.23 |
| 21 | Tex2 | 1.42 | 1.96E-08 | 4.07E-04 | 0.13 | 0.04 | 0.09 | 3.50 |
| 21 | Tcp11l2 | 1.34 | 1.97E-08 | 4.10E-04 | 0.13 | 0.04 | 0.09 | 3.50 |
| 21 | Larp4b | 0.72 | 1.99E-08 | 4.14E-04 | 0.84 | 0.61 | 0.23 | 1.38 |
| 21 | Sidt2 | 1.09 | 2.07E-08 | 4.29E-04 | 0.37 | 0.18 | 0.19 | 2.04 |
| 21 | Fnip1 | 0.87 | 2.08E-08 | 4.33E-04 | 0.46 | 0.24 | 0.22 | 1.94 |
| 21 | Leprotl1 | 0.59 | 2.10E-08 | 4.36E-04 | 0.80 | 0.55 | 0.25 | 1.45 |
| 21 | Sec61b | 0.29 | 2.11E-08 | 4.40E-04 | 0.99 | 0.99 | 0.01 | 1.01 |
| 21 | Atmin | 0.89 | 2.15E-08 | 4.47E-04 | 0.32 | 0.14 | 0.18 | 2.28 |
| 21 | Faap24 | 0.94 | 2.15E-08 | 4.47E-04 | 0.35 | 0.16 | 0.19 | 2.13 |
| 21 | Nr4a2 | 1.45 | 2.17E-08 | 4.51E-04 | 0.25 | 0.10 | 0.15 | 2.42 |
| 21 | Mlxip | 0.66 | 2.17E-08 | 4.52E-04 | 0.36 | 0.16 | 0.20 | 2.23 |
| 21 | Tnfaip8 | 0.44 | 2.20E-08 | 4.57E-04 | 0.94 | 0.76 | 0.18 | 1.24 |
| 21 | Ubald1 | 0.88 | 2.26E-08 | 4.70E-04 | 0.50 | 0.26 | 0.23 | 1.88 |
| 21 | Ubash3b | 0.88 | 2.26E-08 | 4.71E-04 | 0.45 | 0.24 | 0.21 | 1.89 |
| 21 | Eloa | 0.63 | 2.30E-08 | 4.78E-04 | 0.72 | 0.47 | 0.25 | 1.54 |
| 21 | Mrpl57 | 0.54 | 2.34E-08 | 4.87E-04 | 0.90 | 0.78 | 0.13 | 1.16 |
| 21 | Poc1a | 0.65 | 2.35E-08 | 4.89E-04 | 0.44 | 0.21 | 0.23 | 2.06 |
| 21 | Lsm10 | 0.81 | 2.35E-08 | 4.89E-04 | 0.66 | 0.40 | 0.26 | 1.66 |
| 21 | Vps11 | 1.13 | 2.51E-08 | 5.22E-04 | 0.34 | 0.16 | 0.18 | 2.14 |
| 21 | Atp5g1 | 0.33 | 2.55E-08 | 5.31E-04 | 0.99 | 0.99 | 0.01 | 1.01 |
| 21 | Dmxl1 | 0.78 | 2.56E-08 | 5.33E-04 | 0.66 | 0.41 | 0.25 | 1.61 |
| 21 | Imp4 | 0.61 | 2.62E-08 | 5.46E-04 | 0.76 | 0.54 | 0.23 | 1.42 |
| 21 | Slc38a9 | 0.97 | 2.78E-08 | 5.79E-04 | 0.38 | 0.19 | 0.19 | 2.03 |
| 21 | Sec24a | 0.75 | 2.80E-08 | 5.83E-04 | 0.37 | 0.18 | 0.20 | 2.12 |
| 21 | Tmem106c | 0.87 | 2.81E-08 | 5.84E-04 | 0.39 | 0.20 | 0.20 | 2.00 |
| 21 | Hmgcl | 0.73 | 2.82E-08 | 5.87E-04 | 0.65 | 0.41 | 0.25 | 1.60 |
| 21 | Rbm38 | 0.70 | 2.87E-08 | 5.96E-04 | 0.46 | 0.24 | 0.22 | 1.93 |
| 21 | Ethe1 | 0.76 | 3.01E-08 | 6.26E-04 | 0.69 | 0.45 | 0.23 | 1.52 |
| 21 | Flad1 | 0.87 | 3.02E-08 | 6.27E-04 | 0.26 | 0.11 | 0.15 | 2.47 |
| 21 | Zbtb4 | 1.48 | 3.15E-08 | 6.54E-04 | 0.18 | 0.07 | 0.12 | 2.80 |
| 21 | Dnajc16 | 1.22 | 3.34E-08 | 6.94E-04 | 0.17 | 0.06 | 0.11 | 2.95 |
| 21 | Arl4a | 0.72 | 3.38E-08 | 7.03E-04 | 0.41 | 0.20 | 0.21 | 2.02 |
| 21 | Bend4 | 0.99 | 3.39E-08 | 7.05E-04 | 0.27 | 0.12 | 0.16 | 2.35 |
| 21 | Ank | 1.33 | 3.44E-08 | 7.15E-04 | 0.19 | 0.07 | 0.12 | 2.74 |
| 21 | Atxn1 | 1.11 | 3.49E-08 | 7.27E-04 | 0.23 | 0.09 | 0.14 | 2.51 |
| 21 | Cd164 | 0.61 | 3.61E-08 | 7.50E-04 | 0.81 | 0.64 | 0.17 | 1.27 |
| 21 | Atraid | 0.65 | 3.63E-08 | 7.56E-04 | 0.75 | 0.51 | 0.24 | 1.48 |
| 21 | Acaa2 | 0.41 | 3.67E-08 | 7.63E-04 | 0.76 | 0.48 | 0.28 | 1.58 |
| 21 | Chn2 | 1.04 | 3.75E-08 | 7.80E-04 | 0.20 | 0.07 | 0.13 | 2.76 |
| 21 | C1galt1c1 | 1.16 | 3.80E-08 | 7.91E-04 | 0.34 | 0.16 | 0.18 | 2.10 |
| 21 | Mto1 | 0.57 | 3.84E-08 | 7.98E-04 | 0.41 | 0.20 | 0.22 | 2.12 |

**Supplementary Table 1. Marker genes for the original clusters of integrated clustering of E11.5 control and NotchGOF λ-junction cells.**

| | | | | | | | | |
|---|---|---|---|---|---|---|---|---|
| 21 | Snx8 | 1.18 | 4.06E-08 | 8.44E-04 | 0.29 | 0.13 | 0.16 | 2.19 |
| 21 | Lactb2 | 0.86 | 4.09E-08 | 8.50E-04 | 0.37 | 0.18 | 0.20 | 2.11 |
| 21 | Oser1 | 0.72 | 4.18E-08 | 8.70E-04 | 0.68 | 0.46 | 0.22 | 1.47 |
| 21 | Pvr | 1.10 | 4.62E-08 | 9.61E-04 | 0.19 | 0.07 | 0.12 | 2.78 |
| 21 | Armc7 | 1.64 | 4.68E-08 | 9.74E-04 | 0.15 | 0.05 | 0.10 | 3.06 |
| 21 | Ino80c | 0.75 | 4.68E-08 | 9.74E-04 | 0.39 | 0.19 | 0.20 | 2.03 |
| 21 | Zbtb34 | 0.96 | 4.72E-08 | 9.82E-04 | 0.23 | 0.09 | 0.14 | 2.54 |
| 21 | Zfand5 | 0.45 | 4.72E-08 | 9.83E-04 | 0.95 | 0.83 | 0.12 | 1.15 |
| 21 | Bmyc | 0.61 | 4.81E-08 | 1.00E-03 | 0.83 | 0.62 | 0.20 | 1.33 |
| 21 | Rnf149 | 0.80 | 4.88E-08 | 1.01E-03 | 0.43 | 0.22 | 0.21 | 1.96 |
| 21 | Arhgef10l | 0.85 | 4.88E-08 | 1.02E-03 | 0.32 | 0.15 | 0.18 | 2.22 |
| 21 | Spata13 | 1.11 | 4.99E-08 | 1.04E-03 | 0.20 | 0.08 | 0.13 | 2.67 |
| 21 | Cnpy3 | 0.72 | 5.05E-08 | 1.05E-03 | 0.59 | 0.35 | 0.24 | 1.70 |
| 21 | AI987944 | 0.90 | 5.09E-08 | 1.06E-03 | 0.29 | 0.13 | 0.16 | 2.26 |
| 21 | Hectd3 | 1.16 | 5.13E-08 | 1.07E-03 | 0.26 | 0.11 | 0.15 | 2.33 |
| 21 | Rpa1 | 0.61 | 5.29E-08 | 1.10E-03 | 0.66 | 0.42 | 0.25 | 1.58 |
| 21 | Rmdn1 | 0.72 | 5.30E-08 | 1.10E-03 | 0.36 | 0.17 | 0.19 | 2.10 |
| 21 | Ttc7b | 0.70 | 5.51E-08 | 1.15E-03 | 0.35 | 0.16 | 0.19 | 2.20 |
| 21 | Tsta3 | 0.63 | 5.53E-08 | 1.15E-03 | 0.70 | 0.46 | 0.24 | 1.53 |
| 21 | Wdr44 | 1.40 | 5.65E-08 | 1.17E-03 | 0.18 | 0.06 | 0.11 | 2.78 |
| 21 | Prdx1 | 0.47 | 5.77E-08 | 1.20E-03 | 1.00 | 0.99 | 0.01 | 1.01 |
| 21 | Serhl | 0.77 | 5.80E-08 | 1.21E-03 | 0.36 | 0.17 | 0.19 | 2.13 |
| 21 | Stk11 | 0.57 | 5.86E-08 | 1.22E-03 | 0.73 | 0.48 | 0.25 | 1.53 |
| 21 | Metrnl | 1.42 | 5.91E-08 | 1.23E-03 | 0.18 | 0.07 | 0.12 | 2.76 |
| 21 | Casp9 | 0.80 | 6.12E-08 | 1.27E-03 | 0.25 | 0.10 | 0.15 | 2.50 |
| 21 | Zc3h7a | 0.70 | 6.14E-08 | 1.28E-03 | 0.59 | 0.34 | 0.25 | 1.72 |
| 21 | Pias1 | 0.70 | 6.20E-08 | 1.29E-03 | 0.56 | 0.32 | 0.24 | 1.75 |
| 21 | Armc5 | 0.86 | 6.80E-08 | 1.41E-03 | 0.32 | 0.14 | 0.17 | 2.22 |
| 21 | Ppp1r15b | 0.69 | 7.01E-08 | 1.46E-03 | 0.55 | 0.31 | 0.25 | 1.80 |
| 21 | Ikbkb | 0.65 | 7.09E-08 | 1.48E-03 | 0.57 | 0.32 | 0.25 | 1.79 |
| 21 | Mrpl48 | 0.56 | 7.12E-08 | 1.48E-03 | 0.78 | 0.53 | 0.25 | 1.46 |
| 21 | Scp2 | 0.45 | 7.20E-08 | 1.50E-03 | 0.93 | 0.77 | 0.16 | 1.20 |
| 21 | Fyco1 | 0.80 | 7.32E-08 | 1.52E-03 | 0.35 | 0.16 | 0.19 | 2.13 |
| 21 | Cox5a | 0.27 | 7.46E-08 | 1.55E-03 | 1.00 | 0.99 | 0.01 | 1.01 |
| 21 | CAAA01118383.1 | 0.69 | 7.60E-08 | 1.58E-03 | 0.45 | 0.23 | 0.22 | 1.93 |
| 21 | Itpk1 | 0.81 | 7.91E-08 | 1.65E-03 | 0.19 | 0.07 | 0.12 | 2.82 |
| 21 | Hacd2 | 0.68 | 7.94E-08 | 1.65E-03 | 0.62 | 0.37 | 0.24 | 1.65 |
| 21 | Echdc1 | 0.67 | 8.23E-08 | 1.71E-03 | 0.45 | 0.24 | 0.21 | 1.89 |
| 21 | Arrb1 | 1.04 | 8.24E-08 | 1.71E-03 | 0.27 | 0.12 | 0.15 | 2.31 |
| 21 | Psenen | 0.41 | 8.53E-08 | 1.77E-03 | 0.94 | 0.88 | 0.06 | 1.07 |
| 21 | Slc25a10 | 0.90 | 8.65E-08 | 1.80E-03 | 0.44 | 0.24 | 0.21 | 1.87 |
| 21 | Prdx3 | 0.61 | 8.99E-08 | 1.87E-03 | 0.76 | 0.54 | 0.23 | 1.42 |
| 21 | Lyst | 0.97 | 9.00E-08 | 1.87E-03 | 0.28 | 0.12 | 0.16 | 2.28 |
| 21 | Map4k2 | 1.06 | 9.36E-08 | 1.95E-03 | 0.19 | 0.07 | 0.12 | 2.74 |
| 21 | Smim14 | 0.66 | 9.59E-08 | 1.99E-03 | 0.66 | 0.41 | 0.24 | 1.59 |
| 21 | Polr1d | 0.31 | 1.02E-07 | 2.12E-03 | 0.97 | 0.95 | 0.02 | 1.02 |
| 21 | H2-T22 | 1.20 | 1.03E-07 | 2.15E-03 | 0.24 | 0.10 | 0.14 | 2.40 |
| 21 | Ugcg | 0.70 | 1.04E-07 | 2.16E-03 | 0.61 | 0.37 | 0.24 | 1.64 |
| 21 | Zfp622 | 0.60 | 1.08E-07 | 2.25E-03 | 0.66 | 0.42 | 0.25 | 1.59 |
| 21 | Nomo1 | 0.90 | 1.09E-07 | 2.27E-03 | 0.39 | 0.20 | 0.19 | 1.93 |
| 21 | Htra2 | 0.85 | 1.11E-07 | 2.31E-03 | 0.62 | 0.40 | 0.22 | 1.55 |
| 21 | Nfya | 0.56 | 1.16E-07 | 2.41E-03 | 0.56 | 0.32 | 0.24 | 1.76 |
| 21 | Ptrh1 | 1.29 | 1.16E-07 | 2.41E-03 | 0.13 | 0.04 | 0.09 | 3.24 |
| 21 | Kbtbd8 | 1.30 | 1.20E-07 | 2.50E-03 | 0.13 | 0.04 | 0.09 | 3.32 |
| 21 | Eif4ebp1 | 0.49 | 1.24E-07 | 2.57E-03 | 0.87 | 0.69 | 0.18 | 1.26 |
| 21 | Galnt7 | 0.96 | 1.31E-07 | 2.72E-03 | 0.58 | 0.35 | 0.24 | 1.68 |

**Supplementary Table 1. Marker genes for the original clusters of integrated clustering of E11.5 control and NotchGOF λ-junction cells.**

| | | | | | | | | |
|---|---|---|---|---|---|---|---|---|
| 21 | Ncor2 | 0.61 | 1.35E-07 | 2.81E-03 | 0.62 | 0.35 | 0.26 | 1.75 |
| 21 | Cercam | 0.99 | 1.35E-07 | 2.82E-03 | 0.17 | 0.06 | 0.11 | 2.85 |
| 21 | Psma1 | 0.31 | 1.35E-07 | 2.82E-03 | 0.97 | 0.94 | 0.03 | 1.04 |
| 21 | Mon2 | 0.59 | 1.43E-07 | 2.98E-03 | 0.51 | 0.28 | 0.23 | 1.84 |
| 21 | Tmf1 | 0.55 | 1.46E-07 | 3.04E-03 | 0.73 | 0.49 | 0.24 | 1.50 |
| 21 | Rpn2 | 0.40 | 1.46E-07 | 3.04E-03 | 0.95 | 0.79 | 0.16 | 1.21 |
| 21 | Nfs1 | 0.89 | 1.48E-07 | 3.08E-03 | 0.51 | 0.28 | 0.23 | 1.82 |
| 21 | 2900093K20Rik | 1.05 | 1.50E-07 | 3.12E-03 | 0.22 | 0.09 | 0.13 | 2.52 |
| 21 | Tmem243 | 0.78 | 1.51E-07 | 3.13E-03 | 0.41 | 0.21 | 0.20 | 1.96 |
| 21 | Tigar | 1.22 | 1.59E-07 | 3.31E-03 | 0.19 | 0.07 | 0.12 | 2.63 |
| 21 | Rps6 | 0.25 | 1.62E-07 | 3.37E-03 | 1.00 | 1.00 | 0.00 | 1.00 |
| 21 | Rad23a | 0.61 | 1.63E-07 | 3.38E-03 | 0.68 | 0.46 | 0.22 | 1.48 |
| 21 | Ero1lb | 1.67 | 1.66E-07 | 3.45E-03 | 0.11 | 0.03 | 0.08 | 3.62 |
| 21 | Tmem248 | 0.67 | 1.67E-07 | 3.46E-03 | 0.58 | 0.35 | 0.23 | 1.65 |
| 21 | Bag3 | 1.01 | 1.69E-07 | 3.52E-03 | 0.26 | 0.11 | 0.15 | 2.35 |
| 21 | Snx13 | 0.96 | 1.87E-07 | 3.89E-03 | 0.29 | 0.13 | 0.16 | 2.17 |
| 21 | Akip1 | 0.70 | 1.90E-07 | 3.96E-03 | 0.40 | 0.20 | 0.20 | 2.02 |
| 21 | Arhgap6 | 1.50 | 1.93E-07 | 4.02E-03 | 0.18 | 0.07 | 0.11 | 2.69 |
| 21 | Ccdc28b | 0.52 | 1.95E-07 | 4.06E-03 | 0.64 | 0.37 | 0.27 | 1.72 |
| 21 | Faim | 0.54 | 2.02E-07 | 4.20E-03 | 0.72 | 0.46 | 0.26 | 1.57 |
| 21 | Tmc6 | 1.03 | 2.05E-07 | 4.26E-03 | 0.20 | 0.08 | 0.12 | 2.58 |
| 21 | Pet100 | 0.55 | 2.09E-07 | 4.35E-03 | 0.79 | 0.56 | 0.24 | 1.42 |
| 21 | Bcat2 | 0.66 | 2.09E-07 | 4.35E-03 | 0.51 | 0.29 | 0.22 | 1.75 |
| 21 | Slc39a14 | 1.34 | 2.10E-07 | 4.36E-03 | 0.24 | 0.10 | 0.13 | 2.29 |
| 21 | Mrps18b | 0.73 | 2.24E-07 | 4.65E-03 | 0.73 | 0.50 | 0.23 | 1.45 |
| 21 | Vdac2 | 0.31 | 2.24E-07 | 4.66E-03 | 0.98 | 0.96 | 0.02 | 1.02 |
| 21 | Gem | 1.89 | 2.26E-07 | 4.70E-03 | 0.13 | 0.04 | 0.09 | 3.15 |
| 21 | Arid3b | 0.65 | 2.31E-07 | 4.80E-03 | 0.44 | 0.23 | 0.21 | 1.93 |
| 21 | Pacs2 | 0.76 | 2.34E-07 | 4.87E-03 | 0.35 | 0.17 | 0.18 | 2.07 |
| 21 | Acp2 | 0.98 | 2.38E-07 | 4.95E-03 | 0.46 | 0.27 | 0.19 | 1.72 |
| 21 | Thbs1 | 1.38 | 2.38E-07 | 4.95E-03 | 0.27 | 0.12 | 0.15 | 2.24 |
| 21 | Ier5 | 0.43 | 2.38E-07 | 4.95E-03 | 0.86 | 0.65 | 0.21 | 1.32 |
| 21 | Filip1l | 0.67 | 2.40E-07 | 4.99E-03 | 0.47 | 0.26 | 0.21 | 1.82 |
| 21 | Hagh | 0.84 | 2.42E-07 | 5.03E-03 | 0.44 | 0.25 | 0.20 | 1.80 |
| 21 | Irf2bpl | 0.69 | 2.46E-07 | 5.12E-03 | 0.63 | 0.38 | 0.25 | 1.64 |
| 21 | Fkbp1b | 0.90 | 2.52E-07 | 5.23E-03 | 0.30 | 0.14 | 0.16 | 2.12 |
| 21 | Kxd1 | 0.56 | 2.63E-07 | 5.47E-03 | 0.73 | 0.50 | 0.23 | 1.46 |
| 21 | Tspan33 | 0.64 | 2.65E-07 | 5.52E-03 | 0.21 | 0.08 | 0.13 | 2.50 |
| 21 | 2210016F16Rik | 0.66 | 2.67E-07 | 5.54E-03 | 0.57 | 0.35 | 0.22 | 1.63 |
| 21 | Lamtor5 | 0.44 | 2.71E-07 | 5.63E-03 | 0.92 | 0.79 | 0.13 | 1.16 |
| 21 | Pygb | 0.54 | 2.71E-07 | 5.63E-03 | 0.45 | 0.23 | 0.22 | 1.95 |
| 21 | Slc25a32 | 0.88 | 2.77E-07 | 5.77E-03 | 0.22 | 0.09 | 0.13 | 2.49 |
| 21 | Sumf1 | 1.08 | 2.78E-07 | 5.78E-03 | 0.26 | 0.12 | 0.14 | 2.23 |
| 21 | Mrpl33 | 0.55 | 2.78E-07 | 5.78E-03 | 0.94 | 0.87 | 0.06 | 1.07 |
| 21 | Tmem208 | 0.46 | 2.81E-07 | 5.83E-03 | 0.90 | 0.68 | 0.21 | 1.31 |
| 21 | Lrrk1 | 0.91 | 2.83E-07 | 5.89E-03 | 0.32 | 0.15 | 0.16 | 2.09 |
| 21 | Washc5 | 0.60 | 2.83E-07 | 5.89E-03 | 0.55 | 0.32 | 0.23 | 1.72 |
| 21 | Txnrd2 | 0.67 | 2.91E-07 | 6.05E-03 | 0.41 | 0.21 | 0.20 | 1.94 |
| 21 | Tgfbr1 | 0.88 | 2.97E-07 | 6.18E-03 | 0.75 | 0.55 | 0.19 | 1.35 |
| 21 | Zfp217 | 0.80 | 3.01E-07 | 6.27E-03 | 0.39 | 0.20 | 0.19 | 1.97 |
| 21 | Slc25a19 | 0.89 | 3.03E-07 | 6.30E-03 | 0.27 | 0.12 | 0.15 | 2.27 |
| 21 | Nfkbie | 1.58 | 3.06E-07 | 6.36E-03 | 0.13 | 0.04 | 0.09 | 3.15 |
| 21 | Polg2 | 0.95 | 3.09E-07 | 6.42E-03 | 0.23 | 0.10 | 0.13 | 2.38 |
| 21 | Etv6 | 0.74 | 3.09E-07 | 6.42E-03 | 0.48 | 0.28 | 0.20 | 1.71 |
| 21 | Ell2 | 1.08 | 3.10E-07 | 6.45E-03 | 0.22 | 0.09 | 0.13 | 2.44 |
| 21 | Itprid2 | 0.58 | 3.18E-07 | 6.62E-03 | 0.32 | 0.15 | 0.17 | 2.13 |

**Supplementary Table 1. Marker genes for the original clusters of integrated clustering of E11.5 control and NotchGOF λ-junction cells.**

| | | | | | | | | |
|---|---|---|---|---|---|---|---|---|
| 21 | Zfp335 | 1.02 | 3.24E-07 | 6.74E-03 | 0.22 | 0.09 | 0.13 | 2.38 |
| 21 | Gtf2h1 | 0.48 | 3.25E-07 | 6.76E-03 | 0.68 | 0.40 | 0.28 | 1.68 |
| 21 | Amfr | 0.56 | 3.26E-07 | 6.78E-03 | 0.60 | 0.36 | 0.24 | 1.66 |
| 21 | Zswim6 | 0.75 | 3.28E-07 | 6.83E-03 | 0.51 | 0.30 | 0.22 | 1.73 |
| 21 | Ndufaf8 | 0.46 | 3.45E-07 | 7.18E-03 | 0.88 | 0.67 | 0.21 | 1.31 |
| 21 | Lmbrd1 | 0.72 | 3.48E-07 | 7.24E-03 | 0.42 | 0.22 | 0.20 | 1.89 |
| 21 | Ppm1m | 0.64 | 3.65E-07 | 7.60E-03 | 0.46 | 0.25 | 0.22 | 1.87 |
| 21 | Rrm2b | 0.91 | 3.66E-07 | 7.62E-03 | 0.27 | 0.12 | 0.15 | 2.20 |
| 21 | Furin | 0.89 | 3.71E-07 | 7.72E-03 | 0.39 | 0.20 | 0.19 | 1.95 |
| 21 | Mapre2 | 0.55 | 3.78E-07 | 7.85E-03 | 0.65 | 0.40 | 0.25 | 1.62 |
| 21 | Abca2 | 1.49 | 3.89E-07 | 8.10E-03 | 0.11 | 0.03 | 0.08 | 3.50 |
| 21 | Zfyve19 | 0.66 | 3.91E-07 | 8.14E-03 | 0.44 | 0.24 | 0.20 | 1.86 |
| 21 | Atp13a1 | 0.69 | 4.15E-07 | 8.63E-03 | 0.44 | 0.23 | 0.21 | 1.88 |
| 21 | Cluh | 0.76 | 4.16E-07 | 8.65E-03 | 0.42 | 0.23 | 0.19 | 1.85 |
| 21 | Paxx | 0.80 | 4.74E-07 | 9.85E-03 | 0.26 | 0.12 | 0.14 | 2.25 |
| 21 | Gpatch2l | 1.54 | 4.85E-07 | 1.01E-02 | 0.24 | 0.10 | 0.14 | 2.31 |
| 21 | Siae | 0.99 | 4.86E-07 | 1.01E-02 | 0.18 | 0.07 | 0.11 | 2.64 |
| 21 | Surf4 | 0.51 | 5.05E-07 | 1.05E-02 | 0.85 | 0.59 | 0.26 | 1.44 |
| 21 | Smdt1 | 0.35 | 5.17E-07 | 1.08E-02 | 0.99 | 0.94 | 0.04 | 1.05 |
| 21 | Gnptab | 0.68 | 5.32E-07 | 1.11E-02 | 0.32 | 0.15 | 0.17 | 2.11 |
| 21 | Mtpn | 0.40 | 5.40E-07 | 1.12E-02 | 0.93 | 0.81 | 0.12 | 1.15 |
| 21 | Lin54 | 0.71 | 5.42E-07 | 1.13E-02 | 0.43 | 0.24 | 0.20 | 1.85 |
| 21 | Tgoln1 | 0.53 | 5.44E-07 | 1.13E-02 | 0.53 | 0.29 | 0.24 | 1.83 |
| 21 | Nt5c | 0.58 | 5.52E-07 | 1.15E-02 | 0.73 | 0.54 | 0.19 | 1.36 |
| 21 | Nrbf2 | 0.71 | 5.61E-07 | 1.17E-02 | 0.68 | 0.44 | 0.24 | 1.56 |
| 21 | Cnppd1 | 0.60 | 5.68E-07 | 1.18E-02 | 0.41 | 0.21 | 0.20 | 1.96 |
| 21 | Xpr1 | 0.67 | 5.71E-07 | 1.19E-02 | 0.49 | 0.27 | 0.22 | 1.79 |
| 21 | Trim56 | 0.89 | 5.71E-07 | 1.19E-02 | 0.29 | 0.14 | 0.15 | 2.10 |
| 21 | Ndufs8 | 0.36 | 5.75E-07 | 1.20E-02 | 0.97 | 0.84 | 0.13 | 1.15 |
| 21 | Sash1 | 1.04 | 5.76E-07 | 1.20E-02 | 0.60 | 0.40 | 0.20 | 1.50 |
| 21 | Dus1l | 0.53 | 5.79E-07 | 1.20E-02 | 0.64 | 0.40 | 0.24 | 1.61 |
| 21 | Arap2 | 1.05 | 5.87E-07 | 1.22E-02 | 0.15 | 0.06 | 0.10 | 2.80 |
| 21 | Ldha | 0.38 | 5.93E-07 | 1.23E-02 | 0.98 | 0.97 | 0.01 | 1.01 |
| 21 | Tkt | 0.45 | 6.10E-07 | 1.27E-02 | 0.97 | 0.92 | 0.05 | 1.05 |
| 21 | Scoc | 0.86 | 6.15E-07 | 1.28E-02 | 0.78 | 0.63 | 0.15 | 1.24 |
| 21 | Jak2 | 0.65 | 6.21E-07 | 1.29E-02 | 0.32 | 0.16 | 0.17 | 2.08 |
| 21 | Usp38 | 0.70 | 6.29E-07 | 1.31E-02 | 0.41 | 0.21 | 0.20 | 1.93 |
| 21 | Nfkbil1 | 0.59 | 6.45E-07 | 1.34E-02 | 0.50 | 0.28 | 0.23 | 1.81 |
| 21 | Myo18a | 0.49 | 6.55E-07 | 1.36E-02 | 0.45 | 0.24 | 0.21 | 1.91 |
| 21 | Entpd7 | 0.85 | 6.57E-07 | 1.37E-02 | 0.30 | 0.15 | 0.16 | 2.06 |
| 21 | Hs1bp3 | 0.94 | 7.15E-07 | 1.49E-02 | 0.25 | 0.11 | 0.14 | 2.25 |
| 21 | Oxsm | 1.38 | 7.20E-07 | 1.50E-02 | 0.14 | 0.05 | 0.09 | 2.92 |
| 21 | Btaf1 | 0.70 | 7.33E-07 | 1.52E-02 | 0.59 | 0.36 | 0.23 | 1.63 |
| 21 | Slc7a5 | 1.15 | 7.41E-07 | 1.54E-02 | 0.14 | 0.05 | 0.09 | 2.92 |
| 21 | Plekhj1 | 0.42 | 7.45E-07 | 1.55E-02 | 0.91 | 0.77 | 0.14 | 1.18 |
| 21 | Wdr12 | 0.58 | 7.65E-07 | 1.59E-02 | 0.74 | 0.56 | 0.18 | 1.33 |
| 21 | Gpr107 | 0.58 | 7.82E-07 | 1.63E-02 | 0.55 | 0.31 | 0.23 | 1.75 |
| 21 | Acbd5 | 0.74 | 8.10E-07 | 1.68E-02 | 0.57 | 0.36 | 0.21 | 1.59 |
| 21 | Apeh | 0.60 | 8.11E-07 | 1.69E-02 | 0.48 | 0.27 | 0.21 | 1.78 |
| 21 | Snx17 | 0.49 | 8.29E-07 | 1.73E-02 | 0.87 | 0.67 | 0.20 | 1.30 |
| 21 | Ero1l | 1.34 | 8.32E-07 | 1.73E-02 | 0.49 | 0.30 | 0.19 | 1.66 |
| 21 | Sypl | 0.47 | 8.33E-07 | 1.73E-02 | 0.74 | 0.48 | 0.26 | 1.54 |
| 21 | Med12l | 1.33 | 8.39E-07 | 1.74E-02 | 0.13 | 0.05 | 0.09 | 2.96 |
| 21 | Mfsd14b | 0.65 | 8.39E-07 | 1.74E-02 | 0.44 | 0.23 | 0.21 | 1.92 |
| 21 | Gne | 1.10 | 8.60E-07 | 1.79E-02 | 0.20 | 0.08 | 0.12 | 2.48 |
| 21 | Fyn | 0.62 | 8.86E-07 | 1.84E-02 | 0.69 | 0.44 | 0.24 | 1.55 |

**Supplementary Table 1. Marker genes for the original clusters of integrated clustering of E11.5 control and NotchGOF λ-junction cells.**

| | | | | | | | | |
|---|---|---|---|---|---|---|---|---|
| 21 | Acvrl1 | 1.00 | 9.04E-07 | 1.88E-02 | 0.23 | 0.10 | 0.13 | 2.29 |
| 21 | Itgb1bp1 | 0.62 | 9.22E-07 | 1.92E-02 | 0.62 | 0.40 | 0.22 | 1.55 |
| 21 | Bin1 | 0.80 | 9.22E-07 | 1.92E-02 | 0.78 | 0.62 | 0.16 | 1.26 |
| 21 | Rbck1 | 0.54 | 9.42E-07 | 1.96E-02 | 0.60 | 0.38 | 0.22 | 1.57 |
| 21 | Fam13b | 0.59 | 9.47E-07 | 1.97E-02 | 0.52 | 0.30 | 0.22 | 1.75 |
| 21 | Zfyve26 | 0.86 | 9.60E-07 | 2.00E-02 | 0.32 | 0.16 | 0.16 | 2.00 |
| 21 | Ddi2 | 0.52 | 9.91E-07 | 2.06E-02 | 0.65 | 0.41 | 0.24 | 1.59 |
| 21 | Dph5 | 0.67 | 1.04E-06 | 2.17E-02 | 0.42 | 0.23 | 0.19 | 1.85 |
| 21 | Lrp1 | 0.95 | 1.06E-06 | 2.20E-02 | 0.71 | 0.58 | 0.13 | 1.23 |
| 21 | Slc35f6 | 1.34 | 1.10E-06 | 2.28E-02 | 0.24 | 0.11 | 0.13 | 2.22 |
| 21 | Stat5b | 0.99 | 1.10E-06 | 2.29E-02 | 0.27 | 0.13 | 0.15 | 2.13 |
| 21 | Mfsd13a | 1.46 | 1.11E-06 | 2.31E-02 | 0.12 | 0.04 | 0.08 | 3.05 |
| 21 | Vrk3 | 0.55 | 1.12E-06 | 2.33E-02 | 0.58 | 0.34 | 0.24 | 1.70 |
| 21 | Txndc11 | 0.70 | 1.12E-06 | 2.33E-02 | 0.39 | 0.20 | 0.18 | 1.91 |
| 21 | Tatdn2 | 0.75 | 1.12E-06 | 2.33E-02 | 0.32 | 0.15 | 0.16 | 2.06 |
| 21 | Cacul1 | 0.54 | 1.15E-06 | 2.39E-02 | 0.52 | 0.29 | 0.23 | 1.81 |
| 21 | Cep55 | 0.77 | 1.18E-06 | 2.46E-02 | 0.43 | 0.24 | 0.19 | 1.79 |
| 21 | Tpp2 | 0.58 | 1.19E-06 | 2.48E-02 | 0.82 | 0.60 | 0.22 | 1.37 |
| 21 | Ptpa | 0.59 | 1.22E-06 | 2.54E-02 | 0.73 | 0.52 | 0.21 | 1.41 |
| 21 | Tmem30a | 0.50 | 1.25E-06 | 2.60E-02 | 0.73 | 0.51 | 0.22 | 1.42 |
| 21 | Pter | 0.86 | 1.26E-06 | 2.63E-02 | 0.25 | 0.11 | 0.14 | 2.21 |
| 21 | Stx8 | 0.63 | 1.29E-06 | 2.68E-02 | 0.66 | 0.44 | 0.22 | 1.50 |
| 21 | Itgb5 | 0.70 | 1.31E-06 | 2.72E-02 | 0.73 | 0.59 | 0.14 | 1.24 |
| 21 | Hist1h1c | 0.60 | 1.33E-06 | 2.76E-02 | 0.66 | 0.42 | 0.24 | 1.57 |
| 21 | Ubxn8 | 0.57 | 1.36E-06 | 2.82E-02 | 0.39 | 0.20 | 0.19 | 1.96 |
| 21 | Coa4 | 0.55 | 1.36E-06 | 2.82E-02 | 0.48 | 0.27 | 0.22 | 1.82 |
| 21 | Mtmr6 | 0.68 | 1.39E-06 | 2.88E-02 | 0.29 | 0.14 | 0.15 | 2.13 |
| 21 | Zfp597 | 1.00 | 1.40E-06 | 2.91E-02 | 0.19 | 0.08 | 0.11 | 2.49 |
| 21 | Smim3 | 0.62 | 1.41E-06 | 2.94E-02 | 0.36 | 0.19 | 0.18 | 1.97 |
| 21 | Ccdc107 | 0.46 | 1.44E-06 | 3.00E-02 | 0.66 | 0.42 | 0.25 | 1.59 |
| 21 | Sin3b | 0.42 | 1.45E-06 | 3.03E-02 | 0.94 | 0.80 | 0.14 | 1.18 |
| 21 | Prr14 | 0.87 | 1.50E-06 | 3.12E-02 | 0.39 | 0.20 | 0.18 | 1.89 |
| 21 | Uggt1 | 0.58 | 1.51E-06 | 3.14E-02 | 0.57 | 0.36 | 0.22 | 1.61 |
| 21 | Baz2a | 0.47 | 1.53E-06 | 3.17E-02 | 0.64 | 0.39 | 0.25 | 1.65 |
| 21 | Aldh3a2 | 1.06 | 1.53E-06 | 3.18E-02 | 0.20 | 0.08 | 0.12 | 2.42 |
| 21 | Nek9 | 0.46 | 1.53E-06 | 3.19E-02 | 0.52 | 0.29 | 0.23 | 1.78 |
| 21 | Dusp6 | 0.38 | 1.61E-06 | 3.36E-02 | 0.60 | 0.36 | 0.24 | 1.66 |
| 21 | Trib1 | 1.32 | 1.62E-06 | 3.37E-02 | 0.33 | 0.18 | 0.15 | 1.82 |
| 21 | Bpnt1 | 0.73 | 1.63E-06 | 3.40E-02 | 0.29 | 0.14 | 0.15 | 2.11 |
| 21 | Ocrl | 0.60 | 1.70E-06 | 3.53E-02 | 0.48 | 0.27 | 0.21 | 1.77 |
| 21 | Slc31a1 | 0.51 | 1.71E-06 | 3.56E-02 | 0.59 | 0.35 | 0.25 | 1.72 |
| 21 | Tprn | 1.52 | 1.77E-06 | 3.67E-02 | 0.12 | 0.04 | 0.08 | 3.05 |
| 21 | Prkx | 0.76 | 1.77E-06 | 3.69E-02 | 0.32 | 0.15 | 0.16 | 2.05 |
| 21 | Cmtm3 | 0.53 | 1.82E-06 | 3.78E-02 | 0.65 | 0.42 | 0.23 | 1.55 |
| 21 | Smim24 | 1.16 | 1.84E-06 | 3.83E-02 | 0.14 | 0.05 | 0.09 | 2.80 |
| 21 | Cep128 | 0.66 | 1.84E-06 | 3.83E-02 | 0.34 | 0.17 | 0.17 | 1.97 |
| 21 | Gclc | 0.78 | 1.85E-06 | 3.85E-02 | 0.49 | 0.30 | 0.19 | 1.66 |
| 21 | Gm43813 | 1.04 | 1.89E-06 | 3.92E-02 | 0.20 | 0.09 | 0.12 | 2.33 |
| 21 | Scyl3 | 0.73 | 1.91E-06 | 3.97E-02 | 0.28 | 0.13 | 0.15 | 2.11 |
| 21 | Batf3 | 0.79 | 1.99E-06 | 4.14E-02 | 0.32 | 0.15 | 0.16 | 2.05 |
| 21 | Spns1 | 1.00 | 1.99E-06 | 4.15E-02 | 0.60 | 0.41 | 0.19 | 1.47 |
| 21 | Mrtfa | 0.80 | 2.00E-06 | 4.17E-02 | 0.27 | 0.12 | 0.14 | 2.16 |
| 21 | Retreg2 | 0.65 | 2.07E-06 | 4.30E-02 | 0.49 | 0.29 | 0.20 | 1.70 |
| 21 | Yipf3 | 0.50 | 2.14E-06 | 4.46E-02 | 0.66 | 0.42 | 0.25 | 1.60 |
| 21 | Sap30 | 0.49 | 2.15E-06 | 4.47E-02 | 0.85 | 0.65 | 0.20 | 1.31 |
| 21 | Rubcn | 1.05 | 2.15E-06 | 4.48E-02 | 0.20 | 0.08 | 0.11 | 2.39 |

**Supplementary Table 1. Marker genes for the original clusters of integrated clustering of E11.5 control and NotchGOF λ-junction cells.**

| | | | | | | | | |
|---|---|---|---|---|---|---|---|---|
| 21 | Chchd10 | 0.47 | 2.19E-06 | 4.55E-02 | 0.33 | 0.16 | 0.17 | 2.04 |
| 21 | Timm13 | 0.26 | 2.23E-06 | 4.64E-02 | 0.99 | 0.97 | 0.02 | 1.02 |
| 21 | B9d2 | 0.54 | 2.28E-06 | 4.73E-02 | 0.48 | 0.27 | 0.21 | 1.79 |
| 21 | Erp44 | 0.59 | 2.29E-06 | 4.77E-02 | 0.74 | 0.51 | 0.23 | 1.45 |
| 21 | Nudt7 | 1.10 | 2.33E-06 | 4.85E-02 | 0.18 | 0.07 | 0.11 | 2.50 |
| 21 | Pxmp4 | 0.72 | 2.36E-06 | 4.92E-02 | 0.34 | 0.18 | 0.17 | 1.93 |
| 22 | P2rx1 | 9.11 | 0 | 0 | 0.38 | 0.00 | 0.38 | 191.50 |
| 22 | Gp9 | 11.73 | 0 | 0 | 0.38 | 0.00 | 0.38 | 127.67 |
| 22 | Gp1bb | 9.66 | 0 | 0 | 0.38 | 0.00 | 0.38 | 95.75 |
| 22 | Treml1 | 11.69 | 0 | 0 | 0.37 | 0.00 | 0.37 | 185.00 |
| 22 | Rgs18 | 9.53 | 0 | 0 | 0.37 | 0.00 | 0.37 | 123.33 |
| 22 | Tmem40 | 9.78 | 0 | 0 | 0.37 | 0.00 | 0.37 | 123.33 |
| 22 | Sardh | 9.82 | 0 | 0 | 0.37 | 0.00 | 0.37 | 92.50 |
| 22 | Itgb3 | 9.83 | 0 | 0 | 0.37 | 0.01 | 0.36 | 61.67 |
| 22 | Mpl | 10.23 | 0 | 0 | 0.36 | 0.00 | 0.36 | 358.00 |
| 22 | Mrvi1 | 11.02 | 0 | 0 | 0.36 | 0.00 | 0.36 | 358.00 |
| 22 | Ppbp | 13.37 | 0 | 0 | 0.35 | 0.00 | 0.34 | 173.00 |
| 22 | Gp5 | 11.31 | 0 | 0 | 0.35 | 0.00 | 0.34 | 173.00 |
| 22 | F10 | 11.59 | 0 | 0 | 0.33 | 0.00 | 0.33 | 333.00 |
| 22 | F2rl2 | 11.64 | 0 | 0 | 0.33 | 0.00 | 0.33 | 333.00 |
| 22 | Gp1ba | 12.29 | 0 | 0 | 0.32 | 0.00 | 0.32 | #NA |
| 22 | Sla | 7.37 | 0 | 0 | 0.32 | 0.00 | 0.32 | 80.25 |
| 22 | Ly6g6f | 12.33 | 0 | 0 | 0.31 | 0.00 | 0.31 | #NA |
| 22 | F5 | 8.83 | 0 | 0 | 0.31 | 0.00 | 0.31 | 154.50 |
| 22 | Capn3 | 8.85 | 0 | 0 | 0.31 | 0.00 | 0.31 | 154.50 |
| 22 | I830077J02Rik | 7.73 | 0 | 0 | 0.31 | 0.00 | 0.31 | 77.25 |
| 22 | Tubb1 | 11.23 | 0 | 0 | 0.30 | 0.00 | 0.30 | #NA |
| 22 | Slc35d3 | 8.89 | 0 | 0 | 0.30 | 0.00 | 0.29 | 148.00 |
| 22 | Tbxas1 | 7.71 | 0 | 0 | 0.30 | 0.00 | 0.29 | 98.67 |
| 22 | Mpig6b | 7.62 | 0 | 0 | 0.28 | 0.00 | 0.28 | 142.00 |
| 22 | Gp6 | 12.85 | 0 | 0 | 0.27 | 0.00 | 0.27 | #NA |
| 22 | Cd226 | 11.08 | 0 | 0 | 0.27 | 0.00 | 0.27 | 272.00 |
| 22 | Atp2a3 | 7.94 | 0 | 0 | 0.27 | 0.00 | 0.27 | 90.67 |
| 22 | Cmah | 8.08 | 0 | 0 | 0.27 | 0.00 | 0.27 | 90.67 |
| 22 | Tuba8 | 8.96 | 0 | 0 | 0.24 | 0.00 | 0.23 | 235.00 |
| 22 | Gm30411 | 10.30 | 0 | 0 | 0.24 | 0.00 | 0.23 | 235.00 |
| 22 | AI463229 | 9.74 | 0 | 0 | 0.22 | 0.00 | 0.22 | #NA |
| 22 | Il12a | 9.81 | 0 | 0 | 0.22 | 0.00 | 0.22 | 222.00 |
| 22 | 1110028F11Rik | 8.79 | 0 | 0 | 0.22 | 0.00 | 0.22 | 222.00 |
| 22 | Kcnj5 | 8.98 | 0 | 0 | 0.20 | 0.00 | 0.20 | 198.00 |
| 22 | Gm10419 | 10.07 | 0 | 0 | 0.19 | 0.00 | 0.19 | #NA |
| 22 | Serpinb2 | 10.94 | 0 | 0 | 0.17 | 0.00 | 0.17 | #NA |
| 22 | Clec4g | 10.60 | 0 | 0 | 0.15 | 0.00 | 0.15 | #NA |
| 22 | Tnfsf14 | 9.94 | 0 | 0 | 0.15 | 0.00 | 0.15 | #NA |
| 22 | Cass4 | 8.71 | 0 | 0 | 0.14 | 0.00 | 0.14 | #NA |
| 22 | Cc2d2b | 8.59 | 0 | 0 | 0.14 | 0.00 | 0.14 | #NA |
| 22 | Gm16548 | 10.51 | 0 | 0 | 0.12 | 0.00 | 0.12 | #NA |
| 22 | Gm15915 | 8.82 | 0 | 0 | 0.12 | 0.00 | 0.12 | #NA |
| 22 | Glod5 | 8.09 | 0 | 0 | 0.12 | 0.00 | 0.12 | #NA |
| 22 | Sla2 | 8.11 | 1.87E-303 | 3.89E-299 | 0.30 | 0.00 | 0.29 | 74.00 |
| 22 | Ifi47 | 8.43 | 5.95E-295 | 1.24E-290 | 0.22 | 0.00 | 0.22 | 111.00 |
| 22 | Slc14a1 | 7.64 | 5.93E-293 | 1.23E-288 | 0.28 | 0.00 | 0.28 | 71.00 |
| 22 | Gbx2 | 7.84 | 3.41E-279 | 7.09E-275 | 0.14 | 0.00 | 0.14 | 136.00 |
| 22 | Obscn | 6.97 | 3.24E-272 | 6.75E-268 | 0.15 | 0.00 | 0.15 | 148.00 |
| 22 | Ctla2a | 7.80 | 4.91E-269 | 1.02E-264 | 0.56 | 0.02 | 0.54 | 30.89 |
| 22 | Clec1b | 7.53 | 5.73E-268 | 1.19E-263 | 0.53 | 0.02 | 0.52 | 33.19 |

**Supplementary Table 1. Marker genes for the original clusters of integrated clustering of E11.5 control and NotchGOF λ-junction cells.**

| | | | | | | | | |
|---|---|---|---|---|---|---|---|---|
| 22 | Btk | 6.99 | 1.98E-264 | 4.12E-260 | 0.32 | 0.01 | 0.32 | 64.20 |
| 22 | Was | 6.35 | 5.84E-257 | 1.21E-252 | 0.33 | 0.01 | 0.33 | 55.50 |
| 22 | Pkd1l3 | 7.39 | 2.95E-241 | 6.14E-237 | 0.12 | 0.00 | 0.12 | 123.00 |
| 22 | Gm13842 | 7.58 | 3.30E-241 | 6.87E-237 | 0.12 | 0.00 | 0.12 | 123.00 |
| 22 | Itgb2 | 6.25 | 3.80E-235 | 7.91E-231 | 0.26 | 0.00 | 0.26 | 64.75 |
| 22 | Trbc1 | 5.18 | 1.40E-234 | 2.91E-230 | 0.17 | 0.00 | 0.17 | 86.50 |
| 22 | Vwf | 8.17 | 1.89E-233 | 3.92E-229 | 0.35 | 0.01 | 0.34 | 49.43 |
| 22 | Bin2 | 7.77 | 2.12E-228 | 4.41E-224 | 0.38 | 0.01 | 0.37 | 38.30 |
| 22 | Ctla2b | 8.47 | 7.38E-227 | 1.53E-222 | 0.38 | 0.01 | 0.37 | 38.30 |
| 22 | Gm15657 | 7.78 | 1.48E-226 | 3.08E-222 | 0.11 | 0.00 | 0.11 | 111.00 |
| 22 | Plek | 8.50 | 1.14E-219 | 2.37E-215 | 0.41 | 0.01 | 0.40 | 37.00 |
| 22 | Trpc6 | 7.08 | 7.18E-215 | 1.49E-210 | 0.28 | 0.01 | 0.28 | 56.80 |
| 22 | Alox5 | 7.15 | 7.18E-211 | 1.49E-206 | 0.17 | 0.00 | 0.17 | 86.50 |
| 22 | Muc13 | 6.06 | 3.33E-200 | 6.92E-196 | 0.20 | 0.00 | 0.20 | 66.00 |
| 22 | 3300005D01Rik | 7.76 | 4.01E-200 | 8.34E-196 | 0.14 | 0.00 | 0.14 | 136.00 |
| 22 | Lcp2 | 6.44 | 3.84E-196 | 7.99E-192 | 0.30 | 0.01 | 0.29 | 42.29 |
| 22 | Gmfg | 6.12 | 3.61E-194 | 7.50E-190 | 0.56 | 0.03 | 0.53 | 22.24 |
| 22 | Lgals9 | 5.26 | 3.38E-193 | 7.03E-189 | 0.30 | 0.01 | 0.29 | 42.29 |
| 22 | Ubash3a | 6.92 | 1.12E-191 | 2.32E-187 | 0.27 | 0.01 | 0.27 | 45.33 |
| 22 | Cela1 | 6.13 | 3.69E-189 | 7.67E-185 | 0.26 | 0.01 | 0.25 | 51.80 |
| 22 | Srgn | 6.84 | 1.73E-181 | 3.61E-177 | 0.36 | 0.01 | 0.35 | 32.55 |
| 22 | Slpi | 7.89 | 1.75E-174 | 3.64E-170 | 0.16 | 0.00 | 0.16 | 80.00 |
| 22 | F13a1 | 4.83 | 9.91E-174 | 2.06E-169 | 0.22 | 0.00 | 0.22 | 55.50 |
| 22 | F2rl3 | 6.46 | 7.04E-169 | 1.46E-164 | 0.27 | 0.01 | 0.27 | 45.33 |
| 22 | Lyz2 | 4.05 | 1.70E-166 | 3.54E-162 | 0.31 | 0.01 | 0.30 | 38.63 |
| 22 | Klhl6 | 5.86 | 9.21E-159 | 1.91E-154 | 0.43 | 0.02 | 0.41 | 24.00 |
| 22 | Gimap1 | 5.67 | 1.78E-151 | 3.70E-147 | 0.40 | 0.02 | 0.38 | 24.69 |
| 22 | Hcst | 7.16 | 6.20E-149 | 1.29E-144 | 0.17 | 0.00 | 0.17 | 57.67 |
| 22 | Mmrn1 | 6.47 | 1.55E-147 | 3.23E-143 | 0.36 | 0.01 | 0.35 | 27.54 |
| 22 | Ltb4r1 | 6.01 | 1.05E-144 | 2.18E-140 | 0.19 | 0.00 | 0.18 | 61.67 |
| 22 | 5033406O09Rik | 6.43 | 8.93E-141 | 1.86E-136 | 0.32 | 0.01 | 0.31 | 29.18 |
| 22 | Cxcl5 | 7.15 | 7.22E-139 | 1.50E-134 | 0.17 | 0.00 | 0.17 | 57.67 |
| 22 | Il10ra | 5.33 | 5.64E-136 | 1.17E-131 | 0.19 | 0.00 | 0.18 | 46.25 |
| 22 | Lat | 7.08 | 1.96E-135 | 4.07E-131 | 0.36 | 0.02 | 0.34 | 23.87 |
| 22 | Itga2b | 8.47 | 7.82E-135 | 1.63E-130 | 0.38 | 0.02 | 0.37 | 22.53 |
| 22 | D130058E05Rik | 6.16 | 2.10E-132 | 4.37E-128 | 0.15 | 0.00 | 0.15 | 74.00 |
| 22 | Vav1 | 5.68 | 3.32E-131 | 6.91E-127 | 0.24 | 0.01 | 0.23 | 39.17 |
| 22 | Ltb4r2 | 5.45 | 2.57E-123 | 5.35E-119 | 0.15 | 0.00 | 0.15 | 74.00 |
| 22 | Rasgrp2 | 7.14 | 4.24E-119 | 8.82E-115 | 0.59 | 0.05 | 0.54 | 12.10 |
| 22 | Myct1 | 5.55 | 8.66E-117 | 1.80E-112 | 0.33 | 0.02 | 0.32 | 22.20 |
| 22 | Fcer1g | 5.07 | 2.37E-116 | 4.93E-112 | 0.40 | 0.02 | 0.37 | 18.81 |
| 22 | Vamp5 | 5.59 | 1.30E-115 | 2.70E-111 | 0.65 | 0.06 | 0.59 | 10.72 |
| 22 | Treml2 | 6.19 | 2.38E-115 | 4.95E-111 | 0.33 | 0.02 | 0.32 | 22.20 |
| 22 | Lrmp | 5.50 | 1.86E-114 | 3.87E-110 | 0.27 | 0.01 | 0.26 | 27.20 |
| 22 | Btnl9 | 6.59 | 1.78E-112 | 3.71E-108 | 0.11 | 0.00 | 0.11 | 111.00 |
| 22 | Ptpn7 | 5.86 | 1.82E-112 | 3.78E-108 | 0.25 | 0.01 | 0.24 | 30.88 |
| 22 | Cavin2 | 6.94 | 2.28E-111 | 4.74E-107 | 0.38 | 0.02 | 0.36 | 18.24 |
| 22 | Pdgfb | 5.63 | 6.23E-110 | 1.29E-105 | 0.32 | 0.01 | 0.31 | 22.93 |
| 22 | Cd200r1 | 7.06 | 3.42E-105 | 7.12E-101 | 0.12 | 0.00 | 0.12 | 61.50 |
| 22 | Laptm5 | 3.65 | 9.71E-104 | 2.02E-99 | 0.36 | 0.02 | 0.34 | 18.84 |
| 22 | Lrrc32 | 5.69 | 2.63E-102 | 5.48E-98 | 0.20 | 0.01 | 0.19 | 33.00 |
| 22 | Sox18 | 5.85 | 6.29E-102 | 1.31E-97 | 0.36 | 0.02 | 0.34 | 17.90 |
| 22 | Mfsd2b | 6.08 | 2.49E-100 | 5.18E-96 | 0.32 | 0.02 | 0.31 | 20.06 |
| 22 | Slamf1 | 5.56 | 7.63E-97 | 1.59E-92 | 0.27 | 0.01 | 0.26 | 22.67 |
| 22 | Hhex | 5.85 | 5.98E-96 | 1.24E-91 | 0.31 | 0.02 | 0.29 | 20.60 |
| 22 | Gm12999 | 5.78 | 3.63E-93 | 7.55E-89 | 0.16 | 0.00 | 0.16 | 40.00 |

**Supplementary Table 1. Marker genes for the original clusters of integrated clustering of E11.5 control and NotchGOF λ-junction cells.**

| | | | | | | | | |
|---|---|---|---|---|---|---|---|---|
| 22 | Ripor2 | 4.99 | 1.44E-91 | 3.00E-87 | 0.21 | 0.01 | 0.20 | 30.00 |
| 22 | Pf4 | 12.51 | 2.92E-90 | 6.07E-86 | 0.40 | 0.03 | 0.37 | 14.11 |
| 22 | Alox12 | 7.13 | 2.62E-89 | 5.45E-85 | 0.36 | 0.02 | 0.34 | 15.57 |
| 22 | Esam | 4.51 | 4.88E-87 | 1.01E-82 | 0.38 | 0.03 | 0.36 | 14.73 |
| 22 | Cd84 | 5.49 | 7.40E-87 | 1.54E-82 | 0.15 | 0.00 | 0.14 | 37.00 |
| 22 | Cd37 | 4.92 | 1.76E-86 | 3.66E-82 | 0.20 | 0.01 | 0.19 | 28.29 |
| 22 | Lyn | 4.92 | 2.13E-86 | 4.44E-82 | 0.42 | 0.03 | 0.39 | 13.13 |
| 22 | Treml4 | 7.28 | 4.23E-85 | 8.80E-81 | 0.11 | 0.00 | 0.11 | 55.50 |
| 22 | Kcnk6 | 5.41 | 5.84E-85 | 1.21E-80 | 0.22 | 0.01 | 0.21 | 24.67 |
| 22 | Ptk2b | 4.67 | 2.29E-84 | 4.77E-80 | 0.19 | 0.01 | 0.18 | 30.83 |
| 22 | Stx11 | 6.42 | 6.55E-84 | 1.36E-79 | 0.27 | 0.01 | 0.26 | 19.43 |
| 22 | Inpp5d | 4.34 | 7.93E-82 | 1.65E-77 | 0.27 | 0.01 | 0.26 | 19.43 |
| 22 | Gfi1b | 5.43 | 1.02E-81 | 2.12E-77 | 0.31 | 0.02 | 0.29 | 17.17 |
| 22 | Gm37035 | 5.26 | 3.92E-80 | 8.15E-76 | 0.12 | 0.00 | 0.12 | 41.00 |
| 22 | Dapp1 | 5.51 | 1.05E-79 | 2.18E-75 | 0.38 | 0.03 | 0.35 | 13.21 |
| 22 | Rapsn | 4.98 | 5.66E-79 | 1.18E-74 | 0.15 | 0.00 | 0.14 | 37.00 |
| 22 | Ptger3 | 5.04 | 6.00E-78 | 1.25E-73 | 0.25 | 0.01 | 0.24 | 20.58 |
| 22 | Ripor3 | 5.41 | 9.43E-77 | 1.96E-72 | 0.16 | 0.01 | 0.16 | 32.00 |
| 22 | Dok2 | 5.92 | 2.64E-74 | 5.49E-70 | 0.30 | 0.02 | 0.28 | 15.58 |
| 22 | Nbeal2 | 4.76 | 1.05E-73 | 2.17E-69 | 0.20 | 0.01 | 0.19 | 24.75 |
| 22 | Syk | 4.98 | 1.14E-72 | 2.37E-68 | 0.36 | 0.03 | 0.33 | 12.79 |
| 22 | Mctp1 | 4.93 | 1.50E-72 | 3.12E-68 | 0.19 | 0.01 | 0.18 | 26.43 |
| 22 | Abcb4 | 5.61 | 3.29E-71 | 6.85E-67 | 0.12 | 0.00 | 0.12 | 41.00 |
| 22 | Pik3cg | 6.03 | 7.16E-71 | 1.49E-66 | 0.17 | 0.01 | 0.17 | 28.83 |
| 22 | Fam167b | 6.52 | 9.48E-71 | 1.97E-66 | 0.24 | 0.01 | 0.22 | 19.58 |
| 22 | Plcg2 | 4.91 | 2.31E-70 | 4.80E-66 | 0.28 | 0.02 | 0.27 | 15.78 |
| 22 | Sp110 | 4.11 | 5.55E-70 | 1.15E-65 | 0.16 | 0.01 | 0.15 | 26.67 |
| 22 | Ltb | 5.72 | 1.72E-69 | 3.57E-65 | 0.14 | 0.00 | 0.13 | 34.00 |
| 22 | Ache | 4.75 | 2.18E-69 | 4.53E-65 | 0.28 | 0.02 | 0.27 | 15.78 |
| 22 | Tg | 6.30 | 1.30E-68 | 2.70E-64 | 0.12 | 0.00 | 0.12 | 41.00 |
| 22 | Cyth4 | 3.53 | 2.34E-68 | 4.86E-64 | 0.22 | 0.01 | 0.21 | 20.18 |
| 22 | Il21r | 5.70 | 1.77E-67 | 3.68E-63 | 0.12 | 0.00 | 0.12 | 41.00 |
| 22 | Fam78a | 5.52 | 2.21E-67 | 4.59E-63 | 0.24 | 0.01 | 0.22 | 18.08 |
| 22 | Hcls1 | 4.24 | 3.76E-66 | 7.82E-62 | 0.21 | 0.01 | 0.20 | 21.00 |
| 22 | Rhof | 4.72 | 3.67E-63 | 7.63E-59 | 0.26 | 0.02 | 0.24 | 16.19 |
| 22 | Gm13431 | 4.92 | 3.92E-63 | 8.15E-59 | 0.20 | 0.01 | 0.19 | 19.80 |
| 22 | Fxyd5 | 4.43 | 4.49E-63 | 9.34E-59 | 0.35 | 0.03 | 0.32 | 11.93 |
| 22 | Gimap6 | 4.77 | 5.58E-63 | 1.16E-58 | 0.33 | 0.03 | 0.31 | 12.33 |
| 22 | Tec | 5.63 | 9.51E-62 | 1.98E-57 | 0.22 | 0.01 | 0.21 | 18.50 |
| 22 | Rasl10a | 5.54 | 7.32E-61 | 1.52E-56 | 0.12 | 0.00 | 0.12 | 30.75 |
| 22 | Klf2 | 4.72 | 9.73E-61 | 2.02E-56 | 0.38 | 0.04 | 0.35 | 10.35 |
| 22 | Anxa3 | 4.68 | 5.38E-60 | 1.12E-55 | 0.58 | 0.09 | 0.49 | 6.52 |
| 22 | Prune2 | 4.95 | 9.47E-60 | 1.97E-55 | 0.20 | 0.01 | 0.19 | 19.80 |
| 22 | S100a13 | 4.70 | 4.47E-59 | 9.31E-55 | 0.62 | 0.11 | 0.51 | 5.82 |
| 22 | Nfe2 | 5.09 | 5.87E-58 | 1.22E-53 | 0.36 | 0.04 | 0.32 | 10.23 |
| 22 | Tspan32 | 5.85 | 7.04E-58 | 1.46E-53 | 0.37 | 0.04 | 0.33 | 9.74 |
| 22 | Cd55 | 5.66 | 1.03E-57 | 2.15E-53 | 0.30 | 0.02 | 0.27 | 12.33 |
| 22 | Nrgn | 7.65 | 8.60E-57 | 1.79E-52 | 0.41 | 0.05 | 0.36 | 8.66 |
| 22 | Kdr | 3.81 | 3.56E-56 | 7.41E-52 | 0.30 | 0.02 | 0.27 | 12.87 |
| 22 | F630028O10Rik | 5.67 | 4.46E-56 | 9.28E-52 | 0.11 | 0.00 | 0.11 | 37.00 |
| 22 | Tagln2 | 4.71 | 4.38E-55 | 9.12E-51 | 0.86 | 0.30 | 0.57 | 2.90 |
| 22 | Gng11 | 4.63 | 6.83E-55 | 1.42E-50 | 0.90 | 0.31 | 0.59 | 2.87 |
| 22 | Gata1 | 4.93 | 1.11E-54 | 2.30E-50 | 0.33 | 0.03 | 0.30 | 10.41 |
| 22 | Arhgap25 | 4.86 | 3.35E-54 | 6.96E-50 | 0.20 | 0.01 | 0.19 | 18.00 |
| 22 | Rac2 | 2.58 | 4.68E-53 | 9.74E-49 | 0.21 | 0.01 | 0.20 | 16.15 |
| 22 | Rab37 | 3.68 | 5.68E-53 | 1.18E-48 | 0.22 | 0.01 | 0.21 | 15.86 |

**Supplementary Table 1. Marker genes for the original clusters of integrated clustering of E11.5 control and NotchGOF λ-junction cells.**

| | | | | | | | | |
|---|---|---|---|---|---|---|---|---|
| 22 | Kcne3 | 6.39 | 8.55E-53 | 1.78E-48 | 0.35 | 0.04 | 0.31 | 9.89 |
| 22 | Pecam1 | 2.87 | 9.70E-53 | 2.02E-48 | 0.32 | 0.03 | 0.29 | 11.07 |
| 22 | Fermt3 | 6.97 | 2.50E-52 | 5.19E-48 | 0.40 | 0.05 | 0.35 | 8.23 |
| 22 | Unc13d | 4.70 | 3.73E-52 | 7.75E-48 | 0.31 | 0.03 | 0.28 | 11.04 |
| 22 | Lmo2 | 3.74 | 5.57E-52 | 1.16E-47 | 0.64 | 0.13 | 0.52 | 5.10 |
| 22 | Prkcq | 4.92 | 2.19E-49 | 4.56E-45 | 0.33 | 0.04 | 0.30 | 9.51 |
| 22 | Ptgs1 | 4.76 | 2.50E-49 | 5.19E-45 | 0.36 | 0.04 | 0.32 | 8.95 |
| 22 | Gm16104 | 4.42 | 3.92E-49 | 8.15E-45 | 0.37 | 0.04 | 0.33 | 8.81 |
| 22 | Tmsb4x | 3.20 | 2.45E-48 | 5.09E-44 | 0.99 | 0.99 | 0.00 | 1.00 |
| 22 | Cavin1 | 5.35 | 3.37E-48 | 7.00E-44 | 0.25 | 0.02 | 0.23 | 13.00 |
| 22 | Fyb | 5.37 | 4.64E-48 | 9.65E-44 | 0.36 | 0.04 | 0.32 | 8.52 |
| 22 | AI467606 | 5.60 | 1.18E-47 | 2.46E-43 | 0.11 | 0.00 | 0.11 | 27.75 |
| 22 | Samsn1 | 4.46 | 2.05E-47 | 4.26E-43 | 0.21 | 0.01 | 0.20 | 15.00 |
| 22 | Rilp | 4.57 | 2.22E-47 | 4.61E-43 | 0.11 | 0.00 | 0.11 | 27.75 |
| 22 | Myom1 | 4.03 | 1.44E-46 | 3.00E-42 | 0.31 | 0.03 | 0.28 | 9.97 |
| 22 | Gnaz | 5.38 | 7.73E-46 | 1.61E-41 | 0.36 | 0.04 | 0.32 | 8.33 |
| 22 | S100a16 | 5.20 | 8.21E-46 | 1.71E-41 | 0.47 | 0.08 | 0.39 | 6.25 |
| 22 | Emcn | 3.87 | 3.62E-45 | 7.53E-41 | 0.28 | 0.03 | 0.26 | 10.52 |
| 22 | Adgrg1 | 4.44 | 2.75E-44 | 5.72E-40 | 0.35 | 0.04 | 0.30 | 8.24 |
| 22 | Grap | 4.93 | 3.33E-44 | 6.92E-40 | 0.21 | 0.02 | 0.20 | 14.00 |
| 22 | Zc3hav1 | 3.93 | 7.13E-44 | 1.48E-39 | 0.28 | 0.03 | 0.26 | 10.14 |
| 22 | Actb | 1.47 | 1.11E-43 | 2.31E-39 | 1.00 | 1.00 | 0.00 | 1.00 |
| 22 | Dusp3 | 4.26 | 4.17E-43 | 8.68E-39 | 0.40 | 0.06 | 0.34 | 7.05 |
| 22 | Tnni1 | 5.90 | 4.20E-43 | 8.73E-39 | 0.14 | 0.01 | 0.13 | 19.43 |
| 22 | Tnfaip8l2 | 3.22 | 4.26E-43 | 8.86E-39 | 0.14 | 0.01 | 0.13 | 19.43 |
| 22 | Pfn1 | 1.59 | 8.47E-43 | 1.76E-38 | 0.98 | 1.00 | -0.02 | 0.98 |
| 22 | Susd3 | 4.69 | 1.58E-42 | 3.29E-38 | 0.20 | 0.01 | 0.18 | 14.14 |
| 22 | Lyl1 | 4.84 | 6.32E-41 | 1.31E-36 | 0.31 | 0.04 | 0.27 | 8.58 |
| 22 | Lst1 | 2.87 | 2.96E-40 | 6.15E-36 | 0.19 | 0.01 | 0.17 | 14.23 |
| 22 | Rgs10 | 6.14 | 8.84E-40 | 1.84E-35 | 0.47 | 0.09 | 0.38 | 5.33 |
| 22 | Dock8 | 3.57 | 1.67E-39 | 3.48E-35 | 0.30 | 0.03 | 0.26 | 8.97 |
| 22 | 1700003F12Rik | 4.17 | 3.08E-39 | 6.41E-35 | 0.15 | 0.01 | 0.14 | 16.44 |
| 22 | Gucy1b1 | 4.53 | 3.63E-39 | 7.54E-35 | 0.32 | 0.04 | 0.28 | 8.23 |
| 22 | Cdc42ep2 | 4.97 | 4.14E-38 | 8.60E-34 | 0.33 | 0.04 | 0.29 | 7.57 |
| 22 | Rasa3 | 4.56 | 1.14E-37 | 2.37E-33 | 0.43 | 0.08 | 0.36 | 5.68 |
| 22 | Psd4 | 4.02 | 1.25E-37 | 2.60E-33 | 0.12 | 0.01 | 0.12 | 20.50 |
| 22 | Grap2 | 4.28 | 2.57E-37 | 5.34E-33 | 0.26 | 0.03 | 0.23 | 9.59 |
| 22 | Tal1 | 3.75 | 1.37E-36 | 2.85E-32 | 0.35 | 0.05 | 0.30 | 7.21 |
| 22 | Sp140 | 4.10 | 4.77E-36 | 9.93E-32 | 0.16 | 0.01 | 0.15 | 14.55 |
| 22 | Tgfb1 | 5.03 | 4.84E-36 | 1.01E-31 | 0.49 | 0.11 | 0.39 | 4.57 |
| 22 | Rasip1 | 5.04 | 1.50E-35 | 3.11E-31 | 0.32 | 0.04 | 0.28 | 7.30 |
| 22 | Tbxa2r | 4.47 | 1.62E-35 | 3.38E-31 | 0.19 | 0.02 | 0.17 | 12.33 |
| 22 | Myl6 | 2.00 | 2.39E-35 | 4.97E-31 | 0.98 | 0.99 | -0.02 | 0.98 |
| 22 | Arhgap18 | 4.50 | 2.59E-35 | 5.39E-31 | 0.54 | 0.13 | 0.42 | 4.34 |
| 22 | Mef2c | 3.64 | 7.23E-35 | 1.50E-30 | 0.63 | 0.18 | 0.45 | 3.42 |
| 22 | Gpr146 | 4.68 | 4.87E-34 | 1.01E-29 | 0.26 | 0.03 | 0.23 | 8.63 |
| 22 | Cavin3 | 4.86 | 5.24E-34 | 1.09E-29 | 0.53 | 0.12 | 0.41 | 4.32 |
| 22 | Sh3bgrl3 | 2.23 | 5.47E-34 | 1.14E-29 | 0.91 | 0.91 | 0.01 | 1.01 |
| 22 | Gimap5 | 5.49 | 2.59E-33 | 5.38E-29 | 0.11 | 0.01 | 0.11 | 18.50 |
| 22 | Pear1 | 4.61 | 6.13E-33 | 1.28E-28 | 0.26 | 0.03 | 0.23 | 8.63 |
| 22 | Tuba4a | 6.41 | 8.26E-33 | 1.72E-28 | 0.38 | 0.07 | 0.31 | 5.55 |
| 22 | Tnnt1 | 3.99 | 9.50E-33 | 1.98E-28 | 0.35 | 0.05 | 0.29 | 6.53 |
| 22 | Smagp | 5.09 | 1.82E-32 | 3.78E-28 | 0.27 | 0.04 | 0.24 | 7.77 |
| 22 | Tnfrsf13b | 3.43 | 1.86E-32 | 3.86E-28 | 0.14 | 0.01 | 0.13 | 15.11 |
| 22 | Ptpre | 4.24 | 2.05E-32 | 4.26E-28 | 0.28 | 0.04 | 0.25 | 7.68 |
| 22 | Nrros | 2.92 | 2.14E-32 | 4.44E-28 | 0.30 | 0.04 | 0.26 | 7.79 |

**Supplementary Table 1. Marker genes for the original clusters of integrated clustering of E11.5 control and NotchGOF λ-junction cells.**

| | | | | | | | | |
|---|---|---|---|---|---|---|---|---|
| 22 | Reep2 | 3.82 | 5.40E-32 | 1.12E-27 | 0.15 | 0.01 | 0.14 | 13.45 |
| 22 | Gimap4 | 5.48 | 7.40E-32 | 1.54E-27 | 0.19 | 0.02 | 0.17 | 10.88 |
| 22 | Apold1 | 3.87 | 4.07E-31 | 8.47E-27 | 0.20 | 0.02 | 0.18 | 10.42 |
| 22 | Calm1 | 1.47 | 6.79E-31 | 1.41E-26 | 0.98 | 0.99 | -0.02 | 0.98 |
| 22 | S100a1 | 4.23 | 1.33E-30 | 2.76E-26 | 0.37 | 0.07 | 0.31 | 5.69 |
| 22 | Skap2 | 4.31 | 2.16E-29 | 4.48E-25 | 0.33 | 0.06 | 0.28 | 6.05 |
| 22 | Ppp1r14a | 3.80 | 2.85E-29 | 5.93E-25 | 0.38 | 0.07 | 0.31 | 5.32 |
| 22 | Tpm4 | 1.64 | 3.70E-29 | 7.69E-25 | 0.95 | 0.93 | 0.02 | 1.02 |
| 22 | Adcy9 | 3.82 | 3.96E-29 | 8.24E-25 | 0.14 | 0.01 | 0.13 | 13.60 |
| 22 | Arhgap6 | 3.99 | 7.49E-29 | 1.56E-24 | 0.36 | 0.07 | 0.29 | 5.51 |
| 22 | Cited4 | 3.61 | 8.27E-29 | 1.72E-24 | 0.35 | 0.06 | 0.29 | 5.86 |
| 22 | Exoc3l2 | 5.25 | 1.53E-28 | 3.18E-24 | 0.14 | 0.01 | 0.13 | 13.60 |
| 22 | D030007L05Rik | 5.42 | 3.07E-28 | 6.38E-24 | 0.15 | 0.01 | 0.14 | 12.33 |
| 22 | Arhgef3 | 5.18 | 4.73E-28 | 9.83E-24 | 0.19 | 0.02 | 0.17 | 10.28 |
| 22 | AI504432 | 3.79 | 7.44E-28 | 1.55E-23 | 0.28 | 0.04 | 0.24 | 6.76 |
| 22 | Arpc3 | 2.00 | 3.34E-27 | 6.96E-23 | 0.91 | 0.90 | 0.01 | 1.01 |
| 22 | Arpc1b | 3.21 | 4.30E-27 | 8.94E-23 | 0.72 | 0.35 | 0.37 | 2.06 |
| 22 | Rhoj | 3.61 | 4.63E-27 | 9.63E-23 | 0.48 | 0.12 | 0.36 | 3.98 |
| 22 | Cbfa2t3 | 3.37 | 1.52E-26 | 3.16E-22 | 0.32 | 0.06 | 0.27 | 5.84 |
| 22 | Csgalnact1 | 4.72 | 1.70E-26 | 3.54E-22 | 0.12 | 0.01 | 0.11 | 13.67 |
| 22 | Sncg | 5.96 | 2.74E-26 | 5.69E-22 | 0.14 | 0.01 | 0.13 | 12.36 |
| 22 | Fkbp1a | 2.05 | 5.57E-26 | 1.16E-21 | 0.95 | 0.94 | 0.01 | 1.01 |
| 22 | Kyat1 | 3.49 | 7.84E-26 | 1.63E-21 | 0.31 | 0.05 | 0.26 | 5.83 |
| 22 | Proser2 | 3.37 | 1.38E-25 | 2.86E-21 | 0.32 | 0.06 | 0.26 | 5.63 |
| 22 | Rab27b | 5.26 | 2.97E-25 | 6.17E-21 | 0.41 | 0.10 | 0.31 | 4.20 |
| 22 | Actg1 | 1.09 | 1.02E-24 | 2.13E-20 | 1.00 | 1.00 | 0.00 | 1.00 |
| 22 | Arhgap45 | 3.59 | 1.07E-24 | 2.23E-20 | 0.33 | 0.06 | 0.27 | 5.20 |
| 22 | Cfl1 | 0.98 | 1.32E-24 | 2.75E-20 | 0.98 | 1.00 | -0.02 | 0.98 |
| 22 | Vav3 | 3.13 | 1.60E-24 | 3.33E-20 | 0.32 | 0.06 | 0.26 | 5.53 |
| 22 | Cldn5 | 3.37 | 1.83E-24 | 3.80E-20 | 0.19 | 0.02 | 0.16 | 8.81 |
| 22 | Scarf1 | 3.07 | 2.05E-24 | 4.26E-20 | 0.15 | 0.01 | 0.13 | 10.57 |
| 22 | Dusp2 | 4.62 | 4.25E-24 | 8.84E-20 | 0.24 | 0.03 | 0.20 | 7.12 |
| 22 | Tesc | 3.30 | 4.48E-24 | 9.31E-20 | 0.11 | 0.01 | 0.10 | 13.88 |
| 22 | Pou2f2 | 3.59 | 1.67E-23 | 3.48E-19 | 0.21 | 0.03 | 0.18 | 7.78 |
| 22 | Cotl1 | 2.96 | 1.70E-23 | 3.53E-19 | 0.78 | 0.45 | 0.33 | 1.75 |
| 22 | Adgrg3 | 4.51 | 7.33E-23 | 1.52E-18 | 0.11 | 0.01 | 0.10 | 13.88 |
| 22 | Npr1 | 5.03 | 2.62E-21 | 5.45E-17 | 0.15 | 0.02 | 0.13 | 9.25 |
| 22 | Egfl7 | 3.67 | 4.31E-21 | 8.96E-17 | 0.58 | 0.22 | 0.36 | 2.65 |
| 22 | Myo7a | 2.97 | 6.12E-21 | 1.27E-16 | 0.14 | 0.01 | 0.12 | 10.46 |
| 22 | Nfatc1 | 4.58 | 6.62E-21 | 1.38E-16 | 0.22 | 0.03 | 0.19 | 6.73 |
| 22 | Snca | 3.43 | 9.57E-21 | 1.99E-16 | 0.37 | 0.09 | 0.28 | 4.16 |
| 22 | Gsap | 3.07 | 2.05E-20 | 4.26E-16 | 0.24 | 0.04 | 0.20 | 6.35 |
| 22 | Ehbp1l1 | 3.28 | 2.19E-20 | 4.55E-16 | 0.31 | 0.06 | 0.25 | 4.83 |
| 22 | Fbxo6 | 3.64 | 2.71E-20 | 5.63E-16 | 0.31 | 0.06 | 0.25 | 4.90 |
| 22 | Tie1 | 2.51 | 4.55E-20 | 9.46E-16 | 0.15 | 0.02 | 0.13 | 9.25 |
| 22 | Gucy1a1 | 3.19 | 4.55E-20 | 9.46E-16 | 0.33 | 0.08 | 0.26 | 4.38 |
| 22 | Adora2a | 4.99 | 8.06E-20 | 1.68E-15 | 0.12 | 0.01 | 0.11 | 10.25 |
| 22 | Serf2 | 0.81 | 1.79E-19 | 3.72E-15 | 0.95 | 1.00 | -0.05 | 0.95 |
| 22 | Ipo11 | 3.35 | 1.92E-19 | 3.99E-15 | 0.65 | 0.34 | 0.32 | 1.95 |
| 22 | Pstpip2 | 3.58 | 3.28E-19 | 6.82E-15 | 0.12 | 0.01 | 0.11 | 10.25 |
| 22 | Klhl4 | 4.46 | 3.50E-19 | 7.28E-15 | 0.14 | 0.02 | 0.12 | 9.07 |
| 22 | Ptpn18 | 3.33 | 4.64E-19 | 9.66E-15 | 0.33 | 0.08 | 0.26 | 4.32 |
| 22 | Smim5 | 3.16 | 7.90E-19 | 1.64E-14 | 0.19 | 0.03 | 0.16 | 7.12 |
| 22 | Exoc6 | 3.68 | 1.34E-18 | 2.78E-14 | 0.32 | 0.08 | 0.25 | 4.28 |
| 22 | Prkca | 3.96 | 1.81E-18 | 3.77E-14 | 0.33 | 0.08 | 0.25 | 4.01 |
| 22 | Ptprj | 3.55 | 1.88E-18 | 3.90E-14 | 0.22 | 0.04 | 0.18 | 5.84 |

**Supplementary Table 1. Marker genes for the original clusters of integrated clustering of E11.5 control and NotchGOF λ-junction cells.**

| | | | | | | | | |
|---|---|---|---|---|---|---|---|---|
| 22 | Pik3cb | 3.18 | 1.95E-18 | 4.06E-14 | 0.30 | 0.06 | 0.23 | 4.70 |
| 22 | Crip2 | 4.08 | 2.16E-18 | 4.49E-14 | 0.48 | 0.17 | 0.31 | 2.76 |
| 22 | Eif1 | 0.61 | 2.23E-18 | 4.64E-14 | 0.94 | 1.00 | -0.06 | 0.94 |
| 22 | Ncf1 | 2.54 | 3.75E-18 | 7.81E-14 | 0.11 | 0.01 | 0.10 | 11.10 |
| 22 | Ptpn6 | 3.31 | 4.16E-18 | 8.64E-14 | 0.22 | 0.04 | 0.19 | 6.00 |
| 22 | Larp1b | 3.39 | 8.30E-18 | 1.73E-13 | 0.32 | 0.08 | 0.24 | 4.17 |
| 22 | Ostf1 | 2.92 | 8.45E-18 | 1.76E-13 | 0.58 | 0.28 | 0.30 | 2.10 |
| 22 | Paqr7 | 3.66 | 9.91E-18 | 2.06E-13 | 0.11 | 0.01 | 0.10 | 10.09 |
| 22 | Fgd3 | 3.10 | 1.02E-17 | 2.11E-13 | 0.12 | 0.01 | 0.11 | 9.46 |
| 22 | Rpl6 | 0.50 | 1.20E-17 | 2.50E-13 | 0.98 | 1.00 | -0.03 | 0.98 |
| 22 | Slc6a4 | 5.28 | 1.51E-17 | 3.14E-13 | 0.11 | 0.01 | 0.10 | 10.09 |
| 22 | Actr3 | 1.91 | 1.68E-17 | 3.50E-13 | 0.79 | 0.82 | -0.03 | 0.97 |
| 22 | Parvb | 3.81 | 2.48E-17 | 5.15E-13 | 0.41 | 0.13 | 0.28 | 3.16 |
| 22 | Rap1b | 3.90 | 2.58E-17 | 5.37E-13 | 0.75 | 0.76 | -0.01 | 0.99 |
| 22 | Sox17 | 5.35 | 2.67E-17 | 5.55E-13 | 0.12 | 0.01 | 0.11 | 8.79 |
| 22 | Optn | 3.98 | 3.06E-17 | 6.37E-13 | 0.19 | 0.03 | 0.16 | 6.61 |
| 22 | Tmem163 | 2.35 | 7.61E-17 | 1.58E-12 | 0.22 | 0.04 | 0.18 | 5.69 |
| 22 | Cd300a | 2.33 | 8.13E-17 | 1.69E-12 | 0.12 | 0.01 | 0.11 | 8.79 |
| 22 | Nacc2 | 3.07 | 1.02E-16 | 2.12E-12 | 0.21 | 0.04 | 0.17 | 5.83 |
| 22 | Clic1 | 1.59 | 2.29E-16 | 4.75E-12 | 0.80 | 0.93 | -0.13 | 0.87 |
| 22 | Igtp | 4.24 | 2.43E-16 | 5.06E-12 | 0.15 | 0.02 | 0.13 | 7.40 |
| 22 | Eng | 4.38 | 3.95E-16 | 8.22E-12 | 0.22 | 0.04 | 0.18 | 5.29 |
| 22 | Cdc42 | 1.29 | 4.59E-16 | 9.54E-12 | 0.93 | 0.98 | -0.06 | 0.94 |
| 22 | Bmp2k | 3.77 | 6.06E-16 | 1.26E-11 | 0.42 | 0.15 | 0.27 | 2.78 |
| 22 | Ehd4 | 2.90 | 6.35E-16 | 1.32E-11 | 0.47 | 0.18 | 0.29 | 2.66 |
| 22 | Rabgap1l | 5.54 | 8.31E-16 | 1.73E-11 | 0.40 | 0.14 | 0.26 | 2.86 |
| 22 | Slc25a5 | 1.45 | 1.09E-15 | 2.26E-11 | 0.83 | 0.93 | -0.11 | 0.89 |
| 22 | P2ry1 | 3.95 | 1.10E-15 | 2.28E-11 | 0.25 | 0.05 | 0.19 | 4.66 |
| 22 | Elovl7 | 3.03 | 1.29E-15 | 2.68E-11 | 0.17 | 0.03 | 0.15 | 6.41 |
| 22 | Arpc2 | 1.39 | 1.57E-15 | 3.27E-11 | 0.84 | 0.94 | -0.10 | 0.89 |
| 22 | Alox5ap | 2.93 | 1.64E-15 | 3.41E-11 | 0.21 | 0.04 | 0.17 | 5.38 |
| 22 | Arpc5 | 2.08 | 2.46E-15 | 5.11E-11 | 0.77 | 0.76 | 0.00 | 1.01 |
| 22 | Aldoa | 1.15 | 4.51E-15 | 9.38E-11 | 0.88 | 0.95 | -0.08 | 0.92 |
| 22 | Flt3l | 3.34 | 6.70E-15 | 1.39E-10 | 0.11 | 0.01 | 0.10 | 8.54 |
| 22 | Vasp | 2.70 | 7.15E-15 | 1.49E-10 | 0.68 | 0.49 | 0.19 | 1.40 |
| 22 | Arpc4 | 1.61 | 8.05E-15 | 1.67E-10 | 0.79 | 0.83 | -0.04 | 0.96 |
| 22 | Tubb6 | 4.23 | 9.39E-15 | 1.95E-10 | 0.37 | 0.11 | 0.26 | 3.25 |
| 22 | Dennd2d | 2.63 | 9.39E-15 | 1.95E-10 | 0.11 | 0.01 | 0.10 | 8.54 |
| 22 | Thbs1 | 5.97 | 9.47E-15 | 1.97E-10 | 0.36 | 0.12 | 0.24 | 3.01 |
| 22 | Med12l | 2.65 | 1.13E-14 | 2.34E-10 | 0.22 | 0.05 | 0.18 | 4.93 |
| 22 | Trim47 | 3.56 | 1.68E-14 | 3.49E-10 | 0.14 | 0.02 | 0.12 | 7.16 |
| 22 | Rab11a | 2.57 | 2.24E-14 | 4.67E-10 | 0.72 | 0.64 | 0.07 | 1.12 |
| 22 | Taok3 | 3.61 | 4.49E-14 | 9.33E-10 | 0.33 | 0.10 | 0.23 | 3.20 |
| 22 | Abca7 | 3.09 | 4.82E-14 | 1.00E-09 | 0.15 | 0.02 | 0.13 | 6.73 |
| 22 | Jak3 | 4.24 | 5.12E-14 | 1.06E-09 | 0.17 | 0.03 | 0.14 | 5.77 |
| 22 | Mapkapk3 | 2.45 | 5.90E-14 | 1.23E-09 | 0.20 | 0.04 | 0.16 | 5.35 |
| 22 | Tgfb1i1 | 3.18 | 1.05E-13 | 2.18E-09 | 0.38 | 0.14 | 0.25 | 2.82 |
| 22 | Akap5 | 2.14 | 1.12E-13 | 2.33E-09 | 0.21 | 0.04 | 0.17 | 5.12 |
| 22 | Tln1 | 2.64 | 1.52E-13 | 3.15E-09 | 0.72 | 0.74 | -0.03 | 0.96 |
| 22 | Tpm3 | 1.08 | 2.88E-13 | 6.00E-09 | 0.84 | 0.96 | -0.12 | 0.88 |
| 22 | Ecscr | 3.12 | 2.89E-13 | 6.02E-09 | 0.15 | 0.02 | 0.12 | 6.17 |
| 22 | Arhgef15 | 4.94 | 3.45E-13 | 7.16E-09 | 0.16 | 0.03 | 0.13 | 5.52 |
| 22 | Snx3 | 1.65 | 4.19E-13 | 8.72E-09 | 0.82 | 0.89 | -0.08 | 0.91 |
| 22 | Chst12 | 2.84 | 4.79E-13 | 9.96E-09 | 0.30 | 0.08 | 0.21 | 3.52 |
| 22 | Gnai2 | 1.26 | 4.94E-13 | 1.03E-08 | 0.91 | 0.97 | -0.06 | 0.94 |
| 22 | Exoc3l4 | 3.43 | 5.76E-13 | 1.20E-08 | 0.27 | 0.07 | 0.20 | 3.83 |

**Supplementary Table 1. Marker genes for the original clusters of integrated clustering of E11.5 control and NotchGOF λ-junction cells.**

| | | | | | | | | |
|---|---|---|---|---|---|---|---|---|
| 22 | Ocel1 | 3.27 | 6.07E-13 | 1.26E-08 | 0.24 | 0.05 | 0.18 | 4.35 |
| 22 | Tubb2a | 3.23 | 7.17E-13 | 1.49E-08 | 0.54 | 0.30 | 0.25 | 1.82 |
| 22 | Coro1a | 3.87 | 7.57E-13 | 1.57E-08 | 0.38 | 0.15 | 0.24 | 2.59 |
| 22 | Gfod1 | 3.08 | 9.88E-13 | 2.06E-08 | 0.14 | 0.02 | 0.12 | 6.48 |
| 22 | Sptb | 1.90 | 1.40E-12 | 2.90E-08 | 0.22 | 0.05 | 0.17 | 4.53 |
| 22 | Cnn2 | 2.18 | 1.57E-12 | 3.27E-08 | 0.72 | 0.63 | 0.09 | 1.14 |
| 22 | Rap1a | 2.29 | 1.96E-12 | 4.08E-08 | 0.69 | 0.72 | -0.03 | 0.96 |
| 22 | Gngt2 | 4.69 | 2.33E-12 | 4.85E-08 | 0.19 | 0.04 | 0.15 | 4.74 |
| 22 | Myl12a | 1.19 | 2.50E-12 | 5.21E-08 | 0.83 | 0.95 | -0.13 | 0.87 |
| 22 | Eif5a | 0.62 | 2.81E-12 | 5.84E-08 | 0.98 | 1.00 | -0.02 | 0.98 |
| 22 | Dusp10 | 2.66 | 3.68E-12 | 7.65E-08 | 0.19 | 0.04 | 0.15 | 5.00 |
| 22 | Rps2 | 0.43 | 3.84E-12 | 7.98E-08 | 0.99 | 1.00 | -0.01 | 0.99 |
| 22 | Atp8a1 | 2.83 | 4.07E-12 | 8.46E-08 | 0.30 | 0.09 | 0.21 | 3.36 |
| 22 | Ttc39b | 2.10 | 4.28E-12 | 8.90E-08 | 0.26 | 0.07 | 0.19 | 3.92 |
| 22 | Mknk1 | 2.46 | 4.45E-12 | 9.26E-08 | 0.31 | 0.09 | 0.22 | 3.36 |
| 22 | Pls1 | 3.08 | 7.57E-12 | 1.58E-07 | 0.31 | 0.10 | 0.21 | 3.22 |
| 22 | Egr1 | 2.22 | 1.15E-11 | 2.39E-07 | 0.27 | 0.08 | 0.20 | 3.58 |
| 22 | Rpsa | 0.36 | 1.26E-11 | 2.62E-07 | 1.00 | 1.00 | 0.00 | 1.00 |
| 22 | Ehd2 | 4.61 | 1.31E-11 | 2.71E-07 | 0.21 | 0.05 | 0.16 | 4.12 |
| 22 | Scamp5 | 2.55 | 1.31E-11 | 2.72E-07 | 0.20 | 0.04 | 0.16 | 4.60 |
| 22 | Mfap3l | 3.12 | 1.37E-11 | 2.84E-07 | 0.14 | 0.02 | 0.11 | 5.91 |
| 22 | Hsd3b7 | 2.69 | 1.43E-11 | 2.98E-07 | 0.17 | 0.04 | 0.14 | 4.94 |
| 22 | Neat1 | 3.80 | 2.28E-11 | 4.74E-07 | 0.22 | 0.06 | 0.17 | 4.04 |
| 22 | Engase | 3.03 | 2.62E-11 | 5.44E-07 | 0.20 | 0.04 | 0.15 | 4.50 |
| 22 | Epor | 2.56 | 3.69E-11 | 7.68E-07 | 0.27 | 0.08 | 0.20 | 3.53 |
| 22 | Ost4 | 1.15 | 3.75E-11 | 7.81E-07 | 0.80 | 0.95 | -0.15 | 0.84 |
| 22 | Pros1 | 3.11 | 4.51E-11 | 9.38E-07 | 0.37 | 0.15 | 0.23 | 2.55 |
| 22 | St3gal1 | 2.54 | 4.73E-11 | 9.83E-07 | 0.32 | 0.10 | 0.22 | 3.15 |
| 22 | Wdr1 | 2.11 | 5.45E-11 | 1.13E-06 | 0.64 | 0.55 | 0.10 | 1.17 |
| 22 | Bola2 | 1.17 | 6.23E-11 | 1.30E-06 | 0.77 | 0.89 | -0.12 | 0.86 |
| 22 | Junb | 2.70 | 6.32E-11 | 1.31E-06 | 0.33 | 0.12 | 0.22 | 2.85 |
| 22 | Icam2 | 3.45 | 8.46E-11 | 1.76E-06 | 0.12 | 0.02 | 0.10 | 5.86 |
| 22 | Tpst2 | 2.87 | 9.31E-11 | 1.94E-06 | 0.46 | 0.24 | 0.22 | 1.94 |
| 22 | Trappc2 | 2.54 | 9.97E-11 | 2.07E-06 | 0.40 | 0.16 | 0.23 | 2.41 |
| 22 | Cpeb2 | 2.71 | 1.04E-10 | 2.16E-06 | 0.11 | 0.02 | 0.09 | 6.53 |
| 22 | Mob3c | 2.52 | 1.07E-10 | 2.23E-06 | 0.22 | 0.06 | 0.17 | 4.04 |
| 22 | Sntb1 | 3.13 | 1.38E-10 | 2.86E-06 | 0.15 | 0.03 | 0.12 | 5.29 |
| 22 | Nipal3 | 2.14 | 1.65E-10 | 3.43E-06 | 0.19 | 0.04 | 0.14 | 4.51 |
| 22 | Ywhah | 1.37 | 1.82E-10 | 3.78E-06 | 0.80 | 0.91 | -0.11 | 0.88 |
| 22 | Uba7 | 3.23 | 1.84E-10 | 3.82E-06 | 0.31 | 0.11 | 0.20 | 2.86 |
| 22 | Plvap | 2.08 | 1.91E-10 | 3.97E-06 | 0.19 | 0.04 | 0.14 | 4.40 |
| 22 | Rhoa | 0.99 | 2.73E-10 | 5.68E-06 | 0.88 | 0.97 | -0.09 | 0.91 |
| 22 | Cd38 | 1.86 | 2.83E-10 | 5.88E-06 | 0.21 | 0.05 | 0.16 | 4.12 |
| 22 | Rin3 | 3.81 | 3.09E-10 | 6.42E-06 | 0.14 | 0.03 | 0.11 | 5.23 |
| 22 | Larp7 | 0.27 | 3.28E-10 | 6.82E-06 | 0.37 | 0.77 | -0.40 | 0.48 |
| 22 | Rpl28 | 0.41 | 3.65E-10 | 7.59E-06 | 1.00 | 1.00 | 0.00 | 1.00 |
| 22 | Pip4k2a | 2.46 | 3.75E-10 | 7.79E-06 | 0.32 | 0.12 | 0.20 | 2.74 |
| 22 | Sparc | 0.44 | 3.81E-10 | 7.91E-06 | 0.36 | 0.74 | -0.39 | 0.48 |
| 22 | Gnas | 0.85 | 4.04E-10 | 8.41E-06 | 0.99 | 1.00 | -0.01 | 0.99 |
| 22 | Nab1 | 2.87 | 4.58E-10 | 9.53E-06 | 0.35 | 0.13 | 0.22 | 2.66 |
| 22 | Spsb2 | 2.21 | 4.99E-10 | 1.04E-05 | 0.26 | 0.08 | 0.18 | 3.36 |
| 22 | Fth1 | 0.31 | 6.97E-10 | 1.45E-05 | 0.94 | 1.00 | -0.06 | 0.94 |
| 22 | Zfp36 | 2.82 | 7.93E-10 | 1.65E-05 | 0.25 | 0.07 | 0.18 | 3.43 |
| 22 | Ncf2 | 1.44 | 7.96E-10 | 1.65E-05 | 0.11 | 0.02 | 0.09 | 6.17 |
| 22 | Csrp1 | 2.26 | 8.71E-10 | 1.81E-05 | 0.56 | 0.34 | 0.21 | 1.63 |
| 22 | She | 4.76 | 1.02E-09 | 2.12E-05 | 0.12 | 0.02 | 0.10 | 5.35 |

**Supplementary Table 1. Marker genes for the original clusters of integrated clustering of E11.5 control and NotchGOF λ-junction cells.**

| | | | | | | | | |
|---|---|---|---|---|---|---|---|---|
| 22 | Psen2 | 2.08 | 1.07E-09 | 2.23E-05 | 0.26 | 0.08 | 0.18 | 3.36 |
| 22 | Smim1 | 2.87 | 1.43E-09 | 2.98E-05 | 0.37 | 0.17 | 0.20 | 2.20 |
| 22 | Tubb4b | 1.21 | 1.50E-09 | 3.13E-05 | 0.80 | 0.85 | -0.05 | 0.94 |
| 22 | Eif2s2 | 0.95 | 1.60E-09 | 3.33E-05 | 0.93 | 0.99 | -0.06 | 0.94 |
| 22 | Sh3glb1 | 1.47 | 2.08E-09 | 4.34E-05 | 0.77 | 0.84 | -0.07 | 0.91 |
| 22 | Tm6sf1 | 2.59 | 2.09E-09 | 4.35E-05 | 0.21 | 0.06 | 0.15 | 3.75 |
| 22 | Peg13 | 2.63 | 2.27E-09 | 4.73E-05 | 0.20 | 0.05 | 0.15 | 3.96 |
| 22 | Mfng | 2.57 | 2.32E-09 | 4.82E-05 | 0.19 | 0.05 | 0.14 | 4.02 |
| 22 | Tek | 4.83 | 2.44E-09 | 5.08E-05 | 0.11 | 0.02 | 0.09 | 5.84 |
| 22 | Mrpl17 | 1.47 | 2.59E-09 | 5.39E-05 | 0.75 | 0.80 | -0.04 | 0.95 |
| 22 | Stxbp5 | 1.86 | 2.61E-09 | 5.43E-05 | 0.26 | 0.08 | 0.18 | 3.28 |
| 22 | Mgmt | 3.08 | 2.73E-09 | 5.68E-05 | 0.42 | 0.21 | 0.21 | 2.02 |
| 22 | Rplp1 | 0.33 | 2.92E-09 | 6.06E-05 | 0.99 | 1.00 | -0.01 | 0.99 |
| 22 | Gramd1c | 3.48 | 2.97E-09 | 6.17E-05 | 0.16 | 0.04 | 0.12 | 4.32 |
| 22 | H2-D1 | 2.71 | 2.99E-09 | 6.21E-05 | 0.47 | 0.24 | 0.23 | 1.93 |
| 22 | Map4k1 | 2.38 | 3.56E-09 | 7.41E-05 | 0.19 | 0.05 | 0.14 | 4.11 |
| 22 | Cap1 | 2.51 | 3.88E-09 | 8.06E-05 | 0.61 | 0.55 | 0.06 | 1.11 |
| 22 | Ets1 | 3.27 | 4.26E-09 | 8.87E-05 | 0.38 | 0.18 | 0.20 | 2.14 |
| 22 | Rsu1 | 3.71 | 4.41E-09 | 9.18E-05 | 0.57 | 0.49 | 0.08 | 1.17 |
| 22 | Ecm1 | 3.73 | 4.61E-09 | 9.58E-05 | 0.12 | 0.02 | 0.10 | 5.13 |
| 22 | Atp5g1 | 0.78 | 4.64E-09 | 9.64E-05 | 0.88 | 0.99 | -0.11 | 0.89 |
| 22 | Rac1 | 1.34 | 4.92E-09 | 1.02E-04 | 0.72 | 0.90 | -0.19 | 0.79 |
| 22 | Dok3 | 3.29 | 5.33E-09 | 1.11E-04 | 0.11 | 0.02 | 0.09 | 5.55 |
| 22 | Arl5a | 0.36 | 5.37E-09 | 1.12E-04 | 0.25 | 0.60 | -0.36 | 0.41 |
| 22 | Slc2a3 | 2.55 | 6.64E-09 | 1.38E-04 | 0.33 | 0.14 | 0.19 | 2.40 |
| 22 | Jpt1 | 0.87 | 7.67E-09 | 1.60E-04 | 0.84 | 0.95 | -0.11 | 0.89 |
| 22 | Inf2 | 2.39 | 1.02E-08 | 2.13E-04 | 0.19 | 0.05 | 0.14 | 3.78 |
| 22 | Gchfr | 3.02 | 1.10E-08 | 2.28E-04 | 0.30 | 0.11 | 0.19 | 2.72 |
| 22 | Kin | 0.31 | 1.15E-08 | 2.39E-04 | 0.27 | 0.62 | -0.35 | 0.44 |
| 22 | Stxbp2 | 2.56 | 1.18E-08 | 2.45E-04 | 0.33 | 0.14 | 0.20 | 2.41 |
| 22 | Ndufa8 | 1.22 | 1.21E-08 | 2.51E-04 | 0.82 | 0.90 | -0.08 | 0.91 |
| 22 | Shank3 | 2.88 | 1.69E-08 | 3.52E-04 | 0.11 | 0.02 | 0.09 | 5.29 |
| 22 | Ndrg1 | 2.70 | 1.84E-08 | 3.83E-04 | 0.11 | 0.02 | 0.09 | 5.29 |
| 22 | Ehd3 | 2.61 | 1.84E-08 | 3.83E-04 | 0.28 | 0.11 | 0.17 | 2.58 |
| 22 | Ubash3b | 3.42 | 1.90E-08 | 3.96E-04 | 0.42 | 0.24 | 0.18 | 1.76 |
| 22 | Ppia | 0.45 | 2.00E-08 | 4.15E-04 | 1.00 | 1.00 | 0.00 | 1.00 |
| 22 | B3gnt2 | 2.88 | 2.03E-08 | 4.23E-04 | 0.28 | 0.10 | 0.18 | 2.84 |
| 22 | Rpl36al | 0.50 | 2.85E-08 | 5.92E-04 | 0.89 | 1.00 | -0.11 | 0.89 |
| 22 | Arap1 | 1.47 | 3.08E-08 | 6.40E-04 | 0.24 | 0.07 | 0.16 | 3.31 |
| 22 | Afap1l1 | 2.65 | 3.29E-08 | 6.83E-04 | 0.24 | 0.07 | 0.16 | 3.18 |
| 22 | Rps6 | 0.37 | 3.52E-08 | 7.32E-04 | 0.98 | 1.00 | -0.02 | 0.98 |
| 22 | Ptp4a3 | 3.13 | 3.56E-08 | 7.40E-04 | 0.46 | 0.25 | 0.20 | 1.80 |
| 22 | Nmnat3 | 2.61 | 4.80E-08 | 9.99E-04 | 0.17 | 0.05 | 0.13 | 3.76 |
| 22 | Asb1 | 1.92 | 4.93E-08 | 1.02E-03 | 0.22 | 0.07 | 0.15 | 3.22 |
| 22 | Cpeb4 | 2.41 | 5.20E-08 | 1.08E-03 | 0.26 | 0.10 | 0.16 | 2.73 |
| 22 | Dennd2c | 2.40 | 5.44E-08 | 1.13E-03 | 0.21 | 0.06 | 0.15 | 3.28 |
| 22 | Inpp5b | 2.40 | 5.50E-08 | 1.14E-03 | 0.31 | 0.12 | 0.19 | 2.64 |
| 22 | Gas2l1 | 3.41 | 5.89E-08 | 1.22E-03 | 0.38 | 0.20 | 0.19 | 1.93 |
| 22 | Sec14l2 | 1.98 | 7.04E-08 | 1.46E-03 | 0.12 | 0.03 | 0.10 | 4.73 |
| 22 | Pfkp | 2.39 | 7.42E-08 | 1.54E-03 | 0.19 | 0.05 | 0.13 | 3.56 |
| 22 | Fhod1 | 2.93 | 8.87E-08 | 1.84E-03 | 0.25 | 0.09 | 0.16 | 2.81 |
| 22 | Cycs | 0.86 | 8.94E-08 | 1.86E-03 | 0.89 | 0.96 | -0.07 | 0.92 |
| 22 | Gcnt1 | 2.18 | 9.07E-08 | 1.89E-03 | 0.14 | 0.03 | 0.11 | 4.39 |
| 22 | Sf1 | 0.43 | 9.83E-08 | 2.04E-03 | 0.43 | 0.79 | -0.36 | 0.54 |
| 22 | Ppp1r13b | 3.36 | 1.02E-07 | 2.11E-03 | 0.24 | 0.08 | 0.15 | 2.90 |
| 22 | Traf5 | 2.07 | 1.10E-07 | 2.28E-03 | 0.16 | 0.04 | 0.12 | 3.90 |

**Supplementary Table 1. Marker genes for the original clusters of integrated clustering of E11.5 control and NotchGOF λ-junction cells.**

| | | | | | | | | |
|---|---|---|---|---|---|---|---|---|
| 22 | Agpat4 | 0.31 | 1.22E-07 | 2.53E-03 | 0.16 | 0.44 | -0.28 | 0.36 |
| 22 | Lcp1 | 3.29 | 1.22E-07 | 2.54E-03 | 0.15 | 0.04 | 0.11 | 3.89 |
| 22 | Fam43a | 2.69 | 1.26E-07 | 2.62E-03 | 0.19 | 0.05 | 0.13 | 3.49 |
| 22 | Sytl4 | 2.30 | 1.35E-07 | 2.81E-03 | 0.14 | 0.03 | 0.11 | 4.39 |
| 22 | Odc1 | 1.53 | 1.62E-07 | 3.37E-03 | 0.74 | 0.81 | -0.07 | 0.91 |
| 22 | Tbx1 | 4.07 | 1.84E-07 | 3.83E-03 | 0.12 | 0.03 | 0.09 | 4.24 |
| 22 | Rpl19 | 0.29 | 2.02E-07 | 4.21E-03 | 1.00 | 1.00 | 0.00 | 1.00 |
| 22 | Plxnc1 | 2.16 | 2.48E-07 | 5.16E-03 | 0.16 | 0.04 | 0.12 | 3.72 |
| 22 | Igf2r | 0.61 | 3.00E-07 | 6.23E-03 | 0.19 | 0.50 | -0.32 | 0.37 |
| 22 | Cd44 | 1.76 | 3.02E-07 | 6.28E-03 | 0.15 | 0.04 | 0.11 | 4.00 |
| 22 | Grrp1 | 4.55 | 3.04E-07 | 6.33E-03 | 0.19 | 0.06 | 0.13 | 3.14 |
| 22 | Pitpna | 1.63 | 3.25E-07 | 6.77E-03 | 0.70 | 0.71 | 0.00 | 1.00 |
| 22 | Sec61b | 0.55 | 3.29E-07 | 6.83E-03 | 0.86 | 0.99 | -0.12 | 0.88 |
| 22 | Sec11c | 3.64 | 3.48E-07 | 7.23E-03 | 0.44 | 0.30 | 0.14 | 1.47 |
| 22 | Rab27a | 1.64 | 3.95E-07 | 8.22E-03 | 0.17 | 0.05 | 0.13 | 3.60 |
| 22 | Stk10 | 3.43 | 4.88E-07 | 1.01E-02 | 0.12 | 0.03 | 0.09 | 4.24 |
| 22 | Cox6b1 | 0.54 | 4.95E-07 | 1.03E-02 | 0.94 | 1.00 | -0.06 | 0.94 |
| 22 | Smim3 | 3.70 | 5.12E-07 | 1.06E-02 | 0.35 | 0.19 | 0.16 | 1.86 |
| 22 | Rhob | 2.17 | 5.32E-07 | 1.11E-02 | 0.54 | 0.35 | 0.20 | 1.56 |
| 22 | Syngr1 | 1.90 | 5.43E-07 | 1.13E-02 | 0.21 | 0.07 | 0.14 | 3.09 |
| 22 | Gypc | 2.22 | 5.69E-07 | 1.18E-02 | 0.42 | 0.22 | 0.20 | 1.93 |
| 22 | Gmpr | 2.92 | 6.04E-07 | 1.26E-02 | 0.36 | 0.18 | 0.18 | 2.02 |
| 22 | Ksr1 | 2.79 | 6.18E-07 | 1.28E-02 | 0.14 | 0.03 | 0.10 | 4.00 |
| 22 | Dok4 | 3.63 | 6.25E-07 | 1.30E-02 | 0.22 | 0.08 | 0.14 | 2.71 |
| 22 | Timp3 | 3.49 | 6.30E-07 | 1.31E-02 | 0.33 | 0.17 | 0.16 | 1.94 |
| 22 | Marchf2 | 2.59 | 6.32E-07 | 1.31E-02 | 0.40 | 0.21 | 0.18 | 1.85 |
| 22 | Rps8 | 0.27 | 6.51E-07 | 1.35E-02 | 1.00 | 1.00 | 0.00 | 1.00 |
| 22 | Plekho2 | 2.17 | 6.65E-07 | 1.38E-02 | 0.21 | 0.07 | 0.14 | 3.00 |
| 22 | Ralb | 2.56 | 7.06E-07 | 1.47E-02 | 0.61 | 0.43 | 0.17 | 1.40 |
| 22 | Psmg2 | 0.34 | 7.32E-07 | 1.52E-02 | 0.30 | 0.64 | -0.34 | 0.46 |
| 22 | Jup | 2.08 | 7.83E-07 | 1.63E-02 | 0.41 | 0.20 | 0.21 | 2.04 |
| 22 | Mcrip2 | 3.34 | 7.92E-07 | 1.65E-02 | 0.21 | 0.07 | 0.14 | 3.00 |
| 22 | Tnfaip1 | 0.30 | 8.80E-07 | 1.83E-02 | 0.30 | 0.63 | -0.33 | 0.47 |
| 22 | Cul5 | 0.28 | 9.97E-07 | 2.07E-02 | 0.33 | 0.69 | -0.35 | 0.49 |
| 22 | Abracl | 0.95 | 1.03E-06 | 2.13E-02 | 0.77 | 0.93 | -0.16 | 0.83 |
| 22 | Ufsp1 | 2.36 | 1.07E-06 | 2.23E-02 | 0.28 | 0.12 | 0.17 | 2.47 |
| 22 | Stat3 | 2.31 | 1.12E-06 | 2.33E-02 | 0.31 | 0.13 | 0.18 | 2.34 |
| 22 | Golga7 | 0.48 | 1.20E-06 | 2.49E-02 | 0.43 | 0.74 | -0.31 | 0.58 |
| 22 | Cox5b | 0.54 | 1.25E-06 | 2.60E-02 | 0.89 | 0.99 | -0.10 | 0.90 |
| 22 | St3gal6 | 2.08 | 1.27E-06 | 2.64E-02 | 0.27 | 0.11 | 0.16 | 2.41 |
| 22 | Scarb1 | 1.59 | 1.34E-06 | 2.78E-02 | 0.35 | 0.16 | 0.19 | 2.18 |
| 22 | Bst2 | 3.43 | 1.40E-06 | 2.90E-02 | 0.19 | 0.06 | 0.13 | 3.19 |
| 22 | Myh9 | 2.19 | 1.42E-06 | 2.95E-02 | 0.56 | 0.51 | 0.05 | 1.09 |
| 22 | Anxa5 | 1.91 | 1.42E-06 | 2.96E-02 | 0.64 | 0.61 | 0.04 | 1.06 |
| 22 | Pip5k1b | 2.07 | 1.43E-06 | 2.98E-02 | 0.14 | 0.04 | 0.10 | 3.78 |
| 22 | Ostm1 | 3.09 | 1.51E-06 | 3.15E-02 | 0.21 | 0.07 | 0.14 | 2.88 |
| 22 | Mrpl19 | 0.28 | 1.54E-06 | 3.20E-02 | 0.32 | 0.61 | -0.29 | 0.52 |
| 22 | Rbpms2 | 3.40 | 1.55E-06 | 3.22E-02 | 0.37 | 0.23 | 0.14 | 1.64 |
| 22 | Vdac2 | 0.86 | 1.68E-06 | 3.49E-02 | 0.80 | 0.96 | -0.16 | 0.84 |
| 22 | Dok1 | 2.00 | 1.69E-06 | 3.51E-02 | 0.26 | 0.10 | 0.16 | 2.56 |
| 22 | Rab32 | 0.93 | 1.73E-06 | 3.60E-02 | 0.26 | 0.09 | 0.17 | 2.78 |
| 22 | Prr13 | 2.48 | 1.78E-06 | 3.71E-02 | 0.54 | 0.40 | 0.14 | 1.35 |
| 22 | N4bp3 | 4.05 | 1.99E-06 | 4.13E-02 | 0.19 | 0.07 | 0.12 | 2.85 |
| 22 | Clu | 3.32 | 2.00E-06 | 4.17E-02 | 0.37 | 0.21 | 0.16 | 1.77 |
| 22 | Myzap | 4.65 | 2.06E-06 | 4.29E-02 | 0.12 | 0.03 | 0.09 | 3.73 |
| 22 | Vangl1 | 2.05 | 2.14E-06 | 4.45E-02 | 0.32 | 0.16 | 0.17 | 2.07 |

**Supplementary Table 1. Marker genes for the original clusters of integrated clustering of E11.5 control and NotchGOF λ-junction cells.**

| | | | | | | | | |
|---|---|---|---|---|---|---|---|---|
| 23 | Fabp7 | 10.63 | 0 | 0 | 1.00 | 0.00 | 1.00 | 333.33 |
| 23 | Sox10 | 8.39 | 0 | 0 | 1.00 | 0.01 | 0.99 | 125.00 |
| 23 | Foxd3 | 14.29 | 0 | 0 | 0.94 | 0.00 | 0.94 | #NA |
| 23 | Insc | 9.73 | 0 | 0 | 0.89 | 0.00 | 0.89 | 889.00 |
| 23 | Megf10 | 8.35 | 0 | 0 | 0.89 | 0.01 | 0.88 | 148.17 |
| 23 | Gpr17 | 13.53 | 0 | 0 | 0.83 | 0.00 | 0.83 | #NA |
| 23 | Cdh19 | 9.12 | 0 | 0 | 0.78 | 0.00 | 0.78 | 389.00 |
| 23 | Gm12688 | 12.07 | 0 | 0 | 0.72 | 0.00 | 0.72 | #NA |
| 23 | Gfra3 | 10.96 | 0 | 0 | 0.72 | 0.00 | 0.72 | 722.00 |
| 23 | Tmem132e | 7.54 | 0 | 0 | 0.61 | 0.00 | 0.61 | 152.75 |
| 23 | Npy | 10.13 | 0 | 0 | 0.56 | 0.00 | 0.55 | 278.00 |
| 23 | Scrg1 | 12.18 | 0 | 0 | 0.50 | 0.00 | 0.50 | #NA |
| 23 | Klhl30 | 9.68 | 0 | 0 | 0.33 | 0.00 | 0.33 | #NA |
| 23 | Art3 | 10.10 | 0 | 0 | 0.33 | 0.00 | 0.33 | #NA |
| 23 | Tecrl | 9.98 | 0 | 0 | 0.22 | 0.00 | 0.22 | #NA |
| 23 | Gjc3 | 12.06 | 0 | 0 | 0.22 | 0.00 | 0.22 | #NA |
| 23 | Tmprss11f | 10.89 | 0 | 0 | 0.11 | 0.00 | 0.11 | #NA |
| 23 | Ndst3 | 6.86 | 4.93E-307 | 1.03E-302 | 0.83 | 0.01 | 0.83 | 104.13 |
| 23 | 9330188P03Rik | 7.80 | 8.67E-296 | 1.80E-291 | 0.22 | 0.00 | 0.22 | #NA |
| 23 | Btla | 8.89 | 7.65E-275 | 1.59E-270 | 0.22 | 0.00 | 0.22 | 222.00 |
| 23 | Nppb | 8.82 | 8.96E-275 | 1.86E-270 | 0.22 | 0.00 | 0.22 | 222.00 |
| 23 | Ajap1 | 7.04 | 1.71E-272 | 3.55E-268 | 0.72 | 0.01 | 0.72 | 103.14 |
| 23 | Egfl8 | 7.76 | 2.07E-265 | 4.31E-261 | 0.89 | 0.01 | 0.88 | 80.82 |
| 23 | Adgrg6 | 7.08 | 1.61E-231 | 3.35E-227 | 0.78 | 0.01 | 0.77 | 86.44 |
| 23 | Slitrk2 | 6.07 | 4.11E-194 | 8.55E-190 | 0.89 | 0.02 | 0.87 | 59.27 |
| 23 | Dct | 7.60 | 2.18E-192 | 4.53E-188 | 0.22 | 0.00 | 0.22 | 222.00 |
| 23 | Spon1 | 7.50 | 1.06E-190 | 2.21E-186 | 0.83 | 0.01 | 0.82 | 59.50 |
| 23 | Atp10b | 7.07 | 3.84E-176 | 7.99E-172 | 0.33 | 0.00 | 0.33 | 166.50 |
| 23 | 4930452G13Rik | 9.01 | 3.17E-161 | 6.59E-157 | 0.11 | 0.00 | 0.11 | #NA |
| 23 | Camk2b | 6.29 | 4.44E-157 | 9.24E-153 | 0.67 | 0.01 | 0.66 | 66.70 |
| 23 | Fam178b | 7.41 | 1.15E-156 | 2.40E-152 | 0.44 | 0.01 | 0.44 | 88.80 |
| 23 | Phactr1 | 8.02 | 1.69E-153 | 3.52E-149 | 1.00 | 0.03 | 0.97 | 38.46 |
| 23 | Lims2 | 7.18 | 1.78E-150 | 3.69E-146 | 0.39 | 0.00 | 0.39 | 97.25 |
| 23 | Trpv4 | 6.45 | 1.38E-149 | 2.87E-145 | 0.44 | 0.01 | 0.44 | 88.80 |
| 23 | Npr3 | 6.14 | 2.17E-149 | 4.52E-145 | 0.83 | 0.02 | 0.82 | 49.00 |
| 23 | Plxna4 | 5.85 | 1.72E-147 | 3.59E-143 | 0.94 | 0.02 | 0.92 | 41.04 |
| 23 | Adam12 | 6.27 | 2.07E-139 | 4.30E-135 | 0.72 | 0.01 | 0.71 | 51.57 |
| 23 | 2010320O07Rik | 8.36 | 2.37E-138 | 4.93E-134 | 0.11 | 0.00 | 0.11 | #NA |
| 23 | Deptor | 6.39 | 1.86E-136 | 3.86E-132 | 0.56 | 0.01 | 0.55 | 69.50 |
| 23 | Sema3g | 7.64 | 4.63E-133 | 9.64E-129 | 0.33 | 0.00 | 0.33 | 111.00 |
| 23 | Car6 | 7.38 | 9.35E-133 | 1.94E-128 | 0.22 | 0.00 | 0.22 | 222.00 |
| 23 | Vat1l | 6.36 | 3.23E-130 | 6.71E-126 | 0.67 | 0.01 | 0.65 | 51.31 |
| 23 | Afap1l2 | 6.00 | 1.17E-129 | 2.44E-125 | 0.78 | 0.02 | 0.76 | 45.76 |
| 23 | Cd200 | 6.48 | 1.05E-127 | 2.19E-123 | 0.94 | 0.03 | 0.92 | 34.96 |
| 23 | Lgi4 | 6.71 | 4.34E-125 | 9.02E-121 | 0.44 | 0.01 | 0.44 | 74.00 |
| 23 | Tm4sf1 | 7.14 | 6.81E-117 | 1.42E-112 | 0.72 | 0.02 | 0.71 | 42.47 |
| 23 | Casq2 | 7.89 | 1.76E-114 | 3.67E-110 | 0.17 | 0.00 | 0.17 | 167.00 |
| 23 | Gfra2 | 7.03 | 6.46E-111 | 1.34E-106 | 0.39 | 0.01 | 0.38 | 77.80 |
| 23 | Fgd5 | 4.90 | 1.33E-95 | 2.76E-91 | 0.78 | 0.02 | 0.75 | 32.42 |
| 23 | Matn4 | 6.15 | 2.35E-92 | 4.89E-88 | 0.33 | 0.00 | 0.33 | 83.25 |
| 23 | Igsf11 | 6.71 | 8.57E-92 | 1.78E-87 | 0.22 | 0.00 | 0.22 | 111.00 |
| 23 | Tagln | 5.12 | 2.45E-88 | 5.11E-84 | 0.89 | 0.04 | 0.85 | 25.40 |
| 23 | Ednrb | 5.09 | 4.23E-87 | 8.79E-83 | 0.94 | 0.04 | 0.91 | 24.21 |
| 23 | Cryab | 6.26 | 1.43E-82 | 2.98E-78 | 1.00 | 0.05 | 0.95 | 20.41 |
| 23 | Ngfr | 4.88 | 2.04E-82 | 4.24E-78 | 0.94 | 0.04 | 0.90 | 22.48 |
| 23 | Tspan18 | 4.42 | 5.32E-82 | 1.11E-77 | 0.94 | 0.04 | 0.90 | 23.02 |

**Supplementary Table 1. Marker genes for the original clusters of integrated clustering of E11.5 control and NotchGOF λ-junction cells.**

| | | | | | | | | |
|---|---|---|---|---|---|---|---|---|
| 23 | Gm50147 | 8.56 | 3.20E-81 | 6.65E-77 | 0.11 | 0.00 | 0.11 | 111.00 |
| 23 | Hspb2 | 5.35 | 4.40E-80 | 9.15E-76 | 0.83 | 0.03 | 0.80 | 24.50 |
| 23 | Bche | 5.22 | 3.43E-75 | 7.14E-71 | 0.67 | 0.02 | 0.65 | 30.32 |
| 23 | Cmtm5 | 5.60 | 2.97E-71 | 6.17E-67 | 0.39 | 0.01 | 0.38 | 48.63 |
| 23 | Ina | 2.19 | 7.07E-67 | 1.47E-62 | 0.39 | 0.01 | 0.38 | 48.63 |
| 23 | Rassf4 | 5.25 | 8.33E-67 | 1.73E-62 | 0.72 | 0.03 | 0.69 | 24.07 |
| 23 | Rxrg | 4.86 | 3.16E-65 | 6.57E-61 | 0.44 | 0.01 | 0.43 | 40.36 |
| 23 | Rai2 | 6.94 | 2.70E-64 | 5.61E-60 | 0.17 | 0.00 | 0.17 | 83.50 |
| 23 | Gm29865 | 6.14 | 3.65E-62 | 7.59E-58 | 0.22 | 0.00 | 0.22 | 74.00 |
| 23 | Akap6 | 4.63 | 1.02E-60 | 2.12E-56 | 0.33 | 0.01 | 0.33 | 47.57 |
| 23 | Sntb1 | 5.03 | 2.13E-59 | 4.43E-55 | 0.67 | 0.03 | 0.64 | 23.82 |
| 23 | Dlx1 | 4.68 | 1.89E-58 | 3.92E-54 | 0.89 | 0.05 | 0.84 | 17.10 |
| 23 | Gm16316 | 6.16 | 9.80E-57 | 2.04E-52 | 0.22 | 0.00 | 0.22 | 74.00 |
| 23 | Luzp2 | 4.95 | 5.94E-53 | 1.23E-48 | 0.50 | 0.02 | 0.48 | 27.78 |
| 23 | Mcam | 4.37 | 1.16E-52 | 2.42E-48 | 0.78 | 0.04 | 0.74 | 18.09 |
| 23 | Prima1 | 6.82 | 5.97E-51 | 1.24E-46 | 0.17 | 0.00 | 0.17 | 83.50 |
| 23 | Sema6b | 5.77 | 4.60E-49 | 9.56E-45 | 0.33 | 0.01 | 0.32 | 37.00 |
| 23 | Postn | 5.39 | 1.14E-47 | 2.38E-43 | 1.00 | 0.09 | 0.91 | 11.24 |
| 23 | S100a16 | 3.72 | 2.44E-46 | 5.07E-42 | 0.94 | 0.08 | 0.87 | 12.42 |
| 23 | Prss12 | 4.92 | 9.33E-45 | 1.94E-40 | 0.78 | 0.05 | 0.73 | 14.96 |
| 23 | Cavin2 | 4.79 | 1.98E-43 | 4.13E-39 | 0.50 | 0.02 | 0.48 | 22.73 |
| 23 | Akr1cl | 5.63 | 1.57E-42 | 3.27E-38 | 0.33 | 0.01 | 0.32 | 33.30 |
| 23 | Atp1a2 | 5.62 | 2.41E-41 | 5.01E-37 | 0.33 | 0.01 | 0.32 | 33.30 |
| 23 | Plcg2 | 4.82 | 6.85E-40 | 1.42E-35 | 0.44 | 0.02 | 0.43 | 24.67 |
| 23 | Plp1 | 5.96 | 4.89E-39 | 1.02E-34 | 1.00 | 0.12 | 0.88 | 8.47 |
| 23 | Gja3 | 6.23 | 1.52E-38 | 3.17E-34 | 0.17 | 0.00 | 0.16 | 55.67 |
| 23 | Fitm1 | 6.71 | 6.23E-38 | 1.30E-33 | 0.11 | 0.00 | 0.11 | 111.00 |
| 23 | Il1rap | 4.17 | 1.96E-37 | 4.07E-33 | 0.61 | 0.04 | 0.57 | 16.51 |
| 23 | Adam23 | 4.23 | 1.13E-36 | 2.35E-32 | 0.89 | 0.08 | 0.81 | 10.71 |
| 23 | Olfr889 | 6.62 | 1.33E-36 | 2.77E-32 | 0.28 | 0.01 | 0.27 | 34.75 |
| 23 | Chl1 | 4.24 | 3.64E-35 | 7.56E-31 | 0.39 | 0.02 | 0.37 | 24.31 |
| 23 | Shc4 | 4.40 | 4.43E-35 | 9.21E-31 | 0.50 | 0.03 | 0.47 | 19.23 |
| 23 | Rasa3 | 3.74 | 6.98E-35 | 1.45E-30 | 0.83 | 0.08 | 0.76 | 10.82 |
| 23 | Slc6a15 | 4.10 | 1.14E-34 | 2.37E-30 | 0.56 | 0.03 | 0.52 | 16.85 |
| 23 | Slc35f1 | 4.38 | 4.45E-34 | 9.26E-30 | 0.56 | 0.03 | 0.52 | 16.35 |
| 23 | Arhgap22 | 4.52 | 9.25E-33 | 1.92E-28 | 0.33 | 0.01 | 0.32 | 25.62 |
| 23 | Cbr3 | 4.40 | 3.36E-32 | 7.00E-28 | 0.33 | 0.01 | 0.32 | 25.62 |
| 23 | Bcar3 | 3.94 | 7.18E-32 | 1.49E-27 | 0.67 | 0.05 | 0.62 | 13.08 |
| 23 | Cdh10 | 4.34 | 8.08E-32 | 1.68E-27 | 0.50 | 0.03 | 0.47 | 17.24 |
| 23 | Pou3f2 | 5.54 | 5.55E-31 | 1.15E-26 | 0.17 | 0.00 | 0.16 | 55.67 |
| 23 | Tfap2b | 2.19 | 2.57E-30 | 5.35E-26 | 0.89 | 0.09 | 0.80 | 9.88 |
| 23 | Dlx2 | 3.17 | 1.51E-29 | 3.13E-25 | 1.00 | 0.13 | 0.87 | 7.69 |
| 23 | Padi2 | 5.80 | 1.90E-29 | 3.95E-25 | 0.17 | 0.00 | 0.16 | 41.75 |
| 23 | Gpm6b | 4.28 | 2.72E-29 | 5.66E-25 | 0.72 | 0.07 | 0.66 | 10.78 |
| 23 | Stard13 | 4.16 | 3.29E-29 | 6.83E-25 | 0.44 | 0.03 | 0.42 | 17.76 |
| 23 | Xrra1 | 6.63 | 3.42E-29 | 7.11E-25 | 0.11 | 0.00 | 0.11 | 55.50 |
| 23 | Nkd2 | 3.24 | 4.58E-29 | 9.52E-25 | 1.00 | 0.13 | 0.87 | 7.69 |
| 23 | Zfp488 | 6.23 | 2.23E-28 | 4.65E-24 | 0.11 | 0.00 | 0.11 | 55.50 |
| 23 | Cxcl10 | 6.10 | 1.19E-27 | 2.48E-23 | 0.11 | 0.00 | 0.11 | 55.50 |
| 23 | Ndst4 | 6.61 | 5.82E-27 | 1.21E-22 | 0.11 | 0.00 | 0.11 | 55.50 |
| 23 | Hcn1 | 5.87 | 7.53E-27 | 1.57E-22 | 0.28 | 0.01 | 0.27 | 25.27 |
| 23 | Zfp536 | 2.76 | 1.32E-26 | 2.74E-22 | 0.89 | 0.10 | 0.79 | 8.72 |
| 23 | Erbb4 | 6.12 | 3.38E-26 | 7.02E-22 | 0.11 | 0.00 | 0.11 | 55.50 |
| 23 | Gm16104 | 3.28 | 7.77E-26 | 1.62E-21 | 0.56 | 0.04 | 0.51 | 12.93 |
| 23 | Erbb3 | 3.75 | 8.58E-26 | 1.78E-21 | 1.00 | 0.18 | 0.82 | 5.65 |
| 23 | 6030407O03Rik | 6.49 | 1.38E-25 | 2.87E-21 | 0.11 | 0.00 | 0.11 | 55.50 |

**Supplementary Table 1. Marker genes for the original clusters of integrated clustering of E11.5 control and NotchGOF λ-junction cells.**

| | | | | | | | | |
|---|---|---|---|---|---|---|---|---|
| 23 | Mef2c | 3.94 | 1.87E-25 | 3.90E-21 | 1.00 | 0.19 | 0.81 | 5.38 |
| 23 | Akap12 | 4.93 | 1.98E-25 | 4.11E-21 | 1.00 | 0.20 | 0.80 | 5.08 |
| 23 | Gfra1 | 4.46 | 3.40E-25 | 7.07E-21 | 0.56 | 0.05 | 0.51 | 12.09 |
| 23 | Moxd1 | 3.52 | 1.24E-24 | 2.58E-20 | 0.50 | 0.04 | 0.46 | 13.51 |
| 23 | Fam78b | 4.73 | 1.93E-24 | 4.00E-20 | 0.44 | 0.03 | 0.41 | 14.80 |
| 23 | Egflam | 3.22 | 2.11E-24 | 4.39E-20 | 0.72 | 0.08 | 0.65 | 9.63 |
| 23 | Mpz | 6.40 | 2.15E-24 | 4.47E-20 | 0.22 | 0.01 | 0.21 | 27.75 |
| 23 | Aldh1l2 | 4.38 | 2.49E-24 | 5.17E-20 | 0.28 | 0.01 | 0.27 | 23.17 |
| 23 | Lama4 | 3.75 | 1.05E-23 | 2.18E-19 | 0.89 | 0.13 | 0.76 | 6.63 |
| 23 | Gcnt1 | 4.13 | 2.03E-23 | 4.23E-19 | 0.44 | 0.03 | 0.41 | 14.32 |
| 23 | Dhh | 6.98 | 2.48E-23 | 5.15E-19 | 0.11 | 0.00 | 0.11 | 55.50 |
| 23 | Sox8 | 3.82 | 3.03E-23 | 6.30E-19 | 0.61 | 0.06 | 0.55 | 10.36 |
| 23 | Sorbs1 | 3.84 | 3.87E-23 | 8.05E-19 | 0.72 | 0.08 | 0.64 | 8.60 |
| 23 | Crmp1 | 3.40 | 4.04E-23 | 8.41E-19 | 0.83 | 0.12 | 0.71 | 6.83 |
| 23 | Retreg1 | 3.23 | 4.77E-23 | 9.92E-19 | 0.78 | 0.09 | 0.69 | 8.37 |
| 23 | Tafa4 | 6.05 | 9.99E-23 | 2.08E-18 | 0.11 | 0.00 | 0.11 | 55.50 |
| 23 | Gm43707 | 3.77 | 1.75E-22 | 3.65E-18 | 0.44 | 0.03 | 0.41 | 13.88 |
| 23 | Ets1 | 3.22 | 1.26E-21 | 2.62E-17 | 0.94 | 0.18 | 0.77 | 5.27 |
| 23 | Apold1 | 3.75 | 1.58E-21 | 3.28E-17 | 0.33 | 0.02 | 0.31 | 17.53 |
| 23 | Fads6 | 5.42 | 1.74E-21 | 3.61E-17 | 0.17 | 0.01 | 0.16 | 33.40 |
| 23 | Pcdh1 | 3.45 | 2.15E-21 | 4.48E-17 | 0.56 | 0.05 | 0.50 | 10.69 |
| 23 | Csn3 | 5.61 | 3.80E-21 | 7.91E-17 | 0.17 | 0.01 | 0.16 | 33.40 |
| 23 | Dagla | 4.56 | 4.49E-21 | 9.33E-17 | 0.28 | 0.01 | 0.26 | 19.86 |
| 23 | Man1c1 | 3.79 | 4.84E-21 | 1.01E-16 | 0.44 | 0.03 | 0.41 | 13.06 |
| 23 | L1cam | 3.40 | 1.08E-20 | 2.24E-16 | 0.67 | 0.08 | 0.59 | 8.23 |
| 23 | Map7d2 | 2.59 | 1.90E-20 | 3.96E-16 | 0.72 | 0.09 | 0.64 | 8.49 |
| 23 | Cdc42ep1 | 2.94 | 2.15E-20 | 4.48E-16 | 0.61 | 0.06 | 0.55 | 9.70 |
| 23 | Itga4 | 3.77 | 2.27E-20 | 4.72E-16 | 0.94 | 0.20 | 0.75 | 4.82 |
| 23 | Gm4876 | 4.22 | 9.62E-20 | 2.00E-15 | 0.28 | 0.02 | 0.26 | 18.53 |
| 23 | Sphk1 | 4.61 | 1.09E-19 | 2.26E-15 | 0.22 | 0.01 | 0.21 | 22.20 |
| 23 | Pls3 | 4.73 | 1.34E-19 | 2.78E-15 | 1.00 | 0.30 | 0.70 | 3.34 |
| 23 | Col5a3 | 6.01 | 2.22E-19 | 4.63E-15 | 0.11 | 0.00 | 0.11 | 37.00 |
| 23 | Cdh6 | 4.03 | 3.54E-19 | 7.35E-15 | 1.00 | 0.29 | 0.71 | 3.50 |
| 23 | Coro2b | 3.49 | 8.29E-19 | 1.72E-14 | 0.50 | 0.05 | 0.45 | 10.42 |
| 23 | Myl9 | 3.72 | 1.09E-18 | 2.26E-14 | 0.94 | 0.24 | 0.70 | 3.87 |
| 23 | Klhl21 | 3.59 | 1.59E-18 | 3.30E-14 | 0.50 | 0.05 | 0.45 | 10.00 |
| 23 | Fxyd5 | 2.69 | 1.81E-18 | 3.77E-14 | 0.39 | 0.03 | 0.36 | 12.97 |
| 23 | Mamdc2 | 5.84 | 2.30E-18 | 4.78E-14 | 0.11 | 0.00 | 0.11 | 37.00 |
| 23 | Ptprm | 3.41 | 4.04E-18 | 8.40E-14 | 0.50 | 0.05 | 0.45 | 10.00 |
| 23 | Mmp15 | 3.16 | 4.52E-18 | 9.40E-14 | 0.67 | 0.09 | 0.58 | 7.49 |
| 23 | Kitl | 3.19 | 8.81E-18 | 1.83E-13 | 0.61 | 0.08 | 0.54 | 8.04 |
| 23 | Lpar3 | 5.94 | 1.81E-17 | 3.75E-13 | 0.11 | 0.00 | 0.11 | 37.00 |
| 23 | Fhdc1 | 3.73 | 2.00E-17 | 4.16E-13 | 0.44 | 0.04 | 0.40 | 10.57 |
| 23 | Gas7 | 3.29 | 2.46E-17 | 5.12E-13 | 0.50 | 0.05 | 0.45 | 9.43 |
| 23 | Tspan7 | 3.01 | 2.72E-17 | 5.66E-13 | 1.00 | 0.28 | 0.72 | 3.56 |
| 23 | Timp3 | 3.26 | 5.09E-17 | 1.06E-12 | 0.83 | 0.17 | 0.66 | 4.84 |
| 23 | Slc1a2 | 5.81 | 6.84E-17 | 1.42E-12 | 0.11 | 0.00 | 0.11 | 37.00 |
| 23 | Plvap | 1.83 | 6.92E-17 | 1.44E-12 | 0.44 | 0.04 | 0.40 | 10.57 |
| 23 | Kctd12 | 2.95 | 9.81E-17 | 2.04E-12 | 1.00 | 0.30 | 0.70 | 3.36 |
| 23 | Arhgef26 | 2.76 | 1.05E-16 | 2.18E-12 | 0.78 | 0.13 | 0.65 | 6.17 |
| 23 | Lbh | 2.67 | 1.09E-16 | 2.27E-12 | 0.94 | 0.20 | 0.74 | 4.67 |
| 23 | Gm26688 | 3.93 | 1.43E-16 | 2.98E-12 | 0.28 | 0.02 | 0.26 | 15.44 |
| 23 | Cav1 | 4.16 | 1.70E-16 | 3.55E-12 | 0.22 | 0.01 | 0.21 | 18.50 |
| 23 | Zadh2 | 2.73 | 2.42E-16 | 5.03E-12 | 0.72 | 0.11 | 0.62 | 6.75 |
| 23 | Rhob | 3.37 | 2.47E-16 | 5.14E-12 | 1.00 | 0.35 | 0.65 | 2.88 |
| 23 | Adamts1 | 3.06 | 3.07E-16 | 6.38E-12 | 1.00 | 0.28 | 0.73 | 3.64 |

**Supplementary Table 1. Marker genes for the original clusters of integrated clustering of E11.5 control and NotchGOF λ-junction cells.**

| | | | | | | | | |
|---|---|---|---|---|---|---|---|---|
| 23 | Fads3 | 3.60 | 3.78E-16 | 7.87E-12 | 0.50 | 0.06 | 0.44 | 8.77 |
| 23 | Cthrc1 | 3.54 | 7.84E-16 | 1.63E-11 | 0.89 | 0.19 | 0.70 | 4.58 |
| 23 | Gldn | 4.19 | 1.51E-15 | 3.14E-11 | 0.22 | 0.01 | 0.21 | 18.50 |
| 23 | Dock2 | 4.44 | 1.54E-15 | 3.21E-11 | 0.17 | 0.01 | 0.16 | 23.86 |
| 23 | Heyl | 3.17 | 1.56E-15 | 3.24E-11 | 0.50 | 0.06 | 0.44 | 8.62 |
| 23 | Dpysl3 | 3.83 | 1.82E-15 | 3.79E-11 | 1.00 | 0.45 | 0.55 | 2.21 |
| 23 | Sema3c | 2.21 | 2.96E-15 | 6.17E-11 | 0.67 | 0.09 | 0.57 | 7.10 |
| 23 | Gpr37 | 5.66 | 3.53E-15 | 7.34E-11 | 0.11 | 0.00 | 0.11 | 37.00 |
| 23 | Nrp1 | 2.63 | 4.85E-15 | 1.01E-10 | 0.89 | 0.21 | 0.68 | 4.23 |
| 23 | Gabrb3 | 3.88 | 4.98E-15 | 1.04E-10 | 0.22 | 0.01 | 0.21 | 17.08 |
| 23 | Plxnd1 | 2.42 | 5.37E-15 | 1.12E-10 | 0.50 | 0.06 | 0.44 | 8.47 |
| 23 | Xylt1 | 3.56 | 6.29E-15 | 1.31E-10 | 0.50 | 0.06 | 0.44 | 8.20 |
| 23 | Ptprj | 4.04 | 7.96E-15 | 1.65E-10 | 0.39 | 0.04 | 0.35 | 10.24 |
| 23 | Nacc2 | 3.12 | 9.99E-15 | 2.08E-10 | 0.39 | 0.04 | 0.35 | 10.51 |
| 23 | Anxa1 | 3.28 | 1.15E-14 | 2.40E-10 | 0.33 | 0.03 | 0.31 | 11.89 |
| 23 | Cotl1 | 3.00 | 1.30E-14 | 2.71E-10 | 1.00 | 0.45 | 0.55 | 2.24 |
| 23 | Vgll3 | 3.95 | 1.32E-14 | 2.74E-10 | 0.33 | 0.03 | 0.30 | 11.48 |
| 23 | Synm | 4.44 | 1.45E-14 | 3.02E-10 | 0.28 | 0.02 | 0.26 | 13.24 |
| 23 | Lpcat2 | 3.51 | 1.47E-14 | 3.07E-10 | 0.39 | 0.04 | 0.35 | 9.97 |
| 23 | Cadm1 | 2.81 | 1.67E-14 | 3.48E-10 | 1.00 | 0.39 | 0.61 | 2.57 |
| 23 | Reep1 | 3.11 | 1.75E-14 | 3.64E-10 | 0.44 | 0.05 | 0.40 | 9.25 |
| 23 | Gm16701.1 | 5.19 | 2.40E-14 | 5.00E-10 | 0.11 | 0.00 | 0.11 | 37.00 |
| 23 | Ahr | 2.96 | 2.56E-14 | 5.33E-10 | 0.61 | 0.09 | 0.52 | 6.87 |
| 23 | Metrn | 2.88 | 3.45E-14 | 7.18E-10 | 1.00 | 0.52 | 0.49 | 1.94 |
| 23 | Gamt | 2.83 | 4.68E-14 | 9.74E-10 | 0.61 | 0.09 | 0.52 | 6.57 |
| 23 | Tlr3 | 5.14 | 4.70E-14 | 9.78E-10 | 0.17 | 0.01 | 0.16 | 20.88 |
| 23 | Mxra7 | 2.73 | 5.73E-14 | 1.19E-09 | 0.89 | 0.23 | 0.66 | 3.80 |
| 23 | Afap1l1 | 2.44 | 8.46E-14 | 1.76E-09 | 0.56 | 0.07 | 0.48 | 7.51 |
| 23 | Lipo4 | 5.81 | 1.14E-13 | 2.37E-09 | 0.11 | 0.00 | 0.11 | 27.75 |
| 23 | Ccl6 | 4.07 | 1.21E-13 | 2.52E-09 | 0.11 | 0.00 | 0.11 | 27.75 |
| 23 | Zfyve28 | 5.75 | 1.64E-13 | 3.42E-09 | 0.11 | 0.00 | 0.11 | 27.75 |
| 23 | Pdlim1 | 2.23 | 1.94E-13 | 4.04E-09 | 0.94 | 0.29 | 0.65 | 3.23 |
| 23 | Arpc1b | 2.68 | 2.14E-13 | 4.45E-09 | 0.94 | 0.35 | 0.60 | 2.71 |
| 23 | Htra3 | 4.27 | 2.20E-13 | 4.58E-09 | 0.22 | 0.01 | 0.21 | 15.86 |
| 23 | Map3k1 | 2.42 | 3.20E-13 | 6.66E-09 | 0.94 | 0.26 | 0.68 | 3.59 |
| 23 | Atp6v0e2 | 3.09 | 3.57E-13 | 7.42E-09 | 0.44 | 0.05 | 0.39 | 8.22 |
| 23 | Zeb2 | 2.81 | 3.57E-13 | 7.43E-09 | 1.00 | 0.40 | 0.60 | 2.49 |
| 23 | Rhbdf1 | 2.99 | 4.11E-13 | 8.54E-09 | 0.61 | 0.10 | 0.51 | 6.05 |
| 23 | Dab1 | 1.68 | 4.28E-13 | 8.91E-09 | 0.83 | 0.16 | 0.68 | 5.37 |
| 23 | Dtx4 | 3.06 | 4.69E-13 | 9.75E-09 | 0.39 | 0.04 | 0.35 | 9.49 |
| 23 | Foxn2 | 2.20 | 5.31E-13 | 1.10E-08 | 0.83 | 0.16 | 0.67 | 5.08 |
| 23 | Dbi | 1.80 | 6.53E-13 | 1.36E-08 | 1.00 | 0.98 | 0.02 | 1.02 |
| 23 | Anxa6 | 2.17 | 7.00E-13 | 1.46E-08 | 0.83 | 0.17 | 0.66 | 4.79 |
| 23 | Csrp1 | 2.68 | 8.00E-13 | 1.66E-08 | 0.94 | 0.34 | 0.60 | 2.76 |
| 23 | Slitrk6 | 2.37 | 8.85E-13 | 1.84E-08 | 1.00 | 0.33 | 0.68 | 3.08 |
| 23 | Lamc1 | 2.27 | 9.09E-13 | 1.89E-08 | 0.94 | 0.26 | 0.68 | 3.62 |
| 23 | Klhdc1 | 3.62 | 9.10E-13 | 1.89E-08 | 0.28 | 0.02 | 0.26 | 12.09 |
| 23 | Megf9 | 2.67 | 1.03E-12 | 2.14E-08 | 0.83 | 0.19 | 0.64 | 4.38 |
| 23 | Nnmt | 5.33 | 1.57E-12 | 3.27E-08 | 0.11 | 0.00 | 0.11 | 27.75 |
| 23 | Gm46620 | 5.25 | 1.69E-12 | 3.52E-08 | 0.11 | 0.00 | 0.11 | 27.75 |
| 23 | Fam210b | 2.13 | 1.98E-12 | 4.12E-08 | 0.83 | 0.20 | 0.63 | 4.12 |
| 23 | Kcna1 | 5.47 | 2.18E-12 | 4.54E-08 | 0.11 | 0.00 | 0.11 | 27.75 |
| 23 | Dnajc1 | 2.30 | 2.21E-12 | 4.59E-08 | 1.00 | 0.56 | 0.44 | 1.79 |
| 23 | Enc1 | 2.44 | 4.74E-12 | 9.87E-08 | 0.83 | 0.22 | 0.61 | 3.75 |
| 23 | Sdc3 | 2.04 | 6.77E-12 | 1.41E-07 | 0.89 | 0.21 | 0.68 | 4.34 |
| 23 | Col5a2 | 2.68 | 7.96E-12 | 1.65E-07 | 1.00 | 0.53 | 0.47 | 1.89 |

**Supplementary Table 1. Marker genes for the original clusters of integrated clustering of E11.5 control and NotchGOF λ-junction cells.**

| | | | | | | | | |
|---|---|---|---|---|---|---|---|---|
| 23 | Atg4a | 2.56 | 8.28E-12 | 1.72E-07 | 0.56 | 0.09 | 0.47 | 6.39 |
| 23 | Gm43560 | 3.44 | 8.41E-12 | 1.75E-07 | 0.39 | 0.05 | 0.34 | 8.28 |
| 23 | Kcnk5 | 3.59 | 9.99E-12 | 2.08E-07 | 0.33 | 0.04 | 0.30 | 9.51 |
| 23 | Klhl13 | 2.18 | 1.09E-11 | 2.27E-07 | 0.94 | 0.27 | 0.68 | 3.51 |
| 23 | Sema3d | 1.91 | 1.09E-11 | 2.27E-07 | 0.67 | 0.12 | 0.55 | 5.61 |
| 23 | Selenok-ps1 | 4.57 | 1.12E-11 | 2.32E-07 | 0.17 | 0.01 | 0.16 | 16.70 |
| 23 | Dock11 | 2.70 | 1.12E-11 | 2.34E-07 | 0.83 | 0.23 | 0.60 | 3.64 |
| 23 | Lmo4 | 1.99 | 1.18E-11 | 2.45E-07 | 1.00 | 0.81 | 0.19 | 1.23 |
| 23 | Gdpd5 | 3.17 | 1.19E-11 | 2.47E-07 | 0.39 | 0.05 | 0.34 | 8.28 |
| 23 | Vim | 2.25 | 1.24E-11 | 2.57E-07 | 1.00 | 0.84 | 0.16 | 1.19 |
| 23 | Fbxo7 | 2.55 | 1.52E-11 | 3.17E-07 | 0.67 | 0.14 | 0.53 | 4.90 |
| 23 | Anxa5 | 2.33 | 1.67E-11 | 3.47E-07 | 1.00 | 0.61 | 0.39 | 1.65 |
| 23 | Sparc | 2.38 | 1.89E-11 | 3.93E-07 | 1.00 | 0.74 | 0.26 | 1.35 |
| 23 | Tnik | 2.77 | 2.09E-11 | 4.34E-07 | 0.83 | 0.25 | 0.59 | 3.39 |
| 23 | Egfl7 | 1.50 | 2.23E-11 | 4.63E-07 | 0.89 | 0.22 | 0.67 | 4.04 |
| 23 | Arhgap20 | 4.34 | 2.36E-11 | 4.90E-07 | 0.17 | 0.01 | 0.16 | 16.70 |
| 23 | Paqr4 | 3.01 | 2.52E-11 | 5.23E-07 | 0.33 | 0.04 | 0.30 | 9.25 |
| 23 | Btbd3 | 2.53 | 2.61E-11 | 5.44E-07 | 0.72 | 0.16 | 0.56 | 4.57 |
| 23 | Ptprz1 | 2.53 | 2.71E-11 | 5.63E-07 | 0.61 | 0.11 | 0.50 | 5.71 |
| 23 | Kcna6 | 3.26 | 3.28E-11 | 6.82E-07 | 0.33 | 0.04 | 0.30 | 9.25 |
| 23 | Hs2st1 | 2.65 | 3.31E-11 | 6.89E-07 | 0.94 | 0.57 | 0.38 | 1.67 |
| 23 | Cdh2 | 2.05 | 3.60E-11 | 7.49E-07 | 1.00 | 0.42 | 0.58 | 2.36 |
| 23 | Gimap9 | 2.34 | 3.64E-11 | 7.56E-07 | 0.61 | 0.11 | 0.50 | 5.66 |
| 23 | Celsr2 | 2.59 | 3.80E-11 | 7.89E-07 | 0.56 | 0.09 | 0.46 | 5.98 |
| 23 | Nes | 2.32 | 3.86E-11 | 8.02E-07 | 1.00 | 0.43 | 0.57 | 2.35 |
| 23 | Celf2 | 2.06 | 4.33E-11 | 9.01E-07 | 0.89 | 0.26 | 0.63 | 3.46 |
| 23 | Gnb4 | 2.27 | 4.82E-11 | 1.00E-06 | 0.89 | 0.30 | 0.59 | 2.95 |
| 23 | Tspan4 | 2.25 | 5.02E-11 | 1.04E-06 | 0.67 | 0.13 | 0.54 | 5.21 |
| 23 | Gm19412 | 3.12 | 5.43E-11 | 1.13E-06 | 0.33 | 0.04 | 0.30 | 9.00 |
| 23 | Khdrbs2 | 2.98 | 5.62E-11 | 1.17E-06 | 0.33 | 0.04 | 0.30 | 9.00 |
| 23 | Kifc3 | 2.46 | 5.88E-11 | 1.22E-06 | 0.50 | 0.08 | 0.43 | 6.67 |
| 23 | Itga6 | 2.17 | 5.90E-11 | 1.23E-06 | 0.94 | 0.31 | 0.63 | 3.01 |
| 23 | Tmc6 | 2.53 | 6.03E-11 | 1.25E-06 | 0.50 | 0.08 | 0.42 | 6.58 |
| 23 | Nab1 | 2.24 | 8.20E-11 | 1.70E-06 | 0.67 | 0.13 | 0.54 | 5.09 |
| 23 | Pdzrn3 | 2.38 | 9.94E-11 | 2.07E-06 | 0.78 | 0.17 | 0.60 | 4.47 |
| 23 | Cpne2 | 3.22 | 1.04E-10 | 2.17E-06 | 0.39 | 0.05 | 0.34 | 7.48 |
| 23 | Qpct | 3.26 | 1.19E-10 | 2.48E-06 | 0.33 | 0.04 | 0.30 | 8.76 |
| 23 | Hvcn1 | 2.64 | 1.20E-10 | 2.49E-06 | 0.50 | 0.08 | 0.42 | 6.33 |
| 23 | Grid2 | 3.43 | 1.23E-10 | 2.55E-06 | 0.22 | 0.02 | 0.20 | 12.33 |
| 23 | Gm42670 | 3.75 | 1.24E-10 | 2.57E-06 | 0.22 | 0.02 | 0.20 | 12.33 |
| 23 | Fxyd1 | 2.50 | 1.25E-10 | 2.60E-06 | 0.44 | 0.06 | 0.38 | 7.05 |
| 23 | Chst5 | 3.83 | 1.29E-10 | 2.68E-06 | 0.22 | 0.02 | 0.20 | 12.33 |
| 23 | Acsl4 | 2.73 | 1.31E-10 | 2.73E-06 | 0.67 | 0.14 | 0.53 | 4.73 |
| 23 | Ankrd29 | 2.42 | 1.37E-10 | 2.85E-06 | 0.56 | 0.09 | 0.46 | 5.98 |
| 23 | Gap43 | 2.40 | 1.57E-10 | 3.26E-06 | 0.94 | 0.42 | 0.52 | 2.25 |
| 23 | Jdp2 | 3.06 | 1.63E-10 | 3.38E-06 | 0.33 | 0.04 | 0.30 | 8.76 |
| 23 | Myo18a | 2.41 | 1.65E-10 | 3.43E-06 | 0.83 | 0.24 | 0.60 | 3.53 |
| 23 | Gm47644 | 3.66 | 1.78E-10 | 3.70E-06 | 0.28 | 0.03 | 0.25 | 9.93 |
| 23 | Emb | 2.08 | 1.88E-10 | 3.91E-06 | 1.00 | 0.41 | 0.59 | 2.45 |
| 23 | Cst3 | 1.30 | 2.18E-10 | 4.53E-06 | 1.00 | 0.92 | 0.08 | 1.09 |
| 23 | Lrmp | 3.77 | 2.66E-10 | 5.53E-06 | 0.17 | 0.01 | 0.16 | 15.18 |
| 23 | Pxdc1 | 3.22 | 2.68E-10 | 5.57E-06 | 0.56 | 0.11 | 0.45 | 5.30 |
| 23 | Tagln2 | 2.93 | 2.91E-10 | 6.04E-06 | 0.83 | 0.30 | 0.53 | 2.79 |
| 23 | Nlgn3 | 3.30 | 3.08E-10 | 6.41E-06 | 0.39 | 0.05 | 0.34 | 7.48 |
| 23 | Pros1 | 2.60 | 3.33E-10 | 6.93E-06 | 0.67 | 0.15 | 0.52 | 4.60 |
| 23 | 1700025G04Rik | 1.93 | 3.51E-10 | 7.31E-06 | 1.00 | 0.34 | 0.66 | 2.95 |

**Supplementary Table 1. Marker genes for the original clusters of integrated clustering of E11.5 control and NotchGOF λ-junction cells.**

| | | | | | | | | |
|---|---|---|---|---|---|---|---|---|
| 23 | Il3ra | 2.82 | 3.54E-10 | 7.36E-06 | 0.39 | 0.05 | 0.34 | 7.48 |
| 23 | Pdia5 | 2.34 | 3.60E-10 | 7.48E-06 | 0.61 | 0.12 | 0.49 | 5.22 |
| 23 | Ttyh2 | 3.28 | 3.68E-10 | 7.64E-06 | 0.28 | 0.03 | 0.25 | 9.93 |
| 23 | Alcam | 2.50 | 4.31E-10 | 8.95E-06 | 0.67 | 0.15 | 0.52 | 4.57 |
| 23 | Tmsb4x | 1.16 | 4.47E-10 | 9.29E-06 | 1.00 | 0.99 | 0.01 | 1.01 |
| 23 | Olfm2 | 2.56 | 4.55E-10 | 9.47E-06 | 0.56 | 0.11 | 0.45 | 5.25 |
| 23 | Lipa | 2.77 | 4.72E-10 | 9.82E-06 | 0.78 | 0.22 | 0.56 | 3.55 |
| 23 | Chrdl2 | 3.96 | 7.48E-10 | 1.56E-05 | 0.17 | 0.01 | 0.16 | 15.18 |
| 23 | Ubl3 | 2.27 | 1.20E-09 | 2.50E-05 | 0.89 | 0.37 | 0.52 | 2.39 |
| 23 | Fign | 2.55 | 1.26E-09 | 2.63E-05 | 0.83 | 0.29 | 0.54 | 2.83 |
| 23 | Actg1 | 0.81 | 1.40E-09 | 2.91E-05 | 1.00 | 1.00 | 0.00 | 1.00 |
| 23 | Epha5 | 2.60 | 1.56E-09 | 3.25E-05 | 0.44 | 0.07 | 0.37 | 6.25 |
| 23 | Mid1ip1 | 2.48 | 1.84E-09 | 3.83E-05 | 0.78 | 0.22 | 0.56 | 3.54 |
| 23 | Tshz1 | 2.68 | 2.28E-09 | 4.73E-05 | 0.83 | 0.32 | 0.51 | 2.61 |
| 23 | Chn2 | 2.56 | 2.31E-09 | 4.81E-05 | 0.44 | 0.07 | 0.37 | 6.17 |
| 23 | Clcn5 | 2.14 | 2.54E-09 | 5.28E-05 | 0.94 | 0.39 | 0.55 | 2.41 |
| 23 | Pon2 | 2.09 | 2.69E-09 | 5.60E-05 | 0.72 | 0.18 | 0.54 | 3.97 |
| 23 | Map1a | 2.72 | 2.76E-09 | 5.74E-05 | 0.50 | 0.09 | 0.41 | 5.43 |
| 23 | Dusp3 | 2.55 | 2.90E-09 | 6.04E-05 | 0.39 | 0.06 | 0.33 | 6.82 |
| 23 | Aard | 2.79 | 3.46E-09 | 7.20E-05 | 0.44 | 0.07 | 0.37 | 6.08 |
| 23 | Kif19a | 3.05 | 3.70E-09 | 7.71E-05 | 0.17 | 0.01 | 0.16 | 13.92 |
| 23 | Epb41l4a | 2.33 | 3.73E-09 | 7.77E-05 | 0.56 | 0.11 | 0.45 | 5.20 |
| 23 | Gpsm3 | 2.21 | 3.80E-09 | 7.91E-05 | 0.50 | 0.09 | 0.41 | 5.75 |
| 23 | Fam49a | 2.96 | 3.99E-09 | 8.30E-05 | 0.33 | 0.05 | 0.29 | 7.40 |
| 23 | Trf | 2.51 | 4.18E-09 | 8.68E-05 | 0.33 | 0.04 | 0.29 | 7.74 |
| 23 | Igsf9b | 2.57 | 4.27E-09 | 8.89E-05 | 0.39 | 0.06 | 0.33 | 6.82 |
| 23 | Calm1 | 0.98 | 4.81E-09 | 1.00E-04 | 1.00 | 0.99 | 0.01 | 1.01 |
| 23 | Wdfy1 | 2.46 | 5.20E-09 | 1.08E-04 | 0.83 | 0.35 | 0.48 | 2.37 |
| 23 | Pxn | 2.45 | 5.23E-09 | 1.09E-04 | 0.72 | 0.19 | 0.54 | 3.88 |
| 23 | Cytl1 | 5.08 | 5.41E-09 | 1.13E-04 | 0.11 | 0.01 | 0.11 | 18.50 |
| 23 | Camkv | 4.22 | 5.58E-09 | 1.16E-04 | 0.17 | 0.01 | 0.15 | 12.85 |
| 23 | Ppp1r14c | 3.64 | 6.13E-09 | 1.27E-04 | 0.22 | 0.02 | 0.20 | 10.09 |
| 23 | Fgd1 | 1.73 | 7.42E-09 | 1.54E-04 | 0.94 | 0.32 | 0.62 | 2.92 |
| 23 | Sort1 | 2.00 | 7.51E-09 | 1.56E-04 | 0.78 | 0.21 | 0.57 | 3.80 |
| 23 | Map3k11 | 2.56 | 7.75E-09 | 1.61E-04 | 0.56 | 0.12 | 0.44 | 4.75 |
| 23 | Tubb6 | 2.15 | 8.07E-09 | 1.68E-04 | 0.56 | 0.12 | 0.44 | 4.83 |
| 23 | Mest | 1.56 | 8.42E-09 | 1.75E-04 | 1.00 | 0.91 | 0.09 | 1.10 |
| 23 | Fam89a | 2.35 | 9.24E-09 | 1.92E-04 | 0.44 | 0.08 | 0.37 | 5.92 |
| 23 | Fam111a | 2.30 | 9.24E-09 | 1.92E-04 | 0.83 | 0.33 | 0.51 | 2.56 |
| 23 | Coro1c | 1.94 | 9.30E-09 | 1.93E-04 | 0.94 | 0.35 | 0.59 | 2.70 |
| 23 | Cela1 | 4.26 | 1.16E-08 | 2.41E-04 | 0.11 | 0.01 | 0.11 | 18.50 |
| 23 | Emp3 | 2.27 | 1.21E-08 | 2.52E-04 | 0.61 | 0.14 | 0.47 | 4.46 |
| 23 | Cobll1 | 2.75 | 1.21E-08 | 2.53E-04 | 0.28 | 0.03 | 0.25 | 8.42 |
| 23 | Slc7a1 | 2.75 | 1.23E-08 | 2.56E-04 | 0.44 | 0.08 | 0.37 | 5.62 |
| 23 | Nid1 | 1.73 | 1.26E-08 | 2.62E-04 | 1.00 | 0.35 | 0.65 | 2.89 |
| 23 | Des | 2.84 | 1.52E-08 | 3.16E-04 | 0.33 | 0.05 | 0.29 | 7.09 |
| 23 | Sdsl | 2.46 | 1.69E-08 | 3.51E-04 | 0.39 | 0.06 | 0.33 | 6.48 |
| 23 | Pfn1 | 0.72 | 1.82E-08 | 3.78E-04 | 1.00 | 1.00 | 0.00 | 1.00 |
| 23 | Cnp | 2.47 | 1.83E-08 | 3.80E-04 | 0.61 | 0.16 | 0.45 | 3.89 |
| 23 | AC109619.1 | 4.88 | 1.84E-08 | 3.82E-04 | 0.11 | 0.01 | 0.11 | 18.50 |
| 23 | Tuba1a | 0.92 | 1.90E-08 | 3.95E-04 | 1.00 | 0.98 | 0.02 | 1.02 |
| 23 | Ptpn13 | 1.92 | 2.15E-08 | 4.48E-04 | 0.89 | 0.41 | 0.48 | 2.19 |
| 23 | Psat1 | 1.80 | 2.26E-08 | 4.69E-04 | 0.89 | 0.45 | 0.44 | 1.96 |
| 23 | Cavin3 | 2.11 | 2.29E-08 | 4.76E-04 | 0.56 | 0.12 | 0.43 | 4.48 |
| 23 | Selenof | 0.95 | 2.46E-08 | 5.12E-04 | 1.00 | 0.96 | 0.04 | 1.05 |
| 23 | Ppp1r18 | 2.03 | 2.50E-08 | 5.20E-04 | 0.78 | 0.25 | 0.53 | 3.09 |

**Supplementary Table 1. Marker genes for the original clusters of integrated clustering of E11.5 control and NotchGOF λ-junction cells.**

| | | | | | | | | |
|---|---|---|---|---|---|---|---|---|
| 23 | Zfp979 | 3.35 | 2.58E-08 | 5.38E-04 | 0.28 | 0.04 | 0.24 | 7.94 |
| 23 | Trim71 | 1.79 | 2.91E-08 | 6.05E-04 | 0.78 | 0.21 | 0.57 | 3.69 |
| 23 | Ppic | 1.44 | 3.05E-08 | 6.35E-04 | 1.00 | 0.71 | 0.29 | 1.41 |
| 23 | Ss18l2 | 1.85 | 3.07E-08 | 6.39E-04 | 0.89 | 0.52 | 0.37 | 1.72 |
| 23 | Gnai1 | 1.86 | 3.29E-08 | 6.85E-04 | 0.67 | 0.15 | 0.51 | 4.33 |
| 23 | Grik2 | 3.47 | 3.29E-08 | 6.85E-04 | 0.22 | 0.02 | 0.20 | 9.65 |
| 23 | Trim6 | 3.70 | 3.71E-08 | 7.72E-04 | 0.22 | 0.02 | 0.20 | 9.25 |
| 23 | Plekha4 | 4.93 | 3.72E-08 | 7.74E-04 | 0.11 | 0.01 | 0.11 | 18.50 |
| 23 | Tnfaip8l1 | 2.22 | 3.90E-08 | 8.11E-04 | 0.44 | 0.08 | 0.37 | 5.62 |
| 23 | Adamts12 | 3.19 | 3.93E-08 | 8.18E-04 | 0.28 | 0.04 | 0.24 | 7.94 |
| 23 | Soat1 | 1.99 | 4.07E-08 | 8.47E-04 | 0.94 | 0.44 | 0.50 | 2.14 |
| 23 | Cadm4 | 2.72 | 4.33E-08 | 9.01E-04 | 0.44 | 0.09 | 0.36 | 5.10 |
| 23 | Scap | 1.85 | 4.35E-08 | 9.05E-04 | 0.72 | 0.19 | 0.54 | 3.90 |
| 23 | Itgb1 | 1.39 | 4.62E-08 | 9.60E-04 | 1.00 | 0.85 | 0.15 | 1.17 |
| 23 | Chml | 1.81 | 4.71E-08 | 9.80E-04 | 0.67 | 0.15 | 0.52 | 4.45 |
| 23 | Cd81 | 1.30 | 4.89E-08 | 1.02E-03 | 1.00 | 0.89 | 0.11 | 1.12 |
| 23 | 2900089D17Rik | 4.03 | 4.91E-08 | 1.02E-03 | 0.17 | 0.01 | 0.15 | 11.93 |
| 23 | Metap1d | 1.90 | 4.99E-08 | 1.04E-03 | 0.72 | 0.20 | 0.52 | 3.61 |
| 23 | Sox2 | 0.95 | 5.39E-08 | 1.12E-03 | 0.78 | 0.20 | 0.58 | 3.99 |
| 23 | Flrt1 | 2.95 | 6.02E-08 | 1.25E-03 | 0.22 | 0.02 | 0.20 | 9.25 |
| 23 | Tram2 | 2.39 | 6.08E-08 | 1.26E-03 | 0.50 | 0.10 | 0.40 | 4.85 |
| 23 | Trp53cor1 | 4.22 | 6.62E-08 | 1.38E-03 | 0.11 | 0.01 | 0.10 | 15.86 |
| 23 | 2810430I11Rik | 4.41 | 6.65E-08 | 1.38E-03 | 0.11 | 0.01 | 0.10 | 15.86 |
| 23 | Parp14 | 4.45 | 6.79E-08 | 1.41E-03 | 0.11 | 0.01 | 0.10 | 15.86 |
| 23 | Sppl2b | 2.82 | 6.88E-08 | 1.43E-03 | 0.56 | 0.15 | 0.41 | 3.78 |
| 23 | Prnp | 2.10 | 7.51E-08 | 1.56E-03 | 0.67 | 0.18 | 0.49 | 3.77 |
| 23 | Eif5a | 0.68 | 8.44E-08 | 1.75E-03 | 1.00 | 1.00 | 0.00 | 1.00 |
| 23 | Peg13 | 2.79 | 8.55E-08 | 1.78E-03 | 0.33 | 0.05 | 0.28 | 6.53 |
| 23 | Cep85l | 2.05 | 8.56E-08 | 1.78E-03 | 0.61 | 0.14 | 0.47 | 4.43 |
| 23 | Gramd4 | 2.44 | 8.58E-08 | 1.79E-03 | 0.44 | 0.09 | 0.36 | 5.22 |
| 23 | Mthfd1l | 1.79 | 8.80E-08 | 1.83E-03 | 0.67 | 0.16 | 0.51 | 4.19 |
| 23 | Pea15a | 1.44 | 9.82E-08 | 2.04E-03 | 0.83 | 0.23 | 0.60 | 3.58 |
| 23 | Hsp90aa1 | 0.69 | 1.02E-07 | 2.11E-03 | 1.00 | 1.00 | 0.00 | 1.00 |
| 23 | Odc1 | 1.16 | 1.06E-07 | 2.20E-03 | 1.00 | 0.81 | 0.19 | 1.23 |
| 23 | Rarres2 | 3.29 | 1.09E-07 | 2.26E-03 | 0.17 | 0.01 | 0.15 | 11.93 |
| 23 | Zfp276 | 2.53 | 1.12E-07 | 2.32E-03 | 0.44 | 0.09 | 0.36 | 5.10 |
| 23 | Serpinh1 | 1.12 | 1.14E-07 | 2.36E-03 | 1.00 | 0.94 | 0.06 | 1.06 |
| 23 | Myl6 | 0.73 | 1.18E-07 | 2.45E-03 | 1.00 | 0.99 | 0.01 | 1.01 |
| 23 | Ovca2 | 2.72 | 1.19E-07 | 2.48E-03 | 0.33 | 0.05 | 0.28 | 6.53 |
| 23 | Zbtb38 | 2.58 | 1.26E-07 | 2.62E-03 | 0.56 | 0.14 | 0.42 | 4.06 |
| 23 | Adora1 | 4.33 | 1.37E-07 | 2.84E-03 | 0.11 | 0.01 | 0.10 | 15.86 |
| 23 | Dusp22 | 1.99 | 1.37E-07 | 2.84E-03 | 0.72 | 0.20 | 0.52 | 3.54 |
| 23 | Kif21a | 2.10 | 1.39E-07 | 2.90E-03 | 0.78 | 0.31 | 0.47 | 2.49 |
| 23 | Esyt1 | 3.26 | 1.40E-07 | 2.91E-03 | 0.39 | 0.08 | 0.31 | 5.19 |
| 23 | Fam129b | 1.91 | 1.42E-07 | 2.96E-03 | 0.61 | 0.14 | 0.47 | 4.27 |
| 23 | Slc25a5 | 0.86 | 1.54E-07 | 3.19E-03 | 1.00 | 0.93 | 0.07 | 1.07 |
| 23 | Ctsl | 1.67 | 1.60E-07 | 3.32E-03 | 1.00 | 0.82 | 0.18 | 1.21 |
| 23 | Rin3 | 3.60 | 1.79E-07 | 3.73E-03 | 0.22 | 0.03 | 0.20 | 8.54 |
| 23 | Soga3 | 2.38 | 1.83E-07 | 3.80E-03 | 0.33 | 0.05 | 0.28 | 6.17 |
| 23 | Slc8b1 | 2.84 | 2.01E-07 | 4.18E-03 | 0.22 | 0.03 | 0.20 | 8.88 |
| 23 | Sh3tc2 | 3.60 | 2.12E-07 | 4.40E-03 | 0.22 | 0.03 | 0.20 | 8.54 |
| 23 | Cyp2j9 | 4.56 | 2.22E-07 | 4.61E-03 | 0.11 | 0.01 | 0.10 | 15.86 |
| 23 | Igfbp3 | 1.12 | 2.29E-07 | 4.76E-03 | 0.56 | 0.13 | 0.43 | 4.41 |
| 23 | Rims1 | 3.34 | 2.32E-07 | 4.82E-03 | 0.22 | 0.03 | 0.20 | 8.54 |
| 23 | Cyp2d22 | 2.65 | 2.33E-07 | 4.84E-03 | 0.33 | 0.05 | 0.28 | 6.28 |
| 23 | Ap3m2 | 2.85 | 2.59E-07 | 5.38E-03 | 0.28 | 0.04 | 0.24 | 7.13 |

**Supplementary Table 1. Marker genes for the original clusters of integrated clustering of E11.5 control and NotchGOF λ-junction cells.**

| | | | | | | | | |
|---|---|---|---|---|---|---|---|---|
| 23 | Nudt7 | 2.50 | 2.75E-07 | 5.73E-03 | 0.39 | 0.07 | 0.32 | 5.56 |
| 23 | Cfl1 | 0.61 | 3.12E-07 | 6.49E-03 | 1.00 | 1.00 | 0.00 | 1.00 |
| 23 | Nfic | 1.81 | 3.12E-07 | 6.50E-03 | 0.78 | 0.30 | 0.48 | 2.58 |
| 23 | Ppib | 0.78 | 3.18E-07 | 6.62E-03 | 1.00 | 0.98 | 0.02 | 1.02 |
| 23 | Shank1 | 3.35 | 3.30E-07 | 6.87E-03 | 0.22 | 0.03 | 0.20 | 8.54 |
| 23 | Ywhab | 0.88 | 3.32E-07 | 6.91E-03 | 1.00 | 0.97 | 0.03 | 1.03 |
| 23 | Dusp4 | 1.99 | 3.37E-07 | 7.01E-03 | 0.44 | 0.09 | 0.36 | 5.16 |
| 23 | Gnat2 | 4.54 | 3.46E-07 | 7.19E-03 | 0.11 | 0.01 | 0.10 | 15.86 |
| 23 | Hspd1 | 0.88 | 3.51E-07 | 7.31E-03 | 1.00 | 0.98 | 0.02 | 1.02 |
| 23 | Lrch3 | 2.06 | 3.59E-07 | 7.46E-03 | 0.83 | 0.37 | 0.46 | 2.25 |
| 23 | Serpine2 | 1.47 | 3.66E-07 | 7.60E-03 | 1.00 | 0.71 | 0.29 | 1.42 |
| 23 | Star | 3.24 | 3.69E-07 | 7.68E-03 | 0.22 | 0.03 | 0.20 | 8.22 |
| 23 | Rab8b | 1.43 | 4.10E-07 | 8.53E-03 | 0.83 | 0.24 | 0.59 | 3.43 |
| 23 | Flna | 1.40 | 4.17E-07 | 8.67E-03 | 1.00 | 0.79 | 0.21 | 1.26 |
| 23 | Fabp5 | 0.68 | 4.69E-07 | 9.76E-03 | 1.00 | 0.98 | 0.02 | 1.02 |
| 23 | Snx1 | 1.78 | 5.06E-07 | 1.05E-02 | 0.94 | 0.45 | 0.50 | 2.11 |
| 23 | Sfxn5 | 2.94 | 5.58E-07 | 1.16E-02 | 0.28 | 0.04 | 0.24 | 6.78 |
| 23 | Sh3pxd2a | 1.74 | 5.89E-07 | 1.22E-02 | 0.89 | 0.37 | 0.52 | 2.42 |
| 23 | Rftn1 | 2.47 | 6.03E-07 | 1.25E-02 | 0.28 | 0.04 | 0.24 | 6.95 |
| 23 | Gng5 | 0.50 | 6.12E-07 | 1.27E-02 | 1.00 | 1.00 | 0.00 | 1.00 |
| 23 | Sema3b | 4.12 | 6.46E-07 | 1.34E-02 | 0.17 | 0.02 | 0.15 | 9.82 |
| 23 | Cnn2 | 1.48 | 6.59E-07 | 1.37E-02 | 0.94 | 0.63 | 0.31 | 1.50 |
| 23 | Cdr2 | 1.75 | 6.96E-07 | 1.45E-02 | 0.72 | 0.22 | 0.50 | 3.28 |
| 23 | Sipa1l1 | 1.87 | 7.49E-07 | 1.56E-02 | 0.83 | 0.52 | 0.32 | 1.62 |
| 23 | Gm47061 | 4.44 | 7.89E-07 | 1.64E-02 | 0.11 | 0.01 | 0.10 | 13.88 |
| 23 | Erp29 | 1.16 | 7.91E-07 | 1.65E-02 | 1.00 | 0.88 | 0.12 | 1.14 |
| 23 | Gm48855 | 2.94 | 8.04E-07 | 1.67E-02 | 0.22 | 0.03 | 0.20 | 8.22 |
| 23 | Tgfb1i1 | 1.90 | 8.40E-07 | 1.75E-02 | 0.56 | 0.14 | 0.42 | 4.06 |
| 23 | Sfrp1 | 1.36 | 8.56E-07 | 1.78E-02 | 1.00 | 0.63 | 0.37 | 1.60 |
| 23 | Gm43445 | 1.96 | 8.61E-07 | 1.79E-02 | 0.44 | 0.09 | 0.35 | 4.93 |
| 23 | Cyp2j6 | 2.59 | 8.90E-07 | 1.85E-02 | 0.50 | 0.12 | 0.38 | 4.07 |
| 23 | Lin28a | 4.13 | 9.43E-07 | 1.96E-02 | 0.17 | 0.02 | 0.15 | 9.82 |
| 23 | Ilk | 1.26 | 9.96E-07 | 2.07E-02 | 0.94 | 0.61 | 0.34 | 1.55 |
| 23 | Cltb | 1.65 | 1.02E-06 | 2.11E-02 | 0.67 | 0.18 | 0.48 | 3.63 |
| 23 | Vwa1 | 2.84 | 1.05E-06 | 2.18E-02 | 0.33 | 0.06 | 0.27 | 5.64 |
| 23 | Gm35021 | 4.28 | 1.06E-06 | 2.21E-02 | 0.11 | 0.01 | 0.10 | 13.88 |
| 23 | Tcn2 | 1.61 | 1.09E-06 | 2.27E-02 | 0.67 | 0.18 | 0.49 | 3.75 |
| 23 | Plekho1 | 1.26 | 1.12E-06 | 2.33E-02 | 1.00 | 0.62 | 0.38 | 1.62 |
| 23 | D730045B01Rik | 4.21 | 1.14E-06 | 2.37E-02 | 0.17 | 0.02 | 0.15 | 9.82 |
| 23 | Eng | 2.13 | 1.16E-06 | 2.41E-02 | 0.28 | 0.04 | 0.24 | 6.62 |
| 23 | Pgbd5 | 4.72 | 1.23E-06 | 2.55E-02 | 0.11 | 0.01 | 0.10 | 13.88 |
| 23 | Fam228b | 3.02 | 1.31E-06 | 2.72E-02 | 0.28 | 0.04 | 0.24 | 6.47 |
| 23 | Trub2 | 1.68 | 1.45E-06 | 3.02E-02 | 0.83 | 0.31 | 0.52 | 2.70 |
| 23 | Fxyd3 | 1.56 | 1.48E-06 | 3.07E-02 | 0.44 | 0.09 | 0.35 | 4.77 |
| 23 | Glo1 | 0.91 | 1.55E-06 | 3.23E-02 | 1.00 | 0.85 | 0.15 | 1.18 |
| 23 | Stx7 | 1.54 | 1.55E-06 | 3.23E-02 | 0.83 | 0.31 | 0.52 | 2.67 |
| 23 | Cmtm3 | 1.43 | 1.56E-06 | 3.24E-02 | 0.94 | 0.42 | 0.52 | 2.24 |
| 23 | Msn | 1.60 | 1.56E-06 | 3.25E-02 | 0.83 | 0.52 | 0.32 | 1.62 |
| 23 | Crip2 | 1.21 | 1.56E-06 | 3.25E-02 | 0.61 | 0.17 | 0.44 | 3.51 |
| 23 | C1qtnf12 | 2.45 | 1.59E-06 | 3.30E-02 | 0.56 | 0.17 | 0.39 | 3.33 |
| 23 | Lima1 | 1.19 | 1.78E-06 | 3.70E-02 | 1.00 | 0.74 | 0.26 | 1.35 |
| 23 | Bgn | 1.83 | 1.89E-06 | 3.94E-02 | 0.72 | 0.26 | 0.46 | 2.75 |
| 23 | P2rx4 | 3.32 | 1.90E-06 | 3.95E-02 | 0.22 | 0.03 | 0.19 | 7.40 |
| 23 | Nexn | 2.76 | 1.93E-06 | 4.01E-02 | 0.28 | 0.04 | 0.23 | 6.32 |
| 23 | Marcks | 0.65 | 2.00E-06 | 4.15E-02 | 1.00 | 0.97 | 0.03 | 1.03 |
| 23 | Bag3 | 1.73 | 2.01E-06 | 4.18E-02 | 0.50 | 0.11 | 0.39 | 4.50 |

**Supplementary Table 1. Marker genes for the original clusters of integrated clustering of E11.5 control and NotchGOF λ-junction cells.**

| | | | | | | | | |
|---|---|---|---|---|---|---|---|---|
| 23 | Agrn | 1.66 | 2.09E-06 | 4.35E-02 | 0.83 | 0.39 | 0.44 | 2.13 |
| 23 | Npm1 | 0.44 | 2.22E-06 | 4.61E-02 | 1.00 | 1.00 | 0.00 | 1.00 |
| 23 | Crispld2 | 2.81 | 2.22E-06 | 4.62E-02 | 0.22 | 0.03 | 0.19 | 7.66 |
| 23 | Gstp1 | 1.73 | 2.23E-06 | 4.64E-02 | 0.94 | 0.52 | 0.42 | 1.81 |
| 23 | Snx10 | 2.55 | 2.30E-06 | 4.77E-02 | 0.56 | 0.16 | 0.40 | 3.50 |
| 23 | Pde7b | 2.81 | 2.33E-06 | 4.85E-02 | 0.28 | 0.04 | 0.23 | 6.32 |
| 23 | Nr4a1 | 2.47 | 2.34E-06 | 4.87E-02 | 0.28 | 0.04 | 0.24 | 6.47 |

**Supplementary Table 2. Marker genes for the major cell types of integrated clustering of E11.5 control and NotchGOF λ-junction cells.**

| cluster | gene | avg_log2FC | p_val | p_val_adj | pct.1 | pct.2 | pct1-pct2 | pct1/pct2 |
|---|---|---|---|---|---|---|---|---|
| mesenchyme | Col3a1 | 3.64 | 0 | 0 | 0.96 | 0.22 | 0.74 | 4.31 |
| mesenchyme | Alx1 | 3.67 | 0 | 0 | 0.96 | 0.24 | 0.73 | 4.08 |
| mesenchyme | Col1a2 | 3.44 | 0 | 0 | 0.98 | 0.25 | 0.73 | 3.9 |
| mesenchyme | Cdh11 | 3.44 | 0 | 0 | 0.93 | 0.22 | 0.71 | 4.24 |
| mesenchyme | Prrx2 | 3.47 | 0 | 0 | 0.96 | 0.28 | 0.69 | 3.49 |
| mesenchyme | Clmp | 3.12 | 0 | 0 | 0.86 | 0.18 | 0.68 | 4.8 |
| mesenchyme | Twist1 | 3.6 | 0 | 0 | 0.98 | 0.32 | 0.66 | 3.03 |
| mesenchyme | Lix1 | 3.62 | 0 | 0 | 0.83 | 0.19 | 0.65 | 4.5 |
| mesenchyme | Col26a1 | 3.23 | 0 | 0 | 0.82 | 0.18 | 0.64 | 4.6 |
| mesenchyme | Pth1r | 3.42 | 0 | 0 | 0.77 | 0.15 | 0.62 | 5.12 |
| mesenchyme | Tbx2 | 2.97 | 0 | 0 | 0.84 | 0.23 | 0.62 | 3.69 |
| mesenchyme | Crlf1 | 3.04 | 0 | 0 | 0.84 | 0.23 | 0.61 | 3.69 |
| mesenchyme | Serpinf1 | 3.01 | 0 | 0 | 0.84 | 0.24 | 0.6 | 3.56 |
| mesenchyme | Pdgfra | 3.53 | 0 | 0 | 0.74 | 0.14 | 0.6 | 5.45 |
| mesenchyme | Pcdh18 | 2.85 | 0 | 0 | 0.83 | 0.24 | 0.59 | 3.46 |
| mesenchyme | Prrx1 | 3.21 | 0 | 0 | 0.84 | 0.26 | 0.59 | 3.29 |
| mesenchyme | Dnm3os | 3.48 | 0 | 0 | 0.71 | 0.14 | 0.57 | 5.15 |
| mesenchyme | Ddr2 | 3.44 | 0 | 0 | 0.67 | 0.11 | 0.55 | 5.85 |
| mesenchyme | Tnfaip6 | 3.73 | 0 | 0 | 0.64 | 0.1 | 0.54 | 6.38 |
| mesenchyme | Il11ra1 | 2.61 | 0 | 0 | 0.79 | 0.25 | 0.54 | 3.14 |
| mesenchyme | Lpar1 | 2.57 | 0 | 0 | 0.88 | 0.34 | 0.53 | 2.56 |
| mesenchyme | Mab21l2 | 3.64 | 0 | 0 | 0.67 | 0.14 | 0.53 | 4.77 |
| mesenchyme | Tmem119 | 3.5 | 0 | 0 | 0.62 | 0.11 | 0.52 | 5.83 |
| mesenchyme | Emilin1 | 2.49 | 0 | 0 | 0.7 | 0.18 | 0.52 | 3.85 |
| mesenchyme | Cped1 | 3.4 | 0 | 0 | 0.63 | 0.12 | 0.51 | 5.31 |
| mesenchyme | Fbln2 | 2.89 | 0 | 0 | 0.68 | 0.18 | 0.51 | 3.89 |
| mesenchyme | Runx1t1 | 2.83 | 0 | 0 | 0.69 | 0.19 | 0.5 | 3.67 |
| mesenchyme | Pdgfrb | 3.39 | 0 | 0 | 0.62 | 0.11 | 0.5 | 5.42 |
| mesenchyme | Nbl1 | 3.35 | 0 | 0 | 0.66 | 0.15 | 0.5 | 4.28 |
| mesenchyme | Pcolce | 3.28 | 0 | 0 | 0.62 | 0.12 | 0.5 | 5 |
| mesenchyme | Lgals1 | 2.56 | 0 | 0 | 0.99 | 0.5 | 0.49 | 1.99 |
| mesenchyme | Vcam1 | 3.54 | 0 | 0 | 0.59 | 0.1 | 0.49 | 5.71 |
| mesenchyme | Eva1b | 2.19 | 0 | 0 | 0.7 | 0.21 | 0.48 | 3.26 |
| mesenchyme | Pax3 | 3.33 | 0 | 0 | 0.61 | 0.13 | 0.48 | 4.58 |
| mesenchyme | Dact3 | 2.72 | 0 | 0 | 0.64 | 0.18 | 0.47 | 3.65 |
| mesenchyme | Igf1 | 3.11 | 0 | 0 | 0.63 | 0.16 | 0.46 | 3.85 |
| mesenchyme | Mmp2 | 2.28 | 0 | 0 | 0.67 | 0.21 | 0.45 | 3.11 |
| mesenchyme | Ppfibp2 | 3 | 0 | 0 | 0.58 | 0.13 | 0.45 | 4.61 |
| mesenchyme | Gata2 | 3.24 | 0 | 0 | 0.59 | 0.14 | 0.45 | 4.26 |
| mesenchyme | Sulf2 | 2.01 | 0 | 0 | 0.81 | 0.37 | 0.44 | 2.21 |
| mesenchyme | Rai14 | 2.01 | 0 | 0 | 0.79 | 0.35 | 0.44 | 2.27 |
| mesenchyme | Limch1 | 1.8 | 0 | 0 | 0.82 | 0.38 | 0.44 | 2.16 |
| mesenchyme | Tbx3 | 2.85 | 0 | 0 | 0.58 | 0.14 | 0.44 | 4.08 |
| mesenchyme | Col5a2 | 2.11 | 0 | 0 | 0.67 | 0.24 | 0.43 | 2.85 |
| mesenchyme | Kif26b | 3.25 | 0 | 0 | 0.55 | 0.11 | 0.43 | 4.88 |
| mesenchyme | Rspo2 | 3.67 | 0 | 0 | 0.51 | 0.08 | 0.43 | 6.12 |
| mesenchyme | Snai1 | 3.08 | 0 | 0 | 0.55 | 0.13 | 0.42 | 4.33 |
| mesenchyme | Frem1 | 3.07 | 0 | 0 | 0.54 | 0.12 | 0.42 | 4.52 |
| mesenchyme | Alx4 | 2.76 | 0 | 0 | 0.56 | 0.14 | 0.42 | 3.97 |
| mesenchyme | Crabp1 | 2.61 | 0 | 0 | 0.99 | 0.58 | 0.41 | 1.72 |
| mesenchyme | Hmcn1 | 2.39 | 0 | 0 | 0.67 | 0.26 | 0.41 | 2.59 |
| mesenchyme | Fbn2 | 1.45 | 0 | 0 | 0.87 | 0.46 | 0.41 | 1.89 |
| mesenchyme | Epha7 | 2.22 | 0 | 0 | 0.7 | 0.29 | 0.41 | 2.4 |
| mesenchyme | Scube1 | 3.02 | 0 | 0 | 0.51 | 0.11 | 0.4 | 4.82 |
| mesenchyme | Lima1 | 1.74 | 0 | 0 | 0.87 | 0.47 | 0.4 | 1.85 |

**Supplementary Table 2. Marker genes for the major cell types of integrated clustering of E11.5 control and NotchGOF λ-junction cells.**

| | | | | | | | | |
|---|---|---|---|---|---|---|---|---|
| mesenchyme | Arhgap28 | 2.1 | 0 | 0 | 0.66 | 0.27 | 0.4 | 2.5 |
| mesenchyme | Frzb | 3.48 | 0 | 0 | 0.48 | 0.09 | 0.39 | 5.26 |
| mesenchyme | Lix1l | 2.74 | 0 | 0 | 0.51 | 0.12 | 0.39 | 4.29 |
| mesenchyme | Ednra | 3.66 | 0 | 0 | 0.47 | 0.08 | 0.39 | 5.96 |
| mesenchyme | Fstl1 | 1.88 | 0 | 0 | 0.95 | 0.56 | 0.39 | 1.69 |
| mesenchyme | Serpine2 | 1.96 | 0 | 0 | 0.83 | 0.45 | 0.39 | 1.87 |
| mesenchyme | Fgfr1 | 1.67 | 0 | 0 | 0.78 | 0.4 | 0.38 | 1.96 |
| mesenchyme | Gap43 | 1.21 | 0 | 0 | 0.54 | 0.16 | 0.38 | 3.39 |
| mesenchyme | Epha3 | 3.41 | 0 | 0 | 0.48 | 0.1 | 0.38 | 4.78 |
| mesenchyme | Rspo3 | 3.55 | 0 | 0 | 0.47 | 0.09 | 0.38 | 5.43 |
| mesenchyme | Mme | 2.14 | 0 | 0 | 0.59 | 0.21 | 0.38 | 2.8 |
| mesenchyme | Mfap4 | 1.37 | 0 | 0 | 0.97 | 0.59 | 0.38 | 1.64 |
| mesenchyme | Pax9 | 3.61 | 0 | 0 | 0.49 | 0.11 | 0.38 | 4.58 |
| mesenchyme | Phldb2 | 1.99 | 0 | 0 | 0.66 | 0.28 | 0.38 | 2.35 |
| mesenchyme | Prex2 | 2.33 | 0 | 0 | 0.53 | 0.15 | 0.38 | 3.53 |
| mesenchyme | Inka1 | 1.72 | 0 | 0 | 0.63 | 0.26 | 0.37 | 2.39 |
| mesenchyme | Efnb1 | 1.95 | 0 | 0 | 0.65 | 0.29 | 0.36 | 2.26 |
| mesenchyme | Snai2 | 1.68 | 0 | 0 | 0.58 | 0.22 | 0.36 | 2.63 |
| mesenchyme | Cnn2 | 1.02 | 0 | 0 | 0.75 | 0.39 | 0.36 | 1.94 |
| mesenchyme | Tril | 2.07 | 0 | 0 | 0.66 | 0.31 | 0.36 | 2.17 |
| mesenchyme | Cacna1g | 3.17 | 0 | 0 | 0.44 | 0.09 | 0.35 | 4.83 |
| mesenchyme | Peg10 | 1.77 | 0 | 0 | 0.55 | 0.2 | 0.35 | 2.75 |
| mesenchyme | Oaf | 2.04 | 0 | 0 | 0.57 | 0.22 | 0.35 | 2.56 |
| mesenchyme | Mex3b | 1.53 | 0 | 0 | 0.81 | 0.46 | 0.34 | 1.75 |
| mesenchyme | Col1a1 | 3.66 | 0 | 0 | 0.42 | 0.07 | 0.34 | 5.85 |
| mesenchyme | Alx3 | 3.67 | 0 | 0 | 0.41 | 0.06 | 0.34 | 6.44 |
| mesenchyme | Asph | 1.6 | 0 | 0 | 0.76 | 0.42 | 0.34 | 1.82 |
| mesenchyme | Fli1 | 1.96 | 0 | 0 | 0.48 | 0.14 | 0.34 | 3.44 |
| mesenchyme | Frmd6 | 1.72 | 0 | 0 | 0.6 | 0.26 | 0.34 | 2.34 |
| mesenchyme | Wnt5a | 1.34 | 0 | 0 | 0.75 | 0.41 | 0.34 | 1.83 |
| mesenchyme | Arhgef40 | 1.71 | 0 | 0 | 0.64 | 0.3 | 0.34 | 2.15 |
| mesenchyme | Spin2c | 2.11 | 0 | 0 | 0.53 | 0.19 | 0.34 | 2.8 |
| mesenchyme | Plpp3 | 1.71 | 0 | 0 | 0.77 | 0.43 | 0.34 | 1.78 |
| mesenchyme | Vcan | 1.37 | 0 | 0 | 0.97 | 0.64 | 0.34 | 1.53 |
| mesenchyme | Daam2 | 3.42 | 0 | 0 | 0.41 | 0.07 | 0.34 | 5.96 |
| mesenchyme | Dixdc1 | 2.23 | 0 | 0 | 0.51 | 0.17 | 0.34 | 2.99 |
| mesenchyme | Igsf3 | 1.57 | 0 | 0 | 0.69 | 0.36 | 0.33 | 1.92 |
| mesenchyme | Cald1 | 1.61 | 0 | 0 | 1 | 0.67 | 0.33 | 1.49 |
| mesenchyme | Crym | 2.52 | 0 | 0 | 0.59 | 0.26 | 0.33 | 2.29 |
| mesenchyme | Prdm16 | 2.87 | 0 | 0 | 0.44 | 0.11 | 0.33 | 4.1 |
| mesenchyme | Pcsk5 | 2.36 | 0 | 0 | 0.47 | 0.15 | 0.33 | 3.27 |
| mesenchyme | Rbms3 | 2.05 | 0 | 0 | 0.52 | 0.19 | 0.33 | 2.71 |
| mesenchyme | Nfatc4 | 1.59 | 0 | 0 | 0.66 | 0.33 | 0.33 | 1.99 |
| mesenchyme | Col23a1 | 2.87 | 0 | 0 | 0.42 | 0.09 | 0.33 | 4.55 |
| mesenchyme | Jam3 | 1.46 | 0 | 0 | 0.68 | 0.36 | 0.32 | 1.91 |
| mesenchyme | Msn | 0.58 | 0 | 0 | 0.62 | 0.3 | 0.32 | 2.08 |
| mesenchyme | Pdlim4 | 1.15 | 0 | 0 | 0.73 | 0.41 | 0.32 | 1.78 |
| mesenchyme | Rarb | 2.6 | 0 | 0 | 0.45 | 0.13 | 0.32 | 3.41 |
| mesenchyme | Nid1 | 1.86 | 0 | 0 | 0.45 | 0.13 | 0.32 | 3.48 |
| mesenchyme | Rarg | 1.56 | 0 | 0 | 0.63 | 0.31 | 0.32 | 2.04 |
| mesenchyme | Dnm1 | 2.59 | 0 | 0 | 0.43 | 0.12 | 0.31 | 3.7 |
| mesenchyme | Olfml3 | 3.03 | 0 | 0 | 0.39 | 0.08 | 0.31 | 5.12 |
| mesenchyme | Plekho1 | 1.11 | 0 | 0 | 0.72 | 0.41 | 0.31 | 1.77 |
| mesenchyme | Fbln1 | 1.44 | 0 | 0 | 0.9 | 0.59 | 0.31 | 1.53 |
| mesenchyme | Arhgdib | 1.13 | 0 | 0 | 0.6 | 0.29 | 0.31 | 2.08 |
| mesenchyme | Timp2 | 1.25 | 0 | 0 | 0.77 | 0.46 | 0.31 | 1.68 |

**Supplementary Table 2. Marker genes for the major cell types of integrated clustering of E11.5 control and NotchGOF λ-junction cells.**

| | | | | | | | | |
|---|---|---|---|---|---|---|---|---|
| mesenchyme | Rasl11b | 1.47 | 0 | 0 | 0.59 | 0.28 | 0.31 | 2.12 |
| mesenchyme | Zeb1 | 1.75 | 0 | 0 | 0.45 | 0.14 | 0.31 | 3.22 |
| mesenchyme | Fn1 | 1.59 | 0 | 0 | 0.88 | 0.58 | 0.31 | 1.53 |
| mesenchyme | Wdr86 | 1.44 | 0 | 0 | 0.62 | 0.31 | 0.31 | 1.99 |
| mesenchyme | Hdgfl3 | 1.2 | 0 | 0 | 0.8 | 0.5 | 0.31 | 1.61 |
| mesenchyme | Dchs1 | 1.8 | 0 | 0 | 0.52 | 0.21 | 0.3 | 2.42 |
| mesenchyme | Angptl4 | 2.54 | 0 | 0 | 0.43 | 0.13 | 0.3 | 3.36 |
| mesenchyme | Tpm2 | 1.49 | 0 | 0 | 0.54 | 0.23 | 0.3 | 2.31 |
| mesenchyme | Nkd1 | 2.47 | 0 | 0 | 0.46 | 0.16 | 0.3 | 2.92 |
| mesenchyme | Glipr2 | 2.09 | 0 | 0 | 0.49 | 0.18 | 0.3 | 2.66 |
| mesenchyme | Sh3gl3 | 2.48 | 0 | 0 | 0.43 | 0.13 | 0.3 | 3.34 |
| mesenchyme | Rcn3 | 1.33 | 0 | 0 | 0.83 | 0.53 | 0.3 | 1.57 |
| mesenchyme | Efna2 | 1.2 | 0 | 0 | 0.81 | 0.51 | 0.3 | 1.59 |
| mesenchyme | Ror2 | 2.75 | 0 | 0 | 0.4 | 0.1 | 0.3 | 4 |
| mesenchyme | Igfbp7 | 3.51 | 0 | 0 | 0.36 | 0.06 | 0.3 | 5.77 |
| mesenchyme | Zcchc24 | 2.76 | 0 | 0 | 0.39 | 0.09 | 0.3 | 4.21 |
| mesenchyme | Dab2 | 1.25 | 0 | 0 | 0.45 | 0.16 | 0.3 | 2.9 |
| mesenchyme | Tenm4 | 1.26 | 0 | 0 | 0.71 | 0.41 | 0.3 | 1.71 |
| mesenchyme | Mir100hg | 1.16 | 0 | 0 | 0.86 | 0.56 | 0.29 | 1.52 |
| mesenchyme | Creb5 | 2.56 | 0 | 0 | 0.42 | 0.13 | 0.29 | 3.23 |
| mesenchyme | Dkk1 | 3.69 | 0 | 0 | 0.37 | 0.08 | 0.29 | 4.6 |
| mesenchyme | Tgfbi | 2.8 | 0 | 0 | 0.39 | 0.09 | 0.29 | 4.1 |
| mesenchyme | Tns1 | 2.49 | 0 | 0 | 0.4 | 0.11 | 0.29 | 3.66 |
| mesenchyme | Igfbp4 | 1.58 | 0 | 0 | 1 | 0.71 | 0.29 | 1.41 |
| mesenchyme | Lhx8 | 3.46 | 0 | 0 | 0.37 | 0.08 | 0.29 | 4.82 |
| mesenchyme | Rnf122 | 1.43 | 0 | 0 | 0.63 | 0.34 | 0.29 | 1.85 |
| mesenchyme | Gata3 | 1.89 | 0 | 0 | 0.62 | 0.33 | 0.29 | 1.88 |
| mesenchyme | Fzd2 | 1.21 | 0 | 0 | 0.97 | 0.68 | 0.29 | 1.42 |
| mesenchyme | Dlc1 | 1.71 | 0 | 0 | 0.45 | 0.17 | 0.29 | 2.72 |
| mesenchyme | Cd248 | 3.48 | 0 | 0 | 0.35 | 0.06 | 0.29 | 5.95 |
| mesenchyme | Gsc | 3.32 | 0 | 0 | 0.39 | 0.1 | 0.29 | 3.78 |
| mesenchyme | Actn1 | 1.18 | 0 | 0 | 0.82 | 0.53 | 0.29 | 1.54 |
| mesenchyme | Gas1 | 0.95 | 0 | 0 | 0.79 | 0.51 | 0.28 | 1.55 |
| mesenchyme | S1pr3 | 2.8 | 0 | 0 | 0.37 | 0.09 | 0.28 | 4.28 |
| mesenchyme | Gsto1 | 1.36 | 0 | 0 | 0.74 | 0.46 | 0.28 | 1.61 |
| mesenchyme | Sdc1 | 0.99 | 0 | 0 | 0.79 | 0.51 | 0.28 | 1.54 |
| mesenchyme | Crispld1 | 2.21 | 0 | 0 | 0.42 | 0.14 | 0.28 | 2.99 |
| mesenchyme | Etv4 | 2.21 | 0 | 0 | 0.42 | 0.15 | 0.28 | 2.91 |
| mesenchyme | St6galnac3 | 1.92 | 0 | 0 | 0.48 | 0.21 | 0.28 | 2.32 |
| mesenchyme | Hpgd | 2.5 | 0 | 0 | 0.44 | 0.16 | 0.27 | 2.67 |
| mesenchyme | Car14 | 1.53 | 0 | 0 | 0.51 | 0.24 | 0.27 | 2.14 |
| mesenchyme | Ccne1 | 1.43 | 0 | 0 | 0.59 | 0.32 | 0.27 | 1.85 |
| mesenchyme | Zeb2 | 1.91 | 0 | 0 | 0.49 | 0.22 | 0.27 | 2.24 |
| mesenchyme | Frmd4a | 1.05 | 0 | 0 | 0.76 | 0.49 | 0.27 | 1.55 |
| mesenchyme | Lrp1 | 1.12 | 0 | 0 | 0.67 | 0.4 | 0.27 | 1.67 |
| mesenchyme | Twist2 | 3.51 | 0 | 0 | 0.32 | 0.05 | 0.27 | 6.36 |
| mesenchyme | Ndn | 1.03 | 0 | 0 | 0.83 | 0.56 | 0.27 | 1.48 |
| mesenchyme | Gnb4 | 1.94 | 0 | 0 | 0.39 | 0.12 | 0.27 | 3.23 |
| mesenchyme | Septin6 | 1.21 | 0 | 0 | 0.66 | 0.39 | 0.27 | 1.68 |
| mesenchyme | Rassf3 | 1.83 | 0 | 0 | 0.46 | 0.19 | 0.27 | 2.39 |
| mesenchyme | Satb2 | 3.15 | 0 | 0 | 0.34 | 0.08 | 0.26 | 4.37 |
| mesenchyme | Rnf227 | 1.52 | 0 | 0 | 0.5 | 0.24 | 0.26 | 2.1 |
| mesenchyme | Rab34 | 0.99 | 0 | 0 | 0.8 | 0.54 | 0.26 | 1.49 |
| mesenchyme | Flrt2 | 2.23 | 0 | 0 | 0.54 | 0.28 | 0.26 | 1.95 |
| mesenchyme | Dpep1 | 3.47 | 0 | 0 | 0.31 | 0.05 | 0.26 | 6.24 |
| mesenchyme | Angptl1 | 3.78 | 0 | 0 | 0.31 | 0.05 | 0.26 | 6.55 |

**Supplementary Table 2. Marker genes for the major cell types of integrated clustering of E11.5 control and NotchGOF λ-junction cells.**

| | | | | | | | | |
|---|---|---|---|---|---|---|---|---|
| mesenchyme | Cachd1 | 1.53 | 0 | 0 | 0.61 | 0.35 | 0.26 | 1.76 |
| mesenchyme | Tmem132c | 1.99 | 0 | 0 | 0.41 | 0.15 | 0.26 | 2.79 |
| mesenchyme | Oat | 1.23 | 0 | 0 | 0.77 | 0.5 | 0.26 | 1.52 |
| mesenchyme | Septin11 | 1.33 | 0 | 0 | 0.96 | 0.7 | 0.26 | 1.37 |
| mesenchyme | Limd2 | 1.48 | 0 | 0 | 0.98 | 0.72 | 0.26 | 1.36 |
| mesenchyme | Loxl2 | 1.38 | 0 | 0 | 0.52 | 0.26 | 0.26 | 1.98 |
| mesenchyme | Trim35 | 1.1 | 0 | 0 | 0.79 | 0.53 | 0.26 | 1.49 |
| mesenchyme | Itm2c | 0.91 | 0 | 0 | 0.81 | 0.56 | 0.26 | 1.46 |
| mesenchyme | Dkk2 | 2.42 | 0 | 0 | 0.35 | 0.09 | 0.26 | 3.72 |
| mesenchyme | Ptn | 1.74 | 0 | 0 | 0.98 | 0.73 | 0.26 | 1.35 |
| mesenchyme | Sgcd | 3.44 | 0 | 0 | 0.3 | 0.05 | 0.26 | 6.2 |
| mesenchyme | Dusp7 | 1.27 | 0 | 0 | 0.58 | 0.33 | 0.26 | 1.78 |
| mesenchyme | Gm49708 | 1.37 | 0 | 0 | 0.7 | 0.45 | 0.26 | 1.57 |
| mesenchyme | Vegfc | 2.63 | 0 | 0 | 0.35 | 0.09 | 0.25 | 3.7 |
| mesenchyme | Col9a1 | 2.62 | 0 | 0 | 0.37 | 0.12 | 0.25 | 3.13 |
| mesenchyme | Tmem26 | 2.78 | 0 | 0 | 0.34 | 0.08 | 0.25 | 4.05 |
| mesenchyme | Epha4 | 0.99 | 0 | 0 | 0.73 | 0.48 | 0.25 | 1.52 |
| mesenchyme | Pitpnc1 | 1.18 | 0 | 0 | 0.67 | 0.42 | 0.25 | 1.6 |
| mesenchyme | Synpo | 2.91 | 0 | 0 | 0.32 | 0.07 | 0.25 | 4.52 |
| mesenchyme | Rerg | 2.94 | 0 | 0 | 0.33 | 0.08 | 0.25 | 4.07 |
| mesenchyme | Gjc1 | 0.93 | 0 | 0 | 0.89 | 0.64 | 0.25 | 1.39 |
| mesenchyme | Fndc3c1 | 1.39 | 0 | 0 | 0.55 | 0.3 | 0.25 | 1.82 |
| mesenchyme | Mpped2 | 1.69 | 0 | 0 | 0.93 | 0.68 | 0.25 | 1.36 |
| mesenchyme | Elovl6 | 1 | 0 | 0 | 0.73 | 0.48 | 0.25 | 1.51 |
| mesenchyme | Pax7 | 3.27 | 0 | 0 | 0.32 | 0.07 | 0.25 | 4.53 |
| mesenchyme | Sfrp2 | 1.86 | 0 | 0 | 0.69 | 0.45 | 0.25 | 1.56 |
| mesenchyme | Pja1 | 1.13 | 0 | 0 | 0.76 | 0.51 | 0.25 | 1.48 |
| mesenchyme | Epb41l2 | 0.93 | 0 | 0 | 0.75 | 0.5 | 0.25 | 1.49 |
| mesenchyme | Tlnrd1 | 1.29 | 0 | 0 | 0.58 | 0.34 | 0.25 | 1.73 |
| mesenchyme | Ppp3ca | 1.21 | 0 | 0 | 0.81 | 0.56 | 0.24 | 1.43 |
| mesenchyme | Cdk14 | 1.39 | 0 | 0 | 0.57 | 0.33 | 0.24 | 1.74 |
| mesenchyme | Scarf2 | 2.51 | 0 | 0 | 0.34 | 0.09 | 0.24 | 3.59 |
| mesenchyme | Rest | 0.9 | 0 | 0 | 0.89 | 0.65 | 0.24 | 1.37 |
| mesenchyme | Rftn2 | 2.81 | 0 | 0 | 0.31 | 0.07 | 0.24 | 4.67 |
| mesenchyme | Ndnf | 3.73 | 0 | 0 | 0.29 | 0.05 | 0.24 | 6.26 |
| mesenchyme | Rhoc | 0.64 | 0 | 0 | 0.87 | 0.63 | 0.24 | 1.38 |
| mesenchyme | Cacna1d | 2.76 | 0 | 0 | 0.31 | 0.07 | 0.24 | 4.29 |
| mesenchyme | Eml4 | 1.09 | 0 | 0 | 0.91 | 0.68 | 0.24 | 1.35 |
| mesenchyme | Emilin2 | 2.45 | 0 | 0 | 0.33 | 0.1 | 0.24 | 3.43 |
| mesenchyme | 130024F11Ri | 3.33 | 0 | 0 | 0.3 | 0.06 | 0.24 | 4.93 |
| mesenchyme | Arid5b | 1.01 | 0 | 0 | 0.75 | 0.51 | 0.23 | 1.45 |
| mesenchyme | Reep5 | 1.31 | 0 | 0 | 0.86 | 0.63 | 0.23 | 1.37 |
| mesenchyme | Vegfb | 0.99 | 0 | 0 | 0.74 | 0.5 | 0.23 | 1.46 |
| mesenchyme | Pdlim7 | 0.92 | 0 | 0 | 0.83 | 0.6 | 0.23 | 1.38 |
| mesenchyme | Bin1 | 0.96 | 0 | 0 | 0.7 | 0.47 | 0.23 | 1.49 |
| mesenchyme | Myo1b | 0.78 | 0 | 0 | 0.81 | 0.58 | 0.23 | 1.4 |
| mesenchyme | Colec12 | 2.98 | 0 | 0 | 0.3 | 0.07 | 0.23 | 4.42 |
| mesenchyme | Flna | 0.79 | 0 | 0 | 0.87 | 0.64 | 0.23 | 1.36 |
| mesenchyme | Fxyd6 | 0.9 | 0 | 0 | 0.76 | 0.53 | 0.23 | 1.43 |
| mesenchyme | Tnfaip8 | 1.18 | 0 | 0 | 0.83 | 0.6 | 0.23 | 1.38 |
| mesenchyme | Rev3l | 1.25 | 0 | 0 | 0.66 | 0.44 | 0.23 | 1.52 |
| mesenchyme | Tcf12 | 1.58 | 0 | 0 | 0.95 | 0.72 | 0.22 | 1.31 |
| mesenchyme | Myh10 | 0.87 | 0 | 0 | 0.94 | 0.72 | 0.22 | 1.31 |
| mesenchyme | Mfap2 | 1.33 | 0 | 0 | 0.99 | 0.78 | 0.22 | 1.28 |
| mesenchyme | 610318N02Ri | 1.11 | 0 | 0 | 0.67 | 0.45 | 0.22 | 1.48 |
| mesenchyme | Fam92a | 0.87 | 0 | 0 | 0.83 | 0.61 | 0.22 | 1.36 |

**Supplementary Table 2. Marker genes for the major cell types of integrated clustering of E11.5 control and NotchGOF λ-junction cells.**

| | | | | | | | | |
|---|---|---|---|---|---|---|---|---|
| mesenchyme | Ldhb | 1.04 | 0 | 0 | 0.98 | 0.76 | 0.22 | 1.28 |
| mesenchyme | Ccdc88a | 0.95 | 0 | 0 | 0.88 | 0.67 | 0.22 | 1.33 |
| mesenchyme | Crabp2 | 0.58 | 0 | 0 | 0.99 | 0.77 | 0.22 | 1.28 |
| mesenchyme | Maged2 | 0.8 | 0 | 0 | 0.89 | 0.68 | 0.22 | 1.32 |
| mesenchyme | Sh3bgrl | 0.79 | 0 | 0 | 0.92 | 0.71 | 0.21 | 1.3 |
| mesenchyme | Apip | 1.1 | 0 | 0 | 0.68 | 0.47 | 0.21 | 1.45 |
| mesenchyme | Ptbp2 | 0.82 | 0 | 0 | 0.84 | 0.63 | 0.21 | 1.33 |
| mesenchyme | Gng10 | 0.89 | 0 | 0 | 0.88 | 0.67 | 0.21 | 1.31 |
| mesenchyme | Zcchc3 | 0.86 | 0 | 0 | 0.78 | 0.58 | 0.2 | 1.35 |
| mesenchyme | Rhobtb3 | 0.8 | 0 | 0 | 0.88 | 0.68 | 0.2 | 1.3 |
| mesenchyme | Rbfox2 | 0.8 | 0 | 0 | 0.83 | 0.63 | 0.2 | 1.32 |
| mesenchyme | Oxct1 | 0.71 | 0 | 0 | 0.89 | 0.68 | 0.2 | 1.3 |
| mesenchyme | Yaf2 | 0.95 | 0 | 0 | 0.76 | 0.56 | 0.2 | 1.35 |
| mesenchyme | Dbn1 | 0.71 | 0 | 0 | 0.94 | 0.74 | 0.2 | 1.27 |
| mesenchyme | Acat2 | 0.7 | 0 | 0 | 0.87 | 0.67 | 0.2 | 1.3 |
| mesenchyme | Csrp2 | 0.83 | 0 | 0 | 0.99 | 0.79 | 0.2 | 1.25 |
| mesenchyme | Fscn1 | 0.98 | 0 | 0 | 0.97 | 0.78 | 0.19 | 1.25 |
| mesenchyme | Tead2 | 0.76 | 0 | 0 | 0.96 | 0.77 | 0.19 | 1.24 |
| mesenchyme | Map1lc3a | 0.8 | 0 | 0 | 0.87 | 0.68 | 0.19 | 1.28 |
| mesenchyme | Kifap3 | 0.75 | 0 | 0 | 0.84 | 0.65 | 0.19 | 1.29 |
| mesenchyme | Cdkn1c | 0.94 | 0 | 0 | 0.99 | 0.8 | 0.19 | 1.23 |
| mesenchyme | Trim24 | 0.86 | 0 | 0 | 0.77 | 0.59 | 0.18 | 1.3 |
| mesenchyme | Cd63 | 0.6 | 0 | 0 | 0.97 | 0.79 | 0.18 | 1.22 |
| mesenchyme | Twistnb | 0.97 | 0 | 0 | 0.83 | 0.65 | 0.18 | 1.27 |
| mesenchyme | Arl2bp | 0.78 | 0 | 0 | 0.85 | 0.67 | 0.18 | 1.26 |
| mesenchyme | Adprh | 0.79 | 0 | 0 | 0.84 | 0.67 | 0.17 | 1.26 |
| mesenchyme | Rcn2 | 0.71 | 0 | 0 | 0.97 | 0.8 | 0.17 | 1.22 |
| mesenchyme | Akr7a5 | 0.93 | 0 | 0 | 0.85 | 0.68 | 0.17 | 1.25 |
| mesenchyme | H2afy2 | 1.02 | 0 | 0 | 0.99 | 0.82 | 0.17 | 1.2 |
| mesenchyme | Id2 | 1.08 | 0 | 0 | 0.92 | 0.75 | 0.17 | 1.22 |
| mesenchyme | Dcakd | 0.78 | 0 | 0 | 0.96 | 0.79 | 0.17 | 1.21 |
| mesenchyme | Map4k4 | 0.64 | 0 | 0 | 0.96 | 0.79 | 0.16 | 1.2 |
| mesenchyme | Gm1673 | 0.77 | 0 | 0 | 0.98 | 0.82 | 0.16 | 1.2 |
| mesenchyme | Hnrnph3 | 0.75 | 0 | 0 | 0.91 | 0.75 | 0.16 | 1.22 |
| mesenchyme | Serpinh1 | 0.69 | 0 | 0 | 0.99 | 0.84 | 0.16 | 1.19 |
| mesenchyme | Tpm4 | 1.02 | 0 | 0 | 0.98 | 0.83 | 0.16 | 1.19 |
| mesenchyme | Znrd1 | 0.7 | 0 | 0 | 0.87 | 0.71 | 0.15 | 1.22 |
| mesenchyme | Tut4 | 0.89 | 0 | 0 | 0.97 | 0.82 | 0.15 | 1.19 |
| mesenchyme | Slc38a2 | 0.86 | 0 | 0 | 0.95 | 0.8 | 0.15 | 1.19 |
| mesenchyme | Serf1 | 0.67 | 0 | 0 | 0.97 | 0.82 | 0.15 | 1.18 |
| mesenchyme | Gstm5 | 0.72 | 0 | 0 | 0.93 | 0.78 | 0.15 | 1.19 |
| mesenchyme | Ruvbl1 | 0.75 | 0 | 0 | 0.89 | 0.75 | 0.14 | 1.19 |
| mesenchyme | Nnat | 0.27 | 0 | 0 | 0.99 | 0.85 | 0.14 | 1.17 |
| mesenchyme | Cnn3 | 0.54 | 0 | 0 | 0.98 | 0.84 | 0.14 | 1.17 |
| mesenchyme | Ifitm2 | 0.68 | 0 | 0 | 0.99 | 0.85 | 0.14 | 1.16 |
| mesenchyme | Pclaf | 0.67 | 0 | 0 | 0.93 | 0.8 | 0.14 | 1.17 |
| mesenchyme | Maged1 | 0.58 | 0 | 0 | 0.98 | 0.85 | 0.13 | 1.16 |
| mesenchyme | Baz1b | 0.73 | 0 | 0 | 0.93 | 0.8 | 0.13 | 1.16 |
| mesenchyme | Tsc22d1 | 0.82 | 0 | 0 | 0.98 | 0.84 | 0.13 | 1.16 |
| mesenchyme | Gtf3c6 | 0.69 | 0 | 0 | 0.9 | 0.77 | 0.13 | 1.17 |
| mesenchyme | Npc2 | 0.51 | 0 | 0 | 0.97 | 0.84 | 0.13 | 1.15 |
| mesenchyme | Basp1 | 0.52 | 0 | 0 | 1 | 0.87 | 0.13 | 1.15 |
| mesenchyme | Ssbp3 | 0.97 | 0 | 0 | 0.98 | 0.86 | 0.12 | 1.14 |
| mesenchyme | Bzw2 | 0.58 | 0 | 0 | 0.97 | 0.86 | 0.12 | 1.13 |
| mesenchyme | Ywhah | 0.55 | 0 | 0 | 0.94 | 0.83 | 0.11 | 1.13 |
| mesenchyme | Hdac2 | 0.53 | 0 | 0 | 0.94 | 0.83 | 0.11 | 1.13 |

**Supplementary Table 2. Marker genes for the major cell types of integrated clustering of E11.5 control and NotchGOF λ-junction cells.**

| | | | | | | | | |
|---|---|---|---|---|---|---|---|---|
| mesenchyme | Capzb | 0.65 | 0 | 0 | 0.95 | 0.84 | 0.11 | 1.13 |
| mesenchyme | Pdia6 | 0.6 | 0 | 0 | 0.99 | 0.89 | 0.1 | 1.12 |
| mesenchyme | Eif1b | 0.75 | 0 | 0 | 0.96 | 0.86 | 0.1 | 1.12 |
| mesenchyme | Mettl9 | 0.63 | 0 | 0 | 0.97 | 0.87 | 0.1 | 1.12 |
| mesenchyme | Sox11 | 0.51 | 0 | 0 | 1 | 0.9 | 0.1 | 1.11 |
| mesenchyme | Hsd17b10 | 0.68 | 0 | 0 | 0.94 | 0.84 | 0.1 | 1.12 |
| mesenchyme | Ilf2 | 0.65 | 0 | 0 | 0.98 | 0.88 | 0.1 | 1.11 |
| mesenchyme | Eif4e | 0.58 | 0 | 0 | 0.92 | 0.82 | 0.1 | 1.12 |
| mesenchyme | Ostc | 0.68 | 0 | 0 | 0.98 | 0.88 | 0.1 | 1.11 |
| mesenchyme | Khdrbs1 | 0.68 | 0 | 0 | 0.99 | 0.89 | 0.09 | 1.1 |
| mesenchyme | Nap1l4 | 0.58 | 0 | 0 | 0.96 | 0.87 | 0.09 | 1.11 |
| mesenchyme | Polr2m | 0.53 | 0 | 0 | 0.98 | 0.89 | 0.09 | 1.1 |
| mesenchyme | Ccdc34 | 0.58 | 0 | 0 | 0.95 | 0.86 | 0.09 | 1.1 |
| mesenchyme | Srsf10 | 0.61 | 0 | 0 | 0.97 | 0.89 | 0.09 | 1.1 |
| mesenchyme | Slc25a4 | 0.74 | 0 | 0 | 1 | 0.91 | 0.09 | 1.09 |
| mesenchyme | Lsm3 | 0.61 | 0 | 0 | 0.95 | 0.86 | 0.09 | 1.1 |
| mesenchyme | Arf4 | 0.52 | 0 | 0 | 0.98 | 0.89 | 0.09 | 1.1 |
| mesenchyme | Selenoh | 0.56 | 0 | 0 | 0.99 | 0.91 | 0.08 | 1.09 |
| mesenchyme | Marcks | 0.97 | 0 | 0 | 1 | 0.92 | 0.08 | 1.09 |
| mesenchyme | Ezh2 | 0.5 | 0 | 0 | 0.97 | 0.89 | 0.08 | 1.09 |
| mesenchyme | Lsm5 | 0.52 | 0 | 0 | 0.97 | 0.89 | 0.08 | 1.09 |
| mesenchyme | Tmpo | 0.52 | 0 | 0 | 0.98 | 0.9 | 0.08 | 1.08 |
| mesenchyme | Ptov1 | 0.48 | 0 | 0 | 0.99 | 0.92 | 0.07 | 1.08 |
| mesenchyme | Eid1 | 0.53 | 0 | 0 | 0.99 | 0.92 | 0.07 | 1.08 |
| mesenchyme | Arglu1 | 0.63 | 0 | 0 | 0.99 | 0.91 | 0.07 | 1.08 |
| mesenchyme | Lsm6 | 0.5 | 0 | 0 | 0.96 | 0.89 | 0.07 | 1.08 |
| mesenchyme | Calr | 0.5 | 0 | 0 | 1 | 0.93 | 0.07 | 1.07 |
| mesenchyme | Micos10 | 1.1 | 0 | 0 | 0.99 | 0.92 | 0.07 | 1.07 |
| mesenchyme | Caprin1 | 0.54 | 0 | 0 | 0.98 | 0.92 | 0.07 | 1.07 |
| mesenchyme | Fubp1 | 0.48 | 0 | 0 | 1 | 0.93 | 0.07 | 1.07 |
| mesenchyme | Uqcc2 | 0.51 | 0 | 0 | 0.98 | 0.91 | 0.06 | 1.07 |
| mesenchyme | Septin7 | 0.62 | 0 | 0 | 0.99 | 0.93 | 0.06 | 1.07 |
| mesenchyme | Srsf9 | 0.44 | 0 | 0 | 0.99 | 0.93 | 0.06 | 1.06 |
| mesenchyme | Nasp | 0.66 | 0 | 0 | 1 | 0.94 | 0.06 | 1.06 |
| mesenchyme | Supt16 | 0.51 | 0 | 0 | 0.98 | 0.92 | 0.06 | 1.06 |
| mesenchyme | Hnrnpdl | 0.55 | 0 | 0 | 0.98 | 0.93 | 0.05 | 1.06 |
| mesenchyme | Rbbp4 | 0.49 | 0 | 0 | 0.98 | 0.93 | 0.05 | 1.06 |
| mesenchyme | Cycs | 0.56 | 0 | 0 | 0.98 | 0.93 | 0.05 | 1.06 |
| mesenchyme | Prmt1 | 0.48 | 0 | 0 | 1 | 0.95 | 0.05 | 1.05 |
| mesenchyme | Nme2 | 0.63 | 0 | 0 | 1 | 0.95 | 0.05 | 1.05 |
| mesenchyme | Cbx1 | 0.57 | 0 | 0 | 1 | 0.95 | 0.05 | 1.05 |
| mesenchyme | Ndufc2 | 0.42 | 0 | 0 | 0.99 | 0.94 | 0.05 | 1.05 |
| mesenchyme | Ppib | 0.5 | 0 | 0 | 1 | 0.95 | 0.05 | 1.05 |
| mesenchyme | Hnrnpm | 0.47 | 0 | 0 | 0.99 | 0.95 | 0.05 | 1.05 |
| mesenchyme | Snrpd3 | 0.45 | 0 | 0 | 0.99 | 0.95 | 0.04 | 1.04 |
| mesenchyme | Atp5g3 | 0.43 | 0 | 0 | 0.99 | 0.95 | 0.04 | 1.04 |
| mesenchyme | Cct3 | 0.43 | 0 | 0 | 0.99 | 0.95 | 0.04 | 1.04 |
| mesenchyme | Sfpq | 0.47 | 0 | 0 | 1 | 0.96 | 0.04 | 1.04 |
| mesenchyme | H2afv | 0.44 | 0 | 0 | 1 | 0.96 | 0.04 | 1.04 |
| mesenchyme | Fkbp3 | 0.42 | 0 | 0 | 1 | 0.96 | 0.04 | 1.04 |
| mesenchyme | Nucks1 | 0.51 | 0 | 0 | 0.99 | 0.96 | 0.04 | 1.04 |
| mesenchyme | Mdk | 0.54 | 0 | 0 | 1 | 0.96 | 0.04 | 1.04 |
| mesenchyme | Snrpb | 0.4 | 0 | 0 | 1 | 0.96 | 0.04 | 1.04 |
| mesenchyme | Atp5k | 0.41 | 0 | 0 | 0.99 | 0.95 | 0.04 | 1.04 |
| mesenchyme | Srsf7 | 0.48 | 0 | 0 | 1 | 0.96 | 0.03 | 1.04 |
| mesenchyme | Psma3 | 0.43 | 0 | 0 | 0.99 | 0.96 | 0.03 | 1.03 |

**Supplementary Table 2. Marker genes for the major cell types of integrated clustering of E11.5 control and NotchGOF λ-junction cells.**

| | | | | | | | | |
|---|---|---|---|---|---|---|---|---|
| mesenchyme | Snrpd1 | 0.45 | 0 | 0 | 1 | 0.97 | 0.03 | 1.03 |
| mesenchyme | Ndufb8 | 0.39 | 0 | 0 | 0.99 | 0.97 | 0.03 | 1.03 |
| mesenchyme | Hmgb3 | 0.55 | 0 | 0 | 1 | 0.97 | 0.03 | 1.03 |
| mesenchyme | Nme1 | 0.48 | 0 | 0 | 1 | 0.97 | 0.03 | 1.03 |
| mesenchyme | Hnrnpd | 0.49 | 0 | 0 | 1 | 0.97 | 0.03 | 1.03 |
| mesenchyme | Morf4l1 | 0.38 | 0 | 0 | 1 | 0.98 | 0.02 | 1.02 |
| mesenchyme | Cox7a2 | 0.35 | 0 | 0 | 1 | 0.98 | 0.02 | 1.02 |
| mesenchyme | Hnrnpu | 0.43 | 0 | 0 | 1 | 0.98 | 0.02 | 1.02 |
| mesenchyme | Rbm39 | 0.43 | 0 | 0 | 1 | 0.98 | 0.02 | 1.02 |
| mesenchyme | Ranbp1 | 0.44 | 0 | 0 | 1 | 0.98 | 0.02 | 1.02 |
| mesenchyme | Hnrnpab | 0.38 | 0 | 0 | 1 | 0.98 | 0.02 | 1.02 |
| mesenchyme | Hint1 | 0.44 | 0 | 0 | 1 | 0.98 | 0.02 | 1.02 |
| mesenchyme | Ndufa4 | 0.41 | 0 | 0 | 1 | 0.99 | 0.02 | 1.02 |
| mesenchyme | Atp5j | 0.38 | 0 | 0 | 1 | 0.98 | 0.02 | 1.02 |
| mesenchyme | Ddx5 | 0.37 | 0 | 0 | 1 | 0.99 | 0.01 | 1.01 |
| mesenchyme | Ywhae | 0.33 | 0 | 0 | 1 | 0.99 | 0.01 | 1.01 |
| mesenchyme | Actg1 | 0.4 | 0 | 0 | 1 | 0.99 | 0.01 | 1.01 |
| mesenchyme | Snrpg | 0.34 | 0 | 0 | 1 | 0.99 | 0.01 | 1.01 |
| mesenchyme | Pfn1 | 0.4 | 0 | 0 | 1 | 0.99 | 0.01 | 1.01 |
| mesenchyme | Cox6c | 0.36 | 0 | 0 | 1 | 0.99 | 0.01 | 1.01 |
| mesenchyme | Elob | 0.35 | 0 | 0 | 1 | 0.99 | 0.01 | 1.01 |
| mesenchyme | Srsf3 | 0.43 | 0 | 0 | 1 | 0.99 | 0.01 | 1.01 |
| mesenchyme | Ybx1 | 0.44 | 0 | 0 | 1 | 0.99 | 0.01 | 1.01 |
| mesenchyme | Dynll1 | 0.41 | 0 | 0 | 1 | 0.99 | 0.01 | 1.01 |
| mesenchyme | Ran | 0.34 | 0 | 0 | 1 | 0.99 | 0.01 | 1.01 |
| mesenchyme | Cfl1 | 0.37 | 0 | 0 | 1 | 0.99 | 0.01 | 1.01 |
| mesenchyme | Hnrnpa2b1 | 0.35 | 0 | 0 | 1 | 0.99 | 0.01 | 1.01 |
| mesenchyme | Sumo2 | 0.45 | 0 | 0 | 1 | 0.99 | 0.01 | 1.01 |
| mesenchyme | Tmsb10 | 0.65 | 0 | 0 | 1 | 0.99 | 0.01 | 1.01 |
| mesenchyme | Ubb | 0.41 | 0 | 0 | 1 | 0.99 | 0.01 | 1.01 |
| mesenchyme | Chchd2 | 0.43 | 0 | 0 | 1 | 1 | 0.01 | 1.01 |
| mesenchyme | Tubb5 | 0.41 | 0 | 0 | 1 | 1 | 0.01 | 1.01 |
| mesenchyme | H3f3a | 0.37 | 0 | 0 | 1 | 1 | 0 | 1 |
| mesenchyme | H2afz | 0.62 | 0 | 0 | 1 | 1 | 0 | 1 |
| mesenchyme | Hmgb1 | 0.66 | 0 | 0 | 1 | 1 | 0 | 1 |
| mesenchyme | Ptma | 0.29 | 0 | 0 | 1 | 1 | 0 | 1 |
| mesenchyme | Dnajc8 | 0.46 | 2.41E-307 | 5.01E-303 | 0.96 | 0.91 | 0.06 | 1.06 |
| mesenchyme | Eno3 | 1.61 | 3.02E-306 | 6.27E-302 | 0.46 | 0.2 | 0.25 | 2.26 |
| mesenchyme | Cdk2ap1 | 0.48 | 5.15E-306 | 1.07E-301 | 0.97 | 0.89 | 0.08 | 1.09 |
| mesenchyme | Dnaja1 | 0.39 | 5.58E-306 | 1.16E-301 | 1 | 0.98 | 0.02 | 1.02 |
| mesenchyme | Samd4 | 1.71 | 1.78E-305 | 3.70E-301 | 0.45 | 0.2 | 0.25 | 2.25 |
| mesenchyme | Ift74 | 0.75 | 3.11E-305 | 6.47E-301 | 0.81 | 0.6 | 0.21 | 1.34 |
| mesenchyme | Ctbp2 | 0.86 | 5.61E-302 | 1.17E-297 | 0.75 | 0.55 | 0.2 | 1.36 |
| mesenchyme | Sema5a | 1.74 | 7.01E-301 | 1.46E-296 | 0.46 | 0.21 | 0.25 | 2.2 |
| mesenchyme | Tpd52l1 | 1.95 | 9.19E-301 | 1.91E-296 | 0.41 | 0.17 | 0.24 | 2.44 |
| mesenchyme | Cpxm1 | 1.44 | 9.68E-301 | 2.01E-296 | 0.51 | 0.27 | 0.25 | 1.94 |
| mesenchyme | Stmn1 | 0.29 | 1.76E-299 | 3.67E-295 | 1 | 0.99 | 0.01 | 1.01 |
| mesenchyme | Jun | 1.05 | 3.41E-298 | 7.10E-294 | 0.74 | 0.55 | 0.19 | 1.35 |
| mesenchyme | Rtf1 | 0.51 | 4.54E-298 | 9.45E-294 | 0.95 | 0.86 | 0.09 | 1.11 |
| mesenchyme | Kank4 | 2.96 | 9.51E-298 | 1.98E-293 | 0.27 | 0.05 | 0.22 | 5.04 |
| mesenchyme | Grcc10 | 0.43 | 2.94E-297 | 6.11E-293 | 0.99 | 0.94 | 0.05 | 1.06 |
| mesenchyme | Hmgn1 | 0.33 | 3.45E-297 | 7.17E-293 | 1 | 0.97 | 0.03 | 1.03 |
| mesenchyme | Ldb2 | 1.89 | 4.32E-297 | 8.98E-293 | 0.35 | 0.11 | 0.24 | 3.24 |
| mesenchyme | Itga8 | 3.45 | 8.32E-297 | 1.73E-292 | 0.26 | 0.05 | 0.22 | 5.67 |
| mesenchyme | Skp2 | 0.81 | 4.38E-296 | 9.11E-292 | 0.77 | 0.55 | 0.21 | 1.39 |
| mesenchyme | Top1 | 0.33 | 2.36E-295 | 4.90E-291 | 1 | 0.99 | 0.01 | 1.01 |

**Supplementary Table 2. Marker genes for the major cell types of integrated clustering of E11.5 control and NotchGOF λ-junction cells.**

| | | | | | | | | |
|---|---|---|---|---|---|---|---|---|
| mesenchyme | Mesd | 0.69 | 5.47E-295 | 1.14E-290 | 0.83 | 0.66 | 0.18 | 1.27 |
| mesenchyme | Sec61b | 0.4 | 2.00E-294 | 4.16E-290 | 0.99 | 0.97 | 0.03 | 1.03 |
| mesenchyme | Metrn | 1.22 | 4.44E-293 | 9.24E-289 | 0.59 | 0.37 | 0.22 | 1.59 |
| mesenchyme | Rbm4b | 0.87 | 8.38E-293 | 1.74E-288 | 0.73 | 0.54 | 0.2 | 1.37 |
| mesenchyme | Foxf1 | 3.73 | 2.56E-292 | 5.32E-288 | 0.25 | 0.04 | 0.21 | 6.02 |
| mesenchyme | Atp5h | 0.27 | 2.90E-291 | 6.03E-287 | 1 | 0.99 | 0.01 | 1.01 |
| mesenchyme | Raet1e | 2.64 | 3.68E-291 | 7.65E-287 | 0.3 | 0.08 | 0.22 | 3.8 |
| mesenchyme | H2afy | 0.42 | 8.75E-291 | 1.82E-286 | 0.99 | 0.91 | 0.09 | 1.09 |
| mesenchyme | Dynlrb1 | 0.44 | 1.24E-290 | 2.59E-286 | 0.97 | 0.9 | 0.07 | 1.08 |
| mesenchyme | Hnrnpr | 0.43 | 2.94E-290 | 6.11E-286 | 0.99 | 0.93 | 0.06 | 1.07 |
| mesenchyme | Asxl3 | 1.37 | 6.34E-290 | 1.32E-285 | 0.54 | 0.3 | 0.24 | 1.78 |
| mesenchyme | Bzw1 | 0.4 | 2.40E-288 | 4.99E-284 | 0.99 | 0.95 | 0.04 | 1.04 |
| mesenchyme | Pygo1 | 1.24 | 2.44E-287 | 5.07E-283 | 0.55 | 0.3 | 0.24 | 1.8 |
| mesenchyme | Cbx5 | 0.45 | 1.33E-286 | 2.77E-282 | 0.98 | 0.91 | 0.07 | 1.08 |
| mesenchyme | Scaf11 | 0.5 | 1.78E-286 | 3.70E-282 | 0.96 | 0.87 | 0.09 | 1.1 |
| mesenchyme | Ssbp4 | 0.85 | 1.03E-285 | 2.14E-281 | 0.73 | 0.5 | 0.23 | 1.45 |
| mesenchyme | Zfp326 | 0.59 | 1.92E-285 | 3.98E-281 | 0.91 | 0.76 | 0.15 | 1.19 |
| mesenchyme | Rwdd1 | 0.48 | 2.94E-285 | 6.12E-281 | 0.96 | 0.88 | 0.08 | 1.09 |
| mesenchyme | Dusp9 | 1.38 | 6.57E-285 | 1.37E-280 | 0.49 | 0.25 | 0.24 | 1.95 |
| mesenchyme | Sumo1 | 0.39 | 7.21E-285 | 1.50E-280 | 0.99 | 0.94 | 0.05 | 1.05 |
| mesenchyme | Ttc28 | 0.96 | 1.67E-284 | 3.47E-280 | 0.67 | 0.44 | 0.23 | 1.53 |
| mesenchyme | Adgra2 | 2.37 | 1.73E-284 | 3.61E-280 | 0.31 | 0.09 | 0.22 | 3.59 |
| mesenchyme | Tcf4 | 0.57 | 4.83E-283 | 1.00E-278 | 0.95 | 0.79 | 0.16 | 1.21 |
| mesenchyme | Smarcc1 | 0.44 | 8.50E-283 | 1.77E-278 | 0.99 | 0.92 | 0.07 | 1.07 |
| mesenchyme | Smad7 | 0.79 | 1.20E-282 | 2.49E-278 | 0.77 | 0.55 | 0.22 | 1.4 |
| mesenchyme | Sox5 | 1.5 | 1.66E-282 | 3.45E-278 | 0.48 | 0.23 | 0.24 | 2.05 |
| mesenchyme | Reck | 2.42 | 9.98E-282 | 2.08E-277 | 0.32 | 0.1 | 0.22 | 3.32 |
| mesenchyme | Psma4 | 0.42 | 1.71E-281 | 3.56E-277 | 0.98 | 0.93 | 0.05 | 1.05 |
| mesenchyme | Fzd1 | 1.74 | 5.91E-281 | 1.23E-276 | 0.42 | 0.18 | 0.24 | 2.32 |
| mesenchyme | Marveld1 | 2.46 | 1.80E-280 | 3.75E-276 | 0.31 | 0.09 | 0.22 | 3.49 |
| mesenchyme | Gm8797 | 0.47 | 3.85E-280 | 8.00E-276 | 0.96 | 0.89 | 0.07 | 1.08 |
| mesenchyme | Ogfr | 0.83 | 1.57E-279 | 3.26E-275 | 0.73 | 0.55 | 0.18 | 1.33 |
| mesenchyme | Calm3 | 0.48 | 3.45E-279 | 7.17E-275 | 0.94 | 0.85 | 0.09 | 1.1 |
| mesenchyme | Gpx8 | 0.79 | 1.42E-278 | 2.95E-274 | 0.78 | 0.57 | 0.21 | 1.37 |
| mesenchyme | Hddc2 | 0.76 | 2.02E-278 | 4.20E-274 | 0.81 | 0.66 | 0.16 | 1.24 |
| mesenchyme | Bgn | 2.06 | 1.07E-277 | 2.23E-273 | 0.34 | 0.11 | 0.23 | 3.21 |
| mesenchyme | Ube2r2 | 0.43 | 3.76E-277 | 7.81E-273 | 0.98 | 0.92 | 0.06 | 1.06 |
| mesenchyme | Chchd1 | 0.46 | 1.41E-276 | 2.94E-272 | 0.96 | 0.9 | 0.07 | 1.07 |
| mesenchyme | Hpse2 | 3.55 | 2.66E-276 | 5.53E-272 | 0.24 | 0.04 | 0.2 | 6.56 |
| mesenchyme | Phf5a | 0.42 | 1.23E-275 | 2.57E-271 | 0.97 | 0.91 | 0.06 | 1.07 |
| mesenchyme | Atp5f1 | 0.3 | 6.55E-275 | 1.36E-270 | 1 | 0.98 | 0.02 | 1.02 |
| mesenchyme | Cdc26 | 0.48 | 8.71E-275 | 1.81E-270 | 0.96 | 0.88 | 0.08 | 1.09 |
| mesenchyme | Rbmxl1 | 0.53 | 1.24E-274 | 2.58E-270 | 0.93 | 0.81 | 0.12 | 1.14 |
| mesenchyme | Cks1b | 0.45 | 3.11E-274 | 6.46E-270 | 0.98 | 0.92 | 0.05 | 1.06 |
| mesenchyme | Adgrl3 | 1.39 | 5.48E-274 | 1.14E-269 | 0.52 | 0.29 | 0.23 | 1.81 |
| mesenchyme | Rrm2 | 0.77 | 6.69E-274 | 1.39E-269 | 0.85 | 0.72 | 0.13 | 1.18 |
| mesenchyme | Tcf15 | 1.85 | 1.46E-273 | 3.03E-269 | 0.36 | 0.12 | 0.24 | 2.9 |
| mesenchyme | Zfp637 | 0.78 | 1.44E-272 | 2.99E-268 | 0.78 | 0.63 | 0.16 | 1.25 |
| mesenchyme | Asb4 | 1.36 | 1.58E-272 | 3.29E-268 | 0.38 | 0.13 | 0.25 | 2.9 |
| mesenchyme | Smo | 0.75 | 1.75E-272 | 3.64E-268 | 0.79 | 0.61 | 0.19 | 1.31 |
| mesenchyme | 130114P18Ri | 2.38 | 2.88E-272 | 6.00E-268 | 0.3 | 0.08 | 0.22 | 3.73 |
| mesenchyme | Smarcd1 | 0.61 | 3.14E-272 | 6.53E-268 | 0.89 | 0.74 | 0.15 | 1.2 |
| mesenchyme | Aldh2 | 0.64 | 3.78E-272 | 7.86E-268 | 0.83 | 0.66 | 0.17 | 1.26 |
| mesenchyme | Cnih2 | 1.44 | 5.71E-272 | 1.19E-267 | 0.41 | 0.17 | 0.24 | 2.43 |
| mesenchyme | Fkbp7 | 1.99 | 1.61E-271 | 3.36E-267 | 0.36 | 0.13 | 0.23 | 2.74 |
| mesenchyme | Tcaf1 | 0.8 | 2.39E-271 | 4.98E-267 | 0.74 | 0.53 | 0.21 | 1.4 |

**Supplementary Table 2. Marker genes for the major cell types of integrated clustering of E11.5 control and NotchGOF λ-junction cells.**

| | | | | | | | | |
|---|---|---|---|---|---|---|---|---|
| mesenchyme | Mrpl20 | 0.46 | 1.90E-270 | 3.96E-266 | 0.95 | 0.89 | 0.06 | 1.07 |
| mesenchyme | Tns3 | 1.13 | 1.88E-269 | 3.90E-265 | 0.57 | 0.34 | 0.23 | 1.68 |
| mesenchyme | Dld | 0.7 | 2.20E-269 | 4.57E-265 | 0.81 | 0.68 | 0.14 | 1.2 |
| mesenchyme | Sem1 | 0.29 | 1.98E-268 | 4.11E-264 | 1 | 0.99 | 0.01 | 1.01 |
| mesenchyme | Tceal8 | 0.52 | 2.29E-267 | 4.77E-263 | 0.92 | 0.82 | 0.1 | 1.12 |
| mesenchyme | Septin9 | 0.85 | 2.37E-267 | 4.93E-263 | 0.7 | 0.51 | 0.2 | 1.39 |
| mesenchyme | )610010K14Ri | 0.62 | 2.55E-267 | 5.30E-263 | 0.86 | 0.73 | 0.13 | 1.18 |
| mesenchyme | Gmps | 0.51 | 2.89E-267 | 6.01E-263 | 0.93 | 0.83 | 0.1 | 1.12 |
| mesenchyme | Ndufc1 | 0.39 | 1.07E-266 | 2.22E-262 | 0.98 | 0.94 | 0.04 | 1.04 |
| mesenchyme | Snhg18 | 1.12 | 3.34E-266 | 6.95E-262 | 0.59 | 0.37 | 0.22 | 1.59 |
| mesenchyme | Adamts1 | 2.12 | 6.32E-266 | 1.32E-261 | 0.35 | 0.12 | 0.22 | 2.8 |
| mesenchyme | Pdcd5 | 0.38 | 6.48E-264 | 1.35E-259 | 0.98 | 0.95 | 0.04 | 1.04 |
| mesenchyme | Gpx7 | 1.25 | 5.44E-263 | 1.13E-258 | 0.52 | 0.3 | 0.23 | 1.76 |
| mesenchyme | Eif4a1 | 0.28 | 5.04E-262 | 1.05E-257 | 1 | 0.99 | 0.01 | 1.01 |
| mesenchyme | Nisch | 0.5 | 6.49E-262 | 1.35E-257 | 0.95 | 0.84 | 0.12 | 1.14 |
| mesenchyme | Kank2 | 1.79 | 7.17E-261 | 1.49E-256 | 0.38 | 0.16 | 0.22 | 2.4 |
| mesenchyme | Fap | 2.78 | 1.20E-260 | 2.50E-256 | 0.26 | 0.06 | 0.2 | 4.26 |
| mesenchyme | Gga2 | 0.84 | 1.29E-260 | 2.69E-256 | 0.7 | 0.52 | 0.18 | 1.35 |
| mesenchyme | Cbx3 | 0.37 | 1.45E-260 | 3.02E-256 | 0.99 | 0.96 | 0.03 | 1.03 |
| mesenchyme | Taf1d | 0.53 | 8.58E-260 | 1.78E-255 | 0.92 | 0.83 | 0.1 | 1.12 |
| mesenchyme | Ppic | 0.56 | 9.29E-260 | 1.93E-255 | 0.78 | 0.56 | 0.22 | 1.39 |
| mesenchyme | Tes | 0.79 | 1.04E-258 | 2.16E-254 | 0.76 | 0.56 | 0.19 | 1.34 |
| mesenchyme | Tbcb | 0.5 | 1.05E-258 | 2.19E-254 | 0.92 | 0.81 | 0.12 | 1.14 |
| mesenchyme | Mrpl11 | 0.49 | 1.17E-258 | 2.44E-254 | 0.93 | 0.84 | 0.09 | 1.1 |
| mesenchyme | Lhfpl2 | 1.67 | 2.89E-258 | 6.00E-254 | 0.41 | 0.18 | 0.22 | 2.23 |
| mesenchyme | Prdx4 | 0.51 | 5.09E-258 | 1.06E-253 | 0.93 | 0.81 | 0.13 | 1.15 |
| mesenchyme | Gtf2h5 | 0.42 | 1.35E-257 | 2.81E-253 | 0.96 | 0.91 | 0.05 | 1.06 |
| mesenchyme | Denr | 0.46 | 5.33E-257 | 1.11E-252 | 0.95 | 0.88 | 0.07 | 1.08 |
| mesenchyme | Atp1b3 | 0.77 | 1.31E-256 | 2.73E-252 | 0.77 | 0.61 | 0.16 | 1.25 |
| mesenchyme | Chd4 | 0.44 | 7.70E-256 | 1.60E-251 | 0.98 | 0.91 | 0.08 | 1.08 |
| mesenchyme | mt-Nd1 | 0.33 | 1.64E-254 | 3.40E-250 | 1 | 0.96 | 0.04 | 1.04 |
| mesenchyme | Zcrb1 | 0.46 | 2.47E-254 | 5.14E-250 | 0.95 | 0.89 | 0.06 | 1.07 |
| mesenchyme | Ddx39b | 0.4 | 2.40E-253 | 4.98E-249 | 0.99 | 0.94 | 0.05 | 1.05 |
| mesenchyme | Tcea1 | 0.5 | 1.75E-252 | 3.65E-248 | 0.91 | 0.81 | 0.1 | 1.12 |
| mesenchyme | Lyar | 0.53 | 2.48E-250 | 5.15E-246 | 0.9 | 0.79 | 0.11 | 1.13 |
| mesenchyme | Tpr | 0.45 | 3.45E-250 | 7.17E-246 | 0.99 | 0.95 | 0.04 | 1.05 |
| mesenchyme | Atp5j2 | 0.26 | 4.53E-250 | 9.43E-246 | 1 | 0.98 | 0.01 | 1.01 |
| mesenchyme | U2af1 | 0.38 | 9.16E-250 | 1.91E-245 | 0.98 | 0.94 | 0.04 | 1.05 |
| mesenchyme | Tmem167 | 0.41 | 1.70E-249 | 3.53E-245 | 0.98 | 0.91 | 0.07 | 1.08 |
| mesenchyme | Hook3 | 0.8 | 3.32E-249 | 6.91E-245 | 0.72 | 0.54 | 0.18 | 1.33 |
| mesenchyme | Anp32e | 0.42 | 2.29E-248 | 4.76E-244 | 0.96 | 0.91 | 0.05 | 1.05 |
| mesenchyme | Zc3h15 | 0.4 | 3.81E-248 | 7.93E-244 | 0.97 | 0.92 | 0.05 | 1.06 |
| mesenchyme | Top2b | 0.47 | 1.63E-247 | 3.39E-243 | 0.96 | 0.85 | 0.1 | 1.12 |
| mesenchyme | Rhoq | 1.21 | 2.52E-247 | 5.23E-243 | 0.52 | 0.3 | 0.22 | 1.72 |
| mesenchyme | Bcat1 | 1.61 | 3.82E-247 | 7.94E-243 | 0.38 | 0.16 | 0.22 | 2.34 |
| mesenchyme | Sncaip | 1.77 | 6.11E-247 | 1.27E-242 | 0.37 | 0.16 | 0.22 | 2.38 |
| mesenchyme | Nono | 0.36 | 1.25E-246 | 2.61E-242 | 0.99 | 0.94 | 0.05 | 1.05 |
| mesenchyme | Tcf3 | 0.5 | 1.35E-246 | 2.82E-242 | 0.93 | 0.83 | 0.11 | 1.13 |
| mesenchyme | Cyp51 | 0.79 | 1.27E-245 | 2.65E-241 | 0.75 | 0.57 | 0.18 | 1.32 |
| mesenchyme | Hnrnph1 | 0.41 | 2.03E-244 | 4.23E-240 | 0.99 | 0.91 | 0.07 | 1.08 |
| mesenchyme | Map7d1 | 0.84 | 2.44E-244 | 5.06E-240 | 0.68 | 0.49 | 0.19 | 1.39 |
| mesenchyme | Txn1 | 0.33 | 6.12E-244 | 1.27E-239 | 1 | 0.98 | 0.02 | 1.02 |
| mesenchyme | Cct6a | 0.37 | 6.30E-244 | 1.31E-239 | 0.99 | 0.96 | 0.03 | 1.04 |
| mesenchyme | Rsrc2 | 0.44 | 7.90E-244 | 1.64E-239 | 0.96 | 0.88 | 0.08 | 1.09 |
| mesenchyme | Rbm28 | 0.65 | 1.04E-243 | 2.17E-239 | 0.8 | 0.65 | 0.14 | 1.22 |
| mesenchyme | Adamts19 | 2.03 | 4.17E-243 | 8.67E-239 | 0.32 | 0.11 | 0.21 | 2.97 |

**Supplementary Table 2. Marker genes for the major cell types of integrated clustering of E11.5 control and NotchGOF λ-junction cells.**

| | | | | | | | | |
|---|---|---|---|---|---|---|---|---|
| mesenchyme | Tsen34 | 0.54 | 1.03E-242 | 2.13E-238 | 0.9 | 0.73 | 0.17 | 1.23 |
| mesenchyme | Tma7 | 0.29 | 1.38E-242 | 2.87E-238 | 1 | 0.99 | 0.01 | 1.01 |
| mesenchyme | Ankrd44 | 1.81 | 2.92E-242 | 6.07E-238 | 0.35 | 0.14 | 0.21 | 2.57 |
| mesenchyme | Cfdp1 | 0.42 | 3.35E-242 | 6.97E-238 | 0.96 | 0.89 | 0.08 | 1.08 |
| mesenchyme | Slitrk6 | 1.8 | 3.54E-242 | 7.36E-238 | 0.4 | 0.18 | 0.22 | 2.21 |
| mesenchyme | Matr3 | 0.48 | 6.95E-242 | 1.45E-237 | 0.93 | 0.83 | 0.1 | 1.12 |
| mesenchyme | Hsp90b1 | 0.41 | 6.99E-242 | 1.45E-237 | 1 | 0.93 | 0.06 | 1.07 |
| mesenchyme | Hspa9 | 0.51 | 1.18E-241 | 2.45E-237 | 0.92 | 0.83 | 0.09 | 1.11 |
| mesenchyme | Fat4 | 2 | 1.41E-241 | 2.93E-237 | 0.34 | 0.14 | 0.21 | 2.55 |
| mesenchyme | Dact1 | 1.37 | 1.82E-241 | 3.78E-237 | 0.39 | 0.16 | 0.23 | 2.41 |
| mesenchyme | Cct8 | 0.34 | 2.96E-240 | 6.15E-236 | 0.99 | 0.96 | 0.03 | 1.03 |
| mesenchyme | Ywhaq | 0.34 | 3.78E-240 | 7.86E-236 | 0.99 | 0.96 | 0.03 | 1.03 |
| mesenchyme | Dhx36 | 0.49 | 4.16E-240 | 8.65E-236 | 0.94 | 0.83 | 0.1 | 1.12 |
| mesenchyme | Krtcap2 | 0.42 | 7.23E-240 | 1.50E-235 | 0.97 | 0.87 | 0.1 | 1.11 |
| mesenchyme | Phf6 | 0.7 | 2.03E-239 | 4.21E-235 | 0.77 | 0.62 | 0.15 | 1.24 |
| mesenchyme | Ndufa13 | 0.37 | 2.96E-239 | 6.15E-235 | 0.99 | 0.95 | 0.04 | 1.04 |
| mesenchyme | Kctd15 | 1.26 | 7.97E-239 | 1.66E-234 | 0.49 | 0.28 | 0.21 | 1.77 |
| mesenchyme | Ppp2ca | 0.38 | 9.16E-239 | 1.91E-234 | 0.97 | 0.92 | 0.05 | 1.06 |
| mesenchyme | Polr2i | 0.51 | 9.54E-238 | 1.98E-233 | 0.9 | 0.81 | 0.09 | 1.11 |
| mesenchyme | Ubap2l | 0.42 | 4.90E-236 | 1.02E-231 | 0.96 | 0.89 | 0.08 | 1.08 |
| mesenchyme | Peg12 | 2 | 4.95E-236 | 1.03E-231 | 0.32 | 0.11 | 0.21 | 2.8 |
| mesenchyme | Septin3 | 1.07 | 9.30E-236 | 1.93E-231 | 0.51 | 0.29 | 0.22 | 1.76 |
| mesenchyme | Gspt1 | 0.44 | 1.66E-235 | 3.46E-231 | 0.95 | 0.88 | 0.07 | 1.08 |
| mesenchyme | Timm22 | 0.62 | 2.35E-235 | 4.89E-231 | 0.82 | 0.67 | 0.15 | 1.22 |
| mesenchyme | Mtdh | 0.44 | 5.96E-234 | 1.24E-229 | 0.97 | 0.86 | 0.11 | 1.13 |
| mesenchyme | Tln2 | 0.9 | 1.28E-233 | 2.65E-229 | 0.65 | 0.45 | 0.2 | 1.45 |
| mesenchyme | 430049B03Ri | 1.33 | 1.46E-233 | 3.03E-229 | 0.43 | 0.21 | 0.22 | 2.06 |
| mesenchyme | Hspa5 | 0.4 | 1.98E-233 | 4.13E-229 | 0.99 | 0.92 | 0.07 | 1.08 |
| mesenchyme | Nsg1 | 0.66 | 2.91E-233 | 6.05E-229 | 0.74 | 0.52 | 0.22 | 1.42 |
| mesenchyme | Ube2i | 0.33 | 3.98E-233 | 8.28E-229 | 0.99 | 0.97 | 0.02 | 1.02 |
| mesenchyme | Smarce1 | 0.38 | 1.69E-232 | 3.51E-228 | 0.98 | 0.92 | 0.06 | 1.07 |
| mesenchyme | Cops8 | 0.56 | 1.99E-232 | 4.14E-228 | 0.85 | 0.72 | 0.13 | 1.18 |
| mesenchyme | P3h3 | 1.74 | 5.73E-232 | 1.19E-227 | 0.37 | 0.17 | 0.21 | 2.24 |
| mesenchyme | Foxp2 | 2.4 | 7.43E-231 | 1.54E-226 | 0.41 | 0.22 | 0.19 | 1.88 |
| mesenchyme | Bcl7c | 0.55 | 1.15E-230 | 2.38E-226 | 0.87 | 0.74 | 0.13 | 1.17 |
| mesenchyme | 110004F10Ri | 0.38 | 7.37E-230 | 1.53E-225 | 0.97 | 0.91 | 0.06 | 1.06 |
| mesenchyme | Msx1 | 1.03 | 7.70E-230 | 1.60E-225 | 0.75 | 0.66 | 0.09 | 1.13 |
| mesenchyme | Tnrc6a | 0.61 | 1.44E-228 | 3.00E-224 | 0.82 | 0.69 | 0.13 | 1.19 |
| mesenchyme | Actl6a | 0.49 | 4.25E-228 | 8.84E-224 | 0.91 | 0.8 | 0.1 | 1.13 |
| mesenchyme | Gpc6 | 0.99 | 5.09E-228 | 1.06E-223 | 0.65 | 0.45 | 0.2 | 1.43 |
| mesenchyme | Drap1 | 0.43 | 3.67E-227 | 7.64E-223 | 0.96 | 0.88 | 0.07 | 1.08 |
| mesenchyme | Tspan11 | 2.94 | 1.16E-226 | 2.41E-222 | 0.23 | 0.05 | 0.18 | 4.79 |
| mesenchyme | Tro | 1.16 | 1.31E-226 | 2.72E-222 | 0.49 | 0.28 | 0.21 | 1.76 |
| mesenchyme | Tnrc6c | 0.58 | 1.57E-226 | 3.27E-222 | 0.84 | 0.71 | 0.13 | 1.18 |
| mesenchyme | Tagln3 | 0.8 | 1.13E-225 | 2.34E-221 | 0.33 | 0.12 | 0.21 | 2.7 |
| mesenchyme | Selenof | 0.4 | 6.70E-225 | 1.39E-220 | 0.98 | 0.92 | 0.06 | 1.06 |
| mesenchyme | Atp5md | 0.3 | 1.78E-224 | 3.70E-220 | 0.99 | 0.97 | 0.02 | 1.02 |
| mesenchyme | Pfdn5 | 0.3 | 3.47E-224 | 7.21E-220 | 1 | 0.98 | 0.02 | 1.02 |
| mesenchyme | Nrp2 | 1.67 | 4.45E-224 | 9.26E-220 | 0.44 | 0.23 | 0.21 | 1.92 |
| mesenchyme | Snrpf | 0.3 | 5.95E-224 | 1.24E-219 | 1 | 0.98 | 0.02 | 1.02 |
| mesenchyme | Eif2s2 | 0.31 | 1.63E-222 | 3.40E-218 | 1 | 0.98 | 0.02 | 1.02 |
| mesenchyme | Brd3 | 0.43 | 1.65E-222 | 3.44E-218 | 0.96 | 0.86 | 0.11 | 1.13 |
| mesenchyme | Psmc6 | 0.4 | 5.20E-222 | 1.08E-217 | 0.96 | 0.9 | 0.05 | 1.06 |
| mesenchyme | Idh1 | 0.79 | 1.73E-221 | 3.60E-217 | 0.69 | 0.51 | 0.18 | 1.34 |
| mesenchyme | Fam89b | 0.56 | 2.42E-221 | 5.03E-217 | 0.84 | 0.71 | 0.13 | 1.18 |
| mesenchyme | Zfp644 | 0.51 | 9.64E-221 | 2.00E-216 | 0.9 | 0.78 | 0.13 | 1.16 |

**Supplementary Table 2. Marker genes for the major cell types of integrated clustering of E11.5 control and NotchGOF λ-junction cells.**

| | | | | | | | | |
|---|---|---|---|---|---|---|---|---|
| mesenchyme | Ndufaf2 | 0.64 | 9.97E-221 | 2.07E-216 | 0.78 | 0.65 | 0.13 | 1.21 |
| mesenchyme | Snrpc | 0.38 | 3.51E-220 | 7.29E-216 | 0.97 | 0.91 | 0.05 | 1.06 |
| mesenchyme | Atp5a1 | 0.28 | 1.25E-219 | 2.61E-215 | 1 | 0.98 | 0.02 | 1.02 |
| mesenchyme | Zfp57 | 1.45 | 1.41E-219 | 2.93E-215 | 0.4 | 0.2 | 0.2 | 2.02 |
| mesenchyme | Col6a3 | 2.96 | 1.71E-219 | 3.57E-215 | 0.22 | 0.05 | 0.18 | 4.65 |
| mesenchyme | Edil3 | 2.99 | 2.31E-219 | 4.81E-215 | 0.23 | 0.05 | 0.18 | 4.26 |
| mesenchyme | Cdh6 | 1.66 | 3.99E-219 | 8.29E-215 | 0.35 | 0.14 | 0.21 | 2.45 |
| mesenchyme | Mrpl13 | 0.44 | 5.58E-219 | 1.16E-214 | 0.93 | 0.85 | 0.08 | 1.1 |
| mesenchyme | Col6a1 | 3.45 | 1.93E-218 | 4.02E-214 | 0.2 | 0.03 | 0.17 | 6.03 |
| mesenchyme | Aff3 | 2.17 | 3.16E-218 | 6.57E-214 | 0.28 | 0.09 | 0.19 | 3.05 |
| mesenchyme | Mrpl42 | 0.4 | 3.21E-218 | 6.68E-214 | 0.96 | 0.9 | 0.05 | 1.06 |
| mesenchyme | Fbxl7 | 1.14 | 7.38E-218 | 1.53E-213 | 0.53 | 0.33 | 0.2 | 1.61 |
| mesenchyme | Sf3b1 | 0.35 | 9.34E-218 | 1.94E-213 | 0.99 | 0.95 | 0.04 | 1.04 |
| mesenchyme | Smc6 | 0.51 | 1.74E-217 | 3.63E-213 | 0.9 | 0.81 | 0.09 | 1.11 |
| mesenchyme | Enah | 0.5 | 2.00E-217 | 4.15E-213 | 0.91 | 0.75 | 0.16 | 1.21 |
| mesenchyme | Pmaip1 | 2.39 | 2.25E-217 | 4.68E-213 | 0.31 | 0.12 | 0.19 | 2.66 |
| mesenchyme | B2m | 0.37 | 8.24E-217 | 1.71E-212 | 0.61 | 0.4 | 0.22 | 1.54 |
| mesenchyme | Slc39a6 | 0.86 | 1.40E-216 | 2.91E-212 | 0.63 | 0.44 | 0.2 | 1.45 |
| mesenchyme | Hspe1 | 0.36 | 7.09E-216 | 1.47E-211 | 1 | 0.98 | 0.02 | 1.02 |
| mesenchyme | Hnrnpa1 | 0.29 | 4.91E-215 | 1.02E-210 | 1 | 0.98 | 0.02 | 1.02 |
| mesenchyme | Mns1 | 0.91 | 4.69E-214 | 9.76E-210 | 0.6 | 0.43 | 0.17 | 1.41 |
| mesenchyme | Pdap1 | 0.3 | 8.90E-213 | 1.85E-208 | 1 | 0.98 | 0.02 | 1.02 |
| mesenchyme | Col6a2 | 3.4 | 1.09E-212 | 2.26E-208 | 0.2 | 0.04 | 0.17 | 5.46 |
| mesenchyme | Rbpms2 | 1.67 | 1.22E-212 | 2.55E-208 | 0.29 | 0.1 | 0.19 | 2.89 |
| mesenchyme | Mrfap1 | 0.31 | 1.70E-212 | 3.53E-208 | 1 | 0.97 | 0.03 | 1.03 |
| mesenchyme | S1pr2 | 1.49 | 2.59E-212 | 5.38E-208 | 0.39 | 0.18 | 0.2 | 2.09 |
| mesenchyme | Idh2 | 0.47 | 3.85E-212 | 8.02E-208 | 0.94 | 0.81 | 0.14 | 1.17 |
| mesenchyme | Ddit4l | 2.41 | 4.54E-212 | 9.44E-208 | 0.27 | 0.09 | 0.19 | 3.1 |
| mesenchyme | Iah1 | 0.97 | 8.84E-212 | 1.84E-207 | 0.58 | 0.41 | 0.17 | 1.43 |
| mesenchyme | 330403K07Ri | 0.46 | 1.48E-211 | 3.08E-207 | 0.91 | 0.73 | 0.19 | 1.26 |
| mesenchyme | Elk3 | 0.63 | 1.84E-211 | 3.82E-207 | 0.49 | 0.26 | 0.23 | 1.87 |
| mesenchyme | Psmb1 | 0.3 | 2.70E-211 | 5.61E-207 | 0.99 | 0.97 | 0.03 | 1.03 |
| mesenchyme | Fez1 | 3.13 | 3.91E-211 | 8.12E-207 | 0.2 | 0.04 | 0.17 | 5.58 |
| mesenchyme | Hspd1 | 0.38 | 5.41E-211 | 1.13E-206 | 1 | 0.95 | 0.05 | 1.05 |
| mesenchyme | B3gnt5 | 1.29 | 1.82E-210 | 3.79E-206 | 0.42 | 0.22 | 0.2 | 1.89 |
| mesenchyme | Morf4l2 | 0.39 | 1.99E-210 | 4.14E-206 | 0.96 | 0.88 | 0.09 | 1.1 |
| mesenchyme | Nid2 | 1.76 | 3.30E-210 | 6.86E-206 | 0.3 | 0.11 | 0.19 | 2.76 |
| mesenchyme | Gxylt2 | 1.71 | 5.02E-210 | 1.04E-205 | 0.32 | 0.12 | 0.2 | 2.65 |
| mesenchyme | 110038B12Ri | 0.42 | 5.26E-210 | 1.09E-205 | 0.96 | 0.87 | 0.08 | 1.1 |
| mesenchyme | Kdm2b | 0.9 | 6.04E-210 | 1.26E-205 | 0.62 | 0.45 | 0.18 | 1.39 |
| mesenchyme | Casp8ap2 | 0.6 | 9.70E-210 | 2.02E-205 | 0.84 | 0.7 | 0.14 | 1.19 |
| mesenchyme | Snrpb2 | 0.49 | 1.49E-209 | 3.09E-205 | 0.89 | 0.78 | 0.12 | 1.15 |
| mesenchyme | Cbfb | 0.85 | 1.56E-209 | 3.24E-205 | 0.63 | 0.45 | 0.18 | 1.4 |
| mesenchyme | Mcm4 | 0.69 | 2.00E-209 | 4.15E-205 | 0.79 | 0.63 | 0.16 | 1.26 |
| mesenchyme | Dcp2 | 0.75 | 1.47E-207 | 3.07E-203 | 0.68 | 0.52 | 0.16 | 1.31 |
| mesenchyme | Ndufab1 | 0.32 | 2.14E-207 | 4.45E-203 | 0.99 | 0.95 | 0.04 | 1.04 |
| mesenchyme | Pmp22 | 2.26 | 2.21E-207 | 4.59E-203 | 0.24 | 0.07 | 0.18 | 3.72 |
| mesenchyme | Psmb6 | 0.3 | 2.80E-207 | 5.81E-203 | 0.99 | 0.97 | 0.02 | 1.02 |
| mesenchyme | Psmb4 | 0.35 | 3.03E-207 | 6.31E-203 | 0.98 | 0.94 | 0.04 | 1.04 |
| mesenchyme | Cox5a | 0.3 | 7.01E-207 | 1.46E-202 | 1 | 0.98 | 0.02 | 1.02 |
| mesenchyme | Ndufs5 | 0.36 | 1.08E-206 | 2.25E-202 | 0.97 | 0.92 | 0.04 | 1.05 |
| mesenchyme | Alyref | 0.37 | 2.30E-206 | 4.78E-202 | 0.97 | 0.9 | 0.06 | 1.07 |
| mesenchyme | Sptssa | 0.53 | 3.76E-206 | 7.81E-202 | 0.85 | 0.71 | 0.14 | 1.2 |
| mesenchyme | Rbmx | 0.59 | 7.63E-206 | 1.59E-201 | 0.8 | 0.64 | 0.16 | 1.24 |
| mesenchyme | Clns1a | 0.45 | 1.10E-205 | 2.29E-201 | 0.92 | 0.81 | 0.1 | 1.13 |
| mesenchyme | Cct2 | 0.31 | 1.43E-205 | 2.97E-201 | 0.99 | 0.97 | 0.03 | 1.03 |

**Supplementary Table 2. Marker genes for the major cell types of integrated clustering of E11.5 control and NotchGOF λ-junction cells.**

| | | | | | | | | |
|---|---|---|---|---|---|---|---|---|
| mesenchyme | Psmd4 | 0.35 | 8.07E-205 | 1.68E-200 | 0.97 | 0.93 | 0.04 | 1.04 |
| mesenchyme | Ndufb5 | 0.35 | 1.68E-204 | 3.49E-200 | 0.99 | 0.94 | 0.05 | 1.05 |
| mesenchyme | Tra2a | 0.41 | 2.40E-204 | 4.99E-200 | 0.97 | 0.88 | 0.08 | 1.1 |
| mesenchyme | Rab3il1 | 1.39 | 1.03E-203 | 2.15E-199 | 0.36 | 0.17 | 0.19 | 2.16 |
| mesenchyme | Selenok | 0.42 | 2.21E-203 | 4.59E-199 | 0.94 | 0.87 | 0.07 | 1.08 |
| mesenchyme | Bag1 | 0.4 | 3.41E-203 | 7.09E-199 | 0.94 | 0.88 | 0.06 | 1.07 |
| mesenchyme | Cdc37 | 0.35 | 3.68E-203 | 7.66E-199 | 0.98 | 0.92 | 0.05 | 1.06 |
| mesenchyme | Sfrp1 | 0.62 | 3.83E-203 | 7.97E-199 | 0.71 | 0.46 | 0.25 | 1.53 |
| mesenchyme | Zmat2 | 0.38 | 4.30E-203 | 8.95E-199 | 0.96 | 0.92 | 0.04 | 1.05 |
| mesenchyme | St3gal4 | 1.79 | 5.28E-203 | 1.10E-198 | 0.31 | 0.13 | 0.19 | 2.48 |
| mesenchyme | Dcun1d5 | 0.48 | 7.68E-203 | 1.60E-198 | 0.89 | 0.79 | 0.1 | 1.13 |
| mesenchyme | Immt | 0.46 | 7.96E-203 | 1.66E-198 | 0.91 | 0.83 | 0.08 | 1.09 |
| mesenchyme | St13 | 0.35 | 1.05E-202 | 2.17E-198 | 0.98 | 0.93 | 0.05 | 1.05 |
| mesenchyme | Thrap3 | 0.42 | 1.71E-202 | 3.55E-198 | 0.95 | 0.88 | 0.08 | 1.09 |
| mesenchyme | Nenf | 0.58 | 1.82E-202 | 3.79E-198 | 0.83 | 0.69 | 0.14 | 1.2 |
| mesenchyme | Fbn1 | 1.48 | 2.21E-202 | 4.60E-198 | 0.4 | 0.2 | 0.19 | 1.95 |
| mesenchyme | Dnm3 | 2.56 | 2.56E-202 | 5.32E-198 | 0.22 | 0.05 | 0.17 | 4.36 |
| mesenchyme | Ddx50 | 0.56 | 3.58E-202 | 7.44E-198 | 0.84 | 0.71 | 0.13 | 1.18 |
| mesenchyme | Hmgn3 | 0.53 | 4.84E-202 | 1.01E-197 | 0.81 | 0.63 | 0.18 | 1.28 |
| mesenchyme | Eif3e | 0.33 | 8.43E-202 | 1.75E-197 | 0.99 | 0.96 | 0.03 | 1.03 |
| mesenchyme | Lum | 3.22 | 1.23E-201 | 2.56E-197 | 0.21 | 0.05 | 0.16 | 4.42 |
| mesenchyme | Rrm1 | 0.54 | 1.94E-200 | 4.03E-196 | 0.85 | 0.74 | 0.12 | 1.16 |
| mesenchyme | Anp32b | 0.26 | 3.13E-200 | 6.51E-196 | 1 | 0.98 | 0.02 | 1.02 |
| mesenchyme | Praf2 | 0.86 | 6.47E-200 | 1.35E-195 | 0.59 | 0.41 | 0.18 | 1.45 |
| mesenchyme | Kctd12 | 1.59 | 8.80E-200 | 1.83E-195 | 0.36 | 0.17 | 0.19 | 2.15 |
| mesenchyme | Gpc3 | 0.52 | 9.46E-200 | 1.97E-195 | 0.9 | 0.7 | 0.2 | 1.29 |
| mesenchyme | Cul4b | 0.81 | 1.14E-199 | 2.37E-195 | 0.67 | 0.51 | 0.16 | 1.31 |
| mesenchyme | Traf4 | 0.72 | 2.48E-199 | 5.17E-195 | 0.69 | 0.52 | 0.17 | 1.32 |
| mesenchyme | Foxd2os | 3.29 | 4.30E-199 | 8.94E-195 | 0.2 | 0.04 | 0.16 | 4.72 |
| mesenchyme | Col25a1 | 2.38 | 4.35E-199 | 9.05E-195 | 0.26 | 0.08 | 0.18 | 3.19 |
| mesenchyme | Olfml1 | 3.87 | 5.82E-199 | 1.21E-194 | 0.17 | 0.02 | 0.15 | 8.35 |
| mesenchyme | Dhx9 | 0.44 | 9.73E-199 | 2.02E-194 | 0.93 | 0.84 | 0.09 | 1.11 |
| mesenchyme | Wdr82 | 0.79 | 1.68E-198 | 3.50E-194 | 0.64 | 0.47 | 0.17 | 1.36 |
| mesenchyme | Prpf40a | 0.38 | 4.59E-198 | 9.54E-194 | 0.97 | 0.91 | 0.06 | 1.07 |
| mesenchyme | Eny2 | 0.34 | 4.59E-198 | 9.55E-194 | 0.98 | 0.93 | 0.05 | 1.05 |
| mesenchyme | Slc50a1 | 1.26 | 1.26E-197 | 2.62E-193 | 0.44 | 0.25 | 0.19 | 1.78 |
| mesenchyme | Nudcd2 | 0.43 | 6.03E-197 | 1.26E-192 | 0.92 | 0.84 | 0.08 | 1.1 |
| mesenchyme | Cand2 | 0.98 | 6.12E-197 | 1.27E-192 | 0.55 | 0.37 | 0.18 | 1.49 |
| mesenchyme | Pbdc1 | 0.45 | 1.79E-196 | 3.72E-192 | 0.89 | 0.8 | 0.09 | 1.12 |
| mesenchyme | Echs1 | 0.46 | 1.88E-196 | 3.92E-192 | 0.9 | 0.78 | 0.12 | 1.16 |
| mesenchyme | Adamts10 | 1.48 | 3.68E-196 | 7.66E-192 | 0.39 | 0.2 | 0.19 | 1.94 |
| mesenchyme | Cfl2 | 0.51 | 4.55E-196 | 9.46E-192 | 0.85 | 0.72 | 0.14 | 1.19 |
| mesenchyme | Sf3a3 | 0.45 | 4.98E-196 | 1.04E-191 | 0.89 | 0.79 | 0.1 | 1.12 |
| mesenchyme | Prkar2b | 0.83 | 1.16E-195 | 2.40E-191 | 0.55 | 0.37 | 0.18 | 1.47 |
| mesenchyme | Prickle1 | 1.93 | 1.45E-195 | 3.01E-191 | 0.28 | 0.1 | 0.18 | 2.72 |
| mesenchyme | Pdia3 | 0.36 | 2.67E-195 | 5.56E-191 | 0.99 | 0.9 | 0.09 | 1.1 |
| mesenchyme | Gli3 | 0.85 | 2.99E-195 | 6.22E-191 | 0.64 | 0.47 | 0.17 | 1.37 |
| mesenchyme | Hnrnpk | 0.25 | 8.64E-195 | 1.80E-190 | 1 | 0.99 | 0.01 | 1.01 |
| mesenchyme | Lsm8 | 0.39 | 9.96E-195 | 2.07E-190 | 0.94 | 0.86 | 0.08 | 1.09 |
| mesenchyme | Cacybp | 0.35 | 1.05E-194 | 2.18E-190 | 0.98 | 0.92 | 0.05 | 1.06 |
| mesenchyme | Vasn | 2.01 | 1.11E-194 | 2.30E-190 | 0.28 | 0.1 | 0.18 | 2.82 |
| mesenchyme | Lzts2 | 0.75 | 1.24E-194 | 2.58E-190 | 0.66 | 0.49 | 0.17 | 1.35 |
| mesenchyme | Trim28 | 0.39 | 1.38E-194 | 2.86E-190 | 0.96 | 0.88 | 0.08 | 1.1 |
| mesenchyme | Wnk3 | 1.36 | 5.74E-194 | 1.19E-189 | 0.39 | 0.19 | 0.19 | 1.99 |
| mesenchyme | Cenpv | 0.95 | 6.10E-194 | 1.27E-189 | 0.56 | 0.38 | 0.19 | 1.5 |
| mesenchyme | Sall4 | 2.27 | 6.29E-194 | 1.31E-189 | 0.25 | 0.08 | 0.17 | 3.24 |

**Supplementary Table 2. Marker genes for the major cell types of integrated clustering of E11.5 control and NotchGOF λ-junction cells.**

| | | | | | | | | |
|---|---|---|---|---|---|---|---|---|
| mesenchyme | Kdelr2 | 0.5 | 6.35E-194 | 1.32E-189 | 0.88 | 0.75 | 0.13 | 1.18 |
| mesenchyme | Tra2b | 0.38 | 1.16E-193 | 2.41E-189 | 0.96 | 0.9 | 0.06 | 1.07 |
| mesenchyme | Sarnp | 0.35 | 1.21E-193 | 2.52E-189 | 0.97 | 0.93 | 0.05 | 1.05 |
| mesenchyme | Selenom | 0.32 | 1.56E-193 | 3.24E-189 | 0.59 | 0.39 | 0.19 | 1.49 |
| mesenchyme | Cdk4 | 0.29 | 1.77E-193 | 3.68E-189 | 1 | 0.97 | 0.03 | 1.04 |
| mesenchyme | Rnf7 | 0.34 | 6.07E-193 | 1.26E-188 | 0.98 | 0.93 | 0.04 | 1.05 |
| mesenchyme | Slc35a1 | 1.01 | 1.07E-192 | 2.23E-188 | 0.53 | 0.36 | 0.17 | 1.48 |
| mesenchyme | Ube2k | 0.43 | 1.07E-192 | 2.23E-188 | 0.91 | 0.84 | 0.08 | 1.09 |
| mesenchyme | Vcp | 0.34 | 1.45E-192 | 3.01E-188 | 0.98 | 0.94 | 0.04 | 1.05 |
| mesenchyme | Bex3 | 0.33 | 2.79E-192 | 5.80E-188 | 0.99 | 0.94 | 0.05 | 1.05 |
| mesenchyme | Acin1 | 0.38 | 5.79E-192 | 1.20E-187 | 0.97 | 0.91 | 0.06 | 1.07 |
| mesenchyme | Ctsk | 3.51 | 1.41E-191 | 2.93E-187 | 0.18 | 0.03 | 0.15 | 6.14 |
| mesenchyme | Prpf4b | 0.41 | 1.54E-191 | 3.20E-187 | 0.94 | 0.87 | 0.07 | 1.08 |
| mesenchyme | Bicc1 | 1.85 | 1.53E-190 | 3.19E-186 | 0.29 | 0.11 | 0.18 | 2.58 |
| mesenchyme | Trp53 | 0.4 | 7.05E-190 | 1.47E-185 | 0.95 | 0.85 | 0.09 | 1.11 |
| mesenchyme | Scpep1 | 1.14 | 2.35E-189 | 4.89E-185 | 0.47 | 0.29 | 0.19 | 1.65 |
| mesenchyme | Spats2 | 0.61 | 2.97E-189 | 6.19E-185 | 0.74 | 0.57 | 0.17 | 1.29 |
| mesenchyme | Cops4 | 0.44 | 3.04E-189 | 6.32E-185 | 0.9 | 0.8 | 0.1 | 1.13 |
| mesenchyme | mt-Co3 | 0.26 | 2.43E-188 | 5.06E-184 | 1 | 0.98 | 0.02 | 1.02 |
| mesenchyme | Igf2bp1 | 0.44 | 8.29E-188 | 1.72E-183 | 0.93 | 0.79 | 0.14 | 1.17 |
| mesenchyme | Setbp1 | 1.24 | 2.91E-187 | 6.05E-183 | 0.45 | 0.26 | 0.19 | 1.72 |
| mesenchyme | Ppid | 0.43 | 1.01E-186 | 2.10E-182 | 0.91 | 0.82 | 0.1 | 1.12 |
| mesenchyme | ·930503L19Ri | 1.28 | 2.12E-186 | 4.42E-182 | 0.39 | 0.21 | 0.18 | 1.86 |
| mesenchyme | Srsf2 | 0.29 | 3.78E-186 | 7.86E-182 | 1 | 0.97 | 0.02 | 1.02 |
| mesenchyme | Dtymk | 0.42 | 7.69E-186 | 1.60E-181 | 0.92 | 0.83 | 0.09 | 1.11 |
| mesenchyme | Pakap.1 | 1.04 | 7.87E-186 | 1.64E-181 | 0.48 | 0.28 | 0.19 | 1.68 |
| mesenchyme | Gtf2a2 | 0.43 | 8.37E-186 | 1.74E-181 | 0.91 | 0.83 | 0.08 | 1.09 |
| mesenchyme | Rbpj | 0.61 | 8.86E-186 | 1.84E-181 | 0.75 | 0.61 | 0.14 | 1.23 |
| mesenchyme | Ndufa2 | 0.28 | 1.27E-185 | 2.64E-181 | 0.99 | 0.97 | 0.02 | 1.02 |
| mesenchyme | Sox12 | 0.65 | 1.89E-185 | 3.93E-181 | 0.72 | 0.56 | 0.16 | 1.29 |
| mesenchyme | Thsd7a | 2.59 | 2.86E-185 | 5.95E-181 | 0.2 | 0.05 | 0.15 | 4.42 |
| mesenchyme | Capn6 | 0.74 | 3.31E-185 | 6.88E-181 | 0.61 | 0.41 | 0.21 | 1.5 |
| mesenchyme | 230219D22Ri | 0.44 | 4.24E-185 | 8.81E-181 | 0.91 | 0.8 | 0.12 | 1.14 |
| mesenchyme | Hmgcr | 0.71 | 4.35E-185 | 9.05E-181 | 0.72 | 0.57 | 0.15 | 1.27 |
| mesenchyme | Evi5 | 0.75 | 4.90E-185 | 1.02E-180 | 0.65 | 0.52 | 0.13 | 1.26 |
| mesenchyme | Fibin | 3.7 | 7.71E-185 | 1.60E-180 | 0.17 | 0.03 | 0.14 | 6.11 |
| mesenchyme | Ssr3 | 0.43 | 8.73E-185 | 1.82E-180 | 0.92 | 0.81 | 0.11 | 1.13 |
| mesenchyme | Txndc12 | 0.6 | 1.66E-184 | 3.45E-180 | 0.75 | 0.59 | 0.16 | 1.28 |
| mesenchyme | Lman1 | 0.47 | 2.17E-184 | 4.51E-180 | 0.9 | 0.77 | 0.14 | 1.18 |
| mesenchyme | Ccar1 | 0.41 | 5.14E-184 | 1.07E-179 | 0.95 | 0.87 | 0.08 | 1.09 |
| mesenchyme | Ivns1abp | 0.54 | 2.83E-183 | 5.89E-179 | 0.82 | 0.69 | 0.13 | 1.18 |
| mesenchyme | Hadha | 0.52 | 3.08E-183 | 6.41E-179 | 0.84 | 0.74 | 0.1 | 1.14 |
| mesenchyme | Prrc2c | 0.35 | 3.71E-183 | 7.71E-179 | 0.99 | 0.93 | 0.07 | 1.07 |
| mesenchyme | Alkbh1 | 0.68 | 4.36E-183 | 9.08E-179 | 0.72 | 0.57 | 0.15 | 1.26 |
| mesenchyme | Osbpl1a | 1.17 | 6.54E-183 | 1.36E-178 | 0.45 | 0.27 | 0.18 | 1.67 |
| mesenchyme | Tbx3os1 | 3.2 | 1.09E-182 | 2.26E-178 | 0.19 | 0.04 | 0.15 | 5 |
| mesenchyme | Gcsh | 0.53 | 3.02E-182 | 6.27E-178 | 0.78 | 0.64 | 0.15 | 1.23 |
| mesenchyme | Rbm8a | 0.36 | 5.48E-182 | 1.14E-177 | 0.97 | 0.92 | 0.05 | 1.05 |
| mesenchyme | Sdhb | 0.4 | 6.31E-182 | 1.31E-177 | 0.92 | 0.85 | 0.07 | 1.08 |
| mesenchyme | Thyn1 | 0.57 | 1.73E-181 | 3.60E-177 | 0.78 | 0.64 | 0.14 | 1.22 |
| mesenchyme | Rcbtb2 | 0.53 | 2.85E-181 | 5.93E-177 | 0.81 | 0.66 | 0.14 | 1.21 |
| mesenchyme | Cldnd1 | 0.62 | 3.62E-180 | 7.53E-176 | 0.74 | 0.6 | 0.14 | 1.23 |
| mesenchyme | Nfyb | 0.5 | 4.16E-180 | 8.66E-176 | 0.83 | 0.72 | 0.1 | 1.14 |
| mesenchyme | Arl3 | 0.44 | 6.68E-180 | 1.39E-175 | 0.88 | 0.78 | 0.11 | 1.14 |
| mesenchyme | Psma5 | 0.32 | 9.27E-180 | 1.93E-175 | 0.98 | 0.94 | 0.04 | 1.05 |
| mesenchyme | Usp14 | 0.51 | 1.12E-179 | 2.32E-175 | 0.83 | 0.73 | 0.09 | 1.13 |

**Supplementary Table 2. Marker genes for the major cell types of integrated clustering of E11.5 control and NotchGOF λ-junction cells.**

| | | | | | | | | |
|---|---|---|---|---|---|---|---|---|
| mesenchyme | Taf11 | 0.53 | 1.36E-179 | 2.82E-175 | 0.81 | 0.7 | 0.11 | 1.16 |
| mesenchyme | Gdi2 | 0.34 | 2.08E-179 | 4.32E-175 | 0.97 | 0.9 | 0.07 | 1.07 |
| mesenchyme | Ap3s1 | 0.56 | 2.43E-179 | 5.06E-175 | 0.75 | 0.63 | 0.13 | 1.2 |
| mesenchyme | Mat2a | 0.41 | 9.19E-179 | 1.91E-174 | 0.94 | 0.85 | 0.09 | 1.1 |
| mesenchyme | Prdm16os | 3.54 | 1.12E-178 | 2.33E-174 | 0.17 | 0.03 | 0.14 | 6.35 |
| mesenchyme | Atp5o | 0.26 | 1.22E-178 | 2.54E-174 | 1 | 0.97 | 0.02 | 1.02 |
| mesenchyme | Srsf11 | 0.33 | 2.10E-178 | 4.38E-174 | 0.98 | 0.93 | 0.05 | 1.05 |
| mesenchyme | Eci2 | 0.59 | 7.34E-178 | 1.53E-173 | 0.73 | 0.59 | 0.14 | 1.24 |
| mesenchyme | Pds5b | 0.68 | 1.08E-177 | 2.24E-173 | 0.7 | 0.56 | 0.15 | 1.26 |
| mesenchyme | Adamts20 | 2.84 | 1.09E-177 | 2.26E-173 | 0.2 | 0.05 | 0.15 | 4.24 |
| mesenchyme | Zmiz1 | 0.6 | 1.12E-177 | 2.34E-173 | 0.74 | 0.6 | 0.15 | 1.24 |
| mesenchyme | Adh5 | 0.41 | 1.18E-177 | 2.46E-173 | 0.93 | 0.85 | 0.07 | 1.09 |
| mesenchyme | Fkbp10 | 1.28 | 1.34E-177 | 2.78E-173 | 0.4 | 0.21 | 0.19 | 1.87 |
| mesenchyme | Bach2 | 1.11 | 3.42E-177 | 7.11E-173 | 0.44 | 0.25 | 0.19 | 1.74 |
| mesenchyme | Asap1 | 0.61 | 3.65E-177 | 7.58E-173 | 0.77 | 0.65 | 0.12 | 1.19 |
| mesenchyme | Safb | 0.5 | 5.19E-177 | 1.08E-172 | 0.82 | 0.71 | 0.11 | 1.16 |
| mesenchyme | 610027K06Ri | 2.82 | 7.97E-177 | 1.66E-172 | 0.18 | 0.04 | 0.15 | 4.92 |
| mesenchyme | Unc119 | 0.47 | 1.20E-176 | 2.50E-172 | 0.83 | 0.68 | 0.15 | 1.22 |
| mesenchyme | Srrt | 0.48 | 1.41E-176 | 2.92E-172 | 0.85 | 0.75 | 0.1 | 1.13 |
| mesenchyme | Hmga2 | 0.41 | 2.00E-176 | 4.16E-172 | 0.98 | 0.83 | 0.15 | 1.18 |
| mesenchyme | Ddt | 0.44 | 2.20E-176 | 4.58E-172 | 0.89 | 0.78 | 0.11 | 1.14 |
| mesenchyme | Lama2 | 2.68 | 2.68E-176 | 5.57E-172 | 0.2 | 0.05 | 0.15 | 4.15 |
| mesenchyme | Ssb | 0.29 | 2.88E-176 | 6.00E-172 | 1 | 0.97 | 0.03 | 1.03 |
| mesenchyme | Fndc5 | 2.36 | 4.58E-176 | 9.53E-172 | 0.22 | 0.07 | 0.16 | 3.42 |
| mesenchyme | Rnps1 | 0.35 | 6.45E-176 | 1.34E-171 | 0.97 | 0.91 | 0.06 | 1.07 |
| mesenchyme | Stx18 | 0.93 | 7.73E-176 | 1.61E-171 | 0.57 | 0.42 | 0.15 | 1.36 |
| mesenchyme | Ptpn9 | 0.89 | 1.28E-175 | 2.65E-171 | 0.54 | 0.37 | 0.17 | 1.46 |
| mesenchyme | Mab21l1 | 2.55 | 1.49E-175 | 3.09E-171 | 0.22 | 0.06 | 0.16 | 3.5 |
| mesenchyme | Nudc | 0.32 | 2.01E-175 | 4.19E-171 | 0.98 | 0.93 | 0.05 | 1.05 |
| mesenchyme | Map2k6 | 0.94 | 2.34E-175 | 4.87E-171 | 0.5 | 0.31 | 0.18 | 1.58 |
| mesenchyme | Dgcr6 | 0.57 | 2.51E-175 | 5.23E-171 | 0.77 | 0.64 | 0.13 | 1.2 |
| mesenchyme | Smc3 | 0.35 | 4.20E-175 | 8.75E-171 | 0.97 | 0.91 | 0.06 | 1.07 |
| mesenchyme | Zranb2 | 0.38 | 6.27E-175 | 1.30E-170 | 0.95 | 0.88 | 0.07 | 1.08 |
| mesenchyme | Rfc1 | 0.44 | 7.96E-175 | 1.66E-170 | 0.9 | 0.79 | 0.11 | 1.14 |
| mesenchyme | Dse | 1.52 | 9.35E-175 | 1.95E-170 | 0.33 | 0.15 | 0.18 | 2.17 |
| mesenchyme | Lactb | 1.23 | 2.30E-174 | 4.79E-170 | 0.4 | 0.22 | 0.18 | 1.81 |
| mesenchyme | a | 2.67 | 2.82E-174 | 5.87E-170 | 0.18 | 0.04 | 0.14 | 4.89 |
| mesenchyme | Scx | 2.66 | 3.72E-174 | 7.73E-170 | 0.21 | 0.06 | 0.15 | 3.51 |
| mesenchyme | Pde4d | 1.54 | 4.40E-174 | 9.15E-170 | 0.32 | 0.15 | 0.18 | 2.2 |
| mesenchyme | Lrrc4b | 2.98 | 8.20E-174 | 1.71E-169 | 0.18 | 0.04 | 0.14 | 4.92 |
| mesenchyme | Golm1 | 0.78 | 8.97E-174 | 1.86E-169 | 0.61 | 0.45 | 0.16 | 1.36 |
| mesenchyme | Wnt5b | 1.56 | 9.50E-174 | 1.98E-169 | 0.32 | 0.14 | 0.18 | 2.26 |
| mesenchyme | Smarca5 | 0.31 | 1.11E-173 | 2.30E-169 | 0.99 | 0.95 | 0.04 | 1.04 |
| mesenchyme | Pde3a | 2.74 | 1.15E-173 | 2.39E-169 | 0.18 | 0.04 | 0.14 | 4.94 |
| mesenchyme | Pid1 | 2.17 | 1.22E-173 | 2.53E-169 | 0.25 | 0.09 | 0.16 | 2.92 |
| mesenchyme | Mapre2 | 0.97 | 1.71E-173 | 3.56E-169 | 0.46 | 0.28 | 0.19 | 1.67 |
| mesenchyme | Zfhx4 | 0.72 | 1.74E-173 | 3.62E-169 | 0.79 | 0.64 | 0.14 | 1.22 |
| mesenchyme | Lsm7 | 0.35 | 1.94E-173 | 4.04E-169 | 0.96 | 0.9 | 0.06 | 1.07 |
| mesenchyme | Pdlim3 | 3.33 | 2.15E-173 | 4.47E-169 | 0.2 | 0.05 | 0.15 | 3.96 |
| mesenchyme | Zfp771 | 0.6 | 2.69E-173 | 5.59E-169 | 0.73 | 0.59 | 0.14 | 1.24 |
| mesenchyme | Pgrmc1 | 0.38 | 5.44E-173 | 1.13E-168 | 0.96 | 0.83 | 0.13 | 1.15 |
| mesenchyme | Exosc8 | 0.51 | 6.45E-173 | 1.34E-168 | 0.83 | 0.72 | 0.1 | 1.14 |
| mesenchyme | Hsf2 | 0.91 | 7.08E-173 | 1.47E-168 | 0.54 | 0.37 | 0.17 | 1.47 |
| mesenchyme | Prkar1a | 0.38 | 7.21E-173 | 1.50E-168 | 0.91 | 0.83 | 0.08 | 1.1 |
| mesenchyme | Sgk1 | 1.51 | 9.29E-173 | 1.93E-168 | 0.26 | 0.1 | 0.17 | 2.78 |
| mesenchyme | Runx2 | 2.31 | 4.63E-172 | 9.63E-168 | 0.21 | 0.06 | 0.15 | 3.59 |

**Supplementary Table 2. Marker genes for the major cell types of integrated clustering of E11.5 control and NotchGOF λ-junction cells.**

| | | | | | | | | |
|---|---|---|---|---|---|---|---|---|
| mesenchyme | Nedd8 | 0.26 | 7.92E-172 | 1.65E-167 | 1 | 0.98 | 0.02 | 1.02 |
| mesenchyme | Brd7 | 0.47 | 1.11E-171 | 2.31E-167 | 0.86 | 0.77 | 0.09 | 1.12 |
| mesenchyme | Erc1 | 0.82 | 1.34E-171 | 2.80E-167 | 0.61 | 0.45 | 0.16 | 1.35 |
| mesenchyme | Bcas2 | 0.32 | 1.86E-171 | 3.87E-167 | 0.97 | 0.92 | 0.04 | 1.05 |
| mesenchyme | Rbm26 | 0.44 | 6.37E-171 | 1.33E-166 | 0.87 | 0.76 | 0.12 | 1.15 |
| mesenchyme | Traf3ip2 | 2.01 | 1.39E-170 | 2.89E-166 | 0.28 | 0.11 | 0.16 | 2.48 |
| mesenchyme | Brd9 | 0.76 | 1.41E-170 | 2.93E-166 | 0.6 | 0.45 | 0.16 | 1.35 |
| mesenchyme | Gfpt2 | 2.1 | 2.91E-170 | 6.04E-166 | 0.24 | 0.08 | 0.16 | 3.01 |
| mesenchyme | Ahsa2 | 0.57 | 1.04E-169 | 2.17E-165 | 0.75 | 0.61 | 0.13 | 1.21 |
| mesenchyme | Cox7b | 0.26 | 1.29E-169 | 2.67E-165 | 0.99 | 0.97 | 0.02 | 1.02 |
| mesenchyme | Son | 0.29 | 1.97E-169 | 4.09E-165 | 0.99 | 0.96 | 0.04 | 1.04 |
| mesenchyme | Dzip1 | 1.21 | 2.02E-169 | 4.21E-165 | 0.39 | 0.21 | 0.18 | 1.85 |
| mesenchyme | Psmd8 | 0.33 | 3.83E-169 | 7.97E-165 | 0.97 | 0.92 | 0.04 | 1.05 |
| mesenchyme | Etv1 | 1.31 | 6.27E-169 | 1.30E-164 | 0.36 | 0.18 | 0.18 | 2 |
| mesenchyme | Rab18 | 0.44 | 6.51E-169 | 1.35E-164 | 0.85 | 0.76 | 0.1 | 1.13 |
| mesenchyme | Arhgef25 | 1.17 | 1.38E-168 | 2.87E-164 | 0.43 | 0.25 | 0.18 | 1.7 |
| mesenchyme | 110038F14Ri | 0.52 | 1.59E-168 | 3.31E-164 | 0.77 | 0.64 | 0.13 | 1.21 |
| mesenchyme | AW551984 | 2.97 | 1.73E-168 | 3.59E-164 | 0.17 | 0.03 | 0.14 | 5.39 |
| mesenchyme | Mrpl53 | 0.41 | 9.72E-168 | 2.02E-163 | 0.88 | 0.8 | 0.08 | 1.1 |
| mesenchyme | Mtap | 0.56 | 9.82E-168 | 2.04E-163 | 0.75 | 0.6 | 0.15 | 1.25 |
| mesenchyme | Lap3 | 0.62 | 1.53E-166 | 3.17E-162 | 0.71 | 0.59 | 0.13 | 1.21 |
| mesenchyme | Rasl11a | 1.35 | 3.03E-166 | 6.30E-162 | 0.34 | 0.16 | 0.17 | 2.08 |
| mesenchyme | Mapk1ip1l | 0.55 | 7.42E-166 | 1.54E-161 | 0.76 | 0.63 | 0.13 | 1.2 |
| mesenchyme | Map2k2 | 0.37 | 7.83E-166 | 1.63E-161 | 0.94 | 0.89 | 0.06 | 1.06 |
| mesenchyme | Plat | 1.19 | 8.62E-166 | 1.79E-161 | 0.38 | 0.2 | 0.18 | 1.92 |
| mesenchyme | P4ha2 | 1.43 | 1.31E-165 | 2.73E-161 | 0.33 | 0.16 | 0.17 | 2.07 |
| mesenchyme | Mpzl1 | 0.56 | 1.82E-165 | 3.79E-161 | 0.69 | 0.52 | 0.17 | 1.33 |
| mesenchyme | Strap | 0.4 | 2.02E-165 | 4.21E-161 | 0.92 | 0.85 | 0.07 | 1.08 |
| mesenchyme | Dad1 | 0.35 | 2.74E-165 | 5.70E-161 | 0.96 | 0.89 | 0.07 | 1.08 |
| mesenchyme | Dab2ip | 0.67 | 3.07E-165 | 6.39E-161 | 0.66 | 0.49 | 0.16 | 1.33 |
| mesenchyme | Inpp4b | 2.46 | 3.14E-165 | 6.54E-161 | 0.2 | 0.06 | 0.15 | 3.61 |
| mesenchyme | Slit3 | 2.03 | 3.99E-165 | 8.30E-161 | 0.24 | 0.09 | 0.16 | 2.86 |
| mesenchyme | Fermt2 | 0.52 | 4.36E-165 | 9.07E-161 | 0.78 | 0.6 | 0.18 | 1.3 |
| mesenchyme | Tenm3 | 0.76 | 5.64E-165 | 1.17E-160 | 0.7 | 0.52 | 0.18 | 1.35 |
| mesenchyme | Hnrnpc | 0.28 | 6.80E-165 | 1.41E-160 | 1 | 0.97 | 0.03 | 1.03 |
| mesenchyme | Sned1 | 3.1 | 7.48E-165 | 1.56E-160 | 0.17 | 0.04 | 0.14 | 4.7 |
| mesenchyme | Api5 | 0.48 | 1.16E-164 | 2.42E-160 | 0.83 | 0.73 | 0.1 | 1.13 |
| mesenchyme | Foxl2 | 3.12 | 1.41E-164 | 2.93E-160 | 0.17 | 0.04 | 0.14 | 4.7 |
| mesenchyme | Atp5c1 | 0.27 | 2.44E-164 | 5.07E-160 | 0.99 | 0.97 | 0.03 | 1.03 |
| mesenchyme | Sf3a1 | 0.54 | 2.81E-164 | 5.85E-160 | 0.78 | 0.66 | 0.13 | 1.2 |
| mesenchyme | Epop | 1.07 | 2.93E-164 | 6.10E-160 | 0.44 | 0.27 | 0.17 | 1.64 |
| mesenchyme | Kcnq1ot1 | 0.42 | 6.64E-164 | 1.38E-159 | 0.96 | 0.84 | 0.12 | 1.15 |
| mesenchyme | Add3 | 0.63 | 2.77E-163 | 5.76E-159 | 0.71 | 0.56 | 0.16 | 1.28 |
| mesenchyme | Pxdn | 0.82 | 3.15E-163 | 6.56E-159 | 0.57 | 0.4 | 0.17 | 1.43 |
| mesenchyme | Bmp1 | 1.05 | 3.42E-163 | 7.12E-159 | 0.48 | 0.31 | 0.17 | 1.53 |
| mesenchyme | Ctnnbip1 | 0.44 | 3.81E-162 | 7.92E-158 | 0.86 | 0.72 | 0.14 | 1.19 |
| mesenchyme | Foxc1 | 1.76 | 1.66E-161 | 3.45E-157 | 0.34 | 0.18 | 0.16 | 1.91 |
| mesenchyme | Lsm2 | 0.36 | 1.84E-161 | 3.84E-157 | 0.94 | 0.89 | 0.06 | 1.06 |
| mesenchyme | Hacd4 | 1.26 | 3.04E-161 | 6.33E-157 | 0.31 | 0.14 | 0.17 | 2.26 |
| mesenchyme | Zfp397 | 0.6 | 3.91E-161 | 8.13E-157 | 0.73 | 0.58 | 0.15 | 1.25 |
| mesenchyme | Pttg1 | 0.61 | 9.52E-161 | 1.98E-156 | 0.75 | 0.63 | 0.12 | 1.2 |
| mesenchyme | Mageh1 | 0.74 | 2.72E-160 | 5.66E-156 | 0.62 | 0.45 | 0.17 | 1.37 |
| mesenchyme | Sf3b2 | 0.29 | 2.83E-160 | 5.88E-156 | 0.99 | 0.95 | 0.04 | 1.04 |
| mesenchyme | Cdc42ep4 | 0.87 | 3.18E-160 | 6.61E-156 | 0.5 | 0.33 | 0.17 | 1.51 |
| mesenchyme | Cops6 | 0.41 | 3.38E-160 | 7.03E-156 | 0.9 | 0.82 | 0.08 | 1.09 |
| mesenchyme | As3mt | 1.15 | 3.66E-160 | 7.61E-156 | 0.42 | 0.25 | 0.17 | 1.66 |

**Supplementary Table 2. Marker genes for the major cell types of integrated clustering of E11.5 control and NotchGOF λ-junction cells.**

| | | | | | | | | |
|---|---|---|---|---|---|---|---|---|
| mesenchyme | Ss18 | 0.6 | 3.82E-160 | 7.95E-156 | 0.7 | 0.56 | 0.14 | 1.26 |
| mesenchyme | Coprs | 0.84 | 4.55E-160 | 9.46E-156 | 0.54 | 0.38 | 0.16 | 1.43 |
| mesenchyme | Ptch1 | 1.15 | 5.47E-160 | 1.14E-155 | 0.51 | 0.35 | 0.15 | 1.43 |
| mesenchyme | Ppdpf | 0.29 | 5.91E-160 | 1.23E-155 | 0.98 | 0.92 | 0.06 | 1.07 |
| mesenchyme | Dpysl2 | 0.5 | 1.05E-159 | 2.19E-155 | 0.75 | 0.59 | 0.16 | 1.27 |
| mesenchyme | Rfc3 | 0.49 | 2.39E-159 | 4.97E-155 | 0.81 | 0.69 | 0.12 | 1.18 |
| mesenchyme | Trir | 0.32 | 2.93E-159 | 6.09E-155 | 0.96 | 0.92 | 0.04 | 1.05 |
| mesenchyme | Srek1 | 0.46 | 3.73E-159 | 7.77E-155 | 0.86 | 0.75 | 0.11 | 1.15 |
| mesenchyme | Snrpa1 | 0.35 | 3.95E-159 | 8.22E-155 | 0.95 | 0.88 | 0.07 | 1.08 |
| mesenchyme | Irak1bp1 | 0.76 | 2.60E-158 | 5.41E-154 | 0.6 | 0.44 | 0.16 | 1.37 |
| mesenchyme | Enho | 0.86 | 2.95E-158 | 6.14E-154 | 0.51 | 0.33 | 0.17 | 1.52 |
| mesenchyme | Pla2g4a | 1.53 | 9.75E-158 | 2.03E-153 | 0.31 | 0.14 | 0.17 | 2.19 |
| mesenchyme | Tmem59l | 1.69 | 1.45E-157 | 3.02E-153 | 0.26 | 0.1 | 0.16 | 2.61 |
| mesenchyme | Casd1 | 0.86 | 2.37E-157 | 4.92E-153 | 0.55 | 0.39 | 0.16 | 1.41 |
| mesenchyme | Dhx15 | 0.42 | 3.01E-157 | 6.26E-153 | 0.89 | 0.79 | 0.09 | 1.12 |
| mesenchyme | Angpt1 | 2.5 | 3.46E-157 | 7.19E-153 | 0.19 | 0.05 | 0.14 | 3.69 |
| mesenchyme | Nr2f1 | 1.65 | 5.09E-157 | 1.06E-152 | 0.39 | 0.23 | 0.16 | 1.72 |
| mesenchyme | Aebp1 | 1.91 | 5.74E-157 | 1.19E-152 | 0.26 | 0.1 | 0.15 | 2.5 |
| mesenchyme | Rbm3 | 0.25 | 6.47E-157 | 1.34E-152 | 1 | 0.98 | 0.02 | 1.02 |
| mesenchyme | Chek1 | 0.74 | 6.66E-157 | 1.39E-152 | 0.6 | 0.45 | 0.15 | 1.34 |
| mesenchyme | Dnmt3a | 0.51 | 1.56E-156 | 3.25E-152 | 0.79 | 0.63 | 0.16 | 1.24 |
| mesenchyme | Atad2 | 0.6 | 2.17E-156 | 4.50E-152 | 0.78 | 0.66 | 0.12 | 1.19 |
| mesenchyme | Cep170 | 0.79 | 2.31E-156 | 4.81E-152 | 0.56 | 0.41 | 0.15 | 1.38 |
| mesenchyme | Ric1 | 1.22 | 2.39E-156 | 4.97E-152 | 0.39 | 0.22 | 0.17 | 1.75 |
| mesenchyme | Dnajc15 | 0.48 | 3.15E-156 | 6.56E-152 | 0.81 | 0.71 | 0.11 | 1.15 |
| mesenchyme | Txnl1 | 0.31 | 3.35E-156 | 6.98E-152 | 0.96 | 0.92 | 0.04 | 1.05 |
| mesenchyme | Ptprs | 0.57 | 5.32E-156 | 1.11E-151 | 0.74 | 0.58 | 0.16 | 1.27 |
| mesenchyme | Psmb5 | 0.27 | 5.44E-156 | 1.13E-151 | 0.99 | 0.96 | 0.02 | 1.02 |
| mesenchyme | Ssr1 | 0.51 | 5.48E-156 | 1.14E-151 | 0.8 | 0.69 | 0.11 | 1.15 |
| mesenchyme | Ube2m | 0.3 | 6.76E-156 | 1.41E-151 | 0.97 | 0.94 | 0.03 | 1.04 |
| mesenchyme | Nmral1 | 0.53 | 8.31E-156 | 1.73E-151 | 0.76 | 0.63 | 0.13 | 1.21 |
| mesenchyme | Chn1 | 0.94 | 1.18E-155 | 2.45E-151 | 0.46 | 0.29 | 0.17 | 1.58 |
| mesenchyme | Peg3 | 0.53 | 1.68E-155 | 3.50E-151 | 0.86 | 0.77 | 0.09 | 1.11 |
| mesenchyme | Wwtr1 | 0.9 | 2.07E-155 | 4.31E-151 | 0.49 | 0.32 | 0.17 | 1.53 |
| mesenchyme | Efemp2 | 0.89 | 2.63E-155 | 5.47E-151 | 0.53 | 0.37 | 0.16 | 1.44 |
| mesenchyme | Commd1 | 0.39 | 3.12E-155 | 6.48E-151 | 0.9 | 0.83 | 0.07 | 1.09 |
| mesenchyme | Pnn | 0.34 | 3.35E-155 | 6.97E-151 | 0.97 | 0.91 | 0.06 | 1.07 |
| mesenchyme | Zfp207 | 0.38 | 3.56E-155 | 7.40E-151 | 0.92 | 0.83 | 0.09 | 1.11 |
| mesenchyme | Macf1 | 0.42 | 3.65E-155 | 7.59E-151 | 0.88 | 0.77 | 0.12 | 1.15 |
| mesenchyme | Mmd | 1.08 | 4.39E-155 | 9.13E-151 | 0.41 | 0.24 | 0.17 | 1.72 |
| mesenchyme | Psmc3 | 0.33 | 4.48E-155 | 9.32E-151 | 0.95 | 0.9 | 0.05 | 1.05 |
| mesenchyme | Dnajc10 | 0.58 | 5.45E-155 | 1.13E-150 | 0.73 | 0.59 | 0.14 | 1.23 |
| mesenchyme | Ptges3 | 0.3 | 7.50E-155 | 1.56E-150 | 0.98 | 0.94 | 0.04 | 1.04 |
| mesenchyme | Ilf3 | 0.45 | 9.00E-155 | 1.87E-150 | 0.84 | 0.72 | 0.12 | 1.17 |
| mesenchyme | Ncbp2 | 0.4 | 9.19E-155 | 1.91E-150 | 0.89 | 0.8 | 0.09 | 1.11 |
| mesenchyme | Tnik | 1.25 | 1.19E-154 | 2.47E-150 | 0.3 | 0.13 | 0.17 | 2.29 |
| mesenchyme | Phax | 0.42 | 2.09E-154 | 4.34E-150 | 0.86 | 0.76 | 0.11 | 1.14 |
| mesenchyme | Gnai3 | 0.41 | 2.80E-154 | 5.83E-150 | 0.88 | 0.77 | 0.11 | 1.14 |
| mesenchyme | Zfp422 | 0.42 | 4.46E-154 | 9.28E-150 | 0.89 | 0.76 | 0.13 | 1.17 |
| mesenchyme | Slbp | 0.44 | 6.51E-154 | 1.35E-149 | 0.9 | 0.82 | 0.08 | 1.1 |
| mesenchyme | Farp1 | 0.8 | 7.85E-154 | 1.63E-149 | 0.55 | 0.39 | 0.16 | 1.41 |
| mesenchyme | Eid2 | 0.57 | 3.91E-153 | 8.14E-149 | 0.7 | 0.55 | 0.15 | 1.27 |
| mesenchyme | Gm36988 | 2.52 | 4.05E-153 | 8.42E-149 | 0.19 | 0.05 | 0.14 | 3.72 |
| mesenchyme | Spcs2 | 0.32 | 6.35E-153 | 1.32E-148 | 0.98 | 0.92 | 0.06 | 1.07 |
| mesenchyme | Mllt3 | 0.66 | 7.70E-153 | 1.60E-148 | 0.7 | 0.58 | 0.12 | 1.21 |
| mesenchyme | Aarsd1 | 0.48 | 1.02E-152 | 2.12E-148 | 0.8 | 0.71 | 0.1 | 1.14 |

**Supplementary Table 2. Marker genes for the major cell types of integrated clustering of E11.5 control and NotchGOF λ-junction cells.**

| | | | | | | | | |
|---|---|---|---|---|---|---|---|---|
| mesenchyme | Fam107b | 1.21 | 2.02E-152 | 4.19E-148 | 0.41 | 0.25 | 0.16 | 1.67 |
| mesenchyme | Cntln | 0.82 | 2.63E-152 | 5.47E-148 | 0.54 | 0.38 | 0.17 | 1.44 |
| mesenchyme | Cdk11b | 0.48 | 3.06E-152 | 6.37E-148 | 0.81 | 0.7 | 0.11 | 1.15 |
| mesenchyme | Glul | 0.57 | 4.89E-152 | 1.02E-147 | 0.66 | 0.53 | 0.13 | 1.25 |
| mesenchyme | Fcgrt | 1.59 | 5.27E-152 | 1.10E-147 | 0.19 | 0.05 | 0.14 | 3.78 |
| mesenchyme | Fus | 0.3 | 5.96E-152 | 1.24E-147 | 0.99 | 0.93 | 0.06 | 1.06 |
| mesenchyme | E2f3 | 0.67 | 7.63E-152 | 1.59E-147 | 0.63 | 0.48 | 0.15 | 1.3 |
| mesenchyme | Syde1 | 2.01 | 8.30E-152 | 1.73E-147 | 0.22 | 0.08 | 0.15 | 2.86 |
| mesenchyme | Mtf2 | 0.45 | 1.41E-151 | 2.93E-147 | 0.85 | 0.73 | 0.12 | 1.16 |
| mesenchyme | Sdhd | 0.42 | 1.42E-151 | 2.94E-147 | 0.87 | 0.76 | 0.11 | 1.14 |
| mesenchyme | Ankrd12 | 0.91 | 2.35E-151 | 4.90E-147 | 0.58 | 0.44 | 0.14 | 1.31 |
| mesenchyme | Gna11 | 0.56 | 4.99E-151 | 1.04E-146 | 0.72 | 0.58 | 0.14 | 1.25 |
| mesenchyme | Rsl1d1 | 0.34 | 7.56E-151 | 1.57E-146 | 0.95 | 0.89 | 0.06 | 1.07 |
| mesenchyme | Esd | 0.29 | 1.16E-150 | 2.41E-146 | 0.97 | 0.93 | 0.04 | 1.05 |
| mesenchyme | Shox2 | 2.71 | 3.27E-150 | 6.80E-146 | 0.17 | 0.04 | 0.13 | 4.2 |
| mesenchyme | Nkd2 | 2.61 | 5.45E-150 | 1.13E-145 | 0.17 | 0.04 | 0.13 | 4.12 |
| mesenchyme | Eif4a3 | 0.38 | 1.27E-149 | 2.64E-145 | 0.9 | 0.82 | 0.08 | 1.1 |
| mesenchyme | Foxd2 | 3.42 | 1.57E-149 | 3.27E-145 | 0.15 | 0.03 | 0.12 | 5.1 |
| mesenchyme | Foxd1 | 3.47 | 5.84E-149 | 1.21E-144 | 0.15 | 0.03 | 0.12 | 5.52 |
| mesenchyme | Smarca1 | 0.81 | 8.53E-149 | 1.77E-144 | 0.52 | 0.35 | 0.17 | 1.49 |
| mesenchyme | Banf1 | 0.29 | 1.49E-148 | 3.10E-144 | 0.96 | 0.92 | 0.04 | 1.04 |
| mesenchyme | Ptx3 | 3.27 | 1.64E-148 | 3.40E-144 | 0.15 | 0.03 | 0.12 | 5.21 |
| mesenchyme | Col27a1 | 1.87 | 7.79E-148 | 1.62E-143 | 0.24 | 0.09 | 0.15 | 2.67 |
| mesenchyme | Efnb3 | 1.21 | 9.07E-148 | 1.89E-143 | 0.37 | 0.21 | 0.17 | 1.8 |
| mesenchyme | Cdca7 | 0.49 | 1.08E-147 | 2.25E-143 | 0.77 | 0.61 | 0.16 | 1.26 |
| mesenchyme | Gm30564 | 3.59 | 1.16E-147 | 2.41E-143 | 0.13 | 0.02 | 0.12 | 7.05 |
| mesenchyme | Hp1bp3 | 0.39 | 1.22E-147 | 2.53E-143 | 0.92 | 0.83 | 0.09 | 1.11 |
| mesenchyme | Net1 | 0.75 | 1.45E-147 | 3.02E-143 | 0.63 | 0.51 | 0.12 | 1.23 |
| mesenchyme | Mbd3 | 0.3 | 3.48E-147 | 7.24E-143 | 0.97 | 0.92 | 0.05 | 1.06 |
| mesenchyme | Cdc27 | 0.76 | 4.41E-147 | 9.17E-143 | 0.57 | 0.44 | 0.13 | 1.3 |
| mesenchyme | Rbm25 | 0.3 | 4.63E-147 | 9.62E-143 | 0.99 | 0.94 | 0.05 | 1.05 |
| mesenchyme | E2f6 | 0.77 | 5.89E-147 | 1.22E-142 | 0.58 | 0.43 | 0.15 | 1.34 |
| mesenchyme | Sobp | 1.22 | 2.05E-146 | 4.27E-142 | 0.4 | 0.23 | 0.16 | 1.7 |
| mesenchyme | Mn1 | 1.36 | 2.99E-146 | 6.21E-142 | 0.34 | 0.18 | 0.16 | 1.9 |
| mesenchyme | Rbm5 | 0.46 | 3.31E-146 | 6.89E-142 | 0.86 | 0.76 | 0.1 | 1.13 |
| mesenchyme | Csnk2a1 | 0.29 | 6.01E-146 | 1.25E-141 | 0.97 | 0.93 | 0.04 | 1.04 |
| mesenchyme | Tsc22d3 | 0.7 | 1.01E-145 | 2.10E-141 | 0.58 | 0.43 | 0.16 | 1.36 |
| mesenchyme | Fmr1 | 0.59 | 1.06E-145 | 2.21E-141 | 0.7 | 0.58 | 0.12 | 1.2 |
| mesenchyme | Nfia | 0.84 | 1.83E-145 | 3.81E-141 | 0.53 | 0.36 | 0.17 | 1.47 |
| mesenchyme | Pmepa1 | 0.74 | 4.65E-145 | 9.67E-141 | 0.53 | 0.36 | 0.17 | 1.48 |
| mesenchyme | Trpm7 | 0.56 | 1.10E-144 | 2.28E-140 | 0.73 | 0.61 | 0.12 | 1.19 |
| mesenchyme | Fam171b | 1.37 | 1.25E-144 | 2.60E-140 | 0.3 | 0.14 | 0.16 | 2.14 |
| mesenchyme | Adgra3 | 0.66 | 2.08E-144 | 4.33E-140 | 0.61 | 0.46 | 0.15 | 1.33 |
| mesenchyme | Bdnf | 1.27 | 2.48E-144 | 5.15E-140 | 0.34 | 0.18 | 0.16 | 1.91 |
| mesenchyme | Dync1i2 | 0.35 | 2.62E-144 | 5.44E-140 | 0.92 | 0.83 | 0.09 | 1.11 |
| mesenchyme | Zbed3 | 0.7 | 3.12E-144 | 6.50E-140 | 0.61 | 0.46 | 0.15 | 1.32 |
| mesenchyme | Dbi | 0.27 | 5.93E-144 | 1.23E-139 | 0.99 | 0.94 | 0.05 | 1.05 |
| mesenchyme | Snrnp70 | 0.29 | 2.20E-143 | 4.59E-139 | 0.98 | 0.93 | 0.04 | 1.05 |
| mesenchyme | Grem1 | 3.64 | 2.27E-143 | 4.72E-139 | 0.13 | 0.02 | 0.11 | 6.65 |
| mesenchyme | Zfp354c | 0.97 | 2.35E-143 | 4.89E-139 | 0.44 | 0.28 | 0.16 | 1.56 |
| mesenchyme | Usp3 | 0.49 | 2.56E-143 | 5.32E-139 | 0.77 | 0.65 | 0.12 | 1.19 |
| mesenchyme | Spg20 | 0.72 | 7.98E-143 | 1.66E-138 | 0.59 | 0.44 | 0.14 | 1.32 |
| mesenchyme | Clip3 | 0.69 | 9.30E-143 | 1.93E-138 | 0.57 | 0.41 | 0.17 | 1.41 |
| mesenchyme | Mex3a | 0.38 | 1.15E-142 | 2.40E-138 | 0.9 | 0.75 | 0.16 | 1.21 |
| mesenchyme | Pwwp3a | 0.72 | 4.54E-142 | 9.44E-138 | 0.6 | 0.46 | 0.14 | 1.31 |
| mesenchyme | Ckap4 | 0.43 | 7.47E-142 | 1.55E-137 | 0.85 | 0.69 | 0.16 | 1.23 |

**Supplementary Table 2. Marker genes for the major cell types of integrated clustering of E11.5 control and NotchGOF λ-junction cells.**

| | | | | | | | | |
|---|---|---|---|---|---|---|---|---|
| mesenchyme | Tuba1b | 0.26 | 7.58E-142 | 1.58E-137 | 1 | 0.97 | 0.03 | 1.03 |
| mesenchyme | Foxf2 | 3.7 | 8.64E-142 | 1.80E-137 | 0.13 | 0.02 | 0.11 | 6.6 |
| mesenchyme | Chaf1a | 0.6 | 1.64E-141 | 3.42E-137 | 0.69 | 0.56 | 0.13 | 1.23 |
| mesenchyme | Itgb5 | 0.39 | 2.09E-141 | 4.36E-137 | 0.65 | 0.45 | 0.21 | 1.47 |
| mesenchyme | Igsf10 | 1.56 | 2.20E-141 | 4.57E-137 | 0.29 | 0.14 | 0.15 | 2.13 |
| mesenchyme | Naa50 | 0.44 | 2.32E-141 | 4.82E-137 | 0.82 | 0.73 | 0.09 | 1.12 |
| mesenchyme | Phip | 0.36 | 2.71E-141 | 5.64E-137 | 0.91 | 0.82 | 0.09 | 1.12 |
| mesenchyme | Laptm4b | 0.77 | 2.87E-141 | 5.97E-137 | 0.6 | 0.45 | 0.15 | 1.33 |
| mesenchyme | St3gal2 | 1.5 | 4.56E-141 | 9.49E-137 | 0.3 | 0.15 | 0.15 | 1.97 |
| mesenchyme | Spock3 | 2.46 | 4.62E-141 | 9.61E-137 | 0.17 | 0.05 | 0.13 | 3.82 |
| mesenchyme | Psmd6 | 0.36 | 5.34E-141 | 1.11E-136 | 0.92 | 0.85 | 0.07 | 1.08 |
| mesenchyme | Ndufs2 | 0.33 | 6.30E-141 | 1.31E-136 | 0.95 | 0.89 | 0.06 | 1.07 |
| mesenchyme | Gng2 | 0.53 | 6.87E-141 | 1.43E-136 | 0.63 | 0.45 | 0.18 | 1.39 |
| mesenchyme | Rell1 | 1.07 | 9.07E-141 | 1.89E-136 | 0.4 | 0.24 | 0.16 | 1.65 |
| mesenchyme | Phf20 | 0.63 | 1.16E-140 | 2.42E-136 | 0.65 | 0.51 | 0.14 | 1.27 |
| mesenchyme | Adnp | 0.4 | 1.56E-140 | 3.25E-136 | 0.87 | 0.77 | 0.11 | 1.14 |
| mesenchyme | Slitrk5 | 2.28 | 1.72E-140 | 3.57E-136 | 0.2 | 0.07 | 0.13 | 2.97 |
| mesenchyme | Mrpl28 | 0.37 | 2.24E-140 | 4.66E-136 | 0.91 | 0.84 | 0.07 | 1.08 |
| mesenchyme | Ror1 | 2.57 | 3.09E-140 | 6.42E-136 | 0.16 | 0.04 | 0.12 | 4.18 |
| mesenchyme | Tpm3 | 0.25 | 3.96E-140 | 8.23E-136 | 0.98 | 0.92 | 0.06 | 1.06 |
| mesenchyme | Tm2d2 | 0.51 | 1.84E-139 | 3.83E-135 | 0.77 | 0.64 | 0.13 | 1.2 |
| mesenchyme | Smc2 | 0.43 | 2.64E-139 | 5.48E-135 | 0.89 | 0.81 | 0.08 | 1.1 |
| mesenchyme | Hspa4 | 0.31 | 4.68E-139 | 9.73E-135 | 0.97 | 0.9 | 0.07 | 1.07 |
| mesenchyme | Asf1a | 0.42 | 4.96E-139 | 1.03E-134 | 0.84 | 0.76 | 0.08 | 1.11 |
| mesenchyme | Ppih | 0.42 | 6.12E-139 | 1.27E-134 | 0.82 | 0.73 | 0.09 | 1.13 |
| mesenchyme | Dhx29 | 0.74 | 7.25E-139 | 1.51E-134 | 0.56 | 0.42 | 0.14 | 1.33 |
| mesenchyme | Clint1 | 0.39 | 8.51E-139 | 1.77E-134 | 0.86 | 0.78 | 0.08 | 1.1 |
| mesenchyme | Ctps | 0.64 | 1.45E-138 | 3.01E-134 | 0.63 | 0.5 | 0.13 | 1.25 |
| mesenchyme | Tbx22 | 3.03 | 1.50E-138 | 3.12E-134 | 0.16 | 0.04 | 0.12 | 4.33 |
| mesenchyme | Gcc2 | 0.57 | 2.21E-138 | 4.60E-134 | 0.71 | 0.58 | 0.12 | 1.21 |
| mesenchyme | Atat1 | 0.68 | 2.46E-138 | 5.12E-134 | 0.61 | 0.46 | 0.15 | 1.33 |
| mesenchyme | Barx1 | 1.92 | 6.40E-138 | 1.33E-133 | 0.27 | 0.12 | 0.15 | 2.26 |
| mesenchyme | Ddx1 | 0.4 | 2.09E-137 | 4.35E-133 | 0.86 | 0.78 | 0.09 | 1.11 |
| mesenchyme | Fkbp11 | 1.25 | 2.45E-137 | 5.10E-133 | 0.35 | 0.2 | 0.15 | 1.78 |
| mesenchyme | Lpgat1 | 0.66 | 3.60E-137 | 7.49E-133 | 0.61 | 0.46 | 0.15 | 1.34 |
| mesenchyme | Gng8 | 0.74 | 1.08E-136 | 2.24E-132 | 0.18 | 0.05 | 0.13 | 3.58 |
| mesenchyme | Lpp | 0.65 | 1.61E-136 | 3.35E-132 | 0.61 | 0.45 | 0.16 | 1.36 |
| mesenchyme | Tmem131l | 0.69 | 1.73E-136 | 3.60E-132 | 0.59 | 0.45 | 0.14 | 1.32 |
| mesenchyme | Ppp1cc | 0.3 | 3.66E-136 | 7.61E-132 | 0.96 | 0.9 | 0.06 | 1.06 |
| mesenchyme | Prkra | 0.53 | 3.99E-136 | 8.31E-132 | 0.72 | 0.59 | 0.13 | 1.22 |
| mesenchyme | Sf3b3 | 0.39 | 4.32E-136 | 8.98E-132 | 0.89 | 0.79 | 0.1 | 1.12 |
| mesenchyme | Itga4 | 1.88 | 4.94E-136 | 1.03E-131 | 0.24 | 0.1 | 0.14 | 2.42 |
| mesenchyme | Col13a1 | 3.01 | 6.31E-136 | 1.31E-131 | 0.15 | 0.03 | 0.12 | 4.71 |
| mesenchyme | Hdgfl2 | 0.51 | 6.37E-136 | 1.33E-131 | 0.72 | 0.61 | 0.11 | 1.19 |
| mesenchyme | Manf | 0.38 | 7.29E-136 | 1.52E-131 | 0.92 | 0.84 | 0.09 | 1.1 |
| mesenchyme | Smarcb1 | 0.37 | 9.18E-136 | 1.91E-131 | 0.88 | 0.79 | 0.09 | 1.11 |
| mesenchyme | Zfp266 | 0.42 | 1.06E-135 | 2.21E-131 | 0.84 | 0.71 | 0.13 | 1.19 |
| mesenchyme | Zyx | 0.56 | 1.15E-135 | 2.38E-131 | 0.56 | 0.4 | 0.16 | 1.39 |
| mesenchyme | Orc6 | 0.43 | 1.20E-135 | 2.50E-131 | 0.83 | 0.74 | 0.09 | 1.12 |
| mesenchyme | Erbin | 0.61 | 1.42E-135 | 2.94E-131 | 0.68 | 0.54 | 0.14 | 1.25 |
| mesenchyme | Adora2b | 2.92 | 4.87E-135 | 1.01E-130 | 0.14 | 0.03 | 0.11 | 4.93 |
| mesenchyme | Suclg1 | 0.37 | 5.00E-135 | 1.04E-130 | 0.9 | 0.81 | 0.09 | 1.11 |
| mesenchyme | Mmp16 | 1.54 | 6.18E-135 | 1.29E-130 | 0.29 | 0.14 | 0.15 | 2.07 |
| mesenchyme | Thoc7 | 0.31 | 7.03E-135 | 1.46E-130 | 0.96 | 0.9 | 0.07 | 1.07 |
| mesenchyme | Tardbp | 0.35 | 7.75E-135 | 1.61E-130 | 0.92 | 0.84 | 0.08 | 1.09 |
| mesenchyme | Yipf5 | 0.61 | 8.12E-135 | 1.69E-130 | 0.66 | 0.53 | 0.13 | 1.24 |

**Supplementary Table 2. Marker genes for the major cell types of integrated clustering of E11.5 control and NotchGOF λ-junction cells.**

| | | | | | | | | |
|---|---|---|---|---|---|---|---|---|
| mesenchyme | Anapc16 | 0.44 | 8.55E-135 | 1.78E-130 | 0.79 | 0.69 | 0.11 | 1.15 |
| mesenchyme | Coa3 | 0.33 | 1.25E-134 | 2.60E-130 | 0.92 | 0.86 | 0.07 | 1.08 |
| mesenchyme | Psmd13 | 0.35 | 1.83E-134 | 3.80E-130 | 0.9 | 0.85 | 0.05 | 1.06 |
| mesenchyme | Leprot | 0.68 | 1.89E-134 | 3.93E-130 | 0.57 | 0.42 | 0.15 | 1.35 |
| mesenchyme | Cuta | 0.39 | 2.21E-134 | 4.60E-130 | 0.88 | 0.78 | 0.1 | 1.12 |
| mesenchyme | Smc1a | 0.32 | 3.46E-134 | 7.19E-130 | 0.97 | 0.9 | 0.06 | 1.07 |
| mesenchyme | Cnih1 | 0.4 | 3.80E-134 | 7.89E-130 | 0.86 | 0.76 | 0.1 | 1.13 |
| mesenchyme | Tpgs2 | 0.59 | 3.85E-134 | 8.01E-130 | 0.63 | 0.49 | 0.14 | 1.28 |
| mesenchyme | Ube2e3 | 0.35 | 4.29E-134 | 8.92E-130 | 0.93 | 0.89 | 0.05 | 1.05 |
| mesenchyme | Eloc | 0.28 | 4.39E-134 | 9.14E-130 | 0.96 | 0.92 | 0.04 | 1.04 |
| mesenchyme | Cul5 | 0.53 | 7.87E-134 | 1.64E-129 | 0.72 | 0.61 | 0.11 | 1.18 |
| mesenchyme | Meaf6 | 0.51 | 8.48E-134 | 1.76E-129 | 0.71 | 0.59 | 0.12 | 1.2 |
| mesenchyme | Timm23 | 0.32 | 1.31E-133 | 2.73E-129 | 0.93 | 0.88 | 0.06 | 1.06 |
| mesenchyme | Rfc4 | 0.42 | 2.73E-133 | 5.67E-129 | 0.82 | 0.74 | 0.09 | 1.12 |
| mesenchyme | Zfas1 | 0.39 | 2.99E-133 | 6.22E-129 | 0.85 | 0.76 | 0.09 | 1.12 |
| mesenchyme | Kansl1 | 0.6 | 3.06E-133 | 6.37E-129 | 0.68 | 0.56 | 0.12 | 1.21 |
| mesenchyme | Nolc1 | 0.42 | 3.84E-133 | 7.98E-129 | 0.84 | 0.74 | 0.09 | 1.13 |
| mesenchyme | Ccdc90b | 0.69 | 5.63E-133 | 1.17E-128 | 0.6 | 0.47 | 0.13 | 1.28 |
| mesenchyme | Ube2s | 0.3 | 9.19E-133 | 1.91E-128 | 0.98 | 0.96 | 0.03 | 1.03 |
| mesenchyme | Ebf1 | 1.02 | 9.59E-133 | 2.00E-128 | 0.35 | 0.19 | 0.16 | 1.85 |
| mesenchyme | Tmem100 | 2.14 | 1.08E-132 | 2.24E-128 | 0.19 | 0.06 | 0.13 | 3.19 |
| mesenchyme | Mlf2 | 0.32 | 1.32E-132 | 2.75E-128 | 0.95 | 0.88 | 0.07 | 1.08 |
| mesenchyme | Ik | 0.4 | 1.48E-132 | 3.07E-128 | 0.85 | 0.75 | 0.1 | 1.14 |
| mesenchyme | Eif4g1 | 0.33 | 1.49E-132 | 3.10E-128 | 0.94 | 0.86 | 0.08 | 1.1 |
| mesenchyme | Ckap5 | 0.49 | 1.78E-132 | 3.71E-128 | 0.8 | 0.71 | 0.1 | 1.14 |
| mesenchyme | Brd8 | 0.49 | 2.19E-132 | 4.55E-128 | 0.78 | 0.7 | 0.09 | 1.12 |
| mesenchyme | Dock7 | 0.57 | 4.02E-132 | 8.37E-128 | 0.66 | 0.52 | 0.14 | 1.28 |
| mesenchyme | Ick | 0.83 | 4.10E-132 | 8.52E-128 | 0.49 | 0.34 | 0.15 | 1.42 |
| mesenchyme | Cyp26b1 | 2.45 | 4.80E-132 | 9.98E-128 | 0.17 | 0.05 | 0.12 | 3.69 |
| mesenchyme | Cdc5l | 0.36 | 4.95E-132 | 1.03E-127 | 0.9 | 0.82 | 0.08 | 1.09 |
| mesenchyme | Sec13 | 0.48 | 5.11E-132 | 1.06E-127 | 0.76 | 0.66 | 0.09 | 1.14 |
| mesenchyme | Nedd4 | 0.27 | 5.13E-132 | 1.07E-127 | 1 | 0.95 | 0.05 | 1.05 |
| mesenchyme | Nol11 | 0.49 | 6.25E-132 | 1.30E-127 | 0.75 | 0.64 | 0.11 | 1.17 |
| mesenchyme | Clic4 | 0.53 | 6.99E-132 | 1.45E-127 | 0.71 | 0.61 | 0.11 | 1.17 |
| mesenchyme | Rab2a | 0.32 | 1.24E-131 | 2.58E-127 | 0.94 | 0.86 | 0.09 | 1.1 |
| mesenchyme | Elavl1 | 0.33 | 1.61E-131 | 3.34E-127 | 0.91 | 0.84 | 0.07 | 1.08 |
| mesenchyme | Mdfi | 0.65 | 2.00E-131 | 4.16E-127 | 0.61 | 0.47 | 0.15 | 1.32 |
| mesenchyme | Brms1l | 0.98 | 3.79E-131 | 7.89E-127 | 0.42 | 0.27 | 0.15 | 1.54 |
| mesenchyme | Psmb7 | 0.3 | 5.38E-131 | 1.12E-126 | 0.95 | 0.9 | 0.05 | 1.05 |
| mesenchyme | Rbm22 | 0.51 | 8.37E-131 | 1.74E-126 | 0.73 | 0.63 | 0.1 | 1.16 |
| mesenchyme | Prrg1 | 1.82 | 1.01E-130 | 2.10E-126 | 0.24 | 0.1 | 0.14 | 2.36 |
| mesenchyme | Bccip | 0.4 | 1.03E-130 | 2.15E-126 | 0.83 | 0.75 | 0.08 | 1.11 |
| mesenchyme | Psmd9 | 0.56 | 1.19E-130 | 2.47E-126 | 0.69 | 0.59 | 0.11 | 1.18 |
| mesenchyme | .500009L16Ri | 0.53 | 1.58E-130 | 3.29E-126 | 0.63 | 0.47 | 0.16 | 1.35 |
| mesenchyme | Psmc4 | 0.31 | 2.19E-130 | 4.56E-126 | 0.93 | 0.87 | 0.06 | 1.07 |
| mesenchyme | Nab2 | 1.07 | 2.72E-130 | 5.65E-126 | 0.36 | 0.21 | 0.15 | 1.73 |
| mesenchyme | Serp2 | 1.68 | 3.26E-130 | 6.78E-126 | 0.24 | 0.11 | 0.14 | 2.3 |
| mesenchyme | Soat1 | 0.76 | 3.35E-130 | 6.96E-126 | 0.49 | 0.34 | 0.15 | 1.45 |
| mesenchyme | Ethe1 | 0.79 | 4.02E-130 | 8.36E-126 | 0.5 | 0.35 | 0.15 | 1.41 |
| mesenchyme | Eif5b | 0.27 | 5.71E-130 | 1.19E-125 | 0.98 | 0.94 | 0.04 | 1.04 |
| mesenchyme | AI606473 | 3.43 | 6.80E-130 | 1.42E-125 | 0.12 | 0.02 | 0.1 | 6.47 |
| mesenchyme | Stag1 | 0.48 | 8.40E-130 | 1.75E-125 | 0.79 | 0.69 | 0.1 | 1.14 |
| mesenchyme | Ddx17 | 0.37 | 8.41E-130 | 1.75E-125 | 0.9 | 0.81 | 0.09 | 1.11 |
| mesenchyme | Akr1a1 | 0.27 | 8.76E-130 | 1.82E-125 | 0.98 | 0.92 | 0.06 | 1.07 |
| mesenchyme | Fam193a | 0.6 | 8.77E-130 | 1.82E-125 | 0.66 | 0.54 | 0.12 | 1.22 |
| mesenchyme | Kansl2 | 0.54 | 9.50E-130 | 1.98E-125 | 0.67 | 0.56 | 0.11 | 1.2 |

**Supplementary Table 2. Marker genes for the major cell types of integrated clustering of E11.5 control and NotchGOF λ-junction cells.**

| | | | | | | | | |
|---|---|---|---|---|---|---|---|---|
| mesenchyme | Map1b | 0.36 | 1.16E-129 | 2.41E-125 | 0.68 | 0.53 | 0.14 | 1.27 |
| mesenchyme | Bud31 | 0.39 | 1.67E-129 | 3.48E-125 | 0.85 | 0.78 | 0.07 | 1.09 |
| mesenchyme | Evl | 0.4 | 1.77E-129 | 3.68E-125 | 0.81 | 0.67 | 0.13 | 1.19 |
| mesenchyme | Cdk5rap2 | 0.62 | 1.91E-129 | 3.97E-125 | 0.65 | 0.53 | 0.12 | 1.23 |
| mesenchyme | Hnrnpl | 0.37 | 2.42E-129 | 5.04E-125 | 0.89 | 0.81 | 0.08 | 1.1 |
| mesenchyme | Inppl1 | 0.93 | 3.42E-129 | 7.11E-125 | 0.44 | 0.3 | 0.15 | 1.5 |
| mesenchyme | Tiprl | 0.49 | 7.12E-129 | 1.48E-124 | 0.72 | 0.63 | 0.1 | 1.15 |
| mesenchyme | Dctn4 | 0.58 | 7.47E-129 | 1.55E-124 | 0.68 | 0.56 | 0.11 | 1.2 |
| mesenchyme | Armc1 | 0.48 | 8.84E-129 | 1.84E-124 | 0.76 | 0.67 | 0.08 | 1.12 |
| mesenchyme | Tsn | 0.27 | 1.58E-128 | 3.28E-124 | 0.98 | 0.93 | 0.05 | 1.05 |
| mesenchyme | Dek | 0.29 | 3.22E-128 | 6.70E-124 | 0.98 | 0.93 | 0.05 | 1.06 |
| mesenchyme | Qk | 0.32 | 4.80E-128 | 9.99E-124 | 0.93 | 0.83 | 0.09 | 1.11 |
| mesenchyme | Prps1 | 0.86 | 5.23E-128 | 1.09E-123 | 0.49 | 0.36 | 0.13 | 1.37 |
| mesenchyme | Vps72 | 0.41 | 6.77E-128 | 1.41E-123 | 0.83 | 0.72 | 0.11 | 1.15 |
| mesenchyme | Ptms | 0.29 | 7.40E-128 | 1.54E-123 | 0.99 | 0.88 | 0.11 | 1.12 |
| mesenchyme | Greb1 | 1.09 | 7.89E-128 | 1.64E-123 | 0.35 | 0.19 | 0.15 | 1.8 |
| mesenchyme | Sdc2 | 1 | 1.01E-127 | 2.10E-123 | 0.42 | 0.27 | 0.15 | 1.57 |
| mesenchyme | Lmnb1 | 0.3 | 1.44E-127 | 2.99E-123 | 0.97 | 0.91 | 0.06 | 1.07 |
| mesenchyme | Thoc1 | 0.52 | 1.72E-127 | 3.58E-123 | 0.71 | 0.59 | 0.11 | 1.19 |
| mesenchyme | Uba2 | 0.34 | 2.15E-127 | 4.46E-123 | 0.89 | 0.81 | 0.08 | 1.1 |
| mesenchyme | Zfp428 | 0.48 | 3.71E-127 | 7.72E-123 | 0.73 | 0.6 | 0.13 | 1.22 |
| mesenchyme | Exoc3 | 0.46 | 5.00E-127 | 1.04E-122 | 0.79 | 0.69 | 0.1 | 1.14 |
| mesenchyme | Ccn1 | 0.94 | 7.86E-127 | 1.63E-122 | 0.36 | 0.2 | 0.16 | 1.77 |
| mesenchyme | Arl1 | 0.35 | 9.40E-127 | 1.95E-122 | 0.91 | 0.83 | 0.09 | 1.11 |
| mesenchyme | Hirip3 | 0.54 | 9.53E-127 | 1.98E-122 | 0.73 | 0.61 | 0.12 | 1.19 |
| mesenchyme | Bnc2 | 1.6 | 1.19E-126 | 2.48E-122 | 0.22 | 0.09 | 0.14 | 2.55 |
| mesenchyme | Timeless | 0.68 | 4.09E-126 | 8.51E-122 | 0.59 | 0.45 | 0.14 | 1.32 |
| mesenchyme | Rab13 | 0.72 | 4.71E-126 | 9.81E-122 | 0.48 | 0.32 | 0.16 | 1.49 |
| mesenchyme | Ei24 | 0.54 | 5.21E-126 | 1.08E-121 | 0.74 | 0.63 | 0.1 | 1.16 |
| mesenchyme | Cited1 | 2.3 | 5.27E-126 | 1.10E-121 | 0.16 | 0.04 | 0.12 | 3.74 |
| mesenchyme | Msrb2 | 1.13 | 8.04E-126 | 1.67E-121 | 0.37 | 0.22 | 0.15 | 1.67 |
| mesenchyme | Srsf1 | 0.31 | 9.30E-126 | 1.93E-121 | 0.95 | 0.87 | 0.08 | 1.09 |
| mesenchyme | Mcm7 | 0.38 | 1.14E-125 | 2.37E-121 | 0.89 | 0.81 | 0.09 | 1.11 |
| mesenchyme | Paip2 | 0.27 | 1.25E-125 | 2.60E-121 | 0.99 | 0.95 | 0.04 | 1.04 |
| mesenchyme | Ddhd2 | 0.94 | 1.61E-125 | 3.36E-121 | 0.43 | 0.29 | 0.15 | 1.51 |
| mesenchyme | Dnttip2 | 0.4 | 1.89E-125 | 3.93E-121 | 0.84 | 0.75 | 0.09 | 1.12 |
| mesenchyme | Timm17b | 0.4 | 2.65E-125 | 5.52E-121 | 0.85 | 0.77 | 0.07 | 1.09 |
| mesenchyme | Dicer1 | 0.42 | 2.67E-125 | 5.56E-121 | 0.85 | 0.74 | 0.1 | 1.14 |
| mesenchyme | 230217C12Ri | 1.68 | 4.24E-125 | 8.82E-121 | 0.22 | 0.09 | 0.13 | 2.48 |
| mesenchyme | Sdc3 | 1.56 | 4.46E-125 | 9.27E-121 | 0.25 | 0.11 | 0.14 | 2.21 |
| mesenchyme | Gstp1 | 0.72 | 4.62E-125 | 9.60E-121 | 0.56 | 0.44 | 0.12 | 1.28 |
| mesenchyme | Smim10l1 | 0.6 | 5.26E-125 | 1.09E-120 | 0.66 | 0.52 | 0.14 | 1.27 |
| mesenchyme | Ciao2a | 0.36 | 9.23E-125 | 1.92E-120 | 0.87 | 0.8 | 0.07 | 1.09 |
| mesenchyme | Mcm5 | 0.49 | 1.38E-124 | 2.88E-120 | 0.78 | 0.66 | 0.12 | 1.19 |
| mesenchyme | Slc35b1 | 0.61 | 2.31E-124 | 4.80E-120 | 0.64 | 0.51 | 0.12 | 1.24 |
| mesenchyme | Rpp25l | 0.46 | 2.41E-124 | 5.01E-120 | 0.77 | 0.68 | 0.09 | 1.14 |
| mesenchyme | Aifm1 | 0.62 | 2.90E-124 | 6.03E-120 | 0.64 | 0.53 | 0.11 | 1.21 |
| mesenchyme | Magohb | 0.59 | 3.90E-124 | 8.11E-120 | 0.61 | 0.49 | 0.13 | 1.26 |
| mesenchyme | Luc7l2 | 0.29 | 8.21E-124 | 1.71E-119 | 0.96 | 0.91 | 0.05 | 1.05 |
| mesenchyme | Ip6k2 | 0.6 | 9.16E-124 | 1.91E-119 | 0.6 | 0.47 | 0.14 | 1.29 |
| mesenchyme | Marcksl1 | 0.26 | 9.39E-124 | 1.95E-119 | 1 | 0.93 | 0.07 | 1.07 |
| mesenchyme | Spcs1 | 0.31 | 1.09E-123 | 2.27E-119 | 0.94 | 0.87 | 0.07 | 1.08 |
| mesenchyme | Tcof1 | 0.43 | 2.06E-123 | 4.28E-119 | 0.79 | 0.67 | 0.12 | 1.18 |
| mesenchyme | Mcrip1 | 0.36 | 2.42E-123 | 5.03E-119 | 0.89 | 0.79 | 0.1 | 1.12 |
| mesenchyme | Rrbp1 | 0.27 | 4.78E-123 | 9.94E-119 | 0.94 | 0.8 | 0.14 | 1.18 |
| mesenchyme | Msantd2 | 0.81 | 6.61E-123 | 1.37E-118 | 0.49 | 0.36 | 0.14 | 1.39 |

**Supplementary Table 2. Marker genes for the major cell types of integrated clustering of E11.5 control and NotchGOF λ-junction cells.**

| | | | | | | | | |
|---|---|---|---|---|---|---|---|---|
| mesenchyme | .610008E11Ri | 0.94 | 1.33E-122 | 2.77E-118 | 0.44 | 0.3 | 0.14 | 1.48 |
| mesenchyme | Rpn1 | 0.4 | 1.83E-122 | 3.81E-118 | 0.81 | 0.67 | 0.13 | 1.2 |
| mesenchyme | Tmem45a | 3.52 | 5.59E-122 | 1.16E-117 | 0.12 | 0.02 | 0.1 | 6.44 |
| mesenchyme | Timm8b | 0.26 | 6.49E-122 | 1.35E-117 | 0.96 | 0.93 | 0.03 | 1.03 |
| mesenchyme | Smarca2 | 0.54 | 6.96E-122 | 1.45E-117 | 0.67 | 0.53 | 0.14 | 1.26 |
| mesenchyme | Ndufa10 | 0.34 | 1.02E-121 | 2.12E-117 | 0.92 | 0.86 | 0.06 | 1.07 |
| mesenchyme | Vdac2 | 0.25 | 1.16E-121 | 2.41E-117 | 0.97 | 0.93 | 0.04 | 1.04 |
| mesenchyme | Rbm4 | 0.45 | 1.20E-121 | 2.50E-117 | 0.74 | 0.63 | 0.11 | 1.17 |
| mesenchyme | Numa1 | 0.49 | 1.29E-121 | 2.68E-117 | 0.76 | 0.65 | 0.11 | 1.17 |
| mesenchyme | Svep1 | 3.6 | 1.49E-121 | 3.09E-117 | 0.12 | 0.02 | 0.1 | 6.44 |
| mesenchyme | Robo1 | 0.76 | 2.07E-121 | 4.30E-117 | 0.51 | 0.37 | 0.14 | 1.38 |
| mesenchyme | Ewsr1 | 0.3 | 2.64E-121 | 5.48E-117 | 0.96 | 0.9 | 0.05 | 1.06 |
| mesenchyme | H1fx | 0.28 | 3.07E-121 | 6.39E-117 | 0.97 | 0.84 | 0.14 | 1.16 |
| mesenchyme | Ndufa1 | 0.37 | 4.18E-121 | 8.70E-117 | 0.88 | 0.83 | 0.05 | 1.06 |
| mesenchyme | Atxn7l3b | 0.26 | 4.98E-121 | 1.04E-116 | 1 | 0.94 | 0.05 | 1.06 |
| mesenchyme | Atrx | 0.29 | 1.24E-120 | 2.57E-116 | 0.98 | 0.92 | 0.06 | 1.07 |
| mesenchyme | Polr2k | 0.29 | 1.55E-120 | 3.22E-116 | 0.93 | 0.88 | 0.05 | 1.06 |
| mesenchyme | Nbea | 0.82 | 1.64E-120 | 3.42E-116 | 0.49 | 0.35 | 0.14 | 1.39 |
| mesenchyme | Ctxn1 | 0.37 | 1.89E-120 | 3.92E-116 | 0.83 | 0.69 | 0.14 | 1.21 |
| mesenchyme | Grb10 | 0.31 | 2.44E-120 | 5.06E-116 | 0.97 | 0.87 | 0.1 | 1.12 |
| mesenchyme | Mthfd2l | 0.53 | 2.68E-120 | 5.57E-116 | 0.68 | 0.55 | 0.14 | 1.26 |
| mesenchyme | Zfp462 | 0.44 | 2.93E-120 | 6.08E-116 | 0.78 | 0.63 | 0.15 | 1.24 |
| mesenchyme | Tmem216 | 0.78 | 3.63E-120 | 7.56E-116 | 0.48 | 0.34 | 0.14 | 1.41 |
| mesenchyme | Fendrr | 3.24 | 3.78E-120 | 7.85E-116 | 0.13 | 0.03 | 0.1 | 4.68 |
| mesenchyme | Fip1l1 | 0.37 | 4.64E-120 | 9.65E-116 | 0.86 | 0.79 | 0.07 | 1.09 |
| mesenchyme | Fam110b | 1.15 | 6.14E-120 | 1.28E-115 | 0.36 | 0.21 | 0.14 | 1.68 |
| mesenchyme | Mogat2 | 2.4 | 6.80E-120 | 1.41E-115 | 0.16 | 0.04 | 0.11 | 3.6 |
| mesenchyme | Cct4 | 0.26 | 1.10E-119 | 2.29E-115 | 0.98 | 0.94 | 0.04 | 1.04 |
| mesenchyme | Itpa | 0.42 | 3.16E-119 | 6.57E-115 | 0.8 | 0.71 | 0.09 | 1.13 |
| mesenchyme | Abl1 | 0.71 | 4.93E-119 | 1.03E-114 | 0.54 | 0.41 | 0.13 | 1.33 |
| mesenchyme | Top2a | 0.66 | 5.72E-119 | 1.19E-114 | 0.81 | 0.74 | 0.07 | 1.1 |
| mesenchyme | Dtl | 0.6 | 7.28E-119 | 1.52E-114 | 0.66 | 0.53 | 0.13 | 1.25 |
| mesenchyme | Dpy19l1 | 1.03 | 8.23E-119 | 1.71E-114 | 0.41 | 0.27 | 0.14 | 1.52 |
| mesenchyme | Glud1 | 0.45 | 1.03E-118 | 2.13E-114 | 0.69 | 0.57 | 0.13 | 1.23 |
| mesenchyme | Gm11627 | 2.06 | 1.14E-118 | 2.37E-114 | 0.17 | 0.06 | 0.12 | 3.05 |
| mesenchyme | Mocs2 | 0.56 | 1.31E-118 | 2.72E-114 | 0.64 | 0.53 | 0.12 | 1.22 |
| mesenchyme | Ephb3 | 1.18 | 2.35E-118 | 4.90E-114 | 0.35 | 0.2 | 0.14 | 1.7 |
| mesenchyme | Clmn | 1.21 | 2.97E-118 | 6.17E-114 | 0.32 | 0.17 | 0.14 | 1.81 |
| mesenchyme | Cops2 | 0.37 | 3.46E-118 | 7.20E-114 | 0.82 | 0.76 | 0.07 | 1.09 |
| mesenchyme | Haus7 | 0.77 | 3.98E-118 | 8.27E-114 | 0.51 | 0.38 | 0.12 | 1.32 |
| mesenchyme | Usp34 | 0.37 | 4.44E-118 | 9.23E-114 | 0.88 | 0.78 | 0.09 | 1.12 |
| mesenchyme | Srpk2 | 0.46 | 4.55E-118 | 9.46E-114 | 0.75 | 0.63 | 0.12 | 1.2 |
| mesenchyme | Pex13 | 0.59 | 4.98E-118 | 1.03E-113 | 0.62 | 0.5 | 0.12 | 1.25 |
| mesenchyme | Pfdn6 | 0.3 | 6.16E-118 | 1.28E-113 | 0.92 | 0.87 | 0.05 | 1.06 |
| mesenchyme | Cmtm7 | 0.49 | 6.21E-118 | 1.29E-113 | 0.54 | 0.38 | 0.16 | 1.41 |
| mesenchyme | Pin1 | 0.33 | 7.42E-118 | 1.54E-113 | 0.9 | 0.84 | 0.06 | 1.07 |
| mesenchyme | Tmeff1 | 0.42 | 1.07E-117 | 2.22E-113 | 0.73 | 0.59 | 0.14 | 1.24 |
| mesenchyme | Chpt1 | 0.94 | 1.11E-117 | 2.30E-113 | 0.44 | 0.3 | 0.14 | 1.47 |
| mesenchyme | Utp11 | 0.42 | 1.54E-117 | 3.20E-113 | 0.8 | 0.7 | 0.1 | 1.14 |
| mesenchyme | Clspn | 0.62 | 1.55E-117 | 3.22E-113 | 0.66 | 0.54 | 0.12 | 1.22 |
| mesenchyme | 700081O15Ri | 0.63 | 2.00E-117 | 4.17E-113 | 0.6 | 0.48 | 0.13 | 1.26 |
| mesenchyme | Hacd1 | 0.87 | 2.57E-117 | 5.35E-113 | 0.44 | 0.3 | 0.14 | 1.44 |
| mesenchyme | Pcyt1a | 1.08 | 3.03E-117 | 6.30E-113 | 0.36 | 0.23 | 0.14 | 1.61 |
| mesenchyme | Rangrf | 0.63 | 3.21E-117 | 6.69E-113 | 0.58 | 0.46 | 0.12 | 1.27 |
| mesenchyme | Tmed10 | 0.29 | 3.58E-117 | 7.44E-113 | 0.96 | 0.9 | 0.06 | 1.07 |
| mesenchyme | Ptpn11 | 0.47 | 4.50E-117 | 9.36E-113 | 0.72 | 0.61 | 0.11 | 1.19 |

**Supplementary Table 2. Marker genes for the major cell types of integrated clustering of E11.5 control and NotchGOF λ-junction cells.**

| | | | | | | | | |
|---|---|---|---|---|---|---|---|---|
| mesenchyme | Tln1 | 0.29 | 9.75E-117 | 2.03E-112 | 0.78 | 0.66 | 0.12 | 1.18 |
| mesenchyme | Cygb | 3.39 | 1.25E-116 | 2.60E-112 | 0.12 | 0.02 | 0.1 | 5.85 |
| mesenchyme | Ash2l | 0.38 | 1.48E-116 | 3.09E-112 | 0.8 | 0.71 | 0.1 | 1.13 |
| mesenchyme | Pcnp | 0.33 | 1.67E-116 | 3.48E-112 | 0.9 | 0.84 | 0.06 | 1.07 |
| mesenchyme | Sh3gl1 | 0.53 | 1.84E-116 | 3.82E-112 | 0.66 | 0.54 | 0.12 | 1.22 |
| mesenchyme | Pmpcb | 0.46 | 2.37E-116 | 4.92E-112 | 0.73 | 0.63 | 0.1 | 1.16 |
| mesenchyme | Arhgap42 | 0.85 | 2.82E-116 | 5.87E-112 | 0.42 | 0.27 | 0.15 | 1.55 |
| mesenchyme | Itm2a | 1.28 | 2.86E-116 | 5.95E-112 | 0.58 | 0.48 | 0.1 | 1.22 |
| mesenchyme | Rab31 | 1.36 | 3.25E-116 | 6.76E-112 | 0.27 | 0.13 | 0.14 | 2.01 |
| mesenchyme | Hist1h3c | 0.85 | 4.80E-116 | 9.98E-112 | 0.63 | 0.52 | 0.11 | 1.21 |
| mesenchyme | Sox4 | 0.35 | 4.89E-116 | 1.02E-111 | 0.98 | 0.85 | 0.13 | 1.15 |
| mesenchyme | Pbx2 | 0.38 | 4.92E-116 | 1.02E-111 | 0.84 | 0.74 | 0.1 | 1.14 |
| mesenchyme | Tcerg1 | 0.36 | 6.20E-116 | 1.29E-111 | 0.88 | 0.79 | 0.08 | 1.11 |
| mesenchyme | Cul3 | 0.36 | 7.88E-116 | 1.64E-111 | 0.87 | 0.79 | 0.08 | 1.1 |
| mesenchyme | Mpc1 | 0.54 | 1.08E-115 | 2.25E-111 | 0.66 | 0.56 | 0.1 | 1.18 |
| mesenchyme | Gpatch8 | 0.4 | 1.09E-115 | 2.28E-111 | 0.82 | 0.73 | 0.08 | 1.11 |
| mesenchyme | Dhx8 | 0.58 | 1.20E-115 | 2.50E-111 | 0.63 | 0.51 | 0.12 | 1.23 |
| mesenchyme | Cenpa | 0.54 | 1.45E-115 | 3.02E-111 | 0.82 | 0.73 | 0.09 | 1.12 |
| mesenchyme | U2af2 | 0.35 | 1.50E-115 | 3.11E-111 | 0.87 | 0.79 | 0.08 | 1.1 |
| mesenchyme | H2-Q7 | 2.41 | 1.54E-115 | 3.21E-111 | 0.14 | 0.04 | 0.11 | 3.89 |
| mesenchyme | Srrm1 | 0.26 | 1.79E-115 | 3.72E-111 | 0.98 | 0.93 | 0.05 | 1.05 |
| mesenchyme | Xrn2 | 0.32 | 1.92E-115 | 4.00E-111 | 0.91 | 0.83 | 0.08 | 1.1 |
| mesenchyme | Hspa13 | 0.71 | 2.16E-115 | 4.49E-111 | 0.54 | 0.41 | 0.14 | 1.33 |
| mesenchyme | Mrpl4 | 0.51 | 2.94E-115 | 6.12E-111 | 0.68 | 0.58 | 0.1 | 1.18 |
| mesenchyme | Cacna1c | 2.9 | 3.37E-115 | 7.01E-111 | 0.13 | 0.03 | 0.1 | 4.81 |
| mesenchyme | Zcchc17 | 0.42 | 5.32E-115 | 1.11E-110 | 0.78 | 0.7 | 0.08 | 1.11 |
| mesenchyme | Mrps17 | 0.38 | 8.07E-115 | 1.68E-110 | 0.83 | 0.75 | 0.08 | 1.1 |
| mesenchyme | Lpin2 | 0.63 | 1.19E-114 | 2.47E-110 | 0.59 | 0.46 | 0.13 | 1.27 |
| mesenchyme | Smc5 | 0.54 | 1.80E-114 | 3.74E-110 | 0.68 | 0.56 | 0.11 | 1.2 |
| mesenchyme | Axin2 | 1.06 | 1.85E-114 | 3.85E-110 | 0.39 | 0.24 | 0.14 | 1.58 |
| mesenchyme | Hist1h2ag | 1.12 | 2.78E-114 | 5.79E-110 | 0.36 | 0.23 | 0.13 | 1.59 |
| mesenchyme | Qdpr | 0.5 | 3.53E-114 | 7.34E-110 | 0.58 | 0.45 | 0.12 | 1.27 |
| mesenchyme | Iws1 | 0.48 | 3.54E-114 | 7.37E-110 | 0.74 | 0.63 | 0.1 | 1.16 |
| mesenchyme | Copb1 | 0.42 | 4.24E-114 | 8.81E-110 | 0.81 | 0.71 | 0.1 | 1.14 |
| mesenchyme | Poglut2 | 0.84 | 4.90E-114 | 1.02E-109 | 0.44 | 0.3 | 0.14 | 1.45 |
| mesenchyme | Ddx21 | 0.44 | 5.62E-114 | 1.17E-109 | 0.76 | 0.67 | 0.09 | 1.13 |
| mesenchyme | Dhx30 | 0.58 | 1.33E-113 | 2.77E-109 | 0.65 | 0.53 | 0.12 | 1.23 |
| mesenchyme | Wipi1 | 1 | 1.36E-113 | 2.82E-109 | 0.44 | 0.31 | 0.14 | 1.44 |
| mesenchyme | Arid4b | 0.32 | 1.67E-113 | 3.47E-109 | 0.89 | 0.81 | 0.09 | 1.11 |
| mesenchyme | Bbip1 | 0.45 | 1.78E-113 | 3.71E-109 | 0.73 | 0.65 | 0.08 | 1.13 |
| mesenchyme | Cops9 | 0.28 | 1.94E-113 | 4.05E-109 | 0.94 | 0.89 | 0.05 | 1.05 |
| mesenchyme | Fkbp5 | 0.77 | 2.20E-113 | 4.58E-109 | 0.47 | 0.33 | 0.14 | 1.42 |
| mesenchyme | Slirp | 0.32 | 2.38E-113 | 4.95E-109 | 0.9 | 0.85 | 0.06 | 1.07 |
| mesenchyme | Zfp131 | 0.59 | 2.48E-113 | 5.17E-109 | 0.61 | 0.49 | 0.12 | 1.24 |
| mesenchyme | Rpl7l1 | 0.34 | 3.99E-113 | 8.29E-109 | 0.88 | 0.81 | 0.07 | 1.09 |
| mesenchyme | Ntn3 | 1.81 | 4.55E-113 | 9.47E-109 | 0.2 | 0.08 | 0.12 | 2.55 |
| mesenchyme | Kcp | 2.58 | 6.67E-113 | 1.39E-108 | 0.14 | 0.04 | 0.1 | 3.67 |
| mesenchyme | Shisa4 | 1.01 | 6.69E-113 | 1.39E-108 | 0.38 | 0.24 | 0.14 | 1.59 |
| mesenchyme | Mtfr1l | 0.53 | 7.47E-113 | 1.55E-108 | 0.66 | 0.56 | 0.1 | 1.18 |
| mesenchyme | Tmed1 | 0.5 | 1.00E-112 | 2.08E-108 | 0.71 | 0.58 | 0.13 | 1.23 |
| mesenchyme | Srp9 | 0.29 | 1.23E-112 | 2.56E-108 | 0.94 | 0.88 | 0.06 | 1.07 |
| mesenchyme | Sqle | 0.6 | 1.47E-112 | 3.05E-108 | 0.62 | 0.5 | 0.12 | 1.24 |
| mesenchyme | Tmem126a | 0.44 | 2.40E-112 | 5.00E-108 | 0.76 | 0.65 | 0.1 | 1.16 |
| mesenchyme | Srsf4 | 0.42 | 3.15E-112 | 6.55E-108 | 0.76 | 0.66 | 0.1 | 1.15 |
| mesenchyme | Pam16 | 0.39 | 4.87E-112 | 1.01E-107 | 0.81 | 0.75 | 0.06 | 1.09 |
| mesenchyme | Med4 | 0.43 | 5.35E-112 | 1.11E-107 | 0.74 | 0.64 | 0.1 | 1.16 |

**Supplementary Table 2. Marker genes for the major cell types of integrated clustering of E11.5 control and NotchGOF λ-junction cells.**

| | | | | | | | | |
|---|---|---|---|---|---|---|---|---|
| mesenchyme | Smchd1 | 0.28 | 6.37E-112 | 1.32E-107 | 0.88 | 0.8 | 0.08 | 1.1 |
| mesenchyme | Antxr2 | 1.6 | 6.84E-112 | 1.42E-107 | 0.23 | 0.1 | 0.13 | 2.23 |
| mesenchyme | Hs2st1 | 0.6 | 7.38E-112 | 1.53E-107 | 0.61 | 0.48 | 0.12 | 1.25 |
| mesenchyme | Dctpp1 | 0.28 | 8.81E-112 | 1.83E-107 | 0.95 | 0.9 | 0.04 | 1.05 |
| mesenchyme | Rnf150 | 1.07 | 8.87E-112 | 1.84E-107 | 0.36 | 0.22 | 0.14 | 1.64 |
| mesenchyme | Htatsf1 | 0.43 | 1.17E-111 | 2.43E-107 | 0.78 | 0.68 | 0.09 | 1.14 |
| mesenchyme | Wbp11 | 0.34 | 1.29E-111 | 2.69E-107 | 0.88 | 0.81 | 0.07 | 1.09 |
| mesenchyme | Ssrp1 | 0.28 | 1.43E-111 | 2.97E-107 | 0.95 | 0.88 | 0.07 | 1.08 |
| mesenchyme | Tmsb15b2 | 0.57 | 1.63E-111 | 3.40E-107 | 0.59 | 0.45 | 0.14 | 1.31 |
| mesenchyme | Kif2a | 0.39 | 1.71E-111 | 3.56E-107 | 0.82 | 0.74 | 0.08 | 1.11 |
| mesenchyme | Tmem263 | 0.73 | 1.79E-111 | 3.71E-107 | 0.51 | 0.39 | 0.12 | 1.32 |
| mesenchyme | Dhfr | 0.49 | 2.12E-111 | 4.42E-107 | 0.72 | 0.59 | 0.12 | 1.21 |
| mesenchyme | Snrpa | 0.3 | 2.97E-111 | 6.18E-107 | 0.95 | 0.88 | 0.07 | 1.08 |
| mesenchyme | Itga9 | 1.42 | 3.28E-111 | 6.82E-107 | 0.26 | 0.13 | 0.13 | 2.02 |
| mesenchyme | Adamtsl1 | 2.59 | 3.83E-111 | 7.97E-107 | 0.14 | 0.04 | 0.1 | 3.86 |
| mesenchyme | Lrrc17 | 3.53 | 5.19E-111 | 1.08E-106 | 0.11 | 0.02 | 0.09 | 6.35 |
| mesenchyme | Ptprg | 0.98 | 5.52E-111 | 1.15E-106 | 0.41 | 0.28 | 0.13 | 1.48 |
| mesenchyme | Mrps15 | 0.37 | 6.20E-111 | 1.29E-106 | 0.83 | 0.77 | 0.06 | 1.08 |
| mesenchyme | Chordc1 | 0.41 | 9.49E-111 | 1.97E-106 | 0.77 | 0.69 | 0.09 | 1.13 |
| mesenchyme | Psat1 | 0.7 | 2.41E-110 | 5.00E-106 | 0.5 | 0.36 | 0.14 | 1.38 |
| mesenchyme | Tmem164 | 1.13 | 3.62E-110 | 7.54E-106 | 0.36 | 0.23 | 0.13 | 1.57 |
| mesenchyme | Stra6 | 0.79 | 3.67E-110 | 7.63E-106 | 0.39 | 0.24 | 0.15 | 1.63 |
| mesenchyme | Rnf115 | 0.65 | 3.78E-110 | 7.86E-106 | 0.56 | 0.45 | 0.11 | 1.25 |
| mesenchyme | Fign | 1.08 | 4.50E-110 | 9.37E-106 | 0.34 | 0.2 | 0.14 | 1.7 |
| mesenchyme | Ube3a | 0.36 | 4.92E-110 | 1.02E-105 | 0.87 | 0.78 | 0.09 | 1.11 |
| mesenchyme | Gnpnat1 | 0.72 | 5.49E-110 | 1.14E-105 | 0.47 | 0.34 | 0.13 | 1.4 |
| mesenchyme | Clk1 | 0.49 | 5.83E-110 | 1.21E-105 | 0.71 | 0.62 | 0.1 | 1.15 |
| mesenchyme | Pgm2l1 | 1.24 | 7.18E-110 | 1.49E-105 | 0.4 | 0.28 | 0.12 | 1.43 |
| mesenchyme | Mthfd2 | 0.74 | 7.87E-110 | 1.64E-105 | 0.51 | 0.39 | 0.12 | 1.31 |
| mesenchyme | Tpst1 | 0.99 | 1.03E-109 | 2.14E-105 | 0.38 | 0.25 | 0.14 | 1.55 |
| mesenchyme | Ctcf | 0.33 | 1.19E-109 | 2.48E-105 | 0.91 | 0.84 | 0.07 | 1.08 |
| mesenchyme | Mphosph10 | 0.41 | 1.30E-109 | 2.70E-105 | 0.78 | 0.7 | 0.09 | 1.12 |
| mesenchyme | Dpf1 | 1.8 | 1.72E-109 | 3.57E-105 | 0.2 | 0.08 | 0.12 | 2.44 |
| mesenchyme | Dnajc7 | 0.3 | 2.03E-109 | 4.23E-105 | 0.9 | 0.81 | 0.09 | 1.12 |
| mesenchyme | Nsd1 | 0.33 | 3.00E-109 | 6.24E-105 | 0.9 | 0.81 | 0.1 | 1.12 |
| mesenchyme | Bcl9 | 0.44 | 3.59E-109 | 7.47E-105 | 0.73 | 0.6 | 0.13 | 1.22 |
| mesenchyme | Sav1 | 0.77 | 3.87E-109 | 8.04E-105 | 0.49 | 0.37 | 0.12 | 1.33 |
| mesenchyme | Bmper | 3.46 | 5.68E-109 | 1.18E-104 | 0.11 | 0.02 | 0.09 | 5.79 |
| mesenchyme | Ap1m1 | 0.89 | 5.94E-109 | 1.23E-104 | 0.43 | 0.3 | 0.13 | 1.42 |
| mesenchyme | Zfp334 | 0.93 | 6.18E-109 | 1.28E-104 | 0.38 | 0.24 | 0.14 | 1.57 |
| mesenchyme | Auts2 | 0.35 | 6.45E-109 | 1.34E-104 | 0.92 | 0.77 | 0.15 | 1.19 |
| mesenchyme | Phb | 0.37 | 1.17E-108 | 2.44E-104 | 0.83 | 0.74 | 0.09 | 1.11 |
| mesenchyme | Homer3 | 0.67 | 1.17E-108 | 2.44E-104 | 0.53 | 0.39 | 0.13 | 1.34 |
| mesenchyme | Srprb | 0.68 | 2.66E-108 | 5.52E-104 | 0.54 | 0.42 | 0.12 | 1.28 |
| mesenchyme | Rcc2 | 0.27 | 8.19E-108 | 1.70E-103 | 0.95 | 0.89 | 0.06 | 1.07 |
| mesenchyme | Nrp1 | 1.44 | 8.61E-108 | 1.79E-103 | 0.25 | 0.12 | 0.13 | 2.09 |
| mesenchyme | Fndc4 | 0.8 | 8.98E-108 | 1.87E-103 | 0.46 | 0.32 | 0.13 | 1.41 |
| mesenchyme | Zfp68 | 0.76 | 1.23E-107 | 2.56E-103 | 0.47 | 0.35 | 0.13 | 1.37 |
| mesenchyme | Cops3 | 0.36 | 1.25E-107 | 2.60E-103 | 0.83 | 0.75 | 0.09 | 1.12 |
| mesenchyme | Cdc14b | 0.8 | 1.25E-107 | 2.61E-103 | 0.45 | 0.31 | 0.14 | 1.44 |
| mesenchyme | Ptgr1 | 1.13 | 1.41E-107 | 2.93E-103 | 0.31 | 0.18 | 0.13 | 1.76 |
| mesenchyme | Nbdy | 0.48 | 1.51E-107 | 3.14E-103 | 0.66 | 0.55 | 0.12 | 1.22 |
| mesenchyme | Zfp521 | 1.12 | 1.56E-107 | 3.24E-103 | 0.33 | 0.2 | 0.13 | 1.67 |
| mesenchyme | Flrt3 | 0.8 | 2.60E-107 | 5.40E-103 | 0.49 | 0.35 | 0.14 | 1.4 |
| mesenchyme | Syne1 | 1.93 | 3.11E-107 | 6.47E-103 | 0.17 | 0.06 | 0.11 | 2.8 |
| mesenchyme | Prkacb | 0.61 | 3.43E-107 | 7.12E-103 | 0.57 | 0.45 | 0.12 | 1.27 |

**Supplementary Table 2. Marker genes for the major cell types of integrated clustering of E11.5 control and NotchGOF λ-junction cells.**

| | | | | | | | | |
|---|---|---|---|---|---|---|---|---|
| mesenchyme | Ddah2 | 0.25 | 5.08E-107 | 1.06E-102 | 0.98 | 0.88 | 0.1 | 1.12 |
| mesenchyme | Trmt112 | 0.26 | 7.42E-107 | 1.54E-102 | 0.96 | 0.9 | 0.06 | 1.07 |
| mesenchyme | Plppr3 | 0.79 | 7.57E-107 | 1.58E-102 | 0.43 | 0.29 | 0.14 | 1.49 |
| mesenchyme | Bclaf1 | 0.28 | 7.82E-107 | 1.63E-102 | 0.96 | 0.89 | 0.07 | 1.08 |
| mesenchyme | Clock | 1.02 | 8.60E-107 | 1.79E-102 | 0.36 | 0.22 | 0.14 | 1.61 |
| mesenchyme | Ipo9 | 0.6 | 9.62E-107 | 2.00E-102 | 0.58 | 0.46 | 0.12 | 1.25 |
| mesenchyme | Hypk | 0.33 | 1.64E-106 | 3.41E-102 | 0.86 | 0.81 | 0.05 | 1.07 |
| mesenchyme | Cspp1 | 0.46 | 1.90E-106 | 3.95E-102 | 0.72 | 0.62 | 0.1 | 1.16 |
| mesenchyme | Pnisr | 0.3 | 2.39E-106 | 4.98E-102 | 0.95 | 0.86 | 0.08 | 1.1 |
| mesenchyme | Alkal2 | 3.27 | 2.47E-106 | 5.14E-102 | 0.11 | 0.02 | 0.09 | 5.14 |
| mesenchyme | Gpr45 | 1.53 | 2.86E-106 | 5.94E-102 | 0.22 | 0.1 | 0.12 | 2.21 |
| mesenchyme | Sh3bp5 | 0.37 | 3.80E-106 | 7.89E-102 | 0.56 | 0.4 | 0.16 | 1.4 |
| mesenchyme | Leo1 | 0.5 | 4.03E-106 | 8.38E-102 | 0.67 | 0.56 | 0.1 | 1.18 |
| mesenchyme | 610012G03Ri | 0.3 | 4.40E-106 | 9.16E-102 | 0.89 | 0.81 | 0.08 | 1.09 |
| mesenchyme | Thoc3 | 0.36 | 5.07E-106 | 1.05E-101 | 0.82 | 0.73 | 0.09 | 1.13 |
| mesenchyme | Ptpdc1 | 1.67 | 8.21E-106 | 1.71E-101 | 0.2 | 0.09 | 0.12 | 2.34 |
| mesenchyme | Tmem98 | 0.65 | 8.26E-106 | 1.72E-101 | 0.54 | 0.42 | 0.12 | 1.3 |
| mesenchyme | Vstm2b | 2.35 | 1.34E-105 | 2.79E-101 | 0.13 | 0.03 | 0.1 | 4.13 |
| mesenchyme | Mrpl18 | 0.27 | 1.61E-105 | 3.36E-101 | 0.95 | 0.9 | 0.05 | 1.06 |
| mesenchyme | Ube2l3 | 0.25 | 1.80E-105 | 3.74E-101 | 0.96 | 0.92 | 0.04 | 1.04 |
| mesenchyme | Uhrf2 | 0.45 | 2.53E-105 | 5.26E-101 | 0.76 | 0.68 | 0.09 | 1.13 |
| mesenchyme | Smarcc2 | 0.48 | 2.56E-105 | 5.32E-101 | 0.69 | 0.59 | 0.1 | 1.17 |
| mesenchyme | Ndufb11 | 0.25 | 2.61E-105 | 5.43E-101 | 0.98 | 0.94 | 0.04 | 1.04 |
| mesenchyme | Sephs1 | 0.53 | 3.13E-105 | 6.51E-101 | 0.64 | 0.55 | 0.1 | 1.18 |
| mesenchyme | Armcx2 | 0.57 | 3.16E-105 | 6.58E-101 | 0.58 | 0.44 | 0.14 | 1.31 |
| mesenchyme | Klhdc2 | 0.3 | 8.80E-105 | 1.83E-100 | 0.9 | 0.85 | 0.05 | 1.06 |
| mesenchyme | Bub3 | 0.39 | 9.09E-105 | 1.89E-100 | 0.85 | 0.78 | 0.07 | 1.09 |
| mesenchyme | Thap12 | 0.44 | 1.06E-104 | 2.21E-100 | 0.73 | 0.62 | 0.11 | 1.17 |
| mesenchyme | Gm7324 | 0.49 | 1.73E-104 | 3.60E-100 | 0.65 | 0.55 | 0.11 | 1.19 |
| mesenchyme | Cby1 | 0.62 | 2.40E-104 | 4.98E-100 | 0.56 | 0.44 | 0.12 | 1.27 |
| mesenchyme | Baiap2l2 | 2.14 | 2.61E-104 | 5.42E-100 | 0.15 | 0.05 | 0.1 | 3.24 |
| mesenchyme | Tmem258 | 0.26 | 3.53E-104 | 7.35E-100 | 0.96 | 0.92 | 0.04 | 1.04 |
| mesenchyme | Snx6 | 0.26 | 5.13E-104 | 1.07E-99 | 0.96 | 0.85 | 0.1 | 1.12 |
| mesenchyme | Dock4 | 0.96 | 6.28E-104 | 1.31E-99 | 0.36 | 0.22 | 0.14 | 1.62 |
| mesenchyme | Ddx42 | 0.32 | 7.44E-104 | 1.55E-99 | 0.87 | 0.77 | 0.1 | 1.12 |
| mesenchyme | Lef1 | 0.67 | 1.07E-103 | 2.22E-99 | 0.51 | 0.36 | 0.15 | 1.41 |
| mesenchyme | Tanc1 | 0.7 | 1.10E-103 | 2.29E-99 | 0.49 | 0.36 | 0.14 | 1.38 |
| mesenchyme | Igdcc4 | 2.04 | 1.23E-103 | 2.55E-99 | 0.16 | 0.06 | 0.11 | 2.95 |
| mesenchyme | Impa1 | 0.55 | 1.28E-103 | 2.66E-99 | 0.62 | 0.52 | 0.1 | 1.2 |
| mesenchyme | Ddx24 | 0.28 | 1.58E-103 | 3.28E-99 | 0.93 | 0.88 | 0.05 | 1.05 |
| mesenchyme | Zik1 | 0.9 | 1.65E-103 | 3.44E-99 | 0.38 | 0.25 | 0.14 | 1.54 |
| mesenchyme | Fam133b | 0.38 | 1.84E-103 | 3.83E-99 | 0.81 | 0.72 | 0.09 | 1.12 |
| mesenchyme | \830018L16Ri | 2.14 | 2.02E-103 | 4.21E-99 | 0.16 | 0.06 | 0.11 | 2.91 |
| mesenchyme | Mcm3 | 0.39 | 2.20E-103 | 4.57E-99 | 0.83 | 0.71 | 0.12 | 1.17 |
| mesenchyme | Wipf1 | 1.24 | 2.47E-103 | 5.13E-99 | 0.27 | 0.15 | 0.13 | 1.88 |
| mesenchyme | Emc2 | 0.54 | 2.62E-103 | 5.44E-99 | 0.63 | 0.53 | 0.1 | 1.19 |
| mesenchyme | Abce1 | 0.39 | 4.42E-103 | 9.20E-99 | 0.79 | 0.71 | 0.08 | 1.12 |
| mesenchyme | Islr | 3.18 | 5.42E-103 | 1.13E-98 | 0.11 | 0.02 | 0.09 | 4.87 |
| mesenchyme | Rsl24d1 | 0.36 | 5.95E-103 | 1.24E-98 | 0.76 | 0.67 | 0.09 | 1.14 |
| mesenchyme | Tasor2 | 0.63 | 5.95E-103 | 1.24E-98 | 0.55 | 0.43 | 0.12 | 1.28 |
| mesenchyme | Hax1 | 0.48 | 6.47E-103 | 1.34E-98 | 0.7 | 0.61 | 0.09 | 1.14 |
| mesenchyme | Polr2b | 0.48 | 6.66E-103 | 1.38E-98 | 0.71 | 0.62 | 0.09 | 1.15 |
| mesenchyme | Ergic2 | 0.35 | 8.68E-103 | 1.81E-98 | 0.82 | 0.75 | 0.07 | 1.1 |
| mesenchyme | Zfp61 | 1.23 | 8.89E-103 | 1.85E-98 | 0.28 | 0.15 | 0.13 | 1.83 |
| mesenchyme | 632427E13Ri | 0.47 | 1.03E-102 | 2.14E-98 | 0.69 | 0.57 | 0.12 | 1.21 |
| mesenchyme | Ccdc59 | 0.35 | 1.08E-102 | 2.25E-98 | 0.82 | 0.73 | 0.09 | 1.12 |

**Supplementary Table 2. Marker genes for the major cell types of integrated clustering of E11.5 control and NotchGOF λ-junction cells.**

| | | | | | | | | |
|---|---|---|---|---|---|---|---|---|
| mesenchyme | Jakmip2 | 1.15 | 1.10E-102 | 2.29E-98 | 0.18 | 0.07 | 0.12 | 2.69 |
| mesenchyme | Akap11 | 0.67 | 1.61E-102 | 3.34E-98 | 0.52 | 0.4 | 0.13 | 1.32 |
| mesenchyme | Gusb | 0.74 | 3.38E-102 | 7.03E-98 | 0.35 | 0.22 | 0.14 | 1.63 |
| mesenchyme | Tle4 | 0.61 | 4.15E-102 | 8.64E-98 | 0.62 | 0.51 | 0.11 | 1.21 |
| mesenchyme | Hmg20b | 0.34 | 6.41E-102 | 1.33E-97 | 0.83 | 0.74 | 0.09 | 1.13 |
| mesenchyme | Arhgap5 | 0.32 | 6.43E-102 | 1.34E-97 | 0.9 | 0.78 | 0.13 | 1.16 |
| mesenchyme | Mpdz | 0.59 | 8.47E-102 | 1.76E-97 | 0.57 | 0.44 | 0.13 | 1.3 |
| mesenchyme | Utp3 | 0.28 | 1.07E-101 | 2.23E-97 | 0.9 | 0.83 | 0.07 | 1.09 |
| mesenchyme | Tbc1d16 | 0.67 | 1.29E-101 | 2.68E-97 | 0.49 | 0.36 | 0.13 | 1.36 |
| mesenchyme | Samd14 | 0.72 | 1.36E-101 | 2.83E-97 | 0.45 | 0.33 | 0.12 | 1.38 |
| mesenchyme | Haus3 | 0.64 | 1.48E-101 | 3.08E-97 | 0.53 | 0.43 | 0.11 | 1.25 |
| mesenchyme | Dnmt1 | 0.4 | 1.56E-101 | 3.25E-97 | 0.82 | 0.73 | 0.08 | 1.11 |
| mesenchyme | Dnaja2 | 0.25 | 1.60E-101 | 3.32E-97 | 0.96 | 0.9 | 0.05 | 1.06 |
| mesenchyme | Smndc1 | 0.43 | 2.02E-101 | 4.20E-97 | 0.71 | 0.63 | 0.09 | 1.14 |
| mesenchyme | Dnajc9 | 0.3 | 2.10E-101 | 4.38E-97 | 0.89 | 0.81 | 0.08 | 1.1 |
| mesenchyme | Phactr3 | 3.1 | 2.15E-101 | 4.47E-97 | 0.11 | 0.02 | 0.09 | 5.53 |
| mesenchyme | Sugt1 | 0.28 | 3.02E-101 | 6.27E-97 | 0.9 | 0.82 | 0.08 | 1.09 |
| mesenchyme | Rgs5 | 3.02 | 3.08E-101 | 6.41E-97 | 0.11 | 0.02 | 0.09 | 4.71 |
| mesenchyme | Rbm17 | 0.29 | 4.94E-101 | 1.03E-96 | 0.91 | 0.84 | 0.07 | 1.09 |
| mesenchyme | Ccdc127 | 0.52 | 5.29E-101 | 1.10E-96 | 0.63 | 0.53 | 0.1 | 1.18 |
| mesenchyme | Hnrnpll | 0.4 | 6.32E-101 | 1.31E-96 | 0.76 | 0.65 | 0.1 | 1.16 |
| mesenchyme | Sar1a | 0.39 | 6.45E-101 | 1.34E-96 | 0.79 | 0.72 | 0.07 | 1.1 |
| mesenchyme | Cenpm | 0.54 | 6.66E-101 | 1.38E-96 | 0.63 | 0.54 | 0.1 | 1.18 |
| mesenchyme | Bcl2l12 | 0.62 | 7.23E-101 | 1.50E-96 | 0.54 | 0.42 | 0.12 | 1.29 |
| mesenchyme | Kif3a | 0.48 | 7.71E-101 | 1.60E-96 | 0.66 | 0.54 | 0.12 | 1.22 |
| mesenchyme | Tpmt | 0.84 | 8.04E-101 | 1.67E-96 | 0.41 | 0.29 | 0.13 | 1.45 |
| mesenchyme | Hmgcs1 | 0.36 | 8.88E-101 | 1.85E-96 | 0.81 | 0.7 | 0.11 | 1.16 |
| mesenchyme | Prim1 | 0.41 | 1.01E-100 | 2.10E-96 | 0.71 | 0.6 | 0.11 | 1.17 |
| mesenchyme | Yeats2 | 0.54 | 1.21E-100 | 2.52E-96 | 0.62 | 0.5 | 0.12 | 1.24 |
| mesenchyme | Psmg1 | 0.49 | 1.34E-100 | 2.79E-96 | 0.67 | 0.56 | 0.11 | 1.19 |
| mesenchyme | Sh3kbp1 | 1.24 | 1.45E-100 | 3.02E-96 | 0.26 | 0.14 | 0.12 | 1.87 |
| mesenchyme | Rora | 1.53 | 2.00E-100 | 4.16E-96 | 0.22 | 0.1 | 0.12 | 2.15 |
| mesenchyme | Tyms | 0.35 | 2.24E-100 | 4.66E-96 | 0.86 | 0.78 | 0.08 | 1.1 |
| mesenchyme | Prpf4 | 0.55 | 3.19E-100 | 6.63E-96 | 0.62 | 0.52 | 0.1 | 1.2 |
| mesenchyme | Mapk8 | 0.78 | 3.21E-100 | 6.68E-96 | 0.47 | 0.34 | 0.12 | 1.36 |
| mesenchyme | Gulo | 2.53 | 3.50E-100 | 7.27E-96 | 0.13 | 0.03 | 0.09 | 3.85 |
| mesenchyme | Gtf2e2 | 0.37 | 3.81E-100 | 7.92E-96 | 0.79 | 0.71 | 0.08 | 1.11 |
| mesenchyme | Naa35 | 0.39 | 3.88E-100 | 8.07E-96 | 0.76 | 0.66 | 0.1 | 1.14 |
| mesenchyme | Cand1 | 0.47 | 3.99E-100 | 8.29E-96 | 0.68 | 0.57 | 0.11 | 1.19 |
| mesenchyme | Abhd8 | 0.65 | 4.13E-100 | 8.59E-96 | 0.52 | 0.4 | 0.12 | 1.29 |
| mesenchyme | Rfc2 | 0.35 | 4.42E-100 | 9.19E-96 | 0.83 | 0.74 | 0.09 | 1.13 |
| mesenchyme | Smarca4 | 0.29 | 5.13E-100 | 1.07E-95 | 0.93 | 0.84 | 0.09 | 1.11 |
| mesenchyme | Sgip1 | 2.15 | 5.84E-100 | 1.21E-95 | 0.14 | 0.04 | 0.1 | 3.27 |
| mesenchyme | Cnrip1 | 1.85 | 6.05E-100 | 1.26E-95 | 0.18 | 0.07 | 0.11 | 2.59 |
| mesenchyme | Caml | 0.65 | 8.55E-100 | 1.78E-95 | 0.53 | 0.42 | 0.12 | 1.27 |
| mesenchyme | C1d | 0.32 | 1.71E-99 | 3.56E-95 | 0.87 | 0.79 | 0.08 | 1.1 |
| mesenchyme | Bri3bp | 0.51 | 1.79E-99 | 3.73E-95 | 0.63 | 0.52 | 0.12 | 1.23 |
| mesenchyme | Cdc23 | 0.55 | 2.10E-99 | 4.36E-95 | 0.6 | 0.5 | 0.1 | 1.21 |
| mesenchyme | Akt1 | 0.39 | 2.40E-99 | 5.00E-95 | 0.76 | 0.65 | 0.11 | 1.17 |
| mesenchyme | Bcl2 | 0.83 | 2.61E-99 | 5.43E-95 | 0.4 | 0.26 | 0.13 | 1.51 |
| mesenchyme | Guk1 | 0.37 | 3.20E-99 | 6.66E-95 | 0.8 | 0.74 | 0.07 | 1.09 |
| mesenchyme | Csk | 0.55 | 6.57E-99 | 1.37E-94 | 0.58 | 0.47 | 0.12 | 1.25 |
| mesenchyme | Sdha | 0.39 | 8.56E-99 | 1.78E-94 | 0.8 | 0.72 | 0.08 | 1.12 |
| mesenchyme | Ppp2r2d | 0.41 | 1.13E-98 | 2.35E-94 | 0.71 | 0.62 | 0.09 | 1.15 |
| mesenchyme | Psmc5 | 0.26 | 1.54E-98 | 3.20E-94 | 0.93 | 0.88 | 0.05 | 1.06 |
| mesenchyme | Trappc4 | 0.32 | 1.69E-98 | 3.51E-94 | 0.85 | 0.77 | 0.08 | 1.1 |

**Supplementary Table 2. Marker genes for the major cell types of integrated clustering of E11.5 control and NotchGOF λ-junction cells.**

| | | | | | | | | |
|---|---|---|---|---|---|---|---|---|
| mesenchyme | Lsp1 | 0.58 | 1.75E-98 | 3.64E-94 | 0.2 | 0.08 | 0.12 | 2.4 |
| mesenchyme | Nktr | 0.33 | 1.76E-98 | 3.66E-94 | 0.88 | 0.79 | 0.09 | 1.11 |
| mesenchyme | Casp6 | 0.52 | 1.78E-98 | 3.71E-94 | 0.6 | 0.48 | 0.12 | 1.26 |
| mesenchyme | Wdr6 | 0.44 | 2.18E-98 | 4.53E-94 | 0.71 | 0.58 | 0.13 | 1.22 |
| mesenchyme | Sucla2 | 0.47 | 2.27E-98 | 4.72E-94 | 0.69 | 0.61 | 0.09 | 1.14 |
| mesenchyme | Prkaca | 0.55 | 2.90E-98 | 6.02E-94 | 0.6 | 0.49 | 0.11 | 1.22 |
| mesenchyme | Hip1 | 0.69 | 3.27E-98 | 6.81E-94 | 0.49 | 0.36 | 0.13 | 1.35 |
| mesenchyme | Trappc3 | 0.43 | 3.81E-98 | 7.92E-94 | 0.71 | 0.61 | 0.1 | 1.16 |
| mesenchyme | Ebf3 | 0.81 | 4.31E-98 | 8.97E-94 | 0.21 | 0.1 | 0.12 | 2.24 |
| mesenchyme | Gpbp1 | 0.36 | 7.36E-98 | 1.53E-93 | 0.81 | 0.74 | 0.07 | 1.1 |
| mesenchyme | Nsmce2 | 0.37 | 8.42E-98 | 1.75E-93 | 0.8 | 0.7 | 0.09 | 1.13 |
| mesenchyme | Rsrc1 | 0.41 | 8.68E-98 | 1.81E-93 | 0.77 | 0.69 | 0.09 | 1.12 |
| mesenchyme | Snrnp40 | 0.3 | 9.81E-98 | 2.04E-93 | 0.88 | 0.8 | 0.08 | 1.1 |
| mesenchyme | Nfic | 1 | 1.01E-97 | 2.11E-93 | 0.34 | 0.21 | 0.13 | 1.61 |
| mesenchyme | Lpar4 | 0.81 | 1.61E-97 | 3.34E-93 | 0.37 | 0.23 | 0.14 | 1.59 |
| mesenchyme | Cdc16 | 0.4 | 1.70E-97 | 3.54E-93 | 0.76 | 0.67 | 0.09 | 1.14 |
| mesenchyme | Stard3nl | 0.46 | 1.73E-97 | 3.59E-93 | 0.63 | 0.51 | 0.12 | 1.23 |
| mesenchyme | Usp47 | 0.42 | 2.40E-97 | 4.99E-93 | 0.73 | 0.64 | 0.09 | 1.15 |
| mesenchyme | Kdm4a | 0.75 | 2.51E-97 | 5.22E-93 | 0.45 | 0.33 | 0.12 | 1.37 |
| mesenchyme | Nsfl1c | 0.34 | 4.51E-97 | 9.39E-93 | 0.86 | 0.78 | 0.08 | 1.1 |
| mesenchyme | Csrnp3 | 2.13 | 4.71E-97 | 9.79E-93 | 0.13 | 0.04 | 0.1 | 3.44 |
| mesenchyme | Kcne1l | 1.77 | 6.02E-97 | 1.25E-92 | 0.16 | 0.06 | 0.1 | 2.81 |
| mesenchyme | 930453N24Ri | 0.52 | 7.91E-97 | 1.65E-92 | 0.62 | 0.53 | 0.09 | 1.17 |
| mesenchyme | Pkd2 | 0.86 | 7.93E-97 | 1.65E-92 | 0.43 | 0.31 | 0.12 | 1.39 |
| mesenchyme | Psmd1 | 0.32 | 8.62E-97 | 1.79E-92 | 0.88 | 0.82 | 0.06 | 1.07 |
| mesenchyme | Rflnb | 0.49 | 8.86E-97 | 1.84E-92 | 0.6 | 0.46 | 0.14 | 1.3 |
| mesenchyme | Serinc1 | 0.42 | 9.91E-97 | 2.06E-92 | 0.73 | 0.64 | 0.1 | 1.15 |
| mesenchyme | Pdlim2 | 1.58 | 1.00E-96 | 2.08E-92 | 0.2 | 0.09 | 0.11 | 2.22 |
| mesenchyme | Ccni | 0.35 | 1.21E-96 | 2.52E-92 | 0.83 | 0.75 | 0.07 | 1.1 |
| mesenchyme | Cmpk1 | 0.33 | 1.59E-96 | 3.31E-92 | 0.81 | 0.74 | 0.07 | 1.09 |
| mesenchyme | Abcf1 | 0.29 | 2.41E-96 | 5.00E-92 | 0.92 | 0.86 | 0.06 | 1.07 |
| mesenchyme | Tmem223 | 0.36 | 2.69E-96 | 5.58E-92 | 0.86 | 0.8 | 0.06 | 1.07 |
| mesenchyme | Gm10762 | 0.81 | 2.91E-96 | 6.06E-92 | 0.44 | 0.31 | 0.12 | 1.39 |
| mesenchyme | Zc3h13 | 0.35 | 3.04E-96 | 6.32E-92 | 0.86 | 0.78 | 0.08 | 1.1 |
| mesenchyme | Tax1bp3 | 0.35 | 3.05E-96 | 6.35E-92 | 0.69 | 0.56 | 0.13 | 1.22 |
| mesenchyme | Bod1 | 0.3 | 4.40E-96 | 9.14E-92 | 0.87 | 0.75 | 0.12 | 1.16 |
| mesenchyme | Cib2 | 0.97 | 5.10E-96 | 1.06E-91 | 0.3 | 0.17 | 0.13 | 1.75 |
| mesenchyme | Faap20 | 0.5 | 5.17E-96 | 1.08E-91 | 0.64 | 0.53 | 0.11 | 1.21 |
| mesenchyme | Gm45716 | 0.81 | 5.62E-96 | 1.17E-91 | 0.42 | 0.29 | 0.12 | 1.42 |
| mesenchyme | Ccp110 | 0.61 | 1.36E-95 | 2.84E-91 | 0.47 | 0.35 | 0.12 | 1.35 |
| mesenchyme | Mrps22 | 0.56 | 1.40E-95 | 2.91E-91 | 0.6 | 0.51 | 0.09 | 1.17 |
| mesenchyme | Postn | 2.36 | 1.69E-95 | 3.52E-91 | 0.12 | 0.03 | 0.09 | 4.07 |
| mesenchyme | Aff2 | 3.07 | 1.73E-95 | 3.59E-91 | 0.11 | 0.02 | 0.08 | 4.77 |
| mesenchyme | Ift27 | 0.54 | 1.96E-95 | 4.07E-91 | 0.6 | 0.49 | 0.11 | 1.22 |
| mesenchyme | Klhl22 | 0.92 | 2.08E-95 | 4.32E-91 | 0.38 | 0.25 | 0.13 | 1.49 |
| mesenchyme | Lig1 | 0.37 | 2.48E-95 | 5.16E-91 | 0.82 | 0.72 | 0.1 | 1.14 |
| mesenchyme | Polr2a | 0.36 | 2.92E-95 | 6.06E-91 | 0.82 | 0.75 | 0.08 | 1.1 |
| mesenchyme | Xpo1 | 0.36 | 3.22E-95 | 6.70E-91 | 0.81 | 0.73 | 0.09 | 1.12 |
| mesenchyme | Csrp1 | 0.66 | 3.46E-95 | 7.20E-91 | 0.39 | 0.25 | 0.13 | 1.53 |
| mesenchyme | Ints8 | 0.74 | 3.61E-95 | 7.50E-91 | 0.47 | 0.36 | 0.11 | 1.32 |
| mesenchyme | Rcbtb1 | 0.64 | 3.61E-95 | 7.51E-91 | 0.52 | 0.4 | 0.12 | 1.3 |
| mesenchyme | Mrps7 | 0.35 | 7.18E-95 | 1.49E-90 | 0.8 | 0.73 | 0.07 | 1.1 |
| mesenchyme | Ahi1 | 0.72 | 7.35E-95 | 1.53E-90 | 0.46 | 0.34 | 0.12 | 1.36 |
| mesenchyme | Ddx46 | 0.27 | 7.72E-95 | 1.60E-90 | 0.94 | 0.88 | 0.06 | 1.07 |
| mesenchyme | Tlk2 | 0.49 | 9.41E-95 | 1.96E-90 | 0.64 | 0.53 | 0.11 | 1.2 |
| mesenchyme | Birc5 | 0.39 | 1.00E-94 | 2.09E-90 | 0.83 | 0.76 | 0.07 | 1.09 |

**Supplementary Table 2. Marker genes for the major cell types of integrated clustering of E11.5 control and NotchGOF λ-junction cells.**

| | | | | | | | | |
|---|---|---|---|---|---|---|---|---|
| mesenchyme | Itgb1 | 0.26 | 1.17E-94 | 2.43E-90 | 0.9 | 0.77 | 0.13 | 1.17 |
| mesenchyme | Nt5c3b | 0.41 | 1.19E-94 | 2.48E-90 | 0.72 | 0.6 | 0.12 | 1.2 |
| mesenchyme | Gulp1 | 1.14 | 1.52E-94 | 3.15E-90 | 0.29 | 0.17 | 0.12 | 1.72 |
| mesenchyme | Dctd | 0.68 | 1.89E-94 | 3.94E-90 | 0.51 | 0.39 | 0.12 | 1.3 |
| mesenchyme | Ankrd11 | 0.26 | 1.98E-94 | 4.12E-90 | 0.96 | 0.88 | 0.08 | 1.09 |
| mesenchyme | Cntfr | 1.45 | 2.37E-94 | 4.92E-90 | 0.26 | 0.15 | 0.12 | 1.79 |
| mesenchyme | Syncrip | 0.26 | 2.40E-94 | 5.00E-90 | 0.95 | 0.88 | 0.07 | 1.08 |
| mesenchyme | Ufc1 | 0.32 | 3.35E-94 | 6.97E-90 | 0.86 | 0.78 | 0.08 | 1.1 |
| mesenchyme | Nck1 | 0.56 | 3.77E-94 | 7.83E-90 | 0.57 | 0.47 | 0.11 | 1.23 |
| mesenchyme | Mcc | 1.88 | 3.80E-94 | 7.90E-90 | 0.16 | 0.06 | 0.1 | 2.65 |
| mesenchyme | Cbr1 | 0.29 | 1.05E-93 | 2.19E-89 | 0.79 | 0.7 | 0.08 | 1.12 |
| mesenchyme | Galnt1 | 0.45 | 1.70E-93 | 3.53E-89 | 0.69 | 0.59 | 0.1 | 1.17 |
| mesenchyme | Osr2 | 2.63 | 1.95E-93 | 4.06E-89 | 0.12 | 0.03 | 0.09 | 3.93 |
| mesenchyme | Shd | 1.26 | 2.35E-93 | 4.88E-89 | 0.21 | 0.1 | 0.11 | 2.17 |
| mesenchyme | Hells | 0.29 | 2.39E-93 | 4.96E-89 | 0.83 | 0.67 | 0.16 | 1.24 |
| mesenchyme | Mrpl58 | 0.32 | 2.40E-93 | 5.00E-89 | 0.83 | 0.77 | 0.06 | 1.08 |
| mesenchyme | Cthrc1 | 1.45 | 2.66E-93 | 5.52E-89 | 0.23 | 0.12 | 0.12 | 2 |
| mesenchyme | Rab30 | 1.68 | 3.03E-93 | 6.30E-89 | 0.18 | 0.07 | 0.1 | 2.42 |
| mesenchyme | Ccdc174 | 0.44 | 3.85E-93 | 8.01E-89 | 0.68 | 0.6 | 0.08 | 1.14 |
| mesenchyme | Gmnn | 0.35 | 3.89E-93 | 8.10E-89 | 0.83 | 0.75 | 0.08 | 1.11 |
| mesenchyme | Aco2 | 0.37 | 4.02E-93 | 8.36E-89 | 0.8 | 0.73 | 0.08 | 1.11 |
| mesenchyme | Ivd | 0.75 | 4.41E-93 | 9.18E-89 | 0.45 | 0.33 | 0.12 | 1.37 |
| mesenchyme | Pigx | 0.37 | 5.79E-93 | 1.20E-88 | 0.79 | 0.68 | 0.11 | 1.17 |
| mesenchyme | Csde1 | 0.26 | 7.25E-93 | 1.51E-88 | 0.95 | 0.9 | 0.05 | 1.06 |
| mesenchyme | Dancr | 0.53 | 8.56E-93 | 1.78E-88 | 0.63 | 0.52 | 0.1 | 1.2 |
| mesenchyme | Cdc25a | 0.47 | 1.40E-92 | 2.91E-88 | 0.66 | 0.55 | 0.11 | 1.2 |
| mesenchyme | Ahnak | 0.43 | 1.94E-92 | 4.03E-88 | 0.51 | 0.35 | 0.16 | 1.45 |
| mesenchyme | Egln2 | 0.47 | 1.94E-92 | 4.04E-88 | 0.65 | 0.55 | 0.1 | 1.18 |
| mesenchyme | Zfp638 | 0.34 | 4.17E-92 | 8.68E-88 | 0.84 | 0.75 | 0.09 | 1.12 |
| mesenchyme | Ccdc124 | 0.32 | 4.19E-92 | 8.71E-88 | 0.84 | 0.78 | 0.06 | 1.07 |
| mesenchyme | Eln | 3.03 | 6.44E-92 | 1.34E-87 | 0.1 | 0.02 | 0.08 | 4.52 |
| mesenchyme | Ptbp1 | 0.28 | 1.05E-91 | 2.17E-87 | 0.92 | 0.83 | 0.08 | 1.1 |
| mesenchyme | Gfer | 0.55 | 1.12E-91 | 2.33E-87 | 0.59 | 0.51 | 0.09 | 1.17 |
| mesenchyme | Acads | 0.47 | 1.32E-91 | 2.75E-87 | 0.64 | 0.53 | 0.11 | 1.2 |
| mesenchyme | Stk38 | 0.43 | 1.71E-91 | 3.55E-87 | 0.69 | 0.6 | 0.09 | 1.15 |
| mesenchyme | Rab11b | 0.32 | 1.73E-91 | 3.60E-87 | 0.86 | 0.79 | 0.06 | 1.08 |
| mesenchyme | Yae1d1 | 0.38 | 1.91E-91 | 3.98E-87 | 0.74 | 0.64 | 0.1 | 1.15 |
| mesenchyme | Sox9 | 0.85 | 2.01E-91 | 4.18E-87 | 0.63 | 0.5 | 0.13 | 1.26 |
| mesenchyme | Hadh | 0.37 | 2.04E-91 | 4.24E-87 | 0.81 | 0.72 | 0.09 | 1.13 |
| mesenchyme | Plekha5 | 0.65 | 2.22E-91 | 4.62E-87 | 0.51 | 0.38 | 0.13 | 1.33 |
| mesenchyme | Brd2 | 0.34 | 2.27E-91 | 4.73E-87 | 0.85 | 0.77 | 0.08 | 1.1 |
| mesenchyme | Rbm46 | 2.68 | 3.00E-91 | 6.24E-87 | 0.11 | 0.03 | 0.09 | 4.15 |
| mesenchyme | Umps | 0.47 | 3.05E-91 | 6.35E-87 | 0.64 | 0.54 | 0.1 | 1.18 |
| mesenchyme | Map3k7 | 0.69 | 3.31E-91 | 6.89E-87 | 0.51 | 0.4 | 0.11 | 1.26 |
| mesenchyme | Resf1 | 0.34 | 3.34E-91 | 6.95E-87 | 0.78 | 0.68 | 0.09 | 1.13 |
| mesenchyme | Sec63 | 0.39 | 3.35E-91 | 6.97E-87 | 0.76 | 0.66 | 0.1 | 1.16 |
| mesenchyme | Glrx2 | 0.37 | 3.51E-91 | 7.30E-87 | 0.77 | 0.71 | 0.07 | 1.1 |
| mesenchyme | Phf20l1 | 0.47 | 3.65E-91 | 7.59E-87 | 0.69 | 0.59 | 0.1 | 1.16 |
| mesenchyme | Smim11 | 0.37 | 3.85E-91 | 8.01E-87 | 0.78 | 0.71 | 0.07 | 1.1 |
| mesenchyme | Txlna | 0.44 | 4.32E-91 | 8.98E-87 | 0.69 | 0.59 | 0.1 | 1.16 |
| mesenchyme | Zfp260 | 0.44 | 5.28E-91 | 1.10E-86 | 0.69 | 0.58 | 0.11 | 1.19 |
| mesenchyme | Rtcb | 0.33 | 6.11E-91 | 1.27E-86 | 0.84 | 0.77 | 0.07 | 1.09 |
| mesenchyme | Akr1b3 | 0.42 | 6.24E-91 | 1.30E-86 | 0.71 | 0.61 | 0.1 | 1.16 |
| mesenchyme | Senp6 | 0.3 | 1.08E-90 | 2.25E-86 | 0.91 | 0.82 | 0.08 | 1.1 |
| mesenchyme | Rala | 0.29 | 1.16E-90 | 2.40E-86 | 0.87 | 0.81 | 0.07 | 1.08 |
| mesenchyme | Smoc1 | 2.79 | 1.41E-90 | 2.93E-86 | 0.1 | 0.02 | 0.08 | 4.52 |

**Supplementary Table 2. Marker genes for the major cell types of integrated clustering of E11.5 control and NotchGOF λ-junction cells.**

| | | | | | | | | |
|---|---|---|---|---|---|---|---|---|
| mesenchyme | Atoh8 | 2.28 | 1.42E-90 | 2.95E-86 | 0.12 | 0.03 | 0.09 | 3.64 |
| mesenchyme | Mrpl43 | 0.3 | 1.76E-90 | 3.66E-86 | 0.85 | 0.79 | 0.05 | 1.07 |
| mesenchyme | Ntan1 | 0.53 | 2.29E-90 | 4.77E-86 | 0.7 | 0.61 | 0.09 | 1.15 |
| mesenchyme | Hmox2 | 0.44 | 2.39E-90 | 4.97E-86 | 0.67 | 0.58 | 0.09 | 1.16 |
| mesenchyme | Ndufs8 | 0.31 | 4.01E-90 | 8.34E-86 | 0.87 | 0.79 | 0.07 | 1.09 |
| mesenchyme | Plpbp | 0.76 | 4.24E-90 | 8.82E-86 | 0.43 | 0.32 | 0.11 | 1.36 |
| mesenchyme | Sp3 | 0.48 | 4.53E-90 | 9.41E-86 | 0.66 | 0.57 | 0.09 | 1.15 |
| mesenchyme | Pik3r3 | 0.95 | 5.24E-90 | 1.09E-85 | 0.32 | 0.2 | 0.12 | 1.63 |
| mesenchyme | Srek1ip1 | 0.4 | 6.24E-90 | 1.30E-85 | 0.74 | 0.65 | 0.08 | 1.13 |
| mesenchyme | Dap3 | 0.39 | 6.26E-90 | 1.30E-85 | 0.75 | 0.67 | 0.08 | 1.11 |
| mesenchyme | Atad2b | 0.44 | 9.66E-90 | 2.01E-85 | 0.67 | 0.59 | 0.09 | 1.15 |
| mesenchyme | Tmem29 | 0.56 | 1.01E-89 | 2.11E-85 | 0.55 | 0.46 | 0.09 | 1.21 |
| mesenchyme | Jarid2 | 0.52 | 1.13E-89 | 2.35E-85 | 0.64 | 0.54 | 0.1 | 1.18 |
| mesenchyme | Tsnax | 0.43 | 1.21E-89 | 2.52E-85 | 0.69 | 0.61 | 0.08 | 1.14 |
| mesenchyme | Yif1b | 0.4 | 1.24E-89 | 2.58E-85 | 0.7 | 0.6 | 0.1 | 1.17 |
| mesenchyme | Thap2 | 0.59 | 1.25E-89 | 2.60E-85 | 0.55 | 0.44 | 0.11 | 1.24 |
| mesenchyme | St6galnac4 | 1.25 | 1.60E-89 | 3.33E-85 | 0.24 | 0.12 | 0.11 | 1.93 |
| mesenchyme | Tgs1 | 0.33 | 1.61E-89 | 3.36E-85 | 0.82 | 0.74 | 0.08 | 1.11 |
| mesenchyme | 110017D15Ri | 2.32 | 2.29E-89 | 4.77E-85 | 0.11 | 0.03 | 0.08 | 4.07 |
| mesenchyme | Rxylt1 | 0.66 | 2.55E-89 | 5.29E-85 | 0.51 | 0.4 | 0.11 | 1.28 |
| mesenchyme | Trio | 0.59 | 2.67E-89 | 5.56E-85 | 0.55 | 0.44 | 0.12 | 1.26 |
| mesenchyme | Gfpt1 | 0.78 | 3.88E-89 | 8.07E-85 | 0.42 | 0.3 | 0.12 | 1.39 |
| mesenchyme | Notch2 | 0.78 | 5.68E-89 | 1.18E-84 | 0.41 | 0.28 | 0.13 | 1.46 |
| mesenchyme | Pom121 | 0.49 | 5.81E-89 | 1.21E-84 | 0.63 | 0.54 | 0.09 | 1.17 |
| mesenchyme | G0s2 | 1.55 | 6.43E-89 | 1.34E-84 | 0.19 | 0.08 | 0.11 | 2.3 |
| mesenchyme | Foxp4 | 0.77 | 1.21E-88 | 2.52E-84 | 0.46 | 0.34 | 0.12 | 1.34 |
| mesenchyme | Decr1 | 0.71 | 2.00E-88 | 4.16E-84 | 0.45 | 0.34 | 0.11 | 1.34 |
| mesenchyme | Virma | 0.51 | 2.23E-88 | 4.64E-84 | 0.62 | 0.52 | 0.1 | 1.19 |
| mesenchyme | Mras | 1.1 | 2.55E-88 | 5.31E-84 | 0.25 | 0.13 | 0.12 | 1.89 |
| mesenchyme | Nop56 | 0.3 | 2.82E-88 | 5.86E-84 | 0.89 | 0.82 | 0.08 | 1.09 |
| mesenchyme | 430416N02Ri | 0.69 | 3.45E-88 | 7.18E-84 | 0.5 | 0.39 | 0.11 | 1.28 |
| mesenchyme | Cttnbp2nl | 0.55 | 3.79E-88 | 7.89E-84 | 0.54 | 0.42 | 0.12 | 1.3 |
| mesenchyme | Pcca | 0.97 | 4.08E-88 | 8.49E-84 | 0.33 | 0.22 | 0.12 | 1.54 |
| mesenchyme | Pex5l | 2.75 | 4.09E-88 | 8.50E-84 | 0.12 | 0.03 | 0.09 | 3.5 |
| mesenchyme | Spsb4 | 1.13 | 5.42E-88 | 1.13E-83 | 0.24 | 0.13 | 0.11 | 1.88 |
| mesenchyme | Scd1 | 1.22 | 5.47E-88 | 1.14E-83 | 0.23 | 0.11 | 0.11 | 2.01 |
| mesenchyme | Itpr1 | 1.39 | 5.54E-88 | 1.15E-83 | 0.25 | 0.14 | 0.11 | 1.79 |
| mesenchyme | Mtrex | 0.49 | 6.54E-88 | 1.36E-83 | 0.65 | 0.56 | 0.09 | 1.16 |
| mesenchyme | Psd3 | 1.21 | 7.89E-88 | 1.64E-83 | 0.26 | 0.15 | 0.12 | 1.78 |
| mesenchyme | Ryk | 0.34 | 1.02E-87 | 2.12E-83 | 0.83 | 0.74 | 0.09 | 1.13 |
| mesenchyme | Enpp2 | 1.91 | 1.13E-87 | 2.34E-83 | 0.15 | 0.05 | 0.09 | 2.84 |
| mesenchyme | Fbln5 | 1.56 | 2.98E-87 | 6.20E-83 | 0.18 | 0.08 | 0.1 | 2.36 |
| mesenchyme | Chd1 | 0.33 | 4.23E-87 | 8.80E-83 | 0.82 | 0.72 | 0.1 | 1.13 |
| mesenchyme | Rbm18 | 0.55 | 5.38E-87 | 1.12E-82 | 0.58 | 0.49 | 0.09 | 1.19 |
| mesenchyme | Ddx18 | 0.28 | 6.32E-87 | 1.31E-82 | 0.87 | 0.79 | 0.08 | 1.09 |
| mesenchyme | Cenps | 0.61 | 9.56E-87 | 1.99E-82 | 0.54 | 0.43 | 0.11 | 1.24 |
| mesenchyme | Scoc | 0.42 | 1.33E-86 | 2.76E-82 | 0.66 | 0.57 | 0.09 | 1.16 |
| mesenchyme | Acp1 | 0.25 | 1.36E-86 | 2.83E-82 | 0.95 | 0.89 | 0.05 | 1.06 |
| mesenchyme | Dis3 | 0.53 | 1.70E-86 | 3.54E-82 | 0.59 | 0.48 | 0.1 | 1.21 |
| mesenchyme | Mrpl41 | 0.33 | 2.19E-86 | 4.56E-82 | 0.77 | 0.7 | 0.08 | 1.11 |
| mesenchyme | Adgre5 | 1.72 | 2.62E-86 | 5.46E-82 | 0.16 | 0.06 | 0.1 | 2.55 |
| mesenchyme | Sigmar1 | 0.56 | 3.05E-86 | 6.34E-82 | 0.58 | 0.48 | 0.11 | 1.22 |
| mesenchyme | Mier1 | 0.35 | 4.14E-86 | 8.60E-82 | 0.82 | 0.75 | 0.06 | 1.09 |
| mesenchyme | Mylip | 0.88 | 5.39E-86 | 1.12E-81 | 0.35 | 0.23 | 0.12 | 1.51 |
| mesenchyme | Zfp553 | 0.47 | 5.44E-86 | 1.13E-81 | 0.64 | 0.53 | 0.11 | 1.21 |
| mesenchyme | Phactr2 | 0.78 | 6.21E-86 | 1.29E-81 | 0.49 | 0.38 | 0.11 | 1.3 |

**Supplementary Table 2. Marker genes for the major cell types of integrated clustering of E11.5 control and NotchGOF λ-junction cells.**

| | | | | | | | | |
|---|---|---|---|---|---|---|---|---|
| mesenchyme | Snx18 | 0.75 | 7.22E-86 | 1.50E-81 | 0.42 | 0.3 | 0.12 | 1.39 |
| mesenchyme | Pcolce2 | 1.11 | 7.80E-86 | 1.62E-81 | 0.17 | 0.07 | 0.1 | 2.46 |
| mesenchyme | Raet1d | 2.46 | 1.10E-85 | 2.29E-81 | 0.11 | 0.03 | 0.08 | 3.83 |
| mesenchyme | Ift81 | 0.6 | 1.26E-85 | 2.62E-81 | 0.49 | 0.38 | 0.12 | 1.31 |
| mesenchyme | Orc2 | 0.51 | 1.27E-85 | 2.64E-81 | 0.6 | 0.51 | 0.09 | 1.17 |
| mesenchyme | Casp3 | 0.3 | 1.42E-85 | 2.95E-81 | 0.76 | 0.65 | 0.11 | 1.17 |
| mesenchyme | Rhno1 | 0.4 | 1.51E-85 | 3.15E-81 | 0.7 | 0.61 | 0.08 | 1.14 |
| mesenchyme | Mtx2 | 0.28 | 1.61E-85 | 3.35E-81 | 0.85 | 0.8 | 0.05 | 1.06 |
| mesenchyme | Mrto4 | 0.31 | 1.66E-85 | 3.45E-81 | 0.83 | 0.78 | 0.06 | 1.07 |
| mesenchyme | Kirrel | 1.23 | 1.66E-85 | 3.45E-81 | 0.26 | 0.15 | 0.11 | 1.75 |
| mesenchyme | Plod1 | 1.31 | 1.72E-85 | 3.57E-81 | 0.24 | 0.13 | 0.11 | 1.85 |
| mesenchyme | Emc8 | 0.33 | 2.26E-85 | 4.71E-81 | 0.82 | 0.74 | 0.08 | 1.11 |
| mesenchyme | Zfp9 | 1.47 | 3.52E-85 | 7.33E-81 | 0.2 | 0.09 | 0.1 | 2.11 |
| mesenchyme | Huwe1 | 0.3 | 5.48E-85 | 1.14E-80 | 0.89 | 0.81 | 0.08 | 1.1 |
| mesenchyme | Atad5 | 0.58 | 5.66E-85 | 1.18E-80 | 0.57 | 0.46 | 0.11 | 1.24 |
| mesenchyme | Nradd | 0.55 | 6.14E-85 | 1.28E-80 | 0.57 | 0.45 | 0.11 | 1.25 |
| mesenchyme | Ndufs4 | 0.25 | 6.20E-85 | 1.29E-80 | 0.92 | 0.88 | 0.04 | 1.05 |
| mesenchyme | Lrch3 | 0.77 | 6.23E-85 | 1.30E-80 | 0.41 | 0.29 | 0.12 | 1.4 |
| mesenchyme | Eepd1 | 1.78 | 7.87E-85 | 1.64E-80 | 0.15 | 0.06 | 0.09 | 2.66 |
| mesenchyme | Ppp4r2 | 0.29 | 9.21E-85 | 1.92E-80 | 0.87 | 0.82 | 0.06 | 1.07 |
| mesenchyme | Podxl2 | 0.56 | 1.09E-84 | 2.27E-80 | 0.47 | 0.34 | 0.13 | 1.39 |
| mesenchyme | Rnf144a | 0.66 | 1.34E-84 | 2.78E-80 | 0.48 | 0.36 | 0.12 | 1.32 |
| mesenchyme | Brd4 | 0.32 | 1.55E-84 | 3.21E-80 | 0.83 | 0.75 | 0.08 | 1.11 |
| mesenchyme | Cnep1r1 | 0.56 | 1.58E-84 | 3.28E-80 | 0.57 | 0.48 | 0.09 | 1.19 |
| mesenchyme | Tipin | 0.34 | 1.79E-84 | 3.73E-80 | 0.8 | 0.71 | 0.09 | 1.12 |
| mesenchyme | Vamp3 | 0.49 | 2.27E-84 | 4.73E-80 | 0.6 | 0.51 | 0.09 | 1.18 |
| mesenchyme | Etv5 | 0.79 | 2.40E-84 | 4.98E-80 | 0.36 | 0.23 | 0.12 | 1.53 |
| mesenchyme | Etfb | 0.26 | 2.64E-84 | 5.48E-80 | 0.91 | 0.85 | 0.06 | 1.07 |
| mesenchyme | Apcdd1 | 1.38 | 2.64E-84 | 5.49E-80 | 0.25 | 0.14 | 0.11 | 1.8 |
| mesenchyme | Col5a1 | 1.1 | 2.65E-84 | 5.50E-80 | 0.3 | 0.19 | 0.11 | 1.58 |
| mesenchyme | Vash2 | 0.93 | 4.20E-84 | 8.73E-80 | 0.33 | 0.21 | 0.12 | 1.56 |
| mesenchyme | Pex19 | 0.54 | 4.39E-84 | 9.12E-80 | 0.56 | 0.46 | 0.1 | 1.23 |
| mesenchyme | Gps1 | 0.35 | 4.83E-84 | 1.00E-79 | 0.77 | 0.69 | 0.08 | 1.11 |
| mesenchyme | Gpr180 | 0.97 | 4.84E-84 | 1.01E-79 | 0.34 | 0.22 | 0.12 | 1.52 |
| mesenchyme | Samd1 | 0.74 | 5.91E-84 | 1.23E-79 | 0.44 | 0.34 | 0.11 | 1.32 |
| mesenchyme | Tk1 | 0.56 | 6.32E-84 | 1.32E-79 | 0.58 | 0.48 | 0.1 | 1.21 |
| mesenchyme | Lsm14b | 0.51 | 8.15E-84 | 1.70E-79 | 0.52 | 0.41 | 0.12 | 1.29 |
| mesenchyme | Csnk1g3 | 0.37 | 9.05E-84 | 1.88E-79 | 0.71 | 0.62 | 0.09 | 1.15 |
| mesenchyme | Senp7 | 0.5 | 1.12E-83 | 2.32E-79 | 0.58 | 0.49 | 0.09 | 1.18 |
| mesenchyme | Zc3h18 | 0.38 | 1.13E-83 | 2.35E-79 | 0.75 | 0.65 | 0.1 | 1.15 |
| mesenchyme | Gdf6 | 2.17 | 1.42E-83 | 2.96E-79 | 0.14 | 0.05 | 0.09 | 2.84 |
| mesenchyme | Sdhaf1 | 0.46 | 1.47E-83 | 3.05E-79 | 0.63 | 0.53 | 0.1 | 1.18 |
| mesenchyme | Psrc1 | 0.72 | 1.55E-83 | 3.22E-79 | 0.46 | 0.34 | 0.11 | 1.33 |
| mesenchyme | Ehmt1 | 0.32 | 1.71E-83 | 3.56E-79 | 0.85 | 0.76 | 0.1 | 1.13 |
| mesenchyme | Tiparp | 0.89 | 1.72E-83 | 3.58E-79 | 0.36 | 0.24 | 0.12 | 1.49 |
| mesenchyme | Bms1 | 0.42 | 3.17E-83 | 6.59E-79 | 0.71 | 0.63 | 0.08 | 1.13 |
| mesenchyme | Ttc4 | 0.48 | 3.29E-83 | 6.84E-79 | 0.62 | 0.53 | 0.09 | 1.16 |
| mesenchyme | Asns | 0.71 | 3.37E-83 | 7.01E-79 | 0.48 | 0.37 | 0.1 | 1.28 |
| mesenchyme | Slc9a3r2 | 0.45 | 3.48E-83 | 7.24E-79 | 0.31 | 0.18 | 0.12 | 1.68 |
| mesenchyme | Ciapin1 | 0.33 | 3.73E-83 | 7.76E-79 | 0.76 | 0.69 | 0.07 | 1.11 |
| mesenchyme | Vps4a | 0.58 | 5.96E-83 | 1.24E-78 | 0.54 | 0.45 | 0.09 | 1.19 |
| mesenchyme | Esf1 | 0.28 | 6.46E-83 | 1.34E-78 | 0.89 | 0.83 | 0.06 | 1.08 |
| mesenchyme | Taf1 | 0.49 | 1.35E-82 | 2.81E-78 | 0.61 | 0.52 | 0.1 | 1.18 |
| mesenchyme | Cnpy4 | 0.83 | 1.39E-82 | 2.88E-78 | 0.39 | 0.28 | 0.11 | 1.4 |
| mesenchyme | Hsph1 | 0.43 | 1.40E-82 | 2.91E-78 | 0.71 | 0.63 | 0.08 | 1.13 |
| mesenchyme | Ssbp2 | 0.35 | 1.58E-82 | 3.28E-78 | 0.74 | 0.64 | 0.11 | 1.17 |

**Supplementary Table 2. Marker genes for the major cell types of integrated clustering of E11.5 control and NotchGOF λ-junction cells.**

| | | | | | | | | |
|---|---|---|---|---|---|---|---|---|
| mesenchyme | Znhit6 | 0.48 | 1.72E-82 | 3.57E-78 | 0.63 | 0.55 | 0.09 | 1.16 |
| mesenchyme | Dnajb11 | 0.39 | 1.89E-82 | 3.93E-78 | 0.71 | 0.62 | 0.09 | 1.14 |
| mesenchyme | Boc | 1.29 | 2.28E-82 | 4.75E-78 | 0.24 | 0.13 | 0.11 | 1.83 |
| mesenchyme | Eif4g3 | 0.35 | 2.75E-82 | 5.72E-78 | 0.79 | 0.7 | 0.09 | 1.13 |
| mesenchyme | Tm9sf4 | 0.51 | 5.18E-82 | 1.08E-77 | 0.61 | 0.52 | 0.1 | 1.19 |
| mesenchyme | Dpysl4 | 1.11 | 5.60E-82 | 1.17E-77 | 0.22 | 0.11 | 0.11 | 1.96 |
| mesenchyme | Wsb1 | 0.37 | 5.65E-82 | 1.18E-77 | 0.72 | 0.63 | 0.09 | 1.14 |
| mesenchyme | Inhba | 1.39 | 8.06E-82 | 1.68E-77 | 0.2 | 0.1 | 0.11 | 2.09 |
| mesenchyme | Pafah1b2 | 0.3 | 9.26E-82 | 1.93E-77 | 0.85 | 0.78 | 0.07 | 1.09 |
| mesenchyme | P4hb | 0.27 | 1.01E-81 | 2.09E-77 | 0.92 | 0.81 | 0.11 | 1.13 |
| mesenchyme | 810001G20Ri | 0.78 | 1.04E-81 | 2.17E-77 | 0.43 | 0.32 | 0.11 | 1.35 |
| mesenchyme | Abi2 | 0.64 | 1.38E-81 | 2.87E-77 | 0.48 | 0.37 | 0.1 | 1.28 |
| mesenchyme | Ska2 | 0.52 | 1.67E-81 | 3.47E-77 | 0.57 | 0.48 | 0.09 | 1.19 |
| mesenchyme | Fbxo5 | 0.6 | 2.64E-81 | 5.49E-77 | 0.61 | 0.55 | 0.07 | 1.12 |
| mesenchyme | Dclk1 | 0.95 | 3.15E-81 | 6.56E-77 | 0.29 | 0.18 | 0.11 | 1.64 |
| mesenchyme | Ly6e | 0.35 | 3.57E-81 | 7.42E-77 | 0.49 | 0.37 | 0.12 | 1.34 |
| mesenchyme | Zfp423 | 1.19 | 3.63E-81 | 7.55E-77 | 0.24 | 0.14 | 0.11 | 1.79 |
| mesenchyme | Plod2 | 0.59 | 3.78E-81 | 7.86E-77 | 0.53 | 0.42 | 0.12 | 1.28 |
| mesenchyme | Topbp1 | 0.37 | 4.13E-81 | 8.59E-77 | 0.76 | 0.68 | 0.08 | 1.12 |
| mesenchyme | C1qtnf6 | 1.43 | 4.25E-81 | 8.84E-77 | 0.19 | 0.09 | 0.1 | 2.06 |
| mesenchyme | Zfp322a | 0.44 | 5.97E-81 | 1.24E-76 | 0.67 | 0.58 | 0.09 | 1.16 |
| mesenchyme | Wac | 0.32 | 6.22E-81 | 1.29E-76 | 0.8 | 0.73 | 0.07 | 1.1 |
| mesenchyme | Tmem109 | 0.73 | 6.24E-81 | 1.30E-76 | 0.4 | 0.29 | 0.11 | 1.38 |
| mesenchyme | Sgta | 0.29 | 8.10E-81 | 1.68E-76 | 0.83 | 0.75 | 0.08 | 1.11 |
| mesenchyme | Elp5 | 0.49 | 1.13E-80 | 2.35E-76 | 0.62 | 0.54 | 0.08 | 1.15 |
| mesenchyme | Hsd11b2 | 2.17 | 1.13E-80 | 2.36E-76 | 0.13 | 0.05 | 0.09 | 2.91 |
| mesenchyme | Snhg5 | 0.66 | 1.27E-80 | 2.63E-76 | 0.48 | 0.37 | 0.11 | 1.29 |
| mesenchyme | Cisd2 | 0.34 | 1.35E-80 | 2.80E-76 | 0.8 | 0.72 | 0.07 | 1.1 |
| mesenchyme | Chd9 | 0.51 | 1.45E-80 | 3.02E-76 | 0.57 | 0.46 | 0.11 | 1.23 |
| mesenchyme | Efna4 | 0.4 | 1.47E-80 | 3.06E-76 | 0.66 | 0.54 | 0.12 | 1.22 |
| mesenchyme | Eya4 | 0.95 | 1.90E-80 | 3.95E-76 | 0.33 | 0.21 | 0.12 | 1.56 |
| mesenchyme | Wwp1 | 0.72 | 2.08E-80 | 4.32E-76 | 0.45 | 0.34 | 0.11 | 1.31 |
| mesenchyme | Cdo1 | 0.29 | 2.32E-80 | 4.82E-76 | 0.56 | 0.41 | 0.14 | 1.35 |
| mesenchyme | Ccm2 | 0.71 | 2.34E-80 | 4.87E-76 | 0.43 | 0.33 | 0.1 | 1.32 |
| mesenchyme | Tsen15 | 0.42 | 2.45E-80 | 5.10E-76 | 0.67 | 0.56 | 0.11 | 1.19 |
| mesenchyme | Papss1 | 0.41 | 2.64E-80 | 5.49E-76 | 0.7 | 0.6 | 0.1 | 1.16 |
| mesenchyme | Msantd3 | 0.51 | 2.77E-80 | 5.77E-76 | 0.54 | 0.44 | 0.1 | 1.24 |
| mesenchyme | Lcorl | 0.62 | 2.79E-80 | 5.81E-76 | 0.51 | 0.42 | 0.1 | 1.24 |
| mesenchyme | Tia1 | 0.27 | 4.03E-80 | 8.37E-76 | 0.91 | 0.8 | 0.11 | 1.14 |
| mesenchyme | Tmem44 | 1.12 | 4.43E-80 | 9.21E-76 | 0.25 | 0.14 | 0.11 | 1.77 |
| mesenchyme | Elp1 | 0.41 | 4.44E-80 | 9.22E-76 | 0.68 | 0.58 | 0.1 | 1.17 |
| mesenchyme | Polr3gl | 0.94 | 8.13E-80 | 1.69E-75 | 0.35 | 0.24 | 0.11 | 1.45 |
| mesenchyme | Ppp1r8 | 0.61 | 9.19E-80 | 1.91E-75 | 0.48 | 0.38 | 0.1 | 1.27 |
| mesenchyme | Zfp91 | 0.26 | 1.26E-79 | 2.61E-75 | 0.89 | 0.84 | 0.05 | 1.06 |
| mesenchyme | Rnf128 | 1.87 | 1.42E-79 | 2.95E-75 | 0.18 | 0.08 | 0.1 | 2.19 |
| mesenchyme | Mrpl21 | 0.27 | 1.47E-79 | 3.06E-75 | 0.88 | 0.83 | 0.05 | 1.06 |
| mesenchyme | Fndc3a | 0.37 | 2.73E-79 | 5.67E-75 | 0.72 | 0.61 | 0.11 | 1.18 |
| mesenchyme | Exosc10 | 0.66 | 4.89E-79 | 1.02E-74 | 0.47 | 0.37 | 0.1 | 1.28 |
| mesenchyme | Zfp281 | 0.62 | 5.12E-79 | 1.06E-74 | 0.5 | 0.39 | 0.11 | 1.28 |
| mesenchyme | Ap4m1 | 0.71 | 5.22E-79 | 1.08E-74 | 0.45 | 0.35 | 0.11 | 1.31 |
| mesenchyme | Emsy | 0.45 | 5.40E-79 | 1.12E-74 | 0.65 | 0.56 | 0.1 | 1.17 |
| mesenchyme | Aif1l | 0.66 | 6.30E-79 | 1.31E-74 | 0.42 | 0.3 | 0.11 | 1.38 |
| mesenchyme | Cyp27a1 | 2.31 | 6.40E-79 | 1.33E-74 | 0.11 | 0.03 | 0.08 | 3.44 |
| mesenchyme | Mpp6 | 0.34 | 6.75E-79 | 1.40E-74 | 0.77 | 0.7 | 0.07 | 1.11 |
| mesenchyme | Zc3h14 | 0.34 | 7.83E-79 | 1.63E-74 | 0.79 | 0.72 | 0.08 | 1.11 |
| mesenchyme | Stip1 | 0.26 | 8.24E-79 | 1.71E-74 | 0.9 | 0.83 | 0.07 | 1.09 |

**Supplementary Table 2. Marker genes for the major cell types of integrated clustering of E11.5 control and NotchGOF λ-junction cells.**

| | | | | | | | | |
|---|---|---|---|---|---|---|---|---|
| mesenchyme | Aida | 0.55 | 8.75E-79 | 1.82E-74 | 0.53 | 0.44 | 0.09 | 1.21 |
| mesenchyme | Ephb4 | 0.65 | 1.10E-78 | 2.29E-74 | 0.43 | 0.32 | 0.11 | 1.35 |
| mesenchyme | Wdr75 | 0.48 | 1.19E-78 | 2.47E-74 | 0.61 | 0.53 | 0.08 | 1.16 |
| mesenchyme | Ttyh3 | 0.54 | 1.84E-78 | 3.82E-74 | 0.57 | 0.47 | 0.09 | 1.2 |
| mesenchyme | Gdf11 | 0.79 | 1.97E-78 | 4.10E-74 | 0.36 | 0.24 | 0.12 | 1.48 |
| mesenchyme | Arf2 | 0.69 | 2.68E-78 | 5.57E-74 | 0.42 | 0.32 | 0.11 | 1.33 |
| mesenchyme | Wdr35 | 0.88 | 4.80E-78 | 9.98E-74 | 0.33 | 0.22 | 0.11 | 1.5 |
| mesenchyme | Chtop | 0.3 | 7.42E-78 | 1.54E-73 | 0.82 | 0.74 | 0.08 | 1.1 |
| mesenchyme | Mettl16 | 0.41 | 7.88E-78 | 1.64E-73 | 0.67 | 0.59 | 0.08 | 1.14 |
| mesenchyme | R3hcc1 | 0.5 | 8.65E-78 | 1.80E-73 | 0.59 | 0.49 | 0.1 | 1.2 |
| mesenchyme | Fgd1 | 0.88 | 9.15E-78 | 1.90E-73 | 0.36 | 0.25 | 0.11 | 1.43 |
| mesenchyme | Ripk3 | 1.3 | 9.63E-78 | 2.00E-73 | 0.2 | 0.1 | 0.1 | 2.01 |
| mesenchyme | Ier5l | 0.33 | 1.26E-77 | 2.62E-73 | 0.71 | 0.58 | 0.12 | 1.21 |
| mesenchyme | Rrp8 | 0.52 | 1.26E-77 | 2.63E-73 | 0.56 | 0.47 | 0.09 | 1.19 |
| mesenchyme | Rassf8 | 1.24 | 1.68E-77 | 3.49E-73 | 0.22 | 0.12 | 0.1 | 1.9 |
| mesenchyme | Slc18b1 | 1.67 | 1.74E-77 | 3.62E-73 | 0.15 | 0.06 | 0.09 | 2.41 |
| mesenchyme | Pinx1 | 0.49 | 1.77E-77 | 3.67E-73 | 0.58 | 0.48 | 0.09 | 1.19 |
| mesenchyme | Astn1 | 1.77 | 1.77E-77 | 3.68E-73 | 0.15 | 0.06 | 0.09 | 2.51 |
| mesenchyme | Nynrin | 0.5 | 2.02E-77 | 4.20E-73 | 0.59 | 0.48 | 0.11 | 1.23 |
| mesenchyme | Gfra4 | 1.92 | 2.28E-77 | 4.74E-73 | 0.13 | 0.05 | 0.09 | 2.81 |
| mesenchyme | Tspan31 | 0.33 | 2.39E-77 | 4.98E-73 | 0.77 | 0.66 | 0.11 | 1.16 |
| mesenchyme | Anapc7 | 0.49 | 2.44E-77 | 5.07E-73 | 0.58 | 0.49 | 0.09 | 1.19 |
| mesenchyme | Neo1 | 0.57 | 3.03E-77 | 6.29E-73 | 0.54 | 0.43 | 0.11 | 1.26 |
| mesenchyme | Tmem160 | 0.36 | 3.19E-77 | 6.64E-73 | 0.75 | 0.7 | 0.06 | 1.08 |
| mesenchyme | Plcg1 | 0.7 | 3.51E-77 | 7.29E-73 | 0.44 | 0.33 | 0.11 | 1.33 |
| mesenchyme | Sertad4 | 0.75 | 3.60E-77 | 7.48E-73 | 0.41 | 0.29 | 0.12 | 1.41 |
| mesenchyme | Rnf130 | 0.45 | 4.03E-77 | 8.39E-73 | 0.6 | 0.49 | 0.11 | 1.23 |
| mesenchyme | Mir17hg | 0.86 | 4.48E-77 | 9.32E-73 | 0.3 | 0.19 | 0.11 | 1.57 |
| mesenchyme | Mcfd2 | 0.64 | 6.11E-77 | 1.27E-72 | 0.5 | 0.39 | 0.11 | 1.27 |
| mesenchyme | Mxd3 | 1.08 | 9.36E-77 | 1.95E-72 | 0.28 | 0.17 | 0.1 | 1.6 |
| mesenchyme | Calu | 0.36 | 1.07E-76 | 2.23E-72 | 0.74 | 0.64 | 0.1 | 1.15 |
| mesenchyme | Txnrd1 | 0.28 | 1.19E-76 | 2.48E-72 | 0.89 | 0.82 | 0.07 | 1.08 |
| mesenchyme | Flywch1 | 0.47 | 1.23E-76 | 2.55E-72 | 0.6 | 0.5 | 0.1 | 1.19 |
| mesenchyme | St6galnac6 | 1.48 | 1.35E-76 | 2.81E-72 | 0.19 | 0.1 | 0.1 | 2.01 |
| mesenchyme | Ppm1m | 1.08 | 1.43E-76 | 2.98E-72 | 0.28 | 0.18 | 0.1 | 1.58 |
| mesenchyme | Sirt1 | 0.73 | 1.58E-76 | 3.29E-72 | 0.43 | 0.33 | 0.1 | 1.3 |
| mesenchyme | Fxr1 | 0.25 | 1.62E-76 | 3.36E-72 | 0.87 | 0.81 | 0.06 | 1.08 |
| mesenchyme | Trp53bp1 | 0.46 | 1.70E-76 | 3.54E-72 | 0.6 | 0.5 | 0.1 | 1.21 |
| mesenchyme | Glg1 | 0.38 | 1.83E-76 | 3.81E-72 | 0.74 | 0.64 | 0.1 | 1.16 |
| mesenchyme | Adam19 | 1.29 | 2.34E-76 | 4.87E-72 | 0.19 | 0.09 | 0.1 | 2.1 |
| mesenchyme | Usp1 | 0.3 | 2.46E-76 | 5.13E-72 | 0.84 | 0.75 | 0.09 | 1.11 |
| mesenchyme | Dlst | 0.5 | 2.55E-76 | 5.31E-72 | 0.6 | 0.51 | 0.09 | 1.17 |
| mesenchyme | Vangl2 | 0.45 | 2.62E-76 | 5.45E-72 | 0.61 | 0.49 | 0.12 | 1.24 |
| mesenchyme | Isoc1 | 0.3 | 2.86E-76 | 5.94E-72 | 0.76 | 0.66 | 0.1 | 1.15 |
| mesenchyme | Slf1 | 0.52 | 3.29E-76 | 6.84E-72 | 0.57 | 0.48 | 0.09 | 1.2 |
| mesenchyme | Has2 | 1.18 | 3.67E-76 | 7.63E-72 | 0.23 | 0.12 | 0.1 | 1.84 |
| mesenchyme | Tmem35a | 2.07 | 3.74E-76 | 7.78E-72 | 0.13 | 0.04 | 0.08 | 2.91 |
| mesenchyme | Pin4 | 0.28 | 4.79E-76 | 9.96E-72 | 0.84 | 0.8 | 0.04 | 1.05 |
| mesenchyme | Zc3hc1 | 0.67 | 5.09E-76 | 1.06E-71 | 0.45 | 0.35 | 0.1 | 1.28 |
| mesenchyme | Ppp2r2a | 0.29 | 5.49E-76 | 1.14E-71 | 0.83 | 0.74 | 0.09 | 1.12 |
| mesenchyme | Kbtbd4 | 0.78 | 7.59E-76 | 1.58E-71 | 0.38 | 0.28 | 0.1 | 1.37 |
| mesenchyme | Nsd3 | 0.43 | 8.64E-76 | 1.80E-71 | 0.66 | 0.58 | 0.08 | 1.14 |
| mesenchyme | Gorab | 0.68 | 9.50E-76 | 1.98E-71 | 0.41 | 0.3 | 0.11 | 1.36 |
| mesenchyme | Foxc2 | 1.87 | 1.54E-75 | 3.20E-71 | 0.15 | 0.06 | 0.09 | 2.4 |
| mesenchyme | Topors | 0.4 | 1.56E-75 | 3.25E-71 | 0.69 | 0.6 | 0.09 | 1.15 |
| mesenchyme | Coq7 | 0.38 | 2.03E-75 | 4.22E-71 | 0.7 | 0.61 | 0.09 | 1.15 |

**Supplementary Table 2. Marker genes for the major cell types of integrated clustering of E11.5 control and NotchGOF λ-junction cells.**

| | | | | | | | | |
|---|---|---|---|---|---|---|---|---|
| mesenchyme | Gpat4 | 0.59 | 2.44E-75 | 5.08E-71 | 0.52 | 0.42 | 0.1 | 1.23 |
| mesenchyme | Pkig | 0.26 | 2.89E-75 | 6.02E-71 | 0.81 | 0.74 | 0.06 | 1.08 |
| mesenchyme | Dnah7a | 1.66 | 3.34E-75 | 6.94E-71 | 0.15 | 0.06 | 0.09 | 2.39 |
| mesenchyme | Cbx2 | 0.61 | 3.67E-75 | 7.64E-71 | 0.47 | 0.36 | 0.11 | 1.3 |
| mesenchyme | Mrc2 | 1.04 | 4.15E-75 | 8.62E-71 | 0.25 | 0.15 | 0.11 | 1.73 |
| mesenchyme | Rnf219 | 0.44 | 4.71E-75 | 9.79E-71 | 0.61 | 0.51 | 0.11 | 1.21 |
| mesenchyme | Chst14 | 1.15 | 4.79E-75 | 9.97E-71 | 0.27 | 0.16 | 0.1 | 1.65 |
| mesenchyme | Tmem134 | 0.34 | 5.29E-75 | 1.10E-70 | 0.77 | 0.71 | 0.06 | 1.08 |
| mesenchyme | Anapc4 | 0.51 | 5.49E-75 | 1.14E-70 | 0.56 | 0.47 | 0.09 | 1.2 |
| mesenchyme | Phf23 | 0.41 | 6.34E-75 | 1.32E-70 | 0.66 | 0.57 | 0.09 | 1.16 |
| mesenchyme | L1cam | 1.99 | 6.56E-75 | 1.36E-70 | 0.11 | 0.03 | 0.08 | 3.5 |
| mesenchyme | Anapc1 | 0.46 | 6.76E-75 | 1.41E-70 | 0.61 | 0.52 | 0.09 | 1.17 |
| mesenchyme | Pxdc1 | 2.03 | 7.11E-75 | 1.48E-70 | 0.13 | 0.05 | 0.08 | 2.69 |
| mesenchyme | Pdxp | 0.63 | 8.23E-75 | 1.71E-70 | 0.43 | 0.32 | 0.11 | 1.36 |
| mesenchyme | Arhgap31 | 0.81 | 1.21E-74 | 2.52E-70 | 0.23 | 0.12 | 0.11 | 1.91 |
| mesenchyme | Mrps16 | 0.27 | 1.27E-74 | 2.64E-70 | 0.86 | 0.8 | 0.06 | 1.07 |
| mesenchyme | Dynlt1f | 0.28 | 1.27E-74 | 2.65E-70 | 0.8 | 0.72 | 0.07 | 1.1 |
| mesenchyme | Mrpl35 | 0.33 | 1.29E-74 | 2.68E-70 | 0.75 | 0.69 | 0.06 | 1.08 |
| mesenchyme | Impad1 | 0.33 | 1.36E-74 | 2.83E-70 | 0.76 | 0.64 | 0.11 | 1.18 |
| mesenchyme | Mdp1 | 0.43 | 1.38E-74 | 2.86E-70 | 0.62 | 0.54 | 0.08 | 1.14 |
| mesenchyme | Mad2l2 | 0.64 | 1.38E-74 | 2.87E-70 | 0.45 | 0.35 | 0.1 | 1.28 |
| mesenchyme | Ppil3 | 0.49 | 1.66E-74 | 3.45E-70 | 0.57 | 0.47 | 0.1 | 1.21 |
| mesenchyme | Ntng1 | 1.76 | 1.79E-74 | 3.73E-70 | 0.15 | 0.06 | 0.09 | 2.38 |
| mesenchyme | Gemin7 | 0.35 | 2.03E-74 | 4.22E-70 | 0.72 | 0.67 | 0.06 | 1.08 |
| mesenchyme | Bora | 0.68 | 2.15E-74 | 4.46E-70 | 0.48 | 0.38 | 0.1 | 1.26 |
| mesenchyme | Por | 0.57 | 2.51E-74 | 5.22E-70 | 0.5 | 0.4 | 0.1 | 1.26 |
| mesenchyme | Zmym4 | 0.37 | 2.53E-74 | 5.25E-70 | 0.73 | 0.64 | 0.1 | 1.15 |
| mesenchyme | Dzip3 | 0.43 | 2.69E-74 | 5.60E-70 | 0.64 | 0.54 | 0.1 | 1.19 |
| mesenchyme | Gpt2 | 1.12 | 3.32E-74 | 6.90E-70 | 0.24 | 0.14 | 0.1 | 1.74 |
| mesenchyme | Prpsap1 | 0.36 | 3.74E-74 | 7.78E-70 | 0.71 | 0.63 | 0.09 | 1.14 |
| mesenchyme | Xbp1 | 0.59 | 4.11E-74 | 8.55E-70 | 0.51 | 0.41 | 0.1 | 1.25 |
| mesenchyme | Cetn2 | 0.36 | 4.58E-74 | 9.52E-70 | 0.67 | 0.59 | 0.08 | 1.14 |
| mesenchyme | Kif1b | 0.35 | 5.33E-74 | 1.11E-69 | 0.7 | 0.59 | 0.11 | 1.18 |
| mesenchyme | Antxr1 | 0.72 | 5.62E-74 | 1.17E-69 | 0.35 | 0.24 | 0.11 | 1.48 |
| mesenchyme | Tle2 | 1.62 | 5.74E-74 | 1.19E-69 | 0.15 | 0.06 | 0.09 | 2.4 |
| mesenchyme | Tcim | 0.43 | 5.90E-74 | 1.23E-69 | 0.41 | 0.29 | 0.12 | 1.43 |
| mesenchyme | Arcn1 | 0.4 | 7.07E-74 | 1.47E-69 | 0.68 | 0.6 | 0.08 | 1.13 |
| mesenchyme | Commd8 | 0.38 | 8.27E-74 | 1.72E-69 | 0.69 | 0.62 | 0.08 | 1.13 |
| mesenchyme | Snhg6 | 0.31 | 9.12E-74 | 1.90E-69 | 0.77 | 0.7 | 0.07 | 1.1 |
| mesenchyme | Lyrm2 | 0.4 | 1.94E-73 | 4.03E-69 | 0.66 | 0.57 | 0.09 | 1.15 |
| mesenchyme | Arxes2 | 0.66 | 1.95E-73 | 4.05E-69 | 0.43 | 0.33 | 0.11 | 1.33 |
| mesenchyme | Abat | 1.38 | 2.03E-73 | 4.23E-69 | 0.19 | 0.09 | 0.1 | 2.04 |
| mesenchyme | Cop1 | 0.38 | 2.54E-73 | 5.28E-69 | 0.7 | 0.63 | 0.07 | 1.11 |
| mesenchyme | Rex1bd | 0.3 | 2.55E-73 | 5.29E-69 | 0.88 | 0.83 | 0.05 | 1.06 |
| mesenchyme | Ptges | 1.35 | 2.80E-73 | 5.83E-69 | 0.17 | 0.08 | 0.09 | 2.16 |
| mesenchyme | Mrps11 | 0.33 | 3.73E-73 | 7.77E-69 | 0.73 | 0.65 | 0.07 | 1.11 |
| mesenchyme | Ankrd29 | 1.96 | 4.44E-73 | 9.24E-69 | 0.12 | 0.04 | 0.08 | 2.95 |
| mesenchyme | Rilpl1 | 0.79 | 5.17E-73 | 1.08E-68 | 0.33 | 0.22 | 0.11 | 1.5 |
| mesenchyme | Bub1 | 0.54 | 5.23E-73 | 1.09E-68 | 0.59 | 0.51 | 0.08 | 1.16 |
| mesenchyme | Selenos | 0.45 | 5.54E-73 | 1.15E-68 | 0.64 | 0.55 | 0.09 | 1.16 |
| mesenchyme | Taf7 | 0.38 | 5.87E-73 | 1.22E-68 | 0.72 | 0.65 | 0.06 | 1.09 |
| mesenchyme | Clasp1 | 0.47 | 5.90E-73 | 1.23E-68 | 0.59 | 0.5 | 0.09 | 1.18 |
| mesenchyme | Mrpl50 | 0.32 | 6.90E-73 | 1.44E-68 | 0.76 | 0.68 | 0.08 | 1.11 |
| mesenchyme | Psmc1 | 0.28 | 7.59E-73 | 1.58E-68 | 0.82 | 0.77 | 0.06 | 1.07 |
| mesenchyme | Plekhh2 | 1.38 | 8.60E-73 | 1.79E-68 | 0.19 | 0.09 | 0.09 | 2.03 |
| mesenchyme | Prpf19 | 0.27 | 9.70E-73 | 2.02E-68 | 0.83 | 0.75 | 0.08 | 1.11 |

**Supplementary Table 2. Marker genes for the major cell types of integrated clustering of E11.5 control and NotchGOF λ-junction cells.**

| | | | | | | | | |
|---|---|---|---|---|---|---|---|---|
| mesenchyme | Stau1 | 0.5 | 1.28E-72 | 2.67E-68 | 0.57 | 0.48 | 0.09 | 1.19 |
| mesenchyme | Arhgef9 | 1.21 | 1.68E-72 | 3.49E-68 | 0.22 | 0.12 | 0.1 | 1.83 |
| mesenchyme | Angel2 | 0.44 | 1.91E-72 | 3.96E-68 | 0.62 | 0.53 | 0.09 | 1.16 |
| mesenchyme | Yap1 | 0.36 | 2.06E-72 | 4.29E-68 | 0.65 | 0.55 | 0.11 | 1.2 |
| mesenchyme | Lama4 | 0.73 | 2.46E-72 | 5.11E-68 | 0.16 | 0.07 | 0.09 | 2.34 |
| mesenchyme | Csnk2a2 | 0.3 | 2.61E-72 | 5.43E-68 | 0.81 | 0.74 | 0.07 | 1.09 |
| mesenchyme | Trip13 | 0.48 | 2.63E-72 | 5.47E-68 | 0.57 | 0.48 | 0.09 | 1.19 |
| mesenchyme | Hist1h1b | 1.08 | 2.80E-72 | 5.83E-68 | 0.66 | 0.62 | 0.04 | 1.07 |
| mesenchyme | Marchf5 | 0.26 | 3.43E-72 | 7.14E-68 | 0.89 | 0.85 | 0.04 | 1.04 |
| mesenchyme | Fibp | 0.45 | 3.73E-72 | 7.76E-68 | 0.64 | 0.56 | 0.08 | 1.14 |
| mesenchyme | Chmp4b | 0.26 | 4.31E-72 | 8.96E-68 | 0.87 | 0.81 | 0.06 | 1.08 |
| mesenchyme | Uqcrc1 | 0.27 | 6.63E-72 | 1.38E-67 | 0.9 | 0.83 | 0.07 | 1.08 |
| mesenchyme | Ehmt2 | 0.47 | 8.02E-72 | 1.67E-67 | 0.6 | 0.51 | 0.09 | 1.18 |
| mesenchyme | Kat6b | 0.58 | 1.11E-71 | 2.31E-67 | 0.51 | 0.4 | 0.1 | 1.25 |
| mesenchyme | Wdr61 | 0.29 | 1.23E-71 | 2.55E-67 | 0.79 | 0.73 | 0.06 | 1.08 |
| mesenchyme | Bmerb1 | 1.73 | 1.29E-71 | 2.69E-67 | 0.11 | 0.04 | 0.08 | 3 |
| mesenchyme | Nudcd3 | 0.37 | 1.56E-71 | 3.24E-67 | 0.69 | 0.62 | 0.07 | 1.11 |
| mesenchyme | Mvk | 0.61 | 1.71E-71 | 3.56E-67 | 0.46 | 0.36 | 0.1 | 1.27 |
| mesenchyme | Txndc15 | 0.44 | 1.72E-71 | 3.58E-67 | 0.63 | 0.54 | 0.09 | 1.17 |
| mesenchyme | Prpf31 | 0.3 | 2.32E-71 | 4.82E-67 | 0.78 | 0.7 | 0.08 | 1.11 |
| mesenchyme | Gpsm1 | 0.86 | 3.06E-71 | 6.37E-67 | 0.32 | 0.22 | 0.1 | 1.47 |
| mesenchyme | Lrrc49 | 0.92 | 3.40E-71 | 7.06E-67 | 0.28 | 0.18 | 0.1 | 1.58 |
| mesenchyme | Trim26 | 0.63 | 3.69E-71 | 7.66E-67 | 0.46 | 0.36 | 0.1 | 1.27 |
| mesenchyme | E2f5 | 0.55 | 3.81E-71 | 7.92E-67 | 0.52 | 0.43 | 0.09 | 1.21 |
| mesenchyme | Actr10 | 0.32 | 3.98E-71 | 8.27E-67 | 0.75 | 0.68 | 0.07 | 1.1 |
| mesenchyme | Fdft1 | 0.46 | 1.11E-70 | 2.32E-66 | 0.63 | 0.55 | 0.08 | 1.14 |
| mesenchyme | Tmem132a | 0.35 | 1.13E-70 | 2.35E-66 | 0.67 | 0.57 | 0.11 | 1.19 |
| mesenchyme | Fblim1 | 0.46 | 1.21E-70 | 2.53E-66 | 0.51 | 0.39 | 0.12 | 1.3 |
| mesenchyme | Rnf20 | 0.32 | 1.71E-70 | 3.56E-66 | 0.76 | 0.68 | 0.08 | 1.12 |
| mesenchyme | Taf3 | 0.57 | 1.96E-70 | 4.09E-66 | 0.54 | 0.45 | 0.08 | 1.18 |
| mesenchyme | Cers2 | 0.5 | 2.56E-70 | 5.32E-66 | 0.57 | 0.48 | 0.08 | 1.17 |
| mesenchyme | Arxes1 | 0.7 | 2.95E-70 | 6.14E-66 | 0.41 | 0.3 | 0.11 | 1.35 |
| mesenchyme | Mindy4 | 1.71 | 2.95E-70 | 6.14E-66 | 0.16 | 0.07 | 0.09 | 2.19 |
| mesenchyme | Cblb | 0.72 | 3.17E-70 | 6.60E-66 | 0.38 | 0.27 | 0.11 | 1.39 |
| mesenchyme | Nelfcd | 0.4 | 3.19E-70 | 6.63E-66 | 0.65 | 0.58 | 0.07 | 1.13 |
| mesenchyme | Maea | 0.37 | 3.54E-70 | 7.36E-66 | 0.7 | 0.63 | 0.07 | 1.12 |
| mesenchyme | Midn | 0.32 | 3.89E-70 | 8.09E-66 | 0.76 | 0.66 | 0.1 | 1.15 |
| mesenchyme | Prpf8 | 0.31 | 5.02E-70 | 1.04E-65 | 0.79 | 0.71 | 0.08 | 1.12 |
| mesenchyme | Tomm70a | 0.33 | 7.44E-70 | 1.55E-65 | 0.74 | 0.66 | 0.08 | 1.12 |
| mesenchyme | Hjurp | 0.47 | 7.88E-70 | 1.64E-65 | 0.63 | 0.55 | 0.08 | 1.14 |
| mesenchyme | Snrnp200 | 0.34 | 8.68E-70 | 1.81E-65 | 0.76 | 0.69 | 0.07 | 1.1 |
| mesenchyme | Shf | 0.83 | 8.95E-70 | 1.86E-65 | 0.31 | 0.21 | 0.11 | 1.51 |
| mesenchyme | Kif2c | 0.66 | 9.78E-70 | 2.03E-65 | 0.5 | 0.41 | 0.09 | 1.21 |
| mesenchyme | Abca5 | 1.57 | 9.95E-70 | 2.07E-65 | 0.16 | 0.07 | 0.09 | 2.18 |
| mesenchyme | 833420G17Ri | 0.39 | 1.08E-69 | 2.24E-65 | 0.67 | 0.6 | 0.08 | 1.13 |
| mesenchyme | Ublcp1 | 0.54 | 1.12E-69 | 2.33E-65 | 0.53 | 0.45 | 0.08 | 1.18 |
| mesenchyme | Ap1s1 | 0.29 | 1.27E-69 | 2.64E-65 | 0.79 | 0.72 | 0.07 | 1.09 |
| mesenchyme | Dennd5a | 0.74 | 1.36E-69 | 2.83E-65 | 0.38 | 0.28 | 0.1 | 1.35 |
| mesenchyme | Prpf38a | 0.32 | 1.37E-69 | 2.86E-65 | 0.76 | 0.69 | 0.07 | 1.1 |
| mesenchyme | Mlf1 | 1.01 | 1.63E-69 | 3.40E-65 | 0.24 | 0.14 | 0.1 | 1.7 |
| mesenchyme | Hsd17b12 | 0.32 | 2.00E-69 | 4.16E-65 | 0.75 | 0.65 | 0.1 | 1.15 |
| mesenchyme | Msantd4 | 0.48 | 2.01E-69 | 4.18E-65 | 0.56 | 0.46 | 0.1 | 1.21 |
| mesenchyme | BC005561 | 0.3 | 2.31E-69 | 4.81E-65 | 0.81 | 0.71 | 0.1 | 1.14 |
| mesenchyme | Elp6 | 0.71 | 2.58E-69 | 5.36E-65 | 0.42 | 0.32 | 0.1 | 1.3 |
| mesenchyme | Slc25a24 | 0.64 | 3.58E-69 | 7.44E-65 | 0.38 | 0.28 | 0.11 | 1.39 |
| mesenchyme | Exo1 | 0.6 | 4.22E-69 | 8.79E-65 | 0.49 | 0.39 | 0.1 | 1.26 |

**Supplementary Table 2. Marker genes for the major cell types of integrated clustering of E11.5 control and NotchGOF λ-junction cells.**

| | | | | | | | | |
|---|---|---|---|---|---|---|---|---|
| mesenchyme | Cdc40 | 0.33 | 4.26E-69 | 8.85E-65 | 0.74 | 0.65 | 0.08 | 1.13 |
| mesenchyme | Rnf8 | 0.39 | 4.90E-69 | 1.02E-64 | 0.65 | 0.58 | 0.07 | 1.13 |
| mesenchyme | Setd2 | 0.39 | 5.43E-69 | 1.13E-64 | 0.69 | 0.6 | 0.09 | 1.14 |
| mesenchyme | Rsf1 | 0.27 | 5.71E-69 | 1.19E-64 | 0.85 | 0.76 | 0.09 | 1.12 |
| mesenchyme | '530068E07Ri | 0.32 | 6.08E-69 | 1.26E-64 | 0.77 | 0.68 | 0.09 | 1.13 |
| mesenchyme | Ylpm1 | 0.36 | 7.16E-69 | 1.49E-64 | 0.72 | 0.64 | 0.09 | 1.14 |
| mesenchyme | Ntf3 | 0.85 | 1.05E-68 | 2.18E-64 | 0.33 | 0.22 | 0.11 | 1.48 |
| mesenchyme | Cenpe | 0.57 | 1.08E-68 | 2.26E-64 | 0.68 | 0.61 | 0.07 | 1.11 |
| mesenchyme | Gypc | 1.08 | 1.10E-68 | 2.29E-64 | 0.25 | 0.16 | 0.09 | 1.61 |
| mesenchyme | Wbp1 | 0.28 | 1.14E-68 | 2.37E-64 | 0.8 | 0.72 | 0.08 | 1.11 |
| mesenchyme | Pex3 | 0.86 | 1.15E-68 | 2.38E-64 | 0.35 | 0.25 | 0.1 | 1.4 |
| mesenchyme | Bmpr1a | 0.36 | 1.29E-68 | 2.69E-64 | 0.69 | 0.57 | 0.12 | 1.21 |
| mesenchyme | Plscr1 | 1.04 | 1.42E-68 | 2.96E-64 | 0.27 | 0.17 | 0.1 | 1.59 |
| mesenchyme | Iffo1 | 2.01 | 1.59E-68 | 3.30E-64 | 0.12 | 0.04 | 0.07 | 2.8 |
| mesenchyme | Zfp647 | 0.94 | 1.68E-68 | 3.49E-64 | 0.29 | 0.19 | 0.1 | 1.54 |
| mesenchyme | Terf2 | 0.56 | 1.74E-68 | 3.62E-64 | 0.48 | 0.39 | 0.1 | 1.25 |
| mesenchyme | Peli1 | 0.33 | 1.77E-68 | 3.68E-64 | 0.71 | 0.62 | 0.09 | 1.14 |
| mesenchyme | Polr2g | 0.25 | 1.82E-68 | 3.79E-64 | 0.85 | 0.79 | 0.06 | 1.08 |
| mesenchyme | Glyr1 | 0.29 | 1.93E-68 | 4.01E-64 | 0.77 | 0.68 | 0.09 | 1.14 |
| mesenchyme | Pwp1 | 0.42 | 2.45E-68 | 5.10E-64 | 0.6 | 0.52 | 0.08 | 1.16 |
| mesenchyme | Nme4 | 0.3 | 3.07E-68 | 6.39E-64 | 0.78 | 0.72 | 0.06 | 1.09 |
| mesenchyme | Adgrl2 | 0.32 | 3.09E-68 | 6.42E-64 | 0.73 | 0.61 | 0.11 | 1.18 |
| mesenchyme | Lsm1 | 0.3 | 3.85E-68 | 8.00E-64 | 0.75 | 0.69 | 0.07 | 1.09 |
| mesenchyme | Cwf19l2 | 0.45 | 4.08E-68 | 8.48E-64 | 0.57 | 0.48 | 0.08 | 1.17 |
| mesenchyme | Hdac3 | 0.34 | 4.49E-68 | 9.34E-64 | 0.7 | 0.62 | 0.08 | 1.13 |
| mesenchyme | Pofut2 | 0.53 | 5.19E-68 | 1.08E-63 | 0.51 | 0.42 | 0.09 | 1.22 |
| mesenchyme | Hist1h2bk | 1.1 | 5.44E-68 | 1.13E-63 | 0.26 | 0.16 | 0.09 | 1.58 |
| mesenchyme | Tm9sf3 | 0.26 | 7.02E-68 | 1.46E-63 | 0.89 | 0.81 | 0.09 | 1.11 |
| mesenchyme | !700069I18Ril | 1.6 | 1.13E-67 | 2.36E-63 | 0.16 | 0.07 | 0.09 | 2.16 |
| mesenchyme | Ttc39c | 1.04 | 1.46E-67 | 3.04E-63 | 0.24 | 0.14 | 0.1 | 1.73 |
| mesenchyme | St5 | 0.98 | 1.47E-67 | 3.06E-63 | 0.26 | 0.16 | 0.1 | 1.6 |
| mesenchyme | Pja2 | 0.49 | 1.48E-67 | 3.08E-63 | 0.58 | 0.49 | 0.09 | 1.17 |
| mesenchyme | Med22 | 0.72 | 1.51E-67 | 3.13E-63 | 0.37 | 0.27 | 0.1 | 1.37 |
| mesenchyme | Ifngr2 | 0.55 | 1.81E-67 | 3.77E-63 | 0.41 | 0.3 | 0.11 | 1.36 |
| mesenchyme | Sass6 | 0.51 | 1.88E-67 | 3.92E-63 | 0.53 | 0.45 | 0.08 | 1.18 |
| mesenchyme | Rbm7 | 0.37 | 2.00E-67 | 4.15E-63 | 0.68 | 0.62 | 0.06 | 1.1 |
| mesenchyme | Ube2g1 | 0.32 | 2.69E-67 | 5.60E-63 | 0.64 | 0.55 | 0.09 | 1.17 |
| mesenchyme | Uba5 | 0.42 | 3.91E-67 | 8.13E-63 | 0.62 | 0.54 | 0.08 | 1.14 |
| mesenchyme | Kmt2a | 0.32 | 4.20E-67 | 8.74E-63 | 0.75 | 0.65 | 0.09 | 1.14 |
| mesenchyme | Nuf2 | 0.63 | 4.21E-67 | 8.76E-63 | 0.5 | 0.43 | 0.07 | 1.17 |
| mesenchyme | Rprd1b | 0.48 | 4.70E-67 | 9.77E-63 | 0.56 | 0.48 | 0.08 | 1.17 |
| mesenchyme | Ncoa7 | 0.71 | 7.56E-67 | 1.57E-62 | 0.43 | 0.35 | 0.09 | 1.25 |
| mesenchyme | 930579G24Ri | 0.99 | 7.67E-67 | 1.60E-62 | 0.28 | 0.19 | 0.1 | 1.5 |
| mesenchyme | Dyrk2 | 0.64 | 7.69E-67 | 1.60E-62 | 0.41 | 0.31 | 0.1 | 1.32 |
| mesenchyme | Cdk12 | 0.33 | 1.39E-66 | 2.90E-62 | 0.76 | 0.68 | 0.08 | 1.12 |
| mesenchyme | Msx2 | 0.71 | 1.62E-66 | 3.37E-62 | 0.46 | 0.36 | 0.1 | 1.27 |
| mesenchyme | Mypop | 1.57 | 1.85E-66 | 3.85E-62 | 0.14 | 0.06 | 0.08 | 2.38 |
| mesenchyme | Iglon5 | 1.14 | 1.95E-66 | 4.05E-62 | 0.2 | 0.11 | 0.09 | 1.85 |
| mesenchyme | Timp1 | 1.33 | 2.82E-66 | 5.86E-62 | 0.2 | 0.11 | 0.09 | 1.83 |
| mesenchyme | Cnot3 | 0.34 | 3.21E-66 | 6.67E-62 | 0.73 | 0.65 | 0.08 | 1.13 |
| mesenchyme | Zfp532 | 0.49 | 3.33E-66 | 6.92E-62 | 0.51 | 0.41 | 0.1 | 1.25 |
| mesenchyme | Adamts5 | 2.13 | 3.51E-66 | 7.30E-62 | 0.14 | 0.06 | 0.08 | 2.39 |
| mesenchyme | Cenpw | 0.35 | 3.67E-66 | 7.63E-62 | 0.75 | 0.7 | 0.05 | 1.07 |
| mesenchyme | Ubqln2 | 0.37 | 3.80E-66 | 7.90E-62 | 0.68 | 0.59 | 0.09 | 1.15 |
| mesenchyme | .190005I06Ril | 1.41 | 3.93E-66 | 8.16E-62 | 0.16 | 0.07 | 0.08 | 2.17 |
| mesenchyme | Fam171a1 | 0.75 | 4.44E-66 | 9.24E-62 | 0.36 | 0.26 | 0.1 | 1.37 |

**Supplementary Table 2. Marker genes for the major cell types of integrated clustering of E11.5 control and NotchGOF λ-junction cells.**

| | | | | | | | | |
|---|---|---|---|---|---|---|---|---|
| mesenchyme | Tram1l1 | 1.24 | 4.60E-66 | 9.57E-62 | 0.21 | 0.12 | 0.09 | 1.81 |
| mesenchyme | Spsb1 | 1.35 | 5.02E-66 | 1.04E-61 | 0.18 | 0.09 | 0.09 | 1.94 |
| mesenchyme | Itga5 | 1.01 | 5.21E-66 | 1.08E-61 | 0.25 | 0.15 | 0.1 | 1.65 |
| mesenchyme | Atg5 | 0.44 | 8.50E-66 | 1.77E-61 | 0.6 | 0.52 | 0.08 | 1.15 |
| mesenchyme | Tmem121 | 0.87 | 1.02E-65 | 2.13E-61 | 0.26 | 0.17 | 0.1 | 1.6 |
| mesenchyme | Prrc2a | 0.29 | 1.08E-65 | 2.24E-61 | 0.79 | 0.71 | 0.08 | 1.11 |
| mesenchyme | Srm | 0.31 | 1.08E-65 | 2.25E-61 | 0.71 | 0.62 | 0.09 | 1.14 |
| mesenchyme | Cxxc4 | 0.97 | 1.11E-65 | 2.30E-61 | 0.34 | 0.24 | 0.1 | 1.4 |
| mesenchyme | Car12 | 1.72 | 1.19E-65 | 2.47E-61 | 0.12 | 0.04 | 0.07 | 2.68 |
| mesenchyme | Pgm2 | 0.74 | 1.25E-65 | 2.60E-61 | 0.31 | 0.21 | 0.1 | 1.48 |
| mesenchyme | Golim4 | 0.37 | 1.35E-65 | 2.81E-61 | 0.64 | 0.55 | 0.1 | 1.18 |
| mesenchyme | Mcl1 | 0.26 | 1.37E-65 | 2.85E-61 | 0.84 | 0.79 | 0.05 | 1.06 |
| mesenchyme | Cox20 | 0.38 | 1.38E-65 | 2.87E-61 | 0.69 | 0.63 | 0.06 | 1.1 |
| mesenchyme | Ppp4c | 0.25 | 1.43E-65 | 2.97E-61 | 0.87 | 0.81 | 0.06 | 1.08 |
| mesenchyme | Tiam2 | 1.59 | 1.53E-65 | 3.19E-61 | 0.12 | 0.04 | 0.07 | 2.72 |
| mesenchyme | Tsr2 | 0.61 | 1.60E-65 | 3.33E-61 | 0.43 | 0.33 | 0.1 | 1.3 |
| mesenchyme | Ndufa9 | 0.33 | 1.95E-65 | 4.05E-61 | 0.74 | 0.68 | 0.06 | 1.09 |
| mesenchyme | Dnttip1 | 0.34 | 2.15E-65 | 4.46E-61 | 0.67 | 0.59 | 0.08 | 1.13 |
| mesenchyme | Cul2 | 0.48 | 2.21E-65 | 4.60E-61 | 0.55 | 0.47 | 0.08 | 1.17 |
| mesenchyme | Timm50 | 0.32 | 2.40E-65 | 5.00E-61 | 0.76 | 0.69 | 0.07 | 1.1 |
| mesenchyme | Pitpnb | 0.35 | 2.48E-65 | 5.17E-61 | 0.7 | 0.62 | 0.08 | 1.13 |
| mesenchyme | Cox16 | 0.31 | 3.37E-65 | 7.00E-61 | 0.77 | 0.72 | 0.05 | 1.06 |
| mesenchyme | Fam204a | 0.34 | 3.73E-65 | 7.76E-61 | 0.68 | 0.61 | 0.07 | 1.11 |
| mesenchyme | Dock1 | 0.5 | 3.74E-65 | 7.77E-61 | 0.54 | 0.44 | 0.1 | 1.23 |
| mesenchyme | Ncapd2 | 0.5 | 4.14E-65 | 8.61E-61 | 0.58 | 0.51 | 0.07 | 1.14 |
| mesenchyme | Trappc5 | 0.36 | 4.79E-65 | 9.97E-61 | 0.69 | 0.62 | 0.07 | 1.11 |
| mesenchyme | Ddx23 | 0.4 | 5.22E-65 | 1.08E-60 | 0.62 | 0.54 | 0.08 | 1.15 |
| mesenchyme | Hist1h2ae | 0.83 | 5.87E-65 | 1.22E-60 | 0.72 | 0.66 | 0.07 | 1.1 |
| mesenchyme | Ugdh | 0.5 | 5.98E-65 | 1.24E-60 | 0.5 | 0.4 | 0.1 | 1.25 |
| mesenchyme | Pmf1 | 0.32 | 6.72E-65 | 1.40E-60 | 0.69 | 0.63 | 0.06 | 1.1 |
| mesenchyme | Zrsr2 | 0.27 | 6.85E-65 | 1.42E-60 | 0.79 | 0.71 | 0.08 | 1.12 |
| mesenchyme | )930021J03Ri | 0.45 | 1.33E-64 | 2.76E-60 | 0.58 | 0.49 | 0.09 | 1.19 |
| mesenchyme | Atp2a2 | 0.35 | 1.39E-64 | 2.89E-60 | 0.71 | 0.62 | 0.09 | 1.14 |
| mesenchyme | Stoml2 | 0.26 | 1.87E-64 | 3.89E-60 | 0.86 | 0.81 | 0.05 | 1.06 |
| mesenchyme | Fam118b | 0.63 | 1.90E-64 | 3.95E-60 | 0.4 | 0.31 | 0.1 | 1.31 |
| mesenchyme | Hpf1 | 0.26 | 2.18E-64 | 4.54E-60 | 0.86 | 0.81 | 0.05 | 1.07 |
| mesenchyme | Kdelr1 | 0.27 | 2.45E-64 | 5.09E-60 | 0.86 | 0.78 | 0.08 | 1.1 |
| mesenchyme | Ppp2r3a | 0.72 | 2.98E-64 | 6.19E-60 | 0.39 | 0.29 | 0.1 | 1.35 |
| mesenchyme | Lias | 0.5 | 3.05E-64 | 6.35E-60 | 0.54 | 0.46 | 0.08 | 1.17 |
| mesenchyme | Hist1h2bm | 0.98 | 3.12E-64 | 6.49E-60 | 0.34 | 0.25 | 0.09 | 1.36 |
| mesenchyme | Utp6 | 0.58 | 3.17E-64 | 6.60E-60 | 0.47 | 0.4 | 0.08 | 1.2 |
| mesenchyme | Slain2 | 0.49 | 3.26E-64 | 6.78E-60 | 0.47 | 0.38 | 0.1 | 1.26 |
| mesenchyme | Mertk | 1.16 | 3.42E-64 | 7.11E-60 | 0.19 | 0.1 | 0.09 | 1.9 |
| mesenchyme | Gpn1 | 0.4 | 3.69E-64 | 7.67E-60 | 0.61 | 0.53 | 0.09 | 1.16 |
| mesenchyme | P3h4 | 0.72 | 3.71E-64 | 7.72E-60 | 0.39 | 0.29 | 0.1 | 1.34 |
| mesenchyme | Dusp6 | 0.82 | 4.71E-64 | 9.80E-60 | 0.39 | 0.3 | 0.09 | 1.31 |
| mesenchyme | Znrf1 | 0.36 | 5.23E-64 | 1.09E-59 | 0.63 | 0.54 | 0.08 | 1.15 |
| mesenchyme | Rbm10 | 0.37 | 5.67E-64 | 1.18E-59 | 0.62 | 0.54 | 0.08 | 1.14 |
| mesenchyme | Spin1 | 0.31 | 7.20E-64 | 1.50E-59 | 0.72 | 0.64 | 0.08 | 1.13 |
| mesenchyme | Eefsec | 0.63 | 7.47E-64 | 1.55E-59 | 0.42 | 0.33 | 0.09 | 1.28 |
| mesenchyme | Magee1 | 1.32 | 7.67E-64 | 1.60E-59 | 0.17 | 0.08 | 0.09 | 2.06 |
| mesenchyme | Ccnl2 | 0.31 | 8.53E-64 | 1.77E-59 | 0.73 | 0.66 | 0.07 | 1.11 |
| mesenchyme | Sec23a | 0.73 | 8.84E-64 | 1.84E-59 | 0.37 | 0.28 | 0.09 | 1.34 |
| mesenchyme | 330434G04Ri | 1.86 | 8.86E-64 | 1.84E-59 | 0.11 | 0.04 | 0.07 | 2.71 |
| mesenchyme | Mff | 0.27 | 9.79E-64 | 2.04E-59 | 0.79 | 0.75 | 0.04 | 1.05 |
| mesenchyme | Cyth3 | 0.65 | 1.07E-63 | 2.23E-59 | 0.42 | 0.33 | 0.09 | 1.28 |

**Supplementary Table 2. Marker genes for the major cell types of integrated clustering of E11.5 control and NotchGOF λ-junction cells.**

| | | | | | | | | |
|---|---|---|---|---|---|---|---|---|
| mesenchyme | Czib | 0.46 | 1.10E-63 | 2.28E-59 | 0.56 | 0.48 | 0.08 | 1.16 |
| mesenchyme | Rnf2 | 0.39 | 1.10E-63 | 2.29E-59 | 0.63 | 0.54 | 0.09 | 1.16 |
| mesenchyme | Gxylt1 | 0.54 | 1.23E-63 | 2.56E-59 | 0.5 | 0.4 | 0.09 | 1.23 |
| mesenchyme | Cacna2d1 | 0.82 | 1.34E-63 | 2.79E-59 | 0.15 | 0.07 | 0.08 | 2.26 |
| mesenchyme | Pcna | 0.3 | 1.54E-63 | 3.20E-59 | 0.86 | 0.79 | 0.07 | 1.09 |
| mesenchyme | 930402H24Ri | 0.86 | 1.98E-63 | 4.12E-59 | 0.29 | 0.19 | 0.1 | 1.52 |
| mesenchyme | Spag9 | 0.38 | 2.09E-63 | 4.35E-59 | 0.65 | 0.57 | 0.08 | 1.15 |
| mesenchyme | Clpp | 0.3 | 2.34E-63 | 4.86E-59 | 0.77 | 0.71 | 0.06 | 1.08 |
| mesenchyme | Ensa | 0.38 | 2.95E-63 | 6.14E-59 | 0.64 | 0.58 | 0.07 | 1.11 |
| mesenchyme | Zfp157 | 0.7 | 3.15E-63 | 6.55E-59 | 0.4 | 0.31 | 0.09 | 1.31 |
| mesenchyme | Nvl | 0.42 | 3.21E-63 | 6.67E-59 | 0.61 | 0.53 | 0.08 | 1.15 |
| mesenchyme | Fh1 | 0.31 | 3.26E-63 | 6.77E-59 | 0.74 | 0.68 | 0.06 | 1.09 |
| mesenchyme | Eftud2 | 0.33 | 3.26E-63 | 6.78E-59 | 0.71 | 0.64 | 0.08 | 1.12 |
| mesenchyme | Ddx52 | 0.41 | 4.10E-63 | 8.53E-59 | 0.61 | 0.54 | 0.07 | 1.13 |
| mesenchyme | Ncbp3 | 0.47 | 5.09E-63 | 1.06E-58 | 0.56 | 0.48 | 0.08 | 1.17 |
| mesenchyme | Pop4 | 0.53 | 5.25E-63 | 1.09E-58 | 0.49 | 0.4 | 0.09 | 1.22 |
| mesenchyme | Rab23 | 0.69 | 9.39E-63 | 1.95E-58 | 0.36 | 0.26 | 0.1 | 1.38 |
| mesenchyme | Acadl | 0.5 | 9.82E-63 | 2.04E-58 | 0.56 | 0.49 | 0.07 | 1.15 |
| mesenchyme | Ptpn2 | 0.42 | 1.03E-62 | 2.15E-58 | 0.61 | 0.53 | 0.07 | 1.14 |
| mesenchyme | Kansl1l | 0.86 | 1.07E-62 | 2.22E-58 | 0.3 | 0.2 | 0.1 | 1.49 |
| mesenchyme | R3hdm1 | 0.3 | 1.43E-62 | 2.98E-58 | 0.76 | 0.69 | 0.07 | 1.1 |
| mesenchyme | Ccar2 | 0.53 | 1.94E-62 | 4.04E-58 | 0.5 | 0.4 | 0.09 | 1.23 |
| mesenchyme | Thoc2 | 0.28 | 2.39E-62 | 4.96E-58 | 0.8 | 0.74 | 0.07 | 1.09 |
| mesenchyme | Sec22b | 0.4 | 2.75E-62 | 5.71E-58 | 0.63 | 0.56 | 0.07 | 1.13 |
| mesenchyme | Cep85l | 1.34 | 3.06E-62 | 6.37E-58 | 0.17 | 0.08 | 0.08 | 2.01 |
| mesenchyme | Sfswap | 0.44 | 3.33E-62 | 6.93E-58 | 0.6 | 0.52 | 0.08 | 1.16 |
| mesenchyme | Hsd17b11 | 1.1 | 3.50E-62 | 7.28E-58 | 0.24 | 0.15 | 0.09 | 1.62 |
| mesenchyme | Cryz | 0.66 | 4.08E-62 | 8.48E-58 | 0.41 | 0.31 | 0.1 | 1.31 |
| mesenchyme | Cpsf2 | 0.34 | 4.30E-62 | 8.93E-58 | 0.69 | 0.62 | 0.07 | 1.11 |
| mesenchyme | Pgrmc2 | 0.4 | 4.90E-62 | 1.02E-57 | 0.61 | 0.53 | 0.08 | 1.14 |
| mesenchyme | Csnk1g2 | 0.26 | 6.59E-62 | 1.37E-57 | 0.82 | 0.75 | 0.07 | 1.09 |
| mesenchyme | Tcf7l1 | 0.39 | 7.21E-62 | 1.50E-57 | 0.62 | 0.51 | 0.11 | 1.21 |
| mesenchyme | Mta1 | 0.33 | 7.44E-62 | 1.55E-57 | 0.73 | 0.63 | 0.1 | 1.15 |
| mesenchyme | Ush1c | 1.54 | 1.05E-61 | 2.18E-57 | 0.13 | 0.05 | 0.08 | 2.39 |
| mesenchyme | Ubr5 | 0.37 | 1.15E-61 | 2.40E-57 | 0.69 | 0.62 | 0.07 | 1.12 |
| mesenchyme | Triqk | 1.36 | 1.22E-61 | 2.54E-57 | 0.15 | 0.07 | 0.08 | 2.13 |
| mesenchyme | Ubn1 | 0.3 | 1.30E-61 | 2.71E-57 | 0.79 | 0.73 | 0.06 | 1.08 |
| mesenchyme | Hmgcl | 0.61 | 1.46E-61 | 3.03E-57 | 0.43 | 0.35 | 0.09 | 1.25 |
| mesenchyme | Sacs | 1 | 1.51E-61 | 3.14E-57 | 0.21 | 0.12 | 0.09 | 1.77 |
| mesenchyme | Lnpk | 0.55 | 1.70E-61 | 3.53E-57 | 0.51 | 0.44 | 0.07 | 1.17 |
| mesenchyme | Cenpp | 0.62 | 1.78E-61 | 3.71E-57 | 0.43 | 0.35 | 0.08 | 1.24 |
| mesenchyme | Riok1 | 0.41 | 2.12E-61 | 4.41E-57 | 0.61 | 0.54 | 0.07 | 1.13 |
| mesenchyme | Hist1h2ab | 0.98 | 2.48E-61 | 5.17E-57 | 0.4 | 0.32 | 0.08 | 1.24 |
| mesenchyme | 921531C22Ri | 1.21 | 2.59E-61 | 5.38E-57 | 0.21 | 0.12 | 0.09 | 1.73 |
| mesenchyme | Ccdc25 | 0.35 | 2.76E-61 | 5.75E-57 | 0.67 | 0.6 | 0.06 | 1.11 |
| mesenchyme | Capn2 | 0.48 | 2.90E-61 | 6.04E-57 | 0.49 | 0.39 | 0.1 | 1.26 |
| mesenchyme | Cnot2 | 0.33 | 3.00E-61 | 6.24E-57 | 0.67 | 0.6 | 0.07 | 1.11 |
| mesenchyme | Clip2 | 1.12 | 3.34E-61 | 6.94E-57 | 0.22 | 0.13 | 0.09 | 1.69 |
| mesenchyme | Ino80e | 0.48 | 3.36E-61 | 7.00E-57 | 0.55 | 0.47 | 0.08 | 1.16 |
| mesenchyme | Zfr | 0.29 | 3.53E-61 | 7.35E-57 | 0.79 | 0.73 | 0.06 | 1.08 |
| mesenchyme | Rab36 | 1.42 | 3.54E-61 | 7.37E-57 | 0.15 | 0.07 | 0.08 | 2.16 |
| mesenchyme | Ppp2r1b | 0.62 | 3.76E-61 | 7.81E-57 | 0.43 | 0.34 | 0.09 | 1.26 |
| mesenchyme | Map2k4 | 0.48 | 4.18E-61 | 8.70E-57 | 0.55 | 0.47 | 0.08 | 1.16 |
| mesenchyme | Irx3 | 0.86 | 4.64E-61 | 9.66E-57 | 0.31 | 0.2 | 0.1 | 1.5 |
| mesenchyme | Lrrc59 | 0.26 | 4.72E-61 | 9.81E-57 | 0.82 | 0.74 | 0.08 | 1.1 |
| mesenchyme | Fgfr1op | 0.65 | 5.15E-61 | 1.07E-56 | 0.42 | 0.34 | 0.09 | 1.25 |

**Supplementary Table 2. Marker genes for the major cell types of integrated clustering of E11.5 control and NotchGOF λ-junction cells.**

| | | | | | | | | |
|---|---|---|---|---|---|---|---|---|
| mesenchyme | Angptl2 | 1.09 | 5.29E-61 | 1.10E-56 | 0.22 | 0.13 | 0.09 | 1.71 |
| mesenchyme | Tmod2 | 1.06 | 7.69E-61 | 1.60E-56 | 0.23 | 0.14 | 0.09 | 1.67 |
| mesenchyme | Stk3 | 0.64 | 8.23E-61 | 1.71E-56 | 0.45 | 0.37 | 0.08 | 1.21 |
| mesenchyme | Ube2a | 0.3 | 9.23E-61 | 1.92E-56 | 0.73 | 0.67 | 0.05 | 1.08 |
| mesenchyme | Tex9 | 0.53 | 1.11E-60 | 2.32E-56 | 0.48 | 0.39 | 0.09 | 1.24 |
| mesenchyme | Bud23 | 0.33 | 1.24E-60 | 2.57E-56 | 0.69 | 0.62 | 0.07 | 1.11 |
| mesenchyme | Hsd17b7 | 0.86 | 1.27E-60 | 2.64E-56 | 0.3 | 0.21 | 0.09 | 1.44 |
| mesenchyme | Ubtf | 0.27 | 1.39E-60 | 2.89E-56 | 0.85 | 0.79 | 0.06 | 1.07 |
| mesenchyme | Hist1h3i | 1.11 | 1.68E-60 | 3.48E-56 | 0.27 | 0.18 | 0.09 | 1.49 |
| mesenchyme | Krit1 | 0.39 | 2.04E-60 | 4.24E-56 | 0.65 | 0.59 | 0.06 | 1.1 |
| mesenchyme | Nudt5 | 0.26 | 2.15E-60 | 4.47E-56 | 0.81 | 0.76 | 0.05 | 1.07 |
| mesenchyme | Tnpo3 | 0.34 | 2.38E-60 | 4.96E-56 | 0.7 | 0.63 | 0.07 | 1.11 |
| mesenchyme | Itih5 | 1.24 | 2.42E-60 | 5.03E-56 | 0.15 | 0.07 | 0.08 | 2.11 |
| mesenchyme | Pakap | 1.31 | 2.60E-60 | 5.41E-56 | 0.15 | 0.07 | 0.08 | 2.18 |
| mesenchyme | Hilpda | 0.64 | 3.03E-60 | 6.31E-56 | 0.39 | 0.29 | 0.09 | 1.32 |
| mesenchyme | Hdhd2 | 0.48 | 5.24E-60 | 1.09E-55 | 0.53 | 0.45 | 0.08 | 1.18 |
| mesenchyme | Wdr18 | 0.3 | 5.91E-60 | 1.23E-55 | 0.75 | 0.68 | 0.07 | 1.11 |
| mesenchyme | Ctr9 | 0.29 | 6.76E-60 | 1.41E-55 | 0.71 | 0.63 | 0.08 | 1.12 |
| mesenchyme | Wif1 | 1.98 | 7.10E-60 | 1.48E-55 | 0.12 | 0.05 | 0.07 | 2.54 |
| mesenchyme | Smn1 | 0.29 | 8.48E-60 | 1.76E-55 | 0.75 | 0.68 | 0.07 | 1.1 |
| mesenchyme | Plcb1 | 1.58 | 1.34E-59 | 2.79E-55 | 0.12 | 0.05 | 0.07 | 2.45 |
| mesenchyme | Rtl8a | 0.39 | 1.84E-59 | 3.84E-55 | 0.42 | 0.32 | 0.1 | 1.32 |
| mesenchyme | Ccnh | 0.38 | 1.94E-59 | 4.04E-55 | 0.63 | 0.57 | 0.06 | 1.1 |
| mesenchyme | Gli2 | 0.8 | 2.12E-59 | 4.42E-55 | 0.29 | 0.2 | 0.09 | 1.47 |
| mesenchyme | Gm26648 | 1.76 | 2.13E-59 | 4.43E-55 | 0.11 | 0.04 | 0.07 | 2.58 |
| mesenchyme | Lrch2 | 0.77 | 2.45E-59 | 5.09E-55 | 0.31 | 0.22 | 0.1 | 1.44 |
| mesenchyme | Stn1 | 0.63 | 2.86E-59 | 5.94E-55 | 0.39 | 0.3 | 0.09 | 1.31 |
| mesenchyme | Gpc2 | 0.56 | 3.00E-59 | 6.23E-55 | 0.43 | 0.33 | 0.1 | 1.3 |
| mesenchyme | Lrrcc1 | 0.42 | 3.80E-59 | 7.90E-55 | 0.57 | 0.49 | 0.08 | 1.17 |
| mesenchyme | Ppm1a | 0.27 | 4.16E-59 | 8.66E-55 | 0.74 | 0.68 | 0.05 | 1.08 |
| mesenchyme | Carm1 | 0.39 | 4.86E-59 | 1.01E-54 | 0.58 | 0.5 | 0.08 | 1.16 |
| mesenchyme | Nelfa | 0.4 | 5.20E-59 | 1.08E-54 | 0.57 | 0.49 | 0.08 | 1.16 |
| mesenchyme | Eps8 | 1.13 | 5.41E-59 | 1.12E-54 | 0.19 | 0.11 | 0.09 | 1.81 |
| mesenchyme | Naxe | 0.31 | 5.47E-59 | 1.14E-54 | 0.75 | 0.68 | 0.07 | 1.1 |
| mesenchyme | Ccnl1 | 0.33 | 6.64E-59 | 1.38E-54 | 0.74 | 0.69 | 0.06 | 1.08 |
| mesenchyme | Ilkap | 0.26 | 8.08E-59 | 1.68E-54 | 0.78 | 0.73 | 0.05 | 1.07 |
| mesenchyme | Armcx3 | 0.54 | 9.72E-59 | 2.02E-54 | 0.46 | 0.37 | 0.09 | 1.25 |
| mesenchyme | Stk16 | 0.35 | 9.95E-59 | 2.07E-54 | 0.69 | 0.63 | 0.06 | 1.09 |
| mesenchyme | Rhod | 0.68 | 9.95E-59 | 2.07E-54 | 0.37 | 0.28 | 0.09 | 1.34 |
| mesenchyme | Cse1l | 0.37 | 1.01E-58 | 2.10E-54 | 0.65 | 0.58 | 0.07 | 1.12 |
| mesenchyme | Vcl | 0.51 | 1.09E-58 | 2.27E-54 | 0.43 | 0.33 | 0.09 | 1.28 |
| mesenchyme | Tssc4 | 0.39 | 1.15E-58 | 2.40E-54 | 0.61 | 0.53 | 0.08 | 1.15 |
| mesenchyme | Rad17 | 0.52 | 1.16E-58 | 2.42E-54 | 0.5 | 0.42 | 0.07 | 1.18 |
| mesenchyme | Smad1 | 0.46 | 1.22E-58 | 2.53E-54 | 0.51 | 0.42 | 0.09 | 1.22 |
| mesenchyme | Camk2n1 | 0.4 | 1.22E-58 | 2.54E-54 | 0.22 | 0.13 | 0.09 | 1.71 |
| mesenchyme | Nfe2l2 | 0.56 | 1.27E-58 | 2.65E-54 | 0.47 | 0.4 | 0.08 | 1.2 |
| mesenchyme | Tfap2c | 0.84 | 1.51E-58 | 3.13E-54 | 0.31 | 0.21 | 0.1 | 1.46 |
| mesenchyme | 230038L03Ri | 1.34 | 1.52E-58 | 3.16E-54 | 0.15 | 0.07 | 0.08 | 2.08 |
| mesenchyme | Lysmd2 | 0.93 | 1.76E-58 | 3.67E-54 | 0.25 | 0.16 | 0.09 | 1.56 |
| mesenchyme | Smad4 | 0.27 | 1.84E-58 | 3.83E-54 | 0.8 | 0.71 | 0.09 | 1.12 |
| mesenchyme | Stx12 | 0.4 | 2.22E-58 | 4.63E-54 | 0.55 | 0.47 | 0.08 | 1.18 |
| mesenchyme | Itgb1bp1 | 0.59 | 2.56E-58 | 5.32E-54 | 0.43 | 0.34 | 0.09 | 1.25 |
| mesenchyme | Trim71 | 0.99 | 2.72E-58 | 5.65E-54 | 0.24 | 0.15 | 0.09 | 1.59 |
| mesenchyme | Cdc37l1 | 0.39 | 2.72E-58 | 5.66E-54 | 0.62 | 0.55 | 0.07 | 1.12 |
| mesenchyme | Zkscan3 | 0.33 | 3.55E-58 | 7.39E-54 | 0.7 | 0.62 | 0.08 | 1.13 |
| mesenchyme | Dync2h1 | 0.66 | 4.35E-58 | 9.05E-54 | 0.41 | 0.32 | 0.09 | 1.29 |

**Supplementary Table 2. Marker genes for the major cell types of integrated clustering of E11.5 control and NotchGOF λ-junction cells.**

| | | | | | | | | |
|---|---|---|---|---|---|---|---|---|
| mesenchyme | Mpnd | 0.3 | 4.75E-58 | 9.89E-54 | 0.71 | 0.62 | 0.08 | 1.13 |
| mesenchyme | Eif3j1 | 0.28 | 4.93E-58 | 1.02E-53 | 0.74 | 0.7 | 0.05 | 1.07 |
| mesenchyme | Isy1 | 0.41 | 5.69E-58 | 1.18E-53 | 0.57 | 0.5 | 0.07 | 1.13 |
| mesenchyme | Poldip2 | 0.37 | 6.45E-58 | 1.34E-53 | 0.64 | 0.57 | 0.07 | 1.12 |
| mesenchyme | Ttc14 | 0.31 | 7.04E-58 | 1.46E-53 | 0.76 | 0.69 | 0.07 | 1.1 |
| mesenchyme | Adpgk | 0.76 | 7.28E-58 | 1.51E-53 | 0.34 | 0.25 | 0.09 | 1.36 |
| mesenchyme | Saal1 | 0.53 | 9.93E-58 | 2.06E-53 | 0.49 | 0.41 | 0.08 | 1.19 |
| mesenchyme | Cfap36 | 0.35 | 1.00E-57 | 2.09E-53 | 0.58 | 0.5 | 0.08 | 1.15 |
| mesenchyme | Hsd17b4 | 0.45 | 1.29E-57 | 2.68E-53 | 0.57 | 0.5 | 0.07 | 1.14 |
| mesenchyme | P3h1 | 1.29 | 1.49E-57 | 3.10E-53 | 0.17 | 0.09 | 0.08 | 1.85 |
| mesenchyme | Tm2d1 | 0.42 | 1.52E-57 | 3.17E-53 | 0.54 | 0.46 | 0.08 | 1.16 |
| mesenchyme | Mmp14 | 0.35 | 1.62E-57 | 3.38E-53 | 0.67 | 0.56 | 0.11 | 1.19 |
| mesenchyme | Rap1gds1 | 0.78 | 1.64E-57 | 3.40E-53 | 0.32 | 0.23 | 0.09 | 1.38 |
| mesenchyme | Mapk7 | 0.67 | 1.69E-57 | 3.52E-53 | 0.37 | 0.28 | 0.09 | 1.32 |
| mesenchyme | Pabpc4l | 0.61 | 1.71E-57 | 3.55E-53 | 0.42 | 0.32 | 0.09 | 1.29 |
| mesenchyme | Hist1h3e | 0.7 | 2.15E-57 | 4.47E-53 | 0.45 | 0.38 | 0.08 | 1.2 |
| mesenchyme | Dexi | 0.6 | 2.24E-57 | 4.66E-53 | 0.41 | 0.33 | 0.09 | 1.27 |
| mesenchyme | Tusc3 | 0.42 | 2.80E-57 | 5.82E-53 | 0.59 | 0.52 | 0.07 | 1.14 |
| mesenchyme | Dhrs3 | 0.59 | 3.02E-57 | 6.27E-53 | 0.33 | 0.23 | 0.1 | 1.45 |
| mesenchyme | Ube2q2 | 0.5 | 3.49E-57 | 7.26E-53 | 0.52 | 0.45 | 0.07 | 1.16 |
| mesenchyme | Wdr70 | 0.48 | 3.87E-57 | 8.04E-53 | 0.52 | 0.44 | 0.08 | 1.17 |
| mesenchyme | Zfyve16 | 0.64 | 5.09E-57 | 1.06E-52 | 0.38 | 0.29 | 0.09 | 1.33 |
| mesenchyme | Usf1 | 0.41 | 6.02E-57 | 1.25E-52 | 0.57 | 0.49 | 0.08 | 1.16 |
| mesenchyme | Nudt3 | 0.31 | 6.87E-57 | 1.43E-52 | 0.66 | 0.59 | 0.07 | 1.11 |
| mesenchyme | Rnf157 | 1.28 | 7.01E-57 | 1.46E-52 | 0.18 | 0.1 | 0.08 | 1.83 |
| mesenchyme | Pwwp2a | 0.4 | 7.06E-57 | 1.47E-52 | 0.59 | 0.51 | 0.08 | 1.16 |
| mesenchyme | Polr3d | 0.42 | 7.80E-57 | 1.62E-52 | 0.56 | 0.47 | 0.09 | 1.18 |
| mesenchyme | Mta2 | 0.34 | 8.80E-57 | 1.83E-52 | 0.64 | 0.57 | 0.08 | 1.13 |
| mesenchyme | Rexo1 | 0.39 | 1.20E-56 | 2.49E-52 | 0.64 | 0.57 | 0.07 | 1.13 |
| mesenchyme | Tnrc18 | 0.42 | 1.33E-56 | 2.76E-52 | 0.59 | 0.5 | 0.09 | 1.18 |
| mesenchyme | Ankrd17 | 0.3 | 1.44E-56 | 2.99E-52 | 0.75 | 0.69 | 0.06 | 1.09 |
| mesenchyme | Cwc27 | 0.5 | 1.47E-56 | 3.07E-52 | 0.5 | 0.41 | 0.08 | 1.2 |
| mesenchyme | Ddrgk1 | 0.45 | 1.71E-56 | 3.55E-52 | 0.58 | 0.51 | 0.07 | 1.13 |
| mesenchyme | Aars | 0.51 | 1.72E-56 | 3.58E-52 | 0.49 | 0.42 | 0.07 | 1.17 |
| mesenchyme | Parp2 | 0.4 | 2.01E-56 | 4.19E-52 | 0.6 | 0.52 | 0.07 | 1.14 |
| mesenchyme | Elp3 | 0.45 | 2.27E-56 | 4.72E-52 | 0.53 | 0.44 | 0.08 | 1.19 |
| mesenchyme | Lpl | 0.37 | 2.79E-56 | 5.80E-52 | 0.19 | 0.1 | 0.08 | 1.81 |
| mesenchyme | Hist1h2be | 1.01 | 2.85E-56 | 5.92E-52 | 0.3 | 0.22 | 0.08 | 1.38 |
| mesenchyme | Chchd3 | 0.31 | 3.30E-56 | 6.86E-52 | 0.71 | 0.65 | 0.05 | 1.08 |
| mesenchyme | Camkk2 | 0.89 | 3.44E-56 | 7.16E-52 | 0.26 | 0.18 | 0.09 | 1.5 |
| mesenchyme | Gtf3c1 | 0.39 | 4.70E-56 | 9.76E-52 | 0.6 | 0.52 | 0.08 | 1.16 |
| mesenchyme | Pip4p2 | 0.37 | 4.92E-56 | 1.02E-51 | 0.59 | 0.52 | 0.07 | 1.13 |
| mesenchyme | Mosmo | 0.49 | 6.40E-56 | 1.33E-51 | 0.52 | 0.45 | 0.07 | 1.16 |
| mesenchyme | Gmcl1 | 0.49 | 6.63E-56 | 1.38E-51 | 0.47 | 0.39 | 0.09 | 1.22 |
| mesenchyme | Rara | 0.45 | 8.12E-56 | 1.69E-51 | 0.5 | 0.41 | 0.09 | 1.23 |
| mesenchyme | Spc25 | 0.47 | 8.49E-56 | 1.77E-51 | 0.57 | 0.51 | 0.06 | 1.12 |
| mesenchyme | Fst | 0.7 | 9.50E-56 | 1.98E-51 | 0.15 | 0.07 | 0.08 | 2.1 |
| mesenchyme | Tpx2 | 0.51 | 9.73E-56 | 2.02E-51 | 0.67 | 0.61 | 0.06 | 1.09 |
| mesenchyme | Tdg | 0.28 | 9.91E-56 | 2.06E-51 | 0.69 | 0.62 | 0.07 | 1.12 |
| mesenchyme | Eif4ebp1 | 0.27 | 1.01E-55 | 2.11E-51 | 0.72 | 0.65 | 0.07 | 1.11 |
| mesenchyme | Snx24 | 0.64 | 1.15E-55 | 2.39E-51 | 0.39 | 0.3 | 0.09 | 1.31 |
| mesenchyme | Cep295 | 0.48 | 1.32E-55 | 2.75E-51 | 0.51 | 0.43 | 0.08 | 1.18 |
| mesenchyme | Ppp5c | 0.39 | 1.38E-55 | 2.87E-51 | 0.56 | 0.49 | 0.08 | 1.16 |
| mesenchyme | Hadhb | 0.36 | 1.39E-55 | 2.89E-51 | 0.64 | 0.58 | 0.07 | 1.12 |
| mesenchyme | Rtca | 0.45 | 1.39E-55 | 2.89E-51 | 0.5 | 0.43 | 0.08 | 1.18 |
| mesenchyme | Ikbip | 0.78 | 1.46E-55 | 3.03E-51 | 0.33 | 0.24 | 0.09 | 1.37 |

**Supplementary Table 2. Marker genes for the major cell types of integrated clustering of E11.5 control and NotchGOF λ-junction cells.**

| | | | | | | | | |
|---|---|---|---|---|---|---|---|---|
| mesenchyme | Hccs | 0.7 | 1.56E-55 | 3.25E-51 | 0.35 | 0.26 | 0.09 | 1.34 |
| mesenchyme | Dlg5 | 0.52 | 1.92E-55 | 4.00E-51 | 0.45 | 0.36 | 0.09 | 1.26 |
| mesenchyme | Mtfr1 | 0.47 | 3.28E-55 | 6.82E-51 | 0.54 | 0.48 | 0.06 | 1.13 |
| mesenchyme | Impa2 | 0.76 | 3.70E-55 | 7.70E-51 | 0.33 | 0.24 | 0.09 | 1.37 |
| mesenchyme | Fam168b | 0.28 | 3.88E-55 | 8.07E-51 | 0.71 | 0.65 | 0.07 | 1.1 |
| mesenchyme | Tgfb1i1 | 1.05 | 3.90E-55 | 8.11E-51 | 0.16 | 0.08 | 0.08 | 1.93 |
| mesenchyme | Prim2 | 0.62 | 4.61E-55 | 9.59E-51 | 0.41 | 0.33 | 0.08 | 1.24 |
| mesenchyme | Ift20 | 0.3 | 5.81E-55 | 1.21E-50 | 0.78 | 0.74 | 0.04 | 1.06 |
| mesenchyme | Ptges2 | 0.73 | 6.91E-55 | 1.44E-50 | 0.34 | 0.26 | 0.08 | 1.32 |
| mesenchyme | Nrd1 | 0.26 | 8.83E-55 | 1.84E-50 | 0.82 | 0.78 | 0.04 | 1.05 |
| mesenchyme | Cenpq | 0.45 | 9.00E-55 | 1.87E-50 | 0.58 | 0.53 | 0.05 | 1.1 |
| mesenchyme | Rps19bp1 | 0.3 | 9.61E-55 | 2.00E-50 | 0.73 | 0.68 | 0.05 | 1.08 |
| mesenchyme | Galt | 0.83 | 1.16E-54 | 2.41E-50 | 0.31 | 0.22 | 0.09 | 1.39 |
| mesenchyme | Zcchc8 | 0.48 | 1.19E-54 | 2.48E-50 | 0.5 | 0.43 | 0.07 | 1.17 |
| mesenchyme | Fmnl3 | 0.89 | 1.37E-54 | 2.85E-50 | 0.24 | 0.15 | 0.09 | 1.59 |
| mesenchyme | Rps6ka3 | 0.57 | 1.53E-54 | 3.17E-50 | 0.48 | 0.41 | 0.08 | 1.19 |
| mesenchyme | Gmds | 0.7 | 1.71E-54 | 3.56E-50 | 0.39 | 0.3 | 0.08 | 1.28 |
| mesenchyme | Pnkd | 0.25 | 1.78E-54 | 3.71E-50 | 0.76 | 0.71 | 0.05 | 1.07 |
| mesenchyme | Taf6 | 0.47 | 1.87E-54 | 3.88E-50 | 0.53 | 0.46 | 0.07 | 1.16 |
| mesenchyme | Slc8a1 | 1.21 | 1.92E-54 | 3.99E-50 | 0.15 | 0.07 | 0.08 | 2.04 |
| mesenchyme | Zfp36l2 | 0.38 | 2.43E-54 | 5.06E-50 | 0.46 | 0.37 | 0.09 | 1.24 |
| mesenchyme | Apool | 0.37 | 2.73E-54 | 5.68E-50 | 0.64 | 0.58 | 0.06 | 1.1 |
| mesenchyme | Strn4 | 0.48 | 3.75E-54 | 7.79E-50 | 0.52 | 0.44 | 0.07 | 1.17 |
| mesenchyme | Ankrd26 | 0.55 | 3.80E-54 | 7.89E-50 | 0.46 | 0.38 | 0.08 | 1.22 |
| mesenchyme | Rcc1 | 0.44 | 3.97E-54 | 8.26E-50 | 0.54 | 0.46 | 0.08 | 1.17 |
| mesenchyme | Scara3 | 1.06 | 4.32E-54 | 8.98E-50 | 0.21 | 0.13 | 0.08 | 1.65 |
| mesenchyme | Tubg1 | 0.35 | 4.48E-54 | 9.32E-50 | 0.6 | 0.52 | 0.07 | 1.14 |
| mesenchyme | Zfp580 | 0.45 | 4.78E-54 | 9.93E-50 | 0.47 | 0.38 | 0.09 | 1.24 |
| mesenchyme | Raf1 | 0.37 | 5.53E-54 | 1.15E-49 | 0.59 | 0.52 | 0.07 | 1.14 |
| mesenchyme | Mrpl22 | 0.33 | 7.71E-54 | 1.60E-49 | 0.68 | 0.62 | 0.06 | 1.09 |
| mesenchyme | Sirt2 | 0.43 | 7.86E-54 | 1.63E-49 | 0.53 | 0.47 | 0.06 | 1.14 |
| mesenchyme | Guf1 | 0.59 | 7.87E-54 | 1.64E-49 | 0.44 | 0.36 | 0.08 | 1.23 |
| mesenchyme | Arid1b | 0.5 | 8.14E-54 | 1.69E-49 | 0.51 | 0.44 | 0.07 | 1.17 |
| mesenchyme | Dusp18 | 1.1 | 8.30E-54 | 1.73E-49 | 0.19 | 0.11 | 0.08 | 1.72 |
| mesenchyme | Cmc2 | 0.38 | 9.63E-54 | 2.00E-49 | 0.62 | 0.57 | 0.05 | 1.09 |
| mesenchyme | 700123O20Ri | 0.26 | 1.05E-53 | 2.18E-49 | 0.76 | 0.7 | 0.06 | 1.08 |
| mesenchyme | Bckdhb | 0.6 | 1.13E-53 | 2.34E-49 | 0.39 | 0.31 | 0.09 | 1.28 |
| mesenchyme | Psmc3ip | 0.39 | 1.33E-53 | 2.77E-49 | 0.58 | 0.51 | 0.07 | 1.14 |
| mesenchyme | Gatad1 | 0.26 | 1.36E-53 | 2.84E-49 | 0.82 | 0.77 | 0.04 | 1.06 |
| mesenchyme | Wasf1 | 0.6 | 1.42E-53 | 2.95E-49 | 0.43 | 0.35 | 0.09 | 1.24 |
| mesenchyme | Fopnl | 0.32 | 1.63E-53 | 3.40E-49 | 0.68 | 0.63 | 0.05 | 1.08 |
| mesenchyme | Bag6 | 0.33 | 1.64E-53 | 3.42E-49 | 0.67 | 0.6 | 0.07 | 1.12 |
| mesenchyme | Il6st | 0.8 | 1.90E-53 | 3.96E-49 | 0.31 | 0.23 | 0.09 | 1.38 |
| mesenchyme | ·833439L19Ri | 0.42 | 2.00E-53 | 4.15E-49 | 0.53 | 0.46 | 0.07 | 1.14 |
| mesenchyme | Uba3 | 0.43 | 2.04E-53 | 4.23E-49 | 0.55 | 0.47 | 0.07 | 1.15 |
| mesenchyme | Proser1 | 0.46 | 3.04E-53 | 6.33E-49 | 0.52 | 0.43 | 0.08 | 1.19 |
| mesenchyme | Gabpb1 | 0.48 | 3.65E-53 | 7.59E-49 | 0.49 | 0.42 | 0.07 | 1.17 |
| mesenchyme | Ppp1r12a | 0.32 | 3.66E-53 | 7.61E-49 | 0.75 | 0.69 | 0.06 | 1.08 |
| mesenchyme | Pnpla8 | 0.49 | 3.68E-53 | 7.65E-49 | 0.52 | 0.46 | 0.06 | 1.14 |
| mesenchyme | Smad2 | 0.44 | 3.70E-53 | 7.70E-49 | 0.52 | 0.45 | 0.08 | 1.17 |
| mesenchyme | Me1 | 0.7 | 4.11E-53 | 8.55E-49 | 0.28 | 0.19 | 0.09 | 1.49 |
| mesenchyme | Specc1 | 0.54 | 5.62E-53 | 1.17E-48 | 0.45 | 0.37 | 0.08 | 1.21 |
| mesenchyme | Chsy1 | 0.62 | 5.69E-53 | 1.18E-48 | 0.42 | 0.35 | 0.07 | 1.21 |
| mesenchyme | Zfp639 | 0.47 | 5.82E-53 | 1.21E-48 | 0.49 | 0.41 | 0.08 | 1.2 |
| mesenchyme | Cggbp1 | 0.26 | 6.55E-53 | 1.36E-48 | 0.8 | 0.76 | 0.04 | 1.05 |
| mesenchyme | Zranb1 | 0.35 | 8.17E-53 | 1.70E-48 | 0.63 | 0.57 | 0.06 | 1.11 |

**Supplementary Table 2. Marker genes for the major cell types of integrated clustering of E11.5 control and NotchGOF λ-junction cells.**

| | | | | | | | | |
|---|---|---|---|---|---|---|---|---|
| mesenchyme | Parl | 0.31 | 8.40E-53 | 1.75E-48 | 0.69 | 0.63 | 0.06 | 1.1 |
| mesenchyme | Aamdc | 0.71 | 8.41E-53 | 1.75E-48 | 0.34 | 0.26 | 0.08 | 1.31 |
| mesenchyme | Micu1 | 0.69 | 9.25E-53 | 1.92E-48 | 0.36 | 0.28 | 0.08 | 1.29 |
| mesenchyme | Gins2 | 0.34 | 9.97E-53 | 2.07E-48 | 0.63 | 0.56 | 0.08 | 1.14 |
| mesenchyme | Nup37 | 0.37 | 1.10E-52 | 2.28E-48 | 0.56 | 0.48 | 0.08 | 1.16 |
| mesenchyme | Impact | 0.69 | 1.11E-52 | 2.31E-48 | 0.35 | 0.27 | 0.09 | 1.32 |
| mesenchyme | Pls3 | 0.5 | 1.65E-52 | 3.43E-48 | 0.33 | 0.24 | 0.09 | 1.38 |
| mesenchyme | Mrpl3 | 0.31 | 2.25E-52 | 4.68E-48 | 0.69 | 0.63 | 0.06 | 1.1 |
| mesenchyme | Zscan2 | 0.99 | 2.97E-52 | 6.17E-48 | 0.22 | 0.13 | 0.08 | 1.64 |
| mesenchyme | Dmac1 | 0.26 | 3.06E-52 | 6.37E-48 | 0.78 | 0.72 | 0.06 | 1.08 |
| mesenchyme | Rbm6 | 0.28 | 5.67E-52 | 1.18E-47 | 0.74 | 0.68 | 0.06 | 1.09 |
| mesenchyme | Usp42 | 0.63 | 5.72E-52 | 1.19E-47 | 0.39 | 0.31 | 0.09 | 1.28 |
| mesenchyme | Gng12 | 0.42 | 5.99E-52 | 1.25E-47 | 0.39 | 0.3 | 0.1 | 1.34 |
| mesenchyme | Gstm7 | 0.68 | 6.01E-52 | 1.25E-47 | 0.31 | 0.22 | 0.09 | 1.42 |
| mesenchyme | Prelid3b | 0.3 | 6.47E-52 | 1.35E-47 | 0.67 | 0.61 | 0.06 | 1.09 |
| mesenchyme | Arl6ip6 | 0.3 | 6.57E-52 | 1.37E-47 | 0.7 | 0.63 | 0.07 | 1.11 |
| mesenchyme | Snx17 | 0.31 | 6.89E-52 | 1.43E-47 | 0.69 | 0.64 | 0.05 | 1.08 |
| mesenchyme | Eif4e3 | 0.63 | 7.43E-52 | 1.54E-47 | 0.31 | 0.22 | 0.09 | 1.41 |
| mesenchyme | Kin | 0.33 | 9.04E-52 | 1.88E-47 | 0.64 | 0.58 | 0.06 | 1.1 |
| mesenchyme | Anapc15 | 0.38 | 9.30E-52 | 1.93E-47 | 0.62 | 0.57 | 0.05 | 1.09 |
| mesenchyme | Ube2f | 0.35 | 1.37E-51 | 2.86E-47 | 0.59 | 0.52 | 0.07 | 1.13 |
| mesenchyme | Snrnp48 | 0.28 | 1.41E-51 | 2.94E-47 | 0.72 | 0.66 | 0.06 | 1.09 |
| mesenchyme | Zfp329 | 0.36 | 1.47E-51 | 3.06E-47 | 0.58 | 0.49 | 0.09 | 1.17 |
| mesenchyme | Crtc3 | 0.78 | 1.52E-51 | 3.16E-47 | 0.27 | 0.19 | 0.09 | 1.45 |
| mesenchyme | Aopep | 0.29 | 1.56E-51 | 3.25E-47 | 0.65 | 0.57 | 0.08 | 1.14 |
| mesenchyme | Dipk1b | 0.43 | 2.05E-51 | 4.27E-47 | 0.51 | 0.42 | 0.1 | 1.23 |
| mesenchyme | Gatad2b | 0.34 | 2.06E-51 | 4.28E-47 | 0.66 | 0.6 | 0.06 | 1.1 |
| mesenchyme | Med1 | 0.38 | 2.09E-51 | 4.34E-47 | 0.6 | 0.53 | 0.07 | 1.14 |
| mesenchyme | Elovl5 | 0.7 | 2.25E-51 | 4.68E-47 | 0.33 | 0.24 | 0.09 | 1.36 |
| mesenchyme | Ythdc2 | 0.85 | 2.69E-51 | 5.59E-47 | 0.3 | 0.22 | 0.08 | 1.37 |
| mesenchyme | Ak6 | 0.3 | 3.12E-51 | 6.49E-47 | 0.71 | 0.67 | 0.04 | 1.06 |
| mesenchyme | Polb | 0.33 | 3.14E-51 | 6.52E-47 | 0.61 | 0.55 | 0.07 | 1.12 |
| mesenchyme | Mxra8 | 1.25 | 3.15E-51 | 6.54E-47 | 0.17 | 0.1 | 0.08 | 1.78 |
| mesenchyme | Klf12 | 1.1 | 3.24E-51 | 6.74E-47 | 0.18 | 0.11 | 0.08 | 1.74 |
| mesenchyme | Ash1l | 0.27 | 3.27E-51 | 6.81E-47 | 0.76 | 0.68 | 0.08 | 1.11 |
| mesenchyme | Rnf214 | 0.41 | 3.37E-51 | 7.01E-47 | 0.54 | 0.47 | 0.08 | 1.16 |
| mesenchyme | Zfp24 | 0.35 | 4.17E-51 | 8.68E-47 | 0.61 | 0.54 | 0.07 | 1.12 |
| mesenchyme | Faf2 | 0.31 | 4.44E-51 | 9.24E-47 | 0.69 | 0.62 | 0.07 | 1.12 |
| mesenchyme | Tctex1d2 | 0.51 | 4.74E-51 | 9.87E-47 | 0.46 | 0.38 | 0.08 | 1.22 |
| mesenchyme | Timm9 | 0.34 | 5.00E-51 | 1.04E-46 | 0.64 | 0.58 | 0.06 | 1.1 |
| mesenchyme | Palm | 0.36 | 7.08E-51 | 1.47E-46 | 0.49 | 0.4 | 0.1 | 1.24 |
| mesenchyme | Arl6 | 0.47 | 7.25E-51 | 1.51E-46 | 0.45 | 0.37 | 0.08 | 1.22 |
| mesenchyme | Nit2 | 0.44 | 7.66E-51 | 1.59E-46 | 0.53 | 0.46 | 0.07 | 1.15 |
| mesenchyme | Tnks | 0.4 | 7.68E-51 | 1.60E-46 | 0.57 | 0.49 | 0.08 | 1.16 |
| mesenchyme | Hmgxb4 | 0.32 | 7.80E-51 | 1.62E-46 | 0.65 | 0.58 | 0.07 | 1.12 |
| mesenchyme | Fnbp4 | 0.39 | 8.61E-51 | 1.79E-46 | 0.59 | 0.52 | 0.07 | 1.13 |
| mesenchyme | Pgp | 0.29 | 1.01E-50 | 2.11E-46 | 0.69 | 0.6 | 0.08 | 1.13 |
| mesenchyme | Exoc5 | 0.35 | 1.18E-50 | 2.46E-46 | 0.59 | 0.53 | 0.07 | 1.13 |
| mesenchyme | Taf5 | 0.54 | 1.21E-50 | 2.52E-46 | 0.46 | 0.39 | 0.07 | 1.17 |
| mesenchyme | Dpf2 | 0.29 | 1.40E-50 | 2.90E-46 | 0.68 | 0.61 | 0.07 | 1.11 |
| mesenchyme | Slc27a3 | 0.89 | 1.80E-50 | 3.74E-46 | 0.24 | 0.16 | 0.08 | 1.52 |
| mesenchyme | Ncs1 | 0.62 | 1.94E-50 | 4.03E-46 | 0.34 | 0.25 | 0.09 | 1.35 |
| mesenchyme | Acat1 | 0.44 | 2.03E-50 | 4.23E-46 | 0.49 | 0.41 | 0.09 | 1.21 |
| mesenchyme | N4bp2 | 0.33 | 2.33E-50 | 4.84E-46 | 0.69 | 0.63 | 0.06 | 1.09 |
| mesenchyme | Spata7 | 0.4 | 2.44E-50 | 5.07E-46 | 0.41 | 0.33 | 0.09 | 1.26 |
| mesenchyme | Trp53bp2 | 0.45 | 2.47E-50 | 5.13E-46 | 0.49 | 0.41 | 0.09 | 1.21 |

**Supplementary Table 2. Marker genes for the major cell types of integrated clustering of E11.5 control and NotchGOF λ-junction cells.**

| | | | | | | | | |
|---|---|---|---|---|---|---|---|---|
| mesenchyme | Pheta1 | 0.57 | 2.58E-50 | 5.38E-46 | 0.35 | 0.26 | 0.09 | 1.33 |
| mesenchyme | 2810408I11Rik | 0.71 | 2.59E-50 | 5.39E-46 | 0.33 | 0.25 | 0.08 | 1.33 |
| mesenchyme | Ing1 | 0.28 | 3.25E-50 | 6.77E-46 | 0.75 | 0.69 | 0.06 | 1.09 |
| mesenchyme | Tmem41b | 0.59 | 5.91E-50 | 1.23E-45 | 0.42 | 0.34 | 0.07 | 1.21 |
| mesenchyme | Rpa2 | 0.31 | 6.20E-50 | 1.29E-45 | 0.69 | 0.62 | 0.07 | 1.11 |
| mesenchyme | Fads1 | 0.46 | 6.61E-50 | 1.37E-45 | 0.51 | 0.44 | 0.07 | 1.17 |
| mesenchyme | Gyg | 0.5 | 7.75E-50 | 1.61E-45 | 0.45 | 0.37 | 0.08 | 1.22 |
| mesenchyme | Peak1 | 0.81 | 8.25E-50 | 1.72E-45 | 0.28 | 0.19 | 0.09 | 1.45 |
| mesenchyme | Itgav | 0.62 | 8.32E-50 | 1.73E-45 | 0.31 | 0.22 | 0.09 | 1.42 |
| mesenchyme | Chmp6 | 0.37 | 8.62E-50 | 1.79E-45 | 0.54 | 0.46 | 0.08 | 1.17 |
| mesenchyme | Tgfbr3 | 1.82 | 8.81E-50 | 1.83E-45 | 0.1 | 0.04 | 0.06 | 2.46 |
| mesenchyme | Senp1 | 0.47 | 1.06E-49 | 2.21E-45 | 0.48 | 0.41 | 0.07 | 1.18 |
| mesenchyme | Pcif1 | 0.36 | 1.14E-49 | 2.37E-45 | 0.62 | 0.57 | 0.05 | 1.09 |
| mesenchyme | Cfap97 | 0.49 | 1.31E-49 | 2.73E-45 | 0.48 | 0.4 | 0.08 | 1.19 |
| mesenchyme | Brwd1 | 0.33 | 1.33E-49 | 2.78E-45 | 0.66 | 0.59 | 0.07 | 1.11 |
| mesenchyme | Zfp518b | 0.58 | 1.63E-49 | 3.40E-45 | 0.39 | 0.31 | 0.08 | 1.27 |
| mesenchyme | Trp53i13 | 0.73 | 2.00E-49 | 4.17E-45 | 0.3 | 0.22 | 0.08 | 1.38 |
| mesenchyme | Crot | 0.46 | 2.02E-49 | 4.19E-45 | 0.45 | 0.36 | 0.09 | 1.24 |
| mesenchyme | Pimreg | 0.42 | 2.37E-49 | 4.93E-45 | 0.55 | 0.46 | 0.09 | 1.2 |
| mesenchyme | Rp9 | 0.25 | 2.45E-49 | 5.09E-45 | 0.8 | 0.75 | 0.05 | 1.07 |
| mesenchyme | Rab9 | 0.34 | 3.08E-49 | 6.40E-45 | 0.63 | 0.56 | 0.06 | 1.11 |
| mesenchyme | Nufip2 | 0.39 | 3.35E-49 | 6.97E-45 | 0.57 | 0.5 | 0.07 | 1.14 |
| mesenchyme | Cenph | 0.32 | 3.50E-49 | 7.29E-45 | 0.65 | 0.6 | 0.05 | 1.09 |
| mesenchyme | Fbxo30 | 0.56 | 3.53E-49 | 7.35E-45 | 0.44 | 0.37 | 0.07 | 1.2 |
| mesenchyme | Hnrnph2 | 0.25 | 5.00E-49 | 1.04E-44 | 0.77 | 0.72 | 0.05 | 1.06 |
| mesenchyme | Med21 | 0.27 | 5.47E-49 | 1.14E-44 | 0.76 | 0.7 | 0.06 | 1.08 |
| mesenchyme | Tbpl1 | 0.29 | 6.14E-49 | 1.28E-44 | 0.63 | 0.56 | 0.07 | 1.12 |
| mesenchyme | Mia3 | 0.37 | 6.36E-49 | 1.32E-44 | 0.61 | 0.55 | 0.07 | 1.13 |
| mesenchyme | Zfp280d | 0.31 | 9.29E-49 | 1.93E-44 | 0.69 | 0.61 | 0.07 | 1.12 |
| mesenchyme | Commd2 | 0.28 | 1.11E-48 | 2.31E-44 | 0.71 | 0.66 | 0.05 | 1.08 |
| mesenchyme | Mrps18a | 0.27 | 1.12E-48 | 2.34E-44 | 0.75 | 0.7 | 0.04 | 1.06 |
| mesenchyme | Ech1 | 0.39 | 1.13E-48 | 2.36E-44 | 0.54 | 0.47 | 0.07 | 1.15 |
| mesenchyme | Cyhr1 | 0.34 | 1.20E-48 | 2.50E-44 | 0.65 | 0.6 | 0.05 | 1.08 |
| mesenchyme | Akap8 | 0.3 | 1.25E-48 | 2.60E-44 | 0.68 | 0.62 | 0.06 | 1.09 |
| mesenchyme | Rpap3 | 0.43 | 1.40E-48 | 2.92E-44 | 0.49 | 0.42 | 0.07 | 1.17 |
| mesenchyme | Itch | 0.4 | 1.42E-48 | 2.95E-44 | 0.56 | 0.5 | 0.06 | 1.13 |
| mesenchyme | Slco3a1 | 1.04 | 2.39E-48 | 4.97E-44 | 0.15 | 0.08 | 0.07 | 1.89 |
| mesenchyme | Cox19 | 0.28 | 2.56E-48 | 5.33E-44 | 0.7 | 0.65 | 0.05 | 1.08 |
| mesenchyme | Tvp23a | 1.17 | 3.32E-48 | 6.90E-44 | 0.16 | 0.08 | 0.07 | 1.86 |
| mesenchyme | Me2 | 0.47 | 3.55E-48 | 7.38E-44 | 0.48 | 0.41 | 0.07 | 1.17 |
| mesenchyme | Mrps6 | 0.3 | 4.04E-48 | 8.41E-44 | 0.65 | 0.58 | 0.07 | 1.12 |
| mesenchyme | Bet1 | 0.31 | 4.07E-48 | 8.46E-44 | 0.67 | 0.62 | 0.04 | 1.07 |
| mesenchyme | Trnau1ap | 0.31 | 4.15E-48 | 8.64E-44 | 0.65 | 0.59 | 0.07 | 1.11 |
| mesenchyme | Shcbp1 | 0.49 | 4.18E-48 | 8.69E-44 | 0.49 | 0.42 | 0.07 | 1.16 |
| mesenchyme | Nbas | 0.94 | 4.22E-48 | 8.77E-44 | 0.25 | 0.17 | 0.08 | 1.47 |
| mesenchyme | Ifrd1 | 0.46 | 4.26E-48 | 8.86E-44 | 0.53 | 0.47 | 0.05 | 1.11 |
| mesenchyme | Anxa11 | 1.16 | 4.29E-48 | 8.93E-44 | 0.18 | 0.11 | 0.07 | 1.69 |
| mesenchyme | Akr1e1 | 0.58 | 4.32E-48 | 8.98E-44 | 0.37 | 0.29 | 0.08 | 1.28 |
| mesenchyme | Mrpl19 | 0.32 | 4.45E-48 | 9.25E-44 | 0.63 | 0.57 | 0.06 | 1.1 |
| mesenchyme | Gtf2b | 0.27 | 4.79E-48 | 9.96E-44 | 0.71 | 0.65 | 0.06 | 1.09 |
| mesenchyme | Prkrip1 | 0.42 | 5.06E-48 | 1.05E-43 | 0.51 | 0.46 | 0.06 | 1.13 |
| mesenchyme | Lmnb2 | 0.56 | 6.12E-48 | 1.27E-43 | 0.4 | 0.32 | 0.08 | 1.25 |
| mesenchyme | Prkcsh | 0.27 | 7.84E-48 | 1.63E-43 | 0.75 | 0.67 | 0.08 | 1.12 |
| mesenchyme | Prr12 | 0.47 | 7.94E-48 | 1.65E-43 | 0.47 | 0.39 | 0.08 | 1.2 |
| mesenchyme | Cox10 | 0.6 | 9.26E-48 | 1.93E-43 | 0.36 | 0.28 | 0.08 | 1.28 |
| mesenchyme | Zfand3 | 0.37 | 9.44E-48 | 1.96E-43 | 0.55 | 0.49 | 0.07 | 1.14 |

**Supplementary Table 2. Marker genes for the major cell types of integrated clustering of E11.5 control and NotchGOF λ-junction cells.**

| | | | | | | | | |
|---|---|---|---|---|---|---|---|---|
| mesenchyme | Sin3a | 0.42 | 1.25E-47 | 2.59E-43 | 0.55 | 0.48 | 0.07 | 1.15 |
| mesenchyme | Rasa2 | 0.61 | 1.34E-47 | 2.79E-43 | 0.38 | 0.3 | 0.08 | 1.27 |
| mesenchyme | Aldh7a1 | 0.33 | 1.83E-47 | 3.80E-43 | 0.58 | 0.51 | 0.08 | 1.15 |
| mesenchyme | Hcfc1 | 0.3 | 2.18E-47 | 4.53E-43 | 0.69 | 0.64 | 0.05 | 1.08 |
| mesenchyme | Ndufv1 | 0.26 | 2.40E-47 | 5.00E-43 | 0.77 | 0.73 | 0.04 | 1.06 |
| mesenchyme | 810055G02Ri | 0.94 | 2.62E-47 | 5.44E-43 | 0.24 | 0.16 | 0.08 | 1.48 |
| mesenchyme | Nup50 | 0.33 | 2.68E-47 | 5.57E-43 | 0.63 | 0.58 | 0.05 | 1.09 |
| mesenchyme | Anxa7 | 0.79 | 2.91E-47 | 6.06E-43 | 0.38 | 0.32 | 0.06 | 1.19 |
| mesenchyme | Hist1h1a | 0.78 | 2.97E-47 | 6.18E-43 | 0.44 | 0.39 | 0.06 | 1.15 |
| mesenchyme | 530008M17R | 0.3 | 3.11E-47 | 6.46E-43 | 0.45 | 0.36 | 0.1 | 1.27 |
| mesenchyme | .110059E24Ri | 0.34 | 3.11E-47 | 6.47E-43 | 0.58 | 0.5 | 0.08 | 1.16 |
| mesenchyme | Ube2j2 | 0.26 | 3.15E-47 | 6.55E-43 | 0.72 | 0.68 | 0.04 | 1.06 |
| mesenchyme | Drosha | 0.4 | 3.95E-47 | 8.21E-43 | 0.53 | 0.46 | 0.08 | 1.17 |
| mesenchyme | Hddc3 | 0.3 | 4.78E-47 | 9.94E-43 | 0.6 | 0.53 | 0.07 | 1.13 |
| mesenchyme | Nxpe3 | 1.36 | 4.91E-47 | 1.02E-42 | 0.13 | 0.06 | 0.07 | 2.07 |
| mesenchyme | Tmem97 | 0.42 | 4.94E-47 | 1.03E-42 | 0.49 | 0.41 | 0.08 | 1.2 |
| mesenchyme | Gtf2a1 | 0.42 | 5.02E-47 | 1.04E-42 | 0.52 | 0.45 | 0.07 | 1.16 |
| mesenchyme | Crnkl1 | 0.34 | 5.58E-47 | 1.16E-42 | 0.58 | 0.51 | 0.06 | 1.12 |
| mesenchyme | Ppif | 0.32 | 7.67E-47 | 1.59E-42 | 0.56 | 0.48 | 0.08 | 1.16 |
| mesenchyme | Trim33 | 0.28 | 8.06E-47 | 1.68E-42 | 0.68 | 0.61 | 0.07 | 1.11 |
| mesenchyme | Nicn1 | 0.58 | 8.12E-47 | 1.69E-42 | 0.36 | 0.28 | 0.08 | 1.3 |
| mesenchyme | Ireb2 | 0.36 | 8.25E-47 | 1.72E-42 | 0.57 | 0.5 | 0.07 | 1.14 |
| mesenchyme | Sh3pxd2b | 0.66 | 8.56E-47 | 1.78E-42 | 0.32 | 0.24 | 0.08 | 1.35 |
| mesenchyme | Sde2 | 0.27 | 9.50E-47 | 1.98E-42 | 0.7 | 0.64 | 0.05 | 1.08 |
| mesenchyme | Nkap | 0.34 | 1.20E-46 | 2.49E-42 | 0.62 | 0.56 | 0.06 | 1.1 |
| mesenchyme | Ttk | 0.63 | 1.29E-46 | 2.67E-42 | 0.39 | 0.32 | 0.07 | 1.23 |
| mesenchyme | Cep19 | 0.56 | 1.36E-46 | 2.83E-42 | 0.38 | 0.3 | 0.08 | 1.28 |
| mesenchyme | Drg2 | 0.35 | 1.56E-46 | 3.23E-42 | 0.58 | 0.51 | 0.07 | 1.13 |
| mesenchyme | Hist2h2ac | 0.98 | 1.57E-46 | 3.27E-42 | 0.35 | 0.28 | 0.07 | 1.23 |
| mesenchyme | Supt20 | 0.32 | 1.57E-46 | 3.27E-42 | 0.61 | 0.55 | 0.06 | 1.11 |
| mesenchyme | Tars | 0.33 | 2.39E-46 | 4.98E-42 | 0.62 | 0.57 | 0.06 | 1.1 |
| mesenchyme | Cdyl | 0.59 | 2.46E-46 | 5.11E-42 | 0.41 | 0.34 | 0.07 | 1.21 |
| mesenchyme | Bcas3 | 0.85 | 2.47E-46 | 5.14E-42 | 0.24 | 0.16 | 0.08 | 1.5 |
| mesenchyme | Pdzrn3 | 0.88 | 2.60E-46 | 5.40E-42 | 0.2 | 0.12 | 0.08 | 1.63 |
| mesenchyme | Nsrp1 | 0.27 | 2.77E-46 | 5.76E-42 | 0.72 | 0.67 | 0.05 | 1.07 |
| mesenchyme | Wdhd1 | 0.65 | 3.21E-46 | 6.67E-42 | 0.35 | 0.28 | 0.07 | 1.26 |
| mesenchyme | Zfp30 | 0.5 | 3.76E-46 | 7.82E-42 | 0.46 | 0.38 | 0.08 | 1.22 |
| mesenchyme | Nufip1 | 0.37 | 4.19E-46 | 8.72E-42 | 0.53 | 0.46 | 0.07 | 1.16 |
| mesenchyme | Rad51ap1 | 0.49 | 4.31E-46 | 8.97E-42 | 0.51 | 0.45 | 0.06 | 1.13 |
| mesenchyme | Ghr | 1.14 | 4.46E-46 | 9.27E-42 | 0.16 | 0.09 | 0.07 | 1.76 |
| mesenchyme | Epc2 | 0.33 | 4.50E-46 | 9.37E-42 | 0.64 | 0.57 | 0.07 | 1.12 |
| mesenchyme | Lipt2 | 0.38 | 4.58E-46 | 9.53E-42 | 0.39 | 0.3 | 0.09 | 1.28 |
| mesenchyme | Insig1 | 0.68 | 4.89E-46 | 1.02E-41 | 0.34 | 0.25 | 0.08 | 1.32 |
| mesenchyme | Bloc1s1 | 0.32 | 5.28E-46 | 1.10E-41 | 0.61 | 0.55 | 0.06 | 1.11 |
| mesenchyme | Xpnpep1 | 0.38 | 5.71E-46 | 1.19E-41 | 0.55 | 0.49 | 0.06 | 1.13 |
| mesenchyme | Hist1h2an | 0.85 | 5.73E-46 | 1.19E-41 | 0.33 | 0.26 | 0.07 | 1.27 |
| mesenchyme | Ift46 | 0.41 | 5.98E-46 | 1.24E-41 | 0.52 | 0.44 | 0.08 | 1.17 |
| mesenchyme | Lrp3 | 0.85 | 6.33E-46 | 1.32E-41 | 0.24 | 0.16 | 0.08 | 1.48 |
| mesenchyme | Rprd1a | 0.38 | 6.57E-46 | 1.37E-41 | 0.55 | 0.48 | 0.07 | 1.15 |
| mesenchyme | Ubr7 | 0.43 | 6.89E-46 | 1.43E-41 | 0.5 | 0.42 | 0.08 | 1.18 |
| mesenchyme | Dpm3 | 0.27 | 7.22E-46 | 1.50E-41 | 0.77 | 0.72 | 0.04 | 1.06 |
| mesenchyme | Cdk2 | 0.53 | 7.33E-46 | 1.52E-41 | 0.43 | 0.36 | 0.07 | 1.19 |
| mesenchyme | Simc1 | 1.09 | 8.12E-46 | 1.69E-41 | 0.16 | 0.09 | 0.07 | 1.78 |
| mesenchyme | Ctdspl2 | 0.35 | 8.55E-46 | 1.78E-41 | 0.6 | 0.53 | 0.07 | 1.13 |
| mesenchyme | 130401M01R | 0.31 | 9.45E-46 | 1.97E-41 | 0.61 | 0.53 | 0.08 | 1.15 |
| mesenchyme | Oaz2 | 0.36 | 1.11E-45 | 2.30E-41 | 0.57 | 0.51 | 0.06 | 1.13 |

**Supplementary Table 2. Marker genes for the major cell types of integrated clustering of E11.5 control and NotchGOF λ-junction cells.**

| | | | | | | | | |
|---|---|---|---|---|---|---|---|---|
| mesenchyme | Ice1 | 0.36 | 1.12E-45 | 2.34E-41 | 0.57 | 0.49 | 0.07 | 1.15 |
| mesenchyme | Rgma | 0.76 | 1.19E-45 | 2.48E-41 | 0.27 | 0.19 | 0.08 | 1.41 |
| mesenchyme | Mrpl55 | 0.29 | 1.20E-45 | 2.49E-41 | 0.67 | 0.62 | 0.05 | 1.08 |
| mesenchyme | N6amt1 | 0.44 | 1.21E-45 | 2.52E-41 | 0.49 | 0.42 | 0.07 | 1.18 |
| mesenchyme | Pxylp1 | 1.25 | 1.22E-45 | 2.54E-41 | 0.13 | 0.07 | 0.07 | 2 |
| mesenchyme | Atp7a | 0.81 | 1.25E-45 | 2.59E-41 | 0.26 | 0.19 | 0.08 | 1.42 |
| mesenchyme | Atf2 | 0.37 | 1.35E-45 | 2.80E-41 | 0.56 | 0.49 | 0.07 | 1.14 |
| mesenchyme | B3glct | 0.66 | 1.43E-45 | 2.98E-41 | 0.34 | 0.27 | 0.08 | 1.29 |
| mesenchyme | Crlf3 | 0.47 | 1.46E-45 | 3.04E-41 | 0.47 | 0.41 | 0.07 | 1.16 |
| mesenchyme | Gas2 | 0.8 | 1.49E-45 | 3.11E-41 | 0.29 | 0.21 | 0.08 | 1.38 |
| mesenchyme | Spire1 | 0.64 | 1.55E-45 | 3.23E-41 | 0.37 | 0.3 | 0.07 | 1.23 |
| mesenchyme | Cnot4 | 0.28 | 1.60E-45 | 3.33E-41 | 0.68 | 0.63 | 0.05 | 1.08 |
| mesenchyme | Ndufs1 | 0.36 | 1.72E-45 | 3.58E-41 | 0.61 | 0.56 | 0.05 | 1.09 |
| mesenchyme | Nup35 | 0.44 | 1.86E-45 | 3.87E-41 | 0.49 | 0.44 | 0.06 | 1.13 |
| mesenchyme | Agk | 0.66 | 1.92E-45 | 3.99E-41 | 0.35 | 0.27 | 0.08 | 1.28 |
| mesenchyme | Abraxas2 | 0.39 | 1.93E-45 | 4.02E-41 | 0.54 | 0.48 | 0.07 | 1.14 |
| mesenchyme | Myo9a | 0.57 | 2.08E-45 | 4.33E-41 | 0.4 | 0.32 | 0.08 | 1.25 |
| mesenchyme | Rbms2 | 0.44 | 2.23E-45 | 4.63E-41 | 0.49 | 0.43 | 0.06 | 1.14 |
| mesenchyme | Zfp451 | 0.37 | 2.24E-45 | 4.65E-41 | 0.6 | 0.53 | 0.07 | 1.12 |
| mesenchyme | Acvrl1 | 1.18 | 2.55E-45 | 5.31E-41 | 0.12 | 0.06 | 0.06 | 2.08 |
| mesenchyme | Letm1 | 0.52 | 3.85E-45 | 8.00E-41 | 0.46 | 0.4 | 0.06 | 1.15 |
| mesenchyme | Fam149a | 1.22 | 3.98E-45 | 8.27E-41 | 0.17 | 0.1 | 0.07 | 1.69 |
| mesenchyme | Supt6 | 0.28 | 5.71E-45 | 1.19E-40 | 0.7 | 0.64 | 0.06 | 1.1 |
| mesenchyme | Tet1 | 0.59 | 6.18E-45 | 1.28E-40 | 0.37 | 0.29 | 0.08 | 1.28 |
| mesenchyme | Scamp3 | 0.39 | 6.47E-45 | 1.35E-40 | 0.56 | 0.5 | 0.06 | 1.13 |
| mesenchyme | Gm4258 | 0.37 | 7.70E-45 | 1.60E-40 | 0.52 | 0.44 | 0.08 | 1.18 |
| mesenchyme | 930581F22Ri | 1.57 | 9.30E-45 | 1.93E-40 | 0.11 | 0.05 | 0.06 | 2.15 |
| mesenchyme | Lzic | 0.36 | 1.59E-44 | 3.30E-40 | 0.53 | 0.47 | 0.06 | 1.13 |
| mesenchyme | Zfp830 | 0.4 | 1.66E-44 | 3.45E-40 | 0.5 | 0.44 | 0.06 | 1.15 |
| mesenchyme | Ccdc107 | 0.53 | 1.84E-44 | 3.83E-40 | 0.44 | 0.38 | 0.07 | 1.17 |
| mesenchyme | Ppp1r35 | 0.35 | 2.03E-44 | 4.22E-40 | 0.57 | 0.52 | 0.06 | 1.11 |
| mesenchyme | Mmadhc | 0.3 | 2.07E-44 | 4.31E-40 | 0.64 | 0.59 | 0.05 | 1.08 |
| mesenchyme | Txndc5 | 0.31 | 2.08E-44 | 4.32E-40 | 0.64 | 0.56 | 0.08 | 1.14 |
| mesenchyme | Ncbp1 | 0.31 | 2.14E-44 | 4.46E-40 | 0.62 | 0.56 | 0.06 | 1.11 |
| mesenchyme | Epn1 | 0.31 | 4.33E-44 | 9.01E-40 | 0.64 | 0.6 | 0.04 | 1.07 |
| mesenchyme | 810026B05Ri | 0.33 | 4.56E-44 | 9.49E-40 | 0.62 | 0.56 | 0.06 | 1.11 |
| mesenchyme | Sec61a2 | 0.48 | 5.05E-44 | 1.05E-39 | 0.44 | 0.37 | 0.08 | 1.21 |
| mesenchyme | Lrrc58 | 0.27 | 5.45E-44 | 1.13E-39 | 0.66 | 0.58 | 0.08 | 1.14 |
| mesenchyme | Selenon | 0.57 | 6.26E-44 | 1.30E-39 | 0.4 | 0.32 | 0.07 | 1.23 |
| mesenchyme | Crebbp | 0.26 | 6.76E-44 | 1.41E-39 | 0.77 | 0.71 | 0.06 | 1.08 |
| mesenchyme | Taf13 | 0.37 | 7.84E-44 | 1.63E-39 | 0.54 | 0.47 | 0.06 | 1.14 |
| mesenchyme | Utp14a | 0.35 | 8.07E-44 | 1.68E-39 | 0.59 | 0.54 | 0.06 | 1.11 |
| mesenchyme | Cks2 | 0.32 | 8.86E-44 | 1.84E-39 | 0.85 | 0.81 | 0.04 | 1.05 |
| mesenchyme | Gon4l | 0.27 | 9.16E-44 | 1.90E-39 | 0.71 | 0.65 | 0.07 | 1.1 |
| mesenchyme | Fam193b | 0.45 | 1.01E-43 | 2.10E-39 | 0.48 | 0.42 | 0.07 | 1.16 |
| mesenchyme | Dhodh | 0.66 | 1.02E-43 | 2.12E-39 | 0.32 | 0.24 | 0.08 | 1.33 |
| mesenchyme | Prkd1 | 1.1 | 1.11E-43 | 2.31E-39 | 0.16 | 0.09 | 0.07 | 1.76 |
| mesenchyme | Cntrl | 0.44 | 1.14E-43 | 2.37E-39 | 0.49 | 0.42 | 0.07 | 1.17 |
| mesenchyme | Dipk1a | 0.88 | 1.37E-43 | 2.84E-39 | 0.23 | 0.16 | 0.07 | 1.47 |
| mesenchyme | Cbll1 | 0.37 | 1.46E-43 | 3.04E-39 | 0.54 | 0.47 | 0.06 | 1.14 |
| mesenchyme | Map4 | 0.45 | 1.70E-43 | 3.54E-39 | 0.42 | 0.35 | 0.08 | 1.22 |
| mesenchyme | Asap2 | 0.29 | 1.79E-43 | 3.73E-39 | 0.58 | 0.5 | 0.08 | 1.16 |
| mesenchyme | Nf2 | 0.7 | 1.94E-43 | 4.03E-39 | 0.31 | 0.24 | 0.08 | 1.33 |
| mesenchyme | Stk40 | 0.61 | 2.02E-43 | 4.19E-39 | 0.33 | 0.25 | 0.07 | 1.29 |
| mesenchyme | Tex10 | 0.48 | 2.15E-43 | 4.47E-39 | 0.47 | 0.41 | 0.06 | 1.15 |
| mesenchyme | Fam98b | 0.27 | 2.59E-43 | 5.38E-39 | 0.67 | 0.61 | 0.06 | 1.11 |

**Supplementary Table 2. Marker genes for the major cell types of integrated clustering of E11.5 control and NotchGOF λ-junction cells.**

| | | | | | | | | |
|---|---|---|---|---|---|---|---|---|
| mesenchyme | Dclk2 | 1.41 | 3.13E-43 | 6.50E-39 | 0.14 | 0.07 | 0.06 | 1.85 |
| mesenchyme | Nav1 | 0.31 | 3.24E-43 | 6.74E-39 | 0.57 | 0.48 | 0.09 | 1.19 |
| mesenchyme | Bak1 | 0.47 | 3.37E-43 | 7.01E-39 | 0.44 | 0.36 | 0.08 | 1.21 |
| mesenchyme | Nxt1 | 0.27 | 3.52E-43 | 7.33E-39 | 0.69 | 0.64 | 0.05 | 1.08 |
| mesenchyme | Paxbp1 | 0.3 | 4.44E-43 | 9.23E-39 | 0.65 | 0.59 | 0.06 | 1.09 |
| mesenchyme | Spata5 | 0.61 | 5.04E-43 | 1.05E-38 | 0.36 | 0.29 | 0.07 | 1.25 |
| mesenchyme | Mark3 | 0.28 | 6.38E-43 | 1.33E-38 | 0.68 | 0.62 | 0.06 | 1.09 |
| mesenchyme | Mis12 | 0.39 | 1.04E-42 | 2.17E-38 | 0.51 | 0.45 | 0.06 | 1.14 |
| mesenchyme | Ttc19 | 0.52 | 1.10E-42 | 2.29E-38 | 0.43 | 0.36 | 0.07 | 1.19 |
| mesenchyme | Cldn34c1 | 1.13 | 1.23E-42 | 2.56E-38 | 0.17 | 0.1 | 0.07 | 1.68 |
| mesenchyme | Dennd5b | 0.51 | 1.26E-42 | 2.62E-38 | 0.42 | 0.34 | 0.08 | 1.24 |
| mesenchyme | Kif20a | 0.54 | 1.32E-42 | 2.75E-38 | 0.49 | 0.43 | 0.06 | 1.13 |
| mesenchyme | Gmfb | 0.36 | 1.35E-42 | 2.81E-38 | 0.57 | 0.52 | 0.05 | 1.1 |
| mesenchyme | Blm | 0.38 | 1.40E-42 | 2.91E-38 | 0.54 | 0.48 | 0.06 | 1.12 |
| mesenchyme | Jak1 | 0.27 | 1.43E-42 | 2.96E-38 | 0.63 | 0.56 | 0.07 | 1.13 |
| mesenchyme | Plcl2 | 1.3 | 1.50E-42 | 3.11E-38 | 0.13 | 0.07 | 0.06 | 1.89 |
| mesenchyme | Chd8 | 0.27 | 1.54E-42 | 3.21E-38 | 0.72 | 0.65 | 0.06 | 1.1 |
| mesenchyme | Unc5c | 0.6 | 1.58E-42 | 3.28E-38 | 0.37 | 0.28 | 0.09 | 1.31 |
| mesenchyme | Kctd20 | 0.45 | 1.59E-42 | 3.30E-38 | 0.51 | 0.46 | 0.06 | 1.12 |
| mesenchyme | Sart3 | 0.36 | 1.76E-42 | 3.65E-38 | 0.58 | 0.52 | 0.06 | 1.11 |
| mesenchyme | Eri3 | 0.27 | 1.94E-42 | 4.04E-38 | 0.61 | 0.55 | 0.07 | 1.12 |
| mesenchyme | Pds5a | 0.29 | 1.98E-42 | 4.13E-38 | 0.69 | 0.64 | 0.05 | 1.08 |
| mesenchyme | Trrap | 0.33 | 2.07E-42 | 4.32E-38 | 0.61 | 0.54 | 0.06 | 1.11 |
| mesenchyme | Maip1 | 0.36 | 2.23E-42 | 4.64E-38 | 0.55 | 0.49 | 0.06 | 1.13 |
| mesenchyme | Rnf216 | 0.37 | 2.55E-42 | 5.31E-38 | 0.55 | 0.49 | 0.06 | 1.12 |
| mesenchyme | Smyd2 | 0.48 | 2.66E-42 | 5.53E-38 | 0.44 | 0.36 | 0.07 | 1.2 |
| mesenchyme | Hist1h2ap | 0.67 | 2.66E-42 | 5.54E-38 | 0.76 | 0.74 | 0.03 | 1.03 |
| mesenchyme | Irx5 | 0.67 | 2.78E-42 | 5.79E-38 | 0.31 | 0.22 | 0.09 | 1.38 |
| mesenchyme | Cd320 | 0.34 | 3.14E-42 | 6.53E-38 | 0.59 | 0.51 | 0.07 | 1.14 |
| mesenchyme | Fasn | 0.36 | 3.36E-42 | 6.99E-38 | 0.53 | 0.46 | 0.08 | 1.16 |
| mesenchyme | Nop16 | 0.29 | 3.40E-42 | 7.08E-38 | 0.63 | 0.56 | 0.07 | 1.12 |
| mesenchyme | Tcf7 | 1.08 | 3.76E-42 | 7.82E-38 | 0.17 | 0.1 | 0.07 | 1.68 |
| mesenchyme | Nqo1 | 1.65 | 3.89E-42 | 8.10E-38 | 0.12 | 0.06 | 0.06 | 1.98 |
| mesenchyme | Mllt1 | 0.64 | 5.45E-42 | 1.13E-37 | 0.32 | 0.25 | 0.07 | 1.29 |
| mesenchyme | Dnajb4 | 0.67 | 5.45E-42 | 1.13E-37 | 0.33 | 0.26 | 0.07 | 1.28 |
| mesenchyme | Wdr45b | 0.3 | 5.74E-42 | 1.19E-37 | 0.61 | 0.54 | 0.07 | 1.12 |
| mesenchyme | Cog8 | 0.45 | 5.80E-42 | 1.21E-37 | 0.44 | 0.37 | 0.07 | 1.18 |
| mesenchyme | Pym1 | 0.35 | 6.06E-42 | 1.26E-37 | 0.53 | 0.47 | 0.06 | 1.13 |
| mesenchyme | Ogfrl1 | 0.57 | 6.34E-42 | 1.32E-37 | 0.36 | 0.29 | 0.07 | 1.26 |
| mesenchyme | Gbf1 | 0.51 | 6.86E-42 | 1.43E-37 | 0.46 | 0.4 | 0.06 | 1.15 |
| mesenchyme | Mcm10 | 0.61 | 7.21E-42 | 1.50E-37 | 0.38 | 0.31 | 0.07 | 1.22 |
| mesenchyme | Fam114a1 | 1.09 | 7.35E-42 | 1.53E-37 | 0.19 | 0.12 | 0.07 | 1.58 |
| mesenchyme | Srpr | 0.25 | 7.37E-42 | 1.53E-37 | 0.71 | 0.65 | 0.06 | 1.1 |
| mesenchyme | Casc3 | 0.36 | 8.08E-42 | 1.68E-37 | 0.56 | 0.5 | 0.06 | 1.11 |
| mesenchyme | Pola2 | 0.48 | 8.28E-42 | 1.72E-37 | 0.45 | 0.38 | 0.07 | 1.18 |
| mesenchyme | Ptcd3 | 0.39 | 8.81E-42 | 1.83E-37 | 0.52 | 0.46 | 0.06 | 1.13 |
| mesenchyme | Car13 | 1.26 | 8.91E-42 | 1.85E-37 | 0.12 | 0.06 | 0.06 | 2.02 |
| mesenchyme | Bcl10 | 0.38 | 9.21E-42 | 1.91E-37 | 0.52 | 0.46 | 0.06 | 1.12 |
| mesenchyme | Sgtb | 1.12 | 9.78E-42 | 2.03E-37 | 0.13 | 0.07 | 0.06 | 1.93 |
| mesenchyme | Washc2 | 0.36 | 1.03E-41 | 2.13E-37 | 0.55 | 0.48 | 0.07 | 1.15 |
| mesenchyme | Kars | 0.27 | 1.06E-41 | 2.19E-37 | 0.65 | 0.59 | 0.06 | 1.1 |
| mesenchyme | Rhoj | 0.34 | 1.08E-41 | 2.25E-37 | 0.14 | 0.08 | 0.07 | 1.92 |
| mesenchyme | Copa | 0.27 | 1.16E-41 | 2.42E-37 | 0.67 | 0.61 | 0.06 | 1.1 |
| mesenchyme | Mcrs1 | 0.29 | 1.23E-41 | 2.56E-37 | 0.65 | 0.61 | 0.05 | 1.07 |
| mesenchyme | Med7 | 0.35 | 1.26E-41 | 2.61E-37 | 0.55 | 0.5 | 0.05 | 1.1 |
| mesenchyme | B4galt6 | 0.81 | 1.51E-41 | 3.14E-37 | 0.25 | 0.18 | 0.07 | 1.41 |

**Supplementary Table 2. Marker genes for the major cell types of integrated clustering of E11.5 control and NotchGOF λ-junction cells.**

| | | | | | | | | |
|---|---|---|---|---|---|---|---|---|
| mesenchyme | Tob1 | 0.67 | 1.91E-41 | 3.98E-37 | 0.3 | 0.23 | 0.08 | 1.33 |
| mesenchyme | Llgl1 | 0.42 | 2.31E-41 | 4.81E-37 | 0.48 | 0.42 | 0.07 | 1.16 |
| mesenchyme | Pdhx | 0.64 | 2.36E-41 | 4.91E-37 | 0.33 | 0.26 | 0.07 | 1.26 |
| mesenchyme | Phpt1 | 0.3 | 2.45E-41 | 5.10E-37 | 0.59 | 0.53 | 0.06 | 1.12 |
| mesenchyme | Lsm14a | 0.26 | 3.04E-41 | 6.33E-37 | 0.71 | 0.67 | 0.05 | 1.07 |
| mesenchyme | Dnajc25 | 0.76 | 3.23E-41 | 6.71E-37 | 0.3 | 0.23 | 0.07 | 1.31 |
| mesenchyme | Zfp37 | 0.79 | 3.49E-41 | 7.27E-37 | 0.25 | 0.17 | 0.08 | 1.43 |
| mesenchyme | Elf2 | 0.27 | 4.24E-41 | 8.82E-37 | 0.68 | 0.63 | 0.06 | 1.09 |
| mesenchyme | Farsa | 0.29 | 4.52E-41 | 9.41E-37 | 0.64 | 0.59 | 0.05 | 1.09 |
| mesenchyme | Mapk8ip3 | 0.64 | 4.57E-41 | 9.50E-37 | 0.32 | 0.25 | 0.07 | 1.28 |
| mesenchyme | Dennd2a | 0.65 | 4.83E-41 | 1.00E-36 | 0.31 | 0.24 | 0.07 | 1.31 |
| mesenchyme | Fxn | 0.37 | 5.31E-41 | 1.10E-36 | 0.56 | 0.52 | 0.04 | 1.08 |
| mesenchyme | Zbed5 | 1 | 5.71E-41 | 1.19E-36 | 0.18 | 0.11 | 0.07 | 1.64 |
| mesenchyme | Acadsb | 0.57 | 5.91E-41 | 1.23E-36 | 0.39 | 0.32 | 0.07 | 1.2 |
| mesenchyme | Birc6 | 0.25 | 7.19E-41 | 1.50E-36 | 0.72 | 0.65 | 0.06 | 1.1 |
| mesenchyme | Rbm27 | 0.28 | 7.80E-41 | 1.62E-36 | 0.65 | 0.59 | 0.06 | 1.09 |
| mesenchyme | Tmem39b | 0.52 | 7.80E-41 | 1.62E-36 | 0.38 | 0.31 | 0.07 | 1.24 |
| mesenchyme | Eloa | 0.42 | 8.42E-41 | 1.75E-36 | 0.49 | 0.43 | 0.06 | 1.13 |
| mesenchyme | Pqbp1 | 0.37 | 8.44E-41 | 1.76E-36 | 0.53 | 0.48 | 0.06 | 1.12 |
| mesenchyme | Nol10 | 0.56 | 8.72E-41 | 1.81E-36 | 0.36 | 0.29 | 0.07 | 1.26 |
| mesenchyme | Fzd7 | 0.5 | 9.07E-41 | 1.89E-36 | 0.38 | 0.3 | 0.08 | 1.27 |
| mesenchyme | Crnde | 1.02 | 9.15E-41 | 1.90E-36 | 0.17 | 0.11 | 0.07 | 1.64 |
| mesenchyme | Dctn6 | 0.33 | 1.06E-40 | 2.20E-36 | 0.56 | 0.5 | 0.06 | 1.11 |
| mesenchyme | Evc | 0.99 | 1.08E-40 | 2.25E-36 | 0.17 | 0.11 | 0.07 | 1.63 |
| mesenchyme | Bub1b | 0.38 | 1.10E-40 | 2.28E-36 | 0.58 | 0.52 | 0.06 | 1.12 |
| mesenchyme | Il17rd | 0.26 | 1.25E-40 | 2.61E-36 | 0.54 | 0.43 | 0.11 | 1.25 |
| mesenchyme | Phc1 | 0.49 | 1.46E-40 | 3.03E-36 | 0.39 | 0.32 | 0.08 | 1.24 |
| mesenchyme | Dync2li1 | 0.5 | 1.51E-40 | 3.13E-36 | 0.38 | 0.31 | 0.07 | 1.24 |
| mesenchyme | Slc36a4 | 0.99 | 1.51E-40 | 3.14E-36 | 0.19 | 0.12 | 0.07 | 1.58 |
| mesenchyme | Got1 | 0.4 | 1.54E-40 | 3.21E-36 | 0.48 | 0.41 | 0.07 | 1.16 |
| mesenchyme | Nup98 | 0.4 | 1.57E-40 | 3.27E-36 | 0.5 | 0.44 | 0.06 | 1.14 |
| mesenchyme | Mcts2 | 0.4 | 1.63E-40 | 3.40E-36 | 0.48 | 0.42 | 0.06 | 1.14 |
| mesenchyme | Mex3c | 0.31 | 1.76E-40 | 3.66E-36 | 0.56 | 0.49 | 0.07 | 1.14 |
| mesenchyme | Rcor2 | 0.28 | 1.83E-40 | 3.80E-36 | 0.52 | 0.44 | 0.08 | 1.18 |
| mesenchyme | Acbd3 | 0.3 | 1.85E-40 | 3.85E-36 | 0.64 | 0.58 | 0.06 | 1.1 |
| mesenchyme | Cdc7 | 0.53 | 2.08E-40 | 4.32E-36 | 0.39 | 0.32 | 0.07 | 1.22 |
| mesenchyme | Sertad1 | 0.77 | 2.21E-40 | 4.60E-36 | 0.25 | 0.18 | 0.07 | 1.41 |
| mesenchyme | Kif15 | 0.31 | 2.22E-40 | 4.63E-36 | 0.65 | 0.59 | 0.06 | 1.1 |
| mesenchyme | Akap7 | 0.81 | 2.34E-40 | 4.86E-36 | 0.25 | 0.18 | 0.07 | 1.38 |
| mesenchyme | Cenpc1 | 0.45 | 2.51E-40 | 5.22E-36 | 0.48 | 0.42 | 0.06 | 1.13 |
| mesenchyme | Pspc1 | 0.28 | 2.82E-40 | 5.87E-36 | 0.63 | 0.56 | 0.06 | 1.12 |
| mesenchyme | Gli1 | 1.06 | 3.13E-40 | 6.52E-36 | 0.16 | 0.09 | 0.07 | 1.72 |
| mesenchyme | Rnf168 | 0.39 | 3.31E-40 | 6.88E-36 | 0.55 | 0.48 | 0.07 | 1.13 |
| mesenchyme | Hist1h3d | 1.05 | 5.06E-40 | 1.05E-35 | 0.21 | 0.14 | 0.07 | 1.48 |
| mesenchyme | Cacfd1 | 0.4 | 5.42E-40 | 1.13E-35 | 0.47 | 0.4 | 0.07 | 1.17 |
| mesenchyme | Alkbh8 | 0.48 | 5.66E-40 | 1.18E-35 | 0.44 | 0.38 | 0.06 | 1.16 |
| mesenchyme | Nf1 | 0.52 | 6.00E-40 | 1.25E-35 | 0.42 | 0.36 | 0.06 | 1.18 |
| mesenchyme | Cep83 | 0.35 | 6.14E-40 | 1.28E-35 | 0.56 | 0.52 | 0.04 | 1.09 |
| mesenchyme | Map3k12 | 0.77 | 6.33E-40 | 1.32E-35 | 0.23 | 0.16 | 0.07 | 1.45 |
| mesenchyme | Eva1a | 0.6 | 6.56E-40 | 1.36E-35 | 0.27 | 0.19 | 0.08 | 1.4 |
| mesenchyme | Ttbk2 | 0.92 | 6.68E-40 | 1.39E-35 | 0.2 | 0.14 | 0.07 | 1.51 |
| mesenchyme | Pole | 0.65 | 6.80E-40 | 1.41E-35 | 0.32 | 0.25 | 0.07 | 1.27 |
| mesenchyme | Ccng2 | 0.42 | 7.21E-40 | 1.50E-35 | 0.46 | 0.39 | 0.07 | 1.18 |
| mesenchyme | D10Wsu102e | 0.31 | 7.26E-40 | 1.51E-35 | 0.62 | 0.55 | 0.06 | 1.11 |
| mesenchyme | Fhl3 | 1.25 | 8.21E-40 | 1.71E-35 | 0.13 | 0.07 | 0.06 | 1.9 |
| mesenchyme | Nup54 | 0.34 | 9.10E-40 | 1.89E-35 | 0.58 | 0.53 | 0.05 | 1.1 |

**Supplementary Table 2. Marker genes for the major cell types of integrated clustering of E11.5 control and NotchGOF λ-junction cells.**

| | | | | | | | | |
|---|---|---|---|---|---|---|---|---|
| mesenchyme | Ythdf1 | 0.29 | 9.56E-40 | 1.99E-35 | 0.62 | 0.57 | 0.06 | 1.1 |
| mesenchyme | Ncapd3 | 0.35 | 1.26E-39 | 2.63E-35 | 0.55 | 0.5 | 0.05 | 1.1 |
| mesenchyme | Cxcl12 | 1.19 | 1.94E-39 | 4.03E-35 | 0.2 | 0.13 | 0.07 | 1.52 |
| mesenchyme | Nudt22 | 0.86 | 2.06E-39 | 4.28E-35 | 0.24 | 0.17 | 0.07 | 1.39 |
| mesenchyme | Cdr2l | 0.87 | 2.26E-39 | 4.71E-35 | 0.22 | 0.15 | 0.07 | 1.47 |
| mesenchyme | Stxbp4 | 0.49 | 2.62E-39 | 5.44E-35 | 0.38 | 0.31 | 0.07 | 1.23 |
| mesenchyme | Lrrc20 | 1.04 | 2.82E-39 | 5.86E-35 | 0.17 | 0.11 | 0.07 | 1.6 |
| mesenchyme | Klhdc10 | 0.43 | 2.92E-39 | 6.07E-35 | 0.46 | 0.4 | 0.06 | 1.15 |
| mesenchyme | Socs6 | 0.41 | 2.93E-39 | 6.09E-35 | 0.49 | 0.42 | 0.07 | 1.16 |
| mesenchyme | Utp18 | 0.31 | 3.11E-39 | 6.46E-35 | 0.6 | 0.55 | 0.05 | 1.08 |
| mesenchyme | Kif22 | 0.44 | 3.33E-39 | 6.92E-35 | 0.52 | 0.46 | 0.06 | 1.12 |
| mesenchyme | Fen1 | 0.29 | 3.35E-39 | 6.97E-35 | 0.67 | 0.62 | 0.05 | 1.08 |
| mesenchyme | Zfp449 | 1.09 | 3.53E-39 | 7.35E-35 | 0.14 | 0.08 | 0.06 | 1.76 |
| mesenchyme | Atn1 | 0.37 | 3.55E-39 | 7.38E-35 | 0.47 | 0.4 | 0.07 | 1.18 |
| mesenchyme | Chaf1b | 0.38 | 3.74E-39 | 7.77E-35 | 0.53 | 0.47 | 0.06 | 1.13 |
| mesenchyme | Myof | 1.42 | 3.75E-39 | 7.80E-35 | 0.11 | 0.05 | 0.06 | 2.04 |
| mesenchyme | Ibtk | 0.53 | 4.50E-39 | 9.37E-35 | 0.4 | 0.34 | 0.06 | 1.18 |
| mesenchyme | Ranbp6 | 0.84 | 4.56E-39 | 9.49E-35 | 0.22 | 0.15 | 0.07 | 1.47 |
| mesenchyme | Agpat4 | 0.44 | 4.73E-39 | 9.84E-35 | 0.46 | 0.4 | 0.06 | 1.14 |
| mesenchyme | Mis18bp1 | 0.46 | 5.15E-39 | 1.07E-34 | 0.54 | 0.48 | 0.05 | 1.11 |
| mesenchyme | Tbc1d7 | 0.4 | 5.37E-39 | 1.12E-34 | 0.42 | 0.35 | 0.07 | 1.2 |
| mesenchyme | Fgfrl1 | 0.59 | 5.45E-39 | 1.13E-34 | 0.36 | 0.3 | 0.07 | 1.22 |
| mesenchyme | Gnaq | 0.33 | 5.50E-39 | 1.14E-34 | 0.55 | 0.48 | 0.07 | 1.15 |
| mesenchyme | Cerk | 0.86 | 7.38E-39 | 1.53E-34 | 0.17 | 0.11 | 0.07 | 1.61 |
| mesenchyme | Atxn2l | 0.31 | 7.49E-39 | 1.56E-34 | 0.6 | 0.55 | 0.06 | 1.1 |
| mesenchyme | Bach1 | 0.4 | 7.76E-39 | 1.61E-34 | 0.5 | 0.44 | 0.06 | 1.14 |
| mesenchyme | Jagn1 | 0.32 | 7.92E-39 | 1.65E-34 | 0.6 | 0.54 | 0.05 | 1.1 |
| mesenchyme | Usp39 | 0.28 | 8.02E-39 | 1.67E-34 | 0.6 | 0.56 | 0.04 | 1.08 |
| mesenchyme | Zfp410 | 0.45 | 8.15E-39 | 1.69E-34 | 0.44 | 0.38 | 0.06 | 1.17 |
| mesenchyme | Rab2b | 0.65 | 8.51E-39 | 1.77E-34 | 0.3 | 0.23 | 0.07 | 1.32 |
| mesenchyme | 931406C07Ri | 0.65 | 9.04E-39 | 1.88E-34 | 0.29 | 0.21 | 0.07 | 1.34 |
| mesenchyme | Tubgcp2 | 0.35 | 1.21E-38 | 2.51E-34 | 0.54 | 0.49 | 0.06 | 1.12 |
| mesenchyme | Usp5 | 0.35 | 1.27E-38 | 2.64E-34 | 0.52 | 0.46 | 0.06 | 1.12 |
| mesenchyme | Prdm5 | 0.82 | 1.41E-38 | 2.94E-34 | 0.23 | 0.17 | 0.07 | 1.42 |
| mesenchyme | Btf3l4 | 0.37 | 1.44E-38 | 2.99E-34 | 0.49 | 0.43 | 0.06 | 1.13 |
| mesenchyme | Actr1b | 0.31 | 1.51E-38 | 3.14E-34 | 0.55 | 0.48 | 0.07 | 1.14 |
| mesenchyme | Ppp1r12c | 0.56 | 1.57E-38 | 3.26E-34 | 0.34 | 0.28 | 0.07 | 1.23 |
| mesenchyme | S100pbp | 0.58 | 1.79E-38 | 3.72E-34 | 0.33 | 0.26 | 0.07 | 1.27 |
| mesenchyme | Mad1l1 | 0.5 | 1.80E-38 | 3.75E-34 | 0.38 | 0.31 | 0.07 | 1.22 |
| mesenchyme | Srp68 | 0.33 | 1.97E-38 | 4.10E-34 | 0.55 | 0.49 | 0.06 | 1.11 |
| mesenchyme | Rita1 | 0.83 | 2.32E-38 | 4.83E-34 | 0.22 | 0.15 | 0.07 | 1.45 |
| mesenchyme | Acaca | 0.58 | 2.35E-38 | 4.89E-34 | 0.36 | 0.28 | 0.07 | 1.25 |
| mesenchyme | Fbxo11 | 0.33 | 2.45E-38 | 5.10E-34 | 0.56 | 0.5 | 0.06 | 1.12 |
| mesenchyme | Nsmce3 | 0.3 | 2.51E-38 | 5.22E-34 | 0.62 | 0.58 | 0.05 | 1.08 |
| mesenchyme | Dnlz | 0.25 | 2.77E-38 | 5.76E-34 | 0.67 | 0.63 | 0.04 | 1.07 |
| mesenchyme | Xrcc2 | 1.04 | 2.84E-38 | 5.91E-34 | 0.17 | 0.11 | 0.06 | 1.61 |
| mesenchyme | Cdc42se2 | 0.36 | 3.19E-38 | 6.63E-34 | 0.52 | 0.46 | 0.06 | 1.13 |
| mesenchyme | Grk6 | 0.4 | 3.20E-38 | 6.67E-34 | 0.5 | 0.45 | 0.06 | 1.13 |
| mesenchyme | Armcx6 | 0.86 | 3.21E-38 | 6.68E-34 | 0.23 | 0.16 | 0.07 | 1.43 |
| mesenchyme | Ccdc66 | 0.47 | 3.61E-38 | 7.50E-34 | 0.41 | 0.34 | 0.07 | 1.2 |
| mesenchyme | Pola1 | 0.36 | 4.23E-38 | 8.79E-34 | 0.51 | 0.45 | 0.06 | 1.14 |
| mesenchyme | 110021N24Ri | 1.18 | 4.47E-38 | 9.30E-34 | 0.13 | 0.07 | 0.06 | 1.83 |
| mesenchyme | Ptgfrn | 0.57 | 4.71E-38 | 9.79E-34 | 0.36 | 0.29 | 0.07 | 1.25 |
| mesenchyme | Prxl2b | 0.42 | 5.35E-38 | 1.11E-33 | 0.44 | 0.37 | 0.07 | 1.19 |
| mesenchyme | Smpd4 | 0.5 | 6.26E-38 | 1.30E-33 | 0.41 | 0.35 | 0.06 | 1.17 |
| mesenchyme | Cep192 | 0.45 | 6.53E-38 | 1.36E-33 | 0.46 | 0.41 | 0.06 | 1.15 |

**Supplementary Table 2. Marker genes for the major cell types of integrated clustering of E11.5 control and NotchGOF λ-junction cells.**

| | | | | | | | | |
|---|---|---|---|---|---|---|---|---|
| mesenchyme | Plekhg2 | 0.48 | 6.69E-38 | 1.39E-33 | 0.4 | 0.33 | 0.07 | 1.21 |
| mesenchyme | Heg1 | 0.93 | 6.75E-38 | 1.40E-33 | 0.17 | 0.11 | 0.07 | 1.61 |
| mesenchyme | Nudt9 | 0.34 | 8.48E-38 | 1.76E-33 | 0.5 | 0.44 | 0.06 | 1.14 |
| mesenchyme | Trpc4ap | 0.36 | 9.96E-38 | 2.07E-33 | 0.55 | 0.5 | 0.05 | 1.1 |
| mesenchyme | Gins4 | 0.27 | 1.06E-37 | 2.21E-33 | 0.66 | 0.62 | 0.03 | 1.05 |
| mesenchyme | Zfp101 | 0.54 | 1.30E-37 | 2.70E-33 | 0.39 | 0.32 | 0.07 | 1.21 |
| mesenchyme | Usp37 | 0.41 | 1.31E-37 | 2.73E-33 | 0.47 | 0.41 | 0.06 | 1.14 |
| mesenchyme | Taf2 | 0.43 | 1.74E-37 | 3.62E-33 | 0.47 | 0.41 | 0.06 | 1.14 |
| mesenchyme | Gins3 | 0.7 | 1.85E-37 | 3.84E-33 | 0.28 | 0.21 | 0.07 | 1.33 |
| mesenchyme | Mafb | 0.98 | 1.86E-37 | 3.86E-33 | 0.25 | 0.18 | 0.07 | 1.4 |
| mesenchyme | Far1 | 0.32 | 2.34E-37 | 4.87E-33 | 0.56 | 0.5 | 0.05 | 1.11 |
| mesenchyme | Pfkm | 0.69 | 2.36E-37 | 4.90E-33 | 0.29 | 0.22 | 0.07 | 1.3 |
| mesenchyme | B9d1 | 0.26 | 2.50E-37 | 5.21E-33 | 0.6 | 0.51 | 0.08 | 1.16 |
| mesenchyme | Wdcp | 0.63 | 2.74E-37 | 5.69E-33 | 0.39 | 0.33 | 0.07 | 1.2 |
| mesenchyme | Pias3 | 0.83 | 2.79E-37 | 5.80E-33 | 0.23 | 0.16 | 0.07 | 1.42 |
| mesenchyme | Gtf3c3 | 0.46 | 2.95E-37 | 6.14E-33 | 0.4 | 0.33 | 0.07 | 1.21 |
| mesenchyme | Heatr3 | 0.47 | 3.04E-37 | 6.32E-33 | 0.39 | 0.32 | 0.07 | 1.21 |
| mesenchyme | Mrps10 | 0.29 | 3.06E-37 | 6.36E-33 | 0.6 | 0.56 | 0.04 | 1.08 |
| mesenchyme | Snhg12 | 0.41 | 3.45E-37 | 7.17E-33 | 0.5 | 0.44 | 0.06 | 1.12 |
| mesenchyme | Sccpdh | 0.56 | 3.82E-37 | 7.94E-33 | 0.35 | 0.28 | 0.07 | 1.26 |
| mesenchyme | Nkiras2 | 0.41 | 4.16E-37 | 8.66E-33 | 0.45 | 0.39 | 0.07 | 1.17 |
| mesenchyme | Smyd5 | 0.56 | 4.30E-37 | 8.93E-33 | 0.35 | 0.28 | 0.07 | 1.24 |
| mesenchyme | Smim26 | 0.43 | 4.38E-37 | 9.11E-33 | 0.45 | 0.4 | 0.06 | 1.14 |
| mesenchyme | Pde3b | 0.66 | 4.40E-37 | 9.15E-33 | 0.27 | 0.2 | 0.07 | 1.35 |
| mesenchyme | Pstk | 0.32 | 4.47E-37 | 9.29E-33 | 0.58 | 0.53 | 0.05 | 1.09 |
| mesenchyme | Tial1 | 0.28 | 4.72E-37 | 9.81E-33 | 0.68 | 0.64 | 0.04 | 1.06 |
| mesenchyme | Pbk | 0.36 | 4.99E-37 | 1.04E-32 | 0.56 | 0.51 | 0.05 | 1.1 |
| mesenchyme | Siah1a | 0.33 | 5.30E-37 | 1.10E-32 | 0.51 | 0.45 | 0.06 | 1.14 |
| mesenchyme | Ifit2 | 1.24 | 5.49E-37 | 1.14E-32 | 0.11 | 0.06 | 0.05 | 1.96 |
| mesenchyme | Melk | 0.42 | 7.45E-37 | 1.55E-32 | 0.49 | 0.43 | 0.06 | 1.14 |
| mesenchyme | Eef1akmt1 | 0.26 | 1.16E-36 | 2.42E-32 | 0.67 | 0.61 | 0.06 | 1.09 |
| mesenchyme | Arhgap24 | 1.15 | 1.32E-36 | 2.75E-32 | 0.14 | 0.08 | 0.06 | 1.75 |
| mesenchyme | Hykk | 1.31 | 1.43E-36 | 2.97E-32 | 0.11 | 0.05 | 0.05 | 1.98 |
| mesenchyme | Tle1 | 0.28 | 1.46E-36 | 3.03E-32 | 0.55 | 0.49 | 0.06 | 1.12 |
| mesenchyme | Rsbn1 | 0.3 | 1.85E-36 | 3.85E-32 | 0.58 | 0.51 | 0.07 | 1.13 |
| mesenchyme | Nob1 | 0.33 | 2.12E-36 | 4.41E-32 | 0.53 | 0.48 | 0.05 | 1.11 |
| mesenchyme | Lrp6 | 0.3 | 2.38E-36 | 4.96E-32 | 0.6 | 0.53 | 0.07 | 1.14 |
| mesenchyme | Dnajc30 | 0.41 | 2.46E-36 | 5.11E-32 | 0.47 | 0.41 | 0.06 | 1.13 |
| mesenchyme | Lca5 | 0.75 | 2.75E-36 | 5.72E-32 | 0.24 | 0.17 | 0.07 | 1.4 |
| mesenchyme | Taf12 | 0.31 | 2.81E-36 | 5.85E-32 | 0.6 | 0.56 | 0.04 | 1.07 |
| mesenchyme | Plekha8 | 0.42 | 2.82E-36 | 5.86E-32 | 0.49 | 0.43 | 0.06 | 1.13 |
| mesenchyme | Ints14 | 0.5 | 3.05E-36 | 6.35E-32 | 0.42 | 0.36 | 0.06 | 1.16 |
| mesenchyme | Aven | 0.48 | 3.20E-36 | 6.66E-32 | 0.41 | 0.36 | 0.06 | 1.16 |
| mesenchyme | Ckap2l | 0.53 | 3.48E-36 | 7.24E-32 | 0.5 | 0.46 | 0.04 | 1.1 |
| mesenchyme | Orc3 | 0.46 | 3.69E-36 | 7.68E-32 | 0.41 | 0.35 | 0.06 | 1.16 |
| mesenchyme | Pkdcc | 0.89 | 3.72E-36 | 7.75E-32 | 0.35 | 0.29 | 0.06 | 1.22 |
| mesenchyme | )430038I01Ril | 0.51 | 4.22E-36 | 8.78E-32 | 0.36 | 0.29 | 0.07 | 1.23 |
| mesenchyme | Phf12 | 0.43 | 4.97E-36 | 1.03E-31 | 0.46 | 0.4 | 0.06 | 1.15 |
| mesenchyme | Pip4k2b | 0.8 | 4.99E-36 | 1.04E-31 | 0.24 | 0.18 | 0.07 | 1.38 |
| mesenchyme | Slc4a1ap | 0.49 | 5.01E-36 | 1.04E-31 | 0.4 | 0.34 | 0.06 | 1.18 |
| mesenchyme | Ulk2 | 0.69 | 5.04E-36 | 1.05E-31 | 0.27 | 0.21 | 0.07 | 1.33 |
| mesenchyme | Ndufaf6 | 0.51 | 5.46E-36 | 1.13E-31 | 0.37 | 0.31 | 0.07 | 1.21 |
| mesenchyme | Tbck | 0.64 | 5.92E-36 | 1.23E-31 | 0.31 | 0.24 | 0.07 | 1.28 |
| mesenchyme | Itsn1 | 0.47 | 6.22E-36 | 1.29E-31 | 0.46 | 0.4 | 0.05 | 1.13 |
| mesenchyme | Snx4 | 0.42 | 6.47E-36 | 1.35E-31 | 0.44 | 0.38 | 0.06 | 1.16 |
| mesenchyme | Asb13 | 0.71 | 6.54E-36 | 1.36E-31 | 0.25 | 0.18 | 0.07 | 1.37 |

**Supplementary Table 2. Marker genes for the major cell types of integrated clustering of E11.5 control and NotchGOF λ-junction cells.**

| | | | | | | | | |
|---|---|---|---|---|---|---|---|---|
| mesenchyme | Azin1 | 0.27 | 7.22E-36 | 1.50E-31 | 0.64 | 0.6 | 0.04 | 1.06 |
| mesenchyme | Penk | 1.72 | 7.92E-36 | 1.65E-31 | 0.11 | 0.06 | 0.05 | 1.93 |
| mesenchyme | 310009B15Ri | 0.36 | 7.94E-36 | 1.65E-31 | 0.56 | 0.51 | 0.05 | 1.09 |
| mesenchyme | Pphln1 | 0.46 | 8.32E-36 | 1.73E-31 | 0.42 | 0.36 | 0.06 | 1.16 |
| mesenchyme | Ppp4r1 | 0.5 | 9.82E-36 | 2.04E-31 | 0.42 | 0.36 | 0.06 | 1.17 |
| mesenchyme | Trim59 | 0.28 | 1.14E-35 | 2.37E-31 | 0.73 | 0.7 | 0.03 | 1.04 |
| mesenchyme | Hist1h4h | 0.67 | 1.18E-35 | 2.45E-31 | 0.43 | 0.38 | 0.05 | 1.14 |
| mesenchyme | 130307A14Ri | 0.87 | 1.26E-35 | 2.61E-31 | 0.22 | 0.15 | 0.07 | 1.43 |
| mesenchyme | Msl1 | 0.27 | 1.29E-35 | 2.68E-31 | 0.64 | 0.57 | 0.07 | 1.11 |
| mesenchyme | Itpripl2 | 1.04 | 1.33E-35 | 2.78E-31 | 0.15 | 0.09 | 0.06 | 1.65 |
| mesenchyme | Rcl1 | 0.42 | 1.36E-35 | 2.84E-31 | 0.47 | 0.42 | 0.06 | 1.13 |
| mesenchyme | Arpin | 0.86 | 1.44E-35 | 2.99E-31 | 0.2 | 0.14 | 0.06 | 1.47 |
| mesenchyme | Sesn3 | 0.27 | 1.49E-35 | 3.10E-31 | 0.62 | 0.54 | 0.09 | 1.16 |
| mesenchyme | Fzd10os | 0.75 | 1.72E-35 | 3.57E-31 | 0.23 | 0.16 | 0.07 | 1.44 |
| mesenchyme | Tacc3 | 0.3 | 2.09E-35 | 4.35E-31 | 0.62 | 0.57 | 0.05 | 1.09 |
| mesenchyme | Syce2 | 0.3 | 2.26E-35 | 4.70E-31 | 0.61 | 0.56 | 0.05 | 1.08 |
| mesenchyme | Spindoc | 0.53 | 2.44E-35 | 5.07E-31 | 0.33 | 0.26 | 0.07 | 1.27 |
| mesenchyme | Dhps | 0.37 | 2.52E-35 | 5.24E-31 | 0.52 | 0.47 | 0.05 | 1.1 |
| mesenchyme | Sall1 | 1.42 | 2.61E-35 | 5.42E-31 | 0.11 | 0.06 | 0.05 | 1.9 |
| mesenchyme | Vldlr | 1.12 | 2.73E-35 | 5.67E-31 | 0.14 | 0.08 | 0.06 | 1.73 |
| mesenchyme | Epc1 | 0.39 | 2.77E-35 | 5.76E-31 | 0.49 | 0.44 | 0.05 | 1.11 |
| mesenchyme | Gprasp2 | 0.54 | 2.79E-35 | 5.81E-31 | 0.32 | 0.26 | 0.07 | 1.27 |
| mesenchyme | Cep78 | 0.4 | 2.82E-35 | 5.87E-31 | 0.42 | 0.35 | 0.07 | 1.19 |
| mesenchyme | 110051M20R | 0.41 | 2.97E-35 | 6.17E-31 | 0.37 | 0.3 | 0.07 | 1.23 |
| mesenchyme | Elp2 | 0.32 | 3.04E-35 | 6.33E-31 | 0.56 | 0.51 | 0.06 | 1.11 |
| mesenchyme | Tmem126b | 0.5 | 3.07E-35 | 6.39E-31 | 0.38 | 0.32 | 0.06 | 1.19 |
| mesenchyme | Smg6 | 0.25 | 3.48E-35 | 7.23E-31 | 0.62 | 0.56 | 0.06 | 1.1 |
| mesenchyme | Sart1 | 0.33 | 3.53E-35 | 7.35E-31 | 0.57 | 0.53 | 0.04 | 1.08 |
| mesenchyme | Nfx1 | 0.33 | 3.98E-35 | 8.28E-31 | 0.59 | 0.54 | 0.04 | 1.08 |
| mesenchyme | Aip | 0.43 | 4.16E-35 | 8.65E-31 | 0.43 | 0.38 | 0.05 | 1.14 |
| mesenchyme | Wdr60 | 0.42 | 5.17E-35 | 1.07E-30 | 0.41 | 0.34 | 0.07 | 1.2 |
| mesenchyme | Uggt2 | 0.83 | 5.30E-35 | 1.10E-30 | 0.21 | 0.15 | 0.07 | 1.46 |
| mesenchyme | Kctd2 | 0.48 | 6.53E-35 | 1.36E-30 | 0.39 | 0.33 | 0.06 | 1.19 |
| mesenchyme | Ogdh | 0.38 | 6.63E-35 | 1.38E-30 | 0.54 | 0.5 | 0.04 | 1.08 |
| mesenchyme | Bad | 0.34 | 7.37E-35 | 1.53E-30 | 0.55 | 0.5 | 0.05 | 1.11 |
| mesenchyme | Inpp1 | 0.94 | 7.58E-35 | 1.58E-30 | 0.19 | 0.13 | 0.06 | 1.5 |
| mesenchyme | Casc4 | 0.4 | 7.96E-35 | 1.66E-30 | 0.49 | 0.43 | 0.06 | 1.15 |
| mesenchyme | Tdp2 | 0.26 | 8.12E-35 | 1.69E-30 | 0.59 | 0.53 | 0.05 | 1.1 |
| mesenchyme | Nadk | 0.38 | 8.98E-35 | 1.87E-30 | 0.45 | 0.39 | 0.06 | 1.15 |
| mesenchyme | Zcchc9 | 0.38 | 9.12E-35 | 1.90E-30 | 0.5 | 0.44 | 0.05 | 1.12 |
| mesenchyme | Zdhhc17 | 0.7 | 1.13E-34 | 2.36E-30 | 0.28 | 0.21 | 0.07 | 1.32 |
| mesenchyme | Usp29 | 0.75 | 1.18E-34 | 2.46E-30 | 0.23 | 0.16 | 0.07 | 1.4 |
| mesenchyme | Elovl4 | 0.77 | 1.22E-34 | 2.53E-30 | 0.2 | 0.14 | 0.07 | 1.49 |
| mesenchyme | Zkscan1 | 0.27 | 1.54E-34 | 3.20E-30 | 0.61 | 0.54 | 0.07 | 1.13 |
| mesenchyme | Gpx3 | 0.69 | 1.75E-34 | 3.64E-30 | 0.24 | 0.18 | 0.07 | 1.38 |
| mesenchyme | Qrsl1 | 0.8 | 1.85E-34 | 3.85E-30 | 0.24 | 0.17 | 0.06 | 1.37 |
| mesenchyme | Elp4 | 0.38 | 1.86E-34 | 3.86E-30 | 0.48 | 0.42 | 0.06 | 1.14 |
| mesenchyme | Nutf2-ps1 | 0.28 | 1.86E-34 | 3.87E-30 | 0.55 | 0.49 | 0.06 | 1.11 |
| mesenchyme | Supt3 | 0.47 | 1.88E-34 | 3.91E-30 | 0.41 | 0.36 | 0.06 | 1.16 |
| mesenchyme | Rchy1 | 0.36 | 2.25E-34 | 4.68E-30 | 0.52 | 0.47 | 0.05 | 1.1 |
| mesenchyme | N4bp1 | 0.27 | 2.35E-34 | 4.90E-30 | 0.61 | 0.55 | 0.06 | 1.1 |
| mesenchyme | Vkorc1 | 0.46 | 2.36E-34 | 4.92E-30 | 0.45 | 0.4 | 0.05 | 1.14 |
| mesenchyme | Cth | 0.6 | 2.46E-34 | 5.12E-30 | 0.23 | 0.16 | 0.07 | 1.41 |
| mesenchyme | Nmt1 | 0.26 | 2.53E-34 | 5.27E-30 | 0.63 | 0.59 | 0.04 | 1.07 |
| mesenchyme | Agps | 0.42 | 2.54E-34 | 5.28E-30 | 0.43 | 0.36 | 0.06 | 1.18 |
| mesenchyme | Gtf2h3 | 0.34 | 2.69E-34 | 5.60E-30 | 0.51 | 0.46 | 0.06 | 1.12 |

**Supplementary Table 2. Marker genes for the major cell types of integrated clustering of E11.5 control and NotchGOF λ-junction cells.**

| | | | | | | | | |
|---|---|---|---|---|---|---|---|---|
| mesenchyme | Slc1a3 | 0.97 | 2.78E-34 | 5.78E-30 | 0.25 | 0.18 | 0.06 | 1.35 |
| mesenchyme | Gsdme | 0.65 | 2.83E-34 | 5.88E-30 | 0.26 | 0.2 | 0.07 | 1.33 |
| mesenchyme | Syap1 | 0.48 | 2.90E-34 | 6.04E-30 | 0.38 | 0.31 | 0.07 | 1.21 |
| mesenchyme | Dvl3 | 0.37 | 2.93E-34 | 6.09E-30 | 0.46 | 0.4 | 0.06 | 1.15 |
| mesenchyme | Ptpra | 0.27 | 2.96E-34 | 6.16E-30 | 0.62 | 0.57 | 0.05 | 1.09 |
| mesenchyme | Smad5 | 0.31 | 3.36E-34 | 6.98E-30 | 0.52 | 0.45 | 0.07 | 1.15 |
| mesenchyme | Naa16 | 0.53 | 3.57E-34 | 7.42E-30 | 0.4 | 0.35 | 0.05 | 1.16 |
| mesenchyme | Zfp827 | 0.54 | 3.70E-34 | 7.70E-30 | 0.33 | 0.26 | 0.07 | 1.26 |
| mesenchyme | AI506816 | 0.33 | 3.72E-34 | 7.75E-30 | 0.56 | 0.5 | 0.06 | 1.12 |
| mesenchyme | Rlim | 0.35 | 3.73E-34 | 7.75E-30 | 0.55 | 0.51 | 0.05 | 1.09 |
| mesenchyme | Zfp618 | 0.3 | 3.81E-34 | 7.91E-30 | 0.55 | 0.47 | 0.08 | 1.16 |
| mesenchyme | Dusp23 | 0.71 | 3.85E-34 | 8.01E-30 | 0.2 | 0.13 | 0.07 | 1.51 |
| mesenchyme | G2e3 | 0.46 | 3.89E-34 | 8.09E-30 | 0.49 | 0.44 | 0.05 | 1.1 |
| mesenchyme | Tchp | 0.57 | 3.94E-34 | 8.19E-30 | 0.32 | 0.25 | 0.07 | 1.26 |
| mesenchyme | Pdcd11 | 0.39 | 4.17E-34 | 8.68E-30 | 0.49 | 0.44 | 0.06 | 1.13 |
| mesenchyme | Mterf3 | 0.41 | 4.41E-34 | 9.17E-30 | 0.45 | 0.39 | 0.05 | 1.14 |
| mesenchyme | Ahcy | 0.62 | 4.46E-34 | 9.28E-30 | 0.29 | 0.23 | 0.06 | 1.28 |
| mesenchyme | Ubfd1 | 0.25 | 4.47E-34 | 9.29E-30 | 0.66 | 0.61 | 0.05 | 1.08 |
| mesenchyme | Slc2a4 | 0.92 | 5.06E-34 | 1.05E-29 | 0.19 | 0.13 | 0.06 | 1.47 |
| mesenchyme | Zfp606 | 0.42 | 5.71E-34 | 1.19E-29 | 0.41 | 0.35 | 0.07 | 1.19 |
| mesenchyme | Camk1 | 0.35 | 5.81E-34 | 1.21E-29 | 0.44 | 0.38 | 0.06 | 1.17 |
| mesenchyme | Tox | 0.75 | 6.28E-34 | 1.31E-29 | 0.28 | 0.22 | 0.07 | 1.3 |
| mesenchyme | Eif4enif1 | 0.33 | 8.28E-34 | 1.72E-29 | 0.56 | 0.51 | 0.05 | 1.1 |
| mesenchyme | Mettl6 | 0.35 | 8.67E-34 | 1.80E-29 | 0.52 | 0.47 | 0.05 | 1.11 |
| mesenchyme | Tyw3 | 0.75 | 8.68E-34 | 1.80E-29 | 0.23 | 0.17 | 0.07 | 1.39 |
| mesenchyme | Rabl6 | 0.26 | 1.03E-33 | 2.13E-29 | 0.67 | 0.63 | 0.04 | 1.06 |
| mesenchyme | Nampt | 0.41 | 1.13E-33 | 2.36E-29 | 0.4 | 0.33 | 0.07 | 1.21 |
| mesenchyme | Spry4 | 0.94 | 1.15E-33 | 2.39E-29 | 0.17 | 0.11 | 0.06 | 1.56 |
| mesenchyme | Cops7b | 0.51 | 1.28E-33 | 2.66E-29 | 0.38 | 0.32 | 0.06 | 1.19 |
| mesenchyme | Gde1 | 0.3 | 1.71E-33 | 3.55E-29 | 0.54 | 0.48 | 0.06 | 1.12 |
| mesenchyme | Six4 | 0.37 | 1.73E-33 | 3.59E-29 | 0.49 | 0.41 | 0.07 | 1.18 |
| mesenchyme | Phf2 | 0.55 | 1.73E-33 | 3.61E-29 | 0.33 | 0.26 | 0.07 | 1.25 |
| mesenchyme | 110008P14Ri | 0.38 | 1.96E-33 | 4.09E-29 | 0.49 | 0.44 | 0.05 | 1.11 |
| mesenchyme | Mrps27 | 0.36 | 2.02E-33 | 4.21E-29 | 0.46 | 0.4 | 0.06 | 1.15 |
| mesenchyme | Srfbp1 | 0.45 | 2.07E-33 | 4.30E-29 | 0.42 | 0.36 | 0.06 | 1.16 |
| mesenchyme | Nt5m | 0.5 | 2.12E-33 | 4.42E-29 | 0.37 | 0.31 | 0.06 | 1.2 |
| mesenchyme | Tpp2 | 0.3 | 2.20E-33 | 4.57E-29 | 0.61 | 0.57 | 0.04 | 1.07 |
| mesenchyme | Opa1 | 0.35 | 2.31E-33 | 4.79E-29 | 0.51 | 0.47 | 0.05 | 1.1 |
| mesenchyme | Crebzf | 0.3 | 2.57E-33 | 5.35E-29 | 0.59 | 0.54 | 0.05 | 1.09 |
| mesenchyme | Pde6d | 0.31 | 2.77E-33 | 5.76E-29 | 0.55 | 0.5 | 0.05 | 1.1 |
| mesenchyme | Tab2 | 0.26 | 2.79E-33 | 5.80E-29 | 0.69 | 0.65 | 0.04 | 1.07 |
| mesenchyme | Ndst1 | 0.32 | 3.02E-33 | 6.28E-29 | 0.49 | 0.43 | 0.06 | 1.15 |
| mesenchyme | Dr1 | 0.41 | 3.07E-33 | 6.38E-29 | 0.45 | 0.4 | 0.06 | 1.14 |
| mesenchyme | Ints2 | 0.46 | 3.08E-33 | 6.40E-29 | 0.41 | 0.36 | 0.06 | 1.16 |
| mesenchyme | Nucb1 | 0.36 | 3.19E-33 | 6.63E-29 | 0.53 | 0.49 | 0.05 | 1.09 |
| mesenchyme | Poglut1 | 0.42 | 3.71E-33 | 7.73E-29 | 0.43 | 0.37 | 0.06 | 1.17 |
| mesenchyme | Stxbp1 | 1.02 | 3.95E-33 | 8.21E-29 | 0.11 | 0.06 | 0.05 | 1.84 |
| mesenchyme | Septin8 | 0.75 | 4.10E-33 | 8.53E-29 | 0.24 | 0.18 | 0.06 | 1.34 |
| mesenchyme | Cstf3 | 0.31 | 4.59E-33 | 9.55E-29 | 0.57 | 0.53 | 0.04 | 1.09 |
| mesenchyme | Mgat3 | 0.93 | 4.88E-33 | 1.01E-28 | 0.18 | 0.12 | 0.06 | 1.52 |
| mesenchyme | Fzd4 | 0.76 | 5.16E-33 | 1.07E-28 | 0.27 | 0.21 | 0.06 | 1.3 |
| mesenchyme | Fam216a | 0.28 | 5.31E-33 | 1.10E-28 | 0.55 | 0.49 | 0.06 | 1.13 |
| mesenchyme | Zfp41 | 0.47 | 5.52E-33 | 1.15E-28 | 0.37 | 0.3 | 0.07 | 1.22 |
| mesenchyme | Pold3 | 0.26 | 5.55E-33 | 1.15E-28 | 0.59 | 0.54 | 0.05 | 1.09 |
| mesenchyme | Fam217b | 1.26 | 5.69E-33 | 1.18E-28 | 0.11 | 0.06 | 0.05 | 1.85 |
| mesenchyme | Slc39a7 | 0.33 | 6.03E-33 | 1.25E-28 | 0.55 | 0.5 | 0.05 | 1.11 |

**Supplementary Table 2. Marker genes for the major cell types of integrated clustering of E11.5 control and NotchGOF λ-junction cells.**

| | | | | | | | | |
|---|---|---|---|---|---|---|---|---|
| mesenchyme | :630043F03Ri | 0.94 | 6.22E-33 | 1.29E-28 | 0.17 | 0.11 | 0.06 | 1.56 |
| mesenchyme | Ptgis | 1.2 | 6.31E-33 | 1.31E-28 | 0.13 | 0.08 | 0.05 | 1.7 |
| mesenchyme | Rtkn | 0.71 | 6.38E-33 | 1.33E-28 | 0.24 | 0.17 | 0.07 | 1.38 |
| mesenchyme | Arfgap2 | 0.35 | 6.75E-33 | 1.40E-28 | 0.55 | 0.51 | 0.04 | 1.07 |
| mesenchyme | Zmat5 | 0.35 | 7.17E-33 | 1.49E-28 | 0.5 | 0.45 | 0.05 | 1.11 |
| mesenchyme | Kctd12b | 0.66 | 7.54E-33 | 1.57E-28 | 0.2 | 0.14 | 0.07 | 1.47 |
| mesenchyme | Arl8a | 0.31 | 8.15E-33 | 1.70E-28 | 0.45 | 0.38 | 0.07 | 1.18 |
| mesenchyme | Rhobtb1 | 0.91 | 8.76E-33 | 1.82E-28 | 0.17 | 0.11 | 0.06 | 1.56 |
| mesenchyme | Mxd4 | 0.34 | 8.85E-33 | 1.84E-28 | 0.49 | 0.42 | 0.07 | 1.17 |
| mesenchyme | Bdh2 | 1.19 | 9.22E-33 | 1.92E-28 | 0.11 | 0.06 | 0.05 | 1.84 |
| mesenchyme | Rrp1b | 0.29 | 1.05E-32 | 2.19E-28 | 0.55 | 0.49 | 0.07 | 1.14 |
| mesenchyme | Mrps9 | 0.31 | 1.08E-32 | 2.25E-28 | 0.57 | 0.53 | 0.04 | 1.09 |
| mesenchyme | Bmi1 | 0.38 | 1.28E-32 | 2.66E-28 | 0.43 | 0.37 | 0.06 | 1.17 |
| mesenchyme | Senp2 | 0.3 | 1.29E-32 | 2.68E-28 | 0.58 | 0.55 | 0.04 | 1.07 |
| mesenchyme | Zfp90 | 0.42 | 1.37E-32 | 2.86E-28 | 0.37 | 0.3 | 0.07 | 1.22 |
| mesenchyme | Cenpj | 0.47 | 1.44E-32 | 3.00E-28 | 0.4 | 0.34 | 0.06 | 1.18 |
| mesenchyme | Incenp | 0.4 | 1.51E-32 | 3.13E-28 | 0.6 | 0.58 | 0.03 | 1.05 |
| mesenchyme | 932438A13Ri | 0.53 | 1.56E-32 | 3.24E-28 | 0.37 | 0.31 | 0.06 | 1.19 |
| mesenchyme | Dzip1l | 0.52 | 1.60E-32 | 3.33E-28 | 0.33 | 0.26 | 0.07 | 1.28 |
| mesenchyme | Cep112 | 0.8 | 1.89E-32 | 3.94E-28 | 0.22 | 0.16 | 0.06 | 1.39 |
| mesenchyme | Poc1b | 0.54 | 1.99E-32 | 4.14E-28 | 0.35 | 0.29 | 0.06 | 1.21 |
| mesenchyme | Mfap3 | 0.39 | 2.10E-32 | 4.37E-28 | 0.47 | 0.42 | 0.06 | 1.14 |
| mesenchyme | Clcn5 | 0.44 | 2.80E-32 | 5.82E-28 | 0.41 | 0.35 | 0.07 | 1.19 |
| mesenchyme | Tmem70 | 0.3 | 3.26E-32 | 6.78E-28 | 0.56 | 0.52 | 0.04 | 1.09 |
| mesenchyme | Lrrfip2 | 0.33 | 3.26E-32 | 6.78E-28 | 0.51 | 0.47 | 0.05 | 1.1 |
| mesenchyme | Rufy1 | 0.68 | 3.88E-32 | 8.06E-28 | 0.27 | 0.21 | 0.06 | 1.3 |
| mesenchyme | Pkn1 | 0.49 | 4.00E-32 | 8.32E-28 | 0.38 | 0.32 | 0.06 | 1.2 |
| mesenchyme | Timm21 | 0.3 | 4.02E-32 | 8.36E-28 | 0.56 | 0.51 | 0.05 | 1.09 |
| mesenchyme | Ahctf1 | 0.33 | 4.05E-32 | 8.42E-28 | 0.56 | 0.51 | 0.05 | 1.09 |
| mesenchyme | 110065P20Ri | 0.3 | 4.09E-32 | 8.50E-28 | 0.57 | 0.52 | 0.05 | 1.09 |
| mesenchyme | Srr | 1.06 | 4.29E-32 | 8.92E-28 | 0.13 | 0.08 | 0.05 | 1.67 |
| mesenchyme | Aasdhppt | 0.28 | 4.40E-32 | 9.14E-28 | 0.57 | 0.54 | 0.03 | 1.06 |
| mesenchyme | Upf3a | 0.26 | 5.09E-32 | 1.06E-27 | 0.63 | 0.59 | 0.04 | 1.06 |
| mesenchyme | Zfp846 | 0.67 | 5.25E-32 | 1.09E-27 | 0.27 | 0.21 | 0.06 | 1.3 |
| mesenchyme | Tada1 | 0.44 | 5.33E-32 | 1.11E-27 | 0.4 | 0.34 | 0.06 | 1.17 |
| mesenchyme | Mbip | 0.39 | 5.76E-32 | 1.20E-27 | 0.41 | 0.35 | 0.06 | 1.17 |
| mesenchyme | Rgl1 | 0.62 | 5.85E-32 | 1.22E-27 | 0.17 | 0.11 | 0.06 | 1.55 |
| mesenchyme | Zc3h8 | 0.76 | 5.89E-32 | 1.23E-27 | 0.23 | 0.17 | 0.06 | 1.36 |
| mesenchyme | Fam229b | 1.13 | 6.19E-32 | 1.29E-27 | 0.13 | 0.08 | 0.05 | 1.71 |
| mesenchyme | Ebpl | 0.25 | 6.52E-32 | 1.36E-27 | 0.57 | 0.5 | 0.07 | 1.14 |
| mesenchyme | 310022B05Ri | 0.53 | 6.81E-32 | 1.42E-27 | 0.33 | 0.26 | 0.07 | 1.25 |
| mesenchyme | Zfp286 | 0.59 | 7.67E-32 | 1.60E-27 | 0.27 | 0.2 | 0.07 | 1.32 |
| mesenchyme | Trafd1 | 0.58 | 7.99E-32 | 1.66E-27 | 0.32 | 0.26 | 0.06 | 1.24 |
| mesenchyme | U2af1l4 | 0.3 | 8.10E-32 | 1.68E-27 | 0.53 | 0.49 | 0.04 | 1.09 |
| mesenchyme | Pomgnt1 | 0.48 | 8.14E-32 | 1.69E-27 | 0.35 | 0.29 | 0.06 | 1.21 |
| mesenchyme | Fkbp9 | 0.36 | 9.49E-32 | 1.97E-27 | 0.46 | 0.4 | 0.06 | 1.16 |
| mesenchyme | Zcwpw1 | 0.9 | 9.74E-32 | 2.03E-27 | 0.18 | 0.12 | 0.06 | 1.48 |
| mesenchyme | Ppp1r18 | 0.45 | 1.08E-31 | 2.25E-27 | 0.28 | 0.21 | 0.07 | 1.34 |
| mesenchyme | Mrps30 | 0.26 | 1.21E-31 | 2.51E-27 | 0.62 | 0.58 | 0.04 | 1.06 |
| mesenchyme | Rdh11 | 0.75 | 1.42E-31 | 2.95E-27 | 0.24 | 0.18 | 0.06 | 1.34 |
| mesenchyme | Cnot6l | 0.3 | 1.55E-31 | 3.23E-27 | 0.56 | 0.5 | 0.06 | 1.12 |
| mesenchyme | Suv39h1 | 0.39 | 1.56E-31 | 3.25E-27 | 0.48 | 0.44 | 0.04 | 1.1 |
| mesenchyme | Rnf26 | 0.57 | 1.80E-31 | 3.74E-27 | 0.38 | 0.33 | 0.05 | 1.14 |
| mesenchyme | Spdl1 | 0.44 | 1.86E-31 | 3.87E-27 | 0.45 | 0.4 | 0.05 | 1.12 |
| mesenchyme | Ice2 | 0.67 | 2.16E-31 | 4.48E-27 | 0.27 | 0.21 | 0.06 | 1.3 |
| mesenchyme | Bud13 | 0.53 | 2.18E-31 | 4.53E-27 | 0.31 | 0.24 | 0.06 | 1.26 |

**Supplementary Table 2. Marker genes for the major cell types of integrated clustering of E11.5 control and NotchGOF λ-junction cells.**

| | | | | | | | | |
|---|---|---|---|---|---|---|---|---|
| mesenchyme | Snrnp25 | 0.27 | 2.19E-31 | 4.56E-27 | 0.63 | 0.59 | 0.04 | 1.07 |
| mesenchyme | Ubxn6 | 0.32 | 2.28E-31 | 4.74E-27 | 0.54 | 0.49 | 0.04 | 1.09 |
| mesenchyme | Parg | 0.33 | 2.36E-31 | 4.90E-27 | 0.51 | 0.46 | 0.05 | 1.12 |
| mesenchyme | Slc16a2 | 0.83 | 2.55E-31 | 5.30E-27 | 0.17 | 0.11 | 0.06 | 1.52 |
| mesenchyme | Phf21a | 0.35 | 2.58E-31 | 5.36E-27 | 0.49 | 0.44 | 0.06 | 1.13 |
| mesenchyme | ·110056K07Ri | 0.64 | 2.64E-31 | 5.49E-27 | 0.26 | 0.2 | 0.06 | 1.3 |
| mesenchyme | Adam11 | 1.32 | 2.67E-31 | 5.56E-27 | 0.1 | 0.05 | 0.05 | 1.91 |
| mesenchyme | Ncald | 1.18 | 2.86E-31 | 5.94E-27 | 0.11 | 0.06 | 0.05 | 1.82 |
| mesenchyme | Spast | 0.27 | 3.05E-31 | 6.35E-27 | 0.59 | 0.55 | 0.05 | 1.08 |
| mesenchyme | Rpp30 | 0.31 | 3.19E-31 | 6.64E-27 | 0.57 | 0.54 | 0.04 | 1.07 |
| mesenchyme | Phrf1 | 0.3 | 3.35E-31 | 6.96E-27 | 0.55 | 0.5 | 0.05 | 1.1 |
| mesenchyme | Ccdc32 | 0.5 | 3.60E-31 | 7.48E-27 | 0.33 | 0.27 | 0.06 | 1.23 |
| mesenchyme | Nfyc | 0.28 | 3.71E-31 | 7.73E-27 | 0.61 | 0.57 | 0.04 | 1.07 |
| mesenchyme | Nudt16l1 | 0.29 | 3.94E-31 | 8.20E-27 | 0.54 | 0.48 | 0.06 | 1.11 |
| mesenchyme | Krr1 | 0.3 | 3.98E-31 | 8.28E-27 | 0.53 | 0.48 | 0.05 | 1.1 |
| mesenchyme | Zfp768 | 0.46 | 4.28E-31 | 8.90E-27 | 0.36 | 0.29 | 0.06 | 1.21 |
| mesenchyme | Farsb | 0.28 | 4.51E-31 | 9.38E-27 | 0.58 | 0.54 | 0.04 | 1.07 |
| mesenchyme | Gsn | 0.55 | 5.07E-31 | 1.06E-26 | 0.16 | 0.11 | 0.06 | 1.56 |
| mesenchyme | Unc119b | 0.5 | 5.12E-31 | 1.06E-26 | 0.37 | 0.31 | 0.06 | 1.19 |
| mesenchyme | Aqr | 0.28 | 5.22E-31 | 1.08E-26 | 0.57 | 0.52 | 0.05 | 1.1 |
| mesenchyme | Nckap1 | 0.27 | 5.94E-31 | 1.24E-26 | 0.59 | 0.53 | 0.05 | 1.1 |
| mesenchyme | 530601H04Ri | 0.44 | 6.19E-31 | 1.29E-26 | 0.44 | 0.39 | 0.05 | 1.13 |
| mesenchyme | Bag2 | 0.44 | 6.30E-31 | 1.31E-26 | 0.42 | 0.37 | 0.05 | 1.15 |
| mesenchyme | Ercc6l | 0.47 | 6.44E-31 | 1.34E-26 | 0.38 | 0.32 | 0.06 | 1.17 |
| mesenchyme | Ciz1 | 0.51 | 6.76E-31 | 1.41E-26 | 0.36 | 0.31 | 0.06 | 1.18 |
| mesenchyme | Cd1d1 | 0.88 | 6.88E-31 | 1.43E-26 | 0.19 | 0.13 | 0.06 | 1.47 |
| mesenchyme | Cad | 0.43 | 7.31E-31 | 1.52E-26 | 0.4 | 0.35 | 0.06 | 1.17 |
| mesenchyme | Zc4h2 | 0.76 | 8.03E-31 | 1.67E-26 | 0.21 | 0.15 | 0.06 | 1.4 |
| mesenchyme | Usp28 | 0.67 | 8.52E-31 | 1.77E-26 | 0.25 | 0.18 | 0.06 | 1.34 |
| mesenchyme | Eed | 0.28 | 8.64E-31 | 1.80E-26 | 0.56 | 0.51 | 0.05 | 1.11 |
| mesenchyme | Gprasp1 | 0.35 | 1.02E-30 | 2.11E-26 | 0.51 | 0.45 | 0.06 | 1.12 |
| mesenchyme | Cdk5r1 | 0.84 | 1.39E-30 | 2.89E-26 | 0.13 | 0.07 | 0.05 | 1.73 |
| mesenchyme | Nipsnap2 | 0.34 | 1.47E-30 | 3.05E-26 | 0.47 | 0.42 | 0.05 | 1.12 |
| mesenchyme | ·730409E04Ri | 1.24 | 1.47E-30 | 3.06E-26 | 0.1 | 0.05 | 0.05 | 1.87 |
| mesenchyme | Prpf39 | 0.31 | 1.57E-30 | 3.26E-26 | 0.53 | 0.49 | 0.05 | 1.09 |
| mesenchyme | Stau2 | 0.43 | 1.60E-30 | 3.33E-26 | 0.4 | 0.34 | 0.06 | 1.17 |
| mesenchyme | Cplane1 | 0.46 | 2.43E-30 | 5.05E-26 | 0.37 | 0.31 | 0.06 | 1.19 |
| mesenchyme | Ccl25 | 0.57 | 2.62E-30 | 5.46E-26 | 0.29 | 0.23 | 0.06 | 1.26 |
| mesenchyme | Klhdc8b | 0.9 | 2.90E-30 | 6.03E-26 | 0.18 | 0.12 | 0.06 | 1.47 |
| mesenchyme | Tmem101 | 0.3 | 2.91E-30 | 6.05E-26 | 0.54 | 0.49 | 0.05 | 1.11 |
| mesenchyme | Rnf182 | 0.27 | 3.32E-30 | 6.91E-26 | 0.24 | 0.17 | 0.06 | 1.37 |
| mesenchyme | Cdc25c | 0.55 | 4.73E-30 | 9.84E-26 | 0.37 | 0.31 | 0.05 | 1.17 |
| mesenchyme | Pycrl | 0.28 | 5.06E-30 | 1.05E-25 | 0.57 | 0.52 | 0.05 | 1.1 |
| mesenchyme | Ddx11 | 0.69 | 5.12E-30 | 1.06E-25 | 0.27 | 0.21 | 0.06 | 1.27 |
| mesenchyme | Rnf170 | 0.58 | 5.26E-30 | 1.09E-25 | 0.28 | 0.23 | 0.06 | 1.26 |
| mesenchyme | Megf6 | 0.69 | 5.51E-30 | 1.15E-25 | 0.23 | 0.17 | 0.06 | 1.35 |
| mesenchyme | Zfp248 | 0.88 | 5.77E-30 | 1.20E-25 | 0.17 | 0.11 | 0.06 | 1.49 |
| mesenchyme | Kdm3b | 0.37 | 6.44E-30 | 1.34E-25 | 0.47 | 0.41 | 0.06 | 1.14 |
| mesenchyme | Aifm2 | 1 | 6.65E-30 | 1.38E-25 | 0.12 | 0.07 | 0.05 | 1.72 |
| mesenchyme | Cenpn | 0.49 | 6.75E-30 | 1.40E-25 | 0.4 | 0.35 | 0.05 | 1.14 |
| mesenchyme | Vkorc1l1 | 0.46 | 6.93E-30 | 1.44E-25 | 0.42 | 0.37 | 0.05 | 1.14 |
| mesenchyme | Ado | 0.36 | 6.99E-30 | 1.45E-25 | 0.41 | 0.36 | 0.05 | 1.14 |
| mesenchyme | Tlcd2 | 1.05 | 7.10E-30 | 1.48E-25 | 0.16 | 0.1 | 0.05 | 1.53 |
| mesenchyme | Vopp1 | 0.63 | 7.76E-30 | 1.61E-25 | 0.3 | 0.24 | 0.06 | 1.23 |
| mesenchyme | Uxs1 | 0.58 | 7.85E-30 | 1.63E-25 | 0.33 | 0.28 | 0.05 | 1.2 |
| mesenchyme | Pacsin2 | 0.45 | 8.99E-30 | 1.87E-25 | 0.37 | 0.31 | 0.06 | 1.18 |

**Supplementary Table 2. Marker genes for the major cell types of integrated clustering of E11.5 control and NotchGOF λ-junction cells.**

| | | | | | | | | |
|---|---|---|---|---|---|---|---|---|
| mesenchyme | 010003L21Ri | 0.52 | 9.47E-30 | 1.97E-25 | 0.33 | 0.27 | 0.06 | 1.21 |
| mesenchyme | Atf1 | 0.37 | 9.78E-30 | 2.03E-25 | 0.51 | 0.48 | 0.03 | 1.07 |
| mesenchyme | Cdca2 | 0.48 | 1.03E-29 | 2.15E-25 | 0.45 | 0.4 | 0.05 | 1.12 |
| mesenchyme | Exog | 0.66 | 1.03E-29 | 2.15E-25 | 0.27 | 0.21 | 0.06 | 1.28 |
| mesenchyme | Ubtd2 | 0.47 | 1.21E-29 | 2.51E-25 | 0.35 | 0.28 | 0.06 | 1.22 |
| mesenchyme | Pkd1 | 0.86 | 1.38E-29 | 2.87E-25 | 0.19 | 0.13 | 0.06 | 1.43 |
| mesenchyme | Mrps31 | 0.28 | 1.40E-29 | 2.91E-25 | 0.53 | 0.49 | 0.04 | 1.08 |
| mesenchyme | Cebpd | 0.78 | 1.43E-29 | 2.97E-25 | 0.12 | 0.07 | 0.05 | 1.76 |
| mesenchyme | Sdad1 | 0.33 | 1.45E-29 | 3.01E-25 | 0.49 | 0.44 | 0.05 | 1.12 |
| mesenchyme | Pcgf2 | 0.39 | 1.45E-29 | 3.02E-25 | 0.41 | 0.35 | 0.06 | 1.17 |
| mesenchyme | Zfp189 | 0.77 | 1.52E-29 | 3.16E-25 | 0.13 | 0.08 | 0.05 | 1.68 |
| mesenchyme | Zfp882 | 0.79 | 1.64E-29 | 3.40E-25 | 0.16 | 0.1 | 0.06 | 1.55 |
| mesenchyme | Dcun1d4 | 0.77 | 1.67E-29 | 3.47E-25 | 0.21 | 0.15 | 0.06 | 1.39 |
| mesenchyme | Pwwp3b | 0.49 | 1.69E-29 | 3.51E-25 | 0.34 | 0.27 | 0.07 | 1.24 |
| mesenchyme | Gtf2h2 | 0.45 | 1.73E-29 | 3.59E-25 | 0.4 | 0.35 | 0.05 | 1.15 |
| mesenchyme | Hist1h4d | 0.56 | 1.83E-29 | 3.80E-25 | 0.47 | 0.43 | 0.04 | 1.09 |
| mesenchyme | Prep | 0.46 | 2.05E-29 | 4.27E-25 | 0.41 | 0.36 | 0.05 | 1.15 |
| mesenchyme | 5720427I07Ril | 0.67 | 2.16E-29 | 4.50E-25 | 0.22 | 0.16 | 0.06 | 1.37 |
| mesenchyme | Camsap2 | 0.46 | 2.23E-29 | 4.64E-25 | 0.34 | 0.28 | 0.06 | 1.22 |
| mesenchyme | Fam102b | 0.63 | 2.27E-29 | 4.72E-25 | 0.22 | 0.16 | 0.06 | 1.36 |
| mesenchyme | Coil | 0.4 | 2.30E-29 | 4.79E-25 | 0.41 | 0.35 | 0.06 | 1.16 |
| mesenchyme | Dtx3 | 0.31 | 2.42E-29 | 5.02E-25 | 0.49 | 0.44 | 0.05 | 1.12 |
| mesenchyme | Speg | 0.83 | 2.75E-29 | 5.71E-25 | 0.18 | 0.13 | 0.06 | 1.45 |
| mesenchyme | Rtl8b | 0.38 | 2.88E-29 | 6.00E-25 | 0.36 | 0.29 | 0.06 | 1.22 |
| mesenchyme | Prpf3 | 0.42 | 2.91E-29 | 6.05E-25 | 0.41 | 0.36 | 0.05 | 1.14 |
| mesenchyme | Rybp | 0.37 | 2.95E-29 | 6.14E-25 | 0.44 | 0.39 | 0.05 | 1.14 |
| mesenchyme | Smim15 | 0.26 | 3.15E-29 | 6.54E-25 | 0.59 | 0.56 | 0.03 | 1.06 |
| mesenchyme | Kansl3 | 0.43 | 3.47E-29 | 7.21E-25 | 0.4 | 0.34 | 0.06 | 1.16 |
| mesenchyme | Enkd1 | 0.6 | 3.59E-29 | 7.46E-25 | 0.3 | 0.24 | 0.06 | 1.24 |
| mesenchyme | Gabbr1 | 0.61 | 3.64E-29 | 7.56E-25 | 0.26 | 0.2 | 0.06 | 1.3 |
| mesenchyme | Gm37494 | 0.47 | 3.76E-29 | 7.82E-25 | 0.36 | 0.3 | 0.06 | 1.19 |
| mesenchyme | Zfp871 | 0.28 | 3.79E-29 | 7.88E-25 | 0.6 | 0.55 | 0.05 | 1.09 |
| mesenchyme | Mcm2 | 0.27 | 4.30E-29 | 8.94E-25 | 0.64 | 0.59 | 0.04 | 1.07 |
| mesenchyme | Rnf4 | 0.26 | 5.48E-29 | 1.14E-24 | 0.64 | 0.61 | 0.03 | 1.06 |
| mesenchyme | Nans | 0.26 | 5.63E-29 | 1.17E-24 | 0.61 | 0.59 | 0.02 | 1.04 |
| mesenchyme | Tent2 | 0.35 | 7.24E-29 | 1.51E-24 | 0.5 | 0.46 | 0.04 | 1.09 |
| mesenchyme | Prkar2a | 0.26 | 7.86E-29 | 1.64E-24 | 0.54 | 0.48 | 0.06 | 1.13 |
| mesenchyme | Afap1 | 0.47 | 8.05E-29 | 1.67E-24 | 0.28 | 0.21 | 0.06 | 1.3 |
| mesenchyme | Zup1 | 0.46 | 8.17E-29 | 1.70E-24 | 0.38 | 0.32 | 0.05 | 1.17 |
| mesenchyme | Specc1l | 0.38 | 8.26E-29 | 1.72E-24 | 0.44 | 0.39 | 0.05 | 1.14 |
| mesenchyme | Mfhas1 | 0.76 | 8.62E-29 | 1.79E-24 | 0.21 | 0.16 | 0.06 | 1.37 |
| mesenchyme | Fem1c | 0.38 | 8.82E-29 | 1.83E-24 | 0.4 | 0.34 | 0.06 | 1.16 |
| mesenchyme | Mcmbp | 0.25 | 9.02E-29 | 1.88E-24 | 0.59 | 0.53 | 0.05 | 1.1 |
| mesenchyme | Ankmy2 | 0.52 | 9.16E-29 | 1.91E-24 | 0.31 | 0.25 | 0.06 | 1.24 |
| mesenchyme | Pibf1 | 0.43 | 9.50E-29 | 1.98E-24 | 0.39 | 0.33 | 0.05 | 1.16 |
| mesenchyme | Usp40 | 0.52 | 1.01E-28 | 2.11E-24 | 0.36 | 0.3 | 0.05 | 1.17 |
| mesenchyme | Cc2d2a | 0.68 | 1.10E-28 | 2.30E-24 | 0.23 | 0.17 | 0.06 | 1.34 |
| mesenchyme | Srcap | 0.33 | 1.15E-28 | 2.39E-24 | 0.52 | 0.47 | 0.04 | 1.09 |
| mesenchyme | Edn3 | 0.93 | 1.34E-28 | 2.79E-24 | 0.16 | 0.11 | 0.05 | 1.5 |
| mesenchyme | Nxf1 | 0.32 | 1.38E-28 | 2.86E-24 | 0.5 | 0.46 | 0.04 | 1.09 |
| mesenchyme | Manbal | 0.26 | 1.61E-28 | 3.35E-24 | 0.55 | 0.5 | 0.05 | 1.09 |
| mesenchyme | Ccny | 0.29 | 1.70E-28 | 3.54E-24 | 0.55 | 0.51 | 0.04 | 1.08 |
| mesenchyme | Cxxc1 | 0.35 | 1.74E-28 | 3.62E-24 | 0.46 | 0.42 | 0.05 | 1.11 |
| mesenchyme | Lsg1 | 0.26 | 1.80E-28 | 3.75E-24 | 0.57 | 0.53 | 0.04 | 1.08 |
| mesenchyme | Tada2a | 0.37 | 2.04E-28 | 4.25E-24 | 0.44 | 0.38 | 0.06 | 1.14 |
| mesenchyme | Lhpp | 1.05 | 2.24E-28 | 4.66E-24 | 0.12 | 0.07 | 0.05 | 1.68 |

**Supplementary Table 2. Marker genes for the major cell types of integrated clustering of E11.5 control and NotchGOF λ-junction cells.**

| | | | | | | | | |
|---|---|---|---|---|---|---|---|---|
| mesenchyme | Degs1 | 0.34 | 2.25E-28 | 4.68E-24 | 0.42 | 0.36 | 0.06 | 1.16 |
| mesenchyme | Gmpr2 | 0.37 | 2.37E-28 | 4.94E-24 | 0.44 | 0.39 | 0.05 | 1.13 |
| mesenchyme | Camsap1 | 0.33 | 2.42E-28 | 5.02E-24 | 0.5 | 0.45 | 0.05 | 1.11 |
| mesenchyme | Surf4 | 0.27 | 2.56E-28 | 5.33E-24 | 0.61 | 0.56 | 0.05 | 1.09 |
| mesenchyme | Atp11c | 0.66 | 2.57E-28 | 5.35E-24 | 0.27 | 0.21 | 0.06 | 1.27 |
| mesenchyme | Phlda1 | 0.36 | 2.69E-28 | 5.60E-24 | 0.44 | 0.38 | 0.06 | 1.15 |
| mesenchyme | Slc25a25 | 0.72 | 2.80E-28 | 5.82E-24 | 0.2 | 0.14 | 0.06 | 1.39 |
| mesenchyme | Zfp507 | 0.41 | 2.85E-28 | 5.92E-24 | 0.41 | 0.35 | 0.06 | 1.16 |
| mesenchyme | Ints12 | 0.59 | 2.90E-28 | 6.03E-24 | 0.32 | 0.26 | 0.05 | 1.21 |
| mesenchyme | Kctd5 | 0.27 | 3.15E-28 | 6.55E-24 | 0.55 | 0.5 | 0.05 | 1.1 |
| mesenchyme | Emc4 | 0.26 | 4.47E-28 | 9.29E-24 | 0.61 | 0.56 | 0.05 | 1.09 |
| mesenchyme | Spag5 | 0.57 | 4.53E-28 | 9.42E-24 | 0.32 | 0.27 | 0.05 | 1.2 |
| mesenchyme | Ankrd46 | 0.46 | 4.66E-28 | 9.69E-24 | 0.37 | 0.31 | 0.06 | 1.18 |
| mesenchyme | Fancm | 0.47 | 4.84E-28 | 1.01E-23 | 0.38 | 0.33 | 0.05 | 1.17 |
| mesenchyme | Spout1 | 0.38 | 4.92E-28 | 1.02E-23 | 0.45 | 0.4 | 0.05 | 1.12 |
| mesenchyme | Lrpprc | 0.32 | 4.96E-28 | 1.03E-23 | 0.49 | 0.45 | 0.04 | 1.09 |
| mesenchyme | 933404O12Ri | 0.6 | 5.30E-28 | 1.10E-23 | 0.27 | 0.21 | 0.06 | 1.26 |
| mesenchyme | Mcm3ap | 0.51 | 5.51E-28 | 1.15E-23 | 0.36 | 0.3 | 0.05 | 1.18 |
| mesenchyme | Hist1h3a | 0.81 | 6.29E-28 | 1.31E-23 | 0.21 | 0.16 | 0.05 | 1.34 |
| mesenchyme | Tet2 | 0.55 | 6.42E-28 | 1.33E-23 | 0.27 | 0.21 | 0.06 | 1.29 |
| mesenchyme | Zmym1 | 0.44 | 6.60E-28 | 1.37E-23 | 0.36 | 0.3 | 0.05 | 1.18 |
| mesenchyme | Vhl | 0.65 | 6.83E-28 | 1.42E-23 | 0.26 | 0.2 | 0.06 | 1.28 |
| mesenchyme | Rapgef2 | 0.67 | 7.00E-28 | 1.46E-23 | 0.24 | 0.19 | 0.06 | 1.31 |
| mesenchyme | Clpx | 0.33 | 7.15E-28 | 1.49E-23 | 0.49 | 0.44 | 0.05 | 1.1 |
| mesenchyme | Pole2 | 0.59 | 7.57E-28 | 1.57E-23 | 0.3 | 0.25 | 0.05 | 1.21 |
| mesenchyme | Nif3l1 | 0.42 | 7.95E-28 | 1.65E-23 | 0.38 | 0.32 | 0.05 | 1.17 |
| mesenchyme | Ophn1 | 0.68 | 8.22E-28 | 1.71E-23 | 0.21 | 0.15 | 0.06 | 1.38 |
| mesenchyme | Rrn3 | 0.45 | 8.49E-28 | 1.77E-23 | 0.38 | 0.33 | 0.05 | 1.15 |
| mesenchyme | Asrgl1 | 0.5 | 9.68E-28 | 2.01E-23 | 0.34 | 0.29 | 0.05 | 1.18 |
| mesenchyme | Nedd1 | 0.49 | 9.90E-28 | 2.06E-23 | 0.36 | 0.31 | 0.05 | 1.16 |
| mesenchyme | Ttl | 0.63 | 1.04E-27 | 2.17E-23 | 0.25 | 0.19 | 0.06 | 1.32 |
| mesenchyme | Senp3 | 0.32 | 1.07E-27 | 2.23E-23 | 0.49 | 0.45 | 0.05 | 1.11 |
| mesenchyme | Cmss1 | 0.29 | 1.52E-27 | 3.17E-23 | 0.51 | 0.47 | 0.04 | 1.09 |
| mesenchyme | .700062C07Ri | 0.41 | 1.69E-27 | 3.51E-23 | 0.4 | 0.35 | 0.05 | 1.14 |
| mesenchyme | Itfg2 | 0.49 | 1.71E-27 | 3.55E-23 | 0.36 | 0.31 | 0.05 | 1.17 |
| mesenchyme | Mbtps1 | 0.25 | 2.01E-27 | 4.18E-23 | 0.57 | 0.52 | 0.05 | 1.1 |
| mesenchyme | Nfya | 0.54 | 2.34E-27 | 4.87E-23 | 0.34 | 0.29 | 0.05 | 1.18 |
| mesenchyme | Rab39b | 0.68 | 2.62E-27 | 5.44E-23 | 0.18 | 0.12 | 0.06 | 1.46 |
| mesenchyme | Ccs | 0.36 | 2.81E-27 | 5.85E-23 | 0.47 | 0.42 | 0.04 | 1.1 |
| mesenchyme | Mrpl16 | 0.3 | 3.38E-27 | 7.04E-23 | 0.52 | 0.49 | 0.03 | 1.06 |
| mesenchyme | Gk5 | 1.04 | 3.50E-27 | 7.28E-23 | 0.11 | 0.06 | 0.05 | 1.72 |
| mesenchyme | Haus4 | 0.52 | 3.81E-27 | 7.92E-23 | 0.33 | 0.29 | 0.05 | 1.17 |
| mesenchyme | Rexo4 | 0.27 | 4.07E-27 | 8.46E-23 | 0.47 | 0.42 | 0.05 | 1.11 |
| mesenchyme | .700021F05Ri | 0.35 | 4.19E-27 | 8.72E-23 | 0.48 | 0.44 | 0.04 | 1.09 |
| mesenchyme | Btbd2 | 0.25 | 4.29E-27 | 8.93E-23 | 0.54 | 0.49 | 0.06 | 1.12 |
| mesenchyme | Dscc1 | 0.5 | 4.41E-27 | 9.16E-23 | 0.32 | 0.27 | 0.06 | 1.21 |
| mesenchyme | Tmem138 | 0.38 | 4.56E-27 | 9.49E-23 | 0.41 | 0.35 | 0.06 | 1.16 |
| mesenchyme | Dhx38 | 0.38 | 4.59E-27 | 9.55E-23 | 0.47 | 0.42 | 0.05 | 1.11 |
| mesenchyme | Exosc1 | 0.26 | 4.65E-27 | 9.67E-23 | 0.56 | 0.52 | 0.04 | 1.08 |
| mesenchyme | Crybg3 | 0.62 | 4.86E-27 | 1.01E-22 | 0.23 | 0.17 | 0.06 | 1.33 |
| mesenchyme | Zfp811 | 1.01 | 4.89E-27 | 1.02E-22 | 0.14 | 0.09 | 0.05 | 1.56 |
| mesenchyme | Mrpl1 | 0.31 | 5.20E-27 | 1.08E-22 | 0.47 | 0.43 | 0.04 | 1.09 |
| mesenchyme | Tbc1d10b | 0.45 | 5.53E-27 | 1.15E-22 | 0.37 | 0.32 | 0.05 | 1.16 |
| mesenchyme | Smyd3 | 0.64 | 5.86E-27 | 1.22E-22 | 0.26 | 0.2 | 0.06 | 1.27 |
| mesenchyme | Ncaph | 0.32 | 6.28E-27 | 1.31E-22 | 0.5 | 0.47 | 0.04 | 1.08 |
| mesenchyme | Zfp60 | 0.35 | 6.54E-27 | 1.36E-22 | 0.44 | 0.39 | 0.06 | 1.14 |

**Supplementary Table 2. Marker genes for the major cell types of integrated clustering of E11.5 control and NotchGOF λ-junction cells.**

| | | | | | | | | |
|---|---|---|---|---|---|---|---|---|
| mesenchyme | Ranbp3 | 0.33 | 6.95E-27 | 1.45E-22 | 0.5 | 0.47 | 0.04 | 1.08 |
| mesenchyme | Ckap2 | 0.47 | 7.08E-27 | 1.47E-22 | 0.41 | 0.37 | 0.05 | 1.12 |
| mesenchyme | Sgo2a | 0.43 | 7.30E-27 | 1.52E-22 | 0.5 | 0.45 | 0.04 | 1.09 |
| mesenchyme | Rab22a | 0.31 | 8.01E-27 | 1.67E-22 | 0.49 | 0.45 | 0.04 | 1.09 |
| mesenchyme | Rhobtb2 | 0.98 | 8.44E-27 | 1.76E-22 | 0.14 | 0.09 | 0.05 | 1.56 |
| mesenchyme | Nme7 | 0.36 | 1.08E-26 | 2.25E-22 | 0.41 | 0.36 | 0.05 | 1.14 |
| mesenchyme | Rngtt | 0.42 | 1.22E-26 | 2.53E-22 | 0.41 | 0.37 | 0.04 | 1.12 |
| mesenchyme | Arhgap35 | 0.3 | 1.27E-26 | 2.64E-22 | 0.47 | 0.41 | 0.06 | 1.14 |
| mesenchyme | Chst2 | 0.79 | 1.35E-26 | 2.82E-22 | 0.22 | 0.17 | 0.06 | 1.33 |
| mesenchyme | Tmem184c | 0.43 | 1.38E-26 | 2.87E-22 | 0.37 | 0.31 | 0.06 | 1.19 |
| mesenchyme | Rbsn | 0.55 | 1.41E-26 | 2.94E-22 | 0.29 | 0.23 | 0.05 | 1.23 |
| mesenchyme | Usp21 | 0.6 | 1.43E-26 | 2.97E-22 | 0.27 | 0.21 | 0.06 | 1.26 |
| mesenchyme | Purg | 0.44 | 1.44E-26 | 2.99E-22 | 0.31 | 0.25 | 0.06 | 1.22 |
| mesenchyme | Nup93 | 0.32 | 1.51E-26 | 3.14E-22 | 0.49 | 0.44 | 0.04 | 1.09 |
| mesenchyme | Tmem39a | 0.46 | 1.54E-26 | 3.20E-22 | 0.38 | 0.33 | 0.05 | 1.16 |
| mesenchyme | Ppwd1 | 0.34 | 1.58E-26 | 3.28E-22 | 0.51 | 0.47 | 0.04 | 1.08 |
| mesenchyme | Heatr6 | 0.36 | 1.63E-26 | 3.39E-22 | 0.46 | 0.42 | 0.04 | 1.11 |
| mesenchyme | Ppm1d | 0.44 | 1.77E-26 | 3.67E-22 | 0.4 | 0.36 | 0.04 | 1.11 |
| mesenchyme | Edc4 | 0.44 | 1.85E-26 | 3.85E-22 | 0.35 | 0.3 | 0.05 | 1.18 |
| mesenchyme | Strip1 | 0.49 | 2.02E-26 | 4.20E-22 | 0.36 | 0.31 | 0.05 | 1.17 |
| mesenchyme | 410004B18Ri | 0.26 | 2.29E-26 | 4.76E-22 | 0.53 | 0.49 | 0.04 | 1.09 |
| mesenchyme | Scaf8 | 0.34 | 2.43E-26 | 5.06E-22 | 0.42 | 0.37 | 0.05 | 1.14 |
| mesenchyme | Cmtr1 | 0.48 | 2.44E-26 | 5.08E-22 | 0.35 | 0.31 | 0.05 | 1.16 |
| mesenchyme | Fra10ac1 | 0.32 | 2.59E-26 | 5.39E-22 | 0.49 | 0.44 | 0.05 | 1.11 |
| mesenchyme | Edrf1 | 0.51 | 2.63E-26 | 5.47E-22 | 0.31 | 0.25 | 0.05 | 1.21 |
| mesenchyme | St6gal2 | 0.99 | 2.77E-26 | 5.75E-22 | 0.12 | 0.07 | 0.05 | 1.66 |
| mesenchyme | Utp15 | 0.36 | 2.86E-26 | 5.95E-22 | 0.41 | 0.36 | 0.05 | 1.14 |
| mesenchyme | Arhgef2 | 0.33 | 2.94E-26 | 6.11E-22 | 0.45 | 0.39 | 0.06 | 1.15 |
| mesenchyme | Cdc45 | 0.43 | 3.13E-26 | 6.51E-22 | 0.39 | 0.33 | 0.05 | 1.15 |
| mesenchyme | Xpo4 | 0.39 | 3.39E-26 | 7.05E-22 | 0.45 | 0.41 | 0.04 | 1.1 |
| mesenchyme | Zfp763 | 0.72 | 3.39E-26 | 7.06E-22 | 0.2 | 0.14 | 0.06 | 1.39 |
| mesenchyme | Tex30 | 0.33 | 4.09E-26 | 8.51E-22 | 0.49 | 0.45 | 0.04 | 1.09 |
| mesenchyme | Hgsnat | 0.73 | 4.31E-26 | 8.96E-22 | 0.22 | 0.16 | 0.05 | 1.33 |
| mesenchyme | Ddx25 | 0.77 | 4.51E-26 | 9.37E-22 | 0.16 | 0.11 | 0.05 | 1.5 |
| mesenchyme | Fbxo21 | 0.39 | 4.60E-26 | 9.56E-22 | 0.4 | 0.35 | 0.05 | 1.15 |
| mesenchyme | Nphp1 | 0.4 | 4.79E-26 | 9.96E-22 | 0.32 | 0.26 | 0.06 | 1.21 |
| mesenchyme | Slit2 | 0.63 | 5.32E-26 | 1.11E-21 | 0.39 | 0.34 | 0.05 | 1.15 |
| mesenchyme | Ankle2 | 0.44 | 5.46E-26 | 1.14E-21 | 0.38 | 0.33 | 0.05 | 1.15 |
| mesenchyme | Telo2 | 0.61 | 5.79E-26 | 1.20E-21 | 0.32 | 0.27 | 0.05 | 1.18 |
| mesenchyme | Fam168a | 0.25 | 5.98E-26 | 1.24E-21 | 0.56 | 0.51 | 0.05 | 1.1 |
| mesenchyme | Kif20b | 0.31 | 6.18E-26 | 1.28E-21 | 0.57 | 0.53 | 0.04 | 1.09 |
| mesenchyme | Map1s | 0.43 | 7.01E-26 | 1.46E-21 | 0.39 | 0.34 | 0.05 | 1.15 |
| mesenchyme | Noc3l | 0.38 | 8.19E-26 | 1.70E-21 | 0.42 | 0.37 | 0.05 | 1.14 |
| mesenchyme | Prss23 | 0.62 | 8.43E-26 | 1.75E-21 | 0.33 | 0.28 | 0.05 | 1.19 |
| mesenchyme | B4galt2 | 0.51 | 8.64E-26 | 1.80E-21 | 0.29 | 0.23 | 0.06 | 1.24 |
| mesenchyme | Pogz | 0.43 | 9.53E-26 | 1.98E-21 | 0.39 | 0.35 | 0.05 | 1.14 |
| mesenchyme | Ddx47 | 0.31 | 9.60E-26 | 2.00E-21 | 0.52 | 0.49 | 0.03 | 1.06 |
| mesenchyme | Fut10 | 0.85 | 9.61E-26 | 2.00E-21 | 0.15 | 0.1 | 0.05 | 1.51 |
| mesenchyme | Cttnbp2 | 0.82 | 1.07E-25 | 2.23E-21 | 0.17 | 0.12 | 0.05 | 1.43 |
| mesenchyme | Hibch | 0.55 | 1.13E-25 | 2.36E-21 | 0.3 | 0.24 | 0.05 | 1.21 |
| mesenchyme | Anapc10 | 0.27 | 1.15E-25 | 2.40E-21 | 0.53 | 0.5 | 0.04 | 1.07 |
| mesenchyme | Large1 | 0.67 | 1.17E-25 | 2.43E-21 | 0.21 | 0.16 | 0.05 | 1.34 |
| mesenchyme | Gm10320 | 0.45 | 1.26E-25 | 2.62E-21 | 0.42 | 0.37 | 0.05 | 1.13 |
| mesenchyme | Panx1 | 0.47 | 1.29E-25 | 2.68E-21 | 0.33 | 0.28 | 0.06 | 1.2 |
| mesenchyme | Dmwd | 0.3 | 1.40E-25 | 2.91E-21 | 0.46 | 0.41 | 0.05 | 1.12 |
| mesenchyme | Maml3 | 0.41 | 1.40E-25 | 2.92E-21 | 0.42 | 0.37 | 0.05 | 1.13 |

**Supplementary Table 2. Marker genes for the major cell types of integrated clustering of E11.5 control and NotchGOF λ-junction cells.**

| | | | | | | | | |
|---|---|---|---|---|---|---|---|---|
| mesenchyme | Arl5b | 0.3 | 1.63E-25 | 3.38E-21 | 0.47 | 0.42 | 0.05 | 1.11 |
| mesenchyme | Pcgf1 | 0.77 | 1.73E-25 | 3.59E-21 | 0.19 | 0.14 | 0.05 | 1.36 |
| mesenchyme | Pik3ca | 0.37 | 1.94E-25 | 4.03E-21 | 0.45 | 0.41 | 0.05 | 1.11 |
| mesenchyme | Ptdss2 | 0.35 | 1.95E-25 | 4.06E-21 | 0.52 | 0.49 | 0.03 | 1.05 |
| mesenchyme | Srgap2 | 0.41 | 2.24E-25 | 4.66E-21 | 0.38 | 0.33 | 0.05 | 1.15 |
| mesenchyme | Spata6 | 0.63 | 2.41E-25 | 5.00E-21 | 0.22 | 0.17 | 0.06 | 1.33 |
| mesenchyme | Kif11 | 0.4 | 2.44E-25 | 5.07E-21 | 0.55 | 0.52 | 0.03 | 1.05 |
| mesenchyme | Cdca4 | 0.28 | 2.55E-25 | 5.31E-21 | 0.57 | 0.54 | 0.03 | 1.06 |
| mesenchyme | Thra | 0.4 | 2.69E-25 | 5.60E-21 | 0.44 | 0.39 | 0.04 | 1.11 |
| mesenchyme | Mettl2 | 0.41 | 2.81E-25 | 5.84E-21 | 0.38 | 0.32 | 0.05 | 1.16 |
| mesenchyme | Arl15 | 0.74 | 2.81E-25 | 5.85E-21 | 0.19 | 0.13 | 0.05 | 1.4 |
| mesenchyme | Wee1 | 0.3 | 2.92E-25 | 6.08E-21 | 0.48 | 0.43 | 0.05 | 1.11 |
| mesenchyme | Epm2aip1 | 0.32 | 2.99E-25 | 6.22E-21 | 0.44 | 0.39 | 0.05 | 1.14 |
| mesenchyme | Terf2ip | 0.37 | 3.04E-25 | 6.32E-21 | 0.4 | 0.35 | 0.05 | 1.14 |
| mesenchyme | Rprd2 | 0.35 | 3.14E-25 | 6.52E-21 | 0.45 | 0.41 | 0.04 | 1.1 |
| mesenchyme | Tmem218 | 0.49 | 3.38E-25 | 7.03E-21 | 0.32 | 0.26 | 0.06 | 1.21 |
| mesenchyme | Ppp2r3c | 0.28 | 3.40E-25 | 7.07E-21 | 0.48 | 0.44 | 0.04 | 1.09 |
| mesenchyme | Tmem41a | 0.65 | 3.45E-25 | 7.18E-21 | 0.24 | 0.19 | 0.05 | 1.28 |
| mesenchyme | Polr3f | 0.45 | 3.81E-25 | 7.92E-21 | 0.36 | 0.31 | 0.05 | 1.17 |
| mesenchyme | Dusp12 | 0.36 | 3.87E-25 | 8.04E-21 | 0.43 | 0.38 | 0.05 | 1.13 |
| mesenchyme | Tspan2 | 0.91 | 4.08E-25 | 8.48E-21 | 0.12 | 0.07 | 0.05 | 1.64 |
| mesenchyme | Tle3 | 0.35 | 4.13E-25 | 8.60E-21 | 0.45 | 0.41 | 0.05 | 1.12 |
| mesenchyme | Pus7l | 0.81 | 4.17E-25 | 8.67E-21 | 0.17 | 0.12 | 0.05 | 1.44 |
| mesenchyme | Batf3 | 0.79 | 4.52E-25 | 9.40E-21 | 0.17 | 0.12 | 0.05 | 1.42 |
| mesenchyme | Pcdhga12 | 0.9 | 4.56E-25 | 9.49E-21 | 0.14 | 0.09 | 0.05 | 1.51 |
| mesenchyme | Pdzd4 | 0.8 | 4.83E-25 | 1.00E-20 | 0.17 | 0.12 | 0.05 | 1.44 |
| mesenchyme | Tmem222 | 0.35 | 5.35E-25 | 1.11E-20 | 0.46 | 0.42 | 0.04 | 1.09 |
| mesenchyme | Acap2 | 0.33 | 5.82E-25 | 1.21E-20 | 0.48 | 0.44 | 0.04 | 1.1 |
| mesenchyme | Rilpl2 | 0.48 | 6.02E-25 | 1.25E-20 | 0.3 | 0.24 | 0.06 | 1.23 |
| mesenchyme | Ppat | 0.26 | 6.17E-25 | 1.28E-20 | 0.52 | 0.47 | 0.04 | 1.09 |
| mesenchyme | Anln | 0.47 | 6.79E-25 | 1.41E-20 | 0.4 | 0.36 | 0.04 | 1.11 |
| mesenchyme | Mdfic | 0.85 | 6.87E-25 | 1.43E-20 | 0.24 | 0.19 | 0.05 | 1.26 |
| mesenchyme | Cenpu | 0.51 | 7.02E-25 | 1.46E-20 | 0.33 | 0.28 | 0.05 | 1.16 |
| mesenchyme | Sema4c | 0.49 | 7.08E-25 | 1.47E-20 | 0.29 | 0.24 | 0.06 | 1.23 |
| mesenchyme | 730480H06Ri | 0.4 | 7.59E-25 | 1.58E-20 | 0.36 | 0.31 | 0.05 | 1.17 |
| mesenchyme | Cyp20a1 | 0.62 | 8.24E-25 | 1.71E-20 | 0.27 | 0.22 | 0.05 | 1.22 |
| mesenchyme | Hic2 | 0.47 | 8.93E-25 | 1.86E-20 | 0.31 | 0.26 | 0.05 | 1.2 |
| mesenchyme | Zcchc10 | 0.41 | 9.32E-25 | 1.94E-20 | 0.37 | 0.32 | 0.05 | 1.15 |
| mesenchyme | Gm26724 | 0.86 | 1.01E-24 | 2.10E-20 | 0.17 | 0.12 | 0.05 | 1.42 |
| mesenchyme | Rab4a | 0.38 | 1.01E-24 | 2.11E-20 | 0.37 | 0.32 | 0.06 | 1.17 |
| mesenchyme | Nfil3 | 0.63 | 1.02E-24 | 2.12E-20 | 0.15 | 0.1 | 0.05 | 1.48 |
| mesenchyme | Dlg4 | 0.42 | 1.05E-24 | 2.18E-20 | 0.33 | 0.28 | 0.06 | 1.2 |
| mesenchyme | Ephb6 | 1.17 | 1.14E-24 | 2.37E-20 | 0.1 | 0.06 | 0.04 | 1.71 |
| mesenchyme | Usp2 | 0.9 | 1.18E-24 | 2.45E-20 | 0.16 | 0.11 | 0.05 | 1.45 |
| mesenchyme | Sema6c | 0.73 | 1.47E-24 | 3.06E-20 | 0.18 | 0.13 | 0.05 | 1.41 |
| mesenchyme | L3hypdh | 0.59 | 1.56E-24 | 3.25E-20 | 0.21 | 0.15 | 0.05 | 1.36 |
| mesenchyme | Kpna1 | 0.38 | 1.71E-24 | 3.56E-20 | 0.4 | 0.36 | 0.04 | 1.12 |
| mesenchyme | Gle1 | 0.33 | 2.03E-24 | 4.21E-20 | 0.49 | 0.46 | 0.03 | 1.06 |
| mesenchyme | Plod3 | 0.5 | 2.07E-24 | 4.31E-20 | 0.34 | 0.29 | 0.05 | 1.16 |
| mesenchyme | Msh6 | 0.29 | 2.13E-24 | 4.43E-20 | 0.5 | 0.45 | 0.05 | 1.12 |
| mesenchyme | Eif2ak2 | 0.52 | 2.17E-24 | 4.52E-20 | 0.3 | 0.25 | 0.05 | 1.2 |
| mesenchyme | Rabep1 | 0.25 | 2.21E-24 | 4.60E-20 | 0.55 | 0.5 | 0.05 | 1.09 |
| mesenchyme | Utp20 | 0.41 | 2.56E-24 | 5.32E-20 | 0.35 | 0.3 | 0.05 | 1.17 |
| mesenchyme | Tor1a | 0.28 | 2.79E-24 | 5.81E-20 | 0.53 | 0.49 | 0.04 | 1.08 |
| mesenchyme | Ddx20 | 0.42 | 2.83E-24 | 5.88E-20 | 0.35 | 0.31 | 0.05 | 1.16 |
| mesenchyme | Ercc1 | 0.29 | 2.83E-24 | 5.90E-20 | 0.45 | 0.41 | 0.05 | 1.11 |

**Supplementary Table 2. Marker genes for the major cell types of integrated clustering of E11.5 control and NotchGOF λ-junction cells.**

| | | | | | | | | |
|---|---|---|---|---|---|---|---|---|
| mesenchyme | Lig3 | 0.33 | 3.12E-24 | 6.48E-20 | 0.45 | 0.4 | 0.05 | 1.12 |
| mesenchyme | Zfp821 | 0.28 | 3.14E-24 | 6.53E-20 | 0.47 | 0.42 | 0.05 | 1.12 |
| mesenchyme | Plagl1 | 0.7 | 3.25E-24 | 6.76E-20 | 0.45 | 0.42 | 0.03 | 1.07 |
| mesenchyme | Bmp7 | 0.3 | 3.30E-24 | 6.87E-20 | 0.4 | 0.34 | 0.06 | 1.18 |
| mesenchyme | Vamp4 | 0.34 | 3.43E-24 | 7.14E-20 | 0.45 | 0.4 | 0.04 | 1.1 |
| mesenchyme | Ctu2 | 0.46 | 3.68E-24 | 7.66E-20 | 0.35 | 0.3 | 0.05 | 1.16 |
| mesenchyme | Pcnx | 0.53 | 4.09E-24 | 8.51E-20 | 0.3 | 0.25 | 0.05 | 1.2 |
| mesenchyme | Bcl9l | 0.42 | 4.32E-24 | 8.99E-20 | 0.35 | 0.3 | 0.05 | 1.18 |
| mesenchyme | Wdyhv1 | 0.62 | 4.61E-24 | 9.59E-20 | 0.29 | 0.24 | 0.05 | 1.19 |
| mesenchyme | Tspan9 | 0.57 | 5.50E-24 | 1.14E-19 | 0.21 | 0.16 | 0.05 | 1.34 |
| mesenchyme | Mtx1 | 0.25 | 6.47E-24 | 1.34E-19 | 0.56 | 0.54 | 0.03 | 1.05 |
| mesenchyme | Trerf1 | 0.54 | 6.48E-24 | 1.35E-19 | 0.29 | 0.23 | 0.05 | 1.23 |
| mesenchyme | Dnaaf2 | 0.27 | 7.14E-24 | 1.48E-19 | 0.52 | 0.49 | 0.04 | 1.08 |
| mesenchyme | Lmf1 | 0.63 | 7.85E-24 | 1.63E-19 | 0.24 | 0.19 | 0.05 | 1.27 |
| mesenchyme | Sgcb | 0.63 | 8.44E-24 | 1.76E-19 | 0.23 | 0.17 | 0.05 | 1.3 |
| mesenchyme | Prmt7 | 0.28 | 8.47E-24 | 1.76E-19 | 0.52 | 0.49 | 0.03 | 1.07 |
| mesenchyme | Lnpep | 0.6 | 9.35E-24 | 1.94E-19 | 0.31 | 0.26 | 0.05 | 1.18 |
| mesenchyme | Tmco3 | 0.49 | 9.58E-24 | 1.99E-19 | 0.31 | 0.26 | 0.05 | 1.2 |
| mesenchyme | Kctd17 | 0.38 | 1.07E-23 | 2.22E-19 | 0.29 | 0.24 | 0.06 | 1.24 |
| mesenchyme | Depdc1a | 0.53 | 1.24E-23 | 2.58E-19 | 0.37 | 0.33 | 0.04 | 1.12 |
| mesenchyme | Mtmr11 | 0.82 | 1.29E-23 | 2.69E-19 | 0.12 | 0.07 | 0.05 | 1.62 |
| mesenchyme | Ccnt1 | 0.27 | 1.31E-23 | 2.73E-19 | 0.59 | 0.56 | 0.03 | 1.06 |
| mesenchyme | Sat2 | 0.85 | 1.35E-23 | 2.81E-19 | 0.15 | 0.1 | 0.05 | 1.47 |
| mesenchyme | Klhl12 | 0.53 | 1.48E-23 | 3.09E-19 | 0.29 | 0.24 | 0.05 | 1.21 |
| mesenchyme | Dnpep | 0.26 | 1.63E-23 | 3.38E-19 | 0.52 | 0.48 | 0.04 | 1.08 |
| mesenchyme | D16Ertd472e | 0.42 | 1.73E-23 | 3.59E-19 | 0.29 | 0.23 | 0.06 | 1.24 |
| mesenchyme | Zfp184 | 0.53 | 1.76E-23 | 3.67E-19 | 0.24 | 0.19 | 0.06 | 1.29 |
| mesenchyme | Ptcd2 | 0.31 | 1.84E-23 | 3.83E-19 | 0.46 | 0.42 | 0.04 | 1.1 |
| mesenchyme | Cacna1a | 0.5 | 1.88E-23 | 3.92E-19 | 0.32 | 0.27 | 0.05 | 1.19 |
| mesenchyme | Bap1 | 0.47 | 1.89E-23 | 3.94E-19 | 0.35 | 0.3 | 0.04 | 1.15 |
| mesenchyme | Gcn1 | 0.32 | 2.04E-23 | 4.25E-19 | 0.53 | 0.49 | 0.04 | 1.08 |
| mesenchyme | Smap2 | 0.33 | 2.42E-23 | 5.03E-19 | 0.39 | 0.35 | 0.04 | 1.13 |
| mesenchyme | Rin2 | 0.71 | 2.47E-23 | 5.13E-19 | 0.22 | 0.17 | 0.05 | 1.3 |
| mesenchyme | Ufd1 | 0.31 | 2.63E-23 | 5.46E-19 | 0.54 | 0.52 | 0.02 | 1.04 |
| mesenchyme | Zfp74 | 0.73 | 2.64E-23 | 5.50E-19 | 0.19 | 0.14 | 0.05 | 1.36 |
| mesenchyme | Ube2o | 0.4 | 2.68E-23 | 5.57E-19 | 0.4 | 0.37 | 0.04 | 1.1 |
| mesenchyme | Fam135a | 0.51 | 2.92E-23 | 6.07E-19 | 0.31 | 0.26 | 0.05 | 1.2 |
| mesenchyme | Cstf2t | 0.35 | 3.04E-23 | 6.32E-19 | 0.4 | 0.35 | 0.05 | 1.15 |
| mesenchyme | Crem | 0.54 | 3.15E-23 | 6.54E-19 | 0.2 | 0.15 | 0.05 | 1.36 |
| mesenchyme | Tmx4 | 0.37 | 3.26E-23 | 6.78E-19 | 0.48 | 0.45 | 0.03 | 1.07 |
| mesenchyme | Uimc1 | 0.5 | 4.09E-23 | 8.51E-19 | 0.33 | 0.28 | 0.04 | 1.15 |
| mesenchyme | Surf6 | 0.49 | 4.70E-23 | 9.77E-19 | 0.29 | 0.24 | 0.05 | 1.21 |
| mesenchyme | Fsd1l | 0.46 | 4.92E-23 | 1.02E-18 | 0.29 | 0.24 | 0.05 | 1.22 |
| mesenchyme | Trmt10c | 0.32 | 5.98E-23 | 1.24E-18 | 0.47 | 0.43 | 0.04 | 1.09 |
| mesenchyme | Zdhhc16 | 0.43 | 6.03E-23 | 1.25E-18 | 0.38 | 0.34 | 0.04 | 1.12 |
| mesenchyme | Ptpa | 0.27 | 6.14E-23 | 1.28E-18 | 0.53 | 0.5 | 0.03 | 1.06 |
| mesenchyme | Gfm1 | 0.45 | 6.33E-23 | 1.32E-18 | 0.36 | 0.32 | 0.04 | 1.13 |
| mesenchyme | Aldh18a1 | 0.5 | 7.26E-23 | 1.51E-18 | 0.31 | 0.26 | 0.05 | 1.19 |
| mesenchyme | Trmt11 | 0.56 | 7.28E-23 | 1.51E-18 | 0.31 | 0.26 | 0.05 | 1.17 |
| mesenchyme | Fto | 0.26 | 7.73E-23 | 1.61E-18 | 0.49 | 0.44 | 0.05 | 1.11 |
| mesenchyme | Tor1aip1 | 0.35 | 7.80E-23 | 1.62E-18 | 0.46 | 0.42 | 0.04 | 1.1 |
| mesenchyme | Xrcc6 | 0.32 | 8.71E-23 | 1.81E-18 | 0.45 | 0.41 | 0.04 | 1.1 |
| mesenchyme | Lancl2 | 0.57 | 1.05E-22 | 2.19E-18 | 0.29 | 0.25 | 0.05 | 1.18 |
| mesenchyme | Smpd2 | 0.41 | 1.08E-22 | 2.24E-18 | 0.33 | 0.28 | 0.05 | 1.18 |
| mesenchyme | Qrich1 | 0.28 | 1.12E-22 | 2.32E-18 | 0.53 | 0.5 | 0.04 | 1.07 |
| mesenchyme | Rnf111 | 0.26 | 1.14E-22 | 2.38E-18 | 0.53 | 0.5 | 0.04 | 1.07 |

**Supplementary Table 2. Marker genes for the major cell types of integrated clustering of E11.5 control and NotchGOF λ-junction cells.**

| | | | | | | | | |
|---|---|---|---|---|---|---|---|---|
| mesenchyme | Kif4 | 0.36 | 1.51E-22 | 3.14E-18 | 0.48 | 0.45 | 0.04 | 1.08 |
| mesenchyme | Khdc4 | 0.33 | 1.70E-22 | 3.54E-18 | 0.43 | 0.39 | 0.04 | 1.11 |
| mesenchyme | Gtf2h4 | 0.32 | 1.85E-22 | 3.84E-18 | 0.43 | 0.39 | 0.04 | 1.11 |
| mesenchyme | Ankrd24 | 0.65 | 1.87E-22 | 3.88E-18 | 0.2 | 0.15 | 0.05 | 1.33 |
| mesenchyme | Elmo2 | 0.53 | 2.05E-22 | 4.27E-18 | 0.27 | 0.22 | 0.05 | 1.22 |
| mesenchyme | Zfp454 | 0.89 | 2.11E-22 | 4.38E-18 | 0.14 | 0.09 | 0.05 | 1.48 |
| mesenchyme | Abhd10 | 0.43 | 2.20E-22 | 4.57E-18 | 0.34 | 0.29 | 0.05 | 1.15 |
| mesenchyme | Scfd1 | 0.27 | 2.20E-22 | 4.57E-18 | 0.5 | 0.46 | 0.04 | 1.08 |
| mesenchyme | Gtf3c5 | 0.65 | 2.25E-22 | 4.67E-18 | 0.23 | 0.18 | 0.05 | 1.27 |
| mesenchyme | Tmx3 | 0.26 | 2.29E-22 | 4.76E-18 | 0.54 | 0.5 | 0.04 | 1.08 |
| mesenchyme | Ttc37 | 0.39 | 2.41E-22 | 5.01E-18 | 0.34 | 0.29 | 0.05 | 1.17 |
| mesenchyme | Agtpbp1 | 0.81 | 2.52E-22 | 5.24E-18 | 0.19 | 0.15 | 0.05 | 1.3 |
| mesenchyme | Bloc1s5 | 0.39 | 2.66E-22 | 5.54E-18 | 0.34 | 0.29 | 0.05 | 1.17 |
| mesenchyme | Sh3pxd2a | 0.37 | 2.78E-22 | 5.78E-18 | 0.38 | 0.34 | 0.05 | 1.14 |
| mesenchyme | Nfix | 0.82 | 2.81E-22 | 5.84E-18 | 0.17 | 0.12 | 0.05 | 1.38 |
| mesenchyme | Slc35a4 | 0.34 | 2.88E-22 | 5.99E-18 | 0.46 | 0.43 | 0.03 | 1.08 |
| mesenchyme | Zfp599 | 0.65 | 2.89E-22 | 6.02E-18 | 0.21 | 0.16 | 0.05 | 1.32 |
| mesenchyme | Phf13 | 0.55 | 3.34E-22 | 6.95E-18 | 0.29 | 0.24 | 0.05 | 1.2 |
| mesenchyme | Ubr1 | 0.46 | 3.51E-22 | 7.30E-18 | 0.35 | 0.31 | 0.04 | 1.13 |
| mesenchyme | Ccdc106 | 0.9 | 3.71E-22 | 7.71E-18 | 0.12 | 0.08 | 0.04 | 1.53 |
| mesenchyme | Wdr54 | 0.86 | 3.78E-22 | 7.87E-18 | 0.12 | 0.08 | 0.04 | 1.57 |
| mesenchyme | Kyat3 | 0.74 | 4.12E-22 | 8.56E-18 | 0.15 | 0.11 | 0.05 | 1.44 |
| mesenchyme | Snrnp35 | 0.49 | 4.60E-22 | 9.56E-18 | 0.3 | 0.25 | 0.05 | 1.19 |
| mesenchyme | Cd2bp2 | 0.3 | 4.87E-22 | 1.01E-17 | 0.51 | 0.47 | 0.03 | 1.07 |
| mesenchyme | Acot1 | 0.27 | 6.83E-22 | 1.42E-17 | 0.37 | 0.31 | 0.06 | 1.19 |
| mesenchyme | Tubgcp5 | 0.39 | 6.84E-22 | 1.42E-17 | 0.36 | 0.31 | 0.05 | 1.16 |
| mesenchyme | Lin9 | 0.45 | 7.17E-22 | 1.49E-17 | 0.35 | 0.31 | 0.04 | 1.14 |
| mesenchyme | Fchsd2 | 0.4 | 7.78E-22 | 1.62E-17 | 0.35 | 0.3 | 0.05 | 1.16 |
| mesenchyme | Sec24b | 0.42 | 8.06E-22 | 1.68E-17 | 0.37 | 0.33 | 0.04 | 1.13 |
| mesenchyme | Cyp2j6 | 0.76 | 8.52E-22 | 1.77E-17 | 0.14 | 0.09 | 0.05 | 1.48 |
| mesenchyme | Spg7 | 0.4 | 8.57E-22 | 1.78E-17 | 0.42 | 0.39 | 0.03 | 1.08 |
| mesenchyme | Rspry1 | 0.28 | 8.85E-22 | 1.84E-17 | 0.52 | 0.49 | 0.03 | 1.07 |
| mesenchyme | Man2a2 | 0.57 | 9.07E-22 | 1.89E-17 | 0.27 | 0.23 | 0.05 | 1.21 |
| mesenchyme | Mxra7 | 0.55 | 9.20E-22 | 1.91E-17 | 0.25 | 0.2 | 0.05 | 1.25 |
| mesenchyme | Cep131 | 0.54 | 1.00E-21 | 2.08E-17 | 0.27 | 0.22 | 0.05 | 1.22 |
| mesenchyme | Shisa5 | 0.5 | 1.08E-21 | 2.25E-17 | 0.26 | 0.21 | 0.05 | 1.23 |
| mesenchyme | Ccnj | 0.69 | 1.08E-21 | 2.25E-17 | 0.19 | 0.14 | 0.05 | 1.34 |
| mesenchyme | Wdr20 | 0.55 | 1.12E-21 | 2.33E-17 | 0.3 | 0.26 | 0.04 | 1.17 |
| mesenchyme | Usp19 | 0.26 | 1.30E-21 | 2.70E-17 | 0.55 | 0.51 | 0.03 | 1.06 |
| mesenchyme | Flywch2 | 0.26 | 1.39E-21 | 2.88E-17 | 0.47 | 0.42 | 0.05 | 1.12 |
| mesenchyme | Ccnb2 | 0.27 | 1.39E-21 | 2.89E-17 | 0.61 | 0.57 | 0.05 | 1.08 |
| mesenchyme | Gtf3c4 | 0.38 | 1.44E-21 | 3.00E-17 | 0.38 | 0.33 | 0.05 | 1.14 |
| mesenchyme | Prmt6 | 0.46 | 2.08E-21 | 4.33E-17 | 0.29 | 0.24 | 0.05 | 1.21 |
| mesenchyme | Fam111a | 0.34 | 2.21E-21 | 4.59E-17 | 0.34 | 0.29 | 0.05 | 1.18 |
| mesenchyme | Fignl2 | 0.81 | 2.28E-21 | 4.75E-17 | 0.14 | 0.1 | 0.04 | 1.46 |
| mesenchyme | Spred2 | 0.31 | 2.41E-21 | 5.02E-17 | 0.41 | 0.36 | 0.05 | 1.14 |
| mesenchyme | Fastkd2 | 0.43 | 2.57E-21 | 5.34E-17 | 0.33 | 0.29 | 0.05 | 1.16 |
| mesenchyme | Ccdc167 | 0.31 | 2.70E-21 | 5.61E-17 | 0.46 | 0.44 | 0.03 | 1.06 |
| mesenchyme | Oscp1 | 0.6 | 2.97E-21 | 6.18E-17 | 0.21 | 0.16 | 0.05 | 1.3 |
| mesenchyme | Mir99ahg | 0.28 | 3.26E-21 | 6.79E-17 | 0.5 | 0.45 | 0.05 | 1.11 |
| mesenchyme | Fer | 0.37 | 3.32E-21 | 6.90E-17 | 0.36 | 0.32 | 0.04 | 1.14 |
| mesenchyme | Prdm2 | 0.29 | 3.40E-21 | 7.07E-17 | 0.48 | 0.43 | 0.04 | 1.09 |
| mesenchyme | Lrrc40 | 0.38 | 3.68E-21 | 7.65E-17 | 0.35 | 0.31 | 0.04 | 1.14 |
| mesenchyme | Zfp934 | 0.5 | 4.09E-21 | 8.51E-17 | 0.27 | 0.23 | 0.05 | 1.21 |
| mesenchyme | Atp23 | 0.54 | 4.30E-21 | 8.95E-17 | 0.25 | 0.21 | 0.05 | 1.22 |
| mesenchyme | Plaat3 | 0.61 | 5.03E-21 | 1.05E-16 | 0.24 | 0.19 | 0.05 | 1.25 |

**Supplementary Table 2. Marker genes for the major cell types of integrated clustering of E11.5 control and NotchGOF λ-junction cells.**

| | | | | | | | | |
|---|---|---|---|---|---|---|---|---|
| mesenchyme | Dnajc12 | 0.67 | 5.05E-21 | 1.05E-16 | 0.18 | 0.14 | 0.05 | 1.35 |
| mesenchyme | Galnt11 | 0.36 | 5.17E-21 | 1.08E-16 | 0.36 | 0.31 | 0.05 | 1.16 |
| mesenchyme | Prr11 | 0.65 | 5.25E-21 | 1.09E-16 | 0.21 | 0.16 | 0.05 | 1.28 |
| mesenchyme | Pex14 | 0.38 | 5.60E-21 | 1.16E-16 | 0.36 | 0.31 | 0.05 | 1.15 |
| mesenchyme | Nrk | 0.91 | 6.47E-21 | 1.35E-16 | 0.17 | 0.13 | 0.04 | 1.35 |
| mesenchyme | Kntc1 | 0.66 | 7.16E-21 | 1.49E-16 | 0.21 | 0.16 | 0.05 | 1.28 |
| mesenchyme | Zfp566 | 0.52 | 7.21E-21 | 1.50E-16 | 0.22 | 0.17 | 0.05 | 1.31 |
| mesenchyme | Fhod3 | 1.09 | 7.30E-21 | 1.52E-16 | 0.1 | 0.07 | 0.04 | 1.58 |
| mesenchyme | Ppp2r5d | 0.3 | 7.56E-21 | 1.57E-16 | 0.48 | 0.44 | 0.04 | 1.09 |
| mesenchyme | Dvl2 | 0.32 | 7.73E-21 | 1.61E-16 | 0.4 | 0.35 | 0.05 | 1.13 |
| mesenchyme | Vps50 | 0.4 | 8.72E-21 | 1.81E-16 | 0.33 | 0.29 | 0.05 | 1.16 |
| mesenchyme | Glmn | 0.29 | 8.84E-21 | 1.84E-16 | 0.46 | 0.42 | 0.04 | 1.09 |
| mesenchyme | Sox6 | 1.25 | 1.06E-20 | 2.21E-16 | 0.14 | 0.1 | 0.04 | 1.43 |
| mesenchyme | Sufu | 0.59 | 1.09E-20 | 2.26E-16 | 0.24 | 0.19 | 0.05 | 1.24 |
| mesenchyme | Nln | 0.45 | 1.09E-20 | 2.27E-16 | 0.31 | 0.26 | 0.05 | 1.19 |
| mesenchyme | Dnajc11 | 0.29 | 1.18E-20 | 2.45E-16 | 0.5 | 0.47 | 0.03 | 1.05 |
| mesenchyme | Ikzf4 | 0.71 | 1.29E-20 | 2.68E-16 | 0.16 | 0.12 | 0.05 | 1.38 |
| mesenchyme | Sgo1 | 0.36 | 1.29E-20 | 2.69E-16 | 0.46 | 0.43 | 0.03 | 1.07 |
| mesenchyme | Rbm33 | 0.28 | 1.36E-20 | 2.83E-16 | 0.49 | 0.46 | 0.03 | 1.07 |
| mesenchyme | Alg10b | 0.32 | 1.38E-20 | 2.86E-16 | 0.46 | 0.42 | 0.04 | 1.09 |
| mesenchyme | Ctps2 | 0.42 | 1.45E-20 | 3.01E-16 | 0.3 | 0.26 | 0.05 | 1.18 |
| mesenchyme | Neil3 | 0.66 | 1.48E-20 | 3.08E-16 | 0.25 | 0.2 | 0.04 | 1.22 |
| mesenchyme | Arhgap10 | 0.28 | 1.49E-20 | 3.09E-16 | 0.44 | 0.39 | 0.05 | 1.12 |
| mesenchyme | Apbb2 | 0.44 | 1.54E-20 | 3.20E-16 | 0.32 | 0.27 | 0.05 | 1.18 |
| mesenchyme | Eef2k | 0.53 | 1.56E-20 | 3.24E-16 | 0.26 | 0.22 | 0.05 | 1.22 |
| mesenchyme | Hspa12b | 0.96 | 1.61E-20 | 3.34E-16 | 0.15 | 0.11 | 0.04 | 1.39 |
| mesenchyme | Ap2a1 | 0.45 | 1.69E-20 | 3.52E-16 | 0.34 | 0.3 | 0.04 | 1.13 |
| mesenchyme | Pcx | 0.96 | 1.77E-20 | 3.69E-16 | 0.15 | 0.1 | 0.04 | 1.39 |
| mesenchyme | Ift80 | 0.61 | 1.78E-20 | 3.70E-16 | 0.23 | 0.19 | 0.05 | 1.24 |
| mesenchyme | Ttf2 | 0.4 | 1.87E-20 | 3.89E-16 | 0.33 | 0.29 | 0.04 | 1.15 |
| mesenchyme | Fxr2 | 0.26 | 1.91E-20 | 3.97E-16 | 0.48 | 0.44 | 0.04 | 1.09 |
| mesenchyme | Ccdc28b | 0.35 | 1.96E-20 | 4.07E-16 | 0.39 | 0.35 | 0.04 | 1.12 |
| mesenchyme | Pcf11 | 0.33 | 1.98E-20 | 4.11E-16 | 0.46 | 0.43 | 0.03 | 1.07 |
| mesenchyme | Lmcd1 | 1.02 | 1.99E-20 | 4.15E-16 | 0.1 | 0.06 | 0.04 | 1.59 |
| mesenchyme | Actr1a | 0.26 | 2.07E-20 | 4.30E-16 | 0.48 | 0.44 | 0.04 | 1.08 |
| mesenchyme | Rev1 | 0.33 | 2.07E-20 | 4.31E-16 | 0.37 | 0.32 | 0.04 | 1.14 |
| mesenchyme | Gins1 | 0.33 | 2.08E-20 | 4.32E-16 | 0.4 | 0.37 | 0.03 | 1.09 |
| mesenchyme | Stam | 0.48 | 2.15E-20 | 4.48E-16 | 0.31 | 0.27 | 0.04 | 1.15 |
| mesenchyme | Haspin | 0.53 | 2.20E-20 | 4.57E-16 | 0.3 | 0.26 | 0.04 | 1.17 |
| mesenchyme | Vps9d1 | 0.53 | 2.49E-20 | 5.17E-16 | 0.25 | 0.2 | 0.05 | 1.24 |
| mesenchyme | Trub2 | 0.45 | 2.53E-20 | 5.26E-16 | 0.32 | 0.28 | 0.04 | 1.16 |
| mesenchyme | Haus6 | 0.37 | 3.51E-20 | 7.30E-16 | 0.38 | 0.34 | 0.04 | 1.12 |
| mesenchyme | Adamts7 | 0.99 | 3.53E-20 | 7.35E-16 | 0.11 | 0.07 | 0.04 | 1.55 |
| mesenchyme | Gemin5 | 0.4 | 3.80E-20 | 7.91E-16 | 0.38 | 0.34 | 0.04 | 1.12 |
| mesenchyme | Cog4 | 0.28 | 4.40E-20 | 9.16E-16 | 0.43 | 0.39 | 0.04 | 1.1 |
| mesenchyme | Foxk2 | 0.4 | 4.85E-20 | 1.01E-15 | 0.38 | 0.35 | 0.04 | 1.1 |
| mesenchyme | Zswim4 | 0.49 | 5.07E-20 | 1.05E-15 | 0.25 | 0.2 | 0.05 | 1.25 |
| mesenchyme | 730455P16Ri | 0.43 | 5.27E-20 | 1.10E-15 | 0.32 | 0.28 | 0.04 | 1.16 |
| mesenchyme | Triap1 | 0.26 | 5.96E-20 | 1.24E-15 | 0.52 | 0.5 | 0.03 | 1.05 |
| mesenchyme | Map1a | 0.71 | 6.00E-20 | 1.25E-15 | 0.11 | 0.07 | 0.04 | 1.59 |
| mesenchyme | Cherp | 0.33 | 6.00E-20 | 1.25E-15 | 0.44 | 0.41 | 0.04 | 1.09 |
| mesenchyme | Spry2 | 0.42 | 6.06E-20 | 1.26E-15 | 0.31 | 0.26 | 0.05 | 1.2 |
| mesenchyme | Tm9sf1 | 0.37 | 6.37E-20 | 1.33E-15 | 0.45 | 0.43 | 0.03 | 1.06 |
| mesenchyme | Ppt2 | 0.51 | 6.90E-20 | 1.44E-15 | 0.27 | 0.22 | 0.05 | 1.2 |
| mesenchyme | 430095P16Ri | 1.15 | 7.83E-20 | 1.63E-15 | 0.12 | 0.08 | 0.04 | 1.48 |
| mesenchyme | Wdr48 | 0.27 | 7.92E-20 | 1.65E-15 | 0.46 | 0.43 | 0.04 | 1.09 |

**Supplementary Table 2. Marker genes for the major cell types of integrated clustering of E11.5 control and NotchGOF λ-junction cells.**

| | | | | | | | | |
|---|---|---|---|---|---|---|---|---|
| mesenchyme | Cpne1 | 0.33 | 8.14E-20 | 1.69E-15 | 0.43 | 0.39 | 0.04 | 1.09 |
| mesenchyme | Dnajc24 | 0.25 | 9.09E-20 | 1.89E-15 | 0.49 | 0.45 | 0.04 | 1.08 |
| mesenchyme | Diaph3 | 0.45 | 9.71E-20 | 2.02E-15 | 0.38 | 0.35 | 0.03 | 1.09 |
| mesenchyme | Pcdh10 | 0.81 | 1.06E-19 | 2.21E-15 | 0.12 | 0.08 | 0.04 | 1.52 |
| mesenchyme | Dnajc5 | 0.26 | 1.07E-19 | 2.22E-15 | 0.54 | 0.51 | 0.03 | 1.06 |
| mesenchyme | Commd10 | 0.29 | 1.11E-19 | 2.32E-15 | 0.47 | 0.43 | 0.03 | 1.07 |
| mesenchyme | Atrip | 0.47 | 1.15E-19 | 2.39E-15 | 0.29 | 0.25 | 0.05 | 1.18 |
| mesenchyme | Actr6 | 0.31 | 1.18E-19 | 2.45E-15 | 0.41 | 0.37 | 0.04 | 1.11 |
| mesenchyme | Coq10b | 0.53 | 1.29E-19 | 2.67E-15 | 0.27 | 0.23 | 0.04 | 1.18 |
| mesenchyme | Dido1 | 0.25 | 1.39E-19 | 2.88E-15 | 0.51 | 0.47 | 0.04 | 1.08 |
| mesenchyme | Impdh1 | 0.51 | 1.46E-19 | 3.04E-15 | 0.23 | 0.19 | 0.05 | 1.24 |
| mesenchyme | Afg3l1 | 0.28 | 1.50E-19 | 3.11E-15 | 0.49 | 0.45 | 0.03 | 1.07 |
| mesenchyme | Scyl1 | 0.5 | 1.72E-19 | 3.58E-15 | 0.3 | 0.26 | 0.04 | 1.16 |
| mesenchyme | Msmo1 | 0.27 | 1.76E-19 | 3.67E-15 | 0.51 | 0.47 | 0.04 | 1.08 |
| mesenchyme | Ttc21b | 0.55 | 1.82E-19 | 3.78E-15 | 0.25 | 0.2 | 0.05 | 1.23 |
| mesenchyme | Gmeb1 | 0.34 | 1.87E-19 | 3.90E-15 | 0.42 | 0.39 | 0.04 | 1.09 |
| mesenchyme | Xxylt1 | 0.54 | 2.22E-19 | 4.62E-15 | 0.26 | 0.22 | 0.04 | 1.2 |
| mesenchyme | Pold1 | 0.39 | 2.24E-19 | 4.67E-15 | 0.36 | 0.31 | 0.04 | 1.13 |
| mesenchyme | Mrm2 | 0.55 | 2.31E-19 | 4.81E-15 | 0.22 | 0.18 | 0.05 | 1.25 |
| mesenchyme | Acot2 | 0.76 | 2.51E-19 | 5.21E-15 | 0.15 | 0.11 | 0.04 | 1.4 |
| mesenchyme | Socs1 | 0.65 | 2.92E-19 | 6.06E-15 | 0.17 | 0.13 | 0.04 | 1.34 |
| mesenchyme | Pnpt1 | 0.44 | 2.94E-19 | 6.11E-15 | 0.31 | 0.26 | 0.04 | 1.16 |
| mesenchyme | Ackr3 | 0.31 | 3.03E-19 | 6.30E-15 | 0.43 | 0.39 | 0.04 | 1.1 |
| mesenchyme | Nt5c2 | 0.37 | 3.14E-19 | 6.53E-15 | 0.38 | 0.34 | 0.04 | 1.11 |
| mesenchyme | Hsf1 | 0.4 | 3.40E-19 | 7.07E-15 | 0.36 | 0.32 | 0.04 | 1.11 |
| mesenchyme | Xndc1 | 0.58 | 3.75E-19 | 7.81E-15 | 0.26 | 0.22 | 0.04 | 1.19 |
| mesenchyme | Hist1h2ak | 0.8 | 3.93E-19 | 8.17E-15 | 0.18 | 0.14 | 0.04 | 1.3 |
| mesenchyme | Efl1 | 0.47 | 4.01E-19 | 8.35E-15 | 0.3 | 0.26 | 0.04 | 1.16 |
| mesenchyme | Tbc1d15 | 0.32 | 4.04E-19 | 8.40E-15 | 0.52 | 0.5 | 0.02 | 1.04 |
| mesenchyme | Gas8 | 0.28 | 4.30E-19 | 8.93E-15 | 0.41 | 0.37 | 0.05 | 1.12 |
| mesenchyme | Rab3a | 0.43 | 5.02E-19 | 1.05E-14 | 0.15 | 0.11 | 0.04 | 1.41 |
| mesenchyme | Tgfb3 | 0.88 | 5.13E-19 | 1.07E-14 | 0.11 | 0.07 | 0.04 | 1.54 |
| mesenchyme | Rnf169 | 0.42 | 5.44E-19 | 1.13E-14 | 0.32 | 0.28 | 0.04 | 1.16 |
| mesenchyme | Clk4 | 0.3 | 5.46E-19 | 1.14E-14 | 0.46 | 0.44 | 0.03 | 1.06 |
| mesenchyme | Hbs1l | 0.26 | 5.85E-19 | 1.22E-14 | 0.54 | 0.52 | 0.02 | 1.04 |
| mesenchyme | Ankrd40 | 0.28 | 6.18E-19 | 1.28E-14 | 0.48 | 0.46 | 0.02 | 1.05 |
| mesenchyme | 930095G15Ri | 0.62 | 6.29E-19 | 1.31E-14 | 0.19 | 0.15 | 0.05 | 1.3 |
| mesenchyme | Dcaf15 | 0.41 | 6.48E-19 | 1.35E-14 | 0.36 | 0.32 | 0.04 | 1.11 |
| mesenchyme | Tsga10 | 0.91 | 6.75E-19 | 1.40E-14 | 0.11 | 0.08 | 0.04 | 1.51 |
| mesenchyme | Crkl | 0.37 | 7.02E-19 | 1.46E-14 | 0.39 | 0.35 | 0.04 | 1.1 |
| mesenchyme | Mtpap | 0.32 | 7.45E-19 | 1.55E-14 | 0.38 | 0.35 | 0.04 | 1.11 |
| mesenchyme | Nek9 | 0.45 | 7.46E-19 | 1.55E-14 | 0.31 | 0.26 | 0.04 | 1.16 |
| mesenchyme | Sntb2 | 0.35 | 7.50E-19 | 1.56E-14 | 0.36 | 0.31 | 0.05 | 1.14 |
| mesenchyme | Zfp775 | 0.69 | 7.60E-19 | 1.58E-14 | 0.15 | 0.1 | 0.04 | 1.4 |
| mesenchyme | Gm17092 | 0.55 | 8.43E-19 | 1.75E-14 | 0.26 | 0.22 | 0.04 | 1.19 |
| mesenchyme | Nup205 | 0.4 | 8.46E-19 | 1.76E-14 | 0.36 | 0.32 | 0.04 | 1.12 |
| mesenchyme | Marchf6 | 0.32 | 8.77E-19 | 1.82E-14 | 0.41 | 0.36 | 0.04 | 1.12 |
| mesenchyme | Ncln | 0.38 | 9.06E-19 | 1.88E-14 | 0.39 | 0.35 | 0.04 | 1.11 |
| mesenchyme | Hist1h3b | 0.67 | 9.15E-19 | 1.90E-14 | 0.25 | 0.2 | 0.04 | 1.2 |
| mesenchyme | Scaf4 | 0.34 | 1.02E-18 | 2.12E-14 | 0.43 | 0.39 | 0.04 | 1.1 |
| mesenchyme | Slc7a6 | 0.43 | 1.02E-18 | 2.13E-14 | 0.3 | 0.26 | 0.04 | 1.17 |
| mesenchyme | Clk2 | 0.36 | 1.07E-18 | 2.23E-14 | 0.38 | 0.34 | 0.04 | 1.11 |
| mesenchyme | Hmg20a | 0.3 | 1.12E-18 | 2.33E-14 | 0.44 | 0.4 | 0.04 | 1.09 |
| mesenchyme | Epb41l3 | 0.27 | 1.13E-18 | 2.36E-14 | 0.47 | 0.42 | 0.05 | 1.11 |
| mesenchyme | Ncoa5 | 0.31 | 1.14E-18 | 2.37E-14 | 0.39 | 0.35 | 0.04 | 1.12 |
| mesenchyme | Xpo5 | 0.25 | 1.15E-18 | 2.40E-14 | 0.45 | 0.41 | 0.04 | 1.09 |

**Supplementary Table 2. Marker genes for the major cell types of integrated clustering of E11.5 control and NotchGOF λ-junction cells.**

| | | | | | | | | |
|---|---|---|---|---|---|---|---|---|
| mesenchyme | Rpgr | 0.28 | 1.16E-18 | 2.42E-14 | 0.36 | 0.31 | 0.05 | 1.17 |
| mesenchyme | Spin4 | 0.79 | 1.23E-18 | 2.55E-14 | 0.14 | 0.1 | 0.04 | 1.43 |
| mesenchyme | Stx5a | 0.28 | 1.26E-18 | 2.62E-14 | 0.49 | 0.46 | 0.03 | 1.07 |
| mesenchyme | Necap1 | 0.53 | 1.26E-18 | 2.63E-14 | 0.26 | 0.21 | 0.04 | 1.2 |
| mesenchyme | Sdhaf3 | 0.32 | 1.37E-18 | 2.84E-14 | 0.38 | 0.34 | 0.04 | 1.11 |
| mesenchyme | Hacd3 | 0.28 | 1.50E-18 | 3.11E-14 | 0.44 | 0.41 | 0.04 | 1.09 |
| mesenchyme | Armcx4 | 0.31 | 1.50E-18 | 3.12E-14 | 0.38 | 0.34 | 0.04 | 1.13 |
| mesenchyme | Ercc8 | 0.49 | 1.57E-18 | 3.28E-14 | 0.24 | 0.2 | 0.04 | 1.21 |
| mesenchyme | Snx21 | 0.77 | 1.62E-18 | 3.38E-14 | 0.15 | 0.11 | 0.04 | 1.38 |
| mesenchyme | Prpf40b | 0.52 | 1.92E-18 | 3.99E-14 | 0.25 | 0.21 | 0.04 | 1.2 |
| mesenchyme | Gpaa1 | 0.31 | 2.00E-18 | 4.17E-14 | 0.43 | 0.4 | 0.03 | 1.08 |
| mesenchyme | Glce | 0.56 | 2.18E-18 | 4.54E-14 | 0.2 | 0.15 | 0.04 | 1.29 |
| mesenchyme | Herc2 | 0.36 | 2.35E-18 | 4.89E-14 | 0.42 | 0.39 | 0.03 | 1.07 |
| mesenchyme | Chmp7 | 0.39 | 2.36E-18 | 4.90E-14 | 0.37 | 0.33 | 0.04 | 1.11 |
| mesenchyme | Uspl1 | 0.37 | 2.37E-18 | 4.94E-14 | 0.4 | 0.37 | 0.03 | 1.08 |
| mesenchyme | Ccnyl1 | 0.52 | 2.43E-18 | 5.06E-14 | 0.2 | 0.16 | 0.04 | 1.28 |
| mesenchyme | Irs2 | 0.3 | 3.30E-18 | 6.86E-14 | 0.45 | 0.41 | 0.04 | 1.09 |
| mesenchyme | Ccdc137 | 0.57 | 3.38E-18 | 7.03E-14 | 0.23 | 0.19 | 0.04 | 1.23 |
| mesenchyme | Zswim7 | 0.59 | 3.44E-18 | 7.15E-14 | 0.27 | 0.23 | 0.04 | 1.17 |
| mesenchyme | Pip5k1a | 0.26 | 3.45E-18 | 7.18E-14 | 0.48 | 0.45 | 0.03 | 1.07 |
| mesenchyme | Pggt1b | 0.25 | 3.60E-18 | 7.48E-14 | 0.51 | 0.48 | 0.03 | 1.06 |
| mesenchyme | Ift172 | 0.74 | 4.04E-18 | 8.41E-14 | 0.15 | 0.11 | 0.04 | 1.37 |
| mesenchyme | Mdn1 | 0.31 | 4.28E-18 | 8.91E-14 | 0.43 | 0.38 | 0.04 | 1.11 |
| mesenchyme | Sacm1l | 0.35 | 4.46E-18 | 9.27E-14 | 0.34 | 0.3 | 0.04 | 1.13 |
| mesenchyme | Ints3 | 0.38 | 4.51E-18 | 9.39E-14 | 0.37 | 0.33 | 0.04 | 1.11 |
| mesenchyme | Gpatch2 | 0.29 | 4.71E-18 | 9.80E-14 | 0.43 | 0.39 | 0.04 | 1.11 |
| mesenchyme | Gm3550 | 0.8 | 5.02E-18 | 1.04E-13 | 0.12 | 0.08 | 0.04 | 1.46 |
| mesenchyme | Pif1 | 0.78 | 5.03E-18 | 1.05E-13 | 0.15 | 0.11 | 0.04 | 1.36 |
| mesenchyme | Alkbh7 | 0.28 | 5.24E-18 | 1.09E-13 | 0.41 | 0.38 | 0.03 | 1.08 |
| mesenchyme | Hcfc2 | 0.76 | 5.45E-18 | 1.13E-13 | 0.16 | 0.12 | 0.04 | 1.33 |
| mesenchyme | Depdc5 | 0.59 | 5.69E-18 | 1.18E-13 | 0.22 | 0.18 | 0.04 | 1.23 |
| mesenchyme | Sclt1 | 0.28 | 5.70E-18 | 1.19E-13 | 0.44 | 0.4 | 0.04 | 1.09 |
| mesenchyme | )630045J12Ri | 0.47 | 6.44E-18 | 1.34E-13 | 0.26 | 0.21 | 0.05 | 1.21 |
| mesenchyme | Elmod2 | 0.53 | 6.60E-18 | 1.37E-13 | 0.24 | 0.2 | 0.04 | 1.22 |
| mesenchyme | Pi4k2a | 0.48 | 6.63E-18 | 1.38E-13 | 0.29 | 0.25 | 0.04 | 1.16 |
| mesenchyme | Suv39h2 | 0.27 | 6.66E-18 | 1.39E-13 | 0.43 | 0.39 | 0.04 | 1.11 |
| mesenchyme | Cnst | 0.66 | 7.40E-18 | 1.54E-13 | 0.2 | 0.16 | 0.04 | 1.25 |
| mesenchyme | 031439G07Ri | 0.44 | 7.99E-18 | 1.66E-13 | 0.27 | 0.22 | 0.04 | 1.19 |
| mesenchyme | Terf1 | 0.27 | 8.35E-18 | 1.74E-13 | 0.53 | 0.52 | 0.02 | 1.03 |
| mesenchyme | Nup214 | 0.27 | 8.37E-18 | 1.74E-13 | 0.48 | 0.45 | 0.03 | 1.06 |
| mesenchyme | Mief2 | 0.98 | 8.42E-18 | 1.75E-13 | 0.11 | 0.08 | 0.04 | 1.47 |
| mesenchyme | Slc7a6os | 0.47 | 9.03E-18 | 1.88E-13 | 0.29 | 0.25 | 0.04 | 1.17 |
| mesenchyme | Dlgap5 | 0.31 | 9.88E-18 | 2.05E-13 | 0.5 | 0.47 | 0.03 | 1.06 |
| mesenchyme | Gnai1 | 0.57 | 1.03E-17 | 2.14E-13 | 0.17 | 0.13 | 0.04 | 1.34 |
| mesenchyme | Ormdl3 | 0.31 | 1.06E-17 | 2.21E-13 | 0.47 | 0.45 | 0.02 | 1.04 |
| mesenchyme | Atxn7l3 | 0.3 | 1.19E-17 | 2.47E-13 | 0.42 | 0.38 | 0.04 | 1.1 |
| mesenchyme | 2210408I21Ril | 0.75 | 1.28E-17 | 2.65E-13 | 0.14 | 0.1 | 0.04 | 1.4 |
| mesenchyme | Mrpl39 | 0.27 | 1.31E-17 | 2.73E-13 | 0.47 | 0.45 | 0.02 | 1.05 |
| mesenchyme | Tpgs1 | 0.27 | 1.37E-17 | 2.85E-13 | 0.45 | 0.42 | 0.03 | 1.07 |
| mesenchyme | Stk24 | 0.33 | 1.41E-17 | 2.94E-13 | 0.47 | 0.45 | 0.02 | 1.04 |
| mesenchyme | Cep126 | 0.57 | 1.44E-17 | 3.00E-13 | 0.17 | 0.13 | 0.04 | 1.33 |
| mesenchyme | AI837181 | 0.35 | 1.45E-17 | 3.02E-13 | 0.36 | 0.33 | 0.04 | 1.11 |
| mesenchyme | Hmmr | 0.35 | 1.54E-17 | 3.20E-13 | 0.54 | 0.52 | 0.03 | 1.06 |
| mesenchyme | Zfp560 | 0.35 | 1.64E-17 | 3.40E-13 | 0.38 | 0.35 | 0.03 | 1.08 |
| mesenchyme | Ctc1 | 0.38 | 1.75E-17 | 3.64E-13 | 0.39 | 0.36 | 0.03 | 1.08 |
| mesenchyme | Gm49700 | 0.66 | 1.85E-17 | 3.84E-13 | 0.17 | 0.13 | 0.04 | 1.32 |

**Supplementary Table 2. Marker genes for the major cell types of integrated clustering of E11.5 control and NotchGOF λ-junction cells.**

| | | | | | | | | |
|---|---|---|---|---|---|---|---|---|
| mesenchyme | Swap70 | 0.37 | 1.90E-17 | 3.94E-13 | 0.34 | 0.3 | 0.04 | 1.14 |
| mesenchyme | Sugp2 | 0.56 | 2.43E-17 | 5.06E-13 | 0.21 | 0.17 | 0.04 | 1.25 |
| mesenchyme | Gzf1 | 0.34 | 2.77E-17 | 5.75E-13 | 0.36 | 0.32 | 0.04 | 1.12 |
| mesenchyme | Shc1 | 0.57 | 2.94E-17 | 6.12E-13 | 0.25 | 0.21 | 0.04 | 1.18 |
| mesenchyme | Trps1 | 0.5 | 3.02E-17 | 6.28E-13 | 0.5 | 0.47 | 0.02 | 1.05 |
| mesenchyme | Cdc20 | 0.31 | 3.27E-17 | 6.81E-13 | 0.59 | 0.56 | 0.03 | 1.06 |
| mesenchyme | Kif18a | 0.43 | 3.43E-17 | 7.14E-13 | 0.36 | 0.32 | 0.04 | 1.11 |
| mesenchyme | Zfp239 | 0.43 | 3.66E-17 | 7.61E-13 | 0.25 | 0.21 | 0.05 | 1.22 |
| mesenchyme | Cpt1c | 0.9 | 3.97E-17 | 8.26E-13 | 0.1 | 0.07 | 0.03 | 1.52 |
| mesenchyme | Mtmr2 | 0.3 | 4.19E-17 | 8.72E-13 | 0.39 | 0.36 | 0.04 | 1.1 |
| mesenchyme | 130309D02Ri | 0.36 | 4.26E-17 | 8.85E-13 | 0.37 | 0.34 | 0.03 | 1.09 |
| mesenchyme | Prrc1 | 0.32 | 4.36E-17 | 9.07E-13 | 0.45 | 0.42 | 0.03 | 1.07 |
| mesenchyme | Tnks1bp1 | 0.62 | 4.56E-17 | 9.48E-13 | 0.19 | 0.15 | 0.04 | 1.27 |
| mesenchyme | Mppe1 | 0.73 | 4.73E-17 | 9.83E-13 | 0.15 | 0.11 | 0.04 | 1.35 |
| mesenchyme | Cul7 | 0.35 | 4.78E-17 | 9.95E-13 | 0.38 | 0.34 | 0.04 | 1.1 |
| mesenchyme | Cdc42bpa | 0.35 | 4.79E-17 | 9.97E-13 | 0.39 | 0.36 | 0.03 | 1.09 |
| mesenchyme | Map2k5 | 0.37 | 4.95E-17 | 1.03E-12 | 0.35 | 0.31 | 0.04 | 1.12 |
| mesenchyme | Atm | 0.39 | 5.11E-17 | 1.06E-12 | 0.33 | 0.29 | 0.04 | 1.14 |
| mesenchyme | Ncapg | 0.3 | 5.63E-17 | 1.17E-12 | 0.49 | 0.46 | 0.02 | 1.05 |
| mesenchyme | Cwc25 | 0.27 | 5.86E-17 | 1.22E-12 | 0.46 | 0.43 | 0.03 | 1.07 |
| mesenchyme | Pask | 0.63 | 5.90E-17 | 1.23E-12 | 0.17 | 0.13 | 0.04 | 1.32 |
| mesenchyme | Msl2 | 0.4 | 6.22E-17 | 1.29E-12 | 0.36 | 0.33 | 0.03 | 1.1 |
| mesenchyme | Crtap | 0.53 | 6.76E-17 | 1.41E-12 | 0.29 | 0.25 | 0.04 | 1.15 |
| mesenchyme | Tstd2 | 0.45 | 6.99E-17 | 1.45E-12 | 0.29 | 0.25 | 0.04 | 1.16 |
| mesenchyme | Ccdc102a | 0.33 | 7.02E-17 | 1.46E-12 | 0.29 | 0.25 | 0.04 | 1.17 |
| mesenchyme | Hist1h2ad | 0.85 | 7.11E-17 | 1.48E-12 | 0.11 | 0.08 | 0.04 | 1.45 |
| mesenchyme | Castor2 | 0.8 | 7.44E-17 | 1.55E-12 | 0.15 | 0.11 | 0.04 | 1.36 |
| mesenchyme | Mgat1 | 0.52 | 7.57E-17 | 1.57E-12 | 0.22 | 0.18 | 0.04 | 1.22 |
| mesenchyme | Hist1h2bh | 0.8 | 7.67E-17 | 1.60E-12 | 0.13 | 0.09 | 0.04 | 1.38 |
| mesenchyme | Spryd4 | 0.45 | 7.84E-17 | 1.63E-12 | 0.27 | 0.23 | 0.04 | 1.16 |
| mesenchyme | Ift57 | 0.31 | 8.43E-17 | 1.75E-12 | 0.33 | 0.29 | 0.04 | 1.15 |
| mesenchyme | Eme1 | 0.32 | 8.74E-17 | 1.82E-12 | 0.38 | 0.34 | 0.04 | 1.11 |
| mesenchyme | Efr3b | 0.63 | 8.93E-17 | 1.86E-12 | 0.11 | 0.07 | 0.04 | 1.49 |
| mesenchyme | Gatd3a | 0.27 | 9.83E-17 | 2.04E-12 | 0.41 | 0.38 | 0.03 | 1.08 |
| mesenchyme | Uba7 | 0.9 | 9.98E-17 | 2.08E-12 | 0.12 | 0.09 | 0.03 | 1.4 |
| mesenchyme | Scmh1 | 0.4 | 1.02E-16 | 2.13E-12 | 0.34 | 0.3 | 0.04 | 1.14 |
| mesenchyme | Usp16 | 0.29 | 1.02E-16 | 2.13E-12 | 0.48 | 0.46 | 0.02 | 1.04 |
| mesenchyme | Smim13 | 0.5 | 1.10E-16 | 2.29E-12 | 0.26 | 0.23 | 0.04 | 1.17 |
| mesenchyme | Senp5 | 0.32 | 1.17E-16 | 2.43E-12 | 0.39 | 0.35 | 0.03 | 1.09 |
| mesenchyme | Snhg11 | 0.71 | 1.19E-16 | 2.47E-12 | 0.12 | 0.08 | 0.04 | 1.46 |
| mesenchyme | Clybl | 0.68 | 1.32E-16 | 2.74E-12 | 0.21 | 0.17 | 0.04 | 1.21 |
| mesenchyme | Morc3 | 0.29 | 1.33E-16 | 2.77E-12 | 0.43 | 0.41 | 0.03 | 1.07 |
| mesenchyme | Pitrm1 | 0.26 | 1.36E-16 | 2.83E-12 | 0.54 | 0.53 | 0.01 | 1.02 |
| mesenchyme | Ska1 | 0.31 | 1.38E-16 | 2.88E-12 | 0.43 | 0.4 | 0.03 | 1.07 |
| mesenchyme | Pex5 | 0.5 | 1.46E-16 | 3.03E-12 | 0.26 | 0.22 | 0.04 | 1.18 |
| mesenchyme | Ndc1 | 0.39 | 1.47E-16 | 3.06E-12 | 0.32 | 0.28 | 0.04 | 1.13 |
| mesenchyme | Rnf217 | 0.54 | 1.51E-16 | 3.14E-12 | 0.2 | 0.16 | 0.04 | 1.25 |
| mesenchyme | Sumf2 | 0.3 | 1.59E-16 | 3.30E-12 | 0.42 | 0.39 | 0.04 | 1.09 |
| mesenchyme | Zscan21 | 0.27 | 1.69E-16 | 3.52E-12 | 0.42 | 0.38 | 0.04 | 1.09 |
| mesenchyme | Ccnb1 | 0.27 | 1.94E-16 | 4.04E-12 | 0.59 | 0.56 | 0.03 | 1.05 |
| mesenchyme | Sun2 | 0.67 | 2.10E-16 | 4.38E-12 | 0.19 | 0.15 | 0.04 | 1.26 |
| mesenchyme | Kcnj2 | 0.65 | 2.13E-16 | 4.43E-12 | 0.19 | 0.15 | 0.04 | 1.26 |
| mesenchyme | Ifnar1 | 0.36 | 2.38E-16 | 4.95E-12 | 0.36 | 0.33 | 0.03 | 1.1 |
| mesenchyme | Zfp105 | 0.42 | 2.41E-16 | 5.02E-12 | 0.28 | 0.24 | 0.04 | 1.17 |
| mesenchyme | Nfkb1 | 0.32 | 2.51E-16 | 5.23E-12 | 0.32 | 0.27 | 0.04 | 1.16 |
| mesenchyme | Tfcp2 | 0.37 | 2.52E-16 | 5.23E-12 | 0.29 | 0.25 | 0.04 | 1.16 |

**Supplementary Table 2. Marker genes for the major cell types of integrated clustering of E11.5 control and NotchGOF λ-junction cells.**

| | | | | | | | | |
|---|---|---|---|---|---|---|---|---|
| mesenchyme | Zfp950 | 0.33 | 2.79E-16 | 5.80E-12 | 0.39 | 0.35 | 0.04 | 1.1 |
| mesenchyme | Rpa1 | 0.32 | 2.92E-16 | 6.07E-12 | 0.43 | 0.41 | 0.02 | 1.06 |
| mesenchyme | Capn15 | 0.37 | 3.06E-16 | 6.35E-12 | 0.39 | 0.36 | 0.03 | 1.08 |
| mesenchyme | Rpusd4 | 0.35 | 3.08E-16 | 6.40E-12 | 0.32 | 0.28 | 0.04 | 1.13 |
| mesenchyme | Sbf1 | 0.43 | 3.26E-16 | 6.77E-12 | 0.29 | 0.25 | 0.04 | 1.15 |
| mesenchyme | Slc25a36 | 0.27 | 3.34E-16 | 6.95E-12 | 0.45 | 0.41 | 0.04 | 1.09 |
| mesenchyme | Nlgn2 | 0.51 | 3.36E-16 | 6.99E-12 | 0.25 | 0.21 | 0.04 | 1.18 |
| mesenchyme | Dhx40 | 0.26 | 3.52E-16 | 7.33E-12 | 0.44 | 0.41 | 0.04 | 1.09 |
| mesenchyme | Hist1h2bb | 0.71 | 3.55E-16 | 7.38E-12 | 0.22 | 0.18 | 0.04 | 1.2 |
| mesenchyme | Snip1 | 0.37 | 3.61E-16 | 7.50E-12 | 0.35 | 0.31 | 0.03 | 1.11 |
| mesenchyme | Siah1b | 0.31 | 3.75E-16 | 7.79E-12 | 0.31 | 0.27 | 0.04 | 1.15 |
| mesenchyme | Zbtb10 | 0.55 | 4.11E-16 | 8.56E-12 | 0.21 | 0.17 | 0.04 | 1.24 |
| mesenchyme | Cldn11 | 1.12 | 4.47E-16 | 9.30E-12 | 0.14 | 0.11 | 0.04 | 1.34 |
| mesenchyme | Msh3 | 0.32 | 5.06E-16 | 1.05E-11 | 0.43 | 0.4 | 0.03 | 1.07 |
| mesenchyme | Kif21b | 0.61 | 5.74E-16 | 1.19E-11 | 0.18 | 0.14 | 0.04 | 1.29 |
| mesenchyme | Mfsd10 | 0.36 | 5.80E-16 | 1.21E-11 | 0.36 | 0.33 | 0.04 | 1.11 |
| mesenchyme | Tmem267 | 0.65 | 5.82E-16 | 1.21E-11 | 0.18 | 0.15 | 0.04 | 1.26 |
| mesenchyme | Scaf1 | 0.31 | 6.03E-16 | 1.26E-11 | 0.42 | 0.39 | 0.03 | 1.07 |
| mesenchyme | Zfp770 | 0.4 | 6.08E-16 | 1.26E-11 | 0.3 | 0.26 | 0.04 | 1.14 |
| mesenchyme | Arl16 | 0.32 | 6.20E-16 | 1.29E-11 | 0.37 | 0.34 | 0.03 | 1.1 |
| mesenchyme | Ercc5 | 0.33 | 6.22E-16 | 1.29E-11 | 0.33 | 0.29 | 0.04 | 1.13 |
| mesenchyme | Tbl3 | 0.38 | 6.87E-16 | 1.43E-11 | 0.36 | 0.33 | 0.03 | 1.09 |
| mesenchyme | Bace1 | 0.38 | 7.05E-16 | 1.47E-11 | 0.31 | 0.27 | 0.04 | 1.14 |
| mesenchyme | Zfp62 | 0.28 | 7.62E-16 | 1.59E-11 | 0.45 | 0.41 | 0.03 | 1.07 |
| mesenchyme | Hist1h4c | 0.82 | 7.99E-16 | 1.66E-11 | 0.16 | 0.12 | 0.04 | 1.29 |
| mesenchyme | Zfp146 | 0.34 | 8.02E-16 | 1.67E-11 | 0.35 | 0.31 | 0.04 | 1.12 |
| mesenchyme | St3gal3 | 0.74 | 8.15E-16 | 1.69E-11 | 0.13 | 0.09 | 0.04 | 1.38 |
| mesenchyme | Ndufaf5 | 0.53 | 8.19E-16 | 1.70E-11 | 0.21 | 0.18 | 0.04 | 1.22 |
| mesenchyme | Cip2a | 0.38 | 8.34E-16 | 1.74E-11 | 0.4 | 0.37 | 0.03 | 1.07 |
| mesenchyme | Nup153 | 0.26 | 8.37E-16 | 1.74E-11 | 0.45 | 0.42 | 0.03 | 1.07 |
| mesenchyme | Osbpl6 | 0.73 | 8.70E-16 | 1.81E-11 | 0.16 | 0.12 | 0.04 | 1.31 |
| mesenchyme | St8sia2 | 0.58 | 9.32E-16 | 1.94E-11 | 0.21 | 0.17 | 0.04 | 1.24 |
| mesenchyme | Sgce | 0.34 | 9.67E-16 | 2.01E-11 | 0.34 | 0.3 | 0.04 | 1.13 |
| mesenchyme | Golt1b | 0.27 | 9.78E-16 | 2.03E-11 | 0.47 | 0.44 | 0.02 | 1.05 |
| mesenchyme | Rbm14 | 0.25 | 9.86E-16 | 2.05E-11 | 0.45 | 0.42 | 0.03 | 1.07 |
| mesenchyme | B4gat1 | 0.34 | 1.06E-15 | 2.20E-11 | 0.37 | 0.33 | 0.03 | 1.1 |
| mesenchyme | Trip4 | 0.41 | 1.12E-15 | 2.33E-11 | 0.3 | 0.26 | 0.04 | 1.14 |
| mesenchyme | Osgin2 | 0.57 | 1.16E-15 | 2.41E-11 | 0.2 | 0.16 | 0.04 | 1.23 |
| mesenchyme | Kctd10 | 0.27 | 1.18E-15 | 2.45E-11 | 0.31 | 0.27 | 0.04 | 1.15 |
| mesenchyme | Dip2c | 0.41 | 1.19E-15 | 2.47E-11 | 0.25 | 0.21 | 0.04 | 1.19 |
| mesenchyme | Selenoi | 0.36 | 1.29E-15 | 2.69E-11 | 0.35 | 0.32 | 0.03 | 1.11 |
| mesenchyme | Yes1 | 0.29 | 1.31E-15 | 2.72E-11 | 0.36 | 0.32 | 0.04 | 1.13 |
| mesenchyme | Sorbs3 | 0.63 | 1.36E-15 | 2.82E-11 | 0.15 | 0.11 | 0.04 | 1.32 |
| mesenchyme | 700097O09Ri | 0.29 | 1.36E-15 | 2.84E-11 | 0.36 | 0.33 | 0.03 | 1.09 |
| mesenchyme | Tm7sf3 | 0.51 | 1.38E-15 | 2.87E-11 | 0.24 | 0.2 | 0.04 | 1.18 |
| mesenchyme | Pank3 | 0.32 | 1.46E-15 | 3.04E-11 | 0.39 | 0.36 | 0.03 | 1.09 |
| mesenchyme | Wrap73 | 0.33 | 1.54E-15 | 3.19E-11 | 0.36 | 0.33 | 0.03 | 1.1 |
| mesenchyme | Dtwd2 | 0.7 | 1.55E-15 | 3.23E-11 | 0.13 | 0.1 | 0.04 | 1.37 |
| mesenchyme | Uap1 | 0.34 | 1.57E-15 | 3.27E-11 | 0.33 | 0.3 | 0.03 | 1.11 |
| mesenchyme | Hist1h3f | 0.53 | 1.58E-15 | 3.28E-11 | 0.28 | 0.25 | 0.03 | 1.13 |
| mesenchyme | Hist1h2bl | 0.78 | 1.75E-15 | 3.64E-11 | 0.12 | 0.08 | 0.03 | 1.4 |
| mesenchyme | Gemin2 | 0.32 | 1.87E-15 | 3.88E-11 | 0.41 | 0.38 | 0.03 | 1.07 |
| mesenchyme | Dnajc17 | 0.34 | 1.99E-15 | 4.14E-11 | 0.34 | 0.31 | 0.04 | 1.12 |
| mesenchyme | Tbc1d12 | 0.69 | 2.04E-15 | 4.24E-11 | 0.18 | 0.14 | 0.04 | 1.25 |
| mesenchyme | Rad9a | 0.46 | 2.09E-15 | 4.34E-11 | 0.23 | 0.19 | 0.04 | 1.19 |
| mesenchyme | Fam53b | 0.62 | 2.15E-15 | 4.46E-11 | 0.19 | 0.15 | 0.04 | 1.25 |

**Supplementary Table 2. Marker genes for the major cell types of integrated clustering of E11.5 control and NotchGOF λ-junction cells.**

| | | | | | | | | |
|---|---|---|---|---|---|---|---|---|
| mesenchyme | Acp2 | 0.43 | 2.15E-15 | 4.47E-11 | 0.28 | 0.25 | 0.04 | 1.16 |
| mesenchyme | Cers6 | 0.4 | 2.23E-15 | 4.64E-11 | 0.26 | 0.22 | 0.04 | 1.19 |
| mesenchyme | Amacr | 0.68 | 2.40E-15 | 4.99E-11 | 0.16 | 0.12 | 0.04 | 1.31 |
| mesenchyme | Dhcr7 | 0.3 | 2.53E-15 | 5.26E-11 | 0.34 | 0.31 | 0.04 | 1.12 |
| mesenchyme | Cnr1 | 0.8 | 2.59E-15 | 5.38E-11 | 0.1 | 0.07 | 0.03 | 1.49 |
| mesenchyme | Ttll4 | 0.37 | 2.63E-15 | 5.47E-11 | 0.3 | 0.26 | 0.04 | 1.14 |
| mesenchyme | Hyou1 | 0.43 | 2.78E-15 | 5.79E-11 | 0.3 | 0.27 | 0.04 | 1.13 |
| mesenchyme | Katnal1 | 0.62 | 2.83E-15 | 5.88E-11 | 0.19 | 0.15 | 0.04 | 1.25 |
| mesenchyme | Fam160b1 | 0.4 | 2.89E-15 | 6.01E-11 | 0.3 | 0.26 | 0.04 | 1.14 |
| mesenchyme | Cyth1 | 0.66 | 3.12E-15 | 6.48E-11 | 0.16 | 0.12 | 0.04 | 1.3 |
| mesenchyme | Prkce | 0.72 | 3.16E-15 | 6.56E-11 | 0.17 | 0.13 | 0.04 | 1.28 |
| mesenchyme | Fam13a | 0.84 | 3.43E-15 | 7.13E-11 | 0.11 | 0.07 | 0.03 | 1.45 |
| mesenchyme | Rnpc3 | 0.32 | 3.75E-15 | 7.80E-11 | 0.38 | 0.35 | 0.03 | 1.09 |
| mesenchyme | Nhsl2 | 0.61 | 3.84E-15 | 7.98E-11 | 0.18 | 0.14 | 0.04 | 1.27 |
| mesenchyme | Iqgap2 | 0.35 | 3.89E-15 | 8.10E-11 | 0.38 | 0.34 | 0.03 | 1.1 |
| mesenchyme | Apaf1 | 0.48 | 4.06E-15 | 8.45E-11 | 0.24 | 0.2 | 0.04 | 1.19 |
| mesenchyme | Abt1 | 0.28 | 4.08E-15 | 8.49E-11 | 0.43 | 0.41 | 0.02 | 1.05 |
| mesenchyme | Pias2 | 0.25 | 4.08E-15 | 8.49E-11 | 0.41 | 0.38 | 0.03 | 1.08 |
| mesenchyme | Tmem151b | 0.77 | 4.33E-15 | 9.01E-11 | 0.1 | 0.07 | 0.03 | 1.46 |
| mesenchyme | Dgcr2 | 0.25 | 4.33E-15 | 9.01E-11 | 0.48 | 0.45 | 0.03 | 1.07 |
| mesenchyme | Inpp5a | 0.59 | 4.48E-15 | 9.32E-11 | 0.17 | 0.13 | 0.04 | 1.29 |
| mesenchyme | Pold2 | 0.27 | 4.52E-15 | 9.39E-11 | 0.44 | 0.41 | 0.03 | 1.07 |
| mesenchyme | Abcd4 | 0.59 | 4.58E-15 | 9.53E-11 | 0.17 | 0.13 | 0.04 | 1.29 |
| mesenchyme | Secisbp2l | 0.37 | 4.87E-15 | 1.01E-10 | 0.35 | 0.31 | 0.04 | 1.12 |
| mesenchyme | Adam9 | 0.68 | 5.28E-15 | 1.10E-10 | 0.15 | 0.12 | 0.04 | 1.32 |
| mesenchyme | Tsix | 0.3 | 5.55E-15 | 1.15E-10 | 0.39 | 0.35 | 0.03 | 1.09 |
| mesenchyme | Rab4b | 0.29 | 5.56E-15 | 1.16E-10 | 0.4 | 0.37 | 0.03 | 1.07 |
| mesenchyme | Ccdc138 | 0.51 | 5.70E-15 | 1.18E-10 | 0.21 | 0.17 | 0.04 | 1.22 |
| mesenchyme | Snapc2 | 0.37 | 5.76E-15 | 1.20E-10 | 0.32 | 0.29 | 0.03 | 1.11 |
| mesenchyme | Tspyl4 | 0.37 | 5.81E-15 | 1.21E-10 | 0.25 | 0.21 | 0.04 | 1.19 |
| mesenchyme | Hist1h2bj | 0.54 | 6.11E-15 | 1.27E-10 | 0.26 | 0.23 | 0.03 | 1.14 |
| mesenchyme | Wasf3 | 0.53 | 6.13E-15 | 1.27E-10 | 0.18 | 0.14 | 0.04 | 1.28 |
| mesenchyme | Ykt6 | 0.28 | 6.15E-15 | 1.28E-10 | 0.42 | 0.39 | 0.03 | 1.06 |
| mesenchyme | Exo5 | 0.41 | 6.20E-15 | 1.29E-10 | 0.28 | 0.24 | 0.04 | 1.15 |
| mesenchyme | Tbc1d19 | 0.44 | 6.26E-15 | 1.30E-10 | 0.23 | 0.19 | 0.04 | 1.22 |
| mesenchyme | Klhl26 | 0.54 | 6.57E-15 | 1.37E-10 | 0.19 | 0.15 | 0.04 | 1.25 |
| mesenchyme | Cd276 | 0.32 | 6.84E-15 | 1.42E-10 | 0.34 | 0.3 | 0.04 | 1.13 |
| mesenchyme | Urm1 | 0.27 | 6.88E-15 | 1.43E-10 | 0.39 | 0.36 | 0.03 | 1.09 |
| mesenchyme | Nsdhl | 0.27 | 8.08E-15 | 1.68E-10 | 0.39 | 0.35 | 0.03 | 1.1 |
| mesenchyme | Lrrc8a | 0.44 | 8.10E-15 | 1.68E-10 | 0.29 | 0.25 | 0.04 | 1.14 |
| mesenchyme | Ltn1 | 0.35 | 8.11E-15 | 1.69E-10 | 0.34 | 0.31 | 0.03 | 1.1 |
| mesenchyme | Rpgrip1l | 0.3 | 8.13E-15 | 1.69E-10 | 0.33 | 0.29 | 0.04 | 1.13 |
| mesenchyme | Zfp26 | 0.3 | 8.89E-15 | 1.85E-10 | 0.37 | 0.34 | 0.04 | 1.11 |
| mesenchyme | Wars | 0.3 | 9.42E-15 | 1.96E-10 | 0.42 | 0.4 | 0.02 | 1.06 |
| mesenchyme | Tigd2 | 0.29 | 9.99E-15 | 2.08E-10 | 0.4 | 0.37 | 0.03 | 1.08 |
| mesenchyme | Lipa | 0.37 | 1.00E-14 | 2.09E-10 | 0.23 | 0.19 | 0.04 | 1.21 |
| mesenchyme | Cpq | 0.75 | 1.01E-14 | 2.09E-10 | 0.17 | 0.14 | 0.03 | 1.24 |
| mesenchyme | Glt8d1 | 0.36 | 1.18E-14 | 2.46E-10 | 0.35 | 0.32 | 0.03 | 1.1 |
| mesenchyme | Cnp | 0.57 | 1.28E-14 | 2.67E-10 | 0.17 | 0.13 | 0.04 | 1.27 |
| mesenchyme | Adat2 | 0.44 | 1.46E-14 | 3.04E-10 | 0.31 | 0.28 | 0.03 | 1.11 |
| mesenchyme | Manea | 0.48 | 1.47E-14 | 3.05E-10 | 0.25 | 0.21 | 0.04 | 1.18 |
| mesenchyme | Opa3 | 0.35 | 1.47E-14 | 3.06E-10 | 0.36 | 0.33 | 0.03 | 1.09 |
| mesenchyme | Vcpkmt | 0.52 | 1.52E-14 | 3.16E-10 | 0.22 | 0.19 | 0.04 | 1.19 |
| mesenchyme | Ttc23 | 0.26 | 1.68E-14 | 3.49E-10 | 0.31 | 0.27 | 0.04 | 1.15 |
| mesenchyme | Smarcd3 | 0.29 | 1.77E-14 | 3.68E-10 | 0.29 | 0.25 | 0.04 | 1.17 |
| mesenchyme | Dgcr8 | 0.35 | 1.79E-14 | 3.72E-10 | 0.38 | 0.35 | 0.03 | 1.07 |

**Supplementary Table 2. Marker genes for the major cell types of integrated clustering of E11.5 control and NotchGOF λ-junction cells.**

| | | | | | | | | |
|---|---|---|---|---|---|---|---|---|
| mesenchyme | Zfp689 | 0.76 | 1.91E-14 | 3.97E-10 | 0.11 | 0.08 | 0.03 | 1.42 |
| mesenchyme | Odr4 | 0.4 | 1.99E-14 | 4.13E-10 | 0.34 | 0.31 | 0.03 | 1.1 |
| mesenchyme | Aspm | 0.4 | 2.02E-14 | 4.21E-10 | 0.42 | 0.4 | 0.03 | 1.06 |
| mesenchyme | Plk1 | 0.28 | 2.08E-14 | 4.33E-10 | 0.48 | 0.44 | 0.03 | 1.07 |
| mesenchyme | Nup160 | 0.38 | 2.11E-14 | 4.38E-10 | 0.32 | 0.29 | 0.03 | 1.11 |
| mesenchyme | Coa7 | 0.3 | 2.19E-14 | 4.56E-10 | 0.39 | 0.36 | 0.03 | 1.08 |
| mesenchyme | Akna | 0.67 | 2.23E-14 | 4.64E-10 | 0.12 | 0.09 | 0.03 | 1.39 |
| mesenchyme | Hist1h2bn | 0.8 | 2.32E-14 | 4.83E-10 | 0.16 | 0.13 | 0.03 | 1.26 |
| mesenchyme | Entr1 | 0.26 | 2.40E-14 | 4.99E-10 | 0.44 | 0.42 | 0.02 | 1.05 |
| mesenchyme | Cpsf4 | 0.27 | 2.41E-14 | 5.01E-10 | 0.4 | 0.37 | 0.03 | 1.08 |
| mesenchyme | Stard7 | 0.26 | 2.44E-14 | 5.08E-10 | 0.43 | 0.41 | 0.03 | 1.06 |
| mesenchyme | Limk1 | 0.68 | 2.45E-14 | 5.10E-10 | 0.16 | 0.13 | 0.04 | 1.28 |
| mesenchyme | Rpgrip1 | 0.45 | 2.47E-14 | 5.14E-10 | 0.28 | 0.25 | 0.04 | 1.15 |
| mesenchyme | Cep85 | 0.56 | 2.50E-14 | 5.20E-10 | 0.18 | 0.14 | 0.04 | 1.25 |
| mesenchyme | Pi4kb | 0.54 | 2.58E-14 | 5.37E-10 | 0.21 | 0.17 | 0.04 | 1.21 |
| mesenchyme | Gm42047 | 0.91 | 2.64E-14 | 5.48E-10 | 0.15 | 0.12 | 0.03 | 1.28 |
| mesenchyme | Lrrc41 | 0.35 | 2.85E-14 | 5.93E-10 | 0.39 | 0.37 | 0.02 | 1.07 |
| mesenchyme | Med23 | 0.38 | 3.04E-14 | 6.32E-10 | 0.29 | 0.25 | 0.04 | 1.14 |
| mesenchyme | Pex11b | 0.42 | 3.07E-14 | 6.38E-10 | 0.3 | 0.28 | 0.03 | 1.11 |
| mesenchyme | Mboat7 | 0.34 | 3.20E-14 | 6.65E-10 | 0.34 | 0.31 | 0.03 | 1.11 |
| mesenchyme | Otud7b | 0.28 | 3.21E-14 | 6.67E-10 | 0.4 | 0.37 | 0.03 | 1.08 |
| mesenchyme | Eps15l1 | 0.32 | 3.34E-14 | 6.94E-10 | 0.38 | 0.35 | 0.03 | 1.08 |
| mesenchyme | Lrrc75a | 0.49 | 3.35E-14 | 6.96E-10 | 0.22 | 0.18 | 0.04 | 1.2 |
| mesenchyme | Cenpi | 0.58 | 3.50E-14 | 7.29E-10 | 0.18 | 0.14 | 0.04 | 1.24 |
| mesenchyme | Hus1 | 0.46 | 3.61E-14 | 7.51E-10 | 0.26 | 0.23 | 0.03 | 1.14 |
| mesenchyme | Yju2 | 0.43 | 4.17E-14 | 8.67E-10 | 0.25 | 0.22 | 0.04 | 1.16 |
| mesenchyme | Spidr | 0.4 | 4.25E-14 | 8.85E-10 | 0.32 | 0.29 | 0.03 | 1.1 |
| mesenchyme | 720489N17Ri | 0.61 | 4.49E-14 | 9.34E-10 | 0.19 | 0.15 | 0.04 | 1.24 |
| mesenchyme | Coq3 | 0.5 | 4.52E-14 | 9.39E-10 | 0.2 | 0.16 | 0.04 | 1.23 |
| mesenchyme | Zfp668 | 0.43 | 4.72E-14 | 9.81E-10 | 0.26 | 0.22 | 0.04 | 1.16 |
| mesenchyme | Pik3r2 | 0.31 | 4.96E-14 | 1.03E-09 | 0.33 | 0.29 | 0.04 | 1.13 |
| mesenchyme | Tsc1 | 0.47 | 5.02E-14 | 1.04E-09 | 0.23 | 0.2 | 0.04 | 1.18 |
| mesenchyme | Arhgef10l | 0.57 | 5.63E-14 | 1.17E-09 | 0.16 | 0.12 | 0.04 | 1.29 |
| mesenchyme | Bcat2 | 0.5 | 5.73E-14 | 1.19E-09 | 0.3 | 0.28 | 0.03 | 1.09 |
| mesenchyme | Supt7l | 0.49 | 5.95E-14 | 1.24E-09 | 0.26 | 0.22 | 0.03 | 1.14 |
| mesenchyme | Mdc1 | 0.3 | 6.28E-14 | 1.31E-09 | 0.4 | 0.37 | 0.03 | 1.08 |
| mesenchyme | E2f1 | 0.47 | 6.41E-14 | 1.33E-09 | 0.29 | 0.26 | 0.03 | 1.11 |
| mesenchyme | 610037D02Ri | 0.4 | 6.82E-14 | 1.42E-09 | 0.35 | 0.32 | 0.03 | 1.1 |
| mesenchyme | Magi1 | 0.43 | 6.98E-14 | 1.45E-09 | 0.32 | 0.29 | 0.03 | 1.11 |
| mesenchyme | Msra | 0.62 | 7.19E-14 | 1.50E-09 | 0.18 | 0.14 | 0.03 | 1.24 |
| mesenchyme | Gopc | 0.34 | 7.91E-14 | 1.64E-09 | 0.33 | 0.3 | 0.03 | 1.11 |
| mesenchyme | Dcaf10 | 0.55 | 9.00E-14 | 1.87E-09 | 0.27 | 0.24 | 0.03 | 1.11 |
| mesenchyme | Tube1 | 0.57 | 9.80E-14 | 2.04E-09 | 0.19 | 0.15 | 0.04 | 1.23 |
| mesenchyme | Gstk1 | 0.31 | 1.01E-13 | 2.10E-09 | 0.19 | 0.15 | 0.04 | 1.26 |
| mesenchyme | Evc2 | 0.56 | 1.04E-13 | 2.16E-09 | 0.19 | 0.15 | 0.04 | 1.24 |
| mesenchyme | Ints13 | 0.3 | 1.12E-13 | 2.32E-09 | 0.43 | 0.41 | 0.02 | 1.05 |
| mesenchyme | Dus2 | 0.67 | 1.15E-13 | 2.39E-09 | 0.16 | 0.13 | 0.03 | 1.26 |
| mesenchyme | Nop9 | 0.29 | 1.16E-13 | 2.42E-09 | 0.39 | 0.36 | 0.02 | 1.07 |
| mesenchyme | Ift122 | 0.42 | 1.23E-13 | 2.56E-09 | 0.23 | 0.19 | 0.04 | 1.2 |
| mesenchyme | Arhgap11a | 0.33 | 1.27E-13 | 2.64E-09 | 0.41 | 0.39 | 0.02 | 1.05 |
| mesenchyme | Tmem246 | 0.72 | 1.28E-13 | 2.66E-09 | 0.14 | 0.11 | 0.03 | 1.31 |
| mesenchyme | Tmtc2 | 0.47 | 1.28E-13 | 2.66E-09 | 0.21 | 0.17 | 0.04 | 1.21 |
| mesenchyme | Gm4737 | 0.8 | 1.34E-13 | 2.78E-09 | 0.13 | 0.1 | 0.03 | 1.32 |
| mesenchyme | Cdkn2aip | 0.32 | 1.40E-13 | 2.92E-09 | 0.4 | 0.37 | 0.02 | 1.06 |
| mesenchyme | Mark1 | 0.37 | 1.46E-13 | 3.04E-09 | 0.26 | 0.23 | 0.04 | 1.16 |
| mesenchyme | Sap130 | 0.4 | 1.51E-13 | 3.14E-09 | 0.27 | 0.24 | 0.03 | 1.13 |

**Supplementary Table 2. Marker genes for the major cell types of integrated clustering of E11.5 control and NotchGOF λ-junction cells.**

| | | | | | | | | |
|---|---|---|---|---|---|---|---|---|
| mesenchyme | .610020C07Ri | 0.57 | 1.53E-13 | 3.19E-09 | 0.2 | 0.16 | 0.03 | 1.21 |
| mesenchyme | Ccnk | 0.28 | 1.64E-13 | 3.40E-09 | 0.43 | 0.41 | 0.02 | 1.06 |
| mesenchyme | Brip1 | 0.53 | 1.69E-13 | 3.52E-09 | 0.24 | 0.21 | 0.03 | 1.15 |
| mesenchyme | Axin1 | 0.33 | 1.80E-13 | 3.73E-09 | 0.37 | 0.34 | 0.03 | 1.08 |
| mesenchyme | Hmces | 0.27 | 1.86E-13 | 3.87E-09 | 0.38 | 0.35 | 0.03 | 1.08 |
| mesenchyme | Crat | 0.59 | 1.90E-13 | 3.95E-09 | 0.22 | 0.18 | 0.03 | 1.17 |
| mesenchyme | Map3k3 | 0.57 | 2.09E-13 | 4.35E-09 | 0.17 | 0.13 | 0.03 | 1.26 |
| mesenchyme | Zfp982 | 0.81 | 2.16E-13 | 4.48E-09 | 0.11 | 0.08 | 0.03 | 1.37 |
| mesenchyme | Sipa1l2 | 0.62 | 2.17E-13 | 4.50E-09 | 0.15 | 0.11 | 0.03 | 1.3 |
| mesenchyme | ;930430L01Ri | 0.77 | 2.18E-13 | 4.54E-09 | 0.11 | 0.08 | 0.03 | 1.39 |
| mesenchyme | Rad54l | 0.34 | 2.24E-13 | 4.66E-09 | 0.32 | 0.29 | 0.03 | 1.11 |
| mesenchyme | Tmem168 | 0.42 | 2.36E-13 | 4.92E-09 | 0.29 | 0.26 | 0.03 | 1.13 |
| mesenchyme | Shkbp1 | 0.31 | 2.40E-13 | 4.99E-09 | 0.35 | 0.32 | 0.03 | 1.1 |
| mesenchyme | Alg2 | 0.33 | 2.48E-13 | 5.17E-09 | 0.33 | 0.3 | 0.03 | 1.11 |
| mesenchyme | Dhx57 | 0.33 | 2.53E-13 | 5.26E-09 | 0.3 | 0.27 | 0.04 | 1.14 |
| mesenchyme | Kifc1 | 0.43 | 2.67E-13 | 5.56E-09 | 0.3 | 0.27 | 0.03 | 1.11 |
| mesenchyme | Alyref2 | 0.74 | 2.71E-13 | 5.64E-09 | 0.13 | 0.1 | 0.03 | 1.31 |
| mesenchyme | BC055324 | 0.46 | 2.92E-13 | 6.06E-09 | 0.24 | 0.21 | 0.03 | 1.15 |
| mesenchyme | Cry1 | 0.42 | 3.08E-13 | 6.41E-09 | 0.24 | 0.21 | 0.03 | 1.16 |
| mesenchyme | Hif1an | 0.4 | 3.12E-13 | 6.50E-09 | 0.32 | 0.29 | 0.03 | 1.1 |
| mesenchyme | Prr7 | 0.38 | 3.20E-13 | 6.66E-09 | 0.25 | 0.21 | 0.04 | 1.18 |
| mesenchyme | Camk2g | 0.33 | 3.22E-13 | 6.70E-09 | 0.35 | 0.32 | 0.03 | 1.1 |
| mesenchyme | Tdp1 | 0.34 | 3.49E-13 | 7.25E-09 | 0.32 | 0.29 | 0.03 | 1.09 |
| mesenchyme | Chp1 | 0.28 | 3.57E-13 | 7.43E-09 | 0.4 | 0.38 | 0.02 | 1.05 |
| mesenchyme | Pofut1 | 0.53 | 3.61E-13 | 7.51E-09 | 0.23 | 0.2 | 0.03 | 1.17 |
| mesenchyme | Zbtb45 | 0.46 | 3.77E-13 | 7.84E-09 | 0.23 | 0.2 | 0.03 | 1.17 |
| mesenchyme | 630089N07Ri | 0.72 | 3.83E-13 | 7.96E-09 | 0.15 | 0.12 | 0.03 | 1.28 |
| mesenchyme | Nqo2 | 0.66 | 3.87E-13 | 8.04E-09 | 0.14 | 0.11 | 0.03 | 1.31 |
| mesenchyme | Zfp711 | 0.39 | 4.13E-13 | 8.59E-09 | 0.23 | 0.19 | 0.04 | 1.19 |
| mesenchyme | Fez2 | 0.32 | 4.23E-13 | 8.80E-09 | 0.32 | 0.29 | 0.03 | 1.1 |
| mesenchyme | Mfsd14a | 0.26 | 4.31E-13 | 8.96E-09 | 0.43 | 0.4 | 0.02 | 1.06 |
| mesenchyme | Gm17066 | 0.54 | 4.32E-13 | 8.98E-09 | 0.21 | 0.18 | 0.03 | 1.19 |
| mesenchyme | Snapc3 | 0.32 | 4.65E-13 | 9.68E-09 | 0.34 | 0.31 | 0.03 | 1.09 |
| mesenchyme | Pck2 | 0.62 | 4.74E-13 | 9.86E-09 | 0.22 | 0.19 | 0.03 | 1.16 |
| mesenchyme | Pecr | 0.44 | 4.93E-13 | 1.02E-08 | 0.22 | 0.18 | 0.03 | 1.19 |
| mesenchyme | Hdx | 0.66 | 5.03E-13 | 1.05E-08 | 0.11 | 0.08 | 0.03 | 1.39 |
| mesenchyme | Cept1 | 0.33 | 5.05E-13 | 1.05E-08 | 0.34 | 0.32 | 0.03 | 1.09 |
| mesenchyme | Ypel2 | 0.63 | 5.14E-13 | 1.07E-08 | 0.14 | 0.11 | 0.03 | 1.3 |
| mesenchyme | Dpy19l4 | 0.37 | 5.14E-13 | 1.07E-08 | 0.29 | 0.26 | 0.03 | 1.12 |
| mesenchyme | Mfn1 | 0.34 | 5.23E-13 | 1.09E-08 | 0.3 | 0.27 | 0.03 | 1.12 |
| mesenchyme | Nol9 | 0.32 | 5.34E-13 | 1.11E-08 | 0.33 | 0.3 | 0.03 | 1.1 |
| mesenchyme | Nfatc3 | 0.35 | 5.37E-13 | 1.12E-08 | 0.37 | 0.35 | 0.02 | 1.07 |
| mesenchyme | Lemd2 | 0.37 | 5.86E-13 | 1.22E-08 | 0.33 | 0.3 | 0.03 | 1.09 |
| mesenchyme | Dmap1 | 0.29 | 5.95E-13 | 1.24E-08 | 0.39 | 0.36 | 0.03 | 1.07 |
| mesenchyme | Acd | 0.28 | 6.17E-13 | 1.28E-08 | 0.39 | 0.37 | 0.03 | 1.07 |
| mesenchyme | Samd8 | 0.48 | 6.29E-13 | 1.31E-08 | 0.22 | 0.19 | 0.04 | 1.19 |
| mesenchyme | Pdss2 | 0.5 | 6.44E-13 | 1.34E-08 | 0.24 | 0.21 | 0.03 | 1.14 |
| mesenchyme | Prom1 | 0.25 | 7.04E-13 | 1.47E-08 | 0.42 | 0.39 | 0.04 | 1.09 |
| mesenchyme | Zc3h7a | 0.34 | 7.35E-13 | 1.53E-08 | 0.35 | 0.33 | 0.03 | 1.08 |
| mesenchyme | Zbtb41 | 0.4 | 7.37E-13 | 1.53E-08 | 0.26 | 0.23 | 0.03 | 1.14 |
| mesenchyme | Ppp1r9b | 0.42 | 7.39E-13 | 1.54E-08 | 0.27 | 0.24 | 0.03 | 1.12 |
| mesenchyme | Smg5 | 0.34 | 7.43E-13 | 1.55E-08 | 0.34 | 0.32 | 0.02 | 1.08 |
| mesenchyme | Washc5 | 0.37 | 7.52E-13 | 1.56E-08 | 0.33 | 0.3 | 0.03 | 1.09 |
| mesenchyme | Mvd | 0.29 | 8.14E-13 | 1.69E-08 | 0.31 | 0.28 | 0.04 | 1.13 |
| mesenchyme | Zfp267 | 0.46 | 8.16E-13 | 1.70E-08 | 0.23 | 0.2 | 0.03 | 1.17 |
| mesenchyme | Eml3 | 0.72 | 8.49E-13 | 1.77E-08 | 0.14 | 0.11 | 0.03 | 1.29 |

**Supplementary Table 2. Marker genes for the major cell types of integrated clustering of E11.5 control and NotchGOF λ-junction cells.**

| | | | | | | | | |
|---|---|---|---|---|---|---|---|---|
| mesenchyme | Zfp251 | 0.43 | 9.63E-13 | 2.00E-08 | 0.28 | 0.25 | 0.03 | 1.13 |
| mesenchyme | Mrpl47 | 0.29 | 9.81E-13 | 2.04E-08 | 0.39 | 0.36 | 0.02 | 1.07 |
| mesenchyme | Coq6 | 0.38 | 1.02E-12 | 2.12E-08 | 0.31 | 0.28 | 0.03 | 1.1 |
| mesenchyme | Creld2 | 0.25 | 1.08E-12 | 2.24E-08 | 0.46 | 0.43 | 0.03 | 1.06 |
| mesenchyme | Zfp523 | 0.33 | 1.09E-12 | 2.26E-08 | 0.32 | 0.29 | 0.03 | 1.12 |
| mesenchyme | Setd1b | 0.27 | 1.10E-12 | 2.29E-08 | 0.44 | 0.42 | 0.02 | 1.05 |
| mesenchyme | Usp4 | 0.25 | 1.18E-12 | 2.46E-08 | 0.44 | 0.43 | 0.01 | 1.03 |
| mesenchyme | Dyrk1a | 0.27 | 1.18E-12 | 2.46E-08 | 0.39 | 0.37 | 0.02 | 1.06 |
| mesenchyme | Aurka | 0.37 | 1.20E-12 | 2.50E-08 | 0.46 | 0.44 | 0.02 | 1.04 |
| mesenchyme | Bbs7 | 0.65 | 1.21E-12 | 2.51E-08 | 0.17 | 0.14 | 0.03 | 1.22 |
| mesenchyme | Itprid2 | 0.99 | 1.30E-12 | 2.71E-08 | 0.16 | 0.13 | 0.03 | 1.23 |
| mesenchyme | L3mbtl3 | 0.32 | 1.33E-12 | 2.76E-08 | 0.4 | 0.38 | 0.03 | 1.07 |
| mesenchyme | Sirt6 | 0.28 | 1.33E-12 | 2.77E-08 | 0.35 | 0.32 | 0.03 | 1.09 |
| mesenchyme | Gtpbp10 | 0.29 | 1.35E-12 | 2.81E-08 | 0.32 | 0.29 | 0.03 | 1.11 |
| mesenchyme | Bnip1 | 0.35 | 1.36E-12 | 2.82E-08 | 0.37 | 0.35 | 0.02 | 1.06 |
| mesenchyme | Brca1 | 0.28 | 1.63E-12 | 3.39E-08 | 0.39 | 0.36 | 0.03 | 1.09 |
| mesenchyme | Bloc1s4 | 0.33 | 1.64E-12 | 3.41E-08 | 0.33 | 0.31 | 0.03 | 1.08 |
| mesenchyme | Dtnbp1 | 0.25 | 1.65E-12 | 3.42E-08 | 0.43 | 0.41 | 0.02 | 1.05 |
| mesenchyme | Arhgef17 | 0.27 | 1.71E-12 | 3.55E-08 | 0.37 | 0.34 | 0.04 | 1.1 |
| mesenchyme | Cdc6 | 0.52 | 1.72E-12 | 3.59E-08 | 0.29 | 0.26 | 0.03 | 1.11 |
| mesenchyme | Traip | 0.43 | 1.86E-12 | 3.86E-08 | 0.28 | 0.26 | 0.03 | 1.11 |
| mesenchyme | Wdr5b | 0.8 | 1.87E-12 | 3.90E-08 | 0.1 | 0.07 | 0.03 | 1.4 |
| mesenchyme | Ilvbl | 0.48 | 2.05E-12 | 4.26E-08 | 0.23 | 0.2 | 0.03 | 1.17 |
| mesenchyme | Sco1 | 0.49 | 2.07E-12 | 4.30E-08 | 0.2 | 0.16 | 0.03 | 1.21 |
| mesenchyme | Nup155 | 0.29 | 2.16E-12 | 4.50E-08 | 0.41 | 0.39 | 0.02 | 1.06 |
| mesenchyme | Nupl2 | 0.62 | 2.19E-12 | 4.56E-08 | 0.17 | 0.14 | 0.03 | 1.23 |
| mesenchyme | Gse1 | 0.34 | 2.20E-12 | 4.57E-08 | 0.32 | 0.29 | 0.03 | 1.1 |
| mesenchyme | Elk1 | 0.49 | 2.26E-12 | 4.69E-08 | 0.18 | 0.15 | 0.03 | 1.23 |
| mesenchyme | Ube2g2 | 0.33 | 2.49E-12 | 5.17E-08 | 0.34 | 0.31 | 0.02 | 1.07 |
| mesenchyme | Pms1 | 0.52 | 2.70E-12 | 5.61E-08 | 0.22 | 0.19 | 0.03 | 1.17 |
| mesenchyme | Agbl5 | 0.5 | 2.85E-12 | 5.92E-08 | 0.21 | 0.17 | 0.03 | 1.2 |
| mesenchyme | Rcc1l | 0.36 | 3.10E-12 | 6.45E-08 | 0.25 | 0.22 | 0.03 | 1.15 |
| mesenchyme | Zfp952 | 0.55 | 3.26E-12 | 6.78E-08 | 0.16 | 0.13 | 0.03 | 1.25 |
| mesenchyme | Zranb3 | 0.45 | 3.39E-12 | 7.04E-08 | 0.22 | 0.19 | 0.03 | 1.17 |
| mesenchyme | Nemp1 | 0.62 | 3.97E-12 | 8.25E-08 | 0.14 | 0.11 | 0.03 | 1.28 |
| mesenchyme | Ftsj1 | 0.37 | 4.18E-12 | 8.69E-08 | 0.3 | 0.28 | 0.02 | 1.09 |
| mesenchyme | Brip1os | 0.34 | 4.25E-12 | 8.85E-08 | 0.37 | 0.35 | 0.02 | 1.07 |
| mesenchyme | Hps4 | 0.31 | 4.31E-12 | 8.97E-08 | 0.32 | 0.29 | 0.03 | 1.11 |
| mesenchyme | Rps6ka5 | 0.67 | 4.73E-12 | 9.83E-08 | 0.14 | 0.11 | 0.03 | 1.28 |
| mesenchyme | Steap2 | 0.69 | 5.37E-12 | 1.12E-07 | 0.1 | 0.07 | 0.03 | 1.41 |
| mesenchyme | Alg3 | 0.46 | 5.39E-12 | 1.12E-07 | 0.29 | 0.26 | 0.03 | 1.1 |
| mesenchyme | Ggps1 | 0.27 | 5.45E-12 | 1.13E-07 | 0.43 | 0.41 | 0.02 | 1.04 |
| mesenchyme | Mus81 | 0.48 | 5.84E-12 | 1.21E-07 | 0.19 | 0.15 | 0.03 | 1.21 |
| mesenchyme | Atg4b | 0.29 | 5.99E-12 | 1.25E-07 | 0.37 | 0.35 | 0.02 | 1.06 |
| mesenchyme | Chtf18 | 0.45 | 5.99E-12 | 1.25E-07 | 0.25 | 0.21 | 0.03 | 1.14 |
| mesenchyme | Kat7 | 0.26 | 6.32E-12 | 1.31E-07 | 0.41 | 0.39 | 0.02 | 1.05 |
| mesenchyme | Mipep | 0.43 | 6.36E-12 | 1.32E-07 | 0.27 | 0.24 | 0.03 | 1.11 |
| mesenchyme | Ndufaf1 | 0.34 | 6.42E-12 | 1.33E-07 | 0.31 | 0.28 | 0.03 | 1.1 |
| mesenchyme | Sik2 | 0.71 | 6.47E-12 | 1.35E-07 | 0.17 | 0.14 | 0.03 | 1.21 |
| mesenchyme | Mynn | 0.37 | 6.94E-12 | 1.44E-07 | 0.3 | 0.27 | 0.03 | 1.1 |
| mesenchyme | Bcor | 0.27 | 7.46E-12 | 1.55E-07 | 0.42 | 0.4 | 0.02 | 1.04 |
| mesenchyme | Fam199x | 0.26 | 7.72E-12 | 1.61E-07 | 0.35 | 0.32 | 0.03 | 1.1 |
| mesenchyme | Zswim9 | 0.52 | 7.95E-12 | 1.65E-07 | 0.18 | 0.15 | 0.03 | 1.22 |
| mesenchyme | Mast2 | 0.39 | 8.06E-12 | 1.68E-07 | 0.3 | 0.28 | 0.03 | 1.09 |
| mesenchyme | Ttc26 | 0.75 | 8.21E-12 | 1.71E-07 | 0.12 | 0.09 | 0.03 | 1.32 |
| mesenchyme | Anks3 | 0.51 | 9.40E-12 | 1.95E-07 | 0.21 | 0.18 | 0.03 | 1.17 |

**Supplementary Table 2. Marker genes for the major cell types of integrated clustering of E11.5 control and NotchGOF λ-junction cells.**

| | | | | | | | | |
|---|---|---|---|---|---|---|---|---|
| mesenchyme | Atp6v0a2 | 0.37 | 9.86E-12 | 2.05E-07 | 0.27 | 0.25 | 0.03 | 1.11 |
| mesenchyme | Lrp12 | 0.44 | 1.07E-11 | 2.24E-07 | 0.21 | 0.18 | 0.03 | 1.19 |
| mesenchyme | Zfp280b | 0.56 | 1.14E-11 | 2.36E-07 | 0.17 | 0.14 | 0.03 | 1.22 |
| mesenchyme | Mecom | 0.54 | 1.17E-11 | 2.43E-07 | 0.41 | 0.4 | 0.01 | 1.02 |
| mesenchyme | Taco1 | 0.44 | 1.21E-11 | 2.52E-07 | 0.19 | 0.16 | 0.03 | 1.21 |
| mesenchyme | Slc39a13 | 0.66 | 1.23E-11 | 2.56E-07 | 0.15 | 0.12 | 0.03 | 1.25 |
| mesenchyme | Slc39a14 | 0.55 | 1.24E-11 | 2.57E-07 | 0.12 | 0.09 | 0.03 | 1.35 |
| mesenchyme | Gm49797 | 0.41 | 1.33E-11 | 2.76E-07 | 0.25 | 0.22 | 0.03 | 1.14 |
| mesenchyme | Gpr161 | 0.7 | 1.44E-11 | 3.00E-07 | 0.12 | 0.09 | 0.03 | 1.32 |
| mesenchyme | Trmt13 | 0.41 | 1.45E-11 | 3.02E-07 | 0.3 | 0.27 | 0.03 | 1.1 |
| mesenchyme | Atr | 0.33 | 1.85E-11 | 3.84E-07 | 0.34 | 0.32 | 0.03 | 1.08 |
| mesenchyme | Fbxl5 | 0.29 | 1.86E-11 | 3.87E-07 | 0.37 | 0.35 | 0.02 | 1.06 |
| mesenchyme | Xpc | 0.31 | 2.08E-11 | 4.32E-07 | 0.34 | 0.32 | 0.03 | 1.09 |
| mesenchyme | Smyd4 | 0.53 | 2.14E-11 | 4.45E-07 | 0.19 | 0.16 | 0.03 | 1.19 |
| mesenchyme | Slc5a3 | 0.49 | 2.15E-11 | 4.48E-07 | 0.22 | 0.19 | 0.03 | 1.16 |
| mesenchyme | Zfp275 | 0.42 | 2.21E-11 | 4.59E-07 | 0.23 | 0.2 | 0.03 | 1.15 |
| mesenchyme | Akap8l | 0.28 | 2.27E-11 | 4.73E-07 | 0.38 | 0.36 | 0.02 | 1.06 |
| mesenchyme | Fbh1 | 0.26 | 2.32E-11 | 4.82E-07 | 0.37 | 0.35 | 0.02 | 1.07 |
| mesenchyme | Klf4 | 0.41 | 2.49E-11 | 5.18E-07 | 0.22 | 0.19 | 0.03 | 1.18 |
| mesenchyme | Papolg | 0.5 | 2.55E-11 | 5.31E-07 | 0.2 | 0.17 | 0.03 | 1.17 |
| mesenchyme | Camk2n2 | 0.71 | 2.55E-11 | 5.31E-07 | 0.11 | 0.08 | 0.03 | 1.34 |
| mesenchyme | Pus7 | 0.33 | 2.56E-11 | 5.32E-07 | 0.33 | 0.31 | 0.02 | 1.08 |
| mesenchyme | Aak1 | 0.31 | 3.31E-11 | 6.87E-07 | 0.36 | 0.34 | 0.02 | 1.06 |
| mesenchyme | Epb41l1 | 0.68 | 3.35E-11 | 6.98E-07 | 0.15 | 0.12 | 0.03 | 1.23 |
| mesenchyme | Nup85 | 0.28 | 3.44E-11 | 7.14E-07 | 0.37 | 0.36 | 0.02 | 1.05 |
| mesenchyme | Slc38a6 | 0.54 | 3.55E-11 | 7.39E-07 | 0.16 | 0.13 | 0.03 | 1.24 |
| mesenchyme | Kif1bp | 0.27 | 3.57E-11 | 7.43E-07 | 0.37 | 0.35 | 0.02 | 1.07 |
| mesenchyme | Trim45 | 0.56 | 3.81E-11 | 7.93E-07 | 0.12 | 0.09 | 0.03 | 1.32 |
| mesenchyme | Eif5a2 | 0.69 | 3.92E-11 | 8.15E-07 | 0.13 | 0.1 | 0.03 | 1.28 |
| mesenchyme | Foxn2 | 0.61 | 4.12E-11 | 8.56E-07 | 0.17 | 0.15 | 0.03 | 1.2 |
| mesenchyme | Usp45 | 0.48 | 4.26E-11 | 8.86E-07 | 0.25 | 0.22 | 0.03 | 1.11 |
| mesenchyme | Diaph2 | 0.58 | 4.42E-11 | 9.20E-07 | 0.17 | 0.14 | 0.03 | 1.23 |
| mesenchyme | Mios | 0.37 | 4.44E-11 | 9.23E-07 | 0.29 | 0.26 | 0.03 | 1.1 |
| mesenchyme | Rad1 | 0.31 | 4.52E-11 | 9.41E-07 | 0.33 | 0.3 | 0.03 | 1.09 |
| mesenchyme | Copz2 | 0.32 | 4.64E-11 | 9.64E-07 | 0.27 | 0.24 | 0.03 | 1.13 |
| mesenchyme | Rnft1 | 0.41 | 4.70E-11 | 9.78E-07 | 0.24 | 0.21 | 0.03 | 1.14 |
| mesenchyme | Pgam5 | 0.28 | 4.90E-11 | 1.02E-06 | 0.38 | 0.35 | 0.02 | 1.06 |
| mesenchyme | Apex2 | 0.42 | 5.00E-11 | 1.04E-06 | 0.24 | 0.22 | 0.03 | 1.12 |
| mesenchyme | Thap1 | 0.34 | 5.02E-11 | 1.04E-06 | 0.27 | 0.24 | 0.03 | 1.11 |
| mesenchyme | Csrnp2 | 0.55 | 5.17E-11 | 1.08E-06 | 0.15 | 0.12 | 0.03 | 1.25 |
| mesenchyme | Rapgef1 | 0.36 | 5.22E-11 | 1.09E-06 | 0.28 | 0.25 | 0.03 | 1.11 |
| mesenchyme | Tmem17 | 0.5 | 5.30E-11 | 1.10E-06 | 0.18 | 0.15 | 0.03 | 1.21 |
| mesenchyme | Zfp810 | 0.51 | 5.40E-11 | 1.12E-06 | 0.2 | 0.17 | 0.03 | 1.19 |
| mesenchyme | Tef | 0.5 | 6.01E-11 | 1.25E-06 | 0.13 | 0.1 | 0.03 | 1.31 |
| mesenchyme | Clasp2 | 0.42 | 6.02E-11 | 1.25E-06 | 0.25 | 0.22 | 0.03 | 1.12 |
| mesenchyme | Nck2 | 0.3 | 6.39E-11 | 1.33E-06 | 0.28 | 0.25 | 0.03 | 1.12 |
| mesenchyme | Troap | 0.49 | 6.41E-11 | 1.33E-06 | 0.19 | 0.16 | 0.03 | 1.18 |
| mesenchyme | Fbxl17 | 0.38 | 6.52E-11 | 1.36E-06 | 0.21 | 0.18 | 0.03 | 1.18 |
| mesenchyme | Akt3 | 0.34 | 6.55E-11 | 1.36E-06 | 0.28 | 0.25 | 0.03 | 1.13 |
| mesenchyme | Hdac7 | 0.3 | 6.63E-11 | 1.38E-06 | 0.3 | 0.27 | 0.03 | 1.12 |
| mesenchyme | Tsen2 | 0.3 | 7.09E-11 | 1.48E-06 | 0.28 | 0.25 | 0.03 | 1.12 |
| mesenchyme | Cdk19 | 0.31 | 7.10E-11 | 1.48E-06 | 0.32 | 0.29 | 0.03 | 1.09 |
| mesenchyme | Mccc1 | 0.38 | 7.21E-11 | 1.50E-06 | 0.22 | 0.18 | 0.03 | 1.17 |
| mesenchyme | Nsmf | 0.57 | 7.38E-11 | 1.53E-06 | 0.18 | 0.15 | 0.03 | 1.2 |
| mesenchyme | Zfp746 | 0.34 | 7.59E-11 | 1.58E-06 | 0.33 | 0.3 | 0.03 | 1.08 |
| mesenchyme | Klhdc4 | 0.27 | 7.60E-11 | 1.58E-06 | 0.36 | 0.34 | 0.02 | 1.06 |

**Supplementary Table 2. Marker genes for the major cell types of integrated clustering of E11.5 control and NotchGOF λ-junction cells.**

| | | | | | | | | |
|---|---|---|---|---|---|---|---|---|
| mesenchyme | Creb3l2 | 0.34 | 8.04E-11 | 1.67E-06 | 0.33 | 0.3 | 0.03 | 1.09 |
| mesenchyme | Cox18 | 0.35 | 8.56E-11 | 1.78E-06 | 0.31 | 0.29 | 0.02 | 1.07 |
| mesenchyme | Mpi | 0.47 | 8.66E-11 | 1.80E-06 | 0.21 | 0.18 | 0.03 | 1.15 |
| mesenchyme | Rexo5 | 0.62 | 1.05E-10 | 2.19E-06 | 0.13 | 0.1 | 0.03 | 1.28 |
| mesenchyme | Fbxl15 | 0.5 | 1.08E-10 | 2.24E-06 | 0.2 | 0.18 | 0.03 | 1.16 |
| mesenchyme | Polr1a | 0.3 | 1.09E-10 | 2.27E-06 | 0.37 | 0.35 | 0.02 | 1.06 |
| mesenchyme | Mbd4 | 0.37 | 1.11E-10 | 2.31E-06 | 0.34 | 0.32 | 0.02 | 1.07 |
| mesenchyme | Tubgcp4 | 0.3 | 1.20E-10 | 2.49E-06 | 0.35 | 0.32 | 0.02 | 1.06 |
| mesenchyme | Tmtc4 | 0.46 | 1.29E-10 | 2.68E-06 | 0.3 | 0.27 | 0.03 | 1.09 |
| mesenchyme | Arhgef7 | 0.28 | 1.33E-10 | 2.77E-06 | 0.33 | 0.3 | 0.03 | 1.09 |
| mesenchyme | 933439C10Ri | 0.62 | 1.37E-10 | 2.85E-06 | 0.11 | 0.09 | 0.03 | 1.31 |
| mesenchyme | Rufy2 | 0.34 | 1.40E-10 | 2.92E-06 | 0.27 | 0.24 | 0.03 | 1.13 |
| mesenchyme | Atxn7l1 | 0.58 | 1.54E-10 | 3.19E-06 | 0.18 | 0.15 | 0.03 | 1.19 |
| mesenchyme | Alg6 | 0.33 | 1.54E-10 | 3.21E-06 | 0.31 | 0.29 | 0.02 | 1.08 |
| mesenchyme | Vps16 | 0.37 | 1.63E-10 | 3.40E-06 | 0.3 | 0.28 | 0.02 | 1.08 |
| mesenchyme | Usp33 | 0.32 | 1.69E-10 | 3.52E-06 | 0.34 | 0.32 | 0.02 | 1.06 |
| mesenchyme | Pms2 | 0.29 | 1.79E-10 | 3.72E-06 | 0.33 | 0.3 | 0.02 | 1.07 |
| mesenchyme | Thap3 | 0.31 | 1.81E-10 | 3.77E-06 | 0.32 | 0.29 | 0.02 | 1.08 |
| mesenchyme | Septin2 | 0.44 | 2.08E-10 | 4.32E-06 | 0.19 | 0.16 | 0.03 | 1.19 |
| mesenchyme | Ska3 | 0.33 | 2.08E-10 | 4.33E-06 | 0.29 | 0.27 | 0.03 | 1.09 |
| mesenchyme | Zfp40 | 0.6 | 2.09E-10 | 4.35E-06 | 0.14 | 0.12 | 0.03 | 1.24 |
| mesenchyme | Aars2 | 0.5 | 2.36E-10 | 4.90E-06 | 0.16 | 0.13 | 0.03 | 1.22 |
| mesenchyme | AW554918 | 0.52 | 2.39E-10 | 4.97E-06 | 0.21 | 0.19 | 0.03 | 1.14 |
| mesenchyme | Nsl1 | 0.52 | 2.45E-10 | 5.10E-06 | 0.2 | 0.17 | 0.03 | 1.16 |
| mesenchyme | Zfp930 | 0.49 | 2.46E-10 | 5.12E-06 | 0.18 | 0.15 | 0.03 | 1.2 |
| mesenchyme | Zfp866 | 0.42 | 2.57E-10 | 5.35E-06 | 0.21 | 0.18 | 0.03 | 1.16 |
| mesenchyme | Zbtb37 | 0.32 | 2.96E-10 | 6.16E-06 | 0.3 | 0.27 | 0.03 | 1.11 |
| mesenchyme | Cep72 | 0.44 | 2.99E-10 | 6.22E-06 | 0.16 | 0.13 | 0.03 | 1.21 |
| mesenchyme | Fbxo17 | 0.5 | 3.01E-10 | 6.26E-06 | 0.17 | 0.14 | 0.03 | 1.21 |
| mesenchyme | 933407K13Ri | 0.63 | 3.08E-10 | 6.41E-06 | 0.11 | 0.08 | 0.03 | 1.33 |
| mesenchyme | Clasrp | 0.36 | 3.22E-10 | 6.69E-06 | 0.26 | 0.23 | 0.03 | 1.12 |
| mesenchyme | Actr8 | 0.28 | 3.31E-10 | 6.88E-06 | 0.33 | 0.31 | 0.02 | 1.08 |
| mesenchyme | Rptor | 0.45 | 3.36E-10 | 6.99E-06 | 0.25 | 0.22 | 0.03 | 1.11 |
| mesenchyme | Ric8b | 0.47 | 3.58E-10 | 7.45E-06 | 0.22 | 0.19 | 0.03 | 1.14 |
| mesenchyme | Ctdsp1 | 0.31 | 4.02E-10 | 8.36E-06 | 0.35 | 0.34 | 0.02 | 1.05 |
| mesenchyme | 931428F04Ri | 0.45 | 4.03E-10 | 8.39E-06 | 0.17 | 0.14 | 0.03 | 1.2 |
| mesenchyme | Ankrd54 | 0.31 | 4.69E-10 | 9.76E-06 | 0.34 | 0.32 | 0.02 | 1.06 |
| mesenchyme | Fpgs | 0.48 | 4.77E-10 | 9.92E-06 | 0.2 | 0.17 | 0.03 | 1.16 |
| mesenchyme | Fam57a | 0.52 | 4.78E-10 | 9.93E-06 | 0.15 | 0.12 | 0.03 | 1.23 |
| mesenchyme | Tmem161a | 0.33 | 5.12E-10 | 1.07E-05 | 0.31 | 0.28 | 0.02 | 1.08 |
| mesenchyme | Lztr1 | 0.27 | 5.33E-10 | 1.11E-05 | 0.33 | 0.31 | 0.03 | 1.08 |
| mesenchyme | Ccdc71l | 0.4 | 5.43E-10 | 1.13E-05 | 0.29 | 0.27 | 0.02 | 1.08 |
| mesenchyme | Zcchc14 | 0.29 | 5.58E-10 | 1.16E-05 | 0.33 | 0.31 | 0.03 | 1.08 |
| mesenchyme | Zfp280c | 0.35 | 5.74E-10 | 1.19E-05 | 0.25 | 0.22 | 0.03 | 1.13 |
| mesenchyme | Spaca6 | 0.27 | 5.83E-10 | 1.21E-05 | 0.34 | 0.31 | 0.03 | 1.1 |
| mesenchyme | Rbbp5 | 0.41 | 5.94E-10 | 1.23E-05 | 0.26 | 0.23 | 0.03 | 1.11 |
| mesenchyme | Rnf215 | 0.35 | 5.94E-10 | 1.24E-05 | 0.26 | 0.24 | 0.03 | 1.11 |
| mesenchyme | Pam | 0.56 | 6.28E-10 | 1.31E-05 | 0.16 | 0.13 | 0.03 | 1.21 |
| mesenchyme | Aldh9a1 | 0.35 | 6.32E-10 | 1.31E-05 | 0.32 | 0.31 | 0.02 | 1.05 |
| mesenchyme | Zyg11b | 0.26 | 6.57E-10 | 1.37E-05 | 0.35 | 0.33 | 0.03 | 1.08 |
| mesenchyme | Fbxw11 | 0.33 | 6.71E-10 | 1.40E-05 | 0.34 | 0.32 | 0.02 | 1.06 |
| mesenchyme | Zmym3 | 0.37 | 7.72E-10 | 1.61E-05 | 0.26 | 0.24 | 0.03 | 1.1 |
| mesenchyme | Dusp22 | 0.48 | 8.11E-10 | 1.69E-05 | 0.21 | 0.19 | 0.03 | 1.15 |
| mesenchyme | Arnt | 0.37 | 8.25E-10 | 1.72E-05 | 0.26 | 0.23 | 0.03 | 1.11 |
| mesenchyme | Ubqln4 | 0.25 | 8.56E-10 | 1.78E-05 | 0.38 | 0.36 | 0.02 | 1.05 |
| mesenchyme | Zwilch | 0.32 | 8.68E-10 | 1.81E-05 | 0.32 | 0.3 | 0.02 | 1.07 |

**Supplementary Table 2. Marker genes for the major cell types of integrated clustering of E11.5 control and NotchGOF λ-junction cells.**

| | | | | | | | | |
|---|---|---|---|---|---|---|---|---|
| mesenchyme | 030458C11Ri | 0.3 | 8.74E-10 | 1.82E-05 | 0.34 | 0.32 | 0.02 | 1.07 |
| mesenchyme | Zfp212 | 0.31 | 8.88E-10 | 1.85E-05 | 0.27 | 0.24 | 0.03 | 1.11 |
| mesenchyme | Plekhm2 | 0.44 | 9.22E-10 | 1.92E-05 | 0.21 | 0.18 | 0.03 | 1.15 |
| mesenchyme | Dcp1a | 0.44 | 9.23E-10 | 1.92E-05 | 0.2 | 0.17 | 0.03 | 1.16 |
| mesenchyme | Fam13b | 0.28 | 9.28E-10 | 1.93E-05 | 0.31 | 0.29 | 0.02 | 1.08 |
| mesenchyme | Wdr55 | 0.29 | 9.74E-10 | 2.02E-05 | 0.37 | 0.36 | 0.01 | 1.04 |
| mesenchyme | Zbtb6 | 0.4 | 9.78E-10 | 2.03E-05 | 0.22 | 0.19 | 0.03 | 1.15 |
| mesenchyme | Dna2 | 0.36 | 1.01E-09 | 2.09E-05 | 0.27 | 0.24 | 0.03 | 1.11 |
| mesenchyme | Adcy6 | 0.57 | 1.08E-09 | 2.25E-05 | 0.15 | 0.12 | 0.03 | 1.23 |
| mesenchyme | Cpsf1 | 0.29 | 1.11E-09 | 2.30E-05 | 0.33 | 0.31 | 0.02 | 1.07 |
| mesenchyme | Arnt2 | 0.42 | 1.13E-09 | 2.35E-05 | 0.2 | 0.17 | 0.03 | 1.17 |
| mesenchyme | Rabepk | 0.34 | 1.23E-09 | 2.55E-05 | 0.28 | 0.26 | 0.02 | 1.08 |
| mesenchyme | Slc12a7 | 0.4 | 1.35E-09 | 2.81E-05 | 0.24 | 0.21 | 0.03 | 1.13 |
| mesenchyme | Thap7 | 0.27 | 1.45E-09 | 3.03E-05 | 0.34 | 0.32 | 0.02 | 1.05 |
| mesenchyme | Zfp160 | 0.38 | 1.48E-09 | 3.07E-05 | 0.24 | 0.22 | 0.03 | 1.13 |
| mesenchyme | Adprm | 0.33 | 1.51E-09 | 3.13E-05 | 0.23 | 0.2 | 0.03 | 1.14 |
| mesenchyme | Zdhhc15 | 0.59 | 1.64E-09 | 3.42E-05 | 0.14 | 0.11 | 0.03 | 1.23 |
| mesenchyme | Tmem201 | 0.34 | 1.75E-09 | 3.65E-05 | 0.26 | 0.23 | 0.03 | 1.12 |
| mesenchyme | Prpsap2 | 0.32 | 1.77E-09 | 3.68E-05 | 0.27 | 0.25 | 0.02 | 1.1 |
| mesenchyme | Pskh1 | 0.4 | 1.83E-09 | 3.81E-05 | 0.2 | 0.17 | 0.03 | 1.17 |
| mesenchyme | Parvb | 0.33 | 1.91E-09 | 3.98E-05 | 0.14 | 0.11 | 0.03 | 1.23 |
| mesenchyme | Zfp629 | 0.33 | 1.92E-09 | 3.99E-05 | 0.28 | 0.26 | 0.03 | 1.1 |
| mesenchyme | Upp2 | 0.37 | 1.94E-09 | 4.03E-05 | 0.19 | 0.16 | 0.03 | 1.18 |
| mesenchyme | Ess2 | 0.31 | 2.06E-09 | 4.29E-05 | 0.3 | 0.28 | 0.02 | 1.07 |
| mesenchyme | Csgalnact2 | 0.57 | 2.08E-09 | 4.33E-05 | 0.11 | 0.08 | 0.03 | 1.3 |
| mesenchyme | Entpd7 | 0.58 | 2.11E-09 | 4.39E-05 | 0.16 | 0.13 | 0.03 | 1.2 |
| mesenchyme | Snapc1 | 0.32 | 2.14E-09 | 4.45E-05 | 0.31 | 0.29 | 0.02 | 1.08 |
| mesenchyme | Dhrs7 | 0.59 | 2.18E-09 | 4.53E-05 | 0.14 | 0.11 | 0.03 | 1.23 |
| mesenchyme | Bop1 | 0.25 | 2.27E-09 | 4.71E-05 | 0.38 | 0.36 | 0.02 | 1.04 |
| mesenchyme | Efr3a | 0.32 | 2.36E-09 | 4.92E-05 | 0.36 | 0.35 | 0.01 | 1.04 |
| mesenchyme | Dnajc27 | 0.68 | 2.39E-09 | 4.97E-05 | 0.1 | 0.08 | 0.02 | 1.29 |
| mesenchyme | Ints7 | 0.26 | 2.41E-09 | 5.02E-05 | 0.39 | 0.37 | 0.02 | 1.04 |
| mesenchyme | Ankrd13b | 0.39 | 2.41E-09 | 5.02E-05 | 0.2 | 0.17 | 0.03 | 1.16 |
| mesenchyme | Eea1 | 0.28 | 2.49E-09 | 5.18E-05 | 0.43 | 0.43 | 0 | 1 |
| mesenchyme | Maml1 | 0.4 | 2.70E-09 | 5.62E-05 | 0.23 | 0.2 | 0.03 | 1.14 |
| mesenchyme | Ptpn23 | 0.33 | 2.74E-09 | 5.70E-05 | 0.27 | 0.25 | 0.02 | 1.1 |
| mesenchyme | Stx1a | 0.63 | 2.90E-09 | 6.02E-05 | 0.1 | 0.08 | 0.02 | 1.3 |
| mesenchyme | Ccdc22 | 0.33 | 2.99E-09 | 6.23E-05 | 0.31 | 0.29 | 0.02 | 1.06 |
| mesenchyme | Nphp3 | 0.6 | 3.04E-09 | 6.33E-05 | 0.12 | 0.09 | 0.03 | 1.28 |
| mesenchyme | Srxn1 | 0.73 | 3.10E-09 | 6.45E-05 | 0.12 | 0.1 | 0.02 | 1.25 |
| mesenchyme | Six5 | 0.39 | 3.13E-09 | 6.50E-05 | 0.21 | 0.18 | 0.03 | 1.16 |
| mesenchyme | Pigc | 0.4 | 3.34E-09 | 6.95E-05 | 0.24 | 0.22 | 0.03 | 1.12 |
| mesenchyme | Pgm1 | 0.26 | 3.55E-09 | 7.39E-05 | 0.32 | 0.3 | 0.02 | 1.08 |
| mesenchyme | Fam189b | 0.46 | 3.62E-09 | 7.52E-05 | 0.18 | 0.15 | 0.03 | 1.17 |
| mesenchyme | Aaas | 0.3 | 3.75E-09 | 7.80E-05 | 0.34 | 0.32 | 0.02 | 1.05 |
| mesenchyme | Btaf1 | 0.27 | 3.86E-09 | 8.03E-05 | 0.37 | 0.35 | 0.01 | 1.04 |
| mesenchyme | Cast | 0.26 | 3.96E-09 | 8.23E-05 | 0.25 | 0.23 | 0.03 | 1.12 |
| mesenchyme | Pacs1 | 0.29 | 4.09E-09 | 8.50E-05 | 0.29 | 0.27 | 0.03 | 1.09 |
| mesenchyme | Cers5 | 0.31 | 4.23E-09 | 8.80E-05 | 0.35 | 0.33 | 0.02 | 1.06 |
| mesenchyme | Slc30a5 | 0.32 | 4.37E-09 | 9.08E-05 | 0.32 | 0.3 | 0.02 | 1.06 |
| mesenchyme | Dcun1d3 | 0.49 | 4.45E-09 | 9.26E-05 | 0.16 | 0.13 | 0.03 | 1.2 |
| mesenchyme | Cdadc1 | 0.41 | 4.53E-09 | 9.43E-05 | 0.25 | 0.23 | 0.02 | 1.09 |
| mesenchyme | Gm15446 | 0.63 | 4.71E-09 | 9.79E-05 | 0.12 | 0.1 | 0.03 | 1.26 |
| mesenchyme | Cep135 | 0.3 | 4.97E-09 | 1.03E-04 | 0.3 | 0.28 | 0.02 | 1.08 |
| mesenchyme | Pyroxd1 | 0.3 | 5.13E-09 | 1.07E-04 | 0.25 | 0.23 | 0.03 | 1.12 |
| mesenchyme | Dus4l | 0.47 | 5.76E-09 | 1.20E-04 | 0.17 | 0.14 | 0.03 | 1.19 |

**Supplementary Table 2. Marker genes for the major cell types of integrated clustering of E11.5 control and NotchGOF λ-junction cells.**

| | | | | | | | | |
|---|---|---|---|---|---|---|---|---|
| mesenchyme | Tab1 | 0.4 | 6.01E-09 | 1.25E-04 | 0.23 | 0.2 | 0.03 | 1.12 |
| mesenchyme | Zfp955b | 0.27 | 6.13E-09 | 1.27E-04 | 0.33 | 0.31 | 0.02 | 1.07 |
| mesenchyme | Odf2l | 0.27 | 6.23E-09 | 1.30E-04 | 0.36 | 0.34 | 0.02 | 1.05 |
| mesenchyme | Ift88 | 0.27 | 6.49E-09 | 1.35E-04 | 0.31 | 0.28 | 0.03 | 1.09 |
| mesenchyme | Top3a | 0.46 | 6.59E-09 | 1.37E-04 | 0.18 | 0.15 | 0.03 | 1.17 |
| mesenchyme | Qsox1 | 0.64 | 7.03E-09 | 1.46E-04 | 0.13 | 0.1 | 0.02 | 1.24 |
| mesenchyme | Ftx | 0.6 | 7.21E-09 | 1.50E-04 | 0.15 | 0.12 | 0.03 | 1.2 |
| mesenchyme | Dynlt1c | 0.35 | 7.31E-09 | 1.52E-04 | 0.21 | 0.18 | 0.03 | 1.14 |
| mesenchyme | Mapkbp1 | 0.62 | 7.35E-09 | 1.53E-04 | 0.1 | 0.08 | 0.02 | 1.3 |
| mesenchyme | Gys1 | 0.38 | 7.77E-09 | 1.62E-04 | 0.24 | 0.21 | 0.03 | 1.12 |
| mesenchyme | Rcor3 | 0.39 | 7.89E-09 | 1.64E-04 | 0.24 | 0.22 | 0.02 | 1.11 |
| mesenchyme | Frk | 0.53 | 8.10E-09 | 1.68E-04 | 0.16 | 0.13 | 0.03 | 1.2 |
| mesenchyme | B4galt7 | 0.44 | 8.32E-09 | 1.73E-04 | 0.2 | 0.18 | 0.03 | 1.14 |
| mesenchyme | B3galnt2 | 0.3 | 8.70E-09 | 1.81E-04 | 0.31 | 0.29 | 0.02 | 1.07 |
| mesenchyme | Togaram1 | 0.38 | 8.70E-09 | 1.81E-04 | 0.27 | 0.25 | 0.02 | 1.09 |
| mesenchyme | Klhl11 | 0.39 | 8.83E-09 | 1.84E-04 | 0.19 | 0.17 | 0.03 | 1.15 |
| mesenchyme | Pcsk7 | 0.51 | 8.87E-09 | 1.84E-04 | 0.15 | 0.13 | 0.02 | 1.19 |
| mesenchyme | Zfp976 | 0.48 | 9.55E-09 | 1.99E-04 | 0.14 | 0.11 | 0.03 | 1.22 |
| mesenchyme | Alg11 | 0.39 | 9.71E-09 | 2.02E-04 | 0.25 | 0.23 | 0.02 | 1.1 |
| mesenchyme | Ing3 | 0.27 | 1.12E-08 | 2.32E-04 | 0.37 | 0.35 | 0.02 | 1.05 |
| mesenchyme | Eef1aknmt | 0.37 | 1.27E-08 | 2.63E-04 | 0.24 | 0.21 | 0.02 | 1.11 |
| mesenchyme | Slx4 | 0.61 | 1.28E-08 | 2.65E-04 | 0.14 | 0.11 | 0.02 | 1.21 |
| mesenchyme | Slc16a10 | 0.79 | 1.34E-08 | 2.79E-04 | 0.12 | 0.09 | 0.02 | 1.24 |
| mesenchyme | Hist2h2bb | 0.43 | 1.37E-08 | 2.86E-04 | 0.26 | 0.24 | 0.02 | 1.09 |
| mesenchyme | Zfp932 | 0.36 | 1.39E-08 | 2.89E-04 | 0.25 | 0.23 | 0.02 | 1.1 |
| mesenchyme | Plpp1 | 0.36 | 1.42E-08 | 2.95E-04 | 0.16 | 0.13 | 0.03 | 1.21 |
| mesenchyme | Cep41 | 0.3 | 1.52E-08 | 3.16E-04 | 0.23 | 0.2 | 0.03 | 1.13 |
| mesenchyme | Edem3 | 0.5 | 1.58E-08 | 3.28E-04 | 0.16 | 0.14 | 0.03 | 1.18 |
| mesenchyme | )430019H16Ri | 0.32 | 1.63E-08 | 3.39E-04 | 0.2 | 0.17 | 0.03 | 1.16 |
| mesenchyme | Obsl1 | 0.27 | 1.82E-08 | 3.78E-04 | 0.32 | 0.3 | 0.02 | 1.07 |
| mesenchyme | Pusl1 | 0.29 | 1.97E-08 | 4.10E-04 | 0.27 | 0.24 | 0.03 | 1.1 |
| mesenchyme | Mri1 | 0.37 | 2.09E-08 | 4.35E-04 | 0.28 | 0.26 | 0.02 | 1.06 |
| mesenchyme | Sirt4 | 0.49 | 2.15E-08 | 4.47E-04 | 0.12 | 0.1 | 0.02 | 1.25 |
| mesenchyme | Usb1 | 0.32 | 2.27E-08 | 4.72E-04 | 0.23 | 0.21 | 0.02 | 1.12 |
| mesenchyme | Platr25 | 0.36 | 2.32E-08 | 4.81E-04 | 0.24 | 0.21 | 0.02 | 1.11 |
| mesenchyme | Gm12184 | 0.55 | 2.37E-08 | 4.92E-04 | 0.12 | 0.1 | 0.02 | 1.24 |
| mesenchyme | )610030E20Ri | 0.27 | 2.47E-08 | 5.14E-04 | 0.36 | 0.34 | 0.02 | 1.05 |
| mesenchyme | Mms22l | 0.28 | 2.51E-08 | 5.21E-04 | 0.36 | 0.35 | 0.01 | 1.03 |
| mesenchyme | Pigh | 0.38 | 2.54E-08 | 5.29E-04 | 0.29 | 0.27 | 0.02 | 1.08 |
| mesenchyme | Mbd5 | 0.43 | 2.69E-08 | 5.59E-04 | 0.22 | 0.19 | 0.02 | 1.12 |
| mesenchyme | Stat5b | 0.7 | 2.75E-08 | 5.71E-04 | 0.14 | 0.11 | 0.02 | 1.19 |
| mesenchyme | Gm5914 | 0.45 | 2.89E-08 | 6.02E-04 | 0.14 | 0.12 | 0.02 | 1.2 |
| mesenchyme | Heatr5b | 0.55 | 2.92E-08 | 6.08E-04 | 0.14 | 0.11 | 0.02 | 1.21 |
| mesenchyme | Rint1 | 0.3 | 3.51E-08 | 7.31E-04 | 0.33 | 0.32 | 0.02 | 1.05 |
| mesenchyme | Xrcc3 | 0.45 | 3.52E-08 | 7.32E-04 | 0.11 | 0.09 | 0.02 | 1.26 |
| mesenchyme | Uqcc1 | 0.29 | 3.62E-08 | 7.53E-04 | 0.3 | 0.28 | 0.02 | 1.07 |
| mesenchyme | Rnf19a | 0.42 | 3.82E-08 | 7.95E-04 | 0.23 | 0.21 | 0.02 | 1.11 |
| mesenchyme | Dmac2 | 0.26 | 3.96E-08 | 8.23E-04 | 0.32 | 0.3 | 0.02 | 1.06 |
| mesenchyme | Zfp975 | 0.54 | 4.11E-08 | 8.54E-04 | 0.12 | 0.1 | 0.02 | 1.23 |
| mesenchyme | Rab32 | 0.92 | 4.16E-08 | 8.64E-04 | 0.1 | 0.08 | 0.02 | 1.26 |
| mesenchyme | Ttll12 | 0.29 | 4.16E-08 | 8.65E-04 | 0.33 | 0.31 | 0.01 | 1.05 |
| mesenchyme | Bcap29 | 0.26 | 4.23E-08 | 8.80E-04 | 0.35 | 0.33 | 0.02 | 1.05 |
| mesenchyme | Patl1 | 0.41 | 4.32E-08 | 8.98E-04 | 0.25 | 0.23 | 0.02 | 1.09 |
| mesenchyme | Ddx59 | 0.57 | 4.34E-08 | 9.02E-04 | 0.13 | 0.1 | 0.02 | 1.21 |
| mesenchyme | Zw10 | 0.32 | 4.36E-08 | 9.07E-04 | 0.33 | 0.32 | 0.01 | 1.04 |
| mesenchyme | St8sia4 | 0.31 | 4.45E-08 | 9.25E-04 | 0.1 | 0.08 | 0.02 | 1.31 |

**Supplementary Table 2. Marker genes for the major cell types of integrated clustering of E11.5 control and NotchGOF λ-junction cells.**

| | | | | | | | | |
|---|---|---|---|---|---|---|---|---|
| mesenchyme | Fam118a | 0.48 | 4.48E-08 | 9.31E-04 | 0.18 | 0.15 | 0.02 | 1.16 |
| mesenchyme | Mettl21a | 0.42 | 4.51E-08 | 9.37E-04 | 0.19 | 0.17 | 0.02 | 1.14 |
| mesenchyme | Zfp383 | 0.39 | 4.56E-08 | 9.48E-04 | 0.16 | 0.14 | 0.03 | 1.18 |
| mesenchyme | Qtrt2 | 0.29 | 4.75E-08 | 9.89E-04 | 0.26 | 0.23 | 0.03 | 1.11 |
| mesenchyme | 810009A15Ri | 0.46 | 4.76E-08 | 9.89E-04 | 0.18 | 0.15 | 0.02 | 1.16 |
| mesenchyme | Zfp398 | 0.42 | 4.81E-08 | 9.99E-04 | 0.22 | 0.2 | 0.02 | 1.11 |
| mesenchyme | Gucd1 | 0.48 | 4.92E-08 | 1.02E-03 | 0.19 | 0.17 | 0.02 | 1.13 |
| mesenchyme | Cenpt | 0.33 | 4.95E-08 | 1.03E-03 | 0.26 | 0.24 | 0.02 | 1.08 |
| mesenchyme | Sirt3 | 0.36 | 5.54E-08 | 1.15E-03 | 0.26 | 0.24 | 0.02 | 1.09 |
| mesenchyme | Gtpbp3 | 0.4 | 5.72E-08 | 1.19E-03 | 0.2 | 0.18 | 0.02 | 1.13 |
| mesenchyme | Trmu | 0.4 | 5.78E-08 | 1.20E-03 | 0.22 | 0.19 | 0.02 | 1.12 |
| mesenchyme | Phykpl | 0.31 | 6.02E-08 | 1.25E-03 | 0.26 | 0.24 | 0.02 | 1.1 |
| mesenchyme | Klc2 | 0.45 | 6.20E-08 | 1.29E-03 | 0.17 | 0.15 | 0.02 | 1.15 |
| mesenchyme | Tbc1d24 | 0.6 | 6.26E-08 | 1.30E-03 | 0.11 | 0.09 | 0.02 | 1.26 |
| mesenchyme | Rbm19 | 0.32 | 6.99E-08 | 1.45E-03 | 0.28 | 0.26 | 0.02 | 1.06 |
| mesenchyme | Zscan12 | 0.3 | 7.04E-08 | 1.46E-03 | 0.25 | 0.23 | 0.02 | 1.1 |
| mesenchyme | Sh2b2 | 0.33 | 7.66E-08 | 1.59E-03 | 0.17 | 0.15 | 0.03 | 1.17 |
| mesenchyme | Atrnl1 | 0.42 | 7.83E-08 | 1.63E-03 | 0.21 | 0.19 | 0.02 | 1.12 |
| mesenchyme | Hist1h2br | 0.53 | 8.08E-08 | 1.68E-03 | 0.16 | 0.14 | 0.02 | 1.16 |
| mesenchyme | Lrfn4 | 0.31 | 8.66E-08 | 1.80E-03 | 0.27 | 0.25 | 0.02 | 1.08 |
| mesenchyme | Shisa2 | 0.79 | 8.68E-08 | 1.81E-03 | 0.12 | 0.1 | 0.02 | 1.22 |
| mesenchyme | Srrd | 0.28 | 8.71E-08 | 1.81E-03 | 0.29 | 0.27 | 0.02 | 1.07 |
| mesenchyme | Snx7 | 0.59 | 9.17E-08 | 1.91E-03 | 0.11 | 0.09 | 0.02 | 1.25 |
| mesenchyme | Sorcs2 | 0.43 | 9.55E-08 | 1.99E-03 | 0.12 | 0.09 | 0.02 | 1.26 |
| mesenchyme | Foxred1 | 0.38 | 1.08E-07 | 2.24E-03 | 0.22 | 0.2 | 0.02 | 1.11 |
| mesenchyme | Mtbp | 0.35 | 1.08E-07 | 2.24E-03 | 0.23 | 0.21 | 0.02 | 1.1 |
| mesenchyme | Septin1 | 0.35 | 1.14E-07 | 2.36E-03 | 0.23 | 0.21 | 0.02 | 1.11 |
| mesenchyme | E2f7 | 0.32 | 1.23E-07 | 2.57E-03 | 0.26 | 0.24 | 0.02 | 1.09 |
| mesenchyme | Mmaa | 0.45 | 1.30E-07 | 2.71E-03 | 0.15 | 0.13 | 0.02 | 1.18 |
| mesenchyme | Btd | 0.58 | 1.31E-07 | 2.72E-03 | 0.13 | 0.11 | 0.02 | 1.21 |
| mesenchyme | Faxc | 0.5 | 1.33E-07 | 2.77E-03 | 0.11 | 0.09 | 0.02 | 1.26 |
| mesenchyme | Cep250 | 0.34 | 1.34E-07 | 2.79E-03 | 0.27 | 0.25 | 0.02 | 1.08 |
| mesenchyme | Dpy19l3 | 0.46 | 1.34E-07 | 2.79E-03 | 0.16 | 0.14 | 0.02 | 1.18 |
| mesenchyme | Tmem18 | 0.57 | 1.35E-07 | 2.81E-03 | 0.16 | 0.14 | 0.02 | 1.15 |
| mesenchyme | Soga1 | 0.46 | 1.45E-07 | 3.02E-03 | 0.13 | 0.11 | 0.02 | 1.21 |
| mesenchyme | BC030867 | 0.48 | 1.48E-07 | 3.09E-03 | 0.15 | 0.13 | 0.02 | 1.17 |
| mesenchyme | Fbxw8 | 0.3 | 1.51E-07 | 3.15E-03 | 0.29 | 0.27 | 0.02 | 1.07 |
| mesenchyme | Fancc | 0.6 | 1.52E-07 | 3.15E-03 | 0.11 | 0.09 | 0.02 | 1.24 |
| mesenchyme | Gm1976 | 0.52 | 1.55E-07 | 3.23E-03 | 0.12 | 0.1 | 0.02 | 1.23 |
| mesenchyme | Pdxk | 0.61 | 1.58E-07 | 3.29E-03 | 0.12 | 0.1 | 0.02 | 1.22 |
| mesenchyme | Col9a3 | 0.61 | 1.61E-07 | 3.35E-03 | 0.15 | 0.13 | 0.02 | 1.19 |
| mesenchyme | Sfxn4 | 0.53 | 1.69E-07 | 3.52E-03 | 0.1 | 0.08 | 0.02 | 1.27 |
| mesenchyme | Cntrob | 0.36 | 1.74E-07 | 3.63E-03 | 0.22 | 0.19 | 0.02 | 1.12 |
| mesenchyme | Thumpd2 | 0.38 | 1.79E-07 | 3.72E-03 | 0.17 | 0.14 | 0.02 | 1.16 |
| mesenchyme | Acad9 | 0.38 | 1.85E-07 | 3.85E-03 | 0.26 | 0.24 | 0.02 | 1.08 |
| mesenchyme | Gcfc2 | 0.37 | 1.86E-07 | 3.87E-03 | 0.26 | 0.24 | 0.02 | 1.08 |
| mesenchyme | Fam122a | 0.28 | 1.90E-07 | 3.96E-03 | 0.3 | 0.28 | 0.01 | 1.05 |
| mesenchyme | Slc49a4 | 0.35 | 2.10E-07 | 4.36E-03 | 0.28 | 0.26 | 0.02 | 1.07 |
| mesenchyme | Zfp820 | 0.43 | 2.15E-07 | 4.47E-03 | 0.21 | 0.19 | 0.02 | 1.11 |
| mesenchyme | Evi5l | 0.58 | 2.23E-07 | 4.64E-03 | 0.12 | 0.1 | 0.02 | 1.22 |
| mesenchyme | Eri2 | 0.45 | 2.32E-07 | 4.84E-03 | 0.21 | 0.18 | 0.02 | 1.11 |
| mesenchyme | Pomc | 0.35 | 2.46E-07 | 5.11E-03 | 0.14 | 0.12 | 0.02 | 1.2 |
| mesenchyme | Mrm3 | 0.42 | 2.50E-07 | 5.19E-03 | 0.21 | 0.19 | 0.02 | 1.11 |
| mesenchyme | Ndrg3 | 0.37 | 2.53E-07 | 5.26E-03 | 0.27 | 0.25 | 0.01 | 1.06 |
| mesenchyme | Wwp2 | 0.34 | 2.59E-07 | 5.39E-03 | 0.31 | 0.3 | 0.01 | 1.04 |
| mesenchyme | Kif14 | 0.53 | 2.60E-07 | 5.42E-03 | 0.2 | 0.18 | 0.02 | 1.11 |

**Supplementary Table 2. Marker genes for the major cell types of integrated clustering of E11.5 control and NotchGOF λ-junction cells.**

| | | | | | | | | |
|---|---|---|---|---|---|---|---|---|
| mesenchyme | Mier3 | 0.28 | 2.69E-07 | 5.59E-03 | 0.3 | 0.29 | 0.02 | 1.06 |
| mesenchyme | Arfgap3 | 0.33 | 2.80E-07 | 5.81E-03 | 0.23 | 0.21 | 0.02 | 1.09 |
| mesenchyme | Abr | 0.37 | 3.00E-07 | 6.23E-03 | 0.23 | 0.21 | 0.02 | 1.1 |
| mesenchyme | Extl3 | 0.34 | 3.07E-07 | 6.38E-03 | 0.21 | 0.19 | 0.02 | 1.12 |
| mesenchyme | Zfp617 | 0.32 | 3.20E-07 | 6.66E-03 | 0.31 | 0.29 | 0.02 | 1.05 |
| mesenchyme | Chst15 | 0.58 | 3.26E-07 | 6.78E-03 | 0.18 | 0.16 | 0.02 | 1.13 |
| mesenchyme | Tmem43 | 0.39 | 3.32E-07 | 6.90E-03 | 0.22 | 0.2 | 0.02 | 1.1 |
| mesenchyme | Wdr37 | 0.36 | 3.51E-07 | 7.31E-03 | 0.22 | 0.19 | 0.02 | 1.11 |
| mesenchyme | Phldb1 | 0.38 | 3.63E-07 | 7.56E-03 | 0.21 | 0.19 | 0.02 | 1.11 |
| mesenchyme | Ccsap | 0.45 | 3.70E-07 | 7.69E-03 | 0.12 | 0.1 | 0.02 | 1.21 |
| mesenchyme | Ccdc134 | 0.51 | 3.83E-07 | 7.98E-03 | 0.17 | 0.14 | 0.02 | 1.15 |
| mesenchyme | Morn4 | 0.4 | 3.93E-07 | 8.17E-03 | 0.17 | 0.15 | 0.02 | 1.15 |
| mesenchyme | Foxj3 | 0.32 | 4.13E-07 | 8.60E-03 | 0.29 | 0.28 | 0.01 | 1.05 |
| mesenchyme | Trmt10b | 0.34 | 4.19E-07 | 8.71E-03 | 0.25 | 0.22 | 0.02 | 1.09 |
| mesenchyme | Kat14 | 0.28 | 4.44E-07 | 9.24E-03 | 0.28 | 0.27 | 0.02 | 1.07 |
| mesenchyme | Khnyn | 0.32 | 4.60E-07 | 9.56E-03 | 0.24 | 0.22 | 0.02 | 1.09 |
| mesenchyme | Armcx5 | 0.32 | 4.97E-07 | 1.03E-02 | 0.26 | 0.24 | 0.02 | 1.07 |
| mesenchyme | Ccdc117 | 0.34 | 5.31E-07 | 1.10E-02 | 0.21 | 0.19 | 0.02 | 1.11 |
| mesenchyme | Smarcal1 | 0.32 | 5.37E-07 | 1.12E-02 | 0.25 | 0.24 | 0.01 | 1.06 |
| mesenchyme | 900076A07Ri | 0.3 | 5.41E-07 | 1.13E-02 | 0.31 | 0.3 | 0.02 | 1.05 |
| mesenchyme | Rps6kb2 | 0.31 | 5.49E-07 | 1.14E-02 | 0.29 | 0.28 | 0.01 | 1.04 |
| mesenchyme | Ldlr | 0.36 | 6.02E-07 | 1.25E-02 | 0.25 | 0.23 | 0.02 | 1.08 |
| mesenchyme | Arhgef26 | 0.48 | 6.18E-07 | 1.29E-02 | 0.13 | 0.11 | 0.02 | 1.18 |
| mesenchyme | Rnf40 | 0.3 | 6.19E-07 | 1.29E-02 | 0.27 | 0.25 | 0.02 | 1.07 |
| mesenchyme | Heatr1 | 0.26 | 6.41E-07 | 1.33E-02 | 0.31 | 0.3 | 0.02 | 1.05 |
| mesenchyme | Bicra | 0.48 | 6.85E-07 | 1.42E-02 | 0.16 | 0.14 | 0.02 | 1.16 |
| mesenchyme | Tmem143 | 0.64 | 7.15E-07 | 1.49E-02 | 0.1 | 0.08 | 0.02 | 1.23 |
| mesenchyme | Tbc1d20 | 0.26 | 7.64E-07 | 1.59E-02 | 0.36 | 0.35 | 0.01 | 1.02 |
| mesenchyme | Tedc2 | 0.33 | 8.17E-07 | 1.70E-02 | 0.19 | 0.17 | 0.02 | 1.13 |
| mesenchyme | Lrr1 | 0.35 | 8.27E-07 | 1.72E-02 | 0.27 | 0.26 | 0.01 | 1.05 |
| mesenchyme | Zfp938 | 0.4 | 8.33E-07 | 1.73E-02 | 0.21 | 0.19 | 0.02 | 1.1 |
| mesenchyme | Anks1 | 0.35 | 8.80E-07 | 1.83E-02 | 0.17 | 0.15 | 0.02 | 1.15 |
| mesenchyme | Lss | 0.4 | 9.05E-07 | 1.88E-02 | 0.17 | 0.15 | 0.02 | 1.14 |
| mesenchyme | Azin2 | 0.45 | 9.45E-07 | 1.97E-02 | 0.12 | 0.1 | 0.02 | 1.2 |
| mesenchyme | Tmem50b | 0.43 | 9.46E-07 | 1.97E-02 | 0.19 | 0.17 | 0.02 | 1.12 |
| mesenchyme | Pprc1 | 0.3 | 1.05E-06 | 2.19E-02 | 0.26 | 0.24 | 0.02 | 1.07 |
| mesenchyme | St6gal1 | 0.39 | 1.06E-06 | 2.21E-02 | 0.2 | 0.17 | 0.02 | 1.12 |
| mesenchyme | Acy1 | 0.36 | 1.11E-06 | 2.31E-02 | 0.17 | 0.15 | 0.02 | 1.14 |
| mesenchyme | Mtx3 | 0.46 | 1.11E-06 | 2.31E-02 | 0.14 | 0.12 | 0.02 | 1.17 |
| mesenchyme | Zbtb17 | 0.28 | 1.11E-06 | 2.32E-02 | 0.27 | 0.25 | 0.02 | 1.08 |
| mesenchyme | Dpagt1 | 0.28 | 1.12E-06 | 2.34E-02 | 0.29 | 0.28 | 0.01 | 1.05 |
| mesenchyme | Rxra | 0.4 | 1.13E-06 | 2.34E-02 | 0.2 | 0.18 | 0.02 | 1.11 |
| mesenchyme | Sfxn2 | 0.39 | 1.20E-06 | 2.50E-02 | 0.21 | 0.19 | 0.02 | 1.11 |
| mesenchyme | P4ha3 | 0.48 | 1.20E-06 | 2.50E-02 | 0.17 | 0.15 | 0.02 | 1.14 |
| mesenchyme | Snx30 | 0.36 | 1.23E-06 | 2.56E-02 | 0.16 | 0.14 | 0.02 | 1.15 |
| mesenchyme | Abcc5 | 0.31 | 1.24E-06 | 2.59E-02 | 0.26 | 0.24 | 0.02 | 1.07 |
| mesenchyme | Fastkd3 | 0.37 | 1.25E-06 | 2.61E-02 | 0.17 | 0.15 | 0.02 | 1.13 |
| mesenchyme | Endog | 0.39 | 1.27E-06 | 2.65E-02 | 0.23 | 0.21 | 0.02 | 1.09 |
| mesenchyme | Poc5 | 0.35 | 1.28E-06 | 2.66E-02 | 0.21 | 0.19 | 0.02 | 1.1 |
| mesenchyme | Fnip1 | 0.34 | 1.29E-06 | 2.69E-02 | 0.25 | 0.23 | 0.02 | 1.08 |
| mesenchyme | Dgat1 | 0.39 | 1.31E-06 | 2.73E-02 | 0.2 | 0.18 | 0.02 | 1.11 |
| mesenchyme | Ccdc136 | 0.25 | 1.33E-06 | 2.76E-02 | 0.29 | 0.27 | 0.02 | 1.08 |
| mesenchyme | Wrap53 | 0.32 | 1.41E-06 | 2.93E-02 | 0.23 | 0.22 | 0.02 | 1.08 |
| mesenchyme | Nr4a2 | 0.57 | 1.53E-06 | 3.18E-02 | 0.11 | 0.09 | 0.02 | 1.21 |
| mesenchyme | Lig4 | 0.46 | 1.55E-06 | 3.22E-02 | 0.17 | 0.15 | 0.02 | 1.13 |
| mesenchyme | Asb8 | 0.31 | 1.61E-06 | 3.34E-02 | 0.3 | 0.3 | 0.01 | 1.03 |

**Supplementary Table 2. Marker genes for the major cell types of integrated clustering of E11.5 control and NotchGOF λ-junction cells.**

| | | | | | | | | |
|---|---|---|---|---|---|---|---|---|
| mesenchyme | Invs | 0.54 | 1.64E-06 | 3.41E-02 | 0.14 | 0.12 | 0.02 | 1.17 |
| mesenchyme | Tbc1d13 | 0.5 | 1.73E-06 | 3.60E-02 | 0.15 | 0.13 | 0.02 | 1.16 |
| mesenchyme | Wdr92 | 0.26 | 1.75E-06 | 3.64E-02 | 0.29 | 0.27 | 0.02 | 1.07 |
| mesenchyme | Mul1 | 0.29 | 1.76E-06 | 3.66E-02 | 0.27 | 0.26 | 0.01 | 1.05 |
| mesenchyme | Mlx | 0.25 | 1.78E-06 | 3.70E-02 | 0.31 | 0.3 | 0.01 | 1.04 |
| mesenchyme | Dnajb14 | 0.28 | 1.86E-06 | 3.87E-02 | 0.33 | 0.32 | 0.01 | 1.03 |
| mesenchyme | Ccdc91 | 0.27 | 1.86E-06 | 3.88E-02 | 0.28 | 0.26 | 0.02 | 1.07 |
| mesenchyme | Slc6a8 | 0.33 | 1.88E-06 | 3.91E-02 | 0.2 | 0.18 | 0.02 | 1.12 |
| mesenchyme | Tmem167b | 0.34 | 1.88E-06 | 3.92E-02 | 0.21 | 0.19 | 0.02 | 1.1 |
| mesenchyme | Lama1 | 0.61 | 1.89E-06 | 3.92E-02 | 0.11 | 0.09 | 0.02 | 1.2 |
| mesenchyme | Extl2 | 0.41 | 1.90E-06 | 3.96E-02 | 0.15 | 0.13 | 0.02 | 1.16 |
| mesenchyme | Zfp81 | 0.29 | 2.05E-06 | 4.26E-02 | 0.26 | 0.24 | 0.02 | 1.08 |
| mesenchyme | 430405H02Ri | 0.35 | 2.09E-06 | 4.35E-02 | 0.21 | 0.19 | 0.02 | 1.11 |
| mesenchyme | Slc30a4 | 0.39 | 2.14E-06 | 4.45E-02 | 0.14 | 0.12 | 0.02 | 1.18 |
| mesenchyme | 730063M14R | 0.3 | 2.18E-06 | 4.53E-02 | 0.14 | 0.12 | 0.02 | 1.17 |
| mesenchyme | Zdhhc4 | 0.32 | 2.33E-06 | 4.85E-02 | 0.27 | 0.26 | 0.02 | 1.06 |
| ectoderm | Pdgfa | 5.74 | 0 | 0 | 0.93 | 0.08 | 0.85 | 11.91 |
| ectoderm | Spint2 | 4.72 | 0 | 0 | 0.93 | 0.11 | 0.82 | 8.18 |
| ectoderm | Epcam | 5.54 | 0 | 0 | 0.97 | 0.15 | 0.82 | 6.4 |
| ectoderm | Dsp | 6.4 | 0 | 0 | 0.85 | 0.04 | 0.8 | 19.25 |
| ectoderm | Cldn6 | 6.01 | 0 | 0 | 0.84 | 0.04 | 0.8 | 21.46 |
| ectoderm | Six3 | 5.92 | 0 | 0 | 0.83 | 0.05 | 0.78 | 16.33 |
| ectoderm | Ezr | 4.56 | 0 | 0 | 0.9 | 0.14 | 0.76 | 6.49 |
| ectoderm | Cldn7 | 5.5 | 0 | 0 | 0.82 | 0.06 | 0.76 | 14.05 |
| ectoderm | Krt18 | 5.24 | 0 | 0 | 0.87 | 0.12 | 0.75 | 7.25 |
| ectoderm | Krt8 | 5.21 | 0 | 0 | 0.82 | 0.09 | 0.73 | 9.59 |
| ectoderm | Igfbp5 | 4.5 | 0 | 0 | 0.95 | 0.24 | 0.71 | 3.99 |
| ectoderm | Six3os1 | 5.85 | 0 | 0 | 0.77 | 0.06 | 0.71 | 12.66 |
| ectoderm | Ikzf2 | 4.35 | 0 | 0 | 0.81 | 0.12 | 0.7 | 7.02 |
| ectoderm | Hist3h2ba | 3.46 | 0 | 0 | 0.8 | 0.11 | 0.69 | 7.42 |
| ectoderm | Isl1 | 5.22 | 0 | 0 | 0.74 | 0.06 | 0.69 | 13.49 |
| ectoderm | Tpd52 | 3.39 | 0 | 0 | 0.78 | 0.1 | 0.68 | 7.97 |
| ectoderm | Wnt4 | 6.18 | 0 | 0 | 0.71 | 0.04 | 0.67 | 17.7 |
| ectoderm | Bcam | 5.38 | 0 | 0 | 0.71 | 0.05 | 0.66 | 15.69 |
| ectoderm | Sox2 | 6.1 | 0 | 0 | 0.69 | 0.04 | 0.66 | 19.83 |
| ectoderm | Mpzl2 | 4.97 | 0 | 0 | 0.71 | 0.05 | 0.66 | 14.45 |
| ectoderm | Sfn | 6.52 | 0 | 0 | 0.71 | 0.05 | 0.66 | 14.16 |
| ectoderm | Slc9a3r1 | 4.21 | 0 | 0 | 0.73 | 0.09 | 0.64 | 8.49 |
| ectoderm | Gm16136 | 6.16 | 0 | 0 | 0.65 | 0.02 | 0.63 | 28.3 |
| ectoderm | Pdgfc | 3.65 | 0 | 0 | 0.75 | 0.13 | 0.62 | 5.67 |
| ectoderm | Pitx1 | 5.45 | 0 | 0 | 0.65 | 0.04 | 0.61 | 17.92 |
| ectoderm | Cdh1 | 5.92 | 0 | 0 | 0.63 | 0.02 | 0.61 | 26.33 |
| ectoderm | Col18a1 | 2.97 | 0 | 0 | 0.83 | 0.22 | 0.61 | 3.8 |
| ectoderm | Rab25 | 5.79 | 0 | 0 | 0.62 | 0.02 | 0.6 | 34.22 |
| ectoderm | Esrp1 | 5.39 | 0 | 0 | 0.62 | 0.03 | 0.6 | 24.88 |
| ectoderm | Nav2 | 3.87 | 0 | 0 | 0.69 | 0.1 | 0.59 | 7.03 |
| ectoderm | Acer2 | 2.81 | 0 | 0 | 0.84 | 0.26 | 0.58 | 3.29 |
| ectoderm | Itga6 | 3.11 | 0 | 0 | 0.75 | 0.17 | 0.58 | 4.34 |
| ectoderm | Hey1 | 3.91 | 0 | 0 | 0.74 | 0.16 | 0.58 | 4.55 |
| ectoderm | Cldn3 | 4.98 | 0 | 0 | 0.6 | 0.03 | 0.57 | 23.04 |
| ectoderm | Socs2 | 3.68 | 0 | 0 | 0.77 | 0.21 | 0.56 | 3.68 |
| ectoderm | Fermt1 | 6.99 | 0 | 0 | 0.56 | 0.01 | 0.55 | 55.6 |
| ectoderm | Stard10 | 2.97 | 0 | 0 | 0.71 | 0.17 | 0.55 | 4.22 |
| ectoderm | Ap1m2 | 6.09 | 0 | 0 | 0.56 | 0.01 | 0.54 | 39.71 |
| ectoderm | Kif21a | 2.92 | 0 | 0 | 0.72 | 0.18 | 0.54 | 4.01 |
| ectoderm | Nrarp | 4.62 | 0 | 0 | 0.59 | 0.05 | 0.54 | 11.51 |

**Supplementary Table 2. Marker genes for the major cell types of integrated clustering of E11.5 control and NotchGOF λ-junction cells.**

| | | | | | | | | |
|---|---|---|---|---|---|---|---|---|
| ectoderm | Frem2 | 3.89 | 0 | 0 | 0.6 | 0.07 | 0.53 | 9.06 |
| ectoderm | Ston2 | 4.24 | 0 | 0 | 0.59 | 0.05 | 0.53 | 10.85 |
| ectoderm | Mapk13 | 3.63 | 0 | 0 | 0.59 | 0.08 | 0.52 | 7.82 |
| ectoderm | Perp | 6.61 | 0 | 0 | 0.53 | 0.02 | 0.51 | 29.56 |
| ectoderm | F11r | 3.61 | 0 | 0 | 0.57 | 0.06 | 0.51 | 9.83 |
| ectoderm | Wwc1 | 4.05 | 0 | 0 | 0.57 | 0.06 | 0.51 | 9.63 |
| ectoderm | Fjx1 | 2.92 | 0 | 0 | 0.77 | 0.27 | 0.51 | 2.89 |
| ectoderm | Cd9 | 2.73 | 0 | 0 | 0.68 | 0.18 | 0.51 | 3.85 |
| ectoderm | Lsr | 3.2 | 0 | 0 | 0.59 | 0.09 | 0.5 | 6.93 |
| ectoderm | Nectin4 | 6.82 | 0 | 0 | 0.51 | 0.01 | 0.5 | 46.36 |
| ectoderm | Kcnk1 | 3.75 | 0 | 0 | 0.57 | 0.07 | 0.5 | 8.56 |
| ectoderm | Ccdc88c | 4.42 | 0 | 0 | 0.53 | 0.04 | 0.5 | 14.05 |
| ectoderm | Trp63 | 6.39 | 0 | 0 | 0.51 | 0.02 | 0.49 | 31.88 |
| ectoderm | Bex1 | 3.04 | 0 | 0 | 0.96 | 0.47 | 0.49 | 2.04 |
| ectoderm | Foxg1 | 4.41 | 0 | 0 | 0.51 | 0.03 | 0.49 | 17.72 |
| ectoderm | Sp8 | 5.1 | 0 | 0 | 0.5 | 0.02 | 0.48 | 20.96 |
| ectoderm | Pdlim1 | 3.34 | 0 | 0 | 0.65 | 0.18 | 0.48 | 3.69 |
| ectoderm | Grhl2 | 5.5 | 0 | 0 | 0.49 | 0.02 | 0.47 | 27.06 |
| ectoderm | Jag1 | 3.84 | 0 | 0 | 0.53 | 0.07 | 0.47 | 8.2 |
| ectoderm | Anxa2 | 2.69 | 0 | 0 | 0.77 | 0.3 | 0.47 | 2.54 |
| ectoderm | Aldh1a3 | 5.25 | 0 | 0 | 0.58 | 0.12 | 0.46 | 4.83 |
| ectoderm | Erbb3 | 3 | 0 | 0 | 0.53 | 0.07 | 0.46 | 8.09 |
| ectoderm | Dmrta2 | 4.89 | 0 | 0 | 0.49 | 0.03 | 0.46 | 15.87 |
| ectoderm | Wfdc2 | 3.14 | 0 | 0 | 0.81 | 0.35 | 0.46 | 2.31 |
| ectoderm | Cpm | 4.96 | 0 | 0 | 0.52 | 0.07 | 0.46 | 7.81 |
| ectoderm | Notch1 | 3.18 | 0 | 0 | 0.59 | 0.14 | 0.46 | 4.3 |
| ectoderm | Hspb8 | 5.44 | 0 | 0 | 0.48 | 0.03 | 0.46 | 16.69 |
| ectoderm | Tmem54 | 6.11 | 0 | 0 | 0.47 | 0.01 | 0.45 | 42.27 |
| ectoderm | Pax6 | 5.27 | 0 | 0 | 0.47 | 0.02 | 0.45 | 22.52 |
| ectoderm | Arhgef19 | 4.71 | 0 | 0 | 0.47 | 0.02 | 0.45 | 19.67 |
| ectoderm | Cdh3 | 3 | 0 | 0 | 0.55 | 0.1 | 0.45 | 5.37 |
| ectoderm | Cmtm8 | 2.99 | 0 | 0 | 0.52 | 0.07 | 0.45 | 7.28 |
| ectoderm | Klk8 | 2.6 | 0 | 0 | 0.55 | 0.11 | 0.44 | 5.11 |
| ectoderm | St14 | 5.68 | 0 | 0 | 0.46 | 0.01 | 0.44 | 35.15 |
| ectoderm | Hook1 | 3 | 0 | 0 | 0.54 | 0.1 | 0.44 | 5.6 |
| ectoderm | Krt14 | 6.33 | 0 | 0 | 0.47 | 0.03 | 0.44 | 17.37 |
| ectoderm | Ly6h | 4.07 | 0 | 0 | 0.48 | 0.05 | 0.43 | 10.21 |
| ectoderm | Dsg2 | 4.73 | 0 | 0 | 0.46 | 0.03 | 0.43 | 17.65 |
| ectoderm | Fras1 | 2.51 | 0 | 0 | 0.59 | 0.16 | 0.43 | 3.62 |
| ectoderm | Dab1 | 3.7 | 0 | 0 | 0.48 | 0.05 | 0.43 | 9.21 |
| ectoderm | Samd5 | 2.36 | 0 | 0 | 0.69 | 0.26 | 0.43 | 2.63 |
| ectoderm | Ano1 | 5.08 | 0 | 0 | 0.45 | 0.03 | 0.42 | 15.96 |
| ectoderm | Runx1 | 3.79 | 0 | 0 | 0.51 | 0.09 | 0.42 | 5.44 |
| ectoderm | Crb3 | 5.72 | 0 | 0 | 0.43 | 0.01 | 0.42 | 38.73 |
| ectoderm | Mycl | 2.38 | 0 | 0 | 0.62 | 0.2 | 0.41 | 3.03 |
| ectoderm | Tspan13 | 1.88 | 0 | 0 | 0.73 | 0.31 | 0.41 | 2.32 |
| ectoderm | Ctnnd2 | 4.76 | 0 | 0 | 0.43 | 0.02 | 0.41 | 19.68 |
| ectoderm | Ablim1 | 2.76 | 0 | 0 | 0.54 | 0.13 | 0.41 | 4.2 |
| ectoderm | Otx1 | 5.45 | 0 | 0 | 0.42 | 0.02 | 0.41 | 28.27 |
| ectoderm | Frmd4b | 3.29 | 0 | 0 | 0.48 | 0.07 | 0.41 | 6.73 |
| ectoderm | Cd151 | 2.1 | 0 | 0 | 0.64 | 0.23 | 0.41 | 2.75 |
| ectoderm | Emb | 2.08 | 0 | 0 | 0.71 | 0.31 | 0.4 | 2.29 |
| ectoderm | Mir205hg | 5.1 | 0 | 0 | 0.42 | 0.03 | 0.4 | 16.96 |
| ectoderm | Celf2 | 2.3 | 0 | 0 | 0.56 | 0.16 | 0.4 | 3.47 |
| ectoderm | Tom1l1 | 3.41 | 0 | 0 | 0.46 | 0.07 | 0.4 | 7 |
| ectoderm | Mgst1 | 1.87 | 0 | 0 | 0.68 | 0.28 | 0.39 | 2.4 |

**Supplementary Table 2. Marker genes for the major cell types of integrated clustering of E11.5 control and NotchGOF λ-junction cells.**

| | | | | | | | | |
|---|---|---|---|---|---|---|---|---|
| ectoderm | Pltp | 3.52 | 0 | 0 | 0.46 | 0.07 | 0.39 | 7.05 |
| ectoderm | Cldn4 | 6 | 0 | 0 | 0.4 | 0.01 | 0.39 | 40.2 |
| ectoderm | Map7 | 5.34 | 0 | 0 | 0.41 | 0.01 | 0.39 | 29 |
| ectoderm | Fgf9 | 6.1 | 0 | 0 | 0.41 | 0.01 | 0.39 | 31.15 |
| ectoderm | Lama5 | 4.74 | 0 | 0 | 0.41 | 0.02 | 0.39 | 18.82 |
| ectoderm | Irf6 | 5.77 | 0 | 0 | 0.41 | 0.02 | 0.39 | 25.38 |
| ectoderm | Jup | 2.65 | 0 | 0 | 0.5 | 0.11 | 0.39 | 4.68 |
| ectoderm | Stx3 | 4.14 | 0 | 0 | 0.42 | 0.03 | 0.39 | 12.41 |
| ectoderm | Plp2 | 1.92 | 0 | 0 | 0.67 | 0.29 | 0.39 | 2.34 |
| ectoderm | Ddr1 | 3.63 | 0 | 0 | 0.42 | 0.04 | 0.38 | 10.53 |
| ectoderm | Cadm1 | 2.22 | 0 | 0 | 0.68 | 0.3 | 0.38 | 2.28 |
| ectoderm | Sema3d | 4.55 | 0 | 0 | 0.41 | 0.03 | 0.38 | 15 |
| ectoderm | Tnfaip2 | 4.41 | 0 | 0 | 0.43 | 0.05 | 0.38 | 8.27 |
| ectoderm | Sdc4 | 2.03 | 0 | 0 | 0.63 | 0.26 | 0.38 | 2.47 |
| ectoderm | Spint1 | 5.67 | 0 | 0 | 0.39 | 0.01 | 0.38 | 38.5 |
| ectoderm | Meis2 | 0.85 | 0 | 0 | 0.96 | 0.58 | 0.37 | 1.64 |
| ectoderm | Pla2g12a | 1.96 | 0 | 0 | 0.61 | 0.24 | 0.37 | 2.54 |
| ectoderm | Klf5 | 4 | 0 | 0 | 0.4 | 0.04 | 0.37 | 11.22 |
| ectoderm | Ssx2ip | 1.89 | 0 | 0 | 0.64 | 0.27 | 0.37 | 2.34 |
| ectoderm | Marchf9 | 3.05 | 0 | 0 | 0.43 | 0.06 | 0.37 | 6.98 |
| ectoderm | Otx2 | 4.87 | 0 | 0 | 0.38 | 0.02 | 0.36 | 25.27 |
| ectoderm | Bex4 | 2.18 | 0 | 0 | 0.93 | 0.56 | 0.36 | 1.64 |
| ectoderm | Grb7 | 5.32 | 0 | 0 | 0.37 | 0.01 | 0.36 | 31.08 |
| ectoderm | Ptprf | 2.38 | 0 | 0 | 0.91 | 0.55 | 0.36 | 1.65 |
| ectoderm | Rab15 | 3.5 | 0 | 0 | 0.41 | 0.05 | 0.36 | 8.44 |
| ectoderm | Wnt6 | 6.77 | 0 | 0 | 0.37 | 0.02 | 0.36 | 24.8 |
| ectoderm | Ephb1 | 5.52 | 0 | 0 | 0.37 | 0.01 | 0.36 | 26.43 |
| ectoderm | Lmo7 | 3.37 | 0 | 0 | 0.43 | 0.08 | 0.36 | 5.62 |
| ectoderm | Id4 | 2.39 | 0 | 0 | 0.58 | 0.22 | 0.36 | 2.61 |
| ectoderm | Nebl | 5.09 | 0 | 0 | 0.38 | 0.02 | 0.35 | 15.75 |
| ectoderm | Uchl1 | 1.52 | 0 | 0 | 0.62 | 0.27 | 0.35 | 2.31 |
| ectoderm | Cdkn1a | 4.76 | 0 | 0 | 0.38 | 0.03 | 0.35 | 13.96 |
| ectoderm | Dok5 | 4.35 | 0 | 0 | 0.38 | 0.03 | 0.35 | 11.58 |
| ectoderm | Hes5 | 5.64 | 0 | 0 | 0.38 | 0.03 | 0.35 | 11.58 |
| ectoderm | Prss8 | 5.08 | 0 | 0 | 0.36 | 0.01 | 0.35 | 25.86 |
| ectoderm | Cgn | 4.6 | 0 | 0 | 0.37 | 0.02 | 0.35 | 16.09 |
| ectoderm | Fgd2 | 4.3 | 0 | 0 | 0.36 | 0.02 | 0.35 | 20.17 |
| ectoderm | Khdrbs3 | 2.16 | 0 | 0 | 0.88 | 0.53 | 0.34 | 1.64 |
| ectoderm | Ppp2r2b | 5.21 | 0 | 0 | 0.36 | 0.02 | 0.34 | 23.87 |
| ectoderm | Epb41l5 | 2.06 | 0 | 0 | 0.51 | 0.17 | 0.34 | 2.97 |
| ectoderm | Etl4 | 1.78 | 0 | 0 | 0.57 | 0.23 | 0.34 | 2.46 |
| ectoderm | Slc12a2 | 1.89 | 0 | 0 | 0.69 | 0.36 | 0.33 | 1.94 |
| ectoderm | Espn | 4.18 | 0 | 0 | 0.36 | 0.03 | 0.33 | 12.07 |
| ectoderm | Lad1 | 5.62 | 0 | 0 | 0.34 | 0.01 | 0.33 | 31.09 |
| ectoderm | Igfbp3 | 2.71 | 0 | 0 | 0.38 | 0.05 | 0.33 | 8.17 |
| ectoderm | Efna1 | 2.69 | 0 | 0 | 0.39 | 0.06 | 0.33 | 6.29 |
| ectoderm | Afdn | 1.54 | 0 | 0 | 0.82 | 0.49 | 0.33 | 1.66 |
| ectoderm | Stk26 | 2.15 | 0 | 0 | 0.48 | 0.15 | 0.33 | 3.18 |
| ectoderm | Tubb2b | 1.85 | 0 | 0 | 0.92 | 0.59 | 0.33 | 1.55 |
| ectoderm | Klc3 | 3.07 | 0 | 0 | 0.36 | 0.03 | 0.33 | 11.87 |
| ectoderm | Synrg | 1.99 | 0 | 0 | 0.48 | 0.16 | 0.33 | 3.06 |
| ectoderm | Kdf1 | 5.63 | 0 | 0 | 0.33 | 0.01 | 0.33 | 41.63 |
| ectoderm | Arhgef5 | 2.19 | 0 | 0 | 0.45 | 0.12 | 0.33 | 3.66 |
| ectoderm | Zfp219 | 1.44 | 0 | 0 | 0.65 | 0.32 | 0.33 | 2.01 |
| ectoderm | Ehbp1 | 2.19 | 0 | 0 | 0.49 | 0.16 | 0.33 | 3.01 |
| ectoderm | Celsr1 | 4.15 | 0 | 0 | 0.35 | 0.03 | 0.33 | 13.5 |

**Supplementary Table 2. Marker genes for the major cell types of integrated clustering of E11.5 control and NotchGOF λ-junction cells.**

| | | | | | | | | |
|---|---|---|---|---|---|---|---|---|
| ectoderm | Gkap1 | 1.36 | 0 | 0 | 0.68 | 0.36 | 0.32 | 1.91 |
| ectoderm | Agrn | 1.59 | 0 | 0 | 0.64 | 0.31 | 0.32 | 2.04 |
| ectoderm | Hs6st1 | 2.56 | 0 | 0 | 0.42 | 0.09 | 0.32 | 4.45 |
| ectoderm | Krt5 | 6.56 | 0 | 0 | 0.34 | 0.01 | 0.32 | 24 |
| ectoderm | Dpysl5 | 3.32 | 0 | 0 | 0.35 | 0.03 | 0.32 | 11.39 |
| ectoderm | Cldn9 | 5.05 | 0 | 0 | 0.33 | 0.01 | 0.32 | 27.75 |
| ectoderm | Ormdl2 | 1.64 | 0 | 0 | 0.59 | 0.27 | 0.32 | 2.18 |
| ectoderm | Pcdh7 | 1.14 | 0 | 0 | 0.64 | 0.32 | 0.32 | 2.01 |
| ectoderm | Scube3 | 2.1 | 0 | 0 | 0.45 | 0.13 | 0.32 | 3.52 |
| ectoderm | Hs6st2 | 1.85 | 0 | 0 | 0.53 | 0.21 | 0.32 | 2.53 |
| ectoderm | Pawr | 3.13 | 0 | 0 | 0.37 | 0.06 | 0.31 | 6.71 |
| ectoderm | Ago1 | 1.55 | 0 | 0 | 0.6 | 0.29 | 0.31 | 2.08 |
| ectoderm | Hspb1 | 3.84 | 0 | 0 | 0.39 | 0.08 | 0.31 | 5.17 |
| ectoderm | Nav3 | 3.97 | 0 | 0 | 0.37 | 0.05 | 0.31 | 6.91 |
| ectoderm | Igf2 | 1.78 | 0 | 0 | 0.82 | 0.51 | 0.31 | 1.62 |
| ectoderm | Acvr2a | 1.99 | 0 | 0 | 0.51 | 0.2 | 0.31 | 2.59 |
| ectoderm | Elf1 | 1.86 | 0 | 0 | 0.5 | 0.19 | 0.31 | 2.65 |
| ectoderm | 900026A02Ri | 3.81 | 0 | 0 | 0.34 | 0.03 | 0.31 | 11.33 |
| ectoderm | Arrdc1 | 3.68 | 0 | 0 | 0.35 | 0.04 | 0.31 | 9.38 |
| ectoderm | Dag1 | 1.6 | 0 | 0 | 0.61 | 0.3 | 0.31 | 2.04 |
| ectoderm | Slc2a1 | 1.53 | 0 | 0 | 0.66 | 0.36 | 0.31 | 1.87 |
| ectoderm | Myh9 | 1.56 | 0 | 0 | 0.74 | 0.43 | 0.31 | 1.71 |
| ectoderm | Cystm1 | 3.53 | 0 | 0 | 0.32 | 0.02 | 0.31 | 19 |
| ectoderm | Arl4c | 1.87 | 0 | 0 | 0.56 | 0.26 | 0.31 | 2.19 |
| ectoderm | Lgr4 | 2.21 | 0 | 0 | 0.46 | 0.16 | 0.31 | 2.96 |
| ectoderm | Gm266 | 2.94 | 0 | 0 | 0.54 | 0.24 | 0.3 | 2.27 |
| ectoderm | Acaa2 | 1.7 | 0 | 0 | 0.71 | 0.41 | 0.3 | 1.74 |
| ectoderm | Ntf5 | 4.84 | 0 | 0 | 0.32 | 0.02 | 0.3 | 19.88 |
| ectoderm | Plekha7 | 4.25 | 0 | 0 | 0.32 | 0.02 | 0.3 | 16.1 |
| ectoderm | Ctnnal1 | 1.82 | 0 | 0 | 0.49 | 0.19 | 0.3 | 2.57 |
| ectoderm | Nudt14 | 1.37 | 0 | 0 | 0.63 | 0.33 | 0.3 | 1.9 |
| ectoderm | Zdhhc2 | 3.29 | 0 | 0 | 0.36 | 0.06 | 0.3 | 6.05 |
| ectoderm | Tspan12 | 1.95 | 0 | 0 | 0.44 | 0.14 | 0.3 | 3.11 |
| ectoderm | Crip2 | 1.01 | 0 | 0 | 0.4 | 0.1 | 0.3 | 3.86 |
| ectoderm | Dlx2 | 3.95 | 0 | 0 | 0.35 | 0.06 | 0.29 | 5.97 |
| ectoderm | Zfhx2 | 1.65 | 0 | 0 | 0.48 | 0.18 | 0.29 | 2.61 |
| ectoderm | Plch1 | 3.76 | 0 | 0 | 0.32 | 0.03 | 0.29 | 10.42 |
| ectoderm | Igsf9 | 1.37 | 0 | 0 | 0.63 | 0.34 | 0.29 | 1.87 |
| ectoderm | Dlk1 | 2.15 | 0 | 0 | 0.64 | 0.35 | 0.29 | 1.83 |
| ectoderm | Lratd2 | 3.41 | 0 | 0 | 0.33 | 0.04 | 0.29 | 8.1 |
| ectoderm | Usp25 | 1.61 | 0 | 0 | 0.66 | 0.37 | 0.29 | 1.79 |
| ectoderm | Hist3h2a | 1.82 | 0 | 0 | 0.49 | 0.2 | 0.29 | 2.47 |
| ectoderm | Fzd6 | 4.63 | 0 | 0 | 0.31 | 0.02 | 0.29 | 17.94 |
| ectoderm | Ece1 | 1.89 | 0 | 0 | 0.45 | 0.16 | 0.29 | 2.76 |
| ectoderm | Tmem238 | 2.26 | 0 | 0 | 0.4 | 0.11 | 0.28 | 3.54 |
| ectoderm | Upk3bl | 6.58 | 0 | 0 | 0.29 | 0.01 | 0.28 | 57.8 |
| ectoderm | Rab27b | 3.15 | 0 | 0 | 0.31 | 0.03 | 0.28 | 10.79 |
| ectoderm | Rnh1 | 1.43 | 0 | 0 | 0.77 | 0.49 | 0.28 | 1.58 |
| ectoderm | Rprm | 3.07 | 0 | 0 | 0.38 | 0.09 | 0.28 | 3.99 |
| ectoderm | Dnase2a | 1.95 | 0 | 0 | 0.43 | 0.15 | 0.28 | 2.86 |
| ectoderm | Fzd3 | 1.48 | 0 | 0 | 0.8 | 0.52 | 0.28 | 1.54 |
| ectoderm | Cxadr | 1.21 | 0 | 0 | 0.68 | 0.4 | 0.28 | 1.71 |
| ectoderm | Myh14 | 4.76 | 0 | 0 | 0.3 | 0.01 | 0.28 | 21.07 |
| ectoderm | Ocln | 5.29 | 0 | 0 | 0.29 | 0.01 | 0.28 | 29.1 |
| ectoderm | Cox4i2 | 1.98 | 0 | 0 | 0.48 | 0.2 | 0.28 | 2.41 |
| ectoderm | Ripk4 | 5.71 | 0 | 0 | 0.29 | 0.01 | 0.28 | 41 |

**Supplementary Table 2. Marker genes for the major cell types of integrated clustering of E11.5 control and NotchGOF λ-junction cells.**

| | | | | | | | | |
|---|---|---|---|---|---|---|---|---|
| ectoderm | Igfbp2 | 2.8 | 0 | 0 | 0.97 | 0.69 | 0.28 | 1.41 |
| ectoderm | Myo10 | 1.37 | 0 | 0 | 0.69 | 0.41 | 0.28 | 1.69 |
| ectoderm | Pdk1 | 1.59 | 0 | 0 | 0.47 | 0.19 | 0.28 | 2.46 |
| ectoderm | Chchd10 | 1.97 | 0 | 0 | 0.37 | 0.1 | 0.28 | 3.93 |
| ectoderm | Schip1 | 2.19 | 0 | 0 | 0.37 | 0.1 | 0.28 | 3.92 |
| ectoderm | Gad2 | 5.94 | 0 | 0 | 0.29 | 0.01 | 0.28 | 35.63 |
| ectoderm | Mtln | 1.43 | 0 | 0 | 0.54 | 0.26 | 0.28 | 2.05 |
| ectoderm | Slc2a3 | 3.53 | 0 | 0 | 0.35 | 0.07 | 0.28 | 4.77 |
| ectoderm | Cacna1b | 4.63 | 0 | 0 | 0.29 | 0.02 | 0.27 | 19.27 |
| ectoderm | Cep170b | 2.64 | 0 | 0 | 0.33 | 0.06 | 0.27 | 5.39 |
| ectoderm | Nckap5 | 4.26 | 0 | 0 | 0.3 | 0.03 | 0.27 | 11.46 |
| ectoderm | Cers4 | 2.27 | 0 | 0 | 0.36 | 0.09 | 0.27 | 3.91 |
| ectoderm | Dlx3 | 5.4 | 0 | 0 | 0.28 | 0.01 | 0.27 | 28.1 |
| ectoderm | 110039M20R | 4.14 | 0 | 0 | 0.28 | 0.01 | 0.27 | 23.58 |
| ectoderm | Ndrg2 | 2.59 | 0 | 0 | 0.34 | 0.07 | 0.27 | 4.8 |
| ectoderm | Npnt | 1.67 | 0 | 0 | 0.44 | 0.17 | 0.27 | 2.55 |
| ectoderm | Pkp4 | 1.36 | 0 | 0 | 0.58 | 0.32 | 0.27 | 1.85 |
| ectoderm | Ptpru | 2.21 | 0 | 0 | 0.36 | 0.09 | 0.27 | 3.85 |
| ectoderm | Krt19 | 5.98 | 0 | 0 | 0.28 | 0.01 | 0.27 | 27.8 |
| ectoderm | Dapl1 | 7.1 | 0 | 0 | 0.27 | 0.01 | 0.27 | 45.5 |
| ectoderm | Fxyd3 | 3.96 | 0 | 0 | 0.3 | 0.03 | 0.27 | 10.54 |
| ectoderm | Krtcap3 | 3.02 | 0 | 0 | 0.32 | 0.05 | 0.27 | 6.3 |
| ectoderm | Susd4 | 3.72 | 0 | 0 | 0.29 | 0.03 | 0.27 | 10.46 |
| ectoderm | Apoc1 | 2.99 | 0 | 0 | 0.29 | 0.02 | 0.27 | 13.05 |
| ectoderm | S100a10 | 1.79 | 0 | 0 | 0.75 | 0.49 | 0.26 | 1.53 |
| ectoderm | Fgfr2 | 1.81 | 0 | 0 | 0.48 | 0.22 | 0.26 | 2.22 |
| ectoderm | Capg | 2.34 | 0 | 0 | 0.35 | 0.09 | 0.26 | 3.93 |
| ectoderm | AW011738 | 3.87 | 0 | 0 | 0.29 | 0.03 | 0.26 | 10.67 |
| ectoderm | Slc22a23 | 3.7 | 0 | 0 | 0.29 | 0.03 | 0.26 | 8.88 |
| ectoderm | Tacstd2 | 6.66 | 0 | 0 | 0.27 | 0.01 | 0.26 | 33.38 |
| ectoderm | 310030G06Ri | 5.3 | 0 | 0 | 0.27 | 0.01 | 0.26 | 29.56 |
| ectoderm | Fgf8 | 6.42 | 0 | 0 | 0.28 | 0.02 | 0.26 | 15.28 |
| ectoderm | Elavl2 | 1.56 | 0 | 0 | 0.38 | 0.13 | 0.26 | 3.03 |
| ectoderm | Mbp | 3.23 | 0 | 0 | 0.29 | 0.03 | 0.26 | 8.53 |
| ectoderm | Celsr2 | 3.35 | 0 | 0 | 0.29 | 0.03 | 0.26 | 8.97 |
| ectoderm | Six6 | 4.6 | 0 | 0 | 0.26 | 0.01 | 0.26 | 29.33 |
| ectoderm | Npdc1 | 2.65 | 0 | 0 | 0.3 | 0.05 | 0.25 | 6.64 |
| ectoderm | Palm3 | 2.58 | 0 | 0 | 0.34 | 0.09 | 0.25 | 3.92 |
| ectoderm | Pik3r1 | 1.8 | 0 | 0 | 0.58 | 0.33 | 0.25 | 1.77 |
| ectoderm | Pla2r1 | 4.78 | 0 | 0 | 0.27 | 0.02 | 0.25 | 17.87 |
| ectoderm | Pls1 | 3.14 | 0 | 0 | 0.29 | 0.04 | 0.25 | 8.23 |
| ectoderm | Ppm1l | 1.71 | 0 | 0 | 0.41 | 0.16 | 0.25 | 2.54 |
| ectoderm | Hes1 | 1.42 | 0 | 0 | 0.74 | 0.49 | 0.25 | 1.52 |
| ectoderm | Galm | 2.12 | 0 | 0 | 0.4 | 0.15 | 0.25 | 2.65 |
| ectoderm | Oxtr | 5.78 | 0 | 0 | 0.26 | 0.01 | 0.25 | 32.25 |
| ectoderm | Dtna | 2.05 | 0 | 0 | 0.33 | 0.08 | 0.25 | 3.96 |
| ectoderm | Nectin1 | 1.9 | 0 | 0 | 0.36 | 0.12 | 0.25 | 3.14 |
| ectoderm | Bspry | 5.59 | 0 | 0 | 0.25 | 0.01 | 0.25 | 36.29 |
| ectoderm | Prr5l | 1.95 | 0 | 0 | 0.35 | 0.11 | 0.25 | 3.3 |
| ectoderm | Adamts18 | 4.9 | 0 | 0 | 0.26 | 0.01 | 0.25 | 21.5 |
| ectoderm | Rragd | 1.87 | 0 | 0 | 0.41 | 0.17 | 0.25 | 2.47 |
| ectoderm | Esrp2 | 5.7 | 0 | 0 | 0.25 | 0.01 | 0.25 | 36 |
| ectoderm | Jag2 | 4.15 | 0 | 0 | 0.27 | 0.02 | 0.24 | 11.17 |
| ectoderm | Sorl1 | 2.09 | 0 | 0 | 0.34 | 0.09 | 0.24 | 3.61 |
| ectoderm | Rab11fip1 | 2.68 | 0 | 0 | 0.3 | 0.06 | 0.24 | 5.03 |
| ectoderm | Sptbn2 | 2.46 | 0 | 0 | 0.3 | 0.06 | 0.24 | 5.32 |

**Supplementary Table 2. Marker genes for the major cell types of integrated clustering of E11.5 control and NotchGOF λ-junction cells.**

| | | | | | | | | |
|---|---|---|---|---|---|---|---|---|
| ectoderm | Camsap3 | 3.55 | 0 | 0 | 0.27 | 0.02 | 0.24 | 11.04 |
| ectoderm | Abcg1 | 3.51 | 0 | 0 | 0.27 | 0.03 | 0.24 | 10.19 |
| ectoderm | Pkp3 | 5.2 | 0 | 0 | 0.25 | 0.01 | 0.24 | 24.9 |
| ectoderm | 2610528J11Ri | 5.73 | 0 | 0 | 0.24 | 0.01 | 0.24 | 48.6 |
| ectoderm | Stxbp2 | 2.24 | 0 | 0 | 0.32 | 0.08 | 0.24 | 3.98 |
| ectoderm | Mtus1 | 2.62 | 0 | 0 | 0.29 | 0.05 | 0.24 | 5.84 |
| ectoderm | Ier5 | 1.41 | 0 | 0 | 0.83 | 0.6 | 0.24 | 1.4 |
| ectoderm | Chmp2b | 1.1 | 0 | 0 | 0.74 | 0.51 | 0.24 | 1.46 |
| ectoderm | Prss22 | 6.08 | 0 | 0 | 0.24 | 0.01 | 0.24 | 48 |
| ectoderm | Elf5 | 6.03 | 0 | 0 | 0.24 | 0.01 | 0.23 | 47.8 |
| ectoderm | Cd2ap | 1.35 | 0 | 0 | 0.8 | 0.57 | 0.23 | 1.41 |
| ectoderm | Lurap1l | 4.27 | 0 | 0 | 0.25 | 0.02 | 0.23 | 11.55 |
| ectoderm | Rassf10 | 5.73 | 0 | 0 | 0.24 | 0.01 | 0.23 | 39.5 |
| ectoderm | Plekhg3 | 3.51 | 0 | 0 | 0.26 | 0.03 | 0.23 | 9.52 |
| ectoderm | Smtnl2 | 3.94 | 0 | 0 | 0.25 | 0.02 | 0.23 | 11.86 |
| ectoderm | Lgi2 | 2.92 | 0 | 0 | 0.29 | 0.06 | 0.23 | 4.8 |
| ectoderm | Nectin3 | 1.42 | 0 | 0 | 0.72 | 0.49 | 0.23 | 1.46 |
| ectoderm | Ppfia3 | 2.3 | 0 | 0 | 0.3 | 0.08 | 0.23 | 3.97 |
| ectoderm | Itga3 | 4.42 | 0 | 0 | 0.24 | 0.02 | 0.23 | 16.07 |
| ectoderm | Mtcl1 | 2.21 | 0 | 0 | 0.29 | 0.07 | 0.23 | 4.32 |
| ectoderm | Cpz | 4.19 | 0 | 0 | 0.24 | 0.02 | 0.22 | 14.18 |
| ectoderm | Patj | 2.07 | 0 | 0 | 0.3 | 0.08 | 0.22 | 3.85 |
| ectoderm | Mapk12 | 2.65 | 0 | 0 | 0.28 | 0.06 | 0.22 | 4.56 |
| ectoderm | Cdcp1 | 5.35 | 0 | 0 | 0.23 | 0.01 | 0.22 | 37.67 |
| ectoderm | Dsc2 | 5.72 | 0 | 0 | 0.22 | 0.01 | 0.22 | 36.83 |
| ectoderm | S100a11 | 2.52 | 0 | 0 | 0.92 | 0.71 | 0.21 | 1.3 |
| ectoderm | Bend7 | 3.08 | 0 | 0 | 0.25 | 0.03 | 0.21 | 7.69 |
| ectoderm | Macc1 | 6.01 | 0 | 0 | 0.22 | 0.01 | 0.21 | 43.8 |
| ectoderm | Faah | 5.12 | 0 | 0 | 0.22 | 0.01 | 0.21 | 31.43 |
| ectoderm | Grhl3 | 6.42 | 0 | 0 | 0.22 | 0.01 | 0.21 | 43.4 |
| ectoderm | Rhov | 5.72 | 0 | 0 | 0.22 | 0.01 | 0.21 | 31.14 |
| ectoderm | Gm32585 | 5.9 | 0 | 0 | 0.22 | 0.01 | 0.21 | 43 |
| ectoderm | Lmo2 | 2.18 | 0 | 0 | 0.29 | 0.08 | 0.21 | 3.73 |
| ectoderm | Lypd6b | 3.82 | 0 | 0 | 0.23 | 0.02 | 0.21 | 12 |
| ectoderm | Prr36 | 2.76 | 0 | 0 | 0.25 | 0.04 | 0.21 | 6.5 |
| ectoderm | 0230030E09Ri | 4.98 | 0 | 0 | 0.22 | 0.01 | 0.21 | 30.71 |
| ectoderm | Nrg1 | 2.82 | 0 | 0 | 0.24 | 0.03 | 0.21 | 7.65 |
| ectoderm | Sema3c | 2.96 | 0 | 0 | 0.25 | 0.05 | 0.21 | 5.56 |
| ectoderm | Arhgap27os2 | 3.93 | 0 | 0 | 0.22 | 0.02 | 0.21 | 13.06 |
| ectoderm | Marveld2 | 4.67 | 0 | 0 | 0.22 | 0.01 | 0.21 | 19.64 |
| ectoderm | Lrp2 | 5.65 | 0 | 0 | 0.21 | 0.01 | 0.2 | 35 |
| ectoderm | Krt7 | 6.17 | 0 | 0 | 0.21 | 0 | 0.2 | 51.75 |
| ectoderm | Heyl | 4.79 | 0 | 0 | 0.21 | 0.01 | 0.2 | 21 |
| ectoderm | Pdcd4 | 1.15 | 0 | 0 | 0.79 | 0.59 | 0.2 | 1.34 |
| ectoderm | B3gnt2 | 2.14 | 0 | 0 | 0.25 | 0.05 | 0.2 | 4.83 |
| ectoderm | Tenm2 | 3.48 | 0 | 0 | 0.22 | 0.03 | 0.2 | 8.96 |
| ectoderm | Unc13b | 2.68 | 0 | 0 | 0.24 | 0.04 | 0.2 | 6.53 |
| ectoderm | Wipf3 | 4.23 | 0 | 0 | 0.21 | 0.01 | 0.2 | 16.31 |
| ectoderm | Clic6 | 5.06 | 0 | 0 | 0.21 | 0.01 | 0.2 | 29.43 |
| ectoderm | Tex15 | 2.09 | 0 | 0 | 0.26 | 0.06 | 0.2 | 4.14 |
| ectoderm | Slain1 | 3.97 | 0 | 0 | 0.21 | 0.01 | 0.2 | 16.23 |
| ectoderm | Bicd1 | 2.27 | 0 | 0 | 0.25 | 0.05 | 0.2 | 4.86 |
| ectoderm | Retreg1 | 2.63 | 0 | 0 | 0.24 | 0.05 | 0.2 | 5.28 |
| ectoderm | Arhgef16 | 4.19 | 0 | 0 | 0.21 | 0.01 | 0.2 | 17.42 |
| ectoderm | Dmrt3 | 5.22 | 0 | 0 | 0.2 | 0.01 | 0.2 | 25.38 |
| ectoderm | Pkp2 | 4.86 | 0 | 0 | 0.2 | 0.01 | 0.19 | 25.13 |

**Supplementary Table 2. Marker genes for the major cell types of integrated clustering of E11.5 control and NotchGOF λ-junction cells.**

| | | | | | | | | |
|---|---|---|---|---|---|---|---|---|
| ectoderm | Zfp750 | 6.35 | 0 | 0 | 0.2 | 0 | 0.19 | 49 |
| ectoderm | Large2 | 2.66 | 0 | 0 | 0.23 | 0.04 | 0.19 | 5.63 |
| ectoderm | Sparc | 0.91 | 0 | 0 | 0.89 | 0.7 | 0.19 | 1.27 |
| ectoderm | Samd10 | 2.45 | 0 | 0 | 0.24 | 0.05 | 0.19 | 5.11 |
| ectoderm | Svip | 2.67 | 0 | 0 | 0.24 | 0.05 | 0.19 | 5.02 |
| ectoderm | Syngr1 | 3.18 | 0 | 0 | 0.21 | 0.02 | 0.19 | 9.22 |
| ectoderm | Lmo1 | 3.06 | 0 | 0 | 0.22 | 0.03 | 0.19 | 7.03 |
| ectoderm | 130071C03Ri | 4.69 | 0 | 0 | 0.19 | 0.01 | 0.19 | 24.25 |
| ectoderm | Syt5 | 3.9 | 0 | 0 | 0.2 | 0.02 | 0.18 | 13.27 |
| ectoderm | Eml5 | 2.44 | 0 | 0 | 0.23 | 0.04 | 0.18 | 5.38 |
| ectoderm | Trim29 | 6.37 | 0 | 0 | 0.19 | 0 | 0.18 | 46.5 |
| ectoderm | Tnfrsf12a | 2.88 | 0 | 0 | 0.22 | 0.03 | 0.18 | 6.32 |
| ectoderm | Tjp3 | 5.31 | 0 | 0 | 0.19 | 0.01 | 0.18 | 31 |
| ectoderm | Cyb561 | 3.94 | 0 | 0 | 0.19 | 0.01 | 0.18 | 14.85 |
| ectoderm | Jph4 | 4.97 | 0 | 0 | 0.19 | 0.01 | 0.18 | 16 |
| ectoderm | Rnf43 | 5.15 | 0 | 0 | 0.19 | 0.01 | 0.18 | 26.57 |
| ectoderm | Il17re | 5.96 | 0 | 0 | 0.18 | 0 | 0.18 | 60.33 |
| ectoderm | Hip1r | 2.82 | 0 | 0 | 0.21 | 0.03 | 0.18 | 6.21 |
| ectoderm | Kremen2 | 6.46 | 0 | 0 | 0.18 | 0 | 0.18 | 45.25 |
| ectoderm | Rab38 | 3.42 | 0 | 0 | 0.2 | 0.03 | 0.18 | 7.52 |
| ectoderm | Rims2 | 3.23 | 0 | 0 | 0.19 | 0.02 | 0.18 | 11.29 |
| ectoderm | Epha1 | 3.59 | 0 | 0 | 0.19 | 0.02 | 0.17 | 10.67 |
| ectoderm | Bnipl | 5.71 | 0 | 0 | 0.18 | 0 | 0.17 | 44.25 |
| ectoderm | Hap1 | 3.93 | 0 | 0 | 0.18 | 0.01 | 0.17 | 18.2 |
| ectoderm | Nrcam | 4.12 | 0 | 0 | 0.18 | 0.01 | 0.17 | 16.55 |
| ectoderm | Cryab | 4.53 | 0 | 0 | 0.18 | 0.01 | 0.17 | 22.25 |
| ectoderm | Gjb2 | 5.87 | 0 | 0 | 0.18 | 0.01 | 0.17 | 29.33 |
| ectoderm | Shtn1 | 3.12 | 0 | 0 | 0.18 | 0.01 | 0.17 | 15.08 |
| ectoderm | Ermn | 6.46 | 0 | 0 | 0.17 | 0 | 0.17 | 43 |
| ectoderm | Cux1 | 0.92 | 0 | 0 | 0.83 | 0.66 | 0.17 | 1.26 |
| ectoderm | Foxe1 | 7.03 | 0 | 0 | 0.17 | 0.01 | 0.17 | 28.83 |
| ectoderm | Llgl2 | 4.46 | 0 | 0 | 0.18 | 0.01 | 0.17 | 17.6 |
| ectoderm | Ces1d | 5.79 | 0 | 0 | 0.17 | 0 | 0.17 | 42.25 |
| ectoderm | Wnt7b | 6.56 | 0 | 0 | 0.17 | 0 | 0.17 | 56 |
| ectoderm | Scg5 | 3.06 | 0 | 0 | 0.18 | 0.02 | 0.16 | 11.93 |
| ectoderm | Nr2e1 | 5.73 | 0 | 0 | 0.17 | 0 | 0.16 | 41.75 |
| ectoderm | Arg2 | 3.65 | 0 | 0 | 0.18 | 0.02 | 0.16 | 11.73 |
| ectoderm | Lamc2 | 5.45 | 0 | 0 | 0.17 | 0.01 | 0.16 | 33 |
| ectoderm | Rnf208 | 2.61 | 0 | 0 | 0.19 | 0.03 | 0.16 | 6.13 |
| ectoderm | Itgb6 | 5.75 | 0 | 0 | 0.16 | 0 | 0.16 | 40.75 |
| ectoderm | Epn3 | 3.94 | 0 | 0 | 0.17 | 0.01 | 0.16 | 14.17 |
| ectoderm | Ppl | 5.8 | 0 | 0 | 0.16 | 0 | 0.16 | 40.5 |
| ectoderm | Fgf17 | 5.63 | 0 | 0 | 0.16 | 0.01 | 0.16 | 27.17 |
| ectoderm | Barx2 | 6.65 | 0 | 0 | 0.16 | 0 | 0.16 | 40 |
| ectoderm | Dmbx1 | 4.69 | 0 | 0 | 0.16 | 0.01 | 0.15 | 20.13 |
| ectoderm | Krt15 | 6.85 | 0 | 0 | 0.16 | 0 | 0.15 | 39.25 |
| ectoderm | Arhgap8 | 4.69 | 0 | 0 | 0.16 | 0.01 | 0.15 | 26.5 |
| ectoderm | Ripply3 | 3.18 | 0 | 0 | 0.18 | 0.03 | 0.15 | 7.12 |
| ectoderm | Nup210 | 3 | 0 | 0 | 0.18 | 0.03 | 0.15 | 7.08 |
| ectoderm | Frmd5 | 3.6 | 0 | 0 | 0.17 | 0.02 | 0.15 | 11.07 |
| ectoderm | St8sia6 | 4.27 | 0 | 0 | 0.16 | 0.01 | 0.15 | 12.62 |
| ectoderm | Pqlc3 | 3.41 | 0 | 0 | 0.17 | 0.02 | 0.15 | 8.4 |
| ectoderm | Lypd2 | 6.34 | 0 | 0 | 0.15 | 0 | 0.15 | 50.33 |
| ectoderm | Rasef | 4.79 | 0 | 0 | 0.15 | 0.01 | 0.15 | 25.5 |
| ectoderm | Marveld3 | 6.01 | 0 | 0 | 0.15 | 0 | 0.15 | 74.5 |
| ectoderm | Trpm3 | 3.97 | 0 | 0 | 0.16 | 0.01 | 0.15 | 13.25 |

**Supplementary Table 2. Marker genes for the major cell types of integrated clustering of E11.5 control and NotchGOF λ-junction cells.**

| | | | | | | | | |
|---|---|---|---|---|---|---|---|---|
| ectoderm | Smim22 | 4.78 | 0 | 0 | 0.15 | 0 | 0.15 | 37.25 |
| ectoderm | Eps8l2 | 4.46 | 0 | 0 | 0.15 | 0.01 | 0.14 | 19 |
| ectoderm | Sox3 | 5.3 | 0 | 0 | 0.15 | 0.01 | 0.14 | 25 |
| ectoderm | Fgf12 | 3.64 | 0 | 0 | 0.16 | 0.02 | 0.14 | 10.53 |
| ectoderm | Ppp1r36 | 3.15 | 0 | 0 | 0.16 | 0.02 | 0.14 | 8.1 |
| ectoderm | Exph5 | 6.31 | 0 | 0 | 0.14 | 0 | 0.14 | 72 |
| ectoderm | Fam107a | 4.04 | 0 | 0 | 0.15 | 0.01 | 0.14 | 13.91 |
| ectoderm | Myo5c | 4.94 | 0 | 0 | 0.15 | 0.01 | 0.14 | 29 |
| ectoderm | Cxcr4 | 2.82 | 0 | 0 | 0.16 | 0.02 | 0.14 | 9.18 |
| ectoderm | Cldn23 | 4.96 | 0 | 0 | 0.14 | 0.01 | 0.14 | 28.8 |
| ectoderm | Prr15l | 6.06 | 0 | 0 | 0.14 | 0 | 0.14 | 47.33 |
| ectoderm | Gjb3 | 5.71 | 0 | 0 | 0.14 | 0 | 0.14 | 70 |
| ectoderm | Fam187b | 5 | 0 | 0 | 0.14 | 0.01 | 0.14 | 24 |
| ectoderm | .700023F06Ri | 3.62 | 0 | 0 | 0.15 | 0.01 | 0.14 | 12.33 |
| ectoderm | Hras | 1.34 | 0 | 0 | 0.91 | 0.77 | 0.14 | 1.18 |
| ectoderm | Ildr1 | 4.43 | 0 | 0 | 0.14 | 0.01 | 0.13 | 23.33 |
| ectoderm | Lmo4 | 1.25 | 0 | 0 | 0.91 | 0.78 | 0.13 | 1.17 |
| ectoderm | Cyb5r3 | 1.23 | 0 | 0 | 0.91 | 0.78 | 0.13 | 1.17 |
| ectoderm | Myo5b | 3.33 | 0 | 0 | 0.15 | 0.01 | 0.13 | 11.15 |
| ectoderm | Cblc | 6.49 | 0 | 0 | 0.13 | 0 | 0.13 | 66.5 |
| ectoderm | Cbln2 | 3.96 | 0 | 0 | 0.14 | 0.01 | 0.13 | 12.45 |
| ectoderm | Hspb2 | 5.36 | 0 | 0 | 0.13 | 0 | 0.13 | 32.25 |
| ectoderm | AI661453 | 4.64 | 0 | 0 | 0.13 | 0.01 | 0.13 | 21.83 |
| ectoderm | Nuak2 | 3.88 | 0 | 0 | 0.13 | 0.01 | 0.12 | 16.5 |
| ectoderm | Nfe2l3 | 4.52 | 0 | 0 | 0.13 | 0.01 | 0.12 | 25.8 |
| ectoderm | .310002F09Ri | 4.21 | 0 | 0 | 0.13 | 0.01 | 0.12 | 16.38 |
| ectoderm | Vwa2 | 4.33 | 0 | 0 | 0.13 | 0.01 | 0.12 | 18.57 |
| ectoderm | Lratd1 | 5.23 | 0 | 0 | 0.13 | 0.01 | 0.12 | 21.17 |
| ectoderm | Prom2 | 5.04 | 0 | 0 | 0.12 | 0 | 0.12 | 31 |
| ectoderm | Kcnh8 | 3.76 | 0 | 0 | 0.13 | 0.01 | 0.12 | 21 |
| ectoderm | Zfp385c | 5.47 | 0 | 0 | 0.12 | 0.01 | 0.12 | 24.8 |
| ectoderm | Wnt3 | 7.98 | 0 | 0 | 0.12 | 0 | 0.12 | 120 |
| ectoderm | Sox21 | 5.28 | 0 | 0 | 0.12 | 0.01 | 0.12 | 24.6 |
| ectoderm | Cldn8 | 6.83 | 0 | 0 | 0.12 | 0 | 0.12 | 60 |
| ectoderm | Col17a1 | 4.73 | 0 | 0 | 0.13 | 0.01 | 0.12 | 15.75 |
| ectoderm | Prr15 | 4.77 | 0 | 0 | 0.12 | 0.01 | 0.12 | 20.5 |
| ectoderm | Sinhcaf | 0.81 | 0 | 0 | 0.93 | 0.81 | 0.12 | 1.14 |
| ectoderm | Ctxn3 | 5.83 | 0 | 0 | 0.12 | 0 | 0.12 | 39.33 |
| ectoderm | Sp6 | 5.42 | 0 | 0 | 0.12 | 0 | 0.11 | 39 |
| ectoderm | 930426D05Ri | 4.68 | 0 | 0 | 0.12 | 0.01 | 0.11 | 23.8 |
| ectoderm | Ovol2 | 6.11 | 0 | 0 | 0.12 | 0 | 0.11 | 57.5 |
| ectoderm | Wnt9b | 6.25 | 0 | 0 | 0.12 | 0 | 0.11 | 29.25 |
| ectoderm | Dennd1c | 4.28 | 0 | 0 | 0.12 | 0.01 | 0.11 | 19.67 |
| ectoderm | Itgb4 | 4.59 | 0 | 0 | 0.12 | 0.01 | 0.11 | 19.5 |
| ectoderm | Gm12446 | 6.68 | 0 | 0 | 0.11 | 0 | 0.11 | 54 |
| ectoderm | Npm3 | 0.95 | 0 | 0 | 0.87 | 0.77 | 0.1 | 1.13 |
| ectoderm | Ktn1 | 0.97 | 0 | 0 | 0.9 | 0.8 | 0.09 | 1.11 |
| ectoderm | Itm2b | 1.74 | 0 | 0 | 0.91 | 0.82 | 0.09 | 1.1 |
| ectoderm | Aplp2 | 0.99 | 0 | 0 | 0.88 | 0.8 | 0.07 | 1.09 |
| ectoderm | H19 | 1.69 | 0 | 0 | 0.89 | 0.83 | 0.06 | 1.07 |
| ectoderm | Cd24a | 0.79 | 0 | 0 | 0.96 | 0.91 | 0.05 | 1.06 |
| ectoderm | Dstn | 1.03 | 0 | 0 | 0.98 | 0.93 | 0.05 | 1.05 |
| ectoderm | Atp6v1f | 0.72 | 0 | 0 | 0.94 | 0.89 | 0.05 | 1.06 |
| ectoderm | Cyb5a | 0.93 | 0 | 0 | 0.96 | 0.91 | 0.04 | 1.05 |
| ectoderm | Pbx1 | 0.7 | 0 | 0 | 0.95 | 0.91 | 0.04 | 1.04 |
| ectoderm | Atox1 | 1.49 | 0 | 0 | 0.98 | 0.94 | 0.04 | 1.04 |

**Supplementary Table 2. Marker genes for the major cell types of integrated clustering of E11.5 control and NotchGOF λ-junction cells.**

| | | | | | | | | |
|---|---|---|---|---|---|---|---|---|
| ectoderm | Siva1 | 1.14 | 0 | 0 | 0.97 | 0.94 | 0.03 | 1.03 |
| ectoderm | Tpi1 | 0.87 | 0 | 0 | 0.98 | 0.95 | 0.03 | 1.03 |
| ectoderm | Pgk1 | 0.71 | 0 | 0 | 0.98 | 0.95 | 0.03 | 1.03 |
| ectoderm | Mdh1 | 0.83 | 0 | 0 | 0.97 | 0.94 | 0.03 | 1.03 |
| ectoderm | Aldoa | 0.57 | 0 | 0 | 0.97 | 0.95 | 0.03 | 1.03 |
| ectoderm | Pgam1 | 0.71 | 0 | 0 | 0.99 | 0.96 | 0.03 | 1.03 |
| ectoderm | Ldha | 0.73 | 0 | 0 | 0.99 | 0.97 | 0.02 | 1.02 |
| ectoderm | Eno1 | 0.63 | 0 | 0 | 0.99 | 0.97 | 0.02 | 1.02 |
| ectoderm | Pkm | 0.57 | 0 | 0 | 1 | 0.98 | 0.02 | 1.02 |
| ectoderm | Tceal9 | 0.48 | 0 | 0 | 0.99 | 0.98 | 0.01 | 1.01 |
| ectoderm | Fabp5 | 1.07 | 0 | 0 | 0.99 | 0.98 | 0.01 | 1.01 |
| ectoderm | Hdgf | 0.6 | 0 | 0 | 0.98 | 0.97 | 0 | 1 |
| ectoderm | Rpl7 | 0.36 | 0 | 0 | 1 | 1 | 0 | 1 |
| ectoderm | Rpl12 | 0.32 | 0 | 0 | 1 | 1 | 0 | 1 |
| ectoderm | Rps21 | 0.27 | 0 | 0 | 1 | 1 | 0 | 1 |
| ectoderm | Gapdh | 0.47 | 0 | 0 | 1 | 1 | 0 | 1 |
| ectoderm | Rps18 | 0.36 | 0 | 0 | 1 | 1 | 0 | 1 |
| ectoderm | Fau | 0.33 | 0 | 0 | 1 | 1 | 0 | 1 |
| ectoderm | Rpl36a | 0.29 | 0 | 0 | 1 | 1 | 0 | 1 |
| ectoderm | Rps27 | 0.39 | 0 | 0 | 1 | 1 | 0 | 1 |
| ectoderm | Rps20 | 0.27 | 0 | 0 | 1 | 1 | 0 | 1 |
| ectoderm | Rplp0 | 0.28 | 0 | 0 | 1 | 1 | 0 | 1 |
| ectoderm | Rpl32 | 0.31 | 0 | 0 | 1 | 1 | 0 | 1 |
| ectoderm | Rpl28 | 0.31 | 0 | 0 | 1 | 1 | 0 | 1 |
| ectoderm | Rps5 | 0.3 | 0 | 0 | 1 | 1 | 0 | 1 |
| ectoderm | Rps19 | 0.4 | 0 | 0 | 1 | 1 | 0 | 1 |
| ectoderm | Rps16 | 0.32 | 0 | 0 | 1 | 1 | 0 | 1 |
| ectoderm | Rpl27a | 0.37 | 0 | 0 | 1 | 1 | 0 | 1 |
| ectoderm | Rps13 | 0.28 | 0 | 0 | 1 | 1 | 0 | 1 |
| ectoderm | Rplp2 | 0.29 | 0 | 0 | 1 | 1 | 0 | 1 |
| ectoderm | Rpl18a | 0.29 | 0 | 0 | 1 | 1 | 0 | 1 |
| ectoderm | Rpl14 | 0.26 | 0 | 0 | 1 | 1 | 0 | 1 |
| ectoderm | Rps26 | 0.34 | 0 | 0 | 1 | 1 | 0 | 1 |
| ectoderm | Rps27a | 0.31 | 0 | 0 | 1 | 1 | 0 | 1 |
| ectoderm | Rpl26 | 0.28 | 0 | 0 | 1 | 1 | 0 | 1 |
| ectoderm | Rpl23 | 0.29 | 0 | 0 | 1 | 1 | 0 | 1 |
| ectoderm | Rpl38 | 0.38 | 0 | 0 | 1 | 1 | 0 | 1 |
| ectoderm | Rps7 | 0.27 | 0 | 0 | 1 | 1 | 0 | 1 |
| ectoderm | Tpt1 | 0.26 | 0 | 0 | 1 | 1 | 0 | 1 |
| ectoderm | Rpl3 | 0.26 | 0 | 0 | 1 | 1 | 0 | 1 |
| ectoderm | Rps2 | 0.28 | 0 | 0 | 1 | 1 | 0 | 1 |
| ectoderm | Rps10 | 0.28 | 0 | 0 | 1 | 1 | 0 | 1 |
| ectoderm | Rpl10a | 0.32 | 0 | 0 | 1 | 1 | 0 | 1 |
| ectoderm | Rps28 | 0.27 | 0 | 0 | 1 | 1 | 0 | 1 |
| ectoderm | Hsp90ab1 | 0.4 | 0 | 0 | 1 | 1 | 0 | 1 |
| ectoderm | Rpl36 | 0.28 | 0 | 0 | 1 | 1 | 0 | 1 |
| ectoderm | Rps14 | 0.29 | 0 | 0 | 1 | 1 | 0 | 1 |
| ectoderm | Rpl17 | 0.37 | 0 | 0 | 1 | 1 | 0 | 1 |
| ectoderm | Rps4x | 0.25 | 0 | 0 | 1 | 1 | 0 | 1 |
| ectoderm | Nr2f2 | 1.06 | 4.89E-307 | 1.02E-302 | 0.59 | 0.3 | 0.29 | 1.99 |
| ectoderm | Gm48435 | 5.43 | 3.35E-306 | 6.97E-302 | 0.1 | 0 | 0.1 | 51 |
| ectoderm | Tnk1 | 3.82 | 1.08E-304 | 2.24E-300 | 0.13 | 0.01 | 0.12 | 14.11 |
| ectoderm | Nrtn | 2.27 | 9.80E-303 | 2.04E-298 | 0.32 | 0.11 | 0.22 | 3.05 |
| ectoderm | Lypd3 | 5.87 | 1.10E-301 | 2.29E-297 | 0.1 | 0 | 0.1 | 50.5 |
| ectoderm | Pdpn | 1.92 | 1.45E-301 | 3.02E-297 | 0.37 | 0.13 | 0.23 | 2.77 |
| ectoderm | Syt1 | 3.61 | 1.01E-300 | 2.10E-296 | 0.13 | 0.01 | 0.12 | 12.7 |

**Supplementary Table 2. Marker genes for the major cell types of integrated clustering of E11.5 control and NotchGOF λ-junction cells.**

| | | | | | | | | |
|---|---|---|---|---|---|---|---|---|
| ectoderm | Gpc4 | 2.08 | 1.33E-300 | 2.77E-296 | 0.3 | 0.09 | 0.21 | 3.32 |
| ectoderm | Frrs1 | 1.6 | 1.86E-300 | 3.88E-296 | 0.41 | 0.16 | 0.25 | 2.54 |
| ectoderm | Fam162a | 0.76 | 4.81E-300 | 1.00E-295 | 0.93 | 0.88 | 0.04 | 1.05 |
| ectoderm | Id1 | 0.79 | 1.28E-299 | 2.67E-295 | 0.94 | 0.84 | 0.1 | 1.12 |
| ectoderm | Pclo | 1.49 | 2.16E-298 | 4.49E-294 | 0.26 | 0.06 | 0.19 | 4.18 |
| ectoderm | Pkp1 | 6.31 | 2.95E-297 | 6.14E-293 | 0.1 | 0 | 0.1 | 33.33 |
| ectoderm | Pcgf5 | 2.5 | 2.97E-294 | 6.18E-290 | 0.22 | 0.05 | 0.17 | 4.62 |
| ectoderm | Kremen1 | 1.96 | 4.23E-294 | 8.79E-290 | 0.31 | 0.1 | 0.21 | 3.24 |
| ectoderm | Dock9 | 2.44 | 1.66E-293 | 3.46E-289 | 0.2 | 0.04 | 0.16 | 5.42 |
| ectoderm | Gldc | 3.45 | 3.59E-293 | 7.47E-289 | 0.14 | 0.01 | 0.12 | 10.38 |
| ectoderm | Cox7a2l | 0.53 | 4.51E-291 | 9.38E-287 | 0.96 | 0.96 | 0.01 | 1.01 |
| ectoderm | Gpi1 | 0.67 | 9.93E-291 | 2.07E-286 | 0.91 | 0.82 | 0.09 | 1.11 |
| ectoderm | Gucy1a1 | 2.89 | 1.66E-290 | 3.45E-286 | 0.2 | 0.04 | 0.16 | 5.18 |
| ectoderm | Tmem178 | 4.05 | 1.10E-287 | 2.28E-283 | 0.12 | 0.01 | 0.11 | 16.43 |
| ectoderm | Myl12b | 0.63 | 1.22E-285 | 2.54E-281 | 0.93 | 0.87 | 0.06 | 1.06 |
| ectoderm | Metrnl | 2.66 | 2.12E-285 | 4.41E-281 | 0.18 | 0.03 | 0.15 | 5.97 |
| ectoderm | Foxp1 | 0.61 | 2.85E-283 | 5.92E-279 | 0.84 | 0.53 | 0.32 | 1.6 |
| ectoderm | Plekhg1 | 1.99 | 1.24E-282 | 2.57E-278 | 0.29 | 0.09 | 0.2 | 3.28 |
| ectoderm | Cacnb3 | 1.35 | 5.21E-282 | 1.08E-277 | 0.47 | 0.21 | 0.26 | 2.24 |
| ectoderm | Cadps2 | 3.09 | 1.11E-281 | 2.31E-277 | 0.12 | 0.01 | 0.11 | 12.2 |
| ectoderm | Acvr2b | 0.86 | 3.18E-281 | 6.61E-277 | 0.79 | 0.6 | 0.19 | 1.31 |
| ectoderm | Igsf9b | 2.94 | 1.42E-280 | 2.94E-276 | 0.16 | 0.02 | 0.14 | 6.75 |
| ectoderm | Mkrn1 | 1.06 | 5.84E-280 | 1.22E-275 | 0.63 | 0.36 | 0.27 | 1.73 |
| ectoderm | Kif13b | 1.87 | 2.16E-277 | 4.49E-273 | 0.32 | 0.11 | 0.21 | 2.89 |
| ectoderm | Tfap2b | 3.54 | 4.44E-276 | 9.23E-272 | 0.21 | 0.05 | 0.16 | 4.2 |
| ectoderm | Tafa2 | 3.67 | 6.55E-275 | 1.36E-270 | 0.13 | 0.01 | 0.12 | 10 |
| ectoderm | Apc | 1.09 | 1.38E-273 | 2.88E-269 | 0.75 | 0.58 | 0.17 | 1.29 |
| ectoderm | Rpl5 | 0.25 | 2.49E-273 | 5.17E-269 | 1 | 1 | 0 | 1 |
| ectoderm | Cldn1 | 5.18 | 3.10E-273 | 6.46E-269 | 0.1 | 0.01 | 0.1 | 20.4 |
| ectoderm | Ppfibp1 | 1.27 | 3.98E-272 | 8.27E-268 | 0.62 | 0.37 | 0.24 | 1.66 |
| ectoderm | Knop1 | 0.58 | 3.11E-270 | 6.46E-266 | 0.94 | 0.89 | 0.05 | 1.05 |
| ectoderm | Plch2 | 4.25 | 1.37E-266 | 2.84E-262 | 0.1 | 0.01 | 0.1 | 17.33 |
| ectoderm | Cdh2 | 1.51 | 2.17E-266 | 4.52E-262 | 0.59 | 0.37 | 0.21 | 1.57 |
| ectoderm | Ttc9 | 3.47 | 1.88E-264 | 3.92E-260 | 0.13 | 0.01 | 0.11 | 9.77 |
| ectoderm | Slc15a2 | 3.33 | 2.12E-264 | 4.41E-260 | 0.14 | 0.02 | 0.12 | 7.53 |
| ectoderm | Ide | 1.43 | 2.35E-262 | 4.89E-258 | 0.45 | 0.21 | 0.24 | 2.12 |
| ectoderm | Mir9-3hg | 4.97 | 1.03E-261 | 2.14E-257 | 0.1 | 0.01 | 0.1 | 20 |
| ectoderm | Sh3gl2 | 3.28 | 1.78E-261 | 3.71E-257 | 0.13 | 0.01 | 0.11 | 9.62 |
| ectoderm | Prxl2a | 1.2 | 4.14E-260 | 8.60E-256 | 0.57 | 0.32 | 0.25 | 1.79 |
| ectoderm | Tspan7 | 1.59 | 1.49E-259 | 3.10E-255 | 0.46 | 0.22 | 0.24 | 2.07 |
| ectoderm | H2afj | 0.65 | 1.13E-258 | 2.35E-254 | 0.89 | 0.79 | 0.11 | 1.13 |
| ectoderm | Armc10 | 0.97 | 1.34E-258 | 2.78E-254 | 0.71 | 0.47 | 0.24 | 1.51 |
| ectoderm | Prag1 | 1.89 | 3.32E-258 | 6.91E-254 | 0.29 | 0.09 | 0.2 | 3.1 |
| ectoderm | Pak3 | 1.86 | 4.25E-258 | 8.83E-254 | 0.29 | 0.09 | 0.2 | 3.14 |
| ectoderm | Nol7 | 0.59 | 1.87E-257 | 3.90E-253 | 0.91 | 0.87 | 0.04 | 1.05 |
| ectoderm | Mtmr7 | 2.8 | 1.41E-256 | 2.94E-252 | 0.15 | 0.02 | 0.13 | 7.05 |
| ectoderm | Prps2 | 1.85 | 8.46E-255 | 1.76E-250 | 0.33 | 0.12 | 0.21 | 2.73 |
| ectoderm | Arid3b | 1.55 | 1.35E-254 | 2.80E-250 | 0.4 | 0.18 | 0.23 | 2.3 |
| ectoderm | Ppp1r13l | 2.5 | 1.50E-254 | 3.12E-250 | 0.19 | 0.04 | 0.15 | 4.62 |
| ectoderm | Lrrk1 | 1.71 | 1.58E-254 | 3.28E-250 | 0.31 | 0.1 | 0.2 | 2.96 |
| ectoderm | Eps8l1 | 3.28 | 2.28E-254 | 4.74E-250 | 0.12 | 0.01 | 0.11 | 9.54 |
| ectoderm | Dipk2a | 1.7 | 1.24E-253 | 2.58E-249 | 0.41 | 0.19 | 0.23 | 2.21 |
| ectoderm | Trpm4 | 2.04 | 2.01E-252 | 4.19E-248 | 0.23 | 0.06 | 0.17 | 3.91 |
| ectoderm | Egln3 | 2.38 | 3.44E-252 | 7.16E-248 | 0.19 | 0.04 | 0.15 | 4.63 |
| ectoderm | Map7d2 | 2.28 | 3.53E-251 | 7.35E-247 | 0.2 | 0.05 | 0.16 | 4.32 |
| ectoderm | Pard6b | 2.76 | 1.17E-250 | 2.43E-246 | 0.15 | 0.02 | 0.13 | 6.48 |

**Supplementary Table 2. Marker genes for the major cell types of integrated clustering of E11.5 control and NotchGOF λ-junction cells.**

| | | | | | | | | |
|---|---|---|---|---|---|---|---|---|
| ectoderm | Cdk6 | 1.26 | 1.44E-249 | 2.99E-245 | 0.52 | 0.28 | 0.24 | 1.85 |
| ectoderm | Trmt9b | 2.54 | 2.49E-249 | 5.17E-245 | 0.18 | 0.04 | 0.14 | 5.09 |
| ectoderm | Atp8b1 | 2.81 | 3.57E-248 | 7.43E-244 | 0.13 | 0.02 | 0.11 | 8.53 |
| ectoderm | Sorbs2 | 1.35 | 1.08E-246 | 2.25E-242 | 0.4 | 0.17 | 0.23 | 2.38 |
| ectoderm | Dlx1 | 3.1 | 4.57E-246 | 9.51E-242 | 0.15 | 0.02 | 0.12 | 6.35 |
| ectoderm | Fzd8 | 1.89 | 9.38E-246 | 1.95E-241 | 0.3 | 0.1 | 0.2 | 2.91 |
| ectoderm | Acsl5 | 1.7 | 1.59E-244 | 3.30E-240 | 0.31 | 0.11 | 0.2 | 2.8 |
| ectoderm | Atp6v0e2 | 2.47 | 4.74E-244 | 9.86E-240 | 0.15 | 0.02 | 0.13 | 6.21 |
| ectoderm | Cdca7l | 1.3 | 9.44E-243 | 1.96E-238 | 0.49 | 0.25 | 0.24 | 1.95 |
| ectoderm | Adamts17 | 3.41 | 5.23E-242 | 1.09E-237 | 0.12 | 0.01 | 0.1 | 9.67 |
| ectoderm | Aig1 | 1.22 | 9.35E-242 | 1.95E-237 | 0.52 | 0.27 | 0.25 | 1.9 |
| ectoderm | Ctnnb1 | 0.45 | 1.11E-240 | 2.31E-236 | 0.97 | 0.97 | 0.01 | 1.01 |
| ectoderm | Cntnap2 | 2.89 | 2.22E-240 | 4.62E-236 | 0.14 | 0.02 | 0.12 | 6.26 |
| ectoderm | Rdh10 | 2.19 | 4.05E-240 | 8.43E-236 | 0.3 | 0.11 | 0.19 | 2.72 |
| ectoderm | Tbc1d30 | 3.65 | 1.41E-239 | 2.94E-235 | 0.1 | 0.01 | 0.1 | 13 |
| ectoderm | Srd5a1 | 1.76 | 5.06E-239 | 1.05E-234 | 0.34 | 0.14 | 0.21 | 2.52 |
| ectoderm | F3 | 2.23 | 1.07E-238 | 2.22E-234 | 0.23 | 0.06 | 0.17 | 3.62 |
| ectoderm | Dlx6os1 | 3.91 | 8.00E-235 | 1.66E-230 | 0.12 | 0.02 | 0.11 | 7.69 |
| ectoderm | Nr2f6 | 0.88 | 1.65E-234 | 3.43E-230 | 0.74 | 0.56 | 0.19 | 1.33 |
| ectoderm | Mid1ip1 | 1.83 | 4.05E-234 | 8.43E-230 | 0.38 | 0.17 | 0.21 | 2.27 |
| ectoderm | Egfr | 1.95 | 6.39E-234 | 1.33E-229 | 0.23 | 0.06 | 0.16 | 3.6 |
| ectoderm | Gpm6a | 2.96 | 1.55E-232 | 3.21E-228 | 0.14 | 0.02 | 0.12 | 6.57 |
| ectoderm | Proser2 | 2.87 | 1.77E-232 | 3.67E-228 | 0.15 | 0.03 | 0.12 | 5.43 |
| ectoderm | Cpne5 | 2.83 | 2.68E-232 | 5.57E-228 | 0.15 | 0.02 | 0.12 | 6.04 |
| ectoderm | Pomp | 0.39 | 9.21E-232 | 1.92E-227 | 0.98 | 0.98 | 0 | 1 |
| ectoderm | Vamp8 | 0.83 | 1.44E-231 | 3.00E-227 | 0.77 | 0.56 | 0.2 | 1.36 |
| ectoderm | 330021F23Ri | 3.37 | 1.63E-231 | 3.39E-227 | 0.1 | 0.01 | 0.09 | 11.44 |
| ectoderm | Exoc3l4 | 2.03 | 1.73E-230 | 3.59E-226 | 0.18 | 0.04 | 0.14 | 4.66 |
| ectoderm | Ntm | 2.11 | 3.96E-230 | 8.24E-226 | 0.23 | 0.07 | 0.17 | 3.43 |
| ectoderm | Hpn | 3.1 | 4.21E-230 | 8.76E-226 | 0.12 | 0.01 | 0.1 | 8.43 |
| ectoderm | Ifi30 | 1.08 | 1.46E-229 | 3.04E-225 | 0.48 | 0.24 | 0.24 | 2 |
| ectoderm | H2-Q4 | 1.89 | 1.96E-229 | 4.08E-225 | 0.24 | 0.07 | 0.17 | 3.46 |
| ectoderm | Slc35g1 | 2.61 | 3.91E-229 | 8.13E-225 | 0.15 | 0.03 | 0.12 | 5.43 |
| ectoderm | Sox13 | 2.17 | 2.19E-228 | 4.56E-224 | 0.2 | 0.05 | 0.15 | 3.9 |
| ectoderm | Tead1 | 0.9 | 8.09E-227 | 1.68E-222 | 0.74 | 0.56 | 0.18 | 1.32 |
| ectoderm | Prkci | 1.36 | 1.84E-225 | 3.82E-221 | 0.44 | 0.21 | 0.23 | 2.07 |
| ectoderm | Dlx5 | 1.49 | 3.14E-225 | 6.52E-221 | 0.39 | 0.17 | 0.22 | 2.34 |
| ectoderm | Gipc2 | 2.62 | 1.35E-224 | 2.81E-220 | 0.16 | 0.03 | 0.13 | 4.91 |
| ectoderm | Fhdc1 | 3.03 | 2.74E-224 | 5.70E-220 | 0.12 | 0.02 | 0.11 | 7.24 |
| ectoderm | Ak4 | 1.92 | 1.91E-223 | 3.98E-219 | 0.23 | 0.07 | 0.16 | 3.45 |
| ectoderm | Slc25a23 | 1.47 | 3.71E-222 | 7.71E-218 | 0.31 | 0.12 | 0.2 | 2.71 |
| ectoderm | Snx10 | 1.58 | 2.02E-221 | 4.20E-217 | 0.31 | 0.11 | 0.19 | 2.7 |
| ectoderm | Snn | 1.16 | 9.78E-221 | 2.04E-216 | 0.6 | 0.38 | 0.22 | 1.57 |
| ectoderm | Acot7 | 1.31 | 1.49E-220 | 3.10E-216 | 0.4 | 0.18 | 0.22 | 2.2 |
| ectoderm | Cdc42se1 | 1.03 | 2.54E-220 | 5.29E-216 | 0.6 | 0.37 | 0.23 | 1.62 |
| ectoderm | Nabp2 | 0.8 | 1.21E-218 | 2.51E-214 | 0.77 | 0.62 | 0.15 | 1.25 |
| ectoderm | Evpl | 3.1 | 3.26E-218 | 6.77E-214 | 0.12 | 0.01 | 0.1 | 8.21 |
| ectoderm | Ppp1r1a | 1.62 | 6.56E-216 | 1.36E-211 | 0.31 | 0.12 | 0.19 | 2.58 |
| ectoderm | Tafa5 | 1.72 | 2.45E-215 | 5.10E-211 | 0.25 | 0.08 | 0.17 | 3.06 |
| ectoderm | Trim2 | 1.11 | 2.41E-214 | 5.00E-210 | 0.58 | 0.36 | 0.22 | 1.6 |
| ectoderm | Iqgap1 | 0.99 | 9.82E-214 | 2.04E-209 | 0.64 | 0.42 | 0.22 | 1.52 |
| ectoderm | Crim1 | 1.49 | 2.76E-213 | 5.73E-209 | 0.35 | 0.15 | 0.2 | 2.34 |
| ectoderm | Lrrc1 | 1.58 | 6.28E-213 | 1.31E-208 | 0.29 | 0.11 | 0.18 | 2.73 |
| ectoderm | Zfp467 | 1.19 | 6.39E-213 | 1.33E-208 | 0.47 | 0.24 | 0.23 | 1.97 |
| ectoderm | Lamc1 | 1.21 | 2.13E-212 | 4.43E-208 | 0.43 | 0.21 | 0.22 | 2.08 |
| ectoderm | Pfkl | 0.79 | 2.61E-212 | 5.43E-208 | 0.77 | 0.61 | 0.17 | 1.27 |

**Supplementary Table 2. Marker genes for the major cell types of integrated clustering of E11.5 control and NotchGOF λ-junction cells.**

| | | | | | | | | |
|---|---|---|---|---|---|---|---|---|
| ectoderm | Plec | 1.32 | 6.89E-212 | 1.43E-207 | 0.44 | 0.22 | 0.22 | 1.99 |
| ectoderm | Limk2 | 1.66 | 1.11E-211 | 2.30E-207 | 0.35 | 0.15 | 0.2 | 2.31 |
| ectoderm | Tst | 1.86 | 3.21E-211 | 6.69E-207 | 0.26 | 0.09 | 0.17 | 2.95 |
| ectoderm | Srcin1 | 2.59 | 4.81E-211 | 1.00E-206 | 0.13 | 0.02 | 0.11 | 6.24 |
| ectoderm | Tox2 | 3.04 | 1.09E-210 | 2.27E-206 | 0.11 | 0.02 | 0.1 | 7.53 |
| ectoderm | Hspg2 | 1.08 | 3.15E-210 | 6.55E-206 | 0.39 | 0.18 | 0.22 | 2.23 |
| ectoderm | Gsta4 | 0.76 | 8.62E-210 | 1.79E-205 | 0.83 | 0.73 | 0.1 | 1.14 |
| ectoderm | Cobll1 | 3.6 | 8.86E-210 | 1.84E-205 | 0.1 | 0.01 | 0.09 | 9.27 |
| ectoderm | H2-T23 | 2.93 | 9.03E-210 | 1.88E-205 | 0.14 | 0.03 | 0.12 | 5.26 |
| ectoderm | Prkch | 2 | 5.55E-208 | 1.15E-203 | 0.2 | 0.05 | 0.14 | 3.72 |
| ectoderm | Satb1 | 1.05 | 7.41E-208 | 1.54E-203 | 0.56 | 0.33 | 0.23 | 1.69 |
| ectoderm | Hspa4l | 1.8 | 8.90E-208 | 1.85E-203 | 0.23 | 0.07 | 0.16 | 3.15 |
| ectoderm | Hes6 | 0.72 | 5.76E-207 | 1.20E-202 | 0.51 | 0.28 | 0.23 | 1.81 |
| ectoderm | Clu | 2.21 | 7.08E-207 | 1.47E-202 | 0.36 | 0.16 | 0.19 | 2.18 |
| ectoderm | Cobl | 1.84 | 1.06E-204 | 2.21E-200 | 0.17 | 0.04 | 0.13 | 4.36 |
| ectoderm | Tjp2 | 1.25 | 3.37E-204 | 7.01E-200 | 0.48 | 0.26 | 0.22 | 1.83 |
| ectoderm | Gltp | 1.06 | 2.62E-203 | 5.45E-199 | 0.56 | 0.34 | 0.22 | 1.63 |
| ectoderm | Slc35f1 | 3.13 | 5.65E-202 | 1.17E-197 | 0.1 | 0.01 | 0.09 | 8.58 |
| ectoderm | Sft2d2 | 1.55 | 9.96E-202 | 2.07E-197 | 0.29 | 0.11 | 0.18 | 2.65 |
| ectoderm | Mpp7 | 1.11 | 2.62E-201 | 5.44E-197 | 0.54 | 0.32 | 0.22 | 1.68 |
| ectoderm | Cebpb | 1.62 | 3.27E-201 | 6.80E-197 | 0.25 | 0.08 | 0.16 | 2.99 |
| ectoderm | Plxnb1 | 1.24 | 6.04E-201 | 1.26E-196 | 0.43 | 0.21 | 0.22 | 2.02 |
| ectoderm | Eef2 | 0.28 | 6.96E-201 | 1.45E-196 | 0.99 | 1 | -0.01 | 0.99 |
| ectoderm | Phlpp1 | 1.69 | 1.21E-200 | 2.52E-196 | 0.26 | 0.09 | 0.17 | 2.82 |
| ectoderm | Rtn3 | 0.67 | 5.16E-200 | 1.07E-195 | 0.84 | 0.77 | 0.07 | 1.09 |
| ectoderm | Tpbg | 1.13 | 2.37E-198 | 4.94E-194 | 0.48 | 0.25 | 0.23 | 1.89 |
| ectoderm | Aktip | 1.23 | 2.54E-198 | 5.29E-194 | 0.46 | 0.24 | 0.22 | 1.92 |
| ectoderm | Irf2 | 1.3 | 2.98E-198 | 6.20E-194 | 0.42 | 0.21 | 0.21 | 2.02 |
| ectoderm | Trpc1 | 1.85 | 8.56E-198 | 1.78E-193 | 0.23 | 0.08 | 0.16 | 3.07 |
| ectoderm | Pcdh17 | 1.5 | 8.90E-198 | 1.85E-193 | 0.36 | 0.16 | 0.2 | 2.25 |
| ectoderm | Gsap | 2.98 | 9.31E-198 | 1.94E-193 | 0.11 | 0.02 | 0.1 | 7.33 |
| ectoderm | Elf3 | 2.13 | 1.46E-197 | 3.04E-193 | 0.17 | 0.04 | 0.13 | 4.05 |
| ectoderm | Gm9922 | 2.82 | 1.10E-196 | 2.29E-192 | 0.12 | 0.02 | 0.1 | 6.76 |
| ectoderm | Trp53i11 | 1.02 | 1.18E-196 | 2.45E-192 | 0.39 | 0.18 | 0.21 | 2.19 |
| ectoderm | Ccdc68 | 2.63 | 1.29E-195 | 2.67E-191 | 0.14 | 0.03 | 0.11 | 5.11 |
| ectoderm | Ptpn13 | 1.18 | 2.89E-195 | 6.02E-191 | 0.57 | 0.36 | 0.21 | 1.6 |
| ectoderm | Tcf7l2 | 0.93 | 2.97E-195 | 6.19E-191 | 0.73 | 0.57 | 0.16 | 1.27 |
| ectoderm | Pura | 0.94 | 4.53E-195 | 9.42E-191 | 0.63 | 0.42 | 0.21 | 1.49 |
| ectoderm | Fam214a | 1.85 | 5.19E-195 | 1.08E-190 | 0.21 | 0.07 | 0.15 | 3.29 |
| ectoderm | Dynll2 | 0.58 | 5.58E-195 | 1.16E-190 | 0.89 | 0.83 | 0.06 | 1.07 |
| ectoderm | Rassf7 | 1.36 | 7.67E-194 | 1.59E-189 | 0.35 | 0.15 | 0.2 | 2.28 |
| ectoderm | Pdgfrl | 1.47 | 2.51E-193 | 5.23E-189 | 0.33 | 0.14 | 0.19 | 2.36 |
| ectoderm | Dgka | 2.16 | 5.36E-192 | 1.12E-187 | 0.15 | 0.03 | 0.12 | 4.58 |
| ectoderm | Zfhx3 | 0.38 | 5.42E-192 | 1.13E-187 | 0.91 | 0.78 | 0.13 | 1.17 |
| ectoderm | Erich1 | 1.38 | 8.95E-192 | 1.86E-187 | 0.39 | 0.19 | 0.2 | 2.06 |
| ectoderm | Lgalsl | 1.49 | 1.12E-189 | 2.33E-185 | 0.47 | 0.27 | 0.21 | 1.78 |
| ectoderm | Jade1 | 1.48 | 1.37E-189 | 2.85E-185 | 0.31 | 0.13 | 0.18 | 2.4 |
| ectoderm | Kdm5a | 0.77 | 5.04E-189 | 1.05E-184 | 0.78 | 0.64 | 0.14 | 1.22 |
| ectoderm | Creld1 | 1.53 | 1.06E-188 | 2.20E-184 | 0.26 | 0.09 | 0.17 | 2.81 |
| ectoderm | Tctex1d1 | 3.76 | 1.34E-188 | 2.79E-184 | 0.1 | 0.01 | 0.09 | 7.43 |
| ectoderm | Soga3 | 1.5 | 2.24E-188 | 4.65E-184 | 0.14 | 0.03 | 0.11 | 5.07 |
| ectoderm | Dazap2 | 0.8 | 2.97E-188 | 6.17E-184 | 0.71 | 0.53 | 0.18 | 1.34 |
| ectoderm | Mfsd6 | 2.84 | 8.23E-188 | 1.71E-183 | 0.1 | 0.01 | 0.09 | 7.36 |
| ectoderm | Tfap2a | 1.74 | 2.86E-187 | 5.95E-183 | 0.36 | 0.17 | 0.19 | 2.09 |
| ectoderm | Mettl27 | 1.51 | 4.24E-187 | 8.83E-183 | 0.27 | 0.1 | 0.17 | 2.68 |
| ectoderm | Cryl1 | 1.55 | 1.08E-186 | 2.25E-182 | 0.28 | 0.11 | 0.17 | 2.58 |

**Supplementary Table 2. Marker genes for the major cell types of integrated clustering of E11.5 control and NotchGOF λ-junction cells.**

| | | | | | | | | |
|---|---|---|---|---|---|---|---|---|
| ectoderm | Mgat4a | 1.82 | 1.86E-186 | 3.86E-182 | 0.2 | 0.06 | 0.14 | 3.35 |
| ectoderm | Pnck | 1.95 | 6.68E-186 | 1.39E-181 | 0.17 | 0.04 | 0.13 | 4 |
| ectoderm | Errfi1 | 1.52 | 1.44E-184 | 3.00E-180 | 0.32 | 0.14 | 0.18 | 2.31 |
| ectoderm | .810037I17Ril | 0.58 | 1.53E-184 | 3.19E-180 | 0.89 | 0.82 | 0.07 | 1.08 |
| ectoderm | Nedd9 | 1.54 | 2.32E-184 | 4.82E-180 | 0.31 | 0.13 | 0.18 | 2.4 |
| ectoderm | Clip4 | 2.4 | 2.82E-184 | 5.86E-180 | 0.12 | 0.02 | 0.1 | 5.55 |
| ectoderm | Ifitm3 | 1.81 | 3.13E-184 | 6.52E-180 | 0.44 | 0.25 | 0.19 | 1.73 |
| ectoderm | Rbp1 | 0.89 | 6.95E-184 | 1.45E-179 | 0.94 | 0.9 | 0.04 | 1.04 |
| ectoderm | Sostdc1 | 2.45 | 7.12E-183 | 1.48E-178 | 0.27 | 0.11 | 0.16 | 2.5 |
| ectoderm | Zmat4 | 2.14 | 2.18E-181 | 4.53E-177 | 0.14 | 0.03 | 0.11 | 4.63 |
| ectoderm | Arap2 | 2.4 | 6.84E-181 | 1.42E-176 | 0.14 | 0.03 | 0.11 | 4.76 |
| ectoderm | Rab3ip | 1.1 | 1.01E-180 | 2.11E-176 | 0.47 | 0.26 | 0.21 | 1.82 |
| ectoderm | Smim10l2a | 2.02 | 2.30E-180 | 4.79E-176 | 0.17 | 0.04 | 0.12 | 3.93 |
| ectoderm | Firre | 1.88 | 1.00E-179 | 2.08E-175 | 0.19 | 0.06 | 0.14 | 3.39 |
| ectoderm | Spock2 | 2.99 | 1.38E-179 | 2.86E-175 | 0.11 | 0.02 | 0.09 | 6.47 |
| ectoderm | Lrrfip1 | 1.06 | 2.05E-179 | 4.27E-175 | 0.5 | 0.29 | 0.21 | 1.72 |
| ectoderm | Abca3 | 2.03 | 3.49E-179 | 7.26E-175 | 0.15 | 0.04 | 0.12 | 4.16 |
| ectoderm | Rbms1 | 0.64 | 4.29E-179 | 8.92E-175 | 0.84 | 0.77 | 0.08 | 1.1 |
| ectoderm | Baiap2l1 | 2.09 | 7.36E-179 | 1.53E-174 | 0.15 | 0.04 | 0.12 | 4.31 |
| ectoderm | Ccdc171 | 1.86 | 1.35E-178 | 2.80E-174 | 0.2 | 0.06 | 0.14 | 3.21 |
| ectoderm | 700033N17Ri | 1.44 | 9.15E-178 | 1.90E-173 | 0.24 | 0.08 | 0.16 | 2.9 |
| ectoderm | Bcl11b | 0.89 | 1.14E-177 | 2.38E-173 | 0.71 | 0.55 | 0.16 | 1.28 |
| ectoderm | Ajuba | 1.29 | 2.87E-177 | 5.97E-173 | 0.43 | 0.23 | 0.2 | 1.85 |
| ectoderm | Prkcz | 1.58 | 7.35E-177 | 1.53E-172 | 0.22 | 0.07 | 0.15 | 3.03 |
| ectoderm | Jpt1 | 0.47 | 8.70E-177 | 1.81E-172 | 0.96 | 0.94 | 0.02 | 1.02 |
| ectoderm | Bnip3l | 0.43 | 1.52E-176 | 3.16E-172 | 0.94 | 0.93 | 0.02 | 1.02 |
| ectoderm | Ralgps1 | 2.51 | 5.18E-176 | 1.08E-171 | 0.12 | 0.02 | 0.1 | 5.62 |
| ectoderm | Alad | 0.33 | 1.31E-174 | 2.73E-170 | 0.76 | 0.6 | 0.16 | 1.27 |
| ectoderm | Kank1 | 1.41 | 1.45E-174 | 3.03E-170 | 0.29 | 0.12 | 0.17 | 2.46 |
| ectoderm | Syngr2 | 1.41 | 9.69E-174 | 2.02E-169 | 0.27 | 0.1 | 0.17 | 2.59 |
| ectoderm | Aftph | 1.24 | 2.60E-173 | 5.40E-169 | 0.4 | 0.21 | 0.2 | 1.95 |
| ectoderm | Churc1 | 0.88 | 5.02E-173 | 1.04E-168 | 0.72 | 0.55 | 0.17 | 1.31 |
| ectoderm | Dlx6 | 1.83 | 1.20E-172 | 2.50E-168 | 0.19 | 0.06 | 0.13 | 3.36 |
| ectoderm | Col2a1 | 0.51 | 2.52E-172 | 5.23E-168 | 0.79 | 0.6 | 0.19 | 1.31 |
| ectoderm | Wls | 0.91 | 2.78E-172 | 5.78E-168 | 0.79 | 0.72 | 0.07 | 1.1 |
| ectoderm | Pard3 | 0.76 | 4.92E-172 | 1.02E-167 | 0.71 | 0.54 | 0.17 | 1.32 |
| ectoderm | Serinc2 | 0.89 | 3.47E-171 | 7.21E-167 | 0.56 | 0.34 | 0.22 | 1.66 |
| ectoderm | Nrip1 | 0.78 | 1.85E-170 | 3.85E-166 | 0.81 | 0.74 | 0.07 | 1.1 |
| ectoderm | Sema6a | 1.03 | 2.68E-170 | 5.56E-166 | 0.6 | 0.4 | 0.2 | 1.51 |
| ectoderm | Rbm47 | 2.66 | 4.04E-170 | 8.41E-166 | 0.1 | 0.02 | 0.09 | 6.67 |
| ectoderm | Grip1 | 1.6 | 8.69E-170 | 1.81E-165 | 0.22 | 0.07 | 0.14 | 2.99 |
| ectoderm | Med19 | 0.63 | 1.70E-169 | 3.54E-165 | 0.82 | 0.72 | 0.1 | 1.14 |
| ectoderm | Pcbd1 | 1.54 | 2.66E-169 | 5.53E-165 | 0.24 | 0.09 | 0.15 | 2.75 |
| ectoderm | Pik3cb | 2.18 | 3.89E-169 | 8.10E-165 | 0.15 | 0.04 | 0.11 | 4 |
| ectoderm | Frat2 | 1.31 | 8.86E-169 | 1.84E-164 | 0.38 | 0.19 | 0.19 | 1.98 |
| ectoderm | Tpm1 | 0.75 | 3.75E-168 | 7.80E-164 | 0.93 | 0.9 | 0.03 | 1.03 |
| ectoderm | Gadd45g | 2.94 | 8.43E-168 | 1.75E-163 | 0.34 | 0.18 | 0.16 | 1.91 |
| ectoderm | Hspa2 | 2.43 | 1.06E-166 | 2.20E-162 | 0.14 | 0.03 | 0.1 | 4.25 |
| ectoderm | Adi1 | 1.23 | 1.25E-166 | 2.61E-162 | 0.46 | 0.27 | 0.19 | 1.69 |
| ectoderm | Mpst | 0.93 | 3.13E-165 | 6.50E-161 | 0.57 | 0.37 | 0.2 | 1.53 |
| ectoderm | Nudt18 | 2.03 | 3.39E-165 | 7.06E-161 | 0.15 | 0.04 | 0.11 | 4.03 |
| ectoderm | Ttc3 | 0.38 | 1.15E-164 | 2.40E-160 | 0.98 | 0.95 | 0.02 | 1.03 |
| ectoderm | Ptpn3 | 2.53 | 1.22E-164 | 2.53E-160 | 0.11 | 0.02 | 0.09 | 5.89 |
| ectoderm | Scrn1 | 2.21 | 1.85E-164 | 3.86E-160 | 0.12 | 0.02 | 0.09 | 5.23 |
| ectoderm | Cltb | 1.39 | 2.62E-164 | 5.45E-160 | 0.31 | 0.14 | 0.17 | 2.2 |
| ectoderm | Adgrg1 | 2.14 | 6.21E-164 | 1.29E-159 | 0.11 | 0.02 | 0.09 | 5.33 |

**Supplementary Table 2. Marker genes for the major cell types of integrated clustering of E11.5 control and NotchGOF λ-junction cells.**

| | | | | | | | | |
|---|---|---|---|---|---|---|---|---|
| ectoderm | Klhl23 | 1.01 | 7.50E-164 | 1.56E-159 | 0.52 | 0.33 | 0.2 | 1.61 |
| ectoderm | Skp1a | 0.4 | 6.64E-163 | 1.38E-158 | 0.97 | 0.95 | 0.02 | 1.02 |
| ectoderm | Gtf3a | 0.68 | 1.87E-162 | 3.89E-158 | 0.75 | 0.61 | 0.14 | 1.23 |
| ectoderm | Pnpla2 | 1.76 | 1.91E-162 | 3.98E-158 | 0.19 | 0.06 | 0.13 | 3.17 |
| ectoderm | Kxd1 | 0.83 | 3.14E-162 | 6.53E-158 | 0.65 | 0.46 | 0.19 | 1.41 |
| ectoderm | Chd7 | 0.64 | 3.60E-162 | 7.48E-158 | 0.76 | 0.62 | 0.14 | 1.23 |
| ectoderm | Calm2 | 0.37 | 8.82E-162 | 1.83E-157 | 0.98 | 0.98 | 0.01 | 1.01 |
| ectoderm | Rbbp8 | 0.89 | 5.03E-161 | 1.05E-156 | 0.68 | 0.52 | 0.16 | 1.3 |
| ectoderm | Meg3 | 0.96 | 2.64E-160 | 5.49E-156 | 0.75 | 0.68 | 0.07 | 1.1 |
| ectoderm | Cstb | 0.79 | 6.30E-159 | 1.31E-154 | 0.85 | 0.79 | 0.06 | 1.08 |
| ectoderm | Usp53 | 1.78 | 7.89E-159 | 1.64E-154 | 0.17 | 0.05 | 0.12 | 3.5 |
| ectoderm | Fry | 2.21 | 1.01E-158 | 2.11E-154 | 0.13 | 0.03 | 0.1 | 4.16 |
| ectoderm | Clstn1 | 0.92 | 1.92E-158 | 3.99E-154 | 0.54 | 0.34 | 0.2 | 1.59 |
| ectoderm | Zdhhc21 | 1.01 | 5.44E-158 | 1.13E-153 | 0.44 | 0.24 | 0.2 | 1.8 |
| ectoderm | Ptprz1 | 1.91 | 8.92E-158 | 1.86E-153 | 0.21 | 0.08 | 0.14 | 2.8 |
| ectoderm | Pik3ip1 | 1.73 | 5.41E-157 | 1.13E-152 | 0.18 | 0.06 | 0.13 | 3.17 |
| ectoderm | B3gnt7 | 2.39 | 7.37E-157 | 1.53E-152 | 0.11 | 0.02 | 0.09 | 5.09 |
| ectoderm | Plekhf2 | 1.2 | 1.16E-156 | 2.42E-152 | 0.37 | 0.19 | 0.18 | 1.97 |
| ectoderm | Zfp704 | 0.83 | 1.31E-156 | 2.72E-152 | 0.62 | 0.42 | 0.2 | 1.46 |
| ectoderm | Pitx2 | 2.12 | 6.24E-156 | 1.30E-151 | 0.26 | 0.11 | 0.15 | 2.34 |
| ectoderm | Tjp1 | 0.76 | 3.09E-155 | 6.43E-151 | 0.64 | 0.45 | 0.19 | 1.41 |
| ectoderm | Tmem159 | 1.14 | 8.89E-155 | 1.85E-150 | 0.41 | 0.22 | 0.19 | 1.88 |
| ectoderm | Tmem205 | 1.07 | 1.28E-154 | 2.66E-150 | 0.43 | 0.23 | 0.19 | 1.83 |
| ectoderm | Trappc6a | 0.85 | 4.94E-154 | 1.03E-149 | 0.66 | 0.48 | 0.18 | 1.36 |
| ectoderm | Rb1 | 1.74 | 1.68E-153 | 3.50E-149 | 0.28 | 0.12 | 0.15 | 2.25 |
| ectoderm | Tmem30b | 1.43 | 1.77E-152 | 3.68E-148 | 0.27 | 0.11 | 0.16 | 2.37 |
| ectoderm | Chst11 | 2.1 | 8.82E-152 | 1.83E-147 | 0.16 | 0.05 | 0.11 | 3.4 |
| ectoderm | Cecr2 | 1.07 | 1.06E-151 | 2.20E-147 | 0.36 | 0.18 | 0.18 | 2.01 |
| ectoderm | Gstt3 | 2.19 | 5.38E-151 | 1.12E-146 | 0.12 | 0.03 | 0.1 | 4.52 |
| ectoderm | Syt7 | 2.04 | 1.28E-150 | 2.66E-146 | 0.14 | 0.03 | 0.1 | 4.09 |
| ectoderm | Rhbdf1 | 1.6 | 1.79E-150 | 3.72E-146 | 0.2 | 0.07 | 0.13 | 2.87 |
| ectoderm | Ckmt1 | 0.93 | 2.57E-149 | 5.34E-145 | 0.11 | 0.02 | 0.09 | 5.05 |
| ectoderm | Ldlrap1 | 1.33 | 3.87E-149 | 8.05E-145 | 0.31 | 0.15 | 0.16 | 2.13 |
| ectoderm | Krt1 | 2.18 | 4.56E-149 | 9.49E-145 | 0.12 | 0.03 | 0.1 | 4.28 |
| ectoderm | Golga4 | 0.67 | 2.31E-148 | 4.80E-144 | 0.74 | 0.61 | 0.13 | 1.21 |
| ectoderm | Gsr | 1.01 | 3.32E-148 | 6.91E-144 | 0.48 | 0.29 | 0.19 | 1.65 |
| ectoderm | Ostf1 | 0.85 | 3.37E-148 | 7.01E-144 | 0.42 | 0.23 | 0.19 | 1.82 |
| ectoderm | Stox2 | 0.74 | 2.31E-147 | 4.80E-143 | 0.69 | 0.52 | 0.16 | 1.31 |
| ectoderm | Plekhh1 | 1.8 | 3.30E-147 | 6.86E-143 | 0.15 | 0.04 | 0.11 | 3.7 |
| ectoderm | Vps36 | 0.55 | 3.95E-147 | 8.23E-143 | 0.85 | 0.78 | 0.07 | 1.09 |
| ectoderm | Dnmbp | 1.36 | 6.05E-147 | 1.26E-142 | 0.24 | 0.09 | 0.14 | 2.53 |
| ectoderm | Cdc42ep5 | 0.89 | 1.65E-146 | 3.43E-142 | 0.6 | 0.43 | 0.18 | 1.41 |
| ectoderm | Mettl26 | 0.81 | 2.27E-146 | 4.72E-142 | 0.63 | 0.45 | 0.17 | 1.38 |
| ectoderm | Gcat | 0.83 | 5.09E-146 | 1.06E-141 | 0.62 | 0.44 | 0.18 | 1.41 |
| ectoderm | Fam89a | 1.99 | 3.49E-145 | 7.25E-141 | 0.16 | 0.05 | 0.11 | 3.29 |
| ectoderm | Zbtb18 | 1.45 | 5.46E-144 | 1.14E-139 | 0.21 | 0.08 | 0.13 | 2.7 |
| ectoderm | Jdp2 | 2.89 | 7.62E-144 | 1.58E-139 | 0.1 | 0.02 | 0.08 | 5.26 |
| ectoderm | Wdsub1 | 1.07 | 9.44E-144 | 1.96E-139 | 0.34 | 0.17 | 0.17 | 2.04 |
| ectoderm | Rhob | 1.18 | 1.56E-143 | 3.25E-139 | 0.48 | 0.31 | 0.18 | 1.58 |
| ectoderm | Tshz1 | 0.92 | 6.46E-143 | 1.34E-138 | 0.46 | 0.27 | 0.19 | 1.7 |
| ectoderm | Chd3 | 0.49 | 9.10E-143 | 1.89E-138 | 0.94 | 0.88 | 0.05 | 1.06 |
| ectoderm | H2-K1 | 1.28 | 2.22E-142 | 4.61E-138 | 0.25 | 0.1 | 0.15 | 2.45 |
| ectoderm | Tmem184a | 2.21 | 1.05E-141 | 2.18E-137 | 0.13 | 0.03 | 0.09 | 4.03 |
| ectoderm | Tiam1 | 1.04 | 1.38E-141 | 2.88E-137 | 0.45 | 0.27 | 0.18 | 1.68 |
| ectoderm | Ctdspl | 0.95 | 2.03E-141 | 4.23E-137 | 0.49 | 0.3 | 0.19 | 1.63 |
| ectoderm | Patz1 | 0.74 | 1.47E-140 | 3.05E-136 | 0.65 | 0.48 | 0.17 | 1.35 |

**Supplementary Table 2. Marker genes for the major cell types of integrated clustering of E11.5 control and NotchGOF λ-junction cells.**

| | | | | | | | | |
|---|---|---|---|---|---|---|---|---|
| ectoderm | Kazn | 1.16 | 5.54E-140 | 1.15E-135 | 0.31 | 0.15 | 0.16 | 2.12 |
| ectoderm | Klhl13 | 1.05 | 8.22E-140 | 1.71E-135 | 0.41 | 0.23 | 0.18 | 1.79 |
| ectoderm | Rasal2 | 1.1 | 1.73E-138 | 3.60E-134 | 0.38 | 0.21 | 0.18 | 1.86 |
| ectoderm | Irs1 | 1.26 | 3.12E-138 | 6.50E-134 | 0.3 | 0.14 | 0.16 | 2.13 |
| ectoderm | Pts | 0.83 | 3.42E-138 | 7.11E-134 | 0.55 | 0.36 | 0.19 | 1.52 |
| ectoderm | Rreb1 | 1.11 | 4.65E-138 | 9.67E-134 | 0.34 | 0.17 | 0.17 | 2 |
| ectoderm | Fhl1 | 1.4 | 9.24E-138 | 1.92E-133 | 0.25 | 0.11 | 0.14 | 2.32 |
| ectoderm | Rian | 1.06 | 9.52E-138 | 1.98E-133 | 0.45 | 0.28 | 0.18 | 1.64 |
| ectoderm | Eya2 | 1.25 | 1.49E-137 | 3.10E-133 | 0.36 | 0.19 | 0.17 | 1.87 |
| ectoderm | Pard3b | 1.35 | 2.64E-136 | 5.48E-132 | 0.22 | 0.09 | 0.14 | 2.53 |
| ectoderm | Grb14 | 1.99 | 3.10E-136 | 6.45E-132 | 0.11 | 0.03 | 0.09 | 4.38 |
| ectoderm | Met | 2.32 | 4.09E-135 | 8.51E-131 | 0.11 | 0.02 | 0.08 | 4.61 |
| ectoderm | Npepps | 0.61 | 3.75E-134 | 7.80E-130 | 0.74 | 0.62 | 0.12 | 1.2 |
| ectoderm | Fam173a | 0.76 | 4.64E-134 | 9.65E-130 | 0.63 | 0.47 | 0.16 | 1.35 |
| ectoderm | Adamts9 | 1.93 | 6.80E-134 | 1.41E-129 | 0.26 | 0.12 | 0.14 | 2.21 |
| ectoderm | Thbs1 | 0.66 | 1.42E-133 | 2.95E-129 | 0.22 | 0.09 | 0.14 | 2.55 |
| ectoderm | Tpp1 | 0.93 | 4.20E-133 | 8.74E-129 | 0.4 | 0.22 | 0.18 | 1.79 |
| ectoderm | Mtss1 | 0.98 | 4.90E-133 | 1.02E-128 | 0.44 | 0.25 | 0.18 | 1.73 |
| ectoderm | Nt5c | 0.76 | 5.77E-133 | 1.20E-128 | 0.66 | 0.5 | 0.17 | 1.34 |
| ectoderm | Nab1 | 1.38 | 1.26E-132 | 2.63E-128 | 0.23 | 0.1 | 0.14 | 2.39 |
| ectoderm | Notch3 | 0.91 | 2.38E-132 | 4.95E-128 | 0.54 | 0.37 | 0.17 | 1.47 |
| ectoderm | Ahr | 1.48 | 3.10E-132 | 6.45E-128 | 0.18 | 0.06 | 0.12 | 2.92 |
| ectoderm | Dtx3l | 2.17 | 6.70E-132 | 1.39E-127 | 0.1 | 0.02 | 0.08 | 4.68 |
| ectoderm | Agpat3 | 0.97 | 6.89E-132 | 1.43E-127 | 0.4 | 0.22 | 0.18 | 1.79 |
| ectoderm | Mbnl2 | 0.73 | 8.30E-132 | 1.73E-127 | 0.77 | 0.65 | 0.12 | 1.18 |
| ectoderm | Exosc7 | 0.65 | 8.98E-132 | 1.87E-127 | 0.72 | 0.59 | 0.13 | 1.21 |
| ectoderm | Kdm5b | 0.53 | 9.57E-132 | 1.99E-127 | 0.9 | 0.85 | 0.05 | 1.05 |
| ectoderm | Ptpn4 | 1.24 | 3.49E-131 | 7.25E-127 | 0.27 | 0.12 | 0.15 | 2.19 |
| ectoderm | Rimklb | 0.92 | 1.00E-130 | 2.08E-126 | 0.47 | 0.28 | 0.19 | 1.67 |
| ectoderm | Strbp | 0.55 | 1.28E-129 | 2.66E-125 | 0.79 | 0.68 | 0.11 | 1.15 |
| ectoderm | Xpr1 | 1.13 | 1.29E-129 | 2.68E-125 | 0.4 | 0.23 | 0.17 | 1.73 |
| ectoderm | Thrb | 1.78 | 1.93E-129 | 4.02E-125 | 0.15 | 0.05 | 0.1 | 3.17 |
| ectoderm | Nr3c1 | 1.26 | 2.29E-129 | 4.77E-125 | 0.28 | 0.13 | 0.15 | 2.16 |
| ectoderm | Mcee | 0.9 | 2.38E-129 | 4.95E-125 | 0.57 | 0.4 | 0.17 | 1.43 |
| ectoderm | Cbarp | 1.38 | 2.46E-129 | 5.12E-125 | 0.21 | 0.08 | 0.13 | 2.61 |
| ectoderm | Ephb2 | 0.94 | 2.57E-129 | 5.35E-125 | 0.42 | 0.24 | 0.18 | 1.74 |
| ectoderm | Nt5dc2 | 0.45 | 2.72E-129 | 5.65E-125 | 0.92 | 0.92 | 0 | 1 |
| ectoderm | Txndc17 | 0.54 | 1.27E-128 | 2.64E-124 | 0.93 | 0.92 | 0.02 | 1.02 |
| ectoderm | Pwwp2b | 1.77 | 4.23E-127 | 8.80E-123 | 0.15 | 0.05 | 0.1 | 3.12 |
| ectoderm | Wnk2 | 2.05 | 5.29E-127 | 1.10E-122 | 0.11 | 0.03 | 0.09 | 4.07 |
| ectoderm | Epb41l4a | 1.34 | 8.14E-126 | 1.69E-121 | 0.2 | 0.08 | 0.12 | 2.58 |
| ectoderm | Dap | 0.68 | 1.59E-125 | 3.30E-121 | 0.69 | 0.53 | 0.15 | 1.29 |
| ectoderm | Scd2 | 0.45 | 1.67E-125 | 3.47E-121 | 0.93 | 0.91 | 0.02 | 1.02 |
| ectoderm | Gprc5c | 0.98 | 2.98E-125 | 6.20E-121 | 0.37 | 0.2 | 0.17 | 1.87 |
| ectoderm | Tnfrsf19 | 1.76 | 2.99E-125 | 6.22E-121 | 0.25 | 0.12 | 0.14 | 2.13 |
| ectoderm | Echdc1 | 1.08 | 9.88E-125 | 2.05E-120 | 0.36 | 0.2 | 0.16 | 1.82 |
| ectoderm | Ubald1 | 1.08 | 1.14E-124 | 2.37E-120 | 0.39 | 0.23 | 0.17 | 1.74 |
| ectoderm | Macrod2 | 1.09 | 1.52E-124 | 3.16E-120 | 0.35 | 0.19 | 0.16 | 1.88 |
| ectoderm | Lamb1 | 0.86 | 2.47E-124 | 5.13E-120 | 0.52 | 0.35 | 0.17 | 1.49 |
| ectoderm | Rtn2 | 1.58 | 2.52E-123 | 5.25E-119 | 0.17 | 0.06 | 0.11 | 2.85 |
| ectoderm | Ociad2 | 1.12 | 2.61E-123 | 5.43E-119 | 0.3 | 0.15 | 0.15 | 2.03 |
| ectoderm | Lrrc8b | 1.4 | 7.74E-123 | 1.61E-118 | 0.21 | 0.08 | 0.12 | 2.49 |
| ectoderm | Col11a1 | 0.88 | 8.06E-123 | 1.68E-118 | 0.56 | 0.39 | 0.17 | 1.43 |
| ectoderm | Mxi1 | 0.83 | 1.71E-122 | 3.55E-118 | 0.56 | 0.39 | 0.17 | 1.44 |
| ectoderm | Scp2 | 0.51 | 4.52E-122 | 9.40E-118 | 0.83 | 0.76 | 0.07 | 1.1 |
| ectoderm | Tead3 | 1 | 1.03E-121 | 2.14E-117 | 0.38 | 0.21 | 0.17 | 1.81 |

**Supplementary Table 2. Marker genes for the major cell types of integrated clustering of E11.5 control and NotchGOF λ-junction cells.**

| | | | | | | | | |
|---|---|---|---|---|---|---|---|---|
| ectoderm | Znrf3 | 1.37 | 1.10E-121 | 2.30E-117 | 0.29 | 0.15 | 0.14 | 1.98 |
| ectoderm | Rabgap1 | 0.67 | 1.23E-121 | 2.55E-117 | 0.68 | 0.53 | 0.15 | 1.28 |
| ectoderm | Bdh1 | 1.79 | 1.62E-121 | 3.36E-117 | 0.15 | 0.05 | 0.1 | 3.08 |
| ectoderm | Utrn | 0.74 | 3.15E-121 | 6.55E-117 | 0.71 | 0.59 | 0.12 | 1.2 |
| ectoderm | Pick1 | 0.95 | 6.03E-121 | 1.25E-116 | 0.39 | 0.22 | 0.17 | 1.8 |
| ectoderm | Gnao1 | 1.41 | 2.25E-120 | 4.67E-116 | 0.17 | 0.06 | 0.11 | 2.86 |
| ectoderm | Zdhhc12 | 0.95 | 2.33E-120 | 4.84E-116 | 0.39 | 0.22 | 0.17 | 1.77 |
| ectoderm | Zdbf2 | 0.77 | 2.65E-120 | 5.52E-116 | 0.55 | 0.37 | 0.18 | 1.49 |
| ectoderm | Imp3 | 0.5 | 6.78E-120 | 1.41E-115 | 0.85 | 0.77 | 0.07 | 1.09 |
| ectoderm | Rgs9 | 1.6 | 1.08E-119 | 2.24E-115 | 0.17 | 0.06 | 0.11 | 2.8 |
| ectoderm | Abhd14b | 1.86 | 9.58E-119 | 1.99E-114 | 0.13 | 0.04 | 0.09 | 3.42 |
| ectoderm | Lonrf1 | 1.49 | 9.60E-119 | 2.00E-114 | 0.18 | 0.07 | 0.11 | 2.66 |
| ectoderm | Malat1 | 0.44 | 1.19E-118 | 2.47E-114 | 0.99 | 1 | -0.01 | 0.99 |
| ectoderm | Foxo1 | 1.21 | 1.19E-118 | 2.48E-114 | 0.29 | 0.15 | 0.15 | 1.98 |
| ectoderm | Mtss2 | 1 | 5.40E-118 | 1.12E-113 | 0.4 | 0.23 | 0.17 | 1.71 |
| ectoderm | Sord | 1.54 | 7.98E-118 | 1.66E-113 | 0.25 | 0.12 | 0.13 | 2.14 |
| ectoderm | Hk2 | 1.04 | 1.49E-117 | 3.10E-113 | 0.38 | 0.22 | 0.17 | 1.76 |
| ectoderm | Spats2l | 1.15 | 1.62E-117 | 3.37E-113 | 0.35 | 0.2 | 0.16 | 1.79 |
| ectoderm | AC149090.1 | 0.93 | 5.07E-117 | 1.05E-112 | 0.41 | 0.24 | 0.17 | 1.7 |
| ectoderm | Nipsnap1 | 0.61 | 7.66E-117 | 1.59E-112 | 0.68 | 0.52 | 0.16 | 1.31 |
| ectoderm | Cadm4 | 1.47 | 1.32E-116 | 2.74E-112 | 0.17 | 0.06 | 0.11 | 2.79 |
| ectoderm | Ddx58 | 1.93 | 3.85E-116 | 8.02E-112 | 0.12 | 0.03 | 0.09 | 3.56 |
| ectoderm | Ccdc112 | 0.69 | 4.25E-116 | 8.84E-112 | 0.64 | 0.48 | 0.17 | 1.35 |
| ectoderm | Tmem123 | 0.94 | 4.76E-116 | 9.89E-112 | 0.45 | 0.28 | 0.17 | 1.6 |
| ectoderm | Dusp5 | 1.44 | 6.36E-116 | 1.32E-111 | 0.21 | 0.09 | 0.12 | 2.39 |
| ectoderm | Tgfb2 | 1.41 | 3.75E-115 | 7.81E-111 | 0.58 | 0.46 | 0.12 | 1.27 |
| ectoderm | Baz2b | 0.54 | 3.93E-115 | 8.17E-111 | 0.79 | 0.68 | 0.1 | 1.15 |
| ectoderm | Tedc1 | 1.23 | 7.81E-115 | 1.63E-110 | 0.23 | 0.1 | 0.13 | 2.28 |
| ectoderm | Lrrc8d | 1.04 | 8.69E-115 | 1.81E-110 | 0.31 | 0.16 | 0.15 | 1.94 |
| ectoderm | Bphl | 0.82 | 1.05E-114 | 2.19E-110 | 0.51 | 0.34 | 0.17 | 1.51 |
| ectoderm | Ust | 1.7 | 1.12E-114 | 2.33E-110 | 0.13 | 0.04 | 0.09 | 3.3 |
| ectoderm | Arf1 | 0.29 | 1.27E-114 | 2.64E-110 | 0.97 | 0.98 | 0 | 1 |
| ectoderm | Slc48a1 | 1.14 | 1.35E-114 | 2.81E-110 | 0.27 | 0.14 | 0.14 | 2.03 |
| ectoderm | Pbx4 | 1.4 | 2.00E-114 | 4.15E-110 | 0.17 | 0.06 | 0.11 | 2.76 |
| ectoderm | Uqcr11 | 0.27 | 3.32E-114 | 6.90E-110 | 0.97 | 0.98 | -0.01 | 0.99 |
| ectoderm | Zmiz2 | 0.86 | 8.24E-114 | 1.71E-109 | 0.44 | 0.27 | 0.17 | 1.62 |
| ectoderm | Gstm1 | 1.54 | 7.05E-112 | 1.47E-107 | 0.36 | 0.21 | 0.15 | 1.7 |
| ectoderm | Dcaf4 | 1.08 | 8.03E-112 | 1.67E-107 | 0.36 | 0.21 | 0.15 | 1.75 |
| ectoderm | Baz2a | 0.82 | 9.26E-112 | 1.93E-107 | 0.51 | 0.35 | 0.17 | 1.47 |
| ectoderm | Hltf | 0.9 | 2.53E-111 | 5.27E-107 | 0.44 | 0.28 | 0.16 | 1.58 |
| ectoderm | Vgll4 | 0.5 | 3.52E-111 | 7.33E-107 | 0.84 | 0.76 | 0.08 | 1.1 |
| ectoderm | Mark2 | 0.7 | 4.83E-111 | 1.00E-106 | 0.59 | 0.43 | 0.16 | 1.38 |
| ectoderm | Podxl | 1.69 | 5.59E-111 | 1.16E-106 | 0.1 | 0.03 | 0.08 | 3.92 |
| ectoderm | Ptprk | 0.71 | 1.12E-110 | 2.33E-106 | 0.58 | 0.4 | 0.18 | 1.44 |
| ectoderm | Psap | 0.54 | 1.49E-110 | 3.10E-106 | 0.71 | 0.59 | 0.13 | 1.21 |
| ectoderm | Abhd11 | 0.91 | 4.02E-110 | 8.36E-106 | 0.39 | 0.23 | 0.16 | 1.69 |
| ectoderm | Psip1 | 0.54 | 5.62E-110 | 1.17E-105 | 0.98 | 0.98 | -0.01 | 0.99 |
| ectoderm | Pak2 | 0.45 | 3.93E-109 | 8.17E-105 | 0.88 | 0.82 | 0.06 | 1.07 |
| ectoderm | Phlda3 | 1.06 | 4.42E-109 | 9.20E-105 | 0.52 | 0.38 | 0.14 | 1.38 |
| ectoderm | Vill | 1.39 | 6.19E-109 | 1.29E-104 | 0.16 | 0.06 | 0.1 | 2.78 |
| ectoderm | Cndp2 | 0.84 | 1.44E-108 | 3.00E-104 | 0.49 | 0.34 | 0.16 | 1.47 |
| ectoderm | 010320M18R | 0.86 | 1.78E-108 | 3.71E-104 | 0.47 | 0.31 | 0.17 | 1.54 |
| ectoderm | Cuedc1 | 0.78 | 2.05E-108 | 4.27E-104 | 0.51 | 0.35 | 0.17 | 1.48 |
| ectoderm | Dst | 1.15 | 3.51E-108 | 7.31E-104 | 0.47 | 0.32 | 0.15 | 1.47 |
| ectoderm | Arl6ip5 | 0.78 | 3.80E-108 | 7.90E-104 | 0.45 | 0.28 | 0.17 | 1.61 |
| ectoderm | Nell2 | 1.81 | 4.12E-108 | 8.58E-104 | 0.11 | 0.03 | 0.08 | 3.61 |

**Supplementary Table 2. Marker genes for the major cell types of integrated clustering of E11.5 control and NotchGOF λ-junction cells.**

| | | | | | | | | |
|---|---|---|---|---|---|---|---|---|
| ectoderm | Junb | 1.2 | 4.18E-108 | 8.69E-104 | 0.21 | 0.09 | 0.12 | 2.31 |
| ectoderm | mt-Nd6 | 0.87 | 1.69E-107 | 3.52E-103 | 0.48 | 0.32 | 0.16 | 1.52 |
| ectoderm | Cemip2 | 0.7 | 1.82E-107 | 3.80E-103 | 0.57 | 0.41 | 0.16 | 1.38 |
| ectoderm | Ccdc85c | 1.29 | 1.95E-107 | 4.05E-103 | 0.2 | 0.09 | 0.12 | 2.34 |
| ectoderm | Klf3 | 0.74 | 2.96E-107 | 6.15E-103 | 0.61 | 0.47 | 0.15 | 1.31 |
| ectoderm | Ybx3 | 0.42 | 3.56E-107 | 7.41E-103 | 0.87 | 0.83 | 0.04 | 1.05 |
| ectoderm | Pon2 | 1 | 1.50E-106 | 3.13E-102 | 0.29 | 0.15 | 0.14 | 1.95 |
| ectoderm | Hpcal1 | 0.84 | 1.90E-106 | 3.95E-102 | 0.46 | 0.29 | 0.16 | 1.56 |
| ectoderm | Snx5 | 0.43 | 5.13E-106 | 1.07E-101 | 0.83 | 0.76 | 0.07 | 1.09 |
| ectoderm | Zfp503 | 0.71 | 2.31E-105 | 4.81E-101 | 0.51 | 0.33 | 0.18 | 1.53 |
| ectoderm | Fat1 | 0.68 | 4.17E-105 | 8.68E-101 | 0.63 | 0.48 | 0.15 | 1.32 |
| ectoderm | Pea15a | 1.03 | 4.95E-105 | 1.03E-100 | 0.34 | 0.2 | 0.15 | 1.74 |
| ectoderm | Prr13 | 0.64 | 7.43E-105 | 1.54E-100 | 0.54 | 0.36 | 0.18 | 1.5 |
| ectoderm | Cxxc5 | 0.69 | 1.21E-104 | 2.52E-100 | 0.58 | 0.41 | 0.16 | 1.39 |
| ectoderm | Pak1 | 0.68 | 1.53E-104 | 3.19E-100 | 0.62 | 0.45 | 0.17 | 1.37 |
| ectoderm | .700096K18Ri | 1.29 | 1.61E-104 | 3.35E-100 | 0.18 | 0.07 | 0.11 | 2.53 |
| ectoderm | Reep3 | 0.66 | 1.76E-104 | 3.67E-100 | 0.64 | 0.49 | 0.15 | 1.3 |
| ectoderm | Mcoln3 | 1.67 | 2.16E-104 | 4.50E-100 | 0.12 | 0.04 | 0.08 | 3.31 |
| ectoderm | Emid1 | 1.32 | 5.61E-104 | 1.17E-99 | 0.17 | 0.07 | 0.11 | 2.59 |
| ectoderm | Sort1 | 0.95 | 7.36E-104 | 1.53E-99 | 0.32 | 0.17 | 0.15 | 1.86 |
| ectoderm | Fgd4 | 1.19 | 1.03E-103 | 2.15E-99 | 0.17 | 0.07 | 0.1 | 2.58 |
| ectoderm | Serinc5 | 1.82 | 1.42E-103 | 2.96E-99 | 0.12 | 0.04 | 0.08 | 3.18 |
| ectoderm | Scamp2 | 0.79 | 1.46E-103 | 3.04E-99 | 0.44 | 0.28 | 0.16 | 1.59 |
| ectoderm | Arfgef1 | 0.7 | 2.17E-103 | 4.51E-99 | 0.6 | 0.45 | 0.15 | 1.33 |
| ectoderm | Pygl | 1.67 | 3.89E-103 | 8.08E-99 | 0.18 | 0.07 | 0.11 | 2.44 |
| ectoderm | Nucb2 | 1.09 | 4.56E-103 | 9.48E-99 | 0.26 | 0.13 | 0.13 | 2.05 |
| ectoderm | Prkg1 | 1.37 | 5.25E-103 | 1.09E-98 | 0.26 | 0.13 | 0.13 | 1.98 |
| ectoderm | Dynlt3 | 0.99 | 8.13E-103 | 1.69E-98 | 0.37 | 0.22 | 0.15 | 1.67 |
| ectoderm | Plce1 | 1.09 | 1.14E-102 | 2.37E-98 | 0.34 | 0.19 | 0.15 | 1.75 |
| ectoderm | Zfp612 | 1 | 1.16E-102 | 2.42E-98 | 0.27 | 0.14 | 0.14 | 2 |
| ectoderm | Nedd4l | 1.05 | 1.87E-102 | 3.90E-98 | 0.24 | 0.12 | 0.13 | 2.1 |
| ectoderm | Cdc14a | 1.06 | 2.43E-102 | 5.05E-98 | 0.37 | 0.22 | 0.15 | 1.66 |
| ectoderm | Meis1 | 0.59 | 3.41E-102 | 7.09E-98 | 0.59 | 0.42 | 0.18 | 1.43 |
| ectoderm | Apba2 | 1.58 | 3.71E-102 | 7.71E-98 | 0.13 | 0.04 | 0.09 | 3.1 |
| ectoderm | Dedd2 | 1.5 | 8.55E-102 | 1.78E-97 | 0.15 | 0.05 | 0.09 | 2.75 |
| ectoderm | Enpp5 | 1.87 | 1.07E-101 | 2.22E-97 | 0.1 | 0.03 | 0.07 | 3.7 |
| ectoderm | Plekha1 | 0.99 | 7.90E-101 | 1.64E-96 | 0.31 | 0.17 | 0.14 | 1.86 |
| ectoderm | Fam49a | 2.18 | 1.27E-100 | 2.64E-96 | 0.1 | 0.03 | 0.07 | 3.7 |
| ectoderm | Zfp36l1 | 0.41 | 1.35E-100 | 2.82E-96 | 0.88 | 0.84 | 0.05 | 1.05 |
| ectoderm | Pcm1 | 0.35 | 1.93E-100 | 4.01E-96 | 0.92 | 0.89 | 0.02 | 1.03 |
| ectoderm | Zdhhc13 | 1.24 | 2.94E-100 | 6.12E-96 | 0.23 | 0.11 | 0.12 | 2.09 |
| ectoderm | Vsig10 | 1.72 | 3.18E-100 | 6.61E-96 | 0.12 | 0.04 | 0.08 | 3.29 |
| ectoderm | Slc25a27 | 0.74 | 3.95E-100 | 8.21E-96 | 0.53 | 0.36 | 0.17 | 1.46 |
| ectoderm | Ank3 | 0.64 | 5.30E-100 | 1.10E-95 | 0.61 | 0.45 | 0.16 | 1.35 |
| ectoderm | Chn2 | 1.61 | 5.15E-99 | 1.07E-94 | 0.14 | 0.05 | 0.09 | 2.82 |
| ectoderm | Glrb | 1.75 | 6.92E-99 | 1.44E-94 | 0.11 | 0.03 | 0.08 | 3.44 |
| ectoderm | Cpne3 | 0.63 | 1.06E-98 | 2.20E-94 | 0.64 | 0.49 | 0.15 | 1.31 |
| ectoderm | Ccdc92 | 1.45 | 2.95E-98 | 6.13E-94 | 0.16 | 0.06 | 0.1 | 2.63 |
| ectoderm | Vwa5a | 1.41 | 3.16E-98 | 6.57E-94 | 0.16 | 0.07 | 0.1 | 2.52 |
| ectoderm | Slc44a2 | 0.91 | 4.24E-98 | 8.82E-94 | 0.39 | 0.24 | 0.15 | 1.63 |
| ectoderm | Dpysl3 | 0.94 | 2.00E-97 | 4.15E-93 | 0.55 | 0.42 | 0.13 | 1.31 |
| ectoderm | Ass1 | 1.81 | 2.02E-97 | 4.20E-93 | 0.12 | 0.04 | 0.08 | 3.1 |
| ectoderm | Eef2kmt | 1.06 | 2.34E-97 | 4.86E-93 | 0.29 | 0.15 | 0.13 | 1.88 |
| ectoderm | Ctso | 1.67 | 2.78E-97 | 5.79E-93 | 0.13 | 0.05 | 0.09 | 2.91 |
| ectoderm | Cox14 | 0.42 | 3.22E-97 | 6.70E-93 | 0.85 | 0.79 | 0.06 | 1.08 |
| ectoderm | Pak4 | 0.78 | 4.31E-97 | 8.95E-93 | 0.5 | 0.35 | 0.15 | 1.44 |

**Supplementary Table 2. Marker genes for the major cell types of integrated clustering of E11.5 control and NotchGOF λ-junction cells.**

| | | | | | | | | |
|---|---|---|---|---|---|---|---|---|
| ectoderm | Unc5b | 0.64 | 1.66E-96 | 3.45E-92 | 0.53 | 0.35 | 0.17 | 1.49 |
| ectoderm | Os9 | 0.76 | 4.30E-96 | 8.93E-92 | 0.5 | 0.33 | 0.16 | 1.49 |
| ectoderm | Plagl2 | 0.78 | 7.57E-96 | 1.57E-91 | 0.48 | 0.32 | 0.15 | 1.47 |
| ectoderm | Airn | 0.94 | 1.14E-95 | 2.38E-91 | 0.3 | 0.16 | 0.14 | 1.87 |
| ectoderm | Nme3 | 1.38 | 1.73E-95 | 3.61E-91 | 0.15 | 0.06 | 0.09 | 2.67 |
| ectoderm | Bmyc | 0.57 | 2.56E-95 | 5.32E-91 | 0.73 | 0.59 | 0.14 | 1.23 |
| ectoderm | Grtp1 | 1.17 | 8.00E-95 | 1.66E-90 | 0.23 | 0.11 | 0.12 | 2.04 |
| ectoderm | Slco5a1 | 1.26 | 9.46E-95 | 1.97E-90 | 0.16 | 0.06 | 0.1 | 2.56 |
| ectoderm | Gab1 | 1.04 | 1.09E-94 | 2.26E-90 | 0.29 | 0.16 | 0.13 | 1.84 |
| ectoderm | Nectin2 | 0.66 | 5.64E-94 | 1.17E-89 | 0.59 | 0.45 | 0.14 | 1.32 |
| ectoderm | Id3 | 0.54 | 1.02E-93 | 2.13E-89 | 0.97 | 0.95 | 0.02 | 1.02 |
| ectoderm | Dync1i1 | 2.22 | 1.49E-93 | 3.10E-89 | 0.13 | 0.05 | 0.08 | 2.8 |
| ectoderm | Ebf4 | 1.52 | 2.19E-93 | 4.56E-89 | 0.12 | 0.04 | 0.08 | 3.08 |
| ectoderm | Gnpda2 | 0.92 | 2.69E-93 | 5.60E-89 | 0.4 | 0.25 | 0.15 | 1.59 |
| ectoderm | Tmem63b | 0.75 | 3.29E-93 | 6.83E-89 | 0.46 | 0.3 | 0.16 | 1.53 |
| ectoderm | Efnb2 | 0.48 | 4.31E-93 | 8.97E-89 | 0.58 | 0.41 | 0.18 | 1.43 |
| ectoderm | Akap9 | 0.48 | 6.12E-93 | 1.27E-88 | 0.82 | 0.77 | 0.05 | 1.07 |
| ectoderm | Cmtm4 | 0.86 | 7.72E-93 | 1.60E-88 | 0.37 | 0.22 | 0.15 | 1.66 |
| ectoderm | Ctsh | 1.39 | 2.24E-92 | 4.66E-88 | 0.13 | 0.04 | 0.08 | 2.91 |
| ectoderm | Sdsl | 1.59 | 2.54E-92 | 5.28E-88 | 0.12 | 0.04 | 0.08 | 2.98 |
| ectoderm | Ube2b | 0.3 | 1.01E-91 | 2.11E-87 | 0.93 | 0.93 | 0.01 | 1.01 |
| ectoderm | Pigp | 0.53 | 1.42E-91 | 2.95E-87 | 0.73 | 0.62 | 0.11 | 1.18 |
| ectoderm | Bcar3 | 1.66 | 2.31E-91 | 4.80E-87 | 0.11 | 0.03 | 0.07 | 3.18 |
| ectoderm | Retreg2 | 0.82 | 2.95E-91 | 6.14E-87 | 0.4 | 0.25 | 0.15 | 1.57 |
| ectoderm | Aagab | 1 | 3.14E-91 | 6.53E-87 | 0.27 | 0.15 | 0.13 | 1.88 |
| ectoderm | Plpp2 | 0.85 | 7.08E-91 | 1.47E-86 | 0.38 | 0.23 | 0.15 | 1.64 |
| ectoderm | Nudt11 | 1.01 | 1.02E-90 | 2.12E-86 | 0.28 | 0.15 | 0.13 | 1.88 |
| ectoderm | Gdpd1 | 0.78 | 1.41E-90 | 2.92E-86 | 0.39 | 0.24 | 0.15 | 1.64 |
| ectoderm | Dcaf17 | 0.79 | 1.82E-90 | 3.79E-86 | 0.45 | 0.3 | 0.15 | 1.51 |
| ectoderm | Gm10260 | 0.37 | 2.48E-90 | 5.17E-86 | 0.87 | 0.84 | 0.03 | 1.04 |
| ectoderm | Kit | 1.26 | 2.85E-90 | 5.92E-86 | 0.17 | 0.07 | 0.1 | 2.32 |
| ectoderm | Ctf1 | 0.96 | 3.86E-90 | 8.04E-86 | 0.26 | 0.14 | 0.13 | 1.94 |
| ectoderm | Ell2 | 1.2 | 4.46E-90 | 9.27E-86 | 0.16 | 0.07 | 0.1 | 2.43 |
| ectoderm | Lrrn3 | 1.61 | 4.65E-90 | 9.68E-86 | 0.14 | 0.05 | 0.09 | 2.65 |
| ectoderm | Mien1 | 0.56 | 7.87E-90 | 1.64E-85 | 0.66 | 0.53 | 0.13 | 1.24 |
| ectoderm | Cd47 | 0.63 | 9.43E-90 | 1.96E-85 | 0.56 | 0.41 | 0.15 | 1.37 |
| ectoderm | Gpd2 | 0.88 | 1.25E-89 | 2.60E-85 | 0.34 | 0.2 | 0.14 | 1.71 |
| ectoderm | Smim4 | 0.74 | 5.14E-89 | 1.07E-84 | 0.54 | 0.39 | 0.15 | 1.37 |
| ectoderm | Glcci1 | 1.03 | 7.79E-89 | 1.62E-84 | 0.5 | 0.36 | 0.14 | 1.41 |
| ectoderm | Rnf24 | 0.89 | 2.79E-88 | 5.81E-84 | 0.41 | 0.27 | 0.14 | 1.54 |
| ectoderm | Dbndd2 | 1.34 | 5.18E-88 | 1.08E-83 | 0.15 | 0.06 | 0.09 | 2.51 |
| ectoderm | Cmip | 0.76 | 1.09E-87 | 2.27E-83 | 0.51 | 0.36 | 0.15 | 1.41 |
| ectoderm | Sypl | 0.63 | 1.35E-87 | 2.80E-83 | 0.59 | 0.45 | 0.15 | 1.33 |
| ectoderm | Meis3 | 0.81 | 1.75E-87 | 3.63E-83 | 0.3 | 0.17 | 0.14 | 1.82 |
| ectoderm | St7 | 0.92 | 3.08E-87 | 6.41E-83 | 0.33 | 0.19 | 0.14 | 1.71 |
| ectoderm | Edem2 | 0.93 | 5.36E-87 | 1.11E-82 | 0.29 | 0.16 | 0.13 | 1.8 |
| ectoderm | Pex2 | 0.62 | 5.94E-87 | 1.23E-82 | 0.59 | 0.44 | 0.15 | 1.33 |
| ectoderm | Igdcc3 | 0.99 | 7.44E-87 | 1.55E-82 | 0.34 | 0.21 | 0.14 | 1.65 |
| ectoderm | Hook2 | 0.88 | 7.82E-87 | 1.63E-82 | 0.26 | 0.14 | 0.12 | 1.91 |
| ectoderm | Copg2 | 0.72 | 8.50E-87 | 1.77E-82 | 0.49 | 0.34 | 0.15 | 1.45 |
| ectoderm | Dnajc1 | 0.79 | 1.16E-86 | 2.42E-82 | 0.64 | 0.53 | 0.1 | 1.19 |
| ectoderm | Cd82 | 1.33 | 1.18E-86 | 2.46E-82 | 0.13 | 0.05 | 0.08 | 2.77 |
| ectoderm | Pcbp4 | 0.36 | 1.35E-86 | 2.82E-82 | 0.9 | 0.86 | 0.04 | 1.05 |
| ectoderm | Dhx32 | 1 | 3.66E-86 | 7.61E-82 | 0.32 | 0.19 | 0.13 | 1.7 |
| ectoderm | Kif13a | 0.96 | 8.73E-86 | 1.82E-81 | 0.31 | 0.18 | 0.13 | 1.74 |
| ectoderm | Tnks2 | 0.4 | 1.79E-85 | 3.73E-81 | 0.85 | 0.8 | 0.06 | 1.07 |

**Supplementary Table 2. Marker genes for the major cell types of integrated clustering of E11.5 control and NotchGOF λ-junction cells.**

| | | | | | | | | |
|---|---|---|---|---|---|---|---|---|
| ectoderm | Nlk | 0.72 | 2.84E-85 | 5.91E-81 | 0.38 | 0.24 | 0.14 | 1.6 |
| ectoderm | Daam1 | 0.81 | 5.39E-85 | 1.12E-80 | 0.51 | 0.38 | 0.14 | 1.36 |
| ectoderm | Ccdc181 | 1.35 | 6.58E-85 | 1.37E-80 | 0.13 | 0.05 | 0.08 | 2.76 |
| ectoderm | Parva | 0.56 | 8.87E-85 | 1.85E-80 | 0.65 | 0.52 | 0.13 | 1.26 |
| ectoderm | Pacsin3 | 0.88 | 1.10E-84 | 2.28E-80 | 0.32 | 0.19 | 0.14 | 1.72 |
| ectoderm | Ormdl1 | 0.65 | 1.50E-84 | 3.12E-80 | 0.55 | 0.41 | 0.14 | 1.35 |
| ectoderm | Ric3 | 1.56 | 1.51E-84 | 3.14E-80 | 0.11 | 0.03 | 0.07 | 3.12 |
| ectoderm | Egfl6 | 1.08 | 2.76E-84 | 5.73E-80 | 0.18 | 0.08 | 0.1 | 2.23 |
| ectoderm | Lamtor2 | 0.3 | 7.55E-84 | 1.57E-79 | 0.91 | 0.89 | 0.02 | 1.02 |
| ectoderm | Ptpn14 | 1.11 | 8.68E-84 | 1.81E-79 | 0.24 | 0.13 | 0.12 | 1.92 |
| ectoderm | Tmem189 | 0.79 | 1.69E-83 | 3.52E-79 | 0.41 | 0.27 | 0.14 | 1.53 |
| ectoderm | Larp4b | 0.5 | 1.97E-83 | 4.10E-79 | 0.69 | 0.59 | 0.1 | 1.17 |
| ectoderm | Rcsd1 | 1.22 | 2.69E-83 | 5.60E-79 | 0.17 | 0.07 | 0.09 | 2.31 |
| ectoderm | Tmem53 | 1.24 | 2.80E-83 | 5.83E-79 | 0.16 | 0.07 | 0.09 | 2.39 |
| ectoderm | Srgap3 | 1.06 | 3.31E-83 | 6.88E-79 | 0.22 | 0.11 | 0.11 | 2.02 |
| ectoderm | Lypd6 | 0.9 | 6.11E-83 | 1.27E-78 | 0.21 | 0.1 | 0.11 | 2.09 |
| ectoderm | Pycard | 0.82 | 8.62E-83 | 1.79E-78 | 0.21 | 0.1 | 0.11 | 2.06 |
| ectoderm | Gpr137c | 1.04 | 8.99E-83 | 1.87E-78 | 0.23 | 0.12 | 0.11 | 1.96 |
| ectoderm | Suclg2 | 0.59 | 1.17E-82 | 2.44E-78 | 0.6 | 0.46 | 0.14 | 1.3 |
| ectoderm | Shroom3 | 1 | 1.87E-82 | 3.89E-78 | 0.24 | 0.13 | 0.12 | 1.94 |
| ectoderm | Txnip | 0.82 | 2.11E-82 | 4.39E-78 | 0.45 | 0.31 | 0.14 | 1.46 |
| ectoderm | Nhsl1 | 0.91 | 3.90E-82 | 8.11E-78 | 0.32 | 0.19 | 0.13 | 1.71 |
| ectoderm | Syde2 | 1.25 | 8.33E-82 | 1.73E-77 | 0.14 | 0.05 | 0.08 | 2.56 |
| ectoderm | Kitl | 1.16 | 9.08E-82 | 1.89E-77 | 0.14 | 0.06 | 0.09 | 2.52 |
| ectoderm | Smim20 | 0.67 | 1.21E-81 | 2.52E-77 | 0.52 | 0.37 | 0.15 | 1.4 |
| ectoderm | Eml2 | 1.17 | 1.58E-81 | 3.28E-77 | 0.17 | 0.08 | 0.1 | 2.25 |
| ectoderm | Nudt16 | 1.21 | 1.76E-81 | 3.65E-77 | 0.16 | 0.07 | 0.09 | 2.36 |
| ectoderm | Dmd | 1.16 | 3.03E-81 | 6.29E-77 | 0.16 | 0.07 | 0.09 | 2.35 |
| ectoderm | Ptk7 | 0.76 | 6.23E-81 | 1.30E-76 | 0.48 | 0.33 | 0.15 | 1.44 |
| ectoderm | Sft2d3 | 0.87 | 1.05E-80 | 2.18E-76 | 0.3 | 0.17 | 0.13 | 1.74 |
| ectoderm | E330009J07Ri | 1.12 | 2.16E-80 | 4.48E-76 | 0.2 | 0.1 | 0.1 | 2.07 |
| ectoderm | Trim25 | 1.48 | 2.44E-80 | 5.07E-76 | 0.12 | 0.04 | 0.08 | 2.79 |
| ectoderm | Sec22a | 0.72 | 4.79E-80 | 9.97E-76 | 0.4 | 0.26 | 0.14 | 1.55 |
| ectoderm | Prickle3 | 1.38 | 5.23E-80 | 1.09E-75 | 0.13 | 0.05 | 0.08 | 2.63 |
| ectoderm | Yipf2 | 1.1 | 7.48E-80 | 1.55E-75 | 0.2 | 0.1 | 0.1 | 2.05 |
| ectoderm | Bcl7a | 0.46 | 8.20E-80 | 1.71E-75 | 0.75 | 0.65 | 0.1 | 1.16 |
| ectoderm | Fbxo27 | 1.35 | 1.76E-79 | 3.67E-75 | 0.13 | 0.05 | 0.08 | 2.61 |
| ectoderm | Slc44a1 | 0.8 | 2.05E-79 | 4.26E-75 | 0.34 | 0.21 | 0.14 | 1.66 |
| ectoderm | Igf2bp3 | 0.36 | 2.34E-79 | 4.87E-75 | 0.86 | 0.81 | 0.05 | 1.06 |
| ectoderm | Khk | 1.61 | 2.37E-79 | 4.92E-75 | 0.13 | 0.05 | 0.08 | 2.6 |
| ectoderm | Herc3 | 1.27 | 3.91E-79 | 8.13E-75 | 0.14 | 0.06 | 0.08 | 2.51 |
| ectoderm | Edf1 | 0.26 | 5.78E-79 | 1.20E-74 | 0.95 | 0.96 | -0.01 | 0.99 |
| ectoderm | Bid | 1.09 | 1.17E-78 | 2.44E-74 | 0.2 | 0.1 | 0.1 | 2.06 |
| ectoderm | Endod1 | 1.34 | 1.23E-78 | 2.56E-74 | 0.16 | 0.07 | 0.09 | 2.27 |
| ectoderm | Sh3rf1 | 0.82 | 4.03E-78 | 8.39E-74 | 0.33 | 0.2 | 0.13 | 1.66 |
| ectoderm | Kcnmb4os2 | 1.03 | 4.16E-78 | 8.64E-74 | 0.25 | 0.13 | 0.11 | 1.86 |
| ectoderm | Eya1 | 0.66 | 5.89E-78 | 1.22E-73 | 0.59 | 0.47 | 0.12 | 1.25 |
| ectoderm | Miga1 | 0.69 | 1.10E-77 | 2.28E-73 | 0.41 | 0.27 | 0.14 | 1.54 |
| ectoderm | Saraf | 0.66 | 1.32E-77 | 2.74E-73 | 0.63 | 0.51 | 0.12 | 1.23 |
| ectoderm | Ube2e1 | 0.36 | 1.51E-77 | 3.14E-73 | 0.88 | 0.83 | 0.05 | 1.06 |
| ectoderm | Pnrc1 | 0.42 | 1.51E-77 | 3.15E-73 | 0.89 | 0.86 | 0.03 | 1.03 |
| ectoderm | Actn4 | 0.46 | 1.67E-77 | 3.48E-73 | 0.76 | 0.68 | 0.08 | 1.12 |
| ectoderm | Maf | 0.37 | 6.30E-77 | 1.31E-72 | 0.14 | 0.05 | 0.08 | 2.54 |
| ectoderm | Ppp1r9a | 0.66 | 7.10E-77 | 1.48E-72 | 0.51 | 0.36 | 0.15 | 1.4 |
| ectoderm | Inava | 0.82 | 8.78E-77 | 1.83E-72 | 0.33 | 0.2 | 0.13 | 1.65 |
| ectoderm | Dennd1b | 0.85 | 2.16E-76 | 4.49E-72 | 0.25 | 0.13 | 0.12 | 1.86 |

**Supplementary Table 2. Marker genes for the major cell types of integrated clustering of E11.5 control and NotchGOF λ-junction cells.**

| | | | | | | | | |
|---|---|---|---|---|---|---|---|---|
| ectoderm | Taok1 | 0.52 | 4.50E-76 | 9.35E-72 | 0.68 | 0.58 | 0.1 | 1.18 |
| ectoderm | Amt | 0.98 | 5.47E-76 | 1.14E-71 | 0.23 | 0.12 | 0.11 | 1.92 |
| ectoderm | Osbpl3 | 1.35 | 5.68E-76 | 1.18E-71 | 0.16 | 0.07 | 0.09 | 2.22 |
| ectoderm | Usp46 | 0.51 | 1.09E-75 | 2.26E-71 | 0.72 | 0.6 | 0.12 | 1.19 |
| ectoderm | Tmem200a | 1.39 | 4.14E-75 | 8.60E-71 | 0.11 | 0.04 | 0.07 | 2.75 |
| ectoderm | Klhl8 | 1.4 | 5.15E-75 | 1.07E-70 | 0.16 | 0.07 | 0.09 | 2.21 |
| ectoderm | Rel | 1.38 | 1.47E-74 | 3.07E-70 | 0.16 | 0.07 | 0.09 | 2.23 |
| ectoderm | Fam3c | 0.73 | 1.65E-74 | 3.44E-70 | 0.47 | 0.33 | 0.14 | 1.42 |
| ectoderm | Galnt7 | 0.7 | 1.99E-74 | 4.14E-70 | 0.45 | 0.31 | 0.14 | 1.45 |
| ectoderm | Vwa1 | 1.6 | 2.11E-74 | 4.38E-70 | 0.11 | 0.04 | 0.07 | 2.69 |
| ectoderm | Harbi1 | 0.96 | 4.96E-74 | 1.03E-69 | 0.28 | 0.16 | 0.12 | 1.74 |
| ectoderm | Tet3 | 0.54 | 5.61E-74 | 1.17E-69 | 0.65 | 0.55 | 0.11 | 1.2 |
| ectoderm | Thumpd3 | 0.66 | 6.06E-74 | 1.26E-69 | 0.49 | 0.35 | 0.14 | 1.39 |
| ectoderm | Kctd6 | 0.98 | 1.19E-73 | 2.47E-69 | 0.24 | 0.13 | 0.11 | 1.85 |
| ectoderm | Npas3 | 1.39 | 1.56E-73 | 3.25E-69 | 0.12 | 0.04 | 0.07 | 2.64 |
| ectoderm | Pcdh9 | 1.17 | 1.96E-73 | 4.08E-69 | 0.19 | 0.09 | 0.1 | 2.05 |
| ectoderm | Ptprm | 1.4 | 3.22E-73 | 6.69E-69 | 0.1 | 0.04 | 0.07 | 2.86 |
| ectoderm | Dcbld1 | 0.95 | 5.86E-73 | 1.22E-68 | 0.4 | 0.27 | 0.13 | 1.46 |
| ectoderm | Ift43 | 0.5 | 1.65E-72 | 3.44E-68 | 0.65 | 0.51 | 0.14 | 1.28 |
| ectoderm | Shroom2 | 1.25 | 2.57E-72 | 5.35E-68 | 0.14 | 0.06 | 0.08 | 2.35 |
| ectoderm | Sulf1 | 0.33 | 3.17E-72 | 6.59E-68 | 0.66 | 0.5 | 0.16 | 1.31 |
| ectoderm | Arfip2 | 0.6 | 3.85E-72 | 8.01E-68 | 0.53 | 0.39 | 0.14 | 1.36 |
| ectoderm | Amot | 1.05 | 1.14E-71 | 2.37E-67 | 0.56 | 0.46 | 0.11 | 1.23 |
| ectoderm | Sppl2a | 0.6 | 1.32E-71 | 2.74E-67 | 0.58 | 0.44 | 0.13 | 1.3 |
| ectoderm | Dcxr | 1.27 | 1.82E-71 | 3.78E-67 | 0.16 | 0.08 | 0.09 | 2.16 |
| ectoderm | Rnd3 | 0.52 | 1.85E-71 | 3.85E-67 | 0.73 | 0.62 | 0.11 | 1.18 |
| ectoderm | Map3k20 | 1.15 | 2.14E-71 | 4.45E-67 | 0.19 | 0.1 | 0.1 | 2 |
| ectoderm | Ccpg1os | 0.88 | 4.77E-71 | 9.92E-67 | 0.24 | 0.13 | 0.11 | 1.83 |
| ectoderm | Cdh4 | 1.5 | 5.18E-71 | 1.08E-66 | 0.1 | 0.04 | 0.07 | 2.81 |
| ectoderm | Srebf1 | 0.86 | 9.91E-71 | 2.06E-66 | 0.28 | 0.17 | 0.12 | 1.72 |
| ectoderm | Fam32a | 0.41 | 1.48E-70 | 3.07E-66 | 0.81 | 0.76 | 0.06 | 1.08 |
| ectoderm | Gid4 | 0.7 | 1.61E-70 | 3.35E-66 | 0.44 | 0.31 | 0.13 | 1.43 |
| ectoderm | Plpp6 | 1.22 | 3.49E-70 | 7.26E-66 | 0.13 | 0.05 | 0.08 | 2.5 |
| ectoderm | Cyp39a1 | 0.92 | 6.34E-70 | 1.32E-65 | 0.23 | 0.13 | 0.11 | 1.85 |
| ectoderm | Ccdc115 | 0.58 | 7.18E-70 | 1.49E-65 | 0.59 | 0.46 | 0.13 | 1.28 |
| ectoderm | Maml2 | 1.17 | 1.15E-69 | 2.40E-65 | 0.16 | 0.07 | 0.09 | 2.18 |
| ectoderm | Mboat1 | 1.4 | 2.37E-69 | 4.92E-65 | 0.1 | 0.04 | 0.07 | 2.81 |
| ectoderm | Atp11a | 0.85 | 2.39E-69 | 4.97E-65 | 0.26 | 0.15 | 0.11 | 1.77 |
| ectoderm | Tmem106c | 0.86 | 2.42E-69 | 5.02E-65 | 0.29 | 0.17 | 0.12 | 1.69 |
| ectoderm | Ccdc120 | 1.1 | 3.38E-69 | 7.03E-65 | 0.18 | 0.09 | 0.09 | 2.06 |
| ectoderm | Rassf9 | 0.81 | 3.47E-69 | 7.22E-65 | 0.25 | 0.14 | 0.11 | 1.79 |
| ectoderm | Plxdc2 | 0.72 | 3.68E-69 | 7.65E-65 | 0.35 | 0.22 | 0.13 | 1.6 |
| ectoderm | Cdk2ap2 | 0.55 | 3.75E-69 | 7.81E-65 | 0.56 | 0.43 | 0.13 | 1.31 |
| ectoderm | Wdr31 | 1.22 | 4.60E-69 | 9.56E-65 | 0.15 | 0.07 | 0.08 | 2.2 |
| ectoderm | Carmil1 | 0.99 | 6.66E-69 | 1.38E-64 | 0.23 | 0.12 | 0.1 | 1.85 |
| ectoderm | Ctsc | 0.33 | 7.01E-69 | 1.46E-64 | 0.2 | 0.1 | 0.1 | 1.96 |
| ectoderm | Acly | 0.49 | 7.17E-69 | 1.49E-64 | 0.67 | 0.55 | 0.12 | 1.21 |
| ectoderm | Six2 | 0.4 | 7.35E-69 | 1.53E-64 | 0.56 | 0.42 | 0.15 | 1.35 |
| ectoderm | Psme1 | 0.38 | 1.12E-68 | 2.34E-64 | 0.81 | 0.75 | 0.06 | 1.08 |
| ectoderm | Gnptg | 0.8 | 1.20E-68 | 2.50E-64 | 0.29 | 0.18 | 0.12 | 1.67 |
| ectoderm | Ermp1 | 0.8 | 1.54E-68 | 3.20E-64 | 0.43 | 0.31 | 0.13 | 1.42 |
| ectoderm | Nmt2 | 0.7 | 2.22E-68 | 4.63E-64 | 0.44 | 0.31 | 0.13 | 1.42 |
| ectoderm | Ptprd | 0.53 | 2.57E-68 | 5.35E-64 | 0.55 | 0.41 | 0.14 | 1.34 |
| ectoderm | Sgsm2 | 1.2 | 3.11E-68 | 6.47E-64 | 0.11 | 0.04 | 0.07 | 2.64 |
| ectoderm | Cyb5r4 | 0.6 | 3.92E-68 | 8.15E-64 | 0.5 | 0.37 | 0.13 | 1.36 |
| ectoderm | Pde5a | 0.88 | 4.83E-68 | 1.00E-63 | 0.27 | 0.16 | 0.11 | 1.72 |

**Supplementary Table 2. Marker genes for the major cell types of integrated clustering of E11.5 control and NotchGOF λ-junction cells.**

| | | | | | | | | |
|---|---|---|---|---|---|---|---|---|
| ectoderm | Ankrd49 | 0.59 | 4.93E-68 | 1.03E-63 | 0.53 | 0.41 | 0.13 | 1.31 |
| ectoderm | Ralbp1 | 0.44 | 5.61E-68 | 1.17E-63 | 0.82 | 0.78 | 0.04 | 1.05 |
| ectoderm | Dclre1c | 1.91 | 6.09E-68 | 1.27E-63 | 0.1 | 0.04 | 0.06 | 2.68 |
| ectoderm | Prkcd | 0.76 | 6.44E-68 | 1.34E-63 | 0.31 | 0.19 | 0.12 | 1.66 |
| ectoderm | Pim3 | 0.67 | 7.39E-68 | 1.54E-63 | 0.4 | 0.27 | 0.13 | 1.49 |
| ectoderm | Socs7 | 0.82 | 7.69E-68 | 1.60E-63 | 0.3 | 0.18 | 0.12 | 1.65 |
| ectoderm | Phgdh | 0.34 | 8.79E-68 | 1.83E-63 | 0.86 | 0.81 | 0.04 | 1.06 |
| ectoderm | Rab11fip4 | 0.93 | 9.05E-68 | 1.88E-63 | 0.21 | 0.11 | 0.1 | 1.91 |
| ectoderm | Acp6 | 0.86 | 1.00E-67 | 2.08E-63 | 0.32 | 0.2 | 0.12 | 1.6 |
| ectoderm | Tbc1d10a | 0.92 | 1.02E-67 | 2.11E-63 | 0.29 | 0.17 | 0.11 | 1.66 |
| ectoderm | Agl | 0.76 | 1.60E-67 | 3.32E-63 | 0.35 | 0.22 | 0.12 | 1.55 |
| ectoderm | Gas7 | 1.38 | 1.91E-67 | 3.98E-63 | 0.1 | 0.04 | 0.06 | 2.68 |
| ectoderm | Esco1 | 0.79 | 2.05E-67 | 4.26E-63 | 0.33 | 0.21 | 0.12 | 1.6 |
| ectoderm | Tmem141 | 0.81 | 2.34E-67 | 4.86E-63 | 0.29 | 0.17 | 0.12 | 1.71 |
| ectoderm | Sh3d19 | 0.71 | 2.53E-67 | 5.27E-63 | 0.36 | 0.23 | 0.13 | 1.56 |
| ectoderm | Pacsin1 | 1.01 | 3.72E-67 | 7.73E-63 | 0.17 | 0.08 | 0.09 | 2.07 |
| ectoderm | Kdm3a | 0.65 | 4.26E-67 | 8.87E-63 | 0.47 | 0.34 | 0.13 | 1.39 |
| ectoderm | Alpl | 1.17 | 4.55E-67 | 9.47E-63 | 0.17 | 0.08 | 0.09 | 2.05 |
| ectoderm | Tceal5 | 0.9 | 4.99E-67 | 1.04E-62 | 0.23 | 0.13 | 0.11 | 1.83 |
| ectoderm | Cyfip2 | 0.89 | 1.75E-66 | 3.64E-62 | 0.16 | 0.07 | 0.08 | 2.18 |
| ectoderm | Sertad2 | 0.59 | 1.83E-66 | 3.80E-62 | 0.51 | 0.38 | 0.13 | 1.35 |
| ectoderm | Prkag2 | 0.84 | 3.30E-66 | 6.86E-62 | 0.32 | 0.2 | 0.12 | 1.61 |
| ectoderm | Arhgap18 | 0.36 | 8.13E-66 | 1.69E-61 | 0.2 | 0.1 | 0.1 | 1.95 |
| ectoderm | Acvr1b | 1.11 | 1.30E-65 | 2.69E-61 | 0.18 | 0.09 | 0.09 | 1.98 |
| ectoderm | Ssr4 | 0.3 | 1.38E-65 | 2.86E-61 | 0.86 | 0.83 | 0.03 | 1.03 |
| ectoderm | Magi3 | 0.77 | 2.02E-65 | 4.20E-61 | 0.28 | 0.16 | 0.12 | 1.7 |
| ectoderm | Nomo1 | 0.77 | 2.47E-65 | 5.14E-61 | 0.29 | 0.17 | 0.12 | 1.67 |
| ectoderm | Zcchc18 | 0.69 | 2.79E-65 | 5.81E-61 | 0.44 | 0.31 | 0.13 | 1.43 |
| ectoderm | Dpp3 | 0.48 | 3.61E-65 | 7.50E-61 | 0.66 | 0.55 | 0.11 | 1.19 |
| ectoderm | Aprt | 0.26 | 3.76E-65 | 7.83E-61 | 0.93 | 0.92 | 0.01 | 1.01 |
| ectoderm | Aldh5a1 | 0.71 | 4.35E-65 | 9.05E-61 | 0.34 | 0.21 | 0.12 | 1.58 |
| ectoderm | 700025G04Ri | 0.66 | 4.38E-65 | 9.12E-61 | 0.44 | 0.31 | 0.13 | 1.43 |
| ectoderm | Fmc1 | 0.52 | 4.56E-65 | 9.49E-61 | 0.67 | 0.57 | 0.1 | 1.18 |
| ectoderm | Ext1 | 0.58 | 5.50E-65 | 1.14E-60 | 0.51 | 0.38 | 0.13 | 1.35 |
| ectoderm | Zkscan7 | 1.14 | 5.62E-65 | 1.17E-60 | 0.14 | 0.06 | 0.08 | 2.27 |
| ectoderm | S100a6 | 1.35 | 7.91E-65 | 1.64E-60 | 0.24 | 0.14 | 0.1 | 1.71 |
| ectoderm | Sdr39u1 | 0.91 | 8.41E-65 | 1.75E-60 | 0.24 | 0.13 | 0.1 | 1.78 |
| ectoderm | Stap2 | 1.09 | 1.36E-64 | 2.82E-60 | 0.14 | 0.06 | 0.08 | 2.22 |
| ectoderm | Gpc1 | 0.69 | 2.18E-64 | 4.54E-60 | 0.49 | 0.37 | 0.12 | 1.32 |
| ectoderm | Cxcl14 | 2.66 | 2.69E-64 | 5.60E-60 | 0.22 | 0.13 | 0.09 | 1.67 |
| ectoderm | Lrrc42 | 0.44 | 2.92E-64 | 6.07E-60 | 0.7 | 0.58 | 0.12 | 1.21 |
| ectoderm | Jmy | 0.63 | 3.43E-64 | 7.12E-60 | 0.48 | 0.35 | 0.13 | 1.38 |
| ectoderm | Rnf44 | 0.64 | 4.27E-64 | 8.88E-60 | 0.45 | 0.32 | 0.13 | 1.41 |
| ectoderm | Erbb2 | 0.68 | 4.33E-64 | 9.01E-60 | 0.33 | 0.21 | 0.12 | 1.59 |
| ectoderm | Npepl1 | 0.66 | 7.41E-64 | 1.54E-59 | 0.43 | 0.3 | 0.13 | 1.44 |
| ectoderm | Rnf180 | 1.46 | 7.65E-64 | 1.59E-59 | 0.1 | 0.04 | 0.06 | 2.63 |
| ectoderm | 310039H08Ri | 0.62 | 1.00E-63 | 2.09E-59 | 0.52 | 0.39 | 0.13 | 1.32 |
| ectoderm | Mapk8ip1 | 0.65 | 1.19E-63 | 2.48E-59 | 0.35 | 0.23 | 0.12 | 1.54 |
| ectoderm | Lhfp | 0.93 | 1.30E-63 | 2.71E-59 | 0.32 | 0.21 | 0.11 | 1.53 |
| ectoderm | Atp2b1 | 0.41 | 2.02E-63 | 4.20E-59 | 0.85 | 0.85 | 0 | 1 |
| ectoderm | Cbx6 | 0.71 | 2.07E-63 | 4.31E-59 | 0.37 | 0.24 | 0.13 | 1.53 |
| ectoderm | Clcn3 | 0.59 | 2.59E-63 | 5.39E-59 | 0.48 | 0.35 | 0.13 | 1.36 |
| ectoderm | Taf10 | 0.26 | 2.91E-63 | 6.05E-59 | 0.93 | 0.93 | 0.01 | 1.01 |
| ectoderm | Vmac | 1.04 | 3.22E-63 | 6.71E-59 | 0.16 | 0.08 | 0.08 | 2.11 |
| ectoderm | Wdpcp | 1.16 | 3.91E-63 | 8.13E-59 | 0.14 | 0.06 | 0.08 | 2.2 |
| ectoderm | Pcdh1 | 1.21 | 4.59E-63 | 9.54E-59 | 0.1 | 0.04 | 0.06 | 2.7 |

**Supplementary Table 2. Marker genes for the major cell types of integrated clustering of E11.5 control and NotchGOF λ-junction cells.**

| | | | | | | | | |
|---|---|---|---|---|---|---|---|---|
| ectoderm | Tmc4 | 1.21 | 6.03E-63 | 1.25E-58 | 0.16 | 0.08 | 0.08 | 2.04 |
| ectoderm | Jtb | 0.5 | 1.00E-62 | 2.09E-58 | 0.62 | 0.51 | 0.11 | 1.22 |
| ectoderm | Mllt10 | 0.44 | 1.26E-62 | 2.62E-58 | 0.71 | 0.61 | 0.09 | 1.15 |
| ectoderm | Rida | 0.58 | 2.17E-62 | 4.52E-58 | 0.52 | 0.39 | 0.13 | 1.32 |
| ectoderm | Erf | 0.7 | 2.64E-62 | 5.50E-58 | 0.4 | 0.27 | 0.13 | 1.47 |
| ectoderm | Minar2 | 1.25 | 3.86E-62 | 8.03E-58 | 0.16 | 0.08 | 0.08 | 2.05 |
| ectoderm | Gabarapl2 | 0.32 | 9.18E-62 | 1.91E-57 | 0.86 | 0.81 | 0.04 | 1.05 |
| ectoderm | Setd7 | 0.79 | 1.06E-61 | 2.20E-57 | 0.29 | 0.18 | 0.11 | 1.63 |
| ectoderm | Sik1 | 1.14 | 1.45E-61 | 3.01E-57 | 0.17 | 0.09 | 0.08 | 1.98 |
| ectoderm | Pgap1 | 0.81 | 3.38E-61 | 7.04E-57 | 0.23 | 0.13 | 0.1 | 1.79 |
| ectoderm | Klf10 | 1.08 | 3.62E-61 | 7.53E-57 | 0.18 | 0.09 | 0.09 | 1.97 |
| ectoderm | Thnsl1 | 1.18 | 4.31E-61 | 8.97E-57 | 0.13 | 0.06 | 0.07 | 2.29 |
| ectoderm | Zfp395 | 0.73 | 5.86E-61 | 1.22E-56 | 0.33 | 0.21 | 0.12 | 1.57 |
| ectoderm | Msi1 | 0.62 | 6.89E-61 | 1.43E-56 | 0.47 | 0.34 | 0.13 | 1.38 |
| ectoderm | Letmd1 | 0.74 | 1.12E-60 | 2.33E-56 | 0.31 | 0.2 | 0.12 | 1.59 |
| ectoderm | Ccdc80 | 0.81 | 1.73E-60 | 3.60E-56 | 0.27 | 0.16 | 0.11 | 1.68 |
| ectoderm | Nxn | 0.49 | 2.28E-60 | 4.75E-56 | 0.76 | 0.7 | 0.06 | 1.08 |
| ectoderm | Sema4d | 1.1 | 2.49E-60 | 5.18E-56 | 0.13 | 0.05 | 0.07 | 2.31 |
| ectoderm | Calcoco1 | 0.83 | 2.73E-60 | 5.68E-56 | 0.3 | 0.19 | 0.11 | 1.59 |
| ectoderm | Plcb4 | 0.81 | 3.25E-60 | 6.76E-56 | 0.28 | 0.17 | 0.11 | 1.64 |
| ectoderm | Arf3 | 0.65 | 4.17E-60 | 8.68E-56 | 0.41 | 0.28 | 0.13 | 1.45 |
| ectoderm | Pcdh19 | 1.19 | 4.39E-60 | 9.12E-56 | 0.24 | 0.15 | 0.1 | 1.66 |
| ectoderm | Cd164 | 0.44 | 5.24E-60 | 1.09E-55 | 0.7 | 0.62 | 0.08 | 1.13 |
| ectoderm | Rnf187 | 0.26 | 6.03E-60 | 1.25E-55 | 0.92 | 0.9 | 0.01 | 1.01 |
| ectoderm | Hint2 | 0.45 | 6.38E-60 | 1.33E-55 | 0.68 | 0.59 | 0.09 | 1.16 |
| ectoderm | 430403G16Ri | 1.32 | 9.54E-60 | 1.98E-55 | 0.13 | 0.06 | 0.07 | 2.27 |
| ectoderm | Six1 | 0.42 | 1.67E-59 | 3.47E-55 | 0.69 | 0.59 | 0.1 | 1.18 |
| ectoderm | Mest | 0.52 | 2.91E-59 | 6.06E-55 | 0.91 | 0.91 | -0.01 | 0.99 |
| ectoderm | Zcchc7 | 0.51 | 3.21E-59 | 6.68E-55 | 0.61 | 0.49 | 0.11 | 1.23 |
| ectoderm | Nhs | 0.59 | 5.31E-59 | 1.10E-54 | 0.43 | 0.31 | 0.12 | 1.39 |
| ectoderm | Slc25a17 | 0.49 | 7.33E-59 | 1.52E-54 | 0.69 | 0.6 | 0.09 | 1.16 |
| ectoderm | Ccdc28a | 1.05 | 7.73E-59 | 1.61E-54 | 0.17 | 0.09 | 0.08 | 1.96 |
| ectoderm | Llph | 0.31 | 1.50E-58 | 3.13E-54 | 0.88 | 0.85 | 0.03 | 1.04 |
| ectoderm | Snapin | 0.61 | 1.75E-58 | 3.64E-54 | 0.42 | 0.3 | 0.13 | 1.42 |
| ectoderm | Dhrs4 | 0.45 | 2.10E-58 | 4.37E-54 | 0.7 | 0.62 | 0.08 | 1.14 |
| ectoderm | Zfp7 | 0.84 | 3.07E-58 | 6.39E-54 | 0.22 | 0.12 | 0.1 | 1.78 |
| ectoderm | Dbp | 0.94 | 4.35E-58 | 9.04E-54 | 0.21 | 0.12 | 0.09 | 1.8 |
| ectoderm | Slc7a11 | 1.88 | 5.18E-58 | 1.08E-53 | 0.13 | 0.06 | 0.07 | 2.09 |
| ectoderm | Ccdc18 | 0.68 | 5.42E-58 | 1.13E-53 | 0.4 | 0.28 | 0.12 | 1.43 |
| ectoderm | Rsrp1 | 0.27 | 7.02E-58 | 1.46E-53 | 0.89 | 0.88 | 0.01 | 1.01 |
| ectoderm | Necab2 | 1.08 | 8.09E-58 | 1.68E-53 | 0.28 | 0.18 | 0.1 | 1.54 |
| ectoderm | Itgb8 | 0.84 | 1.66E-57 | 3.44E-53 | 0.2 | 0.11 | 0.09 | 1.84 |
| ectoderm | Sun1 | 0.6 | 1.67E-57 | 3.47E-53 | 0.37 | 0.25 | 0.12 | 1.5 |
| ectoderm | Cd3eap | 0.53 | 1.87E-57 | 3.89E-53 | 0.61 | 0.5 | 0.11 | 1.21 |
| ectoderm | Marchf8 | 0.78 | 2.76E-57 | 5.74E-53 | 0.25 | 0.15 | 0.1 | 1.68 |
| ectoderm | Fam192a | 0.42 | 3.63E-57 | 7.55E-53 | 0.72 | 0.63 | 0.09 | 1.14 |
| ectoderm | Dusp28 | 0.92 | 4.77E-57 | 9.91E-53 | 0.18 | 0.1 | 0.09 | 1.91 |
| ectoderm | Rcan1 | 0.76 | 5.06E-57 | 1.05E-52 | 0.35 | 0.23 | 0.11 | 1.48 |
| ectoderm | Tulp4 | 0.46 | 6.38E-57 | 1.33E-52 | 0.73 | 0.65 | 0.08 | 1.12 |
| ectoderm | Zfp536 | 0.78 | 9.94E-57 | 2.07E-52 | 0.17 | 0.08 | 0.08 | 1.99 |
| ectoderm | Cryzl2 | 0.98 | 1.07E-56 | 2.22E-52 | 0.17 | 0.08 | 0.08 | 1.98 |
| ectoderm | Cyba | 0.26 | 1.89E-56 | 3.94E-52 | 0.56 | 0.42 | 0.14 | 1.33 |
| ectoderm | Olfm1 | 0.66 | 1.92E-56 | 4.00E-52 | 0.32 | 0.2 | 0.11 | 1.56 |
| ectoderm | Bok | 0.62 | 3.93E-56 | 8.17E-52 | 0.28 | 0.18 | 0.11 | 1.62 |
| ectoderm | Pex7 | 0.69 | 4.83E-56 | 1.01E-51 | 0.34 | 0.23 | 0.12 | 1.51 |
| ectoderm | Sema3f | 0.6 | 7.03E-56 | 1.46E-51 | 0.38 | 0.26 | 0.12 | 1.45 |

**Supplementary Table 2. Marker genes for the major cell types of integrated clustering of E11.5 control and NotchGOF λ-junction cells.**

| | | | | | | | | |
|---|---|---|---|---|---|---|---|---|
| ectoderm | Olfm2 | 0.83 | 8.74E-56 | 1.82E-51 | 0.17 | 0.09 | 0.08 | 1.97 |
| ectoderm | Tmem108 | 0.93 | 1.49E-55 | 3.09E-51 | 0.3 | 0.2 | 0.1 | 1.53 |
| ectoderm | Plcd1 | 1.35 | 1.97E-55 | 4.09E-51 | 0.12 | 0.06 | 0.07 | 2.18 |
| ectoderm | Mcf2l | 1.04 | 2.10E-55 | 4.36E-51 | 0.12 | 0.05 | 0.07 | 2.24 |
| ectoderm | Ankrd6 | 1.08 | 2.69E-55 | 5.60E-51 | 0.17 | 0.09 | 0.08 | 1.91 |
| ectoderm | Trak1 | 0.83 | 3.05E-55 | 6.35E-51 | 0.19 | 0.1 | 0.09 | 1.88 |
| ectoderm | Rps27rt | 0.38 | 3.10E-55 | 6.45E-51 | 0.79 | 0.72 | 0.07 | 1.09 |
| ectoderm | Oard1 | 0.43 | 3.61E-55 | 7.50E-51 | 0.72 | 0.66 | 0.06 | 1.1 |
| ectoderm | Gm26917 | 0.61 | 5.71E-55 | 1.19E-50 | 0.62 | 0.54 | 0.09 | 1.16 |
| ectoderm | Tgif1 | 0.51 | 9.96E-55 | 2.07E-50 | 0.7 | 0.61 | 0.09 | 1.14 |
| ectoderm | Gadd45b | 1.21 | 1.02E-54 | 2.11E-50 | 0.21 | 0.12 | 0.09 | 1.71 |
| ectoderm | 230118H07Ri | 0.62 | 1.06E-54 | 2.21E-50 | 0.57 | 0.49 | 0.09 | 1.18 |
| ectoderm | Fgf13 | 0.9 | 1.10E-54 | 2.29E-50 | 0.18 | 0.09 | 0.08 | 1.91 |
| ectoderm | Atp13a2 | 0.64 | 1.14E-54 | 2.38E-50 | 0.35 | 0.23 | 0.12 | 1.5 |
| ectoderm | Wbp2 | 0.66 | 1.34E-54 | 2.79E-50 | 0.36 | 0.25 | 0.11 | 1.45 |
| ectoderm | Nr2c1 | 0.7 | 1.37E-54 | 2.86E-50 | 0.31 | 0.2 | 0.11 | 1.54 |
| ectoderm | Gipc1 | 0.37 | 1.41E-54 | 2.93E-50 | 0.75 | 0.67 | 0.08 | 1.13 |
| ectoderm | Gtf2i | 0.31 | 1.52E-54 | 3.15E-50 | 0.88 | 0.85 | 0.03 | 1.04 |
| ectoderm | Zfp113 | 0.58 | 2.55E-54 | 5.30E-50 | 0.42 | 0.3 | 0.12 | 1.41 |
| ectoderm | Phc2 | 0.4 | 2.62E-54 | 5.45E-50 | 0.75 | 0.67 | 0.08 | 1.11 |
| ectoderm | Kif5c | 0.48 | 2.91E-54 | 6.06E-50 | 0.58 | 0.46 | 0.12 | 1.25 |
| ectoderm | Syne4 | 0.72 | 5.78E-54 | 1.20E-49 | 0.28 | 0.17 | 0.11 | 1.62 |
| ectoderm | Adgrb3 | 1.11 | 6.95E-54 | 1.45E-49 | 0.11 | 0.05 | 0.06 | 2.28 |
| ectoderm | Marchf2 | 0.49 | 1.19E-53 | 2.48E-49 | 0.6 | 0.49 | 0.12 | 1.24 |
| ectoderm | Trim44 | 0.36 | 1.20E-53 | 2.50E-49 | 0.77 | 0.69 | 0.09 | 1.12 |
| ectoderm | Rap2c | 0.55 | 1.34E-53 | 2.78E-49 | 0.67 | 0.58 | 0.09 | 1.16 |
| ectoderm | Slc12a4 | 0.97 | 7.25E-53 | 1.51E-48 | 0.21 | 0.12 | 0.09 | 1.73 |
| ectoderm | 500011B03Ri | 0.36 | 7.50E-53 | 1.56E-48 | 0.73 | 0.65 | 0.08 | 1.13 |
| ectoderm | Cux2 | 0.77 | 7.74E-53 | 1.61E-48 | 0.16 | 0.08 | 0.08 | 1.99 |
| ectoderm | Nes | 0.58 | 8.80E-53 | 1.83E-48 | 0.5 | 0.4 | 0.1 | 1.25 |
| ectoderm | Suco | 0.59 | 1.01E-52 | 2.10E-48 | 0.57 | 0.47 | 0.1 | 1.21 |
| ectoderm | Tfrc | 0.48 | 1.11E-52 | 2.30E-48 | 0.49 | 0.37 | 0.12 | 1.34 |
| ectoderm | Mlec | 0.35 | 1.17E-52 | 2.43E-48 | 0.79 | 0.74 | 0.05 | 1.07 |
| ectoderm | Ist1 | 0.52 | 1.21E-52 | 2.51E-48 | 0.55 | 0.43 | 0.12 | 1.27 |
| ectoderm | Mvb12a | 0.51 | 1.47E-52 | 3.06E-48 | 0.54 | 0.43 | 0.11 | 1.26 |
| ectoderm | Ammecr1 | 0.57 | 1.51E-52 | 3.14E-48 | 0.44 | 0.33 | 0.12 | 1.36 |
| ectoderm | Eno1b | 0.61 | 1.52E-52 | 3.16E-48 | 0.44 | 0.32 | 0.12 | 1.38 |
| ectoderm | Zfp800 | 0.65 | 1.58E-52 | 3.29E-48 | 0.35 | 0.24 | 0.11 | 1.46 |
| ectoderm | Cab39 | 0.5 | 1.71E-52 | 3.56E-48 | 0.65 | 0.56 | 0.09 | 1.16 |
| ectoderm | Acadm | 0.61 | 1.89E-52 | 3.93E-48 | 0.42 | 0.31 | 0.12 | 1.38 |
| ectoderm | Cd59a | 0.9 | 1.91E-52 | 3.97E-48 | 0.3 | 0.2 | 0.1 | 1.52 |
| ectoderm | Tmx2 | 0.52 | 2.51E-52 | 5.22E-48 | 0.57 | 0.47 | 0.11 | 1.23 |
| ectoderm | Ostm1 | 1.08 | 2.79E-52 | 5.79E-48 | 0.12 | 0.06 | 0.07 | 2.16 |
| ectoderm | Sall2 | 0.63 | 3.26E-52 | 6.77E-48 | 0.48 | 0.37 | 0.11 | 1.31 |
| ectoderm | Rnf167 | 0.57 | 3.41E-52 | 7.09E-48 | 0.46 | 0.34 | 0.12 | 1.34 |
| ectoderm | H1f0 | 0.31 | 3.46E-52 | 7.19E-48 | 0.89 | 0.84 | 0.05 | 1.06 |
| ectoderm | 810044D09Ri | 0.98 | 3.71E-52 | 7.71E-48 | 0.17 | 0.09 | 0.08 | 1.85 |
| ectoderm | Dach1 | 0.48 | 4.64E-52 | 9.65E-48 | 0.48 | 0.36 | 0.13 | 1.36 |
| ectoderm | Ddhd1 | 0.85 | 6.70E-52 | 1.39E-47 | 0.24 | 0.15 | 0.09 | 1.63 |
| ectoderm | Sbno2 | 0.89 | 7.58E-52 | 1.58E-47 | 0.18 | 0.1 | 0.08 | 1.84 |
| ectoderm | Cep70 | 0.57 | 1.27E-51 | 2.64E-47 | 0.48 | 0.37 | 0.11 | 1.29 |
| ectoderm | Gm4890 | 1.06 | 1.92E-51 | 3.99E-47 | 0.12 | 0.06 | 0.06 | 2.16 |
| ectoderm | Fat3 | 1.14 | 2.86E-51 | 5.95E-47 | 0.15 | 0.08 | 0.07 | 1.96 |
| ectoderm | Fbxl20 | 0.59 | 4.10E-51 | 8.53E-47 | 0.3 | 0.19 | 0.11 | 1.56 |
| ectoderm | Sh3bp4 | 0.8 | 4.38E-51 | 9.11E-47 | 0.24 | 0.15 | 0.1 | 1.65 |
| ectoderm | Col4a5 | 0.78 | 5.99E-51 | 1.25E-46 | 0.2 | 0.11 | 0.09 | 1.8 |

**Supplementary Table 2. Marker genes for the major cell types of integrated clustering of E11.5 control and NotchGOF λ-junction cells.**

| | | | | | | | | |
|---|---|---|---|---|---|---|---|---|
| ectoderm | Bcl11a | 0.35 | 7.05E-51 | 1.47E-46 | 0.79 | 0.72 | 0.07 | 1.1 |
| ectoderm | Adam22 | 1.11 | 8.27E-51 | 1.72E-46 | 0.13 | 0.06 | 0.07 | 2.1 |
| ectoderm | Ing4 | 0.34 | 1.23E-50 | 2.56E-46 | 0.81 | 0.75 | 0.06 | 1.08 |
| ectoderm | Erc2 | 1.37 | 1.31E-50 | 2.72E-46 | 0.12 | 0.06 | 0.06 | 2.15 |
| ectoderm | Noc2l | 0.41 | 1.42E-50 | 2.96E-46 | 0.68 | 0.59 | 0.08 | 1.14 |
| ectoderm | Ubl3 | 0.5 | 1.64E-50 | 3.41E-46 | 0.46 | 0.34 | 0.12 | 1.34 |
| ectoderm | Dyrk3 | 1.1 | 1.78E-50 | 3.70E-46 | 0.16 | 0.08 | 0.07 | 1.89 |
| ectoderm | Rps6ka6 | 0.49 | 1.98E-50 | 4.12E-46 | 0.55 | 0.43 | 0.12 | 1.28 |
| ectoderm | F8a | 0.94 | 2.25E-50 | 4.69E-46 | 0.14 | 0.07 | 0.07 | 2.03 |
| ectoderm | Rabggta | 0.87 | 2.28E-50 | 4.75E-46 | 0.2 | 0.11 | 0.09 | 1.77 |
| ectoderm | Zswim6 | 0.59 | 2.47E-50 | 5.14E-46 | 0.38 | 0.27 | 0.12 | 1.43 |
| ectoderm | Spop | 0.4 | 2.56E-50 | 5.32E-46 | 0.74 | 0.68 | 0.06 | 1.09 |
| ectoderm | Tmem179b | 0.62 | 3.55E-50 | 7.38E-46 | 0.38 | 0.27 | 0.11 | 1.42 |
| ectoderm | Vezt | 0.53 | 4.63E-50 | 9.63E-46 | 0.41 | 0.3 | 0.12 | 1.4 |
| ectoderm | H2-D1 | 0.52 | 5.00E-50 | 1.04E-45 | 0.33 | 0.22 | 0.11 | 1.5 |
| ectoderm | Fryl | 0.65 | 5.15E-50 | 1.07E-45 | 0.39 | 0.28 | 0.11 | 1.4 |
| ectoderm | Chmp2a | 0.39 | 6.56E-50 | 1.36E-45 | 0.73 | 0.64 | 0.09 | 1.13 |
| ectoderm | Trp53inp1 | 0.87 | 7.61E-50 | 1.58E-45 | 0.26 | 0.16 | 0.1 | 1.58 |
| ectoderm | Atp11b | 0.5 | 7.71E-50 | 1.60E-45 | 0.51 | 0.4 | 0.12 | 1.29 |
| ectoderm | Lmln | 1.19 | 9.33E-50 | 1.94E-45 | 0.1 | 0.04 | 0.06 | 2.38 |
| ectoderm | Hebp2 | 0.9 | 9.44E-50 | 1.96E-45 | 0.32 | 0.22 | 0.1 | 1.47 |
| ectoderm | Zfp217 | 0.65 | 9.84E-50 | 2.05E-45 | 0.28 | 0.18 | 0.1 | 1.57 |
| ectoderm | Esrra | 0.94 | 1.07E-49 | 2.22E-45 | 0.19 | 0.11 | 0.08 | 1.77 |
| ectoderm | Cdkl5 | 1.18 | 1.08E-49 | 2.25E-45 | 0.1 | 0.04 | 0.06 | 2.35 |
| ectoderm | Bckdha | 0.58 | 1.09E-49 | 2.28E-45 | 0.46 | 0.34 | 0.12 | 1.35 |
| ectoderm | Zfp385a | 0.69 | 1.23E-49 | 2.57E-45 | 0.34 | 0.23 | 0.11 | 1.48 |
| ectoderm | Arhgap12 | 0.53 | 1.64E-49 | 3.41E-45 | 0.43 | 0.31 | 0.12 | 1.38 |
| ectoderm | Thtpa | 0.75 | 1.93E-49 | 4.01E-45 | 0.23 | 0.14 | 0.09 | 1.66 |
| ectoderm | Arl6ip1 | 0.27 | 3.01E-49 | 6.25E-45 | 0.91 | 0.88 | 0.03 | 1.03 |
| ectoderm | Zfand1 | 0.57 | 3.42E-49 | 7.11E-45 | 0.44 | 0.32 | 0.11 | 1.35 |
| ectoderm | Acrbp | 0.91 | 3.55E-49 | 7.39E-45 | 0.16 | 0.09 | 0.08 | 1.87 |
| ectoderm | Nsmaf | 0.68 | 5.38E-49 | 1.12E-44 | 0.35 | 0.24 | 0.11 | 1.44 |
| ectoderm | Hmga1 | 0.44 | 5.86E-49 | 1.22E-44 | 0.59 | 0.47 | 0.12 | 1.25 |
| ectoderm | Hunk | 0.45 | 9.10E-49 | 1.89E-44 | 0.59 | 0.47 | 0.12 | 1.25 |
| ectoderm | Smco4 | 0.35 | 9.82E-49 | 2.04E-44 | 0.59 | 0.46 | 0.13 | 1.28 |
| ectoderm | )610010F05Ri | 0.62 | 1.01E-48 | 2.11E-44 | 0.33 | 0.22 | 0.11 | 1.48 |
| ectoderm | Slc35d1 | 0.85 | 1.19E-48 | 2.48E-44 | 0.22 | 0.13 | 0.09 | 1.67 |
| ectoderm | Ankra2 | 0.58 | 1.20E-48 | 2.49E-44 | 0.4 | 0.29 | 0.11 | 1.39 |
| ectoderm | Gm42418 | 0.4 | 1.55E-48 | 3.22E-44 | 1 | 1 | 0 | 1 |
| ectoderm | Zc2hc1a | 0.5 | 1.70E-48 | 3.54E-44 | 0.51 | 0.39 | 0.12 | 1.3 |
| ectoderm | Ccpg1 | 0.8 | 1.78E-48 | 3.71E-44 | 0.23 | 0.14 | 0.09 | 1.64 |
| ectoderm | Slc16a3 | 0.6 | 1.81E-48 | 3.76E-44 | 0.41 | 0.3 | 0.11 | 1.37 |
| ectoderm | Lcor | 0.59 | 2.72E-48 | 5.65E-44 | 0.37 | 0.26 | 0.11 | 1.43 |
| ectoderm | Malt1 | 0.68 | 2.94E-48 | 6.11E-44 | 0.24 | 0.14 | 0.09 | 1.64 |
| ectoderm | 610306M01R | 0.72 | 4.18E-48 | 8.69E-44 | 0.23 | 0.14 | 0.09 | 1.65 |
| ectoderm | Mid1 | 0.65 | 4.19E-48 | 8.71E-44 | 0.33 | 0.22 | 0.11 | 1.49 |
| ectoderm | Slc16a1 | 0.68 | 4.22E-48 | 8.78E-44 | 0.38 | 0.28 | 0.1 | 1.37 |
| ectoderm | Snrpn | 0.45 | 4.85E-48 | 1.01E-43 | 0.56 | 0.44 | 0.11 | 1.26 |
| ectoderm | Cacng4 | 0.66 | 4.94E-48 | 1.03E-43 | 0.34 | 0.23 | 0.11 | 1.45 |
| ectoderm | Tvp23b | 0.49 | 5.63E-48 | 1.17E-43 | 0.5 | 0.38 | 0.12 | 1.31 |
| ectoderm | Map9 | 0.64 | 8.35E-48 | 1.74E-43 | 0.25 | 0.16 | 0.1 | 1.6 |
| ectoderm | Slc25a13 | 0.5 | 9.02E-48 | 1.88E-43 | 0.53 | 0.42 | 0.11 | 1.26 |
| ectoderm | Reps1 | 0.57 | 1.09E-47 | 2.27E-43 | 0.44 | 0.33 | 0.11 | 1.34 |
| ectoderm | Mindy2 | 0.77 | 1.48E-47 | 3.09E-43 | 0.25 | 0.15 | 0.09 | 1.6 |
| ectoderm | Itsn2 | 0.6 | 1.66E-47 | 3.45E-43 | 0.27 | 0.17 | 0.1 | 1.57 |
| ectoderm | Ppip5k2 | 0.45 | 1.73E-47 | 3.60E-43 | 0.54 | 0.43 | 0.11 | 1.25 |

**Supplementary Table 2. Marker genes for the major cell types of integrated clustering of E11.5 control and NotchGOF λ-junction cells.**

| | | | | | | | | |
|---|---|---|---|---|---|---|---|---|
| ectoderm | Exoc6 | 1.11 | 1.92E-47 | 3.99E-43 | 0.12 | 0.06 | 0.06 | 2.05 |
| ectoderm | 3110082I17Ril | 0.6 | 2.66E-47 | 5.54E-43 | 0.38 | 0.27 | 0.11 | 1.42 |
| ectoderm | Mboat2 | 0.54 | 2.71E-47 | 5.64E-43 | 0.44 | 0.32 | 0.12 | 1.36 |
| ectoderm | Dvl1 | 0.79 | 2.95E-47 | 6.14E-43 | 0.2 | 0.11 | 0.08 | 1.73 |
| ectoderm | Plekhb2 | 1.17 | 3.46E-47 | 7.19E-43 | 0.11 | 0.05 | 0.06 | 2.19 |
| ectoderm | Abca1 | 0.78 | 4.11E-47 | 8.54E-43 | 0.12 | 0.06 | 0.06 | 2.07 |
| ectoderm | Prrg3 | 0.78 | 4.13E-47 | 8.59E-43 | 0.17 | 0.09 | 0.08 | 1.82 |
| ectoderm | Znrd1as | 0.89 | 4.20E-47 | 8.74E-43 | 0.19 | 0.11 | 0.08 | 1.73 |
| ectoderm | Tmem47 | 0.78 | 5.73E-47 | 1.19E-42 | 0.45 | 0.34 | 0.11 | 1.31 |
| ectoderm | Sipa1l3 | 0.88 | 5.92E-47 | 1.23E-42 | 0.16 | 0.09 | 0.07 | 1.86 |
| ectoderm | Stk11 | 0.47 | 7.10E-47 | 1.48E-42 | 0.56 | 0.46 | 0.1 | 1.22 |
| ectoderm | Psen2 | 0.89 | 7.14E-47 | 1.48E-42 | 0.13 | 0.06 | 0.07 | 2.05 |
| ectoderm | Asl | 0.72 | 1.05E-46 | 2.18E-42 | 0.22 | 0.13 | 0.09 | 1.67 |
| ectoderm | Rnpep | 0.45 | 1.11E-46 | 2.31E-42 | 0.53 | 0.41 | 0.12 | 1.29 |
| ectoderm | Arvcf | 0.62 | 1.17E-46 | 2.44E-42 | 0.33 | 0.22 | 0.11 | 1.48 |
| ectoderm | Baiap2 | 1.05 | 1.88E-46 | 3.91E-42 | 0.22 | 0.14 | 0.08 | 1.61 |
| ectoderm | Rfxap | 0.49 | 2.95E-46 | 6.13E-42 | 0.6 | 0.51 | 0.09 | 1.18 |
| ectoderm | Afap1l1 | 1.17 | 3.13E-46 | 6.52E-42 | 0.12 | 0.06 | 0.06 | 2.05 |
| ectoderm | Akap12 | 0.6 | 4.34E-46 | 9.02E-42 | 0.27 | 0.17 | 0.1 | 1.55 |
| ectoderm | Msi2 | 0.36 | 4.81E-46 | 1.00E-41 | 0.73 | 0.66 | 0.07 | 1.1 |
| ectoderm | Garem1 | 0.89 | 6.10E-46 | 1.27E-41 | 0.14 | 0.07 | 0.07 | 1.96 |
| ectoderm | Xylt1 | 1.14 | 7.74E-46 | 1.61E-41 | 0.1 | 0.05 | 0.06 | 2.21 |
| ectoderm | Rnpepl1 | 0.58 | 1.19E-45 | 2.48E-41 | 0.34 | 0.23 | 0.11 | 1.46 |
| ectoderm | Tm7sf2 | 0.72 | 1.32E-45 | 2.74E-41 | 0.21 | 0.13 | 0.09 | 1.67 |
| ectoderm | Zc3h12c | 0.97 | 1.54E-45 | 3.20E-41 | 0.11 | 0.05 | 0.06 | 2.18 |
| ectoderm | AU040320 | 0.88 | 1.57E-45 | 3.27E-41 | 0.17 | 0.09 | 0.08 | 1.8 |
| ectoderm | Tamm41 | 0.55 | 2.12E-45 | 4.40E-41 | 0.43 | 0.32 | 0.11 | 1.35 |
| ectoderm | Dcaf6 | 1.01 | 2.61E-45 | 5.43E-41 | 0.13 | 0.07 | 0.07 | 1.97 |
| ectoderm | Ubxn2a | 0.54 | 2.70E-45 | 5.63E-41 | 0.4 | 0.29 | 0.11 | 1.37 |
| ectoderm | Slc37a3 | 0.57 | 3.00E-45 | 6.25E-41 | 0.35 | 0.24 | 0.11 | 1.46 |
| ectoderm | Tmem51 | 0.79 | 4.03E-45 | 8.39E-41 | 0.16 | 0.08 | 0.07 | 1.87 |
| ectoderm | Pigt | 0.48 | 4.19E-45 | 8.72E-41 | 0.51 | 0.4 | 0.11 | 1.27 |
| ectoderm | Kdm6b | 0.45 | 4.62E-45 | 9.60E-41 | 0.71 | 0.64 | 0.08 | 1.12 |
| ectoderm | Kcnk2 | 0.64 | 5.39E-45 | 1.12E-40 | 0.38 | 0.28 | 0.1 | 1.38 |
| ectoderm | Zfp949 | 0.81 | 5.72E-45 | 1.19E-40 | 0.18 | 0.1 | 0.08 | 1.75 |
| ectoderm | Iqsec1 | 0.78 | 7.84E-45 | 1.63E-40 | 0.2 | 0.12 | 0.08 | 1.69 |
| ectoderm | Pole3 | 0.35 | 8.49E-45 | 1.77E-40 | 0.79 | 0.75 | 0.04 | 1.05 |
| ectoderm | Ccdc173 | 0.74 | 1.04E-44 | 2.17E-40 | 0.18 | 0.1 | 0.08 | 1.76 |
| ectoderm | Ddx19b | 0.96 | 1.44E-44 | 2.99E-40 | 0.14 | 0.08 | 0.07 | 1.89 |
| ectoderm | 700037H04Ri | 0.48 | 1.67E-44 | 3.47E-40 | 0.56 | 0.45 | 0.11 | 1.23 |
| ectoderm | Tbc1d9 | 1.09 | 1.98E-44 | 4.12E-40 | 0.12 | 0.06 | 0.06 | 2 |
| ectoderm | Tgfb1 | 0.59 | 3.16E-44 | 6.57E-40 | 0.16 | 0.09 | 0.07 | 1.77 |
| ectoderm | AW146154 | 0.92 | 3.22E-44 | 6.70E-40 | 0.14 | 0.07 | 0.07 | 1.93 |
| ectoderm | Efs | 0.54 | 3.32E-44 | 6.90E-40 | 0.37 | 0.26 | 0.11 | 1.43 |
| ectoderm | Ptbp3 | 0.43 | 3.47E-44 | 7.22E-40 | 0.68 | 0.61 | 0.07 | 1.12 |
| ectoderm | Zfp664 | 0.31 | 3.57E-44 | 7.43E-40 | 0.8 | 0.73 | 0.06 | 1.09 |
| ectoderm | Smpdl3a | 0.8 | 4.44E-44 | 9.24E-40 | 0.19 | 0.11 | 0.08 | 1.7 |
| ectoderm | Ppil1 | 0.31 | 4.80E-44 | 9.99E-40 | 0.83 | 0.81 | 0.02 | 1.03 |
| ectoderm | Micu2 | 0.43 | 4.93E-44 | 1.03E-39 | 0.59 | 0.49 | 0.11 | 1.22 |
| ectoderm | Snhg14 | 0.65 | 5.07E-44 | 1.05E-39 | 0.25 | 0.16 | 0.09 | 1.57 |
| ectoderm | Fmn2 | 1.03 | 5.08E-44 | 1.06E-39 | 0.12 | 0.06 | 0.06 | 2.05 |
| ectoderm | Naa80 | 0.82 | 6.35E-44 | 1.32E-39 | 0.22 | 0.14 | 0.08 | 1.6 |
| ectoderm | Otud4 | 0.52 | 7.78E-44 | 1.62E-39 | 0.39 | 0.28 | 0.11 | 1.38 |
| ectoderm | Serpinb6a | 0.52 | 8.35E-44 | 1.74E-39 | 0.48 | 0.36 | 0.11 | 1.3 |
| ectoderm | Hsdl2 | 0.73 | 8.37E-44 | 1.74E-39 | 0.34 | 0.25 | 0.1 | 1.4 |
| ectoderm | Lyplal1 | 0.65 | 1.09E-43 | 2.27E-39 | 0.32 | 0.22 | 0.1 | 1.45 |

**Supplementary Table 2. Marker genes for the major cell types of integrated clustering of E11.5 control and NotchGOF λ-junction cells.**

| | | | | | | | | |
|---|---|---|---|---|---|---|---|---|
| ectoderm | Rnf6 | 0.52 | 1.25E-43 | 2.59E-39 | 0.44 | 0.33 | 0.11 | 1.34 |
| ectoderm | Tuba1c | 0.65 | 1.36E-43 | 2.84E-39 | 0.51 | 0.42 | 0.1 | 1.23 |
| ectoderm | Ralgapa2 | 0.77 | 1.39E-43 | 2.88E-39 | 0.2 | 0.12 | 0.08 | 1.68 |
| ectoderm | Crebrf | 0.89 | 2.46E-43 | 5.11E-39 | 0.17 | 0.09 | 0.07 | 1.77 |
| ectoderm | Ybx2 | 0.87 | 3.12E-43 | 6.48E-39 | 0.12 | 0.06 | 0.06 | 2.05 |
| ectoderm | Igsf8 | 0.57 | 3.37E-43 | 7.01E-39 | 0.32 | 0.22 | 0.1 | 1.47 |
| ectoderm | Mrps28 | 0.39 | 4.60E-43 | 9.57E-39 | 0.67 | 0.61 | 0.07 | 1.11 |
| ectoderm | Fam114a2 | 0.36 | 5.30E-43 | 1.10E-38 | 0.73 | 0.66 | 0.07 | 1.1 |
| ectoderm | Vps13a | 0.46 | 6.67E-43 | 1.39E-38 | 0.43 | 0.32 | 0.11 | 1.35 |
| ectoderm | Purb | 0.27 | 7.45E-43 | 1.55E-38 | 0.87 | 0.87 | 0 | 1 |
| ectoderm | )030056L22Ri | 0.54 | 8.07E-43 | 1.68E-38 | 0.55 | 0.46 | 0.09 | 1.19 |
| ectoderm | Ankhd1 | 0.47 | 9.06E-43 | 1.88E-38 | 0.57 | 0.47 | 0.1 | 1.22 |
| ectoderm | Pank1 | 0.54 | 1.04E-42 | 2.17E-38 | 0.4 | 0.29 | 0.11 | 1.37 |
| ectoderm | Trim37 | 0.5 | 1.66E-42 | 3.45E-38 | 0.47 | 0.37 | 0.11 | 1.29 |
| ectoderm | Lpar6 | 0.54 | 1.66E-42 | 3.45E-38 | 0.48 | 0.38 | 0.1 | 1.27 |
| ectoderm | Rdm1 | 0.54 | 2.36E-42 | 4.90E-38 | 0.4 | 0.3 | 0.11 | 1.36 |
| ectoderm | Pbxip1 | 0.66 | 2.49E-42 | 5.17E-38 | 0.28 | 0.18 | 0.09 | 1.52 |
| ectoderm | Exosc5 | 0.4 | 2.58E-42 | 5.36E-38 | 0.65 | 0.57 | 0.09 | 1.15 |
| ectoderm | Gstz1 | 0.59 | 3.06E-42 | 6.37E-38 | 0.35 | 0.25 | 0.1 | 1.41 |
| ectoderm | Ascc1 | 0.49 | 3.94E-42 | 8.19E-38 | 0.48 | 0.37 | 0.11 | 1.29 |
| ectoderm | Nr2c2ap | 0.56 | 3.99E-42 | 8.30E-38 | 0.31 | 0.21 | 0.1 | 1.47 |
| ectoderm | Kcnmb4 | 0.63 | 4.41E-42 | 9.18E-38 | 0.4 | 0.3 | 0.1 | 1.35 |
| ectoderm | Diablo | 0.34 | 4.92E-42 | 1.02E-37 | 0.69 | 0.61 | 0.08 | 1.13 |
| ectoderm | Atp9a | 0.6 | 5.52E-42 | 1.15E-37 | 0.25 | 0.16 | 0.09 | 1.57 |
| ectoderm | Vps4b | 0.39 | 5.86E-42 | 1.22E-37 | 0.69 | 0.61 | 0.08 | 1.13 |
| ectoderm | Stk4 | 0.5 | 6.65E-42 | 1.38E-37 | 0.44 | 0.34 | 0.11 | 1.32 |
| ectoderm | Fndc3b | 0.61 | 6.77E-42 | 1.41E-37 | 0.32 | 0.22 | 0.1 | 1.44 |
| ectoderm | Rnf11 | 0.39 | 8.80E-42 | 1.83E-37 | 0.68 | 0.61 | 0.07 | 1.11 |
| ectoderm | Map4k5 | 0.57 | 9.50E-42 | 1.98E-37 | 0.39 | 0.29 | 0.1 | 1.35 |
| ectoderm | Lbh | 0.81 | 1.02E-41 | 2.12E-37 | 0.27 | 0.18 | 0.09 | 1.48 |
| ectoderm | Mapk11 | 1.02 | 1.42E-41 | 2.96E-37 | 0.12 | 0.06 | 0.06 | 1.98 |
| ectoderm | Egln1 | 0.33 | 1.86E-41 | 3.86E-37 | 0.74 | 0.67 | 0.07 | 1.1 |
| ectoderm | Polr3h | 0.49 | 2.24E-41 | 4.65E-37 | 0.46 | 0.36 | 0.11 | 1.3 |
| ectoderm | Kifc3 | 1.09 | 2.27E-41 | 4.72E-37 | 0.12 | 0.06 | 0.06 | 1.97 |
| ectoderm | Apobec3 | 0.77 | 2.39E-41 | 4.97E-37 | 0.13 | 0.07 | 0.06 | 1.93 |
| ectoderm | Marchf7 | 0.33 | 2.83E-41 | 5.89E-37 | 0.76 | 0.72 | 0.05 | 1.06 |
| ectoderm | Ccdc191 | 0.59 | 3.11E-41 | 6.47E-37 | 0.29 | 0.2 | 0.1 | 1.48 |
| ectoderm | Etv3 | 0.7 | 5.40E-41 | 1.12E-36 | 0.29 | 0.2 | 0.09 | 1.45 |
| ectoderm | Atp2c1 | 0.46 | 6.80E-41 | 1.41E-36 | 0.48 | 0.38 | 0.11 | 1.28 |
| ectoderm | Camta1 | 0.37 | 7.17E-41 | 1.49E-36 | 0.68 | 0.6 | 0.07 | 1.12 |
| ectoderm | Stxbp6 | 0.68 | 7.80E-41 | 1.62E-36 | 0.2 | 0.12 | 0.08 | 1.66 |
| ectoderm | Slc45a4 | 0.76 | 1.17E-40 | 2.44E-36 | 0.2 | 0.12 | 0.08 | 1.67 |
| ectoderm | Igf2r | 0.45 | 1.35E-40 | 2.81E-36 | 0.57 | 0.48 | 0.1 | 1.2 |
| ectoderm | Ankzf1 | 0.58 | 1.62E-40 | 3.37E-36 | 0.29 | 0.2 | 0.1 | 1.49 |
| ectoderm | Plek2 | 0.93 | 3.28E-40 | 6.81E-36 | 0.1 | 0.05 | 0.05 | 2.1 |
| ectoderm | Ankrd13c | 0.62 | 3.45E-40 | 7.18E-36 | 0.31 | 0.22 | 0.09 | 1.42 |
| ectoderm | Scai | 0.49 | 4.35E-40 | 9.04E-36 | 0.43 | 0.33 | 0.11 | 1.32 |
| ectoderm | Zfp593 | 0.46 | 5.20E-40 | 1.08E-35 | 0.55 | 0.45 | 0.1 | 1.22 |
| ectoderm | Pgap2 | 0.42 | 5.44E-40 | 1.13E-35 | 0.53 | 0.43 | 0.1 | 1.24 |
| ectoderm | Ino80b | 0.49 | 5.89E-40 | 1.23E-35 | 0.53 | 0.43 | 0.1 | 1.22 |
| ectoderm | Mrpl44 | 0.48 | 6.17E-40 | 1.28E-35 | 0.49 | 0.39 | 0.1 | 1.26 |
| ectoderm | Kdsr | 0.57 | 6.51E-40 | 1.35E-35 | 0.35 | 0.25 | 0.1 | 1.4 |
| ectoderm | Txndc16 | 0.74 | 6.54E-40 | 1.36E-35 | 0.2 | 0.13 | 0.08 | 1.63 |
| ectoderm | Atp6ap2 | 0.48 | 7.13E-40 | 1.48E-35 | 0.5 | 0.4 | 0.1 | 1.25 |
| ectoderm | Mitd1 | 0.57 | 9.37E-40 | 1.95E-35 | 0.37 | 0.27 | 0.1 | 1.36 |
| ectoderm | Dcaf12l1 | 0.71 | 1.22E-39 | 2.53E-35 | 0.17 | 0.1 | 0.07 | 1.73 |

**Supplementary Table 2. Marker genes for the major cell types of integrated clustering of E11.5 control and NotchGOF λ-junction cells.**

| | | | | | | | | |
|---|---|---|---|---|---|---|---|---|
| ectoderm | Wtip | 0.39 | 1.41E-39 | 2.93E-35 | 0.63 | 0.54 | 0.09 | 1.17 |
| ectoderm | Pradc1 | 0.68 | 1.52E-39 | 3.16E-35 | 0.29 | 0.2 | 0.09 | 1.46 |
| ectoderm | 510002D24Ri | 0.45 | 2.55E-39 | 5.30E-35 | 0.48 | 0.37 | 0.11 | 1.28 |
| ectoderm | Gprc5b | 0.82 | 2.96E-39 | 6.17E-35 | 0.15 | 0.08 | 0.07 | 1.79 |
| ectoderm | Mif4gd | 0.64 | 3.66E-39 | 7.61E-35 | 0.23 | 0.15 | 0.08 | 1.55 |
| ectoderm | Nudt10 | 0.84 | 4.43E-39 | 9.22E-35 | 0.17 | 0.1 | 0.07 | 1.7 |
| ectoderm | Ptrh2 | 0.34 | 5.80E-39 | 1.21E-34 | 0.73 | 0.64 | 0.08 | 1.13 |
| ectoderm | C1qtnf12 | 0.75 | 5.93E-39 | 1.23E-34 | 0.23 | 0.15 | 0.08 | 1.55 |
| ectoderm | Isyna1 | 0.35 | 9.62E-39 | 2.00E-34 | 0.72 | 0.67 | 0.05 | 1.08 |
| ectoderm | Nr1h2 | 0.56 | 1.03E-38 | 2.14E-34 | 0.39 | 0.29 | 0.1 | 1.34 |
| ectoderm | Ahcyl2 | 0.57 | 1.05E-38 | 2.18E-34 | 0.37 | 0.27 | 0.1 | 1.36 |
| ectoderm | Borcs8 | 0.42 | 1.41E-38 | 2.93E-34 | 0.62 | 0.53 | 0.09 | 1.17 |
| ectoderm | Bag4 | 0.44 | 1.61E-38 | 3.35E-34 | 0.48 | 0.37 | 0.11 | 1.29 |
| ectoderm | Fahd2a | 0.61 | 2.03E-38 | 4.22E-34 | 0.26 | 0.17 | 0.09 | 1.49 |
| ectoderm | Celf4 | 0.58 | 2.46E-38 | 5.11E-34 | 0.3 | 0.21 | 0.09 | 1.44 |
| ectoderm | 410022M11R | 0.74 | 2.89E-38 | 6.02E-34 | 0.17 | 0.1 | 0.07 | 1.71 |
| ectoderm | Pisd | 0.57 | 3.39E-38 | 7.04E-34 | 0.28 | 0.19 | 0.09 | 1.48 |
| ectoderm | Rac3 | 0.5 | 3.56E-38 | 7.40E-34 | 0.41 | 0.31 | 0.1 | 1.33 |
| ectoderm | Rps6ka1 | 0.51 | 3.66E-38 | 7.61E-34 | 0.32 | 0.22 | 0.1 | 1.44 |
| ectoderm | Smox | 0.62 | 3.88E-38 | 8.07E-34 | 0.31 | 0.22 | 0.09 | 1.42 |
| ectoderm | Mttp | 0.66 | 4.14E-38 | 8.62E-34 | 0.19 | 0.12 | 0.08 | 1.65 |
| ectoderm | Nsun2 | 0.34 | 9.17E-38 | 1.91E-33 | 0.7 | 0.62 | 0.08 | 1.12 |
| ectoderm | Ccdc51 | 0.87 | 9.47E-38 | 1.97E-33 | 0.17 | 0.1 | 0.07 | 1.67 |
| ectoderm | Ahdc1 | 0.6 | 9.65E-38 | 2.01E-33 | 0.22 | 0.14 | 0.08 | 1.58 |
| ectoderm | Gm45051 | 0.85 | 1.10E-37 | 2.29E-33 | 0.12 | 0.06 | 0.06 | 1.92 |
| ectoderm | Nuak1 | 0.72 | 1.15E-37 | 2.39E-33 | 0.27 | 0.19 | 0.09 | 1.46 |
| ectoderm | Paip2b | 0.55 | 1.24E-37 | 2.59E-33 | 0.37 | 0.27 | 0.1 | 1.36 |
| ectoderm | Cep152 | 0.7 | 1.47E-37 | 3.05E-33 | 0.18 | 0.11 | 0.07 | 1.66 |
| ectoderm | Dgkz | 0.57 | 1.58E-37 | 3.29E-33 | 0.36 | 0.26 | 0.1 | 1.36 |
| ectoderm | Elk4 | 0.6 | 1.90E-37 | 3.96E-33 | 0.3 | 0.21 | 0.09 | 1.44 |
| ectoderm | Bclaf3 | 0.44 | 2.85E-37 | 5.93E-33 | 0.42 | 0.31 | 0.11 | 1.34 |
| ectoderm | Alcam | 0.42 | 3.50E-37 | 7.27E-33 | 0.21 | 0.13 | 0.08 | 1.62 |
| ectoderm | Pan3 | 0.5 | 3.89E-37 | 8.08E-33 | 0.41 | 0.31 | 0.1 | 1.33 |
| ectoderm | Tmem38a | 0.79 | 4.11E-37 | 8.56E-33 | 0.15 | 0.09 | 0.07 | 1.75 |
| ectoderm | Map3k11 | 0.71 | 4.77E-37 | 9.92E-33 | 0.17 | 0.1 | 0.07 | 1.7 |
| ectoderm | Zfp651 | 0.59 | 5.16E-37 | 1.07E-32 | 0.22 | 0.14 | 0.08 | 1.6 |
| ectoderm | Insr | 0.56 | 5.48E-37 | 1.14E-32 | 0.31 | 0.22 | 0.09 | 1.43 |
| ectoderm | Kidins220 | 0.54 | 5.78E-37 | 1.20E-32 | 0.35 | 0.26 | 0.1 | 1.37 |
| ectoderm | Krcc1 | 0.4 | 6.11E-37 | 1.27E-32 | 0.63 | 0.56 | 0.08 | 1.13 |
| ectoderm | Akr1b10 | 0.73 | 8.51E-37 | 1.77E-32 | 0.17 | 0.1 | 0.07 | 1.68 |
| ectoderm | Madd | 0.86 | 1.13E-36 | 2.36E-32 | 0.12 | 0.06 | 0.06 | 1.93 |
| ectoderm | Slc39a10 | 0.38 | 1.24E-36 | 2.58E-32 | 0.73 | 0.67 | 0.06 | 1.1 |
| ectoderm | Tspan14 | 0.47 | 1.27E-36 | 2.64E-32 | 0.41 | 0.3 | 0.11 | 1.35 |
| ectoderm | Mmp11 | 0.5 | 1.48E-36 | 3.09E-32 | 0.4 | 0.29 | 0.1 | 1.35 |
| ectoderm | Hibadh | 0.45 | 1.61E-36 | 3.36E-32 | 0.62 | 0.53 | 0.09 | 1.16 |
| ectoderm | Bbx | 0.32 | 1.77E-36 | 3.67E-32 | 0.66 | 0.56 | 0.1 | 1.17 |
| ectoderm | Mllt6 | 0.61 | 2.00E-36 | 4.17E-32 | 0.26 | 0.18 | 0.09 | 1.48 |
| ectoderm | Fam98c | 0.72 | 2.03E-36 | 4.21E-32 | 0.17 | 0.1 | 0.07 | 1.67 |
| ectoderm | Fam120a | 0.42 | 2.27E-36 | 4.72E-32 | 0.53 | 0.44 | 0.09 | 1.22 |
| ectoderm | Pdcd6 | 0.27 | 2.31E-36 | 4.80E-32 | 0.81 | 0.78 | 0.04 | 1.05 |
| ectoderm | Abhd17c | 0.47 | 2.78E-36 | 5.77E-32 | 0.48 | 0.38 | 0.1 | 1.26 |
| ectoderm | Mlycd | 0.63 | 3.03E-36 | 6.31E-32 | 0.2 | 0.13 | 0.08 | 1.61 |
| ectoderm | Zbtb8os | 0.42 | 3.65E-36 | 7.59E-32 | 0.57 | 0.48 | 0.09 | 1.19 |
| ectoderm | Glrx | 0.85 | 3.79E-36 | 7.89E-32 | 0.3 | 0.22 | 0.08 | 1.38 |
| ectoderm | Ero1l | 0.47 | 3.91E-36 | 8.13E-32 | 0.37 | 0.27 | 0.1 | 1.36 |
| ectoderm | Sh3glb2 | 0.47 | 3.93E-36 | 8.17E-32 | 0.4 | 0.3 | 0.1 | 1.34 |

**Supplementary Table 2. Marker genes for the major cell types of integrated clustering of E11.5 control and NotchGOF λ-junction cells.**

| | | | | | | | | |
|---|---|---|---|---|---|---|---|---|
| ectoderm | Cnot7 | 0.34 | 5.10E-36 | 1.06E-31 | 0.73 | 0.66 | 0.06 | 1.1 |
| ectoderm | Ppp1r13b | 0.77 | 5.71E-36 | 1.19E-31 | 0.13 | 0.07 | 0.06 | 1.87 |
| ectoderm | Hbp1 | 0.39 | 6.46E-36 | 1.34E-31 | 0.64 | 0.54 | 0.09 | 1.17 |
| ectoderm | Vps54 | 0.5 | 1.11E-35 | 2.32E-31 | 0.39 | 0.29 | 0.1 | 1.34 |
| ectoderm | Rnf38 | 0.56 | 1.21E-35 | 2.53E-31 | 0.35 | 0.25 | 0.09 | 1.37 |
| ectoderm | Dkk3 | 0.83 | 1.21E-35 | 2.53E-31 | 0.17 | 0.1 | 0.07 | 1.65 |
| ectoderm | Sema3a | 0.68 | 1.28E-35 | 2.66E-31 | 0.48 | 0.38 | 0.1 | 1.26 |
| ectoderm | Tmem245 | 0.46 | 1.29E-35 | 2.68E-31 | 0.48 | 0.38 | 0.1 | 1.25 |
| ectoderm | Col4a2 | 0.3 | 1.29E-35 | 2.68E-31 | 0.5 | 0.4 | 0.1 | 1.25 |
| ectoderm | Cgnl1 | 1.15 | 1.35E-35 | 2.80E-31 | 0.17 | 0.11 | 0.07 | 1.61 |
| ectoderm | Ddb2 | 0.81 | 1.66E-35 | 3.46E-31 | 0.15 | 0.08 | 0.06 | 1.75 |
| ectoderm | Abhd4 | 0.72 | 1.66E-35 | 3.46E-31 | 0.2 | 0.12 | 0.07 | 1.58 |
| ectoderm | Gm9856 | 0.59 | 1.98E-35 | 4.13E-31 | 0.21 | 0.14 | 0.08 | 1.57 |
| ectoderm | Flot2 | 0.58 | 2.22E-35 | 4.62E-31 | 0.28 | 0.19 | 0.09 | 1.44 |
| ectoderm | Dnajc21 | 0.37 | 2.39E-35 | 4.98E-31 | 0.65 | 0.58 | 0.07 | 1.12 |
| ectoderm | Sesn2 | 1.02 | 2.49E-35 | 5.17E-31 | 0.11 | 0.06 | 0.05 | 1.91 |
| ectoderm | Zfp458 | 0.96 | 2.65E-35 | 5.51E-31 | 0.1 | 0.05 | 0.05 | 2 |
| ectoderm | Rtkn2 | 0.82 | 2.76E-35 | 5.74E-31 | 0.12 | 0.06 | 0.06 | 1.88 |
| ectoderm | Tspan5 | 0.49 | 3.06E-35 | 6.37E-31 | 0.34 | 0.25 | 0.1 | 1.39 |
| ectoderm | Tab3 | 0.83 | 3.52E-35 | 7.33E-31 | 0.13 | 0.08 | 0.06 | 1.79 |
| ectoderm | Per3 | 0.69 | 3.72E-35 | 7.74E-31 | 0.15 | 0.09 | 0.06 | 1.74 |
| ectoderm | Zfp568 | 0.59 | 3.82E-35 | 7.95E-31 | 0.34 | 0.25 | 0.09 | 1.36 |
| ectoderm | Sgpl1 | 0.49 | 3.98E-35 | 8.28E-31 | 0.42 | 0.32 | 0.1 | 1.32 |
| ectoderm | Kif16b | 0.62 | 4.03E-35 | 8.37E-31 | 0.21 | 0.14 | 0.08 | 1.56 |
| ectoderm | Chmp1b | 0.49 | 4.72E-35 | 9.83E-31 | 0.43 | 0.33 | 0.1 | 1.29 |
| ectoderm | Arhgef12 | 0.36 | 5.23E-35 | 1.09E-30 | 0.64 | 0.56 | 0.08 | 1.13 |
| ectoderm | Zc3h6 | 0.74 | 7.46E-35 | 1.55E-30 | 0.13 | 0.08 | 0.06 | 1.79 |
| ectoderm | Rcan3 | 0.82 | 8.15E-35 | 1.69E-30 | 0.12 | 0.07 | 0.06 | 1.85 |
| ectoderm | Phtf2 | 0.62 | 8.41E-35 | 1.75E-30 | 0.26 | 0.18 | 0.08 | 1.46 |
| ectoderm | Foxn3 | 0.5 | 1.54E-34 | 3.20E-30 | 0.72 | 0.68 | 0.04 | 1.06 |
| ectoderm | Ly75 | 0.6 | 1.55E-34 | 3.22E-30 | 0.21 | 0.14 | 0.08 | 1.55 |
| ectoderm | Mmgt2 | 0.39 | 1.97E-34 | 4.10E-30 | 0.52 | 0.41 | 0.1 | 1.25 |
| ectoderm | Rffl | 1.1 | 2.50E-34 | 5.21E-30 | 0.11 | 0.06 | 0.05 | 1.85 |
| ectoderm | Hps1 | 0.66 | 2.73E-34 | 5.69E-30 | 0.18 | 0.11 | 0.07 | 1.62 |
| ectoderm | Tmem42 | 0.63 | 2.87E-34 | 5.98E-30 | 0.21 | 0.14 | 0.08 | 1.56 |
| ectoderm | Entpd5 | 0.79 | 2.88E-34 | 5.99E-30 | 0.12 | 0.07 | 0.06 | 1.85 |
| ectoderm | Pmm1 | 0.59 | 3.03E-34 | 6.30E-30 | 0.34 | 0.25 | 0.09 | 1.35 |
| ectoderm | Tshz2 | 0.56 | 3.22E-34 | 6.69E-30 | 0.41 | 0.32 | 0.09 | 1.29 |
| ectoderm | Agap3 | 0.51 | 3.45E-34 | 7.18E-30 | 0.43 | 0.34 | 0.1 | 1.28 |
| ectoderm | Ddi2 | 0.53 | 3.49E-34 | 7.26E-30 | 0.48 | 0.39 | 0.09 | 1.23 |
| ectoderm | Tmed5 | 0.43 | 4.48E-34 | 9.31E-30 | 0.55 | 0.47 | 0.09 | 1.19 |
| ectoderm | Lin28b | 0.48 | 4.55E-34 | 9.46E-30 | 0.28 | 0.19 | 0.09 | 1.47 |
| ectoderm | Irf2bpl | 0.51 | 5.05E-34 | 1.05E-29 | 0.46 | 0.36 | 0.09 | 1.26 |
| ectoderm | E2f8 | 0.68 | 5.57E-34 | 1.16E-29 | 0.27 | 0.19 | 0.08 | 1.43 |
| ectoderm | Fgd6 | 0.7 | 6.02E-34 | 1.25E-29 | 0.22 | 0.15 | 0.08 | 1.52 |
| ectoderm | Bcl2l1 | 0.44 | 7.51E-34 | 1.56E-29 | 0.44 | 0.35 | 0.09 | 1.27 |
| ectoderm | Kantr | 0.57 | 9.84E-34 | 2.05E-29 | 0.23 | 0.15 | 0.08 | 1.52 |
| ectoderm | Luzp1 | 0.3 | 1.04E-33 | 2.16E-29 | 0.81 | 0.76 | 0.05 | 1.06 |
| ectoderm | Smad3 | 0.47 | 1.21E-33 | 2.51E-29 | 0.34 | 0.25 | 0.09 | 1.38 |
| ectoderm | Lrig3 | 0.32 | 1.35E-33 | 2.80E-29 | 0.3 | 0.21 | 0.09 | 1.45 |
| ectoderm | Cib1 | 0.43 | 1.37E-33 | 2.85E-29 | 0.48 | 0.38 | 0.1 | 1.25 |
| ectoderm | Plxnb2 | 0.41 | 1.40E-33 | 2.92E-29 | 0.6 | 0.51 | 0.09 | 1.17 |
| ectoderm | Spred3 | 0.62 | 1.46E-33 | 3.04E-29 | 0.14 | 0.08 | 0.06 | 1.75 |
| ectoderm | .190007I07Ril | 0.56 | 1.56E-33 | 3.24E-29 | 0.35 | 0.26 | 0.09 | 1.35 |
| ectoderm | Cyb561d2 | 0.58 | 2.04E-33 | 4.25E-29 | 0.23 | 0.15 | 0.08 | 1.51 |
| ectoderm | Axl | 0.53 | 2.12E-33 | 4.41E-29 | 0.21 | 0.14 | 0.08 | 1.56 |

**Supplementary Table 2. Marker genes for the major cell types of integrated clustering of E11.5 control and NotchGOF λ-junction cells.**

| | | | | | | | | |
|---|---|---|---|---|---|---|---|---|
| ectoderm | Prr5 | 0.59 | 2.18E-33 | 4.54E-29 | 0.17 | 0.1 | 0.07 | 1.64 |
| ectoderm | Gpbp1l1 | 0.5 | 2.31E-33 | 4.80E-29 | 0.46 | 0.36 | 0.09 | 1.26 |
| ectoderm | Oxsr1 | 0.51 | 2.53E-33 | 5.25E-29 | 0.39 | 0.3 | 0.09 | 1.31 |
| ectoderm | Ankrd37 | 0.68 | 3.30E-33 | 6.85E-29 | 0.22 | 0.14 | 0.07 | 1.52 |
| ectoderm | Ipmk | 0.68 | 3.35E-33 | 6.97E-29 | 0.25 | 0.17 | 0.08 | 1.46 |
| ectoderm | Hmga1b | 0.39 | 4.41E-33 | 9.18E-29 | 0.52 | 0.41 | 0.1 | 1.25 |
| ectoderm | Rhou | 0.43 | 4.53E-33 | 9.41E-29 | 0.49 | 0.39 | 0.1 | 1.25 |
| ectoderm | 110032A03Ri | 0.5 | 6.08E-33 | 1.26E-28 | 0.4 | 0.3 | 0.1 | 1.33 |
| ectoderm | Stxbp3 | 0.58 | 6.32E-33 | 1.32E-28 | 0.3 | 0.22 | 0.09 | 1.39 |
| ectoderm | Usp31 | 0.76 | 6.37E-33 | 1.32E-28 | 0.16 | 0.1 | 0.06 | 1.66 |
| ectoderm | Tmem87b | 0.55 | 6.58E-33 | 1.37E-28 | 0.28 | 0.2 | 0.08 | 1.43 |
| ectoderm | Maf1 | 0.37 | 7.43E-33 | 1.54E-28 | 0.6 | 0.51 | 0.09 | 1.17 |
| ectoderm | Fam241b | 0.59 | 8.39E-33 | 1.74E-28 | 0.23 | 0.15 | 0.08 | 1.52 |
| ectoderm | Mterf1a | 0.82 | 9.91E-33 | 2.06E-28 | 0.13 | 0.07 | 0.06 | 1.79 |
| ectoderm | Ccnd2 | 0.82 | 1.00E-32 | 2.08E-28 | 0.81 | 0.87 | -0.06 | 0.93 |
| ectoderm | Hdhd3 | 0.66 | 1.09E-32 | 2.27E-28 | 0.22 | 0.15 | 0.07 | 1.5 |
| ectoderm | Abcb8 | 0.54 | 1.12E-32 | 2.34E-28 | 0.24 | 0.16 | 0.08 | 1.49 |
| ectoderm | Pla2g6 | 0.87 | 1.22E-32 | 2.53E-28 | 0.12 | 0.07 | 0.05 | 1.78 |
| ectoderm | Hivep2 | 0.65 | 1.24E-32 | 2.57E-28 | 0.12 | 0.06 | 0.05 | 1.84 |
| ectoderm | Erlin2 | 0.45 | 1.28E-32 | 2.67E-28 | 0.4 | 0.3 | 0.1 | 1.31 |
| ectoderm | Shoc2 | 0.41 | 1.53E-32 | 3.18E-28 | 0.53 | 0.44 | 0.09 | 1.21 |
| ectoderm | Zfp53 | 0.87 | 1.72E-32 | 3.58E-28 | 0.13 | 0.08 | 0.06 | 1.75 |
| ectoderm | Rbbp9 | 0.45 | 1.87E-32 | 3.89E-28 | 0.47 | 0.37 | 0.1 | 1.26 |
| ectoderm | Gas2l3 | 0.5 | 2.06E-32 | 4.29E-28 | 0.24 | 0.17 | 0.08 | 1.47 |
| ectoderm | Ppp1r12b | 1.02 | 2.07E-32 | 4.31E-28 | 0.16 | 0.1 | 0.06 | 1.62 |
| ectoderm | Trit1 | 0.63 | 2.73E-32 | 5.68E-28 | 0.26 | 0.18 | 0.08 | 1.44 |
| ectoderm | Nras | 0.3 | 2.88E-32 | 5.99E-28 | 0.74 | 0.68 | 0.06 | 1.09 |
| ectoderm | Bmf | 0.62 | 3.52E-32 | 7.31E-28 | 0.16 | 0.09 | 0.06 | 1.68 |
| ectoderm | Nudt7 | 0.89 | 3.75E-32 | 7.80E-28 | 0.11 | 0.06 | 0.05 | 1.88 |
| ectoderm | Insig2 | 0.49 | 3.93E-32 | 8.16E-28 | 0.34 | 0.25 | 0.09 | 1.36 |
| ectoderm | Tox3 | 0.45 | 4.53E-32 | 9.43E-28 | 0.3 | 0.21 | 0.09 | 1.4 |
| ectoderm | Acyp1 | 0.44 | 5.21E-32 | 1.08E-27 | 0.5 | 0.41 | 0.09 | 1.22 |
| ectoderm | Rnf149 | 0.66 | 5.27E-32 | 1.10E-27 | 0.29 | 0.2 | 0.08 | 1.4 |
| ectoderm | Iqcb1 | 0.32 | 5.85E-32 | 1.22E-27 | 0.6 | 0.5 | 0.1 | 1.19 |
| ectoderm | Dennd2c | 1.27 | 6.47E-32 | 1.35E-27 | 0.1 | 0.05 | 0.05 | 1.89 |
| ectoderm | Myc | 0.73 | 6.55E-32 | 1.36E-27 | 0.22 | 0.15 | 0.07 | 1.49 |
| ectoderm | Tmem14a | 0.71 | 6.58E-32 | 1.37E-27 | 0.15 | 0.09 | 0.06 | 1.69 |
| ectoderm | Slc35a3 | 0.67 | 6.59E-32 | 1.37E-27 | 0.18 | 0.11 | 0.07 | 1.62 |
| ectoderm | Anxa4 | 0.76 | 6.76E-32 | 1.41E-27 | 0.16 | 0.1 | 0.06 | 1.65 |
| ectoderm | Fam129b | 0.7 | 7.15E-32 | 1.49E-27 | 0.2 | 0.13 | 0.07 | 1.54 |
| ectoderm | Rfx3 | 0.54 | 7.96E-32 | 1.66E-27 | 0.28 | 0.19 | 0.08 | 1.44 |
| ectoderm | Hsdl1 | 0.61 | 1.05E-31 | 2.18E-27 | 0.21 | 0.14 | 0.07 | 1.51 |
| ectoderm | 933434E20Ri | 0.39 | 1.19E-31 | 2.47E-27 | 0.56 | 0.48 | 0.09 | 1.19 |
| ectoderm | Tmem19 | 0.56 | 1.24E-31 | 2.58E-27 | 0.26 | 0.18 | 0.08 | 1.46 |
| ectoderm | Grik5 | 0.48 | 1.29E-31 | 2.68E-27 | 0.22 | 0.14 | 0.08 | 1.54 |
| ectoderm | Wdr89 | 0.26 | 2.17E-31 | 4.52E-27 | 0.8 | 0.76 | 0.04 | 1.05 |
| ectoderm | Tmem250-ps | 0.69 | 2.29E-31 | 4.76E-27 | 0.2 | 0.13 | 0.07 | 1.54 |
| ectoderm | Aldh4a1 | 0.78 | 2.52E-31 | 5.24E-27 | 0.14 | 0.08 | 0.06 | 1.7 |
| ectoderm | Fbxo10 | 0.78 | 2.52E-31 | 5.25E-27 | 0.12 | 0.07 | 0.05 | 1.8 |
| ectoderm | Ncstn | 0.45 | 3.32E-31 | 6.90E-27 | 0.36 | 0.27 | 0.09 | 1.34 |
| ectoderm | Raver2 | 0.59 | 3.98E-31 | 8.27E-27 | 0.22 | 0.15 | 0.07 | 1.5 |
| ectoderm | Tgif2 | 0.42 | 4.13E-31 | 8.59E-27 | 0.53 | 0.44 | 0.09 | 1.21 |
| ectoderm | Naf1 | 0.4 | 4.16E-31 | 8.66E-27 | 0.47 | 0.38 | 0.09 | 1.25 |
| ectoderm | Cenpb | 0.34 | 7.50E-31 | 1.56E-26 | 0.7 | 0.65 | 0.05 | 1.07 |
| ectoderm | Sema4b | 0.78 | 8.19E-31 | 1.70E-26 | 0.13 | 0.08 | 0.06 | 1.74 |
| ectoderm | Mplkip | 0.29 | 8.25E-31 | 1.72E-26 | 0.72 | 0.66 | 0.06 | 1.09 |

**Supplementary Table 2. Marker genes for the major cell types of integrated clustering of E11.5 control and NotchGOF λ-junction cells.**

| | | | | | | | | |
|---|---|---|---|---|---|---|---|---|
| ectoderm | Pard6g | 0.34 | 8.51E-31 | 1.77E-26 | 0.66 | 0.58 | 0.08 | 1.14 |
| ectoderm | Wrnip1 | 0.56 | 8.68E-31 | 1.81E-26 | 0.26 | 0.18 | 0.08 | 1.44 |
| ectoderm | Cask | 0.46 | 8.88E-31 | 1.85E-26 | 0.41 | 0.32 | 0.09 | 1.29 |
| ectoderm | Kctd1 | 0.55 | 1.03E-30 | 2.14E-26 | 0.52 | 0.45 | 0.07 | 1.15 |
| ectoderm | Polr3g | 0.56 | 1.10E-30 | 2.28E-26 | 0.34 | 0.26 | 0.08 | 1.32 |
| ectoderm | Rrp7a | 0.37 | 1.23E-30 | 2.56E-26 | 0.55 | 0.46 | 0.09 | 1.2 |
| ectoderm | C2cd3 | 0.42 | 1.24E-30 | 2.58E-26 | 0.38 | 0.29 | 0.09 | 1.33 |
| ectoderm | Arhgap32 | 0.53 | 1.30E-30 | 2.69E-26 | 0.25 | 0.17 | 0.08 | 1.45 |
| ectoderm | Myo1e | 0.51 | 1.34E-30 | 2.78E-26 | 0.3 | 0.22 | 0.09 | 1.4 |
| ectoderm | 700012D14Ri | 0.65 | 1.54E-30 | 3.21E-26 | 0.18 | 0.12 | 0.07 | 1.57 |
| ectoderm | Cdk17 | 0.53 | 2.20E-30 | 4.58E-26 | 0.27 | 0.19 | 0.08 | 1.42 |
| ectoderm | Hlcs | 0.6 | 2.37E-30 | 4.93E-26 | 0.18 | 0.12 | 0.07 | 1.57 |
| ectoderm | Zfp316 | 0.6 | 2.49E-30 | 5.17E-26 | 0.2 | 0.13 | 0.07 | 1.54 |
| ectoderm | Aspscr1 | 0.5 | 2.73E-30 | 5.67E-26 | 0.52 | 0.44 | 0.08 | 1.18 |
| ectoderm | Arhgap29 | 0.76 | 3.31E-30 | 6.88E-26 | 0.42 | 0.35 | 0.07 | 1.2 |
| ectoderm | Pex16 | 0.6 | 3.45E-30 | 7.18E-26 | 0.24 | 0.17 | 0.08 | 1.45 |
| ectoderm | Rtl6 | 0.61 | 3.80E-30 | 7.90E-26 | 0.16 | 0.1 | 0.06 | 1.63 |
| ectoderm | Gpalpp1 | 0.48 | 3.84E-30 | 7.99E-26 | 0.43 | 0.34 | 0.09 | 1.27 |
| ectoderm | Sirt7 | 0.59 | 3.99E-30 | 8.29E-26 | 0.23 | 0.16 | 0.07 | 1.47 |
| ectoderm | Jmjd1c | 0.36 | 4.23E-30 | 8.80E-26 | 0.56 | 0.46 | 0.1 | 1.21 |
| ectoderm | Arhgef28 | 0.75 | 5.39E-30 | 1.12E-25 | 0.15 | 0.09 | 0.06 | 1.66 |
| ectoderm | Rrp12 | 0.61 | 6.63E-30 | 1.38E-25 | 0.26 | 0.18 | 0.08 | 1.42 |
| ectoderm | Emp1 | 0.92 | 6.68E-30 | 1.39E-25 | 0.18 | 0.12 | 0.06 | 1.53 |
| ectoderm | Rnf145 | 0.43 | 8.56E-30 | 1.78E-25 | 0.44 | 0.35 | 0.09 | 1.26 |
| ectoderm | Zadh2 | 0.73 | 1.06E-29 | 2.20E-25 | 0.15 | 0.09 | 0.06 | 1.63 |
| ectoderm | Sp4 | 0.49 | 1.17E-29 | 2.42E-25 | 0.3 | 0.22 | 0.08 | 1.38 |
| ectoderm | Dlg3 | 0.47 | 1.19E-29 | 2.48E-25 | 0.39 | 0.3 | 0.09 | 1.31 |
| ectoderm | Gm20186 | 0.86 | 1.35E-29 | 2.80E-25 | 0.1 | 0.06 | 0.05 | 1.87 |
| ectoderm | Polr2h | 0.25 | 1.54E-29 | 3.20E-25 | 0.81 | 0.78 | 0.03 | 1.04 |
| ectoderm | Dtd1 | 0.41 | 1.86E-29 | 3.88E-25 | 0.49 | 0.39 | 0.09 | 1.24 |
| ectoderm | Prelid2 | 0.47 | 2.03E-29 | 4.22E-25 | 0.44 | 0.35 | 0.09 | 1.25 |
| ectoderm | Zfp652 | 0.34 | 2.36E-29 | 4.91E-25 | 0.58 | 0.49 | 0.09 | 1.17 |
| ectoderm | Map3k1 | 0.36 | 2.44E-29 | 5.07E-25 | 0.33 | 0.24 | 0.09 | 1.37 |
| ectoderm | Tradd | 0.75 | 3.34E-29 | 6.95E-25 | 0.16 | 0.1 | 0.06 | 1.58 |
| ectoderm | Qars | 0.35 | 3.53E-29 | 7.35E-25 | 0.65 | 0.58 | 0.07 | 1.12 |
| ectoderm | Stim2 | 0.5 | 4.22E-29 | 8.79E-25 | 0.3 | 0.22 | 0.08 | 1.38 |
| ectoderm | Ttc5 | 0.32 | 4.49E-29 | 9.33E-25 | 0.61 | 0.52 | 0.09 | 1.17 |
| ectoderm | Csnk1g1 | 0.42 | 4.87E-29 | 1.01E-24 | 0.43 | 0.34 | 0.09 | 1.26 |
| ectoderm | Rab21 | 0.25 | 5.90E-29 | 1.23E-24 | 0.54 | 0.45 | 0.1 | 1.21 |
| ectoderm | Atg101 | 0.38 | 6.66E-29 | 1.38E-24 | 0.56 | 0.47 | 0.08 | 1.18 |
| ectoderm | Rrp36 | 0.46 | 8.22E-29 | 1.71E-24 | 0.39 | 0.3 | 0.09 | 1.31 |
| ectoderm | Sephs2 | 0.42 | 9.28E-29 | 1.93E-24 | 0.49 | 0.4 | 0.08 | 1.2 |
| ectoderm | M6pr | 0.38 | 1.11E-28 | 2.31E-24 | 0.51 | 0.42 | 0.09 | 1.21 |
| ectoderm | Klhl2 | 0.66 | 1.16E-28 | 2.41E-24 | 0.17 | 0.11 | 0.06 | 1.57 |
| ectoderm | Shb | 0.55 | 1.32E-28 | 2.74E-24 | 0.25 | 0.18 | 0.07 | 1.41 |
| ectoderm | Mbnl3 | 0.6 | 1.35E-28 | 2.81E-24 | 0.18 | 0.11 | 0.06 | 1.57 |
| ectoderm | Fam172a | 0.42 | 1.38E-28 | 2.86E-24 | 0.42 | 0.33 | 0.09 | 1.28 |
| ectoderm | Rpia | 0.53 | 1.43E-28 | 2.98E-24 | 0.36 | 0.28 | 0.08 | 1.29 |
| ectoderm | Rabif | 0.47 | 1.66E-28 | 3.46E-24 | 0.31 | 0.23 | 0.08 | 1.35 |
| ectoderm | Tyro3 | 0.51 | 1.77E-28 | 3.69E-24 | 0.23 | 0.16 | 0.07 | 1.47 |
| ectoderm | Klhl7 | 0.37 | 2.27E-28 | 4.72E-24 | 0.58 | 0.5 | 0.09 | 1.17 |
| ectoderm | Tmub1 | 0.48 | 2.35E-28 | 4.90E-24 | 0.26 | 0.18 | 0.08 | 1.43 |
| ectoderm | Pex26 | 0.71 | 2.93E-28 | 6.09E-24 | 0.12 | 0.07 | 0.05 | 1.71 |
| ectoderm | Slc38a10 | 0.38 | 3.42E-28 | 7.11E-24 | 0.41 | 0.32 | 0.09 | 1.29 |
| ectoderm | Hipk2 | 0.31 | 4.28E-28 | 8.91E-24 | 0.68 | 0.64 | 0.04 | 1.07 |
| ectoderm | Ak3 | 0.41 | 4.43E-28 | 9.21E-24 | 0.45 | 0.36 | 0.09 | 1.24 |

**Supplementary Table 2. Marker genes for the major cell types of integrated clustering of E11.5 control and NotchGOF λ-junction cells.**

| | | | | | | | | |
|---|---|---|---|---|---|---|---|---|
| ectoderm | Ngrn | 0.35 | 4.56E-28 | 9.49E-24 | 0.53 | 0.45 | 0.08 | 1.19 |
| ectoderm | Zfp667 | 0.69 | 4.64E-28 | 9.66E-24 | 0.14 | 0.09 | 0.06 | 1.64 |
| ectoderm | Map4k3 | 0.46 | 5.22E-28 | 1.09E-23 | 0.35 | 0.26 | 0.09 | 1.33 |
| ectoderm | Tmem170 | 0.7 | 5.28E-28 | 1.10E-23 | 0.14 | 0.08 | 0.05 | 1.65 |
| ectoderm | Il10rb | 0.8 | 5.36E-28 | 1.11E-23 | 0.13 | 0.07 | 0.05 | 1.7 |
| ectoderm | Tmem192 | 0.56 | 5.70E-28 | 1.19E-23 | 0.24 | 0.17 | 0.07 | 1.44 |
| ectoderm | Dph6 | 0.39 | 6.52E-28 | 1.36E-23 | 0.48 | 0.39 | 0.09 | 1.23 |
| ectoderm | Cope | 0.27 | 7.08E-28 | 1.47E-23 | 0.79 | 0.76 | 0.03 | 1.03 |
| ectoderm | BC005537 | 0.35 | 8.87E-28 | 1.84E-23 | 0.56 | 0.48 | 0.08 | 1.18 |
| ectoderm | Mterf4 | 0.42 | 8.95E-28 | 1.86E-23 | 0.41 | 0.32 | 0.09 | 1.28 |
| ectoderm | Fgfr1op2 | 0.31 | 9.48E-28 | 1.97E-23 | 0.72 | 0.67 | 0.05 | 1.08 |
| ectoderm | Fbxo16 | 0.63 | 1.04E-27 | 2.15E-23 | 0.15 | 0.1 | 0.06 | 1.62 |
| ectoderm | Mcrip2 | 0.77 | 1.19E-27 | 2.48E-23 | 0.11 | 0.06 | 0.05 | 1.78 |
| ectoderm | Rnf114 | 0.4 | 1.36E-27 | 2.83E-23 | 0.54 | 0.46 | 0.08 | 1.18 |
| ectoderm | Cep164 | 0.34 | 1.48E-27 | 3.08E-23 | 0.67 | 0.6 | 0.07 | 1.12 |
| ectoderm | Pgpep1 | 0.51 | 1.64E-27 | 3.41E-23 | 0.35 | 0.27 | 0.08 | 1.3 |
| ectoderm | C1galt1c1 | 0.47 | 1.73E-27 | 3.59E-23 | 0.22 | 0.14 | 0.07 | 1.49 |
| ectoderm | Stxbp5 | 0.68 | 2.06E-27 | 4.28E-23 | 0.12 | 0.07 | 0.05 | 1.74 |
| ectoderm | .210016F16Ri | 0.45 | 2.16E-27 | 4.50E-23 | 0.42 | 0.33 | 0.08 | 1.25 |
| ectoderm | Chmp1a | 0.38 | 2.25E-27 | 4.69E-23 | 0.49 | 0.41 | 0.09 | 1.21 |
| ectoderm | Sdf2 | 0.29 | 2.60E-27 | 5.42E-23 | 0.62 | 0.54 | 0.08 | 1.14 |
| ectoderm | Arhgap21 | 0.53 | 2.76E-27 | 5.75E-23 | 0.28 | 0.2 | 0.08 | 1.37 |
| ectoderm | Frs2 | 0.57 | 3.73E-27 | 7.75E-23 | 0.34 | 0.26 | 0.08 | 1.3 |
| ectoderm | Ap1ar | 0.34 | 4.06E-27 | 8.45E-23 | 0.61 | 0.54 | 0.07 | 1.14 |
| ectoderm | Disp1 | 0.67 | 4.49E-27 | 9.34E-23 | 0.15 | 0.1 | 0.06 | 1.6 |
| ectoderm | Fbxo25 | 0.62 | 5.24E-27 | 1.09E-22 | 0.2 | 0.14 | 0.07 | 1.47 |
| ectoderm | Nsun3 | 0.76 | 5.61E-27 | 1.17E-22 | 0.11 | 0.07 | 0.05 | 1.74 |
| ectoderm | Fbxw7 | 0.42 | 5.70E-27 | 1.19E-22 | 0.45 | 0.36 | 0.09 | 1.23 |
| ectoderm | C1qtnf4 | 0.46 | 6.83E-27 | 1.42E-22 | 0.36 | 0.27 | 0.09 | 1.32 |
| ectoderm | Slc20a2 | 0.45 | 7.15E-27 | 1.49E-22 | 0.3 | 0.22 | 0.08 | 1.37 |
| ectoderm | Rab6a | 0.28 | 7.63E-27 | 1.59E-22 | 0.75 | 0.71 | 0.04 | 1.06 |
| ectoderm | Oga | 0.29 | 8.10E-27 | 1.68E-22 | 0.71 | 0.65 | 0.06 | 1.09 |
| ectoderm | Brms1 | 0.48 | 8.62E-27 | 1.79E-22 | 0.41 | 0.33 | 0.08 | 1.26 |
| ectoderm | Rnf181 | 0.39 | 9.83E-27 | 2.04E-22 | 0.45 | 0.37 | 0.08 | 1.23 |
| ectoderm | Fbxw4 | 0.73 | 1.19E-26 | 2.48E-22 | 0.19 | 0.13 | 0.06 | 1.48 |
| ectoderm | Zfp608 | 0.32 | 1.40E-26 | 2.91E-22 | 0.6 | 0.52 | 0.09 | 1.17 |
| ectoderm | Wfs1 | 0.71 | 1.72E-26 | 3.59E-22 | 0.1 | 0.06 | 0.05 | 1.82 |
| ectoderm | Trmt2b | 0.68 | 1.88E-26 | 3.91E-22 | 0.13 | 0.08 | 0.05 | 1.68 |
| ectoderm | Sirt5 | 0.43 | 2.14E-26 | 4.44E-22 | 0.33 | 0.25 | 0.08 | 1.34 |
| ectoderm | Dcp1b | 0.65 | 2.45E-26 | 5.10E-22 | 0.13 | 0.08 | 0.05 | 1.65 |
| ectoderm | 600012H06Ri | 0.46 | 2.69E-26 | 5.59E-22 | 0.28 | 0.2 | 0.08 | 1.37 |
| ectoderm | Plcb3 | 0.43 | 2.82E-26 | 5.87E-22 | 0.36 | 0.27 | 0.09 | 1.31 |
| ectoderm | Grhpr | 0.43 | 2.86E-26 | 5.95E-22 | 0.5 | 0.42 | 0.08 | 1.19 |
| ectoderm | Zfp809 | 0.84 | 3.07E-26 | 6.38E-22 | 0.11 | 0.06 | 0.05 | 1.75 |
| ectoderm | Ydjc | 0.75 | 3.36E-26 | 6.99E-22 | 0.12 | 0.07 | 0.05 | 1.67 |
| ectoderm | Furin | 0.53 | 3.48E-26 | 7.23E-22 | 0.26 | 0.18 | 0.07 | 1.4 |
| ectoderm | BC052040 | 0.51 | 4.25E-26 | 8.83E-22 | 0.23 | 0.16 | 0.07 | 1.43 |
| ectoderm | Renbp | 0.71 | 4.26E-26 | 8.85E-22 | 0.12 | 0.07 | 0.05 | 1.72 |
| ectoderm | Zfp994 | 0.7 | 4.86E-26 | 1.01E-21 | 0.13 | 0.08 | 0.05 | 1.64 |
| ectoderm | Cmas | 0.37 | 5.42E-26 | 1.13E-21 | 0.64 | 0.58 | 0.06 | 1.1 |
| ectoderm | Ppm1b | 0.34 | 5.54E-26 | 1.15E-21 | 0.6 | 0.53 | 0.08 | 1.14 |
| ectoderm | .110012L19Ri | 0.48 | 5.75E-26 | 1.20E-21 | 0.34 | 0.26 | 0.08 | 1.31 |
| ectoderm | Acad11 | 0.74 | 6.06E-26 | 1.26E-21 | 0.13 | 0.08 | 0.05 | 1.66 |
| ectoderm | Chd2 | 0.3 | 6.41E-26 | 1.33E-21 | 0.67 | 0.61 | 0.06 | 1.1 |
| ectoderm | Ring1 | 0.47 | 6.70E-26 | 1.39E-21 | 0.31 | 0.23 | 0.08 | 1.34 |
| ectoderm | Thbs3 | 0.56 | 7.92E-26 | 1.65E-21 | 0.14 | 0.09 | 0.06 | 1.63 |

**Supplementary Table 2. Marker genes for the major cell types of integrated clustering of E11.5 control and NotchGOF λ-junction cells.**

| | | | | | | | | |
|---|---|---|---|---|---|---|---|---|
| ectoderm | Haghl | 0.43 | 7.95E-26 | 1.65E-21 | 0.42 | 0.33 | 0.09 | 1.26 |
| ectoderm | Prdm10 | 0.7 | 9.50E-26 | 1.98E-21 | 0.16 | 0.1 | 0.06 | 1.56 |
| ectoderm | Slc39a3 | 0.65 | 1.01E-25 | 2.10E-21 | 0.17 | 0.11 | 0.06 | 1.54 |
| ectoderm | Prss36 | 0.7 | 1.04E-25 | 2.16E-21 | 0.13 | 0.08 | 0.05 | 1.65 |
| ectoderm | Senp8 | 0.54 | 1.29E-25 | 2.67E-21 | 0.26 | 0.19 | 0.07 | 1.38 |
| ectoderm | Dtd2 | 0.43 | 1.30E-25 | 2.71E-21 | 0.39 | 0.31 | 0.08 | 1.27 |
| ectoderm | Rpp21 | 0.31 | 1.50E-25 | 3.12E-21 | 0.63 | 0.56 | 0.07 | 1.13 |
| ectoderm | Cfap298 | 0.43 | 1.66E-25 | 3.45E-21 | 0.43 | 0.35 | 0.08 | 1.24 |
| ectoderm | Atp1b1 | 0.39 | 1.70E-25 | 3.54E-21 | 0.13 | 0.08 | 0.05 | 1.67 |
| ectoderm | Dbnl | 0.27 | 1.71E-25 | 3.56E-21 | 0.7 | 0.65 | 0.06 | 1.09 |
| ectoderm | Commd6 | 0.31 | 1.90E-25 | 3.94E-21 | 0.7 | 0.64 | 0.06 | 1.09 |
| ectoderm | Ypel1 | 0.31 | 2.00E-25 | 4.17E-21 | 0.57 | 0.48 | 0.09 | 1.18 |
| ectoderm | Gstt2 | 0.75 | 2.07E-25 | 4.32E-21 | 0.11 | 0.07 | 0.05 | 1.72 |
| ectoderm | Bckdk | 0.35 | 2.27E-25 | 4.73E-21 | 0.55 | 0.47 | 0.08 | 1.16 |
| ectoderm | Hnrnpul2 | 0.25 | 2.40E-25 | 5.00E-21 | 0.79 | 0.76 | 0.03 | 1.03 |
| ectoderm | Slc4a2 | 0.48 | 2.60E-25 | 5.42E-21 | 0.27 | 0.2 | 0.07 | 1.36 |
| ectoderm | Zbtb43 | 0.54 | 2.99E-25 | 6.22E-21 | 0.29 | 0.21 | 0.07 | 1.35 |
| ectoderm | Bola1 | 0.32 | 3.16E-25 | 6.58E-21 | 0.66 | 0.61 | 0.06 | 1.09 |
| ectoderm | Ubn2 | 0.25 | 3.20E-25 | 6.66E-21 | 0.74 | 0.69 | 0.05 | 1.08 |
| ectoderm | Fbxo44 | 0.59 | 3.31E-25 | 6.88E-21 | 0.13 | 0.08 | 0.05 | 1.63 |
| ectoderm | Btbd9 | 0.43 | 3.55E-25 | 7.38E-21 | 0.36 | 0.28 | 0.08 | 1.3 |
| ectoderm | Poglut3 | 0.55 | 3.83E-25 | 7.96E-21 | 0.18 | 0.12 | 0.06 | 1.5 |
| ectoderm | Efcab14 | 0.38 | 3.88E-25 | 8.06E-21 | 0.49 | 0.4 | 0.09 | 1.21 |
| ectoderm | Tlcd1 | 0.69 | 4.01E-25 | 8.35E-21 | 0.13 | 0.08 | 0.05 | 1.64 |
| ectoderm | Fuom | 0.4 | 4.17E-25 | 8.67E-21 | 0.34 | 0.26 | 0.08 | 1.32 |
| ectoderm | Etfa | 0.27 | 4.47E-25 | 9.30E-21 | 0.76 | 0.73 | 0.04 | 1.05 |
| ectoderm | Gramd3 | 0.48 | 5.28E-25 | 1.10E-20 | 0.24 | 0.16 | 0.07 | 1.43 |
| ectoderm | 810402E24Ri | 0.47 | 5.34E-25 | 1.11E-20 | 0.27 | 0.2 | 0.08 | 1.38 |
| ectoderm | Kif26a | 0.49 | 6.23E-25 | 1.30E-20 | 0.16 | 0.1 | 0.06 | 1.56 |
| ectoderm | Cradd | 0.57 | 6.48E-25 | 1.35E-20 | 0.22 | 0.16 | 0.07 | 1.43 |
| ectoderm | Xkr6 | 0.41 | 6.79E-25 | 1.41E-20 | 0.17 | 0.11 | 0.06 | 1.54 |
| ectoderm | Rfk | 0.6 | 7.71E-25 | 1.60E-20 | 0.52 | 0.47 | 0.05 | 1.12 |
| ectoderm | Lyrm9 | 0.57 | 8.90E-25 | 1.85E-20 | 0.22 | 0.16 | 0.07 | 1.42 |
| ectoderm | Nsun4 | 0.5 | 9.27E-25 | 1.93E-20 | 0.21 | 0.14 | 0.07 | 1.46 |
| ectoderm | Spata13 | 0.67 | 1.14E-24 | 2.37E-20 | 0.11 | 0.07 | 0.05 | 1.7 |
| ectoderm | Ncoa2 | 0.4 | 1.21E-24 | 2.51E-20 | 0.36 | 0.27 | 0.08 | 1.31 |
| ectoderm | Dsel | 0.44 | 1.42E-24 | 2.95E-20 | 0.4 | 0.31 | 0.08 | 1.27 |
| ectoderm | Ank2 | 0.39 | 1.74E-24 | 3.62E-20 | 0.3 | 0.23 | 0.08 | 1.33 |
| ectoderm | Bnip3 | 0.58 | 2.16E-24 | 4.48E-20 | 0.47 | 0.4 | 0.07 | 1.16 |
| ectoderm | Nfat5 | 0.26 | 2.18E-24 | 4.53E-20 | 0.63 | 0.56 | 0.07 | 1.12 |
| ectoderm | Slc4a4 | 0.67 | 2.75E-24 | 5.72E-20 | 0.16 | 0.1 | 0.05 | 1.53 |
| ectoderm | Tacc2 | 0.35 | 2.83E-24 | 5.88E-20 | 0.54 | 0.47 | 0.07 | 1.16 |
| ectoderm | Zkscan2 | 0.61 | 3.03E-24 | 6.29E-20 | 0.15 | 0.09 | 0.05 | 1.58 |
| ectoderm | Mccc2 | 0.41 | 3.11E-24 | 6.47E-20 | 0.34 | 0.26 | 0.08 | 1.31 |
| ectoderm | Commd9 | 0.32 | 3.27E-24 | 6.81E-20 | 0.56 | 0.47 | 0.09 | 1.18 |
| ectoderm | Tfb1m | 0.53 | 3.31E-24 | 6.88E-20 | 0.17 | 0.11 | 0.06 | 1.53 |
| ectoderm | Bin3 | 0.44 | 3.74E-24 | 7.78E-20 | 0.35 | 0.27 | 0.08 | 1.29 |
| ectoderm | Mettl23 | 0.28 | 3.77E-24 | 7.85E-20 | 0.67 | 0.61 | 0.06 | 1.1 |
| ectoderm | Lrrc61 | 0.65 | 4.03E-24 | 8.39E-20 | 0.14 | 0.09 | 0.05 | 1.59 |
| ectoderm | Slc3a2 | 0.38 | 4.22E-24 | 8.77E-20 | 0.66 | 0.62 | 0.04 | 1.07 |
| ectoderm | Dph5 | 0.48 | 4.42E-24 | 9.18E-20 | 0.28 | 0.21 | 0.07 | 1.35 |
| ectoderm | Nop53 | 0.25 | 5.25E-24 | 1.09E-19 | 0.76 | 0.73 | 0.03 | 1.05 |
| ectoderm | Galnt2 | 0.37 | 5.26E-24 | 1.09E-19 | 0.52 | 0.45 | 0.08 | 1.18 |
| ectoderm | Atp6v0e | 0.28 | 5.36E-24 | 1.11E-19 | 0.75 | 0.71 | 0.04 | 1.06 |
| ectoderm | Slc12a6 | 0.56 | 5.39E-24 | 1.12E-19 | 0.18 | 0.12 | 0.06 | 1.5 |
| ectoderm | Pus1 | 0.46 | 6.27E-24 | 1.30E-19 | 0.35 | 0.28 | 0.08 | 1.28 |

**Supplementary Table 2. Marker genes for the major cell types of integrated clustering of E11.5 control and NotchGOF λ-junction cells.**

| | | | | | | | | |
|---|---|---|---|---|---|---|---|---|
| ectoderm | Traf6 | 0.55 | 7.80E-24 | 1.62E-19 | 0.19 | 0.13 | 0.06 | 1.46 |
| ectoderm | Trip10 | 0.48 | 9.11E-24 | 1.90E-19 | 0.27 | 0.2 | 0.07 | 1.37 |
| ectoderm | Cc2d1b | 0.54 | 1.01E-23 | 2.11E-19 | 0.23 | 0.16 | 0.07 | 1.42 |
| ectoderm | Dpm2 | 0.3 | 1.16E-23 | 2.42E-19 | 0.63 | 0.56 | 0.08 | 1.14 |
| ectoderm | Lrrc56 | 0.6 | 1.31E-23 | 2.72E-19 | 0.14 | 0.09 | 0.05 | 1.59 |
| ectoderm | Numbl | 0.39 | 1.36E-23 | 2.83E-19 | 0.18 | 0.12 | 0.06 | 1.5 |
| ectoderm | Tmem38b | 0.5 | 1.38E-23 | 2.87E-19 | 0.2 | 0.13 | 0.06 | 1.46 |
| ectoderm | 810408A11Ri | 0.57 | 1.54E-23 | 3.21E-19 | 0.2 | 0.14 | 0.06 | 1.45 |
| ectoderm | Ddit4 | 0.56 | 1.59E-23 | 3.30E-19 | 0.43 | 0.36 | 0.07 | 1.2 |
| ectoderm | Wdtc1 | 0.46 | 1.65E-23 | 3.43E-19 | 0.23 | 0.16 | 0.07 | 1.41 |
| ectoderm | Bbc3 | 0.65 | 1.71E-23 | 3.57E-19 | 0.14 | 0.09 | 0.05 | 1.57 |
| ectoderm | Cdk7 | 0.32 | 1.85E-23 | 3.85E-19 | 0.55 | 0.47 | 0.08 | 1.17 |
| ectoderm | Amer1 | 0.39 | 2.04E-23 | 4.23E-19 | 0.57 | 0.49 | 0.07 | 1.14 |
| ectoderm | Fbxl14 | 0.35 | 2.07E-23 | 4.31E-19 | 0.45 | 0.36 | 0.09 | 1.23 |
| ectoderm | Ears2 | 0.5 | 2.26E-23 | 4.69E-19 | 0.21 | 0.14 | 0.06 | 1.44 |
| ectoderm | Tceal3 | 0.5 | 2.39E-23 | 4.98E-19 | 0.16 | 0.11 | 0.06 | 1.53 |
| ectoderm | Cacul1 | 0.45 | 2.50E-23 | 5.20E-19 | 0.35 | 0.27 | 0.08 | 1.29 |
| ectoderm | Tgoln1 | 0.39 | 2.93E-23 | 6.09E-19 | 0.35 | 0.27 | 0.08 | 1.29 |
| ectoderm | Pex11g | 0.76 | 3.27E-23 | 6.80E-19 | 0.13 | 0.08 | 0.05 | 1.61 |
| ectoderm | Morn2 | 0.46 | 3.83E-23 | 7.96E-19 | 0.39 | 0.31 | 0.08 | 1.26 |
| ectoderm | Agap1 | 0.34 | 4.77E-23 | 9.91E-19 | 0.48 | 0.4 | 0.09 | 1.21 |
| ectoderm | Cacnb2 | 0.67 | 4.83E-23 | 1.00E-18 | 0.11 | 0.07 | 0.05 | 1.69 |
| ectoderm | Pdcd6ip | 0.33 | 4.95E-23 | 1.03E-18 | 0.54 | 0.47 | 0.08 | 1.16 |
| ectoderm | Nit1 | 0.39 | 5.01E-23 | 1.04E-18 | 0.41 | 0.33 | 0.08 | 1.25 |
| ectoderm | Wasl | 0.35 | 5.78E-23 | 1.20E-18 | 0.61 | 0.54 | 0.07 | 1.13 |
| ectoderm | Gabarapl1 | 0.42 | 5.81E-23 | 1.21E-18 | 0.39 | 0.31 | 0.08 | 1.25 |
| ectoderm | Csf2ra | 0.52 | 8.14E-23 | 1.69E-18 | 0.19 | 0.13 | 0.06 | 1.46 |
| ectoderm | 610002M06R | 0.38 | 8.32E-23 | 1.73E-18 | 0.35 | 0.27 | 0.08 | 1.28 |
| ectoderm | Pik3c2a | 0.37 | 8.48E-23 | 1.76E-18 | 0.47 | 0.39 | 0.08 | 1.21 |
| ectoderm | Dapk1 | 0.35 | 8.81E-23 | 1.83E-18 | 0.34 | 0.26 | 0.08 | 1.32 |
| ectoderm | Slc35b3 | 0.64 | 9.61E-23 | 2.00E-18 | 0.27 | 0.2 | 0.07 | 1.33 |
| ectoderm | Plxna2 | 1 | 9.92E-23 | 2.06E-18 | 0.28 | 0.22 | 0.06 | 1.28 |
| ectoderm | Flad1 | 0.67 | 1.08E-22 | 2.25E-18 | 0.15 | 0.09 | 0.05 | 1.54 |
| ectoderm | Cpt1a | 0.62 | 1.13E-22 | 2.35E-18 | 0.14 | 0.09 | 0.05 | 1.56 |
| ectoderm | Snx16 | 0.44 | 1.63E-22 | 3.39E-18 | 0.24 | 0.18 | 0.07 | 1.37 |
| ectoderm | AV356131 | 0.56 | 1.96E-22 | 4.08E-18 | 0.11 | 0.07 | 0.05 | 1.66 |
| ectoderm | Echdc2 | 0.38 | 1.96E-22 | 4.08E-18 | 0.32 | 0.24 | 0.08 | 1.34 |
| ectoderm | Pcmtd1 | 0.37 | 2.09E-22 | 4.34E-18 | 0.53 | 0.45 | 0.08 | 1.17 |
| ectoderm | Golga1 | 0.37 | 2.29E-22 | 4.77E-18 | 0.35 | 0.27 | 0.08 | 1.29 |
| ectoderm | Hif1a | 0.25 | 2.65E-22 | 5.52E-18 | 0.78 | 0.75 | 0.03 | 1.04 |
| ectoderm | Smarcd2 | 0.38 | 2.78E-22 | 5.78E-18 | 0.35 | 0.27 | 0.07 | 1.27 |
| ectoderm | Dnaaf5 | 0.57 | 3.64E-22 | 7.56E-18 | 0.13 | 0.08 | 0.05 | 1.59 |
| ectoderm | Slc25a33 | 0.52 | 3.90E-22 | 8.11E-18 | 0.22 | 0.15 | 0.06 | 1.41 |
| ectoderm | Spry1 | 0.44 | 4.76E-22 | 9.90E-18 | 0.35 | 0.27 | 0.08 | 1.28 |
| ectoderm | Ung | 0.52 | 5.18E-22 | 1.08E-17 | 0.4 | 0.33 | 0.07 | 1.22 |
| ectoderm | Syf2 | 0.27 | 5.29E-22 | 1.10E-17 | 0.7 | 0.64 | 0.05 | 1.08 |
| ectoderm | Naxd | 0.26 | 5.60E-22 | 1.16E-17 | 0.69 | 0.63 | 0.06 | 1.09 |
| ectoderm | Avpi1 | 0.44 | 5.64E-22 | 1.17E-17 | 0.38 | 0.3 | 0.07 | 1.24 |
| ectoderm | Egr1 | 1 | 6.90E-22 | 1.43E-17 | 0.11 | 0.07 | 0.04 | 1.64 |
| ectoderm | Itpripl1 | 0.69 | 6.93E-22 | 1.44E-17 | 0.13 | 0.08 | 0.05 | 1.58 |
| ectoderm | Camkmt | 0.48 | 7.23E-22 | 1.50E-17 | 0.23 | 0.17 | 0.06 | 1.38 |
| ectoderm | Rmnd5a | 0.36 | 7.72E-22 | 1.61E-17 | 0.46 | 0.39 | 0.07 | 1.19 |
| ectoderm | Ikbkg | 0.66 | 7.76E-22 | 1.61E-17 | 0.16 | 0.11 | 0.05 | 1.49 |
| ectoderm | Rsbn1l | 0.28 | 7.80E-22 | 1.62E-17 | 0.67 | 0.63 | 0.05 | 1.08 |
| ectoderm | Rps6ka2 | 0.6 | 8.06E-22 | 1.68E-17 | 0.12 | 0.07 | 0.05 | 1.64 |
| ectoderm | Jrkl | 0.48 | 1.04E-21 | 2.17E-17 | 0.22 | 0.16 | 0.06 | 1.4 |

**Supplementary Table 2. Marker genes for the major cell types of integrated clustering of E11.5 control and NotchGOF λ-junction cells.**

| | | | | | | | | |
|---|---|---|---|---|---|---|---|---|
| ectoderm | Efhd2 | 0.29 | 1.13E-21 | 2.36E-17 | 0.26 | 0.19 | 0.07 | 1.36 |
| ectoderm | Gchfr | 0.56 | 1.26E-21 | 2.61E-17 | 0.15 | 0.1 | 0.05 | 1.53 |
| ectoderm | Fuca2 | 0.3 | 1.27E-21 | 2.65E-17 | 0.3 | 0.22 | 0.08 | 1.35 |
| ectoderm | Lrba | 0.26 | 1.35E-21 | 2.82E-17 | 0.29 | 0.21 | 0.08 | 1.37 |
| ectoderm | Pacs2 | 0.42 | 1.36E-21 | 2.83E-17 | 0.22 | 0.16 | 0.06 | 1.41 |
| ectoderm | Ncor2 | 0.36 | 1.48E-21 | 3.08E-17 | 0.41 | 0.33 | 0.08 | 1.24 |
| ectoderm | B4galnt4 | 0.37 | 1.51E-21 | 3.13E-17 | 0.3 | 0.23 | 0.08 | 1.33 |
| ectoderm | Tusc2 | 0.35 | 1.56E-21 | 3.24E-17 | 0.45 | 0.37 | 0.08 | 1.21 |
| ectoderm | Mink1 | 0.52 | 1.59E-21 | 3.31E-17 | 0.22 | 0.16 | 0.06 | 1.39 |
| ectoderm | Prmt3 | 0.38 | 1.61E-21 | 3.34E-17 | 0.39 | 0.32 | 0.08 | 1.24 |
| ectoderm | Fubp3 | 0.31 | 1.64E-21 | 3.40E-17 | 0.52 | 0.43 | 0.08 | 1.19 |
| ectoderm | Gm11361 | 0.36 | 1.71E-21 | 3.56E-17 | 0.47 | 0.39 | 0.07 | 1.19 |
| ectoderm | Pef1 | 0.35 | 1.90E-21 | 3.96E-17 | 0.42 | 0.34 | 0.08 | 1.23 |
| ectoderm | Slk | 0.29 | 2.12E-21 | 4.41E-17 | 0.59 | 0.52 | 0.07 | 1.14 |
| ectoderm | Ppil2 | 0.3 | 2.13E-21 | 4.43E-17 | 0.65 | 0.59 | 0.06 | 1.1 |
| ectoderm | Pqlc1 | 0.58 | 2.20E-21 | 4.57E-17 | 0.18 | 0.13 | 0.06 | 1.44 |
| ectoderm | 300002E11Ri | 0.42 | 2.22E-21 | 4.61E-17 | 0.38 | 0.31 | 0.08 | 1.25 |
| ectoderm | Lats1 | 0.43 | 2.33E-21 | 4.86E-17 | 0.32 | 0.24 | 0.07 | 1.29 |
| ectoderm | Rmnd5b | 0.47 | 2.37E-21 | 4.93E-17 | 0.22 | 0.16 | 0.06 | 1.39 |
| ectoderm | Larp1b | 0.73 | 2.47E-21 | 5.13E-17 | 0.11 | 0.07 | 0.04 | 1.62 |
| ectoderm | Rad18 | 0.46 | 2.55E-21 | 5.30E-17 | 0.42 | 0.36 | 0.06 | 1.18 |
| ectoderm | Ctsl | 0.39 | 2.75E-21 | 5.71E-17 | 0.83 | 0.82 | 0.01 | 1.01 |
| ectoderm | Ccl27a | 0.38 | 2.97E-21 | 6.18E-17 | 0.36 | 0.28 | 0.08 | 1.28 |
| ectoderm | Cdk5rap3 | 0.34 | 3.03E-21 | 6.30E-17 | 0.5 | 0.41 | 0.09 | 1.21 |
| ectoderm | Zfp318 | 0.25 | 3.16E-21 | 6.57E-17 | 0.54 | 0.45 | 0.08 | 1.19 |
| ectoderm | Nbr1 | 0.35 | 3.25E-21 | 6.76E-17 | 0.45 | 0.37 | 0.08 | 1.21 |
| ectoderm | Aebp2 | 0.32 | 3.28E-21 | 6.81E-17 | 0.48 | 0.4 | 0.08 | 1.21 |
| ectoderm | Uhmk1 | 0.55 | 4.40E-21 | 9.14E-17 | 0.26 | 0.19 | 0.06 | 1.33 |
| ectoderm | Cdk5rap1 | 0.65 | 4.96E-21 | 1.03E-16 | 0.12 | 0.08 | 0.05 | 1.57 |
| ectoderm | Arl2 | 0.26 | 5.11E-21 | 1.06E-16 | 0.69 | 0.64 | 0.05 | 1.08 |
| ectoderm | Yif1a | 0.31 | 5.21E-21 | 1.08E-16 | 0.54 | 0.46 | 0.08 | 1.17 |
| ectoderm | Aptx | 0.38 | 5.43E-21 | 1.13E-16 | 0.31 | 0.24 | 0.07 | 1.3 |
| ectoderm | Dlg1 | 0.37 | 6.03E-21 | 1.25E-16 | 0.43 | 0.36 | 0.07 | 1.2 |
| ectoderm | Rtf2 | 0.25 | 6.34E-21 | 1.32E-16 | 0.7 | 0.65 | 0.05 | 1.07 |
| ectoderm | Ppp2cb | 0.35 | 6.88E-21 | 1.43E-16 | 0.5 | 0.42 | 0.07 | 1.17 |
| ectoderm | Hexb | 0.45 | 7.92E-21 | 1.65E-16 | 0.28 | 0.21 | 0.07 | 1.31 |
| ectoderm | Arl8b | 0.36 | 8.67E-21 | 1.80E-16 | 0.53 | 0.46 | 0.07 | 1.16 |
| ectoderm | Mbd2 | 0.34 | 8.68E-21 | 1.81E-16 | 0.6 | 0.55 | 0.05 | 1.1 |
| ectoderm | Inafm2 | 0.59 | 9.14E-21 | 1.90E-16 | 0.17 | 0.11 | 0.05 | 1.46 |
| ectoderm | Coq2 | 0.33 | 1.01E-20 | 2.09E-16 | 0.52 | 0.44 | 0.08 | 1.17 |
| ectoderm | Rragb | 0.68 | 1.08E-20 | 2.24E-16 | 0.12 | 0.08 | 0.04 | 1.58 |
| ectoderm | Eef1e1 | 0.25 | 1.20E-20 | 2.49E-16 | 0.69 | 0.64 | 0.05 | 1.08 |
| ectoderm | Fyco1 | 0.45 | 1.38E-20 | 2.87E-16 | 0.21 | 0.15 | 0.06 | 1.4 |
| ectoderm | Wdr45 | 0.56 | 1.84E-20 | 3.83E-16 | 0.17 | 0.12 | 0.05 | 1.45 |
| ectoderm | Rere | 0.34 | 1.88E-20 | 3.91E-16 | 0.64 | 0.58 | 0.06 | 1.1 |
| ectoderm | Mgme1 | 0.42 | 1.93E-20 | 4.01E-16 | 0.29 | 0.22 | 0.07 | 1.31 |
| ectoderm | BC004004 | 0.36 | 2.03E-20 | 4.23E-16 | 0.38 | 0.3 | 0.07 | 1.25 |
| ectoderm | Homez | 0.56 | 2.04E-20 | 4.25E-16 | 0.13 | 0.09 | 0.05 | 1.55 |
| ectoderm | Man2c1 | 0.51 | 2.19E-20 | 4.55E-16 | 0.14 | 0.1 | 0.05 | 1.52 |
| ectoderm | Wdfy1 | 0.39 | 2.22E-20 | 4.61E-16 | 0.41 | 0.33 | 0.08 | 1.23 |
| ectoderm | Snx14 | 0.4 | 2.22E-20 | 4.62E-16 | 0.21 | 0.15 | 0.06 | 1.4 |
| ectoderm | Atpaf1 | 0.34 | 2.26E-20 | 4.70E-16 | 0.44 | 0.36 | 0.08 | 1.22 |
| ectoderm | Gm17251 | 0.58 | 2.84E-20 | 5.90E-16 | 0.2 | 0.14 | 0.06 | 1.4 |
| ectoderm | Sil1 | 0.48 | 2.95E-20 | 6.14E-16 | 0.15 | 0.1 | 0.05 | 1.49 |
| ectoderm | Kdm7a | 0.39 | 3.00E-20 | 6.23E-16 | 0.4 | 0.34 | 0.07 | 1.2 |
| ectoderm | Strn | 0.49 | 3.82E-20 | 7.94E-16 | 0.22 | 0.16 | 0.06 | 1.37 |

**Supplementary Table 2. Marker genes for the major cell types of integrated clustering of E11.5 control and NotchGOF λ-junction cells.**

| | | | | | | | | |
|---|---|---|---|---|---|---|---|---|
| ectoderm | AW549877 | 0.39 | 3.94E-20 | 8.19E-16 | 0.26 | 0.2 | 0.07 | 1.34 |
| ectoderm | :030006K11Ri | 0.62 | 4.43E-20 | 9.22E-16 | 0.12 | 0.08 | 0.04 | 1.55 |
| ectoderm | Ctnnd1 | 0.31 | 4.67E-20 | 9.72E-16 | 0.52 | 0.44 | 0.08 | 1.18 |
| ectoderm | Phkg2 | 0.46 | 4.68E-20 | 9.74E-16 | 0.3 | 0.23 | 0.07 | 1.3 |
| ectoderm | Lmtk2 | 0.56 | 4.73E-20 | 9.84E-16 | 0.13 | 0.09 | 0.05 | 1.53 |
| ectoderm | Ofd1 | 0.36 | 4.86E-20 | 1.01E-15 | 0.37 | 0.3 | 0.07 | 1.25 |
| ectoderm | Sidt2 | 0.53 | 5.07E-20 | 1.06E-15 | 0.23 | 0.17 | 0.06 | 1.36 |
| ectoderm | Prtg | 0.42 | 5.08E-20 | 1.06E-15 | 0.22 | 0.16 | 0.06 | 1.4 |
| ectoderm | Etfrf1 | 0.46 | 5.24E-20 | 1.09E-15 | 0.27 | 0.21 | 0.07 | 1.32 |
| ectoderm | Cox6b2 | 0.28 | 5.45E-20 | 1.13E-15 | 0.53 | 0.45 | 0.08 | 1.18 |
| ectoderm | Clcc1 | 0.36 | 5.45E-20 | 1.13E-15 | 0.47 | 0.4 | 0.07 | 1.18 |
| ectoderm | Vrk3 | 0.37 | 5.49E-20 | 1.14E-15 | 0.4 | 0.33 | 0.07 | 1.22 |
| ectoderm | Fntb | 0.43 | 5.86E-20 | 1.22E-15 | 0.28 | 0.21 | 0.07 | 1.32 |
| ectoderm | Tmtc3 | 0.42 | 5.98E-20 | 1.24E-15 | 0.29 | 0.22 | 0.07 | 1.3 |
| ectoderm | Zfp2 | 0.52 | 6.33E-20 | 1.32E-15 | 0.21 | 0.15 | 0.06 | 1.39 |
| ectoderm | 230354K17Ri | 0.33 | 6.53E-20 | 1.36E-15 | 0.42 | 0.34 | 0.08 | 1.22 |
| ectoderm | Abcd3 | 0.31 | 6.89E-20 | 1.43E-15 | 0.48 | 0.41 | 0.08 | 1.19 |
| ectoderm | Parn | 0.43 | 7.23E-20 | 1.50E-15 | 0.32 | 0.25 | 0.07 | 1.28 |
| ectoderm | Tmem120a | 0.35 | 7.46E-20 | 1.55E-15 | 0.36 | 0.28 | 0.08 | 1.27 |
| ectoderm | Tagap1 | 0.6 | 7.86E-20 | 1.63E-15 | 0.19 | 0.13 | 0.06 | 1.42 |
| ectoderm | Nfs1 | 0.45 | 8.45E-20 | 1.76E-15 | 0.33 | 0.27 | 0.07 | 1.26 |
| ectoderm | Fktn | 0.36 | 9.46E-20 | 1.97E-15 | 0.32 | 0.25 | 0.07 | 1.3 |
| ectoderm | Ptar1 | 0.42 | 1.05E-19 | 2.18E-15 | 0.33 | 0.26 | 0.07 | 1.25 |
| ectoderm | Sqstm1 | 0.36 | 1.09E-19 | 2.27E-15 | 0.46 | 0.4 | 0.07 | 1.17 |
| ectoderm | Gm43445 | 0.64 | 1.15E-19 | 2.38E-15 | 0.12 | 0.08 | 0.04 | 1.54 |
| ectoderm | Dstyk | 0.4 | 1.16E-19 | 2.42E-15 | 0.28 | 0.21 | 0.07 | 1.32 |
| ectoderm | Gm17018 | 0.29 | 1.18E-19 | 2.46E-15 | 0.58 | 0.51 | 0.07 | 1.14 |
| ectoderm | Zfp703 | 0.45 | 1.26E-19 | 2.63E-15 | 0.31 | 0.24 | 0.07 | 1.28 |
| ectoderm | Lin7c | 0.32 | 1.39E-19 | 2.90E-15 | 0.55 | 0.48 | 0.07 | 1.15 |
| ectoderm | Traf3 | 0.41 | 1.40E-19 | 2.92E-15 | 0.24 | 0.18 | 0.06 | 1.35 |
| ectoderm | Nxt2 | 0.65 | 1.55E-19 | 3.21E-15 | 0.11 | 0.07 | 0.04 | 1.61 |
| ectoderm | Gm11110 | 0.68 | 1.67E-19 | 3.48E-15 | 0.12 | 0.08 | 0.04 | 1.54 |
| ectoderm | 230307C23Ri | 0.52 | 1.95E-19 | 4.06E-15 | 0.21 | 0.15 | 0.06 | 1.38 |
| ectoderm | Gm26930 | 0.55 | 2.10E-19 | 4.36E-15 | 0.12 | 0.08 | 0.04 | 1.55 |
| ectoderm | Hcfc1r1 | 0.33 | 2.19E-19 | 4.56E-15 | 0.62 | 0.57 | 0.05 | 1.1 |
| ectoderm | Ccdc43 | 0.37 | 2.20E-19 | 4.58E-15 | 0.47 | 0.39 | 0.07 | 1.19 |
| ectoderm | Srgap1 | 0.32 | 2.40E-19 | 4.99E-15 | 0.4 | 0.33 | 0.08 | 1.24 |
| ectoderm | Slc16a13 | 0.62 | 2.46E-19 | 5.11E-15 | 0.13 | 0.08 | 0.05 | 1.54 |
| ectoderm | Mthfd1 | 0.38 | 2.51E-19 | 5.22E-15 | 0.44 | 0.37 | 0.07 | 1.19 |
| ectoderm | Atxn3 | 0.46 | 2.60E-19 | 5.41E-15 | 0.24 | 0.18 | 0.06 | 1.33 |
| ectoderm | Tmem175 | 0.52 | 2.78E-19 | 5.79E-15 | 0.14 | 0.09 | 0.05 | 1.52 |
| ectoderm | 510039O18Ri | 0.39 | 2.87E-19 | 5.97E-15 | 0.29 | 0.22 | 0.07 | 1.31 |
| ectoderm | Amer2 | 0.51 | 3.04E-19 | 6.31E-15 | 0.15 | 0.1 | 0.05 | 1.49 |
| ectoderm | Mlh3 | 0.51 | 3.16E-19 | 6.57E-15 | 0.22 | 0.16 | 0.06 | 1.37 |
| ectoderm | Zfyve27 | 0.54 | 3.24E-19 | 6.75E-15 | 0.17 | 0.12 | 0.05 | 1.44 |
| ectoderm | Mtmr4 | 0.36 | 4.50E-19 | 9.36E-15 | 0.35 | 0.27 | 0.07 | 1.27 |
| ectoderm | Wipf2 | 0.36 | 4.63E-19 | 9.62E-15 | 0.31 | 0.24 | 0.07 | 1.29 |
| ectoderm | Pip4p1 | 0.38 | 4.64E-19 | 9.65E-15 | 0.42 | 0.35 | 0.07 | 1.21 |
| ectoderm | Plxna3 | 0.51 | 4.69E-19 | 9.76E-15 | 0.13 | 0.09 | 0.05 | 1.52 |
| ectoderm | Scamp1 | 0.43 | 5.18E-19 | 1.08E-14 | 0.22 | 0.16 | 0.06 | 1.37 |
| ectoderm | Zfp605 | 0.48 | 5.64E-19 | 1.17E-14 | 0.16 | 0.11 | 0.05 | 1.46 |
| ectoderm | Ccndbp1 | 0.41 | 6.05E-19 | 1.26E-14 | 0.33 | 0.27 | 0.07 | 1.25 |
| ectoderm | Wdr41 | 0.53 | 6.27E-19 | 1.30E-14 | 0.14 | 0.09 | 0.05 | 1.5 |
| ectoderm | Mrps34 | 0.28 | 6.84E-19 | 1.42E-14 | 0.66 | 0.61 | 0.05 | 1.09 |
| ectoderm | Zbtb38 | 0.37 | 7.48E-19 | 1.56E-14 | 0.18 | 0.12 | 0.05 | 1.44 |
| ectoderm | Tefm | 0.45 | 7.87E-19 | 1.64E-14 | 0.24 | 0.18 | 0.06 | 1.35 |

**Supplementary Table 2. Marker genes for the major cell types of integrated clustering of E11.5 control and NotchGOF λ-junction cells.**

| | | | | | | | | |
|---|---|---|---|---|---|---|---|---|
| ectoderm | Grina | 0.27 | 8.27E-19 | 1.72E-14 | 0.31 | 0.23 | 0.07 | 1.31 |
| ectoderm | Abcb7 | 0.43 | 8.33E-19 | 1.73E-14 | 0.25 | 0.19 | 0.06 | 1.33 |
| ectoderm | Ogfod2 | 0.62 | 8.38E-19 | 1.74E-14 | 0.13 | 0.09 | 0.04 | 1.49 |
| ectoderm | Dck | 0.3 | 8.41E-19 | 1.75E-14 | 0.51 | 0.44 | 0.07 | 1.16 |
| ectoderm | Zfp995 | 0.71 | 1.06E-18 | 2.20E-14 | 0.11 | 0.07 | 0.04 | 1.58 |
| ectoderm | Gart | 0.3 | 1.11E-18 | 2.31E-14 | 0.6 | 0.55 | 0.05 | 1.09 |
| ectoderm | Gm5617 | 0.38 | 1.18E-18 | 2.45E-14 | 0.45 | 0.38 | 0.07 | 1.18 |
| ectoderm | Oxld1 | 0.53 | 1.19E-18 | 2.48E-14 | 0.16 | 0.11 | 0.05 | 1.46 |
| ectoderm | Ino80d | 0.33 | 1.27E-18 | 2.64E-14 | 0.4 | 0.33 | 0.07 | 1.23 |
| ectoderm | Coasy | 0.35 | 1.30E-18 | 2.70E-14 | 0.37 | 0.3 | 0.07 | 1.24 |
| ectoderm | Elmsan1 | 0.53 | 1.33E-18 | 2.76E-14 | 0.16 | 0.11 | 0.05 | 1.45 |
| ectoderm | Rad9b | 0.59 | 1.43E-18 | 2.97E-14 | 0.12 | 0.07 | 0.04 | 1.57 |
| ectoderm | Ncoa3 | 0.45 | 1.50E-18 | 3.12E-14 | 0.28 | 0.21 | 0.06 | 1.3 |
| ectoderm | Commd4 | 0.28 | 1.50E-18 | 3.13E-14 | 0.59 | 0.52 | 0.07 | 1.13 |
| ectoderm | Micu3 | 0.4 | 1.60E-18 | 3.33E-14 | 0.22 | 0.16 | 0.06 | 1.37 |
| ectoderm | Ugcg | 0.34 | 1.61E-18 | 3.34E-14 | 0.43 | 0.36 | 0.07 | 1.2 |
| ectoderm | Kif1c | 0.4 | 1.73E-18 | 3.60E-14 | 0.2 | 0.15 | 0.06 | 1.39 |
| ectoderm | Gpr89 | 0.36 | 1.74E-18 | 3.61E-14 | 0.39 | 0.31 | 0.07 | 1.24 |
| ectoderm | Nubp2 | 0.37 | 1.74E-18 | 3.62E-14 | 0.36 | 0.29 | 0.07 | 1.24 |
| ectoderm | Prepl | 0.49 | 1.80E-18 | 3.74E-14 | 0.16 | 0.11 | 0.05 | 1.44 |
| ectoderm | Cited2 | 0.27 | 1.91E-18 | 3.97E-14 | 0.68 | 0.63 | 0.05 | 1.08 |
| ectoderm | Gria3 | 0.52 | 2.27E-18 | 4.71E-14 | 0.21 | 0.15 | 0.06 | 1.37 |
| ectoderm | 2310068J16Ri | 0.61 | 2.29E-18 | 4.76E-14 | 0.11 | 0.07 | 0.04 | 1.61 |
| ectoderm | Asxl1 | 0.27 | 2.47E-18 | 5.14E-14 | 0.55 | 0.48 | 0.07 | 1.14 |
| ectoderm | Haus1 | 0.32 | 2.52E-18 | 5.25E-14 | 0.56 | 0.5 | 0.06 | 1.12 |
| ectoderm | Smim8 | 0.33 | 2.60E-18 | 5.41E-14 | 0.48 | 0.41 | 0.08 | 1.19 |
| ectoderm | Stx6 | 0.3 | 2.61E-18 | 5.44E-14 | 0.56 | 0.49 | 0.07 | 1.14 |
| ectoderm | Dph2 | 0.56 | 2.82E-18 | 5.87E-14 | 0.13 | 0.09 | 0.04 | 1.49 |
| ectoderm | D11Wsu47e | 0.51 | 3.40E-18 | 7.07E-14 | 0.13 | 0.09 | 0.04 | 1.51 |
| ectoderm | Etfbkmt | 0.62 | 3.47E-18 | 7.22E-14 | 0.1 | 0.06 | 0.04 | 1.61 |
| ectoderm | Nmrk1 | 0.53 | 3.88E-18 | 8.07E-14 | 0.12 | 0.07 | 0.04 | 1.55 |
| ectoderm | Kdm4c | 0.47 | 4.70E-18 | 9.78E-14 | 0.15 | 0.1 | 0.05 | 1.46 |
| ectoderm | Atp13a3 | 0.32 | 5.05E-18 | 1.05E-13 | 0.46 | 0.39 | 0.07 | 1.18 |
| ectoderm | Eif4ebp3 | 0.38 | 7.17E-18 | 1.49E-13 | 0.24 | 0.18 | 0.06 | 1.33 |
| ectoderm | Uaca | 0.35 | 7.37E-18 | 1.53E-13 | 0.39 | 0.32 | 0.07 | 1.21 |
| ectoderm | Deaf1 | 0.33 | 7.55E-18 | 1.57E-13 | 0.27 | 0.21 | 0.07 | 1.32 |
| ectoderm | Gramd4 | 0.54 | 7.57E-18 | 1.57E-13 | 0.12 | 0.08 | 0.04 | 1.53 |
| ectoderm | Stbd1 | 0.64 | 7.61E-18 | 1.58E-13 | 0.14 | 0.1 | 0.05 | 1.47 |
| ectoderm | Thoc6 | 0.29 | 7.68E-18 | 1.60E-13 | 0.53 | 0.47 | 0.06 | 1.13 |
| ectoderm | Pcyox1l | 0.36 | 7.87E-18 | 1.64E-13 | 0.25 | 0.19 | 0.06 | 1.34 |
| ectoderm | Nagk | 0.37 | 8.39E-18 | 1.74E-13 | 0.25 | 0.18 | 0.06 | 1.33 |
| ectoderm | Araf | 0.29 | 9.37E-18 | 1.95E-13 | 0.46 | 0.39 | 0.07 | 1.19 |
| ectoderm | 700001O22Ri | 0.58 | 1.04E-17 | 2.16E-13 | 0.1 | 0.06 | 0.04 | 1.6 |
| ectoderm | Glb1 | 0.43 | 1.13E-17 | 2.35E-13 | 0.21 | 0.15 | 0.06 | 1.37 |
| ectoderm | Jade3 | 0.55 | 1.16E-17 | 2.42E-13 | 0.16 | 0.12 | 0.05 | 1.42 |
| ectoderm | Pigv | 0.59 | 1.47E-17 | 3.06E-13 | 0.13 | 0.09 | 0.04 | 1.49 |
| ectoderm | Pygb | 0.33 | 1.49E-17 | 3.11E-13 | 0.28 | 0.22 | 0.07 | 1.3 |
| ectoderm | Parm1 | 0.57 | 1.52E-17 | 3.16E-13 | 0.2 | 0.15 | 0.05 | 1.34 |
| ectoderm | Hif3a | 0.55 | 1.56E-17 | 3.24E-13 | 0.12 | 0.08 | 0.04 | 1.53 |
| ectoderm | Khsrp | 0.27 | 1.66E-17 | 3.45E-13 | 0.61 | 0.55 | 0.06 | 1.11 |
| ectoderm | Hexim1 | 0.41 | 2.06E-17 | 4.28E-13 | 0.34 | 0.27 | 0.06 | 1.24 |
| ectoderm | Zfp992 | 0.52 | 2.08E-17 | 4.32E-13 | 0.2 | 0.15 | 0.05 | 1.35 |
| ectoderm | 830406C13Ri | 0.43 | 2.12E-17 | 4.40E-13 | 0.29 | 0.23 | 0.06 | 1.27 |
| ectoderm | Usp32 | 0.48 | 2.19E-17 | 4.56E-13 | 0.2 | 0.14 | 0.05 | 1.37 |
| ectoderm | Immp2l | 0.57 | 2.31E-17 | 4.80E-13 | 0.12 | 0.08 | 0.04 | 1.53 |
| ectoderm | Sowahc | 0.6 | 2.36E-17 | 4.92E-13 | 0.18 | 0.13 | 0.05 | 1.4 |

**Supplementary Table 2. Marker genes for the major cell types of integrated clustering of E11.5 control and NotchGOF λ-junction cells.**

| | | | | | | | | |
|---|---|---|---|---|---|---|---|---|
| ectoderm | Exd2 | 0.42 | 2.43E-17 | 5.05E-13 | 0.2 | 0.14 | 0.05 | 1.38 |
| ectoderm | Gls | 0.33 | 2.50E-17 | 5.21E-13 | 0.46 | 0.39 | 0.07 | 1.18 |
| ectoderm | Dalrd3 | 0.33 | 2.75E-17 | 5.73E-13 | 0.32 | 0.25 | 0.07 | 1.27 |
| ectoderm | Mnt | 0.47 | 3.03E-17 | 6.31E-13 | 0.22 | 0.16 | 0.06 | 1.34 |
| ectoderm | Trim32 | 0.38 | 3.05E-17 | 6.35E-13 | 0.4 | 0.33 | 0.07 | 1.21 |
| ectoderm | Rap2b | 0.28 | 3.18E-17 | 6.61E-13 | 0.54 | 0.47 | 0.07 | 1.15 |
| ectoderm | Atl2 | 0.4 | 3.47E-17 | 7.21E-13 | 0.3 | 0.24 | 0.06 | 1.26 |
| ectoderm | Ttc32 | 0.32 | 3.54E-17 | 7.36E-13 | 0.31 | 0.24 | 0.07 | 1.27 |
| ectoderm | Rrs1 | 0.27 | 4.01E-17 | 8.34E-13 | 0.65 | 0.6 | 0.05 | 1.09 |
| ectoderm | Pnpla6 | 0.52 | 4.17E-17 | 8.67E-13 | 0.12 | 0.08 | 0.04 | 1.52 |
| ectoderm | Tarbp2 | 0.3 | 4.20E-17 | 8.73E-13 | 0.56 | 0.5 | 0.06 | 1.13 |
| ectoderm | Ap2b1 | 0.27 | 4.25E-17 | 8.83E-13 | 0.54 | 0.48 | 0.07 | 1.14 |
| ectoderm | 610507B11Ri | 0.28 | 5.16E-17 | 1.07E-12 | 0.51 | 0.45 | 0.06 | 1.14 |
| ectoderm | Lars2 | 0.37 | 5.21E-17 | 1.08E-12 | 0.52 | 0.46 | 0.06 | 1.14 |
| ectoderm | Ppp1r21 | 0.45 | 5.37E-17 | 1.12E-12 | 0.26 | 0.2 | 0.06 | 1.3 |
| ectoderm | Abtb1 | 0.55 | 5.72E-17 | 1.19E-12 | 0.14 | 0.1 | 0.04 | 1.44 |
| ectoderm | Myo9b | 0.39 | 6.57E-17 | 1.37E-12 | 0.28 | 0.22 | 0.06 | 1.29 |
| ectoderm | Hscb | 0.4 | 6.65E-17 | 1.38E-12 | 0.37 | 0.31 | 0.07 | 1.21 |
| ectoderm | Gm15222 | 0.58 | 6.95E-17 | 1.44E-12 | 0.15 | 0.11 | 0.05 | 1.42 |
| ectoderm | Zfp687 | 0.29 | 7.75E-17 | 1.61E-12 | 0.25 | 0.18 | 0.06 | 1.34 |
| ectoderm | Hist1h2bc | 0.55 | 7.79E-17 | 1.62E-12 | 0.31 | 0.25 | 0.06 | 1.23 |
| ectoderm | Dusp16 | 0.42 | 9.44E-17 | 1.96E-12 | 0.26 | 0.2 | 0.06 | 1.3 |
| ectoderm | Tmub2 | 0.37 | 9.76E-17 | 2.03E-12 | 0.28 | 0.22 | 0.06 | 1.28 |
| ectoderm | Fbxo46 | 0.51 | 1.02E-16 | 2.13E-12 | 0.16 | 0.12 | 0.05 | 1.41 |
| ectoderm | Mbd6 | 0.38 | 1.11E-16 | 2.31E-12 | 0.29 | 0.22 | 0.06 | 1.29 |
| ectoderm | Zfp692 | 0.5 | 1.16E-16 | 2.42E-12 | 0.14 | 0.1 | 0.04 | 1.45 |
| ectoderm | Haus2 | 0.35 | 1.30E-16 | 2.71E-12 | 0.32 | 0.26 | 0.07 | 1.26 |
| ectoderm | Herpud2 | 0.32 | 1.32E-16 | 2.74E-12 | 0.34 | 0.27 | 0.07 | 1.25 |
| ectoderm | Ssh2 | 0.33 | 1.41E-16 | 2.94E-12 | 0.28 | 0.22 | 0.06 | 1.3 |
| ectoderm | Krba1 | 0.35 | 1.45E-16 | 3.02E-12 | 0.22 | 0.17 | 0.06 | 1.34 |
| ectoderm | Gpank1 | 0.36 | 1.53E-16 | 3.18E-12 | 0.28 | 0.22 | 0.06 | 1.29 |
| ectoderm | Gm9993 | 0.47 | 1.59E-16 | 3.31E-12 | 0.11 | 0.07 | 0.04 | 1.53 |
| ectoderm | Znrf2 | 0.39 | 1.74E-16 | 3.62E-12 | 0.23 | 0.17 | 0.06 | 1.33 |
| ectoderm | Akap1 | 0.34 | 1.76E-16 | 3.67E-12 | 0.23 | 0.18 | 0.06 | 1.32 |
| ectoderm | Zfp622 | 0.34 | 1.78E-16 | 3.71E-12 | 0.47 | 0.4 | 0.06 | 1.16 |
| ectoderm | Zxdb | 0.46 | 1.79E-16 | 3.72E-12 | 0.14 | 0.09 | 0.04 | 1.47 |
| ectoderm | Tbc1d1 | 0.34 | 1.83E-16 | 3.81E-12 | 0.43 | 0.36 | 0.07 | 1.2 |
| ectoderm | Asb7 | 0.51 | 1.88E-16 | 3.92E-12 | 0.25 | 0.2 | 0.06 | 1.29 |
| ectoderm | Pigl | 0.51 | 2.17E-16 | 4.51E-12 | 0.1 | 0.07 | 0.04 | 1.56 |
| ectoderm | Fdx1 | 0.26 | 2.45E-16 | 5.09E-12 | 0.59 | 0.52 | 0.07 | 1.14 |
| ectoderm | Amn1 | 0.32 | 2.56E-16 | 5.33E-12 | 0.49 | 0.43 | 0.07 | 1.16 |
| ectoderm | Ski | 0.29 | 3.03E-16 | 6.30E-12 | 0.47 | 0.4 | 0.07 | 1.18 |
| ectoderm | Pigs | 0.26 | 3.32E-16 | 6.91E-12 | 0.36 | 0.29 | 0.07 | 1.24 |
| ectoderm | Mpp5 | 0.29 | 3.50E-16 | 7.27E-12 | 0.43 | 0.36 | 0.07 | 1.19 |
| ectoderm | Plxna1 | 0.45 | 3.78E-16 | 7.87E-12 | 0.25 | 0.19 | 0.06 | 1.3 |
| ectoderm | Efna5 | 0.3 | 3.87E-16 | 8.06E-12 | 0.52 | 0.46 | 0.07 | 1.15 |
| ectoderm | Trim3 | 0.54 | 4.77E-16 | 9.93E-12 | 0.13 | 0.09 | 0.04 | 1.47 |
| ectoderm | Ppt1 | 0.29 | 5.41E-16 | 1.12E-11 | 0.37 | 0.3 | 0.07 | 1.24 |
| ectoderm | Zcchc2 | 0.42 | 5.47E-16 | 1.14E-11 | 0.13 | 0.09 | 0.04 | 1.48 |
| ectoderm | Adipor2 | 0.34 | 5.49E-16 | 1.14E-11 | 0.4 | 0.33 | 0.07 | 1.2 |
| ectoderm | Gimap9 | 0.61 | 5.65E-16 | 1.18E-11 | 0.14 | 0.1 | 0.04 | 1.41 |
| ectoderm | Tfb2m | 0.3 | 5.73E-16 | 1.19E-11 | 0.38 | 0.31 | 0.07 | 1.22 |
| ectoderm | Pex12 | 0.56 | 5.74E-16 | 1.19E-11 | 0.13 | 0.09 | 0.04 | 1.47 |
| ectoderm | Fgfbp3 | 0.46 | 6.12E-16 | 1.27E-11 | 0.13 | 0.09 | 0.04 | 1.46 |
| ectoderm | Top3b | 0.32 | 6.61E-16 | 1.38E-11 | 0.39 | 0.33 | 0.07 | 1.21 |
| ectoderm | Ninl | 0.37 | 6.70E-16 | 1.39E-11 | 0.29 | 0.23 | 0.06 | 1.26 |

**Supplementary Table 2. Marker genes for the major cell types of integrated clustering of E11.5 control and NotchGOF λ-junction cells.**

| | | | | | | | | |
|---|---|---|---|---|---|---|---|---|
| ectoderm | Atg13 | 0.45 | 6.85E-16 | 1.42E-11 | 0.19 | 0.14 | 0.05 | 1.36 |
| ectoderm | Src | 0.32 | 7.11E-16 | 1.48E-11 | 0.28 | 0.22 | 0.06 | 1.29 |
| ectoderm | Usp20 | 0.48 | 7.58E-16 | 1.58E-11 | 0.11 | 0.07 | 0.04 | 1.53 |
| ectoderm | B3galt6 | 0.29 | 8.31E-16 | 1.73E-11 | 0.36 | 0.29 | 0.07 | 1.23 |
| ectoderm | Pold4 | 0.37 | 9.16E-16 | 1.91E-11 | 0.23 | 0.17 | 0.06 | 1.32 |
| ectoderm | Galc | 0.42 | 1.02E-15 | 2.12E-11 | 0.22 | 0.16 | 0.05 | 1.32 |
| ectoderm | Ift52 | 0.31 | 1.19E-15 | 2.48E-11 | 0.45 | 0.38 | 0.07 | 1.17 |
| ectoderm | Atmin | 0.49 | 1.20E-15 | 2.51E-11 | 0.18 | 0.13 | 0.05 | 1.38 |
| ectoderm | Ttc13 | 0.32 | 1.25E-15 | 2.61E-11 | 0.28 | 0.22 | 0.06 | 1.28 |
| ectoderm | Rxrb | 0.25 | 1.27E-15 | 2.64E-11 | 0.47 | 0.4 | 0.08 | 1.19 |
| ectoderm | Med17 | 0.31 | 1.36E-15 | 2.83E-11 | 0.39 | 0.32 | 0.07 | 1.22 |
| ectoderm | Grk5 | 0.46 | 1.38E-15 | 2.87E-11 | 0.16 | 0.11 | 0.05 | 1.41 |
| ectoderm | Lmbrd1 | 0.3 | 1.41E-15 | 2.94E-11 | 0.27 | 0.21 | 0.06 | 1.29 |
| ectoderm | Lamb2 | 0.36 | 1.60E-15 | 3.34E-11 | 0.17 | 0.12 | 0.05 | 1.4 |
| ectoderm | Gm3839 | 0.44 | 1.74E-15 | 3.63E-11 | 0.21 | 0.15 | 0.05 | 1.34 |
| ectoderm | Gm6710 | 0.59 | 1.80E-15 | 3.74E-11 | 0.13 | 0.09 | 0.04 | 1.46 |
| ectoderm | Uevld | 0.53 | 1.90E-15 | 3.94E-11 | 0.16 | 0.12 | 0.04 | 1.38 |
| ectoderm | Exoc4 | 0.25 | 2.25E-15 | 4.69E-11 | 0.46 | 0.39 | 0.07 | 1.18 |
| ectoderm | Scap | 0.36 | 2.40E-15 | 4.99E-11 | 0.23 | 0.17 | 0.06 | 1.33 |
| ectoderm | Aggf1 | 0.25 | 2.46E-15 | 5.12E-11 | 0.63 | 0.58 | 0.05 | 1.09 |
| ectoderm | Htatip2 | 0.79 | 2.61E-15 | 5.44E-11 | 0.1 | 0.07 | 0.03 | 1.49 |
| ectoderm | Atg4d | 0.42 | 2.72E-15 | 5.65E-11 | 0.25 | 0.19 | 0.05 | 1.28 |
| ectoderm | .600002K03Ri | 0.6 | 3.50E-15 | 7.28E-11 | 0.15 | 0.11 | 0.04 | 1.4 |
| ectoderm | Osbpl8 | 0.45 | 3.75E-15 | 7.80E-11 | 0.28 | 0.22 | 0.06 | 1.25 |
| ectoderm | Chfr | 0.39 | 3.81E-15 | 7.92E-11 | 0.3 | 0.24 | 0.06 | 1.24 |
| ectoderm | Lypla2 | 0.32 | 3.82E-15 | 7.94E-11 | 0.41 | 0.34 | 0.07 | 1.19 |
| ectoderm | Btg3 | 0.35 | 3.83E-15 | 7.96E-11 | 0.37 | 0.31 | 0.07 | 1.21 |
| ectoderm | Wdr77 | 0.26 | 4.10E-15 | 8.52E-11 | 0.58 | 0.52 | 0.06 | 1.11 |
| ectoderm | Slc35a5 | 0.47 | 4.23E-15 | 8.80E-11 | 0.13 | 0.09 | 0.04 | 1.45 |
| ectoderm | Pex6 | 0.34 | 4.47E-15 | 9.29E-11 | 0.34 | 0.28 | 0.07 | 1.23 |
| ectoderm | Ammecr1l | 0.33 | 5.49E-15 | 1.14E-10 | 0.35 | 0.28 | 0.07 | 1.23 |
| ectoderm | Ogfod3 | 0.43 | 6.05E-15 | 1.26E-10 | 0.18 | 0.13 | 0.05 | 1.37 |
| ectoderm | Fam117a | 0.38 | 6.37E-15 | 1.33E-10 | 0.28 | 0.22 | 0.06 | 1.25 |
| ectoderm | Atp6v1d | 0.25 | 7.17E-15 | 1.49E-10 | 0.67 | 0.63 | 0.04 | 1.06 |
| ectoderm | Jpt2 | 0.25 | 8.41E-15 | 1.75E-10 | 0.5 | 0.43 | 0.07 | 1.16 |
| ectoderm | Zfp282 | 0.38 | 8.76E-15 | 1.82E-10 | 0.31 | 0.25 | 0.06 | 1.24 |
| ectoderm | Nog | 0.34 | 8.83E-15 | 1.84E-10 | 0.13 | 0.09 | 0.04 | 1.45 |
| ectoderm | Letm2 | 0.37 | 8.88E-15 | 1.85E-10 | 0.2 | 0.15 | 0.05 | 1.34 |
| ectoderm | Med13 | 0.31 | 8.98E-15 | 1.87E-10 | 0.6 | 0.54 | 0.05 | 1.1 |
| ectoderm | Stk39 | 0.35 | 9.14E-15 | 1.90E-10 | 0.31 | 0.25 | 0.06 | 1.24 |
| ectoderm | Parp8 | 0.44 | 9.49E-15 | 1.97E-10 | 0.18 | 0.13 | 0.05 | 1.37 |
| ectoderm | Wiz | 0.27 | 1.04E-14 | 2.17E-10 | 0.47 | 0.4 | 0.06 | 1.16 |
| ectoderm | Ppp2r3d | 0.33 | 1.05E-14 | 2.18E-10 | 0.27 | 0.21 | 0.06 | 1.28 |
| ectoderm | Abhd13 | 0.37 | 1.18E-14 | 2.46E-10 | 0.23 | 0.18 | 0.06 | 1.31 |
| ectoderm | Ap4s1 | 0.31 | 1.41E-14 | 2.94E-10 | 0.47 | 0.41 | 0.06 | 1.15 |
| ectoderm | Fcf1 | 0.25 | 1.63E-14 | 3.38E-10 | 0.7 | 0.66 | 0.04 | 1.05 |
| ectoderm | Unc50 | 0.25 | 1.63E-14 | 3.39E-10 | 0.6 | 0.55 | 0.05 | 1.09 |
| ectoderm | Tdrd3 | 0.29 | 1.66E-14 | 3.46E-10 | 0.39 | 0.33 | 0.07 | 1.21 |
| ectoderm | Lmbr1l | 0.4 | 1.72E-14 | 3.58E-10 | 0.22 | 0.17 | 0.05 | 1.31 |
| ectoderm | Dennd4c | 0.4 | 1.81E-14 | 3.77E-10 | 0.21 | 0.16 | 0.05 | 1.32 |
| ectoderm | Bmt2 | 0.3 | 1.85E-14 | 3.84E-10 | 0.37 | 0.3 | 0.07 | 1.21 |
| ectoderm | Atxn7 | 0.34 | 1.86E-14 | 3.87E-10 | 0.26 | 0.2 | 0.06 | 1.28 |
| ectoderm | Rabgef1 | 0.48 | 2.06E-14 | 4.29E-10 | 0.2 | 0.15 | 0.05 | 1.31 |
| ectoderm | Tm2d3 | 0.25 | 2.08E-14 | 4.32E-10 | 0.55 | 0.49 | 0.06 | 1.13 |
| ectoderm | Sapcd2 | 0.36 | 2.12E-14 | 4.40E-10 | 0.3 | 0.24 | 0.06 | 1.23 |
| ectoderm | Dcaf12 | 0.32 | 2.49E-14 | 5.18E-10 | 0.39 | 0.33 | 0.06 | 1.18 |

**Supplementary Table 2. Marker genes for the major cell types of integrated clustering of E11.5 control and NotchGOF λ-junction cells.**

| | | | | | | | | |
|---|---|---|---|---|---|---|---|---|
| ectoderm | Otulin | 0.29 | 2.57E-14 | 5.34E-10 | 0.46 | 0.4 | 0.06 | 1.15 |
| ectoderm | Zfp646 | 0.28 | 2.83E-14 | 5.88E-10 | 0.46 | 0.4 | 0.07 | 1.17 |
| ectoderm | Tada3 | 0.3 | 3.18E-14 | 6.61E-10 | 0.41 | 0.35 | 0.06 | 1.18 |
| ectoderm | Lmbr1 | 0.62 | 3.36E-14 | 6.99E-10 | 0.12 | 0.08 | 0.04 | 1.46 |
| ectoderm | Mettl5 | 0.31 | 3.55E-14 | 7.39E-10 | 0.36 | 0.3 | 0.06 | 1.2 |
| ectoderm | Dtnb | 0.3 | 4.03E-14 | 8.39E-10 | 0.32 | 0.26 | 0.06 | 1.23 |
| ectoderm | Caskin2 | 0.4 | 4.11E-14 | 8.56E-10 | 0.15 | 0.1 | 0.04 | 1.4 |
| ectoderm | Zfp937 | 0.42 | 4.91E-14 | 1.02E-09 | 0.13 | 0.09 | 0.04 | 1.43 |
| ectoderm | Pip4k2c | 0.36 | 4.96E-14 | 1.03E-09 | 0.22 | 0.17 | 0.05 | 1.29 |
| ectoderm | Ncoa1 | 0.32 | 5.35E-14 | 1.11E-09 | 0.21 | 0.16 | 0.05 | 1.32 |
| ectoderm | Scaper | 0.26 | 5.51E-14 | 1.15E-09 | 0.34 | 0.28 | 0.06 | 1.23 |
| ectoderm | ·930523C07Ri | 0.66 | 5.67E-14 | 1.18E-09 | 0.1 | 0.07 | 0.03 | 1.49 |
| ectoderm | Zfp367 | 0.41 | 5.74E-14 | 1.19E-09 | 0.18 | 0.13 | 0.05 | 1.34 |
| ectoderm | Chchd5 | 0.28 | 6.47E-14 | 1.34E-09 | 0.51 | 0.45 | 0.06 | 1.13 |
| ectoderm | Lztfl1 | 0.27 | 7.90E-14 | 1.64E-09 | 0.54 | 0.48 | 0.07 | 1.14 |
| ectoderm | Glis2 | 0.3 | 8.18E-14 | 1.70E-09 | 0.24 | 0.18 | 0.06 | 1.3 |
| ectoderm | Ptgr2 | 0.48 | 8.86E-14 | 1.84E-09 | 0.19 | 0.14 | 0.05 | 1.32 |
| ectoderm | Reep4 | 0.36 | 8.88E-14 | 1.85E-09 | 0.37 | 0.32 | 0.06 | 1.18 |
| ectoderm | Col4a3bp | 0.3 | 8.92E-14 | 1.86E-09 | 0.37 | 0.31 | 0.06 | 1.2 |
| ectoderm | Mettl14 | 0.29 | 9.24E-14 | 1.92E-09 | 0.32 | 0.26 | 0.06 | 1.24 |
| ectoderm | Mmp15 | 0.37 | 9.92E-14 | 2.06E-09 | 0.12 | 0.08 | 0.04 | 1.46 |
| ectoderm | Rras2 | 0.34 | 1.04E-13 | 2.17E-09 | 0.32 | 0.26 | 0.06 | 1.23 |
| ectoderm | Zmpste24 | 0.31 | 1.18E-13 | 2.46E-09 | 0.36 | 0.3 | 0.06 | 1.2 |
| ectoderm | Asxl2 | 0.27 | 1.30E-13 | 2.71E-09 | 0.46 | 0.4 | 0.06 | 1.15 |
| ectoderm | Avl9 | 0.28 | 1.40E-13 | 2.91E-09 | 0.4 | 0.33 | 0.06 | 1.19 |
| ectoderm | Ube2e2 | 0.27 | 1.40E-13 | 2.91E-09 | 0.43 | 0.37 | 0.07 | 1.19 |
| ectoderm | L2hgdh | 0.4 | 1.64E-13 | 3.42E-09 | 0.25 | 0.2 | 0.05 | 1.27 |
| ectoderm | Ano10 | 0.33 | 1.71E-13 | 3.56E-09 | 0.16 | 0.12 | 0.05 | 1.38 |
| ectoderm | Ercc4 | 0.59 | 1.88E-13 | 3.90E-09 | 0.12 | 0.08 | 0.04 | 1.44 |
| ectoderm | Lpar2 | 0.32 | 2.22E-13 | 4.62E-09 | 0.1 | 0.07 | 0.03 | 1.49 |
| ectoderm | Xpa | 0.3 | 2.31E-13 | 4.80E-09 | 0.49 | 0.43 | 0.06 | 1.14 |
| ectoderm | Fuz | 0.4 | 2.44E-13 | 5.07E-09 | 0.14 | 0.1 | 0.04 | 1.4 |
| ectoderm | Babam2 | 0.33 | 2.73E-13 | 5.67E-09 | 0.39 | 0.34 | 0.06 | 1.17 |
| ectoderm | Trmt1l | 0.27 | 3.06E-13 | 6.36E-09 | 0.31 | 0.25 | 0.06 | 1.23 |
| ectoderm | Fbxl3 | 0.29 | 3.06E-13 | 6.36E-09 | 0.3 | 0.24 | 0.06 | 1.24 |
| ectoderm | Chd1l | 0.31 | 3.08E-13 | 6.41E-09 | 0.24 | 0.19 | 0.05 | 1.27 |
| ectoderm | Bmp4 | 0.38 | 3.57E-13 | 7.43E-09 | 0.55 | 0.51 | 0.05 | 1.09 |
| ectoderm | Pi4k2b | 0.34 | 3.64E-13 | 7.57E-09 | 0.33 | 0.27 | 0.06 | 1.21 |
| ectoderm | Wdr4 | 0.33 | 4.08E-13 | 8.49E-09 | 0.31 | 0.25 | 0.06 | 1.23 |
| ectoderm | Ddx10 | 0.26 | 4.33E-13 | 9.01E-09 | 0.53 | 0.47 | 0.06 | 1.12 |
| ectoderm | Cdon | 0.28 | 4.47E-13 | 9.31E-09 | 0.26 | 0.21 | 0.06 | 1.26 |
| ectoderm | Tmem127 | 0.31 | 4.49E-13 | 9.35E-09 | 0.23 | 0.18 | 0.05 | 1.29 |
| ectoderm | Zbtb8a | 0.49 | 4.60E-13 | 9.57E-09 | 0.1 | 0.07 | 0.03 | 1.47 |
| ectoderm | Nek7 | 0.27 | 4.61E-13 | 9.59E-09 | 0.34 | 0.28 | 0.06 | 1.22 |
| ectoderm | Skida1 | 0.37 | 4.72E-13 | 9.82E-09 | 0.19 | 0.15 | 0.05 | 1.32 |
| ectoderm | Cdc73 | 0.25 | 4.90E-13 | 1.02E-08 | 0.53 | 0.46 | 0.06 | 1.13 |
| ectoderm | Rpp38 | 0.44 | 5.07E-13 | 1.06E-08 | 0.21 | 0.16 | 0.05 | 1.29 |
| ectoderm | Nudt8 | 0.37 | 5.47E-13 | 1.14E-08 | 0.26 | 0.21 | 0.05 | 1.24 |
| ectoderm | Aasdh | 0.48 | 5.69E-13 | 1.18E-08 | 0.12 | 0.09 | 0.04 | 1.42 |
| ectoderm | Hyal1 | 0.51 | 5.94E-13 | 1.24E-08 | 0.12 | 0.08 | 0.04 | 1.43 |
| ectoderm | Mocs3 | 0.47 | 5.94E-13 | 1.24E-08 | 0.11 | 0.08 | 0.04 | 1.46 |
| ectoderm | Pink1 | 0.39 | 6.15E-13 | 1.28E-08 | 0.26 | 0.21 | 0.05 | 1.24 |
| ectoderm | Rmdn1 | 0.38 | 6.26E-13 | 1.30E-08 | 0.21 | 0.16 | 0.05 | 1.29 |
| ectoderm | Dbt | 0.31 | 6.32E-13 | 1.31E-08 | 0.34 | 0.28 | 0.06 | 1.2 |
| ectoderm | Vps37d | 0.31 | 6.37E-13 | 1.32E-08 | 0.34 | 0.28 | 0.06 | 1.21 |
| ectoderm | Ldlrad3 | 0.3 | 6.64E-13 | 1.38E-08 | 0.36 | 0.3 | 0.06 | 1.2 |

**Supplementary Table 2. Marker genes for the major cell types of integrated clustering of E11.5 control and NotchGOF λ-junction cells.**

| | | | | | | | | |
|---|---|---|---|---|---|---|---|---|
| ectoderm | Gcc1 | 0.49 | 6.90E-13 | 1.43E-08 | 0.1 | 0.07 | 0.03 | 1.47 |
| ectoderm | Hgh1 | 0.32 | 8.23E-13 | 1.71E-08 | 0.17 | 0.12 | 0.04 | 1.35 |
| ectoderm | Wwc2 | 0.27 | 8.38E-13 | 1.74E-08 | 0.61 | 0.57 | 0.04 | 1.08 |
| ectoderm | Slc35e3 | 0.36 | 8.69E-13 | 1.81E-08 | 0.17 | 0.12 | 0.04 | 1.35 |
| ectoderm | Etaa1 | 0.29 | 9.03E-13 | 1.88E-08 | 0.58 | 0.52 | 0.05 | 1.1 |
| ectoderm | Wdfy2 | 0.36 | 9.16E-13 | 1.91E-08 | 0.2 | 0.15 | 0.05 | 1.3 |
| ectoderm | Fkbp1b | 0.33 | 9.55E-13 | 1.99E-08 | 0.18 | 0.13 | 0.04 | 1.33 |
| ectoderm | Zbtb9 | 0.42 | 9.84E-13 | 2.05E-08 | 0.14 | 0.1 | 0.04 | 1.39 |
| ectoderm | Rgs3 | 0.37 | 1.11E-12 | 2.30E-08 | 0.22 | 0.17 | 0.05 | 1.29 |
| ectoderm | Pomk | 0.42 | 1.18E-12 | 2.46E-08 | 0.12 | 0.08 | 0.04 | 1.43 |
| ectoderm | Casp8 | 0.3 | 1.24E-12 | 2.58E-08 | 0.22 | 0.17 | 0.05 | 1.3 |
| ectoderm | Btbd3 | 0.35 | 1.42E-12 | 2.96E-08 | 0.19 | 0.15 | 0.05 | 1.31 |
| ectoderm | Zfp654 | 0.34 | 1.46E-12 | 3.05E-08 | 0.18 | 0.14 | 0.04 | 1.32 |
| ectoderm | Rpe | 0.32 | 1.55E-12 | 3.23E-08 | 0.38 | 0.32 | 0.06 | 1.19 |
| ectoderm | Slc7a1 | 0.81 | 1.68E-12 | 3.50E-08 | 0.1 | 0.07 | 0.03 | 1.43 |
| ectoderm | Prkx | 0.38 | 1.69E-12 | 3.52E-08 | 0.19 | 0.14 | 0.05 | 1.31 |
| ectoderm | Pdcl | 0.27 | 1.81E-12 | 3.76E-08 | 0.49 | 0.42 | 0.06 | 1.15 |
| ectoderm | Trappc12 | 0.29 | 1.90E-12 | 3.95E-08 | 0.25 | 0.2 | 0.05 | 1.27 |
| ectoderm | AI987944 | 0.42 | 2.28E-12 | 4.75E-08 | 0.16 | 0.12 | 0.04 | 1.34 |
| ectoderm | Fbxw9 | 0.34 | 2.33E-12 | 4.86E-08 | 0.23 | 0.18 | 0.05 | 1.28 |
| ectoderm | Med14 | 0.29 | 2.57E-12 | 5.34E-08 | 0.31 | 0.25 | 0.05 | 1.21 |
| ectoderm | Otud5 | 0.26 | 2.74E-12 | 5.71E-08 | 0.42 | 0.36 | 0.06 | 1.16 |
| ectoderm | Ptpn21 | 0.38 | 2.76E-12 | 5.74E-08 | 0.15 | 0.11 | 0.04 | 1.36 |
| ectoderm | Atp6v1b2 | 0.26 | 2.79E-12 | 5.79E-08 | 0.44 | 0.37 | 0.06 | 1.17 |
| ectoderm | Trip6 | 0.26 | 2.89E-12 | 6.01E-08 | 0.52 | 0.45 | 0.07 | 1.14 |
| ectoderm | Psph | 0.31 | 2.96E-12 | 6.16E-08 | 0.33 | 0.27 | 0.06 | 1.21 |
| ectoderm | Tnpo2 | 0.3 | 3.03E-12 | 6.30E-08 | 0.28 | 0.23 | 0.05 | 1.24 |
| ectoderm | Dyrk1b | 0.35 | 3.03E-12 | 6.30E-08 | 0.18 | 0.13 | 0.04 | 1.33 |
| ectoderm | Taf5l | 0.29 | 3.10E-12 | 6.45E-08 | 0.32 | 0.27 | 0.06 | 1.21 |
| ectoderm | Coa4 | 0.3 | 3.12E-12 | 6.49E-08 | 0.31 | 0.25 | 0.06 | 1.22 |
| ectoderm | Zfp867 | 0.44 | 3.15E-12 | 6.55E-08 | 0.15 | 0.11 | 0.04 | 1.36 |
| ectoderm | Neu1 | 0.32 | 3.25E-12 | 6.76E-08 | 0.16 | 0.12 | 0.04 | 1.34 |
| ectoderm | 500015A07Ri | 0.4 | 3.75E-12 | 7.79E-08 | 0.16 | 0.12 | 0.04 | 1.34 |
| ectoderm | Sos1 | 0.26 | 3.84E-12 | 7.99E-08 | 0.4 | 0.34 | 0.06 | 1.18 |
| ectoderm | Arhgef18 | 0.4 | 3.92E-12 | 8.15E-08 | 0.17 | 0.13 | 0.04 | 1.33 |
| ectoderm | Cybc1 | 0.36 | 3.97E-12 | 8.26E-08 | 0.26 | 0.21 | 0.05 | 1.24 |
| ectoderm | Bcl2l11 | 0.34 | 4.89E-12 | 1.02E-07 | 0.4 | 0.34 | 0.06 | 1.18 |
| ectoderm | Mib1 | 0.26 | 4.95E-12 | 1.03E-07 | 0.51 | 0.46 | 0.05 | 1.12 |
| ectoderm | Prorp | 0.46 | 5.05E-12 | 1.05E-07 | 0.17 | 0.13 | 0.04 | 1.32 |
| ectoderm | Zbtb7a | 0.33 | 5.06E-12 | 1.05E-07 | 0.38 | 0.32 | 0.06 | 1.17 |
| ectoderm | Gphn | 0.29 | 5.21E-12 | 1.08E-07 | 0.23 | 0.18 | 0.05 | 1.29 |
| ectoderm | Scarb1 | 0.33 | 5.41E-12 | 1.12E-07 | 0.19 | 0.15 | 0.05 | 1.31 |
| ectoderm | Zbtb34 | 0.53 | 5.46E-12 | 1.14E-07 | 0.12 | 0.08 | 0.03 | 1.4 |
| ectoderm | Tom1 | 0.31 | 5.90E-12 | 1.23E-07 | 0.14 | 0.1 | 0.04 | 1.37 |
| ectoderm | Heca | 0.29 | 6.06E-12 | 1.26E-07 | 0.27 | 0.22 | 0.05 | 1.23 |
| ectoderm | Xpot | 0.31 | 6.27E-12 | 1.30E-07 | 0.46 | 0.4 | 0.06 | 1.15 |
| ectoderm | Rhot2 | 0.33 | 6.32E-12 | 1.31E-07 | 0.22 | 0.17 | 0.05 | 1.28 |
| ectoderm | Lmbrd2 | 0.3 | 6.53E-12 | 1.36E-07 | 0.22 | 0.17 | 0.05 | 1.27 |
| ectoderm | Braf | 0.29 | 6.65E-12 | 1.38E-07 | 0.38 | 0.32 | 0.06 | 1.17 |
| ectoderm | Mfge8 | 0.4 | 6.67E-12 | 1.39E-07 | 0.56 | 0.51 | 0.04 | 1.08 |
| ectoderm | Gba | 0.36 | 7.93E-12 | 1.65E-07 | 0.12 | 0.09 | 0.04 | 1.4 |
| ectoderm | Gm4924 | 0.41 | 8.02E-12 | 1.67E-07 | 0.17 | 0.13 | 0.04 | 1.32 |
| ectoderm | 310015A10Ri | 0.37 | 8.07E-12 | 1.68E-07 | 0.1 | 0.07 | 0.03 | 1.47 |
| ectoderm | Slf2 | 0.29 | 8.22E-12 | 1.71E-07 | 0.37 | 0.31 | 0.06 | 1.19 |
| ectoderm | Prr14l | 0.26 | 8.34E-12 | 1.73E-07 | 0.28 | 0.23 | 0.05 | 1.24 |
| ectoderm | Vav2 | 0.4 | 8.37E-12 | 1.74E-07 | 0.11 | 0.08 | 0.03 | 1.42 |

**Supplementary Table 2. Marker genes for the major cell types of integrated clustering of E11.5 control and NotchGOF λ-junction cells.**

| | | | | | | | | |
|---|---|---|---|---|---|---|---|---|
| ectoderm | Ulk4 | 0.43 | 8.79E-12 | 1.83E-07 | 0.23 | 0.19 | 0.05 | 1.24 |
| ectoderm | Gmppa | 0.31 | 9.04E-12 | 1.88E-07 | 0.21 | 0.16 | 0.05 | 1.29 |
| ectoderm | \130010J15Ri | 0.4 | 9.41E-12 | 1.96E-07 | 0.19 | 0.14 | 0.04 | 1.3 |
| ectoderm | Zswim8 | 0.3 | 9.43E-12 | 1.96E-07 | 0.38 | 0.33 | 0.06 | 1.17 |
| ectoderm | Coq4 | 0.28 | 9.80E-12 | 2.04E-07 | 0.38 | 0.32 | 0.06 | 1.18 |
| ectoderm | Rars2 | 0.33 | 9.97E-12 | 2.07E-07 | 0.38 | 0.32 | 0.06 | 1.18 |
| ectoderm | 732440D04Ri | 0.4 | 1.09E-11 | 2.27E-07 | 0.13 | 0.09 | 0.04 | 1.39 |
| ectoderm | Zfpl1 | 0.26 | 1.11E-11 | 2.31E-07 | 0.47 | 0.41 | 0.06 | 1.14 |
| ectoderm | Nol8 | 0.27 | 1.12E-11 | 2.33E-07 | 0.46 | 0.41 | 0.06 | 1.14 |
| ectoderm | Qtrt1 | 0.28 | 1.20E-11 | 2.50E-07 | 0.29 | 0.23 | 0.06 | 1.24 |
| ectoderm | Fam214b | 0.47 | 1.35E-11 | 2.81E-07 | 0.12 | 0.09 | 0.04 | 1.39 |
| ectoderm | Lin37 | 0.29 | 1.38E-11 | 2.87E-07 | 0.28 | 0.23 | 0.05 | 1.22 |
| ectoderm | Hdac6 | 0.34 | 1.40E-11 | 2.90E-07 | 0.28 | 0.23 | 0.05 | 1.23 |
| ectoderm | Slc25a26 | 0.41 | 1.43E-11 | 2.97E-07 | 0.12 | 0.08 | 0.03 | 1.4 |
| ectoderm | Parp11 | 0.3 | 1.53E-11 | 3.19E-07 | 0.2 | 0.15 | 0.05 | 1.3 |
| ectoderm | Adck1 | 0.28 | 1.56E-11 | 3.23E-07 | 0.2 | 0.15 | 0.05 | 1.3 |
| ectoderm | Szrd1 | 0.25 | 1.59E-11 | 3.30E-07 | 0.55 | 0.5 | 0.05 | 1.09 |
| ectoderm | 2310011J03Ri | 0.27 | 1.60E-11 | 3.32E-07 | 0.53 | 0.47 | 0.06 | 1.12 |
| ectoderm | Stim1 | 0.39 | 1.61E-11 | 3.34E-07 | 0.18 | 0.14 | 0.04 | 1.3 |
| ectoderm | Txlng | 0.31 | 1.85E-11 | 3.86E-07 | 0.26 | 0.21 | 0.05 | 1.24 |
| ectoderm | Agpat2 | 0.29 | 1.88E-11 | 3.91E-07 | 0.2 | 0.16 | 0.05 | 1.29 |
| ectoderm | Nfkbib | 0.3 | 1.97E-11 | 4.11E-07 | 0.28 | 0.23 | 0.05 | 1.22 |
| ectoderm | Whamm | 0.4 | 2.12E-11 | 4.42E-07 | 0.13 | 0.09 | 0.03 | 1.37 |
| ectoderm | Socs4 | 0.25 | 2.31E-11 | 4.81E-07 | 0.39 | 0.33 | 0.06 | 1.18 |
| ectoderm | B3galnt1 | 0.41 | 2.32E-11 | 4.82E-07 | 0.17 | 0.13 | 0.04 | 1.3 |
| ectoderm | Zfp626 | 0.33 | 2.59E-11 | 5.39E-07 | 0.17 | 0.13 | 0.04 | 1.32 |
| ectoderm | Msto1 | 0.31 | 2.84E-11 | 5.91E-07 | 0.26 | 0.21 | 0.05 | 1.23 |
| ectoderm | Ascc2 | 0.25 | 2.86E-11 | 5.95E-07 | 0.3 | 0.25 | 0.06 | 1.22 |
| ectoderm | Gm10130 | 0.39 | 2.94E-11 | 6.11E-07 | 0.12 | 0.09 | 0.03 | 1.39 |
| ectoderm | Zbtb33 | 0.32 | 2.97E-11 | 6.18E-07 | 0.31 | 0.25 | 0.06 | 1.22 |
| ectoderm | Ap5m1 | 0.26 | 3.26E-11 | 6.78E-07 | 0.28 | 0.23 | 0.05 | 1.23 |
| ectoderm | Gnptab | 0.27 | 3.72E-11 | 7.74E-07 | 0.18 | 0.14 | 0.04 | 1.3 |
| ectoderm | Adat1 | 0.4 | 3.95E-11 | 8.21E-07 | 0.17 | 0.13 | 0.04 | 1.3 |
| ectoderm | Mapk14 | 0.25 | 3.96E-11 | 8.24E-07 | 0.37 | 0.32 | 0.06 | 1.18 |
| ectoderm | Stx4a | 0.29 | 4.10E-11 | 8.52E-07 | 0.36 | 0.31 | 0.06 | 1.19 |
| ectoderm | Arhgap23 | 0.51 | 4.13E-11 | 8.59E-07 | 0.12 | 0.09 | 0.03 | 1.36 |
| ectoderm | Prxl2c | 0.35 | 4.40E-11 | 9.15E-07 | 0.33 | 0.28 | 0.05 | 1.19 |
| ectoderm | Zfp672 | 0.35 | 4.46E-11 | 9.28E-07 | 0.2 | 0.16 | 0.04 | 1.27 |
| ectoderm | Lrp10 | 0.26 | 4.89E-11 | 1.02E-06 | 0.21 | 0.16 | 0.05 | 1.28 |
| ectoderm | Poll | 0.27 | 5.19E-11 | 1.08E-06 | 0.21 | 0.16 | 0.05 | 1.28 |
| ectoderm | Mecr | 0.26 | 5.20E-11 | 1.08E-06 | 0.35 | 0.3 | 0.06 | 1.19 |
| ectoderm | Fam126a | 0.39 | 5.23E-11 | 1.09E-06 | 0.29 | 0.24 | 0.05 | 1.2 |
| ectoderm | Fam126b | 0.3 | 5.36E-11 | 1.11E-06 | 0.23 | 0.18 | 0.05 | 1.27 |
| ectoderm | Zfp729a | 0.36 | 5.50E-11 | 1.14E-06 | 0.18 | 0.14 | 0.04 | 1.29 |
| ectoderm | Gpn2 | 0.32 | 5.86E-11 | 1.22E-06 | 0.24 | 0.19 | 0.05 | 1.25 |
| ectoderm | Cmtm6 | 0.41 | 6.24E-11 | 1.30E-06 | 0.16 | 0.12 | 0.04 | 1.3 |
| ectoderm | Rasa3 | 0.34 | 6.67E-11 | 1.39E-06 | 0.1 | 0.07 | 0.03 | 1.41 |
| ectoderm | Ube2w | 0.28 | 6.91E-11 | 1.44E-06 | 0.39 | 0.34 | 0.05 | 1.16 |
| ectoderm | Vps13b | 0.34 | 7.18E-11 | 1.49E-06 | 0.22 | 0.17 | 0.05 | 1.26 |
| ectoderm | Zxdc | 0.41 | 7.92E-11 | 1.65E-06 | 0.13 | 0.1 | 0.03 | 1.35 |
| ectoderm | Kdm1b | 0.44 | 8.28E-11 | 1.72E-06 | 0.13 | 0.09 | 0.03 | 1.36 |
| ectoderm | Tnip1 | 0.33 | 8.52E-11 | 1.77E-06 | 0.13 | 0.1 | 0.03 | 1.35 |
| ectoderm | Gm29170 | 0.46 | 8.87E-11 | 1.85E-06 | 0.1 | 0.07 | 0.03 | 1.42 |
| ectoderm | Ccng1 | 0.33 | 9.23E-11 | 1.92E-06 | 0.24 | 0.19 | 0.05 | 1.25 |
| ectoderm | Alkbh6 | 0.25 | 9.30E-11 | 1.93E-06 | 0.49 | 0.43 | 0.06 | 1.14 |
| ectoderm | Pou2f1 | 0.28 | 9.67E-11 | 2.01E-06 | 0.4 | 0.35 | 0.06 | 1.16 |

**Supplementary Table 2. Marker genes for the major cell types of integrated clustering of E11.5 control and NotchGOF λ-junction cells.**

| | | | | | | | | |
|---|---|---|---|---|---|---|---|---|
| ectoderm | .700008J07Ri | 0.31 | 1.03E-10 | 2.14E-06 | 0.2 | 0.15 | 0.04 | 1.29 |
| ectoderm | Tmem161b | 0.35 | 1.13E-10 | 2.36E-06 | 0.19 | 0.15 | 0.04 | 1.28 |
| ectoderm | Cpe | 0.33 | 1.23E-10 | 2.56E-06 | 0.46 | 0.41 | 0.05 | 1.13 |
| ectoderm | Nanp | 0.27 | 1.32E-10 | 2.74E-06 | 0.43 | 0.37 | 0.06 | 1.15 |
| ectoderm | Ube2t | 0.36 | 1.37E-10 | 2.84E-06 | 0.37 | 0.32 | 0.05 | 1.15 |
| ectoderm | Bbs2 | 0.35 | 1.37E-10 | 2.85E-06 | 0.12 | 0.09 | 0.03 | 1.38 |
| ectoderm | Mgat5 | 0.27 | 1.44E-10 | 3.00E-06 | 0.3 | 0.25 | 0.05 | 1.21 |
| ectoderm | Shmt2 | 0.27 | 1.54E-10 | 3.20E-06 | 0.38 | 0.33 | 0.05 | 1.16 |
| ectoderm | Gnl3l | 0.26 | 1.55E-10 | 3.23E-06 | 0.37 | 0.31 | 0.06 | 1.18 |
| ectoderm | Acot8 | 0.4 | 1.73E-10 | 3.60E-06 | 0.18 | 0.14 | 0.04 | 1.27 |
| ectoderm | AW209491 | 0.27 | 1.74E-10 | 3.62E-06 | 0.24 | 0.2 | 0.05 | 1.23 |
| ectoderm | Uxt | 0.29 | 1.78E-10 | 3.70E-06 | 0.42 | 0.37 | 0.05 | 1.14 |
| ectoderm | Mob2 | 0.3 | 2.07E-10 | 4.31E-06 | 0.45 | 0.39 | 0.06 | 1.14 |
| ectoderm | Apmap | 0.3 | 2.07E-10 | 4.31E-06 | 0.22 | 0.18 | 0.05 | 1.25 |
| ectoderm | Mdm2 | 0.26 | 2.19E-10 | 4.55E-06 | 0.48 | 0.42 | 0.06 | 1.13 |
| ectoderm | Chchd4 | 0.26 | 2.19E-10 | 4.56E-06 | 0.52 | 0.47 | 0.05 | 1.11 |
| ectoderm | Relch | 0.33 | 2.27E-10 | 4.72E-06 | 0.27 | 0.22 | 0.05 | 1.21 |
| ectoderm | Armh4 | 0.25 | 2.35E-10 | 4.88E-06 | 0.18 | 0.14 | 0.04 | 1.29 |
| ectoderm | Blcap | 0.28 | 2.38E-10 | 4.94E-06 | 0.36 | 0.3 | 0.05 | 1.18 |
| ectoderm | Zfp317 | 0.33 | 2.38E-10 | 4.96E-06 | 0.24 | 0.19 | 0.05 | 1.24 |
| ectoderm | Wwox | 0.31 | 2.78E-10 | 5.78E-06 | 0.13 | 0.1 | 0.03 | 1.35 |
| ectoderm | Pter | 0.46 | 2.92E-10 | 6.07E-06 | 0.14 | 0.11 | 0.03 | 1.32 |
| ectoderm | Dars2 | 0.32 | 2.92E-10 | 6.07E-06 | 0.24 | 0.19 | 0.05 | 1.24 |
| ectoderm | Ikbkb | 0.27 | 2.93E-10 | 6.09E-06 | 0.36 | 0.3 | 0.06 | 1.18 |
| ectoderm | Zfp229 | 0.34 | 3.00E-10 | 6.24E-06 | 0.12 | 0.08 | 0.03 | 1.38 |
| ectoderm | Pcmtd2 | 0.28 | 3.14E-10 | 6.53E-06 | 0.42 | 0.36 | 0.05 | 1.14 |
| ectoderm | Pik3c3 | 0.26 | 3.26E-10 | 6.78E-06 | 0.21 | 0.17 | 0.04 | 1.26 |
| ectoderm | Zpr1 | 0.26 | 3.34E-10 | 6.96E-06 | 0.47 | 0.42 | 0.06 | 1.13 |
| ectoderm | Mroh1 | 0.36 | 3.40E-10 | 7.06E-06 | 0.1 | 0.07 | 0.03 | 1.41 |
| ectoderm | Spice1 | 0.25 | 3.57E-10 | 7.42E-06 | 0.3 | 0.25 | 0.05 | 1.21 |
| ectoderm | Atp2b4 | 0.33 | 3.74E-10 | 7.77E-06 | 0.24 | 0.19 | 0.04 | 1.22 |
| ectoderm | Ppp6r1 | 0.31 | 4.17E-10 | 8.67E-06 | 0.29 | 0.24 | 0.05 | 1.2 |
| ectoderm | 130311K13Ri | 0.43 | 4.28E-10 | 8.90E-06 | 0.13 | 0.1 | 0.03 | 1.32 |
| ectoderm | Miip | 0.37 | 4.41E-10 | 9.17E-06 | 0.25 | 0.21 | 0.04 | 1.21 |
| ectoderm | Rad51d | 0.34 | 4.42E-10 | 9.19E-06 | 0.16 | 0.13 | 0.04 | 1.29 |
| ectoderm | Cnnm3 | 0.28 | 5.05E-10 | 1.05E-05 | 0.21 | 0.17 | 0.04 | 1.26 |
| ectoderm | Mthfsd | 0.25 | 5.35E-10 | 1.11E-05 | 0.29 | 0.24 | 0.05 | 1.21 |
| ectoderm | )130020L05Ri | 0.35 | 5.43E-10 | 1.13E-05 | 0.17 | 0.13 | 0.04 | 1.29 |
| ectoderm | Jmjd4 | 0.39 | 5.93E-10 | 1.23E-05 | 0.15 | 0.11 | 0.04 | 1.32 |
| ectoderm | Uvrag | 0.28 | 5.93E-10 | 1.23E-05 | 0.26 | 0.22 | 0.05 | 1.21 |
| ectoderm | Recql5 | 0.35 | 6.07E-10 | 1.26E-05 | 0.11 | 0.08 | 0.03 | 1.38 |
| ectoderm | Ndor1 | 0.36 | 6.13E-10 | 1.28E-05 | 0.13 | 0.1 | 0.03 | 1.34 |
| ectoderm | Fsd1 | 0.27 | 6.27E-10 | 1.30E-05 | 0.28 | 0.23 | 0.05 | 1.22 |
| ectoderm | Inpp5k | 0.4 | 6.52E-10 | 1.36E-05 | 0.14 | 0.11 | 0.03 | 1.31 |
| ectoderm | Ube2h | 0.3 | 7.08E-10 | 1.47E-05 | 0.5 | 0.46 | 0.04 | 1.1 |
| ectoderm | Gm19426 | 0.37 | 7.53E-10 | 1.57E-05 | 0.15 | 0.12 | 0.04 | 1.31 |
| ectoderm | Plp1 | 0.26 | 7.79E-10 | 1.62E-05 | 0.15 | 0.11 | 0.04 | 1.32 |
| ectoderm | Slc30a1 | 0.5 | 9.56E-10 | 1.99E-05 | 0.11 | 0.08 | 0.03 | 1.36 |
| ectoderm | Mtg2 | 0.29 | 9.74E-10 | 2.03E-05 | 0.3 | 0.25 | 0.05 | 1.19 |
| ectoderm | Ccdc6 | 0.27 | 9.76E-10 | 2.03E-05 | 0.32 | 0.27 | 0.05 | 1.19 |
| ectoderm | 630072M18R | 0.38 | 1.04E-09 | 2.16E-05 | 0.13 | 0.1 | 0.03 | 1.34 |
| ectoderm | Atxn1 | 0.75 | 1.07E-09 | 2.23E-05 | 0.12 | 0.09 | 0.03 | 1.34 |
| ectoderm | Klhl24 | 0.26 | 1.09E-09 | 2.28E-05 | 0.63 | 0.59 | 0.04 | 1.06 |
| ectoderm | Dhdds | 0.27 | 1.27E-09 | 2.65E-05 | 0.33 | 0.28 | 0.05 | 1.18 |
| ectoderm | Wdr47 | 0.44 | 1.33E-09 | 2.76E-05 | 0.12 | 0.09 | 0.03 | 1.34 |
| ectoderm | Repin1 | 0.26 | 1.33E-09 | 2.77E-05 | 0.16 | 0.13 | 0.04 | 1.3 |

**Supplementary Table 2. Marker genes for the major cell types of integrated clustering of E11.5 control and NotchGOF λ-junction cells.**

| | | | | | | | | |
|---|---|---|---|---|---|---|---|---|
| ectoderm | Wdr24 | 0.37 | 1.36E-09 | 2.83E-05 | 0.14 | 0.11 | 0.03 | 1.31 |
| ectoderm | Kmt2d | 0.29 | 1.46E-09 | 3.03E-05 | 0.29 | 0.24 | 0.05 | 1.2 |
| ectoderm | Adk | 0.27 | 1.48E-09 | 3.08E-05 | 0.63 | 0.6 | 0.03 | 1.05 |
| ectoderm | Arhgap39 | 0.27 | 1.58E-09 | 3.28E-05 | 0.21 | 0.17 | 0.04 | 1.25 |
| ectoderm | Cldn12 | 0.36 | 1.67E-09 | 3.46E-05 | 0.14 | 0.1 | 0.03 | 1.32 |
| ectoderm | Slc29a3 | 0.32 | 1.73E-09 | 3.60E-05 | 0.13 | 0.1 | 0.03 | 1.33 |
| ectoderm | Lekr1 | 0.29 | 1.74E-09 | 3.61E-05 | 0.14 | 0.11 | 0.03 | 1.31 |
| ectoderm | Usp24 | 0.25 | 1.76E-09 | 3.67E-05 | 0.35 | 0.3 | 0.05 | 1.18 |
| ectoderm | Card19 | 0.39 | 1.87E-09 | 3.89E-05 | 0.29 | 0.24 | 0.05 | 1.19 |
| ectoderm | Tti1 | 0.28 | 1.87E-09 | 3.89E-05 | 0.2 | 0.16 | 0.04 | 1.25 |
| ectoderm | Mpv17l2 | 0.31 | 2.00E-09 | 4.16E-05 | 0.26 | 0.22 | 0.05 | 1.21 |
| ectoderm | Gemin8 | 0.3 | 2.21E-09 | 4.59E-05 | 0.23 | 0.18 | 0.04 | 1.24 |
| ectoderm | Nr1h5 | 0.28 | 2.22E-09 | 4.62E-05 | 0.16 | 0.13 | 0.04 | 1.29 |
| ectoderm | Arid3a | 0.27 | 2.26E-09 | 4.70E-05 | 0.37 | 0.32 | 0.05 | 1.15 |
| ectoderm | Tk2 | 0.39 | 2.29E-09 | 4.76E-05 | 0.12 | 0.09 | 0.03 | 1.32 |
| ectoderm | Arfip1 | 0.29 | 2.40E-09 | 4.99E-05 | 0.22 | 0.18 | 0.04 | 1.24 |
| ectoderm | Dennd6b | 0.29 | 2.46E-09 | 5.11E-05 | 0.12 | 0.09 | 0.03 | 1.34 |
| ectoderm | Slc4a3 | 0.26 | 2.54E-09 | 5.29E-05 | 0.17 | 0.13 | 0.04 | 1.29 |
| ectoderm | Intu | 0.33 | 2.59E-09 | 5.38E-05 | 0.13 | 0.1 | 0.03 | 1.33 |
| ectoderm | Cbr4 | 0.26 | 2.66E-09 | 5.53E-05 | 0.27 | 0.22 | 0.05 | 1.21 |
| ectoderm | 330018D20Ri | 0.34 | 3.53E-09 | 7.34E-05 | 0.22 | 0.18 | 0.04 | 1.23 |
| ectoderm | Rela | 0.27 | 3.54E-09 | 7.36E-05 | 0.28 | 0.24 | 0.05 | 1.2 |
| ectoderm | Dolpp1 | 0.32 | 3.77E-09 | 7.84E-05 | 0.17 | 0.13 | 0.04 | 1.28 |
| ectoderm | Scnm1 | 0.28 | 3.81E-09 | 7.92E-05 | 0.46 | 0.41 | 0.05 | 1.13 |
| ectoderm | Rab10os | 0.28 | 3.87E-09 | 8.04E-05 | 0.22 | 0.18 | 0.04 | 1.23 |
| ectoderm | Comtd1 | 0.32 | 4.41E-09 | 9.17E-05 | 0.17 | 0.13 | 0.04 | 1.26 |
| ectoderm | Polr3e | 0.29 | 4.54E-09 | 9.45E-05 | 0.25 | 0.21 | 0.04 | 1.21 |
| ectoderm | Taf1c | 0.37 | 4.74E-09 | 9.86E-05 | 0.15 | 0.11 | 0.03 | 1.29 |
| ectoderm | Zbtb5 | 0.26 | 4.92E-09 | 1.02E-04 | 0.34 | 0.29 | 0.05 | 1.18 |
| ectoderm | Tekt1 | 0.37 | 5.61E-09 | 1.17E-04 | 0.11 | 0.09 | 0.03 | 1.34 |
| ectoderm | 810403D21Ri | 0.34 | 6.20E-09 | 1.29E-04 | 0.17 | 0.14 | 0.04 | 1.26 |
| ectoderm | Pex11a | 0.32 | 6.33E-09 | 1.32E-04 | 0.14 | 0.1 | 0.03 | 1.31 |
| ectoderm | Zfp64 | 0.29 | 6.62E-09 | 1.38E-04 | 0.35 | 0.3 | 0.05 | 1.17 |
| ectoderm | Syt11 | 0.45 | 7.44E-09 | 1.55E-04 | 0.34 | 0.3 | 0.04 | 1.13 |
| ectoderm | Slx4ip | 0.26 | 7.93E-09 | 1.65E-04 | 0.29 | 0.24 | 0.05 | 1.19 |
| ectoderm | Rnase4 | 0.31 | 8.25E-09 | 1.72E-04 | 0.12 | 0.09 | 0.03 | 1.31 |
| ectoderm | Zfp65 | 0.31 | 8.53E-09 | 1.77E-04 | 0.21 | 0.17 | 0.04 | 1.23 |
| ectoderm | Zfp943 | 0.39 | 8.70E-09 | 1.81E-04 | 0.18 | 0.14 | 0.04 | 1.25 |
| ectoderm | Jmjd7 | 0.26 | 8.88E-09 | 1.85E-04 | 0.2 | 0.16 | 0.04 | 1.25 |
| ectoderm | Hint3 | 0.31 | 9.08E-09 | 1.89E-04 | 0.27 | 0.23 | 0.04 | 1.19 |
| ectoderm | Slc27a1 | 0.35 | 1.04E-08 | 2.17E-04 | 0.11 | 0.08 | 0.03 | 1.35 |
| ectoderm | Ercc6 | 0.28 | 1.08E-08 | 2.24E-04 | 0.19 | 0.15 | 0.04 | 1.25 |
| ectoderm | Slc29a4 | 0.31 | 1.23E-08 | 2.56E-04 | 0.13 | 0.1 | 0.03 | 1.31 |
| ectoderm | Atg14 | 0.33 | 1.24E-08 | 2.58E-04 | 0.27 | 0.23 | 0.04 | 1.19 |
| ectoderm | Klf13 | 0.29 | 1.55E-08 | 3.23E-04 | 0.41 | 0.37 | 0.04 | 1.11 |
| ectoderm | Inpp5f | 0.3 | 1.59E-08 | 3.32E-04 | 0.27 | 0.23 | 0.04 | 1.18 |
| ectoderm | Slc35a2 | 0.28 | 1.64E-08 | 3.40E-04 | 0.18 | 0.14 | 0.04 | 1.25 |
| ectoderm | Trmt61a | 0.31 | 1.65E-08 | 3.43E-04 | 0.18 | 0.14 | 0.03 | 1.24 |
| ectoderm | Zbtb21 | 0.32 | 1.72E-08 | 3.58E-04 | 0.16 | 0.13 | 0.04 | 1.27 |
| ectoderm | Txnrd3 | 0.29 | 1.73E-08 | 3.60E-04 | 0.14 | 0.11 | 0.03 | 1.29 |
| ectoderm | 2310061I04Ril | 0.26 | 1.79E-08 | 3.73E-04 | 0.22 | 0.18 | 0.04 | 1.22 |
| ectoderm | Naa30 | 0.27 | 2.18E-08 | 4.53E-04 | 0.27 | 0.22 | 0.04 | 1.19 |
| ectoderm | Aph1b | 0.27 | 2.51E-08 | 5.22E-04 | 0.11 | 0.09 | 0.03 | 1.33 |
| ectoderm | Cds2 | 0.27 | 2.80E-08 | 5.82E-04 | 0.21 | 0.17 | 0.04 | 1.22 |
| ectoderm | Mfsd1 | 0.28 | 2.82E-08 | 5.86E-04 | 0.4 | 0.35 | 0.04 | 1.12 |
| ectoderm | Apba3 | 0.29 | 3.16E-08 | 6.57E-04 | 0.18 | 0.14 | 0.04 | 1.24 |

**Supplementary Table 2. Marker genes for the major cell types of integrated clustering of E11.5 control and NotchGOF λ-junction cells.**

| | | | | | | | | |
|---|---|---|---|---|---|---|---|---|
| ectoderm | Zfp991 | 0.31 | 3.96E-08 | 8.23E-04 | 0.11 | 0.08 | 0.03 | 1.33 |
| ectoderm | Uckl1 | 0.3 | 4.37E-08 | 9.08E-04 | 0.14 | 0.11 | 0.03 | 1.28 |
| ectoderm | Arrdc3 | 0.28 | 4.50E-08 | 9.35E-04 | 0.24 | 0.2 | 0.04 | 1.2 |
| ectoderm | Tbcel | 0.27 | 5.93E-08 | 1.23E-03 | 0.25 | 0.21 | 0.04 | 1.2 |
| ectoderm | Slc33a1 | 0.28 | 6.36E-08 | 1.32E-03 | 0.14 | 0.11 | 0.03 | 1.28 |
| ectoderm | Ciao3 | 0.28 | 6.94E-08 | 1.44E-03 | 0.2 | 0.16 | 0.04 | 1.23 |
| ectoderm | Tmem87a | 0.28 | 7.34E-08 | 1.53E-03 | 0.23 | 0.19 | 0.04 | 1.21 |
| ectoderm | Fam160b2 | 0.25 | 8.44E-08 | 1.75E-03 | 0.17 | 0.13 | 0.03 | 1.25 |
| ectoderm | Zfp777 | 0.29 | 9.07E-08 | 1.89E-03 | 0.22 | 0.18 | 0.04 | 1.22 |
| ectoderm | Tut1 | 0.27 | 9.39E-08 | 1.95E-03 | 0.17 | 0.14 | 0.03 | 1.24 |
| ectoderm | Coq10a | 0.27 | 1.09E-07 | 2.26E-03 | 0.1 | 0.08 | 0.03 | 1.32 |
| ectoderm | Zfp974 | 0.38 | 1.16E-07 | 2.41E-03 | 0.11 | 0.08 | 0.03 | 1.31 |
| ectoderm | Cln6 | 0.31 | 1.19E-07 | 2.48E-03 | 0.12 | 0.09 | 0.03 | 1.3 |
| ectoderm | Noct | 0.34 | 1.25E-07 | 2.61E-03 | 0.22 | 0.18 | 0.04 | 1.2 |
| ectoderm | Hinfp | 0.31 | 1.34E-07 | 2.78E-03 | 0.18 | 0.15 | 0.04 | 1.24 |
| ectoderm | Qsox2 | 0.28 | 1.35E-07 | 2.80E-03 | 0.16 | 0.13 | 0.03 | 1.26 |
| ectoderm | Cnksr3 | 0.26 | 1.46E-07 | 3.04E-03 | 0.13 | 0.1 | 0.03 | 1.29 |
| ectoderm | .130014A01Ri | 0.25 | 1.55E-07 | 3.22E-03 | 0.1 | 0.08 | 0.03 | 1.33 |
| ectoderm | Tmco6 | 0.27 | 1.75E-07 | 3.64E-03 | 0.15 | 0.12 | 0.03 | 1.25 |
| ectoderm | Ubox5 | 0.33 | 1.98E-07 | 4.11E-03 | 0.12 | 0.09 | 0.03 | 1.3 |
| ectoderm | Eipr1 | 0.26 | 2.09E-07 | 4.35E-03 | 0.27 | 0.23 | 0.04 | 1.17 |
| ectoderm | Ect2 | 0.25 | 2.22E-07 | 4.61E-03 | 0.44 | 0.4 | 0.03 | 1.08 |
| ectoderm | Mkks | 0.26 | 2.23E-07 | 4.65E-03 | 0.2 | 0.16 | 0.04 | 1.22 |
| ectoderm | Tpra1 | 0.27 | 2.43E-07 | 5.06E-03 | 0.15 | 0.12 | 0.03 | 1.25 |
| ectoderm | Chtf8 | 0.33 | 2.55E-07 | 5.30E-03 | 0.15 | 0.12 | 0.03 | 1.26 |
| ectoderm | Bloc1s3 | 0.44 | 2.60E-07 | 5.40E-03 | 0.11 | 0.08 | 0.03 | 1.3 |
| ectoderm | Pptc7 | 0.26 | 3.22E-07 | 6.71E-03 | 0.22 | 0.18 | 0.04 | 1.2 |
| ectoderm | Taf4 | 0.29 | 3.40E-07 | 7.08E-03 | 0.16 | 0.13 | 0.03 | 1.24 |
| ectoderm | Pfkfb3 | 0.38 | 3.50E-07 | 7.27E-03 | 0.13 | 0.1 | 0.03 | 1.27 |
| ectoderm | Aldh6a1 | 0.31 | 4.28E-07 | 8.90E-03 | 0.13 | 0.11 | 0.03 | 1.27 |
| ectoderm | Fkbpl | 0.34 | 5.56E-07 | 1.16E-02 | 0.17 | 0.14 | 0.03 | 1.22 |
| ectoderm | Rad51b | 0.28 | 6.25E-07 | 1.30E-02 | 0.14 | 0.11 | 0.03 | 1.25 |
| ectoderm | Cep55 | 0.28 | 6.41E-07 | 1.33E-02 | 0.27 | 0.23 | 0.04 | 1.16 |
| ectoderm | Clba1 | 0.33 | 6.48E-07 | 1.35E-02 | 0.1 | 0.08 | 0.02 | 1.3 |
| ectoderm | Arrb2 | 0.29 | 6.73E-07 | 1.40E-02 | 0.2 | 0.16 | 0.03 | 1.2 |
| ectoderm | Polg2 | 0.33 | 7.63E-07 | 1.59E-02 | 0.12 | 0.09 | 0.03 | 1.27 |
| ectoderm | Atg10 | 0.3 | 7.82E-07 | 1.63E-02 | 0.17 | 0.14 | 0.03 | 1.21 |
| ectoderm | Lifr | 0.32 | 8.37E-07 | 1.74E-02 | 0.13 | 0.1 | 0.03 | 1.26 |
| ectoderm | Primpol | 0.31 | 8.88E-07 | 1.85E-02 | 0.13 | 0.11 | 0.03 | 1.26 |
| ectoderm | Bsdc1 | 0.25 | 1.28E-06 | 2.66E-02 | 0.25 | 0.21 | 0.04 | 1.18 |
| ectoderm | Rbm45 | 0.28 | 1.38E-06 | 2.88E-02 | 0.19 | 0.16 | 0.03 | 1.21 |
| ectoderm | Aplf | 0.32 | 1.44E-06 | 2.99E-02 | 0.13 | 0.1 | 0.03 | 1.25 |
| ectoderm | Zfp524 | 0.45 | 1.45E-06 | 3.01E-02 | 0.12 | 0.1 | 0.02 | 1.24 |
| ectoderm | Tbc1d22b | 0.25 | 1.57E-06 | 3.27E-02 | 0.13 | 0.1 | 0.03 | 1.28 |
| ectoderm | Fam234a | 0.3 | 1.68E-06 | 3.49E-02 | 0.17 | 0.14 | 0.03 | 1.22 |
| ectoderm | Ipp | 0.28 | 1.73E-06 | 3.60E-02 | 0.17 | 0.14 | 0.03 | 1.22 |
| ectoderm | Acox1 | 0.26 | 1.81E-06 | 3.76E-02 | 0.17 | 0.14 | 0.03 | 1.22 |
| ectoderm | Rpap2 | 0.29 | 1.94E-06 | 4.04E-02 | 0.26 | 0.23 | 0.04 | 1.16 |
| ectoderm | Setd6 | 0.33 | 2.05E-06 | 4.27E-02 | 0.15 | 0.12 | 0.03 | 1.23 |
| ectoderm | Gbe1 | 0.26 | 2.26E-06 | 4.70E-02 | 0.12 | 0.1 | 0.03 | 1.26 |
| red blood cells | Gypa | 9.27 | 0 | 0 | 0.99 | 0.01 | 0.98 | 89.64 |
| red blood cells | Slc4a1 | 7.87 | 0 | 0 | 1 | 0.02 | 0.98 | 41.63 |
| red blood cells | Alas2 | 9.06 | 0 | 0 | 0.99 | 0.02 | 0.98 | 52.32 |
| red blood cells | Rhd | 8.87 | 0 | 0 | 0.98 | 0.01 | 0.97 | 98.4 |
| red blood cells | Hemgn | 8.75 | 0 | 0 | 0.98 | 0.02 | 0.96 | 57.59 |
| red blood cells | Ermap | 8.59 | 0 | 0 | 0.96 | 0.01 | 0.96 | 160.5 |

**Supplementary Table 2. Marker genes for the major cell types of integrated clustering of E11.5 control and NotchGOF λ-junction cells.**

| | | | | | | | | |
|---|---|---|---|---|---|---|---|---|
| red blood cells | Trim10 | 9.2 | 0 | 0 | 0.96 | 0 | 0.95 | 238.75 |
| red blood cells | Cited4 | 6.75 | 0 | 0 | 0.97 | 0.02 | 0.95 | 46.05 |
| red blood cells | Klf1 | 6.91 | 0 | 0 | 0.96 | 0.02 | 0.94 | 53.22 |
| red blood cells | Acp5 | 6.16 | 0 | 0 | 0.98 | 0.05 | 0.93 | 18.92 |
| red blood cells | Pdzk1ip1 | 8.39 | 0 | 0 | 0.93 | 0.01 | 0.92 | 103.11 |
| red blood cells | Car2 | 7.55 | 0 | 0 | 1 | 0.08 | 0.92 | 12.33 |
| red blood cells | Icam4 | 8.45 | 0 | 0 | 0.92 | 0.01 | 0.92 | 153.83 |
| red blood cells | Tspan33 | 5.86 | 0 | 0 | 0.96 | 0.05 | 0.91 | 20 |
| red blood cells | Ncf4 | 7.52 | 0 | 0 | 0.92 | 0.01 | 0.91 | 92 |
| red blood cells | Rhag | 9 | 0 | 0 | 0.91 | 0 | 0.91 | 227.75 |
| red blood cells | Spta1 | 8.45 | 0 | 0 | 0.91 | 0.01 | 0.9 | 181.2 |
| red blood cells | Mt2 | 5.89 | 0 | 0 | 0.98 | 0.1 | 0.89 | 10.33 |
| red blood cells | Snca | 4.7 | 0 | 0 | 0.94 | 0.06 | 0.88 | 17.05 |
| red blood cells | Sptb | 6.86 | 0 | 0 | 0.89 | 0.01 | 0.88 | 63.86 |
| red blood cells | Tal1 | 6.57 | 0 | 0 | 0.89 | 0.01 | 0.88 | 63.64 |
| red blood cells | Ube2l6 | 4.46 | 0 | 0 | 0.97 | 0.1 | 0.87 | 9.68 |
| red blood cells | Kel | 7.61 | 0 | 0 | 0.87 | 0.01 | 0.86 | 123.57 |
| red blood cells | Pnpo | 4.5 | 0 | 0 | 0.97 | 0.14 | 0.83 | 7.03 |
| red blood cells | Add2 | 4.49 | 0 | 0 | 0.88 | 0.06 | 0.82 | 15.39 |
| red blood cells | Smim1 | 4.61 | 0 | 0 | 0.95 | 0.14 | 0.82 | 7.07 |
| red blood cells | Epb42 | 8.32 | 0 | 0 | 0.82 | 0 | 0.81 | 204.5 |
| red blood cells | Slc39a8 | 3.83 | 0 | 0 | 0.95 | 0.14 | 0.81 | 6.83 |
| red blood cells | Fam210b | 3.99 | 0 | 0 | 0.98 | 0.17 | 0.81 | 5.79 |
| red blood cells | Cldn13 | 8.82 | 0 | 0 | 0.81 | 0 | 0.81 | 270.33 |
| red blood cells | Hsd3b6 | 9.11 | 0 | 0 | 0.81 | 0 | 0.8 | 201.75 |
| red blood cells | Tuba4a | 3.81 | 0 | 0 | 0.84 | 0.04 | 0.8 | 22.7 |
| red blood cells | Slc38a5 | 8.21 | 0 | 0 | 0.8 | 0 | 0.8 | 200.5 |
| red blood cells | Hbb-bs | 9.2 | 0 | 0 | 0.99 | 0.21 | 0.79 | 4.77 |
| red blood cells | Slc25a37 | 5.52 | 0 | 0 | 1 | 0.21 | 0.79 | 4.74 |
| red blood cells | Gmpr | 3.7 | 0 | 0 | 0.92 | 0.15 | 0.78 | 6.32 |
| red blood cells | Marchf2 | 3.36 | 0 | 0 | 0.96 | 0.18 | 0.78 | 5.22 |
| red blood cells | Cd59a | 3.1 | 0 | 0 | 0.96 | 0.19 | 0.77 | 5.02 |
| red blood cells | Tspan32 | 5.73 | 0 | 0 | 0.77 | 0.01 | 0.77 | 96.75 |
| red blood cells | Chchd10 | 3.07 | 0 | 0 | 0.9 | 0.13 | 0.77 | 6.85 |
| red blood cells | Mt1 | 6.02 | 0 | 0 | 0.99 | 0.24 | 0.76 | 4.18 |
| red blood cells | Hbq1b | 8.86 | 0 | 0 | 0.75 | 0 | 0.75 | 748 |
| red blood cells | Epor | 4.24 | 0 | 0 | 0.79 | 0.05 | 0.75 | 16.87 |
| red blood cells | Fech | 4.01 | 0 | 0 | 0.99 | 0.25 | 0.75 | 4.04 |
| red blood cells | Ptp4a3 | 3.12 | 0 | 0 | 0.97 | 0.23 | 0.74 | 4.31 |
| red blood cells | Hebp1 | 4 | 0 | 0 | 0.98 | 0.24 | 0.74 | 4.08 |
| red blood cells | Hbb-bt | 9.11 | 0 | 0 | 1 | 0.27 | 0.73 | 3.65 |
| red blood cells | Atp8a1 | 3.73 | 0 | 0 | 0.78 | 0.06 | 0.72 | 13.24 |
| red blood cells | Abcb10 | 3.08 | 0 | 0 | 0.92 | 0.2 | 0.72 | 4.61 |
| red blood cells | Ell2 | 3.93 | 0 | 0 | 0.78 | 0.06 | 0.72 | 12.82 |
| red blood cells | .300017J02Ri | 7.53 | 0 | 0 | 0.72 | 0 | 0.72 | 180.5 |
| red blood cells | Tent5c | 6.08 | 0 | 0 | 0.73 | 0.01 | 0.72 | 56.15 |
| red blood cells | Plek2 | 4.7 | 0 | 0 | 0.75 | 0.03 | 0.72 | 22.73 |
| red blood cells | Slc30a10 | 5.74 | 0 | 0 | 0.73 | 0.01 | 0.72 | 56.08 |
| red blood cells | Gata1 | 6.73 | 0 | 0 | 0.72 | 0 | 0.71 | 179.25 |
| red blood cells | Cfap57 | 8.81 | 0 | 0 | 0.71 | 0 | 0.71 | 237.33 |
| red blood cells | Gm16867 | 5.41 | 0 | 0 | 0.73 | 0.02 | 0.71 | 36.45 |
| red blood cells | Rbm38 | 2.71 | 0 | 0 | 0.92 | 0.21 | 0.71 | 4.38 |
| red blood cells | Kcnn4 | 7.37 | 0 | 0 | 0.71 | 0 | 0.7 | 177 |
| red blood cells | Cpox | 4.27 | 0 | 0 | 0.99 | 0.29 | 0.7 | 3.39 |
| red blood cells | Ank1 | 6.71 | 0 | 0 | 0.7 | 0.01 | 0.69 | 99.71 |
| red blood cells | Nfe2 | 5.8 | 0 | 0 | 0.7 | 0.01 | 0.69 | 87 |

**Supplementary Table 2. Marker genes for the major cell types of integrated clustering of E11.5 control and NotchGOF λ-junction cells.**

| | | | | | | | | |
|---|---|---|---|---|---|---|---|---|
| red blood cells | Uros | 2.76 | 0 | 0 | 0.83 | 0.14 | 0.68 | 5.73 |
| red blood cells | Lmx1a | 8.01 | 0 | 0 | 0.68 | 0 | 0.68 | 227 |
| red blood cells | Ctse | 7.73 | 0 | 0 | 0.68 | 0 | 0.67 | 225.33 |
| red blood cells | Grina | 2.29 | 0 | 0 | 0.9 | 0.22 | 0.67 | 4 |
| red blood cells | Blvrb | 5.01 | 0 | 0 | 0.99 | 0.32 | 0.67 | 3.11 |
| red blood cells | Creg1 | 3.38 | 0 | 0 | 0.99 | 0.33 | 0.66 | 3.01 |
| red blood cells | Ucp2 | 3.02 | 0 | 0 | 0.99 | 0.33 | 0.65 | 2.97 |
| red blood cells | Rfesd | 2.77 | 0 | 0 | 0.93 | 0.28 | 0.65 | 3.3 |
| red blood cells | Asb17os | 8.92 | 0 | 0 | 0.65 | 0 | 0.65 | 649 |
| red blood cells | Tmod1 | 4.2 | 0 | 0 | 0.68 | 0.04 | 0.65 | 18.46 |
| red blood cells | Dmtn | 3.99 | 0 | 0 | 0.68 | 0.04 | 0.64 | 16.66 |
| red blood cells | Ncoa4 | 2.26 | 0 | 0 | 0.93 | 0.29 | 0.63 | 3.16 |
| red blood cells | Serinc3 | 2.66 | 0 | 0 | 0.97 | 0.34 | 0.63 | 2.87 |
| red blood cells | Chac2 | 2.97 | 0 | 0 | 0.74 | 0.11 | 0.63 | 6.78 |
| red blood cells | St3gal5 | 2.21 | 0 | 0 | 0.85 | 0.22 | 0.63 | 3.89 |
| red blood cells | Grap2 | 7.42 | 0 | 0 | 0.63 | 0 | 0.62 | 209 |
| red blood cells | Gclm | 3.55 | 0 | 0 | 0.99 | 0.37 | 0.62 | 2.7 |
| red blood cells | Aqp3 | 5.3 | 0 | 0 | 0.63 | 0.01 | 0.62 | 52.67 |
| red blood cells | Tspo | 2.25 | 0 | 0 | 0.94 | 0.33 | 0.62 | 2.88 |
| red blood cells | Fam241a | 2.84 | 0 | 0 | 0.71 | 0.1 | 0.61 | 7.34 |
| red blood cells | Cpeb4 | 3.34 | 0 | 0 | 0.68 | 0.07 | 0.61 | 9.52 |
| red blood cells | Zfpm1 | 2.02 | 0 | 0 | 0.9 | 0.29 | 0.61 | 3.08 |
| red blood cells | Hbb-bh1 | 9.52 | 0 | 0 | 1 | 0.4 | 0.6 | 2.52 |
| red blood cells | Slc25a21 | 3.48 | 0 | 0 | 0.63 | 0.04 | 0.59 | 14.41 |
| red blood cells | Josd2 | 3.14 | 0 | 0 | 0.98 | 0.4 | 0.58 | 2.47 |
| red blood cells | Ifrd2 | 2.05 | 0 | 0 | 0.89 | 0.31 | 0.58 | 2.88 |
| red blood cells | Slc43a1 | 2.33 | 0 | 0 | 0.71 | 0.14 | 0.57 | 5.14 |
| red blood cells | Vangl1 | 2.38 | 0 | 0 | 0.7 | 0.13 | 0.57 | 5.29 |
| red blood cells | Dhrs11 | 4.34 | 0 | 0 | 0.59 | 0.02 | 0.56 | 24.38 |
| red blood cells | Xk | 5.34 | 0 | 0 | 0.57 | 0.01 | 0.56 | 44 |
| red blood cells | Prr13 | 2.05 | 0 | 0 | 0.94 | 0.38 | 0.56 | 2.47 |
| red blood cells | Tmem238 | 1.89 | 0 | 0 | 0.71 | 0.16 | 0.55 | 4.48 |
| red blood cells | Lyrm9 | 2.07 | 0 | 0 | 0.7 | 0.15 | 0.55 | 4.63 |
| red blood cells | Arrb1 | 2.71 | 0 | 0 | 0.64 | 0.09 | 0.55 | 6.85 |
| red blood cells | Bmp2k | 2.3 | 0 | 0 | 0.68 | 0.13 | 0.55 | 5.22 |
| red blood cells | Lyl1 | 4.74 | 0 | 0 | 0.56 | 0.02 | 0.55 | 37.6 |
| red blood cells | Bpgm | 3.08 | 0 | 0 | 0.98 | 0.44 | 0.54 | 2.24 |
| red blood cells | Gm2061 | 8.12 | 0 | 0 | 0.54 | 0 | 0.54 | 271 |
| red blood cells | Tinagl1 | 3.51 | 0 | 0 | 0.58 | 0.04 | 0.54 | 15.18 |
| red blood cells | Gm867 | 8.32 | 0 | 0 | 0.54 | 0 | 0.54 | 268.5 |
| red blood cells | Reln | 4.53 | 0 | 0 | 0.53 | 0.02 | 0.51 | 29.22 |
| red blood cells | Mgst3 | 4.44 | 0 | 0 | 0.99 | 0.49 | 0.5 | 2.03 |
| red blood cells | Pkhd1l1 | 8.21 | 0 | 0 | 0.5 | 0 | 0.5 | 250.5 |
| red blood cells | Cox6b2 | 2.09 | 0 | 0 | 0.95 | 0.45 | 0.5 | 2.11 |
| red blood cells | Fbxo6 | 3.09 | 0 | 0 | 0.54 | 0.04 | 0.49 | 12.2 |
| red blood cells | Gpr146 | 4.87 | 0 | 0 | 0.5 | 0.01 | 0.49 | 45.09 |
| red blood cells | Rec114 | 5.8 | 0 | 0 | 0.49 | 0.01 | 0.48 | 70 |
| red blood cells | Dapk2 | 3.69 | 0 | 0 | 0.51 | 0.03 | 0.48 | 17.55 |
| red blood cells | Tmem184a | 3.58 | 0 | 0 | 0.51 | 0.04 | 0.48 | 14.6 |
| red blood cells | Pklr | 7.91 | 0 | 0 | 0.46 | 0 | 0.46 | 462 |
| red blood cells | Svip | 2.1 | 0 | 0 | 0.53 | 0.07 | 0.46 | 7.2 |
| red blood cells | Htatip2 | 2.75 | 0 | 0 | 0.51 | 0.06 | 0.45 | 9.11 |
| red blood cells | Steap3 | 5.89 | 0 | 0 | 0.46 | 0.01 | 0.45 | 75.83 |
| red blood cells | Ikzf1 | 4.77 | 0 | 0 | 0.46 | 0.01 | 0.45 | 45.9 |
| red blood cells | Urod | 2.63 | 0 | 0 | 0.99 | 0.55 | 0.44 | 1.8 |
| red blood cells | Aqp8 | 8.92 | 0 | 0 | 0.44 | 0 | 0.44 | 438 |

**Supplementary Table 2. Marker genes for the major cell types of integrated clustering of E11.5 control and NotchGOF λ-junction cells.**

| | | | | | | | | |
|---|---|---|---|---|---|---|---|---|
| red blood cells | Gm32051 | 8.63 | 0 | 0 | 0.43 | 0 | 0.43 | 434 |
| red blood cells | Mylk3 | 5.04 | 0 | 0 | 0.44 | 0.01 | 0.43 | 48.78 |
| red blood cells | Slc6a9 | 3.58 | 0 | 0 | 0.45 | 0.03 | 0.42 | 15.59 |
| red blood cells | Hmbs | 3.4 | 0 | 0 | 0.99 | 0.57 | 0.42 | 1.74 |
| red blood cells | Gfi1b | 6.58 | 0 | 0 | 0.42 | 0 | 0.41 | 138.33 |
| red blood cells | Cmpk2 | 3.9 | 0 | 0 | 0.43 | 0.02 | 0.41 | 20.38 |
| red blood cells | Gm14275 | 9.06 | 0 | 0 | 0.4 | 0 | 0.4 | 399 |
| red blood cells | Nmnat3 | 3.29 | 0 | 0 | 0.43 | 0.03 | 0.4 | 13.84 |
| red blood cells | Redrum | 5.34 | 0 | 0 | 0.4 | 0.01 | 0.39 | 57.29 |
| red blood cells | Slc22a4 | 6.7 | 0 | 0 | 0.4 | 0 | 0.39 | 131.67 |
| red blood cells | Pheta2 | 2.7 | 0 | 0 | 0.44 | 0.05 | 0.39 | 9.52 |
| red blood cells | 030468B19Ri | 9.14 | 0 | 0 | 0.39 | 0 | 0.39 | 392 |
| red blood cells | Larp1b | 2.32 | 0 | 0 | 0.45 | 0.06 | 0.39 | 7.11 |
| red blood cells | Slfn14 | 8.33 | 0 | 0 | 0.38 | 0 | 0.38 | #NA |
| red blood cells | Alad | 3.34 | 0 | 0 | 1 | 0.62 | 0.37 | 1.6 |
| red blood cells | Ccdc141 | 5.22 | 0 | 0 | 0.38 | 0.01 | 0.37 | 38 |
| red blood cells | Asb17 | 7.32 | 0 | 0 | 0.37 | 0 | 0.36 | 182.5 |
| red blood cells | Cd59b | 3.09 | 0 | 0 | 0.39 | 0.03 | 0.36 | 12.45 |
| red blood cells | Gm38619.1 | 8.33 | 0 | 0 | 0.36 | 0 | 0.36 | 356 |
| red blood cells | Cdkn2c | 3.39 | 0 | 0 | 0.38 | 0.03 | 0.35 | 14.62 |
| red blood cells | Hba-a2 | 8.45 | 0 | 0 | 1 | 0.65 | 0.35 | 1.55 |
| red blood cells | Ifi35 | 3.91 | 0 | 0 | 0.37 | 0.02 | 0.35 | 21.82 |
| red blood cells | Zc3hav1 | 4.38 | 0 | 0 | 0.37 | 0.01 | 0.35 | 26.14 |
| red blood cells | Ttc39a | 2.44 | 0 | 0 | 0.4 | 0.05 | 0.35 | 8.61 |
| red blood cells | Spns2 | 3.98 | 0 | 0 | 0.37 | 0.02 | 0.35 | 24.33 |
| red blood cells | Rnf212 | 8.45 | 0 | 0 | 0.34 | 0 | 0.34 | 340 |
| red blood cells | Ache | 5.11 | 0 | 0 | 0.34 | 0.01 | 0.33 | 55.83 |
| red blood cells | Adra2b | 7.82 | 0 | 0 | 0.33 | 0 | 0.33 | 328 |
| red blood cells | Ccdc92b | 6.54 | 0 | 0 | 0.33 | 0 | 0.33 | 109.33 |
| red blood cells | Mrap | 8 | 0 | 0 | 0.32 | 0 | 0.32 | 324 |
| red blood cells | Hba-a1 | 9.7 | 0 | 0 | 1 | 0.68 | 0.32 | 1.46 |
| red blood cells | Gm43154 | 7.65 | 0 | 0 | 0.31 | 0 | 0.31 | 314 |
| red blood cells | Tmem86b | 3.73 | 0 | 0 | 0.33 | 0.02 | 0.31 | 18.28 |
| red blood cells | Gm20161 | 9.24 | 0 | 0 | 0.3 | 0 | 0.3 | #NA |
| red blood cells | 430401H09Ri | 7.46 | 0 | 0 | 0.3 | 0 | 0.3 | 302 |
| red blood cells | Wnk4 | 2.97 | 0 | 0 | 0.32 | 0.02 | 0.3 | 13.42 |
| red blood cells | Mfsd2b | 4.94 | 0 | 0 | 0.3 | 0.01 | 0.3 | 60.6 |
| red blood cells | Acss1 | 3.97 | 0 | 0 | 0.3 | 0.01 | 0.28 | 21.29 |
| red blood cells | Treml2 | 4.66 | 0 | 0 | 0.28 | 0.01 | 0.27 | 55.4 |
| red blood cells | Art4 | 3.21 | 0 | 0 | 0.29 | 0.02 | 0.27 | 15.94 |
| red blood cells | Trib3 | 4.12 | 0 | 0 | 0.28 | 0.01 | 0.27 | 23.08 |
| red blood cells | Igtp | 3.69 | 0 | 0 | 0.26 | 0.01 | 0.25 | 25.9 |
| red blood cells | Ypel4 | 3.54 | 0 | 0 | 0.27 | 0.02 | 0.25 | 13.95 |
| red blood cells | Tmem56 | 3.9 | 0 | 0 | 0.26 | 0.01 | 0.25 | 19.92 |
| red blood cells | Gm9895 | 5.47 | 0 | 0 | 0.24 | 0 | 0.24 | 81 |
| red blood cells | Gm10390 | 7.33 | 0 | 0 | 0.24 | 0 | 0.24 | 239 |
| red blood cells | Smim5 | 3.06 | 0 | 0 | 0.25 | 0.02 | 0.24 | 14.82 |
| red blood cells | Popdc2 | 5.22 | 0 | 0 | 0.24 | 0 | 0.23 | 78.67 |
| red blood cells | Nudt4 | 3.13 | 0 | 0 | 1 | 0.76 | 0.23 | 1.3 |
| red blood cells | Cox17 | 1.53 | 0 | 0 | 0.98 | 0.76 | 0.23 | 1.3 |
| red blood cells | Flt3l | 5.19 | 0 | 0 | 0.23 | 0 | 0.22 | 56.5 |
| red blood cells | Ccrl2 | 4.9 | 0 | 0 | 0.22 | 0 | 0.21 | 54.5 |
| red blood cells | Arsg | 3.85 | 0 | 0 | 0.21 | 0.01 | 0.21 | 26.75 |
| red blood cells | Ackr1 | 4.07 | 0 | 0 | 0.21 | 0.01 | 0.2 | 26.13 |
| red blood cells | Gm38619 | 9.23 | 0 | 0 | 0.2 | 0 | 0.2 | #NA |
| red blood cells | Abcg4 | 3.49 | 0 | 0 | 0.21 | 0.01 | 0.2 | 18.82 |

**Supplementary Table 2. Marker genes for the major cell types of integrated clustering of E11.5 control and NotchGOF λ-junction cells.**

| | | | | | | | | |
|---|---|---|---|---|---|---|---|---|
| red blood cells | Gm10603 | 5.31 | 0 | 0 | 0.19 | 0 | 0.19 | 48.25 |
| red blood cells | Lrrc39 | 6.05 | 0 | 0 | 0.18 | 0 | 0.18 | 92 |
| red blood cells | Hopx | 3.43 | 0 | 0 | 0.18 | 0.01 | 0.17 | 22.75 |
| red blood cells | 930455G09Ri | 3.83 | 0 | 0 | 0.18 | 0.01 | 0.17 | 25.57 |
| red blood cells | Tgm2 | 3.46 | 0 | 0 | 0.18 | 0.01 | 0.17 | 19.56 |
| red blood cells | Csf2rb | 4.39 | 0 | 0 | 0.17 | 0.01 | 0.17 | 34.2 |
| red blood cells | Adrb2 | 3.68 | 0 | 0 | 0.17 | 0.01 | 0.16 | 24.29 |
| red blood cells | 810053B23Ri | 8.09 | 0 | 0 | 0.16 | 0 | 0.16 | #NA |
| red blood cells | Nxpe2 | 5.72 | 0 | 0 | 0.15 | 0 | 0.15 | 76.5 |
| red blood cells | Apol8 | 5.4 | 0 | 0 | 0.15 | 0 | 0.15 | 74.5 |
| red blood cells | Lefty1 | 4.19 | 0 | 0 | 0.15 | 0.01 | 0.14 | 29.4 |
| red blood cells | H2-T24 | 5.11 | 0 | 0 | 0.14 | 0 | 0.14 | 70.5 |
| red blood cells | Gpr150 | 5.99 | 0 | 0 | 0.13 | 0 | 0.13 | 127 |
| red blood cells | Il1bos | 8.62 | 0 | 0 | 0.12 | 0 | 0.12 | #NA |
| red blood cells | Hba-x | 9.47 | 0 | 0 | 1 | 0.89 | 0.11 | 1.13 |
| red blood cells | Cyp4b1 | 8.27 | 0 | 0 | 0.11 | 0 | 0.11 | #NA |
| red blood cells | Glrx5 | 1.92 | 0 | 0 | 0.99 | 0.88 | 0.11 | 1.12 |
| red blood cells | Pdia2 | 6.99 | 0 | 0 | 0.11 | 0 | 0.11 | #NA |
| red blood cells | Mst1 | 5.95 | 0 | 0 | 0.1 | 0 | 0.1 | 103 |
| red blood cells | Tmem14c | 1.93 | 0 | 0 | 0.99 | 0.92 | 0.07 | 1.07 |
| red blood cells | Gpx1 | 3.43 | 0 | 0 | 1 | 0.95 | 0.05 | 1.05 |
| red blood cells | Hbb-y | 9.38 | 0 | 0 | 1 | 0.98 | 0.02 | 1.02 |
| red blood cells | Fth1 | 3.25 | 0 | 0 | 1 | 1 | 0 | 1 |
| red blood cells | Ftl1 | 0.74 | 0 | 0 | 1 | 1 | 0 | 1 |
| red blood cells | Prdx2 | 1.04 | 0 | 0 | 1 | 1 | 0 | 1 |
| red blood cells | Lmo2 | 0.92 | 2.81E-302 | 5.85E-298 | 0.6 | 0.11 | 0.49 | 5.48 |
| red blood cells | Samd11 | 4.32 | 2.18E-298 | 4.54E-294 | 0.14 | 0 | 0.13 | 33.75 |
| red blood cells | St3gal6 | 2.12 | 2.32E-298 | 4.82E-294 | 0.54 | 0.1 | 0.44 | 5.63 |
| red blood cells | E2f2 | 2.19 | 4.18E-298 | 8.69E-294 | 0.53 | 0.1 | 0.44 | 5.61 |
| red blood cells | Rgs10 | 1.46 | 6.76E-296 | 1.41E-291 | 0.47 | 0.07 | 0.39 | 6.4 |
| red blood cells | Ankrd9 | 1.95 | 1.73E-288 | 3.60E-284 | 0.56 | 0.11 | 0.46 | 5.29 |
| red blood cells | 930523C07Ri | 2.58 | 3.21E-288 | 6.67E-284 | 0.42 | 0.06 | 0.35 | 6.71 |
| red blood cells | Acyp2 | 2.11 | 1.54E-285 | 3.20E-281 | 0.51 | 0.09 | 0.42 | 5.62 |
| red blood cells | Atp1b2 | 3.37 | 3.31E-285 | 6.89E-281 | 0.19 | 0.01 | 0.18 | 17 |
| red blood cells | Ubac1 | 1.71 | 1.18E-283 | 2.45E-279 | 0.95 | 0.56 | 0.4 | 1.71 |
| red blood cells | Fam117a | 1.9 | 8.50E-273 | 1.77E-268 | 0.77 | 0.22 | 0.55 | 3.58 |
| red blood cells | Mkrn1 | 1.7 | 1.58E-272 | 3.28E-268 | 0.94 | 0.41 | 0.53 | 2.3 |
| red blood cells | Ccnd3 | 1.22 | 1.46E-270 | 3.03E-266 | 0.99 | 0.85 | 0.14 | 1.17 |
| red blood cells | Btg2 | 1.46 | 1.22E-269 | 2.54E-265 | 0.83 | 0.25 | 0.57 | 3.27 |
| red blood cells | Txnrd2 | 1.82 | 5.54E-269 | 1.15E-264 | 0.73 | 0.19 | 0.54 | 3.82 |
| red blood cells | Marchf3 | 4.57 | 2.19E-268 | 4.56E-264 | 0.1 | 0 | 0.1 | 52 |
| red blood cells | Pigq | 1.89 | 7.93E-265 | 1.65E-260 | 0.77 | 0.22 | 0.56 | 3.56 |
| red blood cells | Tmcc2 | 2.52 | 4.23E-264 | 8.79E-260 | 0.43 | 0.07 | 0.36 | 6.09 |
| red blood cells | Ubash3a | 4.47 | 6.19E-264 | 1.29E-259 | 0.1 | 0 | 0.1 | 34.67 |
| red blood cells | Gcnt1 | 3.34 | 6.49E-259 | 1.35E-254 | 0.24 | 0.02 | 0.22 | 10.57 |
| red blood cells | Slc40a1 | 1.82 | 1.08E-257 | 2.25E-253 | 0.35 | 0.05 | 0.3 | 7.31 |
| red blood cells | Ppox | 1.77 | 7.80E-257 | 1.62E-252 | 0.76 | 0.22 | 0.54 | 3.44 |
| red blood cells | Dnaja4 | 2.31 | 6.72E-252 | 1.40E-247 | 0.32 | 0.04 | 0.28 | 7.69 |
| red blood cells | Nt5c3 | 1.78 | 1.45E-249 | 3.01E-245 | 0.9 | 0.38 | 0.52 | 2.37 |
| red blood cells | Isca1 | 1.71 | 4.13E-248 | 8.58E-244 | 0.85 | 0.29 | 0.56 | 2.91 |
| red blood cells | Pip5k1b | 2.47 | 7.77E-248 | 1.62E-243 | 0.26 | 0.03 | 0.23 | 9.48 |
| red blood cells | Tfr2 | 3.14 | 2.62E-243 | 5.45E-239 | 0.16 | 0.01 | 0.15 | 17.22 |
| red blood cells | Pim1 | 1.73 | 6.86E-243 | 1.43E-238 | 0.93 | 0.45 | 0.48 | 2.06 |
| red blood cells | Fads3 | 2.15 | 1.17E-242 | 2.43E-238 | 0.33 | 0.05 | 0.29 | 7.22 |
| red blood cells | Stom | 2.63 | 7.39E-242 | 1.54E-237 | 0.29 | 0.04 | 0.26 | 8.08 |
| red blood cells | Cdc42ep3 | 1.28 | 5.23E-240 | 1.09E-235 | 0.63 | 0.15 | 0.49 | 4.34 |

**Supplementary Table 2. Marker genes for the major cell types of integrated clustering of E11.5 control and NotchGOF λ-junction cells.**

| | | | | | | | | |
|---|---|---|---|---|---|---|---|---|
| red blood cells | Tfrc | 1.89 | 7.80E-239 | 1.62E-234 | 0.89 | 0.38 | 0.51 | 2.35 |
| red blood cells | Fam78a | 3.61 | 1.71E-236 | 3.56E-232 | 0.15 | 0.01 | 0.14 | 18.13 |
| red blood cells | Nrtn | 1.45 | 9.88E-232 | 2.06E-227 | 0.6 | 0.14 | 0.46 | 4.33 |
| red blood cells | Prdx3 | 1.49 | 8.30E-231 | 1.73E-226 | 0.95 | 0.52 | 0.43 | 1.83 |
| red blood cells | Spaca9 | 2.46 | 5.36E-230 | 1.11E-225 | 0.24 | 0.03 | 0.22 | 9.6 |
| red blood cells | Gm42418 | 0.73 | 1.70E-229 | 3.53E-225 | 1 | 1 | 0 | 1 |
| red blood cells | Exoc6 | 1.95 | 1.69E-223 | 3.51E-219 | 0.38 | 0.06 | 0.31 | 5.98 |
| red blood cells | C1qtnf4 | 1.51 | 1.39E-221 | 2.90E-217 | 0.81 | 0.27 | 0.54 | 3.03 |
| red blood cells | Pfkfb1 | 2.32 | 3.68E-221 | 7.65E-217 | 0.27 | 0.03 | 0.24 | 7.97 |
| red blood cells | 430431A17Ri | 3.71 | 1.09E-219 | 2.26E-215 | 0.11 | 0.01 | 0.11 | 22.2 |
| red blood cells | Slc29a1 | 1.51 | 1.14E-215 | 2.36E-211 | 0.5 | 0.11 | 0.39 | 4.65 |
| red blood cells | Slc16a6 | 2.37 | 8.96E-215 | 1.86E-210 | 0.31 | 0.04 | 0.26 | 6.98 |
| red blood cells | 030028A08Ri | 2.13 | 4.24E-213 | 8.82E-209 | 0.33 | 0.05 | 0.28 | 6.54 |
| red blood cells | Abcg2 | 1.54 | 1.71E-208 | 3.56E-204 | 0.84 | 0.33 | 0.51 | 2.55 |
| red blood cells | Cltb | 1.59 | 2.71E-206 | 5.63E-202 | 0.62 | 0.17 | 0.46 | 3.75 |
| red blood cells | Prokr1 | 3.15 | 6.46E-203 | 1.34E-198 | 0.14 | 0.01 | 0.13 | 15.44 |
| red blood cells | Asb1 | 1.89 | 9.88E-202 | 2.05E-197 | 0.35 | 0.06 | 0.29 | 5.95 |
| red blood cells | Tango2 | 1.68 | 2.33E-196 | 4.84E-192 | 0.56 | 0.14 | 0.42 | 3.98 |
| red blood cells | Dnajb3 | 1.52 | 2.10E-194 | 4.37E-190 | 0.55 | 0.13 | 0.42 | 4.12 |
| red blood cells | Lockd | 1.37 | 1.50E-192 | 3.12E-188 | 0.86 | 0.34 | 0.52 | 2.5 |
| red blood cells | Ppp1r15a | 1.44 | 7.14E-191 | 1.48E-186 | 0.74 | 0.23 | 0.51 | 3.19 |
| red blood cells | 033406O09Ri | 3.21 | 1.31E-189 | 2.72E-185 | 0.13 | 0.01 | 0.12 | 15.63 |
| red blood cells | Fbxo7 | 1.63 | 1.86E-189 | 3.87E-185 | 0.51 | 0.12 | 0.39 | 4.25 |
| red blood cells | Arhgap23 | 1.73 | 2.55E-188 | 5.30E-184 | 0.41 | 0.08 | 0.33 | 4.92 |
| red blood cells | Slc20a1 | 1.69 | 2.48E-187 | 5.17E-183 | 0.82 | 0.3 | 0.52 | 2.71 |
| red blood cells | Endod1 | 1.63 | 1.56E-184 | 3.25E-180 | 0.4 | 0.08 | 0.32 | 5.01 |
| red blood cells | Ssx2ip | 1.22 | 9.98E-183 | 2.08E-178 | 0.85 | 0.34 | 0.51 | 2.49 |
| red blood cells | Ocel1 | 1.95 | 1.81E-181 | 3.76E-177 | 0.29 | 0.05 | 0.24 | 6.38 |
| red blood cells | Slc43a3 | 1.48 | 1.83E-181 | 3.80E-177 | 0.77 | 0.27 | 0.5 | 2.88 |
| red blood cells | Cdc25b | 1.69 | 2.30E-180 | 4.77E-176 | 0.62 | 0.19 | 0.43 | 3.29 |
| red blood cells | Cdr2 | 1.54 | 1.94E-178 | 4.04E-174 | 0.66 | 0.2 | 0.46 | 3.27 |
| red blood cells | Bola3 | 1.03 | 2.51E-178 | 5.22E-174 | 0.98 | 0.74 | 0.24 | 1.32 |
| red blood cells | Lmna | 1.48 | 4.83E-177 | 1.00E-172 | 0.72 | 0.26 | 0.47 | 2.82 |
| red blood cells | Mindy2 | 1.4 | 7.14E-176 | 1.49E-171 | 0.58 | 0.16 | 0.42 | 3.64 |
| red blood cells | Hsd17b14 | 3.09 | 1.02E-169 | 2.12E-165 | 0.13 | 0.01 | 0.12 | 13.4 |
| red blood cells | Ehbp1l1 | 1.84 | 3.79E-168 | 7.89E-164 | 0.31 | 0.06 | 0.25 | 5.62 |
| red blood cells | Tuft1 | 2.15 | 1.17E-167 | 2.43E-163 | 0.23 | 0.03 | 0.2 | 7.29 |
| red blood cells | Ehd1 | 1.2 | 1.41E-166 | 2.94E-162 | 0.94 | 0.51 | 0.43 | 1.84 |
| red blood cells | Trak2 | 1.65 | 4.33E-161 | 9.00E-157 | 0.48 | 0.12 | 0.36 | 3.91 |
| red blood cells | Sgk3 | 1.43 | 1.55E-160 | 3.21E-156 | 0.51 | 0.14 | 0.38 | 3.76 |
| red blood cells | H2afj | 0.86 | 7.19E-160 | 1.50E-155 | 0.98 | 0.81 | 0.18 | 1.22 |
| red blood cells | Fzd5 | 2.44 | 7.84E-157 | 1.63E-152 | 0.16 | 0.02 | 0.14 | 9.47 |
| red blood cells | Cdkn3 | 1.38 | 8.72E-157 | 1.81E-152 | 0.73 | 0.28 | 0.45 | 2.63 |
| red blood cells | St3gal1 | 1.57 | 1.06E-154 | 2.20E-150 | 0.4 | 0.09 | 0.31 | 4.44 |
| red blood cells | Rexo2 | 1.1 | 7.84E-154 | 1.63E-149 | 0.94 | 0.58 | 0.36 | 1.62 |
| red blood cells | Apeh | 1.36 | 3.48E-153 | 7.23E-149 | 0.71 | 0.26 | 0.46 | 2.8 |
| red blood cells | Pde10a | 1.88 | 8.99E-153 | 1.87E-148 | 0.24 | 0.04 | 0.2 | 6.43 |
| red blood cells | Hesx1 | 3.09 | 1.94E-152 | 4.04E-148 | 0.1 | 0.01 | 0.1 | 14.86 |
| red blood cells | Gadd45a | 1.09 | 2.94E-152 | 6.11E-148 | 0.69 | 0.23 | 0.46 | 3 |
| red blood cells | Slc16a10 | 1.32 | 2.50E-147 | 5.21E-143 | 0.41 | 0.1 | 0.31 | 4.23 |
| red blood cells | Slc48a1 | 1.37 | 7.72E-147 | 1.61E-142 | 0.54 | 0.15 | 0.38 | 3.5 |
| red blood cells | Unc13d | 2.24 | 2.21E-145 | 4.59E-141 | 0.18 | 0.02 | 0.16 | 7.91 |
| red blood cells | Slc25a39 | 0.96 | 1.28E-144 | 2.67E-140 | 0.97 | 0.7 | 0.27 | 1.39 |
| red blood cells | Dyrk3 | 1.47 | 5.01E-142 | 1.04E-137 | 0.38 | 0.09 | 0.29 | 4.27 |
| red blood cells | Brpf3 | 1.65 | 1.23E-140 | 2.56E-136 | 0.44 | 0.11 | 0.32 | 3.89 |
| red blood cells | Slc41a3 | 2.28 | 8.44E-140 | 1.75E-135 | 0.16 | 0.02 | 0.14 | 8.53 |

**Supplementary Table 2. Marker genes for the major cell types of integrated clustering of E11.5 control and NotchGOF λ-junction cells.**

| | | | | | | | | |
|---|---|---|---|---|---|---|---|---|
| red blood cells | Taldo1 | 0.74 | 1.24E-135 | 2.58E-131 | 0.99 | 0.89 | 0.1 | 1.11 |
| red blood cells | Pon2 | 0.99 | 1.22E-134 | 2.53E-130 | 0.56 | 0.17 | 0.39 | 3.36 |
| red blood cells | Uba7 | 1.43 | 4.87E-130 | 1.01E-125 | 0.39 | 0.1 | 0.29 | 4.02 |
| red blood cells | Cd55 | 2.06 | 1.03E-128 | 2.14E-124 | 0.16 | 0.02 | 0.14 | 8.21 |
| red blood cells | Sh3yl1 | 1.29 | 3.89E-127 | 8.09E-123 | 0.57 | 0.18 | 0.39 | 3.16 |
| red blood cells | Tubb4b | 0.89 | 9.73E-123 | 2.02E-118 | 0.97 | 0.85 | 0.13 | 1.15 |
| red blood cells | Capn1 | 1.63 | 8.42E-122 | 1.75E-117 | 0.23 | 0.04 | 0.19 | 5.44 |
| red blood cells | Mylpf | 1.45 | 1.02E-117 | 2.12E-113 | 0.3 | 0.07 | 0.23 | 4.45 |
| red blood cells | Cercam | 1.38 | 4.72E-115 | 9.83E-111 | 0.25 | 0.05 | 0.2 | 4.98 |
| red blood cells | Ppp1r9b | 1.16 | 6.60E-115 | 1.37E-110 | 0.66 | 0.24 | 0.42 | 2.74 |
| red blood cells | Dusp8 | 1.69 | 2.44E-112 | 5.08E-108 | 0.13 | 0.02 | 0.11 | 8.47 |
| red blood cells | Fermt3 | 0.54 | 3.76E-111 | 7.82E-107 | 0.22 | 0.04 | 0.18 | 5.29 |
| red blood cells | Reep6 | 1.56 | 4.80E-111 | 9.98E-107 | 0.23 | 0.04 | 0.19 | 5.2 |
| red blood cells | S100a13 | 0.77 | 1.65E-110 | 3.44E-106 | 0.37 | 0.1 | 0.27 | 3.81 |
| red blood cells | Arl4a | 0.97 | 4.44E-109 | 9.23E-105 | 0.56 | 0.19 | 0.37 | 2.93 |
| red blood cells | Fam110c | 2.62 | 6.19E-109 | 1.29E-104 | 0.1 | 0.01 | 0.09 | 11.11 |
| red blood cells | Slc30a1 | 1.44 | 1.61E-108 | 3.35E-104 | 0.31 | 0.08 | 0.23 | 4.11 |
| red blood cells | i430095P16Ri | 1.2 | 2.20E-108 | 4.58E-104 | 0.36 | 0.1 | 0.27 | 3.79 |
| red blood cells | Hagh | 0.97 | 8.86E-106 | 1.84E-101 | 0.63 | 0.23 | 0.4 | 2.74 |
| red blood cells | Katnb1 | 1.07 | 2.43E-104 | 5.06E-100 | 0.58 | 0.21 | 0.37 | 2.79 |
| red blood cells | Vamp8 | 0.84 | 6.18E-104 | 1.29E-99 | 0.92 | 0.6 | 0.32 | 1.54 |
| red blood cells | Rnf123 | 1.18 | 2.27E-103 | 4.71E-99 | 0.39 | 0.11 | 0.28 | 3.5 |
| red blood cells | Dnase1l1 | 1.44 | 2.53E-103 | 5.26E-99 | 0.21 | 0.04 | 0.17 | 5.17 |
| red blood cells | Arid3a | 1.02 | 2.94E-103 | 6.11E-99 | 0.75 | 0.32 | 0.43 | 2.36 |
| red blood cells | Mbnl1 | 1.03 | 6.52E-102 | 1.36E-97 | 0.87 | 0.46 | 0.41 | 1.9 |
| red blood cells | Mki67 | 0.93 | 6.71E-102 | 1.40E-97 | 0.96 | 0.71 | 0.25 | 1.35 |
| red blood cells | Fos | 0.62 | 5.65E-101 | 1.17E-96 | 0.42 | 0.13 | 0.29 | 3.26 |
| red blood cells | Hscb | 0.93 | 9.89E-101 | 2.06E-96 | 0.74 | 0.31 | 0.43 | 2.4 |
| red blood cells | Tmc6 | 1.31 | 2.20E-100 | 4.57E-96 | 0.28 | 0.07 | 0.22 | 4.16 |
| red blood cells | Itpka | 2.16 | 4.25E-99 | 8.84E-95 | 0.13 | 0.02 | 0.11 | 7.59 |
| red blood cells | Gypc | 1.01 | 1.39E-98 | 2.89E-94 | 0.58 | 0.2 | 0.37 | 2.83 |
| red blood cells | Nudt19 | 0.99 | 4.49E-95 | 9.34E-91 | 0.7 | 0.31 | 0.39 | 2.27 |
| red blood cells | Atpif1 | 0.36 | 1.41E-93 | 2.94E-89 | 1 | 1 | 0 | 1 |
| red blood cells | 600006K01Ri | 1.73 | 1.88E-92 | 3.92E-88 | 0.13 | 0.02 | 0.11 | 6.95 |
| red blood cells | Tom1l1 | 0.71 | 4.88E-92 | 1.02E-87 | 0.46 | 0.15 | 0.31 | 3.08 |
| red blood cells | Rbks | 1.57 | 1.81E-91 | 3.76E-87 | 0.17 | 0.03 | 0.14 | 5.69 |
| red blood cells | Ankle1 | 1.3 | 1.21E-87 | 2.52E-83 | 0.31 | 0.09 | 0.22 | 3.61 |
| red blood cells | Rffl | 1.28 | 3.21E-87 | 6.68E-83 | 0.26 | 0.06 | 0.2 | 4.05 |
| red blood cells | Ift140 | 0.99 | 3.38E-87 | 7.04E-83 | 0.45 | 0.15 | 0.3 | 2.97 |
| red blood cells | Skap2 | 0.93 | 8.59E-87 | 1.79E-82 | 0.22 | 0.05 | 0.17 | 4.47 |
| red blood cells | S100a1 | 1.08 | 3.43E-83 | 7.14E-79 | 0.24 | 0.06 | 0.18 | 4.07 |
| red blood cells | Ifi30 | 0.7 | 1.24E-82 | 2.58E-78 | 0.66 | 0.28 | 0.38 | 2.36 |
| red blood cells | Nipa1 | 0.84 | 5.16E-81 | 1.07E-76 | 0.41 | 0.13 | 0.27 | 3.07 |
| red blood cells | 700037H04Ri | 0.79 | 7.54E-80 | 1.57E-75 | 0.85 | 0.46 | 0.39 | 1.84 |
| red blood cells | Dennd2c | 1.12 | 7.81E-80 | 1.62E-75 | 0.23 | 0.06 | 0.18 | 4.03 |
| red blood cells | Tuba1c | 0.96 | 2.06E-79 | 4.29E-75 | 0.76 | 0.43 | 0.34 | 1.78 |
| red blood cells | Ybx3 | 0.58 | 5.51E-78 | 1.15E-73 | 0.98 | 0.83 | 0.15 | 1.17 |
| red blood cells | Mpst | 0.78 | 9.80E-77 | 2.04E-72 | 0.8 | 0.4 | 0.39 | 1.97 |
| red blood cells | Mafk | 1.48 | 2.31E-74 | 4.81E-70 | 0.17 | 0.04 | 0.13 | 4.8 |
| red blood cells | Pop5 | 0.67 | 4.80E-74 | 9.99E-70 | 0.93 | 0.64 | 0.29 | 1.44 |
| red blood cells | Rad23a | 0.82 | 7.74E-73 | 1.61E-68 | 0.85 | 0.44 | 0.4 | 1.91 |
| red blood cells | Gch1 | 0.91 | 1.31E-72 | 2.73E-68 | 0.26 | 0.07 | 0.19 | 3.66 |
| red blood cells | Slc25a38 | 0.92 | 2.73E-72 | 5.67E-68 | 0.6 | 0.26 | 0.34 | 2.31 |
| red blood cells | Hspa4l | 0.81 | 3.85E-72 | 8.01E-68 | 0.33 | 0.1 | 0.22 | 3.14 |
| red blood cells | Arf5 | 0.34 | 1.08E-71 | 2.25E-67 | 1 | 0.98 | 0.01 | 1.01 |
| red blood cells | Pycr2 | 0.7 | 3.94E-71 | 8.19E-67 | 0.91 | 0.6 | 0.31 | 1.52 |

**Supplementary Table 2. Marker genes for the major cell types of integrated clustering of E11.5 control and NotchGOF λ-junction cells.**

| | | | | | | | | |
|---|---|---|---|---|---|---|---|---|
| red blood cells | Nadk2 | 0.79 | 1.37E-70 | 2.85E-66 | 0.71 | 0.34 | 0.37 | 2.1 |
| red blood cells | Nars | 0.59 | 4.65E-69 | 9.68E-65 | 0.96 | 0.8 | 0.17 | 1.21 |
| red blood cells | Adcy7 | 1.83 | 1.97E-68 | 4.10E-64 | 0.12 | 0.02 | 0.1 | 5.9 |
| red blood cells | Ftl1-ps1 | 0.69 | 7.75E-68 | 1.61E-63 | 0.79 | 0.39 | 0.4 | 2.01 |
| red blood cells | Tmem179b | 0.68 | 8.97E-67 | 1.87E-62 | 0.64 | 0.28 | 0.36 | 2.26 |
| red blood cells | Optn | 1.48 | 9.01E-67 | 1.87E-62 | 0.13 | 0.03 | 0.11 | 5.24 |
| red blood cells | Cep76 | 0.82 | 2.63E-66 | 5.47E-62 | 0.42 | 0.15 | 0.27 | 2.73 |
| red blood cells | Stx2 | 0.78 | 7.04E-65 | 1.46E-60 | 0.51 | 0.21 | 0.3 | 2.43 |
| red blood cells | Aff1 | 1.3 | 1.35E-64 | 2.80E-60 | 0.23 | 0.07 | 0.17 | 3.55 |
| red blood cells | Rab3il1 | 0.61 | 2.18E-64 | 4.53E-60 | 0.65 | 0.28 | 0.36 | 2.28 |
| red blood cells | Zfp367 | 0.92 | 2.37E-64 | 4.94E-60 | 0.38 | 0.14 | 0.24 | 2.76 |
| red blood cells | Pabpc4 | 0.73 | 1.83E-63 | 3.80E-59 | 0.93 | 0.64 | 0.28 | 1.44 |
| red blood cells | Smox | 0.68 | 2.10E-63 | 4.37E-59 | 0.55 | 0.23 | 0.32 | 2.39 |
| red blood cells | Cdkn2d | 0.98 | 3.80E-63 | 7.91E-59 | 0.67 | 0.37 | 0.3 | 1.81 |
| red blood cells | H2-T23 | 0.99 | 5.03E-63 | 1.05E-58 | 0.19 | 0.05 | 0.14 | 3.92 |
| red blood cells | Iba57 | 1.35 | 6.20E-62 | 1.29E-57 | 0.15 | 0.03 | 0.12 | 4.59 |
| red blood cells | Tatdn3 | 0.86 | 4.43E-60 | 9.21E-56 | 0.3 | 0.1 | 0.2 | 3.01 |
| red blood cells | Pcgf5 | 0.9 | 6.61E-60 | 1.37E-55 | 0.26 | 0.08 | 0.18 | 3.21 |
| red blood cells | Nup210 | 1.47 | 1.16E-59 | 2.40E-55 | 0.2 | 0.06 | 0.15 | 3.64 |
| red blood cells | Frrs1 | 0.76 | 1.81E-59 | 3.77E-55 | 0.5 | 0.21 | 0.29 | 2.36 |
| red blood cells | Spsb2 | 0.78 | 1.85E-59 | 3.86E-55 | 0.24 | 0.07 | 0.17 | 3.41 |
| red blood cells | Hist1h4i | 0.75 | 2.01E-59 | 4.17E-55 | 0.81 | 0.45 | 0.36 | 1.81 |
| red blood cells | Nqo1 | 0.86 | 3.57E-59 | 7.43E-55 | 0.28 | 0.09 | 0.19 | 3.09 |
| red blood cells | Fbxo9 | 0.64 | 5.91E-59 | 1.23E-54 | 0.62 | 0.28 | 0.34 | 2.22 |
| red blood cells | Pir | 0.82 | 3.34E-58 | 6.95E-54 | 0.33 | 0.12 | 0.22 | 2.81 |
| red blood cells | Stradb | 0.77 | 4.02E-58 | 8.36E-54 | 0.45 | 0.18 | 0.27 | 2.49 |
| red blood cells | Yod1 | 0.8 | 2.29E-57 | 4.75E-53 | 0.37 | 0.14 | 0.23 | 2.69 |
| red blood cells | Pdk2 | 0.86 | 3.79E-56 | 7.87E-52 | 0.25 | 0.08 | 0.17 | 3.21 |
| red blood cells | Rmc1 | 0.86 | 3.87E-56 | 8.04E-52 | 0.24 | 0.08 | 0.17 | 3.21 |
| red blood cells | Asns | 0.7 | 4.48E-56 | 9.32E-52 | 0.77 | 0.43 | 0.34 | 1.8 |
| red blood cells | Capn5 | 1.64 | 1.15E-55 | 2.39E-51 | 0.12 | 0.02 | 0.1 | 4.96 |
| red blood cells | Fam174b | 0.73 | 4.13E-55 | 8.58E-51 | 0.3 | 0.11 | 0.2 | 2.83 |
| red blood cells | Smim4 | 0.58 | 5.39E-55 | 1.12E-50 | 0.78 | 0.41 | 0.37 | 1.88 |
| red blood cells | Tarsl2 | 0.73 | 1.08E-54 | 2.25E-50 | 0.27 | 0.09 | 0.18 | 3.02 |
| red blood cells | Glrx | 0.47 | 1.46E-53 | 3.03E-49 | 0.52 | 0.23 | 0.29 | 2.31 |
| red blood cells | Akip1 | 0.58 | 1.56E-53 | 3.25E-49 | 0.46 | 0.19 | 0.27 | 2.42 |
| red blood cells | Foxo3 | 0.95 | 4.76E-53 | 9.91E-49 | 0.33 | 0.12 | 0.21 | 2.71 |
| red blood cells | Slc11a2 | 0.68 | 7.23E-53 | 1.50E-48 | 0.41 | 0.16 | 0.25 | 2.52 |
| red blood cells | Hspa12b | 0.64 | 1.30E-52 | 2.70E-48 | 0.34 | 0.13 | 0.21 | 2.71 |
| red blood cells | Skiv2l | 0.82 | 2.40E-52 | 4.99E-48 | 0.53 | 0.23 | 0.3 | 2.26 |
| red blood cells | Ank | 0.98 | 5.88E-52 | 1.22E-47 | 0.21 | 0.06 | 0.15 | 3.31 |
| red blood cells | Gm35082 | 1.43 | 1.09E-50 | 2.26E-46 | 0.12 | 0.03 | 0.09 | 4.64 |
| red blood cells | Dleu2 | 0.71 | 7.84E-50 | 1.63E-45 | 0.53 | 0.24 | 0.29 | 2.17 |
| red blood cells | Sord | 0.52 | 8.22E-49 | 1.71E-44 | 0.35 | 0.14 | 0.22 | 2.56 |
| red blood cells | Khk | 0.66 | 1.07E-48 | 2.23E-44 | 0.2 | 0.06 | 0.14 | 3.31 |
| red blood cells | Ogfrl1 | 0.59 | 1.58E-48 | 3.29E-44 | 0.66 | 0.33 | 0.33 | 2.02 |
| red blood cells | Abcb6 | 0.86 | 1.72E-48 | 3.58E-44 | 0.3 | 0.11 | 0.19 | 2.67 |
| red blood cells | Psme3 | 0.59 | 8.52E-48 | 1.77E-43 | 0.83 | 0.49 | 0.35 | 1.72 |
| red blood cells | Eif2ak1 | 0.58 | 1.06E-47 | 2.20E-43 | 0.76 | 0.41 | 0.35 | 1.86 |
| red blood cells | Narf | 0.52 | 1.90E-46 | 3.94E-42 | 0.74 | 0.41 | 0.33 | 1.82 |
| red blood cells | Abhd5 | 0.5 | 3.29E-46 | 6.84E-42 | 0.44 | 0.19 | 0.25 | 2.34 |
| red blood cells | Zfand2b | 0.5 | 4.93E-46 | 1.02E-41 | 0.54 | 0.25 | 0.29 | 2.18 |
| red blood cells | Rap1a | 0.45 | 1.04E-45 | 2.16E-41 | 0.93 | 0.71 | 0.22 | 1.32 |
| red blood cells | Tmem120b | 0.83 | 2.26E-45 | 4.69E-41 | 0.19 | 0.06 | 0.13 | 3.26 |
| red blood cells | Mthfd2 | 0.55 | 2.82E-45 | 5.86E-41 | 0.8 | 0.46 | 0.34 | 1.73 |
| red blood cells | Stxbp2 | 0.4 | 1.04E-44 | 2.16E-40 | 0.33 | 0.13 | 0.2 | 2.55 |

**Supplementary Table 2. Marker genes for the major cell types of integrated clustering of E11.5 control and NotchGOF λ-junction cells.**

| | | | | | | | | |
|---|---|---|---|---|---|---|---|---|
| red blood cells | Tnfrsf21 | 0.69 | 3.69E-44 | 7.67E-40 | 0.34 | 0.14 | 0.2 | 2.48 |
| red blood cells | Ctsb | 0.29 | 5.87E-44 | 1.22E-39 | 0.96 | 0.74 | 0.22 | 1.29 |
| red blood cells | Fryl | 0.59 | 2.01E-43 | 4.18E-39 | 0.59 | 0.29 | 0.3 | 2.03 |
| red blood cells | Bcor | 0.64 | 6.94E-43 | 1.44E-38 | 0.72 | 0.4 | 0.32 | 1.78 |
| red blood cells | Ubl4a | 0.5 | 7.14E-43 | 1.48E-38 | 0.89 | 0.56 | 0.32 | 1.57 |
| red blood cells | Pabpc1 | 0.31 | 4.65E-41 | 9.67E-37 | 1 | 0.99 | 0.01 | 1.01 |
| red blood cells | Clk3 | 0.57 | 1.36E-40 | 2.84E-36 | 0.69 | 0.37 | 0.32 | 1.86 |
| red blood cells | Taok3 | 0.48 | 1.81E-40 | 3.77E-36 | 0.26 | 0.1 | 0.16 | 2.65 |
| red blood cells | Mvb12b | 0.46 | 3.90E-40 | 8.12E-36 | 0.47 | 0.22 | 0.25 | 2.16 |
| red blood cells | Zfp524 | 0.55 | 1.14E-39 | 2.37E-35 | 0.26 | 0.1 | 0.16 | 2.64 |
| red blood cells | Ap1g2 | 0.89 | 1.81E-39 | 3.76E-35 | 0.14 | 0.04 | 0.1 | 3.51 |
| red blood cells | Neat1 | 0.71 | 2.64E-39 | 5.49E-35 | 0.17 | 0.05 | 0.11 | 3.24 |
| red blood cells | E2f8 | 0.46 | 8.40E-39 | 1.75E-34 | 0.43 | 0.2 | 0.23 | 2.19 |
| red blood cells | Podxl | 0.69 | 2.45E-38 | 5.09E-34 | 0.14 | 0.04 | 0.1 | 3.5 |
| red blood cells | Il10rb | 0.48 | 3.17E-38 | 6.59E-34 | 0.22 | 0.08 | 0.14 | 2.77 |
| red blood cells | Rgcc | 0.61 | 4.63E-38 | 9.64E-34 | 0.23 | 0.09 | 0.15 | 2.68 |
| red blood cells | Dedd2 | 0.75 | 7.31E-38 | 1.52E-33 | 0.2 | 0.07 | 0.13 | 2.89 |
| red blood cells | Stat5a | 1.24 | 7.37E-38 | 1.53E-33 | 0.11 | 0.03 | 0.08 | 4.04 |
| red blood cells | Irak2 | 0.69 | 1.06E-37 | 2.20E-33 | 0.16 | 0.05 | 0.11 | 3.2 |
| red blood cells | Atg4d | 0.49 | 5.79E-37 | 1.20E-32 | 0.43 | 0.2 | 0.23 | 2.17 |
| red blood cells | Chst10 | 0.75 | 8.03E-37 | 1.67E-32 | 0.21 | 0.08 | 0.14 | 2.74 |
| red blood cells | Retreg2 | 0.42 | 1.96E-36 | 4.07E-32 | 0.56 | 0.28 | 0.28 | 2.02 |
| red blood cells | Atp2b4 | 0.5 | 2.35E-36 | 4.88E-32 | 0.42 | 0.19 | 0.23 | 2.18 |
| red blood cells | Ier3 | 0.87 | 3.35E-36 | 6.96E-32 | 0.2 | 0.07 | 0.13 | 2.81 |
| red blood cells | Snx15 | 0.41 | 8.27E-36 | 1.72E-31 | 0.58 | 0.3 | 0.28 | 1.96 |
| red blood cells | Tlcd1 | 0.79 | 4.60E-35 | 9.57E-31 | 0.22 | 0.09 | 0.14 | 2.64 |
| red blood cells | Ccdc82 | 0.46 | 5.83E-35 | 1.21E-30 | 0.28 | 0.11 | 0.16 | 2.44 |
| red blood cells | Ptgr2 | 0.35 | 1.34E-34 | 2.79E-30 | 0.33 | 0.14 | 0.19 | 2.32 |
| red blood cells | Plin2 | 0.47 | 2.33E-34 | 4.85E-30 | 0.81 | 0.52 | 0.29 | 1.55 |
| red blood cells | Ccndbp1 | 0.26 | 3.30E-34 | 6.87E-30 | 0.55 | 0.27 | 0.28 | 2.04 |
| red blood cells | D1Ertd622e | 0.54 | 2.18E-33 | 4.54E-29 | 0.29 | 0.13 | 0.17 | 2.34 |
| red blood cells | Mad2l1bp | 0.33 | 2.31E-33 | 4.81E-29 | 0.47 | 0.23 | 0.25 | 2.08 |
| red blood cells | Selenbp1 | 0.41 | 6.71E-33 | 1.40E-28 | 0.21 | 0.08 | 0.13 | 2.63 |
| red blood cells | Fbxo34 | 0.42 | 6.81E-33 | 1.42E-28 | 0.56 | 0.29 | 0.27 | 1.94 |
| red blood cells | Synj1 | 0.82 | 1.42E-32 | 2.95E-28 | 0.25 | 0.1 | 0.15 | 2.44 |
| red blood cells | .110008P14Ri | 0.38 | 3.35E-32 | 6.96E-28 | 0.78 | 0.46 | 0.33 | 1.72 |
| red blood cells | Tapbp | 0.56 | 3.70E-32 | 7.69E-28 | 0.21 | 0.08 | 0.13 | 2.63 |
| red blood cells | Agfg2 | 0.37 | 7.99E-32 | 1.66E-27 | 0.43 | 0.21 | 0.22 | 2.09 |
| red blood cells | Afg3l2 | 0.36 | 1.07E-31 | 2.22E-27 | 0.63 | 0.33 | 0.3 | 1.9 |
| red blood cells | Wbp2 | 0.31 | 1.37E-31 | 2.85E-27 | 0.54 | 0.27 | 0.27 | 2 |
| red blood cells | Dusp1 | 0.53 | 2.09E-31 | 4.34E-27 | 0.54 | 0.28 | 0.26 | 1.9 |
| red blood cells | Cacna2d3 | 0.33 | 2.13E-31 | 4.42E-27 | 0.2 | 0.07 | 0.12 | 2.68 |
| red blood cells | Mbd2 | 0.42 | 5.43E-31 | 1.13E-26 | 0.86 | 0.55 | 0.31 | 1.56 |
| red blood cells | Zbtb7a | 0.4 | 6.72E-31 | 1.40E-26 | 0.6 | 0.32 | 0.28 | 1.86 |
| red blood cells | Cmas | 0.4 | 4.82E-30 | 1.00E-25 | 0.87 | 0.59 | 0.28 | 1.48 |
| red blood cells | Hist1h2bj | 0.46 | 9.66E-30 | 2.01E-25 | 0.47 | 0.24 | 0.23 | 1.97 |
| red blood cells | Piezo1 | 0.53 | 2.04E-29 | 4.23E-25 | 0.27 | 0.12 | 0.15 | 2.29 |
| red blood cells | Eipr1 | 0.34 | 4.08E-29 | 8.48E-25 | 0.46 | 0.23 | 0.23 | 2 |
| red blood cells | Tacc1 | 0.34 | 4.17E-29 | 8.66E-25 | 0.58 | 0.31 | 0.27 | 1.88 |
| red blood cells | Pip4k2a | 0.52 | 6.30E-29 | 1.31E-24 | 0.26 | 0.11 | 0.15 | 2.33 |
| red blood cells | Micall2 | 0.61 | 1.92E-28 | 4.00E-24 | 0.17 | 0.06 | 0.11 | 2.67 |
| red blood cells | Noct | 0.37 | 2.10E-28 | 4.38E-24 | 0.38 | 0.18 | 0.2 | 2.08 |
| red blood cells | Napepld | 0.54 | 3.28E-28 | 6.82E-24 | 0.16 | 0.06 | 0.1 | 2.77 |
| red blood cells | Adipor1 | 0.37 | 3.62E-28 | 7.54E-24 | 0.86 | 0.55 | 0.31 | 1.55 |
| red blood cells | Otub2 | 0.75 | 2.54E-26 | 5.29E-22 | 0.13 | 0.05 | 0.09 | 2.85 |
| red blood cells | Hipk1 | 0.37 | 3.35E-26 | 6.96E-22 | 0.65 | 0.36 | 0.29 | 1.81 |

**Supplementary Table 2. Marker genes for the major cell types of integrated clustering of E11.5 control and NotchGOF λ-junction cells.**

| | | | | | | | | |
|---|---|---|---|---|---|---|---|---|
| red blood cells | Clcn3 | 0.53 | 1.54E-25 | 3.21E-21 | 0.67 | 0.37 | 0.3 | 1.8 |
| red blood cells | Fam104a | 0.28 | 1.96E-25 | 4.07E-21 | 0.97 | 0.88 | 0.09 | 1.11 |
| red blood cells | Bcat2 | 0.29 | 2.09E-25 | 4.34E-21 | 0.53 | 0.28 | 0.25 | 1.87 |
| red blood cells | Hif3a | 0.45 | 4.10E-25 | 8.54E-21 | 0.2 | 0.08 | 0.12 | 2.41 |
| red blood cells | Wdr81 | 0.72 | 4.23E-25 | 8.80E-21 | 0.13 | 0.05 | 0.09 | 2.81 |
| red blood cells | Adgrg1 | 0.68 | 5.55E-25 | 1.15E-20 | 0.12 | 0.04 | 0.08 | 2.95 |
| red blood cells | Rab8b | 0.35 | 9.16E-25 | 1.91E-20 | 0.45 | 0.24 | 0.22 | 1.92 |
| red blood cells | Slc23a2 | 0.64 | 2.00E-24 | 4.16E-20 | 0.18 | 0.07 | 0.11 | 2.42 |
| red blood cells | Pnpla7 | 1.29 | 3.71E-24 | 7.72E-20 | 0.11 | 0.03 | 0.07 | 3.09 |
| red blood cells | Ogfod2 | 0.33 | 8.72E-24 | 1.81E-19 | 0.22 | 0.1 | 0.12 | 2.29 |
| red blood cells | Sod2 | 0.3 | 1.24E-23 | 2.57E-19 | 0.88 | 0.62 | 0.27 | 1.44 |
| red blood cells | 600002K03Ri | 0.31 | 1.43E-23 | 2.97E-19 | 0.24 | 0.11 | 0.13 | 2.19 |
| red blood cells | Slc7a5 | 0.6 | 1.83E-23 | 3.81E-19 | 0.13 | 0.05 | 0.08 | 2.8 |
| red blood cells | Myc | 0.31 | 2.57E-22 | 5.35E-18 | 0.31 | 0.16 | 0.16 | 1.98 |
| red blood cells | Emc9 | 0.51 | 4.35E-22 | 9.04E-18 | 0.12 | 0.05 | 0.08 | 2.76 |
| red blood cells | Polh | 0.47 | 7.76E-22 | 1.61E-17 | 0.19 | 0.09 | 0.11 | 2.28 |
| red blood cells | Pik3cb | 0.46 | 8.11E-22 | 1.69E-17 | 0.15 | 0.06 | 0.09 | 2.5 |
| red blood cells | Jak2 | 0.29 | 2.53E-21 | 5.27E-17 | 0.3 | 0.15 | 0.15 | 2.01 |
| red blood cells | Grtp1 | 0.27 | 2.92E-21 | 6.07E-17 | 0.28 | 0.14 | 0.14 | 2.06 |
| red blood cells | Sppl2b | 0.32 | 9.42E-21 | 1.96E-16 | 0.29 | 0.14 | 0.15 | 2.02 |
| red blood cells | Snrnp25 | 0.37 | 1.61E-20 | 3.34E-16 | 0.83 | 0.61 | 0.23 | 1.38 |
| red blood cells | Ninl | 0.43 | 1.98E-20 | 4.12E-16 | 0.43 | 0.23 | 0.2 | 1.84 |
| red blood cells | Traf5 | 0.58 | 2.65E-20 | 5.51E-16 | 0.11 | 0.04 | 0.07 | 2.77 |
| red blood cells | Marchf8 | 0.37 | 3.81E-20 | 7.92E-16 | 0.32 | 0.17 | 0.15 | 1.93 |
| red blood cells | Cbfa2t3 | 0.81 | 1.33E-19 | 2.76E-15 | 0.13 | 0.05 | 0.08 | 2.47 |
| red blood cells | Slc7a1 | 0.34 | 1.46E-19 | 3.03E-15 | 0.17 | 0.08 | 0.1 | 2.26 |
| red blood cells | Hipk2 | 0.35 | 4.64E-19 | 9.65E-15 | 0.87 | 0.64 | 0.24 | 1.37 |
| red blood cells | Ddit3 | 0.32 | 6.52E-19 | 1.36E-14 | 0.37 | 0.2 | 0.17 | 1.85 |
| red blood cells | Man1a | 0.76 | 1.65E-18 | 3.44E-14 | 0.14 | 0.06 | 0.08 | 2.39 |
| red blood cells | Flvcr1 | 0.48 | 2.76E-18 | 5.73E-14 | 0.2 | 0.09 | 0.1 | 2.11 |
| red blood cells | Kat2b | 0.63 | 2.86E-18 | 5.96E-14 | 0.11 | 0.04 | 0.07 | 2.59 |
| red blood cells | Arrdc2 | 0.79 | 3.20E-18 | 6.66E-14 | 0.11 | 0.04 | 0.07 | 2.58 |
| red blood cells | Eif2b3 | 0.31 | 4.64E-18 | 9.64E-14 | 0.72 | 0.47 | 0.25 | 1.53 |
| red blood cells | Ghitm | 0.28 | 5.00E-18 | 1.04E-13 | 0.89 | 0.66 | 0.22 | 1.34 |
| red blood cells | Pet100 | 0.3 | 1.25E-17 | 2.60E-13 | 0.82 | 0.55 | 0.27 | 1.5 |
| red blood cells | Gm4013 | 0.44 | 1.42E-17 | 2.96E-13 | 0.12 | 0.05 | 0.07 | 2.51 |
| red blood cells | Rhbdd1 | 0.4 | 3.96E-16 | 8.23E-12 | 0.12 | 0.05 | 0.07 | 2.38 |
| red blood cells | Mrm1 | 0.58 | 1.86E-15 | 3.87E-11 | 0.1 | 0.04 | 0.06 | 2.46 |
| red blood cells | Znfx1 | 0.4 | 2.40E-15 | 4.98E-11 | 0.13 | 0.06 | 0.07 | 2.28 |
| red blood cells | Ddc | 0.44 | 2.80E-15 | 5.83E-11 | 0.12 | 0.05 | 0.07 | 2.31 |
| red blood cells | Cish | 0.5 | 3.03E-15 | 6.29E-11 | 0.14 | 0.06 | 0.07 | 2.21 |
| red blood cells | Fam234b | 0.33 | 9.51E-13 | 1.98E-08 | 0.12 | 0.06 | 0.06 | 2.14 |
| red blood cells | Ehd3 | 0.28 | 3.72E-11 | 7.74E-07 | 0.2 | 0.11 | 0.09 | 1.83 |
| red blood cells | Spata13 | 0.7 | 6.61E-11 | 1.37E-06 | 0.14 | 0.08 | 0.07 | 1.89 |
| red blood cells | Ranbp10 | 0.34 | 1.45E-10 | 3.01E-06 | 0.47 | 0.29 | 0.17 | 1.59 |
| red blood cells | Ap1b1 | 0.32 | 6.07E-09 | 1.26E-04 | 0.38 | 0.24 | 0.14 | 1.58 |
| endothelium | Vamp5 | 6.91 | 0 | 0 | 0.92 | 0.05 | 0.87 | 19.57 |
| endothelium | Esam | 8.77 | 0 | 0 | 0.86 | 0.01 | 0.85 | 77.82 |
| endothelium | Ctla2a | 10.08 | 0 | 0 | 0.84 | 0 | 0.84 | 210.75 |
| endothelium | Pecam1 | 8.42 | 0 | 0 | 0.85 | 0.01 | 0.84 | 60.79 |
| endothelium | Kdr | 9.7 | 0 | 0 | 0.85 | 0.01 | 0.84 | 93.89 |
| endothelium | Clec1b | 10.44 | 0 | 0 | 0.84 | 0 | 0.84 | 418.5 |
| endothelium | Emcn | 9.79 | 0 | 0 | 0.85 | 0.01 | 0.83 | 70.42 |
| endothelium | Cldn5 | 10.28 | 0 | 0 | 0.82 | 0.01 | 0.82 | 137.33 |
| endothelium | Rasip1 | 6.95 | 0 | 0 | 0.85 | 0.03 | 0.82 | 28.17 |
| endothelium | Gimap6 | 7.77 | 0 | 0 | 0.82 | 0.01 | 0.81 | 63.38 |

**Supplementary Table 2. Marker genes for the major cell types of integrated clustering of E11.5 control and NotchGOF λ-junction cells.**

| | | | | | | | | |
|---|---|---|---|---|---|---|---|---|
| endothelium | S100a16 | 6.57 | 0 | 0 | 0.87 | 0.06 | 0.81 | 14.06 |
| endothelium | Sox18 | 9.25 | 0 | 0 | 0.82 | 0.01 | 0.81 | 136 |
| endothelium | Anxa3 | 5.4 | 0 | 0 | 0.89 | 0.08 | 0.81 | 11.64 |
| endothelium | Ecscr | 9.31 | 0 | 0 | 0.81 | 0.01 | 0.8 | 90.33 |
| endothelium | Plvap | 8.1 | 0 | 0 | 0.82 | 0.03 | 0.8 | 30.52 |
| endothelium | Cdh5 | 9.67 | 0 | 0 | 0.8 | 0.01 | 0.8 | 100.38 |
| endothelium | Icam2 | 9.45 | 0 | 0 | 0.8 | 0.01 | 0.79 | 133.33 |
| endothelium | Gmfg | 5.94 | 0 | 0 | 0.81 | 0.01 | 0.79 | 67.08 |
| endothelium | Fxyd5 | 6.22 | 0 | 0 | 0.8 | 0.02 | 0.79 | 53.53 |
| endothelium | Cd93 | 9.83 | 0 | 0 | 0.79 | 0 | 0.79 | 197.25 |
| endothelium | S100a13 | 4.57 | 0 | 0 | 0.88 | 0.09 | 0.78 | 9.33 |
| endothelium | Aplnr | 9.77 | 0 | 0 | 0.78 | 0.01 | 0.78 | 156.8 |
| endothelium | Lmo2 | 3.86 | 0 | 0 | 0.89 | 0.11 | 0.77 | 7.79 |
| endothelium | Eng | 6.48 | 0 | 0 | 0.8 | 0.03 | 0.77 | 28.57 |
| endothelium | Tgfb1 | 4.69 | 0 | 0 | 0.86 | 0.1 | 0.77 | 9.09 |
| endothelium | Col15a1 | 8.7 | 0 | 0 | 0.78 | 0.01 | 0.77 | 97 |
| endothelium | Flt1 | 9.26 | 0 | 0 | 0.77 | 0.01 | 0.77 | 128.83 |
| endothelium | Tspan18 | 6.3 | 0 | 0 | 0.79 | 0.03 | 0.76 | 29.15 |
| endothelium | Plxnd1 | 6.69 | 0 | 0 | 0.81 | 0.05 | 0.76 | 17.89 |
| endothelium | Arhgap18 | 4.85 | 0 | 0 | 0.87 | 0.11 | 0.76 | 7.72 |
| endothelium | Rasgrp2 | 6.58 | 0 | 0 | 0.8 | 0.04 | 0.76 | 21.49 |
| endothelium | Tmem255a | 7.32 | 0 | 0 | 0.77 | 0.02 | 0.76 | 51.53 |
| endothelium | Gngt2 | 6.27 | 0 | 0 | 0.78 | 0.03 | 0.76 | 31.24 |
| endothelium | Cavin3 | 5.26 | 0 | 0 | 0.86 | 0.11 | 0.75 | 7.74 |
| endothelium | Mef2c | 4.06 | 0 | 0 | 0.91 | 0.17 | 0.74 | 5.3 |
| endothelium | Gimap1 | 8.56 | 0 | 0 | 0.74 | 0 | 0.74 | 247 |
| endothelium | Pde8a | 7.14 | 0 | 0 | 0.74 | 0.01 | 0.73 | 73.9 |
| endothelium | Lcp1 | 4.78 | 0 | 0 | 0.75 | 0.03 | 0.73 | 30.08 |
| endothelium | Mfng | 4.23 | 0 | 0 | 0.76 | 0.03 | 0.72 | 22.94 |
| endothelium | Crip2 | 5.18 | 0 | 0 | 0.88 | 0.16 | 0.72 | 5.48 |
| endothelium | Gimap4 | 9.22 | 0 | 0 | 0.73 | 0 | 0.72 | 181.25 |
| endothelium | Tie1 | 9.08 | 0 | 0 | 0.72 | 0 | 0.72 | 241 |
| endothelium | Rhoj | 4 | 0 | 0 | 0.83 | 0.11 | 0.72 | 7.66 |
| endothelium | Cd34 | 5.7 | 0 | 0 | 0.84 | 0.12 | 0.72 | 6.96 |
| endothelium | Smagp | 6.04 | 0 | 0 | 0.74 | 0.02 | 0.71 | 33.45 |
| endothelium | Adgrf5 | 9.56 | 0 | 0 | 0.71 | 0 | 0.71 | 237.33 |
| endothelium | Myct1 | 8.91 | 0 | 0 | 0.71 | 0 | 0.7 | 235.67 |
| endothelium | Ehd2 | 5.51 | 0 | 0 | 0.74 | 0.04 | 0.7 | 19 |
| endothelium | Egfl7 | 5.45 | 0 | 0 | 0.91 | 0.21 | 0.7 | 4.36 |
| endothelium | Dok4 | 4.97 | 0 | 0 | 0.77 | 0.07 | 0.7 | 10.97 |
| endothelium | Tfpi | 3.98 | 0 | 0 | 0.8 | 0.1 | 0.7 | 7.95 |
| endothelium | Ets1 | 4.64 | 0 | 0 | 0.86 | 0.17 | 0.69 | 5.14 |
| endothelium | Klhl4 | 9.29 | 0 | 0 | 0.69 | 0 | 0.69 | 346.5 |
| endothelium | Tubb6 | 4.53 | 0 | 0 | 0.79 | 0.1 | 0.69 | 7.76 |
| endothelium | She | 6.82 | 0 | 0 | 0.7 | 0.01 | 0.69 | 63.55 |
| endothelium | N4bp3 | 5.13 | 0 | 0 | 0.74 | 0.05 | 0.69 | 14.21 |
| endothelium | Flt4 | 9.11 | 0 | 0 | 0.69 | 0 | 0.68 | 228.33 |
| endothelium | Mmrn1 | 9.54 | 0 | 0 | 0.68 | 0 | 0.68 | 340 |
| endothelium | Rasgrp3 | 8.12 | 0 | 0 | 0.68 | 0.01 | 0.68 | 136.6 |
| endothelium | Tagln2 | 4.66 | 0 | 0 | 0.96 | 0.29 | 0.68 | 3.36 |
| endothelium | Lyn | 5.19 | 0 | 0 | 0.7 | 0.02 | 0.68 | 33.14 |
| endothelium | Ramp2 | 5.56 | 0 | 0 | 0.85 | 0.17 | 0.67 | 4.87 |
| endothelium | Kcne3 | 7.69 | 0 | 0 | 0.7 | 0.02 | 0.67 | 29 |
| endothelium | Gng11 | 5.58 | 0 | 0 | 0.97 | 0.3 | 0.67 | 3.21 |
| endothelium | Afap1l1 | 4.11 | 0 | 0 | 0.73 | 0.06 | 0.67 | 11.79 |
| endothelium | Tmem88 | 5.01 | 0 | 0 | 0.72 | 0.05 | 0.67 | 14.94 |

**Supplementary Table 2. Marker genes for the major cell types of integrated clustering of E11.5 control and NotchGOF λ-junction cells.**

| | | | | | | | | |
|---|---|---|---|---|---|---|---|---|
| endothelium | Arhgef15 | 6.06 | 0 | 0 | 0.68 | 0.02 | 0.66 | 40 |
| endothelium | Sema6d | 4.82 | 0 | 0 | 0.72 | 0.05 | 0.66 | 13.75 |
| endothelium | Gm16104 | 4.99 | 0 | 0 | 0.69 | 0.03 | 0.66 | 22.35 |
| endothelium | Timp3 | 4.58 | 0 | 0 | 0.82 | 0.16 | 0.66 | 5.13 |
| endothelium | Myzap | 5.87 | 0 | 0 | 0.68 | 0.02 | 0.66 | 33.85 |
| endothelium | Mmrn2 | 5.28 | 0 | 0 | 0.7 | 0.04 | 0.66 | 17.4 |
| endothelium | Klhl6 | 6.59 | 0 | 0 | 0.66 | 0.01 | 0.65 | 94.43 |
| endothelium | Dysf | 5.58 | 0 | 0 | 0.68 | 0.03 | 0.65 | 26.04 |
| endothelium | Stab1 | 4.91 | 0 | 0 | 0.66 | 0.01 | 0.65 | 60.09 |
| endothelium | Ptpn18 | 3.64 | 0 | 0 | 0.72 | 0.07 | 0.65 | 10.83 |
| endothelium | Klf2 | 6.66 | 0 | 0 | 0.68 | 0.03 | 0.65 | 25.96 |
| endothelium | Slc29a1 | 4.32 | 0 | 0 | 0.76 | 0.11 | 0.65 | 6.82 |
| endothelium | Thsd1 | 7.23 | 0 | 0 | 0.65 | 0.01 | 0.64 | 108 |
| endothelium | Arap3 | 5.07 | 0 | 0 | 0.67 | 0.03 | 0.63 | 20.21 |
| endothelium | Kank3 | 4.52 | 0 | 0 | 0.69 | 0.06 | 0.63 | 12.51 |
| endothelium | Dusp3 | 4.69 | 0 | 0 | 0.68 | 0.05 | 0.63 | 15 |
| endothelium | Tek | 7.54 | 0 | 0 | 0.64 | 0.01 | 0.63 | 79.63 |
| endothelium | Cavin1 | 7.07 | 0 | 0 | 0.64 | 0.01 | 0.63 | 79.63 |
| endothelium | )030007L05Ri | 10.54 | 0 | 0 | 0.63 | 0 | 0.63 | 629 |
| endothelium | Gpr183 | 6.63 | 0 | 0 | 0.63 | 0.01 | 0.63 | 105.33 |
| endothelium | Tbx1 | 5.81 | 0 | 0 | 0.64 | 0.02 | 0.63 | 37.82 |
| endothelium | Tm6sf1 | 4.33 | 0 | 0 | 0.67 | 0.04 | 0.63 | 15.2 |
| endothelium | Kitl | 4.32 | 0 | 0 | 0.69 | 0.07 | 0.62 | 10.54 |
| endothelium | Fam167b | 9.98 | 0 | 0 | 0.62 | 0 | 0.62 | 621 |
| endothelium | Dusp2 | 5.39 | 0 | 0 | 0.64 | 0.02 | 0.62 | 29.09 |
| endothelium | Sparcl1 | 6.86 | 0 | 0 | 0.64 | 0.02 | 0.62 | 37.35 |
| endothelium | Pde2a | 8.33 | 0 | 0 | 0.62 | 0 | 0.62 | 309.5 |
| endothelium | Mcam | 5.07 | 0 | 0 | 0.65 | 0.03 | 0.62 | 20.25 |
| endothelium | Adgrl4 | 8.49 | 0 | 0 | 0.62 | 0 | 0.61 | 205.33 |
| endothelium | Scarf1 | 8.13 | 0 | 0 | 0.62 | 0 | 0.61 | 205.33 |
| endothelium | Hapln1 | 5.32 | 0 | 0 | 0.64 | 0.03 | 0.61 | 18.82 |
| endothelium | S1pr1 | 4.44 | 0 | 0 | 0.65 | 0.05 | 0.6 | 12.52 |
| endothelium | Grrp1 | 4.75 | 0 | 0 | 0.65 | 0.05 | 0.6 | 13.16 |
| endothelium | Pdgfb | 7.6 | 0 | 0 | 0.6 | 0 | 0.6 | 150 |
| endothelium | Gchfr | 3.44 | 0 | 0 | 0.69 | 0.1 | 0.6 | 7.07 |
| endothelium | Apold1 | 6.95 | 0 | 0 | 0.6 | 0.01 | 0.59 | 75 |
| endothelium | Grap | 7.37 | 0 | 0 | 0.6 | 0.01 | 0.59 | 119.4 |
| endothelium | Tmem204 | 7.71 | 0 | 0 | 0.58 | 0 | 0.58 | 194.67 |
| endothelium | Adam15 | 3.62 | 0 | 0 | 0.67 | 0.09 | 0.58 | 7.67 |
| endothelium | Sox17 | 7.62 | 0 | 0 | 0.58 | 0 | 0.58 | 193 |
| endothelium | Fgd5 | 5.92 | 0 | 0 | 0.59 | 0.01 | 0.58 | 45.31 |
| endothelium | Exoc3l4 | 3.8 | 0 | 0 | 0.62 | 0.06 | 0.56 | 10.18 |
| endothelium | .610203C22Ri | 8.02 | 0 | 0 | 0.56 | 0 | 0.56 | 187.67 |
| endothelium | Pcdh12 | 8.58 | 0 | 0 | 0.56 | 0 | 0.56 | 557 |
| endothelium | Robo4 | 10.14 | 0 | 0 | 0.56 | 0 | 0.55 | 555 |
| endothelium | Cd38 | 4.09 | 0 | 0 | 0.6 | 0.04 | 0.55 | 14.51 |
| endothelium | Ptprb | 7.83 | 0 | 0 | 0.56 | 0 | 0.55 | 185 |
| endothelium | Cyyr1 | 4.49 | 0 | 0 | 0.6 | 0.04 | 0.55 | 13.84 |
| endothelium | Nrros | 4 | 0 | 0 | 0.58 | 0.03 | 0.55 | 19.86 |
| endothelium | Erg | 5.23 | 0 | 0 | 0.57 | 0.02 | 0.55 | 31.39 |
| endothelium | Podxl | 4.92 | 0 | 0 | 0.58 | 0.03 | 0.55 | 17.03 |
| endothelium | Ppp1r13b | 3.65 | 0 | 0 | 0.61 | 0.07 | 0.54 | 8.63 |
| endothelium | Npr1 | 6.94 | 0 | 0 | 0.54 | 0.01 | 0.54 | 90.67 |
| endothelium | Epas1 | 7.85 | 0 | 0 | 0.54 | 0 | 0.54 | 180.33 |
| endothelium | Bst2 | 3.91 | 0 | 0 | 0.58 | 0.05 | 0.54 | 11.92 |
| endothelium | Cdc42ep2 | 4.71 | 0 | 0 | 0.56 | 0.04 | 0.52 | 15.91 |

**Supplementary Table 2. Marker genes for the major cell types of integrated clustering of E11.5 control and NotchGOF λ-junction cells.**

| | | | | | | | | |
|---|---|---|---|---|---|---|---|---|
| endothelium | Clec14a | 4.19 | 0 | 0 | 0.57 | 0.05 | 0.52 | 12.56 |
| endothelium | Pcdh1 | 4 | 0 | 0 | 0.56 | 0.04 | 0.52 | 13.09 |
| endothelium | Bcl6b | 6.42 | 0 | 0 | 0.53 | 0.01 | 0.52 | 58.67 |
| endothelium | Abi3 | 6.31 | 0 | 0 | 0.53 | 0.01 | 0.52 | 75 |
| endothelium | Sncg | 8.87 | 0 | 0 | 0.52 | 0 | 0.51 | 257.5 |
| endothelium | Nos3 | 6.77 | 0 | 0 | 0.51 | 0.01 | 0.51 | 85.33 |
| endothelium | Upp1 | 7.67 | 0 | 0 | 0.51 | 0 | 0.51 | 127.25 |
| endothelium | Itga2 | 7.15 | 0 | 0 | 0.51 | 0.01 | 0.5 | 101.8 |
| endothelium | Ushbp1 | 7.84 | 0 | 0 | 0.5 | 0 | 0.5 | 252 |
| endothelium | Samsn1 | 6.09 | 0 | 0 | 0.51 | 0.01 | 0.5 | 84.5 |
| endothelium | Vav3 | 3.51 | 0 | 0 | 0.55 | 0.05 | 0.5 | 10.98 |
| endothelium | Fam43a | 3.61 | 0 | 0 | 0.53 | 0.05 | 0.49 | 11.84 |
| endothelium | Adora2a | 7.48 | 0 | 0 | 0.49 | 0 | 0.49 | 163.67 |
| endothelium | Hhex | 6.42 | 0 | 0 | 0.49 | 0.01 | 0.49 | 70.43 |
| endothelium | Icam1 | 6.17 | 0 | 0 | 0.49 | 0.01 | 0.48 | 54.78 |
| endothelium | Cd40 | 4.82 | 0 | 0 | 0.5 | 0.02 | 0.48 | 29.35 |
| endothelium | Ptprm | 4.07 | 0 | 0 | 0.52 | 0.04 | 0.48 | 12.38 |
| endothelium | Arhgap27 | 4.79 | 0 | 0 | 0.5 | 0.02 | 0.48 | 21.78 |
| endothelium | Exoc3l | 8.35 | 0 | 0 | 0.48 | 0 | 0.48 | 238.5 |
| endothelium | Elmo1 | 3.86 | 0 | 0 | 0.51 | 0.04 | 0.47 | 13.47 |
| endothelium | Calcrl | 4.31 | 0 | 0 | 0.5 | 0.03 | 0.47 | 19.19 |
| endothelium | Cd109 | 3.68 | 0 | 0 | 0.52 | 0.05 | 0.47 | 11.49 |
| endothelium | Cda | 4.89 | 0 | 0 | 0.47 | 0.02 | 0.45 | 23.35 |
| endothelium | Cav2 | 5.26 | 0 | 0 | 0.45 | 0.01 | 0.44 | 34.69 |
| endothelium | Plac1 | 5.08 | 0 | 0 | 0.45 | 0.02 | 0.44 | 28.19 |
| endothelium | Itga1 | 4.24 | 0 | 0 | 0.46 | 0.03 | 0.43 | 15.3 |
| endothelium | Ccm2l | 8.41 | 0 | 0 | 0.43 | 0 | 0.43 | 429 |
| endothelium | Bik | 5.2 | 0 | 0 | 0.44 | 0.02 | 0.43 | 24.61 |
| endothelium | Exoc3l2 | 7.46 | 0 | 0 | 0.42 | 0 | 0.42 | 141.33 |
| endothelium | Lat | 6.51 | 0 | 0 | 0.43 | 0.01 | 0.42 | 53.63 |
| endothelium | Sox7 | 9.48 | 0 | 0 | 0.42 | 0 | 0.42 | 421 |
| endothelium | Rapgef3 | 6.57 | 0 | 0 | 0.42 | 0.01 | 0.42 | 84.8 |
| endothelium | Gdpd5 | 3.59 | 0 | 0 | 0.46 | 0.04 | 0.42 | 11.69 |
| endothelium | Ctla2b | 7.91 | 0 | 0 | 0.42 | 0 | 0.42 | 139.67 |
| endothelium | Rin3 | 4.62 | 0 | 0 | 0.43 | 0.02 | 0.41 | 24 |
| endothelium | Ggta1 | 3.66 | 0 | 0 | 0.45 | 0.04 | 0.41 | 12.11 |
| endothelium | Cav1 | 7.06 | 0 | 0 | 0.41 | 0 | 0.41 | 103.25 |
| endothelium | 130074G19Ri | 8.22 | 0 | 0 | 0.41 | 0 | 0.41 | 408 |
| endothelium | Spns2 | 4.62 | 0 | 0 | 0.42 | 0.02 | 0.4 | 19.05 |
| endothelium | Nova2 | 4.41 | 0 | 0 | 0.42 | 0.02 | 0.4 | 20.8 |
| endothelium | Enpp3 | 4.18 | 0 | 0 | 0.42 | 0.03 | 0.39 | 15.59 |
| endothelium | Adgrg3 | 8.22 | 0 | 0 | 0.39 | 0 | 0.39 | 392 |
| endothelium | Hlx | 6.76 | 0 | 0 | 0.39 | 0 | 0.39 | 97.25 |
| endothelium | Cracr2b | 4.02 | 0 | 0 | 0.41 | 0.03 | 0.38 | 14.17 |
| endothelium | Atf3 | 3.82 | 0 | 0 | 0.4 | 0.02 | 0.38 | 19.85 |
| endothelium | Ppp1r16b | 8.43 | 0 | 0 | 0.38 | 0 | 0.38 | 189.5 |
| endothelium | Inpp5d | 4.65 | 0 | 0 | 0.38 | 0.01 | 0.38 | 48 |
| endothelium | Lbp | 3.84 | 0 | 0 | 0.4 | 0.02 | 0.38 | 16.67 |
| endothelium | Jak3 | 4.2 | 0 | 0 | 0.4 | 0.02 | 0.37 | 17.26 |
| endothelium | Lyl1 | 4.26 | 0 | 0 | 0.4 | 0.03 | 0.37 | 13.43 |
| endothelium | Shank3 | 4.54 | 0 | 0 | 0.38 | 0.01 | 0.37 | 27.21 |
| endothelium | Gm13889 | 5.81 | 0 | 0 | 0.37 | 0.01 | 0.37 | 53.29 |
| endothelium | Pear1 | 3.98 | 0 | 0 | 0.39 | 0.02 | 0.37 | 16.21 |
| endothelium | Pon3 | 3.62 | 0 | 0 | 0.39 | 0.03 | 0.36 | 14.88 |
| endothelium | Zfpm2 | 6.49 | 0 | 0 | 0.36 | 0.01 | 0.36 | 72.6 |
| endothelium | Cd200 | 3.91 | 0 | 0 | 0.38 | 0.02 | 0.36 | 18.05 |

**Supplementary Table 2. Marker genes for the major cell types of integrated clustering of E11.5 control and NotchGOF λ-junction cells.**

| | | | | | | | | |
|---|---|---|---|---|---|---|---|---|
| endothelium | Adam12 | 5.84 | 0 | 0 | 0.37 | 0.01 | 0.36 | 45.63 |
| endothelium | Tm4sf1 | 5.46 | 0 | 0 | 0.37 | 0.01 | 0.35 | 33.18 |
| endothelium | Laptm5 | 2.85 | 0 | 0 | 0.37 | 0.01 | 0.35 | 26.29 |
| endothelium | Tbxa2r | 5.45 | 0 | 0 | 0.36 | 0.01 | 0.35 | 40 |
| endothelium | Card10 | 4.07 | 0 | 0 | 0.36 | 0.02 | 0.35 | 22.69 |
| endothelium | Abcc9 | 6.78 | 0 | 0 | 0.34 | 0 | 0.34 | 114.67 |
| endothelium | Sqor | 4.41 | 0 | 0 | 0.34 | 0.01 | 0.33 | 24.57 |
| endothelium | Dll4 | 8.47 | 0 | 0 | 0.32 | 0 | 0.32 | 323 |
| endothelium | Apln | 4.61 | 0 | 0 | 0.34 | 0.02 | 0.32 | 17.05 |
| endothelium | Rgs7bp | 4.09 | 0 | 0 | 0.33 | 0.02 | 0.32 | 18.5 |
| endothelium | Elf4 | 4 | 0 | 0 | 0.33 | 0.02 | 0.31 | 17.42 |
| endothelium | Thbd | 5.47 | 0 | 0 | 0.32 | 0.01 | 0.31 | 52.83 |
| endothelium | Car8 | 4.09 | 0 | 0 | 0.32 | 0.02 | 0.31 | 21.53 |
| endothelium | F2rl3 | 7.32 | 0 | 0 | 0.31 | 0 | 0.31 | 154.5 |
| endothelium | Csgalnact1 | 6.29 | 0 | 0 | 0.3 | 0 | 0.3 | 76 |
| endothelium | Gm32688 | 4.99 | 0 | 0 | 0.3 | 0.01 | 0.29 | 30.4 |
| endothelium | Gimap5 | 8.13 | 0 | 0 | 0.29 | 0 | 0.29 | 293 |
| endothelium | Prl3b1 | 10.76 | 0 | 0 | 0.29 | 0 | 0.29 | #NA |
| endothelium | Trpc6 | 7.18 | 0 | 0 | 0.29 | 0 | 0.29 | 291 |
| endothelium | Trim47 | 4.35 | 0 | 0 | 0.3 | 0.01 | 0.28 | 21.14 |
| endothelium | Gm9917 | 4.96 | 0 | 0 | 0.29 | 0.01 | 0.28 | 31.67 |
| endothelium | Rapgef4 | 5.13 | 0 | 0 | 0.28 | 0.01 | 0.28 | 35.38 |
| endothelium | Clec1a | 8.68 | 0 | 0 | 0.26 | 0 | 0.26 | 264 |
| endothelium | Zfp366 | 7.91 | 0 | 0 | 0.25 | 0 | 0.25 | 253 |
| endothelium | Fam78a | 5.28 | 0 | 0 | 0.26 | 0.01 | 0.25 | 28.44 |
| endothelium | Sh3tc1 | 6.27 | 0 | 0 | 0.25 | 0 | 0.25 | 82.67 |
| endothelium | Hcls1 | 4.13 | 0 | 0 | 0.24 | 0.01 | 0.23 | 34.29 |
| endothelium | Prnd | 8.61 | 0 | 0 | 0.23 | 0 | 0.23 | 232 |
| endothelium | Arhgap25 | 4.58 | 0 | 0 | 0.24 | 0.01 | 0.23 | 33.86 |
| endothelium | Gpihbp1 | 8.92 | 0 | 0 | 0.23 | 0 | 0.23 | #NA |
| endothelium | 033406O09Ri | 5.59 | 0 | 0 | 0.24 | 0.01 | 0.23 | 29.38 |
| endothelium | Prdm1 | 4.7 | 0 | 0 | 0.23 | 0.01 | 0.23 | 33.14 |
| endothelium | Depp1 | 9.93 | 0 | 0 | 0.22 | 0 | 0.22 | #NA |
| endothelium | Ptpn7 | 5.08 | 0 | 0 | 0.22 | 0.01 | 0.22 | 44.8 |
| endothelium | Pik3r6 | 5.97 | 0 | 0 | 0.21 | 0 | 0.2 | 68.33 |
| endothelium | Sardh | 8.49 | 0 | 0 | 0.19 | 0 | 0.19 | 96 |
| endothelium | 230102K24Ri | 8.75 | 0 | 0 | 0.19 | 0 | 0.19 | #NA |
| endothelium | Ppm1j | 5.86 | 0 | 0 | 0.19 | 0.01 | 0.18 | 37.4 |
| endothelium | Rsad2 | 6.38 | 0 | 0 | 0.17 | 0 | 0.17 | 86.5 |
| endothelium | Madcam1 | 8.43 | 0 | 0 | 0.17 | 0 | 0.17 | #NA |
| endothelium | Gimap8 | 6.17 | 0 | 0 | 0.16 | 0 | 0.16 | 163 |
| endothelium | Adamtsl2 | 7.01 | 0 | 0 | 0.16 | 0 | 0.16 | 157 |
| endothelium | Tfpi2 | 7.28 | 0 | 0 | 0.15 | 0 | 0.15 | #NA |
| endothelium | Unc45b | 6.72 | 0 | 0 | 0.15 | 0 | 0.15 | 74.5 |
| endothelium | Nos2 | 9.92 | 0 | 0 | 0.14 | 0 | 0.14 | #NA |
| endothelium | AU021092 | 7.55 | 0 | 0 | 0.13 | 0 | 0.13 | 133 |
| endothelium | Gpr182 | 9.31 | 0 | 0 | 0.13 | 0 | 0.13 | #NA |
| endothelium | Gm13056 | 8 | 0 | 0 | 0.13 | 0 | 0.13 | #NA |
| endothelium | Kcnq1 | 7.01 | 0 | 0 | 0.13 | 0 | 0.12 | 125 |
| endothelium | Esm1 | 10.62 | 0 | 0 | 0.12 | 0 | 0.12 | #NA |
| endothelium | Mmp25 | 5.82 | 0 | 0 | 0.13 | 0 | 0.12 | 62.5 |
| endothelium | Gm38414 | 6.01 | 0 | 0 | 0.12 | 0 | 0.12 | 117 |
| endothelium | Sema3g | 6.16 | 0 | 0 | 0.12 | 0 | 0.11 | 115 |
| endothelium | 330037F02Ri | 8.44 | 0 | 0 | 0.11 | 0 | 0.11 | #NA |
| endothelium | Gm38832 | 7.73 | 0 | 0 | 0.11 | 0 | 0.11 | 107 |
| endothelium | Cdc42ep1 | 3.6 | 2.09E-305 | 4.34E-301 | 0.53 | 0.05 | 0.47 | 9.78 |

**Supplementary Table 2. Marker genes for the major cell types of integrated clustering of E11.5 control and NotchGOF λ-junction cells.**

| | | | | | | | | |
|---|---|---|---|---|---|---|---|---|
| endothelium | Ptgs1 | 3.76 | 3.43E-303 | 7.13E-299 | 0.42 | 0.03 | 0.39 | 12.47 |
| endothelium | Pon2 | 3.4 | 5.91E-303 | 1.23E-298 | 0.84 | 0.17 | 0.67 | 4.91 |
| endothelium | BC028528 | 3.47 | 1.31E-297 | 2.72E-293 | 0.58 | 0.07 | 0.51 | 8.6 |
| endothelium | Colgalt2 | 4.64 | 2.52E-290 | 5.23E-286 | 0.15 | 0 | 0.15 | 50.67 |
| endothelium | Junb | 3.35 | 1.00E-289 | 2.08E-285 | 0.7 | 0.11 | 0.59 | 6.59 |
| endothelium | Gm33370 | 6.49 | 3.14E-288 | 6.54E-284 | 0.11 | 0 | 0.11 | 107 |
| endothelium | Gm17745 | 6.27 | 3.39E-288 | 7.04E-284 | 0.1 | 0 | 0.1 | 104 |
| endothelium | Ripply3 | 3.2 | 9.24E-288 | 1.92E-283 | 0.51 | 0.05 | 0.46 | 9.48 |
| endothelium | Emp1 | 3.52 | 4.04E-287 | 8.40E-283 | 0.72 | 0.12 | 0.6 | 5.98 |
| endothelium | Lama4 | 3.5 | 5.22E-284 | 1.09E-279 | 0.72 | 0.12 | 0.6 | 5.88 |
| endothelium | Rasa3 | 3.65 | 7.53E-284 | 1.57E-279 | 0.57 | 0.07 | 0.5 | 8.35 |
| endothelium | Tnnt2 | 6.17 | 4.46E-282 | 9.28E-278 | 0.12 | 0 | 0.12 | 61.5 |
| endothelium | Bace2 | 4.56 | 1.24E-278 | 2.59E-274 | 0.2 | 0.01 | 0.19 | 27.86 |
| endothelium | Ifi47 | 7.09 | 1.46E-278 | 3.04E-274 | 0.1 | 0 | 0.1 | 101 |
| endothelium | Sipa1 | 3.47 | 1.33E-277 | 2.76E-273 | 0.8 | 0.17 | 0.63 | 4.8 |
| endothelium | Srgn | 4.84 | 2.20E-276 | 4.58E-272 | 0.21 | 0.01 | 0.2 | 26.38 |
| endothelium | Tgfbr2 | 3.37 | 7.04E-275 | 1.46E-270 | 0.59 | 0.08 | 0.51 | 7.72 |
| endothelium | Samd9l | 5.46 | 5.11E-273 | 1.06E-268 | 0.12 | 0 | 0.12 | 60 |
| endothelium | Gja4 | 6.41 | 1.06E-268 | 2.19E-264 | 0.17 | 0.01 | 0.16 | 33.6 |
| endothelium | Procr | 4.02 | 1.26E-268 | 2.62E-264 | 0.25 | 0.01 | 0.24 | 19.31 |
| endothelium | B3gnt2 | 3.39 | 1.26E-268 | 2.62E-264 | 0.63 | 0.09 | 0.54 | 6.91 |
| endothelium | Dusp27 | 4.79 | 3.82E-267 | 7.94E-263 | 0.17 | 0.01 | 0.16 | 33.6 |
| endothelium | Cbfa2t3 | 3.52 | 6.26E-266 | 1.30E-261 | 0.46 | 0.05 | 0.42 | 9.67 |
| endothelium | Ecm1 | 3.96 | 4.57E-263 | 9.50E-259 | 0.3 | 0.02 | 0.28 | 15.74 |
| endothelium | Prkch | 3.12 | 1.60E-262 | 3.34E-258 | 0.59 | 0.08 | 0.51 | 7.43 |
| endothelium | Filip1 | 3.46 | 6.81E-257 | 1.42E-252 | 0.37 | 0.03 | 0.34 | 11.97 |
| endothelium | Man1c1 | 3.62 | 1.26E-256 | 2.63E-252 | 0.36 | 0.03 | 0.33 | 12.75 |
| endothelium | Notch4 | 3.3 | 3.08E-255 | 6.41E-251 | 0.47 | 0.05 | 0.42 | 9.2 |
| endothelium | Kcnk6 | 4.81 | 1.66E-254 | 3.45E-250 | 0.18 | 0.01 | 0.18 | 30.17 |
| endothelium | Stk10 | 3.8 | 5.89E-254 | 1.22E-249 | 0.33 | 0.02 | 0.3 | 13.54 |
| endothelium | Crmp1 | 2.99 | 2.98E-253 | 6.20E-249 | 0.67 | 0.11 | 0.56 | 6 |
| endothelium | Fam89a | 3.35 | 2.17E-250 | 4.52E-246 | 0.53 | 0.07 | 0.46 | 8 |
| endothelium | Pald1 | 3.26 | 4.07E-250 | 8.47E-246 | 0.56 | 0.08 | 0.49 | 7.47 |
| endothelium | Eogt | 3.42 | 3.43E-249 | 7.14E-245 | 0.4 | 0.04 | 0.36 | 10.73 |
| endothelium | Efna1 | 3.16 | 3.03E-247 | 6.31E-243 | 0.72 | 0.13 | 0.59 | 5.47 |
| endothelium | Rcsd1 | 3.18 | 4.32E-247 | 8.98E-243 | 0.59 | 0.09 | 0.51 | 6.96 |
| endothelium | Jcad | 3.62 | 5.50E-246 | 1.14E-241 | 0.46 | 0.05 | 0.41 | 9.12 |
| endothelium | Tnfaip8l1 | 3.48 | 9.71E-246 | 2.02E-241 | 0.53 | 0.07 | 0.46 | 7.61 |
| endothelium | Scn1b | 3.92 | 1.01E-244 | 2.10E-240 | 0.29 | 0.02 | 0.27 | 14.65 |
| endothelium | Epha2 | 3.91 | 3.71E-244 | 7.73E-240 | 0.34 | 0.03 | 0.32 | 12.29 |
| endothelium | BC049762 | 5.24 | 5.09E-244 | 1.06E-239 | 0.12 | 0 | 0.12 | 41 |
| endothelium | Arpc1b | 3.16 | 8.00E-244 | 1.66E-239 | 0.92 | 0.34 | 0.59 | 2.74 |
| endothelium | Plau | 5.01 | 2.09E-241 | 4.35E-237 | 0.18 | 0.01 | 0.17 | 25.86 |
| endothelium | Arhgef3 | 4.27 | 3.94E-240 | 8.20E-236 | 0.25 | 0.01 | 0.24 | 17.93 |
| endothelium | Rapgef5 | 3.19 | 3.50E-238 | 7.28E-234 | 0.65 | 0.11 | 0.54 | 5.92 |
| endothelium | Sh2d3c | 3.24 | 6.91E-238 | 1.44E-233 | 0.6 | 0.09 | 0.51 | 6.52 |
| endothelium | Hecw2 | 3.24 | 1.00E-237 | 2.09E-233 | 0.51 | 0.07 | 0.45 | 7.88 |
| endothelium | Dock6 | 3.13 | 2.41E-235 | 5.01E-231 | 0.61 | 0.1 | 0.52 | 6.26 |
| endothelium | Nfatc1 | 3.89 | 7.59E-234 | 1.58E-229 | 0.34 | 0.03 | 0.31 | 12 |
| endothelium | Mmp15 | 3.09 | 9.86E-234 | 2.05E-229 | 0.56 | 0.08 | 0.48 | 6.95 |
| endothelium | Bok | 3.35 | 2.43E-228 | 5.05E-224 | 0.78 | 0.19 | 0.59 | 4.1 |
| endothelium | Ostf1 | 3.03 | 1.40E-227 | 2.91E-223 | 0.86 | 0.27 | 0.6 | 3.25 |
| endothelium | Sema6b | 4.38 | 6.02E-225 | 1.25E-220 | 0.16 | 0.01 | 0.16 | 27.17 |
| endothelium | Zc3hav1 | 3.7 | 6.78E-225 | 1.41E-220 | 0.3 | 0.02 | 0.28 | 13.09 |
| endothelium | Ipo11 | 3.15 | 6.83E-225 | 1.42E-220 | 0.9 | 0.33 | 0.58 | 2.78 |
| endothelium | Tmsb4x | 2.78 | 8.21E-225 | 1.71E-220 | 1 | 0.99 | 0.01 | 1.01 |

**Supplementary Table 2. Marker genes for the major cell types of integrated clustering of E11.5 control and NotchGOF λ-junction cells.**

| | | | | | | | | |
|---|---|---|---|---|---|---|---|---|
| endothelium | Cgnl1 | 3.14 | 1.85E-224 | 3.85E-220 | 0.65 | 0.11 | 0.53 | 5.66 |
| endothelium | Lrrc8c | 2.94 | 3.03E-224 | 6.30E-220 | 0.66 | 0.12 | 0.54 | 5.63 |
| endothelium | Igfbp3 | 3.91 | 4.18E-222 | 8.70E-218 | 0.64 | 0.12 | 0.52 | 5.44 |
| endothelium | Cmtm8 | 2.88 | 9.73E-222 | 2.02E-217 | 0.76 | 0.17 | 0.59 | 4.47 |
| endothelium | Slc31a2 | 3.08 | 1.21E-221 | 2.51E-217 | 0.5 | 0.07 | 0.43 | 7.68 |
| endothelium | Cotl1 | 3.08 | 8.89E-221 | 1.85E-216 | 0.95 | 0.44 | 0.51 | 2.17 |
| endothelium | H2-D1 | 3.09 | 1.10E-220 | 2.28E-216 | 0.84 | 0.23 | 0.6 | 3.6 |
| endothelium | Dnah2os | 5.68 | 2.66E-218 | 5.54E-214 | 0.1 | 0 | 0.1 | 50.5 |
| endothelium | 430548M08R | 4.02 | 5.92E-218 | 1.23E-213 | 0.21 | 0.01 | 0.2 | 19.36 |
| endothelium | Lrrc32 | 5.4 | 7.71E-218 | 1.60E-213 | 0.14 | 0 | 0.13 | 34 |
| endothelium | Ramp3 | 3.62 | 1.04E-217 | 2.16E-213 | 0.32 | 0.03 | 0.29 | 12.12 |
| endothelium | Dock9 | 3.08 | 3.81E-217 | 7.92E-213 | 0.5 | 0.07 | 0.43 | 7.52 |
| endothelium | Acvrl1 | 2.85 | 1.40E-216 | 2.92E-212 | 0.59 | 0.09 | 0.5 | 6.33 |
| endothelium | Anxa6 | 3.2 | 6.83E-216 | 1.42E-211 | 0.73 | 0.16 | 0.57 | 4.47 |
| endothelium | Ulbp1 | 3.81 | 3.56E-215 | 7.41E-211 | 0.28 | 0.02 | 0.26 | 13.75 |
| endothelium | Rassf2 | 3.57 | 3.74E-214 | 7.77E-210 | 0.26 | 0.02 | 0.25 | 14.67 |
| endothelium | Klf7 | 2.83 | 2.44E-210 | 5.08E-206 | 0.81 | 0.22 | 0.59 | 3.68 |
| endothelium | Cyth4 | 2.83 | 3.27E-210 | 6.81E-206 | 0.18 | 0.01 | 0.18 | 20.44 |
| endothelium | Hspg2 | 3.35 | 6.34E-209 | 1.32E-204 | 0.77 | 0.22 | 0.55 | 3.54 |
| endothelium | F11r | 2.46 | 1.80E-206 | 3.75E-202 | 0.77 | 0.17 | 0.59 | 4.45 |
| endothelium | Ednrb | 2.99 | 5.66E-204 | 1.18E-199 | 0.35 | 0.03 | 0.32 | 10.26 |
| endothelium | Fkbp1a | 2.05 | 1.03E-203 | 2.15E-199 | 0.99 | 0.94 | 0.05 | 1.05 |
| endothelium | Orai1 | 2.77 | 1.98E-203 | 4.12E-199 | 0.73 | 0.16 | 0.57 | 4.56 |
| endothelium | Asb4 | 2.71 | 4.84E-203 | 1.01E-198 | 0.87 | 0.29 | 0.58 | 3.02 |
| endothelium | Il16 | 4.19 | 5.66E-203 | 1.18E-198 | 0.14 | 0.01 | 0.14 | 28.8 |
| endothelium | Syk | 3.68 | 6.04E-202 | 1.26E-197 | 0.29 | 0.02 | 0.27 | 12.13 |
| endothelium | Trpc3 | 3.28 | 1.52E-201 | 3.16E-197 | 0.32 | 0.03 | 0.29 | 11.03 |
| endothelium | Tlr3 | 4.41 | 3.77E-201 | 7.83E-197 | 0.15 | 0.01 | 0.14 | 29.4 |
| endothelium | Shroom2 | 2.93 | 4.01E-199 | 8.34E-195 | 0.5 | 0.07 | 0.43 | 6.99 |
| endothelium | Bvht | 3.38 | 3.12E-198 | 6.50E-194 | 0.42 | 0.05 | 0.37 | 8.25 |
| endothelium | Coro2b | 3.14 | 1.25E-196 | 2.60E-192 | 0.38 | 0.04 | 0.34 | 9.02 |
| endothelium | Fam124b | 4.6 | 6.05E-195 | 1.26E-190 | 0.1 | 0 | 0.1 | 50.5 |
| endothelium | Calm1 | 1.4 | 1.15E-193 | 2.40E-189 | 0.99 | 0.99 | 0 | 1 |
| endothelium | Slc25a45 | 4.21 | 7.66E-193 | 1.59E-188 | 0.18 | 0.01 | 0.17 | 19.56 |
| endothelium | Ptp4a3 | 3.29 | 1.00E-192 | 2.09E-188 | 0.8 | 0.24 | 0.56 | 3.29 |
| endothelium | Actb | 1.1 | 1.12E-192 | 2.33E-188 | 1 | 1 | 0 | 1 |
| endothelium | Tinagl1 | 3.41 | 2.66E-192 | 5.54E-188 | 0.42 | 0.05 | 0.36 | 7.85 |
| endothelium | Sh3bgrl3 | 1.85 | 1.66E-191 | 3.46E-187 | 0.98 | 0.91 | 0.07 | 1.08 |
| endothelium | Gm36028 | 4.83 | 1.12E-190 | 2.33E-186 | 0.1 | 0 | 0.1 | 52 |
| endothelium | Tbkbp1 | 2.97 | 7.88E-187 | 1.64E-182 | 0.44 | 0.06 | 0.38 | 7.63 |
| endothelium | Prcp | 2.83 | 3.39E-185 | 7.05E-181 | 0.8 | 0.25 | 0.55 | 3.23 |
| endothelium | Gnai2 | 1.8 | 8.18E-185 | 1.70E-180 | 0.98 | 0.97 | 0.01 | 1.01 |
| endothelium | Tspan4 | 2.66 | 1.72E-184 | 3.58E-180 | 0.62 | 0.12 | 0.5 | 5.2 |
| endothelium | Gm13431 | 4.25 | 1.80E-184 | 3.73E-180 | 0.16 | 0.01 | 0.15 | 22.86 |
| endothelium | Cdkl1 | 3.04 | 5.83E-184 | 1.21E-179 | 0.32 | 0.03 | 0.28 | 10.16 |
| endothelium | Fes | 3.25 | 6.30E-184 | 1.31E-179 | 0.35 | 0.04 | 0.31 | 9.38 |
| endothelium | Arpc3 | 1.84 | 1.95E-183 | 4.06E-179 | 0.98 | 0.9 | 0.08 | 1.09 |
| endothelium | Arhgap31 | 2.91 | 2.36E-183 | 4.91E-179 | 0.72 | 0.18 | 0.54 | 3.97 |
| endothelium | Akap12 | 2.92 | 7.80E-182 | 1.62E-177 | 0.73 | 0.19 | 0.54 | 3.91 |
| endothelium | Tead4 | 3.83 | 9.21E-181 | 1.92E-176 | 0.2 | 0.01 | 0.19 | 16.42 |
| endothelium | Pfn1 | 1.13 | 7.94E-180 | 1.65E-175 | 1 | 1 | 0 | 1 |
| endothelium | Jup | 2.45 | 1.10E-177 | 2.30E-173 | 0.75 | 0.19 | 0.56 | 3.96 |
| endothelium | Map3k11 | 2.65 | 9.68E-177 | 2.01E-172 | 0.58 | 0.11 | 0.47 | 5.36 |
| endothelium | Myl6 | 1.37 | 9.96E-177 | 2.07E-172 | 0.99 | 0.99 | 0 | 1 |
| endothelium | Clic1 | 1.81 | 1.81E-176 | 3.77E-172 | 0.96 | 0.93 | 0.03 | 1.03 |
| endothelium | Tal1 | 2.71 | 4.16E-176 | 8.66E-172 | 0.37 | 0.04 | 0.33 | 8.56 |

**Supplementary Table 2. Marker genes for the major cell types of integrated clustering of E11.5 control and NotchGOF λ-junction cells.**

| | | | | | | | | |
|---|---|---|---|---|---|---|---|---|
| endothelium | Cmbl | 3.76 | 1.20E-173 | 2.49E-169 | 0.21 | 0.01 | 0.19 | 14.64 |
| endothelium | Ets2 | 2.97 | 2.90E-173 | 6.03E-169 | 0.78 | 0.25 | 0.53 | 3.15 |
| endothelium | Slfn5 | 3.92 | 5.93E-173 | 1.23E-168 | 0.15 | 0.01 | 0.15 | 21.71 |
| endothelium | Ripor1 | 2.69 | 6.95E-173 | 1.44E-168 | 0.5 | 0.08 | 0.42 | 6.15 |
| endothelium | Pcdh17 | 2.81 | 1.14E-172 | 2.36E-168 | 0.73 | 0.2 | 0.53 | 3.66 |
| endothelium | Pea15a | 2.65 | 1.58E-172 | 3.29E-168 | 0.78 | 0.22 | 0.55 | 3.46 |
| endothelium | Ppp1r14a | 2.55 | 3.28E-172 | 6.81E-168 | 0.45 | 0.07 | 0.39 | 6.83 |
| endothelium | Rnf125 | 3.43 | 6.01E-172 | 1.25E-167 | 0.2 | 0.01 | 0.18 | 16.25 |
| endothelium | Skap2 | 3.17 | 1.36E-171 | 2.83E-167 | 0.39 | 0.05 | 0.34 | 7.74 |
| endothelium | Vim | 3.13 | 4.31E-170 | 8.96E-166 | 0.95 | 0.84 | 0.11 | 1.13 |
| endothelium | Tubb2a | 2.89 | 7.46E-169 | 1.55E-164 | 0.83 | 0.29 | 0.54 | 2.88 |
| endothelium | Tmem173 | 2.99 | 1.45E-167 | 3.01E-163 | 0.39 | 0.05 | 0.34 | 7.59 |
| endothelium | Ccdc85b | 2.82 | 1.63E-167 | 3.38E-163 | 0.92 | 0.58 | 0.34 | 1.58 |
| endothelium | Slc7a7 | 3.66 | 6.44E-166 | 1.34E-161 | 0.21 | 0.02 | 0.19 | 13.87 |
| endothelium | Slc9a3r2 | 3.11 | 8.34E-166 | 1.74E-161 | 0.78 | 0.26 | 0.52 | 3.03 |
| endothelium | Ralb | 2.65 | 5.04E-165 | 1.05E-160 | 0.9 | 0.43 | 0.47 | 2.11 |
| endothelium | Pfkp | 2.96 | 5.17E-165 | 1.07E-160 | 0.37 | 0.05 | 0.32 | 7.83 |
| endothelium | Msn | 3 | 6.02E-165 | 1.25E-160 | 0.89 | 0.51 | 0.38 | 1.74 |
| endothelium | Parvb | 2.94 | 4.66E-164 | 9.68E-160 | 0.6 | 0.12 | 0.47 | 4.92 |
| endothelium | Nrp1 | 2.21 | 1.83E-163 | 3.80E-159 | 0.75 | 0.2 | 0.55 | 3.76 |
| endothelium | Hapln3 | 3.65 | 8.69E-162 | 1.81E-157 | 0.2 | 0.01 | 0.18 | 14.07 |
| endothelium | Ica1 | 2.51 | 1.37E-160 | 2.85E-156 | 0.57 | 0.11 | 0.46 | 5.03 |
| endothelium | Slc43a2 | 2.73 | 6.07E-159 | 1.26E-154 | 0.32 | 0.04 | 0.28 | 8.89 |
| endothelium | Rras | 2.43 | 2.02E-156 | 4.20E-152 | 0.74 | 0.21 | 0.53 | 3.53 |
| endothelium | Plaur | 4.33 | 3.13E-156 | 6.51E-152 | 0.14 | 0.01 | 0.13 | 19.86 |
| endothelium | Piezo1 | 2.62 | 3.14E-156 | 6.52E-152 | 0.57 | 0.12 | 0.45 | 4.85 |
| endothelium | Sh2b3 | 2.52 | 4.88E-156 | 1.02E-151 | 0.44 | 0.07 | 0.37 | 6.42 |
| endothelium | Gimap9 | 2.5 | 1.49E-155 | 3.09E-151 | 0.54 | 0.1 | 0.44 | 5.31 |
| endothelium | Zfp69 | 3.27 | 3.13E-155 | 6.51E-151 | 0.31 | 0.04 | 0.27 | 8.83 |
| endothelium | Cmklr1 | 4.98 | 6.11E-155 | 1.27E-150 | 0.11 | 0 | 0.11 | 27.25 |
| endothelium | Hid1 | 3.64 | 3.24E-154 | 6.74E-150 | 0.17 | 0.01 | 0.16 | 16.8 |
| endothelium | Gm47664 | 3.61 | 1.01E-153 | 2.09E-149 | 0.17 | 0.01 | 0.16 | 16.8 |
| endothelium | Gnas | 1.12 | 1.03E-153 | 2.13E-149 | 1 | 1 | 0 | 1 |
| endothelium | Myo6 | 2.4 | 1.44E-153 | 3.00E-149 | 0.69 | 0.18 | 0.51 | 3.84 |
| endothelium | Gas6 | 2.88 | 2.04E-153 | 4.24E-149 | 0.25 | 0.02 | 0.23 | 10.54 |
| endothelium | Card6 | 3.42 | 2.65E-153 | 5.52E-149 | 0.18 | 0.01 | 0.17 | 14.92 |
| endothelium | Arhgap29 | 2.56 | 7.30E-153 | 1.52E-148 | 0.85 | 0.36 | 0.49 | 2.36 |
| endothelium | Itga6 | 2.13 | 1.00E-152 | 2.08E-148 | 0.85 | 0.3 | 0.55 | 2.81 |
| endothelium | Abcg2 | 2.59 | 1.08E-152 | 2.25E-148 | 0.83 | 0.34 | 0.49 | 2.44 |
| endothelium | Tpm4 | 1.32 | 5.40E-152 | 1.12E-147 | 0.99 | 0.93 | 0.05 | 1.06 |
| endothelium | Pkn3 | 3.11 | 9.46E-152 | 1.97E-147 | 0.34 | 0.04 | 0.3 | 7.93 |
| endothelium | Piezo2 | 2.2 | 1.58E-151 | 3.29E-147 | 0.76 | 0.22 | 0.54 | 3.46 |
| endothelium | Trim25 | 2.85 | 6.35E-151 | 1.32E-146 | 0.38 | 0.05 | 0.33 | 7.06 |
| endothelium | Hbegf | 2.84 | 3.09E-150 | 6.43E-146 | 0.45 | 0.08 | 0.38 | 5.88 |
| endothelium | Anxa2 | 2.25 | 7.87E-150 | 1.64E-145 | 0.88 | 0.41 | 0.48 | 2.16 |
| endothelium | Lxn | 2.73 | 1.14E-149 | 2.37E-145 | 0.84 | 0.36 | 0.48 | 2.36 |
| endothelium | Vwf | 6.23 | 2.05E-149 | 4.26E-145 | 0.13 | 0.01 | 0.13 | 22.17 |
| endothelium | Tpst2 | 2.53 | 1.90E-148 | 3.96E-144 | 0.75 | 0.23 | 0.52 | 3.28 |
| endothelium | Cd81 | 2.01 | 3.04E-148 | 6.33E-144 | 0.93 | 0.89 | 0.04 | 1.04 |
| endothelium | Gprc5a | 3.71 | 1.21E-147 | 2.52E-143 | 0.18 | 0.01 | 0.17 | 13.92 |
| endothelium | Rhob | 2.43 | 1.99E-146 | 4.14E-142 | 0.85 | 0.34 | 0.51 | 2.51 |
| endothelium | Col4a2 | 3.22 | 2.92E-146 | 6.08E-142 | 0.81 | 0.42 | 0.39 | 1.92 |
| endothelium | Map4k2 | 2.64 | 1.17E-145 | 2.43E-141 | 0.41 | 0.06 | 0.35 | 6.42 |
| endothelium | Arpc5 | 1.78 | 2.58E-144 | 5.37E-140 | 0.94 | 0.76 | 0.18 | 1.24 |
| endothelium | Rftn1 | 2.69 | 2.63E-144 | 5.47E-140 | 0.3 | 0.04 | 0.26 | 8.54 |
| endothelium | 110032F04Ri | 3.89 | 3.12E-144 | 6.49E-140 | 0.22 | 0.02 | 0.2 | 11.53 |

**Supplementary Table 2. Marker genes for the major cell types of integrated clustering of E11.5 control and NotchGOF λ-junction cells.**

| | | | | | | | | |
|---|---|---|---|---|---|---|---|---|
| endothelium | Il2rg | 4.38 | 6.38E-144 | 1.33E-139 | 0.11 | 0 | 0.11 | 27.25 |
| endothelium | Col18a1 | 2.43 | 6.10E-143 | 1.27E-138 | 0.84 | 0.36 | 0.48 | 2.35 |
| endothelium | Tgfb1i1 | 2.6 | 2.38E-142 | 4.96E-138 | 0.58 | 0.13 | 0.45 | 4.54 |
| endothelium | Sacs | 2.5 | 5.39E-140 | 1.12E-135 | 0.65 | 0.17 | 0.48 | 3.86 |
| endothelium | Rab11a | 2.12 | 1.07E-139 | 2.22E-135 | 0.92 | 0.64 | 0.28 | 1.44 |
| endothelium | Myo18a | 2.4 | 1.72E-139 | 3.58E-135 | 0.73 | 0.23 | 0.5 | 3.22 |
| endothelium | Elk3 | 2.71 | 2.02E-139 | 4.21E-135 | 0.81 | 0.41 | 0.4 | 1.96 |
| endothelium | Cfap45 | 4.03 | 2.93E-139 | 6.10E-135 | 0.13 | 0.01 | 0.13 | 19 |
| endothelium | Vasp | 2.26 | 3.76E-139 | 7.82E-135 | 0.89 | 0.48 | 0.42 | 1.87 |
| endothelium | Csrp1 | 2.21 | 9.70E-139 | 2.02E-134 | 0.84 | 0.33 | 0.5 | 2.51 |
| endothelium | Klk8 | 2.16 | 7.92E-138 | 1.65E-133 | 0.73 | 0.21 | 0.52 | 3.5 |
| endothelium | Zbtb46 | 2.92 | 3.64E-137 | 7.58E-133 | 0.3 | 0.04 | 0.26 | 8.31 |
| endothelium | Ppp1r2 | 1.85 | 3.83E-136 | 7.98E-132 | 0.93 | 0.75 | 0.18 | 1.25 |
| endothelium | Cfl1 | 0.75 | 2.00E-135 | 4.16E-131 | 1 | 1 | 0 | 1 |
| endothelium | Ehd4 | 2.23 | 3.79E-135 | 7.89E-131 | 0.65 | 0.17 | 0.48 | 3.86 |
| endothelium | Tgm2 | 4.3 | 3.85E-135 | 8.00E-131 | 0.17 | 0.01 | 0.16 | 14.25 |
| endothelium | Hsd3b7 | 2.88 | 1.81E-134 | 3.77E-130 | 0.27 | 0.03 | 0.24 | 8.68 |
| endothelium | St8sia1 | 2.97 | 2.09E-134 | 4.35E-130 | 0.25 | 0.03 | 0.23 | 9.65 |
| endothelium | Hebp1 | 1.87 | 5.53E-134 | 1.15E-129 | 0.79 | 0.26 | 0.53 | 3.04 |
| endothelium | Slc25a5 | 1.32 | 6.04E-134 | 1.26E-129 | 0.96 | 0.93 | 0.03 | 1.03 |
| endothelium | Ttc9 | 2.78 | 8.00E-134 | 1.66E-129 | 0.29 | 0.04 | 0.26 | 8.14 |
| endothelium | Ppic | 1.99 | 6.20E-132 | 1.29E-127 | 0.9 | 0.71 | 0.2 | 1.28 |
| endothelium | Npdc1 | 2.16 | 2.47E-131 | 5.13E-127 | 0.5 | 0.1 | 0.4 | 4.99 |
| endothelium | Serf2 | 0.7 | 2.65E-131 | 5.51E-127 | 0.99 | 1 | -0.01 | 0.99 |
| endothelium | Itsn2 | 2.3 | 5.25E-131 | 1.09E-126 | 0.67 | 0.18 | 0.49 | 3.66 |
| endothelium | Peg13 | 2.69 | 1.18E-130 | 2.46E-126 | 0.33 | 0.05 | 0.28 | 7.13 |
| endothelium | Ndrg1 | 3.13 | 1.00E-129 | 2.09E-125 | 0.2 | 0.02 | 0.18 | 11.11 |
| endothelium | Actg1 | 0.79 | 2.04E-128 | 4.25E-124 | 1 | 1 | 0 | 1 |
| endothelium | Cebpd | 2.57 | 3.19E-128 | 6.64E-124 | 0.47 | 0.09 | 0.38 | 5.08 |
| endothelium | Dgkh | 2.52 | 4.79E-128 | 9.97E-124 | 0.45 | 0.08 | 0.37 | 5.4 |
| endothelium | Sigirr | 2.92 | 7.67E-128 | 1.60E-123 | 0.23 | 0.02 | 0.21 | 9.96 |
| endothelium | St8sia4 | 2.49 | 1.87E-127 | 3.88E-123 | 0.45 | 0.09 | 0.37 | 5.21 |
| endothelium | Fyn | 2.13 | 5.95E-127 | 1.24E-122 | 0.85 | 0.44 | 0.41 | 1.95 |
| endothelium | Myo7a | 3.26 | 6.81E-127 | 1.42E-122 | 0.16 | 0.01 | 0.15 | 14.27 |
| endothelium | Mef2a | 2.23 | 8.57E-127 | 1.78E-122 | 0.83 | 0.38 | 0.45 | 2.16 |
| endothelium | Cdc42 | 1 | 2.04E-126 | 4.23E-122 | 0.98 | 0.98 | -0.01 | 0.99 |
| endothelium | Plpp1 | 2.43 | 4.92E-126 | 1.02E-121 | 0.57 | 0.14 | 0.43 | 4.09 |
| endothelium | Snx3 | 1.35 | 5.94E-126 | 1.24E-121 | 0.96 | 0.89 | 0.06 | 1.07 |
| endothelium | Cmtm3 | 2.25 | 2.17E-125 | 4.51E-121 | 0.84 | 0.41 | 0.43 | 2.03 |
| endothelium | Mest | 1.9 | 2.40E-125 | 4.99E-121 | 0.92 | 0.91 | 0.01 | 1.01 |
| endothelium | Pde4b | 3.09 | 4.72E-125 | 9.82E-121 | 0.29 | 0.04 | 0.25 | 7.51 |
| endothelium | Smtn | 2.4 | 1.69E-124 | 3.52E-120 | 0.57 | 0.14 | 0.43 | 4.17 |
| endothelium | Cnn2 | 1.78 | 7.38E-123 | 1.53E-118 | 0.93 | 0.63 | 0.3 | 1.48 |
| endothelium | Col4a1 | 3.29 | 1.77E-122 | 3.68E-118 | 0.83 | 0.62 | 0.22 | 1.35 |
| endothelium | Rgl1 | 2.45 | 9.30E-122 | 1.94E-117 | 0.58 | 0.15 | 0.43 | 3.97 |
| endothelium | Mtss1 | 2.09 | 2.19E-120 | 4.55E-116 | 0.77 | 0.29 | 0.48 | 2.68 |
| endothelium | Notch1 | 1.68 | 7.42E-120 | 1.54E-115 | 0.75 | 0.24 | 0.51 | 3.11 |
| endothelium | Adcy4 | 2.47 | 8.92E-120 | 1.86E-115 | 0.36 | 0.06 | 0.3 | 6.15 |
| endothelium | Zfp979 | 2.76 | 9.20E-120 | 1.91E-115 | 0.26 | 0.03 | 0.23 | 8.26 |
| endothelium | Disc1 | 2.9 | 4.47E-119 | 9.29E-115 | 0.18 | 0.02 | 0.16 | 11.93 |
| endothelium | Cd9 | 2.84 | 1.21E-118 | 2.52E-114 | 0.79 | 0.29 | 0.5 | 2.71 |
| endothelium | Tspan17 | 2.93 | 2.35E-118 | 4.88E-114 | 0.25 | 0.03 | 0.22 | 8.37 |
| endothelium | Rcan3 | 2.53 | 5.33E-118 | 1.11E-113 | 0.4 | 0.07 | 0.33 | 5.6 |
| endothelium | Plk2 | 2.33 | 6.36E-118 | 1.32E-113 | 0.79 | 0.37 | 0.43 | 2.16 |
| endothelium | Tns2 | 2.53 | 2.36E-117 | 4.91E-113 | 0.33 | 0.05 | 0.28 | 6.56 |
| endothelium | Cx3cl1 | 3.44 | 2.44E-116 | 5.07E-112 | 0.13 | 0.01 | 0.12 | 14.78 |

**Supplementary Table 2. Marker genes for the major cell types of integrated clustering of E11.5 control and NotchGOF λ-junction cells.**

| | | | | | | | | |
|---|---|---|---|---|---|---|---|---|
| endothelium | Nes | 2.35 | 1.20E-115 | 2.50E-111 | 0.81 | 0.42 | 0.39 | 1.92 |
| endothelium | Nlrc3 | 3.22 | 1.65E-115 | 3.44E-111 | 0.18 | 0.02 | 0.17 | 10.82 |
| endothelium | Myo1c | 2.36 | 3.37E-115 | 7.02E-111 | 0.45 | 0.09 | 0.36 | 4.87 |
| endothelium | Tspan9 | 2.17 | 7.86E-115 | 1.63E-110 | 0.65 | 0.19 | 0.46 | 3.43 |
| endothelium | Sgk1 | 2.35 | 1.28E-114 | 2.66E-110 | 0.66 | 0.2 | 0.46 | 3.26 |
| endothelium | Arpc4 | 1.37 | 2.80E-114 | 5.82E-110 | 0.94 | 0.82 | 0.12 | 1.14 |
| endothelium | Vsir | 3.59 | 1.05E-112 | 2.18E-108 | 0.16 | 0.01 | 0.15 | 12.54 |
| endothelium | Pik3cg | 4.55 | 2.54E-112 | 5.29E-108 | 0.1 | 0.01 | 0.1 | 20.8 |
| endothelium | Tmsb10 | 0.8 | 2.86E-112 | 5.94E-108 | 1 | 1 | 0 | 1 |
| endothelium | Abcg1 | 2.02 | 6.46E-112 | 1.34E-107 | 0.41 | 0.08 | 0.34 | 5.29 |
| endothelium | Nab1 | 2.19 | 6.68E-112 | 1.39E-107 | 0.53 | 0.12 | 0.41 | 4.29 |
| endothelium | Anxa5 | 1.81 | 1.74E-111 | 3.63E-107 | 0.9 | 0.6 | 0.3 | 1.49 |
| endothelium | Grasp | 1.96 | 2.81E-111 | 5.84E-107 | 0.65 | 0.19 | 0.46 | 3.47 |
| endothelium | Sh3glb1 | 1.35 | 3.67E-111 | 7.63E-107 | 0.94 | 0.84 | 0.11 | 1.13 |
| endothelium | Bmp2k | 2.57 | 4.66E-111 | 9.69E-107 | 0.56 | 0.15 | 0.42 | 3.88 |
| endothelium | Plagl1 | 1.87 | 4.90E-111 | 1.02E-106 | 0.84 | 0.43 | 0.41 | 1.96 |
| endothelium | Gmpr | 2.17 | 5.92E-111 | 1.23E-106 | 0.62 | 0.17 | 0.45 | 3.66 |
| endothelium | Pnp | 1.68 | 1.19E-110 | 2.48E-106 | 0.91 | 0.66 | 0.25 | 1.38 |
| endothelium | Ehbp1l1 | 2.6 | 1.37E-110 | 2.85E-106 | 0.35 | 0.06 | 0.29 | 5.82 |
| endothelium | Sptbn1 | 1.9 | 1.48E-110 | 3.08E-106 | 0.88 | 0.76 | 0.12 | 1.16 |
| endothelium | Ripk1 | 2.4 | 4.65E-110 | 9.66E-106 | 0.51 | 0.12 | 0.39 | 4.28 |
| endothelium | Slco3a1 | 2.46 | 2.52E-109 | 5.24E-105 | 0.51 | 0.12 | 0.38 | 4.12 |
| endothelium | Prex1 | 2.29 | 6.26E-109 | 1.30E-104 | 0.63 | 0.19 | 0.44 | 3.36 |
| endothelium | Actr3 | 1.38 | 1.25E-108 | 2.59E-104 | 0.94 | 0.82 | 0.13 | 1.16 |
| endothelium | Sparc | 2.28 | 2.06E-108 | 4.29E-104 | 0.86 | 0.74 | 0.12 | 1.16 |
| endothelium | Sp140 | 3.38 | 2.16E-108 | 4.48E-104 | 0.13 | 0.01 | 0.12 | 14.78 |
| endothelium | Acot9 | 2.12 | 3.22E-108 | 6.71E-104 | 0.58 | 0.15 | 0.43 | 3.81 |
| endothelium | Ndufa8 | 1.23 | 1.05E-107 | 2.19E-103 | 0.94 | 0.9 | 0.05 | 1.05 |
| endothelium | Rhoa | 0.94 | 1.87E-107 | 3.88E-103 | 0.97 | 0.97 | 0.01 | 1.01 |
| endothelium | Pitpna | 1.53 | 3.71E-107 | 7.72E-103 | 0.92 | 0.7 | 0.22 | 1.31 |
| endothelium | Rap1a | 1.66 | 4.05E-107 | 8.42E-103 | 0.92 | 0.71 | 0.2 | 1.28 |
| endothelium | Fam13c | 2.58 | 5.25E-107 | 1.09E-102 | 0.36 | 0.07 | 0.3 | 5.54 |
| endothelium | Snx9 | 1.91 | 1.45E-106 | 3.01E-102 | 0.77 | 0.31 | 0.47 | 2.52 |
| endothelium | Mgat4b | 2.06 | 2.56E-106 | 5.32E-102 | 0.81 | 0.39 | 0.42 | 2.07 |
| endothelium | Itpkb | 2.06 | 3.55E-106 | 7.37E-102 | 0.63 | 0.19 | 0.44 | 3.38 |
| endothelium | Tjp1 | 2.11 | 4.34E-106 | 9.03E-102 | 0.82 | 0.49 | 0.33 | 1.68 |
| endothelium | Trf | 2.11 | 9.39E-105 | 1.95E-100 | 0.28 | 0.04 | 0.24 | 7.05 |
| endothelium | Abhd17a | 1.8 | 1.00E-104 | 2.08E-100 | 0.89 | 0.7 | 0.18 | 1.26 |
| endothelium | Arpc2 | 1.05 | 1.93E-104 | 4.01E-100 | 0.96 | 0.94 | 0.02 | 1.02 |
| endothelium | Itm2b | 1.31 | 3.52E-104 | 7.31E-100 | 0.91 | 0.84 | 0.06 | 1.08 |
| endothelium | Lfng | 4.05 | 7.70E-104 | 1.60E-99 | 0.17 | 0.02 | 0.16 | 10.69 |
| endothelium | Tacc1 | 2.09 | 1.78E-103 | 3.70E-99 | 0.77 | 0.31 | 0.46 | 2.49 |
| endothelium | Arl4a | 1.99 | 2.17E-103 | 4.51E-99 | 0.65 | 0.2 | 0.45 | 3.3 |
| endothelium | Rbpms | 1.89 | 3.62E-103 | 7.53E-99 | 0.85 | 0.54 | 0.31 | 1.57 |
| endothelium | Ppp1r18 | 1.92 | 4.21E-103 | 8.76E-99 | 0.71 | 0.24 | 0.47 | 2.92 |
| endothelium | Trim16 | 2.9 | 5.40E-103 | 1.12E-98 | 0.19 | 0.02 | 0.17 | 9.6 |
| endothelium | Nid1 | 2.05 | 2.08E-102 | 4.32E-98 | 0.77 | 0.34 | 0.43 | 2.27 |
| endothelium | Il4ra | 2.78 | 2.94E-102 | 6.12E-98 | 0.17 | 0.02 | 0.15 | 11 |
| endothelium | Ece1 | 2.04 | 3.81E-102 | 7.93E-98 | 0.66 | 0.22 | 0.44 | 2.95 |
| endothelium | Selenop | 1.71 | 1.41E-101 | 2.92E-97 | 0.61 | 0.18 | 0.43 | 3.37 |
| endothelium | Fhod1 | 2.41 | 1.42E-101 | 2.95E-97 | 0.41 | 0.08 | 0.32 | 4.88 |
| endothelium | Tpm3 | 0.97 | 1.69E-101 | 3.52E-97 | 0.96 | 0.96 | 0 | 1 |
| endothelium | Lamc1 | 1.99 | 2.27E-100 | 4.71E-96 | 0.71 | 0.25 | 0.45 | 2.79 |
| endothelium | Svil | 1.89 | 3.52E-100 | 7.32E-96 | 0.8 | 0.38 | 0.42 | 2.12 |
| endothelium | Gng12 | 1.94 | 3.61E-100 | 7.51E-96 | 0.77 | 0.36 | 0.42 | 2.18 |
| endothelium | S100a10 | 1.64 | 3.87E-99 | 8.06E-95 | 0.89 | 0.55 | 0.34 | 1.62 |

**Supplementary Table 2. Marker genes for the major cell types of integrated clustering of E11.5 control and NotchGOF λ-junction cells.**

| | | | | | | | | |
|---|---|---|---|---|---|---|---|---|
| endothelium | Rps2 | 0.49 | 7.97E-99 | 1.66E-94 | 1 | 1 | 0 | 1 |
| endothelium | Eif1 | 0.5 | 2.07E-98 | 4.31E-94 | 0.99 | 1 | -0.01 | 0.99 |
| endothelium | Tmem37 | 2.24 | 2.72E-98 | 5.65E-94 | 0.39 | 0.08 | 0.31 | 4.9 |
| endothelium | Ddah1 | 2.18 | 1.35E-97 | 2.80E-93 | 0.52 | 0.13 | 0.38 | 3.84 |
| endothelium | Gnb2 | 0.88 | 1.70E-97 | 3.54E-93 | 0.96 | 0.96 | -0.01 | 0.99 |
| endothelium | Dgkz | 1.85 | 6.05E-97 | 1.26E-92 | 0.74 | 0.28 | 0.46 | 2.65 |
| endothelium | Tln1 | 1.67 | 7.09E-97 | 1.47E-92 | 0.93 | 0.74 | 0.19 | 1.26 |
| endothelium | Pgm2 | 1.99 | 1.06E-96 | 2.21E-92 | 0.73 | 0.27 | 0.46 | 2.73 |
| endothelium | 900026A02Ri | 1.86 | 1.10E-96 | 2.29E-92 | 0.45 | 0.1 | 0.35 | 4.49 |
| endothelium | Inf2 | 2.48 | 1.49E-96 | 3.11E-92 | 0.29 | 0.05 | 0.24 | 6.33 |
| endothelium | Slc2a1 | 2.06 | 1.92E-95 | 4.00E-91 | 0.81 | 0.42 | 0.39 | 1.93 |
| endothelium | Mid2 | 2.99 | 5.22E-95 | 1.09E-90 | 0.21 | 0.03 | 0.18 | 8.2 |
| endothelium | Galnt12 | 2.88 | 1.30E-94 | 2.71E-90 | 0.19 | 0.02 | 0.17 | 9 |
| endothelium | Rsu1 | 2.4 | 1.37E-94 | 2.84E-90 | 0.86 | 0.48 | 0.38 | 1.79 |
| endothelium | Elovl1 | 1.95 | 2.76E-94 | 5.75E-90 | 0.69 | 0.25 | 0.44 | 2.73 |
| endothelium | Kctd12b | 2.13 | 1.08E-93 | 2.25E-89 | 0.57 | 0.17 | 0.4 | 3.29 |
| endothelium | Degs1 | 1.96 | 2.64E-93 | 5.50E-89 | 0.79 | 0.39 | 0.4 | 2.01 |
| endothelium | Parp4 | 3.19 | 5.43E-93 | 1.13E-88 | 0.17 | 0.02 | 0.15 | 9.71 |
| endothelium | Schip1 | 2.01 | 2.45E-92 | 5.09E-88 | 0.56 | 0.16 | 0.41 | 3.61 |
| endothelium | Frmd4b | 1.56 | 6.01E-92 | 1.25E-87 | 0.57 | 0.16 | 0.41 | 3.52 |
| endothelium | Aprt | 1.12 | 2.04E-91 | 4.24E-87 | 0.96 | 0.92 | 0.03 | 1.04 |
| endothelium | Rap1b | 2.33 | 2.17E-91 | 4.51E-87 | 0.92 | 0.76 | 0.16 | 1.22 |
| endothelium | Tcf15 | 1.85 | 1.18E-90 | 2.45E-86 | 0.72 | 0.28 | 0.44 | 2.61 |
| endothelium | Rgs3 | 2.14 | 1.18E-90 | 2.46E-86 | 0.58 | 0.18 | 0.4 | 3.27 |
| endothelium | Myh9 | 1.87 | 4.19E-90 | 8.72E-86 | 0.84 | 0.5 | 0.34 | 1.68 |
| endothelium | Lrp10 | 1.97 | 9.11E-90 | 1.90E-85 | 0.56 | 0.17 | 0.4 | 3.39 |
| endothelium | D5Ertd605e | 3.65 | 1.89E-89 | 3.92E-85 | 0.1 | 0.01 | 0.1 | 14.86 |
| endothelium | Chst1 | 2.91 | 4.02E-89 | 8.36E-85 | 0.27 | 0.04 | 0.22 | 6.21 |
| endothelium | Abca1 | 1.73 | 1.57E-88 | 3.28E-84 | 0.34 | 0.07 | 0.27 | 5.09 |
| endothelium | Maged2 | 1.54 | 4.46E-88 | 9.29E-84 | 0.94 | 0.82 | 0.12 | 1.14 |
| endothelium | Ywhah | 1.04 | 1.06E-87 | 2.20E-83 | 0.96 | 0.91 | 0.05 | 1.05 |
| endothelium | B2m | 1.51 | 1.12E-87 | 2.33E-83 | 0.86 | 0.54 | 0.33 | 1.61 |
| endothelium | Maff | 2.42 | 3.85E-87 | 8.01E-83 | 0.26 | 0.04 | 0.22 | 6.21 |
| endothelium | Myl12a | 0.92 | 8.63E-87 | 1.79E-82 | 0.96 | 0.95 | 0.01 | 1.01 |
| endothelium | Dapk2 | 2.83 | 8.77E-87 | 1.82E-82 | 0.27 | 0.05 | 0.22 | 5.93 |
| endothelium | Galnt18 | 2.68 | 1.73E-86 | 3.59E-82 | 0.22 | 0.03 | 0.19 | 7.06 |
| endothelium | Adam19 | 1.87 | 1.26E-85 | 2.62E-81 | 0.53 | 0.15 | 0.38 | 3.52 |
| endothelium | Ldb2 | 1.88 | 1.45E-85 | 3.01E-81 | 0.68 | 0.27 | 0.42 | 2.56 |
| endothelium | Mpp1 | 1.86 | 1.61E-85 | 3.34E-81 | 0.65 | 0.23 | 0.42 | 2.8 |
| endothelium | Pdlim5 | 1.78 | 3.13E-85 | 6.52E-81 | 0.64 | 0.22 | 0.42 | 2.94 |
| endothelium | Gng5 | 0.56 | 3.26E-85 | 6.78E-81 | 0.98 | 1 | -0.01 | 0.99 |
| endothelium | Stard8 | 1.87 | 5.47E-85 | 1.14E-80 | 0.38 | 0.08 | 0.3 | 4.54 |
| endothelium | Gm26862 | 3.01 | 9.78E-85 | 2.03E-80 | 0.12 | 0.01 | 0.11 | 12.78 |
| endothelium | Coro1c | 1.83 | 1.66E-84 | 3.46E-80 | 0.75 | 0.34 | 0.41 | 2.18 |
| endothelium | Ubtd1 | 1.91 | 2.31E-84 | 4.80E-80 | 0.52 | 0.14 | 0.37 | 3.59 |
| endothelium | Ppm1f | 2.04 | 2.69E-84 | 5.60E-80 | 0.42 | 0.1 | 0.32 | 4.16 |
| endothelium | Zeb1 | 1.85 | 2.21E-83 | 4.59E-79 | 0.74 | 0.34 | 0.4 | 2.16 |
| endothelium | Tdrd7 | 3.02 | 2.95E-83 | 6.14E-79 | 0.2 | 0.03 | 0.17 | 7.8 |
| endothelium | Sh3bp5 | 1.95 | 5.31E-83 | 1.10E-78 | 0.8 | 0.5 | 0.3 | 1.6 |
| endothelium | Paqr4 | 2.44 | 1.11E-82 | 2.31E-78 | 0.22 | 0.03 | 0.19 | 6.91 |
| endothelium | Dram2 | 1.72 | 1.29E-82 | 2.69E-78 | 0.61 | 0.2 | 0.41 | 3.08 |
| endothelium | Vat1 | 1.98 | 1.31E-82 | 2.73E-78 | 0.48 | 0.13 | 0.35 | 3.66 |
| endothelium | Pold4 | 1.84 | 1.49E-82 | 3.11E-78 | 0.58 | 0.18 | 0.4 | 3.23 |
| endothelium | Acsl5 | 1.9 | 5.02E-82 | 1.04E-77 | 0.52 | 0.15 | 0.37 | 3.42 |
| endothelium | Kif26a | 1.9 | 1.00E-81 | 2.08E-77 | 0.44 | 0.11 | 0.33 | 3.99 |
| endothelium | Wdr1 | 1.54 | 1.57E-81 | 3.26E-77 | 0.86 | 0.54 | 0.32 | 1.59 |

**Supplementary Table 2. Marker genes for the major cell types of integrated clustering of E11.5 control and NotchGOF λ-junction cells.**

| | | | | | | | | |
|---|---|---|---|---|---|---|---|---|
| endothelium | Fli1 | 1.62 | 2.22E-81 | 4.61E-77 | 0.78 | 0.37 | 0.42 | 2.15 |
| endothelium | Tcim | 2.17 | 2.38E-81 | 4.96E-77 | 0.74 | 0.36 | 0.38 | 2.05 |
| endothelium | Fzd4 | 1.6 | 2.90E-81 | 6.04E-77 | 0.68 | 0.25 | 0.43 | 2.75 |
| endothelium | Hprt | 1.73 | 3.23E-81 | 6.72E-77 | 0.81 | 0.43 | 0.38 | 1.9 |
| endothelium | H19 | 1.21 | 4.77E-81 | 9.92E-77 | 0.93 | 0.84 | 0.09 | 1.1 |
| endothelium | Als2cl | 3.18 | 5.58E-81 | 1.16E-76 | 0.11 | 0.01 | 0.1 | 12.44 |
| endothelium | Ppp2r5a | 1.91 | 1.71E-80 | 3.55E-76 | 0.72 | 0.31 | 0.4 | 2.28 |
| endothelium | Slc7a1 | 1.81 | 3.44E-80 | 7.15E-76 | 0.35 | 0.07 | 0.28 | 4.72 |
| endothelium | H2-K1 | 1.89 | 3.59E-80 | 7.47E-76 | 0.47 | 0.13 | 0.34 | 3.66 |
| endothelium | Kcna5 | 3.15 | 5.20E-80 | 1.08E-75 | 0.16 | 0.02 | 0.14 | 9.12 |
| endothelium | Frmd8 | 1.88 | 9.63E-80 | 2.00E-75 | 0.6 | 0.2 | 0.4 | 3.02 |
| endothelium | Abcc4 | 2.87 | 1.09E-79 | 2.26E-75 | 0.16 | 0.02 | 0.14 | 8.58 |
| endothelium | Ly6h | 1.43 | 2.73E-79 | 5.67E-75 | 0.51 | 0.15 | 0.37 | 3.51 |
| endothelium | Crip1 | 1.83 | 4.06E-79 | 8.44E-75 | 0.77 | 0.4 | 0.38 | 1.94 |
| endothelium | Sntb1 | 2.58 | 6.04E-79 | 1.26E-74 | 0.19 | 0.03 | 0.17 | 7.38 |
| endothelium | Rinl | 2.94 | 7.49E-79 | 1.56E-74 | 0.15 | 0.02 | 0.13 | 9.8 |
| endothelium | Lifr | 1.98 | 1.25E-78 | 2.60E-74 | 0.42 | 0.1 | 0.31 | 4 |
| endothelium | Caskin2 | 2.12 | 1.87E-78 | 3.89E-74 | 0.42 | 0.11 | 0.32 | 3.93 |
| endothelium | Fmnl3 | 1.88 | 2.05E-78 | 4.26E-74 | 0.59 | 0.2 | 0.39 | 2.93 |
| endothelium | Itga5 | 1.74 | 4.56E-78 | 9.49E-74 | 0.62 | 0.21 | 0.41 | 2.93 |
| endothelium | Rasd1 | 2.38 | 4.69E-78 | 9.76E-74 | 0.24 | 0.04 | 0.2 | 6.15 |
| endothelium | Itga2b | 6.37 | 8.36E-78 | 1.74E-73 | 0.15 | 0.02 | 0.13 | 9.19 |
| endothelium | Ccdc82 | 1.93 | 1.47E-77 | 3.06E-73 | 0.44 | 0.11 | 0.32 | 3.85 |
| endothelium | Pam | 1.62 | 2.79E-77 | 5.79E-73 | 0.51 | 0.15 | 0.36 | 3.47 |
| endothelium | Fam129b | 1.86 | 2.96E-77 | 6.16E-73 | 0.49 | 0.14 | 0.35 | 3.56 |
| endothelium | Ywhab | 0.74 | 5.13E-77 | 1.07E-72 | 0.96 | 0.97 | -0.01 | 0.99 |
| endothelium | Pros1 | 2.19 | 6.21E-77 | 1.29E-72 | 0.48 | 0.14 | 0.34 | 3.47 |
| endothelium | Nhp2 | 1.05 | 6.71E-77 | 1.40E-72 | 0.93 | 0.88 | 0.05 | 1.05 |
| endothelium | Gpx3 | 1.69 | 8.69E-77 | 1.81E-72 | 0.62 | 0.21 | 0.4 | 2.91 |
| endothelium | Uaca | 1.71 | 1.14E-76 | 2.37E-72 | 0.72 | 0.33 | 0.39 | 2.18 |
| endothelium | Cabp1 | 3.12 | 1.83E-76 | 3.80E-72 | 0.13 | 0.01 | 0.11 | 10.42 |
| endothelium | Fam102b | 1.84 | 2.73E-76 | 5.69E-72 | 0.58 | 0.19 | 0.39 | 3.01 |
| endothelium | Matrchf2 | 1.63 | 6.15E-76 | 1.28E-71 | 0.84 | 0.51 | 0.33 | 1.64 |
| endothelium | 930012K11Ri | 2.12 | 9.70E-76 | 2.02E-71 | 0.3 | 0.06 | 0.24 | 5.1 |
| endothelium | Pitpnm2 | 1.81 | 1.43E-75 | 2.98E-71 | 0.36 | 0.08 | 0.28 | 4.44 |
| endothelium | Itgb1 | 1.17 | 1.53E-75 | 3.18E-71 | 0.89 | 0.85 | 0.03 | 1.04 |
| endothelium | 931406P16Ri | 1.7 | 3.01E-75 | 6.26E-71 | 0.74 | 0.35 | 0.39 | 2.12 |
| endothelium | Scarb1 | 1.8 | 3.28E-75 | 6.83E-71 | 0.5 | 0.15 | 0.35 | 3.29 |
| endothelium | St6galnac4 | 1.7 | 5.81E-75 | 1.21E-70 | 0.58 | 0.19 | 0.39 | 3.04 |
| endothelium | Igf2 | 1.72 | 6.01E-75 | 1.25E-70 | 0.84 | 0.58 | 0.26 | 1.44 |
| endothelium | Klhl36 | 1.97 | 6.72E-75 | 1.40E-70 | 0.31 | 0.06 | 0.25 | 4.9 |
| endothelium | Dlc1 | 1.61 | 1.45E-74 | 3.03E-70 | 0.74 | 0.36 | 0.38 | 2.07 |
| endothelium | Prr5l | 1.72 | 1.65E-74 | 3.44E-70 | 0.52 | 0.16 | 0.36 | 3.27 |
| endothelium | Cd79b | 2.53 | 2.25E-74 | 4.69E-70 | 0.12 | 0.01 | 0.11 | 10.25 |
| endothelium | Pomp | 0.78 | 2.51E-74 | 5.21E-70 | 0.97 | 0.98 | -0.01 | 0.99 |
| endothelium | Klf10 | 1.87 | 4.18E-74 | 8.70E-70 | 0.41 | 0.1 | 0.3 | 3.92 |
| endothelium | Heg1 | 1.95 | 5.09E-74 | 1.06E-69 | 0.49 | 0.15 | 0.35 | 3.36 |
| endothelium | Spata13 | 2.51 | 6.73E-74 | 1.40E-69 | 0.33 | 0.07 | 0.26 | 4.6 |
| endothelium | Magi3 | 1.75 | 8.49E-74 | 1.77E-69 | 0.56 | 0.19 | 0.37 | 3 |
| endothelium | Bgn | 1.6 | 1.18E-73 | 2.45E-69 | 0.65 | 0.26 | 0.39 | 2.53 |
| endothelium | Slc43a3 | 1.71 | 1.66E-73 | 3.45E-69 | 0.68 | 0.28 | 0.4 | 2.42 |
| endothelium | Rhoc | 1.29 | 2.15E-73 | 4.48E-69 | 0.93 | 0.79 | 0.14 | 1.18 |
| endothelium | Klf6 | 1.91 | 4.83E-73 | 1.00E-68 | 0.61 | 0.23 | 0.38 | 2.63 |
| endothelium | Cd151 | 1.42 | 7.62E-73 | 1.58E-68 | 0.76 | 0.32 | 0.43 | 2.34 |
| endothelium | Prkd2 | 2.22 | 7.95E-73 | 1.65E-68 | 0.4 | 0.1 | 0.3 | 3.93 |
| endothelium | Chst7 | 2.98 | 1.82E-72 | 3.78E-68 | 0.14 | 0.02 | 0.12 | 9.07 |

**Supplementary Table 2. Marker genes for the major cell types of integrated clustering of E11.5 control and NotchGOF λ-junction cells.**

| | | | | | | | | |
|---|---|---|---|---|---|---|---|---|
| endothelium | Gpr146 | 3.33 | 2.13E-72 | 4.43E-68 | 0.19 | 0.03 | 0.16 | 6.75 |
| endothelium | Nrarp | 1.29 | 2.91E-72 | 6.06E-68 | 0.57 | 0.17 | 0.39 | 3.26 |
| endothelium | Rac1 | 0.95 | 5.62E-72 | 1.17E-67 | 0.93 | 0.9 | 0.03 | 1.03 |
| endothelium | Rasal2 | 1.58 | 1.12E-71 | 2.34E-67 | 0.63 | 0.24 | 0.39 | 2.6 |
| endothelium | Stim1 | 1.81 | 1.16E-71 | 2.41E-67 | 0.48 | 0.14 | 0.34 | 3.46 |
| endothelium | Fcer1g | 2.9 | 7.18E-71 | 1.49E-66 | 0.16 | 0.02 | 0.14 | 7.85 |
| endothelium | Kcnj15 | 2.57 | 1.58E-70 | 3.29E-66 | 0.14 | 0.02 | 0.13 | 8.81 |
| endothelium | Ifi35 | 2.9 | 2.12E-70 | 4.41E-66 | 0.19 | 0.03 | 0.16 | 6.75 |
| endothelium | Ppfibp1 | 1.52 | 2.99E-70 | 6.21E-66 | 0.78 | 0.43 | 0.35 | 1.83 |
| endothelium | Prkce | 1.62 | 3.26E-70 | 6.79E-66 | 0.5 | 0.15 | 0.35 | 3.37 |
| endothelium | Ocel1 | 2.38 | 4.08E-70 | 8.48E-66 | 0.26 | 0.05 | 0.21 | 5.18 |
| endothelium | Arhgef5 | 1.67 | 4.81E-70 | 1.00E-65 | 0.56 | 0.2 | 0.37 | 2.87 |
| endothelium | Cap1 | 1.64 | 7.35E-70 | 1.53E-65 | 0.84 | 0.54 | 0.3 | 1.55 |
| endothelium | Ripk3 | 1.6 | 1.72E-69 | 3.57E-65 | 0.52 | 0.16 | 0.36 | 3.25 |
| endothelium | Nexmif | 2.58 | 3.19E-69 | 6.64E-65 | 0.2 | 0.03 | 0.17 | 6.34 |
| endothelium | Rpl28 | 0.32 | 4.70E-69 | 9.78E-65 | 1 | 1 | 0 | 1 |
| endothelium | Tax1bp3 | 1.45 | 4.87E-69 | 1.01E-64 | 0.86 | 0.64 | 0.22 | 1.34 |
| endothelium | Lrmp | 3.72 | 6.51E-69 | 1.35E-64 | 0.1 | 0.01 | 0.1 | 11.56 |
| endothelium | Peg10 | 1.59 | 9.16E-69 | 1.91E-64 | 0.76 | 0.43 | 0.33 | 1.76 |
| endothelium | 430049B03Ri | 1.72 | 9.47E-69 | 1.97E-64 | 0.71 | 0.35 | 0.36 | 2.05 |
| endothelium | Stard4 | 1.64 | 2.30E-68 | 4.79E-64 | 0.68 | 0.27 | 0.4 | 2.48 |
| endothelium | Pxn | 1.58 | 5.86E-68 | 1.22E-63 | 0.54 | 0.18 | 0.36 | 3.02 |
| endothelium | Gng2 | 1.43 | 7.31E-68 | 1.52E-63 | 0.84 | 0.57 | 0.28 | 1.49 |
| endothelium | Pik3c2b | 1.99 | 1.00E-67 | 2.08E-63 | 0.3 | 0.07 | 0.24 | 4.54 |
| endothelium | Nid2 | 1.74 | 1.39E-67 | 2.88E-63 | 0.61 | 0.23 | 0.38 | 2.63 |
| endothelium | Rpsa | 0.31 | 1.49E-67 | 3.11E-63 | 1 | 1 | 0 | 1 |
| endothelium | Slc6a6 | 1.8 | 3.57E-67 | 7.42E-63 | 0.72 | 0.35 | 0.38 | 2.08 |
| endothelium | Mapk12 | 1.6 | 1.02E-66 | 2.12E-62 | 0.41 | 0.11 | 0.3 | 3.71 |
| endothelium | Ppia | 0.38 | 1.77E-66 | 3.68E-62 | 1 | 1 | 0 | 1 |
| endothelium | Lipt2 | 2.08 | 1.84E-66 | 3.83E-62 | 0.69 | 0.36 | 0.34 | 1.95 |
| endothelium | Plekha1 | 1.72 | 2.71E-66 | 5.65E-62 | 0.55 | 0.2 | 0.36 | 2.83 |
| endothelium | Pdia5 | 1.94 | 3.47E-66 | 7.22E-62 | 0.41 | 0.11 | 0.29 | 3.62 |
| endothelium | Tecpr1 | 1.78 | 4.00E-66 | 8.32E-62 | 0.45 | 0.13 | 0.32 | 3.37 |
| endothelium | Cpne8 | 1.7 | 5.17E-66 | 1.08E-61 | 0.33 | 0.08 | 0.25 | 4.33 |
| endothelium | Hdac7 | 1.59 | 7.95E-66 | 1.65E-61 | 0.66 | 0.28 | 0.38 | 2.37 |
| endothelium | Gale | 1.74 | 9.68E-66 | 2.01E-61 | 0.43 | 0.12 | 0.31 | 3.53 |
| endothelium | Lpar4 | 1.74 | 9.84E-66 | 2.05E-61 | 0.69 | 0.32 | 0.37 | 2.16 |
| endothelium | Tnfrsf1a | 1.67 | 1.26E-65 | 2.61E-61 | 0.68 | 0.3 | 0.38 | 2.28 |
| endothelium | Lrrfip1 | 1.55 | 1.75E-65 | 3.65E-61 | 0.73 | 0.34 | 0.4 | 2.19 |
| endothelium | Ctbs | 1.88 | 1.95E-65 | 4.05E-61 | 0.36 | 0.09 | 0.27 | 3.97 |
| endothelium | Twf2 | 1.49 | 1.42E-64 | 2.95E-60 | 0.8 | 0.42 | 0.38 | 1.9 |
| endothelium | Bin2 | 5.56 | 2.22E-64 | 4.62E-60 | 0.1 | 0.01 | 0.09 | 11.22 |
| endothelium | Lst1 | 1.46 | 3.39E-64 | 7.04E-60 | 0.12 | 0.01 | 0.1 | 9.58 |
| endothelium | Mef2d | 1.78 | 8.58E-64 | 1.79E-59 | 0.45 | 0.13 | 0.32 | 3.37 |
| endothelium | Baiap2 | 1.52 | 1.65E-63 | 3.44E-59 | 0.48 | 0.15 | 0.33 | 3.24 |
| endothelium | Sec11c | 2.26 | 2.62E-63 | 5.44E-59 | 0.67 | 0.3 | 0.38 | 2.28 |
| endothelium | Mxra7 | 1.43 | 3.29E-63 | 6.84E-59 | 0.61 | 0.23 | 0.38 | 2.67 |
| endothelium | Map4k5 | 1.72 | 4.88E-63 | 1.01E-58 | 0.66 | 0.3 | 0.35 | 2.17 |
| endothelium | Dock11 | 1.72 | 8.30E-63 | 1.73E-58 | 0.58 | 0.22 | 0.35 | 2.58 |
| endothelium | Nckap5l | 1.94 | 9.49E-63 | 1.97E-58 | 0.46 | 0.14 | 0.32 | 3.3 |
| endothelium | Fam78b | 2.3 | 2.29E-62 | 4.76E-58 | 0.18 | 0.03 | 0.15 | 6.39 |
| endothelium | Kctd17 | 1.5 | 2.70E-62 | 5.62E-58 | 0.65 | 0.27 | 0.38 | 2.41 |
| endothelium | Gnb4 | 1.55 | 3.48E-62 | 7.24E-58 | 0.69 | 0.29 | 0.39 | 2.33 |
| endothelium | Slco2a1 | 2.49 | 5.99E-62 | 1.25E-57 | 0.16 | 0.02 | 0.14 | 6.96 |
| endothelium | Arhgef28 | 1.86 | 6.17E-62 | 1.28E-57 | 0.37 | 0.1 | 0.27 | 3.79 |
| endothelium | Hfe | 2.54 | 7.47E-62 | 1.55E-57 | 0.12 | 0.01 | 0.11 | 9.23 |

**Supplementary Table 2. Marker genes for the major cell types of integrated clustering of E11.5 control and NotchGOF λ-junction cells.**

| | | | | | | | | |
|---|---|---|---|---|---|---|---|---|
| endothelium | Tnfaip2 | 0.62 | 7.66E-62 | 1.59E-57 | 0.48 | 0.14 | 0.34 | 3.44 |
| endothelium | Wasf2 | 1.17 | 1.11E-61 | 2.31E-57 | 0.89 | 0.71 | 0.18 | 1.25 |
| endothelium | Fhl2 | 2.08 | 1.67E-61 | 3.48E-57 | 0.33 | 0.08 | 0.25 | 3.96 |
| endothelium | Dock8 | 2.21 | 3.17E-61 | 6.59E-57 | 0.19 | 0.03 | 0.16 | 6.1 |
| endothelium | Reep1 | 2.11 | 5.96E-61 | 1.24E-56 | 0.23 | 0.05 | 0.19 | 5.16 |
| endothelium | Prickle3 | 1.77 | 7.08E-61 | 1.47E-56 | 0.28 | 0.06 | 0.22 | 4.49 |
| endothelium | Casp7 | 1.61 | 1.46E-60 | 3.04E-56 | 0.49 | 0.17 | 0.33 | 2.99 |
| endothelium | Rab13 | 1.49 | 2.21E-60 | 4.60E-56 | 0.78 | 0.42 | 0.36 | 1.86 |
| endothelium | Casp8 | 1.51 | 2.54E-60 | 5.27E-56 | 0.52 | 0.18 | 0.34 | 2.89 |
| endothelium | Arid3a | 1.42 | 2.71E-60 | 5.65E-56 | 0.7 | 0.33 | 0.37 | 2.12 |
| endothelium | Cpne2 | 1.92 | 3.77E-60 | 7.84E-56 | 0.24 | 0.05 | 0.2 | 5.06 |
| endothelium | Tspan15 | 1.62 | 4.28E-60 | 8.90E-56 | 0.53 | 0.19 | 0.34 | 2.84 |
| endothelium | Odc1 | 1.52 | 4.81E-60 | 1.00E-55 | 0.89 | 0.81 | 0.08 | 1.1 |
| endothelium | Cd55 | 3.8 | 4.85E-60 | 1.01E-55 | 0.16 | 0.02 | 0.13 | 7.05 |
| endothelium | Marchf2 | 1.64 | 5.86E-60 | 1.22E-55 | 0.57 | 0.21 | 0.36 | 2.75 |
| endothelium | Lamb1 | 1.4 | 8.61E-60 | 1.79E-55 | 0.73 | 0.38 | 0.35 | 1.91 |
| endothelium | Actn4 | 1.18 | 1.63E-59 | 3.39E-55 | 0.86 | 0.7 | 0.16 | 1.23 |
| endothelium | Serpinh1 | 0.8 | 1.68E-59 | 3.48E-55 | 0.89 | 0.94 | -0.06 | 0.94 |
| endothelium | Plec | 1.46 | 1.95E-59 | 4.05E-55 | 0.64 | 0.27 | 0.37 | 2.39 |
| endothelium | Sri | 1.1 | 4.42E-59 | 9.20E-55 | 0.89 | 0.77 | 0.12 | 1.15 |
| endothelium | Ost4 | 0.75 | 4.97E-59 | 1.03E-54 | 0.96 | 0.95 | 0 | 1 |
| endothelium | Nr4a1 | 2.42 | 5.31E-59 | 1.10E-54 | 0.21 | 0.04 | 0.17 | 5.33 |
| endothelium | Ctsz | 1.39 | 9.14E-59 | 1.90E-54 | 0.84 | 0.57 | 0.27 | 1.47 |
| endothelium | Rap2a | 1.53 | 1.04E-58 | 2.17E-54 | 0.69 | 0.32 | 0.38 | 2.18 |
| endothelium | Rrbp1 | 1.04 | 1.11E-58 | 2.30E-54 | 0.86 | 0.9 | -0.04 | 0.96 |
| endothelium | Snrk | 1.5 | 2.52E-58 | 5.23E-54 | 0.62 | 0.26 | 0.37 | 2.43 |
| endothelium | Arhgdia | 0.81 | 2.90E-58 | 6.02E-54 | 0.92 | 0.88 | 0.03 | 1.04 |
| endothelium | Actr2 | 1.27 | 3.04E-58 | 6.33E-54 | 0.86 | 0.63 | 0.23 | 1.37 |
| endothelium | Nol4l | 1.55 | 3.27E-58 | 6.80E-54 | 0.49 | 0.16 | 0.33 | 3.05 |
| endothelium | Irgm1 | 2.73 | 3.35E-58 | 6.97E-54 | 0.11 | 0.01 | 0.1 | 9.08 |
| endothelium | Ssh2 | 1.36 | 4.14E-58 | 8.61E-54 | 0.59 | 0.23 | 0.36 | 2.62 |
| endothelium | Itpr2 | 1.85 | 5.04E-58 | 1.05E-53 | 0.23 | 0.04 | 0.18 | 5.16 |
| endothelium | Smpd5 | 2.65 | 7.11E-58 | 1.48E-53 | 0.1 | 0.01 | 0.09 | 10.1 |
| endothelium | Gpsm3 | 2.02 | 1.06E-57 | 2.20E-53 | 0.33 | 0.08 | 0.25 | 3.95 |
| endothelium | Arl6ip5 | 1.33 | 1.18E-57 | 2.45E-53 | 0.7 | 0.32 | 0.39 | 2.23 |
| endothelium | Spns1 | 1.83 | 1.26E-57 | 2.61E-53 | 0.74 | 0.41 | 0.34 | 1.83 |
| endothelium | Arhgef1 | 1.5 | 1.70E-57 | 3.53E-53 | 0.62 | 0.26 | 0.36 | 2.36 |
| endothelium | Kifc3 | 1.8 | 2.90E-57 | 6.03E-53 | 0.3 | 0.07 | 0.23 | 4.21 |
| endothelium | Exoc6 | 2.49 | 3.65E-57 | 7.59E-53 | 0.29 | 0.07 | 0.22 | 4.04 |
| endothelium | Arsa | 1.73 | 4.03E-57 | 8.39E-53 | 0.28 | 0.06 | 0.22 | 4.38 |
| endothelium | Ap1s2 | 1.8 | 5.87E-57 | 1.22E-52 | 0.42 | 0.13 | 0.29 | 3.22 |
| endothelium | Myadm | 1.8 | 1.36E-56 | 2.84E-52 | 0.41 | 0.12 | 0.28 | 3.35 |
| endothelium | Wscd1 | 2.28 | 1.72E-56 | 3.57E-52 | 0.16 | 0.02 | 0.13 | 6.54 |
| endothelium | Peak1 | 1.45 | 2.21E-56 | 4.60E-52 | 0.6 | 0.24 | 0.36 | 2.5 |
| endothelium | Cyba | 1.69 | 4.42E-56 | 9.20E-52 | 0.77 | 0.45 | 0.32 | 1.72 |
| endothelium | Stat3 | 1.73 | 6.07E-56 | 1.26E-51 | 0.42 | 0.13 | 0.29 | 3.25 |
| endothelium | Bnip2 | 1.05 | 7.15E-56 | 1.49E-51 | 0.85 | 0.68 | 0.17 | 1.24 |
| endothelium | Cmtm6 | 1.54 | 9.45E-56 | 1.96E-51 | 0.42 | 0.13 | 0.29 | 3.3 |
| endothelium | Wsb2 | 1.51 | 1.01E-55 | 2.09E-51 | 0.68 | 0.31 | 0.37 | 2.21 |
| endothelium | Ccser2 | 1.48 | 1.22E-55 | 2.54E-51 | 0.63 | 0.27 | 0.36 | 2.37 |
| endothelium | Cdc42se1 | 1.32 | 1.75E-55 | 3.64E-51 | 0.77 | 0.42 | 0.35 | 1.84 |
| endothelium | Sorbs1 | 1.6 | 1.83E-55 | 3.82E-51 | 0.32 | 0.08 | 0.24 | 3.96 |
| endothelium | Rgl2 | 1.44 | 2.34E-55 | 4.88E-51 | 0.54 | 0.2 | 0.34 | 2.72 |
| endothelium | Stx2 | 1.43 | 2.45E-55 | 5.10E-51 | 0.57 | 0.22 | 0.35 | 2.62 |
| endothelium | Sptan1 | 1.4 | 2.61E-55 | 5.42E-51 | 0.81 | 0.56 | 0.25 | 1.44 |
| endothelium | Ebf3 | 1.14 | 2.88E-55 | 5.99E-51 | 0.5 | 0.17 | 0.33 | 2.93 |

**Supplementary Table 2. Marker genes for the major cell types of integrated clustering of E11.5 control and NotchGOF λ-junction cells.**

| | | | | | | | | |
|---|---|---|---|---|---|---|---|---|
| endothelium | Adgre5 | 1.47 | 5.24E-55 | 1.09E-50 | 0.41 | 0.12 | 0.28 | 3.32 |
| endothelium | Pxdc1 | 1.41 | 9.56E-55 | 1.99E-50 | 0.36 | 0.1 | 0.26 | 3.59 |
| endothelium | Usp43 | 2.69 | 1.22E-54 | 2.54E-50 | 0.12 | 0.02 | 0.11 | 8.2 |
| endothelium | Fam102a | 1.9 | 2.06E-54 | 4.29E-50 | 0.44 | 0.15 | 0.29 | 2.99 |
| endothelium | Ap2s1 | 0.86 | 2.22E-54 | 4.61E-50 | 0.93 | 0.89 | 0.04 | 1.05 |
| endothelium | Ptpn1 | 1.29 | 7.02E-54 | 1.46E-49 | 0.83 | 0.52 | 0.31 | 1.59 |
| endothelium | Git1 | 1.5 | 8.07E-54 | 1.68E-49 | 0.61 | 0.26 | 0.35 | 2.38 |
| endothelium | Dock4 | 1.48 | 8.47E-54 | 1.76E-49 | 0.66 | 0.31 | 0.35 | 2.13 |
| endothelium | Tmc6 | 1.88 | 1.13E-53 | 2.36E-49 | 0.29 | 0.07 | 0.22 | 4.04 |
| endothelium | Nmi | 1.69 | 1.18E-53 | 2.46E-49 | 0.33 | 0.09 | 0.24 | 3.73 |
| endothelium | Nt5c | 1.2 | 1.42E-53 | 2.95E-49 | 0.82 | 0.53 | 0.29 | 1.54 |
| endothelium | Cdipt | 1.44 | 2.26E-53 | 4.69E-49 | 0.72 | 0.35 | 0.36 | 2.03 |
| endothelium | Cdc42bpb | 1.39 | 3.56E-53 | 7.41E-49 | 0.71 | 0.35 | 0.36 | 2.04 |
| endothelium | Rab8b | 1.61 | 6.75E-53 | 1.40E-48 | 0.59 | 0.24 | 0.35 | 2.49 |
| endothelium | Wipf1 | 1.33 | 9.30E-53 | 1.93E-48 | 0.58 | 0.23 | 0.36 | 2.58 |
| endothelium | Tmbim1 | 1.68 | 9.89E-53 | 2.06E-48 | 0.34 | 0.09 | 0.25 | 3.61 |
| endothelium | Srgap1 | 1.43 | 1.46E-52 | 3.04E-48 | 0.68 | 0.34 | 0.34 | 2 |
| endothelium | Sdcbp | 1.1 | 2.41E-52 | 5.01E-48 | 0.88 | 0.69 | 0.19 | 1.27 |
| endothelium | Rgs10 | 4.07 | 5.62E-52 | 1.17E-47 | 0.31 | 0.09 | 0.22 | 3.64 |
| endothelium | Vangl1 | 1.7 | 1.29E-51 | 2.68E-47 | 0.44 | 0.15 | 0.28 | 2.88 |
| endothelium | Hspa12b | 1.7 | 1.29E-51 | 2.69E-47 | 0.41 | 0.13 | 0.28 | 3.14 |
| endothelium | Mgmt | 1.89 | 1.82E-51 | 3.79E-47 | 0.53 | 0.2 | 0.33 | 2.62 |
| endothelium | Mapk3 | 1.39 | 2.46E-51 | 5.12E-47 | 0.76 | 0.45 | 0.31 | 1.7 |
| endothelium | Mrpl17 | 1.01 | 2.86E-51 | 5.96E-47 | 0.9 | 0.79 | 0.11 | 1.14 |
| endothelium | Btg2 | 1.11 | 3.98E-51 | 8.27E-47 | 0.62 | 0.27 | 0.36 | 2.32 |
| endothelium | Sptlc2 | 1.31 | 4.34E-51 | 9.03E-47 | 0.68 | 0.32 | 0.36 | 2.11 |
| endothelium | Vps29 | 0.94 | 5.16E-51 | 1.07E-46 | 0.91 | 0.78 | 0.13 | 1.17 |
| endothelium | Crem | 1.63 | 6.14E-51 | 1.28E-46 | 0.49 | 0.18 | 0.31 | 2.79 |
| endothelium | Lrp11 | 2.04 | 8.95E-51 | 1.86E-46 | 0.18 | 0.03 | 0.15 | 5.48 |
| endothelium | Prr5 | 1.78 | 1.03E-50 | 2.14E-46 | 0.38 | 0.12 | 0.27 | 3.31 |
| endothelium | Krtcap2 | 0.68 | 1.31E-50 | 2.72E-46 | 0.92 | 0.94 | -0.02 | 0.98 |
| endothelium | Bag3 | 1.52 | 1.48E-50 | 3.07E-46 | 0.37 | 0.11 | 0.26 | 3.41 |
| endothelium | Plekhg5 | 1.58 | 1.63E-50 | 3.39E-46 | 0.34 | 0.1 | 0.24 | 3.57 |
| endothelium | Palm | 1.34 | 2.25E-50 | 4.68E-46 | 0.74 | 0.46 | 0.29 | 1.63 |
| endothelium | Lpar6 | 1.36 | 2.48E-50 | 5.15E-46 | 0.73 | 0.4 | 0.33 | 1.81 |
| endothelium | Bola2 | 0.77 | 8.38E-50 | 1.74E-45 | 0.93 | 0.88 | 0.05 | 1.05 |
| endothelium | Dapp1 | 3.54 | 2.23E-49 | 4.64E-45 | 0.16 | 0.03 | 0.13 | 5.71 |
| endothelium | Ybx3 | 0.86 | 2.47E-49 | 5.13E-45 | 0.92 | 0.84 | 0.08 | 1.09 |
| endothelium | Tmem120b | 1.66 | 3.21E-49 | 6.68E-45 | 0.25 | 0.06 | 0.19 | 4.28 |
| endothelium | Yes1 | 1.45 | 4.79E-49 | 9.96E-45 | 0.68 | 0.34 | 0.33 | 1.96 |
| endothelium | Ltbr | 1.36 | 8.01E-49 | 1.67E-44 | 0.57 | 0.23 | 0.34 | 2.51 |
| endothelium | Pim3 | 1.42 | 1.25E-48 | 2.60E-44 | 0.63 | 0.29 | 0.33 | 2.14 |
| endothelium | Eva1b | 1.14 | 1.33E-48 | 2.78E-44 | 0.81 | 0.54 | 0.28 | 1.51 |
| endothelium | Fermt2 | 1.05 | 2.32E-48 | 4.83E-44 | 0.85 | 0.72 | 0.13 | 1.18 |
| endothelium | Foxo1 | 1.25 | 3.60E-48 | 7.48E-44 | 0.48 | 0.18 | 0.31 | 2.71 |
| endothelium | Stim2 | 1.33 | 3.80E-48 | 7.90E-44 | 0.57 | 0.23 | 0.34 | 2.47 |
| endothelium | Ccnyl1 | 1.62 | 4.66E-48 | 9.70E-44 | 0.48 | 0.18 | 0.31 | 2.71 |
| endothelium | Acy3 | 2.41 | 5.43E-48 | 1.13E-43 | 0.1 | 0.01 | 0.09 | 8 |
| endothelium | Sypl | 1.16 | 5.61E-48 | 1.17E-43 | 0.78 | 0.48 | 0.3 | 1.62 |
| endothelium | Hivep3 | 2.1 | 6.38E-48 | 1.33E-43 | 0.13 | 0.02 | 0.11 | 6.89 |
| endothelium | Nfic | 1.14 | 6.81E-48 | 1.42E-43 | 0.66 | 0.3 | 0.37 | 2.24 |
| endothelium | Aim2 | 2.4 | 7.95E-48 | 1.65E-43 | 0.25 | 0.06 | 0.19 | 4.13 |
| endothelium | Peli2 | 1.26 | 9.60E-48 | 2.00E-43 | 0.5 | 0.18 | 0.32 | 2.78 |
| endothelium | Btbd3 | 1.68 | 1.57E-47 | 3.26E-43 | 0.44 | 0.15 | 0.28 | 2.84 |
| endothelium | Fam111a | 1.3 | 2.35E-47 | 4.88E-43 | 0.66 | 0.32 | 0.34 | 2.07 |
| endothelium | Syngr2 | 1.3 | 2.53E-47 | 5.27E-43 | 0.42 | 0.14 | 0.28 | 2.99 |

**Supplementary Table 2. Marker genes for the major cell types of integrated clustering of E11.5 control and NotchGOF λ-junction cells.**

| | | | | | | | | |
|---|---|---|---|---|---|---|---|---|
| endothelium | Stxbp1 | 1.74 | 3.44E-47 | 7.16E-43 | 0.32 | 0.09 | 0.23 | 3.55 |
| endothelium | Eif4ebp1 | 1.08 | 4.57E-47 | 9.50E-43 | 0.86 | 0.69 | 0.16 | 1.24 |
| endothelium | Gamt | 1.39 | 4.94E-47 | 1.03E-42 | 0.32 | 0.09 | 0.23 | 3.56 |
| endothelium | Vamp8 | 1.11 | 7.27E-47 | 1.51E-42 | 0.86 | 0.61 | 0.25 | 1.41 |
| endothelium | Afdn | 1.04 | 7.77E-47 | 1.62E-42 | 0.81 | 0.57 | 0.24 | 1.43 |
| endothelium | Asah2 | 1.73 | 9.26E-47 | 1.93E-42 | 0.21 | 0.04 | 0.16 | 4.73 |
| endothelium | Commd7 | 1.03 | 1.04E-46 | 2.17E-42 | 0.83 | 0.57 | 0.26 | 1.46 |
| endothelium | Prelid1 | 0.72 | 1.12E-46 | 2.34E-42 | 0.94 | 0.93 | 0.01 | 1.01 |
| endothelium | Spata6 | 1.37 | 1.21E-46 | 2.52E-42 | 0.51 | 0.2 | 0.31 | 2.58 |
| endothelium | Cdc42ep3 | 1.37 | 1.23E-46 | 2.56E-42 | 0.45 | 0.16 | 0.29 | 2.78 |
| endothelium | Creg1 | 1.15 | 1.26E-46 | 2.63E-42 | 0.7 | 0.35 | 0.36 | 2.02 |
| endothelium | Esyt1 | 1.52 | 1.31E-46 | 2.72E-42 | 0.28 | 0.07 | 0.2 | 3.87 |
| endothelium | Strn | 1.51 | 3.26E-46 | 6.78E-42 | 0.47 | 0.17 | 0.3 | 2.73 |
| endothelium | Nr2f2 | 1.09 | 6.89E-46 | 1.43E-41 | 0.69 | 0.36 | 0.33 | 1.91 |
| endothelium | Lnx2 | 1.78 | 7.93E-46 | 1.65E-41 | 0.24 | 0.06 | 0.18 | 4.16 |
| endothelium | Nme2 | 0.39 | 9.35E-46 | 1.94E-41 | 0.99 | 0.98 | 0.01 | 1.01 |
| endothelium | Irf2 | 1.31 | 1.10E-45 | 2.28E-41 | 0.59 | 0.25 | 0.34 | 2.34 |
| endothelium | Nav3 | 0.86 | 1.16E-45 | 2.42E-41 | 0.39 | 0.12 | 0.27 | 3.14 |
| endothelium | Gm43260 | 1.93 | 1.53E-45 | 3.18E-41 | 0.17 | 0.03 | 0.14 | 5.24 |
| endothelium | Manba | 2.76 | 1.60E-45 | 3.33E-41 | 0.11 | 0.02 | 0.1 | 7.47 |
| endothelium | Trib1 | 1.5 | 2.56E-45 | 5.32E-41 | 0.47 | 0.18 | 0.3 | 2.67 |
| endothelium | Mindy2 | 1.48 | 3.59E-45 | 7.46E-41 | 0.46 | 0.17 | 0.29 | 2.68 |
| endothelium | Jak1 | 1.11 | 4.39E-45 | 9.12E-41 | 0.83 | 0.6 | 0.23 | 1.38 |
| endothelium | Rps3 | 0.26 | 9.85E-45 | 2.05E-40 | 1 | 1 | 0 | 1 |
| endothelium | Triobp | 1.5 | 1.19E-44 | 2.48E-40 | 0.64 | 0.31 | 0.34 | 2.09 |
| endothelium | Nqo2 | 1.42 | 1.61E-44 | 3.35E-40 | 0.38 | 0.13 | 0.26 | 3.05 |
| endothelium | Zfp36 | 1.87 | 2.54E-44 | 5.29E-40 | 0.26 | 0.07 | 0.19 | 3.78 |
| endothelium | Vwa1 | 1.92 | 2.57E-44 | 5.35E-40 | 0.23 | 0.06 | 0.18 | 4.14 |
| endothelium | Fez2 | 1.28 | 2.61E-44 | 5.43E-40 | 0.65 | 0.31 | 0.35 | 2.13 |
| endothelium | Smap2 | 1.36 | 3.46E-44 | 7.19E-40 | 0.7 | 0.37 | 0.34 | 1.91 |
| endothelium | Nrp2 | 0.94 | 4.00E-44 | 8.31E-40 | 0.72 | 0.37 | 0.35 | 1.96 |
| endothelium | Arhgap23 | 1.88 | 4.56E-44 | 9.47E-40 | 0.31 | 0.09 | 0.22 | 3.36 |
| endothelium | Pip4k2a | 1.65 | 4.71E-44 | 9.80E-40 | 0.35 | 0.11 | 0.24 | 3.12 |
| endothelium | Arrb1 | 1.46 | 7.41E-44 | 1.54E-39 | 0.35 | 0.11 | 0.24 | 3.14 |
| endothelium | Rtl8a | 1.41 | 7.94E-44 | 1.65E-39 | 0.72 | 0.39 | 0.33 | 1.86 |
| endothelium | Necap2 | 1.33 | 8.11E-44 | 1.69E-39 | 0.73 | 0.39 | 0.34 | 1.86 |
| endothelium | Rab12 | 1.2 | 8.17E-44 | 1.70E-39 | 0.77 | 0.48 | 0.29 | 1.6 |
| endothelium | F2r | 1.19 | 8.40E-44 | 1.75E-39 | 0.82 | 0.57 | 0.25 | 1.45 |
| endothelium | Man1a | 1.4 | 9.13E-44 | 1.90E-39 | 0.24 | 0.06 | 0.18 | 4.07 |
| endothelium | Prrg3 | 1.54 | 1.05E-43 | 2.19E-39 | 0.34 | 0.11 | 0.24 | 3.19 |
| endothelium | Ldlrap1 | 1.41 | 1.27E-43 | 2.65E-39 | 0.48 | 0.18 | 0.3 | 2.66 |
| endothelium | Chst2 | 1.28 | 1.81E-43 | 3.77E-39 | 0.5 | 0.2 | 0.3 | 2.49 |
| endothelium | Hexa | 1.37 | 2.03E-43 | 4.22E-39 | 0.65 | 0.31 | 0.34 | 2.09 |
| endothelium | Tmem63a | 2.19 | 3.63E-43 | 7.55E-39 | 0.14 | 0.02 | 0.12 | 5.88 |
| endothelium | Cyp20a1 | 1.16 | 3.73E-43 | 7.77E-39 | 0.59 | 0.25 | 0.34 | 2.35 |
| endothelium | Tmed5 | 1.1 | 4.57E-43 | 9.51E-39 | 0.77 | 0.48 | 0.29 | 1.59 |
| endothelium | Mydgf | 0.86 | 5.88E-43 | 1.22E-38 | 0.88 | 0.73 | 0.14 | 1.19 |
| endothelium | Gap43 | 1.27 | 6.60E-43 | 1.37E-38 | 0.72 | 0.41 | 0.31 | 1.74 |
| endothelium | Rps6ka1 | 1.32 | 6.76E-43 | 1.41E-38 | 0.57 | 0.24 | 0.33 | 2.37 |
| endothelium | Slc43a1 | 1.21 | 1.18E-42 | 2.46E-38 | 0.44 | 0.16 | 0.28 | 2.79 |
| endothelium | Uvrag | 1.29 | 1.32E-42 | 2.74E-38 | 0.53 | 0.22 | 0.31 | 2.38 |
| endothelium | Mpzl1 | 1.17 | 1.51E-42 | 3.14E-38 | 0.8 | 0.63 | 0.17 | 1.26 |
| endothelium | Iqgap1 | 1.09 | 1.52E-42 | 3.17E-38 | 0.76 | 0.47 | 0.29 | 1.61 |
| endothelium | Ptpn12 | 1.34 | 1.64E-42 | 3.41E-38 | 0.67 | 0.35 | 0.32 | 1.91 |
| endothelium | Lgals8 | 1.81 | 1.69E-42 | 3.52E-38 | 0.29 | 0.08 | 0.21 | 3.49 |
| endothelium | Sox13 | 1.37 | 1.86E-42 | 3.87E-38 | 0.3 | 0.09 | 0.21 | 3.48 |

**Supplementary Table 2. Marker genes for the major cell types of integrated clustering of E11.5 control and NotchGOF λ-junction cells.**

| | | | | | | | | |
|---|---|---|---|---|---|---|---|---|
| endothelium | D1Ertd622e | 1.52 | 1.98E-42 | 4.12E-38 | 0.38 | 0.13 | 0.25 | 2.96 |
| endothelium | Hspa8 | 0.5 | 2.28E-42 | 4.74E-38 | 0.99 | 1 | -0.01 | 0.99 |
| endothelium | Rgcc | 1.81 | 2.76E-42 | 5.73E-38 | 0.3 | 0.09 | 0.21 | 3.38 |
| endothelium | Npr3 | 2.14 | 3.06E-42 | 6.35E-38 | 0.11 | 0.02 | 0.1 | 7 |
| endothelium | Cdk17 | 1.33 | 3.28E-42 | 6.81E-38 | 0.5 | 0.2 | 0.3 | 2.46 |
| endothelium | Ifngr1 | 1.2 | 3.52E-42 | 7.31E-38 | 0.56 | 0.24 | 0.32 | 2.31 |
| endothelium | Il10rb | 1.51 | 4.45E-42 | 9.25E-38 | 0.29 | 0.08 | 0.21 | 3.47 |
| endothelium | Tspan14 | 1.16 | 5.32E-42 | 1.11E-37 | 0.66 | 0.32 | 0.34 | 2.06 |
| endothelium | Egln3 | 1.92 | 6.24E-42 | 1.30E-37 | 0.27 | 0.07 | 0.19 | 3.66 |
| endothelium | Rgs2 | 1.29 | 6.48E-42 | 1.35E-37 | 0.65 | 0.34 | 0.31 | 1.92 |
| endothelium | Rhog | 1.28 | 7.15E-42 | 1.49E-37 | 0.68 | 0.35 | 0.33 | 1.94 |
| endothelium | Plod1 | 1.2 | 7.19E-42 | 1.50E-37 | 0.5 | 0.2 | 0.3 | 2.55 |
| endothelium | Map3k20 | 1.65 | 9.77E-42 | 2.03E-37 | 0.35 | 0.11 | 0.24 | 3.09 |
| endothelium | Pqlc1 | 1.33 | 1.22E-41 | 2.54E-37 | 0.39 | 0.13 | 0.26 | 2.9 |
| endothelium | Cast | 1.41 | 1.70E-41 | 3.53E-37 | 0.56 | 0.24 | 0.32 | 2.32 |
| endothelium | Mfge8 | 1.11 | 1.74E-41 | 3.61E-37 | 0.77 | 0.52 | 0.25 | 1.49 |
| endothelium | Rock2 | 0.96 | 1.95E-41 | 4.05E-37 | 0.84 | 0.66 | 0.18 | 1.27 |
| endothelium | Dusp6 | 1.02 | 2.56E-41 | 5.32E-37 | 0.72 | 0.36 | 0.36 | 2 |
| endothelium | Cavin2 | 4.79 | 3.68E-41 | 7.66E-37 | 0.12 | 0.02 | 0.1 | 6.15 |
| endothelium | Apobec3 | 1.25 | 5.33E-41 | 1.11E-36 | 0.29 | 0.08 | 0.2 | 3.52 |
| endothelium | Slc38a10 | 1.16 | 6.25E-41 | 1.30E-36 | 0.67 | 0.33 | 0.34 | 2.02 |
| endothelium | Ndel1 | 1.2 | 6.26E-41 | 1.30E-36 | 0.65 | 0.31 | 0.34 | 2.08 |
| endothelium | Tstd3 | 1.34 | 6.58E-41 | 1.37E-36 | 0.5 | 0.21 | 0.3 | 2.46 |
| endothelium | Prdx1 | 0.42 | 7.24E-41 | 1.51E-36 | 0.98 | 0.99 | -0.01 | 0.99 |
| endothelium | Prnp | 1.21 | 1.20E-40 | 2.50E-36 | 0.45 | 0.17 | 0.28 | 2.62 |
| endothelium | Atp8b2 | 1.26 | 1.22E-40 | 2.55E-36 | 0.29 | 0.08 | 0.21 | 3.51 |
| endothelium | Kctd10 | 1.29 | 1.34E-40 | 2.78E-36 | 0.62 | 0.29 | 0.33 | 2.13 |
| endothelium | Phactr2 | 1.03 | 1.37E-40 | 2.85E-36 | 0.77 | 0.45 | 0.32 | 1.71 |
| endothelium | Usp12 | 1.41 | 1.41E-40 | 2.93E-36 | 0.5 | 0.2 | 0.3 | 2.49 |
| endothelium | Prkar1a | 0.68 | 1.80E-40 | 3.74E-36 | 0.93 | 0.88 | 0.05 | 1.06 |
| endothelium | Ksr1 | 2.29 | 1.89E-40 | 3.92E-36 | 0.16 | 0.03 | 0.13 | 5 |
| endothelium | Glud1 | 1.03 | 1.91E-40 | 3.96E-36 | 0.83 | 0.65 | 0.18 | 1.28 |
| endothelium | Ptbp3 | 0.97 | 2.07E-40 | 4.31E-36 | 0.83 | 0.62 | 0.21 | 1.34 |
| endothelium | Kit | 1.74 | 2.13E-40 | 4.42E-36 | 0.3 | 0.09 | 0.21 | 3.23 |
| endothelium | Endod1 | 1.47 | 2.47E-40 | 5.14E-36 | 0.29 | 0.09 | 0.21 | 3.33 |
| endothelium | Pelo | 1.3 | 2.89E-40 | 6.02E-36 | 0.69 | 0.39 | 0.3 | 1.77 |
| endothelium | Xbp1 | 1.14 | 4.50E-40 | 9.35E-36 | 0.76 | 0.47 | 0.29 | 1.61 |
| endothelium | Ptpre | 2.54 | 5.18E-40 | 1.08E-35 | 0.17 | 0.04 | 0.13 | 4.8 |
| endothelium | Hras | 0.79 | 6.75E-40 | 1.40E-35 | 0.9 | 0.8 | 0.1 | 1.12 |
| endothelium | Nrgn | 5.49 | 8.12E-40 | 1.69E-35 | 0.19 | 0.05 | 0.15 | 4.27 |
| endothelium | Chmp2a | 0.89 | 1.37E-39 | 2.86E-35 | 0.84 | 0.66 | 0.18 | 1.27 |
| endothelium | Pls3 | 1.42 | 1.48E-39 | 3.08E-35 | 0.6 | 0.29 | 0.31 | 2.05 |
| endothelium | Cdyl2 | 1.58 | 1.54E-39 | 3.20E-35 | 0.21 | 0.05 | 0.16 | 4.16 |
| endothelium | Arhgef7 | 1.25 | 1.62E-39 | 3.36E-35 | 0.62 | 0.31 | 0.31 | 1.99 |
| endothelium | Clta | 0.42 | 1.84E-39 | 3.83E-35 | 0.94 | 0.98 | -0.04 | 0.96 |
| endothelium | Mgat1 | 1.51 | 2.22E-39 | 4.63E-35 | 0.49 | 0.2 | 0.29 | 2.42 |
| endothelium | Tuba1a | 0.5 | 3.26E-39 | 6.78E-35 | 0.99 | 0.98 | 0.01 | 1.01 |
| endothelium | Prag1 | 1.38 | 3.45E-39 | 7.18E-35 | 0.39 | 0.14 | 0.25 | 2.85 |
| endothelium | Gypc | 1.42 | 4.35E-39 | 9.05E-35 | 0.51 | 0.21 | 0.3 | 2.38 |
| endothelium | Gne | 1.46 | 5.50E-39 | 1.14E-34 | 0.27 | 0.08 | 0.19 | 3.49 |
| endothelium | Acot7 | 1.15 | 6.23E-39 | 1.30E-34 | 0.55 | 0.23 | 0.32 | 2.37 |
| endothelium | Mgat2 | 1.06 | 7.95E-39 | 1.65E-34 | 0.74 | 0.46 | 0.28 | 1.61 |
| endothelium | Git2 | 1.21 | 8.37E-39 | 1.74E-34 | 0.66 | 0.33 | 0.33 | 1.97 |
| endothelium | Lrrc58 | 0.94 | 1.05E-38 | 2.19E-34 | 0.82 | 0.63 | 0.2 | 1.31 |
| endothelium | Flnb | 1.23 | 1.25E-38 | 2.60E-34 | 0.6 | 0.29 | 0.3 | 2.04 |
| endothelium | Relb | 1.88 | 1.35E-38 | 2.81E-34 | 0.16 | 0.03 | 0.13 | 4.85 |

**Supplementary Table 2. Marker genes for the major cell types of integrated clustering of E11.5 control and NotchGOF λ-junction cells.**

| | | | | | | | | |
|---|---|---|---|---|---|---|---|---|
| endothelium | Chst15 | 0.99 | 2.00E-38 | 4.16E-34 | 0.45 | 0.17 | 0.28 | 2.68 |
| endothelium | Itpripl2 | 1.14 | 2.29E-38 | 4.75E-34 | 0.38 | 0.13 | 0.25 | 2.94 |
| endothelium | Iqcg | 1.76 | 2.82E-38 | 5.87E-34 | 0.17 | 0.04 | 0.13 | 4.71 |
| endothelium | Mylk | 1.45 | 3.03E-38 | 6.29E-34 | 0.44 | 0.17 | 0.27 | 2.62 |
| endothelium | Zfp948 | 1.57 | 3.44E-38 | 7.16E-34 | 0.3 | 0.09 | 0.21 | 3.29 |
| endothelium | .730020E08Ri | 1.59 | 3.72E-38 | 7.73E-34 | 0.18 | 0.04 | 0.14 | 4.51 |
| endothelium | Shc1 | 1.21 | 4.00E-38 | 8.32E-34 | 0.54 | 0.23 | 0.31 | 2.31 |
| endothelium | Gabarapl1 | 1.45 | 4.14E-38 | 8.60E-34 | 0.65 | 0.33 | 0.32 | 1.98 |
| endothelium | Vash1 | 2.58 | 4.39E-38 | 9.14E-34 | 0.11 | 0.02 | 0.09 | 6.22 |
| endothelium | Pdzd2 | 1.84 | 4.44E-38 | 9.23E-34 | 0.16 | 0.03 | 0.13 | 4.91 |
| endothelium | Hmgn5 | 1.1 | 4.46E-38 | 9.28E-34 | 0.87 | 0.71 | 0.16 | 1.22 |
| endothelium | Klhl3 | 1.94 | 4.63E-38 | 9.64E-34 | 0.13 | 0.02 | 0.11 | 5.54 |
| endothelium | Pde10a | 1.69 | 4.73E-38 | 9.84E-34 | 0.18 | 0.04 | 0.14 | 4.38 |
| endothelium | Ccser1 | 2.59 | 6.30E-38 | 1.31E-33 | 0.15 | 0.03 | 0.12 | 4.9 |
| endothelium | Ppargc1b | 2.08 | 6.65E-38 | 1.38E-33 | 0.14 | 0.03 | 0.11 | 5.22 |
| endothelium | Il3ra | 1.63 | 7.03E-38 | 1.46E-33 | 0.2 | 0.05 | 0.15 | 4.14 |
| endothelium | Med10 | 0.96 | 7.09E-38 | 1.48E-33 | 0.86 | 0.72 | 0.14 | 1.19 |
| endothelium | Dcbld1 | 1 | 8.30E-38 | 1.73E-33 | 0.62 | 0.3 | 0.33 | 2.11 |
| endothelium | Klf13 | 1.06 | 1.15E-37 | 2.40E-33 | 0.69 | 0.37 | 0.32 | 1.86 |
| endothelium | Entpd1 | 1.74 | 1.30E-37 | 2.71E-33 | 0.22 | 0.06 | 0.17 | 3.8 |
| endothelium | Capns1 | 0.81 | 1.36E-37 | 2.83E-33 | 0.9 | 0.8 | 0.1 | 1.12 |
| endothelium | Zfp710 | 1.13 | 1.49E-37 | 3.10E-33 | 0.45 | 0.18 | 0.27 | 2.55 |
| endothelium | Plekha3 | 1.25 | 1.54E-37 | 3.21E-33 | 0.55 | 0.25 | 0.31 | 2.23 |
| endothelium | Cnrip1 | 0.98 | 1.68E-37 | 3.49E-33 | 0.39 | 0.14 | 0.26 | 2.88 |
| endothelium | Sft2d1 | 1.19 | 3.65E-37 | 7.60E-33 | 0.52 | 0.22 | 0.3 | 2.35 |
| endothelium | Mvb12b | 1.31 | 4.27E-37 | 8.88E-33 | 0.52 | 0.22 | 0.29 | 2.3 |
| endothelium | Gm43573 | 2.04 | 8.13E-37 | 1.69E-32 | 0.14 | 0.03 | 0.12 | 4.97 |
| endothelium | Tspan6 | 1.02 | 8.46E-37 | 1.76E-32 | 0.83 | 0.74 | 0.09 | 1.12 |
| endothelium | Pakap.1 | 1.15 | 9.76E-37 | 2.03E-32 | 0.71 | 0.41 | 0.3 | 1.73 |
| endothelium | Zyx | 1.69 | 1.00E-36 | 2.09E-32 | 0.78 | 0.51 | 0.27 | 1.53 |
| endothelium | Cr1l | 1.04 | 1.11E-36 | 2.31E-32 | 0.63 | 0.32 | 0.31 | 2 |
| endothelium | Sertad1 | 1.41 | 1.39E-36 | 2.89E-32 | 0.51 | 0.22 | 0.29 | 2.33 |
| endothelium | Anxa4 | 1.36 | 1.48E-36 | 3.08E-32 | 0.33 | 0.11 | 0.22 | 3.04 |
| endothelium | Itga9 | 1.18 | 1.68E-36 | 3.50E-32 | 0.5 | 0.21 | 0.29 | 2.39 |
| endothelium | Tspan13 | 1.09 | 1.71E-36 | 3.55E-32 | 0.71 | 0.41 | 0.3 | 1.73 |
| endothelium | Gdi2 | 0.52 | 1.71E-36 | 3.56E-32 | 0.95 | 0.95 | 0 | 1 |
| endothelium | Lrrk2 | 1.82 | 1.74E-36 | 3.62E-32 | 0.15 | 0.03 | 0.12 | 4.97 |
| endothelium | Slc23a2 | 1.55 | 2.11E-36 | 4.39E-32 | 0.26 | 0.07 | 0.18 | 3.46 |
| endothelium | St3gal5 | 1.1 | 2.18E-36 | 4.54E-32 | 0.53 | 0.24 | 0.3 | 2.25 |
| endothelium | St5 | 1.34 | 2.21E-36 | 4.59E-32 | 0.53 | 0.22 | 0.3 | 2.35 |
| endothelium | Mtus1 | 1.26 | 4.00E-36 | 8.31E-32 | 0.32 | 0.1 | 0.21 | 3.06 |
| endothelium | Tspan2 | 1.26 | 4.11E-36 | 8.56E-32 | 0.3 | 0.1 | 0.21 | 3.17 |
| endothelium | Swap70 | 1.29 | 4.69E-36 | 9.76E-32 | 0.62 | 0.32 | 0.3 | 1.93 |
| endothelium | Gmds | 1.08 | 5.75E-36 | 1.20E-31 | 0.67 | 0.36 | 0.32 | 1.89 |
| endothelium | Lhx6 | 1.44 | 8.75E-36 | 1.82E-31 | 0.22 | 0.06 | 0.16 | 3.73 |
| endothelium | Lrrc8b | 1.35 | 1.03E-35 | 2.14E-31 | 0.33 | 0.11 | 0.22 | 2.98 |
| endothelium | Npepl1 | 1.08 | 1.06E-35 | 2.20E-31 | 0.66 | 0.32 | 0.34 | 2.03 |
| endothelium | Mob3c | 1.74 | 1.46E-35 | 3.03E-31 | 0.21 | 0.05 | 0.15 | 3.87 |
| endothelium | Stat1 | 1.79 | 1.58E-35 | 3.29E-31 | 0.3 | 0.1 | 0.2 | 3.12 |
| endothelium | Crlf2 | 1.44 | 1.64E-35 | 3.42E-31 | 0.24 | 0.07 | 0.17 | 3.59 |
| endothelium | Trappc2 | 1.58 | 1.72E-35 | 3.58E-31 | 0.41 | 0.16 | 0.25 | 2.55 |
| endothelium | Neu1 | 1.24 | 1.74E-35 | 3.63E-31 | 0.36 | 0.13 | 0.23 | 2.81 |
| endothelium | Cd47 | 1.33 | 1.86E-35 | 3.87E-31 | 0.75 | 0.44 | 0.31 | 1.71 |
| endothelium | Nectin2 | 1.01 | 2.38E-35 | 4.94E-31 | 0.74 | 0.48 | 0.27 | 1.57 |
| endothelium | St3gal1 | 1.62 | 2.46E-35 | 5.11E-31 | 0.3 | 0.1 | 0.2 | 3.04 |
| endothelium | Ssu72 | 0.71 | 2.62E-35 | 5.46E-31 | 0.91 | 0.84 | 0.07 | 1.09 |

**Supplementary Table 2. Marker genes for the major cell types of integrated clustering of E11.5 control and NotchGOF λ-junction cells.**

| | | | | | | | | |
|---|---|---|---|---|---|---|---|---|
| endothelium | Nipsnap3b | 1.02 | 3.63E-35 | 7.56E-31 | 0.74 | 0.43 | 0.31 | 1.71 |
| endothelium | Gm42702 | 1.79 | 3.68E-35 | 7.65E-31 | 0.16 | 0.04 | 0.13 | 4.53 |
| endothelium | Chst12 | 1.7 | 4.14E-35 | 8.60E-31 | 0.27 | 0.08 | 0.19 | 3.26 |
| endothelium | Gm10638 | 1.8 | 8.76E-35 | 1.82E-30 | 0.2 | 0.05 | 0.15 | 3.94 |
| endothelium | Cpd | 1.11 | 9.49E-35 | 1.97E-30 | 0.65 | 0.33 | 0.31 | 1.94 |
| endothelium | Pf4 | 10.3 | 1.01E-34 | 2.11E-30 | 0.13 | 0.03 | 0.11 | 4.93 |
| endothelium | Emilin1 | 0.91 | 1.03E-34 | 2.15E-30 | 0.79 | 0.53 | 0.27 | 1.51 |
| endothelium | Plp2 | 1.12 | 1.06E-34 | 2.21E-30 | 0.68 | 0.38 | 0.3 | 1.8 |
| endothelium | Ctnnb1 | 0.46 | 1.40E-34 | 2.92E-30 | 0.95 | 0.97 | -0.01 | 0.99 |
| endothelium | Sorbs3 | 1.19 | 1.44E-34 | 2.99E-30 | 0.37 | 0.13 | 0.24 | 2.79 |
| endothelium | Tmbim6 | 0.75 | 1.53E-34 | 3.17E-30 | 0.88 | 0.84 | 0.04 | 1.05 |
| endothelium | Cd59a | 1.23 | 1.54E-34 | 3.20E-30 | 0.51 | 0.22 | 0.3 | 2.36 |
| endothelium | Homer3 | 0.99 | 1.79E-34 | 3.73E-30 | 0.76 | 0.48 | 0.28 | 1.58 |
| endothelium | Stard13 | 1.87 | 1.89E-34 | 3.92E-30 | 0.13 | 0.02 | 0.1 | 5.43 |
| endothelium | Mogat2 | 1.25 | 1.94E-34 | 4.04E-30 | 0.33 | 0.12 | 0.22 | 2.9 |
| endothelium | Chmp3 | 1.09 | 2.33E-34 | 4.85E-30 | 0.75 | 0.45 | 0.29 | 1.65 |
| endothelium | Tnnt1 | 2.33 | 2.38E-34 | 4.95E-30 | 0.2 | 0.05 | 0.15 | 3.79 |
| endothelium | Rbms1 | 0.77 | 2.51E-34 | 5.21E-30 | 0.87 | 0.78 | 0.09 | 1.12 |
| endothelium | Dhrs7 | 1.21 | 2.89E-34 | 6.01E-30 | 0.35 | 0.12 | 0.23 | 2.84 |
| endothelium | Hmox1 | 0.91 | 2.99E-34 | 6.21E-30 | 0.46 | 0.19 | 0.28 | 2.48 |
| endothelium | Plcb4 | 1.11 | 3.09E-34 | 6.43E-30 | 0.47 | 0.19 | 0.28 | 2.43 |
| endothelium | Eif5a | 0.38 | 3.44E-34 | 7.16E-30 | 0.99 | 1 | -0.01 | 0.99 |
| endothelium | Renbp | 1.48 | 3.88E-34 | 8.07E-30 | 0.25 | 0.08 | 0.18 | 3.37 |
| endothelium | Rexo2 | 0.98 | 4.15E-34 | 8.62E-30 | 0.8 | 0.59 | 0.21 | 1.35 |
| endothelium | Acyp2 | 1.25 | 4.50E-34 | 9.37E-30 | 0.31 | 0.1 | 0.21 | 3 |
| endothelium | Dennd2c | 1.63 | 5.01E-34 | 1.04E-29 | 0.22 | 0.06 | 0.16 | 3.56 |
| endothelium | B4galt5 | 1.39 | 6.25E-34 | 1.30E-29 | 0.31 | 0.1 | 0.2 | 2.98 |
| endothelium | Siah2 | 1.33 | 7.54E-34 | 1.57E-29 | 0.25 | 0.07 | 0.18 | 3.4 |
| endothelium | Casd1 | 0.91 | 8.62E-34 | 1.79E-29 | 0.78 | 0.5 | 0.28 | 1.57 |
| endothelium | Gm42670 | 2.26 | 1.26E-33 | 2.62E-29 | 0.1 | 0.02 | 0.08 | 5.94 |
| endothelium | Capza2 | 0.62 | 1.46E-33 | 3.03E-29 | 0.92 | 0.86 | 0.06 | 1.07 |
| endothelium | Egr1 | 1.21 | 1.57E-33 | 3.27E-29 | 0.25 | 0.07 | 0.17 | 3.36 |
| endothelium | Gab2 | 1.62 | 2.08E-33 | 4.32E-29 | 0.18 | 0.04 | 0.14 | 4.07 |
| endothelium | Zfp36l2 | 0.96 | 2.61E-33 | 5.43E-29 | 0.73 | 0.43 | 0.3 | 1.7 |
| endothelium | Gnai1 | 1.08 | 2.69E-33 | 5.59E-29 | 0.39 | 0.15 | 0.24 | 2.61 |
| endothelium | Rabac1 | 0.94 | 2.94E-33 | 6.11E-29 | 0.8 | 0.55 | 0.25 | 1.45 |
| endothelium | Mthfd1l | 1.25 | 3.43E-33 | 7.13E-29 | 0.4 | 0.16 | 0.25 | 2.58 |
| endothelium | Rit1 | 1.21 | 3.52E-33 | 7.32E-29 | 0.53 | 0.25 | 0.28 | 2.12 |
| endothelium | Ptk2 | 1.05 | 3.75E-33 | 7.80E-29 | 0.78 | 0.53 | 0.25 | 1.46 |
| endothelium | Ccdc126 | 1.55 | 4.10E-33 | 8.53E-29 | 0.27 | 0.09 | 0.19 | 3.2 |
| endothelium | Gatm | 1.29 | 5.46E-33 | 1.14E-28 | 0.17 | 0.04 | 0.13 | 4.23 |
| endothelium | Adamts9 | 0.83 | 5.66E-33 | 1.18E-28 | 0.39 | 0.15 | 0.24 | 2.63 |
| endothelium | 200002D01Ri | 1.91 | 8.09E-33 | 1.68E-28 | 0.16 | 0.04 | 0.12 | 4.43 |
| endothelium | Itpr3 | 1.63 | 1.07E-32 | 2.22E-28 | 0.19 | 0.05 | 0.14 | 3.86 |
| endothelium | Nin | 1.1 | 1.25E-32 | 2.59E-28 | 0.55 | 0.27 | 0.29 | 2.07 |
| endothelium | Def6 | 1.97 | 1.26E-32 | 2.63E-28 | 0.16 | 0.04 | 0.13 | 4.29 |
| endothelium | Dnajc3 | 0.94 | 1.28E-32 | 2.66E-28 | 0.77 | 0.52 | 0.25 | 1.49 |
| endothelium | Xpnpep1 | 1.03 | 1.29E-32 | 2.68E-28 | 0.78 | 0.52 | 0.26 | 1.49 |
| endothelium | Aplp2 | 0.77 | 1.63E-32 | 3.38E-28 | 0.84 | 0.82 | 0.02 | 1.02 |
| endothelium | Cers4 | 1.06 | 1.88E-32 | 3.92E-28 | 0.4 | 0.16 | 0.25 | 2.58 |
| endothelium | Atox1 | 0.43 | 1.89E-32 | 3.93E-28 | 0.96 | 0.95 | 0.01 | 1.01 |
| endothelium | Trp53i11 | 1.14 | 2.27E-32 | 4.71E-28 | 0.5 | 0.22 | 0.27 | 2.22 |
| endothelium | Dmtn | 1.64 | 2.29E-32 | 4.77E-28 | 0.22 | 0.06 | 0.16 | 3.45 |
| endothelium | Ubald2 | 0.88 | 2.29E-32 | 4.77E-28 | 0.85 | 0.69 | 0.16 | 1.23 |
| endothelium | Tbc1d4 | 1.29 | 2.85E-32 | 5.93E-28 | 0.32 | 0.11 | 0.21 | 2.86 |
| endothelium | Zfp715 | 1.33 | 3.28E-32 | 6.83E-28 | 0.26 | 0.08 | 0.18 | 3.22 |

**Supplementary Table 2. Marker genes for the major cell types of integrated clustering of E11.5 control and NotchGOF λ-junction cells.**

| | | | | | | | | |
|---|---|---|---|---|---|---|---|---|
| endothelium | Ier2 | 0.8 | 7.26E-32 | 1.51E-27 | 0.88 | 0.79 | 0.09 | 1.11 |
| endothelium | Rnasek | 1.27 | 8.17E-32 | 1.70E-27 | 0.86 | 0.72 | 0.15 | 1.2 |
| endothelium | Tlnrd1 | 0.94 | 8.54E-32 | 1.78E-27 | 0.77 | 0.5 | 0.28 | 1.55 |
| endothelium | Gpi1 | 0.67 | 1.32E-31 | 2.74E-27 | 0.93 | 0.84 | 0.09 | 1.11 |
| endothelium | Mark4 | 1.24 | 1.43E-31 | 2.98E-27 | 0.48 | 0.21 | 0.27 | 2.29 |
| endothelium | Cdr2l | 1.08 | 1.55E-31 | 3.23E-27 | 0.45 | 0.19 | 0.26 | 2.37 |
| endothelium | Vdac2 | 0.53 | 2.18E-31 | 4.53E-27 | 0.95 | 0.96 | -0.01 | 0.99 |
| endothelium | Acsf2 | 1.17 | 2.32E-31 | 4.83E-27 | 0.36 | 0.14 | 0.23 | 2.63 |
| endothelium | Gm17018 | 0.99 | 2.69E-31 | 5.59E-27 | 0.77 | 0.52 | 0.25 | 1.47 |
| endothelium | Jag2 | 0.82 | 2.70E-31 | 5.63E-27 | 0.26 | 0.08 | 0.18 | 3.24 |
| endothelium | Iffo2 | 0.97 | 2.88E-31 | 6.00E-27 | 0.36 | 0.13 | 0.23 | 2.73 |
| endothelium | Fam219a | 1.78 | 3.17E-31 | 6.59E-27 | 0.14 | 0.03 | 0.11 | 4.5 |
| endothelium | Atl1 | 1.44 | 3.35E-31 | 6.96E-27 | 0.18 | 0.05 | 0.13 | 3.85 |
| endothelium | Npl | 1 | 3.38E-31 | 7.03E-27 | 0.29 | 0.1 | 0.19 | 3 |
| endothelium | Gm26632 | 1.73 | 4.86E-31 | 1.01E-26 | 0.13 | 0.03 | 0.1 | 4.85 |
| endothelium | Dnm2 | 1.02 | 5.57E-31 | 1.16E-26 | 0.69 | 0.38 | 0.31 | 1.8 |
| endothelium | Ssr4 | 0.77 | 7.24E-31 | 1.51E-26 | 0.87 | 0.84 | 0.03 | 1.03 |
| endothelium | Abca3 | 1.74 | 8.02E-31 | 1.67E-26 | 0.22 | 0.06 | 0.15 | 3.43 |
| endothelium | Tmem123 | 1.05 | 8.76E-31 | 1.82E-26 | 0.62 | 0.32 | 0.3 | 1.95 |
| endothelium | Wwtr1 | 1.11 | 9.24E-31 | 1.92E-26 | 0.68 | 0.43 | 0.25 | 1.59 |
| endothelium | Acadl | 0.88 | 1.15E-30 | 2.39E-26 | 0.79 | 0.53 | 0.26 | 1.48 |
| endothelium | Stap2 | 2 | 1.46E-30 | 3.03E-26 | 0.25 | 0.08 | 0.17 | 3.18 |
| endothelium | Hpgd | 0.32 | 1.61E-30 | 3.34E-26 | 0.7 | 0.34 | 0.36 | 2.04 |
| endothelium | Mmd | 1.11 | 1.76E-30 | 3.66E-26 | 0.64 | 0.35 | 0.29 | 1.85 |
| endothelium | Slc41a3 | 2 | 1.78E-30 | 3.71E-26 | 0.12 | 0.02 | 0.09 | 5.09 |
| endothelium | Tbrg1 | 0.68 | 1.96E-30 | 4.07E-26 | 0.89 | 0.83 | 0.06 | 1.07 |
| endothelium | Mast4 | 1.04 | 2.40E-30 | 4.98E-26 | 0.72 | 0.45 | 0.27 | 1.61 |
| endothelium | Hivep2 | 1.37 | 2.80E-30 | 5.83E-26 | 0.24 | 0.07 | 0.17 | 3.24 |
| endothelium | Capn2 | 1.1 | 3.34E-30 | 6.94E-26 | 0.73 | 0.45 | 0.28 | 1.62 |
| endothelium | Ccdc12 | 1.03 | 4.98E-30 | 1.04E-25 | 0.85 | 0.7 | 0.15 | 1.21 |
| endothelium | Cltb | 1.1 | 5.23E-30 | 1.09E-25 | 0.43 | 0.18 | 0.25 | 2.38 |
| endothelium | Fzd10 | 1.18 | 5.93E-30 | 1.23E-25 | 0.35 | 0.14 | 0.21 | 2.57 |
| endothelium | Gas7 | 1.02 | 5.93E-30 | 1.23E-25 | 0.19 | 0.05 | 0.14 | 3.71 |
| endothelium | Polr1d | 0.51 | 6.38E-30 | 1.33E-25 | 0.94 | 0.95 | -0.01 | 0.99 |
| endothelium | Fbxo6 | 2.13 | 8.91E-30 | 1.85E-25 | 0.21 | 0.06 | 0.15 | 3.35 |
| endothelium | Mbd2 | 0.98 | 1.08E-29 | 2.25E-25 | 0.81 | 0.56 | 0.25 | 1.46 |
| endothelium | S100a1 | 2.44 | 1.09E-29 | 2.27E-25 | 0.21 | 0.06 | 0.15 | 3.35 |
| endothelium | Prodh | 1.13 | 1.24E-29 | 2.58E-25 | 0.3 | 0.11 | 0.2 | 2.84 |
| endothelium | Nfkb1 | 1.14 | 1.42E-29 | 2.94E-25 | 0.57 | 0.3 | 0.27 | 1.92 |
| endothelium | Raph1 | 1.06 | 1.53E-29 | 3.19E-25 | 0.68 | 0.42 | 0.26 | 1.61 |
| endothelium | St3gal6 | 1.41 | 1.53E-29 | 3.19E-25 | 0.3 | 0.11 | 0.19 | 2.74 |
| endothelium | Ltv1 | 1.09 | 1.74E-29 | 3.61E-25 | 0.72 | 0.46 | 0.26 | 1.56 |
| endothelium | Myc | 1.2 | 1.99E-29 | 4.14E-25 | 0.39 | 0.16 | 0.23 | 2.43 |
| endothelium | Gmppb | 1.18 | 2.46E-29 | 5.11E-25 | 0.44 | 0.19 | 0.25 | 2.34 |
| endothelium | Inka1 | 1.04 | 2.49E-29 | 5.17E-25 | 0.78 | 0.51 | 0.28 | 1.54 |
| endothelium | Lims1 | 1.4 | 2.77E-29 | 5.76E-25 | 0.78 | 0.57 | 0.21 | 1.38 |
| endothelium | Ube2a | 0.9 | 2.83E-29 | 5.89E-25 | 0.85 | 0.71 | 0.14 | 1.2 |
| endothelium | Sh3tc2 | 2.05 | 2.87E-29 | 5.97E-25 | 0.12 | 0.03 | 0.1 | 4.8 |
| endothelium | Arf1 | 0.39 | 3.04E-29 | 6.32E-25 | 0.95 | 0.98 | -0.03 | 0.97 |
| endothelium | Sgms1 | 0.84 | 3.69E-29 | 7.67E-25 | 0.41 | 0.17 | 0.25 | 2.47 |
| endothelium | Tmem184b | 1.17 | 4.05E-29 | 8.43E-25 | 0.43 | 0.18 | 0.25 | 2.35 |
| endothelium | Shroom4 | 1.24 | 4.61E-29 | 9.60E-25 | 0.26 | 0.09 | 0.18 | 3.03 |
| endothelium | Slc30a1 | 1.27 | 4.73E-29 | 9.84E-25 | 0.25 | 0.08 | 0.17 | 3.1 |
| endothelium | Rpl36al | 0.32 | 4.88E-29 | 1.01E-24 | 0.97 | 1 | -0.02 | 0.98 |
| endothelium | Rmc1 | 1.36 | 5.67E-29 | 1.18E-24 | 0.25 | 0.08 | 0.17 | 3.1 |
| endothelium | Pdcd6ip | 0.89 | 6.19E-29 | 1.29E-24 | 0.73 | 0.48 | 0.25 | 1.51 |

**Supplementary Table 2. Marker genes for the major cell types of integrated clustering of E11.5 control and NotchGOF λ-junction cells.**

| | | | | | | | | |
|---|---|---|---|---|---|---|---|---|
| endothelium | Dad1 | 0.5 | 6.33E-29 | 1.32E-24 | 0.9 | 0.94 | -0.04 | 0.96 |
| endothelium | Septin4 | 2.12 | 8.37E-29 | 1.74E-24 | 0.11 | 0.02 | 0.09 | 5.19 |
| endothelium | Ldlrad4 | 1.65 | 8.50E-29 | 1.77E-24 | 0.14 | 0.03 | 0.11 | 4.39 |
| endothelium | Plekho1 | 0.92 | 1.16E-28 | 2.41E-24 | 0.81 | 0.61 | 0.19 | 1.32 |
| endothelium | Rreb1 | 1.1 | 1.39E-28 | 2.90E-24 | 0.46 | 0.21 | 0.26 | 2.26 |
| endothelium | Hipk3 | 1.06 | 1.82E-28 | 3.78E-24 | 0.38 | 0.15 | 0.23 | 2.52 |
| endothelium | Slk | 0.89 | 2.32E-28 | 4.83E-24 | 0.77 | 0.53 | 0.24 | 1.45 |
| endothelium | Acsl1 | 1.03 | 2.37E-28 | 4.92E-24 | 0.29 | 0.1 | 0.19 | 2.85 |
| endothelium | Capza1 | 0.81 | 2.92E-28 | 6.08E-24 | 0.82 | 0.64 | 0.18 | 1.28 |
| endothelium | Rab5c | 0.81 | 3.43E-28 | 7.13E-24 | 0.81 | 0.65 | 0.16 | 1.25 |
| endothelium | Nek6 | 1.32 | 4.02E-28 | 8.37E-24 | 0.58 | 0.31 | 0.27 | 1.89 |
| endothelium | Slc8b1 | 2.27 | 5.22E-28 | 1.09E-23 | 0.12 | 0.02 | 0.09 | 4.79 |
| endothelium | Rhbdf1 | 1.05 | 5.36E-28 | 1.11E-23 | 0.28 | 0.1 | 0.19 | 2.89 |
| endothelium | Txn1 | 0.33 | 5.86E-28 | 1.22E-23 | 0.96 | 0.99 | -0.03 | 0.97 |
| endothelium | Ninj1 | 0.97 | 6.13E-28 | 1.27E-23 | 0.44 | 0.19 | 0.25 | 2.3 |
| endothelium | Kbtbd11 | 1.75 | 7.09E-28 | 1.48E-23 | 0.13 | 0.03 | 0.1 | 4.46 |
| endothelium | Rilpl1 | 1.23 | 7.18E-28 | 1.49E-23 | 0.56 | 0.29 | 0.28 | 1.97 |
| endothelium | Lck | 1.76 | 8.23E-28 | 1.71E-23 | 0.21 | 0.07 | 0.15 | 3.28 |
| endothelium | Grpel1 | 0.67 | 8.27E-28 | 1.72E-23 | 0.88 | 0.83 | 0.05 | 1.06 |
| endothelium | Abhd5 | 1.03 | 1.01E-27 | 2.09E-23 | 0.44 | 0.2 | 0.24 | 2.24 |
| endothelium | Tcf4 | 0.62 | 1.07E-27 | 2.23E-23 | 0.89 | 0.9 | -0.01 | 0.98 |
| endothelium | Engase | 1.78 | 1.11E-27 | 2.30E-23 | 0.16 | 0.04 | 0.12 | 3.88 |
| endothelium | Ccdc125 | 1.34 | 1.23E-27 | 2.55E-23 | 0.2 | 0.06 | 0.14 | 3.38 |
| endothelium | Tprgl | 0.87 | 1.26E-27 | 2.62E-23 | 0.79 | 0.6 | 0.19 | 1.31 |
| endothelium | Ssbp4 | 0.82 | 1.38E-27 | 2.86E-23 | 0.84 | 0.65 | 0.19 | 1.29 |
| endothelium | Adam10 | 0.9 | 1.41E-27 | 2.93E-23 | 0.76 | 0.53 | 0.23 | 1.42 |
| endothelium | Carhsp1 | 0.89 | 1.85E-27 | 3.85E-23 | 0.81 | 0.59 | 0.22 | 1.37 |
| endothelium | Fkbp10 | 0.87 | 1.98E-27 | 4.11E-23 | 0.62 | 0.33 | 0.28 | 1.85 |
| endothelium | Fscn1 | 0.57 | 2.01E-27 | 4.18E-23 | 0.9 | 0.91 | -0.02 | 0.98 |
| endothelium | Myd88 | 1.24 | 2.15E-27 | 4.46E-23 | 0.18 | 0.05 | 0.13 | 3.54 |
| endothelium | Gch1 | 1.33 | 2.89E-27 | 6.00E-23 | 0.23 | 0.08 | 0.16 | 3.09 |
| endothelium | Zbtb38 | 1.01 | 2.89E-27 | 6.01E-23 | 0.34 | 0.13 | 0.21 | 2.59 |
| endothelium | Ophn1 | 0.96 | 3.18E-27 | 6.61E-23 | 0.44 | 0.19 | 0.25 | 2.31 |
| endothelium | Enc1 | 0.99 | 3.43E-27 | 7.14E-23 | 0.48 | 0.22 | 0.26 | 2.19 |
| endothelium | Tmed3 | 0.77 | 3.90E-27 | 8.12E-23 | 0.73 | 0.44 | 0.29 | 1.65 |
| endothelium | Desi2 | 0.84 | 4.21E-27 | 8.75E-23 | 0.7 | 0.44 | 0.26 | 1.59 |
| endothelium | Ccnd3 | 0.73 | 4.90E-27 | 1.02E-22 | 0.91 | 0.85 | 0.06 | 1.06 |
| endothelium | Iqsec1 | 1.13 | 6.04E-27 | 1.26E-22 | 0.34 | 0.14 | 0.21 | 2.55 |
| endothelium | Enox2 | 1.19 | 6.67E-27 | 1.39E-22 | 0.22 | 0.07 | 0.15 | 3.22 |
| endothelium | Ddx41 | 0.97 | 6.72E-27 | 1.40E-22 | 0.78 | 0.56 | 0.22 | 1.39 |
| endothelium | Calm2 | 0.4 | 7.07E-27 | 1.47E-22 | 0.96 | 0.98 | -0.02 | 0.98 |
| endothelium | Vcl | 1.38 | 7.77E-27 | 1.62E-22 | 0.67 | 0.39 | 0.28 | 1.71 |
| endothelium | Cd302 | 0.94 | 9.52E-27 | 1.98E-22 | 0.56 | 0.28 | 0.28 | 1.99 |
| endothelium | Rdx | 0.7 | 1.14E-26 | 2.37E-22 | 0.92 | 0.96 | -0.05 | 0.95 |
| endothelium | Kti12 | 0.98 | 1.27E-26 | 2.65E-22 | 0.69 | 0.44 | 0.25 | 1.57 |
| endothelium | Rmnd5b | 1.09 | 1.31E-26 | 2.72E-22 | 0.4 | 0.17 | 0.23 | 2.36 |
| endothelium | C2cd2 | 1.38 | 1.38E-26 | 2.86E-22 | 0.28 | 0.1 | 0.18 | 2.77 |
| endothelium | Hk1 | 1.09 | 1.39E-26 | 2.90E-22 | 0.51 | 0.25 | 0.26 | 2.06 |
| endothelium | Ifitm3 | 0.69 | 1.60E-26 | 3.33E-22 | 0.59 | 0.29 | 0.29 | 2 |
| endothelium | Numb | 0.89 | 1.68E-26 | 3.50E-22 | 0.59 | 0.32 | 0.28 | 1.88 |
| endothelium | Atp5g1 | 0.43 | 2.18E-26 | 4.53E-22 | 0.97 | 0.99 | -0.02 | 0.98 |
| endothelium | Ass1 | 1.43 | 2.80E-26 | 5.83E-22 | 0.19 | 0.06 | 0.14 | 3.37 |
| endothelium | Mospd1 | 0.96 | 2.82E-26 | 5.86E-22 | 0.51 | 0.24 | 0.27 | 2.09 |
| endothelium | Tnfaip1 | 0.89 | 2.89E-26 | 6.01E-22 | 0.79 | 0.62 | 0.17 | 1.27 |
| endothelium | Fam3c | 0.91 | 3.68E-26 | 7.65E-22 | 0.65 | 0.36 | 0.29 | 1.8 |
| endothelium | Rwdd2a | 1.83 | 3.83E-26 | 7.97E-22 | 0.1 | 0.02 | 0.08 | 4.95 |

**Supplementary Table 2. Marker genes for the major cell types of integrated clustering of E11.5 control and NotchGOF λ-junction cells.**

| | | | | | | | | |
|---|---|---|---|---|---|---|---|---|
| endothelium | Map3k3 | 1.32 | 3.85E-26 | 8.00E-22 | 0.37 | 0.15 | 0.21 | 2.4 |
| endothelium | Myo10 | 0.94 | 4.28E-26 | 8.91E-22 | 0.7 | 0.47 | 0.23 | 1.5 |
| endothelium | Rps6ka2 | 1.3 | 4.59E-26 | 9.54E-22 | 0.24 | 0.08 | 0.16 | 3.01 |
| endothelium | Bahcc1 | 1.07 | 4.68E-26 | 9.74E-22 | 0.52 | 0.27 | 0.25 | 1.93 |
| endothelium | Anxa11 | 1.29 | 4.76E-26 | 9.90E-22 | 0.37 | 0.15 | 0.22 | 2.47 |
| endothelium | Eef1akmt4 | 0.96 | 4.99E-26 | 1.04E-21 | 0.49 | 0.23 | 0.26 | 2.12 |
| endothelium | Ampd2 | 0.76 | 5.30E-26 | 1.10E-21 | 0.38 | 0.16 | 0.23 | 2.48 |
| endothelium | Ppp1ca | 0.36 | 5.71E-26 | 1.19E-21 | 0.95 | 0.98 | -0.04 | 0.96 |
| endothelium | Naa20 | 0.85 | 6.03E-26 | 1.25E-21 | 0.85 | 0.72 | 0.13 | 1.18 |
| endothelium | Msrb3 | 1.26 | 6.20E-26 | 1.29E-21 | 0.3 | 0.11 | 0.19 | 2.67 |
| endothelium | Bambi | 0.69 | 8.03E-26 | 1.67E-21 | 0.82 | 0.64 | 0.18 | 1.29 |
| endothelium | Ydjc | 1.27 | 8.88E-26 | 1.85E-21 | 0.24 | 0.08 | 0.16 | 2.93 |
| endothelium | Tubb4b | 0.66 | 9.79E-26 | 2.04E-21 | 0.93 | 0.85 | 0.09 | 1.1 |
| endothelium | Fnbp1 | 1.16 | 1.21E-25 | 2.53E-21 | 0.49 | 0.23 | 0.26 | 2.11 |
| endothelium | Slc12a2 | 0.64 | 1.25E-25 | 2.60E-21 | 0.71 | 0.43 | 0.28 | 1.65 |
| endothelium | Prex2 | 0.97 | 1.31E-25 | 2.73E-21 | 0.67 | 0.4 | 0.27 | 1.67 |
| endothelium | Adarb1 | 1.65 | 1.47E-25 | 3.06E-21 | 0.21 | 0.07 | 0.14 | 3.09 |
| endothelium | Slc44a2 | 0.81 | 1.47E-25 | 3.07E-21 | 0.55 | 0.27 | 0.28 | 2.01 |
| endothelium | Ctnna1 | 0.67 | 1.55E-25 | 3.22E-21 | 0.89 | 0.85 | 0.05 | 1.06 |
| endothelium | Cerk | 1.18 | 2.05E-25 | 4.26E-21 | 0.36 | 0.15 | 0.21 | 2.45 |
| endothelium | Stx4a | 1.07 | 2.32E-25 | 4.82E-21 | 0.58 | 0.31 | 0.27 | 1.86 |
| endothelium | Ctnnd1 | 0.85 | 2.46E-25 | 5.12E-21 | 0.71 | 0.46 | 0.25 | 1.54 |
| endothelium | Tmem241 | 1.55 | 2.52E-25 | 5.23E-21 | 0.15 | 0.04 | 0.11 | 3.87 |
| endothelium | Psen2 | 1.23 | 2.66E-25 | 5.52E-21 | 0.23 | 0.08 | 0.15 | 3.03 |
| endothelium | Rell1 | 1.01 | 2.87E-25 | 5.97E-21 | 0.62 | 0.35 | 0.27 | 1.78 |
| endothelium | Septin7 | 0.45 | 2.87E-25 | 5.97E-21 | 0.97 | 0.97 | 0 | 1 |
| endothelium | C1qtnf6 | 0.95 | 3.84E-25 | 7.98E-21 | 0.38 | 0.16 | 0.22 | 2.38 |
| endothelium | AI504432 | 2.1 | 4.12E-25 | 8.57E-21 | 0.15 | 0.04 | 0.11 | 3.71 |
| endothelium | Rpl4 | 0.27 | 4.36E-25 | 9.07E-21 | 0.99 | 1 | -0.01 | 0.99 |
| endothelium | Taok2 | 0.97 | 5.06E-25 | 1.05E-20 | 0.58 | 0.31 | 0.27 | 1.88 |
| endothelium | Impdh1 | 1.22 | 5.64E-25 | 1.17E-20 | 0.46 | 0.21 | 0.25 | 2.17 |
| endothelium | Qk | 0.52 | 6.69E-25 | 1.39E-20 | 0.9 | 0.9 | 0 | 1 |
| endothelium | Dusp1 | 1.14 | 7.19E-25 | 1.50E-20 | 0.54 | 0.29 | 0.25 | 1.85 |
| endothelium | Dpagt1 | 1.09 | 7.89E-25 | 1.64E-20 | 0.55 | 0.28 | 0.27 | 1.95 |
| endothelium | Rbm43 | 0.99 | 8.27E-25 | 1.72E-20 | 0.29 | 0.11 | 0.18 | 2.72 |
| endothelium | Ahr | 0.8 | 8.31E-25 | 1.73E-20 | 0.25 | 0.09 | 0.16 | 2.89 |
| endothelium | Dnaja1 | 0.36 | 8.54E-25 | 1.78E-20 | 0.98 | 0.99 | -0.02 | 0.98 |
| endothelium | Cdc42bpa | 0.83 | 1.04E-24 | 2.16E-20 | 0.65 | 0.37 | 0.27 | 1.72 |
| endothelium | Nhsl2 | 0.94 | 1.20E-24 | 2.50E-20 | 0.37 | 0.16 | 0.21 | 2.33 |
| endothelium | Gmip | 1.43 | 1.47E-24 | 3.06E-20 | 0.18 | 0.05 | 0.13 | 3.35 |
| endothelium | 610012G03Ri | 0.61 | 1.50E-24 | 3.13E-20 | 0.92 | 0.86 | 0.05 | 1.06 |
| endothelium | Txnip | 0.94 | 1.55E-24 | 3.23E-20 | 0.62 | 0.34 | 0.28 | 1.84 |
| endothelium | Fam53b | 0.91 | 1.88E-24 | 3.92E-20 | 0.4 | 0.18 | 0.23 | 2.29 |
| endothelium | Aldoa | 0.54 | 1.93E-24 | 4.02E-20 | 0.97 | 0.95 | 0.02 | 1.02 |
| endothelium | Dennd5b | 0.92 | 2.30E-24 | 4.79E-20 | 0.65 | 0.39 | 0.26 | 1.68 |
| endothelium | Pygb | 1.1 | 2.32E-24 | 4.82E-20 | 0.48 | 0.23 | 0.25 | 2.1 |
| endothelium | Tram1 | 0.89 | 2.64E-24 | 5.50E-20 | 0.73 | 0.5 | 0.23 | 1.45 |
| endothelium | Fry | 1.15 | 2.96E-24 | 6.16E-20 | 0.18 | 0.05 | 0.13 | 3.35 |
| endothelium | Bcar1 | 1.21 | 3.25E-24 | 6.76E-20 | 0.49 | 0.24 | 0.25 | 2.02 |
| endothelium | Comt | 0.76 | 3.92E-24 | 8.15E-20 | 0.77 | 0.53 | 0.23 | 1.43 |
| endothelium | Nomo1 | 1.34 | 4.02E-24 | 8.37E-20 | 0.42 | 0.2 | 0.22 | 2.14 |
| endothelium | Gm43259 | 1.79 | 4.89E-24 | 1.02E-19 | 0.11 | 0.02 | 0.08 | 4.65 |
| endothelium | Irak4 | 1.39 | 5.12E-24 | 1.07E-19 | 0.16 | 0.05 | 0.12 | 3.54 |
| endothelium | Litaf | 0.89 | 5.72E-24 | 1.19E-19 | 0.74 | 0.5 | 0.24 | 1.49 |
| endothelium | Ap2m1 | 0.58 | 6.23E-24 | 1.30E-19 | 0.91 | 0.86 | 0.05 | 1.06 |
| endothelium | Nras | 0.73 | 6.37E-24 | 1.32E-19 | 0.85 | 0.69 | 0.16 | 1.23 |

**Supplementary Table 2. Marker genes for the major cell types of integrated clustering of E11.5 control and NotchGOF λ-junction cells.**

| | | | | | | | | |
|---|---|---|---|---|---|---|---|---|
| endothelium | Ttc7b | 1.15 | 6.70E-24 | 1.39E-19 | 0.36 | 0.16 | 0.21 | 2.33 |
| endothelium | 330009N23Ri | 1.42 | 6.77E-24 | 1.41E-19 | 0.11 | 0.02 | 0.08 | 4.65 |
| endothelium | Fam57a | 1.2 | 7.46E-24 | 1.55E-19 | 0.33 | 0.14 | 0.19 | 2.38 |
| endothelium | Blvra | 1.16 | 1.29E-23 | 2.67E-19 | 0.42 | 0.2 | 0.22 | 2.1 |
| endothelium | Tgfbr3 | 1.03 | 1.42E-23 | 2.95E-19 | 0.23 | 0.08 | 0.15 | 2.9 |
| endothelium | Rab2a | 0.49 | 1.65E-23 | 3.44E-19 | 0.92 | 0.92 | 0.01 | 1.01 |
| endothelium | Wdfy1 | 0.83 | 1.75E-23 | 3.64E-19 | 0.63 | 0.35 | 0.29 | 1.82 |
| endothelium | Rab1a | 0.7 | 1.85E-23 | 3.85E-19 | 0.87 | 0.71 | 0.16 | 1.22 |
| endothelium | Ufm1 | 0.63 | 2.12E-23 | 4.40E-19 | 0.84 | 0.69 | 0.15 | 1.22 |
| endothelium | Inpp5a | 1.2 | 2.38E-23 | 4.95E-19 | 0.35 | 0.15 | 0.2 | 2.36 |
| endothelium | Map4 | 0.97 | 2.60E-23 | 5.41E-19 | 0.66 | 0.39 | 0.27 | 1.68 |
| endothelium | Ephb4 | 0.95 | 2.98E-23 | 6.19E-19 | 0.65 | 0.39 | 0.26 | 1.65 |
| endothelium | Cpeb4 | 1.38 | 3.37E-23 | 7.00E-19 | 0.25 | 0.09 | 0.16 | 2.67 |
| endothelium | Nagpa | 0.86 | 3.76E-23 | 7.81E-19 | 0.27 | 0.1 | 0.17 | 2.72 |
| endothelium | Arhgap1 | 0.97 | 3.94E-23 | 8.19E-19 | 0.51 | 0.26 | 0.25 | 1.97 |
| endothelium | Cox5b | 0.33 | 4.30E-23 | 8.94E-19 | 0.97 | 0.99 | -0.01 | 0.99 |
| endothelium | Fam171a2 | 0.85 | 4.37E-23 | 9.09E-19 | 0.68 | 0.45 | 0.23 | 1.51 |
| endothelium | Gpd1l | 1.27 | 4.96E-23 | 1.03E-18 | 0.21 | 0.07 | 0.14 | 2.97 |
| endothelium | Nckipsd | 1.1 | 5.02E-23 | 1.04E-18 | 0.2 | 0.07 | 0.14 | 3.08 |
| endothelium | Bicd2 | 0.96 | 5.25E-23 | 1.09E-18 | 0.56 | 0.29 | 0.27 | 1.95 |
| endothelium | Fosl2 | 1.36 | 5.53E-23 | 1.15E-18 | 0.12 | 0.03 | 0.09 | 4.14 |
| endothelium | Emc7 | 0.86 | 5.69E-23 | 1.18E-18 | 0.73 | 0.49 | 0.23 | 1.47 |
| endothelium | Snx7 | 1.02 | 6.20E-23 | 1.29E-18 | 0.27 | 0.1 | 0.17 | 2.66 |
| endothelium | Acsl4 | 0.97 | 6.25E-23 | 1.30E-18 | 0.33 | 0.14 | 0.19 | 2.4 |
| endothelium | Ccnd1 | 0.63 | 8.35E-23 | 1.74E-18 | 0.84 | 0.8 | 0.04 | 1.05 |
| endothelium | Cds2 | 1.02 | 8.81E-23 | 1.83E-18 | 0.4 | 0.18 | 0.22 | 2.25 |
| endothelium | Irak2 | 1.4 | 8.91E-23 | 1.85E-18 | 0.17 | 0.05 | 0.12 | 3.29 |
| endothelium | Ckb | 0.66 | 9.66E-23 | 2.01E-18 | 0.59 | 0.33 | 0.26 | 1.79 |
| endothelium | Capg | 0.75 | 1.17E-22 | 2.44E-18 | 0.35 | 0.15 | 0.2 | 2.36 |
| endothelium | 933415A04Ri | 1.13 | 1.20E-22 | 2.50E-18 | 0.21 | 0.07 | 0.14 | 2.97 |
| endothelium | Anxa1 | 1.75 | 1.24E-22 | 2.58E-18 | 0.11 | 0.03 | 0.09 | 4.15 |
| endothelium | Lats2 | 1.07 | 1.28E-22 | 2.67E-18 | 0.41 | 0.18 | 0.22 | 2.21 |
| endothelium | Nrbp1 | 0.86 | 1.51E-22 | 3.14E-18 | 0.77 | 0.56 | 0.22 | 1.39 |
| endothelium | Ifnar2 | 0.89 | 1.58E-22 | 3.29E-18 | 0.58 | 0.33 | 0.26 | 1.79 |
| endothelium | Ptrhd1 | 0.9 | 1.79E-22 | 3.73E-18 | 0.61 | 0.36 | 0.26 | 1.72 |
| endothelium | Rtl8b | 0.86 | 1.97E-22 | 4.10E-18 | 0.59 | 0.33 | 0.26 | 1.79 |
| endothelium | Uchl5 | 0.77 | 2.07E-22 | 4.32E-18 | 0.75 | 0.54 | 0.21 | 1.39 |
| endothelium | Rnf19a | 0.91 | 2.42E-22 | 5.03E-18 | 0.45 | 0.22 | 0.23 | 2.07 |
| endothelium | Cox5a | 0.32 | 2.60E-22 | 5.41E-18 | 0.98 | 0.99 | -0.01 | 0.99 |
| endothelium | Atp6v0e | 0.75 | 2.72E-22 | 5.65E-18 | 0.83 | 0.72 | 0.12 | 1.16 |
| endothelium | Ctso | 1.07 | 2.89E-22 | 6.01E-18 | 0.2 | 0.06 | 0.13 | 3.05 |
| endothelium | Pdcd10 | 0.8 | 2.98E-22 | 6.19E-18 | 0.83 | 0.72 | 0.11 | 1.15 |
| endothelium | Pla2g6 | 1.05 | 3.20E-22 | 6.65E-18 | 0.22 | 0.08 | 0.15 | 2.84 |
| endothelium | Sec14l2 | 1.7 | 3.85E-22 | 8.01E-18 | 0.11 | 0.03 | 0.08 | 4.28 |
| endothelium | Pik3cb | 1.7 | 4.09E-22 | 8.51E-18 | 0.19 | 0.06 | 0.13 | 3.07 |
| endothelium | Riox1 | 0.9 | 4.41E-22 | 9.17E-18 | 0.53 | 0.28 | 0.25 | 1.9 |
| endothelium | Pak2 | 0.57 | 4.66E-22 | 9.70E-18 | 0.88 | 0.84 | 0.05 | 1.06 |
| endothelium | Dusp23 | 1.07 | 5.40E-22 | 1.12E-17 | 0.38 | 0.17 | 0.21 | 2.24 |
| endothelium | Adgrg1 | 2.55 | 5.53E-22 | 1.15E-17 | 0.14 | 0.04 | 0.1 | 3.44 |
| endothelium | Hps3 | 1.14 | 6.20E-22 | 1.29E-17 | 0.23 | 0.08 | 0.15 | 2.84 |
| endothelium | Rabep2 | 0.89 | 6.85E-22 | 1.42E-17 | 0.41 | 0.19 | 0.22 | 2.16 |
| endothelium | Naa38 | 0.6 | 7.13E-22 | 1.48E-17 | 0.85 | 0.74 | 0.11 | 1.14 |
| endothelium | Yaf2 | 0.74 | 7.67E-22 | 1.59E-17 | 0.86 | 0.7 | 0.16 | 1.24 |
| endothelium | Stx7 | 0.94 | 7.70E-22 | 1.60E-17 | 0.56 | 0.31 | 0.25 | 1.82 |
| endothelium | Rassf1 | 1.06 | 7.83E-22 | 1.63E-17 | 0.65 | 0.4 | 0.25 | 1.64 |
| endothelium | Plcg1 | 0.79 | 7.87E-22 | 1.64E-17 | 0.65 | 0.4 | 0.25 | 1.64 |

**Supplementary Table 2. Marker genes for the major cell types of integrated clustering of E11.5 control and NotchGOF λ-junction cells.**

| | | | | | | | | |
|---|---|---|---|---|---|---|---|---|
| endothelium | Pcbp3 | 1.01 | 8.13E-22 | 1.69E-17 | 0.38 | 0.17 | 0.21 | 2.21 |
| endothelium | Top1mt | 1.4 | 8.33E-22 | 1.73E-17 | 0.17 | 0.06 | 0.12 | 3.15 |
| endothelium | Snx4 | 0.78 | 9.58E-22 | 1.99E-17 | 0.69 | 0.42 | 0.27 | 1.65 |
| endothelium | Cycs | 0.46 | 1.01E-21 | 2.10E-17 | 0.97 | 0.96 | 0.01 | 1.01 |
| endothelium | Optn | 2.37 | 1.02E-21 | 2.13E-17 | 0.11 | 0.03 | 0.08 | 4 |
| endothelium | Edf1 | 0.35 | 1.08E-21 | 2.25E-17 | 0.94 | 0.96 | -0.02 | 0.98 |
| endothelium | Btbd10 | 0.81 | 1.22E-21 | 2.54E-17 | 0.49 | 0.25 | 0.25 | 2.01 |
| endothelium | Cdr2 | 1.23 | 1.60E-21 | 3.32E-17 | 0.44 | 0.22 | 0.22 | 2.04 |
| endothelium | Sat1 | 0.77 | 1.61E-21 | 3.35E-17 | 0.57 | 0.3 | 0.26 | 1.87 |
| endothelium | Orai3 | 1.17 | 1.61E-21 | 3.36E-17 | 0.28 | 0.11 | 0.17 | 2.5 |
| endothelium | Lta4h | 0.84 | 1.65E-21 | 3.42E-17 | 0.69 | 0.46 | 0.22 | 1.48 |
| endothelium | Man2a1 | 0.92 | 2.29E-21 | 4.75E-17 | 0.47 | 0.23 | 0.24 | 2.03 |
| endothelium | Eps15 | 0.75 | 2.37E-21 | 4.92E-17 | 0.56 | 0.3 | 0.26 | 1.86 |
| endothelium | Slc35d2 | 1.43 | 2.39E-21 | 4.97E-17 | 0.11 | 0.03 | 0.08 | 4.28 |
| endothelium | Tmod3 | 0.78 | 2.54E-21 | 5.29E-17 | 0.81 | 0.64 | 0.17 | 1.27 |
| endothelium | Stc1 | 1.59 | 2.62E-21 | 5.44E-17 | 0.2 | 0.07 | 0.13 | 2.9 |
| endothelium | Hspa4l | 1.22 | 3.24E-21 | 6.74E-17 | 0.27 | 0.11 | 0.16 | 2.47 |
| endothelium | Srgap2 | 0.79 | 3.58E-21 | 7.45E-17 | 0.61 | 0.36 | 0.26 | 1.73 |
| endothelium | Ppib | 0.31 | 3.64E-21 | 7.56E-17 | 0.93 | 0.98 | -0.05 | 0.94 |
| endothelium | Rab35 | 0.83 | 3.92E-21 | 8.16E-17 | 0.62 | 0.36 | 0.26 | 1.71 |
| endothelium | Tsc22d4 | 0.67 | 4.18E-21 | 8.69E-17 | 0.8 | 0.61 | 0.19 | 1.3 |
| endothelium | Dynlt3 | 1.01 | 4.28E-21 | 8.90E-17 | 0.49 | 0.25 | 0.24 | 1.93 |
| endothelium | Nme1 | 0.34 | 4.30E-21 | 8.95E-17 | 0.97 | 0.99 | -0.02 | 0.98 |
| endothelium | Map3k14 | 2.08 | 4.47E-21 | 9.30E-17 | 0.13 | 0.04 | 0.1 | 3.64 |
| endothelium | Rap1gds1 | 0.8 | 4.50E-21 | 9.36E-17 | 0.55 | 0.29 | 0.26 | 1.91 |
| endothelium | Fam122b | 1.18 | 4.96E-21 | 1.03E-16 | 0.22 | 0.08 | 0.14 | 2.73 |
| endothelium | Cables1 | 1.26 | 5.08E-21 | 1.06E-16 | 0.23 | 0.08 | 0.15 | 2.73 |
| endothelium | B4galt1 | 0.89 | 5.31E-21 | 1.10E-16 | 0.45 | 0.23 | 0.23 | 2 |
| endothelium | Gna12 | 0.9 | 6.06E-21 | 1.26E-16 | 0.56 | 0.31 | 0.25 | 1.82 |
| endothelium | Whamm | 1.13 | 6.14E-21 | 1.28E-16 | 0.25 | 0.1 | 0.16 | 2.58 |
| endothelium | Stat6 | 1.04 | 6.58E-21 | 1.37E-16 | 0.17 | 0.05 | 0.12 | 3.17 |
| endothelium | Pcp4l1 | 1.09 | 8.33E-21 | 1.73E-16 | 0.1 | 0.02 | 0.08 | 4.21 |
| endothelium | Mapre1 | 0.51 | 9.06E-21 | 1.88E-16 | 0.93 | 0.91 | 0.02 | 1.03 |
| endothelium | Ppp1r14b | 0.31 | 9.68E-21 | 2.01E-16 | 0.96 | 0.99 | -0.03 | 0.97 |
| endothelium | Utp11 | 0.56 | 9.99E-21 | 2.08E-16 | 0.86 | 0.77 | 0.09 | 1.12 |
| endothelium | Nagk | 0.97 | 1.03E-20 | 2.15E-16 | 0.41 | 0.2 | 0.21 | 2.08 |
| endothelium | Plcb1 | 1.41 | 1.11E-20 | 2.31E-16 | 0.25 | 0.09 | 0.15 | 2.61 |
| endothelium | Igfbp4 | 0.43 | 1.23E-20 | 2.55E-16 | 0.91 | 0.9 | 0 | 1 |
| endothelium | Tmem14c | 0.82 | 1.32E-20 | 2.75E-16 | 0.89 | 0.92 | -0.03 | 0.97 |
| endothelium | Nedd9 | 0.8 | 1.58E-20 | 3.29E-16 | 0.36 | 0.17 | 0.2 | 2.19 |
| endothelium | Cers5 | 0.66 | 1.80E-20 | 3.74E-16 | 0.61 | 0.34 | 0.27 | 1.8 |
| endothelium | Ctsd | 0.65 | 2.28E-20 | 4.74E-16 | 0.78 | 0.61 | 0.18 | 1.29 |
| endothelium | Reep3 | 0.76 | 2.33E-20 | 4.85E-16 | 0.76 | 0.52 | 0.24 | 1.47 |
| endothelium | Tgds | 0.85 | 2.39E-20 | 4.98E-16 | 0.41 | 0.2 | 0.22 | 2.1 |
| endothelium | Babam1 | 0.72 | 2.48E-20 | 5.17E-16 | 0.79 | 0.58 | 0.21 | 1.36 |
| endothelium | Cish | 1.09 | 2.65E-20 | 5.51E-16 | 0.18 | 0.06 | 0.12 | 2.97 |
| endothelium | Kdelr2 | 0.5 | 3.17E-20 | 6.59E-16 | 0.85 | 0.83 | 0.02 | 1.02 |
| endothelium | Arf6 | 0.89 | 3.47E-20 | 7.21E-16 | 0.64 | 0.39 | 0.25 | 1.63 |
| endothelium | Alad | 0.47 | 3.83E-20 | 7.96E-16 | 0.82 | 0.63 | 0.18 | 1.29 |
| endothelium | Stk25 | 0.86 | 3.91E-20 | 8.14E-16 | 0.71 | 0.45 | 0.26 | 1.58 |
| endothelium | Zdhhc12 | 0.75 | 4.29E-20 | 8.92E-16 | 0.5 | 0.26 | 0.24 | 1.93 |
| endothelium | Colgalt1 | 0.82 | 4.58E-20 | 9.53E-16 | 0.71 | 0.47 | 0.24 | 1.5 |
| endothelium | Rheb | 0.53 | 5.01E-20 | 1.04E-15 | 0.93 | 0.87 | 0.07 | 1.08 |
| endothelium | Mcur1 | 1.23 | 5.52E-20 | 1.15E-15 | 0.42 | 0.21 | 0.21 | 1.97 |
| endothelium | Spag9 | 0.72 | 5.81E-20 | 1.21E-15 | 0.79 | 0.62 | 0.17 | 1.27 |
| endothelium | Ccnjl | 0.81 | 6.78E-20 | 1.41E-15 | 0.66 | 0.41 | 0.25 | 1.6 |

**Supplementary Table 2. Marker genes for the major cell types of integrated clustering of E11.5 control and NotchGOF λ-junction cells.**

| | | | | | | | | |
|---|---|---|---|---|---|---|---|---|
| endothelium | Lman2 | 0.67 | 7.44E-20 | 1.55E-15 | 0.78 | 0.62 | 0.17 | 1.27 |
| endothelium | Ucp2 | 0.29 | 8.41E-20 | 1.75E-15 | 0.64 | 0.35 | 0.29 | 1.81 |
| endothelium | Hmg20b | 0.71 | 8.49E-20 | 1.77E-15 | 0.89 | 0.8 | 0.09 | 1.11 |
| endothelium | Pacc1 | 0.92 | 8.58E-20 | 1.79E-15 | 0.24 | 0.09 | 0.15 | 2.61 |
| endothelium | Pkn1 | 0.7 | 8.77E-20 | 1.82E-15 | 0.61 | 0.35 | 0.26 | 1.73 |
| endothelium | B4galt7 | 0.69 | 9.09E-20 | 1.89E-15 | 0.41 | 0.19 | 0.22 | 2.13 |
| endothelium | Ran | 0.28 | 9.97E-20 | 2.07E-15 | 0.98 | 1 | -0.01 | 0.99 |
| endothelium | Mri1 | 0.72 | 1.25E-19 | 2.60E-15 | 0.52 | 0.27 | 0.25 | 1.91 |
| endothelium | Slc9a9 | 1.37 | 1.26E-19 | 2.63E-15 | 0.13 | 0.04 | 0.09 | 3.54 |
| endothelium | Cltc | 0.63 | 1.34E-19 | 2.78E-15 | 0.85 | 0.75 | 0.1 | 1.13 |
| endothelium | Shisa5 | 0.57 | 1.49E-19 | 3.10E-15 | 0.48 | 0.24 | 0.24 | 2 |
| endothelium | Lamp1 | 0.53 | 1.57E-19 | 3.27E-15 | 0.87 | 0.86 | 0.01 | 1.01 |
| endothelium | Rnpep | 0.83 | 1.60E-19 | 3.32E-15 | 0.66 | 0.43 | 0.23 | 1.52 |
| endothelium | Haus4 | 0.84 | 1.93E-19 | 4.00E-15 | 0.56 | 0.31 | 0.24 | 1.78 |
| endothelium | Tifa | 1.5 | 1.97E-19 | 4.10E-15 | 0.11 | 0.03 | 0.08 | 3.89 |
| endothelium | Nfe2l2 | 0.73 | 2.05E-19 | 4.27E-15 | 0.7 | 0.44 | 0.26 | 1.59 |
| endothelium | Dnajb11 | 0.71 | 2.09E-19 | 4.35E-15 | 0.82 | 0.68 | 0.14 | 1.2 |
| endothelium | Tnip1 | 1.54 | 2.30E-19 | 4.78E-15 | 0.25 | 0.1 | 0.15 | 2.5 |
| endothelium | Gls | 0.79 | 2.37E-19 | 4.92E-15 | 0.64 | 0.41 | 0.24 | 1.58 |
| endothelium | Srp9 | 0.47 | 2.50E-19 | 5.21E-15 | 0.91 | 0.92 | -0.01 | 0.99 |
| endothelium | Rab5if | 0.48 | 2.67E-19 | 5.56E-15 | 0.89 | 0.91 | -0.02 | 0.97 |
| endothelium | Saraf | 0.6 | 3.36E-19 | 6.98E-15 | 0.76 | 0.54 | 0.22 | 1.41 |
| endothelium | Ube2d3 | 0.37 | 3.46E-19 | 7.20E-15 | 0.96 | 0.97 | -0.01 | 0.99 |
| endothelium | Acat1 | 0.72 | 3.47E-19 | 7.21E-15 | 0.7 | 0.46 | 0.24 | 1.51 |
| endothelium | Srm | 0.81 | 3.62E-19 | 7.53E-15 | 0.82 | 0.68 | 0.15 | 1.22 |
| endothelium | Fkbp7 | 0.66 | 3.62E-19 | 7.54E-15 | 0.52 | 0.28 | 0.24 | 1.87 |
| endothelium | Cers2 | 0.63 | 4.23E-19 | 8.80E-15 | 0.75 | 0.54 | 0.22 | 1.4 |
| endothelium | Zdhhc9 | 1 | 4.40E-19 | 9.16E-15 | 0.23 | 0.09 | 0.14 | 2.6 |
| endothelium | Zfpm1 | 0.97 | 5.07E-19 | 1.06E-14 | 0.54 | 0.31 | 0.23 | 1.74 |
| endothelium | Trak2 | 0.92 | 5.85E-19 | 1.22E-14 | 0.31 | 0.13 | 0.17 | 2.29 |
| endothelium | Cgrrf1 | 0.8 | 7.56E-19 | 1.57E-14 | 0.49 | 0.25 | 0.24 | 1.93 |
| endothelium | Atp6v0a2 | 0.83 | 9.61E-19 | 2.00E-14 | 0.49 | 0.26 | 0.23 | 1.88 |
| endothelium | Alg1 | 0.8 | 1.03E-18 | 2.14E-14 | 0.21 | 0.08 | 0.13 | 2.74 |
| endothelium | Rrm2b | 0.99 | 1.04E-18 | 2.16E-14 | 0.29 | 0.12 | 0.17 | 2.36 |
| endothelium | Rps6ka3 | 0.79 | 1.17E-18 | 2.44E-14 | 0.7 | 0.45 | 0.25 | 1.54 |
| endothelium | Amotl1 | 0.6 | 1.36E-18 | 2.82E-14 | 0.78 | 0.69 | 0.09 | 1.13 |
| endothelium | Fundc2 | 0.57 | 1.48E-18 | 3.07E-14 | 0.86 | 0.77 | 0.09 | 1.12 |
| endothelium | Mrps18b | 0.89 | 1.69E-18 | 3.51E-14 | 0.72 | 0.5 | 0.23 | 1.45 |
| endothelium | Ccdc102a | 0.83 | 1.81E-18 | 3.77E-14 | 0.51 | 0.27 | 0.24 | 1.86 |
| endothelium | Aph1a | 0.69 | 1.82E-18 | 3.79E-14 | 0.74 | 0.5 | 0.24 | 1.48 |
| endothelium | Btbd19 | 1.42 | 2.15E-18 | 4.47E-14 | 0.1 | 0.03 | 0.08 | 3.85 |
| endothelium | Gm43328 | 1.59 | 2.20E-18 | 4.57E-14 | 0.1 | 0.03 | 0.08 | 3.85 |
| endothelium | Plaat3 | 0.96 | 2.23E-18 | 4.64E-14 | 0.44 | 0.22 | 0.22 | 1.99 |
| endothelium | Nudt9 | 0.76 | 2.66E-18 | 5.53E-14 | 0.71 | 0.48 | 0.23 | 1.48 |
| endothelium | Smdt1 | 0.39 | 2.84E-18 | 5.91E-14 | 0.93 | 0.94 | -0.01 | 0.98 |
| endothelium | Cflar | 0.69 | 2.91E-18 | 6.05E-14 | 0.55 | 0.31 | 0.24 | 1.78 |
| endothelium | Mtmr6 | 1 | 2.91E-18 | 6.06E-14 | 0.3 | 0.13 | 0.17 | 2.28 |
| endothelium | Fignl2 | 0.83 | 3.39E-18 | 7.05E-14 | 0.29 | 0.12 | 0.16 | 2.32 |
| endothelium | Bcat2 | 0.66 | 3.56E-18 | 7.40E-14 | 0.53 | 0.29 | 0.24 | 1.84 |
| endothelium | Uqcrb | 0.31 | 3.83E-18 | 7.96E-14 | 0.96 | 0.98 | -0.03 | 0.97 |
| endothelium | Apbb2 | 0.8 | 4.31E-18 | 8.97E-14 | 0.53 | 0.3 | 0.23 | 1.76 |
| endothelium | Fam107b | 0.58 | 4.43E-18 | 9.22E-14 | 0.62 | 0.35 | 0.27 | 1.76 |
| endothelium | Slc39a8 | 0.52 | 4.58E-18 | 9.52E-14 | 0.37 | 0.17 | 0.2 | 2.17 |
| endothelium | Snap23 | 0.85 | 6.42E-18 | 1.33E-13 | 0.77 | 0.56 | 0.22 | 1.39 |
| endothelium | Mars2 | 1.11 | 7.38E-18 | 1.54E-13 | 0.19 | 0.07 | 0.12 | 2.74 |
| endothelium | Traf5 | 1.48 | 7.53E-18 | 1.57E-13 | 0.13 | 0.04 | 0.09 | 3.28 |

**Supplementary Table 2. Marker genes for the major cell types of integrated clustering of E11.5 control and NotchGOF λ-junction cells.**

| | | | | | | | | |
|---|---|---|---|---|---|---|---|---|
| endothelium | Zdhhc7 | 0.86 | 7.79E-18 | 1.62E-13 | 0.3 | 0.14 | 0.17 | 2.25 |
| endothelium | Ly96 | 1.05 | 8.84E-18 | 1.84E-13 | 0.17 | 0.06 | 0.11 | 2.95 |
| endothelium | Layn | 1.61 | 1.02E-17 | 2.13E-13 | 0.1 | 0.03 | 0.08 | 3.71 |
| endothelium | Timp1 | 0.69 | 1.04E-17 | 2.16E-13 | 0.36 | 0.17 | 0.19 | 2.14 |
| endothelium | Akt1 | 0.6 | 1.11E-17 | 2.30E-13 | 0.85 | 0.72 | 0.13 | 1.18 |
| endothelium | Eif2s2 | 0.41 | 1.16E-17 | 2.41E-13 | 0.98 | 0.99 | -0.01 | 0.99 |
| endothelium | Fkbp15 | 0.72 | 1.20E-17 | 2.50E-13 | 0.3 | 0.13 | 0.17 | 2.28 |
| endothelium | Yrdc | 0.86 | 1.22E-17 | 2.53E-13 | 0.66 | 0.41 | 0.24 | 1.58 |
| endothelium | Bcl2 | 0.96 | 1.33E-17 | 2.76E-13 | 0.57 | 0.35 | 0.22 | 1.64 |
| endothelium | Manf | 0.48 | 1.43E-17 | 2.97E-13 | 0.88 | 0.9 | -0.01 | 0.98 |
| endothelium | Dubr | 1.77 | 1.43E-17 | 2.97E-13 | 0.12 | 0.04 | 0.09 | 3.42 |
| endothelium | Sash1 | 0.88 | 1.51E-17 | 3.14E-13 | 0.63 | 0.4 | 0.24 | 1.59 |
| endothelium | Eef2kmt | 0.69 | 1.65E-17 | 3.44E-13 | 0.38 | 0.18 | 0.2 | 2.08 |
| endothelium | Vps26c | 0.74 | 1.67E-17 | 3.47E-13 | 0.44 | 0.22 | 0.22 | 1.98 |
| endothelium | Gab1 | 0.99 | 1.78E-17 | 3.69E-13 | 0.38 | 0.19 | 0.19 | 2.02 |
| endothelium | Sik3 | 0.69 | 1.79E-17 | 3.72E-13 | 0.53 | 0.28 | 0.25 | 1.87 |
| endothelium | Rbfa | 1.37 | 1.94E-17 | 4.03E-13 | 0.53 | 0.31 | 0.22 | 1.73 |
| endothelium | Twf1 | 0.65 | 2.07E-17 | 4.31E-13 | 0.72 | 0.49 | 0.23 | 1.48 |
| endothelium | Edem1 | 0.97 | 2.08E-17 | 4.32E-13 | 0.45 | 0.24 | 0.21 | 1.87 |
| endothelium | Scamp2 | 0.66 | 2.15E-17 | 4.46E-13 | 0.56 | 0.31 | 0.25 | 1.8 |
| endothelium | Nfkbie | 1.4 | 2.15E-17 | 4.48E-13 | 0.13 | 0.04 | 0.09 | 3.28 |
| endothelium | Dync1li1 | 0.66 | 2.29E-17 | 4.77E-13 | 0.73 | 0.52 | 0.21 | 1.41 |
| endothelium | Fosb | 1.56 | 2.87E-17 | 5.98E-13 | 0.11 | 0.03 | 0.08 | 3.5 |
| endothelium | Fam241a | 0.92 | 3.00E-17 | 6.24E-13 | 0.27 | 0.12 | 0.15 | 2.31 |
| endothelium | Ilk | 1.17 | 3.43E-17 | 7.14E-13 | 0.8 | 0.61 | 0.19 | 1.32 |
| endothelium | Brk1 | 0.42 | 3.46E-17 | 7.20E-13 | 0.92 | 0.92 | 0 | 1 |
| endothelium | Tspo | 1.16 | 3.56E-17 | 7.41E-13 | 0.58 | 0.35 | 0.23 | 1.66 |
| endothelium | Ubash3b | 1.72 | 3.69E-17 | 7.67E-13 | 0.44 | 0.24 | 0.2 | 1.86 |
| endothelium | Trdmt1 | 1.04 | 3.70E-17 | 7.69E-13 | 0.16 | 0.05 | 0.1 | 2.96 |
| endothelium | Aacs | 0.67 | 3.87E-17 | 8.05E-13 | 0.26 | 0.11 | 0.15 | 2.38 |
| endothelium | Ktn1 | 0.52 | 3.92E-17 | 8.14E-13 | 0.84 | 0.83 | 0.01 | 1.01 |
| endothelium | BC029722 | 1.2 | 4.40E-17 | 9.14E-13 | 0.26 | 0.11 | 0.15 | 2.34 |
| endothelium | Tram2 | 0.78 | 5.76E-17 | 1.20E-12 | 0.25 | 0.1 | 0.14 | 2.43 |
| endothelium | Serpinb6a | 0.68 | 5.77E-17 | 1.20E-12 | 0.63 | 0.39 | 0.24 | 1.63 |
| endothelium | Chmp6 | 0.76 | 6.91E-17 | 1.44E-12 | 0.7 | 0.51 | 0.2 | 1.39 |
| endothelium | Mrpl57 | 0.58 | 7.01E-17 | 1.46E-12 | 0.86 | 0.78 | 0.09 | 1.11 |
| endothelium | Tpcn1 | 0.89 | 7.45E-17 | 1.55E-12 | 0.22 | 0.09 | 0.13 | 2.48 |
| endothelium | Nudt16 | 0.95 | 7.47E-17 | 1.55E-12 | 0.22 | 0.09 | 0.13 | 2.54 |
| endothelium | Map3k1 | 0.82 | 1.17E-16 | 2.44E-12 | 0.48 | 0.26 | 0.22 | 1.83 |
| endothelium | Cd2ap | 0.61 | 1.22E-16 | 2.54E-12 | 0.79 | 0.62 | 0.17 | 1.28 |
| endothelium | Galns | 1.25 | 1.24E-16 | 2.58E-12 | 0.11 | 0.03 | 0.08 | 3.5 |
| endothelium | Trim3 | 0.93 | 1.30E-16 | 2.70E-12 | 0.23 | 0.09 | 0.14 | 2.47 |
| endothelium | C2cd2l | 1.17 | 1.35E-16 | 2.81E-12 | 0.15 | 0.05 | 0.1 | 2.92 |
| endothelium | Nr2c2ap | 0.89 | 1.40E-16 | 2.91E-12 | 0.44 | 0.23 | 0.21 | 1.92 |
| endothelium | Klhl2 | 1.23 | 1.43E-16 | 2.98E-12 | 0.27 | 0.12 | 0.15 | 2.25 |
| endothelium | Cptp | 0.85 | 1.46E-16 | 3.03E-12 | 0.29 | 0.13 | 0.16 | 2.2 |
| endothelium | Dnajc13 | 0.66 | 1.59E-16 | 3.30E-12 | 0.61 | 0.37 | 0.23 | 1.63 |
| endothelium | Idh2 | 0.41 | 1.62E-16 | 3.36E-12 | 0.91 | 0.9 | 0.01 | 1.01 |
| endothelium | Plekhg1 | 0.76 | 2.11E-16 | 4.38E-12 | 0.3 | 0.14 | 0.16 | 2.19 |
| endothelium | Tbc1d2b | 1.18 | 2.12E-16 | 4.41E-12 | 0.21 | 0.08 | 0.13 | 2.54 |
| endothelium | Zswim6 | 0.64 | 2.16E-16 | 4.49E-12 | 0.53 | 0.29 | 0.24 | 1.81 |
| endothelium | Tnfrsf21 | 0.82 | 2.22E-16 | 4.62E-12 | 0.3 | 0.14 | 0.16 | 2.17 |
| endothelium | Pdia6 | 0.34 | 2.31E-16 | 4.81E-12 | 0.9 | 0.96 | -0.06 | 0.94 |
| endothelium | Micall2 | 1.19 | 2.36E-16 | 4.91E-12 | 0.18 | 0.07 | 0.11 | 2.71 |
| endothelium | Vamp3 | 0.72 | 2.42E-16 | 5.04E-12 | 0.78 | 0.57 | 0.22 | 1.38 |
| endothelium | Wbp2 | 0.68 | 2.50E-16 | 5.20E-12 | 0.49 | 0.27 | 0.22 | 1.79 |

**Supplementary Table 2. Marker genes for the major cell types of integrated clustering of E11.5 control and NotchGOF λ-junction cells.**

| | | | | | | | | |
|---|---|---|---|---|---|---|---|---|
| endothelium | Add2 | 0.66 | 2.54E-16 | 5.29E-12 | 0.22 | 0.09 | 0.13 | 2.49 |
| endothelium | Lrp5 | 0.6 | 3.07E-16 | 6.39E-12 | 0.36 | 0.17 | 0.19 | 2.07 |
| endothelium | Sirt7 | 1.02 | 3.53E-16 | 7.35E-12 | 0.34 | 0.17 | 0.17 | 2.02 |
| endothelium | Erp29 | 0.36 | 3.64E-16 | 7.57E-12 | 0.88 | 0.88 | 0 | 1 |
| endothelium | Coq10b | 0.64 | 3.86E-16 | 8.03E-12 | 0.47 | 0.25 | 0.22 | 1.86 |
| endothelium | Lrrc8a | 0.72 | 3.94E-16 | 8.20E-12 | 0.49 | 0.27 | 0.22 | 1.83 |
| endothelium | Prkab1 | 0.89 | 3.99E-16 | 8.29E-12 | 0.55 | 0.33 | 0.22 | 1.68 |
| endothelium | Efnb2 | 0.67 | 4.01E-16 | 8.33E-12 | 0.67 | 0.45 | 0.23 | 1.5 |
| endothelium | Exoc5 | 0.65 | 4.08E-16 | 8.48E-12 | 0.76 | 0.57 | 0.19 | 1.34 |
| endothelium | Spryd3 | 1.05 | 4.10E-16 | 8.52E-12 | 0.19 | 0.07 | 0.12 | 2.6 |
| endothelium | Rrp7a | 0.99 | 4.26E-16 | 8.87E-12 | 0.69 | 0.47 | 0.21 | 1.45 |
| endothelium | Txndc17 | 0.32 | 4.26E-16 | 8.87E-12 | 0.92 | 0.92 | 0 | 1 |
| endothelium | Herc6 | 1.02 | 4.41E-16 | 9.17E-12 | 0.15 | 0.05 | 0.1 | 2.92 |
| endothelium | Tmem109 | 0.83 | 4.48E-16 | 9.31E-12 | 0.59 | 0.36 | 0.23 | 1.63 |
| endothelium | Slc25a30 | 0.94 | 4.66E-16 | 9.70E-12 | 0.31 | 0.15 | 0.16 | 2.12 |
| endothelium | Mcrip2 | 2.15 | 4.99E-16 | 1.04E-11 | 0.18 | 0.07 | 0.11 | 2.63 |
| endothelium | Hdac10 | 1.11 | 5.10E-16 | 1.06E-11 | 0.12 | 0.04 | 0.09 | 3.24 |
| endothelium | Pxk | 0.72 | 5.21E-16 | 1.08E-11 | 0.34 | 0.16 | 0.18 | 2.08 |
| endothelium | Lrrc59 | 0.49 | 5.28E-16 | 1.10E-11 | 0.85 | 0.79 | 0.06 | 1.07 |
| endothelium | Tradd | 0.95 | 5.78E-16 | 1.20E-11 | 0.26 | 0.11 | 0.15 | 2.27 |
| endothelium | Ddrgk1 | 0.7 | 5.79E-16 | 1.20E-11 | 0.74 | 0.55 | 0.19 | 1.35 |
| endothelium | Polg | 0.85 | 6.04E-16 | 1.26E-11 | 0.43 | 0.22 | 0.2 | 1.91 |
| endothelium | Gmpr2 | 0.75 | 6.32E-16 | 1.31E-11 | 0.64 | 0.41 | 0.22 | 1.53 |
| endothelium | Psat1 | 0.61 | 6.36E-16 | 1.32E-11 | 0.68 | 0.45 | 0.23 | 1.51 |
| endothelium | Fyb | 3.2 | 6.40E-16 | 1.33E-11 | 0.13 | 0.04 | 0.08 | 3.05 |
| endothelium | Plekho2 | 1.12 | 6.50E-16 | 1.35E-11 | 0.18 | 0.07 | 0.11 | 2.62 |
| endothelium | Nans | 0.57 | 6.98E-16 | 1.45E-11 | 0.76 | 0.6 | 0.16 | 1.26 |
| endothelium | Pwwp2b | 0.83 | 7.82E-16 | 1.63E-11 | 0.19 | 0.07 | 0.12 | 2.63 |
| endothelium | Airn | 0.84 | 8.16E-16 | 1.70E-11 | 0.38 | 0.19 | 0.19 | 2.02 |
| endothelium | Chn2 | 1.06 | 8.22E-16 | 1.71E-11 | 0.18 | 0.07 | 0.11 | 2.63 |
| endothelium | Cyfip1 | 0.56 | 9.26E-16 | 1.93E-11 | 0.82 | 0.7 | 0.12 | 1.17 |
| endothelium | Fhl3 | 1.03 | 1.02E-15 | 2.13E-11 | 0.24 | 0.11 | 0.14 | 2.31 |
| endothelium | Bbc3 | 1.42 | 1.07E-15 | 2.22E-11 | 0.24 | 0.1 | 0.14 | 2.35 |
| endothelium | Smpdl3b | 1.11 | 1.07E-15 | 2.23E-11 | 0.18 | 0.07 | 0.11 | 2.67 |
| endothelium | Sub1 | 0.3 | 1.08E-15 | 2.24E-11 | 0.98 | 0.99 | -0.02 | 0.98 |
| endothelium | Ubxn8 | 0.72 | 1.08E-15 | 2.24E-11 | 0.38 | 0.19 | 0.19 | 1.95 |
| endothelium | Adgra2 | 0.82 | 1.12E-15 | 2.33E-11 | 0.44 | 0.23 | 0.21 | 1.89 |
| endothelium | Prelid2 | 0.64 | 1.19E-15 | 2.47E-11 | 0.6 | 0.37 | 0.23 | 1.63 |
| endothelium | Ap1s1 | 0.54 | 1.25E-15 | 2.59E-11 | 0.85 | 0.76 | 0.09 | 1.11 |
| endothelium | Nlk | 0.69 | 1.33E-15 | 2.76E-11 | 0.48 | 0.27 | 0.21 | 1.78 |
| endothelium | Cryzl2 | 0.84 | 1.41E-15 | 2.93E-11 | 0.24 | 0.1 | 0.14 | 2.35 |
| endothelium | Rel | 0.78 | 1.44E-15 | 2.98E-11 | 0.22 | 0.09 | 0.13 | 2.43 |
| endothelium | Bcl10 | 0.71 | 1.54E-15 | 3.20E-11 | 0.72 | 0.49 | 0.23 | 1.46 |
| endothelium | Mapk11 | 0.92 | 1.77E-15 | 3.67E-11 | 0.18 | 0.07 | 0.11 | 2.59 |
| endothelium | 010204K13Ri | 0.94 | 1.85E-15 | 3.84E-11 | 0.45 | 0.25 | 0.2 | 1.78 |
| endothelium | Slc20a1 | 0.56 | 1.88E-15 | 3.91E-11 | 0.55 | 0.32 | 0.23 | 1.73 |
| endothelium | Cdk1 | 0.71 | 1.95E-15 | 4.06E-11 | 0.82 | 0.67 | 0.15 | 1.23 |
| endothelium | Ndfip2 | 0.7 | 2.04E-15 | 4.25E-11 | 0.73 | 0.51 | 0.22 | 1.42 |
| endothelium | Mindy3 | 0.89 | 2.09E-15 | 4.35E-11 | 0.49 | 0.28 | 0.21 | 1.76 |
| endothelium | Tpm3-rs7 | 0.92 | 2.11E-15 | 4.38E-11 | 0.25 | 0.11 | 0.14 | 2.3 |
| endothelium | Arf5 | 0.27 | 2.25E-15 | 4.67E-11 | 0.93 | 0.99 | -0.06 | 0.94 |
| endothelium | Poglut3 | 0.74 | 2.27E-15 | 4.73E-11 | 0.29 | 0.13 | 0.16 | 2.17 |
| endothelium | Mcl1 | 0.47 | 2.30E-15 | 4.78E-11 | 0.86 | 0.82 | 0.04 | 1.05 |
| endothelium | 610010K14Ri | 0.43 | 2.33E-15 | 4.85E-11 | 0.87 | 0.81 | 0.06 | 1.07 |
| endothelium | Cdc37 | 0.33 | 2.47E-15 | 5.14E-11 | 0.94 | 0.96 | -0.02 | 0.98 |
| endothelium | Inpp1 | 0.71 | 2.50E-15 | 5.20E-11 | 0.34 | 0.16 | 0.18 | 2.07 |

**Supplementary Table 2. Marker genes for the major cell types of integrated clustering of E11.5 control and NotchGOF λ-junction cells.**

| | | | | | | | | |
|---|---|---|---|---|---|---|---|---|
| endothelium | Gclm | 0.7 | 2.55E-15 | 5.31E-11 | 0.63 | 0.39 | 0.25 | 1.64 |
| endothelium | Jpt1 | 0.4 | 2.63E-15 | 5.47E-11 | 0.96 | 0.94 | 0.01 | 1.01 |
| endothelium | Leprot | 0.77 | 2.88E-15 | 6.00E-11 | 0.71 | 0.52 | 0.19 | 1.37 |
| endothelium | Tmbim4 | 0.85 | 2.89E-15 | 6.01E-11 | 0.7 | 0.49 | 0.2 | 1.41 |
| endothelium | Uckl1 | 1.2 | 3.26E-15 | 6.78E-11 | 0.26 | 0.12 | 0.14 | 2.23 |
| endothelium | Parp9 | 1.32 | 3.30E-15 | 6.86E-11 | 0.1 | 0.03 | 0.07 | 3.35 |
| endothelium | Prr13 | 1.12 | 3.62E-15 | 7.52E-11 | 0.64 | 0.4 | 0.24 | 1.6 |
| endothelium | Arhgap6 | 2.11 | 3.81E-15 | 7.93E-11 | 0.17 | 0.06 | 0.1 | 2.58 |
| endothelium | Arpc5l | 0.5 | 3.96E-15 | 8.24E-11 | 0.89 | 0.82 | 0.07 | 1.09 |
| endothelium | Itpr1 | 0.65 | 3.98E-15 | 8.27E-11 | 0.4 | 0.21 | 0.19 | 1.9 |
| endothelium | Mitd1 | 1.24 | 4.02E-15 | 8.36E-11 | 0.51 | 0.3 | 0.22 | 1.74 |
| endothelium | Uqcr10 | 0.33 | 4.53E-15 | 9.41E-11 | 0.97 | 0.98 | -0.02 | 0.98 |
| endothelium | Edem2 | 0.69 | 4.85E-15 | 1.01E-10 | 0.38 | 0.19 | 0.19 | 1.97 |
| endothelium | Ntan1 | 0.54 | 5.25E-15 | 1.09E-10 | 0.81 | 0.67 | 0.15 | 1.22 |
| endothelium | Ddx39 | 0.6 | 5.41E-15 | 1.12E-10 | 0.86 | 0.79 | 0.07 | 1.08 |
| endothelium | Clic4 | 0.55 | 6.53E-15 | 1.36E-10 | 0.82 | 0.68 | 0.14 | 1.21 |
| endothelium | Vdac1 | 0.33 | 6.58E-15 | 1.37E-10 | 0.93 | 0.95 | -0.02 | 0.98 |
| endothelium | Pgp | 0.56 | 6.69E-15 | 1.39E-10 | 0.81 | 0.66 | 0.16 | 1.24 |
| endothelium | Dnmbp | 1.08 | 7.07E-15 | 1.47E-10 | 0.28 | 0.13 | 0.15 | 2.2 |
| endothelium | Macrod2 | 0.68 | 7.50E-15 | 1.56E-10 | 0.42 | 0.22 | 0.2 | 1.9 |
| endothelium | Crtap | 0.59 | 9.00E-15 | 1.87E-10 | 0.49 | 0.27 | 0.21 | 1.78 |
| endothelium | Prkaa1 | 0.78 | 9.68E-15 | 2.01E-10 | 0.32 | 0.15 | 0.16 | 2.06 |
| endothelium | Plpp2 | 0.68 | 1.01E-14 | 2.10E-10 | 0.46 | 0.26 | 0.2 | 1.77 |
| endothelium | Inpp4b | 0.56 | 1.02E-14 | 2.13E-10 | 0.32 | 0.15 | 0.17 | 2.13 |
| endothelium | Cox18 | 0.67 | 1.06E-14 | 2.21E-10 | 0.52 | 0.3 | 0.22 | 1.73 |
| endothelium | Mtmr12 | 0.8 | 1.09E-14 | 2.27E-10 | 0.25 | 0.11 | 0.14 | 2.24 |
| endothelium | Plekhh3 | 1.43 | 1.16E-14 | 2.41E-10 | 0.11 | 0.04 | 0.08 | 3.2 |
| endothelium | Selenof | 0.33 | 1.18E-14 | 2.46E-10 | 0.91 | 0.96 | -0.04 | 0.95 |
| endothelium | Serhl | 0.65 | 1.24E-14 | 2.57E-10 | 0.34 | 0.17 | 0.17 | 2.04 |
| endothelium | Pttg1ip | 0.9 | 1.28E-14 | 2.66E-10 | 0.62 | 0.39 | 0.23 | 1.59 |
| endothelium | Dennd4a | 1.11 | 1.30E-14 | 2.70E-10 | 0.2 | 0.08 | 0.12 | 2.43 |
| endothelium | Cln5 | 0.65 | 1.42E-14 | 2.95E-10 | 0.41 | 0.21 | 0.19 | 1.92 |
| endothelium | Crlf3 | 0.69 | 1.52E-14 | 3.15E-10 | 0.66 | 0.45 | 0.21 | 1.47 |
| endothelium | Gramd4 | 1.42 | 1.59E-14 | 3.30E-10 | 0.2 | 0.08 | 0.12 | 2.41 |
| endothelium | Mfsd13a | 1.03 | 1.62E-14 | 3.38E-10 | 0.12 | 0.04 | 0.08 | 3.08 |
| endothelium | Mvp | 1 | 2.06E-14 | 4.27E-10 | 0.16 | 0.06 | 0.1 | 2.67 |
| endothelium | Arhgef2 | 0.6 | 2.14E-14 | 4.45E-10 | 0.64 | 0.42 | 0.22 | 1.51 |
| endothelium | Tcn2 | 0.73 | 2.21E-14 | 4.60E-10 | 0.35 | 0.18 | 0.17 | 1.98 |
| endothelium | Gem | 0.98 | 2.30E-14 | 4.78E-10 | 0.12 | 0.04 | 0.08 | 3.08 |
| endothelium | Igf2r | 0.77 | 2.42E-14 | 5.03E-10 | 0.71 | 0.5 | 0.21 | 1.43 |
| endothelium | Sdf2l1 | 0.71 | 2.52E-14 | 5.25E-10 | 0.68 | 0.47 | 0.21 | 1.44 |
| endothelium | Mfsd5 | 0.68 | 2.64E-14 | 5.48E-10 | 0.43 | 0.24 | 0.19 | 1.81 |
| endothelium | Dph5 | 0.75 | 2.70E-14 | 5.61E-10 | 0.41 | 0.23 | 0.19 | 1.83 |
| endothelium | Grb2 | 0.55 | 2.75E-14 | 5.71E-10 | 0.81 | 0.64 | 0.18 | 1.28 |
| endothelium | Card19 | 0.77 | 2.77E-14 | 5.76E-10 | 0.46 | 0.25 | 0.21 | 1.82 |
| endothelium | Nfkbil1 | 0.8 | 2.96E-14 | 6.16E-10 | 0.48 | 0.28 | 0.21 | 1.75 |
| endothelium | Tnk2 | 0.67 | 3.02E-14 | 6.28E-10 | 0.3 | 0.14 | 0.15 | 2.08 |
| endothelium | Stx12 | 0.65 | 3.17E-14 | 6.59E-10 | 0.72 | 0.52 | 0.21 | 1.4 |
| endothelium | Jmjd1c | 0.52 | 3.19E-14 | 6.63E-10 | 0.71 | 0.48 | 0.23 | 1.47 |
| endothelium | Ggh | 0.62 | 3.20E-14 | 6.65E-10 | 0.45 | 0.25 | 0.2 | 1.8 |
| endothelium | Yif1b | 0.73 | 3.29E-14 | 6.84E-10 | 0.8 | 0.67 | 0.13 | 1.19 |
| endothelium | 810006K23Ri | 0.59 | 3.40E-14 | 7.08E-10 | 0.58 | 0.35 | 0.23 | 1.66 |
| endothelium | Ankrd39 | 0.82 | 3.60E-14 | 7.49E-10 | 0.32 | 0.16 | 0.16 | 2.05 |
| endothelium | Diaph2 | 0.66 | 4.60E-14 | 9.56E-10 | 0.31 | 0.15 | 0.16 | 2.04 |
| endothelium | Rgs19 | 0.67 | 4.91E-14 | 1.02E-09 | 0.66 | 0.45 | 0.21 | 1.47 |
| endothelium | Txnrd2 | 0.62 | 5.82E-14 | 1.21E-09 | 0.4 | 0.21 | 0.19 | 1.9 |

**Supplementary Table 2. Marker genes for the major cell types of integrated clustering of E11.5 control and NotchGOF λ-junction cells.**

| | | | | | | | | |
|---|---|---|---|---|---|---|---|---|
| endothelium | Rpp21 | 0.57 | 6.36E-14 | 1.32E-09 | 0.74 | 0.57 | 0.17 | 1.3 |
| endothelium | Acap2 | 0.6 | 6.77E-14 | 1.41E-09 | 0.69 | 0.46 | 0.22 | 1.49 |
| endothelium | Gars | 0.52 | 6.95E-14 | 1.45E-09 | 0.82 | 0.67 | 0.15 | 1.22 |
| endothelium | Gnb1 | 0.32 | 8.10E-14 | 1.68E-09 | 0.93 | 0.96 | -0.03 | 0.97 |
| endothelium | Srp19 | 0.41 | 8.29E-14 | 1.72E-09 | 0.91 | 0.88 | 0.03 | 1.03 |
| endothelium | Creld1 | 0.66 | 8.94E-14 | 1.86E-09 | 0.27 | 0.13 | 0.14 | 2.13 |
| endothelium | Cyb5a | 0.37 | 9.67E-14 | 2.01E-09 | 0.92 | 0.92 | -0.01 | 0.99 |
| endothelium | Asl | 0.61 | 9.69E-14 | 2.01E-09 | 0.3 | 0.15 | 0.16 | 2.07 |
| endothelium | Szrd1 | 0.56 | 1.03E-13 | 2.13E-09 | 0.72 | 0.51 | 0.21 | 1.42 |
| endothelium | Mrpl36 | 0.47 | 1.03E-13 | 2.14E-09 | 0.82 | 0.72 | 0.11 | 1.15 |
| endothelium | Ccdc32 | 0.64 | 1.03E-13 | 2.15E-09 | 0.52 | 0.31 | 0.21 | 1.69 |
| endothelium | Pde4d | 0.65 | 1.07E-13 | 2.22E-09 | 0.46 | 0.26 | 0.2 | 1.75 |
| endothelium | Grk2 | 0.8 | 1.14E-13 | 2.37E-09 | 0.54 | 0.33 | 0.22 | 1.66 |
| endothelium | Acap3 | 0.74 | 1.18E-13 | 2.44E-09 | 0.18 | 0.07 | 0.11 | 2.48 |
| endothelium | Surf4 | 0.53 | 1.19E-13 | 2.48E-09 | 0.78 | 0.59 | 0.2 | 1.34 |
| endothelium | Tmem150a | 0.93 | 1.51E-13 | 3.15E-09 | 0.16 | 0.06 | 0.1 | 2.58 |
| endothelium | Gorasp2 | 0.58 | 1.62E-13 | 3.38E-09 | 0.73 | 0.53 | 0.19 | 1.36 |
| endothelium | Alg3 | 0.61 | 1.79E-13 | 3.73E-09 | 0.49 | 0.28 | 0.21 | 1.77 |
| endothelium | Tmem181a | 0.64 | 1.79E-13 | 3.73E-09 | 0.23 | 0.1 | 0.13 | 2.27 |
| endothelium | Mtln | 0.58 | 1.89E-13 | 3.93E-09 | 0.54 | 0.33 | 0.21 | 1.65 |
| endothelium | Nfkbib | 0.99 | 1.91E-13 | 3.98E-09 | 0.43 | 0.24 | 0.19 | 1.78 |
| endothelium | 610507B11Ri | 0.65 | 1.92E-13 | 3.99E-09 | 0.68 | 0.46 | 0.21 | 1.46 |
| endothelium | Tbc1d9b | 0.67 | 2.20E-13 | 4.58E-09 | 0.4 | 0.21 | 0.19 | 1.88 |
| endothelium | Zbtb1 | 0.63 | 2.24E-13 | 4.66E-09 | 0.58 | 0.35 | 0.23 | 1.65 |
| endothelium | Man2b1 | 0.39 | 2.28E-13 | 4.74E-09 | 0.42 | 0.23 | 0.2 | 1.86 |
| endothelium | Mcc | 0.79 | 2.29E-13 | 4.77E-09 | 0.27 | 0.13 | 0.14 | 2.07 |
| endothelium | Naa60 | 0.73 | 2.33E-13 | 4.85E-09 | 0.45 | 0.25 | 0.2 | 1.82 |
| endothelium | Gusb | 0.46 | 2.75E-13 | 5.71E-09 | 0.53 | 0.31 | 0.22 | 1.73 |
| endothelium | Sec24d | 0.64 | 2.77E-13 | 5.76E-09 | 0.2 | 0.08 | 0.11 | 2.41 |
| endothelium | Faap24 | 0.78 | 2.79E-13 | 5.81E-09 | 0.32 | 0.16 | 0.16 | 1.96 |
| endothelium | Mrtfb | 1.01 | 2.82E-13 | 5.87E-09 | 0.44 | 0.24 | 0.19 | 1.8 |
| endothelium | Emp3 | 1.12 | 3.31E-13 | 6.88E-09 | 0.28 | 0.14 | 0.14 | 2.04 |
| endothelium | Abl2 | 0.76 | 3.39E-13 | 7.05E-09 | 0.36 | 0.19 | 0.17 | 1.91 |
| endothelium | Mrrf | 0.53 | 3.47E-13 | 7.22E-09 | 0.57 | 0.36 | 0.21 | 1.59 |
| endothelium | Lmf2 | 0.59 | 3.54E-13 | 7.35E-09 | 0.36 | 0.19 | 0.17 | 1.93 |
| endothelium | Mrps16 | 0.54 | 4.03E-13 | 8.39E-09 | 0.88 | 0.84 | 0.04 | 1.05 |
| endothelium | Klhl8 | 1.36 | 4.05E-13 | 8.43E-09 | 0.21 | 0.09 | 0.12 | 2.29 |
| endothelium | Pgs1 | 0.68 | 4.06E-13 | 8.45E-09 | 0.5 | 0.3 | 0.21 | 1.7 |
| endothelium | Atp2b4 | 0.81 | 4.21E-13 | 8.76E-09 | 0.37 | 0.2 | 0.17 | 1.86 |
| endothelium | Efhd2 | 0.44 | 4.29E-13 | 8.93E-09 | 0.38 | 0.2 | 0.18 | 1.87 |
| endothelium | Atf7 | 0.6 | 4.45E-13 | 9.26E-09 | 0.38 | 0.2 | 0.18 | 1.92 |
| endothelium | Ubc | 0.34 | 4.46E-13 | 9.27E-09 | 0.96 | 0.98 | -0.01 | 0.99 |
| endothelium | Crtc2 | 0.74 | 4.46E-13 | 9.27E-09 | 0.25 | 0.12 | 0.13 | 2.16 |
| endothelium | Prkar2b | 0.9 | 4.48E-13 | 9.31E-09 | 0.33 | 0.5 | -0.17 | 0.66 |
| endothelium | Tro | 0.69 | 4.54E-13 | 9.44E-09 | 0.64 | 0.42 | 0.22 | 1.52 |
| endothelium | Pfkfb3 | 0.96 | 4.61E-13 | 9.60E-09 | 0.23 | 0.1 | 0.13 | 2.2 |
| endothelium | Arap2 | 0.98 | 4.69E-13 | 9.75E-09 | 0.14 | 0.05 | 0.09 | 2.67 |
| endothelium | Ppp1r15a | 0.71 | 4.74E-13 | 9.86E-09 | 0.43 | 0.25 | 0.18 | 1.74 |
| endothelium | Bop1 | 0.62 | 5.27E-13 | 1.10E-08 | 0.59 | 0.37 | 0.22 | 1.61 |
| endothelium | Prdx6 | 0.3 | 5.38E-13 | 1.12E-08 | 0.95 | 0.97 | -0.02 | 0.98 |
| endothelium | Grk5 | 0.99 | 5.63E-13 | 1.17E-08 | 0.25 | 0.12 | 0.13 | 2.13 |
| endothelium | Nck1 | 0.78 | 5.69E-13 | 1.18E-08 | 0.72 | 0.54 | 0.19 | 1.35 |
| endothelium | Tgoln1 | 0.61 | 5.78E-13 | 1.20E-08 | 0.49 | 0.29 | 0.2 | 1.7 |
| endothelium | Filip1l | 0.72 | 5.81E-13 | 1.21E-08 | 0.45 | 0.26 | 0.19 | 1.75 |
| endothelium | Xaf1 | 0.7 | 5.99E-13 | 1.24E-08 | 0.15 | 0.06 | 0.09 | 2.57 |
| endothelium | Pisd | 0.61 | 6.46E-13 | 1.34E-08 | 0.38 | 0.21 | 0.17 | 1.84 |

**Supplementary Table 2. Marker genes for the major cell types of integrated clustering of E11.5 control and NotchGOF λ-junction cells.**

| | | | | | | | | |
|---|---|---|---|---|---|---|---|---|
| endothelium | Lpcat1 | 0.63 | 6.63E-13 | 1.38E-08 | 0.66 | 0.47 | 0.19 | 1.41 |
| endothelium | Tbc1d12 | 0.91 | 6.88E-13 | 1.43E-08 | 0.32 | 0.17 | 0.16 | 1.96 |
| endothelium | Prdx3 | 0.72 | 7.30E-13 | 1.52E-08 | 0.73 | 0.53 | 0.19 | 1.36 |
| endothelium | Ralgapa2 | 0.81 | 7.63E-13 | 1.59E-08 | 0.28 | 0.14 | 0.14 | 2.06 |
| endothelium | Arrdc1 | 0.66 | 8.08E-13 | 1.68E-08 | 0.24 | 0.11 | 0.13 | 2.18 |
| endothelium | Lpp | 0.51 | 8.39E-13 | 1.75E-08 | 0.71 | 0.56 | 0.15 | 1.27 |
| endothelium | Rb1 | 0.54 | 8.41E-13 | 1.75E-08 | 0.31 | 0.16 | 0.15 | 1.97 |
| endothelium | Cenpx | 0.44 | 9.08E-13 | 1.89E-08 | 0.85 | 0.79 | 0.06 | 1.08 |
| endothelium | Rab21 | 0.7 | 9.26E-13 | 1.93E-08 | 0.66 | 0.46 | 0.2 | 1.43 |
| endothelium | Prkab2 | 0.54 | 1.12E-12 | 2.33E-08 | 0.26 | 0.13 | 0.14 | 2.11 |
| endothelium | Mknk1 | 1.11 | 1.21E-12 | 2.53E-08 | 0.2 | 0.09 | 0.11 | 2.23 |
| endothelium | Cfap298 | 0.58 | 1.23E-12 | 2.57E-08 | 0.58 | 0.37 | 0.22 | 1.59 |
| endothelium | Rer1 | 0.39 | 1.28E-12 | 2.67E-08 | 0.86 | 0.84 | 0.02 | 1.03 |
| endothelium | Pigk | 0.6 | 1.29E-12 | 2.68E-08 | 0.49 | 0.29 | 0.2 | 1.68 |
| endothelium | Gipc1 | 0.56 | 1.29E-12 | 2.69E-08 | 0.83 | 0.69 | 0.14 | 1.21 |
| endothelium | 630072M18R | 1.26 | 1.30E-12 | 2.70E-08 | 0.23 | 0.1 | 0.12 | 2.18 |
| endothelium | Tbc1d1 | 0.63 | 1.31E-12 | 2.73E-08 | 0.57 | 0.37 | 0.2 | 1.54 |
| endothelium | Stx6 | 0.58 | 1.47E-12 | 3.05E-08 | 0.7 | 0.5 | 0.2 | 1.4 |
| endothelium | Tsen15 | 0.58 | 1.51E-12 | 3.14E-08 | 0.78 | 0.63 | 0.15 | 1.23 |
| endothelium | Armcx4 | 0.58 | 1.53E-12 | 3.18E-08 | 0.57 | 0.36 | 0.21 | 1.58 |
| endothelium | Gemin7 | 0.64 | 1.57E-12 | 3.27E-08 | 0.82 | 0.7 | 0.12 | 1.17 |
| endothelium | Mapre2 | 0.54 | 1.59E-12 | 3.30E-08 | 0.61 | 0.4 | 0.21 | 1.52 |
| endothelium | Hsd17b11 | 0.57 | 1.65E-12 | 3.42E-08 | 0.38 | 0.2 | 0.18 | 1.88 |
| endothelium | Xrcc4 | 0.73 | 1.65E-12 | 3.42E-08 | 0.34 | 0.18 | 0.16 | 1.87 |
| endothelium | Septin10 | 0.81 | 1.69E-12 | 3.52E-08 | 0.57 | 0.37 | 0.2 | 1.54 |
| endothelium | Atg3 | 0.54 | 1.70E-12 | 3.54E-08 | 0.66 | 0.45 | 0.22 | 1.49 |
| endothelium | Pla2g15 | 1.04 | 1.73E-12 | 3.60E-08 | 0.17 | 0.07 | 0.1 | 2.46 |
| endothelium | Ppp1r12c | 0.85 | 1.98E-12 | 4.12E-08 | 0.52 | 0.32 | 0.2 | 1.62 |
| endothelium | Slc16a6 | 1.09 | 2.13E-12 | 4.44E-08 | 0.14 | 0.05 | 0.09 | 2.62 |
| endothelium | Pdia3 | 0.28 | 2.14E-12 | 4.45E-08 | 0.89 | 0.96 | -0.07 | 0.93 |
| endothelium | Map1lc3b | 0.37 | 2.25E-12 | 4.68E-08 | 0.93 | 0.93 | 0 | 1 |
| endothelium | Ric8a | 0.54 | 2.43E-12 | 5.05E-08 | 0.44 | 0.25 | 0.19 | 1.76 |
| endothelium | Snap29 | 0.63 | 2.69E-12 | 5.60E-08 | 0.36 | 0.19 | 0.17 | 1.87 |
| endothelium | Abhd16a | 0.84 | 2.75E-12 | 5.72E-08 | 0.47 | 0.27 | 0.2 | 1.72 |
| endothelium | Fam120a | 0.61 | 2.88E-12 | 6.00E-08 | 0.68 | 0.46 | 0.22 | 1.48 |
| endothelium | Pnpla2 | 0.98 | 2.94E-12 | 6.10E-08 | 0.2 | 0.09 | 0.11 | 2.24 |
| endothelium | Cep55 | 0.87 | 2.96E-12 | 6.16E-08 | 0.41 | 0.24 | 0.17 | 1.69 |
| endothelium | Imp4 | 0.54 | 3.03E-12 | 6.31E-08 | 0.72 | 0.54 | 0.19 | 1.35 |
| endothelium | Crybg3 | 0.61 | 3.08E-12 | 6.41E-08 | 0.38 | 0.21 | 0.17 | 1.82 |
| endothelium | Tomm34 | 0.44 | 3.25E-12 | 6.76E-08 | 0.64 | 0.41 | 0.23 | 1.55 |
| endothelium | Acot8 | 0.77 | 3.36E-12 | 6.99E-08 | 0.29 | 0.15 | 0.15 | 1.98 |
| endothelium | Cmpk2 | 1.48 | 3.39E-12 | 7.05E-08 | 0.11 | 0.04 | 0.07 | 2.97 |
| endothelium | Atf6 | 0.62 | 3.63E-12 | 7.56E-08 | 0.28 | 0.14 | 0.14 | 2.01 |
| endothelium | .510009E07Ri | 0.81 | 3.71E-12 | 7.72E-08 | 0.16 | 0.06 | 0.09 | 2.5 |
| endothelium | F8 | 1.12 | 3.75E-12 | 7.80E-08 | 0.11 | 0.04 | 0.07 | 2.87 |
| endothelium | Hexb | 0.32 | 3.79E-12 | 7.88E-08 | 0.41 | 0.22 | 0.19 | 1.83 |
| endothelium | Psma6 | 0.36 | 3.84E-12 | 7.99E-08 | 0.93 | 0.95 | -0.01 | 0.99 |
| endothelium | Irf2bp2 | 0.62 | 3.85E-12 | 8.02E-08 | 0.57 | 0.35 | 0.22 | 1.63 |
| endothelium | Ap3m2 | 1.01 | 3.90E-12 | 8.12E-08 | 0.11 | 0.04 | 0.07 | 2.87 |
| endothelium | Stard3 | 0.57 | 4.01E-12 | 8.33E-08 | 0.42 | 0.23 | 0.19 | 1.79 |
| endothelium | Trmt10a | 0.51 | 4.14E-12 | 8.60E-08 | 0.55 | 0.34 | 0.21 | 1.63 |
| endothelium | Cyth1 | 0.79 | 4.14E-12 | 8.61E-08 | 0.29 | 0.14 | 0.14 | 2 |
| endothelium | Pigl | 0.9 | 4.16E-12 | 8.65E-08 | 0.17 | 0.07 | 0.1 | 2.37 |
| endothelium | Jph1 | 0.71 | 4.24E-12 | 8.83E-08 | 0.17 | 0.07 | 0.1 | 2.4 |
| endothelium | Zfp961 | 0.75 | 4.28E-12 | 8.90E-08 | 0.33 | 0.17 | 0.16 | 1.91 |
| endothelium | Akip1 | 0.61 | 4.29E-12 | 8.93E-08 | 0.36 | 0.2 | 0.17 | 1.85 |

**Supplementary Table 2. Marker genes for the major cell types of integrated clustering of E11.5 control and NotchGOF λ-junction cells.**

| | | | | | | | | |
|---|---|---|---|---|---|---|---|---|
| endothelium | Tjp2 | 0.45 | 4.35E-12 | 9.04E-08 | 0.52 | 0.31 | 0.21 | 1.67 |
| endothelium | Fermt3 | 4.76 | 4.37E-12 | 9.10E-08 | 0.12 | 0.05 | 0.08 | 2.56 |
| endothelium | Brms1 | 0.58 | 4.44E-12 | 9.24E-08 | 0.56 | 0.34 | 0.21 | 1.62 |
| endothelium | Ints6l | 0.61 | 4.48E-12 | 9.32E-08 | 0.33 | 0.17 | 0.16 | 1.92 |
| endothelium | Creld2 | 0.48 | 4.49E-12 | 9.34E-08 | 0.66 | 0.44 | 0.21 | 1.48 |
| endothelium | Slc16a13 | 0.73 | 4.56E-12 | 9.48E-08 | 0.2 | 0.09 | 0.11 | 2.21 |
| endothelium | Creb3 | 0.74 | 4.76E-12 | 9.91E-08 | 0.35 | 0.19 | 0.16 | 1.86 |
| endothelium | Lsm11 | 0.57 | 5.10E-12 | 1.06E-07 | 0.36 | 0.19 | 0.17 | 1.86 |
| endothelium | Nxpe4 | 1 | 5.92E-12 | 1.23E-07 | 0.12 | 0.04 | 0.08 | 2.79 |
| endothelium | Timm10b | 0.49 | 6.69E-12 | 1.39E-07 | 0.87 | 0.85 | 0.03 | 1.03 |
| endothelium | Slc22a15 | 0.63 | 6.84E-12 | 1.42E-07 | 0.21 | 0.09 | 0.11 | 2.21 |
| endothelium | Larp1b | 1.65 | 7.08E-12 | 1.47E-07 | 0.17 | 0.08 | 0.1 | 2.28 |
| endothelium | Cmtm7 | 0.49 | 7.24E-12 | 1.50E-07 | 0.68 | 0.48 | 0.2 | 1.41 |
| endothelium | Lysmd2 | 0.74 | 7.35E-12 | 1.53E-07 | 0.39 | 0.22 | 0.18 | 1.82 |
| endothelium | Ankrd37 | 0.84 | 7.41E-12 | 1.54E-07 | 0.3 | 0.16 | 0.14 | 1.89 |
| endothelium | Pemt | 0.91 | 7.67E-12 | 1.59E-07 | 0.12 | 0.04 | 0.08 | 2.79 |
| endothelium | Akirin1 | 0.52 | 7.79E-12 | 1.62E-07 | 0.75 | 0.57 | 0.18 | 1.31 |
| endothelium | Jmjd6 | 0.7 | 7.93E-12 | 1.65E-07 | 0.72 | 0.52 | 0.2 | 1.38 |
| endothelium | Myo1b | 0.51 | 8.25E-12 | 1.72E-07 | 0.79 | 0.73 | 0.06 | 1.08 |
| endothelium | Tmc4 | 0.55 | 8.50E-12 | 1.77E-07 | 0.21 | 0.1 | 0.12 | 2.2 |
| endothelium | Ubl3 | 0.61 | 8.61E-12 | 1.79E-07 | 0.58 | 0.37 | 0.21 | 1.57 |
| endothelium | Uck2 | 0.6 | 1.01E-11 | 2.11E-07 | 0.72 | 0.52 | 0.2 | 1.37 |
| endothelium | Slc1a5 | 0.81 | 1.05E-11 | 2.19E-07 | 0.36 | 0.2 | 0.16 | 1.79 |
| endothelium | Nav1 | 0.5 | 1.09E-11 | 2.27E-07 | 0.72 | 0.54 | 0.18 | 1.34 |
| endothelium | Gm26532 | 0.88 | 1.10E-11 | 2.29E-07 | 0.11 | 0.04 | 0.07 | 2.87 |
| endothelium | Elf2 | 0.59 | 1.11E-11 | 2.31E-07 | 0.79 | 0.66 | 0.12 | 1.19 |
| endothelium | Gm12166 | 0.62 | 1.16E-11 | 2.41E-07 | 0.26 | 0.13 | 0.13 | 2.02 |
| endothelium | Faap100 | 0.73 | 1.20E-11 | 2.49E-07 | 0.42 | 0.24 | 0.18 | 1.73 |
| endothelium | Uqcrq | 0.26 | 1.21E-11 | 2.51E-07 | 0.95 | 0.97 | -0.02 | 0.98 |
| endothelium | Sbf2 | 0.58 | 1.21E-11 | 2.52E-07 | 0.53 | 0.34 | 0.19 | 1.55 |
| endothelium | Fam160b1 | 0.59 | 1.31E-11 | 2.72E-07 | 0.47 | 0.28 | 0.19 | 1.67 |
| endothelium | Ppp1r15b | 0.52 | 1.33E-11 | 2.76E-07 | 0.51 | 0.31 | 0.2 | 1.66 |
| endothelium | Isca1 | 0.62 | 1.36E-11 | 2.83E-07 | 0.51 | 0.31 | 0.2 | 1.64 |
| endothelium | Septin2 | 0.58 | 1.37E-11 | 2.85E-07 | 0.33 | 0.17 | 0.16 | 1.9 |
| endothelium | Ssna1 | 0.5 | 1.46E-11 | 3.04E-07 | 0.88 | 0.85 | 0.03 | 1.04 |
| endothelium | Plpp5 | 0.43 | 1.54E-11 | 3.20E-07 | 0.36 | 0.19 | 0.17 | 1.88 |
| endothelium | Stxbp5 | 1.07 | 1.55E-11 | 3.23E-07 | 0.18 | 0.08 | 0.1 | 2.29 |
| endothelium | Ski | 0.54 | 1.69E-11 | 3.52E-07 | 0.62 | 0.41 | 0.21 | 1.52 |
| endothelium | Armc6 | 0.49 | 1.82E-11 | 3.78E-07 | 0.28 | 0.14 | 0.14 | 2 |
| endothelium | Sft2d2 | 0.68 | 1.85E-11 | 3.84E-07 | 0.29 | 0.15 | 0.14 | 1.91 |
| endothelium | Cog8 | 0.7 | 1.91E-11 | 3.98E-07 | 0.64 | 0.42 | 0.22 | 1.53 |
| endothelium | .600002K03Ri | 0.56 | 2.02E-11 | 4.21E-07 | 0.24 | 0.11 | 0.12 | 2.08 |
| endothelium | Mdga2 | 0.84 | 2.04E-11 | 4.23E-07 | 0.17 | 0.07 | 0.1 | 2.36 |
| endothelium | Cox17 | 0.65 | 2.05E-11 | 4.26E-07 | 0.86 | 0.77 | 0.1 | 1.13 |
| endothelium | Mpdu1 | 0.55 | 2.10E-11 | 4.38E-07 | 0.68 | 0.5 | 0.18 | 1.35 |
| endothelium | Kbtbd2 | 0.55 | 2.11E-11 | 4.39E-07 | 0.54 | 0.33 | 0.2 | 1.61 |
| endothelium | Taf9b | 0.98 | 2.18E-11 | 4.54E-07 | 0.11 | 0.04 | 0.07 | 2.79 |
| endothelium | Pus1 | 0.54 | 2.19E-11 | 4.56E-07 | 0.48 | 0.29 | 0.19 | 1.66 |
| endothelium | Cxxc5 | 0.41 | 2.21E-11 | 4.60E-07 | 0.67 | 0.45 | 0.22 | 1.5 |
| endothelium | Lipa | 0.45 | 2.30E-11 | 4.79E-07 | 0.39 | 0.22 | 0.18 | 1.81 |
| endothelium | Sgsm3 | 0.87 | 2.32E-11 | 4.82E-07 | 0.21 | 0.1 | 0.11 | 2.11 |
| endothelium | Gsr | 0.58 | 2.40E-11 | 4.99E-07 | 0.53 | 0.33 | 0.2 | 1.59 |
| endothelium | Gosr1 | 0.66 | 2.44E-11 | 5.08E-07 | 0.32 | 0.17 | 0.15 | 1.89 |
| endothelium | Ccn2 | 1.26 | 2.51E-11 | 5.22E-07 | 0.1 | 0.04 | 0.07 | 2.89 |
| endothelium | Dok1 | 1 | 2.52E-11 | 5.25E-07 | 0.21 | 0.1 | 0.11 | 2.11 |
| endothelium | Plxnc1 | 1.23 | 2.54E-11 | 5.28E-07 | 0.12 | 0.04 | 0.07 | 2.74 |

**Supplementary Table 2. Marker genes for the major cell types of integrated clustering of E11.5 control and NotchGOF λ-junction cells.**

| | | | | | | | | |
|---|---|---|---|---|---|---|---|---|
| endothelium | Adipor1 | 0.62 | 2.58E-11 | 5.37E-07 | 0.73 | 0.56 | 0.17 | 1.3 |
| endothelium | Mtmr11 | 0.86 | 2.63E-11 | 5.47E-07 | 0.21 | 0.1 | 0.11 | 2.11 |
| endothelium | Rnf141 | 0.62 | 2.67E-11 | 5.56E-07 | 0.25 | 0.12 | 0.13 | 2.03 |
| endothelium | Hip1 | 0.61 | 2.81E-11 | 5.85E-07 | 0.63 | 0.45 | 0.19 | 1.42 |
| endothelium | 510039O18Ri | 0.5 | 2.88E-11 | 5.99E-07 | 0.41 | 0.23 | 0.18 | 1.77 |
| endothelium | Znrf2 | 0.54 | 2.88E-11 | 5.99E-07 | 0.34 | 0.18 | 0.16 | 1.86 |
| endothelium | Cttnbp2 | 0.66 | 2.91E-11 | 6.05E-07 | 0.3 | 0.16 | 0.14 | 1.91 |
| endothelium | Map4k4 | 0.34 | 2.91E-11 | 6.05E-07 | 0.86 | 0.9 | -0.05 | 0.95 |
| endothelium | Agfg1 | 0.63 | 3.00E-11 | 6.25E-07 | 0.76 | 0.59 | 0.17 | 1.28 |
| endothelium | Slc41a1 | 0.53 | 3.04E-11 | 6.33E-07 | 0.28 | 0.14 | 0.14 | 1.97 |
| endothelium | Fam98c | 0.7 | 3.10E-11 | 6.45E-07 | 0.24 | 0.12 | 0.12 | 2.04 |
| endothelium | Tatdn3 | 0.98 | 3.14E-11 | 6.53E-07 | 0.22 | 0.11 | 0.11 | 2.09 |
| endothelium | Pip5k1c | 0.54 | 3.16E-11 | 6.57E-07 | 0.41 | 0.23 | 0.18 | 1.79 |
| endothelium | Gramd1a | 0.43 | 3.21E-11 | 6.68E-07 | 0.43 | 0.24 | 0.19 | 1.79 |
| endothelium | Shkbp1 | 0.76 | 3.29E-11 | 6.85E-07 | 0.53 | 0.34 | 0.19 | 1.57 |
| endothelium | Cfap20 | 0.48 | 3.33E-11 | 6.92E-07 | 0.82 | 0.75 | 0.08 | 1.1 |
| endothelium | Alox5ap | 1.29 | 3.42E-11 | 7.10E-07 | 0.11 | 0.04 | 0.07 | 2.82 |
| endothelium | Immp2l | 0.95 | 3.57E-11 | 7.42E-07 | 0.18 | 0.08 | 0.1 | 2.22 |
| endothelium | Acadm | 0.57 | 3.57E-11 | 7.43E-07 | 0.53 | 0.33 | 0.2 | 1.6 |
| endothelium | Sil1 | 0.55 | 3.71E-11 | 7.72E-07 | 0.23 | 0.11 | 0.12 | 2.07 |
| endothelium | Slc9a1 | 0.61 | 3.78E-11 | 7.87E-07 | 0.23 | 0.11 | 0.12 | 2.1 |
| endothelium | Spr | 0.62 | 4.01E-11 | 8.33E-07 | 0.69 | 0.5 | 0.19 | 1.38 |
| endothelium | Emc2 | 0.64 | 4.10E-11 | 8.53E-07 | 0.77 | 0.59 | 0.18 | 1.31 |
| endothelium | Lrch1 | 0.7 | 4.15E-11 | 8.62E-07 | 0.18 | 0.08 | 0.1 | 2.27 |
| endothelium | Esyt2 | 0.56 | 4.16E-11 | 8.65E-07 | 0.37 | 0.21 | 0.16 | 1.76 |
| endothelium | Katnbl1 | 0.6 | 4.33E-11 | 9.01E-07 | 0.55 | 0.35 | 0.2 | 1.57 |
| endothelium | Nucb2 | 0.53 | 4.57E-11 | 9.50E-07 | 0.3 | 0.15 | 0.15 | 1.94 |
| endothelium | Tiam1 | 0.43 | 5.39E-11 | 1.12E-06 | 0.51 | 0.31 | 0.2 | 1.64 |
| endothelium | Sin3b | 0.49 | 6.42E-11 | 1.34E-06 | 0.88 | 0.8 | 0.08 | 1.09 |
| endothelium | Fbxl12 | 0.44 | 6.48E-11 | 1.35E-06 | 0.36 | 0.2 | 0.16 | 1.83 |
| endothelium | Uqcc2 | 0.29 | 6.49E-11 | 1.35E-06 | 0.94 | 0.96 | -0.02 | 0.98 |
| endothelium | Flna | 0.5 | 6.52E-11 | 1.36E-06 | 0.84 | 0.79 | 0.04 | 1.06 |
| endothelium | Baz1a | 0.43 | 6.65E-11 | 1.38E-06 | 0.74 | 0.56 | 0.18 | 1.33 |
| endothelium | Septin1 | 0.71 | 6.74E-11 | 1.40E-06 | 0.39 | 0.22 | 0.17 | 1.75 |
| endothelium | Zfp984 | 0.69 | 6.81E-11 | 1.42E-06 | 0.17 | 0.07 | 0.1 | 2.3 |
| endothelium | Akap13 | 0.54 | 6.85E-11 | 1.42E-06 | 0.69 | 0.5 | 0.19 | 1.38 |
| endothelium | Plekhm1 | 1.13 | 6.93E-11 | 1.44E-06 | 0.14 | 0.06 | 0.08 | 2.4 |
| endothelium | 4921524J17Ri | 0.59 | 7.22E-11 | 1.50E-06 | 0.66 | 0.49 | 0.17 | 1.35 |
| endothelium | Ptpn4 | 0.72 | 7.52E-11 | 1.56E-06 | 0.29 | 0.16 | 0.14 | 1.85 |
| endothelium | Mad2l1bp | 0.69 | 7.57E-11 | 1.57E-06 | 0.41 | 0.23 | 0.17 | 1.73 |
| endothelium | Btbd1 | 0.52 | 7.66E-11 | 1.59E-06 | 0.8 | 0.63 | 0.17 | 1.26 |
| endothelium | Ankrd50 | 0.58 | 7.79E-11 | 1.62E-06 | 0.4 | 0.23 | 0.17 | 1.73 |
| endothelium | Epn2 | 0.66 | 7.85E-11 | 1.63E-06 | 0.46 | 0.28 | 0.18 | 1.66 |
| endothelium | Icmt | 0.58 | 7.94E-11 | 1.65E-06 | 0.25 | 0.13 | 0.13 | 1.99 |
| endothelium | Pigq | 0.57 | 8.07E-11 | 1.68E-06 | 0.41 | 0.24 | 0.18 | 1.74 |
| endothelium | Vps28 | 0.5 | 8.10E-11 | 1.68E-06 | 0.77 | 0.62 | 0.15 | 1.24 |
| endothelium | Nmb | 0.83 | 8.11E-11 | 1.69E-06 | 0.17 | 0.08 | 0.1 | 2.25 |
| endothelium | Ginm1 | 0.46 | 8.20E-11 | 1.71E-06 | 0.48 | 0.29 | 0.19 | 1.67 |
| endothelium | Atl3 | 0.49 | 8.27E-11 | 1.72E-06 | 0.39 | 0.22 | 0.17 | 1.77 |
| endothelium | Elk4 | 0.5 | 9.02E-11 | 1.87E-06 | 0.4 | 0.23 | 0.17 | 1.76 |
| endothelium | Fancg | 1.07 | 9.14E-11 | 1.90E-06 | 0.13 | 0.05 | 0.08 | 2.52 |
| endothelium | Trmt61a | 0.62 | 9.22E-11 | 1.92E-06 | 0.28 | 0.15 | 0.14 | 1.93 |
| endothelium | Zdhhc20 | 0.47 | 9.47E-11 | 1.97E-06 | 0.76 | 0.58 | 0.18 | 1.31 |
| endothelium | Sco2 | 0.5 | 9.58E-11 | 1.99E-06 | 0.34 | 0.19 | 0.15 | 1.81 |
| endothelium | Zfp524 | 0.75 | 1.10E-10 | 2.28E-06 | 0.21 | 0.1 | 0.11 | 2.05 |
| endothelium | Sap18 | 0.38 | 1.17E-10 | 2.42E-06 | 0.9 | 0.85 | 0.05 | 1.06 |

**Supplementary Table 2. Marker genes for the major cell types of integrated clustering of E11.5 control and NotchGOF λ-junction cells.**

| | | | | | | | | |
|---|---|---|---|---|---|---|---|---|
| endothelium | Map7d1 | 0.49 | 1.25E-10 | 2.60E-06 | 0.75 | 0.62 | 0.14 | 1.22 |
| endothelium | Amdhd2 | 0.49 | 1.27E-10 | 2.64E-06 | 0.42 | 0.24 | 0.18 | 1.74 |
| endothelium | Wdr45b | 0.53 | 1.31E-10 | 2.72E-06 | 0.76 | 0.58 | 0.17 | 1.3 |
| endothelium | Zfp560 | 0.55 | 1.33E-10 | 2.76E-06 | 0.57 | 0.37 | 0.2 | 1.55 |
| endothelium | Dgke | 0.68 | 1.34E-10 | 2.78E-06 | 0.16 | 0.07 | 0.09 | 2.33 |
| endothelium | Dnmt3b | 0.46 | 1.36E-10 | 2.82E-06 | 0.38 | 0.21 | 0.16 | 1.77 |
| endothelium | Sugt1 | 0.43 | 1.37E-10 | 2.85E-06 | 0.92 | 0.87 | 0.05 | 1.06 |
| endothelium | Spcs3 | 0.5 | 1.41E-10 | 2.94E-06 | 0.58 | 0.38 | 0.2 | 1.54 |
| endothelium | Abracl | 0.36 | 1.41E-10 | 2.94E-06 | 0.95 | 0.93 | 0.02 | 1.02 |
| endothelium | Slc25a24 | 0.57 | 1.46E-10 | 3.04E-06 | 0.55 | 0.35 | 0.2 | 1.58 |
| endothelium | Bag1 | 0.33 | 1.54E-10 | 3.20E-06 | 0.92 | 0.92 | 0 | 1 |
| endothelium | Nus1 | 0.53 | 1.60E-10 | 3.32E-06 | 0.67 | 0.48 | 0.18 | 1.38 |
| endothelium | Ogfod2 | 0.55 | 1.62E-10 | 3.37E-06 | 0.21 | 0.1 | 0.11 | 2.12 |
| endothelium | Dgkd | 0.72 | 1.63E-10 | 3.40E-06 | 0.4 | 0.23 | 0.16 | 1.69 |
| endothelium | 230307C23Ri | 0.72 | 1.64E-10 | 3.42E-06 | 0.31 | 0.16 | 0.14 | 1.88 |
| endothelium | Xylt1 | 0.71 | 1.67E-10 | 3.47E-06 | 0.14 | 0.06 | 0.08 | 2.4 |
| endothelium | Limch1 | 0.42 | 1.71E-10 | 3.55E-06 | 0.78 | 0.68 | 0.1 | 1.15 |
| endothelium | Ppp2cb | 0.53 | 1.76E-10 | 3.66E-06 | 0.64 | 0.44 | 0.2 | 1.45 |
| endothelium | Glb1 | 0.44 | 1.79E-10 | 3.73E-06 | 0.3 | 0.16 | 0.14 | 1.88 |
| endothelium | Fam49b | 0.45 | 1.80E-10 | 3.74E-06 | 0.78 | 0.65 | 0.13 | 1.19 |
| endothelium | Rab1b | 0.44 | 1.81E-10 | 3.76E-06 | 0.73 | 0.53 | 0.2 | 1.38 |
| endothelium | Txnl1 | 0.32 | 2.01E-10 | 4.18E-06 | 0.94 | 0.95 | -0.01 | 0.99 |
| endothelium | Lin28b | 0.63 | 2.04E-10 | 4.23E-06 | 0.36 | 0.21 | 0.16 | 1.76 |
| endothelium | Czib | 0.62 | 2.06E-10 | 4.28E-06 | 0.7 | 0.53 | 0.17 | 1.33 |
| endothelium | Arap1 | 0.98 | 2.06E-10 | 4.29E-06 | 0.16 | 0.07 | 0.09 | 2.29 |
| endothelium | Pcbd1 | 0.52 | 2.37E-10 | 4.93E-06 | 0.24 | 0.12 | 0.12 | 2.01 |
| endothelium | Mtcl1 | 0.56 | 2.40E-10 | 4.98E-06 | 0.24 | 0.12 | 0.12 | 1.98 |
| endothelium | Spty2d1 | 0.54 | 2.41E-10 | 5.00E-06 | 0.42 | 0.25 | 0.17 | 1.7 |
| endothelium | Heatr5a | 0.5 | 2.49E-10 | 5.18E-06 | 0.33 | 0.19 | 0.15 | 1.79 |
| endothelium | Bysl | 0.75 | 2.60E-10 | 5.40E-06 | 0.53 | 0.34 | 0.2 | 1.58 |
| endothelium | Add1 | 0.53 | 2.61E-10 | 5.43E-06 | 0.79 | 0.65 | 0.14 | 1.22 |
| endothelium | Sav1 | 0.48 | 2.80E-10 | 5.83E-06 | 0.66 | 0.45 | 0.21 | 1.47 |
| endothelium | Rpn2 | 0.34 | 2.86E-10 | 5.96E-06 | 0.84 | 0.79 | 0.05 | 1.06 |
| endothelium | Mcu | 0.67 | 2.96E-10 | 6.16E-06 | 0.29 | 0.15 | 0.13 | 1.88 |
| endothelium | Pigt | 0.43 | 2.97E-10 | 6.17E-06 | 0.62 | 0.43 | 0.19 | 1.45 |
| endothelium | Fkbp9 | 0.52 | 2.97E-10 | 6.18E-06 | 0.65 | 0.44 | 0.21 | 1.49 |
| endothelium | Cnbp | 0.29 | 3.22E-10 | 6.69E-06 | 0.96 | 0.98 | -0.02 | 0.98 |
| endothelium | Coa3 | 0.36 | 3.28E-10 | 6.83E-06 | 0.9 | 0.9 | 0 | 1 |
| endothelium | Hmox2 | 0.48 | 3.32E-10 | 6.91E-06 | 0.76 | 0.64 | 0.12 | 1.19 |
| endothelium | Socs5 | 0.47 | 3.47E-10 | 7.22E-06 | 0.27 | 0.14 | 0.13 | 1.94 |
| endothelium | Ttc32 | 0.54 | 3.62E-10 | 7.53E-06 | 0.43 | 0.25 | 0.18 | 1.7 |
| endothelium | Plpp7 | 0.86 | 3.71E-10 | 7.71E-06 | 0.1 | 0.04 | 0.06 | 2.73 |
| endothelium | Mtbp | 0.56 | 3.72E-10 | 7.73E-06 | 0.38 | 0.22 | 0.16 | 1.74 |
| endothelium | Sharpin | 0.42 | 3.76E-10 | 7.82E-06 | 0.31 | 0.17 | 0.15 | 1.87 |
| endothelium | Ppt2 | 0.47 | 4.09E-10 | 8.51E-06 | 0.43 | 0.25 | 0.18 | 1.7 |
| endothelium | Flii | 0.51 | 4.37E-10 | 9.09E-06 | 0.56 | 0.37 | 0.2 | 1.53 |
| endothelium | Gyg | 0.53 | 4.44E-10 | 9.23E-06 | 0.62 | 0.42 | 0.2 | 1.48 |
| endothelium | Ppp1r11 | 0.44 | 4.57E-10 | 9.49E-06 | 0.89 | 0.81 | 0.08 | 1.09 |
| endothelium | Sec14l1 | 0.44 | 4.62E-10 | 9.61E-06 | 0.52 | 0.32 | 0.2 | 1.6 |
| endothelium | Urm1 | 0.5 | 4.68E-10 | 9.73E-06 | 0.57 | 0.37 | 0.2 | 1.54 |
| endothelium | Tex261 | 0.49 | 4.79E-10 | 9.97E-06 | 0.76 | 0.6 | 0.16 | 1.26 |
| endothelium | Tspan12 | 0.45 | 4.85E-10 | 1.01E-05 | 0.37 | 0.21 | 0.16 | 1.78 |
| endothelium | Ccdc85c | 1.01 | 4.86E-10 | 1.01E-05 | 0.22 | 0.11 | 0.11 | 1.98 |
| endothelium | Golm1 | 0.49 | 5.25E-10 | 1.09E-05 | 0.7 | 0.55 | 0.15 | 1.27 |
| endothelium | Afg1l | 1.22 | 5.26E-10 | 1.09E-05 | 0.12 | 0.05 | 0.07 | 2.56 |
| endothelium | Crim1 | 0.33 | 5.33E-10 | 1.11E-05 | 0.35 | 0.2 | 0.16 | 1.79 |

**Supplementary Table 2. Marker genes for the major cell types of integrated clustering of E11.5 control and NotchGOF λ-junction cells.**

| | | | | | | | | |
|---|---|---|---|---|---|---|---|---|
| endothelium | Marchf8 | 0.34 | 5.39E-10 | 1.12E-05 | 0.32 | 0.17 | 0.15 | 1.86 |
| endothelium | Hadhb | 0.44 | 5.56E-10 | 1.16E-05 | 0.75 | 0.62 | 0.13 | 1.21 |
| endothelium | Gipc2 | 0.54 | 5.63E-10 | 1.17E-05 | 0.15 | 0.06 | 0.08 | 2.33 |
| endothelium | Eif3b | 0.42 | 5.66E-10 | 1.18E-05 | 0.86 | 0.82 | 0.04 | 1.05 |
| endothelium | Gpr180 | 0.42 | 5.67E-10 | 1.18E-05 | 0.49 | 0.3 | 0.2 | 1.66 |
| endothelium | .930006K02Ri | 1.1 | 5.80E-10 | 1.21E-05 | 0.11 | 0.04 | 0.07 | 2.55 |
| endothelium | Mafk | 1.11 | 6.55E-10 | 1.36E-05 | 0.1 | 0.04 | 0.07 | 2.67 |
| endothelium | Fars2 | 0.41 | 6.57E-10 | 1.37E-05 | 0.4 | 0.23 | 0.17 | 1.72 |
| endothelium | Mvb12a | 0.57 | 7.13E-10 | 1.48E-05 | 0.66 | 0.46 | 0.2 | 1.45 |
| endothelium | Dazap2 | 0.45 | 7.60E-10 | 1.58E-05 | 0.74 | 0.57 | 0.17 | 1.3 |
| endothelium | Ccny | 0.55 | 7.68E-10 | 1.60E-05 | 0.7 | 0.54 | 0.16 | 1.31 |
| endothelium | Prkx | 0.62 | 7.80E-10 | 1.62E-05 | 0.29 | 0.15 | 0.13 | 1.86 |
| endothelium | AI413582 | 1.13 | 8.65E-10 | 1.80E-05 | 0.19 | 0.09 | 0.1 | 2.11 |
| endothelium | Bak1 | 0.58 | 9.02E-10 | 1.88E-05 | 0.61 | 0.41 | 0.2 | 1.49 |
| endothelium | Mcm3 | 0.45 | 9.16E-10 | 1.91E-05 | 0.86 | 0.79 | 0.07 | 1.09 |
| endothelium | Galnt4 | 0.96 | 9.20E-10 | 1.91E-05 | 0.11 | 0.04 | 0.07 | 2.61 |
| endothelium | Tyk2 | 0.66 | 9.32E-10 | 1.94E-05 | 0.14 | 0.06 | 0.08 | 2.39 |
| endothelium | Ebp | 0.64 | 9.78E-10 | 2.03E-05 | 0.6 | 0.4 | 0.19 | 1.48 |
| endothelium | Slc39a11 | 0.71 | 1.00E-09 | 2.08E-05 | 0.16 | 0.07 | 0.09 | 2.28 |
| endothelium | Kif1c | 0.66 | 1.01E-09 | 2.09E-05 | 0.29 | 0.16 | 0.13 | 1.85 |
| endothelium | Zfp287 | 0.56 | 1.02E-09 | 2.12E-05 | 0.16 | 0.07 | 0.09 | 2.23 |
| endothelium | Itga3 | 0.5 | 1.06E-09 | 2.21E-05 | 0.16 | 0.07 | 0.09 | 2.25 |
| endothelium | Tcta | 0.69 | 1.06E-09 | 2.21E-05 | 0.14 | 0.06 | 0.08 | 2.31 |
| endothelium | Vps33a | 0.77 | 1.15E-09 | 2.39E-05 | 0.26 | 0.14 | 0.12 | 1.89 |
| endothelium | Il13ra1 | 0.74 | 1.16E-09 | 2.42E-05 | 0.12 | 0.05 | 0.07 | 2.56 |
| endothelium | Atg4a | 0.86 | 1.18E-09 | 2.46E-05 | 0.18 | 0.09 | 0.1 | 2.13 |
| endothelium | Rnf14 | 0.47 | 1.21E-09 | 2.52E-05 | 0.55 | 0.35 | 0.2 | 1.57 |
| endothelium | Spindoc | 0.45 | 1.25E-09 | 2.60E-05 | 0.49 | 0.3 | 0.18 | 1.6 |
| endothelium | Mlx | 0.59 | 1.25E-09 | 2.61E-05 | 0.49 | 0.3 | 0.18 | 1.6 |
| endothelium | Tomm5 | 0.36 | 1.26E-09 | 2.62E-05 | 0.9 | 0.9 | 0 | 1 |
| endothelium | Rbm38 | 1.12 | 1.31E-09 | 2.72E-05 | 0.38 | 0.24 | 0.15 | 1.63 |
| endothelium | Vsig10 | 0.7 | 1.33E-09 | 2.77E-05 | 0.13 | 0.05 | 0.08 | 2.42 |
| endothelium | Sbds | 0.43 | 1.36E-09 | 2.82E-05 | 0.7 | 0.52 | 0.18 | 1.35 |
| endothelium | Synj1 | 0.65 | 1.42E-09 | 2.95E-05 | 0.21 | 0.11 | 0.11 | 1.99 |
| endothelium | Fnip2 | 0.72 | 1.43E-09 | 2.97E-05 | 0.2 | 0.1 | 0.1 | 2.03 |
| endothelium | Vapb | 0.57 | 1.55E-09 | 3.22E-05 | 0.75 | 0.61 | 0.15 | 1.24 |
| endothelium | Slc12a6 | 0.69 | 1.58E-09 | 3.28E-05 | 0.25 | 0.13 | 0.12 | 1.89 |
| endothelium | Nln | 0.55 | 1.61E-09 | 3.36E-05 | 0.47 | 0.29 | 0.18 | 1.63 |
| endothelium | Btg3 | 0.58 | 1.75E-09 | 3.63E-05 | 0.5 | 0.32 | 0.18 | 1.56 |
| endothelium | Sh3pxd2b | 0.46 | 1.75E-09 | 3.64E-05 | 0.47 | 0.29 | 0.18 | 1.63 |
| endothelium | Xiap | 0.45 | 1.77E-09 | 3.67E-05 | 0.67 | 0.49 | 0.18 | 1.36 |
| endothelium | Arhgef12 | 0.49 | 1.78E-09 | 3.70E-05 | 0.74 | 0.58 | 0.16 | 1.28 |
| endothelium | Atp6v0a1 | 0.57 | 1.79E-09 | 3.72E-05 | 0.34 | 0.2 | 0.14 | 1.72 |
| endothelium | Fdx2 | 0.55 | 1.82E-09 | 3.78E-05 | 0.76 | 0.59 | 0.17 | 1.28 |
| endothelium | Txndc11 | 0.43 | 1.90E-09 | 3.95E-05 | 0.35 | 0.2 | 0.15 | 1.76 |
| endothelium | Tusc2 | 0.59 | 1.98E-09 | 4.12E-05 | 0.58 | 0.39 | 0.19 | 1.49 |
| endothelium | Map4k1 | 1.12 | 2.03E-09 | 4.22E-05 | 0.11 | 0.05 | 0.07 | 2.49 |
| endothelium | Slc31a1 | 0.48 | 2.09E-09 | 4.35E-05 | 0.53 | 0.35 | 0.19 | 1.54 |
| endothelium | Selenom | 0.33 | 2.17E-09 | 4.52E-05 | 0.7 | 0.52 | 0.18 | 1.34 |
| endothelium | Ech1 | 0.64 | 2.19E-09 | 4.56E-05 | 0.69 | 0.51 | 0.18 | 1.34 |
| endothelium | Cib1 | 0.54 | 2.22E-09 | 4.61E-05 | 0.6 | 0.4 | 0.2 | 1.49 |
| endothelium | Trappc6a | 0.53 | 2.24E-09 | 4.66E-05 | 0.71 | 0.52 | 0.19 | 1.36 |
| endothelium | Pcgf5 | 0.53 | 2.30E-09 | 4.77E-05 | 0.18 | 0.09 | 0.09 | 2.08 |
| endothelium | Rab43 | 1.08 | 2.31E-09 | 4.81E-05 | 0.15 | 0.07 | 0.08 | 2.26 |
| endothelium | Fbxw11 | 0.45 | 2.34E-09 | 4.87E-05 | 0.52 | 0.33 | 0.19 | 1.57 |
| endothelium | Ebna1bp2 | 0.43 | 2.35E-09 | 4.89E-05 | 0.84 | 0.8 | 0.04 | 1.05 |

**Supplementary Table 2. Marker genes for the major cell types of integrated clustering of E11.5 control and NotchGOF λ-junction cells.**

| | | | | | | | | |
|---|---|---|---|---|---|---|---|---|
| endothelium | Nts | 0.54 | 2.35E-09 | 4.89E-05 | 0.13 | 0.06 | 0.08 | 2.34 |
| endothelium | Mink1 | 0.39 | 2.39E-09 | 4.98E-05 | 0.31 | 0.17 | 0.14 | 1.84 |
| endothelium | Rab29 | 0.94 | 2.40E-09 | 4.99E-05 | 0.12 | 0.05 | 0.07 | 2.41 |
| endothelium | Cyb561d2 | 0.69 | 2.42E-09 | 5.02E-05 | 0.3 | 0.17 | 0.13 | 1.78 |
| endothelium | Pacsin2 | 0.46 | 2.45E-09 | 5.10E-05 | 0.54 | 0.35 | 0.19 | 1.55 |
| endothelium | Picalm | 0.4 | 2.54E-09 | 5.29E-05 | 0.75 | 0.62 | 0.13 | 1.21 |
| endothelium | Ttpal | 0.66 | 2.82E-09 | 5.86E-05 | 0.22 | 0.11 | 0.11 | 1.94 |
| endothelium | Fads3 | 0.75 | 2.84E-09 | 5.90E-05 | 0.13 | 0.06 | 0.08 | 2.34 |
| endothelium | Zfp994 | 0.57 | 2.84E-09 | 5.91E-05 | 0.19 | 0.09 | 0.1 | 2.05 |
| endothelium | Prkca | 2.03 | 2.87E-09 | 5.98E-05 | 0.17 | 0.08 | 0.09 | 2.02 |
| endothelium | Ppp6c | 0.45 | 2.96E-09 | 6.15E-05 | 0.65 | 0.47 | 0.19 | 1.4 |
| endothelium | .810037I17Ril | 0.45 | 2.97E-09 | 6.17E-05 | 0.87 | 0.84 | 0.04 | 1.04 |
| endothelium | Ier3 | 0.66 | 3.05E-09 | 6.35E-05 | 0.16 | 0.08 | 0.09 | 2.14 |
| endothelium | Ncoa3 | 0.65 | 3.09E-09 | 6.42E-05 | 0.37 | 0.23 | 0.15 | 1.65 |
| endothelium | Stard9 | 0.62 | 3.13E-09 | 6.52E-05 | 0.15 | 0.07 | 0.08 | 2.22 |
| endothelium | Ccm2 | 0.57 | 3.14E-09 | 6.54E-05 | 0.6 | 0.4 | 0.21 | 1.52 |
| endothelium | Stt3b | 0.45 | 3.23E-09 | 6.71E-05 | 0.69 | 0.51 | 0.19 | 1.37 |
| endothelium | Serinc5 | 0.92 | 3.24E-09 | 6.73E-05 | 0.13 | 0.06 | 0.08 | 2.34 |
| endothelium | Aldh3a2 | 0.87 | 3.73E-09 | 7.77E-05 | 0.17 | 0.08 | 0.09 | 2.14 |
| endothelium | Tmem256 | 0.27 | 3.85E-09 | 8.01E-05 | 0.9 | 0.89 | 0.01 | 1.01 |
| endothelium | Maml2 | 0.42 | 4.02E-09 | 8.35E-05 | 0.19 | 0.09 | 0.1 | 2.06 |
| endothelium | Ankrd13a | 0.43 | 4.07E-09 | 8.46E-05 | 0.46 | 0.29 | 0.18 | 1.61 |
| endothelium | Fam98a | 0.46 | 4.10E-09 | 8.52E-05 | 0.51 | 0.33 | 0.19 | 1.57 |
| endothelium | Rhobtb1 | 0.38 | 4.10E-09 | 8.53E-05 | 0.27 | 0.15 | 0.13 | 1.86 |
| endothelium | Serinc3 | 0.31 | 4.19E-09 | 8.72E-05 | 0.54 | 0.36 | 0.18 | 1.5 |
| endothelium | Inpp5b | 1.06 | 4.21E-09 | 8.77E-05 | 0.22 | 0.12 | 0.11 | 1.92 |
| endothelium | Dip2a | 0.68 | 4.26E-09 | 8.86E-05 | 0.2 | 0.1 | 0.1 | 2.01 |
| endothelium | Pcyox1l | 0.34 | 4.31E-09 | 8.97E-05 | 0.36 | 0.2 | 0.15 | 1.77 |
| endothelium | Taok3 | 1.82 | 4.38E-09 | 9.12E-05 | 0.2 | 0.1 | 0.09 | 1.88 |
| endothelium | Pvr | 0.7 | 4.57E-09 | 9.51E-05 | 0.15 | 0.07 | 0.08 | 2.22 |
| endothelium | L3hypdh | 0.44 | 4.88E-09 | 1.02E-04 | 0.33 | 0.19 | 0.15 | 1.78 |
| endothelium | Coro1b | 0.61 | 4.91E-09 | 1.02E-04 | 0.15 | 0.07 | 0.08 | 2.2 |
| endothelium | Lix1l | 0.41 | 4.98E-09 | 1.04E-04 | 0.58 | 0.38 | 0.2 | 1.54 |
| endothelium | Ero1l | 0.33 | 5.05E-09 | 1.05E-04 | 0.47 | 0.29 | 0.18 | 1.61 |
| endothelium | Zfp28 | 0.45 | 5.15E-09 | 1.07E-04 | 0.17 | 0.08 | 0.09 | 2.14 |
| endothelium | Etfb | 0.47 | 5.83E-09 | 1.21E-04 | 0.9 | 0.89 | 0.01 | 1.01 |
| endothelium | Spry4 | 0.76 | 6.36E-09 | 1.32E-04 | 0.26 | 0.14 | 0.12 | 1.83 |
| endothelium | Map2k1 | 0.44 | 6.70E-09 | 1.39E-04 | 0.53 | 0.34 | 0.19 | 1.54 |
| endothelium | Slc35e1 | 0.43 | 6.85E-09 | 1.42E-04 | 0.3 | 0.17 | 0.13 | 1.77 |
| endothelium | Fuca1 | 0.33 | 7.41E-09 | 1.54E-04 | 0.55 | 0.35 | 0.2 | 1.56 |
| endothelium | Ndufs5 | 0.29 | 7.70E-09 | 1.60E-04 | 0.94 | 0.95 | -0.01 | 0.99 |
| endothelium | Ralgps2 | 0.51 | 7.92E-09 | 1.65E-04 | 0.81 | 0.71 | 0.1 | 1.14 |
| endothelium | 2810408I11Ril | 0.46 | 7.93E-09 | 1.65E-04 | 0.48 | 0.3 | 0.18 | 1.59 |
| endothelium | Gabarapl2 | 0.32 | 8.18E-09 | 1.70E-04 | 0.87 | 0.82 | 0.05 | 1.06 |
| endothelium | Fut4 | 0.7 | 8.32E-09 | 1.73E-04 | 0.14 | 0.06 | 0.08 | 2.27 |
| endothelium | Sh3glb2 | 0.41 | 8.55E-09 | 1.78E-04 | 0.51 | 0.32 | 0.19 | 1.61 |
| endothelium | Actr1a | 0.49 | 8.80E-09 | 1.83E-04 | 0.64 | 0.46 | 0.18 | 1.39 |
| endothelium | Galk1 | 0.45 | 8.93E-09 | 1.86E-04 | 0.81 | 0.71 | 0.1 | 1.13 |
| endothelium | Dapk3 | 0.6 | 9.36E-09 | 1.95E-04 | 0.64 | 0.47 | 0.17 | 1.36 |
| endothelium | Ube2g2 | 0.69 | 9.53E-09 | 1.98E-04 | 0.5 | 0.33 | 0.18 | 1.55 |
| endothelium | Echdc1 | 0.47 | 9.59E-09 | 1.99E-04 | 0.4 | 0.24 | 0.16 | 1.67 |
| endothelium | Gpr108 | 0.46 | 9.72E-09 | 2.02E-04 | 0.29 | 0.16 | 0.13 | 1.8 |
| endothelium | Adamts4 | 0.51 | 9.77E-09 | 2.03E-04 | 0.13 | 0.06 | 0.08 | 2.29 |
| endothelium | Zdhhc18 | 0.41 | 9.94E-09 | 2.07E-04 | 0.27 | 0.15 | 0.13 | 1.86 |
| endothelium | Acox1 | 0.43 | 9.95E-09 | 2.07E-04 | 0.26 | 0.14 | 0.12 | 1.85 |
| endothelium | Gpatch3 | 0.58 | 9.95E-09 | 2.07E-04 | 0.15 | 0.07 | 0.08 | 2.19 |

**Supplementary Table 2. Marker genes for the major cell types of integrated clustering of E11.5 control and NotchGOF λ-junction cells.**

| | | | | | | | | |
|---|---|---|---|---|---|---|---|---|
| endothelium | Atp1b3 | 0.36 | 1.00E-08 | 2.09E-04 | 0.82 | 0.72 | 0.11 | 1.15 |
| endothelium | Capzb | 0.3 | 1.02E-08 | 2.12E-04 | 0.92 | 0.92 | 0.01 | 1.01 |
| endothelium | Tcea2 | 1.59 | 1.13E-08 | 2.36E-04 | 0.11 | 0.05 | 0.07 | 2.43 |
| endothelium | Slc25a25 | 0.69 | 1.17E-08 | 2.43E-04 | 0.31 | 0.18 | 0.13 | 1.73 |
| endothelium | Baz2a | 0.39 | 1.18E-08 | 2.46E-04 | 0.58 | 0.39 | 0.19 | 1.5 |
| endothelium | Pter | 0.56 | 1.18E-08 | 2.46E-04 | 0.22 | 0.11 | 0.11 | 1.94 |
| endothelium | Plod3 | 0.33 | 1.19E-08 | 2.47E-04 | 0.52 | 0.32 | 0.19 | 1.6 |
| endothelium | Cul7 | 0.63 | 1.21E-08 | 2.51E-04 | 0.56 | 0.36 | 0.2 | 1.55 |
| endothelium | Arfgef2 | 1.32 | 1.27E-08 | 2.65E-04 | 0.18 | 0.09 | 0.09 | 2.02 |
| endothelium | Micall1 | 0.9 | 1.28E-08 | 2.67E-04 | 0.22 | 0.12 | 0.1 | 1.87 |
| endothelium | Trip10 | 0.51 | 1.30E-08 | 2.70E-04 | 0.36 | 0.22 | 0.14 | 1.65 |
| endothelium | Cemip2 | 0.31 | 1.31E-08 | 2.72E-04 | 0.62 | 0.45 | 0.18 | 1.39 |
| endothelium | Srd5a3 | 0.52 | 1.31E-08 | 2.72E-04 | 0.36 | 0.21 | 0.15 | 1.69 |
| endothelium | Rnf13 | 0.31 | 1.32E-08 | 2.74E-04 | 0.41 | 0.24 | 0.17 | 1.69 |
| endothelium | Cbl | 0.41 | 1.34E-08 | 2.79E-04 | 0.65 | 0.47 | 0.18 | 1.39 |
| endothelium | Riok3 | 0.5 | 1.36E-08 | 2.83E-04 | 0.68 | 0.5 | 0.18 | 1.36 |
| endothelium | Rnf31 | 0.56 | 1.40E-08 | 2.90E-04 | 0.15 | 0.07 | 0.08 | 2.19 |
| endothelium | Gm11772 | 1.22 | 1.43E-08 | 2.97E-04 | 0.11 | 0.05 | 0.07 | 2.43 |
| endothelium | Aen | 0.58 | 1.57E-08 | 3.26E-04 | 0.58 | 0.39 | 0.19 | 1.47 |
| endothelium | Sipa1l2 | 0.39 | 1.59E-08 | 3.32E-04 | 0.25 | 0.13 | 0.12 | 1.86 |
| endothelium | Dera | 0.52 | 1.60E-08 | 3.33E-04 | 0.78 | 0.66 | 0.12 | 1.17 |
| endothelium | Dennd5a | 0.45 | 1.62E-08 | 3.37E-04 | 0.53 | 0.35 | 0.18 | 1.53 |
| endothelium | Amfr | 0.44 | 1.67E-08 | 3.47E-04 | 0.56 | 0.36 | 0.2 | 1.55 |
| endothelium | Gbe1 | 0.48 | 1.73E-08 | 3.60E-04 | 0.2 | 0.1 | 0.1 | 1.98 |
| endothelium | Tomm40l | 0.5 | 1.76E-08 | 3.66E-04 | 0.29 | 0.16 | 0.13 | 1.78 |
| endothelium | Cyc1 | 0.28 | 1.79E-08 | 3.72E-04 | 0.9 | 0.9 | 0 | 1 |
| endothelium | Scyl1 | 0.4 | 1.86E-08 | 3.87E-04 | 0.45 | 0.28 | 0.17 | 1.59 |
| endothelium | Inpp5k | 0.48 | 2.05E-08 | 4.27E-04 | 0.21 | 0.11 | 0.1 | 1.92 |
| endothelium | Hdlbp | 0.31 | 2.08E-08 | 4.32E-04 | 0.87 | 0.86 | 0.01 | 1.01 |
| endothelium | Anxa7 | 0.4 | 2.10E-08 | 4.37E-04 | 0.56 | 0.36 | 0.2 | 1.55 |
| endothelium | Mob3a | 0.6 | 2.12E-08 | 4.40E-04 | 0.28 | 0.16 | 0.12 | 1.77 |
| endothelium | Dcaf6 | 0.51 | 2.14E-08 | 4.46E-04 | 0.17 | 0.08 | 0.09 | 2.05 |
| endothelium | Tbrg4 | 0.59 | 2.19E-08 | 4.55E-04 | 0.32 | 0.19 | 0.13 | 1.7 |
| endothelium | Tmem144 | 0.73 | 2.21E-08 | 4.61E-04 | 0.11 | 0.05 | 0.07 | 2.38 |
| endothelium | Retreg1 | 0.4 | 2.24E-08 | 4.65E-04 | 0.18 | 0.09 | 0.09 | 2 |
| endothelium | Tigar | 0.6 | 2.29E-08 | 4.77E-04 | 0.15 | 0.07 | 0.08 | 2.14 |
| endothelium | Zmpste24 | 0.36 | 2.35E-08 | 4.88E-04 | 0.49 | 0.31 | 0.18 | 1.57 |
| endothelium | Mrps6 | 0.63 | 2.35E-08 | 4.88E-04 | 0.78 | 0.63 | 0.15 | 1.24 |
| endothelium | Arrb2 | 0.68 | 2.35E-08 | 4.89E-04 | 0.29 | 0.17 | 0.12 | 1.72 |
| endothelium | Cox7a2l | 0.25 | 2.35E-08 | 4.90E-04 | 0.93 | 0.96 | -0.02 | 0.97 |
| endothelium | Entr1 | 0.56 | 2.47E-08 | 5.14E-04 | 0.63 | 0.43 | 0.2 | 1.46 |
| endothelium | Ano6 | 0.48 | 2.50E-08 | 5.21E-04 | 0.4 | 0.25 | 0.16 | 1.64 |
| endothelium | Rybp | 0.47 | 2.51E-08 | 5.23E-04 | 0.61 | 0.42 | 0.19 | 1.45 |
| endothelium | Cd276 | 0.38 | 2.52E-08 | 5.25E-04 | 0.51 | 0.33 | 0.18 | 1.55 |
| endothelium | Klhl21 | 0.7 | 2.58E-08 | 5.36E-04 | 0.12 | 0.05 | 0.07 | 2.35 |
| endothelium | Bcap31 | 0.45 | 2.58E-08 | 5.37E-04 | 0.82 | 0.75 | 0.07 | 1.1 |
| endothelium | Agpat5 | 0.63 | 2.68E-08 | 5.58E-04 | 0.62 | 0.45 | 0.17 | 1.39 |
| endothelium | Tmem219 | 0.34 | 2.74E-08 | 5.70E-04 | 0.28 | 0.15 | 0.12 | 1.8 |
| endothelium | Zfp335 | 0.36 | 2.77E-08 | 5.76E-04 | 0.19 | 0.09 | 0.09 | 2.01 |
| endothelium | Med27 | 0.5 | 2.83E-08 | 5.88E-04 | 0.41 | 0.25 | 0.16 | 1.63 |
| endothelium | Tmem170 | 0.31 | 2.86E-08 | 5.96E-04 | 0.19 | 0.09 | 0.1 | 2.01 |
| endothelium | Ppid | 0.37 | 2.91E-08 | 6.06E-04 | 0.89 | 0.88 | 0.01 | 1.01 |
| endothelium | Polk | 0.63 | 2.94E-08 | 6.11E-04 | 0.23 | 0.12 | 0.11 | 1.89 |
| endothelium | Ciapin1 | 0.36 | 2.98E-08 | 6.19E-04 | 0.82 | 0.74 | 0.08 | 1.11 |
| endothelium | Gpatch4 | 0.58 | 3.12E-08 | 6.49E-04 | 0.69 | 0.55 | 0.14 | 1.26 |
| endothelium | Snx2 | 0.46 | 3.12E-08 | 6.49E-04 | 0.74 | 0.58 | 0.17 | 1.29 |

**Supplementary Table 2. Marker genes for the major cell types of integrated clustering of E11.5 control and NotchGOF λ-junction cells.**

| | | | | | | | | |
|---|---|---|---|---|---|---|---|---|
| endothelium | Pcbd2 | 0.54 | 3.20E-08 | 6.65E-04 | 0.61 | 0.43 | 0.18 | 1.42 |
| endothelium | Fam149a | 0.36 | 3.38E-08 | 7.03E-04 | 0.27 | 0.15 | 0.12 | 1.84 |
| endothelium | Neurl2 | 0.79 | 3.39E-08 | 7.04E-04 | 0.14 | 0.07 | 0.08 | 2.15 |
| endothelium | Gm49066 | 0.75 | 3.49E-08 | 7.25E-04 | 0.11 | 0.04 | 0.06 | 2.43 |
| endothelium | Hmbox1 | 0.45 | 3.53E-08 | 7.34E-04 | 0.5 | 0.33 | 0.17 | 1.53 |
| endothelium | Agtrap | 0.47 | 3.53E-08 | 7.35E-04 | 0.26 | 0.15 | 0.11 | 1.78 |
| endothelium | Rab31 | 0.5 | 3.64E-08 | 7.57E-04 | 0.37 | 0.22 | 0.15 | 1.67 |
| endothelium | Fpgs | 0.52 | 3.67E-08 | 7.63E-04 | 0.32 | 0.19 | 0.13 | 1.71 |
| endothelium | Tyw5 | 0.41 | 3.68E-08 | 7.66E-04 | 0.37 | 0.22 | 0.15 | 1.66 |
| endothelium | Pdcl3 | 0.43 | 3.75E-08 | 7.81E-04 | 0.81 | 0.71 | 0.1 | 1.15 |
| endothelium | Frg1 | 0.51 | 3.79E-08 | 7.89E-04 | 0.82 | 0.74 | 0.08 | 1.11 |
| endothelium | Yars | 0.53 | 3.80E-08 | 7.90E-04 | 0.54 | 0.37 | 0.17 | 1.47 |
| endothelium | Sh3bp4 | 0.53 | 3.82E-08 | 7.94E-04 | 0.29 | 0.17 | 0.12 | 1.72 |
| endothelium | Gstk1 | 0.54 | 4.05E-08 | 8.42E-04 | 0.3 | 0.18 | 0.13 | 1.73 |
| endothelium | Zfp800 | 0.45 | 4.09E-08 | 8.50E-04 | 0.43 | 0.27 | 0.17 | 1.63 |
| endothelium | St6galnac6 | 0.51 | 4.15E-08 | 8.63E-04 | 0.28 | 0.16 | 0.12 | 1.77 |
| endothelium | Limd1 | 0.38 | 4.26E-08 | 8.85E-04 | 0.69 | 0.51 | 0.18 | 1.34 |
| endothelium | Entpd6 | 0.43 | 4.27E-08 | 8.88E-04 | 0.26 | 0.15 | 0.12 | 1.79 |
| endothelium | Pop7 | 0.44 | 4.33E-08 | 9.00E-04 | 0.68 | 0.52 | 0.16 | 1.3 |
| endothelium | Dock1 | 0.48 | 4.36E-08 | 9.06E-04 | 0.67 | 0.5 | 0.17 | 1.34 |
| endothelium | Cstf2 | 0.41 | 4.38E-08 | 9.12E-04 | 0.53 | 0.34 | 0.18 | 1.54 |
| endothelium | Cars | 0.55 | 4.46E-08 | 9.27E-04 | 0.46 | 0.3 | 0.17 | 1.55 |
| endothelium | Mtmr9 | 0.27 | 4.70E-08 | 9.77E-04 | 0.26 | 0.15 | 0.12 | 1.82 |
| endothelium | Slc37a4 | 0.41 | 4.78E-08 | 9.93E-04 | 0.13 | 0.06 | 0.07 | 2.22 |
| endothelium | Ano10 | 0.42 | 4.93E-08 | 1.02E-03 | 0.24 | 0.13 | 0.11 | 1.85 |
| endothelium | Srf | 0.79 | 5.26E-08 | 1.09E-03 | 0.24 | 0.13 | 0.11 | 1.82 |
| endothelium | Etv6 | 0.47 | 5.37E-08 | 1.12E-03 | 0.43 | 0.28 | 0.15 | 1.54 |
| endothelium | Nat10 | 0.52 | 5.48E-08 | 1.14E-03 | 0.32 | 0.19 | 0.13 | 1.67 |
| endothelium | Tango2 | 0.71 | 5.61E-08 | 1.17E-03 | 0.27 | 0.16 | 0.12 | 1.75 |
| endothelium | Camkk2 | 0.51 | 6.04E-08 | 1.26E-03 | 0.38 | 0.23 | 0.15 | 1.66 |
| endothelium | Malt1 | 0.55 | 6.07E-08 | 1.26E-03 | 0.29 | 0.16 | 0.12 | 1.76 |
| endothelium | Nudt14 | 0.35 | 6.20E-08 | 1.29E-03 | 0.59 | 0.4 | 0.19 | 1.48 |
| endothelium | Eea1 | 0.33 | 6.70E-08 | 1.39E-03 | 0.61 | 0.43 | 0.18 | 1.42 |
| endothelium | Slc25a32 | 0.45 | 6.96E-08 | 1.45E-03 | 0.17 | 0.09 | 0.09 | 2.01 |
| endothelium | Nfkb2 | 0.83 | 7.27E-08 | 1.51E-03 | 0.14 | 0.07 | 0.07 | 2.1 |
| endothelium | Ifrd2 | 0.55 | 7.34E-08 | 1.53E-03 | 0.5 | 0.33 | 0.18 | 1.53 |
| endothelium | Zdhhc17 | 0.35 | 7.39E-08 | 1.54E-03 | 0.41 | 0.25 | 0.16 | 1.62 |
| endothelium | Phf7 | 0.51 | 7.42E-08 | 1.54E-03 | 0.26 | 0.15 | 0.11 | 1.79 |
| endothelium | Wdr11 | 0.31 | 7.74E-08 | 1.61E-03 | 0.39 | 0.24 | 0.16 | 1.66 |
| endothelium | Tollip | 0.43 | 7.74E-08 | 1.61E-03 | 0.28 | 0.16 | 0.12 | 1.74 |
| endothelium | Svbp | 0.32 | 7.84E-08 | 1.63E-03 | 0.89 | 0.87 | 0.02 | 1.02 |
| endothelium | Tubd1 | 0.46 | 8.00E-08 | 1.66E-03 | 0.19 | 0.1 | 0.09 | 1.95 |
| endothelium | Pde12 | 0.48 | 8.06E-08 | 1.68E-03 | 0.44 | 0.28 | 0.16 | 1.57 |
| endothelium | Prps2 | 0.28 | 8.11E-08 | 1.69E-03 | 0.29 | 0.17 | 0.12 | 1.73 |
| endothelium | Lypla2 | 0.38 | 8.23E-08 | 1.71E-03 | 0.54 | 0.36 | 0.18 | 1.51 |
| endothelium | Klhl5 | 0.64 | 9.07E-08 | 1.89E-03 | 0.3 | 0.17 | 0.13 | 1.74 |
| endothelium | Armcx1 | 0.43 | 9.26E-08 | 1.93E-03 | 0.52 | 0.34 | 0.18 | 1.53 |
| endothelium | Mfhas1 | 0.42 | 9.27E-08 | 1.93E-03 | 0.33 | 0.19 | 0.14 | 1.72 |
| endothelium | Hif3a | 0.66 | 9.70E-08 | 2.02E-03 | 0.17 | 0.09 | 0.08 | 1.97 |
| endothelium | Arxes1 | 0.3 | 9.70E-08 | 2.02E-03 | 0.54 | 0.37 | 0.17 | 1.46 |
| endothelium | Tmem30a | 0.4 | 1.00E-07 | 2.08E-03 | 0.68 | 0.51 | 0.17 | 1.33 |
| endothelium | Ammecr1 | 0.45 | 1.00E-07 | 2.09E-03 | 0.53 | 0.35 | 0.18 | 1.5 |
| endothelium | Sgpp1 | 0.38 | 1.01E-07 | 2.09E-03 | 0.2 | 0.1 | 0.1 | 1.93 |
| endothelium | Eif6 | 0.37 | 1.03E-07 | 2.15E-03 | 0.87 | 0.87 | 0 | 1 |
| endothelium | Trim71 | 0.37 | 1.04E-07 | 2.16E-03 | 0.35 | 0.21 | 0.14 | 1.66 |
| endothelium | Ptgr2 | 0.31 | 1.04E-07 | 2.17E-03 | 0.26 | 0.15 | 0.12 | 1.77 |

**Supplementary Table 2. Marker genes for the major cell types of integrated clustering of E11.5 control and NotchGOF λ-junction cells.**

| | | | | | | | | |
|---|---|---|---|---|---|---|---|---|
| endothelium | Cnn3 | 0.28 | 1.05E-07 | 2.18E-03 | 0.92 | 0.94 | -0.02 | 0.98 |
| endothelium | Foxn2 | 0.66 | 1.07E-07 | 2.22E-03 | 0.28 | 0.16 | 0.11 | 1.7 |
| endothelium | Fkbp14 | 0.33 | 1.10E-07 | 2.29E-03 | 0.25 | 0.14 | 0.11 | 1.8 |
| endothelium | Gas2l3 | 0.51 | 1.12E-07 | 2.32E-03 | 0.3 | 0.18 | 0.12 | 1.65 |
| endothelium | Snx25 | 0.57 | 1.12E-07 | 2.33E-03 | 0.14 | 0.06 | 0.07 | 2.13 |
| endothelium | Acox3 | 0.64 | 1.15E-07 | 2.39E-03 | 0.15 | 0.07 | 0.08 | 2.07 |
| endothelium | Wwc2 | 0.33 | 1.16E-07 | 2.42E-03 | 0.73 | 0.58 | 0.16 | 1.27 |
| endothelium | Klf9 | 1.41 | 1.18E-07 | 2.44E-03 | 0.11 | 0.05 | 0.06 | 2.24 |
| endothelium | Otud4 | 0.3 | 1.18E-07 | 2.46E-03 | 0.48 | 0.31 | 0.17 | 1.55 |
| endothelium | Cacul1 | 0.31 | 1.20E-07 | 2.50E-03 | 0.45 | 0.28 | 0.17 | 1.6 |
| endothelium | Itgb1bp1 | 0.51 | 1.23E-07 | 2.55E-03 | 0.57 | 0.39 | 0.18 | 1.45 |
| endothelium | Babam2 | 0.49 | 1.23E-07 | 2.56E-03 | 0.51 | 0.35 | 0.17 | 1.48 |
| endothelium | Fhl1 | 0.96 | 1.24E-07 | 2.58E-03 | 0.25 | 0.14 | 0.1 | 1.73 |
| endothelium | Sdccag8 | 0.43 | 1.26E-07 | 2.61E-03 | 0.23 | 0.13 | 0.1 | 1.83 |
| endothelium | Bpnt1 | 0.26 | 1.27E-07 | 2.64E-03 | 0.25 | 0.14 | 0.11 | 1.81 |
| endothelium | Sgf29 | 0.52 | 1.29E-07 | 2.68E-03 | 0.49 | 0.32 | 0.17 | 1.52 |
| endothelium | Paox | 0.28 | 1.34E-07 | 2.79E-03 | 0.28 | 0.16 | 0.12 | 1.78 |
| endothelium | Trabd | 0.57 | 1.36E-07 | 2.83E-03 | 0.57 | 0.42 | 0.15 | 1.36 |
| endothelium | Sbno2 | 0.59 | 1.36E-07 | 2.83E-03 | 0.21 | 0.11 | 0.1 | 1.85 |
| endothelium | Tcirg1 | 0.77 | 1.46E-07 | 3.05E-03 | 0.17 | 0.09 | 0.09 | 1.97 |
| endothelium | Gga3 | 0.59 | 1.53E-07 | 3.17E-03 | 0.25 | 0.14 | 0.11 | 1.76 |
| endothelium | Ttc13 | 0.35 | 1.53E-07 | 3.18E-03 | 0.37 | 0.23 | 0.14 | 1.63 |
| endothelium | Zfp36l1 | 0.31 | 1.54E-07 | 3.21E-03 | 0.84 | 0.85 | -0.01 | 0.98 |
| endothelium | Plcl1 | 0.46 | 1.54E-07 | 3.21E-03 | 0.13 | 0.06 | 0.07 | 2.15 |
| endothelium | Prelid3b | 0.43 | 1.57E-07 | 3.27E-03 | 0.77 | 0.65 | 0.12 | 1.18 |
| endothelium | Eif4e2 | 0.36 | 1.63E-07 | 3.38E-03 | 0.89 | 0.86 | 0.03 | 1.04 |
| endothelium | Esco1 | 0.41 | 1.64E-07 | 3.41E-03 | 0.38 | 0.23 | 0.15 | 1.63 |
| endothelium | Smarcd2 | 0.52 | 1.69E-07 | 3.52E-03 | 0.45 | 0.29 | 0.16 | 1.57 |
| endothelium | Desi1 | 0.49 | 1.73E-07 | 3.60E-03 | 0.55 | 0.38 | 0.17 | 1.45 |
| endothelium | Dnajc2 | 0.31 | 1.76E-07 | 3.65E-03 | 0.88 | 0.86 | 0.02 | 1.02 |
| endothelium | Man2c1 | 0.87 | 1.79E-07 | 3.71E-03 | 0.2 | 0.11 | 0.09 | 1.88 |
| endothelium | Septin8 | 0.34 | 1.82E-07 | 3.78E-03 | 0.36 | 0.22 | 0.14 | 1.65 |
| endothelium | Eif4h | 0.27 | 1.85E-07 | 3.84E-03 | 0.91 | 0.93 | -0.02 | 0.98 |
| endothelium | Atp6v0b | 0.3 | 1.86E-07 | 3.88E-03 | 0.82 | 0.74 | 0.08 | 1.11 |
| endothelium | Sh3kbp1 | 0.37 | 1.96E-07 | 4.08E-03 | 0.37 | 0.22 | 0.15 | 1.66 |
| endothelium | Morc4 | 0.32 | 2.11E-07 | 4.38E-03 | 0.26 | 0.15 | 0.11 | 1.79 |
| endothelium | Tm2d3 | 0.5 | 2.13E-07 | 4.43E-03 | 0.68 | 0.5 | 0.18 | 1.35 |
| endothelium | Nt5dc1 | 0.46 | 2.14E-07 | 4.45E-03 | 0.2 | 0.11 | 0.09 | 1.87 |
| endothelium | Mrps11 | 0.5 | 2.14E-07 | 4.46E-03 | 0.81 | 0.7 | 0.11 | 1.16 |
| endothelium | Samm50 | 0.38 | 2.15E-07 | 4.46E-03 | 0.81 | 0.72 | 0.09 | 1.12 |
| endothelium | Nckap1 | 0.39 | 2.18E-07 | 4.54E-03 | 0.73 | 0.56 | 0.16 | 1.29 |
| endothelium | Tmcc1 | 0.32 | 2.28E-07 | 4.74E-03 | 0.62 | 0.44 | 0.18 | 1.42 |
| endothelium | Ptdss1 | 0.34 | 2.31E-07 | 4.81E-03 | 0.33 | 0.2 | 0.13 | 1.66 |
| endothelium | Epn1 | 0.36 | 2.34E-07 | 4.87E-03 | 0.77 | 0.62 | 0.14 | 1.23 |
| endothelium | Alg14 | 0.52 | 2.42E-07 | 5.03E-03 | 0.47 | 0.31 | 0.16 | 1.51 |
| endothelium | Cit | 0.32 | 2.48E-07 | 5.17E-03 | 0.55 | 0.37 | 0.18 | 1.47 |
| endothelium | Tnfsf12 | 0.45 | 2.54E-07 | 5.29E-03 | 0.14 | 0.07 | 0.07 | 2.06 |
| endothelium | Hint3 | 0.71 | 2.55E-07 | 5.30E-03 | 0.38 | 0.24 | 0.14 | 1.61 |
| endothelium | Ifngr2 | 0.52 | 2.71E-07 | 5.64E-03 | 0.56 | 0.37 | 0.19 | 1.5 |
| endothelium | Ccdc62 | 0.75 | 2.79E-07 | 5.80E-03 | 0.14 | 0.07 | 0.07 | 2.04 |
| endothelium | Eif2s1 | 0.25 | 2.85E-07 | 5.92E-03 | 0.88 | 0.9 | -0.03 | 0.97 |
| endothelium | Lbr | 0.45 | 2.89E-07 | 6.00E-03 | 0.66 | 0.5 | 0.16 | 1.32 |
| endothelium | Emc6 | 0.34 | 2.93E-07 | 6.10E-03 | 0.88 | 0.83 | 0.04 | 1.05 |
| endothelium | Rasa1 | 0.4 | 2.95E-07 | 6.13E-03 | 0.76 | 0.61 | 0.15 | 1.25 |
| endothelium | Rapgef2 | 0.55 | 3.49E-07 | 7.26E-03 | 0.36 | 0.22 | 0.13 | 1.61 |
| endothelium | Smurf1 | 0.56 | 3.56E-07 | 7.40E-03 | 0.25 | 0.14 | 0.11 | 1.77 |

**Supplementary Table 2. Marker genes for the major cell types of integrated clustering of E11.5 control and NotchGOF λ-junction cells.**

| | | | | | | | | |
|---|---|---|---|---|---|---|---|---|
| endothelium | Coa5 | 0.77 | 3.56E-07 | 7.41E-03 | 0.41 | 0.27 | 0.14 | 1.53 |
| endothelium | Qsox1 | 0.32 | 3.58E-07 | 7.44E-03 | 0.21 | 0.12 | 0.1 | 1.82 |
| endothelium | Atg16l1 | 0.33 | 3.62E-07 | 7.54E-03 | 0.34 | 0.2 | 0.13 | 1.66 |
| endothelium | Sec11a | 0.26 | 3.63E-07 | 7.55E-03 | 0.88 | 0.9 | -0.02 | 0.98 |
| endothelium | Zyg11b | 0.34 | 3.68E-07 | 7.66E-03 | 0.51 | 0.34 | 0.17 | 1.5 |
| endothelium | Pofut2 | 0.42 | 3.69E-07 | 7.68E-03 | 0.65 | 0.48 | 0.17 | 1.35 |
| endothelium | Trp53inp2 | 0.47 | 3.80E-07 | 7.90E-03 | 0.18 | 0.1 | 0.09 | 1.89 |
| endothelium | Ranbp9 | 0.54 | 3.85E-07 | 8.00E-03 | 0.51 | 0.34 | 0.17 | 1.51 |
| endothelium | Rwdd2b | 0.5 | 3.87E-07 | 8.04E-03 | 0.21 | 0.12 | 0.1 | 1.83 |
| endothelium | Nox4 | 0.41 | 3.89E-07 | 8.09E-03 | 0.16 | 0.08 | 0.08 | 1.98 |
| endothelium | Mrtfa | 0.61 | 3.89E-07 | 8.10E-03 | 0.22 | 0.12 | 0.09 | 1.77 |
| endothelium | Hyal2 | 0.35 | 3.91E-07 | 8.13E-03 | 0.71 | 0.53 | 0.18 | 1.34 |
| endothelium | Mtpn | 0.28 | 3.92E-07 | 8.16E-03 | 0.84 | 0.81 | 0.03 | 1.04 |
| endothelium | Yipf6 | 0.32 | 3.98E-07 | 8.28E-03 | 0.32 | 0.19 | 0.13 | 1.7 |
| endothelium | Atp9a | 0.43 | 4.01E-07 | 8.35E-03 | 0.3 | 0.18 | 0.12 | 1.7 |
| endothelium | Clcn4 | 0.28 | 4.01E-07 | 8.35E-03 | 0.47 | 0.3 | 0.17 | 1.55 |
| endothelium | Dnase1l1 | 0.89 | 4.07E-07 | 8.47E-03 | 0.1 | 0.05 | 0.06 | 2.26 |
| endothelium | Zbtb11 | 0.37 | 4.09E-07 | 8.51E-03 | 0.37 | 0.23 | 0.14 | 1.61 |
| endothelium | Plcb3 | 0.26 | 4.17E-07 | 8.68E-03 | 0.45 | 0.29 | 0.16 | 1.56 |
| endothelium | Atp6v0d1 | 0.42 | 4.23E-07 | 8.80E-03 | 0.69 | 0.54 | 0.16 | 1.29 |
| endothelium | Brox | 0.72 | 4.26E-07 | 8.85E-03 | 0.37 | 0.22 | 0.14 | 1.64 |
| endothelium | Mtg1 | 0.31 | 4.31E-07 | 8.96E-03 | 0.26 | 0.15 | 0.11 | 1.75 |
| endothelium | Taldo1 | 0.38 | 4.32E-07 | 8.99E-03 | 0.91 | 0.89 | 0.02 | 1.02 |
| endothelium | Slc25a26 | 0.6 | 4.41E-07 | 9.16E-03 | 0.17 | 0.09 | 0.08 | 1.92 |
| endothelium | Nop2 | 0.37 | 4.41E-07 | 9.18E-03 | 0.5 | 0.33 | 0.17 | 1.51 |
| endothelium | Calm3 | 0.25 | 4.44E-07 | 9.23E-03 | 0.89 | 0.91 | -0.02 | 0.98 |
| endothelium | Fam174a | 0.3 | 4.46E-07 | 9.28E-03 | 0.48 | 0.31 | 0.17 | 1.55 |
| endothelium | Gcnt2 | 1.21 | 4.49E-07 | 9.34E-03 | 0.1 | 0.05 | 0.06 | 2.21 |
| endothelium | Luzp1 | 0.4 | 4.54E-07 | 9.43E-03 | 0.82 | 0.77 | 0.05 | 1.06 |
| endothelium | Utrn | 0.28 | 4.54E-07 | 9.43E-03 | 0.76 | 0.61 | 0.15 | 1.25 |
| endothelium | Ncoa4 | 0.63 | 4.99E-07 | 1.04E-02 | 0.48 | 0.32 | 0.16 | 1.51 |
| endothelium | Klf16 | 0.51 | 5.00E-07 | 1.04E-02 | 0.16 | 0.08 | 0.08 | 1.99 |
| endothelium | Traf4 | 0.5 | 5.02E-07 | 1.04E-02 | 0.75 | 0.63 | 0.12 | 1.19 |
| endothelium | Fbxl17 | 0.5 | 5.31E-07 | 1.10E-02 | 0.33 | 0.2 | 0.13 | 1.66 |
| endothelium | Pxmp2 | 0.72 | 5.32E-07 | 1.11E-02 | 0.15 | 0.07 | 0.07 | 2.01 |
| endothelium | Cenpt | 0.45 | 5.33E-07 | 1.11E-02 | 0.4 | 0.25 | 0.15 | 1.6 |
| endothelium | Pard3b | 0.73 | 5.37E-07 | 1.12E-02 | 0.21 | 0.12 | 0.09 | 1.79 |
| endothelium | Zdhhc15 | 0.28 | 5.42E-07 | 1.13E-02 | 0.23 | 0.13 | 0.1 | 1.8 |
| endothelium | Usp14 | 0.41 | 5.51E-07 | 1.15E-02 | 0.87 | 0.79 | 0.08 | 1.09 |
| endothelium | Tpk1 | 0.4 | 5.54E-07 | 1.15E-02 | 0.12 | 0.06 | 0.07 | 2.16 |
| endothelium | Gnpda2 | 0.35 | 5.59E-07 | 1.16E-02 | 0.44 | 0.28 | 0.16 | 1.57 |
| endothelium | Dmwd | 0.48 | 5.67E-07 | 1.18E-02 | 0.59 | 0.45 | 0.14 | 1.32 |
| endothelium | Antxr2 | 0.28 | 5.79E-07 | 1.20E-02 | 0.31 | 0.19 | 0.12 | 1.67 |
| endothelium | Pop5 | 0.4 | 5.80E-07 | 1.21E-02 | 0.76 | 0.65 | 0.11 | 1.16 |
| endothelium | Ranbp2 | 0.46 | 6.09E-07 | 1.27E-02 | 0.78 | 0.66 | 0.12 | 1.18 |
| endothelium | Tpgs1 | 0.49 | 6.24E-07 | 1.30E-02 | 0.6 | 0.43 | 0.17 | 1.38 |
| endothelium | Tmem86a | 0.6 | 6.41E-07 | 1.33E-02 | 0.19 | 0.1 | 0.09 | 1.85 |
| endothelium | Nsf | 0.54 | 6.86E-07 | 1.43E-02 | 0.49 | 0.34 | 0.15 | 1.46 |
| endothelium | Phldb1 | 0.25 | 6.95E-07 | 1.45E-02 | 0.33 | 0.2 | 0.13 | 1.66 |
| endothelium | Rab10os | 0.39 | 6.95E-07 | 1.45E-02 | 0.3 | 0.18 | 0.12 | 1.65 |
| endothelium | Sgce | 0.6 | 6.99E-07 | 1.45E-02 | 0.48 | 0.32 | 0.15 | 1.47 |
| endothelium | Pgm3 | 0.51 | 7.24E-07 | 1.51E-02 | 0.44 | 0.29 | 0.15 | 1.53 |
| endothelium | Them4 | 0.48 | 7.79E-07 | 1.62E-02 | 0.22 | 0.13 | 0.1 | 1.75 |
| endothelium | Dph2 | 0.41 | 7.91E-07 | 1.64E-02 | 0.18 | 0.1 | 0.09 | 1.86 |
| endothelium | Hacd1 | 0.6 | 7.99E-07 | 1.66E-02 | 0.56 | 0.39 | 0.17 | 1.42 |
| endothelium | Ndufa5 | 0.27 | 8.03E-07 | 1.67E-02 | 0.91 | 0.92 | -0.01 | 0.99 |

**Supplementary Table 2. Marker genes for the major cell types of integrated clustering of E11.5 control and NotchGOF λ-junction cells.**

| | | | | | | | | |
|---|---|---|---|---|---|---|---|---|
| endothelium | AA01118383 | 0.3 | 8.20E-07 | 1.71E-02 | 0.37 | 0.23 | 0.14 | 1.59 |
| endothelium | Dusp19 | 0.32 | 8.54E-07 | 1.78E-02 | 0.3 | 0.18 | 0.12 | 1.68 |
| endothelium | Ddi2 | 0.49 | 8.59E-07 | 1.79E-02 | 0.56 | 0.41 | 0.15 | 1.36 |
| endothelium | Gas2l1 | 1.61 | 8.66E-07 | 1.80E-02 | 0.31 | 0.2 | 0.11 | 1.56 |
| endothelium | Neat1 | 1.81 | 9.11E-07 | 1.90E-02 | 0.12 | 0.06 | 0.06 | 2.09 |
| endothelium | Esrra | 0.36 | 9.29E-07 | 1.93E-02 | 0.22 | 0.12 | 0.1 | 1.78 |
| endothelium | Tdrkh | 0.79 | 9.31E-07 | 1.94E-02 | 0.13 | 0.06 | 0.07 | 2.08 |
| endothelium | Dcaf12 | 0.31 | 9.31E-07 | 1.94E-02 | 0.51 | 0.34 | 0.17 | 1.5 |
| endothelium | Atp6v1a | 0.34 | 9.54E-07 | 1.98E-02 | 0.69 | 0.53 | 0.17 | 1.32 |
| endothelium | Stam2 | 0.28 | 9.83E-07 | 2.04E-02 | 0.48 | 0.31 | 0.17 | 1.54 |
| endothelium | Exoc8 | 0.55 | 9.94E-07 | 2.07E-02 | 0.18 | 0.1 | 0.08 | 1.86 |
| endothelium | Vps37b | 0.37 | 1.00E-06 | 2.08E-02 | 0.45 | 0.29 | 0.16 | 1.54 |
| endothelium | Ropn1l | 0.44 | 1.01E-06 | 2.10E-02 | 0.13 | 0.06 | 0.07 | 2.08 |
| endothelium | Slc35a3 | 0.29 | 1.05E-06 | 2.17E-02 | 0.22 | 0.12 | 0.1 | 1.78 |
| endothelium | Tomm22 | 0.26 | 1.06E-06 | 2.19E-02 | 0.89 | 0.9 | -0.01 | 0.99 |
| endothelium | Ticam1 | 0.57 | 1.17E-06 | 2.43E-02 | 0.11 | 0.05 | 0.06 | 2.18 |
| endothelium | Madd | 0.68 | 1.18E-06 | 2.46E-02 | 0.14 | 0.07 | 0.07 | 1.93 |
| endothelium | Uba1 | 0.27 | 1.39E-06 | 2.88E-02 | 0.84 | 0.76 | 0.08 | 1.1 |
| endothelium | Smad1 | 0.46 | 1.39E-06 | 2.90E-02 | 0.63 | 0.48 | 0.15 | 1.32 |
| endothelium | Slc16a3 | 0.29 | 1.40E-06 | 2.91E-02 | 0.48 | 0.33 | 0.16 | 1.48 |
| endothelium | Zfp597 | 0.98 | 1.41E-06 | 2.94E-02 | 0.15 | 0.08 | 0.07 | 1.93 |
| endothelium | Psmb10 | 0.61 | 1.44E-06 | 2.99E-02 | 0.36 | 0.23 | 0.13 | 1.57 |
| endothelium | Cables2 | 0.43 | 1.45E-06 | 3.02E-02 | 0.3 | 0.18 | 0.12 | 1.63 |
| endothelium | Atg7 | 0.27 | 1.47E-06 | 3.05E-02 | 0.22 | 0.13 | 0.1 | 1.77 |
| endothelium | Rab7 | 0.36 | 1.52E-06 | 3.16E-02 | 0.79 | 0.69 | 0.1 | 1.15 |
| endothelium | Ttll1 | 0.74 | 1.55E-06 | 3.23E-02 | 0.26 | 0.16 | 0.11 | 1.66 |
| endothelium | L3mbtl2 | 0.28 | 1.62E-06 | 3.37E-02 | 0.31 | 0.19 | 0.12 | 1.65 |
| endothelium | Arhgap17 | 0.29 | 1.67E-06 | 3.47E-02 | 0.54 | 0.37 | 0.17 | 1.46 |
| endothelium | Mknk2 | 0.29 | 1.77E-06 | 3.69E-02 | 0.59 | 0.41 | 0.18 | 1.44 |
| endothelium | Lmtk2 | 0.61 | 1.79E-06 | 3.72E-02 | 0.18 | 0.1 | 0.08 | 1.86 |
| endothelium | Slmap | 0.57 | 1.80E-06 | 3.74E-02 | 0.48 | 0.33 | 0.15 | 1.47 |
| endothelium | Pcsk7 | 0.69 | 1.80E-06 | 3.74E-02 | 0.25 | 0.14 | 0.1 | 1.73 |
| endothelium | Mcts1 | 0.39 | 1.86E-06 | 3.86E-02 | 0.65 | 0.47 | 0.18 | 1.38 |
| endothelium | Mak16 | 0.38 | 1.89E-06 | 3.93E-02 | 0.78 | 0.66 | 0.11 | 1.17 |
| endothelium | Borcs6 | 0.53 | 1.92E-06 | 3.99E-02 | 0.25 | 0.15 | 0.1 | 1.71 |
| endothelium | Smim26 | 0.5 | 1.96E-06 | 4.07E-02 | 0.59 | 0.43 | 0.16 | 1.37 |
| endothelium | Psmc4 | 0.27 | 1.98E-06 | 4.11E-02 | 0.92 | 0.91 | 0.01 | 1.01 |
| endothelium | Rad51b | 0.62 | 2.00E-06 | 4.16E-02 | 0.2 | 0.12 | 0.09 | 1.77 |
| endothelium | Pcyt1a | 0.33 | 2.05E-06 | 4.27E-02 | 0.48 | 0.32 | 0.16 | 1.51 |
| endothelium | Cpt1a | 0.47 | 2.11E-06 | 4.40E-02 | 0.18 | 0.1 | 0.08 | 1.82 |
| endothelium | Rab4b | 0.4 | 2.15E-06 | 4.46E-02 | 0.55 | 0.39 | 0.16 | 1.42 |
| endothelium | Rabif | 0.53 | 2.16E-06 | 4.49E-02 | 0.4 | 0.25 | 0.15 | 1.59 |
| endothelium | Asah1 | 0.42 | 2.19E-06 | 4.56E-02 | 0.54 | 0.37 | 0.17 | 1.46 |
| endothelium | Chml | 0.46 | 2.23E-06 | 4.64E-02 | 0.25 | 0.15 | 0.1 | 1.7 |
| endothelium | Acvr1 | 0.36 | 2.24E-06 | 4.67E-02 | 0.31 | 0.19 | 0.12 | 1.62 |
| endothelium | Tmem248 | 0.35 | 2.26E-06 | 4.70E-02 | 0.51 | 0.35 | 0.16 | 1.45 |
| endothelium | Cttnbp2nl | 0.55 | 2.27E-06 | 4.73E-02 | 0.64 | 0.5 | 0.14 | 1.29 |
| endothelium | Agpat3 | 0.3 | 2.32E-06 | 4.82E-02 | 0.41 | 0.26 | 0.14 | 1.53 |
| endothelium | Bach1 | 0.41 | 2.36E-06 | 4.91E-02 | 0.64 | 0.48 | 0.17 | 1.35 |
| olfactory neurons | Lhx2 | 10.28 | 0 | 0 | 0.88 | 0.02 | 0.86 | 55 |
| olfactory neurons | Insm1 | 9.09 | 0 | 0 | 0.86 | 0.01 | 0.85 | 85.5 |
| olfactory neurons | Elavl4 | 8.32 | 0 | 0 | 0.87 | 0.06 | 0.81 | 14.95 |
| olfactory neurons | Nhlh2 | 10.84 | 0 | 0 | 0.81 | 0 | 0.8 | 269 |
| olfactory neurons | Neurod1 | 11.75 | 0 | 0 | 0.8 | 0 | 0.8 | 267 |
| olfactory neurons | Igfbpl1 | 10.67 | 0 | 0 | 0.8 | 0.01 | 0.8 | 133.5 |
| olfactory neurons | Elavl3 | 10.25 | 0 | 0 | 0.8 | 0.01 | 0.79 | 99.38 |

**Supplementary Table 2. Marker genes for the major cell types of integrated clustering of E11.5 control and NotchGOF λ-junction cells.**

| | | | | | | | | |
|---|---|---|---|---|---|---|---|---|
| olfactory neurons | Ebf2 | 6.67 | 0 | 0 | 0.81 | 0.04 | 0.78 | 23.23 |
| olfactory neurons | Ppp1r14a | 6.64 | 0 | 0 | 0.84 | 0.07 | 0.77 | 12.49 |
| olfactory neurons | Srrm4 | 10.32 | 0 | 0 | 0.77 | 0 | 0.76 | 255 |
| olfactory neurons | Mfng | 6.75 | 0 | 0 | 0.8 | 0.04 | 0.76 | 20.03 |
| olfactory neurons | Neurog1 | 11.14 | 0 | 0 | 0.75 | 0 | 0.75 | 373.5 |
| olfactory neurons | Wdr66 | 7.77 | 0 | 0 | 0.7 | 0.01 | 0.69 | 49.93 |
| olfactory neurons | Nhlh1 | 11.43 | 0 | 0 | 0.66 | 0 | 0.66 | 657 |
| olfactory neurons | Soga3 | 6 | 0 | 0 | 0.7 | 0.05 | 0.65 | 14.56 |
| olfactory neurons | Fezf1 | 6.92 | 0 | 0 | 0.63 | 0.02 | 0.61 | 33.32 |
| olfactory neurons | Btbd17 | 9.11 | 0 | 0 | 0.62 | 0.01 | 0.61 | 103.33 |
| olfactory neurons | Dcx | 6.64 | 0 | 0 | 0.63 | 0.02 | 0.61 | 27.52 |
| olfactory neurons | Miat | 7.31 | 0 | 0 | 0.6 | 0.01 | 0.59 | 60.2 |
| olfactory neurons | Ckmt1 | 6.7 | 0 | 0 | 0.63 | 0.04 | 0.59 | 16.08 |
| olfactory neurons | Sowaha | 10.59 | 0 | 0 | 0.59 | 0 | 0.59 | 295 |
| olfactory neurons | Stmn3 | 7.43 | 0 | 0 | 0.61 | 0.03 | 0.58 | 19 |
| olfactory neurons | Syp | 7.04 | 0 | 0 | 0.58 | 0.01 | 0.57 | 41.71 |
| olfactory neurons | Kif19a | 7.67 | 0 | 0 | 0.56 | 0.01 | 0.55 | 70 |
| olfactory neurons | Cdhr1 | 9.99 | 0 | 0 | 0.54 | 0 | 0.54 | 180.67 |
| olfactory neurons | St18 | 11.39 | 0 | 0 | 0.52 | 0 | 0.52 | 518 |
| olfactory neurons | Rbm24 | 7.24 | 0 | 0 | 0.52 | 0.01 | 0.51 | 43.67 |
| olfactory neurons | Syt13 | 7.04 | 0 | 0 | 0.51 | 0.01 | 0.5 | 51.2 |
| olfactory neurons | Chga | 8.62 | 0 | 0 | 0.51 | 0.01 | 0.5 | 56.22 |
| olfactory neurons | Ina | 10.9 | 0 | 0 | 0.5 | 0.01 | 0.5 | 100 |
| olfactory neurons | Coro2a | 6.1 | 0 | 0 | 0.51 | 0.02 | 0.49 | 29.76 |
| olfactory neurons | Thsd7b | 6 | 0 | 0 | 0.49 | 0.02 | 0.47 | 24.7 |
| olfactory neurons | Uncx | 6.8 | 0 | 0 | 0.49 | 0.02 | 0.47 | 23.52 |
| olfactory neurons | Nrxn1 | 6.27 | 0 | 0 | 0.49 | 0.02 | 0.47 | 21.48 |
| olfactory neurons | Slc17a6 | 11.56 | 0 | 0 | 0.46 | 0 | 0.46 | #NA |
| olfactory neurons | Dcc | 7.62 | 0 | 0 | 0.47 | 0.01 | 0.46 | 58.75 |
| olfactory neurons | Gdap1l1 | 6.99 | 0 | 0 | 0.47 | 0.01 | 0.46 | 33.57 |
| olfactory neurons | Olig2 | 11.82 | 0 | 0 | 0.44 | 0 | 0.44 | #NA |
| olfactory neurons | Celf3 | 8.96 | 0 | 0 | 0.43 | 0 | 0.43 | 217 |
| olfactory neurons | Foxo6 | 7.05 | 0 | 0 | 0.44 | 0.01 | 0.43 | 40 |
| olfactory neurons | Brsk2 | 6.38 | 0 | 0 | 0.42 | 0.01 | 0.41 | 42.2 |
| olfactory neurons | Myt1l | 10.99 | 0 | 0 | 0.4 | 0 | 0.4 | 398 |
| olfactory neurons | Atp8b4 | 9.83 | 0 | 0 | 0.4 | 0 | 0.4 | 199 |
| olfactory neurons | Scg3 | 8.16 | 0 | 0 | 0.4 | 0.01 | 0.39 | 79.6 |
| olfactory neurons | Ttc9b | 8.31 | 0 | 0 | 0.38 | 0 | 0.38 | 95 |
| olfactory neurons | Syt4 | 6.8 | 0 | 0 | 0.37 | 0.01 | 0.36 | 61.17 |
| olfactory neurons | Inka2 | 6.64 | 0 | 0 | 0.36 | 0.01 | 0.35 | 45.13 |
| olfactory neurons | Scn9a | 6.49 | 0 | 0 | 0.36 | 0.01 | 0.35 | 40.11 |
| olfactory neurons | Shisal1 | 7.06 | 0 | 0 | 0.36 | 0.01 | 0.35 | 59.17 |
| olfactory neurons | Fstl5 | 8.25 | 0 | 0 | 0.34 | 0.01 | 0.34 | 42.88 |
| olfactory neurons | Dscaml1 | 9.46 | 0 | 0 | 0.32 | 0 | 0.32 | 319 |
| olfactory neurons | Nxph4 | 8.54 | 0 | 0 | 0.32 | 0 | 0.32 | 159.5 |
| olfactory neurons | Pcdh8 | 7.16 | 0 | 0 | 0.32 | 0.01 | 0.31 | 53.17 |
| olfactory neurons | Akap6 | 8.15 | 0 | 0 | 0.3 | 0.01 | 0.3 | 60.2 |
| olfactory neurons | Olig1 | 9.83 | 0 | 0 | 0.29 | 0 | 0.29 | 289 |
| olfactory neurons | Rgs20 | 7.71 | 0 | 0 | 0.27 | 0 | 0.27 | 90.33 |
| olfactory neurons | Clvs1 | 8.15 | 0 | 0 | 0.27 | 0 | 0.27 | 90.33 |
| olfactory neurons | Igsf21 | 9.22 | 0 | 0 | 0.27 | 0 | 0.26 | 265 |
| olfactory neurons | Stmn4 | 8.17 | 0 | 0 | 0.27 | 0 | 0.26 | 66.25 |
| olfactory neurons | Dner | 6.49 | 0 | 0 | 0.26 | 0 | 0.26 | 64.75 |
| olfactory neurons | Mast1 | 6.52 | 0 | 0 | 0.26 | 0 | 0.26 | 64.75 |
| olfactory neurons | Nol4 | 6.87 | 0 | 0 | 0.26 | 0.01 | 0.25 | 51.8 |
| olfactory neurons | Myt1 | 9.91 | 0 | 0 | 0.24 | 0 | 0.24 | #NA |

**Supplementary Table 2. Marker genes for the major cell types of integrated clustering of E11.5 control and NotchGOF λ-junction cells.**

| | | | | | | | | |
|---|---|---|---|---|---|---|---|---|
| olfactory neurons | Chgb | 8.24 | 0 | 0 | 0.23 | 0 | 0.23 | 76.33 |
| olfactory neurons | Slc17a7 | 7.8 | 0 | 0 | 0.22 | 0 | 0.22 | 223 |
| olfactory neurons | Smim18 | 10.63 | 0 | 0 | 0.22 | 0 | 0.22 | #NA |
| olfactory neurons | Pcsk2 | 8.89 | 0 | 0 | 0.22 | 0 | 0.22 | 217 |
| olfactory neurons | Actl6b | 11.39 | 0 | 0 | 0.21 | 0 | 0.21 | #NA |
| olfactory neurons | 730403I07Ri | 9.53 | 0 | 0 | 0.21 | 0 | 0.21 | 211 |
| olfactory neurons | Pirt | 11.07 | 0 | 0 | 0.19 | 0 | 0.19 | #NA |
| olfactory neurons | 230051N06Ri | 10.58 | 0 | 0 | 0.18 | 0 | 0.18 | #NA |
| olfactory neurons | Chrna3 | 8.51 | 0 | 0 | 0.18 | 0 | 0.17 | 175 |
| olfactory neurons | Gfy | 10.3 | 0 | 0 | 0.17 | 0 | 0.17 | #NA |
| olfactory neurons | Ppp1r17 | 10.39 | 0 | 0 | 0.16 | 0 | 0.16 | 157 |
| olfactory neurons | Gm3234 | 7.97 | 0 | 0 | 0.16 | 0 | 0.16 | 157 |
| olfactory neurons | Tmprss6 | 8.87 | 0 | 0 | 0.16 | 0 | 0.16 | 157 |
| olfactory neurons | Gda | 8.64 | 0 | 0 | 0.16 | 0 | 0.16 | 157 |
| olfactory neurons | Rd3 | 9.94 | 0 | 0 | 0.15 | 0 | 0.15 | #NA |
| olfactory neurons | Gm27197 | 10.5 | 0 | 0 | 0.13 | 0 | 0.13 | #NA |
| olfactory neurons | Gm31819 | 9.65 | 0 | 0 | 0.11 | 0 | 0.11 | #NA |
| olfactory neurons | Eef1a2 | 8.96 | 0 | 0 | 0.1 | 0 | 0.1 | #NA |
| olfactory neurons | Kcnk10 | 8.89 | 0 | 0 | 0.1 | 0 | 0.1 | #NA |
| olfactory neurons | Islr2 | 7.8 | 1.83E-295 | 3.80E-291 | 0.16 | 0 | 0.16 | 81.5 |
| olfactory neurons | Srrm3 | 6.19 | 6.35E-292 | 1.32E-287 | 0.28 | 0.01 | 0.27 | 39.57 |
| olfactory neurons | Rnf112 | 6.49 | 1.99E-288 | 4.14E-284 | 0.19 | 0 | 0.19 | 64.33 |
| olfactory neurons | Syt2 | 6.89 | 5.08E-285 | 1.06E-280 | 0.22 | 0 | 0.21 | 54.25 |
| olfactory neurons | Mreg | 6.29 | 1.01E-284 | 2.10E-280 | 0.34 | 0.01 | 0.33 | 30.64 |
| olfactory neurons | Gm17322 | 8.4 | 5.90E-283 | 1.23E-278 | 0.16 | 0 | 0.16 | 78.5 |
| olfactory neurons | Svop | 8.11 | 4.12E-278 | 8.58E-274 | 0.11 | 0 | 0.11 | 108 |
| olfactory neurons | Tesc | 8.4 | 1.34E-277 | 2.79E-273 | 0.26 | 0.01 | 0.25 | 43.17 |
| olfactory neurons | Map6 | 6.6 | 2.41E-277 | 5.01E-273 | 0.34 | 0.01 | 0.33 | 28.08 |
| olfactory neurons | Gm13589 | 7.14 | 4.54E-277 | 9.44E-273 | 0.19 | 0 | 0.18 | 62.33 |
| olfactory neurons | Ccdc187 | 8 | 8.96E-266 | 1.86E-261 | 0.1 | 0 | 0.1 | 102 |
| olfactory neurons | Nsg2 | 6.49 | 4.21E-265 | 8.75E-261 | 0.33 | 0.01 | 0.32 | 27.58 |
| olfactory neurons | Spock1 | 7.09 | 2.52E-259 | 5.25E-255 | 0.19 | 0 | 0.19 | 64.33 |
| olfactory neurons | Sez6l2 | 7.12 | 2.95E-257 | 6.13E-253 | 0.18 | 0 | 0.17 | 58.33 |
| olfactory neurons | Rundc3a | 5.62 | 4.59E-257 | 9.54E-253 | 0.5 | 0.03 | 0.47 | 17.24 |
| olfactory neurons | Ntrk1 | 8.4 | 2.10E-252 | 4.36E-248 | 0.11 | 0 | 0.11 | 108 |
| olfactory neurons | Scrt1 | 6.45 | 3.89E-252 | 8.09E-248 | 0.23 | 0.01 | 0.22 | 45.8 |
| olfactory neurons | Lingo2 | 6.94 | 1.50E-247 | 3.13E-243 | 0.22 | 0.01 | 0.22 | 44.6 |
| olfactory neurons | Ndrg4 | 5.44 | 3.53E-242 | 7.35E-238 | 0.49 | 0.03 | 0.46 | 15.94 |
| olfactory neurons | Cpeb1 | 8.69 | 5.27E-231 | 1.10E-226 | 0.11 | 0 | 0.11 | 108 |
| olfactory neurons | Cntn2 | 6.16 | 3.52E-229 | 7.33E-225 | 0.34 | 0.02 | 0.33 | 22.87 |
| olfactory neurons | Tubb3 | 7.61 | 1.66E-226 | 3.44E-222 | 0.98 | 0.17 | 0.81 | 5.64 |
| olfactory neurons | Nrn1 | 7.47 | 1.21E-222 | 2.52E-218 | 0.22 | 0.01 | 0.22 | 37.17 |
| olfactory neurons | Prmt8 | 8.25 | 4.62E-220 | 9.62E-216 | 0.11 | 0 | 0.11 | 114 |
| olfactory neurons | Asic1 | 6.28 | 8.79E-219 | 1.83E-214 | 0.18 | 0 | 0.17 | 58.33 |
| olfactory neurons | Brinp2 | 6.13 | 3.44E-216 | 7.16E-212 | 0.19 | 0 | 0.19 | 48.25 |
| olfactory neurons | Scg5 | 4.74 | 3.91E-212 | 8.14E-208 | 0.58 | 0.05 | 0.53 | 11.68 |
| olfactory neurons | 930020B18Ri | 5.86 | 4.56E-212 | 9.48E-208 | 0.36 | 0.02 | 0.34 | 19.72 |
| olfactory neurons | Fam57b | 6.41 | 6.21E-212 | 1.29E-207 | 0.2 | 0.01 | 0.19 | 39.8 |
| olfactory neurons | Shisa7 | 6 | 8.99E-210 | 1.87E-205 | 0.21 | 0.01 | 0.21 | 35.17 |
| olfactory neurons | Vwa5b2 | 5.76 | 1.77E-205 | 3.68E-201 | 0.17 | 0 | 0.17 | 56.33 |
| olfactory neurons | Gdap1 | 5.29 | 8.28E-205 | 1.72E-200 | 0.36 | 0.02 | 0.34 | 19 |
| olfactory neurons | Sct | 7.64 | 1.55E-200 | 3.22E-196 | 0.42 | 0.03 | 0.39 | 16 |
| olfactory neurons | Stmn2 | 6.83 | 1.48E-199 | 3.09E-195 | 0.75 | 0.1 | 0.65 | 7.78 |
| olfactory neurons | Cntn1 | 6.22 | 4.98E-187 | 1.04E-182 | 0.21 | 0.01 | 0.2 | 34.17 |
| olfactory neurons | Ap3b2 | 5.43 | 1.62E-185 | 3.37E-181 | 0.34 | 0.02 | 0.32 | 18.72 |
| olfactory neurons | Dll1 | 6.66 | 3.42E-185 | 7.10E-181 | 0.16 | 0 | 0.15 | 52.33 |

**Supplementary Table 2. Marker genes for the major cell types of integrated clustering of E11.5 control and NotchGOF λ-junction cells.**

| | | | | | | | | |
|---|---|---|---|---|---|---|---|---|
| olfactory neurons | Myo7a | 5.83 | 2.41E-181 | 5.00E-177 | 0.27 | 0.01 | 0.26 | 22.58 |
| olfactory neurons | Gm2694 | 5.52 | 3.67E-179 | 7.63E-175 | 0.33 | 0.02 | 0.31 | 19.12 |
| olfactory neurons | Tuba4a | 4.49 | 2.06E-178 | 4.28E-174 | 0.61 | 0.07 | 0.54 | 9.35 |
| olfactory neurons | Snap25 | 6.09 | 2.86E-178 | 5.94E-174 | 0.23 | 0.01 | 0.22 | 28.63 |
| olfactory neurons | Arhgdig | 7.11 | 6.09E-174 | 1.27E-169 | 0.64 | 0.08 | 0.56 | 8.19 |
| olfactory neurons | 930578M01R | 5.24 | 5.09E-173 | 1.06E-168 | 0.37 | 0.02 | 0.35 | 15.54 |
| olfactory neurons | Gm16364 | 7.74 | 1.21E-171 | 2.51E-167 | 0.1 | 0 | 0.1 | 102 |
| olfactory neurons | Lrrc10b | 6 | 2.90E-170 | 6.04E-166 | 0.17 | 0 | 0.17 | 42.25 |
| olfactory neurons | Dll3 | 5.2 | 8.65E-157 | 1.80E-152 | 0.29 | 0.02 | 0.27 | 18.06 |
| olfactory neurons | Pnma2 | 5.48 | 2.43E-152 | 5.06E-148 | 0.24 | 0.01 | 0.23 | 23.5 |
| olfactory neurons | Lamp5 | 5.96 | 9.41E-152 | 1.96E-147 | 0.18 | 0.01 | 0.18 | 30.17 |
| olfactory neurons | Mapk10 | 6.04 | 4.10E-149 | 8.53E-145 | 0.18 | 0.01 | 0.18 | 30.17 |
| olfactory neurons | Atp1a3 | 6.56 | 2.20E-142 | 4.59E-138 | 0.13 | 0 | 0.12 | 42.33 |
| olfactory neurons | Gpx2 | 6.9 | 3.46E-141 | 7.19E-137 | 0.1 | 0 | 0.1 | 51 |
| olfactory neurons | Emx2 | 5.07 | 6.42E-140 | 1.34E-135 | 0.48 | 0.05 | 0.43 | 9.52 |
| olfactory neurons | Dpysl5 | 4.06 | 1.63E-137 | 3.40E-133 | 0.67 | 0.11 | 0.56 | 6.37 |
| olfactory neurons | Apba1 | 6.84 | 1.13E-135 | 2.35E-131 | 0.1 | 0 | 0.1 | 51 |
| olfactory neurons | Rgs11 | 5.17 | 1.88E-135 | 3.90E-131 | 0.23 | 0.01 | 0.22 | 20.82 |
| olfactory neurons | Mapk8ip2 | 4.78 | 7.20E-135 | 1.50E-130 | 0.4 | 0.04 | 0.37 | 10.92 |
| olfactory neurons | Rap1gap2 | 5.22 | 6.75E-133 | 1.40E-128 | 0.25 | 0.01 | 0.24 | 18.07 |
| olfactory neurons | Celf6 | 6.77 | 3.93E-132 | 8.16E-128 | 0.11 | 0 | 0.11 | 54 |
| olfactory neurons | Kcnmb2 | 5.59 | 2.04E-130 | 4.24E-126 | 0.3 | 0.02 | 0.28 | 14.33 |
| olfactory neurons | Rtn1 | 5.66 | 5.78E-130 | 1.20E-125 | 0.89 | 0.25 | 0.64 | 3.51 |
| olfactory neurons | Klc3 | 4.57 | 3.88E-129 | 8.07E-125 | 0.65 | 0.11 | 0.55 | 6.2 |
| olfactory neurons | Celsr3 | 5.83 | 1.98E-128 | 4.13E-124 | 0.13 | 0 | 0.12 | 42.33 |
| olfactory neurons | Nrg1 | 4.06 | 1.84E-127 | 3.84E-123 | 0.57 | 0.08 | 0.49 | 7.35 |
| olfactory neurons | Dkkl1 | 5.32 | 2.92E-127 | 6.08E-123 | 0.19 | 0.01 | 0.18 | 23.38 |
| olfactory neurons | Gng3 | 5.62 | 1.24E-126 | 2.58E-122 | 0.52 | 0.07 | 0.46 | 7.59 |
| olfactory neurons | Galnt17 | 5.09 | 3.91E-126 | 8.14E-122 | 0.19 | 0.01 | 0.19 | 24.13 |
| olfactory neurons | Kcnh8 | 4.47 | 2.69E-124 | 5.59E-120 | 0.37 | 0.03 | 0.34 | 11.47 |
| olfactory neurons | Cxcr4 | 4.32 | 4.04E-123 | 8.41E-119 | 0.44 | 0.05 | 0.39 | 9.36 |
| olfactory neurons | Dusp8 | 5.63 | 3.88E-121 | 8.08E-117 | 0.27 | 0.02 | 0.25 | 15.59 |
| olfactory neurons | Ell3 | 4.75 | 7.41E-121 | 1.54E-116 | 0.34 | 0.03 | 0.31 | 11.83 |
| olfactory neurons | Gm13648 | 4.91 | 2.02E-119 | 4.20E-115 | 0.31 | 0.02 | 0.28 | 12.79 |
| olfactory neurons | Shtn1 | 4.86 | 6.97E-119 | 1.45E-114 | 0.44 | 0.05 | 0.39 | 8.8 |
| olfactory neurons | Adgrg1 | 4.04 | 2.27E-118 | 4.72E-114 | 0.4 | 0.04 | 0.36 | 9.95 |
| olfactory neurons | Trp53i11 | 4.07 | 1.24E-116 | 2.57E-112 | 0.83 | 0.22 | 0.61 | 3.73 |
| olfactory neurons | Cplx2 | 4.95 | 1.04E-115 | 2.15E-111 | 0.74 | 0.17 | 0.57 | 4.44 |
| olfactory neurons | Crmp1 | 4.62 | 3.47E-114 | 7.22E-110 | 0.65 | 0.12 | 0.53 | 5.47 |
| olfactory neurons | Ccdc184 | 6.88 | 1.22E-111 | 2.54E-107 | 0.11 | 0 | 0.11 | 38 |
| olfactory neurons | Elavl2 | 3.89 | 1.54E-109 | 3.20E-105 | 0.76 | 0.18 | 0.58 | 4.15 |
| olfactory neurons | Tpd52 | 3.27 | 2.00E-108 | 4.15E-104 | 0.88 | 0.26 | 0.62 | 3.38 |
| olfactory neurons | Rem2 | 4.7 | 1.47E-107 | 3.06E-103 | 0.47 | 0.06 | 0.41 | 7.58 |
| olfactory neurons | Cadps | 5.08 | 6.84E-107 | 1.42E-102 | 0.28 | 0.02 | 0.26 | 13.19 |
| olfactory neurons | Hes6 | 5.35 | 1.64E-106 | 3.40E-102 | 0.89 | 0.33 | 0.56 | 2.68 |
| olfactory neurons | Fgd4 | 3.84 | 2.21E-106 | 4.59E-102 | 0.55 | 0.09 | 0.47 | 6.37 |
| olfactory neurons | Tmcc3 | 4.24 | 1.46E-103 | 3.04E-99 | 0.34 | 0.03 | 0.3 | 10.21 |
| olfactory neurons | Cotl1 | 4.3 | 2.45E-103 | 5.10E-99 | 0.95 | 0.44 | 0.5 | 2.14 |
| olfactory neurons | P2rx3 | 5.6 | 4.07E-103 | 8.46E-99 | 0.13 | 0 | 0.12 | 31.75 |
| olfactory neurons | Cystm1 | 4.47 | 7.74E-103 | 1.61E-98 | 0.54 | 0.09 | 0.45 | 6.16 |
| olfactory neurons | Gm14226 | 5.45 | 2.29E-102 | 4.77E-98 | 0.12 | 0 | 0.12 | 30 |
| olfactory neurons | Sult4a1 | 4.46 | 1.09E-100 | 2.27E-96 | 0.35 | 0.04 | 0.31 | 9.69 |
| olfactory neurons | Art5 | 4.66 | 1.46E-100 | 3.03E-96 | 0.2 | 0.01 | 0.19 | 18.09 |
| olfactory neurons | Cadm3 | 4.77 | 2.09E-100 | 4.35E-96 | 0.25 | 0.02 | 0.23 | 13.32 |
| olfactory neurons | Tmsb4x | 2.62 | 1.61E-99 | 3.36E-95 | 1 | 0.99 | 0.01 | 1.01 |
| olfactory neurons | Spock2 | 3.98 | 2.41E-99 | 5.02E-95 | 0.36 | 0.04 | 0.32 | 9.59 |

**Supplementary Table 2. Marker genes for the major cell types of integrated clustering of E11.5 control and NotchGOF λ-junction cells.**

| | | | | | | | | |
|---|---|---|---|---|---|---|---|---|
| olfactory neurons | Fam110a | 4.13 | 4.31E-98 | 8.95E-94 | 0.82 | 0.26 | 0.56 | 3.17 |
| olfactory neurons | Pclo | 3.98 | 3.36E-97 | 6.99E-93 | 0.57 | 0.11 | 0.47 | 5.45 |
| olfactory neurons | Atp1b1 | 4.73 | 8.95E-96 | 1.86E-91 | 0.51 | 0.09 | 0.43 | 6.02 |
| olfactory neurons | Gfra1 | 4.26 | 1.08E-94 | 2.25E-90 | 0.37 | 0.04 | 0.33 | 8.48 |
| olfactory neurons | Dusp26 | 5.5 | 6.55E-94 | 1.36E-89 | 0.15 | 0.01 | 0.14 | 24.17 |
| olfactory neurons | Gng8 | 4.69 | 1.16E-93 | 2.42E-89 | 0.62 | 0.13 | 0.49 | 4.66 |
| olfactory neurons | Uchl1 | 4.15 | 7.09E-92 | 1.47E-87 | 0.89 | 0.35 | 0.54 | 2.54 |
| olfactory neurons | Hspa4l | 3.69 | 1.39E-91 | 2.89E-87 | 0.57 | 0.11 | 0.46 | 5.25 |
| olfactory neurons | Necab3 | 6.65 | 2.06E-90 | 4.28E-86 | 0.15 | 0.01 | 0.14 | 20.71 |
| olfactory neurons | 330009N23Ri | 4.78 | 4.56E-90 | 9.48E-86 | 0.27 | 0.02 | 0.24 | 11.52 |
| olfactory neurons | Gnrh1 | 9.9 | 1.40E-89 | 2.91E-85 | 0.13 | 0.01 | 0.13 | 22.17 |
| olfactory neurons | Gm26871 | 4.97 | 1.43E-89 | 2.97E-85 | 0.18 | 0.01 | 0.17 | 17.5 |
| olfactory neurons | Foxg1 | 3.28 | 2.56E-89 | 5.32E-85 | 0.65 | 0.14 | 0.5 | 4.51 |
| olfactory neurons | Camk2n1 | 4.16 | 3.30E-89 | 6.86E-85 | 0.69 | 0.19 | 0.51 | 3.71 |
| olfactory neurons | Nfasc | 4.53 | 2.28E-88 | 4.75E-84 | 0.27 | 0.03 | 0.25 | 10.84 |
| olfactory neurons | Nacad | 5.29 | 2.90E-88 | 6.02E-84 | 0.15 | 0.01 | 0.14 | 21.57 |
| olfactory neurons | Eya2 | 3.4 | 3.18E-88 | 6.61E-84 | 0.78 | 0.23 | 0.55 | 3.4 |
| olfactory neurons | Cbfa2t3 | 3.74 | 2.42E-87 | 5.02E-83 | 0.4 | 0.05 | 0.35 | 7.51 |
| olfactory neurons | Sgpp2 | 3.93 | 6.00E-87 | 1.25E-82 | 0.3 | 0.03 | 0.27 | 10.03 |
| olfactory neurons | 330159F19Ri | 4.08 | 7.14E-87 | 1.48E-82 | 0.27 | 0.02 | 0.24 | 11.04 |
| olfactory neurons | Unc13a | 4.63 | 2.17E-86 | 4.50E-82 | 0.18 | 0.01 | 0.17 | 16.45 |
| olfactory neurons | Pcsk1n | 6.66 | 8.96E-86 | 1.86E-81 | 0.19 | 0.01 | 0.18 | 15.58 |
| olfactory neurons | Serping1 | 4.02 | 2.13E-85 | 4.42E-81 | 0.4 | 0.06 | 0.35 | 7.21 |
| olfactory neurons | Tubb2b | 3.34 | 2.29E-85 | 4.76E-81 | 0.97 | 0.67 | 0.3 | 1.45 |
| olfactory neurons | Trim36 | 4.25 | 6.96E-84 | 1.45E-79 | 0.28 | 0.03 | 0.26 | 10.11 |
| olfactory neurons | Rab6b | 5.15 | 1.19E-82 | 2.48E-78 | 0.33 | 0.04 | 0.29 | 8.28 |
| olfactory neurons | Pnmal1 | 5.43 | 4.15E-82 | 8.64E-78 | 0.2 | 0.01 | 0.19 | 14.21 |
| olfactory neurons | Cep152 | 3.58 | 6.23E-82 | 1.30E-77 | 0.57 | 0.12 | 0.45 | 4.69 |
| olfactory neurons | Psip1 | 1.7 | 1.17E-81 | 2.43E-77 | 0.98 | 0.98 | 0 | 1 |
| olfactory neurons | Grasp | 4.3 | 1.32E-81 | 2.74E-77 | 0.68 | 0.19 | 0.48 | 3.52 |
| olfactory neurons | Hist3h2ba | 3.37 | 2.45E-81 | 5.09E-77 | 0.83 | 0.27 | 0.56 | 3.06 |
| olfactory neurons | Gnas | 1.6 | 4.99E-81 | 1.04E-76 | 1 | 1 | 0 | 1 |
| olfactory neurons | Snca | 4.11 | 1.42E-80 | 2.95E-76 | 0.49 | 0.09 | 0.41 | 5.68 |
| olfactory neurons | Cyfip2 | 4.12 | 5.83E-80 | 1.21E-75 | 0.48 | 0.09 | 0.39 | 5.48 |
| olfactory neurons | Gadd45a | 4.36 | 6.72E-80 | 1.40E-75 | 0.74 | 0.24 | 0.49 | 3.02 |
| olfactory neurons | 110039M20R | 3.88 | 2.30E-79 | 4.79E-75 | 0.45 | 0.08 | 0.38 | 6.03 |
| olfactory neurons | Gprin1 | 4.19 | 2.76E-79 | 5.75E-75 | 0.25 | 0.02 | 0.22 | 10.74 |
| olfactory neurons | Kif5a | 4.35 | 3.21E-79 | 6.67E-75 | 0.35 | 0.05 | 0.3 | 7.76 |
| olfactory neurons | Plk3 | 4.82 | 1.72E-78 | 3.57E-74 | 0.24 | 0.02 | 0.22 | 10.95 |
| olfactory neurons | Map1b | 3.9 | 1.93E-78 | 4.01E-74 | 0.95 | 0.63 | 0.32 | 1.51 |
| olfactory neurons | Nova2 | 4.33 | 2.18E-78 | 4.53E-74 | 0.26 | 0.03 | 0.23 | 10.36 |
| olfactory neurons | Btg2 | 3.96 | 4.21E-78 | 8.76E-74 | 0.77 | 0.27 | 0.5 | 2.85 |
| olfactory neurons | Ssc5d | 4.87 | 1.33E-77 | 2.76E-73 | 0.15 | 0.01 | 0.14 | 16.78 |
| olfactory neurons | Dpysl3 | 3.44 | 5.89E-77 | 1.23E-72 | 0.89 | 0.45 | 0.44 | 1.98 |
| olfactory neurons | Fam169b | 5.14 | 1.05E-76 | 2.19E-72 | 0.1 | 0 | 0.1 | 25.5 |
| olfactory neurons | Prnp | 3.62 | 1.16E-75 | 2.42E-71 | 0.65 | 0.17 | 0.48 | 3.76 |
| olfactory neurons | Apc2 | 4.58 | 1.24E-74 | 2.58E-70 | 0.24 | 0.02 | 0.21 | 10.68 |
| olfactory neurons | Tuba1a | 3.06 | 1.42E-74 | 2.95E-70 | 0.99 | 0.98 | 0 | 1 |
| olfactory neurons | Stmn1 | 1.45 | 9.19E-74 | 1.91E-69 | 0.99 | 1 | 0 | 1 |
| olfactory neurons | Flrt1 | 4.27 | 1.12E-73 | 2.34E-69 | 0.24 | 0.02 | 0.21 | 10.68 |
| olfactory neurons | Dusp4 | 3.7 | 7.05E-73 | 1.47E-68 | 0.46 | 0.08 | 0.38 | 5.52 |
| olfactory neurons | Scg2 | 4.08 | 9.15E-73 | 1.90E-68 | 0.15 | 0.01 | 0.14 | 16.78 |
| olfactory neurons | Cadps2 | 4.34 | 1.14E-72 | 2.38E-68 | 0.3 | 0.04 | 0.26 | 8.43 |
| olfactory neurons | Map2 | 3.63 | 1.23E-72 | 2.57E-68 | 0.69 | 0.21 | 0.47 | 3.21 |
| olfactory neurons | Ebf3 | 4.04 | 1.87E-72 | 3.90E-68 | 0.63 | 0.17 | 0.46 | 3.7 |
| olfactory neurons | Riiad1 | 4.91 | 1.32E-71 | 2.74E-67 | 0.15 | 0.01 | 0.14 | 16.11 |

**Supplementary Table 2. Marker genes for the major cell types of integrated clustering of E11.5 control and NotchGOF λ-junction cells.**

| | | | | | | | | |
|---|---|---|---|---|---|---|---|---|
| olfactory neurons | Tmem163 | 4.99 | 8.21E-71 | 1.71E-66 | 0.3 | 0.04 | 0.26 | 7.92 |
| olfactory neurons | Mapt | 5.08 | 1.71E-70 | 3.55E-66 | 0.31 | 0.04 | 0.27 | 7.63 |
| olfactory neurons | Rab3c | 5.02 | 1.68E-69 | 3.49E-65 | 0.16 | 0.01 | 0.15 | 15.7 |
| olfactory neurons | Syt1 | 4.12 | 2.80E-69 | 5.82E-65 | 0.3 | 0.04 | 0.26 | 8.19 |
| olfactory neurons | Kif1a | 4.1 | 4.39E-69 | 9.12E-65 | 0.37 | 0.06 | 0.32 | 6.43 |
| olfactory neurons | Ckb | 3.75 | 8.91E-68 | 1.85E-63 | 0.78 | 0.33 | 0.45 | 2.36 |
| olfactory neurons | Rims2 | 3.86 | 1.11E-67 | 2.30E-63 | 0.37 | 0.06 | 0.31 | 6.44 |
| olfactory neurons | Gadd45g | 3.38 | 2.33E-67 | 4.84E-63 | 0.68 | 0.21 | 0.47 | 3.24 |
| olfactory neurons | Mllt11 | 4.27 | 2.86E-67 | 5.95E-63 | 0.72 | 0.26 | 0.47 | 2.8 |
| olfactory neurons | Cldn3 | 2.89 | 4.58E-67 | 9.52E-63 | 0.63 | 0.16 | 0.47 | 3.91 |
| olfactory neurons | Ripor2 | 5.1 | 5.23E-66 | 1.09E-61 | 0.13 | 0.01 | 0.12 | 18.14 |
| olfactory neurons | Arfgef3 | 5.05 | 8.07E-66 | 1.68E-61 | 0.19 | 0.02 | 0.17 | 11.69 |
| olfactory neurons | Phactr1 | 4.22 | 1.40E-65 | 2.92E-61 | 0.24 | 0.03 | 0.21 | 9.4 |
| olfactory neurons | Ebf1 | 3.33 | 2.75E-65 | 5.73E-61 | 0.75 | 0.3 | 0.46 | 2.55 |
| olfactory neurons | Rassf4 | 4.26 | 3.70E-65 | 7.70E-61 | 0.25 | 0.03 | 0.23 | 9.04 |
| olfactory neurons | Cerkl | 4.31 | 1.14E-64 | 2.37E-60 | 0.14 | 0.01 | 0.13 | 15.44 |
| olfactory neurons | Tagln3 | 4.16 | 1.97E-64 | 4.11E-60 | 0.72 | 0.26 | 0.46 | 2.75 |
| olfactory neurons | Cdyl2 | 4.55 | 8.88E-64 | 1.85E-59 | 0.33 | 0.05 | 0.28 | 6.49 |
| olfactory neurons | Cadm1 | 2.93 | 3.16E-63 | 6.57E-59 | 0.8 | 0.39 | 0.42 | 2.08 |
| olfactory neurons | Lmo1 | 3.74 | 5.56E-63 | 1.16E-58 | 0.41 | 0.07 | 0.34 | 5.62 |
| olfactory neurons | Trim9 | 5.07 | 5.60E-63 | 1.16E-58 | 0.1 | 0.01 | 0.1 | 20.4 |
| olfactory neurons | Klf7 | 3.78 | 6.49E-62 | 1.35E-57 | 0.67 | 0.23 | 0.44 | 2.95 |
| olfactory neurons | Tox3 | 3.19 | 7.89E-62 | 1.64E-57 | 0.68 | 0.23 | 0.45 | 2.96 |
| olfactory neurons | Atf5 | 4.45 | 7.89E-62 | 1.64E-57 | 0.62 | 0.19 | 0.43 | 3.21 |
| olfactory neurons | Nkain4 | 4.62 | 2.32E-61 | 4.83E-57 | 0.22 | 0.02 | 0.2 | 9.29 |
| olfactory neurons | 500004A13Ri | 4.19 | 5.51E-61 | 1.15E-56 | 0.39 | 0.07 | 0.32 | 5.51 |
| olfactory neurons | Cacna2d1 | 3.97 | 7.55E-61 | 1.57E-56 | 0.49 | 0.12 | 0.38 | 4.22 |
| olfactory neurons | Hecw1 | 3.72 | 1.00E-60 | 2.08E-56 | 0.24 | 0.03 | 0.21 | 9.04 |
| olfactory neurons | Ucp2 | 3.16 | 1.65E-60 | 3.43E-56 | 0.8 | 0.36 | 0.45 | 2.26 |
| olfactory neurons | Nrsn1 | 5.26 | 2.13E-60 | 4.43E-56 | 0.15 | 0.01 | 0.14 | 14.5 |
| olfactory neurons | Hid1 | 4.6 | 3.19E-60 | 6.63E-56 | 0.16 | 0.01 | 0.15 | 13.08 |
| olfactory neurons | Fam20c | 3.95 | 5.83E-60 | 1.21E-55 | 0.21 | 0.02 | 0.19 | 9.59 |
| olfactory neurons | Basp1 | 1.93 | 6.02E-60 | 1.25E-55 | 0.96 | 0.96 | 0.01 | 1.01 |
| olfactory neurons | Efna3 | 4.74 | 7.59E-60 | 1.58E-55 | 0.35 | 0.06 | 0.29 | 5.92 |
| olfactory neurons | Numbl | 3.29 | 2.46E-59 | 5.13E-55 | 0.52 | 0.13 | 0.39 | 4.03 |
| olfactory neurons | Zc3h12c | 3.36 | 8.37E-59 | 1.74E-54 | 0.36 | 0.06 | 0.29 | 5.82 |
| olfactory neurons | Npdc1 | 3.23 | 1.07E-58 | 2.22E-54 | 0.47 | 0.1 | 0.37 | 4.52 |
| olfactory neurons | Ksr2 | 4.71 | 1.10E-58 | 2.29E-54 | 0.15 | 0.01 | 0.13 | 13.18 |
| olfactory neurons | Lrp11 | 3.6 | 1.80E-58 | 3.75E-54 | 0.27 | 0.03 | 0.23 | 7.79 |
| olfactory neurons | Serpini1 | 3.44 | 3.92E-58 | 8.15E-54 | 0.36 | 0.06 | 0.29 | 5.82 |
| olfactory neurons | Ndufa4l2 | 4.25 | 3.93E-57 | 8.18E-53 | 0.23 | 0.03 | 0.2 | 8.48 |
| olfactory neurons | Mical2 | 3.64 | 1.15E-56 | 2.40E-52 | 0.22 | 0.03 | 0.2 | 8.92 |
| olfactory neurons | Nrxn2 | 3.81 | 1.64E-56 | 3.41E-52 | 0.3 | 0.05 | 0.26 | 6.54 |
| olfactory neurons | AC109619.1 | 4.71 | 5.93E-56 | 1.23E-51 | 0.1 | 0.01 | 0.1 | 20.4 |
| olfactory neurons | Smpd3 | 4.01 | 6.03E-56 | 1.25E-51 | 0.15 | 0.01 | 0.14 | 12.58 |
| olfactory neurons | Cemip2 | 3 | 2.51E-55 | 5.23E-51 | 0.79 | 0.45 | 0.34 | 1.76 |
| olfactory neurons | Emx2os | 3.87 | 3.90E-55 | 8.10E-51 | 0.18 | 0.02 | 0.16 | 10.94 |
| olfactory neurons | Map1a | 3.21 | 4.53E-55 | 9.42E-51 | 0.42 | 0.09 | 0.33 | 4.69 |
| olfactory neurons | Aplp1 | 3.55 | 6.79E-55 | 1.41E-50 | 0.62 | 0.21 | 0.41 | 2.95 |
| olfactory neurons | Fam124a | 4.29 | 6.90E-55 | 1.43E-50 | 0.14 | 0.01 | 0.13 | 13.9 |
| olfactory neurons | Gjb5 | 5.24 | 2.43E-54 | 5.06E-50 | 0.11 | 0.01 | 0.11 | 16.29 |
| olfactory neurons | Crip2 | 2.61 | 1.03E-53 | 2.15E-49 | 0.6 | 0.17 | 0.43 | 3.49 |
| olfactory neurons | Ptms | 1.56 | 1.42E-53 | 2.95E-49 | 0.95 | 0.95 | -0.01 | 0.99 |
| olfactory neurons | Art1 | 5.47 | 5.46E-53 | 1.14E-48 | 0.12 | 0.01 | 0.11 | 15 |
| olfactory neurons | Spint2 | 1.96 | 7.88E-53 | 1.64E-48 | 0.81 | 0.31 | 0.5 | 2.6 |
| olfactory neurons | Rgs16 | 3.99 | 2.20E-52 | 4.57E-48 | 0.29 | 0.05 | 0.24 | 6.42 |

**Supplementary Table 2. Marker genes for the major cell types of integrated clustering of E11.5 control and NotchGOF λ-junction cells.**

| | | | | | | | | |
|---|---|---|---|---|---|---|---|---|
| olfactory neurons | Slc27a2 | 4.16 | 7.94E-52 | 1.65E-47 | 0.16 | 0.01 | 0.14 | 11.21 |
| olfactory neurons | Sel1l3 | 4.06 | 9.97E-52 | 2.07E-47 | 0.18 | 0.02 | 0.16 | 10.29 |
| olfactory neurons | Kdm7a | 2.58 | 1.52E-51 | 3.17E-47 | 0.75 | 0.35 | 0.41 | 2.16 |
| olfactory neurons | Atl1 | 3.29 | 2.03E-51 | 4.23E-47 | 0.3 | 0.05 | 0.25 | 6.28 |
| olfactory neurons | Hpca | 5.58 | 2.43E-51 | 5.04E-47 | 0.11 | 0.01 | 0.1 | 15.43 |
| olfactory neurons | Chrnb2 | 4.47 | 6.46E-51 | 1.34E-46 | 0.11 | 0.01 | 0.1 | 15.43 |
| olfactory neurons | Snap91 | 3.48 | 1.36E-50 | 2.83E-46 | 0.22 | 0.03 | 0.19 | 8.35 |
| olfactory neurons | Nnat | 2.69 | 2.40E-50 | 5.00E-46 | 0.97 | 0.94 | 0.03 | 1.03 |
| olfactory neurons | Ada | 3.52 | 1.00E-49 | 2.08E-45 | 0.38 | 0.08 | 0.3 | 4.81 |
| olfactory neurons | Mtcl1 | 2.51 | 1.13E-48 | 2.34E-44 | 0.48 | 0.12 | 0.36 | 3.97 |
| olfactory neurons | Apba2 | 3.56 | 1.57E-48 | 3.27E-44 | 0.33 | 0.06 | 0.27 | 5.42 |
| olfactory neurons | Gnao1 | 3.56 | 5.70E-48 | 1.19E-43 | 0.38 | 0.08 | 0.3 | 4.63 |
| olfactory neurons | Col9a2 | 4.16 | 8.36E-48 | 1.74E-43 | 0.42 | 0.1 | 0.32 | 4.14 |
| olfactory neurons | Robo2 | 2.92 | 8.97E-48 | 1.86E-43 | 0.58 | 0.19 | 0.39 | 3.04 |
| olfactory neurons | Mxi1 | 2.51 | 2.13E-47 | 4.43E-43 | 0.79 | 0.43 | 0.36 | 1.85 |
| olfactory neurons | Phyhipl | 3.46 | 2.49E-47 | 5.17E-43 | 0.25 | 0.04 | 0.22 | 6.66 |
| olfactory neurons | Pcbp4 | 1.58 | 2.70E-47 | 5.62E-43 | 0.92 | 0.87 | 0.05 | 1.06 |
| olfactory neurons | Myo5b | 3.3 | 5.04E-47 | 1.05E-42 | 0.27 | 0.04 | 0.23 | 6.3 |
| olfactory neurons | Bcl2l1 | 2.43 | 2.31E-46 | 4.80E-42 | 0.75 | 0.37 | 0.38 | 2.02 |
| olfactory neurons | Fam78b | 4.06 | 3.84E-46 | 7.99E-42 | 0.22 | 0.03 | 0.19 | 7.48 |
| olfactory neurons | Tenm1 | 4.07 | 7.63E-46 | 1.59E-41 | 0.14 | 0.01 | 0.13 | 11.58 |
| olfactory neurons | Map9 | 2.82 | 8.16E-46 | 1.70E-41 | 0.55 | 0.18 | 0.37 | 3.08 |
| olfactory neurons | Tmem198 | 4.94 | 1.30E-45 | 2.70E-41 | 0.1 | 0.01 | 0.1 | 14.57 |
| olfactory neurons | 630019K06Ri | 3.33 | 1.81E-45 | 3.77E-41 | 0.4 | 0.1 | 0.31 | 4.25 |
| olfactory neurons | Kif5c | 2.8 | 6.47E-45 | 1.35E-40 | 0.8 | 0.49 | 0.32 | 1.65 |
| olfactory neurons | Ddr1 | 2.51 | 9.36E-45 | 1.95E-40 | 0.48 | 0.13 | 0.35 | 3.71 |
| olfactory neurons | Svip | 3.23 | 1.19E-44 | 2.47E-40 | 0.39 | 0.09 | 0.3 | 4.31 |
| olfactory neurons | Parp6 | 3.1 | 1.40E-44 | 2.91E-40 | 0.43 | 0.11 | 0.32 | 3.88 |
| olfactory neurons | Selenom | 3.85 | 4.01E-44 | 8.33E-40 | 0.78 | 0.52 | 0.26 | 1.5 |
| olfactory neurons | Plekha6 | 4.23 | 5.08E-44 | 1.06E-39 | 0.16 | 0.02 | 0.14 | 9.81 |
| olfactory neurons | Nlk | 2.86 | 5.49E-44 | 1.14E-39 | 0.64 | 0.27 | 0.37 | 2.35 |
| olfactory neurons | Nuak2 | 3.54 | 6.26E-44 | 1.30E-39 | 0.24 | 0.04 | 0.2 | 6.51 |
| olfactory neurons | Calm2 | 1.14 | 8.03E-44 | 1.67E-39 | 0.96 | 0.98 | -0.02 | 0.98 |
| olfactory neurons | Sptbn2 | 2.94 | 2.14E-43 | 4.44E-39 | 0.43 | 0.11 | 0.32 | 3.88 |
| olfactory neurons | Rnf182 | 3.73 | 5.06E-43 | 1.05E-38 | 0.57 | 0.21 | 0.35 | 2.67 |
| olfactory neurons | Ptma | 0.76 | 1.02E-42 | 2.13E-38 | 1 | 1 | 0 | 1 |
| olfactory neurons | Plk4 | 2.75 | 6.37E-42 | 1.33E-37 | 0.79 | 0.55 | 0.24 | 1.43 |
| olfactory neurons | Stk33 | 4.92 | 9.06E-42 | 1.88E-37 | 0.11 | 0.01 | 0.11 | 12.67 |
| olfactory neurons | Abcd2 | 3.05 | 1.54E-41 | 3.20E-37 | 0.34 | 0.07 | 0.27 | 4.68 |
| olfactory neurons | Zbtb18 | 3 | 1.54E-41 | 3.21E-37 | 0.42 | 0.11 | 0.31 | 3.82 |
| olfactory neurons | Mturn | 3.13 | 1.65E-41 | 3.42E-37 | 0.3 | 0.06 | 0.24 | 5.1 |
| olfactory neurons | Jpt1 | 1.35 | 2.01E-41 | 4.18E-37 | 0.94 | 0.94 | 0 | 1 |
| olfactory neurons | Amotl2 | 2.51 | 3.65E-41 | 7.60E-37 | 0.69 | 0.34 | 0.35 | 2.04 |
| olfactory neurons | Rgmb | 3.19 | 7.09E-41 | 1.47E-36 | 0.45 | 0.12 | 0.32 | 3.6 |
| olfactory neurons | Pak3 | 2.86 | 8.04E-41 | 1.67E-36 | 0.46 | 0.14 | 0.33 | 3.41 |
| olfactory neurons | Rgs4 | 4.38 | 1.29E-40 | 2.68E-36 | 0.23 | 0.04 | 0.19 | 6.36 |
| olfactory neurons | Amer2 | 2.69 | 2.28E-40 | 4.74E-36 | 0.42 | 0.11 | 0.31 | 3.92 |
| olfactory neurons | Mbp | 3.03 | 2.63E-40 | 5.47E-36 | 0.39 | 0.09 | 0.29 | 4.11 |
| olfactory neurons | Gal | 3.16 | 3.77E-40 | 7.84E-36 | 0.13 | 0.01 | 0.12 | 10.23 |
| olfactory neurons | Plcxd3 | 4.8 | 7.70E-40 | 1.60E-35 | 0.11 | 0.01 | 0.1 | 12 |
| olfactory neurons | Col16a1 | 4.8 | 1.12E-39 | 2.33E-35 | 0.27 | 0.05 | 0.22 | 5.52 |
| olfactory neurons | Ndufaf3 | 2.61 | 1.61E-39 | 3.34E-35 | 0.72 | 0.42 | 0.31 | 1.74 |
| olfactory neurons | Rbfox3 | 4.18 | 2.98E-39 | 6.20E-35 | 0.11 | 0.01 | 0.1 | 12 |
| olfactory neurons | Eif4e3 | 2.47 | 3.33E-39 | 6.93E-35 | 0.64 | 0.28 | 0.36 | 2.3 |
| olfactory neurons | Ctnna2 | 3.23 | 3.88E-39 | 8.07E-35 | 0.27 | 0.05 | 0.22 | 5.52 |
| olfactory neurons | Ptpn6 | 3.14 | 5.46E-39 | 1.13E-34 | 0.23 | 0.04 | 0.19 | 6.36 |

**Supplementary Table 2. Marker genes for the major cell types of integrated clustering of E11.5 control and NotchGOF λ-junction cells.**

| | | | | | | | | |
|---|---|---|---|---|---|---|---|---|
| olfactory neurons | Sult2b1 | 3.89 | 6.74E-39 | 1.40E-34 | 0.15 | 0.02 | 0.13 | 8.88 |
| olfactory neurons | Fam107a | 2.71 | 1.21E-38 | 2.51E-34 | 0.25 | 0.04 | 0.21 | 5.75 |
| olfactory neurons | Ssx2ip | 2.29 | 1.87E-38 | 3.89E-34 | 0.72 | 0.36 | 0.36 | 1.99 |
| olfactory neurons | Olfm1 | 2.8 | 4.31E-38 | 8.96E-34 | 0.57 | 0.23 | 0.35 | 2.53 |
| olfactory neurons | Bcl7a | 1.86 | 6.98E-38 | 1.45E-33 | 0.83 | 0.67 | 0.16 | 1.24 |
| olfactory neurons | Fam219a | 3.37 | 1.13E-37 | 2.35E-33 | 0.21 | 0.03 | 0.18 | 6.39 |
| olfactory neurons | Sox11 | 0.9 | 1.28E-37 | 2.66E-33 | 0.95 | 0.97 | -0.02 | 0.98 |
| olfactory neurons | Kcnn2 | 3.94 | 1.36E-37 | 2.82E-33 | 0.12 | 0.01 | 0.11 | 10.91 |
| olfactory neurons | Cyth2 | 1.87 | 1.47E-37 | 3.06E-33 | 0.83 | 0.67 | 0.16 | 1.23 |
| olfactory neurons | Cecr2 | 2.31 | 1.50E-37 | 3.12E-33 | 0.58 | 0.22 | 0.36 | 2.6 |
| olfactory neurons | Cldn9 | 2.44 | 1.75E-37 | 3.63E-33 | 0.37 | 0.09 | 0.28 | 4.17 |
| olfactory neurons | Slc43a2 | 3.17 | 1.93E-37 | 4.02E-33 | 0.24 | 0.04 | 0.2 | 5.88 |
| olfactory neurons | Fam131a | 3.71 | 2.06E-37 | 4.28E-33 | 0.16 | 0.02 | 0.14 | 8.15 |
| olfactory neurons | 530008M17R | 2.36 | 2.15E-37 | 4.48E-33 | 0.73 | 0.42 | 0.31 | 1.74 |
| olfactory neurons | Nat14 | 3.2 | 2.76E-37 | 5.74E-33 | 0.24 | 0.04 | 0.2 | 5.74 |
| olfactory neurons | Baz1a | 2.81 | 3.33E-37 | 6.92E-33 | 0.8 | 0.56 | 0.24 | 1.43 |
| olfactory neurons | Wdr47 | 2.66 | 7.03E-37 | 1.46E-32 | 0.37 | 0.09 | 0.28 | 3.97 |
| olfactory neurons | Gabarapl2 | 1.49 | 1.26E-36 | 2.63E-32 | 0.88 | 0.82 | 0.06 | 1.07 |
| olfactory neurons | Wscd1 | 4.03 | 1.55E-36 | 3.23E-32 | 0.18 | 0.03 | 0.16 | 7.24 |
| olfactory neurons | Smarcd2 | 2.72 | 3.93E-36 | 8.18E-32 | 0.63 | 0.29 | 0.34 | 2.17 |
| olfactory neurons | Jag2 | 2.35 | 1.20E-35 | 2.50E-31 | 0.34 | 0.08 | 0.26 | 4.23 |
| olfactory neurons | Rhbdl3 | 3.9 | 1.88E-35 | 3.91E-31 | 0.11 | 0.01 | 0.1 | 10.8 |
| olfactory neurons | Ascl1 | 4.41 | 2.46E-35 | 5.11E-31 | 0.13 | 0.01 | 0.11 | 9.77 |
| olfactory neurons | Ank2 | 3.64 | 1.20E-34 | 2.50E-30 | 0.56 | 0.25 | 0.32 | 2.29 |
| olfactory neurons | Syt11 | 3.03 | 2.15E-34 | 4.48E-30 | 0.63 | 0.31 | 0.32 | 2.06 |
| olfactory neurons | Dlx6 | 3.38 | 2.98E-34 | 6.20E-30 | 0.34 | 0.09 | 0.26 | 3.99 |
| olfactory neurons | Ccser2 | 2.35 | 3.68E-34 | 7.64E-30 | 0.6 | 0.27 | 0.33 | 2.24 |
| olfactory neurons | Nol4l | 2.55 | 4.29E-34 | 8.91E-30 | 0.48 | 0.17 | 0.32 | 2.92 |
| olfactory neurons | Trnp1 | 4.11 | 5.48E-34 | 1.14E-29 | 0.18 | 0.03 | 0.15 | 7 |
| olfactory neurons | Rtn2 | 2.85 | 5.73E-34 | 1.19E-29 | 0.34 | 0.08 | 0.25 | 4.06 |
| olfactory neurons | Otx2 | 2.8 | 7.86E-34 | 1.63E-29 | 0.38 | 0.1 | 0.28 | 3.76 |
| olfactory neurons | Lipg | 3.3 | 9.91E-34 | 2.06E-29 | 0.21 | 0.04 | 0.18 | 5.86 |
| olfactory neurons | Tekt2 | 4.15 | 1.21E-33 | 2.51E-29 | 0.12 | 0.01 | 0.11 | 9.23 |
| olfactory neurons | Krt8 | 1.59 | 1.33E-33 | 2.77E-29 | 0.68 | 0.26 | 0.42 | 2.62 |
| olfactory neurons | Mkrn1 | 1.99 | 2.35E-33 | 4.89E-29 | 0.74 | 0.43 | 0.32 | 1.74 |
| olfactory neurons | Jakmip2 | 2.95 | 2.81E-33 | 5.84E-29 | 0.43 | 0.14 | 0.29 | 3.01 |
| olfactory neurons | Mapk8ip1 | 2.65 | 2.91E-33 | 6.05E-29 | 0.58 | 0.26 | 0.33 | 2.27 |
| olfactory neurons | Grik2 | 3.62 | 4.14E-33 | 8.60E-29 | 0.16 | 0.02 | 0.14 | 7.41 |
| olfactory neurons | Actb | 0.61 | 5.11E-33 | 1.06E-28 | 1 | 1 | 0 | 1 |
| olfactory neurons | Map1lc3b | 1.31 | 9.99E-33 | 2.08E-28 | 0.9 | 0.93 | -0.02 | 0.98 |
| olfactory neurons | Syt5 | 3.1 | 1.62E-32 | 3.36E-28 | 0.27 | 0.06 | 0.21 | 4.67 |
| olfactory neurons | Mbd2 | 2.04 | 1.65E-32 | 3.44E-28 | 0.79 | 0.56 | 0.23 | 1.41 |
| olfactory neurons | Trim62 | 2.91 | 1.87E-32 | 3.89E-28 | 0.36 | 0.1 | 0.26 | 3.61 |
| olfactory neurons | Pcdh20 | 3.84 | 3.39E-32 | 7.05E-28 | 0.13 | 0.01 | 0.11 | 9.07 |
| olfactory neurons | Rab3b | 3.93 | 3.90E-32 | 8.10E-28 | 0.27 | 0.06 | 0.21 | 4.44 |
| olfactory neurons | Cdkn1a | 2.14 | 4.38E-32 | 9.11E-28 | 0.4 | 0.11 | 0.29 | 3.62 |
| olfactory neurons | Mdm1 | 2.81 | 5.52E-32 | 1.15E-27 | 0.6 | 0.3 | 0.3 | 2.02 |
| olfactory neurons | Hpcal1 | 2.31 | 5.65E-32 | 1.17E-27 | 0.66 | 0.33 | 0.33 | 1.98 |
| olfactory neurons | Trp53inp2 | 2.66 | 6.54E-32 | 1.36E-27 | 0.36 | 0.1 | 0.26 | 3.7 |
| olfactory neurons | Onecut2 | 4.5 | 1.10E-31 | 2.29E-27 | 0.15 | 0.02 | 0.13 | 7.55 |
| olfactory neurons | Ildr1 | 3.09 | 1.31E-31 | 2.72E-27 | 0.21 | 0.04 | 0.17 | 5.7 |
| olfactory neurons | Prr36 | 2.45 | 1.94E-31 | 4.04E-27 | 0.34 | 0.09 | 0.25 | 3.87 |
| olfactory neurons | Abcg1 | 2.77 | 2.72E-31 | 5.66E-27 | 0.33 | 0.08 | 0.24 | 3.96 |
| olfactory neurons | .330084C13Ri | 3.72 | 2.89E-31 | 6.02E-27 | 0.1 | 0.01 | 0.09 | 10.2 |
| olfactory neurons | Rab3a | 3.15 | 4.09E-31 | 8.50E-27 | 0.42 | 0.14 | 0.28 | 3.08 |
| olfactory neurons | Scrn1 | 3.79 | 5.69E-31 | 1.18E-26 | 0.22 | 0.04 | 0.18 | 5.19 |

**Supplementary Table 2. Marker genes for the major cell types of integrated clustering of E11.5 control and NotchGOF λ-junction cells.**

| | | | | | | | | |
|---|---|---|---|---|---|---|---|---|
| olfactory neurons | Dbndd1 | 2.99 | 6.20E-31 | 1.29E-26 | 0.25 | 0.05 | 0.2 | 4.69 |
| olfactory neurons | Samd10 | 2.63 | 9.55E-31 | 1.99E-26 | 0.34 | 0.09 | 0.25 | 3.81 |
| olfactory neurons | Eid1 | 0.97 | 1.87E-30 | 3.90E-26 | 0.94 | 0.97 | -0.03 | 0.97 |
| olfactory neurons | Ccdc88c | 2.01 | 2.49E-30 | 5.19E-26 | 0.46 | 0.16 | 0.31 | 2.96 |
| olfactory neurons | Shb | 2.55 | 6.37E-30 | 1.33E-25 | 0.5 | 0.2 | 0.3 | 2.55 |
| olfactory neurons | Epcam | 0.95 | 7.20E-30 | 1.50E-25 | 0.78 | 0.35 | 0.44 | 2.26 |
| olfactory neurons | Gdi1 | 2.26 | 9.08E-30 | 1.89E-25 | 0.7 | 0.42 | 0.28 | 1.67 |
| olfactory neurons | Tomm34 | 2.03 | 9.56E-30 | 1.99E-25 | 0.69 | 0.42 | 0.28 | 1.67 |
| olfactory neurons | Ube2b | 1.11 | 1.28E-29 | 2.67E-25 | 0.91 | 0.93 | -0.02 | 0.98 |
| olfactory neurons | Tfeb | 3.36 | 2.15E-29 | 4.48E-25 | 0.16 | 0.02 | 0.13 | 6.83 |
| olfactory neurons | Sp8 | 1.82 | 2.26E-29 | 4.70E-25 | 0.44 | 0.14 | 0.3 | 3.17 |
| olfactory neurons | Slc26a10 | 3.28 | 2.32E-29 | 4.82E-25 | 0.13 | 0.02 | 0.11 | 8.47 |
| olfactory neurons | Cspg5 | 3.88 | 2.96E-29 | 6.16E-25 | 0.14 | 0.02 | 0.12 | 7.32 |
| olfactory neurons | Gdpd1 | 2.49 | 3.53E-29 | 7.35E-25 | 0.58 | 0.27 | 0.31 | 2.15 |
| olfactory neurons | Zmat4 | 3.67 | 4.69E-29 | 9.76E-25 | 0.25 | 0.06 | 0.19 | 4.49 |
| olfactory neurons | Ccdc96 | 3.28 | 5.59E-29 | 1.16E-24 | 0.15 | 0.02 | 0.13 | 6.86 |
| olfactory neurons | Celf5 | 3.59 | 9.91E-29 | 2.06E-24 | 0.15 | 0.02 | 0.13 | 6.86 |
| olfactory neurons | Tdrkh | 2.91 | 9.96E-29 | 2.07E-24 | 0.26 | 0.06 | 0.2 | 4.32 |
| olfactory neurons | Tmcc2 | 2.42 | 1.07E-28 | 2.23E-24 | 0.31 | 0.08 | 0.23 | 3.82 |
| olfactory neurons | Hspa12a | 3.14 | 1.29E-28 | 2.68E-24 | 0.18 | 0.03 | 0.15 | 6.03 |
| olfactory neurons | Dlx6os1 | 3.16 | 1.81E-28 | 3.75E-24 | 0.21 | 0.04 | 0.17 | 5.28 |
| olfactory neurons | Rcor2 | 1.83 | 1.82E-28 | 3.78E-24 | 0.72 | 0.49 | 0.23 | 1.48 |
| olfactory neurons | Prex1 | 2.33 | 2.09E-28 | 4.34E-24 | 0.49 | 0.19 | 0.3 | 2.53 |
| olfactory neurons | Mcf2l | 2.87 | 3.31E-28 | 6.89E-24 | 0.28 | 0.07 | 0.21 | 4.07 |
| olfactory neurons | Pitx1 | 1.69 | 3.91E-28 | 8.13E-24 | 0.51 | 0.18 | 0.33 | 2.81 |
| olfactory neurons | Gse1 | 2.12 | 5.44E-28 | 1.13E-23 | 0.6 | 0.3 | 0.29 | 1.97 |
| olfactory neurons | Zfhx2 | 1.91 | 6.07E-28 | 1.26E-23 | 0.57 | 0.25 | 0.32 | 2.25 |
| olfactory neurons | Stox1 | 4.03 | 7.41E-28 | 1.54E-23 | 0.16 | 0.03 | 0.14 | 6.27 |
| olfactory neurons | Hap1 | 3.31 | 7.82E-28 | 1.63E-23 | 0.24 | 0.05 | 0.18 | 4.61 |
| olfactory neurons | Pnmal2 | 3.03 | 9.53E-28 | 1.98E-23 | 0.27 | 0.06 | 0.2 | 4.14 |
| olfactory neurons | Nefm | 4.38 | 9.73E-28 | 2.02E-23 | 0.16 | 0.03 | 0.14 | 6.27 |
| olfactory neurons | Lrrn3 | 2.71 | 1.53E-27 | 3.18E-23 | 0.28 | 0.07 | 0.21 | 3.99 |
| olfactory neurons | Fyn | 1.76 | 7.96E-27 | 1.66E-22 | 0.72 | 0.44 | 0.28 | 1.64 |
| olfactory neurons | Ddit4 | 2.21 | 8.25E-27 | 1.72E-22 | 0.68 | 0.38 | 0.3 | 1.8 |
| olfactory neurons | Stxbp1 | 2.62 | 8.57E-27 | 1.78E-22 | 0.33 | 0.09 | 0.23 | 3.46 |
| olfactory neurons | Kif21a | 1.99 | 8.71E-27 | 1.81E-22 | 0.63 | 0.31 | 0.32 | 2.02 |
| olfactory neurons | Ralgds | 2.18 | 8.87E-27 | 1.85E-22 | 0.52 | 0.22 | 0.3 | 2.35 |
| olfactory neurons | Srgap1 | 2.04 | 1.23E-26 | 2.56E-22 | 0.63 | 0.34 | 0.29 | 1.86 |
| olfactory neurons | Gm19938 | 3.31 | 1.25E-26 | 2.61E-22 | 0.15 | 0.02 | 0.13 | 6.57 |
| olfactory neurons | Gck | 3.37 | 1.35E-26 | 2.81E-22 | 0.16 | 0.03 | 0.13 | 6.28 |
| olfactory neurons | Rnf5 | 1.74 | 1.72E-26 | 3.57E-22 | 0.81 | 0.65 | 0.16 | 1.25 |
| olfactory neurons | Calm1 | 1.17 | 1.95E-26 | 4.06E-22 | 0.95 | 0.99 | -0.05 | 0.95 |
| olfactory neurons | Lzts1 | 2.85 | 3.28E-26 | 6.82E-22 | 0.2 | 0.04 | 0.16 | 5.1 |
| olfactory neurons | Mycl | 2.06 | 4.02E-26 | 8.36E-22 | 0.6 | 0.3 | 0.29 | 1.97 |
| olfactory neurons | Eif1 | 0.49 | 4.93E-26 | 1.03E-21 | 0.98 | 1 | -0.02 | 0.98 |
| olfactory neurons | Marchf9 | 2.24 | 6.52E-26 | 1.36E-21 | 0.42 | 0.15 | 0.27 | 2.85 |
| olfactory neurons | Adgrb2 | 2.76 | 6.58E-26 | 1.37E-21 | 0.21 | 0.04 | 0.17 | 4.91 |
| olfactory neurons | Mybl1 | 3.48 | 7.08E-26 | 1.47E-21 | 0.25 | 0.06 | 0.19 | 4.19 |
| olfactory neurons | Lpcat2 | 2.91 | 9.36E-26 | 1.95E-21 | 0.19 | 0.04 | 0.16 | 5.08 |
| olfactory neurons | Fgf12 | 2.95 | 1.02E-25 | 2.12E-21 | 0.22 | 0.05 | 0.18 | 4.65 |
| olfactory neurons | Ddx5 | 0.53 | 1.24E-25 | 2.58E-21 | 0.99 | 1 | -0.01 | 0.99 |
| olfactory neurons | Acer2 | 1.61 | 1.55E-25 | 3.22E-21 | 0.71 | 0.4 | 0.32 | 1.8 |
| olfactory neurons | Ppp1r14b | 0.63 | 3.01E-25 | 6.27E-21 | 0.95 | 0.99 | -0.04 | 0.96 |
| olfactory neurons | Neurl1a | 2.46 | 3.12E-25 | 6.49E-21 | 0.26 | 0.06 | 0.2 | 4.05 |
| olfactory neurons | Elmo1 | 3.53 | 3.78E-25 | 7.85E-21 | 0.21 | 0.05 | 0.17 | 4.69 |
| olfactory neurons | Rasgef1b | 3.08 | 3.78E-25 | 7.85E-21 | 0.22 | 0.05 | 0.17 | 4.62 |

**Supplementary Table 2. Marker genes for the major cell types of integrated clustering of E11.5 control and NotchGOF λ-junction cells.**

| | | | | | | | | |
|---|---|---|---|---|---|---|---|---|
| olfactory neurons | Gabrb3 | 3.45 | 3.90E-25 | 8.12E-21 | 0.1 | 0.01 | 0.09 | 8.5 |
| olfactory neurons | Kdm4b | 2.22 | 5.45E-25 | 1.13E-20 | 0.44 | 0.17 | 0.27 | 2.65 |
| olfactory neurons | Stil | 2.53 | 5.66E-25 | 1.18E-20 | 0.45 | 0.19 | 0.26 | 2.34 |
| olfactory neurons | Kdm1a | 0.96 | 7.00E-25 | 1.46E-20 | 0.88 | 0.92 | -0.04 | 0.95 |
| olfactory neurons | Mfsd6 | 2.82 | 7.94E-25 | 1.65E-20 | 0.18 | 0.03 | 0.15 | 5.32 |
| olfactory neurons | Casp3 | 1.67 | 1.02E-24 | 2.12E-20 | 0.82 | 0.72 | 0.1 | 1.13 |
| olfactory neurons | Slco5a1 | 2.55 | 1.11E-24 | 2.30E-20 | 0.3 | 0.08 | 0.21 | 3.6 |
| olfactory neurons | Rb1 | 2.06 | 1.28E-24 | 2.65E-20 | 0.43 | 0.16 | 0.28 | 2.73 |
| olfactory neurons | Arpc5 | 1.4 | 1.55E-24 | 3.22E-20 | 0.84 | 0.76 | 0.08 | 1.11 |
| olfactory neurons | Chd7 | 1.62 | 1.67E-24 | 3.47E-20 | 0.77 | 0.66 | 0.11 | 1.17 |
| olfactory neurons | Rufy3 | 3.31 | 1.72E-24 | 3.58E-20 | 0.49 | 0.23 | 0.26 | 2.15 |
| olfactory neurons | Actg1 | 0.64 | 1.78E-24 | 3.71E-20 | 0.99 | 1 | -0.01 | 0.99 |
| olfactory neurons | Prdx2 | 0.56 | 2.23E-24 | 4.63E-20 | 1 | 1 | 0 | 1 |
| olfactory neurons | Tle1 | 1.79 | 2.43E-24 | 5.05E-20 | 0.77 | 0.53 | 0.23 | 1.44 |
| olfactory neurons | Cdh2 | 1.35 | 2.84E-24 | 5.91E-20 | 0.71 | 0.42 | 0.29 | 1.68 |
| olfactory neurons | Synrg | 1.96 | 3.39E-24 | 7.04E-20 | 0.52 | 0.24 | 0.29 | 2.23 |
| olfactory neurons | Olfm2 | 2.47 | 3.50E-24 | 7.27E-20 | 0.34 | 0.1 | 0.23 | 3.24 |
| olfactory neurons | Tshz1 | 2.41 | 4.25E-24 | 8.85E-20 | 0.58 | 0.32 | 0.27 | 1.84 |
| olfactory neurons | Tsg101 | 1.21 | 4.82E-24 | 1.00E-19 | 0.87 | 0.85 | 0.02 | 1.03 |
| olfactory neurons | Atp6v1g1 | 0.92 | 5.37E-24 | 1.12E-19 | 0.9 | 0.97 | -0.06 | 0.94 |
| olfactory neurons | Nab1 | 2.09 | 5.73E-24 | 1.19E-19 | 0.38 | 0.13 | 0.25 | 2.95 |
| olfactory neurons | Napb | 2.71 | 6.15E-24 | 1.28E-19 | 0.23 | 0.05 | 0.18 | 4.24 |
| olfactory neurons | Sertad2 | 1.87 | 1.15E-23 | 2.39E-19 | 0.66 | 0.41 | 0.26 | 1.63 |
| olfactory neurons | Nedd4l | 2.31 | 1.24E-23 | 2.59E-19 | 0.39 | 0.14 | 0.25 | 2.72 |
| olfactory neurons | Rapgef5 | 2.52 | 1.71E-23 | 3.55E-19 | 0.36 | 0.12 | 0.24 | 3.03 |
| olfactory neurons | Spsb4 | 2.17 | 2.03E-23 | 4.23E-19 | 0.48 | 0.21 | 0.28 | 2.34 |
| olfactory neurons | Dctn3 | 1.37 | 2.17E-23 | 4.52E-19 | 0.81 | 0.79 | 0.03 | 1.04 |
| olfactory neurons | 700088E04Ri | 2.86 | 2.41E-23 | 5.02E-19 | 0.22 | 0.05 | 0.17 | 4.21 |
| olfactory neurons | Klf13 | 2.02 | 8.50E-23 | 1.77E-18 | 0.61 | 0.37 | 0.24 | 1.65 |
| olfactory neurons | Pdrg1 | 2.09 | 9.05E-23 | 1.88E-18 | 0.74 | 0.54 | 0.2 | 1.37 |
| olfactory neurons | Saraf | 2.07 | 1.87E-22 | 3.88E-18 | 0.71 | 0.54 | 0.17 | 1.31 |
| olfactory neurons | Bex2 | 1.05 | 2.18E-22 | 4.53E-18 | 0.91 | 0.85 | 0.06 | 1.07 |
| olfactory neurons | Fdps | 1.26 | 2.18E-22 | 4.53E-18 | 0.86 | 0.82 | 0.04 | 1.05 |
| olfactory neurons | Prox1 | 3.37 | 2.49E-22 | 5.19E-18 | 0.13 | 0.02 | 0.11 | 6.05 |
| olfactory neurons | Sh3bgrl | 1.16 | 5.30E-22 | 1.10E-17 | 0.87 | 0.86 | 0.02 | 1.02 |
| olfactory neurons | Ube2s | 1.12 | 5.68E-22 | 1.18E-17 | 0.94 | 0.97 | -0.03 | 0.97 |
| olfactory neurons | Fry | 2.67 | 6.90E-22 | 1.43E-17 | 0.22 | 0.06 | 0.17 | 4.05 |
| olfactory neurons | Kcnma1 | 4.18 | 7.51E-22 | 1.56E-17 | 0.11 | 0.02 | 0.09 | 7.2 |
| olfactory neurons | Rnf165 | 2 | 8.43E-22 | 1.75E-17 | 0.51 | 0.25 | 0.27 | 2.08 |
| olfactory neurons | 230030E09Ri | 2.74 | 1.01E-21 | 2.11E-17 | 0.23 | 0.06 | 0.17 | 4.02 |
| olfactory neurons | Kif5b | 0.81 | 1.71E-21 | 3.55E-17 | 0.91 | 0.94 | -0.03 | 0.97 |
| olfactory neurons | Atp6v0b | 1.4 | 1.97E-21 | 4.09E-17 | 0.8 | 0.74 | 0.06 | 1.09 |
| olfactory neurons | Gm5468 | 3.43 | 2.13E-21 | 4.43E-17 | 0.1 | 0.01 | 0.09 | 7.29 |
| olfactory neurons | Add2 | 2.39 | 2.74E-21 | 5.70E-17 | 0.29 | 0.09 | 0.2 | 3.25 |
| olfactory neurons | 310040G24Ri | 3.76 | 3.45E-21 | 7.18E-17 | 0.16 | 0.03 | 0.13 | 4.94 |
| olfactory neurons | Atp8a1 | 2.68 | 5.12E-21 | 1.07E-16 | 0.28 | 0.09 | 0.2 | 3.25 |
| olfactory neurons | Espn | 2.27 | 6.63E-21 | 1.38E-16 | 0.33 | 0.11 | 0.22 | 3.04 |
| olfactory neurons | Ttll7 | 2.83 | 7.75E-21 | 1.61E-16 | 0.25 | 0.07 | 0.18 | 3.47 |
| olfactory neurons | Slain1 | 2.78 | 8.23E-21 | 1.71E-16 | 0.23 | 0.06 | 0.17 | 3.82 |
| olfactory neurons | Rab39b | 2.19 | 8.37E-21 | 1.74E-16 | 0.4 | 0.16 | 0.25 | 2.56 |
| olfactory neurons | Ubb | 0.47 | 1.07E-20 | 2.22E-16 | 0.99 | 1 | -0.01 | 0.99 |
| olfactory neurons | Ccdc92 | 2.54 | 1.27E-20 | 2.65E-16 | 0.27 | 0.08 | 0.19 | 3.35 |
| olfactory neurons | Ppp2r5b | 2.25 | 1.32E-20 | 2.75E-16 | 0.29 | 0.09 | 0.2 | 3.25 |
| olfactory neurons | Mien1 | 1.51 | 1.43E-20 | 2.98E-16 | 0.74 | 0.56 | 0.17 | 1.3 |
| olfactory neurons | Ncam1 | 3.45 | 1.72E-20 | 3.57E-16 | 0.58 | 0.36 | 0.22 | 1.61 |
| olfactory neurons | Ap3m2 | 2.75 | 1.98E-20 | 4.11E-16 | 0.18 | 0.04 | 0.14 | 4.61 |

**Supplementary Table 2. Marker genes for the major cell types of integrated clustering of E11.5 control and NotchGOF λ-junction cells.**

| | | | | | | | | |
|---|---|---|---|---|---|---|---|---|
| olfactory neurons | Enc1 | 2.61 | 2.08E-20 | 4.32E-16 | 0.46 | 0.22 | 0.24 | 2.1 |
| olfactory neurons | Mfap4 | 1.32 | 2.23E-20 | 4.63E-16 | 0.87 | 0.85 | 0.02 | 1.02 |
| olfactory neurons | Tmem200a | 2.89 | 2.35E-20 | 4.90E-16 | 0.22 | 0.06 | 0.16 | 3.88 |
| olfactory neurons | Cend1 | 2.96 | 2.42E-20 | 5.04E-16 | 0.22 | 0.06 | 0.16 | 3.78 |
| olfactory neurons | Rprm | 1.55 | 3.23E-20 | 6.71E-16 | 0.42 | 0.16 | 0.26 | 2.58 |
| olfactory neurons | [illegible]130071C03Ri | 2.51 | 3.25E-20 | 6.76E-16 | 0.21 | 0.05 | 0.16 | 4.06 |
| olfactory neurons | Bicd1 | 2.54 | 3.85E-20 | 8.00E-16 | 0.3 | 0.1 | 0.2 | 3.07 |
| olfactory neurons | Tcta | 2.63 | 4.60E-20 | 9.56E-16 | 0.23 | 0.06 | 0.17 | 3.75 |
| olfactory neurons | Eif4g2 | 0.49 | 5.06E-20 | 1.05E-15 | 0.93 | 0.99 | -0.05 | 0.94 |
| olfactory neurons | Cmpk1 | 1.14 | 5.20E-20 | 1.08E-15 | 0.83 | 0.79 | 0.04 | 1.05 |
| olfactory neurons | Lsr | 1.59 | 5.79E-20 | 1.21E-15 | 0.48 | 0.21 | 0.27 | 2.31 |
| olfactory neurons | Nabp1 | 2.87 | 5.94E-20 | 1.24E-15 | 0.15 | 0.03 | 0.12 | 5.03 |
| olfactory neurons | Tspan13 | 1.65 | 6.09E-20 | 1.27E-15 | 0.66 | 0.41 | 0.25 | 1.61 |
| olfactory neurons | Phf21b | 3.2 | 1.00E-19 | 2.09E-15 | 0.18 | 0.04 | 0.14 | 4.41 |
| olfactory neurons | Dubr | 2.76 | 1.31E-19 | 2.73E-15 | 0.17 | 0.04 | 0.13 | 4.57 |
| olfactory neurons | Ccsap | 2.58 | 1.42E-19 | 2.96E-15 | 0.32 | 0.11 | 0.21 | 2.87 |
| olfactory neurons | Gpm6b | 3 | 1.78E-19 | 3.70E-15 | 0.24 | 0.07 | 0.17 | 3.51 |
| olfactory neurons | [illegible]610010F05Ri | 1.82 | 1.79E-19 | 3.72E-15 | 0.5 | 0.25 | 0.25 | 2.02 |
| olfactory neurons | Stx1a | 2.29 | 3.12E-19 | 6.49E-15 | 0.29 | 0.09 | 0.2 | 3.07 |
| olfactory neurons | Ralyl | 4.17 | 4.56E-19 | 9.48E-15 | 0.14 | 0.03 | 0.11 | 5.35 |
| olfactory neurons | Arid3b | 1.86 | 4.62E-19 | 9.61E-15 | 0.48 | 0.23 | 0.25 | 2.11 |
| olfactory neurons | Afap1 | 2.2 | 4.73E-19 | 9.85E-15 | 0.49 | 0.26 | 0.24 | 1.94 |
| olfactory neurons | H3f3b | 0.41 | 7.75E-19 | 1.61E-14 | 1 | 1 | 0 | 1 |
| olfactory neurons | Rab3d | 3.25 | 8.63E-19 | 1.80E-14 | 0.12 | 0.02 | 0.1 | 5.71 |
| olfactory neurons | Apbb1 | 2.1 | 9.23E-19 | 1.92E-14 | 0.55 | 0.32 | 0.23 | 1.71 |
| olfactory neurons | Ccp110 | 2.02 | 1.04E-18 | 2.16E-14 | 0.63 | 0.43 | 0.2 | 1.48 |
| olfactory neurons | Hist3h2a | 2.28 | 1.05E-18 | 2.18E-14 | 0.51 | 0.27 | 0.25 | 1.93 |
| olfactory neurons | Dpysl2 | 1.4 | 1.29E-18 | 2.69E-14 | 0.78 | 0.7 | 0.09 | 1.12 |
| olfactory neurons | BC031181 | 1.3 | 1.33E-18 | 2.78E-14 | 0.8 | 0.74 | 0.06 | 1.08 |
| olfactory neurons | Ccpg1 | 1.96 | 1.34E-18 | 2.78E-14 | 0.4 | 0.16 | 0.24 | 2.46 |
| olfactory neurons | Ackr1 | 3.57 | 1.59E-18 | 3.31E-14 | 0.1 | 0.02 | 0.09 | 6.38 |
| olfactory neurons | Dbn1 | 0.99 | 1.69E-18 | 3.51E-14 | 0.83 | 0.88 | -0.06 | 0.94 |
| olfactory neurons | Pou2f2 | 3.33 | 2.13E-18 | 4.43E-14 | 0.14 | 0.03 | 0.11 | 5.15 |
| olfactory neurons | Mtmr7 | 2.57 | 2.53E-18 | 5.27E-14 | 0.2 | 0.05 | 0.15 | 3.9 |
| olfactory neurons | Dpysl4 | 2.07 | 2.70E-18 | 5.62E-14 | 0.41 | 0.18 | 0.23 | 2.25 |
| olfactory neurons | Baz2b | 1.36 | 2.76E-18 | 5.75E-14 | 0.79 | 0.71 | 0.08 | 1.12 |
| olfactory neurons | Dstn | 0.73 | 3.43E-18 | 7.13E-14 | 0.91 | 0.94 | -0.03 | 0.97 |
| olfactory neurons | Mospd1 | 2.11 | 4.01E-18 | 8.33E-14 | 0.49 | 0.25 | 0.24 | 1.98 |
| olfactory neurons | Ralgps1 | 2.48 | 4.18E-18 | 8.69E-14 | 0.18 | 0.04 | 0.14 | 4.21 |
| olfactory neurons | Ddah2 | 0.89 | 4.84E-18 | 1.01E-13 | 0.93 | 0.95 | -0.02 | 0.98 |
| olfactory neurons | Scn3a | 3.32 | 5.01E-18 | 1.04E-13 | 0.21 | 0.06 | 0.15 | 3.66 |
| olfactory neurons | Tmem176b | 1.49 | 5.41E-18 | 1.13E-13 | 0.72 | 0.57 | 0.15 | 1.27 |
| olfactory neurons | Frrs1l | 2.85 | 6.72E-18 | 1.40E-13 | 0.15 | 0.03 | 0.12 | 4.83 |
| olfactory neurons | Tubb5 | 0.5 | 7.02E-18 | 1.46E-13 | 1 | 1 | 0 | 1 |
| olfactory neurons | Tecpr1 | 1.88 | 7.70E-18 | 1.60E-13 | 0.36 | 0.14 | 0.22 | 2.57 |
| olfactory neurons | Actr3b | 2.92 | 9.92E-18 | 2.06E-13 | 0.14 | 0.03 | 0.11 | 4.96 |
| olfactory neurons | Ttll1 | 2.29 | 9.97E-18 | 2.07E-13 | 0.39 | 0.16 | 0.23 | 2.43 |
| olfactory neurons | Akap17b | 2.68 | 1.27E-17 | 2.64E-13 | 0.32 | 0.12 | 0.2 | 2.64 |
| olfactory neurons | Riok3 | 1.51 | 1.33E-17 | 2.77E-13 | 0.68 | 0.5 | 0.18 | 1.35 |
| olfactory neurons | Pdgfa | 1.08 | 1.74E-17 | 3.63E-13 | 0.59 | 0.28 | 0.31 | 2.08 |
| olfactory neurons | Aldh1b1 | 3.08 | 1.76E-17 | 3.67E-13 | 0.11 | 0.02 | 0.09 | 6 |
| olfactory neurons | Smarca5 | 0.61 | 2.12E-17 | 4.41E-13 | 0.93 | 0.98 | -0.05 | 0.95 |
| olfactory neurons | Zfp386 | 1.81 | 3.28E-17 | 6.83E-13 | 0.65 | 0.41 | 0.23 | 1.56 |
| olfactory neurons | Pcp4l1 | 2.91 | 3.42E-17 | 7.10E-13 | 0.13 | 0.02 | 0.1 | 5.29 |
| olfactory neurons | Malat1 | 1.13 | 6.58E-17 | 1.37E-12 | 0.92 | 1 | -0.07 | 0.93 |
| olfactory neurons | Ypel3 | 1.71 | 7.11E-17 | 1.48E-12 | 0.71 | 0.57 | 0.14 | 1.25 |

**Supplementary Table 2. Marker genes for the major cell types of integrated clustering of E11.5 control and NotchGOF λ-junction cells.**

| | | | | | | | | |
|---|---|---|---|---|---|---|---|---|
| olfactory neurons | Fabp5 | 0.51 | 7.13E-17 | 1.48E-12 | 0.95 | 0.98 | -0.03 | 0.97 |
| olfactory neurons | Gm10544 | 3.04 | 7.69E-17 | 1.60E-12 | 0.15 | 0.03 | 0.11 | 4.68 |
| olfactory neurons | Cldn7 | 0.75 | 1.05E-16 | 2.19E-12 | 0.55 | 0.24 | 0.31 | 2.28 |
| olfactory neurons | Cox6a1 | 0.68 | 1.11E-16 | 2.31E-12 | 0.93 | 0.96 | -0.03 | 0.97 |
| olfactory neurons | Carhsp1 | 1.52 | 1.20E-16 | 2.50E-12 | 0.72 | 0.59 | 0.12 | 1.21 |
| olfactory neurons | Rsph9 | 2.29 | 1.22E-16 | 2.54E-12 | 0.3 | 0.11 | 0.19 | 2.76 |
| olfactory neurons | Tppp3 | 4.41 | 1.58E-16 | 3.28E-12 | 0.16 | 0.04 | 0.12 | 4.08 |
| olfactory neurons | Ywhag | 1.52 | 1.98E-16 | 4.11E-12 | 0.73 | 0.63 | 0.1 | 1.15 |
| olfactory neurons | Racgap1 | 1.49 | 2.05E-16 | 4.25E-12 | 0.71 | 0.53 | 0.19 | 1.35 |
| olfactory neurons | Amigo1 | 2.82 | 2.09E-16 | 4.36E-12 | 0.11 | 0.02 | 0.09 | 5.43 |
| olfactory neurons | Cxxc5 | 1.74 | 2.39E-16 | 4.96E-12 | 0.65 | 0.45 | 0.2 | 1.44 |
| olfactory neurons | Ago1 | 1.56 | 2.45E-16 | 5.09E-12 | 0.58 | 0.36 | 0.22 | 1.6 |
| olfactory neurons | Atp8a2 | 3.09 | 2.79E-16 | 5.81E-12 | 0.12 | 0.02 | 0.1 | 5.22 |
| olfactory neurons | Gm49066 | 2.4 | 2.80E-16 | 5.82E-12 | 0.18 | 0.04 | 0.13 | 3.98 |
| olfactory neurons | Trim24 | 1.34 | 3.30E-16 | 6.86E-12 | 0.77 | 0.71 | 0.05 | 1.07 |
| olfactory neurons | Sdk1 | 2.86 | 3.99E-16 | 8.30E-12 | 0.25 | 0.08 | 0.17 | 3.01 |
| olfactory neurons | Pim3 | 2.54 | 4.02E-16 | 8.36E-12 | 0.51 | 0.3 | 0.22 | 1.72 |
| olfactory neurons | Pacsin1 | 1.94 | 4.13E-16 | 8.58E-12 | 0.29 | 0.1 | 0.19 | 2.81 |
| olfactory neurons | Unc13b | 2.04 | 4.57E-16 | 9.50E-12 | 0.25 | 0.08 | 0.17 | 3.01 |
| olfactory neurons | Cep120 | 1.72 | 5.06E-16 | 1.05E-11 | 0.46 | 0.24 | 0.23 | 1.96 |
| olfactory neurons | Gm30015 | 4.2 | 5.50E-16 | 1.14E-11 | 0.1 | 0.02 | 0.08 | 5.67 |
| olfactory neurons | Dnajb6 | 0.96 | 5.99E-16 | 1.25E-11 | 0.88 | 0.89 | -0.01 | 0.99 |
| olfactory neurons | Cep97 | 2.34 | 6.13E-16 | 1.28E-11 | 0.41 | 0.21 | 0.21 | 2 |
| olfactory neurons | Selenok | 0.99 | 6.15E-16 | 1.28E-11 | 0.86 | 0.92 | -0.06 | 0.93 |
| olfactory neurons | Pick1 | 1.87 | 6.48E-16 | 1.35E-11 | 0.49 | 0.26 | 0.23 | 1.91 |
| olfactory neurons | Gm4651 | 2.5 | 7.87E-16 | 1.64E-11 | 0.13 | 0.03 | 0.1 | 4.88 |
| olfactory neurons | Pou3f1 | 3.08 | 7.95E-16 | 1.65E-11 | 0.13 | 0.03 | 0.1 | 4.88 |
| olfactory neurons | Cd55 | 2.56 | 8.92E-16 | 1.86E-11 | 0.12 | 0.02 | 0.1 | 5 |
| olfactory neurons | Rmst | 2.27 | 9.73E-16 | 2.02E-11 | 0.24 | 0.08 | 0.16 | 3.13 |
| olfactory neurons | Rnf32 | 2.81 | 9.77E-16 | 2.03E-11 | 0.12 | 0.02 | 0.1 | 5 |
| olfactory neurons | Msi2 | 1.11 | 1.22E-15 | 2.53E-11 | 0.78 | 0.68 | 0.1 | 1.14 |
| olfactory neurons | Dtna | 1.97 | 1.44E-15 | 3.00E-11 | 0.34 | 0.14 | 0.2 | 2.4 |
| olfactory neurons | Ache | 3.98 | 1.48E-15 | 3.07E-11 | 0.1 | 0.02 | 0.08 | 5.67 |
| olfactory neurons | Tmem108 | 1.84 | 1.64E-15 | 3.42E-11 | 0.44 | 0.22 | 0.22 | 2.02 |
| olfactory neurons | Sh2d3c | 2.06 | 1.65E-15 | 3.44E-11 | 0.28 | 0.1 | 0.18 | 2.77 |
| olfactory neurons | Kalrn | 1.99 | 1.84E-15 | 3.83E-11 | 0.4 | 0.2 | 0.21 | 2.05 |
| olfactory neurons | Fmn2 | 2.35 | 1.88E-15 | 3.92E-11 | 0.22 | 0.07 | 0.15 | 3.19 |
| olfactory neurons | Iqgap1 | 1.28 | 2.01E-15 | 4.18E-11 | 0.7 | 0.47 | 0.23 | 1.48 |
| olfactory neurons | Srp14 | 0.76 | 2.10E-15 | 4.36E-11 | 0.9 | 0.95 | -0.05 | 0.95 |
| olfactory neurons | Selenow | 0.65 | 2.94E-15 | 6.12E-11 | 0.95 | 0.99 | -0.04 | 0.96 |
| olfactory neurons | Kdm5b | 0.84 | 2.96E-15 | 6.15E-11 | 0.86 | 0.86 | 0 | 1 |
| olfactory neurons | Atp6v0e2 | 3.18 | 3.27E-15 | 6.80E-11 | 0.19 | 0.05 | 0.13 | 3.53 |
| olfactory neurons | Prkar1a | 1.05 | 3.45E-15 | 7.17E-11 | 0.88 | 0.88 | 0 | 1 |
| olfactory neurons | Jund | 0.87 | 3.58E-15 | 7.45E-11 | 0.89 | 0.94 | -0.05 | 0.94 |
| olfactory neurons | Dlx5 | 2.16 | 3.90E-15 | 8.11E-11 | 0.44 | 0.22 | 0.22 | 2.03 |
| olfactory neurons | Rtn3 | 0.93 | 4.00E-15 | 8.32E-11 | 0.81 | 0.79 | 0.02 | 1.03 |
| olfactory neurons | Tubb4a | 3.36 | 4.02E-15 | 8.37E-11 | 0.15 | 0.04 | 0.11 | 4.08 |
| olfactory neurons | Coro1a | 1.95 | 4.70E-15 | 9.78E-11 | 0.35 | 0.15 | 0.2 | 2.36 |
| olfactory neurons | Ubald2 | 1.16 | 5.47E-15 | 1.14E-10 | 0.78 | 0.69 | 0.09 | 1.13 |
| olfactory neurons | Septin3 | 1.55 | 6.48E-15 | 1.35E-10 | 0.61 | 0.44 | 0.17 | 1.39 |
| olfactory neurons | Cep170 | 1.34 | 7.54E-15 | 1.57E-10 | 0.68 | 0.51 | 0.17 | 1.33 |
| olfactory neurons | Gm15417 | 2.82 | 7.93E-15 | 1.65E-10 | 0.12 | 0.03 | 0.1 | 4.8 |
| olfactory neurons | Chfr | 1.62 | 8.88E-15 | 1.85E-10 | 0.48 | 0.26 | 0.22 | 1.86 |
| olfactory neurons | Gpi1 | 1.06 | 9.07E-15 | 1.89E-10 | 0.86 | 0.84 | 0.01 | 1.02 |
| olfactory neurons | Syt7 | 3.04 | 1.02E-14 | 2.13E-10 | 0.19 | 0.06 | 0.14 | 3.39 |
| olfactory neurons | Nfe2l3 | 2.43 | 1.15E-14 | 2.40E-10 | 0.15 | 0.04 | 0.11 | 4.14 |

**Supplementary Table 2. Marker genes for the major cell types of integrated clustering of E11.5 control and NotchGOF λ-junction cells.**

| | | | | | | | | |
|---|---|---|---|---|---|---|---|---|
| olfactory neurons | Ank3 | 2.01 | 1.23E-14 | 2.56E-10 | 0.65 | 0.49 | 0.16 | 1.33 |
| olfactory neurons | Maml3 | 1.32 | 1.35E-14 | 2.80E-10 | 0.61 | 0.4 | 0.21 | 1.53 |
| olfactory neurons | Tgfbr1 | 1.83 | 1.50E-14 | 3.13E-10 | 0.68 | 0.55 | 0.12 | 1.22 |
| olfactory neurons | Pkig | 1.06 | 1.62E-14 | 3.37E-10 | 0.86 | 0.79 | 0.07 | 1.09 |
| olfactory neurons | Rgs17 | 2.65 | 1.84E-14 | 3.83E-10 | 0.15 | 0.04 | 0.11 | 4.14 |
| olfactory neurons | Map7d2 | 2.08 | 2.01E-14 | 4.17E-10 | 0.24 | 0.08 | 0.16 | 2.87 |
| olfactory neurons | Gm43260 | 2.48 | 2.06E-14 | 4.29E-10 | 0.15 | 0.04 | 0.11 | 4.14 |
| olfactory neurons | Hsp90aa1 | 0.4 | 2.14E-14 | 4.45E-10 | 0.97 | 1 | -0.03 | 0.97 |
| olfactory neurons | Ctxn1 | 1.26 | 2.38E-14 | 4.95E-10 | 0.8 | 0.78 | 0.01 | 1.02 |
| olfactory neurons | Fbxl16 | 3.3 | 2.46E-14 | 5.12E-10 | 0.13 | 0.03 | 0.1 | 4.54 |
| olfactory neurons | Fgf15 | 3.67 | 2.58E-14 | 5.37E-10 | 0.12 | 0.03 | 0.09 | 4.62 |
| olfactory neurons | Marveld2 | 1.97 | 2.74E-14 | 5.71E-10 | 0.2 | 0.06 | 0.14 | 3.32 |
| olfactory neurons | Ccdc171 | 2.08 | 2.77E-14 | 5.75E-10 | 0.27 | 0.1 | 0.17 | 2.79 |
| olfactory neurons | Marchf8 | 1.88 | 2.81E-14 | 5.85E-10 | 0.37 | 0.17 | 0.2 | 2.16 |
| olfactory neurons | B3gnt4 | 2.56 | 3.11E-14 | 6.46E-10 | 0.16 | 0.04 | 0.12 | 3.93 |
| olfactory neurons | Mtch1 | 0.68 | 3.40E-14 | 7.07E-10 | 0.93 | 0.96 | -0.02 | 0.98 |
| olfactory neurons | Nefl | 4.36 | 4.22E-14 | 8.77E-10 | 0.13 | 0.03 | 0.1 | 4.29 |
| olfactory neurons | Igf1r | 1.19 | 5.16E-14 | 1.07E-09 | 0.79 | 0.77 | 0.02 | 1.02 |
| olfactory neurons | Hook1 | 1.45 | 5.55E-14 | 1.15E-09 | 0.43 | 0.2 | 0.23 | 2.12 |
| olfactory neurons | Snx32 | 2.82 | 6.08E-14 | 1.26E-09 | 0.11 | 0.02 | 0.09 | 4.75 |
| olfactory neurons | Zcchc18 | 1.66 | 6.59E-14 | 1.37E-09 | 0.54 | 0.34 | 0.2 | 1.6 |
| olfactory neurons | Gldc | 2.59 | 7.12E-14 | 1.48E-09 | 0.16 | 0.04 | 0.12 | 3.83 |
| olfactory neurons | Gap43 | 3.34 | 7.99E-14 | 1.66E-09 | 0.58 | 0.42 | 0.16 | 1.38 |
| olfactory neurons | Lmo2 | 2.35 | 8.36E-14 | 1.74E-09 | 0.31 | 0.13 | 0.19 | 2.46 |
| olfactory neurons | Baz2a | 1.23 | 8.83E-14 | 1.84E-09 | 0.61 | 0.39 | 0.22 | 1.58 |
| olfactory neurons | Klc1 | 1.97 | 9.12E-14 | 1.90E-09 | 0.63 | 0.47 | 0.16 | 1.33 |
| olfactory neurons | Six3 | 0.87 | 1.04E-13 | 2.16E-09 | 0.51 | 0.24 | 0.27 | 2.11 |
| olfactory neurons | Msi1 | 1.45 | 1.11E-13 | 2.32E-09 | 0.58 | 0.37 | 0.21 | 1.55 |
| olfactory neurons | Ift43 | 1.66 | 1.33E-13 | 2.76E-09 | 0.71 | 0.55 | 0.16 | 1.29 |
| olfactory neurons | Prr15 | 2.18 | 1.50E-13 | 3.11E-09 | 0.14 | 0.03 | 0.11 | 4.09 |
| olfactory neurons | Ssh2 | 1.54 | 1.50E-13 | 3.11E-09 | 0.43 | 0.23 | 0.2 | 1.89 |
| olfactory neurons | Rtn4 | 0.81 | 1.83E-13 | 3.80E-09 | 0.83 | 0.94 | -0.11 | 0.88 |
| olfactory neurons | Gpr137c | 1.53 | 1.84E-13 | 3.83E-09 | 0.34 | 0.15 | 0.19 | 2.32 |
| olfactory neurons | Gfod1 | 2.99 | 1.85E-13 | 3.85E-09 | 0.1 | 0.02 | 0.08 | 4.86 |
| olfactory neurons | Tubb2a | 1.89 | 2.22E-13 | 4.61E-09 | 0.5 | 0.3 | 0.2 | 1.68 |
| olfactory neurons | Dusp14 | 3.12 | 2.22E-13 | 4.61E-09 | 0.18 | 0.05 | 0.12 | 3.43 |
| olfactory neurons | Lcorl | 1.58 | 2.73E-13 | 5.69E-09 | 0.64 | 0.48 | 0.16 | 1.33 |
| olfactory neurons | Cpe | 1.5 | 2.88E-13 | 5.98E-09 | 0.6 | 0.42 | 0.18 | 1.43 |
| olfactory neurons | Tulp4 | 1.11 | 3.41E-13 | 7.09E-09 | 0.77 | 0.67 | 0.1 | 1.15 |
| olfactory neurons | Nrcam | 2.56 | 3.82E-13 | 7.95E-09 | 0.18 | 0.05 | 0.12 | 3.37 |
| olfactory neurons | C9orf72 | 2.48 | 4.05E-13 | 8.42E-09 | 0.28 | 0.11 | 0.17 | 2.47 |
| olfactory neurons | Csnk1e | 0.94 | 4.19E-13 | 8.72E-09 | 0.84 | 0.84 | -0.01 | 0.99 |
| olfactory neurons | Rac3 | 1.95 | 4.27E-13 | 8.88E-09 | 0.51 | 0.33 | 0.18 | 1.54 |
| olfactory neurons | Dmxl2 | 2.19 | 4.41E-13 | 9.18E-09 | 0.22 | 0.07 | 0.14 | 2.93 |
| olfactory neurons | Rala | 0.82 | 4.66E-13 | 9.69E-09 | 0.84 | 0.85 | -0.02 | 0.98 |
| olfactory neurons | Arhgap33 | 1.74 | 4.79E-13 | 9.97E-09 | 0.31 | 0.13 | 0.18 | 2.34 |
| olfactory neurons | Dynll1 | 0.51 | 4.85E-13 | 1.01E-08 | 0.99 | 1 | -0.01 | 0.99 |
| olfactory neurons | Acap3 | 2.01 | 5.13E-13 | 1.07E-08 | 0.22 | 0.07 | 0.14 | 2.93 |
| olfactory neurons | C2cd2l | 2.49 | 5.28E-13 | 1.10E-08 | 0.18 | 0.05 | 0.12 | 3.3 |
| olfactory neurons | Tanc2 | 1.6 | 5.50E-13 | 1.14E-08 | 0.43 | 0.23 | 0.2 | 1.87 |
| olfactory neurons | Pabpc1l | 2.76 | 5.81E-13 | 1.21E-08 | 0.12 | 0.03 | 0.09 | 4.29 |
| olfactory neurons | Ankrd37 | 1.62 | 5.93E-13 | 1.23E-08 | 0.35 | 0.16 | 0.19 | 2.21 |
| olfactory neurons | Ptprf | 0.76 | 6.28E-13 | 1.31E-08 | 0.8 | 0.64 | 0.16 | 1.24 |
| olfactory neurons | Them6 | 2.93 | 6.36E-13 | 1.32E-08 | 0.1 | 0.02 | 0.08 | 4.86 |
| olfactory neurons | Cep63 | 1.43 | 7.06E-13 | 1.47E-08 | 0.52 | 0.32 | 0.19 | 1.6 |
| olfactory neurons | Six1 | 1.11 | 7.82E-13 | 1.63E-08 | 0.75 | 0.61 | 0.14 | 1.23 |

**Supplementary Table 2. Marker genes for the major cell types of integrated clustering of E11.5 control and NotchGOF λ-junction cells.**

| | | | | | | | | |
|---|---|---|---|---|---|---|---|---|
| olfactory neurons | Zfp667 | 2.12 | 7.91E-13 | 1.64E-08 | 0.26 | 0.1 | 0.16 | 2.62 |
| olfactory neurons | Tnfrsf12a | 1.79 | 9.94E-13 | 2.07E-08 | 0.22 | 0.08 | 0.15 | 2.9 |
| olfactory neurons | Ccpg1os | 1.79 | 1.13E-12 | 2.34E-08 | 0.34 | 0.16 | 0.19 | 2.17 |
| olfactory neurons | Mapk11 | 2.12 | 1.18E-12 | 2.46E-08 | 0.21 | 0.07 | 0.14 | 2.93 |
| olfactory neurons | Nectin1 | 1.79 | 1.22E-12 | 2.54E-08 | 0.36 | 0.18 | 0.19 | 2.06 |
| olfactory neurons | Meg3 | 1.89 | 1.23E-12 | 2.57E-08 | 0.74 | 0.69 | 0.04 | 1.06 |
| olfactory neurons | Ttc5 | 1.17 | 1.31E-12 | 2.73E-08 | 0.69 | 0.54 | 0.14 | 1.26 |
| olfactory neurons | Kctd13 | 1.7 | 1.44E-12 | 2.99E-08 | 0.33 | 0.15 | 0.18 | 2.16 |
| olfactory neurons | Crabp2 | 0.65 | 1.48E-12 | 3.08E-08 | 0.95 | 0.92 | 0.03 | 1.03 |
| olfactory neurons | Cpeb4 | 2.09 | 1.59E-12 | 3.31E-08 | 0.25 | 0.1 | 0.15 | 2.6 |
| olfactory neurons | Ncoa1 | 2.16 | 1.65E-12 | 3.42E-08 | 0.35 | 0.17 | 0.18 | 2.03 |
| olfactory neurons | Bcl11b | 1.89 | 1.82E-12 | 3.79E-08 | 0.63 | 0.59 | 0.04 | 1.06 |
| olfactory neurons | Parm1 | 2.26 | 1.97E-12 | 4.09E-08 | 0.34 | 0.16 | 0.18 | 2.12 |
| olfactory neurons | Rian | 1.89 | 2.06E-12 | 4.29E-08 | 0.51 | 0.32 | 0.19 | 1.61 |
| olfactory neurons | Eml1 | 1.78 | 2.25E-12 | 4.68E-08 | 0.35 | 0.17 | 0.18 | 2.09 |
| olfactory neurons | Chchd10 | 1.64 | 2.28E-12 | 4.75E-08 | 0.35 | 0.16 | 0.19 | 2.17 |
| olfactory neurons | Fam187b | 1.89 | 2.32E-12 | 4.84E-08 | 0.15 | 0.04 | 0.11 | 3.72 |
| olfactory neurons | Card10 | 2.9 | 2.80E-12 | 5.81E-08 | 0.1 | 0.02 | 0.08 | 4.64 |
| olfactory neurons | Rab33a | 3.06 | 3.40E-12 | 7.07E-08 | 0.13 | 0.03 | 0.1 | 4.1 |
| olfactory neurons | Myo1b | 0.91 | 3.45E-12 | 7.18E-08 | 0.8 | 0.73 | 0.06 | 1.09 |
| olfactory neurons | Ric3 | 2.17 | 3.47E-12 | 7.23E-08 | 0.17 | 0.05 | 0.12 | 3.31 |
| olfactory neurons | Cox7a1 | 3.7 | 3.59E-12 | 7.47E-08 | 0.1 | 0.02 | 0.08 | 4.64 |
| olfactory neurons | Mex3a | 0.83 | 3.66E-12 | 7.62E-08 | 0.83 | 0.85 | -0.03 | 0.97 |
| olfactory neurons | Nav1 | 1.21 | 3.78E-12 | 7.86E-08 | 0.66 | 0.54 | 0.12 | 1.22 |
| olfactory neurons | Wfs1 | 1.81 | 3.85E-12 | 8.02E-08 | 0.2 | 0.07 | 0.13 | 3.02 |
| olfactory neurons | Rnasek | 1 | 4.07E-12 | 8.47E-08 | 0.76 | 0.72 | 0.04 | 1.05 |
| olfactory neurons | Phyh | 1.61 | 4.99E-12 | 1.04E-07 | 0.48 | 0.29 | 0.19 | 1.65 |
| olfactory neurons | Plch1 | 1.51 | 6.16E-12 | 1.28E-07 | 0.26 | 0.1 | 0.16 | 2.56 |
| olfactory neurons | Psma7 | 0.55 | 6.32E-12 | 1.31E-07 | 0.96 | 0.99 | -0.03 | 0.97 |
| olfactory neurons | Gabarap | 0.61 | 6.78E-12 | 1.41E-07 | 0.91 | 0.98 | -0.07 | 0.93 |
| olfactory neurons | Prmt2 | 1.7 | 7.43E-12 | 1.54E-07 | 0.48 | 0.31 | 0.17 | 1.55 |
| olfactory neurons | Ralgapa2 | 1.79 | 7.51E-12 | 1.56E-07 | 0.3 | 0.14 | 0.16 | 2.18 |
| olfactory neurons | Gm5860 | 3.19 | 9.22E-12 | 1.92E-07 | 0.13 | 0.03 | 0.09 | 3.85 |
| olfactory neurons | Tex15 | 4.24 | 9.54E-12 | 1.98E-07 | 0.26 | 0.11 | 0.15 | 2.35 |
| olfactory neurons | Usp29 | 1.52 | 9.81E-12 | 2.04E-07 | 0.4 | 0.21 | 0.19 | 1.93 |
| olfactory neurons | Mapre3 | 2.43 | 1.06E-11 | 2.20E-07 | 0.27 | 0.11 | 0.16 | 2.38 |
| olfactory neurons | Gm42702 | 2.25 | 1.06E-11 | 2.20E-07 | 0.14 | 0.04 | 0.1 | 3.66 |
| olfactory neurons | Tmem178 | 2.89 | 1.13E-11 | 2.34E-07 | 0.13 | 0.03 | 0.09 | 3.85 |
| olfactory neurons | Snrpn | 1.78 | 1.14E-11 | 2.36E-07 | 0.63 | 0.47 | 0.16 | 1.34 |
| olfactory neurons | Dync1i2 | 0.73 | 1.37E-11 | 2.84E-07 | 0.84 | 0.89 | -0.05 | 0.94 |
| olfactory neurons | Lrrc42 | 1.51 | 1.47E-11 | 3.05E-07 | 0.68 | 0.61 | 0.07 | 1.11 |
| olfactory neurons | Lman2l | 1.52 | 1.62E-11 | 3.37E-07 | 0.53 | 0.35 | 0.18 | 1.53 |
| olfactory neurons | Atp6v1e1 | 1.09 | 1.89E-11 | 3.93E-07 | 0.72 | 0.68 | 0.04 | 1.07 |
| olfactory neurons | Unc45a | 1.8 | 2.13E-11 | 4.43E-07 | 0.45 | 0.27 | 0.18 | 1.67 |
| olfactory neurons | Jade1 | 1.98 | 2.57E-11 | 5.35E-07 | 0.34 | 0.17 | 0.17 | 1.98 |
| olfactory neurons | Tmeff1 | 1.11 | 2.67E-11 | 5.56E-07 | 0.74 | 0.69 | 0.05 | 1.07 |
| olfactory neurons | Arpc3 | 0.7 | 2.74E-11 | 5.70E-07 | 0.87 | 0.9 | -0.03 | 0.97 |
| olfactory neurons | Kit | 2.58 | 2.82E-11 | 5.86E-07 | 0.24 | 0.1 | 0.15 | 2.51 |
| olfactory neurons | Prkx | 1.69 | 2.84E-11 | 5.90E-07 | 0.32 | 0.15 | 0.17 | 2.07 |
| olfactory neurons | Fkbp1a | 0.56 | 3.06E-11 | 6.37E-07 | 0.89 | 0.94 | -0.05 | 0.94 |
| olfactory neurons | Atp6v0d1 | 1.31 | 3.19E-11 | 6.64E-07 | 0.69 | 0.54 | 0.15 | 1.28 |
| olfactory neurons | Akna | 1.96 | 3.22E-11 | 6.69E-07 | 0.26 | 0.11 | 0.15 | 2.44 |
| olfactory neurons | Frmd5 | 2.03 | 3.40E-11 | 7.07E-07 | 0.16 | 0.05 | 0.11 | 3.2 |
| olfactory neurons | Dnaja1 | 0.38 | 3.42E-11 | 7.11E-07 | 0.93 | 0.99 | -0.06 | 0.94 |
| olfactory neurons | Gamt | 2.85 | 3.45E-11 | 7.18E-07 | 0.24 | 0.09 | 0.14 | 2.55 |
| olfactory neurons | 930452B06Ri | 2.46 | 3.67E-11 | 7.63E-07 | 0.28 | 0.12 | 0.16 | 2.28 |

**Supplementary Table 2. Marker genes for the major cell types of integrated clustering of E11.5 control and NotchGOF λ-junction cells.**

| | | | | | | | | |
|---|---|---|---|---|---|---|---|---|
| olfactory neurons | Mgst3 | 1.45 | 3.74E-11 | 7.78E-07 | 0.63 | 0.51 | 0.13 | 1.25 |
| olfactory neurons | Tnik | 1.77 | 3.75E-11 | 7.81E-07 | 0.42 | 0.25 | 0.17 | 1.7 |
| olfactory neurons | Dda1 | 1.32 | 4.15E-11 | 8.63E-07 | 0.69 | 0.57 | 0.12 | 1.21 |
| olfactory neurons | 810455O05Ri | 2.74 | 4.71E-11 | 9.79E-07 | 0.1 | 0.02 | 0.08 | 4.25 |
| olfactory neurons | Rnd2 | 2.26 | 4.77E-11 | 9.92E-07 | 0.23 | 0.09 | 0.14 | 2.6 |
| olfactory neurons | Snta1 | 2.23 | 5.13E-11 | 1.07E-06 | 0.16 | 0.05 | 0.11 | 3.2 |
| olfactory neurons | Hist2h2be | 2.57 | 5.21E-11 | 1.08E-06 | 0.13 | 0.04 | 0.1 | 3.59 |
| olfactory neurons | Tnfrsf21 | 1.78 | 5.66E-11 | 1.18E-06 | 0.31 | 0.14 | 0.17 | 2.16 |
| olfactory neurons | Strbp | 1.06 | 5.67E-11 | 1.18E-06 | 0.75 | 0.71 | 0.04 | 1.06 |
| olfactory neurons | Jup | 1.17 | 5.71E-11 | 1.19E-06 | 0.4 | 0.2 | 0.2 | 2 |
| olfactory neurons | .900093K20Ri | 1.99 | 5.91E-11 | 1.23E-06 | 0.22 | 0.09 | 0.14 | 2.59 |
| olfactory neurons | Tmcc1 | 1.24 | 6.07E-11 | 1.26E-06 | 0.6 | 0.44 | 0.16 | 1.37 |
| olfactory neurons | Cyb5r4 | 1.4 | 6.23E-11 | 1.29E-06 | 0.57 | 0.4 | 0.17 | 1.43 |
| olfactory neurons | Kras | 0.87 | 7.34E-11 | 1.53E-06 | 0.82 | 0.85 | -0.03 | 0.97 |
| olfactory neurons | Fam122b | 1.94 | 7.57E-11 | 1.57E-06 | 0.22 | 0.08 | 0.13 | 2.61 |
| olfactory neurons | Ezr | 0.71 | 8.04E-11 | 1.67E-06 | 0.57 | 0.32 | 0.25 | 1.76 |
| olfactory neurons | Daam1 | 1.33 | 8.57E-11 | 1.78E-06 | 0.58 | 0.41 | 0.18 | 1.43 |
| olfactory neurons | Mob3b | 1.8 | 9.02E-11 | 1.88E-06 | 0.23 | 0.09 | 0.14 | 2.54 |
| olfactory neurons | Cyb561 | 2.3 | 1.00E-10 | 2.08E-06 | 0.17 | 0.06 | 0.11 | 3.02 |
| olfactory neurons | Dram2 | 1.43 | 1.02E-10 | 2.12E-06 | 0.39 | 0.21 | 0.19 | 1.91 |
| olfactory neurons | Lurap1l | 1.71 | 1.21E-10 | 2.51E-06 | 0.21 | 0.08 | 0.13 | 2.74 |
| olfactory neurons | Myef2 | 0.88 | 1.21E-10 | 2.53E-06 | 0.75 | 0.69 | 0.06 | 1.09 |
| olfactory neurons | Cdkn2d | 1.71 | 1.29E-10 | 2.69E-06 | 0.53 | 0.38 | 0.15 | 1.38 |
| olfactory neurons | Kcnn1 | 2.18 | 1.33E-10 | 2.77E-06 | 0.15 | 0.04 | 0.1 | 3.37 |
| olfactory neurons | Susd6 | 1.68 | 1.35E-10 | 2.80E-06 | 0.37 | 0.2 | 0.17 | 1.86 |
| olfactory neurons | Lbh | 1.97 | 1.49E-10 | 3.09E-06 | 0.37 | 0.2 | 0.17 | 1.85 |
| olfactory neurons | .700023F06Ri | 2.63 | 1.52E-10 | 3.17E-06 | 0.15 | 0.04 | 0.1 | 3.3 |
| olfactory neurons | Arpc1a | 0.73 | 1.54E-10 | 3.20E-06 | 0.85 | 0.88 | -0.04 | 0.96 |
| olfactory neurons | Gdf11 | 1.44 | 1.55E-10 | 3.22E-06 | 0.49 | 0.32 | 0.17 | 1.53 |
| olfactory neurons | Twf1 | 1.92 | 1.58E-10 | 3.29E-06 | 0.61 | 0.49 | 0.12 | 1.24 |
| olfactory neurons | Ubl5 | 0.53 | 1.59E-10 | 3.30E-06 | 0.93 | 0.98 | -0.05 | 0.95 |
| olfactory neurons | Cdc42ep3 | 1.57 | 1.66E-10 | 3.44E-06 | 0.34 | 0.17 | 0.17 | 2.04 |
| olfactory neurons | Rundc3b | 2.4 | 1.75E-10 | 3.64E-06 | 0.14 | 0.04 | 0.1 | 3.39 |
| olfactory neurons | Kif3c | 1.82 | 1.89E-10 | 3.93E-06 | 0.27 | 0.12 | 0.15 | 2.25 |
| olfactory neurons | Gm42372 | 2.14 | 1.90E-10 | 3.94E-06 | 0.16 | 0.05 | 0.11 | 3.14 |
| olfactory neurons | Dctn1 | 1.46 | 2.30E-10 | 4.78E-06 | 0.62 | 0.52 | 0.1 | 1.19 |
| olfactory neurons | Dnajc6 | 2.54 | 2.38E-10 | 4.94E-06 | 0.1 | 0.03 | 0.08 | 4.08 |
| olfactory neurons | Camsap3 | 2.08 | 2.49E-10 | 5.17E-06 | 0.21 | 0.08 | 0.13 | 2.57 |
| olfactory neurons | Srcin1 | 2.03 | 2.56E-10 | 5.33E-06 | 0.15 | 0.05 | 0.1 | 3.21 |
| olfactory neurons | Trp53inp1 | 1.42 | 2.64E-10 | 5.48E-06 | 0.37 | 0.19 | 0.19 | 2.01 |
| olfactory neurons | Fam222a | 2.97 | 2.81E-10 | 5.85E-06 | 0.11 | 0.03 | 0.08 | 3.68 |
| olfactory neurons | Vamp2 | 1.8 | 3.29E-10 | 6.85E-06 | 0.45 | 0.3 | 0.15 | 1.52 |
| olfactory neurons | Zfp664 | 0.85 | 3.30E-10 | 6.86E-06 | 0.78 | 0.75 | 0.03 | 1.04 |
| olfactory neurons | Sbk1 | 1.19 | 3.34E-10 | 6.95E-06 | 0.62 | 0.52 | 0.1 | 1.18 |
| olfactory neurons | Id4 | 2.64 | 3.71E-10 | 7.71E-06 | 0.47 | 0.31 | 0.16 | 1.53 |
| olfactory neurons | Ppp1r11 | 0.84 | 3.78E-10 | 7.87E-06 | 0.83 | 0.81 | 0.02 | 1.03 |
| olfactory neurons | Gm2464 | 3.29 | 3.84E-10 | 7.98E-06 | 0.11 | 0.03 | 0.08 | 3.86 |
| olfactory neurons | Celsr2 | 1.7 | 3.90E-10 | 8.11E-06 | 0.23 | 0.09 | 0.14 | 2.46 |
| olfactory neurons | Cxxc4 | 1.79 | 4.14E-10 | 8.62E-06 | 0.46 | 0.31 | 0.15 | 1.5 |
| olfactory neurons | Cdk5r1 | 2.17 | 4.17E-10 | 8.68E-06 | 0.25 | 0.11 | 0.14 | 2.29 |
| olfactory neurons | Map2k1 | 1.64 | 4.43E-10 | 9.20E-06 | 0.5 | 0.35 | 0.15 | 1.45 |
| olfactory neurons | Dynll2 | 0.86 | 4.69E-10 | 9.76E-06 | 0.81 | 0.84 | -0.04 | 0.96 |
| olfactory neurons | Kcnh2 | 1.95 | 5.12E-10 | 1.06E-05 | 0.15 | 0.05 | 0.1 | 3.22 |
| olfactory neurons | Sdccag8 | 2.01 | 5.41E-10 | 1.13E-05 | 0.28 | 0.13 | 0.15 | 2.2 |
| olfactory neurons | Ciao2b | 0.96 | 5.53E-10 | 1.15E-05 | 0.75 | 0.73 | 0.02 | 1.02 |
| olfactory neurons | Map3k13 | 3.11 | 5.73E-10 | 1.19E-05 | 0.12 | 0.03 | 0.09 | 3.53 |

**Supplementary Table 2. Marker genes for the major cell types of integrated clustering of E11.5 control and NotchGOF λ-junction cells.**

| | | | | | | | | |
|---|---|---|---|---|---|---|---|---|
| olfactory neurons | Rbfox2 | 0.88 | 5.93E-10 | 1.23E-05 | 0.78 | 0.77 | 0.02 | 1.02 |
| olfactory neurons | Gng2 | 1.2 | 6.52E-10 | 1.36E-05 | 0.67 | 0.57 | 0.1 | 1.17 |
| olfactory neurons | Armh4 | 2.47 | 7.75E-10 | 1.61E-05 | 0.3 | 0.15 | 0.15 | 1.97 |
| olfactory neurons | Abca3 | 1.87 | 8.09E-10 | 1.68E-05 | 0.18 | 0.07 | 0.12 | 2.78 |
| olfactory neurons | Nbea | 1.2 | 8.54E-10 | 1.78E-05 | 0.59 | 0.44 | 0.15 | 1.33 |
| olfactory neurons | Ift22 | 1.27 | 8.74E-10 | 1.82E-05 | 0.51 | 0.35 | 0.15 | 1.43 |
| olfactory neurons | Cib1 | 1.36 | 9.60E-10 | 2.00E-05 | 0.55 | 0.4 | 0.15 | 1.38 |
| olfactory neurons | Dync1li1 | 1.21 | 1.00E-09 | 2.08E-05 | 0.64 | 0.52 | 0.12 | 1.23 |
| olfactory neurons | Pbx3 | 1.23 | 1.14E-09 | 2.37E-05 | 0.65 | 0.53 | 0.12 | 1.23 |
| olfactory neurons | Plxnc1 | 2.02 | 1.23E-09 | 2.57E-05 | 0.14 | 0.04 | 0.1 | 3.23 |
| olfactory neurons | 700037H04Ri | 1.59 | 1.27E-09 | 2.64E-05 | 0.62 | 0.48 | 0.14 | 1.3 |
| olfactory neurons | Plppr2 | 2.11 | 1.36E-09 | 2.83E-05 | 0.18 | 0.06 | 0.11 | 2.78 |
| olfactory neurons | Rcan1 | 2.08 | 1.40E-09 | 2.90E-05 | 0.42 | 0.26 | 0.16 | 1.62 |
| olfactory neurons | Peg13 | 2.11 | 1.58E-09 | 3.29E-05 | 0.15 | 0.05 | 0.1 | 3.02 |
| olfactory neurons | Gnaz | 2.32 | 1.60E-09 | 3.32E-05 | 0.14 | 0.04 | 0.1 | 3.16 |
| olfactory neurons | Arl8a | 1.69 | 1.67E-09 | 3.47E-05 | 0.54 | 0.43 | 0.11 | 1.26 |
| olfactory neurons | Zc2hc1a | 1.26 | 1.67E-09 | 3.48E-05 | 0.57 | 0.42 | 0.15 | 1.37 |
| olfactory neurons | Zbtb20 | 2.04 | 1.71E-09 | 3.56E-05 | 0.44 | 0.29 | 0.15 | 1.5 |
| olfactory neurons | Hspa2 | 2.07 | 1.75E-09 | 3.64E-05 | 0.16 | 0.06 | 0.11 | 2.91 |
| olfactory neurons | Akr1b8 | 1.78 | 1.75E-09 | 3.64E-05 | 0.2 | 0.08 | 0.12 | 2.58 |
| olfactory neurons | Mindy2 | 1.73 | 1.80E-09 | 3.74E-05 | 0.33 | 0.18 | 0.15 | 1.86 |
| olfactory neurons | Crtc3 | 1.75 | 1.81E-09 | 3.77E-05 | 0.4 | 0.25 | 0.15 | 1.62 |
| olfactory neurons | H13 | 0.92 | 1.84E-09 | 3.82E-05 | 0.72 | 0.63 | 0.08 | 1.13 |
| olfactory neurons | Rab6a | 1.14 | 1.95E-09 | 4.06E-05 | 0.74 | 0.72 | 0.03 | 1.03 |
| olfactory neurons | Clta | 0.55 | 2.03E-09 | 4.22E-05 | 0.94 | 0.98 | -0.04 | 0.96 |
| olfactory neurons | Gnptg | 1.65 | 2.26E-09 | 4.69E-05 | 0.37 | 0.2 | 0.17 | 1.82 |
| olfactory neurons | Cfap36 | 1.32 | 2.32E-09 | 4.82E-05 | 0.69 | 0.55 | 0.14 | 1.25 |
| olfactory neurons | Arg2 | 2 | 2.41E-09 | 5.01E-05 | 0.16 | 0.05 | 0.1 | 2.96 |
| olfactory neurons | Cfl1 | 0.31 | 2.44E-09 | 5.08E-05 | 0.99 | 1 | -0.01 | 0.99 |
| olfactory neurons | Ttc39b | 2 | 2.45E-09 | 5.09E-05 | 0.18 | 0.07 | 0.12 | 2.74 |
| olfactory neurons | Ccdc28b | 1.35 | 2.53E-09 | 5.25E-05 | 0.53 | 0.38 | 0.16 | 1.41 |
| olfactory neurons | Eml5 | 1.89 | 2.61E-09 | 5.44E-05 | 0.21 | 0.09 | 0.13 | 2.45 |
| olfactory neurons | Mapk8 | 1.05 | 2.78E-09 | 5.78E-05 | 0.59 | 0.43 | 0.17 | 1.39 |
| olfactory neurons | Atp6v1h | 1.04 | 2.88E-09 | 5.99E-05 | 0.54 | 0.36 | 0.17 | 1.48 |
| olfactory neurons | Rasal2 | 1.81 | 2.92E-09 | 6.08E-05 | 0.4 | 0.25 | 0.16 | 1.62 |
| olfactory neurons | Dedd2 | 1.9 | 3.00E-09 | 6.23E-05 | 0.19 | 0.08 | 0.12 | 2.57 |
| olfactory neurons | Stx7 | 2.22 | 3.07E-09 | 6.38E-05 | 0.45 | 0.31 | 0.14 | 1.45 |
| olfactory neurons | 830012L14Ri | 2.28 | 3.08E-09 | 6.40E-05 | 0.16 | 0.06 | 0.1 | 2.85 |
| olfactory neurons | Mterf2 | 1.51 | 3.27E-09 | 6.81E-05 | 0.37 | 0.21 | 0.16 | 1.77 |
| olfactory neurons | Smim22 | 3.07 | 3.43E-09 | 7.14E-05 | 0.13 | 0.04 | 0.09 | 3.26 |
| olfactory neurons | Palm | 1.46 | 3.52E-09 | 7.33E-05 | 0.58 | 0.46 | 0.12 | 1.25 |
| olfactory neurons | Clcn2 | 2.06 | 4.03E-09 | 8.38E-05 | 0.2 | 0.08 | 0.12 | 2.55 |
| olfactory neurons | Akirin1 | 1.31 | 4.14E-09 | 8.61E-05 | 0.65 | 0.58 | 0.07 | 1.12 |
| olfactory neurons | Slc24a5 | 1.08 | 4.54E-09 | 9.43E-05 | 0.57 | 0.4 | 0.16 | 1.4 |
| olfactory neurons | Gde1 | 1.17 | 4.67E-09 | 9.70E-05 | 0.63 | 0.52 | 0.11 | 1.2 |
| olfactory neurons | Bdh1 | 2.23 | 4.69E-09 | 9.76E-05 | 0.19 | 0.07 | 0.12 | 2.6 |
| olfactory neurons | Ndufb1-ps | 0.63 | 4.81E-09 | 9.99E-05 | 0.89 | 0.94 | -0.05 | 0.95 |
| olfactory neurons | Lcor | 1.26 | 4.83E-09 | 1.00E-04 | 0.45 | 0.28 | 0.16 | 1.57 |
| olfactory neurons | Garnl3 | 2.31 | 5.09E-09 | 1.06E-04 | 0.11 | 0.03 | 0.08 | 3.6 |
| olfactory neurons | Dmrta2 | 1.58 | 5.18E-09 | 1.08E-04 | 0.29 | 0.14 | 0.15 | 2.04 |
| olfactory neurons | Hspa8 | 0.32 | 5.24E-09 | 1.09E-04 | 0.98 | 1 | -0.02 | 0.98 |
| olfactory neurons | Cep250 | 1.4 | 5.27E-09 | 1.10E-04 | 0.42 | 0.26 | 0.16 | 1.6 |
| olfactory neurons | Mgat5b | 2.04 | 5.29E-09 | 1.10E-04 | 0.13 | 0.04 | 0.09 | 3.26 |
| olfactory neurons | Mapk3 | 1.32 | 5.57E-09 | 1.16E-04 | 0.58 | 0.45 | 0.13 | 1.28 |
| olfactory neurons | Ciapin1 | 0.92 | 6.30E-09 | 1.31E-04 | 0.78 | 0.74 | 0.05 | 1.06 |
| olfactory neurons | Pfdn4 | 0.99 | 6.30E-09 | 1.31E-04 | 0.81 | 0.75 | 0.05 | 1.07 |

**Supplementary Table 2. Marker genes for the major cell types of integrated clustering of E11.5 control and NotchGOF λ-junction cells.**

| | | | | | | | | |
|---|---|---|---|---|---|---|---|---|
| olfactory neurons | Wbp2 | 1.55 | 6.55E-09 | 1.36E-04 | 0.43 | 0.28 | 0.16 | 1.57 |
| olfactory neurons | Ppp2cb | 1.79 | 7.03E-09 | 1.46E-04 | 0.57 | 0.44 | 0.13 | 1.3 |
| olfactory neurons | Mycbp2 | 0.92 | 7.22E-09 | 1.50E-04 | 0.73 | 0.7 | 0.03 | 1.04 |
| olfactory neurons | Lpin1 | 1.78 | 7.31E-09 | 1.52E-04 | 0.18 | 0.07 | 0.11 | 2.65 |
| olfactory neurons | Enox1 | 2.27 | 7.35E-09 | 1.53E-04 | 0.19 | 0.08 | 0.12 | 2.51 |
| olfactory neurons | Cnih2 | 2.01 | 7.36E-09 | 1.53E-04 | 0.46 | 0.33 | 0.14 | 1.42 |
| olfactory neurons | Esrp1 | 0.75 | 7.37E-09 | 1.53E-04 | 0.35 | 0.17 | 0.18 | 2.07 |
| olfactory neurons | Wnk2 | 1.81 | 8.05E-09 | 1.68E-04 | 0.15 | 0.05 | 0.1 | 3.02 |
| olfactory neurons | Akap9 | 0.8 | 8.23E-09 | 1.71E-04 | 0.77 | 0.78 | -0.01 | 0.98 |
| olfactory neurons | Gm36839 | 2.24 | 8.33E-09 | 1.73E-04 | 0.11 | 0.03 | 0.08 | 3.35 |
| olfactory neurons | Micu3 | 2.3 | 8.37E-09 | 1.74E-04 | 0.33 | 0.17 | 0.15 | 1.87 |
| olfactory neurons | Vps28 | 1.14 | 8.47E-09 | 1.76E-04 | 0.71 | 0.62 | 0.09 | 1.14 |
| olfactory neurons | Cacnb3 | 1.04 | 8.63E-09 | 1.79E-04 | 0.45 | 0.27 | 0.18 | 1.65 |
| olfactory neurons | Vav2 | 1.76 | 8.95E-09 | 1.86E-04 | 0.21 | 0.09 | 0.12 | 2.41 |
| olfactory neurons | Zdhhc12 | 1.65 | 1.06E-08 | 2.20E-04 | 0.42 | 0.26 | 0.16 | 1.62 |
| olfactory neurons | Dynlt1f | 0.91 | 1.08E-08 | 2.24E-04 | 0.8 | 0.77 | 0.02 | 1.03 |
| olfactory neurons | Rsrp1 | 0.73 | 1.15E-08 | 2.39E-04 | 0.83 | 0.89 | -0.06 | 0.94 |
| olfactory neurons | Mdm2 | 0.94 | 1.20E-08 | 2.50E-04 | 0.59 | 0.44 | 0.15 | 1.35 |
| olfactory neurons | Hmgcll1 | 2.14 | 1.33E-08 | 2.76E-04 | 0.11 | 0.03 | 0.08 | 3.35 |
| olfactory neurons | Trappc2l | 1.21 | 1.40E-08 | 2.90E-04 | 0.7 | 0.62 | 0.08 | 1.13 |
| olfactory neurons | Ywhah | 0.79 | 1.40E-08 | 2.91E-04 | 0.88 | 0.91 | -0.03 | 0.97 |
| olfactory neurons | Med12l | 3.27 | 1.40E-08 | 2.91E-04 | 0.13 | 0.05 | 0.09 | 2.96 |
| olfactory neurons | Tspyl4 | 1.82 | 1.41E-08 | 2.92E-04 | 0.38 | 0.24 | 0.14 | 1.59 |
| olfactory neurons | Fgd2 | 2.02 | 1.44E-08 | 2.99E-04 | 0.23 | 0.1 | 0.13 | 2.27 |
| olfactory neurons | Btbd10 | 1.57 | 1.45E-08 | 3.01E-04 | 0.4 | 0.25 | 0.16 | 1.63 |
| olfactory neurons | Zdhhc2 | 1.27 | 1.47E-08 | 3.06E-04 | 0.28 | 0.13 | 0.15 | 2.13 |
| olfactory neurons | Hdac2 | 0.66 | 1.57E-08 | 3.27E-04 | 0.86 | 0.9 | -0.05 | 0.95 |
| olfactory neurons | Celf4 | 1.72 | 1.59E-08 | 3.30E-04 | 0.37 | 0.23 | 0.15 | 1.65 |
| olfactory neurons | Myl6 | 0.41 | 1.63E-08 | 3.40E-04 | 0.92 | 0.99 | -0.08 | 0.92 |
| olfactory neurons | Ccndbp1 | 1.76 | 1.64E-08 | 3.42E-04 | 0.45 | 0.28 | 0.17 | 1.59 |
| olfactory neurons | Grcc10 | 0.65 | 1.76E-08 | 3.65E-04 | 0.93 | 0.98 | -0.04 | 0.96 |
| olfactory neurons | Nuak1 | 1.61 | 1.79E-08 | 3.72E-04 | 0.36 | 0.2 | 0.15 | 1.74 |
| olfactory neurons | AI413582 | 1.96 | 1.80E-08 | 3.74E-04 | 0.21 | 0.09 | 0.12 | 2.29 |
| olfactory neurons | Enho | 1.19 | 1.85E-08 | 3.84E-04 | 0.58 | 0.45 | 0.13 | 1.28 |
| olfactory neurons | Ank | 2.14 | 1.92E-08 | 3.99E-04 | 0.18 | 0.07 | 0.11 | 2.54 |
| olfactory neurons | Atp6v1b2 | 1.39 | 1.94E-08 | 4.04E-04 | 0.55 | 0.39 | 0.16 | 1.42 |
| olfactory neurons | Pak7 | 2.24 | 1.96E-08 | 4.09E-04 | 0.14 | 0.05 | 0.09 | 2.9 |
| olfactory neurons | Cebpg | 1.21 | 2.10E-08 | 4.36E-04 | 0.65 | 0.55 | 0.1 | 1.19 |
| olfactory neurons | Katnal2 | 1.89 | 2.21E-08 | 4.60E-04 | 0.17 | 0.06 | 0.11 | 2.68 |
| olfactory neurons | Fgf9 | 0.66 | 2.24E-08 | 4.65E-04 | 0.25 | 0.11 | 0.14 | 2.31 |
| olfactory neurons | Rnd3 | 1.08 | 2.24E-08 | 4.66E-04 | 0.7 | 0.64 | 0.06 | 1.09 |
| olfactory neurons | Trak1 | 1.49 | 2.29E-08 | 4.77E-04 | 0.25 | 0.12 | 0.13 | 2.13 |
| olfactory neurons | Igfbp2 | 0.5 | 2.30E-08 | 4.78E-04 | 0.78 | 0.76 | 0.03 | 1.04 |
| olfactory neurons | Chmp1b | 1.34 | 2.35E-08 | 4.89E-04 | 0.51 | 0.35 | 0.15 | 1.44 |
| olfactory neurons | Carmil3 | 2.25 | 2.47E-08 | 5.14E-04 | 0.15 | 0.06 | 0.1 | 2.75 |
| olfactory neurons | Ankrd13b | 1.49 | 2.48E-08 | 5.16E-04 | 0.34 | 0.19 | 0.15 | 1.8 |
| olfactory neurons | Snx11 | 1.66 | 2.51E-08 | 5.23E-04 | 0.3 | 0.16 | 0.14 | 1.91 |
| olfactory neurons | Dennd1a | 1.22 | 2.53E-08 | 5.26E-04 | 0.36 | 0.21 | 0.15 | 1.73 |
| olfactory neurons | Pax6 | 1.33 | 2.66E-08 | 5.53E-04 | 0.27 | 0.13 | 0.14 | 2.08 |
| olfactory neurons | Wsb2 | 1.35 | 3.13E-08 | 6.50E-04 | 0.46 | 0.31 | 0.15 | 1.47 |
| olfactory neurons | Smim20 | 1.37 | 3.18E-08 | 6.61E-04 | 0.55 | 0.41 | 0.15 | 1.36 |
| olfactory neurons | Acrbp | 1.83 | 3.19E-08 | 6.64E-04 | 0.23 | 0.1 | 0.13 | 2.2 |
| olfactory neurons | Rc3h2 | 1.15 | 3.34E-08 | 6.94E-04 | 0.63 | 0.53 | 0.11 | 1.2 |
| olfactory neurons | Pls1 | 2.17 | 3.35E-08 | 6.96E-04 | 0.22 | 0.1 | 0.12 | 2.26 |
| olfactory neurons | Mdh2 | 0.64 | 3.42E-08 | 7.11E-04 | 0.89 | 0.95 | -0.07 | 0.93 |
| olfactory neurons | Gm42418 | 0.29 | 3.56E-08 | 7.40E-04 | 0.99 | 1 | 0 | 1 |

**Supplementary Table 2. Marker genes for the major cell types of integrated clustering of E11.5 control and NotchGOF λ-junction cells.**

| | | | | | | | | |
|---|---|---|---|---|---|---|---|---|
| olfactory neurons | B3gnt2 | 2.13 | 3.58E-08 | 7.44E-04 | 0.22 | 0.1 | 0.12 | 2.17 |
| olfactory neurons | Ap2m1 | 0.66 | 3.72E-08 | 7.74E-04 | 0.82 | 0.86 | -0.04 | 0.95 |
| olfactory neurons | Ptprz1 | 2.07 | 3.84E-08 | 8.00E-04 | 0.23 | 0.11 | 0.12 | 2.14 |
| olfactory neurons | Psmc5 | 0.53 | 4.89E-08 | 1.02E-03 | 0.85 | 0.92 | -0.07 | 0.93 |
| olfactory neurons | Snn | 0.93 | 5.16E-08 | 1.07E-03 | 0.58 | 0.44 | 0.15 | 1.34 |
| olfactory neurons | Zfp428 | 0.85 | 5.25E-08 | 1.09E-03 | 0.74 | 0.69 | 0.04 | 1.06 |
| olfactory neurons | Ttc3 | 0.44 | 5.95E-08 | 1.24E-03 | 0.86 | 0.96 | -0.1 | 0.9 |
| olfactory neurons | Mtus2 | 2.4 | 6.00E-08 | 1.25E-03 | 0.12 | 0.04 | 0.08 | 3 |
| olfactory neurons | Sirt7 | 1.73 | 6.01E-08 | 1.25E-03 | 0.31 | 0.17 | 0.14 | 1.82 |
| olfactory neurons | Fbxo32 | 2.01 | 6.17E-08 | 1.28E-03 | 0.1 | 0.03 | 0.07 | 3.4 |
| olfactory neurons | Dennd1b | 1.54 | 6.18E-08 | 1.28E-03 | 0.3 | 0.16 | 0.14 | 1.88 |
| olfactory neurons | Khdrbs2 | 1.9 | 6.42E-08 | 1.33E-03 | 0.11 | 0.04 | 0.08 | 3.17 |
| olfactory neurons | Slc25a33 | 1.65 | 6.61E-08 | 1.37E-03 | 0.3 | 0.17 | 0.13 | 1.8 |
| olfactory neurons | Psmc2 | 0.65 | 6.75E-08 | 1.40E-03 | 0.82 | 0.88 | -0.06 | 0.93 |
| olfactory neurons | Smchd1 | 1.59 | 6.81E-08 | 1.42E-03 | 0.79 | 0.86 | -0.07 | 0.92 |
| olfactory neurons | Fzd3 | 0.85 | 6.84E-08 | 1.42E-03 | 0.68 | 0.59 | 0.09 | 1.15 |
| olfactory neurons | Chmp5 | 0.67 | 6.87E-08 | 1.43E-03 | 0.8 | 0.8 | 0 | 1 |
| olfactory neurons | Gsk3b | 0.76 | 6.87E-08 | 1.43E-03 | 0.82 | 0.89 | -0.07 | 0.92 |
| olfactory neurons | Hsdl1 | 1.72 | 7.35E-08 | 1.53E-03 | 0.29 | 0.15 | 0.14 | 1.88 |
| olfactory neurons | Esyt1 | 1.84 | 7.48E-08 | 1.56E-03 | 0.18 | 0.07 | 0.11 | 2.45 |
| olfactory neurons | Ntm | 1.87 | 7.58E-08 | 1.58E-03 | 0.23 | 0.11 | 0.12 | 2.14 |
| olfactory neurons | Cog1 | 1.61 | 7.89E-08 | 1.64E-03 | 0.31 | 0.17 | 0.14 | 1.83 |
| olfactory neurons | Hnrnpa2b1 | 0.33 | 8.02E-08 | 1.67E-03 | 0.98 | 1 | -0.02 | 0.98 |
| olfactory neurons | Rb1cc1 | 0.8 | 8.20E-08 | 1.70E-03 | 0.72 | 0.66 | 0.07 | 1.1 |
| olfactory neurons | Amn1 | 1.04 | 9.05E-08 | 1.88E-03 | 0.59 | 0.44 | 0.15 | 1.34 |
| olfactory neurons | Rab3ip | 1.44 | 9.28E-08 | 1.93E-03 | 0.45 | 0.31 | 0.14 | 1.46 |
| olfactory neurons | .700096K18Ri | 1.72 | 9.63E-08 | 2.00E-03 | 0.21 | 0.1 | 0.12 | 2.22 |
| olfactory neurons | Chchd6 | 1.17 | 9.73E-08 | 2.02E-03 | 0.64 | 0.54 | 0.1 | 1.19 |
| olfactory neurons | Katnal1 | 1.56 | 9.93E-08 | 2.06E-03 | 0.32 | 0.17 | 0.15 | 1.83 |
| olfactory neurons | Chn2 | 1.69 | 1.01E-07 | 2.09E-03 | 0.18 | 0.07 | 0.1 | 2.46 |
| olfactory neurons | Isl1 | 1.31 | 1.10E-07 | 2.28E-03 | 0.4 | 0.22 | 0.18 | 1.8 |
| olfactory neurons | Six6 | 3.34 | 1.10E-07 | 2.29E-03 | 0.17 | 0.07 | 0.1 | 2.38 |
| olfactory neurons | Ypel5 | 1.13 | 1.16E-07 | 2.42E-03 | 0.42 | 0.25 | 0.17 | 1.66 |
| olfactory neurons | Capn10 | 1.12 | 1.17E-07 | 2.44E-03 | 0.37 | 0.22 | 0.15 | 1.69 |
| olfactory neurons | L1cam | 2.12 | 1.17E-07 | 2.44E-03 | 0.19 | 0.08 | 0.11 | 2.34 |
| olfactory neurons | Cmpk2 | 2.11 | 1.19E-07 | 2.47E-03 | 0.11 | 0.04 | 0.08 | 3.08 |
| olfactory neurons | Rtl8a | 1.72 | 1.22E-07 | 2.55E-03 | 0.51 | 0.39 | 0.12 | 1.31 |
| olfactory neurons | Jarid2 | 0.97 | 1.24E-07 | 2.58E-03 | 0.66 | 0.61 | 0.05 | 1.09 |
| olfactory neurons | Dynlrb1 | 0.65 | 1.30E-07 | 2.70E-03 | 0.91 | 0.95 | -0.03 | 0.96 |
| olfactory neurons | Wwc1 | 0.99 | 1.43E-07 | 2.98E-03 | 0.34 | 0.18 | 0.16 | 1.85 |
| olfactory neurons | Nebl | 1.01 | 1.60E-07 | 3.32E-03 | 0.24 | 0.11 | 0.13 | 2.16 |
| olfactory neurons | Klhl29 | 1.8 | 1.62E-07 | 3.37E-03 | 0.13 | 0.05 | 0.09 | 2.83 |
| olfactory neurons | Leprotl1 | 0.79 | 1.63E-07 | 3.39E-03 | 0.65 | 0.55 | 0.1 | 1.17 |
| olfactory neurons | Dclk1 | 1.76 | 1.66E-07 | 3.45E-03 | 0.39 | 0.26 | 0.13 | 1.51 |
| olfactory neurons | Dst | 1.35 | 1.71E-07 | 3.55E-03 | 0.48 | 0.36 | 0.13 | 1.35 |
| olfactory neurons | Evl | 0.94 | 1.86E-07 | 3.88E-03 | 0.78 | 0.76 | 0.01 | 1.02 |
| olfactory neurons | Acvr2a | 1.01 | 1.90E-07 | 3.94E-03 | 0.43 | 0.27 | 0.16 | 1.59 |
| olfactory neurons | Septin5 | 2.01 | 1.93E-07 | 4.02E-03 | 0.16 | 0.07 | 0.1 | 2.43 |
| olfactory neurons | Sort1 | 1.67 | 1.97E-07 | 4.10E-03 | 0.34 | 0.2 | 0.14 | 1.68 |
| olfactory neurons | Epb41 | 0.94 | 1.98E-07 | 4.11E-03 | 0.74 | 0.76 | -0.02 | 0.98 |
| olfactory neurons | Dyrk1b | 1.22 | 2.14E-07 | 4.46E-03 | 0.28 | 0.14 | 0.13 | 1.94 |
| olfactory neurons | Cdh4 | 2.29 | 2.25E-07 | 4.67E-03 | 0.14 | 0.05 | 0.09 | 2.73 |
| olfactory neurons | Chmp2a | 0.92 | 2.26E-07 | 4.71E-03 | 0.72 | 0.66 | 0.06 | 1.09 |
| olfactory neurons | Csnk1g3 | 0.91 | 2.39E-07 | 4.97E-03 | 0.75 | 0.68 | 0.06 | 1.09 |
| olfactory neurons | Krtcap3 | 1.38 | 2.39E-07 | 4.97E-03 | 0.24 | 0.11 | 0.12 | 2.06 |
| olfactory neurons | Fam204a | 1.01 | 2.41E-07 | 5.01E-03 | 0.73 | 0.65 | 0.08 | 1.12 |

**Supplementary Table 2. Marker genes for the major cell types of integrated clustering of E11.5 control and NotchGOF λ-junction cells.**

| | | | | | | | | |
|---|---|---|---|---|---|---|---|---|
| olfactory neurons | Akap5 | 2.03 | 2.46E-07 | 5.11E-03 | 0.12 | 0.04 | 0.08 | 2.93 |
| olfactory neurons | Kbtbd8 | 2.01 | 2.51E-07 | 5.22E-03 | 0.11 | 0.04 | 0.08 | 3 |
| olfactory neurons | Afdn | 0.74 | 2.70E-07 | 5.62E-03 | 0.71 | 0.57 | 0.13 | 1.23 |
| olfactory neurons | Vasp | 1.2 | 2.75E-07 | 5.73E-03 | 0.58 | 0.49 | 0.1 | 1.2 |
| olfactory neurons | Fam227a | 1.87 | 2.84E-07 | 5.91E-03 | 0.14 | 0.05 | 0.09 | 2.73 |
| olfactory neurons | Dmrt3 | 1.45 | 3.24E-07 | 6.75E-03 | 0.15 | 0.05 | 0.09 | 2.69 |
| olfactory neurons | Dach1 | 1.19 | 3.34E-07 | 6.95E-03 | 0.53 | 0.39 | 0.15 | 1.38 |
| olfactory neurons | Zswim5 | 2.22 | 3.42E-07 | 7.11E-03 | 0.1 | 0.03 | 0.07 | 3.09 |
| olfactory neurons | Bex1 | 0.35 | 3.46E-07 | 7.19E-03 | 0.77 | 0.59 | 0.17 | 1.29 |
| olfactory neurons | Dctn2 | 0.9 | 4.03E-07 | 8.37E-03 | 0.8 | 0.83 | -0.03 | 0.96 |
| olfactory neurons | Pik3ip1 | 1.9 | 4.14E-07 | 8.62E-03 | 0.19 | 0.09 | 0.11 | 2.19 |
| olfactory neurons | Chn1 | 1.42 | 4.39E-07 | 9.12E-03 | 0.52 | 0.4 | 0.12 | 1.29 |
| olfactory neurons | Ufm1 | 0.93 | 4.58E-07 | 9.52E-03 | 0.72 | 0.69 | 0.03 | 1.04 |
| olfactory neurons | Ggt7 | 2.04 | 4.60E-07 | 9.57E-03 | 0.1 | 0.03 | 0.07 | 3.09 |
| olfactory neurons | Nin | 1.29 | 4.77E-07 | 9.92E-03 | 0.41 | 0.27 | 0.14 | 1.51 |
| olfactory neurons | Peli1 | 0.83 | 4.86E-07 | 1.01E-02 | 0.71 | 0.68 | 0.03 | 1.05 |
| olfactory neurons | Cep290 | 1.03 | 4.94E-07 | 1.03E-02 | 0.61 | 0.51 | 0.09 | 1.18 |
| olfactory neurons | Ly6h | 1.98 | 5.06E-07 | 1.05E-02 | 0.28 | 0.15 | 0.13 | 1.82 |
| olfactory neurons | Nell2 | 2.81 | 5.13E-07 | 1.07E-02 | 0.13 | 0.05 | 0.08 | 2.66 |
| olfactory neurons | Gipc2 | 1.67 | 5.14E-07 | 1.07E-02 | 0.16 | 0.06 | 0.09 | 2.45 |
| olfactory neurons | 110006O24Ri | 2.83 | 5.16E-07 | 1.07E-02 | 0.11 | 0.04 | 0.08 | 2.92 |
| olfactory neurons | Ssbp3 | 0.47 | 5.40E-07 | 1.12E-02 | 0.86 | 0.94 | -0.08 | 0.92 |
| olfactory neurons | Gls | 1.03 | 5.48E-07 | 1.14E-02 | 0.52 | 0.41 | 0.12 | 1.28 |
| olfactory neurons | Znrf1 | 0.98 | 5.80E-07 | 1.21E-02 | 0.68 | 0.6 | 0.08 | 1.13 |
| olfactory neurons | Madd | 1.84 | 5.95E-07 | 1.24E-02 | 0.17 | 0.07 | 0.1 | 2.32 |
| olfactory neurons | 700097O09Ri | 1.43 | 5.99E-07 | 1.25E-02 | 0.46 | 0.35 | 0.11 | 1.33 |
| olfactory neurons | Pcdh17 | 1.54 | 6.10E-07 | 1.27E-02 | 0.34 | 0.21 | 0.14 | 1.66 |
| olfactory neurons | Kcne1l | 2.23 | 6.65E-07 | 1.38E-02 | 0.24 | 0.13 | 0.12 | 1.91 |
| olfactory neurons | Rbm18 | 0.97 | 6.92E-07 | 1.44E-02 | 0.64 | 0.55 | 0.09 | 1.16 |
| olfactory neurons | Minar2 | 1.86 | 6.92E-07 | 1.44E-02 | 0.21 | 0.1 | 0.11 | 2.14 |
| olfactory neurons | BC005537 | 0.91 | 7.08E-07 | 1.47E-02 | 0.59 | 0.5 | 0.09 | 1.18 |
| olfactory neurons | Lrrn1 | 1.69 | 7.31E-07 | 1.52E-02 | 0.13 | 0.05 | 0.08 | 2.76 |
| olfactory neurons | Rnf121 | 1.16 | 7.44E-07 | 1.55E-02 | 0.49 | 0.37 | 0.12 | 1.33 |
| olfactory neurons | Slc43a1 | 1.36 | 7.53E-07 | 1.57E-02 | 0.29 | 0.16 | 0.13 | 1.81 |
| olfactory neurons | Psen2 | 2.41 | 8.00E-07 | 1.66E-02 | 0.18 | 0.08 | 0.1 | 2.27 |
| olfactory neurons | Stx2 | 1.38 | 8.11E-07 | 1.69E-02 | 0.34 | 0.22 | 0.12 | 1.55 |
| olfactory neurons | Ncstn | 1.3 | 8.51E-07 | 1.77E-02 | 0.42 | 0.29 | 0.13 | 1.45 |
| olfactory neurons | Prr13 | 1.06 | 8.58E-07 | 1.79E-02 | 0.54 | 0.4 | 0.14 | 1.35 |
| olfactory neurons | Hmgn5 | 1.05 | 8.59E-07 | 1.79E-02 | 0.71 | 0.71 | -0.01 | 0.99 |
| olfactory neurons | Synj1 | 2 | 8.77E-07 | 1.82E-02 | 0.22 | 0.11 | 0.11 | 2.01 |
| olfactory neurons | Slc9a3r1 | 0.62 | 8.98E-07 | 1.87E-02 | 0.42 | 0.24 | 0.17 | 1.72 |
| olfactory neurons | Brsk1 | 1.72 | 9.13E-07 | 1.90E-02 | 0.18 | 0.08 | 0.1 | 2.23 |
| olfactory neurons | Gkap1 | 1.2 | 9.21E-07 | 1.92E-02 | 0.57 | 0.44 | 0.14 | 1.31 |
| olfactory neurons | Meak7 | 2.17 | 9.29E-07 | 1.93E-02 | 0.15 | 0.06 | 0.09 | 2.54 |
| olfactory neurons | Tnnt1 | 1.86 | 9.55E-07 | 1.99E-02 | 0.14 | 0.05 | 0.09 | 2.57 |
| olfactory neurons | Cep70 | 0.91 | 9.61E-07 | 2.00E-02 | 0.54 | 0.4 | 0.14 | 1.35 |
| olfactory neurons | Rnf38 | 1.17 | 1.09E-06 | 2.26E-02 | 0.4 | 0.27 | 0.13 | 1.47 |
| olfactory neurons | Ell2 | 1.62 | 1.10E-06 | 2.29E-02 | 0.19 | 0.09 | 0.1 | 2.17 |
| olfactory neurons | Atp6v0e | 0.78 | 1.14E-06 | 2.38E-02 | 0.7 | 0.72 | -0.02 | 0.97 |
| olfactory neurons | Trappc6b | 0.89 | 1.16E-06 | 2.41E-02 | 0.63 | 0.54 | 0.09 | 1.17 |
| olfactory neurons | Pard6a | 1.76 | 1.28E-06 | 2.65E-02 | 0.29 | 0.17 | 0.12 | 1.73 |
| olfactory neurons | Thnsl1 | 1.62 | 1.29E-06 | 2.68E-02 | 0.17 | 0.07 | 0.1 | 2.32 |
| olfactory neurons | Smarcc2 | 0.89 | 1.37E-06 | 2.84E-02 | 0.71 | 0.65 | 0.06 | 1.09 |
| olfactory neurons | Hs6st1 | 1.27 | 1.59E-06 | 3.31E-02 | 0.3 | 0.17 | 0.13 | 1.75 |
| olfactory neurons | Calm3 | 0.7 | 1.60E-06 | 3.32E-02 | 0.87 | 0.91 | -0.05 | 0.95 |
| olfactory neurons | Meis3 | 2.06 | 1.61E-06 | 3.35E-02 | 0.33 | 0.2 | 0.13 | 1.63 |

**Supplementary Table 2. Marker genes for the major cell types of integrated clustering of E11.5 control and NotchGOF λ-junction cells.**

| | | | | | | | | |
|---|---|---|---|---|---|---|---|---|
| olfactory neurons | Fam8a1 | 1.38 | 1.65E-06 | 3.43E-02 | 0.39 | 0.27 | 0.12 | 1.46 |
| olfactory neurons | Ooep | 2.44 | 1.65E-06 | 3.44E-02 | 0.11 | 0.04 | 0.07 | 2.84 |
| olfactory neurons | Reep1 | 1.73 | 1.67E-06 | 3.47E-02 | 0.13 | 0.05 | 0.08 | 2.65 |
| olfactory neurons | Ndfip2 | 0.99 | 1.74E-06 | 3.62E-02 | 0.63 | 0.51 | 0.12 | 1.22 |
| olfactory neurons | Dysf | 1.59 | 1.75E-06 | 3.63E-02 | 0.11 | 0.04 | 0.07 | 2.84 |
| olfactory neurons | Kctd10 | 1.3 | 1.75E-06 | 3.63E-02 | 0.43 | 0.3 | 0.14 | 1.46 |
| olfactory neurons | Spa17 | 1.58 | 1.76E-06 | 3.65E-02 | 0.28 | 0.15 | 0.12 | 1.81 |
| olfactory neurons | Mpc1 | 1.08 | 1.78E-06 | 3.70E-02 | 0.68 | 0.63 | 0.06 | 1.09 |
| olfactory neurons | Atp6v1f | 0.55 | 1.81E-06 | 3.77E-02 | 0.9 | 0.9 | -0.01 | 0.99 |
| olfactory neurons | Morn4 | 1.37 | 1.88E-06 | 3.92E-02 | 0.29 | 0.16 | 0.13 | 1.76 |
| olfactory neurons | Map3k12 | 1.14 | 1.95E-06 | 4.05E-02 | 0.34 | 0.21 | 0.13 | 1.61 |
| olfactory neurons | Rab15 | 1.27 | 1.96E-06 | 4.09E-02 | 0.25 | 0.13 | 0.12 | 1.89 |
| olfactory neurons | Atp6v1d | 0.85 | 2.08E-06 | 4.33E-02 | 0.68 | 0.64 | 0.04 | 1.06 |
| olfactory neurons | Arl6 | 1.37 | 2.15E-06 | 4.48E-02 | 0.54 | 0.42 | 0.12 | 1.28 |
| olfactory neurons | 110008P14Ri | 1.19 | 2.24E-06 | 4.65E-02 | 0.56 | 0.47 | 0.09 | 1.2 |
| olfactory neurons | Prepl | 1.17 | 2.28E-06 | 4.73E-02 | 0.24 | 0.13 | 0.12 | 1.93 |
| olfactory neurons | Whamm | 1.45 | 2.38E-06 | 4.95E-02 | 0.21 | 0.1 | 0.11 | 2.05 |
| other bloold cells | Tyrobp | 12.38 | 0 | 0 | 1 | 0.01 | 1 | 200 |
| other bloold cells | Rac2 | 10.45 | 0 | 0 | 1 | 0.01 | 0.99 | 166.67 |
| other bloold cells | Fcer1g | 9.28 | 0 | 0 | 0.99 | 0.02 | 0.98 | 66.2 |
| other bloold cells | Laptm5 | 9 | 0 | 0 | 0.99 | 0.01 | 0.97 | 75.85 |
| other bloold cells | Spi1 | 11.82 | 0 | 0 | 0.96 | 0 | 0.96 | 479 |
| other bloold cells | Cyth4 | 9.28 | 0 | 0 | 0.96 | 0.01 | 0.95 | 191.6 |
| other bloold cells | Lcp1 | 7.11 | 0 | 0 | 0.98 | 0.03 | 0.95 | 30.59 |
| other bloold cells | Cd52 | 12.36 | 0 | 0 | 0.94 | 0 | 0.94 | 944 |
| other bloold cells | Gmfg | 6.53 | 0 | 0 | 0.96 | 0.02 | 0.94 | 45.62 |
| other bloold cells | Lst1 | 9.49 | 0 | 0 | 0.94 | 0.01 | 0.94 | 134.86 |
| other bloold cells | Ncf2 | 8.22 | 0 | 0 | 0.94 | 0.01 | 0.93 | 78.67 |
| other bloold cells | Ptprc | 11.48 | 0 | 0 | 0.92 | 0 | 0.92 | 923 |
| other bloold cells | Ptpn6 | 6.5 | 0 | 0 | 0.95 | 0.03 | 0.92 | 30.68 |
| other bloold cells | Cd53 | 11.63 | 0 | 0 | 0.92 | 0 | 0.92 | 916 |
| other bloold cells | Ptpn18 | 6.32 | 0 | 0 | 0.99 | 0.07 | 0.91 | 13.69 |
| other bloold cells | Fermt3 | 5.6 | 0 | 0 | 0.95 | 0.04 | 0.91 | 22.64 |
| other bloold cells | Bin2 | 6.86 | 0 | 0 | 0.89 | 0.01 | 0.88 | 177.6 |
| other bloold cells | Fyb | 6.87 | 0 | 0 | 0.92 | 0.04 | 0.88 | 25.44 |
| other bloold cells | Nrros | 6.02 | 0 | 0 | 0.9 | 0.03 | 0.87 | 27.33 |
| other bloold cells | Apbb1ip | 8.25 | 0 | 0 | 0.87 | 0.01 | 0.87 | 109.25 |
| other bloold cells | Tm6sf1 | 5.32 | 0 | 0 | 0.92 | 0.05 | 0.87 | 18.32 |
| other bloold cells | Fcgr3 | 11.84 | 0 | 0 | 0.87 | 0 | 0.87 | 433.5 |
| other bloold cells | Evi2a | 9.92 | 0 | 0 | 0.87 | 0 | 0.87 | 433.5 |
| other bloold cells | Cx3cr1 | 11.25 | 0 | 0 | 0.87 | 0 | 0.86 | 216.75 |
| other bloold cells | Lyn | 6.15 | 0 | 0 | 0.89 | 0.03 | 0.86 | 32.89 |
| other bloold cells | Pou2f2 | 6.69 | 0 | 0 | 0.88 | 0.02 | 0.86 | 40.05 |
| other bloold cells | Arhgap30 | 9.86 | 0 | 0 | 0.86 | 0 | 0.86 | 430 |
| other bloold cells | Csf1r | 11.74 | 0 | 0 | 0.86 | 0 | 0.86 | 430 |
| other bloold cells | Tnfaip8l2 | 10.95 | 0 | 0 | 0.85 | 0 | 0.85 | 853 |
| other bloold cells | Ly86 | 11.9 | 0 | 0 | 0.85 | 0 | 0.85 | 426.5 |
| other bloold cells | Rgs10 | 5.42 | 0 | 0 | 0.93 | 0.08 | 0.85 | 11.2 |
| other bloold cells | Psmb8 | 11.24 | 0 | 0 | 0.85 | 0 | 0.85 | 846 |
| other bloold cells | Inpp5d | 7.6 | 0 | 0 | 0.85 | 0.01 | 0.84 | 94 |
| other bloold cells | Fxyd5 | 5.75 | 0 | 0 | 0.86 | 0.02 | 0.84 | 35.83 |
| other bloold cells | Nckap1l | 8.63 | 0 | 0 | 0.84 | 0 | 0.84 | 209.75 |
| other bloold cells | Alox5ap | 6.05 | 0 | 0 | 0.87 | 0.03 | 0.83 | 25.5 |
| other bloold cells | Pld4 | 7.44 | 0 | 0 | 0.85 | 0.02 | 0.83 | 56.4 |
| other bloold cells | Unc93b1 | 6.28 | 0 | 0 | 0.87 | 0.04 | 0.83 | 20.33 |
| other bloold cells | Ctss | 12.45 | 0 | 0 | 0.83 | 0 | 0.82 | 825 |

**Supplementary Table 2. Marker genes for the major cell types of integrated clustering of E11.5 control and NotchGOF λ-junction cells.**

| | | | | | | | | |
|---|---|---|---|---|---|---|---|---|
| other bloold cells | Lyz2 | 10.19 | 0 | 0 | 0.83 | 0 | 0.82 | 206.25 |
| other bloold cells | Srgn | 7.73 | 0 | 0 | 0.83 | 0.01 | 0.82 | 137.5 |
| other bloold cells | Plek | 5.91 | 0 | 0 | 0.83 | 0.01 | 0.82 | 117.86 |
| other bloold cells | Ighm | 12.37 | 0 | 0 | 0.82 | 0 | 0.82 | 818 |
| other bloold cells | Zfp36 | 5.82 | 0 | 0 | 0.88 | 0.07 | 0.81 | 13.15 |
| other bloold cells | Irf8 | 7.32 | 0 | 0 | 0.83 | 0.02 | 0.81 | 48.53 |
| other bloold cells | Hcls1 | 8.34 | 0 | 0 | 0.81 | 0.01 | 0.81 | 162.2 |
| other bloold cells | Ncf1 | 8.69 | 0 | 0 | 0.8 | 0.01 | 0.8 | 160.8 |
| other bloold cells | Ms4a6c | 10.93 | 0 | 0 | 0.8 | 0 | 0.79 | 199.25 |
| other bloold cells | Skap2 | 5.08 | 0 | 0 | 0.84 | 0.05 | 0.79 | 16.78 |
| other bloold cells | Dock8 | 5.45 | 0 | 0 | 0.82 | 0.03 | 0.79 | 28.21 |
| other bloold cells | Was | 8.48 | 0 | 0 | 0.79 | 0 | 0.79 | 395 |
| other bloold cells | Lgals9 | 9.3 | 0 | 0 | 0.78 | 0 | 0.78 | 391.5 |
| other bloold cells | Otulinl | 4.72 | 0 | 0 | 0.83 | 0.05 | 0.78 | 15.7 |
| other bloold cells | Ptgs1 | 5.44 | 0 | 0 | 0.81 | 0.04 | 0.78 | 22.53 |
| other bloold cells | Lcp2 | 8.14 | 0 | 0 | 0.78 | 0 | 0.77 | 388 |
| other bloold cells | Hpgds | 6.79 | 0 | 0 | 0.8 | 0.03 | 0.77 | 24.36 |
| other bloold cells | Aif1 | 11.26 | 0 | 0 | 0.77 | 0 | 0.77 | 384.5 |
| other bloold cells | Vav1 | 8.87 | 0 | 0 | 0.77 | 0 | 0.77 | 384.5 |
| other bloold cells | Rassf4 | 5.82 | 0 | 0 | 0.79 | 0.03 | 0.77 | 31.6 |
| other bloold cells | Plcg2 | 6.53 | 0 | 0 | 0.78 | 0.01 | 0.76 | 59.69 |
| other bloold cells | Myo1f | 11.7 | 0 | 0 | 0.76 | 0 | 0.76 | #NA |
| other bloold cells | Mpeg1 | 11.67 | 0 | 0 | 0.76 | 0 | 0.75 | 755 |
| other bloold cells | Cd68 | 6.07 | 0 | 0 | 0.78 | 0.04 | 0.75 | 21.75 |
| other bloold cells | Trem2 | 11.91 | 0 | 0 | 0.75 | 0 | 0.75 | 748 |
| other bloold cells | Ccl3 | 11 | 0 | 0 | 0.75 | 0 | 0.75 | 249.33 |
| other bloold cells | Bst2 | 4.82 | 0 | 0 | 0.8 | 0.05 | 0.74 | 14.76 |
| other bloold cells | Fcgr1 | 12.89 | 0 | 0 | 0.73 | 0 | 0.73 | #NA |
| other bloold cells | Klhl6 | 5.8 | 0 | 0 | 0.75 | 0.02 | 0.73 | 49.87 |
| other bloold cells | Ms4a6b | 11.74 | 0 | 0 | 0.73 | 0 | 0.73 | 734 |
| other bloold cells | C3ar1 | 11.25 | 0 | 0 | 0.73 | 0 | 0.73 | 367 |
| other bloold cells | Dock2 | 9.16 | 0 | 0 | 0.73 | 0 | 0.73 | 367 |
| other bloold cells | Fes | 4.84 | 0 | 0 | 0.77 | 0.04 | 0.73 | 20.24 |
| other bloold cells | C1qb | 12.65 | 0 | 0 | 0.73 | 0.01 | 0.73 | 146.8 |
| other bloold cells | Cybb | 12.11 | 0 | 0 | 0.73 | 0 | 0.73 | 727 |
| other bloold cells | Cd300a | 7.73 | 0 | 0 | 0.73 | 0.01 | 0.73 | 81.56 |
| other bloold cells | Neat1 | 5.7 | 0 | 0 | 0.78 | 0.05 | 0.73 | 15.22 |
| other bloold cells | Ncf4 | 3.66 | 0 | 0 | 0.76 | 0.04 | 0.72 | 18.14 |
| other bloold cells | Cd79b | 7.28 | 0 | 0 | 0.73 | 0.01 | 0.72 | 90.88 |
| other bloold cells | Ikzf1 | 6.06 | 0 | 0 | 0.74 | 0.02 | 0.72 | 32.22 |
| other bloold cells | C1qc | 12.45 | 0 | 0 | 0.72 | 0 | 0.72 | 180 |
| other bloold cells | Milr1 | 7.41 | 0 | 0 | 0.72 | 0.01 | 0.71 | 102.86 |
| other bloold cells | Gatm | 6.58 | 0 | 0 | 0.75 | 0.04 | 0.71 | 20.78 |
| other bloold cells | C1qa | 12.93 | 0 | 0 | 0.71 | 0 | 0.71 | 237.67 |
| other bloold cells | Hck | 8.75 | 0 | 0 | 0.71 | 0 | 0.7 | 235.33 |
| other bloold cells | Slfn2 | 7.19 | 0 | 0 | 0.71 | 0.01 | 0.7 | 59.42 |
| other bloold cells | Fcrls | 11.82 | 0 | 0 | 0.7 | 0 | 0.7 | 349.5 |
| other bloold cells | BC035044 | 10.85 | 0 | 0 | 0.69 | 0 | 0.69 | 692 |
| other bloold cells | Cfh | 9.83 | 0 | 0 | 0.69 | 0 | 0.69 | 346 |
| other bloold cells | Gngt2 | 4.36 | 0 | 0 | 0.72 | 0.03 | 0.69 | 21.18 |
| other bloold cells | Gpr34 | 12.1 | 0 | 0 | 0.69 | 0 | 0.69 | #NA |
| other bloold cells | Adgre1 | 10.87 | 0 | 0 | 0.69 | 0 | 0.68 | 685 |
| other bloold cells | Ms4a6d | 11.48 | 0 | 0 | 0.69 | 0 | 0.68 | 685 |
| other bloold cells | P2ry6 | 9.79 | 0 | 0 | 0.69 | 0 | 0.68 | 342.5 |
| other bloold cells | Traf3ip3 | 10.41 | 0 | 0 | 0.68 | 0 | 0.68 | 678 |
| other bloold cells | Btk | 7.74 | 0 | 0 | 0.68 | 0 | 0.68 | 339 |

**Supplementary Table 2. Marker genes for the major cell types of integrated clustering of E11.5 control and NotchGOF λ-junction cells.**

| | | | | | | | | |
|---|---|---|---|---|---|---|---|---|
| other bloold cells | Hhex | 5.82 | 0 | 0 | 0.68 | 0.01 | 0.67 | 56.5 |
| other bloold cells | Psmb9 | 6.66 | 0 | 0 | 0.67 | 0.01 | 0.66 | 74.56 |
| other bloold cells | Nfam1 | 7.76 | 0 | 0 | 0.66 | 0.01 | 0.66 | 110.67 |
| other bloold cells | Pik3ap1 | 6.33 | 0 | 0 | 0.67 | 0.01 | 0.66 | 47.93 |
| other bloold cells | Stab1 | 7.4 | 0 | 0 | 0.67 | 0.02 | 0.65 | 37.28 |
| other bloold cells | Irf5 | 8.14 | 0 | 0 | 0.65 | 0 | 0.65 | 216.67 |
| other bloold cells | Vsir | 6.5 | 0 | 0 | 0.66 | 0.01 | 0.65 | 54.75 |
| other bloold cells | Lpcat2 | 4.98 | 0 | 0 | 0.68 | 0.03 | 0.64 | 19.94 |
| other bloold cells | Il6ra | 10.18 | 0 | 0 | 0.64 | 0 | 0.64 | 643 |
| other bloold cells | Ramp1 | 7.86 | 0 | 0 | 0.64 | 0.01 | 0.64 | 128.6 |
| other bloold cells | Mrc1 | 11.33 | 0 | 0 | 0.64 | 0 | 0.63 | 318 |
| other bloold cells | Lrmp | 6.97 | 0 | 0 | 0.64 | 0.01 | 0.63 | 106 |
| other bloold cells | Adcy7 | 5.46 | 0 | 0 | 0.65 | 0.02 | 0.63 | 32.5 |
| other bloold cells | Sp140 | 6.93 | 0 | 0 | 0.64 | 0.01 | 0.63 | 90.86 |
| other bloold cells | Fmnl1 | 8.78 | 0 | 0 | 0.63 | 0 | 0.63 | 314.5 |
| other bloold cells | Ccr1 | 11.79 | 0 | 0 | 0.62 | 0 | 0.62 | #NA |
| other bloold cells | Arhgef6 | 6.09 | 0 | 0 | 0.64 | 0.01 | 0.62 | 45.43 |
| other bloold cells | Cd300c2 | 10.48 | 0 | 0 | 0.62 | 0 | 0.62 | 622 |
| other bloold cells | Glipr1 | 9.94 | 0 | 0 | 0.62 | 0 | 0.62 | 311 |
| other bloold cells | Tnfrsf13b | 7.24 | 0 | 0 | 0.62 | 0.01 | 0.62 | 124.4 |
| other bloold cells | Selplg | 9.96 | 0 | 0 | 0.61 | 0 | 0.61 | 608 |
| other bloold cells | Trim30a | 10.31 | 0 | 0 | 0.6 | 0 | 0.6 | 601 |
| other bloold cells | Slc7a8 | 8 | 0 | 0 | 0.6 | 0 | 0.6 | 200.33 |
| other bloold cells | Uap1l1 | 5.03 | 0 | 0 | 0.63 | 0.03 | 0.6 | 19.66 |
| other bloold cells | Il4ra | 6.45 | 0 | 0 | 0.61 | 0.01 | 0.59 | 43.43 |
| other bloold cells | Sash3 | 11.23 | 0 | 0 | 0.59 | 0 | 0.59 | #NA |
| other bloold cells | Itgam | 12.17 | 0 | 0 | 0.59 | 0 | 0.59 | #NA |
| other bloold cells | Cd86 | 11.35 | 0 | 0 | 0.59 | 0 | 0.59 | #NA |
| other bloold cells | Apobec1 | 8.09 | 0 | 0 | 0.59 | 0.01 | 0.58 | 97.83 |
| other bloold cells | Cd33 | 11.7 | 0 | 0 | 0.58 | 0 | 0.58 | #NA |
| other bloold cells | Lrrc25 | 11.58 | 0 | 0 | 0.58 | 0 | 0.58 | #NA |
| other bloold cells | Gpr183 | 5.73 | 0 | 0 | 0.59 | 0.01 | 0.58 | 42.43 |
| other bloold cells | Igsf6 | 8.01 | 0 | 0 | 0.58 | 0 | 0.58 | 193.33 |
| other bloold cells | Syk | 4.9 | 0 | 0 | 0.6 | 0.03 | 0.58 | 24.04 |
| other bloold cells | Abi3 | 5.56 | 0 | 0 | 0.59 | 0.01 | 0.57 | 45.15 |
| other bloold cells | Tifab | 11.28 | 0 | 0 | 0.57 | 0 | 0.57 | #NA |
| other bloold cells | Ifi204 | 10.41 | 0 | 0 | 0.57 | 0 | 0.57 | 573 |
| other bloold cells | Snx20 | 5.89 | 0 | 0 | 0.59 | 0.02 | 0.57 | 39.13 |
| other bloold cells | Il17ra | 5.41 | 0 | 0 | 0.59 | 0.02 | 0.57 | 25.83 |
| other bloold cells | Samsn1 | 6.36 | 0 | 0 | 0.58 | 0.01 | 0.57 | 52.73 |
| other bloold cells | Csf2rb | 6.91 | 0 | 0 | 0.57 | 0.01 | 0.57 | 71.63 |
| other bloold cells | Parvg | 8.88 | 0 | 0 | 0.57 | 0 | 0.57 | 566 |
| other bloold cells | Sp110 | 8.97 | 0 | 0 | 0.57 | 0 | 0.56 | 283 |
| other bloold cells | Ptpn7 | 7.35 | 0 | 0 | 0.57 | 0.01 | 0.56 | 113.2 |
| other bloold cells | Ifi202b | 7.27 | 0 | 0 | 0.57 | 0.01 | 0.56 | 94.33 |
| other bloold cells | Ccl9 | 11.39 | 0 | 0 | 0.56 | 0 | 0.56 | #NA |
| other bloold cells | Rhoh | 9.13 | 0 | 0 | 0.56 | 0 | 0.56 | 559 |
| other bloold cells | Sla | 7.45 | 0 | 0 | 0.56 | 0 | 0.56 | 279.5 |
| other bloold cells | Zc3hav1 | 5.06 | 0 | 0 | 0.58 | 0.03 | 0.56 | 23.2 |
| other bloold cells | P2rx4 | 4.78 | 0 | 0 | 0.58 | 0.03 | 0.55 | 22.31 |
| other bloold cells | Gpr65 | 11.23 | 0 | 0 | 0.55 | 0 | 0.55 | #NA |
| other bloold cells | Slc15a3 | 6.37 | 0 | 0 | 0.57 | 0.01 | 0.55 | 40.43 |
| other bloold cells | Csf3r | 10.39 | 0 | 0 | 0.55 | 0 | 0.55 | 552 |
| other bloold cells | Pik3cd | 5.57 | 0 | 0 | 0.57 | 0.02 | 0.55 | 37.73 |
| other bloold cells | Tnfrsf11a | 7.86 | 0 | 0 | 0.55 | 0 | 0.55 | 138 |
| other bloold cells | Cd14 | 5.82 | 0 | 0 | 0.56 | 0.02 | 0.54 | 27.95 |

**Supplementary Table 2. Marker genes for the major cell types of integrated clustering of E11.5 control and NotchGOF λ-junction cells.**

| | | | | | | | | |
|---|---|---|---|---|---|---|---|---|
| other bloold cells | Myo1g | 10.8 | 0 | 0 | 0.53 | 0 | 0.53 | #NA |
| other bloold cells | Slc25a45 | 6.05 | 0 | 0 | 0.53 | 0.01 | 0.52 | 66.38 |
| other bloold cells | Ppfia4 | 5.08 | 0 | 0 | 0.55 | 0.02 | 0.52 | 22.71 |
| other bloold cells | Tfec | 5.83 | 0 | 0 | 0.53 | 0.01 | 0.52 | 44.25 |
| other bloold cells | Gbp7 | 5.78 | 0 | 0 | 0.53 | 0.02 | 0.51 | 31.24 |
| other bloold cells | Icam1 | 5.03 | 0 | 0 | 0.52 | 0.01 | 0.51 | 37.43 |
| other bloold cells | 930111J21Rik | 11.07 | 0 | 0 | 0.51 | 0 | 0.51 | #NA |
| other bloold cells | Ccr5 | 9.6 | 0 | 0 | 0.51 | 0 | 0.51 | 510 |
| other bloold cells | Tnfrsf1b | 8.03 | 0 | 0 | 0.51 | 0 | 0.51 | 170 |
| other bloold cells | Ifi207 | 11.83 | 0 | 0 | 0.5 | 0 | 0.5 | #NA |
| other bloold cells | Adap2 | 11.16 | 0 | 0 | 0.5 | 0 | 0.5 | #NA |
| other bloold cells | Rgs1 | 10.14 | 0 | 0 | 0.5 | 0 | 0.5 | 503 |
| other bloold cells | Slco2b1 | 9.08 | 0 | 0 | 0.5 | 0 | 0.5 | 251.5 |
| other bloold cells | Dok2 | 5.22 | 0 | 0 | 0.52 | 0.02 | 0.5 | 32.31 |
| other bloold cells | Cela1 | 7.83 | 0 | 0 | 0.5 | 0 | 0.5 | 251.5 |
| other bloold cells | Camk1d | 5.87 | 0 | 0 | 0.51 | 0.01 | 0.5 | 51 |
| other bloold cells | Tnni2 | 8.46 | 0 | 0 | 0.5 | 0 | 0.5 | 167.67 |
| other bloold cells | Pik3cg | 7.31 | 0 | 0 | 0.5 | 0 | 0.5 | 167.67 |
| other bloold cells | Clec4a2 | 11.76 | 0 | 0 | 0.5 | 0 | 0.5 | #NA |
| other bloold cells | Itgb2 | 8.31 | 0 | 0 | 0.5 | 0 | 0.5 | 497 |
| other bloold cells | Ccl4 | 11.25 | 0 | 0 | 0.5 | 0 | 0.5 | 248.5 |
| other bloold cells | Slc11a1 | 11.75 | 0 | 0 | 0.49 | 0 | 0.49 | #NA |
| other bloold cells | Cd84 | 9.18 | 0 | 0 | 0.49 | 0 | 0.49 | 490 |
| other bloold cells | Casp1 | 10.48 | 0 | 0 | 0.48 | 0 | 0.48 | #NA |
| other bloold cells | Bcl2a1b | 12.24 | 0 | 0 | 0.48 | 0 | 0.48 | #NA |
| other bloold cells | Cd180 | 10.83 | 0 | 0 | 0.48 | 0 | 0.48 | #NA |
| other bloold cells | Tmem273 | 13.08 | 0 | 0 | 0.48 | 0 | 0.48 | #NA |
| other bloold cells | P2ry12 | 9.66 | 0 | 0 | 0.48 | 0 | 0.48 | 483 |
| other bloold cells | Wdfy4 | 10.1 | 0 | 0 | 0.48 | 0 | 0.48 | #NA |
| other bloold cells | Arhgap25 | 6.13 | 0 | 0 | 0.48 | 0.01 | 0.48 | 60.38 |
| other bloold cells | Lair1 | 10.56 | 0 | 0 | 0.48 | 0 | 0.48 | 476 |
| other bloold cells | F13a1 | 9.85 | 0 | 0 | 0.48 | 0 | 0.48 | 476 |
| other bloold cells | Arid5a | 5.13 | 0 | 0 | 0.48 | 0.01 | 0.46 | 36.62 |
| other bloold cells | Arl11 | 9.79 | 0 | 0 | 0.45 | 0 | 0.45 | 448 |
| other bloold cells | Tmem8 | 5.53 | 0 | 0 | 0.46 | 0.02 | 0.45 | 28.88 |
| other bloold cells | Arhgap4 | 5.65 | 0 | 0 | 0.46 | 0.01 | 0.44 | 41.36 |
| other bloold cells | Hfe | 5.88 | 0 | 0 | 0.46 | 0.01 | 0.44 | 37.92 |
| other bloold cells | Cytip | 11.9 | 0 | 0 | 0.44 | 0 | 0.44 | #NA |
| other bloold cells | Cryba4 | 10.31 | 0 | 0 | 0.44 | 0 | 0.44 | 441 |
| other bloold cells | Il10ra | 9.1 | 0 | 0 | 0.43 | 0 | 0.43 | 434 |
| other bloold cells | Ccl6 | 10.99 | 0 | 0 | 0.43 | 0 | 0.43 | 434 |
| other bloold cells | Clec12a | 8.44 | 0 | 0 | 0.43 | 0 | 0.43 | 144.67 |
| other bloold cells | 630001G21Ri | 5.32 | 0 | 0 | 0.44 | 0.01 | 0.43 | 40.09 |
| other bloold cells | P2ry13 | 10.98 | 0 | 0 | 0.43 | 0 | 0.43 | #NA |
| other bloold cells | Parp14 | 7.2 | 0 | 0 | 0.43 | 0 | 0.42 | 106.75 |
| other bloold cells | Tlr7 | 10.51 | 0 | 0 | 0.42 | 0 | 0.42 | #NA |
| other bloold cells | 630028O10Ri | 8.87 | 0 | 0 | 0.42 | 0 | 0.42 | 420 |
| other bloold cells | Rassf5 | 8.48 | 0 | 0 | 0.41 | 0 | 0.41 | 413 |
| other bloold cells | Hlx | 5.61 | 0 | 0 | 0.42 | 0.01 | 0.41 | 52.5 |
| other bloold cells | Lilrb4a | 10.92 | 0 | 0 | 0.41 | 0 | 0.41 | 413 |
| other bloold cells | Rapsn | 7.76 | 0 | 0 | 0.41 | 0 | 0.41 | 206.5 |
| other bloold cells | Gna15 | 6.45 | 0 | 0 | 0.41 | 0.01 | 0.41 | 82.6 |
| other bloold cells | Ang | 5.15 | 0 | 0 | 0.42 | 0.01 | 0.41 | 32.31 |
| other bloold cells | Themis2 | 8.74 | 0 | 0 | 0.41 | 0 | 0.41 | 406 |
| other bloold cells | Il21r | 8.52 | 0 | 0 | 0.41 | 0 | 0.41 | 406 |
| other bloold cells | 830077J02Ril | 6.73 | 0 | 0 | 0.41 | 0 | 0.4 | 203 |

**Supplementary Table 2. Marker genes for the major cell types of integrated clustering of E11.5 control and NotchGOF λ-junction cells.**

| | | | | | | | | |
|---|---|---|---|---|---|---|---|---|
| other bloold cells | Psd4 | 6.82 | 0 | 0 | 0.41 | 0 | 0.4 | 101.5 |
| other bloold cells | Arhgap15 | 10.57 | 0 | 0 | 0.4 | 0 | 0.4 | #NA |
| other bloold cells | Ccl24 | 12.2 | 0 | 0 | 0.4 | 0 | 0.4 | #NA |
| other bloold cells | Abca9 | 7.12 | 0 | 0 | 0.4 | 0 | 0.4 | 99.75 |
| other bloold cells | Susd3 | 5.14 | 0 | 0 | 0.41 | 0.01 | 0.39 | 33.83 |
| other bloold cells | Pilra | 10.95 | 0 | 0 | 0.39 | 0 | 0.39 | #NA |
| other bloold cells | Nupr1 | 7.9 | 0 | 0 | 0.39 | 0 | 0.39 | 196 |
| other bloold cells | Il16 | 6.94 | 0 | 0 | 0.39 | 0.01 | 0.39 | 78.4 |
| other bloold cells | Adap1 | 5.96 | 0 | 0 | 0.39 | 0.01 | 0.39 | 65.33 |
| other bloold cells | AB124611 | 8.85 | 0 | 0 | 0.39 | 0 | 0.38 | 385 |
| other bloold cells | Trpv2 | 6.19 | 0 | 0 | 0.38 | 0.01 | 0.37 | 75.6 |
| other bloold cells | Ripor2 | 5.66 | 0 | 0 | 0.38 | 0.01 | 0.37 | 75.6 |
| other bloold cells | Ptafr | 10.85 | 0 | 0 | 0.37 | 0 | 0.37 | #NA |
| other bloold cells | Rgs18 | 5.3 | 0 | 0 | 0.37 | 0 | 0.37 | 185.5 |
| other bloold cells | Irgm1 | 5.32 | 0 | 0 | 0.38 | 0.01 | 0.37 | 34.36 |
| other bloold cells | Siglech | 12.5 | 0 | 0 | 0.36 | 0 | 0.36 | #NA |
| other bloold cells | Ms4a7 | 11.81 | 0 | 0 | 0.36 | 0 | 0.36 | #NA |
| other bloold cells | Rasal3 | 8.71 | 0 | 0 | 0.36 | 0 | 0.36 | 364 |
| other bloold cells | Cysltr1 | 5.57 | 0 | 0 | 0.37 | 0.01 | 0.36 | 46.38 |
| other bloold cells | Ptk2b | 6.25 | 0 | 0 | 0.36 | 0 | 0.36 | 91 |
| other bloold cells | Cd37 | 6.97 | 0 | 0 | 0.36 | 0.01 | 0.36 | 72.8 |
| other bloold cells | Clec5a | 10.8 | 0 | 0 | 0.36 | 0 | 0.36 | #NA |
| other bloold cells | C5ar1 | 11.18 | 0 | 0 | 0.36 | 0 | 0.36 | #NA |
| other bloold cells | Pkib | 9.42 | 0 | 0 | 0.36 | 0 | 0.36 | 357 |
| other bloold cells | Ctla2b | 4.75 | 0 | 0 | 0.36 | 0.01 | 0.36 | 40.44 |
| other bloold cells | Ccdc88b | 6.43 | 0 | 0 | 0.35 | 0 | 0.35 | 87.5 |
| other bloold cells | Cd48 | 11.18 | 0 | 0 | 0.34 | 0 | 0.34 | #NA |
| other bloold cells | Epsti1 | 11.21 | 0 | 0 | 0.34 | 0 | 0.34 | #NA |
| other bloold cells | Cd74 | 6.46 | 0 | 0 | 0.34 | 0 | 0.34 | 85.75 |
| other bloold cells | 310001H17Ri | 7.76 | 0 | 0 | 0.34 | 0 | 0.33 | 168 |
| other bloold cells | Neurl3 | 6.96 | 0 | 0 | 0.34 | 0 | 0.33 | 112 |
| other bloold cells | Hk3 | 10.78 | 0 | 0 | 0.32 | 0 | 0.32 | #NA |
| other bloold cells | Evi2 | 6.93 | 0 | 0 | 0.32 | 0 | 0.32 | 161 |
| other bloold cells | Asb2 | 10.2 | 0 | 0 | 0.32 | 0 | 0.32 | #NA |
| other bloold cells | Clec4a1 | 9.55 | 0 | 0 | 0.32 | 0 | 0.31 | 315 |
| other bloold cells | Il18 | 6.26 | 0 | 0 | 0.32 | 0.01 | 0.31 | 63 |
| other bloold cells | Il2rg | 6.43 | 0 | 0 | 0.31 | 0 | 0.3 | 77 |
| other bloold cells | Pstpip1 | 5.85 | 0 | 0 | 0.31 | 0.01 | 0.3 | 51.33 |
| other bloold cells | Tlr13 | 11.04 | 0 | 0 | 0.3 | 0 | 0.3 | #NA |
| other bloold cells | Hcst | 7.13 | 0 | 0 | 0.3 | 0 | 0.3 | 301 |
| other bloold cells | Ptpro | 6.85 | 0 | 0 | 0.3 | 0 | 0.3 | 150.5 |
| other bloold cells | Prkcb | 6.82 | 0 | 0 | 0.3 | 0 | 0.3 | 100.33 |
| other bloold cells | Rnase6 | 7.69 | 0 | 0 | 0.3 | 0 | 0.3 | 100.33 |
| other bloold cells | Lgals3 | 8.2 | 0 | 0 | 0.3 | 0 | 0.3 | 75.25 |
| other bloold cells | Nlrp3 | 11.74 | 0 | 0 | 0.29 | 0 | 0.29 | #NA |
| other bloold cells | Serpinb1a | 9.36 | 0 | 0 | 0.29 | 0 | 0.29 | 294 |
| other bloold cells | Plcb2 | 6.85 | 0 | 0 | 0.29 | 0 | 0.29 | 147 |
| other bloold cells | Tap2 | 5.58 | 0 | 0 | 0.29 | 0.01 | 0.29 | 49 |
| other bloold cells | Ifi211 | 11 | 0 | 0 | 0.29 | 0 | 0.29 | #NA |
| other bloold cells | 430427O19Ri | 9.33 | 0 | 0 | 0.29 | 0 | 0.29 | #NA |
| other bloold cells | Marchf1 | 8.18 | 0 | 0 | 0.29 | 0 | 0.29 | 287 |
| other bloold cells | Tbxas1 | 6.34 | 0 | 0 | 0.29 | 0 | 0.28 | 95.67 |
| other bloold cells | Fcgr4 | 10.47 | 0 | 0 | 0.28 | 0 | 0.28 | #NA |
| other bloold cells | Mndal | 12.19 | 0 | 0 | 0.28 | 0 | 0.28 | #NA |
| other bloold cells | Tlr1 | 9.95 | 0 | 0 | 0.28 | 0 | 0.28 | #NA |
| other bloold cells | Napsa | 8.04 | 0 | 0 | 0.28 | 0 | 0.28 | 140 |

**Supplementary Table 2. Marker genes for the major cell types of integrated clustering of E11.5 control and NotchGOF λ-junction cells.**

| | | | | | | | | |
|---|---|---|---|---|---|---|---|---|
| other bloold cells | Sp100 | 9.9 | 0 | 0 | 0.27 | 0 | 0.27 | #NA |
| other bloold cells | Cd300lf | 8.22 | 0 | 0 | 0.27 | 0 | 0.27 | 273 |
| other bloold cells | Ticam2 | 6.33 | 0 | 0 | 0.27 | 0 | 0.27 | 91 |
| other bloold cells | Stap1 | 6.63 | 0 | 0 | 0.27 | 0 | 0.27 | 68.25 |
| other bloold cells | Apoc2 | 11.55 | 0 | 0 | 0.27 | 0 | 0.27 | #NA |
| other bloold cells | Cd200r1 | 7.87 | 0 | 0 | 0.27 | 0 | 0.27 | 266 |
| other bloold cells | Itgal | 6.28 | 0 | 0 | 0.27 | 0 | 0.26 | 66.5 |
| other bloold cells | Gm44751 | 5.82 | 0 | 0 | 0.27 | 0.01 | 0.26 | 53.2 |
| other bloold cells | Aoah | 9.84 | 0 | 0 | 0.26 | 0 | 0.26 | #NA |
| other bloold cells | Ltc4s | 7.9 | 0 | 0 | 0.26 | 0 | 0.26 | 129.5 |
| other bloold cells | AI467606 | 6.86 | 0 | 0 | 0.26 | 0 | 0.26 | 86.33 |
| other bloold cells | AI662270 | 6.81 | 0 | 0 | 0.26 | 0 | 0.26 | 86.33 |
| other bloold cells | Bank1 | 9.86 | 0 | 0 | 0.25 | 0 | 0.25 | #NA |
| other bloold cells | Clec4a3 | 10.65 | 0 | 0 | 0.25 | 0 | 0.25 | #NA |
| other bloold cells | Phf11b | 12.21 | 0 | 0 | 0.25 | 0 | 0.25 | #NA |
| other bloold cells | Gbp3 | 6.8 | 0 | 0 | 0.25 | 0 | 0.25 | 126 |
| other bloold cells | P2rx7 | 9.33 | 0 | 0 | 0.25 | 0 | 0.25 | #NA |
| other bloold cells | Hp | 10.55 | 0 | 0 | 0.25 | 0 | 0.25 | #NA |
| other bloold cells | Wfdc17 | 12.12 | 0 | 0 | 0.25 | 0 | 0.25 | #NA |
| other bloold cells | Naip2 | 8.17 | 0 | 0 | 0.25 | 0 | 0.24 | 245 |
| other bloold cells | Fgl2 | 7.04 | 0 | 0 | 0.25 | 0 | 0.24 | 122.5 |
| other bloold cells | P2rx1 | 5.45 | 0 | 0 | 0.25 | 0 | 0.24 | 122.5 |
| other bloold cells | Alox5 | 7.82 | 0 | 0 | 0.24 | 0 | 0.24 | 238 |
| other bloold cells | Ptprcap | 8.87 | 0 | 0 | 0.24 | 0 | 0.24 | 119 |
| other bloold cells | Lpxn | 6.89 | 0 | 0 | 0.24 | 0 | 0.24 | 119 |
| other bloold cells | Gpr171 | 8.29 | 0 | 0 | 0.23 | 0 | 0.23 | 231 |
| other bloold cells | Clec10a | 9.41 | 0 | 0 | 0.23 | 0 | 0.23 | 231 |
| other bloold cells | Ccl12 | 10.51 | 0 | 0 | 0.23 | 0 | 0.23 | 231 |
| other bloold cells | Mpo | 10.54 | 0 | 0 | 0.23 | 0 | 0.23 | 77 |
| other bloold cells | Cd69 | 10.58 | 0 | 0 | 0.22 | 0 | 0.22 | #NA |
| other bloold cells | Bex6 | 11.63 | 0 | 0 | 0.22 | 0 | 0.22 | #NA |
| other bloold cells | Rab7b | 7.24 | 0 | 0 | 0.22 | 0 | 0.22 | 112 |
| other bloold cells | Atp2a3 | 6.03 | 0 | 0 | 0.22 | 0 | 0.22 | 112 |
| other bloold cells | Ltb | 7.24 | 0 | 0 | 0.22 | 0 | 0.22 | 74.67 |
| other bloold cells | Clec4n | 11.7 | 0 | 0 | 0.22 | 0 | 0.22 | #NA |
| other bloold cells | Tagap | 8.77 | 0 | 0 | 0.22 | 0 | 0.22 | #NA |
| other bloold cells | Gm26740 | 7.02 | 0 | 0 | 0.22 | 0 | 0.22 | 217 |
| other bloold cells | Tlr4 | 6.86 | 0 | 0 | 0.22 | 0 | 0.22 | 108.5 |
| other bloold cells | Ccr2 | 11.94 | 0 | 0 | 0.21 | 0 | 0.21 | #NA |
| other bloold cells | Tespa1 | 10.69 | 0 | 0 | 0.21 | 0 | 0.21 | #NA |
| other bloold cells | Card9 | 8.14 | 0 | 0 | 0.21 | 0 | 0.21 | 210 |
| other bloold cells | Cmah | 5.85 | 0 | 0 | 0.21 | 0 | 0.21 | 105 |
| other bloold cells | Abcc3 | 7.18 | 0 | 0 | 0.21 | 0 | 0.21 | 70 |
| other bloold cells | Cd28 | 10.8 | 0 | 0 | 0.2 | 0 | 0.2 | #NA |
| other bloold cells | Tmem156 | 9.93 | 0 | 0 | 0.2 | 0 | 0.2 | #NA |
| other bloold cells | Gpr141 | 10.5 | 0 | 0 | 0.2 | 0 | 0.2 | #NA |
| other bloold cells | Mcoln2 | 8.05 | 0 | 0 | 0.2 | 0 | 0.2 | 203 |
| other bloold cells | Mir142hg | 7.59 | 0 | 0 | 0.2 | 0 | 0.2 | 203 |
| other bloold cells | Ifi209 | 11.88 | 0 | 0 | 0.2 | 0 | 0.2 | #NA |
| other bloold cells | Msr1 | 9.32 | 0 | 0 | 0.2 | 0 | 0.2 | #NA |
| other bloold cells | Cfb | 9.75 | 0 | 0 | 0.2 | 0 | 0.2 | #NA |
| other bloold cells | Tnf | 8.83 | 0 | 0 | 0.2 | 0 | 0.2 | #NA |
| other bloold cells | Ifi203 | 10.12 | 0 | 0 | 0.19 | 0 | 0.19 | #NA |
| other bloold cells | Ms4a4c | 11.32 | 0 | 0 | 0.19 | 0 | 0.19 | #NA |
| other bloold cells | Crybb1 | 9.51 | 0 | 0 | 0.18 | 0 | 0.18 | #NA |
| other bloold cells | Ccl2 | 9.91 | 0 | 0 | 0.18 | 0 | 0.18 | #NA |

**Supplementary Table 2. Marker genes for the major cell types of integrated clustering of E11.5 control and NotchGOF λ-junction cells.**

| | | | | | | | | |
|---|---|---|---|---|---|---|---|---|
| other bloold cells | Serpina3g | 8.84 | 0 | 0 | 0.18 | 0 | 0.18 | 182 |
| other bloold cells | Ly9 | 10.79 | 0 | 0 | 0.18 | 0 | 0.18 | #NA |
| other bloold cells | Siglece | 7.98 | 0 | 0 | 0.18 | 0 | 0.18 | #NA |
| other bloold cells | Tpbgl | 9.64 | 0 | 0 | 0.18 | 0 | 0.18 | #NA |
| other bloold cells | Il1b | 8.75 | 0 | 0 | 0.18 | 0 | 0.17 | 175 |
| other bloold cells | Kcnk12 | 7.37 | 0 | 0 | 0.17 | 0 | 0.17 | 168 |
| other bloold cells | Fcgr2b | 9.26 | 0 | 0 | 0.16 | 0 | 0.16 | #NA |
| other bloold cells | Trim30d | 9.38 | 0 | 0 | 0.16 | 0 | 0.16 | #NA |
| other bloold cells | Slamf9 | 8.32 | 0 | 0 | 0.16 | 0 | 0.16 | 161 |
| other bloold cells | Gpr84 | 8 | 0 | 0 | 0.16 | 0 | 0.16 | 161 |
| other bloold cells | Spn | 9.36 | 0 | 0 | 0.15 | 0 | 0.15 | #NA |
| other bloold cells | Cyp4f18 | 8.16 | 0 | 0 | 0.15 | 0 | 0.15 | #NA |
| other bloold cells | Lilr4b | 10.64 | 0 | 0 | 0.15 | 0 | 0.15 | #NA |
| other bloold cells | Oas1a | 7.49 | 0 | 0 | 0.15 | 0 | 0.15 | 154 |
| other bloold cells | Trbc1 | 9.25 | 0 | 0 | 0.15 | 0 | 0.15 | 154 |
| other bloold cells | Havcr2 | 8.21 | 0 | 0 | 0.15 | 0 | 0.15 | 154 |
| other bloold cells | Slfn8 | 7.03 | 0 | 0 | 0.15 | 0 | 0.15 | 154 |
| other bloold cells | Gm38843 | 9.39 | 0 | 0 | 0.15 | 0 | 0.15 | #NA |
| other bloold cells | Abcg3 | 8.17 | 0 | 0 | 0.15 | 0 | 0.15 | 147 |
| other bloold cells | Cst7 | 9.33 | 0 | 0 | 0.14 | 0 | 0.14 | #NA |
| other bloold cells | Gsg1 | 7.33 | 0 | 0 | 0.14 | 0 | 0.14 | 140 |
| other bloold cells | 600010M07R | 9.76 | 0 | 0 | 0.13 | 0 | 0.13 | #NA |
| other bloold cells | Il7r | 10.54 | 0 | 0 | 0.13 | 0 | 0.13 | #NA |
| other bloold cells | Cd300ld | 8.33 | 0 | 0 | 0.13 | 0 | 0.13 | 133 |
| other bloold cells | Gm5547 | 8.53 | 0 | 0 | 0.13 | 0 | 0.13 | #NA |
| other bloold cells | Gdf3 | 10.12 | 0 | 0 | 0.13 | 0 | 0.13 | #NA |
| other bloold cells | Bcl2a1d | 10.38 | 0 | 0 | 0.13 | 0 | 0.13 | #NA |
| other bloold cells | Gm5086 | 9.68 | 0 | 0 | 0.13 | 0 | 0.13 | #NA |
| other bloold cells | Pglyrp2 | 10.66 | 0 | 0 | 0.13 | 0 | 0.13 | #NA |
| other bloold cells | 130050O18Ri | 7.91 | 0 | 0 | 0.13 | 0 | 0.13 | 126 |
| other bloold cells | Scimp | 10.72 | 0 | 0 | 0.12 | 0 | 0.12 | #NA |
| other bloold cells | Adap2os | 9.12 | 0 | 0 | 0.12 | 0 | 0.12 | #NA |
| other bloold cells | Flt3 | 8.41 | 0 | 0 | 0.11 | 0 | 0.11 | #NA |
| other bloold cells | 010013B24Ri | 8.91 | 0 | 0 | 0.11 | 0 | 0.11 | #NA |
| other bloold cells | Gapt | 10.25 | 0 | 0 | 0.11 | 0 | 0.11 | #NA |
| other bloold cells | Ebi3 | 4.95 | 4.72E-308 | 9.83E-304 | 0.43 | 0.01 | 0.41 | 30.5 |
| other bloold cells | Elmo1 | 4.25 | 1.91E-306 | 3.97E-302 | 0.7 | 0.04 | 0.66 | 16.64 |
| other bloold cells | Tlr2 | 4.87 | 1.63E-300 | 3.39E-296 | 0.51 | 0.02 | 0.49 | 23.18 |
| other bloold cells | Ptpn22 | 6.51 | 4.33E-298 | 9.01E-294 | 0.15 | 0 | 0.15 | 147 |
| other bloold cells | Smagp | 4.44 | 1.07E-295 | 2.23E-291 | 0.6 | 0.03 | 0.57 | 19.39 |
| other bloold cells | Parp10 | 6.39 | 3.82E-295 | 7.94E-291 | 0.19 | 0 | 0.19 | 94.5 |
| other bloold cells | Slc7a7 | 5.25 | 2.69E-293 | 5.59E-289 | 0.43 | 0.02 | 0.41 | 28.47 |
| other bloold cells | Gas6 | 6.32 | 4.41E-290 | 9.16E-286 | 0.52 | 0.03 | 0.5 | 20.96 |
| other bloold cells | Ptpre | 4.85 | 5.69E-290 | 1.18E-285 | 0.61 | 0.03 | 0.57 | 17.88 |
| other bloold cells | Klf2 | 4.53 | 3.63E-289 | 7.54E-285 | 0.62 | 0.03 | 0.58 | 18.09 |
| other bloold cells | Hdc | 9.64 | 4.44E-288 | 9.24E-284 | 0.12 | 0 | 0.12 | 119 |
| other bloold cells | Gm44067 | 7.66 | 4.69E-288 | 9.76E-284 | 0.12 | 0 | 0.12 | 119 |
| other bloold cells | Cd44 | 4.6 | 2.30E-287 | 4.79E-283 | 0.61 | 0.03 | 0.57 | 17.88 |
| other bloold cells | Tifa | 4.92 | 2.71E-286 | 5.64E-282 | 0.54 | 0.03 | 0.51 | 20.69 |
| other bloold cells | Tmem106a | 5.61 | 1.96E-277 | 4.07E-273 | 0.33 | 0.01 | 0.32 | 36.56 |
| other bloold cells | Gpr160 | 5.08 | 3.18E-273 | 6.62E-269 | 0.33 | 0.01 | 0.32 | 36.56 |
| other bloold cells | Spns3 | 7.54 | 1.52E-269 | 3.16E-265 | 0.11 | 0 | 0.1 | 105 |
| other bloold cells | Dapp1 | 4.47 | 1.59E-268 | 3.31E-264 | 0.53 | 0.03 | 0.5 | 19.67 |
| other bloold cells | Coro2a | 4.82 | 9.18E-267 | 1.91E-262 | 0.43 | 0.02 | 0.42 | 24.11 |
| other bloold cells | Paqr7 | 4.96 | 3.06E-263 | 6.36E-259 | 0.32 | 0.01 | 0.31 | 35 |
| other bloold cells | Plac8 | 7.8 | 1.29E-262 | 2.69E-258 | 0.43 | 0.02 | 0.41 | 23.72 |

**Supplementary Table 2. Marker genes for the major cell types of integrated clustering of E11.5 control and NotchGOF λ-junction cells.**

| | | | | | | | | |
|---|---|---|---|---|---|---|---|---|
| other bloold cells | Csf2rb2 | 7.55 | 3.57E-262 | 7.42E-258 | 0.13 | 0 | 0.13 | 133 |
| other bloold cells | Myo7a | 4.43 | 1.19E-260 | 2.49E-256 | 0.35 | 0.01 | 0.34 | 31.82 |
| other bloold cells | Slc46a3 | 5.44 | 1.94E-257 | 4.04E-253 | 0.33 | 0.01 | 0.32 | 32.9 |
| other bloold cells | Ctsc | 6.22 | 9.57E-256 | 1.99E-251 | 0.97 | 0.12 | 0.85 | 8.1 |
| other bloold cells | Egr1 | 4.65 | 1.58E-255 | 3.29E-251 | 0.8 | 0.07 | 0.73 | 11.32 |
| other bloold cells | Spp1 | 10.79 | 2.23E-253 | 4.64E-249 | 0.14 | 0 | 0.14 | 140 |
| other bloold cells | Maf | 6.52 | 7.37E-253 | 1.53E-248 | 0.77 | 0.07 | 0.7 | 11.14 |
| other bloold cells | Pycard | 5.27 | 1.51E-252 | 3.13E-248 | 0.97 | 0.12 | 0.85 | 7.9 |
| other bloold cells | Grap | 4.34 | 2.91E-249 | 6.06E-245 | 0.37 | 0.01 | 0.36 | 28.54 |
| other bloold cells | Tgfb1 | 4.07 | 3.18E-249 | 6.62E-245 | 0.94 | 0.1 | 0.83 | 9.1 |
| other bloold cells | Atf3 | 5.71 | 2.12E-246 | 4.40E-242 | 0.48 | 0.02 | 0.45 | 19.83 |
| other bloold cells | Arhgap9 | 4.33 | 1.33E-244 | 2.77E-240 | 0.57 | 0.04 | 0.54 | 16.37 |
| other bloold cells | Lgals3bp | 5.87 | 3.17E-244 | 6.60E-240 | 0.26 | 0.01 | 0.25 | 43.17 |
| other bloold cells | Gpsm3 | 4.28 | 8.70E-244 | 1.81E-239 | 0.85 | 0.08 | 0.76 | 10.32 |
| other bloold cells | Abcb1b | 5.39 | 1.33E-243 | 2.76E-239 | 0.24 | 0.01 | 0.23 | 47.6 |
| other bloold cells | Nfkbid | 5.12 | 4.37E-243 | 9.08E-239 | 0.28 | 0.01 | 0.27 | 35 |
| other bloold cells | Coro1a | 6.28 | 1.19E-241 | 2.48E-237 | 1 | 0.14 | 0.86 | 6.99 |
| other bloold cells | Tec | 4.81 | 4.94E-240 | 1.03E-235 | 0.33 | 0.01 | 0.32 | 29.91 |
| other bloold cells | Plxna4os1 | 6.76 | 9.83E-233 | 2.04E-228 | 0.14 | 0 | 0.14 | 70 |
| other bloold cells | Ehbp1l1 | 4.23 | 3.33E-232 | 6.92E-228 | 0.72 | 0.06 | 0.66 | 11.8 |
| other bloold cells | Ifi35 | 4.55 | 3.73E-231 | 7.75E-227 | 0.5 | 0.03 | 0.47 | 17.75 |
| other bloold cells | Ifi27 | 5.48 | 6.39E-230 | 1.33E-225 | 0.29 | 0.01 | 0.28 | 31.89 |
| other bloold cells | Gas7 | 4.82 | 1.78E-228 | 3.71E-224 | 0.64 | 0.05 | 0.59 | 13.12 |
| other bloold cells | Pirb | 6.05 | 3.78E-228 | 7.87E-224 | 0.18 | 0 | 0.18 | 60.67 |
| other bloold cells | Arhgap45 | 3.85 | 4.18E-228 | 8.68E-224 | 0.72 | 0.06 | 0.66 | 12 |
| other bloold cells | Lyl1 | 4.48 | 1.74E-227 | 3.63E-223 | 0.54 | 0.03 | 0.51 | 16.3 |
| other bloold cells | Trf | 5.73 | 1.22E-226 | 2.53E-222 | 0.58 | 0.04 | 0.54 | 14.5 |
| other bloold cells | Anxa3 | 4.27 | 1.24E-226 | 2.57E-222 | 0.83 | 0.09 | 0.74 | 9.59 |
| other bloold cells | S100a13 | 3.99 | 1.34E-224 | 2.79E-220 | 0.9 | 0.1 | 0.79 | 8.69 |
| other bloold cells | Abhd15 | 6.54 | 2.50E-220 | 5.19E-216 | 0.12 | 0 | 0.12 | 119 |
| other bloold cells | Rasgrp3 | 3.97 | 2.62E-220 | 5.45E-216 | 0.36 | 0.02 | 0.35 | 24.27 |
| other bloold cells | Irf7 | 7.05 | 5.17E-220 | 1.07E-215 | 0.15 | 0 | 0.15 | 77 |
| other bloold cells | Serpinb8 | 5.14 | 1.24E-218 | 2.58E-214 | 0.2 | 0 | 0.2 | 50.75 |
| other bloold cells | Slc22a3 | 8.05 | 3.48E-218 | 7.23E-214 | 0.11 | 0 | 0.11 | 112 |
| other bloold cells | Ptprj | 4.43 | 7.38E-218 | 1.53E-213 | 0.54 | 0.04 | 0.5 | 15.37 |
| other bloold cells | Npl | 5.21 | 2.43E-217 | 5.05E-213 | 0.84 | 0.09 | 0.75 | 8.93 |
| other bloold cells | Atp1a3 | 5.43 | 9.12E-217 | 1.90E-212 | 0.17 | 0 | 0.17 | 56 |
| other bloold cells | Haao | 6.48 | 1.89E-216 | 3.93E-212 | 0.11 | 0 | 0.1 | 105 |
| other bloold cells | Slc43a2 | 4.48 | 8.63E-215 | 1.79E-210 | 0.56 | 0.04 | 0.52 | 14.71 |
| other bloold cells | Lat2 | 4.9 | 1.60E-214 | 3.33E-210 | 0.34 | 0.01 | 0.33 | 24.5 |
| other bloold cells | Parp9 | 4.11 | 2.32E-213 | 4.83E-209 | 0.49 | 0.03 | 0.46 | 16.9 |
| other bloold cells | Dok3 | 4.63 | 4.76E-213 | 9.91E-209 | 0.39 | 0.02 | 0.37 | 22.65 |
| other bloold cells | Treml2 | 4.82 | 6.52E-211 | 1.36E-206 | 0.34 | 0.01 | 0.33 | 24.5 |
| other bloold cells | Elf4 | 4.39 | 7.08E-211 | 1.47E-206 | 0.43 | 0.02 | 0.41 | 19.41 |
| other bloold cells | Cebpb | 4.89 | 1.53E-208 | 3.17E-204 | 0.89 | 0.12 | 0.77 | 7.66 |
| other bloold cells | Fam129a | 6.12 | 3.12E-206 | 6.48E-202 | 0.2 | 0 | 0.19 | 49 |
| other bloold cells | Pik3r6 | 5.57 | 3.64E-206 | 7.58E-202 | 0.21 | 0.01 | 0.21 | 42 |
| other bloold cells | Cd83 | 5.22 | 4.78E-206 | 9.95E-202 | 0.4 | 0.02 | 0.38 | 19.95 |
| other bloold cells | D5Ertd605e | 4.92 | 8.30E-205 | 1.73E-200 | 0.25 | 0.01 | 0.24 | 35 |
| other bloold cells | Tlr6 | 6.29 | 3.32E-204 | 6.90E-200 | 0.12 | 0 | 0.12 | 119 |
| other bloold cells | Sall3 | 6.32 | 8.93E-204 | 1.86E-199 | 0.14 | 0 | 0.14 | 70 |
| other bloold cells | Kynu | 6.23 | 1.05E-203 | 2.18E-199 | 0.14 | 0 | 0.14 | 70 |
| other bloold cells | Zeb2os | 4.33 | 1.38E-201 | 2.87E-197 | 0.58 | 0.04 | 0.54 | 13.18 |
| other bloold cells | Rnase4 | 5.38 | 1.86E-198 | 3.88E-194 | 0.79 | 0.09 | 0.7 | 8.49 |
| other bloold cells | Sirpa | 4.48 | 2.79E-198 | 5.79E-194 | 0.88 | 0.11 | 0.77 | 7.73 |
| other bloold cells | Trim14 | 6.5 | 5.25E-197 | 1.09E-192 | 0.11 | 0 | 0.1 | 105 |

**Supplementary Table 2. Marker genes for the major cell types of integrated clustering of E11.5 control and NotchGOF λ-junction cells.**

| | | | | | | | | |
|---|---|---|---|---|---|---|---|---|
| other bloold cells | Junb | 4.13 | 4.25E-195 | 8.83E-191 | 0.87 | 0.11 | 0.76 | 7.74 |
| other bloold cells | Ctsh | 4.87 | 5.68E-195 | 1.18E-190 | 0.64 | 0.06 | 0.58 | 10.9 |
| other bloold cells | Stk10 | 4.23 | 1.73E-194 | 3.59E-190 | 0.45 | 0.03 | 0.42 | 17.23 |
| other bloold cells | Pparg | 6.11 | 6.71E-194 | 1.40E-189 | 0.13 | 0 | 0.13 | 66.5 |
| other bloold cells | Aim2 | 3.59 | 1.09E-193 | 2.26E-189 | 0.67 | 0.06 | 0.61 | 11.18 |
| other bloold cells | Smim40 | 6.17 | 8.88E-192 | 1.85E-187 | 0.16 | 0 | 0.16 | 53.67 |
| other bloold cells | Kcnk6 | 4.99 | 1.65E-191 | 3.43E-187 | 0.25 | 0.01 | 0.24 | 31.5 |
| other bloold cells | Emp3 | 4.3 | 1.56E-190 | 3.24E-186 | 0.9 | 0.13 | 0.77 | 6.83 |
| other bloold cells | Fos | 5.54 | 3.46E-189 | 7.19E-185 | 0.9 | 0.14 | 0.76 | 6.63 |
| other bloold cells | Gm2a | 3.73 | 7.27E-189 | 1.51E-184 | 0.71 | 0.07 | 0.64 | 10.23 |
| other bloold cells | Pf4 | 1.24 | 3.83E-187 | 7.96E-183 | 0.43 | 0.03 | 0.41 | 16.69 |
| other bloold cells | Ifi27l2a | 6.85 | 2.19E-185 | 4.56E-181 | 0.21 | 0.01 | 0.2 | 35 |
| other bloold cells | Taok3 | 3.85 | 2.48E-185 | 5.16E-181 | 0.82 | 0.1 | 0.72 | 8.18 |
| other bloold cells | Cgas | 4.03 | 2.68E-184 | 5.56E-180 | 0.4 | 0.02 | 0.38 | 18.14 |
| other bloold cells | Cmklr1 | 5.62 | 3.46E-183 | 7.19E-179 | 0.19 | 0.01 | 0.18 | 37.8 |
| other bloold cells | Dusp2 | 3.82 | 1.14E-182 | 2.36E-178 | 0.47 | 0.03 | 0.44 | 15.13 |
| other bloold cells | Jak3 | 3.9 | 2.74E-180 | 5.69E-176 | 0.44 | 0.03 | 0.41 | 16.33 |
| other bloold cells | Man1c1 | 4.16 | 5.98E-179 | 1.24E-174 | 0.47 | 0.03 | 0.44 | 15.13 |
| other bloold cells | Ttc7 | 3.95 | 1.95E-178 | 4.06E-174 | 0.47 | 0.03 | 0.44 | 14.66 |
| other bloold cells | H2-K1 | 3.88 | 1.49E-176 | 3.11E-172 | 0.9 | 0.13 | 0.77 | 6.88 |
| other bloold cells | Plxnc1 | 3.95 | 5.18E-176 | 1.08E-171 | 0.52 | 0.04 | 0.48 | 13.1 |
| other bloold cells | Slc16a6 | 3.97 | 9.17E-174 | 1.91E-169 | 0.58 | 0.05 | 0.53 | 11.37 |
| other bloold cells | Erap1 | 3.84 | 7.07E-173 | 1.47E-168 | 0.48 | 0.04 | 0.45 | 13.8 |
| other bloold cells | Arsb | 4.16 | 6.08E-170 | 1.26E-165 | 0.63 | 0.06 | 0.57 | 9.98 |
| other bloold cells | Rgs14 | 4.93 | 2.61E-169 | 5.42E-165 | 0.16 | 0 | 0.16 | 53.67 |
| other bloold cells | B4galnt1 | 4.13 | 1.14E-167 | 2.38E-163 | 0.52 | 0.04 | 0.48 | 12.19 |
| other bloold cells | Batf | 3.7 | 3.25E-167 | 6.76E-163 | 0.46 | 0.03 | 0.43 | 14.44 |
| other bloold cells | Rinl | 4.3 | 5.39E-165 | 1.12E-160 | 0.32 | 0.02 | 0.31 | 20.13 |
| other bloold cells | Rasa4 | 4.33 | 7.08E-165 | 1.47E-160 | 0.31 | 0.01 | 0.29 | 22 |
| other bloold cells | Mef2c | 4.61 | 2.58E-164 | 5.36E-160 | 0.94 | 0.18 | 0.76 | 5.18 |
| other bloold cells | Sh3tc1 | 5.23 | 7.63E-164 | 1.59E-159 | 0.2 | 0.01 | 0.2 | 33.83 |
| other bloold cells | Tent5a | 4.49 | 1.49E-163 | 3.10E-159 | 0.48 | 0.04 | 0.45 | 13.05 |
| other bloold cells | Pip4k2a | 3.64 | 6.17E-163 | 1.28E-158 | 0.81 | 0.11 | 0.7 | 7.24 |
| other bloold cells | 230029C05Ri | 4.73 | 7.77E-162 | 1.62E-157 | 0.39 | 0.02 | 0.37 | 16.33 |
| other bloold cells | Abca1 | 5.26 | 1.69E-160 | 3.51E-156 | 0.63 | 0.07 | 0.56 | 9.25 |
| other bloold cells | Apobec3 | 3.64 | 1.15E-159 | 2.40E-155 | 0.69 | 0.08 | 0.61 | 8.65 |
| other bloold cells | Def6 | 3.73 | 4.96E-159 | 1.03E-154 | 0.48 | 0.04 | 0.45 | 13.05 |
| other bloold cells | Ier3 | 4.45 | 9.48E-159 | 1.97E-154 | 0.65 | 0.07 | 0.58 | 8.9 |
| other bloold cells | Cbfa2t3 | 4.15 | 1.11E-158 | 2.31E-154 | 0.56 | 0.05 | 0.51 | 10.75 |
| other bloold cells | Anxa1 | 3.77 | 1.66E-157 | 3.46E-153 | 0.4 | 0.03 | 0.37 | 15.35 |
| other bloold cells | Tbc1d10c | 5.49 | 5.50E-157 | 1.14E-152 | 0.19 | 0.01 | 0.18 | 37.8 |
| other bloold cells | Rnf213 | 4.43 | 1.32E-156 | 2.74E-152 | 0.4 | 0.03 | 0.37 | 15.35 |
| other bloold cells | Ehd4 | 4.71 | 5.98E-154 | 1.24E-149 | 0.91 | 0.17 | 0.74 | 5.28 |
| other bloold cells | Xlr | 6.76 | 6.39E-154 | 1.33E-149 | 0.13 | 0 | 0.12 | 63 |
| other bloold cells | Apoc1 | 5.15 | 1.04E-153 | 2.17E-149 | 0.68 | 0.08 | 0.6 | 8.27 |
| other bloold cells | Renbp | 3.55 | 2.40E-153 | 4.99E-149 | 0.66 | 0.07 | 0.59 | 8.97 |
| other bloold cells | Clec2d | 4.12 | 3.54E-152 | 7.36E-148 | 0.36 | 0.02 | 0.34 | 17 |
| other bloold cells | Parp12 | 5.19 | 1.01E-151 | 2.09E-147 | 0.21 | 0.01 | 0.2 | 30 |
| other bloold cells | Dock10 | 3.49 | 7.37E-150 | 1.53E-145 | 0.69 | 0.08 | 0.6 | 8.35 |
| other bloold cells | Efhd2 | 4.51 | 1.63E-149 | 3.39E-145 | 0.94 | 0.2 | 0.74 | 4.71 |
| other bloold cells | Slc37a2 | 4.82 | 6.73E-147 | 1.40E-142 | 0.32 | 0.02 | 0.3 | 17.89 |
| other bloold cells | Lsp1 | 4.78 | 3.47E-146 | 7.22E-142 | 0.85 | 0.16 | 0.69 | 5.46 |
| other bloold cells | Tmem140 | 4.63 | 2.72E-143 | 5.65E-139 | 0.24 | 0.01 | 0.23 | 23.8 |
| other bloold cells | Samd9l | 6.01 | 2.12E-142 | 4.41E-138 | 0.14 | 0 | 0.14 | 46.67 |
| other bloold cells | St8sia4 | 3.7 | 3.39E-142 | 7.06E-138 | 0.69 | 0.09 | 0.6 | 7.69 |
| other bloold cells | Acap1 | 6.03 | 3.20E-141 | 6.65E-137 | 0.15 | 0 | 0.14 | 36.75 |

**Supplementary Table 2. Marker genes for the major cell types of integrated clustering of E11.5 control and NotchGOF λ-junction cells.**

| | | | | | | | | |
|---|---|---|---|---|---|---|---|---|
| other bloold cells | Tgfbr2 | 3.57 | 1.29E-140 | 2.69E-136 | 0.66 | 0.08 | 0.58 | 8.2 |
| other bloold cells | Trim47 | 4.6 | 1.33E-140 | 2.77E-136 | 0.31 | 0.02 | 0.29 | 18.12 |
| other bloold cells | Il18rap | 4.42 | 1.16E-139 | 2.42E-135 | 0.27 | 0.01 | 0.25 | 20.46 |
| other bloold cells | P2ry10b | 6.17 | 2.13E-139 | 4.43E-135 | 0.12 | 0 | 0.12 | 59.5 |
| other bloold cells | Phf11d | 6.37 | 1.79E-138 | 3.72E-134 | 0.11 | 0 | 0.1 | 52.5 |
| other bloold cells | Pstpip2 | 4.44 | 4.61E-138 | 9.60E-134 | 0.25 | 0.01 | 0.23 | 22.27 |
| other bloold cells | Man2b1 | 3.69 | 6.19E-138 | 1.29E-133 | 0.96 | 0.23 | 0.73 | 4.24 |
| other bloold cells | Trim25 | 3.44 | 9.22E-137 | 1.92E-132 | 0.55 | 0.06 | 0.5 | 9.68 |
| other bloold cells | Tmem173 | 3.57 | 1.37E-136 | 2.86E-132 | 0.54 | 0.05 | 0.48 | 9.96 |
| other bloold cells | Ccdc152 | 5.75 | 1.55E-135 | 3.22E-131 | 0.16 | 0.01 | 0.16 | 32.2 |
| other bloold cells | Rin3 | 3.69 | 1.98E-135 | 4.12E-131 | 0.36 | 0.02 | 0.33 | 14.88 |
| other bloold cells | Selenop | 5.37 | 3.83E-135 | 7.97E-131 | 0.9 | 0.19 | 0.72 | 4.88 |
| other bloold cells | Arap1 | 3.52 | 6.80E-135 | 1.41E-130 | 0.59 | 0.07 | 0.53 | 8.74 |
| other bloold cells | Plxnd1 | 2.24 | 1.27E-134 | 2.64E-130 | 0.55 | 0.06 | 0.5 | 10.04 |
| other bloold cells | Man1a | 3.73 | 6.33E-133 | 1.32E-128 | 0.55 | 0.06 | 0.49 | 9.36 |
| other bloold cells | Ostf1 | 3.72 | 6.56E-133 | 1.36E-128 | 0.98 | 0.27 | 0.71 | 3.6 |
| other bloold cells | Gpr146 | 4.12 | 5.68E-131 | 1.18E-126 | 0.38 | 0.03 | 0.35 | 13.5 |
| other bloold cells | Sema4d | 3.55 | 9.36E-131 | 1.95E-126 | 0.59 | 0.07 | 0.52 | 8.63 |
| other bloold cells | Manba | 4.45 | 3.62E-130 | 7.52E-126 | 0.28 | 0.02 | 0.27 | 18.67 |
| other bloold cells | Ifi47 | 5.44 | 1.34E-129 | 2.79E-125 | 0.11 | 0 | 0.11 | 56 |
| other bloold cells | Ocel1 | 3.83 | 1.39E-129 | 2.88E-125 | 0.51 | 0.05 | 0.46 | 9.81 |
| other bloold cells | Blnk | 4.4 | 2.34E-127 | 4.87E-123 | 0.25 | 0.01 | 0.24 | 19.38 |
| other bloold cells | Apobr | 4.43 | 9.46E-127 | 1.97E-122 | 0.25 | 0.01 | 0.23 | 20.42 |
| other bloold cells | Arpc1b | 4.78 | 1.58E-126 | 3.29E-122 | 0.99 | 0.34 | 0.65 | 2.89 |
| other bloold cells | Syngr2 | 3.42 | 2.86E-126 | 5.94E-122 | 0.79 | 0.14 | 0.65 | 5.64 |
| other bloold cells | Cebpa | 4.07 | 2.88E-126 | 5.99E-122 | 0.36 | 0.03 | 0.33 | 13.73 |
| other bloold cells | Pnpla7 | 3.65 | 2.96E-126 | 6.15E-122 | 0.41 | 0.03 | 0.38 | 12.15 |
| other bloold cells | Myd88 | 3.59 | 4.84E-126 | 1.01E-121 | 0.5 | 0.05 | 0.45 | 9.86 |
| other bloold cells | Plekha2 | 3.62 | 6.48E-124 | 1.35E-119 | 0.47 | 0.05 | 0.42 | 10.42 |
| other bloold cells | Bmp2k | 3.34 | 1.81E-123 | 3.76E-119 | 0.81 | 0.15 | 0.66 | 5.48 |
| other bloold cells | Pygm | 5.41 | 3.79E-122 | 7.88E-118 | 0.13 | 0 | 0.13 | 44.33 |
| other bloold cells | Tor3a | 3.56 | 8.56E-122 | 1.78E-117 | 0.43 | 0.04 | 0.39 | 11.24 |
| other bloold cells | Capn3 | 4.6 | 7.28E-119 | 1.51E-114 | 0.11 | 0 | 0.11 | 56 |
| other bloold cells | Amz1 | 4.24 | 2.61E-118 | 5.42E-114 | 0.21 | 0.01 | 0.2 | 23.33 |
| other bloold cells | Runx1 | 2.56 | 4.03E-118 | 8.38E-114 | 0.92 | 0.19 | 0.73 | 4.82 |
| other bloold cells | Il10rb | 3.7 | 5.79E-118 | 1.21E-113 | 0.61 | 0.08 | 0.53 | 7.33 |
| other bloold cells | Ifih1 | 4.46 | 8.64E-118 | 1.80E-113 | 0.22 | 0.01 | 0.21 | 21.7 |
| other bloold cells | Fosb | 3.83 | 3.09E-117 | 6.43E-113 | 0.38 | 0.03 | 0.35 | 12.19 |
| other bloold cells | Morrbid | 4.86 | 9.10E-117 | 1.89E-112 | 0.14 | 0 | 0.14 | 35 |
| other bloold cells | Map3k8 | 4.03 | 1.59E-116 | 3.30E-112 | 0.3 | 0.02 | 0.28 | 15.05 |
| other bloold cells | Rrad | 4.42 | 2.47E-116 | 5.14E-112 | 0.25 | 0.01 | 0.23 | 18.85 |
| other bloold cells | Rassf2 | 4.05 | 1.29E-114 | 2.68E-110 | 0.31 | 0.02 | 0.29 | 14.67 |
| other bloold cells | Ggta1 | 3.6 | 2.71E-113 | 5.64E-109 | 0.43 | 0.04 | 0.39 | 10.33 |
| other bloold cells | Lmo2 | 2.44 | 3.67E-113 | 7.64E-109 | 0.76 | 0.12 | 0.63 | 6.09 |
| other bloold cells | Serinc3 | 3.8 | 5.72E-112 | 1.19E-107 | 0.98 | 0.36 | 0.62 | 2.73 |
| other bloold cells | Hivep3 | 4.1 | 2.40E-111 | 4.98E-107 | 0.29 | 0.02 | 0.27 | 14.7 |
| other bloold cells | Slc16a7 | 4.33 | 4.27E-111 | 8.89E-107 | 0.31 | 0.02 | 0.29 | 14 |
| other bloold cells | Rnasel | 3.51 | 1.69E-110 | 3.52E-106 | 0.6 | 0.08 | 0.52 | 7.24 |
| other bloold cells | Hexb | 4.03 | 6.02E-110 | 1.25E-105 | 0.89 | 0.22 | 0.67 | 4 |
| other bloold cells | Ifngr1 | 3.32 | 2.42E-109 | 5.04E-105 | 0.92 | 0.24 | 0.68 | 3.8 |
| other bloold cells | Stxbp2 | 2.86 | 1.01E-108 | 2.09E-104 | 0.76 | 0.13 | 0.62 | 5.63 |
| other bloold cells | Pik3r5 | 4.62 | 1.06E-108 | 2.21E-104 | 0.18 | 0.01 | 0.17 | 25 |
| other bloold cells | Sat1 | 4.34 | 1.22E-108 | 2.53E-104 | 0.96 | 0.3 | 0.65 | 3.15 |
| other bloold cells | Gmip | 3.29 | 3.90E-108 | 8.11E-104 | 0.48 | 0.05 | 0.43 | 9.11 |
| other bloold cells | Stat1 | 3.17 | 5.25E-108 | 1.09E-103 | 0.64 | 0.1 | 0.54 | 6.69 |
| other bloold cells | Gsdmd | 3.84 | 1.98E-107 | 4.12E-103 | 0.32 | 0.02 | 0.3 | 13.42 |

**Supplementary Table 2. Marker genes for the major cell types of integrated clustering of E11.5 control and NotchGOF λ-junction cells.**

| | | | | | | | | |
|---|---|---|---|---|---|---|---|---|
| other bloold cells | Dclre1c | 3.13 | 2.44E-107 | 5.08E-103 | 0.47 | 0.05 | 0.42 | 9.2 |
| other bloold cells | H2-D1 | 3.05 | 3.37E-107 | 7.02E-103 | 0.92 | 0.24 | 0.68 | 3.83 |
| other bloold cells | Rab32 | 3.07 | 6.88E-107 | 1.43E-102 | 0.62 | 0.09 | 0.53 | 6.91 |
| other bloold cells | Rbm47 | 3.81 | 9.65E-107 | 2.01E-102 | 0.37 | 0.03 | 0.34 | 11.24 |
| other bloold cells | Rcsd1 | 2.86 | 1.22E-106 | 2.53E-102 | 0.63 | 0.09 | 0.54 | 6.91 |
| other bloold cells | AI413582 | 3.1 | 1.56E-106 | 3.25E-102 | 0.62 | 0.09 | 0.53 | 6.91 |
| other bloold cells | Gsn | 3.42 | 4.06E-106 | 8.45E-102 | 0.75 | 0.14 | 0.61 | 5.3 |
| other bloold cells | Cyba | 3.97 | 7.29E-106 | 1.52E-101 | 0.99 | 0.45 | 0.54 | 2.21 |
| other bloold cells | Adrb2 | 5.24 | 8.96E-106 | 1.86E-101 | 0.22 | 0.01 | 0.21 | 18.67 |
| other bloold cells | Psmb10 | 3.03 | 9.32E-106 | 1.94E-101 | 0.9 | 0.23 | 0.67 | 3.93 |
| other bloold cells | Reps2 | 3.79 | 2.26E-105 | 4.71E-101 | 0.34 | 0.03 | 0.32 | 12.25 |
| other bloold cells | Tspo | 3.91 | 2.88E-105 | 5.99E-101 | 0.97 | 0.35 | 0.62 | 2.79 |
| other bloold cells | Dram1 | 4.83 | 5.29E-105 | 1.10E-100 | 0.13 | 0 | 0.13 | 33.25 |
| other bloold cells | Tpd52 | 3.08 | 5.90E-105 | 1.23E-100 | 0.93 | 0.26 | 0.67 | 3.58 |
| other bloold cells | Celf2 | 2.75 | 4.12E-104 | 8.58E-100 | 0.93 | 0.25 | 0.68 | 3.68 |
| other bloold cells | Rtp4 | 4.66 | 8.61E-104 | 1.79E-99 | 0.2 | 0.01 | 0.19 | 21.78 |
| other bloold cells | Nfatc1 | 3.68 | 9.96E-104 | 2.07E-99 | 0.36 | 0.03 | 0.33 | 11.52 |
| other bloold cells | Pla2g15 | 3.22 | 2.72E-103 | 5.66E-99 | 0.52 | 0.07 | 0.46 | 7.94 |
| other bloold cells | Nostrin | 5.75 | 6.83E-103 | 1.42E-98 | 0.11 | 0 | 0.1 | 52.5 |
| other bloold cells | Tmem37 | 3.59 | 9.90E-103 | 2.06E-98 | 0.58 | 0.08 | 0.5 | 6.99 |
| other bloold cells | Plekho2 | 3.34 | 1.45E-102 | 3.03E-98 | 0.52 | 0.07 | 0.46 | 7.71 |
| other bloold cells | Phyhd1 | 3.95 | 2.34E-102 | 4.87E-98 | 0.29 | 0.02 | 0.27 | 13.36 |
| other bloold cells | Nagpa | 3.02 | 4.18E-102 | 8.69E-98 | 0.63 | 0.1 | 0.53 | 6.48 |
| other bloold cells | Stat6 | 3.06 | 1.99E-101 | 4.14E-97 | 0.47 | 0.05 | 0.42 | 8.85 |
| other bloold cells | 610203C22Ri | 3.93 | 2.24E-101 | 4.66E-97 | 0.22 | 0.01 | 0.21 | 18.08 |
| other bloold cells | Guca1a | 3.45 | 2.79E-101 | 5.80E-97 | 0.36 | 0.03 | 0.33 | 11.03 |
| other bloold cells | Sec11c | 3.06 | 3.90E-101 | 8.10E-97 | 0.92 | 0.3 | 0.63 | 3.1 |
| other bloold cells | Cotl1 | 3.22 | 6.56E-100 | 1.36E-95 | 1 | 0.44 | 0.56 | 2.26 |
| other bloold cells | Gm36738 | 3.82 | 2.36E-99 | 4.90E-95 | 0.22 | 0.01 | 0.21 | 18.08 |
| other bloold cells | Zbtb7b | 3.97 | 2.54E-99 | 5.28E-95 | 0.27 | 0.02 | 0.25 | 14.37 |
| other bloold cells | B2m | 3.75 | 2.11E-98 | 4.40E-94 | 1 | 0.54 | 0.46 | 1.86 |
| other bloold cells | Sh2b3 | 3.11 | 3.45E-98 | 7.17E-94 | 0.54 | 0.07 | 0.47 | 7.37 |
| other bloold cells | Gusb | 3.21 | 3.45E-98 | 7.18E-94 | 0.92 | 0.3 | 0.62 | 3.04 |
| other bloold cells | Nr4a1 | 3.74 | 4.47E-98 | 9.30E-94 | 0.4 | 0.04 | 0.36 | 9.73 |
| other bloold cells | Khk | 3.14 | 9.71E-98 | 2.02E-93 | 0.5 | 0.06 | 0.44 | 7.86 |
| other bloold cells | Anxa6 | 2.78 | 1.27E-97 | 2.65E-93 | 0.8 | 0.17 | 0.63 | 4.69 |
| other bloold cells | Arhgdib | 3.27 | 2.46E-97 | 5.12E-93 | 0.99 | 0.5 | 0.5 | 1.99 |
| other bloold cells | Tmem229b | 3.51 | 8.30E-97 | 1.73E-92 | 0.3 | 0.02 | 0.28 | 13.09 |
| other bloold cells | Pon3 | 3.5 | 1.32E-96 | 2.75E-92 | 0.34 | 0.03 | 0.31 | 11.06 |
| other bloold cells | Adam15 | 3.02 | 2.29E-96 | 4.76E-92 | 0.61 | 0.1 | 0.51 | 6.4 |
| other bloold cells | Ckb | 4.27 | 2.79E-96 | 5.81E-92 | 0.94 | 0.33 | 0.61 | 2.86 |
| other bloold cells | Fgd2 | 3.01 | 4.98E-96 | 1.04E-91 | 0.62 | 0.1 | 0.52 | 6.28 |
| other bloold cells | Nabp1 | 3.81 | 5.28E-96 | 1.10E-91 | 0.33 | 0.03 | 0.3 | 11.34 |
| other bloold cells | Ubash3a | 4.8 | 1.96E-95 | 4.07E-91 | 0.15 | 0.01 | 0.14 | 24.5 |
| other bloold cells | Hexa | 3.76 | 6.13E-95 | 1.27E-90 | 0.92 | 0.31 | 0.6 | 2.93 |
| other bloold cells | Mvp | 3.26 | 7.87E-95 | 1.64E-90 | 0.48 | 0.06 | 0.42 | 8.07 |
| other bloold cells | Ucp2 | 3.04 | 2.60E-94 | 5.41E-90 | 0.97 | 0.35 | 0.61 | 2.73 |
| other bloold cells | Sipa1 | 2.6 | 8.35E-94 | 1.74E-89 | 0.8 | 0.17 | 0.63 | 4.65 |
| other bloold cells | Hebp1 | 2.74 | 5.08E-93 | 1.06E-88 | 0.93 | 0.26 | 0.67 | 3.52 |
| other bloold cells | Stard8 | 3.64 | 8.30E-93 | 1.73E-88 | 0.56 | 0.09 | 0.47 | 6.5 |
| other bloold cells | Daglb | 3.05 | 2.56E-92 | 5.33E-88 | 0.63 | 0.11 | 0.52 | 5.93 |
| other bloold cells | Arrb2 | 2.81 | 2.87E-92 | 5.96E-88 | 0.78 | 0.17 | 0.61 | 4.62 |
| other bloold cells | Klf6 | 3.44 | 5.04E-92 | 1.05E-87 | 0.85 | 0.23 | 0.62 | 3.68 |
| other bloold cells | Ninj1 | 3.12 | 6.29E-92 | 1.31E-87 | 0.8 | 0.19 | 0.62 | 4.25 |
| other bloold cells | Abcd1 | 3.06 | 7.98E-92 | 1.66E-87 | 0.48 | 0.06 | 0.42 | 7.79 |
| other bloold cells | Lpl | 4.65 | 1.30E-91 | 2.71E-87 | 0.71 | 0.16 | 0.55 | 4.5 |

**Supplementary Table 2. Marker genes for the major cell types of integrated clustering of E11.5 control and NotchGOF λ-junction cells.**

| | | | | | | | | |
|---|---|---|---|---|---|---|---|---|
| other bloold cells | Ap1g2 | 3.14 | 2.12E-91 | 4.41E-87 | 0.39 | 0.04 | 0.35 | 9.63 |
| other bloold cells | Gimap1 | 3.5 | 5.81E-91 | 1.21E-86 | 0.24 | 0.02 | 0.22 | 14.88 |
| other bloold cells | Ifnar2 | 2.9 | 8.06E-91 | 1.68E-86 | 0.94 | 0.33 | 0.62 | 2.88 |
| other bloold cells | Casp8 | 2.68 | 9.26E-91 | 1.93E-86 | 0.8 | 0.18 | 0.62 | 4.4 |
| other bloold cells | Enpp1 | 4.91 | 6.88E-90 | 1.43E-85 | 0.14 | 0.01 | 0.14 | 28 |
| other bloold cells | Tmsb4x | 3.78 | 1.28E-89 | 2.66E-85 | 1 | 0.99 | 0.01 | 1.01 |
| other bloold cells | Stat3 | 2.73 | 6.59E-88 | 1.37E-83 | 0.69 | 0.13 | 0.56 | 5.31 |
| other bloold cells | Tfeb | 3.43 | 6.95E-88 | 1.45E-83 | 0.28 | 0.02 | 0.26 | 12.73 |
| other bloold cells | Mt1 | 2.98 | 9.78E-88 | 2.03E-83 | 0.9 | 0.27 | 0.64 | 3.4 |
| other bloold cells | E030030I06Ril | 4.97 | 1.14E-87 | 2.36E-83 | 0.13 | 0 | 0.12 | 31.5 |
| other bloold cells | Rasa3 | 3.19 | 1.77E-87 | 3.68E-83 | 0.52 | 0.08 | 0.44 | 6.89 |
| other bloold cells | Padi2 | 5.96 | 4.03E-87 | 8.39E-83 | 0.11 | 0 | 0.1 | 35 |
| other bloold cells | Creg1 | 2.88 | 1.71E-86 | 3.55E-82 | 0.94 | 0.35 | 0.59 | 2.68 |
| other bloold cells | Sh3bgrl3 | 2.21 | 1.76E-86 | 3.67E-82 | 1 | 0.91 | 0.09 | 1.1 |
| other bloold cells | Fam78a | 4.34 | 2.62E-86 | 5.44E-82 | 0.2 | 0.01 | 0.19 | 16.92 |
| other bloold cells | Tagln2 | 3.2 | 1.45E-85 | 3.02E-81 | 0.88 | 0.3 | 0.59 | 2.98 |
| other bloold cells | Lrmda | 4.06 | 1.49E-85 | 3.09E-81 | 0.18 | 0.01 | 0.17 | 19.44 |
| other bloold cells | Rasgrp2 | 2.66 | 1.85E-85 | 3.84E-81 | 0.41 | 0.05 | 0.36 | 8.43 |
| other bloold cells | Nceh1 | 3.44 | 3.84E-85 | 7.98E-81 | 0.41 | 0.05 | 0.36 | 8.26 |
| other bloold cells | Naaa | 3.73 | 5.40E-85 | 1.12E-80 | 0.27 | 0.02 | 0.25 | 12.67 |
| other bloold cells | Ifi30 | 3.82 | 1.53E-84 | 3.19E-80 | 0.87 | 0.29 | 0.58 | 2.99 |
| other bloold cells | Serpinb6b | 3.57 | 2.33E-84 | 4.84E-80 | 0.27 | 0.02 | 0.25 | 12.67 |
| other bloold cells | Acss1 | 3.52 | 2.50E-84 | 5.19E-80 | 0.28 | 0.02 | 0.26 | 12.17 |
| other bloold cells | Egr3 | 5.08 | 1.33E-83 | 2.77E-79 | 0.11 | 0 | 0.1 | 35 |
| other bloold cells | Ppp1r18 | 2.57 | 1.45E-83 | 3.02E-79 | 0.88 | 0.25 | 0.63 | 3.55 |
| other bloold cells | Mtus1 | 3.12 | 2.66E-83 | 5.52E-79 | 0.59 | 0.1 | 0.48 | 5.7 |
| other bloold cells | Arpc2 | 1.89 | 3.75E-83 | 7.81E-79 | 1 | 0.94 | 0.06 | 1.06 |
| other bloold cells | Lacc1 | 3.21 | 6.48E-83 | 1.35E-78 | 0.54 | 0.09 | 0.45 | 6.18 |
| other bloold cells | Apoe | 7.13 | 3.75E-82 | 7.79E-78 | 0.94 | 0.43 | 0.51 | 2.18 |
| other bloold cells | Sqor | 4.02 | 5.28E-82 | 1.10E-77 | 0.25 | 0.02 | 0.23 | 12.89 |
| other bloold cells | Galns | 3.33 | 8.46E-81 | 1.76E-76 | 0.32 | 0.03 | 0.29 | 10.06 |
| other bloold cells | Ctsz | 3.38 | 1.02E-80 | 2.12E-76 | 0.99 | 0.58 | 0.41 | 1.71 |
| other bloold cells | Tmem9b | 2.58 | 1.63E-80 | 3.39E-76 | 0.69 | 0.14 | 0.55 | 5.04 |
| other bloold cells | Rgs7bp | 3.48 | 1.87E-80 | 3.88E-76 | 0.27 | 0.02 | 0.24 | 12.09 |
| other bloold cells | Gbp9 | 4.72 | 2.39E-80 | 4.97E-76 | 0.11 | 0 | 0.1 | 35 |
| other bloold cells | Tbc1d8 | 3.17 | 5.77E-80 | 1.20E-75 | 0.32 | 0.03 | 0.28 | 10.16 |
| other bloold cells | Rnf144b | 4.53 | 4.48E-79 | 9.32E-75 | 0.15 | 0.01 | 0.14 | 21 |
| other bloold cells | Arpc5 | 2.37 | 5.51E-79 | 1.15E-74 | 0.99 | 0.76 | 0.23 | 1.3 |
| other bloold cells | Mctp1 | 4.25 | 8.34E-79 | 1.74E-74 | 0.15 | 0.01 | 0.14 | 21 |
| other bloold cells | Mbnl1 | 2.65 | 1.34E-78 | 2.78E-74 | 0.97 | 0.47 | 0.5 | 2.07 |
| other bloold cells | Cst3 | 3.22 | 1.78E-78 | 3.70E-74 | 1 | 0.92 | 0.08 | 1.09 |
| other bloold cells | Tmcc3 | 3.35 | 3.38E-78 | 7.02E-74 | 0.32 | 0.03 | 0.29 | 9.76 |
| other bloold cells | Prkcd | 2.76 | 6.09E-78 | 1.27E-73 | 0.8 | 0.21 | 0.59 | 3.8 |
| other bloold cells | Ly6e | 3.25 | 9.30E-78 | 1.93E-73 | 0.94 | 0.45 | 0.49 | 2.09 |
| other bloold cells | Tapbp | 2.84 | 1.02E-77 | 2.12E-73 | 0.51 | 0.08 | 0.43 | 6.38 |
| other bloold cells | Arid3a | 2.64 | 2.37E-77 | 4.93E-73 | 0.9 | 0.33 | 0.57 | 2.73 |
| other bloold cells | S100a1 | 2.99 | 5.01E-77 | 1.04E-72 | 0.45 | 0.06 | 0.39 | 7.11 |
| other bloold cells | Cebpd | 2.95 | 5.18E-77 | 1.08E-72 | 0.56 | 0.1 | 0.46 | 5.76 |
| other bloold cells | Ulbp1 | 3.43 | 8.33E-77 | 1.73E-72 | 0.27 | 0.02 | 0.24 | 11.57 |
| other bloold cells | Nbeal2 | 4.1 | 9.76E-77 | 2.03E-72 | 0.15 | 0.01 | 0.15 | 19.25 |
| other bloold cells | Btg2 | 2.98 | 1.30E-76 | 2.70E-72 | 0.85 | 0.27 | 0.58 | 3.12 |
| other bloold cells | Hprt | 2.54 | 1.32E-76 | 2.74E-72 | 0.94 | 0.43 | 0.5 | 2.16 |
| other bloold cells | Cmtm7 | 2.85 | 1.41E-76 | 2.94E-72 | 0.97 | 0.48 | 0.48 | 2 |
| other bloold cells | Capg | 2.96 | 1.54E-76 | 3.21E-72 | 0.69 | 0.15 | 0.54 | 4.6 |
| other bloold cells | Gpx1 | 1.77 | 1.69E-76 | 3.51E-72 | 1 | 0.95 | 0.05 | 1.05 |
| other bloold cells | Fcgrt | 3.73 | 3.23E-76 | 6.72E-72 | 0.65 | 0.14 | 0.51 | 4.61 |

**Supplementary Table 2. Marker genes for the major cell types of integrated clustering of E11.5 control and NotchGOF λ-junction cells.**

| | | | | | | | | |
|---|---|---|---|---|---|---|---|---|
| other bloold cells | Gcnt1 | 3.62 | 3.23E-76 | 6.72E-72 | 0.3 | 0.03 | 0.27 | 10.03 |
| other bloold cells | Tacc1 | 2.42 | 3.41E-76 | 7.09E-72 | 0.9 | 0.32 | 0.58 | 2.84 |
| other bloold cells | Ctsd | 4.5 | 4.31E-76 | 8.95E-72 | 0.99 | 0.61 | 0.39 | 1.64 |
| other bloold cells | Fam111a | 2.71 | 4.94E-76 | 1.03E-71 | 0.9 | 0.32 | 0.57 | 2.77 |
| other bloold cells | Tnip2 | 3.11 | 5.09E-76 | 1.06E-71 | 0.29 | 0.03 | 0.26 | 10.63 |
| other bloold cells | Actr3 | 1.98 | 5.60E-76 | 1.16E-71 | 0.99 | 0.82 | 0.18 | 1.22 |
| other bloold cells | Atp13a2 | 2.52 | 7.95E-76 | 1.65E-71 | 0.86 | 0.26 | 0.61 | 3.37 |
| other bloold cells | Pld1 | 3.62 | 1.07E-75 | 2.22E-71 | 0.24 | 0.02 | 0.22 | 12.53 |
| other bloold cells | Ccrl2 | 4.83 | 1.79E-75 | 3.73E-71 | 0.18 | 0.01 | 0.17 | 16.55 |
| other bloold cells | Erp29 | 2.07 | 1.93E-75 | 4.02E-71 | 0.99 | 0.88 | 0.11 | 1.13 |
| other bloold cells | Rhog | 2.56 | 2.27E-75 | 4.72E-71 | 0.93 | 0.35 | 0.58 | 2.66 |
| other bloold cells | Arpc4 | 1.82 | 4.05E-75 | 8.42E-71 | 1 | 0.83 | 0.18 | 1.21 |
| other bloold cells | Mknk1 | 2.67 | 9.58E-75 | 1.99E-70 | 0.54 | 0.09 | 0.45 | 5.98 |
| other bloold cells | Lgmn | 5.05 | 1.08E-74 | 2.25E-70 | 0.8 | 0.29 | 0.52 | 2.82 |
| other bloold cells | Actb | 1.5 | 1.28E-74 | 2.67E-70 | 1 | 1 | 0 | 1 |
| other bloold cells | Tubb2a | 2.9 | 1.44E-74 | 2.99E-70 | 0.9 | 0.3 | 0.6 | 3.03 |
| other bloold cells | Rps6ka1 | 2.58 | 3.89E-74 | 8.10E-70 | 0.82 | 0.24 | 0.58 | 3.41 |
| other bloold cells | Clic1 | 1.71 | 6.42E-74 | 1.33E-69 | 1 | 0.93 | 0.07 | 1.08 |
| other bloold cells | Kctd12 | 2.62 | 8.30E-74 | 1.73E-69 | 0.89 | 0.3 | 0.59 | 3.01 |
| other bloold cells | Hmox1 | 4.94 | 1.37E-73 | 2.86E-69 | 0.72 | 0.19 | 0.53 | 3.85 |
| other bloold cells | Tnnt1 | 2.72 | 6.42E-73 | 1.33E-68 | 0.4 | 0.05 | 0.35 | 7.67 |
| other bloold cells | Rftn1 | 3.03 | 7.78E-73 | 1.62E-68 | 0.34 | 0.04 | 0.3 | 8.84 |
| other bloold cells | Pdgfb | 3.64 | 7.85E-73 | 1.63E-68 | 0.2 | 0.01 | 0.19 | 14.5 |
| other bloold cells | Clta | 1.71 | 1.83E-72 | 3.81E-68 | 1 | 0.98 | 0.02 | 1.02 |
| other bloold cells | Exoc6 | 2.82 | 2.36E-72 | 4.90E-68 | 0.47 | 0.07 | 0.4 | 6.42 |
| other bloold cells | Fth1 | 2.31 | 5.36E-72 | 1.11E-67 | 1 | 1 | 0 | 1 |
| other bloold cells | Pold4 | 2.45 | 8.05E-72 | 1.67E-67 | 0.75 | 0.18 | 0.56 | 4.07 |
| other bloold cells | Tmem104 | 3.46 | 1.77E-71 | 3.69E-67 | 0.27 | 0.03 | 0.24 | 10.64 |
| other bloold cells | Itpr2 | 3.11 | 2.88E-71 | 6.00E-67 | 0.36 | 0.05 | 0.32 | 8.09 |
| other bloold cells | Mpp1 | 2.87 | 4.73E-71 | 9.83E-67 | 0.8 | 0.24 | 0.56 | 3.36 |
| other bloold cells | Ednrb | 3.65 | 5.91E-71 | 1.23E-66 | 0.33 | 0.04 | 0.29 | 8.66 |
| other bloold cells | Irak4 | 2.72 | 1.69E-70 | 3.51E-66 | 0.37 | 0.05 | 0.33 | 8.07 |
| other bloold cells | Arap3 | 2.8 | 1.75E-70 | 3.63E-66 | 0.36 | 0.04 | 0.31 | 8.3 |
| other bloold cells | Pfn1 | 1.17 | 3.34E-70 | 6.95E-66 | 0.99 | 1 | 0 | 1 |
| other bloold cells | Gnai2 | 1.48 | 3.97E-70 | 8.26E-66 | 1 | 0.97 | 0.03 | 1.03 |
| other bloold cells | Itm2b | 2.83 | 4.84E-70 | 1.01E-65 | 1 | 0.84 | 0.16 | 1.19 |
| other bloold cells | Ctsa | 2.83 | 5.89E-70 | 1.23E-65 | 0.94 | 0.44 | 0.51 | 2.17 |
| other bloold cells | Iqgap1 | 2.25 | 1.54E-69 | 3.21E-65 | 0.97 | 0.47 | 0.49 | 2.05 |
| other bloold cells | Slc9a9 | 3.4 | 1.81E-69 | 3.75E-65 | 0.32 | 0.04 | 0.29 | 8.7 |
| other bloold cells | Acsl5 | 2.59 | 3.85E-69 | 8.00E-65 | 0.68 | 0.16 | 0.52 | 4.35 |
| other bloold cells | Atp8a1 | 2.69 | 5.48E-69 | 1.14E-64 | 0.5 | 0.09 | 0.42 | 5.92 |
| other bloold cells | Vamp8 | 2.14 | 5.65E-69 | 1.18E-64 | 0.97 | 0.61 | 0.36 | 1.59 |
| other bloold cells | Scamp2 | 2.43 | 5.71E-69 | 1.19E-64 | 0.89 | 0.31 | 0.58 | 2.85 |
| other bloold cells | Slfn5 | 5.18 | 1.24E-68 | 2.59E-64 | 0.15 | 0.01 | 0.15 | 17.11 |
| other bloold cells | Aldh3b1 | 4.58 | 1.27E-68 | 2.65E-64 | 0.11 | 0 | 0.1 | 26.25 |
| other bloold cells | Orai1 | 2.35 | 3.43E-68 | 7.13E-64 | 0.71 | 0.17 | 0.54 | 4.28 |
| other bloold cells | Arpc3 | 1.57 | 6.03E-68 | 1.25E-63 | 0.99 | 0.9 | 0.09 | 1.1 |
| other bloold cells | Gm16096 | 4.68 | 6.27E-68 | 1.30E-63 | 0.11 | 0 | 0.1 | 26.25 |
| other bloold cells | Icos | 4.1 | 2.87E-67 | 5.97E-63 | 0.15 | 0.01 | 0.14 | 18.38 |
| other bloold cells | Car9 | 3.97 | 3.96E-67 | 8.23E-63 | 0.15 | 0.01 | 0.14 | 18.38 |
| other bloold cells | Gpr157 | 3.98 | 8.23E-67 | 1.71E-62 | 0.18 | 0.01 | 0.16 | 14.58 |
| other bloold cells | Msn | 2.15 | 1.08E-66 | 2.24E-62 | 0.98 | 0.51 | 0.47 | 1.91 |
| other bloold cells | Entpd1 | 3.27 | 2.50E-66 | 5.20E-62 | 0.41 | 0.06 | 0.35 | 6.88 |
| other bloold cells | Ptgs2 | 3.75 | 2.66E-66 | 5.53E-62 | 0.13 | 0.01 | 0.12 | 21 |
| other bloold cells | Dusp3 | 2.65 | 5.12E-66 | 1.07E-61 | 0.39 | 0.06 | 0.34 | 7.13 |
| other bloold cells | Scamp5 | 3.29 | 6.20E-66 | 1.29E-61 | 0.34 | 0.04 | 0.29 | 8 |

**Supplementary Table 2. Marker genes for the major cell types of integrated clustering of E11.5 control and NotchGOF λ-junction cells.**

| | | | | | | | | |
|---|---|---|---|---|---|---|---|---|
| other bloold cells | Capza2 | 1.72 | 8.42E-66 | 1.75E-61 | 1 | 0.86 | 0.14 | 1.16 |
| other bloold cells | Rel | 2.45 | 1.23E-65 | 2.55E-61 | 0.5 | 0.09 | 0.42 | 5.72 |
| other bloold cells | Bhlhe41 | 4.07 | 1.39E-65 | 2.90E-61 | 0.18 | 0.01 | 0.16 | 14.58 |
| other bloold cells | Atp6v0b | 1.95 | 1.71E-65 | 3.55E-61 | 0.98 | 0.74 | 0.24 | 1.33 |
| other bloold cells | Susd1 | 3.83 | 1.97E-65 | 4.09E-61 | 0.19 | 0.01 | 0.18 | 13.5 |
| other bloold cells | Vwa5a | 2.71 | 4.37E-65 | 9.09E-61 | 0.49 | 0.09 | 0.4 | 5.7 |
| other bloold cells | Rgs2 | 2.55 | 5.62E-65 | 1.17E-60 | 0.89 | 0.34 | 0.55 | 2.6 |
| other bloold cells | Tln1 | 1.73 | 1.22E-64 | 2.54E-60 | 0.99 | 0.74 | 0.25 | 1.33 |
| other bloold cells | Trim12a | 3.08 | 1.48E-64 | 3.08E-60 | 0.24 | 0.02 | 0.22 | 10.82 |
| other bloold cells | Pon2 | 2.03 | 1.03E-63 | 2.14E-59 | 0.72 | 0.18 | 0.54 | 4.02 |
| other bloold cells | Psap | 2.97 | 1.11E-63 | 2.31E-59 | 0.97 | 0.62 | 0.35 | 1.56 |
| other bloold cells | Nfatc2 | 3.91 | 1.39E-63 | 2.89E-59 | 0.19 | 0.01 | 0.18 | 13.5 |
| other bloold cells | Camk2d | 2.6 | 2.36E-63 | 4.92E-59 | 0.71 | 0.19 | 0.52 | 3.68 |
| other bloold cells | Itsn2 | 2.29 | 7.59E-63 | 1.58E-58 | 0.71 | 0.19 | 0.52 | 3.77 |
| other bloold cells | Grn | 3.86 | 1.34E-62 | 2.80E-58 | 0.84 | 0.35 | 0.49 | 2.4 |
| other bloold cells | Zc3h12a | 4.04 | 2.23E-62 | 4.63E-58 | 0.15 | 0.01 | 0.14 | 15.4 |
| other bloold cells | P2ry14 | 3.57 | 2.69E-62 | 5.59E-58 | 0.24 | 0.02 | 0.22 | 10.35 |
| other bloold cells | Elf1 | 2.21 | 2.96E-62 | 6.16E-58 | 0.82 | 0.26 | 0.56 | 3.13 |
| other bloold cells | Nup210 | 2.76 | 3.00E-62 | 6.23E-58 | 0.4 | 0.06 | 0.34 | 6.65 |
| other bloold cells | Fkbp15 | 2.57 | 4.34E-62 | 9.02E-58 | 0.6 | 0.13 | 0.47 | 4.59 |
| other bloold cells | N4bp2l1 | 2.87 | 4.60E-62 | 9.57E-58 | 0.29 | 0.03 | 0.26 | 8.91 |
| other bloold cells | H2afj | 1.62 | 9.13E-62 | 1.90E-57 | 0.99 | 0.81 | 0.18 | 1.22 |
| other bloold cells | Plaur | 4.17 | 4.45E-61 | 9.25E-57 | 0.14 | 0.01 | 0.13 | 17.5 |
| other bloold cells | Psme2b | 4.01 | 4.89E-61 | 1.02E-56 | 0.13 | 0.01 | 0.13 | 19 |
| other bloold cells | Tpp1 | 2.71 | 5.89E-61 | 1.23E-56 | 0.8 | 0.26 | 0.53 | 3.03 |
| other bloold cells | Arl6ip5 | 2.15 | 6.67E-61 | 1.39E-56 | 0.87 | 0.32 | 0.55 | 2.73 |
| other bloold cells | Disc1 | 3.42 | 6.70E-61 | 1.39E-56 | 0.2 | 0.02 | 0.19 | 11.94 |
| other bloold cells | Edem1 | 2.33 | 6.97E-61 | 1.45E-56 | 0.79 | 0.24 | 0.55 | 3.29 |
| other bloold cells | Cpeb2 | 3.51 | 8.01E-61 | 1.66E-56 | 0.2 | 0.02 | 0.18 | 12.25 |
| other bloold cells | Slc48a1 | 2.58 | 8.34E-61 | 1.74E-56 | 0.66 | 0.17 | 0.49 | 3.98 |
| other bloold cells | Abcd2 | 2.74 | 9.59E-61 | 1.99E-56 | 0.43 | 0.07 | 0.36 | 6.03 |
| other bloold cells | Gm13212 | 3.05 | 1.25E-60 | 2.60E-56 | 0.26 | 0.03 | 0.23 | 9.59 |
| other bloold cells | Blvra | 2.32 | 1.25E-60 | 2.61E-56 | 0.73 | 0.2 | 0.53 | 3.6 |
| other bloold cells | Myo5a | 2.45 | 2.22E-60 | 4.62E-56 | 0.87 | 0.32 | 0.56 | 2.74 |
| other bloold cells | Ptger3 | 3.35 | 3.24E-60 | 6.73E-56 | 0.17 | 0.01 | 0.16 | 14 |
| other bloold cells | Itga6 | 1.97 | 3.89E-60 | 8.09E-56 | 0.91 | 0.31 | 0.6 | 2.93 |
| other bloold cells | Sh3kbp1 | 2.25 | 3.91E-60 | 8.14E-56 | 0.77 | 0.22 | 0.55 | 3.48 |
| other bloold cells | Fam49b | 1.74 | 4.72E-60 | 9.83E-56 | 0.99 | 0.65 | 0.33 | 1.51 |
| other bloold cells | Klk8 | 2.34 | 5.38E-60 | 1.12E-55 | 0.74 | 0.21 | 0.53 | 3.48 |
| other bloold cells | Rogdi | 2.51 | 5.51E-60 | 1.15E-55 | 0.76 | 0.25 | 0.52 | 3.1 |
| other bloold cells | Madd | 2.5 | 5.92E-60 | 1.23E-55 | 0.43 | 0.07 | 0.36 | 6.11 |
| other bloold cells | Glrx | 2.32 | 1.14E-59 | 2.38E-55 | 0.76 | 0.23 | 0.53 | 3.27 |
| other bloold cells | Stk17b | 2.75 | 1.84E-59 | 3.82E-55 | 0.58 | 0.13 | 0.45 | 4.36 |
| other bloold cells | Ssr4 | 1.46 | 2.16E-59 | 4.49E-55 | 0.99 | 0.84 | 0.15 | 1.18 |
| other bloold cells | Rab27a | 2.94 | 2.84E-59 | 5.91E-55 | 0.34 | 0.05 | 0.3 | 7.3 |
| other bloold cells | Dtx3l | 3.15 | 4.71E-59 | 9.80E-55 | 0.32 | 0.04 | 0.28 | 7.88 |
| other bloold cells | Tcirg1 | 2.63 | 5.02E-59 | 1.04E-54 | 0.48 | 0.09 | 0.39 | 5.47 |
| other bloold cells | Rap1b | 1.75 | 5.94E-59 | 1.24E-54 | 0.99 | 0.76 | 0.23 | 1.3 |
| other bloold cells | Tep1 | 2.75 | 7.25E-59 | 1.51E-54 | 0.38 | 0.06 | 0.32 | 6.63 |
| other bloold cells | Gba | 2.54 | 8.17E-59 | 1.70E-54 | 0.5 | 0.09 | 0.4 | 5.29 |
| other bloold cells | Zeb2 | 2.26 | 9.62E-59 | 2.00E-54 | 0.92 | 0.4 | 0.52 | 2.3 |
| other bloold cells | Pygl | 2.21 | 1.51E-58 | 3.14E-54 | 0.51 | 0.1 | 0.41 | 5.31 |
| other bloold cells | Slc8b1 | 3.05 | 1.55E-58 | 3.23E-54 | 0.24 | 0.02 | 0.21 | 9.92 |
| other bloold cells | Csf2ra | 2.46 | 1.76E-58 | 3.66E-54 | 0.6 | 0.14 | 0.46 | 4.26 |
| other bloold cells | Rap1a | 1.65 | 2.55E-58 | 5.31E-54 | 0.99 | 0.72 | 0.27 | 1.38 |
| other bloold cells | 600014C10Ri | 3.05 | 6.84E-58 | 1.42E-53 | 0.54 | 0.11 | 0.43 | 4.76 |

**Supplementary Table 2. Marker genes for the major cell types of integrated clustering of E11.5 control and NotchGOF λ-junction cells.**

| | | | | | | | | |
|---|---|---|---|---|---|---|---|---|
| other bloold cells | Lamp1 | 2.03 | 1.16E-57 | 2.42E-53 | 0.98 | 0.86 | 0.12 | 1.14 |
| other bloold cells | Tmc8 | 5.55 | 2.46E-57 | 5.12E-53 | 0.12 | 0.01 | 0.11 | 19.83 |
| other bloold cells | Runx3 | 4.27 | 2.71E-57 | 5.63E-53 | 0.2 | 0.02 | 0.18 | 11.53 |
| other bloold cells | Lrch4 | 2.51 | 5.00E-57 | 1.04E-52 | 0.64 | 0.16 | 0.48 | 3.98 |
| other bloold cells | Snx5 | 1.78 | 5.11E-57 | 1.06E-52 | 0.98 | 0.78 | 0.2 | 1.26 |
| other bloold cells | Prps2 | 2.11 | 5.60E-57 | 1.16E-52 | 0.66 | 0.17 | 0.5 | 3.98 |
| other bloold cells | Plin2 | 3.23 | 5.87E-57 | 1.22E-52 | 0.94 | 0.53 | 0.41 | 1.77 |
| other bloold cells | Rnf130 | 1.76 | 6.05E-57 | 1.26E-52 | 0.97 | 0.56 | 0.41 | 1.73 |
| other bloold cells | Slc8a1 | 2.45 | 9.91E-57 | 2.06E-52 | 0.56 | 0.12 | 0.44 | 4.62 |
| other bloold cells | Twf2 | 1.97 | 1.91E-56 | 3.97E-52 | 0.9 | 0.42 | 0.47 | 2.12 |
| other bloold cells | .330040F15Ri | 3.68 | 2.61E-56 | 5.43E-52 | 0.15 | 0.01 | 0.14 | 14.7 |
| other bloold cells | Dusp27 | 4.16 | 2.85E-56 | 5.93E-52 | 0.13 | 0.01 | 0.12 | 18 |
| other bloold cells | BC028528 | 2.42 | 4.03E-56 | 8.37E-52 | 0.43 | 0.08 | 0.36 | 5.79 |
| other bloold cells | Spaca9 | 3.19 | 8.17E-56 | 1.70E-51 | 0.27 | 0.03 | 0.24 | 8.53 |
| other bloold cells | Stx7 | 2.1 | 8.31E-56 | 1.73E-51 | 0.86 | 0.31 | 0.55 | 2.78 |
| other bloold cells | Grb2 | 1.72 | 1.19E-55 | 2.48E-51 | 0.99 | 0.64 | 0.35 | 1.55 |
| other bloold cells | Rnpep | 2.11 | 1.29E-55 | 2.69E-51 | 0.92 | 0.43 | 0.48 | 2.11 |
| other bloold cells | Hacd4 | 2.65 | 1.36E-55 | 2.84E-51 | 0.75 | 0.25 | 0.5 | 3.03 |
| other bloold cells | Flt3l | 4.26 | 2.53E-55 | 5.25E-51 | 0.16 | 0.01 | 0.15 | 13.42 |
| other bloold cells | Atp6v0e | 1.51 | 2.75E-55 | 5.71E-51 | 0.98 | 0.72 | 0.26 | 1.37 |
| other bloold cells | Rab8b | 2.16 | 8.22E-55 | 1.71E-50 | 0.77 | 0.24 | 0.53 | 3.2 |
| other bloold cells | Cd47 | 2.36 | 8.97E-55 | 1.87E-50 | 0.92 | 0.44 | 0.48 | 2.09 |
| other bloold cells | H2-Oa | 4.24 | 2.06E-54 | 4.27E-50 | 0.12 | 0.01 | 0.11 | 17 |
| other bloold cells | Tpcn1 | 2.36 | 2.41E-54 | 5.02E-50 | 0.47 | 0.09 | 0.38 | 5.33 |
| other bloold cells | Fgd4 | 2.5 | 3.13E-54 | 6.51E-50 | 0.46 | 0.09 | 0.37 | 5.25 |
| other bloold cells | Igtp | 3.73 | 3.45E-54 | 7.18E-50 | 0.2 | 0.02 | 0.18 | 10.68 |
| other bloold cells | Tap1 | 3.06 | 4.12E-54 | 8.57E-50 | 0.25 | 0.03 | 0.22 | 8.69 |
| other bloold cells | Grk2 | 1.93 | 9.26E-54 | 1.93E-49 | 0.85 | 0.33 | 0.52 | 2.6 |
| other bloold cells | Actr2 | 1.69 | 1.08E-53 | 2.24E-49 | 0.97 | 0.63 | 0.34 | 1.55 |
| other bloold cells | Ptbp3 | 1.59 | 1.20E-53 | 2.49E-49 | 0.97 | 0.62 | 0.35 | 1.57 |
| other bloold cells | S100a4 | 4.03 | 1.30E-53 | 2.71E-49 | 0.17 | 0.01 | 0.16 | 12.92 |
| other bloold cells | Lipa | 2.57 | 1.95E-53 | 4.05E-49 | 0.7 | 0.22 | 0.48 | 3.24 |
| other bloold cells | Sigirr | 2.88 | 2.07E-53 | 4.31E-49 | 0.24 | 0.03 | 0.21 | 9.15 |
| other bloold cells | Ripk1 | 2.38 | 2.89E-53 | 6.01E-49 | 0.55 | 0.12 | 0.42 | 4.43 |
| other bloold cells | Rab3il1 | 2.65 | 3.64E-53 | 7.58E-49 | 0.78 | 0.3 | 0.48 | 2.63 |
| other bloold cells | Gamt | 2.41 | 3.82E-53 | 7.94E-49 | 0.47 | 0.09 | 0.38 | 5.21 |
| other bloold cells | Pecam1 | 1.49 | 4.21E-53 | 8.75E-49 | 0.25 | 0.03 | 0.22 | 9 |
| other bloold cells | Cd36 | 4.04 | 7.53E-53 | 1.57E-48 | 0.15 | 0.01 | 0.14 | 14 |
| other bloold cells | Sh3bp1 | 1.94 | 1.23E-52 | 2.55E-48 | 0.8 | 0.29 | 0.52 | 2.81 |
| other bloold cells | Asah1 | 2.4 | 1.39E-52 | 2.88E-48 | 0.86 | 0.37 | 0.49 | 2.34 |
| other bloold cells | Parp4 | 3.3 | 1.41E-52 | 2.93E-48 | 0.2 | 0.02 | 0.18 | 10.89 |
| other bloold cells | Samhd1 | 2.11 | 2.47E-52 | 5.13E-48 | 0.88 | 0.39 | 0.5 | 2.28 |
| other bloold cells | Dram2 | 2.29 | 3.00E-52 | 6.23E-48 | 0.69 | 0.2 | 0.49 | 3.41 |
| other bloold cells | Ssh2 | 2.29 | 3.59E-52 | 7.46E-48 | 0.73 | 0.23 | 0.51 | 3.22 |
| other bloold cells | B3gnt8 | 3.53 | 3.76E-52 | 7.81E-48 | 0.18 | 0.02 | 0.16 | 11.67 |
| other bloold cells | Tfpi | 1.94 | 4.26E-52 | 8.87E-48 | 0.52 | 0.11 | 0.41 | 4.76 |
| other bloold cells | Abca3 | 2.64 | 4.44E-52 | 9.23E-48 | 0.38 | 0.06 | 0.32 | 6 |
| other bloold cells | Tspan4 | 2.23 | 6.93E-52 | 1.44E-47 | 0.55 | 0.13 | 0.43 | 4.42 |
| other bloold cells | Cap1 | 1.94 | 6.94E-52 | 1.44E-47 | 0.93 | 0.54 | 0.39 | 1.72 |
| other bloold cells | Psme1 | 1.5 | 1.12E-51 | 2.33E-47 | 0.98 | 0.76 | 0.22 | 1.29 |
| other bloold cells | Neu1 | 2.25 | 1.13E-51 | 2.34E-47 | 0.56 | 0.13 | 0.43 | 4.33 |
| other bloold cells | Tpst2 | 2.12 | 1.87E-51 | 3.90E-47 | 0.74 | 0.23 | 0.51 | 3.17 |
| other bloold cells | Abhd12 | 2.71 | 3.69E-51 | 7.67E-47 | 0.79 | 0.31 | 0.48 | 2.55 |
| other bloold cells | Anxa4 | 2.64 | 4.14E-51 | 8.61E-47 | 0.5 | 0.11 | 0.39 | 4.6 |
| other bloold cells | Slc25a5 | 1.26 | 8.51E-51 | 1.77E-46 | 1 | 0.93 | 0.07 | 1.07 |
| other bloold cells | Snx2 | 2.34 | 1.24E-50 | 2.57E-46 | 0.93 | 0.58 | 0.35 | 1.61 |

**Supplementary Table 2. Marker genes for the major cell types of integrated clustering of E11.5 control and NotchGOF λ-junction cells.**

| | | | | | | | | |
|---|---|---|---|---|---|---|---|---|
| other bloold cells | Icam2 | 1.07 | 1.63E-50 | 3.38E-46 | 0.2 | 0.02 | 0.18 | 10.15 |
| other bloold cells | Mfng | 1.45 | 4.43E-50 | 9.21E-46 | 0.32 | 0.05 | 0.27 | 7 |
| other bloold cells | Ntpcr | 2.36 | 1.14E-49 | 2.37E-45 | 0.71 | 0.23 | 0.48 | 3.14 |
| other bloold cells | Cyp4f13 | 3.74 | 1.17E-49 | 2.44E-45 | 0.15 | 0.01 | 0.14 | 13.36 |
| other bloold cells | Rplp1 | 0.58 | 1.64E-49 | 3.41E-45 | 1 | 1 | 0 | 1 |
| other bloold cells | Coro7 | 2.15 | 3.89E-49 | 8.10E-45 | 0.49 | 0.1 | 0.39 | 4.71 |
| other bloold cells | Arhgap27 | 2.66 | 3.91E-49 | 8.12E-45 | 0.25 | 0.03 | 0.22 | 8.13 |
| other bloold cells | Hps3 | 2.55 | 4.17E-49 | 8.68E-45 | 0.42 | 0.08 | 0.34 | 5.19 |
| other bloold cells | Ccdc69 | 3.64 | 4.99E-49 | 1.04E-44 | 0.11 | 0.01 | 0.1 | 17.5 |
| other bloold cells | Pros1 | 2.14 | 7.98E-49 | 1.66E-44 | 0.58 | 0.14 | 0.44 | 4.06 |
| other bloold cells | Tmem63a | 3.27 | 9.35E-49 | 1.94E-44 | 0.22 | 0.03 | 0.2 | 8.96 |
| other bloold cells | Tdrd7 | 3.02 | 4.03E-48 | 8.39E-44 | 0.23 | 0.03 | 0.2 | 8.56 |
| other bloold cells | Lgals8 | 2.32 | 4.92E-48 | 1.02E-43 | 0.43 | 0.09 | 0.35 | 5.11 |
| other bloold cells | Irf9 | 2.49 | 7.21E-48 | 1.50E-43 | 0.37 | 0.06 | 0.31 | 5.8 |
| other bloold cells | Rtn4rl1 | 4.49 | 7.76E-48 | 1.61E-43 | 0.11 | 0.01 | 0.11 | 16 |
| other bloold cells | Tal1 | 2.29 | 8.40E-48 | 1.75E-43 | 0.32 | 0.05 | 0.27 | 6.7 |
| other bloold cells | Gpx3 | 2.5 | 8.89E-48 | 1.85E-43 | 0.69 | 0.22 | 0.47 | 3.17 |
| other bloold cells | Ncoa3 | 1.94 | 1.85E-47 | 3.86E-43 | 0.71 | 0.23 | 0.49 | 3.17 |
| other bloold cells | Prex1 | 1.93 | 2.10E-47 | 4.36E-43 | 0.67 | 0.19 | 0.48 | 3.49 |
| other bloold cells | Cib1 | 1.79 | 2.21E-47 | 4.59E-43 | 0.87 | 0.4 | 0.47 | 2.17 |
| other bloold cells | Gab3 | 3.72 | 2.55E-47 | 5.30E-43 | 0.13 | 0.01 | 0.12 | 14 |
| other bloold cells | Ctsb | 4.03 | 3.03E-47 | 6.30E-43 | 0.95 | 0.75 | 0.2 | 1.27 |
| other bloold cells | Man2b2 | 3.52 | 4.63E-47 | 9.63E-43 | 0.17 | 0.02 | 0.15 | 11.2 |
| other bloold cells | Capn1 | 2.84 | 5.50E-47 | 1.14E-42 | 0.32 | 0.05 | 0.27 | 6.43 |
| other bloold cells | Arhgap17 | 1.97 | 5.82E-47 | 1.21E-42 | 0.86 | 0.37 | 0.49 | 2.33 |
| other bloold cells | Ptp4a3 | 1.99 | 5.98E-47 | 1.24E-42 | 0.75 | 0.25 | 0.5 | 2.98 |
| other bloold cells | Mir22hg | 3.94 | 1.06E-46 | 2.20E-42 | 0.15 | 0.01 | 0.14 | 12.25 |
| other bloold cells | Nfkbiz | 2.77 | 1.19E-46 | 2.48E-42 | 0.38 | 0.07 | 0.31 | 5.4 |
| other bloold cells | Itgb7 | 4.72 | 1.55E-46 | 3.23E-42 | 0.12 | 0.01 | 0.11 | 14.88 |
| other bloold cells | Mapkapk3 | 2.82 | 2.46E-46 | 5.11E-42 | 0.27 | 0.04 | 0.23 | 7.39 |
| other bloold cells | Mfsd12 | 2.98 | 3.51E-46 | 7.30E-42 | 0.34 | 0.06 | 0.28 | 5.89 |
| other bloold cells | Zfp36l2 | 1.97 | 3.57E-46 | 7.42E-42 | 0.87 | 0.43 | 0.44 | 2.02 |
| other bloold cells | Sgk3 | 2.01 | 3.58E-46 | 7.44E-42 | 0.58 | 0.15 | 0.43 | 3.92 |
| other bloold cells | Kat2b | 2.6 | 3.65E-46 | 7.59E-42 | 0.3 | 0.05 | 0.26 | 6.69 |
| other bloold cells | Vav3 | 2.48 | 4.23E-46 | 8.80E-42 | 0.34 | 0.06 | 0.29 | 6.02 |
| other bloold cells | Slc6a6 | 1.95 | 4.30E-46 | 8.94E-42 | 0.86 | 0.35 | 0.51 | 2.45 |
| other bloold cells | Pdxk | 2.11 | 6.65E-46 | 1.38E-41 | 0.5 | 0.11 | 0.39 | 4.41 |
| other bloold cells | Scarb2 | 2.42 | 7.25E-46 | 1.51E-41 | 0.62 | 0.18 | 0.44 | 3.42 |
| other bloold cells | Elovl1 | 1.95 | 1.07E-45 | 2.23E-41 | 0.73 | 0.26 | 0.48 | 2.86 |
| other bloold cells | Eng | 1.61 | 1.12E-45 | 2.33E-41 | 0.29 | 0.04 | 0.25 | 7 |
| other bloold cells | Serp1 | 1.88 | 1.30E-45 | 2.70E-41 | 0.97 | 0.76 | 0.22 | 1.29 |
| other bloold cells | Fau | 0.53 | 1.35E-45 | 2.80E-41 | 1 | 1 | 0 | 1 |
| other bloold cells | Shisa5 | 2.01 | 1.69E-45 | 3.52E-41 | 0.73 | 0.24 | 0.5 | 3.07 |
| other bloold cells | Ttyh2 | 3.22 | 1.94E-45 | 4.03E-41 | 0.22 | 0.03 | 0.2 | 8.3 |
| other bloold cells | Tmed5 | 1.63 | 3.75E-45 | 7.79E-41 | 0.93 | 0.48 | 0.45 | 1.92 |
| other bloold cells | Aldoa | 1.09 | 4.77E-45 | 9.92E-41 | 0.99 | 0.95 | 0.04 | 1.04 |
| other bloold cells | Ap1s2 | 2.28 | 5.46E-45 | 1.14E-40 | 0.53 | 0.13 | 0.4 | 3.99 |
| other bloold cells | Slc38a7 | 2.64 | 6.06E-45 | 1.26E-40 | 0.41 | 0.08 | 0.33 | 5.04 |
| other bloold cells | Syngr1 | 3.22 | 6.54E-45 | 1.36E-40 | 0.36 | 0.07 | 0.3 | 5.43 |
| other bloold cells | Gm47664 | 3.55 | 7.40E-45 | 1.54E-40 | 0.15 | 0.01 | 0.14 | 12.25 |
| other bloold cells | Picalm | 2.04 | 7.45E-45 | 1.55E-40 | 0.96 | 0.62 | 0.34 | 1.56 |
| other bloold cells | Cxcl16 | 5.13 | 7.68E-45 | 1.60E-40 | 0.11 | 0.01 | 0.11 | 16 |
| other bloold cells | Calm1 | 0.82 | 1.28E-44 | 2.67E-40 | 1 | 0.99 | 0.01 | 1.01 |
| other bloold cells | Crlf2 | 2.23 | 1.75E-44 | 3.63E-40 | 0.37 | 0.07 | 0.3 | 5.54 |
| other bloold cells | Ly96 | 2.54 | 1.81E-44 | 3.77E-40 | 0.34 | 0.06 | 0.28 | 5.89 |
| other bloold cells | Dpp7 | 2.94 | 2.88E-44 | 5.98E-40 | 0.38 | 0.07 | 0.31 | 5.32 |

**Supplementary Table 2. Marker genes for the major cell types of integrated clustering of E11.5 control and NotchGOF λ-junction cells.**

| | | | | | | | | |
|---|---|---|---|---|---|---|---|---|
| other bloold cells | Tubb6 | 2.07 | 3.48E-44 | 7.24E-40 | 0.49 | 0.11 | 0.38 | 4.34 |
| other bloold cells | Tsc22d4 | 1.52 | 3.75E-44 | 7.80E-40 | 0.97 | 0.61 | 0.35 | 1.58 |
| other bloold cells | Apold1 | 2.69 | 6.18E-44 | 1.29E-39 | 0.18 | 0.02 | 0.16 | 10.11 |
| other bloold cells | Gna12 | 2.06 | 6.41E-44 | 1.33E-39 | 0.78 | 0.31 | 0.48 | 2.56 |
| other bloold cells | Prr5 | 2.42 | 7.97E-44 | 1.66E-39 | 0.49 | 0.12 | 0.37 | 4.15 |
| other bloold cells | Myl6 | 0.92 | 9.50E-44 | 1.98E-39 | 0.99 | 0.99 | 0 | 1 |
| other bloold cells | Aup1 | 1.6 | 9.95E-44 | 2.07E-39 | 0.91 | 0.44 | 0.47 | 2.05 |
| other bloold cells | Rab5if | 1.07 | 1.28E-43 | 2.66E-39 | 0.99 | 0.91 | 0.08 | 1.08 |
| other bloold cells | Xaf1 | 1.53 | 1.48E-43 | 3.08E-39 | 0.34 | 0.06 | 0.28 | 5.89 |
| other bloold cells | Hdac9 | 2.49 | 2.18E-43 | 4.53E-39 | 0.39 | 0.07 | 0.31 | 5.2 |
| other bloold cells | Atp6v1b2 | 2.03 | 2.68E-43 | 5.58E-39 | 0.85 | 0.39 | 0.47 | 2.22 |
| other bloold cells | Akna | 2.07 | 2.84E-43 | 5.90E-39 | 0.47 | 0.11 | 0.36 | 4.47 |
| other bloold cells | )610040J01Ri | 2.59 | 3.00E-43 | 6.24E-39 | 0.23 | 0.03 | 0.2 | 7.97 |
| other bloold cells | Arhgap18 | 1.62 | 3.02E-43 | 6.28E-39 | 0.52 | 0.12 | 0.39 | 4.17 |
| other bloold cells | S1pr1 | 2.88 | 3.84E-43 | 7.99E-39 | 0.34 | 0.06 | 0.28 | 5.53 |
| other bloold cells | Snta1 | 2.61 | 3.86E-43 | 8.02E-39 | 0.31 | 0.05 | 0.26 | 6.16 |
| other bloold cells | Naglu | 2.95 | 3.87E-43 | 8.06E-39 | 0.23 | 0.03 | 0.2 | 7.7 |
| other bloold cells | Mgat4a | 2.21 | 4.06E-43 | 8.45E-39 | 0.43 | 0.09 | 0.34 | 4.72 |
| other bloold cells | Cfl1 | 0.67 | 4.36E-43 | 9.07E-39 | 1 | 1 | 0 | 1 |
| other bloold cells | Gm15417 | 2.82 | 4.58E-43 | 9.53E-39 | 0.21 | 0.03 | 0.19 | 8.4 |
| other bloold cells | Akap13 | 1.65 | 5.23E-43 | 1.09E-38 | 0.92 | 0.5 | 0.42 | 1.83 |
| other bloold cells | Isg15 | 4.57 | 6.90E-43 | 1.44E-38 | 0.13 | 0.01 | 0.12 | 12.09 |
| other bloold cells | Gpi1 | 1.2 | 9.49E-43 | 1.97E-38 | 0.98 | 0.84 | 0.14 | 1.17 |
| other bloold cells | Itga4 | 1.78 | 1.45E-42 | 3.01E-38 | 0.66 | 0.19 | 0.46 | 3.4 |
| other bloold cells | Relt | 3.11 | 1.48E-42 | 3.07E-38 | 0.23 | 0.03 | 0.2 | 7.7 |
| other bloold cells | Wdr1 | 1.73 | 1.61E-42 | 3.35E-38 | 0.9 | 0.55 | 0.36 | 1.66 |
| other bloold cells | Ptpn1 | 1.72 | 1.82E-42 | 3.79E-38 | 0.92 | 0.52 | 0.4 | 1.76 |
| other bloold cells | Prr13 | 1.68 | 2.32E-42 | 4.82E-38 | 0.9 | 0.4 | 0.5 | 2.24 |
| other bloold cells | Irf2bp2 | 1.75 | 2.49E-42 | 5.18E-38 | 0.83 | 0.35 | 0.49 | 2.4 |
| other bloold cells | Myo1c | 2.17 | 3.16E-42 | 6.58E-38 | 0.44 | 0.1 | 0.35 | 4.59 |
| other bloold cells | Pnp | 1.55 | 3.46E-42 | 7.20E-38 | 0.97 | 0.66 | 0.31 | 1.47 |
| other bloold cells | Rhob | 2.28 | 3.72E-42 | 7.73E-38 | 0.8 | 0.35 | 0.45 | 2.31 |
| other bloold cells | Tspan32 | 2.68 | 4.96E-42 | 1.03E-37 | 0.26 | 0.04 | 0.22 | 7 |
| other bloold cells | Sdf2l1 | 1.89 | 6.83E-42 | 1.42E-37 | 0.87 | 0.47 | 0.41 | 1.86 |
| other bloold cells | Acp5 | 1.53 | 1.22E-41 | 2.54E-37 | 0.43 | 0.09 | 0.34 | 4.85 |
| other bloold cells | Cd93 | 1.21 | 1.47E-41 | 3.06E-37 | 0.18 | 0.02 | 0.16 | 9.72 |
| other bloold cells | Gltp | 1.79 | 1.55E-41 | 3.23E-37 | 0.87 | 0.39 | 0.47 | 2.2 |
| other bloold cells | Emb | 2.06 | 1.80E-41 | 3.74E-37 | 0.84 | 0.41 | 0.43 | 2.07 |
| other bloold cells | Htatip2 | 2.48 | 2.31E-41 | 4.80E-37 | 0.37 | 0.07 | 0.3 | 5.08 |
| other bloold cells | Dcxr | 2.13 | 2.72E-41 | 5.66E-37 | 0.43 | 0.09 | 0.34 | 4.62 |
| other bloold cells | Atf6 | 2.11 | 3.21E-41 | 6.68E-37 | 0.53 | 0.14 | 0.39 | 3.82 |
| other bloold cells | Map3k5 | 2.72 | 3.56E-41 | 7.40E-37 | 0.25 | 0.03 | 0.21 | 7.21 |
| other bloold cells | Cryzl2 | 2.16 | 4.13E-41 | 8.60E-37 | 0.45 | 0.1 | 0.35 | 4.39 |
| other bloold cells | Cdc42 | 0.82 | 4.25E-41 | 8.84E-37 | 0.99 | 0.98 | 0.01 | 1.01 |
| other bloold cells | Cd9 | 1.42 | 4.41E-41 | 9.18E-37 | 0.81 | 0.3 | 0.51 | 2.73 |
| other bloold cells | Zfp710 | 2.19 | 5.27E-41 | 1.10E-36 | 0.6 | 0.18 | 0.42 | 3.36 |
| other bloold cells | Map3k11 | 2.13 | 6.36E-41 | 1.32E-36 | 0.48 | 0.11 | 0.36 | 4.18 |
| other bloold cells | Mfsd1 | 1.81 | 7.59E-41 | 1.58E-36 | 0.83 | 0.36 | 0.47 | 2.31 |
| other bloold cells | Plekhb2 | 2.56 | 8.26E-41 | 1.72E-36 | 0.33 | 0.06 | 0.27 | 5.48 |
| other bloold cells | Slc29a1 | 1.77 | 8.86E-41 | 1.84E-36 | 0.5 | 0.12 | 0.38 | 4.11 |
| other bloold cells | Znrf2 | 2.02 | 9.51E-41 | 1.98E-36 | 0.61 | 0.18 | 0.43 | 3.32 |
| other bloold cells | Ncoa4 | 1.99 | 1.79E-40 | 3.72E-36 | 0.77 | 0.32 | 0.45 | 2.44 |
| other bloold cells | Ppm1h | 2.33 | 3.20E-40 | 6.65E-36 | 0.39 | 0.08 | 0.31 | 4.78 |
| other bloold cells | Abca7 | 3.12 | 3.31E-40 | 6.88E-36 | 0.19 | 0.02 | 0.17 | 8.59 |
| other bloold cells | Vamp5 | 1.9 | 3.43E-40 | 7.13E-36 | 0.34 | 0.06 | 0.27 | 5.42 |
| other bloold cells | Irf1 | 2.33 | 3.86E-40 | 8.03E-36 | 0.26 | 0.04 | 0.22 | 6.82 |

**Supplementary Table 2. Marker genes for the major cell types of integrated clustering of E11.5 control and NotchGOF λ-junction cells.**

| | | | | | | | | |
|---|---|---|---|---|---|---|---|---|
| other bloold cells | Il3ra | 2.41 | 4.83E-40 | 1.01E-35 | 0.3 | 0.05 | 0.25 | 6.02 |
| other bloold cells | Mcur1 | 1.75 | 7.47E-40 | 1.55E-35 | 0.68 | 0.22 | 0.46 | 3.15 |
| other bloold cells | Dab2 | 3.68 | 8.81E-40 | 1.83E-35 | 0.71 | 0.35 | 0.36 | 2.02 |
| other bloold cells | Cltc | 1.73 | 1.29E-39 | 2.69E-35 | 0.94 | 0.75 | 0.2 | 1.26 |
| other bloold cells | Esyt1 | 2.23 | 1.49E-39 | 3.10E-35 | 0.37 | 0.07 | 0.3 | 5.08 |
| other bloold cells | Dtx4 | 2.82 | 5.33E-39 | 1.11E-34 | 0.26 | 0.04 | 0.22 | 6.48 |
| other bloold cells | Rps29 | 0.4 | 6.97E-39 | 1.45E-34 | 1 | 1 | 0 | 1 |
| other bloold cells | Rab29 | 2.71 | 9.16E-39 | 1.91E-34 | 0.29 | 0.05 | 0.24 | 5.88 |
| other bloold cells | Slc16a10 | 2.16 | 1.14E-38 | 2.38E-34 | 0.45 | 0.11 | 0.34 | 4.19 |
| other bloold cells | Fnbp1 | 1.84 | 1.22E-38 | 2.53E-34 | 0.68 | 0.24 | 0.44 | 2.89 |
| other bloold cells | Prdx5 | 1.4 | 1.58E-38 | 3.29E-34 | 0.99 | 0.8 | 0.19 | 1.24 |
| other bloold cells | Gns | 2.33 | 2.36E-38 | 4.90E-34 | 0.69 | 0.25 | 0.43 | 2.72 |
| other bloold cells | Prdm1 | 3.79 | 3.09E-38 | 6.43E-34 | 0.13 | 0.01 | 0.12 | 11.45 |
| other bloold cells | Ap5b1 | 3.65 | 3.98E-38 | 8.27E-34 | 0.11 | 0.01 | 0.1 | 14 |
| other bloold cells | Slc9a3r1 | 1.24 | 4.01E-38 | 8.35E-34 | 0.75 | 0.24 | 0.51 | 3.12 |
| other bloold cells | 310040G24Ri | 2.33 | 4.21E-38 | 8.76E-34 | 0.23 | 0.03 | 0.2 | 7.22 |
| other bloold cells | Rreb1 | 1.83 | 4.42E-38 | 9.19E-34 | 0.64 | 0.21 | 0.44 | 3.11 |
| other bloold cells | Sdcbp | 1.81 | 5.67E-38 | 1.18E-33 | 0.94 | 0.69 | 0.25 | 1.36 |
| other bloold cells | Cdc42se1 | 1.53 | 6.08E-38 | 1.27E-33 | 0.91 | 0.42 | 0.49 | 2.14 |
| other bloold cells | Lamp2 | 1.74 | 6.85E-38 | 1.42E-33 | 0.92 | 0.56 | 0.35 | 1.62 |
| other bloold cells | Cln6 | 2.04 | 7.19E-38 | 1.50E-33 | 0.43 | 0.1 | 0.33 | 4.4 |
| other bloold cells | Tmem179b | 1.54 | 8.82E-38 | 1.83E-33 | 0.78 | 0.29 | 0.48 | 2.65 |
| other bloold cells | Rbfa | 1.86 | 1.12E-37 | 2.34E-33 | 0.74 | 0.31 | 0.43 | 2.41 |
| other bloold cells | Txnip | 1.83 | 1.52E-37 | 3.16E-33 | 0.8 | 0.34 | 0.46 | 2.34 |
| other bloold cells | Sirt7 | 1.87 | 1.82E-37 | 3.79E-33 | 0.57 | 0.17 | 0.4 | 3.37 |
| other bloold cells | Snx30 | 2.03 | 3.17E-37 | 6.58E-33 | 0.53 | 0.15 | 0.38 | 3.56 |
| other bloold cells | Mindy2 | 1.96 | 3.61E-37 | 7.50E-33 | 0.58 | 0.17 | 0.41 | 3.33 |
| other bloold cells | D1Ertd622e | 2.06 | 4.42E-37 | 9.19E-33 | 0.49 | 0.13 | 0.36 | 3.8 |
| other bloold cells | Asl | 1.79 | 4.67E-37 | 9.71E-33 | 0.54 | 0.15 | 0.39 | 3.66 |
| other bloold cells | Tmem176a | 2.23 | 4.87E-37 | 1.01E-32 | 0.83 | 0.41 | 0.42 | 2.04 |
| other bloold cells | Exoc3l2 | 3.29 | 5.58E-37 | 1.16E-32 | 0.12 | 0.01 | 0.11 | 11.9 |
| other bloold cells | Cox7a1 | 2.66 | 1.02E-36 | 2.11E-32 | 0.18 | 0.02 | 0.16 | 8.27 |
| other bloold cells | Itpripl2 | 1.93 | 1.52E-36 | 3.16E-32 | 0.5 | 0.13 | 0.37 | 3.82 |
| other bloold cells | Nfkb1 | 1.8 | 1.75E-36 | 3.64E-32 | 0.76 | 0.3 | 0.46 | 2.55 |
| other bloold cells | Slc40a1 | 4.23 | 2.49E-36 | 5.18E-32 | 0.3 | 0.06 | 0.24 | 5.19 |
| other bloold cells | Ppp2r5a | 1.52 | 3.48E-36 | 7.24E-32 | 0.79 | 0.32 | 0.47 | 2.48 |
| other bloold cells | Irf2 | 1.72 | 4.74E-36 | 9.86E-32 | 0.69 | 0.25 | 0.44 | 2.72 |
| other bloold cells | Acer3 | 1.97 | 5.24E-36 | 1.09E-31 | 0.55 | 0.16 | 0.38 | 3.39 |
| other bloold cells | Rnaset2a | 1.49 | 5.28E-36 | 1.10E-31 | 0.92 | 0.52 | 0.4 | 1.77 |
| other bloold cells | Agtrap | 2.01 | 7.49E-36 | 1.56E-31 | 0.52 | 0.15 | 0.37 | 3.54 |
| other bloold cells | Atp6v0c | 1.26 | 7.89E-36 | 1.64E-31 | 1 | 0.96 | 0.04 | 1.04 |
| other bloold cells | Tmem219 | 1.94 | 8.69E-36 | 1.81E-31 | 0.53 | 0.15 | 0.38 | 3.45 |
| other bloold cells | Chst12 | 1.9 | 1.04E-35 | 2.16E-31 | 0.39 | 0.08 | 0.3 | 4.64 |
| other bloold cells | Myh9 | 1.31 | 1.54E-35 | 3.20E-31 | 0.93 | 0.51 | 0.43 | 1.84 |
| other bloold cells | Serf2 | 0.59 | 1.55E-35 | 3.23E-31 | 1 | 1 | 0 | 1 |
| other bloold cells | Wdr26 | 1.37 | 1.59E-35 | 3.30E-31 | 0.94 | 0.71 | 0.23 | 1.33 |
| other bloold cells | Frmd4b | 1.76 | 1.66E-35 | 3.45E-31 | 0.57 | 0.17 | 0.4 | 3.37 |
| other bloold cells | Dgkz | 1.81 | 1.68E-35 | 3.49E-31 | 0.73 | 0.28 | 0.45 | 2.58 |
| other bloold cells | Retreg1 | 2.17 | 2.35E-35 | 4.89E-31 | 0.4 | 0.09 | 0.31 | 4.34 |
| other bloold cells | Bscl2 | 1.49 | 2.38E-35 | 4.95E-31 | 0.65 | 0.21 | 0.44 | 3.11 |
| other bloold cells | Blvrb | 1.6 | 2.59E-35 | 5.38E-31 | 0.8 | 0.34 | 0.46 | 2.34 |
| other bloold cells | Pxn | 1.7 | 2.90E-35 | 6.03E-31 | 0.6 | 0.18 | 0.42 | 3.27 |
| other bloold cells | Engase | 2.5 | 3.11E-35 | 6.48E-31 | 0.26 | 0.04 | 0.22 | 6.02 |
| other bloold cells | Fgd3 | 3.19 | 3.14E-35 | 6.53E-31 | 0.13 | 0.01 | 0.12 | 10.23 |
| other bloold cells | Nrgn | 3.88 | 3.39E-35 | 7.04E-31 | 0.27 | 0.05 | 0.22 | 5.66 |
| other bloold cells | Rnasek | 1.25 | 3.57E-35 | 7.43E-31 | 0.96 | 0.72 | 0.24 | 1.33 |

**Supplementary Table 2. Marker genes for the major cell types of integrated clustering of E11.5 control and NotchGOF λ-junction cells.**

| | | | | | | | | |
|---|---|---|---|---|---|---|---|---|
| other bloold cells | Mroh1 | 2.23 | 3.63E-35 | 7.55E-31 | 0.36 | 0.08 | 0.28 | 4.7 |
| other bloold cells | Cndp2 | 2.12 | 4.41E-35 | 9.18E-31 | 0.79 | 0.37 | 0.42 | 2.14 |
| other bloold cells | Slc3a2 | 1.42 | 4.85E-35 | 1.01E-30 | 0.94 | 0.62 | 0.32 | 1.52 |
| other bloold cells | Lfng | 2.58 | 5.29E-35 | 1.10E-30 | 0.16 | 0.02 | 0.14 | 8.94 |
| other bloold cells | Ftl1 | 1.9 | 8.69E-35 | 1.81E-30 | 1 | 1 | 0 | 1 |
| other bloold cells | Pkn1 | 1.52 | 1.05E-34 | 2.18E-30 | 0.83 | 0.35 | 0.48 | 2.35 |
| other bloold cells | Snx3 | 1.03 | 1.06E-34 | 2.21E-30 | 0.99 | 0.89 | 0.1 | 1.11 |
| other bloold cells | G6pdx | 1.79 | 1.14E-34 | 2.38E-30 | 0.66 | 0.24 | 0.42 | 2.74 |
| other bloold cells | Git2 | 1.54 | 1.17E-34 | 2.43E-30 | 0.8 | 0.34 | 0.46 | 2.36 |
| other bloold cells | Rgl1 | 2.16 | 1.18E-34 | 2.45E-30 | 0.51 | 0.15 | 0.36 | 3.38 |
| other bloold cells | Zfp992 | 1.81 | 1.31E-34 | 2.72E-30 | 0.54 | 0.16 | 0.38 | 3.45 |
| other bloold cells | Cyb561a3 | 2.42 | 1.73E-34 | 3.61E-30 | 0.3 | 0.06 | 0.24 | 5.19 |
| other bloold cells | Nmi | 2.14 | 1.97E-34 | 4.10E-30 | 0.39 | 0.09 | 0.3 | 4.36 |
| other bloold cells | Adam17 | 1.8 | 2.29E-34 | 4.77E-30 | 0.79 | 0.33 | 0.46 | 2.37 |
| other bloold cells | Tmem86a | 3.01 | 2.63E-34 | 5.46E-30 | 0.41 | 0.1 | 0.3 | 3.9 |
| other bloold cells | Synj1 | 2.44 | 2.64E-34 | 5.50E-30 | 0.43 | 0.11 | 0.32 | 3.99 |
| other bloold cells | Hsd3b7 | 2.53 | 3.08E-34 | 6.40E-30 | 0.22 | 0.03 | 0.19 | 6.59 |
| other bloold cells | Gabarap | 1.14 | 3.10E-34 | 6.45E-30 | 1 | 0.98 | 0.02 | 1.02 |
| other bloold cells | Slc12a9 | 2.27 | 3.36E-34 | 6.98E-30 | 0.36 | 0.08 | 0.28 | 4.55 |
| other bloold cells | Tmem144 | 2.72 | 4.62E-34 | 9.61E-30 | 0.27 | 0.05 | 0.22 | 5.66 |
| other bloold cells | Rab43 | 2.28 | 5.48E-34 | 1.14E-29 | 0.32 | 0.07 | 0.26 | 4.95 |
| other bloold cells | Adssl1 | 2.5 | 8.79E-34 | 1.83E-29 | 0.23 | 0.04 | 0.2 | 6.42 |
| other bloold cells | Frrs1 | 1.79 | 9.19E-34 | 1.91E-29 | 0.63 | 0.22 | 0.41 | 2.86 |
| other bloold cells | Slc6a13 | 3.69 | 1.10E-33 | 2.28E-29 | 0.15 | 0.02 | 0.13 | 9.19 |
| other bloold cells | Mitf | 2.96 | 1.11E-33 | 2.31E-29 | 0.17 | 0.02 | 0.15 | 8.4 |
| other bloold cells | Lrp10 | 1.68 | 1.13E-33 | 2.35E-29 | 0.57 | 0.17 | 0.4 | 3.33 |
| other bloold cells | Necap2 | 1.61 | 1.16E-33 | 2.41E-29 | 0.82 | 0.39 | 0.43 | 2.09 |
| other bloold cells | Cyfip1 | 1.71 | 2.43E-33 | 5.06E-29 | 0.94 | 0.7 | 0.24 | 1.35 |
| other bloold cells | Tradd | 1.83 | 3.07E-33 | 6.38E-29 | 0.45 | 0.11 | 0.33 | 3.93 |
| other bloold cells | Mob3c | 2.27 | 3.42E-33 | 7.12E-29 | 0.29 | 0.05 | 0.23 | 5.31 |
| other bloold cells | Slc36a1 | 2.19 | 3.75E-33 | 7.80E-29 | 0.29 | 0.05 | 0.23 | 5.42 |
| other bloold cells | Map4k1 | 2.15 | 4.58E-33 | 9.53E-29 | 0.26 | 0.04 | 0.22 | 5.89 |
| other bloold cells | Rps12 | 0.44 | 4.96E-33 | 1.03E-28 | 1 | 1 | 0 | 1 |
| other bloold cells | Ubl3 | 1.51 | 5.04E-33 | 1.05E-28 | 0.8 | 0.37 | 0.43 | 2.17 |
| other bloold cells | Chmp2a | 1.19 | 7.83E-33 | 1.63E-28 | 0.95 | 0.66 | 0.29 | 1.44 |
| other bloold cells | Ifngr2 | 1.7 | 8.75E-33 | 1.82E-28 | 0.78 | 0.37 | 0.41 | 2.11 |
| other bloold cells | Pitpna | 1.21 | 8.93E-33 | 1.86E-28 | 0.94 | 0.7 | 0.23 | 1.33 |
| other bloold cells | Plekho1 | 1.42 | 9.64E-33 | 2.00E-28 | 0.94 | 0.62 | 0.33 | 1.53 |
| other bloold cells | Pag1 | 2.1 | 1.15E-32 | 2.39E-28 | 0.26 | 0.05 | 0.21 | 5.76 |
| other bloold cells | Pla2g4a | 1.51 | 1.28E-32 | 2.66E-28 | 0.68 | 0.25 | 0.43 | 2.72 |
| other bloold cells | Lrrfip1 | 1.54 | 2.13E-32 | 4.44E-28 | 0.76 | 0.34 | 0.42 | 2.25 |
| other bloold cells | Dleu2 | 1.95 | 2.48E-32 | 5.17E-28 | 0.65 | 0.25 | 0.4 | 2.58 |
| other bloold cells | Fosl2 | 2.53 | 2.59E-32 | 5.38E-28 | 0.2 | 0.03 | 0.17 | 6.77 |
| other bloold cells | Rac1 | 0.92 | 2.70E-32 | 5.62E-28 | 0.99 | 0.9 | 0.08 | 1.09 |
| other bloold cells | Arhgdia | 0.96 | 3.32E-32 | 6.90E-28 | 0.99 | 0.88 | 0.11 | 1.12 |
| other bloold cells | Wdr81 | 2.48 | 4.05E-32 | 8.43E-28 | 0.27 | 0.05 | 0.22 | 5.43 |
| other bloold cells | Pqlc2 | 2.47 | 5.27E-32 | 1.10E-27 | 0.41 | 0.1 | 0.3 | 3.94 |
| other bloold cells | Pnpla2 | 1.92 | 5.50E-32 | 1.14E-27 | 0.38 | 0.09 | 0.29 | 4.3 |
| other bloold cells | Ndufv3 | 0.99 | 7.07E-32 | 1.47E-27 | 0.98 | 0.84 | 0.14 | 1.17 |
| other bloold cells | Vps13c | 1.84 | 7.28E-32 | 1.51E-27 | 0.58 | 0.2 | 0.39 | 2.97 |
| other bloold cells | H2-T23 | 2.44 | 7.36E-32 | 1.53E-27 | 0.28 | 0.05 | 0.23 | 5.28 |
| other bloold cells | Adgre5 | 2.09 | 1.04E-31 | 2.16E-27 | 0.46 | 0.13 | 0.33 | 3.64 |
| other bloold cells | Slc7a1 | 2.22 | 1.27E-31 | 2.64E-27 | 0.34 | 0.08 | 0.27 | 4.4 |
| other bloold cells | Ap1b1 | 1.8 | 1.49E-31 | 3.10E-27 | 0.64 | 0.24 | 0.4 | 2.64 |
| other bloold cells | Lamtor4 | 1.3 | 1.62E-31 | 3.38E-27 | 0.93 | 0.66 | 0.27 | 1.41 |
| other bloold cells | Rap1gap2 | 2.37 | 1.71E-31 | 3.56E-27 | 0.14 | 0.02 | 0.13 | 9.33 |

**Supplementary Table 2. Marker genes for the major cell types of integrated clustering of E11.5 control and NotchGOF λ-junction cells.**

| | | | | | | | | |
|---|---|---|---|---|---|---|---|---|
| other bloold cells | M6pr | 1.5 | 1.74E-31 | 3.61E-27 | 0.85 | 0.44 | 0.41 | 1.92 |
| other bloold cells | Dennd4a | 1.94 | 1.95E-31 | 4.06E-27 | 0.36 | 0.08 | 0.28 | 4.41 |
| other bloold cells | Nhlrc3 | 1.87 | 2.14E-31 | 4.45E-27 | 0.46 | 0.12 | 0.33 | 3.73 |
| other bloold cells | Dazap2 | 1.3 | 2.45E-31 | 5.09E-27 | 0.91 | 0.57 | 0.34 | 1.59 |
| other bloold cells | Stxbp5 | 1.9 | 2.45E-31 | 5.10E-27 | 0.35 | 0.08 | 0.27 | 4.49 |
| other bloold cells | Vasp | 1.34 | 2.77E-31 | 5.77E-27 | 0.88 | 0.48 | 0.4 | 1.83 |
| other bloold cells | Tmbim1 | 2.04 | 3.09E-31 | 6.42E-27 | 0.39 | 0.1 | 0.3 | 4.08 |
| other bloold cells | Cryl1 | 1.85 | 3.11E-31 | 6.46E-27 | 0.5 | 0.15 | 0.35 | 3.34 |
| other bloold cells | Acox3 | 2.07 | 3.16E-31 | 6.58E-27 | 0.33 | 0.07 | 0.26 | 4.63 |
| other bloold cells | Tmem170b | 2.18 | 3.22E-31 | 6.70E-27 | 0.31 | 0.06 | 0.24 | 4.81 |
| other bloold cells | Cd34 | 1.8 | 3.74E-31 | 7.78E-27 | 0.46 | 0.13 | 0.32 | 3.47 |
| other bloold cells | Cdkn2c | 2.79 | 3.99E-31 | 8.31E-27 | 0.23 | 0.04 | 0.19 | 5.92 |
| other bloold cells | Cln3 | 2.11 | 5.03E-31 | 1.05E-26 | 0.48 | 0.14 | 0.34 | 3.43 |
| other bloold cells | Nrp1 | 1.23 | 7.12E-31 | 1.48E-26 | 0.62 | 0.21 | 0.41 | 2.99 |
| other bloold cells | Rhbdf2 | 2.63 | 9.38E-31 | 1.95E-26 | 0.2 | 0.03 | 0.17 | 6.55 |
| other bloold cells | Sh3glb1 | 1.09 | 1.47E-30 | 3.06E-26 | 0.97 | 0.84 | 0.13 | 1.16 |
| other bloold cells | Gdi2 | 0.8 | 1.49E-30 | 3.09E-26 | 0.99 | 0.95 | 0.04 | 1.04 |
| other bloold cells | Glb1 | 1.56 | 1.68E-30 | 3.50E-26 | 0.54 | 0.16 | 0.38 | 3.32 |
| other bloold cells | Atp6v0d1 | 1.34 | 2.04E-30 | 4.25E-26 | 0.9 | 0.54 | 0.37 | 1.69 |
| other bloold cells | Rnf141 | 2.16 | 2.12E-30 | 4.41E-26 | 0.44 | 0.12 | 0.32 | 3.61 |
| other bloold cells | Edem2 | 1.62 | 2.62E-30 | 5.44E-26 | 0.58 | 0.19 | 0.39 | 3.02 |
| other bloold cells | Sesn1 | 1.89 | 3.22E-30 | 6.70E-26 | 0.61 | 0.23 | 0.38 | 2.7 |
| other bloold cells | Ftl1-ps1 | 1.92 | 3.46E-30 | 7.20E-26 | 0.78 | 0.41 | 0.37 | 1.91 |
| other bloold cells | Slc12a6 | 1.91 | 3.79E-30 | 7.88E-26 | 0.46 | 0.13 | 0.33 | 3.47 |
| other bloold cells | Ctla2a | 1.3 | 4.55E-30 | 9.46E-26 | 0.15 | 0.02 | 0.14 | 8.11 |
| other bloold cells | Pdcd6ip | 1.3 | 5.96E-30 | 1.24E-25 | 0.87 | 0.48 | 0.38 | 1.79 |
| other bloold cells | Capzb | 0.86 | 6.07E-30 | 1.26E-25 | 0.99 | 0.92 | 0.07 | 1.08 |
| other bloold cells | Rnf13 | 1.68 | 6.54E-30 | 1.36E-25 | 0.64 | 0.24 | 0.4 | 2.67 |
| other bloold cells | Pcgf5 | 1.75 | 7.38E-30 | 1.53E-25 | 0.36 | 0.09 | 0.28 | 4.23 |
| other bloold cells | Rsu1 | 1.38 | 9.10E-30 | 1.89E-25 | 0.89 | 0.48 | 0.41 | 1.84 |
| other bloold cells | Wasf2 | 1.22 | 9.62E-30 | 2.00E-25 | 0.96 | 0.71 | 0.25 | 1.35 |
| other bloold cells | Znfx1 | 2.22 | 9.66E-30 | 2.01E-25 | 0.29 | 0.06 | 0.23 | 4.86 |
| other bloold cells | Capza1 | 1.14 | 1.20E-29 | 2.49E-25 | 0.92 | 0.65 | 0.28 | 1.43 |
| other bloold cells | Heatr5a | 1.65 | 1.30E-29 | 2.69E-25 | 0.56 | 0.19 | 0.37 | 3.01 |
| other bloold cells | Slc29a3 | 1.89 | 1.50E-29 | 3.12E-25 | 0.39 | 0.1 | 0.29 | 3.92 |
| other bloold cells | Mapk3 | 1.38 | 1.94E-29 | 4.04E-25 | 0.83 | 0.45 | 0.38 | 1.86 |
| other bloold cells | Diaph2 | 1.72 | 2.47E-29 | 5.14E-25 | 0.5 | 0.15 | 0.35 | 3.29 |
| other bloold cells | H6pd | 2.47 | 3.19E-29 | 6.63E-25 | 0.2 | 0.03 | 0.17 | 6.53 |
| other bloold cells | Sptlc2 | 1.57 | 3.60E-29 | 7.49E-25 | 0.76 | 0.33 | 0.44 | 2.33 |
| other bloold cells | Emc9 | 2.35 | 3.67E-29 | 7.64E-25 | 0.25 | 0.05 | 0.21 | 5.36 |
| other bloold cells | Glmp | 1.25 | 3.70E-29 | 7.70E-25 | 0.91 | 0.47 | 0.44 | 1.95 |
| other bloold cells | Arrdc1 | 1.52 | 4.02E-29 | 8.36E-25 | 0.42 | 0.11 | 0.31 | 3.82 |
| other bloold cells | Tfe3 | 1.8 | 4.72E-29 | 9.82E-25 | 0.54 | 0.18 | 0.36 | 2.99 |
| other bloold cells | Nudt16 | 1.9 | 8.54E-29 | 1.78E-24 | 0.36 | 0.09 | 0.27 | 4.15 |
| other bloold cells | Mob1a | 1.35 | 1.11E-28 | 2.31E-24 | 0.85 | 0.45 | 0.39 | 1.87 |
| other bloold cells | Tmem192 | 1.7 | 1.13E-28 | 2.34E-24 | 0.55 | 0.18 | 0.36 | 2.98 |
| other bloold cells | Nfe2l2 | 1.58 | 1.13E-28 | 2.36E-24 | 0.84 | 0.45 | 0.39 | 1.88 |
| other bloold cells | Vps18 | 1.71 | 1.23E-28 | 2.56E-24 | 0.46 | 0.14 | 0.33 | 3.4 |
| other bloold cells | Ccdc82 | 1.74 | 1.45E-28 | 3.02E-24 | 0.43 | 0.12 | 0.31 | 3.65 |
| other bloold cells | Stk4 | 1.51 | 1.97E-28 | 4.10E-24 | 0.76 | 0.36 | 0.4 | 2.1 |
| other bloold cells | Ostm1 | 2.14 | 2.10E-28 | 4.36E-24 | 0.32 | 0.07 | 0.24 | 4.38 |
| other bloold cells | Npc2 | 0.95 | 3.15E-28 | 6.55E-24 | 0.99 | 0.93 | 0.06 | 1.07 |
| other bloold cells | Vps8 | 2.13 | 3.38E-28 | 7.03E-24 | 0.27 | 0.06 | 0.22 | 4.96 |
| other bloold cells | Calhm6 | 2.86 | 4.61E-28 | 9.59E-24 | 0.17 | 0.02 | 0.14 | 7 |
| other bloold cells | Cenpx | 1 | 4.89E-28 | 1.02E-23 | 0.95 | 0.79 | 0.17 | 1.21 |
| other bloold cells | Scarb1 | 1.75 | 4.94E-28 | 1.03E-23 | 0.49 | 0.16 | 0.33 | 3.12 |

**Supplementary Table 2. Marker genes for the major cell types of integrated clustering of E11.5 control and NotchGOF λ-junction cells.**

| | | | | | | | | |
|---|---|---|---|---|---|---|---|---|
| other bloold cells | Tuba1c | 1.25 | 5.49E-28 | 1.14E-23 | 0.84 | 0.44 | 0.4 | 1.92 |
| other bloold cells | Camk1 | 1.57 | 6.75E-28 | 1.40E-23 | 0.82 | 0.42 | 0.4 | 1.94 |
| other bloold cells | Dhx58 | 3.2 | 9.26E-28 | 1.93E-23 | 0.11 | 0.01 | 0.1 | 10.5 |
| other bloold cells | Ost4 | 0.65 | 1.32E-27 | 2.74E-23 | 0.98 | 0.95 | 0.03 | 1.03 |
| other bloold cells | Aff1 | 2.06 | 1.33E-27 | 2.77E-23 | 0.31 | 0.07 | 0.24 | 4.4 |
| other bloold cells | Nlrc3 | 2.78 | 1.47E-27 | 3.06E-23 | 0.15 | 0.02 | 0.13 | 7.74 |
| other bloold cells | Cd302 | 1.59 | 1.50E-27 | 3.12E-23 | 0.67 | 0.29 | 0.39 | 2.35 |
| other bloold cells | Gnb2 | 0.65 | 1.60E-27 | 3.32E-23 | 0.99 | 0.96 | 0.02 | 1.02 |
| other bloold cells | Gab2 | 2.35 | 1.72E-27 | 3.58E-23 | 0.24 | 0.05 | 0.19 | 5.29 |
| other bloold cells | Coa5 | 1.4 | 1.79E-27 | 3.72E-23 | 0.68 | 0.27 | 0.41 | 2.55 |
| other bloold cells | Fli1 | 1.21 | 1.80E-27 | 3.73E-23 | 0.82 | 0.37 | 0.45 | 2.21 |
| other bloold cells | Dnase2a | 1.57 | 1.82E-27 | 3.78E-23 | 0.6 | 0.22 | 0.38 | 2.77 |
| other bloold cells | Stat5a | 2.56 | 2.05E-27 | 4.27E-23 | 0.18 | 0.03 | 0.15 | 6.5 |
| other bloold cells | Eps15 | 1.5 | 2.10E-27 | 4.37E-23 | 0.7 | 0.31 | 0.39 | 2.29 |
| other bloold cells | Atp6v1f | 0.79 | 2.22E-27 | 4.63E-23 | 1 | 0.9 | 0.1 | 1.11 |
| other bloold cells | Rnf135 | 2.24 | 2.49E-27 | 5.18E-23 | 0.21 | 0.04 | 0.17 | 5.83 |
| other bloold cells | Calhm2 | 2.01 | 2.62E-27 | 5.46E-23 | 0.21 | 0.04 | 0.18 | 6 |
| other bloold cells | Stim1 | 1.49 | 2.65E-27 | 5.52E-23 | 0.48 | 0.14 | 0.33 | 3.35 |
| other bloold cells | Lamtor1 | 0.98 | 4.02E-27 | 8.37E-23 | 0.97 | 0.86 | 0.1 | 1.12 |
| other bloold cells | Gpd1l | 1.74 | 5.18E-27 | 1.08E-22 | 0.32 | 0.07 | 0.24 | 4.38 |
| other bloold cells | Atp6ap2 | 1.33 | 5.40E-27 | 1.12E-22 | 0.84 | 0.43 | 0.41 | 1.97 |
| other bloold cells | 033406O09Ri | 3.01 | 5.68E-27 | 1.18E-22 | 0.11 | 0.01 | 0.1 | 9.33 |
| other bloold cells | Tubgcp5 | 1.71 | 5.83E-27 | 1.21E-22 | 0.75 | 0.34 | 0.4 | 2.17 |
| other bloold cells | Fam174a | 1.44 | 5.87E-27 | 1.22E-22 | 0.72 | 0.31 | 0.41 | 2.35 |
| other bloold cells | St6galnac4 | 1.56 | 5.94E-27 | 1.23E-22 | 0.56 | 0.2 | 0.36 | 2.85 |
| other bloold cells | Zfp120 | 1.85 | 5.96E-27 | 1.24E-22 | 0.45 | 0.13 | 0.32 | 3.37 |
| other bloold cells | Pgd | 1.26 | 6.26E-27 | 1.30E-22 | 0.89 | 0.59 | 0.3 | 1.52 |
| other bloold cells | Rp2 | 1.93 | 7.08E-27 | 1.47E-22 | 0.5 | 0.17 | 0.33 | 2.91 |
| other bloold cells | Tcn2 | 1.94 | 7.22E-27 | 1.50E-22 | 0.51 | 0.18 | 0.33 | 2.9 |
| other bloold cells | Aagab | 1.56 | 8.50E-27 | 1.77E-22 | 0.53 | 0.17 | 0.36 | 3.05 |
| other bloold cells | Ptgr2 | 1.75 | 1.04E-26 | 2.16E-22 | 0.47 | 0.15 | 0.32 | 3.17 |
| other bloold cells | Matk | 2.45 | 1.21E-26 | 2.52E-22 | 0.18 | 0.03 | 0.15 | 6.48 |
| other bloold cells | Glud1 | 1.03 | 1.44E-26 | 3.00E-22 | 0.92 | 0.65 | 0.27 | 1.41 |
| other bloold cells | Bid | 1.75 | 1.64E-26 | 3.41E-22 | 0.42 | 0.12 | 0.3 | 3.5 |
| other bloold cells | Cmtm6 | 1.66 | 1.65E-26 | 3.44E-22 | 0.44 | 0.13 | 0.31 | 3.39 |
| other bloold cells | Nek7 | 1.5 | 1.72E-26 | 3.58E-22 | 0.67 | 0.29 | 0.38 | 2.28 |
| other bloold cells | Il15 | 2.57 | 1.89E-26 | 3.93E-22 | 0.16 | 0.02 | 0.14 | 7 |
| other bloold cells | Rab5c | 1.23 | 1.96E-26 | 4.07E-22 | 0.94 | 0.65 | 0.3 | 1.46 |
| other bloold cells | Pld2 | 2.69 | 2.14E-26 | 4.46E-22 | 0.15 | 0.02 | 0.13 | 7.33 |
| other bloold cells | Pea15a | 1.67 | 2.40E-26 | 4.99E-22 | 0.6 | 0.23 | 0.37 | 2.6 |
| other bloold cells | Ppp1r3b | 2.23 | 2.42E-26 | 5.04E-22 | 0.22 | 0.04 | 0.18 | 5.46 |
| other bloold cells | Cfp | 2.48 | 2.46E-26 | 5.11E-22 | 0.23 | 0.04 | 0.19 | 5.25 |
| other bloold cells | Akr1b10 | 1.66 | 2.93E-26 | 6.09E-22 | 0.41 | 0.12 | 0.3 | 3.56 |
| other bloold cells | Ada | 1.82 | 2.98E-26 | 6.20E-22 | 0.33 | 0.08 | 0.25 | 4.11 |
| other bloold cells | Arhgef1 | 1.42 | 3.25E-26 | 6.76E-22 | 0.66 | 0.27 | 0.39 | 2.47 |
| other bloold cells | Comtd1 | 1.65 | 3.49E-26 | 7.26E-22 | 0.46 | 0.14 | 0.32 | 3.25 |
| other bloold cells | Mthfs | 1.88 | 3.57E-26 | 7.43E-22 | 0.46 | 0.14 | 0.32 | 3.3 |
| other bloold cells | Acsl1 | 1.7 | 3.68E-26 | 7.64E-22 | 0.39 | 0.1 | 0.28 | 3.74 |
| other bloold cells | Rpl10a | 0.37 | 3.96E-26 | 8.23E-22 | 0.99 | 1 | -0.01 | 0.99 |
| other bloold cells | Gnpda1 | 2 | 4.87E-26 | 1.01E-21 | 0.52 | 0.18 | 0.33 | 2.83 |
| other bloold cells | Tor1aip1 | 1.35 | 5.09E-26 | 1.06E-21 | 0.85 | 0.44 | 0.41 | 1.94 |
| other bloold cells | Psme2 | 1.01 | 5.20E-26 | 1.08E-21 | 0.96 | 0.76 | 0.2 | 1.26 |
| other bloold cells | Washc4 | 1.47 | 6.13E-26 | 1.28E-21 | 0.72 | 0.31 | 0.41 | 2.29 |
| other bloold cells | Atp6ap1 | 1.28 | 7.03E-26 | 1.46E-21 | 0.76 | 0.35 | 0.41 | 2.15 |
| other bloold cells | Cox17 | 0.88 | 7.38E-26 | 1.53E-21 | 0.98 | 0.77 | 0.21 | 1.28 |
| other bloold cells | Gla | 2.23 | 8.53E-26 | 1.77E-21 | 0.38 | 0.1 | 0.28 | 3.71 |

**Supplementary Table 2. Marker genes for the major cell types of integrated clustering of E11.5 control and NotchGOF λ-junction cells.**

| | | | | | | | | |
|---|---|---|---|---|---|---|---|---|
| other bloold cells | Eif2ak2 | 1.41 | 8.87E-26 | 1.85E-21 | 0.67 | 0.28 | 0.39 | 2.42 |
| other bloold cells | Cnp | 1.61 | 1.08E-25 | 2.25E-21 | 0.48 | 0.16 | 0.33 | 3.12 |
| other bloold cells | Pfkp | 2.04 | 1.13E-25 | 2.35E-21 | 0.25 | 0.05 | 0.2 | 4.94 |
| other bloold cells | Il13ra1 | 2.31 | 1.21E-25 | 2.52E-21 | 0.23 | 0.05 | 0.19 | 5.13 |
| other bloold cells | Slc35c1 | 2.2 | 1.24E-25 | 2.58E-21 | 0.22 | 0.04 | 0.18 | 5.43 |
| other bloold cells | Rpl35a | 0.31 | 1.40E-25 | 2.92E-21 | 0.99 | 1 | -0.01 | 0.99 |
| other bloold cells | Smap2 | 1.31 | 1.50E-25 | 3.12E-21 | 0.77 | 0.37 | 0.4 | 2.07 |
| other bloold cells | Csk | 1.15 | 1.51E-25 | 3.15E-21 | 0.87 | 0.54 | 0.33 | 1.61 |
| other bloold cells | Mcl1 | 0.96 | 1.65E-25 | 3.44E-21 | 0.99 | 0.82 | 0.17 | 1.21 |
| other bloold cells | Dhrs3 | 1.66 | 1.69E-25 | 3.52E-21 | 0.66 | 0.29 | 0.36 | 2.23 |
| other bloold cells | Tpm2 | 1.99 | 2.32E-25 | 4.82E-21 | 0.78 | 0.44 | 0.35 | 1.8 |
| other bloold cells | Swap70 | 1.34 | 2.33E-25 | 4.84E-21 | 0.72 | 0.33 | 0.39 | 2.2 |
| other bloold cells | Marchf2 | 1.25 | 2.41E-25 | 5.01E-21 | 0.59 | 0.21 | 0.37 | 2.76 |
| other bloold cells | Colgalt1 | 1.22 | 2.55E-25 | 5.31E-21 | 0.87 | 0.47 | 0.39 | 1.83 |
| other bloold cells | Aph1c | 2.4 | 2.57E-25 | 5.35E-21 | 0.21 | 0.04 | 0.17 | 5.53 |
| other bloold cells | Zfp979 | 2.24 | 2.61E-25 | 5.44E-21 | 0.2 | 0.03 | 0.16 | 5.76 |
| other bloold cells | Nr3c1 | 1.59 | 2.74E-25 | 5.69E-21 | 0.49 | 0.16 | 0.33 | 3.02 |
| other bloold cells | Ormdl2 | 1.22 | 3.04E-25 | 6.32E-21 | 0.75 | 0.35 | 0.4 | 2.14 |
| other bloold cells | Vps33a | 1.64 | 3.37E-25 | 7.01E-21 | 0.45 | 0.14 | 0.31 | 3.25 |
| other bloold cells | Ints6l | 2 | 3.43E-25 | 7.13E-21 | 0.48 | 0.17 | 0.31 | 2.84 |
| other bloold cells | Mef2a | 1.46 | 3.58E-25 | 7.44E-21 | 0.78 | 0.39 | 0.39 | 2.01 |
| other bloold cells | Cpt1a | 1.93 | 3.79E-25 | 7.89E-21 | 0.36 | 0.1 | 0.26 | 3.64 |
| other bloold cells | Acat1 | 1.2 | 4.45E-25 | 9.26E-21 | 0.85 | 0.46 | 0.38 | 1.82 |
| other bloold cells | Ubtd1 | 1.56 | 4.63E-25 | 9.63E-21 | 0.47 | 0.15 | 0.32 | 3.15 |
| other bloold cells | Gdpd3 | 3.98 | 5.07E-25 | 1.05E-20 | 0.12 | 0.01 | 0.11 | 8.5 |
| other bloold cells | Taldo1 | 0.84 | 5.18E-25 | 1.08E-20 | 0.96 | 0.89 | 0.07 | 1.08 |
| other bloold cells | Dnase1l1 | 2.37 | 6.82E-25 | 1.42E-20 | 0.23 | 0.05 | 0.19 | 5.02 |
| other bloold cells | Prcp | 1.49 | 6.86E-25 | 1.43E-20 | 0.64 | 0.26 | 0.38 | 2.48 |
| other bloold cells | Dennd4b | 2.33 | 7.32E-25 | 1.52E-20 | 0.24 | 0.05 | 0.19 | 4.86 |
| other bloold cells | Cdkn1a | 0.82 | 8.76E-25 | 1.82E-20 | 0.4 | 0.11 | 0.29 | 3.59 |
| other bloold cells | BC005537 | 1.38 | 9.10E-25 | 1.89E-20 | 0.82 | 0.5 | 0.32 | 1.65 |
| other bloold cells | Fuca1 | 1.4 | 9.12E-25 | 1.90E-20 | 0.75 | 0.35 | 0.39 | 2.11 |
| other bloold cells | Slc2a5 | 2.81 | 9.66E-25 | 2.01E-20 | 0.13 | 0.02 | 0.11 | 7.88 |
| other bloold cells | Hpse | 2.78 | 9.91E-25 | 2.06E-20 | 0.13 | 0.02 | 0.12 | 7.82 |
| other bloold cells | Litaf | 1.67 | 1.08E-24 | 2.24E-20 | 0.85 | 0.5 | 0.34 | 1.68 |
| other bloold cells | Cltb | 1.52 | 1.24E-24 | 2.58E-20 | 0.52 | 0.18 | 0.34 | 2.88 |
| other bloold cells | B4galt1 | 1.45 | 1.26E-24 | 2.61E-20 | 0.6 | 0.23 | 0.37 | 2.65 |
| other bloold cells | H2-DMa | 2.2 | 1.34E-24 | 2.78E-20 | 0.25 | 0.05 | 0.2 | 4.67 |
| other bloold cells | Rhoa | 0.75 | 1.66E-24 | 3.45E-20 | 1 | 0.97 | 0.03 | 1.04 |
| other bloold cells | Unc13d | 2.33 | 1.90E-24 | 3.96E-20 | 0.18 | 0.03 | 0.15 | 6.03 |
| other bloold cells | Fbxo6 | 2.31 | 1.94E-24 | 4.03E-20 | 0.27 | 0.06 | 0.21 | 4.33 |
| other bloold cells | Dnm2 | 1.27 | 2.15E-24 | 4.47E-20 | 0.8 | 0.39 | 0.41 | 2.06 |
| other bloold cells | Kif21b | 1.54 | 2.34E-24 | 4.87E-20 | 0.49 | 0.16 | 0.33 | 3.02 |
| other bloold cells | Lpcat3 | 1.31 | 2.87E-24 | 5.97E-20 | 0.71 | 0.31 | 0.4 | 2.31 |
| other bloold cells | Jmjd1c | 1.32 | 3.08E-24 | 6.40E-20 | 0.85 | 0.48 | 0.36 | 1.76 |
| other bloold cells | Osbpl11 | 1.54 | 3.40E-24 | 7.07E-20 | 0.64 | 0.28 | 0.36 | 2.3 |
| other bloold cells | Nlrx1 | 2.63 | 3.59E-24 | 7.47E-20 | 0.12 | 0.01 | 0.11 | 8.5 |
| other bloold cells | Rnaset2b | 1.42 | 3.80E-24 | 7.90E-20 | 0.57 | 0.21 | 0.36 | 2.68 |
| other bloold cells | Ptrhd1 | 1.28 | 4.20E-24 | 8.73E-20 | 0.76 | 0.36 | 0.4 | 2.1 |
| other bloold cells | Ccdc12 | 0.9 | 4.27E-24 | 8.87E-20 | 0.94 | 0.71 | 0.23 | 1.33 |
| other bloold cells | Arf3 | 1.34 | 5.07E-24 | 1.06E-19 | 0.71 | 0.31 | 0.4 | 2.29 |
| other bloold cells | Mcrip2 | 1.93 | 5.22E-24 | 1.09E-19 | 0.29 | 0.07 | 0.22 | 4.16 |
| other bloold cells | Etfb | 0.79 | 5.26E-24 | 1.09E-19 | 0.98 | 0.89 | 0.09 | 1.1 |
| other bloold cells | Shfl | 2.73 | 5.66E-24 | 1.18E-19 | 0.12 | 0.02 | 0.1 | 7.93 |
| other bloold cells | Slc45a4 | 1.74 | 6.02E-24 | 1.25E-19 | 0.43 | 0.13 | 0.29 | 3.19 |
| other bloold cells | Fcho1 | 3.07 | 6.11E-24 | 1.27E-19 | 0.13 | 0.02 | 0.12 | 7.39 |

**Supplementary Table 2. Marker genes for the major cell types of integrated clustering of E11.5 control and NotchGOF λ-junction cells.**

| | | | | | | | | |
|---|---|---|---|---|---|---|---|---|
| other bloold cells | Fuca2 | 1.42 | 6.56E-24 | 1.36E-19 | 0.59 | 0.24 | 0.36 | 2.53 |
| other bloold cells | Tmbim4 | 1.18 | 6.91E-24 | 1.44E-19 | 0.83 | 0.49 | 0.34 | 1.68 |
| other bloold cells | Iqgap2 | 1.33 | 7.05E-24 | 1.47E-19 | 0.73 | 0.36 | 0.37 | 2.02 |
| other bloold cells | Ppp6r1 | 1.37 | 7.90E-24 | 1.64E-19 | 0.62 | 0.25 | 0.37 | 2.46 |
| other bloold cells | Jdp2 | 2 | 8.00E-24 | 1.66E-19 | 0.2 | 0.04 | 0.17 | 5.49 |
| other bloold cells | Wbp2 | 1.32 | 1.02E-23 | 2.12E-19 | 0.65 | 0.28 | 0.37 | 2.36 |
| other bloold cells | Piezo1 | 1.66 | 1.25E-23 | 2.59E-19 | 0.41 | 0.12 | 0.28 | 3.3 |
| other bloold cells | Tpm3 | 0.59 | 1.35E-23 | 2.80E-19 | 0.99 | 0.96 | 0.04 | 1.04 |
| other bloold cells | Dgkd | 1.59 | 1.37E-23 | 2.85E-19 | 0.57 | 0.24 | 0.34 | 2.44 |
| other bloold cells | Rpl36al | 0.44 | 1.47E-23 | 3.05E-19 | 0.99 | 1 | 0 | 1 |
| other bloold cells | AU020206 | 1.39 | 1.49E-23 | 3.10E-19 | 0.59 | 0.22 | 0.37 | 2.66 |
| other bloold cells | 031439G07Ri | 1.53 | 1.51E-23 | 3.13E-19 | 0.59 | 0.25 | 0.34 | 2.38 |
| other bloold cells | Pde4b | 2.16 | 1.58E-23 | 3.28E-19 | 0.22 | 0.04 | 0.17 | 5.05 |
| other bloold cells | Brk1 | 0.72 | 1.59E-23 | 3.31E-19 | 0.99 | 0.92 | 0.07 | 1.08 |
| other bloold cells | Rps11 | 0.26 | 1.64E-23 | 3.42E-19 | 1 | 1 | 0 | 1 |
| other bloold cells | Tbc1d22a | 1.78 | 1.88E-23 | 3.92E-19 | 0.3 | 0.07 | 0.23 | 4.07 |
| other bloold cells | Gng2 | 0.9 | 1.92E-23 | 3.99E-19 | 0.93 | 0.57 | 0.36 | 1.63 |
| other bloold cells | Tmem50a | 0.93 | 2.00E-23 | 4.17E-19 | 0.99 | 0.81 | 0.19 | 1.23 |
| other bloold cells | Dok1 | 1.77 | 2.18E-23 | 4.53E-19 | 0.36 | 0.1 | 0.26 | 3.57 |
| other bloold cells | Fam98c | 1.64 | 2.24E-23 | 4.66E-19 | 0.39 | 0.12 | 0.28 | 3.41 |
| other bloold cells | Psen1 | 1.51 | 2.58E-23 | 5.37E-19 | 0.62 | 0.25 | 0.37 | 2.49 |
| other bloold cells | Mef2d | 1.65 | 2.60E-23 | 5.41E-19 | 0.43 | 0.14 | 0.3 | 3.14 |
| other bloold cells | Sord | 1.46 | 2.76E-23 | 5.74E-19 | 0.45 | 0.15 | 0.3 | 3.09 |
| other bloold cells | Ankrd13a | 1.37 | 2.98E-23 | 6.19E-19 | 0.66 | 0.29 | 0.38 | 2.3 |
| other bloold cells | Fam20c | 3.92 | 3.02E-23 | 6.28E-19 | 0.15 | 0.02 | 0.13 | 6.68 |
| other bloold cells | Ciao2a | 0.84 | 3.07E-23 | 6.38E-19 | 0.95 | 0.85 | 0.1 | 1.12 |
| other bloold cells | Gng12 | 1.56 | 3.44E-23 | 7.16E-19 | 0.72 | 0.36 | 0.36 | 2 |
| other bloold cells | Plcl2 | 1.6 | 3.48E-23 | 7.24E-19 | 0.39 | 0.11 | 0.27 | 3.44 |
| other bloold cells | 700017B05Ri | 1.65 | 3.54E-23 | 7.37E-19 | 0.51 | 0.18 | 0.33 | 2.85 |
| other bloold cells | Rilpl2 | 1.33 | 4.05E-23 | 8.41E-19 | 0.67 | 0.28 | 0.4 | 2.43 |
| other bloold cells | Nptn | 1.32 | 4.41E-23 | 9.17E-19 | 0.8 | 0.45 | 0.34 | 1.76 |
| other bloold cells | Orai2 | 2.22 | 5.03E-23 | 1.05E-18 | 0.22 | 0.04 | 0.17 | 4.93 |
| other bloold cells | Cyb561d2 | 1.52 | 5.21E-23 | 1.08E-18 | 0.49 | 0.17 | 0.32 | 2.9 |
| other bloold cells | Rpl39 | 0.3 | 5.36E-23 | 1.12E-18 | 1 | 1 | 0 | 1 |
| other bloold cells | Frmd8 | 1.39 | 5.51E-23 | 1.15E-18 | 0.55 | 0.2 | 0.34 | 2.7 |
| other bloold cells | Mgat2 | 1.35 | 5.59E-23 | 1.16E-18 | 0.8 | 0.46 | 0.34 | 1.74 |
| other bloold cells | Rab39 | 3 | 5.89E-23 | 1.22E-18 | 0.11 | 0.01 | 0.09 | 8.75 |
| other bloold cells | Sbno2 | 1.62 | 6.91E-23 | 1.44E-18 | 0.39 | 0.11 | 0.27 | 3.38 |
| other bloold cells | Psen2 | 1.87 | 8.71E-23 | 1.81E-18 | 0.3 | 0.08 | 0.22 | 3.91 |
| other bloold cells | Lgals1 | 1.41 | 8.83E-23 | 1.84E-18 | 0.98 | 0.83 | 0.15 | 1.18 |
| other bloold cells | Ogfrl1 | 1.55 | 1.35E-22 | 2.80E-18 | 0.67 | 0.34 | 0.34 | 2 |
| other bloold cells | Zdhhc14 | 2.91 | 1.71E-22 | 3.56E-18 | 0.16 | 0.03 | 0.13 | 5.96 |
| other bloold cells | Stom | 2.22 | 1.95E-22 | 4.05E-18 | 0.22 | 0.05 | 0.17 | 4.82 |
| other bloold cells | Pik3cb | 1.77 | 1.99E-22 | 4.13E-18 | 0.27 | 0.06 | 0.2 | 4.29 |
| other bloold cells | Fnip2 | 2.14 | 2.00E-22 | 4.16E-18 | 0.34 | 0.1 | 0.24 | 3.43 |
| other bloold cells | Msrb1 | 1.46 | 2.03E-22 | 4.22E-18 | 0.83 | 0.49 | 0.34 | 1.69 |
| other bloold cells | Zyx | 1.15 | 2.08E-22 | 4.33E-18 | 0.84 | 0.51 | 0.33 | 1.65 |
| other bloold cells | Ralb | 1.2 | 2.52E-22 | 5.24E-18 | 0.8 | 0.43 | 0.37 | 1.85 |
| other bloold cells | Plk3 | 2.68 | 3.24E-22 | 6.75E-18 | 0.15 | 0.02 | 0.12 | 6.39 |
| other bloold cells | Rnh1 | 0.89 | 3.31E-22 | 6.89E-18 | 0.9 | 0.55 | 0.34 | 1.62 |
| other bloold cells | Tnfrsf1a | 1.32 | 3.39E-22 | 7.05E-18 | 0.69 | 0.3 | 0.39 | 2.3 |
| other bloold cells | Herc6 | 1.84 | 3.41E-22 | 7.09E-18 | 0.24 | 0.05 | 0.19 | 4.67 |
| other bloold cells | Ripk3 | 1.43 | 3.66E-22 | 7.61E-18 | 0.48 | 0.16 | 0.32 | 2.95 |
| other bloold cells | Bcl3 | 2.19 | 3.69E-22 | 7.67E-18 | 0.27 | 0.07 | 0.21 | 4.14 |
| other bloold cells | Hspa4l | 1.31 | 3.85E-22 | 8.02E-18 | 0.38 | 0.11 | 0.27 | 3.41 |
| other bloold cells | Slc15a4 | 1.6 | 5.07E-22 | 1.05E-17 | 0.44 | 0.15 | 0.3 | 3.02 |

**Supplementary Table 2. Marker genes for the major cell types of integrated clustering of E11.5 control and NotchGOF λ-junction cells.**

| | | | | | | | | |
|---|---|---|---|---|---|---|---|---|
| other bloold cells | Arfgef2 | 1.56 | 6.45E-22 | 1.34E-17 | 0.34 | 0.09 | 0.25 | 3.69 |
| other bloold cells | Ap2s1 | 0.9 | 6.47E-22 | 1.35E-17 | 0.99 | 0.89 | 0.11 | 1.12 |
| other bloold cells | Cln5 | 1.38 | 6.52E-22 | 1.36E-17 | 0.55 | 0.21 | 0.34 | 2.59 |
| other bloold cells | Sft2d1 | 1.43 | 6.64E-22 | 1.38E-17 | 0.55 | 0.22 | 0.33 | 2.49 |
| other bloold cells | Glul | 1.69 | 7.80E-22 | 1.62E-17 | 0.87 | 0.62 | 0.25 | 1.41 |
| other bloold cells | Atp6v1a | 1.24 | 9.28E-22 | 1.93E-17 | 0.88 | 0.53 | 0.36 | 1.67 |
| other bloold cells | Npepl1 | 1.13 | 9.28E-22 | 1.93E-17 | 0.71 | 0.33 | 0.39 | 2.18 |
| other bloold cells | H2-M3 | 2.86 | 9.36E-22 | 1.95E-17 | 0.13 | 0.02 | 0.11 | 7 |
| other bloold cells | Cpeb4 | 1.65 | 1.06E-21 | 2.21E-17 | 0.34 | 0.09 | 0.24 | 3.57 |
| other bloold cells | Tmem14c | 0.56 | 1.14E-21 | 2.38E-17 | 0.99 | 0.92 | 0.06 | 1.07 |
| other bloold cells | Rin2 | 1.79 | 1.19E-21 | 2.47E-17 | 0.51 | 0.2 | 0.31 | 2.54 |
| other bloold cells | Fmn1 | 2.83 | 1.25E-21 | 2.59E-17 | 0.13 | 0.02 | 0.11 | 7 |
| other bloold cells | Psat1 | 1.23 | 1.38E-21 | 2.87E-17 | 0.78 | 0.45 | 0.33 | 1.72 |
| other bloold cells | Ophn1 | 1.84 | 1.49E-21 | 3.10E-17 | 0.5 | 0.19 | 0.31 | 2.6 |
| other bloold cells | Tram1 | 1.15 | 1.58E-21 | 3.28E-17 | 0.83 | 0.51 | 0.32 | 1.63 |
| other bloold cells | Orai3 | 1.58 | 1.65E-21 | 3.42E-17 | 0.38 | 0.11 | 0.26 | 3.32 |
| other bloold cells | Derl2 | 1.22 | 1.79E-21 | 3.72E-17 | 0.74 | 0.38 | 0.37 | 1.97 |
| other bloold cells | Mapkapk2 | 1.11 | 1.98E-21 | 4.12E-17 | 0.85 | 0.55 | 0.3 | 1.55 |
| other bloold cells | mt-Co1 | 0.44 | 2.07E-21 | 4.31E-17 | 1 | 0.99 | 0.01 | 1.01 |
| other bloold cells | Nmt1 | 1.07 | 2.13E-21 | 4.42E-17 | 0.9 | 0.61 | 0.29 | 1.47 |
| other bloold cells | Amdhd2 | 1.45 | 2.41E-21 | 5.02E-17 | 0.59 | 0.25 | 0.35 | 2.42 |
| other bloold cells | Eef2kmt | 1.37 | 2.68E-21 | 5.58E-17 | 0.5 | 0.18 | 0.32 | 2.76 |
| other bloold cells | Ppp1r21 | 1.4 | 2.72E-21 | 5.66E-17 | 0.54 | 0.21 | 0.33 | 2.55 |
| other bloold cells | Rab7 | 1 | 2.87E-21 | 5.97E-17 | 0.96 | 0.69 | 0.27 | 1.39 |
| other bloold cells | Ctbs | 1.84 | 2.97E-21 | 6.18E-17 | 0.33 | 0.09 | 0.24 | 3.5 |
| other bloold cells | Cox4i1 | 0.38 | 3.14E-21 | 6.52E-17 | 1 | 1 | 0 | 1 |
| other bloold cells | Tmem176b | 1.68 | 3.22E-21 | 6.69E-17 | 0.81 | 0.57 | 0.24 | 1.43 |
| other bloold cells | Agpat5 | 1.25 | 3.51E-21 | 7.31E-17 | 0.78 | 0.45 | 0.34 | 1.76 |
| other bloold cells | Morc3 | 1.16 | 3.54E-21 | 7.37E-17 | 0.79 | 0.42 | 0.37 | 1.87 |
| other bloold cells | Pip4p1 | 1.26 | 3.82E-21 | 7.95E-17 | 0.72 | 0.36 | 0.36 | 2 |
| other bloold cells | Zzef1 | 1.55 | 3.94E-21 | 8.18E-17 | 0.48 | 0.17 | 0.31 | 2.8 |
| other bloold cells | Lactb | 1.33 | 4.01E-21 | 8.34E-17 | 0.69 | 0.34 | 0.35 | 2.04 |
| other bloold cells | Ccdc22 | 1.16 | 4.05E-21 | 8.43E-17 | 0.69 | 0.3 | 0.39 | 2.32 |
| other bloold cells | D6Wsu163e | 1.72 | 4.15E-21 | 8.63E-17 | 0.34 | 0.1 | 0.24 | 3.54 |
| other bloold cells | Wipf1 | 1.35 | 4.33E-21 | 9.00E-17 | 0.57 | 0.23 | 0.34 | 2.49 |
| other bloold cells | Jak1 | 0.98 | 4.61E-21 | 9.60E-17 | 0.93 | 0.61 | 0.32 | 1.53 |
| other bloold cells | Ldlrad4 | 2.09 | 4.62E-21 | 9.61E-17 | 0.18 | 0.03 | 0.14 | 5.47 |
| other bloold cells | Paqr4 | 2.46 | 4.66E-21 | 9.69E-17 | 0.18 | 0.04 | 0.15 | 5.2 |
| other bloold cells | B3gnt2 | 1.5 | 4.72E-21 | 9.81E-17 | 0.34 | 0.1 | 0.24 | 3.46 |
| other bloold cells | Cstb | 1.3 | 5.71E-21 | 1.19E-16 | 0.93 | 0.8 | 0.13 | 1.16 |
| other bloold cells | Rbm43 | 1.56 | 6.73E-21 | 1.40E-16 | 0.36 | 0.11 | 0.26 | 3.37 |
| other bloold cells | Pomp | 0.54 | 6.79E-21 | 1.41E-16 | 0.99 | 0.98 | 0.02 | 1.02 |
| other bloold cells | Rgs19 | 1.11 | 7.50E-21 | 1.56E-16 | 0.8 | 0.45 | 0.35 | 1.78 |
| other bloold cells | Abr | 1.19 | 8.48E-21 | 1.76E-16 | 0.57 | 0.22 | 0.36 | 2.64 |
| other bloold cells | Nubp2 | 1.21 | 8.81E-21 | 1.83E-16 | 0.67 | 0.31 | 0.37 | 2.19 |
| other bloold cells | Qsox1 | 1.51 | 9.74E-21 | 2.03E-16 | 0.38 | 0.12 | 0.26 | 3.26 |
| other bloold cells | Chmp3 | 1.05 | 1.08E-20 | 2.24E-16 | 0.83 | 0.46 | 0.37 | 1.81 |
| other bloold cells | Spcs3 | 1.12 | 1.14E-20 | 2.36E-16 | 0.76 | 0.38 | 0.38 | 2.02 |
| other bloold cells | Myo9b | 1.26 | 1.16E-20 | 2.41E-16 | 0.58 | 0.23 | 0.35 | 2.52 |
| other bloold cells | Tbc1d14 | 1.45 | 1.23E-20 | 2.55E-16 | 0.51 | 0.19 | 0.32 | 2.64 |
| other bloold cells | Sfxn5 | 2.28 | 1.23E-20 | 2.56E-16 | 0.2 | 0.04 | 0.16 | 4.9 |
| other bloold cells | Tbc1d9 | 1.98 | 1.38E-20 | 2.87E-16 | 0.29 | 0.08 | 0.21 | 3.78 |
| other bloold cells | Arf6 | 1.14 | 1.49E-20 | 3.10E-16 | 0.75 | 0.4 | 0.35 | 1.89 |
| other bloold cells | Dse | 1.46 | 1.63E-20 | 3.40E-16 | 0.59 | 0.27 | 0.33 | 2.22 |
| other bloold cells | Cln8 | 1.69 | 1.69E-20 | 3.51E-16 | 0.52 | 0.21 | 0.32 | 2.56 |
| other bloold cells | Stx4a | 1.3 | 1.78E-20 | 3.71E-16 | 0.69 | 0.32 | 0.37 | 2.17 |

**Supplementary Table 2. Marker genes for the major cell types of integrated clustering of E11.5 control and NotchGOF λ-junction cells.**

| | | | | | | | | |
|---|---|---|---|---|---|---|---|---|
| other bloold cells | Akr1a1 | 0.77 | 2.07E-20 | 4.30E-16 | 1 | 0.96 | 0.04 | 1.04 |
| other bloold cells | Myadm | 1.54 | 2.12E-20 | 4.41E-16 | 0.39 | 0.13 | 0.27 | 3.14 |
| other bloold cells | Vps29 | 0.81 | 2.47E-20 | 5.14E-16 | 0.94 | 0.78 | 0.17 | 1.21 |
| other bloold cells | 510039O18Ri | 1.3 | 2.81E-20 | 5.84E-16 | 0.57 | 0.23 | 0.33 | 2.43 |
| other bloold cells | Bola2 | 0.72 | 3.10E-20 | 6.46E-16 | 0.97 | 0.88 | 0.08 | 1.09 |
| other bloold cells | Snapc5 | 0.97 | 3.59E-20 | 7.47E-16 | 0.86 | 0.55 | 0.31 | 1.56 |
| other bloold cells | B4galt5 | 1.67 | 3.62E-20 | 7.53E-16 | 0.35 | 0.11 | 0.25 | 3.33 |
| other bloold cells | Lman2 | 0.99 | 4.03E-20 | 8.39E-16 | 0.86 | 0.62 | 0.24 | 1.39 |
| other bloold cells | Ech1 | 1 | 4.26E-20 | 8.87E-16 | 0.86 | 0.51 | 0.35 | 1.68 |
| other bloold cells | Pde3b | 1.32 | 4.73E-20 | 9.85E-16 | 0.59 | 0.24 | 0.35 | 2.44 |
| other bloold cells | Atxn3 | 1.23 | 5.93E-20 | 1.23E-15 | 0.5 | 0.19 | 0.31 | 2.66 |
| other bloold cells | Rpl10 | 0.32 | 6.26E-20 | 1.30E-15 | 1 | 1 | 0 | 1 |
| other bloold cells | Cebpzos | 1.36 | 6.43E-20 | 1.34E-15 | 0.87 | 0.57 | 0.3 | 1.53 |
| other bloold cells | Bsn | 2.1 | 6.65E-20 | 1.38E-15 | 0.18 | 0.03 | 0.14 | 5.3 |
| other bloold cells | Herpud1 | 1.3 | 6.73E-20 | 1.40E-15 | 0.6 | 0.25 | 0.35 | 2.41 |
| other bloold cells | Actg1 | 0.44 | 8.61E-20 | 1.79E-15 | 1 | 1 | 0 | 1 |
| other bloold cells | Pign | 1.26 | 8.66E-20 | 1.80E-15 | 0.43 | 0.14 | 0.29 | 3.01 |
| other bloold cells | Mertk | 1.9 | 8.67E-20 | 1.80E-15 | 0.43 | 0.16 | 0.27 | 2.7 |
| other bloold cells | Mob3a | 1.41 | 8.95E-20 | 1.86E-15 | 0.46 | 0.16 | 0.3 | 2.88 |
| other bloold cells | Emilin2 | 1.55 | 9.21E-20 | 1.92E-15 | 0.58 | 0.26 | 0.33 | 2.27 |
| other bloold cells | Acadm | 1.28 | 1.05E-19 | 2.19E-15 | 0.67 | 0.33 | 0.34 | 2.02 |
| other bloold cells | Dynlt3 | 1.19 | 1.08E-19 | 2.25E-15 | 0.61 | 0.26 | 0.35 | 2.38 |
| other bloold cells | Rab3d | 2.33 | 1.08E-19 | 2.25E-15 | 0.13 | 0.02 | 0.11 | 6.33 |
| other bloold cells | Abi1 | 1.18 | 1.25E-19 | 2.60E-15 | 0.76 | 0.39 | 0.37 | 1.96 |
| other bloold cells | Ppp1r13b | 1.58 | 1.36E-19 | 2.83E-15 | 0.29 | 0.08 | 0.21 | 3.68 |
| other bloold cells | Derl1 | 1.09 | 1.42E-19 | 2.96E-15 | 0.8 | 0.49 | 0.32 | 1.65 |
| other bloold cells | Hgf | 2.67 | 1.52E-19 | 3.16E-15 | 0.13 | 0.02 | 0.11 | 6.63 |
| other bloold cells | Pts | 1.13 | 1.66E-19 | 3.44E-15 | 0.78 | 0.41 | 0.38 | 1.93 |
| other bloold cells | Fig4 | 1.76 | 2.07E-19 | 4.30E-15 | 0.28 | 0.08 | 0.21 | 3.73 |
| other bloold cells | Atg3 | 1.16 | 2.19E-19 | 4.55E-15 | 0.78 | 0.45 | 0.33 | 1.74 |
| other bloold cells | 610012G03Ri | 0.84 | 2.54E-19 | 5.28E-15 | 0.99 | 0.86 | 0.12 | 1.14 |
| other bloold cells | Tnfrsf21 | 1.51 | 2.58E-19 | 5.36E-15 | 0.41 | 0.14 | 0.27 | 2.93 |
| other bloold cells | Cdk6 | 1.43 | 2.74E-19 | 5.69E-15 | 0.68 | 0.34 | 0.34 | 1.99 |
| other bloold cells | Trappc6a | 0.97 | 2.77E-19 | 5.77E-15 | 0.89 | 0.52 | 0.37 | 1.7 |
| other bloold cells | Gimap6 | 1.57 | 3.04E-19 | 6.31E-15 | 0.15 | 0.03 | 0.13 | 5.7 |
| other bloold cells | Ap5z1 | 1.74 | 3.17E-19 | 6.60E-15 | 0.22 | 0.05 | 0.17 | 4.31 |
| other bloold cells | Malt1 | 1.46 | 3.53E-19 | 7.34E-15 | 0.45 | 0.17 | 0.28 | 2.72 |
| other bloold cells | Dnajb14 | 1.35 | 4.09E-19 | 8.52E-15 | 0.67 | 0.32 | 0.35 | 2.09 |
| other bloold cells | Card19 | 1.07 | 4.20E-19 | 8.74E-15 | 0.6 | 0.25 | 0.35 | 2.39 |
| other bloold cells | Plgrkt | 1.18 | 4.26E-19 | 8.87E-15 | 0.81 | 0.48 | 0.33 | 1.7 |
| other bloold cells | Hps1 | 1.38 | 5.99E-19 | 1.25E-14 | 0.39 | 0.13 | 0.26 | 3.06 |
| other bloold cells | Osbpl8 | 1.24 | 6.16E-19 | 1.28E-14 | 0.56 | 0.23 | 0.33 | 2.41 |
| other bloold cells | Commd4 | 0.96 | 6.46E-19 | 1.34E-14 | 0.87 | 0.53 | 0.34 | 1.64 |
| other bloold cells | Cds1 | 1.86 | 7.20E-19 | 1.50E-14 | 0.19 | 0.04 | 0.15 | 4.85 |
| other bloold cells | Sys1 | 0.82 | 7.21E-19 | 1.50E-14 | 0.94 | 0.7 | 0.23 | 1.33 |
| other bloold cells | Plod1 | 1.51 | 8.11E-19 | 1.69E-14 | 0.5 | 0.2 | 0.3 | 2.49 |
| other bloold cells | Tapbpl | 2.64 | 8.32E-19 | 1.73E-14 | 0.11 | 0.02 | 0.1 | 7 |
| other bloold cells | Nfe2 | 2.12 | 9.11E-19 | 1.89E-14 | 0.18 | 0.04 | 0.14 | 5 |
| other bloold cells | Ebp | 1.22 | 9.55E-19 | 1.99E-14 | 0.73 | 0.4 | 0.32 | 1.8 |
| other bloold cells | 610507B11Ri | 1.06 | 9.61E-19 | 2.00E-14 | 0.81 | 0.46 | 0.35 | 1.75 |
| other bloold cells | Slc41a3 | 2.73 | 9.73E-19 | 2.02E-14 | 0.14 | 0.02 | 0.12 | 5.83 |
| other bloold cells | Wwp2 | 1.37 | 9.80E-19 | 2.04E-14 | 0.64 | 0.31 | 0.33 | 2.09 |
| other bloold cells | Fam173a | 1.03 | 9.82E-19 | 2.04E-14 | 0.84 | 0.5 | 0.34 | 1.66 |
| other bloold cells | Uba7 | 1.5 | 9.87E-19 | 2.05E-14 | 0.34 | 0.11 | 0.24 | 3.21 |
| other bloold cells | Ppt1 | 1.29 | 1.14E-18 | 2.37E-14 | 0.65 | 0.31 | 0.34 | 2.07 |
| other bloold cells | Dennd1c | 1.9 | 1.20E-18 | 2.49E-14 | 0.17 | 0.03 | 0.14 | 5.09 |

**Supplementary Table 2. Marker genes for the major cell types of integrated clustering of E11.5 control and NotchGOF λ-junction cells.**

| | | | | | | | | |
|---|---|---|---|---|---|---|---|---|
| other bloold cells | Timm10b | 0.69 | 1.33E-18 | 2.77E-14 | 0.96 | 0.85 | 0.11 | 1.13 |
| other bloold cells | Tie1 | 1.53 | 1.39E-18 | 2.90E-14 | 0.11 | 0.02 | 0.1 | 7 |
| other bloold cells | Mgat1 | 1.36 | 1.54E-18 | 3.21E-14 | 0.51 | 0.21 | 0.3 | 2.46 |
| other bloold cells | Vac14 | 1.54 | 1.60E-18 | 3.32E-14 | 0.52 | 0.22 | 0.31 | 2.4 |
| other bloold cells | Uchl5 | 1.13 | 1.62E-18 | 3.37E-14 | 0.79 | 0.54 | 0.25 | 1.46 |
| other bloold cells | Os9 | 0.97 | 1.66E-18 | 3.45E-14 | 0.75 | 0.37 | 0.38 | 2.02 |
| other bloold cells | Ctse | 3.17 | 1.70E-18 | 3.54E-14 | 0.15 | 0.03 | 0.13 | 5.31 |
| other bloold cells | Agps | 1.15 | 1.75E-18 | 3.65E-14 | 0.76 | 0.41 | 0.36 | 1.88 |
| other bloold cells | Use1 | 0.94 | 2.02E-18 | 4.21E-14 | 0.91 | 0.65 | 0.26 | 1.4 |
| other bloold cells | Acot9 | 1.34 | 2.41E-18 | 5.01E-14 | 0.43 | 0.16 | 0.28 | 2.76 |
| other bloold cells | Ppard | 1.62 | 2.45E-18 | 5.09E-14 | 0.33 | 0.1 | 0.23 | 3.26 |
| other bloold cells | Slc31a2 | 1.95 | 2.62E-18 | 5.46E-14 | 0.27 | 0.07 | 0.19 | 3.69 |
| other bloold cells | Rpsa | 0.33 | 2.66E-18 | 5.53E-14 | 1 | 1 | 0 | 1 |
| other bloold cells | Ptpra | 1.13 | 2.66E-18 | 5.53E-14 | 0.87 | 0.6 | 0.26 | 1.44 |
| other bloold cells | Serpinb6a | 1.98 | 2.78E-18 | 5.79E-14 | 0.71 | 0.39 | 0.32 | 1.81 |
| other bloold cells | Slc35c2 | 1.15 | 2.93E-18 | 6.10E-14 | 0.68 | 0.33 | 0.35 | 2.07 |
| other bloold cells | Tuba4a | 1.85 | 3.31E-18 | 6.87E-14 | 0.25 | 0.07 | 0.18 | 3.65 |
| other bloold cells | Tango2 | 1.33 | 3.54E-18 | 7.37E-14 | 0.43 | 0.16 | 0.27 | 2.74 |
| other bloold cells | Gdap2 | 1.4 | 3.61E-18 | 7.51E-14 | 0.36 | 0.11 | 0.24 | 3.13 |
| other bloold cells | Dhrs7 | 1.37 | 3.66E-18 | 7.60E-14 | 0.39 | 0.13 | 0.26 | 3.03 |
| other bloold cells | Sharpin | 1.23 | 3.82E-18 | 7.95E-14 | 0.46 | 0.17 | 0.29 | 2.71 |
| other bloold cells | Flot1 | 1.1 | 4.96E-18 | 1.03E-13 | 0.64 | 0.3 | 0.34 | 2.12 |
| other bloold cells | Trim12c | 1.47 | 5.05E-18 | 1.05E-13 | 0.46 | 0.18 | 0.29 | 2.61 |
| other bloold cells | ·931406C07Ri | 1.34 | 5.08E-18 | 1.06E-13 | 0.58 | 0.26 | 0.32 | 2.24 |
| other bloold cells | Enox2 | 1.8 | 5.40E-18 | 1.12E-13 | 0.26 | 0.07 | 0.19 | 3.7 |
| other bloold cells | Pnpo | 0.79 | 5.98E-18 | 1.24E-13 | 0.47 | 0.17 | 0.3 | 2.76 |
| other bloold cells | Sorl1 | 1.48 | 8.72E-18 | 1.81E-13 | 0.41 | 0.15 | 0.26 | 2.74 |
| other bloold cells | Tmbim6 | 0.71 | 9.24E-18 | 1.92E-13 | 0.97 | 0.84 | 0.13 | 1.15 |
| other bloold cells | Relch | 1.25 | 1.07E-17 | 2.22E-13 | 0.55 | 0.23 | 0.32 | 2.37 |
| other bloold cells | Ptpmt1 | 0.93 | 1.13E-17 | 2.34E-13 | 0.85 | 0.56 | 0.29 | 1.52 |
| other bloold cells | Snap29 | 1.43 | 1.21E-17 | 2.53E-13 | 0.48 | 0.19 | 0.29 | 2.49 |
| other bloold cells | Nagk | 1.53 | 1.23E-17 | 2.56E-13 | 0.48 | 0.2 | 0.29 | 2.45 |
| other bloold cells | Atp6v1e1 | 0.86 | 1.24E-17 | 2.59E-13 | 0.9 | 0.68 | 0.23 | 1.33 |
| other bloold cells | Snx14 | 1.22 | 1.26E-17 | 2.63E-13 | 0.45 | 0.16 | 0.29 | 2.75 |
| other bloold cells | Capns1 | 0.79 | 1.29E-17 | 2.68E-13 | 0.97 | 0.8 | 0.16 | 1.2 |
| other bloold cells | Atp6v0a1 | 1.51 | 1.41E-17 | 2.93E-13 | 0.49 | 0.2 | 0.29 | 2.49 |
| other bloold cells | Tmco1 | 0.84 | 1.47E-17 | 3.06E-13 | 0.92 | 0.7 | 0.23 | 1.32 |
| other bloold cells | Gmpr | 1.01 | 1.48E-17 | 3.07E-13 | 0.47 | 0.18 | 0.29 | 2.68 |
| other bloold cells | Phka2 | 1.55 | 1.50E-17 | 3.13E-13 | 0.37 | 0.13 | 0.24 | 2.92 |
| other bloold cells | Alg1 | 1.73 | 1.57E-17 | 3.26E-13 | 0.27 | 0.08 | 0.2 | 3.55 |
| other bloold cells | Mpv17l2 | 1.23 | 1.57E-17 | 3.27E-13 | 0.54 | 0.23 | 0.31 | 2.38 |
| other bloold cells | Lrpap1 | 0.93 | 1.93E-17 | 4.02E-13 | 0.85 | 0.49 | 0.36 | 1.73 |
| other bloold cells | Ap2m1 | 0.78 | 1.96E-17 | 4.07E-13 | 0.97 | 0.86 | 0.11 | 1.13 |
| other bloold cells | Mien1 | 0.85 | 2.24E-17 | 4.66E-13 | 0.88 | 0.56 | 0.32 | 1.56 |
| other bloold cells | .510009E07Ri | 1.85 | 2.54E-17 | 5.29E-13 | 0.24 | 0.06 | 0.18 | 3.84 |
| other bloold cells | Trappc2l | 0.88 | 2.57E-17 | 5.35E-13 | 0.9 | 0.62 | 0.28 | 1.45 |
| other bloold cells | Lyplal1 | 1.19 | 2.75E-17 | 5.72E-13 | 0.57 | 0.25 | 0.32 | 2.3 |
| other bloold cells | Mafk | 1.86 | 2.83E-17 | 5.89E-13 | 0.18 | 0.04 | 0.14 | 4.55 |
| other bloold cells | Mif4gd | 1.34 | 2.85E-17 | 5.94E-13 | 0.45 | 0.17 | 0.28 | 2.67 |
| other bloold cells | Trim21 | 2.29 | 2.90E-17 | 6.04E-13 | 0.16 | 0.03 | 0.13 | 4.88 |
| other bloold cells | Galk2 | 1.06 | 3.07E-17 | 6.38E-13 | 0.61 | 0.27 | 0.34 | 2.23 |
| other bloold cells | Atp6v1c1 | 1.23 | 3.19E-17 | 6.63E-13 | 0.69 | 0.36 | 0.33 | 1.92 |
| other bloold cells | Gls | 1.21 | 3.27E-17 | 6.80E-13 | 0.73 | 0.41 | 0.32 | 1.78 |
| other bloold cells | Flii | 1.16 | 3.34E-17 | 6.94E-13 | 0.73 | 0.37 | 0.36 | 1.99 |
| other bloold cells | Efr3a | 0.95 | 3.42E-17 | 7.12E-13 | 0.73 | 0.36 | 0.37 | 2.04 |
| other bloold cells | Mid1ip1 | 1.39 | 3.64E-17 | 7.57E-13 | 0.51 | 0.22 | 0.29 | 2.34 |

**Supplementary Table 2. Marker genes for the major cell types of integrated clustering of E11.5 control and NotchGOF λ-junction cells.**

| | | | | | | | | |
|---|---|---|---|---|---|---|---|---|
| other bloold cells | Ifnar1 | 1.12 | 3.74E-17 | 7.77E-13 | 0.69 | 0.35 | 0.35 | 1.99 |
| other bloold cells | Arl4c | 1.36 | 3.94E-17 | 8.20E-13 | 0.64 | 0.33 | 0.31 | 1.95 |
| other bloold cells | Tbkbp1 | 1.33 | 3.95E-17 | 8.21E-13 | 0.25 | 0.06 | 0.18 | 3.89 |
| other bloold cells | Vav2 | 1.58 | 4.21E-17 | 8.76E-13 | 0.29 | 0.08 | 0.2 | 3.42 |
| other bloold cells | St3gal6 | 2.01 | 4.29E-17 | 8.92E-13 | 0.33 | 0.11 | 0.22 | 2.94 |
| other bloold cells | Fam53b | 1.34 | 4.49E-17 | 9.35E-13 | 0.46 | 0.18 | 0.28 | 2.57 |
| other bloold cells | Npr1 | 2.09 | 4.94E-17 | 1.03E-12 | 0.11 | 0.02 | 0.09 | 7 |
| other bloold cells | Grina | 1.13 | 5.09E-17 | 1.06E-12 | 0.57 | 0.25 | 0.32 | 2.3 |
| other bloold cells | Tyk2 | 2 | 5.56E-17 | 1.16E-12 | 0.22 | 0.06 | 0.17 | 3.86 |
| other bloold cells | Sri | 0.82 | 6.16E-17 | 1.28E-12 | 0.97 | 0.77 | 0.2 | 1.26 |
| other bloold cells | Twf1 | 1.16 | 7.26E-17 | 1.51E-12 | 0.8 | 0.49 | 0.31 | 1.63 |
| other bloold cells | Ap2a2 | 1.23 | 7.36E-17 | 1.53E-12 | 0.83 | 0.54 | 0.29 | 1.54 |
| other bloold cells | Cast | 1.19 | 8.37E-17 | 1.74E-12 | 0.56 | 0.24 | 0.32 | 2.3 |
| other bloold cells | Ddx58 | 1.84 | 8.97E-17 | 1.86E-12 | 0.22 | 0.05 | 0.16 | 4.02 |
| other bloold cells | Mt2 | 1.23 | 1.01E-16 | 2.11E-12 | 0.36 | 0.13 | 0.24 | 2.82 |
| other bloold cells | Layn | 2.05 | 1.07E-16 | 2.23E-12 | 0.15 | 0.03 | 0.12 | 5.25 |
| other bloold cells | Pak2 | 0.63 | 1.14E-16 | 2.38E-12 | 0.99 | 0.84 | 0.15 | 1.18 |
| other bloold cells | Mthfsl | 1.14 | 1.15E-16 | 2.38E-12 | 0.55 | 0.23 | 0.32 | 2.39 |
| other bloold cells | Acsl4 | 1.28 | 1.23E-16 | 2.55E-12 | 0.39 | 0.14 | 0.25 | 2.8 |
| other bloold cells | Hsd17b11 | 1.12 | 1.29E-16 | 2.67E-12 | 0.51 | 0.21 | 0.31 | 2.49 |
| other bloold cells | Otulin | 0.96 | 1.37E-16 | 2.84E-12 | 0.73 | 0.41 | 0.32 | 1.79 |
| other bloold cells | Fabp3 | 2.59 | 1.39E-16 | 2.89E-12 | 0.14 | 0.03 | 0.11 | 5.19 |
| other bloold cells | Wdr41 | 1.51 | 1.46E-16 | 3.03E-12 | 0.32 | 0.1 | 0.22 | 3.16 |
| other bloold cells | Mfsd11 | 1.52 | 1.49E-16 | 3.10E-12 | 0.41 | 0.15 | 0.25 | 2.64 |
| other bloold cells | Akr1b3 | 0.77 | 1.51E-16 | 3.14E-12 | 0.89 | 0.68 | 0.21 | 1.31 |
| other bloold cells | Washc2 | 0.9 | 1.83E-16 | 3.81E-12 | 0.83 | 0.52 | 0.31 | 1.6 |
| other bloold cells | Flt1 | 0.72 | 1.85E-16 | 3.85E-12 | 0.12 | 0.02 | 0.1 | 5.95 |
| other bloold cells | Abhd6 | 1.36 | 1.98E-16 | 4.11E-12 | 0.44 | 0.17 | 0.27 | 2.63 |
| other bloold cells | Ppcdc | 1.37 | 1.99E-16 | 4.13E-12 | 0.38 | 0.13 | 0.24 | 2.82 |
| other bloold cells | Hk2 | 1.18 | 1.99E-16 | 4.14E-12 | 0.57 | 0.25 | 0.31 | 2.23 |
| other bloold cells | Coro1c | 1.11 | 2.04E-16 | 4.24E-12 | 0.68 | 0.35 | 0.33 | 1.95 |
| other bloold cells | Cerk | 1.62 | 2.13E-16 | 4.42E-12 | 0.4 | 0.15 | 0.25 | 2.68 |
| other bloold cells | Fchsd2 | 1.22 | 2.17E-16 | 4.52E-12 | 0.66 | 0.33 | 0.33 | 1.99 |
| other bloold cells | Fam49a | 1.9 | 2.19E-16 | 4.55E-12 | 0.19 | 0.04 | 0.15 | 4.3 |
| other bloold cells | Rab1a | 0.86 | 2.24E-16 | 4.67E-12 | 0.94 | 0.71 | 0.23 | 1.33 |
| other bloold cells | Tnfaip3 | 2.84 | 2.26E-16 | 4.69E-12 | 0.11 | 0.02 | 0.09 | 6.56 |
| other bloold cells | Ssbp4 | 0.79 | 2.27E-16 | 4.73E-12 | 0.92 | 0.65 | 0.27 | 1.41 |
| other bloold cells | Haus4 | 1 | 2.44E-16 | 5.08E-12 | 0.66 | 0.32 | 0.34 | 2.09 |
| other bloold cells | Birc3 | 1.94 | 2.49E-16 | 5.17E-12 | 0.19 | 0.04 | 0.15 | 4.3 |
| other bloold cells | Emid1 | 1.82 | 2.50E-16 | 5.21E-12 | 0.29 | 0.09 | 0.2 | 3.23 |
| other bloold cells | Arhgef3 | 2.46 | 2.54E-16 | 5.28E-12 | 0.11 | 0.02 | 0.09 | 6.22 |
| other bloold cells | Crlf3 | 0.98 | 2.55E-16 | 5.30E-12 | 0.79 | 0.45 | 0.34 | 1.75 |
| other bloold cells | Nfic | 1.17 | 2.64E-16 | 5.48E-12 | 0.62 | 0.3 | 0.32 | 2.08 |
| other bloold cells | Arl5c | 2.84 | 2.95E-16 | 6.13E-12 | 0.11 | 0.02 | 0.09 | 6.56 |
| other bloold cells | Emc7 | 0.9 | 3.05E-16 | 6.34E-12 | 0.8 | 0.5 | 0.3 | 1.61 |
| other bloold cells | Sfmbt2 | 2.1 | 3.07E-16 | 6.38E-12 | 0.14 | 0.03 | 0.11 | 5.19 |
| other bloold cells | Rps20 | 0.27 | 3.27E-16 | 6.81E-12 | 1 | 1 | 0 | 1 |
| other bloold cells | Ndufb1-ps | 0.57 | 3.47E-16 | 7.22E-12 | 0.98 | 0.94 | 0.04 | 1.04 |
| other bloold cells | Calm3 | 0.57 | 3.50E-16 | 7.27E-12 | 0.99 | 0.91 | 0.08 | 1.09 |
| other bloold cells | Synrg | 0.93 | 3.54E-16 | 7.37E-12 | 0.55 | 0.24 | 0.32 | 2.35 |
| other bloold cells | Arhgap31 | 1.29 | 3.92E-16 | 8.14E-12 | 0.47 | 0.19 | 0.28 | 2.49 |
| other bloold cells | Rpl12 | 0.26 | 3.92E-16 | 8.15E-12 | 1 | 1 | 0 | 1 |
| other bloold cells | Uvrag | 1.06 | 4.03E-16 | 8.39E-12 | 0.54 | 0.23 | 0.31 | 2.37 |
| other bloold cells | Baz1a | 0.92 | 4.95E-16 | 1.03E-11 | 0.82 | 0.56 | 0.26 | 1.47 |
| other bloold cells | Fez2 | 1.24 | 4.96E-16 | 1.03E-11 | 0.65 | 0.31 | 0.34 | 2.1 |
| other bloold cells | Pnkp | 1.18 | 5.36E-16 | 1.11E-11 | 0.52 | 0.22 | 0.3 | 2.33 |

**Supplementary Table 2. Marker genes for the major cell types of integrated clustering of E11.5 control and NotchGOF λ-junction cells.**

| | | | | | | | | |
|---|---|---|---|---|---|---|---|---|
| other bloold cells | Helz2 | 2.51 | 5.56E-16 | 1.16E-11 | 0.11 | 0.02 | 0.09 | 6.56 |
| other bloold cells | Fbxw4 | 1.13 | 6.19E-16 | 1.29E-11 | 0.4 | 0.14 | 0.26 | 2.79 |
| other bloold cells | AC149090.1 | 1.13 | 6.46E-16 | 1.34E-11 | 0.61 | 0.28 | 0.33 | 2.16 |
| other bloold cells | Mast3 | 1.37 | 6.76E-16 | 1.41E-11 | 0.33 | 0.11 | 0.22 | 3.02 |
| other bloold cells | Uqcrq | 0.46 | 7.76E-16 | 1.61E-11 | 1 | 0.97 | 0.03 | 1.03 |
| other bloold cells | Cracr2b | 2.13 | 8.35E-16 | 1.74E-11 | 0.16 | 0.04 | 0.13 | 4.6 |
| other bloold cells | Fam102b | 1.31 | 8.62E-16 | 1.79E-11 | 0.48 | 0.2 | 0.28 | 2.38 |
| other bloold cells | Pacc1 | 1.87 | 8.94E-16 | 1.86E-11 | 0.3 | 0.09 | 0.21 | 3.2 |
| other bloold cells | Rela | 1.14 | 9.22E-16 | 1.92E-11 | 0.54 | 0.25 | 0.29 | 2.2 |
| other bloold cells | Gak | 1.22 | 1.02E-15 | 2.11E-11 | 0.52 | 0.23 | 0.3 | 2.31 |
| other bloold cells | Atp1b3 | 0.79 | 1.04E-15 | 2.17E-11 | 0.93 | 0.72 | 0.21 | 1.3 |
| other bloold cells | Foxo3 | 1.16 | 1.12E-15 | 2.34E-11 | 0.36 | 0.13 | 0.24 | 2.89 |
| other bloold cells | Rbl2 | 1.39 | 1.24E-15 | 2.58E-11 | 0.29 | 0.09 | 0.2 | 3.26 |
| other bloold cells | Ampd3 | 2.65 | 1.30E-15 | 2.71E-11 | 0.11 | 0.02 | 0.09 | 6.18 |
| other bloold cells | Mtr | 1.39 | 1.34E-15 | 2.79E-11 | 0.5 | 0.23 | 0.27 | 2.18 |
| other bloold cells | Clptm1 | 0.99 | 1.50E-15 | 3.11E-11 | 0.76 | 0.45 | 0.32 | 1.71 |
| other bloold cells | Foxj2 | 1.65 | 1.53E-15 | 3.19E-11 | 0.25 | 0.07 | 0.18 | 3.55 |
| other bloold cells | Rhof | 2.26 | 1.56E-15 | 3.25E-11 | 0.11 | 0.02 | 0.09 | 6.18 |
| other bloold cells | Klhl5 | 1.14 | 1.64E-15 | 3.41E-11 | 0.44 | 0.17 | 0.27 | 2.58 |
| other bloold cells | Pald1 | 1.54 | 1.77E-15 | 3.69E-11 | 0.27 | 0.08 | 0.19 | 3.29 |
| other bloold cells | Mtss1 | 1.1 | 1.96E-15 | 4.08E-11 | 0.62 | 0.3 | 0.32 | 2.08 |
| other bloold cells | Nqo2 | 1.29 | 2.07E-15 | 4.30E-11 | 0.36 | 0.13 | 0.24 | 2.82 |
| other bloold cells | Ydjc | 1.68 | 2.21E-15 | 4.60E-11 | 0.27 | 0.08 | 0.19 | 3.29 |
| other bloold cells | .600002K03Ri | 1.34 | 2.21E-15 | 4.61E-11 | 0.34 | 0.12 | 0.22 | 2.92 |
| other bloold cells | Cflar | 1 | 2.42E-15 | 5.03E-11 | 0.63 | 0.31 | 0.32 | 2.03 |
| other bloold cells | Slc2a3 | 0.56 | 2.43E-15 | 5.05E-11 | 0.39 | 0.14 | 0.25 | 2.79 |
| other bloold cells | Nudt18 | 1.65 | 2.58E-15 | 5.38E-11 | 0.22 | 0.06 | 0.16 | 3.67 |
| other bloold cells | Ccpg1 | 1.2 | 2.72E-15 | 5.65E-11 | 0.42 | 0.16 | 0.26 | 2.59 |
| other bloold cells | Slc38a6 | 1.58 | 2.91E-15 | 6.05E-11 | 0.39 | 0.15 | 0.24 | 2.57 |
| other bloold cells | Iqsec2 | 2.22 | 3.23E-15 | 6.71E-11 | 0.13 | 0.02 | 0.1 | 5.48 |
| other bloold cells | Ranbp2 | 0.77 | 3.24E-15 | 6.75E-11 | 0.9 | 0.66 | 0.25 | 1.38 |
| other bloold cells | Trappc8 | 1.12 | 3.96E-15 | 8.25E-11 | 0.54 | 0.24 | 0.3 | 2.21 |
| other bloold cells | Ptger4 | 2.11 | 4.11E-15 | 8.54E-11 | 0.13 | 0.03 | 0.11 | 5.12 |
| other bloold cells | Inpp5b | 1.34 | 4.43E-15 | 9.21E-11 | 0.34 | 0.12 | 0.22 | 2.9 |
| other bloold cells | Ankrd44 | 1.09 | 4.55E-15 | 9.46E-11 | 0.61 | 0.28 | 0.33 | 2.18 |
| other bloold cells | Adam9 | 1.67 | 4.62E-15 | 9.61E-11 | 0.36 | 0.14 | 0.22 | 2.6 |
| other bloold cells | Rab10os | 1.2 | 4.62E-15 | 9.61E-11 | 0.45 | 0.18 | 0.26 | 2.43 |
| other bloold cells | Pcbd2 | 0.84 | 5.02E-15 | 1.04E-10 | 0.78 | 0.43 | 0.35 | 1.8 |
| other bloold cells | Snx18 | 1.12 | 5.05E-15 | 1.05E-10 | 0.69 | 0.38 | 0.32 | 1.84 |
| other bloold cells | Eml3 | 1.29 | 5.17E-15 | 1.08E-10 | 0.35 | 0.12 | 0.23 | 2.82 |
| other bloold cells | Heatr1 | 1.16 | 5.22E-15 | 1.09E-10 | 0.62 | 0.31 | 0.31 | 2.01 |
| other bloold cells | Inpp5k | 1.51 | 5.32E-15 | 1.11E-10 | 0.32 | 0.11 | 0.21 | 2.88 |
| other bloold cells | Nab1 | 1.19 | 5.35E-15 | 1.11E-10 | 0.36 | 0.13 | 0.24 | 2.82 |
| other bloold cells | Aldh2 | 0.79 | 5.38E-15 | 1.12E-10 | 0.92 | 0.78 | 0.15 | 1.19 |
| other bloold cells | Atp6v1h | 1.06 | 5.68E-15 | 1.18E-10 | 0.68 | 0.36 | 0.32 | 1.88 |
| other bloold cells | Ubc | 0.56 | 5.77E-15 | 1.20E-10 | 1 | 0.98 | 0.03 | 1.03 |
| other bloold cells | Lyrm9 | 1.05 | 5.79E-15 | 1.20E-10 | 0.44 | 0.17 | 0.27 | 2.56 |
| other bloold cells | Smarcd2 | 0.92 | 5.89E-15 | 1.23E-10 | 0.62 | 0.29 | 0.33 | 2.12 |
| other bloold cells | Bcap29 | 0.92 | 5.90E-15 | 1.23E-10 | 0.69 | 0.34 | 0.34 | 2 |
| other bloold cells | Arhgap19 | 1.05 | 5.94E-15 | 1.23E-10 | 0.59 | 0.29 | 0.31 | 2.07 |
| other bloold cells | Map3k1 | 1.03 | 6.35E-15 | 1.32E-10 | 0.57 | 0.26 | 0.31 | 2.17 |
| other bloold cells | Tmem141 | 1.28 | 7.13E-15 | 1.48E-10 | 0.46 | 0.19 | 0.27 | 2.38 |
| other bloold cells | Cbl | 0.83 | 7.15E-15 | 1.49E-10 | 0.81 | 0.47 | 0.34 | 1.73 |
| other bloold cells | Man2c1 | 1.41 | 7.74E-15 | 1.61E-10 | 0.32 | 0.11 | 0.21 | 2.97 |
| other bloold cells | Rbks | 1.83 | 7.91E-15 | 1.65E-10 | 0.15 | 0.03 | 0.12 | 4.67 |
| other bloold cells | Erlin1 | 1.19 | 8.29E-15 | 1.72E-10 | 0.4 | 0.15 | 0.25 | 2.63 |

**Supplementary Table 2. Marker genes for the major cell types of integrated clustering of E11.5 control and NotchGOF λ-junction cells.**

| | | | | | | | | |
|---|---|---|---|---|---|---|---|---|
| other bloold cells | Itpkb | 1.17 | 8.31E-15 | 1.73E-10 | 0.47 | 0.19 | 0.28 | 2.43 |
| other bloold cells | Gm42670 | 2.36 | 8.59E-15 | 1.79E-10 | 0.11 | 0.02 | 0.09 | 6.18 |
| other bloold cells | Slc38a10 | 1.06 | 8.76E-15 | 1.82E-10 | 0.66 | 0.34 | 0.33 | 1.98 |
| other bloold cells | Prtn3 | 4.4 | 9.29E-15 | 1.93E-10 | 0.34 | 0.14 | 0.2 | 2.45 |
| other bloold cells | Rab11a | 0.81 | 9.45E-15 | 1.97E-10 | 0.9 | 0.64 | 0.26 | 1.41 |
| other bloold cells | Dnajc3 | 0.9 | 9.69E-15 | 2.02E-10 | 0.83 | 0.52 | 0.31 | 1.59 |
| other bloold cells | Slc35d2 | 1.79 | 1.02E-14 | 2.12E-10 | 0.13 | 0.03 | 0.11 | 5.12 |
| other bloold cells | Emc2 | 0.72 | 1.05E-14 | 2.18E-10 | 0.88 | 0.59 | 0.29 | 1.49 |
| other bloold cells | Map3k14 | 1.97 | 1.14E-14 | 2.36E-10 | 0.16 | 0.04 | 0.12 | 4.35 |
| other bloold cells | Gaa | 1.31 | 1.15E-14 | 2.39E-10 | 0.45 | 0.18 | 0.26 | 2.43 |
| other bloold cells | Isoc2b | 2.18 | 1.25E-14 | 2.60E-10 | 0.13 | 0.02 | 0.1 | 5.25 |
| other bloold cells | Stard3 | 1.12 | 1.30E-14 | 2.69E-10 | 0.52 | 0.24 | 0.29 | 2.23 |
| other bloold cells | Fbxl8 | 2.05 | 1.30E-14 | 2.71E-10 | 0.12 | 0.02 | 0.1 | 5.41 |
| other bloold cells | Map3k3 | 1.21 | 1.33E-14 | 2.77E-10 | 0.4 | 0.15 | 0.25 | 2.59 |
| other bloold cells | Tet2 | 1.09 | 1.46E-14 | 3.03E-10 | 0.55 | 0.25 | 0.3 | 2.19 |
| other bloold cells | Fxyd2 | 2.89 | 1.51E-14 | 3.15E-10 | 0.11 | 0.02 | 0.09 | 5.6 |
| other bloold cells | Foxn2 | 1.23 | 1.61E-14 | 3.34E-10 | 0.41 | 0.16 | 0.25 | 2.55 |
| other bloold cells | Mpc2 | 0.75 | 1.61E-14 | 3.36E-10 | 0.93 | 0.79 | 0.14 | 1.18 |
| other bloold cells | Trim65 | 1.85 | 1.71E-14 | 3.55E-10 | 0.16 | 0.04 | 0.12 | 4.35 |
| other bloold cells | Selenbp1 | 1.31 | 1.73E-14 | 3.59E-10 | 0.27 | 0.09 | 0.19 | 3.21 |
| other bloold cells | Dtnbp1 | 0.91 | 1.85E-14 | 3.85E-10 | 0.75 | 0.42 | 0.33 | 1.79 |
| other bloold cells | Ssu72 | 0.58 | 1.88E-14 | 3.92E-10 | 0.96 | 0.84 | 0.12 | 1.15 |
| other bloold cells | Idua | 2.09 | 1.89E-14 | 3.93E-10 | 0.15 | 0.03 | 0.12 | 4.53 |
| other bloold cells | Ap3b1 | 0.9 | 1.99E-14 | 4.14E-10 | 0.79 | 0.47 | 0.32 | 1.68 |
| other bloold cells | Arl6ip1 | 0.41 | 2.13E-14 | 4.44E-10 | 0.99 | 0.89 | 0.1 | 1.12 |
| other bloold cells | Mthfd1l | 1.2 | 2.16E-14 | 4.50E-10 | 0.41 | 0.16 | 0.25 | 2.57 |
| other bloold cells | Gyg | 1.08 | 2.29E-14 | 4.76E-10 | 0.73 | 0.42 | 0.31 | 1.74 |
| other bloold cells | Dnajc13 | 1.1 | 2.34E-14 | 4.88E-10 | 0.67 | 0.38 | 0.3 | 1.78 |
| other bloold cells | Klf13 | 1.04 | 2.36E-14 | 4.91E-10 | 0.71 | 0.37 | 0.34 | 1.92 |
| other bloold cells | Cdk2ap2 | 0.84 | 2.36E-14 | 4.92E-10 | 0.78 | 0.46 | 0.32 | 1.71 |
| other bloold cells | Slc17a5 | 1.19 | 2.39E-14 | 4.97E-10 | 0.44 | 0.18 | 0.26 | 2.41 |
| other bloold cells | Sft2d2 | 1.35 | 2.44E-14 | 5.07E-10 | 0.39 | 0.15 | 0.24 | 2.61 |
| other bloold cells | Ten1 | 0.78 | 2.55E-14 | 5.30E-10 | 0.86 | 0.57 | 0.3 | 1.52 |
| other bloold cells | Parp8 | 1.26 | 2.76E-14 | 5.74E-10 | 0.37 | 0.14 | 0.23 | 2.71 |
| other bloold cells | Sppl2a | 0.87 | 2.79E-14 | 5.80E-10 | 0.83 | 0.47 | 0.36 | 1.76 |
| other bloold cells | Aftph | 1.08 | 2.90E-14 | 6.03E-10 | 0.54 | 0.25 | 0.29 | 2.14 |
| other bloold cells | Taf10 | 0.45 | 3.14E-14 | 6.53E-10 | 0.99 | 0.93 | 0.06 | 1.06 |
| other bloold cells | Tmem268 | 1.56 | 3.21E-14 | 6.68E-10 | 0.27 | 0.08 | 0.18 | 3.17 |
| other bloold cells | Mafb | 1.46 | 3.30E-14 | 6.86E-10 | 0.49 | 0.22 | 0.27 | 2.22 |
| other bloold cells | Shmt2 | 1.14 | 3.76E-14 | 7.82E-10 | 0.62 | 0.34 | 0.28 | 1.83 |
| other bloold cells | Siah2 | 1.29 | 3.77E-14 | 7.83E-10 | 0.25 | 0.08 | 0.18 | 3.36 |
| other bloold cells | Rb1 | 0.96 | 3.77E-14 | 7.84E-10 | 0.41 | 0.16 | 0.25 | 2.55 |
| other bloold cells | Zfp385a | 0.99 | 3.92E-14 | 8.16E-10 | 0.54 | 0.25 | 0.28 | 2.12 |
| other bloold cells | Aldh9a1 | 1.12 | 4.05E-14 | 8.43E-10 | 0.62 | 0.32 | 0.31 | 1.97 |
| other bloold cells | Mtmr1 | 1.63 | 4.17E-14 | 8.66E-10 | 0.24 | 0.07 | 0.17 | 3.35 |
| other bloold cells | Fhod1 | 1.5 | 4.35E-14 | 9.04E-10 | 0.27 | 0.09 | 0.19 | 3.14 |
| other bloold cells | Zbtb7a | 1.01 | 4.43E-14 | 9.21E-10 | 0.63 | 0.33 | 0.3 | 1.9 |
| other bloold cells | Bnip2 | 0.74 | 5.00E-14 | 1.04E-09 | 0.92 | 0.69 | 0.23 | 1.34 |
| other bloold cells | Sh2d3c | 1.21 | 5.15E-14 | 1.07E-09 | 0.3 | 0.1 | 0.2 | 3.01 |
| other bloold cells | Edf1 | 0.39 | 5.18E-14 | 1.08E-09 | 0.99 | 0.96 | 0.03 | 1.03 |
| other bloold cells | Pepd | 0.91 | 5.68E-14 | 1.18E-09 | 0.76 | 0.42 | 0.34 | 1.81 |
| other bloold cells | Gcnt2 | 1.99 | 5.70E-14 | 1.19E-09 | 0.18 | 0.05 | 0.14 | 3.87 |
| other bloold cells | Galc | 1.6 | 5.96E-14 | 1.24E-09 | 0.42 | 0.18 | 0.25 | 2.4 |
| other bloold cells | Spty2d1 | 0.99 | 6.00E-14 | 1.25E-09 | 0.54 | 0.25 | 0.29 | 2.14 |
| other bloold cells | Ctso | 1.82 | 6.73E-14 | 1.40E-09 | 0.22 | 0.07 | 0.16 | 3.45 |
| other bloold cells | Plxnb2 | 1.01 | 6.84E-14 | 1.42E-09 | 0.74 | 0.53 | 0.21 | 1.39 |

**Supplementary Table 2. Marker genes for the major cell types of integrated clustering of E11.5 control and NotchGOF λ-junction cells.**

| | | | | | | | | |
|---|---|---|---|---|---|---|---|---|
| other bloold cells | Ankfy1 | 1.01 | 7.29E-14 | 1.52E-09 | 0.59 | 0.28 | 0.31 | 2.08 |
| other bloold cells | Creb3 | 1.1 | 9.15E-14 | 1.90E-09 | 0.45 | 0.19 | 0.26 | 2.37 |
| other bloold cells | Vmp1 | 0.83 | 9.45E-14 | 1.97E-09 | 0.87 | 0.59 | 0.28 | 1.48 |
| other bloold cells | Tmem175 | 1.19 | 1.03E-13 | 2.13E-09 | 0.3 | 0.1 | 0.2 | 2.98 |
| other bloold cells | Vat1 | 1.47 | 1.08E-13 | 2.25E-09 | 0.36 | 0.14 | 0.22 | 2.63 |
| other bloold cells | Rnf31 | 1.33 | 1.09E-13 | 2.28E-09 | 0.23 | 0.07 | 0.16 | 3.45 |
| other bloold cells | Comt | 0.88 | 1.12E-13 | 2.34E-09 | 0.83 | 0.54 | 0.29 | 1.54 |
| other bloold cells | Calr | 0.65 | 1.15E-13 | 2.38E-09 | 0.99 | 0.97 | 0.02 | 1.02 |
| other bloold cells | Agpat2 | 1 | 1.16E-13 | 2.41E-09 | 0.42 | 0.17 | 0.25 | 2.53 |
| other bloold cells | Hist1h2bc | 1.57 | 1.21E-13 | 2.53E-09 | 0.52 | 0.26 | 0.26 | 1.99 |
| other bloold cells | Ghitm | 0.72 | 1.22E-13 | 2.53E-09 | 0.9 | 0.67 | 0.23 | 1.35 |
| other bloold cells | Oplah | 2.44 | 1.31E-13 | 2.71E-09 | 0.11 | 0.02 | 0.09 | 5.33 |
| other bloold cells | Gpr108 | 1.17 | 1.35E-13 | 2.81E-09 | 0.4 | 0.16 | 0.24 | 2.49 |
| other bloold cells | Cd82 | 1.98 | 1.39E-13 | 2.89E-09 | 0.22 | 0.07 | 0.16 | 3.39 |
| other bloold cells | Ilk | 0.86 | 1.39E-13 | 2.90E-09 | 0.89 | 0.61 | 0.28 | 1.46 |
| other bloold cells | Tdrkh | 1.72 | 1.41E-13 | 2.93E-09 | 0.21 | 0.06 | 0.15 | 3.5 |
| other bloold cells | Atg7 | 1.23 | 1.44E-13 | 3.00E-09 | 0.34 | 0.13 | 0.22 | 2.74 |
| other bloold cells | Acaa1a | 1.01 | 1.51E-13 | 3.13E-09 | 0.75 | 0.46 | 0.29 | 1.63 |
| other bloold cells | Arhgap1 | 0.9 | 1.53E-13 | 3.19E-09 | 0.56 | 0.26 | 0.3 | 2.16 |
| other bloold cells | Ufsp2 | 0.8 | 1.57E-13 | 3.26E-09 | 0.8 | 0.49 | 0.31 | 1.62 |
| other bloold cells | Nsf | 0.99 | 1.59E-13 | 3.30E-09 | 0.64 | 0.34 | 0.31 | 1.91 |
| other bloold cells | Ldlrap1 | 1.33 | 1.62E-13 | 3.37E-09 | 0.43 | 0.18 | 0.25 | 2.37 |
| other bloold cells | Gm26532 | 1.88 | 1.91E-13 | 3.97E-09 | 0.16 | 0.04 | 0.12 | 4.13 |
| other bloold cells | Mov10 | 0.87 | 1.92E-13 | 4.00E-09 | 0.47 | 0.19 | 0.28 | 2.42 |
| other bloold cells | Myc | 1.51 | 2.09E-13 | 4.35E-09 | 0.39 | 0.16 | 0.23 | 2.4 |
| other bloold cells | Iscu | 0.78 | 2.15E-13 | 4.46E-09 | 0.9 | 0.64 | 0.25 | 1.39 |
| other bloold cells | Tmem256 | 0.63 | 2.16E-13 | 4.50E-09 | 0.99 | 0.89 | 0.1 | 1.11 |
| other bloold cells | Hpgd | 0.73 | 2.25E-13 | 4.67E-09 | 0.67 | 0.35 | 0.32 | 1.93 |
| other bloold cells | Paox | 1.33 | 2.26E-13 | 4.69E-09 | 0.39 | 0.16 | 0.23 | 2.48 |
| other bloold cells | Egfl7 | 0.75 | 2.28E-13 | 4.74E-09 | 0.48 | 0.22 | 0.26 | 2.17 |
| other bloold cells | Slc23a2 | 1.63 | 2.49E-13 | 5.18E-09 | 0.25 | 0.08 | 0.17 | 3.18 |
| other bloold cells | Atox1 | 0.29 | 2.50E-13 | 5.19E-09 | 1 | 0.95 | 0.05 | 1.05 |
| other bloold cells | Rnf17 | 1.48 | 2.60E-13 | 5.40E-09 | 0.15 | 0.03 | 0.11 | 4.45 |
| other bloold cells | Pacsin2 | 0.96 | 2.63E-13 | 5.47E-09 | 0.65 | 0.35 | 0.3 | 1.87 |
| other bloold cells | Fam107b | 0.85 | 2.68E-13 | 5.57E-09 | 0.65 | 0.36 | 0.3 | 1.83 |
| other bloold cells | Ppp1r9b | 1.14 | 2.68E-13 | 5.57E-09 | 0.53 | 0.26 | 0.28 | 2.08 |
| other bloold cells | Mcts1 | 0.86 | 2.71E-13 | 5.64E-09 | 0.75 | 0.47 | 0.28 | 1.58 |
| other bloold cells | Limd2 | 0.55 | 2.75E-13 | 5.72E-09 | 0.99 | 0.9 | 0.09 | 1.1 |
| other bloold cells | Prelid1 | 0.49 | 2.89E-13 | 6.00E-09 | 0.99 | 0.93 | 0.07 | 1.07 |
| other bloold cells | Tbc1d5 | 1.07 | 3.05E-13 | 6.34E-09 | 0.47 | 0.21 | 0.26 | 2.29 |
| other bloold cells | Arid4a | 0.82 | 3.05E-13 | 6.34E-09 | 0.78 | 0.47 | 0.31 | 1.65 |
| other bloold cells | Arl8b | 0.88 | 3.19E-13 | 6.64E-09 | 0.76 | 0.47 | 0.29 | 1.61 |
| other bloold cells | Zfp984 | 1.65 | 3.28E-13 | 6.82E-09 | 0.24 | 0.07 | 0.17 | 3.26 |
| other bloold cells | Myb | 2.49 | 3.37E-13 | 7.00E-09 | 0.25 | 0.08 | 0.17 | 3 |
| other bloold cells | Ncstn | 1.03 | 3.60E-13 | 7.48E-09 | 0.58 | 0.29 | 0.29 | 1.99 |
| other bloold cells | Vrk2 | 0.88 | 3.64E-13 | 7.57E-09 | 0.51 | 0.23 | 0.28 | 2.26 |
| other bloold cells | Apmap | 1.38 | 3.68E-13 | 7.66E-09 | 0.44 | 0.19 | 0.25 | 2.35 |
| other bloold cells | Myom1 | 1.71 | 3.71E-13 | 7.71E-09 | 0.14 | 0.03 | 0.11 | 4.52 |
| other bloold cells | Epg5 | 1.5 | 3.92E-13 | 8.15E-09 | 0.17 | 0.04 | 0.13 | 4 |
| other bloold cells | Susd6 | 1.16 | 3.98E-13 | 8.27E-09 | 0.46 | 0.2 | 0.26 | 2.31 |
| other bloold cells | Arsa | 1.47 | 4.14E-13 | 8.60E-09 | 0.22 | 0.07 | 0.16 | 3.34 |
| other bloold cells | Fkbp2 | 0.73 | 4.34E-13 | 9.02E-09 | 0.91 | 0.65 | 0.26 | 1.39 |
| other bloold cells | Epn1 | 0.87 | 4.65E-13 | 9.67E-09 | 0.87 | 0.62 | 0.25 | 1.4 |
| other bloold cells | Ptges3l | 1.49 | 4.93E-13 | 1.03E-08 | 0.21 | 0.06 | 0.15 | 3.5 |
| other bloold cells | Ripor1 | 1.55 | 4.98E-13 | 1.04E-08 | 0.27 | 0.09 | 0.18 | 2.99 |
| other bloold cells | Rffl | 1.55 | 5.03E-13 | 1.05E-08 | 0.23 | 0.07 | 0.16 | 3.25 |

**Supplementary Table 2. Marker genes for the major cell types of integrated clustering of E11.5 control and NotchGOF λ-junction cells.**

| | | | | | | | | |
|---|---|---|---|---|---|---|---|---|
| other bloold cells | Mavs | 1.53 | 5.11E-13 | 1.06E-08 | 0.27 | 0.09 | 0.18 | 2.99 |
| other bloold cells | Ctsl | 1.24 | 5.28E-13 | 1.10E-08 | 0.93 | 0.82 | 0.11 | 1.13 |
| other bloold cells | Shtn1 | 1.6 | 5.37E-13 | 1.12E-08 | 0.19 | 0.05 | 0.14 | 3.63 |
| other bloold cells | Wdfy2 | 1.15 | 5.40E-13 | 1.12E-08 | 0.4 | 0.16 | 0.24 | 2.48 |
| other bloold cells | Smg9 | 1.09 | 5.74E-13 | 1.19E-08 | 0.36 | 0.14 | 0.23 | 2.62 |
| other bloold cells | Hltf | 0.83 | 5.93E-13 | 1.23E-08 | 0.64 | 0.32 | 0.32 | 2 |
| other bloold cells | Depdc7 | 1.84 | 6.02E-13 | 1.25E-08 | 0.14 | 0.03 | 0.11 | 4.52 |
| other bloold cells | Jun | 0.84 | 6.39E-13 | 1.33E-08 | 0.9 | 0.67 | 0.23 | 1.34 |
| other bloold cells | Pbxip1 | 1.04 | 6.84E-13 | 1.42E-08 | 0.47 | 0.2 | 0.27 | 2.32 |
| other bloold cells | Bach1 | 0.97 | 6.93E-13 | 1.44E-08 | 0.79 | 0.48 | 0.31 | 1.66 |
| other bloold cells | Pgap1 | 1.31 | 7.02E-13 | 1.46E-08 | 0.37 | 0.15 | 0.22 | 2.46 |
| other bloold cells | Smcr8 | 1.72 | 7.29E-13 | 1.52E-08 | 0.25 | 0.08 | 0.17 | 3.07 |
| other bloold cells | Ppp1ca | 0.36 | 7.46E-13 | 1.55E-08 | 1 | 0.98 | 0.02 | 1.02 |
| other bloold cells | Ecpas | 0.85 | 7.69E-13 | 1.60E-08 | 0.75 | 0.42 | 0.33 | 1.77 |
| other bloold cells | Adrb1 | 2.18 | 8.32E-13 | 1.73E-08 | 0.11 | 0.02 | 0.09 | 5.25 |
| other bloold cells | Nfkb2 | 1.73 | 8.41E-13 | 1.75E-08 | 0.22 | 0.07 | 0.16 | 3.34 |
| other bloold cells | Mapk14 | 0.98 | 8.56E-13 | 1.78E-08 | 0.62 | 0.33 | 0.29 | 1.88 |
| other bloold cells | Commd7 | 0.71 | 9.11E-13 | 1.89E-08 | 0.84 | 0.57 | 0.27 | 1.47 |
| other bloold cells | Dusp22 | 0.99 | 9.29E-13 | 1.93E-08 | 0.46 | 0.2 | 0.26 | 2.28 |
| other bloold cells | Eea1 | 0.79 | 9.46E-13 | 1.97E-08 | 0.78 | 0.43 | 0.35 | 1.82 |
| other bloold cells | Irak1 | 0.84 | 9.64E-13 | 2.00E-08 | 0.75 | 0.44 | 0.31 | 1.7 |
| other bloold cells | Casp7 | 1.09 | 9.88E-13 | 2.05E-08 | 0.41 | 0.17 | 0.24 | 2.44 |
| other bloold cells | Acot8 | 1.07 | 1.04E-12 | 2.16E-08 | 0.38 | 0.15 | 0.23 | 2.54 |
| other bloold cells | Crat | 1.09 | 1.06E-12 | 2.21E-08 | 0.46 | 0.2 | 0.25 | 2.23 |
| other bloold cells | Tbc1d4 | 1.62 | 1.08E-12 | 2.24E-08 | 0.31 | 0.11 | 0.19 | 2.7 |
| other bloold cells | Wnk1 | 0.73 | 1.10E-12 | 2.28E-08 | 0.96 | 0.78 | 0.18 | 1.23 |
| other bloold cells | Vps26a | 0.96 | 1.14E-12 | 2.37E-08 | 0.73 | 0.43 | 0.3 | 1.7 |
| other bloold cells | Cyld | 1.15 | 1.15E-12 | 2.38E-08 | 0.48 | 0.22 | 0.26 | 2.16 |
| other bloold cells | Ciao3 | 1.13 | 1.15E-12 | 2.40E-08 | 0.4 | 0.17 | 0.23 | 2.4 |
| other bloold cells | Rmc1 | 1.26 | 1.17E-12 | 2.44E-08 | 0.25 | 0.08 | 0.17 | 3.11 |
| other bloold cells | Rras | 1.14 | 1.19E-12 | 2.48E-08 | 0.48 | 0.22 | 0.26 | 2.18 |
| other bloold cells | Vps35l | 1.08 | 1.25E-12 | 2.61E-08 | 0.54 | 0.26 | 0.28 | 2.07 |
| other bloold cells | Trpc4ap | 0.85 | 1.30E-12 | 2.70E-08 | 0.82 | 0.53 | 0.29 | 1.54 |
| other bloold cells | Tmem189 | 0.89 | 1.32E-12 | 2.74E-08 | 0.62 | 0.3 | 0.31 | 2.04 |
| other bloold cells | Adgrg1 | 2.89 | 1.43E-12 | 2.98E-08 | 0.16 | 0.04 | 0.12 | 3.83 |
| other bloold cells | Marchf2 | 0.73 | 1.44E-12 | 3.00E-08 | 0.82 | 0.51 | 0.3 | 1.59 |
| other bloold cells | Snx10 | 1.16 | 1.75E-12 | 3.63E-08 | 0.39 | 0.16 | 0.23 | 2.44 |
| other bloold cells | Slc30a5 | 0.94 | 1.79E-12 | 3.72E-08 | 0.61 | 0.31 | 0.3 | 1.95 |
| other bloold cells | Lamtor2 | 0.55 | 1.98E-12 | 4.11E-08 | 0.97 | 0.89 | 0.07 | 1.08 |
| other bloold cells | Rnft1 | 0.94 | 2.11E-12 | 4.38E-08 | 0.5 | 0.23 | 0.27 | 2.21 |
| other bloold cells | Map11 | 1.31 | 2.12E-12 | 4.41E-08 | 0.29 | 0.11 | 0.19 | 2.8 |
| other bloold cells | Atp6v0a2 | 0.97 | 2.24E-12 | 4.67E-08 | 0.55 | 0.26 | 0.29 | 2.11 |
| other bloold cells | Rab11fip5 | 2.02 | 2.44E-12 | 5.08E-08 | 0.14 | 0.03 | 0.11 | 4.24 |
| other bloold cells | Ppp1r15a | 1.09 | 2.57E-12 | 5.35E-08 | 0.51 | 0.25 | 0.26 | 2.04 |
| other bloold cells | Adipor1 | 0.76 | 2.62E-12 | 5.45E-08 | 0.87 | 0.56 | 0.31 | 1.55 |
| other bloold cells | Etv3 | 1.07 | 2.71E-12 | 5.65E-08 | 0.49 | 0.22 | 0.27 | 2.2 |
| other bloold cells | Raph1 | 1.01 | 2.94E-12 | 6.10E-08 | 0.73 | 0.42 | 0.31 | 1.72 |
| other bloold cells | Bcap31 | 0.56 | 2.95E-12 | 6.14E-08 | 0.95 | 0.75 | 0.2 | 1.27 |
| other bloold cells | Usp12 | 0.97 | 3.14E-12 | 6.52E-08 | 0.46 | 0.21 | 0.26 | 2.25 |
| other bloold cells | Agfg2 | 0.86 | 3.20E-12 | 6.66E-08 | 0.48 | 0.21 | 0.26 | 2.25 |
| other bloold cells | Cars2 | 1.06 | 3.37E-12 | 7.01E-08 | 0.42 | 0.18 | 0.24 | 2.37 |
| other bloold cells | Rap2a | 1 | 3.49E-12 | 7.27E-08 | 0.61 | 0.32 | 0.29 | 1.88 |
| other bloold cells | Ifitm3 | 2.04 | 3.76E-12 | 7.83E-08 | 0.54 | 0.3 | 0.24 | 1.81 |
| other bloold cells | Vim | 0.71 | 4.14E-12 | 8.61E-08 | 0.99 | 0.84 | 0.15 | 1.18 |
| other bloold cells | Ggh | 0.95 | 4.26E-12 | 8.85E-08 | 0.54 | 0.25 | 0.28 | 2.12 |
| other bloold cells | Exosc5 | 0.86 | 4.31E-12 | 8.97E-08 | 0.8 | 0.58 | 0.22 | 1.38 |

**Supplementary Table 2. Marker genes for the major cell types of integrated clustering of E11.5 control and NotchGOF λ-junction cells.**

| | | | | | | | | |
|---|---|---|---|---|---|---|---|---|
| other bloold cells | Adar | 1.19 | 4.46E-12 | 9.28E-08 | 0.37 | 0.15 | 0.22 | 2.44 |
| other bloold cells | Chmp4b | 0.57 | 4.98E-12 | 1.04E-07 | 0.97 | 0.85 | 0.12 | 1.14 |
| other bloold cells | Cept1 | 0.84 | 5.05E-12 | 1.05E-07 | 0.62 | 0.33 | 0.29 | 1.87 |
| other bloold cells | Tox4 | 0.76 | 5.34E-12 | 1.11E-07 | 0.81 | 0.52 | 0.29 | 1.56 |
| other bloold cells | Dip2b | 0.91 | 5.41E-12 | 1.13E-07 | 0.64 | 0.34 | 0.29 | 1.85 |
| other bloold cells | Hes6 | 0.51 | 5.93E-12 | 1.23E-07 | 0.62 | 0.33 | 0.29 | 1.87 |
| other bloold cells | Steap3 | 2.54 | 6.20E-12 | 1.29E-07 | 0.11 | 0.02 | 0.09 | 4.67 |
| other bloold cells | Sprtn | 1.34 | 6.46E-12 | 1.34E-07 | 0.26 | 0.09 | 0.17 | 2.94 |
| other bloold cells | Lnpep | 0.79 | 6.63E-12 | 1.38E-07 | 0.59 | 0.29 | 0.3 | 2.04 |
| other bloold cells | Cpd | 1.39 | 6.68E-12 | 1.39E-07 | 0.59 | 0.34 | 0.26 | 1.76 |
| other bloold cells | Chid1 | 0.92 | 6.71E-12 | 1.40E-07 | 0.53 | 0.25 | 0.28 | 2.09 |
| other bloold cells | Irf3 | 0.81 | 7.00E-12 | 1.46E-07 | 0.63 | 0.33 | 0.3 | 1.91 |
| other bloold cells | Fbrs | 0.85 | 7.07E-12 | 1.47E-07 | 0.46 | 0.2 | 0.26 | 2.3 |
| other bloold cells | Flcn | 1.35 | 7.21E-12 | 1.50E-07 | 0.3 | 0.11 | 0.19 | 2.66 |
| other bloold cells | Bahd1 | 1.64 | 7.36E-12 | 1.53E-07 | 0.16 | 0.04 | 0.12 | 3.83 |
| other bloold cells | Ogfod2 | 1.17 | 8.36E-12 | 1.74E-07 | 0.28 | 0.1 | 0.18 | 2.83 |
| other bloold cells | Nrm | 0.95 | 8.38E-12 | 1.74E-07 | 0.64 | 0.35 | 0.28 | 1.8 |
| other bloold cells | Acox1 | 1.43 | 8.48E-12 | 1.76E-07 | 0.35 | 0.14 | 0.21 | 2.43 |
| other bloold cells | Zbtb38 | 1.13 | 8.55E-12 | 1.78E-07 | 0.34 | 0.14 | 0.21 | 2.52 |
| other bloold cells | Xbp1 | 0.84 | 9.20E-12 | 1.91E-07 | 0.77 | 0.48 | 0.29 | 1.61 |
| other bloold cells | Klf7 | 1 | 9.46E-12 | 1.97E-07 | 0.48 | 0.23 | 0.25 | 2.11 |
| other bloold cells | Ecm1 | 2.41 | 9.54E-12 | 1.98E-07 | 0.11 | 0.02 | 0.09 | 4.67 |
| other bloold cells | Usp8 | 0.8 | 1.11E-11 | 2.30E-07 | 0.78 | 0.46 | 0.32 | 1.68 |
| other bloold cells | Tmem87b | 1.03 | 1.11E-11 | 2.31E-07 | 0.47 | 0.21 | 0.26 | 2.2 |
| other bloold cells | Pkig | 0.56 | 1.13E-11 | 2.35E-07 | 0.94 | 0.79 | 0.16 | 1.2 |
| other bloold cells | Tbc1d2b | 1.46 | 1.21E-11 | 2.51E-07 | 0.25 | 0.08 | 0.16 | 2.92 |
| other bloold cells | Krcc1 | 0.7 | 1.22E-11 | 2.53E-07 | 0.85 | 0.58 | 0.28 | 1.48 |
| other bloold cells | Bad | 0.72 | 1.28E-11 | 2.66E-07 | 0.81 | 0.53 | 0.28 | 1.54 |
| other bloold cells | Galnt12 | 2.24 | 1.30E-11 | 2.70E-07 | 0.11 | 0.02 | 0.09 | 4.67 |
| other bloold cells | Soat1 | 1.24 | 1.30E-11 | 2.71E-07 | 0.69 | 0.44 | 0.24 | 1.55 |
| other bloold cells | Auh | 0.9 | 1.36E-11 | 2.82E-07 | 0.55 | 0.28 | 0.27 | 1.95 |
| other bloold cells | Cyfip2 | 1.28 | 1.37E-11 | 2.85E-07 | 0.26 | 0.09 | 0.17 | 2.88 |
| other bloold cells | Hras | 0.5 | 1.43E-11 | 2.97E-07 | 0.94 | 0.8 | 0.14 | 1.18 |
| other bloold cells | Tmed3 | 0.79 | 1.56E-11 | 3.25E-07 | 0.72 | 0.45 | 0.27 | 1.61 |
| other bloold cells | BC004004 | 0.94 | 1.64E-11 | 3.41E-07 | 0.59 | 0.32 | 0.27 | 1.85 |
| other bloold cells | Dennd1b | 1.05 | 1.74E-11 | 3.61E-07 | 0.39 | 0.16 | 0.23 | 2.42 |
| other bloold cells | Zdhhc9 | 1.3 | 1.77E-11 | 3.68E-07 | 0.26 | 0.09 | 0.17 | 2.91 |
| other bloold cells | Gm26782 | 1.63 | 1.82E-11 | 3.78E-07 | 0.21 | 0.07 | 0.14 | 3.18 |
| other bloold cells | Slk | 0.69 | 1.82E-11 | 3.80E-07 | 0.8 | 0.53 | 0.27 | 1.51 |
| other bloold cells | Retsat | 1.56 | 1.83E-11 | 3.81E-07 | 0.18 | 0.05 | 0.13 | 3.43 |
| other bloold cells | Hspa8 | 0.38 | 1.87E-11 | 3.89E-07 | 1 | 1 | 0 | 1 |
| other bloold cells | Nin | 0.88 | 1.89E-11 | 3.93E-07 | 0.54 | 0.27 | 0.27 | 1.99 |
| other bloold cells | Hivep2 | 1.42 | 1.90E-11 | 3.95E-07 | 0.23 | 0.08 | 0.16 | 3.04 |
| other bloold cells | Uckl1 | 0.88 | 2.03E-11 | 4.22E-07 | 0.32 | 0.12 | 0.2 | 2.69 |
| other bloold cells | Nuak2 | 1.56 | 2.03E-11 | 4.23E-07 | 0.15 | 0.04 | 0.11 | 3.97 |
| other bloold cells | .810037I17Ril | 0.55 | 2.14E-11 | 4.45E-07 | 0.94 | 0.84 | 0.1 | 1.12 |
| other bloold cells | Rexo2 | 0.74 | 2.17E-11 | 4.52E-07 | 0.81 | 0.59 | 0.22 | 1.37 |
| other bloold cells | Traf5 | 1.8 | 2.19E-11 | 4.56E-07 | 0.15 | 0.04 | 0.11 | 3.76 |
| other bloold cells | Reep5 | 0.48 | 2.43E-11 | 5.05E-07 | 0.97 | 0.78 | 0.19 | 1.24 |
| other bloold cells | Vps13d | 0.97 | 2.48E-11 | 5.15E-07 | 0.35 | 0.14 | 0.21 | 2.52 |
| other bloold cells | Cr1l | 0.77 | 2.63E-11 | 5.46E-07 | 0.6 | 0.32 | 0.28 | 1.88 |
| other bloold cells | Npc1 | 1.03 | 2.71E-11 | 5.64E-07 | 0.53 | 0.26 | 0.27 | 2.05 |
| other bloold cells | Cox5b | 0.34 | 2.72E-11 | 5.66E-07 | 0.99 | 0.99 | 0.01 | 1.01 |
| other bloold cells | Cybc1 | 0.95 | 2.77E-11 | 5.76E-07 | 0.48 | 0.22 | 0.25 | 2.14 |
| other bloold cells | Nr2c2ap | 0.98 | 2.79E-11 | 5.80E-07 | 0.48 | 0.23 | 0.25 | 2.06 |
| other bloold cells | Dusp1 | 1.69 | 2.81E-11 | 5.85E-07 | 0.52 | 0.29 | 0.22 | 1.76 |

**Supplementary Table 2. Marker genes for the major cell types of integrated clustering of E11.5 control and NotchGOF λ-junction cells.**

| | | | | | | | | |
|---|---|---|---|---|---|---|---|---|
| other bloold cells | Rmnd5b | 1.37 | 2.92E-11 | 6.08E-07 | 0.39 | 0.17 | 0.22 | 2.31 |
| other bloold cells | Mbd2 | 0.64 | 3.06E-11 | 6.36E-07 | 0.83 | 0.56 | 0.27 | 1.49 |
| other bloold cells | Map2k1 | 0.97 | 3.07E-11 | 6.38E-07 | 0.6 | 0.35 | 0.26 | 1.74 |
| other bloold cells | Ier2 | 0.64 | 3.15E-11 | 6.55E-07 | 0.94 | 0.79 | 0.15 | 1.19 |
| other bloold cells | BC029722 | 1.02 | 3.25E-11 | 6.76E-07 | 0.31 | 0.12 | 0.19 | 2.68 |
| other bloold cells | Jtb | 0.78 | 3.33E-11 | 6.92E-07 | 0.8 | 0.53 | 0.26 | 1.5 |
| other bloold cells | Tmem181a | 1.15 | 3.55E-11 | 7.38E-07 | 0.28 | 0.1 | 0.18 | 2.75 |
| other bloold cells | Bcl10 | 0.85 | 3.62E-11 | 7.52E-07 | 0.76 | 0.5 | 0.27 | 1.53 |
| other bloold cells | Oma1 | 0.99 | 3.64E-11 | 7.57E-07 | 0.41 | 0.18 | 0.23 | 2.27 |
| other bloold cells | Prorsd1 | 0.8 | 4.07E-11 | 8.46E-07 | 0.67 | 0.38 | 0.29 | 1.76 |
| other bloold cells | Qprt | 1.47 | 4.07E-11 | 8.46E-07 | 0.16 | 0.04 | 0.12 | 3.66 |
| other bloold cells | Naprt | 2.02 | 4.20E-11 | 8.74E-07 | 0.11 | 0.03 | 0.09 | 4.48 |
| other bloold cells | Rrbp1 | 0.5 | 4.22E-11 | 8.77E-07 | 0.99 | 0.9 | 0.1 | 1.11 |
| other bloold cells | Nipsnap3b | 0.86 | 4.26E-11 | 8.86E-07 | 0.7 | 0.44 | 0.26 | 1.61 |
| other bloold cells | Tnip1 | 1.2 | 4.38E-11 | 9.10E-07 | 0.28 | 0.1 | 0.18 | 2.72 |
| other bloold cells | Pigt | 0.82 | 4.42E-11 | 9.20E-07 | 0.69 | 0.43 | 0.26 | 1.62 |
| other bloold cells | Zfp800 | 0.91 | 4.68E-11 | 9.73E-07 | 0.53 | 0.27 | 0.27 | 2 |
| other bloold cells | Tob2 | 0.98 | 4.79E-11 | 9.96E-07 | 0.5 | 0.24 | 0.26 | 2.07 |
| other bloold cells | Itpripl1 | 1.37 | 4.84E-11 | 1.01E-06 | 0.26 | 0.09 | 0.17 | 2.76 |
| other bloold cells | Rnf166 | 0.85 | 4.88E-11 | 1.01E-06 | 0.53 | 0.26 | 0.27 | 2.04 |
| other bloold cells | Mospd3 | 1.16 | 5.13E-11 | 1.07E-06 | 0.39 | 0.17 | 0.22 | 2.28 |
| other bloold cells | Gpt | 1.91 | 5.55E-11 | 1.15E-06 | 0.15 | 0.04 | 0.11 | 3.67 |
| other bloold cells | Tbk1 | 1.06 | 5.66E-11 | 1.18E-06 | 0.54 | 0.28 | 0.26 | 1.91 |
| other bloold cells | Mrpl36 | 0.69 | 6.06E-11 | 1.26E-06 | 0.92 | 0.72 | 0.2 | 1.28 |
| other bloold cells | Csrnp1 | 2 | 6.69E-11 | 1.39E-06 | 0.13 | 0.03 | 0.1 | 4.03 |
| other bloold cells | Slc35b2 | 0.73 | 6.74E-11 | 1.40E-06 | 0.72 | 0.41 | 0.31 | 1.74 |
| other bloold cells | Spns2 | 2.92 | 6.98E-11 | 1.45E-06 | 0.12 | 0.03 | 0.09 | 4.25 |
| other bloold cells | Rab31 | 0.96 | 7.11E-11 | 1.48E-06 | 0.47 | 0.22 | 0.25 | 2.09 |
| other bloold cells | Ltbr | 0.9 | 7.89E-11 | 1.64E-06 | 0.48 | 0.23 | 0.25 | 2.1 |
| other bloold cells | Elmsan1 | 1.12 | 8.03E-11 | 1.67E-06 | 0.31 | 0.12 | 0.19 | 2.59 |
| other bloold cells | Rit1 | 0.93 | 8.25E-11 | 1.72E-06 | 0.52 | 0.25 | 0.26 | 2.04 |
| other bloold cells | Snx9 | 0.87 | 8.37E-11 | 1.74E-06 | 0.59 | 0.31 | 0.27 | 1.87 |
| other bloold cells | Tbc1d13 | 0.97 | 8.42E-11 | 1.75E-06 | 0.34 | 0.14 | 0.2 | 2.47 |
| other bloold cells | Sqstm1 | 0.8 | 8.54E-11 | 1.78E-06 | 0.71 | 0.41 | 0.3 | 1.73 |
| other bloold cells | Anxa5 | 1 | 8.66E-11 | 1.80E-06 | 0.83 | 0.61 | 0.23 | 1.38 |
| other bloold cells | Usp24 | 0.91 | 8.86E-11 | 1.84E-06 | 0.57 | 0.31 | 0.27 | 1.87 |
| other bloold cells | Iqsec1 | 1.15 | 8.91E-11 | 1.85E-06 | 0.34 | 0.14 | 0.2 | 2.43 |
| other bloold cells | Coq8b | 0.99 | 9.00E-11 | 1.87E-06 | 0.32 | 0.12 | 0.19 | 2.58 |
| other bloold cells | Rap1gds1 | 0.96 | 9.22E-11 | 1.92E-06 | 0.56 | 0.29 | 0.27 | 1.91 |
| other bloold cells | Max | 0.8 | 9.39E-11 | 1.95E-06 | 0.69 | 0.4 | 0.28 | 1.7 |
| other bloold cells | Itgav | 1.07 | 9.90E-11 | 2.06E-06 | 0.53 | 0.27 | 0.26 | 1.94 |
| other bloold cells | Gclm | 0.75 | 1.03E-10 | 2.14E-06 | 0.69 | 0.39 | 0.3 | 1.78 |
| other bloold cells | Akap10 | 1.2 | 1.08E-10 | 2.25E-06 | 0.32 | 0.13 | 0.19 | 2.44 |
| other bloold cells | Slc39a1 | 0.72 | 1.09E-10 | 2.26E-06 | 0.53 | 0.26 | 0.28 | 2.07 |
| other bloold cells | Pim2 | 2.11 | 1.10E-10 | 2.29E-06 | 0.11 | 0.03 | 0.09 | 4.48 |
| other bloold cells | Myl12b | 0.52 | 1.28E-10 | 2.65E-06 | 0.98 | 0.89 | 0.09 | 1.1 |
| other bloold cells | Nans | 0.65 | 1.32E-10 | 2.74E-06 | 0.86 | 0.6 | 0.26 | 1.42 |
| other bloold cells | Atp2b1 | 0.7 | 1.35E-10 | 2.81E-06 | 0.96 | 0.85 | 0.11 | 1.13 |
| other bloold cells | Lin28b | 0.93 | 1.44E-10 | 3.01E-06 | 0.43 | 0.21 | 0.23 | 2.1 |
| other bloold cells | Atp13a3 | 1.05 | 1.45E-10 | 3.01E-06 | 0.68 | 0.41 | 0.27 | 1.66 |
| other bloold cells | Dna2 | 0.83 | 1.47E-10 | 3.06E-06 | 0.52 | 0.26 | 0.26 | 2 |
| other bloold cells | Alkbh3 | 0.86 | 1.48E-10 | 3.08E-06 | 0.43 | 0.2 | 0.24 | 2.18 |
| other bloold cells | Ubl4a | 0.76 | 1.51E-10 | 3.14E-06 | 0.8 | 0.58 | 0.22 | 1.39 |
| other bloold cells | Commd8 | 0.62 | 1.52E-10 | 3.15E-06 | 0.88 | 0.67 | 0.21 | 1.32 |
| other bloold cells | Pfkfb3 | 1.38 | 1.52E-10 | 3.16E-06 | 0.27 | 0.11 | 0.17 | 2.6 |
| other bloold cells | Sgpp1 | 1.05 | 1.61E-10 | 3.34E-06 | 0.28 | 0.1 | 0.18 | 2.72 |

**Supplementary Table 2. Marker genes for the major cell types of integrated clustering of E11.5 control and NotchGOF λ-junction cells.**

| | | | | | | | | |
|---|---|---|---|---|---|---|---|---|
| other bloold cells | Mfsd10 | 0.86 | 1.61E-10 | 3.34E-06 | 0.6 | 0.35 | 0.25 | 1.72 |
| other bloold cells | Idh3g | 0.6 | 1.62E-10 | 3.36E-06 | 0.86 | 0.66 | 0.2 | 1.31 |
| other bloold cells | Spryd7 | 0.9 | 1.63E-10 | 3.39E-06 | 0.36 | 0.15 | 0.21 | 2.38 |
| other bloold cells | Pxk | 0.8 | 1.63E-10 | 3.40E-06 | 0.39 | 0.17 | 0.23 | 2.36 |
| other bloold cells | Mydgf | 0.56 | 1.76E-10 | 3.66E-06 | 0.94 | 0.73 | 0.2 | 1.28 |
| other bloold cells | Eef1akmt4 | 0.88 | 1.81E-10 | 3.76E-06 | 0.48 | 0.23 | 0.25 | 2.07 |
| other bloold cells | Mlec | 0.7 | 1.82E-10 | 3.79E-06 | 0.94 | 0.75 | 0.19 | 1.26 |
| other bloold cells | Slc25a26 | 1.06 | 1.83E-10 | 3.80E-06 | 0.25 | 0.09 | 0.16 | 2.83 |
| other bloold cells | Tm9sf2 | 0.61 | 1.90E-10 | 3.95E-06 | 0.89 | 0.62 | 0.27 | 1.44 |
| other bloold cells | Anxa11 | 0.69 | 1.93E-10 | 4.02E-06 | 0.37 | 0.15 | 0.22 | 2.41 |
| other bloold cells | Rufy1 | 0.93 | 1.98E-10 | 4.11E-06 | 0.5 | 0.25 | 0.25 | 2.02 |
| other bloold cells | Tmem35b | 1.27 | 2.15E-10 | 4.48E-06 | 0.18 | 0.06 | 0.13 | 3.31 |
| other bloold cells | Irak2 | 1.68 | 2.17E-10 | 4.52E-06 | 0.18 | 0.05 | 0.12 | 3.3 |
| other bloold cells | Tmem123 | 0.7 | 2.18E-10 | 4.53E-06 | 0.62 | 0.32 | 0.29 | 1.92 |
| other bloold cells | Hacd1 | 0.85 | 2.21E-10 | 4.60E-06 | 0.65 | 0.39 | 0.26 | 1.65 |
| other bloold cells | Coro1b | 1.31 | 2.25E-10 | 4.69E-06 | 0.21 | 0.07 | 0.14 | 3.04 |
| other bloold cells | Colca2 | 1.49 | 2.32E-10 | 4.82E-06 | 0.14 | 0.04 | 0.1 | 3.78 |
| other bloold cells | Plbd2 | 0.99 | 2.36E-10 | 4.90E-06 | 0.48 | 0.24 | 0.24 | 1.98 |
| other bloold cells | Ttll3 | 2.42 | 2.41E-10 | 5.02E-06 | 0.12 | 0.03 | 0.09 | 4.1 |
| other bloold cells | Vps52 | 0.92 | 2.44E-10 | 5.07E-06 | 0.5 | 0.25 | 0.25 | 1.99 |
| other bloold cells | Zfp991 | 1.28 | 2.48E-10 | 5.15E-06 | 0.25 | 0.09 | 0.16 | 2.78 |
| other bloold cells | Selenot | 0.64 | 2.48E-10 | 5.17E-06 | 0.84 | 0.54 | 0.3 | 1.55 |
| other bloold cells | Impa2 | 0.87 | 2.54E-10 | 5.29E-06 | 0.55 | 0.3 | 0.25 | 1.85 |
| other bloold cells | ·833438C02Ri | 1.64 | 2.59E-10 | 5.38E-06 | 0.13 | 0.03 | 0.1 | 4.06 |
| other bloold cells | Ccdc115 | 0.7 | 2.59E-10 | 5.38E-06 | 0.73 | 0.49 | 0.25 | 1.5 |
| other bloold cells | Sap25 | 1.72 | 2.69E-10 | 5.59E-06 | 0.13 | 0.03 | 0.1 | 3.91 |
| other bloold cells | Tmem250-ps | 0.98 | 2.70E-10 | 5.62E-06 | 0.34 | 0.14 | 0.2 | 2.4 |
| other bloold cells | Fam120a | 0.81 | 2.78E-10 | 5.78E-06 | 0.71 | 0.46 | 0.26 | 1.56 |
| other bloold cells | Dnajc5 | 0.71 | 3.21E-10 | 6.68E-06 | 0.83 | 0.52 | 0.3 | 1.58 |
| other bloold cells | Atp5l | 0.29 | 3.21E-10 | 6.68E-06 | 0.99 | 0.99 | 0 | 1 |
| other bloold cells | Snf8 | 0.58 | 3.42E-10 | 7.11E-06 | 0.89 | 0.75 | 0.14 | 1.18 |
| other bloold cells | Gna13 | 0.83 | 3.44E-10 | 7.16E-06 | 0.69 | 0.43 | 0.26 | 1.61 |
| other bloold cells | Gm12166 | 1.02 | 3.55E-10 | 7.38E-06 | 0.32 | 0.13 | 0.19 | 2.44 |
| other bloold cells | Dipk1a | 0.87 | 3.87E-10 | 8.04E-06 | 0.44 | 0.2 | 0.24 | 2.16 |
| other bloold cells | Ccng1 | 0.79 | 3.97E-10 | 8.25E-06 | 0.43 | 0.2 | 0.23 | 2.16 |
| other bloold cells | 900026A02Ri | 1.18 | 4.12E-10 | 8.57E-06 | 0.27 | 0.1 | 0.17 | 2.63 |
| other bloold cells | C2cd2l | 1.32 | 4.13E-10 | 8.58E-06 | 0.18 | 0.05 | 0.12 | 3.3 |
| other bloold cells | Cox8a | 0.25 | 4.19E-10 | 8.72E-06 | 0.99 | 1 | 0 | 1 |
| other bloold cells | Man2a1 | 0.93 | 4.23E-10 | 8.79E-06 | 0.47 | 0.23 | 0.24 | 2.02 |
| other bloold cells | Zmpste24 | 0.82 | 4.23E-10 | 8.79E-06 | 0.59 | 0.32 | 0.27 | 1.86 |
| other bloold cells | Brms1 | 0.83 | 4.25E-10 | 8.84E-06 | 0.62 | 0.35 | 0.27 | 1.78 |
| other bloold cells | Tmem135 | 0.93 | 4.90E-10 | 1.02E-05 | 0.32 | 0.13 | 0.19 | 2.46 |
| other bloold cells | Calcrl | 1.6 | 5.00E-10 | 1.04E-05 | 0.13 | 0.04 | 0.1 | 3.8 |
| other bloold cells | ·830406C13Ri | 0.81 | 5.14E-10 | 1.07E-05 | 0.5 | 0.24 | 0.26 | 2.05 |
| other bloold cells | Lifr | 1.08 | 5.40E-10 | 1.12E-05 | 0.28 | 0.11 | 0.17 | 2.57 |
| other bloold cells | Necap1 | 0.97 | 5.82E-10 | 1.21E-05 | 0.48 | 0.24 | 0.24 | 2.02 |
| other bloold cells | Ykt6 | 0.75 | 5.89E-10 | 1.22E-05 | 0.69 | 0.41 | 0.28 | 1.7 |
| other bloold cells | Klf10 | 1.06 | 6.00E-10 | 1.25E-05 | 0.28 | 0.11 | 0.17 | 2.57 |
| other bloold cells | Helb | 0.93 | 6.02E-10 | 1.25E-05 | 0.32 | 0.13 | 0.19 | 2.44 |
| other bloold cells | Ctsf | 1.3 | 6.12E-10 | 1.27E-05 | 0.27 | 0.11 | 0.17 | 2.58 |
| other bloold cells | Rragc | 0.89 | 6.25E-10 | 1.30E-05 | 0.74 | 0.45 | 0.29 | 1.63 |
| other bloold cells | Aph1a | 0.69 | 6.28E-10 | 1.31E-05 | 0.79 | 0.5 | 0.29 | 1.57 |
| other bloold cells | Scn1b | 2.24 | 6.52E-10 | 1.36E-05 | 0.11 | 0.02 | 0.08 | 4.38 |
| other bloold cells | Hmox2 | 0.66 | 6.75E-10 | 1.40E-05 | 0.88 | 0.64 | 0.24 | 1.37 |
| other bloold cells | Slc35a3 | 0.94 | 6.84E-10 | 1.42E-05 | 0.31 | 0.12 | 0.19 | 2.5 |
| other bloold cells | Apaf1 | 0.83 | 6.96E-10 | 1.45E-05 | 0.47 | 0.23 | 0.24 | 2.08 |

**Supplementary Table 2. Marker genes for the major cell types of integrated clustering of E11.5 control and NotchGOF λ-junction cells.**

| | | | | | | | | |
|---|---|---|---|---|---|---|---|---|
| other bloold cells | Ywhag | 0.76 | 7.02E-10 | 1.46E-05 | 0.82 | 0.63 | 0.19 | 1.29 |
| other bloold cells | Acap2 | 0.7 | 7.30E-10 | 1.52E-05 | 0.75 | 0.46 | 0.29 | 1.62 |
| other bloold cells | Sh3bp2 | 1.35 | 7.54E-10 | 1.57E-05 | 0.22 | 0.08 | 0.15 | 2.84 |
| other bloold cells | Map3k20 | 1.41 | 7.55E-10 | 1.57E-05 | 0.29 | 0.12 | 0.18 | 2.51 |
| other bloold cells | Anxa7 | 0.72 | 7.60E-10 | 1.58E-05 | 0.63 | 0.36 | 0.27 | 1.74 |
| other bloold cells | Abracl | 0.36 | 8.07E-10 | 1.68E-05 | 0.99 | 0.93 | 0.06 | 1.06 |
| other bloold cells | Speg | 1.43 | 8.39E-10 | 1.75E-05 | 0.36 | 0.16 | 0.19 | 2.18 |
| other bloold cells | Nmrk1 | 1.29 | 8.48E-10 | 1.76E-05 | 0.23 | 0.08 | 0.15 | 2.78 |
| other bloold cells | Esrra | 1.03 | 9.33E-10 | 1.94E-05 | 0.31 | 0.13 | 0.18 | 2.46 |
| other bloold cells | Cdk18 | 2.01 | 9.44E-10 | 1.96E-05 | 0.11 | 0.02 | 0.08 | 4.38 |
| other bloold cells | Srd5a3 | 0.87 | 9.69E-10 | 2.01E-05 | 0.44 | 0.21 | 0.23 | 2.08 |
| other bloold cells | Dock11 | 0.83 | 1.05E-09 | 2.19E-05 | 0.47 | 0.23 | 0.24 | 2.06 |
| other bloold cells | Vps35 | 0.66 | 1.06E-09 | 2.21E-05 | 0.91 | 0.76 | 0.15 | 1.19 |
| other bloold cells | Vti1b | 0.57 | 1.10E-09 | 2.29E-05 | 0.83 | 0.52 | 0.31 | 1.59 |
| other bloold cells | Kit | 1.37 | 1.11E-09 | 2.31E-05 | 0.25 | 0.1 | 0.16 | 2.6 |
| other bloold cells | Otud1 | 1.91 | 1.11E-09 | 2.31E-05 | 0.12 | 0.03 | 0.09 | 3.97 |
| other bloold cells | Cnn2 | 0.61 | 1.22E-09 | 2.53E-05 | 0.87 | 0.63 | 0.25 | 1.39 |
| other bloold cells | Prr14l | 0.69 | 1.24E-09 | 2.58E-05 | 0.49 | 0.24 | 0.25 | 2.03 |
| other bloold cells | Pqlc3 | 1.4 | 1.25E-09 | 2.61E-05 | 0.18 | 0.06 | 0.12 | 3.18 |
| other bloold cells | Igsf8 | 0.78 | 1.28E-09 | 2.66E-05 | 0.49 | 0.24 | 0.25 | 2.01 |
| other bloold cells | Rock2 | 0.73 | 1.32E-09 | 2.75E-05 | 0.87 | 0.66 | 0.2 | 1.31 |
| other bloold cells | Uqcr10 | 0.32 | 1.37E-09 | 2.84E-05 | 0.99 | 0.98 | 0.01 | 1.01 |
| other bloold cells | Vps28 | 0.59 | 1.37E-09 | 2.85E-05 | 0.89 | 0.62 | 0.27 | 1.43 |
| other bloold cells | Tbc1d9b | 0.88 | 1.38E-09 | 2.87E-05 | 0.44 | 0.22 | 0.23 | 2.05 |
| other bloold cells | Usp2 | 1.17 | 1.47E-09 | 3.06E-05 | 0.33 | 0.14 | 0.19 | 2.33 |
| other bloold cells | Pigk | 0.77 | 1.58E-09 | 3.29E-05 | 0.55 | 0.3 | 0.26 | 1.86 |
| other bloold cells | Zfp524 | 1.13 | 1.62E-09 | 3.38E-05 | 0.27 | 0.1 | 0.16 | 2.56 |
| other bloold cells | Sun2 | 1.09 | 1.70E-09 | 3.53E-05 | 0.37 | 0.17 | 0.2 | 2.13 |
| other bloold cells | Snx1 | 0.69 | 1.75E-09 | 3.64E-05 | 0.72 | 0.45 | 0.27 | 1.61 |
| other bloold cells | Sgsh | 2.05 | 1.78E-09 | 3.71E-05 | 0.11 | 0.03 | 0.08 | 4.2 |
| other bloold cells | Klhl18 | 1.19 | 1.80E-09 | 3.75E-05 | 0.23 | 0.08 | 0.15 | 2.75 |
| other bloold cells | Gm20707 | 1.58 | 1.80E-09 | 3.75E-05 | 0.13 | 0.04 | 0.1 | 3.69 |
| other bloold cells | Stxbp3 | 0.85 | 1.86E-09 | 3.87E-05 | 0.48 | 0.24 | 0.24 | 2.02 |
| other bloold cells | Naga | 1.07 | 1.88E-09 | 3.92E-05 | 0.29 | 0.12 | 0.17 | 2.45 |
| other bloold cells | Atg10 | 0.78 | 1.92E-09 | 3.99E-05 | 0.35 | 0.15 | 0.2 | 2.35 |
| other bloold cells | Nipa2 | 0.7 | 2.01E-09 | 4.19E-05 | 0.79 | 0.54 | 0.25 | 1.46 |
| other bloold cells | Slc26a2 | 1.09 | 2.06E-09 | 4.28E-05 | 0.35 | 0.16 | 0.19 | 2.24 |
| other bloold cells | Abcg1 | 1.49 | 2.08E-09 | 4.32E-05 | 0.22 | 0.08 | 0.14 | 2.7 |
| other bloold cells | Gm43328 | 2.02 | 2.16E-09 | 4.50E-05 | 0.11 | 0.03 | 0.08 | 4 |
| other bloold cells | Htt | 0.75 | 2.20E-09 | 4.57E-05 | 0.57 | 0.3 | 0.27 | 1.89 |
| other bloold cells | Slc25a12 | 1.05 | 2.25E-09 | 4.67E-05 | 0.38 | 0.18 | 0.2 | 2.14 |
| other bloold cells | Ube2f | 0.57 | 2.42E-09 | 5.04E-05 | 0.81 | 0.56 | 0.25 | 1.44 |
| other bloold cells | Neurl2 | 1.53 | 2.46E-09 | 5.13E-05 | 0.2 | 0.07 | 0.13 | 2.93 |
| other bloold cells | Fkbp8 | 0.56 | 2.53E-09 | 5.25E-05 | 0.98 | 0.82 | 0.16 | 1.19 |
| other bloold cells | Pex7 | 0.83 | 2.55E-09 | 5.30E-05 | 0.49 | 0.25 | 0.24 | 1.93 |
| other bloold cells | Mbp | 0.8 | 2.73E-09 | 5.68E-05 | 0.25 | 0.1 | 0.16 | 2.65 |
| other bloold cells | Pgk1 | 0.43 | 2.76E-09 | 5.75E-05 | 0.98 | 0.96 | 0.02 | 1.03 |
| other bloold cells | Ttc32 | 0.81 | 2.84E-09 | 5.90E-05 | 0.5 | 0.26 | 0.24 | 1.94 |
| other bloold cells | AU040320 | 1.1 | 2.86E-09 | 5.94E-05 | 0.27 | 0.11 | 0.16 | 2.46 |
| other bloold cells | Atp1a1 | 0.63 | 2.88E-09 | 5.98E-05 | 0.87 | 0.63 | 0.23 | 1.37 |
| other bloold cells | Cdkn3 | 1.06 | 2.94E-09 | 6.12E-05 | 0.52 | 0.29 | 0.22 | 1.76 |
| other bloold cells | Fancg | 1.36 | 2.98E-09 | 6.19E-05 | 0.17 | 0.05 | 0.12 | 3.17 |
| other bloold cells | Churc1 | 0.59 | 2.98E-09 | 6.20E-05 | 0.85 | 0.59 | 0.27 | 1.45 |
| other bloold cells | Ppp1r12c | 0.72 | 2.99E-09 | 6.21E-05 | 0.58 | 0.32 | 0.26 | 1.82 |
| other bloold cells | Zmiz2 | 0.77 | 3.15E-09 | 6.54E-05 | 0.59 | 0.31 | 0.28 | 1.89 |
| other bloold cells | Alkbh2 | 0.91 | 3.16E-09 | 6.58E-05 | 0.37 | 0.17 | 0.21 | 2.23 |

**Supplementary Table 2. Marker genes for the major cell types of integrated clustering of E11.5 control and NotchGOF λ-junction cells.**

| | | | | | | | | |
|---|---|---|---|---|---|---|---|---|
| other bloold cells | Tgfbrap1 | 1.22 | 3.36E-09 | 6.98E-05 | 0.29 | 0.12 | 0.17 | 2.37 |
| other bloold cells | Sel1l | 0.7 | 3.47E-09 | 7.21E-05 | 0.64 | 0.37 | 0.27 | 1.74 |
| other bloold cells | Sgpl1 | 1.04 | 3.49E-09 | 7.25E-05 | 0.55 | 0.34 | 0.21 | 1.6 |
| other bloold cells | Slc30a1 | 1.44 | 3.49E-09 | 7.26E-05 | 0.22 | 0.08 | 0.14 | 2.67 |
| other bloold cells | Ccdc86 | 0.84 | 3.61E-09 | 7.50E-05 | 0.8 | 0.55 | 0.25 | 1.45 |
| other bloold cells | Arhgef2 | 0.73 | 3.90E-09 | 8.11E-05 | 0.68 | 0.43 | 0.25 | 1.6 |
| other bloold cells | Ide | 0.77 | 3.90E-09 | 8.12E-05 | 0.5 | 0.27 | 0.23 | 1.87 |
| other bloold cells | Ptgr1 | 0.85 | 4.00E-09 | 8.32E-05 | 0.51 | 0.26 | 0.25 | 1.93 |
| other bloold cells | Ncln | 0.68 | 4.08E-09 | 8.50E-05 | 0.64 | 0.37 | 0.26 | 1.71 |
| other bloold cells | Plekhf2 | 0.67 | 4.09E-09 | 8.50E-05 | 0.47 | 0.23 | 0.24 | 2.03 |
| other bloold cells | 230307C23Ri | 0.83 | 4.37E-09 | 9.09E-05 | 0.36 | 0.16 | 0.2 | 2.22 |
| other bloold cells | Rsrp1 | 0.49 | 4.54E-09 | 9.45E-05 | 0.97 | 0.89 | 0.08 | 1.09 |
| other bloold cells | Zbtb18 | 1.08 | 4.56E-09 | 9.48E-05 | 0.27 | 0.11 | 0.16 | 2.48 |
| other bloold cells | Trafd1 | 0.74 | 4.87E-09 | 1.01E-04 | 0.55 | 0.3 | 0.25 | 1.85 |
| other bloold cells | Gmppa | 0.89 | 4.87E-09 | 1.01E-04 | 0.37 | 0.17 | 0.2 | 2.17 |
| other bloold cells | Axl | 1.66 | 4.94E-09 | 1.03E-04 | 0.32 | 0.15 | 0.17 | 2.12 |
| other bloold cells | Mrps26 | 0.68 | 5.37E-09 | 1.12E-04 | 0.78 | 0.65 | 0.13 | 1.2 |
| other bloold cells | Wwp1 | 0.79 | 5.45E-09 | 1.13E-04 | 0.69 | 0.41 | 0.27 | 1.66 |
| other bloold cells | Mknk2 | 0.72 | 5.49E-09 | 1.14E-04 | 0.69 | 0.41 | 0.27 | 1.67 |
| other bloold cells | Specc1 | 0.66 | 5.51E-09 | 1.15E-04 | 0.7 | 0.42 | 0.28 | 1.65 |
| other bloold cells | Hipk3 | 0.91 | 5.54E-09 | 1.15E-04 | 0.34 | 0.15 | 0.19 | 2.26 |
| other bloold cells | Hsdl1 | 1 | 5.58E-09 | 1.16E-04 | 0.34 | 0.15 | 0.19 | 2.23 |
| other bloold cells | Ccdc50 | 0.62 | 5.99E-09 | 1.25E-04 | 0.9 | 0.76 | 0.14 | 1.19 |
| other bloold cells | Rabac1 | 0.64 | 5.99E-09 | 1.25E-04 | 0.79 | 0.55 | 0.24 | 1.43 |
| other bloold cells | Pcyox1 | 0.85 | 6.05E-09 | 1.26E-04 | 0.54 | 0.3 | 0.24 | 1.82 |
| other bloold cells | Brox | 0.87 | 6.09E-09 | 1.27E-04 | 0.45 | 0.22 | 0.22 | 2 |
| other bloold cells | Ppargc1b | 1.69 | 6.17E-09 | 1.28E-04 | 0.11 | 0.03 | 0.08 | 3.86 |
| other bloold cells | B3galt4 | 1.59 | 6.27E-09 | 1.30E-04 | 0.14 | 0.04 | 0.1 | 3.41 |
| other bloold cells | Neurl1a | 1.23 | 6.40E-09 | 1.33E-04 | 0.19 | 0.07 | 0.12 | 2.91 |
| other bloold cells | Smim4 | 0.75 | 6.41E-09 | 1.33E-04 | 0.68 | 0.43 | 0.25 | 1.59 |
| other bloold cells | Camta2 | 1.39 | 6.57E-09 | 1.37E-04 | 0.17 | 0.06 | 0.11 | 3.05 |
| other bloold cells | Usp15 | 0.63 | 6.72E-09 | 1.40E-04 | 0.8 | 0.57 | 0.23 | 1.4 |
| other bloold cells | Vps26c | 0.86 | 6.87E-09 | 1.43E-04 | 0.45 | 0.23 | 0.22 | 1.97 |
| other bloold cells | Stx2 | 0.89 | 7.73E-09 | 1.61E-04 | 0.44 | 0.22 | 0.22 | 2 |
| other bloold cells | Hps5 | 0.84 | 7.76E-09 | 1.61E-04 | 0.33 | 0.14 | 0.19 | 2.32 |
| other bloold cells | Cyp4v3 | 1.78 | 8.01E-09 | 1.67E-04 | 0.13 | 0.04 | 0.09 | 3.6 |
| other bloold cells | Ap3m2 | 1.81 | 8.18E-09 | 1.70E-04 | 0.13 | 0.04 | 0.1 | 3.5 |
| other bloold cells | Trmt2b | 1.34 | 8.29E-09 | 1.72E-04 | 0.23 | 0.09 | 0.14 | 2.6 |
| other bloold cells | Atxn7l1 | 0.79 | 8.42E-09 | 1.75E-04 | 0.36 | 0.17 | 0.2 | 2.21 |
| other bloold cells | Mogs | 0.75 | 8.43E-09 | 1.75E-04 | 0.59 | 0.35 | 0.25 | 1.71 |
| other bloold cells | Scly | 0.96 | 8.47E-09 | 1.76E-04 | 0.27 | 0.11 | 0.16 | 2.48 |
| other bloold cells | Pop5 | 0.64 | 8.50E-09 | 1.77E-04 | 0.83 | 0.65 | 0.18 | 1.28 |
| other bloold cells | Mxd4 | 0.65 | 8.54E-09 | 1.78E-04 | 0.73 | 0.47 | 0.26 | 1.55 |
| other bloold cells | Spsb2 | 1.31 | 8.72E-09 | 1.81E-04 | 0.21 | 0.08 | 0.13 | 2.76 |
| other bloold cells | Hdac5 | 0.78 | 8.86E-09 | 1.84E-04 | 0.38 | 0.17 | 0.2 | 2.17 |
| other bloold cells | Nat9 | 0.79 | 9.14E-09 | 1.90E-04 | 0.42 | 0.2 | 0.22 | 2.06 |
| other bloold cells | Adam10 | 0.65 | 9.16E-09 | 1.91E-04 | 0.8 | 0.54 | 0.27 | 1.5 |
| other bloold cells | Hpn | 1.67 | 9.33E-09 | 1.94E-04 | 0.13 | 0.04 | 0.1 | 3.5 |
| other bloold cells | Zfp592 | 0.95 | 9.50E-09 | 1.98E-04 | 0.34 | 0.15 | 0.19 | 2.23 |
| other bloold cells | Rabgap1l | 1 | 9.56E-09 | 1.99E-04 | 0.31 | 0.14 | 0.17 | 2.23 |
| other bloold cells | Crebrf | 1.17 | 9.66E-09 | 2.01E-04 | 0.27 | 0.11 | 0.16 | 2.4 |
| other bloold cells | Pomt1 | 0.97 | 1.04E-08 | 2.16E-04 | 0.29 | 0.12 | 0.17 | 2.39 |
| other bloold cells | Jup | 0.54 | 1.05E-08 | 2.18E-04 | 0.42 | 0.2 | 0.22 | 2.11 |
| other bloold cells | Atl3 | 0.79 | 1.08E-08 | 2.26E-04 | 0.44 | 0.22 | 0.22 | 1.99 |
| other bloold cells | Wdr91 | 0.98 | 1.12E-08 | 2.32E-04 | 0.26 | 0.1 | 0.16 | 2.51 |
| other bloold cells | Degs1 | 0.74 | 1.12E-08 | 2.34E-04 | 0.66 | 0.4 | 0.27 | 1.67 |

**Supplementary Table 2. Marker genes for the major cell types of integrated clustering of E11.5 control and NotchGOF λ-junction cells.**

| | | | | | | | | |
|---|---|---|---|---|---|---|---|---|
| other bloold cells | Ccdc93 | 0.76 | 1.18E-08 | 2.45E-04 | 0.37 | 0.17 | 0.2 | 2.2 |
| other bloold cells | Acvr1b | 0.97 | 1.21E-08 | 2.52E-04 | 0.27 | 0.11 | 0.16 | 2.42 |
| other bloold cells | Stard9 | 1.52 | 1.36E-08 | 2.82E-04 | 0.19 | 0.07 | 0.12 | 2.82 |
| other bloold cells | Chkb | 0.7 | 1.36E-08 | 2.83E-04 | 0.47 | 0.24 | 0.23 | 2 |
| other bloold cells | Anapc15 | 0.55 | 1.36E-08 | 2.83E-04 | 0.84 | 0.6 | 0.24 | 1.39 |
| other bloold cells | Clptm1l | 0.67 | 1.44E-08 | 3.00E-04 | 0.72 | 0.48 | 0.24 | 1.49 |
| other bloold cells | Kidins220 | 0.71 | 1.50E-08 | 3.11E-04 | 0.52 | 0.28 | 0.24 | 1.87 |
| other bloold cells | Alg5 | 0.64 | 1.51E-08 | 3.14E-04 | 0.64 | 0.37 | 0.27 | 1.74 |
| other bloold cells | Vill | 1.23 | 1.51E-08 | 3.15E-04 | 0.22 | 0.08 | 0.14 | 2.65 |
| other bloold cells | Dusp5 | 1.35 | 1.55E-08 | 3.21E-04 | 0.27 | 0.12 | 0.16 | 2.33 |
| other bloold cells | Rnpepl1 | 0.75 | 1.59E-08 | 3.31E-04 | 0.48 | 0.26 | 0.23 | 1.89 |
| other bloold cells | Sh2b2 | 0.97 | 1.62E-08 | 3.36E-04 | 0.34 | 0.16 | 0.18 | 2.13 |
| other bloold cells | Tmem97 | 0.76 | 1.76E-08 | 3.66E-04 | 0.69 | 0.46 | 0.23 | 1.49 |
| other bloold cells | Pim1 | 0.68 | 1.79E-08 | 3.71E-04 | 0.69 | 0.47 | 0.22 | 1.47 |
| other bloold cells | Dpm3 | 0.45 | 1.92E-08 | 4.00E-04 | 0.92 | 0.75 | 0.17 | 1.23 |
| other bloold cells | Tex2 | 1.42 | 1.96E-08 | 4.07E-04 | 0.13 | 0.04 | 0.09 | 3.5 |
| other bloold cells | Tcp11l2 | 1.34 | 1.97E-08 | 4.10E-04 | 0.13 | 0.04 | 0.09 | 3.5 |
| other bloold cells | Larp4b | 0.72 | 1.99E-08 | 4.14E-04 | 0.84 | 0.61 | 0.23 | 1.38 |
| other bloold cells | Sidt2 | 1.09 | 2.07E-08 | 4.29E-04 | 0.37 | 0.18 | 0.19 | 2.04 |
| other bloold cells | Fnip1 | 0.87 | 2.08E-08 | 4.33E-04 | 0.46 | 0.24 | 0.22 | 1.94 |
| other bloold cells | Leprotl1 | 0.59 | 2.10E-08 | 4.36E-04 | 0.8 | 0.55 | 0.25 | 1.45 |
| other bloold cells | Sec61b | 0.29 | 2.11E-08 | 4.40E-04 | 0.99 | 0.99 | 0.01 | 1.01 |
| other bloold cells | Atmin | 0.89 | 2.15E-08 | 4.47E-04 | 0.32 | 0.14 | 0.18 | 2.28 |
| other bloold cells | Faap24 | 0.94 | 2.15E-08 | 4.47E-04 | 0.35 | 0.16 | 0.19 | 2.13 |
| other bloold cells | Nr4a2 | 1.45 | 2.17E-08 | 4.51E-04 | 0.25 | 0.1 | 0.15 | 2.42 |
| other bloold cells | Mlxip | 0.66 | 2.17E-08 | 4.52E-04 | 0.36 | 0.16 | 0.2 | 2.23 |
| other bloold cells | Tnfaip8 | 0.44 | 2.20E-08 | 4.57E-04 | 0.94 | 0.76 | 0.18 | 1.24 |
| other bloold cells | Ubald1 | 0.88 | 2.26E-08 | 4.70E-04 | 0.5 | 0.26 | 0.23 | 1.88 |
| other bloold cells | Ubash3b | 0.88 | 2.26E-08 | 4.71E-04 | 0.45 | 0.24 | 0.21 | 1.89 |
| other bloold cells | Eloa | 0.63 | 2.30E-08 | 4.78E-04 | 0.72 | 0.47 | 0.25 | 1.54 |
| other bloold cells | Mrpl57 | 0.54 | 2.34E-08 | 4.87E-04 | 0.9 | 0.78 | 0.13 | 1.16 |
| other bloold cells | Poc1a | 0.65 | 2.35E-08 | 4.89E-04 | 0.44 | 0.21 | 0.23 | 2.06 |
| other bloold cells | Lsm10 | 0.81 | 2.35E-08 | 4.89E-04 | 0.66 | 0.4 | 0.26 | 1.66 |
| other bloold cells | Vps11 | 1.13 | 2.51E-08 | 5.22E-04 | 0.34 | 0.16 | 0.18 | 2.14 |
| other bloold cells | Atp5g1 | 0.33 | 2.55E-08 | 5.31E-04 | 0.99 | 0.99 | 0.01 | 1.01 |
| other bloold cells | Dmxl1 | 0.78 | 2.56E-08 | 5.33E-04 | 0.66 | 0.41 | 0.25 | 1.61 |
| other bloold cells | Imp4 | 0.61 | 2.62E-08 | 5.46E-04 | 0.76 | 0.54 | 0.23 | 1.42 |
| other bloold cells | Slc38a9 | 0.97 | 2.78E-08 | 5.79E-04 | 0.38 | 0.19 | 0.19 | 2.03 |
| other bloold cells | Sec24a | 0.75 | 2.80E-08 | 5.83E-04 | 0.37 | 0.18 | 0.2 | 2.12 |
| other bloold cells | Tmem106c | 0.87 | 2.81E-08 | 5.84E-04 | 0.39 | 0.2 | 0.2 | 2 |
| other bloold cells | Hmgcl | 0.73 | 2.82E-08 | 5.87E-04 | 0.65 | 0.41 | 0.25 | 1.6 |
| other bloold cells | Rbm38 | 0.7 | 2.87E-08 | 5.96E-04 | 0.46 | 0.24 | 0.22 | 1.93 |
| other bloold cells | Ethe1 | 0.76 | 3.01E-08 | 6.26E-04 | 0.69 | 0.45 | 0.23 | 1.52 |
| other bloold cells | Flad1 | 0.87 | 3.02E-08 | 6.27E-04 | 0.26 | 0.11 | 0.15 | 2.47 |
| other bloold cells | Zbtb4 | 1.48 | 3.15E-08 | 6.54E-04 | 0.18 | 0.07 | 0.12 | 2.8 |
| other bloold cells | Dnajc16 | 1.22 | 3.34E-08 | 6.94E-04 | 0.17 | 0.06 | 0.11 | 2.95 |
| other bloold cells | Arl4a | 0.72 | 3.38E-08 | 7.03E-04 | 0.41 | 0.2 | 0.21 | 2.02 |
| other bloold cells | Bend4 | 0.99 | 3.39E-08 | 7.05E-04 | 0.27 | 0.12 | 0.16 | 2.35 |
| other bloold cells | Ank | 1.33 | 3.44E-08 | 7.15E-04 | 0.19 | 0.07 | 0.12 | 2.74 |
| other bloold cells | Atxn1 | 1.11 | 3.49E-08 | 7.27E-04 | 0.23 | 0.09 | 0.14 | 2.51 |
| other bloold cells | Cd164 | 0.61 | 3.61E-08 | 7.50E-04 | 0.81 | 0.64 | 0.17 | 1.27 |
| other bloold cells | Atraid | 0.65 | 3.63E-08 | 7.56E-04 | 0.75 | 0.51 | 0.24 | 1.48 |
| other bloold cells | Acaa2 | 0.41 | 3.67E-08 | 7.63E-04 | 0.76 | 0.48 | 0.28 | 1.58 |
| other bloold cells | Chn2 | 1.04 | 3.75E-08 | 7.80E-04 | 0.2 | 0.07 | 0.13 | 2.76 |
| other bloold cells | C1galt1c1 | 1.16 | 3.80E-08 | 7.91E-04 | 0.34 | 0.16 | 0.18 | 2.1 |
| other bloold cells | Mto1 | 0.57 | 3.84E-08 | 7.98E-04 | 0.41 | 0.2 | 0.22 | 2.12 |

**Supplementary Table 2. Marker genes for the major cell types of integrated clustering of E11.5 control and NotchGOF λ-junction cells.**

| | | | | | | | | |
|---|---|---|---|---|---|---|---|---|
| other bloold cells | Snx8 | 1.18 | 4.06E-08 | 8.44E-04 | 0.29 | 0.13 | 0.16 | 2.19 |
| other bloold cells | Lactb2 | 0.86 | 4.09E-08 | 8.50E-04 | 0.37 | 0.18 | 0.2 | 2.11 |
| other bloold cells | Oser1 | 0.72 | 4.18E-08 | 8.70E-04 | 0.68 | 0.46 | 0.22 | 1.47 |
| other bloold cells | Pvr | 1.1 | 4.62E-08 | 9.61E-04 | 0.19 | 0.07 | 0.12 | 2.78 |
| other bloold cells | Armc7 | 1.64 | 4.68E-08 | 9.74E-04 | 0.15 | 0.05 | 0.1 | 3.06 |
| other bloold cells | Ino80c | 0.75 | 4.68E-08 | 9.74E-04 | 0.39 | 0.19 | 0.2 | 2.03 |
| other bloold cells | Zbtb34 | 0.96 | 4.72E-08 | 9.82E-04 | 0.23 | 0.09 | 0.14 | 2.54 |
| other bloold cells | Zfand5 | 0.45 | 4.72E-08 | 9.83E-04 | 0.95 | 0.83 | 0.12 | 1.15 |
| other bloold cells | Bmyc | 0.61 | 4.81E-08 | 1.00E-03 | 0.83 | 0.62 | 0.2 | 1.33 |
| other bloold cells | Rnf149 | 0.8 | 4.88E-08 | 1.01E-03 | 0.43 | 0.22 | 0.21 | 1.96 |
| other bloold cells | Arhgef10l | 0.85 | 4.88E-08 | 1.02E-03 | 0.32 | 0.15 | 0.18 | 2.22 |
| other bloold cells | Spata13 | 1.11 | 4.99E-08 | 1.04E-03 | 0.2 | 0.08 | 0.13 | 2.67 |
| other bloold cells | Cnpy3 | 0.72 | 5.05E-08 | 1.05E-03 | 0.59 | 0.35 | 0.24 | 1.7 |
| other bloold cells | AI987944 | 0.9 | 5.09E-08 | 1.06E-03 | 0.29 | 0.13 | 0.16 | 2.26 |
| other bloold cells | Hectd3 | 1.16 | 5.13E-08 | 1.07E-03 | 0.26 | 0.11 | 0.15 | 2.33 |
| other bloold cells | Rpa1 | 0.61 | 5.29E-08 | 1.10E-03 | 0.66 | 0.42 | 0.25 | 1.58 |
| other bloold cells | Rmdn1 | 0.72 | 5.30E-08 | 1.10E-03 | 0.36 | 0.17 | 0.19 | 2.1 |
| other bloold cells | Ttc7b | 0.7 | 5.51E-08 | 1.15E-03 | 0.35 | 0.16 | 0.19 | 2.2 |
| other bloold cells | Tsta3 | 0.63 | 5.53E-08 | 1.15E-03 | 0.7 | 0.46 | 0.24 | 1.53 |
| other bloold cells | Wdr44 | 1.4 | 5.65E-08 | 1.17E-03 | 0.18 | 0.06 | 0.11 | 2.78 |
| other bloold cells | Prdx1 | 0.47 | 5.77E-08 | 1.20E-03 | 1 | 0.99 | 0.01 | 1.01 |
| other bloold cells | Serhl | 0.77 | 5.80E-08 | 1.21E-03 | 0.36 | 0.17 | 0.19 | 2.13 |
| other bloold cells | Stk11 | 0.57 | 5.86E-08 | 1.22E-03 | 0.73 | 0.48 | 0.25 | 1.53 |
| other bloold cells | Metrnl | 1.42 | 5.91E-08 | 1.23E-03 | 0.18 | 0.07 | 0.12 | 2.76 |
| other bloold cells | Casp9 | 0.8 | 6.12E-08 | 1.27E-03 | 0.25 | 0.1 | 0.15 | 2.5 |
| other bloold cells | Zc3h7a | 0.7 | 6.14E-08 | 1.28E-03 | 0.59 | 0.34 | 0.25 | 1.72 |
| other bloold cells | Pias1 | 0.7 | 6.20E-08 | 1.29E-03 | 0.56 | 0.32 | 0.24 | 1.75 |
| other bloold cells | Armc5 | 0.86 | 6.80E-08 | 1.41E-03 | 0.32 | 0.14 | 0.17 | 2.22 |
| other bloold cells | Ppp1r15b | 0.69 | 7.01E-08 | 1.46E-03 | 0.55 | 0.31 | 0.25 | 1.8 |
| other bloold cells | Ikbkb | 0.65 | 7.09E-08 | 1.48E-03 | 0.57 | 0.32 | 0.25 | 1.79 |
| other bloold cells | Mrpl48 | 0.56 | 7.12E-08 | 1.48E-03 | 0.78 | 0.53 | 0.25 | 1.46 |
| other bloold cells | Scp2 | 0.45 | 7.20E-08 | 1.50E-03 | 0.93 | 0.77 | 0.16 | 1.2 |
| other bloold cells | Fyco1 | 0.8 | 7.32E-08 | 1.52E-03 | 0.35 | 0.16 | 0.19 | 2.13 |
| other bloold cells | Cox5a | 0.27 | 7.46E-08 | 1.55E-03 | 1 | 0.99 | 0.01 | 1.01 |
| other bloold cells | AAA01118383 | 0.69 | 7.60E-08 | 1.58E-03 | 0.45 | 0.23 | 0.22 | 1.93 |
| other bloold cells | Itpk1 | 0.81 | 7.91E-08 | 1.65E-03 | 0.19 | 0.07 | 0.12 | 2.82 |
| other bloold cells | Hacd2 | 0.68 | 7.94E-08 | 1.65E-03 | 0.62 | 0.37 | 0.24 | 1.65 |
| other bloold cells | Echdc1 | 0.67 | 8.23E-08 | 1.71E-03 | 0.45 | 0.24 | 0.21 | 1.89 |
| other bloold cells | Arrb1 | 1.04 | 8.24E-08 | 1.71E-03 | 0.27 | 0.12 | 0.15 | 2.31 |
| other bloold cells | Psenen | 0.41 | 8.53E-08 | 1.77E-03 | 0.94 | 0.88 | 0.06 | 1.07 |
| other bloold cells | Slc25a10 | 0.9 | 8.65E-08 | 1.80E-03 | 0.44 | 0.24 | 0.21 | 1.87 |
| other bloold cells | Prdx3 | 0.61 | 8.99E-08 | 1.87E-03 | 0.76 | 0.54 | 0.23 | 1.42 |
| other bloold cells | Lyst | 0.97 | 9.00E-08 | 1.87E-03 | 0.28 | 0.12 | 0.16 | 2.28 |
| other bloold cells | Map4k2 | 1.06 | 9.36E-08 | 1.95E-03 | 0.19 | 0.07 | 0.12 | 2.74 |
| other bloold cells | Smim14 | 0.66 | 9.59E-08 | 1.99E-03 | 0.66 | 0.41 | 0.24 | 1.59 |
| other bloold cells | Polr1d | 0.31 | 1.02E-07 | 2.12E-03 | 0.97 | 0.95 | 0.02 | 1.02 |
| other bloold cells | H2-T22 | 1.2 | 1.03E-07 | 2.15E-03 | 0.24 | 0.1 | 0.14 | 2.4 |
| other bloold cells | Ugcg | 0.7 | 1.04E-07 | 2.16E-03 | 0.61 | 0.37 | 0.24 | 1.64 |
| other bloold cells | Zfp622 | 0.6 | 1.08E-07 | 2.25E-03 | 0.66 | 0.42 | 0.25 | 1.59 |
| other bloold cells | Nomo1 | 0.9 | 1.09E-07 | 2.27E-03 | 0.39 | 0.2 | 0.19 | 1.93 |
| other bloold cells | Htra2 | 0.85 | 1.11E-07 | 2.31E-03 | 0.62 | 0.4 | 0.22 | 1.55 |
| other bloold cells | Nfya | 0.56 | 1.16E-07 | 2.41E-03 | 0.56 | 0.32 | 0.24 | 1.76 |
| other bloold cells | Ptrh1 | 1.29 | 1.16E-07 | 2.41E-03 | 0.13 | 0.04 | 0.09 | 3.24 |
| other bloold cells | Kbtbd8 | 1.3 | 1.20E-07 | 2.50E-03 | 0.13 | 0.04 | 0.09 | 3.32 |
| other bloold cells | Eif4ebp1 | 0.49 | 1.24E-07 | 2.57E-03 | 0.87 | 0.69 | 0.18 | 1.26 |
| other bloold cells | Galnt7 | 0.96 | 1.31E-07 | 2.72E-03 | 0.58 | 0.35 | 0.24 | 1.68 |

**Supplementary Table 2. Marker genes for the major cell types of integrated clustering of E11.5 control and NotchGOF λ-junction cells.**

| | | | | | | | | |
|---|---|---|---|---|---|---|---|---|
| other bloold cells | Ncor2 | 0.61 | 1.35E-07 | 2.81E-03 | 0.62 | 0.35 | 0.26 | 1.75 |
| other bloold cells | Cercam | 0.99 | 1.35E-07 | 2.82E-03 | 0.17 | 0.06 | 0.11 | 2.85 |
| other bloold cells | Psma1 | 0.31 | 1.35E-07 | 2.82E-03 | 0.97 | 0.94 | 0.03 | 1.04 |
| other bloold cells | Mon2 | 0.59 | 1.43E-07 | 2.98E-03 | 0.51 | 0.28 | 0.23 | 1.84 |
| other bloold cells | Tmf1 | 0.55 | 1.46E-07 | 3.04E-03 | 0.73 | 0.49 | 0.24 | 1.5 |
| other bloold cells | Rpn2 | 0.4 | 1.46E-07 | 3.04E-03 | 0.95 | 0.79 | 0.16 | 1.21 |
| other bloold cells | Nfs1 | 0.89 | 1.48E-07 | 3.08E-03 | 0.51 | 0.28 | 0.23 | 1.82 |
| other bloold cells | 900093K20Ri | 1.05 | 1.50E-07 | 3.12E-03 | 0.22 | 0.09 | 0.13 | 2.52 |
| other bloold cells | Tmem243 | 0.78 | 1.51E-07 | 3.13E-03 | 0.41 | 0.21 | 0.2 | 1.96 |
| other bloold cells | Tigar | 1.22 | 1.59E-07 | 3.31E-03 | 0.19 | 0.07 | 0.12 | 2.63 |
| other bloold cells | Rps6 | 0.25 | 1.62E-07 | 3.37E-03 | 1 | 1 | 0 | 1 |
| other bloold cells | Rad23a | 0.61 | 1.63E-07 | 3.38E-03 | 0.68 | 0.46 | 0.22 | 1.48 |
| other bloold cells | Ero1lb | 1.67 | 1.66E-07 | 3.45E-03 | 0.11 | 0.03 | 0.08 | 3.62 |
| other bloold cells | Tmem248 | 0.67 | 1.67E-07 | 3.46E-03 | 0.58 | 0.35 | 0.23 | 1.65 |
| other bloold cells | Bag3 | 1.01 | 1.69E-07 | 3.52E-03 | 0.26 | 0.11 | 0.15 | 2.35 |
| other bloold cells | Snx13 | 0.96 | 1.87E-07 | 3.89E-03 | 0.29 | 0.13 | 0.16 | 2.17 |
| other bloold cells | Akip1 | 0.7 | 1.90E-07 | 3.96E-03 | 0.4 | 0.2 | 0.2 | 2.02 |
| other bloold cells | Arhgap6 | 1.5 | 1.93E-07 | 4.02E-03 | 0.18 | 0.07 | 0.11 | 2.69 |
| other bloold cells | Ccdc28b | 0.52 | 1.95E-07 | 4.06E-03 | 0.64 | 0.37 | 0.27 | 1.72 |
| other bloold cells | Faim | 0.54 | 2.02E-07 | 4.20E-03 | 0.72 | 0.46 | 0.26 | 1.57 |
| other bloold cells | Tmc6 | 1.03 | 2.05E-07 | 4.26E-03 | 0.2 | 0.08 | 0.12 | 2.58 |
| other bloold cells | Pet100 | 0.55 | 2.09E-07 | 4.35E-03 | 0.79 | 0.56 | 0.24 | 1.42 |
| other bloold cells | Bcat2 | 0.66 | 2.09E-07 | 4.35E-03 | 0.51 | 0.29 | 0.22 | 1.75 |
| other bloold cells | Slc39a14 | 1.34 | 2.10E-07 | 4.36E-03 | 0.24 | 0.1 | 0.13 | 2.29 |
| other bloold cells | Mrps18b | 0.73 | 2.24E-07 | 4.65E-03 | 0.73 | 0.5 | 0.23 | 1.45 |
| other bloold cells | Vdac2 | 0.31 | 2.24E-07 | 4.66E-03 | 0.98 | 0.96 | 0.02 | 1.02 |
| other bloold cells | Gem | 1.89 | 2.26E-07 | 4.70E-03 | 0.13 | 0.04 | 0.09 | 3.15 |
| other bloold cells | Arid3b | 0.65 | 2.31E-07 | 4.80E-03 | 0.44 | 0.23 | 0.21 | 1.93 |
| other bloold cells | Pacs2 | 0.76 | 2.34E-07 | 4.87E-03 | 0.35 | 0.17 | 0.18 | 2.07 |
| other bloold cells | Acp2 | 0.98 | 2.38E-07 | 4.95E-03 | 0.46 | 0.27 | 0.19 | 1.72 |
| other bloold cells | Thbs1 | 1.38 | 2.38E-07 | 4.95E-03 | 0.27 | 0.12 | 0.15 | 2.24 |
| other bloold cells | Ier5 | 0.43 | 2.38E-07 | 4.95E-03 | 0.86 | 0.65 | 0.21 | 1.32 |
| other bloold cells | Filip1l | 0.67 | 2.40E-07 | 4.99E-03 | 0.47 | 0.26 | 0.21 | 1.82 |
| other bloold cells | Hagh | 0.84 | 2.42E-07 | 5.03E-03 | 0.44 | 0.25 | 0.2 | 1.8 |
| other bloold cells | Irf2bpl | 0.69 | 2.46E-07 | 5.12E-03 | 0.63 | 0.38 | 0.25 | 1.64 |
| other bloold cells | Fkbp1b | 0.9 | 2.52E-07 | 5.23E-03 | 0.3 | 0.14 | 0.16 | 2.12 |
| other bloold cells | Kxd1 | 0.56 | 2.63E-07 | 5.47E-03 | 0.73 | 0.5 | 0.23 | 1.46 |
| other bloold cells | Tspan33 | 0.64 | 2.65E-07 | 5.52E-03 | 0.21 | 0.08 | 0.13 | 2.5 |
| other bloold cells | 210016F16Ri | 0.66 | 2.67E-07 | 5.54E-03 | 0.57 | 0.35 | 0.22 | 1.63 |
| other bloold cells | Lamtor5 | 0.44 | 2.71E-07 | 5.63E-03 | 0.92 | 0.79 | 0.13 | 1.16 |
| other bloold cells | Pygb | 0.54 | 2.71E-07 | 5.63E-03 | 0.45 | 0.23 | 0.22 | 1.95 |
| other bloold cells | Slc25a32 | 0.88 | 2.77E-07 | 5.77E-03 | 0.22 | 0.09 | 0.13 | 2.49 |
| other bloold cells | Sumf1 | 1.08 | 2.78E-07 | 5.78E-03 | 0.26 | 0.12 | 0.14 | 2.23 |
| other bloold cells | Mrpl33 | 0.55 | 2.78E-07 | 5.78E-03 | 0.94 | 0.87 | 0.06 | 1.07 |
| other bloold cells | Tmem208 | 0.46 | 2.81E-07 | 5.83E-03 | 0.9 | 0.68 | 0.21 | 1.31 |
| other bloold cells | Lrrk1 | 0.91 | 2.83E-07 | 5.89E-03 | 0.32 | 0.15 | 0.16 | 2.09 |
| other bloold cells | Washc5 | 0.6 | 2.83E-07 | 5.89E-03 | 0.55 | 0.32 | 0.23 | 1.72 |
| other bloold cells | Txnrd2 | 0.67 | 2.91E-07 | 6.05E-03 | 0.41 | 0.21 | 0.2 | 1.94 |
| other bloold cells | Tgfbr1 | 0.88 | 2.97E-07 | 6.18E-03 | 0.75 | 0.55 | 0.19 | 1.35 |
| other bloold cells | Zfp217 | 0.8 | 3.01E-07 | 6.27E-03 | 0.39 | 0.2 | 0.19 | 1.97 |
| other bloold cells | Slc25a19 | 0.89 | 3.03E-07 | 6.30E-03 | 0.27 | 0.12 | 0.15 | 2.27 |
| other bloold cells | Nfkbie | 1.58 | 3.06E-07 | 6.36E-03 | 0.13 | 0.04 | 0.09 | 3.15 |
| other bloold cells | Polg2 | 0.95 | 3.09E-07 | 6.42E-03 | 0.23 | 0.1 | 0.13 | 2.38 |
| other bloold cells | Etv6 | 0.74 | 3.09E-07 | 6.42E-03 | 0.48 | 0.28 | 0.2 | 1.71 |
| other bloold cells | Ell2 | 1.08 | 3.10E-07 | 6.45E-03 | 0.22 | 0.09 | 0.13 | 2.44 |
| other bloold cells | Itprid2 | 0.58 | 3.18E-07 | 6.62E-03 | 0.32 | 0.15 | 0.17 | 2.13 |

**Supplementary Table 2. Marker genes for the major cell types of integrated clustering of E11.5 control and NotchGOF λ-junction cells.**

| | | | | | | | | |
|---|---|---|---|---|---|---|---|---|
| other bloold cells | Zfp335 | 1.02 | 3.24E-07 | 6.74E-03 | 0.22 | 0.09 | 0.13 | 2.38 |
| other bloold cells | Gtf2h1 | 0.48 | 3.25E-07 | 6.76E-03 | 0.68 | 0.4 | 0.28 | 1.68 |
| other bloold cells | Amfr | 0.56 | 3.26E-07 | 6.78E-03 | 0.6 | 0.36 | 0.24 | 1.66 |
| other bloold cells | Zswim6 | 0.75 | 3.28E-07 | 6.83E-03 | 0.51 | 0.3 | 0.22 | 1.73 |
| other bloold cells | Ndufaf8 | 0.46 | 3.45E-07 | 7.18E-03 | 0.88 | 0.67 | 0.21 | 1.31 |
| other bloold cells | Lmbrd1 | 0.72 | 3.48E-07 | 7.24E-03 | 0.42 | 0.22 | 0.2 | 1.89 |
| other bloold cells | Ppm1m | 0.64 | 3.65E-07 | 7.60E-03 | 0.46 | 0.25 | 0.22 | 1.87 |
| other bloold cells | Rrm2b | 0.91 | 3.66E-07 | 7.62E-03 | 0.27 | 0.12 | 0.15 | 2.2 |
| other bloold cells | Furin | 0.89 | 3.71E-07 | 7.72E-03 | 0.39 | 0.2 | 0.19 | 1.95 |
| other bloold cells | Mapre2 | 0.55 | 3.78E-07 | 7.85E-03 | 0.65 | 0.4 | 0.25 | 1.62 |
| other bloold cells | Abca2 | 1.49 | 3.89E-07 | 8.10E-03 | 0.11 | 0.03 | 0.08 | 3.5 |
| other bloold cells | Zfyve19 | 0.66 | 3.91E-07 | 8.14E-03 | 0.44 | 0.24 | 0.2 | 1.86 |
| other bloold cells | Atp13a1 | 0.69 | 4.15E-07 | 8.63E-03 | 0.44 | 0.23 | 0.21 | 1.88 |
| other bloold cells | Cluh | 0.76 | 4.16E-07 | 8.65E-03 | 0.42 | 0.23 | 0.19 | 1.85 |
| other bloold cells | Paxx | 0.8 | 4.74E-07 | 9.85E-03 | 0.26 | 0.12 | 0.14 | 2.25 |
| other bloold cells | Gpatch2l | 1.54 | 4.85E-07 | 1.01E-02 | 0.24 | 0.1 | 0.14 | 2.31 |
| other bloold cells | Siae | 0.99 | 4.86E-07 | 1.01E-02 | 0.18 | 0.07 | 0.11 | 2.64 |
| other bloold cells | Surf4 | 0.51 | 5.05E-07 | 1.05E-02 | 0.85 | 0.59 | 0.26 | 1.44 |
| other bloold cells | Smdt1 | 0.35 | 5.17E-07 | 1.08E-02 | 0.99 | 0.94 | 0.04 | 1.05 |
| other bloold cells | Gnptab | 0.68 | 5.32E-07 | 1.11E-02 | 0.32 | 0.15 | 0.17 | 2.11 |
| other bloold cells | Mtpn | 0.4 | 5.40E-07 | 1.12E-02 | 0.93 | 0.81 | 0.12 | 1.15 |
| other bloold cells | Lin54 | 0.71 | 5.42E-07 | 1.13E-02 | 0.43 | 0.24 | 0.2 | 1.85 |
| other bloold cells | Tgoln1 | 0.53 | 5.44E-07 | 1.13E-02 | 0.53 | 0.29 | 0.24 | 1.83 |
| other bloold cells | Nt5c | 0.58 | 5.52E-07 | 1.15E-02 | 0.73 | 0.54 | 0.19 | 1.36 |
| other bloold cells | Nrbf2 | 0.71 | 5.61E-07 | 1.17E-02 | 0.68 | 0.44 | 0.24 | 1.56 |
| other bloold cells | Cnppd1 | 0.6 | 5.68E-07 | 1.18E-02 | 0.41 | 0.21 | 0.2 | 1.96 |
| other bloold cells | Xpr1 | 0.67 | 5.71E-07 | 1.19E-02 | 0.49 | 0.27 | 0.22 | 1.79 |
| other bloold cells | Trim56 | 0.89 | 5.71E-07 | 1.19E-02 | 0.29 | 0.14 | 0.15 | 2.1 |
| other bloold cells | Ndufs8 | 0.36 | 5.75E-07 | 1.20E-02 | 0.97 | 0.84 | 0.13 | 1.15 |
| other bloold cells | Sash1 | 1.04 | 5.76E-07 | 1.20E-02 | 0.6 | 0.4 | 0.2 | 1.5 |
| other bloold cells | Dus1l | 0.53 | 5.79E-07 | 1.20E-02 | 0.64 | 0.4 | 0.24 | 1.61 |
| other bloold cells | Arap2 | 1.05 | 5.87E-07 | 1.22E-02 | 0.15 | 0.06 | 0.1 | 2.8 |
| other bloold cells | Ldha | 0.38 | 5.93E-07 | 1.23E-02 | 0.98 | 0.97 | 0.01 | 1.01 |
| other bloold cells | Tkt | 0.45 | 6.10E-07 | 1.27E-02 | 0.97 | 0.92 | 0.05 | 1.05 |
| other bloold cells | Scoc | 0.86 | 6.15E-07 | 1.28E-02 | 0.78 | 0.63 | 0.15 | 1.24 |
| other bloold cells | Jak2 | 0.65 | 6.21E-07 | 1.29E-02 | 0.32 | 0.16 | 0.17 | 2.08 |
| other bloold cells | Usp38 | 0.7 | 6.29E-07 | 1.31E-02 | 0.41 | 0.21 | 0.2 | 1.93 |
| other bloold cells | Nfkbil1 | 0.59 | 6.45E-07 | 1.34E-02 | 0.5 | 0.28 | 0.23 | 1.81 |
| other bloold cells | Myo18a | 0.49 | 6.55E-07 | 1.36E-02 | 0.45 | 0.24 | 0.21 | 1.91 |
| other bloold cells | Entpd7 | 0.85 | 6.57E-07 | 1.37E-02 | 0.3 | 0.15 | 0.16 | 2.06 |
| other bloold cells | Hs1bp3 | 0.94 | 7.15E-07 | 1.49E-02 | 0.25 | 0.11 | 0.14 | 2.25 |
| other bloold cells | Oxsm | 1.38 | 7.20E-07 | 1.50E-02 | 0.14 | 0.05 | 0.09 | 2.92 |
| other bloold cells | Btaf1 | 0.7 | 7.33E-07 | 1.52E-02 | 0.59 | 0.36 | 0.23 | 1.63 |
| other bloold cells | Slc7a5 | 1.15 | 7.41E-07 | 1.54E-02 | 0.14 | 0.05 | 0.09 | 2.92 |
| other bloold cells | Plekhj1 | 0.42 | 7.45E-07 | 1.55E-02 | 0.91 | 0.77 | 0.14 | 1.18 |
| other bloold cells | Wdr12 | 0.58 | 7.65E-07 | 1.59E-02 | 0.74 | 0.56 | 0.18 | 1.33 |
| other bloold cells | Gpr107 | 0.58 | 7.82E-07 | 1.63E-02 | 0.55 | 0.31 | 0.23 | 1.75 |
| other bloold cells | Acbd5 | 0.74 | 8.10E-07 | 1.68E-02 | 0.57 | 0.36 | 0.21 | 1.59 |
| other bloold cells | Apeh | 0.6 | 8.11E-07 | 1.69E-02 | 0.48 | 0.27 | 0.21 | 1.78 |
| other bloold cells | Snx17 | 0.49 | 8.29E-07 | 1.73E-02 | 0.87 | 0.67 | 0.2 | 1.3 |
| other bloold cells | Ero1l | 1.34 | 8.32E-07 | 1.73E-02 | 0.49 | 0.3 | 0.19 | 1.66 |
| other bloold cells | Sypl | 0.47 | 8.33E-07 | 1.73E-02 | 0.74 | 0.48 | 0.26 | 1.54 |
| other bloold cells | Med12l | 1.33 | 8.39E-07 | 1.74E-02 | 0.13 | 0.05 | 0.09 | 2.96 |
| other bloold cells | Mfsd14b | 0.65 | 8.39E-07 | 1.74E-02 | 0.44 | 0.23 | 0.21 | 1.92 |
| other bloold cells | Gne | 1.1 | 8.60E-07 | 1.79E-02 | 0.2 | 0.08 | 0.12 | 2.48 |
| other bloold cells | Fyn | 0.62 | 8.86E-07 | 1.84E-02 | 0.69 | 0.44 | 0.24 | 1.55 |

**Supplementary Table 2. Marker genes for the major cell types of integrated clustering of E11.5 control and NotchGOF λ-junction cells.**

| | | | | | | | | |
|---|---|---|---|---|---|---|---|---|
| other bloold cells | Acvrl1 | 1 | 9.04E-07 | 1.88E-02 | 0.23 | 0.1 | 0.13 | 2.29 |
| other bloold cells | Itgb1bp1 | 0.62 | 9.22E-07 | 1.92E-02 | 0.62 | 0.4 | 0.22 | 1.55 |
| other bloold cells | Bin1 | 0.8 | 9.22E-07 | 1.92E-02 | 0.78 | 0.62 | 0.16 | 1.26 |
| other bloold cells | Rbck1 | 0.54 | 9.42E-07 | 1.96E-02 | 0.6 | 0.38 | 0.22 | 1.57 |
| other bloold cells | Fam13b | 0.59 | 9.47E-07 | 1.97E-02 | 0.52 | 0.3 | 0.22 | 1.75 |
| other bloold cells | Zfyve26 | 0.86 | 9.60E-07 | 2.00E-02 | 0.32 | 0.16 | 0.16 | 2 |
| other bloold cells | Ddi2 | 0.52 | 9.91E-07 | 2.06E-02 | 0.65 | 0.41 | 0.24 | 1.59 |
| other bloold cells | Dph5 | 0.67 | 1.04E-06 | 2.17E-02 | 0.42 | 0.23 | 0.19 | 1.85 |
| other bloold cells | Lrp1 | 0.95 | 1.06E-06 | 2.20E-02 | 0.71 | 0.58 | 0.13 | 1.23 |
| other bloold cells | Slc35f6 | 1.34 | 1.10E-06 | 2.28E-02 | 0.24 | 0.11 | 0.13 | 2.22 |
| other bloold cells | Stat5b | 0.99 | 1.10E-06 | 2.29E-02 | 0.27 | 0.13 | 0.15 | 2.13 |
| other bloold cells | Mfsd13a | 1.46 | 1.11E-06 | 2.31E-02 | 0.12 | 0.04 | 0.08 | 3.05 |
| other bloold cells | Vrk3 | 0.55 | 1.12E-06 | 2.33E-02 | 0.58 | 0.34 | 0.24 | 1.7 |
| other bloold cells | Txndc11 | 0.7 | 1.12E-06 | 2.33E-02 | 0.39 | 0.2 | 0.18 | 1.91 |
| other bloold cells | Tatdn2 | 0.75 | 1.12E-06 | 2.33E-02 | 0.32 | 0.15 | 0.16 | 2.06 |
| other bloold cells | Cacul1 | 0.54 | 1.15E-06 | 2.39E-02 | 0.52 | 0.29 | 0.23 | 1.81 |
| other bloold cells | Cep55 | 0.77 | 1.18E-06 | 2.46E-02 | 0.43 | 0.24 | 0.19 | 1.79 |
| other bloold cells | Tpp2 | 0.58 | 1.19E-06 | 2.48E-02 | 0.82 | 0.6 | 0.22 | 1.37 |
| other bloold cells | Ptpa | 0.59 | 1.22E-06 | 2.54E-02 | 0.73 | 0.52 | 0.21 | 1.41 |
| other bloold cells | Tmem30a | 0.5 | 1.25E-06 | 2.60E-02 | 0.73 | 0.51 | 0.22 | 1.42 |
| other bloold cells | Pter | 0.86 | 1.26E-06 | 2.63E-02 | 0.25 | 0.11 | 0.14 | 2.21 |
| other bloold cells | Stx8 | 0.63 | 1.29E-06 | 2.68E-02 | 0.66 | 0.44 | 0.22 | 1.5 |
| other bloold cells | Itgb5 | 0.7 | 1.31E-06 | 2.72E-02 | 0.73 | 0.59 | 0.14 | 1.24 |
| other bloold cells | Hist1h1c | 0.6 | 1.33E-06 | 2.76E-02 | 0.66 | 0.42 | 0.24 | 1.57 |
| other bloold cells | Ubxn8 | 0.57 | 1.36E-06 | 2.82E-02 | 0.39 | 0.2 | 0.19 | 1.96 |
| other bloold cells | Coa4 | 0.55 | 1.36E-06 | 2.82E-02 | 0.48 | 0.27 | 0.22 | 1.82 |
| other bloold cells | Mtmr6 | 0.68 | 1.39E-06 | 2.88E-02 | 0.29 | 0.14 | 0.15 | 2.13 |
| other bloold cells | Zfp597 | 1 | 1.40E-06 | 2.91E-02 | 0.19 | 0.08 | 0.11 | 2.49 |
| other bloold cells | Smim3 | 0.62 | 1.41E-06 | 2.94E-02 | 0.36 | 0.19 | 0.18 | 1.97 |
| other bloold cells | Ccdc107 | 0.46 | 1.44E-06 | 3.00E-02 | 0.66 | 0.42 | 0.25 | 1.59 |
| other bloold cells | Sin3b | 0.42 | 1.45E-06 | 3.03E-02 | 0.94 | 0.8 | 0.14 | 1.18 |
| other bloold cells | Prr14 | 0.87 | 1.50E-06 | 3.12E-02 | 0.39 | 0.2 | 0.18 | 1.89 |
| other bloold cells | Uggt1 | 0.58 | 1.51E-06 | 3.14E-02 | 0.57 | 0.36 | 0.22 | 1.61 |
| other bloold cells | Baz2a | 0.47 | 1.53E-06 | 3.17E-02 | 0.64 | 0.39 | 0.25 | 1.65 |
| other bloold cells | Aldh3a2 | 1.06 | 1.53E-06 | 3.18E-02 | 0.2 | 0.08 | 0.12 | 2.42 |
| other bloold cells | Nek9 | 0.46 | 1.53E-06 | 3.19E-02 | 0.52 | 0.29 | 0.23 | 1.78 |
| other bloold cells | Dusp6 | 0.38 | 1.61E-06 | 3.36E-02 | 0.6 | 0.36 | 0.24 | 1.66 |
| other bloold cells | Trib1 | 1.32 | 1.62E-06 | 3.37E-02 | 0.33 | 0.18 | 0.15 | 1.82 |
| other bloold cells | Bpnt1 | 0.73 | 1.63E-06 | 3.40E-02 | 0.29 | 0.14 | 0.15 | 2.11 |
| other bloold cells | Ocrl | 0.6 | 1.70E-06 | 3.53E-02 | 0.48 | 0.27 | 0.21 | 1.77 |
| other bloold cells | Slc31a1 | 0.51 | 1.71E-06 | 3.56E-02 | 0.59 | 0.35 | 0.25 | 1.72 |
| other bloold cells | Tprn | 1.52 | 1.77E-06 | 3.67E-02 | 0.12 | 0.04 | 0.08 | 3.05 |
| other bloold cells | Prkx | 0.76 | 1.77E-06 | 3.69E-02 | 0.32 | 0.15 | 0.16 | 2.05 |
| other bloold cells | Cmtm3 | 0.53 | 1.82E-06 | 3.78E-02 | 0.65 | 0.42 | 0.23 | 1.55 |
| other bloold cells | Smim24 | 1.16 | 1.84E-06 | 3.83E-02 | 0.14 | 0.05 | 0.09 | 2.8 |
| other bloold cells | Cep128 | 0.66 | 1.84E-06 | 3.83E-02 | 0.34 | 0.17 | 0.17 | 1.97 |
| other bloold cells | Gclc | 0.78 | 1.85E-06 | 3.85E-02 | 0.49 | 0.3 | 0.19 | 1.66 |
| other bloold cells | Gm43813 | 1.04 | 1.89E-06 | 3.92E-02 | 0.2 | 0.09 | 0.12 | 2.33 |
| other bloold cells | Scyl3 | 0.73 | 1.91E-06 | 3.97E-02 | 0.28 | 0.13 | 0.15 | 2.11 |
| other bloold cells | Batf3 | 0.79 | 1.99E-06 | 4.14E-02 | 0.32 | 0.15 | 0.16 | 2.05 |
| other bloold cells | Spns1 | 1 | 1.99E-06 | 4.15E-02 | 0.6 | 0.41 | 0.19 | 1.47 |
| other bloold cells | Mrtfa | 0.8 | 2.00E-06 | 4.17E-02 | 0.27 | 0.12 | 0.14 | 2.16 |
| other bloold cells | Retreg2 | 0.65 | 2.07E-06 | 4.30E-02 | 0.49 | 0.29 | 0.2 | 1.7 |
| other bloold cells | Yipf3 | 0.5 | 2.14E-06 | 4.46E-02 | 0.66 | 0.42 | 0.25 | 1.6 |
| other bloold cells | Sap30 | 0.49 | 2.15E-06 | 4.47E-02 | 0.85 | 0.65 | 0.2 | 1.31 |
| other bloold cells | Rubcn | 1.05 | 2.15E-06 | 4.48E-02 | 0.2 | 0.08 | 0.11 | 2.39 |

**Supplementary Table 2. Marker genes for the major cell types of integrated clustering of E11.5 control and NotchGOF λ-junction cells.**

| | | | | | | | | |
|---|---|---|---|---|---|---|---|---|
| other bloold cells | Chchd10 | 0.47 | 2.19E-06 | 4.55E-02 | 0.33 | 0.16 | 0.17 | 2.04 |
| other bloold cells | Timm13 | 0.26 | 2.23E-06 | 4.64E-02 | 0.99 | 0.97 | 0.02 | 1.02 |
| other bloold cells | B9d2 | 0.54 | 2.28E-06 | 4.73E-02 | 0.48 | 0.27 | 0.21 | 1.79 |
| other bloold cells | Erp44 | 0.59 | 2.29E-06 | 4.77E-02 | 0.74 | 0.51 | 0.23 | 1.45 |
| other bloold cells | Nudt7 | 1.1 | 2.33E-06 | 4.85E-02 | 0.18 | 0.07 | 0.11 | 2.5 |
| other bloold cells | Pxmp4 | 0.72 | 2.36E-06 | 4.92E-02 | 0.34 | 0.18 | 0.17 | 1.93 |
| schwann cell progenitors | Fabp7 | 10.63 | 0 | 0 | 1 | 0 | 1 | 333.33 |
| schwann cell progenitors | Sox10 | 8.39 | 0 | 0 | 1 | 0.01 | 0.99 | 125 |
| schwann cell progenitors | Foxd3 | 14.29 | 0 | 0 | 0.94 | 0 | 0.94 | #NA |
| schwann cell progenitors | Insc | 9.73 | 0 | 0 | 0.89 | 0 | 0.89 | 889 |
| schwann cell progenitors | Megf10 | 8.35 | 0 | 0 | 0.89 | 0.01 | 0.88 | 148.17 |
| schwann cell progenitors | Gpr17 | 13.53 | 0 | 0 | 0.83 | 0 | 0.83 | #NA |
| schwann cell progenitors | Cdh19 | 9.12 | 0 | 0 | 0.78 | 0 | 0.78 | 389 |
| schwann cell progenitors | Gm12688 | 12.07 | 0 | 0 | 0.72 | 0 | 0.72 | #NA |
| schwann cell progenitors | Gfra3 | 10.96 | 0 | 0 | 0.72 | 0 | 0.72 | 722 |
| schwann cell progenitors | Tmem132e | 7.54 | 0 | 0 | 0.61 | 0 | 0.61 | 152.75 |
| schwann cell progenitors | Npy | 10.13 | 0 | 0 | 0.56 | 0 | 0.55 | 278 |
| schwann cell progenitors | Scrg1 | 12.18 | 0 | 0 | 0.5 | 0 | 0.5 | #NA |
| schwann cell progenitors | Klhl30 | 9.68 | 0 | 0 | 0.33 | 0 | 0.33 | #NA |
| schwann cell progenitors | Art3 | 10.1 | 0 | 0 | 0.33 | 0 | 0.33 | #NA |
| schwann cell progenitors | Tecrl | 9.98 | 0 | 0 | 0.22 | 0 | 0.22 | #NA |
| schwann cell progenitors | Gjc3 | 12.06 | 0 | 0 | 0.22 | 0 | 0.22 | #NA |
| schwann cell progenitors | Tmprss11f | 10.89 | 0 | 0 | 0.11 | 0 | 0.11 | #NA |
| schwann cell progenitors | Ndst3 | 6.86 | 4.93E-307 | 1.03E-302 | 0.83 | 0.01 | 0.83 | 104.13 |
| schwann cell progenitors | 330188P03Ri | 7.8 | 8.67E-296 | 1.80E-291 | 0.22 | 0 | 0.22 | #NA |
| schwann cell progenitors | Btla | 8.89 | 7.65E-275 | 1.59E-270 | 0.22 | 0 | 0.22 | 222 |
| schwann cell progenitors | Nppb | 8.82 | 8.96E-275 | 1.86E-270 | 0.22 | 0 | 0.22 | 222 |
| schwann cell progenitors | Ajap1 | 7.04 | 1.71E-272 | 3.55E-268 | 0.72 | 0.01 | 0.72 | 103.14 |
| schwann cell progenitors | Egfl8 | 7.76 | 2.07E-265 | 4.31E-261 | 0.89 | 0.01 | 0.88 | 80.82 |
| schwann cell progenitors | Adgrg6 | 7.08 | 1.61E-231 | 3.35E-227 | 0.78 | 0.01 | 0.77 | 86.44 |
| schwann cell progenitors | Slitrk2 | 6.07 | 4.11E-194 | 8.55E-190 | 0.89 | 0.02 | 0.87 | 59.27 |
| schwann cell progenitors | Dct | 7.6 | 2.18E-192 | 4.53E-188 | 0.22 | 0 | 0.22 | 222 |
| schwann cell progenitors | Spon1 | 7.5 | 1.06E-190 | 2.21E-186 | 0.83 | 0.01 | 0.82 | 59.5 |
| schwann cell progenitors | Atp10b | 7.07 | 3.84E-176 | 7.99E-172 | 0.33 | 0 | 0.33 | 166.5 |
| schwann cell progenitors | 930452G13Ri | 9.01 | 3.17E-161 | 6.59E-157 | 0.11 | 0 | 0.11 | #NA |
| schwann cell progenitors | Camk2b | 6.29 | 4.44E-157 | 9.24E-153 | 0.67 | 0.01 | 0.66 | 66.7 |
| schwann cell progenitors | Fam178b | 7.41 | 1.15E-156 | 2.40E-152 | 0.44 | 0.01 | 0.44 | 88.8 |
| schwann cell progenitors | Phactr1 | 8.02 | 1.69E-153 | 3.52E-149 | 1 | 0.03 | 0.97 | 38.46 |
| schwann cell progenitors | Lims2 | 7.18 | 1.78E-150 | 3.69E-146 | 0.39 | 0 | 0.39 | 97.25 |
| schwann cell progenitors | Trpv4 | 6.45 | 1.38E-149 | 2.87E-145 | 0.44 | 0.01 | 0.44 | 88.8 |
| schwann cell progenitors | Npr3 | 6.14 | 2.17E-149 | 4.52E-145 | 0.83 | 0.02 | 0.82 | 49 |
| schwann cell progenitors | Plxna4 | 5.85 | 1.72E-147 | 3.59E-143 | 0.94 | 0.02 | 0.92 | 41.04 |
| schwann cell progenitors | Adam12 | 6.27 | 2.07E-139 | 4.30E-135 | 0.72 | 0.01 | 0.71 | 51.57 |
| schwann cell progenitors | 010320O07Ri | 8.36 | 2.37E-138 | 4.93E-134 | 0.11 | 0 | 0.11 | #NA |
| schwann cell progenitors | Deptor | 6.39 | 1.86E-136 | 3.86E-132 | 0.56 | 0.01 | 0.55 | 69.5 |
| schwann cell progenitors | Sema3g | 7.64 | 4.63E-133 | 9.64E-129 | 0.33 | 0 | 0.33 | 111 |
| schwann cell progenitors | Car6 | 7.38 | 9.35E-133 | 1.94E-128 | 0.22 | 0 | 0.22 | 222 |
| schwann cell progenitors | Vat1l | 6.36 | 3.23E-130 | 6.71E-126 | 0.67 | 0.01 | 0.65 | 51.31 |
| schwann cell progenitors | Afap1l2 | 6 | 1.17E-129 | 2.44E-125 | 0.78 | 0.02 | 0.76 | 45.76 |
| schwann cell progenitors | Cd200 | 6.48 | 1.05E-127 | 2.19E-123 | 0.94 | 0.03 | 0.92 | 34.96 |
| schwann cell progenitors | Lgi4 | 6.71 | 4.34E-125 | 9.02E-121 | 0.44 | 0.01 | 0.44 | 74 |
| schwann cell progenitors | Tm4sf1 | 7.14 | 6.81E-117 | 1.42E-112 | 0.72 | 0.02 | 0.71 | 42.47 |
| schwann cell progenitors | Casq2 | 7.89 | 1.76E-114 | 3.67E-110 | 0.17 | 0 | 0.17 | 167 |
| schwann cell progenitors | Gfra2 | 7.03 | 6.46E-111 | 1.34E-106 | 0.39 | 0.01 | 0.38 | 77.8 |
| schwann cell progenitors | Fgd5 | 4.9 | 1.33E-95 | 2.76E-91 | 0.78 | 0.02 | 0.75 | 32.42 |
| schwann cell progenitors | Matn4 | 6.15 | 2.35E-92 | 4.89E-88 | 0.33 | 0 | 0.33 | 83.25 |

**Supplementary Table 2. Marker genes for the major cell types of integrated clustering of E11.5 control and NotchGOF λ-junction cells.**

| | | | | | | | | |
|---|---|---|---|---|---|---|---|---|
| schwann cell progenitors | Igsf11 | 6.71 | 8.57E-92 | 1.78E-87 | 0.22 | 0 | 0.22 | 111 |
| schwann cell progenitors | Tagln | 5.12 | 2.45E-88 | 5.11E-84 | 0.89 | 0.04 | 0.85 | 25.4 |
| schwann cell progenitors | Ednrb | 5.09 | 4.23E-87 | 8.79E-83 | 0.94 | 0.04 | 0.91 | 24.21 |
| schwann cell progenitors | Cryab | 6.26 | 1.43E-82 | 2.98E-78 | 1 | 0.05 | 0.95 | 20.41 |
| schwann cell progenitors | Ngfr | 4.88 | 2.04E-82 | 4.24E-78 | 0.94 | 0.04 | 0.9 | 22.48 |
| schwann cell progenitors | Tspan18 | 4.42 | 5.32E-82 | 1.11E-77 | 0.94 | 0.04 | 0.9 | 23.02 |
| schwann cell progenitors | Gm50147 | 8.56 | 3.20E-81 | 6.65E-77 | 0.11 | 0 | 0.11 | 111 |
| schwann cell progenitors | Hspb2 | 5.35 | 4.40E-80 | 9.15E-76 | 0.83 | 0.03 | 0.8 | 24.5 |
| schwann cell progenitors | Bche | 5.22 | 3.43E-75 | 7.14E-71 | 0.67 | 0.02 | 0.65 | 30.32 |
| schwann cell progenitors | Cmtm5 | 5.6 | 2.97E-71 | 6.17E-67 | 0.39 | 0.01 | 0.38 | 48.63 |
| schwann cell progenitors | Ina | 2.19 | 7.07E-67 | 1.47E-62 | 0.39 | 0.01 | 0.38 | 48.63 |
| schwann cell progenitors | Rassf4 | 5.25 | 8.33E-67 | 1.73E-62 | 0.72 | 0.03 | 0.69 | 24.07 |
| schwann cell progenitors | Rxrg | 4.86 | 3.16E-65 | 6.57E-61 | 0.44 | 0.01 | 0.43 | 40.36 |
| schwann cell progenitors | Rai2 | 6.94 | 2.70E-64 | 5.61E-60 | 0.17 | 0 | 0.17 | 83.5 |
| schwann cell progenitors | Gm29865 | 6.14 | 3.65E-62 | 7.59E-58 | 0.22 | 0 | 0.22 | 74 |
| schwann cell progenitors | Akap6 | 4.63 | 1.02E-60 | 2.12E-56 | 0.33 | 0.01 | 0.33 | 47.57 |
| schwann cell progenitors | Sntb1 | 5.03 | 2.13E-59 | 4.43E-55 | 0.67 | 0.03 | 0.64 | 23.82 |
| schwann cell progenitors | Dlx1 | 4.68 | 1.89E-58 | 3.92E-54 | 0.89 | 0.05 | 0.84 | 17.1 |
| schwann cell progenitors | Gm16316 | 6.16 | 9.80E-57 | 2.04E-52 | 0.22 | 0 | 0.22 | 74 |
| schwann cell progenitors | Luzp2 | 4.95 | 5.94E-53 | 1.23E-48 | 0.5 | 0.02 | 0.48 | 27.78 |
| schwann cell progenitors | Mcam | 4.37 | 1.16E-52 | 2.42E-48 | 0.78 | 0.04 | 0.74 | 18.09 |
| schwann cell progenitors | Prima1 | 6.82 | 5.97E-51 | 1.24E-46 | 0.17 | 0 | 0.17 | 83.5 |
| schwann cell progenitors | Sema6b | 5.77 | 4.60E-49 | 9.56E-45 | 0.33 | 0.01 | 0.32 | 37 |
| schwann cell progenitors | Postn | 5.39 | 1.14E-47 | 2.38E-43 | 1 | 0.09 | 0.91 | 11.24 |
| schwann cell progenitors | S100a16 | 3.72 | 2.44E-46 | 5.07E-42 | 0.94 | 0.08 | 0.87 | 12.42 |
| schwann cell progenitors | Prss12 | 4.92 | 9.33E-45 | 1.94E-40 | 0.78 | 0.05 | 0.73 | 14.96 |
| schwann cell progenitors | Cavin2 | 4.79 | 1.98E-43 | 4.13E-39 | 0.5 | 0.02 | 0.48 | 22.73 |
| schwann cell progenitors | Akr1cl | 5.63 | 1.57E-42 | 3.27E-38 | 0.33 | 0.01 | 0.32 | 33.3 |
| schwann cell progenitors | Atp1a2 | 5.62 | 2.41E-41 | 5.01E-37 | 0.33 | 0.01 | 0.32 | 33.3 |
| schwann cell progenitors | Plcg2 | 4.82 | 6.85E-40 | 1.42E-35 | 0.44 | 0.02 | 0.43 | 24.67 |
| schwann cell progenitors | Plp1 | 5.96 | 4.89E-39 | 1.02E-34 | 1 | 0.12 | 0.88 | 8.47 |
| schwann cell progenitors | Gja3 | 6.23 | 1.52E-38 | 3.17E-34 | 0.17 | 0 | 0.16 | 55.67 |
| schwann cell progenitors | Fitm1 | 6.71 | 6.23E-38 | 1.30E-33 | 0.11 | 0 | 0.11 | 111 |
| schwann cell progenitors | Il1rap | 4.17 | 1.96E-37 | 4.07E-33 | 0.61 | 0.04 | 0.57 | 16.51 |
| schwann cell progenitors | Adam23 | 4.23 | 1.13E-36 | 2.35E-32 | 0.89 | 0.08 | 0.81 | 10.71 |
| schwann cell progenitors | Olfr889 | 6.62 | 1.33E-36 | 2.77E-32 | 0.28 | 0.01 | 0.27 | 34.75 |
| schwann cell progenitors | Chl1 | 4.24 | 3.64E-35 | 7.56E-31 | 0.39 | 0.02 | 0.37 | 24.31 |
| schwann cell progenitors | Shc4 | 4.4 | 4.43E-35 | 9.21E-31 | 0.5 | 0.03 | 0.47 | 19.23 |
| schwann cell progenitors | Rasa3 | 3.74 | 6.98E-35 | 1.45E-30 | 0.83 | 0.08 | 0.76 | 10.82 |
| schwann cell progenitors | Slc6a15 | 4.1 | 1.14E-34 | 2.37E-30 | 0.56 | 0.03 | 0.52 | 16.85 |
| schwann cell progenitors | Slc35f1 | 4.38 | 4.45E-34 | 9.26E-30 | 0.56 | 0.03 | 0.52 | 16.35 |
| schwann cell progenitors | Arhgap22 | 4.52 | 9.25E-33 | 1.92E-28 | 0.33 | 0.01 | 0.32 | 25.62 |
| schwann cell progenitors | Cbr3 | 4.4 | 3.36E-32 | 7.00E-28 | 0.33 | 0.01 | 0.32 | 25.62 |
| schwann cell progenitors | Bcar3 | 3.94 | 7.18E-32 | 1.49E-27 | 0.67 | 0.05 | 0.62 | 13.08 |
| schwann cell progenitors | Cdh10 | 4.34 | 8.08E-32 | 1.68E-27 | 0.5 | 0.03 | 0.47 | 17.24 |
| schwann cell progenitors | Pou3f2 | 5.54 | 5.55E-31 | 1.15E-26 | 0.17 | 0 | 0.16 | 55.67 |
| schwann cell progenitors | Tfap2b | 2.19 | 2.57E-30 | 5.35E-26 | 0.89 | 0.09 | 0.8 | 9.88 |
| schwann cell progenitors | Dlx2 | 3.17 | 1.51E-29 | 3.13E-25 | 1 | 0.13 | 0.87 | 7.69 |
| schwann cell progenitors | Padi2 | 5.8 | 1.90E-29 | 3.95E-25 | 0.17 | 0 | 0.16 | 41.75 |
| schwann cell progenitors | Gpm6b | 4.28 | 2.72E-29 | 5.66E-25 | 0.72 | 0.07 | 0.66 | 10.78 |
| schwann cell progenitors | Stard13 | 4.16 | 3.29E-29 | 6.83E-25 | 0.44 | 0.03 | 0.42 | 17.76 |
| schwann cell progenitors | Xrra1 | 6.63 | 3.42E-29 | 7.11E-25 | 0.11 | 0 | 0.11 | 55.5 |
| schwann cell progenitors | Nkd2 | 3.24 | 4.58E-29 | 9.52E-25 | 1 | 0.13 | 0.87 | 7.69 |
| schwann cell progenitors | Zfp488 | 6.23 | 2.23E-28 | 4.65E-24 | 0.11 | 0 | 0.11 | 55.5 |
| schwann cell progenitors | Cxcl10 | 6.1 | 1.19E-27 | 2.48E-23 | 0.11 | 0 | 0.11 | 55.5 |
| schwann cell progenitors | Ndst4 | 6.61 | 5.82E-27 | 1.21E-22 | 0.11 | 0 | 0.11 | 55.5 |

**Supplementary Table 2. Marker genes for the major cell types of integrated clustering of E11.5 control and NotchGOF λ-junction cells.**

| | | | | | | | | |
|---|---|---|---|---|---|---|---|---|
| schwann cell progenitors | Hcn1 | 5.87 | 7.53E-27 | 1.57E-22 | 0.28 | 0.01 | 0.27 | 25.27 |
| schwann cell progenitors | Zfp536 | 2.76 | 1.32E-26 | 2.74E-22 | 0.89 | 0.1 | 0.79 | 8.72 |
| schwann cell progenitors | Erbb4 | 6.12 | 3.38E-26 | 7.02E-22 | 0.11 | 0 | 0.11 | 55.5 |
| schwann cell progenitors | Gm16104 | 3.28 | 7.77E-26 | 1.62E-21 | 0.56 | 0.04 | 0.51 | 12.93 |
| schwann cell progenitors | Erbb3 | 3.75 | 8.58E-26 | 1.78E-21 | 1 | 0.18 | 0.82 | 5.65 |
| schwann cell progenitors | 030407O03Ri | 6.49 | 1.38E-25 | 2.87E-21 | 0.11 | 0 | 0.11 | 55.5 |
| schwann cell progenitors | Mef2c | 3.94 | 1.87E-25 | 3.90E-21 | 1 | 0.19 | 0.81 | 5.38 |
| schwann cell progenitors | Akap12 | 4.93 | 1.98E-25 | 4.11E-21 | 1 | 0.2 | 0.8 | 5.08 |
| schwann cell progenitors | Gfra1 | 4.46 | 3.40E-25 | 7.07E-21 | 0.56 | 0.05 | 0.51 | 12.09 |
| schwann cell progenitors | Moxd1 | 3.52 | 1.24E-24 | 2.58E-20 | 0.5 | 0.04 | 0.46 | 13.51 |
| schwann cell progenitors | Fam78b | 4.73 | 1.93E-24 | 4.00E-20 | 0.44 | 0.03 | 0.41 | 14.8 |
| schwann cell progenitors | Egflam | 3.22 | 2.11E-24 | 4.39E-20 | 0.72 | 0.08 | 0.65 | 9.63 |
| schwann cell progenitors | Mpz | 6.4 | 2.15E-24 | 4.47E-20 | 0.22 | 0.01 | 0.21 | 27.75 |
| schwann cell progenitors | Aldh1l2 | 4.38 | 2.49E-24 | 5.17E-20 | 0.28 | 0.01 | 0.27 | 23.17 |
| schwann cell progenitors | Lama4 | 3.75 | 1.05E-23 | 2.18E-19 | 0.89 | 0.13 | 0.76 | 6.63 |
| schwann cell progenitors | Gcnt1 | 4.13 | 2.03E-23 | 4.23E-19 | 0.44 | 0.03 | 0.41 | 14.32 |
| schwann cell progenitors | Dhh | 6.98 | 2.48E-23 | 5.15E-19 | 0.11 | 0 | 0.11 | 55.5 |
| schwann cell progenitors | Sox8 | 3.82 | 3.03E-23 | 6.30E-19 | 0.61 | 0.06 | 0.55 | 10.36 |
| schwann cell progenitors | Sorbs1 | 3.84 | 3.87E-23 | 8.05E-19 | 0.72 | 0.08 | 0.64 | 8.6 |
| schwann cell progenitors | Crmp1 | 3.4 | 4.04E-23 | 8.41E-19 | 0.83 | 0.12 | 0.71 | 6.83 |
| schwann cell progenitors | Retreg1 | 3.23 | 4.77E-23 | 9.92E-19 | 0.78 | 0.09 | 0.69 | 8.37 |
| schwann cell progenitors | Tafa4 | 6.05 | 9.99E-23 | 2.08E-18 | 0.11 | 0 | 0.11 | 55.5 |
| schwann cell progenitors | Gm43707 | 3.77 | 1.75E-22 | 3.65E-18 | 0.44 | 0.03 | 0.41 | 13.88 |
| schwann cell progenitors | Ets1 | 3.22 | 1.26E-21 | 2.62E-17 | 0.94 | 0.18 | 0.77 | 5.27 |
| schwann cell progenitors | Apold1 | 3.75 | 1.58E-21 | 3.28E-17 | 0.33 | 0.02 | 0.31 | 17.53 |
| schwann cell progenitors | Fads6 | 5.42 | 1.74E-21 | 3.61E-17 | 0.17 | 0.01 | 0.16 | 33.4 |
| schwann cell progenitors | Pcdh1 | 3.45 | 2.15E-21 | 4.48E-17 | 0.56 | 0.05 | 0.5 | 10.69 |
| schwann cell progenitors | Csn3 | 5.61 | 3.80E-21 | 7.91E-17 | 0.17 | 0.01 | 0.16 | 33.4 |
| schwann cell progenitors | Dagla | 4.56 | 4.49E-21 | 9.33E-17 | 0.28 | 0.01 | 0.26 | 19.86 |
| schwann cell progenitors | Man1c1 | 3.79 | 4.84E-21 | 1.01E-16 | 0.44 | 0.03 | 0.41 | 13.06 |
| schwann cell progenitors | L1cam | 3.4 | 1.08E-20 | 2.24E-16 | 0.67 | 0.08 | 0.59 | 8.23 |
| schwann cell progenitors | Map7d2 | 2.59 | 1.90E-20 | 3.96E-16 | 0.72 | 0.09 | 0.64 | 8.49 |
| schwann cell progenitors | Cdc42ep1 | 2.94 | 2.15E-20 | 4.48E-16 | 0.61 | 0.06 | 0.55 | 9.7 |
| schwann cell progenitors | Itga4 | 3.77 | 2.27E-20 | 4.72E-16 | 0.94 | 0.2 | 0.75 | 4.82 |
| schwann cell progenitors | Gm4876 | 4.22 | 9.62E-20 | 2.00E-15 | 0.28 | 0.02 | 0.26 | 18.53 |
| schwann cell progenitors | Sphk1 | 4.61 | 1.09E-19 | 2.26E-15 | 0.22 | 0.01 | 0.21 | 22.2 |
| schwann cell progenitors | Pls3 | 4.73 | 1.34E-19 | 2.78E-15 | 1 | 0.3 | 0.7 | 3.34 |
| schwann cell progenitors | Col5a3 | 6.01 | 2.22E-19 | 4.63E-15 | 0.11 | 0 | 0.11 | 37 |
| schwann cell progenitors | Cdh6 | 4.03 | 3.54E-19 | 7.35E-15 | 1 | 0.29 | 0.71 | 3.5 |
| schwann cell progenitors | Coro2b | 3.49 | 8.29E-19 | 1.72E-14 | 0.5 | 0.05 | 0.45 | 10.42 |
| schwann cell progenitors | Myl9 | 3.72 | 1.09E-18 | 2.26E-14 | 0.94 | 0.24 | 0.7 | 3.87 |
| schwann cell progenitors | Klhl21 | 3.59 | 1.59E-18 | 3.30E-14 | 0.5 | 0.05 | 0.45 | 10 |
| schwann cell progenitors | Fxyd5 | 2.69 | 1.81E-18 | 3.77E-14 | 0.39 | 0.03 | 0.36 | 12.97 |
| schwann cell progenitors | Mamdc2 | 5.84 | 2.30E-18 | 4.78E-14 | 0.11 | 0 | 0.11 | 37 |
| schwann cell progenitors | Ptprm | 3.41 | 4.04E-18 | 8.40E-14 | 0.5 | 0.05 | 0.45 | 10 |
| schwann cell progenitors | Mmp15 | 3.16 | 4.52E-18 | 9.40E-14 | 0.67 | 0.09 | 0.58 | 7.49 |
| schwann cell progenitors | Kitl | 3.19 | 8.81E-18 | 1.83E-13 | 0.61 | 0.08 | 0.54 | 8.04 |
| schwann cell progenitors | Lpar3 | 5.94 | 1.81E-17 | 3.75E-13 | 0.11 | 0 | 0.11 | 37 |
| schwann cell progenitors | Fhdc1 | 3.73 | 2.00E-17 | 4.16E-13 | 0.44 | 0.04 | 0.4 | 10.57 |
| schwann cell progenitors | Gas7 | 3.29 | 2.46E-17 | 5.12E-13 | 0.5 | 0.05 | 0.45 | 9.43 |
| schwann cell progenitors | Tspan7 | 3.01 | 2.72E-17 | 5.66E-13 | 1 | 0.28 | 0.72 | 3.56 |
| schwann cell progenitors | Timp3 | 3.26 | 5.09E-17 | 1.06E-12 | 0.83 | 0.17 | 0.66 | 4.84 |
| schwann cell progenitors | Slc1a2 | 5.81 | 6.84E-17 | 1.42E-12 | 0.11 | 0 | 0.11 | 37 |
| schwann cell progenitors | Plvap | 1.83 | 6.92E-17 | 1.44E-12 | 0.44 | 0.04 | 0.4 | 10.57 |
| schwann cell progenitors | Kctd12 | 2.95 | 9.81E-17 | 2.04E-12 | 1 | 0.3 | 0.7 | 3.36 |
| schwann cell progenitors | Arhgef26 | 2.76 | 1.05E-16 | 2.18E-12 | 0.78 | 0.13 | 0.65 | 6.17 |

**Supplementary Table 2. Marker genes for the major cell types of integrated clustering of E11.5 control and NotchGOF λ-junction cells.**

| | | | | | | | | |
|---|---|---|---|---|---|---|---|---|
| schwann cell progenitors | Lbh | 2.67 | 1.09E-16 | 2.27E-12 | 0.94 | 0.2 | 0.74 | 4.67 |
| schwann cell progenitors | Gm26688 | 3.93 | 1.43E-16 | 2.98E-12 | 0.28 | 0.02 | 0.26 | 15.44 |
| schwann cell progenitors | Cav1 | 4.16 | 1.70E-16 | 3.55E-12 | 0.22 | 0.01 | 0.21 | 18.5 |
| schwann cell progenitors | Zadh2 | 2.73 | 2.42E-16 | 5.03E-12 | 0.72 | 0.11 | 0.62 | 6.75 |
| schwann cell progenitors | Rhob | 3.37 | 2.47E-16 | 5.14E-12 | 1 | 0.35 | 0.65 | 2.88 |
| schwann cell progenitors | Adamts1 | 3.06 | 3.07E-16 | 6.38E-12 | 1 | 0.28 | 0.73 | 3.64 |
| schwann cell progenitors | Fads3 | 3.6 | 3.78E-16 | 7.87E-12 | 0.5 | 0.06 | 0.44 | 8.77 |
| schwann cell progenitors | Cthrc1 | 3.54 | 7.84E-16 | 1.63E-11 | 0.89 | 0.19 | 0.7 | 4.58 |
| schwann cell progenitors | Gldn | 4.19 | 1.51E-15 | 3.14E-11 | 0.22 | 0.01 | 0.21 | 18.5 |
| schwann cell progenitors | Dock2 | 4.44 | 1.54E-15 | 3.21E-11 | 0.17 | 0.01 | 0.16 | 23.86 |
| schwann cell progenitors | Heyl | 3.17 | 1.56E-15 | 3.24E-11 | 0.5 | 0.06 | 0.44 | 8.62 |
| schwann cell progenitors | Dpysl3 | 3.83 | 1.82E-15 | 3.79E-11 | 1 | 0.45 | 0.55 | 2.21 |
| schwann cell progenitors | Sema3c | 2.21 | 2.96E-15 | 6.17E-11 | 0.67 | 0.09 | 0.57 | 7.1 |
| schwann cell progenitors | Gpr37 | 5.66 | 3.53E-15 | 7.34E-11 | 0.11 | 0 | 0.11 | 37 |
| schwann cell progenitors | Nrp1 | 2.63 | 4.85E-15 | 1.01E-10 | 0.89 | 0.21 | 0.68 | 4.23 |
| schwann cell progenitors | Gabrb3 | 3.88 | 4.98E-15 | 1.04E-10 | 0.22 | 0.01 | 0.21 | 17.08 |
| schwann cell progenitors | Plxnd1 | 2.42 | 5.37E-15 | 1.12E-10 | 0.5 | 0.06 | 0.44 | 8.47 |
| schwann cell progenitors | Xylt1 | 3.56 | 6.29E-15 | 1.31E-10 | 0.5 | 0.06 | 0.44 | 8.2 |
| schwann cell progenitors | Ptprj | 4.04 | 7.96E-15 | 1.65E-10 | 0.39 | 0.04 | 0.35 | 10.24 |
| schwann cell progenitors | Nacc2 | 3.12 | 9.99E-15 | 2.08E-10 | 0.39 | 0.04 | 0.35 | 10.51 |
| schwann cell progenitors | Anxa1 | 3.28 | 1.15E-14 | 2.40E-10 | 0.33 | 0.03 | 0.31 | 11.89 |
| schwann cell progenitors | Cotl1 | 3 | 1.30E-14 | 2.71E-10 | 1 | 0.45 | 0.55 | 2.24 |
| schwann cell progenitors | Vgll3 | 3.95 | 1.32E-14 | 2.74E-10 | 0.33 | 0.03 | 0.3 | 11.48 |
| schwann cell progenitors | Synm | 4.44 | 1.45E-14 | 3.02E-10 | 0.28 | 0.02 | 0.26 | 13.24 |
| schwann cell progenitors | Lpcat2 | 3.51 | 1.47E-14 | 3.07E-10 | 0.39 | 0.04 | 0.35 | 9.97 |
| schwann cell progenitors | Cadm1 | 2.81 | 1.67E-14 | 3.48E-10 | 1 | 0.39 | 0.61 | 2.57 |
| schwann cell progenitors | Reep1 | 3.11 | 1.75E-14 | 3.64E-10 | 0.44 | 0.05 | 0.4 | 9.25 |
| schwann cell progenitors | Gm16701.1 | 5.19 | 2.40E-14 | 5.00E-10 | 0.11 | 0 | 0.11 | 37 |
| schwann cell progenitors | Ahr | 2.96 | 2.56E-14 | 5.33E-10 | 0.61 | 0.09 | 0.52 | 6.87 |
| schwann cell progenitors | Metrn | 2.88 | 3.45E-14 | 7.18E-10 | 1 | 0.52 | 0.49 | 1.94 |
| schwann cell progenitors | Gamt | 2.83 | 4.68E-14 | 9.74E-10 | 0.61 | 0.09 | 0.52 | 6.57 |
| schwann cell progenitors | Tlr3 | 5.14 | 4.70E-14 | 9.78E-10 | 0.17 | 0.01 | 0.16 | 20.88 |
| schwann cell progenitors | Mxra7 | 2.73 | 5.73E-14 | 1.19E-09 | 0.89 | 0.23 | 0.66 | 3.8 |
| schwann cell progenitors | Afap1l1 | 2.44 | 8.46E-14 | 1.76E-09 | 0.56 | 0.07 | 0.48 | 7.51 |
| schwann cell progenitors | Lipo4 | 5.81 | 1.14E-13 | 2.37E-09 | 0.11 | 0 | 0.11 | 27.75 |
| schwann cell progenitors | Ccl6 | 4.07 | 1.21E-13 | 2.52E-09 | 0.11 | 0 | 0.11 | 27.75 |
| schwann cell progenitors | Zfyve28 | 5.75 | 1.64E-13 | 3.42E-09 | 0.11 | 0 | 0.11 | 27.75 |
| schwann cell progenitors | Pdlim1 | 2.23 | 1.94E-13 | 4.04E-09 | 0.94 | 0.29 | 0.65 | 3.23 |
| schwann cell progenitors | Arpc1b | 2.68 | 2.14E-13 | 4.45E-09 | 0.94 | 0.35 | 0.6 | 2.71 |
| schwann cell progenitors | Htra3 | 4.27 | 2.20E-13 | 4.58E-09 | 0.22 | 0.01 | 0.21 | 15.86 |
| schwann cell progenitors | Map3k1 | 2.42 | 3.20E-13 | 6.66E-09 | 0.94 | 0.26 | 0.68 | 3.59 |
| schwann cell progenitors | Atp6v0e2 | 3.09 | 3.57E-13 | 7.42E-09 | 0.44 | 0.05 | 0.39 | 8.22 |
| schwann cell progenitors | Zeb2 | 2.81 | 3.57E-13 | 7.43E-09 | 1 | 0.4 | 0.6 | 2.49 |
| schwann cell progenitors | Rhbdf1 | 2.99 | 4.11E-13 | 8.54E-09 | 0.61 | 0.1 | 0.51 | 6.05 |
| schwann cell progenitors | Dab1 | 1.68 | 4.28E-13 | 8.91E-09 | 0.83 | 0.16 | 0.68 | 5.37 |
| schwann cell progenitors | Dtx4 | 3.06 | 4.69E-13 | 9.75E-09 | 0.39 | 0.04 | 0.35 | 9.49 |
| schwann cell progenitors | Foxn2 | 2.2 | 5.31E-13 | 1.10E-08 | 0.83 | 0.16 | 0.67 | 5.08 |
| schwann cell progenitors | Dbi | 1.8 | 6.53E-13 | 1.36E-08 | 1 | 0.98 | 0.02 | 1.02 |
| schwann cell progenitors | Anxa6 | 2.17 | 7.00E-13 | 1.46E-08 | 0.83 | 0.17 | 0.66 | 4.79 |
| schwann cell progenitors | Csrp1 | 2.68 | 8.00E-13 | 1.66E-08 | 0.94 | 0.34 | 0.6 | 2.76 |
| schwann cell progenitors | Slitrk6 | 2.37 | 8.85E-13 | 1.84E-08 | 1 | 0.33 | 0.68 | 3.08 |
| schwann cell progenitors | Lamc1 | 2.27 | 9.09E-13 | 1.89E-08 | 0.94 | 0.26 | 0.68 | 3.62 |
| schwann cell progenitors | Klhdc1 | 3.62 | 9.10E-13 | 1.89E-08 | 0.28 | 0.02 | 0.26 | 12.09 |
| schwann cell progenitors | Megf9 | 2.67 | 1.03E-12 | 2.14E-08 | 0.83 | 0.19 | 0.64 | 4.38 |
| schwann cell progenitors | Nnmt | 5.33 | 1.57E-12 | 3.27E-08 | 0.11 | 0 | 0.11 | 27.75 |
| schwann cell progenitors | Gm46620 | 5.25 | 1.69E-12 | 3.52E-08 | 0.11 | 0 | 0.11 | 27.75 |

**Supplementary Table 2. Marker genes for the major cell types of integrated clustering of E11.5 control and NotchGOF λ-junction cells.**

| | | | | | | | | |
|---|---|---|---|---|---|---|---|---|
| schwann cell progenitors | Fam210b | 2.13 | 1.98E-12 | 4.12E-08 | 0.83 | 0.2 | 0.63 | 4.12 |
| schwann cell progenitors | Kcna1 | 5.47 | 2.18E-12 | 4.54E-08 | 0.11 | 0 | 0.11 | 27.75 |
| schwann cell progenitors | Dnajc1 | 2.3 | 2.21E-12 | 4.59E-08 | 1 | 0.56 | 0.44 | 1.79 |
| schwann cell progenitors | Enc1 | 2.44 | 4.74E-12 | 9.87E-08 | 0.83 | 0.22 | 0.61 | 3.75 |
| schwann cell progenitors | Sdc3 | 2.04 | 6.77E-12 | 1.41E-07 | 0.89 | 0.21 | 0.68 | 4.34 |
| schwann cell progenitors | Col5a2 | 2.68 | 7.96E-12 | 1.65E-07 | 1 | 0.53 | 0.47 | 1.89 |
| schwann cell progenitors | Atg4a | 2.56 | 8.28E-12 | 1.72E-07 | 0.56 | 0.09 | 0.47 | 6.39 |
| schwann cell progenitors | Gm43560 | 3.44 | 8.41E-12 | 1.75E-07 | 0.39 | 0.05 | 0.34 | 8.28 |
| schwann cell progenitors | Kcnk5 | 3.59 | 9.99E-12 | 2.08E-07 | 0.33 | 0.04 | 0.3 | 9.51 |
| schwann cell progenitors | Klhl13 | 2.18 | 1.09E-11 | 2.27E-07 | 0.94 | 0.27 | 0.68 | 3.51 |
| schwann cell progenitors | Sema3d | 1.91 | 1.09E-11 | 2.27E-07 | 0.67 | 0.12 | 0.55 | 5.61 |
| schwann cell progenitors | Selenok-ps1 | 4.57 | 1.12E-11 | 2.32E-07 | 0.17 | 0.01 | 0.16 | 16.7 |
| schwann cell progenitors | Dock11 | 2.7 | 1.12E-11 | 2.34E-07 | 0.83 | 0.23 | 0.6 | 3.64 |
| schwann cell progenitors | Lmo4 | 1.99 | 1.18E-11 | 2.45E-07 | 1 | 0.81 | 0.19 | 1.23 |
| schwann cell progenitors | Gdpd5 | 3.17 | 1.19E-11 | 2.47E-07 | 0.39 | 0.05 | 0.34 | 8.28 |
| schwann cell progenitors | Vim | 2.25 | 1.24E-11 | 2.57E-07 | 1 | 0.84 | 0.16 | 1.19 |
| schwann cell progenitors | Fbxo7 | 2.55 | 1.52E-11 | 3.17E-07 | 0.67 | 0.14 | 0.53 | 4.9 |
| schwann cell progenitors | Anxa5 | 2.33 | 1.67E-11 | 3.47E-07 | 1 | 0.61 | 0.39 | 1.65 |
| schwann cell progenitors | Sparc | 2.38 | 1.89E-11 | 3.93E-07 | 1 | 0.74 | 0.26 | 1.35 |
| schwann cell progenitors | Tnik | 2.77 | 2.09E-11 | 4.34E-07 | 0.83 | 0.25 | 0.59 | 3.39 |
| schwann cell progenitors | Egfl7 | 1.5 | 2.23E-11 | 4.63E-07 | 0.89 | 0.22 | 0.67 | 4.04 |
| schwann cell progenitors | Arhgap20 | 4.34 | 2.36E-11 | 4.90E-07 | 0.17 | 0.01 | 0.16 | 16.7 |
| schwann cell progenitors | Paqr4 | 3.01 | 2.52E-11 | 5.23E-07 | 0.33 | 0.04 | 0.3 | 9.25 |
| schwann cell progenitors | Btbd3 | 2.53 | 2.61E-11 | 5.44E-07 | 0.72 | 0.16 | 0.56 | 4.57 |
| schwann cell progenitors | Ptprz1 | 2.53 | 2.71E-11 | 5.63E-07 | 0.61 | 0.11 | 0.5 | 5.71 |
| schwann cell progenitors | Kcna6 | 3.26 | 3.28E-11 | 6.82E-07 | 0.33 | 0.04 | 0.3 | 9.25 |
| schwann cell progenitors | Hs2st1 | 2.65 | 3.31E-11 | 6.89E-07 | 0.94 | 0.57 | 0.38 | 1.67 |
| schwann cell progenitors | Cdh2 | 2.05 | 3.60E-11 | 7.49E-07 | 1 | 0.42 | 0.58 | 2.36 |
| schwann cell progenitors | Gimap9 | 2.34 | 3.64E-11 | 7.56E-07 | 0.61 | 0.11 | 0.5 | 5.66 |
| schwann cell progenitors | Celsr2 | 2.59 | 3.80E-11 | 7.89E-07 | 0.56 | 0.09 | 0.46 | 5.98 |
| schwann cell progenitors | Nes | 2.32 | 3.86E-11 | 8.02E-07 | 1 | 0.43 | 0.57 | 2.35 |
| schwann cell progenitors | Celf2 | 2.06 | 4.33E-11 | 9.01E-07 | 0.89 | 0.26 | 0.63 | 3.46 |
| schwann cell progenitors | Gnb4 | 2.27 | 4.82E-11 | 1.00E-06 | 0.89 | 0.3 | 0.59 | 2.95 |
| schwann cell progenitors | Tspan4 | 2.25 | 5.02E-11 | 1.04E-06 | 0.67 | 0.13 | 0.54 | 5.21 |
| schwann cell progenitors | Gm19412 | 3.12 | 5.43E-11 | 1.13E-06 | 0.33 | 0.04 | 0.3 | 9 |
| schwann cell progenitors | Khdrbs2 | 2.98 | 5.62E-11 | 1.17E-06 | 0.33 | 0.04 | 0.3 | 9 |
| schwann cell progenitors | Kifc3 | 2.46 | 5.88E-11 | 1.22E-06 | 0.5 | 0.08 | 0.43 | 6.67 |
| schwann cell progenitors | Itga6 | 2.17 | 5.90E-11 | 1.23E-06 | 0.94 | 0.31 | 0.63 | 3.01 |
| schwann cell progenitors | Tmc6 | 2.53 | 6.03E-11 | 1.25E-06 | 0.5 | 0.08 | 0.42 | 6.58 |
| schwann cell progenitors | Nab1 | 2.24 | 8.20E-11 | 1.70E-06 | 0.67 | 0.13 | 0.54 | 5.09 |
| schwann cell progenitors | Pdzrn3 | 2.38 | 9.94E-11 | 2.07E-06 | 0.78 | 0.17 | 0.6 | 4.47 |
| schwann cell progenitors | Cpne2 | 3.22 | 1.04E-10 | 2.17E-06 | 0.39 | 0.05 | 0.34 | 7.48 |
| schwann cell progenitors | Qpct | 3.26 | 1.19E-10 | 2.48E-06 | 0.33 | 0.04 | 0.3 | 8.76 |
| schwann cell progenitors | Hvcn1 | 2.64 | 1.20E-10 | 2.49E-06 | 0.5 | 0.08 | 0.42 | 6.33 |
| schwann cell progenitors | Grid2 | 3.43 | 1.23E-10 | 2.55E-06 | 0.22 | 0.02 | 0.2 | 12.33 |
| schwann cell progenitors | Gm42670 | 3.75 | 1.24E-10 | 2.57E-06 | 0.22 | 0.02 | 0.2 | 12.33 |
| schwann cell progenitors | Fxyd1 | 2.5 | 1.25E-10 | 2.60E-06 | 0.44 | 0.06 | 0.38 | 7.05 |
| schwann cell progenitors | Chst5 | 3.83 | 1.29E-10 | 2.68E-06 | 0.22 | 0.02 | 0.2 | 12.33 |
| schwann cell progenitors | Acsl4 | 2.73 | 1.31E-10 | 2.73E-06 | 0.67 | 0.14 | 0.53 | 4.73 |
| schwann cell progenitors | Ankrd29 | 2.42 | 1.37E-10 | 2.85E-06 | 0.56 | 0.09 | 0.46 | 5.98 |
| schwann cell progenitors | Gap43 | 2.4 | 1.57E-10 | 3.26E-06 | 0.94 | 0.42 | 0.52 | 2.25 |
| schwann cell progenitors | Jdp2 | 3.06 | 1.63E-10 | 3.38E-06 | 0.33 | 0.04 | 0.3 | 8.76 |
| schwann cell progenitors | Myo18a | 2.41 | 1.65E-10 | 3.43E-06 | 0.83 | 0.24 | 0.6 | 3.53 |
| schwann cell progenitors | Gm47644 | 3.66 | 1.78E-10 | 3.70E-06 | 0.28 | 0.03 | 0.25 | 9.93 |
| schwann cell progenitors | Emb | 2.08 | 1.88E-10 | 3.91E-06 | 1 | 0.41 | 0.59 | 2.45 |
| schwann cell progenitors | Cst3 | 1.3 | 2.18E-10 | 4.53E-06 | 1 | 0.92 | 0.08 | 1.09 |

**Supplementary Table 2. Marker genes for the major cell types of integrated clustering of E11.5 control and NotchGOF λ-junction cells.**

| | | | | | | | | |
|---|---|---|---|---|---|---|---|---|
| schwann cell progenitors | Lrmp | 3.77 | 2.66E-10 | 5.53E-06 | 0.17 | 0.01 | 0.16 | 15.18 |
| schwann cell progenitors | Pxdc1 | 3.22 | 2.68E-10 | 5.57E-06 | 0.56 | 0.11 | 0.45 | 5.3 |
| schwann cell progenitors | Tagln2 | 2.93 | 2.91E-10 | 6.04E-06 | 0.83 | 0.3 | 0.53 | 2.79 |
| schwann cell progenitors | Nlgn3 | 3.3 | 3.08E-10 | 6.41E-06 | 0.39 | 0.05 | 0.34 | 7.48 |
| schwann cell progenitors | Pros1 | 2.6 | 3.33E-10 | 6.93E-06 | 0.67 | 0.15 | 0.52 | 4.6 |
| schwann cell progenitors | 700025G04Ri | 1.93 | 3.51E-10 | 7.31E-06 | 1 | 0.34 | 0.66 | 2.95 |
| schwann cell progenitors | Il3ra | 2.82 | 3.54E-10 | 7.36E-06 | 0.39 | 0.05 | 0.34 | 7.48 |
| schwann cell progenitors | Pdia5 | 2.34 | 3.60E-10 | 7.48E-06 | 0.61 | 0.12 | 0.49 | 5.22 |
| schwann cell progenitors | Ttyh2 | 3.28 | 3.68E-10 | 7.64E-06 | 0.28 | 0.03 | 0.25 | 9.93 |
| schwann cell progenitors | Alcam | 2.5 | 4.31E-10 | 8.95E-06 | 0.67 | 0.15 | 0.52 | 4.57 |
| schwann cell progenitors | Tmsb4x | 1.16 | 4.47E-10 | 9.29E-06 | 1 | 0.99 | 0.01 | 1.01 |
| schwann cell progenitors | Olfm2 | 2.56 | 4.55E-10 | 9.47E-06 | 0.56 | 0.11 | 0.45 | 5.25 |
| schwann cell progenitors | Lipa | 2.77 | 4.72E-10 | 9.82E-06 | 0.78 | 0.22 | 0.56 | 3.55 |
| schwann cell progenitors | Chrdl2 | 3.96 | 7.48E-10 | 1.56E-05 | 0.17 | 0.01 | 0.16 | 15.18 |
| schwann cell progenitors | Ubl3 | 2.27 | 1.20E-09 | 2.50E-05 | 0.89 | 0.37 | 0.52 | 2.39 |
| schwann cell progenitors | Fign | 2.55 | 1.26E-09 | 2.63E-05 | 0.83 | 0.29 | 0.54 | 2.83 |
| schwann cell progenitors | Actg1 | 0.81 | 1.40E-09 | 2.91E-05 | 1 | 1 | 0 | 1 |
| schwann cell progenitors | Epha5 | 2.6 | 1.56E-09 | 3.25E-05 | 0.44 | 0.07 | 0.37 | 6.25 |
| schwann cell progenitors | Mid1ip1 | 2.48 | 1.84E-09 | 3.83E-05 | 0.78 | 0.22 | 0.56 | 3.54 |
| schwann cell progenitors | Tshz1 | 2.68 | 2.28E-09 | 4.73E-05 | 0.83 | 0.32 | 0.51 | 2.61 |
| schwann cell progenitors | Chn2 | 2.56 | 2.31E-09 | 4.81E-05 | 0.44 | 0.07 | 0.37 | 6.17 |
| schwann cell progenitors | Clcn5 | 2.14 | 2.54E-09 | 5.28E-05 | 0.94 | 0.39 | 0.55 | 2.41 |
| schwann cell progenitors | Pon2 | 2.09 | 2.69E-09 | 5.60E-05 | 0.72 | 0.18 | 0.54 | 3.97 |
| schwann cell progenitors | Map1a | 2.72 | 2.76E-09 | 5.74E-05 | 0.5 | 0.09 | 0.41 | 5.43 |
| schwann cell progenitors | Dusp3 | 2.55 | 2.90E-09 | 6.04E-05 | 0.39 | 0.06 | 0.33 | 6.82 |
| schwann cell progenitors | Aard | 2.79 | 3.46E-09 | 7.20E-05 | 0.44 | 0.07 | 0.37 | 6.08 |
| schwann cell progenitors | Kif19a | 3.05 | 3.70E-09 | 7.71E-05 | 0.17 | 0.01 | 0.16 | 13.92 |
| schwann cell progenitors | Epb41l4a | 2.33 | 3.73E-09 | 7.77E-05 | 0.56 | 0.11 | 0.45 | 5.2 |
| schwann cell progenitors | Gpsm3 | 2.21 | 3.80E-09 | 7.91E-05 | 0.5 | 0.09 | 0.41 | 5.75 |
| schwann cell progenitors | Fam49a | 2.96 | 3.99E-09 | 8.30E-05 | 0.33 | 0.05 | 0.29 | 7.4 |
| schwann cell progenitors | Trf | 2.51 | 4.18E-09 | 8.68E-05 | 0.33 | 0.04 | 0.29 | 7.74 |
| schwann cell progenitors | Igsf9b | 2.57 | 4.27E-09 | 8.89E-05 | 0.39 | 0.06 | 0.33 | 6.82 |
| schwann cell progenitors | Calm1 | 0.98 | 4.81E-09 | 1.00E-04 | 1 | 0.99 | 0.01 | 1.01 |
| schwann cell progenitors | Wdfy1 | 2.46 | 5.20E-09 | 1.08E-04 | 0.83 | 0.35 | 0.48 | 2.37 |
| schwann cell progenitors | Pxn | 2.45 | 5.23E-09 | 1.09E-04 | 0.72 | 0.19 | 0.54 | 3.88 |
| schwann cell progenitors | Cytl1 | 5.08 | 5.41E-09 | 1.13E-04 | 0.11 | 0.01 | 0.11 | 18.5 |
| schwann cell progenitors | Camkv | 4.22 | 5.58E-09 | 1.16E-04 | 0.17 | 0.01 | 0.15 | 12.85 |
| schwann cell progenitors | Ppp1r14c | 3.64 | 6.13E-09 | 1.27E-04 | 0.22 | 0.02 | 0.2 | 10.09 |
| schwann cell progenitors | Fgd1 | 1.73 | 7.42E-09 | 1.54E-04 | 0.94 | 0.32 | 0.62 | 2.92 |
| schwann cell progenitors | Sort1 | 2 | 7.51E-09 | 1.56E-04 | 0.78 | 0.21 | 0.57 | 3.8 |
| schwann cell progenitors | Map3k11 | 2.56 | 7.75E-09 | 1.61E-04 | 0.56 | 0.12 | 0.44 | 4.75 |
| schwann cell progenitors | Tubb6 | 2.15 | 8.07E-09 | 1.68E-04 | 0.56 | 0.12 | 0.44 | 4.83 |
| schwann cell progenitors | Mest | 1.56 | 8.42E-09 | 1.75E-04 | 1 | 0.91 | 0.09 | 1.1 |
| schwann cell progenitors | Fam89a | 2.35 | 9.24E-09 | 1.92E-04 | 0.44 | 0.08 | 0.37 | 5.92 |
| schwann cell progenitors | Fam111a | 2.3 | 9.24E-09 | 1.92E-04 | 0.83 | 0.33 | 0.51 | 2.56 |
| schwann cell progenitors | Coro1c | 1.94 | 9.30E-09 | 1.93E-04 | 0.94 | 0.35 | 0.59 | 2.7 |
| schwann cell progenitors | Cela1 | 4.26 | 1.16E-08 | 2.41E-04 | 0.11 | 0.01 | 0.11 | 18.5 |
| schwann cell progenitors | Emp3 | 2.27 | 1.21E-08 | 2.52E-04 | 0.61 | 0.14 | 0.47 | 4.46 |
| schwann cell progenitors | Cobll1 | 2.75 | 1.21E-08 | 2.53E-04 | 0.28 | 0.03 | 0.25 | 8.42 |
| schwann cell progenitors | Slc7a1 | 2.75 | 1.23E-08 | 2.56E-04 | 0.44 | 0.08 | 0.37 | 5.62 |
| schwann cell progenitors | Nid1 | 1.73 | 1.26E-08 | 2.62E-04 | 1 | 0.35 | 0.65 | 2.89 |
| schwann cell progenitors | Des | 2.84 | 1.52E-08 | 3.16E-04 | 0.33 | 0.05 | 0.29 | 7.09 |
| schwann cell progenitors | Sdsl | 2.46 | 1.69E-08 | 3.51E-04 | 0.39 | 0.06 | 0.33 | 6.48 |
| schwann cell progenitors | Pfn1 | 0.72 | 1.82E-08 | 3.78E-04 | 1 | 1 | 0 | 1 |
| schwann cell progenitors | Cnp | 2.47 | 1.83E-08 | 3.80E-04 | 0.61 | 0.16 | 0.45 | 3.89 |
| schwann cell progenitors | AC109619.1 | 4.88 | 1.84E-08 | 3.82E-04 | 0.11 | 0.01 | 0.11 | 18.5 |

**Supplementary Table 2. Marker genes for the major cell types of integrated clustering of E11.5 control and NotchGOF λ-junction cells.**

| | | | | | | | | |
|---|---|---|---|---|---|---|---|---|
| schwann cell progenitors | Tuba1a | 0.92 | 1.90E-08 | 3.95E-04 | 1 | 0.98 | 0.02 | 1.02 |
| schwann cell progenitors | Ptpn13 | 1.92 | 2.15E-08 | 4.48E-04 | 0.89 | 0.41 | 0.48 | 2.19 |
| schwann cell progenitors | Psat1 | 1.8 | 2.26E-08 | 4.69E-04 | 0.89 | 0.45 | 0.44 | 1.96 |
| schwann cell progenitors | Cavin3 | 2.11 | 2.29E-08 | 4.76E-04 | 0.56 | 0.12 | 0.43 | 4.48 |
| schwann cell progenitors | Selenof | 0.95 | 2.46E-08 | 5.12E-04 | 1 | 0.96 | 0.04 | 1.05 |
| schwann cell progenitors | Ppp1r18 | 2.03 | 2.50E-08 | 5.20E-04 | 0.78 | 0.25 | 0.53 | 3.09 |
| schwann cell progenitors | Zfp979 | 3.35 | 2.58E-08 | 5.38E-04 | 0.28 | 0.04 | 0.24 | 7.94 |
| schwann cell progenitors | Trim71 | 1.79 | 2.91E-08 | 6.05E-04 | 0.78 | 0.21 | 0.57 | 3.69 |
| schwann cell progenitors | Ppic | 1.44 | 3.05E-08 | 6.35E-04 | 1 | 0.71 | 0.29 | 1.41 |
| schwann cell progenitors | Ss18l2 | 1.85 | 3.07E-08 | 6.39E-04 | 0.89 | 0.52 | 0.37 | 1.72 |
| schwann cell progenitors | Gnai1 | 1.86 | 3.29E-08 | 6.85E-04 | 0.67 | 0.15 | 0.51 | 4.33 |
| schwann cell progenitors | Grik2 | 3.47 | 3.29E-08 | 6.85E-04 | 0.22 | 0.02 | 0.2 | 9.65 |
| schwann cell progenitors | Trim6 | 3.7 | 3.71E-08 | 7.72E-04 | 0.22 | 0.02 | 0.2 | 9.25 |
| schwann cell progenitors | Plekha4 | 4.93 | 3.72E-08 | 7.74E-04 | 0.11 | 0.01 | 0.11 | 18.5 |
| schwann cell progenitors | Tnfaip8l1 | 2.22 | 3.90E-08 | 8.11E-04 | 0.44 | 0.08 | 0.37 | 5.62 |
| schwann cell progenitors | Adamts12 | 3.19 | 3.93E-08 | 8.18E-04 | 0.28 | 0.04 | 0.24 | 7.94 |
| schwann cell progenitors | Soat1 | 1.99 | 4.07E-08 | 8.47E-04 | 0.94 | 0.44 | 0.5 | 2.14 |
| schwann cell progenitors | Cadm4 | 2.72 | 4.33E-08 | 9.01E-04 | 0.44 | 0.09 | 0.36 | 5.1 |
| schwann cell progenitors | Scap | 1.85 | 4.35E-08 | 9.05E-04 | 0.72 | 0.19 | 0.54 | 3.9 |
| schwann cell progenitors | Itgb1 | 1.39 | 4.62E-08 | 9.60E-04 | 1 | 0.85 | 0.15 | 1.17 |
| schwann cell progenitors | Chml | 1.81 | 4.71E-08 | 9.80E-04 | 0.67 | 0.15 | 0.52 | 4.45 |
| schwann cell progenitors | Cd81 | 1.3 | 4.89E-08 | 1.02E-03 | 1 | 0.89 | 0.11 | 1.12 |
| schwann cell progenitors | 900089D17Ri | 4.03 | 4.91E-08 | 1.02E-03 | 0.17 | 0.01 | 0.15 | 11.93 |
| schwann cell progenitors | Metap1d | 1.9 | 4.99E-08 | 1.04E-03 | 0.72 | 0.2 | 0.52 | 3.61 |
| schwann cell progenitors | Sox2 | 0.95 | 5.39E-08 | 1.12E-03 | 0.78 | 0.2 | 0.58 | 3.99 |
| schwann cell progenitors | Flrt1 | 2.95 | 6.02E-08 | 1.25E-03 | 0.22 | 0.02 | 0.2 | 9.25 |
| schwann cell progenitors | Tram2 | 2.39 | 6.08E-08 | 1.26E-03 | 0.5 | 0.1 | 0.4 | 4.85 |
| schwann cell progenitors | Trp53cor1 | 4.22 | 6.62E-08 | 1.38E-03 | 0.11 | 0.01 | 0.1 | 15.86 |
| schwann cell progenitors | 2810430I11Ril | 4.41 | 6.65E-08 | 1.38E-03 | 0.11 | 0.01 | 0.1 | 15.86 |
| schwann cell progenitors | Parp14 | 4.45 | 6.79E-08 | 1.41E-03 | 0.11 | 0.01 | 0.1 | 15.86 |
| schwann cell progenitors | Sppl2b | 2.82 | 6.88E-08 | 1.43E-03 | 0.56 | 0.15 | 0.41 | 3.78 |
| schwann cell progenitors | Prnp | 2.1 | 7.51E-08 | 1.56E-03 | 0.67 | 0.18 | 0.49 | 3.77 |
| schwann cell progenitors | Eif5a | 0.68 | 8.44E-08 | 1.75E-03 | 1 | 1 | 0 | 1 |
| schwann cell progenitors | Peg13 | 2.79 | 8.55E-08 | 1.78E-03 | 0.33 | 0.05 | 0.28 | 6.53 |
| schwann cell progenitors | Cep85l | 2.05 | 8.56E-08 | 1.78E-03 | 0.61 | 0.14 | 0.47 | 4.43 |
| schwann cell progenitors | Gramd4 | 2.44 | 8.58E-08 | 1.79E-03 | 0.44 | 0.09 | 0.36 | 5.22 |
| schwann cell progenitors | Mthfd1l | 1.79 | 8.80E-08 | 1.83E-03 | 0.67 | 0.16 | 0.51 | 4.19 |
| schwann cell progenitors | Pea15a | 1.44 | 9.82E-08 | 2.04E-03 | 0.83 | 0.23 | 0.6 | 3.58 |
| schwann cell progenitors | Hsp90aa1 | 0.69 | 1.02E-07 | 2.11E-03 | 1 | 1 | 0 | 1 |
| schwann cell progenitors | Odc1 | 1.16 | 1.06E-07 | 2.20E-03 | 1 | 0.81 | 0.19 | 1.23 |
| schwann cell progenitors | Rarres2 | 3.29 | 1.09E-07 | 2.26E-03 | 0.17 | 0.01 | 0.15 | 11.93 |
| schwann cell progenitors | Zfp276 | 2.53 | 1.12E-07 | 2.32E-03 | 0.44 | 0.09 | 0.36 | 5.1 |
| schwann cell progenitors | Serpinh1 | 1.12 | 1.14E-07 | 2.36E-03 | 1 | 0.94 | 0.06 | 1.06 |
| schwann cell progenitors | Myl6 | 0.73 | 1.18E-07 | 2.45E-03 | 1 | 0.99 | 0.01 | 1.01 |
| schwann cell progenitors | Ovca2 | 2.72 | 1.19E-07 | 2.48E-03 | 0.33 | 0.05 | 0.28 | 6.53 |
| schwann cell progenitors | Zbtb38 | 2.58 | 1.26E-07 | 2.62E-03 | 0.56 | 0.14 | 0.42 | 4.06 |
| schwann cell progenitors | Adora1 | 4.33 | 1.37E-07 | 2.84E-03 | 0.11 | 0.01 | 0.1 | 15.86 |
| schwann cell progenitors | Dusp22 | 1.99 | 1.37E-07 | 2.84E-03 | 0.72 | 0.2 | 0.52 | 3.54 |
| schwann cell progenitors | Kif21a | 2.1 | 1.39E-07 | 2.90E-03 | 0.78 | 0.31 | 0.47 | 2.49 |
| schwann cell progenitors | Esyt1 | 3.26 | 1.40E-07 | 2.91E-03 | 0.39 | 0.08 | 0.31 | 5.19 |
| schwann cell progenitors | Fam129b | 1.91 | 1.42E-07 | 2.96E-03 | 0.61 | 0.14 | 0.47 | 4.27 |
| schwann cell progenitors | Slc25a5 | 0.86 | 1.54E-07 | 3.19E-03 | 1 | 0.93 | 0.07 | 1.07 |
| schwann cell progenitors | Ctsl | 1.67 | 1.60E-07 | 3.32E-03 | 1 | 0.82 | 0.18 | 1.21 |
| schwann cell progenitors | Rin3 | 3.6 | 1.79E-07 | 3.73E-03 | 0.22 | 0.03 | 0.2 | 8.54 |
| schwann cell progenitors | Soga3 | 2.38 | 1.83E-07 | 3.80E-03 | 0.33 | 0.05 | 0.28 | 6.17 |
| schwann cell progenitors | Slc8b1 | 2.84 | 2.01E-07 | 4.18E-03 | 0.22 | 0.03 | 0.2 | 8.88 |

**Supplementary Table 2. Marker genes for the major cell types of integrated clustering of E11.5 control and NotchGOF λ-junction cells.**

| | | | | | | | | |
|---|---|---|---|---|---|---|---|---|
| schwann cell progenitors | Sh3tc2 | 3.6 | 2.12E-07 | 4.40E-03 | 0.22 | 0.03 | 0.2 | 8.54 |
| schwann cell progenitors | Cyp2j9 | 4.56 | 2.22E-07 | 4.61E-03 | 0.11 | 0.01 | 0.1 | 15.86 |
| schwann cell progenitors | Igfbp3 | 1.12 | 2.29E-07 | 4.76E-03 | 0.56 | 0.13 | 0.43 | 4.41 |
| schwann cell progenitors | Rims1 | 3.34 | 2.32E-07 | 4.82E-03 | 0.22 | 0.03 | 0.2 | 8.54 |
| schwann cell progenitors | Cyp2d22 | 2.65 | 2.33E-07 | 4.84E-03 | 0.33 | 0.05 | 0.28 | 6.28 |
| schwann cell progenitors | Ap3m2 | 2.85 | 2.59E-07 | 5.38E-03 | 0.28 | 0.04 | 0.24 | 7.13 |
| schwann cell progenitors | Nudt7 | 2.5 | 2.75E-07 | 5.73E-03 | 0.39 | 0.07 | 0.32 | 5.56 |
| schwann cell progenitors | Cfl1 | 0.61 | 3.12E-07 | 6.49E-03 | 1 | 1 | 0 | 1 |
| schwann cell progenitors | Nfic | 1.81 | 3.12E-07 | 6.50E-03 | 0.78 | 0.3 | 0.48 | 2.58 |
| schwann cell progenitors | Ppib | 0.78 | 3.18E-07 | 6.62E-03 | 1 | 0.98 | 0.02 | 1.02 |
| schwann cell progenitors | Shank1 | 3.35 | 3.30E-07 | 6.87E-03 | 0.22 | 0.03 | 0.2 | 8.54 |
| schwann cell progenitors | Ywhab | 0.88 | 3.32E-07 | 6.91E-03 | 1 | 0.97 | 0.03 | 1.03 |
| schwann cell progenitors | Dusp4 | 1.99 | 3.37E-07 | 7.01E-03 | 0.44 | 0.09 | 0.36 | 5.16 |
| schwann cell progenitors | Gnat2 | 4.54 | 3.46E-07 | 7.19E-03 | 0.11 | 0.01 | 0.1 | 15.86 |
| schwann cell progenitors | Hspd1 | 0.88 | 3.51E-07 | 7.31E-03 | 1 | 0.98 | 0.02 | 1.02 |
| schwann cell progenitors | Lrch3 | 2.06 | 3.59E-07 | 7.46E-03 | 0.83 | 0.37 | 0.46 | 2.25 |
| schwann cell progenitors | Serpine2 | 1.47 | 3.66E-07 | 7.60E-03 | 1 | 0.71 | 0.29 | 1.42 |
| schwann cell progenitors | Star | 3.24 | 3.69E-07 | 7.68E-03 | 0.22 | 0.03 | 0.2 | 8.22 |
| schwann cell progenitors | Rab8b | 1.43 | 4.10E-07 | 8.53E-03 | 0.83 | 0.24 | 0.59 | 3.43 |
| schwann cell progenitors | Flna | 1.4 | 4.17E-07 | 8.67E-03 | 1 | 0.79 | 0.21 | 1.26 |
| schwann cell progenitors | Fabp5 | 0.68 | 4.69E-07 | 9.76E-03 | 1 | 0.98 | 0.02 | 1.02 |
| schwann cell progenitors | Snx1 | 1.78 | 5.06E-07 | 1.05E-02 | 0.94 | 0.45 | 0.5 | 2.11 |
| schwann cell progenitors | Sfxn5 | 2.94 | 5.58E-07 | 1.16E-02 | 0.28 | 0.04 | 0.24 | 6.78 |
| schwann cell progenitors | Sh3pxd2a | 1.74 | 5.89E-07 | 1.22E-02 | 0.89 | 0.37 | 0.52 | 2.42 |
| schwann cell progenitors | Rftn1 | 2.47 | 6.03E-07 | 1.25E-02 | 0.28 | 0.04 | 0.24 | 6.95 |
| schwann cell progenitors | Gng5 | 0.5 | 6.12E-07 | 1.27E-02 | 1 | 1 | 0 | 1 |
| schwann cell progenitors | Sema3b | 4.12 | 6.46E-07 | 1.34E-02 | 0.17 | 0.02 | 0.15 | 9.82 |
| schwann cell progenitors | Cnn2 | 1.48 | 6.59E-07 | 1.37E-02 | 0.94 | 0.63 | 0.31 | 1.5 |
| schwann cell progenitors | Cdr2 | 1.75 | 6.96E-07 | 1.45E-02 | 0.72 | 0.22 | 0.5 | 3.28 |
| schwann cell progenitors | Sipa1l1 | 1.87 | 7.49E-07 | 1.56E-02 | 0.83 | 0.52 | 0.32 | 1.62 |
| schwann cell progenitors | Gm47061 | 4.44 | 7.89E-07 | 1.64E-02 | 0.11 | 0.01 | 0.1 | 13.88 |
| schwann cell progenitors | Erp29 | 1.16 | 7.91E-07 | 1.65E-02 | 1 | 0.88 | 0.12 | 1.14 |
| schwann cell progenitors | Gm48855 | 2.94 | 8.04E-07 | 1.67E-02 | 0.22 | 0.03 | 0.2 | 8.22 |
| schwann cell progenitors | Tgfb1i1 | 1.9 | 8.40E-07 | 1.75E-02 | 0.56 | 0.14 | 0.42 | 4.06 |
| schwann cell progenitors | Sfrp1 | 1.36 | 8.56E-07 | 1.78E-02 | 1 | 0.63 | 0.37 | 1.6 |
| schwann cell progenitors | Gm43445 | 1.96 | 8.61E-07 | 1.79E-02 | 0.44 | 0.09 | 0.35 | 4.93 |
| schwann cell progenitors | Cyp2j6 | 2.59 | 8.90E-07 | 1.85E-02 | 0.5 | 0.12 | 0.38 | 4.07 |
| schwann cell progenitors | Lin28a | 4.13 | 9.43E-07 | 1.96E-02 | 0.17 | 0.02 | 0.15 | 9.82 |
| schwann cell progenitors | Ilk | 1.26 | 9.96E-07 | 2.07E-02 | 0.94 | 0.61 | 0.34 | 1.55 |
| schwann cell progenitors | Cltb | 1.65 | 1.02E-06 | 2.11E-02 | 0.67 | 0.18 | 0.48 | 3.63 |
| schwann cell progenitors | Vwa1 | 2.84 | 1.05E-06 | 2.18E-02 | 0.33 | 0.06 | 0.27 | 5.64 |
| schwann cell progenitors | Gm35021 | 4.28 | 1.06E-06 | 2.21E-02 | 0.11 | 0.01 | 0.1 | 13.88 |
| schwann cell progenitors | Tcn2 | 1.61 | 1.09E-06 | 2.27E-02 | 0.67 | 0.18 | 0.49 | 3.75 |
| schwann cell progenitors | Plekho1 | 1.26 | 1.12E-06 | 2.33E-02 | 1 | 0.62 | 0.38 | 1.62 |
| schwann cell progenitors | 730045B01Ri | 4.21 | 1.14E-06 | 2.37E-02 | 0.17 | 0.02 | 0.15 | 9.82 |
| schwann cell progenitors | Eng | 2.13 | 1.16E-06 | 2.41E-02 | 0.28 | 0.04 | 0.24 | 6.62 |
| schwann cell progenitors | Pgbd5 | 4.72 | 1.23E-06 | 2.55E-02 | 0.11 | 0.01 | 0.1 | 13.88 |
| schwann cell progenitors | Fam228b | 3.02 | 1.31E-06 | 2.72E-02 | 0.28 | 0.04 | 0.24 | 6.47 |
| schwann cell progenitors | Trub2 | 1.68 | 1.45E-06 | 3.02E-02 | 0.83 | 0.31 | 0.52 | 2.7 |
| schwann cell progenitors | Fxyd3 | 1.56 | 1.48E-06 | 3.07E-02 | 0.44 | 0.09 | 0.35 | 4.77 |
| schwann cell progenitors | Glo1 | 0.91 | 1.55E-06 | 3.23E-02 | 1 | 0.85 | 0.15 | 1.18 |
| schwann cell progenitors | Stx7 | 1.54 | 1.55E-06 | 3.23E-02 | 0.83 | 0.31 | 0.52 | 2.67 |
| schwann cell progenitors | Cmtm3 | 1.43 | 1.56E-06 | 3.24E-02 | 0.94 | 0.42 | 0.52 | 2.24 |
| schwann cell progenitors | Msn | 1.6 | 1.56E-06 | 3.25E-02 | 0.83 | 0.52 | 0.32 | 1.62 |
| schwann cell progenitors | Crip2 | 1.21 | 1.56E-06 | 3.25E-02 | 0.61 | 0.17 | 0.44 | 3.51 |
| schwann cell progenitors | C1qtnf12 | 2.45 | 1.59E-06 | 3.30E-02 | 0.56 | 0.17 | 0.39 | 3.33 |

**Supplementary Table 2. Marker genes for the major cell types of integrated clustering of E11.5 control and NotchGOF λ-junction cells.**

| | | | | | | | | |
|---|---|---|---|---|---|---|---|---|
| schwann cell progenitors | Lima1 | 1.19 | 1.78E-06 | 3.70E-02 | 1 | 0.74 | 0.26 | 1.35 |
| schwann cell progenitors | Bgn | 1.83 | 1.89E-06 | 3.94E-02 | 0.72 | 0.26 | 0.46 | 2.75 |
| schwann cell progenitors | P2rx4 | 3.32 | 1.90E-06 | 3.95E-02 | 0.22 | 0.03 | 0.19 | 7.4 |
| schwann cell progenitors | Nexn | 2.76 | 1.93E-06 | 4.01E-02 | 0.28 | 0.04 | 0.23 | 6.32 |
| schwann cell progenitors | Marcks | 0.65 | 2.00E-06 | 4.15E-02 | 1 | 0.97 | 0.03 | 1.03 |
| schwann cell progenitors | Bag3 | 1.73 | 2.01E-06 | 4.18E-02 | 0.5 | 0.11 | 0.39 | 4.5 |
| schwann cell progenitors | Agrn | 1.66 | 2.09E-06 | 4.35E-02 | 0.83 | 0.39 | 0.44 | 2.13 |
| schwann cell progenitors | Npm1 | 0.44 | 2.22E-06 | 4.61E-02 | 1 | 1 | 0 | 1 |
| schwann cell progenitors | Crispld2 | 2.81 | 2.22E-06 | 4.62E-02 | 0.22 | 0.03 | 0.19 | 7.66 |
| schwann cell progenitors | Gstp1 | 1.73 | 2.23E-06 | 4.64E-02 | 0.94 | 0.52 | 0.42 | 1.81 |
| schwann cell progenitors | Snx10 | 2.55 | 2.30E-06 | 4.77E-02 | 0.56 | 0.16 | 0.4 | 3.5 |
| schwann cell progenitors | Pde7b | 2.81 | 2.33E-06 | 4.85E-02 | 0.28 | 0.04 | 0.23 | 6.32 |
| schwann cell progenitors | Nr4a1 | 2.47 | 2.34E-06 | 4.87E-02 | 0.28 | 0.04 | 0.24 | 6.47 |

**Supplementary Table 3. Marker genes for the original clusters of integrated clustering of E11.5 control and NotchGOF λ-junction ectoderm cells.**

| cluster | gene | avg_log2FC | p_val | p_val_adj | pct.1 | pct.2 | pct1-pct2 | pct1/pct2 |
|---|---|---|---|---|---|---|---|---|
| 0 | Aldh1a3 | 2.42 | 1.47E-208 | 3.15E-204 | 0.96 | 0.5 | 0.46 | 1.93 |
| 0 | Stra6 | 2.42 | 4.81E-157 | 1.03E-152 | 0.68 | 0.19 | 0.49 | 3.63 |
| 0 | Six2 | 1.5 | 5.99E-114 | 1.28E-109 | 0.87 | 0.49 | 0.39 | 1.8 |
| 0 | Lhfp | 1.78 | 8.92E-101 | 1.91E-96 | 0.67 | 0.25 | 0.42 | 2.66 |
| 0 | Ebf1 | 1.11 | 5.37E-99 | 1.15E-94 | 0.42 | 0.09 | 0.33 | 4.65 |
| 0 | Unc13c | 2.99 | 3.89E-96 | 8.30E-92 | 0.28 | 0.04 | 0.24 | 7.67 |
| 0 | Mfap4 | 1.19 | 1.57E-93 | 3.35E-89 | 0.93 | 0.62 | 0.31 | 1.5 |
| 0 | D230030E09Rik | 2.39 | 3.70E-89 | 7.90E-85 | 0.52 | 0.17 | 0.35 | 3.01 |
| 0 | Eya2 | 1.25 | 1.10E-78 | 2.35E-74 | 0.73 | 0.31 | 0.41 | 2.32 |
| 0 | Crabp2 | 0.78 | 2.43E-78 | 5.20E-74 | 0.99 | 0.83 | 0.17 | 1.2 |
| 0 | Pcp4 | 3.47 | 1.26E-72 | 2.69E-68 | 0.19 | 0.02 | 0.17 | 8.77 |
| 0 | Eno3 | 2.2 | 6.06E-72 | 1.29E-67 | 0.44 | 0.14 | 0.3 | 3.14 |
| 0 | Tmeff2 | 3.74 | 6.76E-72 | 1.44E-67 | 0.19 | 0.02 | 0.17 | 8.59 |
| 0 | Mgp | 3.77 | 2.46E-71 | 5.24E-67 | 0.23 | 0.04 | 0.19 | 6.36 |
| 0 | Cadm1 | 0.91 | 6.57E-67 | 1.40E-62 | 0.92 | 0.68 | 0.23 | 1.34 |
| 0 | Cthrc1 | 2.47 | 1.84E-65 | 3.93E-61 | 0.31 | 0.07 | 0.23 | 4.21 |
| 0 | Nrg1 | 1.42 | 3.42E-63 | 7.30E-59 | 0.55 | 0.22 | 0.33 | 2.48 |
| 0 | Nt5dc2 | 0.65 | 2.94E-62 | 6.28E-58 | 0.98 | 0.91 | 0.07 | 1.08 |
| 0 | Six3os1 | 0.86 | 9.96E-62 | 2.13E-57 | 0.93 | 0.74 | 0.19 | 1.26 |
| 0 | Fgf12 | 1.77 | 1.23E-59 | 2.63E-55 | 0.39 | 0.13 | 0.27 | 3.1 |
| 0 | Meis2 | 0.68 | 8.20E-58 | 1.75E-53 | 0.99 | 0.95 | 0.04 | 1.05 |
| 0 | Zfhx3 | 0.79 | 1.30E-56 | 2.79E-52 | 0.98 | 0.91 | 0.07 | 1.08 |
| 0 | Lmcd1 | 3.1 | 6.53E-56 | 1.40E-51 | 0.2 | 0.03 | 0.16 | 5.91 |
| 0 | Fgf9 | 1.03 | 1.45E-55 | 3.10E-51 | 0.72 | 0.38 | 0.34 | 1.89 |
| 0 | Meis1 | 0.97 | 7.45E-55 | 1.59E-50 | 0.81 | 0.55 | 0.26 | 1.46 |
| 0 | Pcdh19 | 1.38 | 6.49E-54 | 1.39E-49 | 0.49 | 0.2 | 0.29 | 2.47 |
| 0 | Maob | 2.52 | 2.76E-49 | 5.89E-45 | 0.19 | 0.04 | 0.16 | 5.31 |
| 0 | Tnnt1 | 2.15 | 3.45E-49 | 7.37E-45 | 0.25 | 0.06 | 0.19 | 4.05 |
| 0 | Gm49708 | 1.45 | 7.94E-49 | 1.70E-44 | 0.71 | 0.46 | 0.26 | 1.56 |
| 0 | Dmrt3 | 1.46 | 8.58E-49 | 1.83E-44 | 0.44 | 0.17 | 0.27 | 2.54 |
| 0 | Grb10 | 0.66 | 1.71E-48 | 3.65E-44 | 0.98 | 0.92 | 0.06 | 1.06 |
| 0 | Nfib | 0.73 | 9.11E-47 | 1.95E-42 | 0.92 | 0.77 | 0.15 | 1.2 |
| 0 | Ccdc171 | 1.5 | 1.93E-46 | 4.13E-42 | 0.44 | 0.18 | 0.26 | 2.42 |
| 0 | Gxylt2 | 1.73 | 2.25E-46 | 4.80E-42 | 0.27 | 0.07 | 0.2 | 3.67 |
| 0 | Car14 | 1.4 | 2.75E-46 | 5.87E-42 | 0.46 | 0.2 | 0.27 | 2.33 |
| 0 | Foxc2 | 2.81 | 3.76E-45 | 8.04E-41 | 0.16 | 0.03 | 0.13 | 5.79 |
| 0 | Pcbp4 | 0.5 | 5.12E-45 | 1.09E-40 | 0.98 | 0.9 | 0.08 | 1.09 |
| 0 | Kitl | 1.73 | 1.47E-44 | 3.14E-40 | 0.32 | 0.11 | 0.21 | 2.96 |
| 0 | Rragd | 0.91 | 3.61E-44 | 7.72E-40 | 0.68 | 0.37 | 0.31 | 1.82 |
| 0 | Gsta4 | 0.68 | 6.98E-44 | 1.49E-39 | 0.93 | 0.79 | 0.14 | 1.18 |
| 0 | Nrxn3 | 2.12 | 1.56E-43 | 3.34E-39 | 0.27 | 0.08 | 0.19 | 3.44 |
| 0 | Six3 | 0.64 | 3.96E-43 | 8.46E-39 | 0.97 | 0.82 | 0.15 | 1.18 |
| 0 | Ddx3y | 2.38 | 7.56E-42 | 1.61E-37 | 0.15 | 0.02 | 0.12 | 6.13 |
| 0 | Scn9a | 2.08 | 3.80E-41 | 8.12E-37 | 0.15 | 0.03 | 0.13 | 5.7 |
| 0 | Ltbp1 | 1.34 | 1.16E-40 | 2.47E-36 | 0.46 | 0.21 | 0.25 | 2.16 |
| 0 | Eif2s3y | 2.05 | 4.26E-40 | 9.11E-36 | 0.16 | 0.03 | 0.13 | 5.23 |
| 0 | Grk5 | 1.59 | 4.57E-40 | 9.76E-36 | 0.33 | 0.12 | 0.21 | 2.74 |
| 0 | Il17rd | 0.85 | 5.69E-40 | 1.22E-35 | 0.75 | 0.46 | 0.29 | 1.62 |
| 0 | Cldn9 | 0.97 | 1.13E-38 | 2.41E-34 | 0.59 | 0.31 | 0.28 | 1.88 |
| 0 | Ptchd1 | 1.79 | 1.15E-38 | 2.45E-34 | 0.22 | 0.06 | 0.16 | 3.76 |
| 0 | Fam107a | 1.31 | 1.16E-38 | 2.47E-34 | 0.36 | 0.13 | 0.22 | 2.65 |
| 0 | Zfp608 | 0.89 | 5.83E-38 | 1.24E-33 | 0.79 | 0.58 | 0.2 | 1.35 |
| 0 | Tbc1d1 | 1.03 | 9.62E-38 | 2.05E-33 | 0.64 | 0.38 | 0.25 | 1.66 |
| 0 | Pitx1 | 0.88 | 1.16E-37 | 2.48E-33 | 0.84 | 0.63 | 0.21 | 1.34 |
| 0 | Rbms3 | 1.64 | 1.35E-37 | 2.88E-33 | 0.33 | 0.13 | 0.2 | 2.6 |
| 0 | Egr1 | 2.24 | 2.41E-36 | 5.14E-32 | 0.25 | 0.08 | 0.17 | 3.07 |

**Supplementary Table 3. Marker genes for the original clusters of integrated clustering of E11.5 control and NotchGOF λ-junction ectoderm cells.**

| | | | | | | | | |
|---|---|---|---|---|---|---|---|---|
| 0 | Ubash3b | 1.23 | 1.67E-35 | 3.57E-31 | 0.45 | 0.21 | 0.24 | 2.11 |
| 0 | Psip1 | 0.54 | 2.12E-34 | 4.52E-30 | 1 | 0.98 | 0.02 | 1.02 |
| 0 | Tril | 1.31 | 1.85E-33 | 3.95E-29 | 0.5 | 0.27 | 0.23 | 1.88 |
| 0 | Chd3 | 0.49 | 4.38E-33 | 9.35E-29 | 0.99 | 0.93 | 0.06 | 1.06 |
| 0 | Hdgf | 0.33 | 8.40E-33 | 1.79E-28 | 1 | 0.98 | 0.01 | 1.01 |
| 0 | Gsn | 2.02 | 1.11E-32 | 2.37E-28 | 0.2 | 0.06 | 0.14 | 3.38 |
| 0 | Fgf17 | 1.42 | 3.24E-32 | 6.93E-28 | 0.33 | 0.14 | 0.2 | 2.4 |
| 0 | Nrip1 | 0.59 | 7.84E-32 | 1.67E-27 | 0.94 | 0.8 | 0.14 | 1.18 |
| 0 | Adarb2 | 2.52 | 1.03E-31 | 2.20E-27 | 0.15 | 0.03 | 0.11 | 4.45 |
| 0 | Flrt3 | 0.97 | 1.18E-31 | 2.53E-27 | 0.6 | 0.37 | 0.23 | 1.62 |
| 0 | Ascl1 | 1.8 | 2.38E-31 | 5.08E-27 | 0.1 | 0.02 | 0.09 | 6.5 |
| 0 | Foxg1 | 0.64 | 5.21E-31 | 1.11E-26 | 0.77 | 0.5 | 0.27 | 1.53 |
| 0 | Nfia | 0.92 | 8.57E-31 | 1.83E-26 | 0.58 | 0.34 | 0.24 | 1.7 |
| 0 | Kdm5d | 2.41 | 3.27E-30 | 6.98E-26 | 0.11 | 0.02 | 0.09 | 6.24 |
| 0 | Skp1a | 0.4 | 3.31E-30 | 7.08E-26 | 0.99 | 0.97 | 0.02 | 1.02 |
| 0 | Limch1 | 1.28 | 3.35E-30 | 7.15E-26 | 0.52 | 0.32 | 0.21 | 1.66 |
| 0 | Dmrta2 | 0.7 | 4.12E-29 | 8.81E-25 | 0.73 | 0.47 | 0.26 | 1.55 |
| 0 | Espn | 1.01 | 1.68E-28 | 3.58E-24 | 0.56 | 0.34 | 0.22 | 1.63 |
| 0 | Uchl1 | 0.32 | 2.01E-28 | 4.29E-24 | 0.82 | 0.62 | 0.2 | 1.33 |
| 0 | Hmcn1 | 1.15 | 2.30E-28 | 4.91E-24 | 0.38 | 0.18 | 0.2 | 2.13 |
| 0 | Mpped2 | 0.75 | 2.61E-28 | 5.57E-24 | 0.88 | 0.77 | 0.11 | 1.15 |
| 0 | Tmem176b | 0.74 | 3.16E-28 | 6.76E-24 | 0.77 | 0.58 | 0.19 | 1.33 |
| 0 | Homer2 | 1.04 | 3.46E-28 | 7.40E-24 | 0.38 | 0.18 | 0.2 | 2.11 |
| 0 | Asb7 | 1.39 | 1.64E-27 | 3.49E-23 | 0.42 | 0.22 | 0.2 | 1.9 |
| 0 | Tsc22d2 | 0.87 | 2.55E-27 | 5.45E-23 | 0.63 | 0.4 | 0.23 | 1.56 |
| 0 | Bcl11a | 0.51 | 3.02E-27 | 6.45E-23 | 0.91 | 0.78 | 0.13 | 1.17 |
| 0 | P4ha3 | 1.24 | 3.43E-27 | 7.33E-23 | 0.32 | 0.14 | 0.18 | 2.29 |
| 0 | Fgf16 | 1.78 | 7.68E-27 | 1.64E-22 | 0.18 | 0.05 | 0.12 | 3.32 |
| 0 | Id2 | 0.53 | 1.50E-26 | 3.20E-22 | 0.94 | 0.81 | 0.12 | 1.15 |
| 0 | Tshz2 | 0.93 | 3.48E-26 | 7.44E-22 | 0.59 | 0.37 | 0.22 | 1.61 |
| 0 | Dusp5 | 1.05 | 1.05E-25 | 2.25E-21 | 0.37 | 0.18 | 0.19 | 2.08 |
| 0 | Six1 | 0.47 | 2.50E-25 | 5.35E-21 | 0.9 | 0.65 | 0.25 | 1.38 |
| 0 | Sp8 | 0.65 | 3.88E-25 | 8.29E-21 | 0.72 | 0.49 | 0.23 | 1.47 |
| 0 | Sncaip | 1.32 | 4.05E-25 | 8.64E-21 | 0.29 | 0.12 | 0.16 | 2.32 |
| 0 | Hspa8 | 0.29 | 4.25E-25 | 9.08E-21 | 1 | 1 | 0 | 1 |
| 0 | Rdh10 | 1.07 | 7.70E-25 | 1.64E-20 | 0.47 | 0.28 | 0.2 | 1.71 |
| 0 | Gse1 | 0.81 | 1.10E-24 | 2.34E-20 | 0.45 | 0.24 | 0.21 | 1.86 |
| 0 | Cacna2d3 | 1.63 | 1.11E-24 | 2.37E-20 | 0.18 | 0.06 | 0.12 | 3.07 |
| 0 | Sema3c | 1 | 1.31E-24 | 2.81E-20 | 0.43 | 0.23 | 0.2 | 1.86 |
| 0 | Olfm2 | 1.01 | 1.37E-24 | 2.92E-20 | 0.33 | 0.15 | 0.18 | 2.17 |
| 0 | Card19 | 1.05 | 2.02E-24 | 4.30E-20 | 0.44 | 0.25 | 0.19 | 1.78 |
| 0 | Ntn1 | 1.41 | 2.45E-24 | 5.24E-20 | 0.21 | 0.07 | 0.13 | 2.89 |
| 0 | Zfhx4 | 0.54 | 4.94E-24 | 1.05E-19 | 0.89 | 0.75 | 0.14 | 1.18 |
| 0 | Crlf1 | 1.72 | 6.67E-24 | 1.43E-19 | 0.26 | 0.11 | 0.15 | 2.35 |
| 0 | Slit2 | 0.7 | 7.90E-24 | 1.69E-19 | 0.58 | 0.37 | 0.21 | 1.58 |
| 0 | Tle5 | 0.33 | 9.10E-24 | 1.94E-19 | 0.98 | 0.96 | 0.03 | 1.03 |
| 0 | Lypd6b | 1.13 | 1.17E-23 | 2.49E-19 | 0.39 | 0.21 | 0.18 | 1.87 |
| 0 | Ctxn3 | 1.61 | 1.39E-23 | 2.97E-19 | 0.25 | 0.1 | 0.15 | 2.39 |
| 0 | Syndig1 | 1.51 | 3.35E-23 | 7.15E-19 | 0.13 | 0.04 | 0.1 | 3.8 |
| 0 | Fat3 | 1.46 | 4.09E-23 | 8.73E-19 | 0.29 | 0.13 | 0.15 | 2.18 |
| 0 | Gas1 | 0.52 | 4.66E-23 | 9.95E-19 | 0.75 | 0.54 | 0.21 | 1.39 |
| 0 | Gchfr | 1.11 | 5.11E-23 | 1.09E-18 | 0.28 | 0.12 | 0.16 | 2.32 |
| 0 | Smim4 | 0.62 | 1.04E-22 | 2.23E-18 | 0.72 | 0.51 | 0.21 | 1.42 |
| 0 | Cenpv | 0.7 | 1.74E-22 | 3.72E-18 | 0.57 | 0.35 | 0.22 | 1.62 |
| 0 | Prr15 | 1.18 | 4.82E-22 | 1.03E-17 | 0.26 | 0.11 | 0.15 | 2.37 |
| 0 | St3gal1 | 1.94 | 1.07E-21 | 2.29E-17 | 0.12 | 0.03 | 0.09 | 3.87 |
| 0 | Phgdh | 0.43 | 4.10E-21 | 8.75E-17 | 0.93 | 0.82 | 0.1 | 1.13 |

**Supplementary Table 3. Marker genes for the original clusters of integrated clustering of E11.5 control and NotchGOF λ-junction ectoderm cells.**

| | | | | | | | | |
|---|---|---|---|---|---|---|---|---|
| 0 | Clu | 0.92 | 5.34E-21 | 1.14E-16 | 0.52 | 0.32 | 0.2 | 1.6 |
| 0 | Zfp536 | 1.2 | 5.54E-21 | 1.18E-16 | 0.3 | 0.14 | 0.15 | 2.06 |
| 0 | Ralgapa2 | 1.08 | 5.75E-21 | 1.23E-16 | 0.36 | 0.19 | 0.17 | 1.92 |
| 0 | Mmp16 | 1.26 | 5.91E-21 | 1.26E-16 | 0.26 | 0.12 | 0.14 | 2.23 |
| 0 | Eya1 | 0.53 | 7.85E-21 | 1.68E-16 | 0.79 | 0.56 | 0.23 | 1.4 |
| 0 | Cpne8 | 1.6 | 1.07E-20 | 2.28E-16 | 0.19 | 0.07 | 0.12 | 2.7 |
| 0 | Ddc | 1.61 | 2.11E-20 | 4.52E-16 | 0.19 | 0.07 | 0.12 | 2.63 |
| 0 | Prss22 | 1.01 | 2.49E-20 | 5.32E-16 | 0.4 | 0.22 | 0.18 | 1.79 |
| 0 | Has3 | 2.06 | 2.83E-20 | 6.05E-16 | 0.1 | 0.02 | 0.08 | 4.21 |
| 0 | Tmem108 | 0.94 | 3.07E-20 | 6.56E-16 | 0.47 | 0.28 | 0.19 | 1.67 |
| 0 | Cd24a | 0.3 | 3.49E-20 | 7.45E-16 | 0.99 | 0.96 | 0.03 | 1.03 |
| 0 | Nnat | 0.39 | 3.65E-20 | 7.79E-16 | 0.99 | 0.95 | 0.04 | 1.04 |
| 0 | Podn | 1.97 | 4.83E-20 | 1.03E-15 | 0.13 | 0.04 | 0.09 | 3.28 |
| 0 | Stk32a | 1.99 | 6.03E-20 | 1.29E-15 | 0.1 | 0.03 | 0.08 | 4.08 |
| 0 | Galnt7 | 0.7 | 7.68E-20 | 1.64E-15 | 0.62 | 0.43 | 0.19 | 1.43 |
| 0 | Trim2 | 0.52 | 9.74E-20 | 2.08E-15 | 0.77 | 0.56 | 0.21 | 1.37 |
| 0 | Adamts5 | 1.9 | 1.79E-19 | 3.83E-15 | 0.12 | 0.03 | 0.09 | 3.5 |
| 0 | Slco5a1 | 1.07 | 2.55E-19 | 5.45E-15 | 0.3 | 0.15 | 0.15 | 2.01 |
| 0 | Azin1 | 0.55 | 7.96E-19 | 1.70E-14 | 0.71 | 0.53 | 0.18 | 1.34 |
| 0 | Tcf4 | 0.4 | 8.60E-19 | 1.84E-14 | 0.94 | 0.88 | 0.06 | 1.07 |
| 0 | Lmod1 | 1.57 | 8.70E-19 | 1.86E-14 | 0.14 | 0.05 | 0.1 | 3.02 |
| 0 | 3110039M20Rik | 0.56 | 1.01E-18 | 2.16E-14 | 0.47 | 0.28 | 0.2 | 1.7 |
| 0 | Ppa1 | 0.45 | 1.84E-18 | 3.92E-14 | 0.92 | 0.87 | 0.05 | 1.06 |
| 0 | Lrrn1 | 1.72 | 2.14E-18 | 4.57E-14 | 0.13 | 0.04 | 0.09 | 3.29 |
| 0 | Nhsl2 | 1.11 | 2.34E-18 | 4.99E-14 | 0.24 | 0.11 | 0.13 | 2.19 |
| 0 | Arhgef28 | 1 | 2.61E-18 | 5.58E-14 | 0.27 | 0.13 | 0.14 | 2.11 |
| 0 | Hs6st2 | 0.63 | 2.75E-18 | 5.87E-14 | 0.7 | 0.52 | 0.18 | 1.36 |
| 0 | Lhfpl2 | 1.05 | 3.43E-18 | 7.33E-14 | 0.28 | 0.14 | 0.14 | 2.02 |
| 0 | Tmem59l | 1.27 | 3.47E-18 | 7.40E-14 | 0.19 | 0.07 | 0.11 | 2.54 |
| 0 | Sobp | 0.88 | 3.48E-18 | 7.43E-14 | 0.39 | 0.22 | 0.17 | 1.75 |
| 0 | Lrig3 | 0.81 | 8.08E-18 | 1.73E-13 | 0.45 | 0.27 | 0.18 | 1.65 |
| 0 | Abr | 0.92 | 8.99E-18 | 1.92E-13 | 0.36 | 0.2 | 0.16 | 1.83 |
| 0 | Cdon | 0.94 | 9.52E-18 | 2.03E-13 | 0.4 | 0.24 | 0.16 | 1.68 |
| 0 | Eya4 | 0.83 | 1.16E-17 | 2.48E-13 | 0.36 | 0.2 | 0.16 | 1.83 |
| 0 | Met | 1.27 | 1.71E-17 | 3.66E-13 | 0.21 | 0.09 | 0.12 | 2.27 |
| 0 | Fos | 2.24 | 2.05E-17 | 4.37E-13 | 0.22 | 0.1 | 0.12 | 2.14 |
| 0 | Chmp2b | 0.47 | 2.63E-17 | 5.62E-13 | 0.86 | 0.72 | 0.14 | 1.2 |
| 0 | Hif3a | 1 | 3.68E-17 | 7.86E-13 | 0.22 | 0.1 | 0.13 | 2.26 |
| 0 | Sox12 | 0.51 | 7.02E-17 | 1.50E-12 | 0.75 | 0.58 | 0.17 | 1.3 |
| 0 | Ddhd2 | 0.61 | 1.52E-16 | 3.25E-12 | 0.43 | 0.25 | 0.18 | 1.71 |
| 0 | Ppargc1a | 1.59 | 1.89E-16 | 4.04E-12 | 0.12 | 0.04 | 0.08 | 3.08 |
| 0 | Tmod1 | 1.81 | 2.19E-16 | 4.68E-12 | 0.11 | 0.04 | 0.08 | 3.23 |
| 0 | Sesn3 | 0.53 | 2.82E-16 | 6.03E-12 | 0.75 | 0.6 | 0.15 | 1.25 |
| 0 | Pik3ap1 | 1.53 | 3.34E-16 | 7.13E-12 | 0.12 | 0.04 | 0.08 | 3.03 |
| 0 | Mex3b | 0.54 | 3.77E-16 | 8.05E-12 | 0.63 | 0.44 | 0.18 | 1.41 |
| 0 | Selenbp1 | 1.48 | 3.87E-16 | 8.28E-12 | 0.13 | 0.04 | 0.08 | 2.98 |
| 0 | Slc27a3 | 1.01 | 4.09E-16 | 8.73E-12 | 0.29 | 0.15 | 0.14 | 1.93 |
| 0 | Col11a1 | 0.62 | 5.55E-16 | 1.19E-11 | 0.71 | 0.55 | 0.16 | 1.3 |
| 0 | Dsel | 0.63 | 5.80E-16 | 1.24E-11 | 0.54 | 0.36 | 0.18 | 1.49 |
| 0 | Thbs1 | 0.89 | 6.78E-16 | 1.45E-11 | 0.35 | 0.2 | 0.15 | 1.74 |
| 0 | Pla2r1 | 0.49 | 7.46E-16 | 1.59E-11 | 0.43 | 0.26 | 0.18 | 1.69 |
| 0 | Prim1 | 0.66 | 7.84E-16 | 1.68E-11 | 0.7 | 0.56 | 0.14 | 1.26 |
| 0 | Emb | 0.28 | 1.09E-15 | 2.33E-11 | 0.89 | 0.71 | 0.17 | 1.24 |
| 0 | Isl1 | 0.4 | 1.38E-15 | 2.95E-11 | 0.89 | 0.73 | 0.16 | 1.22 |
| 0 | Serpinh1 | 0.29 | 2.06E-15 | 4.39E-11 | 0.99 | 0.95 | 0.04 | 1.04 |
| 0 | Cox20 | 0.52 | 2.24E-15 | 4.78E-11 | 0.72 | 0.57 | 0.15 | 1.27 |
| 0 | Jade1 | 0.77 | 2.24E-15 | 4.79E-11 | 0.47 | 0.3 | 0.16 | 1.54 |

**Supplementary Table 3. Marker genes for the original clusters of integrated clustering of E11.5 control and NotchGOF λ-junction ectoderm cells.**

| | | | | | | | | |
|---|---|---|---|---|---|---|---|---|
| 0 | Gli3 | 0.48 | 2.42E-15 | 5.17E-11 | 0.7 | 0.5 | 0.2 | 1.4 |
| 0 | Bnip3 | 0.52 | 2.43E-15 | 5.19E-11 | 0.62 | 0.43 | 0.19 | 1.44 |
| 0 | Cacnb2 | 1.16 | 2.50E-15 | 5.35E-11 | 0.21 | 0.1 | 0.11 | 2.14 |
| 0 | Mov10 | 0.81 | 3.00E-15 | 6.41E-11 | 0.34 | 0.2 | 0.15 | 1.75 |
| 0 | Cd47 | 0.6 | 3.10E-15 | 6.63E-11 | 0.71 | 0.55 | 0.16 | 1.29 |
| 0 | 2010300C02Rik | 1.29 | 3.53E-15 | 7.55E-11 | 0.18 | 0.08 | 0.1 | 2.29 |
| 0 | Ctnna2 | 1.14 | 3.56E-15 | 7.60E-11 | 0.16 | 0.06 | 0.1 | 2.5 |
| 0 | Rcc2 | 0.27 | 4.84E-15 | 1.03E-10 | 0.97 | 0.93 | 0.04 | 1.04 |
| 0 | Tmem47 | 0.43 | 5.47E-15 | 1.17E-10 | 0.6 | 0.43 | 0.17 | 1.4 |
| 0 | Cdh2 | 0.27 | 7.80E-15 | 1.67E-10 | 0.82 | 0.58 | 0.24 | 1.42 |
| 0 | Nog | 1.19 | 8.18E-15 | 1.75E-10 | 0.21 | 0.1 | 0.11 | 2.09 |
| 0 | Epha7 | 0.92 | 1.27E-14 | 2.71E-10 | 0.42 | 0.27 | 0.15 | 1.55 |
| 0 | Prkd1 | 1.25 | 1.64E-14 | 3.51E-10 | 0.17 | 0.07 | 0.1 | 2.37 |
| 0 | Atp2b1 | 0.36 | 2.27E-14 | 4.84E-10 | 0.96 | 0.88 | 0.08 | 1.09 |
| 0 | Tent5a | 1.4 | 2.83E-14 | 6.05E-10 | 0.14 | 0.05 | 0.08 | 2.65 |
| 0 | Bach2 | 0.73 | 3.14E-14 | 6.71E-10 | 0.38 | 0.23 | 0.15 | 1.67 |
| 0 | Tead2 | 0.31 | 3.59E-14 | 7.67E-10 | 0.94 | 0.86 | 0.09 | 1.1 |
| 0 | Slc44a3 | 1.6 | 3.62E-14 | 7.74E-10 | 0.1 | 0.03 | 0.07 | 3.06 |
| 0 | Hdgfl3 | 0.51 | 4.12E-14 | 8.80E-10 | 0.67 | 0.52 | 0.16 | 1.31 |
| 0 | Samd1 | 0.68 | 5.90E-14 | 1.26E-09 | 0.45 | 0.3 | 0.16 | 1.53 |
| 0 | Ccn1 | 1.52 | 6.22E-14 | 1.33E-09 | 0.3 | 0.18 | 0.12 | 1.68 |
| 0 | Rest | 0.42 | 7.32E-14 | 1.56E-09 | 0.81 | 0.66 | 0.14 | 1.21 |
| 0 | Vgll3 | 1.18 | 8.16E-14 | 1.74E-09 | 0.19 | 0.09 | 0.1 | 2.19 |
| 0 | Myc | 0.93 | 8.26E-14 | 1.76E-09 | 0.33 | 0.2 | 0.13 | 1.68 |
| 0 | Pax6 | 0.44 | 8.51E-14 | 1.82E-09 | 0.66 | 0.46 | 0.2 | 1.42 |
| 0 | Sec62 | 0.4 | 8.98E-14 | 1.92E-09 | 0.96 | 0.92 | 0.04 | 1.04 |
| 0 | Ces1d | 0.91 | 1.10E-13 | 2.34E-09 | 0.29 | 0.16 | 0.13 | 1.8 |
| 0 | Tmem30b | 0.65 | 1.14E-13 | 2.44E-09 | 0.41 | 0.25 | 0.16 | 1.62 |
| 0 | Tmem39b | 0.6 | 1.27E-13 | 2.71E-09 | 0.46 | 0.3 | 0.16 | 1.53 |
| 0 | Kif26b | 1.81 | 1.40E-13 | 2.99E-09 | 0.12 | 0.04 | 0.07 | 2.76 |
| 0 | Capn6 | 0.42 | 2.88E-13 | 6.16E-09 | 0.61 | 0.43 | 0.18 | 1.43 |
| 0 | Msrb3 | 1.1 | 3.27E-13 | 6.99E-09 | 0.21 | 0.11 | 0.11 | 2 |
| 0 | Kit | 0.77 | 3.55E-13 | 7.58E-09 | 0.29 | 0.16 | 0.13 | 1.79 |
| 0 | Nek7 | 0.76 | 4.31E-13 | 9.21E-09 | 0.47 | 0.31 | 0.16 | 1.5 |
| 0 | Zfp703 | 0.62 | 4.60E-13 | 9.83E-09 | 0.44 | 0.28 | 0.16 | 1.56 |
| 0 | Zfp827 | 0.64 | 5.85E-13 | 1.25E-08 | 0.42 | 0.27 | 0.15 | 1.57 |
| 0 | Zfp467 | 0.53 | 5.96E-13 | 1.27E-08 | 0.61 | 0.45 | 0.16 | 1.36 |
| 0 | Plekho1 | 0.55 | 6.28E-13 | 1.34E-08 | 0.51 | 0.34 | 0.17 | 1.5 |
| 0 | Gad2 | 0.78 | 6.42E-13 | 1.37E-08 | 0.43 | 0.27 | 0.15 | 1.56 |
| 0 | Rrp12 | 0.65 | 7.88E-13 | 1.68E-08 | 0.37 | 0.23 | 0.14 | 1.6 |
| 0 | Apoe | 0.55 | 1.11E-12 | 2.37E-08 | 0.63 | 0.48 | 0.15 | 1.32 |
| 0 | Dhx32 | 0.61 | 1.11E-12 | 2.37E-08 | 0.45 | 0.3 | 0.15 | 1.5 |
| 0 | Mir100hg | 0.44 | 1.19E-12 | 2.54E-08 | 0.77 | 0.61 | 0.16 | 1.25 |
| 0 | Cdv3 | 0.31 | 1.22E-12 | 2.60E-08 | 0.97 | 0.94 | 0.03 | 1.03 |
| 0 | Serpine2 | 0.54 | 1.57E-12 | 3.35E-08 | 0.61 | 0.45 | 0.17 | 1.37 |
| 0 | Mecom | 0.27 | 1.86E-12 | 3.98E-08 | 0.62 | 0.43 | 0.19 | 1.44 |
| 0 | Mfap2 | 0.29 | 1.96E-12 | 4.19E-08 | 0.96 | 0.89 | 0.07 | 1.07 |
| 0 | Ociad2 | 0.63 | 1.98E-12 | 4.23E-08 | 0.44 | 0.28 | 0.16 | 1.55 |
| 0 | Add3 | 0.52 | 2.06E-12 | 4.39E-08 | 0.71 | 0.58 | 0.13 | 1.22 |
| 0 | Parva | 0.42 | 2.38E-12 | 5.09E-08 | 0.75 | 0.63 | 0.13 | 1.2 |
| 0 | Pax3 | 1.45 | 2.64E-12 | 5.65E-08 | 0.13 | 0.05 | 0.08 | 2.52 |
| 0 | Atp8b1 | 0.97 | 3.26E-12 | 6.97E-08 | 0.23 | 0.12 | 0.11 | 1.89 |
| 0 | Maoa | 0.59 | 3.47E-12 | 7.42E-08 | 0.41 | 0.27 | 0.14 | 1.54 |
| 0 | Plat | 0.61 | 4.19E-12 | 8.94E-08 | 0.31 | 0.18 | 0.13 | 1.74 |
| 0 | Dtna | 0.53 | 5.90E-12 | 1.26E-07 | 0.49 | 0.33 | 0.16 | 1.48 |
| 0 | Mllt3 | 0.48 | 6.11E-12 | 1.30E-07 | 0.69 | 0.54 | 0.15 | 1.29 |
| 0 | Glrx2 | 0.36 | 6.19E-12 | 1.32E-07 | 0.79 | 0.67 | 0.12 | 1.17 |

**Supplementary Table 3. Marker genes for the original clusters of integrated clustering of E11.5 control and NotchGOF λ-junction ectoderm cells.**

| | | | | | | | | |
|---|---|---|---|---|---|---|---|---|
| 0 | Man2a1 | 0.71 | 6.62E-12 | 1.41E-07 | 0.36 | 0.23 | 0.14 | 1.59 |
| 0 | Rab27a | 1.65 | 7.20E-12 | 1.54E-07 | 0.11 | 0.05 | 0.07 | 2.53 |
| 0 | Cnn3 | 0.25 | 7.71E-12 | 1.65E-07 | 0.97 | 0.93 | 0.05 | 1.05 |
| 0 | Ly6h | 0.71 | 8.69E-12 | 1.86E-07 | 0.61 | 0.48 | 0.13 | 1.26 |
| 0 | Nans | 0.41 | 9.00E-12 | 1.92E-07 | 0.67 | 0.51 | 0.16 | 1.31 |
| 0 | Frat2 | 0.46 | 9.48E-12 | 2.03E-07 | 0.53 | 0.36 | 0.17 | 1.47 |
| 0 | Tmem176a | 0.48 | 1.49E-11 | 3.19E-07 | 0.56 | 0.4 | 0.16 | 1.39 |
| 0 | Cxxc4 | 0.41 | 1.55E-11 | 3.32E-07 | 0.38 | 0.24 | 0.14 | 1.6 |
| 0 | F3 | 1.24 | 1.85E-11 | 3.96E-07 | 0.34 | 0.22 | 0.12 | 1.56 |
| 0 | Lgr5 | 1.12 | 2.48E-11 | 5.30E-07 | 0.34 | 0.23 | 0.11 | 1.48 |
| 0 | Bphl | 0.44 | 2.75E-11 | 5.88E-07 | 0.65 | 0.49 | 0.16 | 1.32 |
| 0 | Ntf3 | 0.71 | 2.82E-11 | 6.02E-07 | 0.37 | 0.23 | 0.13 | 1.57 |
| 0 | Tanc2 | 0.75 | 2.96E-11 | 6.33E-07 | 0.33 | 0.21 | 0.12 | 1.59 |
| 0 | Msl3 | 0.5 | 3.05E-11 | 6.52E-07 | 0.57 | 0.42 | 0.15 | 1.37 |
| 0 | Otx1 | 0.41 | 3.14E-11 | 6.71E-07 | 0.58 | 0.42 | 0.16 | 1.38 |
| 0 | Kcnd2 | 1.19 | 3.54E-11 | 7.55E-07 | 0.14 | 0.06 | 0.08 | 2.26 |
| 0 | Pdgfc | 0.26 | 4.75E-11 | 1.01E-06 | 0.89 | 0.74 | 0.14 | 1.19 |
| 0 | Galm | 0.46 | 5.98E-11 | 1.28E-06 | 0.53 | 0.38 | 0.15 | 1.38 |
| 0 | Fam171b | 0.62 | 8.39E-11 | 1.79E-06 | 0.23 | 0.12 | 0.11 | 1.88 |
| 0 | Psmg4 | 0.33 | 1.11E-10 | 2.38E-06 | 0.83 | 0.74 | 0.09 | 1.13 |
| 0 | Rnf121 | 0.52 | 1.20E-10 | 2.56E-06 | 0.55 | 0.41 | 0.14 | 1.34 |
| 0 | Bnipl | 0.78 | 1.26E-10 | 2.68E-06 | 0.29 | 0.17 | 0.12 | 1.68 |
| 0 | Mtus1 | 0.52 | 1.34E-10 | 2.86E-06 | 0.42 | 0.28 | 0.14 | 1.51 |
| 0 | Sulf1 | 0.57 | 1.41E-10 | 3.01E-06 | 0.77 | 0.65 | 0.12 | 1.18 |
| 0 | Chn2 | 0.86 | 1.41E-10 | 3.01E-06 | 0.23 | 0.13 | 0.1 | 1.78 |
| 0 | Snx24 | 0.52 | 1.53E-10 | 3.27E-06 | 0.44 | 0.3 | 0.14 | 1.48 |
| 0 | Urod | 0.48 | 1.72E-10 | 3.67E-06 | 0.61 | 0.47 | 0.14 | 1.31 |
| 0 | Cox19 | 0.3 | 1.76E-10 | 3.75E-06 | 0.78 | 0.64 | 0.14 | 1.22 |
| 0 | Krtcap3 | 0.58 | 1.80E-10 | 3.85E-06 | 0.47 | 0.31 | 0.15 | 1.48 |
| 0 | Myl9 | 0.9 | 2.00E-10 | 4.27E-06 | 0.29 | 0.18 | 0.12 | 1.64 |
| 0 | Pak3 | 0.61 | 2.26E-10 | 4.83E-06 | 0.42 | 0.29 | 0.13 | 1.45 |
| 0 | 2700069I18Rik | 1.28 | 2.29E-10 | 4.90E-06 | 0.13 | 0.06 | 0.07 | 2.24 |
| 0 | Nmrk1 | 0.93 | 2.50E-10 | 5.34E-06 | 0.19 | 0.1 | 0.09 | 1.89 |
| 0 | Gabra4 | 1.36 | 2.55E-10 | 5.45E-06 | 0.1 | 0.04 | 0.06 | 2.49 |
| 0 | St3gal5 | 0.55 | 2.61E-10 | 5.57E-06 | 0.35 | 0.23 | 0.13 | 1.56 |
| 0 | Mxd4 | 0.39 | 2.68E-10 | 5.73E-06 | 0.55 | 0.4 | 0.15 | 1.37 |
| 0 | Ube2j1 | 0.49 | 2.85E-10 | 6.08E-06 | 0.39 | 0.26 | 0.14 | 1.53 |
| 0 | 2700033N17Rik | 0.76 | 3.48E-10 | 7.44E-06 | 0.34 | 0.22 | 0.12 | 1.54 |
| 0 | Gm32585 | 0.75 | 3.61E-10 | 7.72E-06 | 0.32 | 0.2 | 0.12 | 1.58 |
| 0 | Dhcr24 | 0.59 | 3.89E-10 | 8.30E-06 | 0.48 | 0.34 | 0.14 | 1.42 |
| 0 | Cdh11 | 0.81 | 4.55E-10 | 9.72E-06 | 0.21 | 0.11 | 0.09 | 1.82 |
| 0 | Vldlr | 1.2 | 5.05E-10 | 1.08E-05 | 0.14 | 0.07 | 0.07 | 2.09 |
| 0 | Pcdh7 | 0.47 | 5.27E-10 | 1.13E-05 | 0.76 | 0.64 | 0.12 | 1.19 |
| 0 | Samd5 | 0.37 | 5.35E-10 | 1.14E-05 | 0.79 | 0.67 | 0.12 | 1.18 |
| 0 | Fbln7 | 1.22 | 5.57E-10 | 1.19E-05 | 0.1 | 0.04 | 0.06 | 2.48 |
| 0 | Slc36a1 | 0.94 | 5.57E-10 | 1.19E-05 | 0.15 | 0.07 | 0.08 | 2.08 |
| 0 | Man1a | 1.07 | 6.46E-10 | 1.38E-05 | 0.11 | 0.04 | 0.06 | 2.47 |
| 0 | Nbea | 0.47 | 6.49E-10 | 1.39E-05 | 0.5 | 0.35 | 0.14 | 1.41 |
| 0 | Sparc | 0.41 | 7.04E-10 | 1.50E-05 | 0.94 | 0.88 | 0.05 | 1.06 |
| 0 | Ctbp2 | 0.38 | 7.04E-10 | 1.50E-05 | 0.71 | 0.57 | 0.14 | 1.24 |
| 0 | Gata3 | 0.87 | 7.56E-10 | 1.61E-05 | 0.44 | 0.33 | 0.11 | 1.33 |
| 0 | Rasgef1b | 0.91 | 8.16E-10 | 1.74E-05 | 0.17 | 0.09 | 0.08 | 1.97 |
| 0 | Dmtn | 0.84 | 8.72E-10 | 1.86E-05 | 0.18 | 0.09 | 0.09 | 1.96 |
| 0 | Tead1 | 0.33 | 9.75E-10 | 2.08E-05 | 0.87 | 0.75 | 0.11 | 1.15 |
| 0 | Twist1 | 0.63 | 1.11E-09 | 2.38E-05 | 0.32 | 0.21 | 0.12 | 1.58 |
| 0 | Zswim6 | 0.53 | 1.16E-09 | 2.47E-05 | 0.5 | 0.36 | 0.14 | 1.38 |
| 0 | Rhobtb3 | 0.28 | 1.20E-09 | 2.57E-05 | 0.86 | 0.75 | 0.1 | 1.14 |

**Supplementary Table 3. Marker genes for the original clusters of integrated clustering of E11.5 control and NotchGOF λ-junction ectoderm cells.**

| | | | | | | | | |
|---|---|---|---|---|---|---|---|---|
| 0 | Errfi1 | 0.61 | 1.23E-09 | 2.63E-05 | 0.43 | 0.3 | 0.13 | 1.44 |
| 0 | Tmem160 | 0.33 | 1.28E-09 | 2.74E-05 | 0.78 | 0.67 | 0.11 | 1.17 |
| 0 | Jag2 | 0.51 | 1.28E-09 | 2.74E-05 | 0.4 | 0.27 | 0.13 | 1.46 |
| 0 | Lmo4 | 0.35 | 1.34E-09 | 2.86E-05 | 0.97 | 0.91 | 0.06 | 1.07 |
| 0 | Kctd12 | 1.29 | 1.42E-09 | 3.04E-05 | 0.18 | 0.1 | 0.09 | 1.87 |
| 0 | Fzd2 | 0.25 | 1.49E-09 | 3.19E-05 | 0.9 | 0.81 | 0.1 | 1.12 |
| 0 | Aldh7a1 | 0.4 | 1.71E-09 | 3.64E-05 | 0.67 | 0.54 | 0.13 | 1.25 |
| 0 | Mir17hg | 0.71 | 1.71E-09 | 3.65E-05 | 0.28 | 0.17 | 0.11 | 1.64 |
| 0 | Slc4a7 | 0.42 | 1.80E-09 | 3.85E-05 | 0.69 | 0.54 | 0.15 | 1.27 |
| 0 | Abcg1 | 0.47 | 1.88E-09 | 4.01E-05 | 0.4 | 0.28 | 0.13 | 1.47 |
| 0 | Nab2 | 0.79 | 2.04E-09 | 4.36E-05 | 0.3 | 0.19 | 0.11 | 1.57 |
| 0 | Slc7a6 | 0.61 | 2.05E-09 | 4.38E-05 | 0.37 | 0.25 | 0.12 | 1.47 |
| 0 | Impad1 | 0.31 | 2.17E-09 | 4.64E-05 | 0.84 | 0.73 | 0.11 | 1.16 |
| 0 | D430041D05Rik | 1.26 | 2.23E-09 | 4.76E-05 | 0.11 | 0.05 | 0.06 | 2.32 |
| 0 | Otud7b | 0.51 | 2.28E-09 | 4.87E-05 | 0.49 | 0.36 | 0.13 | 1.37 |
| 0 | Crim1 | 0.6 | 2.37E-09 | 5.06E-05 | 0.47 | 0.35 | 0.13 | 1.37 |
| 0 | Tut4 | 0.27 | 2.58E-09 | 5.51E-05 | 0.95 | 0.9 | 0.05 | 1.06 |
| 0 | Junb | 0.87 | 2.73E-09 | 5.83E-05 | 0.31 | 0.2 | 0.1 | 1.51 |
| 0 | Six4 | 0.42 | 2.76E-09 | 5.91E-05 | 0.61 | 0.47 | 0.15 | 1.31 |
| 0 | Ankrd6 | 0.94 | 3.19E-09 | 6.81E-05 | 0.25 | 0.16 | 0.09 | 1.6 |
| 0 | Cul1 | 0.29 | 3.23E-09 | 6.91E-05 | 0.9 | 0.85 | 0.05 | 1.06 |
| 0 | Ssc4d | 0.79 | 3.25E-09 | 6.94E-05 | 0.17 | 0.09 | 0.08 | 1.92 |
| 0 | Ppfia3 | 0.52 | 3.28E-09 | 7.00E-05 | 0.41 | 0.28 | 0.13 | 1.44 |
| 0 | Scpep1 | 0.47 | 3.68E-09 | 7.86E-05 | 0.4 | 0.28 | 0.13 | 1.47 |
| 0 | Rnf24 | 0.46 | 3.79E-09 | 8.10E-05 | 0.53 | 0.39 | 0.14 | 1.36 |
| 0 | Ccng1 | 0.54 | 4.60E-09 | 9.82E-05 | 0.33 | 0.22 | 0.12 | 1.54 |
| 0 | Akr1b3 | 0.31 | 4.71E-09 | 1.01E-04 | 0.73 | 0.59 | 0.14 | 1.24 |
| 0 | Acvr2a | 0.42 | 4.85E-09 | 1.04E-04 | 0.63 | 0.52 | 0.11 | 1.21 |
| 0 | Efemp2 | 0.42 | 5.06E-09 | 1.08E-04 | 0.51 | 0.37 | 0.14 | 1.38 |
| 0 | Cldn11 | 0.98 | 5.29E-09 | 1.13E-04 | 0.18 | 0.1 | 0.08 | 1.79 |
| 0 | Tbl1x | 0.26 | 6.74E-09 | 1.44E-04 | 0.89 | 0.8 | 0.09 | 1.11 |
| 0 | Tdrkh | 0.85 | 6.92E-09 | 1.48E-04 | 0.17 | 0.09 | 0.08 | 1.9 |
| 0 | Kcnk2 | 0.79 | 7.07E-09 | 1.51E-04 | 0.47 | 0.36 | 0.11 | 1.3 |
| 0 | Cryl1 | 0.6 | 8.87E-09 | 1.89E-04 | 0.37 | 0.26 | 0.11 | 1.44 |
| 0 | Dmrta2os | 0.83 | 9.70E-09 | 2.07E-04 | 0.15 | 0.08 | 0.08 | 1.97 |
| 0 | Tox3 | 0.36 | 9.96E-09 | 2.13E-04 | 0.42 | 0.3 | 0.12 | 1.42 |
| 0 | Frat1 | 0.66 | 1.15E-08 | 2.46E-04 | 0.16 | 0.08 | 0.08 | 1.93 |
| 0 | Epop | 0.52 | 1.49E-08 | 3.18E-04 | 0.36 | 0.24 | 0.12 | 1.48 |
| 0 | Shroom3 | 0.55 | 1.54E-08 | 3.30E-04 | 0.35 | 0.23 | 0.11 | 1.49 |
| 0 | Btg2 | 0.47 | 1.56E-08 | 3.33E-04 | 0.41 | 0.3 | 0.11 | 1.38 |
| 0 | Tnrc18 | 0.31 | 1.73E-08 | 3.69E-04 | 0.69 | 0.54 | 0.15 | 1.27 |
| 0 | Wipf3 | 0.59 | 2.21E-08 | 4.72E-04 | 0.31 | 0.21 | 0.11 | 1.51 |
| 0 | Cachd1 | 0.44 | 2.23E-08 | 4.76E-04 | 0.45 | 0.32 | 0.13 | 1.4 |
| 0 | Arhgef2 | 0.46 | 2.42E-08 | 5.16E-04 | 0.51 | 0.38 | 0.13 | 1.34 |
| 0 | Serpinf1 | 0.59 | 2.42E-08 | 5.17E-04 | 0.25 | 0.16 | 0.1 | 1.62 |
| 0 | Rprm | 0.38 | 2.43E-08 | 5.18E-04 | 0.49 | 0.36 | 0.13 | 1.35 |
| 0 | Tjap1 | 0.49 | 2.48E-08 | 5.29E-04 | 0.46 | 0.33 | 0.13 | 1.38 |
| 0 | Raver2 | 0.59 | 2.82E-08 | 6.02E-04 | 0.3 | 0.2 | 0.11 | 1.53 |
| 0 | Fbxl7 | 0.43 | 2.82E-08 | 6.03E-04 | 0.45 | 0.32 | 0.13 | 1.4 |
| 0 | Pogk | 0.38 | 2.90E-08 | 6.20E-04 | 0.67 | 0.53 | 0.14 | 1.26 |
| 0 | Tjp3 | 0.57 | 2.95E-08 | 6.29E-04 | 0.28 | 0.18 | 0.1 | 1.57 |
| 0 | Cyp4f16 | 0.8 | 3.31E-08 | 7.06E-04 | 0.16 | 0.09 | 0.08 | 1.86 |
| 0 | Mab21l2 | 0.76 | 3.37E-08 | 7.20E-04 | 0.11 | 0.05 | 0.06 | 2.18 |
| 0 | Sult4a1 | 0.72 | 3.56E-08 | 7.60E-04 | 0.14 | 0.07 | 0.07 | 1.95 |
| 0 | Schip1 | 0.42 | 3.74E-08 | 7.99E-04 | 0.49 | 0.36 | 0.12 | 1.34 |
| 0 | Psat1 | 0.52 | 3.96E-08 | 8.47E-04 | 0.42 | 0.31 | 0.12 | 1.37 |
| 0 | Eci2 | 0.28 | 4.13E-08 | 8.83E-04 | 0.72 | 0.57 | 0.15 | 1.27 |

**Supplementary Table 3. Marker genes for the original clusters of integrated clustering of E11.5 control and NotchGOF λ-junction ectoderm cells.**

| | | | | | | | | |
|---|---|---|---|---|---|---|---|---|
| 0 | Foxn3 | 0.27 | 4.43E-08 | 9.46E-04 | 0.81 | 0.7 | 0.11 | 1.15 |
| 0 | Cpt1c | 0.87 | 4.78E-08 | 1.02E-03 | 0.12 | 0.06 | 0.06 | 2.05 |
| 0 | Fads2 | 0.45 | 5.03E-08 | 1.07E-03 | 0.57 | 0.44 | 0.12 | 1.27 |
| 0 | Lpar4 | 0.72 | 5.16E-08 | 1.10E-03 | 0.29 | 0.19 | 0.1 | 1.55 |
| 0 | Acsl3 | 0.28 | 5.34E-08 | 1.14E-03 | 0.87 | 0.77 | 0.1 | 1.13 |
| 0 | Rbbp8 | 0.34 | 5.35E-08 | 1.14E-03 | 0.79 | 0.66 | 0.13 | 1.19 |
| 0 | Tmem41b | 0.42 | 5.62E-08 | 1.20E-03 | 0.45 | 0.32 | 0.13 | 1.4 |
| 0 | Ttc39b | 1.19 | 5.74E-08 | 1.23E-03 | 0.12 | 0.06 | 0.06 | 2.07 |
| 0 | Gpm6a | 0.58 | 5.75E-08 | 1.23E-03 | 0.22 | 0.13 | 0.09 | 1.67 |
| 0 | Utp14a | 0.37 | 6.21E-08 | 1.33E-03 | 0.64 | 0.52 | 0.13 | 1.24 |
| 0 | Usp49 | 0.59 | 6.66E-08 | 1.42E-03 | 0.18 | 0.1 | 0.08 | 1.81 |
| 0 | Apip | 0.36 | 8.10E-08 | 1.73E-03 | 0.52 | 0.38 | 0.13 | 1.34 |
| 0 | Dmrt2 | 1.11 | 8.21E-08 | 1.75E-03 | 0.16 | 0.09 | 0.07 | 1.78 |
| 0 | Ccdc28b | 0.42 | 8.36E-08 | 1.79E-03 | 0.46 | 0.34 | 0.12 | 1.36 |
| 0 | Bmi1 | 0.36 | 8.46E-08 | 1.81E-03 | 0.51 | 0.38 | 0.13 | 1.34 |
| 0 | Supt20 | 0.35 | 8.72E-08 | 1.86E-03 | 0.68 | 0.57 | 0.11 | 1.19 |
| 0 | Mrps36 | 0.26 | 8.77E-08 | 1.87E-03 | 0.86 | 0.79 | 0.07 | 1.09 |
| 0 | Plcb4 | 0.48 | 9.24E-08 | 1.97E-03 | 0.38 | 0.26 | 0.11 | 1.43 |
| 0 | Avl9 | 0.45 | 9.92E-08 | 2.12E-03 | 0.51 | 0.39 | 0.12 | 1.32 |
| 0 | Ebpl | 0.43 | 1.04E-07 | 2.23E-03 | 0.65 | 0.54 | 0.11 | 1.2 |
| 0 | Mthfd1 | 0.42 | 1.07E-07 | 2.29E-03 | 0.55 | 0.41 | 0.13 | 1.32 |
| 0 | Gli2 | 0.43 | 1.25E-07 | 2.67E-03 | 0.31 | 0.2 | 0.11 | 1.54 |
| 0 | St3gal2 | 0.76 | 1.26E-07 | 2.68E-03 | 0.17 | 0.1 | 0.08 | 1.78 |
| 0 | Lnpep | 0.5 | 1.39E-07 | 2.96E-03 | 0.35 | 0.24 | 0.11 | 1.46 |
| 0 | Gulp1 | 0.61 | 1.43E-07 | 3.05E-03 | 0.23 | 0.14 | 0.09 | 1.61 |
| 0 | Cxcr4 | 0.56 | 1.47E-07 | 3.13E-03 | 0.26 | 0.17 | 0.09 | 1.54 |
| 0 | Myo6 | 0.7 | 1.55E-07 | 3.32E-03 | 0.3 | 0.2 | 0.1 | 1.5 |
| 0 | Noct | 0.37 | 1.57E-07 | 3.35E-03 | 0.3 | 0.2 | 0.11 | 1.56 |
| 0 | Farp1 | 0.37 | 1.63E-07 | 3.48E-03 | 0.53 | 0.4 | 0.13 | 1.32 |
| 0 | Tbl1xr1 | 0.43 | 1.65E-07 | 3.53E-03 | 0.53 | 0.42 | 0.11 | 1.27 |
| 0 | Rab3ip | 0.37 | 1.71E-07 | 3.66E-03 | 0.59 | 0.46 | 0.13 | 1.28 |
| 0 | Tedc1 | 0.57 | 1.73E-07 | 3.69E-03 | 0.32 | 0.22 | 0.1 | 1.46 |
| 0 | Plppr3 | 0.41 | 1.78E-07 | 3.80E-03 | 0.42 | 0.3 | 0.12 | 1.4 |
| 0 | Prss8 | 0.4 | 1.81E-07 | 3.86E-03 | 0.49 | 0.37 | 0.13 | 1.34 |
| 0 | Zfp516 | 0.4 | 1.88E-07 | 4.01E-03 | 0.62 | 0.5 | 0.12 | 1.23 |
| 0 | Upk3bl | 0.29 | 1.92E-07 | 4.09E-03 | 0.4 | 0.29 | 0.12 | 1.41 |
| 0 | Fbxo44 | 0.59 | 1.93E-07 | 4.12E-03 | 0.2 | 0.12 | 0.08 | 1.69 |
| 0 | Nfkbiz | 0.92 | 1.96E-07 | 4.18E-03 | 0.14 | 0.08 | 0.07 | 1.86 |
| 0 | Epha5 | 0.38 | 1.99E-07 | 4.26E-03 | 0.15 | 0.08 | 0.07 | 1.86 |
| 0 | Ica1 | 0.5 | 2.01E-07 | 4.29E-03 | 0.21 | 0.13 | 0.09 | 1.68 |
| 0 | Scnm1 | 0.41 | 2.19E-07 | 4.68E-03 | 0.57 | 0.44 | 0.13 | 1.3 |
| 0 | Zfp511 | 0.4 | 2.38E-07 | 5.08E-03 | 0.48 | 0.36 | 0.12 | 1.34 |
| 0 | Taf1d | 0.26 | 2.63E-07 | 5.62E-03 | 0.9 | 0.86 | 0.04 | 1.05 |
| 0 | Cic | 0.41 | 2.64E-07 | 5.64E-03 | 0.5 | 0.38 | 0.12 | 1.31 |
| 0 | Mrpl4 | 0.29 | 2.66E-07 | 5.68E-03 | 0.67 | 0.54 | 0.12 | 1.23 |
| 0 | Lipa | 0.56 | 2.84E-07 | 6.06E-03 | 0.25 | 0.16 | 0.09 | 1.57 |
| 0 | Clic5 | 1.01 | 3.10E-07 | 6.63E-03 | 0.12 | 0.06 | 0.06 | 1.97 |
| 0 | Cpe | 0.36 | 3.26E-07 | 6.97E-03 | 0.59 | 0.45 | 0.13 | 1.29 |
| 0 | Prdx3 | 0.34 | 3.37E-07 | 7.21E-03 | 0.63 | 0.51 | 0.12 | 1.23 |
| 0 | Rev1 | 0.34 | 3.67E-07 | 7.83E-03 | 0.43 | 0.31 | 0.12 | 1.4 |
| 0 | Ctdsp2 | 0.3 | 3.70E-07 | 7.91E-03 | 0.7 | 0.57 | 0.14 | 1.24 |
| 0 | Dipk1b | 0.32 | 3.91E-07 | 8.36E-03 | 0.61 | 0.48 | 0.13 | 1.26 |
| 0 | Bbip1 | 0.25 | 3.95E-07 | 8.44E-03 | 0.72 | 0.6 | 0.12 | 1.2 |
| 0 | Utp14b | 0.39 | 4.14E-07 | 8.84E-03 | 0.51 | 0.39 | 0.12 | 1.3 |
| 0 | Usp2 | 0.74 | 4.37E-07 | 9.33E-03 | 0.17 | 0.1 | 0.07 | 1.73 |
| 0 | Efnb2 | 0.42 | 4.45E-07 | 9.50E-03 | 0.69 | 0.59 | 0.11 | 1.18 |
| 0 | Pip5k1c | 0.49 | 4.61E-07 | 9.84E-03 | 0.29 | 0.19 | 0.1 | 1.51 |

**Supplementary Table 3. Marker genes for the original clusters of integrated clustering of E11.5 control and NotchGOF λ-junction ectoderm cells.**

| | | | | | | | | |
|---|---|---|---|---|---|---|---|---|
| 0 | Syde2 | 0.61 | 4.80E-07 | 1.03E-02 | 0.21 | 0.13 | 0.08 | 1.64 |
| 0 | Ptp4a3 | 0.57 | 5.08E-07 | 1.09E-02 | 0.23 | 0.15 | 0.09 | 1.6 |
| 0 | Kif26a | 0.61 | 5.23E-07 | 1.12E-02 | 0.22 | 0.14 | 0.08 | 1.61 |
| 0 | Nampt | 0.37 | 5.31E-07 | 1.13E-02 | 0.43 | 0.32 | 0.12 | 1.37 |
| 0 | Elp4 | 0.33 | 5.32E-07 | 1.14E-02 | 0.56 | 0.43 | 0.12 | 1.28 |
| 0 | Itm2a | 0.4 | 5.33E-07 | 1.14E-02 | 0.63 | 0.51 | 0.12 | 1.23 |
| 0 | Wdr6 | 0.28 | 5.55E-07 | 1.19E-02 | 0.75 | 0.64 | 0.11 | 1.18 |
| 0 | Gpc3 | 0.44 | 6.16E-07 | 1.32E-02 | 0.9 | 0.83 | 0.06 | 1.08 |
| 0 | Ostf1 | 0.4 | 6.17E-07 | 1.32E-02 | 0.52 | 0.4 | 0.11 | 1.28 |
| 0 | Ube2q2 | 0.38 | 6.63E-07 | 1.42E-02 | 0.56 | 0.44 | 0.12 | 1.28 |
| 0 | Ank | 0.65 | 6.78E-07 | 1.45E-02 | 0.14 | 0.08 | 0.06 | 1.84 |
| 0 | Hist1h2an | 0.52 | 7.05E-07 | 1.51E-02 | 0.3 | 0.21 | 0.1 | 1.46 |
| 0 | Ggact | 0.74 | 7.11E-07 | 1.52E-02 | 0.13 | 0.07 | 0.06 | 1.87 |
| 0 | Mef2c | 0.86 | 7.28E-07 | 1.56E-02 | 0.18 | 0.11 | 0.07 | 1.66 |
| 0 | Mllt6 | 0.47 | 7.71E-07 | 1.65E-02 | 0.34 | 0.24 | 0.1 | 1.42 |
| 0 | Sp4 | 0.45 | 7.98E-07 | 1.70E-02 | 0.39 | 0.29 | 0.11 | 1.38 |
| 0 | Nol12 | 0.31 | 8.24E-07 | 1.76E-02 | 0.65 | 0.54 | 0.12 | 1.21 |
| 0 | Gm16141 | 0.56 | 9.27E-07 | 1.98E-02 | 0.11 | 0.05 | 0.06 | 2.04 |
| 0 | Rgs9 | 0.66 | 9.44E-07 | 2.02E-02 | 0.24 | 0.16 | 0.09 | 1.54 |
| 0 | Bicd1 | 0.33 | 9.49E-07 | 2.03E-02 | 0.36 | 0.25 | 0.11 | 1.44 |
| 0 | Lad1 | 0.43 | 9.55E-07 | 2.04E-02 | 0.45 | 0.33 | 0.12 | 1.34 |
| 0 | Sertad1 | 0.73 | 9.98E-07 | 2.13E-02 | 0.18 | 0.11 | 0.07 | 1.66 |
| 0 | Socs3 | 0.54 | 1.05E-06 | 2.24E-02 | 0.25 | 0.16 | 0.09 | 1.54 |
| 0 | Fnip1 | 0.47 | 1.08E-06 | 2.31E-02 | 0.32 | 0.22 | 0.1 | 1.47 |
| 0 | Pik3r3 | 0.43 | 1.13E-06 | 2.42E-02 | 0.29 | 0.19 | 0.09 | 1.48 |
| 0 | Aven | 0.33 | 1.19E-06 | 2.53E-02 | 0.43 | 0.31 | 0.12 | 1.39 |
| 0 | Ing3 | 0.37 | 1.46E-06 | 3.11E-02 | 0.46 | 0.35 | 0.11 | 1.32 |
| 0 | Tob1 | 0.46 | 1.47E-06 | 3.13E-02 | 0.33 | 0.23 | 0.1 | 1.42 |
| 0 | Ntn3 | 0.93 | 1.49E-06 | 3.18E-02 | 0.11 | 0.06 | 0.05 | 1.9 |
| 0 | 9-Sep | 0.33 | 1.49E-06 | 3.18E-02 | 0.64 | 0.5 | 0.14 | 1.28 |
| 0 | Cd1d1 | 0.73 | 1.50E-06 | 3.20E-02 | 0.18 | 0.11 | 0.07 | 1.64 |
| 0 | Plin3 | 0.33 | 1.65E-06 | 3.53E-02 | 0.62 | 0.52 | 0.1 | 1.2 |
| 0 | Has2 | 0.65 | 1.69E-06 | 3.61E-02 | 0.19 | 0.12 | 0.07 | 1.6 |
| 0 | Hist1h2bg | 0.89 | 1.75E-06 | 3.74E-02 | 0.13 | 0.07 | 0.06 | 1.82 |
| 0 | Cstf2t | 0.5 | 1.79E-06 | 3.83E-02 | 0.46 | 0.36 | 0.1 | 1.29 |
| 0 | Limd2 | 0.25 | 1.84E-06 | 3.94E-02 | 0.83 | 0.76 | 0.07 | 1.1 |
| 0 | Ypel1 | 0.27 | 2.03E-06 | 4.33E-02 | 0.67 | 0.55 | 0.12 | 1.21 |
| 0 | Immp2l | 0.61 | 2.03E-06 | 4.35E-02 | 0.17 | 0.1 | 0.07 | 1.67 |
| 0 | Nudt14 | 0.27 | 2.05E-06 | 4.37E-02 | 0.74 | 0.62 | 0.11 | 1.18 |
| 0 | Cpz | 0.5 | 2.08E-06 | 4.45E-02 | 0.33 | 0.24 | 0.09 | 1.38 |
| 0 | Nr2e1 | 0.45 | 2.10E-06 | 4.49E-02 | 0.24 | 0.16 | 0.09 | 1.53 |
| 0 | Tbce | 0.41 | 2.21E-06 | 4.73E-02 | 0.36 | 0.26 | 0.1 | 1.39 |
| 0 | Necab2 | 0.59 | 2.26E-06 | 4.83E-02 | 0.36 | 0.28 | 0.09 | 1.32 |
| 0 | Tapt1 | 0.67 | 2.29E-06 | 4.88E-02 | 0.23 | 0.15 | 0.08 | 1.52 |
| 0 | Dchs1 | 0.47 | 2.30E-06 | 4.90E-02 | 0.26 | 0.17 | 0.09 | 1.51 |
| 1 | Oxtr | 2.61 | 1.12E-197 | 2.40E-193 | 0.77 | 0.19 | 0.58 | 3.98 |
| 1 | Cpm | 1.88 | 5.11E-172 | 1.09E-167 | 0.96 | 0.47 | 0.49 | 2.03 |
| 1 | Gstm1 | 2.72 | 3.13E-131 | 6.68E-127 | 0.73 | 0.29 | 0.43 | 2.48 |
| 1 | Gm266 | 1.63 | 3.43E-129 | 7.32E-125 | 0.92 | 0.49 | 0.44 | 1.9 |
| 1 | Tnfaip2 | 1.51 | 1.91E-124 | 4.08E-120 | 0.88 | 0.38 | 0.5 | 2.33 |
| 1 | Clu | 2.43 | 1.53E-120 | 3.27E-116 | 0.74 | 0.29 | 0.44 | 2.5 |
| 1 | Rbp1 | 1.21 | 9.89E-118 | 2.11E-113 | 1 | 0.91 | 0.09 | 1.1 |
| 1 | Aldh1a3 | 1.14 | 7.71E-104 | 1.65E-99 | 0.95 | 0.51 | 0.44 | 1.85 |
| 1 | Ctxn3 | 2.46 | 2.55E-102 | 5.44E-98 | 0.41 | 0.08 | 0.33 | 5.02 |
| 1 | Fxyd7 | 3.07 | 4.06E-98 | 8.67E-94 | 0.31 | 0.05 | 0.27 | 6.8 |
| 1 | Ano1 | 1.52 | 5.26E-96 | 1.12E-91 | 0.82 | 0.42 | 0.4 | 1.97 |
| 1 | Cldn11 | 2.61 | 1.86E-92 | 3.97E-88 | 0.38 | 0.07 | 0.3 | 5.08 |

**Supplementary Table 3. Marker genes for the original clusters of integrated clustering of E11.5 control and NotchGOF λ-junction ectoderm cells.**

| | | | | | | | | |
|---|---|---|---|---|---|---|---|---|
| 1 | Six3 | 0.95 | 5.04E-90 | 1.08E-85 | 0.99 | 0.82 | 0.17 | 1.21 |
| 1 | Hey1 | 0.95 | 7.32E-87 | 1.56E-82 | 0.98 | 0.72 | 0.26 | 1.36 |
| 1 | Samd5 | 1.05 | 3.99E-82 | 8.53E-78 | 0.97 | 0.65 | 0.32 | 1.49 |
| 1 | Khdrbs3 | 0.96 | 4.28E-82 | 9.13E-78 | 0.98 | 0.87 | 0.11 | 1.13 |
| 1 | Gpc1 | 1.33 | 4.00E-79 | 8.54E-75 | 0.82 | 0.46 | 0.36 | 1.78 |
| 1 | Six6 | 1.49 | 3.04E-77 | 6.49E-73 | 0.62 | 0.23 | 0.39 | 2.66 |
| 1 | Krt18 | 0.83 | 7.00E-77 | 1.49E-72 | 0.99 | 0.85 | 0.14 | 1.16 |
| 1 | Crabp2 | 0.79 | 1.34E-75 | 2.85E-71 | 0.99 | 0.83 | 0.17 | 1.2 |
| 1 | Prkg1 | 1.55 | 1.71E-72 | 3.65E-68 | 0.57 | 0.21 | 0.36 | 2.75 |
| 1 | Itih5 | 2.73 | 8.64E-72 | 1.85E-67 | 0.27 | 0.05 | 0.22 | 5.68 |
| 1 | Heyl | 1.8 | 1.32E-70 | 2.82E-66 | 0.51 | 0.18 | 0.34 | 2.92 |
| 1 | Pitx1 | 1.17 | 4.00E-70 | 8.54E-66 | 0.89 | 0.62 | 0.26 | 1.42 |
| 1 | Nuak1 | 1.6 | 6.13E-70 | 1.31E-65 | 0.58 | 0.23 | 0.35 | 2.53 |
| 1 | Eya1 | 1.02 | 1.33E-68 | 2.85E-64 | 0.89 | 0.55 | 0.34 | 1.61 |
| 1 | Pax6 | 1.17 | 1.07E-67 | 2.29E-63 | 0.81 | 0.44 | 0.37 | 1.84 |
| 1 | Id4 | 1.26 | 7.63E-67 | 1.63E-62 | 0.83 | 0.54 | 0.29 | 1.54 |
| 1 | Itga6 | 0.91 | 4.02E-66 | 8.60E-62 | 0.94 | 0.77 | 0.18 | 1.23 |
| 1 | Slfn2 | 3.09 | 5.36E-65 | 1.14E-60 | 0.2 | 0.03 | 0.17 | 7.8 |
| 1 | Six3os1 | 0.75 | 6.26E-65 | 1.34E-60 | 0.99 | 0.74 | 0.25 | 1.34 |
| 1 | Mgst1 | 1.05 | 7.58E-65 | 1.62E-60 | 0.92 | 0.66 | 0.26 | 1.4 |
| 1 | Nrarp | 0.94 | 4.11E-64 | 8.77E-60 | 0.9 | 0.57 | 0.33 | 1.58 |
| 1 | Ccdc80 | 1.47 | 5.83E-64 | 1.25E-59 | 0.56 | 0.22 | 0.34 | 2.53 |
| 1 | Six2 | 1.07 | 7.65E-64 | 1.63E-59 | 0.82 | 0.5 | 0.32 | 1.64 |
| 1 | Gsap | 2.17 | 6.96E-62 | 1.49E-57 | 0.33 | 0.08 | 0.25 | 3.95 |
| 1 | Crym | 1.68 | 1.97E-61 | 4.21E-57 | 0.52 | 0.2 | 0.32 | 2.58 |
| 1 | Tmem108 | 1.55 | 2.84E-61 | 6.07E-57 | 0.59 | 0.26 | 0.33 | 2.25 |
| 1 | Hes5 | 1.02 | 1.95E-59 | 4.17E-55 | 0.73 | 0.34 | 0.39 | 2.13 |
| 1 | Krt8 | 0.66 | 5.60E-59 | 1.20E-54 | 0.98 | 0.8 | 0.19 | 1.23 |
| 1 | Foxg1 | 0.98 | 1.42E-58 | 3.04E-54 | 0.82 | 0.5 | 0.33 | 1.65 |
| 1 | Tubb2b | 0.64 | 1.94E-58 | 4.14E-54 | 0.98 | 0.92 | 0.06 | 1.07 |
| 1 | Ephb1 | 1.25 | 2.78E-58 | 5.93E-54 | 0.68 | 0.34 | 0.34 | 1.99 |
| 1 | Adi1 | 1.17 | 8.26E-57 | 1.76E-52 | 0.74 | 0.42 | 0.32 | 1.76 |
| 1 | Tshz2 | 1.15 | 9.55E-56 | 2.04E-51 | 0.7 | 0.35 | 0.34 | 1.97 |
| 1 | Anxa5 | 1.05 | 3.47E-54 | 7.41E-50 | 0.85 | 0.6 | 0.25 | 1.42 |
| 1 | Dab1 | 1.02 | 4.88E-54 | 1.04E-49 | 0.79 | 0.45 | 0.34 | 1.75 |
| 1 | Fbn2 | 1.17 | 5.61E-54 | 1.20E-49 | 0.74 | 0.43 | 0.31 | 1.73 |
| 1 | Gucy1a1 | 1.28 | 1.28E-53 | 2.74E-49 | 0.45 | 0.16 | 0.3 | 2.87 |
| 1 | 3110039M20Rik | 1.18 | 3.72E-53 | 7.94E-49 | 0.59 | 0.27 | 0.33 | 2.23 |
| 1 | Galnt14 | 2.04 | 8.19E-53 | 1.75E-48 | 0.28 | 0.07 | 0.21 | 4.13 |
| 1 | Dmrta2 | 0.93 | 8.29E-53 | 1.77E-48 | 0.81 | 0.47 | 0.34 | 1.74 |
| 1 | Slc22a23 | 1.32 | 8.68E-53 | 1.85E-48 | 0.59 | 0.27 | 0.32 | 2.18 |
| 1 | Ces1d | 1.75 | 1.00E-51 | 2.14E-47 | 0.41 | 0.14 | 0.27 | 2.85 |
| 1 | S1pr3 | 2.8 | 5.91E-51 | 1.26E-46 | 0.19 | 0.03 | 0.16 | 6.03 |
| 1 | Cyb5a | 0.47 | 6.00E-51 | 1.28E-46 | 0.99 | 0.96 | 0.03 | 1.03 |
| 1 | Six1 | 0.74 | 1.46E-50 | 3.12E-46 | 0.93 | 0.65 | 0.28 | 1.43 |
| 1 | Il7 | 2.77 | 2.53E-50 | 5.41E-46 | 0.19 | 0.03 | 0.16 | 5.91 |
| 1 | Mest | 0.51 | 4.09E-50 | 8.73E-46 | 0.99 | 0.9 | 0.1 | 1.11 |
| 1 | Mef2c | 1.74 | 6.30E-50 | 1.34E-45 | 0.32 | 0.09 | 0.23 | 3.51 |
| 1 | Pbx1 | 0.49 | 1.78E-49 | 3.80E-45 | 0.99 | 0.97 | 0.03 | 1.03 |
| 1 | Fgf10 | 1.96 | 5.27E-49 | 1.13E-44 | 0.32 | 0.09 | 0.22 | 3.41 |
| 1 | Slc12a2 | 0.84 | 3.11E-48 | 6.65E-44 | 0.91 | 0.69 | 0.22 | 1.32 |
| 1 | Dmbx1 | 1.66 | 3.80E-48 | 8.11E-44 | 0.38 | 0.13 | 0.25 | 2.92 |
| 1 | Slc9a3r1 | 0.75 | 5.84E-48 | 1.25E-43 | 0.92 | 0.73 | 0.19 | 1.27 |
| 1 | Cdh2 | 0.79 | 1.17E-47 | 2.50E-43 | 0.9 | 0.57 | 0.33 | 1.59 |
| 1 | Fap | 3.09 | 1.29E-47 | 2.75E-43 | 0.15 | 0.02 | 0.13 | 7.63 |
| 1 | Dlk1 | 0.33 | 4.31E-46 | 9.21E-42 | 0.91 | 0.6 | 0.31 | 1.52 |
| 1 | Ncam1 | 0.38 | 1.33E-44 | 2.84E-40 | 0.63 | 0.29 | 0.34 | 2.15 |

**Supplementary Table 3. Marker genes for the original clusters of integrated clustering of E11.5 control and NotchGOF λ-junction ectoderm cells.**

| | | | | | | | | |
|---|---|---|---|---|---|---|---|---|
| 1 | Gsta4 | 0.64 | 7.22E-44 | 1.54E-39 | 0.95 | 0.79 | 0.16 | 1.2 |
| 1 | Sparc | 0.53 | 1.50E-43 | 3.20E-39 | 0.99 | 0.88 | 0.11 | 1.13 |
| 1 | Exoc3l4 | 1.49 | 2.40E-43 | 5.13E-39 | 0.4 | 0.15 | 0.25 | 2.65 |
| 1 | Pacsin1 | 1.51 | 2.05E-42 | 4.38E-38 | 0.39 | 0.14 | 0.24 | 2.71 |
| 1 | Zfhx4 | 0.65 | 1.01E-40 | 2.17E-36 | 0.93 | 0.75 | 0.18 | 1.24 |
| 1 | Nt5dc2 | 0.46 | 1.97E-39 | 4.21E-35 | 0.98 | 0.91 | 0.07 | 1.08 |
| 1 | Zmat4 | 1.38 | 3.32E-39 | 7.09E-35 | 0.35 | 0.12 | 0.22 | 2.8 |
| 1 | Faah | 1.29 | 8.58E-39 | 1.83E-34 | 0.45 | 0.2 | 0.26 | 2.32 |
| 1 | Lhfpl2 | 1.56 | 9.05E-39 | 1.93E-34 | 0.35 | 0.13 | 0.22 | 2.67 |
| 1 | C130071C03Rik | 1.21 | 2.02E-38 | 4.32E-34 | 0.42 | 0.18 | 0.25 | 2.41 |
| 1 | Igfbp3 | 0.55 | 2.84E-38 | 6.07E-34 | 0.65 | 0.35 | 0.3 | 1.84 |
| 1 | Col11a1 | 0.82 | 5.14E-38 | 1.10E-33 | 0.79 | 0.54 | 0.26 | 1.47 |
| 1 | Fat4 | 1.59 | 1.56E-37 | 3.33E-33 | 0.28 | 0.09 | 0.19 | 3.18 |
| 1 | D230030E09Rik | 1.03 | 2.57E-37 | 5.50E-33 | 0.44 | 0.19 | 0.25 | 2.33 |
| 1 | Chst2 | 1.52 | 6.00E-37 | 1.28E-32 | 0.34 | 0.12 | 0.22 | 2.74 |
| 1 | Dsel | 1.12 | 1.37E-36 | 2.92E-32 | 0.62 | 0.35 | 0.27 | 1.76 |
| 1 | Vim | 0.64 | 7.91E-36 | 1.69E-31 | 0.91 | 0.69 | 0.21 | 1.31 |
| 1 | Plet1 | 2.33 | 1.93E-35 | 4.13E-31 | 0.16 | 0.03 | 0.13 | 4.85 |
| 1 | Fras1 | 0.79 | 2.21E-35 | 4.72E-31 | 0.81 | 0.59 | 0.21 | 1.36 |
| 1 | Rcsd1 | 1.31 | 3.33E-35 | 7.12E-31 | 0.36 | 0.14 | 0.22 | 2.61 |
| 1 | Tkt | 0.49 | 5.72E-35 | 1.22E-30 | 0.97 | 0.87 | 0.1 | 1.11 |
| 1 | Svip | 1.17 | 7.31E-35 | 1.56E-30 | 0.48 | 0.23 | 0.25 | 2.08 |
| 1 | Isl1 | 0.54 | 8.74E-35 | 1.87E-30 | 0.94 | 0.73 | 0.21 | 1.29 |
| 1 | 4930426D05Rik | 1.67 | 2.72E-34 | 5.81E-30 | 0.29 | 0.1 | 0.19 | 2.87 |
| 1 | Mdk | 0.27 | 1.35E-33 | 2.89E-29 | 1 | 0.99 | 0.01 | 1.01 |
| 1 | Hmga2 | 0.44 | 2.94E-33 | 6.27E-29 | 0.99 | 0.92 | 0.07 | 1.08 |
| 1 | Muc1 | 1.91 | 7.97E-33 | 1.70E-28 | 0.21 | 0.06 | 0.15 | 3.51 |
| 1 | Necab2 | 1.14 | 1.07E-32 | 2.29E-28 | 0.5 | 0.26 | 0.25 | 1.97 |
| 1 | Lypd6 | 1.21 | 1.28E-32 | 2.73E-28 | 0.41 | 0.18 | 0.23 | 2.29 |
| 1 | Hmgn2 | 0.37 | 3.55E-32 | 7.58E-28 | 0.99 | 0.98 | 0.02 | 1.02 |
| 1 | AW011738 | 1.12 | 1.06E-31 | 2.27E-27 | 0.52 | 0.27 | 0.25 | 1.94 |
| 1 | Samd4 | 1.33 | 1.75E-31 | 3.74E-27 | 0.35 | 0.14 | 0.21 | 2.42 |
| 1 | B230118H07Rik | 0.8 | 2.37E-31 | 5.07E-27 | 0.76 | 0.54 | 0.22 | 1.4 |
| 1 | Plch1 | 0.96 | 3.15E-31 | 6.72E-27 | 0.56 | 0.31 | 0.25 | 1.8 |
| 1 | Myh14 | 1.04 | 4.39E-31 | 9.37E-27 | 0.52 | 0.27 | 0.25 | 1.92 |
| 1 | Calr | 0.4 | 6.16E-31 | 1.32E-26 | 0.99 | 0.97 | 0.02 | 1.02 |
| 1 | Prrx1 | 1.55 | 1.39E-30 | 2.98E-26 | 0.37 | 0.17 | 0.2 | 2.2 |
| 1 | Tmem200a | 1.46 | 1.90E-30 | 4.06E-26 | 0.28 | 0.1 | 0.18 | 2.79 |
| 1 | Lypd6b | 1.02 | 3.15E-30 | 6.73E-26 | 0.44 | 0.21 | 0.24 | 2.15 |
| 1 | Igf2r | 0.82 | 3.58E-30 | 7.65E-26 | 0.76 | 0.56 | 0.21 | 1.37 |
| 1 | Ubash3b | 1.05 | 4.90E-30 | 1.05E-25 | 0.45 | 0.22 | 0.23 | 2.07 |
| 1 | Rhov | 0.82 | 5.09E-30 | 1.09E-25 | 0.44 | 0.2 | 0.24 | 2.23 |
| 1 | Fam214a | 1.18 | 6.98E-30 | 1.49E-25 | 0.41 | 0.19 | 0.22 | 2.18 |
| 1 | 2010300C02Rik | 1.74 | 1.69E-29 | 3.62E-25 | 0.23 | 0.07 | 0.15 | 3.13 |
| 1 | Atp2b1 | 0.49 | 2.98E-29 | 6.37E-25 | 0.97 | 0.88 | 0.09 | 1.1 |
| 1 | Cldn3 | 0.58 | 5.45E-29 | 1.16E-24 | 0.85 | 0.61 | 0.24 | 1.4 |
| 1 | Gnai2 | 0.32 | 1.20E-28 | 2.57E-24 | 1 | 0.97 | 0.03 | 1.03 |
| 1 | Fdps | 0.6 | 2.95E-28 | 6.29E-24 | 0.91 | 0.8 | 0.11 | 1.14 |
| 1 | Fxyd1 | 1.94 | 3.10E-28 | 6.62E-24 | 0.13 | 0.03 | 0.11 | 4.62 |
| 1 | Notch3 | 0.71 | 3.28E-28 | 7.01E-24 | 0.75 | 0.51 | 0.24 | 1.47 |
| 1 | Gldn | 2.7 | 4.00E-28 | 8.55E-24 | 0.12 | 0.02 | 0.09 | 5.23 |
| 1 | Podn | 1.96 | 1.74E-27 | 3.73E-23 | 0.15 | 0.04 | 0.12 | 4.03 |
| 1 | Ppp1r1a | 0.9 | 2.92E-27 | 6.23E-23 | 0.52 | 0.29 | 0.24 | 1.83 |
| 1 | Lypd2 | 1.27 | 3.15E-27 | 6.72E-23 | 0.32 | 0.14 | 0.19 | 2.35 |
| 1 | Sorbs2 | 0.88 | 4.29E-27 | 9.16E-23 | 0.61 | 0.38 | 0.24 | 1.63 |
| 1 | Arhgap8 | 1.28 | 9.10E-27 | 1.94E-22 | 0.33 | 0.14 | 0.19 | 2.33 |
| 1 | Cdk6 | 0.74 | 1.49E-26 | 3.19E-22 | 0.71 | 0.5 | 0.21 | 1.43 |

**Supplementary Table 3. Marker genes for the original clusters of integrated clustering of E11.5 control and NotchGOF λ-junction ectoderm cells.**

| | | | | | | | | |
|---|---|---|---|---|---|---|---|---|
| 1 | Cyb5r3 | 0.42 | 2.31E-26 | 4.94E-22 | 0.98 | 0.92 | 0.06 | 1.06 |
| 1 | Eps8l1 | 1.27 | 2.79E-26 | 5.96E-22 | 0.28 | 0.11 | 0.17 | 2.59 |
| 1 | Atox1 | 0.47 | 4.26E-26 | 9.09E-22 | 0.99 | 0.98 | 0.01 | 1.01 |
| 1 | BC048679 | 2.38 | 4.41E-26 | 9.41E-22 | 0.11 | 0.02 | 0.09 | 5.05 |
| 1 | Thrb | 1.08 | 2.67E-25 | 5.70E-21 | 0.31 | 0.13 | 0.18 | 2.4 |
| 1 | Camk1 | 0.88 | 3.19E-25 | 6.80E-21 | 0.6 | 0.36 | 0.24 | 1.65 |
| 1 | Lamc1 | 0.87 | 3.26E-25 | 6.97E-21 | 0.63 | 0.41 | 0.22 | 1.53 |
| 1 | Maged1 | 0.38 | 3.80E-25 | 8.12E-21 | 0.98 | 0.94 | 0.04 | 1.04 |
| 1 | Rhoc | 0.64 | 4.19E-25 | 8.96E-21 | 0.8 | 0.63 | 0.17 | 1.27 |
| 1 | Il17rd | 0.67 | 7.55E-25 | 1.61E-20 | 0.72 | 0.47 | 0.25 | 1.52 |
| 1 | Gata3 | 0.7 | 8.04E-25 | 1.72E-20 | 0.56 | 0.31 | 0.25 | 1.79 |
| 1 | Gjb2 | 0.74 | 2.43E-24 | 5.19E-20 | 0.35 | 0.16 | 0.19 | 2.22 |
| 1 | Ppp2r2b | 0.68 | 2.71E-24 | 5.79E-20 | 0.59 | 0.35 | 0.24 | 1.68 |
| 1 | Dusp5 | 1.09 | 3.83E-24 | 8.17E-20 | 0.38 | 0.18 | 0.19 | 2.07 |
| 1 | H1fx | 0.42 | 3.87E-24 | 8.27E-20 | 0.96 | 0.94 | 0.03 | 1.03 |
| 1 | Alad | 0.57 | 5.30E-24 | 1.13E-19 | 0.87 | 0.73 | 0.14 | 1.2 |
| 1 | 9230105E05Rik | 2.01 | 6.23E-24 | 1.33E-19 | 0.11 | 0.02 | 0.09 | 4.71 |
| 1 | Pkdcc | 0.88 | 1.35E-23 | 2.89E-19 | 0.51 | 0.29 | 0.22 | 1.74 |
| 1 | Adarb2 | 1.74 | 1.73E-23 | 3.69E-19 | 0.14 | 0.04 | 0.1 | 3.86 |
| 1 | Tuba1b | 0.37 | 1.85E-23 | 3.95E-19 | 0.99 | 0.97 | 0.03 | 1.03 |
| 1 | Rasgrp2 | 2.03 | 2.64E-23 | 5.63E-19 | 0.14 | 0.04 | 0.1 | 3.76 |
| 1 | Tmem176a | 0.8 | 2.82E-23 | 6.02E-19 | 0.61 | 0.4 | 0.22 | 1.55 |
| 1 | Sall2 | 0.72 | 4.60E-23 | 9.83E-19 | 0.67 | 0.45 | 0.22 | 1.5 |
| 1 | Tril | 0.89 | 5.24E-23 | 1.12E-18 | 0.49 | 0.27 | 0.22 | 1.79 |
| 1 | Frmd4a | 0.6 | 5.41E-23 | 1.16E-18 | 0.71 | 0.46 | 0.25 | 1.53 |
| 1 | Ntng1 | 1.73 | 8.16E-23 | 1.74E-18 | 0.16 | 0.05 | 0.11 | 3.28 |
| 1 | Notch2 | 0.91 | 1.12E-22 | 2.39E-18 | 0.49 | 0.28 | 0.21 | 1.75 |
| 1 | Nfix | 1.43 | 1.45E-22 | 3.10E-18 | 0.21 | 0.08 | 0.13 | 2.77 |
| 1 | Lama1 | 1.47 | 1.81E-22 | 3.87E-18 | 0.23 | 0.09 | 0.14 | 2.64 |
| 1 | Bspry | 1.03 | 2.09E-22 | 4.46E-18 | 0.44 | 0.24 | 0.2 | 1.82 |
| 1 | Ttyh1 | 1.39 | 1.43E-21 | 3.06E-17 | 0.2 | 0.07 | 0.13 | 2.76 |
| 1 | App | 0.43 | 1.83E-21 | 3.91E-17 | 0.94 | 0.84 | 0.1 | 1.12 |
| 1 | Alkbh7 | 0.81 | 1.95E-21 | 4.17E-17 | 0.56 | 0.36 | 0.2 | 1.57 |
| 1 | Isyna1 | 0.52 | 2.72E-21 | 5.81E-17 | 0.84 | 0.69 | 0.15 | 1.21 |
| 1 | Aebp1 | 1.68 | 7.60E-21 | 1.62E-16 | 0.2 | 0.07 | 0.13 | 2.74 |
| 1 | Sod1 | 0.46 | 1.83E-20 | 3.91E-16 | 0.89 | 0.76 | 0.14 | 1.18 |
| 1 | P4ha3 | 1.15 | 2.19E-20 | 4.67E-16 | 0.31 | 0.14 | 0.16 | 2.12 |
| 1 | Ppfibp1 | 0.57 | 3.08E-20 | 6.59E-16 | 0.77 | 0.59 | 0.18 | 1.3 |
| 1 | Naxe | 0.49 | 3.73E-20 | 7.96E-16 | 0.83 | 0.69 | 0.14 | 1.2 |
| 1 | Id1 | 0.43 | 3.77E-20 | 8.04E-16 | 0.98 | 0.92 | 0.05 | 1.06 |
| 1 | Sapcd1 | 1.52 | 4.75E-20 | 1.01E-15 | 0.18 | 0.06 | 0.12 | 2.83 |
| 1 | Apc | 0.39 | 5.90E-20 | 1.26E-15 | 0.91 | 0.77 | 0.14 | 1.18 |
| 1 | Arrdc1 | 0.79 | 6.26E-20 | 1.34E-15 | 0.54 | 0.33 | 0.21 | 1.62 |
| 1 | Rcn1 | 0.4 | 6.40E-20 | 1.37E-15 | 0.94 | 0.84 | 0.1 | 1.12 |
| 1 | Maf | 0.84 | 9.40E-20 | 2.01E-15 | 0.28 | 0.12 | 0.16 | 2.27 |
| 1 | S100a16 | 0.86 | 1.17E-19 | 2.49E-15 | 0.29 | 0.13 | 0.16 | 2.21 |
| 1 | Hes1 | 0.43 | 1.36E-19 | 2.91E-15 | 0.9 | 0.71 | 0.19 | 1.27 |
| 1 | Ust | 1.17 | 2.39E-19 | 5.11E-15 | 0.26 | 0.12 | 0.15 | 2.27 |
| 1 | Iqgap2 | 0.71 | 2.92E-19 | 6.24E-15 | 0.52 | 0.31 | 0.21 | 1.69 |
| 1 | Pcdh19 | 0.67 | 3.17E-19 | 6.78E-15 | 0.41 | 0.22 | 0.2 | 1.91 |
| 1 | Nes | 0.49 | 4.67E-19 | 9.98E-15 | 0.7 | 0.47 | 0.22 | 1.47 |
| 1 | Mtx2 | 0.42 | 5.19E-19 | 1.11E-14 | 0.9 | 0.8 | 0.1 | 1.12 |
| 1 | Rian | 0.47 | 5.49E-19 | 1.17E-14 | 0.65 | 0.44 | 0.21 | 1.47 |
| 1 | Egfl6 | 1.01 | 5.79E-19 | 1.24E-14 | 0.33 | 0.17 | 0.16 | 1.99 |
| 1 | Ntm | 0.54 | 6.96E-19 | 1.49E-14 | 0.42 | 0.22 | 0.2 | 1.88 |
| 1 | Ddah2 | 0.26 | 9.28E-19 | 1.98E-14 | 0.99 | 0.97 | 0.02 | 1.02 |
| 1 | Adam22 | 1.15 | 1.69E-18 | 3.61E-14 | 0.25 | 0.11 | 0.14 | 2.28 |

**Supplementary Table 3. Marker genes for the original clusters of integrated clustering of E11.5 control and NotchGOF λ-junction ectoderm cells.**

| | | | | | | | | |
|---|---|---|---|---|---|---|---|---|
| 1 | Nr2e1 | 1 | 1.93E-18 | 4.11E-14 | 0.31 | 0.15 | 0.16 | 2.03 |
| 1 | Sfrp1 | 0.27 | 2.49E-18 | 5.33E-14 | 0.75 | 0.49 | 0.26 | 1.53 |
| 1 | Prss8 | 0.69 | 3.32E-18 | 7.09E-14 | 0.56 | 0.36 | 0.2 | 1.56 |
| 1 | Lrrfip1 | 0.66 | 4.22E-18 | 9.01E-14 | 0.67 | 0.48 | 0.2 | 1.41 |
| 1 | Snx24 | 0.77 | 7.20E-18 | 1.54E-13 | 0.49 | 0.3 | 0.19 | 1.65 |
| 1 | Nhs | 0.6 | 7.76E-18 | 1.66E-13 | 0.62 | 0.41 | 0.2 | 1.49 |
| 1 | Cops6 | 0.41 | 8.46E-18 | 1.81E-13 | 0.9 | 0.8 | 0.1 | 1.13 |
| 1 | Gm48435 | 1.17 | 9.89E-18 | 2.11E-13 | 0.22 | 0.09 | 0.13 | 2.43 |
| 1 | Nfia | 0.64 | 1.45E-17 | 3.10E-13 | 0.55 | 0.35 | 0.2 | 1.59 |
| 1 | Rad18 | 0.68 | 1.55E-17 | 3.32E-13 | 0.57 | 0.39 | 0.19 | 1.48 |
| 1 | Mapk13 | 0.53 | 2.15E-17 | 4.60E-13 | 0.76 | 0.59 | 0.17 | 1.29 |
| 1 | Gnao1 | 0.77 | 2.26E-17 | 4.82E-13 | 0.32 | 0.16 | 0.16 | 1.98 |
| 1 | Shroom3 | 0.8 | 2.32E-17 | 4.95E-13 | 0.4 | 0.22 | 0.18 | 1.8 |
| 1 | Pwwp3b | 0.88 | 2.36E-17 | 5.03E-13 | 0.45 | 0.27 | 0.18 | 1.66 |
| 1 | Pard3b | 0.86 | 2.93E-17 | 6.27E-13 | 0.38 | 0.21 | 0.18 | 1.85 |
| 1 | Grhl1 | 1.42 | 2.94E-17 | 6.29E-13 | 0.19 | 0.07 | 0.11 | 2.53 |
| 1 | Scara3 | 1.1 | 3.08E-17 | 6.58E-13 | 0.25 | 0.12 | 0.14 | 2.17 |
| 1 | Bex4 | 0.28 | 3.58E-17 | 7.66E-13 | 0.98 | 0.93 | 0.06 | 1.06 |
| 1 | Gstt3 | 0.97 | 3.90E-17 | 8.33E-13 | 0.24 | 0.1 | 0.13 | 2.3 |
| 1 | Mpped2 | 0.4 | 4.13E-17 | 8.81E-13 | 0.88 | 0.77 | 0.11 | 1.15 |
| 1 | Mecom | 0.32 | 4.24E-17 | 9.07E-13 | 0.65 | 0.43 | 0.23 | 1.53 |
| 1 | Tmem200c | 1.16 | 7.05E-17 | 1.51E-12 | 0.19 | 0.08 | 0.12 | 2.51 |
| 1 | Mapk12 | 0.68 | 7.26E-17 | 1.55E-12 | 0.46 | 0.27 | 0.19 | 1.72 |
| 1 | Tceal3 | 1.03 | 8.84E-17 | 1.89E-12 | 0.29 | 0.14 | 0.15 | 2.03 |
| 1 | Sp8 | 0.46 | 9.05E-17 | 1.93E-12 | 0.71 | 0.5 | 0.22 | 1.44 |
| 1 | Palm3 | 0.65 | 1.15E-16 | 2.45E-12 | 0.52 | 0.33 | 0.19 | 1.58 |
| 1 | Alx4 | 1.26 | 1.32E-16 | 2.82E-12 | 0.18 | 0.07 | 0.11 | 2.55 |
| 1 | Fgf15 | 1.17 | 1.33E-16 | 2.85E-12 | 0.21 | 0.09 | 0.12 | 2.3 |
| 1 | Hsp90b1 | 0.3 | 1.91E-16 | 4.08E-12 | 1 | 0.98 | 0.02 | 1.02 |
| 1 | Myl7 | 1.08 | 2.09E-16 | 4.46E-12 | 0.17 | 0.06 | 0.1 | 2.68 |
| 1 | Adgrl2 | 0.45 | 2.10E-16 | 4.48E-12 | 0.81 | 0.69 | 0.12 | 1.17 |
| 1 | Ssbp4 | 0.58 | 2.35E-16 | 5.01E-12 | 0.65 | 0.47 | 0.18 | 1.39 |
| 1 | Tst | 0.83 | 2.54E-16 | 5.42E-12 | 0.41 | 0.24 | 0.17 | 1.72 |
| 1 | Ckap4 | 0.4 | 2.61E-16 | 5.58E-12 | 0.88 | 0.78 | 0.1 | 1.13 |
| 1 | Snx3 | 0.3 | 2.62E-16 | 5.60E-12 | 0.92 | 0.86 | 0.06 | 1.07 |
| 1 | Snn | 0.59 | 2.67E-16 | 5.70E-12 | 0.75 | 0.59 | 0.15 | 1.26 |
| 1 | Ap3b2 | 0.88 | 2.69E-16 | 5.75E-12 | 0.17 | 0.06 | 0.11 | 2.66 |
| 1 | Erbb3 | 0.48 | 2.88E-16 | 6.15E-12 | 0.72 | 0.52 | 0.2 | 1.38 |
| 1 | Rnh1 | 0.42 | 3.51E-16 | 7.50E-12 | 0.87 | 0.77 | 0.1 | 1.13 |
| 1 | Ssbp2 | 0.44 | 4.00E-16 | 8.55E-12 | 0.82 | 0.68 | 0.14 | 1.21 |
| 1 | Cldn9 | 0.55 | 4.38E-16 | 9.35E-12 | 0.53 | 0.33 | 0.2 | 1.62 |
| 1 | Muc4 | 1.39 | 5.54E-16 | 1.18E-11 | 0.13 | 0.04 | 0.09 | 3.02 |
| 1 | Etfa | 0.41 | 5.58E-16 | 1.19E-11 | 0.85 | 0.75 | 0.11 | 1.14 |
| 1 | Dtna | 0.65 | 8.09E-16 | 1.73E-11 | 0.52 | 0.33 | 0.19 | 1.59 |
| 1 | Rprm | 0.49 | 1.03E-15 | 2.20E-11 | 0.55 | 0.36 | 0.19 | 1.54 |
| 1 | Kazn | 0.72 | 1.06E-15 | 2.26E-11 | 0.46 | 0.29 | 0.17 | 1.59 |
| 1 | Mdfic | 1.02 | 1.11E-15 | 2.37E-11 | 0.32 | 0.17 | 0.15 | 1.85 |
| 1 | Klhl23 | 0.49 | 1.12E-15 | 2.39E-11 | 0.7 | 0.5 | 0.21 | 1.41 |
| 1 | Dach1 | 0.48 | 1.30E-15 | 2.78E-11 | 0.67 | 0.47 | 0.21 | 1.44 |
| 1 | Pik3ap1 | 1.75 | 1.41E-15 | 3.01E-11 | 0.12 | 0.04 | 0.08 | 3.02 |
| 1 | Prtg | 0.74 | 1.44E-15 | 3.07E-11 | 0.36 | 0.19 | 0.16 | 1.85 |
| 1 | A330048O09Rik | 1.32 | 1.45E-15 | 3.10E-11 | 0.15 | 0.05 | 0.09 | 2.79 |
| 1 | Ccnb2 | 0.7 | 1.50E-15 | 3.21E-11 | 0.64 | 0.48 | 0.15 | 1.32 |
| 1 | Kcnd2 | 1.47 | 1.54E-15 | 3.29E-11 | 0.16 | 0.06 | 0.1 | 2.6 |
| 1 | Tm7sf2 | 0.87 | 1.63E-15 | 3.48E-11 | 0.36 | 0.2 | 0.16 | 1.78 |
| 1 | Spry4 | 1.36 | 1.71E-15 | 3.66E-11 | 0.18 | 0.08 | 0.11 | 2.41 |
| 1 | Chchd10 | 0.65 | 1.73E-15 | 3.69E-11 | 0.55 | 0.37 | 0.18 | 1.49 |

**Supplementary Table 3. Marker genes for the original clusters of integrated clustering of E11.5 control and NotchGOF λ-junction ectoderm cells.**

| | | | | | | | | |
|---|---|---|---|---|---|---|---|---|
| 1 | Foxp2 | 0.84 | 1.84E-15 | 3.93E-11 | 0.35 | 0.19 | 0.16 | 1.82 |
| 1 | Trib1 | 1.01 | 1.86E-15 | 3.97E-11 | 0.31 | 0.16 | 0.14 | 1.87 |
| 1 | Hs6st2 | 0.53 | 2.08E-15 | 4.44E-11 | 0.7 | 0.52 | 0.18 | 1.34 |
| 1 | Id3 | 0.37 | 2.18E-15 | 4.66E-11 | 0.99 | 0.96 | 0.03 | 1.03 |
| 1 | Lgmn | 0.66 | 4.13E-15 | 8.83E-11 | 0.51 | 0.33 | 0.18 | 1.53 |
| 1 | Rab25 | 0.44 | 4.41E-15 | 9.41E-11 | 0.77 | 0.61 | 0.16 | 1.26 |
| 1 | Lamc2 | 0.71 | 5.78E-15 | 1.23E-10 | 0.31 | 0.16 | 0.15 | 1.94 |
| 1 | Ndufa12 | 0.26 | 5.82E-15 | 1.24E-10 | 0.99 | 0.94 | 0.04 | 1.04 |
| 1 | Tmem176b | 0.35 | 6.62E-15 | 1.42E-10 | 0.76 | 0.58 | 0.18 | 1.3 |
| 1 | Tubb4b | 0.49 | 7.38E-15 | 1.58E-10 | 0.9 | 0.8 | 0.1 | 1.13 |
| 1 | Piezo1 | 0.95 | 8.58E-15 | 1.83E-10 | 0.27 | 0.13 | 0.13 | 2.02 |
| 1 | Pde1b | 1.24 | 9.01E-15 | 1.92E-10 | 0.16 | 0.06 | 0.1 | 2.6 |
| 1 | Scd2 | 0.25 | 9.55E-15 | 2.04E-10 | 0.99 | 0.96 | 0.03 | 1.03 |
| 1 | Maged2 | 0.4 | 1.04E-14 | 2.22E-10 | 0.81 | 0.69 | 0.12 | 1.17 |
| 1 | Gucy1b1 | 1.32 | 1.06E-14 | 2.27E-10 | 0.17 | 0.07 | 0.1 | 2.44 |
| 1 | Crip1 | 0.46 | 1.26E-14 | 2.69E-10 | 0.57 | 0.38 | 0.19 | 1.51 |
| 1 | Mapk8ip1 | 0.53 | 2.12E-14 | 4.52E-10 | 0.51 | 0.35 | 0.17 | 1.49 |
| 1 | Galm | 0.4 | 2.26E-14 | 4.83E-10 | 0.58 | 0.38 | 0.2 | 1.54 |
| 1 | Ppfia3 | 0.76 | 2.27E-14 | 4.85E-10 | 0.45 | 0.28 | 0.17 | 1.59 |
| 1 | Misp | 1.31 | 2.67E-14 | 5.70E-10 | 0.15 | 0.06 | 0.09 | 2.55 |
| 1 | Kcna5 | 1.22 | 3.50E-14 | 7.49E-10 | 0.13 | 0.05 | 0.09 | 2.81 |
| 1 | Cgn | 0.58 | 4.47E-14 | 9.56E-10 | 0.55 | 0.37 | 0.18 | 1.49 |
| 1 | Pea15a | 0.45 | 4.66E-14 | 9.96E-10 | 0.51 | 0.32 | 0.19 | 1.6 |
| 1 | P2rx2 | 1.17 | 5.80E-14 | 1.24E-09 | 0.16 | 0.07 | 0.1 | 2.46 |
| 1 | B3galnt1 | 0.65 | 6.27E-14 | 1.34E-09 | 0.29 | 0.15 | 0.14 | 1.95 |
| 1 | Mgat4a | 0.72 | 7.00E-14 | 1.50E-09 | 0.33 | 0.19 | 0.15 | 1.79 |
| 1 | Gstm7 | 0.71 | 7.12E-14 | 1.52E-09 | 0.38 | 0.22 | 0.16 | 1.7 |
| 1 | Gas2 | 0.85 | 7.98E-14 | 1.70E-09 | 0.34 | 0.2 | 0.15 | 1.74 |
| 1 | Bnipl | 0.78 | 8.38E-14 | 1.79E-09 | 0.31 | 0.17 | 0.14 | 1.83 |
| 1 | Igdcc3 | 0.46 | 8.57E-14 | 1.83E-09 | 0.51 | 0.32 | 0.19 | 1.58 |
| 1 | Tram2 | 1.32 | 8.62E-14 | 1.84E-09 | 0.17 | 0.07 | 0.1 | 2.36 |
| 1 | Dtx1 | 1.08 | 9.21E-14 | 1.97E-09 | 0.13 | 0.05 | 0.08 | 2.78 |
| 1 | Itga4 | 1.44 | 1.17E-13 | 2.50E-09 | 0.13 | 0.05 | 0.08 | 2.72 |
| 1 | Rhou | 0.55 | 1.24E-13 | 2.64E-09 | 0.63 | 0.46 | 0.16 | 1.35 |
| 1 | Bex2 | 0.31 | 1.40E-13 | 2.99E-09 | 0.94 | 0.87 | 0.07 | 1.08 |
| 1 | Tex15 | 0.26 | 1.55E-13 | 3.30E-09 | 0.42 | 0.25 | 0.17 | 1.68 |
| 1 | Mrpl14 | 0.29 | 1.98E-13 | 4.22E-09 | 0.93 | 0.87 | 0.07 | 1.08 |
| 1 | Tekt1 | 0.85 | 2.27E-13 | 4.85E-09 | 0.21 | 0.1 | 0.11 | 2.17 |
| 1 | Etfb | 0.3 | 2.44E-13 | 5.21E-09 | 0.93 | 0.84 | 0.1 | 1.11 |
| 1 | Xylt1 | 0.89 | 2.53E-13 | 5.40E-09 | 0.2 | 0.09 | 0.11 | 2.17 |
| 1 | Slc25a23 | 0.64 | 2.81E-13 | 5.99E-09 | 0.47 | 0.3 | 0.17 | 1.57 |
| 1 | Add3 | 0.48 | 2.81E-13 | 6.01E-09 | 0.73 | 0.58 | 0.16 | 1.27 |
| 1 | Dcps | 0.46 | 2.84E-13 | 6.07E-09 | 0.76 | 0.6 | 0.16 | 1.26 |
| 1 | Sema3f | 0.57 | 5.22E-13 | 1.11E-08 | 0.54 | 0.37 | 0.17 | 1.46 |
| 1 | Smarcd3 | 0.67 | 5.80E-13 | 1.24E-08 | 0.42 | 0.27 | 0.16 | 1.58 |
| 1 | Cpd | 0.65 | 6.78E-13 | 1.45E-08 | 0.52 | 0.36 | 0.16 | 1.44 |
| 1 | Ssrp1 | 0.26 | 6.82E-13 | 1.46E-08 | 0.95 | 0.9 | 0.05 | 1.05 |
| 1 | Cox4i2 | 0.34 | 6.83E-13 | 1.46E-08 | 0.65 | 0.46 | 0.19 | 1.41 |
| 1 | Msmo1 | 0.38 | 8.02E-13 | 1.71E-08 | 0.65 | 0.48 | 0.18 | 1.37 |
| 1 | Elf3 | 0.81 | 8.07E-13 | 1.72E-08 | 0.3 | 0.16 | 0.14 | 1.84 |
| 1 | Dnajc1 | 0.4 | 8.23E-13 | 1.76E-08 | 0.77 | 0.64 | 0.13 | 1.21 |
| 1 | Pop5 | 0.39 | 8.81E-13 | 1.88E-08 | 0.75 | 0.59 | 0.16 | 1.27 |
| 1 | Hmgb2 | 0.3 | 1.11E-12 | 2.36E-08 | 0.99 | 0.97 | 0.02 | 1.02 |
| 1 | Dhrs4 | 0.45 | 1.13E-12 | 2.42E-08 | 0.81 | 0.69 | 0.13 | 1.19 |
| 1 | Fbn1 | 0.84 | 1.19E-12 | 2.54E-08 | 0.33 | 0.19 | 0.14 | 1.73 |
| 1 | Mir99ahg | 0.38 | 1.28E-12 | 2.73E-08 | 0.68 | 0.52 | 0.17 | 1.32 |
| 1 | Scg5 | 0.56 | 1.30E-12 | 2.77E-08 | 0.33 | 0.19 | 0.14 | 1.74 |

**Supplementary Table 3. Marker genes for the original clusters of integrated clustering of E11.5 control and NotchGOF λ-junction ectoderm cells.**

| | | | | | | | | |
|---|---|---|---|---|---|---|---|---|
| 1 | Cdc42ep5 | 0.47 | 1.34E-12 | 2.87E-08 | 0.72 | 0.58 | 0.15 | 1.25 |
| 1 | Spata24 | 0.68 | 1.56E-12 | 3.33E-08 | 0.43 | 0.28 | 0.15 | 1.54 |
| 1 | Tcirg1 | 1.18 | 1.82E-12 | 3.89E-08 | 0.19 | 0.09 | 0.1 | 2.15 |
| 1 | Zdbf2 | 0.47 | 2.03E-12 | 4.34E-08 | 0.69 | 0.53 | 0.16 | 1.3 |
| 1 | Phgdh | 0.28 | 2.47E-12 | 5.28E-08 | 0.94 | 0.82 | 0.12 | 1.15 |
| 1 | Has2 | 0.75 | 3.05E-12 | 6.51E-08 | 0.23 | 0.12 | 0.12 | 1.99 |
| 1 | Endod1 | 0.94 | 3.72E-12 | 7.94E-08 | 0.27 | 0.15 | 0.12 | 1.82 |
| 1 | Rab11fip1 | 0.57 | 4.01E-12 | 8.56E-08 | 0.45 | 0.29 | 0.16 | 1.55 |
| 1 | Kcnb1 | 1.11 | 4.09E-12 | 8.73E-08 | 0.17 | 0.08 | 0.09 | 2.21 |
| 1 | Ier5l | 0.39 | 4.61E-12 | 9.85E-08 | 0.78 | 0.64 | 0.14 | 1.22 |
| 1 | Snx2 | 0.44 | 4.78E-12 | 1.02E-07 | 0.7 | 0.55 | 0.15 | 1.26 |
| 1 | Ccdc8 | 0.47 | 5.19E-12 | 1.11E-07 | 0.53 | 0.36 | 0.16 | 1.45 |
| 1 | Grk5 | 0.96 | 5.26E-12 | 1.12E-07 | 0.25 | 0.14 | 0.12 | 1.84 |
| 1 | Camk2d | 0.52 | 6.62E-12 | 1.41E-07 | 0.38 | 0.23 | 0.15 | 1.67 |
| 1 | Nab2 | 0.74 | 7.36E-12 | 1.57E-07 | 0.32 | 0.19 | 0.13 | 1.71 |
| 1 | Lifr | 0.99 | 8.34E-12 | 1.78E-07 | 0.22 | 0.11 | 0.11 | 1.94 |
| 1 | Garem1 | 0.87 | 8.41E-12 | 1.80E-07 | 0.24 | 0.13 | 0.11 | 1.9 |
| 1 | Smtnl2 | 0.53 | 8.66E-12 | 1.85E-07 | 0.4 | 0.24 | 0.15 | 1.63 |
| 1 | Ccnd3 | 0.3 | 8.86E-12 | 1.89E-07 | 0.9 | 0.79 | 0.11 | 1.14 |
| 1 | Derl3 | 1.33 | 1.05E-11 | 2.25E-07 | 0.12 | 0.05 | 0.07 | 2.54 |
| 1 | Ddx56 | 0.53 | 1.09E-11 | 2.32E-07 | 0.52 | 0.35 | 0.17 | 1.48 |
| 1 | Arhgap27os2 | 0.71 | 1.13E-11 | 2.42E-07 | 0.35 | 0.21 | 0.14 | 1.66 |
| 1 | Frk | 0.77 | 1.20E-11 | 2.57E-07 | 0.25 | 0.13 | 0.12 | 1.91 |
| 1 | Dclk1 | 0.63 | 1.45E-11 | 3.10E-07 | 0.31 | 0.18 | 0.13 | 1.74 |
| 1 | Gchfr | 0.91 | 1.51E-11 | 3.22E-07 | 0.24 | 0.13 | 0.11 | 1.86 |
| 1 | Mcm6 | 0.4 | 1.52E-11 | 3.26E-07 | 0.8 | 0.65 | 0.15 | 1.23 |
| 1 | Slitrk6 | 0.6 | 1.58E-11 | 3.37E-07 | 0.28 | 0.16 | 0.13 | 1.8 |
| 1 | Gmpr | 0.78 | 1.61E-11 | 3.45E-07 | 0.31 | 0.18 | 0.13 | 1.72 |
| 1 | Taf6l | 0.62 | 1.74E-11 | 3.73E-07 | 0.41 | 0.26 | 0.15 | 1.57 |
| 1 | Emid1 | 0.59 | 1.75E-11 | 3.74E-07 | 0.29 | 0.16 | 0.13 | 1.81 |
| 1 | Smim1 | 0.65 | 1.84E-11 | 3.94E-07 | 0.39 | 0.24 | 0.14 | 1.6 |
| 1 | Col2a1 | 0.27 | 1.97E-11 | 4.20E-07 | 0.9 | 0.8 | 0.1 | 1.12 |
| 1 | Slc16a1 | 0.59 | 1.99E-11 | 4.24E-07 | 0.51 | 0.37 | 0.14 | 1.39 |
| 1 | Ppp1r2 | 0.37 | 2.08E-11 | 4.44E-07 | 0.85 | 0.75 | 0.1 | 1.14 |
| 1 | Ttc12 | 1.16 | 2.23E-11 | 4.76E-07 | 0.15 | 0.06 | 0.08 | 2.3 |
| 1 | Spred3 | 0.72 | 2.25E-11 | 4.80E-07 | 0.24 | 0.13 | 0.11 | 1.89 |
| 1 | Laptm4b | 0.5 | 2.29E-11 | 4.90E-07 | 0.63 | 0.47 | 0.16 | 1.33 |
| 1 | Btbd3 | 0.71 | 2.30E-11 | 4.91E-07 | 0.31 | 0.18 | 0.13 | 1.73 |
| 1 | Cope | 0.29 | 2.91E-11 | 6.22E-07 | 0.88 | 0.78 | 0.1 | 1.13 |
| 1 | Fzd2 | 0.29 | 3.02E-11 | 6.46E-07 | 0.91 | 0.81 | 0.11 | 1.13 |
| 1 | Larp7 | 0.35 | 3.11E-11 | 6.64E-07 | 0.85 | 0.77 | 0.08 | 1.1 |
| 1 | 2310033P09Rik | 0.38 | 3.15E-11 | 6.72E-07 | 0.76 | 0.61 | 0.15 | 1.24 |
| 1 | Zfp467 | 0.52 | 3.17E-11 | 6.76E-07 | 0.61 | 0.45 | 0.16 | 1.35 |
| 1 | Fjx1 | 0.3 | 3.33E-11 | 7.10E-07 | 0.91 | 0.78 | 0.14 | 1.17 |
| 1 | Tuba1c | 0.59 | 3.79E-11 | 8.09E-07 | 0.63 | 0.48 | 0.15 | 1.31 |
| 1 | Lamb2 | 0.73 | 3.79E-11 | 8.09E-07 | 0.28 | 0.15 | 0.12 | 1.8 |
| 1 | Myo5c | 0.69 | 3.98E-11 | 8.50E-07 | 0.25 | 0.14 | 0.12 | 1.84 |
| 1 | Cryab | 0.3 | 4.26E-11 | 9.09E-07 | 0.3 | 0.16 | 0.13 | 1.82 |
| 1 | Tcim | 0.75 | 4.36E-11 | 9.31E-07 | 0.39 | 0.25 | 0.14 | 1.53 |
| 1 | Ruvbl2 | 0.38 | 4.61E-11 | 9.84E-07 | 0.79 | 0.65 | 0.14 | 1.21 |
| 1 | Ccnd1 | 0.4 | 4.64E-11 | 9.90E-07 | 0.79 | 0.69 | 0.1 | 1.14 |
| 1 | Acsl3 | 0.31 | 4.65E-11 | 9.92E-07 | 0.89 | 0.77 | 0.12 | 1.16 |
| 1 | Cndp2 | 0.45 | 4.71E-11 | 1.01E-06 | 0.62 | 0.48 | 0.15 | 1.31 |
| 1 | Uqcrc1 | 0.29 | 4.86E-11 | 1.04E-06 | 0.9 | 0.82 | 0.07 | 1.09 |
| 1 | Itm2a | 0.45 | 4.96E-11 | 1.06E-06 | 0.66 | 0.5 | 0.16 | 1.31 |
| 1 | Stra6 | 0.39 | 5.26E-11 | 1.12E-06 | 0.4 | 0.24 | 0.16 | 1.65 |
| 1 | Gipc1 | 0.34 | 5.89E-11 | 1.26E-06 | 0.83 | 0.74 | 0.09 | 1.12 |

**Supplementary Table 3. Marker genes for the original clusters of integrated clustering of E11.5 control and NotchGOF λ-junction ectoderm cells.**

| | | | | | | | | |
|---|---|---|---|---|---|---|---|---|
| 1 | Srebf1 | 0.62 | 6.47E-11 | 1.38E-06 | 0.42 | 0.28 | 0.15 | 1.53 |
| 1 | Sdc2 | 0.45 | 7.41E-11 | 1.58E-06 | 0.44 | 0.29 | 0.16 | 1.55 |
| 1 | Efemp2 | 0.61 | 8.19E-11 | 1.75E-06 | 0.53 | 0.37 | 0.16 | 1.41 |
| 1 | Tmem159 | 0.52 | 8.43E-11 | 1.80E-06 | 0.55 | 0.39 | 0.15 | 1.39 |
| 1 | Nup210 | 0.7 | 8.58E-11 | 1.83E-06 | 0.3 | 0.17 | 0.12 | 1.71 |
| 1 | Tom1 | 0.73 | 8.91E-11 | 1.90E-06 | 0.24 | 0.13 | 0.11 | 1.83 |
| 1 | Ric3 | 0.98 | 9.99E-11 | 2.13E-06 | 0.2 | 0.1 | 0.1 | 1.96 |
| 1 | Dmrt3 | 0.55 | 1.00E-10 | 2.14E-06 | 0.33 | 0.2 | 0.13 | 1.68 |
| 1 | Gria3 | 0.7 | 1.01E-10 | 2.16E-06 | 0.31 | 0.19 | 0.13 | 1.69 |
| 1 | Eci1 | 0.39 | 1.35E-10 | 2.89E-06 | 0.67 | 0.51 | 0.16 | 1.31 |
| 1 | Cenpb | 0.39 | 1.36E-10 | 2.91E-06 | 0.81 | 0.69 | 0.11 | 1.16 |
| 1 | Cetn3 | 0.27 | 1.39E-10 | 2.97E-06 | 0.94 | 0.89 | 0.06 | 1.06 |
| 1 | Tbc1d17 | 0.52 | 1.43E-10 | 3.05E-06 | 0.46 | 0.31 | 0.15 | 1.5 |
| 1 | Irf2bpl | 0.45 | 1.46E-10 | 3.11E-06 | 0.59 | 0.44 | 0.15 | 1.35 |
| 1 | Celsr1 | 0.49 | 1.51E-10 | 3.22E-06 | 0.52 | 0.36 | 0.16 | 1.44 |
| 1 | Prxl2b | 0.51 | 1.61E-10 | 3.43E-06 | 0.53 | 0.37 | 0.16 | 1.43 |
| 1 | Spin4 | 1.04 | 1.68E-10 | 3.58E-06 | 0.18 | 0.09 | 0.09 | 2.03 |
| 1 | Six4 | 0.44 | 1.68E-10 | 3.59E-06 | 0.61 | 0.47 | 0.14 | 1.3 |
| 1 | Epha5 | 0.83 | 1.69E-10 | 3.61E-06 | 0.17 | 0.08 | 0.09 | 2.07 |
| 1 | Tyms | 0.33 | 1.83E-10 | 3.91E-06 | 0.85 | 0.74 | 0.11 | 1.15 |
| 1 | Fech | 0.53 | 1.97E-10 | 4.22E-06 | 0.42 | 0.28 | 0.15 | 1.52 |
| 1 | Rbm47 | 1.01 | 2.15E-10 | 4.60E-06 | 0.18 | 0.09 | 0.09 | 1.99 |
| 1 | Speg | 0.83 | 2.16E-10 | 4.61E-06 | 0.22 | 0.12 | 0.1 | 1.87 |
| 1 | Timeless | 0.42 | 2.18E-10 | 4.66E-06 | 0.6 | 0.44 | 0.15 | 1.34 |
| 1 | Prr5 | 0.72 | 2.22E-10 | 4.75E-06 | 0.27 | 0.15 | 0.12 | 1.76 |
| 1 | Nsmce4a | 0.28 | 2.29E-10 | 4.89E-06 | 0.91 | 0.83 | 0.08 | 1.1 |
| 1 | Rpia | 0.53 | 2.30E-10 | 4.91E-06 | 0.49 | 0.34 | 0.15 | 1.46 |
| 1 | Tenm2 | 0.71 | 2.68E-10 | 5.73E-06 | 0.34 | 0.22 | 0.13 | 1.6 |
| 1 | Snx17 | 0.36 | 2.79E-10 | 5.95E-06 | 0.75 | 0.62 | 0.13 | 1.2 |
| 1 | Pnck | 0.71 | 3.35E-10 | 7.16E-06 | 0.27 | 0.15 | 0.12 | 1.76 |
| 1 | Cd47 | 0.41 | 3.59E-10 | 7.66E-06 | 0.7 | 0.56 | 0.14 | 1.25 |
| 1 | Nr2f6 | 0.34 | 3.70E-10 | 7.91E-06 | 0.85 | 0.73 | 0.11 | 1.15 |
| 1 | Zfpm1 | 0.68 | 3.76E-10 | 8.03E-06 | 0.31 | 0.19 | 0.12 | 1.65 |
| 1 | Prrx2 | 0.59 | 3.85E-10 | 8.23E-06 | 0.3 | 0.18 | 0.12 | 1.66 |
| 1 | Sntb1 | 1.22 | 4.79E-10 | 1.02E-05 | 0.14 | 0.06 | 0.08 | 2.21 |
| 1 | Fdft1 | 0.43 | 4.95E-10 | 1.06E-05 | 0.67 | 0.54 | 0.13 | 1.24 |
| 1 | Abhd14b | 0.8 | 5.08E-10 | 1.08E-05 | 0.22 | 0.12 | 0.1 | 1.86 |
| 1 | Gins2 | 0.4 | 5.08E-10 | 1.08E-05 | 0.7 | 0.54 | 0.16 | 1.29 |
| 1 | Hist1h3a | 0.73 | 5.39E-10 | 1.15E-05 | 0.22 | 0.12 | 0.1 | 1.88 |
| 1 | Mettl26 | 0.35 | 5.74E-10 | 1.23E-05 | 0.76 | 0.62 | 0.14 | 1.22 |
| 1 | Med13l | 0.45 | 5.77E-10 | 1.23E-05 | 0.63 | 0.5 | 0.13 | 1.25 |
| 1 | Arsb | 1.26 | 5.85E-10 | 1.25E-05 | 0.12 | 0.05 | 0.07 | 2.45 |
| 1 | Jag1 | 0.31 | 6.66E-10 | 1.42E-05 | 0.73 | 0.54 | 0.19 | 1.35 |
| 1 | Cmtm8 | 0.41 | 7.00E-10 | 1.50E-05 | 0.66 | 0.52 | 0.15 | 1.28 |
| 1 | Dhcr7 | 0.46 | 7.13E-10 | 1.52E-05 | 0.47 | 0.32 | 0.15 | 1.49 |
| 1 | Sh3pxd2a | 0.63 | 7.22E-10 | 1.54E-05 | 0.45 | 0.32 | 0.13 | 1.41 |
| 1 | Cenpm | 0.4 | 7.40E-10 | 1.58E-05 | 0.63 | 0.49 | 0.14 | 1.29 |
| 1 | Chka | 0.35 | 8.03E-10 | 1.71E-05 | 0.68 | 0.51 | 0.17 | 1.32 |
| 1 | Gstp1 | 0.43 | 8.03E-10 | 1.71E-05 | 0.54 | 0.38 | 0.16 | 1.42 |
| 1 | Hmgn5 | 0.28 | 8.04E-10 | 1.72E-05 | 0.84 | 0.72 | 0.13 | 1.17 |
| 1 | Airn | 0.63 | 8.33E-10 | 1.78E-05 | 0.41 | 0.28 | 0.13 | 1.47 |
| 1 | Prr36 | 0.58 | 9.04E-10 | 1.93E-05 | 0.39 | 0.25 | 0.14 | 1.54 |
| 1 | Trpm4 | 0.5 | 9.57E-10 | 2.04E-05 | 0.35 | 0.22 | 0.14 | 1.62 |
| 1 | Insig1 | 0.69 | 9.69E-10 | 2.07E-05 | 0.37 | 0.24 | 0.13 | 1.54 |
| 1 | Cbln2 | 0.6 | 1.01E-09 | 2.16E-05 | 0.24 | 0.13 | 0.11 | 1.82 |
| 1 | Dnah7a | 1.19 | 1.05E-09 | 2.25E-05 | 0.13 | 0.06 | 0.07 | 2.29 |
| 1 | Zmat1 | 1.12 | 1.10E-09 | 2.34E-05 | 0.11 | 0.05 | 0.07 | 2.4 |

**Supplementary Table 3. Marker genes for the original clusters of integrated clustering of E11.5 control and NotchGOF λ-junction ectoderm cells.**

| | | | | | | | | |
|---|---|---|---|---|---|---|---|---|
| 1 | Lrrk1 | 0.54 | 1.13E-09 | 2.41E-05 | 0.44 | 0.29 | 0.14 | 1.49 |
| 1 | Sfrp2 | 0.57 | 1.15E-09 | 2.46E-05 | 0.66 | 0.5 | 0.16 | 1.32 |
| 1 | Pold2 | 0.47 | 1.24E-09 | 2.65E-05 | 0.55 | 0.4 | 0.15 | 1.38 |
| 1 | Fgf16 | 0.97 | 1.32E-09 | 2.81E-05 | 0.14 | 0.06 | 0.07 | 2.21 |
| 1 | Etv1 | 0.4 | 1.32E-09 | 2.82E-05 | 0.28 | 0.16 | 0.12 | 1.74 |
| 1 | Ramp3 | 1.18 | 1.32E-09 | 2.83E-05 | 0.1 | 0.04 | 0.06 | 2.53 |
| 1 | Srcin1 | 0.75 | 1.45E-09 | 3.11E-05 | 0.23 | 0.13 | 0.1 | 1.8 |
| 1 | Actn1 | 0.39 | 1.46E-09 | 3.12E-05 | 0.7 | 0.56 | 0.14 | 1.25 |
| 1 | Abhd14a | 0.49 | 1.49E-09 | 3.18E-05 | 0.53 | 0.39 | 0.15 | 1.37 |
| 1 | Npepl1 | 0.4 | 1.59E-09 | 3.39E-05 | 0.56 | 0.41 | 0.15 | 1.38 |
| 1 | Blvrb | 0.61 | 1.61E-09 | 3.45E-05 | 0.28 | 0.17 | 0.11 | 1.69 |
| 1 | Wee1 | 0.43 | 1.87E-09 | 3.99E-05 | 0.56 | 0.41 | 0.14 | 1.34 |
| 1 | Adgrb3 | 0.71 | 2.07E-09 | 4.42E-05 | 0.2 | 0.1 | 0.1 | 1.93 |
| 1 | Timp1 | 1.22 | 2.18E-09 | 4.66E-05 | 0.15 | 0.08 | 0.08 | 2.04 |
| 1 | Ahr | 0.44 | 2.21E-09 | 4.71E-05 | 0.28 | 0.17 | 0.12 | 1.7 |
| 1 | Slit3 | 1.04 | 2.24E-09 | 4.78E-05 | 0.14 | 0.06 | 0.07 | 2.18 |
| 1 | Slc9a9 | 0.83 | 2.49E-09 | 5.32E-05 | 0.14 | 0.06 | 0.07 | 2.18 |
| 1 | Tox2 | 0.73 | 2.55E-09 | 5.44E-05 | 0.2 | 0.11 | 0.09 | 1.87 |
| 1 | Glrx | 0.4 | 2.56E-09 | 5.46E-05 | 0.42 | 0.28 | 0.14 | 1.48 |
| 1 | Rims2 | 0.52 | 2.73E-09 | 5.84E-05 | 0.32 | 0.2 | 0.12 | 1.63 |
| 1 | Psme1 | 0.29 | 2.80E-09 | 5.98E-05 | 0.88 | 0.8 | 0.08 | 1.1 |
| 1 | Elf5 | 0.76 | 2.84E-09 | 6.07E-05 | 0.34 | 0.23 | 0.12 | 1.52 |
| 1 | Cd302 | 0.48 | 3.31E-09 | 7.06E-05 | 0.41 | 0.28 | 0.14 | 1.49 |
| 1 | Dap | 0.34 | 3.33E-09 | 7.12E-05 | 0.79 | 0.67 | 0.12 | 1.18 |
| 1 | Nmi | 0.97 | 3.38E-09 | 7.23E-05 | 0.14 | 0.06 | 0.07 | 2.14 |
| 1 | Ndufv1 | 0.34 | 3.42E-09 | 7.30E-05 | 0.8 | 0.7 | 0.1 | 1.14 |
| 1 | Ephb2 | 0.39 | 3.48E-09 | 7.42E-05 | 0.55 | 0.42 | 0.14 | 1.33 |
| 1 | Cntln | 0.46 | 3.49E-09 | 7.47E-05 | 0.52 | 0.38 | 0.14 | 1.35 |
| 1 | Ndrg2 | 0.46 | 3.61E-09 | 7.71E-05 | 0.48 | 0.34 | 0.14 | 1.43 |
| 1 | Rab27a | 1.09 | 3.62E-09 | 7.73E-05 | 0.11 | 0.05 | 0.06 | 2.36 |
| 1 | Fam72a | 1.03 | 3.67E-09 | 7.84E-05 | 0.13 | 0.06 | 0.07 | 2.2 |
| 1 | Pcbd2 | 0.41 | 4.94E-09 | 1.06E-04 | 0.57 | 0.43 | 0.14 | 1.32 |
| 1 | Smc4 | 0.36 | 4.97E-09 | 1.06E-04 | 0.82 | 0.74 | 0.08 | 1.11 |
| 1 | Myo5b | 0.8 | 5.22E-09 | 1.12E-04 | 0.25 | 0.15 | 0.1 | 1.68 |
| 1 | Lamb1 | 0.36 | 5.24E-09 | 1.12E-04 | 0.66 | 0.5 | 0.16 | 1.32 |
| 1 | Pdgfrl | 0.37 | 5.27E-09 | 1.13E-04 | 0.46 | 0.32 | 0.15 | 1.46 |
| 1 | Slc35f1 | 0.9 | 5.43E-09 | 1.16E-04 | 0.18 | 0.1 | 0.09 | 1.87 |
| 1 | Ddt | 0.3 | 5.86E-09 | 1.25E-04 | 0.93 | 0.84 | 0.09 | 1.1 |
| 1 | Gpx3 | 0.75 | 5.91E-09 | 1.26E-04 | 0.21 | 0.12 | 0.1 | 1.8 |
| 1 | Ext2 | 0.4 | 6.24E-09 | 1.33E-04 | 0.57 | 0.42 | 0.15 | 1.35 |
| 1 | Hif3a | 0.91 | 6.42E-09 | 1.37E-04 | 0.2 | 0.11 | 0.09 | 1.84 |
| 1 | Nsfl1c | 0.26 | 6.71E-09 | 1.43E-04 | 0.88 | 0.8 | 0.07 | 1.09 |
| 1 | Pkp4 | 0.29 | 6.78E-09 | 1.45E-04 | 0.72 | 0.57 | 0.15 | 1.27 |
| 1 | Psmg4 | 0.28 | 6.98E-09 | 1.49E-04 | 0.84 | 0.74 | 0.11 | 1.14 |
| 1 | Dync2li1 | 0.46 | 7.27E-09 | 1.55E-04 | 0.47 | 0.33 | 0.14 | 1.43 |
| 1 | Psme2 | 0.28 | 7.35E-09 | 1.57E-04 | 0.84 | 0.75 | 0.09 | 1.11 |
| 1 | Tmem50b | 0.69 | 7.39E-09 | 1.58E-04 | 0.26 | 0.16 | 0.1 | 1.67 |
| 1 | Dcaf12l1 | 0.56 | 7.57E-09 | 1.62E-04 | 0.27 | 0.16 | 0.11 | 1.7 |
| 1 | Nrtn | 0.51 | 7.58E-09 | 1.62E-04 | 0.44 | 0.31 | 0.14 | 1.44 |
| 1 | Arhgef9 | 0.98 | 7.74E-09 | 1.65E-04 | 0.18 | 0.1 | 0.08 | 1.87 |
| 1 | Gnaz | 1 | 7.90E-09 | 1.69E-04 | 0.11 | 0.05 | 0.06 | 2.31 |
| 1 | Oard1 | 0.3 | 8.02E-09 | 1.71E-04 | 0.81 | 0.71 | 0.09 | 1.13 |
| 1 | Ltbp1 | 0.49 | 9.96E-09 | 2.13E-04 | 0.36 | 0.23 | 0.13 | 1.54 |
| 1 | Fuom | 0.47 | 1.03E-08 | 2.21E-04 | 0.45 | 0.31 | 0.14 | 1.46 |
| 1 | Slc4a8 | 0.82 | 1.04E-08 | 2.21E-04 | 0.18 | 0.09 | 0.09 | 1.9 |
| 1 | Eif4ebp1 | 0.32 | 1.06E-08 | 2.26E-04 | 0.75 | 0.62 | 0.14 | 1.23 |
| 1 | Rhbdf1 | 0.67 | 1.09E-08 | 2.33E-04 | 0.3 | 0.19 | 0.11 | 1.59 |

**Supplementary Table 3. Marker genes for the original clusters of integrated clustering of E11.5 control and NotchGOF λ-junction ectoderm cells.**

| | | | | | | | | |
|---|---|---|---|---|---|---|---|---|
| 1 | Mrpl38 | 0.3 | 1.13E-08 | 2.41E-04 | 0.75 | 0.63 | 0.12 | 1.19 |
| 1 | Hmces | 0.56 | 1.14E-08 | 2.43E-04 | 0.46 | 0.33 | 0.12 | 1.37 |
| 1 | Bckdk | 0.36 | 1.18E-08 | 2.52E-04 | 0.66 | 0.53 | 0.13 | 1.25 |
| 1 | Dmd | 0.66 | 1.26E-08 | 2.70E-04 | 0.25 | 0.15 | 0.1 | 1.71 |
| 1 | Birc5 | 0.36 | 1.30E-08 | 2.79E-04 | 0.8 | 0.7 | 0.1 | 1.14 |
| 1 | Pdia4 | 0.26 | 1.41E-08 | 3.01E-04 | 0.89 | 0.8 | 0.09 | 1.11 |
| 1 | Fxn | 0.48 | 1.45E-08 | 3.11E-04 | 0.56 | 0.45 | 0.11 | 1.26 |
| 1 | Nudt4 | 0.27 | 1.51E-08 | 3.23E-04 | 0.87 | 0.77 | 0.1 | 1.13 |
| 1 | Klf3 | 0.31 | 1.64E-08 | 3.50E-04 | 0.73 | 0.59 | 0.13 | 1.22 |
| 1 | Jph4 | 0.77 | 1.66E-08 | 3.54E-04 | 0.3 | 0.19 | 0.11 | 1.56 |
| 1 | Cux2 | 0.65 | 1.71E-08 | 3.64E-04 | 0.25 | 0.15 | 0.1 | 1.7 |
| 1 | Tbc1d30 | 0.76 | 1.75E-08 | 3.73E-04 | 0.19 | 0.1 | 0.09 | 1.85 |
| 1 | Gstt2 | 0.72 | 1.75E-08 | 3.74E-04 | 0.19 | 0.1 | 0.09 | 1.87 |
| 1 | Cox7a1 | 0.98 | 2.00E-08 | 4.27E-04 | 0.11 | 0.05 | 0.06 | 2.27 |
| 1 | Tubb4a | 0.69 | 2.01E-08 | 4.30E-04 | 0.17 | 0.09 | 0.08 | 1.93 |
| 1 | Rgs9 | 0.6 | 2.16E-08 | 4.60E-04 | 0.26 | 0.16 | 0.11 | 1.68 |
| 1 | Wdsub1 | 0.46 | 2.18E-08 | 4.65E-04 | 0.46 | 0.32 | 0.14 | 1.44 |
| 1 | St7 | 0.34 | 2.58E-08 | 5.51E-04 | 0.45 | 0.32 | 0.14 | 1.44 |
| 1 | Gins1 | 0.44 | 2.75E-08 | 5.88E-04 | 0.47 | 0.33 | 0.14 | 1.41 |
| 1 | Cmtm4 | 0.4 | 2.98E-08 | 6.36E-04 | 0.5 | 0.37 | 0.14 | 1.37 |
| 1 | Cgref1 | 0.95 | 3.01E-08 | 6.43E-04 | 0.12 | 0.06 | 0.06 | 2.15 |
| 1 | Tln2 | 0.39 | 3.22E-08 | 6.88E-04 | 0.61 | 0.47 | 0.15 | 1.31 |
| 1 | Mki67 | 0.39 | 3.38E-08 | 7.22E-04 | 0.74 | 0.64 | 0.1 | 1.16 |
| 1 | Dhx32 | 0.42 | 3.64E-08 | 7.77E-04 | 0.43 | 0.3 | 0.13 | 1.44 |
| 1 | Tctex1d2 | 0.41 | 3.69E-08 | 7.88E-04 | 0.53 | 0.39 | 0.14 | 1.36 |
| 1 | Wdr18 | 0.32 | 3.88E-08 | 8.30E-04 | 0.79 | 0.67 | 0.12 | 1.17 |
| 1 | Xkr5 | 0.95 | 3.90E-08 | 8.34E-04 | 0.14 | 0.07 | 0.07 | 1.99 |
| 1 | Gm26917 | 0.27 | 3.91E-08 | 8.36E-04 | 0.75 | 0.63 | 0.12 | 1.19 |
| 1 | Poglut3 | 0.77 | 4.10E-08 | 8.76E-04 | 0.27 | 0.17 | 0.1 | 1.59 |
| 1 | Spc24 | 0.28 | 4.19E-08 | 8.94E-04 | 0.76 | 0.65 | 0.12 | 1.18 |
| 1 | Bcl2l14 | 1.18 | 4.33E-08 | 9.25E-04 | 0.11 | 0.05 | 0.06 | 2.22 |
| 1 | B930095G15Rik | 0.57 | 4.82E-08 | 1.03E-03 | 0.24 | 0.15 | 0.1 | 1.67 |
| 1 | Cep112 | 0.76 | 5.03E-08 | 1.07E-03 | 0.26 | 0.16 | 0.1 | 1.61 |
| 1 | Pbx2 | 0.25 | 5.42E-08 | 1.16E-03 | 0.87 | 0.79 | 0.09 | 1.11 |
| 1 | Arhgef12 | 0.3 | 5.65E-08 | 1.21E-03 | 0.75 | 0.63 | 0.12 | 1.19 |
| 1 | Npas3 | 0.77 | 5.96E-08 | 1.27E-03 | 0.18 | 0.1 | 0.08 | 1.79 |
| 1 | Mettl1 | 0.28 | 6.14E-08 | 1.31E-03 | 0.75 | 0.61 | 0.14 | 1.22 |
| 1 | Triobp | 0.53 | 6.24E-08 | 1.33E-03 | 0.44 | 0.31 | 0.13 | 1.4 |
| 1 | Acaa1a | 0.39 | 6.73E-08 | 1.44E-03 | 0.59 | 0.44 | 0.15 | 1.33 |
| 1 | Mcrip2 | 0.74 | 6.92E-08 | 1.48E-03 | 0.18 | 0.1 | 0.08 | 1.81 |
| 1 | Srgap3 | 0.41 | 7.23E-08 | 1.54E-03 | 0.32 | 0.21 | 0.12 | 1.55 |
| 1 | Tdrkh | 0.63 | 7.66E-08 | 1.64E-03 | 0.17 | 0.09 | 0.08 | 1.89 |
| 1 | Pcna | 0.3 | 7.73E-08 | 1.65E-03 | 0.86 | 0.76 | 0.1 | 1.14 |
| 1 | B4galt5 | 0.81 | 8.06E-08 | 1.72E-03 | 0.2 | 0.11 | 0.08 | 1.74 |
| 1 | Hspa4l | 0.44 | 8.27E-08 | 1.77E-03 | 0.36 | 0.24 | 0.12 | 1.5 |
| 1 | 2310009B15Rik | 0.36 | 8.37E-08 | 1.79E-03 | 0.62 | 0.5 | 0.13 | 1.26 |
| 1 | Myc | 0.41 | 8.56E-08 | 1.83E-03 | 0.31 | 0.2 | 0.11 | 1.56 |
| 1 | Nectin2 | 0.29 | 9.01E-08 | 1.93E-03 | 0.72 | 0.59 | 0.13 | 1.22 |
| 1 | Ldlr | 0.63 | 9.12E-08 | 1.95E-03 | 0.32 | 0.21 | 0.11 | 1.5 |
| 1 | Tmem37 | 1.08 | 9.51E-08 | 2.03E-03 | 0.11 | 0.05 | 0.06 | 2.18 |
| 1 | Cltb | 0.4 | 9.86E-08 | 2.11E-03 | 0.44 | 0.31 | 0.13 | 1.42 |
| 1 | Fut10 | 0.85 | 1.05E-07 | 2.24E-03 | 0.16 | 0.09 | 0.08 | 1.84 |
| 1 | Utp11 | 0.29 | 1.13E-07 | 2.42E-03 | 0.81 | 0.71 | 0.1 | 1.14 |
| 1 | Tyro3 | 0.59 | 1.14E-07 | 2.44E-03 | 0.32 | 0.21 | 0.11 | 1.51 |
| 1 | Atrnl1 | 0.48 | 1.17E-07 | 2.49E-03 | 0.3 | 0.19 | 0.11 | 1.59 |
| 1 | Lmod1 | 1.02 | 1.17E-07 | 2.50E-03 | 0.11 | 0.05 | 0.06 | 2.13 |
| 1 | Adgrl1 | 0.36 | 1.18E-07 | 2.51E-03 | 0.59 | 0.46 | 0.13 | 1.28 |

**Supplementary Table 3. Marker genes for the original clusters of integrated clustering of E11.5 control and NotchGOF λ-junction ectoderm cells.**

| | | | | | | | | |
|---|---|---|---|---|---|---|---|---|
| 1 | Pmf1 | 0.25 | 1.19E-07 | 2.53E-03 | 0.72 | 0.61 | 0.12 | 1.19 |
| 1 | Kif22 | 0.48 | 1.22E-07 | 2.61E-03 | 0.52 | 0.41 | 0.11 | 1.28 |
| 1 | Zmym3 | 0.44 | 1.24E-07 | 2.64E-03 | 0.35 | 0.24 | 0.12 | 1.5 |
| 1 | Mocs2 | 0.31 | 1.35E-07 | 2.88E-03 | 0.64 | 0.51 | 0.13 | 1.25 |
| 1 | Crb3 | 0.4 | 1.39E-07 | 2.98E-03 | 0.55 | 0.43 | 0.12 | 1.29 |
| 1 | Lta4h | 0.39 | 1.40E-07 | 3.00E-03 | 0.59 | 0.46 | 0.13 | 1.27 |
| 1 | Wwc1 | 0.3 | 1.50E-07 | 3.21E-03 | 0.7 | 0.58 | 0.13 | 1.22 |
| 1 | Mboat1 | 0.84 | 1.55E-07 | 3.30E-03 | 0.17 | 0.09 | 0.08 | 1.82 |
| 1 | Tubb2a | 0.32 | 1.62E-07 | 3.46E-03 | 0.48 | 0.35 | 0.13 | 1.37 |
| 1 | Tmem189 | 0.46 | 1.63E-07 | 3.47E-03 | 0.53 | 0.4 | 0.13 | 1.31 |
| 1 | Zc2hc1a | 0.36 | 1.67E-07 | 3.57E-03 | 0.64 | 0.5 | 0.14 | 1.29 |
| 1 | Gsr | 0.38 | 1.70E-07 | 3.64E-03 | 0.61 | 0.47 | 0.15 | 1.32 |
| 1 | Tlcd1 | 0.59 | 1.71E-07 | 3.66E-03 | 0.2 | 0.11 | 0.09 | 1.75 |
| 1 | Cxxc5 | 0.26 | 1.76E-07 | 3.76E-03 | 0.7 | 0.57 | 0.13 | 1.22 |
| 1 | Prr15l | 0.6 | 1.82E-07 | 3.89E-03 | 0.23 | 0.13 | 0.09 | 1.68 |
| 1 | Metrnl | 0.44 | 1.82E-07 | 3.89E-03 | 0.27 | 0.17 | 0.1 | 1.59 |
| 1 | Mpp2 | 0.57 | 1.89E-07 | 4.04E-03 | 0.25 | 0.16 | 0.1 | 1.61 |
| 1 | Ccdc51 | 0.56 | 1.91E-07 | 4.08E-03 | 0.26 | 0.16 | 0.1 | 1.61 |
| 1 | Pgf | 0.9 | 1.95E-07 | 4.16E-03 | 0.16 | 0.09 | 0.07 | 1.82 |
| 1 | Cxcr4 | 0.33 | 1.97E-07 | 4.21E-03 | 0.28 | 0.17 | 0.1 | 1.61 |
| 1 | Hmox2 | 0.32 | 1.98E-07 | 4.22E-03 | 0.68 | 0.56 | 0.12 | 1.21 |
| 1 | Dmrta2os | 0.79 | 2.22E-07 | 4.74E-03 | 0.15 | 0.08 | 0.07 | 1.89 |
| 1 | Pak4 | 0.41 | 2.37E-07 | 5.06E-03 | 0.61 | 0.48 | 0.13 | 1.27 |
| 1 | Mccc2 | 0.49 | 2.45E-07 | 5.23E-03 | 0.45 | 0.33 | 0.12 | 1.38 |
| 1 | P3h3 | 0.61 | 2.54E-07 | 5.42E-03 | 0.23 | 0.14 | 0.09 | 1.64 |
| 1 | Lig1 | 0.27 | 2.58E-07 | 5.52E-03 | 0.78 | 0.68 | 0.1 | 1.15 |
| 1 | Nmral1 | 0.28 | 2.74E-07 | 5.86E-03 | 0.75 | 0.64 | 0.11 | 1.17 |
| 1 | Cyba | 0.29 | 2.94E-07 | 6.28E-03 | 0.66 | 0.55 | 0.11 | 1.21 |
| 1 | Edn3 | 0.73 | 2.97E-07 | 6.34E-03 | 0.19 | 0.11 | 0.08 | 1.76 |
| 1 | Tnks1bp1 | 0.55 | 3.26E-07 | 6.96E-03 | 0.23 | 0.14 | 0.09 | 1.64 |
| 1 | Lrba | 0.42 | 3.46E-07 | 7.38E-03 | 0.39 | 0.27 | 0.12 | 1.45 |
| 1 | Mpp7 | 0.25 | 3.52E-07 | 7.52E-03 | 0.66 | 0.53 | 0.13 | 1.25 |
| 1 | Tmem51 | 0.64 | 3.67E-07 | 7.84E-03 | 0.24 | 0.14 | 0.09 | 1.63 |
| 1 | Lss | 0.55 | 3.96E-07 | 8.46E-03 | 0.24 | 0.15 | 0.09 | 1.62 |
| 1 | Epn3 | 0.58 | 4.08E-07 | 8.72E-03 | 0.26 | 0.17 | 0.1 | 1.58 |
| 1 | Uaca | 0.43 | 4.14E-07 | 8.84E-03 | 0.5 | 0.38 | 0.12 | 1.33 |
| 1 | Zcchc3 | 0.32 | 4.21E-07 | 9.00E-03 | 0.71 | 0.6 | 0.11 | 1.19 |
| 1 | Sgms1 | 0.5 | 4.29E-07 | 9.17E-03 | 0.26 | 0.17 | 0.1 | 1.58 |
| 1 | Dgka | 0.55 | 4.33E-07 | 9.24E-03 | 0.24 | 0.15 | 0.09 | 1.63 |
| 1 | Aldoc | 1.01 | 4.36E-07 | 9.31E-03 | 0.12 | 0.06 | 0.06 | 1.97 |
| 1 | Ankle1 | 0.77 | 4.36E-07 | 9.32E-03 | 0.17 | 0.1 | 0.07 | 1.74 |
| 1 | Hist1h2ag | 0.44 | 4.43E-07 | 9.46E-03 | 0.29 | 0.19 | 0.1 | 1.52 |
| 1 | Dnmt3a | 0.29 | 4.52E-07 | 9.65E-03 | 0.81 | 0.71 | 0.11 | 1.15 |
| 1 | Txndc15 | 0.28 | 4.63E-07 | 9.89E-03 | 0.69 | 0.56 | 0.13 | 1.22 |
| 1 | Aimp2 | 0.31 | 4.64E-07 | 9.91E-03 | 0.65 | 0.53 | 0.12 | 1.22 |
| 1 | Bud23 | 0.29 | 4.66E-07 | 9.96E-03 | 0.72 | 0.62 | 0.1 | 1.17 |
| 1 | Pck2 | 0.56 | 4.81E-07 | 1.03E-02 | 0.25 | 0.16 | 0.09 | 1.58 |
| 1 | Gamt | 0.49 | 4.83E-07 | 1.03E-02 | 0.14 | 0.08 | 0.07 | 1.88 |
| 1 | Acad9 | 0.4 | 4.86E-07 | 1.04E-02 | 0.34 | 0.23 | 0.11 | 1.48 |
| 1 | Mfsd13a | 0.83 | 4.94E-07 | 1.06E-02 | 0.15 | 0.08 | 0.07 | 1.86 |
| 1 | Cdkn3 | 0.54 | 5.11E-07 | 1.09E-02 | 0.34 | 0.24 | 0.1 | 1.43 |
| 1 | Smpdl3a | 0.55 | 5.15E-07 | 1.10E-02 | 0.28 | 0.18 | 0.1 | 1.57 |
| 1 | Exph5 | 0.57 | 5.21E-07 | 1.11E-02 | 0.23 | 0.14 | 0.09 | 1.64 |
| 1 | Sapcd2 | 0.51 | 5.49E-07 | 1.17E-02 | 0.38 | 0.28 | 0.11 | 1.4 |
| 1 | Ptprd | 0.31 | 6.00E-07 | 1.28E-02 | 0.67 | 0.55 | 0.13 | 1.23 |
| 1 | Pspc1 | 0.31 | 6.20E-07 | 1.32E-02 | 0.72 | 0.6 | 0.12 | 1.21 |
| 1 | Anxa6 | 0.51 | 6.29E-07 | 1.34E-02 | 0.2 | 0.12 | 0.08 | 1.7 |

**Supplementary Table 3. Marker genes for the original clusters of integrated clustering of E11.5 control and NotchGOF λ-junction ectoderm cells.**

| | | | | | | | | |
|---|---|---|---|---|---|---|---|---|
| 1 | Nhsl2 | 0.75 | 6.34E-07 | 1.35E-02 | 0.2 | 0.12 | 0.08 | 1.68 |
| 1 | Disp1 | 0.8 | 6.37E-07 | 1.36E-02 | 0.23 | 0.15 | 0.09 | 1.59 |
| 1 | Grhpr | 0.31 | 6.43E-07 | 1.37E-02 | 0.61 | 0.48 | 0.13 | 1.28 |
| 1 | Ptgr1 | 0.54 | 6.64E-07 | 1.42E-02 | 0.24 | 0.15 | 0.09 | 1.61 |
| 1 | Kazald1 | 0.53 | 6.83E-07 | 1.46E-02 | 0.14 | 0.08 | 0.07 | 1.91 |
| 1 | Cisd3 | 0.87 | 6.89E-07 | 1.47E-02 | 0.13 | 0.07 | 0.06 | 1.94 |
| 1 | Ubtd1 | 0.49 | 6.94E-07 | 1.48E-02 | 0.21 | 0.13 | 0.09 | 1.67 |
| 1 | Ubac2 | 0.32 | 6.96E-07 | 1.49E-02 | 0.58 | 0.45 | 0.13 | 1.28 |
| 1 | Hist2h2be | 0.71 | 6.98E-07 | 1.49E-02 | 0.13 | 0.07 | 0.06 | 1.94 |
| 1 | Il17re | 0.43 | 7.15E-07 | 1.53E-02 | 0.27 | 0.18 | 0.1 | 1.56 |
| 1 | Amer1 | 0.35 | 7.38E-07 | 1.58E-02 | 0.69 | 0.57 | 0.12 | 1.2 |
| 1 | Zfp536 | 0.61 | 7.56E-07 | 1.61E-02 | 0.24 | 0.15 | 0.09 | 1.58 |
| 1 | Nbn | 0.35 | 7.70E-07 | 1.64E-02 | 0.44 | 0.32 | 0.12 | 1.37 |
| 1 | Tom1l1 | 0.33 | 7.75E-07 | 1.66E-02 | 0.6 | 0.47 | 0.13 | 1.29 |
| 1 | Ston2 | 0.31 | 7.87E-07 | 1.68E-02 | 0.7 | 0.59 | 0.1 | 1.18 |
| 1 | Aig1 | 0.26 | 7.98E-07 | 1.70E-02 | 0.68 | 0.53 | 0.15 | 1.28 |
| 1 | Srgap2 | 0.49 | 8.19E-07 | 1.75E-02 | 0.41 | 0.3 | 0.11 | 1.37 |
| 1 | Fam20c | 0.72 | 8.29E-07 | 1.77E-02 | 0.14 | 0.08 | 0.07 | 1.88 |
| 1 | Gm26793 | 0.76 | 8.55E-07 | 1.83E-02 | 0.13 | 0.07 | 0.06 | 1.91 |
| 1 | Pdia5 | 0.6 | 8.66E-07 | 1.85E-02 | 0.2 | 0.12 | 0.08 | 1.68 |
| 1 | Rassf10 | 0.46 | 8.72E-07 | 1.86E-02 | 0.34 | 0.24 | 0.11 | 1.46 |
| 1 | Grtp1 | 0.45 | 8.72E-07 | 1.86E-02 | 0.33 | 0.22 | 0.11 | 1.48 |
| 1 | Slc22a17 | 0.37 | 8.84E-07 | 1.89E-02 | 0.51 | 0.39 | 0.13 | 1.33 |
| 1 | St8sia6 | 0.47 | 8.91E-07 | 1.90E-02 | 0.26 | 0.16 | 0.09 | 1.58 |
| 1 | Syngr1 | 0.47 | 9.05E-07 | 1.93E-02 | 0.31 | 0.21 | 0.1 | 1.48 |
| 1 | Hist1h2bb | 0.44 | 9.45E-07 | 2.02E-02 | 0.24 | 0.15 | 0.09 | 1.61 |
| 1 | Prrg3 | 0.51 | 9.71E-07 | 2.07E-02 | 0.25 | 0.16 | 0.09 | 1.58 |
| 1 | Ctnnd2 | 0.31 | 1.00E-06 | 2.14E-02 | 0.57 | 0.44 | 0.13 | 1.29 |
| 1 | Plk1 | 0.53 | 1.00E-06 | 2.14E-02 | 0.49 | 0.39 | 0.1 | 1.25 |
| 1 | Farp1 | 0.3 | 1.04E-06 | 2.22E-02 | 0.53 | 0.4 | 0.13 | 1.32 |
| 1 | Supt3 | 0.36 | 1.04E-06 | 2.23E-02 | 0.48 | 0.36 | 0.12 | 1.35 |
| 1 | Twf2 | 0.4 | 1.05E-06 | 2.23E-02 | 0.46 | 0.35 | 0.12 | 1.34 |
| 1 | Hist1h3e | 0.44 | 1.07E-06 | 2.29E-02 | 0.44 | 0.34 | 0.1 | 1.3 |
| 1 | Slc2a4 | 0.58 | 1.10E-06 | 2.35E-02 | 0.19 | 0.11 | 0.08 | 1.72 |
| 1 | Tnfaip8l1 | 0.91 | 1.20E-06 | 2.56E-02 | 0.11 | 0.05 | 0.06 | 2.06 |
| 1 | Socs3 | 0.46 | 1.21E-06 | 2.59E-02 | 0.26 | 0.16 | 0.09 | 1.57 |
| 1 | Phlpp1 | 0.47 | 1.26E-06 | 2.70E-02 | 0.36 | 0.25 | 0.11 | 1.44 |
| 1 | Drg1 | 0.28 | 1.30E-06 | 2.78E-02 | 0.73 | 0.6 | 0.12 | 1.2 |
| 1 | Mtx1 | 0.32 | 1.36E-06 | 2.90E-02 | 0.63 | 0.51 | 0.13 | 1.25 |
| 1 | Dlg1 | 0.37 | 1.39E-06 | 2.98E-02 | 0.54 | 0.41 | 0.13 | 1.3 |
| 1 | E2f5 | 0.31 | 1.40E-06 | 2.99E-02 | 0.53 | 0.41 | 0.12 | 1.3 |
| 1 | Tmem88 | 0.83 | 1.44E-06 | 3.07E-02 | 0.14 | 0.07 | 0.06 | 1.85 |
| 1 | Sobp | 0.29 | 1.46E-06 | 3.12E-02 | 0.34 | 0.23 | 0.11 | 1.48 |
| 1 | Nsmce1 | 0.3 | 1.60E-06 | 3.41E-02 | 0.76 | 0.67 | 0.09 | 1.13 |
| 1 | Serpinf1 | 0.48 | 1.60E-06 | 3.42E-02 | 0.25 | 0.16 | 0.09 | 1.57 |
| 1 | Oscp1 | 0.46 | 1.63E-06 | 3.47E-02 | 0.27 | 0.17 | 0.09 | 1.52 |
| 1 | Zfyve21 | 0.3 | 1.65E-06 | 3.53E-02 | 0.54 | 0.41 | 0.13 | 1.31 |
| 1 | Cdc20 | 0.45 | 1.75E-06 | 3.73E-02 | 0.62 | 0.52 | 0.1 | 1.19 |
| 1 | Cdca3 | 0.37 | 1.75E-06 | 3.74E-02 | 0.68 | 0.58 | 0.1 | 1.16 |
| 1 | Abr | 0.47 | 1.77E-06 | 3.78E-02 | 0.3 | 0.21 | 0.1 | 1.46 |
| 1 | Gas2l3 | 0.53 | 1.80E-06 | 3.84E-02 | 0.32 | 0.23 | 0.1 | 1.42 |
| 1 | Fam83d | 0.76 | 1.97E-06 | 4.21E-02 | 0.26 | 0.18 | 0.08 | 1.46 |
| 1 | Pi4ka | 0.42 | 2.12E-06 | 4.52E-02 | 0.28 | 0.18 | 0.1 | 1.52 |
| 1 | Klhl42 | 0.59 | 2.12E-06 | 4.53E-02 | 0.16 | 0.09 | 0.07 | 1.76 |
| 1 | Tapt1 | 0.59 | 2.14E-06 | 4.56E-02 | 0.23 | 0.15 | 0.09 | 1.57 |
| 1 | BC055324 | 0.48 | 2.17E-06 | 4.64E-02 | 0.3 | 0.2 | 0.1 | 1.49 |
| 1 | Nek1 | 0.4 | 2.19E-06 | 4.68E-02 | 0.44 | 0.33 | 0.11 | 1.35 |

**Supplementary Table 3. Marker genes for the original clusters of integrated clustering of E11.5 control and NotchGOF λ-junction ectoderm cells.**

| | | | | | | | | |
|---|---|---|---|---|---|---|---|---|
| 1 | Cdk10 | 0.48 | 2.19E-06 | 4.68E-02 | 0.36 | 0.26 | 0.1 | 1.39 |
| 1 | Pkp2 | 0.37 | 2.30E-06 | 4.92E-02 | 0.3 | 0.2 | 0.1 | 1.51 |
| 2 | Sfn | 1.86 | 2.36E-154 | 5.04E-150 | 0.98 | 0.66 | 0.32 | 1.48 |
| 2 | Grhl3 | 2.59 | 8.67E-142 | 1.85E-137 | 0.63 | 0.16 | 0.47 | 3.95 |
| 2 | Zfp750 | 2.46 | 4.62E-132 | 9.88E-128 | 0.6 | 0.15 | 0.45 | 4.1 |
| 2 | Nectin4 | 1.74 | 6.68E-126 | 1.43E-121 | 0.9 | 0.48 | 0.42 | 1.87 |
| 2 | Tctex1d1 | 3.22 | 1.41E-124 | 3.01E-120 | 0.4 | 0.06 | 0.34 | 6.5 |
| 2 | Notch1 | 1.91 | 1.53E-117 | 3.26E-113 | 0.92 | 0.59 | 0.34 | 1.58 |
| 2 | Ppl | 3.01 | 1.28E-116 | 2.73E-112 | 0.5 | 0.12 | 0.39 | 4.24 |
| 2 | Gm16136 | 1.63 | 7.30E-111 | 1.56E-106 | 0.92 | 0.62 | 0.3 | 1.49 |
| 2 | Tgm1 | 3.97 | 1.75E-104 | 3.73E-100 | 0.25 | 0.02 | 0.23 | 10.54 |
| 2 | Cd9 | 1.52 | 3.33E-101 | 7.12E-97 | 0.92 | 0.69 | 0.23 | 1.33 |
| 2 | Nav3 | 1.66 | 1.72E-97 | 3.67E-93 | 0.76 | 0.33 | 0.43 | 2.29 |
| 2 | Tacstd2 | 2.87 | 3.16E-96 | 6.74E-92 | 0.62 | 0.23 | 0.39 | 2.71 |
| 2 | Cstb | 1.31 | 4.09E-95 | 8.74E-91 | 0.97 | 0.83 | 0.13 | 1.16 |
| 2 | Hspb8 | 1.53 | 1.09E-90 | 2.34E-86 | 0.83 | 0.44 | 0.39 | 1.87 |
| 2 | S100a11 | 1.05 | 3.06E-90 | 6.53E-86 | 1 | 0.9 | 0.1 | 1.11 |
| 2 | Ovol1 | 3.13 | 1.79E-89 | 3.82E-85 | 0.3 | 0.05 | 0.26 | 6.33 |
| 2 | Dab1 | 1.43 | 3.79E-86 | 8.09E-82 | 0.82 | 0.45 | 0.38 | 1.85 |
| 2 | Smagp | 3.25 | 3.51E-85 | 7.50E-81 | 0.26 | 0.03 | 0.22 | 7.73 |
| 2 | Hebp2 | 2.1 | 5.63E-85 | 1.20E-80 | 0.63 | 0.27 | 0.36 | 2.36 |
| 2 | Pik3r1 | 1.13 | 2.63E-84 | 5.62E-80 | 0.9 | 0.56 | 0.34 | 1.62 |
| 2 | Irf6 | 1.43 | 3.02E-84 | 6.45E-80 | 0.77 | 0.37 | 0.41 | 2.11 |
| 2 | Dstn | 0.73 | 7.30E-77 | 1.56E-72 | 0.99 | 0.98 | 0.01 | 1.01 |
| 2 | S100a10 | 1.37 | 1.18E-74 | 2.52E-70 | 0.93 | 0.72 | 0.2 | 1.28 |
| 2 | Msx1 | 1.18 | 4.11E-74 | 8.78E-70 | 0.92 | 0.71 | 0.22 | 1.3 |
| 2 | Id3 | 0.94 | 1.46E-73 | 3.11E-69 | 1 | 0.96 | 0.04 | 1.04 |
| 2 | Snx31 | 3.52 | 7.04E-72 | 1.50E-67 | 0.19 | 0.02 | 0.17 | 9.55 |
| 2 | Gabrp | 5.49 | 5.10E-68 | 1.09E-63 | 0.12 | 0.01 | 0.12 | 20.5 |
| 2 | Gsta3 | 2.87 | 1.14E-67 | 2.43E-63 | 0.23 | 0.03 | 0.19 | 6.85 |
| 2 | Cyb5a | 0.66 | 1.64E-67 | 3.50E-63 | 0.99 | 0.96 | 0.03 | 1.03 |
| 2 | Slc2a3 | 2.32 | 4.28E-66 | 9.13E-62 | 0.64 | 0.31 | 0.33 | 2.05 |
| 2 | Acaa2 | 1.12 | 8.97E-65 | 1.92E-60 | 0.91 | 0.69 | 0.22 | 1.32 |
| 2 | Tmsb10 | 0.45 | 5.89E-64 | 1.26E-59 | 1 | 1 | 0 | 1 |
| 2 | Cpm | 1.28 | 2.04E-63 | 4.36E-59 | 0.81 | 0.5 | 0.32 | 1.64 |
| 2 | Rbp1 | 1.29 | 6.22E-63 | 1.33E-58 | 0.96 | 0.92 | 0.05 | 1.05 |
| 2 | Arhgap29 | 1.75 | 7.22E-63 | 1.54E-58 | 0.7 | 0.36 | 0.33 | 1.91 |
| 2 | Krt5 | 1.86 | 1.29E-62 | 2.75E-58 | 0.64 | 0.3 | 0.34 | 2.15 |
| 2 | Cldn6 | 0.74 | 3.45E-62 | 7.37E-58 | 0.98 | 0.86 | 0.12 | 1.14 |
| 2 | Lypd3 | 2.35 | 4.16E-61 | 8.88E-57 | 0.32 | 0.08 | 0.24 | 4.09 |
| 2 | Thrb | 2.02 | 4.40E-61 | 9.39E-57 | 0.39 | 0.12 | 0.27 | 3.34 |
| 2 | Rhov | 1.82 | 2.63E-60 | 5.62E-56 | 0.5 | 0.19 | 0.31 | 2.63 |
| 2 | Cacna1b | 1.44 | 4.37E-59 | 9.33E-55 | 0.6 | 0.27 | 0.33 | 2.24 |
| 2 | Emp1 | 2.52 | 2.53E-58 | 5.40E-54 | 0.41 | 0.14 | 0.27 | 2.92 |
| 2 | Laptm4a | 0.67 | 2.16E-57 | 4.61E-53 | 0.99 | 0.96 | 0.03 | 1.03 |
| 2 | Fabp5 | 0.57 | 7.22E-57 | 1.54E-52 | 1 | 0.99 | 0.01 | 1.01 |
| 2 | Rhcg | 2.57 | 9.73E-57 | 2.08E-52 | 0.18 | 0.02 | 0.15 | 7.38 |
| 2 | Sox9 | 1.12 | 1.43E-56 | 3.06E-52 | 0.84 | 0.54 | 0.3 | 1.55 |
| 2 | Cox4i2 | 1.06 | 1.62E-55 | 3.46E-51 | 0.77 | 0.45 | 0.32 | 1.72 |
| 2 | Pdlim1 | 1.14 | 4.92E-55 | 1.05E-50 | 0.87 | 0.62 | 0.25 | 1.41 |
| 2 | Hexb | 1.72 | 1.58E-54 | 3.38E-50 | 0.54 | 0.24 | 0.3 | 2.27 |
| 2 | Ddx58 | 2.02 | 7.09E-54 | 1.52E-49 | 0.33 | 0.09 | 0.24 | 3.65 |
| 2 | Ikzf2 | 0.8 | 7.29E-54 | 1.56E-49 | 0.94 | 0.83 | 0.11 | 1.14 |
| 2 | Msx2 | 1.06 | 2.16E-53 | 4.60E-49 | 0.71 | 0.36 | 0.35 | 1.97 |
| 2 | Jup | 1.14 | 2.24E-53 | 4.78E-49 | 0.77 | 0.49 | 0.28 | 1.58 |
| 2 | Gucy1a1 | 1.85 | 3.87E-53 | 8.28E-49 | 0.44 | 0.16 | 0.27 | 2.69 |
| 2 | Arf1 | 0.5 | 5.40E-53 | 1.15E-48 | 0.99 | 0.98 | 0.01 | 1.01 |

**Supplementary Table 3. Marker genes for the original clusters of integrated clustering of E11.5 control and NotchGOF λ-junction ectoderm cells.**

| | | | | | | | | |
|---|---|---|---|---|---|---|---|---|
| 2 | Pkp1 | 1.63 | 5.78E-53 | 1.23E-48 | 0.3 | 0.08 | 0.23 | 4.03 |
| 2 | Samd5 | 0.93 | 2.62E-52 | 5.60E-48 | 0.91 | 0.66 | 0.25 | 1.38 |
| 2 | Eaf2 | 2.35 | 8.89E-52 | 1.90E-47 | 0.26 | 0.06 | 0.2 | 4.24 |
| 2 | Palm3 | 1.36 | 1.21E-51 | 2.59E-47 | 0.62 | 0.31 | 0.31 | 1.98 |
| 2 | Fjx1 | 0.8 | 3.40E-51 | 7.27E-47 | 0.93 | 0.78 | 0.16 | 1.2 |
| 2 | Myl12b | 0.62 | 5.31E-51 | 1.13E-46 | 0.98 | 0.93 | 0.04 | 1.04 |
| 2 | Atox1 | 0.54 | 7.12E-51 | 1.52E-46 | 0.99 | 0.98 | 0.02 | 1.02 |
| 2 | Cldn7 | 0.85 | 2.04E-50 | 4.35E-46 | 0.95 | 0.83 | 0.12 | 1.14 |
| 2 | Anxa2 | 0.9 | 3.91E-50 | 8.36E-46 | 0.96 | 0.73 | 0.23 | 1.32 |
| 2 | Bspry | 1.46 | 4.18E-50 | 8.93E-46 | 0.52 | 0.23 | 0.29 | 2.27 |
| 2 | Gja1 | 1.12 | 6.67E-50 | 1.42E-45 | 0.87 | 0.68 | 0.19 | 1.27 |
| 2 | Rgs2 | 2.04 | 1.50E-49 | 3.20E-45 | 0.54 | 0.26 | 0.27 | 2.05 |
| 2 | Ppp2r2b | 1.37 | 2.43E-49 | 5.19E-45 | 0.65 | 0.34 | 0.31 | 1.91 |
| 2 | Alad | 0.85 | 3.05E-49 | 6.51E-45 | 0.91 | 0.73 | 0.18 | 1.25 |
| 2 | Gas1 | 1.38 | 3.46E-49 | 7.40E-45 | 0.77 | 0.54 | 0.23 | 1.41 |
| 2 | St6galnac5 | 3.06 | 8.75E-48 | 1.87E-43 | 0.17 | 0.03 | 0.14 | 6.42 |
| 2 | Cmas | 0.99 | 1.08E-47 | 2.30E-43 | 0.84 | 0.63 | 0.21 | 1.33 |
| 2 | Hras | 0.61 | 2.19E-47 | 4.67E-43 | 0.96 | 0.9 | 0.06 | 1.07 |
| 2 | Smtnl2 | 1.5 | 2.51E-47 | 5.36E-43 | 0.51 | 0.23 | 0.28 | 2.26 |
| 2 | Nectin3 | 0.81 | 6.67E-47 | 1.42E-42 | 0.9 | 0.72 | 0.18 | 1.25 |
| 2 | Klhl23 | 1.09 | 8.48E-47 | 1.81E-42 | 0.75 | 0.49 | 0.26 | 1.53 |
| 2 | D7Ertd443e | 1.86 | 1.11E-46 | 2.38E-42 | 0.28 | 0.08 | 0.21 | 3.71 |
| 2 | Tspan7 | 1.06 | 1.77E-46 | 3.78E-42 | 0.72 | 0.44 | 0.28 | 1.65 |
| 2 | Cldn4 | 1.38 | 2.90E-46 | 6.20E-42 | 0.66 | 0.39 | 0.27 | 1.7 |
| 2 | Gjb3 | 1.69 | 5.92E-46 | 1.26E-41 | 0.36 | 0.12 | 0.24 | 3 |
| 2 | Rhob | 1.15 | 5.17E-45 | 1.10E-40 | 0.71 | 0.45 | 0.26 | 1.59 |
| 2 | Dlx3 | 1.13 | 6.92E-45 | 1.48E-40 | 0.55 | 0.25 | 0.3 | 2.22 |
| 2 | Rab15 | 1.15 | 8.52E-45 | 1.82E-40 | 0.67 | 0.39 | 0.28 | 1.72 |
| 2 | Rab11fip1 | 1.35 | 2.21E-44 | 4.72E-40 | 0.55 | 0.28 | 0.28 | 2 |
| 2 | Wnt4 | 0.69 | 3.14E-44 | 6.71E-40 | 0.93 | 0.68 | 0.25 | 1.36 |
| 2 | Il17re | 1.49 | 3.22E-44 | 6.88E-40 | 0.41 | 0.15 | 0.26 | 2.68 |
| 2 | Efna1 | 1.09 | 5.12E-44 | 1.09E-39 | 0.67 | 0.37 | 0.3 | 1.82 |
| 2 | Cldn8 | 1.94 | 5.38E-44 | 1.15E-39 | 0.32 | 0.1 | 0.22 | 3.17 |
| 2 | Penk | 3.38 | 5.45E-44 | 1.16E-39 | 0.19 | 0.04 | 0.15 | 5 |
| 2 | Anxa9 | 2.93 | 1.56E-43 | 3.34E-39 | 0.15 | 0.02 | 0.13 | 6.73 |
| 2 | Cdh1 | 0.94 | 1.64E-43 | 3.51E-39 | 0.84 | 0.63 | 0.21 | 1.33 |
| 2 | Aplp2 | 0.6 | 8.31E-43 | 1.77E-38 | 0.98 | 0.91 | 0.07 | 1.08 |
| 2 | Kif13b | 1.28 | 1.59E-42 | 3.41E-38 | 0.58 | 0.3 | 0.27 | 1.9 |
| 2 | Rab27b | 1.18 | 4.11E-42 | 8.78E-38 | 0.58 | 0.29 | 0.29 | 2.03 |
| 2 | Dsp | 0.73 | 7.62E-42 | 1.63E-37 | 0.97 | 0.86 | 0.11 | 1.13 |
| 2 | Plcd1 | 2.05 | 7.93E-42 | 1.69E-37 | 0.3 | 0.1 | 0.2 | 3.08 |
| 2 | Cd24a | 0.53 | 9.21E-42 | 1.97E-37 | 0.99 | 0.96 | 0.04 | 1.04 |
| 2 | Hs3st2 | 3.59 | 1.88E-41 | 4.02E-37 | 0.1 | 0.01 | 0.09 | 11.22 |
| 2 | Dennd2c | 1.97 | 2.42E-41 | 5.16E-37 | 0.27 | 0.08 | 0.19 | 3.48 |
| 2 | Kcnk1 | 0.99 | 3.34E-41 | 7.14E-37 | 0.79 | 0.56 | 0.23 | 1.42 |
| 2 | Lurap1l | 1.3 | 3.68E-41 | 7.87E-37 | 0.5 | 0.23 | 0.27 | 2.15 |
| 2 | Id1 | 0.56 | 3.90E-41 | 8.33E-37 | 0.99 | 0.92 | 0.06 | 1.07 |
| 2 | Hes1 | 0.87 | 5.08E-41 | 1.08E-36 | 0.87 | 0.71 | 0.16 | 1.22 |
| 2 | Cldn3 | 0.82 | 9.91E-41 | 2.12E-36 | 0.86 | 0.61 | 0.25 | 1.42 |
| 2 | Cacng4 | 1.15 | 2.71E-40 | 5.80E-36 | 0.59 | 0.31 | 0.28 | 1.9 |
| 2 | Mtss2 | 1.16 | 4.03E-40 | 8.60E-36 | 0.63 | 0.36 | 0.27 | 1.73 |
| 2 | Proser2 | 1.55 | 1.11E-39 | 2.38E-35 | 0.36 | 0.13 | 0.23 | 2.72 |
| 2 | Plce1 | 1.26 | 1.30E-39 | 2.78E-35 | 0.57 | 0.3 | 0.27 | 1.89 |
| 2 | Rbpms | 1.16 | 1.34E-39 | 2.86E-35 | 0.69 | 0.46 | 0.23 | 1.5 |
| 2 | Ablim1 | 0.95 | 1.43E-39 | 3.06E-35 | 0.77 | 0.52 | 0.24 | 1.47 |
| 2 | Itm2b | 0.38 | 4.12E-39 | 8.79E-35 | 0.99 | 0.93 | 0.06 | 1.06 |
| 2 | Rrbp1 | 0.69 | 8.21E-39 | 1.75E-34 | 0.95 | 0.88 | 0.07 | 1.08 |

**Supplementary Table 3. Marker genes for the original clusters of integrated clustering of E11.5 control and NotchGOF λ-junction ectoderm cells.**

| | | | | | | | | |
|---|---|---|---|---|---|---|---|---|
| 2 | Rcsd1 | 1.66 | 1.29E-38 | 2.75E-34 | 0.36 | 0.14 | 0.22 | 2.6 |
| 2 | Plk2 | 1.13 | 1.95E-38 | 4.16E-34 | 0.6 | 0.33 | 0.27 | 1.83 |
| 2 | Wls | 0.58 | 8.46E-38 | 1.81E-33 | 0.95 | 0.78 | 0.17 | 1.22 |
| 2 | Bmp4 | 0.81 | 4.91E-37 | 1.05E-32 | 0.77 | 0.5 | 0.27 | 1.53 |
| 2 | Ahr | 1.45 | 1.63E-36 | 3.49E-32 | 0.38 | 0.15 | 0.23 | 2.48 |
| 2 | Pdzrn3 | 1.73 | 1.72E-36 | 3.68E-32 | 0.29 | 0.1 | 0.2 | 3 |
| 2 | Clcc1 | 1.08 | 2.06E-36 | 4.40E-32 | 0.67 | 0.45 | 0.23 | 1.51 |
| 2 | Cndp1 | 2.54 | 2.71E-36 | 5.79E-32 | 0.14 | 0.03 | 0.12 | 5.54 |
| 2 | Ppp1r13l | 1.44 | 8.29E-36 | 1.77E-31 | 0.4 | 0.17 | 0.23 | 2.37 |
| 2 | Apoc1 | 1.32 | 2.30E-35 | 4.91E-31 | 0.51 | 0.26 | 0.25 | 1.95 |
| 2 | Ptpn13 | 0.94 | 1.49E-34 | 3.17E-30 | 0.77 | 0.55 | 0.22 | 1.41 |
| 2 | Arg1 | 2.56 | 5.41E-34 | 1.15E-29 | 0.17 | 0.04 | 0.13 | 4.46 |
| 2 | Slit3 | 2 | 9.22E-34 | 1.97E-29 | 0.2 | 0.05 | 0.15 | 3.77 |
| 2 | Kdm5b | 0.56 | 9.68E-34 | 2.07E-29 | 0.96 | 0.92 | 0.04 | 1.05 |
| 2 | Mpzl2 | 0.76 | 1.39E-33 | 2.97E-29 | 0.87 | 0.72 | 0.15 | 1.21 |
| 2 | Tpm1 | 0.56 | 1.40E-33 | 2.99E-29 | 0.98 | 0.93 | 0.05 | 1.05 |
| 2 | Nebl | 1.5 | 1.95E-33 | 4.17E-29 | 0.6 | 0.37 | 0.22 | 1.6 |
| 2 | Gria3 | 1.35 | 2.28E-33 | 4.88E-29 | 0.4 | 0.17 | 0.22 | 2.29 |
| 2 | Ctsh | 1.73 | 3.80E-33 | 8.11E-29 | 0.29 | 0.1 | 0.18 | 2.77 |
| 2 | Ostf1 | 1.01 | 6.21E-33 | 1.33E-28 | 0.63 | 0.39 | 0.24 | 1.6 |
| 2 | Evpl | 1.72 | 7.23E-33 | 1.55E-28 | 0.28 | 0.1 | 0.18 | 2.86 |
| 2 | Tgfb2 | 0.53 | 8.53E-33 | 1.82E-28 | 0.81 | 0.55 | 0.26 | 1.48 |
| 2 | Mir9-3hg | 1.46 | 1.32E-32 | 2.83E-28 | 0.26 | 0.08 | 0.18 | 3.13 |
| 2 | Trim2 | 0.86 | 1.37E-32 | 2.93E-28 | 0.76 | 0.57 | 0.19 | 1.33 |
| 2 | Adi1 | 1.07 | 1.66E-32 | 3.54E-28 | 0.65 | 0.44 | 0.21 | 1.49 |
| 2 | Sh2d4a | 2.28 | 1.90E-32 | 4.07E-28 | 0.15 | 0.03 | 0.12 | 4.75 |
| 2 | Ier5 | 0.56 | 2.78E-32 | 5.94E-28 | 0.94 | 0.83 | 0.11 | 1.14 |
| 2 | Esrp2 | 1.3 | 2.93E-32 | 6.25E-28 | 0.46 | 0.23 | 0.23 | 1.97 |
| 2 | Bambi | 0.35 | 1.32E-31 | 2.82E-27 | 0.73 | 0.45 | 0.28 | 1.61 |
| 2 | Ppm1l | 0.99 | 1.46E-31 | 3.12E-27 | 0.63 | 0.4 | 0.23 | 1.56 |
| 2 | Tap1 | 1.81 | 1.57E-31 | 3.35E-27 | 0.2 | 0.06 | 0.15 | 3.61 |
| 2 | Pdcd4 | 0.71 | 1.91E-31 | 4.08E-27 | 0.92 | 0.77 | 0.15 | 1.19 |
| 2 | Dlx2 | 0.73 | 2.76E-31 | 5.90E-27 | 0.59 | 0.31 | 0.28 | 1.89 |
| 2 | Ywhaz | 0.38 | 2.81E-31 | 5.99E-27 | 0.99 | 0.97 | 0.01 | 1.01 |
| 2 | Marveld2 | 1.31 | 5.26E-31 | 1.12E-26 | 0.43 | 0.2 | 0.23 | 2.13 |
| 2 | Cyb5r3 | 0.45 | 5.27E-31 | 1.13E-26 | 0.97 | 0.92 | 0.05 | 1.05 |
| 2 | Cdkn2b | 0.63 | 3.37E-30 | 7.19E-26 | 0.25 | 0.08 | 0.17 | 3.14 |
| 2 | Bicdl2 | 1.98 | 3.89E-30 | 8.30E-26 | 0.16 | 0.04 | 0.12 | 4.13 |
| 2 | Erbb3 | 0.79 | 4.16E-30 | 8.88E-26 | 0.74 | 0.51 | 0.23 | 1.44 |
| 2 | Ptprf | 0.48 | 4.25E-30 | 9.08E-26 | 0.99 | 0.96 | 0.03 | 1.04 |
| 2 | Kcnj15 | 1.99 | 1.17E-29 | 2.51E-25 | 0.17 | 0.04 | 0.13 | 3.89 |
| 2 | Platr21 | 1.72 | 1.50E-29 | 3.20E-25 | 0.25 | 0.08 | 0.16 | 2.95 |
| 2 | Perp | 0.73 | 1.60E-29 | 3.43E-25 | 0.78 | 0.51 | 0.27 | 1.53 |
| 2 | Klf5 | 0.95 | 3.02E-29 | 6.45E-25 | 0.62 | 0.39 | 0.24 | 1.61 |
| 2 | Gas7 | 1.61 | 3.51E-29 | 7.51E-25 | 0.25 | 0.09 | 0.16 | 2.91 |
| 2 | Rcn1 | 0.55 | 3.59E-29 | 7.67E-25 | 0.93 | 0.84 | 0.09 | 1.11 |
| 2 | Rnf24 | 0.89 | 4.77E-29 | 1.02E-24 | 0.61 | 0.38 | 0.23 | 1.6 |
| 2 | Fermt2 | 0.75 | 5.46E-29 | 1.17E-24 | 0.8 | 0.65 | 0.16 | 1.24 |
| 2 | Gstt3 | 1.66 | 7.01E-29 | 1.50E-24 | 0.27 | 0.1 | 0.17 | 2.7 |
| 2 | Spint1 | 1.03 | 7.83E-29 | 1.67E-24 | 0.6 | 0.38 | 0.22 | 1.58 |
| 2 | Gimap9 | 1.39 | 1.22E-28 | 2.61E-24 | 0.29 | 0.11 | 0.18 | 2.6 |
| 2 | Gjb2 | 1.71 | 1.55E-28 | 3.32E-24 | 0.35 | 0.16 | 0.19 | 2.19 |
| 2 | Creb3l2 | 1.15 | 1.70E-28 | 3.63E-24 | 0.52 | 0.3 | 0.22 | 1.73 |
| 2 | Tafa5 | 1.17 | 2.26E-28 | 4.83E-24 | 0.44 | 0.22 | 0.21 | 1.96 |
| 2 | Dcaf4 | 1.04 | 2.61E-28 | 5.57E-24 | 0.56 | 0.32 | 0.24 | 1.73 |
| 2 | Ptprz1 | 1.07 | 2.68E-28 | 5.72E-24 | 0.41 | 0.2 | 0.22 | 2.09 |
| 2 | Lad1 | 1.02 | 3.15E-28 | 6.72E-24 | 0.55 | 0.32 | 0.23 | 1.73 |

**Supplementary Table 3. Marker genes for the original clusters of integrated clustering of E11.5 control and NotchGOF λ-junction ectoderm cells.**

| | | | | | | | | |
|---|---|---|---|---|---|---|---|---|
| 2 | Suco | 0.96 | 3.74E-28 | 7.99E-24 | 0.72 | 0.56 | 0.16 | 1.28 |
| 2 | Ralbp1 | 0.61 | 9.19E-28 | 1.96E-23 | 0.9 | 0.82 | 0.09 | 1.11 |
| 2 | Fgfr3 | 1.94 | 1.50E-27 | 3.21E-23 | 0.15 | 0.04 | 0.12 | 4.11 |
| 2 | Kctd1 | 0.79 | 1.57E-27 | 3.35E-23 | 0.69 | 0.48 | 0.21 | 1.44 |
| 2 | Slc15a2 | 1.28 | 1.64E-27 | 3.51E-23 | 0.31 | 0.13 | 0.19 | 2.46 |
| 2 | Pbx1 | 0.42 | 2.22E-27 | 4.74E-23 | 0.98 | 0.97 | 0.02 | 1.02 |
| 2 | Cers3 | 2.03 | 3.82E-27 | 8.16E-23 | 0.14 | 0.03 | 0.11 | 4.24 |
| 2 | Hmx1 | 2.53 | 4.28E-27 | 9.13E-23 | 0.1 | 0.02 | 0.09 | 5.72 |
| 2 | Six3os1 | 0.55 | 8.58E-27 | 1.83E-22 | 0.9 | 0.75 | 0.15 | 1.2 |
| 2 | Tfap2a | 0.58 | 1.22E-26 | 2.61E-22 | 0.57 | 0.32 | 0.26 | 1.82 |
| 2 | Jag1 | 0.84 | 1.71E-26 | 3.66E-22 | 0.73 | 0.54 | 0.19 | 1.36 |
| 2 | Cd2ap | 0.54 | 2.01E-26 | 4.29E-22 | 0.9 | 0.79 | 0.11 | 1.13 |
| 2 | Nrip1 | 0.57 | 2.02E-26 | 4.31E-22 | 0.92 | 0.8 | 0.12 | 1.15 |
| 2 | Cldn23 | 1.51 | 4.63E-26 | 9.89E-22 | 0.31 | 0.14 | 0.18 | 2.3 |
| 2 | Ahnak | 1.04 | 4.77E-26 | 1.02E-21 | 0.56 | 0.34 | 0.23 | 1.67 |
| 2 | Gadd45b | 1.04 | 6.97E-26 | 1.49E-21 | 0.39 | 0.18 | 0.21 | 2.15 |
| 2 | Dcbld1 | 0.85 | 7.08E-26 | 1.51E-21 | 0.6 | 0.37 | 0.23 | 1.62 |
| 2 | Nrtn | 1.14 | 7.71E-26 | 1.65E-21 | 0.5 | 0.3 | 0.2 | 1.67 |
| 2 | B3galnt1 | 1.31 | 1.08E-25 | 2.31E-21 | 0.33 | 0.14 | 0.18 | 2.27 |
| 2 | Ifitm3 | 0.6 | 1.15E-25 | 2.45E-21 | 0.65 | 0.4 | 0.26 | 1.65 |
| 2 | Me1 | 1.12 | 1.51E-25 | 3.22E-21 | 0.36 | 0.17 | 0.2 | 2.15 |
| 2 | Gab1 | 1.01 | 1.51E-25 | 3.23E-21 | 0.47 | 0.27 | 0.21 | 1.78 |
| 2 | Wwc1 | 0.79 | 2.00E-25 | 4.27E-21 | 0.75 | 0.57 | 0.18 | 1.32 |
| 2 | Ezr | 0.46 | 2.04E-25 | 4.36E-21 | 0.97 | 0.9 | 0.07 | 1.07 |
| 2 | Vcan | 0.8 | 3.97E-25 | 8.48E-21 | 0.84 | 0.74 | 0.1 | 1.14 |
| 2 | Sdc1 | 0.9 | 7.61E-25 | 1.63E-20 | 0.75 | 0.57 | 0.18 | 1.31 |
| 2 | Elovl7 | 1.5 | 7.71E-25 | 1.65E-20 | 0.23 | 0.08 | 0.15 | 2.83 |
| 2 | Esrp1 | 0.76 | 8.61E-25 | 1.84E-20 | 0.77 | 0.63 | 0.15 | 1.23 |
| 2 | Runx1 | 0.31 | 8.91E-25 | 1.90E-20 | 0.74 | 0.49 | 0.26 | 1.52 |
| 2 | Cab39 | 0.76 | 9.73E-25 | 2.08E-20 | 0.77 | 0.64 | 0.13 | 1.2 |
| 2 | Ripply3 | 1.08 | 1.01E-24 | 2.15E-20 | 0.35 | 0.16 | 0.19 | 2.19 |
| 2 | Phldb3 | 1.55 | 1.02E-24 | 2.17E-20 | 0.21 | 0.07 | 0.14 | 2.99 |
| 2 | Stx19 | 2.44 | 1.22E-24 | 2.61E-20 | 0.12 | 0.02 | 0.09 | 4.79 |
| 2 | Papss1 | 0.63 | 1.75E-24 | 3.74E-20 | 0.8 | 0.65 | 0.15 | 1.23 |
| 2 | Capg | 0.9 | 2.00E-24 | 4.28E-20 | 0.55 | 0.33 | 0.22 | 1.66 |
| 2 | Prkch | 1.15 | 2.05E-24 | 4.37E-20 | 0.37 | 0.18 | 0.19 | 2.04 |
| 2 | Stk39 | 0.98 | 2.89E-24 | 6.17E-20 | 0.49 | 0.28 | 0.21 | 1.74 |
| 2 | Plekhg1 | 1.13 | 3.03E-24 | 6.47E-20 | 0.47 | 0.27 | 0.2 | 1.73 |
| 2 | Bex1 | 0.35 | 4.19E-24 | 8.95E-20 | 0.98 | 0.96 | 0.02 | 1.02 |
| 2 | Rfk | 0.79 | 4.80E-24 | 1.02E-19 | 0.69 | 0.5 | 0.2 | 1.39 |
| 2 | Nxn | 0.54 | 5.48E-24 | 1.17E-19 | 0.86 | 0.74 | 0.12 | 1.16 |
| 2 | Elf5 | 1.1 | 5.60E-24 | 1.20E-19 | 0.41 | 0.22 | 0.2 | 1.9 |
| 2 | Tfap2c | 0.86 | 5.76E-24 | 1.23E-19 | 0.41 | 0.2 | 0.21 | 2.07 |
| 2 | Cdc42ep5 | 0.73 | 6.18E-24 | 1.32E-19 | 0.73 | 0.58 | 0.16 | 1.27 |
| 2 | Smim5 | 1.72 | 6.48E-24 | 1.38E-19 | 0.17 | 0.05 | 0.12 | 3.38 |
| 2 | H19 | 0.56 | 6.60E-24 | 1.41E-19 | 0.97 | 0.87 | 0.1 | 1.12 |
| 2 | Limk2 | 1.24 | 6.79E-24 | 1.45E-19 | 0.52 | 0.33 | 0.18 | 1.55 |
| 2 | Trmt9b | 1.21 | 6.88E-24 | 1.47E-19 | 0.34 | 0.16 | 0.18 | 2.15 |
| 2 | Pycard | 1.26 | 7.71E-24 | 1.65E-19 | 0.38 | 0.19 | 0.19 | 1.96 |
| 2 | Cnn2 | 1 | 1.27E-23 | 2.72E-19 | 0.49 | 0.29 | 0.21 | 1.71 |
| 2 | Ermp1 | 0.82 | 1.43E-23 | 3.05E-19 | 0.62 | 0.41 | 0.21 | 1.52 |
| 2 | Ppfibp1 | 0.69 | 1.85E-23 | 3.95E-19 | 0.77 | 0.6 | 0.17 | 1.29 |
| 2 | Fermt1 | 0.52 | 2.67E-23 | 5.71E-19 | 0.79 | 0.53 | 0.26 | 1.5 |
| 2 | Enc1 | 0.81 | 3.31E-23 | 7.08E-19 | 0.43 | 0.23 | 0.2 | 1.87 |
| 2 | Ddah2 | 0.33 | 6.48E-23 | 1.38E-18 | 0.99 | 0.97 | 0.02 | 1.02 |
| 2 | Nfkbia | 0.77 | 1.38E-22 | 2.96E-18 | 0.66 | 0.47 | 0.19 | 1.4 |
| 2 | Rnh1 | 0.53 | 1.63E-22 | 3.48E-18 | 0.86 | 0.77 | 0.1 | 1.12 |

**Supplementary Table 3. Marker genes for the original clusters of integrated clustering of E11.5 control and NotchGOF λ-junction ectoderm cells.**

| | | | | | | | | |
|---|---|---|---|---|---|---|---|---|
| 2 | Prr5l | 0.91 | 1.67E-22 | 3.56E-18 | 0.53 | 0.33 | 0.2 | 1.59 |
| 2 | Ptgr1 | 1.38 | 2.81E-22 | 6.00E-18 | 0.3 | 0.14 | 0.16 | 2.16 |
| 2 | Gipc1 | 0.57 | 3.17E-22 | 6.77E-18 | 0.86 | 0.74 | 0.12 | 1.17 |
| 2 | H2-T23 | 1.27 | 3.38E-22 | 7.22E-18 | 0.3 | 0.13 | 0.17 | 2.26 |
| 2 | Metrnl | 1.5 | 3.54E-22 | 7.56E-18 | 0.33 | 0.16 | 0.17 | 2.06 |
| 2 | Iqgap1 | 0.65 | 5.29E-22 | 1.13E-17 | 0.82 | 0.67 | 0.15 | 1.23 |
| 2 | Wnt5a | 0.54 | 6.18E-22 | 1.32E-17 | 0.63 | 0.41 | 0.22 | 1.54 |
| 2 | Mgst1 | 0.6 | 7.47E-22 | 1.60E-17 | 0.84 | 0.68 | 0.16 | 1.24 |
| 2 | Lhfp | 0.79 | 8.06E-22 | 1.72E-17 | 0.5 | 0.29 | 0.21 | 1.73 |
| 2 | Mpp7 | 0.76 | 8.32E-22 | 1.78E-17 | 0.71 | 0.53 | 0.18 | 1.34 |
| 2 | Unc5b | 0.72 | 1.31E-21 | 2.80E-17 | 0.7 | 0.51 | 0.19 | 1.37 |
| 2 | Cdkn1a | 0.27 | 1.38E-21 | 2.94E-17 | 0.59 | 0.36 | 0.23 | 1.62 |
| 2 | A730020E08Rik | 1.5 | 1.56E-21 | 3.32E-17 | 0.2 | 0.07 | 0.13 | 2.78 |
| 2 | Dhrs3 | 1.08 | 2.15E-21 | 4.60E-17 | 0.37 | 0.19 | 0.18 | 1.94 |
| 2 | Mycl | 0.58 | 2.99E-21 | 6.40E-17 | 0.78 | 0.62 | 0.16 | 1.26 |
| 2 | Slc38a4 | 0.98 | 6.68E-21 | 1.43E-16 | 0.45 | 0.26 | 0.19 | 1.75 |
| 2 | Ocln | 0.93 | 7.61E-21 | 1.63E-16 | 0.48 | 0.28 | 0.19 | 1.69 |
| 2 | Elf1 | 0.72 | 7.96E-21 | 1.70E-16 | 0.67 | 0.48 | 0.19 | 1.4 |
| 2 | Erbb2 | 1.02 | 9.92E-21 | 2.12E-16 | 0.49 | 0.3 | 0.18 | 1.6 |
| 2 | Cebpb | 1.02 | 1.02E-20 | 2.17E-16 | 0.41 | 0.23 | 0.18 | 1.81 |
| 2 | Sh3bp5 | 0.83 | 1.14E-20 | 2.43E-16 | 0.56 | 0.36 | 0.2 | 1.54 |
| 2 | Dok5 | 0.7 | 1.19E-20 | 2.55E-16 | 0.57 | 0.35 | 0.22 | 1.62 |
| 2 | Pcdh1 | 1.47 | 1.52E-20 | 3.25E-16 | 0.22 | 0.09 | 0.14 | 2.55 |
| 2 | Rnd3 | 0.46 | 1.78E-20 | 3.79E-16 | 0.87 | 0.72 | 0.15 | 1.21 |
| 2 | Krt7 | 1.27 | 2.29E-20 | 4.90E-16 | 0.36 | 0.19 | 0.17 | 1.89 |
| 2 | Fam117a | 0.97 | 2.54E-20 | 5.43E-16 | 0.44 | 0.26 | 0.19 | 1.73 |
| 2 | Nrarp | 0.59 | 3.23E-20 | 6.89E-16 | 0.79 | 0.58 | 0.21 | 1.35 |
| 2 | Ank3 | 0.54 | 3.72E-20 | 7.95E-16 | 0.76 | 0.62 | 0.14 | 1.23 |
| 2 | S100a16 | 1.22 | 4.23E-20 | 9.03E-16 | 0.29 | 0.13 | 0.16 | 2.18 |
| 2 | Dusp2 | 1.9 | 8.17E-20 | 1.74E-15 | 0.12 | 0.03 | 0.09 | 3.9 |
| 2 | Anp32a | 0.25 | 1.02E-19 | 2.17E-15 | 1 | 0.99 | 0.01 | 1.01 |
| 2 | Lrrfip1 | 0.71 | 1.29E-19 | 2.75E-15 | 0.66 | 0.48 | 0.18 | 1.37 |
| 2 | Mdh1 | 0.34 | 1.40E-19 | 2.99E-15 | 0.99 | 0.98 | 0.01 | 1.01 |
| 2 | Sh3bgrl3 | 0.38 | 1.50E-19 | 3.20E-15 | 0.96 | 0.91 | 0.05 | 1.06 |
| 2 | Lmo7 | 1.03 | 1.63E-19 | 3.47E-15 | 0.6 | 0.43 | 0.16 | 1.37 |
| 2 | Macc1 | 1.05 | 1.76E-19 | 3.75E-15 | 0.38 | 0.2 | 0.17 | 1.85 |
| 2 | Prxl2a | 0.61 | 1.85E-19 | 3.95E-15 | 0.74 | 0.57 | 0.17 | 1.3 |
| 2 | Dsg2 | 0.67 | 2.63E-19 | 5.62E-15 | 0.65 | 0.47 | 0.18 | 1.39 |
| 2 | Arl4c | 0.52 | 3.93E-19 | 8.40E-15 | 0.74 | 0.56 | 0.18 | 1.32 |
| 2 | Isl2 | 1.08 | 5.18E-19 | 1.11E-14 | 0.28 | 0.13 | 0.15 | 2.21 |
| 2 | Fech | 0.97 | 7.24E-19 | 1.55E-14 | 0.45 | 0.27 | 0.17 | 1.64 |
| 2 | Tbx2 | 1 | 8.23E-19 | 1.76E-14 | 0.28 | 0.13 | 0.15 | 2.16 |
| 2 | Anxa1 | 1.72 | 1.32E-18 | 2.81E-14 | 0.15 | 0.05 | 0.1 | 2.98 |
| 2 | Sorl1 | 0.86 | 2.48E-18 | 5.29E-14 | 0.51 | 0.34 | 0.18 | 1.53 |
| 2 | Dbi | 0.3 | 2.79E-18 | 5.96E-14 | 0.99 | 0.97 | 0.03 | 1.03 |
| 2 | Etv3 | 0.85 | 3.41E-18 | 7.28E-14 | 0.45 | 0.27 | 0.18 | 1.65 |
| 2 | Pygl | 1.29 | 3.82E-18 | 8.16E-14 | 0.32 | 0.16 | 0.15 | 1.93 |
| 2 | Myh9 | 0.5 | 4.27E-18 | 9.13E-14 | 0.86 | 0.74 | 0.12 | 1.16 |
| 2 | Plp2 | 0.56 | 5.90E-18 | 1.26E-13 | 0.8 | 0.69 | 0.12 | 1.17 |
| 2 | Psap | 0.51 | 7.08E-18 | 1.51E-13 | 0.85 | 0.73 | 0.12 | 1.16 |
| 2 | Gata3 | 0.7 | 8.79E-18 | 1.88E-13 | 0.51 | 0.32 | 0.19 | 1.59 |
| 2 | Hibadh | 0.65 | 1.17E-17 | 2.50E-13 | 0.74 | 0.6 | 0.14 | 1.23 |
| 2 | Igsf9b | 1.11 | 1.34E-17 | 2.87E-13 | 0.3 | 0.15 | 0.15 | 1.99 |
| 2 | Epn3 | 1.1 | 1.62E-17 | 3.45E-13 | 0.31 | 0.16 | 0.15 | 1.96 |
| 2 | Cblc | 1.24 | 1.82E-17 | 3.89E-13 | 0.26 | 0.12 | 0.14 | 2.14 |
| 2 | Tor3a | 1.36 | 2.05E-17 | 4.37E-13 | 0.19 | 0.07 | 0.11 | 2.57 |
| 2 | Epha4 | 0.36 | 2.67E-17 | 5.71E-13 | 0.73 | 0.54 | 0.19 | 1.35 |

**Supplementary Table 3. Marker genes for the original clusters of integrated clustering of E11.5 control and NotchGOF λ-junction ectoderm cells.**

| | | | | | | | | |
|---|---|---|---|---|---|---|---|---|
| 2 | Cux1 | 0.4 | 2.72E-17 | 5.80E-13 | 0.91 | 0.83 | 0.08 | 1.09 |
| 2 | Nxph3 | 1.37 | 3.23E-17 | 6.91E-13 | 0.15 | 0.05 | 0.1 | 2.96 |
| 2 | Prdx5 | 0.72 | 3.49E-17 | 7.45E-13 | 0.87 | 0.8 | 0.08 | 1.09 |
| 2 | Bcl11b | 0.34 | 3.53E-17 | 7.55E-13 | 0.85 | 0.7 | 0.15 | 1.21 |
| 2 | Ier5l | 0.56 | 4.19E-17 | 8.95E-13 | 0.79 | 0.64 | 0.15 | 1.24 |
| 2 | Dlgap4 | 0.44 | 8.77E-17 | 1.87E-12 | 0.81 | 0.68 | 0.13 | 1.18 |
| 2 | Smpdl3a | 0.98 | 1.06E-16 | 2.26E-12 | 0.32 | 0.17 | 0.16 | 1.92 |
| 2 | Eml5 | 0.93 | 1.09E-16 | 2.33E-12 | 0.38 | 0.22 | 0.17 | 1.77 |
| 2 | AW011738 | 0.79 | 1.11E-16 | 2.36E-12 | 0.45 | 0.28 | 0.17 | 1.62 |
| 2 | Cebpa | 1.34 | 1.15E-16 | 2.46E-12 | 0.18 | 0.07 | 0.11 | 2.55 |
| 2 | Bcam | 0.39 | 1.25E-16 | 2.67E-12 | 0.87 | 0.71 | 0.17 | 1.23 |
| 2 | Sinhcaf | 0.33 | 1.37E-16 | 2.93E-12 | 0.97 | 0.94 | 0.03 | 1.04 |
| 2 | Crabp1 | 0.36 | 1.49E-16 | 3.18E-12 | 0.69 | 0.55 | 0.14 | 1.26 |
| 2 | Crybg1 | 1.57 | 1.51E-16 | 3.24E-12 | 0.13 | 0.04 | 0.09 | 3.21 |
| 2 | Sash1 | 0.69 | 1.82E-16 | 3.90E-12 | 0.58 | 0.4 | 0.18 | 1.45 |
| 2 | Npm3 | 0.39 | 2.20E-16 | 4.70E-12 | 0.95 | 0.87 | 0.08 | 1.1 |
| 2 | Sh3tc2 | 1.67 | 2.23E-16 | 4.76E-12 | 0.13 | 0.05 | 0.09 | 2.98 |
| 2 | Churc1 | 0.55 | 2.49E-16 | 5.33E-12 | 0.82 | 0.72 | 0.11 | 1.15 |
| 2 | Tgif1 | 0.46 | 4.69E-16 | 1.00E-11 | 0.8 | 0.67 | 0.13 | 1.19 |
| 2 | AI661453 | 1.06 | 5.32E-16 | 1.14E-11 | 0.26 | 0.12 | 0.14 | 2.13 |
| 2 | Gpc3 | 0.25 | 6.95E-16 | 1.49E-11 | 0.95 | 0.83 | 0.13 | 1.15 |
| 2 | Gtf2i | 0.35 | 7.61E-16 | 1.63E-11 | 0.95 | 0.9 | 0.05 | 1.06 |
| 2 | Tbc1d9 | 1.04 | 1.11E-15 | 2.37E-11 | 0.24 | 0.11 | 0.13 | 2.15 |
| 2 | Frmd4b | 0.49 | 1.11E-15 | 2.37E-11 | 0.66 | 0.48 | 0.18 | 1.39 |
| 2 | Camsap3 | 0.79 | 1.14E-15 | 2.44E-11 | 0.43 | 0.26 | 0.17 | 1.63 |
| 2 | Selenoh | 0.27 | 1.38E-15 | 2.95E-11 | 0.98 | 0.95 | 0.03 | 1.03 |
| 2 | Axin2 | 0.61 | 1.63E-15 | 3.47E-11 | 0.41 | 0.24 | 0.17 | 1.7 |
| 2 | Exph5 | 1.06 | 1.88E-15 | 4.02E-11 | 0.27 | 0.14 | 0.13 | 1.99 |
| 2 | Naaladl2 | 1.49 | 2.14E-15 | 4.57E-11 | 0.12 | 0.04 | 0.08 | 3.19 |
| 2 | Adcy1 | 1.48 | 2.18E-15 | 4.66E-11 | 0.14 | 0.05 | 0.09 | 2.77 |
| 2 | Etl4 | 0.49 | 2.46E-15 | 5.26E-11 | 0.73 | 0.56 | 0.17 | 1.3 |
| 2 | Junb | 0.77 | 2.55E-15 | 5.45E-11 | 0.35 | 0.2 | 0.16 | 1.8 |
| 2 | Nab1 | 0.68 | 2.85E-15 | 6.08E-11 | 0.39 | 0.22 | 0.17 | 1.75 |
| 2 | Hey1 | 0.39 | 3.03E-15 | 6.47E-11 | 0.9 | 0.73 | 0.17 | 1.23 |
| 2 | Syt7 | 0.98 | 3.94E-15 | 8.42E-11 | 0.26 | 0.13 | 0.13 | 2.01 |
| 2 | Fzd3 | 0.4 | 3.95E-15 | 8.43E-11 | 0.92 | 0.84 | 0.08 | 1.1 |
| 2 | Celf4 | 0.77 | 4.02E-15 | 8.58E-11 | 0.45 | 0.28 | 0.16 | 1.58 |
| 2 | Hspa2 | 0.95 | 4.04E-15 | 8.62E-11 | 0.26 | 0.13 | 0.13 | 2.04 |
| 2 | Mindy2 | 0.78 | 4.31E-15 | 9.20E-11 | 0.4 | 0.23 | 0.16 | 1.7 |
| 2 | Rab25 | 0.51 | 4.66E-15 | 9.96E-11 | 0.77 | 0.61 | 0.15 | 1.25 |
| 2 | Fzd10os | 0.94 | 6.05E-15 | 1.29E-10 | 0.27 | 0.14 | 0.14 | 1.99 |
| 2 | Cldn1 | 0.69 | 6.30E-15 | 1.35E-10 | 0.21 | 0.09 | 0.12 | 2.31 |
| 2 | Gldc | 1.06 | 6.79E-15 | 1.45E-10 | 0.26 | 0.13 | 0.13 | 1.99 |
| 2 | Meis2 | 0.28 | 7.21E-15 | 1.54E-10 | 0.98 | 0.95 | 0.03 | 1.03 |
| 2 | Mark2 | 0.56 | 7.28E-15 | 1.56E-10 | 0.7 | 0.57 | 0.13 | 1.22 |
| 2 | Hspb1 | 1.27 | 9.35E-15 | 2.00E-10 | 0.52 | 0.37 | 0.15 | 1.41 |
| 2 | Tiam1 | 0.7 | 1.03E-14 | 2.20E-10 | 0.6 | 0.44 | 0.16 | 1.36 |
| 2 | Frem2 | 0.5 | 1.22E-14 | 2.61E-10 | 0.77 | 0.61 | 0.16 | 1.26 |
| 2 | Oxct1 | 0.45 | 1.28E-14 | 2.74E-10 | 0.86 | 0.77 | 0.09 | 1.12 |
| 2 | Fhdc1 | 0.99 | 1.58E-14 | 3.38E-10 | 0.24 | 0.11 | 0.12 | 2.08 |
| 2 | Rbms1 | 0.34 | 1.76E-14 | 3.77E-10 | 0.9 | 0.84 | 0.05 | 1.06 |
| 2 | Mtss1 | 0.75 | 2.04E-14 | 4.36E-10 | 0.56 | 0.42 | 0.14 | 1.34 |
| 2 | Igf2r | 0.49 | 2.45E-14 | 5.24E-10 | 0.72 | 0.56 | 0.15 | 1.27 |
| 2 | Satb1 | 0.51 | 2.59E-14 | 5.53E-10 | 0.69 | 0.55 | 0.14 | 1.25 |
| 2 | Hspa1b | 1.63 | 2.67E-14 | 5.71E-10 | 0.11 | 0.03 | 0.07 | 3.18 |
| 2 | App | 0.33 | 2.96E-14 | 6.32E-10 | 0.91 | 0.85 | 0.06 | 1.07 |
| 2 | Mgll | 1.35 | 3.15E-14 | 6.74E-10 | 0.15 | 0.06 | 0.09 | 2.57 |

**Supplementary Table 3. Marker genes for the original clusters of integrated clustering of E11.5 control and NotchGOF λ-junction ectoderm cells.**

| | | | | | | | | |
|---|---|---|---|---|---|---|---|---|
| 2 | Pros1 | 1.11 | 4.34E-14 | 9.28E-10 | 0.2 | 0.09 | 0.11 | 2.19 |
| 2 | Cd36 | 1.22 | 4.36E-14 | 9.31E-10 | 0.11 | 0.04 | 0.08 | 3.08 |
| 2 | Podxl2 | 0.59 | 4.43E-14 | 9.47E-10 | 0.53 | 0.37 | 0.16 | 1.44 |
| 2 | Crot | 0.73 | 4.89E-14 | 1.04E-09 | 0.51 | 0.35 | 0.15 | 1.44 |
| 2 | Lgalsl | 0.49 | 5.37E-14 | 1.15E-09 | 0.62 | 0.45 | 0.17 | 1.37 |
| 2 | Arl8b | 0.59 | 6.26E-14 | 1.34E-09 | 0.66 | 0.51 | 0.15 | 1.29 |
| 2 | Zfp503 | 0.45 | 8.00E-14 | 1.71E-09 | 0.66 | 0.49 | 0.18 | 1.36 |
| 2 | Rhou | 0.66 | 8.22E-14 | 1.76E-09 | 0.61 | 0.47 | 0.14 | 1.3 |
| 2 | Mmp11 | 0.57 | 9.89E-14 | 2.11E-09 | 0.54 | 0.37 | 0.17 | 1.44 |
| 2 | Sp6 | 0.81 | 1.06E-13 | 2.25E-09 | 0.23 | 0.11 | 0.12 | 2.11 |
| 2 | Faah | 0.81 | 1.07E-13 | 2.28E-09 | 0.35 | 0.21 | 0.14 | 1.67 |
| 2 | Tmem54 | 0.65 | 1.13E-13 | 2.41E-09 | 0.6 | 0.46 | 0.14 | 1.29 |
| 2 | Tsc22d4 | 0.56 | 1.15E-13 | 2.46E-09 | 0.72 | 0.61 | 0.1 | 1.17 |
| 2 | Lamp2 | 0.56 | 1.20E-13 | 2.57E-09 | 0.72 | 0.59 | 0.13 | 1.21 |
| 2 | Ccdc18 | 0.73 | 1.20E-13 | 2.57E-09 | 0.51 | 0.37 | 0.15 | 1.4 |
| 2 | Prr15l | 1.21 | 1.31E-13 | 2.79E-09 | 0.25 | 0.13 | 0.12 | 1.92 |
| 2 | Myzap | 1.46 | 1.68E-13 | 3.60E-09 | 0.11 | 0.04 | 0.07 | 3 |
| 2 | Myo10 | 0.42 | 1.92E-13 | 4.10E-09 | 0.8 | 0.69 | 0.11 | 1.16 |
| 2 | Tpd52 | 0.32 | 2.19E-13 | 4.68E-09 | 0.9 | 0.8 | 0.09 | 1.11 |
| 2 | Eps8l2 | 0.81 | 2.22E-13 | 4.75E-09 | 0.27 | 0.14 | 0.13 | 1.93 |
| 2 | Gm2464 | 1.22 | 2.29E-13 | 4.90E-09 | 0.16 | 0.06 | 0.09 | 2.44 |
| 2 | Plscr3 | 0.54 | 2.79E-13 | 5.97E-09 | 0.69 | 0.52 | 0.17 | 1.32 |
| 2 | Pdlim4 | 0.5 | 3.16E-13 | 6.74E-09 | 0.59 | 0.42 | 0.17 | 1.41 |
| 2 | Tmem123 | 0.75 | 3.30E-13 | 7.05E-09 | 0.57 | 0.44 | 0.14 | 1.31 |
| 2 | St14 | 0.53 | 3.59E-13 | 7.66E-09 | 0.62 | 0.46 | 0.15 | 1.33 |
| 2 | Slc7a11 | 0.54 | 4.52E-13 | 9.66E-09 | 0.24 | 0.12 | 0.12 | 2.04 |
| 2 | Ctsc | 0.97 | 4.66E-13 | 9.95E-09 | 0.32 | 0.19 | 0.13 | 1.71 |
| 2 | Irx5 | 0.53 | 5.34E-13 | 1.14E-08 | 0.38 | 0.23 | 0.16 | 1.68 |
| 2 | Cd44 | 1.22 | 5.50E-13 | 1.18E-08 | 0.12 | 0.04 | 0.08 | 2.83 |
| 2 | Prelid2 | 0.58 | 6.08E-13 | 1.30E-08 | 0.56 | 0.41 | 0.15 | 1.37 |
| 2 | Adrb1 | 1.15 | 6.48E-13 | 1.38E-08 | 0.14 | 0.06 | 0.09 | 2.55 |
| 2 | Nt5dc2 | 0.29 | 6.49E-13 | 1.39E-08 | 0.97 | 0.91 | 0.05 | 1.06 |
| 2 | Zdhhc13 | 0.81 | 6.69E-13 | 1.43E-08 | 0.37 | 0.23 | 0.14 | 1.63 |
| 2 | Rab21 | 0.55 | 7.35E-13 | 1.57E-08 | 0.67 | 0.52 | 0.14 | 1.27 |
| 2 | Mid1ip1 | 0.79 | 7.44E-13 | 1.59E-08 | 0.51 | 0.37 | 0.15 | 1.4 |
| 2 | Tjp2 | 0.64 | 7.68E-13 | 1.64E-08 | 0.59 | 0.46 | 0.13 | 1.28 |
| 2 | F11r | 0.44 | 8.17E-13 | 1.75E-08 | 0.74 | 0.59 | 0.15 | 1.25 |
| 2 | Acbd4 | 1.19 | 1.02E-12 | 2.18E-08 | 0.17 | 0.08 | 0.1 | 2.25 |
| 2 | Epcam | 0.27 | 1.10E-12 | 2.34E-08 | 1 | 0.98 | 0.02 | 1.02 |
| 2 | Casp8 | 0.84 | 1.14E-12 | 2.44E-08 | 0.34 | 0.2 | 0.14 | 1.68 |
| 2 | Cxxc5 | 0.34 | 1.21E-12 | 2.58E-08 | 0.72 | 0.57 | 0.15 | 1.27 |
| 2 | Stard10 | 0.42 | 1.36E-12 | 2.90E-08 | 0.82 | 0.71 | 0.11 | 1.15 |
| 2 | Ankrd22 | 1.62 | 1.58E-12 | 3.36E-08 | 0.14 | 0.06 | 0.08 | 2.47 |
| 2 | Tmem79 | 1.18 | 1.64E-12 | 3.50E-08 | 0.16 | 0.07 | 0.09 | 2.33 |
| 2 | Nav2 | 0.37 | 1.71E-12 | 3.65E-08 | 0.82 | 0.69 | 0.13 | 1.18 |
| 2 | St7 | 0.63 | 1.82E-12 | 3.90E-08 | 0.48 | 0.31 | 0.16 | 1.52 |
| 2 | Tshz1 | 0.46 | 2.03E-12 | 4.34E-08 | 0.6 | 0.46 | 0.14 | 1.3 |
| 2 | Utrn | 0.37 | 2.21E-12 | 4.72E-08 | 0.83 | 0.71 | 0.12 | 1.17 |
| 2 | Tacc2 | 0.49 | 2.22E-12 | 4.74E-08 | 0.66 | 0.52 | 0.14 | 1.27 |
| 2 | Sult2b1 | 1.29 | 2.32E-12 | 4.96E-08 | 0.14 | 0.06 | 0.08 | 2.45 |
| 2 | Aopep | 0.39 | 2.32E-12 | 4.97E-08 | 0.74 | 0.6 | 0.14 | 1.23 |
| 2 | Slc44a1 | 0.68 | 2.38E-12 | 5.09E-08 | 0.48 | 0.33 | 0.14 | 1.44 |
| 2 | Pcdh9 | 0.77 | 2.39E-12 | 5.12E-08 | 0.3 | 0.17 | 0.13 | 1.77 |
| 2 | Slc16a12 | 1.18 | 2.40E-12 | 5.13E-08 | 0.11 | 0.04 | 0.07 | 2.87 |
| 2 | Fam49a | 0.8 | 2.76E-12 | 5.90E-08 | 0.19 | 0.09 | 0.1 | 2.17 |
| 2 | Hdhd3 | 0.81 | 2.81E-12 | 6.01E-08 | 0.34 | 0.2 | 0.14 | 1.69 |
| 2 | Ntn4 | 1.08 | 3.18E-12 | 6.79E-08 | 0.13 | 0.05 | 0.08 | 2.59 |

**Supplementary Table 3. Marker genes for the original clusters of integrated clustering of E11.5 control and NotchGOF λ-junction ectoderm cells.**

| | | | | | | | | |
|---|---|---|---|---|---|---|---|---|
| 2 | Lima1 | 0.52 | 3.23E-12 | 6.90E-08 | 0.61 | 0.47 | 0.15 | 1.32 |
| 2 | Dsc2 | 0.79 | 3.42E-12 | 7.31E-08 | 0.35 | 0.22 | 0.14 | 1.62 |
| 2 | Tead1 | 0.41 | 3.44E-12 | 7.35E-08 | 0.84 | 0.76 | 0.08 | 1.11 |
| 2 | Khdrbs3 | 0.41 | 3.72E-12 | 7.94E-08 | 0.91 | 0.88 | 0.04 | 1.04 |
| 2 | Nsg1 | 0.39 | 4.05E-12 | 8.66E-08 | 0.74 | 0.59 | 0.15 | 1.25 |
| 2 | Ddi2 | 0.63 | 4.08E-12 | 8.71E-08 | 0.59 | 0.46 | 0.13 | 1.28 |
| 2 | Tpm4 | 0.31 | 4.21E-12 | 9.00E-08 | 0.92 | 0.87 | 0.06 | 1.06 |
| 2 | Tcn2 | 0.92 | 4.26E-12 | 9.10E-08 | 0.33 | 0.19 | 0.14 | 1.7 |
| 2 | Nectin2 | 0.5 | 4.51E-12 | 9.63E-08 | 0.7 | 0.6 | 0.1 | 1.17 |
| 2 | Zfp185 | 1.07 | 5.35E-12 | 1.14E-07 | 0.16 | 0.07 | 0.09 | 2.29 |
| 2 | Prag1 | 0.6 | 5.78E-12 | 1.24E-07 | 0.43 | 0.28 | 0.15 | 1.55 |
| 2 | Slc44a2 | 0.61 | 5.92E-12 | 1.26E-07 | 0.54 | 0.38 | 0.16 | 1.41 |
| 2 | Kremen1 | 0.69 | 6.23E-12 | 1.33E-07 | 0.45 | 0.3 | 0.15 | 1.5 |
| 2 | Rtkn | 0.99 | 6.50E-12 | 1.39E-07 | 0.27 | 0.15 | 0.12 | 1.82 |
| 2 | Tmcc3 | 0.85 | 7.03E-12 | 1.50E-07 | 0.2 | 0.1 | 0.1 | 2.08 |
| 2 | Zfhx3 | 0.25 | 7.34E-12 | 1.57E-07 | 0.98 | 0.92 | 0.06 | 1.07 |
| 2 | Kbtbd11 | 1.59 | 7.67E-12 | 1.64E-07 | 0.11 | 0.04 | 0.07 | 2.69 |
| 2 | Ldlrap1 | 0.6 | 7.85E-12 | 1.68E-07 | 0.45 | 0.3 | 0.15 | 1.52 |
| 2 | Adamts9 | 0.57 | 8.28E-12 | 1.77E-07 | 0.38 | 0.24 | 0.14 | 1.57 |
| 2 | Unc5c | 0.38 | 9.73E-12 | 2.08E-07 | 0.49 | 0.32 | 0.17 | 1.54 |
| 2 | Sh3bp1 | 0.64 | 1.08E-11 | 2.30E-07 | 0.42 | 0.28 | 0.14 | 1.51 |
| 2 | Cdc42se1 | 0.51 | 1.11E-11 | 2.38E-07 | 0.7 | 0.6 | 0.11 | 1.18 |
| 2 | Plekhg3 | 0.71 | 1.31E-11 | 2.79E-07 | 0.39 | 0.25 | 0.14 | 1.55 |
| 2 | Itm2c | 0.41 | 1.35E-11 | 2.88E-07 | 0.72 | 0.58 | 0.14 | 1.24 |
| 2 | Id4 | 0.54 | 1.45E-11 | 3.10E-07 | 0.65 | 0.57 | 0.08 | 1.15 |
| 2 | Pik3cb | 0.93 | 1.45E-11 | 3.10E-07 | 0.26 | 0.14 | 0.12 | 1.83 |
| 2 | Trappc6a | 0.36 | 1.49E-11 | 3.18E-07 | 0.79 | 0.65 | 0.14 | 1.21 |
| 2 | Ubald1 | 0.54 | 1.52E-11 | 3.25E-07 | 0.52 | 0.37 | 0.15 | 1.39 |
| 2 | Acap2 | 0.55 | 1.53E-11 | 3.26E-07 | 0.57 | 0.42 | 0.15 | 1.36 |
| 2 | Six3 | 0.27 | 1.53E-11 | 3.27E-07 | 0.91 | 0.83 | 0.08 | 1.09 |
| 2 | Eppk1 | 1.04 | 1.58E-11 | 3.37E-07 | 0.17 | 0.08 | 0.09 | 2.2 |
| 2 | Smad6 | 0.76 | 1.74E-11 | 3.72E-07 | 0.39 | 0.25 | 0.14 | 1.56 |
| 2 | 2610002M06Rik | 0.65 | 1.81E-11 | 3.87E-07 | 0.47 | 0.33 | 0.14 | 1.41 |
| 2 | Dgka | 0.89 | 1.97E-11 | 4.22E-07 | 0.26 | 0.14 | 0.12 | 1.8 |
| 2 | Lamtor2 | 0.26 | 2.47E-11 | 5.28E-07 | 0.95 | 0.91 | 0.03 | 1.04 |
| 2 | Efnb2 | 0.4 | 2.47E-11 | 5.28E-07 | 0.72 | 0.58 | 0.14 | 1.23 |
| 2 | Sox14 | 1.39 | 2.69E-11 | 5.75E-07 | 0.11 | 0.04 | 0.07 | 2.63 |
| 2 | Cacna2d3 | 0.99 | 2.87E-11 | 6.13E-07 | 0.15 | 0.07 | 0.09 | 2.26 |
| 2 | Casz1 | 1.4 | 2.92E-11 | 6.23E-07 | 0.13 | 0.06 | 0.08 | 2.39 |
| 2 | A130010J15Rik | 0.73 | 2.93E-11 | 6.26E-07 | 0.29 | 0.17 | 0.13 | 1.77 |
| 2 | Sypl | 0.44 | 3.06E-11 | 6.53E-07 | 0.71 | 0.59 | 0.12 | 1.21 |
| 2 | Cited2 | 0.4 | 3.12E-11 | 6.66E-07 | 0.76 | 0.68 | 0.09 | 1.13 |
| 2 | Itgb6 | 0.77 | 3.14E-11 | 6.71E-07 | 0.27 | 0.15 | 0.12 | 1.78 |
| 2 | Plxdc2 | 0.63 | 3.18E-11 | 6.80E-07 | 0.47 | 0.33 | 0.14 | 1.41 |
| 2 | Ralgps2 | 0.41 | 3.31E-11 | 7.08E-07 | 0.8 | 0.72 | 0.08 | 1.11 |
| 2 | Ckap4 | 0.35 | 3.80E-11 | 8.11E-07 | 0.86 | 0.78 | 0.08 | 1.11 |
| 2 | Col18a1 | 0.26 | 3.94E-11 | 8.42E-07 | 0.94 | 0.85 | 0.09 | 1.1 |
| 2 | Tnfaip8 | 0.51 | 4.19E-11 | 8.94E-07 | 0.66 | 0.56 | 0.1 | 1.19 |
| 2 | Bdh1 | 0.93 | 4.42E-11 | 9.44E-07 | 0.25 | 0.14 | 0.11 | 1.78 |
| 2 | A630072M18Rik | 0.95 | 4.54E-11 | 9.70E-07 | 0.22 | 0.12 | 0.1 | 1.9 |
| 2 | Epb41 | 0.31 | 4.88E-11 | 1.04E-06 | 0.82 | 0.74 | 0.09 | 1.12 |
| 2 | Fxyd3 | 0.46 | 5.06E-11 | 1.08E-06 | 0.45 | 0.29 | 0.16 | 1.54 |
| 2 | Rap2c | 0.4 | 5.14E-11 | 1.10E-06 | 0.76 | 0.65 | 0.11 | 1.17 |
| 2 | Spty2d1 | 0.81 | 5.88E-11 | 1.26E-06 | 0.39 | 0.26 | 0.13 | 1.51 |
| 2 | Rps6ka6 | 0.49 | 6.02E-11 | 1.28E-06 | 0.65 | 0.52 | 0.14 | 1.26 |
| 2 | Mfsd1 | 0.7 | 6.29E-11 | 1.34E-06 | 0.53 | 0.39 | 0.14 | 1.37 |
| 2 | Ptbp3 | 0.32 | 7.35E-11 | 1.57E-06 | 0.77 | 0.68 | 0.09 | 1.13 |

**Supplementary Table 3. Marker genes for the original clusters of integrated clustering of E11.5 control and NotchGOF λ-junction ectoderm cells.**

| | | | | | | | | |
|---|---|---|---|---|---|---|---|---|
| 2 | Nhsl1 | 0.67 | 7.46E-11 | 1.59E-06 | 0.45 | 0.31 | 0.14 | 1.45 |
| 2 | Elmsan1 | 0.93 | 7.85E-11 | 1.68E-06 | 0.25 | 0.14 | 0.11 | 1.8 |
| 2 | Cds1 | 1.21 | 8.10E-11 | 1.73E-06 | 0.15 | 0.07 | 0.08 | 2.17 |
| 2 | Plxna1 | 0.79 | 8.14E-11 | 1.74E-06 | 0.36 | 0.24 | 0.13 | 1.55 |
| 2 | Cited4 | 1.45 | 9.10E-11 | 1.94E-06 | 0.12 | 0.05 | 0.07 | 2.46 |
| 2 | Adgrb3 | 0.86 | 9.35E-11 | 2.00E-06 | 0.2 | 0.1 | 0.1 | 1.96 |
| 2 | Irf2 | 0.58 | 9.35E-11 | 2.00E-06 | 0.54 | 0.41 | 0.13 | 1.33 |
| 2 | Med13l | 0.52 | 1.19E-10 | 2.54E-06 | 0.62 | 0.5 | 0.12 | 1.23 |
| 2 | Tet3 | 0.42 | 1.21E-10 | 2.58E-06 | 0.77 | 0.66 | 0.11 | 1.17 |
| 2 | Tbc1d10a | 0.67 | 1.27E-10 | 2.72E-06 | 0.4 | 0.27 | 0.14 | 1.51 |
| 2 | Cryl1 | 0.64 | 1.31E-10 | 2.81E-06 | 0.4 | 0.26 | 0.14 | 1.53 |
| 2 | Nab2 | 0.62 | 1.37E-10 | 2.94E-06 | 0.31 | 0.19 | 0.12 | 1.65 |
| 2 | Atp2b1 | 0.33 | 1.38E-10 | 2.95E-06 | 0.92 | 0.88 | 0.04 | 1.04 |
| 2 | Nipal2 | 0.89 | 1.43E-10 | 3.06E-06 | 0.18 | 0.09 | 0.09 | 2.06 |
| 2 | Arl5a | 0.45 | 1.44E-10 | 3.07E-06 | 0.71 | 0.62 | 0.09 | 1.15 |
| 2 | Pwwp2b | 0.89 | 1.53E-10 | 3.26E-06 | 0.25 | 0.14 | 0.11 | 1.79 |
| 2 | Cnn3 | 0.27 | 1.53E-10 | 3.28E-06 | 0.97 | 0.93 | 0.04 | 1.04 |
| 2 | Pqlc1 | 0.97 | 1.54E-10 | 3.30E-06 | 0.28 | 0.16 | 0.12 | 1.71 |
| 2 | Zdhhc5 | 0.66 | 1.57E-10 | 3.36E-06 | 0.45 | 0.32 | 0.13 | 1.41 |
| 2 | Mgat5 | 0.57 | 1.57E-10 | 3.36E-06 | 0.42 | 0.28 | 0.14 | 1.5 |
| 2 | Mbtd1 | 0.37 | 1.64E-10 | 3.50E-06 | 0.85 | 0.76 | 0.09 | 1.12 |
| 2 | Stx4a | 0.56 | 1.71E-10 | 3.66E-06 | 0.47 | 0.33 | 0.14 | 1.42 |
| 2 | Lratd2 | 0.49 | 1.93E-10 | 4.13E-06 | 0.47 | 0.32 | 0.14 | 1.44 |
| 2 | Tjp1 | 0.44 | 2.12E-10 | 4.52E-06 | 0.72 | 0.63 | 0.1 | 1.15 |
| 2 | Dlx1 | 0.7 | 2.34E-10 | 5.01E-06 | 0.25 | 0.14 | 0.11 | 1.83 |
| 2 | Svil | 0.6 | 2.48E-10 | 5.31E-06 | 0.54 | 0.41 | 0.12 | 1.3 |
| 2 | Prom2 | 1.04 | 2.61E-10 | 5.58E-06 | 0.22 | 0.12 | 0.1 | 1.83 |
| 2 | Ptges | 1.17 | 3.06E-10 | 6.54E-06 | 0.15 | 0.07 | 0.08 | 2.18 |
| 2 | Ece1 | 0.43 | 3.83E-10 | 8.17E-06 | 0.6 | 0.45 | 0.15 | 1.33 |
| 2 | Lamb1 | 0.4 | 3.90E-10 | 8.34E-06 | 0.65 | 0.5 | 0.15 | 1.3 |
| 2 | Acp5 | 0.88 | 3.92E-10 | 8.38E-06 | 0.14 | 0.06 | 0.08 | 2.25 |
| 2 | Tnik | 0.87 | 4.15E-10 | 8.88E-06 | 0.19 | 0.1 | 0.09 | 1.97 |
| 2 | Notch3 | 0.36 | 4.28E-10 | 9.14E-06 | 0.68 | 0.53 | 0.15 | 1.29 |
| 2 | Plxnb2 | 0.48 | 4.83E-10 | 1.03E-05 | 0.71 | 0.61 | 0.1 | 1.16 |
| 2 | 1810037I17Rik | 0.3 | 4.84E-10 | 1.03E-05 | 0.93 | 0.89 | 0.04 | 1.04 |
| 2 | Alx4 | 0.94 | 5.35E-10 | 1.14E-05 | 0.16 | 0.08 | 0.08 | 2.11 |
| 2 | Spats2 | 0.39 | 5.39E-10 | 1.15E-05 | 0.72 | 0.61 | 0.11 | 1.17 |
| 2 | Ppfibp2 | 1.13 | 5.66E-10 | 1.21E-05 | 0.13 | 0.06 | 0.07 | 2.28 |
| 2 | Tmem159 | 0.62 | 5.76E-10 | 1.23E-05 | 0.52 | 0.4 | 0.13 | 1.32 |
| 2 | Cit | 0.49 | 6.19E-10 | 1.32E-05 | 0.53 | 0.39 | 0.14 | 1.35 |
| 2 | Chmp4c | 1.15 | 7.44E-10 | 1.59E-05 | 0.13 | 0.06 | 0.07 | 2.24 |
| 2 | Skil | 0.36 | 7.88E-10 | 1.68E-05 | 0.69 | 0.56 | 0.14 | 1.25 |
| 2 | Arl1 | 0.27 | 7.93E-10 | 1.69E-05 | 0.91 | 0.88 | 0.02 | 1.02 |
| 2 | Elovl1 | 0.63 | 8.03E-10 | 1.71E-05 | 0.39 | 0.26 | 0.13 | 1.51 |
| 2 | Foxp1 | 0.25 | 8.67E-10 | 1.85E-05 | 0.95 | 0.85 | 0.09 | 1.11 |
| 2 | Tspan9 | 0.71 | 9.02E-10 | 1.93E-05 | 0.22 | 0.12 | 0.1 | 1.85 |
| 2 | Klf6 | 0.7 | 9.03E-10 | 1.93E-05 | 0.35 | 0.23 | 0.12 | 1.54 |
| 2 | 0610009E02Rik | 1.34 | 9.81E-10 | 2.10E-05 | 0.15 | 0.07 | 0.08 | 2.11 |
| 2 | Akap17b | 0.71 | 1.07E-09 | 2.28E-05 | 0.24 | 0.14 | 0.1 | 1.74 |
| 2 | Hacd4 | 0.82 | 1.07E-09 | 2.28E-05 | 0.18 | 0.09 | 0.09 | 1.97 |
| 2 | Nsun4 | 0.66 | 1.09E-09 | 2.33E-05 | 0.31 | 0.19 | 0.12 | 1.61 |
| 2 | Gm35040 | 1.03 | 1.13E-09 | 2.41E-05 | 0.12 | 0.05 | 0.07 | 2.37 |
| 2 | Lipg | 0.94 | 1.13E-09 | 2.42E-05 | 0.19 | 0.1 | 0.09 | 1.93 |
| 2 | Necap2 | 0.56 | 1.14E-09 | 2.44E-05 | 0.51 | 0.37 | 0.14 | 1.37 |
| 2 | Zfpm1 | 0.71 | 1.30E-09 | 2.77E-05 | 0.3 | 0.19 | 0.11 | 1.61 |
| 2 | Dact1 | 0.56 | 1.31E-09 | 2.80E-05 | 0.24 | 0.14 | 0.11 | 1.79 |
| 2 | Reep3 | 0.44 | 1.32E-09 | 2.81E-05 | 0.74 | 0.64 | 0.09 | 1.14 |

**Supplementary Table 3. Marker genes for the original clusters of integrated clustering of E11.5 control and NotchGOF λ-junction ectoderm cells.**

| | | | | | | | | |
|---|---|---|---|---|---|---|---|---|
| 2 | Ddr1 | 0.43 | 1.35E-09 | 2.89E-05 | 0.59 | 0.44 | 0.15 | 1.33 |
| 2 | Ston2 | 0.41 | 1.47E-09 | 3.13E-05 | 0.72 | 0.59 | 0.13 | 1.23 |
| 2 | Frk | 0.83 | 1.47E-09 | 3.13E-05 | 0.23 | 0.13 | 0.1 | 1.78 |
| 2 | Skap2 | 1.34 | 1.67E-09 | 3.56E-05 | 0.15 | 0.07 | 0.08 | 2.05 |
| 2 | Tcf7 | 0.91 | 1.68E-09 | 3.58E-05 | 0.17 | 0.09 | 0.09 | 1.97 |
| 2 | Ank2 | 0.4 | 1.76E-09 | 3.75E-05 | 0.43 | 0.31 | 0.13 | 1.41 |
| 2 | Dazap2 | 0.37 | 1.77E-09 | 3.77E-05 | 0.79 | 0.71 | 0.08 | 1.11 |
| 2 | Mpzl3 | 1.28 | 1.83E-09 | 3.91E-05 | 0.11 | 0.05 | 0.06 | 2.42 |
| 2 | Bcl7a | 0.26 | 2.20E-09 | 4.70E-05 | 0.84 | 0.76 | 0.08 | 1.11 |
| 2 | Cttnbp2nl | 0.5 | 2.45E-09 | 5.24E-05 | 0.54 | 0.42 | 0.13 | 1.31 |
| 2 | Sptlc3 | 0.83 | 2.50E-09 | 5.34E-05 | 0.14 | 0.07 | 0.08 | 2.15 |
| 2 | Ccser1 | 1.05 | 3.00E-09 | 6.41E-05 | 0.15 | 0.07 | 0.08 | 2.09 |
| 2 | Mlec | 0.28 | 3.06E-09 | 6.54E-05 | 0.87 | 0.8 | 0.07 | 1.09 |
| 2 | Irx3 | 0.37 | 3.21E-09 | 6.86E-05 | 0.32 | 0.2 | 0.12 | 1.63 |
| 2 | Add3 | 0.4 | 3.29E-09 | 7.03E-05 | 0.7 | 0.58 | 0.12 | 1.21 |
| 2 | Ppp1r36 | 0.72 | 3.49E-09 | 7.46E-05 | 0.26 | 0.15 | 0.11 | 1.72 |
| 2 | Acot1 | 0.52 | 3.53E-09 | 7.54E-05 | 0.46 | 0.34 | 0.12 | 1.36 |
| 2 | Klf4 | 0.69 | 3.70E-09 | 7.91E-05 | 0.29 | 0.18 | 0.11 | 1.61 |
| 2 | Apcdd1 | 0.64 | 3.93E-09 | 8.40E-05 | 0.23 | 0.13 | 0.1 | 1.77 |
| 2 | Cyth3 | 0.59 | 4.27E-09 | 9.12E-05 | 0.43 | 0.31 | 0.12 | 1.4 |
| 2 | Klc3 | 0.26 | 4.47E-09 | 9.54E-05 | 0.52 | 0.38 | 0.14 | 1.37 |
| 2 | Atp11b | 0.47 | 5.40E-09 | 1.15E-04 | 0.64 | 0.52 | 0.12 | 1.24 |
| 2 | Cndp2 | 0.49 | 5.58E-09 | 1.19E-04 | 0.59 | 0.48 | 0.11 | 1.24 |
| 2 | Pfkl | 0.34 | 5.80E-09 | 1.24E-04 | 0.84 | 0.76 | 0.08 | 1.11 |
| 2 | Etfa | 0.36 | 6.21E-09 | 1.33E-04 | 0.83 | 0.75 | 0.08 | 1.11 |
| 2 | Dach1 | 0.41 | 6.95E-09 | 1.48E-04 | 0.6 | 0.48 | 0.12 | 1.25 |
| 2 | Iqcb1 | 0.38 | 7.45E-09 | 1.59E-04 | 0.7 | 0.59 | 0.11 | 1.18 |
| 2 | Aktip | 0.47 | 7.50E-09 | 1.60E-04 | 0.58 | 0.45 | 0.13 | 1.28 |
| 2 | Sgpl1 | 0.51 | 7.83E-09 | 1.67E-04 | 0.54 | 0.41 | 0.13 | 1.32 |
| 2 | Fzd7 | 0.61 | 8.50E-09 | 1.82E-04 | 0.46 | 0.34 | 0.12 | 1.35 |
| 2 | Dusp16 | 0.64 | 8.54E-09 | 1.82E-04 | 0.36 | 0.24 | 0.12 | 1.49 |
| 2 | Adam23 | 1.21 | 9.32E-09 | 1.99E-04 | 0.12 | 0.06 | 0.07 | 2.2 |
| 2 | Insr | 0.55 | 9.78E-09 | 2.09E-04 | 0.43 | 0.3 | 0.13 | 1.44 |
| 2 | Ncaph2 | 0.37 | 1.04E-08 | 2.23E-04 | 0.71 | 0.6 | 0.11 | 1.18 |
| 2 | Mafb | 0.67 | 1.05E-08 | 2.24E-04 | 0.25 | 0.15 | 0.1 | 1.67 |
| 2 | Efna5 | 0.44 | 1.08E-08 | 2.31E-04 | 0.63 | 0.51 | 0.12 | 1.23 |
| 2 | Ccdc88c | 0.43 | 1.12E-08 | 2.40E-04 | 0.67 | 0.56 | 0.11 | 1.19 |
| 2 | Ccnjl | 0.47 | 1.14E-08 | 2.43E-04 | 0.55 | 0.43 | 0.12 | 1.29 |
| 2 | Klf3 | 0.36 | 1.26E-08 | 2.69E-04 | 0.7 | 0.6 | 0.11 | 1.18 |
| 2 | Adgrl3 | 0.59 | 1.32E-08 | 2.81E-04 | 0.43 | 0.3 | 0.12 | 1.41 |
| 2 | Tmed3 | 0.44 | 1.41E-08 | 3.01E-04 | 0.61 | 0.49 | 0.12 | 1.26 |
| 2 | Mertk | 0.9 | 1.54E-08 | 3.28E-04 | 0.17 | 0.09 | 0.08 | 1.92 |
| 2 | Myh14 | 0.49 | 1.78E-08 | 3.80E-04 | 0.41 | 0.29 | 0.12 | 1.43 |
| 2 | Hint2 | 0.3 | 1.99E-08 | 4.26E-04 | 0.77 | 0.68 | 0.08 | 1.12 |
| 2 | Bnipl | 0.79 | 2.05E-08 | 4.38E-04 | 0.28 | 0.17 | 0.1 | 1.59 |
| 2 | Grb7 | 0.49 | 2.19E-08 | 4.68E-04 | 0.5 | 0.37 | 0.13 | 1.34 |
| 2 | Calml3 | 1.08 | 2.24E-08 | 4.78E-04 | 0.1 | 0.04 | 0.06 | 2.34 |
| 2 | Trpm3 | 0.62 | 2.28E-08 | 4.87E-04 | 0.26 | 0.16 | 0.1 | 1.66 |
| 2 | Dap | 0.35 | 2.58E-08 | 5.51E-04 | 0.76 | 0.67 | 0.08 | 1.12 |
| 2 | Ripk4 | 0.46 | 2.61E-08 | 5.57E-04 | 0.41 | 0.29 | 0.12 | 1.43 |
| 2 | Heyl | 0.6 | 2.71E-08 | 5.80E-04 | 0.32 | 0.21 | 0.11 | 1.53 |
| 2 | Rer1 | 0.26 | 2.76E-08 | 5.89E-04 | 0.91 | 0.84 | 0.06 | 1.08 |
| 2 | Atp6v1d | 0.33 | 2.79E-08 | 5.95E-04 | 0.76 | 0.66 | 0.09 | 1.14 |
| 2 | Barx2 | 1.16 | 2.95E-08 | 6.30E-04 | 0.25 | 0.16 | 0.09 | 1.59 |
| 2 | Arl6ip1 | 0.3 | 3.02E-08 | 6.45E-04 | 0.97 | 0.94 | 0.03 | 1.03 |
| 2 | Dipk2a | 0.35 | 3.16E-08 | 6.75E-04 | 0.54 | 0.41 | 0.13 | 1.32 |
| 2 | Serinc5 | 0.77 | 3.35E-08 | 7.16E-04 | 0.2 | 0.11 | 0.09 | 1.77 |

**Supplementary Table 3. Marker genes for the original clusters of integrated clustering of E11.5 control and NotchGOF λ-junction ectoderm cells.**

| | | | | | | | | |
|---|---|---|---|---|---|---|---|---|
| 2 | Mbd2 | 0.26 | 3.39E-08 | 7.23E-04 | 0.71 | 0.6 | 0.11 | 1.18 |
| 2 | Spg21 | 0.32 | 3.39E-08 | 7.23E-04 | 0.75 | 0.65 | 0.09 | 1.14 |
| 2 | Cux2 | 0.86 | 3.44E-08 | 7.34E-04 | 0.24 | 0.15 | 0.09 | 1.63 |
| 2 | Foxc1 | 0.93 | 3.59E-08 | 7.67E-04 | 0.23 | 0.13 | 0.09 | 1.69 |
| 2 | Zbtb7a | 0.54 | 3.64E-08 | 7.77E-04 | 0.48 | 0.36 | 0.12 | 1.33 |
| 2 | Hspg2 | 0.5 | 3.70E-08 | 7.91E-04 | 0.5 | 0.38 | 0.12 | 1.31 |
| 2 | Foxo1 | 0.49 | 3.77E-08 | 8.05E-04 | 0.4 | 0.28 | 0.12 | 1.43 |
| 2 | Slc2a1 | 0.27 | 3.87E-08 | 8.27E-04 | 0.8 | 0.68 | 0.12 | 1.17 |
| 2 | Ndufa4l2 | 0.69 | 3.92E-08 | 8.37E-04 | 0.17 | 0.09 | 0.08 | 1.86 |
| 2 | Atp2c2 | 1.11 | 3.92E-08 | 8.37E-04 | 0.11 | 0.05 | 0.06 | 2.22 |
| 2 | Zfp282 | 0.53 | 3.95E-08 | 8.43E-04 | 0.42 | 0.29 | 0.13 | 1.43 |
| 2 | Bcap31 | 0.31 | 3.98E-08 | 8.50E-04 | 0.86 | 0.79 | 0.07 | 1.08 |
| 2 | Fat1 | 0.3 | 4.40E-08 | 9.39E-04 | 0.72 | 0.63 | 0.09 | 1.14 |
| 2 | Plec | 0.34 | 4.54E-08 | 9.71E-04 | 0.57 | 0.44 | 0.14 | 1.31 |
| 2 | Afdn | 0.27 | 4.55E-08 | 9.71E-04 | 0.88 | 0.86 | 0.03 | 1.03 |
| 2 | Renbp | 0.65 | 4.62E-08 | 9.87E-04 | 0.19 | 0.1 | 0.09 | 1.84 |
| 2 | Phlpp1 | 0.48 | 5.17E-08 | 1.10E-03 | 0.37 | 0.25 | 0.12 | 1.46 |
| 2 | Pag1 | 1.03 | 5.18E-08 | 1.11E-03 | 0.11 | 0.05 | 0.06 | 2.22 |
| 2 | Trps1 | 0.3 | 5.44E-08 | 1.16E-03 | 0.63 | 0.52 | 0.11 | 1.21 |
| 2 | Mylk | 0.77 | 5.84E-08 | 1.25E-03 | 0.29 | 0.19 | 0.1 | 1.51 |
| 2 | Cast | 0.72 | 6.03E-08 | 1.29E-03 | 0.28 | 0.18 | 0.1 | 1.55 |
| 2 | Gramd3 | 0.64 | 6.13E-08 | 1.31E-03 | 0.32 | 0.22 | 0.1 | 1.48 |
| 2 | H2-D1 | 0.78 | 6.22E-08 | 1.33E-03 | 0.43 | 0.32 | 0.11 | 1.33 |
| 2 | Tril | 0.39 | 6.29E-08 | 1.34E-03 | 0.41 | 0.29 | 0.12 | 1.44 |
| 2 | Sun2 | 0.92 | 6.64E-08 | 1.42E-03 | 0.21 | 0.13 | 0.09 | 1.68 |
| 2 | Atp2b4 | 0.51 | 7.00E-08 | 1.49E-03 | 0.34 | 0.22 | 0.12 | 1.52 |
| 2 | Itgb4 | 0.65 | 7.06E-08 | 1.51E-03 | 0.2 | 0.11 | 0.09 | 1.77 |
| 2 | Tmbim1 | 0.97 | 7.31E-08 | 1.56E-03 | 0.17 | 0.09 | 0.08 | 1.84 |
| 2 | Arrb2 | 0.81 | 7.51E-08 | 1.60E-03 | 0.28 | 0.18 | 0.1 | 1.53 |
| 2 | Vps36 | 0.27 | 7.61E-08 | 1.62E-03 | 0.9 | 0.85 | 0.05 | 1.06 |
| 2 | Pon2 | 0.35 | 8.88E-08 | 1.90E-03 | 0.41 | 0.28 | 0.13 | 1.45 |
| 2 | Cmtm3 | 0.46 | 8.89E-08 | 1.90E-03 | 0.48 | 0.35 | 0.13 | 1.37 |
| 2 | Rsbn1l | 0.29 | 8.97E-08 | 1.92E-03 | 0.75 | 0.67 | 0.08 | 1.11 |
| 2 | Pdgfrl | 0.46 | 9.06E-08 | 1.94E-03 | 0.44 | 0.32 | 0.12 | 1.38 |
| 2 | Sema3f | 0.48 | 9.08E-08 | 1.94E-03 | 0.5 | 0.37 | 0.13 | 1.34 |
| 2 | Ptk7 | 0.41 | 9.27E-08 | 1.98E-03 | 0.59 | 0.48 | 0.11 | 1.23 |
| 2 | Ctdspl | 0.4 | 9.31E-08 | 1.99E-03 | 0.59 | 0.48 | 0.12 | 1.24 |
| 2 | Rab11a | 0.35 | 9.45E-08 | 2.02E-03 | 0.76 | 0.68 | 0.08 | 1.12 |
| 2 | Phactr2 | 0.53 | 9.77E-08 | 2.09E-03 | 0.47 | 0.36 | 0.11 | 1.3 |
| 2 | Sh3d19 | 0.45 | 1.00E-07 | 2.14E-03 | 0.45 | 0.33 | 0.12 | 1.35 |
| 2 | Cpne3 | 0.37 | 1.09E-07 | 2.32E-03 | 0.72 | 0.63 | 0.09 | 1.14 |
| 2 | Crlf3 | 0.46 | 1.23E-07 | 2.63E-03 | 0.42 | 0.31 | 0.12 | 1.38 |
| 2 | Sbno2 | 0.69 | 1.38E-07 | 2.95E-03 | 0.27 | 0.17 | 0.1 | 1.56 |
| 2 | Alcam | 0.28 | 1.39E-07 | 2.97E-03 | 0.3 | 0.19 | 0.11 | 1.57 |
| 2 | Sppl2a | 0.48 | 1.40E-07 | 2.98E-03 | 0.67 | 0.58 | 0.09 | 1.15 |
| 2 | Igf2os | 0.81 | 1.41E-07 | 3.02E-03 | 0.13 | 0.07 | 0.07 | 2 |
| 2 | Mprip | 0.3 | 1.55E-07 | 3.32E-03 | 0.81 | 0.73 | 0.08 | 1.11 |
| 2 | Arfgef1 | 0.38 | 1.63E-07 | 3.49E-03 | 0.69 | 0.6 | 0.1 | 1.16 |
| 2 | Wiz | 0.4 | 1.64E-07 | 3.50E-03 | 0.56 | 0.46 | 0.1 | 1.22 |
| 2 | Fgd6 | 0.61 | 1.66E-07 | 3.55E-03 | 0.32 | 0.21 | 0.11 | 1.52 |
| 2 | Prrx1 | 0.25 | 1.74E-07 | 3.72E-03 | 0.29 | 0.18 | 0.11 | 1.59 |
| 2 | Gucy1b1 | 1.12 | 1.77E-07 | 3.77E-03 | 0.14 | 0.07 | 0.07 | 1.92 |
| 2 | Pkp4 | 0.3 | 1.80E-07 | 3.85E-03 | 0.67 | 0.57 | 0.1 | 1.17 |
| 2 | Rnf43 | 0.61 | 2.04E-07 | 4.37E-03 | 0.28 | 0.18 | 0.1 | 1.55 |
| 2 | Krt1 | 0.72 | 2.13E-07 | 4.54E-03 | 0.2 | 0.12 | 0.08 | 1.7 |
| 2 | Ptpn14 | 0.54 | 2.23E-07 | 4.76E-03 | 0.33 | 0.22 | 0.11 | 1.51 |
| 2 | 2700033N17Rik | 0.43 | 2.27E-07 | 4.84E-03 | 0.34 | 0.23 | 0.11 | 1.49 |

**Supplementary Table 3. Marker genes for the original clusters of integrated clustering of E11.5 control and NotchGOF λ-junction ectoderm cells.**

| | | | | | | | | |
|---|---|---|---|---|---|---|---|---|
| 2 | Chst5 | 0.88 | 2.28E-07 | 4.86E-03 | 0.12 | 0.06 | 0.06 | 2.09 |
| 2 | Rhbdf1 | 0.64 | 2.33E-07 | 4.98E-03 | 0.29 | 0.19 | 0.1 | 1.51 |
| 2 | Galnt18 | 0.48 | 2.41E-07 | 5.15E-03 | 0.17 | 0.09 | 0.08 | 1.86 |
| 2 | Abhd11 | 0.4 | 2.44E-07 | 5.21E-03 | 0.51 | 0.39 | 0.12 | 1.31 |
| 2 | Plod2 | 0.41 | 2.56E-07 | 5.47E-03 | 0.58 | 0.47 | 0.11 | 1.24 |
| 2 | Reep4 | 0.48 | 2.58E-07 | 5.51E-03 | 0.47 | 0.36 | 0.11 | 1.3 |
| 2 | Col4a3bp | 0.44 | 2.77E-07 | 5.91E-03 | 0.46 | 0.35 | 0.11 | 1.32 |
| 2 | Heca | 0.52 | 2.92E-07 | 6.24E-03 | 0.37 | 0.26 | 0.11 | 1.43 |
| 2 | Reep6 | 0.97 | 2.94E-07 | 6.28E-03 | 0.15 | 0.08 | 0.07 | 1.87 |
| 2 | Piezo2 | 0.25 | 3.03E-07 | 6.46E-03 | 0.26 | 0.16 | 0.1 | 1.61 |
| 2 | Chmp2a | 0.27 | 3.15E-07 | 6.73E-03 | 0.81 | 0.73 | 0.08 | 1.11 |
| 2 | Ltbr | 0.67 | 3.33E-07 | 7.11E-03 | 0.3 | 0.2 | 0.1 | 1.51 |
| 2 | Lrrk1 | 0.56 | 3.38E-07 | 7.23E-03 | 0.4 | 0.3 | 0.1 | 1.34 |
| 2 | Pfn2 | 0.46 | 3.49E-07 | 7.45E-03 | 0.77 | 0.7 | 0.07 | 1.11 |
| 2 | Acsl5 | 0.51 | 3.51E-07 | 7.51E-03 | 0.42 | 0.31 | 0.11 | 1.35 |
| 2 | Fam89a | 0.49 | 3.53E-07 | 7.54E-03 | 0.25 | 0.15 | 0.1 | 1.63 |
| 2 | Fam83h | 0.82 | 3.57E-07 | 7.62E-03 | 0.16 | 0.09 | 0.07 | 1.86 |
| 2 | Arl13b | 0.46 | 3.73E-07 | 7.96E-03 | 0.51 | 0.39 | 0.12 | 1.3 |
| 2 | Arhgdib | 0.47 | 3.87E-07 | 8.28E-03 | 0.3 | 0.2 | 0.1 | 1.5 |
| 2 | Leprot | 0.42 | 4.05E-07 | 8.65E-03 | 0.5 | 0.39 | 0.11 | 1.28 |
| 2 | Rabgap1 | 0.26 | 4.07E-07 | 8.69E-03 | 0.77 | 0.68 | 0.09 | 1.13 |
| 2 | Nfe2l3 | 0.65 | 4.28E-07 | 9.14E-03 | 0.22 | 0.13 | 0.09 | 1.65 |
| 2 | Tax1bp3 | 0.31 | 4.53E-07 | 9.69E-03 | 0.66 | 0.55 | 0.11 | 1.2 |
| 2 | Nmrk1 | 0.91 | 4.55E-07 | 9.72E-03 | 0.18 | 0.11 | 0.08 | 1.71 |
| 2 | Pak1 | 0.36 | 4.58E-07 | 9.79E-03 | 0.7 | 0.62 | 0.08 | 1.13 |
| 2 | Gjb6 | 0.85 | 4.60E-07 | 9.82E-03 | 0.12 | 0.06 | 0.06 | 2.05 |
| 2 | Zfp704 | 0.31 | 4.63E-07 | 9.89E-03 | 0.7 | 0.62 | 0.08 | 1.14 |
| 2 | Gm28112 | 0.86 | 4.93E-07 | 1.05E-02 | 0.15 | 0.08 | 0.07 | 1.87 |
| 2 | Rab38 | 0.66 | 5.05E-07 | 1.08E-02 | 0.29 | 0.19 | 0.1 | 1.49 |
| 2 | Map7 | 0.42 | 5.08E-07 | 1.08E-02 | 0.53 | 0.42 | 0.11 | 1.26 |
| 2 | Marveld3 | 0.71 | 5.27E-07 | 1.13E-02 | 0.24 | 0.15 | 0.09 | 1.6 |
| 2 | Fkbp1b | 0.54 | 5.47E-07 | 1.17E-02 | 0.25 | 0.16 | 0.09 | 1.57 |
| 2 | Aass | 0.81 | 5.53E-07 | 1.18E-02 | 0.13 | 0.07 | 0.06 | 1.94 |
| 2 | Braf | 0.53 | 5.89E-07 | 1.26E-02 | 0.47 | 0.37 | 0.11 | 1.29 |
| 2 | Ten1 | 0.34 | 6.04E-07 | 1.29E-02 | 0.7 | 0.6 | 0.1 | 1.17 |
| 2 | Hunk | 0.35 | 6.12E-07 | 1.31E-02 | 0.67 | 0.58 | 0.09 | 1.16 |
| 2 | Jak1 | 0.34 | 6.12E-07 | 1.31E-02 | 0.64 | 0.55 | 0.1 | 1.18 |
| 2 | Gm9908 | 0.69 | 6.16E-07 | 1.32E-02 | 0.13 | 0.07 | 0.06 | 1.94 |
| 2 | Mxi1 | 0.31 | 6.46E-07 | 1.38E-02 | 0.66 | 0.56 | 0.09 | 1.16 |
| 2 | Ass1 | 0.76 | 6.52E-07 | 1.39E-02 | 0.19 | 0.11 | 0.08 | 1.71 |
| 2 | Ehd1 | 0.49 | 6.69E-07 | 1.43E-02 | 0.55 | 0.46 | 0.1 | 1.21 |
| 2 | Rmnd5a | 0.36 | 6.79E-07 | 1.45E-02 | 0.56 | 0.44 | 0.12 | 1.27 |
| 2 | Ndrg2 | 0.5 | 7.84E-07 | 1.68E-02 | 0.45 | 0.34 | 0.11 | 1.31 |
| 2 | Rarg | 0.51 | 8.05E-07 | 1.72E-02 | 0.39 | 0.28 | 0.11 | 1.38 |
| 2 | A430018G15Rik | 0.87 | 8.30E-07 | 1.77E-02 | 0.14 | 0.08 | 0.07 | 1.85 |
| 2 | Arhgap32 | 0.52 | 8.38E-07 | 1.79E-02 | 0.34 | 0.24 | 0.1 | 1.44 |
| 2 | Grhl1 | 0.79 | 8.76E-07 | 1.87E-02 | 0.15 | 0.08 | 0.07 | 1.85 |
| 2 | Trim59 | 0.31 | 8.87E-07 | 1.90E-02 | 0.78 | 0.71 | 0.07 | 1.1 |
| 2 | Plaat3 | 0.92 | 8.90E-07 | 1.90E-02 | 0.24 | 0.16 | 0.08 | 1.5 |
| 2 | Krcc1 | 0.32 | 9.38E-07 | 2.00E-02 | 0.71 | 0.62 | 0.08 | 1.13 |
| 2 | Arhgap23 | 0.74 | 9.66E-07 | 2.06E-02 | 0.18 | 0.11 | 0.08 | 1.71 |
| 2 | Kifc3 | 0.64 | 9.71E-07 | 2.07E-02 | 0.19 | 0.11 | 0.08 | 1.71 |
| 2 | Mtln | 0.39 | 1.03E-06 | 2.19E-02 | 0.64 | 0.54 | 0.1 | 1.19 |
| 2 | Tmod3 | 0.28 | 1.04E-06 | 2.21E-02 | 0.74 | 0.65 | 0.09 | 1.13 |
| 2 | Ctsd | 0.34 | 1.06E-06 | 2.27E-02 | 0.77 | 0.7 | 0.08 | 1.11 |
| 2 | Prtg | 0.73 | 1.09E-06 | 2.32E-02 | 0.29 | 0.2 | 0.09 | 1.45 |
| 2 | Pold4 | 0.61 | 1.14E-06 | 2.43E-02 | 0.31 | 0.22 | 0.1 | 1.44 |

**Supplementary Table 3. Marker genes for the original clusters of integrated clustering of E11.5 control and NotchGOF λ-junction ectoderm cells.**

| | | | | | | | | |
|---|---|---|---|---|---|---|---|---|
| 2 | Ttc7b | 0.65 | 1.15E-06 | 2.45E-02 | 0.24 | 0.15 | 0.08 | 1.56 |
| 2 | Fam129b | 0.52 | 1.19E-06 | 2.55E-02 | 0.27 | 0.18 | 0.09 | 1.51 |
| 2 | Eps8 | 0.78 | 1.21E-06 | 2.59E-02 | 0.17 | 0.1 | 0.07 | 1.74 |
| 2 | C1qtnf12 | 0.67 | 1.26E-06 | 2.70E-02 | 0.31 | 0.22 | 0.09 | 1.43 |
| 2 | Cdk6 | 0.26 | 1.35E-06 | 2.89E-02 | 0.64 | 0.51 | 0.14 | 1.27 |
| 2 | Fam114a1 | 0.81 | 1.39E-06 | 2.97E-02 | 0.17 | 0.1 | 0.07 | 1.71 |
| 2 | C2cd2 | 0.72 | 1.41E-06 | 3.00E-02 | 0.16 | 0.09 | 0.07 | 1.76 |
| 2 | Rnf149 | 0.58 | 1.41E-06 | 3.00E-02 | 0.38 | 0.28 | 0.1 | 1.35 |
| 2 | Add1 | 0.29 | 1.47E-06 | 3.15E-02 | 0.77 | 0.67 | 0.1 | 1.15 |
| 2 | Arhgef17 | 0.43 | 1.56E-06 | 3.34E-02 | 0.48 | 0.38 | 0.1 | 1.26 |
| 2 | Lmna | 0.54 | 1.58E-06 | 3.37E-02 | 0.27 | 0.18 | 0.09 | 1.5 |
| 2 | Copg2 | 0.33 | 1.63E-06 | 3.49E-02 | 0.59 | 0.48 | 0.11 | 1.23 |
| 2 | Bcl9 | 0.29 | 1.70E-06 | 3.63E-02 | 0.74 | 0.68 | 0.06 | 1.1 |
| 2 | Tmprss13 | 1.11 | 1.74E-06 | 3.71E-02 | 0.11 | 0.06 | 0.06 | 1.98 |
| 2 | Map2k3 | 0.34 | 1.74E-06 | 3.72E-02 | 0.52 | 0.42 | 0.1 | 1.25 |
| 2 | Mdfic | 0.56 | 1.75E-06 | 3.73E-02 | 0.27 | 0.18 | 0.09 | 1.51 |
| 2 | Bcl2l14 | 1.01 | 1.77E-06 | 3.78E-02 | 0.1 | 0.05 | 0.05 | 2.06 |
| 2 | Ralgps1 | 0.57 | 1.81E-06 | 3.86E-02 | 0.2 | 0.12 | 0.08 | 1.65 |
| 2 | Rdm1 | 0.42 | 1.82E-06 | 3.88E-02 | 0.48 | 0.38 | 0.1 | 1.27 |
| 2 | Slc25a27 | 0.38 | 1.83E-06 | 3.90E-02 | 0.62 | 0.54 | 0.08 | 1.15 |
| 2 | Ifngr2 | 0.4 | 1.86E-06 | 3.96E-02 | 0.37 | 0.27 | 0.11 | 1.39 |
| 2 | Rusc1 | 0.63 | 1.88E-06 | 4.01E-02 | 0.23 | 0.15 | 0.08 | 1.56 |
| 2 | Hexim1 | 0.51 | 1.88E-06 | 4.02E-02 | 0.42 | 0.32 | 0.09 | 1.29 |
| 2 | Vps4b | 0.29 | 1.95E-06 | 4.17E-02 | 0.77 | 0.68 | 0.08 | 1.12 |
| 2 | Tmem50b | 0.75 | 1.97E-06 | 4.21E-02 | 0.24 | 0.16 | 0.08 | 1.52 |
| 2 | Igsf10 | 0.74 | 2.14E-06 | 4.56E-02 | 0.19 | 0.11 | 0.07 | 1.67 |
| 2 | Hip1r | 0.66 | 2.14E-06 | 4.56E-02 | 0.29 | 0.21 | 0.09 | 1.43 |
| 2 | Tspan5 | 0.43 | 2.18E-06 | 4.66E-02 | 0.44 | 0.33 | 0.1 | 1.31 |
| 2 | 2310022B05Rik | 0.49 | 2.18E-06 | 4.66E-02 | 0.36 | 0.26 | 0.11 | 1.41 |
| 2 | Trp53bp2 | 0.39 | 2.29E-06 | 4.89E-02 | 0.53 | 0.44 | 0.09 | 1.2 |
| 4 | Cdkl1 | 4.94 | 0 | 0 | 0.63 | 0.03 | 0.6 | 18.62 |
| 4 | Uncx | 2.32 | 8.61E-235 | 1.84E-230 | 0.55 | 0.04 | 0.5 | 12.72 |
| 4 | St8sia6 | 3.17 | 3.51E-222 | 7.51E-218 | 0.71 | 0.11 | 0.6 | 6.42 |
| 4 | Sox3 | 3.05 | 1.13E-214 | 2.41E-210 | 0.66 | 0.09 | 0.57 | 7.24 |
| 4 | Hes5 | 2.55 | 2.02E-210 | 4.31E-206 | 0.97 | 0.32 | 0.65 | 3 |
| 4 | Cbln2 | 3.06 | 2.89E-201 | 6.18E-197 | 0.62 | 0.09 | 0.54 | 7.31 |
| 4 | Gm266 | 2.32 | 1.20E-199 | 2.57E-195 | 1 | 0.49 | 0.51 | 2.03 |
| 4 | Tnfaip2 | 2.43 | 2.13E-196 | 4.56E-192 | 0.96 | 0.38 | 0.58 | 2.51 |
| 4 | Sox21 | 2.97 | 1.90E-178 | 4.05E-174 | 0.56 | 0.08 | 0.48 | 7.34 |
| 4 | Prr15 | 3.25 | 1.66E-176 | 3.54E-172 | 0.55 | 0.08 | 0.48 | 7.1 |
| 4 | Zdhhc2 | 2.5 | 9.68E-171 | 2.07E-166 | 0.87 | 0.32 | 0.55 | 2.72 |
| 4 | Pcdh19 | 2.51 | 1.00E-160 | 2.14E-156 | 0.74 | 0.18 | 0.56 | 4.06 |
| 4 | Ckb | 2.22 | 3.10E-160 | 6.63E-156 | 0.86 | 0.29 | 0.56 | 2.93 |
| 4 | Kazald1 | 3.5 | 9.60E-156 | 2.05E-151 | 0.42 | 0.04 | 0.38 | 9.57 |
| 4 | Tmsb4x | 1.37 | 2.40E-149 | 5.13E-145 | 1 | 1 | 0 | 1 |
| 4 | Cdh4 | 3.1 | 1.23E-147 | 2.63E-143 | 0.47 | 0.06 | 0.41 | 7.34 |
| 4 | Tex15 | 2.04 | 1.76E-134 | 3.76E-130 | 0.74 | 0.22 | 0.52 | 3.35 |
| 4 | Psip1 | 0.95 | 3.88E-129 | 8.28E-125 | 1 | 0.98 | 0.02 | 1.02 |
| 4 | Prss12 | 4.06 | 5.30E-127 | 1.13E-122 | 0.28 | 0.02 | 0.26 | 14 |
| 4 | Rps2 | 0.48 | 6.14E-127 | 1.31E-122 | 1 | 1 | 0 | 1 |
| 4 | Plat | 2.61 | 1.06E-126 | 2.27E-122 | 0.61 | 0.15 | 0.47 | 4.15 |
| 4 | Wfdc2 | 1.25 | 1.25E-124 | 2.67E-120 | 0.99 | 0.8 | 0.19 | 1.24 |
| 4 | Hsp90aa1 | 0.69 | 2.20E-123 | 4.70E-119 | 1 | 1 | 0 | 1 |
| 4 | Cdh2 | 1.45 | 2.52E-122 | 5.38E-118 | 0.97 | 0.57 | 0.41 | 1.71 |
| 4 | Scrn1 | 2.46 | 9.54E-120 | 2.04E-115 | 0.48 | 0.08 | 0.4 | 5.77 |
| 4 | Gnai2 | 0.84 | 1.23E-118 | 2.62E-114 | 1 | 0.97 | 0.02 | 1.02 |
| 4 | Rims4 | 3.66 | 2.18E-117 | 4.65E-113 | 0.3 | 0.03 | 0.27 | 11 |

**Supplementary Table 3. Marker genes for the original clusters of integrated clustering of E11.5 control and NotchGOF λ-junction ectoderm cells.**

| | | | | | | | | |
|---|---|---|---|---|---|---|---|---|
| 4 | Pea15a | 1.87 | 2.51E-117 | 5.37E-113 | 0.79 | 0.29 | 0.51 | 2.77 |
| 4 | Sfrp1 | 1.59 | 2.43E-116 | 5.18E-112 | 0.92 | 0.47 | 0.45 | 1.94 |
| 4 | Vim | 1.28 | 6.04E-116 | 1.29E-111 | 0.96 | 0.69 | 0.27 | 1.4 |
| 4 | Ccnd1 | 1.38 | 1.12E-115 | 2.38E-111 | 0.97 | 0.68 | 0.3 | 1.44 |
| 4 | Sorbs2 | 1.79 | 8.74E-114 | 1.87E-109 | 0.84 | 0.35 | 0.49 | 2.38 |
| 4 | Cenpv | 1.79 | 4.69E-113 | 1.00E-108 | 0.83 | 0.33 | 0.5 | 2.49 |
| 4 | Cyp4v3 | 2.95 | 8.10E-113 | 1.73E-108 | 0.4 | 0.06 | 0.34 | 6.59 |
| 4 | Ncam1 | 1.38 | 2.91E-111 | 6.21E-107 | 0.8 | 0.28 | 0.52 | 2.82 |
| 4 | Hspd1 | 0.82 | 1.03E-110 | 2.21E-106 | 1 | 0.98 | 0.02 | 1.02 |
| 4 | Crip2 | 1.62 | 1.79E-109 | 3.83E-105 | 0.85 | 0.37 | 0.48 | 2.29 |
| 4 | Mif | 0.61 | 8.07E-109 | 1.72E-104 | 1 | 1 | 0 | 1 |
| 4 | Fam189a2 | 3.01 | 4.13E-108 | 8.83E-104 | 0.37 | 0.05 | 0.32 | 7.06 |
| 4 | Rpl41 | 0.35 | 4.62E-108 | 9.87E-104 | 1 | 1 | 0 | 1 |
| 4 | Eif5a | 0.62 | 1.70E-107 | 3.64E-103 | 1 | 0.99 | 0.01 | 1.01 |
| 4 | Rgcc | 2.92 | 1.78E-107 | 3.81E-103 | 0.42 | 0.07 | 0.35 | 5.81 |
| 4 | Tubb2b | 0.88 | 3.30E-107 | 7.05E-103 | 1 | 0.92 | 0.09 | 1.09 |
| 4 | Socs2 | 1.17 | 8.44E-106 | 1.80E-101 | 0.99 | 0.76 | 0.23 | 1.31 |
| 4 | Sox1 | 3.46 | 8.55E-101 | 1.83E-96 | 0.25 | 0.02 | 0.23 | 11.81 |
| 4 | Crb2 | 2.63 | 2.85E-100 | 6.08E-96 | 0.37 | 0.06 | 0.31 | 6.51 |
| 4 | Rplp1 | 0.4 | 2.30E-99 | 4.91E-95 | 1 | 1 | 0 | 1 |
| 4 | Ppa1 | 1.01 | 5.26E-99 | 1.12E-94 | 0.99 | 0.86 | 0.14 | 1.16 |
| 4 | Rprm | 1.38 | 1.03E-95 | 2.19E-91 | 0.83 | 0.33 | 0.51 | 2.53 |
| 4 | Epb41l2 | 1.78 | 5.43E-95 | 1.16E-90 | 0.85 | 0.46 | 0.4 | 1.87 |
| 4 | Stbd1 | 2.35 | 1.04E-94 | 2.23E-90 | 0.47 | 0.1 | 0.37 | 4.54 |
| 4 | Eya4 | 1.9 | 1.81E-94 | 3.86E-90 | 0.61 | 0.18 | 0.43 | 3.47 |
| 4 | Tagln2 | 2.12 | 2.13E-94 | 4.55E-90 | 0.67 | 0.23 | 0.43 | 2.86 |
| 4 | Slitrk5 | 3.52 | 1.67E-93 | 3.57E-89 | 0.25 | 0.03 | 0.23 | 10.08 |
| 4 | Apc | 1.29 | 5.89E-92 | 1.26E-87 | 0.97 | 0.77 | 0.2 | 1.26 |
| 4 | Dmrta2 | 1.36 | 1.16E-91 | 2.47E-87 | 0.9 | 0.46 | 0.44 | 1.94 |
| 4 | Mapk12 | 1.67 | 4.83E-91 | 1.03E-86 | 0.7 | 0.24 | 0.46 | 2.88 |
| 4 | Nr2e1 | 1.87 | 1.06E-90 | 2.25E-86 | 0.53 | 0.13 | 0.4 | 4.11 |
| 4 | Jph4 | 2.17 | 2.31E-90 | 4.93E-86 | 0.56 | 0.16 | 0.4 | 3.52 |
| 4 | Snhg4 | 1.78 | 1.88E-89 | 4.01E-85 | 0.81 | 0.39 | 0.42 | 2.1 |
| 4 | Dut | 0.92 | 1.93E-89 | 4.11E-85 | 1 | 0.9 | 0.1 | 1.11 |
| 4 | Galm | 1.57 | 4.03E-89 | 8.60E-85 | 0.81 | 0.36 | 0.46 | 2.28 |
| 4 | Pls1 | 1.86 | 5.05E-87 | 1.08E-82 | 0.68 | 0.26 | 0.42 | 2.63 |
| 4 | Fut9 | 2.54 | 1.39E-86 | 2.98E-82 | 0.37 | 0.07 | 0.31 | 5.42 |
| 4 | Otx2 | 1.39 | 1.40E-86 | 3.00E-82 | 0.8 | 0.36 | 0.45 | 2.26 |
| 4 | Slc27a2 | 3.24 | 2.63E-86 | 5.62E-82 | 0.28 | 0.04 | 0.24 | 7.89 |
| 4 | Rragd | 1.47 | 3.31E-86 | 7.07E-82 | 0.81 | 0.37 | 0.44 | 2.19 |
| 4 | Ran | 0.56 | 1.08E-84 | 2.31E-80 | 1 | 0.99 | 0.01 | 1.01 |
| 4 | Hspe1 | 0.56 | 1.64E-84 | 3.50E-80 | 1 | 0.99 | 0.01 | 1.01 |
| 4 | Gm10076 | 0.38 | 4.21E-82 | 8.98E-78 | 1 | 1 | 0 | 1 |
| 4 | Egfr | 1.93 | 1.40E-81 | 2.99E-77 | 0.59 | 0.19 | 0.4 | 3.1 |
| 4 | Fzd9 | 3.34 | 6.39E-81 | 1.36E-76 | 0.25 | 0.03 | 0.22 | 8.13 |
| 4 | Ncl | 0.65 | 1.41E-80 | 3.01E-76 | 1 | 0.99 | 0.01 | 1.01 |
| 4 | Gpm6a | 2.09 | 2.16E-80 | 4.61E-76 | 0.45 | 0.11 | 0.35 | 4.19 |
| 4 | Tpi1 | 0.69 | 3.37E-80 | 7.19E-76 | 1 | 0.98 | 0.02 | 1.02 |
| 4 | Ybx1 | 0.45 | 1.79E-78 | 3.83E-74 | 1 | 1 | 0 | 1 |
| 4 | Gpc1 | 1.28 | 5.30E-78 | 1.13E-73 | 0.86 | 0.46 | 0.4 | 1.87 |
| 4 | Cadm1 | 1.03 | 8.35E-77 | 1.78E-72 | 0.96 | 0.69 | 0.27 | 1.39 |
| 4 | Il17rd | 1.28 | 2.31E-76 | 4.94E-72 | 0.86 | 0.46 | 0.4 | 1.88 |
| 4 | Cacybp | 0.74 | 5.45E-76 | 1.16E-71 | 1 | 0.93 | 0.07 | 1.07 |
| 4 | H2afz | 0.51 | 1.09E-75 | 2.33E-71 | 1 | 1 | 0 | 1 |
| 4 | Nipal1 | 3.47 | 4.93E-75 | 1.05E-70 | 0.2 | 0.02 | 0.18 | 10.15 |
| 4 | Tshz3 | 2.19 | 1.05E-74 | 2.25E-70 | 0.4 | 0.09 | 0.31 | 4.49 |
| 4 | Shisa3 | 3.3 | 1.89E-74 | 4.04E-70 | 0.21 | 0.02 | 0.19 | 9.81 |

**Supplementary Table 3. Marker genes for the original clusters of integrated clustering of E11.5 control and NotchGOF λ-junction ectoderm cells.**

| | | | | | | | | |
|---|---|---|---|---|---|---|---|---|
| 4 | C130071C03Rik | 1.83 | 1.38E-73 | 2.94E-69 | 0.54 | 0.17 | 0.37 | 3.23 |
| 4 | Slc22a23 | 1.53 | 1.54E-73 | 3.28E-69 | 0.69 | 0.27 | 0.42 | 2.56 |
| 4 | Nrcam | 1.96 | 5.21E-72 | 1.11E-67 | 0.52 | 0.16 | 0.36 | 3.28 |
| 4 | Rfx4 | 5.49 | 6.69E-72 | 1.43E-67 | 0.11 | 0 | 0.11 | 53.5 |
| 4 | Lyar | 0.93 | 1.06E-71 | 2.27E-67 | 0.97 | 0.79 | 0.18 | 1.23 |
| 4 | Marcks | 0.63 | 1.09E-71 | 2.32E-67 | 1 | 0.99 | 0.01 | 1.01 |
| 4 | Ranbp1 | 0.56 | 1.27E-71 | 2.72E-67 | 1 | 0.98 | 0.02 | 1.02 |
| 4 | Mpp6 | 1.16 | 1.80E-71 | 3.85E-67 | 0.91 | 0.63 | 0.28 | 1.45 |
| 4 | Tkt | 0.79 | 4.72E-71 | 1.01E-66 | 1 | 0.87 | 0.13 | 1.14 |
| 4 | Stox1 | 2.3 | 5.81E-70 | 1.24E-65 | 0.31 | 0.06 | 0.25 | 5.46 |
| 4 | Pcdh11x | 1.83 | 7.17E-70 | 1.53E-65 | 0.4 | 0.1 | 0.31 | 4.21 |
| 4 | Slc1a3 | 1.86 | 8.43E-70 | 1.80E-65 | 0.53 | 0.17 | 0.37 | 3.21 |
| 4 | Slitrk6 | 1.98 | 2.65E-69 | 5.67E-65 | 0.48 | 0.14 | 0.34 | 3.47 |
| 4 | Heg1 | 2.44 | 6.71E-69 | 1.43E-64 | 0.29 | 0.05 | 0.24 | 5.65 |
| 4 | Trpc1 | 1.76 | 1.61E-68 | 3.44E-64 | 0.58 | 0.21 | 0.37 | 2.81 |
| 4 | Rhov | 1.49 | 2.30E-68 | 4.91E-64 | 0.58 | 0.19 | 0.39 | 3.03 |
| 4 | Usp25 | 1.1 | 2.51E-68 | 5.35E-64 | 0.9 | 0.64 | 0.26 | 1.41 |
| 4 | Cdc14a | 1.53 | 2.81E-68 | 6.00E-64 | 0.72 | 0.33 | 0.39 | 2.17 |
| 4 | Espn | 1.28 | 5.01E-68 | 1.07E-63 | 0.75 | 0.33 | 0.42 | 2.26 |
| 4 | Mt1 | 1.27 | 7.18E-68 | 1.53E-63 | 0.84 | 0.47 | 0.37 | 1.78 |
| 4 | Necab2 | 1.57 | 1.63E-67 | 3.47E-63 | 0.64 | 0.25 | 0.39 | 2.58 |
| 4 | Zbtb7c | 3.15 | 5.82E-67 | 1.24E-62 | 0.19 | 0.02 | 0.17 | 10.28 |
| 4 | Itga6 | 0.92 | 7.86E-67 | 1.68E-62 | 0.97 | 0.77 | 0.21 | 1.27 |
| 4 | Pax6 | 1.2 | 2.47E-66 | 5.27E-62 | 0.85 | 0.45 | 0.4 | 1.88 |
| 4 | Tmem121 | 1.8 | 1.79E-65 | 3.83E-61 | 0.48 | 0.15 | 0.34 | 3.29 |
| 4 | Syt11 | 1.13 | 2.21E-65 | 4.72E-61 | 0.7 | 0.3 | 0.4 | 2.31 |
| 4 | Nme1 | 0.54 | 2.78E-65 | 5.95E-61 | 1 | 0.98 | 0.02 | 1.02 |
| 4 | Crym | 1.42 | 3.63E-65 | 7.76E-61 | 0.58 | 0.2 | 0.38 | 2.92 |
| 4 | Ybx3 | 0.78 | 7.16E-65 | 1.53E-60 | 0.97 | 0.86 | 0.11 | 1.12 |
| 4 | Ddah1 | 1.87 | 1.38E-64 | 2.95E-60 | 0.44 | 0.12 | 0.31 | 3.55 |
| 4 | Cd83 | 2.92 | 1.46E-64 | 3.11E-60 | 0.2 | 0.03 | 0.18 | 8.12 |
| 4 | Fam126a | 1.5 | 3.17E-64 | 6.77E-60 | 0.62 | 0.25 | 0.38 | 2.54 |
| 4 | Nr2f1 | 1.88 | 4.20E-64 | 8.97E-60 | 0.52 | 0.18 | 0.35 | 2.94 |
| 4 | Cxcl12 | 1.81 | 2.00E-63 | 4.26E-59 | 0.41 | 0.11 | 0.3 | 3.84 |
| 4 | Nrarp | 0.99 | 3.49E-63 | 7.46E-59 | 0.94 | 0.57 | 0.37 | 1.65 |
| 4 | Pkdcc | 1.36 | 5.47E-62 | 1.17E-57 | 0.66 | 0.28 | 0.38 | 2.37 |
| 4 | Nap1l1 | 0.43 | 1.84E-61 | 3.94E-57 | 1 | 0.99 | 0 | 1 |
| 4 | Set | 0.4 | 2.28E-61 | 4.87E-57 | 1 | 1 | 0 | 1 |
| 4 | Ppp1r1a | 1.35 | 4.12E-61 | 8.80E-57 | 0.66 | 0.28 | 0.39 | 2.41 |
| 4 | Snn | 1.24 | 4.68E-61 | 1.00E-56 | 0.86 | 0.58 | 0.28 | 1.48 |
| 4 | Dek | 0.66 | 6.17E-61 | 1.32E-56 | 1 | 0.92 | 0.08 | 1.09 |
| 4 | Fgf20 | 3.63 | 1.07E-60 | 2.28E-56 | 0.15 | 0.01 | 0.14 | 11 |
| 4 | Phb2 | 0.62 | 2.01E-60 | 4.30E-56 | 1 | 0.96 | 0.04 | 1.04 |
| 4 | Kcnb1 | 2.34 | 2.95E-60 | 6.30E-56 | 0.3 | 0.06 | 0.24 | 4.75 |
| 4 | Fzd4 | 1.73 | 8.78E-60 | 1.88E-55 | 0.47 | 0.15 | 0.32 | 3.14 |
| 4 | Ddx39 | 0.88 | 1.15E-59 | 2.45E-55 | 0.94 | 0.74 | 0.21 | 1.28 |
| 4 | Prr7 | 1.62 | 2.60E-59 | 5.56E-55 | 0.53 | 0.19 | 0.35 | 2.85 |
| 4 | Camk2n2 | 2.25 | 4.27E-59 | 9.13E-55 | 0.29 | 0.06 | 0.23 | 5 |
| 4 | Usp51 | 2.22 | 6.45E-59 | 1.38E-54 | 0.26 | 0.05 | 0.21 | 5.67 |
| 4 | Tuba1b | 0.58 | 2.83E-57 | 6.05E-53 | 1 | 0.97 | 0.03 | 1.03 |
| 4 | Prkg1 | 1.46 | 2.92E-57 | 6.23E-53 | 0.58 | 0.22 | 0.36 | 2.64 |
| 4 | 11-Sep | 1.05 | 3.63E-57 | 7.76E-53 | 0.91 | 0.71 | 0.2 | 1.28 |
| 4 | Sox2 | 0.76 | 7.28E-57 | 1.56E-52 | 0.97 | 0.67 | 0.31 | 1.46 |
| 4 | Frmd3 | 2.28 | 1.23E-56 | 2.63E-52 | 0.24 | 0.04 | 0.2 | 5.79 |
| 4 | Myc | 1.58 | 6.95E-55 | 1.49E-50 | 0.51 | 0.18 | 0.33 | 2.81 |
| 4 | Tyro3 | 1.62 | 2.78E-54 | 5.95E-50 | 0.51 | 0.19 | 0.33 | 2.74 |
| 4 | Mt2 | 1.36 | 3.76E-54 | 8.03E-50 | 0.59 | 0.24 | 0.36 | 2.49 |

**Supplementary Table 3. Marker genes for the original clusters of integrated clustering of E11.5 control and NotchGOF λ-junction ectoderm cells.**

| | | | | | | | | |
|---|---|---|---|---|---|---|---|---|
| 4 | 4930426D05Rik | 1.85 | 9.31E-54 | 1.99E-49 | 0.36 | 0.1 | 0.26 | 3.67 |
| 4 | Tspan12 | 1.19 | 1.15E-53 | 2.46E-49 | 0.76 | 0.42 | 0.34 | 1.82 |
| 4 | Igdcc3 | 1.18 | 1.37E-53 | 2.92E-49 | 0.69 | 0.3 | 0.39 | 2.28 |
| 4 | Fgf12 | 1.73 | 1.76E-53 | 3.76E-49 | 0.43 | 0.14 | 0.29 | 3.17 |
| 4 | Faah | 1.47 | 2.06E-53 | 4.40E-49 | 0.53 | 0.19 | 0.34 | 2.75 |
| 4 | Lsm5 | 0.63 | 6.47E-53 | 1.38E-48 | 0.98 | 0.89 | 0.08 | 1.09 |
| 4 | Cntfr | 1.57 | 8.67E-53 | 1.85E-48 | 0.4 | 0.12 | 0.28 | 3.38 |
| 4 | Rassf10 | 1.55 | 3.66E-52 | 7.81E-48 | 0.54 | 0.21 | 0.33 | 2.52 |
| 4 | Dlk1 | 0.27 | 2.04E-51 | 4.36E-47 | 0.96 | 0.6 | 0.36 | 1.59 |
| 4 | Ap3b2 | 1.97 | 2.16E-51 | 4.62E-47 | 0.26 | 0.06 | 0.21 | 4.71 |
| 4 | Idi1 | 1.11 | 3.41E-51 | 7.29E-47 | 0.87 | 0.56 | 0.31 | 1.55 |
| 4 | Celsr1 | 1.3 | 5.81E-51 | 1.24E-46 | 0.69 | 0.34 | 0.35 | 2.02 |
| 4 | Eef1d | 0.47 | 7.15E-51 | 1.53E-46 | 1 | 0.97 | 0.02 | 1.02 |
| 4 | Nme2 | 0.39 | 1.07E-50 | 2.28E-46 | 1 | 0.99 | 0.01 | 1.01 |
| 4 | Mapk11 | 1.76 | 1.78E-50 | 3.81E-46 | 0.35 | 0.1 | 0.26 | 3.64 |
| 4 | B230118H07Rik | 1.02 | 1.00E-49 | 2.14E-45 | 0.83 | 0.54 | 0.29 | 1.53 |
| 4 | Hells | 0.87 | 2.27E-49 | 4.85E-45 | 0.93 | 0.7 | 0.23 | 1.33 |
| 4 | Tmem238 | 1.16 | 2.79E-49 | 5.96E-45 | 0.72 | 0.37 | 0.35 | 1.93 |
| 4 | Txn1 | 0.44 | 4.23E-49 | 9.03E-45 | 1 | 0.98 | 0.02 | 1.02 |
| 4 | Snhg3 | 1.31 | 7.71E-49 | 1.65E-44 | 0.77 | 0.48 | 0.3 | 1.62 |
| 4 | Six1 | 0.73 | 1.35E-48 | 2.88E-44 | 0.98 | 0.65 | 0.32 | 1.49 |
| 4 | Hacd1 | 1.36 | 1.63E-48 | 3.49E-44 | 0.54 | 0.22 | 0.32 | 2.48 |
| 4 | Ptges3 | 0.52 | 1.64E-48 | 3.51E-44 | 1 | 0.95 | 0.05 | 1.05 |
| 4 | Manf | 0.7 | 2.66E-48 | 5.69E-44 | 0.97 | 0.88 | 0.1 | 1.11 |
| 4 | Klhl14 | 2.56 | 3.37E-48 | 7.20E-44 | 0.21 | 0.04 | 0.17 | 5.53 |
| 4 | Dmrt3 | 1.47 | 5.70E-48 | 1.22E-43 | 0.49 | 0.18 | 0.31 | 2.7 |
| 4 | Nhs | 1.13 | 7.63E-48 | 1.63E-43 | 0.74 | 0.4 | 0.33 | 1.82 |
| 4 | Cox5a | 0.42 | 1.44E-47 | 3.08E-43 | 1 | 0.99 | 0.02 | 1.02 |
| 4 | Nhp2 | 0.64 | 2.48E-47 | 5.31E-43 | 0.97 | 0.89 | 0.08 | 1.09 |
| 4 | Hnrnpdl | 0.58 | 4.14E-47 | 8.84E-43 | 0.99 | 0.95 | 0.04 | 1.05 |
| 4 | Higd1a | 0.71 | 4.41E-47 | 9.43E-43 | 0.96 | 0.84 | 0.12 | 1.14 |
| 4 | Rps27l | 0.42 | 7.93E-47 | 1.69E-42 | 1 | 0.99 | 0.01 | 1.01 |
| 4 | Usp1 | 0.75 | 8.04E-47 | 1.72E-42 | 0.95 | 0.73 | 0.22 | 1.3 |
| 4 | Cmtm7 | 1.13 | 1.05E-46 | 2.25E-42 | 0.68 | 0.33 | 0.35 | 2.05 |
| 4 | Frmd4a | 1.02 | 1.08E-46 | 2.32E-42 | 0.79 | 0.46 | 0.33 | 1.71 |
| 4 | Map7d2 | 1.44 | 1.64E-46 | 3.51E-42 | 0.49 | 0.18 | 0.3 | 2.65 |
| 4 | Tubb4b | 0.66 | 2.65E-46 | 5.67E-42 | 0.99 | 0.79 | 0.2 | 1.25 |
| 4 | Fam107a | 1.55 | 2.85E-46 | 6.09E-42 | 0.41 | 0.14 | 0.28 | 3.02 |
| 4 | Crabp2 | 0.62 | 5.15E-46 | 1.10E-41 | 0.99 | 0.83 | 0.16 | 1.19 |
| 4 | Pde1c | 1.84 | 1.39E-45 | 2.97E-41 | 0.31 | 0.08 | 0.23 | 3.76 |
| 4 | Eif1ax | 0.53 | 1.41E-45 | 3.00E-41 | 0.98 | 0.93 | 0.04 | 1.05 |
| 4 | Mab21l1 | 2.49 | 2.29E-45 | 4.90E-41 | 0.16 | 0.02 | 0.14 | 6.83 |
| 4 | Tmem97 | 1.17 | 3.03E-45 | 6.48E-41 | 0.71 | 0.37 | 0.34 | 1.92 |
| 4 | Plekha5 | 1.14 | 3.25E-45 | 6.94E-41 | 0.71 | 0.4 | 0.31 | 1.79 |
| 4 | Fam181b | 1.68 | 6.49E-45 | 1.39E-40 | 0.34 | 0.1 | 0.24 | 3.49 |
| 4 | Ly75 | 1.39 | 1.79E-44 | 3.82E-40 | 0.48 | 0.18 | 0.3 | 2.66 |
| 4 | Slc16a1 | 1.1 | 1.82E-44 | 3.89E-40 | 0.69 | 0.35 | 0.34 | 1.98 |
| 4 | Iqgap2 | 1.12 | 1.94E-44 | 4.14E-40 | 0.63 | 0.3 | 0.34 | 2.14 |
| 4 | Nes | 1.05 | 3.27E-44 | 6.99E-40 | 0.77 | 0.47 | 0.3 | 1.64 |
| 4 | Dmrta2os | 2.04 | 5.10E-44 | 1.09E-39 | 0.27 | 0.07 | 0.2 | 4.01 |
| 4 | Gm17167 | 2.87 | 7.95E-44 | 1.70E-39 | 0.14 | 0.02 | 0.12 | 8.5 |
| 4 | Hmgn2 | 0.44 | 9.44E-44 | 2.02E-39 | 0.99 | 0.98 | 0.02 | 1.02 |
| 4 | Exoc3l4 | 1.59 | 1.18E-43 | 2.51E-39 | 0.43 | 0.16 | 0.28 | 2.77 |
| 4 | Synm | 2.25 | 1.25E-43 | 2.68E-39 | 0.22 | 0.05 | 0.18 | 4.77 |
| 4 | Fkbp3 | 0.41 | 1.28E-43 | 2.73E-39 | 1 | 0.98 | 0.02 | 1.02 |
| 4 | Eva1a | 1.24 | 1.41E-43 | 3.02E-39 | 0.45 | 0.16 | 0.29 | 2.78 |
| 4 | Slc25a5 | 0.59 | 1.48E-43 | 3.17E-39 | 0.98 | 0.9 | 0.08 | 1.09 |

**Supplementary Table 3. Marker genes for the original clusters of integrated clustering of E11.5 control and NotchGOF λ-junction ectoderm cells.**

| | | | | | | | | |
|---|---|---|---|---|---|---|---|---|
| 4 | Pgp | 1.07 | 2.80E-43 | 5.97E-39 | 0.79 | 0.49 | 0.3 | 1.61 |
| 4 | Grtp1 | 1.43 | 3.36E-43 | 7.17E-39 | 0.5 | 0.2 | 0.3 | 2.48 |
| 4 | Psat1 | 1.19 | 1.12E-42 | 2.39E-38 | 0.61 | 0.29 | 0.32 | 2.11 |
| 4 | Selenoh | 0.47 | 1.25E-42 | 2.66E-38 | 1 | 0.95 | 0.05 | 1.05 |
| 4 | Lhfpl4 | 2.14 | 1.26E-42 | 2.69E-38 | 0.2 | 0.04 | 0.16 | 5.3 |
| 4 | Pclaf | 0.64 | 2.86E-42 | 6.12E-38 | 0.99 | 0.77 | 0.22 | 1.28 |
| 4 | Msmo1 | 1.16 | 3.58E-42 | 7.65E-38 | 0.77 | 0.47 | 0.3 | 1.64 |
| 4 | Cct3 | 0.44 | 3.76E-42 | 8.03E-38 | 0.99 | 0.97 | 0.02 | 1.02 |
| 4 | Kpnb1 | 0.68 | 4.08E-42 | 8.72E-38 | 0.94 | 0.8 | 0.15 | 1.18 |
| 4 | Ddx21 | 0.84 | 4.83E-42 | 1.03E-37 | 0.89 | 0.63 | 0.26 | 1.4 |
| 4 | Ccna2 | 0.89 | 5.37E-42 | 1.15E-37 | 0.84 | 0.53 | 0.32 | 1.6 |
| 4 | Npas3 | 1.9 | 5.70E-42 | 1.22E-37 | 0.31 | 0.09 | 0.22 | 3.46 |
| 4 | Rhoc | 0.81 | 1.00E-41 | 2.14E-37 | 0.9 | 0.62 | 0.28 | 1.44 |
| 4 | Hsd17b11 | 1.85 | 1.61E-41 | 3.43E-37 | 0.32 | 0.09 | 0.22 | 3.39 |
| 4 | Nudc | 0.47 | 1.61E-41 | 3.44E-37 | 0.99 | 0.95 | 0.05 | 1.05 |
| 4 | Pa2g4 | 0.56 | 1.64E-41 | 3.50E-37 | 0.98 | 0.88 | 0.1 | 1.12 |
| 4 | Hist1h4i | 1.07 | 2.10E-41 | 4.48E-37 | 0.72 | 0.4 | 0.31 | 1.78 |
| 4 | Bmpr1b | 2.19 | 2.76E-41 | 5.90E-37 | 0.25 | 0.06 | 0.19 | 4.13 |
| 4 | Rrp12 | 1.31 | 3.74E-41 | 7.99E-37 | 0.51 | 0.22 | 0.29 | 2.33 |
| 4 | Hmcn1 | 1.46 | 5.39E-41 | 1.15E-36 | 0.46 | 0.18 | 0.28 | 2.56 |
| 4 | Vdac3 | 0.55 | 2.76E-40 | 5.89E-36 | 0.98 | 0.93 | 0.06 | 1.06 |
| 4 | Gm4675 | 2.3 | 3.07E-40 | 6.57E-36 | 0.16 | 0.03 | 0.14 | 6.19 |
| 4 | Cyp26a1 | 1.88 | 3.46E-40 | 7.39E-36 | 0.23 | 0.05 | 0.18 | 4.36 |
| 4 | Shisa2 | 1.54 | 4.72E-40 | 1.01E-35 | 0.31 | 0.09 | 0.22 | 3.48 |
| 4 | Cldn9 | 1.06 | 5.49E-40 | 1.17E-35 | 0.65 | 0.32 | 0.33 | 2.04 |
| 4 | Gsta4 | 0.63 | 6.01E-40 | 1.28E-35 | 0.98 | 0.79 | 0.19 | 1.24 |
| 4 | Srm | 0.91 | 6.69E-40 | 1.43E-35 | 0.83 | 0.59 | 0.24 | 1.41 |
| 4 | Fgfr1 | 1.09 | 6.71E-40 | 1.43E-35 | 0.68 | 0.38 | 0.3 | 1.8 |
| 4 | Actn1 | 0.89 | 6.85E-40 | 1.46E-35 | 0.82 | 0.55 | 0.28 | 1.5 |
| 4 | Tmem37 | 2.31 | 9.63E-40 | 2.06E-35 | 0.2 | 0.04 | 0.16 | 4.79 |
| 4 | Timm13 | 0.47 | 1.82E-39 | 3.89E-35 | 0.99 | 0.96 | 0.03 | 1.03 |
| 4 | Svopl | 2.55 | 3.43E-39 | 7.33E-35 | 0.13 | 0.02 | 0.12 | 7.39 |
| 4 | Gnao1 | 1.19 | 4.06E-39 | 8.68E-35 | 0.42 | 0.16 | 0.27 | 2.73 |
| 4 | Elavl2 | 0.78 | 5.32E-39 | 1.14E-34 | 0.72 | 0.38 | 0.34 | 1.89 |
| 4 | Hspa5 | 0.51 | 5.70E-39 | 1.22E-34 | 1 | 0.97 | 0.03 | 1.03 |
| 4 | Cycs | 0.51 | 7.23E-39 | 1.55E-34 | 0.98 | 0.92 | 0.06 | 1.07 |
| 4 | Eya1 | 0.8 | 9.77E-39 | 2.09E-34 | 0.86 | 0.56 | 0.3 | 1.54 |
| 4 | Gm19938 | 1.56 | 2.31E-38 | 4.94E-34 | 0.28 | 0.08 | 0.2 | 3.59 |
| 4 | Pappa | 2.19 | 4.03E-38 | 8.60E-34 | 0.18 | 0.03 | 0.14 | 5.3 |
| 4 | Tipin | 0.72 | 4.27E-38 | 9.13E-34 | 0.93 | 0.66 | 0.27 | 1.4 |
| 4 | Stc2 | 2.35 | 5.40E-38 | 1.15E-33 | 0.18 | 0.03 | 0.14 | 5.24 |
| 4 | Fkbp4 | 0.41 | 1.01E-37 | 2.17E-33 | 1 | 0.97 | 0.02 | 1.02 |
| 4 | Gipc2 | 1.4 | 1.07E-37 | 2.29E-33 | 0.4 | 0.14 | 0.25 | 2.74 |
| 4 | Cdk2ap1 | 0.47 | 1.31E-37 | 2.79E-33 | 0.99 | 0.92 | 0.06 | 1.07 |
| 4 | Banf1 | 0.49 | 2.90E-37 | 6.20E-33 | 0.98 | 0.92 | 0.06 | 1.07 |
| 4 | Uqcrq | 0.44 | 3.46E-37 | 7.39E-33 | 0.99 | 0.96 | 0.04 | 1.04 |
| 4 | Tcf4 | 0.58 | 5.76E-37 | 1.23E-32 | 0.97 | 0.88 | 0.09 | 1.1 |
| 4 | Rpl31 | 0.27 | 6.04E-37 | 1.29E-32 | 1 | 1 | 0 | 1 |
| 4 | Impdh2 | 0.58 | 7.08E-37 | 1.51E-32 | 0.98 | 0.84 | 0.14 | 1.16 |
| 4 | Rsl1d1 | 0.5 | 7.44E-37 | 1.59E-32 | 0.99 | 0.92 | 0.06 | 1.07 |
| 4 | Rasef | 1.34 | 7.53E-37 | 1.61E-32 | 0.38 | 0.13 | 0.24 | 2.85 |
| 4 | Shc3 | 2.15 | 7.56E-37 | 1.62E-32 | 0.19 | 0.04 | 0.15 | 4.92 |
| 4 | Plpp3 | 1 | 1.13E-36 | 2.42E-32 | 0.71 | 0.41 | 0.3 | 1.75 |
| 4 | Sobp | 1.18 | 1.33E-36 | 2.85E-32 | 0.5 | 0.22 | 0.29 | 2.32 |
| 4 | Mrpl12 | 0.55 | 1.41E-36 | 3.01E-32 | 0.96 | 0.88 | 0.09 | 1.1 |
| 4 | Tenm2 | 1.27 | 1.46E-36 | 3.11E-32 | 0.48 | 0.2 | 0.28 | 2.36 |
| 4 | Pbdc1 | 0.63 | 1.56E-36 | 3.34E-32 | 0.94 | 0.8 | 0.14 | 1.18 |

**Supplementary Table 3. Marker genes for the original clusters of integrated clustering of E11.5 control and NotchGOF λ-junction ectoderm cells.**

| | | | | | | | | |
|---|---|---|---|---|---|---|---|---|
| 4 | Ldha | 0.4 | 2.28E-36 | 4.88E-32 | 1 | 0.99 | 0.02 | 1.02 |
| 4 | Fam13c | 1.93 | 2.39E-36 | 5.10E-32 | 0.23 | 0.06 | 0.17 | 4.02 |
| 4 | Gcnt2 | 1.92 | 3.13E-36 | 6.68E-32 | 0.24 | 0.06 | 0.18 | 3.87 |
| 4 | Snrpf | 0.31 | 3.37E-36 | 7.20E-32 | 1 | 0.99 | 0.01 | 1.01 |
| 4 | Heyl | 1.1 | 3.49E-36 | 7.46E-32 | 0.47 | 0.19 | 0.28 | 2.5 |
| 4 | Bend4 | 1.56 | 3.77E-36 | 8.06E-32 | 0.34 | 0.11 | 0.22 | 2.95 |
| 4 | Ank | 1.77 | 3.95E-36 | 8.44E-32 | 0.25 | 0.07 | 0.18 | 3.77 |
| 4 | Oxtr | 0.85 | 4.30E-36 | 9.19E-32 | 0.55 | 0.24 | 0.32 | 2.35 |
| 4 | Tspan13 | 0.7 | 4.71E-36 | 1.01E-31 | 0.93 | 0.75 | 0.18 | 1.24 |
| 4 | C130026L21Rik | 2.12 | 5.38E-36 | 1.15E-31 | 0.17 | 0.03 | 0.14 | 5.18 |
| 4 | Filip1 | 1.85 | 5.39E-36 | 1.15E-31 | 0.23 | 0.06 | 0.17 | 3.9 |
| 4 | Timm8a1 | 0.81 | 6.53E-36 | 1.40E-31 | 0.85 | 0.63 | 0.22 | 1.35 |
| 4 | Rassf9 | 1.07 | 7.15E-36 | 1.53E-31 | 0.5 | 0.21 | 0.29 | 2.35 |
| 4 | Cdca8 | 0.69 | 7.57E-36 | 1.62E-31 | 0.94 | 0.67 | 0.27 | 1.4 |
| 4 | Cct6a | 0.38 | 8.87E-36 | 1.89E-31 | 1 | 0.98 | 0.01 | 1.01 |
| 4 | Ubash3b | 1.19 | 1.01E-35 | 2.17E-31 | 0.5 | 0.22 | 0.28 | 2.28 |
| 4 | Thbs3 | 1.46 | 1.13E-35 | 2.41E-31 | 0.34 | 0.12 | 0.22 | 2.95 |
| 4 | Slc35d1 | 1.22 | 1.56E-35 | 3.34E-31 | 0.47 | 0.2 | 0.27 | 2.37 |
| 4 | Slc1a4 | 1.73 | 1.93E-35 | 4.12E-31 | 0.27 | 0.08 | 0.19 | 3.41 |
| 4 | Anp32b | 0.36 | 3.12E-35 | 6.67E-31 | 1 | 0.99 | 0.01 | 1.01 |
| 4 | Rangap1 | 0.7 | 3.18E-35 | 6.80E-31 | 0.91 | 0.68 | 0.23 | 1.34 |
| 4 | Flrt3 | 0.86 | 4.61E-35 | 9.85E-31 | 0.69 | 0.37 | 0.32 | 1.86 |
| 4 | Hif1a | 0.67 | 6.04E-35 | 1.29E-30 | 0.9 | 0.77 | 0.13 | 1.17 |
| 4 | Srsf2 | 0.37 | 6.49E-35 | 1.39E-30 | 1 | 0.98 | 0.02 | 1.02 |
| 4 | Ipo5 | 0.74 | 9.37E-35 | 2.00E-30 | 0.89 | 0.66 | 0.22 | 1.34 |
| 4 | Npm1 | 0.26 | 1.21E-34 | 2.57E-30 | 1 | 1 | 0 | 1 |
| 4 | Sypl2 | 2.28 | 1.84E-34 | 3.93E-30 | 0.15 | 0.03 | 0.13 | 5.63 |
| 4 | Gar1 | 0.66 | 2.88E-34 | 6.14E-30 | 0.93 | 0.73 | 0.2 | 1.27 |
| 4 | Emid1 | 1.43 | 4.43E-34 | 9.46E-30 | 0.39 | 0.15 | 0.24 | 2.57 |
| 4 | Fbl | 0.5 | 5.55E-34 | 1.19E-29 | 0.97 | 0.9 | 0.07 | 1.08 |
| 4 | Plch1 | 1.03 | 7.96E-34 | 1.70E-29 | 0.61 | 0.31 | 0.3 | 1.96 |
| 4 | Gapdh | 0.28 | 1.05E-33 | 2.25E-29 | 1 | 1 | 0 | 1 |
| 4 | Dclk1 | 1.1 | 1.08E-33 | 2.31E-29 | 0.42 | 0.17 | 0.25 | 2.52 |
| 4 | Gmnn | 0.63 | 1.27E-33 | 2.72E-29 | 0.96 | 0.7 | 0.26 | 1.37 |
| 4 | Atp7b | 1.96 | 1.93E-33 | 4.11E-29 | 0.23 | 0.06 | 0.17 | 3.72 |
| 4 | Ggt7 | 2.05 | 2.43E-33 | 5.20E-29 | 0.2 | 0.05 | 0.15 | 4.23 |
| 4 | Mecom | 0.58 | 2.49E-33 | 5.33E-29 | 0.74 | 0.42 | 0.32 | 1.76 |
| 4 | Pdia6 | 0.48 | 2.67E-33 | 5.70E-29 | 1 | 0.95 | 0.05 | 1.05 |
| 4 | Shisa4 | 1.09 | 2.95E-33 | 6.29E-29 | 0.49 | 0.21 | 0.28 | 2.31 |
| 4 | Ttyh1 | 1.83 | 2.95E-33 | 6.30E-29 | 0.25 | 0.07 | 0.18 | 3.47 |
| 4 | Rpia | 1.02 | 2.99E-33 | 6.38E-29 | 0.62 | 0.33 | 0.29 | 1.9 |
| 4 | Por | 1.08 | 3.70E-33 | 7.91E-29 | 0.67 | 0.4 | 0.27 | 1.69 |
| 4 | Rpl22l1 | 0.28 | 3.75E-33 | 8.01E-29 | 1 | 0.99 | 0.01 | 1.01 |
| 4 | Snrpd2 | 0.31 | 3.98E-33 | 8.50E-29 | 1 | 0.99 | 0.01 | 1.01 |
| 4 | C1qbp | 0.5 | 4.39E-33 | 9.37E-29 | 0.97 | 0.91 | 0.06 | 1.07 |
| 4 | Nuak2 | 1.41 | 4.77E-33 | 1.02E-28 | 0.34 | 0.12 | 0.22 | 2.8 |
| 4 | Snx3 | 0.55 | 5.75E-33 | 1.23E-28 | 0.96 | 0.86 | 0.1 | 1.12 |
| 4 | Wwtr1 | 1.05 | 1.34E-32 | 2.86E-28 | 0.55 | 0.27 | 0.28 | 2.05 |
| 4 | Rtn2 | 1.22 | 1.52E-32 | 3.26E-28 | 0.4 | 0.16 | 0.24 | 2.51 |
| 4 | Psmg4 | 0.66 | 1.64E-32 | 3.50E-28 | 0.91 | 0.73 | 0.18 | 1.24 |
| 4 | Bag2 | 1.06 | 1.81E-32 | 3.87E-28 | 0.6 | 0.32 | 0.28 | 1.88 |
| 4 | Phf21b | 1.82 | 2.12E-32 | 4.52E-28 | 0.22 | 0.06 | 0.16 | 3.84 |
| 4 | Rrm2 | 0.63 | 2.69E-32 | 5.76E-28 | 0.9 | 0.65 | 0.25 | 1.39 |
| 4 | Cyp51 | 0.86 | 5.24E-32 | 1.12E-27 | 0.83 | 0.59 | 0.25 | 1.42 |
| 4 | Mapk8ip2 | 1.31 | 5.57E-32 | 1.19E-27 | 0.28 | 0.09 | 0.19 | 3.16 |
| 4 | B3gnt7 | 1.47 | 6.30E-32 | 1.35E-27 | 0.29 | 0.09 | 0.2 | 3.09 |
| 4 | Frat2 | 0.97 | 8.36E-32 | 1.79E-27 | 0.64 | 0.35 | 0.28 | 1.81 |

**Supplementary Table 3. Marker genes for the original clusters of integrated clustering of E11.5 control and NotchGOF λ-junction ectoderm cells.**

| | | | | | | | | |
|---|---|---|---|---|---|---|---|---|
| 4 | Slc7a5 | 1.72 | 8.73E-32 | 1.86E-27 | 0.23 | 0.07 | 0.17 | 3.6 |
| 4 | Ostc | 0.47 | 9.51E-32 | 2.03E-27 | 0.97 | 0.92 | 0.05 | 1.05 |
| 4 | Zmat4 | 1.29 | 9.69E-32 | 2.07E-27 | 0.35 | 0.13 | 0.22 | 2.7 |
| 4 | Hnrnpu | 0.33 | 1.23E-31 | 2.63E-27 | 1 | 0.99 | 0.01 | 1.01 |
| 4 | Fgf15 | 1.63 | 1.57E-31 | 3.35E-27 | 0.28 | 0.09 | 0.19 | 3.17 |
| 4 | Ccdc34 | 0.49 | 2.07E-31 | 4.41E-27 | 0.97 | 0.85 | 0.12 | 1.14 |
| 4 | Dkc1 | 0.66 | 2.15E-31 | 4.60E-27 | 0.92 | 0.75 | 0.17 | 1.22 |
| 4 | Col9a2 | 0.77 | 3.04E-31 | 6.48E-27 | 0.32 | 0.11 | 0.21 | 2.85 |
| 4 | Rragb | 1.7 | 4.53E-31 | 9.67E-27 | 0.29 | 0.1 | 0.19 | 2.91 |
| 4 | Eed | 0.81 | 4.59E-31 | 9.80E-27 | 0.76 | 0.5 | 0.26 | 1.52 |
| 4 | Hnrnpd | 0.35 | 5.05E-31 | 1.08E-26 | 1 | 0.99 | 0.01 | 1.01 |
| 4 | Scg5 | 0.93 | 5.74E-31 | 1.23E-26 | 0.44 | 0.18 | 0.26 | 2.4 |
| 4 | Lsm3 | 0.46 | 6.20E-31 | 1.32E-26 | 0.97 | 0.88 | 0.1 | 1.11 |
| 4 | Acsl6 | 1.89 | 6.58E-31 | 1.41E-26 | 0.17 | 0.04 | 0.13 | 4.44 |
| 4 | Mns1 | 0.91 | 8.58E-31 | 1.83E-26 | 0.67 | 0.37 | 0.29 | 1.79 |
| 4 | Trpm3 | 1.33 | 9.29E-31 | 1.98E-26 | 0.37 | 0.15 | 0.23 | 2.56 |
| 4 | Tox2 | 1.5 | 9.93E-31 | 2.12E-26 | 0.29 | 0.1 | 0.2 | 2.97 |
| 4 | Tst | 1.22 | 1.32E-30 | 2.81E-26 | 0.49 | 0.23 | 0.26 | 2.09 |
| 4 | Gnl3 | 0.66 | 1.53E-30 | 3.27E-26 | 0.89 | 0.72 | 0.18 | 1.25 |
| 4 | Eif2s2 | 0.35 | 2.83E-30 | 6.04E-26 | 1 | 0.99 | 0.01 | 1.01 |
| 4 | Cad | 0.97 | 2.98E-30 | 6.36E-26 | 0.59 | 0.32 | 0.28 | 1.87 |
| 4 | Rnf152 | 1.97 | 2.98E-30 | 6.36E-26 | 0.15 | 0.03 | 0.12 | 4.9 |
| 4 | Ermn | 1.13 | 3.93E-30 | 8.39E-26 | 0.38 | 0.15 | 0.23 | 2.56 |
| 4 | Prps2 | 0.95 | 4.16E-30 | 8.89E-26 | 0.58 | 0.3 | 0.28 | 1.96 |
| 4 | Fgf13 | 1.24 | 4.40E-30 | 9.39E-26 | 0.38 | 0.15 | 0.23 | 2.48 |
| 4 | Ccdc177 | 1.6 | 4.46E-30 | 9.52E-26 | 0.19 | 0.05 | 0.14 | 4.11 |
| 4 | Abcc4 | 2.09 | 4.75E-30 | 1.02E-25 | 0.15 | 0.03 | 0.12 | 4.81 |
| 4 | Dhx32 | 0.88 | 5.40E-30 | 1.15E-25 | 0.58 | 0.29 | 0.29 | 2.02 |
| 4 | Tmem229b | 2.06 | 8.85E-30 | 1.89E-25 | 0.19 | 0.05 | 0.14 | 3.92 |
| 4 | Fam149a | 1.72 | 1.03E-29 | 2.19E-25 | 0.23 | 0.07 | 0.17 | 3.39 |
| 4 | Bok | 0.99 | 1.14E-29 | 2.45E-25 | 0.54 | 0.26 | 0.28 | 2.06 |
| 4 | Tcim | 0.99 | 1.52E-29 | 3.25E-25 | 0.51 | 0.24 | 0.26 | 2.09 |
| 4 | Ano1 | 0.73 | 2.62E-29 | 5.60E-25 | 0.75 | 0.43 | 0.32 | 1.73 |
| 4 | A830082K12Rik | 2.04 | 2.76E-29 | 5.89E-25 | 0.19 | 0.05 | 0.14 | 4.02 |
| 4 | Tsen15 | 0.77 | 3.90E-29 | 8.33E-25 | 0.81 | 0.56 | 0.25 | 1.44 |
| 4 | Mdk | 0.27 | 4.16E-29 | 8.89E-25 | 1 | 0.99 | 0.01 | 1.01 |
| 4 | Tmem216 | 0.89 | 5.83E-29 | 1.25E-24 | 0.62 | 0.33 | 0.29 | 1.86 |
| 4 | Snrpa1 | 0.48 | 6.07E-29 | 1.30E-24 | 0.97 | 0.9 | 0.08 | 1.08 |
| 4 | Mn1 | 1.25 | 6.45E-29 | 1.38E-24 | 0.36 | 0.14 | 0.22 | 2.57 |
| 4 | Ttc39b | 1.61 | 6.64E-29 | 1.42E-24 | 0.19 | 0.05 | 0.14 | 3.92 |
| 4 | Gm11627 | 1.97 | 6.93E-29 | 1.48E-24 | 0.16 | 0.04 | 0.13 | 4.47 |
| 4 | Hmgb2 | 0.47 | 7.87E-29 | 1.68E-24 | 1 | 0.97 | 0.03 | 1.03 |
| 4 | Nectin3 | 0.62 | 8.39E-29 | 1.79E-24 | 0.9 | 0.72 | 0.18 | 1.25 |
| 4 | Srgap3 | 1.16 | 1.02E-28 | 2.17E-24 | 0.44 | 0.2 | 0.24 | 2.19 |
| 4 | Prdx1 | 0.31 | 1.03E-28 | 2.19E-24 | 1 | 0.99 | 0.01 | 1.01 |
| 4 | Rcc1 | 0.84 | 1.16E-28 | 2.48E-24 | 0.71 | 0.44 | 0.27 | 1.61 |
| 4 | Calr | 0.42 | 1.32E-28 | 2.81E-24 | 1 | 0.97 | 0.03 | 1.03 |
| 4 | Psmb3 | 0.36 | 1.43E-28 | 3.05E-24 | 0.99 | 0.97 | 0.02 | 1.02 |
| 4 | Nop58 | 0.46 | 1.49E-28 | 3.19E-24 | 0.97 | 0.89 | 0.09 | 1.1 |
| 4 | Eef1g | 0.25 | 2.23E-28 | 4.77E-24 | 1 | 1 | 0.01 | 1.01 |
| 4 | Dhcr7 | 1.11 | 2.36E-28 | 5.05E-24 | 0.57 | 0.31 | 0.26 | 1.85 |
| 4 | Mcm7 | 0.56 | 2.48E-28 | 5.29E-24 | 0.96 | 0.8 | 0.15 | 1.19 |
| 4 | Pop5 | 0.72 | 2.70E-28 | 5.78E-24 | 0.83 | 0.58 | 0.25 | 1.42 |
| 4 | Ptgis | 1.66 | 2.93E-28 | 6.26E-24 | 0.22 | 0.06 | 0.16 | 3.44 |
| 4 | Gpm6b | 1.23 | 3.79E-28 | 8.10E-24 | 0.26 | 0.08 | 0.17 | 3.15 |
| 4 | D030056L22Rik | 0.77 | 3.97E-28 | 8.48E-24 | 0.78 | 0.52 | 0.26 | 1.49 |
| 4 | Pwwp3b | 1.04 | 4.87E-28 | 1.04E-23 | 0.53 | 0.27 | 0.26 | 1.98 |

**Supplementary Table 3. Marker genes for the original clusters of integrated clustering of E11.5 control and NotchGOF λ-junction ectoderm cells.**

| | | | | | | | | |
|---|---|---|---|---|---|---|---|---|
| 4 | Dnajc9 | 0.51 | 5.14E-28 | 1.10E-23 | 0.97 | 0.8 | 0.17 | 1.22 |
| 4 | Mcm3 | 0.69 | 5.52E-28 | 1.18E-23 | 0.89 | 0.67 | 0.22 | 1.33 |
| 4 | Pcna | 0.59 | 5.60E-28 | 1.20E-23 | 0.95 | 0.75 | 0.2 | 1.26 |
| 4 | Cdk6 | 0.8 | 7.12E-28 | 1.52E-23 | 0.76 | 0.5 | 0.27 | 1.54 |
| 4 | Polr2f | 0.39 | 8.52E-28 | 1.82E-23 | 0.99 | 0.95 | 0.04 | 1.04 |
| 4 | Ntn1 | 1.59 | 8.65E-28 | 1.85E-23 | 0.24 | 0.07 | 0.16 | 3.19 |
| 4 | Psen2 | 1.41 | 1.06E-27 | 2.25E-23 | 0.3 | 0.11 | 0.19 | 2.67 |
| 4 | Bccip | 0.63 | 1.18E-27 | 2.53E-23 | 0.91 | 0.73 | 0.18 | 1.24 |
| 4 | Pradc1 | 1 | 1.18E-27 | 2.53E-23 | 0.54 | 0.27 | 0.27 | 1.98 |
| 4 | Iqcg | 1.66 | 1.80E-27 | 3.84E-23 | 0.18 | 0.05 | 0.13 | 3.96 |
| 4 | Mapk13 | 0.64 | 1.96E-27 | 4.19E-23 | 0.83 | 0.59 | 0.24 | 1.42 |
| 4 | Mcm5 | 0.65 | 1.97E-27 | 4.20E-23 | 0.87 | 0.61 | 0.26 | 1.42 |
| 4 | Hsp90b1 | 0.34 | 3.32E-27 | 7.10E-23 | 1 | 0.98 | 0.02 | 1.02 |
| 4 | Sod1 | 0.61 | 3.63E-27 | 7.75E-23 | 0.92 | 0.76 | 0.16 | 1.21 |
| 4 | Pawr | 0.86 | 3.89E-27 | 8.30E-23 | 0.62 | 0.35 | 0.27 | 1.79 |
| 4 | Ephb1 | 0.77 | 4.93E-27 | 1.05E-22 | 0.66 | 0.35 | 0.3 | 1.86 |
| 4 | H2-K1 | 1.15 | 5.48E-27 | 1.17E-22 | 0.47 | 0.23 | 0.25 | 2.08 |
| 4 | Mcrip2 | 1.47 | 5.56E-27 | 1.19E-22 | 0.26 | 0.09 | 0.17 | 2.9 |
| 4 | Soga3 | 0.54 | 6.18E-27 | 1.32E-22 | 0.37 | 0.15 | 0.22 | 2.49 |
| 4 | Pmm1 | 0.98 | 7.25E-27 | 1.55E-22 | 0.58 | 0.32 | 0.26 | 1.83 |
| 4 | Cdc42ep4 | 0.9 | 7.27E-27 | 1.55E-22 | 0.58 | 0.31 | 0.27 | 1.85 |
| 4 | Bola3 | 0.65 | 7.31E-27 | 1.56E-22 | 0.84 | 0.62 | 0.22 | 1.36 |
| 4 | Mapk8ip1 | 0.7 | 8.00E-27 | 1.71E-22 | 0.62 | 0.34 | 0.29 | 1.86 |
| 4 | Pkm | 0.27 | 9.84E-27 | 2.10E-22 | 1 | 1 | 0 | 1 |
| 4 | Cul1 | 0.5 | 1.26E-26 | 2.70E-22 | 0.94 | 0.85 | 0.1 | 1.12 |
| 4 | Birc5 | 0.56 | 1.74E-26 | 3.71E-22 | 0.93 | 0.69 | 0.24 | 1.35 |
| 4 | Tacc2 | 0.84 | 2.05E-26 | 4.39E-22 | 0.74 | 0.51 | 0.23 | 1.44 |
| 4 | Maf | 0.76 | 2.58E-26 | 5.51E-22 | 0.32 | 0.12 | 0.2 | 2.66 |
| 4 | Fscn1 | 0.47 | 3.34E-26 | 7.13E-22 | 0.94 | 0.85 | 0.09 | 1.1 |
| 4 | Rad51 | 0.71 | 4.29E-26 | 9.17E-22 | 0.77 | 0.48 | 0.29 | 1.61 |
| 4 | Ppp1r14b | 0.25 | 4.94E-26 | 1.06E-21 | 1 | 0.99 | 0.01 | 1.01 |
| 4 | Cmtm8 | 0.7 | 6.16E-26 | 1.31E-21 | 0.79 | 0.5 | 0.29 | 1.57 |
| 4 | Nasp | 0.36 | 6.71E-26 | 1.43E-21 | 1 | 0.96 | 0.04 | 1.04 |
| 4 | Emg1 | 0.42 | 7.72E-26 | 1.65E-21 | 0.98 | 0.9 | 0.08 | 1.09 |
| 4 | Dipk1c | 2.48 | 8.78E-26 | 1.88E-21 | 0.12 | 0.02 | 0.1 | 5.04 |
| 4 | Cdt1 | 0.89 | 8.81E-26 | 1.88E-21 | 0.63 | 0.37 | 0.26 | 1.69 |
| 4 | Ccdc171 | 1.06 | 1.05E-25 | 2.24E-21 | 0.43 | 0.2 | 0.23 | 2.18 |
| 4 | B4galt5 | 1.22 | 1.15E-25 | 2.46E-21 | 0.29 | 0.1 | 0.18 | 2.74 |
| 4 | Adgrg2 | 2.12 | 1.20E-25 | 2.56E-21 | 0.11 | 0.02 | 0.09 | 5.83 |
| 4 | Ldlr | 1.1 | 1.35E-25 | 2.89E-21 | 0.43 | 0.2 | 0.23 | 2.12 |
| 4 | Dtl | 0.73 | 2.66E-25 | 5.68E-21 | 0.76 | 0.51 | 0.25 | 1.48 |
| 4 | Alyref | 0.4 | 2.89E-25 | 6.17E-21 | 0.98 | 0.91 | 0.07 | 1.08 |
| 4 | Kif15 | 0.6 | 3.02E-25 | 6.45E-21 | 0.84 | 0.56 | 0.28 | 1.5 |
| 4 | Dhfr | 0.69 | 3.55E-25 | 7.58E-21 | 0.83 | 0.56 | 0.27 | 1.48 |
| 4 | Lsm6 | 0.41 | 4.38E-25 | 9.36E-21 | 0.97 | 0.88 | 0.09 | 1.11 |
| 4 | Dctpp1 | 0.41 | 4.76E-25 | 1.02E-20 | 0.98 | 0.9 | 0.09 | 1.09 |
| 4 | Fzd2 | 0.53 | 5.35E-25 | 1.14E-20 | 0.93 | 0.81 | 0.13 | 1.16 |
| 4 | Mocs2 | 0.75 | 5.40E-25 | 1.15E-20 | 0.76 | 0.5 | 0.26 | 1.51 |
| 4 | Sfrp2 | 0.67 | 6.63E-25 | 1.42E-20 | 0.74 | 0.49 | 0.25 | 1.52 |
| 4 | Mfsd2a | 1.74 | 7.03E-25 | 1.50E-20 | 0.19 | 0.05 | 0.13 | 3.53 |
| 4 | Galnt16 | 1.58 | 7.15E-25 | 1.53E-20 | 0.18 | 0.05 | 0.13 | 3.67 |
| 4 | Chst2 | 1.32 | 7.23E-25 | 1.54E-20 | 0.33 | 0.13 | 0.19 | 2.46 |
| 4 | 2810408I11Rik | 1.07 | 7.78E-25 | 1.66E-20 | 0.4 | 0.18 | 0.22 | 2.19 |
| 4 | Olfm1 | 0.71 | 8.13E-25 | 1.74E-20 | 0.57 | 0.31 | 0.27 | 1.88 |
| 4 | Ide | 0.78 | 8.16E-25 | 1.74E-20 | 0.69 | 0.43 | 0.26 | 1.59 |
| 4 | Hspa8 | 0.34 | 1.22E-24 | 2.60E-20 | 1 | 1 | 0 | 1 |
| 4 | Hey2 | 1.25 | 1.29E-24 | 2.75E-20 | 0.23 | 0.08 | 0.16 | 3.08 |

**Supplementary Table 3. Marker genes for the original clusters of integrated clustering of E11.5 control and NotchGOF λ-junction ectoderm cells.**

| | | | | | | | | |
|---|---|---|---|---|---|---|---|---|
| 4 | Pantr1 | 2.14 | 1.35E-24 | 2.89E-20 | 0.1 | 0.02 | 0.09 | 5.72 |
| 4 | Fads1 | 0.82 | 1.77E-24 | 3.78E-20 | 0.66 | 0.41 | 0.25 | 1.62 |
| 4 | Rnaseh2c | 0.45 | 2.04E-24 | 4.36E-20 | 0.97 | 0.84 | 0.12 | 1.15 |
| 4 | Tspan4 | 1.46 | 2.13E-24 | 4.56E-20 | 0.29 | 0.11 | 0.18 | 2.59 |
| 4 | Prc1 | 0.76 | 2.28E-24 | 4.88E-20 | 0.76 | 0.49 | 0.27 | 1.54 |
| 4 | Acss2 | 1.59 | 2.40E-24 | 5.14E-20 | 0.22 | 0.08 | 0.15 | 2.99 |
| 4 | Rtn3 | 0.45 | 2.43E-24 | 5.20E-20 | 0.96 | 0.87 | 0.09 | 1.1 |
| 4 | Lrrc8c | 1.34 | 2.70E-24 | 5.76E-20 | 0.23 | 0.08 | 0.16 | 3.04 |
| 4 | Tspan33 | 1.17 | 2.81E-24 | 5.99E-20 | 0.32 | 0.13 | 0.19 | 2.52 |
| 4 | Elof1 | 0.44 | 2.82E-24 | 6.01E-20 | 0.95 | 0.87 | 0.08 | 1.09 |
| 4 | Sdf2l1 | 0.74 | 3.52E-24 | 7.51E-20 | 0.74 | 0.47 | 0.27 | 1.56 |
| 4 | Nop10 | 0.36 | 3.60E-24 | 7.70E-20 | 0.98 | 0.94 | 0.04 | 1.05 |
| 4 | Spc24 | 0.56 | 4.48E-24 | 9.57E-20 | 0.89 | 0.64 | 0.25 | 1.4 |
| 4 | Snu13 | 0.33 | 4.86E-24 | 1.04E-19 | 0.99 | 0.97 | 0.02 | 1.02 |
| 4 | Hk1 | 0.81 | 5.34E-24 | 1.14E-19 | 0.42 | 0.19 | 0.23 | 2.23 |
| 4 | Ppm1g | 0.56 | 5.96E-24 | 1.27E-19 | 0.9 | 0.76 | 0.14 | 1.19 |
| 4 | Sqle | 0.67 | 5.98E-24 | 1.28E-19 | 0.78 | 0.51 | 0.27 | 1.53 |
| 4 | AW011738 | 0.76 | 6.03E-24 | 1.29E-19 | 0.54 | 0.27 | 0.27 | 1.98 |
| 4 | Aprt | 0.39 | 6.67E-24 | 1.43E-19 | 0.98 | 0.93 | 0.05 | 1.05 |
| 4 | Hspa4l | 0.85 | 8.55E-24 | 1.83E-19 | 0.46 | 0.23 | 0.23 | 2 |
| 4 | Plppr1 | 1.33 | 9.59E-24 | 2.05E-19 | 0.24 | 0.08 | 0.16 | 2.9 |
| 4 | Bex4 | 0.39 | 1.03E-23 | 2.21E-19 | 0.99 | 0.93 | 0.07 | 1.07 |
| 4 | Stip1 | 0.43 | 1.29E-23 | 2.75E-19 | 0.96 | 0.85 | 0.11 | 1.12 |
| 4 | Tfdp1 | 0.63 | 1.30E-23 | 2.78E-19 | 0.86 | 0.64 | 0.23 | 1.35 |
| 4 | Nabp2 | 0.51 | 1.42E-23 | 3.03E-19 | 0.91 | 0.77 | 0.15 | 1.19 |
| 4 | Dpy30 | 0.39 | 1.48E-23 | 3.16E-19 | 0.97 | 0.89 | 0.08 | 1.09 |
| 4 | Nolc1 | 0.53 | 1.52E-23 | 3.25E-19 | 0.89 | 0.75 | 0.13 | 1.18 |
| 4 | Atl1 | 1.29 | 1.68E-23 | 3.58E-19 | 0.2 | 0.06 | 0.14 | 3.32 |
| 4 | Rad18 | 0.73 | 1.95E-23 | 4.16E-19 | 0.65 | 0.38 | 0.27 | 1.69 |
| 4 | Sall2 | 0.75 | 2.10E-23 | 4.48E-19 | 0.71 | 0.45 | 0.26 | 1.58 |
| 4 | E2f8 | 0.87 | 2.26E-23 | 4.83E-19 | 0.48 | 0.24 | 0.24 | 2.01 |
| 4 | Cnbp | 0.31 | 2.33E-23 | 4.99E-19 | 1 | 0.98 | 0.02 | 1.02 |
| 4 | Aen | 0.85 | 2.34E-23 | 4.99E-19 | 0.61 | 0.36 | 0.25 | 1.69 |
| 4 | Sigmar1 | 0.74 | 2.53E-23 | 5.40E-19 | 0.71 | 0.48 | 0.22 | 1.46 |
| 4 | Msi1 | 0.73 | 3.46E-23 | 7.38E-19 | 0.7 | 0.46 | 0.24 | 1.53 |
| 4 | Muc1 | 1.41 | 3.47E-23 | 7.42E-19 | 0.21 | 0.07 | 0.14 | 3.12 |
| 4 | Snrpb | 0.31 | 3.49E-23 | 7.46E-19 | 0.99 | 0.97 | 0.02 | 1.02 |
| 4 | Calm3 | 0.4 | 3.60E-23 | 7.69E-19 | 0.96 | 0.83 | 0.13 | 1.16 |
| 4 | Hip1 | 0.79 | 4.13E-23 | 8.83E-19 | 0.59 | 0.33 | 0.26 | 1.78 |
| 4 | Ydjc | 1.44 | 4.39E-23 | 9.38E-19 | 0.28 | 0.11 | 0.17 | 2.58 |
| 4 | Nsdhl | 0.9 | 4.66E-23 | 9.96E-19 | 0.59 | 0.35 | 0.24 | 1.7 |
| 4 | Insig1 | 1 | 6.93E-23 | 1.48E-18 | 0.46 | 0.23 | 0.22 | 1.96 |
| 4 | Pde3b | 1.03 | 9.64E-23 | 2.06E-18 | 0.39 | 0.18 | 0.21 | 2.17 |
| 4 | Hmgcs1 | 0.58 | 1.22E-22 | 2.61E-18 | 0.9 | 0.74 | 0.15 | 1.2 |
| 4 | Mmp16 | 1.27 | 1.29E-22 | 2.75E-18 | 0.29 | 0.12 | 0.18 | 2.49 |
| 4 | Auts2 | 0.43 | 1.60E-22 | 3.42E-18 | 0.98 | 0.9 | 0.08 | 1.09 |
| 4 | Cenpa | 0.47 | 1.68E-22 | 3.58E-18 | 0.91 | 0.66 | 0.25 | 1.37 |
| 4 | Mrpl42 | 0.39 | 1.68E-22 | 3.58E-18 | 0.97 | 0.9 | 0.07 | 1.08 |
| 4 | Gpatch4 | 0.66 | 1.76E-22 | 3.77E-18 | 0.77 | 0.52 | 0.25 | 1.48 |
| 4 | Ccne1 | 0.94 | 1.88E-22 | 4.01E-18 | 0.5 | 0.27 | 0.23 | 1.87 |
| 4 | Srsf7 | 0.28 | 2.06E-22 | 4.40E-18 | 1 | 0.98 | 0.02 | 1.02 |
| 4 | Polr2l | 0.53 | 2.08E-22 | 4.45E-18 | 0.9 | 0.76 | 0.14 | 1.18 |
| 4 | Bop1 | 0.86 | 2.25E-22 | 4.81E-18 | 0.55 | 0.32 | 0.24 | 1.75 |
| 4 | Psmd13 | 0.43 | 2.79E-22 | 5.96E-18 | 0.94 | 0.84 | 0.1 | 1.12 |
| 4 | Tceal3 | 1.21 | 2.91E-22 | 6.21E-18 | 0.33 | 0.14 | 0.19 | 2.34 |
| 4 | Rab5if | 0.4 | 3.26E-22 | 6.96E-18 | 0.96 | 0.88 | 0.08 | 1.09 |
| 4 | Mtr | 1.1 | 3.40E-22 | 7.26E-18 | 0.4 | 0.19 | 0.21 | 2.07 |

**Supplementary Table 3. Marker genes for the original clusters of integrated clustering of E11.5 control and NotchGOF λ-junction ectoderm cells.**

| | | | | | | | | |
|---|---|---|---|---|---|---|---|---|
| 4 | Lig1 | 0.47 | 3.74E-22 | 7.98E-18 | 0.92 | 0.66 | 0.26 | 1.39 |
| 4 | Syngr1 | 0.96 | 3.79E-22 | 8.09E-18 | 0.42 | 0.2 | 0.22 | 2.08 |
| 4 | Clspn | 0.55 | 4.91E-22 | 1.05E-17 | 0.77 | 0.49 | 0.28 | 1.57 |
| 4 | Slc9a3r1 | 0.54 | 4.98E-22 | 1.06E-17 | 0.88 | 0.74 | 0.14 | 1.2 |
| 4 | Lsm2 | 0.38 | 5.08E-22 | 1.09E-17 | 0.98 | 0.88 | 0.1 | 1.11 |
| 4 | Dpysl4 | 1.13 | 5.30E-22 | 1.13E-17 | 0.26 | 0.1 | 0.16 | 2.65 |
| 4 | Tcof1 | 0.54 | 5.44E-22 | 1.16E-17 | 0.88 | 0.68 | 0.2 | 1.29 |
| 4 | H2afx | 0.39 | 5.69E-22 | 1.22E-17 | 0.99 | 0.89 | 0.1 | 1.11 |
| 4 | Plagl1 | 0.59 | 6.51E-22 | 1.39E-17 | 0.67 | 0.4 | 0.27 | 1.67 |
| 4 | Tubb2a | 0.77 | 7.23E-22 | 1.54E-17 | 0.58 | 0.34 | 0.24 | 1.7 |
| 4 | Wrb | 0.87 | 7.24E-22 | 1.55E-17 | 0.48 | 0.25 | 0.23 | 1.93 |
| 4 | Gcsh | 0.59 | 8.24E-22 | 1.76E-17 | 0.84 | 0.64 | 0.2 | 1.31 |
| 4 | Trib2 | 0.97 | 8.38E-22 | 1.79E-17 | 0.39 | 0.18 | 0.21 | 2.16 |
| 4 | Nop56 | 0.47 | 8.85E-22 | 1.89E-17 | 0.94 | 0.83 | 0.11 | 1.13 |
| 4 | 2610318N02Rik | 0.71 | 8.99E-22 | 1.92E-17 | 0.66 | 0.4 | 0.26 | 1.64 |
| 4 | Rwdd1 | 0.41 | 9.47E-22 | 2.02E-17 | 0.97 | 0.89 | 0.08 | 1.09 |
| 4 | Mtfp1 | 1.3 | 9.53E-22 | 2.04E-17 | 0.2 | 0.06 | 0.13 | 3.11 |
| 4 | Abce1 | 0.59 | 1.06E-21 | 2.27E-17 | 0.86 | 0.7 | 0.17 | 1.24 |
| 4 | Dnajc15 | 0.51 | 1.15E-21 | 2.45E-17 | 0.86 | 0.71 | 0.16 | 1.22 |
| 4 | Sfpq | 0.32 | 1.27E-21 | 2.71E-17 | 1 | 0.98 | 0.02 | 1.02 |
| 4 | Aarsd1 | 0.57 | 1.29E-21 | 2.75E-17 | 0.88 | 0.66 | 0.22 | 1.33 |
| 4 | Phf5a | 0.37 | 1.32E-21 | 2.82E-17 | 0.98 | 0.92 | 0.06 | 1.06 |
| 4 | A830018L16Rik | 2.03 | 1.40E-21 | 2.99E-17 | 0.14 | 0.04 | 0.11 | 3.86 |
| 4 | Larp1b | 1.41 | 1.52E-21 | 3.25E-17 | 0.26 | 0.1 | 0.16 | 2.56 |
| 4 | Ssrp1 | 0.4 | 1.68E-21 | 3.59E-17 | 0.98 | 0.9 | 0.08 | 1.09 |
| 4 | Krt8 | 0.3 | 1.68E-21 | 3.60E-17 | 1 | 0.8 | 0.19 | 1.24 |
| 4 | Agpat5 | 0.86 | 1.95E-21 | 4.16E-17 | 0.67 | 0.45 | 0.23 | 1.5 |
| 4 | Mir670hg | 1.27 | 2.09E-21 | 4.46E-17 | 0.15 | 0.04 | 0.11 | 3.8 |
| 4 | Ppid | 0.47 | 2.13E-21 | 4.56E-17 | 0.94 | 0.84 | 0.1 | 1.12 |
| 4 | Cenpf | 0.49 | 2.20E-21 | 4.69E-17 | 0.89 | 0.64 | 0.25 | 1.38 |
| 4 | Uchl3 | 0.45 | 2.35E-21 | 5.03E-17 | 0.94 | 0.81 | 0.13 | 1.16 |
| 4 | Pgam1 | 0.26 | 2.36E-21 | 5.05E-17 | 1 | 0.99 | 0.01 | 1.01 |
| 4 | Klhl13 | 0.72 | 2.39E-21 | 5.10E-17 | 0.63 | 0.38 | 0.25 | 1.66 |
| 4 | Pgk1 | 0.34 | 2.94E-21 | 6.28E-17 | 1 | 0.98 | 0.02 | 1.02 |
| 4 | Exo1 | 0.69 | 3.25E-21 | 6.94E-17 | 0.62 | 0.37 | 0.26 | 1.7 |
| 4 | Cdv3 | 0.36 | 3.36E-21 | 7.18E-17 | 0.99 | 0.94 | 0.05 | 1.05 |
| 4 | Eif3b | 0.46 | 3.46E-21 | 7.40E-17 | 0.93 | 0.8 | 0.12 | 1.15 |
| 4 | Schip1 | 0.83 | 3.74E-21 | 7.99E-17 | 0.59 | 0.36 | 0.24 | 1.66 |
| 4 | Asf1b | 0.8 | 4.31E-21 | 9.22E-17 | 0.58 | 0.34 | 0.24 | 1.71 |
| 4 | Umps | 0.64 | 5.42E-21 | 1.16E-16 | 0.76 | 0.51 | 0.24 | 1.47 |
| 4 | Map4k5 | 0.76 | 5.85E-21 | 1.25E-16 | 0.59 | 0.36 | 0.24 | 1.66 |
| 4 | Mcm6 | 0.56 | 6.26E-21 | 1.34E-16 | 0.86 | 0.65 | 0.21 | 1.32 |
| 4 | Ruvbl1 | 0.51 | 8.17E-21 | 1.74E-16 | 0.91 | 0.75 | 0.17 | 1.22 |
| 4 | Spata13 | 1.35 | 9.32E-21 | 1.99E-16 | 0.25 | 0.1 | 0.15 | 2.55 |
| 4 | Cybrd1 | 1.34 | 9.97E-21 | 2.13E-16 | 0.22 | 0.08 | 0.14 | 2.82 |
| 4 | Rrm1 | 0.49 | 1.00E-20 | 2.14E-16 | 0.9 | 0.69 | 0.21 | 1.3 |
| 4 | Boc | 1.16 | 1.23E-20 | 2.62E-16 | 0.28 | 0.11 | 0.17 | 2.49 |
| 4 | Acat2 | 0.51 | 1.27E-20 | 2.72E-16 | 0.89 | 0.71 | 0.18 | 1.25 |
| 4 | Gde1 | 0.65 | 1.27E-20 | 2.72E-16 | 0.72 | 0.48 | 0.24 | 1.51 |
| 4 | Cdca3 | 0.51 | 1.31E-20 | 2.81E-16 | 0.81 | 0.57 | 0.25 | 1.43 |
| 4 | Cisd2 | 0.52 | 1.33E-20 | 2.84E-16 | 0.9 | 0.73 | 0.17 | 1.23 |
| 4 | Cct8 | 0.3 | 1.36E-20 | 2.91E-16 | 0.99 | 0.97 | 0.02 | 1.02 |
| 4 | Nars | 0.5 | 1.62E-20 | 3.47E-16 | 0.89 | 0.75 | 0.14 | 1.18 |
| 4 | Enkd1 | 0.93 | 1.75E-20 | 3.74E-16 | 0.45 | 0.24 | 0.21 | 1.91 |
| 4 | Atp5g1 | 0.28 | 1.95E-20 | 4.17E-16 | 1 | 0.97 | 0.02 | 1.02 |
| 4 | Jam3 | 0.82 | 2.06E-20 | 4.40E-16 | 0.57 | 0.34 | 0.23 | 1.67 |
| 4 | Krt18 | 0.31 | 2.17E-20 | 4.64E-16 | 0.99 | 0.85 | 0.14 | 1.16 |

**Supplementary Table 3. Marker genes for the original clusters of integrated clustering of E11.5 control and NotchGOF λ-junction ectoderm cells.**

| | | | | | | | | |
|---|---|---|---|---|---|---|---|---|
| 4 | Mki67 | 0.47 | 2.26E-20 | 4.82E-16 | 0.9 | 0.62 | 0.27 | 1.44 |
| 4 | Fdps | 0.52 | 2.42E-20 | 5.16E-16 | 0.94 | 0.8 | 0.13 | 1.16 |
| 4 | 2010204K13Rik | 0.84 | 2.50E-20 | 5.33E-16 | 0.46 | 0.24 | 0.22 | 1.94 |
| 4 | Supt16 | 0.35 | 2.53E-20 | 5.41E-16 | 0.99 | 0.95 | 0.04 | 1.04 |
| 4 | Selenof | 0.34 | 2.92E-20 | 6.23E-16 | 0.98 | 0.94 | 0.04 | 1.04 |
| 4 | Fgfrl1 | 0.82 | 2.99E-20 | 6.39E-16 | 0.52 | 0.28 | 0.23 | 1.83 |
| 4 | Mrps24 | 0.45 | 3.41E-20 | 7.28E-16 | 0.94 | 0.82 | 0.12 | 1.14 |
| 4 | Uqcc2 | 0.38 | 3.60E-20 | 7.68E-16 | 0.97 | 0.91 | 0.06 | 1.07 |
| 4 | Mrpl28 | 0.41 | 3.87E-20 | 8.28E-16 | 0.95 | 0.83 | 0.12 | 1.14 |
| 4 | Hspa14 | 0.61 | 4.45E-20 | 9.51E-16 | 0.81 | 0.61 | 0.21 | 1.34 |
| 4 | Emc8 | 0.5 | 4.48E-20 | 9.56E-16 | 0.9 | 0.76 | 0.14 | 1.18 |
| 4 | Uhrf1 | 0.56 | 4.49E-20 | 9.59E-16 | 0.85 | 0.62 | 0.23 | 1.38 |
| 4 | Myo5c | 1.08 | 4.84E-20 | 1.03E-15 | 0.31 | 0.13 | 0.17 | 2.3 |
| 4 | Cct5 | 0.27 | 4.84E-20 | 1.03E-15 | 1 | 0.98 | 0.02 | 1.02 |
| 4 | Phgdh | 0.4 | 5.20E-20 | 1.11E-15 | 0.96 | 0.82 | 0.14 | 1.17 |
| 4 | Pld1 | 1.83 | 5.26E-20 | 1.12E-15 | 0.12 | 0.03 | 0.09 | 4 |
| 4 | Tk1 | 0.64 | 6.17E-20 | 1.32E-15 | 0.69 | 0.43 | 0.26 | 1.6 |
| 4 | Tomm40 | 0.48 | 6.49E-20 | 1.39E-15 | 0.9 | 0.8 | 0.11 | 1.13 |
| 4 | Aunip | 0.95 | 6.77E-20 | 1.45E-15 | 0.41 | 0.22 | 0.2 | 1.92 |
| 4 | Hmgcr | 0.6 | 6.77E-20 | 1.45E-15 | 0.8 | 0.59 | 0.21 | 1.36 |
| 4 | Nfix | 1.22 | 6.98E-20 | 1.49E-15 | 0.22 | 0.08 | 0.14 | 2.79 |
| 4 | Hist1h2ap | 0.58 | 7.04E-20 | 1.50E-15 | 0.9 | 0.67 | 0.22 | 1.33 |
| 4 | Naa50 | 0.5 | 7.18E-20 | 1.53E-15 | 0.88 | 0.7 | 0.18 | 1.25 |
| 4 | Gpx4 | 0.27 | 7.92E-20 | 1.69E-15 | 1 | 0.98 | 0.02 | 1.02 |
| 4 | Myo6 | 0.97 | 8.20E-20 | 1.75E-15 | 0.39 | 0.2 | 0.2 | 2.01 |
| 4 | Ccnb1 | 0.48 | 8.52E-20 | 1.82E-15 | 0.76 | 0.5 | 0.26 | 1.52 |
| 4 | Tfrc | 0.63 | 8.69E-20 | 1.86E-15 | 0.72 | 0.49 | 0.23 | 1.46 |
| 4 | Sdhd | 0.5 | 1.02E-19 | 2.18E-15 | 0.92 | 0.76 | 0.16 | 1.21 |
| 4 | Cinp | 0.73 | 1.03E-19 | 2.21E-15 | 0.67 | 0.43 | 0.24 | 1.56 |
| 4 | Hey1 | 0.33 | 1.19E-19 | 2.54E-15 | 0.97 | 0.72 | 0.25 | 1.35 |
| 4 | Nsd2 | 0.54 | 1.19E-19 | 2.55E-15 | 0.9 | 0.73 | 0.16 | 1.22 |
| 4 | Cks2 | 0.46 | 1.28E-19 | 2.73E-15 | 0.93 | 0.76 | 0.17 | 1.22 |
| 4 | Arhgef26 | 1.22 | 1.45E-19 | 3.09E-15 | 0.26 | 0.1 | 0.15 | 2.48 |
| 4 | Pfdn6 | 0.36 | 1.48E-19 | 3.16E-15 | 0.95 | 0.86 | 0.08 | 1.1 |
| 4 | B9d1 | 0.58 | 2.28E-19 | 4.87E-15 | 0.78 | 0.57 | 0.21 | 1.36 |
| 4 | Gsr | 0.65 | 2.35E-19 | 5.02E-15 | 0.71 | 0.46 | 0.25 | 1.54 |
| 4 | Ipo7 | 0.57 | 2.41E-19 | 5.14E-15 | 0.79 | 0.59 | 0.19 | 1.32 |
| 4 | Exosc8 | 0.5 | 2.42E-19 | 5.17E-15 | 0.89 | 0.71 | 0.18 | 1.25 |
| 4 | Pnn | 0.34 | 2.52E-19 | 5.38E-15 | 0.98 | 0.95 | 0.03 | 1.04 |
| 4 | Ywhah | 0.35 | 2.55E-19 | 5.46E-15 | 0.94 | 0.79 | 0.15 | 1.19 |
| 4 | Lockd | 0.71 | 2.80E-19 | 5.98E-15 | 0.55 | 0.32 | 0.24 | 1.74 |
| 4 | Gm47283 | 0.41 | 2.83E-19 | 6.05E-15 | 0.98 | 0.91 | 0.07 | 1.08 |
| 4 | Sfxn1 | 0.53 | 2.88E-19 | 6.14E-15 | 0.88 | 0.69 | 0.19 | 1.27 |
| 4 | Creld2 | 0.78 | 3.27E-19 | 6.98E-15 | 0.65 | 0.43 | 0.22 | 1.51 |
| 4 | Lap3 | 0.6 | 3.38E-19 | 7.22E-15 | 0.76 | 0.55 | 0.22 | 1.39 |
| 4 | Dtymk | 0.45 | 3.60E-19 | 7.69E-15 | 0.95 | 0.82 | 0.13 | 1.15 |
| 4 | Pdlim7 | 0.47 | 4.10E-19 | 8.75E-15 | 0.81 | 0.6 | 0.21 | 1.34 |
| 4 | Akap12 | 0.85 | 4.40E-19 | 9.40E-15 | 0.46 | 0.26 | 0.21 | 1.8 |
| 4 | Ildr2 | 1.58 | 4.73E-19 | 1.01E-14 | 0.15 | 0.04 | 0.11 | 3.41 |
| 4 | Metrn | 0.79 | 4.84E-19 | 1.03E-14 | 0.54 | 0.32 | 0.22 | 1.67 |
| 4 | Bex2 | 0.32 | 4.87E-19 | 1.04E-14 | 0.97 | 0.87 | 0.1 | 1.12 |
| 4 | Gas1 | 0.42 | 5.05E-19 | 1.08E-14 | 0.8 | 0.55 | 0.25 | 1.46 |
| 4 | Ppp1cc | 0.33 | 5.06E-19 | 1.08E-14 | 0.98 | 0.92 | 0.06 | 1.06 |
| 4 | Erg28 | 0.55 | 5.63E-19 | 1.20E-14 | 0.85 | 0.68 | 0.17 | 1.26 |
| 4 | Taf10 | 0.32 | 6.95E-19 | 1.48E-14 | 0.98 | 0.94 | 0.05 | 1.05 |
| 4 | Psmd14 | 0.45 | 7.01E-19 | 1.50E-14 | 0.9 | 0.77 | 0.14 | 1.18 |
| 4 | Melk | 0.62 | 7.41E-19 | 1.58E-14 | 0.66 | 0.42 | 0.24 | 1.59 |

**Supplementary Table 3. Marker genes for the original clusters of integrated clustering of E11.5 control and NotchGOF λ-junction ectoderm cells.**

| | | | | | | | | |
|---|---|---|---|---|---|---|---|---|
| 4 | Atp5g3 | 0.28 | 7.45E-19 | 1.59E-14 | 1 | 0.95 | 0.04 | 1.05 |
| 4 | Krtcap2 | 0.34 | 7.85E-19 | 1.68E-14 | 0.99 | 0.91 | 0.08 | 1.09 |
| 4 | Nsmce4a | 0.42 | 8.73E-19 | 1.87E-14 | 0.96 | 0.83 | 0.14 | 1.16 |
| 4 | Irs1 | 0.82 | 8.76E-19 | 1.87E-14 | 0.51 | 0.29 | 0.22 | 1.77 |
| 4 | Ezr | 0.34 | 9.25E-19 | 1.98E-14 | 0.97 | 0.9 | 0.07 | 1.08 |
| 4 | Ndufa12 | 0.31 | 9.58E-19 | 2.05E-14 | 0.99 | 0.94 | 0.04 | 1.04 |
| 4 | Atp2a2 | 0.49 | 9.84E-19 | 2.10E-14 | 0.85 | 0.65 | 0.19 | 1.3 |
| 4 | Wdr76 | 0.72 | 1.12E-18 | 2.39E-14 | 0.56 | 0.33 | 0.23 | 1.7 |
| 4 | Kif26b | 1.44 | 1.22E-18 | 2.61E-14 | 0.15 | 0.04 | 0.1 | 3.45 |
| 4 | Dmbx1 | 1.14 | 1.27E-18 | 2.70E-14 | 0.31 | 0.15 | 0.17 | 2.14 |
| 4 | Pus7 | 0.73 | 1.27E-18 | 2.71E-14 | 0.51 | 0.28 | 0.23 | 1.8 |
| 4 | Rad51ap1 | 0.6 | 1.38E-18 | 2.96E-14 | 0.64 | 0.38 | 0.25 | 1.66 |
| 4 | Syce2 | 0.53 | 1.42E-18 | 3.03E-14 | 0.76 | 0.51 | 0.25 | 1.48 |
| 4 | Raver2 | 0.9 | 1.45E-18 | 3.10E-14 | 0.38 | 0.19 | 0.19 | 1.97 |
| 4 | Gart | 0.56 | 1.55E-18 | 3.30E-14 | 0.77 | 0.58 | 0.19 | 1.33 |
| 4 | Fasn | 0.67 | 1.77E-18 | 3.77E-14 | 0.67 | 0.45 | 0.21 | 1.47 |
| 4 | Fkbp1a | 0.3 | 1.77E-18 | 3.78E-14 | 0.99 | 0.94 | 0.05 | 1.06 |
| 4 | Dnajc2 | 0.37 | 1.86E-18 | 3.97E-14 | 0.97 | 0.87 | 0.1 | 1.11 |
| 4 | Hat1 | 0.46 | 1.93E-18 | 4.12E-14 | 0.9 | 0.74 | 0.16 | 1.21 |
| 4 | Pgm1 | 0.8 | 1.95E-18 | 4.17E-14 | 0.5 | 0.29 | 0.22 | 1.76 |
| 4 | Nuf2 | 0.7 | 2.11E-18 | 4.51E-14 | 0.6 | 0.36 | 0.24 | 1.65 |
| 4 | Eif4e | 0.4 | 2.22E-18 | 4.75E-14 | 0.95 | 0.82 | 0.13 | 1.15 |
| 4 | Dnph1 | 0.59 | 2.29E-18 | 4.90E-14 | 0.74 | 0.51 | 0.23 | 1.44 |
| 4 | Rgma | 0.85 | 2.31E-18 | 4.94E-14 | 0.38 | 0.19 | 0.19 | 2.01 |
| 4 | Hmmr | 0.55 | 2.51E-18 | 5.36E-14 | 0.69 | 0.44 | 0.25 | 1.57 |
| 4 | Vars | 0.61 | 2.55E-18 | 5.45E-14 | 0.76 | 0.58 | 0.18 | 1.31 |
| 4 | Kitl | 0.73 | 2.55E-18 | 5.45E-14 | 0.29 | 0.12 | 0.17 | 2.36 |
| 4 | Gfer | 0.68 | 2.70E-18 | 5.78E-14 | 0.63 | 0.42 | 0.21 | 1.51 |
| 4 | Slc3a2 | 0.51 | 2.87E-18 | 6.13E-14 | 0.84 | 0.66 | 0.18 | 1.27 |
| 4 | Tmem30b | 0.63 | 2.99E-18 | 6.40E-14 | 0.47 | 0.25 | 0.22 | 1.87 |
| 4 | Ndufa13 | 0.28 | 3.52E-18 | 7.51E-14 | 0.99 | 0.97 | 0.02 | 1.02 |
| 4 | Smc4 | 0.43 | 3.61E-18 | 7.71E-14 | 0.93 | 0.73 | 0.2 | 1.28 |
| 4 | U2af1 | 0.29 | 3.66E-18 | 7.82E-14 | 0.99 | 0.96 | 0.03 | 1.03 |
| 4 | Lsm7 | 0.33 | 3.77E-18 | 8.06E-14 | 0.98 | 0.92 | 0.06 | 1.07 |
| 4 | Ift27 | 0.62 | 4.68E-18 | 1.00E-13 | 0.73 | 0.5 | 0.23 | 1.47 |
| 4 | Dnajb11 | 0.57 | 4.95E-18 | 1.06E-13 | 0.81 | 0.61 | 0.2 | 1.32 |
| 4 | Cldn3 | 0.38 | 5.12E-18 | 1.09E-13 | 0.87 | 0.61 | 0.27 | 1.44 |
| 4 | Gpd2 | 0.7 | 5.23E-18 | 1.12E-13 | 0.54 | 0.32 | 0.22 | 1.71 |
| 4 | Lin28b | 0.82 | 5.47E-18 | 1.17E-13 | 0.46 | 0.26 | 0.21 | 1.81 |
| 4 | Phtf1 | 0.74 | 5.61E-18 | 1.20E-13 | 0.64 | 0.41 | 0.23 | 1.57 |
| 4 | Rassf1 | 0.8 | 6.04E-18 | 1.29E-13 | 0.57 | 0.36 | 0.21 | 1.57 |
| 4 | Cenpm | 0.58 | 6.32E-18 | 1.35E-13 | 0.72 | 0.48 | 0.24 | 1.49 |
| 4 | Pabpc4 | 0.55 | 6.83E-18 | 1.46E-13 | 0.82 | 0.62 | 0.2 | 1.32 |
| 4 | Pimreg | 0.56 | 6.87E-18 | 1.47E-13 | 0.68 | 0.45 | 0.24 | 1.53 |
| 4 | Atad2 | 0.43 | 6.88E-18 | 1.47E-13 | 0.87 | 0.62 | 0.25 | 1.4 |
| 4 | Rpl7l1 | 0.42 | 7.15E-18 | 1.53E-13 | 0.94 | 0.82 | 0.12 | 1.15 |
| 4 | Thop1 | 0.59 | 7.24E-18 | 1.55E-13 | 0.71 | 0.48 | 0.22 | 1.46 |
| 4 | Zeb2 | 1.23 | 7.96E-18 | 1.70E-13 | 0.19 | 0.07 | 0.12 | 2.78 |
| 4 | Cdca7l | 0.59 | 8.14E-18 | 1.74E-13 | 0.69 | 0.46 | 0.23 | 1.5 |
| 4 | Zdbf2 | 0.66 | 8.14E-18 | 1.74E-13 | 0.73 | 0.53 | 0.2 | 1.38 |
| 4 | Plscr1 | 1 | 8.67E-18 | 1.85E-13 | 0.31 | 0.14 | 0.17 | 2.19 |
| 4 | Ftsj3 | 0.6 | 8.83E-18 | 1.89E-13 | 0.77 | 0.58 | 0.19 | 1.34 |
| 4 | Wdr77 | 0.57 | 9.38E-18 | 2.00E-13 | 0.76 | 0.55 | 0.22 | 1.4 |
| 4 | Dyrk3 | 0.98 | 9.55E-18 | 2.04E-13 | 0.32 | 0.15 | 0.17 | 2.15 |
| 4 | Pin1 | 0.35 | 1.06E-17 | 2.26E-13 | 0.94 | 0.85 | 0.09 | 1.11 |
| 4 | Chchd1 | 0.34 | 1.09E-17 | 2.33E-13 | 0.97 | 0.9 | 0.06 | 1.07 |
| 4 | Scd1 | 1.47 | 1.11E-17 | 2.36E-13 | 0.25 | 0.11 | 0.14 | 2.33 |

**Supplementary Table 3. Marker genes for the original clusters of integrated clustering of E11.5 control and NotchGOF λ-junction ectoderm cells.**

| | | | | | | | | |
|---|---|---|---|---|---|---|---|---|
| 4 | Slc16a3 | 0.78 | 1.19E-17 | 2.53E-13 | 0.61 | 0.4 | 0.21 | 1.52 |
| 4 | Pold3 | 0.6 | 1.20E-17 | 2.56E-13 | 0.74 | 0.51 | 0.23 | 1.46 |
| 4 | Doc2a | 1.76 | 1.26E-17 | 2.68E-13 | 0.13 | 0.04 | 0.09 | 3.54 |
| 4 | Cars | 0.75 | 1.28E-17 | 2.73E-13 | 0.49 | 0.28 | 0.21 | 1.77 |
| 4 | Hnrnpll | 0.45 | 1.29E-17 | 2.76E-13 | 0.85 | 0.67 | 0.17 | 1.26 |
| 4 | Iars | 0.67 | 1.31E-17 | 2.79E-13 | 0.65 | 0.44 | 0.22 | 1.5 |
| 4 | Vegfb | 0.58 | 1.44E-17 | 3.09E-13 | 0.74 | 0.52 | 0.23 | 1.44 |
| 4 | B830012L14Rik | 1.11 | 1.46E-17 | 3.12E-13 | 0.22 | 0.08 | 0.13 | 2.58 |
| 4 | Clstn1 | 0.62 | 1.71E-17 | 3.66E-13 | 0.75 | 0.54 | 0.21 | 1.39 |
| 4 | Trim37 | 0.61 | 1.72E-17 | 3.68E-13 | 0.67 | 0.46 | 0.22 | 1.48 |
| 4 | Slc35f1 | 1.27 | 1.78E-17 | 3.79E-13 | 0.23 | 0.09 | 0.14 | 2.46 |
| 4 | Tma16 | 0.57 | 1.90E-17 | 4.05E-13 | 0.74 | 0.55 | 0.2 | 1.36 |
| 4 | Brsk2 | 1.27 | 1.96E-17 | 4.18E-13 | 0.14 | 0.04 | 0.1 | 3.45 |
| 4 | Gsdme | 0.92 | 1.96E-17 | 4.20E-13 | 0.36 | 0.18 | 0.18 | 2.03 |
| 4 | Ccdc124 | 0.44 | 1.98E-17 | 4.22E-13 | 0.9 | 0.75 | 0.15 | 1.21 |
| 4 | Knstrn | 0.47 | 2.08E-17 | 4.45E-13 | 0.8 | 0.56 | 0.24 | 1.43 |
| 4 | Eif2s1 | 0.36 | 2.39E-17 | 5.11E-13 | 0.97 | 0.87 | 0.09 | 1.11 |
| 4 | Pdk3 | 0.67 | 2.64E-17 | 5.65E-13 | 0.62 | 0.4 | 0.22 | 1.56 |
| 4 | Rangrf | 0.66 | 2.79E-17 | 5.96E-13 | 0.65 | 0.44 | 0.21 | 1.48 |
| 4 | Mphosph10 | 0.47 | 2.98E-17 | 6.37E-13 | 0.85 | 0.7 | 0.15 | 1.22 |
| 4 | Dhcr24 | 0.72 | 3.08E-17 | 6.57E-13 | 0.54 | 0.33 | 0.21 | 1.63 |
| 4 | Rims2 | 0.77 | 3.71E-17 | 7.92E-13 | 0.37 | 0.19 | 0.18 | 1.95 |
| 4 | St8sia4 | 1.52 | 3.80E-17 | 8.12E-13 | 0.11 | 0.03 | 0.08 | 4.04 |
| 4 | Prrx1 | 1.06 | 4.03E-17 | 8.61E-13 | 0.35 | 0.18 | 0.17 | 1.96 |
| 4 | Dynll2 | 0.34 | 4.34E-17 | 9.26E-13 | 0.97 | 0.89 | 0.08 | 1.09 |
| 4 | Phyhipl | 1.14 | 4.44E-17 | 9.48E-13 | 0.23 | 0.09 | 0.14 | 2.49 |
| 4 | Ebna1bp2 | 0.39 | 4.75E-17 | 1.02E-12 | 0.92 | 0.82 | 0.1 | 1.12 |
| 4 | Cmss1 | 0.67 | 4.98E-17 | 1.06E-12 | 0.63 | 0.42 | 0.21 | 1.49 |
| 4 | Dusp8 | 0.84 | 5.16E-17 | 1.10E-12 | 0.14 | 0.04 | 0.1 | 3.33 |
| 4 | Ctps | 0.66 | 5.61E-17 | 1.20E-12 | 0.69 | 0.47 | 0.22 | 1.47 |
| 4 | Mrto4 | 0.41 | 5.75E-17 | 1.23E-12 | 0.89 | 0.78 | 0.11 | 1.14 |
| 4 | Mthfd2 | 0.7 | 6.21E-17 | 1.33E-12 | 0.47 | 0.27 | 0.21 | 1.78 |
| 4 | Knl1 | 0.59 | 6.79E-17 | 1.45E-12 | 0.69 | 0.45 | 0.24 | 1.52 |
| 4 | Mllt3 | 0.54 | 6.99E-17 | 1.49E-12 | 0.74 | 0.54 | 0.21 | 1.39 |
| 4 | Cyb5b | 0.59 | 7.57E-17 | 1.62E-12 | 0.72 | 0.51 | 0.21 | 1.42 |
| 4 | Pde1b | 1.36 | 7.67E-17 | 1.64E-12 | 0.17 | 0.06 | 0.11 | 2.84 |
| 4 | Desi2 | 0.63 | 8.07E-17 | 1.72E-12 | 0.62 | 0.4 | 0.22 | 1.55 |
| 4 | Nop16 | 0.59 | 8.12E-17 | 1.74E-12 | 0.74 | 0.56 | 0.18 | 1.32 |
| 4 | Cmc2 | 0.57 | 9.57E-17 | 2.04E-12 | 0.72 | 0.5 | 0.22 | 1.43 |
| 4 | Hdlbp | 0.42 | 1.10E-16 | 2.34E-12 | 0.94 | 0.83 | 0.11 | 1.13 |
| 4 | Ung | 0.67 | 1.21E-16 | 2.59E-12 | 0.57 | 0.35 | 0.22 | 1.61 |
| 4 | Mcm4 | 0.5 | 1.37E-16 | 2.92E-12 | 0.81 | 0.6 | 0.21 | 1.35 |
| 4 | Ugt8a | 1.6 | 1.39E-16 | 2.96E-12 | 0.15 | 0.05 | 0.1 | 3.15 |
| 4 | Cdc34 | 0.41 | 1.39E-16 | 2.97E-12 | 0.93 | 0.78 | 0.15 | 1.19 |
| 4 | Slc1a5 | 0.88 | 1.50E-16 | 3.20E-12 | 0.36 | 0.19 | 0.18 | 1.94 |
| 4 | Tpst1 | 0.86 | 1.57E-16 | 3.35E-12 | 0.42 | 0.23 | 0.19 | 1.83 |
| 4 | Nuak1 | 0.58 | 1.61E-16 | 3.44E-12 | 0.47 | 0.25 | 0.21 | 1.85 |
| 4 | Clip1 | 0.62 | 1.73E-16 | 3.70E-12 | 0.72 | 0.53 | 0.2 | 1.37 |
| 4 | Nde1 | 0.49 | 1.88E-16 | 4.02E-12 | 0.77 | 0.58 | 0.19 | 1.34 |
| 4 | AI506816 | 0.68 | 1.99E-16 | 4.25E-12 | 0.69 | 0.49 | 0.21 | 1.43 |
| 4 | Hmbs | 0.58 | 2.22E-16 | 4.73E-12 | 0.72 | 0.5 | 0.21 | 1.42 |
| 4 | Cd302 | 0.75 | 2.61E-16 | 5.58E-12 | 0.48 | 0.27 | 0.21 | 1.77 |
| 4 | Cltb | 0.63 | 2.68E-16 | 5.72E-12 | 0.52 | 0.3 | 0.22 | 1.74 |
| 4 | Oscp1 | 0.94 | 2.93E-16 | 6.26E-12 | 0.34 | 0.17 | 0.17 | 2 |
| 4 | Me2 | 0.69 | 3.13E-16 | 6.68E-12 | 0.58 | 0.37 | 0.21 | 1.56 |
| 4 | Glrx5 | 0.35 | 3.27E-16 | 6.98E-12 | 0.94 | 0.87 | 0.08 | 1.09 |
| 4 | Rpa2 | 0.52 | 3.28E-16 | 7.01E-12 | 0.76 | 0.59 | 0.17 | 1.29 |

**Supplementary Table 3. Marker genes for the original clusters of integrated clustering of E11.5 control and NotchGOF λ-junction ectoderm cells.**

| | | | | | | | | |
|---|---|---|---|---|---|---|---|---|
| 4 | Lpl | 1.39 | 3.30E-16 | 7.05E-12 | 0.18 | 0.07 | 0.11 | 2.68 |
| 4 | Ccnb2 | 0.38 | 3.39E-16 | 7.24E-12 | 0.75 | 0.47 | 0.28 | 1.58 |
| 4 | Gpr19 | 1.42 | 3.45E-16 | 7.37E-12 | 0.18 | 0.06 | 0.11 | 2.73 |
| 4 | Rian | 0.45 | 3.67E-16 | 7.85E-12 | 0.69 | 0.44 | 0.24 | 1.55 |
| 4 | Them6 | 1.65 | 3.72E-16 | 7.94E-12 | 0.12 | 0.03 | 0.09 | 3.5 |
| 4 | Smpd3 | 1.87 | 3.92E-16 | 8.37E-12 | 0.1 | 0.03 | 0.08 | 4 |
| 4 | Epb41l4a | 0.76 | 4.47E-16 | 9.54E-12 | 0.36 | 0.18 | 0.18 | 1.98 |
| 4 | Gm42047 | 1.54 | 4.52E-16 | 9.66E-12 | 0.2 | 0.08 | 0.12 | 2.55 |
| 4 | Isyna1 | 0.44 | 5.41E-16 | 1.16E-11 | 0.88 | 0.69 | 0.18 | 1.26 |
| 4 | Eif1b | 0.39 | 5.62E-16 | 1.20E-11 | 0.95 | 0.85 | 0.1 | 1.11 |
| 4 | Lbr | 0.62 | 5.74E-16 | 1.23E-11 | 0.72 | 0.52 | 0.2 | 1.39 |
| 4 | Rab3c | 1.4 | 6.02E-16 | 1.29E-11 | 0.12 | 0.04 | 0.09 | 3.44 |
| 4 | Bub1 | 0.5 | 6.61E-16 | 1.41E-11 | 0.7 | 0.47 | 0.22 | 1.47 |
| 4 | Golm1 | 0.65 | 6.85E-16 | 1.46E-11 | 0.64 | 0.42 | 0.22 | 1.51 |
| 4 | Pkig | 0.42 | 7.00E-16 | 1.50E-11 | 0.88 | 0.74 | 0.14 | 1.19 |
| 4 | 1500004A13Rik | 0.54 | 7.27E-16 | 1.55E-11 | 0.2 | 0.08 | 0.12 | 2.57 |
| 4 | Nav1 | 0.55 | 8.37E-16 | 1.79E-11 | 0.7 | 0.49 | 0.21 | 1.43 |
| 4 | Fen1 | 0.49 | 8.85E-16 | 1.89E-11 | 0.79 | 0.57 | 0.22 | 1.39 |
| 4 | Mxd3 | 1.15 | 8.88E-16 | 1.90E-11 | 0.22 | 0.09 | 0.13 | 2.41 |
| 4 | Kctd6 | 0.68 | 1.02E-15 | 2.17E-11 | 0.42 | 0.23 | 0.2 | 1.88 |
| 4 | Slbp | 0.4 | 1.15E-15 | 2.45E-11 | 0.95 | 0.8 | 0.15 | 1.19 |
| 4 | Lrrc59 | 0.42 | 1.21E-15 | 2.58E-11 | 0.89 | 0.74 | 0.14 | 1.19 |
| 4 | L2hgdh | 0.93 | 1.21E-15 | 2.58E-11 | 0.41 | 0.23 | 0.18 | 1.8 |
| 4 | Lrrc4 | 1.39 | 1.32E-15 | 2.82E-11 | 0.13 | 0.04 | 0.09 | 3.28 |
| 4 | Dpy19l1 | 0.75 | 1.35E-15 | 2.88E-11 | 0.41 | 0.22 | 0.19 | 1.84 |
| 4 | H2-D1 | 0.72 | 1.39E-15 | 2.98E-11 | 0.52 | 0.31 | 0.21 | 1.66 |
| 4 | Jag1 | 0.39 | 1.42E-15 | 3.03E-11 | 0.78 | 0.54 | 0.25 | 1.46 |
| 4 | Snrnp40 | 0.36 | 1.43E-15 | 3.05E-11 | 0.92 | 0.82 | 0.1 | 1.13 |
| 4 | Hsph1 | 0.51 | 1.45E-15 | 3.09E-11 | 0.8 | 0.63 | 0.16 | 1.26 |
| 4 | Lrig3 | 0.67 | 1.54E-15 | 3.29E-11 | 0.48 | 0.28 | 0.2 | 1.73 |
| 4 | Snx10 | 0.66 | 1.56E-15 | 3.34E-11 | 0.51 | 0.29 | 0.22 | 1.74 |
| 4 | Pak3 | 0.58 | 1.59E-15 | 3.39E-11 | 0.49 | 0.29 | 0.21 | 1.72 |
| 4 | Rrp1b | 0.59 | 1.72E-15 | 3.68E-11 | 0.69 | 0.5 | 0.19 | 1.38 |
| 4 | Zfp706 | 0.28 | 1.76E-15 | 3.75E-11 | 0.98 | 0.92 | 0.06 | 1.06 |
| 4 | Sall1 | 1.25 | 1.81E-15 | 3.87E-11 | 0.14 | 0.05 | 0.09 | 3.04 |
| 4 | Racgap1 | 0.48 | 1.92E-15 | 4.10E-11 | 0.72 | 0.5 | 0.23 | 1.45 |
| 4 | Rilpl2 | 0.73 | 1.95E-15 | 4.17E-11 | 0.4 | 0.21 | 0.19 | 1.87 |
| 4 | Ndufab1 | 0.26 | 1.97E-15 | 4.20E-11 | 1 | 0.96 | 0.03 | 1.04 |
| 4 | 4933404O12Rik | 0.73 | 1.97E-15 | 4.21E-11 | 0.34 | 0.17 | 0.17 | 1.98 |
| 4 | Aldoa | 0.28 | 2.01E-15 | 4.28E-11 | 0.99 | 0.97 | 0.01 | 1.01 |
| 4 | Map1a | 0.94 | 2.28E-15 | 4.87E-11 | 0.15 | 0.05 | 0.1 | 3 |
| 4 | Maneal | 1.58 | 2.33E-15 | 4.97E-11 | 0.13 | 0.04 | 0.09 | 3.17 |
| 4 | Cenpq | 0.57 | 2.53E-15 | 5.40E-11 | 0.68 | 0.48 | 0.2 | 1.43 |
| 4 | Eif4g1 | 0.35 | 2.61E-15 | 5.57E-11 | 0.96 | 0.9 | 0.05 | 1.06 |
| 4 | Stxbp6 | 0.86 | 2.71E-15 | 5.80E-11 | 0.35 | 0.18 | 0.17 | 1.97 |
| 4 | Phlpp1 | 0.84 | 3.04E-15 | 6.49E-11 | 0.43 | 0.25 | 0.18 | 1.74 |
| 4 | Ckap2 | 0.69 | 3.39E-15 | 7.24E-11 | 0.52 | 0.32 | 0.2 | 1.61 |
| 4 | Mgat4b | 0.59 | 3.66E-15 | 7.82E-11 | 0.64 | 0.43 | 0.21 | 1.5 |
| 4 | Psmc2 | 0.34 | 3.89E-15 | 8.31E-11 | 0.94 | 0.85 | 0.09 | 1.1 |
| 4 | Ppan | 0.6 | 4.12E-15 | 8.80E-11 | 0.67 | 0.46 | 0.21 | 1.46 |
| 4 | Fam136a | 0.43 | 4.23E-15 | 9.03E-11 | 0.93 | 0.8 | 0.13 | 1.17 |
| 4 | Hist1h1b | 0.47 | 4.33E-15 | 9.26E-11 | 0.79 | 0.55 | 0.24 | 1.44 |
| 4 | Ric3 | 0.93 | 4.41E-15 | 9.43E-11 | 0.23 | 0.1 | 0.13 | 2.32 |
| 4 | Rfc4 | 0.43 | 4.51E-15 | 9.64E-11 | 0.88 | 0.72 | 0.17 | 1.23 |
| 4 | Mtdh | 0.34 | 5.18E-15 | 1.11E-10 | 0.98 | 0.93 | 0.05 | 1.05 |
| 4 | Rpn2 | 0.38 | 5.20E-15 | 1.11E-10 | 0.9 | 0.77 | 0.13 | 1.16 |
| 4 | Lrp11 | 1.09 | 5.22E-15 | 1.11E-10 | 0.18 | 0.07 | 0.11 | 2.69 |

**Supplementary Table 3. Marker genes for the original clusters of integrated clustering of E11.5 control and NotchGOF λ-junction ectoderm cells.**

| | | | | | | | | |
|---|---|---|---|---|---|---|---|---|
| 4 | Mybbp1a | 0.41 | 5.34E-15 | 1.14E-10 | 0.88 | 0.78 | 0.1 | 1.13 |
| 4 | Gspt1 | 0.33 | 6.52E-15 | 1.39E-10 | 0.96 | 0.89 | 0.07 | 1.08 |
| 4 | Mrps28 | 0.5 | 6.63E-15 | 1.42E-10 | 0.83 | 0.66 | 0.17 | 1.26 |
| 4 | Timm9 | 0.51 | 6.71E-15 | 1.43E-10 | 0.77 | 0.57 | 0.2 | 1.36 |
| 4 | Plk1 | 0.58 | 7.75E-15 | 1.66E-10 | 0.59 | 0.39 | 0.21 | 1.54 |
| 4 | Fam57a | 1.02 | 8.19E-15 | 1.75E-10 | 0.23 | 0.1 | 0.13 | 2.31 |
| 4 | Nip7 | 0.42 | 8.84E-15 | 1.89E-10 | 0.86 | 0.73 | 0.13 | 1.18 |
| 4 | Notch3 | 0.5 | 9.74E-15 | 2.08E-10 | 0.74 | 0.52 | 0.22 | 1.42 |
| 4 | Orc2 | 0.52 | 9.82E-15 | 2.10E-10 | 0.67 | 0.48 | 0.2 | 1.41 |
| 4 | Pcgf5 | 0.83 | 1.01E-14 | 2.15E-10 | 0.38 | 0.21 | 0.17 | 1.84 |
| 4 | Rif1 | 0.36 | 1.01E-14 | 2.16E-10 | 0.93 | 0.81 | 0.12 | 1.15 |
| 4 | G3bp1 | 0.25 | 1.04E-14 | 2.23E-10 | 1 | 0.97 | 0.02 | 1.02 |
| 4 | Chchd6 | 0.52 | 1.07E-14 | 2.30E-10 | 0.71 | 0.51 | 0.2 | 1.39 |
| 4 | Chek1 | 0.63 | 1.09E-14 | 2.33E-10 | 0.62 | 0.43 | 0.2 | 1.46 |
| 4 | Fndc3c1 | 0.67 | 1.13E-14 | 2.41E-10 | 0.48 | 0.28 | 0.2 | 1.69 |
| 4 | Pip4k2a | 1.21 | 1.26E-14 | 2.68E-10 | 0.22 | 0.09 | 0.12 | 2.33 |
| 4 | Atf4 | 0.45 | 1.27E-14 | 2.72E-10 | 0.83 | 0.66 | 0.17 | 1.25 |
| 4 | Mpp2 | 0.9 | 1.30E-14 | 2.77E-10 | 0.31 | 0.15 | 0.15 | 2 |
| 4 | Ppat | 0.61 | 1.34E-14 | 2.87E-10 | 0.65 | 0.43 | 0.21 | 1.49 |
| 4 | Hprt | 0.6 | 1.39E-14 | 2.96E-10 | 0.66 | 0.45 | 0.2 | 1.45 |
| 4 | Tspan3 | 0.29 | 1.45E-14 | 3.10E-10 | 0.99 | 0.95 | 0.04 | 1.04 |
| 4 | Mars | 0.71 | 1.45E-14 | 3.10E-10 | 0.51 | 0.32 | 0.19 | 1.61 |
| 4 | Sapcd2 | 0.61 | 1.51E-14 | 3.23E-10 | 0.46 | 0.27 | 0.19 | 1.72 |
| 4 | Fgfr1op | 0.7 | 1.52E-14 | 3.25E-10 | 0.52 | 0.32 | 0.2 | 1.63 |
| 4 | Pak1ip1 | 0.46 | 1.54E-14 | 3.29E-10 | 0.82 | 0.67 | 0.15 | 1.22 |
| 4 | Scarb1 | 0.74 | 1.58E-14 | 3.38E-10 | 0.34 | 0.18 | 0.17 | 1.95 |
| 4 | Esco2 | 0.47 | 1.58E-14 | 3.38E-10 | 0.62 | 0.39 | 0.23 | 1.58 |
| 4 | Heatr1 | 0.76 | 1.64E-14 | 3.50E-10 | 0.47 | 0.28 | 0.19 | 1.67 |
| 4 | Akr1b3 | 0.49 | 1.72E-14 | 3.68E-10 | 0.78 | 0.59 | 0.19 | 1.33 |
| 4 | Xylb | 1.36 | 1.82E-14 | 3.90E-10 | 0.15 | 0.06 | 0.1 | 2.75 |
| 4 | Elovl5 | 0.69 | 1.99E-14 | 4.25E-10 | 0.41 | 0.23 | 0.18 | 1.81 |
| 4 | Jmjd6 | 0.49 | 2.25E-14 | 4.80E-10 | 0.69 | 0.49 | 0.21 | 1.42 |
| 4 | Mark1 | 0.71 | 2.32E-14 | 4.96E-10 | 0.42 | 0.24 | 0.18 | 1.75 |
| 4 | Aacs | 1.15 | 2.34E-14 | 5.00E-10 | 0.22 | 0.1 | 0.12 | 2.29 |
| 4 | Bub1b | 0.51 | 2.35E-14 | 5.02E-10 | 0.68 | 0.48 | 0.2 | 1.43 |
| 4 | Tnfrsf21 | 1 | 2.41E-14 | 5.14E-10 | 0.21 | 0.09 | 0.12 | 2.36 |
| 4 | Tmx2 | 0.49 | 2.45E-14 | 5.24E-10 | 0.78 | 0.57 | 0.21 | 1.37 |
| 4 | Prdx3 | 0.54 | 2.54E-14 | 5.42E-10 | 0.72 | 0.5 | 0.22 | 1.44 |
| 4 | Nxt1 | 0.46 | 2.60E-14 | 5.54E-10 | 0.81 | 0.61 | 0.2 | 1.33 |
| 4 | Naf1 | 0.57 | 2.60E-14 | 5.56E-10 | 0.64 | 0.43 | 0.21 | 1.48 |
| 4 | Epb41l4b | 0.95 | 2.62E-14 | 5.59E-10 | 0.21 | 0.09 | 0.12 | 2.39 |
| 4 | Ak4 | 0.73 | 2.67E-14 | 5.69E-10 | 0.39 | 0.21 | 0.18 | 1.83 |
| 4 | Lpcat1 | 0.57 | 2.79E-14 | 5.97E-10 | 0.65 | 0.44 | 0.21 | 1.47 |
| 4 | Hs6st2 | 0.4 | 2.97E-14 | 6.35E-10 | 0.74 | 0.52 | 0.22 | 1.43 |
| 4 | Crip1 | 0.39 | 3.05E-14 | 6.52E-10 | 0.61 | 0.38 | 0.23 | 1.59 |
| 4 | Eif3j1 | 0.46 | 3.17E-14 | 6.77E-10 | 0.82 | 0.65 | 0.16 | 1.25 |
| 4 | Pbk | 0.47 | 3.25E-14 | 6.94E-10 | 0.69 | 0.46 | 0.23 | 1.49 |
| 4 | Chchd4 | 0.54 | 3.30E-14 | 7.05E-10 | 0.7 | 0.49 | 0.21 | 1.43 |
| 4 | Gcat | 0.46 | 3.50E-14 | 7.48E-10 | 0.8 | 0.59 | 0.21 | 1.35 |
| 4 | E2f2 | 1.01 | 3.62E-14 | 7.74E-10 | 0.24 | 0.11 | 0.13 | 2.2 |
| 4 | Nol7 | 0.27 | 3.69E-14 | 7.88E-10 | 0.97 | 0.92 | 0.05 | 1.05 |
| 4 | Atad5 | 0.55 | 3.93E-14 | 8.40E-10 | 0.65 | 0.44 | 0.21 | 1.48 |
| 4 | Bysl | 0.67 | 4.09E-14 | 8.74E-10 | 0.46 | 0.27 | 0.19 | 1.71 |
| 4 | Cep55 | 0.74 | 4.09E-14 | 8.75E-10 | 0.43 | 0.25 | 0.18 | 1.72 |
| 4 | C530008M17Rik | 0.39 | 4.20E-14 | 8.97E-10 | 0.59 | 0.38 | 0.21 | 1.54 |
| 4 | Pole2 | 0.72 | 4.20E-14 | 8.98E-10 | 0.36 | 0.19 | 0.17 | 1.88 |
| 4 | Baiap2l1 | 0.86 | 4.28E-14 | 9.13E-10 | 0.29 | 0.14 | 0.15 | 2.04 |

**Supplementary Table 3. Marker genes for the original clusters of integrated clustering of E11.5 control and NotchGOF λ-junction ectoderm cells.**

| | | | | | | | | |
|---|---|---|---|---|---|---|---|---|
| 4 | Tra2b | 0.29 | 4.32E-14 | 9.23E-10 | 0.98 | 0.92 | 0.06 | 1.07 |
| 4 | Fndc4 | 0.65 | 4.45E-14 | 9.50E-10 | 0.53 | 0.32 | 0.21 | 1.65 |
| 4 | Flvcr1 | 1.01 | 4.63E-14 | 9.90E-10 | 0.19 | 0.07 | 0.11 | 2.53 |
| 4 | Mcm10 | 0.65 | 4.73E-14 | 1.01E-09 | 0.46 | 0.27 | 0.19 | 1.7 |
| 4 | Speg | 0.98 | 5.22E-14 | 1.11E-09 | 0.25 | 0.12 | 0.13 | 2.16 |
| 4 | Mageh1 | 0.56 | 5.79E-14 | 1.24E-09 | 0.68 | 0.5 | 0.18 | 1.36 |
| 4 | Pla2g12a | 0.44 | 5.88E-14 | 1.26E-09 | 0.8 | 0.61 | 0.19 | 1.31 |
| 4 | Klf13 | 0.43 | 5.90E-14 | 1.26E-09 | 0.61 | 0.39 | 0.22 | 1.55 |
| 4 | Chaf1a | 0.47 | 5.92E-14 | 1.26E-09 | 0.77 | 0.54 | 0.23 | 1.42 |
| 4 | Grwd1 | 0.59 | 6.06E-14 | 1.29E-09 | 0.62 | 0.42 | 0.2 | 1.47 |
| 4 | Carmil3 | 1.03 | 6.34E-14 | 1.35E-09 | 0.19 | 0.08 | 0.11 | 2.43 |
| 4 | Fam102b | 1.14 | 6.45E-14 | 1.38E-09 | 0.23 | 0.1 | 0.12 | 2.2 |
| 4 | D930020B18Rik | 0.88 | 6.74E-14 | 1.44E-09 | 0.15 | 0.05 | 0.1 | 2.83 |
| 4 | Dgcr6 | 0.42 | 6.93E-14 | 1.48E-09 | 0.84 | 0.65 | 0.18 | 1.28 |
| 4 | 2900026A02Rik | 0.53 | 6.96E-14 | 1.49E-09 | 0.54 | 0.34 | 0.21 | 1.62 |
| 4 | Fam13a | 1.21 | 7.06E-14 | 1.51E-09 | 0.17 | 0.07 | 0.11 | 2.59 |
| 4 | Tenm4 | 0.43 | 7.40E-14 | 1.58E-09 | 0.68 | 0.44 | 0.23 | 1.52 |
| 4 | Tpm3 | 0.27 | 7.70E-14 | 1.65E-09 | 0.99 | 0.94 | 0.04 | 1.05 |
| 4 | Homer1 | 0.7 | 7.97E-14 | 1.70E-09 | 0.45 | 0.27 | 0.18 | 1.69 |
| 4 | Mrps18b | 0.53 | 7.97E-14 | 1.70E-09 | 0.68 | 0.48 | 0.21 | 1.43 |
| 4 | Znrd2 | 0.38 | 8.17E-14 | 1.74E-09 | 0.89 | 0.76 | 0.13 | 1.17 |
| 4 | Rtkn2 | 1.1 | 8.25E-14 | 1.76E-09 | 0.23 | 0.11 | 0.13 | 2.17 |
| 4 | Akap1 | 0.72 | 8.98E-14 | 1.92E-09 | 0.4 | 0.22 | 0.17 | 1.77 |
| 4 | Skp2 | 0.52 | 1.01E-13 | 2.16E-09 | 0.77 | 0.57 | 0.2 | 1.36 |
| 4 | Rundc3a | 0.47 | 1.03E-13 | 2.20E-09 | 0.2 | 0.08 | 0.12 | 2.43 |
| 4 | Pcyt1a | 0.8 | 1.07E-13 | 2.28E-09 | 0.32 | 0.17 | 0.16 | 1.93 |
| 4 | Arl2bp | 0.4 | 1.07E-13 | 2.28E-09 | 0.86 | 0.68 | 0.18 | 1.27 |
| 4 | Mycbp | 0.59 | 1.12E-13 | 2.39E-09 | 0.67 | 0.46 | 0.21 | 1.47 |
| 4 | Mrps26 | 0.46 | 1.15E-13 | 2.46E-09 | 0.78 | 0.59 | 0.19 | 1.32 |
| 4 | Rcbtb1 | 0.62 | 1.19E-13 | 2.53E-09 | 0.59 | 0.4 | 0.2 | 1.5 |
| 4 | Tpx2 | 0.42 | 1.22E-13 | 2.61E-09 | 0.76 | 0.55 | 0.21 | 1.39 |
| 4 | Tmem107 | 0.66 | 1.32E-13 | 2.82E-09 | 0.5 | 0.31 | 0.2 | 1.64 |
| 4 | Rxylt1 | 0.58 | 1.38E-13 | 2.94E-09 | 0.6 | 0.38 | 0.21 | 1.55 |
| 4 | Lama1 | 0.97 | 1.46E-13 | 3.12E-09 | 0.21 | 0.09 | 0.12 | 2.28 |
| 4 | Tex30 | 0.58 | 1.49E-13 | 3.18E-09 | 0.62 | 0.41 | 0.2 | 1.49 |
| 4 | Mcm2 | 0.5 | 1.49E-13 | 3.19E-09 | 0.75 | 0.54 | 0.21 | 1.38 |
| 4 | Alpl | 0.7 | 1.50E-13 | 3.21E-09 | 0.31 | 0.16 | 0.15 | 1.96 |
| 4 | Grhl1 | 1.15 | 1.50E-13 | 3.21E-09 | 0.19 | 0.08 | 0.11 | 2.43 |
| 4 | Traf7 | 0.5 | 1.65E-13 | 3.52E-09 | 0.72 | 0.52 | 0.2 | 1.39 |
| 4 | Sdk1 | 0.67 | 1.65E-13 | 3.53E-09 | 0.2 | 0.08 | 0.12 | 2.42 |
| 4 | Timm23 | 0.31 | 1.65E-13 | 3.53E-09 | 0.96 | 0.89 | 0.07 | 1.08 |
| 4 | Hgh1 | 0.8 | 1.68E-13 | 3.59E-09 | 0.29 | 0.15 | 0.15 | 2 |
| 4 | Kif22 | 0.49 | 1.85E-13 | 3.96E-09 | 0.61 | 0.4 | 0.21 | 1.53 |
| 4 | Gm15417 | 1.1 | 1.97E-13 | 4.22E-09 | 0.17 | 0.06 | 0.1 | 2.59 |
| 4 | Plekhj1 | 0.38 | 2.03E-13 | 4.34E-09 | 0.88 | 0.75 | 0.13 | 1.18 |
| 4 | Rpn1 | 0.45 | 2.05E-13 | 4.37E-09 | 0.87 | 0.72 | 0.16 | 1.22 |
| 4 | Dnajc21 | 0.44 | 2.14E-13 | 4.58E-09 | 0.8 | 0.64 | 0.16 | 1.25 |
| 4 | Cenpu | 0.67 | 2.21E-13 | 4.71E-09 | 0.42 | 0.24 | 0.18 | 1.73 |
| 4 | Cep89 | 0.73 | 2.21E-13 | 4.72E-09 | 0.51 | 0.34 | 0.17 | 1.51 |
| 4 | Jpt2 | 0.55 | 2.26E-13 | 4.82E-09 | 0.68 | 0.48 | 0.21 | 1.43 |
| 4 | Mrps25 | 0.44 | 2.29E-13 | 4.89E-09 | 0.85 | 0.7 | 0.15 | 1.21 |
| 4 | Hirip3 | 0.48 | 2.33E-13 | 4.97E-09 | 0.81 | 0.61 | 0.2 | 1.32 |
| 4 | Usp39 | 0.5 | 2.54E-13 | 5.43E-09 | 0.72 | 0.53 | 0.18 | 1.34 |
| 4 | Polr1b | 0.94 | 2.81E-13 | 6.00E-09 | 0.3 | 0.16 | 0.15 | 1.93 |
| 4 | Maz | 0.34 | 3.15E-13 | 6.73E-09 | 0.92 | 0.79 | 0.12 | 1.16 |
| 4 | Ltv1 | 0.52 | 3.25E-13 | 6.95E-09 | 0.66 | 0.47 | 0.19 | 1.4 |
| 4 | Parp1 | 0.32 | 3.29E-13 | 7.03E-09 | 0.95 | 0.87 | 0.08 | 1.09 |

**Supplementary Table 3. Marker genes for the original clusters of integrated clustering of E11.5 control and NotchGOF λ-junction ectoderm cells.**

| | | | | | | | | |
|---|---|---|---|---|---|---|---|---|
| 4 | Peg12 | 1.1 | 3.68E-13 | 7.86E-09 | 0.17 | 0.07 | 0.1 | 2.55 |
| 4 | Ube2c | 0.3 | 3.71E-13 | 7.93E-09 | 0.84 | 0.65 | 0.19 | 1.3 |
| 4 | Pmvk | 0.55 | 3.94E-13 | 8.41E-09 | 0.69 | 0.51 | 0.18 | 1.35 |
| 4 | Dph3 | 0.41 | 4.09E-13 | 8.75E-09 | 0.87 | 0.74 | 0.13 | 1.18 |
| 4 | Mad2l1 | 0.44 | 4.15E-13 | 8.88E-09 | 0.74 | 0.52 | 0.22 | 1.41 |
| 4 | Prdm16 | 1.01 | 4.34E-13 | 9.28E-09 | 0.13 | 0.05 | 0.09 | 2.89 |
| 4 | Tmem251 | 0.77 | 4.42E-13 | 9.44E-09 | 0.4 | 0.24 | 0.16 | 1.68 |
| 4 | Rab30 | 1.25 | 4.54E-13 | 9.70E-09 | 0.14 | 0.05 | 0.09 | 2.76 |
| 4 | Gpam | 0.98 | 4.72E-13 | 1.01E-08 | 0.19 | 0.08 | 0.11 | 2.37 |
| 4 | Tekt1 | 0.96 | 4.96E-13 | 1.06E-08 | 0.22 | 0.1 | 0.12 | 2.21 |
| 4 | Las1l | 0.52 | 5.23E-13 | 1.12E-08 | 0.69 | 0.5 | 0.2 | 1.4 |
| 4 | Nup93 | 0.58 | 5.24E-13 | 1.12E-08 | 0.62 | 0.41 | 0.21 | 1.5 |
| 4 | Tmpo | 0.28 | 5.32E-13 | 1.14E-08 | 0.98 | 0.9 | 0.08 | 1.09 |
| 4 | Dnajc19 | 0.36 | 5.35E-13 | 1.14E-08 | 0.9 | 0.76 | 0.13 | 1.18 |
| 4 | Ska3 | 0.63 | 5.50E-13 | 1.17E-08 | 0.41 | 0.24 | 0.17 | 1.73 |
| 4 | Sfmbt1 | 0.8 | 5.54E-13 | 1.18E-08 | 0.31 | 0.17 | 0.15 | 1.88 |
| 4 | Wars | 0.6 | 5.60E-13 | 1.20E-08 | 0.58 | 0.37 | 0.2 | 1.54 |
| 4 | Nrm | 0.6 | 5.94E-13 | 1.27E-08 | 0.56 | 0.36 | 0.2 | 1.54 |
| 4 | Sav1 | 0.7 | 6.00E-13 | 1.28E-08 | 0.49 | 0.31 | 0.18 | 1.6 |
| 4 | Rfc5 | 0.43 | 6.20E-13 | 1.32E-08 | 0.76 | 0.57 | 0.2 | 1.34 |
| 4 | Atic | 0.43 | 6.24E-13 | 1.33E-08 | 0.84 | 0.67 | 0.16 | 1.24 |
| 4 | Eif4ebp1 | 0.45 | 6.31E-13 | 1.35E-08 | 0.8 | 0.61 | 0.19 | 1.31 |
| 4 | Rars | 0.37 | 6.82E-13 | 1.46E-08 | 0.86 | 0.71 | 0.15 | 1.22 |
| 4 | Cd82 | 0.97 | 6.99E-13 | 1.49E-08 | 0.25 | 0.12 | 0.13 | 2.07 |
| 4 | Cdk1 | 0.33 | 7.40E-13 | 1.58E-08 | 0.8 | 0.58 | 0.22 | 1.38 |
| 4 | Pola1 | 0.45 | 7.81E-13 | 1.67E-08 | 0.64 | 0.42 | 0.21 | 1.5 |
| 4 | Aldh18a1 | 0.66 | 8.84E-13 | 1.89E-08 | 0.41 | 0.24 | 0.17 | 1.7 |
| 4 | Noa1 | 0.6 | 9.32E-13 | 1.99E-08 | 0.49 | 0.3 | 0.18 | 1.61 |
| 4 | Ola1 | 0.36 | 9.41E-13 | 2.01E-08 | 0.92 | 0.82 | 0.1 | 1.12 |
| 4 | Sh3gl2 | 0.96 | 9.42E-13 | 2.01E-08 | 0.25 | 0.12 | 0.13 | 2.08 |
| 4 | Pkia | 0.72 | 9.58E-13 | 2.05E-08 | 0.34 | 0.19 | 0.15 | 1.81 |
| 4 | Hist1h3c | 0.56 | 9.66E-13 | 2.06E-08 | 0.65 | 0.45 | 0.2 | 1.44 |
| 4 | Cby1 | 0.45 | 9.80E-13 | 2.09E-08 | 0.65 | 0.44 | 0.21 | 1.47 |
| 4 | Rbmx | 0.37 | 9.94E-13 | 2.12E-08 | 0.84 | 0.69 | 0.16 | 1.23 |
| 4 | Cetn2 | 0.47 | 1.03E-12 | 2.19E-08 | 0.75 | 0.58 | 0.17 | 1.28 |
| 4 | Eif1ad | 0.43 | 1.06E-12 | 2.26E-08 | 0.79 | 0.62 | 0.17 | 1.27 |
| 4 | Kif23 | 0.44 | 1.15E-12 | 2.47E-08 | 0.61 | 0.4 | 0.21 | 1.52 |
| 4 | Fbn2 | 0.3 | 1.19E-12 | 2.54E-08 | 0.68 | 0.45 | 0.23 | 1.52 |
| 4 | Chd7 | 0.32 | 1.23E-12 | 2.62E-08 | 0.89 | 0.78 | 0.11 | 1.14 |
| 4 | Minpp1 | 0.5 | 1.28E-12 | 2.74E-08 | 0.68 | 0.48 | 0.2 | 1.42 |
| 4 | Hist1h4d | 0.58 | 1.35E-12 | 2.88E-08 | 0.57 | 0.37 | 0.19 | 1.52 |
| 4 | Cep135 | 0.59 | 1.48E-12 | 3.16E-08 | 0.44 | 0.26 | 0.18 | 1.69 |
| 4 | Gm42372 | 1.24 | 1.52E-12 | 3.24E-08 | 0.11 | 0.04 | 0.08 | 3.14 |
| 4 | P4hb | 0.32 | 1.55E-12 | 3.31E-08 | 0.97 | 0.89 | 0.08 | 1.09 |
| 4 | Lmnb1 | 0.26 | 1.57E-12 | 3.36E-08 | 0.97 | 0.93 | 0.05 | 1.05 |
| 4 | Ppih | 0.35 | 1.63E-12 | 3.48E-08 | 0.91 | 0.73 | 0.18 | 1.24 |
| 4 | Znrd1 | 0.42 | 1.76E-12 | 3.76E-08 | 0.86 | 0.72 | 0.14 | 1.19 |
| 4 | Donson | 0.75 | 1.76E-12 | 3.77E-08 | 0.36 | 0.21 | 0.16 | 1.76 |
| 4 | Scn8a | 1.08 | 1.78E-12 | 3.79E-08 | 0.16 | 0.07 | 0.1 | 2.48 |
| 4 | Bbip1 | 0.43 | 1.87E-12 | 4.00E-08 | 0.8 | 0.59 | 0.21 | 1.35 |
| 4 | Nqo2 | 0.97 | 2.06E-12 | 4.39E-08 | 0.18 | 0.07 | 0.1 | 2.43 |
| 4 | Acsl4 | 0.94 | 2.10E-12 | 4.49E-08 | 0.24 | 0.12 | 0.12 | 2.04 |
| 4 | Fam98a | 0.6 | 2.15E-12 | 4.58E-08 | 0.51 | 0.33 | 0.19 | 1.57 |
| 4 | Fgfbp3 | 0.95 | 2.21E-12 | 4.72E-08 | 0.24 | 0.12 | 0.12 | 2.04 |
| 4 | Samhd1 | 0.53 | 2.22E-12 | 4.74E-08 | 0.59 | 0.39 | 0.2 | 1.51 |
| 4 | Cdc20 | 0.38 | 2.31E-12 | 4.94E-08 | 0.7 | 0.51 | 0.19 | 1.38 |
| 4 | Pomt1 | 1.07 | 2.42E-12 | 5.17E-08 | 0.22 | 0.11 | 0.12 | 2.11 |

**Supplementary Table 3. Marker genes for the original clusters of integrated clustering of E11.5 control and NotchGOF λ-junction ectoderm cells.**

| | | | | | | | | |
|---|---|---|---|---|---|---|---|---|
| 4 | Eprs | 0.29 | 2.51E-12 | 5.36E-08 | 0.97 | 0.91 | 0.06 | 1.06 |
| 4 | E2f5 | 0.52 | 2.74E-12 | 5.86E-08 | 0.61 | 0.41 | 0.2 | 1.49 |
| 4 | Hist1h2ae | 0.42 | 3.00E-12 | 6.41E-08 | 0.77 | 0.58 | 0.19 | 1.32 |
| 4 | Kbtbd11 | 1.47 | 3.01E-12 | 6.44E-08 | 0.12 | 0.04 | 0.08 | 2.88 |
| 4 | Tor2a | 0.54 | 3.14E-12 | 6.70E-08 | 0.58 | 0.4 | 0.19 | 1.47 |
| 4 | Top2a | 0.28 | 3.20E-12 | 6.85E-08 | 0.91 | 0.67 | 0.23 | 1.35 |
| 4 | Tctex1d2 | 0.64 | 3.23E-12 | 6.91E-08 | 0.57 | 0.39 | 0.18 | 1.47 |
| 4 | Vill | 0.88 | 3.56E-12 | 7.60E-08 | 0.28 | 0.15 | 0.14 | 1.93 |
| 4 | Ccdc30 | 0.85 | 3.66E-12 | 7.82E-08 | 0.2 | 0.09 | 0.11 | 2.26 |
| 4 | Mogs | 0.56 | 3.73E-12 | 7.97E-08 | 0.54 | 0.36 | 0.19 | 1.52 |
| 4 | Rfc1 | 0.34 | 3.86E-12 | 8.24E-08 | 0.94 | 0.83 | 0.11 | 1.13 |
| 4 | Srrt | 0.34 | 3.98E-12 | 8.50E-08 | 0.9 | 0.76 | 0.14 | 1.19 |
| 4 | Usp7 | 0.41 | 4.53E-12 | 9.68E-08 | 0.83 | 0.66 | 0.16 | 1.24 |
| 4 | Polr1e | 0.62 | 4.55E-12 | 9.73E-08 | 0.41 | 0.24 | 0.17 | 1.68 |
| 4 | Mrpl51 | 0.29 | 4.59E-12 | 9.80E-08 | 0.94 | 0.83 | 0.11 | 1.13 |
| 4 | Bcl7c | 0.36 | 4.99E-12 | 1.07E-07 | 0.9 | 0.77 | 0.13 | 1.16 |
| 4 | Taldo1 | 0.27 | 5.31E-12 | 1.13E-07 | 0.95 | 0.87 | 0.08 | 1.1 |
| 4 | Ifrd2 | 0.65 | 5.54E-12 | 1.18E-07 | 0.45 | 0.28 | 0.17 | 1.62 |
| 4 | Dipk1b | 0.46 | 5.62E-12 | 1.20E-07 | 0.69 | 0.48 | 0.21 | 1.44 |
| 4 | Wdr46 | 0.51 | 5.63E-12 | 1.20E-07 | 0.65 | 0.45 | 0.2 | 1.44 |
| 4 | Ipo4 | 0.66 | 5.73E-12 | 1.22E-07 | 0.5 | 0.32 | 0.18 | 1.57 |
| 4 | Mat2a | 0.31 | 6.13E-12 | 1.31E-07 | 0.96 | 0.89 | 0.07 | 1.08 |
| 4 | Zswim5 | 1.01 | 6.25E-12 | 1.34E-07 | 0.19 | 0.09 | 0.11 | 2.26 |
| 4 | Dtx4 | 1.34 | 6.78E-12 | 1.45E-07 | 0.14 | 0.05 | 0.09 | 2.59 |
| 4 | Slc5a6 | 1.12 | 6.78E-12 | 1.45E-07 | 0.15 | 0.06 | 0.09 | 2.59 |
| 4 | Thyn1 | 0.39 | 6.87E-12 | 1.47E-07 | 0.82 | 0.64 | 0.17 | 1.27 |
| 4 | Wipf3 | 0.61 | 7.09E-12 | 1.51E-07 | 0.37 | 0.2 | 0.16 | 1.8 |
| 4 | Ahsa1 | 0.37 | 7.24E-12 | 1.55E-07 | 0.86 | 0.71 | 0.15 | 1.21 |
| 4 | Slc2a1 | 0.4 | 7.28E-12 | 1.56E-07 | 0.81 | 0.68 | 0.13 | 1.2 |
| 4 | Gramd1b | 1.78 | 7.88E-12 | 1.68E-07 | 0.1 | 0.03 | 0.07 | 3.12 |
| 4 | mt-Nd6 | 0.5 | 8.32E-12 | 1.78E-07 | 0.67 | 0.49 | 0.18 | 1.36 |
| 4 | Exosc7 | 0.38 | 8.49E-12 | 1.81E-07 | 0.87 | 0.7 | 0.17 | 1.25 |
| 4 | Msn | 0.66 | 8.52E-12 | 1.82E-07 | 0.27 | 0.14 | 0.13 | 1.97 |
| 4 | Hspa4 | 0.26 | 8.79E-12 | 1.88E-07 | 0.98 | 0.94 | 0.04 | 1.04 |
| 4 | Syncrip | 0.26 | 9.01E-12 | 1.93E-07 | 0.97 | 0.92 | 0.05 | 1.06 |
| 4 | Grhpr | 0.53 | 9.68E-12 | 2.07E-07 | 0.66 | 0.48 | 0.18 | 1.37 |
| 4 | Rad54l | 0.6 | 1.04E-11 | 2.22E-07 | 0.46 | 0.28 | 0.18 | 1.63 |
| 4 | Pdrg1 | 0.29 | 1.05E-11 | 2.25E-07 | 0.75 | 0.56 | 0.18 | 1.33 |
| 4 | Hspe1-rs1 | 0.81 | 1.06E-11 | 2.25E-07 | 0.25 | 0.12 | 0.12 | 2.02 |
| 4 | Sac3d1 | 0.51 | 1.06E-11 | 2.26E-07 | 0.64 | 0.46 | 0.18 | 1.4 |
| 4 | Snx24 | 0.62 | 1.07E-11 | 2.28E-07 | 0.48 | 0.3 | 0.17 | 1.57 |
| 4 | Cenpp | 0.63 | 1.09E-11 | 2.32E-07 | 0.49 | 0.32 | 0.17 | 1.54 |
| 4 | Ivns1abp | 0.37 | 1.11E-11 | 2.36E-07 | 0.86 | 0.7 | 0.16 | 1.23 |
| 4 | Arg2 | 0.81 | 1.11E-11 | 2.38E-07 | 0.31 | 0.17 | 0.14 | 1.84 |
| 4 | Ect2 | 0.45 | 1.13E-11 | 2.41E-07 | 0.6 | 0.4 | 0.2 | 1.5 |
| 4 | Strap | 0.29 | 1.13E-11 | 2.42E-07 | 0.95 | 0.86 | 0.09 | 1.1 |
| 4 | Nusap1 | 0.38 | 1.18E-11 | 2.52E-07 | 0.65 | 0.45 | 0.2 | 1.45 |
| 4 | Atp1b3 | 0.46 | 1.20E-11 | 2.57E-07 | 0.75 | 0.59 | 0.17 | 1.28 |
| 4 | Trip13 | 0.43 | 1.23E-11 | 2.63E-07 | 0.66 | 0.46 | 0.2 | 1.44 |
| 4 | Etf1 | 0.33 | 1.25E-11 | 2.66E-07 | 0.91 | 0.77 | 0.14 | 1.18 |
| 4 | Klhl7 | 0.46 | 1.25E-11 | 2.66E-07 | 0.74 | 0.57 | 0.17 | 1.31 |
| 4 | Reep1 | 1.34 | 1.29E-11 | 2.75E-07 | 0.13 | 0.05 | 0.08 | 2.74 |
| 4 | Atp1a1 | 0.36 | 1.31E-11 | 2.79E-07 | 0.84 | 0.67 | 0.17 | 1.26 |
| 4 | Dnaaf2 | 0.52 | 1.35E-11 | 2.89E-07 | 0.67 | 0.49 | 0.19 | 1.38 |
| 4 | Ska1 | 0.44 | 1.38E-11 | 2.95E-07 | 0.52 | 0.33 | 0.19 | 1.59 |
| 4 | Cnih1 | 0.35 | 1.40E-11 | 2.99E-07 | 0.92 | 0.8 | 0.12 | 1.15 |
| 4 | Tmem158 | 0.75 | 1.47E-11 | 3.15E-07 | 0.27 | 0.14 | 0.13 | 1.94 |

**Supplementary Table 3. Marker genes for the original clusters of integrated clustering of E11.5 control and NotchGOF λ-junction ectoderm cells.**

| | | | | | | | | |
|---|---|---|---|---|---|---|---|---|
| 4 | Fkbp2 | 0.43 | 1.51E-11 | 3.23E-07 | 0.8 | 0.62 | 0.18 | 1.29 |
| 4 | Mirg | 0.95 | 1.54E-11 | 3.28E-07 | 0.16 | 0.07 | 0.1 | 2.41 |
| 4 | Nck2 | 0.63 | 1.61E-11 | 3.44E-07 | 0.42 | 0.25 | 0.17 | 1.67 |
| 4 | E2f4 | 0.46 | 1.63E-11 | 3.49E-07 | 0.72 | 0.55 | 0.17 | 1.31 |
| 4 | Mvd | 0.58 | 1.64E-11 | 3.50E-07 | 0.45 | 0.28 | 0.17 | 1.62 |
| 4 | Tmem147 | 0.42 | 1.65E-11 | 3.53E-07 | 0.83 | 0.67 | 0.15 | 1.23 |
| 4 | H2-Ke6 | 0.62 | 1.68E-11 | 3.59E-07 | 0.41 | 0.24 | 0.16 | 1.67 |
| 4 | Morn2 | 0.49 | 1.69E-11 | 3.62E-07 | 0.57 | 0.37 | 0.2 | 1.53 |
| 4 | Trmt61a | 0.75 | 1.71E-11 | 3.66E-07 | 0.29 | 0.16 | 0.14 | 1.86 |
| 4 | Lrrn1 | 1.44 | 1.74E-11 | 3.71E-07 | 0.12 | 0.04 | 0.08 | 2.77 |
| 4 | Wee1 | 0.51 | 1.82E-11 | 3.88E-07 | 0.59 | 0.41 | 0.18 | 1.43 |
| 4 | Snhg8 | 0.32 | 1.83E-11 | 3.92E-07 | 0.89 | 0.77 | 0.12 | 1.16 |
| 4 | Cmtm4 | 0.55 | 1.89E-11 | 4.03E-07 | 0.55 | 0.36 | 0.19 | 1.51 |
| 4 | Elovl2 | 0.87 | 1.93E-11 | 4.11E-07 | 0.22 | 0.11 | 0.12 | 2.06 |
| 4 | Psmd6 | 0.28 | 2.01E-11 | 4.29E-07 | 0.95 | 0.85 | 0.1 | 1.12 |
| 4 | Med22 | 0.65 | 2.02E-11 | 4.32E-07 | 0.43 | 0.27 | 0.16 | 1.59 |
| 4 | Cdca5 | 0.54 | 2.21E-11 | 4.72E-07 | 0.49 | 0.31 | 0.18 | 1.57 |
| 4 | Ercc6l | 0.55 | 2.27E-11 | 4.85E-07 | 0.46 | 0.29 | 0.17 | 1.6 |
| 4 | Ddx27 | 0.37 | 2.36E-11 | 5.03E-07 | 0.83 | 0.67 | 0.15 | 1.23 |
| 4 | Ggct | 0.75 | 2.36E-11 | 5.03E-07 | 0.27 | 0.14 | 0.13 | 1.96 |
| 4 | Mak16 | 0.44 | 2.39E-11 | 5.11E-07 | 0.79 | 0.64 | 0.15 | 1.24 |
| 4 | Cyp39a1 | 0.65 | 2.39E-11 | 5.11E-07 | 0.37 | 0.22 | 0.15 | 1.7 |
| 4 | Pno1 | 0.43 | 2.49E-11 | 5.32E-07 | 0.76 | 0.6 | 0.16 | 1.27 |
| 4 | Rerg | 1.3 | 2.50E-11 | 5.34E-07 | 0.12 | 0.04 | 0.08 | 2.79 |
| 4 | Fam217b | 1.43 | 2.50E-11 | 5.34E-07 | 0.1 | 0.03 | 0.07 | 3.03 |
| 4 | Got2 | 0.34 | 2.57E-11 | 5.48E-07 | 0.9 | 0.78 | 0.12 | 1.15 |
| 4 | Gatm | 1.04 | 2.65E-11 | 5.65E-07 | 0.15 | 0.06 | 0.09 | 2.44 |
| 4 | Prmt7 | 0.48 | 2.67E-11 | 5.69E-07 | 0.65 | 0.46 | 0.19 | 1.4 |
| 4 | Hdgfl3 | 0.4 | 2.86E-11 | 6.11E-07 | 0.71 | 0.52 | 0.19 | 1.36 |
| 4 | Abr | 0.68 | 2.88E-11 | 6.15E-07 | 0.35 | 0.2 | 0.15 | 1.73 |
| 4 | Slc29a2 | 0.99 | 2.91E-11 | 6.22E-07 | 0.21 | 0.1 | 0.11 | 2.08 |
| 4 | Tuba1c | 0.33 | 3.24E-11 | 6.92E-07 | 0.66 | 0.48 | 0.18 | 1.38 |
| 4 | Baz1b | 0.31 | 3.29E-11 | 7.03E-07 | 0.93 | 0.83 | 0.1 | 1.12 |
| 4 | Kdelr2 | 0.38 | 3.34E-11 | 7.14E-07 | 0.91 | 0.77 | 0.14 | 1.19 |
| 4 | Oma1 | 0.62 | 3.62E-11 | 7.74E-07 | 0.34 | 0.19 | 0.15 | 1.78 |
| 4 | Exosc3 | 0.4 | 3.69E-11 | 7.88E-07 | 0.85 | 0.68 | 0.17 | 1.24 |
| 4 | Rrs1 | 0.44 | 3.80E-11 | 8.12E-07 | 0.78 | 0.63 | 0.15 | 1.24 |
| 4 | Gm48226 | 0.86 | 3.89E-11 | 8.31E-07 | 0.33 | 0.18 | 0.14 | 1.78 |
| 4 | Ccdc86 | 0.44 | 3.94E-11 | 8.41E-07 | 0.75 | 0.55 | 0.19 | 1.35 |
| 4 | Asns | 0.63 | 3.98E-11 | 8.50E-07 | 0.43 | 0.26 | 0.16 | 1.62 |
| 4 | Lrp8 | 1.04 | 3.99E-11 | 8.53E-07 | 0.18 | 0.08 | 0.1 | 2.22 |
| 4 | Gjc1 | 0.37 | 4.15E-11 | 8.86E-07 | 0.87 | 0.72 | 0.15 | 1.21 |
| 4 | Prom1 | 0.5 | 4.15E-11 | 8.87E-07 | 0.61 | 0.44 | 0.17 | 1.39 |
| 4 | Hyou1 | 0.67 | 4.17E-11 | 8.92E-07 | 0.43 | 0.27 | 0.16 | 1.62 |
| 4 | Nsun2 | 0.36 | 4.20E-11 | 8.97E-07 | 0.84 | 0.68 | 0.16 | 1.24 |
| 4 | Cdc45 | 0.55 | 4.29E-11 | 9.17E-07 | 0.47 | 0.3 | 0.17 | 1.57 |
| 4 | Timm10b | 0.31 | 4.41E-11 | 9.42E-07 | 0.91 | 0.82 | 0.09 | 1.11 |
| 4 | Rnf141 | 0.67 | 4.43E-11 | 9.47E-07 | 0.26 | 0.13 | 0.13 | 1.97 |
| 4 | Timm10 | 0.45 | 4.51E-11 | 9.64E-07 | 0.73 | 0.57 | 0.17 | 1.3 |
| 4 | 9530077C05Rik | 1.19 | 4.55E-11 | 9.71E-07 | 0.11 | 0.04 | 0.07 | 2.89 |
| 4 | Gclc | 0.63 | 4.62E-11 | 9.87E-07 | 0.43 | 0.26 | 0.16 | 1.62 |
| 4 | 3110021N24Rik | 1.11 | 4.67E-11 | 9.98E-07 | 0.16 | 0.07 | 0.09 | 2.34 |
| 4 | Rccd1 | 0.79 | 4.71E-11 | 1.01E-06 | 0.26 | 0.14 | 0.12 | 1.92 |
| 4 | Mesd | 0.39 | 4.87E-11 | 1.04E-06 | 0.86 | 0.7 | 0.17 | 1.24 |
| 4 | Cenpw | 0.35 | 4.89E-11 | 1.04E-06 | 0.83 | 0.64 | 0.18 | 1.29 |
| 4 | Ruvbl2 | 0.41 | 4.89E-11 | 1.04E-06 | 0.81 | 0.65 | 0.16 | 1.24 |
| 4 | Tsr2 | 0.54 | 4.97E-11 | 1.06E-06 | 0.51 | 0.32 | 0.18 | 1.57 |

**Supplementary Table 3. Marker genes for the original clusters of integrated clustering of E11.5 control and NotchGOF λ-junction ectoderm cells.**

| | | | | | | | | |
|---|---|---|---|---|---|---|---|---|
| 4 | Nup210 | 0.7 | 5.07E-11 | 1.08E-06 | 0.31 | 0.18 | 0.14 | 1.79 |
| 4 | Smim11 | 0.34 | 5.09E-11 | 1.09E-06 | 0.85 | 0.7 | 0.15 | 1.22 |
| 4 | Tmem126a | 0.41 | 5.14E-11 | 1.10E-06 | 0.8 | 0.67 | 0.13 | 1.2 |
| 4 | Txndc11 | 0.72 | 5.32E-11 | 1.14E-06 | 0.33 | 0.19 | 0.15 | 1.78 |
| 4 | Nphp1 | 0.51 | 5.37E-11 | 1.15E-06 | 0.44 | 0.27 | 0.17 | 1.61 |
| 4 | Ppp2r5a | 0.51 | 5.58E-11 | 1.19E-06 | 0.47 | 0.29 | 0.18 | 1.6 |
| 4 | Wdfy1 | 0.45 | 6.03E-11 | 1.29E-06 | 0.57 | 0.39 | 0.18 | 1.47 |
| 4 | Zfp91 | 0.29 | 6.04E-11 | 1.29E-06 | 0.94 | 0.85 | 0.09 | 1.1 |
| 4 | Prpf19 | 0.32 | 6.36E-11 | 1.36E-06 | 0.89 | 0.77 | 0.12 | 1.16 |
| 4 | Gmps | 0.29 | 6.41E-11 | 1.37E-06 | 0.94 | 0.86 | 0.09 | 1.1 |
| 4 | Creb5 | 1.03 | 6.51E-11 | 1.39E-06 | 0.16 | 0.07 | 0.09 | 2.34 |
| 4 | Wdr12 | 0.44 | 6.76E-11 | 1.44E-06 | 0.71 | 0.53 | 0.18 | 1.34 |
| 4 | Dis3 | 0.44 | 7.22E-11 | 1.54E-06 | 0.66 | 0.48 | 0.18 | 1.37 |
| 4 | Klhdc4 | 0.49 | 7.51E-11 | 1.60E-06 | 0.48 | 0.3 | 0.18 | 1.6 |
| 4 | Hist1h2ab | 0.51 | 7.57E-11 | 1.62E-06 | 0.43 | 0.26 | 0.17 | 1.64 |
| 4 | Ddx18 | 0.31 | 7.68E-11 | 1.64E-06 | 0.92 | 0.82 | 0.1 | 1.12 |
| 4 | Mapt | 0.64 | 7.75E-11 | 1.65E-06 | 0.11 | 0.04 | 0.07 | 2.92 |
| 4 | Cse1l | 0.43 | 7.83E-11 | 1.67E-06 | 0.75 | 0.58 | 0.18 | 1.31 |
| 4 | Gins2 | 0.41 | 7.97E-11 | 1.70E-06 | 0.73 | 0.54 | 0.19 | 1.35 |
| 4 | Gars | 0.42 | 8.23E-11 | 1.76E-06 | 0.79 | 0.65 | 0.15 | 1.23 |
| 4 | Myh10 | 0.32 | 8.42E-11 | 1.80E-06 | 0.9 | 0.79 | 0.11 | 1.13 |
| 4 | Slc7a1 | 0.73 | 9.33E-11 | 1.99E-06 | 0.2 | 0.09 | 0.11 | 2.14 |
| 4 | Kmt5a | 0.33 | 9.43E-11 | 2.01E-06 | 0.93 | 0.82 | 0.11 | 1.14 |
| 4 | Pitpnc1 | 0.61 | 9.57E-11 | 2.04E-06 | 0.56 | 0.39 | 0.16 | 1.41 |
| 4 | Bri3bp | 0.43 | 1.00E-10 | 2.14E-06 | 0.71 | 0.54 | 0.17 | 1.31 |
| 4 | Nup37 | 0.43 | 1.01E-10 | 2.16E-06 | 0.67 | 0.49 | 0.18 | 1.37 |
| 4 | Fkbp14 | 0.66 | 1.03E-10 | 2.20E-06 | 0.25 | 0.13 | 0.12 | 1.98 |
| 4 | Tmem237 | 0.51 | 1.06E-10 | 2.27E-06 | 0.58 | 0.41 | 0.17 | 1.42 |
| 4 | Shmt1 | 0.48 | 1.08E-10 | 2.30E-06 | 0.52 | 0.33 | 0.19 | 1.56 |
| 4 | Rexo2 | 0.43 | 1.09E-10 | 2.32E-06 | 0.71 | 0.56 | 0.15 | 1.27 |
| 4 | Hvcn1 | 1.47 | 1.09E-10 | 2.33E-06 | 0.1 | 0.03 | 0.07 | 2.94 |
| 4 | Nudt9 | 0.44 | 1.11E-10 | 2.36E-06 | 0.59 | 0.41 | 0.19 | 1.45 |
| 4 | Rrp15 | 0.36 | 1.12E-10 | 2.39E-06 | 0.82 | 0.65 | 0.16 | 1.25 |
| 4 | Gsap | 0.89 | 1.13E-10 | 2.41E-06 | 0.21 | 0.1 | 0.11 | 2.07 |
| 4 | Lsm12 | 0.32 | 1.13E-10 | 2.42E-06 | 0.93 | 0.82 | 0.11 | 1.14 |
| 4 | Nudcd2 | 0.29 | 1.14E-10 | 2.43E-06 | 0.92 | 0.85 | 0.07 | 1.08 |
| 4 | Tex9 | 0.48 | 1.14E-10 | 2.43E-06 | 0.61 | 0.42 | 0.19 | 1.45 |
| 4 | Ccp110 | 0.34 | 1.18E-10 | 2.52E-06 | 0.51 | 0.34 | 0.17 | 1.51 |
| 4 | Timm50 | 0.35 | 1.22E-10 | 2.61E-06 | 0.84 | 0.69 | 0.16 | 1.23 |
| 4 | Ormdl2 | 0.46 | 1.27E-10 | 2.72E-06 | 0.77 | 0.6 | 0.17 | 1.28 |
| 4 | Urb2 | 0.95 | 1.31E-10 | 2.81E-06 | 0.18 | 0.08 | 0.1 | 2.19 |
| 4 | Pop1 | 0.75 | 1.33E-10 | 2.85E-06 | 0.34 | 0.2 | 0.14 | 1.7 |
| 4 | Fbxo5 | 0.34 | 1.34E-10 | 2.87E-06 | 0.69 | 0.49 | 0.2 | 1.41 |
| 4 | Noc2l | 0.4 | 1.35E-10 | 2.88E-06 | 0.81 | 0.66 | 0.14 | 1.22 |
| 4 | Eif1a | 0.37 | 1.37E-10 | 2.92E-06 | 0.83 | 0.68 | 0.15 | 1.23 |
| 4 | Arpc5l | 0.3 | 1.47E-10 | 3.14E-06 | 0.9 | 0.79 | 0.12 | 1.15 |
| 4 | Arfip2 | 0.47 | 1.51E-10 | 3.22E-06 | 0.68 | 0.51 | 0.17 | 1.34 |
| 4 | Larp1 | 0.39 | 1.56E-10 | 3.33E-06 | 0.77 | 0.61 | 0.16 | 1.27 |
| 4 | Cep83 | 0.49 | 1.57E-10 | 3.36E-06 | 0.63 | 0.47 | 0.17 | 1.35 |
| 4 | Ust | 0.88 | 1.60E-10 | 3.42E-06 | 0.24 | 0.12 | 0.12 | 1.93 |
| 4 | Mybl2 | 0.84 | 1.61E-10 | 3.44E-06 | 0.33 | 0.2 | 0.13 | 1.67 |
| 4 | Hook1 | 0.41 | 1.63E-10 | 3.49E-06 | 0.73 | 0.55 | 0.18 | 1.33 |
| 4 | Ncapg | 0.48 | 1.75E-10 | 3.73E-06 | 0.58 | 0.41 | 0.17 | 1.43 |
| 4 | S1pr1 | 1.33 | 1.77E-10 | 3.78E-06 | 0.12 | 0.04 | 0.07 | 2.66 |
| 4 | Psmb10 | 0.62 | 1.78E-10 | 3.81E-06 | 0.4 | 0.23 | 0.16 | 1.7 |
| 4 | Tomm5 | 0.27 | 1.78E-10 | 3.81E-06 | 0.94 | 0.9 | 0.04 | 1.05 |
| 4 | B4galt4 | 0.54 | 1.80E-10 | 3.85E-06 | 0.41 | 0.25 | 0.16 | 1.66 |

**Supplementary Table 3. Marker genes for the original clusters of integrated clustering of E11.5 control and NotchGOF λ-junction ectoderm cells.**

| | | | | | | | | |
|---|---|---|---|---|---|---|---|---|
| 4 | Gas6 | 1.32 | 1.93E-10 | 4.12E-06 | 0.11 | 0.04 | 0.07 | 2.84 |
| 4 | Utp18 | 0.46 | 1.94E-10 | 4.15E-06 | 0.71 | 0.53 | 0.18 | 1.34 |
| 4 | Kif24 | 1.08 | 2.02E-10 | 4.32E-06 | 0.18 | 0.08 | 0.1 | 2.17 |
| 4 | Orc6 | 0.32 | 2.04E-10 | 4.36E-06 | 0.87 | 0.72 | 0.16 | 1.22 |
| 4 | Kti12 | 0.55 | 2.07E-10 | 4.41E-06 | 0.58 | 0.42 | 0.16 | 1.39 |
| 4 | Tmem120a | 0.55 | 2.19E-10 | 4.68E-06 | 0.51 | 0.34 | 0.17 | 1.5 |
| 4 | Nek6 | 0.6 | 2.36E-10 | 5.03E-06 | 0.45 | 0.3 | 0.16 | 1.52 |
| 4 | Iscu | 0.41 | 2.52E-10 | 5.38E-06 | 0.77 | 0.58 | 0.19 | 1.33 |
| 4 | Rpa1 | 0.55 | 2.66E-10 | 5.68E-06 | 0.52 | 0.35 | 0.16 | 1.46 |
| 4 | Wdr18 | 0.36 | 2.77E-10 | 5.92E-06 | 0.83 | 0.67 | 0.16 | 1.24 |
| 4 | Cfap410 | 0.73 | 2.78E-10 | 5.93E-06 | 0.35 | 0.21 | 0.14 | 1.66 |
| 4 | Ift22 | 0.47 | 2.85E-10 | 6.09E-06 | 0.56 | 0.37 | 0.19 | 1.51 |
| 4 | Snhg12 | 0.5 | 2.91E-10 | 6.22E-06 | 0.61 | 0.43 | 0.18 | 1.42 |
| 4 | Rnf165 | 0.41 | 3.04E-10 | 6.49E-06 | 0.43 | 0.26 | 0.17 | 1.66 |
| 4 | Ccdc85c | 0.74 | 3.10E-10 | 6.63E-06 | 0.33 | 0.19 | 0.14 | 1.7 |
| 4 | Naa10 | 0.37 | 3.23E-10 | 6.90E-06 | 0.84 | 0.69 | 0.16 | 1.23 |
| 4 | Hist1h2bh | 1.02 | 3.47E-10 | 7.42E-06 | 0.16 | 0.07 | 0.09 | 2.27 |
| 4 | Dlgap5 | 0.45 | 3.51E-10 | 7.51E-06 | 0.59 | 0.42 | 0.17 | 1.4 |
| 4 | Rasgrp2 | 1.28 | 3.55E-10 | 7.58E-06 | 0.11 | 0.04 | 0.07 | 2.65 |
| 4 | Sel1l | 0.55 | 3.70E-10 | 7.91E-06 | 0.57 | 0.4 | 0.18 | 1.44 |
| 4 | Misp | 1.14 | 3.82E-10 | 8.16E-06 | 0.15 | 0.06 | 0.08 | 2.33 |
| 4 | Rasl11a | 0.6 | 3.96E-10 | 8.46E-06 | 0.28 | 0.15 | 0.13 | 1.84 |
| 4 | Tsr1 | 0.49 | 4.45E-10 | 9.51E-06 | 0.53 | 0.37 | 0.16 | 1.43 |
| 4 | Ncbp1 | 0.39 | 4.57E-10 | 9.76E-06 | 0.75 | 0.56 | 0.19 | 1.34 |
| 4 | Zbtb10 | 0.67 | 4.65E-10 | 9.94E-06 | 0.29 | 0.16 | 0.13 | 1.82 |
| 4 | Dhx29 | 0.48 | 4.66E-10 | 9.95E-06 | 0.58 | 0.41 | 0.17 | 1.41 |
| 4 | Hist1h1a | 0.49 | 4.80E-10 | 1.03E-05 | 0.49 | 0.33 | 0.16 | 1.5 |
| 4 | Chrna7 | 1.22 | 5.05E-10 | 1.08E-05 | 0.1 | 0.04 | 0.07 | 2.78 |
| 4 | Yars2 | 0.45 | 5.09E-10 | 1.09E-05 | 0.58 | 0.4 | 0.18 | 1.46 |
| 4 | Cpd | 0.42 | 5.11E-10 | 1.09E-05 | 0.53 | 0.36 | 0.17 | 1.48 |
| 4 | Nadk2 | 0.55 | 5.14E-10 | 1.10E-05 | 0.44 | 0.28 | 0.16 | 1.58 |
| 4 | Psmd1 | 0.28 | 5.17E-10 | 1.11E-05 | 0.92 | 0.82 | 0.1 | 1.12 |
| 4 | Mthfd1l | 0.75 | 5.29E-10 | 1.13E-05 | 0.25 | 0.13 | 0.12 | 1.88 |
| 4 | Plekhh1 | 0.77 | 5.74E-10 | 1.23E-05 | 0.26 | 0.14 | 0.12 | 1.84 |
| 4 | Sec11c | 0.62 | 5.79E-10 | 1.24E-05 | 0.29 | 0.16 | 0.13 | 1.79 |
| 4 | Rab25 | 0.29 | 5.88E-10 | 1.26E-05 | 0.81 | 0.61 | 0.2 | 1.33 |
| 4 | Gm26793 | 1.05 | 5.94E-10 | 1.27E-05 | 0.15 | 0.07 | 0.09 | 2.26 |
| 4 | Rsph9 | 0.65 | 6.13E-10 | 1.31E-05 | 0.22 | 0.11 | 0.11 | 2.03 |
| 4 | Sf3a1 | 0.38 | 6.17E-10 | 1.32E-05 | 0.82 | 0.68 | 0.14 | 1.21 |
| 4 | AU020206 | 0.68 | 6.36E-10 | 1.36E-05 | 0.34 | 0.2 | 0.14 | 1.69 |
| 4 | Hus1 | 0.58 | 6.45E-10 | 1.38E-05 | 0.36 | 0.21 | 0.14 | 1.67 |
| 4 | Trak1 | 0.77 | 6.54E-10 | 1.40E-05 | 0.31 | 0.18 | 0.13 | 1.73 |
| 4 | Lactb | 0.65 | 6.75E-10 | 1.44E-05 | 0.29 | 0.16 | 0.13 | 1.79 |
| 4 | Cfap36 | 0.3 | 6.87E-10 | 1.47E-05 | 0.67 | 0.48 | 0.19 | 1.4 |
| 4 | Nop2 | 0.58 | 6.88E-10 | 1.47E-05 | 0.48 | 0.32 | 0.16 | 1.51 |
| 4 | Myg1 | 0.41 | 6.92E-10 | 1.48E-05 | 0.75 | 0.58 | 0.17 | 1.29 |
| 4 | Gemin6 | 0.48 | 7.03E-10 | 1.50E-05 | 0.57 | 0.41 | 0.17 | 1.41 |
| 4 | Top1mt | 1.17 | 7.10E-10 | 1.52E-05 | 0.14 | 0.06 | 0.08 | 2.34 |
| 4 | 4930503L19Rik | 0.57 | 7.12E-10 | 1.52E-05 | 0.29 | 0.16 | 0.13 | 1.81 |
| 4 | Kif20b | 0.33 | 7.18E-10 | 1.53E-05 | 0.7 | 0.5 | 0.2 | 1.4 |
| 4 | Orc1 | 0.68 | 7.23E-10 | 1.54E-05 | 0.22 | 0.11 | 0.11 | 1.99 |
| 4 | Cdc6 | 0.61 | 7.28E-10 | 1.56E-05 | 0.34 | 0.2 | 0.14 | 1.67 |
| 4 | Apba2 | 0.58 | 7.33E-10 | 1.57E-05 | 0.24 | 0.13 | 0.12 | 1.9 |
| 4 | Cdk18 | 1.16 | 7.58E-10 | 1.62E-05 | 0.11 | 0.04 | 0.07 | 2.68 |
| 4 | Pck2 | 0.6 | 7.85E-10 | 1.68E-05 | 0.29 | 0.16 | 0.13 | 1.83 |
| 4 | Slc8a2 | 1.13 | 8.24E-10 | 1.76E-05 | 0.12 | 0.05 | 0.07 | 2.52 |
| 4 | Plpbp | 0.51 | 8.43E-10 | 1.80E-05 | 0.46 | 0.29 | 0.17 | 1.58 |

**Supplementary Table 3. Marker genes for the original clusters of integrated clustering of E11.5 control and NotchGOF λ-junction ectoderm cells.**

| | | | | | | | | |
|---|---|---|---|---|---|---|---|---|
| 4 | Nup50 | 0.38 | 8.74E-10 | 1.87E-05 | 0.72 | 0.56 | 0.16 | 1.29 |
| 4 | Sulf2 | 0.53 | 9.04E-10 | 1.93E-05 | 0.48 | 0.32 | 0.16 | 1.48 |
| 4 | Fanca | 1.07 | 9.13E-10 | 1.95E-05 | 0.19 | 0.09 | 0.1 | 2.06 |
| 4 | Glo1 | 0.26 | 9.16E-10 | 1.96E-05 | 0.92 | 0.82 | 0.1 | 1.13 |
| 4 | Wdhd1 | 0.58 | 9.23E-10 | 1.97E-05 | 0.38 | 0.23 | 0.15 | 1.63 |
| 4 | Ufsp1 | 0.77 | 9.50E-10 | 2.03E-05 | 0.21 | 0.1 | 0.11 | 2.04 |
| 4 | Impa2 | 0.59 | 9.58E-10 | 2.05E-05 | 0.34 | 0.2 | 0.14 | 1.7 |
| 4 | Pinx1 | 0.42 | 9.66E-10 | 2.06E-05 | 0.65 | 0.48 | 0.17 | 1.35 |
| 4 | Oard1 | 0.34 | 1.01E-09 | 2.15E-05 | 0.86 | 0.71 | 0.15 | 1.2 |
| 4 | Anapc15 | 0.44 | 1.02E-09 | 2.19E-05 | 0.7 | 0.52 | 0.18 | 1.35 |
| 4 | Ccnyl1 | 0.74 | 1.05E-09 | 2.24E-05 | 0.23 | 0.12 | 0.11 | 1.93 |
| 4 | Hpf1 | 0.34 | 1.05E-09 | 2.24E-05 | 0.92 | 0.78 | 0.14 | 1.17 |
| 4 | Bola1 | 0.36 | 1.09E-09 | 2.33E-05 | 0.79 | 0.66 | 0.13 | 1.19 |
| 4 | Pvt1 | 1.3 | 1.13E-09 | 2.41E-05 | 0.11 | 0.04 | 0.07 | 2.62 |
| 4 | Lmln | 1 | 1.19E-09 | 2.53E-05 | 0.19 | 0.1 | 0.1 | 2.02 |
| 4 | Vrk1 | 0.33 | 1.22E-09 | 2.61E-05 | 0.75 | 0.56 | 0.19 | 1.34 |
| 4 | Aig1 | 0.4 | 1.23E-09 | 2.63E-05 | 0.69 | 0.53 | 0.16 | 1.3 |
| 4 | Gphn | 0.59 | 1.24E-09 | 2.65E-05 | 0.34 | 0.21 | 0.14 | 1.67 |
| 4 | Ccdc25 | 0.42 | 1.25E-09 | 2.67E-05 | 0.76 | 0.6 | 0.17 | 1.28 |
| 4 | Rab9 | 0.41 | 1.26E-09 | 2.68E-05 | 0.74 | 0.55 | 0.19 | 1.33 |
| 4 | Zcchc2 | 0.78 | 1.26E-09 | 2.69E-05 | 0.22 | 0.11 | 0.11 | 1.95 |
| 4 | Incenp | 0.33 | 1.30E-09 | 2.78E-05 | 0.71 | 0.52 | 0.19 | 1.36 |
| 4 | Pigh | 0.49 | 1.34E-09 | 2.87E-05 | 0.44 | 0.28 | 0.16 | 1.57 |
| 4 | Hap1 | 0.71 | 1.36E-09 | 2.91E-05 | 0.32 | 0.19 | 0.13 | 1.7 |
| 4 | Pacsin3 | 0.51 | 1.41E-09 | 3.02E-05 | 0.46 | 0.3 | 0.16 | 1.53 |
| 4 | Lss | 0.88 | 1.42E-09 | 3.03E-05 | 0.26 | 0.15 | 0.11 | 1.77 |
| 4 | Suclg1 | 0.27 | 1.50E-09 | 3.20E-05 | 0.92 | 0.83 | 0.1 | 1.12 |
| 4 | Kpna2 | 0.56 | 1.51E-09 | 3.23E-05 | 0.51 | 0.36 | 0.16 | 1.44 |
| 4 | Ecpas | 0.51 | 1.53E-09 | 3.27E-05 | 0.56 | 0.4 | 0.17 | 1.42 |
| 4 | Endod1 | 0.81 | 1.54E-09 | 3.29E-05 | 0.27 | 0.15 | 0.12 | 1.79 |
| 4 | Zdhhc23 | 1.37 | 1.55E-09 | 3.31E-05 | 0.1 | 0.04 | 0.07 | 2.71 |
| 4 | Fam216a | 0.38 | 1.57E-09 | 3.35E-05 | 0.68 | 0.52 | 0.17 | 1.32 |
| 4 | Pdzd4 | 0.7 | 1.58E-09 | 3.37E-05 | 0.22 | 0.11 | 0.11 | 1.96 |
| 4 | Sys1 | 0.3 | 1.61E-09 | 3.44E-05 | 0.82 | 0.7 | 0.11 | 1.16 |
| 4 | A330009N23Rik | 0.85 | 1.61E-09 | 3.44E-05 | 0.1 | 0.04 | 0.07 | 2.71 |
| 4 | Ndc80 | 0.34 | 1.63E-09 | 3.48E-05 | 0.55 | 0.37 | 0.18 | 1.49 |
| 4 | Sox6 | 0.57 | 1.64E-09 | 3.50E-05 | 0.19 | 0.09 | 0.1 | 2.09 |
| 4 | Lrrc8b | 0.58 | 1.71E-09 | 3.66E-05 | 0.34 | 0.2 | 0.14 | 1.68 |
| 4 | Ssx2ip | 0.34 | 1.72E-09 | 3.68E-05 | 0.8 | 0.65 | 0.16 | 1.25 |
| 4 | Rapgef5 | 0.5 | 1.81E-09 | 3.86E-05 | 0.29 | 0.16 | 0.12 | 1.77 |
| 4 | Mpi | 0.67 | 1.85E-09 | 3.95E-05 | 0.29 | 0.16 | 0.12 | 1.75 |
| 4 | Atp6v0e2 | 0.47 | 2.05E-09 | 4.38E-05 | 0.27 | 0.15 | 0.12 | 1.83 |
| 4 | Blm | 0.35 | 2.06E-09 | 4.41E-05 | 0.63 | 0.45 | 0.19 | 1.41 |
| 4 | 3110082I17Rik | 0.51 | 2.18E-09 | 4.65E-05 | 0.54 | 0.37 | 0.17 | 1.47 |
| 4 | Hist1h3i | 0.96 | 2.18E-09 | 4.67E-05 | 0.25 | 0.14 | 0.11 | 1.77 |
| 4 | Ndufaf4 | 0.34 | 2.23E-09 | 4.76E-05 | 0.73 | 0.57 | 0.17 | 1.29 |
| 4 | Prmt5 | 0.35 | 2.29E-09 | 4.90E-05 | 0.77 | 0.6 | 0.17 | 1.28 |
| 4 | Gmpr | 0.74 | 2.32E-09 | 4.96E-05 | 0.31 | 0.18 | 0.13 | 1.71 |
| 4 | Med25 | 0.43 | 2.33E-09 | 4.97E-05 | 0.65 | 0.49 | 0.16 | 1.33 |
| 4 | Mrpl57 | 0.3 | 2.36E-09 | 5.04E-05 | 0.86 | 0.73 | 0.13 | 1.18 |
| 4 | Mrpl38 | 0.4 | 2.44E-09 | 5.21E-05 | 0.79 | 0.63 | 0.16 | 1.25 |
| 4 | P4ha3 | 0.68 | 2.47E-09 | 5.29E-05 | 0.27 | 0.15 | 0.12 | 1.78 |
| 4 | Sipa1 | 0.82 | 2.55E-09 | 5.44E-05 | 0.15 | 0.07 | 0.09 | 2.23 |
| 4 | Hectd2 | 1.01 | 2.62E-09 | 5.59E-05 | 0.13 | 0.05 | 0.07 | 2.42 |
| 4 | Ssr1 | 0.38 | 2.63E-09 | 5.61E-05 | 0.83 | 0.69 | 0.14 | 1.2 |
| 4 | Grsf1 | 0.39 | 2.64E-09 | 5.64E-05 | 0.76 | 0.62 | 0.14 | 1.22 |
| 4 | Arl6 | 0.46 | 2.66E-09 | 5.69E-05 | 0.53 | 0.37 | 0.16 | 1.42 |

**Supplementary Table 3. Marker genes for the original clusters of integrated clustering of E11.5 control and NotchGOF λ-junction ectoderm cells.**

| | | | | | | | | |
|---|---|---|---|---|---|---|---|---|
| 4 | Brca1 | 0.38 | 2.69E-09 | 5.74E-05 | 0.52 | 0.35 | 0.18 | 1.51 |
| 4 | Fzd8 | 0.29 | 2.84E-09 | 6.07E-05 | 0.46 | 0.29 | 0.17 | 1.59 |
| 4 | Bicd1 | 0.45 | 2.87E-09 | 6.14E-05 | 0.4 | 0.25 | 0.15 | 1.58 |
| 4 | Lbhd1 | 0.94 | 2.94E-09 | 6.28E-05 | 0.15 | 0.07 | 0.08 | 2.26 |
| 4 | Rbm38 | 0.56 | 2.97E-09 | 6.34E-05 | 0.38 | 0.24 | 0.14 | 1.59 |
| 4 | Rap2a | 0.55 | 3.09E-09 | 6.60E-05 | 0.44 | 0.29 | 0.15 | 1.52 |
| 4 | Nfe2l3 | 0.77 | 3.09E-09 | 6.61E-05 | 0.24 | 0.13 | 0.11 | 1.85 |
| 4 | Sgo1 | 0.32 | 3.10E-09 | 6.63E-05 | 0.56 | 0.38 | 0.18 | 1.46 |
| 4 | Serp1 | 0.33 | 3.23E-09 | 6.89E-05 | 0.85 | 0.73 | 0.13 | 1.18 |
| 4 | Abcf2 | 0.34 | 3.30E-09 | 7.06E-05 | 0.8 | 0.63 | 0.17 | 1.27 |
| 4 | Dus1l | 0.37 | 3.34E-09 | 7.14E-05 | 0.56 | 0.38 | 0.18 | 1.46 |
| 4 | Bub3 | 0.33 | 3.37E-09 | 7.20E-05 | 0.88 | 0.77 | 0.11 | 1.14 |
| 4 | Lmnb2 | 0.45 | 3.42E-09 | 7.30E-05 | 0.47 | 0.31 | 0.16 | 1.51 |
| 4 | Thap12 | 0.31 | 3.44E-09 | 7.35E-05 | 0.81 | 0.65 | 0.16 | 1.24 |
| 4 | Grb14 | 0.63 | 3.45E-09 | 7.37E-05 | 0.21 | 0.11 | 0.1 | 1.96 |
| 4 | Zmynd19 | 0.41 | 3.59E-09 | 7.66E-05 | 0.64 | 0.46 | 0.18 | 1.38 |
| 4 | Kif4 | 0.35 | 3.74E-09 | 7.99E-05 | 0.59 | 0.41 | 0.18 | 1.45 |
| 4 | Ubr7 | 0.5 | 3.79E-09 | 8.10E-05 | 0.58 | 0.41 | 0.17 | 1.41 |
| 4 | Yaf2 | 0.34 | 3.80E-09 | 8.11E-05 | 0.69 | 0.51 | 0.17 | 1.34 |
| 4 | Rpp30 | 0.41 | 3.87E-09 | 8.27E-05 | 0.67 | 0.51 | 0.17 | 1.33 |
| 4 | Cdon | 0.55 | 3.88E-09 | 8.29E-05 | 0.39 | 0.25 | 0.14 | 1.58 |
| 4 | Cisd1 | 0.31 | 3.94E-09 | 8.42E-05 | 0.9 | 0.78 | 0.12 | 1.16 |
| 4 | Mylk | 0.47 | 4.01E-09 | 8.57E-05 | 0.33 | 0.19 | 0.13 | 1.7 |
| 4 | Cds1 | 0.84 | 4.03E-09 | 8.61E-05 | 0.16 | 0.07 | 0.09 | 2.18 |
| 4 | Amd1 | 0.41 | 4.09E-09 | 8.74E-05 | 0.68 | 0.5 | 0.17 | 1.35 |
| 4 | Nin | 0.53 | 4.16E-09 | 8.88E-05 | 0.42 | 0.27 | 0.15 | 1.57 |
| 4 | Gen1 | 0.51 | 4.27E-09 | 9.11E-05 | 0.38 | 0.24 | 0.14 | 1.6 |
| 4 | Gm9008 | 0.96 | 4.28E-09 | 9.15E-05 | 0.18 | 0.09 | 0.09 | 2.02 |
| 4 | Cep19 | 0.46 | 4.31E-09 | 9.20E-05 | 0.47 | 0.31 | 0.16 | 1.53 |
| 4 | Slc19a1 | 0.73 | 4.31E-09 | 9.22E-05 | 0.25 | 0.13 | 0.11 | 1.84 |
| 4 | Taf1d | 0.28 | 4.33E-09 | 9.24E-05 | 0.95 | 0.85 | 0.09 | 1.11 |
| 4 | Dgcr8 | 0.51 | 4.45E-09 | 9.52E-05 | 0.5 | 0.34 | 0.16 | 1.46 |
| 4 | Stoml2 | 0.31 | 4.47E-09 | 9.56E-05 | 0.91 | 0.81 | 0.1 | 1.13 |
| 4 | Wdr4 | 0.45 | 4.67E-09 | 9.97E-05 | 0.44 | 0.28 | 0.16 | 1.55 |
| 4 | Rrp8 | 0.36 | 4.77E-09 | 1.02E-04 | 0.64 | 0.46 | 0.18 | 1.39 |
| 4 | Uchl5 | 0.34 | 4.80E-09 | 1.03E-04 | 0.7 | 0.51 | 0.19 | 1.36 |
| 4 | Ptprm | 0.66 | 4.92E-09 | 1.05E-04 | 0.19 | 0.09 | 0.1 | 2.04 |
| 4 | Gyg | 0.47 | 4.97E-09 | 1.06E-04 | 0.49 | 0.33 | 0.16 | 1.47 |
| 4 | Uck2 | 0.39 | 5.09E-09 | 1.09E-04 | 0.7 | 0.54 | 0.16 | 1.29 |
| 4 | Haus3 | 0.42 | 5.18E-09 | 1.11E-04 | 0.56 | 0.39 | 0.17 | 1.44 |
| 4 | Jade1 | 0.39 | 5.33E-09 | 1.14E-04 | 0.47 | 0.31 | 0.16 | 1.53 |
| 4 | Nup85 | 0.55 | 5.76E-09 | 1.23E-04 | 0.48 | 0.32 | 0.16 | 1.48 |
| 4 | Sdc2 | 0.61 | 5.87E-09 | 1.25E-04 | 0.44 | 0.29 | 0.15 | 1.52 |
| 4 | Ncln | 0.5 | 5.90E-09 | 1.26E-04 | 0.5 | 0.33 | 0.17 | 1.5 |
| 4 | Inppl1 | 0.59 | 5.93E-09 | 1.27E-04 | 0.41 | 0.26 | 0.15 | 1.55 |
| 4 | Rrp7a | 0.41 | 6.10E-09 | 1.30E-04 | 0.69 | 0.52 | 0.17 | 1.32 |
| 4 | Haus1 | 0.43 | 6.20E-09 | 1.32E-04 | 0.7 | 0.53 | 0.16 | 1.31 |
| 4 | Focad | 1.28 | 6.42E-09 | 1.37E-04 | 0.12 | 0.05 | 0.07 | 2.38 |
| 4 | Txndc5 | 0.36 | 6.50E-09 | 1.39E-04 | 0.75 | 0.61 | 0.15 | 1.24 |
| 4 | Tmem14c | 0.26 | 6.57E-09 | 1.40E-04 | 0.95 | 0.88 | 0.07 | 1.08 |
| 4 | Pwp1 | 0.41 | 6.75E-09 | 1.44E-04 | 0.67 | 0.52 | 0.15 | 1.28 |
| 4 | Xpo4 | 0.39 | 6.84E-09 | 1.46E-04 | 0.57 | 0.4 | 0.17 | 1.43 |
| 4 | Dhrs4 | 0.31 | 6.97E-09 | 1.49E-04 | 0.83 | 0.69 | 0.14 | 1.2 |
| 4 | Nfs1 | 0.48 | 7.04E-09 | 1.50E-04 | 0.48 | 0.32 | 0.17 | 1.52 |
| 4 | Brca2 | 0.38 | 7.04E-09 | 1.50E-04 | 0.64 | 0.45 | 0.19 | 1.41 |
| 4 | Ampd2 | 0.69 | 7.37E-09 | 1.57E-04 | 0.26 | 0.14 | 0.11 | 1.8 |
| 4 | Eefsec | 0.49 | 7.53E-09 | 1.61E-04 | 0.46 | 0.31 | 0.16 | 1.51 |

**Supplementary Table 3. Marker genes for the original clusters of integrated clustering of E11.5 control and NotchGOF λ-junction ectoderm cells.**

| | | | | | | | | |
|---|---|---|---|---|---|---|---|---|
| 4 | Polr2h | 0.29 | 7.98E-09 | 1.70E-04 | 0.91 | 0.81 | 0.11 | 1.13 |
| 4 | Slf1 | 0.36 | 8.06E-09 | 1.72E-04 | 0.66 | 0.48 | 0.18 | 1.38 |
| 4 | Fmc1 | 0.31 | 8.50E-09 | 1.82E-04 | 0.8 | 0.66 | 0.14 | 1.22 |
| 4 | Hist1h2bm | 0.72 | 8.57E-09 | 1.83E-04 | 0.33 | 0.21 | 0.12 | 1.6 |
| 4 | Mrpl16 | 0.39 | 8.74E-09 | 1.87E-04 | 0.62 | 0.45 | 0.17 | 1.38 |
| 4 | Tbrg4 | 0.55 | 8.87E-09 | 1.90E-04 | 0.3 | 0.17 | 0.13 | 1.72 |
| 4 | Dph2 | 0.72 | 9.07E-09 | 1.94E-04 | 0.23 | 0.12 | 0.11 | 1.86 |
| 4 | Ehd3 | 0.86 | 9.19E-09 | 1.96E-04 | 0.15 | 0.07 | 0.08 | 2.2 |
| 4 | Grk5 | 0.53 | 9.26E-09 | 1.98E-04 | 0.26 | 0.14 | 0.12 | 1.82 |
| 4 | Opn3 | 0.98 | 9.55E-09 | 2.04E-04 | 0.19 | 0.1 | 0.09 | 1.97 |
| 4 | Rtca | 0.43 | 9.58E-09 | 2.05E-04 | 0.57 | 0.41 | 0.16 | 1.38 |
| 4 | Rad23a | 0.41 | 9.64E-09 | 2.06E-04 | 0.65 | 0.48 | 0.17 | 1.35 |
| 4 | Efnb1 | 0.48 | 9.65E-09 | 2.06E-04 | 0.43 | 0.28 | 0.15 | 1.54 |
| 4 | Ddx1 | 0.26 | 9.78E-09 | 2.09E-04 | 0.91 | 0.8 | 0.12 | 1.15 |
| 4 | Zcchc3 | 0.38 | 1.04E-08 | 2.22E-04 | 0.76 | 0.6 | 0.16 | 1.26 |
| 4 | Nek2 | 0.38 | 1.04E-08 | 2.23E-04 | 0.36 | 0.23 | 0.14 | 1.61 |
| 4 | Kcnk2 | 0.46 | 1.06E-08 | 2.27E-04 | 0.51 | 0.36 | 0.15 | 1.41 |
| 4 | Ints1 | 0.45 | 1.09E-08 | 2.32E-04 | 0.41 | 0.27 | 0.15 | 1.56 |
| 4 | Chaf1b | 0.4 | 1.09E-08 | 2.34E-04 | 0.62 | 0.46 | 0.16 | 1.36 |
| 4 | Ociad1 | 0.3 | 1.10E-08 | 2.34E-04 | 0.85 | 0.7 | 0.15 | 1.22 |
| 4 | Eme1 | 0.43 | 1.16E-08 | 2.47E-04 | 0.5 | 0.34 | 0.16 | 1.47 |
| 4 | Gdap1 | 0.49 | 1.16E-08 | 2.48E-04 | 0.12 | 0.05 | 0.07 | 2.42 |
| 4 | Nap1l3 | 0.93 | 1.19E-08 | 2.54E-04 | 0.15 | 0.07 | 0.08 | 2.2 |
| 4 | A430005L14Rik | 0.41 | 1.22E-08 | 2.61E-04 | 0.64 | 0.48 | 0.15 | 1.32 |
| 4 | Mcmbp | 0.38 | 1.27E-08 | 2.71E-04 | 0.71 | 0.55 | 0.17 | 1.31 |
| 4 | 9530059O14Rik | 0.94 | 1.29E-08 | 2.75E-04 | 0.11 | 0.04 | 0.06 | 2.49 |
| 4 | Actr5 | 0.58 | 1.32E-08 | 2.82E-04 | 0.31 | 0.19 | 0.13 | 1.67 |
| 4 | Haspin | 0.59 | 1.34E-08 | 2.87E-04 | 0.34 | 0.21 | 0.13 | 1.62 |
| 4 | Chsy1 | 0.47 | 1.37E-08 | 2.93E-04 | 0.48 | 0.32 | 0.16 | 1.5 |
| 4 | Acer3 | 0.6 | 1.42E-08 | 3.02E-04 | 0.29 | 0.17 | 0.12 | 1.71 |
| 4 | Prkar2a | 0.42 | 1.42E-08 | 3.03E-04 | 0.64 | 0.5 | 0.14 | 1.27 |
| 4 | Pprc1 | 0.57 | 1.43E-08 | 3.06E-04 | 0.35 | 0.22 | 0.13 | 1.61 |
| 4 | Sptssa | 0.29 | 1.45E-08 | 3.10E-04 | 0.87 | 0.75 | 0.13 | 1.17 |
| 4 | Rap1a | 0.34 | 1.48E-08 | 3.16E-04 | 0.77 | 0.63 | 0.14 | 1.21 |
| 4 | Syngr2 | 0.62 | 1.49E-08 | 3.19E-04 | 0.4 | 0.27 | 0.13 | 1.5 |
| 4 | Setd6 | 0.66 | 1.51E-08 | 3.22E-04 | 0.24 | 0.13 | 0.11 | 1.82 |
| 4 | Pola2 | 0.49 | 1.54E-08 | 3.28E-04 | 0.5 | 0.35 | 0.16 | 1.46 |
| 4 | Pdcd11 | 0.42 | 1.57E-08 | 3.36E-04 | 0.6 | 0.44 | 0.16 | 1.36 |
| 4 | Oaf | 0.65 | 1.58E-08 | 3.38E-04 | 0.25 | 0.14 | 0.11 | 1.79 |
| 4 | Neil3 | 0.62 | 1.59E-08 | 3.39E-04 | 0.26 | 0.15 | 0.11 | 1.76 |
| 4 | Casp8ap2 | 0.29 | 1.60E-08 | 3.43E-04 | 0.87 | 0.72 | 0.15 | 1.2 |
| 4 | Cnpy3 | 0.41 | 1.61E-08 | 3.44E-04 | 0.52 | 0.35 | 0.17 | 1.49 |
| 4 | Chchd3 | 0.3 | 1.68E-08 | 3.58E-04 | 0.77 | 0.62 | 0.15 | 1.25 |
| 4 | Suv39h1 | 0.42 | 1.68E-08 | 3.59E-04 | 0.57 | 0.4 | 0.18 | 1.44 |
| 4 | Ift74 | 0.32 | 1.78E-08 | 3.81E-04 | 0.83 | 0.66 | 0.17 | 1.25 |
| 4 | Gbe1 | 0.76 | 1.80E-08 | 3.85E-04 | 0.21 | 0.11 | 0.1 | 1.87 |
| 4 | Dclk2 | 1 | 1.82E-08 | 3.89E-04 | 0.1 | 0.04 | 0.06 | 2.58 |
| 4 | Ckap5 | 0.26 | 1.83E-08 | 3.91E-04 | 0.84 | 0.72 | 0.13 | 1.17 |
| 4 | Dnmt1 | 0.3 | 1.90E-08 | 4.05E-04 | 0.85 | 0.72 | 0.13 | 1.18 |
| 4 | Irak1bp1 | 0.38 | 1.91E-08 | 4.07E-04 | 0.63 | 0.48 | 0.16 | 1.33 |
| 4 | Dpysl5 | 0.26 | 1.94E-08 | 4.15E-04 | 0.54 | 0.37 | 0.17 | 1.45 |
| 4 | Rad21 | 0.27 | 1.97E-08 | 4.21E-04 | 0.88 | 0.78 | 0.1 | 1.12 |
| 4 | Pja1 | 0.39 | 1.99E-08 | 4.24E-04 | 0.66 | 0.51 | 0.15 | 1.29 |
| 4 | Rab11fip4 | 0.57 | 1.99E-08 | 4.25E-04 | 0.32 | 0.19 | 0.12 | 1.63 |
| 4 | Prpsap2 | 0.46 | 2.01E-08 | 4.29E-04 | 0.39 | 0.24 | 0.14 | 1.59 |
| 4 | Rad23b | 0.3 | 2.01E-08 | 4.29E-04 | 0.89 | 0.78 | 0.12 | 1.15 |
| 4 | Kif11 | 0.32 | 2.03E-08 | 4.33E-04 | 0.65 | 0.48 | 0.18 | 1.37 |

**Supplementary Table 3. Marker genes for the original clusters of integrated clustering of E11.5 control and NotchGOF λ-junction ectoderm cells.**

| | | | | | | | | |
|---|---|---|---|---|---|---|---|---|
| 4 | Ptpa | 0.41 | 2.06E-08 | 4.40E-04 | 0.61 | 0.45 | 0.16 | 1.36 |
| 4 | Nfkbiz | 0.82 | 2.15E-08 | 4.59E-04 | 0.16 | 0.08 | 0.08 | 2.06 |
| 4 | Wdfy2 | 0.44 | 2.16E-08 | 4.62E-04 | 0.32 | 0.19 | 0.13 | 1.69 |
| 4 | Arf2 | 0.48 | 2.19E-08 | 4.67E-04 | 0.43 | 0.29 | 0.14 | 1.49 |
| 4 | Eif3j2 | 0.48 | 2.20E-08 | 4.70E-04 | 0.47 | 0.32 | 0.15 | 1.48 |
| 4 | Maoa | 0.47 | 2.23E-08 | 4.76E-04 | 0.42 | 0.27 | 0.15 | 1.55 |
| 4 | Slc35f2 | 1.07 | 2.24E-08 | 4.79E-04 | 0.12 | 0.05 | 0.07 | 2.34 |
| 4 | Ccdc181 | 0.56 | 2.24E-08 | 4.79E-04 | 0.22 | 0.12 | 0.1 | 1.87 |
| 4 | 1110038B12Rik | 0.25 | 2.40E-08 | 5.13E-04 | 0.96 | 0.92 | 0.04 | 1.05 |
| 4 | Rab26os | 0.58 | 2.51E-08 | 5.36E-04 | 0.33 | 0.2 | 0.13 | 1.64 |
| 4 | Prrg2 | 0.45 | 2.52E-08 | 5.38E-04 | 0.39 | 0.25 | 0.14 | 1.58 |
| 4 | Hist1h3f | 0.79 | 2.57E-08 | 5.50E-04 | 0.32 | 0.2 | 0.12 | 1.57 |
| 4 | Crat | 0.63 | 2.61E-08 | 5.57E-04 | 0.24 | 0.13 | 0.11 | 1.82 |
| 4 | Sgsm1 | 0.94 | 2.61E-08 | 5.58E-04 | 0.16 | 0.08 | 0.08 | 2.05 |
| 4 | Noc4l | 0.4 | 2.64E-08 | 5.64E-04 | 0.59 | 0.43 | 0.16 | 1.38 |
| 4 | Lrr1 | 0.57 | 2.75E-08 | 5.88E-04 | 0.33 | 0.21 | 0.12 | 1.58 |
| 4 | Shroom3 | 0.48 | 2.86E-08 | 6.10E-04 | 0.37 | 0.23 | 0.14 | 1.58 |
| 4 | Pcyt2 | 0.51 | 2.92E-08 | 6.23E-04 | 0.34 | 0.21 | 0.13 | 1.62 |
| 4 | Recql4 | 0.96 | 3.01E-08 | 6.42E-04 | 0.15 | 0.07 | 0.08 | 2.13 |
| 4 | Xbp1 | 0.37 | 3.02E-08 | 6.46E-04 | 0.54 | 0.37 | 0.17 | 1.45 |
| 4 | Mdn1 | 0.41 | 3.04E-08 | 6.49E-04 | 0.54 | 0.39 | 0.16 | 1.4 |
| 4 | Gnl1 | 0.38 | 3.07E-08 | 6.55E-04 | 0.64 | 0.47 | 0.17 | 1.36 |
| 4 | Gnptg | 0.48 | 3.09E-08 | 6.60E-04 | 0.44 | 0.3 | 0.15 | 1.5 |
| 4 | Prodh | 0.69 | 3.10E-08 | 6.62E-04 | 0.21 | 0.11 | 0.1 | 1.89 |
| 4 | Ncapg2 | 0.37 | 3.13E-08 | 6.68E-04 | 0.53 | 0.37 | 0.16 | 1.43 |
| 4 | Fignl1 | 0.29 | 3.27E-08 | 6.98E-04 | 0.65 | 0.46 | 0.19 | 1.41 |
| 4 | Ncapd3 | 0.33 | 3.35E-08 | 7.15E-04 | 0.65 | 0.47 | 0.18 | 1.38 |
| 4 | Zdhhc21 | 0.43 | 3.40E-08 | 7.27E-04 | 0.58 | 0.43 | 0.14 | 1.32 |
| 4 | Nfasc | 0.66 | 3.46E-08 | 7.38E-04 | 0.18 | 0.09 | 0.09 | 1.96 |
| 4 | Ccne2 | 0.7 | 3.49E-08 | 7.45E-04 | 0.28 | 0.17 | 0.12 | 1.68 |
| 4 | Srebf2 | 0.39 | 3.67E-08 | 7.84E-04 | 0.68 | 0.52 | 0.16 | 1.31 |
| 4 | Six4 | 0.41 | 3.72E-08 | 7.95E-04 | 0.65 | 0.47 | 0.18 | 1.38 |
| 4 | Topbp1 | 0.27 | 3.78E-08 | 8.08E-04 | 0.84 | 0.67 | 0.17 | 1.26 |
| 4 | Aimp2 | 0.37 | 3.89E-08 | 8.32E-04 | 0.68 | 0.53 | 0.15 | 1.29 |
| 4 | Epha5 | 0.55 | 3.93E-08 | 8.40E-04 | 0.17 | 0.08 | 0.08 | 2 |
| 4 | Stard4 | 0.63 | 3.95E-08 | 8.44E-04 | 0.35 | 0.23 | 0.12 | 1.53 |
| 4 | Zdhhc6 | 0.32 | 3.96E-08 | 8.47E-04 | 0.64 | 0.47 | 0.17 | 1.37 |
| 4 | Kbtbd8 | 1.18 | 4.04E-08 | 8.63E-04 | 0.14 | 0.06 | 0.07 | 2.16 |
| 4 | Mapre2 | 0.41 | 4.17E-08 | 8.92E-04 | 0.36 | 0.23 | 0.13 | 1.59 |
| 4 | Larp4 | 0.4 | 4.20E-08 | 8.96E-04 | 0.74 | 0.61 | 0.14 | 1.23 |
| 4 | Wdr43 | 0.35 | 4.32E-08 | 9.24E-04 | 0.85 | 0.7 | 0.15 | 1.21 |
| 4 | Coq4 | 0.4 | 4.38E-08 | 9.36E-04 | 0.51 | 0.36 | 0.15 | 1.41 |
| 4 | Cpsf4 | 0.36 | 4.38E-08 | 9.36E-04 | 0.52 | 0.36 | 0.17 | 1.46 |
| 4 | Pxylp1 | 0.9 | 4.41E-08 | 9.43E-04 | 0.13 | 0.06 | 0.07 | 2.22 |
| 4 | Wrap53 | 0.55 | 4.47E-08 | 9.56E-04 | 0.33 | 0.2 | 0.13 | 1.64 |
| 4 | Rbpms2 | 1.07 | 4.53E-08 | 9.67E-04 | 0.12 | 0.06 | 0.07 | 2.25 |
| 4 | Stk16 | 0.36 | 4.54E-08 | 9.70E-04 | 0.77 | 0.62 | 0.15 | 1.24 |
| 4 | Mettl1 | 0.3 | 4.71E-08 | 1.01E-03 | 0.79 | 0.61 | 0.18 | 1.29 |
| 4 | Rab39b | 0.49 | 4.75E-08 | 1.01E-03 | 0.23 | 0.13 | 0.1 | 1.82 |
| 4 | Rbm47 | 0.75 | 4.80E-08 | 1.03E-03 | 0.18 | 0.1 | 0.09 | 1.92 |
| 4 | Cabin1 | 0.36 | 4.89E-08 | 1.04E-03 | 0.63 | 0.45 | 0.18 | 1.41 |
| 4 | Xpot | 0.4 | 4.92E-08 | 1.05E-03 | 0.59 | 0.43 | 0.16 | 1.37 |
| 4 | Rpap3 | 0.35 | 4.94E-08 | 1.05E-03 | 0.58 | 0.41 | 0.17 | 1.41 |
| 4 | Rbm10 | 0.4 | 4.96E-08 | 1.06E-03 | 0.69 | 0.54 | 0.15 | 1.28 |
| 4 | Traip | 0.51 | 4.96E-08 | 1.06E-03 | 0.37 | 0.24 | 0.13 | 1.56 |
| 4 | Poc1b | 0.5 | 4.99E-08 | 1.07E-03 | 0.43 | 0.29 | 0.14 | 1.49 |
| 4 | Daam1 | 0.42 | 5.15E-08 | 1.10E-03 | 0.67 | 0.52 | 0.15 | 1.29 |

**Supplementary Table 3. Marker genes for the original clusters of integrated clustering of E11.5 control and NotchGOF λ-junction ectoderm cells.**

| | | | | | | | | |
|---|---|---|---|---|---|---|---|---|
| 4 | Ykt6 | 0.38 | 5.18E-08 | 1.11E-03 | 0.52 | 0.36 | 0.16 | 1.46 |
| 4 | Szrd1 | 0.35 | 5.32E-08 | 1.14E-03 | 0.69 | 0.53 | 0.16 | 1.31 |
| 4 | Itgb8 | 0.37 | 5.38E-08 | 1.15E-03 | 0.32 | 0.19 | 0.13 | 1.67 |
| 4 | Zc3h7a | 0.47 | 5.39E-08 | 1.15E-03 | 0.44 | 0.29 | 0.15 | 1.5 |
| 4 | Exosc1 | 0.38 | 5.41E-08 | 1.16E-03 | 0.68 | 0.52 | 0.16 | 1.31 |
| 4 | Ssbp4 | 0.33 | 5.42E-08 | 1.16E-03 | 0.65 | 0.47 | 0.18 | 1.39 |
| 4 | Lsm1 | 0.29 | 5.47E-08 | 1.17E-03 | 0.82 | 0.7 | 0.12 | 1.18 |
| 4 | Aars | 0.42 | 5.47E-08 | 1.17E-03 | 0.54 | 0.38 | 0.16 | 1.43 |
| 4 | Prmt3 | 0.45 | 5.49E-08 | 1.17E-03 | 0.52 | 0.37 | 0.15 | 1.41 |
| 4 | Ephb2 | 0.39 | 5.68E-08 | 1.21E-03 | 0.6 | 0.41 | 0.18 | 1.44 |
| 4 | Psmc1 | 0.27 | 6.08E-08 | 1.30E-03 | 0.88 | 0.76 | 0.12 | 1.16 |
| 4 | Tsku | 1.07 | 6.15E-08 | 1.31E-03 | 0.11 | 0.04 | 0.06 | 2.39 |
| 4 | Tank | 0.42 | 6.16E-08 | 1.32E-03 | 0.4 | 0.26 | 0.14 | 1.54 |
| 4 | Crcp | 0.37 | 6.28E-08 | 1.34E-03 | 0.51 | 0.35 | 0.16 | 1.46 |
| 4 | Rpp25l | 0.3 | 6.33E-08 | 1.35E-03 | 0.83 | 0.7 | 0.13 | 1.19 |
| 4 | Bend7 | 0.45 | 6.38E-08 | 1.36E-03 | 0.39 | 0.25 | 0.14 | 1.56 |
| 4 | Galnt1 | 0.34 | 6.39E-08 | 1.37E-03 | 0.75 | 0.59 | 0.16 | 1.27 |
| 4 | Cenps | 0.48 | 6.61E-08 | 1.41E-03 | 0.57 | 0.42 | 0.16 | 1.38 |
| 4 | Rbfa | 0.36 | 6.64E-08 | 1.42E-03 | 0.5 | 0.33 | 0.16 | 1.48 |
| 4 | Urb1 | 0.6 | 6.71E-08 | 1.43E-03 | 0.22 | 0.12 | 0.1 | 1.82 |
| 4 | Phb | 0.28 | 6.88E-08 | 1.47E-03 | 0.88 | 0.75 | 0.13 | 1.17 |
| 4 | Praf2 | 0.35 | 6.99E-08 | 1.49E-03 | 0.57 | 0.41 | 0.16 | 1.4 |
| 4 | Btaf1 | 0.45 | 6.99E-08 | 1.49E-03 | 0.47 | 0.33 | 0.15 | 1.44 |
| 4 | Mical2 | 0.84 | 7.09E-08 | 1.51E-03 | 0.12 | 0.05 | 0.07 | 2.29 |
| 4 | Tor1a | 0.3 | 7.12E-08 | 1.52E-03 | 0.67 | 0.5 | 0.17 | 1.35 |
| 4 | Cfap298 | 0.38 | 7.16E-08 | 1.53E-03 | 0.58 | 0.41 | 0.17 | 1.41 |
| 4 | Sap18 | 0.26 | 7.32E-08 | 1.56E-03 | 0.93 | 0.81 | 0.12 | 1.15 |
| 4 | Rbm14 | 0.42 | 7.45E-08 | 1.59E-03 | 0.56 | 0.41 | 0.15 | 1.37 |
| 4 | Fam169a | 0.8 | 7.45E-08 | 1.59E-03 | 0.24 | 0.14 | 0.1 | 1.73 |
| 4 | Mrpl19 | 0.33 | 7.45E-08 | 1.59E-03 | 0.72 | 0.59 | 0.14 | 1.23 |
| 4 | Tbc1d4 | 0.69 | 7.50E-08 | 1.60E-03 | 0.17 | 0.09 | 0.09 | 1.97 |
| 4 | Hist1h1e | 0.3 | 7.64E-08 | 1.63E-03 | 0.77 | 0.61 | 0.16 | 1.26 |
| 4 | Sgms2 | 0.99 | 7.78E-08 | 1.66E-03 | 0.12 | 0.05 | 0.07 | 2.25 |
| 4 | Gspt2 | 0.69 | 7.86E-08 | 1.68E-03 | 0.27 | 0.16 | 0.11 | 1.66 |
| 4 | Adarb1 | 0.77 | 8.02E-08 | 1.71E-03 | 0.14 | 0.07 | 0.08 | 2.13 |
| 4 | Haus6 | 0.41 | 8.39E-08 | 1.79E-03 | 0.47 | 0.32 | 0.15 | 1.46 |
| 4 | Cd3eap | 0.36 | 8.43E-08 | 1.80E-03 | 0.73 | 0.59 | 0.14 | 1.24 |
| 4 | Setbp1 | 0.54 | 8.53E-08 | 1.82E-03 | 0.4 | 0.27 | 0.13 | 1.48 |
| 4 | Gpd1l | 0.84 | 8.63E-08 | 1.84E-03 | 0.15 | 0.07 | 0.08 | 2.07 |
| 4 | Ndufv3 | 0.26 | 8.63E-08 | 1.84E-03 | 0.89 | 0.82 | 0.08 | 1.1 |
| 4 | Elovl6 | 0.39 | 8.69E-08 | 1.86E-03 | 0.67 | 0.51 | 0.16 | 1.31 |
| 4 | Cip2a | 0.34 | 9.00E-08 | 1.92E-03 | 0.51 | 0.35 | 0.16 | 1.45 |
| 4 | Aurkb | 0.29 | 9.00E-08 | 1.92E-03 | 0.61 | 0.43 | 0.18 | 1.42 |
| 4 | Cdk2ap2 | 0.36 | 9.04E-08 | 1.93E-03 | 0.71 | 0.55 | 0.16 | 1.29 |
| 4 | Galk1 | 0.3 | 9.12E-08 | 1.95E-03 | 0.83 | 0.71 | 0.13 | 1.18 |
| 4 | Rrp9 | 0.42 | 9.27E-08 | 1.98E-03 | 0.48 | 0.33 | 0.15 | 1.45 |
| 4 | Lonp1 | 0.33 | 9.49E-08 | 2.03E-03 | 0.7 | 0.52 | 0.18 | 1.34 |
| 4 | Mis12 | 0.36 | 9.55E-08 | 2.04E-03 | 0.6 | 0.43 | 0.16 | 1.38 |
| 4 | Flywch1 | 0.31 | 9.79E-08 | 2.09E-03 | 0.69 | 0.52 | 0.16 | 1.31 |
| 4 | Ptpn11 | 0.3 | 1.00E-07 | 2.14E-03 | 0.77 | 0.62 | 0.15 | 1.24 |
| 4 | 2810001G20Rik | 0.35 | 1.00E-07 | 2.14E-03 | 0.47 | 0.31 | 0.15 | 1.49 |
| 4 | Gnptab | 0.42 | 1.01E-07 | 2.17E-03 | 0.3 | 0.18 | 0.12 | 1.66 |
| 4 | Cptp | 0.7 | 1.05E-07 | 2.23E-03 | 0.22 | 0.12 | 0.1 | 1.79 |
| 4 | Tm9sf1 | 0.38 | 1.07E-07 | 2.28E-03 | 0.59 | 0.43 | 0.16 | 1.39 |
| 4 | Iqgap3 | 0.57 | 1.11E-07 | 2.37E-03 | 0.33 | 0.21 | 0.12 | 1.58 |
| 4 | Fam92a | 0.31 | 1.13E-07 | 2.41E-03 | 0.78 | 0.65 | 0.13 | 1.2 |
| 4 | Pih1d2 | 0.72 | 1.16E-07 | 2.47E-03 | 0.15 | 0.07 | 0.08 | 2.05 |

**Supplementary Table 3. Marker genes for the original clusters of integrated clustering of E11.5 control and NotchGOF λ-junction ectoderm cells.**

| | | | | | | | | |
|---|---|---|---|---|---|---|---|---|
| 4 | Smn1 | 0.27 | 1.19E-07 | 2.54E-03 | 0.82 | 0.7 | 0.12 | 1.17 |
| 4 | Sirpa | 0.82 | 1.19E-07 | 2.54E-03 | 0.17 | 0.09 | 0.08 | 1.92 |
| 4 | Polr1a | 0.4 | 1.23E-07 | 2.62E-03 | 0.51 | 0.35 | 0.16 | 1.46 |
| 4 | Dusp18 | 0.79 | 1.23E-07 | 2.63E-03 | 0.19 | 0.1 | 0.09 | 1.85 |
| 4 | Faap24 | 0.57 | 1.24E-07 | 2.65E-03 | 0.26 | 0.15 | 0.11 | 1.71 |
| 4 | B4galt2 | 0.44 | 1.24E-07 | 2.65E-03 | 0.37 | 0.23 | 0.13 | 1.58 |
| 4 | Slc25a30 | 0.5 | 1.25E-07 | 2.67E-03 | 0.26 | 0.15 | 0.11 | 1.7 |
| 4 | Dusp5 | 0.52 | 1.26E-07 | 2.68E-03 | 0.32 | 0.2 | 0.12 | 1.62 |
| 4 | Polr2b | 0.31 | 1.27E-07 | 2.71E-03 | 0.76 | 0.64 | 0.12 | 1.18 |
| 4 | Shmt2 | 0.41 | 1.28E-07 | 2.74E-03 | 0.52 | 0.36 | 0.16 | 1.44 |
| 4 | Tmed5 | 0.33 | 1.29E-07 | 2.75E-03 | 0.7 | 0.55 | 0.15 | 1.28 |
| 4 | Adrm1 | 0.33 | 1.34E-07 | 2.86E-03 | 0.76 | 0.64 | 0.12 | 1.18 |
| 4 | Nadk | 0.38 | 1.34E-07 | 2.87E-03 | 0.53 | 0.37 | 0.16 | 1.44 |
| 4 | Ogdh | 0.36 | 1.43E-07 | 3.04E-03 | 0.61 | 0.44 | 0.17 | 1.38 |
| 4 | E2f1 | 0.52 | 1.44E-07 | 3.08E-03 | 0.32 | 0.2 | 0.12 | 1.59 |
| 4 | Lpar1 | 0.47 | 1.46E-07 | 3.13E-03 | 0.47 | 0.32 | 0.15 | 1.45 |
| 4 | Sppl2b | 0.46 | 1.54E-07 | 3.30E-03 | 0.25 | 0.14 | 0.11 | 1.76 |
| 4 | Ccdc12 | 0.26 | 1.56E-07 | 3.34E-03 | 0.86 | 0.73 | 0.13 | 1.18 |
| 4 | Fads2 | 0.4 | 1.59E-07 | 3.39E-03 | 0.6 | 0.45 | 0.15 | 1.34 |
| 4 | Kpna4 | 0.32 | 1.62E-07 | 3.46E-03 | 0.72 | 0.6 | 0.13 | 1.21 |
| 4 | Dvl1 | 0.53 | 1.63E-07 | 3.49E-03 | 0.3 | 0.19 | 0.12 | 1.63 |
| 4 | Afg3l1 | 0.42 | 1.66E-07 | 3.54E-03 | 0.61 | 0.45 | 0.16 | 1.35 |
| 4 | Acsl5 | 0.42 | 1.67E-07 | 3.57E-03 | 0.45 | 0.31 | 0.14 | 1.45 |
| 4 | 2700038G22Rik | 0.54 | 1.69E-07 | 3.61E-03 | 0.39 | 0.26 | 0.13 | 1.49 |
| 4 | Pold1 | 0.53 | 1.70E-07 | 3.63E-03 | 0.43 | 0.3 | 0.13 | 1.45 |
| 4 | Enoph1 | 0.47 | 1.77E-07 | 3.79E-03 | 0.39 | 0.26 | 0.13 | 1.51 |
| 4 | Stard7 | 0.35 | 1.80E-07 | 3.85E-03 | 0.55 | 0.4 | 0.15 | 1.38 |
| 4 | Sar1b | 0.27 | 1.85E-07 | 3.96E-03 | 0.83 | 0.74 | 0.09 | 1.12 |
| 4 | Abhd14a | 0.32 | 1.92E-07 | 4.11E-03 | 0.55 | 0.39 | 0.16 | 1.41 |
| 4 | Gps1 | 0.25 | 1.98E-07 | 4.23E-03 | 0.83 | 0.69 | 0.14 | 1.2 |
| 4 | D030055H07Rik | 0.87 | 1.99E-07 | 4.25E-03 | 0.13 | 0.06 | 0.07 | 2.11 |
| 4 | Mrpl50 | 0.26 | 2.00E-07 | 4.28E-03 | 0.82 | 0.7 | 0.12 | 1.18 |
| 4 | Micu3 | 0.52 | 2.09E-07 | 4.47E-03 | 0.33 | 0.21 | 0.12 | 1.55 |
| 4 | Rab28 | 0.39 | 2.12E-07 | 4.53E-03 | 0.65 | 0.5 | 0.14 | 1.29 |
| 4 | Tmod2 | 0.58 | 2.13E-07 | 4.54E-03 | 0.21 | 0.12 | 0.09 | 1.78 |
| 4 | Rpp14 | 0.31 | 2.19E-07 | 4.67E-03 | 0.64 | 0.47 | 0.17 | 1.36 |
| 4 | Snrnp25 | 0.33 | 2.27E-07 | 4.84E-03 | 0.7 | 0.52 | 0.18 | 1.35 |
| 4 | Vac14 | 0.55 | 2.28E-07 | 4.87E-03 | 0.32 | 0.21 | 0.12 | 1.56 |
| 4 | Fxn | 0.39 | 2.31E-07 | 4.93E-03 | 0.61 | 0.44 | 0.17 | 1.37 |
| 4 | Dnmt3b | 0.39 | 2.31E-07 | 4.94E-03 | 0.35 | 0.22 | 0.13 | 1.58 |
| 4 | Bphl | 0.31 | 2.32E-07 | 4.95E-03 | 0.68 | 0.5 | 0.18 | 1.36 |
| 4 | Slfn9 | 0.36 | 2.35E-07 | 5.02E-03 | 0.36 | 0.23 | 0.13 | 1.56 |
| 4 | Abcb6 | 0.8 | 2.40E-07 | 5.12E-03 | 0.19 | 0.1 | 0.08 | 1.82 |
| 4 | Tnfrsf12a | 0.5 | 2.40E-07 | 5.13E-03 | 0.34 | 0.22 | 0.12 | 1.54 |
| 4 | Myorg | 1.01 | 2.42E-07 | 5.18E-03 | 0.11 | 0.05 | 0.06 | 2.29 |
| 4 | Ipo11 | 0.43 | 2.43E-07 | 5.20E-03 | 0.47 | 0.33 | 0.14 | 1.43 |
| 4 | Gins1 | 0.45 | 2.43E-07 | 5.20E-03 | 0.47 | 0.34 | 0.14 | 1.41 |
| 4 | Letm1 | 0.35 | 2.48E-07 | 5.30E-03 | 0.53 | 0.37 | 0.16 | 1.42 |
| 4 | Eftud2 | 0.28 | 2.49E-07 | 5.31E-03 | 0.8 | 0.65 | 0.15 | 1.24 |
| 4 | A930004D18Rik | 0.66 | 2.52E-07 | 5.38E-03 | 0.17 | 0.09 | 0.08 | 1.93 |
| 4 | Derl2 | 0.39 | 2.52E-07 | 5.38E-03 | 0.54 | 0.38 | 0.15 | 1.4 |
| 4 | Dleu2 | 0.45 | 2.54E-07 | 5.42E-03 | 0.31 | 0.2 | 0.12 | 1.59 |
| 4 | Naa40 | 0.35 | 2.55E-07 | 5.46E-03 | 0.54 | 0.39 | 0.15 | 1.4 |
| 4 | Trappc13 | 0.33 | 2.56E-07 | 5.46E-03 | 0.61 | 0.44 | 0.17 | 1.39 |
| 4 | Smarcd3 | 0.41 | 2.63E-07 | 5.62E-03 | 0.41 | 0.27 | 0.14 | 1.51 |
| 4 | Prpf4 | 0.36 | 2.64E-07 | 5.64E-03 | 0.68 | 0.51 | 0.17 | 1.34 |
| 4 | Abca2 | 0.8 | 2.68E-07 | 5.73E-03 | 0.15 | 0.07 | 0.07 | 2.04 |

**Supplementary Table 3. Marker genes for the original clusters of integrated clustering of E11.5 control and NotchGOF λ-junction ectoderm cells.**

| | | | | | | | | |
|---|---|---|---|---|---|---|---|---|
| 4 | Ncapd2 | 0.32 | 2.70E-07 | 5.78E-03 | 0.64 | 0.47 | 0.17 | 1.36 |
| 4 | Rhpn1 | 1.15 | 2.73E-07 | 5.83E-03 | 0.11 | 0.05 | 0.06 | 2.23 |
| 4 | Kars | 0.35 | 2.81E-07 | 6.00E-03 | 0.75 | 0.58 | 0.17 | 1.29 |
| 4 | Alg13 | 0.49 | 2.85E-07 | 6.09E-03 | 0.37 | 0.25 | 0.13 | 1.5 |
| 4 | Cpsf2 | 0.34 | 2.89E-07 | 6.17E-03 | 0.76 | 0.61 | 0.15 | 1.25 |
| 4 | Apoo | 0.31 | 2.94E-07 | 6.29E-03 | 0.64 | 0.47 | 0.16 | 1.35 |
| 4 | Cdc7 | 0.35 | 2.95E-07 | 6.30E-03 | 0.44 | 0.3 | 0.14 | 1.47 |
| 4 | Pgd | 0.31 | 2.96E-07 | 6.33E-03 | 0.72 | 0.55 | 0.17 | 1.3 |
| 4 | Notch4 | 0.98 | 2.97E-07 | 6.34E-03 | 0.15 | 0.07 | 0.07 | 2.01 |
| 4 | Rcl1 | 0.46 | 2.97E-07 | 6.35E-03 | 0.55 | 0.4 | 0.15 | 1.36 |
| 4 | Ahcy | 0.61 | 3.05E-07 | 6.51E-03 | 0.33 | 0.21 | 0.11 | 1.54 |
| 4 | Cep170b | 0.39 | 3.15E-07 | 6.74E-03 | 0.47 | 0.32 | 0.15 | 1.46 |
| 4 | Kremen1 | 0.31 | 3.16E-07 | 6.75E-03 | 0.45 | 0.31 | 0.14 | 1.47 |
| 4 | Srcin1 | 0.51 | 3.16E-07 | 6.75E-03 | 0.23 | 0.13 | 0.1 | 1.75 |
| 4 | Gemin5 | 0.51 | 3.20E-07 | 6.83E-03 | 0.47 | 0.33 | 0.14 | 1.43 |
| 4 | Gins3 | 0.48 | 3.34E-07 | 7.13E-03 | 0.3 | 0.19 | 0.11 | 1.6 |
| 4 | Pcgf6 | 0.32 | 3.36E-07 | 7.18E-03 | 0.64 | 0.49 | 0.15 | 1.31 |
| 4 | Epdr1 | 0.57 | 3.49E-07 | 7.46E-03 | 0.19 | 0.1 | 0.09 | 1.83 |
| 4 | Mgat2 | 0.35 | 3.64E-07 | 7.78E-03 | 0.61 | 0.45 | 0.16 | 1.37 |
| 4 | Osgepl1 | 0.36 | 3.65E-07 | 7.80E-03 | 0.3 | 0.18 | 0.12 | 1.65 |
| 4 | Oat | 0.35 | 3.66E-07 | 7.82E-03 | 0.62 | 0.46 | 0.16 | 1.35 |
| 4 | Gla | 0.79 | 3.72E-07 | 7.95E-03 | 0.19 | 0.1 | 0.08 | 1.82 |
| 4 | Nol11 | 0.32 | 3.77E-07 | 8.06E-03 | 0.76 | 0.65 | 0.11 | 1.17 |
| 4 | Lin54 | 0.42 | 3.79E-07 | 8.10E-03 | 0.34 | 0.22 | 0.12 | 1.58 |
| 4 | Pask | 0.62 | 3.79E-07 | 8.11E-03 | 0.21 | 0.12 | 0.09 | 1.76 |
| 4 | Pfkl | 0.28 | 3.81E-07 | 8.13E-03 | 0.86 | 0.76 | 0.1 | 1.13 |
| 4 | Psmg3 | 0.28 | 3.84E-07 | 8.19E-03 | 0.7 | 0.54 | 0.16 | 1.29 |
| 4 | Nudt5 | 0.26 | 3.89E-07 | 8.32E-03 | 0.87 | 0.75 | 0.11 | 1.15 |
| 4 | Gtf2f1 | 0.35 | 3.93E-07 | 8.41E-03 | 0.69 | 0.55 | 0.14 | 1.25 |
| 4 | Prpf8 | 0.27 | 4.00E-07 | 8.54E-03 | 0.87 | 0.74 | 0.13 | 1.17 |
| 4 | Snhg14 | 0.39 | 4.09E-07 | 8.73E-03 | 0.37 | 0.24 | 0.13 | 1.54 |
| 4 | Dusp16 | 0.36 | 4.09E-07 | 8.74E-03 | 0.37 | 0.24 | 0.13 | 1.54 |
| 4 | Kpna3 | 0.38 | 4.11E-07 | 8.78E-03 | 0.71 | 0.6 | 0.11 | 1.18 |
| 4 | Suv39h2 | 0.37 | 4.16E-07 | 8.90E-03 | 0.54 | 0.4 | 0.15 | 1.37 |
| 4 | Tatdn2 | 0.61 | 4.17E-07 | 8.91E-03 | 0.24 | 0.14 | 0.1 | 1.69 |
| 4 | Capn15 | 0.41 | 4.19E-07 | 8.95E-03 | 0.49 | 0.35 | 0.15 | 1.42 |
| 4 | Entpd5 | 0.58 | 4.55E-07 | 9.72E-03 | 0.21 | 0.12 | 0.09 | 1.79 |
| 4 | Efnb3 | 0.47 | 4.75E-07 | 1.01E-02 | 0.33 | 0.21 | 0.12 | 1.59 |
| 4 | Rcc1l | 0.57 | 4.77E-07 | 1.02E-02 | 0.32 | 0.21 | 0.11 | 1.54 |
| 4 | Tctn2 | 0.73 | 5.01E-07 | 1.07E-02 | 0.19 | 0.11 | 0.09 | 1.81 |
| 4 | Farsa | 0.3 | 5.08E-07 | 1.08E-02 | 0.73 | 0.57 | 0.17 | 1.3 |
| 4 | Ccdc8 | 0.41 | 5.09E-07 | 1.09E-02 | 0.51 | 0.37 | 0.15 | 1.4 |
| 4 | Zfp41 | 0.47 | 5.25E-07 | 1.12E-02 | 0.44 | 0.31 | 0.13 | 1.42 |
| 4 | Endog | 0.58 | 5.53E-07 | 1.18E-02 | 0.29 | 0.18 | 0.11 | 1.62 |
| 4 | Aurka | 0.25 | 5.56E-07 | 1.19E-02 | 0.53 | 0.38 | 0.16 | 1.42 |
| 4 | Cox10 | 0.4 | 5.82E-07 | 1.24E-02 | 0.39 | 0.26 | 0.13 | 1.51 |
| 4 | Fbln7 | 0.95 | 5.85E-07 | 1.25E-02 | 0.1 | 0.05 | 0.06 | 2.29 |
| 4 | Trip10 | 0.4 | 6.17E-07 | 1.32E-02 | 0.37 | 0.24 | 0.13 | 1.53 |
| 4 | Gmds | 0.37 | 6.24E-07 | 1.33E-02 | 0.4 | 0.27 | 0.13 | 1.49 |
| 4 | Mrpl37 | 0.42 | 6.24E-07 | 1.33E-02 | 0.53 | 0.38 | 0.15 | 1.4 |
| 4 | Eif2b4 | 0.32 | 6.39E-07 | 1.37E-02 | 0.47 | 0.32 | 0.15 | 1.46 |
| 4 | Adgrg1 | 0.37 | 6.50E-07 | 1.39E-02 | 0.22 | 0.13 | 0.1 | 1.75 |
| 4 | C030037D09Rik | 0.93 | 6.59E-07 | 1.41E-02 | 0.11 | 0.05 | 0.06 | 2.15 |
| 4 | Katnal2 | 0.75 | 6.59E-07 | 1.41E-02 | 0.13 | 0.06 | 0.07 | 2.05 |
| 4 | Men1 | 0.37 | 6.59E-07 | 1.41E-02 | 0.54 | 0.39 | 0.15 | 1.4 |
| 4 | Slc43a1 | 0.64 | 6.64E-07 | 1.42E-02 | 0.18 | 0.1 | 0.08 | 1.84 |
| 4 | Coprs | 0.41 | 6.69E-07 | 1.43E-02 | 0.51 | 0.36 | 0.15 | 1.42 |

**Supplementary Table 3. Marker genes for the original clusters of integrated clustering of E11.5 control and NotchGOF λ-junction ectoderm cells.**

| | | | | | | | | |
|---|---|---|---|---|---|---|---|---|
| 4 | Ppfia3 | 0.43 | 6.79E-07 | 1.45E-02 | 0.42 | 0.29 | 0.13 | 1.47 |
| 4 | Psrc1 | 0.26 | 6.82E-07 | 1.46E-02 | 0.48 | 0.33 | 0.15 | 1.45 |
| 4 | Spin4 | 0.78 | 6.88E-07 | 1.47E-02 | 0.17 | 0.09 | 0.08 | 1.87 |
| 4 | Nifk | 0.28 | 7.10E-07 | 1.52E-02 | 0.81 | 0.68 | 0.13 | 1.18 |
| 4 | Mxra7 | 0.53 | 7.26E-07 | 1.55E-02 | 0.28 | 0.17 | 0.11 | 1.62 |
| 4 | Txnrd2 | 0.68 | 7.29E-07 | 1.56E-02 | 0.24 | 0.15 | 0.1 | 1.65 |
| 4 | Pitpna | 0.26 | 7.29E-07 | 1.56E-02 | 0.79 | 0.67 | 0.12 | 1.18 |
| 4 | Hist2h2bb | 0.59 | 7.65E-07 | 1.63E-02 | 0.31 | 0.2 | 0.11 | 1.54 |
| 4 | Chml | 0.47 | 7.75E-07 | 1.66E-02 | 0.25 | 0.15 | 0.1 | 1.68 |
| 4 | Prkag2 | 0.3 | 7.75E-07 | 1.66E-02 | 0.45 | 0.31 | 0.15 | 1.48 |
| 4 | Mta1 | 0.32 | 7.78E-07 | 1.66E-02 | 0.8 | 0.68 | 0.11 | 1.17 |
| 4 | Dnajb2 | 0.4 | 7.86E-07 | 1.68E-02 | 0.44 | 0.29 | 0.14 | 1.49 |
| 4 | Paxip1 | 0.44 | 8.04E-07 | 1.72E-02 | 0.47 | 0.34 | 0.13 | 1.38 |
| 4 | Nup35 | 0.35 | 8.22E-07 | 1.76E-02 | 0.54 | 0.4 | 0.14 | 1.34 |
| 4 | Pdpn | 0.31 | 8.37E-07 | 1.79E-02 | 0.5 | 0.35 | 0.15 | 1.42 |
| 4 | Tmem189 | 0.39 | 8.43E-07 | 1.80E-02 | 0.54 | 0.41 | 0.14 | 1.34 |
| 4 | Nol9 | 0.41 | 8.43E-07 | 1.80E-02 | 0.41 | 0.28 | 0.13 | 1.46 |
| 4 | Exosc2 | 0.3 | 8.47E-07 | 1.81E-02 | 0.61 | 0.45 | 0.16 | 1.37 |
| 4 | Trim27 | 0.26 | 8.81E-07 | 1.88E-02 | 0.84 | 0.71 | 0.13 | 1.18 |
| 4 | Wdr31 | 0.66 | 8.82E-07 | 1.88E-02 | 0.24 | 0.15 | 0.1 | 1.65 |
| 4 | Tspyl5 | 0.69 | 9.05E-07 | 1.93E-02 | 0.15 | 0.08 | 0.07 | 1.96 |
| 4 | Aifm1 | 0.32 | 9.09E-07 | 1.94E-02 | 0.69 | 0.52 | 0.17 | 1.33 |
| 4 | Paqr4 | 0.71 | 9.20E-07 | 1.97E-02 | 0.14 | 0.07 | 0.07 | 2 |
| 4 | Aspm | 0.34 | 9.28E-07 | 1.98E-02 | 0.5 | 0.37 | 0.13 | 1.35 |
| 4 | Trappc5 | 0.3 | 9.31E-07 | 1.99E-02 | 0.73 | 0.6 | 0.14 | 1.23 |
| 4 | Tex261 | 0.31 | 9.64E-07 | 2.06E-02 | 0.74 | 0.58 | 0.16 | 1.27 |
| 4 | 9-Mar | 0.32 | 9.70E-07 | 2.07E-02 | 0.6 | 0.44 | 0.16 | 1.36 |
| 4 | Qrsl1 | 0.59 | 9.74E-07 | 2.08E-02 | 0.26 | 0.15 | 0.1 | 1.66 |
| 4 | Kif18a | 0.34 | 9.89E-07 | 2.11E-02 | 0.43 | 0.3 | 0.14 | 1.46 |
| 4 | Wdr75 | 0.35 | 9.95E-07 | 2.13E-02 | 0.65 | 0.51 | 0.14 | 1.28 |
| 4 | Mis18a | 0.34 | 1.03E-06 | 2.20E-02 | 0.6 | 0.44 | 0.17 | 1.38 |
| 4 | Prim1 | 0.26 | 1.04E-06 | 2.22E-02 | 0.73 | 0.56 | 0.18 | 1.32 |
| 4 | E2f3 | 0.37 | 1.04E-06 | 2.22E-02 | 0.64 | 0.5 | 0.14 | 1.27 |
| 4 | Mcrs1 | 0.29 | 1.06E-06 | 2.27E-02 | 0.73 | 0.6 | 0.13 | 1.22 |
| 4 | Ube2g2 | 0.42 | 1.06E-06 | 2.27E-02 | 0.41 | 0.28 | 0.13 | 1.46 |
| 4 | Rnf121 | 0.36 | 1.08E-06 | 2.32E-02 | 0.56 | 0.42 | 0.15 | 1.35 |
| 4 | Agfg2 | 0.55 | 1.11E-06 | 2.38E-02 | 0.3 | 0.2 | 0.11 | 1.54 |
| 4 | Dixdc1 | 0.52 | 1.13E-06 | 2.40E-02 | 0.23 | 0.14 | 0.1 | 1.68 |
| 4 | Tm7sf2 | 0.5 | 1.16E-06 | 2.48E-02 | 0.32 | 0.21 | 0.11 | 1.54 |
| 4 | Mok | 0.7 | 1.17E-06 | 2.49E-02 | 0.12 | 0.06 | 0.06 | 2.09 |
| 4 | Txndc15 | 0.31 | 1.20E-06 | 2.57E-02 | 0.74 | 0.56 | 0.18 | 1.32 |
| 4 | Cgn | 0.31 | 1.21E-06 | 2.58E-02 | 0.52 | 0.38 | 0.15 | 1.39 |
| 4 | Hspbap1 | 0.39 | 1.21E-06 | 2.58E-02 | 0.32 | 0.21 | 0.12 | 1.56 |
| 4 | Chpf | 0.5 | 1.22E-06 | 2.62E-02 | 0.28 | 0.17 | 0.11 | 1.61 |
| 4 | Arl6ip6 | 0.27 | 1.26E-06 | 2.69E-02 | 0.79 | 0.65 | 0.14 | 1.21 |
| 4 | Arxes2 | 0.33 | 1.28E-06 | 2.74E-02 | 0.47 | 0.33 | 0.14 | 1.41 |
| 4 | Amfr | 0.32 | 1.28E-06 | 2.74E-02 | 0.55 | 0.39 | 0.16 | 1.4 |
| 4 | Ralgapb | 0.41 | 1.29E-06 | 2.75E-02 | 0.28 | 0.17 | 0.11 | 1.63 |
| 4 | Bpnt1 | 0.35 | 1.29E-06 | 2.76E-02 | 0.24 | 0.14 | 0.1 | 1.69 |
| 4 | Prkar2b | 0.39 | 1.30E-06 | 2.78E-02 | 0.43 | 0.3 | 0.13 | 1.43 |
| 4 | Tmem219 | 0.38 | 1.32E-06 | 2.81E-02 | 0.27 | 0.16 | 0.11 | 1.65 |
| 4 | G6pc3 | 0.52 | 1.35E-06 | 2.88E-02 | 0.38 | 0.27 | 0.12 | 1.45 |
| 4 | Msh6 | 0.3 | 1.38E-06 | 2.96E-02 | 0.6 | 0.46 | 0.14 | 1.31 |
| 4 | Poglut2 | 0.33 | 1.39E-06 | 2.96E-02 | 0.44 | 0.29 | 0.15 | 1.5 |
| 4 | Ado | 0.42 | 1.41E-06 | 3.01E-02 | 0.47 | 0.34 | 0.13 | 1.38 |
| 4 | Nbn | 0.48 | 1.42E-06 | 3.03E-02 | 0.45 | 0.32 | 0.13 | 1.42 |
| 4 | Dna2 | 0.49 | 1.44E-06 | 3.08E-02 | 0.33 | 0.22 | 0.11 | 1.51 |

**Supplementary Table 3. Marker genes for the original clusters of integrated clustering of E11.5 control and NotchGOF λ-junction ectoderm cells.**

| | | | | | | | | |
|---|---|---|---|---|---|---|---|---|
| 4 | Atp11c | 0.43 | 1.56E-06 | 3.33E-02 | 0.31 | 0.19 | 0.11 | 1.59 |
| 4 | Naa20 | 0.26 | 1.57E-06 | 3.36E-02 | 0.79 | 0.69 | 0.1 | 1.15 |
| 4 | Pdss2 | 0.46 | 1.57E-06 | 3.36E-02 | 0.31 | 0.2 | 0.11 | 1.56 |
| 4 | Flnb | 0.38 | 1.58E-06 | 3.37E-02 | 0.43 | 0.3 | 0.13 | 1.45 |
| 4 | Asxl3 | 0.31 | 1.64E-06 | 3.50E-02 | 0.44 | 0.31 | 0.14 | 1.45 |
| 4 | Ercc8 | 0.47 | 1.67E-06 | 3.58E-02 | 0.29 | 0.18 | 0.1 | 1.57 |
| 4 | Hist1h3b | 0.57 | 1.68E-06 | 3.58E-02 | 0.26 | 0.17 | 0.1 | 1.59 |
| 4 | Ncoa6 | 0.35 | 1.68E-06 | 3.59E-02 | 0.5 | 0.36 | 0.14 | 1.4 |
| 4 | Mef2c | 0.48 | 1.72E-06 | 3.67E-02 | 0.2 | 0.11 | 0.08 | 1.73 |
| 4 | Tchp | 0.3 | 1.73E-06 | 3.69E-02 | 0.38 | 0.25 | 0.13 | 1.52 |
| 4 | Cenpi | 0.74 | 1.73E-06 | 3.70E-02 | 0.21 | 0.12 | 0.09 | 1.69 |
| 4 | Hlcs | 0.53 | 1.74E-06 | 3.72E-02 | 0.28 | 0.18 | 0.1 | 1.57 |
| 4 | Ccdc85b | 0.29 | 1.78E-06 | 3.79E-02 | 0.68 | 0.53 | 0.15 | 1.29 |
| 4 | Sypl | 0.29 | 1.79E-06 | 3.83E-02 | 0.74 | 0.59 | 0.15 | 1.26 |
| 4 | Kcna5 | 1.13 | 1.80E-06 | 3.84E-02 | 0.11 | 0.05 | 0.06 | 2.12 |
| 4 | Nf1 | 0.31 | 1.80E-06 | 3.85E-02 | 0.51 | 0.36 | 0.15 | 1.41 |
| 4 | Nacc1 | 0.31 | 1.82E-06 | 3.90E-02 | 0.54 | 0.4 | 0.14 | 1.35 |
| 4 | Fbln1 | 0.27 | 1.83E-06 | 3.90E-02 | 0.78 | 0.64 | 0.14 | 1.22 |
| 4 | Tcf19 | 0.73 | 1.84E-06 | 3.92E-02 | 0.26 | 0.17 | 0.09 | 1.56 |
| 4 | L3mbtl2 | 0.55 | 1.84E-06 | 3.93E-02 | 0.29 | 0.19 | 0.1 | 1.55 |
| 4 | Tdp1 | 0.42 | 1.84E-06 | 3.94E-02 | 0.4 | 0.28 | 0.13 | 1.46 |
| 4 | Tusc2 | 0.31 | 1.87E-06 | 4.00E-02 | 0.6 | 0.44 | 0.16 | 1.36 |
| 4 | Ppp1r35 | 0.32 | 1.92E-06 | 4.11E-02 | 0.64 | 0.49 | 0.15 | 1.29 |
| 4 | Nsun5 | 0.53 | 1.93E-06 | 4.12E-02 | 0.27 | 0.17 | 0.1 | 1.6 |
| 4 | Lpin1 | 0.86 | 1.98E-06 | 4.22E-02 | 0.16 | 0.08 | 0.07 | 1.87 |
| 4 | Abca3 | 0.42 | 1.98E-06 | 4.23E-02 | 0.26 | 0.16 | 0.1 | 1.65 |
| 4 | Zfp961 | 0.47 | 2.02E-06 | 4.31E-02 | 0.26 | 0.16 | 0.1 | 1.63 |
| 4 | Casp9 | 0.59 | 2.05E-06 | 4.37E-02 | 0.18 | 0.1 | 0.08 | 1.8 |
| 4 | Bcl2 | 0.5 | 2.13E-06 | 4.55E-02 | 0.35 | 0.24 | 0.11 | 1.46 |
| 4 | Hist1h2bp | 1.02 | 2.15E-06 | 4.60E-02 | 0.1 | 0.05 | 0.06 | 2.15 |
| 4 | Spcs3 | 0.37 | 2.16E-06 | 4.61E-02 | 0.51 | 0.38 | 0.14 | 1.36 |
| 4 | 5031439G07Rik | 0.47 | 2.16E-06 | 4.62E-02 | 0.32 | 0.21 | 0.11 | 1.54 |
| 4 | Polq | 0.71 | 2.19E-06 | 4.68E-02 | 0.16 | 0.09 | 0.07 | 1.85 |
| 4 | Ppnr | 0.64 | 2.19E-06 | 4.69E-02 | 0.11 | 0.05 | 0.06 | 2.19 |
| 4 | Ccdc134 | 0.51 | 2.22E-06 | 4.74E-02 | 0.23 | 0.14 | 0.09 | 1.67 |
| 4 | Msrb2 | 0.34 | 2.23E-06 | 4.77E-02 | 0.32 | 0.2 | 0.12 | 1.58 |
| 4 | Fam118a | 0.52 | 2.23E-06 | 4.77E-02 | 0.24 | 0.15 | 0.09 | 1.63 |
| 4 | Arhgap19 | 0.32 | 2.27E-06 | 4.85E-02 | 0.39 | 0.26 | 0.13 | 1.5 |
| 4 | Foxm1 | 0.37 | 2.30E-06 | 4.91E-02 | 0.36 | 0.24 | 0.12 | 1.52 |
| 4 | Trim71 | 0.48 | 2.30E-06 | 4.92E-02 | 0.22 | 0.13 | 0.09 | 1.7 |
| 4 | Spdl1 | 0.38 | 2.31E-06 | 4.93E-02 | 0.52 | 0.37 | 0.15 | 1.39 |
| 5 | Pitx2 | 3.67 | 1.96E-303 | 4.18E-299 | 0.93 | 0.18 | 0.75 | 5.25 |
| 5 | Shh | 4.09 | 5.57E-146 | 1.19E-141 | 0.35 | 0.03 | 0.32 | 12.03 |
| 5 | Unc5c | 2.21 | 9.80E-130 | 2.09E-125 | 0.81 | 0.28 | 0.53 | 2.87 |
| 5 | Epha4 | 1.87 | 1.08E-112 | 2.31E-108 | 0.91 | 0.52 | 0.39 | 1.76 |
| 5 | Krt14 | 1.71 | 4.71E-112 | 1.01E-107 | 0.93 | 0.41 | 0.52 | 2.25 |
| 5 | Foxe1 | 1.53 | 2.50E-109 | 5.35E-105 | 0.59 | 0.13 | 0.46 | 4.56 |
| 5 | Chl1 | 3.44 | 1.71E-108 | 3.64E-104 | 0.3 | 0.03 | 0.27 | 9.5 |
| 5 | Ifitm3 | 1.97 | 2.82E-106 | 6.02E-102 | 0.85 | 0.38 | 0.47 | 2.24 |
| 5 | Fgf9 | 1.89 | 8.04E-106 | 1.72E-101 | 0.83 | 0.38 | 0.45 | 2.18 |
| 5 | Col14a1 | 4.06 | 1.44E-102 | 3.07E-98 | 0.23 | 0.02 | 0.22 | 13.76 |
| 5 | Igfbp4 | 1.34 | 7.70E-100 | 1.65E-95 | 0.96 | 0.72 | 0.24 | 1.34 |
| 5 | Phlda3 | 1.73 | 2.02E-98 | 4.32E-94 | 0.87 | 0.46 | 0.41 | 1.88 |
| 5 | Tac2 | 3.09 | 4.58E-98 | 9.77E-94 | 0.23 | 0.02 | 0.22 | 13 |
| 5 | Csrp2 | 1.52 | 1.04E-96 | 2.23E-92 | 0.96 | 0.83 | 0.14 | 1.16 |
| 5 | Dsc3 | 3.58 | 5.92E-90 | 1.26E-85 | 0.31 | 0.04 | 0.26 | 7 |
| 5 | Fgf8 | 1.2 | 1.75E-87 | 3.73E-83 | 0.69 | 0.23 | 0.46 | 2.97 |

**Supplementary Table 3. Marker genes for the original clusters of integrated clustering of E11.5 control and NotchGOF λ-junction ectoderm cells.**

| | | | | | | | | |
|---|---|---|---|---|---|---|---|---|
| 5 | Cldn10 | 3.59 | 9.62E-86 | 2.05E-81 | 0.31 | 0.05 | 0.26 | 6.39 |
| 5 | Pcolce2 | 2.53 | 5.29E-85 | 1.13E-80 | 0.29 | 0.04 | 0.25 | 7.17 |
| 5 | Ccnd2 | 1.19 | 1.38E-83 | 2.94E-79 | 0.99 | 0.76 | 0.22 | 1.29 |
| 5 | Fabp5 | 0.71 | 8.50E-83 | 1.82E-78 | 1 | 0.99 | 0.01 | 1.01 |
| 5 | Ifitm1 | 2.46 | 1.34E-77 | 2.87E-73 | 0.53 | 0.16 | 0.36 | 3.22 |
| 5 | Ptch1 | 1.94 | 2.14E-76 | 4.57E-72 | 0.7 | 0.32 | 0.38 | 2.2 |
| 5 | Capn6 | 1.6 | 2.49E-74 | 5.31E-70 | 0.78 | 0.42 | 0.37 | 1.88 |
| 5 | Cd59a | 2.02 | 5.44E-74 | 1.16E-69 | 0.65 | 0.27 | 0.38 | 2.44 |
| 5 | Fxyd3 | 1.4 | 9.29E-71 | 1.98E-66 | 0.69 | 0.27 | 0.42 | 2.56 |
| 5 | Siva1 | 0.81 | 5.77E-70 | 1.23E-65 | 1 | 0.97 | 0.03 | 1.03 |
| 5 | Dapl1 | 1.71 | 3.71E-68 | 7.93E-64 | 0.62 | 0.23 | 0.39 | 2.66 |
| 5 | Irx1 | 2.48 | 2.21E-66 | 4.73E-62 | 0.29 | 0.05 | 0.24 | 5.56 |
| 5 | Rpl23 | 0.29 | 2.63E-66 | 5.61E-62 | 1 | 1 | 0 | 1 |
| 5 | Ascl4 | 3.78 | 2.81E-64 | 6.00E-60 | 0.17 | 0.02 | 0.15 | 11.07 |
| 5 | Anxa8 | 2.04 | 2.46E-61 | 5.24E-57 | 0.33 | 0.07 | 0.26 | 4.58 |
| 5 | Isl1 | 0.95 | 1.28E-60 | 2.74E-56 | 0.97 | 0.73 | 0.23 | 1.32 |
| 5 | Rflnb | 1.35 | 2.25E-60 | 4.80E-56 | 0.8 | 0.49 | 0.31 | 1.64 |
| 5 | Kctd1 | 1.3 | 2.94E-58 | 6.28E-54 | 0.8 | 0.47 | 0.33 | 1.71 |
| 5 | Trp63 | 0.89 | 1.77E-57 | 3.78E-53 | 0.88 | 0.48 | 0.41 | 1.86 |
| 5 | Rps20 | 0.27 | 2.07E-57 | 4.43E-53 | 1 | 1 | 0 | 1 |
| 5 | Emp3 | 2.5 | 1.54E-56 | 3.29E-52 | 0.26 | 0.05 | 0.21 | 5.31 |
| 5 | Runx1 | 0.76 | 1.99E-56 | 4.25E-52 | 0.86 | 0.48 | 0.37 | 1.77 |
| 5 | Trim29 | 1.71 | 3.59E-54 | 7.68E-50 | 0.47 | 0.16 | 0.31 | 2.92 |
| 5 | Lef1 | 1.31 | 1.39E-53 | 2.96E-49 | 0.7 | 0.35 | 0.36 | 2.03 |
| 5 | Dlx2 | 1.14 | 1.42E-53 | 3.03E-49 | 0.69 | 0.31 | 0.39 | 2.25 |
| 5 | Gm12446 | 1.65 | 2.90E-53 | 6.20E-49 | 0.35 | 0.09 | 0.26 | 3.99 |
| 5 | Rpl34 | 0.26 | 6.69E-53 | 1.43E-48 | 1 | 1 | 0 | 1 |
| 5 | Spred1 | 1.28 | 6.96E-53 | 1.49E-48 | 0.77 | 0.48 | 0.3 | 1.62 |
| 5 | Amot | 1.29 | 9.48E-52 | 2.03E-47 | 0.82 | 0.55 | 0.27 | 1.49 |
| 5 | Pltp | 1.04 | 2.00E-50 | 4.28E-46 | 0.8 | 0.43 | 0.37 | 1.86 |
| 5 | 1500009L16Rik | 1.22 | 5.18E-50 | 1.11E-45 | 0.77 | 0.5 | 0.27 | 1.54 |
| 5 | Sult5a1 | 2.28 | 1.40E-49 | 2.98E-45 | 0.28 | 0.07 | 0.22 | 4.24 |
| 5 | Npm3 | 0.75 | 2.30E-49 | 4.92E-45 | 0.98 | 0.87 | 0.12 | 1.13 |
| 5 | Pkp4 | 1.1 | 5.23E-49 | 1.12E-44 | 0.8 | 0.56 | 0.24 | 1.43 |
| 5 | Rxfp1 | 2.29 | 9.55E-49 | 2.04E-44 | 0.21 | 0.04 | 0.17 | 5.75 |
| 5 | Pdgfa | 0.63 | 1.34E-48 | 2.86E-44 | 0.99 | 0.93 | 0.06 | 1.06 |
| 5 | Perp | 0.7 | 7.88E-46 | 1.68E-41 | 0.91 | 0.5 | 0.41 | 1.81 |
| 5 | Zfp385c | 1.67 | 1.42E-44 | 3.03E-40 | 0.34 | 0.1 | 0.24 | 3.42 |
| 5 | Fermt1 | 0.83 | 1.95E-43 | 4.16E-39 | 0.87 | 0.52 | 0.34 | 1.65 |
| 5 | Lmo2 | 1 | 5.23E-43 | 1.12E-38 | 0.59 | 0.27 | 0.33 | 2.23 |
| 5 | Ntm | 1.69 | 1.35E-42 | 2.89E-38 | 0.5 | 0.22 | 0.28 | 2.29 |
| 5 | Fbln2 | 2.13 | 1.90E-42 | 4.07E-38 | 0.3 | 0.09 | 0.22 | 3.5 |
| 5 | Itgb5 | 0.93 | 2.00E-42 | 4.27E-38 | 0.79 | 0.47 | 0.32 | 1.68 |
| 5 | Rps27 | 0.27 | 3.25E-42 | 6.94E-38 | 1 | 1 | 0 | 1 |
| 5 | Ctnnb1 | 0.42 | 4.48E-42 | 9.57E-38 | 1 | 0.99 | 0.01 | 1.01 |
| 5 | Rpl12 | 0.26 | 6.95E-42 | 1.48E-37 | 1 | 1 | 0 | 1 |
| 5 | Ptch2 | 2.67 | 2.05E-41 | 4.38E-37 | 0.22 | 0.05 | 0.17 | 4.48 |
| 5 | Ly6g6e | 2.15 | 8.53E-41 | 1.82E-36 | 0.23 | 0.05 | 0.18 | 4.4 |
| 5 | Palmd | 1.95 | 1.76E-40 | 3.77E-36 | 0.21 | 0.04 | 0.16 | 5 |
| 5 | Kif13a | 1.43 | 5.19E-40 | 1.11E-35 | 0.56 | 0.28 | 0.28 | 1.99 |
| 5 | Npm1 | 0.27 | 7.95E-40 | 1.70E-35 | 1 | 1 | 0 | 1 |
| 5 | H19 | 0.48 | 7.32E-39 | 1.56E-34 | 1 | 0.87 | 0.13 | 1.15 |
| 5 | Rab38 | 1.44 | 1.49E-38 | 3.18E-34 | 0.44 | 0.18 | 0.26 | 2.49 |
| 5 | Bmp7 | 1.12 | 1.66E-38 | 3.54E-34 | 0.67 | 0.38 | 0.29 | 1.76 |
| 5 | Eef1b2 | 0.28 | 3.29E-38 | 7.03E-34 | 1 | 1 | 0 | 1 |
| 5 | Emb | 0.98 | 6.15E-38 | 1.31E-33 | 0.87 | 0.72 | 0.15 | 1.2 |
| 5 | Vwa1 | 1.71 | 7.36E-38 | 1.57E-33 | 0.31 | 0.09 | 0.21 | 3.29 |

**Supplementary Table 3. Marker genes for the original clusters of integrated clustering of E11.5 control and NotchGOF λ-junction ectoderm cells.**

| | | | | | | | | |
|---|---|---|---|---|---|---|---|---|
| 5 | Serpinb6b | 1.93 | 2.19E-37 | 4.67E-33 | 0.21 | 0.05 | 0.16 | 4.47 |
| 5 | Itm2b | 0.5 | 3.58E-36 | 7.66E-32 | 0.99 | 0.93 | 0.06 | 1.06 |
| 5 | Sema3d | 0.92 | 6.65E-36 | 1.42E-31 | 0.69 | 0.39 | 0.3 | 1.77 |
| 5 | Cnksr2 | 1.19 | 1.40E-35 | 3.00E-31 | 0.59 | 0.31 | 0.28 | 1.91 |
| 5 | Pabpc1 | 0.39 | 4.52E-35 | 9.65E-31 | 0.99 | 0.99 | 0.01 | 1.01 |
| 5 | Spats2l | 1.22 | 3.15E-34 | 6.73E-30 | 0.59 | 0.31 | 0.28 | 1.88 |
| 5 | Pdlim4 | 1.01 | 9.84E-34 | 2.10E-29 | 0.68 | 0.42 | 0.27 | 1.64 |
| 5 | Gli1 | 1.97 | 1.76E-33 | 3.75E-29 | 0.25 | 0.07 | 0.18 | 3.44 |
| 5 | Ifitm2 | 0.5 | 1.87E-33 | 3.98E-29 | 0.99 | 0.93 | 0.06 | 1.07 |
| 5 | Jph4 | 1.3 | 1.95E-33 | 4.16E-29 | 0.43 | 0.18 | 0.25 | 2.42 |
| 5 | Krt17 | 2.2 | 3.03E-33 | 6.46E-29 | 0.23 | 0.07 | 0.17 | 3.6 |
| 5 | Gpc3 | 0.6 | 3.25E-33 | 6.94E-29 | 0.97 | 0.83 | 0.14 | 1.17 |
| 5 | Agl | 1.32 | 7.29E-33 | 1.56E-28 | 0.57 | 0.33 | 0.25 | 1.76 |
| 5 | Cald1 | 0.8 | 1.10E-32 | 2.35E-28 | 0.9 | 0.68 | 0.22 | 1.33 |
| 5 | Dsg2 | 1.06 | 2.13E-32 | 4.55E-28 | 0.72 | 0.46 | 0.26 | 1.55 |
| 5 | Vcan | 0.55 | 7.35E-32 | 1.57E-27 | 0.94 | 0.73 | 0.21 | 1.29 |
| 5 | Tnfrsf19 | 1.41 | 1.32E-31 | 2.81E-27 | 0.48 | 0.23 | 0.25 | 2.12 |
| 5 | P3h2 | 2.99 | 3.10E-31 | 6.61E-27 | 0.12 | 0.02 | 0.1 | 6.56 |
| 5 | Dsp | 0.48 | 4.21E-31 | 8.99E-27 | 0.99 | 0.86 | 0.14 | 1.16 |
| 5 | Dlx1 | 1.37 | 6.20E-31 | 1.33E-26 | 0.34 | 0.13 | 0.22 | 2.69 |
| 5 | Acaa2 | 0.72 | 8.32E-31 | 1.78E-26 | 0.88 | 0.7 | 0.18 | 1.26 |
| 5 | Cdca7 | 0.8 | 3.83E-30 | 8.18E-26 | 0.84 | 0.63 | 0.22 | 1.35 |
| 5 | Fign | 1.3 | 5.56E-30 | 1.19E-25 | 0.4 | 0.17 | 0.23 | 2.32 |
| 5 | Robo1 | 0.9 | 6.62E-30 | 1.41E-25 | 0.65 | 0.38 | 0.27 | 1.71 |
| 5 | Gja1 | 0.63 | 1.72E-29 | 3.68E-25 | 0.88 | 0.69 | 0.19 | 1.27 |
| 5 | Lpar6 | 0.92 | 3.89E-29 | 8.30E-25 | 0.69 | 0.46 | 0.24 | 1.52 |
| 5 | Ets2 | 1.17 | 4.82E-29 | 1.03E-24 | 0.47 | 0.23 | 0.24 | 2.04 |
| 5 | Hspb2 | 2.06 | 1.21E-28 | 2.59E-24 | 0.3 | 0.11 | 0.19 | 2.69 |
| 5 | Barx1 | 1.23 | 1.97E-28 | 4.20E-24 | 0.27 | 0.09 | 0.18 | 3.07 |
| 5 | Dst | 0.9 | 2.55E-28 | 5.45E-24 | 0.69 | 0.46 | 0.23 | 1.5 |
| 5 | Lgr6 | 2.4 | 1.04E-27 | 2.23E-23 | 0.13 | 0.02 | 0.1 | 5.33 |
| 5 | S100a10 | 0.37 | 1.24E-27 | 2.65E-23 | 0.94 | 0.73 | 0.21 | 1.29 |
| 5 | Serpinb9 | 2.52 | 6.72E-27 | 1.44E-22 | 0.12 | 0.02 | 0.1 | 5.62 |
| 5 | Osbpl3 | 1.35 | 1.28E-26 | 2.74E-22 | 0.34 | 0.14 | 0.2 | 2.43 |
| 5 | Copz2 | 1.22 | 2.88E-26 | 6.15E-22 | 0.45 | 0.23 | 0.22 | 1.96 |
| 5 | Nkain3 | 2.14 | 4.21E-26 | 8.99E-22 | 0.19 | 0.05 | 0.14 | 3.52 |
| 5 | Filip1l | 1.1 | 5.92E-26 | 1.26E-21 | 0.48 | 0.25 | 0.23 | 1.93 |
| 5 | Hmga2 | 0.45 | 6.16E-26 | 1.32E-21 | 0.99 | 0.92 | 0.08 | 1.08 |
| 5 | Cdh6 | 1.37 | 6.50E-26 | 1.39E-21 | 0.29 | 0.11 | 0.18 | 2.65 |
| 5 | Nrxn3 | 1.58 | 6.60E-26 | 1.41E-21 | 0.25 | 0.09 | 0.17 | 2.88 |
| 5 | Apcdd1 | 1.51 | 2.42E-25 | 5.17E-21 | 0.31 | 0.13 | 0.19 | 2.49 |
| 5 | Runx2 | 2.41 | 1.25E-24 | 2.68E-20 | 0.15 | 0.03 | 0.11 | 4.26 |
| 5 | Epha7 | 1.16 | 1.75E-24 | 3.75E-20 | 0.48 | 0.27 | 0.21 | 1.8 |
| 5 | Peg3 | 0.41 | 2.64E-24 | 5.65E-20 | 0.94 | 0.82 | 0.13 | 1.15 |
| 5 | Kctd12b | 1.69 | 4.49E-24 | 9.60E-20 | 0.25 | 0.09 | 0.16 | 2.79 |
| 5 | Tns3 | 1.03 | 1.35E-23 | 2.88E-19 | 0.54 | 0.32 | 0.22 | 1.67 |
| 5 | Prss23 | 1.03 | 2.50E-23 | 5.35E-19 | 0.5 | 0.27 | 0.22 | 1.81 |
| 5 | Hpcal1 | 0.75 | 2.60E-23 | 5.56E-19 | 0.68 | 0.45 | 0.23 | 1.51 |
| 5 | Nt5c | 0.63 | 5.90E-23 | 1.26E-18 | 0.82 | 0.65 | 0.17 | 1.26 |
| 5 | Dmrt2 | 1.37 | 5.92E-23 | 1.26E-18 | 0.24 | 0.08 | 0.16 | 2.85 |
| 5 | Slc39a8 | 1.63 | 9.29E-23 | 1.99E-18 | 0.24 | 0.08 | 0.15 | 2.81 |
| 5 | Lgi2 | 0.97 | 1.13E-22 | 2.41E-18 | 0.5 | 0.28 | 0.22 | 1.8 |
| 5 | Sdc4 | 0.65 | 1.38E-22 | 2.94E-18 | 0.82 | 0.63 | 0.19 | 1.3 |
| 5 | Ece1 | 0.96 | 1.78E-22 | 3.80E-18 | 0.64 | 0.45 | 0.19 | 1.42 |
| 5 | Notum | 1.55 | 3.49E-22 | 7.46E-18 | 0.19 | 0.06 | 0.13 | 3.32 |
| 5 | Slitrk6 | 1.35 | 3.65E-22 | 7.80E-18 | 0.34 | 0.16 | 0.18 | 2.17 |
| 5 | Aqp3 | 2.21 | 4.01E-22 | 8.57E-18 | 0.13 | 0.03 | 0.1 | 4.16 |

**Supplementary Table 3. Marker genes for the original clusters of integrated clustering of E11.5 control and NotchGOF λ-junction ectoderm cells.**

| | | | | | | | | |
|---|---|---|---|---|---|---|---|---|
| 5 | Tgfb1 | 0.89 | 7.21E-22 | 1.54E-17 | 0.34 | 0.15 | 0.19 | 2.3 |
| 5 | Slc4a4 | 1.3 | 7.30E-22 | 1.56E-17 | 0.31 | 0.13 | 0.18 | 2.32 |
| 5 | Lgr5 | 0.8 | 8.32E-22 | 1.78E-17 | 0.44 | 0.22 | 0.22 | 2.01 |
| 5 | Slc7a10 | 2.04 | 8.73E-22 | 1.86E-17 | 0.14 | 0.04 | 0.11 | 3.94 |
| 5 | Pitx1 | 0.35 | 1.07E-21 | 2.29E-17 | 0.91 | 0.63 | 0.28 | 1.44 |
| 5 | Ankrd6 | 1.41 | 3.57E-21 | 7.62E-17 | 0.33 | 0.15 | 0.18 | 2.14 |
| 5 | Dsc2 | 1.38 | 5.60E-21 | 1.20E-16 | 0.41 | 0.22 | 0.19 | 1.87 |
| 5 | Plce1 | 0.86 | 8.77E-21 | 1.87E-16 | 0.52 | 0.31 | 0.21 | 1.68 |
| 5 | Lpin2 | 0.87 | 1.16E-20 | 2.47E-16 | 0.63 | 0.45 | 0.19 | 1.42 |
| 5 | Pik3r1 | 0.49 | 1.22E-20 | 2.60E-16 | 0.81 | 0.58 | 0.23 | 1.4 |
| 5 | Cltrn | 2.25 | 1.40E-20 | 2.98E-16 | 0.12 | 0.03 | 0.09 | 4.29 |
| 5 | Ppp1r12b | 1.41 | 2.81E-20 | 6.00E-16 | 0.32 | 0.15 | 0.17 | 2.18 |
| 5 | Pdgfc | 0.46 | 3.03E-20 | 6.48E-16 | 0.9 | 0.75 | 0.15 | 1.2 |
| 5 | Bcl11a | 0.56 | 6.59E-20 | 1.41E-15 | 0.92 | 0.79 | 0.13 | 1.17 |
| 5 | Hspb1 | 0.36 | 1.13E-19 | 2.41E-15 | 0.62 | 0.36 | 0.25 | 1.7 |
| 5 | Dok5 | 0.71 | 1.16E-19 | 2.47E-15 | 0.59 | 0.36 | 0.24 | 1.67 |
| 5 | Greb1 | 1.35 | 1.50E-19 | 3.21E-15 | 0.35 | 0.18 | 0.17 | 1.96 |
| 5 | Tpd52l1 | 1.27 | 4.65E-19 | 9.94E-15 | 0.3 | 0.14 | 0.16 | 2.15 |
| 5 | Snai2 | 0.9 | 9.40E-19 | 2.01E-14 | 0.35 | 0.16 | 0.19 | 2.13 |
| 5 | Btg1 | 0.65 | 9.55E-19 | 2.04E-14 | 0.84 | 0.74 | 0.1 | 1.13 |
| 5 | Col4a2 | 0.59 | 1.11E-18 | 2.36E-14 | 0.71 | 0.48 | 0.23 | 1.47 |
| 5 | Amotl1 | 0.45 | 1.97E-18 | 4.21E-14 | 0.85 | 0.68 | 0.16 | 1.24 |
| 5 | Etl4 | 0.62 | 3.36E-18 | 7.18E-14 | 0.75 | 0.56 | 0.19 | 1.33 |
| 5 | Capg | 0.8 | 3.71E-18 | 7.92E-14 | 0.55 | 0.34 | 0.21 | 1.61 |
| 5 | Ugdh | 0.77 | 3.85E-18 | 8.23E-14 | 0.61 | 0.42 | 0.19 | 1.44 |
| 5 | Ablim1 | 0.59 | 4.06E-18 | 8.66E-14 | 0.73 | 0.53 | 0.19 | 1.36 |
| 5 | Hdac8 | 1.08 | 4.85E-18 | 1.04E-13 | 0.4 | 0.22 | 0.18 | 1.83 |
| 5 | Bag3 | 1.59 | 5.95E-18 | 1.27E-13 | 0.18 | 0.06 | 0.12 | 2.86 |
| 5 | Klf5 | 0.73 | 8.89E-18 | 1.90E-13 | 0.61 | 0.39 | 0.21 | 1.54 |
| 5 | Ppp3ca | 0.74 | 9.35E-18 | 2.00E-13 | 0.7 | 0.53 | 0.17 | 1.31 |
| 5 | Dmtn | 1.45 | 9.71E-18 | 2.07E-13 | 0.22 | 0.09 | 0.13 | 2.49 |
| 5 | Cd109 | 1.57 | 1.28E-17 | 2.74E-13 | 0.14 | 0.04 | 0.1 | 3.46 |
| 5 | Bcam | 0.43 | 1.70E-17 | 3.64E-13 | 0.9 | 0.71 | 0.19 | 1.27 |
| 5 | Anxa2 | 0.3 | 5.89E-17 | 1.26E-12 | 0.96 | 0.74 | 0.22 | 1.3 |
| 5 | Gask1b | 1.7 | 6.81E-17 | 1.45E-12 | 0.17 | 0.06 | 0.11 | 2.91 |
| 5 | Mycl | 0.46 | 7.54E-17 | 1.61E-12 | 0.8 | 0.63 | 0.18 | 1.28 |
| 5 | Nell2 | 1.19 | 8.10E-17 | 1.73E-12 | 0.25 | 0.11 | 0.14 | 2.32 |
| 5 | Adgra3 | 0.82 | 1.20E-16 | 2.56E-12 | 0.63 | 0.48 | 0.15 | 1.31 |
| 5 | Slc35g1 | 1.24 | 1.26E-16 | 2.69E-12 | 0.29 | 0.14 | 0.15 | 2.04 |
| 5 | Stk26 | 0.65 | 1.62E-16 | 3.46E-12 | 0.64 | 0.46 | 0.19 | 1.41 |
| 5 | Pard6g | 0.54 | 2.09E-16 | 4.46E-12 | 0.78 | 0.63 | 0.15 | 1.23 |
| 5 | Camk4 | 1.24 | 2.22E-16 | 4.75E-12 | 0.13 | 0.04 | 0.09 | 3.33 |
| 5 | Tiam1 | 0.77 | 3.35E-16 | 7.15E-12 | 0.61 | 0.44 | 0.17 | 1.38 |
| 5 | Edaradd | 1.4 | 3.87E-16 | 8.28E-12 | 0.19 | 0.07 | 0.12 | 2.66 |
| 5 | Mrpl23 | 0.34 | 5.36E-16 | 1.14E-11 | 0.97 | 0.92 | 0.05 | 1.05 |
| 5 | Mycn | 0.73 | 6.35E-16 | 1.36E-11 | 0.52 | 0.34 | 0.18 | 1.54 |
| 5 | Nrp2 | 0.97 | 6.63E-16 | 1.42E-11 | 0.32 | 0.16 | 0.16 | 1.99 |
| 5 | Kif1b | 0.58 | 7.54E-16 | 1.61E-11 | 0.76 | 0.64 | 0.12 | 1.19 |
| 5 | Gm2694 | 1.39 | 8.66E-16 | 1.85E-11 | 0.17 | 0.06 | 0.11 | 2.71 |
| 5 | Lmo4 | 0.31 | 8.75E-16 | 1.87E-11 | 0.97 | 0.91 | 0.06 | 1.07 |
| 5 | Cox7a2l | 0.28 | 9.18E-16 | 1.96E-11 | 1 | 0.98 | 0.02 | 1.02 |
| 5 | Mir205hg | 0.56 | 1.14E-15 | 2.44E-11 | 0.64 | 0.42 | 0.22 | 1.51 |
| 5 | Rab34 | 0.54 | 1.17E-15 | 2.49E-11 | 0.74 | 0.61 | 0.14 | 1.22 |
| 5 | Ralgps2 | 0.57 | 1.40E-15 | 3.00E-11 | 0.84 | 0.72 | 0.12 | 1.17 |
| 5 | Tpgs2 | 0.79 | 1.54E-15 | 3.28E-11 | 0.65 | 0.49 | 0.16 | 1.32 |
| 5 | 2-Mar | 0.56 | 2.23E-15 | 4.76E-11 | 0.75 | 0.59 | 0.15 | 1.26 |
| 5 | Rest | 0.45 | 3.01E-15 | 6.42E-11 | 0.83 | 0.67 | 0.17 | 1.25 |

**Supplementary Table 3. Marker genes for the original clusters of integrated clustering of E11.5 control and NotchGOF λ-junction ectoderm cells.**

| | | | | | | | | |
|---|---|---|---|---|---|---|---|---|
| 5 | B130024G19Rik | 1.75 | 3.13E-15 | 6.69E-11 | 0.12 | 0.04 | 0.09 | 3.32 |
| 5 | Polr3g | 0.79 | 3.36E-15 | 7.17E-11 | 0.49 | 0.32 | 0.17 | 1.55 |
| 5 | Nampt | 0.82 | 5.66E-15 | 1.21E-10 | 0.49 | 0.32 | 0.17 | 1.55 |
| 5 | Arid3b | 0.64 | 5.72E-15 | 1.22E-10 | 0.57 | 0.4 | 0.17 | 1.44 |
| 5 | Srd5a1 | 0.87 | 7.26E-15 | 1.55E-10 | 0.5 | 0.32 | 0.17 | 1.53 |
| 5 | Serpine2 | 0.75 | 8.68E-15 | 1.85E-10 | 0.61 | 0.45 | 0.16 | 1.35 |
| 5 | Slc39a10 | 0.49 | 1.13E-14 | 2.42E-10 | 0.84 | 0.75 | 0.1 | 1.13 |
| 5 | Frem2 | 0.46 | 1.22E-14 | 2.61E-10 | 0.79 | 0.61 | 0.18 | 1.29 |
| 5 | Mfge8 | 0.36 | 1.59E-14 | 3.39E-10 | 0.73 | 0.54 | 0.19 | 1.35 |
| 5 | D630045J12Rik | 0.88 | 1.63E-14 | 3.48E-10 | 0.39 | 0.23 | 0.16 | 1.72 |
| 5 | Samd5 | 0.41 | 2.04E-14 | 4.36E-10 | 0.85 | 0.67 | 0.18 | 1.27 |
| 5 | Cd44 | 1.79 | 2.71E-14 | 5.79E-10 | 0.13 | 0.04 | 0.09 | 3.02 |
| 5 | Cbr2 | 1.64 | 3.73E-14 | 7.96E-10 | 0.11 | 0.03 | 0.08 | 3.48 |
| 5 | Plcd3 | 1.62 | 4.38E-14 | 9.35E-10 | 0.14 | 0.05 | 0.09 | 2.91 |
| 5 | Vwa2 | 1.22 | 4.50E-14 | 9.60E-10 | 0.25 | 0.12 | 0.13 | 2.07 |
| 5 | Nfkb1 | 0.86 | 5.02E-14 | 1.07E-09 | 0.4 | 0.23 | 0.17 | 1.71 |
| 5 | Pnrc1 | 0.33 | 5.69E-14 | 1.22E-09 | 0.94 | 0.9 | 0.04 | 1.04 |
| 5 | D16Ertd472e | 0.81 | 7.76E-14 | 1.66E-09 | 0.39 | 0.23 | 0.16 | 1.73 |
| 5 | Ntf5 | 0.7 | 8.99E-14 | 1.92E-09 | 0.48 | 0.3 | 0.17 | 1.57 |
| 5 | Zfp703 | 0.92 | 9.05E-14 | 1.93E-09 | 0.44 | 0.28 | 0.16 | 1.56 |
| 5 | Tmem256 | 0.34 | 1.30E-13 | 2.78E-09 | 0.93 | 0.87 | 0.06 | 1.07 |
| 5 | Msx2 | 0.53 | 1.63E-13 | 3.48E-09 | 0.59 | 0.39 | 0.2 | 1.53 |
| 5 | Npnt | 0.52 | 1.64E-13 | 3.50E-09 | 0.62 | 0.43 | 0.19 | 1.44 |
| 5 | Serpinb6a | 0.59 | 1.71E-13 | 3.65E-09 | 0.62 | 0.47 | 0.15 | 1.33 |
| 5 | Ahnak | 0.47 | 1.80E-13 | 3.86E-09 | 0.54 | 0.34 | 0.2 | 1.58 |
| 5 | Spred2 | 0.63 | 1.82E-13 | 3.89E-09 | 0.54 | 0.37 | 0.18 | 1.48 |
| 5 | Clta | 0.35 | 1.91E-13 | 4.09E-09 | 0.99 | 0.98 | 0.01 | 1.01 |
| 5 | Kremen2 | 0.72 | 2.31E-13 | 4.93E-09 | 0.32 | 0.17 | 0.15 | 1.89 |
| 5 | Col5a2 | 0.98 | 2.58E-13 | 5.51E-09 | 0.29 | 0.15 | 0.14 | 1.91 |
| 5 | Kcnmb4 | 0.64 | 2.97E-13 | 6.34E-09 | 0.57 | 0.39 | 0.18 | 1.46 |
| 5 | Tmtc1 | 1.5 | 3.72E-13 | 7.94E-09 | 0.15 | 0.06 | 0.09 | 2.67 |
| 5 | Celf2 | 0.62 | 4.70E-13 | 1.00E-08 | 0.71 | 0.57 | 0.14 | 1.25 |
| 5 | Usp46 | 0.4 | 5.76E-13 | 1.23E-08 | 0.84 | 0.71 | 0.13 | 1.19 |
| 5 | Btrc | 0.81 | 5.99E-13 | 1.28E-08 | 0.48 | 0.32 | 0.16 | 1.5 |
| 5 | Pam | 1.78 | 7.35E-13 | 1.57E-08 | 0.21 | 0.1 | 0.11 | 2.16 |
| 5 | Hspb8 | 0.35 | 1.02E-12 | 2.18E-08 | 0.67 | 0.47 | 0.2 | 1.43 |
| 5 | Nr2f2 | 0.54 | 1.23E-12 | 2.63E-08 | 0.71 | 0.58 | 0.14 | 1.23 |
| 5 | Rgs12 | 0.42 | 1.63E-12 | 3.49E-08 | 0.63 | 0.45 | 0.19 | 1.41 |
| 5 | 1600014C10Rik | 1.24 | 1.69E-12 | 3.60E-08 | 0.23 | 0.12 | 0.12 | 2.02 |
| 5 | Itprid2 | 1.24 | 1.89E-12 | 4.03E-08 | 0.23 | 0.11 | 0.12 | 2.03 |
| 5 | Cox6b2 | 0.62 | 2.20E-12 | 4.69E-08 | 0.64 | 0.5 | 0.14 | 1.28 |
| 5 | Sp6 | 1.1 | 2.96E-12 | 6.32E-08 | 0.23 | 0.11 | 0.12 | 2.06 |
| 5 | Ackr3 | 0.46 | 3.73E-12 | 7.97E-08 | 0.54 | 0.36 | 0.18 | 1.48 |
| 5 | Mbnl2 | 0.33 | 4.84E-12 | 1.03E-07 | 0.89 | 0.78 | 0.11 | 1.15 |
| 5 | Plec | 0.57 | 5.63E-12 | 1.20E-07 | 0.6 | 0.44 | 0.16 | 1.37 |
| 5 | Aprt | 0.32 | 6.27E-12 | 1.34E-07 | 0.98 | 0.93 | 0.05 | 1.06 |
| 5 | Fam83f | 1.42 | 6.56E-12 | 1.40E-07 | 0.11 | 0.03 | 0.07 | 3.12 |
| 5 | Armh4 | 1.2 | 7.77E-12 | 1.66E-07 | 0.3 | 0.18 | 0.12 | 1.71 |
| 5 | Igf2 | 0.3 | 7.96E-12 | 1.70E-07 | 0.9 | 0.81 | 0.1 | 1.12 |
| 5 | Tbx3 | 1.11 | 8.37E-12 | 1.79E-07 | 0.18 | 0.08 | 0.1 | 2.32 |
| 5 | Bmp4 | 0.52 | 1.58E-11 | 3.38E-07 | 0.71 | 0.52 | 0.19 | 1.36 |
| 5 | Irx2 | 1.48 | 1.89E-11 | 4.04E-07 | 0.13 | 0.05 | 0.08 | 2.67 |
| 5 | Minar2 | 1.06 | 1.98E-11 | 4.24E-07 | 0.28 | 0.15 | 0.13 | 1.86 |
| 5 | Hs3st3b1 | 1.7 | 2.36E-11 | 5.04E-07 | 0.11 | 0.04 | 0.07 | 2.85 |
| 5 | Jag2 | 0.55 | 2.92E-11 | 6.23E-07 | 0.44 | 0.27 | 0.16 | 1.6 |
| 5 | Bdh2 | 1.45 | 4.45E-11 | 9.51E-07 | 0.14 | 0.05 | 0.08 | 2.5 |
| 5 | Dusp23 | 1.12 | 4.63E-11 | 9.90E-07 | 0.21 | 0.11 | 0.11 | 2 |

**Supplementary Table 3. Marker genes for the original clusters of integrated clustering of E11.5 control and NotchGOF λ-junction ectoderm cells.**

| | | | | | | | | |
|---|---|---|---|---|---|---|---|---|
| 5 | Klk8 | 0.55 | 5.71E-11 | 1.22E-06 | 0.7 | 0.55 | 0.15 | 1.27 |
| 5 | Cers4 | 0.67 | 6.54E-11 | 1.40E-06 | 0.51 | 0.37 | 0.14 | 1.39 |
| 5 | Frmd6 | 0.8 | 8.86E-11 | 1.89E-06 | 0.36 | 0.22 | 0.14 | 1.62 |
| 5 | Krt5 | 0.35 | 1.03E-10 | 2.21E-06 | 0.49 | 0.33 | 0.16 | 1.49 |
| 5 | Inpp5f | 0.79 | 1.08E-10 | 2.31E-06 | 0.39 | 0.25 | 0.14 | 1.54 |
| 5 | Gnb5 | 0.85 | 1.16E-10 | 2.47E-06 | 0.38 | 0.24 | 0.13 | 1.55 |
| 5 | Osbpl6 | 1.07 | 1.44E-10 | 3.08E-06 | 0.24 | 0.13 | 0.11 | 1.85 |
| 5 | Efcab1 | 1.09 | 1.48E-10 | 3.17E-06 | 0.19 | 0.09 | 0.1 | 2.14 |
| 5 | Prkci | 0.58 | 1.55E-10 | 3.32E-06 | 0.57 | 0.43 | 0.15 | 1.34 |
| 5 | Pid1 | 1.57 | 1.72E-10 | 3.67E-06 | 0.14 | 0.06 | 0.08 | 2.45 |
| 5 | Sesn1 | 0.84 | 1.82E-10 | 3.88E-06 | 0.33 | 0.2 | 0.13 | 1.64 |
| 5 | Nav2 | 0.37 | 1.88E-10 | 4.01E-06 | 0.82 | 0.69 | 0.13 | 1.18 |
| 5 | Xpa | 0.56 | 1.88E-10 | 4.01E-06 | 0.61 | 0.47 | 0.14 | 1.3 |
| 5 | Col18a1 | 0.29 | 2.08E-10 | 4.45E-06 | 0.93 | 0.85 | 0.08 | 1.09 |
| 5 | Pde10a | 1.47 | 2.66E-10 | 5.69E-06 | 0.11 | 0.04 | 0.07 | 2.77 |
| 5 | Col9a3 | 0.76 | 3.03E-10 | 6.47E-06 | 0.26 | 0.14 | 0.12 | 1.86 |
| 5 | Spon2 | 1.36 | 3.26E-10 | 6.95E-06 | 0.13 | 0.05 | 0.08 | 2.51 |
| 5 | Trp53 | 0.29 | 3.31E-10 | 7.08E-06 | 0.93 | 0.89 | 0.05 | 1.05 |
| 5 | 2610301B20Rik | 0.86 | 3.37E-10 | 7.19E-06 | 0.41 | 0.29 | 0.12 | 1.41 |
| 5 | Cemip2 | 0.33 | 3.38E-10 | 7.23E-06 | 0.72 | 0.59 | 0.13 | 1.21 |
| 5 | Micu2 | 0.48 | 3.76E-10 | 8.03E-06 | 0.7 | 0.58 | 0.12 | 1.2 |
| 5 | Cpe | 0.52 | 3.76E-10 | 8.04E-06 | 0.6 | 0.46 | 0.14 | 1.31 |
| 5 | Rpl9-ps6 | 0.32 | 4.10E-10 | 8.76E-06 | 0.88 | 0.8 | 0.08 | 1.1 |
| 5 | Plek2 | 1.36 | 4.63E-10 | 9.90E-06 | 0.19 | 0.1 | 0.09 | 1.98 |
| 5 | Acad8 | 0.75 | 5.11E-10 | 1.09E-05 | 0.44 | 0.32 | 0.13 | 1.4 |
| 5 | Etv1 | 1.05 | 6.72E-10 | 1.44E-05 | 0.28 | 0.16 | 0.11 | 1.69 |
| 5 | Yap1 | 0.42 | 7.40E-10 | 1.58E-05 | 0.72 | 0.6 | 0.12 | 1.2 |
| 5 | Plxnc1 | 1.14 | 7.98E-10 | 1.70E-05 | 0.16 | 0.07 | 0.09 | 2.21 |
| 5 | Slc12a4 | 0.63 | 8.11E-10 | 1.73E-05 | 0.33 | 0.2 | 0.13 | 1.66 |
| 5 | Mdga1 | 1.47 | 8.13E-10 | 1.74E-05 | 0.11 | 0.04 | 0.07 | 2.63 |
| 5 | Ext1 | 0.55 | 8.33E-10 | 1.78E-05 | 0.63 | 0.51 | 0.12 | 1.24 |
| 5 | Adamts18 | 0.48 | 8.76E-10 | 1.87E-05 | 0.41 | 0.26 | 0.15 | 1.58 |
| 5 | 2410006H16Rik | 0.44 | 9.37E-10 | 2.00E-05 | 0.92 | 0.86 | 0.07 | 1.08 |
| 5 | Zfp36l2 | 0.54 | 9.52E-10 | 2.03E-05 | 0.38 | 0.24 | 0.14 | 1.58 |
| 5 | Lmo7 | 0.62 | 9.86E-10 | 2.11E-05 | 0.57 | 0.44 | 0.13 | 1.29 |
| 5 | Srpk1 | 0.32 | 9.88E-10 | 2.11E-05 | 0.9 | 0.83 | 0.07 | 1.08 |
| 5 | Chst11 | 0.86 | 1.04E-09 | 2.22E-05 | 0.27 | 0.15 | 0.12 | 1.76 |
| 5 | Tmod3 | 0.38 | 1.47E-09 | 3.14E-05 | 0.75 | 0.65 | 0.09 | 1.14 |
| 5 | Rps27rt | 0.36 | 1.53E-09 | 3.27E-05 | 0.85 | 0.78 | 0.08 | 1.1 |
| 5 | Golim4 | 0.46 | 1.66E-09 | 3.55E-05 | 0.71 | 0.6 | 0.11 | 1.18 |
| 5 | Dse | 1.01 | 1.80E-09 | 3.85E-05 | 0.2 | 0.1 | 0.1 | 1.94 |
| 5 | Flot2 | 0.7 | 1.96E-09 | 4.18E-05 | 0.39 | 0.27 | 0.12 | 1.46 |
| 5 | Bdnf | 0.82 | 2.18E-09 | 4.65E-05 | 0.29 | 0.17 | 0.12 | 1.68 |
| 5 | Mdm1 | 0.53 | 2.94E-09 | 6.28E-05 | 0.43 | 0.29 | 0.14 | 1.47 |
| 5 | Pthlh | 0.84 | 3.33E-09 | 7.12E-05 | 0.12 | 0.05 | 0.07 | 2.52 |
| 5 | Gas7 | 1.02 | 3.83E-09 | 8.18E-05 | 0.19 | 0.1 | 0.09 | 1.98 |
| 5 | Zfp9 | 1.15 | 4.06E-09 | 8.66E-05 | 0.14 | 0.06 | 0.08 | 2.26 |
| 5 | 9330162012Rik | 1.52 | 4.16E-09 | 8.89E-05 | 0.1 | 0.04 | 0.06 | 2.66 |
| 5 | Col27a1 | 1.39 | 4.62E-09 | 9.88E-05 | 0.14 | 0.06 | 0.08 | 2.25 |
| 5 | Rbm43 | 1.06 | 4.93E-09 | 1.05E-04 | 0.18 | 0.09 | 0.09 | 1.97 |
| 5 | Trim8 | 0.63 | 5.03E-09 | 1.07E-04 | 0.45 | 0.32 | 0.13 | 1.41 |
| 5 | Gm9922 | 1.3 | 5.41E-09 | 1.16E-04 | 0.2 | 0.11 | 0.09 | 1.85 |
| 5 | Eef2k | 0.82 | 5.46E-09 | 1.17E-04 | 0.28 | 0.17 | 0.12 | 1.68 |
| 5 | Baiap2 | 0.56 | 6.04E-09 | 1.29E-04 | 0.34 | 0.21 | 0.13 | 1.62 |
| 5 | Inka1 | 0.73 | 7.18E-09 | 1.53E-04 | 0.16 | 0.07 | 0.08 | 2.15 |
| 5 | 1700007K13Rik | 1.1 | 8.28E-09 | 1.77E-04 | 0.12 | 0.05 | 0.07 | 2.4 |
| 5 | Snhg1 | 0.28 | 8.43E-09 | 1.80E-04 | 0.96 | 0.92 | 0.04 | 1.05 |

**Supplementary Table 3. Marker genes for the original clusters of integrated clustering of E11.5 control and NotchGOF λ-junction ectoderm cells.**

| | | | | | | | | |
|---|---|---|---|---|---|---|---|---|
| 5 | Zfp503 | 0.4 | 8.55E-09 | 1.83E-04 | 0.64 | 0.49 | 0.15 | 1.3 |
| 5 | Cpne5 | 1.06 | 8.66E-09 | 1.85E-04 | 0.24 | 0.14 | 0.1 | 1.69 |
| 5 | Frmpd1 | 1.48 | 9.27E-09 | 1.98E-04 | 0.1 | 0.04 | 0.06 | 2.59 |
| 5 | Dbp | 0.82 | 1.01E-08 | 2.16E-04 | 0.31 | 0.19 | 0.11 | 1.59 |
| 5 | Tmc7 | 1.02 | 1.01E-08 | 2.17E-04 | 0.13 | 0.06 | 0.08 | 2.29 |
| 5 | Cuedc1 | 0.43 | 1.02E-08 | 2.17E-04 | 0.62 | 0.51 | 0.12 | 1.23 |
| 5 | Hprt | 0.48 | 1.18E-08 | 2.52E-04 | 0.58 | 0.46 | 0.12 | 1.25 |
| 5 | Bard1 | 0.84 | 1.34E-08 | 2.86E-04 | 0.28 | 0.18 | 0.11 | 1.62 |
| 5 | Ptprm | 1.15 | 1.41E-08 | 3.02E-04 | 0.18 | 0.1 | 0.09 | 1.91 |
| 5 | Cdk5r1 | 0.98 | 1.75E-08 | 3.74E-04 | 0.14 | 0.06 | 0.08 | 2.21 |
| 5 | Synm | 1.33 | 1.83E-08 | 3.91E-04 | 0.13 | 0.06 | 0.07 | 2.2 |
| 5 | Ldlrap1 | 0.48 | 1.87E-08 | 4.00E-04 | 0.44 | 0.3 | 0.14 | 1.45 |
| 5 | N4bp3 | 1.31 | 1.95E-08 | 4.16E-04 | 0.15 | 0.07 | 0.08 | 2.07 |
| 5 | Sertad4 | 0.53 | 2.37E-08 | 5.06E-04 | 0.45 | 0.32 | 0.13 | 1.42 |
| 5 | Chchd5 | 0.46 | 2.79E-08 | 5.96E-04 | 0.62 | 0.49 | 0.13 | 1.27 |
| 5 | Vwa5a | 0.88 | 3.70E-08 | 7.91E-04 | 0.26 | 0.16 | 0.1 | 1.66 |
| 5 | Sgk3 | 0.74 | 3.71E-08 | 7.92E-04 | 0.21 | 0.12 | 0.1 | 1.83 |
| 5 | Pcdh10 | 1.02 | 4.40E-08 | 9.39E-04 | 0.16 | 0.08 | 0.08 | 2.01 |
| 5 | Arhgdib | 0.51 | 4.52E-08 | 9.65E-04 | 0.32 | 0.2 | 0.12 | 1.6 |
| 5 | Celsr2 | 0.6 | 5.41E-08 | 1.15E-03 | 0.41 | 0.29 | 0.11 | 1.39 |
| 5 | 10-Sep | 0.53 | 5.57E-08 | 1.19E-03 | 0.48 | 0.36 | 0.13 | 1.36 |
| 5 | Bach2 | 0.59 | 6.16E-08 | 1.32E-03 | 0.35 | 0.24 | 0.12 | 1.49 |
| 5 | Gga2 | 0.41 | 6.36E-08 | 1.36E-03 | 0.63 | 0.5 | 0.13 | 1.26 |
| 5 | Sik1 | 0.64 | 9.11E-08 | 1.95E-03 | 0.26 | 0.15 | 0.1 | 1.68 |
| 5 | Isoc1 | 0.3 | 1.04E-07 | 2.22E-03 | 0.77 | 0.7 | 0.08 | 1.11 |
| 5 | Cgnl1 | 0.37 | 1.23E-07 | 2.63E-03 | 0.27 | 0.16 | 0.11 | 1.67 |
| 5 | Pdzrn3 | 0.74 | 1.30E-07 | 2.79E-03 | 0.21 | 0.11 | 0.09 | 1.8 |
| 5 | Chst15 | 0.72 | 1.35E-07 | 2.89E-03 | 0.23 | 0.13 | 0.1 | 1.75 |
| 5 | Irs2 | 0.47 | 1.40E-07 | 2.99E-03 | 0.55 | 0.43 | 0.12 | 1.28 |
| 5 | Dkk3 | 0.82 | 1.48E-07 | 3.17E-03 | 0.25 | 0.16 | 0.1 | 1.61 |
| 5 | Dgkh | 1.29 | 1.55E-07 | 3.30E-03 | 0.1 | 0.04 | 0.06 | 2.36 |
| 5 | Rap2c | 0.31 | 1.65E-07 | 3.53E-03 | 0.75 | 0.66 | 0.09 | 1.14 |
| 5 | Dzip1 | 0.65 | 1.74E-07 | 3.72E-03 | 0.31 | 0.2 | 0.1 | 1.51 |
| 5 | Col4a1 | 0.34 | 1.89E-07 | 4.04E-03 | 0.81 | 0.67 | 0.14 | 1.21 |
| 5 | Prkd3 | 0.51 | 2.17E-07 | 4.64E-03 | 0.63 | 0.54 | 0.09 | 1.17 |
| 5 | Parvb | 1.5 | 2.17E-07 | 4.64E-03 | 0.1 | 0.05 | 0.06 | 2.31 |
| 5 | Pcsk6 | 1.38 | 2.17E-07 | 4.64E-03 | 0.1 | 0.04 | 0.06 | 2.3 |
| 5 | Sall4 | 1.45 | 2.41E-07 | 5.14E-03 | 0.1 | 0.04 | 0.06 | 2.35 |
| 5 | Pqlc3 | 0.73 | 2.58E-07 | 5.51E-03 | 0.27 | 0.16 | 0.1 | 1.62 |
| 5 | Arhgap42 | 0.6 | 2.60E-07 | 5.56E-03 | 0.38 | 0.27 | 0.11 | 1.42 |
| 5 | Ptpn13 | 0.29 | 2.82E-07 | 6.01E-03 | 0.69 | 0.56 | 0.13 | 1.23 |
| 5 | Rac3 | 0.35 | 2.92E-07 | 6.23E-03 | 0.53 | 0.4 | 0.12 | 1.3 |
| 5 | Adamts17 | 1.09 | 2.99E-07 | 6.39E-03 | 0.19 | 0.11 | 0.08 | 1.74 |
| 5 | Ddx25 | 0.9 | 3.03E-07 | 6.48E-03 | 0.19 | 0.11 | 0.08 | 1.79 |
| 5 | Plagl1 | 0.37 | 3.30E-07 | 7.04E-03 | 0.54 | 0.42 | 0.12 | 1.29 |
| 5 | Arhgap24 | 0.95 | 3.52E-07 | 7.52E-03 | 0.14 | 0.07 | 0.07 | 2.03 |
| 5 | Luzp1 | 0.26 | 3.65E-07 | 7.79E-03 | 0.88 | 0.8 | 0.08 | 1.1 |
| 5 | Sh3bp1 | 0.62 | 3.79E-07 | 8.09E-03 | 0.4 | 0.29 | 0.12 | 1.4 |
| 5 | Arhgap10 | 0.49 | 4.07E-07 | 8.69E-03 | 0.55 | 0.42 | 0.13 | 1.3 |
| 5 | Trim44 | 0.25 | 4.18E-07 | 8.92E-03 | 0.84 | 0.78 | 0.06 | 1.07 |
| 5 | Foxo1 | 0.54 | 4.58E-07 | 9.78E-03 | 0.39 | 0.28 | 0.11 | 1.38 |
| 5 | Jade2 | 1.53 | 5.10E-07 | 1.09E-02 | 0.1 | 0.05 | 0.06 | 2.24 |
| 5 | Sorcs2 | 0.56 | 5.16E-07 | 1.10E-02 | 0.17 | 0.09 | 0.08 | 1.88 |
| 5 | Slc25a17 | 0.39 | 5.17E-07 | 1.10E-02 | 0.79 | 0.7 | 0.08 | 1.12 |
| 5 | Fam120a | 0.43 | 5.27E-07 | 1.12E-02 | 0.62 | 0.52 | 0.1 | 1.2 |
| 5 | Agap1 | 0.46 | 5.28E-07 | 1.13E-02 | 0.57 | 0.47 | 0.1 | 1.21 |
| 5 | Mbd2 | 0.31 | 5.68E-07 | 1.21E-02 | 0.7 | 0.61 | 0.09 | 1.15 |

**Supplementary Table 3. Marker genes for the original clusters of integrated clustering of E11.5 control and NotchGOF λ-junction ectoderm cells.**

| | | | | | | | | |
|---|---|---|---|---|---|---|---|---|
| 5 | Glul | 0.34 | 5.82E-07 | 1.24E-02 | 0.58 | 0.45 | 0.13 | 1.28 |
| 5 | Ccdc148 | 1.35 | 6.22E-07 | 1.33E-02 | 0.11 | 0.05 | 0.06 | 2.16 |
| 5 | Rnf217 | 0.72 | 6.51E-07 | 1.39E-02 | 0.25 | 0.16 | 0.09 | 1.6 |
| 5 | Fgfr1op2 | 0.35 | 6.83E-07 | 1.46E-02 | 0.82 | 0.72 | 0.1 | 1.14 |
| 5 | Pnldc1 | 1.01 | 7.10E-07 | 1.52E-02 | 0.11 | 0.05 | 0.06 | 2.2 |
| 5 | Slc25a12 | 0.87 | 7.25E-07 | 1.55E-02 | 0.24 | 0.15 | 0.09 | 1.6 |
| 5 | Cox4i2 | 0.31 | 7.43E-07 | 1.59E-02 | 0.61 | 0.47 | 0.13 | 1.28 |
| 5 | Crabp1 | 0.3 | 8.15E-07 | 1.74E-02 | 0.65 | 0.56 | 0.1 | 1.17 |
| 5 | Tulp3 | 0.63 | 9.09E-07 | 1.94E-02 | 0.34 | 0.24 | 0.1 | 1.43 |
| 5 | Exosc2 | 0.42 | 9.37E-07 | 2.00E-02 | 0.56 | 0.46 | 0.11 | 1.23 |
| 5 | Rdm1 | 0.42 | 1.02E-06 | 2.17E-02 | 0.49 | 0.38 | 0.12 | 1.31 |
| 5 | Peg12 | 0.99 | 1.11E-06 | 2.36E-02 | 0.14 | 0.07 | 0.07 | 1.96 |
| 5 | Cdc42ep3 | 0.51 | 1.12E-06 | 2.40E-02 | 0.27 | 0.17 | 0.1 | 1.58 |
| 5 | S100a6 | 0.38 | 1.15E-06 | 2.46E-02 | 0.34 | 0.23 | 0.11 | 1.47 |
| 5 | Cnn2 | 0.37 | 1.18E-06 | 2.53E-02 | 0.42 | 0.3 | 0.12 | 1.39 |
| 5 | Hs3st1 | 0.92 | 1.21E-06 | 2.58E-02 | 0.13 | 0.07 | 0.07 | 1.99 |
| 5 | Ptpru | 0.43 | 1.32E-06 | 2.82E-02 | 0.48 | 0.37 | 0.11 | 1.3 |
| 5 | Col17a1 | 0.39 | 1.33E-06 | 2.85E-02 | 0.21 | 0.12 | 0.09 | 1.72 |
| 5 | Tgif1 | 0.3 | 1.41E-06 | 3.01E-02 | 0.78 | 0.68 | 0.1 | 1.15 |
| 5 | Man2a1 | 0.64 | 1.45E-06 | 3.10E-02 | 0.34 | 0.24 | 0.1 | 1.41 |
| 5 | F11r | 0.32 | 1.49E-06 | 3.18E-02 | 0.72 | 0.6 | 0.12 | 1.19 |
| 5 | Map3k20 | 0.81 | 1.69E-06 | 3.61E-02 | 0.27 | 0.18 | 0.09 | 1.5 |
| 5 | Cyfip1 | 0.3 | 1.75E-06 | 3.74E-02 | 0.81 | 0.73 | 0.08 | 1.11 |
| 5 | Enho | 0.41 | 1.84E-06 | 3.93E-02 | 0.44 | 0.33 | 0.11 | 1.34 |
| 5 | Sox6 | 0.97 | 1.94E-06 | 4.14E-02 | 0.17 | 0.1 | 0.07 | 1.78 |
| 6 | Dlx6os1 | 4.51 | 5.10E-285 | 1.09E-280 | 0.73 | 0.07 | 0.66 | 10.15 |
| 6 | Fezf2 | 7.83 | 1.51E-184 | 3.22E-180 | 0.26 | 0 | 0.26 | 86.67 |
| 6 | Dlk1 | 3.11 | 8.46E-173 | 1.81E-168 | 0.99 | 0.61 | 0.38 | 1.62 |
| 6 | Flrt2 | 3.92 | 4.33E-162 | 9.24E-158 | 0.77 | 0.21 | 0.57 | 3.75 |
| 6 | Dlx5 | 2.81 | 2.47E-161 | 5.29E-157 | 0.91 | 0.35 | 0.56 | 2.62 |
| 6 | Lgr4 | 2.52 | 3.12E-139 | 6.66E-135 | 0.91 | 0.44 | 0.47 | 2.07 |
| 6 | Car2 | 3.43 | 6.49E-135 | 1.39E-130 | 0.6 | 0.12 | 0.48 | 4.92 |
| 6 | Dync1i1 | 3.37 | 4.00E-133 | 8.54E-129 | 0.54 | 0.09 | 0.45 | 5.87 |
| 6 | Mest | 1.78 | 1.01E-116 | 2.16E-112 | 0.99 | 0.9 | 0.09 | 1.1 |
| 6 | Sox2 | 1.75 | 2.69E-108 | 5.74E-104 | 0.98 | 0.68 | 0.31 | 1.45 |
| 6 | Slitrk1 | 5.6 | 6.13E-108 | 1.31E-103 | 0.19 | 0.01 | 0.19 | 27.71 |
| 6 | Mecom | 2.44 | 2.10E-105 | 4.49E-101 | 0.84 | 0.42 | 0.42 | 2 |
| 6 | Mir9-3hg | 3.2 | 2.43E-95 | 5.20E-91 | 0.42 | 0.08 | 0.35 | 5.65 |
| 6 | Helt | 8.78 | 3.84E-94 | 8.21E-90 | 0.13 | 0 | 0.12 | 125 |
| 6 | Tmem47 | 2.36 | 1.31E-91 | 2.79E-87 | 0.8 | 0.42 | 0.38 | 1.89 |
| 6 | Fezf1 | 1.84 | 3.32E-88 | 7.09E-84 | 0.41 | 0.07 | 0.34 | 5.93 |
| 6 | Meg3 | 1.56 | 5.44E-85 | 1.16E-80 | 0.97 | 0.75 | 0.22 | 1.3 |
| 6 | Mir670hg | 3.39 | 3.07E-78 | 6.57E-74 | 0.27 | 0.03 | 0.24 | 8.15 |
| 6 | Six3os1 | 1.28 | 1.24E-75 | 2.64E-71 | 0.96 | 0.75 | 0.21 | 1.29 |
| 6 | Slit2 | 2.22 | 3.41E-75 | 7.28E-71 | 0.75 | 0.36 | 0.38 | 2.05 |
| 6 | Tgfb2 | 1.04 | 3.86E-75 | 8.24E-71 | 0.95 | 0.55 | 0.39 | 1.71 |
| 6 | Sem1 | 0.72 | 3.64E-74 | 7.78E-70 | 1 | 0.99 | 0.01 | 1.01 |
| 6 | Rprm | 2.23 | 1.34E-73 | 2.86E-69 | 0.75 | 0.35 | 0.4 | 2.13 |
| 6 | Six3 | 1.12 | 1.48E-73 | 3.16E-69 | 0.99 | 0.83 | 0.16 | 1.19 |
| 6 | Syt1 | 2.34 | 1.63E-73 | 3.47E-69 | 0.47 | 0.12 | 0.35 | 4.02 |
| 6 | Mme | 2.66 | 2.63E-70 | 5.62E-66 | 0.51 | 0.15 | 0.36 | 3.31 |
| 6 | Dlx6 | 2.32 | 1.20E-68 | 2.57E-64 | 0.53 | 0.17 | 0.36 | 3.12 |
| 6 | Ermn | 2.27 | 4.20E-63 | 8.97E-59 | 0.49 | 0.14 | 0.34 | 3.38 |
| 6 | Sfrp1 | 1.67 | 2.32E-62 | 4.95E-58 | 0.85 | 0.49 | 0.35 | 1.71 |
| 6 | Pcdh11x | 2.62 | 4.75E-62 | 1.02E-57 | 0.41 | 0.1 | 0.3 | 3.9 |
| 6 | Gnai2 | 0.72 | 2.22E-61 | 4.75E-57 | 0.99 | 0.97 | 0.02 | 1.02 |
| 6 | Fzd8 | 2.08 | 1.08E-60 | 2.30E-56 | 0.64 | 0.28 | 0.36 | 2.32 |

**Supplementary Table 3. Marker genes for the original clusters of integrated clustering of E11.5 control and NotchGOF λ-junction ectoderm cells.**

| | | | | | | | | |
|---|---|---|---|---|---|---|---|---|
| 6 | Marcks | 0.71 | 3.64E-58 | 7.78E-54 | 1 | 0.99 | 0.01 | 1.01 |
| 6 | Col2a1 | 1.1 | 4.35E-56 | 9.30E-52 | 0.93 | 0.8 | 0.13 | 1.16 |
| 6 | Syt11 | 1.85 | 5.85E-56 | 1.25E-51 | 0.66 | 0.32 | 0.35 | 2.08 |
| 6 | Crabp2 | 0.89 | 8.97E-56 | 1.92E-51 | 0.99 | 0.84 | 0.15 | 1.18 |
| 6 | Cdh2 | 1.22 | 1.76E-53 | 3.75E-49 | 0.88 | 0.59 | 0.29 | 1.49 |
| 6 | Gna14 | 4 | 6.36E-53 | 1.36E-48 | 0.14 | 0.01 | 0.13 | 11.92 |
| 6 | Gad2 | 1.49 | 7.85E-51 | 1.68E-46 | 0.61 | 0.27 | 0.34 | 2.29 |
| 6 | Gpm6a | 2.24 | 2.76E-49 | 5.91E-45 | 0.4 | 0.12 | 0.28 | 3.33 |
| 6 | Hey1 | 1.06 | 1.97E-48 | 4.22E-44 | 0.91 | 0.74 | 0.18 | 1.24 |
| 6 | Sp8 | 1.27 | 3.09E-48 | 6.59E-44 | 0.82 | 0.5 | 0.32 | 1.64 |
| 6 | Rian | 1.6 | 1.58E-47 | 3.38E-43 | 0.73 | 0.44 | 0.29 | 1.65 |
| 6 | Lrrn3 | 2.52 | 5.24E-47 | 1.12E-42 | 0.4 | 0.13 | 0.27 | 3.08 |
| 6 | Six6 | 1.43 | 5.58E-44 | 1.19E-39 | 0.59 | 0.26 | 0.33 | 2.31 |
| 6 | Negr1 | 2.91 | 2.03E-42 | 4.34E-38 | 0.19 | 0.03 | 0.16 | 6.27 |
| 6 | Otx2 | 1.48 | 4.96E-42 | 1.06E-37 | 0.69 | 0.38 | 0.32 | 1.84 |
| 6 | Id2 | 1 | 6.34E-42 | 1.35E-37 | 0.96 | 0.82 | 0.14 | 1.16 |
| 6 | C230057A21Rik | 3.01 | 1.06E-40 | 2.26E-36 | 0.19 | 0.03 | 0.15 | 5.97 |
| 6 | Igfbp2 | 0.7 | 7.15E-40 | 1.53E-35 | 1 | 0.98 | 0.01 | 1.01 |
| 6 | Dock10 | 3.08 | 4.02E-39 | 8.60E-35 | 0.21 | 0.04 | 0.17 | 5.02 |
| 6 | Wfdc2 | 0.84 | 2.35E-37 | 5.03E-33 | 0.95 | 0.81 | 0.14 | 1.17 |
| 6 | Mapk12 | 1.54 | 1.16E-36 | 2.49E-32 | 0.55 | 0.27 | 0.28 | 2.03 |
| 6 | Gm35040 | 2.58 | 1.32E-36 | 2.82E-32 | 0.22 | 0.05 | 0.17 | 4.67 |
| 6 | Psip1 | 0.58 | 1.13E-35 | 2.40E-31 | 0.99 | 0.98 | 0.01 | 1.01 |
| 6 | Rtn3 | 0.71 | 4.90E-35 | 1.05E-30 | 0.95 | 0.87 | 0.08 | 1.09 |
| 6 | 1500004A13Rik | 1.98 | 7.69E-35 | 1.64E-30 | 0.28 | 0.08 | 0.2 | 3.67 |
| 6 | Stmn3 | 0.77 | 1.70E-34 | 3.62E-30 | 0.32 | 0.1 | 0.22 | 3.22 |
| 6 | Galm | 1.38 | 4.94E-34 | 1.06E-29 | 0.65 | 0.38 | 0.27 | 1.71 |
| 6 | Sox3 | 2.09 | 7.50E-34 | 1.60E-29 | 0.36 | 0.13 | 0.23 | 2.74 |
| 6 | Crip1 | 1.3 | 3.55E-33 | 7.59E-29 | 0.64 | 0.38 | 0.26 | 1.67 |
| 6 | Slc25a13 | 1.07 | 3.93E-33 | 8.39E-29 | 0.72 | 0.51 | 0.21 | 1.41 |
| 6 | Dusp6 | 1.71 | 8.08E-32 | 1.73E-27 | 0.49 | 0.25 | 0.25 | 2 |
| 6 | Six1 | 0.79 | 1.54E-31 | 3.29E-27 | 0.91 | 0.67 | 0.24 | 1.36 |
| 6 | Etv5 | 1.84 | 2.98E-31 | 6.37E-27 | 0.45 | 0.21 | 0.24 | 2.12 |
| 6 | Nr2e1 | 1.82 | 3.46E-31 | 7.39E-27 | 0.39 | 0.15 | 0.23 | 2.55 |
| 6 | Sall2 | 1.23 | 4.24E-31 | 9.05E-27 | 0.68 | 0.46 | 0.23 | 1.49 |
| 6 | Vstm2b | 3.25 | 1.28E-30 | 2.73E-26 | 0.1 | 0.01 | 0.09 | 8.42 |
| 6 | Hes5 | 1.26 | 1.98E-30 | 4.23E-26 | 0.65 | 0.37 | 0.27 | 1.74 |
| 6 | Wdfy1 | 1.34 | 2.41E-30 | 5.15E-26 | 0.62 | 0.39 | 0.24 | 1.61 |
| 6 | Dpysl3 | 1.1 | 4.79E-30 | 1.02E-25 | 0.76 | 0.56 | 0.2 | 1.36 |
| 6 | Nnat | 0.37 | 6.37E-30 | 1.36E-25 | 0.99 | 0.95 | 0.04 | 1.04 |
| 6 | Foxg1 | 0.92 | 9.87E-30 | 2.11E-25 | 0.79 | 0.52 | 0.27 | 1.51 |
| 6 | Sox2ot | 2.91 | 1.40E-29 | 3.00E-25 | 0.22 | 0.06 | 0.16 | 3.76 |
| 6 | Ssx2ip | 0.87 | 1.85E-29 | 3.96E-25 | 0.82 | 0.65 | 0.17 | 1.26 |
| 6 | Gpx4 | 0.46 | 1.86E-29 | 3.97E-25 | 0.99 | 0.98 | 0.01 | 1.01 |
| 6 | Id3 | 0.68 | 2.30E-28 | 4.91E-24 | 0.98 | 0.96 | 0.01 | 1.01 |
| 6 | Gpc4 | 1.39 | 2.42E-28 | 5.18E-24 | 0.54 | 0.29 | 0.25 | 1.86 |
| 6 | Lmo4 | 0.73 | 3.88E-28 | 8.28E-24 | 0.98 | 0.91 | 0.07 | 1.08 |
| 6 | Bpgm | 1.38 | 3.97E-28 | 8.48E-24 | 0.62 | 0.41 | 0.21 | 1.51 |
| 6 | Fbn2 | 1.16 | 1.41E-27 | 3.01E-23 | 0.69 | 0.45 | 0.24 | 1.53 |
| 6 | Id4 | 0.77 | 2.89E-27 | 6.17E-23 | 0.79 | 0.56 | 0.23 | 1.42 |
| 6 | Mir99ahg | 1.26 | 4.09E-27 | 8.74E-23 | 0.71 | 0.52 | 0.19 | 1.35 |
| 6 | Ndfip1 | 0.68 | 5.51E-27 | 1.18E-22 | 0.9 | 0.83 | 0.07 | 1.09 |
| 6 | Rragd | 1.19 | 8.60E-27 | 1.84E-22 | 0.64 | 0.4 | 0.24 | 1.61 |
| 6 | Fzd2 | 0.74 | 2.54E-26 | 5.42E-22 | 0.9 | 0.81 | 0.09 | 1.11 |
| 6 | Bag2 | 1.27 | 1.75E-25 | 3.75E-21 | 0.56 | 0.33 | 0.22 | 1.67 |
| 6 | Nckap5 | 1.31 | 5.16E-25 | 1.10E-20 | 0.53 | 0.29 | 0.24 | 1.81 |
| 6 | Peg3 | 0.91 | 1.40E-24 | 2.98E-20 | 0.91 | 0.82 | 0.09 | 1.11 |

**Supplementary Table 3. Marker genes for the original clusters of integrated clustering of E11.5 control and NotchGOF λ-junction ectoderm cells.**

| | | | | | | | | |
|---|---|---|---|---|---|---|---|---|
| 6 | Krt8 | 0.49 | 4.26E-24 | 9.11E-20 | 0.97 | 0.81 | 0.17 | 1.2 |
| 6 | Fam126a | 1.51 | 6.67E-24 | 1.42E-19 | 0.48 | 0.27 | 0.21 | 1.78 |
| 6 | Auts2 | 0.63 | 1.34E-23 | 2.87E-19 | 0.94 | 0.9 | 0.04 | 1.04 |
| 6 | Socs2 | 0.55 | 3.04E-23 | 6.49E-19 | 0.93 | 0.77 | 0.16 | 1.2 |
| 6 | Espn | 1.17 | 3.37E-23 | 7.21E-19 | 0.58 | 0.36 | 0.23 | 1.63 |
| 6 | Nes | 0.95 | 4.59E-23 | 9.81E-19 | 0.69 | 0.49 | 0.21 | 1.43 |
| 6 | Sox21 | 1.91 | 9.96E-23 | 2.13E-18 | 0.29 | 0.11 | 0.18 | 2.59 |
| 6 | Mfap2 | 0.59 | 1.53E-22 | 3.26E-18 | 0.95 | 0.9 | 0.05 | 1.05 |
| 6 | Pea15a | 1.2 | 1.79E-22 | 3.82E-18 | 0.54 | 0.32 | 0.22 | 1.67 |
| 6 | Mapk11 | 1.74 | 2.58E-22 | 5.51E-18 | 0.28 | 0.11 | 0.17 | 2.58 |
| 6 | Fgf17 | 2.2 | 3.55E-22 | 7.59E-18 | 0.34 | 0.15 | 0.19 | 2.22 |
| 6 | Eya4 | 1.43 | 5.15E-22 | 1.10E-17 | 0.41 | 0.21 | 0.2 | 2 |
| 6 | Tenm3 | 0.93 | 5.27E-22 | 1.13E-17 | 0.74 | 0.61 | 0.13 | 1.22 |
| 6 | Plch1 | 1.21 | 3.93E-21 | 8.40E-17 | 0.53 | 0.33 | 0.2 | 1.62 |
| 6 | Fgfr1 | 1.12 | 5.28E-21 | 1.13E-16 | 0.58 | 0.39 | 0.19 | 1.48 |
| 6 | Tox2 | 1.82 | 3.34E-20 | 7.13E-16 | 0.27 | 0.11 | 0.16 | 2.49 |
| 6 | Rhoc | 0.79 | 4.13E-20 | 8.82E-16 | 0.79 | 0.64 | 0.14 | 1.22 |
| 6 | Abcg1 | 1.43 | 4.47E-20 | 9.54E-16 | 0.47 | 0.28 | 0.19 | 1.68 |
| 6 | Lmo3 | 2.45 | 4.99E-20 | 1.07E-15 | 0.15 | 0.04 | 0.11 | 3.71 |
| 6 | Prdm16 | 2.32 | 1.13E-19 | 2.42E-15 | 0.16 | 0.05 | 0.12 | 3.5 |
| 6 | Pdlim7 | 0.87 | 2.63E-19 | 5.63E-15 | 0.74 | 0.62 | 0.13 | 1.2 |
| 6 | Phlpp1 | 1.28 | 3.15E-19 | 6.73E-15 | 0.44 | 0.25 | 0.19 | 1.75 |
| 6 | Galnt2 | 0.97 | 5.33E-19 | 1.14E-14 | 0.68 | 0.53 | 0.16 | 1.3 |
| 6 | Scg5 | 1.31 | 8.05E-19 | 1.72E-14 | 0.38 | 0.2 | 0.18 | 1.94 |
| 6 | St8sia4 | 3.12 | 1.07E-18 | 2.29E-14 | 0.12 | 0.03 | 0.09 | 4.3 |
| 6 | Adrb1 | 2.44 | 1.16E-18 | 2.49E-14 | 0.18 | 0.06 | 0.12 | 3.14 |
| 6 | Dnajc1 | 0.72 | 1.56E-18 | 3.32E-14 | 0.77 | 0.64 | 0.13 | 1.2 |
| 6 | Peg10 | 1.67 | 2.09E-18 | 4.47E-14 | 0.25 | 0.1 | 0.15 | 2.56 |
| 6 | Ascl1 | 3.72 | 2.75E-18 | 5.88E-14 | 0.1 | 0.02 | 0.08 | 4.73 |
| 6 | Emb | 0.88 | 5.11E-18 | 1.09E-13 | 0.82 | 0.73 | 0.09 | 1.12 |
| 6 | C130071C03Rik | 1.44 | 9.24E-18 | 1.97E-13 | 0.37 | 0.19 | 0.17 | 1.89 |
| 6 | Fam3c | 1.04 | 1.60E-17 | 3.42E-13 | 0.61 | 0.46 | 0.15 | 1.33 |
| 6 | Slc6a15 | 1.98 | 1.61E-17 | 3.44E-13 | 0.2 | 0.07 | 0.13 | 2.78 |
| 6 | Top1 | 0.3 | 2.41E-17 | 5.15E-13 | 0.99 | 0.99 | 0 | 1 |
| 6 | Klhl7 | 0.83 | 3.88E-17 | 8.29E-13 | 0.71 | 0.57 | 0.14 | 1.24 |
| 6 | Rnf182 | 0.81 | 3.90E-17 | 8.34E-13 | 0.36 | 0.18 | 0.18 | 1.97 |
| 6 | Mpped1 | 2.25 | 4.25E-17 | 9.07E-13 | 0.15 | 0.04 | 0.11 | 3.39 |
| 6 | Phlda1 | 1.13 | 4.69E-17 | 1.00E-12 | 0.55 | 0.37 | 0.18 | 1.48 |
| 6 | Bex2 | 0.5 | 7.39E-17 | 1.58E-12 | 0.9 | 0.88 | 0.03 | 1.03 |
| 6 | Nog | 1.79 | 7.58E-17 | 1.62E-12 | 0.25 | 0.11 | 0.15 | 2.37 |
| 6 | Car14 | 1.29 | 1.05E-16 | 2.25E-12 | 0.4 | 0.22 | 0.17 | 1.78 |
| 6 | Pcdh17 | 1.41 | 1.10E-16 | 2.35E-12 | 0.53 | 0.36 | 0.17 | 1.47 |
| 6 | Frmd3 | 2.08 | 1.12E-16 | 2.39E-12 | 0.17 | 0.06 | 0.11 | 3.04 |
| 6 | Mip | 2.25 | 1.72E-16 | 3.68E-12 | 0.14 | 0.04 | 0.1 | 3.41 |
| 6 | Fkbp4 | 0.31 | 2.51E-16 | 5.36E-12 | 0.99 | 0.98 | 0.01 | 1.01 |
| 6 | Kif21a | 0.57 | 2.73E-16 | 5.83E-12 | 0.81 | 0.74 | 0.06 | 1.08 |
| 6 | Nuak2 | 1.43 | 2.79E-16 | 5.97E-12 | 0.29 | 0.13 | 0.16 | 2.19 |
| 6 | C530008M17Rik | 1.04 | 3.32E-16 | 7.09E-12 | 0.56 | 0.39 | 0.17 | 1.45 |
| 6 | Gulp1 | 1.51 | 6.07E-16 | 1.30E-11 | 0.3 | 0.14 | 0.15 | 2.07 |
| 6 | Gas5 | 0.3 | 6.77E-16 | 1.45E-11 | 1 | 0.99 | 0.01 | 1.01 |
| 6 | Epcam | 0.36 | 8.68E-16 | 1.85E-11 | 1 | 0.98 | 0.01 | 1.01 |
| 6 | Serpinh1 | 0.37 | 9.43E-16 | 2.01E-11 | 0.99 | 0.95 | 0.03 | 1.03 |
| 6 | Pcbp4 | 0.38 | 1.18E-15 | 2.52E-11 | 0.95 | 0.91 | 0.04 | 1.04 |
| 6 | Tcf7l2 | 0.63 | 1.21E-15 | 2.58E-11 | 0.82 | 0.72 | 0.09 | 1.13 |
| 6 | Spint2 | 0.34 | 1.40E-15 | 2.99E-11 | 0.98 | 0.97 | 0.02 | 1.02 |
| 6 | Kif5c | 0.67 | 1.45E-15 | 3.09E-11 | 0.7 | 0.59 | 0.11 | 1.19 |
| 6 | Npdc1 | 1.03 | 1.70E-15 | 3.62E-11 | 0.49 | 0.32 | 0.17 | 1.55 |

**Supplementary Table 3. Marker genes for the original clusters of integrated clustering of E11.5 control and NotchGOF λ-junction ectoderm cells.**

| | | | | | | | | |
|---|---|---|---|---|---|---|---|---|
| 6 | Btg2 | 1.27 | 2.13E-15 | 4.55E-11 | 0.48 | 0.3 | 0.18 | 1.59 |
| 6 | Khdrbs3 | 0.39 | 2.32E-15 | 4.96E-11 | 0.93 | 0.88 | 0.05 | 1.05 |
| 6 | Pbx1 | 0.36 | 2.96E-15 | 6.33E-11 | 0.98 | 0.97 | 0.01 | 1.01 |
| 6 | Lmo1 | 0.97 | 3.15E-15 | 6.73E-11 | 0.39 | 0.22 | 0.17 | 1.79 |
| 6 | Glo1 | 0.51 | 3.58E-15 | 7.64E-11 | 0.88 | 0.82 | 0.06 | 1.08 |
| 6 | Sgce | 1.14 | 3.92E-15 | 8.37E-11 | 0.47 | 0.31 | 0.16 | 1.51 |
| 6 | Sall1 | 2.3 | 5.72E-15 | 1.22E-10 | 0.15 | 0.05 | 0.1 | 3.11 |
| 6 | Irf2bpl | 0.92 | 1.03E-14 | 2.20E-10 | 0.59 | 0.44 | 0.15 | 1.33 |
| 6 | Cldn7 | 0.36 | 1.80E-14 | 3.85E-10 | 0.95 | 0.84 | 0.11 | 1.13 |
| 6 | Pgrmc1 | 0.46 | 2.71E-14 | 5.80E-10 | 0.95 | 0.94 | 0.01 | 1.01 |
| 6 | Stc1 | 1.92 | 5.75E-14 | 1.23E-09 | 0.19 | 0.08 | 0.11 | 2.51 |
| 6 | Pls1 | 0.93 | 6.84E-14 | 1.46E-09 | 0.45 | 0.29 | 0.16 | 1.56 |
| 6 | Pip4p2 | 0.84 | 8.05E-14 | 1.72E-09 | 0.63 | 0.52 | 0.12 | 1.23 |
| 6 | Sorbs2 | 0.82 | 1.05E-13 | 2.25E-09 | 0.56 | 0.39 | 0.17 | 1.43 |
| 6 | Ugcg | 0.88 | 1.08E-13 | 2.31E-09 | 0.56 | 0.42 | 0.14 | 1.34 |
| 6 | Mirg | 1.82 | 1.08E-13 | 2.32E-09 | 0.18 | 0.07 | 0.11 | 2.59 |
| 6 | Wasl | 0.74 | 1.25E-13 | 2.67E-09 | 0.7 | 0.6 | 0.11 | 1.18 |
| 6 | Dmrta1 | 1.72 | 1.41E-13 | 3.01E-09 | 0.2 | 0.08 | 0.12 | 2.43 |
| 6 | Sox5 | 1.25 | 1.62E-13 | 3.45E-09 | 0.38 | 0.23 | 0.15 | 1.68 |
| 6 | Nrcam | 1.19 | 1.68E-13 | 3.59E-09 | 0.34 | 0.19 | 0.15 | 1.81 |
| 6 | Rassf9 | 1.16 | 1.84E-13 | 3.92E-09 | 0.39 | 0.23 | 0.16 | 1.7 |
| 6 | Atp2b1 | 0.4 | 1.91E-13 | 4.07E-09 | 0.94 | 0.88 | 0.06 | 1.07 |
| 6 | Shtn1 | 1.25 | 2.17E-13 | 4.63E-09 | 0.34 | 0.19 | 0.15 | 1.79 |
| 6 | Shisa4 | 1.02 | 2.26E-13 | 4.83E-09 | 0.39 | 0.23 | 0.16 | 1.69 |
| 6 | Prrg2 | 1.32 | 2.32E-13 | 4.95E-09 | 0.4 | 0.25 | 0.15 | 1.59 |
| 6 | Gpm6b | 1.46 | 5.48E-13 | 1.17E-08 | 0.21 | 0.09 | 0.12 | 2.32 |
| 6 | Fbln5 | 1.94 | 5.57E-13 | 1.19E-08 | 0.16 | 0.06 | 0.1 | 2.65 |
| 6 | Syndig1 | 2.62 | 5.85E-13 | 1.25E-08 | 0.13 | 0.04 | 0.09 | 3.05 |
| 6 | Fgf3 | 2.31 | 5.94E-13 | 1.27E-08 | 0.1 | 0.03 | 0.07 | 3.61 |
| 6 | Aig1 | 0.8 | 7.53E-13 | 1.61E-08 | 0.64 | 0.54 | 0.1 | 1.19 |
| 6 | Bmp4 | 0.45 | 1.14E-12 | 2.43E-08 | 0.7 | 0.52 | 0.18 | 1.34 |
| 6 | Pnp | 0.66 | 1.18E-12 | 2.52E-08 | 0.73 | 0.65 | 0.08 | 1.13 |
| 6 | Krt18 | 0.37 | 1.43E-12 | 3.05E-08 | 0.96 | 0.86 | 0.1 | 1.12 |
| 6 | Tox3 | 0.87 | 1.44E-12 | 3.08E-08 | 0.46 | 0.3 | 0.16 | 1.54 |
| 6 | Crip2 | 0.64 | 1.67E-12 | 3.56E-08 | 0.58 | 0.41 | 0.17 | 1.4 |
| 6 | Saraf | 0.68 | 1.84E-12 | 3.93E-08 | 0.73 | 0.66 | 0.08 | 1.12 |
| 6 | Tle5 | 0.34 | 1.99E-12 | 4.26E-08 | 0.99 | 0.96 | 0.03 | 1.03 |
| 6 | Elavl2 | 0.82 | 2.01E-12 | 4.30E-08 | 0.55 | 0.4 | 0.15 | 1.36 |
| 6 | Adgrl2 | 0.61 | 2.26E-12 | 4.84E-08 | 0.77 | 0.7 | 0.07 | 1.1 |
| 6 | Tspan3 | 0.35 | 3.41E-12 | 7.29E-08 | 0.96 | 0.96 | 0 | 1 |
| 6 | Rab28 | 0.73 | 3.98E-12 | 8.49E-08 | 0.62 | 0.51 | 0.11 | 1.21 |
| 6 | Klf13 | 0.75 | 4.54E-12 | 9.70E-08 | 0.55 | 0.4 | 0.15 | 1.36 |
| 6 | Cldn3 | 0.43 | 4.64E-12 | 9.92E-08 | 0.8 | 0.62 | 0.18 | 1.28 |
| 6 | Shisa2 | 1.64 | 5.65E-12 | 1.21E-07 | 0.22 | 0.1 | 0.12 | 2.14 |
| 6 | Serinc2 | 0.66 | 6.07E-12 | 1.30E-07 | 0.68 | 0.57 | 0.12 | 1.21 |
| 6 | C130026L21Rik | 2.09 | 6.10E-12 | 1.30E-07 | 0.12 | 0.04 | 0.08 | 2.98 |
| 6 | Ncald | 2.18 | 7.29E-12 | 1.56E-07 | 0.13 | 0.04 | 0.08 | 2.91 |
| 6 | Hmgn2 | 0.28 | 9.03E-12 | 1.93E-07 | 0.99 | 0.98 | 0.01 | 1.01 |
| 6 | Epha5 | 1.74 | 9.38E-12 | 2.00E-07 | 0.19 | 0.08 | 0.11 | 2.27 |
| 6 | Tnnt1 | 1.62 | 9.50E-12 | 2.03E-07 | 0.19 | 0.08 | 0.11 | 2.34 |
| 6 | Plcl1 | 1.95 | 9.89E-12 | 2.11E-07 | 0.12 | 0.04 | 0.08 | 3.05 |
| 6 | Cldn9 | 0.89 | 1.84E-11 | 3.92E-07 | 0.49 | 0.34 | 0.15 | 1.43 |
| 6 | Serpinf1 | 1.4 | 1.92E-11 | 4.11E-07 | 0.29 | 0.16 | 0.13 | 1.81 |
| 6 | Cxcl12 | 1.49 | 3.38E-11 | 7.23E-07 | 0.25 | 0.13 | 0.12 | 1.92 |
| 6 | Furin | 1.2 | 3.57E-11 | 7.63E-07 | 0.37 | 0.24 | 0.13 | 1.54 |
| 6 | Plin3 | 0.76 | 3.61E-11 | 7.71E-07 | 0.63 | 0.52 | 0.11 | 1.21 |
| 6 | Jund | 0.37 | 4.02E-11 | 8.59E-07 | 0.96 | 0.92 | 0.04 | 1.04 |

**Supplementary Table 3. Marker genes for the original clusters of integrated clustering of E11.5 control and NotchGOF λ-junction ectoderm cells.**

| | | | | | | | | |
|---|---|---|---|---|---|---|---|---|
| 6 | Adam10 | 0.77 | 4.16E-11 | 8.88E-07 | 0.66 | 0.56 | 0.09 | 1.16 |
| 6 | Rcn2 | 0.42 | 4.42E-11 | 9.43E-07 | 0.93 | 0.91 | 0.03 | 1.03 |
| 6 | Ncam1 | 0.7 | 5.68E-11 | 1.21E-06 | 0.46 | 0.33 | 0.14 | 1.42 |
| 6 | Rcn1 | 0.45 | 5.77E-11 | 1.23E-06 | 0.89 | 0.85 | 0.04 | 1.05 |
| 6 | Prr7 | 1.16 | 6.18E-11 | 1.32E-06 | 0.34 | 0.21 | 0.13 | 1.61 |
| 6 | Id1 | 0.39 | 6.47E-11 | 1.38E-06 | 0.96 | 0.93 | 0.04 | 1.04 |
| 6 | Nudt4 | 0.38 | 8.52E-11 | 1.82E-06 | 0.84 | 0.78 | 0.07 | 1.09 |
| 6 | Mettl26 | 0.62 | 8.60E-11 | 1.84E-06 | 0.72 | 0.63 | 0.09 | 1.15 |
| 6 | Fam110b | 1.37 | 1.06E-10 | 2.27E-06 | 0.33 | 0.2 | 0.12 | 1.61 |
| 6 | Cadps2 | 1.43 | 1.18E-10 | 2.52E-06 | 0.25 | 0.13 | 0.12 | 1.89 |
| 6 | Nfia | 0.83 | 1.21E-10 | 2.58E-06 | 0.5 | 0.36 | 0.14 | 1.38 |
| 6 | Sypl | 0.7 | 1.23E-10 | 2.62E-06 | 0.69 | 0.6 | 0.1 | 1.16 |
| 6 | Eya1 | 0.56 | 1.44E-10 | 3.07E-06 | 0.71 | 0.58 | 0.12 | 1.21 |
| 6 | Dsel | 0.93 | 1.52E-10 | 3.25E-06 | 0.51 | 0.38 | 0.13 | 1.35 |
| 6 | Fzd4 | 1.32 | 1.91E-10 | 4.09E-06 | 0.3 | 0.18 | 0.12 | 1.7 |
| 6 | Cnih1 | 0.47 | 1.97E-10 | 4.20E-06 | 0.85 | 0.81 | 0.05 | 1.06 |
| 6 | Sord | 1.54 | 2.58E-10 | 5.52E-06 | 0.35 | 0.23 | 0.12 | 1.52 |
| 6 | Jam3 | 0.85 | 3.68E-10 | 7.86E-06 | 0.48 | 0.36 | 0.12 | 1.34 |
| 6 | Slc25a5 | 0.32 | 4.40E-10 | 9.41E-06 | 0.95 | 0.91 | 0.04 | 1.04 |
| 6 | Sms | 0.36 | 5.71E-10 | 1.22E-05 | 0.89 | 0.86 | 0.03 | 1.04 |
| 6 | Ssbp2 | 0.57 | 6.36E-10 | 1.36E-05 | 0.77 | 0.69 | 0.08 | 1.11 |
| 6 | Rhcg | 2.35 | 6.46E-10 | 1.38E-05 | 0.11 | 0.04 | 0.07 | 2.82 |
| 6 | Msi1 | 0.72 | 7.93E-10 | 1.69E-05 | 0.58 | 0.47 | 0.1 | 1.22 |
| 6 | Krtcap2 | 0.35 | 8.88E-10 | 1.90E-05 | 0.94 | 0.92 | 0.02 | 1.02 |
| 6 | Tpm3 | 0.35 | 1.05E-09 | 2.25E-05 | 0.97 | 0.95 | 0.02 | 1.02 |
| 6 | Dmd | 1.32 | 1.53E-09 | 3.27E-05 | 0.26 | 0.15 | 0.11 | 1.74 |
| 6 | Arxes1 | 0.98 | 1.55E-09 | 3.31E-05 | 0.43 | 0.31 | 0.12 | 1.39 |
| 6 | Fgfbp3 | 1.44 | 2.47E-09 | 5.28E-05 | 0.23 | 0.12 | 0.11 | 1.86 |
| 6 | Ralyl | 1.43 | 2.94E-09 | 6.27E-05 | 0.15 | 0.07 | 0.08 | 2.29 |
| 6 | Klhl5 | 1.22 | 3.26E-09 | 6.96E-05 | 0.28 | 0.16 | 0.12 | 1.72 |
| 6 | Gipc2 | 1.31 | 3.55E-09 | 7.58E-05 | 0.27 | 0.16 | 0.11 | 1.68 |
| 6 | Upp2 | 1.38 | 4.15E-09 | 8.87E-05 | 0.26 | 0.15 | 0.11 | 1.72 |
| 6 | Cxxc5 | 0.62 | 4.27E-09 | 9.13E-05 | 0.67 | 0.58 | 0.09 | 1.15 |
| 6 | Col9a1 | 1.34 | 4.28E-09 | 9.15E-05 | 0.18 | 0.09 | 0.09 | 2.07 |
| 6 | Mgst1 | 0.45 | 4.43E-09 | 9.46E-05 | 0.76 | 0.69 | 0.07 | 1.1 |
| 6 | Akr1a1 | 0.31 | 4.48E-09 | 9.57E-05 | 0.95 | 0.96 | -0.01 | 0.99 |
| 6 | Rasef | 1.38 | 5.02E-09 | 1.07E-04 | 0.26 | 0.15 | 0.11 | 1.72 |
| 6 | Mdfi | 0.68 | 5.15E-09 | 1.10E-04 | 0.6 | 0.49 | 0.11 | 1.22 |
| 6 | Enox1 | 1.74 | 5.22E-09 | 1.12E-04 | 0.12 | 0.05 | 0.07 | 2.53 |
| 6 | Srgap3 | 0.98 | 5.34E-09 | 1.14E-04 | 0.33 | 0.21 | 0.12 | 1.56 |
| 6 | Mir100hg | 0.71 | 5.74E-09 | 1.23E-04 | 0.68 | 0.63 | 0.06 | 1.09 |
| 6 | Fndc3c1 | 0.88 | 6.40E-09 | 1.37E-04 | 0.41 | 0.29 | 0.12 | 1.41 |
| 6 | Aplp2 | 0.4 | 7.16E-09 | 1.53E-04 | 0.94 | 0.92 | 0.02 | 1.02 |
| 6 | Etv4 | 1.53 | 7.33E-09 | 1.57E-04 | 0.19 | 0.1 | 0.1 | 1.96 |
| 6 | Mageh1 | 0.73 | 8.42E-09 | 1.80E-04 | 0.6 | 0.51 | 0.09 | 1.17 |
| 6 | Usp51 | 1.75 | 9.02E-09 | 1.93E-04 | 0.14 | 0.06 | 0.08 | 2.3 |
| 6 | Tle1 | 0.58 | 9.21E-09 | 1.97E-04 | 0.62 | 0.55 | 0.08 | 1.14 |
| 6 | Dpysl4 | 1.31 | 1.00E-08 | 2.14E-04 | 0.2 | 0.11 | 0.1 | 1.92 |
| 6 | Fscn1 | 0.31 | 1.03E-08 | 2.21E-04 | 0.88 | 0.86 | 0.02 | 1.02 |
| 6 | Slc8a1 | 2.13 | 1.05E-08 | 2.25E-04 | 0.12 | 0.05 | 0.07 | 2.48 |
| 6 | Fam214a | 1.16 | 1.25E-08 | 2.68E-04 | 0.32 | 0.21 | 0.11 | 1.53 |
| 6 | Tm9sf3 | 0.36 | 1.30E-08 | 2.78E-04 | 0.9 | 0.87 | 0.03 | 1.03 |
| 6 | Pawr | 0.8 | 1.45E-08 | 3.10E-04 | 0.48 | 0.37 | 0.11 | 1.31 |
| 6 | Hey2 | 1.81 | 1.45E-08 | 3.10E-04 | 0.17 | 0.09 | 0.09 | 2 |
| 6 | Adgrv1 | 1.67 | 1.63E-08 | 3.49E-04 | 0.13 | 0.05 | 0.07 | 2.37 |
| 6 | Dtna | 0.83 | 1.77E-08 | 3.78E-04 | 0.45 | 0.34 | 0.11 | 1.31 |
| 6 | Gad1 | 1.44 | 2.10E-08 | 4.49E-04 | 0.13 | 0.05 | 0.08 | 2.42 |

**Supplementary Table 3. Marker genes for the original clusters of integrated clustering of E11.5 control and NotchGOF λ-junction ectoderm cells.**

| | | | | | | | | |
|---|---|---|---|---|---|---|---|---|
| 6 | B830012L14Rik | 1.63 | 2.14E-08 | 4.56E-04 | 0.18 | 0.09 | 0.09 | 1.97 |
| 6 | Fam171b | 1.35 | 2.15E-08 | 4.60E-04 | 0.23 | 0.13 | 0.1 | 1.79 |
| 6 | Cd63 | 0.35 | 2.21E-08 | 4.71E-04 | 0.93 | 0.92 | 0.01 | 1.01 |
| 6 | Xpr1 | 0.89 | 2.22E-08 | 4.74E-04 | 0.5 | 0.42 | 0.09 | 1.21 |
| 6 | Syt5 | 0.97 | 2.34E-08 | 4.99E-04 | 0.33 | 0.21 | 0.12 | 1.56 |
| 6 | Ppa1 | 0.26 | 2.44E-08 | 5.20E-04 | 0.93 | 0.87 | 0.07 | 1.08 |
| 6 | Nfix | 1.72 | 2.65E-08 | 5.66E-04 | 0.17 | 0.09 | 0.09 | 2 |
| 6 | Txndc15 | 0.53 | 2.66E-08 | 5.68E-04 | 0.65 | 0.57 | 0.08 | 1.15 |
| 6 | Cobl | 1.09 | 2.90E-08 | 6.20E-04 | 0.28 | 0.17 | 0.11 | 1.64 |
| 6 | Dhx32 | 1 | 2.94E-08 | 6.27E-04 | 0.41 | 0.31 | 0.1 | 1.31 |
| 6 | Ckb | 1.18 | 2.96E-08 | 6.32E-04 | 0.47 | 0.34 | 0.13 | 1.38 |
| 6 | Nhs | 0.6 | 3.78E-08 | 8.07E-04 | 0.54 | 0.43 | 0.11 | 1.26 |
| 6 | Tmem121 | 1.02 | 4.10E-08 | 8.76E-04 | 0.28 | 0.17 | 0.11 | 1.62 |
| 6 | Tox | 0.98 | 4.95E-08 | 1.06E-03 | 0.34 | 0.23 | 0.11 | 1.47 |
| 6 | Hsd17b12 | 0.43 | 5.23E-08 | 1.12E-03 | 0.76 | 0.7 | 0.06 | 1.09 |
| 6 | Mpped2 | 0.42 | 5.93E-08 | 1.27E-03 | 0.83 | 0.78 | 0.05 | 1.06 |
| 6 | Gpd2 | 0.86 | 6.99E-08 | 1.49E-03 | 0.43 | 0.33 | 0.1 | 1.3 |
| 6 | Arxes2 | 0.92 | 7.74E-08 | 1.65E-03 | 0.44 | 0.34 | 0.1 | 1.29 |
| 6 | Eno3 | 1.06 | 8.07E-08 | 1.72E-03 | 0.28 | 0.18 | 0.11 | 1.61 |
| 6 | Calm3 | 0.32 | 8.59E-08 | 1.84E-03 | 0.88 | 0.84 | 0.04 | 1.05 |
| 6 | Ndn | 0.48 | 9.54E-08 | 2.04E-03 | 0.67 | 0.61 | 0.06 | 1.09 |
| 6 | Vangl2 | 0.59 | 1.10E-07 | 2.35E-03 | 0.65 | 0.57 | 0.08 | 1.13 |
| 6 | Nav3 | 0.56 | 1.21E-07 | 2.58E-03 | 0.51 | 0.38 | 0.13 | 1.35 |
| 6 | Fam102b | 1.49 | 1.26E-07 | 2.69E-03 | 0.2 | 0.11 | 0.09 | 1.81 |
| 6 | Nebl | 0.38 | 1.46E-07 | 3.11E-03 | 0.52 | 0.39 | 0.13 | 1.32 |
| 6 | Cyp51 | 0.53 | 1.72E-07 | 3.68E-03 | 0.68 | 0.61 | 0.07 | 1.12 |
| 6 | Kcnd2 | 1.47 | 2.38E-07 | 5.08E-03 | 0.14 | 0.07 | 0.07 | 2.12 |
| 6 | Olfm2 | 1.07 | 2.63E-07 | 5.62E-03 | 0.27 | 0.17 | 0.1 | 1.57 |
| 6 | Mgst3 | 0.56 | 3.19E-07 | 6.82E-03 | 0.56 | 0.49 | 0.07 | 1.15 |
| 6 | Praf2 | 0.7 | 3.26E-07 | 6.97E-03 | 0.51 | 0.42 | 0.09 | 1.21 |
| 6 | Pbx3 | 0.5 | 3.49E-07 | 7.45E-03 | 0.65 | 0.58 | 0.06 | 1.11 |
| 6 | Tmx2 | 0.55 | 3.57E-07 | 7.62E-03 | 0.66 | 0.59 | 0.07 | 1.12 |
| 6 | Tmx4 | 0.83 | 4.32E-07 | 9.22E-03 | 0.54 | 0.47 | 0.07 | 1.16 |
| 6 | Slc1a4 | 1.31 | 5.06E-07 | 1.08E-02 | 0.17 | 0.09 | 0.08 | 1.87 |
| 6 | Cystm1 | 0.58 | 5.15E-07 | 1.10E-02 | 0.45 | 0.35 | 0.1 | 1.28 |
| 6 | S100a6 | 0.43 | 6.54E-07 | 1.40E-02 | 0.35 | 0.23 | 0.12 | 1.5 |
| 6 | 4930523C07Rik | 1.06 | 6.75E-07 | 1.44E-02 | 0.18 | 0.1 | 0.08 | 1.88 |
| 6 | Rtn2 | 0.92 | 6.79E-07 | 1.45E-02 | 0.28 | 0.18 | 0.1 | 1.55 |
| 6 | Tceal3 | 1.1 | 7.29E-07 | 1.56E-02 | 0.25 | 0.15 | 0.09 | 1.61 |
| 6 | Pdia6 | 0.28 | 7.55E-07 | 1.61E-02 | 0.95 | 0.95 | 0 | 1 |
| 6 | Sdc2 | 0.85 | 7.93E-07 | 1.69E-02 | 0.39 | 0.3 | 0.09 | 1.31 |
| 6 | Ssr3 | 0.36 | 8.05E-07 | 1.72E-02 | 0.87 | 0.87 | 0 | 1 |
| 6 | Palld | 0.42 | 9.02E-07 | 1.93E-02 | 0.71 | 0.64 | 0.07 | 1.11 |
| 6 | Trim35 | 0.57 | 9.68E-07 | 2.07E-02 | 0.63 | 0.55 | 0.08 | 1.14 |
| 6 | Nbdy | 0.56 | 9.89E-07 | 2.11E-02 | 0.61 | 0.57 | 0.04 | 1.08 |
| 6 | Cd9 | 0.3 | 1.08E-06 | 2.31E-02 | 0.8 | 0.71 | 0.09 | 1.13 |
| 6 | Sdhd | 0.34 | 1.12E-06 | 2.40E-02 | 0.79 | 0.78 | 0.02 | 1.02 |
| 6 | A930004D18Rik | 1.29 | 1.21E-06 | 2.59E-02 | 0.17 | 0.09 | 0.08 | 1.88 |
| 6 | Plod2 | 0.54 | 1.24E-06 | 2.65E-02 | 0.56 | 0.48 | 0.08 | 1.17 |
| 6 | Thbs3 | 1.17 | 1.25E-06 | 2.67E-02 | 0.22 | 0.13 | 0.09 | 1.65 |
| 6 | Iglon5 | 1.32 | 1.31E-06 | 2.80E-02 | 0.19 | 0.11 | 0.08 | 1.74 |
| 6 | Cby1 | 0.59 | 1.45E-06 | 3.10E-02 | 0.53 | 0.46 | 0.08 | 1.17 |
| 6 | Dnajc21 | 0.39 | 1.46E-06 | 3.12E-02 | 0.71 | 0.65 | 0.06 | 1.09 |
| 6 | Glrx2 | 0.51 | 1.49E-06 | 3.18E-02 | 0.71 | 0.69 | 0.02 | 1.03 |
| 6 | Cpt1a | 1.1 | 1.51E-06 | 3.22E-02 | 0.22 | 0.13 | 0.09 | 1.65 |
| 6 | Mllt3 | 0.53 | 1.57E-06 | 3.35E-02 | 0.62 | 0.55 | 0.07 | 1.13 |
| 6 | Pax6 | 0.39 | 1.58E-06 | 3.37E-02 | 0.6 | 0.48 | 0.12 | 1.24 |

**Supplementary Table 3. Marker genes for the original clusters of integrated clustering of E11.5 control and NotchGOF λ-junction ectoderm cells.**

| | | | | | | | | |
|---|---|---|---|---|---|---|---|---|
| 6 | Sod1 | 0.33 | 1.60E-06 | 3.42E-02 | 0.82 | 0.77 | 0.05 | 1.06 |
| 6 | Pmepa1 | 0.76 | 1.65E-06 | 3.53E-02 | 0.46 | 0.36 | 0.1 | 1.26 |
| 6 | Azin1 | 0.51 | 1.68E-06 | 3.59E-02 | 0.62 | 0.55 | 0.07 | 1.12 |
| 6 | Tmem238 | 0.63 | 1.94E-06 | 4.13E-02 | 0.5 | 0.4 | 0.1 | 1.24 |
| 6 | Dach1 | 0.52 | 2.11E-06 | 4.51E-02 | 0.56 | 0.49 | 0.07 | 1.15 |
| 6 | Rrbp1 | 0.35 | 2.18E-06 | 4.65E-02 | 0.92 | 0.89 | 0.03 | 1.04 |
| 6 | Ppp1r1a | 0.84 | 2.31E-06 | 4.94E-02 | 0.4 | 0.31 | 0.09 | 1.28 |
| 7 | Wnt3 | 4.3 | 0 | 0 | 0.83 | 0.07 | 0.77 | 12.42 |
| 7 | Rprml | 6.5 | 0 | 0 | 0.51 | 0.01 | 0.5 | 46.27 |
| 7 | Cxcl14 | 3.78 | 4.17E-268 | 8.90E-264 | 0.96 | 0.15 | 0.81 | 6.34 |
| 7 | Tfap2b | 3.61 | 2.00E-265 | 4.27E-261 | 0.94 | 0.14 | 0.8 | 6.6 |
| 7 | Wnt9b | 3.59 | 2.56E-236 | 5.46E-232 | 0.72 | 0.07 | 0.65 | 9.72 |
| 7 | Sostdc1 | 4.14 | 7.65E-217 | 1.63E-212 | 0.92 | 0.21 | 0.71 | 4.37 |
| 7 | Pcp4l1 | 3.81 | 2.55E-200 | 5.44E-196 | 0.55 | 0.05 | 0.51 | 12.24 |
| 7 | Wnt7b | 3.75 | 3.90E-192 | 8.33E-188 | 0.75 | 0.13 | 0.63 | 6.03 |
| 7 | Krt15 | 3.96 | 1.73E-185 | 3.71E-181 | 0.73 | 0.11 | 0.61 | 6.42 |
| 7 | Wnt6 | 2.95 | 1.38E-181 | 2.95E-177 | 0.98 | 0.33 | 0.65 | 2.97 |
| 7 | Kremen2 | 3.31 | 7.19E-179 | 1.54E-174 | 0.77 | 0.14 | 0.63 | 5.55 |
| 7 | Wnt10a | 5.33 | 7.49E-176 | 1.60E-171 | 0.31 | 0.01 | 0.3 | 33.89 |
| 7 | Piezo2 | 3.08 | 7.71E-171 | 1.65E-166 | 0.75 | 0.13 | 0.62 | 5.76 |
| 7 | Irx5 | 3.04 | 1.59E-170 | 3.40E-166 | 0.86 | 0.2 | 0.66 | 4.3 |
| 7 | Npr3 | 4.52 | 1.31E-162 | 2.79E-158 | 0.39 | 0.03 | 0.36 | 15.56 |
| 7 | Plxna2 | 2.95 | 4.20E-152 | 8.96E-148 | 0.85 | 0.23 | 0.63 | 3.79 |
| 7 | Irx3 | 3.15 | 2.54E-149 | 5.43E-145 | 0.76 | 0.17 | 0.59 | 4.43 |
| 7 | Irx4 | 6.59 | 4.10E-148 | 8.75E-144 | 0.21 | 0 | 0.21 | 71.33 |
| 7 | C130021I20Rik | 5.27 | 3.56E-140 | 7.59E-136 | 0.26 | 0.01 | 0.25 | 28.89 |
| 7 | Fth1 | 1.24 | 1.14E-139 | 2.44E-135 | 1 | 1 | 0 | 1 |
| 7 | Tspan2 | 4.19 | 2.81E-138 | 6.00E-134 | 0.36 | 0.03 | 0.33 | 13.77 |
| 7 | Gria2 | 3.96 | 1.20E-134 | 2.56E-130 | 0.47 | 0.06 | 0.42 | 8.46 |
| 7 | Edaradd | 3.51 | 1.08E-128 | 2.30E-124 | 0.45 | 0.05 | 0.4 | 8.55 |
| 7 | Fgfr2 | 2.3 | 2.30E-123 | 4.92E-119 | 0.95 | 0.44 | 0.51 | 2.15 |
| 7 | Snai2 | 2.53 | 6.76E-119 | 1.44E-114 | 0.68 | 0.15 | 0.53 | 4.67 |
| 7 | Tnfrsf19 | 2.5 | 9.36E-116 | 2.00E-111 | 0.76 | 0.21 | 0.55 | 3.58 |
| 7 | Lgals7 | 3.9 | 3.86E-112 | 8.25E-108 | 0.34 | 0.03 | 0.31 | 11.23 |
| 7 | Ctsl | 1.63 | 9.98E-112 | 2.13E-107 | 0.98 | 0.86 | 0.13 | 1.15 |
| 7 | Rnase4 | 3.09 | 1.52E-109 | 3.25E-105 | 0.52 | 0.09 | 0.43 | 5.86 |
| 7 | Crnde | 3.33 | 3.93E-107 | 8.39E-103 | 0.46 | 0.07 | 0.39 | 6.71 |
| 7 | Tfap2a | 2.29 | 1.10E-103 | 2.35E-99 | 0.84 | 0.31 | 0.53 | 2.7 |
| 7 | Sema3a | 2.58 | 1.70E-103 | 3.63E-99 | 0.88 | 0.46 | 0.43 | 1.94 |
| 7 | Hapln1 | 5.65 | 2.38E-103 | 5.09E-99 | 0.19 | 0.01 | 0.19 | 27.57 |
| 7 | Tmsb10 | 0.73 | 2.24E-94 | 4.79E-90 | 1 | 1 | 0 | 1 |
| 7 | Cobll1 | 2.83 | 1.13E-93 | 2.41E-89 | 0.45 | 0.08 | 0.38 | 5.96 |
| 7 | Nr2e3 | 5.75 | 4.38E-93 | 9.36E-89 | 0.14 | 0 | 0.14 | 68.5 |
| 7 | Lmx1b | 4.37 | 7.22E-92 | 1.54E-87 | 0.21 | 0.01 | 0.2 | 18.82 |
| 7 | Cdkn1c | 1.7 | 9.17E-91 | 1.96E-86 | 0.99 | 0.88 | 0.12 | 1.13 |
| 7 | Col17a1 | 2.57 | 2.93E-90 | 6.25E-86 | 0.52 | 0.1 | 0.41 | 5.06 |
| 7 | Pdlim1 | 1.51 | 3.31E-82 | 7.07E-78 | 0.95 | 0.63 | 0.32 | 1.51 |
| 7 | Trp63 | 1.61 | 4.79E-81 | 1.02E-76 | 0.96 | 0.48 | 0.48 | 1.98 |
| 7 | Mir205hg | 1.87 | 1.82E-80 | 3.89E-76 | 0.86 | 0.41 | 0.45 | 2.08 |
| 7 | Txndc17 | 1.33 | 2.44E-80 | 5.22E-76 | 0.97 | 0.94 | 0.03 | 1.03 |
| 7 | Robo2 | 2.4 | 5.98E-77 | 1.28E-72 | 0.65 | 0.22 | 0.43 | 2.99 |
| 7 | Atxn10 | 1.15 | 1.06E-73 | 2.27E-69 | 0.98 | 0.91 | 0.07 | 1.07 |
| 7 | Ccnd2 | 1.26 | 1.53E-73 | 3.28E-69 | 1 | 0.77 | 0.23 | 1.29 |
| 7 | Pdgfa | 1.1 | 7.43E-73 | 1.59E-68 | 1 | 0.93 | 0.07 | 1.07 |
| 7 | Timp3 | 3.35 | 8.65E-72 | 1.85E-67 | 0.33 | 0.05 | 0.28 | 6.39 |
| 7 | Lmo1 | 2.44 | 2.19E-71 | 4.68E-67 | 0.62 | 0.2 | 0.41 | 3.03 |
| 7 | Crabp1 | 1.98 | 5.56E-70 | 1.19E-65 | 0.9 | 0.54 | 0.36 | 1.67 |

**Supplementary Table 3. Marker genes for the original clusters of integrated clustering of E11.5 control and NotchGOF λ-junction ectoderm cells.**

| | | | | | | | | |
|---|---|---|---|---|---|---|---|---|
| 7 | Itgb5 | 1.65 | 4.34E-69 | 9.28E-65 | 0.88 | 0.47 | 0.4 | 1.85 |
| 7 | Cd24a | 0.86 | 4.77E-65 | 1.02E-60 | 0.99 | 0.96 | 0.03 | 1.04 |
| 7 | Tspan18 | 3.42 | 1.78E-64 | 3.79E-60 | 0.27 | 0.04 | 0.23 | 7.42 |
| 7 | Bnc1 | 3.08 | 3.34E-62 | 7.13E-58 | 0.23 | 0.03 | 0.2 | 8.77 |
| 7 | Jag2 | 1.78 | 4.31E-62 | 9.20E-58 | 0.7 | 0.26 | 0.44 | 2.69 |
| 7 | Perp | 1.37 | 9.28E-62 | 1.98E-57 | 0.93 | 0.52 | 0.42 | 1.81 |
| 7 | Hras | 0.95 | 7.91E-61 | 1.69E-56 | 0.98 | 0.9 | 0.08 | 1.08 |
| 7 | Jdp2 | 2.59 | 9.21E-61 | 1.97E-56 | 0.38 | 0.08 | 0.3 | 4.75 |
| 7 | Brinp1 | 3.28 | 1.09E-60 | 2.32E-56 | 0.23 | 0.03 | 0.2 | 8.65 |
| 7 | Tnik | 2.09 | 5.87E-60 | 1.25E-55 | 0.4 | 0.09 | 0.31 | 4.65 |
| 7 | Ldhb | 1.01 | 4.04E-59 | 8.63E-55 | 0.95 | 0.87 | 0.09 | 1.1 |
| 7 | Meis1 | 1.52 | 1.20E-58 | 2.56E-54 | 0.89 | 0.57 | 0.32 | 1.57 |
| 7 | Wls | 1.16 | 3.26E-58 | 6.96E-54 | 0.97 | 0.79 | 0.18 | 1.23 |
| 7 | Cald1 | 1 | 6.02E-58 | 1.29E-53 | 0.97 | 0.68 | 0.29 | 1.42 |
| 7 | Fermt1 | 1.34 | 2.56E-57 | 5.46E-53 | 0.89 | 0.54 | 0.35 | 1.66 |
| 7 | Insyn2b | 2.65 | 3.35E-56 | 7.15E-52 | 0.31 | 0.06 | 0.25 | 5.55 |
| 7 | Pdgfc | 1.18 | 2.55E-55 | 5.44E-51 | 0.92 | 0.75 | 0.17 | 1.22 |
| 7 | Nrep | 1.11 | 5.21E-55 | 1.11E-50 | 0.95 | 0.8 | 0.15 | 1.19 |
| 7 | Itgav | 2.14 | 1.63E-54 | 3.47E-50 | 0.54 | 0.19 | 0.36 | 2.91 |
| 7 | Msx2 | 1.63 | 1.39E-53 | 2.98E-49 | 0.77 | 0.38 | 0.39 | 2.02 |
| 7 | Plk2 | 1.87 | 1.42E-53 | 3.03E-49 | 0.71 | 0.34 | 0.37 | 2.09 |
| 7 | Nbl1 | 2.79 | 3.80E-53 | 8.11E-49 | 0.32 | 0.07 | 0.25 | 4.91 |
| 7 | Igfbp2 | 0.88 | 1.80E-52 | 3.85E-48 | 1 | 0.98 | 0.02 | 1.02 |
| 7 | Ube2e3 | 1.02 | 9.30E-52 | 1.99E-47 | 0.97 | 0.9 | 0.07 | 1.07 |
| 7 | Pthlh | 3.37 | 1.34E-51 | 2.86E-47 | 0.25 | 0.04 | 0.21 | 6.31 |
| 7 | Wnt7a | 4.89 | 1.86E-50 | 3.98E-46 | 0.11 | 0.01 | 0.1 | 21.8 |
| 7 | Artn | 3.49 | 2.06E-50 | 4.40E-46 | 0.16 | 0.02 | 0.15 | 10.06 |
| 7 | Igfbp4 | 1.11 | 6.06E-50 | 1.29E-45 | 0.94 | 0.73 | 0.22 | 1.3 |
| 7 | Sgk3 | 2.38 | 1.30E-49 | 2.78E-45 | 0.4 | 0.11 | 0.29 | 3.77 |
| 7 | Wnt10b | 3.9 | 1.53E-49 | 3.27E-45 | 0.15 | 0.01 | 0.13 | 11.31 |
| 7 | Gstk1 | 2.32 | 4.67E-49 | 9.98E-45 | 0.43 | 0.12 | 0.31 | 3.48 |
| 7 | Chst11 | 2.12 | 4.99E-49 | 1.06E-44 | 0.46 | 0.14 | 0.32 | 3.22 |
| 7 | Sema3c | 2.13 | 5.62E-49 | 1.20E-44 | 0.58 | 0.23 | 0.35 | 2.5 |
| 7 | Dnajc1 | 1.42 | 6.58E-49 | 1.41E-44 | 0.86 | 0.64 | 0.22 | 1.35 |
| 7 | Art4 | 2.92 | 3.19E-48 | 6.81E-44 | 0.23 | 0.04 | 0.2 | 6.44 |
| 7 | Col5a2 | 2.09 | 9.19E-48 | 1.96E-43 | 0.46 | 0.14 | 0.31 | 3.21 |
| 7 | Plbd1 | 2.93 | 4.02E-47 | 8.58E-43 | 0.2 | 0.03 | 0.17 | 7.41 |
| 7 | Tmem132c | 2.64 | 1.23E-46 | 2.63E-42 | 0.39 | 0.11 | 0.28 | 3.54 |
| 7 | Galnt18 | 2.47 | 1.12E-44 | 2.39E-40 | 0.33 | 0.08 | 0.25 | 4.06 |
| 7 | Cnr1 | 2.65 | 1.23E-44 | 2.63E-40 | 0.27 | 0.06 | 0.22 | 4.89 |
| 7 | Wnt4 | 1.3 | 2.99E-44 | 6.38E-40 | 0.9 | 0.7 | 0.2 | 1.28 |
| 7 | Sox14 | 2.75 | 9.70E-43 | 2.07E-38 | 0.23 | 0.04 | 0.19 | 5.92 |
| 7 | Fzd10 | 2.41 | 4.47E-42 | 9.55E-38 | 0.39 | 0.12 | 0.27 | 3.25 |
| 7 | Col18a1 | 1.02 | 6.15E-42 | 1.31E-37 | 0.93 | 0.86 | 0.07 | 1.09 |
| 7 | Tmem178 | 2.25 | 1.17E-41 | 2.51E-37 | 0.37 | 0.1 | 0.26 | 3.58 |
| 7 | Trp73 | 3.06 | 1.58E-41 | 3.38E-37 | 0.16 | 0.02 | 0.14 | 8.05 |
| 7 | Kitl | 2.4 | 2.36E-41 | 5.05E-37 | 0.39 | 0.12 | 0.27 | 3.25 |
| 7 | Pgam1 | 0.5 | 4.36E-41 | 9.30E-37 | 1 | 0.99 | 0.01 | 1.01 |
| 7 | Sox11 | 0.71 | 1.28E-40 | 2.74E-36 | 1 | 0.99 | 0.01 | 1.01 |
| 7 | Slc7a11 | 1.5 | 3.27E-40 | 6.99E-36 | 0.39 | 0.12 | 0.28 | 3.42 |
| 7 | Zfp36l2 | 2.11 | 6.62E-40 | 1.41E-35 | 0.53 | 0.23 | 0.3 | 2.31 |
| 7 | Arf5 | 0.51 | 8.04E-38 | 1.72E-33 | 1 | 0.99 | 0.01 | 1.01 |
| 7 | Bnc2 | 2.52 | 8.97E-38 | 1.92E-33 | 0.27 | 0.06 | 0.21 | 4.29 |
| 7 | Arhgdib | 1.84 | 1.19E-37 | 2.54E-33 | 0.49 | 0.19 | 0.3 | 2.54 |
| 7 | Ngf | 2.61 | 1.27E-37 | 2.70E-33 | 0.18 | 0.03 | 0.15 | 6.63 |
| 7 | Lef1 | 0.99 | 1.17E-36 | 2.50E-32 | 0.72 | 0.36 | 0.37 | 2.03 |
| 7 | Krt14 | 0.96 | 2.91E-36 | 6.22E-32 | 0.81 | 0.44 | 0.37 | 1.84 |

**Supplementary Table 3. Marker genes for the original clusters of integrated clustering of E11.5 control and NotchGOF λ-junction ectoderm cells.**

| | | | | | | | | |
|---|---|---|---|---|---|---|---|---|
| 7 | Hspb1 | 1.09 | 8.98E-36 | 1.92E-31 | 0.71 | 0.37 | 0.34 | 1.93 |
| 7 | Etaa1os | 2.47 | 1.22E-35 | 2.60E-31 | 0.28 | 0.07 | 0.21 | 3.89 |
| 7 | Gm42418 | 1.06 | 1.90E-35 | 4.06E-31 | 1 | 1 | 0 | 1 |
| 7 | Cdh3 | 1.25 | 3.59E-35 | 7.67E-31 | 0.8 | 0.56 | 0.24 | 1.43 |
| 7 | Tfap2c | 1.66 | 3.69E-35 | 7.87E-31 | 0.5 | 0.2 | 0.29 | 2.44 |
| 7 | Ifitm2 | 0.68 | 4.52E-35 | 9.66E-31 | 0.98 | 0.93 | 0.05 | 1.06 |
| 7 | Dlk2 | 3.09 | 7.76E-35 | 1.66E-30 | 0.15 | 0.02 | 0.13 | 7.35 |
| 7 | Htra1 | 2.27 | 8.32E-35 | 1.78E-30 | 0.24 | 0.05 | 0.19 | 4.51 |
| 7 | Ccdc68 | 2.05 | 8.34E-35 | 1.78E-30 | 0.37 | 0.12 | 0.25 | 3.1 |
| 7 | Tbx3 | 2.19 | 1.19E-34 | 2.55E-30 | 0.28 | 0.07 | 0.21 | 3.96 |
| 7 | 2610204G07Rik | 3.04 | 1.30E-34 | 2.77E-30 | 0.15 | 0.02 | 0.13 | 7 |
| 7 | P4ha1 | 1.3 | 2.39E-34 | 5.12E-30 | 0.76 | 0.51 | 0.25 | 1.49 |
| 7 | Selenow | 0.43 | 4.08E-34 | 8.72E-30 | 1 | 0.99 | 0.01 | 1.01 |
| 7 | Slc35b3 | 1.7 | 2.21E-33 | 4.71E-29 | 0.53 | 0.25 | 0.29 | 2.17 |
| 7 | Sema3e | 2.68 | 3.20E-33 | 6.84E-29 | 0.15 | 0.02 | 0.13 | 7 |
| 7 | Prps2 | 1.69 | 3.37E-33 | 7.20E-29 | 0.59 | 0.31 | 0.28 | 1.92 |
| 7 | Mir100hg | 1.13 | 5.62E-33 | 1.20E-28 | 0.84 | 0.62 | 0.23 | 1.36 |
| 7 | Gm32585 | 1.46 | 1.12E-32 | 2.38E-28 | 0.49 | 0.2 | 0.29 | 2.46 |
| 7 | Pltp | 1.14 | 1.60E-32 | 3.42E-28 | 0.76 | 0.45 | 0.31 | 1.7 |
| 7 | Chchd2 | 0.32 | 1.91E-32 | 4.07E-28 | 1 | 1 | 0 | 1 |
| 7 | Stk26 | 1.12 | 5.01E-32 | 1.07E-27 | 0.73 | 0.45 | 0.28 | 1.61 |
| 7 | Scn3a | 2.34 | 5.54E-32 | 1.18E-27 | 0.13 | 0.02 | 0.12 | 7.39 |
| 7 | Ajuba | 1.28 | 8.13E-32 | 1.74E-27 | 0.68 | 0.41 | 0.28 | 1.67 |
| 7 | Luzp1 | 0.81 | 1.57E-31 | 3.36E-27 | 0.92 | 0.8 | 0.12 | 1.16 |
| 7 | Adgrl2 | 1.02 | 1.98E-31 | 4.23E-27 | 0.89 | 0.69 | 0.2 | 1.29 |
| 7 | Filip1l | 1.58 | 2.15E-31 | 4.59E-27 | 0.53 | 0.25 | 0.28 | 2.12 |
| 7 | Npnt | 1.21 | 4.99E-31 | 1.07E-26 | 0.71 | 0.43 | 0.27 | 1.64 |
| 7 | Cth | 1.98 | 1.03E-30 | 2.20E-26 | 0.39 | 0.15 | 0.24 | 2.61 |
| 7 | Phlda2 | 3.08 | 2.91E-30 | 6.21E-26 | 0.19 | 0.04 | 0.15 | 4.85 |
| 7 | Hpca | 2.26 | 3.92E-30 | 8.37E-26 | 0.14 | 0.02 | 0.12 | 6.85 |
| 7 | Slc1a3 | 1.76 | 3.99E-30 | 8.52E-26 | 0.44 | 0.19 | 0.26 | 2.38 |
| 7 | Pla2r1 | 1.63 | 4.12E-30 | 8.79E-26 | 0.54 | 0.26 | 0.28 | 2.07 |
| 7 | Chst15 | 1.97 | 5.95E-30 | 1.27E-25 | 0.35 | 0.13 | 0.23 | 2.83 |
| 7 | Sema5a | 1.72 | 9.26E-30 | 1.98E-25 | 0.45 | 0.19 | 0.26 | 2.4 |
| 7 | Adk | 1.08 | 9.85E-30 | 2.10E-25 | 0.79 | 0.61 | 0.18 | 1.3 |
| 7 | Dock8 | 1.94 | 1.18E-29 | 2.53E-25 | 0.28 | 0.08 | 0.2 | 3.46 |
| 7 | Tgfbi | 3.17 | 1.01E-28 | 2.16E-24 | 0.13 | 0.02 | 0.11 | 6.84 |
| 7 | Ctnnal1 | 1.38 | 1.09E-28 | 2.33E-24 | 0.69 | 0.48 | 0.22 | 1.45 |
| 7 | Dkk4 | 2.43 | 2.53E-28 | 5.41E-24 | 0.14 | 0.02 | 0.12 | 6.23 |
| 7 | Cbr2 | 2.47 | 3.91E-28 | 8.35E-24 | 0.16 | 0.03 | 0.13 | 5.37 |
| 7 | Fam162a | 0.66 | 6.22E-28 | 1.33E-23 | 0.95 | 0.93 | 0.03 | 1.03 |
| 7 | Cib2 | 1.47 | 1.27E-27 | 2.72E-23 | 0.41 | 0.16 | 0.25 | 2.52 |
| 7 | Nav2 | 0.81 | 2.13E-27 | 4.55E-23 | 0.87 | 0.69 | 0.18 | 1.26 |
| 7 | Blvra | 1.8 | 2.21E-27 | 4.72E-23 | 0.34 | 0.12 | 0.22 | 2.81 |
| 7 | Ntf5 | 1.19 | 2.96E-27 | 6.31E-23 | 0.58 | 0.3 | 0.28 | 1.94 |
| 7 | Rxfp1 | 2.71 | 8.23E-27 | 1.76E-22 | 0.19 | 0.04 | 0.15 | 4.4 |
| 7 | Col4a2 | 1.11 | 1.12E-26 | 2.40E-22 | 0.73 | 0.49 | 0.24 | 1.49 |
| 7 | Parm1 | 1.23 | 1.80E-26 | 3.85E-22 | 0.45 | 0.19 | 0.26 | 2.37 |
| 7 | Pla2g4a | 1.9 | 3.10E-26 | 6.63E-22 | 0.29 | 0.09 | 0.2 | 3.16 |
| 7 | Bgn | 2.78 | 6.26E-26 | 1.34E-21 | 0.14 | 0.03 | 0.12 | 5.54 |
| 7 | Rest | 0.8 | 8.52E-26 | 1.82E-21 | 0.88 | 0.67 | 0.21 | 1.32 |
| 7 | Stmn1 | 0.31 | 9.75E-26 | 2.08E-21 | 1 | 1 | 0 | 1 |
| 7 | Gdf6 | 2.48 | 1.10E-25 | 2.35E-21 | 0.15 | 0.03 | 0.12 | 5.13 |
| 7 | Meis2 | 0.51 | 1.16E-25 | 2.47E-21 | 0.98 | 0.95 | 0.03 | 1.03 |
| 7 | Gadd45g | 0.37 | 1.70E-25 | 3.63E-21 | 0.64 | 0.32 | 0.32 | 1.97 |
| 7 | Cox6c | 0.33 | 2.26E-25 | 4.82E-21 | 1 | 1 | 0.01 | 1.01 |
| 7 | Man1a | 2.56 | 2.78E-25 | 5.94E-21 | 0.18 | 0.04 | 0.14 | 4.33 |

**Supplementary Table 3. Marker genes for the original clusters of integrated clustering of E11.5 control and NotchGOF λ-junction ectoderm cells.**

| | | | | | | | | |
|---|---|---|---|---|---|---|---|---|
| 7 | Pcyt1b | 1.8 | 3.51E-25 | 7.50E-21 | 0.31 | 0.11 | 0.2 | 2.86 |
| 7 | Dag1 | 1.03 | 3.58E-25 | 7.64E-21 | 0.79 | 0.62 | 0.16 | 1.26 |
| 7 | Plxna4 | 2.4 | 3.77E-25 | 8.05E-21 | 0.19 | 0.05 | 0.15 | 4.11 |
| 7 | Hmgn3 | 0.79 | 6.07E-25 | 1.30E-20 | 0.87 | 0.71 | 0.16 | 1.23 |
| 7 | H19 | 0.46 | 8.91E-25 | 1.90E-20 | 0.98 | 0.87 | 0.11 | 1.12 |
| 7 | Arhgef3 | 3.29 | 9.10E-25 | 1.94E-20 | 0.11 | 0.02 | 0.09 | 6.81 |
| 7 | Ahnak | 1.06 | 9.61E-25 | 2.05E-20 | 0.63 | 0.34 | 0.28 | 1.83 |
| 7 | Sox6 | 2.06 | 1.15E-24 | 2.45E-20 | 0.27 | 0.09 | 0.18 | 3.04 |
| 7 | Pdpn | 1.18 | 1.47E-24 | 3.14E-20 | 0.61 | 0.35 | 0.26 | 1.75 |
| 7 | Cnih2 | 1.38 | 1.66E-24 | 3.55E-20 | 0.36 | 0.14 | 0.22 | 2.52 |
| 7 | Megf6 | 1.66 | 4.04E-24 | 8.62E-20 | 0.39 | 0.17 | 0.23 | 2.34 |
| 7 | Dlx2 | 0.89 | 4.73E-24 | 1.01E-19 | 0.62 | 0.33 | 0.3 | 1.9 |
| 7 | Atpif1 | 0.35 | 5.88E-24 | 1.26E-19 | 1 | 0.99 | 0 | 1 |
| 7 | Aldh2 | 0.88 | 2.25E-23 | 4.80E-19 | 0.81 | 0.69 | 0.12 | 1.17 |
| 7 | Dok5 | 1.1 | 2.29E-23 | 4.89E-19 | 0.63 | 0.36 | 0.26 | 1.73 |
| 7 | Vcan | 0.75 | 3.83E-23 | 8.19E-19 | 0.91 | 0.74 | 0.17 | 1.23 |
| 7 | Ttc28 | 1.11 | 3.95E-23 | 8.44E-19 | 0.65 | 0.44 | 0.22 | 1.5 |
| 7 | Nxn | 0.65 | 4.02E-23 | 8.59E-19 | 0.9 | 0.75 | 0.15 | 1.2 |
| 7 | Klhl13 | 1.18 | 4.89E-23 | 1.04E-18 | 0.61 | 0.39 | 0.23 | 1.58 |
| 7 | Btg1 | 0.86 | 7.07E-23 | 1.51E-18 | 0.85 | 0.74 | 0.11 | 1.15 |
| 7 | Dbi | 0.44 | 1.62E-22 | 3.46E-18 | 0.99 | 0.97 | 0.02 | 1.02 |
| 7 | Pard6g | 0.75 | 2.48E-22 | 5.29E-18 | 0.82 | 0.64 | 0.18 | 1.28 |
| 7 | Sox12 | 0.98 | 2.84E-22 | 6.07E-18 | 0.76 | 0.59 | 0.17 | 1.29 |
| 7 | Trps1 | 1.11 | 2.86E-22 | 6.10E-18 | 0.71 | 0.52 | 0.19 | 1.36 |
| 7 | Rap2c | 0.79 | 5.58E-22 | 1.19E-17 | 0.81 | 0.66 | 0.16 | 1.24 |
| 7 | Dst | 0.98 | 6.28E-22 | 1.34E-17 | 0.72 | 0.47 | 0.25 | 1.53 |
| 7 | Anxa2 | 0.57 | 6.90E-22 | 1.47E-17 | 0.95 | 0.75 | 0.2 | 1.27 |
| 7 | Mycn | 1.29 | 7.75E-22 | 1.66E-17 | 0.57 | 0.34 | 0.23 | 1.68 |
| 7 | Arhgap22 | 2.4 | 8.28E-22 | 1.77E-17 | 0.16 | 0.04 | 0.12 | 4.39 |
| 7 | Pcdh18 | 1.62 | 1.24E-21 | 2.64E-17 | 0.38 | 0.17 | 0.21 | 2.24 |
| 7 | Rnd3 | 0.68 | 1.43E-21 | 3.06E-17 | 0.88 | 0.73 | 0.16 | 1.22 |
| 7 | Bzw2 | 0.47 | 1.58E-21 | 3.37E-17 | 0.97 | 0.94 | 0.03 | 1.03 |
| 7 | Ackr3 | 1.27 | 1.77E-21 | 3.79E-17 | 0.58 | 0.37 | 0.22 | 1.59 |
| 7 | Eif2s3y | 2.21 | 2.57E-21 | 5.50E-17 | 0.17 | 0.04 | 0.13 | 4.1 |
| 7 | Fry | 1.57 | 6.98E-21 | 1.49E-16 | 0.33 | 0.13 | 0.2 | 2.48 |
| 7 | Col12a1 | 2.03 | 8.35E-21 | 1.78E-16 | 0.21 | 0.06 | 0.15 | 3.34 |
| 7 | Astn2 | 2.47 | 8.50E-21 | 1.82E-16 | 0.12 | 0.02 | 0.1 | 5.13 |
| 7 | Gng5 | 0.26 | 9.31E-21 | 1.99E-16 | 0.99 | 0.99 | 0 | 1 |
| 7 | Antxr1 | 1.33 | 1.07E-20 | 2.29E-16 | 0.46 | 0.24 | 0.22 | 1.91 |
| 7 | Rbbp8 | 0.8 | 1.14E-20 | 2.43E-16 | 0.83 | 0.67 | 0.16 | 1.24 |
| 7 | Gjb6 | 2.5 | 1.73E-20 | 3.70E-16 | 0.19 | 0.06 | 0.14 | 3.45 |
| 7 | Bambi | 0.57 | 2.82E-20 | 6.02E-16 | 0.69 | 0.48 | 0.22 | 1.45 |
| 7 | Lratd2 | 1.16 | 3.30E-20 | 7.04E-16 | 0.55 | 0.32 | 0.23 | 1.7 |
| 7 | Fmod | 2.58 | 3.60E-20 | 7.69E-16 | 0.11 | 0.02 | 0.09 | 5.74 |
| 7 | Kcnq1ot1 | 0.65 | 4.45E-20 | 9.51E-16 | 0.96 | 0.94 | 0.02 | 1.02 |
| 7 | Btbd11 | 2.43 | 5.87E-20 | 1.25E-15 | 0.14 | 0.03 | 0.11 | 4.57 |
| 7 | Dusp9 | 1.4 | 6.10E-20 | 1.30E-15 | 0.44 | 0.23 | 0.21 | 1.93 |
| 7 | Dusp7 | 1.34 | 7.63E-20 | 1.63E-15 | 0.51 | 0.3 | 0.21 | 1.71 |
| 7 | Cd59a | 1.01 | 1.04E-19 | 2.22E-15 | 0.53 | 0.29 | 0.24 | 1.84 |
| 7 | Slc39a10 | 0.78 | 2.17E-19 | 4.64E-15 | 0.87 | 0.75 | 0.12 | 1.16 |
| 7 | Sulf2 | 1.12 | 2.23E-19 | 4.76E-15 | 0.55 | 0.32 | 0.23 | 1.7 |
| 7 | Ehbp1 | 1.06 | 2.41E-19 | 5.15E-15 | 0.67 | 0.48 | 0.19 | 1.38 |
| 7 | Gjb2 | 1.69 | 2.78E-19 | 5.93E-15 | 0.37 | 0.17 | 0.2 | 2.16 |
| 7 | Afap1l1 | 1.57 | 3.18E-19 | 6.78E-15 | 0.28 | 0.11 | 0.17 | 2.58 |
| 7 | Nrp2 | 1.28 | 5.02E-19 | 1.07E-14 | 0.37 | 0.16 | 0.2 | 2.24 |
| 7 | Aqp4 | 2.68 | 6.09E-19 | 1.30E-14 | 0.13 | 0.03 | 0.1 | 4.48 |
| 7 | Ror1 | 2.55 | 7.03E-19 | 1.50E-14 | 0.11 | 0.02 | 0.09 | 5.19 |

**Supplementary Table 3. Marker genes for the original clusters of integrated clustering of E11.5 control and NotchGOF λ-junction ectoderm cells.**

| | | | | | | | | |
|---|---|---|---|---|---|---|---|---|
| 7 | Atxn7l3b | 0.32 | 7.60E-19 | 1.62E-14 | 0.99 | 0.99 | 0 | 1 |
| 7 | Nell2 | 1.58 | 8.60E-19 | 1.84E-14 | 0.28 | 0.11 | 0.17 | 2.55 |
| 7 | Sfr1 | 0.37 | 1.41E-18 | 3.01E-14 | 0.99 | 0.98 | 0.02 | 1.02 |
| 7 | Ednrb | 2.57 | 1.51E-18 | 3.22E-14 | 0.14 | 0.03 | 0.11 | 4.28 |
| 7 | Zfp503 | 0.82 | 1.90E-18 | 4.06E-14 | 0.71 | 0.49 | 0.22 | 1.45 |
| 7 | Efcab1 | 1.64 | 2.36E-18 | 5.05E-14 | 0.25 | 0.09 | 0.16 | 2.83 |
| 7 | Cox6b2 | 0.95 | 2.64E-18 | 5.63E-14 | 0.67 | 0.5 | 0.17 | 1.35 |
| 7 | Tpm2 | 1.42 | 3.10E-18 | 6.62E-14 | 0.37 | 0.18 | 0.19 | 2.09 |
| 7 | Arpc2 | 0.4 | 7.41E-18 | 1.58E-13 | 0.96 | 0.96 | 0 | 1 |
| 7 | Emp1 | 1.21 | 7.56E-18 | 1.61E-13 | 0.36 | 0.16 | 0.19 | 2.18 |
| 7 | Znrf3 | 1.17 | 1.19E-17 | 2.55E-13 | 0.49 | 0.28 | 0.21 | 1.76 |
| 7 | Ucp2 | 0.74 | 1.52E-17 | 3.26E-13 | 0.66 | 0.47 | 0.2 | 1.43 |
| 7 | Sh3bgrl | 0.55 | 2.18E-17 | 4.65E-13 | 0.87 | 0.75 | 0.12 | 1.16 |
| 7 | Uty | 2.28 | 2.25E-17 | 4.80E-13 | 0.12 | 0.03 | 0.09 | 4.64 |
| 7 | Bnip3 | 0.79 | 2.47E-17 | 5.28E-13 | 0.67 | 0.44 | 0.23 | 1.51 |
| 7 | Cpe | 0.86 | 2.73E-17 | 5.84E-13 | 0.66 | 0.46 | 0.2 | 1.43 |
| 7 | AY036118 | 1 | 3.49E-17 | 7.46E-13 | 0.78 | 0.64 | 0.15 | 1.23 |
| 7 | Mbnl2 | 0.55 | 3.92E-17 | 8.36E-13 | 0.91 | 0.78 | 0.13 | 1.17 |
| 7 | Pid1 | 1.85 | 5.54E-17 | 1.18E-12 | 0.18 | 0.06 | 0.12 | 3.25 |
| 7 | Zfp462 | 0.63 | 8.75E-17 | 1.87E-12 | 0.84 | 0.73 | 0.11 | 1.14 |
| 7 | Snhg11 | 1.75 | 8.83E-17 | 1.89E-12 | 0.21 | 0.07 | 0.14 | 2.96 |
| 7 | Spats2l | 0.85 | 9.81E-17 | 2.10E-12 | 0.56 | 0.32 | 0.23 | 1.72 |
| 7 | Gprc5b | 1.6 | 1.48E-16 | 3.16E-12 | 0.31 | 0.14 | 0.17 | 2.24 |
| 7 | Nudt11 | 1.34 | 1.58E-16 | 3.37E-12 | 0.46 | 0.27 | 0.19 | 1.7 |
| 7 | Prdx6 | 0.45 | 1.63E-16 | 3.49E-12 | 0.96 | 0.95 | 0.01 | 1.01 |
| 7 | Fbln1 | 0.76 | 1.71E-16 | 3.65E-12 | 0.79 | 0.65 | 0.14 | 1.22 |
| 7 | Pakap | 2.02 | 1.84E-16 | 3.94E-12 | 0.19 | 0.06 | 0.13 | 3.1 |
| 7 | Ddit4 | 0.89 | 1.85E-16 | 3.95E-12 | 0.62 | 0.43 | 0.19 | 1.44 |
| 7 | Igf2 | 0.46 | 1.97E-16 | 4.21E-12 | 0.93 | 0.81 | 0.12 | 1.15 |
| 7 | Adamts18 | 1.1 | 2.18E-16 | 4.65E-12 | 0.47 | 0.26 | 0.21 | 1.83 |
| 7 | Khdrbs1 | 0.38 | 2.35E-16 | 5.02E-12 | 0.97 | 0.94 | 0.02 | 1.02 |
| 7 | Mbnl1 | 1.16 | 2.42E-16 | 5.17E-12 | 0.6 | 0.42 | 0.18 | 1.44 |
| 7 | Ccdc126 | 1.77 | 2.74E-16 | 5.84E-12 | 0.23 | 0.08 | 0.14 | 2.71 |
| 7 | Ptn | 0.58 | 3.33E-16 | 7.12E-12 | 0.92 | 0.86 | 0.06 | 1.07 |
| 7 | Gpx1 | 0.39 | 3.73E-16 | 7.97E-12 | 0.97 | 0.95 | 0.02 | 1.02 |
| 7 | Tmem123 | 0.92 | 3.82E-16 | 8.16E-12 | 0.63 | 0.44 | 0.19 | 1.43 |
| 7 | Gstm5 | 0.6 | 3.96E-16 | 8.45E-12 | 0.91 | 0.82 | 0.09 | 1.1 |
| 7 | Rtn4ip1 | 1.77 | 3.96E-16 | 8.46E-12 | 0.21 | 0.08 | 0.14 | 2.78 |
| 7 | Taf4b | 2.25 | 4.08E-16 | 8.73E-12 | 0.15 | 0.04 | 0.11 | 3.51 |
| 7 | Nudt10 | 1.33 | 4.65E-16 | 9.93E-12 | 0.34 | 0.16 | 0.18 | 2.12 |
| 7 | Ankrd29 | 2.71 | 4.95E-16 | 1.06E-11 | 0.11 | 0.02 | 0.08 | 4.57 |
| 7 | Bcam | 0.54 | 5.76E-16 | 1.23E-11 | 0.89 | 0.72 | 0.17 | 1.24 |
| 7 | Micos10 | 0.4 | 6.07E-16 | 1.30E-11 | 0.94 | 0.92 | 0.02 | 1.02 |
| 7 | Ddx3y | 2.33 | 6.69E-16 | 1.43E-11 | 0.13 | 0.04 | 0.1 | 3.8 |
| 7 | Serpinb6a | 0.86 | 7.34E-16 | 1.57E-11 | 0.64 | 0.47 | 0.17 | 1.36 |
| 7 | Castor1 | 2.06 | 1.02E-15 | 2.19E-11 | 0.16 | 0.05 | 0.11 | 3.36 |
| 7 | Gnai1 | 1.61 | 1.11E-15 | 2.37E-11 | 0.25 | 0.1 | 0.15 | 2.49 |
| 7 | Gng11 | 1.55 | 1.38E-15 | 2.96E-11 | 0.26 | 0.11 | 0.15 | 2.39 |
| 7 | Tes | 0.78 | 1.48E-15 | 3.16E-11 | 0.75 | 0.61 | 0.14 | 1.23 |
| 7 | Calml3 | 2.24 | 1.56E-15 | 3.33E-11 | 0.15 | 0.04 | 0.11 | 3.43 |
| 7 | Calm2 | 0.29 | 1.67E-15 | 3.57E-11 | 1 | 0.99 | 0.01 | 1.01 |
| 7 | Pcdh7 | 0.72 | 1.68E-15 | 3.58E-11 | 0.78 | 0.65 | 0.13 | 1.21 |
| 7 | Prkar2b | 1.17 | 2.80E-15 | 5.98E-11 | 0.48 | 0.3 | 0.18 | 1.61 |
| 7 | Col4a1 | 0.72 | 2.91E-15 | 6.23E-11 | 0.81 | 0.68 | 0.14 | 1.2 |
| 7 | Ndufaf4 | 0.9 | 3.31E-15 | 7.06E-11 | 0.7 | 0.58 | 0.12 | 1.21 |
| 7 | Edf1 | 0.36 | 5.06E-15 | 1.08E-10 | 0.98 | 0.96 | 0.01 | 1.01 |
| 7 | Fundc2 | 0.66 | 6.27E-15 | 1.34E-10 | 0.79 | 0.69 | 0.1 | 1.14 |

**Supplementary Table 3. Marker genes for the original clusters of integrated clustering of E11.5 control and NotchGOF λ-junction ectoderm cells.**

| | | | | | | | | |
|---|---|---|---|---|---|---|---|---|
| 7 | Lbh | 0.94 | 6.44E-15 | 1.38E-10 | 0.46 | 0.26 | 0.2 | 1.77 |
| 7 | Arl4a | 1.66 | 7.20E-15 | 1.54E-10 | 0.31 | 0.15 | 0.16 | 2.04 |
| 7 | Dzip1 | 1.28 | 1.09E-14 | 2.34E-10 | 0.37 | 0.2 | 0.17 | 1.86 |
| 7 | Gpx8 | 0.71 | 1.33E-14 | 2.84E-10 | 0.78 | 0.65 | 0.13 | 1.21 |
| 7 | Gga2 | 0.77 | 1.80E-14 | 3.84E-10 | 0.66 | 0.5 | 0.16 | 1.32 |
| 7 | Kdm5d | 2.15 | 1.93E-14 | 4.12E-10 | 0.11 | 0.03 | 0.08 | 4.2 |
| 7 | Nckap5 | 1.11 | 2.45E-14 | 5.22E-10 | 0.48 | 0.3 | 0.18 | 1.6 |
| 7 | Erf | 0.94 | 3.21E-14 | 6.86E-10 | 0.56 | 0.38 | 0.19 | 1.49 |
| 7 | Rapgefl1 | 1.73 | 3.55E-14 | 7.57E-10 | 0.19 | 0.07 | 0.12 | 2.82 |
| 7 | Greb1 | 1.37 | 3.56E-14 | 7.61E-10 | 0.35 | 0.18 | 0.17 | 1.91 |
| 7 | Pfn1 | 0.28 | 3.94E-14 | 8.42E-10 | 1 | 0.99 | 0.01 | 1.01 |
| 7 | Serpinh1 | 0.39 | 4.39E-14 | 9.37E-10 | 0.98 | 0.95 | 0.03 | 1.03 |
| 7 | Foxp1 | 0.47 | 4.79E-14 | 1.02E-09 | 0.92 | 0.86 | 0.06 | 1.07 |
| 7 | Rarg | 1.13 | 5.95E-14 | 1.27E-09 | 0.46 | 0.28 | 0.18 | 1.63 |
| 7 | Gm16141 | 1.98 | 6.25E-14 | 1.33E-09 | 0.16 | 0.05 | 0.11 | 3.04 |
| 7 | Cep164 | 0.64 | 7.86E-14 | 1.68E-09 | 0.79 | 0.66 | 0.13 | 1.19 |
| 7 | Sorcs2 | 1.42 | 1.46E-13 | 3.12E-09 | 0.23 | 0.09 | 0.13 | 2.47 |
| 7 | Ywhah | 0.43 | 1.68E-13 | 3.59E-09 | 0.86 | 0.8 | 0.06 | 1.08 |
| 7 | Tasor2 | 0.84 | 2.04E-13 | 4.36E-09 | 0.61 | 0.44 | 0.17 | 1.39 |
| 7 | Arhgap24 | 1.75 | 2.40E-13 | 5.13E-09 | 0.19 | 0.07 | 0.12 | 2.7 |
| 7 | Lgals1 | 0.86 | 2.54E-13 | 5.42E-09 | 0.52 | 0.35 | 0.17 | 1.47 |
| 7 | Ddr1 | 0.75 | 3.32E-13 | 7.09E-09 | 0.63 | 0.45 | 0.19 | 1.41 |
| 7 | Itm2a | 0.66 | 3.38E-13 | 7.23E-09 | 0.67 | 0.51 | 0.16 | 1.3 |
| 7 | Trp53 | 0.48 | 3.80E-13 | 8.11E-09 | 0.93 | 0.89 | 0.05 | 1.05 |
| 7 | Susd4 | 1.09 | 3.81E-13 | 8.14E-09 | 0.47 | 0.3 | 0.17 | 1.58 |
| 7 | Barx2 | 1.39 | 3.85E-13 | 8.23E-09 | 0.31 | 0.16 | 0.16 | 2 |
| 7 | Bag1 | 0.43 | 4.02E-13 | 8.58E-09 | 0.9 | 0.87 | 0.03 | 1.03 |
| 7 | Gas1 | 0.71 | 4.62E-13 | 9.87E-09 | 0.73 | 0.56 | 0.17 | 1.3 |
| 7 | Cd109 | 1.95 | 4.68E-13 | 1.00E-08 | 0.14 | 0.04 | 0.1 | 3.26 |
| 7 | Lfng | 1.97 | 5.26E-13 | 1.12E-08 | 0.14 | 0.04 | 0.1 | 3.26 |
| 7 | Prickle1 | 1.76 | 7.98E-13 | 1.71E-08 | 0.17 | 0.06 | 0.11 | 2.85 |
| 7 | D16Ertd472e | 1.06 | 1.09E-12 | 2.32E-08 | 0.4 | 0.23 | 0.16 | 1.71 |
| 7 | Eml4 | 0.57 | 1.10E-12 | 2.36E-08 | 0.8 | 0.71 | 0.09 | 1.13 |
| 7 | Nt5c | 0.63 | 1.24E-12 | 2.65E-08 | 0.77 | 0.66 | 0.11 | 1.16 |
| 7 | Mlec | 0.6 | 1.32E-12 | 2.82E-08 | 0.86 | 0.81 | 0.05 | 1.07 |
| 7 | Otx1 | 1 | 1.42E-12 | 3.04E-08 | 0.57 | 0.43 | 0.14 | 1.32 |
| 7 | Avpi1 | 0.96 | 1.89E-12 | 4.03E-08 | 0.51 | 0.35 | 0.16 | 1.45 |
| 7 | H2afy2 | 0.35 | 1.92E-12 | 4.10E-08 | 0.97 | 0.95 | 0.03 | 1.03 |
| 7 | Smarcad1 | 0.52 | 2.19E-12 | 4.67E-08 | 0.83 | 0.77 | 0.06 | 1.08 |
| 7 | Sema4g | 1.39 | 2.43E-12 | 5.18E-08 | 0.28 | 0.14 | 0.14 | 2.05 |
| 7 | Abracl | 0.33 | 2.64E-12 | 5.64E-08 | 0.93 | 0.93 | 0 | 1 |
| 7 | Malat1 | 0.35 | 2.65E-12 | 5.66E-08 | 0.99 | 1 | 0 | 1 |
| 7 | Itm2c | 0.68 | 2.74E-12 | 5.84E-08 | 0.71 | 0.59 | 0.12 | 1.2 |
| 7 | Ier5 | 0.51 | 2.81E-12 | 6.00E-08 | 0.92 | 0.84 | 0.09 | 1.1 |
| 7 | Pde10a | 1.81 | 3.31E-12 | 7.07E-08 | 0.13 | 0.04 | 0.09 | 3.25 |
| 7 | Laptm4a | 0.34 | 3.50E-12 | 7.47E-08 | 0.98 | 0.96 | 0.02 | 1.03 |
| 7 | Cers6 | 1.27 | 4.10E-12 | 8.76E-08 | 0.38 | 0.23 | 0.16 | 1.68 |
| 7 | Helb | 1.32 | 4.17E-12 | 8.91E-08 | 0.27 | 0.13 | 0.14 | 2.09 |
| 7 | Cisd1 | 0.47 | 5.38E-12 | 1.15E-07 | 0.86 | 0.78 | 0.08 | 1.1 |
| 7 | Arf4 | 0.36 | 6.67E-12 | 1.43E-07 | 0.98 | 0.94 | 0.03 | 1.04 |
| 7 | Tiparp | 1.2 | 7.30E-12 | 1.56E-07 | 0.38 | 0.22 | 0.15 | 1.7 |
| 7 | Commd3 | 0.43 | 7.53E-12 | 1.61E-07 | 0.87 | 0.82 | 0.05 | 1.06 |
| 7 | Smarcc1 | 0.34 | 7.60E-12 | 1.62E-07 | 0.98 | 0.97 | 0.01 | 1.01 |
| 7 | Sdc1 | 0.57 | 7.84E-12 | 1.68E-07 | 0.73 | 0.58 | 0.14 | 1.25 |
| 7 | Arhgap42 | 0.87 | 7.86E-12 | 1.68E-07 | 0.44 | 0.27 | 0.17 | 1.65 |
| 7 | Smad7 | 0.59 | 9.05E-12 | 1.93E-07 | 0.76 | 0.6 | 0.16 | 1.27 |
| 7 | Dlx1 | 1.02 | 9.28E-12 | 1.98E-07 | 0.29 | 0.14 | 0.15 | 2.07 |

**Supplementary Table 3. Marker genes for the original clusters of integrated clustering of E11.5 control and NotchGOF λ-junction ectoderm cells.**

| | | | | | | | | |
|---|---|---|---|---|---|---|---|---|
| 7 | Crppa | 1.6 | 1.09E-11 | 2.34E-07 | 0.17 | 0.06 | 0.11 | 2.67 |
| 7 | Mast4 | 0.78 | 1.31E-11 | 2.80E-07 | 0.63 | 0.49 | 0.14 | 1.28 |
| 7 | Rab38 | 0.94 | 1.36E-11 | 2.91E-07 | 0.35 | 0.19 | 0.16 | 1.82 |
| 7 | Mmp2 | 1.49 | 1.47E-11 | 3.14E-07 | 0.24 | 0.11 | 0.13 | 2.14 |
| 7 | Flna | 0.62 | 1.50E-11 | 3.20E-07 | 0.77 | 0.69 | 0.09 | 1.12 |
| 7 | Pxn | 1.38 | 1.60E-11 | 3.43E-07 | 0.31 | 0.17 | 0.14 | 1.85 |
| 7 | Bri3 | 0.32 | 1.78E-11 | 3.81E-07 | 0.98 | 0.97 | 0.01 | 1.01 |
| 7 | mt-Co1 | 0.26 | 2.05E-11 | 4.38E-07 | 1 | 0.99 | 0.01 | 1.01 |
| 7 | Eef2k | 0.97 | 2.84E-11 | 6.07E-07 | 0.32 | 0.17 | 0.15 | 1.9 |
| 7 | Cachd1 | 0.88 | 3.84E-11 | 8.20E-07 | 0.5 | 0.33 | 0.17 | 1.51 |
| 7 | Hmga2 | 0.43 | 4.31E-11 | 9.21E-07 | 0.97 | 0.92 | 0.04 | 1.05 |
| 7 | Ogfrl1 | 0.99 | 4.68E-11 | 1.00E-06 | 0.32 | 0.18 | 0.15 | 1.85 |
| 7 | Lpp | 0.77 | 4.81E-11 | 1.03E-06 | 0.6 | 0.45 | 0.15 | 1.34 |
| 7 | Smad5 | 0.77 | 7.54E-11 | 1.61E-06 | 0.6 | 0.47 | 0.13 | 1.28 |
| 7 | Scmh1 | 0.94 | 8.01E-11 | 1.71E-06 | 0.46 | 0.31 | 0.15 | 1.49 |
| 7 | Oxct1 | 0.62 | 8.27E-11 | 1.77E-06 | 0.82 | 0.77 | 0.05 | 1.06 |
| 7 | Ankrd45 | 1.95 | 1.02E-10 | 2.18E-06 | 0.12 | 0.04 | 0.08 | 3 |
| 7 | Mfge8 | 0.54 | 1.17E-10 | 2.51E-06 | 0.7 | 0.55 | 0.15 | 1.27 |
| 7 | Pkp3 | 0.94 | 1.19E-10 | 2.55E-06 | 0.4 | 0.24 | 0.15 | 1.63 |
| 7 | Gpc2 | 0.82 | 1.25E-10 | 2.67E-06 | 0.52 | 0.36 | 0.16 | 1.45 |
| 7 | Cntnap2 | 1.54 | 1.52E-10 | 3.24E-06 | 0.27 | 0.14 | 0.13 | 1.88 |
| 7 | Scoc | 0.71 | 1.61E-10 | 3.44E-06 | 0.66 | 0.55 | 0.1 | 1.18 |
| 7 | Itpripl2 | 1.83 | 1.80E-10 | 3.85E-06 | 0.13 | 0.04 | 0.08 | 2.93 |
| 7 | Col4a5 | 1.41 | 2.13E-10 | 4.55E-06 | 0.32 | 0.18 | 0.13 | 1.74 |
| 7 | Slc16a13 | 1.2 | 2.16E-10 | 4.61E-06 | 0.25 | 0.12 | 0.13 | 2.08 |
| 7 | Mylip | 0.97 | 2.26E-10 | 4.83E-06 | 0.34 | 0.2 | 0.15 | 1.74 |
| 7 | Frmd4b | 0.72 | 2.35E-10 | 5.01E-06 | 0.64 | 0.49 | 0.15 | 1.3 |
| 7 | Itga3 | 1.14 | 2.50E-10 | 5.35E-06 | 0.39 | 0.25 | 0.14 | 1.56 |
| 7 | 6330403K07Rik | 0.43 | 2.68E-10 | 5.72E-06 | 0.9 | 0.85 | 0.05 | 1.06 |
| 7 | Dancr | 0.72 | 2.92E-10 | 6.25E-06 | 0.68 | 0.55 | 0.14 | 1.25 |
| 7 | Mbd2 | 0.45 | 3.06E-10 | 6.55E-06 | 0.72 | 0.61 | 0.11 | 1.18 |
| 7 | Vwa1 | 1.22 | 3.29E-10 | 7.04E-06 | 0.23 | 0.11 | 0.12 | 2.13 |
| 7 | Tanc2 | 1 | 3.37E-10 | 7.21E-06 | 0.37 | 0.22 | 0.15 | 1.67 |
| 7 | Txnrd1 | 0.37 | 3.41E-10 | 7.28E-06 | 0.92 | 0.87 | 0.05 | 1.06 |
| 7 | Osbpl3 | 1.36 | 3.74E-10 | 7.98E-06 | 0.28 | 0.15 | 0.13 | 1.86 |
| 7 | Ugdh | 0.81 | 3.86E-10 | 8.24E-06 | 0.55 | 0.43 | 0.12 | 1.27 |
| 7 | Efnb2 | 0.63 | 4.13E-10 | 8.82E-06 | 0.72 | 0.59 | 0.13 | 1.22 |
| 7 | Pik3r3 | 1.15 | 4.56E-10 | 9.73E-06 | 0.33 | 0.2 | 0.13 | 1.68 |
| 7 | Fubp1 | 0.28 | 4.66E-10 | 9.95E-06 | 0.99 | 0.98 | 0.01 | 1.01 |
| 7 | Yap1 | 0.59 | 5.12E-10 | 1.09E-05 | 0.72 | 0.61 | 0.11 | 1.18 |
| 7 | Dapl1 | 0.74 | 6.04E-10 | 1.29E-05 | 0.44 | 0.26 | 0.17 | 1.66 |
| 7 | Adamts17 | 1.35 | 6.65E-10 | 1.42E-05 | 0.23 | 0.11 | 0.12 | 2.05 |
| 7 | Tm2d2 | 0.53 | 7.38E-10 | 1.58E-05 | 0.76 | 0.68 | 0.08 | 1.12 |
| 7 | 1110008P14Rik | 0.8 | 9.05E-10 | 1.93E-05 | 0.5 | 0.36 | 0.14 | 1.39 |
| 7 | Lars2 | 0.65 | 9.40E-10 | 2.01E-05 | 0.68 | 0.52 | 0.16 | 1.31 |
| 7 | Eef1e1 | 0.52 | 9.42E-10 | 2.01E-05 | 0.74 | 0.68 | 0.06 | 1.1 |
| 7 | Ccdc102a | 1.01 | 9.85E-10 | 2.10E-05 | 0.37 | 0.23 | 0.14 | 1.61 |
| 7 | Rac3 | 0.84 | 1.04E-09 | 2.22E-05 | 0.54 | 0.41 | 0.14 | 1.34 |
| 7 | Col5a1 | 1.17 | 1.08E-09 | 2.31E-05 | 0.31 | 0.18 | 0.13 | 1.74 |
| 7 | Rbbp4 | 0.29 | 1.09E-09 | 2.34E-05 | 0.98 | 0.94 | 0.03 | 1.04 |
| 7 | Adamts19 | 1.58 | 1.23E-09 | 2.62E-05 | 0.19 | 0.08 | 0.1 | 2.24 |
| 7 | Lin28a | 1.95 | 1.25E-09 | 2.68E-05 | 0.11 | 0.03 | 0.07 | 3.09 |
| 7 | Cacng4 | 0.69 | 1.28E-09 | 2.74E-05 | 0.5 | 0.34 | 0.16 | 1.47 |
| 7 | Ppic | 0.66 | 1.31E-09 | 2.80E-05 | 0.7 | 0.59 | 0.11 | 1.19 |
| 7 | Lpin2 | 0.71 | 1.69E-09 | 3.60E-05 | 0.58 | 0.46 | 0.13 | 1.27 |
| 7 | Pikfyve | 1.08 | 1.90E-09 | 4.06E-05 | 0.37 | 0.24 | 0.13 | 1.55 |
| 7 | B230219D22Rik | 0.33 | 2.26E-09 | 4.83E-05 | 0.92 | 0.86 | 0.06 | 1.06 |

**Supplementary Table 3. Marker genes for the original clusters of integrated clustering of E11.5 control and NotchGOF λ-junction ectoderm cells.**

| | | | | | | | | |
|---|---|---|---|---|---|---|---|---|
| 7 | Mtf2 | 0.42 | 2.40E-09 | 5.12E-05 | 0.84 | 0.78 | 0.07 | 1.09 |
| 7 | Ipmk | 1.03 | 3.04E-09 | 6.50E-05 | 0.37 | 0.23 | 0.13 | 1.56 |
| 7 | Ldb1 | 0.46 | 3.08E-09 | 6.57E-05 | 0.74 | 0.64 | 0.11 | 1.17 |
| 7 | Cpxm1 | 0.93 | 3.12E-09 | 6.66E-05 | 0.4 | 0.26 | 0.14 | 1.53 |
| 7 | Rbis | 0.35 | 3.12E-09 | 6.66E-05 | 0.89 | 0.82 | 0.07 | 1.08 |
| 7 | Sytl1 | 1.79 | 3.35E-09 | 7.15E-05 | 0.11 | 0.04 | 0.07 | 2.95 |
| 7 | Slc2a1 | 0.59 | 3.39E-09 | 7.23E-05 | 0.79 | 0.69 | 0.11 | 1.16 |
| 7 | Ddhd2 | 0.81 | 3.43E-09 | 7.33E-05 | 0.41 | 0.27 | 0.14 | 1.53 |
| 7 | Pnrc1 | 0.31 | 3.46E-09 | 7.38E-05 | 0.96 | 0.9 | 0.05 | 1.06 |
| 7 | Comtd1 | 1.2 | 3.65E-09 | 7.80E-05 | 0.28 | 0.16 | 0.13 | 1.79 |
| 7 | Fign | 0.98 | 3.80E-09 | 8.13E-05 | 0.32 | 0.18 | 0.14 | 1.74 |
| 7 | Jarid2 | 0.7 | 3.85E-09 | 8.22E-05 | 0.64 | 0.55 | 0.09 | 1.17 |
| 7 | Zbtb34 | 1.42 | 4.18E-09 | 8.94E-05 | 0.21 | 0.1 | 0.11 | 2.06 |
| 7 | Knop1 | 0.26 | 4.37E-09 | 9.34E-05 | 0.96 | 0.95 | 0.01 | 1.01 |
| 7 | Tmem132a | 0.59 | 4.66E-09 | 9.95E-05 | 0.75 | 0.66 | 0.09 | 1.13 |
| 7 | Slc26a2 | 1.19 | 4.78E-09 | 1.02E-04 | 0.29 | 0.17 | 0.12 | 1.75 |
| 7 | Klhl29 | 1.77 | 4.87E-09 | 1.04E-04 | 0.11 | 0.04 | 0.07 | 2.87 |
| 7 | Csde1 | 0.31 | 5.31E-09 | 1.13E-04 | 0.95 | 0.93 | 0.02 | 1.02 |
| 7 | Purb | 0.39 | 6.64E-09 | 1.42E-04 | 0.91 | 0.89 | 0.02 | 1.03 |
| 7 | Cmtm3 | 0.76 | 7.14E-09 | 1.53E-04 | 0.5 | 0.36 | 0.14 | 1.39 |
| 7 | Jak1 | 0.55 | 8.44E-09 | 1.80E-04 | 0.66 | 0.55 | 0.11 | 1.19 |
| 7 | Cst3 | 0.32 | 8.90E-09 | 1.90E-04 | 0.95 | 0.93 | 0.02 | 1.02 |
| 7 | Ptpn14 | 0.93 | 9.11E-09 | 1.95E-04 | 0.36 | 0.22 | 0.14 | 1.61 |
| 7 | Trim32 | 0.88 | 9.60E-09 | 2.05E-04 | 0.5 | 0.37 | 0.13 | 1.34 |
| 7 | Lrrc4 | 1.86 | 1.07E-08 | 2.29E-04 | 0.12 | 0.05 | 0.07 | 2.64 |
| 7 | Slc7a10 | 1.53 | 1.12E-08 | 2.38E-04 | 0.12 | 0.04 | 0.07 | 2.76 |
| 7 | Tmem167 | 0.31 | 1.14E-08 | 2.43E-04 | 0.94 | 0.94 | 0 | 1 |
| 7 | Myo1b | 0.44 | 1.14E-08 | 2.43E-04 | 0.74 | 0.62 | 0.12 | 1.2 |
| 7 | Gpc3 | 0.33 | 1.16E-08 | 2.49E-04 | 0.93 | 0.84 | 0.09 | 1.11 |
| 7 | Sord | 0.91 | 1.20E-08 | 2.56E-04 | 0.36 | 0.23 | 0.13 | 1.57 |
| 7 | Ctr9 | 0.5 | 1.30E-08 | 2.77E-04 | 0.74 | 0.68 | 0.06 | 1.1 |
| 7 | Tdrp | 1.38 | 1.41E-08 | 3.01E-04 | 0.19 | 0.09 | 0.1 | 2.08 |
| 7 | Ahcyl2 | 0.66 | 1.41E-08 | 3.01E-04 | 0.5 | 0.37 | 0.13 | 1.36 |
| 7 | Stxbp4 | 0.82 | 1.43E-08 | 3.05E-04 | 0.46 | 0.33 | 0.13 | 1.4 |
| 7 | Kank1 | 1.14 | 1.45E-08 | 3.10E-04 | 0.41 | 0.29 | 0.12 | 1.43 |
| 7 | Polr3g | 0.76 | 1.54E-08 | 3.29E-04 | 0.46 | 0.32 | 0.13 | 1.41 |
| 7 | Spats2 | 0.54 | 1.68E-08 | 3.59E-04 | 0.71 | 0.62 | 0.09 | 1.14 |
| 7 | Acyp1 | 0.63 | 1.89E-08 | 4.04E-04 | 0.63 | 0.49 | 0.13 | 1.27 |
| 7 | Cdh6 | 1.19 | 2.36E-08 | 5.03E-04 | 0.24 | 0.12 | 0.11 | 1.93 |
| 7 | Ccnd3 | 0.42 | 2.41E-08 | 5.14E-04 | 0.88 | 0.8 | 0.08 | 1.1 |
| 7 | Net1 | 0.64 | 2.42E-08 | 5.18E-04 | 0.62 | 0.5 | 0.12 | 1.24 |
| 7 | Ctsb | 0.48 | 2.43E-08 | 5.19E-04 | 0.73 | 0.64 | 0.08 | 1.13 |
| 7 | Chp1 | 0.72 | 2.47E-08 | 5.28E-04 | 0.47 | 0.35 | 0.13 | 1.37 |
| 7 | Ndufaf2 | 0.48 | 2.53E-08 | 5.41E-04 | 0.73 | 0.66 | 0.07 | 1.1 |
| 7 | Rab12 | 0.67 | 2.55E-08 | 5.44E-04 | 0.58 | 0.46 | 0.12 | 1.27 |
| 7 | Apoe | 0.64 | 2.55E-08 | 5.45E-04 | 0.61 | 0.49 | 0.12 | 1.25 |
| 7 | Tulp4 | 0.42 | 2.87E-08 | 6.12E-04 | 0.86 | 0.74 | 0.11 | 1.15 |
| 7 | Slc25a33 | 0.85 | 2.98E-08 | 6.38E-04 | 0.33 | 0.2 | 0.13 | 1.63 |
| 7 | Trp53i11 | 0.39 | 3.11E-08 | 6.65E-04 | 0.55 | 0.42 | 0.14 | 1.33 |
| 7 | St6gal1 | 1.03 | 3.50E-08 | 7.47E-04 | 0.3 | 0.18 | 0.12 | 1.67 |
| 7 | Sh3bp5 | 0.62 | 3.75E-08 | 8.01E-04 | 0.51 | 0.38 | 0.13 | 1.35 |
| 7 | Zfp57 | 1.16 | 4.30E-08 | 9.19E-04 | 0.28 | 0.17 | 0.11 | 1.68 |
| 7 | Cnst | 1.17 | 4.36E-08 | 9.31E-04 | 0.25 | 0.14 | 0.11 | 1.78 |
| 7 | Tgif1 | 0.44 | 4.85E-08 | 1.04E-03 | 0.77 | 0.68 | 0.09 | 1.13 |
| 7 | Tecr | 0.26 | 4.90E-08 | 1.05E-03 | 0.99 | 0.98 | 0.01 | 1.01 |
| 7 | Selenos | 0.54 | 5.34E-08 | 1.14E-03 | 0.69 | 0.58 | 0.1 | 1.18 |
| 7 | Card10 | 1.78 | 5.72E-08 | 1.22E-03 | 0.12 | 0.05 | 0.07 | 2.58 |

**Supplementary Table 3. Marker genes for the original clusters of integrated clustering of E11.5 control and NotchGOF λ-junction ectoderm cells.**

| | | | | | | | | |
|---|---|---|---|---|---|---|---|---|
| 7 | Jakmip2 | 1.05 | 5.91E-08 | 1.26E-03 | 0.14 | 0.06 | 0.08 | 2.45 |
| 7 | Tpd52l1 | 1.03 | 6.33E-08 | 1.35E-03 | 0.26 | 0.15 | 0.11 | 1.77 |
| 7 | Stard8 | 1.9 | 6.53E-08 | 1.39E-03 | 0.11 | 0.04 | 0.07 | 2.69 |
| 7 | Ccdc50 | 0.39 | 6.55E-08 | 1.40E-03 | 0.82 | 0.76 | 0.06 | 1.08 |
| 7 | Cep85l | 1.79 | 6.71E-08 | 1.43E-03 | 0.12 | 0.05 | 0.07 | 2.46 |
| 7 | Osbpl1a | 0.79 | 6.78E-08 | 1.45E-03 | 0.37 | 0.24 | 0.13 | 1.55 |
| 7 | Zfp90 | 0.73 | 6.79E-08 | 1.45E-03 | 0.45 | 0.32 | 0.13 | 1.4 |
| 7 | Mat2b | 0.52 | 7.69E-08 | 1.64E-03 | 0.69 | 0.59 | 0.1 | 1.18 |
| 7 | Tinagl1 | 1.19 | 8.10E-08 | 1.73E-03 | 0.18 | 0.09 | 0.09 | 2.07 |
| 7 | Etaa1 | 0.69 | 8.87E-08 | 1.89E-03 | 0.63 | 0.55 | 0.07 | 1.13 |
| 7 | Txn2 | 0.31 | 8.96E-08 | 1.91E-03 | 0.9 | 0.84 | 0.06 | 1.07 |
| 7 | Plin3 | 0.53 | 9.34E-08 | 2.00E-03 | 0.65 | 0.52 | 0.13 | 1.24 |
| 7 | Palmd | 1.24 | 9.36E-08 | 2.00E-03 | 0.13 | 0.05 | 0.08 | 2.45 |
| 7 | Cited2 | 0.43 | 1.10E-07 | 2.35E-03 | 0.8 | 0.68 | 0.12 | 1.18 |
| 7 | Asap1 | 0.51 | 1.14E-07 | 2.43E-03 | 0.72 | 0.64 | 0.08 | 1.13 |
| 7 | Col4a6 | 1.72 | 1.34E-07 | 2.87E-03 | 0.13 | 0.05 | 0.08 | 2.41 |
| 7 | Nme4 | 0.37 | 1.38E-07 | 2.96E-03 | 0.79 | 0.73 | 0.06 | 1.08 |
| 7 | Ppdpf | 0.26 | 1.40E-07 | 3.00E-03 | 0.97 | 0.97 | 0 | 1 |
| 7 | Aass | 1.51 | 1.44E-07 | 3.08E-03 | 0.15 | 0.07 | 0.08 | 2.19 |
| 7 | Baz1a | 0.31 | 1.44E-07 | 3.08E-03 | 0.66 | 0.58 | 0.08 | 1.13 |
| 7 | Tspan6 | 0.41 | 1.47E-07 | 3.15E-03 | 0.81 | 0.75 | 0.07 | 1.09 |
| 7 | Mark4 | 0.88 | 1.54E-07 | 3.30E-03 | 0.32 | 0.2 | 0.12 | 1.58 |
| 7 | Gimap9 | 0.99 | 1.64E-07 | 3.50E-03 | 0.24 | 0.13 | 0.11 | 1.84 |
| 7 | Sdhaf4 | 0.44 | 1.77E-07 | 3.78E-03 | 0.74 | 0.67 | 0.07 | 1.11 |
| 7 | Caprin1 | 0.26 | 1.82E-07 | 3.89E-03 | 0.97 | 0.96 | 0.01 | 1.01 |
| 7 | Ndufa1 | 0.37 | 1.95E-07 | 4.16E-03 | 0.87 | 0.83 | 0.04 | 1.05 |
| 7 | Lsm14b | 0.59 | 2.01E-07 | 4.29E-03 | 0.54 | 0.42 | 0.12 | 1.27 |
| 7 | Hddc2 | 0.46 | 2.08E-07 | 4.44E-03 | 0.74 | 0.7 | 0.04 | 1.06 |
| 7 | Dlg5 | 0.83 | 2.12E-07 | 4.52E-03 | 0.49 | 0.39 | 0.1 | 1.27 |
| 7 | Bmi1 | 0.62 | 2.15E-07 | 4.59E-03 | 0.51 | 0.39 | 0.12 | 1.3 |
| 7 | Snap47 | 0.39 | 2.16E-07 | 4.61E-03 | 0.76 | 0.68 | 0.08 | 1.12 |
| 7 | Pak2 | 0.27 | 2.27E-07 | 4.85E-03 | 0.93 | 0.89 | 0.04 | 1.05 |
| 7 | Znrf1 | 0.56 | 2.31E-07 | 4.93E-03 | 0.64 | 0.56 | 0.07 | 1.13 |
| 7 | Hadha | 0.42 | 2.51E-07 | 5.36E-03 | 0.82 | 0.74 | 0.08 | 1.1 |
| 7 | Shf | 0.88 | 2.53E-07 | 5.40E-03 | 0.32 | 0.2 | 0.12 | 1.6 |
| 7 | Fam83b | 1.37 | 3.10E-07 | 6.62E-03 | 0.14 | 0.07 | 0.08 | 2.22 |
| 7 | Atf7ip | 0.47 | 3.22E-07 | 6.88E-03 | 0.72 | 0.64 | 0.08 | 1.13 |
| 7 | Psph | 0.88 | 3.32E-07 | 7.10E-03 | 0.42 | 0.31 | 0.11 | 1.36 |
| 7 | Tnrc6c | 0.36 | 3.58E-07 | 7.65E-03 | 0.83 | 0.75 | 0.08 | 1.1 |
| 7 | Ttc23 | 0.77 | 3.63E-07 | 7.76E-03 | 0.41 | 0.29 | 0.13 | 1.44 |
| 7 | Negr1 | 1.64 | 3.70E-07 | 7.91E-03 | 0.1 | 0.04 | 0.06 | 2.62 |
| 7 | Dlx3 | 0.72 | 3.77E-07 | 8.06E-03 | 0.41 | 0.28 | 0.13 | 1.46 |
| 7 | Tro | 0.77 | 3.79E-07 | 8.09E-03 | 0.37 | 0.25 | 0.12 | 1.49 |
| 7 | Ilrun | 0.55 | 4.33E-07 | 9.24E-03 | 0.58 | 0.47 | 0.11 | 1.24 |
| 7 | Arl2bp | 0.48 | 4.52E-07 | 9.67E-03 | 0.79 | 0.7 | 0.1 | 1.14 |
| 7 | Fmr1 | 0.51 | 4.54E-07 | 9.71E-03 | 0.68 | 0.59 | 0.09 | 1.15 |
| 7 | Igf1 | 1.82 | 4.74E-07 | 1.01E-02 | 0.12 | 0.05 | 0.07 | 2.42 |
| 7 | Npc2 | 0.3 | 4.80E-07 | 1.02E-02 | 0.94 | 0.92 | 0.02 | 1.02 |
| 7 | Unc13c | 1.35 | 4.82E-07 | 1.03E-02 | 0.14 | 0.07 | 0.08 | 2.18 |
| 7 | Zfp521 | 0.89 | 4.94E-07 | 1.06E-02 | 0.29 | 0.18 | 0.11 | 1.59 |
| 7 | Ccdc88a | 0.38 | 5.04E-07 | 1.08E-02 | 0.77 | 0.7 | 0.07 | 1.1 |
| 7 | Slc39a6 | 0.56 | 5.45E-07 | 1.16E-02 | 0.58 | 0.48 | 0.1 | 1.21 |
| 7 | Ppp2r2a | 0.34 | 5.50E-07 | 1.17E-02 | 0.84 | 0.78 | 0.06 | 1.08 |
| 7 | Glud1 | 0.56 | 5.51E-07 | 1.18E-02 | 0.65 | 0.56 | 0.09 | 1.16 |
| 7 | Reep5 | 0.49 | 5.74E-07 | 1.23E-02 | 0.65 | 0.56 | 0.1 | 1.18 |
| 7 | Nox4 | 1.36 | 5.89E-07 | 1.26E-02 | 0.17 | 0.09 | 0.09 | 2.02 |
| 7 | Galk1 | 0.47 | 6.17E-07 | 1.32E-02 | 0.76 | 0.72 | 0.04 | 1.06 |

**Supplementary Table 3. Marker genes for the original clusters of integrated clustering of E11.5 control and NotchGOF λ-junction ectoderm cells.**

| | | | | | | | | |
|---|---|---|---|---|---|---|---|---|
| 7 | Capns1 | 0.34 | 6.35E-07 | 1.36E-02 | 0.8 | 0.75 | 0.05 | 1.07 |
| 7 | Mllt10 | 0.36 | 6.54E-07 | 1.40E-02 | 0.78 | 0.72 | 0.06 | 1.09 |
| 7 | Fryl | 0.7 | 6.55E-07 | 1.40E-02 | 0.5 | 0.38 | 0.11 | 1.29 |
| 7 | Large2 | 1 | 6.80E-07 | 1.45E-02 | 0.34 | 0.23 | 0.11 | 1.48 |
| 7 | Hdac2 | 0.34 | 6.87E-07 | 1.47E-02 | 0.89 | 0.89 | 0.01 | 1.01 |
| 7 | Bc1 | 1.2 | 7.61E-07 | 1.62E-02 | 0.25 | 0.15 | 0.1 | 1.68 |
| 7 | Gosr2 | 0.76 | 8.03E-07 | 1.71E-02 | 0.44 | 0.33 | 0.11 | 1.33 |
| 7 | Ddx6 | 0.3 | 8.26E-07 | 1.76E-02 | 0.88 | 0.86 | 0.02 | 1.02 |
| 7 | Bdnf | 0.93 | 8.29E-07 | 1.77E-02 | 0.29 | 0.18 | 0.11 | 1.61 |
| 7 | Dynlrb1 | 0.27 | 8.54E-07 | 1.82E-02 | 0.94 | 0.94 | 0 | 1 |
| 7 | Dennd2c | 0.98 | 8.77E-07 | 1.87E-02 | 0.19 | 0.1 | 0.09 | 1.96 |
| 7 | A330076H08Rik | 1.86 | 8.98E-07 | 1.92E-02 | 0.11 | 0.04 | 0.06 | 2.5 |
| 7 | Mipep | 0.92 | 9.65E-07 | 2.06E-02 | 0.35 | 0.24 | 0.12 | 1.49 |
| 7 | Flywch2 | 0.52 | 1.02E-06 | 2.17E-02 | 0.6 | 0.48 | 0.12 | 1.24 |
| 7 | Fzd3 | 0.29 | 1.03E-06 | 2.20E-02 | 0.89 | 0.84 | 0.04 | 1.05 |
| 7 | Sh2b2 | 0.89 | 1.03E-06 | 2.21E-02 | 0.24 | 0.14 | 0.11 | 1.77 |
| 7 | Slc4a2 | 0.76 | 1.03E-06 | 2.21E-02 | 0.39 | 0.27 | 0.12 | 1.44 |
| 7 | Pakap.1 | 0.82 | 1.16E-06 | 2.47E-02 | 0.37 | 0.26 | 0.11 | 1.44 |
| 7 | Ap4s1 | 0.6 | 1.27E-06 | 2.71E-02 | 0.55 | 0.45 | 0.1 | 1.22 |
| 7 | Smarca2 | 0.52 | 1.29E-06 | 2.76E-02 | 0.68 | 0.58 | 0.1 | 1.18 |
| 7 | Mapre2 | 0.62 | 1.30E-06 | 2.79E-02 | 0.35 | 0.23 | 0.12 | 1.5 |
| 7 | Ifitm3 | 0.45 | 1.30E-06 | 2.79E-02 | 0.57 | 0.42 | 0.15 | 1.37 |
| 7 | Smo | 0.4 | 1.34E-06 | 2.87E-02 | 0.77 | 0.69 | 0.08 | 1.12 |
| 7 | Zfp677 | 1.35 | 1.36E-06 | 2.90E-02 | 0.11 | 0.05 | 0.07 | 2.38 |
| 7 | Slc7a1 | 1.59 | 1.36E-06 | 2.91E-02 | 0.18 | 0.1 | 0.08 | 1.82 |
| 7 | Apbb2 | 0.91 | 1.40E-06 | 2.98E-02 | 0.38 | 0.28 | 0.1 | 1.36 |
| 7 | Usp14 | 0.42 | 1.42E-06 | 3.04E-02 | 0.76 | 0.72 | 0.03 | 1.05 |
| 7 | Rell1 | 0.95 | 1.42E-06 | 3.04E-02 | 0.3 | 0.2 | 0.1 | 1.51 |
| 7 | Tln2 | 0.6 | 1.44E-06 | 3.07E-02 | 0.58 | 0.48 | 0.1 | 1.21 |
| 7 | Gm1976 | 1.31 | 1.59E-06 | 3.39E-02 | 0.19 | 0.1 | 0.09 | 1.86 |
| 7 | Dcp2 | 0.48 | 1.63E-06 | 3.48E-02 | 0.61 | 0.51 | 0.1 | 1.19 |
| 7 | Atp2c2 | 1.33 | 1.72E-06 | 3.66E-02 | 0.12 | 0.05 | 0.07 | 2.29 |
| 7 | Clns1a | 0.31 | 2.07E-06 | 4.42E-02 | 0.88 | 0.85 | 0.02 | 1.03 |
| 7 | B130024G19Rik | 1.75 | 2.14E-06 | 4.57E-02 | 0.1 | 0.04 | 0.06 | 2.43 |
| 7 | Top2b | 0.28 | 2.15E-06 | 4.60E-02 | 0.94 | 0.93 | 0.01 | 1.02 |
| 7 | Ssbp2 | 0.45 | 2.19E-06 | 4.68E-02 | 0.75 | 0.69 | 0.06 | 1.08 |
| 7 | Ctdspl | 0.56 | 2.21E-06 | 4.72E-02 | 0.58 | 0.49 | 0.1 | 1.2 |
| 7 | Arhgap10 | 0.62 | 2.34E-06 | 4.99E-02 | 0.52 | 0.43 | 0.1 | 1.23 |
| 8 | Foxe1 | 4.3 | 0 | 0 | 0.96 | 0.12 | 0.84 | 7.89 |
| 8 | Gm12446 | 4.61 | 0 | 0 | 0.86 | 0.06 | 0.8 | 14.05 |
| 8 | Barx1 | 4.78 | 3.80E-276 | 8.11E-272 | 0.74 | 0.06 | 0.67 | 11.87 |
| 8 | Cgnl1 | 3.88 | 3.01E-208 | 6.42E-204 | 0.8 | 0.13 | 0.68 | 6.38 |
| 8 | Krt19 | 3.24 | 3.94E-151 | 8.42E-147 | 0.88 | 0.25 | 0.63 | 3.57 |
| 8 | Cryab | 3.77 | 1.57E-146 | 3.34E-142 | 0.71 | 0.14 | 0.57 | 4.95 |
| 8 | Foxq1 | 4.61 | 8.12E-146 | 1.73E-141 | 0.31 | 0.01 | 0.3 | 22.07 |
| 8 | S100a6 | 3.19 | 8.87E-137 | 1.89E-132 | 0.79 | 0.2 | 0.59 | 3.9 |
| 8 | Csrp2 | 2.48 | 6.15E-133 | 1.31E-128 | 1 | 0.83 | 0.17 | 1.21 |
| 8 | Foxa1 | 4.68 | 2.73E-126 | 5.83E-122 | 0.29 | 0.02 | 0.27 | 19.13 |
| 8 | H19 | 2.13 | 2.30E-125 | 4.92E-121 | 1 | 0.87 | 0.13 | 1.15 |
| 8 | Cdo1 | 2.55 | 3.25E-122 | 6.95E-118 | 0.94 | 0.44 | 0.5 | 2.14 |
| 8 | Tbx1 | 3.89 | 6.18E-121 | 1.32E-116 | 0.35 | 0.03 | 0.32 | 12.39 |
| 8 | Upk2 | 3.85 | 1.74E-120 | 3.72E-116 | 0.4 | 0.04 | 0.36 | 10 |
| 8 | Amot | 2.42 | 1.73E-118 | 3.70E-114 | 0.96 | 0.55 | 0.41 | 1.74 |
| 8 | Barx2 | 2.51 | 3.42E-113 | 7.31E-109 | 0.66 | 0.13 | 0.53 | 4.98 |
| 8 | Upk3bl | 2.6 | 6.80E-113 | 1.45E-108 | 0.83 | 0.27 | 0.56 | 3.11 |
| 8 | Krt8 | 1.58 | 4.96E-110 | 1.06E-105 | 1 | 0.81 | 0.19 | 1.24 |
| 8 | Dmrt2 | 2.9 | 7.77E-109 | 1.66E-104 | 0.49 | 0.07 | 0.42 | 6.86 |

**Supplementary Table 3. Marker genes for the original clusters of integrated clustering of E11.5 control and NotchGOF λ-junction ectoderm cells.**

| | | | | | | | | |
|---|---|---|---|---|---|---|---|---|
| 8 | Nr2f2 | 2.09 | 3.58E-104 | 7.65E-100 | 0.94 | 0.57 | 0.38 | 1.67 |
| 8 | Plcd1 | 2.71 | 2.71E-92 | 5.78E-88 | 0.51 | 0.1 | 0.42 | 5.34 |
| 8 | Anxa1 | 3.42 | 4.74E-92 | 1.01E-87 | 0.36 | 0.04 | 0.32 | 8.33 |
| 8 | Siva1 | 1.17 | 9.97E-91 | 2.13E-86 | 1 | 0.97 | 0.03 | 1.03 |
| 8 | Atox1 | 1.07 | 7.54E-90 | 1.61E-85 | 1 | 0.98 | 0.02 | 1.02 |
| 8 | Krt7 | 2.44 | 3.95E-87 | 8.45E-83 | 0.66 | 0.18 | 0.48 | 3.63 |
| 8 | Anxa2 | 1.4 | 1.17E-83 | 2.50E-79 | 1 | 0.74 | 0.25 | 1.34 |
| 8 | Tacstd2 | 1.69 | 5.00E-83 | 1.07E-78 | 0.76 | 0.24 | 0.52 | 3.12 |
| 8 | Pon2 | 2.23 | 1.40E-82 | 2.99E-78 | 0.75 | 0.26 | 0.48 | 2.83 |
| 8 | Glrx | 2.35 | 4.68E-81 | 1.00E-76 | 0.74 | 0.27 | 0.47 | 2.74 |
| 8 | Nrk | 2.89 | 1.07E-79 | 2.28E-75 | 0.49 | 0.1 | 0.38 | 4.68 |
| 8 | Igf2 | 1.6 | 2.59E-79 | 5.53E-75 | 0.93 | 0.81 | 0.12 | 1.14 |
| 8 | Pitx2 | 1.39 | 4.77E-79 | 1.02E-74 | 0.76 | 0.22 | 0.54 | 3.45 |
| 8 | Slc39a8 | 2.8 | 1.03E-77 | 2.19E-73 | 0.43 | 0.08 | 0.35 | 5.53 |
| 8 | Fzd5 | 3.62 | 1.80E-77 | 3.84E-73 | 0.26 | 0.03 | 0.24 | 9.78 |
| 8 | Isoc1 | 1.6 | 4.08E-76 | 8.71E-72 | 0.94 | 0.69 | 0.25 | 1.36 |
| 8 | Krt18 | 1.19 | 3.52E-74 | 7.53E-70 | 1 | 0.86 | 0.14 | 1.16 |
| 8 | S100a10 | 1.27 | 1.29E-69 | 2.75E-65 | 1 | 0.73 | 0.26 | 1.36 |
| 8 | Hspb2 | 2.54 | 2.38E-69 | 5.08E-65 | 0.47 | 0.11 | 0.37 | 4.45 |
| 8 | Tpm1 | 1.07 | 2.58E-69 | 5.52E-65 | 1 | 0.93 | 0.07 | 1.07 |
| 8 | Lypd2 | 2.29 | 4.13E-65 | 8.81E-61 | 0.52 | 0.14 | 0.38 | 3.8 |
| 8 | Arhgap29 | 1.6 | 5.09E-65 | 1.09E-60 | 0.82 | 0.38 | 0.45 | 2.19 |
| 8 | Igdcc3 | 1.83 | 5.88E-65 | 1.26E-60 | 0.76 | 0.32 | 0.44 | 2.39 |
| 8 | Grhl3 | 1.88 | 8.83E-63 | 1.89E-58 | 0.62 | 0.19 | 0.43 | 3.21 |
| 8 | Cldn8 | 2.27 | 4.10E-62 | 8.76E-58 | 0.45 | 0.11 | 0.35 | 4.31 |
| 8 | Arl4c | 1.8 | 9.55E-62 | 2.04E-57 | 0.87 | 0.56 | 0.31 | 1.55 |
| 8 | Plagl1 | 1.88 | 2.24E-60 | 4.79E-56 | 0.79 | 0.41 | 0.38 | 1.94 |
| 8 | Nrtn | 1.92 | 6.06E-60 | 1.29E-55 | 0.72 | 0.3 | 0.42 | 2.43 |
| 8 | Dpp6 | 3.83 | 1.76E-59 | 3.77E-55 | 0.15 | 0.01 | 0.14 | 16.33 |
| 8 | Slc38a4 | 1.93 | 1.97E-59 | 4.21E-55 | 0.67 | 0.25 | 0.42 | 2.65 |
| 8 | Tcf7l2 | 1.4 | 9.86E-59 | 2.11E-54 | 0.93 | 0.72 | 0.21 | 1.29 |
| 8 | Adamts18 | 1.73 | 2.74E-57 | 5.85E-53 | 0.66 | 0.25 | 0.42 | 2.71 |
| 8 | Pof1b | 3.12 | 1.10E-55 | 2.36E-51 | 0.25 | 0.03 | 0.21 | 7.21 |
| 8 | Gask1b | 2.45 | 2.70E-55 | 5.77E-51 | 0.3 | 0.05 | 0.25 | 5.92 |
| 8 | Lmod2 | 4.64 | 8.41E-55 | 1.80E-50 | 0.11 | 0 | 0.11 | 27.25 |
| 8 | Lamc2 | 2.26 | 9.29E-55 | 1.99E-50 | 0.51 | 0.15 | 0.35 | 3.31 |
| 8 | Evpl | 2.3 | 1.07E-53 | 2.29E-49 | 0.42 | 0.1 | 0.32 | 4.15 |
| 8 | Sytl2 | 4.13 | 1.42E-53 | 3.04E-49 | 0.14 | 0.01 | 0.13 | 15.11 |
| 8 | Fgfbp1 | 3.74 | 2.19E-53 | 4.67E-49 | 0.17 | 0.02 | 0.15 | 11.07 |
| 8 | Gm16136 | 1.19 | 4.90E-53 | 1.05E-48 | 0.94 | 0.63 | 0.31 | 1.49 |
| 8 | Wfdc2 | 1.18 | 9.84E-53 | 2.10E-48 | 0.99 | 0.81 | 0.18 | 1.22 |
| 8 | Mt1 | 1.64 | 1.09E-51 | 2.33E-47 | 0.85 | 0.48 | 0.37 | 1.75 |
| 8 | Mt2 | 1.74 | 1.98E-51 | 4.24E-47 | 0.65 | 0.25 | 0.4 | 2.6 |
| 8 | Isl2 | 2.3 | 7.86E-51 | 1.68E-46 | 0.45 | 0.12 | 0.32 | 3.62 |
| 8 | Klf5 | 1.49 | 9.90E-51 | 2.12E-46 | 0.77 | 0.39 | 0.38 | 1.98 |
| 8 | Fxyd3 | 1.69 | 7.39E-50 | 1.58E-45 | 0.69 | 0.29 | 0.41 | 2.42 |
| 8 | Cald1 | 0.94 | 2.31E-49 | 4.94E-45 | 0.97 | 0.69 | 0.28 | 1.41 |
| 8 | Krt15 | 1.77 | 3.60E-49 | 7.70E-45 | 0.47 | 0.14 | 0.34 | 3.5 |
| 8 | Ahnak | 1.52 | 6.61E-49 | 1.41E-44 | 0.73 | 0.34 | 0.39 | 2.17 |
| 8 | Rab27b | 1.59 | 1.49E-48 | 3.19E-44 | 0.69 | 0.3 | 0.39 | 2.31 |
| 8 | Gpc3 | 0.84 | 4.54E-48 | 9.70E-44 | 1 | 0.83 | 0.17 | 1.2 |
| 8 | Mycl | 1.16 | 7.44E-48 | 1.59E-43 | 0.92 | 0.63 | 0.3 | 1.47 |
| 8 | Npnt | 1.49 | 1.73E-47 | 3.69E-43 | 0.8 | 0.43 | 0.37 | 1.87 |
| 8 | Fam210b | 2.23 | 9.45E-47 | 2.02E-42 | 0.53 | 0.2 | 0.34 | 2.73 |
| 8 | Nudt4 | 1.13 | 1.27E-46 | 2.70E-42 | 0.94 | 0.77 | 0.17 | 1.22 |
| 8 | Palm3 | 1.64 | 2.87E-46 | 6.13E-42 | 0.7 | 0.33 | 0.37 | 2.12 |
| 8 | Anxa8 | 2.31 | 4.07E-46 | 8.69E-42 | 0.35 | 0.08 | 0.27 | 4.33 |

**Supplementary Table 3. Marker genes for the original clusters of integrated clustering of E11.5 control and NotchGOF λ-junction ectoderm cells.**

| | | | | | | | | |
|---|---|---|---|---|---|---|---|---|
| 8 | Arg1 | 2.84 | 1.11E-45 | 2.37E-41 | 0.24 | 0.04 | 0.2 | 6.05 |
| 8 | Cstb | 0.86 | 9.33E-45 | 1.99E-40 | 0.99 | 0.84 | 0.15 | 1.18 |
| 8 | Cfc1 | 3.17 | 3.24E-44 | 6.92E-40 | 0.16 | 0.02 | 0.14 | 9.29 |
| 8 | Ccnd2 | 0.94 | 1.28E-42 | 2.74E-38 | 0.99 | 0.77 | 0.21 | 1.27 |
| 8 | Anxa3 | 2.4 | 2.63E-42 | 5.62E-38 | 0.28 | 0.06 | 0.22 | 4.91 |
| 8 | Sfn | 0.66 | 1.23E-41 | 2.63E-37 | 0.99 | 0.68 | 0.31 | 1.45 |
| 8 | Mmp11 | 1.49 | 3.33E-41 | 7.11E-37 | 0.73 | 0.37 | 0.35 | 1.95 |
| 8 | Lcp1 | 3.42 | 5.18E-41 | 1.11E-36 | 0.11 | 0.01 | 0.1 | 13.63 |
| 8 | Afap1l1 | 2.17 | 2.56E-40 | 5.46E-36 | 0.37 | 0.1 | 0.27 | 3.6 |
| 8 | Tmod3 | 1.27 | 2.84E-39 | 6.06E-35 | 0.87 | 0.65 | 0.22 | 1.34 |
| 8 | Ccdc68 | 1.81 | 3.52E-39 | 7.51E-35 | 0.4 | 0.12 | 0.29 | 3.39 |
| 8 | Unc5c | 1.15 | 6.25E-39 | 1.33E-34 | 0.69 | 0.31 | 0.38 | 2.23 |
| 8 | Cldn23 | 1.93 | 2.99E-38 | 6.40E-34 | 0.42 | 0.14 | 0.28 | 3.04 |
| 8 | Shh | 1.96 | 4.42E-38 | 9.43E-34 | 0.25 | 0.05 | 0.2 | 5.08 |
| 8 | 2200002D01Rik | 2.52 | 4.71E-38 | 1.01E-33 | 0.26 | 0.06 | 0.21 | 4.73 |
| 8 | Dsp | 0.82 | 7.17E-38 | 1.53E-33 | 0.98 | 0.86 | 0.12 | 1.14 |
| 8 | Igfbp5 | 0.74 | 1.12E-37 | 2.39E-33 | 1 | 0.96 | 0.04 | 1.04 |
| 8 | Hspb1 | 1.12 | 2.67E-37 | 5.69E-33 | 0.74 | 0.36 | 0.38 | 2.04 |
| 8 | 2610016A17Rik | 2.99 | 1.04E-36 | 2.22E-32 | 0.19 | 0.03 | 0.16 | 6.3 |
| 8 | Arid3a | 1.42 | 4.19E-36 | 8.95E-32 | 0.67 | 0.34 | 0.33 | 1.96 |
| 8 | Utrn | 0.98 | 4.19E-36 | 8.95E-32 | 0.93 | 0.71 | 0.22 | 1.31 |
| 8 | Ralbp1 | 0.84 | 6.78E-36 | 1.45E-31 | 0.96 | 0.82 | 0.14 | 1.17 |
| 8 | Ghr | 2.24 | 9.42E-36 | 2.01E-31 | 0.28 | 0.07 | 0.22 | 4.16 |
| 8 | Foxc1 | 1.88 | 1.47E-35 | 3.14E-31 | 0.4 | 0.13 | 0.27 | 3.09 |
| 8 | Sema3c | 1.47 | 1.54E-35 | 3.28E-31 | 0.56 | 0.24 | 0.33 | 2.38 |
| 8 | Zbtb42 | 2.62 | 2.27E-34 | 4.86E-30 | 0.19 | 0.03 | 0.16 | 5.91 |
| 8 | Calm1 | 0.42 | 3.84E-34 | 8.20E-30 | 1 | 0.99 | 0.01 | 1.01 |
| 8 | Fgfr3 | 2.51 | 1.30E-33 | 2.78E-29 | 0.21 | 0.04 | 0.17 | 5.15 |
| 8 | Perp | 0.78 | 2.52E-33 | 5.37E-29 | 0.91 | 0.52 | 0.39 | 1.74 |
| 8 | Ogfrl1 | 1.85 | 1.32E-32 | 2.81E-28 | 0.44 | 0.17 | 0.28 | 2.65 |
| 8 | Mfge8 | 1.27 | 1.91E-32 | 4.07E-28 | 0.81 | 0.54 | 0.27 | 1.49 |
| 8 | B230208H11Rik | 2.41 | 2.28E-31 | 4.86E-27 | 0.22 | 0.05 | 0.17 | 4.76 |
| 8 | Hebp1 | 1.45 | 5.32E-31 | 1.14E-26 | 0.51 | 0.22 | 0.29 | 2.37 |
| 8 | Pmp22 | 3.02 | 8.80E-31 | 1.88E-26 | 0.14 | 0.02 | 0.12 | 7.16 |
| 8 | Grb10 | 0.63 | 1.12E-29 | 2.39E-25 | 1 | 0.93 | 0.07 | 1.08 |
| 8 | Srd5a1 | 1.29 | 1.89E-29 | 4.03E-25 | 0.62 | 0.32 | 0.3 | 1.92 |
| 8 | Tinagl1 | 2.05 | 3.10E-29 | 6.63E-25 | 0.29 | 0.08 | 0.21 | 3.54 |
| 8 | A730020E08Rik | 2.01 | 3.16E-29 | 6.76E-25 | 0.28 | 0.08 | 0.2 | 3.67 |
| 8 | Slc2a3 | 0.93 | 3.85E-29 | 8.23E-25 | 0.65 | 0.34 | 0.32 | 1.94 |
| 8 | Aldh3b2 | 2.76 | 4.20E-29 | 8.97E-25 | 0.16 | 0.03 | 0.13 | 5.79 |
| 8 | Ppl | 1.21 | 9.75E-29 | 2.08E-24 | 0.42 | 0.15 | 0.27 | 2.77 |
| 8 | Cmas | 0.95 | 1.54E-28 | 3.28E-24 | 0.84 | 0.64 | 0.2 | 1.31 |
| 8 | Cers4 | 1.36 | 1.68E-28 | 3.58E-24 | 0.66 | 0.36 | 0.3 | 1.82 |
| 8 | Col5a2 | 1.51 | 2.02E-28 | 4.31E-24 | 0.41 | 0.15 | 0.26 | 2.78 |
| 8 | Alcam | 1.36 | 6.34E-28 | 1.35E-23 | 0.46 | 0.19 | 0.28 | 2.48 |
| 8 | Igf2bp3 | 0.67 | 6.37E-28 | 1.36E-23 | 0.95 | 0.86 | 0.09 | 1.1 |
| 8 | Fignl2 | 1.91 | 1.15E-27 | 2.45E-23 | 0.25 | 0.07 | 0.19 | 3.78 |
| 8 | Arf1 | 0.4 | 1.52E-27 | 3.24E-23 | 0.99 | 0.98 | 0.01 | 1.01 |
| 8 | Iqgap1 | 0.87 | 1.54E-27 | 3.29E-23 | 0.85 | 0.67 | 0.17 | 1.26 |
| 8 | Cd9 | 0.73 | 1.60E-27 | 3.42E-23 | 0.92 | 0.7 | 0.21 | 1.3 |
| 8 | Ppia | 0.26 | 2.50E-27 | 5.33E-23 | 1 | 1 | 0 | 1 |
| 8 | Myh14 | 1.35 | 2.61E-27 | 5.58E-23 | 0.56 | 0.29 | 0.28 | 1.97 |
| 8 | Prom2 | 1.77 | 7.27E-27 | 1.55E-22 | 0.34 | 0.12 | 0.23 | 2.96 |
| 8 | Il17re | 1.63 | 1.18E-26 | 2.53E-22 | 0.42 | 0.17 | 0.25 | 2.47 |
| 8 | Ppp1r13b | 1.69 | 2.19E-26 | 4.69E-22 | 0.34 | 0.12 | 0.23 | 2.91 |
| 8 | Prss22 | 1.24 | 3.46E-26 | 7.39E-22 | 0.51 | 0.23 | 0.28 | 2.24 |
| 8 | Platr21 | 1.91 | 1.36E-25 | 2.91E-21 | 0.29 | 0.09 | 0.2 | 3.2 |

**Supplementary Table 3. Marker genes for the original clusters of integrated clustering of E11.5 control and NotchGOF λ-junction ectoderm cells.**

| | | | | | | | | |
|---|---|---|---|---|---|---|---|---|
| 8 | Hsdl2 | 1.3 | 7.56E-25 | 1.62E-20 | 0.59 | 0.33 | 0.26 | 1.78 |
| 8 | Alox12 | 2.08 | 8.08E-25 | 1.73E-20 | 0.25 | 0.07 | 0.18 | 3.47 |
| 8 | Smtnl2 | 1.38 | 8.40E-25 | 1.79E-20 | 0.51 | 0.25 | 0.26 | 2.08 |
| 8 | Pwwp2b | 1.49 | 1.55E-24 | 3.32E-20 | 0.37 | 0.14 | 0.24 | 2.71 |
| 8 | Ptpn13 | 0.82 | 2.55E-24 | 5.45E-20 | 0.81 | 0.56 | 0.25 | 1.44 |
| 8 | Wwc2 | 0.93 | 2.82E-24 | 6.02E-20 | 0.82 | 0.59 | 0.22 | 1.37 |
| 8 | Hsd17b14 | 2.9 | 3.26E-24 | 6.97E-20 | 0.14 | 0.02 | 0.11 | 5.67 |
| 8 | Pcdh1 | 1.79 | 5.40E-24 | 1.15E-19 | 0.29 | 0.09 | 0.2 | 3.12 |
| 8 | Fstl1 | 0.9 | 6.82E-24 | 1.46E-19 | 0.79 | 0.6 | 0.2 | 1.33 |
| 8 | Cyb5r3 | 0.52 | 1.09E-23 | 2.32E-19 | 0.97 | 0.93 | 0.04 | 1.04 |
| 8 | Foxp1 | 0.58 | 1.37E-23 | 2.94E-19 | 0.96 | 0.86 | 0.1 | 1.11 |
| 8 | Nectin4 | 0.83 | 1.61E-23 | 3.43E-19 | 0.8 | 0.52 | 0.28 | 1.54 |
| 8 | Esrp2 | 1.23 | 1.84E-23 | 3.92E-19 | 0.52 | 0.25 | 0.27 | 2.11 |
| 8 | Ctsh | 1.63 | 2.93E-23 | 6.26E-19 | 0.32 | 0.11 | 0.21 | 2.82 |
| 8 | Tek | 2.72 | 4.74E-23 | 1.01E-18 | 0.12 | 0.02 | 0.1 | 6.16 |
| 8 | Rassf8 | 2.04 | 4.84E-23 | 1.03E-18 | 0.29 | 0.1 | 0.19 | 2.93 |
| 8 | Mpp7 | 0.99 | 5.70E-23 | 1.22E-18 | 0.74 | 0.54 | 0.2 | 1.38 |
| 8 | Mansc4 | 2.18 | 8.90E-23 | 1.90E-18 | 0.17 | 0.04 | 0.13 | 4.47 |
| 8 | Limk2 | 1.08 | 9.83E-23 | 2.10E-18 | 0.6 | 0.34 | 0.26 | 1.78 |
| 8 | Phactr2 | 1.17 | 1.50E-22 | 3.21E-18 | 0.6 | 0.36 | 0.24 | 1.67 |
| 8 | Stx19 | 2.33 | 3.11E-22 | 6.64E-18 | 0.14 | 0.03 | 0.12 | 5.11 |
| 8 | Cdh1 | 0.76 | 9.74E-22 | 2.08E-17 | 0.83 | 0.65 | 0.18 | 1.28 |
| 8 | Chst5 | 2.12 | 1.43E-21 | 3.05E-17 | 0.2 | 0.06 | 0.15 | 3.71 |
| 8 | Fam83f | 2.26 | 1.43E-21 | 3.06E-17 | 0.16 | 0.03 | 0.12 | 4.56 |
| 8 | Gpc4 | 1.18 | 1.96E-21 | 4.18E-17 | 0.55 | 0.3 | 0.25 | 1.85 |
| 8 | Eci1 | 0.91 | 2.38E-21 | 5.09E-17 | 0.75 | 0.52 | 0.23 | 1.44 |
| 8 | Shisa4 | 1.24 | 2.42E-21 | 5.16E-17 | 0.48 | 0.23 | 0.25 | 2.1 |
| 8 | Rbms1 | 0.62 | 3.58E-21 | 7.64E-17 | 0.93 | 0.84 | 0.08 | 1.1 |
| 8 | Idh2 | 0.56 | 4.33E-21 | 9.26E-17 | 0.96 | 0.87 | 0.09 | 1.11 |
| 8 | Nr3c1 | 1.24 | 4.69E-21 | 1.00E-16 | 0.51 | 0.26 | 0.25 | 1.95 |
| 8 | Elf5 | 1.28 | 7.16E-21 | 1.53E-16 | 0.47 | 0.23 | 0.24 | 2.07 |
| 8 | Epb41l5 | 0.92 | 1.46E-20 | 3.13E-16 | 0.73 | 0.51 | 0.22 | 1.44 |
| 8 | Cers3 | 2.15 | 2.68E-20 | 5.72E-16 | 0.17 | 0.04 | 0.13 | 4.15 |
| 8 | 2310030G06Rik | 1.29 | 2.71E-20 | 5.79E-16 | 0.49 | 0.26 | 0.23 | 1.89 |
| 8 | Gjb3 | 1.28 | 2.86E-20 | 6.10E-16 | 0.35 | 0.14 | 0.21 | 2.56 |
| 8 | Junb | 1.3 | 3.65E-20 | 7.80E-16 | 0.44 | 0.2 | 0.24 | 2.18 |
| 8 | Dstn | 0.35 | 4.19E-20 | 8.96E-16 | 0.99 | 0.98 | 0.01 | 1.01 |
| 8 | S100a11 | 0.44 | 4.82E-20 | 1.03E-15 | 1 | 0.9 | 0.1 | 1.11 |
| 8 | Ldhb | 0.48 | 6.08E-20 | 1.30E-15 | 0.98 | 0.87 | 0.12 | 1.13 |
| 8 | Ccnd3 | 0.61 | 6.48E-20 | 1.38E-15 | 0.92 | 0.8 | 0.12 | 1.15 |
| 8 | Sh3tc2 | 2.02 | 7.06E-20 | 1.51E-15 | 0.18 | 0.05 | 0.13 | 3.85 |
| 8 | Fhl1 | 1.32 | 9.85E-20 | 2.10E-15 | 0.47 | 0.24 | 0.23 | 1.95 |
| 8 | Spsb1 | 1.73 | 9.97E-20 | 2.13E-15 | 0.21 | 0.06 | 0.15 | 3.41 |
| 8 | Cldn7 | 0.61 | 3.13E-19 | 6.69E-15 | 0.97 | 0.84 | 0.14 | 1.16 |
| 8 | Eps8l2 | 1.62 | 3.42E-19 | 7.31E-15 | 0.34 | 0.15 | 0.2 | 2.35 |
| 8 | Ier2 | 0.64 | 6.17E-19 | 1.32E-14 | 0.9 | 0.79 | 0.12 | 1.15 |
| 8 | Fign | 1.49 | 6.93E-19 | 1.48E-14 | 0.39 | 0.18 | 0.21 | 2.15 |
| 8 | Sfr1 | 0.39 | 7.39E-19 | 1.58E-14 | 1 | 0.98 | 0.02 | 1.02 |
| 8 | Sema3d | 0.82 | 8.38E-19 | 1.79E-14 | 0.67 | 0.41 | 0.26 | 1.65 |
| 8 | Rai14 | 1.28 | 9.38E-19 | 2.00E-14 | 0.51 | 0.29 | 0.22 | 1.77 |
| 8 | Jph1 | 1.87 | 1.04E-18 | 2.22E-14 | 0.25 | 0.09 | 0.16 | 2.86 |
| 8 | Fhl2 | 2 | 1.32E-18 | 2.83E-14 | 0.19 | 0.05 | 0.13 | 3.56 |
| 8 | Fabp5 | 0.31 | 1.37E-18 | 2.93E-14 | 1 | 0.99 | 0.01 | 1.01 |
| 8 | Dmrta2 | 0.87 | 1.39E-18 | 2.98E-14 | 0.74 | 0.5 | 0.25 | 1.5 |
| 8 | S100a16 | 1.73 | 1.55E-18 | 3.32E-14 | 0.33 | 0.14 | 0.19 | 2.39 |
| 8 | Lypd6 | 1.4 | 2.05E-18 | 4.38E-14 | 0.41 | 0.19 | 0.22 | 2.11 |
| 8 | B130024G19Rik | 2.09 | 2.35E-18 | 5.01E-14 | 0.16 | 0.04 | 0.12 | 4.08 |

**Supplementary Table 3. Marker genes for the original clusters of integrated clustering of E11.5 control and NotchGOF λ-junction ectoderm cells.**

| | | | | | | | | |
|---|---|---|---|---|---|---|---|---|
| 8 | Lgi2 | 1.11 | 2.39E-18 | 5.11E-14 | 0.52 | 0.28 | 0.24 | 1.83 |
| 8 | Fuom | 1.05 | 2.45E-18 | 5.24E-14 | 0.55 | 0.31 | 0.24 | 1.75 |
| 8 | F3 | 1.34 | 2.64E-18 | 5.64E-14 | 0.43 | 0.22 | 0.22 | 1.99 |
| 8 | Spats2l | 0.89 | 2.71E-18 | 5.80E-14 | 0.57 | 0.33 | 0.25 | 1.75 |
| 8 | F11r | 0.76 | 3.57E-18 | 7.64E-14 | 0.8 | 0.6 | 0.21 | 1.34 |
| 8 | Rflnb | 0.88 | 4.05E-18 | 8.65E-14 | 0.74 | 0.51 | 0.23 | 1.46 |
| 8 | Net1 | 0.97 | 4.41E-18 | 9.43E-14 | 0.69 | 0.49 | 0.2 | 1.4 |
| 8 | Arhgap24 | 1.6 | 6.01E-18 | 1.28E-13 | 0.22 | 0.07 | 0.15 | 3.16 |
| 8 | Mapkapk2 | 0.81 | 7.54E-18 | 1.61E-13 | 0.73 | 0.54 | 0.19 | 1.36 |
| 8 | Negr1 | 2.07 | 1.04E-17 | 2.23E-13 | 0.15 | 0.04 | 0.11 | 4.08 |
| 8 | Capg | 0.99 | 1.25E-17 | 2.67E-13 | 0.58 | 0.34 | 0.24 | 1.69 |
| 8 | Macc1 | 1.2 | 2.48E-17 | 5.29E-13 | 0.42 | 0.21 | 0.21 | 1.99 |
| 8 | Dusp9 | 1.43 | 4.92E-17 | 1.05E-12 | 0.43 | 0.23 | 0.21 | 1.9 |
| 8 | Scaper | 1.41 | 5.48E-17 | 1.17E-12 | 0.54 | 0.33 | 0.21 | 1.63 |
| 8 | Pik3ip1 | 1.01 | 1.01E-16 | 2.17E-12 | 0.4 | 0.18 | 0.21 | 2.18 |
| 8 | Arap2 | 1.49 | 1.26E-16 | 2.69E-12 | 0.31 | 0.13 | 0.18 | 2.34 |
| 8 | Gca | 1.95 | 1.29E-16 | 2.76E-12 | 0.21 | 0.07 | 0.14 | 2.93 |
| 8 | Pfn2 | 0.69 | 1.30E-16 | 2.77E-12 | 0.84 | 0.7 | 0.15 | 1.21 |
| 8 | Pdpn | 1 | 2.15E-16 | 4.60E-12 | 0.58 | 0.36 | 0.22 | 1.63 |
| 8 | Egfl6 | 1.38 | 3.64E-16 | 7.77E-12 | 0.37 | 0.17 | 0.19 | 2.1 |
| 8 | Ctdspl | 0.74 | 4.94E-16 | 1.06E-11 | 0.7 | 0.48 | 0.23 | 1.47 |
| 8 | Lifr | 1.39 | 4.95E-16 | 1.06E-11 | 0.28 | 0.12 | 0.17 | 2.42 |
| 8 | Ptprf | 0.42 | 6.27E-16 | 1.34E-11 | 0.99 | 0.96 | 0.03 | 1.03 |
| 8 | Hs6st2 | 0.73 | 8.91E-16 | 1.90E-11 | 0.75 | 0.53 | 0.22 | 1.41 |
| 8 | Lmnb1 | 0.47 | 8.95E-16 | 1.91E-11 | 0.99 | 0.93 | 0.06 | 1.07 |
| 8 | Egln3 | 1.26 | 1.22E-15 | 2.61E-11 | 0.37 | 0.18 | 0.19 | 2.05 |
| 8 | C430049B03Rik | 1.5 | 1.37E-15 | 2.94E-11 | 0.31 | 0.14 | 0.17 | 2.22 |
| 8 | Nebl | 0.58 | 2.17E-15 | 4.63E-11 | 0.63 | 0.39 | 0.24 | 1.62 |
| 8 | Sox2 | 0.42 | 2.57E-15 | 5.49E-11 | 0.95 | 0.69 | 0.26 | 1.38 |
| 8 | Fos | 1.35 | 3.04E-15 | 6.49E-11 | 0.27 | 0.11 | 0.16 | 2.48 |
| 8 | Pon3 | 2.34 | 3.05E-15 | 6.51E-11 | 0.11 | 0.02 | 0.08 | 4.61 |
| 8 | Rxfp1 | 1.94 | 3.73E-15 | 7.96E-11 | 0.16 | 0.05 | 0.11 | 3.43 |
| 8 | Gadd45b | 1.14 | 5.82E-15 | 1.24E-10 | 0.39 | 0.2 | 0.19 | 1.99 |
| 8 | Jade3 | 1.23 | 5.91E-15 | 1.26E-10 | 0.34 | 0.16 | 0.18 | 2.17 |
| 8 | Dcaf4 | 0.97 | 6.10E-15 | 1.30E-10 | 0.54 | 0.34 | 0.2 | 1.59 |
| 8 | Phldb3 | 1.64 | 6.16E-15 | 1.32E-10 | 0.22 | 0.08 | 0.14 | 2.76 |
| 8 | Plch2 | 1.52 | 6.70E-15 | 1.43E-10 | 0.25 | 0.1 | 0.15 | 2.5 |
| 8 | Dlc1 | 1.55 | 7.20E-15 | 1.54E-10 | 0.22 | 0.08 | 0.14 | 2.79 |
| 8 | Lima1 | 0.74 | 7.76E-15 | 1.66E-10 | 0.68 | 0.47 | 0.21 | 1.45 |
| 8 | Zswim8 | 0.88 | 8.27E-15 | 1.77E-10 | 0.58 | 0.37 | 0.21 | 1.57 |
| 8 | Fosb | 1.51 | 8.29E-15 | 1.77E-10 | 0.11 | 0.02 | 0.08 | 4.61 |
| 8 | Myl12a | 0.36 | 8.31E-15 | 1.78E-10 | 0.98 | 0.93 | 0.04 | 1.05 |
| 8 | Serpinb6a | 0.74 | 1.66E-14 | 3.55E-10 | 0.68 | 0.47 | 0.21 | 1.45 |
| 8 | Wnt4 | 0.32 | 1.72E-14 | 3.68E-10 | 0.94 | 0.7 | 0.24 | 1.35 |
| 8 | Dpysl3 | 0.34 | 1.88E-14 | 4.02E-10 | 0.8 | 0.56 | 0.25 | 1.45 |
| 8 | Bcl9l | 0.94 | 1.99E-14 | 4.24E-10 | 0.53 | 0.32 | 0.21 | 1.66 |
| 8 | Zfp568 | 1.03 | 2.25E-14 | 4.80E-10 | 0.52 | 0.33 | 0.2 | 1.6 |
| 8 | Tcf7l1 | 0.75 | 2.40E-14 | 5.13E-10 | 0.77 | 0.56 | 0.2 | 1.36 |
| 8 | Ndufa11 | 0.35 | 2.41E-14 | 5.14E-10 | 0.99 | 0.95 | 0.04 | 1.04 |
| 8 | Tceal9 | 0.27 | 2.61E-14 | 5.57E-10 | 1 | 0.99 | 0.01 | 1.01 |
| 8 | Bmp7 | 0.76 | 3.04E-14 | 6.50E-10 | 0.6 | 0.4 | 0.21 | 1.52 |
| 8 | Eif1a | 0.69 | 3.76E-14 | 8.03E-10 | 0.82 | 0.68 | 0.14 | 1.2 |
| 8 | Pfdn1 | 0.36 | 3.81E-14 | 8.13E-10 | 0.98 | 0.94 | 0.04 | 1.04 |
| 8 | Rest | 0.56 | 4.95E-14 | 1.06E-09 | 0.88 | 0.67 | 0.21 | 1.31 |
| 8 | Sos1 | 0.83 | 5.15E-14 | 1.10E-09 | 0.59 | 0.39 | 0.2 | 1.51 |
| 8 | Crip1 | 0.84 | 5.69E-14 | 1.21E-09 | 0.59 | 0.39 | 0.2 | 1.51 |
| 8 | Rbbp7 | 0.51 | 5.75E-14 | 1.23E-09 | 0.93 | 0.85 | 0.08 | 1.1 |

**Supplementary Table 3. Marker genes for the original clusters of integrated clustering of E11.5 control and NotchGOF λ-junction ectoderm cells.**

| | | | | | | | | |
|---|---|---|---|---|---|---|---|---|
| 8 | Cgn | 0.86 | 6.46E-14 | 1.38E-09 | 0.57 | 0.38 | 0.19 | 1.5 |
| 8 | Stard3nl | 0.79 | 9.30E-14 | 1.99E-09 | 0.7 | 0.53 | 0.17 | 1.33 |
| 8 | Stard10 | 0.62 | 9.77E-14 | 2.09E-09 | 0.86 | 0.72 | 0.14 | 1.19 |
| 8 | Endod1 | 1.24 | 9.79E-14 | 2.09E-09 | 0.32 | 0.15 | 0.17 | 2.11 |
| 8 | Lyrm9 | 1.14 | 1.01E-13 | 2.15E-09 | 0.39 | 0.21 | 0.18 | 1.83 |
| 8 | Klf6 | 1.02 | 1.09E-13 | 2.33E-09 | 0.43 | 0.23 | 0.2 | 1.87 |
| 8 | Foxp4 | 0.79 | 1.13E-13 | 2.41E-09 | 0.56 | 0.36 | 0.2 | 1.55 |
| 8 | Ppp1r14c | 2.17 | 1.24E-13 | 2.64E-09 | 0.14 | 0.04 | 0.1 | 3.41 |
| 8 | Parp1 | 0.42 | 1.60E-13 | 3.42E-09 | 0.94 | 0.88 | 0.06 | 1.07 |
| 8 | Erbb2 | 0.86 | 1.74E-13 | 3.72E-09 | 0.53 | 0.31 | 0.21 | 1.68 |
| 8 | 4930523C07Rik | 1.67 | 1.75E-13 | 3.74E-09 | 0.23 | 0.09 | 0.14 | 2.47 |
| 8 | Plet1 | 2.06 | 2.40E-13 | 5.14E-09 | 0.14 | 0.04 | 0.1 | 3.33 |
| 8 | Ece1 | 0.77 | 2.58E-13 | 5.50E-09 | 0.65 | 0.45 | 0.2 | 1.44 |
| 8 | Acadl | 0.75 | 2.85E-13 | 6.08E-09 | 0.62 | 0.42 | 0.21 | 1.49 |
| 8 | Lrrk1 | 0.96 | 3.68E-13 | 7.86E-09 | 0.49 | 0.3 | 0.2 | 1.66 |
| 8 | Tjp2 | 0.75 | 4.20E-13 | 8.98E-09 | 0.66 | 0.47 | 0.19 | 1.42 |
| 8 | Pdlim1 | 0.32 | 4.31E-13 | 9.21E-09 | 0.84 | 0.64 | 0.21 | 1.32 |
| 8 | Sh3bgrl3 | 0.37 | 4.42E-13 | 9.44E-09 | 0.98 | 0.91 | 0.07 | 1.07 |
| 8 | Ank3 | 0.59 | 4.53E-13 | 9.68E-09 | 0.79 | 0.63 | 0.17 | 1.27 |
| 8 | Fgfr2 | 0.51 | 5.35E-13 | 1.14E-08 | 0.7 | 0.46 | 0.24 | 1.52 |
| 8 | Ccn2 | 1.74 | 6.17E-13 | 1.32E-08 | 0.21 | 0.08 | 0.13 | 2.51 |
| 8 | Gls | 0.92 | 7.24E-13 | 1.55E-08 | 0.62 | 0.47 | 0.15 | 1.33 |
| 8 | Itprid2 | 1.36 | 8.97E-13 | 1.92E-08 | 0.26 | 0.12 | 0.15 | 2.26 |
| 8 | Tbrg1 | 0.47 | 9.12E-13 | 1.95E-08 | 0.93 | 0.84 | 0.08 | 1.1 |
| 8 | Ywhab | 0.32 | 1.22E-12 | 2.60E-08 | 0.99 | 0.96 | 0.03 | 1.03 |
| 8 | Lacc1 | 1.74 | 1.24E-12 | 2.64E-08 | 0.2 | 0.08 | 0.12 | 2.61 |
| 8 | Itgb1 | 0.41 | 1.39E-12 | 2.96E-08 | 0.91 | 0.85 | 0.07 | 1.08 |
| 8 | Kank1 | 0.88 | 1.51E-12 | 3.23E-08 | 0.47 | 0.28 | 0.19 | 1.68 |
| 8 | Mbnl3 | 1.17 | 1.69E-12 | 3.61E-08 | 0.33 | 0.16 | 0.16 | 2.01 |
| 8 | Gsto1 | 0.84 | 1.97E-12 | 4.22E-08 | 0.58 | 0.4 | 0.18 | 1.45 |
| 8 | Rassf9 | 1.08 | 2.03E-12 | 4.34E-08 | 0.41 | 0.23 | 0.18 | 1.76 |
| 8 | Krcc1 | 0.6 | 2.66E-12 | 5.68E-08 | 0.79 | 0.62 | 0.16 | 1.26 |
| 8 | Sumo3 | 0.39 | 2.91E-12 | 6.21E-08 | 0.95 | 0.91 | 0.04 | 1.05 |
| 8 | Bcam | 0.43 | 3.35E-12 | 7.15E-08 | 0.9 | 0.72 | 0.18 | 1.25 |
| 8 | Anp32b | 0.26 | 3.93E-12 | 8.39E-08 | 1 | 0.99 | 0.01 | 1.01 |
| 8 | Rassf10 | 0.86 | 4.68E-12 | 9.99E-08 | 0.43 | 0.24 | 0.19 | 1.8 |
| 8 | Cmtm8 | 0.69 | 5.16E-12 | 1.10E-07 | 0.69 | 0.52 | 0.16 | 1.31 |
| 8 | Ppp2r2b | 0.69 | 5.23E-12 | 1.12E-07 | 0.57 | 0.37 | 0.2 | 1.53 |
| 8 | Spint2 | 0.29 | 6.46E-12 | 1.38E-07 | 0.99 | 0.97 | 0.02 | 1.02 |
| 8 | Aplp2 | 0.3 | 8.34E-12 | 1.78E-07 | 0.95 | 0.92 | 0.03 | 1.04 |
| 8 | Ubl4a | 0.68 | 9.67E-12 | 2.07E-07 | 0.72 | 0.57 | 0.15 | 1.26 |
| 8 | Vps36 | 0.47 | 9.91E-12 | 2.12E-07 | 0.95 | 0.85 | 0.1 | 1.12 |
| 8 | Elf1 | 0.6 | 1.00E-11 | 2.14E-07 | 0.68 | 0.5 | 0.18 | 1.37 |
| 8 | Kctd11 | 1.42 | 1.12E-11 | 2.40E-07 | 0.2 | 0.08 | 0.12 | 2.55 |
| 8 | Ubald1 | 0.74 | 1.21E-11 | 2.58E-07 | 0.57 | 0.38 | 0.19 | 1.5 |
| 8 | 1810058I24Rik | 0.3 | 1.25E-11 | 2.67E-07 | 0.98 | 0.93 | 0.05 | 1.05 |
| 8 | Lims1 | 0.68 | 1.27E-11 | 2.71E-07 | 0.68 | 0.52 | 0.16 | 1.31 |
| 8 | 4930556M19Rik | 2.04 | 1.30E-11 | 2.77E-07 | 0.11 | 0.03 | 0.08 | 3.63 |
| 8 | Myo1c | 1.62 | 1.35E-11 | 2.87E-07 | 0.2 | 0.08 | 0.12 | 2.48 |
| 8 | St7 | 0.9 | 1.48E-11 | 3.15E-07 | 0.5 | 0.32 | 0.18 | 1.56 |
| 8 | Tmem256 | 0.36 | 1.62E-11 | 3.46E-07 | 0.94 | 0.88 | 0.06 | 1.07 |
| 8 | Cystm1 | 0.73 | 1.82E-11 | 3.89E-07 | 0.53 | 0.35 | 0.18 | 1.5 |
| 8 | Pdcd10 | 0.5 | 1.92E-11 | 4.11E-07 | 0.81 | 0.7 | 0.11 | 1.16 |
| 8 | Sri | 0.49 | 1.94E-11 | 4.14E-07 | 0.82 | 0.71 | 0.11 | 1.15 |
| 8 | Hpcal1 | 0.6 | 1.95E-11 | 4.16E-07 | 0.65 | 0.46 | 0.19 | 1.42 |
| 8 | Wnt7b | 0.49 | 2.38E-11 | 5.08E-07 | 0.33 | 0.16 | 0.17 | 2.09 |
| 8 | Amotl1 | 0.47 | 2.38E-11 | 5.08E-07 | 0.83 | 0.69 | 0.14 | 1.2 |

**Supplementary Table 3. Marker genes for the original clusters of integrated clustering of E11.5 control and NotchGOF λ-junction ectoderm cells.**

| | | | | | | | | |
|---|---|---|---|---|---|---|---|---|
| 8 | Akt1 | 0.61 | 2.68E-11 | 5.72E-07 | 0.77 | 0.67 | 0.1 | 1.16 |
| 8 | Ptpn14 | 0.92 | 2.74E-11 | 5.86E-07 | 0.4 | 0.22 | 0.18 | 1.79 |
| 8 | Adgrg6 | 1.95 | 2.84E-11 | 6.08E-07 | 0.1 | 0.03 | 0.08 | 3.78 |
| 8 | Arhgef16 | 1.1 | 2.89E-11 | 6.17E-07 | 0.37 | 0.21 | 0.16 | 1.77 |
| 8 | Slc15a2 | 1.1 | 3.71E-11 | 7.93E-07 | 0.29 | 0.14 | 0.15 | 2.06 |
| 8 | Etfa | 0.46 | 3.75E-11 | 8.02E-07 | 0.86 | 0.75 | 0.11 | 1.15 |
| 8 | Odc1 | 0.45 | 3.98E-11 | 8.50E-07 | 0.92 | 0.83 | 0.09 | 1.11 |
| 8 | Sntb1 | 1.63 | 5.35E-11 | 1.14E-06 | 0.17 | 0.07 | 0.11 | 2.62 |
| 8 | Ptch1 | 0.68 | 5.48E-11 | 1.17E-06 | 0.53 | 0.35 | 0.19 | 1.53 |
| 8 | Lrp2 | 0.98 | 6.51E-11 | 1.39E-06 | 0.38 | 0.21 | 0.16 | 1.76 |
| 8 | Fkbp5 | 0.89 | 6.84E-11 | 1.46E-06 | 0.48 | 0.31 | 0.17 | 1.54 |
| 8 | Cldn3 | 0.51 | 7.85E-11 | 1.68E-06 | 0.82 | 0.62 | 0.2 | 1.32 |
| 8 | Prr15l | 1.21 | 8.14E-11 | 1.74E-06 | 0.28 | 0.14 | 0.14 | 2.02 |
| 8 | Myh9 | 0.56 | 9.45E-11 | 2.02E-06 | 0.84 | 0.75 | 0.09 | 1.12 |
| 8 | Rnf128 | 1.66 | 1.04E-10 | 2.22E-06 | 0.17 | 0.07 | 0.11 | 2.56 |
| 8 | Cdca7 | 0.5 | 1.05E-10 | 2.24E-06 | 0.81 | 0.64 | 0.17 | 1.27 |
| 8 | Srrm1 | 0.3 | 1.07E-10 | 2.29E-06 | 0.98 | 0.97 | 0.02 | 1.02 |
| 8 | Atp6v1d | 0.55 | 1.14E-10 | 2.43E-06 | 0.79 | 0.67 | 0.12 | 1.18 |
| 8 | Gab1 | 0.81 | 1.15E-10 | 2.46E-06 | 0.46 | 0.28 | 0.18 | 1.63 |
| 8 | Ctnna1 | 0.38 | 1.37E-10 | 2.94E-06 | 0.93 | 0.87 | 0.06 | 1.07 |
| 8 | Vmac | 1.04 | 1.38E-10 | 2.94E-06 | 0.3 | 0.15 | 0.15 | 1.95 |
| 8 | Tcf15 | 1.96 | 1.48E-10 | 3.16E-06 | 0.12 | 0.04 | 0.08 | 3.18 |
| 8 | Tap1 | 1.62 | 1.53E-10 | 3.27E-06 | 0.17 | 0.07 | 0.11 | 2.56 |
| 8 | Hacd2 | 0.8 | 1.55E-10 | 3.30E-06 | 0.55 | 0.39 | 0.16 | 1.41 |
| 8 | Hnrnpa3 | 0.26 | 1.64E-10 | 3.50E-06 | 1 | 1 | 0.01 | 1.01 |
| 8 | Cnst | 1.07 | 1.67E-10 | 3.57E-06 | 0.28 | 0.14 | 0.14 | 2.02 |
| 8 | Eppk1 | 1.4 | 1.67E-10 | 3.57E-06 | 0.19 | 0.08 | 0.11 | 2.43 |
| 8 | Ndufb3 | 0.34 | 2.14E-10 | 4.57E-06 | 0.95 | 0.9 | 0.05 | 1.06 |
| 8 | Cldn4 | 0.63 | 2.42E-10 | 5.16E-06 | 0.57 | 0.41 | 0.16 | 1.4 |
| 8 | Mgat3 | 1.3 | 2.50E-10 | 5.34E-06 | 0.25 | 0.12 | 0.13 | 2.11 |
| 8 | Ugt8a | 1.61 | 2.55E-10 | 5.44E-06 | 0.14 | 0.05 | 0.09 | 2.8 |
| 8 | Anapc13 | 0.38 | 2.74E-10 | 5.85E-06 | 0.93 | 0.84 | 0.09 | 1.1 |
| 8 | Prrg3 | 1.11 | 2.76E-10 | 5.89E-06 | 0.31 | 0.16 | 0.15 | 1.9 |
| 8 | Kcnk1 | 0.51 | 2.80E-10 | 5.99E-06 | 0.72 | 0.58 | 0.14 | 1.25 |
| 8 | Fignl1 | 0.74 | 3.24E-10 | 6.92E-06 | 0.61 | 0.47 | 0.14 | 1.29 |
| 8 | Anxa5 | 0.47 | 3.33E-10 | 7.12E-06 | 0.79 | 0.62 | 0.17 | 1.28 |
| 8 | Jup | 0.58 | 3.55E-10 | 7.58E-06 | 0.67 | 0.51 | 0.16 | 1.32 |
| 8 | Snx31 | 1.8 | 3.99E-10 | 8.53E-06 | 0.12 | 0.04 | 0.08 | 3.16 |
| 8 | Ranbp1 | 0.26 | 4.01E-10 | 8.56E-06 | 1 | 0.99 | 0.02 | 1.02 |
| 8 | Id1 | 0.34 | 4.35E-10 | 9.30E-06 | 0.99 | 0.93 | 0.06 | 1.07 |
| 8 | Sdc4 | 0.48 | 4.43E-10 | 9.45E-06 | 0.79 | 0.64 | 0.15 | 1.23 |
| 8 | Sptlc3 | 1.09 | 4.48E-10 | 9.57E-06 | 0.18 | 0.07 | 0.11 | 2.53 |
| 8 | Cbx6 | 0.8 | 4.56E-10 | 9.75E-06 | 0.52 | 0.36 | 0.16 | 1.43 |
| 8 | Klc3 | 0.47 | 4.70E-10 | 1.00E-05 | 0.57 | 0.39 | 0.18 | 1.47 |
| 8 | Nr2f6 | 0.47 | 5.18E-10 | 1.11E-05 | 0.86 | 0.74 | 0.12 | 1.16 |
| 8 | Esrp1 | 0.48 | 6.16E-10 | 1.32E-05 | 0.77 | 0.64 | 0.14 | 1.22 |
| 8 | Cxxc4 | 0.74 | 6.32E-10 | 1.35E-05 | 0.4 | 0.25 | 0.16 | 1.64 |
| 8 | Tmem109 | 0.86 | 8.06E-10 | 1.72E-05 | 0.41 | 0.25 | 0.16 | 1.62 |
| 8 | Cebpa | 1.41 | 9.25E-10 | 1.98E-05 | 0.19 | 0.08 | 0.11 | 2.37 |
| 8 | Smarca5 | 0.25 | 9.33E-10 | 1.99E-05 | 0.98 | 0.97 | 0.01 | 1.01 |
| 8 | Erbb3 | 0.51 | 9.47E-10 | 2.02E-05 | 0.71 | 0.53 | 0.17 | 1.33 |
| 8 | Shisa2 | 1.19 | 9.62E-10 | 2.06E-05 | 0.22 | 0.1 | 0.12 | 2.14 |
| 8 | Larp7 | 0.42 | 9.67E-10 | 2.07E-05 | 0.88 | 0.77 | 0.11 | 1.14 |
| 8 | Nab2 | 1.04 | 1.10E-09 | 2.34E-05 | 0.34 | 0.2 | 0.15 | 1.74 |
| 8 | Fermt1 | 0.38 | 1.12E-09 | 2.40E-05 | 0.79 | 0.54 | 0.25 | 1.45 |
| 8 | Tln1 | 0.49 | 1.17E-09 | 2.50E-05 | 0.77 | 0.65 | 0.12 | 1.18 |
| 8 | Efna1 | 0.58 | 1.50E-09 | 3.20E-05 | 0.57 | 0.39 | 0.18 | 1.46 |

**Supplementary Table 3. Marker genes for the original clusters of integrated clustering of E11.5 control and NotchGOF λ-junction ectoderm cells.**

| | | | | | | | | |
|---|---|---|---|---|---|---|---|---|
| 8 | Slfn9 | 0.85 | 1.74E-09 | 3.71E-05 | 0.4 | 0.24 | 0.16 | 1.66 |
| 8 | Ifitm2 | 0.27 | 2.21E-09 | 4.73E-05 | 0.98 | 0.93 | 0.05 | 1.06 |
| 8 | Rab15 | 0.59 | 2.39E-09 | 5.11E-05 | 0.6 | 0.41 | 0.19 | 1.45 |
| 8 | Lmna | 0.88 | 2.59E-09 | 5.53E-05 | 0.33 | 0.18 | 0.15 | 1.81 |
| 8 | Amer1 | 0.51 | 2.65E-09 | 5.65E-05 | 0.72 | 0.58 | 0.14 | 1.25 |
| 8 | Usp43 | 1.53 | 2.85E-09 | 6.09E-05 | 0.14 | 0.05 | 0.09 | 2.69 |
| 8 | Isl1 | 0.31 | 2.93E-09 | 6.26E-05 | 0.92 | 0.75 | 0.18 | 1.23 |
| 8 | Hip1r | 0.96 | 3.00E-09 | 6.42E-05 | 0.36 | 0.21 | 0.15 | 1.71 |
| 8 | Slc44a2 | 0.69 | 3.01E-09 | 6.42E-05 | 0.54 | 0.39 | 0.15 | 1.39 |
| 8 | Hmg20b | 0.46 | 3.32E-09 | 7.09E-05 | 0.85 | 0.76 | 0.09 | 1.12 |
| 8 | Cxcl12 | 1.3 | 3.41E-09 | 7.29E-05 | 0.25 | 0.13 | 0.12 | 1.95 |
| 8 | Purb | 0.34 | 4.47E-09 | 9.55E-05 | 0.96 | 0.89 | 0.07 | 1.08 |
| 8 | Flywch2 | 0.65 | 4.64E-09 | 9.91E-05 | 0.62 | 0.48 | 0.14 | 1.28 |
| 8 | Aqp11 | 1.68 | 4.71E-09 | 1.01E-04 | 0.14 | 0.05 | 0.09 | 2.64 |
| 8 | Kras | 0.34 | 5.78E-09 | 1.24E-04 | 0.91 | 0.83 | 0.08 | 1.09 |
| 8 | Brwd3 | 0.62 | 5.98E-09 | 1.28E-04 | 0.65 | 0.5 | 0.14 | 1.29 |
| 8 | Stx3 | 0.63 | 6.06E-09 | 1.30E-04 | 0.59 | 0.43 | 0.16 | 1.38 |
| 8 | Dek | 0.31 | 6.18E-09 | 1.32E-04 | 0.99 | 0.93 | 0.07 | 1.07 |
| 8 | Cdc42ep5 | 0.49 | 6.80E-09 | 1.45E-04 | 0.73 | 0.59 | 0.15 | 1.25 |
| 8 | Hes1 | 0.35 | 7.60E-09 | 1.62E-04 | 0.86 | 0.72 | 0.13 | 1.19 |
| 8 | Lrrfip1 | 0.68 | 7.90E-09 | 1.69E-04 | 0.62 | 0.5 | 0.12 | 1.24 |
| 8 | Sh2d4a | 1.54 | 9.04E-09 | 1.93E-04 | 0.12 | 0.04 | 0.08 | 2.81 |
| 8 | Arhgap23 | 1.04 | 1.04E-08 | 2.22E-04 | 0.23 | 0.11 | 0.12 | 2.07 |
| 8 | Syngr1 | 0.93 | 1.05E-08 | 2.23E-04 | 0.36 | 0.22 | 0.14 | 1.67 |
| 8 | Emp1 | 0.46 | 1.09E-08 | 2.32E-04 | 0.31 | 0.17 | 0.15 | 1.86 |
| 8 | Tnk1 | 1.08 | 1.10E-08 | 2.35E-04 | 0.25 | 0.13 | 0.12 | 1.95 |
| 8 | Bmp8b | 1.83 | 1.15E-08 | 2.45E-04 | 0.1 | 0.03 | 0.07 | 3.09 |
| 8 | Ptprm | 1.42 | 1.27E-08 | 2.70E-04 | 0.2 | 0.1 | 0.11 | 2.08 |
| 8 | Pdcd4 | 0.31 | 1.33E-08 | 2.83E-04 | 0.91 | 0.78 | 0.13 | 1.17 |
| 8 | Col4a1 | 0.39 | 1.34E-08 | 2.86E-04 | 0.8 | 0.68 | 0.12 | 1.18 |
| 8 | Bex4 | 0.29 | 1.50E-08 | 3.20E-04 | 0.98 | 0.93 | 0.05 | 1.05 |
| 8 | Egr1 | 0.79 | 1.61E-08 | 3.44E-04 | 0.21 | 0.1 | 0.11 | 2.15 |
| 8 | Serpini1 | 1.2 | 1.62E-08 | 3.45E-04 | 0.12 | 0.04 | 0.08 | 2.81 |
| 8 | S100a1 | 1.48 | 1.63E-08 | 3.49E-04 | 0.15 | 0.06 | 0.09 | 2.44 |
| 8 | Aktip | 0.59 | 1.65E-08 | 3.53E-04 | 0.62 | 0.46 | 0.16 | 1.34 |
| 8 | Myzap | 1.59 | 1.66E-08 | 3.55E-04 | 0.12 | 0.04 | 0.08 | 2.85 |
| 8 | Tmem79 | 1.52 | 1.76E-08 | 3.76E-04 | 0.17 | 0.08 | 0.1 | 2.27 |
| 8 | Cntnap2 | 1.06 | 1.95E-08 | 4.16E-04 | 0.26 | 0.14 | 0.12 | 1.85 |
| 8 | Ccser1 | 1.23 | 2.03E-08 | 4.33E-04 | 0.17 | 0.07 | 0.1 | 2.33 |
| 8 | Rnh1 | 0.36 | 2.15E-08 | 4.58E-04 | 0.9 | 0.77 | 0.13 | 1.16 |
| 8 | Tuft1 | 1.14 | 2.26E-08 | 4.82E-04 | 0.17 | 0.07 | 0.1 | 2.33 |
| 8 | Dock9 | 0.95 | 3.00E-08 | 6.40E-04 | 0.33 | 0.19 | 0.13 | 1.68 |
| 8 | Dap | 0.49 | 3.29E-08 | 7.02E-04 | 0.78 | 0.68 | 0.1 | 1.15 |
| 8 | Dag1 | 0.5 | 3.31E-08 | 7.08E-04 | 0.73 | 0.63 | 0.1 | 1.16 |
| 8 | H2afj | 0.3 | 4.19E-08 | 8.94E-04 | 0.96 | 0.9 | 0.06 | 1.07 |
| 8 | Pkp4 | 0.55 | 4.19E-08 | 8.94E-04 | 0.7 | 0.58 | 0.12 | 1.21 |
| 8 | Llgl2 | 0.86 | 4.40E-08 | 9.40E-04 | 0.31 | 0.18 | 0.13 | 1.74 |
| 8 | Car13 | 1.76 | 4.51E-08 | 9.64E-04 | 0.12 | 0.04 | 0.07 | 2.66 |
| 8 | Inf2 | 1.53 | 4.53E-08 | 9.68E-04 | 0.11 | 0.04 | 0.07 | 2.79 |
| 8 | Impdh1 | 0.97 | 4.54E-08 | 9.71E-04 | 0.29 | 0.17 | 0.12 | 1.73 |
| 8 | Cldn6 | 0.29 | 4.57E-08 | 9.76E-04 | 0.96 | 0.87 | 0.09 | 1.1 |
| 8 | Rab14 | 0.25 | 5.13E-08 | 1.10E-03 | 0.94 | 0.93 | 0.01 | 1.01 |
| 8 | Mark2 | 0.49 | 5.38E-08 | 1.15E-03 | 0.72 | 0.58 | 0.14 | 1.25 |
| 8 | Irf6 | 0.27 | 5.79E-08 | 1.24E-03 | 0.62 | 0.4 | 0.21 | 1.52 |
| 8 | Erbin | 0.47 | 5.83E-08 | 1.25E-03 | 0.71 | 0.56 | 0.14 | 1.25 |
| 8 | Tmem54 | 0.63 | 6.87E-08 | 1.47E-03 | 0.61 | 0.47 | 0.14 | 1.29 |
| 8 | Pcbd1 | 0.88 | 7.37E-08 | 1.57E-03 | 0.36 | 0.23 | 0.13 | 1.58 |

**Supplementary Table 3. Marker genes for the original clusters of integrated clustering of E11.5 control and NotchGOF λ-junction ectoderm cells.**

| | | | | | | | | |
|---|---|---|---|---|---|---|---|---|
| 8 | Lypd3 | 1.02 | 7.63E-08 | 1.63E-03 | 0.21 | 0.1 | 0.11 | 2.06 |
| 8 | Morc4 | 0.99 | 7.83E-08 | 1.67E-03 | 0.26 | 0.14 | 0.12 | 1.84 |
| 8 | Tfap2c | 0.69 | 7.89E-08 | 1.69E-03 | 0.36 | 0.21 | 0.14 | 1.66 |
| 8 | Prkci | 0.59 | 8.03E-08 | 1.72E-03 | 0.57 | 0.43 | 0.14 | 1.33 |
| 8 | Nabp1 | 1.52 | 8.06E-08 | 1.72E-03 | 0.14 | 0.06 | 0.08 | 2.41 |
| 8 | Xpa | 0.57 | 8.44E-08 | 1.80E-03 | 0.63 | 0.47 | 0.16 | 1.33 |
| 8 | Tead1 | 0.37 | 9.03E-08 | 1.93E-03 | 0.86 | 0.76 | 0.1 | 1.12 |
| 8 | Coro1c | 0.66 | 9.60E-08 | 2.05E-03 | 0.46 | 0.32 | 0.14 | 1.43 |
| 8 | Mafb | 0.71 | 1.00E-07 | 2.15E-03 | 0.28 | 0.15 | 0.13 | 1.84 |
| 8 | Msh2 | 0.45 | 1.02E-07 | 2.18E-03 | 0.79 | 0.69 | 0.1 | 1.14 |
| 8 | Bcat1 | 1.15 | 1.03E-07 | 2.21E-03 | 0.24 | 0.13 | 0.11 | 1.86 |
| 8 | Reep6 | 1.46 | 1.04E-07 | 2.23E-03 | 0.17 | 0.08 | 0.09 | 2.12 |
| 8 | Etl4 | 0.43 | 1.11E-07 | 2.37E-03 | 0.71 | 0.57 | 0.14 | 1.24 |
| 8 | Tnfaip8 | 0.48 | 1.12E-07 | 2.39E-03 | 0.71 | 0.56 | 0.15 | 1.27 |
| 8 | Plpp2 | 0.59 | 1.34E-07 | 2.85E-03 | 0.53 | 0.37 | 0.15 | 1.42 |
| 8 | Efna5 | 0.5 | 1.35E-07 | 2.88E-03 | 0.65 | 0.52 | 0.13 | 1.25 |
| 8 | Bcl2l11 | 0.48 | 1.45E-07 | 3.09E-03 | 0.53 | 0.38 | 0.15 | 1.4 |
| 8 | Nynrin | 0.54 | 1.45E-07 | 3.10E-03 | 0.66 | 0.53 | 0.13 | 1.25 |
| 8 | Capn1 | 1.61 | 1.54E-07 | 3.29E-03 | 0.12 | 0.05 | 0.07 | 2.6 |
| 8 | Fam3c | 0.55 | 1.56E-07 | 3.33E-03 | 0.59 | 0.47 | 0.12 | 1.27 |
| 8 | Adamts15 | 1.53 | 1.66E-07 | 3.54E-03 | 0.11 | 0.04 | 0.07 | 2.66 |
| 8 | Synj2 | 1.72 | 1.67E-07 | 3.56E-03 | 0.13 | 0.05 | 0.08 | 2.5 |
| 8 | Creb3l2 | 0.57 | 1.96E-07 | 4.19E-03 | 0.47 | 0.32 | 0.15 | 1.48 |
| 8 | Shroom4 | 1.32 | 2.00E-07 | 4.27E-03 | 0.14 | 0.06 | 0.08 | 2.37 |
| 8 | Ajuba | 0.56 | 2.05E-07 | 4.39E-03 | 0.57 | 0.42 | 0.15 | 1.35 |
| 8 | Reep5 | 0.43 | 2.09E-07 | 4.46E-03 | 0.69 | 0.55 | 0.13 | 1.24 |
| 8 | Naa38 | 0.38 | 2.10E-07 | 4.49E-03 | 0.85 | 0.76 | 0.1 | 1.13 |
| 8 | Arhgap8 | 1.07 | 2.12E-07 | 4.54E-03 | 0.27 | 0.16 | 0.11 | 1.72 |
| 8 | Tagln2 | 0.7 | 2.31E-07 | 4.93E-03 | 0.41 | 0.27 | 0.14 | 1.51 |
| 8 | Mdc1 | 0.62 | 2.57E-07 | 5.49E-03 | 0.51 | 0.37 | 0.14 | 1.39 |
| 8 | Nbeal1 | 0.51 | 2.80E-07 | 5.98E-03 | 0.62 | 0.48 | 0.14 | 1.3 |
| 8 | Cyb5r1 | 0.69 | 2.84E-07 | 6.06E-03 | 0.42 | 0.28 | 0.14 | 1.49 |
| 8 | Tjp1 | 0.44 | 2.86E-07 | 6.12E-03 | 0.74 | 0.63 | 0.11 | 1.17 |
| 8 | Sh3bgrl2 | 1.43 | 2.95E-07 | 6.30E-03 | 0.16 | 0.08 | 0.09 | 2.13 |
| 8 | Arhgap18 | 0.82 | 2.97E-07 | 6.35E-03 | 0.33 | 0.2 | 0.13 | 1.65 |
| 8 | Epb41l4b | 1.43 | 3.10E-07 | 6.62E-03 | 0.19 | 0.1 | 0.1 | 2 |
| 8 | Capn2 | 0.62 | 3.15E-07 | 6.72E-03 | 0.54 | 0.39 | 0.15 | 1.37 |
| 8 | Zfp711 | 0.76 | 3.34E-07 | 7.13E-03 | 0.33 | 0.19 | 0.13 | 1.68 |
| 8 | Fam83h | 1.27 | 3.47E-07 | 7.41E-03 | 0.18 | 0.09 | 0.09 | 2.06 |
| 8 | Mprip | 0.36 | 3.55E-07 | 7.59E-03 | 0.8 | 0.73 | 0.07 | 1.1 |
| 8 | Tk1 | 0.5 | 3.57E-07 | 7.62E-03 | 0.6 | 0.45 | 0.15 | 1.35 |
| 8 | Cryl1 | 0.78 | 3.72E-07 | 7.95E-03 | 0.4 | 0.27 | 0.13 | 1.5 |
| 8 | Dnmt1 | 0.38 | 4.00E-07 | 8.54E-03 | 0.83 | 0.73 | 0.1 | 1.14 |
| 8 | Fam241a | 1.12 | 4.28E-07 | 9.15E-03 | 0.2 | 0.11 | 0.1 | 1.94 |
| 8 | Thrap3 | 0.3 | 4.32E-07 | 9.23E-03 | 0.96 | 0.92 | 0.04 | 1.04 |
| 8 | Mme | 0.68 | 4.47E-07 | 9.55E-03 | 0.3 | 0.18 | 0.12 | 1.69 |
| 8 | Snx5 | 0.28 | 4.59E-07 | 9.80E-03 | 0.89 | 0.82 | 0.07 | 1.09 |
| 8 | Sestd1 | 0.58 | 4.78E-07 | 1.02E-02 | 0.59 | 0.46 | 0.13 | 1.28 |
| 8 | Tmed2 | 0.25 | 4.80E-07 | 1.02E-02 | 0.96 | 0.92 | 0.04 | 1.04 |
| 8 | Cenpb | 0.39 | 4.96E-07 | 1.06E-02 | 0.8 | 0.7 | 0.1 | 1.15 |
| 8 | Plagl2 | 0.61 | 5.17E-07 | 1.10E-02 | 0.59 | 0.47 | 0.12 | 1.25 |
| 8 | Jarid2 | 0.45 | 5.34E-07 | 1.14E-02 | 0.66 | 0.54 | 0.11 | 1.21 |
| 8 | Fzd1 | 0.91 | 5.57E-07 | 1.19E-02 | 0.29 | 0.17 | 0.12 | 1.7 |
| 8 | Plcl1 | 1.12 | 6.12E-07 | 1.31E-02 | 0.11 | 0.04 | 0.07 | 2.65 |
| 8 | Stk26 | 0.46 | 6.32E-07 | 1.35E-02 | 0.63 | 0.46 | 0.16 | 1.35 |
| 8 | Six6 | 0.59 | 6.37E-07 | 1.36E-02 | 0.42 | 0.27 | 0.14 | 1.51 |
| 8 | Trim29 | 0.62 | 6.72E-07 | 1.44E-02 | 0.32 | 0.18 | 0.13 | 1.72 |

**Supplementary Table 3. Marker genes for the original clusters of integrated clustering of E11.5 control and NotchGOF λ-junction ectoderm cells.**

| | | | | | | | | |
|---|---|---|---|---|---|---|---|---|
| 8 | 9130017K11Rik | 1.8 | 6.78E-07 | 1.45E-02 | 0.1 | 0.04 | 0.06 | 2.62 |
| 8 | Tgm1 | 1.11 | 6.93E-07 | 1.48E-02 | 0.12 | 0.05 | 0.07 | 2.47 |
| 8 | Colgalt1 | 0.56 | 7.02E-07 | 1.50E-02 | 0.59 | 0.45 | 0.13 | 1.29 |
| 8 | Tbc1d4 | 1.62 | 7.30E-07 | 1.56E-02 | 0.18 | 0.09 | 0.09 | 1.95 |
| 8 | Capn6 | 0.43 | 8.02E-07 | 1.71E-02 | 0.59 | 0.44 | 0.15 | 1.34 |
| 8 | Ccdc18 | 0.69 | 8.91E-07 | 1.90E-02 | 0.51 | 0.38 | 0.13 | 1.34 |
| 8 | Fn1 | 0.38 | 9.21E-07 | 1.97E-02 | 0.7 | 0.58 | 0.12 | 1.2 |
| 8 | Mcm5 | 0.38 | 9.42E-07 | 2.01E-02 | 0.77 | 0.63 | 0.15 | 1.23 |
| 8 | Timeless | 0.57 | 9.74E-07 | 2.08E-02 | 0.59 | 0.46 | 0.13 | 1.28 |
| 8 | Smagp | 1.34 | 9.76E-07 | 2.08E-02 | 0.13 | 0.06 | 0.08 | 2.32 |
| 8 | Timp2 | 0.54 | 1.00E-06 | 2.14E-02 | 0.64 | 0.51 | 0.14 | 1.27 |
| 8 | Ssna1 | 0.29 | 1.15E-06 | 2.46E-02 | 0.89 | 0.82 | 0.07 | 1.09 |
| 8 | Pik3cb | 1 | 1.23E-06 | 2.62E-02 | 0.25 | 0.15 | 0.11 | 1.72 |
| 8 | Hadh | 0.32 | 1.25E-06 | 2.66E-02 | 0.85 | 0.76 | 0.09 | 1.12 |
| 8 | Atic | 0.36 | 1.25E-06 | 2.68E-02 | 0.82 | 0.68 | 0.14 | 1.2 |
| 8 | Tcim | 0.89 | 1.27E-06 | 2.70E-02 | 0.39 | 0.26 | 0.12 | 1.46 |
| 8 | Shtn1 | 0.58 | 1.28E-06 | 2.73E-02 | 0.31 | 0.2 | 0.12 | 1.59 |
| 8 | Elovl6 | 0.49 | 1.29E-06 | 2.75E-02 | 0.64 | 0.52 | 0.12 | 1.24 |
| 8 | Hdhd3 | 0.83 | 1.31E-06 | 2.80E-02 | 0.33 | 0.21 | 0.12 | 1.55 |
| 8 | Msn | 0.89 | 1.38E-06 | 2.94E-02 | 0.25 | 0.14 | 0.1 | 1.73 |
| 8 | Nrarp | 0.29 | 1.55E-06 | 3.32E-02 | 0.77 | 0.6 | 0.18 | 1.29 |
| 8 | Spc24 | 0.4 | 1.57E-06 | 3.35E-02 | 0.8 | 0.65 | 0.14 | 1.22 |
| 8 | Wsb1 | 0.38 | 1.59E-06 | 3.40E-02 | 0.76 | 0.66 | 0.1 | 1.16 |
| 8 | St14 | 0.6 | 1.64E-06 | 3.49E-02 | 0.59 | 0.47 | 0.11 | 1.23 |
| 8 | Ifit2 | 1.34 | 1.64E-06 | 3.50E-02 | 0.11 | 0.05 | 0.07 | 2.4 |
| 8 | Sh3rf1 | 0.57 | 1.66E-06 | 3.55E-02 | 0.46 | 0.32 | 0.14 | 1.43 |
| 8 | Ccdc112 | 0.4 | 1.75E-06 | 3.73E-02 | 0.75 | 0.64 | 0.11 | 1.17 |
| 8 | Fzd8 | 0.49 | 1.79E-06 | 3.83E-02 | 0.43 | 0.3 | 0.14 | 1.47 |
| 8 | Orc1 | 0.99 | 1.91E-06 | 4.07E-02 | 0.21 | 0.12 | 0.1 | 1.83 |
| 8 | Map3k20 | 0.83 | 1.93E-06 | 4.13E-02 | 0.29 | 0.18 | 0.11 | 1.62 |
| 8 | Metrnl | 0.75 | 1.96E-06 | 4.19E-02 | 0.29 | 0.17 | 0.12 | 1.68 |
| 8 | Pla2g4a | 0.92 | 1.97E-06 | 4.22E-02 | 0.19 | 0.1 | 0.09 | 1.94 |
| 8 | Ahdc1 | 0.7 | 1.98E-06 | 4.22E-02 | 0.34 | 0.22 | 0.12 | 1.54 |
| 8 | Sh3d19 | 0.52 | 2.18E-06 | 4.65E-02 | 0.48 | 0.34 | 0.15 | 1.43 |
| 8 | Fkbp9 | 0.57 | 2.28E-06 | 4.87E-02 | 0.54 | 0.42 | 0.12 | 1.28 |
| 9 | Fgf8 | 3.79 | 5.37E-150 | 1.15E-145 | 0.88 | 0.24 | 0.63 | 3.62 |
| 9 | Cdkn2b | 4.19 | 4.87E-142 | 1.04E-137 | 0.55 | 0.07 | 0.48 | 7.73 |
| 9 | Wnt5a | 2.96 | 1.19E-132 | 2.54E-128 | 0.95 | 0.4 | 0.55 | 2.38 |
| 9 | Gad1 | 3.71 | 6.40E-129 | 1.37E-124 | 0.4 | 0.04 | 0.37 | 11.22 |
| 9 | Cldn1 | 3.75 | 5.18E-126 | 1.11E-121 | 0.54 | 0.08 | 0.46 | 6.97 |
| 9 | Cdkn1a | 3.41 | 3.63E-122 | 7.75E-118 | 0.91 | 0.36 | 0.55 | 2.54 |
| 9 | Itm2b | 2.47 | 9.75E-121 | 2.08E-116 | 1 | 0.93 | 0.07 | 1.07 |
| 9 | Runx1 | 2.91 | 1.47E-119 | 3.14E-115 | 0.96 | 0.49 | 0.47 | 1.96 |
| 9 | Gcg | 6.74 | 4.80E-118 | 1.03E-113 | 0.19 | 0 | 0.19 | 48 |
| 9 | Inhba | 3.84 | 3.46E-114 | 7.39E-110 | 0.48 | 0.07 | 0.41 | 7.24 |
| 9 | Fst | 4.85 | 5.94E-114 | 1.27E-109 | 0.4 | 0.04 | 0.36 | 9.18 |
| 9 | Msx1 | 2.25 | 6.25E-108 | 1.33E-103 | 0.99 | 0.72 | 0.28 | 1.38 |
| 9 | Tgfb1 | 2.81 | 1.67E-105 | 3.57E-101 | 0.64 | 0.14 | 0.5 | 4.64 |
| 9 | Cdkn2a | 4.96 | 2.32E-102 | 4.95E-98 | 0.27 | 0.02 | 0.25 | 15.06 |
| 9 | Gadd45g | 3.34 | 2.90E-99 | 6.19E-95 | 0.81 | 0.32 | 0.5 | 2.58 |
| 9 | Fabp5 | 1.53 | 6.22E-97 | 1.33E-92 | 1 | 0.99 | 0.01 | 1.01 |
| 9 | Notum | 3.54 | 9.39E-95 | 2.01E-90 | 0.38 | 0.05 | 0.34 | 8 |
| 9 | Pkp1 | 3.05 | 1.85E-92 | 3.95E-88 | 0.47 | 0.08 | 0.39 | 5.96 |
| 9 | Bambi | 3.13 | 1.68E-90 | 3.58E-86 | 0.88 | 0.46 | 0.41 | 1.89 |
| 9 | Gja1 | 2.03 | 7.33E-89 | 1.57E-84 | 0.97 | 0.69 | 0.28 | 1.41 |
| 9 | Tgfb2 | 2.67 | 7.62E-89 | 1.63E-84 | 0.94 | 0.56 | 0.38 | 1.67 |
| 9 | Zfp385c | 2.97 | 6.71E-85 | 1.43E-80 | 0.5 | 0.1 | 0.4 | 4.98 |

**Supplementary Table 3. Marker genes for the original clusters of integrated clustering of E11.5 control and NotchGOF λ-junction ectoderm cells.**

| | | | | | | | | |
|---|---|---|---|---|---|---|---|---|
| 9 | Lmo2 | 2.39 | 1.35E-84 | 2.89E-80 | 0.76 | 0.27 | 0.49 | 2.81 |
| 9 | Sfn | 1.81 | 5.12E-84 | 1.09E-79 | 0.99 | 0.69 | 0.31 | 1.45 |
| 9 | Sp5 | 4.52 | 6.34E-84 | 1.35E-79 | 0.26 | 0.02 | 0.24 | 11.77 |
| 9 | Lratd1 | 3.05 | 9.60E-82 | 2.05E-77 | 0.51 | 0.11 | 0.4 | 4.77 |
| 9 | Lgalsl | 2.6 | 5.70E-80 | 1.22E-75 | 0.84 | 0.45 | 0.4 | 1.89 |
| 9 | Igfbp5 | 1.67 | 2.53E-79 | 5.40E-75 | 1 | 0.96 | 0.04 | 1.04 |
| 9 | Dlx2 | 2.12 | 7.10E-78 | 1.52E-73 | 0.82 | 0.32 | 0.51 | 2.6 |
| 9 | Aopep | 1.94 | 5.60E-76 | 1.20E-71 | 0.92 | 0.6 | 0.32 | 1.54 |
| 9 | Atxn1 | 3.35 | 3.15E-75 | 6.72E-71 | 0.45 | 0.09 | 0.36 | 4.91 |
| 9 | Gpr87 | 4.24 | 3.25E-75 | 6.95E-71 | 0.24 | 0.02 | 0.22 | 11.38 |
| 9 | Gal | 5.71 | 1.90E-74 | 4.06E-70 | 0.23 | 0.02 | 0.21 | 11.55 |
| 9 | Foxn3 | 1.73 | 5.89E-74 | 1.26E-69 | 0.93 | 0.7 | 0.22 | 1.31 |
| 9 | Glcci1 | 2.3 | 6.11E-74 | 1.31E-69 | 0.86 | 0.48 | 0.38 | 1.78 |
| 9 | S100a11 | 1.1 | 9.75E-71 | 2.08E-66 | 1 | 0.9 | 0.1 | 1.11 |
| 9 | Ambn | 7.85 | 7.27E-69 | 1.55E-64 | 0.11 | 0 | 0.11 | 55 |
| 9 | Pik3r1 | 1.82 | 1.41E-68 | 3.01E-64 | 0.92 | 0.58 | 0.35 | 1.6 |
| 9 | Adamts9 | 2.74 | 3.30E-66 | 7.05E-62 | 0.64 | 0.24 | 0.41 | 2.74 |
| 9 | Krt14 | 1.97 | 9.29E-66 | 1.98E-61 | 0.88 | 0.44 | 0.44 | 2 |
| 9 | Lef1 | 2.21 | 9.99E-66 | 2.13E-61 | 0.77 | 0.36 | 0.41 | 2.16 |
| 9 | Trp63 | 1.51 | 4.35E-65 | 9.29E-61 | 0.93 | 0.49 | 0.44 | 1.91 |
| 9 | Tspan7 | 1.9 | 2.22E-64 | 4.74E-60 | 0.82 | 0.45 | 0.37 | 1.81 |
| 9 | Rb1 | 2.28 | 6.48E-64 | 1.38E-59 | 0.68 | 0.27 | 0.42 | 2.55 |
| 9 | Cd2ap | 1.4 | 8.34E-64 | 1.78E-59 | 0.96 | 0.8 | 0.16 | 1.21 |
| 9 | Tafa2 | 2.71 | 5.57E-60 | 1.19E-55 | 0.45 | 0.11 | 0.34 | 4.14 |
| 9 | Baiap2 | 2.45 | 1.98E-58 | 4.23E-54 | 0.57 | 0.2 | 0.38 | 2.92 |
| 9 | Krt5 | 2.07 | 2.12E-58 | 4.53E-54 | 0.73 | 0.32 | 0.42 | 2.32 |
| 9 | Spock2 | 3.47 | 6.53E-58 | 1.40E-53 | 0.44 | 0.11 | 0.33 | 3.99 |
| 9 | Sox11 | 0.96 | 2.16E-57 | 4.62E-53 | 1 | 0.99 | 0 | 1 |
| 9 | Wnt6 | 1.32 | 3.85E-57 | 8.23E-53 | 0.84 | 0.35 | 0.49 | 2.43 |
| 9 | Clta | 0.89 | 2.22E-56 | 4.75E-52 | 0.99 | 0.98 | 0.01 | 1.01 |
| 9 | Eef1a1 | 0.38 | 9.36E-56 | 2.00E-51 | 1 | 1 | 0 | 1 |
| 9 | Diras2 | 2.99 | 7.52E-55 | 1.61E-50 | 0.36 | 0.07 | 0.28 | 4.82 |
| 9 | Epha4 | 1.51 | 1.49E-54 | 3.17E-50 | 0.87 | 0.54 | 0.33 | 1.61 |
| 9 | Krt28 | 6.43 | 2.24E-53 | 4.78E-49 | 0.11 | 0 | 0.1 | 26.5 |
| 9 | Tmtc1 | 2.81 | 2.81E-53 | 6.00E-49 | 0.29 | 0.05 | 0.24 | 5.92 |
| 9 | Rfk | 1.61 | 6.03E-52 | 1.29E-47 | 0.85 | 0.5 | 0.35 | 1.69 |
| 9 | Fam49a | 2.8 | 3.27E-51 | 6.99E-47 | 0.37 | 0.09 | 0.28 | 4.34 |
| 9 | Sat1 | 2.25 | 2.50E-50 | 5.34E-46 | 0.64 | 0.28 | 0.36 | 2.26 |
| 9 | Ly6h | 1.8 | 8.47E-50 | 1.81E-45 | 0.8 | 0.48 | 0.32 | 1.67 |
| 9 | Frmd6 | 2.22 | 1.08E-49 | 2.31E-45 | 0.57 | 0.21 | 0.36 | 2.67 |
| 9 | Dcbld1 | 1.85 | 2.53E-49 | 5.39E-45 | 0.72 | 0.38 | 0.35 | 1.93 |
| 9 | Palld | 1.49 | 2.93E-49 | 6.26E-45 | 0.85 | 0.63 | 0.22 | 1.34 |
| 9 | Arid5b | 1.62 | 5.48E-47 | 1.17E-42 | 0.81 | 0.54 | 0.27 | 1.5 |
| 9 | Rpl7 | 0.37 | 6.77E-47 | 1.45E-42 | 1 | 1 | 0 | 1 |
| 9 | Rasl11b | 1.93 | 8.42E-47 | 1.80E-42 | 0.62 | 0.26 | 0.37 | 2.42 |
| 9 | Dkk4 | 4.05 | 2.50E-46 | 5.34E-42 | 0.18 | 0.02 | 0.16 | 8.8 |
| 9 | Nav2 | 1.22 | 7.41E-46 | 1.58E-41 | 0.9 | 0.69 | 0.21 | 1.3 |
| 9 | 5033430I15Rik | 3 | 8.14E-46 | 1.74E-41 | 0.31 | 0.07 | 0.25 | 4.55 |
| 9 | Ldlrad4 | 2.97 | 4.48E-45 | 9.58E-41 | 0.24 | 0.04 | 0.2 | 6.13 |
| 9 | Dlx1 | 1.96 | 3.76E-44 | 8.04E-40 | 0.44 | 0.13 | 0.31 | 3.43 |
| 9 | Lurap1l | 2.04 | 5.18E-43 | 1.11E-38 | 0.58 | 0.24 | 0.34 | 2.38 |
| 9 | Slc16a10 | 4.29 | 1.22E-42 | 2.61E-38 | 0.16 | 0.02 | 0.14 | 8.94 |
| 9 | Pnldc1 | 2.86 | 1.72E-41 | 3.67E-37 | 0.24 | 0.04 | 0.2 | 5.56 |
| 9 | Dsp | 0.99 | 4.23E-41 | 9.04E-37 | 0.98 | 0.86 | 0.11 | 1.13 |
| 9 | Insyn2b | 2.49 | 6.74E-41 | 1.44E-36 | 0.28 | 0.06 | 0.22 | 4.86 |
| 9 | Smad7 | 1.26 | 1.21E-40 | 2.58E-36 | 0.86 | 0.6 | 0.26 | 1.44 |
| 9 | Plce1 | 1.87 | 1.59E-40 | 3.39E-36 | 0.64 | 0.31 | 0.33 | 2.05 |

**Supplementary Table 3. Marker genes for the original clusters of integrated clustering of E11.5 control and NotchGOF λ-junction ectoderm cells.**

| | | | | | | | | |
|---|---|---|---|---|---|---|---|---|
| 9 | Tagln | 2.9 | 2.47E-40 | 5.29E-36 | 0.23 | 0.04 | 0.19 | 5.68 |
| 9 | Kdm5b | 0.83 | 5.44E-40 | 1.16E-35 | 0.97 | 0.92 | 0.05 | 1.05 |
| 9 | Apcdd1 | 2.27 | 7.30E-40 | 1.56E-35 | 0.41 | 0.13 | 0.29 | 3.27 |
| 9 | Ptprz1 | 2.05 | 7.66E-40 | 1.64E-35 | 0.52 | 0.2 | 0.32 | 2.56 |
| 9 | Ptn | 1.41 | 9.02E-40 | 1.93E-35 | 0.94 | 0.86 | 0.08 | 1.09 |
| 9 | Retreg1 | 1.93 | 9.54E-40 | 2.04E-35 | 0.55 | 0.24 | 0.32 | 2.33 |
| 9 | Npm3 | 0.98 | 1.79E-39 | 3.83E-35 | 0.92 | 0.87 | 0.05 | 1.05 |
| 9 | Ermp1 | 1.45 | 2.43E-39 | 5.18E-35 | 0.73 | 0.41 | 0.32 | 1.77 |
| 9 | Fam89a | 2.04 | 2.43E-39 | 5.18E-35 | 0.44 | 0.15 | 0.3 | 3.06 |
| 9 | Parm1 | 1.88 | 3.62E-39 | 7.74E-35 | 0.5 | 0.19 | 0.31 | 2.64 |
| 9 | Aspscr1 | 1.54 | 4.08E-39 | 8.72E-35 | 0.76 | 0.5 | 0.25 | 1.5 |
| 9 | Amtn | 4.21 | 4.78E-39 | 1.02E-34 | 0.1 | 0.01 | 0.1 | 14.57 |
| 9 | Malat1 | 0.68 | 4.90E-39 | 1.05E-34 | 1 | 1 | 0 | 1 |
| 9 | Pmepa1 | 1.84 | 7.69E-39 | 1.64E-34 | 0.66 | 0.35 | 0.3 | 1.87 |
| 9 | Me1 | 2.11 | 9.61E-39 | 2.05E-34 | 0.48 | 0.17 | 0.31 | 2.77 |
| 9 | Mbnl2 | 1.03 | 2.40E-38 | 5.12E-34 | 0.93 | 0.78 | 0.15 | 1.19 |
| 9 | Amotl1 | 1.21 | 3.73E-38 | 7.97E-34 | 0.87 | 0.69 | 0.18 | 1.26 |
| 9 | Erc2 | 2.61 | 5.95E-38 | 1.27E-33 | 0.36 | 0.1 | 0.26 | 3.47 |
| 9 | Tpm1 | 1.06 | 6.56E-38 | 1.40E-33 | 0.97 | 0.93 | 0.04 | 1.05 |
| 9 | Peg3 | 1.21 | 9.59E-38 | 2.05E-33 | 0.93 | 0.82 | 0.11 | 1.14 |
| 9 | Saraf | 1.04 | 9.95E-38 | 2.13E-33 | 0.89 | 0.65 | 0.25 | 1.38 |
| 9 | Cbr3 | 3.81 | 1.30E-37 | 2.77E-33 | 0.13 | 0.01 | 0.11 | 10.42 |
| 9 | Hspb8 | 1.31 | 1.42E-37 | 3.04E-33 | 0.79 | 0.47 | 0.32 | 1.68 |
| 9 | Cux1 | 0.91 | 2.68E-37 | 5.73E-33 | 0.94 | 0.84 | 0.1 | 1.12 |
| 9 | Abracl | 0.79 | 1.14E-36 | 2.44E-32 | 0.96 | 0.92 | 0.04 | 1.04 |
| 9 | Perp | 0.97 | 2.12E-36 | 4.54E-32 | 0.9 | 0.52 | 0.38 | 1.72 |
| 9 | Zfhx3 | 0.74 | 2.31E-36 | 4.94E-32 | 0.99 | 0.92 | 0.07 | 1.08 |
| 9 | Proser2 | 2.02 | 2.50E-36 | 5.34E-32 | 0.42 | 0.14 | 0.28 | 2.99 |
| 9 | Pdcd4 | 1.06 | 3.02E-36 | 6.46E-32 | 0.93 | 0.78 | 0.15 | 1.19 |
| 9 | Nrip1 | 1.09 | 7.82E-36 | 1.67E-31 | 0.94 | 0.81 | 0.13 | 1.16 |
| 9 | Trp53inp1 | 1.82 | 1.12E-35 | 2.39E-31 | 0.55 | 0.24 | 0.31 | 2.27 |
| 9 | Pdgfrl | 1.71 | 2.61E-35 | 5.58E-31 | 0.62 | 0.32 | 0.31 | 1.97 |
| 9 | Pcdh9 | 1.91 | 4.38E-35 | 9.37E-31 | 0.46 | 0.17 | 0.29 | 2.7 |
| 9 | Rps27 | 0.31 | 1.02E-34 | 2.19E-30 | 1 | 1 | 0 | 1 |
| 9 | Cbr1 | 1.2 | 1.36E-34 | 2.90E-30 | 0.89 | 0.72 | 0.17 | 1.24 |
| 9 | Zfp503 | 1.58 | 2.72E-34 | 5.81E-30 | 0.75 | 0.49 | 0.25 | 1.51 |
| 9 | H3f3b | 0.39 | 3.03E-34 | 6.48E-30 | 1 | 1 | 0 | 1 |
| 9 | Dact1 | 2.52 | 4.04E-34 | 8.64E-30 | 0.39 | 0.13 | 0.26 | 2.94 |
| 9 | Hras | 0.65 | 4.82E-34 | 1.03E-29 | 0.98 | 0.9 | 0.08 | 1.09 |
| 9 | Rpl18a | 0.27 | 5.37E-34 | 1.15E-29 | 1 | 1 | 0 | 1 |
| 9 | 6330403K07Rik | 0.95 | 6.70E-34 | 1.43E-29 | 0.93 | 0.85 | 0.09 | 1.1 |
| 9 | Klf4 | 2.15 | 7.07E-34 | 1.51E-29 | 0.46 | 0.18 | 0.28 | 2.54 |
| 9 | Skil | 1.35 | 7.87E-34 | 1.68E-29 | 0.79 | 0.56 | 0.23 | 1.41 |
| 9 | Eya2 | 1.18 | 8.03E-34 | 1.71E-29 | 0.68 | 0.35 | 0.33 | 1.93 |
| 9 | Fau | 0.26 | 8.46E-34 | 1.81E-29 | 1 | 1 | 0 | 1 |
| 9 | Ptbp3 | 1.26 | 1.12E-33 | 2.40E-29 | 0.83 | 0.68 | 0.14 | 1.21 |
| 9 | Ctnnb1 | 0.59 | 1.64E-33 | 3.50E-29 | 0.99 | 0.99 | 0.01 | 1.01 |
| 9 | Fxyd3 | 1.4 | 2.78E-33 | 5.93E-29 | 0.63 | 0.29 | 0.34 | 2.17 |
| 9 | Cald1 | 1.28 | 7.32E-33 | 1.56E-28 | 0.91 | 0.69 | 0.22 | 1.31 |
| 9 | Gad2 | 1.87 | 7.72E-33 | 1.65E-28 | 0.57 | 0.28 | 0.29 | 2.06 |
| 9 | Itgb5 | 1.1 | 1.23E-32 | 2.62E-28 | 0.78 | 0.48 | 0.3 | 1.61 |
| 9 | Ier5 | 0.87 | 4.15E-32 | 8.87E-28 | 0.95 | 0.84 | 0.12 | 1.14 |
| 9 | Irf6 | 1.23 | 4.84E-32 | 1.03E-27 | 0.72 | 0.4 | 0.33 | 1.82 |
| 9 | Pcdh17 | 1.61 | 5.41E-32 | 1.16E-27 | 0.66 | 0.36 | 0.3 | 1.85 |
| 9 | Vcan | 1.18 | 1.25E-31 | 2.67E-27 | 0.92 | 0.74 | 0.18 | 1.24 |
| 9 | Pnrc1 | 0.74 | 1.29E-31 | 2.75E-27 | 0.97 | 0.9 | 0.06 | 1.07 |
| 9 | Klhl24 | 1.08 | 1.96E-31 | 4.18E-27 | 0.85 | 0.63 | 0.22 | 1.36 |

**Supplementary Table 3. Marker genes for the original clusters of integrated clustering of E11.5 control and NotchGOF λ-junction ectoderm cells.**

| | | | | | | | | |
|---|---|---|---|---|---|---|---|---|
| 9 | Gtf2i | 0.75 | 2.13E-31 | 4.55E-27 | 0.97 | 0.9 | 0.07 | 1.08 |
| 9 | Acot1 | 1.68 | 2.67E-31 | 5.71E-27 | 0.61 | 0.33 | 0.28 | 1.84 |
| 9 | Ndrg1 | 2.93 | 5.59E-31 | 1.19E-26 | 0.19 | 0.04 | 0.15 | 5.37 |
| 9 | Stac | 3.86 | 8.87E-31 | 1.90E-26 | 0.11 | 0.01 | 0.1 | 9.64 |
| 9 | Myo10 | 1.12 | 1.31E-30 | 2.80E-26 | 0.84 | 0.7 | 0.14 | 1.2 |
| 9 | Calcoco1 | 1.55 | 1.34E-30 | 2.87E-26 | 0.57 | 0.28 | 0.29 | 2.04 |
| 9 | Kcnj2 | 2.13 | 1.45E-30 | 3.10E-26 | 0.39 | 0.14 | 0.25 | 2.78 |
| 9 | Socs2 | 1.19 | 1.89E-30 | 4.03E-26 | 0.89 | 0.78 | 0.12 | 1.15 |
| 9 | Fgf9 | 1.55 | 2.27E-30 | 4.85E-26 | 0.69 | 0.41 | 0.27 | 1.67 |
| 9 | Map1lc3b | 0.65 | 3.41E-30 | 7.28E-26 | 0.95 | 0.93 | 0.02 | 1.02 |
| 9 | Gabarap | 0.59 | 3.93E-30 | 8.39E-26 | 0.99 | 0.97 | 0.02 | 1.02 |
| 9 | Phlda3 | 1.35 | 7.97E-30 | 1.70E-25 | 0.75 | 0.49 | 0.26 | 1.54 |
| 9 | Chd3 | 0.68 | 1.36E-29 | 2.92E-25 | 0.98 | 0.94 | 0.04 | 1.05 |
| 9 | Kdm6b | 1.08 | 3.27E-29 | 6.98E-25 | 0.86 | 0.72 | 0.14 | 1.19 |
| 9 | Ypel3 | 1.04 | 5.00E-29 | 1.07E-24 | 0.78 | 0.56 | 0.22 | 1.39 |
| 9 | Sh3bp5 | 1.52 | 5.14E-29 | 1.10E-24 | 0.63 | 0.37 | 0.25 | 1.68 |
| 9 | Dclre1c | 2.5 | 8.56E-29 | 1.83E-24 | 0.3 | 0.09 | 0.21 | 3.31 |
| 9 | Irx1 | 2.04 | 1.23E-28 | 2.62E-24 | 0.26 | 0.06 | 0.19 | 3.98 |
| 9 | Tiam2 | 3.13 | 1.38E-28 | 2.95E-24 | 0.16 | 0.03 | 0.13 | 5.75 |
| 9 | Syt1 | 1.76 | 1.49E-28 | 3.19E-24 | 0.38 | 0.13 | 0.25 | 2.91 |
| 9 | Tgif1 | 1.05 | 1.92E-28 | 4.11E-24 | 0.83 | 0.68 | 0.16 | 1.23 |
| 9 | Cmip | 1.27 | 9.20E-28 | 1.97E-23 | 0.72 | 0.5 | 0.22 | 1.44 |
| 9 | Cdh6 | 1.95 | 2.41E-27 | 5.16E-23 | 0.35 | 0.12 | 0.23 | 2.97 |
| 9 | Jun | 1.61 | 3.50E-27 | 7.47E-23 | 0.74 | 0.53 | 0.21 | 1.39 |
| 9 | Dkk2 | 3.39 | 4.62E-27 | 9.88E-23 | 0.2 | 0.04 | 0.15 | 4.45 |
| 9 | Ly6g6e | 2.25 | 4.69E-27 | 1.00E-22 | 0.24 | 0.06 | 0.18 | 3.98 |
| 9 | Nxph3 | 2.17 | 5.54E-27 | 1.18E-22 | 0.22 | 0.05 | 0.17 | 4.23 |
| 9 | Kcnmb4os2 | 1.68 | 6.18E-27 | 1.32E-22 | 0.49 | 0.23 | 0.26 | 2.15 |
| 9 | Ndufa4l2 | 2.21 | 7.25E-27 | 1.55E-22 | 0.29 | 0.09 | 0.2 | 3.3 |
| 9 | Npc2 | 0.66 | 1.21E-26 | 2.58E-22 | 0.97 | 0.92 | 0.05 | 1.06 |
| 9 | Pcsk5 | 2.07 | 1.73E-26 | 3.69E-22 | 0.31 | 0.1 | 0.21 | 3.05 |
| 9 | Znrf3 | 1.61 | 2.93E-26 | 6.26E-22 | 0.53 | 0.28 | 0.26 | 1.92 |
| 9 | Dapl1 | 1.73 | 4.52E-26 | 9.65E-22 | 0.53 | 0.26 | 0.28 | 2.08 |
| 9 | Rgs12 | 1.36 | 5.64E-26 | 1.20E-21 | 0.67 | 0.45 | 0.22 | 1.48 |
| 9 | Fnbp1 | 1.89 | 6.67E-26 | 1.42E-21 | 0.44 | 0.2 | 0.24 | 2.2 |
| 9 | Minar2 | 1.83 | 8.30E-26 | 1.77E-21 | 0.38 | 0.15 | 0.23 | 2.57 |
| 9 | Mtss1 | 1.25 | 1.06E-25 | 2.27E-21 | 0.67 | 0.42 | 0.25 | 1.58 |
| 9 | Bnc1 | 2.82 | 1.29E-25 | 2.74E-21 | 0.17 | 0.03 | 0.13 | 5.16 |
| 9 | Csnk1e | 0.72 | 1.63E-25 | 3.47E-21 | 0.94 | 0.87 | 0.07 | 1.08 |
| 9 | Irx2 | 2.44 | 1.63E-25 | 3.49E-21 | 0.2 | 0.05 | 0.15 | 4.35 |
| 9 | Sox9 | 1.5 | 1.96E-25 | 4.18E-21 | 0.78 | 0.57 | 0.21 | 1.37 |
| 9 | Aplp2 | 0.73 | 2.15E-25 | 4.60E-21 | 0.96 | 0.92 | 0.04 | 1.04 |
| 9 | Actg1 | 0.41 | 2.99E-25 | 6.39E-21 | 1 | 1 | 0 | 1 |
| 9 | Pag1 | 2.5 | 3.39E-25 | 7.23E-21 | 0.2 | 0.05 | 0.15 | 4.26 |
| 9 | Igfbp2 | 0.77 | 6.50E-25 | 1.39E-20 | 1 | 0.99 | 0.01 | 1.01 |
| 9 | Gxylt2 | 2.38 | 6.80E-25 | 1.45E-20 | 0.28 | 0.09 | 0.19 | 3.12 |
| 9 | Hs3st1 | 2.57 | 9.95E-25 | 2.13E-20 | 0.23 | 0.06 | 0.17 | 3.67 |
| 9 | Neat1 | 2.23 | 2.05E-24 | 4.38E-20 | 0.25 | 0.07 | 0.18 | 3.53 |
| 9 | Igfbp4 | 1.28 | 2.55E-24 | 5.44E-20 | 0.88 | 0.73 | 0.14 | 1.19 |
| 9 | Gadd45b | 1.69 | 5.22E-24 | 1.11E-19 | 0.44 | 0.19 | 0.24 | 2.25 |
| 9 | Mid1ip1 | 1.86 | 6.16E-24 | 1.32E-19 | 0.59 | 0.37 | 0.22 | 1.6 |
| 9 | Trps1 | 1.29 | 9.01E-24 | 1.92E-19 | 0.7 | 0.53 | 0.18 | 1.33 |
| 9 | Gnpda2 | 1.48 | 9.09E-24 | 1.94E-19 | 0.6 | 0.39 | 0.21 | 1.54 |
| 9 | Sema6a | 1.18 | 1.10E-23 | 2.36E-19 | 0.78 | 0.61 | 0.18 | 1.29 |
| 9 | Prss23 | 2.02 | 1.34E-23 | 2.86E-19 | 0.52 | 0.28 | 0.24 | 1.85 |
| 9 | Med13 | 1.29 | 1.92E-23 | 4.10E-19 | 0.75 | 0.59 | 0.16 | 1.28 |
| 9 | Frmd4b | 1.12 | 2.02E-23 | 4.32E-19 | 0.71 | 0.49 | 0.22 | 1.45 |

**Supplementary Table 3. Marker genes for the original clusters of integrated clustering of E11.5 control and NotchGOF λ-junction ectoderm cells.**

| | | | | | | | | |
|---|---|---|---|---|---|---|---|---|
| 9 | Eef2 | 0.33 | 2.04E-23 | 4.36E-19 | 1 | 1 | 0 | 1 |
| 9 | Sp6 | 1.99 | 2.05E-23 | 4.38E-19 | 0.31 | 0.11 | 0.2 | 2.83 |
| 9 | Fat3 | 1.94 | 2.17E-23 | 4.63E-19 | 0.35 | 0.14 | 0.21 | 2.52 |
| 9 | Arpp21 | 3.08 | 3.26E-23 | 6.97E-19 | 0.13 | 0.02 | 0.11 | 5.78 |
| 9 | Bdnf | 2.05 | 3.53E-23 | 7.54E-19 | 0.4 | 0.17 | 0.22 | 2.3 |
| 9 | Fbxw4 | 1.74 | 3.60E-23 | 7.68E-19 | 0.41 | 0.18 | 0.23 | 2.28 |
| 9 | Rdh10 | 1.95 | 4.34E-23 | 9.26E-19 | 0.53 | 0.29 | 0.24 | 1.83 |
| 9 | Tmem178 | 1.76 | 4.60E-23 | 9.82E-19 | 0.31 | 0.11 | 0.2 | 2.9 |
| 9 | Ube2h | 1.18 | 5.35E-23 | 1.14E-18 | 0.69 | 0.5 | 0.2 | 1.39 |
| 9 | Fhl2 | 2.53 | 6.75E-23 | 1.44E-18 | 0.2 | 0.05 | 0.15 | 3.92 |
| 9 | Mylip | 1.7 | 1.14E-22 | 2.43E-18 | 0.42 | 0.19 | 0.23 | 2.2 |
| 9 | Gpr4 | 2.6 | 1.18E-22 | 2.52E-18 | 0.11 | 0.02 | 0.1 | 6.33 |
| 9 | Trim16 | 2.84 | 1.21E-22 | 2.58E-18 | 0.17 | 0.04 | 0.13 | 4.58 |
| 9 | 1810058I24Rik | 0.62 | 1.49E-22 | 3.19E-18 | 0.95 | 0.93 | 0.02 | 1.02 |
| 9 | Fblim1 | 1.26 | 1.97E-22 | 4.20E-18 | 0.62 | 0.4 | 0.22 | 1.55 |
| 9 | Zfand5 | 0.93 | 2.00E-22 | 4.28E-18 | 0.89 | 0.83 | 0.07 | 1.08 |
| 9 | Scnn1a | 3.02 | 2.64E-22 | 5.65E-18 | 0.12 | 0.02 | 0.1 | 6.21 |
| 9 | Fosl2 | 1.88 | 3.17E-22 | 6.76E-18 | 0.26 | 0.08 | 0.18 | 3.31 |
| 9 | Mir100hg | 0.86 | 3.67E-22 | 7.83E-18 | 0.82 | 0.62 | 0.2 | 1.32 |
| 9 | Robo1 | 1.35 | 4.62E-22 | 9.86E-18 | 0.61 | 0.39 | 0.22 | 1.56 |
| 9 | Dlx3 | 1.41 | 5.13E-22 | 1.10E-17 | 0.51 | 0.27 | 0.24 | 1.89 |
| 9 | Btrc | 1.83 | 1.17E-21 | 2.50E-17 | 0.53 | 0.32 | 0.21 | 1.66 |
| 9 | Chst3 | 2.42 | 1.31E-21 | 2.80E-17 | 0.18 | 0.05 | 0.14 | 4 |
| 9 | Ablim1 | 0.86 | 1.73E-21 | 3.70E-17 | 0.76 | 0.54 | 0.22 | 1.41 |
| 9 | Mfsd1 | 1.22 | 1.91E-21 | 4.08E-17 | 0.62 | 0.39 | 0.23 | 1.58 |
| 9 | Serinc5 | 1.91 | 2.04E-21 | 4.35E-17 | 0.31 | 0.11 | 0.19 | 2.73 |
| 9 | Bcl11b | 0.83 | 2.93E-21 | 6.25E-17 | 0.86 | 0.71 | 0.15 | 1.2 |
| 9 | Tmem59 | 0.56 | 3.44E-21 | 7.35E-17 | 0.96 | 0.89 | 0.07 | 1.08 |
| 9 | Lmo4 | 0.62 | 3.72E-21 | 7.95E-17 | 0.95 | 0.91 | 0.04 | 1.04 |
| 9 | Mfap4 | 0.67 | 5.54E-21 | 1.18E-16 | 0.84 | 0.65 | 0.2 | 1.3 |
| 9 | Slc39a14 | 2.22 | 5.77E-21 | 1.23E-16 | 0.22 | 0.07 | 0.16 | 3.34 |
| 9 | Filip1l | 1.58 | 6.72E-21 | 1.44E-16 | 0.49 | 0.26 | 0.23 | 1.89 |
| 9 | Cited4 | 2.23 | 8.13E-21 | 1.74E-16 | 0.19 | 0.05 | 0.14 | 3.84 |
| 9 | Msx2 | 1.12 | 8.53E-21 | 1.82E-16 | 0.63 | 0.39 | 0.24 | 1.61 |
| 9 | Nab1 | 1.43 | 1.02E-20 | 2.18E-16 | 0.46 | 0.23 | 0.23 | 1.99 |
| 9 | Kcnmb4 | 1.24 | 1.22E-20 | 2.61E-16 | 0.6 | 0.39 | 0.21 | 1.54 |
| 9 | Tctex1d1 | 1.59 | 1.50E-20 | 3.20E-16 | 0.28 | 0.1 | 0.18 | 2.93 |
| 9 | Gm49708 | 1.27 | 1.68E-20 | 3.60E-16 | 0.68 | 0.48 | 0.2 | 1.42 |
| 9 | Cox7a2l | 0.42 | 2.11E-20 | 4.51E-16 | 0.99 | 0.98 | 0.01 | 1.01 |
| 9 | Nedd9 | 1.41 | 2.83E-20 | 6.05E-16 | 0.52 | 0.3 | 0.22 | 1.74 |
| 9 | Bmp4 | 1.42 | 5.50E-20 | 1.18E-15 | 0.72 | 0.53 | 0.19 | 1.37 |
| 9 | Slc25a17 | 0.72 | 5.81E-20 | 1.24E-15 | 0.85 | 0.7 | 0.14 | 1.2 |
| 9 | Sptlc3 | 2.29 | 6.16E-20 | 1.32E-15 | 0.22 | 0.07 | 0.15 | 3.28 |
| 9 | Col9a3 | 1.67 | 7.70E-20 | 1.64E-15 | 0.34 | 0.14 | 0.2 | 2.44 |
| 9 | Tnfrsf1a | 1.44 | 9.81E-20 | 2.10E-15 | 0.53 | 0.31 | 0.21 | 1.67 |
| 9 | Rel | 1.72 | 1.14E-19 | 2.43E-15 | 0.35 | 0.15 | 0.2 | 2.35 |
| 9 | Ddhd1 | 1.49 | 1.34E-19 | 2.87E-15 | 0.45 | 0.23 | 0.22 | 1.94 |
| 9 | Ppp1r13l | 1.61 | 1.49E-19 | 3.19E-15 | 0.39 | 0.19 | 0.21 | 2.11 |
| 9 | Ddi2 | 1.13 | 2.29E-19 | 4.90E-15 | 0.65 | 0.47 | 0.18 | 1.39 |
| 9 | Notch1 | 0.96 | 4.03E-19 | 8.61E-15 | 0.75 | 0.62 | 0.12 | 1.2 |
| 9 | Ell2 | 1.79 | 4.22E-19 | 9.01E-15 | 0.36 | 0.17 | 0.2 | 2.17 |
| 9 | Itm2c | 0.85 | 4.55E-19 | 9.73E-15 | 0.75 | 0.59 | 0.16 | 1.27 |
| 9 | Kif21a | 0.81 | 5.72E-19 | 1.22E-14 | 0.84 | 0.74 | 0.1 | 1.13 |
| 9 | S100a10 | 0.55 | 5.94E-19 | 1.27E-14 | 0.92 | 0.74 | 0.18 | 1.24 |
| 9 | Kdm5a | 0.74 | 8.22E-19 | 1.76E-14 | 0.88 | 0.8 | 0.08 | 1.1 |
| 9 | Abca1 | 1.76 | 8.82E-19 | 1.88E-14 | 0.29 | 0.11 | 0.18 | 2.63 |
| 9 | Map1lc3a | 0.76 | 1.76E-18 | 3.76E-14 | 0.84 | 0.73 | 0.11 | 1.15 |

**Supplementary Table 3. Marker genes for the original clusters of integrated clustering of E11.5 control and NotchGOF λ-junction ectoderm cells.**

| | | | | | | | | |
|---|---|---|---|---|---|---|---|---|
| 9 | Mir205hg | 0.94 | 2.07E-18 | 4.43E-14 | 0.68 | 0.43 | 0.25 | 1.57 |
| 9 | Pla2r1 | 1.28 | 2.64E-18 | 5.63E-14 | 0.49 | 0.27 | 0.22 | 1.84 |
| 9 | Txnip | 1.15 | 2.67E-18 | 5.71E-14 | 0.64 | 0.44 | 0.2 | 1.45 |
| 9 | Cd59a | 1.2 | 2.83E-18 | 6.04E-14 | 0.52 | 0.29 | 0.23 | 1.78 |
| 9 | Bcl2l11 | 1.52 | 2.93E-18 | 6.26E-14 | 0.57 | 0.38 | 0.19 | 1.5 |
| 9 | Rere | 0.99 | 3.55E-18 | 7.58E-14 | 0.76 | 0.64 | 0.12 | 1.19 |
| 9 | Pcmtd1 | 0.99 | 4.53E-18 | 9.67E-14 | 0.69 | 0.51 | 0.18 | 1.35 |
| 9 | Nxn | 0.69 | 5.61E-18 | 1.20E-13 | 0.89 | 0.75 | 0.15 | 1.2 |
| 9 | Plp2 | 0.82 | 5.92E-18 | 1.26E-13 | 0.81 | 0.69 | 0.11 | 1.16 |
| 9 | Foxo1 | 1.31 | 8.28E-18 | 1.77E-13 | 0.49 | 0.28 | 0.21 | 1.74 |
| 9 | Tns4 | 2.68 | 8.72E-18 | 1.86E-13 | 0.12 | 0.03 | 0.1 | 4.88 |
| 9 | Wls | 0.54 | 1.04E-17 | 2.23E-13 | 0.95 | 0.79 | 0.15 | 1.19 |
| 9 | Rnf149 | 1.58 | 1.33E-17 | 2.84E-13 | 0.47 | 0.28 | 0.19 | 1.7 |
| 9 | Atp6v1g1 | 0.42 | 1.38E-17 | 2.95E-13 | 0.98 | 0.97 | 0.01 | 1.01 |
| 9 | B9d2 | 1.39 | 1.48E-17 | 3.15E-13 | 0.47 | 0.27 | 0.21 | 1.77 |
| 9 | Eif3f | 0.28 | 1.78E-17 | 3.80E-13 | 1 | 0.99 | 0.01 | 1.01 |
| 9 | Wipi1 | 1.34 | 2.38E-17 | 5.08E-13 | 0.5 | 0.3 | 0.21 | 1.7 |
| 9 | Bmpr2 | 0.87 | 2.75E-17 | 5.88E-13 | 0.75 | 0.58 | 0.17 | 1.3 |
| 9 | Cnksr2 | 1.25 | 2.79E-17 | 5.96E-13 | 0.53 | 0.33 | 0.21 | 1.63 |
| 9 | Ogt | 0.56 | 2.85E-17 | 6.10E-13 | 0.91 | 0.8 | 0.11 | 1.14 |
| 9 | Gnas | 0.29 | 6.60E-17 | 1.41E-12 | 1 | 1 | 0 | 1 |
| 9 | Cpne8 | 2.04 | 1.01E-16 | 2.16E-12 | 0.22 | 0.08 | 0.15 | 2.91 |
| 9 | Bnip3 | 1.22 | 1.03E-16 | 2.19E-12 | 0.63 | 0.45 | 0.18 | 1.4 |
| 9 | Rab6a | 0.78 | 1.16E-16 | 2.48E-12 | 0.84 | 0.76 | 0.08 | 1.11 |
| 9 | Supt4a | 0.44 | 1.22E-16 | 2.62E-12 | 0.94 | 0.91 | 0.04 | 1.04 |
| 9 | Nectin3 | 0.77 | 1.35E-16 | 2.87E-12 | 0.85 | 0.73 | 0.12 | 1.16 |
| 9 | Prickle1 | 2.22 | 1.59E-16 | 3.40E-12 | 0.19 | 0.06 | 0.13 | 3.19 |
| 9 | Kctd1 | 0.77 | 2.20E-16 | 4.70E-12 | 0.7 | 0.49 | 0.21 | 1.44 |
| 9 | Afdn | 0.57 | 2.34E-16 | 4.99E-12 | 0.9 | 0.86 | 0.04 | 1.05 |
| 9 | Tmem140 | 2.68 | 3.35E-16 | 7.16E-12 | 0.11 | 0.02 | 0.09 | 5.05 |
| 9 | Arl4c | 0.9 | 4.05E-16 | 8.66E-12 | 0.74 | 0.57 | 0.17 | 1.3 |
| 9 | Lamp2 | 0.71 | 4.89E-16 | 1.04E-11 | 0.76 | 0.6 | 0.16 | 1.27 |
| 9 | Duoxa1 | 2.74 | 4.90E-16 | 1.05E-11 | 0.11 | 0.02 | 0.09 | 4.78 |
| 9 | Plec | 1.03 | 5.61E-16 | 1.20E-11 | 0.62 | 0.44 | 0.18 | 1.4 |
| 9 | Enpp2 | 3.15 | 5.77E-16 | 1.23E-11 | 0.13 | 0.03 | 0.1 | 4.17 |
| 9 | Chmp5 | 0.61 | 6.55E-16 | 1.40E-11 | 0.89 | 0.79 | 0.1 | 1.12 |
| 9 | Snai2 | 1.26 | 6.92E-16 | 1.48E-11 | 0.37 | 0.17 | 0.2 | 2.16 |
| 9 | Rhob | 1 | 9.02E-16 | 1.93E-11 | 0.66 | 0.47 | 0.18 | 1.39 |
| 9 | Pygl | 1.5 | 1.17E-15 | 2.49E-11 | 0.35 | 0.17 | 0.18 | 2.06 |
| 9 | Tnks2 | 0.55 | 1.23E-15 | 2.62E-11 | 0.89 | 0.88 | 0.01 | 1.02 |
| 9 | Mafg | 0.98 | 1.37E-15 | 2.92E-11 | 0.68 | 0.53 | 0.15 | 1.28 |
| 9 | Ppfibp1 | 0.89 | 1.77E-15 | 3.78E-11 | 0.75 | 0.61 | 0.14 | 1.23 |
| 9 | Klhl8 | 1.53 | 1.86E-15 | 3.97E-11 | 0.32 | 0.15 | 0.17 | 2.16 |
| 9 | Nectin4 | 0.71 | 1.88E-15 | 4.01E-11 | 0.73 | 0.52 | 0.21 | 1.39 |
| 9 | Ifngr2 | 1.36 | 1.90E-15 | 4.05E-11 | 0.46 | 0.27 | 0.19 | 1.71 |
| 9 | Cdh3 | 0.75 | 2.04E-15 | 4.36E-11 | 0.73 | 0.56 | 0.16 | 1.29 |
| 9 | Ppp1r12b | 1.31 | 2.20E-15 | 4.69E-11 | 0.33 | 0.15 | 0.18 | 2.15 |
| 9 | Ptk7 | 0.95 | 2.22E-15 | 4.74E-11 | 0.66 | 0.48 | 0.17 | 1.36 |
| 9 | Hs3st6 | 1.94 | 2.25E-15 | 4.80E-11 | 0.26 | 0.11 | 0.15 | 2.44 |
| 9 | Vgll4 | 0.69 | 2.37E-15 | 5.06E-11 | 0.92 | 0.85 | 0.07 | 1.08 |
| 9 | Khdrbs2 | 2.32 | 2.67E-15 | 5.70E-11 | 0.13 | 0.03 | 0.1 | 3.91 |
| 9 | Zfp704 | 0.84 | 3.13E-15 | 6.69E-11 | 0.73 | 0.62 | 0.11 | 1.18 |
| 9 | Ptger4 | 2.93 | 4.74E-15 | 1.01E-10 | 0.11 | 0.02 | 0.08 | 4.61 |
| 9 | Dst | 1.03 | 4.78E-15 | 1.02E-10 | 0.68 | 0.47 | 0.2 | 1.42 |
| 9 | Akirin2 | 0.66 | 5.47E-15 | 1.17E-10 | 0.88 | 0.81 | 0.07 | 1.08 |
| 9 | Cited2 | 0.89 | 7.19E-15 | 1.54E-10 | 0.79 | 0.68 | 0.11 | 1.16 |
| 9 | Bcl9 | 0.72 | 9.61E-15 | 2.05E-10 | 0.79 | 0.68 | 0.11 | 1.16 |

**Supplementary Table 3. Marker genes for the original clusters of integrated clustering of E11.5 control and NotchGOF λ-junction ectoderm cells.**

| | | | | | | | | |
|---|---|---|---|---|---|---|---|---|
| 9 | Ptpn13 | 0.77 | 1.08E-14 | 2.31E-10 | 0.73 | 0.57 | 0.17 | 1.3 |
| 9 | Axin2 | 1.57 | 1.29E-14 | 2.76E-10 | 0.43 | 0.25 | 0.18 | 1.7 |
| 9 | Mtmr11 | 2.19 | 1.39E-14 | 2.96E-10 | 0.18 | 0.06 | 0.12 | 3.03 |
| 9 | Gm42418 | 0.33 | 1.53E-14 | 3.26E-10 | 1 | 1 | 0 | 1 |
| 9 | Sema3d | 1.19 | 1.87E-14 | 4.00E-10 | 0.6 | 0.41 | 0.19 | 1.47 |
| 9 | Vwa5a | 1.47 | 1.89E-14 | 4.04E-10 | 0.33 | 0.16 | 0.17 | 2.1 |
| 9 | Cnn2 | 1.6 | 2.39E-14 | 5.10E-10 | 0.49 | 0.3 | 0.19 | 1.61 |
| 9 | Atxn7l3b | 0.29 | 2.64E-14 | 5.65E-10 | 1 | 0.99 | 0.01 | 1.01 |
| 9 | Cst3 | 0.4 | 2.71E-14 | 5.79E-10 | 0.97 | 0.93 | 0.04 | 1.05 |
| 9 | Peli1 | 0.78 | 2.90E-14 | 6.20E-10 | 0.78 | 0.65 | 0.13 | 1.2 |
| 9 | Slc7a11 | 1.43 | 3.11E-14 | 6.64E-10 | 0.29 | 0.12 | 0.17 | 2.34 |
| 9 | Jup | 0.8 | 3.44E-14 | 7.34E-10 | 0.68 | 0.51 | 0.16 | 1.32 |
| 9 | Map4k4 | 0.53 | 3.48E-14 | 7.43E-10 | 0.89 | 0.89 | 0.01 | 1.01 |
| 9 | Plod2 | 1.04 | 3.65E-14 | 7.80E-10 | 0.65 | 0.47 | 0.18 | 1.37 |
| 9 | Igfbp3 | 1.94 | 4.86E-14 | 1.04E-09 | 0.54 | 0.38 | 0.16 | 1.41 |
| 9 | Pgpep1 | 1.26 | 4.92E-14 | 1.05E-09 | 0.51 | 0.34 | 0.17 | 1.51 |
| 9 | Serf1 | 0.52 | 4.98E-14 | 1.06E-09 | 0.94 | 0.93 | 0.01 | 1.01 |
| 9 | Lmo7 | 1.06 | 6.35E-14 | 1.36E-09 | 0.6 | 0.45 | 0.16 | 1.35 |
| 9 | Churc1 | 0.81 | 6.70E-14 | 1.43E-09 | 0.81 | 0.72 | 0.08 | 1.12 |
| 9 | Cdkn1c | 0.9 | 6.93E-14 | 1.48E-09 | 0.91 | 0.88 | 0.03 | 1.04 |
| 9 | Col17a1 | 1.12 | 8.02E-14 | 1.71E-09 | 0.29 | 0.12 | 0.17 | 2.36 |
| 9 | Tmem30b | 1.54 | 1.11E-13 | 2.37E-09 | 0.43 | 0.27 | 0.17 | 1.62 |
| 9 | Palmd | 2.28 | 1.16E-13 | 2.47E-09 | 0.16 | 0.05 | 0.11 | 3.16 |
| 9 | Socs7 | 1.17 | 1.18E-13 | 2.53E-09 | 0.47 | 0.29 | 0.18 | 1.61 |
| 9 | Xpr1 | 1.19 | 1.25E-13 | 2.66E-09 | 0.58 | 0.41 | 0.17 | 1.41 |
| 9 | Sertad4 | 1.1 | 1.25E-13 | 2.67E-09 | 0.49 | 0.32 | 0.18 | 1.55 |
| 9 | Lbh | 1.25 | 2.08E-13 | 4.45E-09 | 0.44 | 0.26 | 0.17 | 1.67 |
| 9 | Kdm1a | 0.38 | 2.64E-13 | 5.64E-09 | 0.97 | 0.94 | 0.03 | 1.04 |
| 9 | Ypel5 | 1.23 | 2.82E-13 | 6.01E-09 | 0.43 | 0.25 | 0.18 | 1.71 |
| 9 | Atp6v0e | 0.62 | 2.85E-13 | 6.09E-09 | 0.82 | 0.77 | 0.04 | 1.06 |
| 9 | Rgs2 | 1.26 | 3.07E-13 | 6.56E-09 | 0.47 | 0.29 | 0.19 | 1.65 |
| 9 | Myl12a | 0.47 | 4.38E-13 | 9.35E-09 | 0.97 | 0.94 | 0.03 | 1.04 |
| 9 | Sulf1 | 0.74 | 4.49E-13 | 9.58E-09 | 0.79 | 0.66 | 0.13 | 1.2 |
| 9 | Fbxo32 | 2.17 | 6.03E-13 | 1.29E-08 | 0.18 | 0.07 | 0.12 | 2.77 |
| 9 | Cd151 | 0.66 | 6.34E-13 | 1.35E-08 | 0.79 | 0.66 | 0.13 | 1.2 |
| 9 | Itgb6 | 1.35 | 6.71E-13 | 1.43E-08 | 0.32 | 0.16 | 0.16 | 2.03 |
| 9 | Tfap2a | 0.87 | 6.79E-13 | 1.45E-08 | 0.53 | 0.34 | 0.2 | 1.59 |
| 9 | Lrrk2 | 2.3 | 8.11E-13 | 1.73E-08 | 0.12 | 0.03 | 0.09 | 3.81 |
| 9 | Eppk1 | 2 | 8.92E-13 | 1.91E-08 | 0.2 | 0.08 | 0.13 | 2.58 |
| 9 | Spsb1 | 2.48 | 8.95E-13 | 1.91E-08 | 0.18 | 0.06 | 0.11 | 2.79 |
| 9 | Tulp4 | 0.59 | 9.49E-13 | 2.03E-08 | 0.84 | 0.75 | 0.09 | 1.12 |
| 9 | Slc16a7 | 2.44 | 1.27E-12 | 2.72E-08 | 0.14 | 0.04 | 0.1 | 3.2 |
| 9 | Col23a1 | 2.47 | 1.49E-12 | 3.17E-08 | 0.11 | 0.03 | 0.08 | 3.8 |
| 9 | Nsg1 | 0.72 | 1.49E-12 | 3.19E-08 | 0.72 | 0.61 | 0.12 | 1.19 |
| 9 | Col4a2 | 0.78 | 1.60E-12 | 3.42E-08 | 0.66 | 0.5 | 0.17 | 1.33 |
| 9 | Adgrl3 | 1.33 | 1.73E-12 | 3.69E-08 | 0.48 | 0.31 | 0.17 | 1.55 |
| 9 | Rab38 | 1.34 | 1.78E-12 | 3.81E-08 | 0.37 | 0.19 | 0.17 | 1.88 |
| 9 | Inpp5f | 1.43 | 1.90E-12 | 4.06E-08 | 0.42 | 0.26 | 0.16 | 1.63 |
| 9 | Fzd10os | 1.21 | 2.15E-12 | 4.60E-08 | 0.31 | 0.15 | 0.16 | 2.1 |
| 9 | Kidins220 | 1.01 | 2.68E-12 | 5.72E-08 | 0.52 | 0.36 | 0.16 | 1.44 |
| 9 | Usp22 | 0.6 | 2.92E-12 | 6.24E-08 | 0.78 | 0.66 | 0.13 | 1.19 |
| 9 | Cpm | 0.62 | 2.98E-12 | 6.37E-08 | 0.73 | 0.52 | 0.21 | 1.39 |
| 9 | Rb1cc1 | 0.64 | 3.08E-12 | 6.58E-08 | 0.78 | 0.7 | 0.08 | 1.12 |
| 9 | Ttc3 | 0.36 | 3.50E-12 | 7.48E-08 | 1 | 0.99 | 0.01 | 1.01 |
| 9 | Lrrc8d | 1.16 | 3.76E-12 | 8.02E-08 | 0.48 | 0.32 | 0.16 | 1.5 |
| 9 | Pdgfc | 0.56 | 3.90E-12 | 8.33E-08 | 0.85 | 0.76 | 0.09 | 1.12 |
| 9 | Kdm7a | 0.96 | 4.03E-12 | 8.61E-08 | 0.57 | 0.41 | 0.16 | 1.39 |

**Supplementary Table 3. Marker genes for the original clusters of integrated clustering of E11.5 control and NotchGOF λ-junction ectoderm cells.**

| | | | | | | | | |
|---|---|---|---|---|---|---|---|---|
| 9 | Pcmtd2 | 0.83 | 4.20E-12 | 8.98E-08 | 0.57 | 0.41 | 0.16 | 1.39 |
| 9 | Mapk6 | 0.73 | 4.22E-12 | 9.00E-08 | 0.68 | 0.54 | 0.14 | 1.26 |
| 9 | Trp53inp2 | 1.76 | 4.90E-12 | 1.05E-07 | 0.19 | 0.07 | 0.12 | 2.59 |
| 9 | Thsd7b | 2 | 5.00E-12 | 1.07E-07 | 0.17 | 0.06 | 0.11 | 2.89 |
| 9 | Ddr1 | 0.76 | 5.15E-12 | 1.10E-07 | 0.61 | 0.45 | 0.16 | 1.36 |
| 9 | Gpr107 | 1.07 | 6.15E-12 | 1.31E-07 | 0.49 | 0.34 | 0.15 | 1.45 |
| 9 | Vezf1 | 0.49 | 7.02E-12 | 1.50E-07 | 0.87 | 0.82 | 0.05 | 1.06 |
| 9 | Cebpb | 1.64 | 8.14E-12 | 1.74E-07 | 0.39 | 0.24 | 0.15 | 1.65 |
| 9 | Adamts18 | 1.24 | 8.25E-12 | 1.76E-07 | 0.43 | 0.26 | 0.17 | 1.63 |
| 9 | Pdlim4 | 0.86 | 9.12E-12 | 1.95E-07 | 0.58 | 0.43 | 0.15 | 1.35 |
| 9 | Klc3 | 0.71 | 1.04E-11 | 2.21E-07 | 0.55 | 0.39 | 0.16 | 1.41 |
| 9 | Fzd6 | 1.06 | 1.04E-11 | 2.23E-07 | 0.48 | 0.32 | 0.16 | 1.5 |
| 9 | D7Ertd443e | 1.56 | 1.12E-11 | 2.40E-07 | 0.22 | 0.09 | 0.13 | 2.38 |
| 9 | Sesn1 | 1.43 | 1.28E-11 | 2.72E-07 | 0.36 | 0.2 | 0.16 | 1.78 |
| 9 | Tns3 | 1.09 | 1.30E-11 | 2.78E-07 | 0.48 | 0.33 | 0.15 | 1.44 |
| 9 | Jag1 | 0.77 | 1.35E-11 | 2.89E-07 | 0.71 | 0.55 | 0.16 | 1.29 |
| 9 | Dync1i1 | 1.43 | 1.56E-11 | 3.33E-07 | 0.26 | 0.12 | 0.14 | 2.17 |
| 9 | Chmp1b | 0.84 | 1.65E-11 | 3.52E-07 | 0.57 | 0.43 | 0.15 | 1.35 |
| 9 | Osbpl8 | 1.15 | 1.68E-11 | 3.59E-07 | 0.43 | 0.26 | 0.16 | 1.62 |
| 9 | Cdca7 | 0.63 | 1.85E-11 | 3.95E-07 | 0.78 | 0.64 | 0.14 | 1.21 |
| 9 | Sh3glb1 | 0.49 | 1.95E-11 | 4.16E-07 | 0.84 | 0.79 | 0.05 | 1.07 |
| 9 | Myo1b | 0.61 | 2.11E-11 | 4.50E-07 | 0.76 | 0.62 | 0.14 | 1.22 |
| 9 | Ddah2 | 0.33 | 2.55E-11 | 5.46E-07 | 0.99 | 0.97 | 0.02 | 1.02 |
| 9 | Ifitm1 | 1.05 | 2.56E-11 | 5.47E-07 | 0.35 | 0.19 | 0.16 | 1.85 |
| 9 | Alpl | 1.55 | 2.70E-11 | 5.77E-07 | 0.31 | 0.16 | 0.15 | 1.89 |
| 9 | Prag1 | 1.17 | 2.83E-11 | 6.04E-07 | 0.45 | 0.29 | 0.16 | 1.56 |
| 9 | Rora | 1.76 | 2.90E-11 | 6.19E-07 | 0.21 | 0.09 | 0.12 | 2.36 |
| 9 | Cpxm1 | 1.1 | 3.18E-11 | 6.79E-07 | 0.42 | 0.26 | 0.16 | 1.61 |
| 9 | Fmnl2 | 1 | 3.66E-11 | 7.82E-07 | 0.6 | 0.47 | 0.13 | 1.28 |
| 9 | Bcl11a | 0.59 | 3.80E-11 | 8.13E-07 | 0.86 | 0.8 | 0.07 | 1.09 |
| 9 | Hbp1 | 0.62 | 3.92E-11 | 8.36E-07 | 0.77 | 0.63 | 0.14 | 1.22 |
| 9 | Pax7 | 2.4 | 4.35E-11 | 9.29E-07 | 0.11 | 0.03 | 0.08 | 3.66 |
| 9 | Bckdha | 0.87 | 4.59E-11 | 9.81E-07 | 0.61 | 0.46 | 0.15 | 1.34 |
| 9 | Aff4 | 0.71 | 5.00E-11 | 1.07E-06 | 0.75 | 0.66 | 0.09 | 1.14 |
| 9 | Rap2c | 0.68 | 5.23E-11 | 1.12E-06 | 0.75 | 0.66 | 0.09 | 1.13 |
| 9 | Usp46 | 0.65 | 5.26E-11 | 1.12E-06 | 0.78 | 0.72 | 0.06 | 1.09 |
| 9 | Ctsl | 0.44 | 5.34E-11 | 1.14E-06 | 0.92 | 0.86 | 0.06 | 1.07 |
| 9 | Irs2 | 1 | 6.21E-11 | 1.33E-06 | 0.57 | 0.44 | 0.13 | 1.3 |
| 9 | Gm15222 | 1.38 | 6.42E-11 | 1.37E-06 | 0.29 | 0.14 | 0.14 | 1.99 |
| 9 | Rsrp1 | 0.4 | 6.87E-11 | 1.47E-06 | 0.94 | 0.92 | 0.03 | 1.03 |
| 9 | Jmy | 0.85 | 7.21E-11 | 1.54E-06 | 0.6 | 0.48 | 0.13 | 1.27 |
| 9 | Zfas1 | 0.48 | 8.68E-11 | 1.85E-06 | 0.89 | 0.8 | 0.09 | 1.11 |
| 9 | Creb3l2 | 1.01 | 9.86E-11 | 2.11E-06 | 0.48 | 0.32 | 0.16 | 1.52 |
| 9 | Ddb1 | 0.44 | 1.49E-10 | 3.18E-06 | 0.92 | 0.89 | 0.03 | 1.03 |
| 9 | Macf1 | 0.57 | 1.52E-10 | 3.24E-06 | 0.88 | 0.83 | 0.05 | 1.06 |
| 9 | Nhsl1 | 0.96 | 1.63E-10 | 3.49E-06 | 0.47 | 0.32 | 0.15 | 1.47 |
| 9 | Lpp | 0.89 | 1.78E-10 | 3.80E-06 | 0.59 | 0.45 | 0.14 | 1.31 |
| 9 | Dnase2a | 0.9 | 1.93E-10 | 4.13E-06 | 0.58 | 0.43 | 0.14 | 1.33 |
| 9 | Nfil3 | 2 | 2.10E-10 | 4.49E-06 | 0.19 | 0.08 | 0.11 | 2.34 |
| 9 | Dusp2 | 2.11 | 2.35E-10 | 5.01E-06 | 0.12 | 0.04 | 0.08 | 3.28 |
| 9 | Arl5b | 0.76 | 2.46E-10 | 5.26E-06 | 0.6 | 0.45 | 0.15 | 1.33 |
| 9 | Dlgap4 | 0.64 | 2.50E-10 | 5.35E-06 | 0.79 | 0.69 | 0.1 | 1.14 |
| 9 | Ndufa6 | 0.41 | 2.68E-10 | 5.73E-06 | 0.97 | 0.94 | 0.03 | 1.03 |
| 9 | Satb1 | 0.73 | 3.28E-10 | 7.01E-06 | 0.68 | 0.57 | 0.11 | 1.2 |
| 9 | Crebrf | 1.13 | 3.29E-10 | 7.03E-06 | 0.31 | 0.17 | 0.15 | 1.88 |
| 9 | Usp3 | 0.54 | 3.33E-10 | 7.11E-06 | 0.78 | 0.71 | 0.07 | 1.09 |
| 9 | Frem2 | 0.56 | 3.38E-10 | 7.22E-06 | 0.75 | 0.62 | 0.13 | 1.21 |

**Supplementary Table 3. Marker genes for the original clusters of integrated clustering of E11.5 control and NotchGOF λ-junction ectoderm cells.**

| | | | | | | | | |
|---|---|---|---|---|---|---|---|---|
| 9 | Slc16a2 | 1.63 | 3.42E-10 | 7.30E-06 | 0.24 | 0.12 | 0.13 | 2.06 |
| 9 | Hspb1 | 0.6 | 3.43E-10 | 7.32E-06 | 0.57 | 0.38 | 0.19 | 1.5 |
| 9 | Fermt1 | 0.56 | 4.10E-10 | 8.75E-06 | 0.73 | 0.55 | 0.19 | 1.34 |
| 9 | Gm4258 | 0.89 | 4.33E-10 | 9.26E-06 | 0.62 | 0.51 | 0.11 | 1.23 |
| 9 | Sgpl1 | 0.8 | 5.53E-10 | 1.18E-05 | 0.55 | 0.41 | 0.14 | 1.34 |
| 9 | Zdhhc20 | 0.66 | 5.83E-10 | 1.25E-05 | 0.71 | 0.61 | 0.11 | 1.17 |
| 9 | Hes1 | 0.74 | 6.84E-10 | 1.46E-05 | 0.79 | 0.73 | 0.06 | 1.09 |
| 9 | Cnpy2 | 0.37 | 6.95E-10 | 1.49E-05 | 0.96 | 0.95 | 0.01 | 1.01 |
| 9 | Pik3ip1 | 0.94 | 6.99E-10 | 1.49E-05 | 0.33 | 0.19 | 0.15 | 1.78 |
| 9 | Btg1 | 0.67 | 7.71E-10 | 1.65E-05 | 0.8 | 0.75 | 0.05 | 1.07 |
| 9 | Dpysl3 | 0.34 | 9.04E-10 | 1.93E-05 | 0.72 | 0.56 | 0.16 | 1.28 |
| 9 | Unc5c | 0.73 | 1.04E-09 | 2.22E-05 | 0.49 | 0.33 | 0.17 | 1.52 |
| 9 | Vwa1 | 1.55 | 1.19E-09 | 2.55E-05 | 0.23 | 0.11 | 0.12 | 2.1 |
| 9 | Mafb | 1.4 | 1.20E-09 | 2.55E-05 | 0.29 | 0.15 | 0.13 | 1.88 |
| 9 | Dok5 | 0.79 | 1.33E-09 | 2.83E-05 | 0.54 | 0.37 | 0.17 | 1.46 |
| 9 | Ago1 | 0.61 | 1.45E-09 | 3.10E-05 | 0.71 | 0.62 | 0.09 | 1.15 |
| 9 | Zfp750 | 1.14 | 1.48E-09 | 3.16E-05 | 0.34 | 0.19 | 0.14 | 1.74 |
| 9 | Dusp23 | 1.41 | 1.48E-09 | 3.16E-05 | 0.23 | 0.11 | 0.12 | 2.08 |
| 9 | Cdc14a | 1.02 | 1.50E-09 | 3.20E-05 | 0.5 | 0.37 | 0.14 | 1.37 |
| 9 | Rest | 0.49 | 1.50E-09 | 3.21E-05 | 0.8 | 0.68 | 0.12 | 1.18 |
| 9 | Dhrs3 | 1.36 | 1.71E-09 | 3.64E-05 | 0.34 | 0.21 | 0.13 | 1.65 |
| 9 | Erich1 | 0.78 | 1.72E-09 | 3.67E-05 | 0.53 | 0.39 | 0.14 | 1.36 |
| 9 | Serinc1 | 0.55 | 1.83E-09 | 3.92E-05 | 0.75 | 0.66 | 0.09 | 1.13 |
| 9 | Prxl2a | 0.65 | 1.84E-09 | 3.93E-05 | 0.68 | 0.58 | 0.1 | 1.16 |
| 9 | Gdi1 | 0.64 | 2.08E-09 | 4.45E-05 | 0.59 | 0.45 | 0.14 | 1.31 |
| 9 | Adam19 | 2.11 | 2.13E-09 | 4.56E-05 | 0.1 | 0.03 | 0.07 | 3.29 |
| 9 | Nradd | 0.66 | 2.29E-09 | 4.89E-05 | 0.61 | 0.49 | 0.12 | 1.24 |
| 9 | Mob4 | 0.44 | 2.37E-09 | 5.06E-05 | 0.85 | 0.81 | 0.04 | 1.05 |
| 9 | Unc119 | 0.54 | 2.38E-09 | 5.08E-05 | 0.84 | 0.76 | 0.08 | 1.11 |
| 9 | Unc5b | 0.72 | 2.53E-09 | 5.39E-05 | 0.64 | 0.53 | 0.11 | 1.22 |
| 9 | Tbx2 | 1.62 | 2.64E-09 | 5.63E-05 | 0.26 | 0.14 | 0.12 | 1.87 |
| 9 | Tax1bp1 | 0.37 | 2.71E-09 | 5.79E-05 | 0.95 | 0.92 | 0.02 | 1.03 |
| 9 | Atp6v0b | 0.47 | 2.77E-09 | 5.93E-05 | 0.84 | 0.77 | 0.07 | 1.09 |
| 9 | Baz2b | 0.47 | 2.86E-09 | 6.12E-05 | 0.87 | 0.81 | 0.06 | 1.07 |
| 9 | Abcg1 | 0.75 | 2.90E-09 | 6.19E-05 | 0.44 | 0.28 | 0.15 | 1.53 |
| 9 | Mex3a | 0.38 | 2.90E-09 | 6.19E-05 | 0.93 | 0.87 | 0.06 | 1.07 |
| 9 | Btbd7 | 0.57 | 2.99E-09 | 6.40E-05 | 0.78 | 0.71 | 0.07 | 1.09 |
| 9 | Heph | 0.92 | 3.17E-09 | 6.78E-05 | 0.41 | 0.26 | 0.15 | 1.58 |
| 9 | Map3k20 | 1.3 | 3.21E-09 | 6.86E-05 | 0.31 | 0.18 | 0.13 | 1.71 |
| 9 | Alcam | 1.53 | 3.37E-09 | 7.20E-05 | 0.33 | 0.2 | 0.14 | 1.69 |
| 9 | Atp6v0d1 | 0.65 | 3.88E-09 | 8.28E-05 | 0.65 | 0.56 | 0.09 | 1.17 |
| 9 | Ubald2 | 0.52 | 4.01E-09 | 8.56E-05 | 0.78 | 0.71 | 0.07 | 1.1 |
| 9 | Ptprk | 0.59 | 4.15E-09 | 8.87E-05 | 0.69 | 0.59 | 0.1 | 1.17 |
| 9 | Rrbp1 | 0.46 | 4.21E-09 | 8.99E-05 | 0.93 | 0.89 | 0.03 | 1.04 |
| 9 | Trip12 | 0.53 | 4.22E-09 | 9.00E-05 | 0.77 | 0.71 | 0.06 | 1.09 |
| 9 | Myh9 | 0.56 | 4.45E-09 | 9.50E-05 | 0.81 | 0.75 | 0.06 | 1.08 |
| 9 | Rnaset2a | 0.94 | 4.75E-09 | 1.01E-04 | 0.61 | 0.53 | 0.08 | 1.16 |
| 9 | Ppdpf | 0.27 | 4.97E-09 | 1.06E-04 | 0.98 | 0.97 | 0.01 | 1.01 |
| 9 | Stx18 | 0.67 | 5.81E-09 | 1.24E-04 | 0.54 | 0.41 | 0.13 | 1.31 |
| 9 | Hspa2 | 1.18 | 5.99E-09 | 1.28E-04 | 0.26 | 0.13 | 0.13 | 1.96 |
| 9 | Tanc2 | 1.21 | 6.00E-09 | 1.28E-04 | 0.36 | 0.22 | 0.14 | 1.63 |
| 9 | Nsa2 | 0.26 | 6.03E-09 | 1.29E-04 | 0.97 | 0.97 | 0 | 1 |
| 9 | Renbp | 1.53 | 6.76E-09 | 1.44E-04 | 0.22 | 0.11 | 0.11 | 2.02 |
| 9 | Ankhd1 | 0.64 | 7.75E-09 | 1.66E-04 | 0.67 | 0.58 | 0.1 | 1.16 |
| 9 | Prrc2b | 0.44 | 7.79E-09 | 1.67E-04 | 0.85 | 0.79 | 0.06 | 1.08 |
| 9 | Itgb4 | 1.23 | 8.00E-09 | 1.71E-04 | 0.24 | 0.12 | 0.12 | 2.01 |
| 9 | Grina | 1.12 | 8.22E-09 | 1.75E-04 | 0.43 | 0.3 | 0.13 | 1.42 |

**Supplementary Table 3. Marker genes for the original clusters of integrated clustering of E11.5 control and NotchGOF λ-junction ectoderm cells.**

| | | | | | | | | |
|---|---|---|---|---|---|---|---|---|
| 9 | Ulk1 | 0.99 | 8.29E-09 | 1.77E-04 | 0.4 | 0.27 | 0.13 | 1.49 |
| 9 | 2610035D17Rik | 1.87 | 9.39E-09 | 2.01E-04 | 0.13 | 0.05 | 0.08 | 2.72 |
| 9 | Hsdl2 | 1.06 | 9.77E-09 | 2.09E-04 | 0.46 | 0.34 | 0.12 | 1.36 |
| 9 | Ube2b | 0.31 | 9.78E-09 | 2.09E-04 | 0.94 | 0.95 | -0.01 | 0.99 |
| 9 | Dkk3 | 1.32 | 9.95E-09 | 2.13E-04 | 0.28 | 0.16 | 0.12 | 1.77 |
| 9 | Tacc1 | 1.29 | 1.03E-08 | 2.20E-04 | 0.31 | 0.19 | 0.13 | 1.67 |
| 9 | Nfic | 1.31 | 1.18E-08 | 2.51E-04 | 0.28 | 0.16 | 0.12 | 1.75 |
| 9 | Oga | 0.48 | 1.24E-08 | 2.65E-04 | 0.8 | 0.72 | 0.08 | 1.11 |
| 9 | Sbk1 | 0.58 | 1.37E-08 | 2.92E-04 | 0.67 | 0.56 | 0.11 | 1.19 |
| 9 | Klk8 | 0.6 | 1.39E-08 | 2.96E-04 | 0.66 | 0.56 | 0.11 | 1.19 |
| 9 | Mbtd1 | 0.43 | 1.44E-08 | 3.07E-04 | 0.84 | 0.77 | 0.07 | 1.09 |
| 9 | Arl8a | 0.71 | 1.44E-08 | 3.08E-04 | 0.52 | 0.38 | 0.14 | 1.37 |
| 9 | Itga11 | 1.95 | 1.52E-08 | 3.24E-04 | 0.1 | 0.03 | 0.07 | 3.09 |
| 9 | E130102H24Rik | 1.98 | 1.53E-08 | 3.28E-04 | 0.11 | 0.04 | 0.07 | 2.97 |
| 9 | Wnt5b | 1.47 | 1.63E-08 | 3.48E-04 | 0.25 | 0.13 | 0.12 | 1.89 |
| 9 | Sdc4 | 0.58 | 1.71E-08 | 3.65E-04 | 0.73 | 0.65 | 0.08 | 1.12 |
| 9 | Glmp | 0.65 | 1.79E-08 | 3.83E-04 | 0.61 | 0.49 | 0.12 | 1.25 |
| 9 | 2410006H16Rik | 0.43 | 1.81E-08 | 3.86E-04 | 0.9 | 0.86 | 0.04 | 1.05 |
| 9 | Camk2n1 | 0.65 | 1.88E-08 | 4.02E-04 | 0.23 | 0.12 | 0.12 | 2.01 |
| 9 | Mapkbp1 | 1.67 | 1.93E-08 | 4.13E-04 | 0.17 | 0.07 | 0.09 | 2.32 |
| 9 | Pax3 | 1.75 | 1.96E-08 | 4.19E-04 | 0.14 | 0.06 | 0.09 | 2.52 |
| 9 | Tet2 | 0.95 | 1.99E-08 | 4.24E-04 | 0.34 | 0.21 | 0.13 | 1.63 |
| 9 | Meg3 | 0.64 | 2.03E-08 | 4.34E-04 | 0.83 | 0.76 | 0.07 | 1.09 |
| 9 | Wdr26 | 0.53 | 2.38E-08 | 5.09E-04 | 0.81 | 0.75 | 0.06 | 1.08 |
| 9 | Capg | 0.81 | 2.52E-08 | 5.38E-04 | 0.48 | 0.35 | 0.13 | 1.36 |
| 9 | Sbspon | 2.18 | 2.66E-08 | 5.68E-04 | 0.11 | 0.04 | 0.07 | 2.94 |
| 9 | Pkp3 | 0.79 | 2.86E-08 | 6.12E-04 | 0.39 | 0.24 | 0.14 | 1.59 |
| 9 | Pdlim5 | 1.35 | 2.88E-08 | 6.14E-04 | 0.29 | 0.17 | 0.12 | 1.7 |
| 9 | Pak2 | 0.4 | 2.94E-08 | 6.29E-04 | 0.91 | 0.89 | 0.02 | 1.02 |
| 9 | Amn1 | 0.64 | 3.09E-08 | 6.59E-04 | 0.61 | 0.5 | 0.11 | 1.22 |
| 9 | Myl9 | 1.63 | 3.10E-08 | 6.63E-04 | 0.31 | 0.19 | 0.12 | 1.65 |
| 9 | Cdc42ep3 | 1.31 | 3.11E-08 | 6.64E-04 | 0.3 | 0.17 | 0.13 | 1.74 |
| 9 | Sesn2 | 1.5 | 3.21E-08 | 6.85E-04 | 0.21 | 0.11 | 0.11 | 2.02 |
| 9 | Spry1 | 1.3 | 3.23E-08 | 6.89E-04 | 0.44 | 0.32 | 0.12 | 1.38 |
| 9 | Kxd1 | 0.49 | 3.24E-08 | 6.91E-04 | 0.73 | 0.63 | 0.1 | 1.16 |
| 9 | Cacna1b | 0.93 | 3.29E-08 | 7.03E-04 | 0.43 | 0.3 | 0.13 | 1.43 |
| 9 | Siva1 | 0.4 | 3.37E-08 | 7.20E-04 | 0.98 | 0.97 | 0.01 | 1.01 |
| 9 | Tmtc2 | 0.94 | 4.03E-08 | 8.62E-04 | 0.31 | 0.18 | 0.13 | 1.74 |
| 9 | Arid3b | 0.66 | 4.08E-08 | 8.71E-04 | 0.55 | 0.41 | 0.14 | 1.35 |
| 9 | Sqstm1 | 0.61 | 4.26E-08 | 9.11E-04 | 0.58 | 0.46 | 0.13 | 1.28 |
| 9 | Qk | 0.38 | 4.29E-08 | 9.16E-04 | 0.91 | 0.87 | 0.03 | 1.04 |
| 9 | Ccdc28a | 1.01 | 4.34E-08 | 9.27E-04 | 0.28 | 0.16 | 0.12 | 1.79 |
| 9 | Mgat5 | 0.84 | 4.54E-08 | 9.69E-04 | 0.43 | 0.29 | 0.14 | 1.48 |
| 9 | Snhg6 | 0.48 | 4.56E-08 | 9.75E-04 | 0.78 | 0.73 | 0.05 | 1.07 |
| 9 | Cd81 | 0.34 | 4.68E-08 | 1.00E-03 | 0.94 | 0.92 | 0.02 | 1.02 |
| 9 | Bmp7 | 0.79 | 4.69E-08 | 1.00E-03 | 0.53 | 0.4 | 0.13 | 1.32 |
| 9 | Slc30a4 | 1.55 | 4.70E-08 | 1.00E-03 | 0.22 | 0.11 | 0.11 | 1.95 |
| 9 | Bag3 | 1.72 | 4.82E-08 | 1.03E-03 | 0.16 | 0.07 | 0.09 | 2.27 |
| 9 | Plin2 | 0.88 | 5.06E-08 | 1.08E-03 | 0.49 | 0.36 | 0.13 | 1.36 |
| 9 | Mtus1 | 0.93 | 6.33E-08 | 1.35E-03 | 0.42 | 0.29 | 0.13 | 1.43 |
| 9 | Rnf167 | 0.63 | 6.50E-08 | 1.39E-03 | 0.58 | 0.47 | 0.12 | 1.25 |
| 9 | Hey1 | 0.47 | 6.57E-08 | 1.40E-03 | 0.86 | 0.74 | 0.12 | 1.16 |
| 9 | Slc12a4 | 1.05 | 6.59E-08 | 1.41E-03 | 0.33 | 0.2 | 0.13 | 1.64 |
| 9 | Ccng2 | 0.78 | 7.12E-08 | 1.52E-03 | 0.51 | 0.39 | 0.12 | 1.31 |
| 9 | Etv3 | 0.9 | 7.90E-08 | 1.69E-03 | 0.41 | 0.29 | 0.13 | 1.45 |
| 9 | Tulp1 | 1.4 | 8.10E-08 | 1.73E-03 | 0.2 | 0.1 | 0.1 | 1.98 |
| 9 | Skp1a | 0.26 | 8.27E-08 | 1.77E-03 | 0.99 | 0.97 | 0.02 | 1.02 |

**Supplementary Table 3. Marker genes for the original clusters of integrated clustering of E11.5 control and NotchGOF λ-junction ectoderm cells.**

| | | | | | | | | |
|---|---|---|---|---|---|---|---|---|
| 9 | Atp6v1e1 | 0.52 | 8.39E-08 | 1.79E-03 | 0.74 | 0.69 | 0.05 | 1.08 |
| 9 | Rtn4 | 0.29 | 8.50E-08 | 1.82E-03 | 0.98 | 0.96 | 0.03 | 1.03 |
| 9 | Zkscan3 | 0.45 | 9.50E-08 | 2.03E-03 | 0.77 | 0.67 | 0.1 | 1.15 |
| 9 | Tiam1 | 0.69 | 9.68E-08 | 2.07E-03 | 0.57 | 0.45 | 0.12 | 1.26 |
| 9 | A430035B10Rik | 1.59 | 9.72E-08 | 2.08E-03 | 0.11 | 0.04 | 0.07 | 2.79 |
| 9 | Filip1 | 1.97 | 1.01E-07 | 2.16E-03 | 0.16 | 0.07 | 0.09 | 2.21 |
| 9 | Gng12 | 1.03 | 1.02E-07 | 2.18E-03 | 0.35 | 0.23 | 0.12 | 1.52 |
| 9 | Kremen1 | 1.07 | 1.04E-07 | 2.23E-03 | 0.44 | 0.31 | 0.12 | 1.39 |
| 9 | Elk3 | 1.1 | 1.05E-07 | 2.24E-03 | 0.31 | 0.19 | 0.12 | 1.64 |
| 9 | C1qtnf12 | 1.36 | 1.06E-07 | 2.26E-03 | 0.34 | 0.22 | 0.11 | 1.5 |
| 9 | Clip3 | 0.73 | 1.07E-07 | 2.29E-03 | 0.55 | 0.46 | 0.1 | 1.21 |
| 9 | Smarce1 | 0.26 | 1.34E-07 | 2.85E-03 | 0.98 | 0.97 | 0 | 1 |
| 9 | Rnf24 | 0.79 | 1.34E-07 | 2.86E-03 | 0.53 | 0.41 | 0.13 | 1.32 |
| 9 | Pura | 0.51 | 1.42E-07 | 3.04E-03 | 0.72 | 0.64 | 0.08 | 1.13 |
| 9 | Ppp1r36 | 1.06 | 1.47E-07 | 3.13E-03 | 0.28 | 0.16 | 0.12 | 1.77 |
| 9 | Fgd6 | 1 | 1.55E-07 | 3.32E-03 | 0.34 | 0.22 | 0.12 | 1.57 |
| 9 | Zmiz1 | 0.62 | 1.62E-07 | 3.46E-03 | 0.7 | 0.62 | 0.08 | 1.12 |
| 9 | Tmem184b | 1.14 | 1.68E-07 | 3.58E-03 | 0.32 | 0.2 | 0.11 | 1.56 |
| 9 | Lpin2 | 0.78 | 1.74E-07 | 3.71E-03 | 0.55 | 0.46 | 0.09 | 1.19 |
| 9 | Sox14 | 1.5 | 1.74E-07 | 3.72E-03 | 0.12 | 0.05 | 0.08 | 2.6 |
| 9 | Ehbp1 | 0.68 | 1.77E-07 | 3.78E-03 | 0.6 | 0.49 | 0.11 | 1.23 |
| 9 | Rgl1 | 1.69 | 2.00E-07 | 4.28E-03 | 0.14 | 0.06 | 0.08 | 2.36 |
| 9 | Mxd4 | 0.62 | 2.04E-07 | 4.35E-03 | 0.54 | 0.41 | 0.13 | 1.31 |
| 9 | Ass1 | 1.1 | 2.20E-07 | 4.71E-03 | 0.22 | 0.12 | 0.11 | 1.91 |
| 9 | Mbd2 | 0.48 | 2.39E-07 | 5.11E-03 | 0.69 | 0.61 | 0.08 | 1.12 |
| 9 | Bach1 | 0.69 | 2.40E-07 | 5.13E-03 | 0.53 | 0.42 | 0.11 | 1.26 |
| 9 | Nfe2l3 | 1.22 | 2.43E-07 | 5.19E-03 | 0.24 | 0.14 | 0.11 | 1.8 |
| 9 | Kcnj15 | 1.6 | 2.62E-07 | 5.59E-03 | 0.13 | 0.06 | 0.08 | 2.42 |
| 9 | Macrod2 | 1.22 | 2.70E-07 | 5.76E-03 | 0.46 | 0.35 | 0.11 | 1.32 |
| 9 | Ahr | 1.52 | 2.73E-07 | 5.82E-03 | 0.28 | 0.18 | 0.11 | 1.61 |
| 9 | Lrp1 | 0.71 | 2.77E-07 | 5.92E-03 | 0.54 | 0.43 | 0.11 | 1.25 |
| 9 | Txndc17 | 0.31 | 2.77E-07 | 5.93E-03 | 0.95 | 0.95 | 0 | 1 |
| 9 | F11r | 0.49 | 2.82E-07 | 6.02E-03 | 0.69 | 0.61 | 0.09 | 1.14 |
| 9 | Htra1 | 1.69 | 3.02E-07 | 6.44E-03 | 0.14 | 0.06 | 0.08 | 2.31 |
| 9 | Fzd3 | 0.29 | 3.36E-07 | 7.17E-03 | 0.89 | 0.84 | 0.05 | 1.05 |
| 9 | Slc25a27 | 0.55 | 3.46E-07 | 7.39E-03 | 0.64 | 0.54 | 0.09 | 1.17 |
| 9 | Trappc6a | 0.47 | 3.47E-07 | 7.42E-03 | 0.72 | 0.66 | 0.06 | 1.09 |
| 9 | Plekhg3 | 1.08 | 3.48E-07 | 7.43E-03 | 0.38 | 0.27 | 0.12 | 1.43 |
| 9 | Dusp11 | 0.46 | 3.53E-07 | 7.55E-03 | 0.73 | 0.66 | 0.07 | 1.1 |
| 9 | Herc3 | 1.48 | 3.57E-07 | 7.63E-03 | 0.24 | 0.14 | 0.1 | 1.76 |
| 9 | Sinhcaf | 0.27 | 3.68E-07 | 7.86E-03 | 0.95 | 0.94 | 0.01 | 1.01 |
| 9 | Rara | 0.73 | 3.74E-07 | 7.99E-03 | 0.55 | 0.45 | 0.1 | 1.22 |
| 9 | Nfat5 | 0.43 | 4.10E-07 | 8.75E-03 | 0.72 | 0.63 | 0.09 | 1.14 |
| 9 | Gpbp1l1 | 0.65 | 4.14E-07 | 8.85E-03 | 0.56 | 0.45 | 0.11 | 1.25 |
| 9 | Nkd1 | 1.5 | 4.23E-07 | 9.04E-03 | 0.22 | 0.12 | 0.1 | 1.82 |
| 9 | Tspan5 | 0.93 | 4.24E-07 | 9.05E-03 | 0.44 | 0.34 | 0.1 | 1.3 |
| 9 | Znrf1 | 0.48 | 4.42E-07 | 9.44E-03 | 0.66 | 0.56 | 0.1 | 1.17 |
| 9 | Kmt2e | 0.38 | 4.43E-07 | 9.47E-03 | 0.88 | 0.86 | 0.02 | 1.02 |
| 9 | Anxa4 | 1.08 | 4.49E-07 | 9.59E-03 | 0.27 | 0.16 | 0.11 | 1.72 |
| 9 | Dok1 | 1.51 | 4.57E-07 | 9.76E-03 | 0.2 | 0.11 | 0.1 | 1.91 |
| 9 | Cpeb4 | 1.53 | 4.60E-07 | 9.83E-03 | 0.15 | 0.07 | 0.08 | 2.19 |
| 9 | Trappc2l | 0.48 | 4.77E-07 | 1.02E-02 | 0.68 | 0.61 | 0.07 | 1.12 |
| 9 | Mylk | 1.17 | 4.82E-07 | 1.03E-02 | 0.31 | 0.2 | 0.11 | 1.55 |
| 9 | Rap2b | 0.71 | 4.89E-07 | 1.04E-02 | 0.61 | 0.53 | 0.08 | 1.15 |
| 9 | Cuedc1 | 0.63 | 5.34E-07 | 1.14E-02 | 0.61 | 0.51 | 0.1 | 1.2 |
| 9 | Ubxn6 | 0.63 | 5.64E-07 | 1.21E-02 | 0.59 | 0.49 | 0.1 | 1.2 |
| 9 | Neil2 | 1.8 | 6.17E-07 | 1.32E-02 | 0.1 | 0.04 | 0.06 | 2.68 |

**Supplementary Table 3. Marker genes for the original clusters of integrated clustering of E11.5 control and NotchGOF λ-junction ectoderm cells.**

| | | | | | | | | |
|---|---|---|---|---|---|---|---|---|
| 9 | Ankrd12 | 0.66 | 6.89E-07 | 1.47E-02 | 0.53 | 0.42 | 0.11 | 1.26 |
| 9 | Arhgef19 | 0.68 | 6.98E-07 | 1.49E-02 | 0.58 | 0.49 | 0.1 | 1.2 |
| 9 | Lhfp | 0.86 | 7.07E-07 | 1.51E-02 | 0.44 | 0.31 | 0.13 | 1.42 |
| 9 | Akap9 | 0.31 | 7.39E-07 | 1.58E-02 | 0.9 | 0.87 | 0.03 | 1.03 |
| 9 | Arhgap42 | 0.84 | 7.46E-07 | 1.59E-02 | 0.39 | 0.27 | 0.12 | 1.43 |
| 9 | Bcam | 0.46 | 7.72E-07 | 1.65E-02 | 0.82 | 0.72 | 0.1 | 1.14 |
| 9 | Crlf1 | 1.42 | 7.91E-07 | 1.69E-02 | 0.22 | 0.13 | 0.1 | 1.78 |
| 9 | Hook3 | 0.54 | 8.06E-07 | 1.72E-02 | 0.64 | 0.53 | 0.11 | 1.2 |
| 9 | Eps15 | 0.78 | 8.38E-07 | 1.79E-02 | 0.44 | 0.32 | 0.11 | 1.34 |
| 9 | Metap2 | 0.28 | 8.59E-07 | 1.84E-02 | 0.98 | 0.97 | 0.01 | 1.01 |
| 9 | Tcf4 | 0.43 | 8.97E-07 | 1.92E-02 | 0.91 | 0.89 | 0.03 | 1.03 |
| 9 | Ddx17 | 0.36 | 9.06E-07 | 1.94E-02 | 0.89 | 0.87 | 0.02 | 1.02 |
| 9 | Rps27rt | 0.4 | 9.55E-07 | 2.04E-02 | 0.84 | 0.78 | 0.06 | 1.08 |
| 9 | Rasgef1b | 1.22 | 9.62E-07 | 2.06E-02 | 0.18 | 0.09 | 0.09 | 1.98 |
| 9 | Sppl3 | 0.57 | 9.91E-07 | 2.12E-02 | 0.71 | 0.64 | 0.06 | 1.1 |
| 9 | Sema6c | 1.3 | 1.02E-06 | 2.19E-02 | 0.22 | 0.13 | 0.1 | 1.79 |
| 9 | Castor1 | 1.52 | 1.03E-06 | 2.19E-02 | 0.12 | 0.05 | 0.07 | 2.44 |
| 9 | Elf1 | 0.54 | 1.07E-06 | 2.30E-02 | 0.61 | 0.5 | 0.11 | 1.22 |
| 9 | Gpcpd1 | 1.02 | 1.12E-06 | 2.39E-02 | 0.32 | 0.21 | 0.11 | 1.53 |
| 9 | Gpr153 | 1.5 | 1.13E-06 | 2.42E-02 | 0.11 | 0.04 | 0.07 | 2.59 |
| 9 | Sik1 | 1.03 | 1.14E-06 | 2.44E-02 | 0.27 | 0.16 | 0.11 | 1.69 |
| 9 | Nfatc4 | 0.89 | 1.16E-06 | 2.47E-02 | 0.44 | 0.34 | 0.1 | 1.3 |
| 9 | Mrps33 | 0.27 | 1.19E-06 | 2.53E-02 | 0.95 | 0.94 | 0.01 | 1.01 |
| 9 | Glul | 0.64 | 1.19E-06 | 2.54E-02 | 0.57 | 0.46 | 0.11 | 1.23 |
| 9 | Rflnb | 0.64 | 1.34E-06 | 2.86E-02 | 0.62 | 0.52 | 0.1 | 1.2 |
| 9 | Kansl1l | 1.04 | 1.36E-06 | 2.91E-02 | 0.31 | 0.2 | 0.11 | 1.56 |
| 9 | Pink1 | 0.87 | 1.42E-06 | 3.03E-02 | 0.36 | 0.25 | 0.11 | 1.44 |
| 9 | Efna1 | 0.56 | 1.45E-06 | 3.10E-02 | 0.52 | 0.4 | 0.12 | 1.3 |
| 9 | Smad5 | 0.63 | 1.61E-06 | 3.44E-02 | 0.57 | 0.47 | 0.1 | 1.22 |
| 9 | Ctnnd1 | 0.63 | 1.62E-06 | 3.45E-02 | 0.6 | 0.52 | 0.08 | 1.16 |
| 9 | Cdk19 | 0.84 | 1.64E-06 | 3.50E-02 | 0.4 | 0.3 | 0.11 | 1.36 |
| 9 | Ebpl | 0.54 | 1.76E-06 | 3.75E-02 | 0.64 | 0.55 | 0.09 | 1.17 |
| 9 | Heca | 0.77 | 1.77E-06 | 3.79E-02 | 0.38 | 0.26 | 0.12 | 1.46 |
| 9 | Kdm3a | 0.65 | 1.80E-06 | 3.85E-02 | 0.56 | 0.47 | 0.09 | 1.2 |
| 9 | Cttnbp2nl | 0.77 | 1.87E-06 | 4.00E-02 | 0.51 | 0.43 | 0.08 | 1.19 |
| 9 | Stim1 | 1.03 | 1.88E-06 | 4.01E-02 | 0.28 | 0.17 | 0.11 | 1.64 |
| 9 | Rbm5 | 0.37 | 1.90E-06 | 4.06E-02 | 0.84 | 0.81 | 0.03 | 1.04 |
| 9 | Lrrc58 | 0.47 | 1.91E-06 | 4.07E-02 | 0.68 | 0.61 | 0.07 | 1.11 |
| 9 | Cir1 | 0.49 | 2.00E-06 | 4.28E-02 | 0.71 | 0.64 | 0.07 | 1.11 |
| 9 | Hmox1 | 1.26 | 2.01E-06 | 4.29E-02 | 0.17 | 0.09 | 0.08 | 1.99 |
| 9 | Pard6g | 0.53 | 2.03E-06 | 4.34E-02 | 0.69 | 0.65 | 0.05 | 1.07 |
| 9 | H2-T23 | 1.15 | 2.09E-06 | 4.47E-02 | 0.25 | 0.15 | 0.1 | 1.69 |
| 9 | Numb | 1.03 | 2.15E-06 | 4.59E-02 | 0.42 | 0.32 | 0.1 | 1.31 |
| 9 | Gabarapl2 | 0.34 | 2.19E-06 | 4.67E-02 | 0.88 | 0.87 | 0.01 | 1.01 |
| 9 | Zfp637 | 0.48 | 2.27E-06 | 4.85E-02 | 0.71 | 0.65 | 0.06 | 1.09 |
| 9 | 4930402H24Rik | 1.01 | 2.30E-06 | 4.91E-02 | 0.26 | 0.16 | 0.1 | 1.65 |
| 10 | Cxcl14 | 3.23 | 7.94E-176 | 1.70E-171 | 0.95 | 0.17 | 0.78 | 5.55 |
| 10 | Tfap2b | 2.87 | 4.31E-155 | 9.20E-151 | 0.91 | 0.16 | 0.75 | 5.54 |
| 10 | Wnt3 | 2.89 | 1.11E-146 | 2.36E-142 | 0.71 | 0.09 | 0.62 | 7.76 |
| 10 | Slc7a11 | 3.71 | 8.21E-123 | 1.75E-118 | 0.66 | 0.11 | 0.55 | 6.06 |
| 10 | Wnt9b | 3.26 | 1.14E-119 | 2.43E-115 | 0.63 | 0.09 | 0.54 | 6.69 |
| 10 | Piezo2 | 2.94 | 1.37E-105 | 2.93E-101 | 0.72 | 0.15 | 0.57 | 4.87 |
| 10 | Plxna2 | 2.74 | 2.21E-97 | 4.72E-93 | 0.82 | 0.24 | 0.58 | 3.38 |
| 10 | Pcp4l1 | 3.19 | 2.53E-95 | 5.41E-91 | 0.47 | 0.06 | 0.41 | 7.74 |
| 10 | Wnt6 | 2.04 | 2.19E-91 | 4.68E-87 | 0.96 | 0.35 | 0.61 | 2.76 |
| 10 | Col17a1 | 3 | 7.95E-73 | 1.70E-68 | 0.53 | 0.11 | 0.42 | 4.8 |
| 10 | Sostdc1 | 1.89 | 9.44E-70 | 2.02E-65 | 0.76 | 0.24 | 0.52 | 3.2 |

**Supplementary Table 3. Marker genes for the original clusters of integrated clustering of E11.5 control and NotchGOF λ-junction ectoderm cells.**

| | | | | | | | | |
|---|---|---|---|---|---|---|---|---|
| 10 | Ccnd2 | 1.74 | 7.75E-66 | 1.66E-61 | 1 | 0.78 | 0.22 | 1.28 |
| 10 | Irx5 | 2.33 | 4.74E-63 | 1.01E-58 | 0.69 | 0.23 | 0.47 | 3.07 |
| 10 | Tfap2a | 2.19 | 7.11E-62 | 1.52E-57 | 0.78 | 0.33 | 0.45 | 2.38 |
| 10 | Perp | 1.62 | 2.04E-58 | 4.35E-54 | 0.96 | 0.52 | 0.44 | 1.84 |
| 10 | Crabp1 | 1.99 | 2.71E-57 | 5.79E-53 | 0.9 | 0.55 | 0.35 | 1.64 |
| 10 | Bambi | 1.75 | 3.27E-56 | 6.99E-52 | 0.87 | 0.47 | 0.4 | 1.86 |
| 10 | Cox4i2 | 1.97 | 3.33E-53 | 7.11E-49 | 0.83 | 0.47 | 0.36 | 1.77 |
| 10 | Pdlim1 | 1.48 | 1.21E-49 | 2.58E-45 | 0.92 | 0.64 | 0.29 | 1.45 |
| 10 | Kremen2 | 2.14 | 1.24E-49 | 2.65E-45 | 0.56 | 0.17 | 0.39 | 3.38 |
| 10 | Tnfrsf19 | 2 | 6.68E-48 | 1.43E-43 | 0.64 | 0.23 | 0.41 | 2.77 |
| 10 | Trp63 | 1.53 | 3.25E-47 | 6.95E-43 | 0.89 | 0.5 | 0.4 | 1.79 |
| 10 | Irf6 | 1.97 | 1.37E-46 | 2.92E-42 | 0.77 | 0.4 | 0.37 | 1.92 |
| 10 | Hras | 1.01 | 4.79E-45 | 1.02E-40 | 0.98 | 0.91 | 0.07 | 1.08 |
| 10 | Wls | 1.12 | 3.25E-43 | 6.93E-39 | 0.97 | 0.79 | 0.18 | 1.22 |
| 10 | Itgb5 | 1.65 | 8.38E-43 | 1.79E-38 | 0.83 | 0.49 | 0.34 | 1.7 |
| 10 | Col12a1 | 3.05 | 6.64E-41 | 1.42E-36 | 0.31 | 0.06 | 0.25 | 4.92 |
| 10 | Lef1 | 1.61 | 9.46E-41 | 2.02E-36 | 0.76 | 0.36 | 0.4 | 2.1 |
| 10 | Malat1 | 0.86 | 1.19E-40 | 2.53E-36 | 1 | 1 | 0 | 1 |
| 10 | Ctsl | 1.17 | 1.27E-40 | 2.72E-36 | 0.96 | 0.86 | 0.11 | 1.12 |
| 10 | Dlx2 | 1.67 | 5.51E-40 | 1.18E-35 | 0.73 | 0.33 | 0.4 | 2.22 |
| 10 | Nav3 | 1.9 | 4.16E-38 | 8.89E-34 | 0.72 | 0.37 | 0.35 | 1.94 |
| 10 | Xist | 0.83 | 6.71E-38 | 1.43E-33 | 0.99 | 0.91 | 0.08 | 1.09 |
| 10 | Slc2a3 | 1.84 | 4.33E-37 | 9.25E-33 | 0.69 | 0.34 | 0.35 | 2.01 |
| 10 | Dock8 | 2.59 | 5.48E-37 | 1.17E-32 | 0.35 | 0.08 | 0.26 | 4.16 |
| 10 | Irx3 | 1.95 | 7.02E-37 | 1.50E-32 | 0.54 | 0.2 | 0.35 | 2.77 |
| 10 | Wnt4 | 1.3 | 6.98E-36 | 1.49E-31 | 0.92 | 0.7 | 0.22 | 1.31 |
| 10 | Nebl | 1.82 | 1.83E-35 | 3.90E-31 | 0.71 | 0.39 | 0.32 | 1.83 |
| 10 | Btg1 | 1.58 | 3.83E-35 | 8.19E-31 | 0.87 | 0.75 | 0.12 | 1.17 |
| 10 | Irx2 | 2.91 | 9.42E-35 | 2.01E-30 | 0.25 | 0.05 | 0.2 | 5.41 |
| 10 | Snai2 | 2.09 | 3.11E-34 | 6.64E-30 | 0.49 | 0.17 | 0.32 | 2.9 |
| 10 | Sox11 | 0.86 | 9.61E-34 | 2.05E-29 | 1 | 0.99 | 0.01 | 1.01 |
| 10 | D7Ertd443e | 2.66 | 1.53E-33 | 3.28E-29 | 0.35 | 0.09 | 0.26 | 3.83 |
| 10 | Mbnl2 | 1.13 | 3.30E-33 | 7.04E-29 | 0.9 | 0.78 | 0.11 | 1.15 |
| 10 | Fzd3 | 1.17 | 4.79E-33 | 1.02E-28 | 0.95 | 0.84 | 0.11 | 1.13 |
| 10 | Laptm4a | 0.77 | 1.13E-32 | 2.41E-28 | 1 | 0.96 | 0.04 | 1.04 |
| 10 | Fjx1 | 1.12 | 1.18E-32 | 2.52E-28 | 0.93 | 0.79 | 0.14 | 1.18 |
| 10 | Wnt7b | 1.83 | 1.87E-32 | 3.99E-28 | 0.46 | 0.16 | 0.31 | 2.98 |
| 10 | Serpinb6b | 3.07 | 2.35E-32 | 5.02E-28 | 0.26 | 0.05 | 0.21 | 4.89 |
| 10 | Msx1 | 1.29 | 4.37E-32 | 9.34E-28 | 0.93 | 0.72 | 0.21 | 1.29 |
| 10 | Rftn1 | 3.27 | 5.01E-32 | 1.07E-27 | 0.22 | 0.04 | 0.18 | 5.59 |
| 10 | Pik3r1 | 1.43 | 1.08E-31 | 2.31E-27 | 0.83 | 0.59 | 0.24 | 1.41 |
| 10 | Fth1 | 0.58 | 1.12E-31 | 2.39E-27 | 1 | 1 | 0.01 | 1.01 |
| 10 | H19 | 0.96 | 4.97E-31 | 1.06E-26 | 0.99 | 0.88 | 0.12 | 1.13 |
| 10 | Jdp2 | 2.51 | 9.04E-31 | 1.93E-26 | 0.33 | 0.09 | 0.24 | 3.71 |
| 10 | Nrep | 0.94 | 1.30E-30 | 2.77E-26 | 0.93 | 0.8 | 0.13 | 1.16 |
| 10 | Fermt1 | 1.19 | 2.62E-30 | 5.60E-26 | 0.83 | 0.55 | 0.28 | 1.51 |
| 10 | Tpt1 | 0.27 | 2.67E-30 | 5.70E-26 | 1 | 1 | 0 | 1 |
| 10 | Htra1 | 2.6 | 3.10E-30 | 6.62E-26 | 0.26 | 0.06 | 0.2 | 4.63 |
| 10 | Nckap5 | 1.92 | 6.71E-30 | 1.43E-25 | 0.59 | 0.3 | 0.3 | 1.99 |
| 10 | mt-Co1 | 0.48 | 1.09E-29 | 2.32E-25 | 1 | 0.99 | 0.01 | 1.01 |
| 10 | mt-Co2 | 0.39 | 1.30E-29 | 2.77E-25 | 1 | 1 | 0 | 1 |
| 10 | Gria2 | 2.28 | 1.58E-29 | 3.37E-25 | 0.3 | 0.08 | 0.22 | 3.99 |
| 10 | Itm2b | 0.92 | 2.66E-29 | 5.69E-25 | 0.99 | 0.94 | 0.05 | 1.05 |
| 10 | Sorcs2 | 2.61 | 4.48E-29 | 9.56E-25 | 0.32 | 0.09 | 0.23 | 3.6 |
| 10 | Hspb1 | 1.39 | 6.26E-29 | 1.34E-24 | 0.71 | 0.37 | 0.33 | 1.89 |
| 10 | Krt14 | 1.18 | 8.24E-29 | 1.76E-24 | 0.79 | 0.45 | 0.34 | 1.75 |
| 10 | Gm42418 | 0.57 | 9.20E-29 | 1.97E-24 | 1 | 1 | 0 | 1 |

**Supplementary Table 3. Marker genes for the original clusters of integrated clustering of E11.5 control and NotchGOF λ-junction ectoderm cells.**

| | | | | | | | | |
|---|---|---|---|---|---|---|---|---|
| 10 | Chst11 | 2.06 | 4.59E-28 | 9.80E-24 | 0.42 | 0.15 | 0.27 | 2.77 |
| 10 | mt-Co3 | 0.4 | 6.16E-28 | 1.32E-23 | 1 | 1 | 0 | 1 |
| 10 | Rnd3 | 1.1 | 1.27E-27 | 2.71E-23 | 0.88 | 0.73 | 0.15 | 1.2 |
| 10 | Ehbp1 | 1.37 | 1.90E-27 | 4.07E-23 | 0.72 | 0.48 | 0.23 | 1.48 |
| 10 | Rps27 | 0.34 | 2.79E-27 | 5.97E-23 | 1 | 1 | 0 | 1 |
| 10 | Acaa2 | 1.09 | 1.22E-26 | 2.60E-22 | 0.85 | 0.71 | 0.14 | 1.19 |
| 10 | Fam89a | 2.04 | 1.72E-26 | 3.67E-22 | 0.41 | 0.15 | 0.26 | 2.72 |
| 10 | Foxp1 | 0.92 | 1.30E-25 | 2.78E-21 | 0.94 | 0.86 | 0.08 | 1.09 |
| 10 | Cdh6 | 2.34 | 1.92E-25 | 4.10E-21 | 0.36 | 0.12 | 0.24 | 3.05 |
| 10 | Dok5 | 1.4 | 2.29E-25 | 4.90E-21 | 0.66 | 0.37 | 0.29 | 1.79 |
| 10 | Rpl23 | 0.25 | 2.33E-25 | 4.98E-21 | 1 | 1 | 0 | 1 |
| 10 | Cobll1 | 2.53 | 3.71E-25 | 7.92E-21 | 0.31 | 0.09 | 0.22 | 3.33 |
| 10 | Cd59a | 1.62 | 4.15E-25 | 8.86E-21 | 0.58 | 0.29 | 0.29 | 1.98 |
| 10 | Nav2 | 1.2 | 5.04E-25 | 1.08E-20 | 0.86 | 0.7 | 0.16 | 1.23 |
| 10 | Gja1 | 1.14 | 5.89E-25 | 1.26E-20 | 0.87 | 0.7 | 0.17 | 1.24 |
| 10 | Col4a2 | 1.46 | 5.91E-25 | 1.26E-20 | 0.7 | 0.5 | 0.2 | 1.41 |
| 10 | Cdkn1c | 0.95 | 7.33E-25 | 1.57E-20 | 0.95 | 0.88 | 0.07 | 1.08 |
| 10 | Slc3a2 | 1.19 | 7.79E-25 | 1.66E-20 | 0.82 | 0.67 | 0.16 | 1.23 |
| 10 | Sdc1 | 1.43 | 9.41E-25 | 2.01E-20 | 0.75 | 0.58 | 0.16 | 1.28 |
| 10 | Insyn2b | 2.27 | 1.53E-23 | 3.26E-19 | 0.25 | 0.06 | 0.19 | 3.89 |
| 10 | Igfbp4 | 1.16 | 2.14E-23 | 4.57E-19 | 0.85 | 0.74 | 0.11 | 1.15 |
| 10 | Cited4 | 2.48 | 3.91E-23 | 8.36E-19 | 0.22 | 0.05 | 0.17 | 4.27 |
| 10 | Igfbp5 | 0.75 | 4.80E-23 | 1.02E-18 | 0.99 | 0.96 | 0.03 | 1.03 |
| 10 | Nsg1 | 1.11 | 5.38E-23 | 1.15E-18 | 0.8 | 0.6 | 0.2 | 1.33 |
| 10 | Fgfr2 | 1.21 | 5.70E-23 | 1.22E-18 | 0.72 | 0.47 | 0.26 | 1.55 |
| 10 | Pcdh9 | 2.01 | 6.12E-23 | 1.31E-18 | 0.42 | 0.18 | 0.24 | 2.38 |
| 10 | Adk | 1.13 | 7.68E-23 | 1.64E-18 | 0.76 | 0.61 | 0.15 | 1.25 |
| 10 | Txndc17 | 0.82 | 2.05E-22 | 4.37E-18 | 0.93 | 0.95 | -0.01 | 0.99 |
| 10 | Dlx1 | 2.1 | 2.17E-22 | 4.63E-18 | 0.37 | 0.14 | 0.23 | 2.69 |
| 10 | Frmd4b | 1.35 | 2.28E-22 | 4.87E-18 | 0.69 | 0.49 | 0.2 | 1.41 |
| 10 | Ahnak | 1.39 | 6.75E-22 | 1.44E-17 | 0.62 | 0.35 | 0.27 | 1.78 |
| 10 | Ngf | 2.74 | 1.81E-21 | 3.87E-17 | 0.16 | 0.03 | 0.13 | 5.06 |
| 10 | Krt5 | 1.48 | 2.21E-21 | 4.73E-17 | 0.6 | 0.33 | 0.28 | 1.84 |
| 10 | Mir205hg | 1.18 | 2.38E-21 | 5.08E-17 | 0.69 | 0.43 | 0.25 | 1.58 |
| 10 | Col18a1 | 0.77 | 3.95E-21 | 8.43E-17 | 0.9 | 0.86 | 0.05 | 1.05 |
| 10 | Stk26 | 1.42 | 6.70E-21 | 1.43E-16 | 0.68 | 0.46 | 0.21 | 1.45 |
| 10 | Dab1 | 1.17 | 1.12E-20 | 2.39E-16 | 0.73 | 0.48 | 0.24 | 1.5 |
| 10 | Fau | 0.26 | 1.25E-20 | 2.66E-16 | 1 | 1 | 0 | 1 |
| 10 | Lratd2 | 1.77 | 1.90E-20 | 4.06E-16 | 0.55 | 0.33 | 0.22 | 1.66 |
| 10 | Plxna4 | 2.68 | 2.37E-20 | 5.06E-16 | 0.2 | 0.05 | 0.15 | 4.06 |
| 10 | Msx2 | 1.23 | 2.90E-20 | 6.20E-16 | 0.66 | 0.39 | 0.26 | 1.67 |
| 10 | Gm32585 | 1.66 | 1.12E-19 | 2.39E-15 | 0.45 | 0.21 | 0.24 | 2.16 |
| 10 | Pygl | 1.9 | 1.73E-19 | 3.70E-15 | 0.4 | 0.17 | 0.22 | 2.3 |
| 10 | Gabarapl1 | 1.49 | 3.58E-19 | 7.66E-15 | 0.58 | 0.39 | 0.19 | 1.49 |
| 10 | Cdh3 | 1.21 | 4.10E-19 | 8.75E-15 | 0.72 | 0.57 | 0.15 | 1.26 |
| 10 | Bnc2 | 2.25 | 4.51E-19 | 9.62E-15 | 0.24 | 0.07 | 0.17 | 3.41 |
| 10 | Nrp2 | 1.82 | 5.81E-19 | 1.24E-14 | 0.39 | 0.17 | 0.22 | 2.33 |
| 10 | Plbd1 | 2.91 | 6.90E-19 | 1.47E-14 | 0.16 | 0.03 | 0.12 | 4.76 |
| 10 | Cald1 | 0.6 | 7.42E-19 | 1.59E-14 | 0.9 | 0.69 | 0.21 | 1.3 |
| 10 | Jag2 | 1.55 | 7.94E-19 | 1.70E-14 | 0.51 | 0.28 | 0.24 | 1.85 |
| 10 | Tmsb10 | 0.3 | 2.74E-18 | 5.86E-14 | 1 | 1 | 0 | 1 |
| 10 | Sema5a | 1.98 | 2.85E-18 | 6.09E-14 | 0.41 | 0.19 | 0.21 | 2.09 |
| 10 | Cox7a2l | 0.52 | 2.91E-18 | 6.21E-14 | 1 | 0.98 | 0.02 | 1.02 |
| 10 | Igfbp2 | 0.51 | 4.35E-18 | 9.29E-14 | 1 | 0.99 | 0.01 | 1.01 |
| 10 | Dapl1 | 1.18 | 5.41E-18 | 1.16E-13 | 0.52 | 0.26 | 0.26 | 1.98 |
| 10 | Art4 | 2.68 | 6.00E-18 | 1.28E-13 | 0.18 | 0.04 | 0.13 | 4.05 |
| 10 | Dsp | 0.75 | 9.77E-18 | 2.09E-13 | 0.97 | 0.87 | 0.11 | 1.12 |

**Supplementary Table 3. Marker genes for the original clusters of integrated clustering of E11.5 control and NotchGOF λ-junction ectoderm cells.**

| | | | | | | | | |
|---|---|---|---|---|---|---|---|---|
| 10 | Gstk1 | 1.87 | 1.63E-17 | 3.49E-13 | 0.34 | 0.14 | 0.2 | 2.48 |
| 10 | Irx1 | 2.13 | 1.94E-17 | 4.15E-13 | 0.23 | 0.07 | 0.16 | 3.3 |
| 10 | Lgals7 | 1.95 | 2.70E-17 | 5.76E-13 | 0.18 | 0.05 | 0.14 | 3.98 |
| 10 | Ednrb | 2.71 | 3.22E-17 | 6.88E-13 | 0.15 | 0.03 | 0.12 | 4.47 |
| 10 | mt-Atp6 | 0.31 | 4.74E-17 | 1.01E-12 | 1 | 1 | 0 | 1 |
| 10 | Fabp5 | 0.42 | 5.18E-17 | 1.11E-12 | 1 | 0.99 | 0.01 | 1.01 |
| 10 | Pthlh | 2.08 | 6.03E-17 | 1.29E-12 | 0.18 | 0.05 | 0.14 | 3.89 |
| 10 | Abca1 | 1.97 | 7.81E-17 | 1.67E-12 | 0.3 | 0.11 | 0.19 | 2.62 |
| 10 | Cib2 | 1.85 | 8.67E-17 | 1.85E-12 | 0.38 | 0.17 | 0.21 | 2.2 |
| 10 | Plk2 | 1.27 | 1.01E-16 | 2.16E-12 | 0.58 | 0.35 | 0.23 | 1.64 |
| 10 | Ablim1 | 1.06 | 2.57E-16 | 5.48E-12 | 0.69 | 0.55 | 0.14 | 1.26 |
| 10 | mt-Nd4 | 0.31 | 3.21E-16 | 6.86E-12 | 1 | 0.99 | 0.01 | 1.01 |
| 10 | Ccdc3 | 2.68 | 3.51E-16 | 7.50E-12 | 0.14 | 0.03 | 0.11 | 4.44 |
| 10 | Eva1a | 1.81 | 3.97E-16 | 8.48E-12 | 0.38 | 0.18 | 0.2 | 2.08 |
| 10 | Clta | 0.55 | 4.10E-16 | 8.77E-12 | 0.98 | 0.98 | 0 | 1 |
| 10 | Arhgap42 | 1.63 | 4.42E-16 | 9.45E-12 | 0.47 | 0.27 | 0.2 | 1.74 |
| 10 | Znrf3 | 1.58 | 5.98E-16 | 1.28E-11 | 0.48 | 0.28 | 0.2 | 1.7 |
| 10 | Ackr3 | 1.53 | 7.69E-16 | 1.64E-11 | 0.55 | 0.37 | 0.18 | 1.48 |
| 10 | Baiap2 | 1.66 | 2.59E-15 | 5.54E-11 | 0.41 | 0.21 | 0.2 | 1.96 |
| 10 | Vcan | 0.5 | 3.79E-15 | 8.10E-11 | 0.91 | 0.74 | 0.17 | 1.23 |
| 10 | Pnrc1 | 0.63 | 4.60E-15 | 9.82E-11 | 0.94 | 0.91 | 0.04 | 1.04 |
| 10 | Mylip | 1.68 | 4.89E-15 | 1.04E-10 | 0.39 | 0.2 | 0.19 | 1.96 |
| 10 | Otx1 | 1.22 | 5.06E-15 | 1.08E-10 | 0.6 | 0.44 | 0.16 | 1.38 |
| 10 | Arid5b | 1.06 | 5.46E-15 | 1.17E-10 | 0.7 | 0.55 | 0.15 | 1.27 |
| 10 | Trp73 | 2.61 | 7.22E-15 | 1.54E-10 | 0.12 | 0.03 | 0.1 | 4.88 |
| 10 | Npc2 | 0.59 | 9.36E-15 | 2.00E-10 | 0.94 | 0.92 | 0.02 | 1.02 |
| 10 | Notch1 | 0.86 | 1.22E-14 | 2.61E-10 | 0.75 | 0.62 | 0.13 | 1.21 |
| 10 | Npnt | 1.11 | 1.86E-14 | 3.97E-10 | 0.64 | 0.44 | 0.19 | 1.44 |
| 10 | Ntf5 | 1.32 | 1.87E-14 | 4.00E-10 | 0.52 | 0.31 | 0.21 | 1.67 |
| 10 | Ptpn13 | 0.84 | 2.90E-14 | 6.19E-10 | 0.74 | 0.57 | 0.17 | 1.29 |
| 10 | Ikzf2 | 0.76 | 3.00E-14 | 6.41E-10 | 0.9 | 0.84 | 0.07 | 1.08 |
| 10 | Itgav | 1.66 | 3.48E-14 | 7.42E-10 | 0.4 | 0.2 | 0.19 | 1.95 |
| 10 | Bdh1 | 1.72 | 6.60E-14 | 1.41E-09 | 0.33 | 0.15 | 0.18 | 2.2 |
| 10 | Tmem132a | 0.88 | 6.62E-14 | 1.41E-09 | 0.74 | 0.66 | 0.07 | 1.11 |
| 10 | Zfp36l2 | 1.52 | 6.65E-14 | 1.42E-09 | 0.43 | 0.24 | 0.19 | 1.77 |
| 10 | Antxr1 | 1.65 | 1.05E-13 | 2.25E-09 | 0.43 | 0.25 | 0.18 | 1.7 |
| 10 | Sfn | 0.28 | 1.19E-13 | 2.55E-09 | 0.94 | 0.69 | 0.25 | 1.36 |
| 10 | Dkk2 | 1.84 | 1.41E-13 | 3.01E-09 | 0.17 | 0.05 | 0.12 | 3.5 |
| 10 | Sgk3 | 1.85 | 1.60E-13 | 3.43E-09 | 0.28 | 0.12 | 0.17 | 2.41 |
| 10 | Igf2 | 0.64 | 1.66E-13 | 3.55E-09 | 0.88 | 0.81 | 0.07 | 1.09 |
| 10 | AY036118 | 0.81 | 1.94E-13 | 4.14E-09 | 0.76 | 0.64 | 0.12 | 1.19 |
| 10 | Ttc28 | 1.15 | 2.69E-13 | 5.75E-09 | 0.6 | 0.44 | 0.16 | 1.37 |
| 10 | mt-Cytb | 0.27 | 3.06E-13 | 6.53E-09 | 1 | 0.99 | 0.01 | 1.01 |
| 10 | Dst | 1.03 | 5.35E-13 | 1.14E-08 | 0.64 | 0.48 | 0.16 | 1.34 |
| 10 | Ajuba | 1.09 | 6.81E-13 | 1.45E-08 | 0.6 | 0.42 | 0.18 | 1.43 |
| 10 | Cldn1 | 1.53 | 7.34E-13 | 1.57E-08 | 0.25 | 0.1 | 0.16 | 2.57 |
| 10 | Ier5 | 0.59 | 8.87E-13 | 1.90E-08 | 0.91 | 0.84 | 0.07 | 1.08 |
| 10 | Rb1 | 1.28 | 8.94E-13 | 1.91E-08 | 0.47 | 0.29 | 0.19 | 1.66 |
| 10 | Timp3 | 2.42 | 9.56E-13 | 2.04E-08 | 0.19 | 0.06 | 0.13 | 3.02 |
| 10 | Dcbld1 | 1.28 | 1.04E-12 | 2.23E-08 | 0.55 | 0.39 | 0.16 | 1.42 |
| 10 | Fam162a | 0.49 | 1.24E-12 | 2.64E-08 | 0.94 | 0.93 | 0.02 | 1.02 |
| 10 | Chst15 | 1.78 | 1.46E-12 | 3.11E-08 | 0.29 | 0.13 | 0.16 | 2.19 |
| 10 | 1810058I24Rik | 0.48 | 2.22E-12 | 4.74E-08 | 0.96 | 0.93 | 0.03 | 1.03 |
| 10 | Kctd1 | 0.94 | 4.18E-12 | 8.94E-08 | 0.64 | 0.5 | 0.15 | 1.29 |
| 10 | Kcnq1ot1 | 0.64 | 4.33E-12 | 9.25E-08 | 0.92 | 0.94 | -0.02 | 0.98 |
| 10 | Zfp462 | 0.74 | 7.37E-12 | 1.57E-07 | 0.8 | 0.73 | 0.06 | 1.09 |
| 10 | Pdcd4 | 0.59 | 7.42E-12 | 1.58E-07 | 0.88 | 0.79 | 0.09 | 1.12 |

**Supplementary Table 3. Marker genes for the original clusters of integrated clustering of E11.5 control and NotchGOF λ-junction ectoderm cells.**

| | | | | | | | | |
|---|---|---|---|---|---|---|---|---|
| 10 | Itm2c | 1.1 | 8.08E-12 | 1.73E-07 | 0.66 | 0.6 | 0.06 | 1.1 |
| 10 | Hmx1 | 2.44 | 1.11E-11 | 2.37E-07 | 0.11 | 0.03 | 0.08 | 4.28 |
| 10 | Bcam | 0.62 | 1.12E-11 | 2.39E-07 | 0.86 | 0.72 | 0.13 | 1.19 |
| 10 | Pltp | 1.04 | 1.22E-11 | 2.61E-07 | 0.62 | 0.46 | 0.16 | 1.35 |
| 10 | Robo2 | 1.3 | 1.23E-11 | 2.63E-07 | 0.42 | 0.24 | 0.18 | 1.75 |
| 10 | Prdx6 | 0.43 | 1.71E-11 | 3.66E-07 | 0.96 | 0.95 | 0.01 | 1.01 |
| 10 | Tfap2c | 1.57 | 1.91E-11 | 4.07E-07 | 0.39 | 0.22 | 0.17 | 1.8 |
| 10 | Sema3a | 1.12 | 2.18E-11 | 4.66E-07 | 0.61 | 0.48 | 0.13 | 1.27 |
| 10 | Hspb8 | 0.98 | 2.26E-11 | 4.82E-07 | 0.64 | 0.48 | 0.15 | 1.31 |
| 10 | Afdn | 0.62 | 3.95E-11 | 8.44E-07 | 0.89 | 0.86 | 0.03 | 1.04 |
| 10 | Klhl24 | 0.8 | 6.11E-11 | 1.31E-06 | 0.73 | 0.64 | 0.09 | 1.14 |
| 10 | Galnt18 | 1.91 | 6.31E-11 | 1.35E-06 | 0.23 | 0.09 | 0.14 | 2.45 |
| 10 | Dlx3 | 1.24 | 7.18E-11 | 1.53E-06 | 0.45 | 0.28 | 0.17 | 1.62 |
| 10 | Tbx3 | 1.74 | 1.16E-10 | 2.47E-06 | 0.21 | 0.08 | 0.13 | 2.6 |
| 10 | Ldhb | 0.54 | 1.16E-10 | 2.48E-06 | 0.89 | 0.87 | 0.02 | 1.02 |
| 10 | Parm1 | 1.52 | 1.65E-10 | 3.53E-06 | 0.37 | 0.2 | 0.16 | 1.82 |
| 10 | Slc15a2 | 1.64 | 2.22E-10 | 4.74E-06 | 0.29 | 0.14 | 0.15 | 2.04 |
| 10 | Dhrs3 | 1.73 | 2.59E-10 | 5.54E-06 | 0.36 | 0.21 | 0.15 | 1.73 |
| 10 | Stx18 | 1.13 | 2.64E-10 | 5.65E-06 | 0.53 | 0.42 | 0.12 | 1.28 |
| 10 | Adgrl2 | 0.76 | 3.18E-10 | 6.79E-06 | 0.74 | 0.71 | 0.04 | 1.05 |
| 10 | Lratd1 | 1.45 | 3.72E-10 | 7.94E-06 | 0.27 | 0.13 | 0.14 | 2.15 |
| 10 | Gadd45g | 0.72 | 3.95E-10 | 8.44E-06 | 0.53 | 0.34 | 0.2 | 1.58 |
| 10 | Cst3 | 0.51 | 4.66E-10 | 9.95E-06 | 0.94 | 0.93 | 0.02 | 1.02 |
| 10 | Notum | 1.7 | 5.52E-10 | 1.18E-05 | 0.18 | 0.06 | 0.11 | 2.78 |
| 10 | Mfge8 | 0.84 | 5.64E-10 | 1.20E-05 | 0.66 | 0.56 | 0.1 | 1.18 |
| 10 | Ipmk | 1.46 | 5.67E-10 | 1.21E-05 | 0.39 | 0.24 | 0.15 | 1.64 |
| 10 | Cebpb | 1.57 | 6.04E-10 | 1.29E-05 | 0.39 | 0.24 | 0.15 | 1.62 |
| 10 | Tmem123 | 1.02 | 8.75E-10 | 1.87E-05 | 0.55 | 0.45 | 0.1 | 1.23 |
| 10 | Atxn7l3b | 0.27 | 8.91E-10 | 1.90E-05 | 1 | 0.99 | 0 | 1 |
| 10 | Osbpl8 | 1.23 | 9.36E-10 | 2.00E-05 | 0.42 | 0.27 | 0.15 | 1.56 |
| 10 | Rap2c | 0.87 | 1.03E-09 | 2.20E-05 | 0.72 | 0.67 | 0.06 | 1.08 |
| 10 | Slc25a27 | 0.92 | 1.10E-09 | 2.35E-05 | 0.63 | 0.55 | 0.08 | 1.15 |
| 10 | Btbd11 | 2.36 | 1.23E-09 | 2.64E-05 | 0.12 | 0.03 | 0.08 | 3.44 |
| 10 | Cd44 | 2.3 | 1.44E-09 | 3.08E-05 | 0.14 | 0.05 | 0.1 | 3.02 |
| 10 | Glcci1 | 1.12 | 1.47E-09 | 3.14E-05 | 0.6 | 0.5 | 0.1 | 1.19 |
| 10 | Ppp1r3b | 2.36 | 1.50E-09 | 3.21E-05 | 0.15 | 0.05 | 0.1 | 2.94 |
| 10 | Calm2 | 0.33 | 1.53E-09 | 3.27E-05 | 0.98 | 0.99 | -0.01 | 0.99 |
| 10 | Mbnl1 | 1.08 | 1.64E-09 | 3.50E-05 | 0.54 | 0.43 | 0.12 | 1.27 |
| 10 | Mt1 | 1.08 | 2.21E-09 | 4.72E-05 | 0.6 | 0.5 | 0.1 | 1.2 |
| 10 | Eif3f | 0.3 | 2.30E-09 | 4.90E-05 | 0.99 | 0.99 | -0.01 | 0.99 |
| 10 | Card10 | 2.16 | 2.75E-09 | 5.87E-05 | 0.14 | 0.05 | 0.09 | 3.04 |
| 10 | Dnase2a | 1.02 | 3.24E-09 | 6.93E-05 | 0.55 | 0.44 | 0.11 | 1.26 |
| 10 | Amotl1 | 0.57 | 3.42E-09 | 7.29E-05 | 0.78 | 0.7 | 0.09 | 1.12 |
| 10 | Tgfb1 | 1.8 | 3.85E-09 | 8.23E-05 | 0.31 | 0.16 | 0.14 | 1.88 |
| 10 | Tmem178 | 1.6 | 3.96E-09 | 8.45E-05 | 0.24 | 0.11 | 0.13 | 2.14 |
| 10 | Anxa8 | 1.74 | 4.87E-09 | 1.04E-04 | 0.21 | 0.09 | 0.12 | 2.29 |
| 10 | Dusp7 | 1.16 | 5.12E-09 | 1.09E-04 | 0.44 | 0.3 | 0.13 | 1.44 |
| 10 | Ypel3 | 0.81 | 5.17E-09 | 1.10E-04 | 0.66 | 0.57 | 0.09 | 1.15 |
| 10 | Apbb2 | 1.36 | 5.88E-09 | 1.26E-04 | 0.41 | 0.28 | 0.13 | 1.47 |
| 10 | Meis1 | 0.75 | 6.50E-09 | 1.39E-04 | 0.68 | 0.59 | 0.09 | 1.15 |
| 10 | Sowahc | 1.68 | 6.91E-09 | 1.48E-04 | 0.3 | 0.16 | 0.14 | 1.82 |
| 10 | Frmd6 | 1.28 | 7.15E-09 | 1.53E-04 | 0.38 | 0.23 | 0.15 | 1.65 |
| 10 | Gas5 | 0.3 | 7.17E-09 | 1.53E-04 | 1 | 0.99 | 0.01 | 1.01 |
| 10 | Ank3 | 0.63 | 8.12E-09 | 1.73E-04 | 0.74 | 0.63 | 0.1 | 1.16 |
| 10 | Luzp1 | 0.5 | 8.39E-09 | 1.79E-04 | 0.83 | 0.81 | 0.02 | 1.02 |
| 10 | Pard6g | 0.68 | 9.17E-09 | 1.96E-04 | 0.7 | 0.65 | 0.05 | 1.08 |
| 10 | Ifitm2 | 0.35 | 9.29E-09 | 1.98E-04 | 0.95 | 0.93 | 0.02 | 1.02 |

**Supplementary Table 3. Marker genes for the original clusters of integrated clustering of E11.5 control and NotchGOF λ-junction ectoderm cells.**

| | | | | | | | | |
|---|---|---|---|---|---|---|---|---|
| 10 | Cacng4 | 1.04 | 1.04E-08 | 2.23E-04 | 0.47 | 0.34 | 0.13 | 1.38 |
| 10 | Fzd10 | 1.4 | 1.52E-08 | 3.24E-04 | 0.26 | 0.13 | 0.13 | 1.97 |
| 10 | Ctnnb1 | 0.34 | 1.58E-08 | 3.37E-04 | 0.99 | 0.99 | 0 | 1 |
| 10 | Dennd2c | 1.67 | 1.64E-08 | 3.50E-04 | 0.21 | 0.1 | 0.12 | 2.22 |
| 10 | Myo1b | 0.65 | 1.66E-08 | 3.55E-04 | 0.69 | 0.63 | 0.06 | 1.09 |
| 10 | Pde10a | 2.06 | 2.18E-08 | 4.66E-04 | 0.13 | 0.04 | 0.09 | 3.02 |
| 10 | Smad7 | 0.64 | 2.26E-08 | 4.83E-04 | 0.69 | 0.61 | 0.08 | 1.13 |
| 10 | Tnik | 1.35 | 2.38E-08 | 5.09E-04 | 0.22 | 0.1 | 0.12 | 2.19 |
| 10 | Car12 | 2.57 | 2.78E-08 | 5.95E-04 | 0.11 | 0.04 | 0.08 | 3.2 |
| 10 | Ogt | 0.59 | 2.80E-08 | 5.97E-04 | 0.79 | 0.81 | -0.02 | 0.98 |
| 10 | Klk8 | 0.88 | 2.81E-08 | 6.01E-04 | 0.66 | 0.56 | 0.1 | 1.17 |
| 10 | Ctsb | 0.79 | 3.24E-08 | 6.93E-04 | 0.69 | 0.65 | 0.04 | 1.06 |
| 10 | Pcdh18 | 1.51 | 4.27E-08 | 9.11E-04 | 0.31 | 0.18 | 0.13 | 1.74 |
| 10 | Eif3h | 0.27 | 4.82E-08 | 1.03E-03 | 1 | 0.99 | 0.01 | 1.01 |
| 10 | Fzd6 | 1.33 | 4.95E-08 | 1.06E-03 | 0.43 | 0.32 | 0.11 | 1.34 |
| 10 | Rbpms | 0.81 | 5.27E-08 | 1.13E-03 | 0.58 | 0.49 | 0.1 | 1.2 |
| 10 | Dlgap4 | 0.6 | 5.43E-08 | 1.16E-03 | 0.73 | 0.7 | 0.03 | 1.04 |
| 10 | Efna1 | 1.09 | 6.22E-08 | 1.33E-03 | 0.51 | 0.4 | 0.11 | 1.27 |
| 10 | Rest | 0.52 | 6.22E-08 | 1.33E-03 | 0.77 | 0.68 | 0.09 | 1.14 |
| 10 | Trp53inp1 | 1.2 | 1.00E-07 | 2.14E-03 | 0.39 | 0.26 | 0.13 | 1.5 |
| 10 | Lpp | 0.86 | 1.04E-07 | 2.23E-03 | 0.55 | 0.46 | 0.09 | 1.2 |
| 10 | Fundc2 | 0.66 | 1.14E-07 | 2.43E-03 | 0.72 | 0.7 | 0.02 | 1.02 |
| 10 | Pmaip1 | 1.76 | 1.19E-07 | 2.53E-03 | 0.2 | 0.09 | 0.11 | 2.18 |
| 10 | Fblim1 | 0.95 | 1.32E-07 | 2.82E-03 | 0.52 | 0.41 | 0.11 | 1.26 |
| 10 | Crnde | 1.56 | 1.34E-07 | 2.87E-03 | 0.2 | 0.09 | 0.11 | 2.18 |
| 10 | Col4a1 | 0.59 | 1.56E-07 | 3.32E-03 | 0.74 | 0.68 | 0.05 | 1.08 |
| 10 | 10-Sep | 1 | 1.61E-07 | 3.45E-03 | 0.47 | 0.36 | 0.1 | 1.29 |
| 10 | Atp2c2 | 2.02 | 1.74E-07 | 3.72E-03 | 0.14 | 0.05 | 0.09 | 2.63 |
| 10 | Phlda3 | 0.61 | 1.87E-07 | 3.99E-03 | 0.61 | 0.5 | 0.11 | 1.23 |
| 10 | Sh3bp5 | 1.03 | 1.97E-07 | 4.20E-03 | 0.49 | 0.38 | 0.11 | 1.28 |
| 10 | Slc25a17 | 0.52 | 2.24E-07 | 4.78E-03 | 0.75 | 0.71 | 0.04 | 1.05 |
| 10 | Wnt5a | 1.09 | 2.40E-07 | 5.14E-03 | 0.53 | 0.43 | 0.1 | 1.24 |
| 10 | Kit | 1.39 | 2.44E-07 | 5.22E-03 | 0.3 | 0.18 | 0.12 | 1.69 |
| 10 | Cited2 | 0.67 | 2.51E-07 | 5.37E-03 | 0.73 | 0.68 | 0.05 | 1.07 |
| 10 | Col5a2 | 1.27 | 2.73E-07 | 5.82E-03 | 0.28 | 0.16 | 0.13 | 1.8 |
| 10 | Greb1 | 1.35 | 3.20E-07 | 6.83E-03 | 0.31 | 0.19 | 0.12 | 1.63 |
| 10 | Mcee | 0.7 | 4.87E-07 | 1.04E-02 | 0.62 | 0.56 | 0.06 | 1.1 |
| 10 | Trappc6a | 0.67 | 5.10E-07 | 1.09E-02 | 0.69 | 0.67 | 0.02 | 1.04 |
| 10 | Kdm5b | 0.42 | 5.52E-07 | 1.18E-02 | 0.92 | 0.93 | 0 | 1 |
| 10 | Ripk4 | 1.04 | 5.77E-07 | 1.23E-02 | 0.42 | 0.3 | 0.12 | 1.41 |
| 10 | Pkp3 | 1.38 | 6.37E-07 | 1.36E-02 | 0.36 | 0.25 | 0.11 | 1.45 |
| 10 | Sema3d | 0.72 | 6.74E-07 | 1.44E-02 | 0.54 | 0.42 | 0.12 | 1.29 |
| 10 | Evl | 0.5 | 6.77E-07 | 1.45E-02 | 0.77 | 0.73 | 0.04 | 1.05 |
| 10 | Gm16136 | 0.54 | 6.92E-07 | 1.48E-02 | 0.74 | 0.65 | 0.09 | 1.13 |
| 10 | Foxn3 | 0.51 | 7.57E-07 | 1.62E-02 | 0.76 | 0.72 | 0.05 | 1.06 |
| 10 | Fat1 | 0.67 | 7.68E-07 | 1.64E-02 | 0.67 | 0.64 | 0.03 | 1.05 |
| 10 | Ube2e3 | 0.33 | 7.74E-07 | 1.65E-02 | 0.91 | 0.9 | 0.01 | 1.01 |
| 10 | mt-Nd3 | 0.28 | 8.22E-07 | 1.76E-02 | 1 | 0.98 | 0.01 | 1.01 |
| 10 | Mbd2 | 0.61 | 8.33E-07 | 1.78E-02 | 0.66 | 0.61 | 0.05 | 1.08 |
| 10 | Purb | 0.42 | 1.07E-06 | 2.28E-02 | 0.89 | 0.89 | 0 | 1 |
| 10 | Rap2b | 0.69 | 1.13E-06 | 2.42E-02 | 0.59 | 0.53 | 0.06 | 1.12 |
| 10 | Chd6 | 0.47 | 1.26E-06 | 2.69E-02 | 0.77 | 0.76 | 0.01 | 1.02 |
| 10 | Cd2ap | 0.41 | 1.28E-06 | 2.73E-02 | 0.82 | 0.81 | 0.01 | 1.01 |
| 10 | Top2b | 0.29 | 1.42E-06 | 3.04E-02 | 0.89 | 0.93 | -0.04 | 0.96 |
| 10 | Slc35b3 | 1.12 | 1.54E-06 | 3.28E-02 | 0.38 | 0.26 | 0.12 | 1.44 |
| 10 | Ube2g1 | 0.66 | 2.05E-06 | 4.39E-02 | 0.62 | 0.57 | 0.05 | 1.1 |
| 10 | Ctso | 1.61 | 2.06E-06 | 4.41E-02 | 0.23 | 0.13 | 0.1 | 1.79 |

**Supplementary Table 3. Marker genes for the original clusters of integrated clustering of E11.5 control and NotchGOF λ-junction ectoderm cells.**

| | | | | | | | | |
|---|---|---|---|---|---|---|---|---|
| 10 | Vgll4 | 0.49 | 2.12E-06 | 4.54E-02 | 0.83 | 0.86 | -0.03 | 0.97 |
| 10 | Cd109 | 2.11 | 2.18E-06 | 4.65E-02 | 0.12 | 0.05 | 0.07 | 2.6 |
| 11 | Lhx2 | 10.06 | 0 | 0 | 0.93 | 0.02 | 0.91 | 38.71 |
| 11 | Nhlh2 | 11.1 | 0 | 0 | 0.87 | 0 | 0.86 | 216.25 |
| 11 | Igfbpl1 | 9.51 | 0 | 0 | 0.85 | 0.01 | 0.84 | 71 |
| 11 | Ppp1r14a | 7.64 | 0 | 0 | 0.88 | 0.04 | 0.84 | 20.88 |
| 11 | Ebf2 | 9.72 | 0 | 0 | 0.83 | 0 | 0.83 | 208 |
| 11 | Neurod1 | 11.59 | 0 | 0 | 0.83 | 0 | 0.82 | 275.33 |
| 11 | Insm1 | 7.47 | 0 | 0 | 0.85 | 0.02 | 0.82 | 36.74 |
| 11 | Elavl3 | 9.48 | 0 | 0 | 0.83 | 0.01 | 0.82 | 64 |
| 11 | Srrm4 | 9.29 | 0 | 0 | 0.79 | 0.01 | 0.78 | 112.43 |
| 11 | Neurog1 | 8.96 | 0 | 0 | 0.74 | 0.01 | 0.73 | 147 |
| 11 | Nhlh1 | 11.32 | 0 | 0 | 0.7 | 0 | 0.7 | 703 |
| 11 | Gng8 | 8.7 | 0 | 0 | 0.66 | 0.01 | 0.65 | 54.83 |
| 11 | Ebf3 | 7.79 | 0 | 0 | 0.66 | 0.02 | 0.64 | 32.9 |
| 11 | Sowaha | 10.18 | 0 | 0 | 0.63 | 0 | 0.63 | 626 |
| 11 | Btbd17 | 7.71 | 0 | 0 | 0.63 | 0.02 | 0.62 | 39.5 |
| 11 | Miat | 6.47 | 0 | 0 | 0.59 | 0.01 | 0.58 | 54 |
| 11 | St18 | 10.57 | 0 | 0 | 0.55 | 0 | 0.55 | 548 |
| 11 | Cdhr1 | 8.38 | 0 | 0 | 0.56 | 0.01 | 0.55 | 61.67 |
| 11 | Ina | 10.42 | 0 | 0 | 0.54 | 0.01 | 0.53 | 76.43 |
| 11 | Slc17a6 | 11.09 | 0 | 0 | 0.5 | 0 | 0.5 | #NA |
| 11 | Olig2 | 11.15 | 0 | 0 | 0.48 | 0 | 0.48 | 477 |
| 11 | Celf3 | 8.37 | 0 | 0 | 0.46 | 0 | 0.46 | 152.67 |
| 11 | Atp8b4 | 10.14 | 0 | 0 | 0.43 | 0 | 0.43 | 216 |
| 11 | Myt1l | 10.05 | 0 | 0 | 0.43 | 0 | 0.42 | 142 |
| 11 | Wdr66 | 6.38 | 2.39E-277 | 5.11E-273 | 0.72 | 0.04 | 0.69 | 20.66 |
| 11 | Elavl4 | 7.98 | 2.94E-268 | 6.27E-264 | 0.91 | 0.08 | 0.83 | 11.67 |
| 11 | Cbfa2t3 | 7.17 | 1.47E-267 | 3.14E-263 | 0.42 | 0.01 | 0.41 | 83.8 |
| 11 | Tubb3 | 8.09 | 4.71E-261 | 1.01E-256 | 0.99 | 0.11 | 0.89 | 9.29 |
| 11 | Ttc9b | 7.86 | 8.30E-258 | 1.77E-253 | 0.41 | 0.01 | 0.4 | 81.2 |
| 11 | Rbm24 | 6.56 | 1.14E-257 | 2.44E-253 | 0.52 | 0.01 | 0.5 | 36.86 |
| 11 | Dscaml1 | 7.97 | 6.66E-248 | 1.42E-243 | 0.34 | 0 | 0.33 | 167.5 |
| 11 | Kif19a | 6.2 | 5.64E-247 | 1.20E-242 | 0.59 | 0.02 | 0.56 | 25.52 |
| 11 | Nxph4 | 8.1 | 1.76E-239 | 3.76E-235 | 0.34 | 0 | 0.33 | 167.5 |
| 11 | Akap6 | 8.99 | 3.70E-233 | 7.91E-229 | 0.32 | 0 | 0.32 | 161.5 |
| 11 | Syp | 5.95 | 3.17E-232 | 6.78E-228 | 0.62 | 0.03 | 0.59 | 19.97 |
| 11 | Emx2 | 6.68 | 7.90E-228 | 1.69E-223 | 0.5 | 0.02 | 0.48 | 31.06 |
| 11 | Map6 | 7.91 | 3.80E-226 | 8.11E-222 | 0.36 | 0 | 0.35 | 88.75 |
| 11 | Ndrg4 | 6.21 | 3.67E-224 | 7.84E-220 | 0.54 | 0.02 | 0.51 | 25.48 |
| 11 | Syt13 | 5.97 | 1.43E-222 | 3.06E-218 | 0.55 | 0.02 | 0.53 | 23.83 |
| 11 | Dcc | 6.29 | 4.20E-217 | 8.98E-213 | 0.5 | 0.02 | 0.48 | 26.47 |
| 11 | Igsf21 | 8.34 | 8.40E-201 | 1.79E-196 | 0.28 | 0 | 0.28 | 142 |
| 11 | Slc17a7 | 8.67 | 1.23E-195 | 2.62E-191 | 0.24 | 0 | 0.24 | #NA |
| 11 | Scg3 | 6.43 | 6.28E-191 | 1.34E-186 | 0.43 | 0.01 | 0.41 | 30.43 |
| 11 | Rgs20 | 7.78 | 1.05E-190 | 2.24E-186 | 0.29 | 0 | 0.29 | 96.67 |
| 11 | Nrn1 | 10.64 | 2.74E-190 | 5.85E-186 | 0.23 | 0 | 0.23 | #NA |
| 11 | Mfng | 5.85 | 6.54E-190 | 1.40E-185 | 0.79 | 0.09 | 0.71 | 9.13 |
| 11 | Smim18 | 8.98 | 8.79E-185 | 1.88E-180 | 0.23 | 0 | 0.23 | #NA |
| 11 | Olig1 | 7.94 | 3.81E-184 | 8.14E-180 | 0.31 | 0.01 | 0.31 | 62 |
| 11 | Inka2 | 6.18 | 1.15E-181 | 2.45E-177 | 0.37 | 0.01 | 0.36 | 34 |
| 11 | Tagln3 | 6.22 | 1.53E-181 | 3.27E-177 | 0.77 | 0.08 | 0.69 | 9.37 |
| 11 | Myt1 | 8.14 | 1.20E-180 | 2.57E-176 | 0.24 | 0 | 0.24 | 239 |
| 11 | 5730403I07Rik | 8.97 | 4.22E-180 | 9.02E-176 | 0.23 | 0 | 0.23 | 232 |
| 11 | Abcd2 | 6.17 | 1.26E-179 | 2.70E-175 | 0.35 | 0.01 | 0.34 | 43.5 |
| 11 | Clvs1 | 7.54 | 1.32E-177 | 2.83E-173 | 0.29 | 0 | 0.29 | 72.5 |
| 11 | Actl6b | 9.11 | 2.90E-174 | 6.19E-170 | 0.22 | 0 | 0.22 | 219 |

**Supplementary Table 3. Marker genes for the original clusters of integrated clustering of E11.5 control and NotchGOF λ-junction ectoderm cells.**

| | | | | | | | | |
|---|---|---|---|---|---|---|---|---|
| 11 | Fstl5 | 7.5 | 3.16E-173 | 6.75E-169 | 0.37 | 0.01 | 0.36 | 31.17 |
| 11 | Chgb | 8.9 | 4.30E-173 | 9.19E-169 | 0.25 | 0 | 0.24 | 122.5 |
| 11 | Chga | 6.76 | 4.50E-170 | 9.60E-166 | 0.54 | 0.04 | 0.5 | 15.29 |
| 11 | Zeb1 | 5.77 | 1.26E-169 | 2.69E-165 | 0.42 | 0.02 | 0.4 | 24.65 |
| 11 | Dcx | 5.44 | 2.38E-169 | 5.09E-165 | 0.66 | 0.06 | 0.6 | 10.97 |
| 11 | Rgs4 | 9.56 | 7.74E-168 | 1.65E-163 | 0.24 | 0 | 0.24 | 119.5 |
| 11 | Runx1t1 | 5.2 | 6.57E-164 | 1.40E-159 | 0.53 | 0.04 | 0.49 | 15.11 |
| 11 | Pirt | 9.33 | 3.50E-163 | 7.47E-159 | 0.2 | 0 | 0.2 | #NA |
| 11 | Tuba4a | 5.78 | 1.00E-160 | 2.14E-156 | 0.63 | 0.06 | 0.57 | 10.98 |
| 11 | Gap43 | 7.01 | 3.60E-160 | 7.68E-156 | 0.61 | 0.05 | 0.55 | 11.65 |
| 11 | Pcsk2 | 8.24 | 2.66E-157 | 5.67E-153 | 0.23 | 0 | 0.22 | 113 |
| 11 | Stmn4 | 7.12 | 2.56E-153 | 5.46E-149 | 0.29 | 0.01 | 0.28 | 41.43 |
| 11 | A230051N06Rik | 8.62 | 1.28E-152 | 2.73E-148 | 0.19 | 0 | 0.19 | 194 |
| 11 | Camk2n1 | 5.3 | 6.23E-150 | 1.33E-145 | 0.74 | 0.1 | 0.64 | 7.58 |
| 11 | Ebf1 | 5.23 | 6.43E-149 | 1.37E-144 | 0.78 | 0.11 | 0.67 | 6.85 |
| 11 | Gng3 | 6.26 | 1.56E-148 | 3.34E-144 | 0.57 | 0.05 | 0.52 | 11.59 |
| 11 | Ppp1r17 | 11.47 | 4.51E-147 | 9.63E-143 | 0.17 | 0 | 0.17 | #NA |
| 11 | Gfy | 9.49 | 5.58E-147 | 1.19E-142 | 0.18 | 0 | 0.18 | #NA |
| 11 | Gm26871 | 8.94 | 6.02E-147 | 1.29E-142 | 0.18 | 0 | 0.18 | #NA |
| 11 | Chrna3 | 9.54 | 6.02E-147 | 1.29E-142 | 0.18 | 0 | 0.18 | #NA |
| 11 | Tmem163 | 6.36 | 1.06E-145 | 2.25E-141 | 0.32 | 0.01 | 0.31 | 29.36 |
| 11 | Coro2a | 5.14 | 2.78E-145 | 5.95E-141 | 0.54 | 0.04 | 0.5 | 12.32 |
| 11 | Fezf1 | 5.02 | 4.71E-145 | 1.01E-140 | 0.67 | 0.08 | 0.59 | 8.87 |
| 11 | Pnmal1 | 7.78 | 1.23E-142 | 2.63E-138 | 0.21 | 0 | 0.21 | 106.5 |
| 11 | Cacna2d1 | 5.67 | 1.59E-141 | 3.39E-137 | 0.53 | 0.04 | 0.49 | 12.02 |
| 11 | Lamp5 | 7.99 | 9.87E-138 | 2.11E-133 | 0.19 | 0 | 0.19 | 187 |
| 11 | Syt4 | 5.03 | 1.92E-135 | 4.10E-131 | 0.39 | 0.02 | 0.37 | 18.76 |
| 11 | Srrm3 | 6.23 | 6.58E-135 | 1.41E-130 | 0.29 | 0.01 | 0.28 | 32.22 |
| 11 | Art5 | 6.99 | 7.53E-134 | 1.61E-129 | 0.21 | 0 | 0.2 | 103 |
| 11 | Rtn1 | 5.96 | 3.06E-133 | 6.54E-129 | 0.92 | 0.22 | 0.7 | 4.14 |
| 11 | Thsd7b | 4.49 | 3.65E-130 | 7.80E-126 | 0.52 | 0.05 | 0.48 | 11.62 |
| 11 | Dll3 | 5.37 | 3.38E-129 | 7.22E-125 | 0.3 | 0.01 | 0.29 | 29.7 |
| 11 | Rgs16 | 5.69 | 1.02E-128 | 2.19E-124 | 0.28 | 0.01 | 0.27 | 34.63 |
| 11 | Foxo6 | 5.41 | 3.03E-127 | 6.46E-123 | 0.47 | 0.04 | 0.43 | 12.92 |
| 11 | Gdap1l1 | 5.53 | 6.97E-127 | 1.49E-122 | 0.5 | 0.04 | 0.46 | 11.43 |
| 11 | Tmprss6 | 8.21 | 2.81E-126 | 6.01E-122 | 0.17 | 0 | 0.17 | 168 |
| 11 | Rd3 | 9.94 | 2.23E-125 | 4.77E-121 | 0.16 | 0 | 0.16 | #NA |
| 11 | Gda | 7.62 | 8.23E-122 | 1.76E-117 | 0.17 | 0 | 0.17 | 168 |
| 11 | Clmp | 4.91 | 3.44E-121 | 7.35E-117 | 0.6 | 0.07 | 0.53 | 8.11 |
| 11 | Gm27197 | 9.13 | 4.68E-120 | 9.99E-116 | 0.14 | 0 | 0.14 | #NA |
| 11 | Gm13589 | 6.83 | 4.14E-119 | 8.84E-115 | 0.2 | 0 | 0.2 | 66.67 |
| 11 | Islr2 | 8.18 | 4.66E-119 | 9.95E-115 | 0.17 | 0 | 0.17 | 87 |
| 11 | Nol4 | 6.1 | 1.02E-118 | 2.19E-114 | 0.27 | 0.01 | 0.26 | 29.44 |
| 11 | Cend1 | 6.14 | 4.82E-118 | 1.03E-113 | 0.24 | 0.01 | 0.23 | 39.83 |
| 11 | Gfra1 | 5.15 | 3.14E-117 | 6.70E-113 | 0.38 | 0.02 | 0.36 | 15.88 |
| 11 | Vwa5b2 | 6.73 | 8.89E-117 | 1.90E-112 | 0.18 | 0 | 0.18 | 90.5 |
| 11 | Stmn3 | 5.75 | 2.03E-116 | 4.34E-112 | 0.65 | 0.1 | 0.56 | 6.79 |
| 11 | Sct | 7.02 | 3.93E-114 | 8.39E-110 | 0.45 | 0.04 | 0.41 | 11.71 |
| 11 | Scrt1 | 6.18 | 4.01E-113 | 8.56E-109 | 0.25 | 0.01 | 0.24 | 30.63 |
| 11 | Brsk2 | 4.57 | 4.51E-112 | 9.64E-108 | 0.43 | 0.04 | 0.4 | 12.34 |
| 11 | Dner | 5.25 | 1.42E-111 | 3.04E-107 | 0.27 | 0.01 | 0.26 | 26.5 |
| 11 | Syt2 | 6.36 | 2.98E-111 | 6.37E-107 | 0.23 | 0.01 | 0.22 | 37.67 |
| 11 | Shisal1 | 5.07 | 4.90E-111 | 1.05E-106 | 0.36 | 0.02 | 0.33 | 16.14 |
| 11 | Mast1 | 5.09 | 2.11E-109 | 4.52E-105 | 0.28 | 0.01 | 0.27 | 23.08 |
| 11 | Stmn2 | 6 | 2.07E-108 | 4.42E-104 | 0.79 | 0.18 | 0.62 | 4.46 |
| 11 | Col16a1 | 6.75 | 4.66E-107 | 9.95E-103 | 0.25 | 0.01 | 0.24 | 28 |
| 11 | Soga3 | 4.36 | 5.74E-106 | 1.23E-101 | 0.74 | 0.15 | 0.59 | 4.95 |

**Supplementary Table 3. Marker genes for the original clusters of integrated clustering of E11.5 control and NotchGOF λ-junction ectoderm cells.**

| | | | | | | | | |
|---|---|---|---|---|---|---|---|---|
| 11 | Myo7a | 5.27 | 1.18E-105 | 2.53E-101 | 0.27 | 0.01 | 0.26 | 22.58 |
| 11 | Jakmip2 | 4.57 | 1.54E-105 | 3.29E-101 | 0.47 | 0.05 | 0.42 | 10.11 |
| 11 | Nrxn1 | 4.82 | 1.78E-104 | 3.81E-100 | 0.53 | 0.06 | 0.47 | 8.27 |
| 11 | Ckmt1 | 5.27 | 6.12E-104 | 1.31E-99 | 0.67 | 0.12 | 0.55 | 5.64 |
| 11 | Gm31819 | 8.78 | 7.11E-104 | 1.52E-99 | 0.12 | 0 | 0.12 | #NA |
| 11 | Stxbp1 | 4.83 | 1.01E-103 | 2.16E-99 | 0.35 | 0.02 | 0.33 | 15.13 |
| 11 | Cotl1 | 4.59 | 1.61E-103 | 3.43E-99 | 0.98 | 0.47 | 0.51 | 2.08 |
| 11 | Lingo2 | 6.14 | 1.36E-100 | 2.91E-96 | 0.23 | 0.01 | 0.22 | 29 |
| 11 | Spock1 | 6.38 | 9.06E-100 | 1.94E-95 | 0.21 | 0.01 | 0.2 | 34.33 |
| 11 | Rassf4 | 5.78 | 1.35E-99 | 2.89E-95 | 0.27 | 0.01 | 0.26 | 20.85 |
| 11 | Lrrc10b | 6.45 | 4.94E-99 | 1.06E-94 | 0.18 | 0 | 0.18 | 45.25 |
| 11 | Sez6l2 | 6.36 | 7.20E-99 | 1.54E-94 | 0.19 | 0 | 0.18 | 46.75 |
| 11 | Map1a | 4.41 | 1.35E-98 | 2.88E-94 | 0.45 | 0.05 | 0.4 | 9.89 |
| 11 | Ntrk1 | 9.46 | 1.74E-98 | 3.72E-94 | 0.12 | 0 | 0.12 | #NA |
| 11 | Crmp1 | 4.73 | 5.65E-98 | 1.21E-93 | 0.68 | 0.13 | 0.55 | 5.25 |
| 11 | Tesc | 7.47 | 7.75E-98 | 1.65E-93 | 0.28 | 0.02 | 0.27 | 18.93 |
| 11 | Cplx2 | 4.85 | 3.90E-97 | 8.34E-93 | 0.77 | 0.19 | 0.58 | 4.15 |
| 11 | Nsg2 | 5.4 | 6.29E-97 | 1.34E-92 | 0.35 | 0.03 | 0.32 | 13.92 |
| 11 | Rnf112 | 5.62 | 6.41E-97 | 1.37E-92 | 0.2 | 0.01 | 0.19 | 33.33 |
| 11 | Mreg | 5.26 | 7.16E-97 | 1.53E-92 | 0.34 | 0.02 | 0.32 | 14.25 |
| 11 | Atp1a3 | 7.65 | 2.18E-95 | 4.65E-91 | 0.14 | 0 | 0.13 | 135 |
| 11 | Gm17322 | 7.73 | 1.01E-94 | 2.16E-90 | 0.17 | 0 | 0.17 | 56 |
| 11 | Asic1 | 5.84 | 2.52E-94 | 5.39E-90 | 0.19 | 0.01 | 0.18 | 37.4 |
| 11 | Nrxn2 | 5.14 | 2.84E-94 | 6.08E-90 | 0.32 | 0.02 | 0.3 | 14.68 |
| 11 | Ccdc184 | 8.73 | 1.73E-93 | 3.69E-89 | 0.12 | 0 | 0.12 | 123 |
| 11 | Sgpp2 | 4.75 | 2.20E-93 | 4.69E-89 | 0.31 | 0.02 | 0.29 | 15.5 |
| 11 | Kcnk10 | 9.12 | 4.25E-93 | 9.09E-89 | 0.11 | 0 | 0.11 | #NA |
| 11 | Tmsb4x | 2.63 | 1.64E-92 | 3.49E-88 | 1 | 1 | 0 | 1 |
| 11 | Cdyl2 | 5.61 | 3.01E-92 | 6.42E-88 | 0.35 | 0.03 | 0.32 | 12.43 |
| 11 | Gadd45a | 4.72 | 3.56E-92 | 7.61E-88 | 0.76 | 0.19 | 0.56 | 3.95 |
| 11 | Scn9a | 4.58 | 1.24E-91 | 2.65E-87 | 0.37 | 0.03 | 0.34 | 11.33 |
| 11 | Serpini1 | 4.28 | 1.09E-89 | 2.32E-85 | 0.38 | 0.04 | 0.35 | 10.89 |
| 11 | Gm3234 | 6.3 | 1.44E-89 | 3.08E-85 | 0.17 | 0 | 0.16 | 42 |
| 11 | Cerkl | 6.48 | 1.49E-89 | 3.18E-85 | 0.14 | 0 | 0.14 | 71 |
| 11 | Cpeb1 | 8.63 | 3.70E-88 | 7.91E-84 | 0.12 | 0 | 0.12 | 116 |
| 11 | Uncx | 4.93 | 4.70E-88 | 1.00E-83 | 0.53 | 0.08 | 0.45 | 6.61 |
| 11 | Lhx3 | 8.16 | 1.03E-87 | 2.21E-83 | 0.1 | 0 | 0.1 | #NA |
| 11 | Chrna9 | 8.57 | 1.03E-87 | 2.21E-83 | 0.1 | 0 | 0.1 | #NA |
| 11 | Neurod4 | 9.82 | 1.03E-87 | 2.21E-83 | 0.1 | 0 | 0.1 | #NA |
| 11 | Apba1 | 8.67 | 1.10E-87 | 2.34E-83 | 0.11 | 0 | 0.11 | #NA |
| 11 | Ccdc187 | 8.56 | 1.11E-87 | 2.37E-83 | 0.11 | 0 | 0.11 | #NA |
| 11 | Eef1a2 | 8.33 | 1.12E-87 | 2.40E-83 | 0.11 | 0 | 0.11 | #NA |
| 11 | Emx2os | 5.64 | 1.55E-87 | 3.32E-83 | 0.18 | 0.01 | 0.18 | 36.2 |
| 11 | Rundc3a | 4.4 | 5.95E-86 | 1.27E-81 | 0.52 | 0.08 | 0.45 | 6.79 |
| 11 | Pcdh8 | 5.13 | 8.31E-86 | 1.78E-81 | 0.32 | 0.02 | 0.29 | 13.17 |
| 11 | Cadps | 4.94 | 2.02E-85 | 4.31E-81 | 0.3 | 0.02 | 0.28 | 14.14 |
| 11 | Map1b | 4.85 | 3.86E-85 | 8.25E-81 | 0.95 | 0.56 | 0.39 | 1.71 |
| 11 | Fam110a | 4.09 | 1.29E-84 | 2.75E-80 | 0.83 | 0.28 | 0.56 | 3.03 |
| 11 | Dusp4 | 4.25 | 4.24E-84 | 9.06E-80 | 0.48 | 0.06 | 0.41 | 7.45 |
| 11 | Apc2 | 5.32 | 1.07E-83 | 2.29E-79 | 0.25 | 0.01 | 0.24 | 18 |
| 11 | 4930578M01Rik | 4.63 | 4.27E-83 | 9.12E-79 | 0.38 | 0.04 | 0.34 | 9.77 |
| 11 | Trim66 | 8.33 | 2.57E-82 | 5.50E-78 | 0.1 | 0 | 0.1 | #NA |
| 11 | Mapk10 | 5.82 | 3.41E-82 | 7.28E-78 | 0.19 | 0.01 | 0.19 | 27.71 |
| 11 | Stmn1 | 1.64 | 1.07E-81 | 2.28E-77 | 1 | 1 | 0 | 1 |
| 11 | Serping1 | 4.55 | 1.19E-81 | 2.55E-77 | 0.4 | 0.04 | 0.36 | 9.09 |
| 11 | Grasp | 4.46 | 5.62E-81 | 1.20E-76 | 0.68 | 0.17 | 0.52 | 4.05 |
| 11 | Plk3 | 5.53 | 8.72E-81 | 1.86E-76 | 0.24 | 0.01 | 0.23 | 18.38 |

**Supplementary Table 3. Marker genes for the original clusters of integrated clustering of E11.5 control and NotchGOF λ-junction ectoderm cells.**

| | | | | | | | | |
|---|---|---|---|---|---|---|---|---|
| 11 | Rab6b | 5.61 | 1.39E-80 | 2.96E-76 | 0.34 | 0.03 | 0.31 | 10.69 |
| 11 | Tuba1a | 3.37 | 1.18E-78 | 2.53E-74 | 0.99 | 0.98 | 0.02 | 1.02 |
| 11 | Gnas | 1.63 | 2.73E-78 | 5.84E-74 | 1 | 1 | 0 | 1 |
| 11 | Nacad | 5.7 | 8.14E-78 | 1.74E-73 | 0.16 | 0 | 0.15 | 38.75 |
| 11 | Gm14226 | 7.38 | 9.59E-78 | 2.05E-73 | 0.12 | 0 | 0.12 | 61.5 |
| 11 | Basp1 | 2.35 | 3.57E-77 | 7.63E-73 | 0.99 | 0.97 | 0.03 | 1.03 |
| 11 | Cntn1 | 5.41 | 1.60E-76 | 3.42E-72 | 0.22 | 0.01 | 0.21 | 19.91 |
| 11 | Dkkl1 | 5.16 | 7.58E-76 | 1.62E-71 | 0.2 | 0.01 | 0.19 | 22.22 |
| 11 | Galnt17 | 4.93 | 1.04E-75 | 2.22E-71 | 0.21 | 0.01 | 0.2 | 20.6 |
| 11 | Rem2 | 4.47 | 3.50E-75 | 7.48E-71 | 0.5 | 0.08 | 0.42 | 6.29 |
| 11 | Mapt | 5.45 | 3.55E-74 | 7.58E-70 | 0.33 | 0.03 | 0.3 | 10.28 |
| 11 | Gdap1 | 4.19 | 4.24E-73 | 9.05E-69 | 0.38 | 0.05 | 0.34 | 8.47 |
| 11 | Coro1a | 4.83 | 6.29E-71 | 1.34E-66 | 0.36 | 0.04 | 0.32 | 8.8 |
| 11 | Rap1gap2 | 4.84 | 2.81E-70 | 5.99E-66 | 0.26 | 0.02 | 0.24 | 13.58 |
| 11 | Cadm3 | 4.7 | 3.28E-70 | 7.01E-66 | 0.27 | 0.02 | 0.25 | 12.32 |
| 11 | Arhgdig | 5.92 | 1.22E-69 | 2.60E-65 | 0.66 | 0.18 | 0.48 | 3.66 |
| 11 | Kcnh3 | 6.94 | 2.65E-69 | 5.67E-65 | 0.1 | 0 | 0.1 | 103 |
| 11 | Ada | 4.48 | 2.66E-69 | 5.69E-65 | 0.4 | 0.05 | 0.35 | 7.55 |
| 11 | Gm16364 | 7.26 | 1.23E-67 | 2.62E-63 | 0.11 | 0 | 0.11 | 55 |
| 11 | Prnp | 3.51 | 2.19E-67 | 4.68E-63 | 0.66 | 0.17 | 0.49 | 3.85 |
| 11 | Grik2 | 5.25 | 7.03E-67 | 1.50E-62 | 0.17 | 0.01 | 0.17 | 21.75 |
| 11 | Trp53i11 | 3.19 | 8.78E-67 | 1.88E-62 | 0.86 | 0.41 | 0.45 | 2.11 |
| 11 | Kif5a | 4.49 | 1.56E-66 | 3.34E-62 | 0.37 | 0.05 | 0.32 | 7.83 |
| 11 | Selenom | 4.71 | 7.00E-66 | 1.50E-61 | 0.83 | 0.39 | 0.44 | 2.15 |
| 11 | Nefl | 7.77 | 3.57E-65 | 7.64E-61 | 0.14 | 0 | 0.14 | 35.5 |
| 11 | Fam169b | 6.61 | 1.30E-64 | 2.78E-60 | 0.11 | 0 | 0.11 | 55 |
| 11 | Svop | 6.36 | 4.59E-64 | 9.81E-60 | 0.12 | 0 | 0.11 | 58 |
| 11 | Hes6 | 4.28 | 4.80E-64 | 1.03E-59 | 0.87 | 0.52 | 0.35 | 1.67 |
| 11 | Fam57b | 5.02 | 1.99E-63 | 4.25E-59 | 0.21 | 0.01 | 0.2 | 15.21 |
| 11 | Kcnmb2 | 4.87 | 3.36E-63 | 7.18E-59 | 0.32 | 0.04 | 0.28 | 9.03 |
| 11 | Map2 | 3.67 | 4.70E-63 | 1.00E-58 | 0.71 | 0.24 | 0.47 | 2.98 |
| 11 | L1cam | 5.3 | 5.61E-62 | 1.20E-57 | 0.2 | 0.01 | 0.19 | 16.67 |
| 11 | D930020B18Rik | 4.39 | 6.31E-62 | 1.35E-57 | 0.37 | 0.05 | 0.32 | 7.36 |
| 11 | Snca | 4.36 | 9.16E-62 | 1.96E-57 | 0.53 | 0.11 | 0.42 | 4.77 |
| 11 | Ptms | 1.71 | 6.55E-61 | 1.40E-56 | 0.95 | 0.98 | -0.03 | 0.97 |
| 11 | Cnrip1 | 4.82 | 1.44E-60 | 3.07E-56 | 0.25 | 0.02 | 0.23 | 11.45 |
| 11 | Fam78b | 5 | 2.16E-60 | 4.61E-56 | 0.23 | 0.02 | 0.21 | 13.29 |
| 11 | Actb | 1 | 9.41E-60 | 2.01E-55 | 1 | 1 | 0 | 1 |
| 11 | Scg2 | 5.7 | 2.18E-59 | 4.65E-55 | 0.16 | 0.01 | 0.15 | 20.13 |
| 11 | Tubb2b | 2.14 | 3.48E-59 | 7.43E-55 | 0.98 | 0.92 | 0.06 | 1.06 |
| 11 | Fam219a | 4.4 | 4.20E-59 | 8.97E-55 | 0.23 | 0.02 | 0.21 | 12.56 |
| 11 | a | 5.18 | 1.90E-57 | 4.05E-53 | 0.19 | 0.01 | 0.18 | 14.92 |
| 11 | Eif4e3 | 3.14 | 4.60E-57 | 9.82E-53 | 0.66 | 0.21 | 0.45 | 3.21 |
| 11 | Pde2a | 6.47 | 4.94E-57 | 1.05E-52 | 0.1 | 0 | 0.1 | 51.5 |
| 11 | Dpysl3 | 2.73 | 9.41E-57 | 2.01E-52 | 0.9 | 0.56 | 0.35 | 1.62 |
| 11 | Prex1 | 3.43 | 1.05E-56 | 2.23E-52 | 0.5 | 0.11 | 0.39 | 4.61 |
| 11 | Frrs1l | 5.22 | 1.07E-56 | 2.28E-52 | 0.15 | 0.01 | 0.14 | 21.14 |
| 11 | Atp1b1 | 4.15 | 1.50E-56 | 3.20E-52 | 0.54 | 0.13 | 0.41 | 4.04 |
| 11 | Btg2 | 3.28 | 3.75E-56 | 8.00E-52 | 0.76 | 0.3 | 0.46 | 2.54 |
| 11 | Atf5 | 4.46 | 1.20E-55 | 2.56E-51 | 0.65 | 0.21 | 0.44 | 3.12 |
| 11 | Nrsn1 | 5.07 | 2.24E-55 | 4.77E-51 | 0.15 | 0.01 | 0.14 | 21.14 |
| 11 | Scn3a | 4.86 | 5.65E-55 | 1.21E-50 | 0.22 | 0.02 | 0.2 | 12.17 |
| 11 | P2rx3 | 5.36 | 2.00E-54 | 4.28E-50 | 0.14 | 0.01 | 0.13 | 22.5 |
| 11 | Brinp2 | 4.37 | 4.84E-54 | 1.03E-49 | 0.21 | 0.02 | 0.19 | 12.88 |
| 11 | Psip1 | 1.28 | 5.84E-54 | 1.25E-49 | 0.98 | 0.98 | 0 | 1 |
| 11 | Mllt11 | 4.07 | 3.67E-53 | 7.84E-49 | 0.75 | 0.32 | 0.43 | 2.34 |
| 11 | Ptma | 0.87 | 7.37E-53 | 1.57E-48 | 1 | 1 | 0 | 1 |

**Supplementary Table 3. Marker genes for the original clusters of integrated clustering of E11.5 control and NotchGOF λ-junction ectoderm cells.**

| | | | | | | | | |
|---|---|---|---|---|---|---|---|---|
| 11 | Ckb | 3 | 1.92E-52 | 4.10E-48 | 0.77 | 0.34 | 0.44 | 2.3 |
| 11 | Uchl1 | 3.04 | 3.42E-52 | 7.31E-48 | 0.9 | 0.64 | 0.26 | 1.4 |
| 11 | Gprin1 | 4.02 | 4.13E-52 | 8.82E-48 | 0.27 | 0.03 | 0.24 | 9.14 |
| 11 | Pou2f2 | 5.94 | 5.77E-52 | 1.23E-47 | 0.15 | 0.01 | 0.14 | 18.5 |
| 11 | Rnf182 | 4.17 | 8.62E-52 | 1.84E-47 | 0.59 | 0.18 | 0.41 | 3.26 |
| 11 | Tppp3 | 6.42 | 1.04E-51 | 2.22E-47 | 0.17 | 0.01 | 0.16 | 15.27 |
| 11 | Phactr1 | 4.61 | 1.07E-51 | 2.29E-47 | 0.25 | 0.03 | 0.23 | 9.33 |
| 11 | Shisa7 | 4.14 | 1.33E-51 | 2.85E-47 | 0.23 | 0.02 | 0.21 | 10.76 |
| 11 | Ripor2 | 5.09 | 1.40E-51 | 2.99E-47 | 0.14 | 0.01 | 0.13 | 22.5 |
| 11 | Flrt1 | 4.19 | 1.87E-51 | 3.99E-47 | 0.25 | 0.03 | 0.22 | 9.8 |
| 11 | Cntn2 | 4.17 | 4.64E-51 | 9.92E-47 | 0.36 | 0.06 | 0.3 | 6.12 |
| 11 | Pnma2 | 4.3 | 5.56E-51 | 1.19E-46 | 0.25 | 0.03 | 0.23 | 9.33 |
| 11 | Actg1 | 1.04 | 5.84E-51 | 1.25E-46 | 0.99 | 1 | -0.01 | 0.99 |
| 11 | Snap25 | 4.63 | 3.35E-50 | 7.16E-46 | 0.23 | 0.02 | 0.21 | 10.09 |
| 11 | Nnat | 2.82 | 5.22E-50 | 1.11E-45 | 0.96 | 0.95 | 0 | 1 |
| 11 | Riiad1 | 4.88 | 6.02E-50 | 1.29E-45 | 0.16 | 0.01 | 0.15 | 17.22 |
| 11 | Ssc5d | 4.76 | 6.33E-49 | 1.35E-44 | 0.16 | 0.01 | 0.15 | 16.1 |
| 11 | Necab3 | 6.01 | 1.24E-48 | 2.64E-44 | 0.15 | 0.01 | 0.14 | 18.5 |
| 11 | Dysf | 5.15 | 3.73E-48 | 7.97E-44 | 0.12 | 0 | 0.11 | 29 |
| 11 | Aplp1 | 3.49 | 4.78E-48 | 1.02E-43 | 0.65 | 0.23 | 0.41 | 2.78 |
| 11 | Rab3a | 3.82 | 5.42E-48 | 1.16E-43 | 0.44 | 0.09 | 0.35 | 4.67 |
| 11 | Nova2 | 4.02 | 8.38E-48 | 1.79E-43 | 0.27 | 0.03 | 0.24 | 7.97 |
| 11 | Ucp2 | 3.02 | 5.55E-47 | 1.19E-42 | 0.83 | 0.47 | 0.36 | 1.78 |
| 11 | Pcsk1n | 6.1 | 8.50E-47 | 1.81E-42 | 0.2 | 0.02 | 0.18 | 11.11 |
| 11 | Scg5 | 2.84 | 4.25E-46 | 9.09E-42 | 0.61 | 0.19 | 0.41 | 3.12 |
| 11 | Eid1 | 1.33 | 1.40E-45 | 3.00E-41 | 0.94 | 0.97 | -0.04 | 0.96 |
| 11 | Mfap4 | 2.47 | 1.85E-45 | 3.95E-41 | 0.87 | 0.65 | 0.21 | 1.33 |
| 11 | B630019K06Rik | 3.61 | 2.39E-45 | 5.11E-41 | 0.41 | 0.09 | 0.33 | 4.8 |
| 11 | Adgrb2 | 3.89 | 2.51E-45 | 5.37E-41 | 0.21 | 0.02 | 0.19 | 10.14 |
| 11 | Eya2 | 2.54 | 4.45E-45 | 9.51E-41 | 0.78 | 0.36 | 0.42 | 2.18 |
| 11 | Tnik | 3.14 | 4.97E-45 | 1.06E-40 | 0.43 | 0.1 | 0.34 | 4.55 |
| 11 | Kif1a | 3.75 | 5.30E-45 | 1.13E-40 | 0.4 | 0.08 | 0.32 | 4.88 |
| 11 | Plk4 | 2.98 | 5.68E-45 | 1.21E-40 | 0.8 | 0.51 | 0.3 | 1.58 |
| 11 | 9330159F19Rik | 3.47 | 7.40E-45 | 1.58E-40 | 0.28 | 0.04 | 0.25 | 7.28 |
| 11 | Hecw1 | 3.74 | 8.69E-45 | 1.86E-40 | 0.24 | 0.03 | 0.21 | 8.54 |
| 11 | Ap3b2 | 3.62 | 2.57E-44 | 5.50E-40 | 0.36 | 0.07 | 0.29 | 5.39 |
| 11 | Spsb4 | 3.12 | 3.61E-44 | 7.71E-40 | 0.48 | 0.12 | 0.36 | 3.97 |
| 11 | Rgs11 | 3.79 | 4.00E-44 | 8.54E-40 | 0.25 | 0.03 | 0.22 | 7.88 |
| 11 | Cep152 | 2.94 | 4.17E-44 | 8.90E-40 | 0.57 | 0.19 | 0.39 | 3.1 |
| 11 | Gnrh1 | 9.24 | 4.67E-44 | 9.97E-40 | 0.14 | 0.01 | 0.13 | 16.88 |
| 11 | Klf7 | 3.53 | 7.85E-44 | 1.68E-39 | 0.68 | 0.29 | 0.38 | 2.3 |
| 11 | Mapk8ip2 | 3.43 | 1.08E-43 | 2.31E-39 | 0.43 | 0.1 | 0.33 | 4.48 |
| 11 | Fgd4 | 2.82 | 1.21E-43 | 2.58E-39 | 0.58 | 0.18 | 0.4 | 3.23 |
| 11 | Sh2d3c | 3.54 | 2.43E-43 | 5.20E-39 | 0.28 | 0.04 | 0.24 | 7.1 |
| 11 | 1500004A13Rik | 3.85 | 2.80E-43 | 5.99E-39 | 0.39 | 0.08 | 0.31 | 4.84 |
| 11 | Sult4a1 | 3.59 | 3.23E-43 | 6.90E-39 | 0.37 | 0.07 | 0.3 | 5.11 |
| 11 | Parp6 | 3.21 | 3.51E-43 | 7.51E-39 | 0.46 | 0.11 | 0.35 | 4.09 |
| 11 | A330009N23Rik | 4.32 | 4.57E-43 | 9.76E-39 | 0.27 | 0.04 | 0.23 | 7.36 |
| 11 | Add2 | 3.58 | 6.20E-43 | 1.32E-38 | 0.31 | 0.05 | 0.26 | 6.2 |
| 11 | Elavl2 | 2.65 | 8.11E-43 | 1.73E-38 | 0.76 | 0.4 | 0.35 | 1.88 |
| 11 | Cyth2 | 2.1 | 1.18E-42 | 2.51E-38 | 0.85 | 0.64 | 0.21 | 1.33 |
| 11 | Dusp8 | 4.09 | 4.98E-42 | 1.06E-37 | 0.28 | 0.04 | 0.24 | 6.6 |
| 11 | Gm2694 | 3.68 | 7.57E-42 | 1.62E-37 | 0.34 | 0.06 | 0.28 | 5.43 |
| 11 | Ackr1 | 5.48 | 1.00E-41 | 2.14E-37 | 0.1 | 0 | 0.1 | 25.75 |
| 11 | Inka1 | 3.18 | 3.79E-41 | 8.10E-37 | 0.36 | 0.07 | 0.29 | 5.16 |
| 11 | Trim36 | 3.62 | 7.04E-41 | 1.50E-36 | 0.3 | 0.05 | 0.25 | 6.19 |
| 11 | Ddx5 | 0.69 | 2.30E-40 | 4.90E-36 | 0.98 | 1 | -0.02 | 0.98 |

**Supplementary Table 3. Marker genes for the original clusters of integrated clustering of E11.5 control and NotchGOF λ-junction ectoderm cells.**

| | | | | | | | | |
|---|---|---|---|---|---|---|---|---|
| 11 | Trp53inp2 | 3 | 2.96E-40 | 6.32E-36 | 0.36 | 0.07 | 0.29 | 5.16 |
| 11 | Dll1 | 4.59 | 3.46E-40 | 7.40E-36 | 0.13 | 0.01 | 0.12 | 16.13 |
| 11 | Sox11 | 1.06 | 5.92E-40 | 1.27E-35 | 0.94 | 1 | -0.06 | 0.94 |
| 11 | Ndufaf3 | 2.7 | 7.66E-40 | 1.64E-35 | 0.74 | 0.41 | 0.32 | 1.79 |
| 11 | Tmcc2 | 2.97 | 2.79E-39 | 5.95E-35 | 0.32 | 0.06 | 0.26 | 5.64 |
| 11 | Cyfip2 | 3.34 | 5.89E-39 | 1.26E-34 | 0.51 | 0.16 | 0.35 | 3.11 |
| 11 | Elmo1 | 4.3 | 7.55E-39 | 1.61E-34 | 0.22 | 0.03 | 0.19 | 8.11 |
| 11 | Cnih2 | 3.67 | 8.90E-39 | 1.90E-34 | 0.49 | 0.15 | 0.34 | 3.36 |
| 11 | Sh3bgrl | 1.82 | 1.27E-38 | 2.71E-34 | 0.87 | 0.75 | 0.12 | 1.16 |
| 11 | Prdx2 | 0.75 | 1.63E-38 | 3.49E-34 | 1 | 1 | 0 | 1 |
| 11 | Col9a2 | 4.4 | 1.71E-38 | 3.66E-34 | 0.45 | 0.12 | 0.32 | 3.62 |
| 11 | C530008M17Rik | 2.47 | 2.16E-38 | 4.61E-34 | 0.74 | 0.39 | 0.34 | 1.88 |
| 11 | Unc13a | 3.91 | 5.09E-38 | 1.09E-33 | 0.19 | 0.02 | 0.17 | 9.24 |
| 11 | Nol4l | 2.87 | 5.91E-38 | 1.26E-33 | 0.5 | 0.15 | 0.35 | 3.34 |
| 11 | Ubb | 0.7 | 8.91E-38 | 1.90E-33 | 0.99 | 1 | -0.01 | 0.99 |
| 11 | Npr2 | 4.21 | 1.25E-37 | 2.67E-33 | 0.14 | 0.01 | 0.13 | 12.91 |
| 11 | Efna3 | 4.33 | 1.32E-37 | 2.82E-33 | 0.37 | 0.09 | 0.29 | 4.35 |
| 11 | Cfl1 | 0.7 | 2.10E-37 | 4.49E-33 | 0.99 | 1 | -0.01 | 0.99 |
| 11 | Kdm7a | 2.21 | 2.24E-37 | 4.79E-33 | 0.76 | 0.41 | 0.35 | 1.85 |
| 11 | Tubb5 | 0.84 | 3.66E-37 | 7.81E-33 | 1 | 1 | 0 | 1 |
| 11 | Robo2 | 2.76 | 3.82E-37 | 8.16E-33 | 0.6 | 0.23 | 0.37 | 2.59 |
| 11 | Ache | 5.47 | 6.30E-37 | 1.35E-32 | 0.11 | 0.01 | 0.1 | 18.33 |
| 11 | Nkain4 | 4.38 | 7.75E-37 | 1.66E-32 | 0.23 | 0.03 | 0.2 | 7.29 |
| 11 | Tox3 | 2.72 | 7.85E-37 | 1.68E-32 | 0.67 | 0.3 | 0.37 | 2.22 |
| 11 | Celf6 | 5.42 | 1.59E-36 | 3.40E-32 | 0.12 | 0.01 | 0.11 | 16.57 |
| 11 | Akr1b8 | 3.54 | 1.64E-36 | 3.50E-32 | 0.21 | 0.03 | 0.18 | 8.24 |
| 11 | Dpysl4 | 3.09 | 2.96E-36 | 6.32E-32 | 0.41 | 0.1 | 0.3 | 3.98 |
| 11 | Pcbp4 | 1.34 | 4.28E-36 | 9.15E-32 | 0.91 | 0.91 | 0 | 1 |
| 11 | Kcne1l | 4.24 | 8.03E-36 | 1.72E-31 | 0.26 | 0.04 | 0.22 | 6.14 |
| 11 | Lzts1 | 3.49 | 9.52E-36 | 2.03E-31 | 0.21 | 0.03 | 0.18 | 7.92 |
| 11 | Cemip2 | 2.32 | 1.10E-35 | 2.35E-31 | 0.79 | 0.6 | 0.19 | 1.31 |
| 11 | Numbl | 2.91 | 1.39E-35 | 2.96E-31 | 0.53 | 0.18 | 0.35 | 2.94 |
| 11 | AC109619.1 | 4.85 | 5.91E-35 | 1.26E-30 | 0.11 | 0.01 | 0.1 | 15.71 |
| 11 | Celsr3 | 4.5 | 1.18E-34 | 2.51E-30 | 0.12 | 0.01 | 0.11 | 13.67 |
| 11 | Gse1 | 2.5 | 1.59E-34 | 3.39E-30 | 0.61 | 0.26 | 0.35 | 2.35 |
| 11 | Rmst | 3.71 | 3.02E-34 | 6.45E-30 | 0.25 | 0.04 | 0.21 | 6.28 |
| 11 | Rgmb | 3.16 | 5.53E-34 | 1.18E-29 | 0.46 | 0.14 | 0.32 | 3.37 |
| 11 | Fam43a | 4.78 | 7.53E-34 | 1.61E-29 | 0.12 | 0.01 | 0.11 | 13.67 |
| 11 | Rab3b | 4.22 | 1.13E-33 | 2.41E-29 | 0.28 | 0.05 | 0.23 | 5.26 |
| 11 | Pnmal2 | 3.58 | 1.15E-33 | 2.45E-29 | 0.27 | 0.05 | 0.22 | 5.53 |
| 11 | Baz1a | 2.82 | 1.73E-33 | 3.69E-29 | 0.8 | 0.58 | 0.22 | 1.38 |
| 11 | Kif5c | 2.39 | 2.51E-33 | 5.36E-29 | 0.81 | 0.59 | 0.23 | 1.39 |
| 11 | Calm1 | 1.38 | 7.00E-33 | 1.50E-28 | 0.95 | 0.99 | -0.04 | 0.96 |
| 11 | Trim62 | 3.09 | 1.25E-32 | 2.67E-28 | 0.37 | 0.09 | 0.28 | 3.98 |
| 11 | Ppp1r14b | 0.78 | 1.54E-32 | 3.28E-28 | 0.96 | 0.99 | -0.03 | 0.97 |
| 11 | Celf5 | 4.2 | 4.22E-32 | 9.02E-28 | 0.16 | 0.02 | 0.14 | 8.94 |
| 11 | Ppp2r5b | 2.85 | 4.58E-32 | 9.78E-28 | 0.3 | 0.06 | 0.24 | 4.89 |
| 11 | Pclo | 2.77 | 8.68E-32 | 1.86E-27 | 0.6 | 0.28 | 0.32 | 2.16 |
| 11 | Fyn | 2.26 | 1.09E-31 | 2.34E-27 | 0.73 | 0.4 | 0.33 | 1.84 |
| 11 | Stil | 2.9 | 1.53E-31 | 3.26E-27 | 0.48 | 0.17 | 0.31 | 2.84 |
| 11 | Nat14 | 3.21 | 1.98E-31 | 4.23E-27 | 0.26 | 0.05 | 0.21 | 5.49 |
| 11 | Atl1 | 2.83 | 1.99E-31 | 4.24E-27 | 0.31 | 0.07 | 0.24 | 4.7 |
| 11 | Amotl2 | 2.32 | 2.54E-31 | 5.42E-27 | 0.68 | 0.36 | 0.32 | 1.88 |
| 11 | Trnp1 | 4.29 | 3.12E-31 | 6.67E-27 | 0.19 | 0.03 | 0.16 | 7.48 |
| 11 | Ube2s | 1.43 | 3.58E-31 | 7.66E-27 | 0.95 | 0.96 | -0.01 | 0.99 |
| 11 | Marcks | 0.71 | 4.40E-31 | 9.40E-27 | 0.95 | 0.99 | -0.04 | 0.96 |
| 11 | E130114P18Rik | 3.77 | 8.63E-31 | 1.84E-26 | 0.26 | 0.05 | 0.21 | 5.49 |

**Supplementary Table 3. Marker genes for the original clusters of integrated clustering of E11.5 control and NotchGOF λ-junction ectoderm cells.**

| | | | | | | | | |
|---|---|---|---|---|---|---|---|---|
| 11 | Hspa4l | 2.45 | 1.06E-30 | 2.27E-26 | 0.58 | 0.24 | 0.34 | 2.38 |
| 11 | Fam131a | 3.82 | 1.41E-30 | 3.02E-26 | 0.17 | 0.02 | 0.15 | 8.4 |
| 11 | Mturn | 2.99 | 1.85E-30 | 3.96E-26 | 0.31 | 0.07 | 0.24 | 4.49 |
| 11 | Calm2 | 0.91 | 1.96E-30 | 4.18E-26 | 0.95 | 0.99 | -0.04 | 0.96 |
| 11 | Bcl2l1 | 2.07 | 2.25E-30 | 4.80E-26 | 0.74 | 0.45 | 0.29 | 1.64 |
| 11 | Gabarapl2 | 1.32 | 2.98E-30 | 6.36E-26 | 0.89 | 0.87 | 0.02 | 1.02 |
| 11 | 1110017D15Rik | 4.16 | 3.19E-30 | 6.82E-26 | 0.14 | 0.01 | 0.12 | 10.38 |
| 11 | Dynll1 | 0.82 | 4.02E-30 | 8.58E-26 | 0.99 | 1 | 0 | 1 |
| 11 | Dpysl5 | 2.21 | 4.20E-30 | 8.98E-26 | 0.69 | 0.38 | 0.31 | 1.84 |
| 11 | Ccser2 | 2.24 | 5.76E-30 | 1.23E-25 | 0.62 | 0.29 | 0.33 | 2.11 |
| 11 | Dscam | 4.16 | 9.43E-30 | 2.01E-25 | 0.1 | 0.01 | 0.1 | 14.71 |
| 11 | Ywhah | 1.58 | 1.44E-29 | 3.07E-25 | 0.89 | 0.8 | 0.09 | 1.11 |
| 11 | Adgrg1 | 2.51 | 2.24E-29 | 4.79E-25 | 0.42 | 0.13 | 0.29 | 3.33 |
| 11 | Afap1 | 2.65 | 2.97E-29 | 6.35E-25 | 0.53 | 0.21 | 0.32 | 2.57 |
| 11 | 3-Sep | 2.4 | 3.50E-29 | 7.49E-25 | 0.61 | 0.3 | 0.31 | 2.03 |
| 11 | Jpt1 | 1.08 | 7.18E-29 | 1.53E-24 | 0.94 | 0.97 | -0.03 | 0.97 |
| 11 | Mdm1 | 2.78 | 7.20E-29 | 1.54E-24 | 0.6 | 0.29 | 0.31 | 2.05 |
| 11 | Cep170 | 2.09 | 8.20E-29 | 1.75E-24 | 0.68 | 0.39 | 0.3 | 1.77 |
| 11 | Stx1a | 2.93 | 8.50E-29 | 1.82E-24 | 0.3 | 0.07 | 0.23 | 4.39 |
| 11 | Dbn1 | 1.41 | 9.60E-29 | 2.05E-24 | 0.84 | 0.87 | -0.03 | 0.96 |
| 11 | Tmsb10 | 0.98 | 1.09E-28 | 2.33E-24 | 1 | 1 | 0 | 1 |
| 11 | Trim24 | 2.06 | 1.13E-28 | 2.41E-24 | 0.77 | 0.6 | 0.17 | 1.29 |
| 11 | Rufy3 | 3.61 | 1.32E-28 | 2.82E-24 | 0.5 | 0.2 | 0.3 | 2.49 |
| 11 | Nrp1 | 3.69 | 1.54E-28 | 3.29E-24 | 0.23 | 0.04 | 0.19 | 5.79 |
| 11 | Casp3 | 1.89 | 1.62E-28 | 3.47E-24 | 0.83 | 0.7 | 0.13 | 1.19 |
| 11 | Dpysl2 | 1.84 | 1.70E-28 | 3.63E-24 | 0.8 | 0.63 | 0.18 | 1.28 |
| 11 | Cmpk1 | 1.51 | 2.64E-28 | 5.64E-24 | 0.83 | 0.72 | 0.11 | 1.15 |
| 11 | Shox2 | 5.72 | 2.93E-28 | 6.26E-24 | 0.14 | 0.02 | 0.13 | 8.88 |
| 11 | Smoc1 | 3.77 | 3.86E-28 | 8.23E-24 | 0.12 | 0.01 | 0.11 | 11.6 |
| 11 | Snap91 | 3.11 | 4.85E-28 | 1.04E-23 | 0.22 | 0.04 | 0.18 | 5.92 |
| 11 | Tpd52 | 1.48 | 5.50E-28 | 1.17E-23 | 0.88 | 0.81 | 0.07 | 1.09 |
| 11 | Col26a1 | 3.01 | 6.50E-28 | 1.39E-23 | 0.32 | 0.08 | 0.24 | 4.04 |
| 11 | Gpx2 | 4.51 | 9.87E-28 | 2.11E-23 | 0.1 | 0.01 | 0.1 | 12.88 |
| 11 | Myo1b | 1.55 | 1.42E-27 | 3.04E-23 | 0.81 | 0.62 | 0.18 | 1.3 |
| 11 | Klc3 | 2.7 | 1.56E-27 | 3.33E-23 | 0.68 | 0.39 | 0.29 | 1.75 |
| 11 | Neurl1a | 2.78 | 2.53E-27 | 5.40E-23 | 0.27 | 0.06 | 0.21 | 4.75 |
| 11 | Bcl7a | 1.51 | 4.92E-27 | 1.05E-22 | 0.83 | 0.77 | 0.06 | 1.08 |
| 11 | Zc3h12c | 2.6 | 9.38E-27 | 2.00E-22 | 0.37 | 0.11 | 0.26 | 3.4 |
| 11 | Tomm34 | 1.94 | 1.04E-26 | 2.23E-22 | 0.69 | 0.43 | 0.26 | 1.62 |
| 11 | H3f3a | 0.4 | 1.08E-26 | 2.31E-22 | 1 | 1 | 0 | 1 |
| 11 | Nlk | 2.26 | 1.11E-26 | 2.38E-22 | 0.66 | 0.4 | 0.26 | 1.66 |
| 11 | Dnaja1 | 0.67 | 1.31E-26 | 2.79E-22 | 0.94 | 0.99 | -0.05 | 0.95 |
| 11 | Hid1 | 3.69 | 1.37E-26 | 2.92E-22 | 0.17 | 0.02 | 0.15 | 7.3 |
| 11 | Ccsap | 2.98 | 1.46E-26 | 3.12E-22 | 0.32 | 0.08 | 0.24 | 3.89 |
| 11 | Rab39b | 2.57 | 1.82E-26 | 3.89E-22 | 0.41 | 0.13 | 0.28 | 3.22 |
| 11 | Smarcd2 | 2.4 | 1.87E-26 | 3.99E-22 | 0.65 | 0.35 | 0.29 | 1.84 |
| 11 | Nrg1 | 2.29 | 2.06E-26 | 4.40E-22 | 0.57 | 0.26 | 0.32 | 2.22 |
| 11 | Ank2 | 3.19 | 3.26E-26 | 6.96E-22 | 0.59 | 0.31 | 0.28 | 1.91 |
| 11 | Nefm | 5.19 | 4.06E-26 | 8.66E-22 | 0.17 | 0.02 | 0.14 | 7 |
| 11 | Ttll7 | 3.26 | 4.93E-26 | 1.05E-21 | 0.26 | 0.06 | 0.2 | 4.61 |
| 11 | Ssbp3 | 1.17 | 6.09E-26 | 1.30E-21 | 0.86 | 0.89 | -0.03 | 0.96 |
| 11 | Selenok | 1.35 | 1.06E-25 | 2.27E-21 | 0.88 | 0.9 | -0.02 | 0.97 |
| 11 | Map9 | 2.29 | 1.42E-25 | 3.04E-21 | 0.56 | 0.26 | 0.3 | 2.15 |
| 11 | Rbfox3 | 3.85 | 1.80E-25 | 3.85E-21 | 0.12 | 0.01 | 0.11 | 10.55 |
| 11 | Ralgds | 2.2 | 2.68E-25 | 5.72E-21 | 0.54 | 0.23 | 0.31 | 2.32 |
| 11 | Ccp110 | 2.47 | 3.24E-25 | 6.92E-21 | 0.63 | 0.34 | 0.28 | 1.82 |
| 11 | Map1lc3b | 1.15 | 3.47E-25 | 7.42E-21 | 0.89 | 0.94 | -0.05 | 0.95 |

**Supplementary Table 3. Marker genes for the original clusters of integrated clustering of E11.5 control and NotchGOF λ-junction ectoderm cells.**

| | | | | | | | | |
|---|---|---|---|---|---|---|---|---|
| 11 | Eif1 | 0.51 | 4.06E-25 | 8.67E-21 | 0.99 | 1 | -0.01 | 0.99 |
| 11 | Ell3 | 2.93 | 7.88E-25 | 1.68E-20 | 0.36 | 0.11 | 0.25 | 3.38 |
| 11 | Gdi1 | 2.17 | 8.78E-25 | 1.88E-20 | 0.7 | 0.45 | 0.25 | 1.56 |
| 11 | Tle1 | 1.91 | 9.21E-25 | 1.97E-20 | 0.77 | 0.54 | 0.23 | 1.43 |
| 11 | Pex5l | 3.77 | 1.19E-24 | 2.55E-20 | 0.15 | 0.02 | 0.13 | 7.4 |
| 11 | Rcor2 | 1.88 | 1.44E-24 | 3.07E-20 | 0.7 | 0.49 | 0.22 | 1.44 |
| 11 | Gm5468 | 4.26 | 1.47E-24 | 3.14E-20 | 0.1 | 0.01 | 0.09 | 11.44 |
| 11 | Kdm4b | 2.35 | 1.64E-24 | 3.50E-20 | 0.45 | 0.16 | 0.29 | 2.8 |
| 11 | Akna | 2.97 | 2.67E-24 | 5.71E-20 | 0.27 | 0.06 | 0.21 | 4.42 |
| 11 | Prkar1a | 1.36 | 2.80E-24 | 5.99E-20 | 0.88 | 0.85 | 0.03 | 1.04 |
| 11 | Cxcr4 | 2.48 | 3.14E-24 | 6.70E-20 | 0.46 | 0.17 | 0.29 | 2.65 |
| 11 | Amer2 | 2.28 | 3.45E-24 | 7.38E-20 | 0.43 | 0.15 | 0.28 | 2.92 |
| 11 | Kcnma1 | 4.68 | 3.84E-24 | 8.21E-20 | 0.12 | 0.01 | 0.1 | 9.67 |
| 11 | Kdm1a | 1.02 | 4.14E-24 | 8.85E-20 | 0.89 | 0.94 | -0.05 | 0.95 |
| 11 | Thsd7a | 4.99 | 5.03E-24 | 1.07E-19 | 0.12 | 0.01 | 0.1 | 9.67 |
| 11 | Syt11 | 2.56 | 5.37E-24 | 1.15E-19 | 0.61 | 0.34 | 0.28 | 1.82 |
| 11 | Dusp26 | 3.78 | 5.46E-24 | 1.17E-19 | 0.16 | 0.02 | 0.13 | 7.05 |
| 11 | Mbd2 | 1.76 | 5.79E-24 | 1.24E-19 | 0.79 | 0.61 | 0.18 | 1.29 |
| 11 | Wdr47 | 2.23 | 5.98E-24 | 1.28E-19 | 0.39 | 0.12 | 0.27 | 3.2 |
| 11 | Gng2 | 1.96 | 7.28E-24 | 1.56E-19 | 0.68 | 0.43 | 0.26 | 1.6 |
| 11 | Ncam1 | 3.45 | 1.32E-23 | 2.81E-19 | 0.6 | 0.33 | 0.27 | 1.83 |
| 11 | Ftl1 | 0.49 | 1.97E-23 | 4.21E-19 | 1 | 1 | 0 | 1 |
| 11 | Mxi1 | 1.75 | 2.00E-23 | 4.27E-19 | 0.77 | 0.57 | 0.2 | 1.35 |
| 11 | Aff3 | 2.93 | 2.15E-23 | 4.59E-19 | 0.28 | 0.07 | 0.21 | 3.96 |
| 11 | Rhbdl3 | 3.63 | 3.61E-23 | 7.72E-19 | 0.12 | 0.01 | 0.1 | 8.92 |
| 11 | Olfm1 | 2.27 | 5.33E-23 | 1.14E-18 | 0.6 | 0.32 | 0.28 | 1.86 |
| 11 | Dubr | 3.11 | 1.05E-22 | 2.24E-18 | 0.18 | 0.03 | 0.15 | 5.84 |
| 11 | Hspa12a | 3.23 | 1.15E-22 | 2.46E-18 | 0.17 | 0.03 | 0.15 | 6 |
| 11 | Lsamp | 4.35 | 1.63E-22 | 3.48E-18 | 0.12 | 0.01 | 0.1 | 8.92 |
| 11 | Smarca5 | 0.75 | 1.97E-22 | 4.20E-18 | 0.94 | 0.97 | -0.04 | 0.96 |
| 11 | Shtn1 | 2.97 | 2.26E-22 | 4.83E-18 | 0.46 | 0.19 | 0.26 | 2.36 |
| 11 | Bex2 | 1.07 | 2.30E-22 | 4.92E-18 | 0.92 | 0.88 | 0.04 | 1.05 |
| 11 | Cdk5r1 | 2.98 | 2.38E-22 | 5.09E-18 | 0.26 | 0.06 | 0.2 | 4.1 |
| 11 | Tsg101 | 1.19 | 2.46E-22 | 5.25E-18 | 0.87 | 0.86 | 0.01 | 1.01 |
| 11 | Atp6v1g1 | 0.9 | 5.90E-22 | 1.26E-17 | 0.91 | 0.97 | -0.06 | 0.94 |
| 11 | Tmcc3 | 2.62 | 9.81E-22 | 2.10E-17 | 0.33 | 0.1 | 0.23 | 3.26 |
| 11 | Lrp11 | 2.65 | 1.16E-21 | 2.48E-17 | 0.27 | 0.07 | 0.2 | 3.93 |
| 11 | Adam19 | 2.78 | 1.20E-21 | 2.56E-17 | 0.17 | 0.03 | 0.14 | 5.8 |
| 11 | Ddah2 | 0.94 | 1.53E-21 | 3.26E-17 | 0.94 | 0.98 | -0.04 | 0.96 |
| 11 | Ctnna2 | 2.81 | 1.71E-21 | 3.64E-17 | 0.27 | 0.07 | 0.2 | 3.82 |
| 11 | Lcorl | 2.11 | 1.78E-21 | 3.80E-17 | 0.65 | 0.4 | 0.25 | 1.62 |
| 11 | Cpeb4 | 2.74 | 2.64E-21 | 5.64E-17 | 0.27 | 0.07 | 0.2 | 3.96 |
| 11 | Slc43a2 | 2.64 | 2.73E-21 | 5.84E-17 | 0.25 | 0.06 | 0.19 | 4.06 |
| 11 | Trim9 | 3.8 | 2.81E-21 | 6.00E-17 | 0.11 | 0.01 | 0.1 | 9.17 |
| 11 | Spock2 | 2.1 | 2.82E-21 | 6.02E-17 | 0.37 | 0.12 | 0.25 | 3.07 |
| 11 | Rbfox2 | 1.43 | 2.86E-21 | 6.10E-17 | 0.79 | 0.71 | 0.08 | 1.12 |
| 11 | Pkig | 1.33 | 4.56E-21 | 9.74E-17 | 0.87 | 0.75 | 0.12 | 1.15 |
| 11 | Cadm1 | 1.44 | 5.30E-21 | 1.13E-16 | 0.8 | 0.71 | 0.09 | 1.12 |
| 11 | Arpc5 | 1.27 | 7.83E-21 | 1.67E-16 | 0.86 | 0.82 | 0.03 | 1.04 |
| 11 | Tenm1 | 3.46 | 8.61E-21 | 1.84E-16 | 0.14 | 0.02 | 0.12 | 6.76 |
| 11 | Kalrn | 2.33 | 8.65E-21 | 1.85E-16 | 0.43 | 0.17 | 0.26 | 2.49 |
| 11 | Csrnp3 | 3.1 | 8.67E-21 | 1.85E-16 | 0.14 | 0.02 | 0.12 | 7.11 |
| 11 | Dctn3 | 1.27 | 8.88E-21 | 1.90E-16 | 0.82 | 0.81 | 0.01 | 1.01 |
| 11 | Selenow | 0.79 | 1.11E-20 | 2.36E-16 | 0.96 | 0.99 | -0.04 | 0.96 |
| 11 | Caly | 3.67 | 1.48E-20 | 3.15E-16 | 0.11 | 0.01 | 0.1 | 8.46 |
| 11 | Snta1 | 3.05 | 1.59E-20 | 3.39E-16 | 0.16 | 0.03 | 0.13 | 5.75 |
| 11 | Pdrg1 | 2.04 | 1.59E-20 | 3.39E-16 | 0.76 | 0.57 | 0.18 | 1.32 |

**Supplementary Table 3. Marker genes for the original clusters of integrated clustering of E11.5 control and NotchGOF λ-junction ectoderm cells.**

| | | | | | | | | |
|---|---|---|---|---|---|---|---|---|
| 11 | Calm3 | 1.27 | 1.73E-20 | 3.69E-16 | 0.87 | 0.84 | 0.03 | 1.04 |
| 11 | Eif4g2 | 0.51 | 2.70E-20 | 5.77E-16 | 0.94 | 1 | -0.06 | 0.94 |
| 11 | Spns2 | 3.78 | 2.83E-20 | 6.05E-16 | 0.1 | 0.01 | 0.09 | 8.58 |
| 11 | Rnf5 | 1.56 | 3.19E-20 | 6.81E-16 | 0.81 | 0.7 | 0.12 | 1.17 |
| 11 | Acap3 | 2.62 | 3.53E-20 | 7.54E-16 | 0.23 | 0.06 | 0.18 | 4.22 |
| 11 | Stk33 | 4.42 | 3.85E-20 | 8.21E-16 | 0.11 | 0.01 | 0.1 | 8.46 |
| 11 | Hnrnpa2b1 | 0.52 | 4.53E-20 | 9.67E-16 | 0.98 | 1 | -0.02 | 0.98 |
| 11 | Gm13648 | 2.89 | 4.83E-20 | 1.03E-15 | 0.32 | 0.1 | 0.22 | 3.13 |
| 11 | Kif5b | 0.79 | 7.15E-20 | 1.53E-15 | 0.91 | 0.95 | -0.04 | 0.96 |
| 11 | Gadd45g | 1.31 | 7.24E-20 | 1.55E-15 | 0.65 | 0.33 | 0.32 | 1.95 |
| 11 | Mybl1 | 3.64 | 1.07E-19 | 2.29E-15 | 0.25 | 0.07 | 0.19 | 3.82 |
| 11 | Atp8a2 | 3.69 | 1.11E-19 | 2.36E-15 | 0.12 | 0.02 | 0.11 | 7.24 |
| 11 | Shb | 2.19 | 1.12E-19 | 2.39E-15 | 0.52 | 0.25 | 0.26 | 2.05 |
| 11 | Crabp2 | 0.92 | 1.36E-19 | 2.90E-15 | 0.96 | 0.85 | 0.12 | 1.14 |
| 11 | Csrp1 | 2.54 | 1.37E-19 | 2.93E-15 | 0.42 | 0.17 | 0.25 | 2.46 |
| 11 | Napb | 2.66 | 1.40E-19 | 2.98E-15 | 0.24 | 0.06 | 0.18 | 4.05 |
| 11 | Enho | 1.91 | 2.03E-19 | 4.33E-15 | 0.59 | 0.33 | 0.26 | 1.78 |
| 11 | Srgap1 | 1.83 | 2.31E-19 | 4.94E-15 | 0.64 | 0.4 | 0.24 | 1.6 |
| 11 | Mical2 | 2.7 | 2.64E-19 | 5.64E-15 | 0.22 | 0.05 | 0.17 | 4.21 |
| 11 | Riok3 | 1.64 | 2.80E-19 | 5.98E-15 | 0.68 | 0.47 | 0.21 | 1.44 |
| 11 | Chn1 | 2.3 | 4.51E-19 | 9.63E-15 | 0.52 | 0.27 | 0.25 | 1.94 |
| 11 | 2310040G24Rik | 3.95 | 5.45E-19 | 1.16E-14 | 0.17 | 0.03 | 0.14 | 5.25 |
| 11 | Crtc3 | 2.47 | 9.66E-19 | 2.06E-14 | 0.4 | 0.17 | 0.23 | 2.41 |
| 11 | Dynlrb1 | 0.96 | 1.03E-18 | 2.19E-14 | 0.91 | 0.94 | -0.03 | 0.97 |
| 11 | Carhsp1 | 1.67 | 1.14E-18 | 2.44E-14 | 0.73 | 0.58 | 0.15 | 1.26 |
| 11 | Cystm1 | 2.53 | 1.16E-18 | 2.48E-14 | 0.58 | 0.35 | 0.23 | 1.64 |
| 11 | Usp29 | 2.04 | 1.32E-18 | 2.81E-14 | 0.41 | 0.16 | 0.25 | 2.54 |
| 11 | Klf13 | 1.8 | 1.41E-18 | 3.01E-14 | 0.63 | 0.41 | 0.22 | 1.54 |
| 11 | Gdf11 | 1.97 | 1.42E-18 | 3.02E-14 | 0.51 | 0.26 | 0.25 | 1.97 |
| 11 | Chrnb2 | 3.26 | 1.47E-18 | 3.14E-14 | 0.12 | 0.02 | 0.1 | 7.25 |
| 11 | Cox6a1 | 0.76 | 1.52E-18 | 3.25E-14 | 0.93 | 0.95 | -0.02 | 0.97 |
| 11 | Map3k13 | 4.01 | 1.85E-18 | 3.96E-14 | 0.13 | 0.02 | 0.11 | 6.45 |
| 11 | Ddit4 | 1.86 | 2.08E-18 | 4.44E-14 | 0.68 | 0.43 | 0.25 | 1.58 |
| 11 | Fam124a | 3.11 | 2.58E-18 | 5.50E-14 | 0.15 | 0.03 | 0.12 | 5.69 |
| 11 | Nbea | 1.8 | 2.68E-18 | 5.73E-14 | 0.6 | 0.36 | 0.24 | 1.65 |
| 11 | Tekt2 | 3.67 | 3.04E-18 | 6.49E-14 | 0.13 | 0.02 | 0.11 | 6.45 |
| 11 | Kcnh8 | 2.4 | 3.20E-18 | 6.84E-14 | 0.37 | 0.14 | 0.23 | 2.69 |
| 11 | Enc1 | 2.58 | 3.23E-18 | 6.89E-14 | 0.49 | 0.25 | 0.24 | 1.98 |
| 11 | Rnf165 | 1.94 | 3.26E-18 | 6.97E-14 | 0.52 | 0.27 | 0.25 | 1.93 |
| 11 | Gjb5 | 3.8 | 4.51E-18 | 9.62E-14 | 0.12 | 0.02 | 0.11 | 6.83 |
| 11 | Smpd3 | 2.85 | 4.53E-18 | 9.68E-14 | 0.16 | 0.03 | 0.13 | 5.54 |
| 11 | Cdkn2d | 2.26 | 4.65E-18 | 9.94E-14 | 0.55 | 0.32 | 0.23 | 1.73 |
| 11 | Apbb1 | 2.12 | 5.20E-18 | 1.11E-13 | 0.57 | 0.34 | 0.23 | 1.67 |
| 11 | Fkbp1a | 0.84 | 5.63E-18 | 1.20E-13 | 0.9 | 0.94 | -0.05 | 0.95 |
| 11 | Tmem198 | 4.02 | 6.52E-18 | 1.39E-13 | 0.11 | 0.02 | 0.1 | 7.33 |
| 11 | Tnfrsf21 | 2.45 | 6.63E-18 | 1.42E-13 | 0.3 | 0.09 | 0.2 | 3.16 |
| 11 | Fdps | 1.14 | 8.35E-18 | 1.78E-13 | 0.85 | 0.82 | 0.04 | 1.04 |
| 11 | Ube2b | 0.84 | 9.80E-18 | 2.09E-13 | 0.9 | 0.95 | -0.05 | 0.94 |
| 11 | B3gnt4 | 2.79 | 1.02E-17 | 2.18E-13 | 0.17 | 0.03 | 0.14 | 5.09 |
| 11 | Fbxl16 | 3.94 | 1.07E-17 | 2.29E-13 | 0.14 | 0.02 | 0.11 | 5.87 |
| 11 | Enox1 | 3.07 | 1.68E-17 | 3.59E-13 | 0.2 | 0.05 | 0.15 | 4.26 |
| 11 | Ctxn1 | 1.46 | 1.71E-17 | 3.65E-13 | 0.81 | 0.79 | 0.02 | 1.02 |
| 11 | Atp6v0b | 1.28 | 2.22E-17 | 4.74E-13 | 0.8 | 0.77 | 0.03 | 1.04 |
| 11 | Gabarap | 0.77 | 2.57E-17 | 5.49E-13 | 0.91 | 0.97 | -0.06 | 0.94 |
| 11 | Stx7 | 2.78 | 2.67E-17 | 5.70E-13 | 0.48 | 0.24 | 0.24 | 1.97 |
| 11 | Rala | 0.98 | 2.72E-17 | 5.80E-13 | 0.85 | 0.84 | 0 | 1 |
| 11 | Mospd1 | 2.13 | 2.90E-17 | 6.18E-13 | 0.5 | 0.25 | 0.25 | 2 |

**Supplementary Table 3. Marker genes for the original clusters of integrated clustering of E11.5 control and NotchGOF λ-junction ectoderm cells.**

| | | | | | | | | |
|---|---|---|---|---|---|---|---|---|
| 11 | Syngr3 | 3.38 | 3.18E-17 | 6.79E-13 | 0.12 | 0.02 | 0.1 | 6.82 |
| 11 | Ywhag | 1.63 | 3.46E-17 | 7.40E-13 | 0.73 | 0.63 | 0.1 | 1.17 |
| 11 | Dync1i2 | 0.94 | 4.13E-17 | 8.81E-13 | 0.85 | 0.88 | -0.03 | 0.96 |
| 11 | Atp8a1 | 2.64 | 4.32E-17 | 9.23E-13 | 0.3 | 0.1 | 0.2 | 2.97 |
| 11 | Mpc1 | 1.8 | 4.40E-17 | 9.41E-13 | 0.67 | 0.48 | 0.2 | 1.41 |
| 11 | Mapk8ip1 | 2.12 | 4.86E-17 | 1.04E-12 | 0.58 | 0.36 | 0.22 | 1.62 |
| 11 | Tmem176b | 1.48 | 5.24E-17 | 1.12E-12 | 0.74 | 0.6 | 0.13 | 1.22 |
| 11 | Samd14 | 1.99 | 8.58E-17 | 1.83E-12 | 0.5 | 0.26 | 0.24 | 1.91 |
| 11 | Cep120 | 1.9 | 8.87E-17 | 1.89E-12 | 0.47 | 0.22 | 0.24 | 2.09 |
| 11 | B230216N24Rik | 3.61 | 1.05E-16 | 2.24E-12 | 0.1 | 0.01 | 0.09 | 7.36 |
| 11 | Ccdc88a | 1.15 | 1.30E-16 | 2.78E-12 | 0.77 | 0.7 | 0.07 | 1.11 |
| 11 | Rtl8a | 2.35 | 1.45E-16 | 3.10E-12 | 0.54 | 0.3 | 0.24 | 1.79 |
| 11 | Racgap1 | 1.5 | 1.48E-16 | 3.17E-12 | 0.72 | 0.51 | 0.21 | 1.41 |
| 11 | Cox7a2 | 0.51 | 1.62E-16 | 3.46E-12 | 0.94 | 0.99 | -0.05 | 0.95 |
| 11 | Dclk1 | 2.52 | 1.63E-16 | 3.48E-12 | 0.41 | 0.18 | 0.22 | 2.21 |
| 11 | Cxxc4 | 2.58 | 1.85E-16 | 3.96E-12 | 0.47 | 0.25 | 0.22 | 1.88 |
| 11 | Aldh1b1 | 3.03 | 2.27E-16 | 4.85E-12 | 0.12 | 0.02 | 0.1 | 6.44 |
| 11 | Actr3b | 3.16 | 2.31E-16 | 4.92E-12 | 0.14 | 0.03 | 0.12 | 5.26 |
| 11 | Gabrb3 | 2.94 | 2.87E-16 | 6.14E-12 | 0.11 | 0.02 | 0.09 | 6.88 |
| 11 | Gnao1 | 2.46 | 3.46E-16 | 7.40E-12 | 0.39 | 0.18 | 0.22 | 2.25 |
| 11 | Snx32 | 3.16 | 3.86E-16 | 8.24E-12 | 0.12 | 0.02 | 0.1 | 5.86 |
| 11 | Ap3m2 | 2.58 | 3.96E-16 | 8.46E-12 | 0.19 | 0.05 | 0.14 | 4.16 |
| 11 | Shd | 2.81 | 4.38E-16 | 9.35E-12 | 0.27 | 0.09 | 0.18 | 3.12 |
| 11 | 5-Sep | 2.68 | 4.52E-16 | 9.66E-12 | 0.18 | 0.04 | 0.14 | 4.21 |
| 11 | Tdrkh | 2.4 | 5.33E-16 | 1.14E-11 | 0.28 | 0.09 | 0.18 | 2.98 |
| 11 | Mapk8 | 1.49 | 5.98E-16 | 1.28E-11 | 0.57 | 0.34 | 0.24 | 1.71 |
| 11 | Palm | 1.93 | 7.83E-16 | 1.67E-11 | 0.61 | 0.41 | 0.2 | 1.49 |
| 11 | Ypel3 | 1.65 | 8.90E-16 | 1.90E-11 | 0.72 | 0.57 | 0.15 | 1.26 |
| 11 | Lpcat2 | 2.48 | 9.18E-16 | 1.96E-11 | 0.21 | 0.05 | 0.15 | 3.81 |
| 11 | Rsph9 | 2.39 | 1.06E-15 | 2.25E-11 | 0.31 | 0.11 | 0.2 | 2.82 |
| 11 | Hdac2 | 0.93 | 1.09E-15 | 2.33E-11 | 0.85 | 0.89 | -0.03 | 0.96 |
| 11 | Pth1r | 2.47 | 1.56E-15 | 3.33E-11 | 0.2 | 0.05 | 0.15 | 3.85 |
| 11 | Mgst3 | 2.43 | 1.61E-15 | 3.45E-11 | 0.65 | 0.49 | 0.16 | 1.32 |
| 11 | Atp5f1 | 0.48 | 2.86E-15 | 6.12E-11 | 0.95 | 0.99 | -0.04 | 0.96 |
| 11 | Akap17b | 2.48 | 2.96E-15 | 6.32E-11 | 0.35 | 0.14 | 0.21 | 2.47 |
| 11 | Ccdc96 | 2.72 | 3.62E-15 | 7.73E-11 | 0.16 | 0.04 | 0.13 | 4.47 |
| 11 | Gm1673 | 0.81 | 3.63E-15 | 7.75E-11 | 0.88 | 0.93 | -0.05 | 0.94 |
| 11 | Gdpd1 | 1.89 | 3.83E-15 | 8.18E-11 | 0.61 | 0.4 | 0.21 | 1.53 |
| 11 | Hpcal1 | 1.71 | 4.60E-15 | 9.83E-11 | 0.65 | 0.46 | 0.19 | 1.41 |
| 11 | Draxin | 3.3 | 5.06E-15 | 1.08E-10 | 0.1 | 0.02 | 0.09 | 6.44 |
| 11 | 1700088E04Rik | 2.49 | 5.55E-15 | 1.19E-10 | 0.24 | 0.07 | 0.17 | 3.23 |
| 11 | Zfp386 | 1.78 | 6.85E-15 | 1.46E-10 | 0.65 | 0.44 | 0.21 | 1.48 |
| 11 | Nfasc | 2.59 | 7.41E-15 | 1.58E-10 | 0.27 | 0.09 | 0.18 | 2.91 |
| 11 | Maml3 | 1.46 | 7.86E-15 | 1.68E-10 | 0.61 | 0.4 | 0.21 | 1.53 |
| 11 | Tecpr1 | 1.84 | 7.97E-15 | 1.70E-10 | 0.36 | 0.15 | 0.21 | 2.44 |
| 11 | C2cd2l | 2.76 | 8.08E-15 | 1.73E-10 | 0.18 | 0.05 | 0.14 | 4.02 |
| 11 | Eif4a1 | 0.44 | 8.98E-15 | 1.92E-10 | 0.98 | 1 | -0.02 | 0.98 |
| 11 | Sdk1 | 2.92 | 9.03E-15 | 1.93E-10 | 0.26 | 0.09 | 0.17 | 3 |
| 11 | Cecr2 | 1.5 | 9.52E-15 | 2.03E-10 | 0.6 | 0.38 | 0.22 | 1.59 |
| 11 | Pcolce2 | 3.79 | 1.04E-14 | 2.21E-10 | 0.21 | 0.06 | 0.15 | 3.44 |
| 11 | Rab3c | 3.2 | 1.32E-14 | 2.81E-10 | 0.17 | 0.04 | 0.13 | 4.1 |
| 11 | Onecut2 | 3.71 | 1.41E-14 | 3.02E-10 | 0.16 | 0.04 | 0.12 | 4.24 |
| 11 | Art1 | 3.79 | 1.43E-14 | 3.05E-10 | 0.12 | 0.02 | 0.1 | 5.35 |
| 11 | Tfeb | 2.82 | 1.57E-14 | 3.35E-10 | 0.16 | 0.04 | 0.12 | 4.35 |
| 11 | Plppr2 | 2.74 | 1.75E-14 | 3.75E-10 | 0.18 | 0.05 | 0.14 | 3.93 |
| 11 | BC031181 | 1.19 | 1.88E-14 | 4.01E-10 | 0.79 | 0.78 | 0.02 | 1.02 |
| 11 | Vim | 2.08 | 1.91E-14 | 4.08E-10 | 0.77 | 0.72 | 0.05 | 1.07 |

**Supplementary Table 3. Marker genes for the original clusters of integrated clustering of E11.5 control and NotchGOF λ-junction ectoderm cells.**

| | | | | | | | | |
|---|---|---|---|---|---|---|---|---|
| 11 | Dnajb6 | 0.98 | 2.33E-14 | 4.97E-10 | 0.87 | 0.9 | -0.02 | 0.97 |
| 11 | Arhgap33 | 2 | 2.55E-14 | 5.46E-10 | 0.32 | 0.12 | 0.19 | 2.57 |
| 11 | Traf4 | 1.49 | 3.06E-14 | 6.53E-10 | 0.67 | 0.51 | 0.16 | 1.32 |
| 11 | 1110008P14Rik | 1.8 | 3.13E-14 | 6.68E-10 | 0.57 | 0.36 | 0.21 | 1.59 |
| 11 | Ppp3ca | 1.45 | 3.37E-14 | 7.19E-10 | 0.67 | 0.54 | 0.13 | 1.24 |
| 11 | Plekho1 | 1.47 | 3.37E-14 | 7.20E-10 | 0.56 | 0.35 | 0.2 | 1.57 |
| 11 | Dbndd1 | 2.33 | 3.60E-14 | 7.69E-10 | 0.26 | 0.09 | 0.17 | 2.93 |
| 11 | Slc43a1 | 2.09 | 3.72E-14 | 7.95E-10 | 0.28 | 0.1 | 0.18 | 2.81 |
| 11 | Stx2 | 2.02 | 3.76E-14 | 8.03E-10 | 0.36 | 0.16 | 0.2 | 2.23 |
| 11 | Nmnat2 | 4.19 | 3.97E-14 | 8.47E-10 | 0.1 | 0.02 | 0.09 | 6.06 |
| 11 | Scube1 | 2.78 | 4.91E-14 | 1.05E-09 | 0.18 | 0.05 | 0.13 | 3.85 |
| 11 | Ccdc28b | 1.72 | 5.07E-14 | 1.08E-09 | 0.56 | 0.35 | 0.21 | 1.61 |
| 11 | Rapgef5 | 2.37 | 5.30E-14 | 1.13E-09 | 0.37 | 0.17 | 0.2 | 2.22 |
| 11 | Dda1 | 1.49 | 5.63E-14 | 1.20E-09 | 0.7 | 0.56 | 0.15 | 1.26 |
| 11 | Klc1 | 1.98 | 6.13E-14 | 1.31E-09 | 0.65 | 0.49 | 0.16 | 1.32 |
| 11 | Tmeff1 | 1.25 | 6.32E-14 | 1.35E-09 | 0.76 | 0.68 | 0.07 | 1.11 |
| 11 | Dram2 | 1.72 | 6.46E-14 | 1.38E-09 | 0.4 | 0.18 | 0.22 | 2.23 |
| 11 | Ciapin1 | 1.22 | 6.73E-14 | 1.44E-09 | 0.78 | 0.69 | 0.09 | 1.14 |
| 11 | Acsl1 | 2.3 | 1.13E-13 | 2.41E-09 | 0.21 | 0.06 | 0.15 | 3.33 |
| 11 | Gamt | 3.36 | 1.16E-13 | 2.47E-09 | 0.24 | 0.08 | 0.16 | 3.03 |
| 11 | Agtpbp1 | 2.28 | 1.16E-13 | 2.48E-09 | 0.28 | 0.1 | 0.17 | 2.69 |
| 11 | Apba2 | 2.37 | 1.19E-13 | 2.54E-09 | 0.32 | 0.13 | 0.19 | 2.43 |
| 11 | Cfap36 | 1.59 | 1.27E-13 | 2.72E-09 | 0.69 | 0.49 | 0.2 | 1.4 |
| 11 | Plcxd3 | 3.76 | 1.39E-13 | 2.97E-09 | 0.11 | 0.02 | 0.09 | 5.5 |
| 11 | Hsp90aa1 | 0.45 | 1.47E-13 | 3.14E-09 | 0.97 | 1 | -0.03 | 0.97 |
| 11 | Nav1 | 1.35 | 1.47E-13 | 3.14E-09 | 0.67 | 0.51 | 0.16 | 1.31 |
| 11 | Mdh2 | 0.83 | 1.60E-13 | 3.41E-09 | 0.88 | 0.94 | -0.06 | 0.94 |
| 11 | Mex3a | 0.91 | 1.69E-13 | 3.62E-09 | 0.83 | 0.88 | -0.04 | 0.95 |
| 11 | Zmiz1 | 1.32 | 1.82E-13 | 3.89E-09 | 0.73 | 0.62 | 0.11 | 1.17 |
| 11 | Hist3h2ba | 1.59 | 2.00E-13 | 4.28E-09 | 0.85 | 0.83 | 0.02 | 1.03 |
| 11 | A330084C13Rik | 2.75 | 2.11E-13 | 4.50E-09 | 0.11 | 0.02 | 0.09 | 5.5 |
| 11 | Pak7 | 2.77 | 2.12E-13 | 4.54E-09 | 0.15 | 0.03 | 0.11 | 4.35 |
| 11 | Fam89b | 1.17 | 2.18E-13 | 4.66E-09 | 0.8 | 0.75 | 0.05 | 1.06 |
| 11 | Arl8a | 1.95 | 2.28E-13 | 4.86E-09 | 0.56 | 0.38 | 0.18 | 1.46 |
| 11 | Arfgef3 | 3.19 | 2.41E-13 | 5.15E-09 | 0.2 | 0.06 | 0.14 | 3.33 |
| 11 | Ttll1 | 2.21 | 2.93E-13 | 6.26E-09 | 0.39 | 0.19 | 0.21 | 2.13 |
| 11 | 2810455O05Rik | 3.31 | 3.14E-13 | 6.71E-09 | 0.1 | 0.02 | 0.09 | 5.72 |
| 11 | Slc26a10 | 2.44 | 3.39E-13 | 7.25E-09 | 0.14 | 0.03 | 0.11 | 4.66 |
| 11 | Rnasek | 1.06 | 3.43E-13 | 7.34E-09 | 0.77 | 0.73 | 0.04 | 1.06 |
| 11 | Cntfr | 2.33 | 3.58E-13 | 7.65E-09 | 0.33 | 0.14 | 0.19 | 2.32 |
| 11 | Grcc10 | 0.83 | 5.30E-13 | 1.13E-08 | 0.92 | 0.98 | -0.05 | 0.95 |
| 11 | Srp14 | 0.74 | 5.67E-13 | 1.21E-08 | 0.9 | 0.95 | -0.05 | 0.95 |
| 11 | Jarid2 | 1.33 | 6.22E-13 | 1.33E-08 | 0.67 | 0.55 | 0.12 | 1.23 |
| 11 | Phyhipl | 2.14 | 6.65E-13 | 1.42E-08 | 0.27 | 0.1 | 0.17 | 2.74 |
| 11 | Dmxl2 | 2.25 | 7.67E-13 | 1.64E-08 | 0.23 | 0.08 | 0.16 | 3.01 |
| 11 | Map3k12 | 1.68 | 7.90E-13 | 1.69E-08 | 0.36 | 0.16 | 0.2 | 2.25 |
| 11 | Ubald2 | 1.11 | 9.14E-13 | 1.95E-08 | 0.78 | 0.71 | 0.07 | 1.1 |
| 11 | Mapre2 | 2.24 | 9.46E-13 | 2.02E-08 | 0.45 | 0.23 | 0.21 | 1.93 |
| 11 | Gal | 0.98 | 1.02E-12 | 2.18E-08 | 0.14 | 0.03 | 0.11 | 4.5 |
| 11 | Gm42372 | 2.6 | 1.04E-12 | 2.22E-08 | 0.16 | 0.04 | 0.12 | 3.97 |
| 11 | Plekha6 | 2.91 | 1.06E-12 | 2.27E-08 | 0.16 | 0.04 | 0.12 | 3.83 |
| 11 | Astn1 | 2.58 | 1.07E-12 | 2.28E-08 | 0.16 | 0.04 | 0.12 | 3.83 |
| 11 | Tanc2 | 1.59 | 1.08E-12 | 2.31E-08 | 0.43 | 0.22 | 0.21 | 1.96 |
| 11 | Cep250 | 1.74 | 1.11E-12 | 2.36E-08 | 0.44 | 0.24 | 0.2 | 1.85 |
| 11 | Gm10544 | 2.87 | 1.16E-12 | 2.47E-08 | 0.15 | 0.04 | 0.11 | 4.11 |
| 11 | Fscn1 | 0.8 | 1.17E-12 | 2.49E-08 | 0.83 | 0.86 | -0.04 | 0.96 |
| 11 | Psmc5 | 0.71 | 1.28E-12 | 2.72E-08 | 0.85 | 0.9 | -0.04 | 0.95 |

**Supplementary Table 3. Marker genes for the original clusters of integrated clustering of E11.5 control and NotchGOF λ-junction ectoderm cells.**

| | | | | | | | | |
|---|---|---|---|---|---|---|---|---|
| 11 | Gck | 2.73 | 1.31E-12 | 2.79E-08 | 0.16 | 0.04 | 0.12 | 3.83 |
| 11 | Khdrbs1 | 0.61 | 1.43E-12 | 3.05E-08 | 0.89 | 0.95 | -0.06 | 0.94 |
| 11 | Dctn1 | 1.59 | 1.53E-12 | 3.26E-08 | 0.65 | 0.51 | 0.13 | 1.26 |
| 11 | Ap2m1 | 0.86 | 1.73E-12 | 3.69E-08 | 0.83 | 0.83 | 0 | 1 |
| 11 | Zfp428 | 1.2 | 1.78E-12 | 3.80E-08 | 0.75 | 0.67 | 0.08 | 1.11 |
| 11 | Kif3c | 1.97 | 2.61E-12 | 5.57E-08 | 0.28 | 0.11 | 0.17 | 2.54 |
| 11 | Gpm6b | 2.61 | 2.88E-12 | 6.14E-08 | 0.25 | 0.09 | 0.16 | 2.68 |
| 11 | Ptpn6 | 2.73 | 3.13E-12 | 6.69E-08 | 0.23 | 0.08 | 0.15 | 2.97 |
| 11 | Cadps2 | 2.38 | 3.25E-12 | 6.95E-08 | 0.31 | 0.13 | 0.18 | 2.31 |
| 11 | Dact1 | 1.48 | 3.58E-12 | 7.65E-08 | 0.34 | 0.14 | 0.19 | 2.38 |
| 11 | Atp6v1e1 | 1.11 | 4.14E-12 | 8.84E-08 | 0.74 | 0.69 | 0.05 | 1.08 |
| 11 | Unc45a | 1.85 | 4.70E-12 | 1.00E-07 | 0.48 | 0.28 | 0.2 | 1.73 |
| 11 | Kctd13 | 1.72 | 5.81E-12 | 1.24E-07 | 0.34 | 0.16 | 0.19 | 2.19 |
| 11 | Mien1 | 1.12 | 6.79E-12 | 1.45E-07 | 0.75 | 0.68 | 0.07 | 1.1 |
| 11 | Fam20c | 2.24 | 7.87E-12 | 1.68E-07 | 0.23 | 0.08 | 0.15 | 2.9 |
| 11 | Zcrb1 | 0.8 | 8.18E-12 | 1.75E-07 | 0.85 | 0.9 | -0.05 | 0.95 |
| 11 | Rtn4 | 0.74 | 8.37E-12 | 1.79E-07 | 0.83 | 0.96 | -0.13 | 0.86 |
| 11 | Kcnn2 | 2.73 | 8.42E-12 | 1.80E-07 | 0.12 | 0.03 | 0.09 | 4.64 |
| 11 | Gm8797 | 0.81 | 8.69E-12 | 1.86E-07 | 0.88 | 0.93 | -0.04 | 0.96 |
| 11 | Lman2l | 1.58 | 9.56E-12 | 2.04E-07 | 0.55 | 0.36 | 0.19 | 1.54 |
| 11 | Foxg1 | 1.3 | 1.02E-11 | 2.18E-07 | 0.65 | 0.54 | 0.11 | 1.2 |
| 11 | Zbtb18 | 1.93 | 1.09E-11 | 2.33E-07 | 0.41 | 0.23 | 0.19 | 1.83 |
| 11 | Hpca | 3.82 | 1.12E-11 | 2.39E-07 | 0.12 | 0.03 | 0.09 | 4.64 |
| 11 | Crip2 | 1.93 | 1.15E-11 | 2.46E-07 | 0.61 | 0.42 | 0.2 | 1.47 |
| 11 | Evl | 1.16 | 1.17E-11 | 2.49E-07 | 0.78 | 0.73 | 0.05 | 1.07 |
| 11 | Mcf2l | 2.02 | 1.28E-11 | 2.74E-07 | 0.3 | 0.13 | 0.17 | 2.34 |
| 11 | Jund | 0.79 | 1.33E-11 | 2.84E-07 | 0.87 | 0.93 | -0.06 | 0.94 |
| 11 | Cog1 | 1.97 | 1.34E-11 | 2.85E-07 | 0.32 | 0.14 | 0.18 | 2.31 |
| 11 | Sertad2 | 1.26 | 1.42E-11 | 3.04E-07 | 0.66 | 0.52 | 0.14 | 1.26 |
| 11 | Stat3 | 2.11 | 1.71E-11 | 3.66E-07 | 0.25 | 0.09 | 0.15 | 2.69 |
| 11 | Katnal1 | 2.04 | 1.73E-11 | 3.69E-07 | 0.32 | 0.14 | 0.18 | 2.32 |
| 11 | Znrf1 | 1.29 | 1.83E-11 | 3.91E-07 | 0.69 | 0.56 | 0.13 | 1.23 |
| 11 | Oaf | 2.06 | 1.85E-11 | 3.95E-07 | 0.31 | 0.14 | 0.17 | 2.18 |
| 11 | Ksr2 | 2.73 | 1.86E-11 | 3.98E-07 | 0.16 | 0.04 | 0.11 | 3.69 |
| 11 | Tbcb | 0.82 | 2.11E-11 | 4.51E-07 | 0.82 | 0.85 | -0.03 | 0.96 |
| 11 | Hecw2 | 2.7 | 2.35E-11 | 5.02E-07 | 0.14 | 0.03 | 0.1 | 3.97 |
| 11 | Dusp1 | 1.83 | 2.45E-11 | 5.23E-07 | 0.43 | 0.24 | 0.19 | 1.8 |
| 11 | Nrp2 | 2.16 | 2.60E-11 | 5.56E-07 | 0.36 | 0.17 | 0.19 | 2.09 |
| 11 | E2f1 | 1.74 | 2.69E-11 | 5.75E-07 | 0.39 | 0.21 | 0.19 | 1.89 |
| 11 | Cspg5 | 2.95 | 2.76E-11 | 5.90E-07 | 0.14 | 0.04 | 0.11 | 3.84 |
| 11 | Efr3b | 2.51 | 2.80E-11 | 5.98E-07 | 0.19 | 0.06 | 0.13 | 3.08 |
| 11 | Psmc2 | 0.86 | 4.03E-11 | 8.61E-07 | 0.83 | 0.87 | -0.04 | 0.95 |
| 11 | Bmerb1 | 3.03 | 4.05E-11 | 8.64E-07 | 0.13 | 0.03 | 0.1 | 4.16 |
| 11 | Ndufb1-ps | 0.74 | 4.06E-11 | 8.68E-07 | 0.89 | 0.93 | -0.04 | 0.95 |
| 11 | Tgfbr1 | 1.65 | 4.37E-11 | 9.33E-07 | 0.68 | 0.61 | 0.07 | 1.12 |
| 11 | C9orf72 | 2.41 | 4.39E-11 | 9.38E-07 | 0.28 | 0.12 | 0.16 | 2.37 |
| 11 | Saraf | 1.5 | 4.74E-11 | 1.01E-06 | 0.72 | 0.66 | 0.05 | 1.08 |
| 11 | Pgm2l1 | 1.54 | 5.58E-11 | 1.19E-06 | 0.41 | 0.22 | 0.19 | 1.86 |
| 11 | Rbm18 | 1.33 | 5.61E-11 | 1.20E-06 | 0.64 | 0.49 | 0.15 | 1.29 |
| 11 | Ywhaq | 0.41 | 5.97E-11 | 1.27E-06 | 0.92 | 0.99 | -0.07 | 0.93 |
| 11 | Dynlt1f | 1.02 | 6.42E-11 | 1.37E-06 | 0.79 | 0.75 | 0.05 | 1.06 |
| 11 | Smchd1 | 1.82 | 6.43E-11 | 1.37E-06 | 0.79 | 0.84 | -0.04 | 0.95 |
| 11 | Ttc39b | 2.43 | 6.88E-11 | 1.47E-06 | 0.19 | 0.06 | 0.13 | 3.12 |
| 11 | Dync1li1 | 1.37 | 6.98E-11 | 1.49E-06 | 0.64 | 0.5 | 0.14 | 1.27 |
| 11 | Ciao2b | 1.09 | 8.21E-11 | 1.75E-06 | 0.76 | 0.74 | 0.02 | 1.02 |
| 11 | Gulo | 3.22 | 8.92E-11 | 1.91E-06 | 0.1 | 0.02 | 0.08 | 4.9 |
| 11 | 0610010F05Rik | 1.42 | 8.94E-11 | 1.91E-06 | 0.52 | 0.33 | 0.18 | 1.54 |

**Supplementary Table 3. Marker genes for the original clusters of integrated clustering of E11.5 control and NotchGOF λ-junction ectoderm cells.**

| | | | | | | | | |
|---|---|---|---|---|---|---|---|---|
| 11 | Mkrn1 | 1.1 | 1.02E-10 | 2.17E-06 | 0.74 | 0.65 | 0.09 | 1.13 |
| 11 | Dact3 | 2.01 | 1.08E-10 | 2.30E-06 | 0.28 | 0.12 | 0.16 | 2.39 |
| 11 | Lmo1 | 2.13 | 1.09E-10 | 2.32E-06 | 0.42 | 0.23 | 0.19 | 1.85 |
| 11 | Pdlim7 | 1.03 | 1.14E-10 | 2.43E-06 | 0.73 | 0.62 | 0.11 | 1.17 |
| 11 | Cep63 | 1.38 | 1.18E-10 | 2.51E-06 | 0.52 | 0.35 | 0.18 | 1.5 |
| 11 | Pcdh20 | 2.55 | 1.34E-10 | 2.87E-06 | 0.12 | 0.03 | 0.09 | 4.3 |
| 11 | Cby1 | 1.41 | 1.38E-10 | 2.95E-06 | 0.63 | 0.46 | 0.17 | 1.38 |
| 11 | Eml1 | 1.84 | 1.74E-10 | 3.72E-06 | 0.34 | 0.17 | 0.17 | 2.01 |
| 11 | Csnk1e | 0.85 | 1.76E-10 | 3.75E-06 | 0.83 | 0.88 | -0.04 | 0.95 |
| 11 | Atp5md | 0.53 | 1.96E-10 | 4.18E-06 | 0.92 | 0.98 | -0.07 | 0.93 |
| 11 | Ppp1r11 | 0.91 | 1.97E-10 | 4.21E-06 | 0.82 | 0.82 | 0 | 1 |
| 11 | Tmcc1 | 1.22 | 2.10E-10 | 4.48E-06 | 0.62 | 0.46 | 0.16 | 1.35 |
| 11 | Rnf32 | 2.4 | 2.92E-10 | 6.23E-06 | 0.13 | 0.03 | 0.1 | 3.91 |
| 11 | Rims2 | 2.04 | 2.93E-10 | 6.27E-06 | 0.38 | 0.21 | 0.18 | 1.86 |
| 11 | Gm5860 | 3.43 | 3.46E-10 | 7.40E-06 | 0.12 | 0.03 | 0.09 | 3.97 |
| 11 | H3f3b | 0.28 | 3.56E-10 | 7.61E-06 | 1 | 1 | 0 | 1 |
| 11 | Hspa8 | 0.34 | 3.58E-10 | 7.64E-06 | 0.98 | 1 | -0.02 | 0.98 |
| 11 | Ndufa4l2 | 2.36 | 3.71E-10 | 7.92E-06 | 0.24 | 0.1 | 0.14 | 2.52 |
| 11 | Chd7 | 1.02 | 4.10E-10 | 8.76E-06 | 0.75 | 0.79 | -0.04 | 0.95 |
| 11 | Tspyl4 | 2.03 | 4.10E-10 | 8.77E-06 | 0.4 | 0.24 | 0.16 | 1.69 |
| 11 | Smarcc2 | 1.08 | 4.56E-10 | 9.74E-06 | 0.71 | 0.62 | 0.09 | 1.15 |
| 11 | Syt1 | 2.1 | 4.81E-10 | 1.03E-05 | 0.31 | 0.14 | 0.17 | 2.23 |
| 11 | Gnb4 | 2.63 | 4.92E-10 | 1.05E-05 | 0.12 | 0.03 | 0.09 | 3.97 |
| 11 | Cdc42ep3 | 1.72 | 5.01E-10 | 1.07E-05 | 0.35 | 0.17 | 0.18 | 2.02 |
| 11 | Tubb2a | 1.83 | 5.94E-10 | 1.27E-05 | 0.52 | 0.36 | 0.17 | 1.46 |
| 11 | Chd4 | 0.68 | 6.44E-10 | 1.37E-05 | 0.88 | 0.97 | -0.09 | 0.91 |
| 11 | Akirin1 | 1.38 | 6.80E-10 | 1.45E-05 | 0.66 | 0.58 | 0.08 | 1.14 |
| 11 | Trim35 | 1.07 | 6.85E-10 | 1.46E-05 | 0.68 | 0.55 | 0.13 | 1.23 |
| 11 | Dab2ip | 1.33 | 6.88E-10 | 1.47E-05 | 0.63 | 0.53 | 0.1 | 1.19 |
| 11 | Fam122b | 1.88 | 7.20E-10 | 1.54E-05 | 0.23 | 0.09 | 0.14 | 2.63 |
| 11 | Cep97 | 2.01 | 7.93E-10 | 1.69E-05 | 0.39 | 0.23 | 0.16 | 1.71 |
| 11 | Mtcl1 | 1.11 | 8.15E-10 | 1.74E-05 | 0.5 | 0.31 | 0.19 | 1.63 |
| 11 | Npdc1 | 1.61 | 8.84E-10 | 1.89E-05 | 0.49 | 0.32 | 0.17 | 1.51 |
| 11 | Kctd10 | 1.69 | 9.06E-10 | 1.94E-05 | 0.43 | 0.25 | 0.18 | 1.71 |
| 11 | Phf21b | 2.49 | 9.30E-10 | 1.99E-05 | 0.19 | 0.07 | 0.13 | 2.85 |
| 11 | Prmt2 | 1.61 | 1.07E-09 | 2.29E-05 | 0.5 | 0.35 | 0.15 | 1.42 |
| 11 | Bpgm | 1.35 | 1.11E-09 | 2.37E-05 | 0.57 | 0.42 | 0.15 | 1.35 |
| 11 | Tcta | 2.16 | 1.12E-09 | 2.40E-05 | 0.23 | 0.09 | 0.14 | 2.54 |
| 11 | Gm43260 | 2.14 | 1.12E-09 | 2.40E-05 | 0.16 | 0.05 | 0.11 | 3.3 |
| 11 | Fam171b | 2.08 | 1.17E-09 | 2.51E-05 | 0.28 | 0.13 | 0.15 | 2.18 |
| 11 | Cycs | 0.77 | 1.18E-09 | 2.52E-05 | 0.87 | 0.93 | -0.06 | 0.93 |
| 11 | Klhl29 | 2 | 1.27E-09 | 2.72E-05 | 0.14 | 0.04 | 0.1 | 3.55 |
| 11 | Mapre3 | 2.28 | 1.53E-09 | 3.27E-05 | 0.28 | 0.13 | 0.15 | 2.2 |
| 11 | Ywhae | 0.37 | 1.54E-09 | 3.29E-05 | 0.96 | 1 | -0.04 | 0.96 |
| 11 | Arpc3 | 0.67 | 1.58E-09 | 3.37E-05 | 0.87 | 0.92 | -0.05 | 0.95 |
| 11 | Tmem41a | 1.48 | 1.62E-09 | 3.45E-05 | 0.34 | 0.17 | 0.17 | 2.03 |
| 11 | Dennd1a | 1.4 | 1.78E-09 | 3.81E-05 | 0.37 | 0.2 | 0.17 | 1.85 |
| 11 | Arl6 | 1.68 | 2.06E-09 | 4.39E-05 | 0.54 | 0.38 | 0.16 | 1.42 |
| 11 | Tshz1 | 1.71 | 2.06E-09 | 4.40E-05 | 0.59 | 0.48 | 0.11 | 1.23 |
| 11 | Zbtb20 | 2.19 | 2.07E-09 | 4.43E-05 | 0.45 | 0.3 | 0.15 | 1.49 |
| 11 | Clcn2 | 1.66 | 2.16E-09 | 4.62E-05 | 0.21 | 0.08 | 0.13 | 2.75 |
| 11 | Vamp2 | 1.81 | 2.39E-09 | 5.10E-05 | 0.47 | 0.32 | 0.15 | 1.46 |
| 11 | Csnk1g3 | 1.09 | 2.58E-09 | 5.51E-05 | 0.75 | 0.67 | 0.08 | 1.12 |
| 11 | H13 | 0.93 | 2.86E-09 | 6.11E-05 | 0.72 | 0.65 | 0.08 | 1.12 |
| 11 | Olfm2 | 1.81 | 3.06E-09 | 6.53E-05 | 0.34 | 0.18 | 0.17 | 1.95 |
| 11 | Actr10 | 1 | 3.11E-09 | 6.64E-05 | 0.73 | 0.69 | 0.04 | 1.05 |
| 11 | Pik3r3 | 1.96 | 3.16E-09 | 6.76E-05 | 0.37 | 0.2 | 0.17 | 1.85 |

**Supplementary Table 3. Marker genes for the original clusters of integrated clustering of E11.5 control and NotchGOF λ-junction ectoderm cells.**

| | | | | | | | | |
|---|---|---|---|---|---|---|---|---|
| 11 | Atp6v0d1 | 1.15 | 3.27E-09 | 6.99E-05 | 0.7 | 0.56 | 0.14 | 1.25 |
| 11 | Peli1 | 0.94 | 3.30E-09 | 7.04E-05 | 0.72 | 0.66 | 0.06 | 1.09 |
| 11 | Chchd2 | 0.28 | 3.33E-09 | 7.12E-05 | 0.98 | 1 | -0.02 | 0.98 |
| 11 | Ahi1 | 1.35 | 3.36E-09 | 7.17E-05 | 0.51 | 0.33 | 0.18 | 1.53 |
| 11 | Cacna1h | 2.32 | 3.37E-09 | 7.19E-05 | 0.14 | 0.04 | 0.1 | 3.38 |
| 11 | 2700097O09Rik | 1.66 | 3.52E-09 | 7.52E-05 | 0.48 | 0.33 | 0.15 | 1.45 |
| 11 | Kras | 0.86 | 3.67E-09 | 7.83E-05 | 0.81 | 0.84 | -0.03 | 0.97 |
| 11 | 3110039M20Rik | 1.97 | 3.86E-09 | 8.25E-05 | 0.45 | 0.3 | 0.15 | 1.5 |
| 11 | Twf1 | 1.85 | 5.14E-09 | 1.10E-04 | 0.63 | 0.54 | 0.08 | 1.15 |
| 11 | Trappc2l | 1.34 | 5.30E-09 | 1.13E-04 | 0.68 | 0.61 | 0.07 | 1.12 |
| 11 | Pdap1 | 0.52 | 5.38E-09 | 1.15E-04 | 0.93 | 0.99 | -0.06 | 0.94 |
| 11 | Phyh | 1.37 | 5.58E-09 | 1.19E-04 | 0.49 | 0.33 | 0.16 | 1.48 |
| 11 | Gfod1 | 2.6 | 5.84E-09 | 1.25E-04 | 0.11 | 0.03 | 0.08 | 3.93 |
| 11 | Mapk3 | 1.32 | 5.99E-09 | 1.28E-04 | 0.61 | 0.48 | 0.12 | 1.25 |
| 11 | Tle4 | 1.33 | 6.42E-09 | 1.37E-04 | 0.63 | 0.54 | 0.09 | 1.17 |
| 11 | Chchd6 | 1.35 | 6.48E-09 | 1.38E-04 | 0.65 | 0.53 | 0.13 | 1.24 |
| 11 | BC030867 | 2.2 | 6.88E-09 | 1.47E-04 | 0.24 | 0.1 | 0.14 | 2.3 |
| 11 | Pou3f1 | 2.24 | 7.07E-09 | 1.51E-04 | 0.14 | 0.04 | 0.1 | 3.38 |
| 11 | Peg13 | 2.13 | 7.67E-09 | 1.64E-04 | 0.16 | 0.05 | 0.11 | 2.98 |
| 11 | Arhgef2 | 1.27 | 8.27E-09 | 1.77E-04 | 0.53 | 0.39 | 0.14 | 1.36 |
| 11 | Mtch1 | 0.57 | 8.35E-09 | 1.78E-04 | 0.94 | 0.97 | -0.04 | 0.96 |
| 11 | Atf4 | 0.99 | 8.96E-09 | 1.91E-04 | 0.74 | 0.68 | 0.07 | 1.1 |
| 11 | Snx11 | 1.78 | 9.65E-09 | 2.06E-04 | 0.31 | 0.15 | 0.16 | 2.01 |
| 11 | Gpr85 | 2.47 | 9.85E-09 | 2.10E-04 | 0.11 | 0.03 | 0.08 | 3.79 |
| 11 | Cxxc5 | 1.3 | 1.10E-08 | 2.35E-04 | 0.67 | 0.59 | 0.09 | 1.15 |
| 11 | Ndrg3 | 1.85 | 1.12E-08 | 2.40E-04 | 0.39 | 0.22 | 0.17 | 1.74 |
| 11 | Pbx3 | 1.17 | 1.18E-08 | 2.51E-04 | 0.68 | 0.59 | 0.09 | 1.16 |
| 11 | Mterf2 | 1.55 | 1.20E-08 | 2.56E-04 | 0.39 | 0.22 | 0.17 | 1.74 |
| 11 | Ndufa11 | 0.71 | 1.20E-08 | 2.56E-04 | 0.92 | 0.95 | -0.03 | 0.96 |
| 11 | Sdccag8 | 2.02 | 1.26E-08 | 2.69E-04 | 0.28 | 0.14 | 0.15 | 2.1 |
| 11 | Chfr | 1.24 | 1.29E-08 | 2.75E-04 | 0.48 | 0.31 | 0.17 | 1.55 |
| 11 | Lrrn3 | 1.6 | 1.44E-08 | 3.07E-04 | 0.3 | 0.15 | 0.15 | 2.05 |
| 11 | Shisa4 | 1.39 | 1.45E-08 | 3.10E-04 | 0.4 | 0.24 | 0.16 | 1.69 |
| 11 | Vps28 | 1.19 | 1.47E-08 | 3.14E-04 | 0.7 | 0.62 | 0.09 | 1.14 |
| 11 | Podxl2 | 1.49 | 1.53E-08 | 3.27E-04 | 0.52 | 0.38 | 0.14 | 1.37 |
| 11 | Gsk3b | 0.77 | 1.58E-08 | 3.38E-04 | 0.83 | 0.91 | -0.07 | 0.92 |
| 11 | Ankrd13b | 1.7 | 1.60E-08 | 3.41E-04 | 0.34 | 0.18 | 0.15 | 1.85 |
| 11 | H2afy | 0.42 | 1.60E-08 | 3.43E-04 | 0.92 | 0.98 | -0.07 | 0.93 |
| 11 | Clip3 | 0.97 | 1.63E-08 | 3.48E-04 | 0.59 | 0.46 | 0.14 | 1.3 |
| 11 | Dennd2a | 1.51 | 1.66E-08 | 3.54E-04 | 0.41 | 0.24 | 0.17 | 1.72 |
| 11 | Gde1 | 1.23 | 1.77E-08 | 3.79E-04 | 0.61 | 0.5 | 0.11 | 1.21 |
| 11 | Asxl3 | 1.55 | 1.99E-08 | 4.24E-04 | 0.47 | 0.32 | 0.16 | 1.5 |
| 11 | Mob3b | 1.7 | 2.01E-08 | 4.30E-04 | 0.23 | 0.1 | 0.13 | 2.32 |
| 11 | Plppr3 | 1.18 | 2.05E-08 | 4.39E-04 | 0.48 | 0.31 | 0.16 | 1.52 |
| 11 | Kif3a | 1.14 | 2.06E-08 | 4.41E-04 | 0.68 | 0.59 | 0.1 | 1.16 |
| 11 | Dusp18 | 1.74 | 2.19E-08 | 4.67E-04 | 0.24 | 0.11 | 0.13 | 2.28 |
| 11 | Cpe | 1.2 | 2.21E-08 | 4.72E-04 | 0.59 | 0.47 | 0.13 | 1.27 |
| 11 | Fam204a | 1.08 | 2.23E-08 | 4.76E-04 | 0.72 | 0.63 | 0.09 | 1.14 |
| 11 | Lrrc75a | 1.58 | 2.34E-08 | 4.99E-04 | 0.36 | 0.19 | 0.16 | 1.83 |
| 11 | Trappc4 | 0.87 | 2.87E-08 | 6.13E-04 | 0.8 | 0.81 | -0.01 | 0.98 |
| 11 | Idh1 | 1.1 | 3.18E-08 | 6.79E-04 | 0.65 | 0.55 | 0.1 | 1.18 |
| 11 | Gm49066 | 1.8 | 3.21E-08 | 6.85E-04 | 0.18 | 0.07 | 0.11 | 2.66 |
| 11 | Susd6 | 1.59 | 3.38E-08 | 7.23E-04 | 0.37 | 0.21 | 0.16 | 1.74 |
| 11 | Baz2b | 0.91 | 3.58E-08 | 7.65E-04 | 0.79 | 0.81 | -0.02 | 0.98 |
| 11 | Pkn1 | 1.45 | 3.69E-08 | 7.88E-04 | 0.44 | 0.29 | 0.15 | 1.54 |
| 11 | Slc27a2 | 2.11 | 3.79E-08 | 8.10E-04 | 0.16 | 0.06 | 0.1 | 2.82 |
| 11 | Pdxp | 1.29 | 3.81E-08 | 8.15E-04 | 0.5 | 0.34 | 0.16 | 1.47 |

**Supplementary Table 3. Marker genes for the original clusters of integrated clustering of E11.5 control and NotchGOF λ-junction ectoderm cells.**

| | | | | | | | | |
|---|---|---|---|---|---|---|---|---|
| 11 | Gdap2 | 1.74 | 3.97E-08 | 8.47E-04 | 0.23 | 0.1 | 0.13 | 2.31 |
| 11 | Ncoa1 | 1.92 | 4.18E-08 | 8.93E-04 | 0.36 | 0.21 | 0.15 | 1.7 |
| 11 | Psmb7 | 0.61 | 4.94E-08 | 1.05E-03 | 0.86 | 0.91 | -0.05 | 0.94 |
| 11 | Ywhab | 0.51 | 5.00E-08 | 1.07E-03 | 0.91 | 0.97 | -0.06 | 0.94 |
| 11 | Tmem131l | 1.03 | 5.46E-08 | 1.17E-03 | 0.59 | 0.48 | 0.11 | 1.22 |
| 11 | Rc3h2 | 1.11 | 5.59E-08 | 1.19E-03 | 0.66 | 0.56 | 0.1 | 1.18 |
| 11 | Slco3a1 | 2.36 | 5.62E-08 | 1.20E-03 | 0.14 | 0.04 | 0.09 | 3.14 |
| 11 | 2900093K20Rik | 1.82 | 6.20E-08 | 1.32E-03 | 0.23 | 0.1 | 0.13 | 2.23 |
| 11 | Sh3gl3 | 1.91 | 7.07E-08 | 1.51E-03 | 0.21 | 0.09 | 0.12 | 2.39 |
| 11 | Magee1 | 2.01 | 7.73E-08 | 1.65E-03 | 0.18 | 0.07 | 0.11 | 2.62 |
| 11 | Apip | 1.23 | 7.81E-08 | 1.67E-03 | 0.56 | 0.4 | 0.16 | 1.4 |
| 11 | Khdrbs2 | 1.79 | 7.83E-08 | 1.67E-03 | 0.12 | 0.04 | 0.09 | 3.32 |
| 11 | Esyt1 | 1.9 | 7.83E-08 | 1.67E-03 | 0.19 | 0.08 | 0.12 | 2.49 |
| 11 | Zcchc18 | 1.4 | 7.91E-08 | 1.69E-03 | 0.57 | 0.44 | 0.13 | 1.29 |
| 11 | Ndn | 1.63 | 7.96E-08 | 1.70E-03 | 0.67 | 0.61 | 0.06 | 1.1 |
| 11 | Arpc1a | 0.65 | 8.16E-08 | 1.74E-03 | 0.87 | 0.91 | -0.05 | 0.95 |
| 11 | Bach2 | 1.41 | 9.28E-08 | 1.98E-03 | 0.4 | 0.24 | 0.16 | 1.65 |
| 11 | Pak3 | 1.48 | 9.85E-08 | 2.10E-03 | 0.45 | 0.3 | 0.14 | 1.47 |
| 11 | Ankrd13a | 1.53 | 1.01E-07 | 2.16E-03 | 0.41 | 0.27 | 0.15 | 1.55 |
| 11 | Atp6v1h | 0.99 | 1.01E-07 | 2.16E-03 | 0.54 | 0.38 | 0.16 | 1.43 |
| 11 | Ufm1 | 0.97 | 1.01E-07 | 2.16E-03 | 0.72 | 0.68 | 0.04 | 1.06 |
| 11 | Rnd2 | 2.16 | 1.04E-07 | 2.23E-03 | 0.24 | 0.11 | 0.13 | 2.19 |
| 11 | Rtl8b | 1.88 | 1.06E-07 | 2.26E-03 | 0.43 | 0.28 | 0.15 | 1.52 |
| 11 | Igf1r | 0.79 | 1.06E-07 | 2.27E-03 | 0.78 | 0.83 | -0.05 | 0.94 |
| 11 | Ttc5 | 0.92 | 1.07E-07 | 2.28E-03 | 0.69 | 0.63 | 0.06 | 1.1 |
| 11 | Ssh2 | 1.19 | 1.09E-07 | 2.32E-03 | 0.43 | 0.28 | 0.15 | 1.54 |
| 11 | Lrrc42 | 1.25 | 1.11E-07 | 2.38E-03 | 0.7 | 0.71 | -0.01 | 0.98 |
| 11 | Capn10 | 1.23 | 1.17E-07 | 2.49E-03 | 0.37 | 0.22 | 0.16 | 1.71 |
| 11 | Sgip1 | 2.34 | 1.18E-07 | 2.52E-03 | 0.1 | 0.03 | 0.07 | 3.55 |
| 11 | Klf12 | 2.4 | 1.23E-07 | 2.63E-03 | 0.23 | 0.1 | 0.12 | 2.22 |
| 11 | Msi2 | 0.75 | 1.28E-07 | 2.72E-03 | 0.77 | 0.75 | 0.01 | 1.02 |
| 11 | Chmp5 | 0.7 | 1.29E-07 | 2.76E-03 | 0.79 | 0.8 | -0.01 | 0.99 |
| 11 | Prkx | 1.79 | 1.33E-07 | 2.85E-03 | 0.33 | 0.19 | 0.14 | 1.77 |
| 11 | Tmem151b | 1.81 | 1.36E-07 | 2.91E-03 | 0.19 | 0.08 | 0.11 | 2.49 |
| 11 | Rtn2 | 1.67 | 1.43E-07 | 3.05E-03 | 0.33 | 0.18 | 0.15 | 1.81 |
| 11 | Katnal2 | 1.98 | 1.56E-07 | 3.33E-03 | 0.17 | 0.07 | 0.11 | 2.6 |
| 11 | Tmod2 | 1.98 | 1.63E-07 | 3.48E-03 | 0.25 | 0.13 | 0.13 | 2.02 |
| 11 | Sv2a | 2.59 | 1.65E-07 | 3.52E-03 | 0.11 | 0.03 | 0.08 | 3.44 |
| 11 | Smap2 | 1.49 | 1.87E-07 | 4.00E-03 | 0.42 | 0.27 | 0.15 | 1.55 |
| 11 | Bbip1 | 1 | 1.90E-07 | 4.05E-03 | 0.68 | 0.61 | 0.07 | 1.11 |
| 11 | Ube2m | 0.5 | 1.92E-07 | 4.11E-03 | 0.88 | 0.95 | -0.06 | 0.93 |
| 11 | Malat1 | 0.73 | 2.07E-07 | 4.42E-03 | 0.93 | 1 | -0.07 | 0.93 |
| 11 | Bmpr1b | 1.83 | 2.08E-07 | 4.44E-03 | 0.19 | 0.08 | 0.11 | 2.46 |
| 11 | Rhobtb2 | 2.14 | 2.11E-07 | 4.52E-03 | 0.19 | 0.08 | 0.11 | 2.46 |
| 11 | Ift22 | 1.2 | 2.13E-07 | 4.54E-03 | 0.5 | 0.38 | 0.12 | 1.32 |
| 11 | Kif1b | 1.26 | 2.13E-07 | 4.56E-03 | 0.68 | 0.65 | 0.03 | 1.04 |
| 11 | Shc2 | 2.58 | 2.19E-07 | 4.67E-03 | 0.18 | 0.07 | 0.11 | 2.45 |
| 11 | Kcnh2 | 2.1 | 2.24E-07 | 4.78E-03 | 0.14 | 0.05 | 0.09 | 2.84 |
| 11 | 4930452B06Rik | 2.35 | 2.28E-07 | 4.88E-03 | 0.28 | 0.15 | 0.14 | 1.91 |
| 11 | Nedd4l | 1.47 | 2.29E-07 | 4.89E-03 | 0.39 | 0.25 | 0.14 | 1.56 |
| 11 | Gm36839 | 2.16 | 2.33E-07 | 4.98E-03 | 0.12 | 0.04 | 0.08 | 3.15 |
| 11 | Gm4419 | 2.49 | 2.36E-07 | 5.04E-03 | 0.1 | 0.03 | 0.07 | 3.43 |
| 11 | Nelfcd | 0.89 | 2.37E-07 | 5.07E-03 | 0.65 | 0.58 | 0.07 | 1.13 |
| 11 | Epb41 | 1.01 | 2.48E-07 | 5.29E-03 | 0.74 | 0.75 | -0.01 | 0.98 |
| 11 | Ccpg1 | 1.27 | 2.55E-07 | 5.46E-03 | 0.4 | 0.24 | 0.16 | 1.67 |
| 11 | Clstn2 | 2.45 | 2.72E-07 | 5.82E-03 | 0.1 | 0.03 | 0.07 | 3.43 |
| 11 | Rac3 | 1.58 | 2.73E-07 | 5.83E-03 | 0.51 | 0.41 | 0.1 | 1.24 |

**Supplementary Table 3. Marker genes for the original clusters of integrated clustering of E11.5 control and NotchGOF λ-junction ectoderm cells.**

| | | | | | | | | |
|---|---|---|---|---|---|---|---|---|
| 11 | 4930402H24Rik | 1.4 | 2.80E-07 | 5.98E-03 | 0.3 | 0.16 | 0.14 | 1.89 |
| 11 | Cenpj | 1.31 | 2.82E-07 | 6.03E-03 | 0.47 | 0.34 | 0.13 | 1.39 |
| 11 | Sbk1 | 1.08 | 2.83E-07 | 6.04E-03 | 0.62 | 0.57 | 0.05 | 1.09 |
| 11 | Rab3d | 2.36 | 2.87E-07 | 6.13E-03 | 0.13 | 0.04 | 0.09 | 3 |
| 11 | Btbd10 | 1.58 | 2.94E-07 | 6.29E-03 | 0.41 | 0.27 | 0.14 | 1.53 |
| 11 | Pim1 | 1.49 | 3.06E-07 | 6.53E-03 | 0.57 | 0.45 | 0.12 | 1.26 |
| 11 | Pim3 | 2.05 | 3.19E-07 | 6.82E-03 | 0.52 | 0.41 | 0.11 | 1.27 |
| 11 | Prelid3b | 1.02 | 3.22E-07 | 6.89E-03 | 0.67 | 0.59 | 0.08 | 1.14 |
| 11 | Ttbk2 | 1.47 | 3.29E-07 | 7.03E-03 | 0.25 | 0.12 | 0.13 | 2.07 |
| 11 | Ror2 | 1.9 | 3.32E-07 | 7.09E-03 | 0.15 | 0.05 | 0.09 | 2.74 |
| 11 | Ndufb10 | 0.74 | 3.38E-07 | 7.21E-03 | 0.86 | 0.9 | -0.04 | 0.96 |
| 11 | Mindy3 | 1.3 | 3.40E-07 | 7.26E-03 | 0.39 | 0.25 | 0.14 | 1.58 |
| 11 | Nkiras1 | 2.15 | 3.68E-07 | 7.86E-03 | 0.25 | 0.12 | 0.13 | 2.03 |
| 11 | Rasgef1b | 1.92 | 3.86E-07 | 8.24E-03 | 0.21 | 0.1 | 0.12 | 2.24 |
| 11 | 4833439L19Rik | 1.38 | 4.13E-07 | 8.82E-03 | 0.55 | 0.46 | 0.09 | 1.19 |
| 11 | Trappc6b | 0.98 | 4.28E-07 | 9.15E-03 | 0.63 | 0.53 | 0.1 | 1.18 |
| 11 | Smarce1 | 0.34 | 4.29E-07 | 9.17E-03 | 0.91 | 0.98 | -0.06 | 0.93 |
| 11 | Coq10b | 1.33 | 4.40E-07 | 9.41E-03 | 0.32 | 0.18 | 0.14 | 1.79 |
| 11 | Sel1l3 | 1.92 | 4.69E-07 | 1.00E-02 | 0.17 | 0.07 | 0.1 | 2.49 |
| 11 | Mab21l2 | 2.51 | 5.07E-07 | 1.08E-02 | 0.15 | 0.06 | 0.09 | 2.64 |
| 11 | Med22 | 1.57 | 5.39E-07 | 1.15E-02 | 0.43 | 0.28 | 0.14 | 1.51 |
| 11 | Scrn1 | 2.38 | 5.41E-07 | 1.16E-02 | 0.24 | 0.12 | 0.12 | 1.98 |
| 11 | Sema6c | 1.33 | 5.77E-07 | 1.23E-02 | 0.26 | 0.13 | 0.13 | 2.05 |
| 11 | Ccdc34 | 0.69 | 6.21E-07 | 1.33E-02 | 0.85 | 0.86 | -0.02 | 0.98 |
| 11 | Ift43 | 1.35 | 6.40E-07 | 1.37E-02 | 0.72 | 0.67 | 0.05 | 1.08 |
| 11 | Lmna | 1.82 | 6.41E-07 | 1.37E-02 | 0.32 | 0.19 | 0.14 | 1.72 |
| 11 | Svip | 1.61 | 6.84E-07 | 1.46E-02 | 0.4 | 0.26 | 0.14 | 1.54 |
| 11 | Mex3b | 0.9 | 6.86E-07 | 1.46E-02 | 0.59 | 0.47 | 0.12 | 1.26 |
| 11 | Nabp1 | 1.89 | 6.90E-07 | 1.47E-02 | 0.16 | 0.06 | 0.1 | 2.63 |
| 11 | Wsb2 | 1.29 | 7.02E-07 | 1.50E-02 | 0.47 | 0.33 | 0.13 | 1.4 |
| 11 | Pacsin1 | 1.3 | 7.51E-07 | 1.60E-02 | 0.31 | 0.17 | 0.14 | 1.82 |
| 11 | Tmem108 | 1.09 | 7.53E-07 | 1.61E-02 | 0.46 | 0.3 | 0.16 | 1.53 |
| 11 | Atg4a | 2.4 | 7.72E-07 | 1.65E-02 | 0.16 | 0.06 | 0.1 | 2.63 |
| 11 | Rnf14 | 1.2 | 8.01E-07 | 1.71E-02 | 0.47 | 0.34 | 0.13 | 1.39 |
| 11 | Armh4 | 2.23 | 8.16E-07 | 1.74E-02 | 0.3 | 0.18 | 0.12 | 1.66 |
| 11 | Rassf3 | 1.79 | 8.17E-07 | 1.75E-02 | 0.25 | 0.13 | 0.12 | 1.96 |
| 11 | Vbp1 | 0.82 | 8.30E-07 | 1.77E-02 | 0.73 | 0.68 | 0.04 | 1.07 |
| 11 | 8-Mar | 1.32 | 8.49E-07 | 1.81E-02 | 0.39 | 0.25 | 0.14 | 1.57 |
| 11 | Psmd6 | 0.68 | 9.10E-07 | 1.94E-02 | 0.85 | 0.86 | -0.01 | 0.99 |
| 11 | Gnb1 | 0.52 | 9.22E-07 | 1.97E-02 | 0.88 | 0.97 | -0.09 | 0.91 |
| 11 | Akr1a1 | 0.57 | 9.36E-07 | 2.00E-02 | 0.88 | 0.97 | -0.08 | 0.92 |
| 11 | Ypel5 | 1.09 | 9.71E-07 | 2.08E-02 | 0.42 | 0.26 | 0.16 | 1.64 |
| 11 | Lrrc49 | 1.24 | 1.05E-06 | 2.24E-02 | 0.32 | 0.18 | 0.14 | 1.76 |
| 11 | Morn4 | 1.51 | 1.09E-06 | 2.33E-02 | 0.29 | 0.16 | 0.13 | 1.85 |
| 11 | Ubl5 | 0.43 | 1.09E-06 | 2.34E-02 | 0.93 | 0.99 | -0.06 | 0.94 |
| 11 | Spa17 | 1.7 | 1.10E-06 | 2.35E-02 | 0.29 | 0.16 | 0.13 | 1.85 |
| 11 | Mtmr6 | 2.18 | 1.11E-06 | 2.37E-02 | 0.24 | 0.12 | 0.12 | 1.98 |
| 11 | Asb1 | 2.56 | 1.20E-06 | 2.57E-02 | 0.14 | 0.05 | 0.09 | 2.76 |
| 11 | Hbb-y | 2.36 | 1.26E-06 | 2.69E-02 | 0.94 | 0.98 | -0.04 | 0.96 |
| 11 | Map2k1 | 1.62 | 1.28E-06 | 2.73E-02 | 0.5 | 0.4 | 0.1 | 1.24 |
| 11 | Slco5a1 | 1.58 | 1.30E-06 | 2.78E-02 | 0.3 | 0.17 | 0.13 | 1.78 |
| 11 | Glipr2 | 2.27 | 1.35E-06 | 2.88E-02 | 0.17 | 0.07 | 0.1 | 2.4 |
| 11 | Zfp667 | 1.68 | 1.36E-06 | 2.90E-02 | 0.27 | 0.14 | 0.13 | 1.88 |
| 11 | Actr1a | 1.16 | 1.39E-06 | 2.98E-02 | 0.56 | 0.44 | 0.11 | 1.26 |
| 11 | Pfdn4 | 0.91 | 1.40E-06 | 2.98E-02 | 0.79 | 0.78 | 0.02 | 1.02 |
| 11 | 1110006O24Rik | 2.94 | 1.42E-06 | 3.04E-02 | 0.12 | 0.04 | 0.08 | 2.97 |
| 11 | Gnaz | 2.3 | 1.46E-06 | 3.11E-02 | 0.14 | 0.05 | 0.09 | 2.63 |

**Supplementary Table 3. Marker genes for the original clusters of integrated clustering of E11.5 control and NotchGOF λ-junction ectoderm cells.**

| | | | | | | | | |
|---|---|---|---|---|---|---|---|---|
| 11 | Garnl3 | 2.15 | 1.52E-06 | 3.24E-02 | 0.11 | 0.04 | 0.08 | 3.14 |
| 11 | Nab1 | 1.16 | 1.57E-06 | 3.35E-02 | 0.39 | 0.24 | 0.15 | 1.61 |
| 11 | Ahsa2 | 0.87 | 1.61E-06 | 3.45E-02 | 0.68 | 0.63 | 0.04 | 1.07 |
| 11 | Taf13 | 1.13 | 1.65E-06 | 3.53E-02 | 0.6 | 0.49 | 0.11 | 1.22 |
| 11 | Synj1 | 1.78 | 1.70E-06 | 3.64E-02 | 0.23 | 0.12 | 0.12 | 1.98 |
| 11 | Six1 | 0.83 | 1.73E-06 | 3.70E-02 | 0.74 | 0.69 | 0.05 | 1.08 |
| 11 | Purg | 1.8 | 1.83E-06 | 3.92E-02 | 0.4 | 0.27 | 0.13 | 1.47 |
| 11 | Nin | 1.31 | 1.86E-06 | 3.98E-02 | 0.41 | 0.28 | 0.13 | 1.48 |
| 11 | Fam227a | 2.01 | 1.87E-06 | 4.00E-02 | 0.14 | 0.05 | 0.09 | 2.63 |
| 11 | Prtn3 | 1.59 | 1.88E-06 | 4.01E-02 | 0.22 | 0.1 | 0.12 | 2.11 |
| 11 | Cib2 | 1.41 | 1.94E-06 | 4.15E-02 | 0.31 | 0.18 | 0.13 | 1.76 |
| 11 | Frs3 | 3.14 | 2.14E-06 | 4.58E-02 | 0.12 | 0.04 | 0.08 | 2.97 |
| 11 | Wscd1 | 2.27 | 2.34E-06 | 5.00E-02 | 0.19 | 0.08 | 0.1 | 2.23 |

**Supplementary Table 4. Marker genes for the annotated cell populations of integrated clustering of E11.5 control and NotchGOF λ-junction ectoderm cells.**

| cluster | gene | avg_log2FC | p_val | p_val_adj | pct.1 | pct.2 | pct1-pct2 | pct1/pct2 |
|---|---|---|---|---|---|---|---|---|
| NE | Cdh2 | 2.22 | 0 | 0 | 0.89 | 0.37 | 0.52 | 2.44 |
| NE | Crabp2 | 1.68 | 0 | 0 | 0.99 | 0.72 | 0.27 | 1.38 |
| NE | Psip1 | 1.11 | 1.77E-299 | 3.78E-295 | 1.00 | 0.97 | 0.03 | 1.03 |
| NE | Gm266 | 2.79 | 5.78E-274 | 1.23E-269 | 0.79 | 0.33 | 0.46 | 2.41 |
| NE | Sfrp1 | 2.30 | 6.62E-250 | 1.41E-245 | 0.78 | 0.29 | 0.48 | 2.64 |
| NE | Six1 | 1.39 | 2.84E-241 | 6.07E-237 | 0.93 | 0.47 | 0.45 | 1.96 |
| NE | Rprm | 3.35 | 7.51E-231 | 1.60E-226 | 0.63 | 0.16 | 0.47 | 3.95 |
| NE | Tubb2b | 0.81 | 3.23E-214 | 6.90E-210 | 0.98 | 0.88 | 0.10 | 1.11 |
| NE | Pax6 | 1.91 | 4.27E-204 | 9.13E-200 | 0.73 | 0.28 | 0.46 | 2.67 |
| NE | Pcdh19 | 4.17 | 1.73E-202 | 3.69E-198 | 0.46 | 0.05 | 0.41 | 9.18 |
| NE | Mecom | 2.34 | 2.38E-202 | 5.09E-198 | 0.70 | 0.24 | 0.45 | 2.88 |
| NE | Eya1 | 1.59 | 1.79E-200 | 3.82E-196 | 0.82 | 0.39 | 0.43 | 2.08 |
| NE | Il17rd | 1.88 | 5.59E-200 | 1.19E-195 | 0.74 | 0.29 | 0.45 | 2.53 |
| NE | Gsta4 | 1.14 | 1.71E-193 | 3.65E-189 | 0.94 | 0.70 | 0.24 | 1.34 |
| NE | Galm | 2.32 | 1.00E-185 | 2.14E-181 | 0.63 | 0.20 | 0.43 | 3.11 |
| NE | Dlk1 | 2.04 | 1.15E-182 | 2.45E-178 | 0.84 | 0.46 | 0.39 | 1.84 |
| NE | Six2 | 1.78 | 9.71E-178 | 2.07E-173 | 0.75 | 0.36 | 0.39 | 2.08 |
| NE | Gnai2 | 0.75 | 2.81E-177 | 6.00E-173 | 0.99 | 0.96 | 0.03 | 1.03 |
| NE | Nr2e1 | 5.33 | 7.10E-177 | 1.52E-172 | 0.35 | 0.01 | 0.34 | 35.10 |
| NE | Rragd | 2.03 | 2.15E-176 | 4.58E-172 | 0.64 | 0.22 | 0.43 | 2.96 |
| NE | Sox2 | 1.35 | 1.34E-174 | 2.86E-170 | 0.91 | 0.52 | 0.40 | 1.77 |
| NE | Foxg1 | 1.45 | 1.41E-173 | 3.00E-169 | 0.77 | 0.34 | 0.43 | 2.28 |
| NE | Cldn9 | 2.00 | 1.45E-157 | 3.11E-153 | 0.57 | 0.16 | 0.41 | 3.51 |
| NE | Pitx1 | 1.45 | 9.99E-157 | 2.13E-152 | 0.84 | 0.49 | 0.35 | 1.70 |
| NE | Rplp1 | 0.34 | 1.03E-155 | 2.20E-151 | 1.00 | 1.00 | 0.00 | 1.00 |
| NE | Six3 | 1.12 | 1.29E-155 | 2.75E-151 | 0.96 | 0.74 | 0.21 | 1.29 |
| NE | Eya4 | 3.25 | 2.87E-154 | 6.13E-150 | 0.40 | 0.06 | 0.35 | 6.97 |
| NE | Clu | 2.58 | 7.53E-151 | 1.61E-146 | 0.56 | 0.17 | 0.38 | 3.25 |
| NE | Espn | 1.99 | 1.20E-149 | 2.56E-145 | 0.58 | 0.19 | 0.39 | 3.03 |
| NE | Wfdc2 | 0.93 | 1.04E-146 | 2.22E-142 | 0.95 | 0.71 | 0.24 | 1.34 |
| NE | Sorbs2 | 1.99 | 6.36E-144 | 1.36E-139 | 0.60 | 0.23 | 0.37 | 2.61 |
| NE | Cenpv | 1.81 | 1.60E-141 | 3.42E-137 | 0.58 | 0.21 | 0.38 | 2.82 |
| NE | Sox3 | 4.84 | 1.49E-140 | 3.19E-136 | 0.31 | 0.02 | 0.29 | 19.06 |
| NE | Ppa1 | 0.90 | 2.64E-137 | 5.64E-133 | 0.94 | 0.81 | 0.13 | 1.16 |
| NE | Sp8 | 1.30 | 2.00E-134 | 4.28E-130 | 0.74 | 0.33 | 0.41 | 2.22 |
| NE | Hes5 | 2.55 | 1.14E-131 | 2.43E-127 | 0.57 | 0.24 | 0.33 | 2.39 |
| NE | Hsp90aa1 | 0.49 | 1.58E-131 | 3.37E-127 | 1.00 | 1.00 | 0.00 | 1.00 |
| NE | Dmrta2 | 1.41 | 2.10E-129 | 4.48E-125 | 0.70 | 0.35 | 0.35 | 2.02 |
| NE | Mapk12 | 2.20 | 9.17E-129 | 1.96E-124 | 0.47 | 0.13 | 0.34 | 3.62 |
| NE | Aldh1a3 | 1.93 | 2.44E-126 | 5.20E-122 | 0.72 | 0.43 | 0.29 | 1.68 |
| NE | Marcks | 0.59 | 1.23E-125 | 2.63E-121 | 1.00 | 0.98 | 0.02 | 1.02 |
| NE | Krt18 | 0.75 | 2.84E-125 | 6.07E-121 | 0.97 | 0.78 | 0.19 | 1.25 |
| NE | Six3os1 | 1.12 | 1.30E-124 | 2.79E-120 | 0.88 | 0.67 | 0.21 | 1.31 |
| NE | Gpm6a | 4.19 | 2.52E-124 | 5.38E-120 | 0.29 | 0.02 | 0.27 | 15.05 |
| NE | Tkt | 0.77 | 6.75E-122 | 1.44E-117 | 0.95 | 0.83 | 0.12 | 1.15 |
| NE | Dmrt3 | 2.57 | 1.94E-119 | 4.14E-115 | 0.37 | 0.07 | 0.30 | 5.34 |
| NE | Ano1 | 1.44 | 5.28E-119 | 1.13E-114 | 0.66 | 0.30 | 0.37 | 2.24 |
| NE | Pea15a | 1.81 | 7.57E-119 | 1.62E-114 | 0.52 | 0.18 | 0.34 | 2.86 |
| NE | Cadm1 | 0.93 | 1.14E-118 | 2.44E-114 | 0.86 | 0.59 | 0.27 | 1.46 |
| NE | Tnfaip2 | 1.93 | 2.57E-116 | 5.49E-112 | 0.61 | 0.30 | 0.31 | 2.05 |
| NE | Fam107a | 2.99 | 1.35E-114 | 2.89E-110 | 0.31 | 0.04 | 0.27 | 8.16 |
| NE | Plch1 | 1.77 | 7.91E-112 | 1.69E-107 | 0.52 | 0.19 | 0.33 | 2.77 |
| NE | Ubash3b | 2.21 | 1.47E-110 | 3.13E-106 | 0.41 | 0.10 | 0.30 | 3.95 |
| NE | Slc22a23 | 1.91 | 6.78E-110 | 1.45E-105 | 0.49 | 0.16 | 0.32 | 2.98 |

**Supplementary Table 4. Marker genes for the annotated cell populations of integrated clustering of E11.5 control and NotchGOF λ-junction ectoderm cells.**

| | | | | | | | | |
|---|---|---|---|---|---|---|---|---|
| NE | Otx2 | 1.40 | 4.81E-109 | 1.03E-104 | 0.59 | 0.24 | 0.35 | 2.44 |
| NE | Pcdh11x | 4.33 | 2.87E-107 | 6.12E-103 | 0.25 | 0.02 | 0.23 | 13.32 |
| NE | Pcbp4 | 0.56 | 6.89E-107 | 1.47E-102 | 0.97 | 0.86 | 0.11 | 1.12 |
| NE | Plat | 2.73 | 1.01E-104 | 2.15E-100 | 0.34 | 0.07 | 0.27 | 4.97 |
| NE | C130071C03Rik | 2.42 | 1.18E-101 | 2.52E-97 | 0.35 | 0.08 | 0.27 | 4.47 |
| NE | Meg3 | 0.81 | 8.01E-100 | 1.71E-95 | 0.91 | 0.63 | 0.28 | 1.44 |
| NE | Ccnd1 | 0.94 | 2.02E-99 | 4.32E-95 | 0.83 | 0.60 | 0.23 | 1.37 |
| NE | Fzd2 | 0.81 | 2.78E-99 | 5.93E-95 | 0.91 | 0.74 | 0.18 | 1.24 |
| NE | Fgf12 | 2.54 | 3.49E-99 | 7.45E-95 | 0.30 | 0.05 | 0.25 | 6.25 |
| NE | Crym | 1.95 | 1.53E-97 | 3.28E-93 | 0.39 | 0.11 | 0.28 | 3.65 |
| NE | Cbln2 | 3.23 | 2.03E-97 | 4.33E-93 | 0.27 | 0.03 | 0.23 | 8.03 |
| NE | Atp2b1 | 0.68 | 4.76E-97 | 1.02E-92 | 0.96 | 0.82 | 0.14 | 1.17 |
| NE | Fbn2 | 1.23 | 8.66E-97 | 1.85E-92 | 0.65 | 0.31 | 0.34 | 2.09 |
| NE | Dsel | 1.60 | 1.93E-96 | 4.13E-92 | 0.55 | 0.24 | 0.31 | 2.30 |
| NE | Prr15 | 3.63 | 4.98E-96 | 1.06E-91 | 0.25 | 0.03 | 0.22 | 9.88 |
| NE | Nt5dc2 | 0.59 | 8.78E-96 | 1.87E-91 | 0.97 | 0.88 | 0.09 | 1.10 |
| NE | Crip2 | 1.12 | 4.28E-95 | 9.14E-91 | 0.59 | 0.28 | 0.31 | 2.14 |
| NE | Ckb | 1.25 | 2.16E-94 | 4.62E-90 | 0.52 | 0.21 | 0.31 | 2.49 |
| NE | Flrt3 | 1.23 | 2.63E-92 | 5.63E-88 | 0.57 | 0.26 | 0.32 | 2.25 |
| NE | Krt8 | 0.49 | 4.37E-91 | 9.34E-87 | 0.97 | 0.69 | 0.28 | 1.40 |
| NE | 3110039M20Rik | 1.18 | 5.98E-91 | 1.28E-86 | 0.47 | 0.16 | 0.31 | 2.88 |
| NE | Vim | 0.64 | 4.83E-90 | 1.03E-85 | 0.84 | 0.61 | 0.24 | 1.39 |
| NE | Slit2 | 1.64 | 2.29E-89 | 4.88E-85 | 0.55 | 0.26 | 0.30 | 2.16 |
| NE | Sox21 | 3.46 | 8.62E-88 | 1.84E-83 | 0.24 | 0.03 | 0.21 | 8.54 |
| NE | Sobp | 1.78 | 9.67E-88 | 2.07E-83 | 0.39 | 0.12 | 0.28 | 3.35 |
| NE | Mfap4 | 0.59 | 2.96E-87 | 6.33E-83 | 0.81 | 0.52 | 0.29 | 1.56 |
| NE | Stra6 | 1.84 | 4.57E-87 | 9.77E-83 | 0.41 | 0.13 | 0.28 | 3.15 |
| NE | Nes | 1.12 | 7.52E-87 | 1.61E-82 | 0.66 | 0.36 | 0.30 | 1.84 |
| NE | Necab2 | 1.88 | 1.41E-86 | 3.01E-82 | 0.43 | 0.16 | 0.27 | 2.71 |
| NE | Ctxn3 | 3.60 | 4.37E-86 | 9.33E-82 | 0.23 | 0.03 | 0.21 | 8.36 |
| NE | Gpc1 | 1.24 | 5.45E-84 | 1.16E-79 | 0.64 | 0.38 | 0.26 | 1.70 |
| NE | Nfia | 1.25 | 1.45E-83 | 3.10E-79 | 0.53 | 0.23 | 0.30 | 2.29 |
| NE | Sall2 | 1.26 | 4.74E-83 | 1.01E-78 | 0.63 | 0.34 | 0.29 | 1.87 |
| NE | Calr | 0.49 | 2.48E-82 | 5.30E-78 | 0.99 | 0.96 | 0.03 | 1.03 |
| NE | Nap1l1 | 0.35 | 5.26E-82 | 1.12E-77 | 1.00 | 0.99 | 0.01 | 1.01 |
| NE | Prkg1 | 1.60 | 1.40E-81 | 2.99E-77 | 0.40 | 0.13 | 0.27 | 3.11 |
| NE | Hmgn2 | 0.44 | 3.29E-81 | 7.03E-77 | 0.99 | 0.97 | 0.02 | 1.02 |
| NE | Dhx32 | 1.47 | 3.20E-80 | 6.83E-76 | 0.47 | 0.19 | 0.28 | 2.52 |
| NE | D230030E09Rik | 2.20 | 3.64E-80 | 7.77E-76 | 0.35 | 0.11 | 0.25 | 3.30 |
| NE | Ppp1r1a | 1.44 | 1.42E-79 | 3.03E-75 | 0.46 | 0.19 | 0.28 | 2.51 |
| NE | Ran | 0.38 | 2.11E-79 | 4.52E-75 | 1.00 | 0.99 | 0.01 | 1.01 |
| NE | Sod1 | 0.72 | 3.35E-79 | 7.16E-75 | 0.87 | 0.69 | 0.18 | 1.26 |
| NE | St8sia6 | 2.34 | 3.45E-79 | 7.36E-75 | 0.29 | 0.07 | 0.23 | 4.39 |
| NE | Zdhhc2 | 1.64 | 3.48E-79 | 7.44E-75 | 0.52 | 0.26 | 0.26 | 2.03 |
| NE | Fam126a | 1.69 | 4.74E-79 | 1.01E-74 | 0.43 | 0.16 | 0.27 | 2.65 |
| NE | P4ha3 | 2.34 | 1.02E-78 | 2.19E-74 | 0.28 | 0.06 | 0.22 | 4.72 |
| NE | Iqgap2 | 1.39 | 1.25E-78 | 2.68E-74 | 0.48 | 0.20 | 0.29 | 2.44 |
| NE | Mef2c | 3.21 | 5.51E-78 | 1.18E-73 | 0.23 | 0.03 | 0.20 | 7.50 |
| NE | Rhoc | 0.91 | 3.60E-77 | 7.70E-73 | 0.78 | 0.54 | 0.24 | 1.44 |
| NE | Ncam1 | 0.48 | 4.51E-76 | 9.64E-72 | 0.49 | 0.21 | 0.28 | 2.35 |
| NE | Tuba1b | 0.50 | 6.46E-76 | 1.38E-71 | 0.99 | 0.96 | 0.03 | 1.04 |
| NE | 4930426D05Rik | 2.91 | 1.41E-75 | 3.02E-71 | 0.23 | 0.03 | 0.20 | 6.74 |
| NE | Nhs | 1.11 | 3.81E-75 | 8.15E-71 | 0.59 | 0.31 | 0.28 | 1.90 |
| NE | Tex15 | 0.92 | 1.72E-73 | 3.67E-69 | 0.41 | 0.16 | 0.26 | 2.65 |
| NE | Rgcc | 3.21 | 2.17E-73 | 4.63E-69 | 0.20 | 0.02 | 0.18 | 8.50 |

**Supplementary Table 4. Marker genes for the annotated cell populations of integrated clustering of E11.5 control and NotchGOF λ-junction ectoderm cells.**

| | | | | | | | | |
|---|---|---|---|---|---|---|---|---|
| NE | Gstm1 | 1.94 | 2.72E-73 | 5.81E-69 | 0.48 | 0.23 | 0.25 | 2.09 |
| NE | Dusp5 | 1.77 | 2.25E-71 | 4.81E-67 | 0.33 | 0.10 | 0.23 | 3.41 |
| NE | Dut | 0.58 | 3.05E-71 | 6.51E-67 | 0.96 | 0.86 | 0.10 | 1.11 |
| NE | Kazald1 | 4.33 | 1.66E-70 | 3.55E-66 | 0.17 | 0.01 | 0.16 | 16.70 |
| NE | Hey1 | 0.71 | 5.44E-70 | 1.16E-65 | 0.87 | 0.64 | 0.23 | 1.36 |
| NE | Snx3 | 0.55 | 5.73E-70 | 1.22E-65 | 0.92 | 0.83 | 0.10 | 1.12 |
| NE | Mdk | 0.30 | 1.57E-69 | 3.36E-65 | 1.00 | 0.99 | 0.01 | 1.01 |
| NE | Psmg4 | 0.70 | 1.27E-68 | 2.71E-64 | 0.84 | 0.67 | 0.17 | 1.24 |
| NE | Rian | 1.01 | 6.63E-68 | 1.42E-63 | 0.61 | 0.34 | 0.27 | 1.81 |
| NE | Pkdcc | 1.30 | 1.14E-67 | 2.44E-63 | 0.46 | 0.20 | 0.26 | 2.28 |
| NE | Phgdh | 0.55 | 2.00E-67 | 4.27E-63 | 0.92 | 0.77 | 0.15 | 1.20 |
| NE | Ddah1 | 2.19 | 6.09E-67 | 1.30E-62 | 0.26 | 0.06 | 0.20 | 4.23 |
| NE | Ephb1 | 1.14 | 6.47E-67 | 1.38E-62 | 0.53 | 0.26 | 0.27 | 2.05 |
| NE | Tmem108 | 1.59 | 7.01E-67 | 1.50E-62 | 0.43 | 0.19 | 0.24 | 2.26 |
| NE | Zfhx4 | 0.68 | 3.69E-66 | 7.89E-62 | 0.87 | 0.69 | 0.18 | 1.27 |
| NE | Cdkl1 | 3.40 | 4.05E-65 | 8.66E-61 | 0.18 | 0.02 | 0.16 | 8.00 |
| NE | Pls1 | 1.42 | 8.37E-65 | 1.79E-60 | 0.43 | 0.19 | 0.24 | 2.28 |
| NE | Idi1 | 0.91 | 1.61E-64 | 3.43E-60 | 0.73 | 0.48 | 0.24 | 1.51 |
| NE | Tmem121 | 1.71 | 2.47E-64 | 5.28E-60 | 0.29 | 0.08 | 0.21 | 3.53 |
| NE | Syt11 | 0.92 | 2.80E-64 | 5.98E-60 | 0.48 | 0.23 | 0.26 | 2.15 |
| NE | Gapdh | 0.25 | 5.19E-64 | 1.11E-59 | 1.00 | 1.00 | 0.00 | 1.00 |
| NE | Hspa8 | 0.36 | 1.09E-63 | 2.33E-59 | 1.00 | 1.00 | 0.00 | 1.00 |
| NE | Crip1 | 0.95 | 7.90E-63 | 1.69E-58 | 0.55 | 0.28 | 0.27 | 1.97 |
| NE | Scrn1 | 1.68 | 8.35E-63 | 1.78E-58 | 0.22 | 0.04 | 0.18 | 5.39 |
| NE | Dmbx1 | 2.20 | 1.63E-62 | 3.49E-58 | 0.27 | 0.07 | 0.20 | 3.71 |
| NE | Nfib | 0.58 | 1.81E-62 | 3.87E-58 | 0.88 | 0.71 | 0.17 | 1.24 |
| NE | Hspd1 | 0.44 | 3.69E-62 | 7.89E-58 | 1.00 | 0.97 | 0.03 | 1.03 |
| NE | Slc9a3r1 | 0.66 | 6.11E-62 | 1.31E-57 | 0.85 | 0.67 | 0.18 | 1.27 |
| NE | Ccdc171 | 1.72 | 1.85E-61 | 3.94E-57 | 0.34 | 0.12 | 0.22 | 2.86 |
| NE | Fgfr1 | 1.11 | 7.47E-61 | 1.60E-56 | 0.54 | 0.29 | 0.25 | 1.86 |
| NE | Mif | 0.31 | 1.42E-60 | 3.03E-56 | 1.00 | 1.00 | 0.00 | 1.00 |
| NE | Myc | 1.65 | 1.73E-60 | 3.70E-56 | 0.33 | 0.12 | 0.21 | 2.85 |
| NE | Khdrbs3 | 0.53 | 1.90E-60 | 4.06E-56 | 0.95 | 0.82 | 0.13 | 1.16 |
| NE | Dtna | 1.15 | 2.06E-60 | 4.40E-56 | 0.49 | 0.23 | 0.26 | 2.10 |
| NE | Apc | 0.77 | 4.65E-60 | 9.94E-56 | 0.86 | 0.73 | 0.13 | 1.18 |
| NE | Dclk1 | 1.15 | 1.18E-59 | 2.53E-55 | 0.30 | 0.09 | 0.21 | 3.27 |
| NE | Slc16a1 | 1.07 | 3.84E-59 | 8.20E-55 | 0.52 | 0.27 | 0.25 | 1.94 |
| NE | H2afz | 0.33 | 4.29E-59 | 9.16E-55 | 1.00 | 1.00 | 0.00 | 1.00 |
| NE | Nuak1 | 1.22 | 6.83E-59 | 1.46E-54 | 0.40 | 0.16 | 0.23 | 2.44 |
| NE | Cldn3 | 0.51 | 1.02E-58 | 2.19E-54 | 0.79 | 0.50 | 0.30 | 1.60 |
| NE | Grk5 | 1.93 | 2.14E-58 | 4.58E-54 | 0.25 | 0.07 | 0.19 | 3.88 |
| NE | Frat2 | 1.03 | 2.96E-58 | 6.32E-54 | 0.52 | 0.26 | 0.25 | 1.95 |
| NE | Col11a1 | 0.82 | 1.77E-57 | 3.77E-53 | 0.70 | 0.46 | 0.24 | 1.52 |
| NE | Eno3 | 2.00 | 5.19E-57 | 1.11E-52 | 0.29 | 0.09 | 0.20 | 3.12 |
| NE | Msmo1 | 0.97 | 1.42E-56 | 3.04E-52 | 0.63 | 0.39 | 0.24 | 1.63 |
| NE | Rasef | 1.89 | 1.98E-56 | 4.22E-52 | 0.26 | 0.07 | 0.19 | 3.66 |
| NE | Lrrn3 | 1.92 | 1.24E-55 | 2.64E-51 | 0.25 | 0.07 | 0.18 | 3.74 |
| NE | Hnrnpdl | 0.42 | 1.33E-55 | 2.83E-51 | 0.97 | 0.94 | 0.03 | 1.03 |
| NE | B230118H07Rik | 0.88 | 1.48E-55 | 3.15E-51 | 0.68 | 0.47 | 0.21 | 1.44 |
| NE | Tnnt1 | 2.76 | 2.32E-55 | 4.95E-51 | 0.16 | 0.02 | 0.14 | 7.76 |
| NE | Nuak2 | 1.79 | 3.22E-55 | 6.89E-51 | 0.24 | 0.06 | 0.18 | 3.90 |
| NE | Fam189a2 | 2.91 | 6.69E-55 | 1.43E-50 | 0.16 | 0.02 | 0.14 | 7.95 |
| NE | Epb41l2 | 1.13 | 9.57E-55 | 2.04E-50 | 0.62 | 0.39 | 0.22 | 1.57 |
| NE | Fzd8 | 1.39 | 1.34E-54 | 2.85E-50 | 0.42 | 0.20 | 0.23 | 2.13 |
| NE | Bex4 | 0.39 | 4.13E-54 | 8.83E-50 | 0.97 | 0.90 | 0.07 | 1.08 |

**Supplementary Table 4. Marker genes for the annotated cell populations of integrated clustering of E11.5 control and NotchGOF λ-junction ectoderm cells.**

| | | | | | | | | |
|---|---|---|---|---|---|---|---|---|
| NE | Hspa5 | 0.41 | 7.39E-54 | 1.58E-49 | 0.99 | 0.96 | 0.03 | 1.03 |
| NE | Scg5 | 1.04 | 3.22E-53 | 6.89E-49 | 0.32 | 0.11 | 0.21 | 2.82 |
| NE | Rbp1 | 0.27 | 5.64E-53 | 1.20E-48 | 0.98 | 0.88 | 0.10 | 1.12 |
| NE | Crb2 | 2.63 | 7.15E-53 | 1.53E-48 | 0.17 | 0.02 | 0.14 | 6.92 |
| NE | Oxtr | 1.49 | 1.46E-52 | 3.12E-48 | 0.38 | 0.17 | 0.21 | 2.19 |
| NE | Epha5 | 1.94 | 1.65E-52 | 3.52E-48 | 0.17 | 0.03 | 0.14 | 6.46 |
| NE | Ybx1 | 0.26 | 2.15E-52 | 4.59E-48 | 1.00 | 0.99 | 0.01 | 1.01 |
| NE | Zmat4 | 1.35 | 3.06E-52 | 6.53E-48 | 0.25 | 0.07 | 0.18 | 3.62 |
| NE | Fdps | 0.55 | 1.41E-51 | 3.01E-47 | 0.89 | 0.75 | 0.14 | 1.18 |
| NE | Rrp12 | 1.32 | 1.79E-51 | 3.82E-47 | 0.36 | 0.15 | 0.21 | 2.36 |
| NE | Mapk13 | 0.68 | 2.20E-51 | 4.70E-47 | 0.73 | 0.50 | 0.23 | 1.45 |
| NE | Faah | 1.36 | 3.39E-51 | 7.24E-47 | 0.34 | 0.13 | 0.21 | 2.55 |
| NE | Ezr | 0.38 | 4.56E-51 | 9.73E-47 | 0.97 | 0.86 | 0.10 | 1.12 |
| NE | Ermn | 1.68 | 8.27E-51 | 1.77E-46 | 0.27 | 0.09 | 0.18 | 3.09 |
| NE | Lyar | 0.57 | 1.05E-50 | 2.24E-46 | 0.87 | 0.75 | 0.12 | 1.16 |
| NE | Slc25a5 | 0.44 | 1.29E-50 | 2.75E-46 | 0.95 | 0.88 | 0.07 | 1.08 |
| NE | Ces1d | 1.69 | 1.62E-50 | 3.45E-46 | 0.28 | 0.09 | 0.18 | 2.99 |
| NE | Jund | 0.39 | 1.74E-50 | 3.72E-46 | 0.97 | 0.89 | 0.08 | 1.09 |
| NE | Tead2 | 0.44 | 4.64E-50 | 9.92E-46 | 0.94 | 0.81 | 0.14 | 1.17 |
| NE | Muc1 | 2.89 | 4.80E-50 | 1.03E-45 | 0.15 | 0.02 | 0.13 | 7.45 |
| NE | Hsp90b1 | 0.35 | 1.47E-49 | 3.14E-45 | 1.00 | 0.97 | 0.03 | 1.03 |
| NE | Epcam | 0.32 | 1.58E-49 | 3.37E-45 | 1.00 | 0.97 | 0.02 | 1.02 |
| NE | Insig1 | 1.39 | 2.50E-49 | 5.34E-45 | 0.37 | 0.16 | 0.20 | 2.24 |
| NE | Gpx4 | 0.31 | 3.48E-49 | 7.44E-45 | 0.99 | 0.97 | 0.03 | 1.03 |
| NE | Impdh2 | 0.47 | 4.10E-49 | 8.76E-45 | 0.91 | 0.81 | 0.10 | 1.12 |
| NE | Car14 | 1.40 | 2.55E-48 | 5.44E-44 | 0.34 | 0.15 | 0.20 | 2.36 |
| NE | Snx24 | 1.06 | 6.94E-48 | 1.48E-43 | 0.44 | 0.22 | 0.22 | 2.02 |
| NE | Kcnb1 | 2.80 | 1.10E-47 | 2.34E-43 | 0.16 | 0.03 | 0.13 | 5.71 |
| NE | Txn1 | 0.30 | 1.57E-47 | 3.36E-43 | 0.99 | 0.98 | 0.02 | 1.02 |
| NE | Prss8 | 0.92 | 2.29E-47 | 4.90E-43 | 0.51 | 0.28 | 0.23 | 1.84 |
| NE | Col2a1 | 0.55 | 2.85E-47 | 6.08E-43 | 0.90 | 0.74 | 0.15 | 1.21 |
| NE | Grtp1 | 1.22 | 3.22E-47 | 6.88E-43 | 0.34 | 0.14 | 0.20 | 2.43 |
| NE | Ppfia3 | 1.15 | 4.81E-47 | 1.03E-42 | 0.41 | 0.20 | 0.21 | 2.03 |
| NE | Frmd3 | 3.49 | 6.16E-47 | 1.32E-42 | 0.12 | 0.01 | 0.11 | 11.18 |
| NE | Celsr1 | 1.04 | 7.39E-47 | 1.58E-42 | 0.49 | 0.28 | 0.21 | 1.77 |
| NE | Tmem47 | 1.24 | 1.04E-46 | 2.22E-42 | 0.57 | 0.36 | 0.21 | 1.59 |
| NE | Snhg4 | 1.12 | 1.12E-46 | 2.39E-42 | 0.54 | 0.34 | 0.21 | 1.61 |
| NE | Lgr4 | 1.15 | 1.21E-46 | 2.58E-42 | 0.59 | 0.38 | 0.22 | 1.58 |
| NE | Thbs3 | 1.56 | 3.30E-46 | 7.05E-42 | 0.22 | 0.06 | 0.16 | 3.48 |
| NE | Tmem238 | 0.93 | 9.82E-46 | 2.10E-41 | 0.53 | 0.31 | 0.22 | 1.72 |
| NE | Six4 | 0.78 | 1.11E-45 | 2.37E-41 | 0.61 | 0.38 | 0.23 | 1.59 |
| NE | Cdh4 | 2.16 | 2.65E-45 | 5.67E-41 | 0.18 | 0.04 | 0.14 | 4.33 |
| NE | Trpc1 | 1.25 | 3.25E-45 | 6.94E-41 | 0.35 | 0.15 | 0.20 | 2.27 |
| NE | Gsr | 0.79 | 3.91E-45 | 8.36E-41 | 0.61 | 0.38 | 0.23 | 1.61 |
| NE | Isyna1 | 0.56 | 3.92E-45 | 8.38E-41 | 0.80 | 0.64 | 0.17 | 1.26 |
| NE | Fkbp3 | 0.28 | 4.61E-45 | 9.84E-41 | 0.99 | 0.97 | 0.02 | 1.02 |
| NE | Lrig3 | 1.10 | 5.12E-45 | 1.09E-40 | 0.41 | 0.20 | 0.21 | 2.06 |
| NE | Actn1 | 0.71 | 7.24E-45 | 1.55E-40 | 0.68 | 0.48 | 0.20 | 1.41 |
| NE | Cyp4v3 | 2.18 | 8.78E-45 | 1.88E-40 | 0.17 | 0.04 | 0.13 | 4.80 |
| NE | Scd2 | 0.40 | 1.14E-44 | 2.44E-40 | 0.99 | 0.95 | 0.05 | 1.05 |
| NE | Gad2 | 0.76 | 3.16E-44 | 6.75E-40 | 0.41 | 0.19 | 0.22 | 2.14 |
| NE | Fzd9 | 4.06 | 3.70E-44 | 7.91E-40 | 0.11 | 0.01 | 0.10 | 15.43 |
| NE | Mapk11 | 1.81 | 4.53E-44 | 9.68E-40 | 0.20 | 0.06 | 0.15 | 3.67 |
| NE | Dlx6os1 | 2.38 | 5.89E-44 | 1.26E-39 | 0.20 | 0.06 | 0.15 | 3.46 |
| NE | Pop5 | 0.64 | 7.41E-44 | 1.58E-39 | 0.72 | 0.51 | 0.21 | 1.40 |

**Supplementary Table 4. Marker genes for the annotated cell populations of integrated clustering of E11.5 control and NotchGOF λ-junction ectoderm cells.**

| | | | | | | | | |
|---|---|---|---|---|---|---|---|---|
| NE | Ddx39 | 0.56 | 8.02E-44 | 1.71E-39 | 0.83 | 0.69 | 0.14 | 1.20 |
| NE | Shroom3 | 1.16 | 1.23E-43 | 2.63E-39 | 0.35 | 0.16 | 0.19 | 2.24 |
| NE | Serpinf1 | 1.44 | 5.92E-43 | 1.26E-38 | 0.26 | 0.09 | 0.17 | 2.82 |
| NE | Nfix | 2.32 | 1.34E-42 | 2.87E-38 | 0.16 | 0.03 | 0.13 | 4.82 |
| NE | Dlx5 | 1.16 | 1.58E-42 | 3.37E-38 | 0.51 | 0.30 | 0.21 | 1.68 |
| NE | Dlx6 | 1.07 | 2.37E-42 | 5.07E-38 | 0.29 | 0.12 | 0.18 | 2.52 |
| NE | Mapk8ip1 | 0.63 | 2.73E-42 | 5.83E-38 | 0.48 | 0.26 | 0.22 | 1.84 |
| NE | Hmgb2 | 0.42 | 2.91E-42 | 6.21E-38 | 0.99 | 0.95 | 0.03 | 1.04 |
| NE | Dmrta2os | 2.01 | 3.22E-42 | 6.87E-38 | 0.15 | 0.03 | 0.12 | 5.13 |
| NE | Mllt3 | 0.71 | 5.05E-42 | 1.08E-37 | 0.67 | 0.46 | 0.22 | 1.47 |
| NE | Mmp16 | 1.72 | 5.15E-42 | 1.10E-37 | 0.22 | 0.07 | 0.15 | 3.22 |
| NE | Dhcr24 | 0.97 | 7.51E-42 | 1.60E-37 | 0.46 | 0.26 | 0.20 | 1.79 |
| NE | Hs6st2 | 0.67 | 1.44E-41 | 3.07E-37 | 0.66 | 0.44 | 0.22 | 1.50 |
| NE | Eif1ax | 0.34 | 1.63E-41 | 3.49E-37 | 0.96 | 0.92 | 0.04 | 1.05 |
| NE | Tyro3 | 1.30 | 2.06E-41 | 4.41E-37 | 0.32 | 0.14 | 0.18 | 2.31 |
| NE | Tox2 | 1.92 | 2.11E-41 | 4.51E-37 | 0.19 | 0.06 | 0.14 | 3.53 |
| NE | Bag2 | 0.98 | 2.25E-41 | 4.80E-37 | 0.46 | 0.25 | 0.21 | 1.82 |
| NE | Ptgis | 2.37 | 2.33E-41 | 4.97E-37 | 0.14 | 0.03 | 0.12 | 5.72 |
| NE | 2610318N02Rik | 0.80 | 2.67E-41 | 5.70E-37 | 0.55 | 0.33 | 0.22 | 1.66 |
| NE | Kpnb1 | 0.49 | 3.47E-41 | 7.41E-37 | 0.87 | 0.76 | 0.11 | 1.14 |
| NE | Id4 | 0.59 | 4.69E-41 | 1.00E-36 | 0.69 | 0.48 | 0.20 | 1.42 |
| NE | Maged1 | 0.35 | 1.42E-40 | 3.04E-36 | 0.97 | 0.92 | 0.05 | 1.05 |
| NE | Ly75 | 1.15 | 1.48E-40 | 3.16E-36 | 0.31 | 0.13 | 0.18 | 2.41 |
| NE | Usp51 | 2.21 | 1.56E-40 | 3.32E-36 | 0.12 | 0.02 | 0.11 | 7.29 |
| NE | Gsap | 1.89 | 1.82E-40 | 3.89E-36 | 0.19 | 0.05 | 0.14 | 3.67 |
| NE | Irf2bpl | 0.77 | 3.01E-40 | 6.43E-36 | 0.57 | 0.36 | 0.21 | 1.59 |
| NE | Tubb4b | 0.53 | 3.25E-40 | 6.95E-36 | 0.88 | 0.75 | 0.13 | 1.17 |
| NE | Cacybp | 0.38 | 3.86E-40 | 8.24E-36 | 0.96 | 0.91 | 0.05 | 1.05 |
| NE | Mir99ahg | 0.80 | 7.30E-40 | 1.56E-35 | 0.64 | 0.44 | 0.20 | 1.46 |
| NE | Eif5a | 0.27 | 1.04E-39 | 2.22E-35 | 1.00 | 0.99 | 0.00 | 1.00 |
| NE | Fzd4 | 1.36 | 1.08E-39 | 2.31E-35 | 0.27 | 0.11 | 0.17 | 2.53 |
| NE | Pdia6 | 0.35 | 2.27E-39 | 4.85E-35 | 0.97 | 0.93 | 0.04 | 1.04 |
| NE | Tst | 1.12 | 3.68E-39 | 7.86E-35 | 0.36 | 0.17 | 0.18 | 2.06 |
| NE | Dkc1 | 0.49 | 4.01E-39 | 8.58E-35 | 0.84 | 0.70 | 0.14 | 1.20 |
| NE | Klhl14 | 3.27 | 4.47E-39 | 9.54E-35 | 0.11 | 0.01 | 0.10 | 9.73 |
| NE | Hmga2 | 0.32 | 5.55E-39 | 1.19E-34 | 0.98 | 0.88 | 0.10 | 1.11 |
| NE | Pdlim7 | 0.55 | 7.90E-39 | 1.69E-34 | 0.73 | 0.54 | 0.19 | 1.36 |
| NE | Abr | 1.17 | 9.30E-39 | 1.99E-34 | 0.31 | 0.14 | 0.18 | 2.28 |
| NE | Pwwp3b | 1.10 | 1.35E-38 | 2.88E-34 | 0.39 | 0.21 | 0.19 | 1.91 |
| NE | Tm7sf2 | 1.21 | 1.48E-38 | 3.16E-34 | 0.31 | 0.14 | 0.17 | 2.25 |
| NE | Prr7 | 1.13 | 5.27E-38 | 1.13E-33 | 0.32 | 0.14 | 0.18 | 2.25 |
| NE | Cyp51 | 0.71 | 9.83E-38 | 2.10E-33 | 0.71 | 0.53 | 0.17 | 1.33 |
| NE | Ranbp1 | 0.28 | 1.43E-37 | 3.06E-33 | 0.99 | 0.98 | 0.01 | 1.01 |
| NE | Ssx2ip | 0.54 | 2.08E-37 | 4.45E-33 | 0.76 | 0.58 | 0.19 | 1.32 |
| NE | Nrcam | 1.28 | 4.33E-37 | 9.26E-33 | 0.29 | 0.12 | 0.16 | 2.36 |
| NE | Rassf10 | 1.12 | 4.64E-37 | 9.91E-33 | 0.34 | 0.17 | 0.18 | 2.06 |
| NE | Zdbf2 | 0.68 | 7.24E-37 | 1.55E-32 | 0.66 | 0.46 | 0.19 | 1.42 |
| NE | Tmem30b | 0.73 | 9.67E-37 | 2.06E-32 | 0.38 | 0.19 | 0.19 | 2.05 |
| NE | Elavl2 | 0.45 | 1.56E-36 | 3.34E-32 | 0.53 | 0.32 | 0.21 | 1.66 |
| NE | Sfrp2 | 0.72 | 1.88E-36 | 4.02E-32 | 0.62 | 0.42 | 0.20 | 1.48 |
| NE | Mrps24 | 0.41 | 2.86E-36 | 6.10E-32 | 0.89 | 0.78 | 0.11 | 1.15 |
| NE | Tshz2 | 0.95 | 3.42E-36 | 7.30E-32 | 0.50 | 0.31 | 0.18 | 1.59 |
| NE | Kcnd2 | 2.53 | 3.58E-36 | 7.65E-32 | 0.13 | 0.02 | 0.10 | 5.52 |
| NE | Mme | 1.61 | 3.64E-36 | 7.78E-32 | 0.26 | 0.11 | 0.15 | 2.34 |
| NE | Cmtm7 | 0.75 | 4.87E-36 | 1.04E-31 | 0.47 | 0.28 | 0.20 | 1.71 |

**Supplementary Table 4. Marker genes for the annotated cell populations of integrated clustering of E11.5 control and NotchGOF λ-junction ectoderm cells.**

| | | | | | | | | |
|---|---|---|---|---|---|---|---|---|
| NE | Glo1 | 0.42 | 5.28E-36 | 1.13E-31 | 0.88 | 0.78 | 0.11 | 1.14 |
| NE | Pgp | 0.72 | 6.06E-36 | 1.29E-31 | 0.63 | 0.44 | 0.19 | 1.43 |
| NE | Dek | 0.36 | 6.30E-36 | 1.35E-31 | 0.96 | 0.90 | 0.06 | 1.07 |
| NE | Podn | 3.26 | 6.47E-36 | 1.38E-31 | 0.10 | 0.01 | 0.09 | 9.09 |
| NE | Gchfr | 1.51 | 7.33E-36 | 1.56E-31 | 0.22 | 0.08 | 0.14 | 2.82 |
| NE | Tshz3 | 1.62 | 8.41E-36 | 1.80E-31 | 0.19 | 0.06 | 0.13 | 3.24 |
| NE | Lhfp | 0.88 | 9.22E-36 | 1.97E-31 | 0.42 | 0.22 | 0.19 | 1.86 |
| NE | Anp32b | 0.25 | 1.18E-35 | 2.52E-31 | 1.00 | 0.98 | 0.02 | 1.02 |
| NE | Auts2 | 0.40 | 1.41E-35 | 3.02E-31 | 0.93 | 0.88 | 0.05 | 1.06 |
| NE | Nme1 | 0.28 | 1.49E-35 | 3.18E-31 | 0.99 | 0.97 | 0.02 | 1.02 |
| NE | Tmem37 | 2.89 | 1.50E-35 | 3.21E-31 | 0.11 | 0.02 | 0.09 | 7.20 |
| NE | Lhfpl2 | 1.41 | 1.55E-35 | 3.32E-31 | 0.24 | 0.09 | 0.15 | 2.59 |
| NE | Speg | 1.59 | 1.57E-35 | 3.36E-31 | 0.20 | 0.07 | 0.13 | 3.00 |
| NE | Flrt2 | 1.94 | 1.86E-35 | 3.97E-31 | 0.34 | 0.18 | 0.17 | 1.95 |
| NE | Ptges3 | 0.31 | 1.90E-35 | 4.05E-31 | 0.98 | 0.94 | 0.03 | 1.03 |
| NE | Tcim | 1.01 | 2.33E-35 | 4.97E-31 | 0.37 | 0.19 | 0.18 | 1.96 |
| NE | Cmtm4 | 0.76 | 2.54E-35 | 5.42E-31 | 0.49 | 0.29 | 0.20 | 1.69 |
| NE | Calm3 | 0.32 | 2.67E-35 | 5.70E-31 | 0.90 | 0.79 | 0.10 | 1.13 |
| NE | Phb2 | 0.30 | 3.05E-35 | 6.52E-31 | 0.97 | 0.95 | 0.02 | 1.02 |
| NE | Ccna2 | 0.65 | 3.14E-35 | 6.72E-31 | 0.65 | 0.48 | 0.17 | 1.36 |
| NE | Hmcn1 | 1.16 | 3.33E-35 | 7.11E-31 | 0.29 | 0.13 | 0.16 | 2.24 |
| NE | Pclaf | 0.41 | 3.59E-35 | 7.66E-31 | 0.87 | 0.73 | 0.14 | 1.19 |
| NE | Ttyh1 | 2.06 | 4.00E-35 | 8.55E-31 | 0.15 | 0.04 | 0.11 | 3.97 |
| NE | Cd302 | 0.91 | 4.22E-35 | 9.01E-31 | 0.39 | 0.20 | 0.19 | 1.93 |
| NE | Ltbp1 | 1.16 | 4.54E-35 | 9.69E-31 | 0.34 | 0.17 | 0.17 | 1.99 |
| NE | Bbip1 | 0.53 | 5.12E-35 | 1.09E-30 | 0.72 | 0.52 | 0.19 | 1.37 |
| NE | Nrg1 | 0.73 | 5.93E-35 | 1.27E-30 | 0.37 | 0.18 | 0.18 | 2.01 |
| NE | Tubb2a | 0.64 | 6.58E-35 | 1.41E-30 | 0.47 | 0.27 | 0.20 | 1.73 |
| NE | Psat1 | 0.89 | 8.16E-35 | 1.74E-30 | 0.42 | 0.24 | 0.19 | 1.78 |
| NE | Dhcr7 | 0.90 | 9.17E-35 | 1.96E-30 | 0.44 | 0.25 | 0.19 | 1.78 |
| NE | Fgfrl1 | 0.84 | 1.67E-34 | 3.57E-30 | 0.41 | 0.22 | 0.19 | 1.88 |
| NE | Spata13 | 1.69 | 2.18E-34 | 4.65E-30 | 0.18 | 0.06 | 0.13 | 3.25 |
| NE | Rims4 | 2.82 | 2.32E-34 | 4.95E-30 | 0.10 | 0.01 | 0.09 | 7.92 |
| NE | Ndufa13 | 0.26 | 2.51E-34 | 5.37E-30 | 0.98 | 0.96 | 0.02 | 1.02 |
| NE | Cldn11 | 1.96 | 2.82E-34 | 6.02E-30 | 0.18 | 0.06 | 0.12 | 3.18 |
| NE | Wdfy1 | 0.79 | 2.89E-34 | 6.17E-30 | 0.51 | 0.32 | 0.20 | 1.62 |
| NE | Tril | 1.11 | 5.80E-34 | 1.24E-29 | 0.39 | 0.22 | 0.18 | 1.80 |
| NE | Fbl | 0.34 | 6.28E-34 | 1.34E-29 | 0.94 | 0.88 | 0.06 | 1.07 |
| NE | Fkbp4 | 0.26 | 6.88E-34 | 1.47E-29 | 0.99 | 0.97 | 0.02 | 1.02 |
| NE | Stox1 | 1.58 | 7.50E-34 | 1.60E-29 | 0.14 | 0.03 | 0.11 | 4.34 |
| NE | Mpped2 | 0.57 | 1.20E-33 | 2.57E-29 | 0.85 | 0.72 | 0.13 | 1.18 |
| NE | Schip1 | 0.75 | 1.24E-33 | 2.65E-29 | 0.48 | 0.29 | 0.19 | 1.67 |
| NE | Snn | 0.74 | 1.46E-33 | 3.12E-29 | 0.69 | 0.54 | 0.15 | 1.27 |
| NE | Bccip | 0.48 | 1.68E-33 | 3.59E-29 | 0.82 | 0.69 | 0.12 | 1.18 |
| NE | Ust | 1.50 | 2.76E-33 | 5.89E-29 | 0.21 | 0.07 | 0.13 | 2.82 |
| NE | Ssrp1 | 0.33 | 2.84E-33 | 6.08E-29 | 0.94 | 0.87 | 0.06 | 1.07 |
| NE | Cpm | 0.56 | 3.62E-33 | 7.73E-29 | 0.64 | 0.44 | 0.20 | 1.46 |
| NE | Card19 | 1.06 | 6.85E-33 | 1.46E-28 | 0.36 | 0.20 | 0.17 | 1.86 |
| NE | Cdv3 | 0.34 | 7.79E-33 | 1.66E-28 | 0.97 | 0.92 | 0.05 | 1.05 |
| NE | H1fx | 0.33 | 1.38E-32 | 2.94E-28 | 0.97 | 0.91 | 0.06 | 1.07 |
| NE | Cct3 | 0.27 | 1.96E-32 | 4.19E-28 | 0.98 | 0.96 | 0.02 | 1.02 |
| NE | Phlpp1 | 1.01 | 2.02E-32 | 4.32E-28 | 0.36 | 0.19 | 0.17 | 1.90 |
| NE | Map7d2 | 1.06 | 2.59E-32 | 5.53E-28 | 0.30 | 0.14 | 0.16 | 2.13 |
| NE | Rassf9 | 0.97 | 2.83E-32 | 6.04E-28 | 0.33 | 0.16 | 0.17 | 2.02 |
| NE | Zfp467 | 0.71 | 4.63E-32 | 9.90E-28 | 0.57 | 0.38 | 0.20 | 1.52 |

**Supplementary Table 4. Marker genes for the annotated cell populations of integrated clustering of E11.5 control and NotchGOF λ-junction ectoderm cells.**

| | | | | | | | | |
|---|---|---|---|---|---|---|---|---|
| NE | Atp7b | 2.06 | 5.20E-32 | 1.11E-27 | 0.13 | 0.03 | 0.10 | 4.26 |
| NE | Pa2g4 | 0.35 | 7.54E-32 | 1.61E-27 | 0.93 | 0.85 | 0.08 | 1.10 |
| NE | Uqcc2 | 0.31 | 7.99E-32 | 1.71E-27 | 0.95 | 0.88 | 0.07 | 1.08 |
| NE | Ipo5 | 0.54 | 1.03E-31 | 2.19E-27 | 0.76 | 0.62 | 0.15 | 1.23 |
| NE | Ntn1 | 1.94 | 1.03E-31 | 2.20E-27 | 0.15 | 0.04 | 0.11 | 3.70 |
| NE | Has2 | 1.32 | 1.05E-31 | 2.24E-27 | 0.20 | 0.07 | 0.13 | 2.80 |
| NE | Cox20 | 0.58 | 1.32E-31 | 2.82E-27 | 0.69 | 0.51 | 0.18 | 1.34 |
| NE | Acsl3 | 0.40 | 1.48E-31 | 3.17E-27 | 0.87 | 0.71 | 0.15 | 1.22 |
| NE | Tceal3 | 1.42 | 1.51E-31 | 3.23E-27 | 0.23 | 0.10 | 0.14 | 2.44 |
| NE | Hacd1 | 0.85 | 1.78E-31 | 3.80E-27 | 0.34 | 0.17 | 0.17 | 1.95 |
| NE | Prim1 | 0.58 | 2.59E-31 | 5.53E-27 | 0.68 | 0.49 | 0.19 | 1.39 |
| NE | Mcm6 | 0.49 | 2.73E-31 | 5.82E-27 | 0.77 | 0.59 | 0.18 | 1.30 |
| NE | Cpd | 0.82 | 3.50E-31 | 7.47E-27 | 0.48 | 0.29 | 0.18 | 1.62 |
| NE | Tsc22d2 | 0.72 | 5.17E-31 | 1.10E-26 | 0.54 | 0.35 | 0.19 | 1.56 |
| NE | Syngr1 | 0.94 | 6.95E-31 | 1.48E-26 | 0.31 | 0.15 | 0.16 | 2.07 |
| NE | Ncl | 0.29 | 7.11E-31 | 1.52E-26 | 1.00 | 0.99 | 0.01 | 1.01 |
| NE | Dtymk | 0.39 | 7.40E-31 | 1.58E-26 | 0.89 | 0.79 | 0.11 | 1.13 |
| NE | Fgf10 | 1.68 | 9.72E-31 | 2.08E-26 | 0.19 | 0.07 | 0.12 | 2.82 |
| NE | Nsmce4a | 0.38 | 1.07E-30 | 2.29E-26 | 0.90 | 0.79 | 0.11 | 1.14 |
| NE | Uchl3 | 0.36 | 2.06E-30 | 4.41E-26 | 0.88 | 0.77 | 0.11 | 1.14 |
| NE | Sqle | 0.57 | 4.15E-30 | 8.86E-26 | 0.64 | 0.45 | 0.19 | 1.43 |
| NE | Akr1b3 | 0.52 | 4.17E-30 | 8.91E-26 | 0.70 | 0.53 | 0.17 | 1.32 |
| NE | Gnao1 | 0.81 | 4.65E-30 | 9.94E-26 | 0.26 | 0.12 | 0.14 | 2.24 |
| NE | Hist1h4i | 0.72 | 6.76E-30 | 1.44E-25 | 0.53 | 0.35 | 0.17 | 1.49 |
| NE | Itga6 | 0.55 | 7.27E-30 | 1.55E-25 | 0.83 | 0.75 | 0.08 | 1.11 |
| NE | Snhg3 | 0.76 | 7.77E-30 | 1.66E-25 | 0.59 | 0.43 | 0.16 | 1.38 |
| NE | Smarcd3 | 0.84 | 1.02E-29 | 2.17E-25 | 0.37 | 0.21 | 0.17 | 1.80 |
| NE | Rab25 | 0.42 | 1.45E-29 | 3.10E-25 | 0.74 | 0.54 | 0.20 | 1.36 |
| NE | Nav1 | 0.55 | 1.98E-29 | 4.22E-25 | 0.61 | 0.43 | 0.19 | 1.44 |
| NE | Nop58 | 0.33 | 2.04E-29 | 4.35E-25 | 0.93 | 0.87 | 0.06 | 1.07 |
| NE | Fgf13 | 1.18 | 2.09E-29 | 4.47E-25 | 0.25 | 0.11 | 0.14 | 2.23 |
| NE | Tmem176a | 0.66 | 2.09E-29 | 4.47E-25 | 0.52 | 0.34 | 0.19 | 1.56 |
| NE | Snrpa1 | 0.32 | 2.34E-29 | 4.99E-25 | 0.94 | 0.88 | 0.06 | 1.07 |
| NE | Hells | 0.44 | 2.51E-29 | 5.36E-25 | 0.80 | 0.65 | 0.15 | 1.23 |
| NE | Mns1 | 0.71 | 2.59E-29 | 5.54E-25 | 0.50 | 0.33 | 0.17 | 1.52 |
| NE | Dnajc9 | 0.37 | 2.76E-29 | 5.90E-25 | 0.88 | 0.76 | 0.12 | 1.16 |
| NE | Ldlr | 1.06 | 4.24E-29 | 9.06E-25 | 0.31 | 0.16 | 0.15 | 1.95 |
| NE | Tspan12 | 0.71 | 5.83E-29 | 1.25E-24 | 0.54 | 0.37 | 0.17 | 1.46 |
| NE | C1qbp | 0.30 | 6.97E-29 | 1.49E-24 | 0.94 | 0.90 | 0.04 | 1.05 |
| NE | Glrx5 | 0.34 | 7.15E-29 | 1.53E-24 | 0.92 | 0.84 | 0.07 | 1.09 |
| NE | Tcf4 | 0.36 | 8.37E-29 | 1.79E-24 | 0.92 | 0.86 | 0.06 | 1.07 |
| NE | Srgap3 | 0.86 | 8.40E-29 | 1.79E-24 | 0.30 | 0.15 | 0.15 | 2.00 |
| NE | Wipf3 | 0.98 | 1.15E-28 | 2.46E-24 | 0.30 | 0.15 | 0.15 | 2.01 |
| NE | Notch3 | 0.55 | 1.27E-28 | 2.72E-24 | 0.65 | 0.45 | 0.19 | 1.43 |
| NE | Cltb | 0.68 | 1.34E-28 | 2.87E-24 | 0.42 | 0.24 | 0.18 | 1.74 |
| NE | Dmd | 1.29 | 1.45E-28 | 3.09E-24 | 0.23 | 0.10 | 0.13 | 2.34 |
| NE | Txndc15 | 0.52 | 1.57E-28 | 3.34E-24 | 0.67 | 0.49 | 0.19 | 1.38 |
| NE | Ssbp4 | 0.61 | 1.60E-28 | 3.41E-24 | 0.59 | 0.41 | 0.18 | 1.44 |
| NE | Pcdh17 | 0.55 | 1.61E-28 | 3.43E-24 | 0.48 | 0.28 | 0.19 | 1.68 |
| NE | B4galt5 | 1.32 | 1.63E-28 | 3.49E-24 | 0.19 | 0.07 | 0.12 | 2.74 |
| NE | Cmtm8 | 0.53 | 1.71E-28 | 3.66E-24 | 0.64 | 0.44 | 0.20 | 1.45 |
| NE | Ccdc8 | 0.64 | 1.87E-28 | 3.99E-24 | 0.48 | 0.30 | 0.18 | 1.61 |
| NE | Dnajc1 | 0.35 | 1.87E-28 | 4.00E-24 | 0.75 | 0.57 | 0.18 | 1.31 |
| NE | Tipin | 0.41 | 2.54E-28 | 5.42E-24 | 0.78 | 0.62 | 0.16 | 1.26 |
| NE | Npas3 | 1.53 | 3.82E-28 | 8.15E-24 | 0.17 | 0.06 | 0.11 | 2.85 |

**Supplementary Table 4. Marker genes for the annotated cell populations of integrated clustering of E11.5 control and NotchGOF λ-junction ectoderm cells.**

| | | | | | | | | |
|---|---|---|---|---|---|---|---|---|
| NE | Scd1 | 1.58 | 5.03E-28 | 1.08E-23 | 0.18 | 0.07 | 0.11 | 2.71 |
| NE | Spc24 | 0.45 | 6.33E-28 | 1.35E-23 | 0.74 | 0.59 | 0.15 | 1.25 |
| NE | Dipk1b | 0.54 | 6.92E-28 | 1.48E-23 | 0.60 | 0.41 | 0.19 | 1.46 |
| NE | Chchd10 | 0.66 | 8.31E-28 | 1.78E-23 | 0.49 | 0.31 | 0.18 | 1.60 |
| NE | Fads1 | 0.64 | 8.44E-28 | 1.80E-23 | 0.53 | 0.35 | 0.18 | 1.50 |
| NE | Gatm | 1.99 | 1.01E-27 | 2.15E-23 | 0.12 | 0.03 | 0.09 | 4.03 |
| NE | Cxcl12 | 1.22 | 1.46E-27 | 3.11E-23 | 0.20 | 0.08 | 0.12 | 2.49 |
| NE | Ssbp2 | 0.39 | 1.59E-27 | 3.39E-23 | 0.78 | 0.62 | 0.16 | 1.25 |
| NE | Sdhd | 0.40 | 1.82E-27 | 3.88E-23 | 0.84 | 0.72 | 0.12 | 1.17 |
| NE | Rnaseh2c | 0.32 | 1.83E-27 | 3.90E-23 | 0.90 | 0.82 | 0.08 | 1.09 |
| NE | Ccdc80 | 0.83 | 2.82E-27 | 6.03E-23 | 0.35 | 0.19 | 0.16 | 1.83 |
| NE | Srcin1 | 1.15 | 3.09E-27 | 6.59E-23 | 0.21 | 0.08 | 0.12 | 2.48 |
| NE | Rtn3 | 0.36 | 3.13E-27 | 6.69E-23 | 0.93 | 0.84 | 0.09 | 1.11 |
| NE | Rgma | 0.96 | 3.19E-27 | 6.81E-23 | 0.29 | 0.14 | 0.14 | 2.00 |
| NE | Pard3b | 0.82 | 3.27E-27 | 6.98E-23 | 0.31 | 0.16 | 0.15 | 1.97 |
| NE | Slc35f1 | 1.65 | 3.67E-27 | 7.84E-23 | 0.17 | 0.06 | 0.11 | 2.86 |
| NE | Cad | 0.72 | 3.91E-27 | 8.35E-23 | 0.44 | 0.27 | 0.17 | 1.65 |
| NE | Rims2 | 0.75 | 3.93E-27 | 8.40E-23 | 0.29 | 0.14 | 0.15 | 2.02 |
| NE | Plscr1 | 1.10 | 3.97E-27 | 8.48E-23 | 0.23 | 0.10 | 0.13 | 2.31 |
| NE | Manf | 0.36 | 4.29E-27 | 9.16E-23 | 0.92 | 0.85 | 0.07 | 1.08 |
| NE | Emg1 | 0.28 | 5.75E-27 | 1.23E-22 | 0.93 | 0.89 | 0.04 | 1.05 |
| NE | Myo6 | 1.00 | 6.31E-27 | 1.35E-22 | 0.29 | 0.15 | 0.14 | 1.97 |
| NE | Bex2 | 0.29 | 6.92E-27 | 1.48E-22 | 0.92 | 0.85 | 0.07 | 1.09 |
| NE | Hmgcs1 | 0.45 | 7.25E-27 | 1.55E-22 | 0.82 | 0.70 | 0.12 | 1.17 |
| NE | Pbx1 | 0.28 | 7.33E-27 | 1.57E-22 | 0.98 | 0.96 | 0.03 | 1.03 |
| NE | Hmgcr | 0.49 | 8.12E-27 | 1.73E-22 | 0.70 | 0.52 | 0.18 | 1.34 |
| NE | Zcchc3 | 0.53 | 8.19E-27 | 1.75E-22 | 0.70 | 0.54 | 0.16 | 1.29 |
| NE | Nol7 | 0.26 | 1.07E-26 | 2.29E-22 | 0.94 | 0.91 | 0.04 | 1.04 |
| NE | Ptchd1 | 1.64 | 1.10E-26 | 2.36E-22 | 0.13 | 0.04 | 0.09 | 3.54 |
| NE | Gfer | 0.60 | 1.12E-26 | 2.39E-22 | 0.54 | 0.36 | 0.18 | 1.49 |
| NE | Cybrd1 | 1.50 | 1.19E-26 | 2.55E-22 | 0.15 | 0.05 | 0.10 | 3.11 |
| NE | Bicd1 | 0.74 | 1.37E-26 | 2.93E-22 | 0.35 | 0.19 | 0.16 | 1.83 |
| NE | Birc5 | 0.45 | 1.62E-26 | 3.45E-22 | 0.79 | 0.65 | 0.13 | 1.21 |
| NE | Camk2n2 | 1.92 | 1.75E-26 | 3.73E-22 | 0.13 | 0.04 | 0.09 | 3.45 |
| NE | Gpatch4 | 0.51 | 1.90E-26 | 4.06E-22 | 0.64 | 0.47 | 0.17 | 1.37 |
| NE | Fndc3c1 | 0.78 | 2.39E-26 | 5.11E-22 | 0.39 | 0.23 | 0.16 | 1.73 |
| NE | Srm | 0.55 | 2.73E-26 | 5.84E-22 | 0.69 | 0.55 | 0.14 | 1.26 |
| NE | Acat2 | 0.43 | 3.26E-26 | 6.96E-22 | 0.80 | 0.67 | 0.13 | 1.19 |
| NE | Maz | 0.35 | 3.43E-26 | 7.32E-22 | 0.86 | 0.76 | 0.10 | 1.14 |
| NE | Lig1 | 0.36 | 3.71E-26 | 7.92E-22 | 0.79 | 0.61 | 0.18 | 1.29 |
| NE | Cntfr | 0.87 | 4.12E-26 | 8.81E-22 | 0.22 | 0.09 | 0.12 | 2.36 |
| NE | Slc1a4 | 1.34 | 4.26E-26 | 9.09E-22 | 0.15 | 0.05 | 0.10 | 2.96 |
| NE | Cdon | 0.85 | 4.58E-26 | 9.78E-22 | 0.34 | 0.19 | 0.15 | 1.79 |
| NE | Fgf17 | 1.59 | 4.67E-26 | 9.98E-22 | 0.23 | 0.11 | 0.12 | 2.14 |
| NE | Ndufv2 | 0.29 | 5.12E-26 | 1.09E-21 | 0.93 | 0.87 | 0.06 | 1.07 |
| NE | Maob | 2.11 | 5.35E-26 | 1.14E-21 | 0.10 | 0.02 | 0.08 | 4.76 |
| NE | Rragb | 1.57 | 6.95E-26 | 1.48E-21 | 0.18 | 0.07 | 0.11 | 2.56 |
| NE | Misp | 1.81 | 7.65E-26 | 1.63E-21 | 0.12 | 0.03 | 0.09 | 3.81 |
| NE | Tmem97 | 0.68 | 9.41E-26 | 2.01E-21 | 0.49 | 0.33 | 0.17 | 1.51 |
| NE | Lap3 | 0.50 | 1.08E-25 | 2.31E-21 | 0.66 | 0.50 | 0.16 | 1.31 |
| NE | Sparc | 0.38 | 1.11E-25 | 2.38E-21 | 0.93 | 0.85 | 0.08 | 1.10 |
| NE | Fasn | 0.60 | 1.12E-25 | 2.39E-21 | 0.56 | 0.40 | 0.17 | 1.41 |
| NE | Mpp6 | 0.53 | 1.13E-25 | 2.41E-21 | 0.73 | 0.60 | 0.13 | 1.22 |
| NE | Sigmar1 | 0.55 | 1.40E-25 | 2.99E-21 | 0.59 | 0.43 | 0.16 | 1.38 |
| NE | Aig1 | 0.55 | 1.43E-25 | 3.05E-21 | 0.64 | 0.46 | 0.18 | 1.38 |

**Supplementary Table 4. Marker genes for the annotated cell populations of integrated clustering of E11.5 control and NotchGOF λ-junction ectoderm cells.**

| | | | | | | | | |
|---|---|---|---|---|---|---|---|---|
| NE | Usp1 | 0.39 | 1.48E-25 | 3.15E-21 | 0.82 | 0.70 | 0.12 | 1.17 |
| NE | Cdca3 | 0.49 | 1.74E-25 | 3.72E-21 | 0.67 | 0.52 | 0.15 | 1.28 |
| NE | Pcna | 0.40 | 1.80E-25 | 3.84E-21 | 0.85 | 0.71 | 0.14 | 1.20 |
| NE | Mtr | 0.92 | 1.92E-25 | 4.10E-21 | 0.29 | 0.15 | 0.14 | 1.95 |
| NE | Lypd6b | 0.91 | 1.97E-25 | 4.20E-21 | 0.31 | 0.17 | 0.14 | 1.84 |
| NE | Acss2 | 1.57 | 2.05E-25 | 4.38E-21 | 0.14 | 0.05 | 0.10 | 3.07 |
| NE | Stbd1 | 1.29 | 2.15E-25 | 4.58E-21 | 0.21 | 0.09 | 0.12 | 2.33 |
| NE | Tmem200c | 1.48 | 2.16E-25 | 4.61E-21 | 0.14 | 0.05 | 0.10 | 3.06 |
| NE | Ccdc34 | 0.31 | 2.33E-25 | 4.98E-21 | 0.90 | 0.83 | 0.08 | 1.09 |
| NE | Ruvbl1 | 0.37 | 2.97E-25 | 6.34E-21 | 0.83 | 0.70 | 0.13 | 1.18 |
| NE | B9d1 | 0.49 | 3.39E-25 | 7.25E-21 | 0.68 | 0.51 | 0.17 | 1.32 |
| NE | Cmss1 | 0.57 | 3.45E-25 | 7.36E-21 | 0.54 | 0.37 | 0.17 | 1.47 |
| NE | Cxxc4 | 0.36 | 3.83E-25 | 8.18E-21 | 0.34 | 0.18 | 0.16 | 1.88 |
| NE | Nhsl2 | 1.15 | 3.96E-25 | 8.46E-21 | 0.19 | 0.08 | 0.11 | 2.48 |
| NE | Hdgfl3 | 0.42 | 3.96E-25 | 8.46E-21 | 0.64 | 0.45 | 0.18 | 1.40 |
| NE | Rpia | 0.68 | 4.40E-25 | 9.39E-21 | 0.44 | 0.28 | 0.17 | 1.60 |
| NE | Ap3b2 | 0.91 | 8.03E-25 | 1.72E-20 | 0.13 | 0.04 | 0.09 | 3.47 |
| NE | Pmvk | 0.48 | 9.38E-25 | 2.00E-20 | 0.62 | 0.45 | 0.17 | 1.37 |
| NE | H2-K1 | 0.85 | 1.02E-24 | 2.18E-20 | 0.33 | 0.19 | 0.15 | 1.78 |
| NE | H2afx | 0.32 | 1.14E-24 | 2.44E-20 | 0.94 | 0.87 | 0.07 | 1.09 |
| NE | Tsen15 | 0.52 | 1.52E-24 | 3.24E-20 | 0.67 | 0.52 | 0.15 | 1.29 |
| NE | Lsm5 | 0.29 | 1.60E-24 | 3.41E-20 | 0.93 | 0.88 | 0.06 | 1.06 |
| NE | Ttc39b | 1.52 | 1.79E-24 | 3.83E-20 | 0.11 | 0.03 | 0.08 | 4.12 |
| NE | Prdx3 | 0.53 | 1.83E-24 | 3.91E-20 | 0.61 | 0.45 | 0.16 | 1.36 |
| NE | Gas2 | 0.92 | 2.33E-24 | 4.98E-20 | 0.29 | 0.15 | 0.14 | 1.91 |
| NE | Mki67 | 0.46 | 2.65E-24 | 5.67E-20 | 0.74 | 0.58 | 0.17 | 1.29 |
| NE | Gde1 | 0.46 | 2.85E-24 | 6.08E-20 | 0.60 | 0.42 | 0.18 | 1.41 |
| NE | Tmem216 | 0.64 | 3.45E-24 | 7.36E-20 | 0.45 | 0.29 | 0.16 | 1.57 |
| NE | Mettl26 | 0.43 | 4.30E-24 | 9.19E-20 | 0.72 | 0.56 | 0.16 | 1.29 |
| NE | Nabp2 | 0.35 | 4.83E-24 | 1.03E-19 | 0.84 | 0.73 | 0.11 | 1.15 |
| NE | Smc4 | 0.44 | 6.09E-24 | 1.30E-19 | 0.80 | 0.70 | 0.10 | 1.15 |
| NE | Pbdc1 | 0.37 | 6.12E-24 | 1.31E-19 | 0.86 | 0.77 | 0.09 | 1.11 |
| NE | Arxes2 | 0.64 | 6.72E-24 | 1.44E-19 | 0.43 | 0.27 | 0.16 | 1.60 |
| NE | Banf1 | 0.26 | 7.06E-24 | 1.51E-19 | 0.95 | 0.90 | 0.05 | 1.05 |
| NE | Ostc | 0.27 | 7.11E-24 | 1.52E-19 | 0.95 | 0.90 | 0.05 | 1.05 |
| NE | Polr2l | 0.36 | 7.87E-24 | 1.68E-19 | 0.83 | 0.73 | 0.10 | 1.14 |
| NE | Cdca8 | 0.40 | 8.09E-24 | 1.73E-19 | 0.77 | 0.63 | 0.14 | 1.22 |
| NE | Boc | 1.05 | 1.07E-23 | 2.29E-19 | 0.19 | 0.08 | 0.11 | 2.43 |
| NE | Myo5c | 1.10 | 1.11E-23 | 2.38E-19 | 0.22 | 0.10 | 0.12 | 2.19 |
| NE | Hif3a | 1.23 | 1.41E-23 | 3.00E-19 | 0.17 | 0.07 | 0.10 | 2.49 |
| NE | Fgf15 | 1.18 | 1.82E-23 | 3.88E-19 | 0.16 | 0.06 | 0.10 | 2.67 |
| NE | Ank | 1.32 | 1.82E-23 | 3.90E-19 | 0.13 | 0.04 | 0.09 | 3.12 |
| NE | B830012L14Rik | 1.36 | 1.95E-23 | 4.17E-19 | 0.15 | 0.05 | 0.10 | 2.79 |
| NE | Eed | 0.52 | 2.24E-23 | 4.78E-19 | 0.62 | 0.45 | 0.16 | 1.36 |
| NE | Fam149a | 1.49 | 2.65E-23 | 5.66E-19 | 0.14 | 0.05 | 0.09 | 3.00 |
| NE | Heyl | 0.94 | 2.89E-23 | 6.18E-19 | 0.29 | 0.16 | 0.13 | 1.84 |
| NE | Lmod1 | 1.72 | 3.09E-23 | 6.60E-19 | 0.10 | 0.03 | 0.08 | 4.04 |
| NE | Mcrip2 | 1.32 | 3.98E-23 | 8.50E-19 | 0.16 | 0.06 | 0.10 | 2.61 |
| NE | Mest | 0.53 | 4.83E-23 | 1.03E-18 | 0.97 | 0.86 | 0.11 | 1.13 |
| NE | Car2 | 1.53 | 4.93E-23 | 1.05E-18 | 0.22 | 0.11 | 0.11 | 2.01 |
| NE | Msi1 | 0.52 | 5.70E-23 | 1.22E-18 | 0.57 | 0.40 | 0.17 | 1.42 |
| NE | Gpd2 | 0.67 | 5.71E-23 | 1.22E-18 | 0.42 | 0.27 | 0.15 | 1.57 |
| NE | Pak3 | 0.62 | 6.38E-23 | 1.36E-18 | 0.39 | 0.24 | 0.15 | 1.65 |
| NE | Dpysl4 | 0.80 | 6.59E-23 | 1.41E-18 | 0.17 | 0.07 | 0.10 | 2.58 |
| NE | Ppm1g | 0.36 | 7.16E-23 | 1.53E-18 | 0.84 | 0.72 | 0.12 | 1.17 |

**Supplementary Table 4. Marker genes for the annotated cell populations of integrated clustering of E11.5 control and NotchGOF λ-junction ectoderm cells.**

| | | | | | | | | |
|---|---|---|---|---|---|---|---|---|
| NE | Hspa4l | 0.49 | 7.91E-23 | 1.69E-18 | 0.34 | 0.19 | 0.15 | 1.79 |
| NE | Jade1 | 0.66 | 7.93E-23 | 1.69E-18 | 0.41 | 0.25 | 0.16 | 1.62 |
| NE | Hist1h2ae | 0.52 | 8.03E-23 | 1.71E-18 | 0.69 | 0.53 | 0.16 | 1.29 |
| NE | Timm8a1 | 0.45 | 8.14E-23 | 1.74E-18 | 0.72 | 0.59 | 0.13 | 1.22 |
| NE | Clic5 | 1.86 | 1.11E-22 | 2.37E-18 | 0.11 | 0.03 | 0.08 | 3.60 |
| NE | Fam181b | 1.18 | 1.32E-22 | 2.83E-18 | 0.18 | 0.07 | 0.10 | 2.41 |
| NE | Ggt7 | 1.71 | 1.47E-22 | 3.13E-18 | 0.10 | 0.03 | 0.08 | 3.85 |
| NE | Itih5 | 1.61 | 1.55E-22 | 3.31E-18 | 0.12 | 0.04 | 0.08 | 3.22 |
| NE | Wrb | 0.69 | 1.56E-22 | 3.33E-18 | 0.35 | 0.21 | 0.15 | 1.71 |
| NE | Fgf16 | 1.64 | 1.82E-22 | 3.90E-18 | 0.11 | 0.03 | 0.08 | 3.42 |
| NE | Abhd14a | 0.53 | 1.96E-22 | 4.19E-18 | 0.49 | 0.33 | 0.16 | 1.49 |
| NE | Tspan3 | 0.25 | 2.01E-22 | 4.28E-18 | 0.98 | 0.94 | 0.04 | 1.04 |
| NE | Fam13c | 1.56 | 2.09E-22 | 4.47E-18 | 0.12 | 0.04 | 0.08 | 3.24 |
| NE | Pabpc4 | 0.42 | 2.20E-22 | 4.70E-18 | 0.72 | 0.56 | 0.16 | 1.29 |
| NE | Olfm1 | 0.41 | 2.25E-22 | 4.81E-18 | 0.42 | 0.26 | 0.16 | 1.62 |
| NE | Hirip3 | 0.46 | 2.51E-22 | 5.36E-18 | 0.71 | 0.57 | 0.14 | 1.25 |
| NE | Tuba1c | 0.51 | 2.88E-22 | 6.14E-18 | 0.58 | 0.43 | 0.15 | 1.36 |
| NE | Bphl | 0.45 | 3.34E-22 | 7.14E-18 | 0.60 | 0.44 | 0.17 | 1.39 |
| NE | Mfap2 | 0.31 | 3.38E-22 | 7.22E-18 | 0.94 | 0.87 | 0.06 | 1.07 |
| NE | Ift27 | 0.52 | 3.55E-22 | 7.58E-18 | 0.61 | 0.44 | 0.16 | 1.37 |
| NE | Plpp3 | 0.44 | 3.70E-22 | 7.91E-18 | 0.53 | 0.36 | 0.17 | 1.47 |
| NE | Rad18 | 0.50 | 3.77E-22 | 8.05E-18 | 0.50 | 0.33 | 0.17 | 1.51 |
| NE | Fscn1 | 0.27 | 4.03E-22 | 8.60E-18 | 0.90 | 0.83 | 0.07 | 1.09 |
| NE | Cldn6 | 0.25 | 4.10E-22 | 8.76E-18 | 0.94 | 0.82 | 0.12 | 1.15 |
| NE | Nsd2 | 0.38 | 4.83E-22 | 1.03E-17 | 0.82 | 0.69 | 0.13 | 1.18 |
| NE | Ruvbl2 | 0.40 | 5.61E-22 | 1.20E-17 | 0.74 | 0.60 | 0.14 | 1.23 |
| NE | Rangrf | 0.54 | 5.88E-22 | 1.26E-17 | 0.55 | 0.39 | 0.16 | 1.40 |
| NE | Stip1 | 0.30 | 6.37E-22 | 1.36E-17 | 0.90 | 0.83 | 0.07 | 1.09 |
| NE | Thyn1 | 0.41 | 6.62E-22 | 1.41E-17 | 0.74 | 0.59 | 0.15 | 1.24 |
| NE | Ctnna2 | 1.12 | 6.79E-22 | 1.45E-17 | 0.12 | 0.04 | 0.08 | 3.15 |
| NE | Rwdd1 | 0.27 | 8.41E-22 | 1.80E-17 | 0.93 | 0.87 | 0.06 | 1.07 |
| NE | Slc27a2 | 1.79 | 8.51E-22 | 1.82E-17 | 0.10 | 0.03 | 0.07 | 3.85 |
| NE | Sfxn1 | 0.39 | 9.12E-22 | 1.95E-17 | 0.77 | 0.66 | 0.12 | 1.18 |
| NE | Hnrnpll | 0.38 | 1.14E-21 | 2.44E-17 | 0.76 | 0.63 | 0.12 | 1.20 |
| NE | Cks2 | 0.44 | 1.29E-21 | 2.75E-17 | 0.83 | 0.74 | 0.08 | 1.11 |
| NE | Cacnb2 | 1.37 | 1.33E-21 | 2.83E-17 | 0.17 | 0.07 | 0.10 | 2.39 |
| NE | Trim37 | 0.47 | 1.63E-21 | 3.49E-17 | 0.57 | 0.40 | 0.16 | 1.41 |
| NE | Arxes1 | 0.73 | 1.68E-21 | 3.59E-17 | 0.39 | 0.25 | 0.15 | 1.58 |
| NE | Homer2 | 0.45 | 1.83E-21 | 3.91E-17 | 0.27 | 0.15 | 0.13 | 1.86 |
| NE | Baiap2l1 | 0.90 | 1.89E-21 | 4.04E-17 | 0.22 | 0.10 | 0.11 | 2.11 |
| NE | Eif4ebp1 | 0.41 | 2.08E-21 | 4.44E-17 | 0.71 | 0.56 | 0.15 | 1.27 |
| NE | Ccnb2 | 0.50 | 2.38E-21 | 5.09E-17 | 0.59 | 0.43 | 0.16 | 1.39 |
| NE | Rtn2 | 0.71 | 2.70E-21 | 5.78E-17 | 0.25 | 0.13 | 0.12 | 1.96 |
| NE | Ak2 | 0.30 | 2.75E-21 | 5.88E-17 | 0.89 | 0.81 | 0.09 | 1.11 |
| NE | Pradc1 | 0.73 | 2.85E-21 | 6.09E-17 | 0.38 | 0.23 | 0.14 | 1.62 |
| NE | Pkp2 | 0.77 | 3.04E-21 | 6.49E-17 | 0.28 | 0.15 | 0.13 | 1.85 |
| NE | Emb | 0.28 | 3.34E-21 | 7.14E-17 | 0.83 | 0.66 | 0.17 | 1.25 |
| NE | Pde1b | 1.59 | 3.37E-21 | 7.20E-17 | 0.11 | 0.04 | 0.08 | 3.26 |
| NE | Mrpl12 | 0.28 | 3.41E-21 | 7.28E-17 | 0.91 | 0.86 | 0.05 | 1.06 |
| NE | Cul1 | 0.32 | 4.12E-21 | 8.81E-17 | 0.89 | 0.83 | 0.06 | 1.07 |
| NE | Pgm1 | 0.64 | 4.33E-21 | 9.24E-17 | 0.39 | 0.24 | 0.15 | 1.62 |
| NE | Lmnb1 | 0.25 | 4.40E-21 | 9.39E-17 | 0.96 | 0.90 | 0.06 | 1.07 |
| NE | Ormdl2 | 0.42 | 4.74E-21 | 1.01E-16 | 0.70 | 0.54 | 0.16 | 1.30 |
| NE | Rbm47 | 1.21 | 5.44E-21 | 1.16E-16 | 0.15 | 0.06 | 0.09 | 2.55 |
| NE | Sapcd2 | 0.68 | 5.47E-21 | 1.17E-16 | 0.36 | 0.22 | 0.14 | 1.61 |

**Supplementary Table 4. Marker genes for the annotated cell populations of integrated clustering of E11.5 control and NotchGOF λ-junction ectoderm cells.**

| | | | | | | | | |
|---|---|---|---|---|---|---|---|---|
| NE | Tdrkh | 1.12 | 5.66E-21 | 1.21E-16 | 0.15 | 0.06 | 0.09 | 2.60 |
| NE | Dctpp1 | 0.26 | 5.93E-21 | 1.27E-16 | 0.95 | 0.87 | 0.07 | 1.08 |
| NE | Elovl6 | 0.48 | 6.75E-21 | 1.44E-16 | 0.61 | 0.45 | 0.16 | 1.36 |
| NE | Smim11 | 0.35 | 7.09E-21 | 1.51E-16 | 0.78 | 0.66 | 0.12 | 1.18 |
| NE | Raver2 | 0.75 | 7.27E-21 | 1.55E-16 | 0.28 | 0.15 | 0.13 | 1.83 |
| NE | Gins2 | 0.44 | 8.41E-21 | 1.80E-16 | 0.64 | 0.49 | 0.16 | 1.32 |
| NE | Crb3 | 0.51 | 8.44E-21 | 1.80E-16 | 0.53 | 0.37 | 0.17 | 1.46 |
| NE | Atic | 0.34 | 8.73E-21 | 1.86E-16 | 0.77 | 0.62 | 0.16 | 1.25 |
| NE | Tmem39b | 0.59 | 9.10E-21 | 1.94E-16 | 0.40 | 0.26 | 0.15 | 1.58 |
| NE | Bola3 | 0.42 | 9.73E-21 | 2.08E-16 | 0.72 | 0.58 | 0.14 | 1.25 |
| NE | Mov10 | 0.81 | 9.88E-21 | 2.11E-16 | 0.28 | 0.16 | 0.13 | 1.80 |
| NE | Oscp1 | 0.81 | 1.06E-20 | 2.27E-16 | 0.25 | 0.13 | 0.12 | 1.92 |
| NE | Por | 0.62 | 1.21E-20 | 2.59E-16 | 0.51 | 0.36 | 0.15 | 1.43 |
| NE | Dnajc15 | 0.32 | 1.25E-20 | 2.67E-16 | 0.79 | 0.67 | 0.12 | 1.18 |
| NE | Atp6v0e2 | 0.60 | 1.48E-20 | 3.16E-16 | 0.22 | 0.11 | 0.12 | 2.07 |
| NE | Ric3 | 1.09 | 1.49E-20 | 3.19E-16 | 0.17 | 0.07 | 0.10 | 2.37 |
| NE | Rpn2 | 0.34 | 1.56E-20 | 3.33E-16 | 0.84 | 0.74 | 0.10 | 1.14 |
| NE | Tmx2 | 0.43 | 1.67E-20 | 3.58E-16 | 0.68 | 0.51 | 0.17 | 1.32 |
| NE | Mageh1 | 0.47 | 1.79E-20 | 3.82E-16 | 0.60 | 0.44 | 0.16 | 1.35 |
| NE | Creld2 | 0.56 | 1.85E-20 | 3.96E-16 | 0.53 | 0.38 | 0.15 | 1.40 |
| NE | Maf | 0.51 | 2.13E-20 | 4.54E-16 | 0.20 | 0.09 | 0.11 | 2.17 |
| NE | Snhg8 | 0.32 | 2.28E-20 | 4.86E-16 | 0.84 | 0.73 | 0.11 | 1.14 |
| NE | Cthrc1 | 1.36 | 2.28E-20 | 4.87E-16 | 0.16 | 0.07 | 0.09 | 2.40 |
| NE | Acvr2a | 0.39 | 2.38E-20 | 5.09E-16 | 0.62 | 0.46 | 0.16 | 1.36 |
| NE | Klhl7 | 0.44 | 2.63E-20 | 5.62E-16 | 0.67 | 0.51 | 0.16 | 1.31 |
| NE | Ccdc85b | 0.42 | 2.76E-20 | 5.89E-16 | 0.63 | 0.47 | 0.16 | 1.33 |
| NE | Hist1h3c | 0.60 | 2.94E-20 | 6.29E-16 | 0.55 | 0.40 | 0.15 | 1.37 |
| NE | Galnt7 | 0.50 | 3.03E-20 | 6.48E-16 | 0.55 | 0.39 | 0.16 | 1.42 |
| NE | Nsdhl | 0.63 | 3.04E-20 | 6.49E-16 | 0.45 | 0.30 | 0.15 | 1.48 |
| NE | Tyms | 0.35 | 3.46E-20 | 7.39E-16 | 0.82 | 0.69 | 0.13 | 1.19 |
| NE | Lpcat1 | 0.48 | 3.48E-20 | 7.44E-16 | 0.55 | 0.38 | 0.17 | 1.43 |
| NE | Cdt1 | 0.53 | 3.62E-20 | 7.73E-16 | 0.48 | 0.33 | 0.16 | 1.47 |
| NE | Vegfb | 0.43 | 3.82E-20 | 8.17E-16 | 0.63 | 0.47 | 0.16 | 1.34 |
| NE | Mcm7 | 0.31 | 3.92E-20 | 8.38E-16 | 0.87 | 0.77 | 0.10 | 1.13 |
| NE | Hip1 | 0.53 | 4.96E-20 | 1.06E-15 | 0.44 | 0.29 | 0.15 | 1.52 |
| NE | Id2 | 0.43 | 5.12E-20 | 1.09E-15 | 0.88 | 0.79 | 0.08 | 1.11 |
| NE | Lamb2 | 0.88 | 5.13E-20 | 1.09E-15 | 0.23 | 0.12 | 0.11 | 1.97 |
| NE | Gmpr | 0.80 | 5.93E-20 | 1.27E-15 | 0.26 | 0.14 | 0.12 | 1.88 |
| NE | Slc1a3 | 0.76 | 6.73E-20 | 1.44E-15 | 0.27 | 0.15 | 0.12 | 1.83 |
| NE | Cdc20 | 0.51 | 7.13E-20 | 1.52E-15 | 0.60 | 0.46 | 0.14 | 1.31 |
| NE | Anxa5 | 0.46 | 7.95E-20 | 1.70E-15 | 0.70 | 0.57 | 0.13 | 1.24 |
| NE | Rif1 | 0.31 | 8.30E-20 | 1.77E-15 | 0.87 | 0.77 | 0.10 | 1.13 |
| NE | Mirg | 1.31 | 8.76E-20 | 1.87E-15 | 0.12 | 0.04 | 0.08 | 2.93 |
| NE | Spin4 | 1.34 | 9.00E-20 | 1.92E-15 | 0.15 | 0.06 | 0.09 | 2.50 |
| NE | Wdr18 | 0.37 | 9.48E-20 | 2.02E-15 | 0.76 | 0.62 | 0.14 | 1.23 |
| NE | Tspan33 | 0.88 | 1.01E-19 | 2.16E-15 | 0.20 | 0.10 | 0.11 | 2.10 |
| NE | Exoc3l4 | 0.95 | 1.06E-19 | 2.27E-15 | 0.25 | 0.13 | 0.11 | 1.87 |
| NE | Fmc1 | 0.38 | 1.10E-19 | 2.35E-15 | 0.74 | 0.61 | 0.13 | 1.21 |
| NE | Kitl | 0.71 | 1.13E-19 | 2.41E-15 | 0.20 | 0.09 | 0.10 | 2.13 |
| NE | Knstrn | 0.37 | 1.28E-19 | 2.74E-15 | 0.67 | 0.51 | 0.16 | 1.31 |
| NE | Mcm5 | 0.37 | 1.37E-19 | 2.93E-15 | 0.72 | 0.57 | 0.15 | 1.26 |
| NE | Igdcc3 | 0.41 | 1.38E-19 | 2.96E-15 | 0.43 | 0.27 | 0.16 | 1.59 |
| NE | Skp2 | 0.45 | 1.47E-19 | 3.14E-15 | 0.67 | 0.53 | 0.14 | 1.27 |
| NE | Eif1b | 0.28 | 2.02E-19 | 4.32E-15 | 0.91 | 0.82 | 0.08 | 1.10 |
| NE | Tedc1 | 0.71 | 2.44E-19 | 5.20E-15 | 0.30 | 0.17 | 0.13 | 1.72 |

**Supplementary Table 4. Marker genes for the annotated cell populations of integrated clustering of E11.5 control and NotchGOF λ-junction ectoderm cells.**

| | | | | | | | | |
|---|---|---|---|---|---|---|---|---|
| NE | Cd47 | 0.45 | 2.48E-19 | 5.30E-15 | 0.66 | 0.51 | 0.15 | 1.29 |
| NE | Hmbs | 0.46 | 2.74E-19 | 5.86E-15 | 0.60 | 0.46 | 0.15 | 1.32 |
| NE | Slc25a23 | 0.57 | 2.75E-19 | 5.88E-15 | 0.40 | 0.25 | 0.14 | 1.57 |
| NE | Gart | 0.39 | 3.12E-19 | 6.66E-15 | 0.67 | 0.54 | 0.13 | 1.24 |
| NE | Ccnb1 | 0.47 | 3.25E-19 | 6.94E-15 | 0.60 | 0.46 | 0.14 | 1.32 |
| NE | Npepl1 | 0.48 | 3.34E-19 | 7.14E-15 | 0.51 | 0.35 | 0.15 | 1.43 |
| NE | Snx2 | 0.42 | 3.62E-19 | 7.72E-15 | 0.65 | 0.50 | 0.15 | 1.31 |
| NE | 2310033P09Rik | 0.37 | 3.71E-19 | 7.93E-15 | 0.72 | 0.56 | 0.16 | 1.28 |
| NE | Gar1 | 0.35 | 3.80E-19 | 8.12E-15 | 0.81 | 0.70 | 0.11 | 1.15 |
| NE | Hap1 | 0.66 | 4.00E-19 | 8.54E-15 | 0.26 | 0.14 | 0.12 | 1.83 |
| NE | Sdc2 | 0.59 | 4.05E-19 | 8.64E-15 | 0.38 | 0.24 | 0.14 | 1.59 |
| NE | Emc8 | 0.32 | 4.16E-19 | 8.89E-15 | 0.83 | 0.73 | 0.10 | 1.14 |
| NE | Hist1h2ap | 0.33 | 4.67E-19 | 9.97E-15 | 0.78 | 0.63 | 0.15 | 1.24 |
| NE | Pus7 | 0.58 | 4.86E-19 | 1.04E-14 | 0.38 | 0.24 | 0.14 | 1.59 |
| NE | Hk1 | 0.66 | 4.92E-19 | 1.05E-14 | 0.28 | 0.16 | 0.12 | 1.78 |
| NE | Erg28 | 0.36 | 5.43E-19 | 1.16E-14 | 0.76 | 0.64 | 0.12 | 1.19 |
| NE | Chka | 0.42 | 5.47E-19 | 1.17E-14 | 0.62 | 0.46 | 0.16 | 1.35 |
| NE | Ftsj3 | 0.44 | 5.62E-19 | 1.20E-14 | 0.67 | 0.53 | 0.14 | 1.26 |
| NE | Gars | 0.38 | 5.78E-19 | 1.23E-14 | 0.73 | 0.60 | 0.13 | 1.21 |
| NE | Psen2 | 0.85 | 5.87E-19 | 1.25E-14 | 0.19 | 0.09 | 0.10 | 2.11 |
| NE | Mrpl28 | 0.27 | 5.95E-19 | 1.27E-14 | 0.88 | 0.81 | 0.07 | 1.09 |
| NE | Mrps28 | 0.37 | 5.97E-19 | 1.28E-14 | 0.74 | 0.62 | 0.12 | 1.20 |
| NE | Ybx3 | 0.29 | 6.30E-19 | 1.35E-14 | 0.91 | 0.84 | 0.07 | 1.09 |
| NE | Heg1 | 1.18 | 6.61E-19 | 1.41E-14 | 0.12 | 0.04 | 0.08 | 2.83 |
| NE | Nrarp | 0.48 | 7.03E-19 | 1.50E-14 | 0.67 | 0.56 | 0.11 | 1.20 |
| NE | Cenpa | 0.31 | 7.37E-19 | 1.57E-14 | 0.76 | 0.62 | 0.14 | 1.22 |
| NE | Tenm4 | 0.36 | 7.57E-19 | 1.62E-14 | 0.55 | 0.39 | 0.16 | 1.41 |
| NE | Ubr5 | 0.40 | 7.98E-19 | 1.71E-14 | 0.74 | 0.60 | 0.14 | 1.23 |
| NE | Dnph1 | 0.42 | 8.20E-19 | 1.75E-14 | 0.61 | 0.47 | 0.14 | 1.30 |
| NE | Hmces | 0.57 | 8.47E-19 | 1.81E-14 | 0.42 | 0.28 | 0.14 | 1.51 |
| NE | Ube2c | 0.40 | 9.14E-19 | 1.95E-14 | 0.74 | 0.61 | 0.14 | 1.22 |
| NE | Socs3 | 0.82 | 9.53E-19 | 2.04E-14 | 0.23 | 0.12 | 0.11 | 1.90 |
| NE | Iars | 0.51 | 9.72E-19 | 2.08E-14 | 0.54 | 0.39 | 0.15 | 1.38 |
| NE | Kif22 | 0.53 | 1.05E-18 | 2.24E-14 | 0.49 | 0.36 | 0.14 | 1.39 |
| NE | Arhgap8 | 0.88 | 1.49E-18 | 3.19E-14 | 0.22 | 0.12 | 0.11 | 1.91 |
| NE | Noa1 | 0.58 | 1.71E-18 | 3.65E-14 | 0.40 | 0.26 | 0.14 | 1.55 |
| NE | Pla2g12a | 0.36 | 1.92E-18 | 4.09E-14 | 0.71 | 0.55 | 0.17 | 1.30 |
| NE | Gstm7 | 0.60 | 1.92E-18 | 4.09E-14 | 0.31 | 0.18 | 0.13 | 1.70 |
| NE | Myh10 | 0.31 | 2.11E-18 | 4.50E-14 | 0.86 | 0.75 | 0.11 | 1.14 |
| NE | Dhfr | 0.39 | 2.38E-18 | 5.08E-14 | 0.67 | 0.51 | 0.16 | 1.31 |
| NE | Gm48435 | 0.99 | 2.54E-18 | 5.42E-14 | 0.15 | 0.07 | 0.09 | 2.34 |
| NE | Rcc1 | 0.47 | 3.00E-18 | 6.40E-14 | 0.55 | 0.40 | 0.14 | 1.35 |
| NE | Gm42047 | 1.58 | 3.30E-18 | 7.05E-14 | 0.13 | 0.05 | 0.08 | 2.49 |
| NE | Cyb5b | 0.46 | 3.63E-18 | 7.76E-14 | 0.60 | 0.46 | 0.14 | 1.31 |
| NE | Svip | 0.64 | 3.65E-18 | 7.80E-14 | 0.33 | 0.21 | 0.13 | 1.61 |
| NE | Tenm3 | 0.40 | 3.68E-18 | 7.86E-14 | 0.70 | 0.55 | 0.14 | 1.26 |
| NE | Rrm1 | 0.30 | 3.86E-18 | 8.25E-14 | 0.79 | 0.65 | 0.14 | 1.21 |
| NE | Gcsh | 0.34 | 3.96E-18 | 8.45E-14 | 0.73 | 0.60 | 0.13 | 1.22 |
| NE | Akap1 | 0.64 | 4.10E-18 | 8.77E-14 | 0.31 | 0.18 | 0.13 | 1.69 |
| NE | Cask | 0.44 | 4.16E-18 | 8.89E-14 | 0.50 | 0.35 | 0.15 | 1.44 |
| NE | Prrg2 | 0.66 | 4.27E-18 | 9.11E-14 | 0.33 | 0.20 | 0.13 | 1.63 |
| NE | Prrx1 | 1.15 | 4.72E-18 | 1.01E-13 | 0.25 | 0.15 | 0.10 | 1.71 |
| NE | Camk1 | 0.53 | 5.01E-18 | 1.07E-13 | 0.47 | 0.32 | 0.15 | 1.47 |
| NE | Gulp1 | 0.90 | 5.29E-18 | 1.13E-13 | 0.21 | 0.11 | 0.10 | 1.95 |
| NE | Stard4 | 0.71 | 5.51E-18 | 1.18E-13 | 0.30 | 0.18 | 0.12 | 1.66 |

**Supplementary Table 4. Marker genes for the annotated cell populations of integrated clustering of E11.5 control and NotchGOF λ-junction ectoderm cells.**

| | | | | | | | | |
|---|---|---|---|---|---|---|---|---|
| NE | Gcnt2 | 1.22 | 5.53E-18 | 1.18E-13 | 0.12 | 0.04 | 0.08 | 2.73 |
| NE | Scara3 | 0.97 | 6.04E-18 | 1.29E-13 | 0.18 | 0.09 | 0.10 | 2.07 |
| NE | Metrn | 0.53 | 6.60E-18 | 1.41E-13 | 0.42 | 0.28 | 0.14 | 1.49 |
| NE | Creld1 | 0.64 | 7.30E-18 | 1.56E-13 | 0.34 | 0.21 | 0.13 | 1.62 |
| NE | Gdf11 | 0.50 | 7.30E-18 | 1.56E-13 | 0.34 | 0.21 | 0.14 | 1.66 |
| NE | Timm10b | 0.26 | 8.08E-18 | 1.73E-13 | 0.88 | 0.79 | 0.09 | 1.11 |
| NE | Polr2h | 0.30 | 8.48E-18 | 1.81E-13 | 0.87 | 0.77 | 0.10 | 1.12 |
| NE | Arhgef9 | 1.09 | 8.91E-18 | 1.90E-13 | 0.15 | 0.07 | 0.09 | 2.26 |
| NE | Mycbp | 0.47 | 9.29E-18 | 1.98E-13 | 0.56 | 0.41 | 0.16 | 1.39 |
| NE | Cenpm | 0.40 | 9.73E-18 | 2.08E-13 | 0.59 | 0.44 | 0.15 | 1.33 |
| NE | Cdc42ep4 | 0.49 | 9.81E-18 | 2.10E-13 | 0.42 | 0.27 | 0.14 | 1.52 |
| NE | Ccne1 | 0.54 | 9.88E-18 | 2.11E-13 | 0.36 | 0.23 | 0.13 | 1.59 |
| NE | Bend7 | 0.56 | 1.17E-17 | 2.50E-13 | 0.33 | 0.20 | 0.13 | 1.65 |
| NE | Rab28 | 0.46 | 1.17E-17 | 2.51E-13 | 0.60 | 0.45 | 0.15 | 1.32 |
| NE | Lss | 0.94 | 1.33E-17 | 2.85E-13 | 0.21 | 0.11 | 0.10 | 1.90 |
| NE | Tpst1 | 0.66 | 1.38E-17 | 2.94E-13 | 0.32 | 0.19 | 0.13 | 1.66 |
| NE | Nusap1 | 0.40 | 1.48E-17 | 3.16E-13 | 0.55 | 0.40 | 0.15 | 1.37 |
| NE | Plk1 | 0.63 | 1.49E-17 | 3.19E-13 | 0.47 | 0.35 | 0.13 | 1.36 |
| NE | Clstn1 | 0.37 | 1.57E-17 | 3.35E-13 | 0.64 | 0.49 | 0.16 | 1.32 |
| NE | Lgmn | 0.48 | 1.58E-17 | 3.37E-13 | 0.43 | 0.29 | 0.14 | 1.49 |
| NE | Sms | 0.26 | 1.65E-17 | 3.52E-13 | 0.91 | 0.82 | 0.09 | 1.11 |
| NE | Srebf1 | 0.64 | 1.69E-17 | 3.61E-13 | 0.36 | 0.23 | 0.13 | 1.57 |
| NE | Prdx4 | 0.25 | 1.86E-17 | 3.97E-13 | 0.92 | 0.84 | 0.08 | 1.09 |
| NE | Rtkn2 | 1.04 | 1.93E-17 | 4.12E-13 | 0.17 | 0.08 | 0.09 | 2.11 |
| NE | Rad51 | 0.40 | 1.93E-17 | 4.12E-13 | 0.59 | 0.44 | 0.15 | 1.35 |
| NE | Pimreg | 0.42 | 1.95E-17 | 4.17E-13 | 0.55 | 0.40 | 0.14 | 1.35 |
| NE | Mcm3 | 0.32 | 2.07E-17 | 4.43E-13 | 0.77 | 0.63 | 0.14 | 1.23 |
| NE | Jam3 | 0.56 | 2.08E-17 | 4.45E-13 | 0.44 | 0.30 | 0.14 | 1.46 |
| NE | D030056L22Rik | 0.42 | 2.91E-17 | 6.21E-13 | 0.62 | 0.49 | 0.14 | 1.28 |
| NE | Kctd6 | 0.56 | 3.03E-17 | 6.48E-13 | 0.31 | 0.19 | 0.13 | 1.68 |
| NE | Cenpb | 0.34 | 3.19E-17 | 6.81E-13 | 0.78 | 0.65 | 0.13 | 1.20 |
| NE | Bnipl | 0.72 | 3.73E-17 | 7.97E-13 | 0.25 | 0.14 | 0.11 | 1.80 |
| NE | Itm2a | 0.37 | 3.89E-17 | 8.30E-13 | 0.61 | 0.45 | 0.16 | 1.36 |
| NE | Aldh7a1 | 0.39 | 3.93E-17 | 8.40E-13 | 0.64 | 0.49 | 0.15 | 1.31 |
| NE | Aimp2 | 0.40 | 3.94E-17 | 8.42E-13 | 0.62 | 0.47 | 0.15 | 1.31 |
| NE | Ect2 | 0.43 | 3.97E-17 | 8.49E-13 | 0.50 | 0.35 | 0.14 | 1.41 |
| NE | Kif23 | 0.46 | 4.17E-17 | 8.91E-13 | 0.50 | 0.35 | 0.14 | 1.40 |
| NE | Atf4 | 0.33 | 4.20E-17 | 8.98E-13 | 0.75 | 0.62 | 0.13 | 1.22 |
| NE | 2010204K13Rik | 0.53 | 4.41E-17 | 9.42E-13 | 0.33 | 0.20 | 0.13 | 1.64 |
| NE | Slc35d1 | 0.61 | 4.59E-17 | 9.81E-13 | 0.29 | 0.17 | 0.12 | 1.68 |
| NE | Npdc1 | 0.43 | 4.72E-17 | 1.01E-12 | 0.41 | 0.26 | 0.14 | 1.53 |
| NE | Fut9 | 1.24 | 4.77E-17 | 1.02E-12 | 0.15 | 0.06 | 0.08 | 2.27 |
| NE | Thop1 | 0.39 | 4.91E-17 | 1.05E-12 | 0.58 | 0.44 | 0.14 | 1.33 |
| NE | Nog | 1.24 | 4.97E-17 | 1.06E-12 | 0.16 | 0.08 | 0.09 | 2.09 |
| NE | Timm50 | 0.30 | 5.06E-17 | 1.08E-12 | 0.77 | 0.64 | 0.13 | 1.21 |
| NE | Cdk6 | 0.43 | 5.15E-17 | 1.10E-12 | 0.60 | 0.46 | 0.13 | 1.29 |
| NE | Knl1 | 0.44 | 5.74E-17 | 1.23E-12 | 0.55 | 0.41 | 0.14 | 1.35 |
| NE | Kif15 | 0.37 | 5.82E-17 | 1.24E-12 | 0.67 | 0.52 | 0.15 | 1.28 |
| NE | Gipc2 | 0.90 | 6.32E-17 | 1.35E-12 | 0.22 | 0.12 | 0.10 | 1.81 |
| NE | Tctex1d2 | 0.52 | 7.75E-17 | 1.65E-12 | 0.48 | 0.34 | 0.14 | 1.40 |
| NE | Ckap2 | 0.60 | 7.75E-17 | 1.66E-12 | 0.41 | 0.28 | 0.13 | 1.47 |
| NE | Prc1 | 0.45 | 8.93E-17 | 1.91E-12 | 0.59 | 0.46 | 0.13 | 1.28 |
| NE | Ccdc124 | 0.28 | 9.05E-17 | 1.93E-12 | 0.82 | 0.71 | 0.11 | 1.15 |
| NE | Ndfip1 | 0.28 | 9.06E-17 | 1.94E-12 | 0.88 | 0.79 | 0.09 | 1.11 |
| NE | Cadps2 | 0.71 | 9.25E-17 | 1.98E-12 | 0.19 | 0.10 | 0.10 | 1.99 |

**Supplementary Table 4. Marker genes for the annotated cell populations of integrated clustering of E11.5 control and NotchGOF λ-junction ectoderm cells.**

| | | | | | | | | |
|---|---|---|---|---|---|---|---|---|
| NE | Bmpr1b | 1.23 | 9.33E-17 | 1.99E-12 | 0.12 | 0.05 | 0.07 | 2.59 |
| NE | Desi2 | 0.46 | 9.84E-17 | 2.10E-12 | 0.50 | 0.36 | 0.14 | 1.40 |
| NE | Bst2 | 1.21 | 1.02E-16 | 2.18E-12 | 0.14 | 0.06 | 0.08 | 2.31 |
| NE | Smim4 | 0.38 | 1.10E-16 | 2.35E-12 | 0.62 | 0.47 | 0.15 | 1.32 |
| NE | Bcl7c | 0.30 | 1.13E-16 | 2.42E-12 | 0.84 | 0.74 | 0.10 | 1.13 |
| NE | Pcyt1a | 0.65 | 1.21E-16 | 2.59E-12 | 0.24 | 0.13 | 0.11 | 1.80 |
| NE | Pck2 | 0.79 | 1.23E-16 | 2.62E-12 | 0.22 | 0.12 | 0.10 | 1.84 |
| NE | Ccdc12 | 0.25 | 1.26E-16 | 2.68E-12 | 0.80 | 0.69 | 0.11 | 1.16 |
| NE | Gli3 | 0.36 | 1.31E-16 | 2.81E-12 | 0.61 | 0.46 | 0.15 | 1.32 |
| NE | Laptm4b | 0.44 | 1.35E-16 | 2.88E-12 | 0.57 | 0.42 | 0.15 | 1.35 |
| NE | Gpx3 | 0.87 | 1.35E-16 | 2.88E-12 | 0.18 | 0.09 | 0.09 | 2.05 |
| NE | Inppl1 | 0.50 | 1.42E-16 | 3.03E-12 | 0.35 | 0.22 | 0.13 | 1.59 |
| NE | Morn2 | 0.40 | 1.50E-16 | 3.21E-12 | 0.47 | 0.32 | 0.15 | 1.46 |
| NE | Hsd17b11 | 0.95 | 1.51E-16 | 3.23E-12 | 0.16 | 0.08 | 0.09 | 2.13 |
| NE | Ctbp2 | 0.37 | 1.54E-16 | 3.28E-12 | 0.67 | 0.52 | 0.15 | 1.29 |
| NE | Grwd1 | 0.40 | 1.58E-16 | 3.37E-12 | 0.52 | 0.37 | 0.15 | 1.40 |
| NE | Suclg1 | 0.26 | 1.77E-16 | 3.78E-12 | 0.88 | 0.80 | 0.07 | 1.09 |
| NE | Gmnn | 0.28 | 1.78E-16 | 3.81E-12 | 0.79 | 0.66 | 0.13 | 1.20 |
| NE | Ppan | 0.41 | 1.82E-16 | 3.89E-12 | 0.56 | 0.42 | 0.14 | 1.34 |
| NE | Cuta | 0.29 | 1.97E-16 | 4.22E-12 | 0.85 | 0.77 | 0.08 | 1.10 |
| NE | E2f5 | 0.42 | 2.03E-16 | 4.34E-12 | 0.50 | 0.36 | 0.14 | 1.40 |
| NE | Shisa2 | 0.96 | 2.04E-16 | 4.36E-12 | 0.16 | 0.07 | 0.08 | 2.17 |
| NE | Mapk8ip2 | 0.43 | 2.06E-16 | 4.41E-12 | 0.15 | 0.07 | 0.08 | 2.22 |
| NE | Mrpl38 | 0.35 | 2.42E-16 | 5.17E-12 | 0.71 | 0.58 | 0.13 | 1.22 |
| NE | Tekt1 | 0.81 | 2.55E-16 | 5.46E-12 | 0.16 | 0.07 | 0.09 | 2.21 |
| NE | Minpp1 | 0.37 | 2.57E-16 | 5.49E-12 | 0.58 | 0.43 | 0.15 | 1.35 |
| NE | Srebf2 | 0.40 | 2.66E-16 | 5.67E-12 | 0.61 | 0.47 | 0.14 | 1.30 |
| NE | Zfp536 | 0.86 | 2.66E-16 | 5.69E-12 | 0.22 | 0.12 | 0.10 | 1.82 |
| NE | Prr36 | 0.51 | 2.77E-16 | 5.91E-12 | 0.34 | 0.21 | 0.13 | 1.62 |
| NE | Slitrk6 | 0.86 | 2.92E-16 | 6.24E-12 | 0.23 | 0.13 | 0.10 | 1.79 |
| NE | Dnajc21 | 0.34 | 3.01E-16 | 6.43E-12 | 0.72 | 0.60 | 0.13 | 1.22 |
| NE | Ext2 | 0.43 | 3.12E-16 | 6.66E-12 | 0.51 | 0.37 | 0.14 | 1.39 |
| NE | Scarb1 | 0.67 | 3.12E-16 | 6.67E-12 | 0.25 | 0.14 | 0.11 | 1.76 |
| NE | Tpx2 | 0.34 | 3.18E-16 | 6.79E-12 | 0.64 | 0.50 | 0.14 | 1.28 |
| NE | Cops6 | 0.26 | 3.21E-16 | 6.85E-12 | 0.86 | 0.77 | 0.08 | 1.11 |
| NE | Farp1 | 0.38 | 3.26E-16 | 6.96E-12 | 0.50 | 0.35 | 0.15 | 1.42 |
| NE | Cars | 0.49 | 3.40E-16 | 7.27E-12 | 0.37 | 0.24 | 0.13 | 1.54 |
| NE | Mpp2 | 0.64 | 3.42E-16 | 7.31E-12 | 0.22 | 0.12 | 0.10 | 1.83 |
| NE | Dnajc19 | 0.27 | 3.44E-16 | 7.35E-12 | 0.83 | 0.72 | 0.11 | 1.16 |
| NE | Slc4a8 | 0.98 | 3.87E-16 | 8.27E-12 | 0.15 | 0.07 | 0.08 | 2.21 |
| NE | Nup210 | 0.77 | 4.38E-16 | 9.35E-12 | 0.25 | 0.14 | 0.11 | 1.75 |
| NE | Fut10 | 1.04 | 4.67E-16 | 9.97E-12 | 0.14 | 0.06 | 0.08 | 2.26 |
| NE | Vill | 0.83 | 4.70E-16 | 1.00E-11 | 0.21 | 0.12 | 0.10 | 1.84 |
| NE | Hsd17b12 | 0.31 | 4.96E-16 | 1.06E-11 | 0.78 | 0.65 | 0.13 | 1.20 |
| NE | Syne4 | 0.54 | 5.07E-16 | 1.08E-11 | 0.35 | 0.22 | 0.13 | 1.57 |
| NE | Vars | 0.39 | 5.17E-16 | 1.10E-11 | 0.67 | 0.53 | 0.13 | 1.25 |
| NE | Tmem147 | 0.29 | 5.18E-16 | 1.11E-11 | 0.76 | 0.63 | 0.13 | 1.20 |
| NE | Mrps36 | 0.29 | 5.53E-16 | 1.18E-11 | 0.85 | 0.76 | 0.09 | 1.11 |
| NE | Tenm2 | 0.59 | 6.01E-16 | 1.28E-11 | 0.29 | 0.18 | 0.12 | 1.66 |
| NE | Naa10 | 0.31 | 6.31E-16 | 1.35E-11 | 0.77 | 0.65 | 0.12 | 1.18 |
| NE | Mcm2 | 0.37 | 6.61E-16 | 1.41E-11 | 0.63 | 0.50 | 0.14 | 1.27 |
| NE | Aifm1 | 0.36 | 6.62E-16 | 1.41E-11 | 0.62 | 0.47 | 0.15 | 1.32 |
| NE | Mgat4b | 0.41 | 6.91E-16 | 1.48E-11 | 0.52 | 0.38 | 0.14 | 1.38 |
| NE | Ipo4 | 0.55 | 7.03E-16 | 1.50E-11 | 0.41 | 0.28 | 0.13 | 1.47 |
| NE | Timp1 | 1.40 | 7.40E-16 | 1.58E-11 | 0.12 | 0.05 | 0.07 | 2.41 |

**Supplementary Table 4. Marker genes for the annotated cell populations of integrated clustering of E11.5 control and NotchGOF λ-junction ectoderm cells.**

| | | | | | | | | |
|---|---|---|---|---|---|---|---|---|
| NE | Gucy1a1 | 0.57 | 7.48E-16 | 1.60E-11 | 0.25 | 0.15 | 0.11 | 1.73 |
| NE | Dyrk3 | 0.61 | 8.75E-16 | 1.87E-11 | 0.22 | 0.12 | 0.10 | 1.86 |
| NE | Ildr1 | 0.73 | 9.28E-16 | 1.98E-11 | 0.21 | 0.11 | 0.10 | 1.88 |
| NE | Cfap36 | 0.29 | 9.49E-16 | 2.03E-11 | 0.58 | 0.43 | 0.15 | 1.35 |
| NE | Rdh10 | 0.40 | 9.62E-16 | 2.06E-11 | 0.37 | 0.25 | 0.13 | 1.51 |
| NE | Nup93 | 0.44 | 9.71E-16 | 2.07E-11 | 0.51 | 0.37 | 0.14 | 1.37 |
| NE | Synm | 1.37 | 1.11E-15 | 2.36E-11 | 0.10 | 0.04 | 0.06 | 2.78 |
| NE | Clcn5 | 0.51 | 1.14E-15 | 2.44E-11 | 0.46 | 0.32 | 0.14 | 1.44 |
| NE | Mta1 | 0.32 | 1.17E-15 | 2.49E-11 | 0.75 | 0.64 | 0.11 | 1.17 |
| NE | Cenpf | 0.33 | 1.17E-15 | 2.50E-11 | 0.74 | 0.61 | 0.13 | 1.21 |
| NE | Slc22a17 | 0.48 | 1.20E-15 | 2.56E-11 | 0.48 | 0.34 | 0.14 | 1.42 |
| NE | Bop1 | 0.56 | 1.26E-15 | 2.68E-11 | 0.41 | 0.28 | 0.13 | 1.45 |
| NE | Fras1 | 0.36 | 1.31E-15 | 2.79E-11 | 0.69 | 0.56 | 0.13 | 1.24 |
| NE | Mocs2 | 0.38 | 1.36E-15 | 2.90E-11 | 0.60 | 0.46 | 0.14 | 1.30 |
| NE | Phtf1 | 0.51 | 1.36E-15 | 2.90E-11 | 0.50 | 0.37 | 0.14 | 1.37 |
| NE | Tmem189 | 0.50 | 1.41E-15 | 3.01E-11 | 0.49 | 0.36 | 0.14 | 1.39 |
| NE | Slc7a6 | 0.57 | 1.52E-15 | 3.26E-11 | 0.33 | 0.22 | 0.12 | 1.55 |
| NE | Hist1h2ab | 0.55 | 1.53E-15 | 3.26E-11 | 0.34 | 0.22 | 0.12 | 1.55 |
| NE | Rassf1 | 0.53 | 1.58E-15 | 3.37E-11 | 0.45 | 0.33 | 0.13 | 1.38 |
| NE | Sfmbt1 | 0.54 | 1.71E-15 | 3.66E-11 | 0.24 | 0.13 | 0.10 | 1.78 |
| NE | Nop56 | 0.26 | 1.72E-15 | 3.67E-11 | 0.88 | 0.81 | 0.07 | 1.09 |
| NE | Wee1 | 0.45 | 1.75E-15 | 3.73E-11 | 0.50 | 0.37 | 0.13 | 1.35 |
| NE | Tcirg1 | 1.32 | 1.76E-15 | 3.76E-11 | 0.14 | 0.06 | 0.08 | 2.19 |
| NE | Decr1 | 0.47 | 1.80E-15 | 3.85E-11 | 0.41 | 0.27 | 0.13 | 1.49 |
| NE | Tspan4 | 0.84 | 2.02E-15 | 4.32E-11 | 0.18 | 0.09 | 0.09 | 1.98 |
| NE | Umps | 0.34 | 2.11E-15 | 4.50E-11 | 0.62 | 0.47 | 0.15 | 1.31 |
| NE | Rnf208 | 0.67 | 2.33E-15 | 4.98E-11 | 0.25 | 0.15 | 0.10 | 1.70 |
| NE | Slc25a13 | 0.39 | 2.59E-15 | 5.54E-11 | 0.60 | 0.46 | 0.14 | 1.31 |
| NE | Elovl5 | 0.56 | 2.65E-15 | 5.65E-11 | 0.31 | 0.19 | 0.12 | 1.62 |
| NE | Tlcd1 | 0.82 | 2.68E-15 | 5.73E-11 | 0.17 | 0.09 | 0.09 | 2.01 |
| NE | Eci1 | 0.32 | 2.80E-15 | 5.97E-11 | 0.62 | 0.46 | 0.16 | 1.35 |
| NE | Gpm6b | 0.59 | 3.36E-15 | 7.17E-11 | 0.14 | 0.06 | 0.08 | 2.24 |
| NE | Ccdc25 | 0.37 | 3.71E-15 | 7.92E-11 | 0.68 | 0.55 | 0.13 | 1.23 |
| NE | Syce2 | 0.33 | 4.19E-15 | 8.96E-11 | 0.62 | 0.47 | 0.15 | 1.32 |
| NE | Pkia | 0.58 | 5.40E-15 | 1.15E-10 | 0.26 | 0.16 | 0.11 | 1.68 |
| NE | Aebp1 | 1.18 | 5.53E-15 | 1.18E-10 | 0.13 | 0.06 | 0.07 | 2.29 |
| NE | Mettl1 | 0.30 | 5.71E-15 | 1.22E-10 | 0.70 | 0.56 | 0.14 | 1.24 |
| NE | Prom1 | 0.44 | 5.77E-15 | 1.23E-10 | 0.53 | 0.39 | 0.14 | 1.36 |
| NE | Psmc1 | 0.27 | 5.82E-15 | 1.24E-10 | 0.83 | 0.73 | 0.10 | 1.14 |
| NE | Golm1 | 0.37 | 6.53E-15 | 1.39E-10 | 0.52 | 0.38 | 0.14 | 1.37 |
| NE | Bub1b | 0.37 | 6.61E-15 | 1.41E-10 | 0.57 | 0.44 | 0.13 | 1.30 |
| NE | Pgf | 1.01 | 7.10E-15 | 1.52E-10 | 0.14 | 0.06 | 0.08 | 2.21 |
| NE | Ide | 0.41 | 7.15E-15 | 1.53E-10 | 0.53 | 0.40 | 0.13 | 1.33 |
| NE | Pdk3 | 0.43 | 7.18E-15 | 1.53E-10 | 0.49 | 0.36 | 0.13 | 1.36 |
| NE | Chpf | 0.83 | 7.24E-15 | 1.55E-10 | 0.23 | 0.14 | 0.10 | 1.72 |
| NE | Clip1 | 0.41 | 7.26E-15 | 1.55E-10 | 0.62 | 0.48 | 0.13 | 1.27 |
| NE | Plcb4 | 0.55 | 7.27E-15 | 1.55E-10 | 0.34 | 0.22 | 0.12 | 1.52 |
| NE | Znrd1 | 0.28 | 7.53E-15 | 1.61E-10 | 0.80 | 0.68 | 0.12 | 1.18 |
| NE | Snx1 | 0.41 | 7.72E-15 | 1.65E-10 | 0.55 | 0.41 | 0.14 | 1.34 |
| NE | Hmmr | 0.30 | 8.03E-15 | 1.72E-10 | 0.54 | 0.40 | 0.14 | 1.35 |
| NE | Krtcap3 | 0.53 | 8.14E-15 | 1.74E-10 | 0.41 | 0.27 | 0.13 | 1.48 |
| NE | Fam3c | 0.42 | 8.68E-15 | 1.85E-10 | 0.55 | 0.41 | 0.14 | 1.35 |
| NE | B3gnt7 | 0.85 | 9.44E-15 | 2.02E-10 | 0.16 | 0.08 | 0.08 | 2.05 |
| NE | Rgs9 | 0.77 | 9.64E-15 | 2.06E-10 | 0.22 | 0.12 | 0.10 | 1.77 |
| NE | Tspan13 | 0.26 | 9.74E-15 | 2.08E-10 | 0.82 | 0.72 | 0.10 | 1.14 |

**Supplementary Table 4. Marker genes for the annotated cell populations of integrated clustering of E11.5 control and NotchGOF λ-junction ectoderm cells.**

| | | | | | | | | |
|---|---|---|---|---|---|---|---|---|
| NE | Sptbn1 | 0.30 | 9.85E-15 | 2.10E-10 | 0.80 | 0.70 | 0.10 | 1.15 |
| NE | Rxylt1 | 0.42 | 1.04E-14 | 2.22E-10 | 0.48 | 0.34 | 0.14 | 1.42 |
| NE | Bud23 | 0.33 | 1.04E-14 | 2.23E-10 | 0.69 | 0.57 | 0.12 | 1.20 |
| NE | Men1 | 0.48 | 1.06E-14 | 2.27E-10 | 0.48 | 0.34 | 0.14 | 1.40 |
| NE | Nfasc | 0.74 | 1.11E-14 | 2.38E-10 | 0.14 | 0.06 | 0.08 | 2.19 |
| NE | Efemp2 | 0.45 | 1.15E-14 | 2.46E-10 | 0.47 | 0.33 | 0.14 | 1.41 |
| NE | Nsfl1c | 0.26 | 1.24E-14 | 2.66E-10 | 0.86 | 0.77 | 0.09 | 1.11 |
| NE | Dcps | 0.33 | 1.26E-14 | 2.68E-10 | 0.69 | 0.56 | 0.12 | 1.22 |
| NE | Cdc25b | 0.65 | 1.28E-14 | 2.73E-10 | 0.24 | 0.14 | 0.10 | 1.71 |
| NE | Pold3 | 0.36 | 1.36E-14 | 2.91E-10 | 0.60 | 0.47 | 0.14 | 1.29 |
| NE | Lockd | 0.42 | 1.42E-14 | 3.04E-10 | 0.41 | 0.28 | 0.13 | 1.45 |
| NE | Rnf165 | 0.38 | 1.49E-14 | 3.18E-10 | 0.34 | 0.22 | 0.12 | 1.57 |
| NE | Kpna2 | 0.60 | 1.53E-14 | 3.27E-10 | 0.44 | 0.32 | 0.12 | 1.38 |
| NE | Spata24 | 0.45 | 1.58E-14 | 3.38E-10 | 0.37 | 0.24 | 0.13 | 1.52 |
| NE | Efnb3 | 0.53 | 1.60E-14 | 3.43E-10 | 0.28 | 0.17 | 0.11 | 1.66 |
| NE | Tbc1d4 | 0.80 | 1.68E-14 | 3.59E-10 | 0.14 | 0.06 | 0.08 | 2.21 |
| NE | Larp1b | 0.90 | 1.70E-14 | 3.62E-10 | 0.16 | 0.08 | 0.08 | 2.00 |
| NE | Unc13c | 1.12 | 1.75E-14 | 3.74E-10 | 0.11 | 0.04 | 0.06 | 2.56 |
| NE | Anapc15 | 0.38 | 1.76E-14 | 3.75E-10 | 0.61 | 0.47 | 0.14 | 1.30 |
| NE | Zeb2 | 0.83 | 1.77E-14 | 3.78E-10 | 0.12 | 0.05 | 0.07 | 2.36 |
| NE | 2010300C02Rik | 1.20 | 2.07E-14 | 4.42E-10 | 0.13 | 0.06 | 0.07 | 2.20 |
| NE | Fdft1 | 0.45 | 2.11E-14 | 4.51E-10 | 0.62 | 0.51 | 0.11 | 1.21 |
| NE | Fam171b | 0.64 | 2.47E-14 | 5.28E-10 | 0.18 | 0.10 | 0.09 | 1.92 |
| NE | Ing3 | 0.39 | 2.49E-14 | 5.32E-10 | 0.44 | 0.30 | 0.13 | 1.45 |
| NE | Plpbp | 0.52 | 2.50E-14 | 5.34E-10 | 0.37 | 0.25 | 0.12 | 1.49 |
| NE | Lin28b | 0.55 | 2.72E-14 | 5.82E-10 | 0.34 | 0.22 | 0.12 | 1.53 |
| NE | Sncaip | 0.79 | 2.87E-14 | 6.14E-10 | 0.19 | 0.11 | 0.09 | 1.82 |
| NE | Gins1 | 0.46 | 2.93E-14 | 6.27E-10 | 0.42 | 0.29 | 0.13 | 1.43 |
| NE | Cd82 | 0.67 | 2.98E-14 | 6.37E-10 | 0.18 | 0.10 | 0.09 | 1.92 |
| NE | Tubb4a | 0.68 | 3.01E-14 | 6.44E-10 | 0.14 | 0.06 | 0.07 | 2.18 |
| NE | Fam216a | 0.32 | 3.05E-14 | 6.50E-10 | 0.60 | 0.47 | 0.13 | 1.29 |
| NE | Tmem126a | 0.30 | 3.23E-14 | 6.89E-10 | 0.74 | 0.64 | 0.10 | 1.16 |
| NE | Arg2 | 0.58 | 3.25E-14 | 6.95E-10 | 0.24 | 0.14 | 0.10 | 1.73 |
| NE | Ube2j1 | 0.47 | 3.34E-14 | 7.13E-10 | 0.34 | 0.22 | 0.12 | 1.54 |
| NE | Cenpp | 0.50 | 3.49E-14 | 7.46E-10 | 0.40 | 0.28 | 0.12 | 1.44 |
| NE | Rad23b | 0.28 | 3.52E-14 | 7.53E-10 | 0.84 | 0.74 | 0.09 | 1.13 |
| NE | Ptpa | 0.40 | 3.58E-14 | 7.64E-10 | 0.54 | 0.40 | 0.13 | 1.33 |
| NE | Emid1 | 0.63 | 3.59E-14 | 7.67E-10 | 0.23 | 0.13 | 0.10 | 1.73 |
| NE | Prtg | 0.51 | 3.80E-14 | 8.11E-10 | 0.27 | 0.17 | 0.11 | 1.64 |
| NE | Tmbim6 | 0.25 | 3.80E-14 | 8.11E-10 | 0.89 | 0.80 | 0.09 | 1.11 |
| NE | Dtl | 0.33 | 3.85E-14 | 8.23E-10 | 0.61 | 0.48 | 0.13 | 1.27 |
| NE | Pcgf5 | 0.53 | 3.93E-14 | 8.39E-10 | 0.28 | 0.17 | 0.11 | 1.63 |
| NE | Adgrl1 | 0.41 | 4.38E-14 | 9.35E-10 | 0.55 | 0.42 | 0.13 | 1.32 |
| NE | Mrpl58 | 0.26 | 4.55E-14 | 9.72E-10 | 0.84 | 0.74 | 0.11 | 1.14 |
| NE | Tor2a | 0.44 | 4.82E-14 | 1.03E-09 | 0.48 | 0.36 | 0.12 | 1.35 |
| NE | Alkbh7 | 0.40 | 5.01E-14 | 1.07E-09 | 0.45 | 0.32 | 0.13 | 1.41 |
| NE | Gnl3 | 0.29 | 5.55E-14 | 1.19E-09 | 0.79 | 0.69 | 0.10 | 1.15 |
| NE | Mesd | 0.27 | 5.59E-14 | 1.19E-09 | 0.78 | 0.66 | 0.11 | 1.17 |
| NE | Cgn | 0.40 | 5.62E-14 | 1.20E-09 | 0.46 | 0.33 | 0.13 | 1.41 |
| NE | Hist1h1a | 0.47 | 5.67E-14 | 1.21E-09 | 0.41 | 0.29 | 0.12 | 1.43 |
| NE | Slc50a1 | 0.57 | 5.68E-14 | 1.21E-09 | 0.28 | 0.17 | 0.11 | 1.62 |
| NE | Mthfd2 | 0.33 | 6.08E-14 | 1.30E-09 | 0.35 | 0.23 | 0.12 | 1.52 |
| NE | Exo1 | 0.39 | 6.37E-14 | 1.36E-09 | 0.46 | 0.33 | 0.13 | 1.39 |
| NE | Il6st | 0.65 | 6.70E-14 | 1.43E-09 | 0.27 | 0.17 | 0.10 | 1.60 |
| NE | Coprs | 0.45 | 7.04E-14 | 1.50E-09 | 0.45 | 0.32 | 0.13 | 1.41 |

**Supplementary Table 4. Marker genes for the annotated cell populations of integrated clustering of E11.5 control and NotchGOF λ-junction ectoderm cells.**

| | | | | | | | | |
|---|---|---|---|---|---|---|---|---|
| NE | Fads2 | 0.46 | 7.24E-14 | 1.55E-09 | 0.53 | 0.40 | 0.13 | 1.32 |
| NE | Cmc2 | 0.37 | 7.40E-14 | 1.58E-09 | 0.59 | 0.46 | 0.13 | 1.27 |
| NE | Rnf121 | 0.38 | 8.40E-14 | 1.79E-09 | 0.50 | 0.37 | 0.13 | 1.35 |
| NE | Pmm1 | 0.49 | 9.16E-14 | 1.96E-09 | 0.41 | 0.29 | 0.12 | 1.41 |
| NE | Chst2 | 0.82 | 9.47E-14 | 2.02E-09 | 0.20 | 0.11 | 0.09 | 1.79 |
| NE | Timm9 | 0.33 | 9.61E-14 | 2.05E-09 | 0.65 | 0.53 | 0.12 | 1.23 |
| NE | Mtx1 | 0.33 | 1.01E-13 | 2.15E-09 | 0.59 | 0.46 | 0.13 | 1.29 |
| NE | Daam1 | 0.37 | 1.02E-13 | 2.17E-09 | 0.61 | 0.48 | 0.13 | 1.27 |
| NE | Gm19938 | 0.72 | 1.08E-13 | 2.31E-09 | 0.14 | 0.07 | 0.07 | 2.12 |
| NE | B4galt4 | 0.54 | 1.24E-13 | 2.65E-09 | 0.32 | 0.21 | 0.11 | 1.54 |
| NE | Sypl | 0.34 | 1.24E-13 | 2.65E-09 | 0.67 | 0.54 | 0.14 | 1.25 |
| NE | Pepd | 0.43 | 1.31E-13 | 2.79E-09 | 0.51 | 0.38 | 0.13 | 1.34 |
| NE | Gyg | 0.39 | 1.40E-13 | 2.99E-09 | 0.42 | 0.29 | 0.13 | 1.44 |
| NE | Bend4 | 0.84 | 1.47E-13 | 3.13E-09 | 0.18 | 0.10 | 0.08 | 1.84 |
| NE | Melk | 0.35 | 1.54E-13 | 3.29E-09 | 0.51 | 0.38 | 0.13 | 1.35 |
| NE | Polr1e | 0.47 | 1.59E-13 | 3.39E-09 | 0.32 | 0.21 | 0.11 | 1.55 |
| NE | Pde3b | 0.65 | 1.70E-13 | 3.62E-09 | 0.26 | 0.16 | 0.10 | 1.63 |
| NE | Fam102b | 1.02 | 1.82E-13 | 3.89E-09 | 0.16 | 0.08 | 0.08 | 1.93 |
| NE | Fam136a | 0.27 | 1.87E-13 | 4.00E-09 | 0.85 | 0.77 | 0.08 | 1.11 |
| NE | Dync2li1 | 0.42 | 1.94E-13 | 4.13E-09 | 0.41 | 0.29 | 0.13 | 1.45 |
| NE | Megf8 | 0.65 | 2.01E-13 | 4.29E-09 | 0.22 | 0.13 | 0.09 | 1.72 |
| NE | Gpam | 0.73 | 2.25E-13 | 4.81E-09 | 0.13 | 0.06 | 0.07 | 2.20 |
| NE | Mrps25 | 0.29 | 2.27E-13 | 4.84E-09 | 0.77 | 0.67 | 0.11 | 1.16 |
| NE | Pacsin3 | 0.49 | 2.27E-13 | 4.86E-09 | 0.38 | 0.26 | 0.12 | 1.44 |
| NE | Ahsa1 | 0.26 | 2.37E-13 | 5.06E-09 | 0.79 | 0.67 | 0.12 | 1.18 |
| NE | Notch2 | 0.51 | 2.50E-13 | 5.34E-09 | 0.37 | 0.26 | 0.12 | 1.45 |
| NE | Nars | 0.26 | 2.62E-13 | 5.61E-09 | 0.82 | 0.73 | 0.09 | 1.12 |
| NE | Cyba | 0.33 | 2.69E-13 | 5.74E-09 | 0.63 | 0.50 | 0.13 | 1.26 |
| NE | Dnajb11 | 0.32 | 2.71E-13 | 5.79E-09 | 0.70 | 0.58 | 0.12 | 1.21 |
| NE | Exosc3 | 0.28 | 2.75E-13 | 5.88E-09 | 0.77 | 0.64 | 0.12 | 1.19 |
| NE | Epdr1 | 0.84 | 3.40E-13 | 7.25E-09 | 0.15 | 0.08 | 0.07 | 1.96 |
| NE | Mydgf | 0.27 | 3.44E-13 | 7.34E-09 | 0.80 | 0.69 | 0.11 | 1.15 |
| NE | Cisd3 | 0.83 | 3.59E-13 | 7.66E-09 | 0.11 | 0.05 | 0.06 | 2.35 |
| NE | Pawr | 0.44 | 3.61E-13 | 7.70E-09 | 0.44 | 0.32 | 0.12 | 1.38 |
| NE | Cxcr4 | 0.31 | 4.46E-13 | 9.53E-09 | 0.24 | 0.14 | 0.10 | 1.69 |
| NE | Top2a | 0.27 | 4.49E-13 | 9.59E-09 | 0.77 | 0.64 | 0.13 | 1.20 |
| NE | Nup85 | 0.51 | 4.54E-13 | 9.70E-09 | 0.40 | 0.28 | 0.12 | 1.41 |
| NE | Cby1 | 0.31 | 4.60E-13 | 9.83E-09 | 0.53 | 0.40 | 0.13 | 1.33 |
| NE | Cep57 | 0.33 | 4.74E-13 | 1.01E-08 | 0.68 | 0.56 | 0.12 | 1.21 |
| NE | Cdk1 | 0.31 | 4.99E-13 | 1.07E-08 | 0.67 | 0.55 | 0.11 | 1.20 |
| NE | Srrt | 0.26 | 5.00E-13 | 1.07E-08 | 0.83 | 0.73 | 0.11 | 1.14 |
| NE | Triobp | 0.44 | 5.09E-13 | 1.09E-08 | 0.40 | 0.27 | 0.12 | 1.45 |
| NE | Galnt2 | 0.33 | 5.30E-13 | 1.13E-08 | 0.61 | 0.48 | 0.13 | 1.27 |
| NE | Ska1 | 0.38 | 5.35E-13 | 1.14E-08 | 0.41 | 0.29 | 0.12 | 1.41 |
| NE | Scpep1 | 0.45 | 5.39E-13 | 1.15E-08 | 0.36 | 0.24 | 0.12 | 1.49 |
| NE | Kars | 0.32 | 5.42E-13 | 1.16E-08 | 0.66 | 0.54 | 0.12 | 1.23 |
| NE | Hist1h3i | 0.78 | 5.50E-13 | 1.17E-08 | 0.20 | 0.11 | 0.09 | 1.75 |
| NE | Hist1h3d | 0.85 | 5.67E-13 | 1.21E-08 | 0.16 | 0.08 | 0.08 | 1.94 |
| NE | Ddx56 | 0.43 | 5.93E-13 | 1.27E-08 | 0.44 | 0.31 | 0.12 | 1.39 |
| NE | Myo5b | 0.75 | 6.14E-13 | 1.31E-08 | 0.21 | 0.12 | 0.09 | 1.70 |
| NE | Prmt5 | 0.33 | 6.62E-13 | 1.41E-08 | 0.68 | 0.57 | 0.12 | 1.21 |
| NE | Lonp1 | 0.31 | 6.77E-13 | 1.45E-08 | 0.61 | 0.47 | 0.14 | 1.29 |
| NE | Rev1 | 0.45 | 7.33E-13 | 1.57E-08 | 0.39 | 0.27 | 0.12 | 1.44 |
| NE | Rrp9 | 0.44 | 7.48E-13 | 1.60E-08 | 0.41 | 0.29 | 0.12 | 1.41 |
| NE | Ift57 | 0.49 | 8.15E-13 | 1.74E-08 | 0.38 | 0.26 | 0.11 | 1.43 |

**Supplementary Table 4. Marker genes for the annotated cell populations of integrated clustering of E11.5 control and NotchGOF λ-junction ectoderm cells.**

| | | | | | | | | |
|---|---|---|---|---|---|---|---|---|
| NE | Bcl2 | 0.57 | 8.71E-13 | 1.86E-08 | 0.31 | 0.21 | 0.10 | 1.50 |
| NE | Wdr46 | 0.36 | 8.74E-13 | 1.87E-08 | 0.55 | 0.41 | 0.14 | 1.35 |
| NE | Esco2 | 0.38 | 9.33E-13 | 1.99E-08 | 0.48 | 0.36 | 0.12 | 1.34 |
| NE | Rcbtb1 | 0.41 | 9.83E-13 | 2.10E-08 | 0.48 | 0.36 | 0.13 | 1.36 |
| NE | Gas2l3 | 0.60 | 1.10E-12 | 2.36E-08 | 0.29 | 0.19 | 0.10 | 1.52 |
| NE | Tmem176b | 0.25 | 1.11E-12 | 2.37E-08 | 0.67 | 0.55 | 0.12 | 1.23 |
| NE | Lbr | 0.36 | 1.23E-12 | 2.63E-08 | 0.61 | 0.48 | 0.13 | 1.27 |
| NE | Xylt1 | 0.95 | 1.30E-12 | 2.79E-08 | 0.15 | 0.07 | 0.07 | 1.99 |
| NE | Rsph9 | 0.53 | 1.32E-12 | 2.81E-08 | 0.16 | 0.08 | 0.08 | 1.93 |
| NE | Pycrl | 0.33 | 1.32E-12 | 2.81E-08 | 0.60 | 0.48 | 0.13 | 1.26 |
| NE | Kdelr2 | 0.26 | 1.33E-12 | 2.84E-08 | 0.84 | 0.74 | 0.10 | 1.14 |
| NE | Abcg1 | 0.51 | 1.42E-12 | 3.04E-08 | 0.35 | 0.24 | 0.11 | 1.46 |
| NE | Wwtr1 | 0.46 | 1.45E-12 | 3.09E-08 | 0.36 | 0.25 | 0.11 | 1.45 |
| NE | Sox2ot | 1.42 | 1.50E-12 | 3.20E-08 | 0.10 | 0.04 | 0.06 | 2.34 |
| NE | Fam214a | 0.69 | 1.57E-12 | 3.36E-08 | 0.27 | 0.17 | 0.10 | 1.55 |
| NE | Fxn | 0.39 | 1.60E-12 | 3.41E-08 | 0.53 | 0.40 | 0.13 | 1.32 |
| NE | Kirrel | 0.65 | 1.70E-12 | 3.64E-08 | 0.19 | 0.10 | 0.08 | 1.79 |
| NE | Mrpl40 | 0.29 | 1.77E-12 | 3.77E-08 | 0.68 | 0.55 | 0.13 | 1.23 |
| NE | Fxyd7 | 1.26 | 1.78E-12 | 3.80E-08 | 0.11 | 0.05 | 0.06 | 2.17 |
| NE | Slc6a15 | 1.04 | 1.96E-12 | 4.19E-08 | 0.12 | 0.05 | 0.06 | 2.17 |
| NE | Chpt1 | 0.42 | 1.97E-12 | 4.20E-08 | 0.37 | 0.25 | 0.11 | 1.45 |
| NE | Ddc | 1.00 | 1.97E-12 | 4.22E-08 | 0.12 | 0.06 | 0.07 | 2.10 |
| NE | Eftud2 | 0.29 | 2.04E-12 | 4.36E-08 | 0.73 | 0.61 | 0.12 | 1.19 |
| NE | D030055H07Rik | 0.88 | 2.21E-12 | 4.73E-08 | 0.10 | 0.04 | 0.06 | 2.37 |
| NE | Enkd1 | 0.51 | 2.22E-12 | 4.75E-08 | 0.32 | 0.21 | 0.11 | 1.52 |
| NE | Pola1 | 0.32 | 2.24E-12 | 4.78E-08 | 0.52 | 0.39 | 0.13 | 1.34 |
| NE | Oma1 | 0.56 | 2.27E-12 | 4.85E-08 | 0.26 | 0.16 | 0.10 | 1.60 |
| NE | Fbxo27 | 0.73 | 2.35E-12 | 5.02E-08 | 0.16 | 0.09 | 0.08 | 1.86 |
| NE | Gusb | 0.54 | 2.52E-12 | 5.39E-08 | 0.22 | 0.13 | 0.09 | 1.69 |
| NE | Tcof1 | 0.28 | 2.61E-12 | 5.58E-08 | 0.76 | 0.65 | 0.12 | 1.18 |
| NE | Setbp1 | 0.44 | 2.67E-12 | 5.69E-08 | 0.34 | 0.23 | 0.11 | 1.46 |
| NE | Gphn | 0.55 | 2.98E-12 | 6.36E-08 | 0.27 | 0.17 | 0.10 | 1.56 |
| NE | Pomt1 | 0.85 | 3.04E-12 | 6.49E-08 | 0.16 | 0.08 | 0.07 | 1.88 |
| NE | Ttyh3 | 0.34 | 3.06E-12 | 6.53E-08 | 0.53 | 0.40 | 0.13 | 1.32 |
| NE | Ccdc189 | 1.13 | 3.09E-12 | 6.60E-08 | 0.11 | 0.05 | 0.06 | 2.23 |
| NE | Mir17hg | 0.68 | 3.14E-12 | 6.71E-08 | 0.23 | 0.14 | 0.09 | 1.62 |
| NE | Sp4 | 0.40 | 3.22E-12 | 6.87E-08 | 0.36 | 0.25 | 0.11 | 1.46 |
| NE | Lpar1 | 0.42 | 3.49E-12 | 7.46E-08 | 0.40 | 0.28 | 0.12 | 1.42 |
| NE | Ctps | 0.38 | 3.51E-12 | 7.49E-08 | 0.55 | 0.44 | 0.12 | 1.27 |
| NE | Fanca | 1.01 | 3.60E-12 | 7.69E-08 | 0.14 | 0.07 | 0.07 | 1.97 |
| NE | Hey2 | 1.17 | 3.72E-12 | 7.95E-08 | 0.13 | 0.06 | 0.06 | 2.02 |
| NE | Entpd5 | 0.65 | 3.96E-12 | 8.45E-08 | 0.16 | 0.09 | 0.08 | 1.84 |
| NE | Ino80b | 0.33 | 4.10E-12 | 8.76E-08 | 0.59 | 0.47 | 0.12 | 1.26 |
| NE | G6pc3 | 0.44 | 4.25E-12 | 9.08E-08 | 0.34 | 0.23 | 0.11 | 1.48 |
| NE | Pcyt2 | 0.42 | 4.28E-12 | 9.13E-08 | 0.28 | 0.18 | 0.10 | 1.59 |
| NE | Hist1h2be | 0.69 | 4.43E-12 | 9.46E-08 | 0.24 | 0.15 | 0.09 | 1.59 |
| NE | Ncln | 0.44 | 4.71E-12 | 1.01E-07 | 0.41 | 0.30 | 0.12 | 1.40 |
| NE | 4932422M17Rik | 0.76 | 4.86E-12 | 1.04E-07 | 0.14 | 0.07 | 0.07 | 1.96 |
| NE | Atp23 | 0.59 | 4.91E-12 | 1.05E-07 | 0.24 | 0.15 | 0.09 | 1.61 |
| NE | C2cd5 | 0.48 | 5.33E-12 | 1.14E-07 | 0.30 | 0.20 | 0.10 | 1.52 |
| NE | Nfkbiz | 0.89 | 5.46E-12 | 1.17E-07 | 0.12 | 0.06 | 0.06 | 2.11 |
| NE | Fkbp2 | 0.30 | 5.59E-12 | 1.19E-07 | 0.70 | 0.58 | 0.12 | 1.20 |
| NE | Ugcg | 0.41 | 5.81E-12 | 1.24E-07 | 0.50 | 0.37 | 0.12 | 1.33 |
| NE | Azin1 | 0.30 | 6.00E-12 | 1.28E-07 | 0.63 | 0.50 | 0.13 | 1.26 |
| NE | Asf1b | 0.44 | 6.11E-12 | 1.30E-07 | 0.42 | 0.31 | 0.11 | 1.37 |

**Supplementary Table 4. Marker genes for the annotated cell populations of integrated clustering of E11.5 control and NotchGOF λ-junction ectoderm cells.**

| | | | | | | | | |
|---|---|---|---|---|---|---|---|---|
| NE | Gaa | 0.49 | 6.61E-12 | 1.41E-07 | 0.26 | 0.16 | 0.09 | 1.57 |
| NE | Mtfp1 | 0.83 | 6.74E-12 | 1.44E-07 | 0.11 | 0.05 | 0.06 | 2.20 |
| NE | Agpat5 | 0.38 | 6.77E-12 | 1.45E-07 | 0.54 | 0.41 | 0.13 | 1.30 |
| NE | Med25 | 0.33 | 6.80E-12 | 1.45E-07 | 0.57 | 0.45 | 0.12 | 1.27 |
| NE | Trib2 | 0.48 | 6.82E-12 | 1.46E-07 | 0.25 | 0.16 | 0.10 | 1.61 |
| NE | Dpy19l1 | 0.51 | 7.89E-12 | 1.69E-07 | 0.30 | 0.20 | 0.10 | 1.52 |
| NE | Taf6l | 0.45 | 9.11E-12 | 1.95E-07 | 0.33 | 0.23 | 0.11 | 1.47 |
| NE | A430005L14Rik | 0.33 | 9.38E-12 | 2.00E-07 | 0.57 | 0.44 | 0.13 | 1.29 |
| NE | Tacc2 | 0.36 | 9.57E-12 | 2.04E-07 | 0.60 | 0.48 | 0.12 | 1.26 |
| NE | Lpl | 1.01 | 9.81E-12 | 2.10E-07 | 0.11 | 0.05 | 0.06 | 2.15 |
| NE | Pcbd2 | 0.34 | 1.07E-11 | 2.28E-07 | 0.51 | 0.39 | 0.12 | 1.32 |
| NE | Ocln | 0.34 | 1.08E-11 | 2.30E-07 | 0.37 | 0.25 | 0.12 | 1.48 |
| NE | Tapt1 | 0.63 | 1.12E-11 | 2.39E-07 | 0.20 | 0.12 | 0.08 | 1.68 |
| NE | Trmt2a | 0.40 | 1.14E-11 | 2.44E-07 | 0.36 | 0.25 | 0.11 | 1.43 |
| NE | Fbxo44 | 0.63 | 1.16E-11 | 2.48E-07 | 0.17 | 0.10 | 0.08 | 1.80 |
| NE | Tusc2 | 0.36 | 1.19E-11 | 2.53E-07 | 0.53 | 0.40 | 0.13 | 1.32 |
| NE | Cers2 | 0.37 | 1.19E-11 | 2.55E-07 | 0.53 | 0.39 | 0.13 | 1.33 |
| NE | Nr2f6 | 0.26 | 1.22E-11 | 2.60E-07 | 0.80 | 0.70 | 0.10 | 1.14 |
| NE | Ecpas | 0.37 | 1.23E-11 | 2.62E-07 | 0.48 | 0.36 | 0.12 | 1.34 |
| NE | Sgpp1 | 0.90 | 1.24E-11 | 2.64E-07 | 0.14 | 0.08 | 0.07 | 1.91 |
| NE | Hspa14 | 0.31 | 1.25E-11 | 2.67E-07 | 0.69 | 0.58 | 0.11 | 1.19 |
| NE | Wdr76 | 0.39 | 1.31E-11 | 2.79E-07 | 0.42 | 0.30 | 0.12 | 1.40 |
| NE | Hist1h3e | 0.45 | 1.32E-11 | 2.83E-07 | 0.41 | 0.30 | 0.11 | 1.36 |
| NE | Med9 | 0.30 | 1.41E-11 | 3.01E-07 | 0.65 | 0.52 | 0.13 | 1.25 |
| NE | Ung | 0.33 | 1.43E-11 | 3.05E-07 | 0.44 | 0.32 | 0.12 | 1.37 |
| NE | Bysl | 0.48 | 1.55E-11 | 3.32E-07 | 0.35 | 0.24 | 0.11 | 1.44 |
| NE | H2-Ke6 | 0.50 | 1.55E-11 | 3.32E-07 | 0.32 | 0.21 | 0.10 | 1.48 |
| NE | Cfap410 | 0.53 | 1.65E-11 | 3.53E-07 | 0.28 | 0.18 | 0.10 | 1.53 |
| NE | Zswim5 | 0.83 | 1.68E-11 | 3.58E-07 | 0.13 | 0.07 | 0.07 | 1.97 |
| NE | Ppp1r35 | 0.33 | 1.72E-11 | 3.68E-07 | 0.57 | 0.45 | 0.12 | 1.26 |
| NE | Rfc5 | 0.29 | 1.77E-11 | 3.79E-07 | 0.65 | 0.53 | 0.12 | 1.22 |
| NE | Mgat4a | 0.59 | 1.85E-11 | 3.96E-07 | 0.25 | 0.17 | 0.09 | 1.53 |
| NE | Nuf2 | 0.36 | 1.88E-11 | 4.01E-07 | 0.45 | 0.33 | 0.12 | 1.35 |
| NE | Phf21b | 0.73 | 1.95E-11 | 4.17E-07 | 0.10 | 0.05 | 0.06 | 2.24 |
| NE | Plppr1 | 0.82 | 2.39E-11 | 5.11E-07 | 0.13 | 0.07 | 0.07 | 1.96 |
| NE | Unc13b | 0.42 | 2.55E-11 | 5.44E-07 | 0.30 | 0.20 | 0.10 | 1.49 |
| NE | Trip6 | 0.30 | 2.68E-11 | 5.73E-07 | 0.57 | 0.44 | 0.13 | 1.30 |
| NE | Rpn1 | 0.26 | 2.74E-11 | 5.86E-07 | 0.79 | 0.69 | 0.10 | 1.15 |
| NE | Cdc14a | 0.42 | 2.91E-11 | 6.22E-07 | 0.43 | 0.33 | 0.11 | 1.32 |
| NE | Prkd1 | 0.99 | 2.94E-11 | 6.29E-07 | 0.12 | 0.06 | 0.06 | 2.05 |
| NE | Tacc3 | 0.27 | 2.94E-11 | 6.29E-07 | 0.59 | 0.47 | 0.12 | 1.26 |
| NE | Trip10 | 0.45 | 3.06E-11 | 6.54E-07 | 0.31 | 0.21 | 0.10 | 1.49 |
| NE | Zc3h8 | 0.55 | 3.30E-11 | 7.05E-07 | 0.21 | 0.13 | 0.08 | 1.66 |
| NE | Mars | 0.40 | 3.39E-11 | 7.24E-07 | 0.40 | 0.28 | 0.11 | 1.39 |
| NE | Acsl4 | 0.63 | 3.46E-11 | 7.39E-07 | 0.17 | 0.10 | 0.07 | 1.76 |
| NE | Cdca7l | 0.31 | 3.62E-11 | 7.72E-07 | 0.55 | 0.43 | 0.12 | 1.27 |
| NE | Mvd | 0.39 | 3.84E-11 | 8.20E-07 | 0.35 | 0.25 | 0.10 | 1.42 |
| NE | Ccdc88c | 0.27 | 4.02E-11 | 8.59E-07 | 0.65 | 0.51 | 0.14 | 1.26 |
| NE | Utp11 | 0.25 | 4.16E-11 | 8.88E-07 | 0.78 | 0.68 | 0.10 | 1.15 |
| NE | Tnfrsf12a | 0.48 | 4.27E-11 | 9.12E-07 | 0.28 | 0.19 | 0.10 | 1.52 |
| NE | Trpm3 | 0.60 | 4.33E-11 | 9.25E-07 | 0.21 | 0.13 | 0.08 | 1.62 |
| NE | Abca3 | 0.53 | 4.35E-11 | 9.30E-07 | 0.21 | 0.13 | 0.08 | 1.65 |
| NE | Mogs | 0.37 | 4.93E-11 | 1.05E-06 | 0.44 | 0.32 | 0.12 | 1.37 |
| NE | Grik5 | 0.46 | 5.10E-11 | 1.09E-06 | 0.27 | 0.18 | 0.09 | 1.53 |
| NE | Tbc1d1 | 0.39 | 5.30E-11 | 1.13E-06 | 0.48 | 0.37 | 0.11 | 1.30 |

**Supplementary Table 4. Marker genes for the annotated cell populations of integrated clustering of E11.5 control and NotchGOF λ-junction ectoderm cells.**

| | | | | | | | | |
|---|---|---|---|---|---|---|---|---|
| NE | Hist1h2bm | 0.57 | 5.50E-11 | 1.18E-06 | 0.27 | 0.18 | 0.09 | 1.49 |
| NE | B930095G15Rik | 0.55 | 5.65E-11 | 1.21E-06 | 0.20 | 0.12 | 0.08 | 1.67 |
| NE | Gpc4 | 0.38 | 5.78E-11 | 1.24E-06 | 0.37 | 0.26 | 0.11 | 1.44 |
| NE | Ccn1 | 0.80 | 5.79E-11 | 1.24E-06 | 0.24 | 0.16 | 0.08 | 1.51 |
| NE | Ly6e | 0.31 | 5.79E-11 | 1.24E-06 | 0.47 | 0.34 | 0.13 | 1.37 |
| NE | Cep126 | 0.59 | 6.03E-11 | 1.29E-06 | 0.18 | 0.10 | 0.08 | 1.72 |
| NE | Gnptg | 0.38 | 6.08E-11 | 1.30E-06 | 0.37 | 0.26 | 0.11 | 1.42 |
| NE | Mrpl4 | 0.28 | 6.10E-11 | 1.30E-06 | 0.63 | 0.50 | 0.13 | 1.26 |
| NE | Tagln2 | 0.66 | 6.37E-11 | 1.36E-06 | 0.33 | 0.24 | 0.10 | 1.40 |
| NE | Rnf139 | 0.55 | 6.47E-11 | 1.38E-06 | 0.31 | 0.22 | 0.10 | 1.44 |
| NE | 2310015A10Rik | 0.82 | 6.57E-11 | 1.40E-06 | 0.14 | 0.07 | 0.06 | 1.89 |
| NE | Slc29a1 | 0.81 | 7.10E-11 | 1.52E-06 | 0.13 | 0.07 | 0.06 | 1.94 |
| NE | Tmx4 | 0.40 | 7.11E-11 | 1.52E-06 | 0.53 | 0.42 | 0.12 | 1.28 |
| NE | Tmem107 | 0.35 | 7.18E-11 | 1.53E-06 | 0.38 | 0.27 | 0.11 | 1.40 |
| NE | Grhl1 | 0.91 | 7.45E-11 | 1.59E-06 | 0.12 | 0.06 | 0.06 | 1.98 |
| NE | Irs1 | 0.46 | 7.53E-11 | 1.61E-06 | 0.37 | 0.26 | 0.11 | 1.40 |
| NE | Slc19a1 | 0.57 | 7.55E-11 | 1.61E-06 | 0.19 | 0.11 | 0.08 | 1.72 |
| NE | Cipc | 0.48 | 7.83E-11 | 1.67E-06 | 0.28 | 0.18 | 0.10 | 1.55 |
| NE | Nxt1 | 0.27 | 7.98E-11 | 1.70E-06 | 0.69 | 0.57 | 0.12 | 1.21 |
| NE | Camkmt | 0.42 | 8.20E-11 | 1.75E-06 | 0.29 | 0.19 | 0.10 | 1.50 |
| NE | Ctdnep1 | 0.28 | 8.31E-11 | 1.78E-06 | 0.67 | 0.55 | 0.12 | 1.21 |
| NE | Tex30 | 0.33 | 8.46E-11 | 1.81E-06 | 0.50 | 0.38 | 0.12 | 1.32 |
| NE | Slc2a4 | 0.74 | 8.62E-11 | 1.84E-06 | 0.16 | 0.09 | 0.07 | 1.78 |
| NE | Ccdc85c | 0.56 | 8.82E-11 | 1.89E-06 | 0.25 | 0.17 | 0.09 | 1.53 |
| NE | Arfip2 | 0.30 | 8.98E-11 | 1.92E-06 | 0.59 | 0.46 | 0.13 | 1.27 |
| NE | Scrn2 | 0.42 | 9.61E-11 | 2.05E-06 | 0.46 | 0.35 | 0.11 | 1.32 |
| NE | Dgcr2 | 0.36 | 9.66E-11 | 2.06E-06 | 0.55 | 0.43 | 0.12 | 1.28 |
| NE | Ppat | 0.35 | 9.73E-11 | 2.08E-06 | 0.51 | 0.40 | 0.11 | 1.28 |
| NE | Fam169a | 0.65 | 9.91E-11 | 2.12E-06 | 0.19 | 0.11 | 0.08 | 1.67 |
| NE | Wdr6 | 0.26 | 1.02E-10 | 2.18E-06 | 0.72 | 0.60 | 0.12 | 1.20 |
| NE | Gla | 0.77 | 1.03E-10 | 2.21E-06 | 0.15 | 0.08 | 0.07 | 1.81 |
| NE | Dctd | 0.33 | 1.07E-10 | 2.28E-06 | 0.48 | 0.36 | 0.11 | 1.32 |
| NE | Rpp25l | 0.25 | 1.13E-10 | 2.42E-06 | 0.77 | 0.66 | 0.11 | 1.16 |
| NE | Aarsd1 | 0.26 | 1.17E-10 | 2.49E-06 | 0.74 | 0.63 | 0.11 | 1.17 |
| NE | Ska3 | 0.35 | 1.17E-10 | 2.50E-06 | 0.31 | 0.21 | 0.10 | 1.47 |
| NE | St3gal5 | 0.48 | 1.21E-10 | 2.57E-06 | 0.29 | 0.20 | 0.09 | 1.47 |
| NE | Cdca5 | 0.33 | 1.21E-10 | 2.58E-06 | 0.39 | 0.28 | 0.11 | 1.39 |
| NE | March9 | 0.28 | 1.21E-10 | 2.59E-06 | 0.53 | 0.40 | 0.13 | 1.32 |
| NE | Mpi | 0.53 | 1.24E-10 | 2.66E-06 | 0.22 | 0.14 | 0.08 | 1.60 |
| NE | Ggact | 0.82 | 1.29E-10 | 2.75E-06 | 0.11 | 0.05 | 0.06 | 2.06 |
| NE | Ccdc134 | 0.53 | 1.30E-10 | 2.77E-06 | 0.19 | 0.11 | 0.08 | 1.69 |
| NE | Tmem128 | 0.26 | 1.30E-10 | 2.79E-06 | 0.73 | 0.62 | 0.11 | 1.18 |
| NE | Lrrc75a | 0.43 | 1.31E-10 | 2.81E-06 | 0.25 | 0.16 | 0.09 | 1.56 |
| NE | Fat4 | 0.68 | 1.33E-10 | 2.83E-06 | 0.15 | 0.08 | 0.07 | 1.81 |
| NE | Thap3 | 0.33 | 1.33E-10 | 2.84E-06 | 0.36 | 0.25 | 0.11 | 1.44 |
| NE | Gsdme | 0.55 | 1.34E-10 | 2.87E-06 | 0.24 | 0.15 | 0.09 | 1.56 |
| NE | Gpr27 | 0.45 | 1.36E-10 | 2.91E-06 | 0.21 | 0.13 | 0.08 | 1.61 |
| NE | Pspc1 | 0.29 | 1.40E-10 | 3.00E-06 | 0.67 | 0.56 | 0.11 | 1.19 |
| NE | Acsl5 | 0.38 | 1.42E-10 | 3.03E-06 | 0.38 | 0.28 | 0.10 | 1.38 |
| NE | Heatr1 | 0.40 | 1.47E-10 | 3.14E-06 | 0.35 | 0.25 | 0.10 | 1.42 |
| NE | Tmem237 | 0.27 | 1.47E-10 | 3.14E-06 | 0.48 | 0.37 | 0.11 | 1.30 |
| NE | Hist1h3a | 0.68 | 1.50E-10 | 3.20E-06 | 0.17 | 0.10 | 0.07 | 1.76 |
| NE | Tmed1 | 0.27 | 1.52E-10 | 3.24E-06 | 0.72 | 0.60 | 0.11 | 1.19 |
| NE | Farsa | 0.27 | 1.55E-10 | 3.30E-06 | 0.65 | 0.53 | 0.12 | 1.23 |
| NE | Add3 | 0.30 | 1.55E-10 | 3.31E-06 | 0.66 | 0.54 | 0.12 | 1.21 |

**Supplementary Table 4. Marker genes for the annotated cell populations of integrated clustering of E11.5 control and NotchGOF λ-junction ectoderm cells.**

| | | | | | | | | |
|---|---|---|---|---|---|---|---|---|
| NE | Blm | 0.28 | 1.58E-10 | 3.37E-06 | 0.53 | 0.41 | 0.12 | 1.30 |
| NE | Dvl1 | 0.50 | 1.58E-10 | 3.37E-06 | 0.24 | 0.16 | 0.09 | 1.55 |
| NE | Clpb | 0.33 | 1.73E-10 | 3.70E-06 | 0.43 | 0.31 | 0.12 | 1.38 |
| NE | Gm10184 | 0.77 | 1.74E-10 | 3.73E-06 | 0.16 | 0.09 | 0.07 | 1.74 |
| NE | Snx17 | 0.25 | 1.76E-10 | 3.75E-06 | 0.70 | 0.58 | 0.11 | 1.19 |
| NE | Qtrt1 | 0.37 | 1.86E-10 | 3.98E-06 | 0.33 | 0.23 | 0.10 | 1.43 |
| NE | Hist2h2be | 0.81 | 1.86E-10 | 3.98E-06 | 0.10 | 0.05 | 0.06 | 2.12 |
| NE | Incenp | 0.30 | 1.91E-10 | 4.08E-06 | 0.59 | 0.49 | 0.10 | 1.21 |
| NE | Aen | 0.33 | 1.92E-10 | 4.10E-06 | 0.45 | 0.34 | 0.12 | 1.35 |
| NE | Rnf26 | 0.40 | 1.94E-10 | 4.15E-06 | 0.38 | 0.27 | 0.10 | 1.38 |
| NE | Mn1 | 0.66 | 1.98E-10 | 4.23E-06 | 0.21 | 0.13 | 0.08 | 1.61 |
| NE | Slc29a2 | 0.82 | 2.02E-10 | 4.31E-06 | 0.15 | 0.08 | 0.07 | 1.79 |
| NE | Amd1 | 0.29 | 2.15E-10 | 4.59E-06 | 0.58 | 0.46 | 0.12 | 1.26 |
| NE | Galnt14 | 0.71 | 2.15E-10 | 4.60E-06 | 0.13 | 0.07 | 0.06 | 1.90 |
| NE | Chaf1b | 0.32 | 2.16E-10 | 4.62E-06 | 0.54 | 0.42 | 0.12 | 1.28 |
| NE | Crat | 0.68 | 2.25E-10 | 4.80E-06 | 0.18 | 0.11 | 0.07 | 1.68 |
| NE | Myh14 | 0.41 | 2.27E-10 | 4.85E-06 | 0.36 | 0.26 | 0.11 | 1.41 |
| NE | Arhgap19 | 0.33 | 2.28E-10 | 4.86E-06 | 0.33 | 0.22 | 0.10 | 1.47 |
| NE | Ngdn | 0.28 | 2.40E-10 | 5.12E-06 | 0.57 | 0.46 | 0.12 | 1.26 |
| NE | Eps8l1 | 0.74 | 2.44E-10 | 5.21E-06 | 0.16 | 0.10 | 0.07 | 1.71 |
| NE | Cep55 | 0.49 | 2.45E-10 | 5.23E-06 | 0.32 | 0.22 | 0.10 | 1.43 |
| NE | Gmds | 0.38 | 2.61E-10 | 5.59E-06 | 0.34 | 0.24 | 0.10 | 1.44 |
| NE | mt-Nd6 | 0.32 | 2.77E-10 | 5.91E-06 | 0.57 | 0.46 | 0.11 | 1.24 |
| NE | Mrps18b | 0.27 | 2.77E-10 | 5.92E-06 | 0.57 | 0.44 | 0.13 | 1.28 |
| NE | Ctps2 | 0.42 | 2.82E-10 | 6.03E-06 | 0.32 | 0.23 | 0.10 | 1.43 |
| NE | Plod2 | 0.28 | 2.84E-10 | 6.07E-06 | 0.54 | 0.43 | 0.11 | 1.26 |
| NE | Wdr77 | 0.30 | 2.92E-10 | 6.24E-06 | 0.63 | 0.52 | 0.12 | 1.23 |
| NE | Fam110b | 0.56 | 2.94E-10 | 6.28E-06 | 0.26 | 0.17 | 0.09 | 1.50 |
| NE | Setd6 | 0.61 | 2.95E-10 | 6.30E-06 | 0.18 | 0.11 | 0.07 | 1.69 |
| NE | Mccc2 | 0.32 | 3.06E-10 | 6.53E-06 | 0.40 | 0.29 | 0.11 | 1.39 |
| NE | Wdr35 | 0.39 | 3.11E-10 | 6.65E-06 | 0.30 | 0.20 | 0.10 | 1.48 |
| NE | Aven | 0.32 | 3.14E-10 | 6.70E-06 | 0.38 | 0.27 | 0.11 | 1.42 |
| NE | Pgd | 0.27 | 3.25E-10 | 6.94E-06 | 0.64 | 0.51 | 0.12 | 1.24 |
| NE | Kif20b | 0.31 | 3.30E-10 | 7.05E-06 | 0.58 | 0.47 | 0.11 | 1.24 |
| NE | Slc1a5 | 0.45 | 3.33E-10 | 7.11E-06 | 0.25 | 0.17 | 0.09 | 1.51 |
| NE | Nacc1 | 0.36 | 3.37E-10 | 7.21E-06 | 0.48 | 0.37 | 0.11 | 1.30 |
| NE | Xrcc6 | 0.33 | 3.39E-10 | 7.23E-06 | 0.47 | 0.35 | 0.12 | 1.34 |
| NE | Gclc | 0.42 | 3.44E-10 | 7.34E-06 | 0.33 | 0.23 | 0.10 | 1.43 |
| NE | Hlcs | 0.45 | 3.45E-10 | 7.38E-06 | 0.23 | 0.15 | 0.08 | 1.56 |
| NE | Spc25 | 0.30 | 3.45E-10 | 7.38E-06 | 0.53 | 0.42 | 0.11 | 1.26 |
| NE | Nphp1 | 0.33 | 3.71E-10 | 7.91E-06 | 0.34 | 0.25 | 0.10 | 1.40 |
| NE | Adsl | 0.29 | 3.88E-10 | 8.30E-06 | 0.56 | 0.45 | 0.12 | 1.26 |
| NE | Nde1 | 0.27 | 3.93E-10 | 8.40E-06 | 0.65 | 0.55 | 0.11 | 1.19 |
| NE | Jpt2 | 0.28 | 3.93E-10 | 8.40E-06 | 0.56 | 0.44 | 0.12 | 1.28 |
| NE | Endog | 0.55 | 3.93E-10 | 8.40E-06 | 0.24 | 0.15 | 0.08 | 1.55 |
| NE | Acaa1a | 0.32 | 4.30E-10 | 9.19E-06 | 0.52 | 0.41 | 0.12 | 1.29 |
| NE | Hist1h1d | 0.40 | 4.54E-10 | 9.71E-06 | 0.39 | 0.29 | 0.10 | 1.34 |
| NE | Hist1h2bb | 0.57 | 4.62E-10 | 9.86E-06 | 0.20 | 0.13 | 0.08 | 1.59 |
| NE | Gpr19 | 1.00 | 4.83E-10 | 1.03E-05 | 0.10 | 0.05 | 0.05 | 2.04 |
| NE | Cenpu | 0.38 | 4.92E-10 | 1.05E-05 | 0.31 | 0.22 | 0.10 | 1.44 |
| NE | Prpsap2 | 0.37 | 4.93E-10 | 1.05E-05 | 0.31 | 0.21 | 0.10 | 1.45 |
| NE | Twf2 | 0.33 | 5.15E-10 | 1.10E-05 | 0.42 | 0.31 | 0.11 | 1.35 |
| NE | Samd4 | 0.53 | 5.51E-10 | 1.18E-05 | 0.21 | 0.14 | 0.08 | 1.57 |
| NE | Garem1 | 0.64 | 5.62E-10 | 1.20E-05 | 0.18 | 0.11 | 0.07 | 1.66 |
| NE | Cse1l | 0.27 | 5.93E-10 | 1.27E-05 | 0.65 | 0.55 | 0.10 | 1.19 |

**Supplementary Table 4. Marker genes for the annotated cell populations of integrated clustering of E11.5 control and NotchGOF λ-junction ectoderm cells.**

| | | | | | | | | |
|---|---|---|---|---|---|---|---|---|
| NE | Aars | 0.31 | 6.17E-10 | 1.32E-05 | 0.46 | 0.34 | 0.11 | 1.33 |
| NE | Zcchc14 | 0.34 | 6.30E-10 | 1.35E-05 | 0.39 | 0.28 | 0.11 | 1.38 |
| NE | Wdcp | 0.48 | 6.32E-10 | 1.35E-05 | 0.36 | 0.26 | 0.10 | 1.39 |
| NE | Praf2 | 0.26 | 6.46E-10 | 1.38E-05 | 0.49 | 0.37 | 0.12 | 1.32 |
| NE | Wrap53 | 0.44 | 6.57E-10 | 1.40E-05 | 0.26 | 0.18 | 0.09 | 1.51 |
| NE | Gemin5 | 0.38 | 6.74E-10 | 1.44E-05 | 0.40 | 0.29 | 0.11 | 1.36 |
| NE | Cinp | 0.32 | 6.84E-10 | 1.46E-05 | 0.51 | 0.41 | 0.11 | 1.27 |
| NE | Shisa4 | 0.40 | 6.85E-10 | 1.46E-05 | 0.29 | 0.20 | 0.09 | 1.46 |
| NE | Elf3 | 0.56 | 7.11E-10 | 1.52E-05 | 0.22 | 0.14 | 0.08 | 1.55 |
| NE | Bora | 0.44 | 7.27E-10 | 1.55E-05 | 0.44 | 0.33 | 0.10 | 1.31 |
| NE | Me2 | 0.31 | 7.72E-10 | 1.65E-05 | 0.45 | 0.34 | 0.12 | 1.34 |
| NE | Wdhd1 | 0.47 | 7.90E-10 | 1.69E-05 | 0.30 | 0.21 | 0.09 | 1.44 |
| NE | Wdr4 | 0.40 | 8.16E-10 | 1.74E-05 | 0.35 | 0.25 | 0.10 | 1.39 |
| NE | Hist1h4d | 0.36 | 8.64E-10 | 1.84E-05 | 0.45 | 0.34 | 0.11 | 1.32 |
| NE | Syngr2 | 0.36 | 8.89E-10 | 1.90E-05 | 0.33 | 0.23 | 0.10 | 1.43 |
| NE | Fkbp11 | 0.52 | 8.92E-10 | 1.90E-05 | 0.23 | 0.15 | 0.08 | 1.55 |
| NE | Cfap298 | 0.29 | 9.01E-10 | 1.92E-05 | 0.49 | 0.38 | 0.12 | 1.31 |
| NE | Dennd1b | 0.41 | 9.10E-10 | 1.94E-05 | 0.31 | 0.22 | 0.10 | 1.45 |
| NE | Aurka | 0.28 | 9.33E-10 | 1.99E-05 | 0.45 | 0.34 | 0.11 | 1.32 |
| NE | Heatr3 | 0.38 | 9.44E-10 | 2.02E-05 | 0.38 | 0.27 | 0.10 | 1.37 |
| NE | Cntln | 0.28 | 9.47E-10 | 2.02E-05 | 0.46 | 0.35 | 0.11 | 1.32 |
| NE | Acadvl | 0.39 | 9.70E-10 | 2.07E-05 | 0.31 | 0.22 | 0.09 | 1.43 |
| NE | Fbxl14 | 0.30 | 9.91E-10 | 2.12E-05 | 0.51 | 0.40 | 0.12 | 1.29 |
| NE | B4galnt4 | 0.39 | 1.02E-09 | 2.17E-05 | 0.35 | 0.25 | 0.10 | 1.38 |
| NE | Tbc1d30 | 0.64 | 1.02E-09 | 2.17E-05 | 0.15 | 0.08 | 0.06 | 1.79 |
| NE | Bcar3 | 0.75 | 1.03E-09 | 2.20E-05 | 0.14 | 0.08 | 0.06 | 1.76 |
| NE | Chsy1 | 0.26 | 1.04E-09 | 2.21E-05 | 0.40 | 0.28 | 0.11 | 1.40 |
| NE | Keap1 | 0.29 | 1.04E-09 | 2.21E-05 | 0.51 | 0.40 | 0.12 | 1.29 |
| NE | Tob1 | 0.34 | 1.05E-09 | 2.25E-05 | 0.29 | 0.20 | 0.09 | 1.47 |
| NE | Pbk | 0.30 | 1.12E-09 | 2.40E-05 | 0.55 | 0.44 | 0.11 | 1.26 |
| NE | Rpp30 | 0.32 | 1.14E-09 | 2.43E-05 | 0.58 | 0.47 | 0.11 | 1.23 |
| NE | Ubtd1 | 0.37 | 1.16E-09 | 2.47E-05 | 0.18 | 0.11 | 0.07 | 1.68 |
| NE | Ift22 | 0.27 | 1.17E-09 | 2.49E-05 | 0.45 | 0.33 | 0.12 | 1.35 |
| NE | Nbn | 0.34 | 1.19E-09 | 2.54E-05 | 0.39 | 0.29 | 0.10 | 1.35 |
| NE | Myg1 | 0.27 | 1.21E-09 | 2.58E-05 | 0.66 | 0.55 | 0.11 | 1.19 |
| NE | Mxra7 | 0.38 | 1.27E-09 | 2.71E-05 | 0.22 | 0.14 | 0.08 | 1.56 |
| NE | Kti12 | 0.33 | 1.35E-09 | 2.88E-05 | 0.50 | 0.39 | 0.11 | 1.28 |
| NE | Hist2h2bb | 0.51 | 1.36E-09 | 2.90E-05 | 0.26 | 0.18 | 0.08 | 1.47 |
| NE | Ptdss2 | 0.28 | 1.40E-09 | 2.98E-05 | 0.52 | 0.41 | 0.12 | 1.29 |
| NE | Aunip | 0.42 | 1.40E-09 | 3.00E-05 | 0.28 | 0.20 | 0.09 | 1.45 |
| NE | Rbfa | 0.31 | 1.41E-09 | 3.00E-05 | 0.41 | 0.30 | 0.11 | 1.35 |
| NE | Ndufv1 | 0.27 | 1.49E-09 | 3.19E-05 | 0.76 | 0.67 | 0.09 | 1.14 |
| NE | Ankrd9 | 0.60 | 1.50E-09 | 3.21E-05 | 0.17 | 0.10 | 0.07 | 1.67 |
| NE | Fkbp14 | 0.48 | 1.53E-09 | 3.27E-05 | 0.18 | 0.11 | 0.07 | 1.68 |
| NE | Six6 | 0.41 | 1.53E-09 | 3.28E-05 | 0.33 | 0.24 | 0.08 | 1.34 |
| NE | Snapc2 | 0.30 | 1.56E-09 | 3.33E-05 | 0.36 | 0.26 | 0.10 | 1.39 |
| NE | Gm26917 | 0.32 | 1.63E-09 | 3.47E-05 | 0.70 | 0.60 | 0.10 | 1.16 |
| NE | Psrc1 | 0.27 | 1.67E-09 | 3.57E-05 | 0.40 | 0.30 | 0.11 | 1.36 |
| NE | Fam20c | 0.58 | 1.70E-09 | 3.63E-05 | 0.11 | 0.06 | 0.06 | 1.95 |
| NE | Ncapg | 0.35 | 1.71E-09 | 3.65E-05 | 0.48 | 0.37 | 0.11 | 1.29 |
| NE | Dis3 | 0.27 | 1.71E-09 | 3.65E-05 | 0.56 | 0.45 | 0.11 | 1.25 |
| NE | Hspbp1 | 0.28 | 1.73E-09 | 3.69E-05 | 0.49 | 0.38 | 0.11 | 1.30 |
| NE | Nek2 | 0.29 | 1.76E-09 | 3.75E-05 | 0.29 | 0.20 | 0.09 | 1.45 |
| NE | Ddx21 | 0.26 | 1.82E-09 | 3.88E-05 | 0.71 | 0.62 | 0.09 | 1.15 |
| NE | Rnaseh2b | 0.26 | 1.88E-09 | 4.01E-05 | 0.58 | 0.46 | 0.12 | 1.27 |

**Supplementary Table 4. Marker genes for the annotated cell populations of integrated clustering of E11.5 control and NotchGOF λ-junction ectoderm cells.**

| | | | | | | | | |
|---|---|---|---|---|---|---|---|---|
| NE | Stc1 | 0.82 | 1.88E-09 | 4.02E-05 | 0.11 | 0.06 | 0.05 | 1.92 |
| NE | Htra2 | 0.29 | 1.90E-09 | 4.05E-05 | 0.42 | 0.31 | 0.11 | 1.35 |
| NE | Hsph1 | 0.26 | 1.99E-09 | 4.26E-05 | 0.70 | 0.61 | 0.09 | 1.15 |
| NE | Gtf2h4 | 0.29 | 2.02E-09 | 4.32E-05 | 0.46 | 0.34 | 0.12 | 1.35 |
| NE | Fbxo17 | 0.50 | 2.05E-09 | 4.38E-05 | 0.20 | 0.13 | 0.08 | 1.59 |
| NE | Sipa1 | 0.85 | 2.11E-09 | 4.50E-05 | 0.11 | 0.05 | 0.05 | 1.96 |
| NE | Ifrd2 | 0.40 | 2.12E-09 | 4.52E-05 | 0.35 | 0.25 | 0.10 | 1.40 |
| NE | Cetn2 | 0.25 | 2.16E-09 | 4.61E-05 | 0.66 | 0.55 | 0.10 | 1.19 |
| NE | Fgfbp3 | 0.74 | 2.17E-09 | 4.63E-05 | 0.17 | 0.10 | 0.07 | 1.65 |
| NE | Ydjc | 0.49 | 2.22E-09 | 4.75E-05 | 0.16 | 0.09 | 0.07 | 1.73 |
| NE | Rad51b | 0.60 | 2.23E-09 | 4.77E-05 | 0.17 | 0.11 | 0.07 | 1.66 |
| NE | Ppt1 | 0.36 | 2.27E-09 | 4.84E-05 | 0.42 | 0.32 | 0.10 | 1.32 |
| NE | H2-D1 | 0.29 | 2.27E-09 | 4.84E-05 | 0.39 | 0.29 | 0.11 | 1.37 |
| NE | Spred3 | 0.52 | 2.31E-09 | 4.93E-05 | 0.18 | 0.11 | 0.07 | 1.62 |
| NE | Pex10 | 0.51 | 2.33E-09 | 4.97E-05 | 0.21 | 0.14 | 0.08 | 1.56 |
| NE | 2310061I04Rik | 0.36 | 2.43E-09 | 5.19E-05 | 0.26 | 0.18 | 0.09 | 1.48 |
| NE | Rab11fip4 | 0.35 | 2.51E-09 | 5.37E-05 | 0.25 | 0.17 | 0.08 | 1.50 |
| NE | Lrpprc | 0.30 | 2.59E-09 | 5.52E-05 | 0.52 | 0.40 | 0.11 | 1.28 |
| NE | Bpgm | 0.33 | 2.62E-09 | 5.59E-05 | 0.49 | 0.38 | 0.11 | 1.29 |
| NE | Uvrag | 0.32 | 2.64E-09 | 5.63E-05 | 0.31 | 0.21 | 0.09 | 1.43 |
| NE | Dnajb14 | 0.34 | 2.66E-09 | 5.68E-05 | 0.38 | 0.28 | 0.10 | 1.38 |
| NE | Evi5 | 0.29 | 2.67E-09 | 5.70E-05 | 0.53 | 0.42 | 0.11 | 1.26 |
| NE | Nrm | 0.39 | 2.76E-09 | 5.90E-05 | 0.44 | 0.33 | 0.10 | 1.31 |
| NE | Las1l | 0.26 | 2.77E-09 | 5.93E-05 | 0.58 | 0.46 | 0.12 | 1.26 |
| NE | Cenpn | 0.32 | 2.79E-09 | 5.95E-05 | 0.39 | 0.29 | 0.10 | 1.34 |
| NE | Haus4 | 0.34 | 2.81E-09 | 6.01E-05 | 0.27 | 0.18 | 0.09 | 1.47 |
| NE | Amer1 | 0.32 | 3.05E-09 | 6.51E-05 | 0.65 | 0.54 | 0.11 | 1.20 |
| NE | Aurkb | 0.32 | 3.08E-09 | 6.58E-05 | 0.50 | 0.40 | 0.10 | 1.26 |
| NE | Tjp3 | 0.43 | 3.19E-09 | 6.80E-05 | 0.23 | 0.16 | 0.08 | 1.51 |
| NE | Pde1c | 0.55 | 3.20E-09 | 6.83E-05 | 0.14 | 0.08 | 0.06 | 1.78 |
| NE | Dnmt3b | 0.52 | 3.40E-09 | 7.25E-05 | 0.28 | 0.19 | 0.09 | 1.45 |
| NE | Pard6b | 0.48 | 3.47E-09 | 7.42E-05 | 0.20 | 0.12 | 0.07 | 1.60 |
| NE | Txnrd2 | 0.52 | 3.62E-09 | 7.73E-05 | 0.19 | 0.12 | 0.07 | 1.58 |
| NE | Nap1l3 | 0.97 | 3.63E-09 | 7.75E-05 | 0.10 | 0.05 | 0.05 | 1.98 |
| NE | Ntf3 | 0.44 | 3.71E-09 | 7.92E-05 | 0.30 | 0.21 | 0.09 | 1.43 |
| NE | Cdh24 | 0.56 | 3.76E-09 | 8.03E-05 | 0.17 | 0.10 | 0.07 | 1.65 |
| NE | Dmrta1 | 0.66 | 3.83E-09 | 8.18E-05 | 0.12 | 0.06 | 0.06 | 1.88 |
| NE | Bspry | 0.32 | 3.94E-09 | 8.42E-05 | 0.31 | 0.22 | 0.09 | 1.43 |
| NE | Pcbd1 | 0.44 | 3.95E-09 | 8.45E-05 | 0.28 | 0.20 | 0.09 | 1.44 |
| NE | Nol12 | 0.28 | 3.97E-09 | 8.47E-05 | 0.61 | 0.50 | 0.11 | 1.23 |
| NE | Afg3l1 | 0.26 | 4.01E-09 | 8.56E-05 | 0.53 | 0.41 | 0.12 | 1.30 |
| NE | Tsr2 | 0.31 | 4.05E-09 | 8.66E-05 | 0.40 | 0.29 | 0.10 | 1.36 |
| NE | Nop16 | 0.27 | 4.24E-09 | 9.05E-05 | 0.64 | 0.53 | 0.11 | 1.21 |
| NE | Prmt3 | 0.33 | 4.25E-09 | 9.07E-05 | 0.44 | 0.33 | 0.11 | 1.32 |
| NE | Cep135 | 0.33 | 4.39E-09 | 9.38E-05 | 0.33 | 0.23 | 0.09 | 1.40 |
| NE | Vmp1 | 0.29 | 4.41E-09 | 9.42E-05 | 0.65 | 0.54 | 0.11 | 1.20 |
| NE | Hist1h2bh | 0.81 | 4.65E-09 | 9.94E-05 | 0.11 | 0.06 | 0.05 | 1.95 |
| NE | Elovl2 | 0.60 | 4.75E-09 | 1.02E-04 | 0.16 | 0.09 | 0.06 | 1.68 |
| NE | Gucy1b1 | 0.76 | 4.76E-09 | 1.02E-04 | 0.11 | 0.06 | 0.05 | 1.93 |
| NE | Lama1 | 0.71 | 4.77E-09 | 1.02E-04 | 0.14 | 0.08 | 0.06 | 1.77 |
| NE | Hist1h2bj | 0.43 | 4.93E-09 | 1.05E-04 | 0.22 | 0.14 | 0.08 | 1.54 |
| NE | Mrpl47 | 0.28 | 4.95E-09 | 1.06E-04 | 0.42 | 0.32 | 0.11 | 1.34 |
| NE | Sgsm1 | 0.68 | 5.06E-09 | 1.08E-04 | 0.12 | 0.06 | 0.06 | 1.87 |
| NE | Tsr1 | 0.27 | 5.51E-09 | 1.18E-04 | 0.45 | 0.34 | 0.11 | 1.32 |
| NE | Glt8d1 | 0.33 | 5.61E-09 | 1.20E-04 | 0.40 | 0.30 | 0.10 | 1.34 |

**Supplementary Table 4. Marker genes for the annotated cell populations of integrated clustering of E11.5 control and NotchGOF λ-junction ectoderm cells.**

| | | | | | | | | |
|---|---|---|---|---|---|---|---|---|
| NE | Ndufaf6 | 0.42 | 5.68E-09 | 1.21E-04 | 0.36 | 0.26 | 0.10 | 1.38 |
| NE | Rbm14 | 0.33 | 5.77E-09 | 1.23E-04 | 0.49 | 0.38 | 0.11 | 1.29 |
| NE | Mdn1 | 0.33 | 5.82E-09 | 1.24E-04 | 0.46 | 0.35 | 0.11 | 1.30 |
| NE | Sppl2b | 0.43 | 5.89E-09 | 1.26E-04 | 0.19 | 0.12 | 0.07 | 1.62 |
| NE | Mfsd13a | 0.79 | 5.95E-09 | 1.27E-04 | 0.12 | 0.06 | 0.05 | 1.84 |
| NE | Gstp1 | 0.32 | 5.99E-09 | 1.28E-04 | 0.46 | 0.35 | 0.11 | 1.31 |
| NE | Rpa1 | 0.35 | 6.06E-09 | 1.30E-04 | 0.42 | 0.32 | 0.10 | 1.31 |
| NE | Tmem164 | 0.36 | 6.07E-09 | 1.30E-04 | 0.24 | 0.15 | 0.08 | 1.53 |
| NE | Myl7 | 0.49 | 6.32E-09 | 1.35E-04 | 0.10 | 0.05 | 0.05 | 1.98 |
| NE | Zfp367 | 0.41 | 6.35E-09 | 1.36E-04 | 0.21 | 0.14 | 0.08 | 1.55 |
| NE | Herpud1 | 0.28 | 6.58E-09 | 1.40E-04 | 0.32 | 0.22 | 0.09 | 1.42 |
| NE | Zc3h7a | 0.33 | 6.65E-09 | 1.42E-04 | 0.36 | 0.26 | 0.09 | 1.35 |
| NE | Iqce | 0.57 | 6.68E-09 | 1.43E-04 | 0.18 | 0.11 | 0.07 | 1.61 |
| NE | Slc25a30 | 0.36 | 6.74E-09 | 1.44E-04 | 0.20 | 0.13 | 0.07 | 1.57 |
| NE | Rilpl2 | 0.41 | 7.30E-09 | 1.56E-04 | 0.28 | 0.19 | 0.09 | 1.45 |
| NE | Ints1 | 0.31 | 7.56E-09 | 1.61E-04 | 0.33 | 0.24 | 0.09 | 1.38 |
| NE | Epop | 0.44 | 7.97E-09 | 1.70E-04 | 0.31 | 0.22 | 0.09 | 1.40 |
| NE | Tram2 | 0.92 | 7.98E-09 | 1.70E-04 | 0.11 | 0.06 | 0.05 | 1.87 |
| NE | Mcm10 | 0.36 | 8.02E-09 | 1.71E-04 | 0.34 | 0.25 | 0.10 | 1.39 |
| NE | Naf1 | 0.27 | 8.29E-09 | 1.77E-04 | 0.51 | 0.40 | 0.11 | 1.28 |
| NE | Mark1 | 0.42 | 8.52E-09 | 1.82E-04 | 0.31 | 0.22 | 0.09 | 1.39 |
| NE | Nek7 | 0.42 | 8.73E-09 | 1.87E-04 | 0.39 | 0.29 | 0.09 | 1.32 |
| NE | Zcchc2 | 0.58 | 8.90E-09 | 1.90E-04 | 0.16 | 0.10 | 0.06 | 1.66 |
| NE | Ssc4d | 0.65 | 9.01E-09 | 1.92E-04 | 0.13 | 0.07 | 0.06 | 1.77 |
| NE | Bub3 | 0.25 | 9.19E-09 | 1.96E-04 | 0.81 | 0.75 | 0.06 | 1.08 |
| NE | Plod3 | 0.40 | 9.33E-09 | 1.99E-04 | 0.34 | 0.25 | 0.10 | 1.39 |
| NE | Tbl3 | 0.28 | 9.53E-09 | 2.04E-04 | 0.38 | 0.27 | 0.11 | 1.39 |
| NE | Hist1h2br | 0.63 | 9.89E-09 | 2.11E-04 | 0.17 | 0.11 | 0.06 | 1.59 |
| NE | Cep89 | 0.39 | 9.94E-09 | 2.12E-04 | 0.41 | 0.31 | 0.10 | 1.31 |
| NE | Gpr180 | 0.46 | 9.97E-09 | 2.13E-04 | 0.25 | 0.17 | 0.08 | 1.47 |
| NE | P3h3 | 0.45 | 1.03E-08 | 2.19E-04 | 0.19 | 0.12 | 0.07 | 1.58 |
| NE | Polr1a | 0.26 | 1.05E-08 | 2.24E-04 | 0.43 | 0.32 | 0.11 | 1.34 |
| NE | Lrp8 | 0.71 | 1.05E-08 | 2.25E-04 | 0.12 | 0.07 | 0.05 | 1.81 |
| NE | Sel1l | 0.28 | 1.05E-08 | 2.25E-04 | 0.47 | 0.36 | 0.11 | 1.32 |
| NE | Pola2 | 0.30 | 1.09E-08 | 2.32E-04 | 0.41 | 0.31 | 0.10 | 1.32 |
| NE | Grhpr | 0.27 | 1.12E-08 | 2.40E-04 | 0.55 | 0.45 | 0.10 | 1.22 |
| NE | Aldh18a1 | 0.32 | 1.14E-08 | 2.43E-04 | 0.31 | 0.22 | 0.09 | 1.41 |
| NE | Wdsub1 | 0.39 | 1.23E-08 | 2.62E-04 | 0.39 | 0.29 | 0.09 | 1.32 |
| NE | Psmb10 | 0.38 | 1.23E-08 | 2.62E-04 | 0.30 | 0.21 | 0.09 | 1.43 |
| NE | Snx10 | 0.30 | 1.23E-08 | 2.63E-04 | 0.37 | 0.27 | 0.10 | 1.37 |
| NE | Bcl7b | 0.26 | 1.24E-08 | 2.65E-04 | 0.44 | 0.34 | 0.11 | 1.32 |
| NE | Sdf2l1 | 0.30 | 1.24E-08 | 2.66E-04 | 0.56 | 0.45 | 0.11 | 1.24 |
| NE | Gmpr2 | 0.30 | 1.27E-08 | 2.71E-04 | 0.42 | 0.31 | 0.11 | 1.35 |
| NE | Hspa13 | 0.31 | 1.28E-08 | 2.73E-04 | 0.50 | 0.39 | 0.11 | 1.28 |
| NE | AU020206 | 0.37 | 1.35E-08 | 2.89E-04 | 0.26 | 0.18 | 0.09 | 1.48 |
| NE | Chek1 | 0.26 | 1.43E-08 | 3.06E-04 | 0.51 | 0.40 | 0.11 | 1.28 |
| NE | Arrdc1 | 0.39 | 1.43E-08 | 3.06E-04 | 0.41 | 0.31 | 0.10 | 1.30 |
| NE | Aars2 | 0.48 | 1.47E-08 | 3.14E-04 | 0.16 | 0.10 | 0.06 | 1.66 |
| NE | Ccdc181 | 0.44 | 1.47E-08 | 3.14E-04 | 0.17 | 0.10 | 0.07 | 1.64 |
| NE | Hist2h2ac | 0.48 | 1.56E-08 | 3.33E-04 | 0.28 | 0.20 | 0.08 | 1.40 |
| NE | Nle1 | 0.42 | 1.59E-08 | 3.39E-04 | 0.27 | 0.18 | 0.08 | 1.44 |
| NE | Txndc11 | 0.42 | 1.61E-08 | 3.44E-04 | 0.25 | 0.17 | 0.08 | 1.48 |
| NE | Pold2 | 0.29 | 1.63E-08 | 3.49E-04 | 0.47 | 0.37 | 0.10 | 1.28 |
| NE | Eif3j2 | 0.33 | 1.64E-08 | 3.51E-04 | 0.39 | 0.29 | 0.10 | 1.34 |
| NE | Egfr | 0.50 | 1.70E-08 | 3.63E-04 | 0.27 | 0.20 | 0.08 | 1.39 |

**Supplementary Table 4. Marker genes for the annotated cell populations of integrated clustering of E11.5 control and NotchGOF λ-junction ectoderm cells.**

| | | | | | | | | |
|---|---|---|---|---|---|---|---|---|
| NE | Ino80e | 0.29 | 1.71E-08 | 3.65E-04 | 0.52 | 0.41 | 0.11 | 1.27 |
| NE | Ppp2r5d | 0.25 | 1.76E-08 | 3.76E-04 | 0.51 | 0.40 | 0.11 | 1.26 |
| NE | Rab9 | 0.26 | 1.82E-08 | 3.89E-04 | 0.63 | 0.52 | 0.11 | 1.22 |
| NE | Neil3 | 0.48 | 1.82E-08 | 3.90E-04 | 0.20 | 0.13 | 0.07 | 1.55 |
| NE | Hist1h2ak | 0.53 | 1.96E-08 | 4.20E-04 | 0.15 | 0.09 | 0.06 | 1.65 |
| NE | Cptp | 0.42 | 2.10E-08 | 4.49E-04 | 0.17 | 0.10 | 0.06 | 1.61 |
| NE | Ccnf | 0.38 | 2.12E-08 | 4.53E-04 | 0.37 | 0.28 | 0.09 | 1.31 |
| NE | Sept11 | 0.32 | 2.24E-08 | 4.79E-04 | 0.77 | 0.71 | 0.06 | 1.09 |
| NE | Ltbp3 | 0.70 | 2.29E-08 | 4.90E-04 | 0.12 | 0.07 | 0.05 | 1.76 |
| NE | Pole2 | 0.41 | 2.41E-08 | 5.15E-04 | 0.25 | 0.17 | 0.08 | 1.46 |
| NE | Cdca2 | 0.35 | 2.52E-08 | 5.38E-04 | 0.42 | 0.33 | 0.09 | 1.28 |
| NE | Atp2a2 | 0.27 | 2.58E-08 | 5.52E-04 | 0.72 | 0.63 | 0.09 | 1.15 |
| NE | Coq4 | 0.26 | 2.58E-08 | 5.52E-04 | 0.43 | 0.32 | 0.10 | 1.32 |
| NE | Aldh9a1 | 0.37 | 2.58E-08 | 5.52E-04 | 0.30 | 0.21 | 0.09 | 1.41 |
| NE | 3110082I17Rik | 0.28 | 2.64E-08 | 5.65E-04 | 0.44 | 0.34 | 0.10 | 1.31 |
| NE | Ttc12 | 0.65 | 2.67E-08 | 5.69E-04 | 0.10 | 0.05 | 0.05 | 1.92 |
| NE | Pold1 | 0.44 | 2.74E-08 | 5.84E-04 | 0.36 | 0.27 | 0.09 | 1.32 |
| NE | Nudt19 | 0.38 | 2.78E-08 | 5.94E-04 | 0.32 | 0.23 | 0.09 | 1.38 |
| NE | Hist1h3f | 0.44 | 2.86E-08 | 6.10E-04 | 0.25 | 0.18 | 0.08 | 1.43 |
| NE | Pdcd11 | 0.29 | 2.86E-08 | 6.11E-04 | 0.51 | 0.41 | 0.10 | 1.25 |
| NE | Nutf2-ps1 | 0.25 | 2.88E-08 | 6.16E-04 | 0.56 | 0.45 | 0.11 | 1.25 |
| NE | Dna2 | 0.37 | 2.95E-08 | 6.29E-04 | 0.27 | 0.19 | 0.08 | 1.42 |
| NE | Ccdc92 | 0.34 | 3.02E-08 | 6.45E-04 | 0.20 | 0.13 | 0.07 | 1.54 |
| NE | Plekha5 | 0.37 | 3.09E-08 | 6.59E-04 | 0.48 | 0.39 | 0.09 | 1.22 |
| NE | Ncaph | 0.29 | 3.14E-08 | 6.70E-04 | 0.50 | 0.40 | 0.10 | 1.26 |
| NE | Lmln | 0.64 | 3.27E-08 | 6.99E-04 | 0.14 | 0.08 | 0.06 | 1.71 |
| NE | Pip4p2 | 0.31 | 3.32E-08 | 7.09E-04 | 0.58 | 0.48 | 0.10 | 1.21 |
| NE | Iqgap3 | 0.44 | 3.32E-08 | 7.10E-04 | 0.26 | 0.19 | 0.08 | 1.41 |
| NE | Nek1 | 0.28 | 3.38E-08 | 7.22E-04 | 0.39 | 0.29 | 0.10 | 1.34 |
| NE | Haspin | 0.44 | 3.65E-08 | 7.80E-04 | 0.26 | 0.18 | 0.08 | 1.42 |
| NE | Filip1 | 0.49 | 3.83E-08 | 8.18E-04 | 0.10 | 0.06 | 0.05 | 1.86 |
| NE | Nek6 | 0.34 | 3.95E-08 | 8.43E-04 | 0.36 | 0.27 | 0.09 | 1.32 |
| NE | Trim3 | 0.63 | 3.97E-08 | 8.47E-04 | 0.16 | 0.10 | 0.06 | 1.62 |
| NE | Trib1 | 0.43 | 4.17E-08 | 8.92E-04 | 0.22 | 0.15 | 0.07 | 1.49 |
| NE | 1810009A15Rik | 0.42 | 4.25E-08 | 9.09E-04 | 0.19 | 0.12 | 0.07 | 1.57 |
| NE | Rnd2 | 0.48 | 4.31E-08 | 9.21E-04 | 0.15 | 0.09 | 0.06 | 1.67 |
| NE | Tdp1 | 0.34 | 4.35E-08 | 9.30E-04 | 0.34 | 0.25 | 0.09 | 1.35 |
| NE | Gm15417 | 0.77 | 4.49E-08 | 9.59E-04 | 0.10 | 0.05 | 0.05 | 1.89 |
| NE | Nop2 | 0.32 | 4.56E-08 | 9.74E-04 | 0.38 | 0.29 | 0.10 | 1.33 |
| NE | Dbf4 | 0.28 | 4.65E-08 | 9.93E-04 | 0.67 | 0.60 | 0.08 | 1.13 |
| NE | Gtse1 | 0.61 | 4.65E-08 | 9.94E-04 | 0.20 | 0.13 | 0.07 | 1.49 |
| NE | Lrr1 | 0.33 | 4.73E-08 | 1.01E-03 | 0.27 | 0.19 | 0.08 | 1.42 |
| NE | Ccdc30 | 0.43 | 4.73E-08 | 1.01E-03 | 0.13 | 0.08 | 0.06 | 1.73 |
| NE | Aaas | 0.32 | 5.01E-08 | 1.07E-03 | 0.35 | 0.26 | 0.09 | 1.36 |
| NE | Slc7a5 | 0.82 | 5.15E-08 | 1.10E-03 | 0.11 | 0.06 | 0.05 | 1.82 |
| NE | Asns | 0.34 | 5.34E-08 | 1.14E-03 | 0.33 | 0.24 | 0.09 | 1.37 |
| NE | Pdia5 | 0.54 | 5.52E-08 | 1.18E-03 | 0.16 | 0.10 | 0.06 | 1.62 |
| NE | Tomm40l | 0.38 | 5.55E-08 | 1.19E-03 | 0.20 | 0.13 | 0.07 | 1.54 |
| NE | Maoa | 0.27 | 5.62E-08 | 1.20E-03 | 0.33 | 0.25 | 0.09 | 1.36 |
| NE | Hist1h2an | 0.41 | 5.73E-08 | 1.22E-03 | 0.26 | 0.18 | 0.08 | 1.42 |
| NE | Telo2 | 0.32 | 6.19E-08 | 1.32E-03 | 0.30 | 0.21 | 0.09 | 1.41 |
| NE | Nbdy | 0.26 | 6.23E-08 | 1.33E-03 | 0.63 | 0.52 | 0.11 | 1.21 |
| NE | Sirt3 | 0.29 | 6.40E-08 | 1.37E-03 | 0.29 | 0.21 | 0.09 | 1.41 |
| NE | Echdc2 | 0.34 | 6.57E-08 | 1.40E-03 | 0.35 | 0.26 | 0.09 | 1.35 |
| NE | Lpin1 | 0.68 | 7.06E-08 | 1.51E-03 | 0.12 | 0.07 | 0.05 | 1.75 |

**Supplementary Table 4. Marker genes for the annotated cell populations of integrated clustering of E11.5 control and NotchGOF λ-junction ectoderm cells.**

| | | | | | | | | |
|---|---|---|---|---|---|---|---|---|
| NE | Hspbap1 | 0.34 | 7.22E-08 | 1.54E-03 | 0.26 | 0.18 | 0.08 | 1.44 |
| NE | Ppp2r5a | 0.34 | 7.24E-08 | 1.55E-03 | 0.36 | 0.27 | 0.09 | 1.34 |
| NE | Sac3d1 | 0.28 | 7.25E-08 | 1.55E-03 | 0.53 | 0.43 | 0.10 | 1.24 |
| NE | Sgsm2 | 0.50 | 7.27E-08 | 1.55E-03 | 0.15 | 0.09 | 0.06 | 1.65 |
| NE | Creg1 | 0.49 | 7.36E-08 | 1.57E-03 | 0.35 | 0.27 | 0.09 | 1.32 |
| NE | Slc52a2 | 0.53 | 7.39E-08 | 1.58E-03 | 0.17 | 0.11 | 0.06 | 1.56 |
| NE | Slc43a1 | 0.50 | 7.63E-08 | 1.63E-03 | 0.14 | 0.08 | 0.06 | 1.67 |
| NE | Cpq | 0.68 | 7.69E-08 | 1.64E-03 | 0.17 | 0.11 | 0.06 | 1.54 |
| NE | Srgap2 | 0.28 | 8.42E-08 | 1.80E-03 | 0.36 | 0.27 | 0.09 | 1.34 |
| NE | Ralgapa2 | 0.45 | 8.44E-08 | 1.80E-03 | 0.25 | 0.18 | 0.08 | 1.43 |
| NE | Flvcr1 | 0.66 | 8.48E-08 | 1.81E-03 | 0.11 | 0.06 | 0.05 | 1.79 |
| NE | Pigh | 0.27 | 8.81E-08 | 1.88E-03 | 0.35 | 0.25 | 0.09 | 1.37 |
| NE | Crocc | 0.45 | 8.99E-08 | 1.92E-03 | 0.19 | 0.13 | 0.06 | 1.50 |
| NE | Caml | 0.26 | 9.01E-08 | 1.93E-03 | 0.49 | 0.39 | 0.10 | 1.26 |
| NE | Pip4k2b | 0.44 | 9.10E-08 | 1.94E-03 | 0.21 | 0.14 | 0.07 | 1.50 |
| NE | Sgf29 | 0.25 | 9.14E-08 | 1.95E-03 | 0.36 | 0.27 | 0.10 | 1.36 |
| NE | Cdc45 | 0.26 | 9.25E-08 | 1.98E-03 | 0.37 | 0.28 | 0.09 | 1.33 |
| NE | Mthfr | 0.42 | 9.34E-08 | 2.00E-03 | 0.13 | 0.08 | 0.05 | 1.70 |
| NE | Cdc25c | 0.30 | 9.42E-08 | 2.01E-03 | 0.35 | 0.27 | 0.09 | 1.33 |
| NE | Mpdu1 | 0.27 | 9.45E-08 | 2.02E-03 | 0.57 | 0.46 | 0.11 | 1.23 |
| NE | Troap | 0.54 | 9.73E-08 | 2.08E-03 | 0.18 | 0.12 | 0.06 | 1.54 |
| NE | Pcf11 | 0.26 | 1.01E-07 | 2.16E-03 | 0.49 | 0.39 | 0.10 | 1.27 |
| NE | Alg6 | 0.31 | 1.03E-07 | 2.19E-03 | 0.35 | 0.26 | 0.09 | 1.36 |
| NE | Rad54l | 0.32 | 1.05E-07 | 2.25E-03 | 0.34 | 0.26 | 0.09 | 1.34 |
| NE | Plod1 | 0.55 | 1.08E-07 | 2.30E-03 | 0.10 | 0.05 | 0.05 | 1.85 |
| NE | Msra | 0.58 | 1.09E-07 | 2.33E-03 | 0.16 | 0.10 | 0.06 | 1.60 |
| NE | Donson | 0.39 | 1.09E-07 | 2.33E-03 | 0.26 | 0.19 | 0.08 | 1.41 |
| NE | Kbtbd4 | 0.27 | 1.10E-07 | 2.35E-03 | 0.31 | 0.23 | 0.08 | 1.36 |
| NE | Wdr90 | 0.53 | 1.12E-07 | 2.39E-03 | 0.17 | 0.11 | 0.06 | 1.57 |
| NE | Upp2 | 0.55 | 1.14E-07 | 2.43E-03 | 0.19 | 0.13 | 0.06 | 1.48 |
| NE | Sirpa | 0.67 | 1.15E-07 | 2.46E-03 | 0.12 | 0.07 | 0.05 | 1.70 |
| NE | Cnpy3 | 0.27 | 1.19E-07 | 2.54E-03 | 0.42 | 0.32 | 0.10 | 1.32 |
| NE | Rasl11a | 0.25 | 1.29E-07 | 2.76E-03 | 0.20 | 0.13 | 0.07 | 1.54 |
| NE | Wasf3 | 0.41 | 1.31E-07 | 2.80E-03 | 0.20 | 0.13 | 0.07 | 1.52 |
| NE | Lrba | 0.36 | 1.42E-07 | 3.04E-03 | 0.33 | 0.24 | 0.08 | 1.34 |
| NE | Pusl1 | 0.38 | 1.52E-07 | 3.26E-03 | 0.29 | 0.21 | 0.08 | 1.38 |
| NE | Notch4 | 0.75 | 1.54E-07 | 3.28E-03 | 0.10 | 0.06 | 0.05 | 1.79 |
| NE | Wdr31 | 0.45 | 1.58E-07 | 3.37E-03 | 0.19 | 0.13 | 0.07 | 1.52 |
| NE | Ppp1r16a | 0.41 | 1.59E-07 | 3.39E-03 | 0.23 | 0.15 | 0.07 | 1.47 |
| NE | Prkab1 | 0.35 | 1.71E-07 | 3.66E-03 | 0.38 | 0.29 | 0.09 | 1.31 |
| NE | Amt | 0.35 | 1.73E-07 | 3.70E-03 | 0.28 | 0.20 | 0.08 | 1.39 |
| NE | Prps1 | 0.28 | 1.75E-07 | 3.74E-03 | 0.35 | 0.26 | 0.09 | 1.35 |
| NE | C630043F03Rik | 0.62 | 1.75E-07 | 3.75E-03 | 0.14 | 0.09 | 0.05 | 1.60 |
| NE | Foxm1 | 0.40 | 1.76E-07 | 3.75E-03 | 0.29 | 0.21 | 0.08 | 1.37 |
| NE | Adam22 | 0.71 | 1.80E-07 | 3.84E-03 | 0.16 | 0.10 | 0.06 | 1.56 |
| NE | Dlgap5 | 0.28 | 1.88E-07 | 4.01E-03 | 0.49 | 0.39 | 0.10 | 1.25 |
| NE | Kif20a | 0.28 | 1.91E-07 | 4.08E-03 | 0.46 | 0.37 | 0.09 | 1.25 |
| NE | Kmt5c | 0.27 | 1.96E-07 | 4.18E-03 | 0.38 | 0.29 | 0.09 | 1.32 |
| NE | Rab26os | 0.35 | 1.98E-07 | 4.23E-03 | 0.26 | 0.18 | 0.08 | 1.41 |
| NE | Cldn12 | 0.58 | 2.06E-07 | 4.39E-03 | 0.17 | 0.11 | 0.06 | 1.54 |
| NE | Mrnip | 0.39 | 2.10E-07 | 4.49E-03 | 0.21 | 0.14 | 0.07 | 1.46 |
| NE | Xbp1 | 0.25 | 2.12E-07 | 4.53E-03 | 0.44 | 0.34 | 0.09 | 1.27 |
| NE | Gclm | 0.26 | 2.19E-07 | 4.68E-03 | 0.57 | 0.46 | 0.10 | 1.22 |
| NE | Rpain | 0.32 | 2.26E-07 | 4.83E-03 | 0.33 | 0.24 | 0.09 | 1.36 |
| NE | L2hgdh | 0.32 | 2.27E-07 | 4.85E-03 | 0.29 | 0.21 | 0.08 | 1.39 |

**Supplementary Table 4. Marker genes for the annotated cell populations of integrated clustering of E11.5 control and NotchGOF λ-junction ectoderm cells.**

| | | | | | | | | |
|---|---|---|---|---|---|---|---|---|
| NE | Pdss2 | 0.31 | 2.28E-07 | 4.86E-03 | 0.25 | 0.17 | 0.07 | 1.43 |
| NE | Tma16 | 0.25 | 2.30E-07 | 4.91E-03 | 0.62 | 0.52 | 0.09 | 1.18 |
| NE | Ndufaf5 | 0.36 | 2.37E-07 | 5.07E-03 | 0.21 | 0.14 | 0.07 | 1.48 |
| NE | Gm45051 | 0.38 | 2.38E-07 | 5.08E-03 | 0.15 | 0.10 | 0.06 | 1.61 |
| NE | Ankle1 | 0.63 | 2.39E-07 | 5.11E-03 | 0.14 | 0.09 | 0.05 | 1.61 |
| NE | Rdh11 | 0.40 | 2.43E-07 | 5.18E-03 | 0.20 | 0.14 | 0.07 | 1.48 |
| NE | Tbrg4 | 0.32 | 2.45E-07 | 5.24E-03 | 0.23 | 0.16 | 0.07 | 1.45 |
| NE | Utp14b | 0.27 | 2.48E-07 | 5.30E-03 | 0.46 | 0.37 | 0.10 | 1.27 |
| NE | Gpaa1 | 0.31 | 2.49E-07 | 5.32E-03 | 0.46 | 0.37 | 0.09 | 1.24 |
| NE | Gm4258 | 0.26 | 2.51E-07 | 5.36E-03 | 0.57 | 0.47 | 0.10 | 1.22 |
| NE | Hist1h2bl | 0.70 | 2.52E-07 | 5.38E-03 | 0.10 | 0.06 | 0.05 | 1.79 |
| NE | Nr2f1 | 0.41 | 2.54E-07 | 5.42E-03 | 0.25 | 0.18 | 0.07 | 1.40 |
| NE | Ring1 | 0.31 | 2.54E-07 | 5.43E-03 | 0.35 | 0.26 | 0.09 | 1.34 |
| NE | Hspe1-rs1 | 0.37 | 2.59E-07 | 5.53E-03 | 0.17 | 0.11 | 0.06 | 1.57 |
| NE | Scap | 0.31 | 2.71E-07 | 5.79E-03 | 0.26 | 0.19 | 0.08 | 1.41 |
| NE | 9430038I01Rik | 0.32 | 2.77E-07 | 5.91E-03 | 0.34 | 0.25 | 0.09 | 1.34 |
| NE | Lin54 | 0.33 | 2.78E-07 | 5.94E-03 | 0.27 | 0.19 | 0.08 | 1.40 |
| NE | Tm9sf1 | 0.25 | 2.83E-07 | 6.04E-03 | 0.49 | 0.40 | 0.10 | 1.25 |
| NE | Etv5 | 0.54 | 2.91E-07 | 6.22E-03 | 0.27 | 0.20 | 0.07 | 1.35 |
| NE | Cic | 0.29 | 2.95E-07 | 6.30E-03 | 0.45 | 0.35 | 0.10 | 1.29 |
| NE | Msrb2 | 0.31 | 2.99E-07 | 6.39E-03 | 0.25 | 0.18 | 0.08 | 1.43 |
| NE | Hagh | 0.25 | 3.02E-07 | 6.45E-03 | 0.30 | 0.22 | 0.08 | 1.37 |
| NE | Cep19 | 0.26 | 3.02E-07 | 6.46E-03 | 0.37 | 0.28 | 0.09 | 1.33 |
| NE | Cep170b | 0.31 | 3.08E-07 | 6.58E-03 | 0.39 | 0.30 | 0.09 | 1.32 |
| NE | Nme6 | 0.26 | 3.11E-07 | 6.63E-03 | 0.48 | 0.38 | 0.10 | 1.25 |
| NE | Sept9 | 0.30 | 3.13E-07 | 6.68E-03 | 0.58 | 0.47 | 0.11 | 1.23 |
| NE | Hist1h2ag | 0.28 | 3.23E-07 | 6.90E-03 | 0.25 | 0.17 | 0.07 | 1.43 |
| NE | Smg9 | 0.39 | 3.27E-07 | 6.99E-03 | 0.18 | 0.12 | 0.06 | 1.53 |
| NE | Actr5 | 0.40 | 3.39E-07 | 7.25E-03 | 0.24 | 0.17 | 0.07 | 1.43 |
| NE | Yju2 | 0.37 | 3.47E-07 | 7.40E-03 | 0.25 | 0.18 | 0.07 | 1.41 |
| NE | Lifr | 0.57 | 3.53E-07 | 7.55E-03 | 0.16 | 0.10 | 0.06 | 1.56 |
| NE | Gm26793 | 0.71 | 3.57E-07 | 7.62E-03 | 0.10 | 0.06 | 0.04 | 1.79 |
| NE | Dleu2 | 0.34 | 3.64E-07 | 7.78E-03 | 0.25 | 0.18 | 0.07 | 1.40 |
| NE | Pi4ka | 0.28 | 3.80E-07 | 8.12E-03 | 0.23 | 0.16 | 0.07 | 1.44 |
| NE | Rnf141 | 0.33 | 3.85E-07 | 8.23E-03 | 0.18 | 0.12 | 0.06 | 1.52 |
| NE | 3110009E18Rik | 0.28 | 3.92E-07 | 8.38E-03 | 0.26 | 0.18 | 0.08 | 1.43 |
| NE | Snx8 | 0.47 | 3.93E-07 | 8.40E-03 | 0.17 | 0.11 | 0.06 | 1.52 |
| NE | Zswim1 | 0.46 | 4.06E-07 | 8.66E-03 | 0.16 | 0.10 | 0.06 | 1.56 |
| NE | Ttk | 0.34 | 4.25E-07 | 9.08E-03 | 0.34 | 0.26 | 0.08 | 1.32 |
| NE | Tspyl5 | 0.45 | 4.46E-07 | 9.53E-03 | 0.11 | 0.06 | 0.05 | 1.75 |
| NE | Lrwd1 | 0.36 | 4.49E-07 | 9.60E-03 | 0.40 | 0.32 | 0.09 | 1.27 |
| NE | Spag5 | 0.33 | 4.53E-07 | 9.67E-03 | 0.29 | 0.22 | 0.08 | 1.36 |
| NE | C2cd3 | 0.31 | 4.74E-07 | 1.01E-02 | 0.42 | 0.33 | 0.09 | 1.28 |
| NE | Fgfr1op | 0.30 | 4.88E-07 | 1.04E-02 | 0.39 | 0.30 | 0.09 | 1.31 |
| NE | Lrrc8b | 0.48 | 4.89E-07 | 1.04E-02 | 0.25 | 0.19 | 0.07 | 1.37 |
| NE | Cadm4 | 0.40 | 5.10E-07 | 1.09E-02 | 0.21 | 0.15 | 0.07 | 1.45 |
| NE | Elovl4 | 0.30 | 5.34E-07 | 1.14E-02 | 0.19 | 0.12 | 0.06 | 1.50 |
| NE | Mdfic | 0.44 | 5.35E-07 | 1.14E-02 | 0.23 | 0.16 | 0.07 | 1.42 |
| NE | Fcor | 0.68 | 5.39E-07 | 1.15E-02 | 0.11 | 0.07 | 0.05 | 1.69 |
| NE | Ddx49 | 0.27 | 5.65E-07 | 1.21E-02 | 0.41 | 0.32 | 0.09 | 1.27 |
| NE | Lamc1 | 0.27 | 5.89E-07 | 1.26E-02 | 0.49 | 0.40 | 0.09 | 1.23 |
| NE | Ttc13 | 0.29 | 6.15E-07 | 1.31E-02 | 0.32 | 0.24 | 0.08 | 1.33 |
| NE | Mal2 | 0.73 | 6.16E-07 | 1.32E-02 | 0.12 | 0.07 | 0.05 | 1.65 |
| NE | 1810055G02Rik | 0.40 | 6.32E-07 | 1.35E-02 | 0.18 | 0.12 | 0.06 | 1.50 |
| NE | Tnks1bp1 | 0.48 | 6.84E-07 | 1.46E-02 | 0.18 | 0.12 | 0.06 | 1.49 |

**Supplementary Table 4. Marker genes for the annotated cell populations of integrated clustering of E11.5 control and NotchGOF λ-junction ectoderm cells.**

| | | | | | | | | |
|---|---|---|---|---|---|---|---|---|
| NE | Hist1h2bg | 0.70 | 7.09E-07 | 1.51E-02 | 0.11 | 0.06 | 0.04 | 1.72 |
| NE | Pcbp3 | 0.44 | 7.22E-07 | 1.54E-02 | 0.19 | 0.13 | 0.06 | 1.47 |
| NE | Pwwp3a | 0.27 | 8.20E-07 | 1.75E-02 | 0.51 | 0.43 | 0.09 | 1.21 |
| NE | Slc48a1 | 0.29 | 8.24E-07 | 1.76E-02 | 0.32 | 0.24 | 0.08 | 1.35 |
| NE | B4gat1 | 0.26 | 8.28E-07 | 1.77E-02 | 0.41 | 0.32 | 0.09 | 1.30 |
| NE | Fbn1 | 0.39 | 8.63E-07 | 1.84E-02 | 0.24 | 0.17 | 0.07 | 1.40 |
| NE | Ykt6 | 0.26 | 8.79E-07 | 1.88E-02 | 0.43 | 0.33 | 0.10 | 1.29 |
| NE | Cux2 | 0.35 | 8.87E-07 | 1.90E-02 | 0.19 | 0.13 | 0.06 | 1.48 |
| NE | Armcx4 | 0.27 | 9.05E-07 | 1.93E-02 | 0.41 | 0.32 | 0.09 | 1.29 |
| NE | Mfn2 | 0.34 | 9.29E-07 | 1.98E-02 | 0.21 | 0.14 | 0.07 | 1.46 |
| NE | Ranbp6 | 0.49 | 9.72E-07 | 2.08E-02 | 0.17 | 0.11 | 0.06 | 1.53 |
| NE | Ctu2 | 0.29 | 9.74E-07 | 2.08E-02 | 0.35 | 0.27 | 0.08 | 1.32 |
| NE | Amfr | 0.25 | 9.79E-07 | 2.09E-02 | 0.46 | 0.36 | 0.10 | 1.26 |
| NE | E2f2 | 0.38 | 1.01E-06 | 2.17E-02 | 0.15 | 0.10 | 0.05 | 1.55 |
| NE | Ap1m1 | 0.25 | 1.04E-06 | 2.22E-02 | 0.31 | 0.23 | 0.08 | 1.36 |
| NE | Gtf3c4 | 0.29 | 1.08E-06 | 2.31E-02 | 0.39 | 0.31 | 0.08 | 1.26 |
| NE | Gm48226 | 0.37 | 1.08E-06 | 2.31E-02 | 0.24 | 0.17 | 0.07 | 1.42 |
| NE | Bbs7 | 0.50 | 1.10E-06 | 2.34E-02 | 0.15 | 0.10 | 0.05 | 1.56 |
| NE | Rab13 | 0.29 | 1.11E-06 | 2.36E-02 | 0.34 | 0.26 | 0.08 | 1.32 |
| NE | Trak1 | 0.37 | 1.11E-06 | 2.38E-02 | 0.23 | 0.16 | 0.07 | 1.42 |
| NE | Gm10282 | 0.32 | 1.12E-06 | 2.40E-02 | 0.20 | 0.13 | 0.06 | 1.48 |
| NE | Xxylt1 | 0.36 | 1.18E-06 | 2.53E-02 | 0.26 | 0.19 | 0.07 | 1.39 |
| NE | Carmil3 | 0.39 | 1.20E-06 | 2.57E-02 | 0.12 | 0.07 | 0.05 | 1.67 |
| NE | Gpx7 | 0.28 | 1.21E-06 | 2.59E-02 | 0.35 | 0.26 | 0.08 | 1.32 |
| NE | Aacs | 0.43 | 1.25E-06 | 2.67E-02 | 0.14 | 0.09 | 0.05 | 1.58 |
| NE | Zswim6 | 0.26 | 1.30E-06 | 2.77E-02 | 0.43 | 0.34 | 0.10 | 1.28 |
| NE | Nek3 | 0.47 | 1.31E-06 | 2.80E-02 | 0.14 | 0.09 | 0.05 | 1.57 |
| NE | Slc9a3r2 | 0.27 | 1.37E-06 | 2.93E-02 | 0.17 | 0.11 | 0.06 | 1.53 |
| NE | Dync1i1 | 0.91 | 1.47E-06 | 3.15E-02 | 0.16 | 0.11 | 0.05 | 1.45 |
| NE | Dcbld2 | 0.27 | 1.53E-06 | 3.26E-02 | 0.29 | 0.22 | 0.08 | 1.35 |
| NE | Ift80 | 0.35 | 1.53E-06 | 3.28E-02 | 0.22 | 0.16 | 0.07 | 1.41 |
| NE | Acaca | 0.28 | 1.58E-06 | 3.38E-02 | 0.34 | 0.26 | 0.08 | 1.31 |
| NE | Pbx4 | 0.37 | 1.61E-06 | 3.44E-02 | 0.21 | 0.15 | 0.06 | 1.43 |
| NE | Rtel1 | 0.27 | 1.61E-06 | 3.45E-02 | 0.21 | 0.15 | 0.06 | 1.42 |
| NE | Pxk | 0.47 | 1.62E-06 | 3.45E-02 | 0.16 | 0.11 | 0.05 | 1.51 |
| NE | Slc43a3 | 0.50 | 1.62E-06 | 3.46E-02 | 0.18 | 0.13 | 0.06 | 1.45 |
| NE | Kcnk2 | 0.44 | 1.65E-06 | 3.53E-02 | 0.42 | 0.34 | 0.07 | 1.21 |
| NE | Arhgef26 | 0.50 | 1.84E-06 | 3.93E-02 | 0.15 | 0.10 | 0.05 | 1.53 |
| NE | 6720427I07Rik | 0.34 | 1.88E-06 | 4.02E-02 | 0.21 | 0.15 | 0.06 | 1.41 |
| NE | Ggct | 0.33 | 1.90E-06 | 4.07E-02 | 0.18 | 0.12 | 0.06 | 1.49 |
| NE | Uxs1 | 0.34 | 1.91E-06 | 4.07E-02 | 0.31 | 0.23 | 0.07 | 1.31 |
| NE | Itga5 | 0.52 | 1.93E-06 | 4.12E-02 | 0.13 | 0.08 | 0.05 | 1.59 |
| NE | Hist4h4 | 0.55 | 2.01E-06 | 4.28E-02 | 0.10 | 0.06 | 0.04 | 1.72 |
| NE | Gspt2 | 0.33 | 2.01E-06 | 4.29E-02 | 0.21 | 0.15 | 0.06 | 1.43 |
| NE | Dph2 | 0.40 | 2.01E-06 | 4.29E-02 | 0.16 | 0.11 | 0.06 | 1.51 |
| NE | Fpgs | 0.35 | 2.10E-06 | 4.49E-02 | 0.20 | 0.14 | 0.06 | 1.44 |
| NE | Vrk2 | 0.25 | 2.23E-06 | 4.77E-02 | 0.24 | 0.17 | 0.07 | 1.39 |
| NE | Agtpbp1 | 0.39 | 2.24E-06 | 4.78E-02 | 0.14 | 0.09 | 0.05 | 1.58 |
| NE | Man2b1 | 0.30 | 2.26E-06 | 4.83E-02 | 0.28 | 0.21 | 0.07 | 1.34 |
| NE | Tbl2 | 0.39 | 2.33E-06 | 4.98E-02 | 0.15 | 0.10 | 0.05 | 1.54 |
| periderm | Sfn | 1.86 | 2.36E-154 | 5.04E-150 | 0.98 | 0.66 | 0.32 | 1.48 |
| periderm | Grhl3 | 2.59 | 8.67E-142 | 1.85E-137 | 0.63 | 0.16 | 0.47 | 3.95 |
| periderm | Zfp750 | 2.46 | 4.62E-132 | 9.88E-128 | 0.60 | 0.15 | 0.45 | 4.10 |
| periderm | Nectin4 | 1.74 | 6.68E-126 | 1.43E-121 | 0.90 | 0.48 | 0.42 | 1.87 |
| periderm | Tctex1d1 | 3.22 | 1.41E-124 | 3.01E-120 | 0.40 | 0.06 | 0.34 | 6.50 |

**Supplementary Table 4. Marker genes for the annotated cell populations of integrated clustering of E11.5 control and NotchGOF λ-junction ectoderm cells.**

| | | | | | | | | |
|---|---|---|---|---|---|---|---|---|
| periderm | Notch1 | 1.91 | 1.53E-117 | 3.26E-113 | 0.92 | 0.59 | 0.34 | 1.58 |
| periderm | Ppl | 3.01 | 1.28E-116 | 2.73E-112 | 0.50 | 0.12 | 0.39 | 4.24 |
| periderm | Gm16136 | 1.63 | 7.30E-111 | 1.56E-106 | 0.92 | 0.62 | 0.30 | 1.49 |
| periderm | Tgm1 | 3.97 | 1.75E-104 | 3.73E-100 | 0.25 | 0.02 | 0.23 | 10.54 |
| periderm | Cd9 | 1.52 | 3.33E-101 | 7.12E-97 | 0.92 | 0.69 | 0.23 | 1.33 |
| periderm | Nav3 | 1.66 | 1.72E-97 | 3.67E-93 | 0.76 | 0.33 | 0.43 | 2.29 |
| periderm | Tacstd2 | 2.87 | 3.16E-96 | 6.74E-92 | 0.62 | 0.23 | 0.39 | 2.71 |
| periderm | Cstb | 1.31 | 4.09E-95 | 8.74E-91 | 0.97 | 0.83 | 0.13 | 1.16 |
| periderm | Hspb8 | 1.53 | 1.09E-90 | 2.34E-86 | 0.83 | 0.44 | 0.39 | 1.87 |
| periderm | S100a11 | 1.05 | 3.06E-90 | 6.53E-86 | 1.00 | 0.90 | 0.10 | 1.11 |
| periderm | Ovol1 | 3.13 | 1.79E-89 | 3.82E-85 | 0.30 | 0.05 | 0.26 | 6.33 |
| periderm | Dab1 | 1.43 | 3.79E-86 | 8.09E-82 | 0.82 | 0.45 | 0.38 | 1.85 |
| periderm | Smagp | 3.25 | 3.51E-85 | 7.50E-81 | 0.26 | 0.03 | 0.22 | 7.73 |
| periderm | Hebp2 | 2.10 | 5.63E-85 | 1.20E-80 | 0.63 | 0.27 | 0.36 | 2.36 |
| periderm | Pik3r1 | 1.13 | 2.63E-84 | 5.62E-80 | 0.90 | 0.56 | 0.34 | 1.62 |
| periderm | Irf6 | 1.43 | 3.02E-84 | 6.45E-80 | 0.77 | 0.37 | 0.41 | 2.11 |
| periderm | Dstn | 0.73 | 7.30E-77 | 1.56E-72 | 0.99 | 0.98 | 0.01 | 1.01 |
| periderm | S100a10 | 1.37 | 1.18E-74 | 2.52E-70 | 0.93 | 0.72 | 0.20 | 1.28 |
| periderm | Msx1 | 1.18 | 4.11E-74 | 8.78E-70 | 0.92 | 0.71 | 0.22 | 1.30 |
| periderm | Id3 | 0.94 | 1.46E-73 | 3.11E-69 | 1.00 | 0.96 | 0.04 | 1.04 |
| periderm | Snx31 | 3.52 | 7.04E-72 | 1.50E-67 | 0.19 | 0.02 | 0.17 | 9.55 |
| periderm | Gabrp | 5.49 | 5.10E-68 | 1.09E-63 | 0.12 | 0.01 | 0.12 | 20.50 |
| periderm | Gsta3 | 2.87 | 1.14E-67 | 2.43E-63 | 0.23 | 0.03 | 0.19 | 6.85 |
| periderm | Cyb5a | 0.66 | 1.64E-67 | 3.50E-63 | 0.99 | 0.96 | 0.03 | 1.03 |
| periderm | Slc2a3 | 2.32 | 4.28E-66 | 9.13E-62 | 0.64 | 0.31 | 0.33 | 2.05 |
| periderm | Acaa2 | 1.12 | 8.97E-65 | 1.92E-60 | 0.91 | 0.69 | 0.22 | 1.32 |
| periderm | Tmsb10 | 0.45 | 5.89E-64 | 1.26E-59 | 1.00 | 1.00 | 0.00 | 1.00 |
| periderm | Cpm | 1.28 | 2.04E-63 | 4.36E-59 | 0.81 | 0.50 | 0.32 | 1.64 |
| periderm | Rbp1 | 1.29 | 6.22E-63 | 1.33E-58 | 0.96 | 0.92 | 0.05 | 1.05 |
| periderm | Arhgap29 | 1.75 | 7.22E-63 | 1.54E-58 | 0.70 | 0.36 | 0.33 | 1.91 |
| periderm | Krt5 | 1.86 | 1.29E-62 | 2.75E-58 | 0.64 | 0.30 | 0.34 | 2.15 |
| periderm | Cldn6 | 0.74 | 3.45E-62 | 7.37E-58 | 0.98 | 0.86 | 0.12 | 1.14 |
| periderm | Lypd3 | 2.35 | 4.16E-61 | 8.88E-57 | 0.32 | 0.08 | 0.24 | 4.09 |
| periderm | Thrb | 2.02 | 4.40E-61 | 9.39E-57 | 0.39 | 0.12 | 0.27 | 3.34 |
| periderm | Rhov | 1.82 | 2.63E-60 | 5.62E-56 | 0.50 | 0.19 | 0.31 | 2.63 |
| periderm | Cacna1b | 1.44 | 4.37E-59 | 9.33E-55 | 0.60 | 0.27 | 0.33 | 2.24 |
| periderm | Emp1 | 2.52 | 2.53E-58 | 5.40E-54 | 0.41 | 0.14 | 0.27 | 2.92 |
| periderm | Laptm4a | 0.67 | 2.16E-57 | 4.61E-53 | 0.99 | 0.96 | 0.03 | 1.03 |
| periderm | Fabp5 | 0.57 | 7.22E-57 | 1.54E-52 | 1.00 | 0.99 | 0.01 | 1.01 |
| periderm | Rhcg | 2.57 | 9.73E-57 | 2.08E-52 | 0.18 | 0.02 | 0.15 | 7.38 |
| periderm | Sox9 | 1.12 | 1.43E-56 | 3.06E-52 | 0.84 | 0.54 | 0.30 | 1.55 |
| periderm | Cox4i2 | 1.06 | 1.62E-55 | 3.46E-51 | 0.77 | 0.45 | 0.32 | 1.72 |
| periderm | Pdlim1 | 1.14 | 4.92E-55 | 1.05E-50 | 0.87 | 0.62 | 0.25 | 1.41 |
| periderm | Hexb | 1.72 | 1.58E-54 | 3.38E-50 | 0.54 | 0.24 | 0.30 | 2.27 |
| periderm | Ddx58 | 2.02 | 7.09E-54 | 1.52E-49 | 0.33 | 0.09 | 0.24 | 3.65 |
| periderm | Ikzf2 | 0.80 | 7.29E-54 | 1.56E-49 | 0.94 | 0.83 | 0.11 | 1.14 |
| periderm | Msx2 | 1.06 | 2.16E-53 | 4.60E-49 | 0.71 | 0.36 | 0.35 | 1.97 |
| periderm | Jup | 1.14 | 2.24E-53 | 4.78E-49 | 0.77 | 0.49 | 0.28 | 1.58 |
| periderm | Gucy1a1 | 1.85 | 3.87E-53 | 8.28E-49 | 0.44 | 0.16 | 0.27 | 2.69 |
| periderm | Arf1 | 0.50 | 5.40E-53 | 1.15E-48 | 0.99 | 0.98 | 0.01 | 1.01 |
| periderm | Pkp1 | 1.63 | 5.78E-53 | 1.23E-48 | 0.30 | 0.08 | 0.23 | 4.03 |
| periderm | Samd5 | 0.93 | 2.62E-52 | 5.60E-48 | 0.91 | 0.66 | 0.25 | 1.38 |
| periderm | Eaf2 | 2.35 | 8.89E-52 | 1.90E-47 | 0.26 | 0.06 | 0.20 | 4.24 |
| periderm | Palm3 | 1.36 | 1.21E-51 | 2.59E-47 | 0.62 | 0.31 | 0.31 | 1.98 |
| periderm | Fjx1 | 0.80 | 3.40E-51 | 7.27E-47 | 0.93 | 0.78 | 0.16 | 1.20 |

**Supplementary Table 4. Marker genes for the annotated cell populations of integrated clustering of E11.5 control and NotchGOF λ-junction ectoderm cells.**

| | | | | | | | | |
|---|---|---|---|---|---|---|---|---|
| periderm | Myl12b | 0.62 | 5.31E-51 | 1.13E-46 | 0.98 | 0.93 | 0.04 | 1.04 |
| periderm | Atox1 | 0.54 | 7.12E-51 | 1.52E-46 | 0.99 | 0.98 | 0.02 | 1.02 |
| periderm | Cldn7 | 0.85 | 2.04E-50 | 4.35E-46 | 0.95 | 0.83 | 0.12 | 1.14 |
| periderm | Anxa2 | 0.90 | 3.91E-50 | 8.36E-46 | 0.96 | 0.73 | 0.23 | 1.32 |
| periderm | Bspry | 1.46 | 4.18E-50 | 8.93E-46 | 0.52 | 0.23 | 0.29 | 2.27 |
| periderm | Gja1 | 1.12 | 6.67E-50 | 1.42E-45 | 0.87 | 0.68 | 0.19 | 1.27 |
| periderm | Rgs2 | 2.04 | 1.50E-49 | 3.20E-45 | 0.54 | 0.26 | 0.27 | 2.05 |
| periderm | Ppp2r2b | 1.37 | 2.43E-49 | 5.19E-45 | 0.65 | 0.34 | 0.31 | 1.91 |
| periderm | Alad | 0.85 | 3.05E-49 | 6.51E-45 | 0.91 | 0.73 | 0.18 | 1.25 |
| periderm | Gas1 | 1.38 | 3.46E-49 | 7.40E-45 | 0.77 | 0.54 | 0.23 | 1.41 |
| periderm | St6galnac5 | 3.06 | 8.75E-48 | 1.87E-43 | 0.17 | 0.03 | 0.14 | 6.42 |
| periderm | Cmas | 0.99 | 1.08E-47 | 2.30E-43 | 0.84 | 0.63 | 0.21 | 1.33 |
| periderm | Hras | 0.61 | 2.19E-47 | 4.67E-43 | 0.96 | 0.90 | 0.06 | 1.07 |
| periderm | Smtnl2 | 1.50 | 2.51E-47 | 5.36E-43 | 0.51 | 0.23 | 0.28 | 2.26 |
| periderm | Nectin3 | 0.81 | 6.67E-47 | 1.42E-42 | 0.90 | 0.72 | 0.18 | 1.25 |
| periderm | Klhl23 | 1.09 | 8.48E-47 | 1.81E-42 | 0.75 | 0.49 | 0.26 | 1.53 |
| periderm | D7Ertd443e | 1.86 | 1.11E-46 | 2.38E-42 | 0.28 | 0.08 | 0.21 | 3.71 |
| periderm | Tspan7 | 1.06 | 1.77E-46 | 3.78E-42 | 0.72 | 0.44 | 0.28 | 1.65 |
| periderm | Cldn4 | 1.38 | 2.90E-46 | 6.20E-42 | 0.66 | 0.39 | 0.27 | 1.70 |
| periderm | Gjb3 | 1.69 | 5.92E-46 | 1.26E-41 | 0.36 | 0.12 | 0.24 | 3.00 |
| periderm | Rhob | 1.15 | 5.17E-45 | 1.10E-40 | 0.71 | 0.45 | 0.26 | 1.59 |
| periderm | Dlx3 | 1.13 | 6.92E-45 | 1.48E-40 | 0.55 | 0.25 | 0.30 | 2.22 |
| periderm | Rab15 | 1.15 | 8.52E-45 | 1.82E-40 | 0.67 | 0.39 | 0.28 | 1.72 |
| periderm | Rab11fip1 | 1.35 | 2.21E-44 | 4.72E-40 | 0.55 | 0.28 | 0.28 | 2.00 |
| periderm | Wnt4 | 0.69 | 3.14E-44 | 6.71E-40 | 0.93 | 0.68 | 0.25 | 1.36 |
| periderm | Il17re | 1.49 | 3.22E-44 | 6.88E-40 | 0.41 | 0.15 | 0.26 | 2.68 |
| periderm | Efna1 | 1.09 | 5.12E-44 | 1.09E-39 | 0.67 | 0.37 | 0.30 | 1.82 |
| periderm | Cldn8 | 1.94 | 5.38E-44 | 1.15E-39 | 0.32 | 0.10 | 0.22 | 3.17 |
| periderm | Penk | 3.38 | 5.45E-44 | 1.16E-39 | 0.19 | 0.04 | 0.15 | 5.00 |
| periderm | Anxa9 | 2.93 | 1.56E-43 | 3.34E-39 | 0.15 | 0.02 | 0.13 | 6.73 |
| periderm | Cdh1 | 0.94 | 1.64E-43 | 3.51E-39 | 0.84 | 0.63 | 0.21 | 1.33 |
| periderm | Aplp2 | 0.60 | 8.31E-43 | 1.77E-38 | 0.98 | 0.91 | 0.07 | 1.08 |
| periderm | Kif13b | 1.28 | 1.59E-42 | 3.41E-38 | 0.58 | 0.30 | 0.27 | 1.90 |
| periderm | Rab27b | 1.18 | 4.11E-42 | 8.78E-38 | 0.58 | 0.29 | 0.29 | 2.03 |
| periderm | Dsp | 0.73 | 7.62E-42 | 1.63E-37 | 0.97 | 0.86 | 0.11 | 1.13 |
| periderm | Plcd1 | 2.05 | 7.93E-42 | 1.69E-37 | 0.30 | 0.10 | 0.20 | 3.08 |
| periderm | Cd24a | 0.53 | 9.21E-42 | 1.97E-37 | 0.99 | 0.96 | 0.04 | 1.04 |
| periderm | Hs3st2 | 3.59 | 1.88E-41 | 4.02E-37 | 0.10 | 0.01 | 0.09 | 11.22 |
| periderm | Dennd2c | 1.97 | 2.42E-41 | 5.16E-37 | 0.27 | 0.08 | 0.19 | 3.48 |
| periderm | Kcnk1 | 0.99 | 3.34E-41 | 7.14E-37 | 0.79 | 0.56 | 0.23 | 1.42 |
| periderm | Lurap1l | 1.30 | 3.68E-41 | 7.87E-37 | 0.50 | 0.23 | 0.27 | 2.15 |
| periderm | Id1 | 0.56 | 3.90E-41 | 8.33E-37 | 0.99 | 0.92 | 0.06 | 1.07 |
| periderm | Hes1 | 0.87 | 5.08E-41 | 1.08E-36 | 0.87 | 0.71 | 0.16 | 1.22 |
| periderm | Cldn3 | 0.82 | 9.91E-41 | 2.12E-36 | 0.86 | 0.61 | 0.25 | 1.42 |
| periderm | Cacng4 | 1.15 | 2.71E-40 | 5.80E-36 | 0.59 | 0.31 | 0.28 | 1.90 |
| periderm | Mtss2 | 1.16 | 4.03E-40 | 8.60E-36 | 0.63 | 0.36 | 0.27 | 1.73 |
| periderm | Proser2 | 1.55 | 1.11E-39 | 2.38E-35 | 0.36 | 0.13 | 0.23 | 2.72 |
| periderm | Plce1 | 1.26 | 1.30E-39 | 2.78E-35 | 0.57 | 0.30 | 0.27 | 1.89 |
| periderm | Rbpms | 1.16 | 1.34E-39 | 2.86E-35 | 0.69 | 0.46 | 0.23 | 1.50 |
| periderm | Ablim1 | 0.95 | 1.43E-39 | 3.06E-35 | 0.77 | 0.52 | 0.24 | 1.47 |
| periderm | Itm2b | 0.38 | 4.12E-39 | 8.79E-35 | 0.99 | 0.93 | 0.06 | 1.06 |
| periderm | Rrbp1 | 0.69 | 8.21E-39 | 1.75E-34 | 0.95 | 0.88 | 0.07 | 1.08 |
| periderm | Rcsd1 | 1.66 | 1.29E-38 | 2.75E-34 | 0.36 | 0.14 | 0.22 | 2.60 |
| periderm | Plk2 | 1.13 | 1.95E-38 | 4.16E-34 | 0.60 | 0.33 | 0.27 | 1.83 |
| periderm | Wls | 0.58 | 8.46E-38 | 1.81E-33 | 0.95 | 0.78 | 0.17 | 1.22 |

**Supplementary Table 4. Marker genes for the annotated cell populations of integrated clustering of E11.5 control and NotchGOF λ-junction ectoderm cells.**

| | | | | | | | | |
|---|---|---|---|---|---|---|---|---|
| periderm | Bmp4 | 0.81 | 4.91E-37 | 1.05E-32 | 0.77 | 0.50 | 0.27 | 1.53 |
| periderm | Ahr | 1.45 | 1.63E-36 | 3.49E-32 | 0.38 | 0.15 | 0.23 | 2.48 |
| periderm | Pdzrn3 | 1.73 | 1.72E-36 | 3.68E-32 | 0.29 | 0.10 | 0.20 | 3.00 |
| periderm | Clcc1 | 1.08 | 2.06E-36 | 4.40E-32 | 0.67 | 0.45 | 0.23 | 1.51 |
| periderm | Cndp1 | 2.54 | 2.71E-36 | 5.79E-32 | 0.14 | 0.03 | 0.12 | 5.54 |
| periderm | Ppp1r13l | 1.44 | 8.29E-36 | 1.77E-31 | 0.40 | 0.17 | 0.23 | 2.37 |
| periderm | Apoc1 | 1.32 | 2.30E-35 | 4.91E-31 | 0.51 | 0.26 | 0.25 | 1.95 |
| periderm | Ptpn13 | 0.94 | 1.49E-34 | 3.17E-30 | 0.77 | 0.55 | 0.22 | 1.41 |
| periderm | Arg1 | 2.56 | 5.41E-34 | 1.15E-29 | 0.17 | 0.04 | 0.13 | 4.46 |
| periderm | Slit3 | 2.00 | 9.22E-34 | 1.97E-29 | 0.20 | 0.05 | 0.15 | 3.77 |
| periderm | Kdm5b | 0.56 | 9.68E-34 | 2.07E-29 | 0.96 | 0.92 | 0.04 | 1.05 |
| periderm | Mpzl2 | 0.76 | 1.39E-33 | 2.97E-29 | 0.87 | 0.72 | 0.15 | 1.21 |
| periderm | Tpm1 | 0.56 | 1.40E-33 | 2.99E-29 | 0.98 | 0.93 | 0.05 | 1.05 |
| periderm | Nebl | 1.50 | 1.95E-33 | 4.17E-29 | 0.60 | 0.37 | 0.22 | 1.60 |
| periderm | Gria3 | 1.35 | 2.28E-33 | 4.88E-29 | 0.40 | 0.17 | 0.22 | 2.29 |
| periderm | Ctsh | 1.73 | 3.80E-33 | 8.11E-29 | 0.29 | 0.10 | 0.18 | 2.77 |
| periderm | Ostf1 | 1.01 | 6.21E-33 | 1.33E-28 | 0.63 | 0.39 | 0.24 | 1.60 |
| periderm | Evpl | 1.72 | 7.23E-33 | 1.55E-28 | 0.28 | 0.10 | 0.18 | 2.86 |
| periderm | Tgfb2 | 0.53 | 8.53E-33 | 1.82E-28 | 0.81 | 0.55 | 0.26 | 1.48 |
| periderm | Mir9-3hg | 1.46 | 1.32E-32 | 2.83E-28 | 0.26 | 0.08 | 0.18 | 3.13 |
| periderm | Trim2 | 0.86 | 1.37E-32 | 2.93E-28 | 0.76 | 0.57 | 0.19 | 1.33 |
| periderm | Adi1 | 1.07 | 1.66E-32 | 3.54E-28 | 0.65 | 0.44 | 0.21 | 1.49 |
| periderm | Sh2d4a | 2.28 | 1.90E-32 | 4.07E-28 | 0.15 | 0.03 | 0.12 | 4.75 |
| periderm | Ier5 | 0.56 | 2.78E-32 | 5.94E-28 | 0.94 | 0.83 | 0.11 | 1.14 |
| periderm | Esrp2 | 1.30 | 2.93E-32 | 6.25E-28 | 0.46 | 0.23 | 0.23 | 1.97 |
| periderm | Bambi | 0.35 | 1.32E-31 | 2.82E-27 | 0.73 | 0.45 | 0.28 | 1.61 |
| periderm | Ppm1l | 0.99 | 1.46E-31 | 3.12E-27 | 0.63 | 0.40 | 0.23 | 1.56 |
| periderm | Tap1 | 1.81 | 1.57E-31 | 3.35E-27 | 0.20 | 0.06 | 0.15 | 3.61 |
| periderm | Pdcd4 | 0.71 | 1.91E-31 | 4.08E-27 | 0.92 | 0.77 | 0.15 | 1.19 |
| periderm | Dlx2 | 0.73 | 2.76E-31 | 5.90E-27 | 0.59 | 0.31 | 0.28 | 1.89 |
| periderm | Ywhaz | 0.38 | 2.81E-31 | 5.99E-27 | 0.99 | 0.97 | 0.01 | 1.01 |
| periderm | Marveld2 | 1.31 | 5.26E-31 | 1.12E-26 | 0.43 | 0.20 | 0.23 | 2.13 |
| periderm | Cyb5r3 | 0.45 | 5.27E-31 | 1.13E-26 | 0.97 | 0.92 | 0.05 | 1.05 |
| periderm | Cdkn2b | 0.63 | 3.37E-30 | 7.19E-26 | 0.25 | 0.08 | 0.17 | 3.14 |
| periderm | Bicdl2 | 1.98 | 3.89E-30 | 8.30E-26 | 0.16 | 0.04 | 0.12 | 4.13 |
| periderm | Erbb3 | 0.79 | 4.16E-30 | 8.88E-26 | 0.74 | 0.51 | 0.23 | 1.44 |
| periderm | Ptprf | 0.48 | 4.25E-30 | 9.08E-26 | 0.99 | 0.96 | 0.03 | 1.04 |
| periderm | Kcnj15 | 1.99 | 1.17E-29 | 2.51E-25 | 0.17 | 0.04 | 0.13 | 3.89 |
| periderm | Platr21 | 1.72 | 1.50E-29 | 3.20E-25 | 0.25 | 0.08 | 0.16 | 2.95 |
| periderm | Perp | 0.73 | 1.60E-29 | 3.43E-25 | 0.78 | 0.51 | 0.27 | 1.53 |
| periderm | Klf5 | 0.95 | 3.02E-29 | 6.45E-25 | 0.62 | 0.39 | 0.24 | 1.61 |
| periderm | Gas7 | 1.61 | 3.51E-29 | 7.51E-25 | 0.25 | 0.09 | 0.16 | 2.91 |
| periderm | Rcn1 | 0.55 | 3.59E-29 | 7.67E-25 | 0.93 | 0.84 | 0.09 | 1.11 |
| periderm | Rnf24 | 0.89 | 4.77E-29 | 1.02E-24 | 0.61 | 0.38 | 0.23 | 1.60 |
| periderm | Fermt2 | 0.75 | 5.46E-29 | 1.17E-24 | 0.80 | 0.65 | 0.16 | 1.24 |
| periderm | Gstt3 | 1.66 | 7.01E-29 | 1.50E-24 | 0.27 | 0.10 | 0.17 | 2.70 |
| periderm | Spint1 | 1.03 | 7.83E-29 | 1.67E-24 | 0.60 | 0.38 | 0.22 | 1.58 |
| periderm | Gimap9 | 1.39 | 1.22E-28 | 2.61E-24 | 0.29 | 0.11 | 0.18 | 2.60 |
| periderm | Gjb2 | 1.71 | 1.55E-28 | 3.32E-24 | 0.35 | 0.16 | 0.19 | 2.19 |
| periderm | Creb3l2 | 1.15 | 1.70E-28 | 3.63E-24 | 0.52 | 0.30 | 0.22 | 1.73 |
| periderm | Tafa5 | 1.17 | 2.26E-28 | 4.83E-24 | 0.44 | 0.22 | 0.21 | 1.96 |
| periderm | Dcaf4 | 1.04 | 2.61E-28 | 5.57E-24 | 0.56 | 0.32 | 0.24 | 1.73 |
| periderm | Ptprz1 | 1.07 | 2.68E-28 | 5.72E-24 | 0.41 | 0.20 | 0.22 | 2.09 |
| periderm | Lad1 | 1.02 | 3.15E-28 | 6.72E-24 | 0.55 | 0.32 | 0.23 | 1.73 |
| periderm | Suco | 0.96 | 3.74E-28 | 7.99E-24 | 0.72 | 0.56 | 0.16 | 1.28 |

**Supplementary Table 4. Marker genes for the annotated cell populations of integrated clustering of E11.5 control and NotchGOF λ-junction ectoderm cells.**

| | | | | | | | | |
|---|---|---|---|---|---|---|---|---|
| periderm | Ralbp1 | 0.61 | 9.19E-28 | 1.96E-23 | 0.90 | 0.82 | 0.09 | 1.11 |
| periderm | Fgfr3 | 1.94 | 1.50E-27 | 3.21E-23 | 0.15 | 0.04 | 0.12 | 4.11 |
| periderm | Kctd1 | 0.79 | 1.57E-27 | 3.35E-23 | 0.69 | 0.48 | 0.21 | 1.44 |
| periderm | Slc15a2 | 1.28 | 1.64E-27 | 3.51E-23 | 0.31 | 0.13 | 0.19 | 2.46 |
| periderm | Pbx1 | 0.42 | 2.22E-27 | 4.74E-23 | 0.98 | 0.97 | 0.02 | 1.02 |
| periderm | Cers3 | 2.03 | 3.82E-27 | 8.16E-23 | 0.14 | 0.03 | 0.11 | 4.24 |
| periderm | Hmx1 | 2.53 | 4.28E-27 | 9.13E-23 | 0.10 | 0.02 | 0.09 | 5.72 |
| periderm | Six3os1 | 0.55 | 8.58E-27 | 1.83E-22 | 0.90 | 0.75 | 0.15 | 1.20 |
| periderm | Tfap2a | 0.58 | 1.22E-26 | 2.61E-22 | 0.57 | 0.32 | 0.26 | 1.82 |
| periderm | Jag1 | 0.84 | 1.71E-26 | 3.66E-22 | 0.73 | 0.54 | 0.19 | 1.36 |
| periderm | Cd2ap | 0.54 | 2.01E-26 | 4.29E-22 | 0.90 | 0.79 | 0.11 | 1.13 |
| periderm | Nrip1 | 0.57 | 2.02E-26 | 4.31E-22 | 0.92 | 0.80 | 0.12 | 1.15 |
| periderm | Cldn23 | 1.51 | 4.63E-26 | 9.89E-22 | 0.31 | 0.14 | 0.18 | 2.30 |
| periderm | Ahnak | 1.04 | 4.77E-26 | 1.02E-21 | 0.56 | 0.34 | 0.23 | 1.67 |
| periderm | Gadd45b | 1.04 | 6.97E-26 | 1.49E-21 | 0.39 | 0.18 | 0.21 | 2.15 |
| periderm | Dcbld1 | 0.85 | 7.08E-26 | 1.51E-21 | 0.60 | 0.37 | 0.23 | 1.62 |
| periderm | Nrtn | 1.14 | 7.71E-26 | 1.65E-21 | 0.50 | 0.30 | 0.20 | 1.67 |
| periderm | B3galnt1 | 1.31 | 1.08E-25 | 2.31E-21 | 0.33 | 0.14 | 0.18 | 2.27 |
| periderm | Ifitm3 | 0.60 | 1.15E-25 | 2.45E-21 | 0.65 | 0.40 | 0.26 | 1.65 |
| periderm | Me1 | 1.12 | 1.51E-25 | 3.22E-21 | 0.36 | 0.17 | 0.20 | 2.15 |
| periderm | Gab1 | 1.01 | 1.51E-25 | 3.23E-21 | 0.47 | 0.27 | 0.21 | 1.78 |
| periderm | Wwc1 | 0.79 | 2.00E-25 | 4.27E-21 | 0.75 | 0.57 | 0.18 | 1.32 |
| periderm | Ezr | 0.46 | 2.04E-25 | 4.36E-21 | 0.97 | 0.90 | 0.07 | 1.07 |
| periderm | Vcan | 0.80 | 3.97E-25 | 8.48E-21 | 0.84 | 0.74 | 0.10 | 1.14 |
| periderm | Sdc1 | 0.90 | 7.61E-25 | 1.63E-20 | 0.75 | 0.57 | 0.18 | 1.31 |
| periderm | Elovl7 | 1.50 | 7.71E-25 | 1.65E-20 | 0.23 | 0.08 | 0.15 | 2.83 |
| periderm | Esrp1 | 0.76 | 8.61E-25 | 1.84E-20 | 0.77 | 0.63 | 0.15 | 1.23 |
| periderm | Runx1 | 0.31 | 8.91E-25 | 1.90E-20 | 0.74 | 0.49 | 0.26 | 1.52 |
| periderm | Cab39 | 0.76 | 9.73E-25 | 2.08E-20 | 0.77 | 0.64 | 0.13 | 1.20 |
| periderm | Ripply3 | 1.08 | 1.01E-24 | 2.15E-20 | 0.35 | 0.16 | 0.19 | 2.19 |
| periderm | Phldb3 | 1.55 | 1.02E-24 | 2.17E-20 | 0.21 | 0.07 | 0.14 | 2.99 |
| periderm | Stx19 | 2.44 | 1.22E-24 | 2.61E-20 | 0.12 | 0.02 | 0.09 | 4.79 |
| periderm | Papss1 | 0.63 | 1.75E-24 | 3.74E-20 | 0.80 | 0.65 | 0.15 | 1.23 |
| periderm | Capg | 0.90 | 2.00E-24 | 4.28E-20 | 0.55 | 0.33 | 0.22 | 1.66 |
| periderm | Prkch | 1.15 | 2.05E-24 | 4.37E-20 | 0.37 | 0.18 | 0.19 | 2.04 |
| periderm | Stk39 | 0.98 | 2.89E-24 | 6.17E-20 | 0.49 | 0.28 | 0.21 | 1.74 |
| periderm | Plekhg1 | 1.13 | 3.03E-24 | 6.47E-20 | 0.47 | 0.27 | 0.20 | 1.73 |
| periderm | Bex1 | 0.35 | 4.19E-24 | 8.95E-20 | 0.98 | 0.96 | 0.02 | 1.02 |
| periderm | Rfk | 0.79 | 4.80E-24 | 1.02E-19 | 0.69 | 0.50 | 0.20 | 1.39 |
| periderm | Nxn | 0.54 | 5.48E-24 | 1.17E-19 | 0.86 | 0.74 | 0.12 | 1.16 |
| periderm | Elf5 | 1.10 | 5.60E-24 | 1.20E-19 | 0.41 | 0.22 | 0.20 | 1.90 |
| periderm | Tfap2c | 0.86 | 5.76E-24 | 1.23E-19 | 0.41 | 0.20 | 0.21 | 2.07 |
| periderm | Cdc42ep5 | 0.73 | 6.18E-24 | 1.32E-19 | 0.73 | 0.58 | 0.16 | 1.27 |
| periderm | Smim5 | 1.72 | 6.48E-24 | 1.38E-19 | 0.17 | 0.05 | 0.12 | 3.38 |
| periderm | H19 | 0.56 | 6.60E-24 | 1.41E-19 | 0.97 | 0.87 | 0.10 | 1.12 |
| periderm | Limk2 | 1.24 | 6.79E-24 | 1.45E-19 | 0.52 | 0.33 | 0.18 | 1.55 |
| periderm | Trmt9b | 1.21 | 6.88E-24 | 1.47E-19 | 0.34 | 0.16 | 0.18 | 2.15 |
| periderm | Pycard | 1.26 | 7.71E-24 | 1.65E-19 | 0.38 | 0.19 | 0.19 | 1.96 |
| periderm | Cnn2 | 1.00 | 1.27E-23 | 2.72E-19 | 0.49 | 0.29 | 0.21 | 1.71 |
| periderm | Ermp1 | 0.82 | 1.43E-23 | 3.05E-19 | 0.62 | 0.41 | 0.21 | 1.52 |
| periderm | Ppfibp1 | 0.69 | 1.85E-23 | 3.95E-19 | 0.77 | 0.60 | 0.17 | 1.29 |
| periderm | Fermt1 | 0.52 | 2.67E-23 | 5.71E-19 | 0.79 | 0.53 | 0.26 | 1.50 |
| periderm | Enc1 | 0.81 | 3.31E-23 | 7.08E-19 | 0.43 | 0.23 | 0.20 | 1.87 |
| periderm | Ddah2 | 0.33 | 6.48E-23 | 1.38E-18 | 0.99 | 0.97 | 0.02 | 1.02 |
| periderm | Nfkbia | 0.77 | 1.38E-22 | 2.96E-18 | 0.66 | 0.47 | 0.19 | 1.40 |

**Supplementary Table 4. Marker genes for the annotated cell populations of integrated clustering of E11.5 control and NotchGOF λ-junction ectoderm cells.**

| | | | | | | | | |
|---|---|---|---|---|---|---|---|---|
| periderm | Rnh1 | 0.53 | 1.63E-22 | 3.48E-18 | 0.86 | 0.77 | 0.10 | 1.12 |
| periderm | Prr5l | 0.91 | 1.67E-22 | 3.56E-18 | 0.53 | 0.33 | 0.20 | 1.59 |
| periderm | Ptgr1 | 1.38 | 2.81E-22 | 6.00E-18 | 0.30 | 0.14 | 0.16 | 2.16 |
| periderm | Gipc1 | 0.57 | 3.17E-22 | 6.77E-18 | 0.86 | 0.74 | 0.12 | 1.17 |
| periderm | H2-T23 | 1.27 | 3.38E-22 | 7.22E-18 | 0.30 | 0.13 | 0.17 | 2.26 |
| periderm | Metrnl | 1.50 | 3.54E-22 | 7.56E-18 | 0.33 | 0.16 | 0.17 | 2.06 |
| periderm | Iqgap1 | 0.65 | 5.29E-22 | 1.13E-17 | 0.82 | 0.67 | 0.15 | 1.23 |
| periderm | Wnt5a | 0.54 | 6.18E-22 | 1.32E-17 | 0.63 | 0.41 | 0.22 | 1.54 |
| periderm | Mgst1 | 0.60 | 7.47E-22 | 1.60E-17 | 0.84 | 0.68 | 0.16 | 1.24 |
| periderm | Lhfp | 0.79 | 8.06E-22 | 1.72E-17 | 0.50 | 0.29 | 0.21 | 1.73 |
| periderm | Mpp7 | 0.76 | 8.32E-22 | 1.78E-17 | 0.71 | 0.53 | 0.18 | 1.34 |
| periderm | Unc5b | 0.72 | 1.31E-21 | 2.80E-17 | 0.70 | 0.51 | 0.19 | 1.37 |
| periderm | Cdkn1a | 0.27 | 1.38E-21 | 2.94E-17 | 0.59 | 0.36 | 0.23 | 1.62 |
| periderm | A730020E08Rik | 1.50 | 1.56E-21 | 3.32E-17 | 0.20 | 0.07 | 0.13 | 2.78 |
| periderm | Dhrs3 | 1.08 | 2.15E-21 | 4.60E-17 | 0.37 | 0.19 | 0.18 | 1.94 |
| periderm | Mycl | 0.58 | 2.99E-21 | 6.40E-17 | 0.78 | 0.62 | 0.16 | 1.26 |
| periderm | Slc38a4 | 0.98 | 6.68E-21 | 1.43E-16 | 0.45 | 0.26 | 0.19 | 1.75 |
| periderm | Ocln | 0.93 | 7.61E-21 | 1.63E-16 | 0.48 | 0.28 | 0.19 | 1.69 |
| periderm | Elf1 | 0.72 | 7.96E-21 | 1.70E-16 | 0.67 | 0.48 | 0.19 | 1.40 |
| periderm | Erbb2 | 1.02 | 9.92E-21 | 2.12E-16 | 0.49 | 0.30 | 0.18 | 1.60 |
| periderm | Cebpb | 1.02 | 1.02E-20 | 2.17E-16 | 0.41 | 0.23 | 0.18 | 1.81 |
| periderm | Sh3bp5 | 0.83 | 1.14E-20 | 2.43E-16 | 0.56 | 0.36 | 0.20 | 1.54 |
| periderm | Dok5 | 0.70 | 1.19E-20 | 2.55E-16 | 0.57 | 0.35 | 0.22 | 1.62 |
| periderm | Pcdh1 | 1.47 | 1.52E-20 | 3.25E-16 | 0.22 | 0.09 | 0.14 | 2.55 |
| periderm | Rnd3 | 0.46 | 1.78E-20 | 3.79E-16 | 0.87 | 0.72 | 0.15 | 1.21 |
| periderm | Krt7 | 1.27 | 2.29E-20 | 4.90E-16 | 0.36 | 0.19 | 0.17 | 1.89 |
| periderm | Fam117a | 0.97 | 2.54E-20 | 5.43E-16 | 0.44 | 0.26 | 0.19 | 1.73 |
| periderm | Nrarp | 0.59 | 3.23E-20 | 6.89E-16 | 0.79 | 0.58 | 0.21 | 1.35 |
| periderm | Ank3 | 0.54 | 3.72E-20 | 7.95E-16 | 0.76 | 0.62 | 0.14 | 1.23 |
| periderm | S100a16 | 1.22 | 4.23E-20 | 9.03E-16 | 0.29 | 0.13 | 0.16 | 2.18 |
| periderm | Dusp2 | 1.90 | 8.17E-20 | 1.74E-15 | 0.12 | 0.03 | 0.09 | 3.90 |
| periderm | Anp32a | 0.25 | 1.02E-19 | 2.17E-15 | 1.00 | 0.99 | 0.01 | 1.01 |
| periderm | Lrrfip1 | 0.71 | 1.29E-19 | 2.75E-15 | 0.66 | 0.48 | 0.18 | 1.37 |
| periderm | Mdh1 | 0.34 | 1.40E-19 | 2.99E-15 | 0.99 | 0.98 | 0.01 | 1.01 |
| periderm | Sh3bgrl3 | 0.38 | 1.50E-19 | 3.20E-15 | 0.96 | 0.91 | 0.05 | 1.06 |
| periderm | Lmo7 | 1.03 | 1.63E-19 | 3.47E-15 | 0.60 | 0.43 | 0.16 | 1.37 |
| periderm | Macc1 | 1.05 | 1.76E-19 | 3.75E-15 | 0.38 | 0.20 | 0.17 | 1.85 |
| periderm | Prxl2a | 0.61 | 1.85E-19 | 3.95E-15 | 0.74 | 0.57 | 0.17 | 1.30 |
| periderm | Dsg2 | 0.67 | 2.63E-19 | 5.62E-15 | 0.65 | 0.47 | 0.18 | 1.39 |
| periderm | Arl4c | 0.52 | 3.93E-19 | 8.40E-15 | 0.74 | 0.56 | 0.18 | 1.32 |
| periderm | Isl2 | 1.08 | 5.18E-19 | 1.11E-14 | 0.28 | 0.13 | 0.15 | 2.21 |
| periderm | Fech | 0.97 | 7.24E-19 | 1.55E-14 | 0.45 | 0.27 | 0.17 | 1.64 |
| periderm | Tbx2 | 1.00 | 8.23E-19 | 1.76E-14 | 0.28 | 0.13 | 0.15 | 2.16 |
| periderm | Anxa1 | 1.72 | 1.32E-18 | 2.81E-14 | 0.15 | 0.05 | 0.10 | 2.98 |
| periderm | Sorl1 | 0.86 | 2.48E-18 | 5.29E-14 | 0.51 | 0.34 | 0.18 | 1.53 |
| periderm | Dbi | 0.30 | 2.79E-18 | 5.96E-14 | 0.99 | 0.97 | 0.03 | 1.03 |
| periderm | Etv3 | 0.85 | 3.41E-18 | 7.28E-14 | 0.45 | 0.27 | 0.18 | 1.65 |
| periderm | Pygl | 1.29 | 3.82E-18 | 8.16E-14 | 0.32 | 0.16 | 0.15 | 1.93 |
| periderm | Myh9 | 0.50 | 4.27E-18 | 9.13E-14 | 0.86 | 0.74 | 0.12 | 1.16 |
| periderm | Plp2 | 0.56 | 5.90E-18 | 1.26E-13 | 0.80 | 0.69 | 0.12 | 1.17 |
| periderm | Psap | 0.51 | 7.08E-18 | 1.51E-13 | 0.85 | 0.73 | 0.12 | 1.16 |
| periderm | Gata3 | 0.70 | 8.79E-18 | 1.88E-13 | 0.51 | 0.32 | 0.19 | 1.59 |
| periderm | Hibadh | 0.65 | 1.17E-17 | 2.50E-13 | 0.74 | 0.60 | 0.14 | 1.23 |
| periderm | Igsf9b | 1.11 | 1.34E-17 | 2.87E-13 | 0.30 | 0.15 | 0.15 | 1.99 |
| periderm | Epn3 | 1.10 | 1.62E-17 | 3.45E-13 | 0.31 | 0.16 | 0.15 | 1.96 |

**Supplementary Table 4. Marker genes for the annotated cell populations of integrated clustering of E11.5 control and NotchGOF λ-junction ectoderm cells.**

| | | | | | | | | |
|---|---|---|---|---|---|---|---|---|
| periderm | Cblc | 1.24 | 1.82E-17 | 3.89E-13 | 0.26 | 0.12 | 0.14 | 2.14 |
| periderm | Tor3a | 1.36 | 2.05E-17 | 4.37E-13 | 0.19 | 0.07 | 0.11 | 2.57 |
| periderm | Epha4 | 0.36 | 2.67E-17 | 5.71E-13 | 0.73 | 0.54 | 0.19 | 1.35 |
| periderm | Cux1 | 0.40 | 2.72E-17 | 5.80E-13 | 0.91 | 0.83 | 0.08 | 1.09 |
| periderm | Nxph3 | 1.37 | 3.23E-17 | 6.91E-13 | 0.15 | 0.05 | 0.10 | 2.96 |
| periderm | Prdx5 | 0.72 | 3.49E-17 | 7.45E-13 | 0.87 | 0.80 | 0.08 | 1.09 |
| periderm | Bcl11b | 0.34 | 3.53E-17 | 7.55E-13 | 0.85 | 0.70 | 0.15 | 1.21 |
| periderm | Ier5l | 0.56 | 4.19E-17 | 8.95E-13 | 0.79 | 0.64 | 0.15 | 1.24 |
| periderm | Dlgap4 | 0.44 | 8.77E-17 | 1.87E-12 | 0.81 | 0.68 | 0.13 | 1.18 |
| periderm | Smpdl3a | 0.98 | 1.06E-16 | 2.26E-12 | 0.32 | 0.17 | 0.16 | 1.92 |
| periderm | Eml5 | 0.93 | 1.09E-16 | 2.33E-12 | 0.38 | 0.22 | 0.17 | 1.77 |
| periderm | AW011738 | 0.79 | 1.11E-16 | 2.36E-12 | 0.45 | 0.28 | 0.17 | 1.62 |
| periderm | Cebpa | 1.34 | 1.15E-16 | 2.46E-12 | 0.18 | 0.07 | 0.11 | 2.55 |
| periderm | Bcam | 0.39 | 1.25E-16 | 2.67E-12 | 0.87 | 0.71 | 0.17 | 1.23 |
| periderm | Sinhcaf | 0.33 | 1.37E-16 | 2.93E-12 | 0.97 | 0.94 | 0.03 | 1.04 |
| periderm | Crabp1 | 0.36 | 1.49E-16 | 3.18E-12 | 0.69 | 0.55 | 0.14 | 1.26 |
| periderm | Crybg1 | 1.57 | 1.51E-16 | 3.24E-12 | 0.13 | 0.04 | 0.09 | 3.21 |
| periderm | Sash1 | 0.69 | 1.82E-16 | 3.90E-12 | 0.58 | 0.40 | 0.18 | 1.45 |
| periderm | Npm3 | 0.39 | 2.20E-16 | 4.70E-12 | 0.95 | 0.87 | 0.08 | 1.10 |
| periderm | Sh3tc2 | 1.67 | 2.23E-16 | 4.76E-12 | 0.13 | 0.05 | 0.09 | 2.98 |
| periderm | Churc1 | 0.55 | 2.49E-16 | 5.33E-12 | 0.82 | 0.72 | 0.11 | 1.15 |
| periderm | Tgif1 | 0.46 | 4.69E-16 | 1.00E-11 | 0.80 | 0.67 | 0.13 | 1.19 |
| periderm | AI661453 | 1.06 | 5.32E-16 | 1.14E-11 | 0.26 | 0.12 | 0.14 | 2.13 |
| periderm | Gpc3 | 0.25 | 6.95E-16 | 1.49E-11 | 0.95 | 0.83 | 0.13 | 1.15 |
| periderm | Gtf2i | 0.35 | 7.61E-16 | 1.63E-11 | 0.95 | 0.90 | 0.05 | 1.06 |
| periderm | Tbc1d9 | 1.04 | 1.11E-15 | 2.37E-11 | 0.24 | 0.11 | 0.13 | 2.15 |
| periderm | Frmd4b | 0.49 | 1.11E-15 | 2.37E-11 | 0.66 | 0.48 | 0.18 | 1.39 |
| periderm | Camsap3 | 0.79 | 1.14E-15 | 2.44E-11 | 0.43 | 0.26 | 0.17 | 1.63 |
| periderm | Selenoh | 0.27 | 1.38E-15 | 2.95E-11 | 0.98 | 0.95 | 0.03 | 1.03 |
| periderm | Axin2 | 0.61 | 1.63E-15 | 3.47E-11 | 0.41 | 0.24 | 0.17 | 1.70 |
| periderm | Exph5 | 1.06 | 1.88E-15 | 4.02E-11 | 0.27 | 0.14 | 0.13 | 1.99 |
| periderm | Naaladl2 | 1.49 | 2.14E-15 | 4.57E-11 | 0.12 | 0.04 | 0.08 | 3.19 |
| periderm | Adcy1 | 1.48 | 2.18E-15 | 4.66E-11 | 0.14 | 0.05 | 0.09 | 2.77 |
| periderm | Etl4 | 0.49 | 2.46E-15 | 5.26E-11 | 0.73 | 0.56 | 0.17 | 1.30 |
| periderm | Junb | 0.77 | 2.55E-15 | 5.45E-11 | 0.35 | 0.20 | 0.16 | 1.80 |
| periderm | Nab1 | 0.68 | 2.85E-15 | 6.08E-11 | 0.39 | 0.22 | 0.17 | 1.75 |
| periderm | Hey1 | 0.39 | 3.03E-15 | 6.47E-11 | 0.90 | 0.73 | 0.17 | 1.23 |
| periderm | Syt7 | 0.98 | 3.94E-15 | 8.42E-11 | 0.26 | 0.13 | 0.13 | 2.01 |
| periderm | Fzd3 | 0.40 | 3.95E-15 | 8.43E-11 | 0.92 | 0.84 | 0.08 | 1.10 |
| periderm | Celf4 | 0.77 | 4.02E-15 | 8.58E-11 | 0.45 | 0.28 | 0.16 | 1.58 |
| periderm | Hspa2 | 0.95 | 4.04E-15 | 8.62E-11 | 0.26 | 0.13 | 0.13 | 2.04 |
| periderm | Mindy2 | 0.78 | 4.31E-15 | 9.20E-11 | 0.40 | 0.23 | 0.16 | 1.70 |
| periderm | Rab25 | 0.51 | 4.66E-15 | 9.96E-11 | 0.77 | 0.61 | 0.15 | 1.25 |
| periderm | Fzd10os | 0.94 | 6.05E-15 | 1.29E-10 | 0.27 | 0.14 | 0.14 | 1.99 |
| periderm | Cldn1 | 0.69 | 6.30E-15 | 1.35E-10 | 0.21 | 0.09 | 0.12 | 2.31 |
| periderm | Gldc | 1.06 | 6.79E-15 | 1.45E-10 | 0.26 | 0.13 | 0.13 | 1.99 |
| periderm | Meis2 | 0.28 | 7.21E-15 | 1.54E-10 | 0.98 | 0.95 | 0.03 | 1.03 |
| periderm | Mark2 | 0.56 | 7.28E-15 | 1.56E-10 | 0.70 | 0.57 | 0.13 | 1.22 |
| periderm | Hspb1 | 1.27 | 9.35E-15 | 2.00E-10 | 0.52 | 0.37 | 0.15 | 1.41 |
| periderm | Tiam1 | 0.70 | 1.03E-14 | 2.20E-10 | 0.60 | 0.44 | 0.16 | 1.36 |
| periderm | Frem2 | 0.50 | 1.22E-14 | 2.61E-10 | 0.77 | 0.61 | 0.16 | 1.26 |
| periderm | Oxct1 | 0.45 | 1.28E-14 | 2.74E-10 | 0.86 | 0.77 | 0.09 | 1.12 |
| periderm | Fhdc1 | 0.99 | 1.58E-14 | 3.38E-10 | 0.24 | 0.11 | 0.12 | 2.08 |
| periderm | Rbms1 | 0.34 | 1.76E-14 | 3.77E-10 | 0.90 | 0.84 | 0.05 | 1.06 |
| periderm | Mtss1 | 0.75 | 2.04E-14 | 4.36E-10 | 0.56 | 0.42 | 0.14 | 1.34 |

**Supplementary Table 4. Marker genes for the annotated cell populations of integrated clustering of E11.5 control and NotchGOF λ-junction ectoderm cells.**

| | | | | | | | | |
|---|---|---|---|---|---|---|---|---|
| periderm | Igf2r | 0.49 | 2.45E-14 | 5.24E-10 | 0.72 | 0.56 | 0.15 | 1.27 |
| periderm | Satb1 | 0.51 | 2.59E-14 | 5.53E-10 | 0.69 | 0.55 | 0.14 | 1.25 |
| periderm | Hspa1b | 1.63 | 2.67E-14 | 5.71E-10 | 0.11 | 0.03 | 0.07 | 3.18 |
| periderm | App | 0.33 | 2.96E-14 | 6.32E-10 | 0.91 | 0.85 | 0.06 | 1.07 |
| periderm | Mgll | 1.35 | 3.15E-14 | 6.74E-10 | 0.15 | 0.06 | 0.09 | 2.57 |
| periderm | Pros1 | 1.11 | 4.34E-14 | 9.28E-10 | 0.20 | 0.09 | 0.11 | 2.19 |
| periderm | Cd36 | 1.22 | 4.36E-14 | 9.31E-10 | 0.11 | 0.04 | 0.08 | 3.08 |
| periderm | Podxl2 | 0.59 | 4.43E-14 | 9.47E-10 | 0.53 | 0.37 | 0.16 | 1.44 |
| periderm | Crot | 0.73 | 4.89E-14 | 1.04E-09 | 0.51 | 0.35 | 0.15 | 1.44 |
| periderm | Lgalsl | 0.49 | 5.37E-14 | 1.15E-09 | 0.62 | 0.45 | 0.17 | 1.37 |
| periderm | Arl8b | 0.59 | 6.26E-14 | 1.34E-09 | 0.66 | 0.51 | 0.15 | 1.29 |
| periderm | Zfp503 | 0.45 | 8.00E-14 | 1.71E-09 | 0.66 | 0.49 | 0.18 | 1.36 |
| periderm | Rhou | 0.66 | 8.22E-14 | 1.76E-09 | 0.61 | 0.47 | 0.14 | 1.30 |
| periderm | Mmp11 | 0.57 | 9.89E-14 | 2.11E-09 | 0.54 | 0.37 | 0.17 | 1.44 |
| periderm | Sp6 | 0.81 | 1.06E-13 | 2.25E-09 | 0.23 | 0.11 | 0.12 | 2.11 |
| periderm | Faah | 0.81 | 1.07E-13 | 2.28E-09 | 0.35 | 0.21 | 0.14 | 1.67 |
| periderm | Tmem54 | 0.65 | 1.13E-13 | 2.41E-09 | 0.60 | 0.46 | 0.14 | 1.29 |
| periderm | Tsc22d4 | 0.56 | 1.15E-13 | 2.46E-09 | 0.72 | 0.61 | 0.10 | 1.17 |
| periderm | Lamp2 | 0.56 | 1.20E-13 | 2.57E-09 | 0.72 | 0.59 | 0.13 | 1.21 |
| periderm | Ccdc18 | 0.73 | 1.20E-13 | 2.57E-09 | 0.51 | 0.37 | 0.15 | 1.40 |
| periderm | Prr15l | 1.21 | 1.31E-13 | 2.79E-09 | 0.25 | 0.13 | 0.12 | 1.92 |
| periderm | Myzap | 1.46 | 1.68E-13 | 3.60E-09 | 0.11 | 0.04 | 0.07 | 3.00 |
| periderm | Myo10 | 0.42 | 1.92E-13 | 4.10E-09 | 0.80 | 0.69 | 0.11 | 1.16 |
| periderm | Tpd52 | 0.32 | 2.19E-13 | 4.68E-09 | 0.90 | 0.80 | 0.09 | 1.11 |
| periderm | Eps8l2 | 0.81 | 2.22E-13 | 4.75E-09 | 0.27 | 0.14 | 0.13 | 1.93 |
| periderm | Gm2464 | 1.22 | 2.29E-13 | 4.90E-09 | 0.16 | 0.06 | 0.09 | 2.44 |
| periderm | Plscr3 | 0.54 | 2.79E-13 | 5.97E-09 | 0.69 | 0.52 | 0.17 | 1.32 |
| periderm | Pdlim4 | 0.50 | 3.16E-13 | 6.74E-09 | 0.59 | 0.42 | 0.17 | 1.41 |
| periderm | Tmem123 | 0.75 | 3.30E-13 | 7.05E-09 | 0.57 | 0.44 | 0.14 | 1.31 |
| periderm | St14 | 0.53 | 3.59E-13 | 7.66E-09 | 0.62 | 0.46 | 0.15 | 1.33 |
| periderm | Slc7a11 | 0.54 | 4.52E-13 | 9.66E-09 | 0.24 | 0.12 | 0.12 | 2.04 |
| periderm | Ctsc | 0.97 | 4.66E-13 | 9.95E-09 | 0.32 | 0.19 | 0.13 | 1.71 |
| periderm | Irx5 | 0.53 | 5.34E-13 | 1.14E-08 | 0.38 | 0.23 | 0.16 | 1.68 |
| periderm | Cd44 | 1.22 | 5.50E-13 | 1.18E-08 | 0.12 | 0.04 | 0.08 | 2.83 |
| periderm | Prelid2 | 0.58 | 6.08E-13 | 1.30E-08 | 0.56 | 0.41 | 0.15 | 1.37 |
| periderm | Adrb1 | 1.15 | 6.48E-13 | 1.38E-08 | 0.14 | 0.06 | 0.09 | 2.55 |
| periderm | Nt5dc2 | 0.29 | 6.49E-13 | 1.39E-08 | 0.97 | 0.91 | 0.05 | 1.06 |
| periderm | Zdhhc13 | 0.81 | 6.69E-13 | 1.43E-08 | 0.37 | 0.23 | 0.14 | 1.63 |
| periderm | Rab21 | 0.55 | 7.35E-13 | 1.57E-08 | 0.67 | 0.52 | 0.14 | 1.27 |
| periderm | Mid1ip1 | 0.79 | 7.44E-13 | 1.59E-08 | 0.51 | 0.37 | 0.15 | 1.40 |
| periderm | Tjp2 | 0.64 | 7.68E-13 | 1.64E-08 | 0.59 | 0.46 | 0.13 | 1.28 |
| periderm | F11r | 0.44 | 8.17E-13 | 1.75E-08 | 0.74 | 0.59 | 0.15 | 1.25 |
| periderm | Acbd4 | 1.19 | 1.02E-12 | 2.18E-08 | 0.17 | 0.08 | 0.10 | 2.25 |
| periderm | Epcam | 0.27 | 1.10E-12 | 2.34E-08 | 1.00 | 0.98 | 0.02 | 1.02 |
| periderm | Casp8 | 0.84 | 1.14E-12 | 2.44E-08 | 0.34 | 0.20 | 0.14 | 1.68 |
| periderm | Cxxc5 | 0.34 | 1.21E-12 | 2.58E-08 | 0.72 | 0.57 | 0.15 | 1.27 |
| periderm | Stard10 | 0.42 | 1.36E-12 | 2.90E-08 | 0.82 | 0.71 | 0.11 | 1.15 |
| periderm | Ankrd22 | 1.62 | 1.58E-12 | 3.36E-08 | 0.14 | 0.06 | 0.08 | 2.47 |
| periderm | Tmem79 | 1.18 | 1.64E-12 | 3.50E-08 | 0.16 | 0.07 | 0.09 | 2.33 |
| periderm | Nav2 | 0.37 | 1.71E-12 | 3.65E-08 | 0.82 | 0.69 | 0.13 | 1.18 |
| periderm | St7 | 0.63 | 1.82E-12 | 3.90E-08 | 0.48 | 0.31 | 0.16 | 1.52 |
| periderm | Tshz1 | 0.46 | 2.03E-12 | 4.34E-08 | 0.60 | 0.46 | 0.14 | 1.30 |
| periderm | Utrn | 0.37 | 2.21E-12 | 4.72E-08 | 0.83 | 0.71 | 0.12 | 1.17 |
| periderm | Tacc2 | 0.49 | 2.22E-12 | 4.74E-08 | 0.66 | 0.52 | 0.14 | 1.27 |
| periderm | Sult2b1 | 1.29 | 2.32E-12 | 4.96E-08 | 0.14 | 0.06 | 0.08 | 2.45 |

**Supplementary Table 4. Marker genes for the annotated cell populations of integrated clustering of E11.5 control and NotchGOF λ-junction ectoderm cells.**

| | | | | | | | | |
|---|---|---|---|---|---|---|---|---|
| periderm | Aopep | 0.39 | 2.32E-12 | 4.97E-08 | 0.74 | 0.60 | 0.14 | 1.23 |
| periderm | Slc44a1 | 0.68 | 2.38E-12 | 5.09E-08 | 0.48 | 0.33 | 0.14 | 1.44 |
| periderm | Pcdh9 | 0.77 | 2.39E-12 | 5.12E-08 | 0.30 | 0.17 | 0.13 | 1.77 |
| periderm | Slc16a12 | 1.18 | 2.40E-12 | 5.13E-08 | 0.11 | 0.04 | 0.07 | 2.87 |
| periderm | Fam49a | 0.80 | 2.76E-12 | 5.90E-08 | 0.19 | 0.09 | 0.10 | 2.17 |
| periderm | Hdhd3 | 0.81 | 2.81E-12 | 6.01E-08 | 0.34 | 0.20 | 0.14 | 1.69 |
| periderm | Ntn4 | 1.08 | 3.18E-12 | 6.79E-08 | 0.13 | 0.05 | 0.08 | 2.59 |
| periderm | Lima1 | 0.52 | 3.23E-12 | 6.90E-08 | 0.61 | 0.47 | 0.15 | 1.32 |
| periderm | Dsc2 | 0.79 | 3.42E-12 | 7.31E-08 | 0.35 | 0.22 | 0.14 | 1.62 |
| periderm | Tead1 | 0.41 | 3.44E-12 | 7.35E-08 | 0.84 | 0.76 | 0.08 | 1.11 |
| periderm | Khdrbs3 | 0.41 | 3.72E-12 | 7.94E-08 | 0.91 | 0.88 | 0.04 | 1.04 |
| periderm | Nsg1 | 0.39 | 4.05E-12 | 8.66E-08 | 0.74 | 0.59 | 0.15 | 1.25 |
| periderm | Ddi2 | 0.63 | 4.08E-12 | 8.71E-08 | 0.59 | 0.46 | 0.13 | 1.28 |
| periderm | Tpm4 | 0.31 | 4.21E-12 | 9.00E-08 | 0.92 | 0.87 | 0.06 | 1.06 |
| periderm | Tcn2 | 0.92 | 4.26E-12 | 9.10E-08 | 0.33 | 0.19 | 0.14 | 1.70 |
| periderm | Nectin2 | 0.50 | 4.51E-12 | 9.63E-08 | 0.70 | 0.60 | 0.10 | 1.17 |
| periderm | Zfp185 | 1.07 | 5.35E-12 | 1.14E-07 | 0.16 | 0.07 | 0.09 | 2.29 |
| periderm | Prag1 | 0.60 | 5.78E-12 | 1.24E-07 | 0.43 | 0.28 | 0.15 | 1.55 |
| periderm | Slc44a2 | 0.61 | 5.92E-12 | 1.26E-07 | 0.54 | 0.38 | 0.16 | 1.41 |
| periderm | Kremen1 | 0.69 | 6.23E-12 | 1.33E-07 | 0.45 | 0.30 | 0.15 | 1.50 |
| periderm | Rtkn | 0.99 | 6.50E-12 | 1.39E-07 | 0.27 | 0.15 | 0.12 | 1.82 |
| periderm | Tmcc3 | 0.85 | 7.03E-12 | 1.50E-07 | 0.20 | 0.10 | 0.10 | 2.08 |
| periderm | Zfhx3 | 0.25 | 7.34E-12 | 1.57E-07 | 0.98 | 0.92 | 0.06 | 1.07 |
| periderm | Kbtbd11 | 1.59 | 7.67E-12 | 1.64E-07 | 0.11 | 0.04 | 0.07 | 2.69 |
| periderm | Ldlrap1 | 0.60 | 7.85E-12 | 1.68E-07 | 0.45 | 0.30 | 0.15 | 1.52 |
| periderm | Adamts9 | 0.57 | 8.28E-12 | 1.77E-07 | 0.38 | 0.24 | 0.14 | 1.57 |
| periderm | Unc5c | 0.38 | 9.73E-12 | 2.08E-07 | 0.49 | 0.32 | 0.17 | 1.54 |
| periderm | Sh3bp1 | 0.64 | 1.08E-11 | 2.30E-07 | 0.42 | 0.28 | 0.14 | 1.51 |
| periderm | Cdc42se1 | 0.51 | 1.11E-11 | 2.38E-07 | 0.70 | 0.60 | 0.11 | 1.18 |
| periderm | Plekhg3 | 0.71 | 1.31E-11 | 2.79E-07 | 0.39 | 0.25 | 0.14 | 1.55 |
| periderm | Itm2c | 0.41 | 1.35E-11 | 2.88E-07 | 0.72 | 0.58 | 0.14 | 1.24 |
| periderm | Id4 | 0.54 | 1.45E-11 | 3.10E-07 | 0.65 | 0.57 | 0.08 | 1.15 |
| periderm | Pik3cb | 0.93 | 1.45E-11 | 3.10E-07 | 0.26 | 0.14 | 0.12 | 1.83 |
| periderm | Trappc6a | 0.36 | 1.49E-11 | 3.18E-07 | 0.79 | 0.65 | 0.14 | 1.21 |
| periderm | Ubald1 | 0.54 | 1.52E-11 | 3.25E-07 | 0.52 | 0.37 | 0.15 | 1.39 |
| periderm | Acap2 | 0.55 | 1.53E-11 | 3.26E-07 | 0.57 | 0.42 | 0.15 | 1.36 |
| periderm | Six3 | 0.27 | 1.53E-11 | 3.27E-07 | 0.91 | 0.83 | 0.08 | 1.09 |
| periderm | Eppk1 | 1.04 | 1.58E-11 | 3.37E-07 | 0.17 | 0.08 | 0.09 | 2.20 |
| periderm | Smad6 | 0.76 | 1.74E-11 | 3.72E-07 | 0.39 | 0.25 | 0.14 | 1.56 |
| periderm | 2610002M06Rik | 0.65 | 1.81E-11 | 3.87E-07 | 0.47 | 0.33 | 0.14 | 1.41 |
| periderm | Dgka | 0.89 | 1.97E-11 | 4.22E-07 | 0.26 | 0.14 | 0.12 | 1.80 |
| periderm | Lamtor2 | 0.26 | 2.47E-11 | 5.28E-07 | 0.95 | 0.91 | 0.03 | 1.04 |
| periderm | Efnb2 | 0.40 | 2.47E-11 | 5.28E-07 | 0.72 | 0.58 | 0.14 | 1.23 |
| periderm | Sox14 | 1.39 | 2.69E-11 | 5.75E-07 | 0.11 | 0.04 | 0.07 | 2.63 |
| periderm | Cacna2d3 | 0.99 | 2.87E-11 | 6.13E-07 | 0.15 | 0.07 | 0.09 | 2.26 |
| periderm | Casz1 | 1.40 | 2.92E-11 | 6.23E-07 | 0.13 | 0.06 | 0.08 | 2.39 |
| periderm | A130010J15Rik | 0.73 | 2.93E-11 | 6.26E-07 | 0.29 | 0.17 | 0.13 | 1.77 |
| periderm | Sypl | 0.44 | 3.06E-11 | 6.53E-07 | 0.71 | 0.59 | 0.12 | 1.21 |
| periderm | Cited2 | 0.40 | 3.12E-11 | 6.66E-07 | 0.76 | 0.68 | 0.09 | 1.13 |
| periderm | Itgb6 | 0.77 | 3.14E-11 | 6.71E-07 | 0.27 | 0.15 | 0.12 | 1.78 |
| periderm | Plxdc2 | 0.63 | 3.18E-11 | 6.80E-07 | 0.47 | 0.33 | 0.14 | 1.41 |
| periderm | Ralgps2 | 0.41 | 3.31E-11 | 7.08E-07 | 0.80 | 0.72 | 0.08 | 1.11 |
| periderm | Ckap4 | 0.35 | 3.80E-11 | 8.11E-07 | 0.86 | 0.78 | 0.08 | 1.11 |
| periderm | Col18a1 | 0.26 | 3.94E-11 | 8.42E-07 | 0.94 | 0.85 | 0.09 | 1.10 |
| periderm | Tnfaip8 | 0.51 | 4.19E-11 | 8.94E-07 | 0.66 | 0.56 | 0.10 | 1.19 |

**Supplementary Table 4. Marker genes for the annotated cell populations of integrated clustering of E11.5 control and NotchGOF λ-junction ectoderm cells.**

| | | | | | | | | |
|---|---|---|---|---|---|---|---|---|
| periderm | Bdh1 | 0.93 | 4.42E-11 | 9.44E-07 | 0.25 | 0.14 | 0.11 | 1.78 |
| periderm | A630072M18Rik | 0.95 | 4.54E-11 | 9.70E-07 | 0.22 | 0.12 | 0.10 | 1.90 |
| periderm | Epb41 | 0.31 | 4.88E-11 | 1.04E-06 | 0.82 | 0.74 | 0.09 | 1.12 |
| periderm | Fxyd3 | 0.46 | 5.06E-11 | 1.08E-06 | 0.45 | 0.29 | 0.16 | 1.54 |
| periderm | Rap2c | 0.40 | 5.14E-11 | 1.10E-06 | 0.76 | 0.65 | 0.11 | 1.17 |
| periderm | Spty2d1 | 0.81 | 5.88E-11 | 1.26E-06 | 0.39 | 0.26 | 0.13 | 1.51 |
| periderm | Rps6ka6 | 0.49 | 6.02E-11 | 1.28E-06 | 0.65 | 0.52 | 0.14 | 1.26 |
| periderm | Mfsd1 | 0.70 | 6.29E-11 | 1.34E-06 | 0.53 | 0.39 | 0.14 | 1.37 |
| periderm | Ptbp3 | 0.32 | 7.35E-11 | 1.57E-06 | 0.77 | 0.68 | 0.09 | 1.13 |
| periderm | Nhsl1 | 0.67 | 7.46E-11 | 1.59E-06 | 0.45 | 0.31 | 0.14 | 1.45 |
| periderm | Elmsan1 | 0.93 | 7.85E-11 | 1.68E-06 | 0.25 | 0.14 | 0.11 | 1.80 |
| periderm | Cds1 | 1.21 | 8.10E-11 | 1.73E-06 | 0.15 | 0.07 | 0.08 | 2.17 |
| periderm | Plxna1 | 0.79 | 8.14E-11 | 1.74E-06 | 0.36 | 0.24 | 0.13 | 1.55 |
| periderm | Cited4 | 1.45 | 9.10E-11 | 1.94E-06 | 0.12 | 0.05 | 0.07 | 2.46 |
| periderm | Adgrb3 | 0.86 | 9.35E-11 | 2.00E-06 | 0.20 | 0.10 | 0.10 | 1.96 |
| periderm | Irf2 | 0.58 | 9.35E-11 | 2.00E-06 | 0.54 | 0.41 | 0.13 | 1.33 |
| periderm | Med13l | 0.52 | 1.19E-10 | 2.54E-06 | 0.62 | 0.50 | 0.12 | 1.23 |
| periderm | Tet3 | 0.42 | 1.21E-10 | 2.58E-06 | 0.77 | 0.66 | 0.11 | 1.17 |
| periderm | Tbc1d10a | 0.67 | 1.27E-10 | 2.72E-06 | 0.40 | 0.27 | 0.14 | 1.51 |
| periderm | Cryl1 | 0.64 | 1.31E-10 | 2.81E-06 | 0.40 | 0.26 | 0.14 | 1.53 |
| periderm | Nab2 | 0.62 | 1.37E-10 | 2.94E-06 | 0.31 | 0.19 | 0.12 | 1.65 |
| periderm | Atp2b1 | 0.33 | 1.38E-10 | 2.95E-06 | 0.92 | 0.88 | 0.04 | 1.04 |
| periderm | Nipal2 | 0.89 | 1.43E-10 | 3.06E-06 | 0.18 | 0.09 | 0.09 | 2.06 |
| periderm | Arl5a | 0.45 | 1.44E-10 | 3.07E-06 | 0.71 | 0.62 | 0.09 | 1.15 |
| periderm | Pwwp2b | 0.89 | 1.53E-10 | 3.26E-06 | 0.25 | 0.14 | 0.11 | 1.79 |
| periderm | Cnn3 | 0.27 | 1.53E-10 | 3.28E-06 | 0.97 | 0.93 | 0.04 | 1.04 |
| periderm | Pqlc1 | 0.97 | 1.54E-10 | 3.30E-06 | 0.28 | 0.16 | 0.12 | 1.71 |
| periderm | Zdhhc5 | 0.66 | 1.57E-10 | 3.36E-06 | 0.45 | 0.32 | 0.13 | 1.41 |
| periderm | Mgat5 | 0.57 | 1.57E-10 | 3.36E-06 | 0.42 | 0.28 | 0.14 | 1.50 |
| periderm | Mbtd1 | 0.37 | 1.64E-10 | 3.50E-06 | 0.85 | 0.76 | 0.09 | 1.12 |
| periderm | Stx4a | 0.56 | 1.71E-10 | 3.66E-06 | 0.47 | 0.33 | 0.14 | 1.42 |
| periderm | Lratd2 | 0.49 | 1.93E-10 | 4.13E-06 | 0.47 | 0.32 | 0.14 | 1.44 |
| periderm | Tjp1 | 0.44 | 2.12E-10 | 4.52E-06 | 0.72 | 0.63 | 0.10 | 1.15 |
| periderm | Dlx1 | 0.70 | 2.34E-10 | 5.01E-06 | 0.25 | 0.14 | 0.11 | 1.83 |
| periderm | Svil | 0.60 | 2.48E-10 | 5.31E-06 | 0.54 | 0.41 | 0.12 | 1.30 |
| periderm | Prom2 | 1.04 | 2.61E-10 | 5.58E-06 | 0.22 | 0.12 | 0.10 | 1.83 |
| periderm | Ptges | 1.17 | 3.06E-10 | 6.54E-06 | 0.15 | 0.07 | 0.08 | 2.18 |
| periderm | Ece1 | 0.43 | 3.83E-10 | 8.17E-06 | 0.60 | 0.45 | 0.15 | 1.33 |
| periderm | Lamb1 | 0.40 | 3.90E-10 | 8.34E-06 | 0.65 | 0.50 | 0.15 | 1.30 |
| periderm | Acp5 | 0.88 | 3.92E-10 | 8.38E-06 | 0.14 | 0.06 | 0.08 | 2.25 |
| periderm | Tnik | 0.87 | 4.15E-10 | 8.88E-06 | 0.19 | 0.10 | 0.09 | 1.97 |
| periderm | Notch3 | 0.36 | 4.28E-10 | 9.14E-06 | 0.68 | 0.53 | 0.15 | 1.29 |
| periderm | Plxnb2 | 0.48 | 4.83E-10 | 1.03E-05 | 0.71 | 0.61 | 0.10 | 1.16 |
| periderm | 1810037I17Rik | 0.30 | 4.84E-10 | 1.03E-05 | 0.93 | 0.89 | 0.04 | 1.04 |
| periderm | Alx4 | 0.94 | 5.35E-10 | 1.14E-05 | 0.16 | 0.08 | 0.08 | 2.11 |
| periderm | Spats2 | 0.39 | 5.39E-10 | 1.15E-05 | 0.72 | 0.61 | 0.11 | 1.17 |
| periderm | Ppfibp2 | 1.13 | 5.66E-10 | 1.21E-05 | 0.13 | 0.06 | 0.07 | 2.28 |
| periderm | Tmem159 | 0.62 | 5.76E-10 | 1.23E-05 | 0.52 | 0.40 | 0.13 | 1.32 |
| periderm | Cit | 0.49 | 6.19E-10 | 1.32E-05 | 0.53 | 0.39 | 0.14 | 1.35 |
| periderm | Chmp4c | 1.15 | 7.44E-10 | 1.59E-05 | 0.13 | 0.06 | 0.07 | 2.24 |
| periderm | Skil | 0.36 | 7.88E-10 | 1.68E-05 | 0.69 | 0.56 | 0.14 | 1.25 |
| periderm | Arl1 | 0.27 | 7.93E-10 | 1.69E-05 | 0.91 | 0.88 | 0.02 | 1.02 |
| periderm | Elovl1 | 0.63 | 8.03E-10 | 1.71E-05 | 0.39 | 0.26 | 0.13 | 1.51 |
| periderm | Foxp1 | 0.25 | 8.67E-10 | 1.85E-05 | 0.95 | 0.85 | 0.09 | 1.11 |
| periderm | Tspan9 | 0.71 | 9.02E-10 | 1.93E-05 | 0.22 | 0.12 | 0.10 | 1.85 |

**Supplementary Table 4. Marker genes for the annotated cell populations of integrated clustering of E11.5 control and NotchGOF λ-junction ectoderm cells.**

| | | | | | | | | |
|---|---|---|---|---|---|---|---|---|
| periderm | Klf6 | 0.70 | 9.03E-10 | 1.93E-05 | 0.35 | 0.23 | 0.12 | 1.54 |
| periderm | 0610009E02Rik | 1.34 | 9.81E-10 | 2.10E-05 | 0.15 | 0.07 | 0.08 | 2.11 |
| periderm | Akap17b | 0.71 | 1.07E-09 | 2.28E-05 | 0.24 | 0.14 | 0.10 | 1.74 |
| periderm | Hacd4 | 0.82 | 1.07E-09 | 2.28E-05 | 0.18 | 0.09 | 0.09 | 1.97 |
| periderm | Nsun4 | 0.66 | 1.09E-09 | 2.33E-05 | 0.31 | 0.19 | 0.12 | 1.61 |
| periderm | Gm35040 | 1.03 | 1.13E-09 | 2.41E-05 | 0.12 | 0.05 | 0.07 | 2.37 |
| periderm | Lipg | 0.94 | 1.13E-09 | 2.42E-05 | 0.19 | 0.10 | 0.09 | 1.93 |
| periderm | Necap2 | 0.56 | 1.14E-09 | 2.44E-05 | 0.51 | 0.37 | 0.14 | 1.37 |
| periderm | Zfpm1 | 0.71 | 1.30E-09 | 2.77E-05 | 0.30 | 0.19 | 0.11 | 1.61 |
| periderm | Dact1 | 0.56 | 1.31E-09 | 2.80E-05 | 0.24 | 0.14 | 0.11 | 1.79 |
| periderm | Reep3 | 0.44 | 1.32E-09 | 2.81E-05 | 0.74 | 0.64 | 0.09 | 1.14 |
| periderm | Ddr1 | 0.43 | 1.35E-09 | 2.89E-05 | 0.59 | 0.44 | 0.15 | 1.33 |
| periderm | Ston2 | 0.41 | 1.47E-09 | 3.13E-05 | 0.72 | 0.59 | 0.13 | 1.23 |
| periderm | Frk | 0.83 | 1.47E-09 | 3.13E-05 | 0.23 | 0.13 | 0.10 | 1.78 |
| periderm | Skap2 | 1.34 | 1.67E-09 | 3.56E-05 | 0.15 | 0.07 | 0.08 | 2.05 |
| periderm | Tcf7 | 0.91 | 1.68E-09 | 3.58E-05 | 0.17 | 0.09 | 0.09 | 1.97 |
| periderm | Ank2 | 0.40 | 1.76E-09 | 3.75E-05 | 0.43 | 0.31 | 0.13 | 1.41 |
| periderm | Dazap2 | 0.37 | 1.77E-09 | 3.77E-05 | 0.79 | 0.71 | 0.08 | 1.11 |
| periderm | Mpzl3 | 1.28 | 1.83E-09 | 3.91E-05 | 0.11 | 0.05 | 0.06 | 2.42 |
| periderm | Bcl7a | 0.26 | 2.20E-09 | 4.70E-05 | 0.84 | 0.76 | 0.08 | 1.11 |
| periderm | Cttnbp2nl | 0.50 | 2.45E-09 | 5.24E-05 | 0.54 | 0.42 | 0.13 | 1.31 |
| periderm | Sptlc3 | 0.83 | 2.50E-09 | 5.34E-05 | 0.14 | 0.07 | 0.08 | 2.15 |
| periderm | Ccser1 | 1.05 | 3.00E-09 | 6.41E-05 | 0.15 | 0.07 | 0.08 | 2.09 |
| periderm | Mlec | 0.28 | 3.06E-09 | 6.54E-05 | 0.87 | 0.80 | 0.07 | 1.09 |
| periderm | Irx3 | 0.37 | 3.21E-09 | 6.86E-05 | 0.32 | 0.20 | 0.12 | 1.63 |
| periderm | Add3 | 0.40 | 3.29E-09 | 7.03E-05 | 0.70 | 0.58 | 0.12 | 1.21 |
| periderm | Ppp1r36 | 0.72 | 3.49E-09 | 7.46E-05 | 0.26 | 0.15 | 0.11 | 1.72 |
| periderm | Acot1 | 0.52 | 3.53E-09 | 7.54E-05 | 0.46 | 0.34 | 0.12 | 1.36 |
| periderm | Klf4 | 0.69 | 3.70E-09 | 7.91E-05 | 0.29 | 0.18 | 0.11 | 1.61 |
| periderm | Apcdd1 | 0.64 | 3.93E-09 | 8.40E-05 | 0.23 | 0.13 | 0.10 | 1.77 |
| periderm | Cyth3 | 0.59 | 4.27E-09 | 9.12E-05 | 0.43 | 0.31 | 0.12 | 1.40 |
| periderm | Klc3 | 0.26 | 4.47E-09 | 9.54E-05 | 0.52 | 0.38 | 0.14 | 1.37 |
| periderm | Atp11b | 0.47 | 5.40E-09 | 1.15E-04 | 0.64 | 0.52 | 0.12 | 1.24 |
| periderm | Cndp2 | 0.49 | 5.58E-09 | 1.19E-04 | 0.59 | 0.48 | 0.11 | 1.24 |
| periderm | Pfkl | 0.34 | 5.80E-09 | 1.24E-04 | 0.84 | 0.76 | 0.08 | 1.11 |
| periderm | Etfa | 0.36 | 6.21E-09 | 1.33E-04 | 0.83 | 0.75 | 0.08 | 1.11 |
| periderm | Dach1 | 0.41 | 6.95E-09 | 1.48E-04 | 0.60 | 0.48 | 0.12 | 1.25 |
| periderm | Iqcb1 | 0.38 | 7.45E-09 | 1.59E-04 | 0.70 | 0.59 | 0.11 | 1.18 |
| periderm | Aktip | 0.47 | 7.50E-09 | 1.60E-04 | 0.58 | 0.45 | 0.13 | 1.28 |
| periderm | Sgpl1 | 0.51 | 7.83E-09 | 1.67E-04 | 0.54 | 0.41 | 0.13 | 1.32 |
| periderm | Fzd7 | 0.61 | 8.50E-09 | 1.82E-04 | 0.46 | 0.34 | 0.12 | 1.35 |
| periderm | Dusp16 | 0.64 | 8.54E-09 | 1.82E-04 | 0.36 | 0.24 | 0.12 | 1.49 |
| periderm | Adam23 | 1.21 | 9.32E-09 | 1.99E-04 | 0.12 | 0.06 | 0.07 | 2.20 |
| periderm | Insr | 0.55 | 9.78E-09 | 2.09E-04 | 0.43 | 0.30 | 0.13 | 1.44 |
| periderm | Ncaph2 | 0.37 | 1.04E-08 | 2.23E-04 | 0.71 | 0.60 | 0.11 | 1.18 |
| periderm | Mafb | 0.67 | 1.05E-08 | 2.24E-04 | 0.25 | 0.15 | 0.10 | 1.67 |
| periderm | Efna5 | 0.44 | 1.08E-08 | 2.31E-04 | 0.63 | 0.51 | 0.12 | 1.23 |
| periderm | Ccdc88c | 0.43 | 1.12E-08 | 2.40E-04 | 0.67 | 0.56 | 0.11 | 1.19 |
| periderm | Ccnjl | 0.47 | 1.14E-08 | 2.43E-04 | 0.55 | 0.43 | 0.12 | 1.29 |
| periderm | Klf3 | 0.36 | 1.26E-08 | 2.69E-04 | 0.70 | 0.60 | 0.11 | 1.18 |
| periderm | Adgrl3 | 0.59 | 1.32E-08 | 2.81E-04 | 0.43 | 0.30 | 0.12 | 1.41 |
| periderm | Tmed3 | 0.44 | 1.41E-08 | 3.01E-04 | 0.61 | 0.49 | 0.12 | 1.26 |
| periderm | Mertk | 0.90 | 1.54E-08 | 3.28E-04 | 0.17 | 0.09 | 0.08 | 1.92 |
| periderm | Myh14 | 0.49 | 1.78E-08 | 3.80E-04 | 0.41 | 0.29 | 0.12 | 1.43 |
| periderm | Hint2 | 0.30 | 1.99E-08 | 4.26E-04 | 0.77 | 0.68 | 0.08 | 1.12 |

**Supplementary Table 4. Marker genes for the annotated cell populations of integrated clustering of E11.5 control and NotchGOF λ-junction ectoderm cells.**

| | | | | | | | | |
|---|---|---|---|---|---|---|---|---|
| periderm | Bnipl | 0.79 | 2.05E-08 | 4.38E-04 | 0.28 | 0.17 | 0.10 | 1.59 |
| periderm | Grb7 | 0.49 | 2.19E-08 | 4.68E-04 | 0.50 | 0.37 | 0.13 | 1.34 |
| periderm | Calml3 | 1.08 | 2.24E-08 | 4.78E-04 | 0.10 | 0.04 | 0.06 | 2.34 |
| periderm | Trpm3 | 0.62 | 2.28E-08 | 4.87E-04 | 0.26 | 0.16 | 0.10 | 1.66 |
| periderm | Dap | 0.35 | 2.58E-08 | 5.51E-04 | 0.76 | 0.67 | 0.08 | 1.12 |
| periderm | Ripk4 | 0.46 | 2.61E-08 | 5.57E-04 | 0.41 | 0.29 | 0.12 | 1.43 |
| periderm | Heyl | 0.60 | 2.71E-08 | 5.80E-04 | 0.32 | 0.21 | 0.11 | 1.53 |
| periderm | Rer1 | 0.26 | 2.76E-08 | 5.89E-04 | 0.91 | 0.84 | 0.06 | 1.08 |
| periderm | Atp6v1d | 0.33 | 2.79E-08 | 5.95E-04 | 0.76 | 0.66 | 0.09 | 1.14 |
| periderm | Barx2 | 1.16 | 2.95E-08 | 6.30E-04 | 0.25 | 0.16 | 0.09 | 1.59 |
| periderm | Arl6ip1 | 0.30 | 3.02E-08 | 6.45E-04 | 0.97 | 0.94 | 0.03 | 1.03 |
| periderm | Dipk2a | 0.35 | 3.16E-08 | 6.75E-04 | 0.54 | 0.41 | 0.13 | 1.32 |
| periderm | Serinc5 | 0.77 | 3.35E-08 | 7.16E-04 | 0.20 | 0.11 | 0.09 | 1.77 |
| periderm | Mbd2 | 0.26 | 3.39E-08 | 7.23E-04 | 0.71 | 0.60 | 0.11 | 1.18 |
| periderm | Spg21 | 0.32 | 3.39E-08 | 7.23E-04 | 0.75 | 0.65 | 0.09 | 1.14 |
| periderm | Cux2 | 0.86 | 3.44E-08 | 7.34E-04 | 0.24 | 0.15 | 0.09 | 1.63 |
| periderm | Foxc1 | 0.93 | 3.59E-08 | 7.67E-04 | 0.23 | 0.13 | 0.09 | 1.69 |
| periderm | Zbtb7a | 0.54 | 3.64E-08 | 7.77E-04 | 0.48 | 0.36 | 0.12 | 1.33 |
| periderm | Hspg2 | 0.50 | 3.70E-08 | 7.91E-04 | 0.50 | 0.38 | 0.12 | 1.31 |
| periderm | Foxo1 | 0.49 | 3.77E-08 | 8.05E-04 | 0.40 | 0.28 | 0.12 | 1.43 |
| periderm | Slc2a1 | 0.27 | 3.87E-08 | 8.27E-04 | 0.80 | 0.68 | 0.12 | 1.17 |
| periderm | Ndufa4l2 | 0.69 | 3.92E-08 | 8.37E-04 | 0.17 | 0.09 | 0.08 | 1.86 |
| periderm | Atp2c2 | 1.11 | 3.92E-08 | 8.37E-04 | 0.11 | 0.05 | 0.06 | 2.22 |
| periderm | Zfp282 | 0.53 | 3.95E-08 | 8.43E-04 | 0.42 | 0.29 | 0.13 | 1.43 |
| periderm | Bcap31 | 0.31 | 3.98E-08 | 8.50E-04 | 0.86 | 0.79 | 0.07 | 1.08 |
| periderm | Fat1 | 0.30 | 4.40E-08 | 9.39E-04 | 0.72 | 0.63 | 0.09 | 1.14 |
| periderm | Plec | 0.34 | 4.54E-08 | 9.71E-04 | 0.57 | 0.44 | 0.14 | 1.31 |
| periderm | Afdn | 0.27 | 4.55E-08 | 9.71E-04 | 0.88 | 0.86 | 0.03 | 1.03 |
| periderm | Renbp | 0.65 | 4.62E-08 | 9.87E-04 | 0.19 | 0.10 | 0.09 | 1.84 |
| periderm | Phlpp1 | 0.48 | 5.17E-08 | 1.10E-03 | 0.37 | 0.25 | 0.12 | 1.46 |
| periderm | Pag1 | 1.03 | 5.18E-08 | 1.11E-03 | 0.11 | 0.05 | 0.06 | 2.22 |
| periderm | Trps1 | 0.30 | 5.44E-08 | 1.16E-03 | 0.63 | 0.52 | 0.11 | 1.21 |
| periderm | Mylk | 0.77 | 5.84E-08 | 1.25E-03 | 0.29 | 0.19 | 0.10 | 1.51 |
| periderm | Cast | 0.72 | 6.03E-08 | 1.29E-03 | 0.28 | 0.18 | 0.10 | 1.55 |
| periderm | Gramd3 | 0.64 | 6.13E-08 | 1.31E-03 | 0.32 | 0.22 | 0.10 | 1.48 |
| periderm | H2-D1 | 0.78 | 6.22E-08 | 1.33E-03 | 0.43 | 0.32 | 0.11 | 1.33 |
| periderm | Tril | 0.39 | 6.29E-08 | 1.34E-03 | 0.41 | 0.29 | 0.12 | 1.44 |
| periderm | Sun2 | 0.92 | 6.64E-08 | 1.42E-03 | 0.21 | 0.13 | 0.09 | 1.68 |
| periderm | Atp2b4 | 0.51 | 7.00E-08 | 1.49E-03 | 0.34 | 0.22 | 0.12 | 1.52 |
| periderm | Itgb4 | 0.65 | 7.06E-08 | 1.51E-03 | 0.20 | 0.11 | 0.09 | 1.77 |
| periderm | Tmbim1 | 0.97 | 7.31E-08 | 1.56E-03 | 0.17 | 0.09 | 0.08 | 1.84 |
| periderm | Arrb2 | 0.81 | 7.51E-08 | 1.60E-03 | 0.28 | 0.18 | 0.10 | 1.53 |
| periderm | Vps36 | 0.27 | 7.61E-08 | 1.62E-03 | 0.90 | 0.85 | 0.05 | 1.06 |
| periderm | Pon2 | 0.35 | 8.88E-08 | 1.90E-03 | 0.41 | 0.28 | 0.13 | 1.45 |
| periderm | Cmtm3 | 0.46 | 8.89E-08 | 1.90E-03 | 0.48 | 0.35 | 0.13 | 1.37 |
| periderm | Rsbn1l | 0.29 | 8.97E-08 | 1.92E-03 | 0.75 | 0.67 | 0.08 | 1.11 |
| periderm | Pdgfrl | 0.46 | 9.06E-08 | 1.94E-03 | 0.44 | 0.32 | 0.12 | 1.38 |
| periderm | Sema3f | 0.48 | 9.08E-08 | 1.94E-03 | 0.50 | 0.37 | 0.13 | 1.34 |
| periderm | Ptk7 | 0.41 | 9.27E-08 | 1.98E-03 | 0.59 | 0.48 | 0.11 | 1.23 |
| periderm | Ctdspl | 0.40 | 9.31E-08 | 1.99E-03 | 0.59 | 0.48 | 0.12 | 1.24 |
| periderm | Rab11a | 0.35 | 9.45E-08 | 2.02E-03 | 0.76 | 0.68 | 0.08 | 1.12 |
| periderm | Phactr2 | 0.53 | 9.77E-08 | 2.09E-03 | 0.47 | 0.36 | 0.11 | 1.30 |
| periderm | Sh3d19 | 0.45 | 1.00E-07 | 2.14E-03 | 0.45 | 0.33 | 0.12 | 1.35 |
| periderm | Cpne3 | 0.37 | 1.09E-07 | 2.32E-03 | 0.72 | 0.63 | 0.09 | 1.14 |
| periderm | Crlf3 | 0.46 | 1.23E-07 | 2.63E-03 | 0.42 | 0.31 | 0.12 | 1.38 |

**Supplementary Table 4. Marker genes for the annotated cell populations of integrated clustering of E11.5 control and NotchGOF λ-junction ectoderm cells.**

| | | | | | | | | |
|---|---|---|---|---|---|---|---|---|
| periderm | Sbno2 | 0.69 | 1.38E-07 | 2.95E-03 | 0.27 | 0.17 | 0.10 | 1.56 |
| periderm | Alcam | 0.28 | 1.39E-07 | 2.97E-03 | 0.30 | 0.19 | 0.11 | 1.57 |
| periderm | Sppl2a | 0.48 | 1.40E-07 | 2.98E-03 | 0.67 | 0.58 | 0.09 | 1.15 |
| periderm | Igf2os | 0.81 | 1.41E-07 | 3.02E-03 | 0.13 | 0.07 | 0.07 | 2.00 |
| periderm | Mprip | 0.30 | 1.55E-07 | 3.32E-03 | 0.81 | 0.73 | 0.08 | 1.11 |
| periderm | Arfgef1 | 0.38 | 1.63E-07 | 3.49E-03 | 0.69 | 0.60 | 0.10 | 1.16 |
| periderm | Wiz | 0.40 | 1.64E-07 | 3.50E-03 | 0.56 | 0.46 | 0.10 | 1.22 |
| periderm | Fgd6 | 0.61 | 1.66E-07 | 3.55E-03 | 0.32 | 0.21 | 0.11 | 1.52 |
| periderm | Prrx1 | 0.25 | 1.74E-07 | 3.72E-03 | 0.29 | 0.18 | 0.11 | 1.59 |
| periderm | Gucy1b1 | 1.12 | 1.77E-07 | 3.77E-03 | 0.14 | 0.07 | 0.07 | 1.92 |
| periderm | Pkp4 | 0.30 | 1.80E-07 | 3.85E-03 | 0.67 | 0.57 | 0.10 | 1.17 |
| periderm | Rnf43 | 0.61 | 2.04E-07 | 4.37E-03 | 0.28 | 0.18 | 0.10 | 1.55 |
| periderm | Krt1 | 0.72 | 2.13E-07 | 4.54E-03 | 0.20 | 0.12 | 0.08 | 1.70 |
| periderm | Ptpn14 | 0.54 | 2.23E-07 | 4.76E-03 | 0.33 | 0.22 | 0.11 | 1.51 |
| periderm | 2700033N17Rik | 0.43 | 2.27E-07 | 4.84E-03 | 0.34 | 0.23 | 0.11 | 1.49 |
| periderm | Chst5 | 0.88 | 2.28E-07 | 4.86E-03 | 0.12 | 0.06 | 0.06 | 2.09 |
| periderm | Rhbdf1 | 0.64 | 2.33E-07 | 4.98E-03 | 0.29 | 0.19 | 0.10 | 1.51 |
| periderm | Galnt18 | 0.48 | 2.41E-07 | 5.15E-03 | 0.17 | 0.09 | 0.08 | 1.86 |
| periderm | Abhd11 | 0.40 | 2.44E-07 | 5.21E-03 | 0.51 | 0.39 | 0.12 | 1.31 |
| periderm | Plod2 | 0.41 | 2.56E-07 | 5.47E-03 | 0.58 | 0.47 | 0.11 | 1.24 |
| periderm | Reep4 | 0.48 | 2.58E-07 | 5.51E-03 | 0.47 | 0.36 | 0.11 | 1.30 |
| periderm | Col4a3bp | 0.44 | 2.77E-07 | 5.91E-03 | 0.46 | 0.35 | 0.11 | 1.32 |
| periderm | Heca | 0.52 | 2.92E-07 | 6.24E-03 | 0.37 | 0.26 | 0.11 | 1.43 |
| periderm | Reep6 | 0.97 | 2.94E-07 | 6.28E-03 | 0.15 | 0.08 | 0.07 | 1.87 |
| periderm | Piezo2 | 0.25 | 3.03E-07 | 6.46E-03 | 0.26 | 0.16 | 0.10 | 1.61 |
| periderm | Chmp2a | 0.27 | 3.15E-07 | 6.73E-03 | 0.81 | 0.73 | 0.08 | 1.11 |
| periderm | Ltbr | 0.67 | 3.33E-07 | 7.11E-03 | 0.30 | 0.20 | 0.10 | 1.51 |
| periderm | Lrrk1 | 0.56 | 3.38E-07 | 7.23E-03 | 0.40 | 0.30 | 0.10 | 1.34 |
| periderm | Pfn2 | 0.46 | 3.49E-07 | 7.45E-03 | 0.77 | 0.70 | 0.07 | 1.11 |
| periderm | Acsl5 | 0.51 | 3.51E-07 | 7.51E-03 | 0.42 | 0.31 | 0.11 | 1.35 |
| periderm | Fam89a | 0.49 | 3.53E-07 | 7.54E-03 | 0.25 | 0.15 | 0.10 | 1.63 |
| periderm | Fam83h | 0.82 | 3.57E-07 | 7.62E-03 | 0.16 | 0.09 | 0.07 | 1.86 |
| periderm | Arl13b | 0.46 | 3.73E-07 | 7.96E-03 | 0.51 | 0.39 | 0.12 | 1.30 |
| periderm | Arhgdib | 0.47 | 3.87E-07 | 8.28E-03 | 0.30 | 0.20 | 0.10 | 1.50 |
| periderm | Leprot | 0.42 | 4.05E-07 | 8.65E-03 | 0.50 | 0.39 | 0.11 | 1.28 |
| periderm | Rabgap1 | 0.26 | 4.07E-07 | 8.69E-03 | 0.77 | 0.68 | 0.09 | 1.13 |
| periderm | Nfe2l3 | 0.65 | 4.28E-07 | 9.14E-03 | 0.22 | 0.13 | 0.09 | 1.65 |
| periderm | Tax1bp3 | 0.31 | 4.53E-07 | 9.69E-03 | 0.66 | 0.55 | 0.11 | 1.20 |
| periderm | Nmrk1 | 0.91 | 4.55E-07 | 9.72E-03 | 0.18 | 0.11 | 0.08 | 1.71 |
| periderm | Pak1 | 0.36 | 4.58E-07 | 9.79E-03 | 0.70 | 0.62 | 0.08 | 1.13 |
| periderm | Gjb6 | 0.85 | 4.60E-07 | 9.82E-03 | 0.12 | 0.06 | 0.06 | 2.05 |
| periderm | Zfp704 | 0.31 | 4.63E-07 | 9.89E-03 | 0.70 | 0.62 | 0.08 | 1.14 |
| periderm | Gm28112 | 0.86 | 4.93E-07 | 1.05E-02 | 0.15 | 0.08 | 0.07 | 1.87 |
| periderm | Rab38 | 0.66 | 5.05E-07 | 1.08E-02 | 0.29 | 0.19 | 0.10 | 1.49 |
| periderm | Map7 | 0.42 | 5.08E-07 | 1.08E-02 | 0.53 | 0.42 | 0.11 | 1.26 |
| periderm | Marveld3 | 0.71 | 5.27E-07 | 1.13E-02 | 0.24 | 0.15 | 0.09 | 1.60 |
| periderm | Fkbp1b | 0.54 | 5.47E-07 | 1.17E-02 | 0.25 | 0.16 | 0.09 | 1.57 |
| periderm | Aass | 0.81 | 5.53E-07 | 1.18E-02 | 0.13 | 0.07 | 0.06 | 1.94 |
| periderm | Braf | 0.53 | 5.89E-07 | 1.26E-02 | 0.47 | 0.37 | 0.11 | 1.29 |
| periderm | Ten1 | 0.34 | 6.04E-07 | 1.29E-02 | 0.70 | 0.60 | 0.10 | 1.17 |
| periderm | Hunk | 0.35 | 6.12E-07 | 1.31E-02 | 0.67 | 0.58 | 0.09 | 1.16 |
| periderm | Jak1 | 0.34 | 6.12E-07 | 1.31E-02 | 0.64 | 0.55 | 0.10 | 1.18 |
| periderm | Gm9908 | 0.69 | 6.16E-07 | 1.32E-02 | 0.13 | 0.07 | 0.06 | 1.94 |
| periderm | Mxi1 | 0.31 | 6.46E-07 | 1.38E-02 | 0.66 | 0.56 | 0.09 | 1.16 |
| periderm | Ass1 | 0.76 | 6.52E-07 | 1.39E-02 | 0.19 | 0.11 | 0.08 | 1.71 |

**Supplementary Table 4. Marker genes for the annotated cell populations of integrated clustering of E11.5 control and NotchGOF λ-junction ectoderm cells.**

| | | | | | | | | |
|---|---|---|---|---|---|---|---|---|
| periderm | Ehd1 | 0.49 | 6.69E-07 | 1.43E-02 | 0.55 | 0.46 | 0.10 | 1.21 |
| periderm | Rmnd5a | 0.36 | 6.79E-07 | 1.45E-02 | 0.56 | 0.44 | 0.12 | 1.27 |
| periderm | Ndrg2 | 0.50 | 7.84E-07 | 1.68E-02 | 0.45 | 0.34 | 0.11 | 1.31 |
| periderm | Rarg | 0.51 | 8.05E-07 | 1.72E-02 | 0.39 | 0.28 | 0.11 | 1.38 |
| periderm | A430018G15Rik | 0.87 | 8.30E-07 | 1.77E-02 | 0.14 | 0.08 | 0.07 | 1.85 |
| periderm | Arhgap32 | 0.52 | 8.38E-07 | 1.79E-02 | 0.34 | 0.24 | 0.10 | 1.44 |
| periderm | Grhl1 | 0.79 | 8.76E-07 | 1.87E-02 | 0.15 | 0.08 | 0.07 | 1.85 |
| periderm | Trim59 | 0.31 | 8.87E-07 | 1.90E-02 | 0.78 | 0.71 | 0.07 | 1.10 |
| periderm | Plaat3 | 0.92 | 8.90E-07 | 1.90E-02 | 0.24 | 0.16 | 0.08 | 1.50 |
| periderm | Krcc1 | 0.32 | 9.38E-07 | 2.00E-02 | 0.71 | 0.62 | 0.08 | 1.13 |
| periderm | Arhgap23 | 0.74 | 9.66E-07 | 2.06E-02 | 0.18 | 0.11 | 0.08 | 1.71 |
| periderm | Kifc3 | 0.64 | 9.71E-07 | 2.07E-02 | 0.19 | 0.11 | 0.08 | 1.71 |
| periderm | Mtln | 0.39 | 1.03E-06 | 2.19E-02 | 0.64 | 0.54 | 0.10 | 1.19 |
| periderm | Tmod3 | 0.28 | 1.04E-06 | 2.21E-02 | 0.74 | 0.65 | 0.09 | 1.13 |
| periderm | Ctsd | 0.34 | 1.06E-06 | 2.27E-02 | 0.77 | 0.70 | 0.08 | 1.11 |
| periderm | Prtg | 0.73 | 1.09E-06 | 2.32E-02 | 0.29 | 0.20 | 0.09 | 1.45 |
| periderm | Pold4 | 0.61 | 1.14E-06 | 2.43E-02 | 0.31 | 0.22 | 0.10 | 1.44 |
| periderm | Ttc7b | 0.65 | 1.15E-06 | 2.45E-02 | 0.24 | 0.15 | 0.08 | 1.56 |
| periderm | Fam129b | 0.52 | 1.19E-06 | 2.55E-02 | 0.27 | 0.18 | 0.09 | 1.51 |
| periderm | Eps8 | 0.78 | 1.21E-06 | 2.59E-02 | 0.17 | 0.10 | 0.07 | 1.74 |
| periderm | C1qtnf12 | 0.67 | 1.26E-06 | 2.70E-02 | 0.31 | 0.22 | 0.09 | 1.43 |
| periderm | Cdk6 | 0.26 | 1.35E-06 | 2.89E-02 | 0.64 | 0.51 | 0.14 | 1.27 |
| periderm | Fam114a1 | 0.81 | 1.39E-06 | 2.97E-02 | 0.17 | 0.10 | 0.07 | 1.71 |
| periderm | C2cd2 | 0.72 | 1.41E-06 | 3.00E-02 | 0.16 | 0.09 | 0.07 | 1.76 |
| periderm | Rnf149 | 0.58 | 1.41E-06 | 3.00E-02 | 0.38 | 0.28 | 0.10 | 1.35 |
| periderm | Add1 | 0.29 | 1.47E-06 | 3.15E-02 | 0.77 | 0.67 | 0.10 | 1.15 |
| periderm | Arhgef17 | 0.43 | 1.56E-06 | 3.34E-02 | 0.48 | 0.38 | 0.10 | 1.26 |
| periderm | Lmna | 0.54 | 1.58E-06 | 3.37E-02 | 0.27 | 0.18 | 0.09 | 1.50 |
| periderm | Copg2 | 0.33 | 1.63E-06 | 3.49E-02 | 0.59 | 0.48 | 0.11 | 1.23 |
| periderm | Bcl9 | 0.29 | 1.70E-06 | 3.63E-02 | 0.74 | 0.68 | 0.06 | 1.10 |
| periderm | Tmprss13 | 1.11 | 1.74E-06 | 3.71E-02 | 0.11 | 0.06 | 0.06 | 1.98 |
| periderm | Map2k3 | 0.34 | 1.74E-06 | 3.72E-02 | 0.52 | 0.42 | 0.10 | 1.25 |
| periderm | Mdfic | 0.56 | 1.75E-06 | 3.73E-02 | 0.27 | 0.18 | 0.09 | 1.51 |
| periderm | Bcl2l14 | 1.01 | 1.77E-06 | 3.78E-02 | 0.10 | 0.05 | 0.05 | 2.06 |
| periderm | Ralgps1 | 0.57 | 1.81E-06 | 3.86E-02 | 0.20 | 0.12 | 0.08 | 1.65 |
| periderm | Rdm1 | 0.42 | 1.82E-06 | 3.88E-02 | 0.48 | 0.38 | 0.10 | 1.27 |
| periderm | Slc25a27 | 0.38 | 1.83E-06 | 3.90E-02 | 0.62 | 0.54 | 0.08 | 1.15 |
| periderm | Ifngr2 | 0.40 | 1.86E-06 | 3.96E-02 | 0.37 | 0.27 | 0.11 | 1.39 |
| periderm | Rusc1 | 0.63 | 1.88E-06 | 4.01E-02 | 0.23 | 0.15 | 0.08 | 1.56 |
| periderm | Hexim1 | 0.51 | 1.88E-06 | 4.02E-02 | 0.42 | 0.32 | 0.09 | 1.29 |
| periderm | Vps4b | 0.29 | 1.95E-06 | 4.17E-02 | 0.77 | 0.68 | 0.08 | 1.12 |
| periderm | Tmem50b | 0.75 | 1.97E-06 | 4.21E-02 | 0.24 | 0.16 | 0.08 | 1.52 |
| periderm | Igsf10 | 0.74 | 2.14E-06 | 4.56E-02 | 0.19 | 0.11 | 0.07 | 1.67 |
| periderm | Hip1r | 0.66 | 2.14E-06 | 4.56E-02 | 0.29 | 0.21 | 0.09 | 1.43 |
| periderm | Tspan5 | 0.43 | 2.18E-06 | 4.66E-02 | 0.44 | 0.33 | 0.10 | 1.31 |
| periderm | 2310022B05Rik | 0.49 | 2.18E-06 | 4.66E-02 | 0.36 | 0.26 | 0.11 | 1.41 |
| periderm | Trp53bp2 | 0.39 | 2.29E-06 | 4.89E-02 | 0.53 | 0.44 | 0.09 | 1.20 |
| dental | Pitx2 | 3.67 | 1.96E-303 | 4.18E-299 | 0.93 | 0.18 | 0.75 | 5.25 |
| dental | Shh | 4.09 | 5.57E-146 | 1.19E-141 | 0.35 | 0.03 | 0.32 | 12.03 |
| dental | Unc5c | 2.21 | 9.80E-130 | 2.09E-125 | 0.81 | 0.28 | 0.53 | 2.87 |
| dental | Epha4 | 1.87 | 1.08E-112 | 2.31E-108 | 0.91 | 0.52 | 0.39 | 1.76 |
| dental | Krt14 | 1.71 | 4.71E-112 | 1.01E-107 | 0.93 | 0.41 | 0.52 | 2.25 |
| dental | Foxe1 | 1.53 | 2.50E-109 | 5.35E-105 | 0.59 | 0.13 | 0.46 | 4.56 |
| dental | Chl1 | 3.44 | 1.71E-108 | 3.64E-104 | 0.30 | 0.03 | 0.27 | 9.50 |
| dental | Ifitm3 | 1.97 | 2.82E-106 | 6.02E-102 | 0.85 | 0.38 | 0.47 | 2.24 |

**Supplementary Table 4. Marker genes for the annotated cell populations of integrated clustering of E11.5 control and NotchGOF λ-junction ectoderm cells.**

| | | | | | | | | |
|---|---|---|---|---|---|---|---|---|
| dental | Fgf9 | 1.89 | 8.04E-106 | 1.72E-101 | 0.83 | 0.38 | 0.45 | 2.18 |
| dental | Col14a1 | 4.06 | 1.44E-102 | 3.07E-98 | 0.23 | 0.02 | 0.22 | 13.76 |
| dental | Igfbp4 | 1.34 | 7.70E-100 | 1.65E-95 | 0.96 | 0.72 | 0.24 | 1.34 |
| dental | Phlda3 | 1.73 | 2.02E-98 | 4.32E-94 | 0.87 | 0.46 | 0.41 | 1.88 |
| dental | Tac2 | 3.09 | 4.58E-98 | 9.77E-94 | 0.23 | 0.02 | 0.22 | 13.00 |
| dental | Csrp2 | 1.52 | 1.04E-96 | 2.23E-92 | 0.96 | 0.83 | 0.14 | 1.16 |
| dental | Dsc3 | 3.58 | 5.92E-90 | 1.26E-85 | 0.31 | 0.04 | 0.26 | 7.00 |
| dental | Fgf8 | 1.20 | 1.75E-87 | 3.73E-83 | 0.69 | 0.23 | 0.46 | 2.97 |
| dental | Cldn10 | 3.59 | 9.62E-86 | 2.05E-81 | 0.31 | 0.05 | 0.26 | 6.39 |
| dental | Pcolce2 | 2.53 | 5.29E-85 | 1.13E-80 | 0.29 | 0.04 | 0.25 | 7.17 |
| dental | Ccnd2 | 1.19 | 1.38E-83 | 2.94E-79 | 0.99 | 0.76 | 0.22 | 1.29 |
| dental | Fabp5 | 0.71 | 8.50E-83 | 1.82E-78 | 1.00 | 0.99 | 0.01 | 1.01 |
| dental | Ifitm1 | 2.46 | 1.34E-77 | 2.87E-73 | 0.53 | 0.16 | 0.36 | 3.22 |
| dental | Ptch1 | 1.94 | 2.14E-76 | 4.57E-72 | 0.70 | 0.32 | 0.38 | 2.20 |
| dental | Capn6 | 1.60 | 2.49E-74 | 5.31E-70 | 0.78 | 0.42 | 0.37 | 1.88 |
| dental | Cd59a | 2.02 | 5.44E-74 | 1.16E-69 | 0.65 | 0.27 | 0.38 | 2.44 |
| dental | Fxyd3 | 1.40 | 9.29E-71 | 1.98E-66 | 0.69 | 0.27 | 0.42 | 2.56 |
| dental | Siva1 | 0.81 | 5.77E-70 | 1.23E-65 | 1.00 | 0.97 | 0.03 | 1.03 |
| dental | Dapl1 | 1.71 | 3.71E-68 | 7.93E-64 | 0.62 | 0.23 | 0.39 | 2.66 |
| dental | Irx1 | 2.48 | 2.21E-66 | 4.73E-62 | 0.29 | 0.05 | 0.24 | 5.56 |
| dental | Rpl23 | 0.29 | 2.63E-66 | 5.61E-62 | 1.00 | 1.00 | 0.00 | 1.00 |
| dental | Ascl4 | 3.78 | 2.81E-64 | 6.00E-60 | 0.17 | 0.02 | 0.15 | 11.07 |
| dental | Anxa8 | 2.04 | 2.46E-61 | 5.24E-57 | 0.33 | 0.07 | 0.26 | 4.58 |
| dental | Isl1 | 0.95 | 1.28E-60 | 2.74E-56 | 0.97 | 0.73 | 0.23 | 1.32 |
| dental | Rflnb | 1.35 | 2.25E-60 | 4.80E-56 | 0.80 | 0.49 | 0.31 | 1.64 |
| dental | Kctd1 | 1.30 | 2.94E-58 | 6.28E-54 | 0.80 | 0.47 | 0.33 | 1.71 |
| dental | Trp63 | 0.89 | 1.77E-57 | 3.78E-53 | 0.88 | 0.48 | 0.41 | 1.86 |
| dental | Rps20 | 0.27 | 2.07E-57 | 4.43E-53 | 1.00 | 1.00 | 0.00 | 1.00 |
| dental | Emp3 | 2.50 | 1.54E-56 | 3.29E-52 | 0.26 | 0.05 | 0.21 | 5.31 |
| dental | Runx1 | 0.76 | 1.99E-56 | 4.25E-52 | 0.86 | 0.48 | 0.37 | 1.77 |
| dental | Trim29 | 1.71 | 3.59E-54 | 7.68E-50 | 0.47 | 0.16 | 0.31 | 2.92 |
| dental | Lef1 | 1.31 | 1.39E-53 | 2.96E-49 | 0.70 | 0.35 | 0.36 | 2.03 |
| dental | Dlx2 | 1.14 | 1.42E-53 | 3.03E-49 | 0.69 | 0.31 | 0.39 | 2.25 |
| dental | Gm12446 | 1.65 | 2.90E-53 | 6.20E-49 | 0.35 | 0.09 | 0.26 | 3.99 |
| dental | Rpl34 | 0.26 | 6.69E-53 | 1.43E-48 | 1.00 | 1.00 | 0.00 | 1.00 |
| dental | Spred1 | 1.28 | 6.96E-53 | 1.49E-48 | 0.77 | 0.48 | 0.30 | 1.62 |
| dental | Amot | 1.29 | 9.48E-52 | 2.03E-47 | 0.82 | 0.55 | 0.27 | 1.49 |
| dental | Pltp | 1.04 | 2.00E-50 | 4.28E-46 | 0.80 | 0.43 | 0.37 | 1.86 |
| dental | 1500009L16Rik | 1.22 | 5.18E-50 | 1.11E-45 | 0.77 | 0.50 | 0.27 | 1.54 |
| dental | Sult5a1 | 2.28 | 1.40E-49 | 2.98E-45 | 0.28 | 0.07 | 0.22 | 4.24 |
| dental | Npm3 | 0.75 | 2.30E-49 | 4.92E-45 | 0.98 | 0.87 | 0.12 | 1.13 |
| dental | Pkp4 | 1.10 | 5.23E-49 | 1.12E-44 | 0.80 | 0.56 | 0.24 | 1.43 |
| dental | Rxfp1 | 2.29 | 9.55E-49 | 2.04E-44 | 0.21 | 0.04 | 0.17 | 5.75 |
| dental | Pdgfa | 0.63 | 1.34E-48 | 2.86E-44 | 0.99 | 0.93 | 0.06 | 1.06 |
| dental | Perp | 0.70 | 7.88E-46 | 1.68E-41 | 0.91 | 0.50 | 0.41 | 1.81 |
| dental | Zfp385c | 1.67 | 1.42E-44 | 3.03E-40 | 0.34 | 0.10 | 0.24 | 3.42 |
| dental | Fermt1 | 0.83 | 1.95E-43 | 4.16E-39 | 0.87 | 0.52 | 0.34 | 1.65 |
| dental | Lmo2 | 1.00 | 5.23E-43 | 1.12E-38 | 0.59 | 0.27 | 0.33 | 2.23 |
| dental | Ntm | 1.69 | 1.35E-42 | 2.89E-38 | 0.50 | 0.22 | 0.28 | 2.29 |
| dental | Fbln2 | 2.13 | 1.90E-42 | 4.07E-38 | 0.30 | 0.09 | 0.22 | 3.50 |
| dental | Itgb5 | 0.93 | 2.00E-42 | 4.27E-38 | 0.79 | 0.47 | 0.32 | 1.68 |
| dental | Rps27 | 0.27 | 3.25E-42 | 6.94E-38 | 1.00 | 1.00 | 0.00 | 1.00 |
| dental | Ctnnb1 | 0.42 | 4.48E-42 | 9.57E-38 | 1.00 | 0.99 | 0.01 | 1.01 |
| dental | Rpl12 | 0.26 | 6.95E-42 | 1.48E-37 | 1.00 | 1.00 | 0.00 | 1.00 |
| dental | Ptch2 | 2.67 | 2.05E-41 | 4.38E-37 | 0.22 | 0.05 | 0.17 | 4.48 |

**Supplementary Table 4. Marker genes for the annotated cell populations of integrated clustering of E11.5 control and NotchGOF λ-junction ectoderm cells.**

| | | | | | | | | |
|---|---|---|---|---|---|---|---|---|
| dental | Ly6g6e | 2.15 | 8.53E-41 | 1.82E-36 | 0.23 | 0.05 | 0.18 | 4.40 |
| dental | Palmd | 1.95 | 1.76E-40 | 3.77E-36 | 0.21 | 0.04 | 0.16 | 5.00 |
| dental | Kif13a | 1.43 | 5.19E-40 | 1.11E-35 | 0.56 | 0.28 | 0.28 | 1.99 |
| dental | Npm1 | 0.27 | 7.95E-40 | 1.70E-35 | 1.00 | 1.00 | 0.00 | 1.00 |
| dental | H19 | 0.48 | 7.32E-39 | 1.56E-34 | 1.00 | 0.87 | 0.13 | 1.15 |
| dental | Rab38 | 1.44 | 1.49E-38 | 3.18E-34 | 0.44 | 0.18 | 0.26 | 2.49 |
| dental | Bmp7 | 1.12 | 1.66E-38 | 3.54E-34 | 0.67 | 0.38 | 0.29 | 1.76 |
| dental | Eef1b2 | 0.28 | 3.29E-38 | 7.03E-34 | 1.00 | 1.00 | 0.00 | 1.00 |
| dental | Emb | 0.98 | 6.15E-38 | 1.31E-33 | 0.87 | 0.72 | 0.15 | 1.20 |
| dental | Vwa1 | 1.71 | 7.36E-38 | 1.57E-33 | 0.31 | 0.09 | 0.21 | 3.29 |
| dental | Serpinb6b | 1.93 | 2.19E-37 | 4.67E-33 | 0.21 | 0.05 | 0.16 | 4.47 |
| dental | Itm2b | 0.50 | 3.58E-36 | 7.66E-32 | 0.99 | 0.93 | 0.06 | 1.06 |
| dental | Sema3d | 0.92 | 6.65E-36 | 1.42E-31 | 0.69 | 0.39 | 0.30 | 1.77 |
| dental | Cnksr2 | 1.19 | 1.40E-35 | 3.00E-31 | 0.59 | 0.31 | 0.28 | 1.91 |
| dental | Pabpc1 | 0.39 | 4.52E-35 | 9.65E-31 | 0.99 | 0.99 | 0.01 | 1.01 |
| dental | Spats2l | 1.22 | 3.15E-34 | 6.73E-30 | 0.59 | 0.31 | 0.28 | 1.88 |
| dental | Pdlim4 | 1.01 | 9.84E-34 | 2.10E-29 | 0.68 | 0.42 | 0.27 | 1.64 |
| dental | Gli1 | 1.97 | 1.76E-33 | 3.75E-29 | 0.25 | 0.07 | 0.18 | 3.44 |
| dental | Ifitm2 | 0.50 | 1.87E-33 | 3.98E-29 | 0.99 | 0.93 | 0.06 | 1.07 |
| dental | Jph4 | 1.30 | 1.95E-33 | 4.16E-29 | 0.43 | 0.18 | 0.25 | 2.42 |
| dental | Krt17 | 2.20 | 3.03E-33 | 6.46E-29 | 0.23 | 0.07 | 0.17 | 3.60 |
| dental | Gpc3 | 0.60 | 3.25E-33 | 6.94E-29 | 0.97 | 0.83 | 0.14 | 1.17 |
| dental | Agl | 1.32 | 7.29E-33 | 1.56E-28 | 0.57 | 0.33 | 0.25 | 1.76 |
| dental | Cald1 | 0.80 | 1.10E-32 | 2.35E-28 | 0.90 | 0.68 | 0.22 | 1.33 |
| dental | Dsg2 | 1.06 | 2.13E-32 | 4.55E-28 | 0.72 | 0.46 | 0.26 | 1.55 |
| dental | Vcan | 0.55 | 7.35E-32 | 1.57E-27 | 0.94 | 0.73 | 0.21 | 1.29 |
| dental | Tnfrsf19 | 1.41 | 1.32E-31 | 2.81E-27 | 0.48 | 0.23 | 0.25 | 2.12 |
| dental | P3h2 | 2.99 | 3.10E-31 | 6.61E-27 | 0.12 | 0.02 | 0.10 | 6.56 |
| dental | Dsp | 0.48 | 4.21E-31 | 8.99E-27 | 0.99 | 0.86 | 0.14 | 1.16 |
| dental | Dlx1 | 1.37 | 6.20E-31 | 1.33E-26 | 0.34 | 0.13 | 0.22 | 2.69 |
| dental | Acaa2 | 0.72 | 8.32E-31 | 1.78E-26 | 0.88 | 0.70 | 0.18 | 1.26 |
| dental | Cdca7 | 0.80 | 3.83E-30 | 8.18E-26 | 0.84 | 0.63 | 0.22 | 1.35 |
| dental | Fign | 1.30 | 5.56E-30 | 1.19E-25 | 0.40 | 0.17 | 0.23 | 2.32 |
| dental | Robo1 | 0.90 | 6.62E-30 | 1.41E-25 | 0.65 | 0.38 | 0.27 | 1.71 |
| dental | Gja1 | 0.63 | 1.72E-29 | 3.68E-25 | 0.88 | 0.69 | 0.19 | 1.27 |
| dental | Lpar6 | 0.92 | 3.89E-29 | 8.30E-25 | 0.69 | 0.46 | 0.24 | 1.52 |
| dental | Ets2 | 1.17 | 4.82E-29 | 1.03E-24 | 0.47 | 0.23 | 0.24 | 2.04 |
| dental | Hspb2 | 2.06 | 1.21E-28 | 2.59E-24 | 0.30 | 0.11 | 0.19 | 2.69 |
| dental | Barx1 | 1.23 | 1.97E-28 | 4.20E-24 | 0.27 | 0.09 | 0.18 | 3.07 |
| dental | Dst | 0.90 | 2.55E-28 | 5.45E-24 | 0.69 | 0.46 | 0.23 | 1.50 |
| dental | Lgr6 | 2.40 | 1.04E-27 | 2.23E-23 | 0.13 | 0.02 | 0.10 | 5.33 |
| dental | S100a10 | 0.37 | 1.24E-27 | 2.65E-23 | 0.94 | 0.73 | 0.21 | 1.29 |
| dental | Serpinb9 | 2.52 | 6.72E-27 | 1.44E-22 | 0.12 | 0.02 | 0.10 | 5.62 |
| dental | Osbpl3 | 1.35 | 1.28E-26 | 2.74E-22 | 0.34 | 0.14 | 0.20 | 2.43 |
| dental | Copz2 | 1.22 | 2.88E-26 | 6.15E-22 | 0.45 | 0.23 | 0.22 | 1.96 |
| dental | Nkain3 | 2.14 | 4.21E-26 | 8.99E-22 | 0.19 | 0.05 | 0.14 | 3.52 |
| dental | Filip1l | 1.10 | 5.92E-26 | 1.26E-21 | 0.48 | 0.25 | 0.23 | 1.93 |
| dental | Hmga2 | 0.45 | 6.16E-26 | 1.32E-21 | 0.99 | 0.92 | 0.08 | 1.08 |
| dental | Cdh6 | 1.37 | 6.50E-26 | 1.39E-21 | 0.29 | 0.11 | 0.18 | 2.65 |
| dental | Nrxn3 | 1.58 | 6.60E-26 | 1.41E-21 | 0.25 | 0.09 | 0.17 | 2.88 |
| dental | Apcdd1 | 1.51 | 2.42E-25 | 5.17E-21 | 0.31 | 0.13 | 0.19 | 2.49 |
| dental | Runx2 | 2.41 | 1.25E-24 | 2.68E-20 | 0.15 | 0.03 | 0.11 | 4.26 |
| dental | Epha7 | 1.16 | 1.75E-24 | 3.75E-20 | 0.48 | 0.27 | 0.21 | 1.80 |
| dental | Peg3 | 0.41 | 2.64E-24 | 5.65E-20 | 0.94 | 0.82 | 0.13 | 1.15 |
| dental | Kctd12b | 1.69 | 4.49E-24 | 9.60E-20 | 0.25 | 0.09 | 0.16 | 2.79 |

**Supplementary Table 4. Marker genes for the annotated cell populations of integrated clustering of E11.5 control and NotchGOF λ-junction ectoderm cells.**

| | | | | | | | | |
|---|---|---|---|---|---|---|---|---|
| dental | Tns3 | 1.03 | 1.35E-23 | 2.88E-19 | 0.54 | 0.32 | 0.22 | 1.67 |
| dental | Prss23 | 1.03 | 2.50E-23 | 5.35E-19 | 0.50 | 0.27 | 0.22 | 1.81 |
| dental | Hpcal1 | 0.75 | 2.60E-23 | 5.56E-19 | 0.68 | 0.45 | 0.23 | 1.51 |
| dental | Nt5c | 0.63 | 5.90E-23 | 1.26E-18 | 0.82 | 0.65 | 0.17 | 1.26 |
| dental | Dmrt2 | 1.37 | 5.92E-23 | 1.26E-18 | 0.24 | 0.08 | 0.16 | 2.85 |
| dental | Slc39a8 | 1.63 | 9.29E-23 | 1.99E-18 | 0.24 | 0.08 | 0.15 | 2.81 |
| dental | Lgi2 | 0.97 | 1.13E-22 | 2.41E-18 | 0.50 | 0.28 | 0.22 | 1.80 |
| dental | Sdc4 | 0.65 | 1.38E-22 | 2.94E-18 | 0.82 | 0.63 | 0.19 | 1.30 |
| dental | Ece1 | 0.96 | 1.78E-22 | 3.80E-18 | 0.64 | 0.45 | 0.19 | 1.42 |
| dental | Notum | 1.55 | 3.49E-22 | 7.46E-18 | 0.19 | 0.06 | 0.13 | 3.32 |
| dental | Slitrk6 | 1.35 | 3.65E-22 | 7.80E-18 | 0.34 | 0.16 | 0.18 | 2.17 |
| dental | Aqp3 | 2.21 | 4.01E-22 | 8.57E-18 | 0.13 | 0.03 | 0.10 | 4.16 |
| dental | Tgfb1 | 0.89 | 7.21E-22 | 1.54E-17 | 0.34 | 0.15 | 0.19 | 2.30 |
| dental | Slc4a4 | 1.30 | 7.30E-22 | 1.56E-17 | 0.31 | 0.13 | 0.18 | 2.32 |
| dental | Lgr5 | 0.80 | 8.32E-22 | 1.78E-17 | 0.44 | 0.22 | 0.22 | 2.01 |
| dental | Slc7a10 | 2.04 | 8.73E-22 | 1.86E-17 | 0.14 | 0.04 | 0.11 | 3.94 |
| dental | Pitx1 | 0.35 | 1.07E-21 | 2.29E-17 | 0.91 | 0.63 | 0.28 | 1.44 |
| dental | Ankrd6 | 1.41 | 3.57E-21 | 7.62E-17 | 0.33 | 0.15 | 0.18 | 2.14 |
| dental | Dsc2 | 1.38 | 5.60E-21 | 1.20E-16 | 0.41 | 0.22 | 0.19 | 1.87 |
| dental | Plce1 | 0.86 | 8.77E-21 | 1.87E-16 | 0.52 | 0.31 | 0.21 | 1.68 |
| dental | Lpin2 | 0.87 | 1.16E-20 | 2.47E-16 | 0.63 | 0.45 | 0.19 | 1.42 |
| dental | Pik3r1 | 0.49 | 1.22E-20 | 2.60E-16 | 0.81 | 0.58 | 0.23 | 1.40 |
| dental | Cltrn | 2.25 | 1.40E-20 | 2.98E-16 | 0.12 | 0.03 | 0.09 | 4.29 |
| dental | Ppp1r12b | 1.41 | 2.81E-20 | 6.00E-16 | 0.32 | 0.15 | 0.17 | 2.18 |
| dental | Pdgfc | 0.46 | 3.03E-20 | 6.48E-16 | 0.90 | 0.75 | 0.15 | 1.20 |
| dental | Bcl11a | 0.56 | 6.59E-20 | 1.41E-15 | 0.92 | 0.79 | 0.13 | 1.17 |
| dental | Hspb1 | 0.36 | 1.13E-19 | 2.41E-15 | 0.62 | 0.36 | 0.25 | 1.70 |
| dental | Dok5 | 0.71 | 1.16E-19 | 2.47E-15 | 0.59 | 0.36 | 0.24 | 1.67 |
| dental | Greb1 | 1.35 | 1.50E-19 | 3.21E-15 | 0.35 | 0.18 | 0.17 | 1.96 |
| dental | Tpd52l1 | 1.27 | 4.65E-19 | 9.94E-15 | 0.30 | 0.14 | 0.16 | 2.15 |
| dental | Snai2 | 0.90 | 9.40E-19 | 2.01E-14 | 0.35 | 0.16 | 0.19 | 2.13 |
| dental | Btg1 | 0.65 | 9.55E-19 | 2.04E-14 | 0.84 | 0.74 | 0.10 | 1.13 |
| dental | Col4a2 | 0.59 | 1.11E-18 | 2.36E-14 | 0.71 | 0.48 | 0.23 | 1.47 |
| dental | Amotl1 | 0.45 | 1.97E-18 | 4.21E-14 | 0.85 | 0.68 | 0.16 | 1.24 |
| dental | Etl4 | 0.62 | 3.36E-18 | 7.18E-14 | 0.75 | 0.56 | 0.19 | 1.33 |
| dental | Capg | 0.80 | 3.71E-18 | 7.92E-14 | 0.55 | 0.34 | 0.21 | 1.61 |
| dental | Ugdh | 0.77 | 3.85E-18 | 8.23E-14 | 0.61 | 0.42 | 0.19 | 1.44 |
| dental | Ablim1 | 0.59 | 4.06E-18 | 8.66E-14 | 0.73 | 0.53 | 0.19 | 1.36 |
| dental | Hdac8 | 1.08 | 4.85E-18 | 1.04E-13 | 0.40 | 0.22 | 0.18 | 1.83 |
| dental | Bag3 | 1.59 | 5.95E-18 | 1.27E-13 | 0.18 | 0.06 | 0.12 | 2.86 |
| dental | Klf5 | 0.73 | 8.89E-18 | 1.90E-13 | 0.61 | 0.39 | 0.21 | 1.54 |
| dental | Ppp3ca | 0.74 | 9.35E-18 | 2.00E-13 | 0.70 | 0.53 | 0.17 | 1.31 |
| dental | Dmtn | 1.45 | 9.71E-18 | 2.07E-13 | 0.22 | 0.09 | 0.13 | 2.49 |
| dental | Cd109 | 1.57 | 1.28E-17 | 2.74E-13 | 0.14 | 0.04 | 0.10 | 3.46 |
| dental | Bcam | 0.43 | 1.70E-17 | 3.64E-13 | 0.90 | 0.71 | 0.19 | 1.27 |
| dental | Anxa2 | 0.30 | 5.89E-17 | 1.26E-12 | 0.96 | 0.74 | 0.22 | 1.30 |
| dental | Gask1b | 1.70 | 6.81E-17 | 1.45E-12 | 0.17 | 0.06 | 0.11 | 2.91 |
| dental | Mycl | 0.46 | 7.54E-17 | 1.61E-12 | 0.80 | 0.63 | 0.18 | 1.28 |
| dental | Nell2 | 1.19 | 8.10E-17 | 1.73E-12 | 0.25 | 0.11 | 0.14 | 2.32 |
| dental | Adgra3 | 0.82 | 1.20E-16 | 2.56E-12 | 0.63 | 0.48 | 0.15 | 1.31 |
| dental | Slc35g1 | 1.24 | 1.26E-16 | 2.69E-12 | 0.29 | 0.14 | 0.15 | 2.04 |
| dental | Stk26 | 0.65 | 1.62E-16 | 3.46E-12 | 0.64 | 0.46 | 0.19 | 1.41 |
| dental | Pard6g | 0.54 | 2.09E-16 | 4.46E-12 | 0.78 | 0.63 | 0.15 | 1.23 |
| dental | Camk4 | 1.24 | 2.22E-16 | 4.75E-12 | 0.13 | 0.04 | 0.09 | 3.33 |
| dental | Tiam1 | 0.77 | 3.35E-16 | 7.15E-12 | 0.61 | 0.44 | 0.17 | 1.38 |

**Supplementary Table 4. Marker genes for the annotated cell populations of integrated clustering of E11.5 control and NotchGOF λ-junction ectoderm cells.**

| | | | | | | | | |
|---|---|---|---|---|---|---|---|---|
| dental | Edaradd | 1.40 | 3.87E-16 | 8.28E-12 | 0.19 | 0.07 | 0.12 | 2.66 |
| dental | Mrpl23 | 0.34 | 5.36E-16 | 1.14E-11 | 0.97 | 0.92 | 0.05 | 1.05 |
| dental | Mycn | 0.73 | 6.35E-16 | 1.36E-11 | 0.52 | 0.34 | 0.18 | 1.54 |
| dental | Nrp2 | 0.97 | 6.63E-16 | 1.42E-11 | 0.32 | 0.16 | 0.16 | 1.99 |
| dental | Kif1b | 0.58 | 7.54E-16 | 1.61E-11 | 0.76 | 0.64 | 0.12 | 1.19 |
| dental | Gm2694 | 1.39 | 8.66E-16 | 1.85E-11 | 0.17 | 0.06 | 0.11 | 2.71 |
| dental | Lmo4 | 0.31 | 8.75E-16 | 1.87E-11 | 0.97 | 0.91 | 0.06 | 1.07 |
| dental | Cox7a2l | 0.28 | 9.18E-16 | 1.96E-11 | 1.00 | 0.98 | 0.02 | 1.02 |
| dental | Mir205hg | 0.56 | 1.14E-15 | 2.44E-11 | 0.64 | 0.42 | 0.22 | 1.51 |
| dental | Rab34 | 0.54 | 1.17E-15 | 2.49E-11 | 0.74 | 0.61 | 0.14 | 1.22 |
| dental | Ralgps2 | 0.57 | 1.40E-15 | 3.00E-11 | 0.84 | 0.72 | 0.12 | 1.17 |
| dental | Tpgs2 | 0.79 | 1.54E-15 | 3.28E-11 | 0.65 | 0.49 | 0.16 | 1.32 |
| dental | Marc2 | 0.56 | 2.23E-15 | 4.76E-11 | 0.75 | 0.59 | 0.15 | 1.26 |
| dental | Rest | 0.45 | 3.01E-15 | 6.42E-11 | 0.83 | 0.67 | 0.17 | 1.25 |
| dental | B130024G19Rik | 1.75 | 3.13E-15 | 6.69E-11 | 0.12 | 0.04 | 0.09 | 3.32 |
| dental | Polr3g | 0.79 | 3.36E-15 | 7.17E-11 | 0.49 | 0.32 | 0.17 | 1.55 |
| dental | Nampt | 0.82 | 5.66E-15 | 1.21E-10 | 0.49 | 0.32 | 0.17 | 1.55 |
| dental | Arid3b | 0.64 | 5.72E-15 | 1.22E-10 | 0.57 | 0.40 | 0.17 | 1.44 |
| dental | Srd5a1 | 0.87 | 7.26E-15 | 1.55E-10 | 0.50 | 0.32 | 0.17 | 1.53 |
| dental | Serpine2 | 0.75 | 8.68E-15 | 1.85E-10 | 0.61 | 0.45 | 0.16 | 1.35 |
| dental | Slc39a10 | 0.49 | 1.13E-14 | 2.42E-10 | 0.84 | 0.75 | 0.10 | 1.13 |
| dental | Frem2 | 0.46 | 1.22E-14 | 2.61E-10 | 0.79 | 0.61 | 0.18 | 1.29 |
| dental | Mfge8 | 0.36 | 1.59E-14 | 3.39E-10 | 0.73 | 0.54 | 0.19 | 1.35 |
| dental | D630045J12Rik | 0.88 | 1.63E-14 | 3.48E-10 | 0.39 | 0.23 | 0.16 | 1.72 |
| dental | Samd5 | 0.41 | 2.04E-14 | 4.36E-10 | 0.85 | 0.67 | 0.18 | 1.27 |
| dental | Cd44 | 1.79 | 2.71E-14 | 5.79E-10 | 0.13 | 0.04 | 0.09 | 3.02 |
| dental | Cbr2 | 1.64 | 3.73E-14 | 7.96E-10 | 0.11 | 0.03 | 0.08 | 3.48 |
| dental | Plcd3 | 1.62 | 4.38E-14 | 9.35E-10 | 0.14 | 0.05 | 0.09 | 2.91 |
| dental | Vwa2 | 1.22 | 4.50E-14 | 9.60E-10 | 0.25 | 0.12 | 0.13 | 2.07 |
| dental | Nfkb1 | 0.86 | 5.02E-14 | 1.07E-09 | 0.40 | 0.23 | 0.17 | 1.71 |
| dental | Pnrc1 | 0.33 | 5.69E-14 | 1.22E-09 | 0.94 | 0.90 | 0.04 | 1.04 |
| dental | D16Ertd472e | 0.81 | 7.76E-14 | 1.66E-09 | 0.39 | 0.23 | 0.16 | 1.73 |
| dental | Ntf5 | 0.70 | 8.99E-14 | 1.92E-09 | 0.48 | 0.30 | 0.17 | 1.57 |
| dental | Zfp703 | 0.92 | 9.05E-14 | 1.93E-09 | 0.44 | 0.28 | 0.16 | 1.56 |
| dental | Tmem256 | 0.34 | 1.30E-13 | 2.78E-09 | 0.93 | 0.87 | 0.06 | 1.07 |
| dental | Msx2 | 0.53 | 1.63E-13 | 3.48E-09 | 0.59 | 0.39 | 0.20 | 1.53 |
| dental | Npnt | 0.52 | 1.64E-13 | 3.50E-09 | 0.62 | 0.43 | 0.19 | 1.44 |
| dental | Serpinb6a | 0.59 | 1.71E-13 | 3.65E-09 | 0.62 | 0.47 | 0.15 | 1.33 |
| dental | Ahnak | 0.47 | 1.80E-13 | 3.86E-09 | 0.54 | 0.34 | 0.20 | 1.58 |
| dental | Spred2 | 0.63 | 1.82E-13 | 3.89E-09 | 0.54 | 0.37 | 0.18 | 1.48 |
| dental | Clta | 0.35 | 1.91E-13 | 4.09E-09 | 0.99 | 0.98 | 0.01 | 1.01 |
| dental | Kremen2 | 0.72 | 2.31E-13 | 4.93E-09 | 0.32 | 0.17 | 0.15 | 1.89 |
| dental | Col5a2 | 0.98 | 2.58E-13 | 5.51E-09 | 0.29 | 0.15 | 0.14 | 1.91 |
| dental | Kcnmb4 | 0.64 | 2.97E-13 | 6.34E-09 | 0.57 | 0.39 | 0.18 | 1.46 |
| dental | Tmtc1 | 1.50 | 3.72E-13 | 7.94E-09 | 0.15 | 0.06 | 0.09 | 2.67 |
| dental | Celf2 | 0.62 | 4.70E-13 | 1.00E-08 | 0.71 | 0.57 | 0.14 | 1.25 |
| dental | Usp46 | 0.40 | 5.76E-13 | 1.23E-08 | 0.84 | 0.71 | 0.13 | 1.19 |
| dental | Btrc | 0.81 | 5.99E-13 | 1.28E-08 | 0.48 | 0.32 | 0.16 | 1.50 |
| dental | Pam | 1.78 | 7.35E-13 | 1.57E-08 | 0.21 | 0.10 | 0.11 | 2.16 |
| dental | Hspb8 | 0.35 | 1.02E-12 | 2.18E-08 | 0.67 | 0.47 | 0.20 | 1.43 |
| dental | Nr2f2 | 0.54 | 1.23E-12 | 2.63E-08 | 0.71 | 0.58 | 0.14 | 1.23 |
| dental | Rgs12 | 0.42 | 1.63E-12 | 3.49E-08 | 0.63 | 0.45 | 0.19 | 1.41 |
| dental | 1600014C10Rik | 1.24 | 1.69E-12 | 3.60E-08 | 0.23 | 0.12 | 0.12 | 2.02 |
| dental | Itprid2 | 1.24 | 1.89E-12 | 4.03E-08 | 0.23 | 0.11 | 0.12 | 2.03 |
| dental | Cox6b2 | 0.62 | 2.20E-12 | 4.69E-08 | 0.64 | 0.50 | 0.14 | 1.28 |

**Supplementary Table 4. Marker genes for the annotated cell populations of integrated clustering of E11.5 control and NotchGOF λ-junction ectoderm cells.**

| | | | | | | | | |
|---|---|---|---|---|---|---|---|---|
| dental | Sp6 | 1.10 | 2.96E-12 | 6.32E-08 | 0.23 | 0.11 | 0.12 | 2.06 |
| dental | Ackr3 | 0.46 | 3.73E-12 | 7.97E-08 | 0.54 | 0.36 | 0.18 | 1.48 |
| dental | Mbnl2 | 0.33 | 4.84E-12 | 1.03E-07 | 0.89 | 0.78 | 0.11 | 1.15 |
| dental | Plec | 0.57 | 5.63E-12 | 1.20E-07 | 0.60 | 0.44 | 0.16 | 1.37 |
| dental | Aprt | 0.32 | 6.27E-12 | 1.34E-07 | 0.98 | 0.93 | 0.05 | 1.06 |
| dental | Fam83f | 1.42 | 6.56E-12 | 1.40E-07 | 0.11 | 0.03 | 0.07 | 3.12 |
| dental | Armh4 | 1.20 | 7.77E-12 | 1.66E-07 | 0.30 | 0.18 | 0.12 | 1.71 |
| dental | Igf2 | 0.30 | 7.96E-12 | 1.70E-07 | 0.90 | 0.81 | 0.10 | 1.12 |
| dental | Tbx3 | 1.11 | 8.37E-12 | 1.79E-07 | 0.18 | 0.08 | 0.10 | 2.32 |
| dental | Bmp4 | 0.52 | 1.58E-11 | 3.38E-07 | 0.71 | 0.52 | 0.19 | 1.36 |
| dental | Irx2 | 1.48 | 1.89E-11 | 4.04E-07 | 0.13 | 0.05 | 0.08 | 2.67 |
| dental | Minar2 | 1.06 | 1.98E-11 | 4.24E-07 | 0.28 | 0.15 | 0.13 | 1.86 |
| dental | Hs3st3b1 | 1.70 | 2.36E-11 | 5.04E-07 | 0.11 | 0.04 | 0.07 | 2.85 |
| dental | Jag2 | 0.55 | 2.92E-11 | 6.23E-07 | 0.44 | 0.27 | 0.16 | 1.60 |
| dental | Bdh2 | 1.45 | 4.45E-11 | 9.51E-07 | 0.14 | 0.05 | 0.08 | 2.50 |
| dental | Dusp23 | 1.12 | 4.63E-11 | 9.90E-07 | 0.21 | 0.11 | 0.11 | 2.00 |
| dental | Klk8 | 0.55 | 5.71E-11 | 1.22E-06 | 0.70 | 0.55 | 0.15 | 1.27 |
| dental | Cers4 | 0.67 | 6.54E-11 | 1.40E-06 | 0.51 | 0.37 | 0.14 | 1.39 |
| dental | Frmd6 | 0.80 | 8.86E-11 | 1.89E-06 | 0.36 | 0.22 | 0.14 | 1.62 |
| dental | Krt5 | 0.35 | 1.03E-10 | 2.21E-06 | 0.49 | 0.33 | 0.16 | 1.49 |
| dental | Inpp5f | 0.79 | 1.08E-10 | 2.31E-06 | 0.39 | 0.25 | 0.14 | 1.54 |
| dental | Gnb5 | 0.85 | 1.16E-10 | 2.47E-06 | 0.38 | 0.24 | 0.13 | 1.55 |
| dental | Osbpl6 | 1.07 | 1.44E-10 | 3.08E-06 | 0.24 | 0.13 | 0.11 | 1.85 |
| dental | Efcab1 | 1.09 | 1.48E-10 | 3.17E-06 | 0.19 | 0.09 | 0.10 | 2.14 |
| dental | Prkci | 0.58 | 1.55E-10 | 3.32E-06 | 0.57 | 0.43 | 0.15 | 1.34 |
| dental | Pid1 | 1.57 | 1.72E-10 | 3.67E-06 | 0.14 | 0.06 | 0.08 | 2.45 |
| dental | Sesn1 | 0.84 | 1.82E-10 | 3.88E-06 | 0.33 | 0.20 | 0.13 | 1.64 |
| dental | Nav2 | 0.37 | 1.88E-10 | 4.01E-06 | 0.82 | 0.69 | 0.13 | 1.18 |
| dental | Xpa | 0.56 | 1.88E-10 | 4.01E-06 | 0.61 | 0.47 | 0.14 | 1.30 |
| dental | Col18a1 | 0.29 | 2.08E-10 | 4.45E-06 | 0.93 | 0.85 | 0.08 | 1.09 |
| dental | Pde10a | 1.47 | 2.66E-10 | 5.69E-06 | 0.11 | 0.04 | 0.07 | 2.77 |
| dental | Col9a3 | 0.76 | 3.03E-10 | 6.47E-06 | 0.26 | 0.14 | 0.12 | 1.86 |
| dental | Spon2 | 1.36 | 3.26E-10 | 6.95E-06 | 0.13 | 0.05 | 0.08 | 2.51 |
| dental | Trp53 | 0.29 | 3.31E-10 | 7.08E-06 | 0.93 | 0.89 | 0.05 | 1.05 |
| dental | 2610301B20Rik | 0.86 | 3.37E-10 | 7.19E-06 | 0.41 | 0.29 | 0.12 | 1.41 |
| dental | Cemip2 | 0.33 | 3.38E-10 | 7.23E-06 | 0.72 | 0.59 | 0.13 | 1.21 |
| dental | Micu2 | 0.48 | 3.76E-10 | 8.03E-06 | 0.70 | 0.58 | 0.12 | 1.20 |
| dental | Cpe | 0.52 | 3.76E-10 | 8.04E-06 | 0.60 | 0.46 | 0.14 | 1.31 |
| dental | Rpl9-ps6 | 0.32 | 4.10E-10 | 8.76E-06 | 0.88 | 0.80 | 0.08 | 1.10 |
| dental | Plek2 | 1.36 | 4.63E-10 | 9.90E-06 | 0.19 | 0.10 | 0.09 | 1.98 |
| dental | Acad8 | 0.75 | 5.11E-10 | 1.09E-05 | 0.44 | 0.32 | 0.13 | 1.40 |
| dental | Etv1 | 1.05 | 6.72E-10 | 1.44E-05 | 0.28 | 0.16 | 0.11 | 1.69 |
| dental | Yap1 | 0.42 | 7.40E-10 | 1.58E-05 | 0.72 | 0.60 | 0.12 | 1.20 |
| dental | Plxnc1 | 1.14 | 7.98E-10 | 1.70E-05 | 0.16 | 0.07 | 0.09 | 2.21 |
| dental | Slc12a4 | 0.63 | 8.11E-10 | 1.73E-05 | 0.33 | 0.20 | 0.13 | 1.66 |
| dental | Mdga1 | 1.47 | 8.13E-10 | 1.74E-05 | 0.11 | 0.04 | 0.07 | 2.63 |
| dental | Ext1 | 0.55 | 8.33E-10 | 1.78E-05 | 0.63 | 0.51 | 0.12 | 1.24 |
| dental | Adamts18 | 0.48 | 8.76E-10 | 1.87E-05 | 0.41 | 0.26 | 0.15 | 1.58 |
| dental | 2410006H16Rik | 0.44 | 9.37E-10 | 2.00E-05 | 0.92 | 0.86 | 0.07 | 1.08 |
| dental | Zfp36l2 | 0.54 | 9.52E-10 | 2.03E-05 | 0.38 | 0.24 | 0.14 | 1.58 |
| dental | Lmo7 | 0.62 | 9.86E-10 | 2.11E-05 | 0.57 | 0.44 | 0.13 | 1.29 |
| dental | Srpk1 | 0.32 | 9.88E-10 | 2.11E-05 | 0.90 | 0.83 | 0.07 | 1.08 |
| dental | Chst11 | 0.86 | 1.04E-09 | 2.22E-05 | 0.27 | 0.15 | 0.12 | 1.76 |
| dental | Tmod3 | 0.38 | 1.47E-09 | 3.14E-05 | 0.75 | 0.65 | 0.09 | 1.14 |
| dental | Rps27rt | 0.36 | 1.53E-09 | 3.27E-05 | 0.85 | 0.78 | 0.08 | 1.10 |

**Supplementary Table 4. Marker genes for the annotated cell populations of integrated clustering of E11.5 control and NotchGOF λ-junction ectoderm cells.**

| | | | | | | | | |
|---|---|---|---|---|---|---|---|---|
| dental | Golim4 | 0.46 | 1.66E-09 | 3.55E-05 | 0.71 | 0.60 | 0.11 | 1.18 |
| dental | Dse | 1.01 | 1.80E-09 | 3.85E-05 | 0.20 | 0.10 | 0.10 | 1.94 |
| dental | Flot2 | 0.70 | 1.96E-09 | 4.18E-05 | 0.39 | 0.27 | 0.12 | 1.46 |
| dental | Bdnf | 0.82 | 2.18E-09 | 4.65E-05 | 0.29 | 0.17 | 0.12 | 1.68 |
| dental | Mdm1 | 0.53 | 2.94E-09 | 6.28E-05 | 0.43 | 0.29 | 0.14 | 1.47 |
| dental | Pthlh | 0.84 | 3.33E-09 | 7.12E-05 | 0.12 | 0.05 | 0.07 | 2.52 |
| dental | Gas7 | 1.02 | 3.83E-09 | 8.18E-05 | 0.19 | 0.10 | 0.09 | 1.98 |
| dental | Zfp9 | 1.15 | 4.06E-09 | 8.66E-05 | 0.14 | 0.06 | 0.08 | 2.26 |
| dental | 9330162012Rik | 1.52 | 4.16E-09 | 8.89E-05 | 0.10 | 0.04 | 0.06 | 2.66 |
| dental | Col27a1 | 1.39 | 4.62E-09 | 9.88E-05 | 0.14 | 0.06 | 0.08 | 2.25 |
| dental | Rbm43 | 1.06 | 4.93E-09 | 1.05E-04 | 0.18 | 0.09 | 0.09 | 1.97 |
| dental | Trim8 | 0.63 | 5.03E-09 | 1.07E-04 | 0.45 | 0.32 | 0.13 | 1.41 |
| dental | Gm9922 | 1.30 | 5.41E-09 | 1.16E-04 | 0.20 | 0.11 | 0.09 | 1.85 |
| dental | Eef2k | 0.82 | 5.46E-09 | 1.17E-04 | 0.28 | 0.17 | 0.12 | 1.68 |
| dental | Baiap2 | 0.56 | 6.04E-09 | 1.29E-04 | 0.34 | 0.21 | 0.13 | 1.62 |
| dental | Inka1 | 0.73 | 7.18E-09 | 1.53E-04 | 0.16 | 0.07 | 0.08 | 2.15 |
| dental | 1700007K13Rik | 1.10 | 8.28E-09 | 1.77E-04 | 0.12 | 0.05 | 0.07 | 2.40 |
| dental | Snhg1 | 0.28 | 8.43E-09 | 1.80E-04 | 0.96 | 0.92 | 0.04 | 1.05 |
| dental | Zfp503 | 0.40 | 8.55E-09 | 1.83E-04 | 0.64 | 0.49 | 0.15 | 1.30 |
| dental | Cpne5 | 1.06 | 8.66E-09 | 1.85E-04 | 0.24 | 0.14 | 0.10 | 1.69 |
| dental | Frmpd1 | 1.48 | 9.27E-09 | 1.98E-04 | 0.10 | 0.04 | 0.06 | 2.59 |
| dental | Dbp | 0.82 | 1.01E-08 | 2.16E-04 | 0.31 | 0.19 | 0.11 | 1.59 |
| dental | Tmc7 | 1.02 | 1.01E-08 | 2.17E-04 | 0.13 | 0.06 | 0.08 | 2.29 |
| dental | Cuedc1 | 0.43 | 1.02E-08 | 2.17E-04 | 0.62 | 0.51 | 0.12 | 1.23 |
| dental | Hprt | 0.48 | 1.18E-08 | 2.52E-04 | 0.58 | 0.46 | 0.12 | 1.25 |
| dental | Bard1 | 0.84 | 1.34E-08 | 2.86E-04 | 0.28 | 0.18 | 0.11 | 1.62 |
| dental | Ptprm | 1.15 | 1.41E-08 | 3.02E-04 | 0.18 | 0.10 | 0.09 | 1.91 |
| dental | Cdk5r1 | 0.98 | 1.75E-08 | 3.74E-04 | 0.14 | 0.06 | 0.08 | 2.21 |
| dental | Synm | 1.33 | 1.83E-08 | 3.91E-04 | 0.13 | 0.06 | 0.07 | 2.20 |
| dental | Ldlrap1 | 0.48 | 1.87E-08 | 4.00E-04 | 0.44 | 0.30 | 0.14 | 1.45 |
| dental | N4bp3 | 1.31 | 1.95E-08 | 4.16E-04 | 0.15 | 0.07 | 0.08 | 2.07 |
| dental | Sertad4 | 0.53 | 2.37E-08 | 5.06E-04 | 0.45 | 0.32 | 0.13 | 1.42 |
| dental | Chchd5 | 0.46 | 2.79E-08 | 5.96E-04 | 0.62 | 0.49 | 0.13 | 1.27 |
| dental | Vwa5a | 0.88 | 3.70E-08 | 7.91E-04 | 0.26 | 0.16 | 0.10 | 1.66 |
| dental | Sgk3 | 0.74 | 3.71E-08 | 7.92E-04 | 0.21 | 0.12 | 0.10 | 1.83 |
| dental | Pcdh10 | 1.02 | 4.40E-08 | 9.39E-04 | 0.16 | 0.08 | 0.08 | 2.01 |
| dental | Arhgdib | 0.51 | 4.52E-08 | 9.65E-04 | 0.32 | 0.20 | 0.12 | 1.60 |
| dental | Celsr2 | 0.60 | 5.41E-08 | 1.15E-03 | 0.41 | 0.29 | 0.11 | 1.39 |
| dental | Sept10 | 0.53 | 5.57E-08 | 1.19E-03 | 0.48 | 0.36 | 0.13 | 1.36 |
| dental | Bach2 | 0.59 | 6.16E-08 | 1.32E-03 | 0.35 | 0.24 | 0.12 | 1.49 |
| dental | Gga2 | 0.41 | 6.36E-08 | 1.36E-03 | 0.63 | 0.50 | 0.13 | 1.26 |
| dental | Sik1 | 0.64 | 9.11E-08 | 1.95E-03 | 0.26 | 0.15 | 0.10 | 1.68 |
| dental | Isoc1 | 0.30 | 1.04E-07 | 2.22E-03 | 0.77 | 0.70 | 0.08 | 1.11 |
| dental | Cgnl1 | 0.37 | 1.23E-07 | 2.63E-03 | 0.27 | 0.16 | 0.11 | 1.67 |
| dental | Pdzrn3 | 0.74 | 1.30E-07 | 2.79E-03 | 0.21 | 0.11 | 0.09 | 1.80 |
| dental | Chst15 | 0.72 | 1.35E-07 | 2.89E-03 | 0.23 | 0.13 | 0.10 | 1.75 |
| dental | Irs2 | 0.47 | 1.40E-07 | 2.99E-03 | 0.55 | 0.43 | 0.12 | 1.28 |
| dental | Dkk3 | 0.82 | 1.48E-07 | 3.17E-03 | 0.25 | 0.16 | 0.10 | 1.61 |
| dental | Dgkh | 1.29 | 1.55E-07 | 3.30E-03 | 0.10 | 0.04 | 0.06 | 2.36 |
| dental | Rap2c | 0.31 | 1.65E-07 | 3.53E-03 | 0.75 | 0.66 | 0.09 | 1.14 |
| dental | Dzip1 | 0.65 | 1.74E-07 | 3.72E-03 | 0.31 | 0.20 | 0.10 | 1.51 |
| dental | Col4a1 | 0.34 | 1.89E-07 | 4.04E-03 | 0.81 | 0.67 | 0.14 | 1.21 |
| dental | Prkd3 | 0.51 | 2.17E-07 | 4.64E-03 | 0.63 | 0.54 | 0.09 | 1.17 |
| dental | Parvb | 1.50 | 2.17E-07 | 4.64E-03 | 0.10 | 0.05 | 0.06 | 2.31 |
| dental | Pcsk6 | 1.38 | 2.17E-07 | 4.64E-03 | 0.10 | 0.04 | 0.06 | 2.30 |

**Supplementary Table 4. Marker genes for the annotated cell populations of integrated clustering of E11.5 control and NotchGOF λ-junction ectoderm cells.**

| | | | | | | | | |
|---|---|---|---|---|---|---|---|---|
| dental | Sall4 | 1.45 | 2.41E-07 | 5.14E-03 | 0.10 | 0.04 | 0.06 | 2.35 |
| dental | Pqlc3 | 0.73 | 2.58E-07 | 5.51E-03 | 0.27 | 0.16 | 0.10 | 1.62 |
| dental | Arhgap42 | 0.60 | 2.60E-07 | 5.56E-03 | 0.38 | 0.27 | 0.11 | 1.42 |
| dental | Ptpn13 | 0.29 | 2.82E-07 | 6.01E-03 | 0.69 | 0.56 | 0.13 | 1.23 |
| dental | Rac3 | 0.35 | 2.92E-07 | 6.23E-03 | 0.53 | 0.40 | 0.12 | 1.30 |
| dental | Adamts17 | 1.09 | 2.99E-07 | 6.39E-03 | 0.19 | 0.11 | 0.08 | 1.74 |
| dental | Ddx25 | 0.90 | 3.03E-07 | 6.48E-03 | 0.19 | 0.11 | 0.08 | 1.79 |
| dental | Plagl1 | 0.37 | 3.30E-07 | 7.04E-03 | 0.54 | 0.42 | 0.12 | 1.29 |
| dental | Arhgap24 | 0.95 | 3.52E-07 | 7.52E-03 | 0.14 | 0.07 | 0.07 | 2.03 |
| dental | Luzp1 | 0.26 | 3.65E-07 | 7.79E-03 | 0.88 | 0.80 | 0.08 | 1.10 |
| dental | Sh3bp1 | 0.62 | 3.79E-07 | 8.09E-03 | 0.40 | 0.29 | 0.12 | 1.40 |
| dental | Arhgap10 | 0.49 | 4.07E-07 | 8.69E-03 | 0.55 | 0.42 | 0.13 | 1.30 |
| dental | Trim44 | 0.25 | 4.18E-07 | 8.92E-03 | 0.84 | 0.78 | 0.06 | 1.07 |
| dental | Foxo1 | 0.54 | 4.58E-07 | 9.78E-03 | 0.39 | 0.28 | 0.11 | 1.38 |
| dental | Jade2 | 1.53 | 5.10E-07 | 1.09E-02 | 0.10 | 0.05 | 0.06 | 2.24 |
| dental | Sorcs2 | 0.56 | 5.16E-07 | 1.10E-02 | 0.17 | 0.09 | 0.08 | 1.88 |
| dental | Slc25a17 | 0.39 | 5.17E-07 | 1.10E-02 | 0.79 | 0.70 | 0.08 | 1.12 |
| dental | Fam120a | 0.43 | 5.27E-07 | 1.12E-02 | 0.62 | 0.52 | 0.10 | 1.20 |
| dental | Agap1 | 0.46 | 5.28E-07 | 1.13E-02 | 0.57 | 0.47 | 0.10 | 1.21 |
| dental | Mbd2 | 0.31 | 5.68E-07 | 1.21E-02 | 0.70 | 0.61 | 0.09 | 1.15 |
| dental | Glul | 0.34 | 5.82E-07 | 1.24E-02 | 0.58 | 0.45 | 0.13 | 1.28 |
| dental | Ccdc148 | 1.35 | 6.22E-07 | 1.33E-02 | 0.11 | 0.05 | 0.06 | 2.16 |
| dental | Rnf217 | 0.72 | 6.51E-07 | 1.39E-02 | 0.25 | 0.16 | 0.09 | 1.60 |
| dental | Fgfr1op2 | 0.35 | 6.83E-07 | 1.46E-02 | 0.82 | 0.72 | 0.10 | 1.14 |
| dental | Pnldc1 | 1.01 | 7.10E-07 | 1.52E-02 | 0.11 | 0.05 | 0.06 | 2.20 |
| dental | Slc25a12 | 0.87 | 7.25E-07 | 1.55E-02 | 0.24 | 0.15 | 0.09 | 1.60 |
| dental | Cox4i2 | 0.31 | 7.43E-07 | 1.59E-02 | 0.61 | 0.47 | 0.13 | 1.28 |
| dental | Crabp1 | 0.30 | 8.15E-07 | 1.74E-02 | 0.65 | 0.56 | 0.10 | 1.17 |
| dental | Tulp3 | 0.63 | 9.09E-07 | 1.94E-02 | 0.34 | 0.24 | 0.10 | 1.43 |
| dental | Exosc2 | 0.42 | 9.37E-07 | 2.00E-02 | 0.56 | 0.46 | 0.11 | 1.23 |
| dental | Rdm1 | 0.42 | 1.02E-06 | 2.17E-02 | 0.49 | 0.38 | 0.12 | 1.31 |
| dental | Peg12 | 0.99 | 1.11E-06 | 2.36E-02 | 0.14 | 0.07 | 0.07 | 1.96 |
| dental | Cdc42ep3 | 0.51 | 1.12E-06 | 2.40E-02 | 0.27 | 0.17 | 0.10 | 1.58 |
| dental | S100a6 | 0.38 | 1.15E-06 | 2.46E-02 | 0.34 | 0.23 | 0.11 | 1.47 |
| dental | Cnn2 | 0.37 | 1.18E-06 | 2.53E-02 | 0.42 | 0.30 | 0.12 | 1.39 |
| dental | Hs3st1 | 0.92 | 1.21E-06 | 2.58E-02 | 0.13 | 0.07 | 0.07 | 1.99 |
| dental | Ptpru | 0.43 | 1.32E-06 | 2.82E-02 | 0.48 | 0.37 | 0.11 | 1.30 |
| dental | Col17a1 | 0.39 | 1.33E-06 | 2.85E-02 | 0.21 | 0.12 | 0.09 | 1.72 |
| dental | Tgif1 | 0.30 | 1.41E-06 | 3.01E-02 | 0.78 | 0.68 | 0.10 | 1.15 |
| dental | Man2a1 | 0.64 | 1.45E-06 | 3.10E-02 | 0.34 | 0.24 | 0.10 | 1.41 |
| dental | F11r | 0.32 | 1.49E-06 | 3.18E-02 | 0.72 | 0.60 | 0.12 | 1.19 |
| dental | Map3k20 | 0.81 | 1.69E-06 | 3.61E-02 | 0.27 | 0.18 | 0.09 | 1.50 |
| dental | Cyfip1 | 0.30 | 1.75E-06 | 3.74E-02 | 0.81 | 0.73 | 0.08 | 1.11 |
| dental | Enho | 0.41 | 1.84E-06 | 3.93E-02 | 0.44 | 0.33 | 0.11 | 1.34 |
| dental | Sox6 | 0.97 | 1.94E-06 | 4.14E-02 | 0.17 | 0.10 | 0.07 | 1.78 |
| SE_basal | Cxcl14 | 5.29 | 0 | 0 | 0.95 | 0.11 | 0.85 | 9.00 |
| SE_basal | Tfap2b | 4.54 | 0 | 0 | 0.93 | 0.10 | 0.83 | 9.29 |
| SE_basal | Wnt3 | 5.76 | 0 | 0 | 0.78 | 0.03 | 0.75 | 26.00 |
| SE_basal | Wnt9b | 4.99 | 0 | 0 | 0.68 | 0.04 | 0.64 | 16.26 |
| SE_basal | Pcp4l1 | 5.26 | 0 | 0 | 0.52 | 0.02 | 0.50 | 24.71 |
| SE_basal | Sostdc1 | 4.36 | 5.13E-293 | 1.10E-288 | 0.86 | 0.18 | 0.68 | 4.75 |
| SE_basal | Piezo2 | 3.95 | 2.98E-292 | 6.36E-288 | 0.74 | 0.10 | 0.64 | 7.62 |
| SE_basal | Wnt6 | 3.15 | 7.20E-287 | 1.54E-282 | 0.97 | 0.30 | 0.68 | 3.29 |
| SE_basal | Plxna2 | 3.64 | 1.06E-263 | 2.27E-259 | 0.84 | 0.19 | 0.65 | 4.36 |
| SE_basal | Irx5 | 3.42 | 6.01E-241 | 1.28E-236 | 0.79 | 0.17 | 0.62 | 4.57 |

**Supplementary Table 4. Marker genes for the annotated cell populations of integrated clustering of E11.5 control and NotchGOF λ-junction ectoderm cells.**

| | | | | | | | | |
|---|---|---|---|---|---|---|---|---|
| SE_basal | Kremen2 | 3.56 | 3.77E-230 | 8.04E-226 | 0.69 | 0.12 | 0.57 | 5.96 |
| SE_basal | Rprml | 6.55 | 3.59E-217 | 7.67E-213 | 0.33 | 0.01 | 0.32 | 41.25 |
| SE_basal | Wnt7b | 3.85 | 1.67E-214 | 3.56E-210 | 0.64 | 0.11 | 0.53 | 5.99 |
| SE_basal | Irx3 | 3.29 | 5.21E-185 | 1.11E-180 | 0.67 | 0.15 | 0.52 | 4.47 |
| SE_basal | Tfap2a | 2.66 | 4.29E-174 | 9.15E-170 | 0.81 | 0.28 | 0.53 | 2.86 |
| SE_basal | Col17a1 | 3.49 | 1.26E-171 | 2.69E-167 | 0.52 | 0.08 | 0.45 | 6.71 |
| SE_basal | Tnfrsf19 | 2.72 | 1.93E-169 | 4.11E-165 | 0.71 | 0.19 | 0.52 | 3.78 |
| SE_basal | Fth1 | 1.09 | 1.44E-168 | 3.08E-164 | 1.00 | 1.00 | 0.00 | 1.00 |
| SE_basal | Gria2 | 4.43 | 7.77E-161 | 1.66E-156 | 0.40 | 0.04 | 0.36 | 9.57 |
| SE_basal | Ctsl | 1.62 | 3.22E-156 | 6.87E-152 | 0.98 | 0.85 | 0.13 | 1.15 |
| SE_basal | Snai2 | 2.80 | 5.06E-154 | 1.08E-149 | 0.60 | 0.13 | 0.47 | 4.76 |
| SE_basal | Slc7a11 | 3.35 | 6.67E-151 | 1.42E-146 | 0.50 | 0.08 | 0.42 | 5.95 |
| SE_basal | Ccnd2 | 1.66 | 4.72E-146 | 1.01E-141 | 1.00 | 0.76 | 0.24 | 1.31 |
| SE_basal | Fgfr2 | 2.17 | 1.70E-140 | 3.62E-136 | 0.86 | 0.43 | 0.43 | 2.01 |
| SE_basal | Pdlim1 | 1.69 | 2.78E-138 | 5.94E-134 | 0.94 | 0.61 | 0.33 | 1.54 |
| SE_basal | Trp63 | 1.78 | 2.36E-134 | 5.03E-130 | 0.93 | 0.46 | 0.47 | 2.03 |
| SE_basal | Crabp1 | 2.31 | 1.75E-133 | 3.73E-129 | 0.90 | 0.52 | 0.38 | 1.73 |
| SE_basal | Krt15 | 3.49 | 2.54E-127 | 5.42E-123 | 0.51 | 0.11 | 0.40 | 4.66 |
| SE_basal | Perp | 1.66 | 1.92E-125 | 4.09E-121 | 0.95 | 0.49 | 0.46 | 1.93 |
| SE_basal | Lgals7 | 4.07 | 3.19E-121 | 6.81E-117 | 0.27 | 0.02 | 0.25 | 12.45 |
| SE_basal | Cobll1 | 3.35 | 2.97E-118 | 6.34E-114 | 0.39 | 0.06 | 0.33 | 6.25 |
| SE_basal | Itgb5 | 1.87 | 4.07E-117 | 8.69E-113 | 0.86 | 0.45 | 0.40 | 1.89 |
| SE_basal | Cdkn1c | 1.58 | 3.16E-115 | 6.76E-111 | 0.98 | 0.87 | 0.11 | 1.12 |
| SE_basal | Npr3 | 4.12 | 5.12E-111 | 1.09E-106 | 0.26 | 0.02 | 0.24 | 11.77 |
| SE_basal | Hras | 1.08 | 7.55E-111 | 1.61E-106 | 0.98 | 0.90 | 0.08 | 1.09 |
| SE_basal | Wnt10a | 4.84 | 3.54E-108 | 7.57E-104 | 0.19 | 0.01 | 0.18 | 23.88 |
| SE_basal | Tmsb10 | 0.61 | 3.90E-108 | 8.33E-104 | 1.00 | 1.00 | 0.00 | 1.00 |
| SE_basal | Wls | 1.26 | 4.09E-106 | 8.73E-102 | 0.97 | 0.78 | 0.19 | 1.25 |
| SE_basal | C130021I20Rik | 5.40 | 9.37E-104 | 2.00E-99 | 0.18 | 0.01 | 0.17 | 29.33 |
| SE_basal | Sema3a | 2.38 | 8.82E-103 | 1.88E-98 | 0.77 | 0.45 | 0.33 | 1.73 |
| SE_basal | Txndc17 | 1.25 | 2.39E-102 | 5.11E-98 | 0.96 | 0.94 | 0.01 | 1.01 |
| SE_basal | Mir205hg | 1.81 | 6.75E-101 | 1.44E-96 | 0.79 | 0.40 | 0.39 | 1.98 |
| SE_basal | Tspan2 | 3.97 | 6.56E-100 | 1.40E-95 | 0.25 | 0.02 | 0.22 | 10.65 |
| SE_basal | Crnde | 3.25 | 7.35E-97 | 1.57E-92 | 0.35 | 0.06 | 0.29 | 5.75 |
| SE_basal | Jdp2 | 3.11 | 6.83E-95 | 1.46E-90 | 0.36 | 0.07 | 0.29 | 5.49 |
| SE_basal | Rnase4 | 2.88 | 4.45E-91 | 9.50E-87 | 0.39 | 0.08 | 0.31 | 4.73 |
| SE_basal | Fermt1 | 1.42 | 6.43E-91 | 1.37E-86 | 0.86 | 0.52 | 0.35 | 1.67 |
| SE_basal | Nrep | 1.15 | 5.97E-89 | 1.27E-84 | 0.94 | 0.79 | 0.15 | 1.19 |
| SE_basal | Irx4 | 6.20 | 2.43E-88 | 5.20E-84 | 0.13 | 0.00 | 0.13 | 65.50 |
| SE_basal | Edaradd | 3.11 | 4.11E-86 | 8.79E-82 | 0.31 | 0.05 | 0.26 | 6.02 |
| SE_basal | Wnt4 | 1.45 | 2.23E-83 | 4.75E-79 | 0.91 | 0.69 | 0.22 | 1.32 |
| SE_basal | Robo2 | 2.29 | 5.21E-82 | 1.11E-77 | 0.55 | 0.20 | 0.35 | 2.71 |
| SE_basal | Insyn2b | 3.00 | 1.29E-81 | 2.76E-77 | 0.28 | 0.04 | 0.24 | 6.41 |
| SE_basal | Jag2 | 1.91 | 8.51E-81 | 1.82E-76 | 0.62 | 0.24 | 0.38 | 2.55 |
| SE_basal | Chst11 | 2.44 | 5.64E-80 | 1.20E-75 | 0.44 | 0.13 | 0.32 | 3.50 |
| SE_basal | Timp3 | 3.78 | 9.75E-80 | 2.08E-75 | 0.27 | 0.04 | 0.23 | 6.33 |
| SE_basal | Lef1 | 1.42 | 1.52E-79 | 3.24E-75 | 0.74 | 0.33 | 0.41 | 2.21 |
| SE_basal | Sox11 | 0.84 | 3.27E-77 | 6.98E-73 | 1.00 | 0.99 | 0.01 | 1.01 |
| SE_basal | Cald1 | 0.92 | 4.55E-77 | 9.72E-73 | 0.94 | 0.67 | 0.27 | 1.41 |
| SE_basal | Igfbp4 | 1.25 | 2.16E-75 | 4.62E-71 | 0.91 | 0.72 | 0.19 | 1.26 |
| SE_basal | Msx2 | 1.65 | 6.42E-75 | 1.37E-70 | 0.72 | 0.36 | 0.36 | 1.98 |
| SE_basal | Lmx1b | 4.41 | 4.25E-72 | 9.07E-68 | 0.15 | 0.01 | 0.14 | 16.11 |
| SE_basal | Igfbp2 | 0.79 | 5.46E-71 | 1.17E-66 | 1.00 | 0.98 | 0.01 | 1.01 |
| SE_basal | Bambi | 1.29 | 7.45E-71 | 1.59E-66 | 0.77 | 0.45 | 0.31 | 1.69 |
| SE_basal | Hapln1 | 5.86 | 1.18E-70 | 2.52E-66 | 0.13 | 0.01 | 0.12 | 25.40 |

**Supplementary Table 4. Marker genes for the annotated cell populations of integrated clustering of E11.5 control and NotchGOF λ-junction ectoderm cells.**

| | | | | | | | | |
|---|---|---|---|---|---|---|---|---|
| SE_basal | Plk2 | 1.84 | 4.07E-70 | 8.68E-66 | 0.65 | 0.32 | 0.33 | 2.02 |
| SE_basal | Pthlh | 3.57 | 1.25E-68 | 2.66E-64 | 0.22 | 0.03 | 0.19 | 7.33 |
| SE_basal | Atxn10 | 0.93 | 9.36E-68 | 2.00E-63 | 0.93 | 0.91 | 0.03 | 1.03 |
| SE_basal | Dock8 | 2.65 | 1.01E-67 | 2.15E-63 | 0.31 | 0.07 | 0.24 | 4.61 |
| SE_basal | Krt14 | 1.16 | 2.12E-67 | 4.53E-63 | 0.80 | 0.42 | 0.38 | 1.91 |
| SE_basal | Itgav | 2.24 | 2.78E-67 | 5.93E-63 | 0.48 | 0.18 | 0.31 | 2.76 |
| SE_basal | Htra1 | 2.90 | 2.99E-67 | 6.39E-63 | 0.25 | 0.04 | 0.21 | 6.02 |
| SE_basal | Gstk1 | 2.49 | 4.65E-67 | 9.94E-63 | 0.39 | 0.11 | 0.28 | 3.51 |
| SE_basal | Plbd1 | 3.71 | 4.79E-67 | 1.02E-62 | 0.18 | 0.02 | 0.16 | 9.15 |
| SE_basal | Hspb1 | 1.36 | 5.45E-67 | 1.16E-62 | 0.71 | 0.35 | 0.36 | 2.04 |
| SE_basal | Pdgfa | 0.89 | 7.29E-67 | 1.56E-62 | 0.98 | 0.93 | 0.05 | 1.05 |
| SE_basal | Art4 | 3.52 | 7.37E-67 | 1.57E-62 | 0.21 | 0.03 | 0.18 | 7.50 |
| SE_basal | Gm42418 | 0.95 | 1.83E-66 | 3.92E-62 | 1.00 | 1.00 | 0.00 | 1.00 |
| SE_basal | Ldhb | 0.90 | 1.45E-65 | 3.10E-61 | 0.93 | 0.87 | 0.06 | 1.07 |
| SE_basal | Col18a1 | 1.01 | 8.40E-65 | 1.79E-60 | 0.92 | 0.85 | 0.07 | 1.08 |
| SE_basal | Cd24a | 0.68 | 7.01E-64 | 1.50E-59 | 0.99 | 0.96 | 0.03 | 1.03 |
| SE_basal | Dlx2 | 1.40 | 2.59E-63 | 5.52E-59 | 0.66 | 0.30 | 0.36 | 2.19 |
| SE_basal | Meis1 | 1.36 | 4.83E-62 | 1.03E-57 | 0.80 | 0.56 | 0.24 | 1.43 |
| SE_basal | Tnik | 2.05 | 4.91E-62 | 1.05E-57 | 0.33 | 0.08 | 0.25 | 4.21 |
| SE_basal | Sgk3 | 2.52 | 5.78E-62 | 1.24E-57 | 0.35 | 0.10 | 0.26 | 3.69 |
| SE_basal | Ngf | 3.28 | 6.98E-61 | 1.49E-56 | 0.17 | 0.02 | 0.15 | 8.60 |
| SE_basal | Col12a1 | 3.07 | 9.71E-60 | 2.07E-55 | 0.25 | 0.05 | 0.20 | 5.06 |
| SE_basal | Bnc2 | 2.87 | 3.06E-58 | 6.53E-54 | 0.26 | 0.05 | 0.20 | 4.85 |
| SE_basal | Btg1 | 1.33 | 1.11E-57 | 2.37E-53 | 0.86 | 0.74 | 0.12 | 1.17 |
| SE_basal | Trp73 | 3.57 | 1.54E-56 | 3.29E-52 | 0.15 | 0.01 | 0.13 | 10.36 |
| SE_basal | H19 | 0.75 | 2.03E-56 | 4.33E-52 | 0.98 | 0.87 | 0.12 | 1.13 |
| SE_basal | Cdh3 | 1.36 | 1.06E-55 | 2.26E-51 | 0.76 | 0.55 | 0.22 | 1.39 |
| SE_basal | Stk26 | 1.39 | 1.67E-54 | 3.56E-50 | 0.71 | 0.44 | 0.27 | 1.60 |
| SE_basal | Adk | 1.21 | 3.79E-54 | 8.09E-50 | 0.78 | 0.60 | 0.18 | 1.30 |
| SE_basal | Brinp1 | 3.09 | 4.78E-54 | 1.02E-49 | 0.17 | 0.02 | 0.15 | 7.73 |
| SE_basal | Gm32585 | 1.74 | 8.02E-54 | 1.71E-49 | 0.48 | 0.19 | 0.29 | 2.55 |
| SE_basal | Nav2 | 1.08 | 8.51E-54 | 1.82E-49 | 0.87 | 0.68 | 0.18 | 1.27 |
| SE_basal | Zfp36l2 | 2.15 | 9.61E-54 | 2.05E-49 | 0.49 | 0.22 | 0.27 | 2.25 |
| SE_basal | Galnt18 | 2.62 | 1.01E-53 | 2.15E-49 | 0.29 | 0.07 | 0.22 | 3.92 |
| SE_basal | Col4a2 | 1.41 | 6.46E-53 | 1.38E-48 | 0.72 | 0.48 | 0.24 | 1.50 |
| SE_basal | Ube2e3 | 0.83 | 1.63E-52 | 3.47E-48 | 0.94 | 0.90 | 0.05 | 1.05 |
| SE_basal | Col5a2 | 2.02 | 3.53E-50 | 7.54E-46 | 0.39 | 0.13 | 0.25 | 2.90 |
| SE_basal | Wnt10b | 3.94 | 7.53E-50 | 1.61E-45 | 0.12 | 0.01 | 0.11 | 12.89 |
| SE_basal | Pdgfc | 0.97 | 8.29E-50 | 1.77E-45 | 0.86 | 0.75 | 0.11 | 1.14 |
| SE_basal | Dok5 | 1.37 | 1.62E-49 | 3.45E-45 | 0.64 | 0.35 | 0.29 | 1.85 |
| SE_basal | Rnd3 | 0.95 | 1.89E-49 | 4.03E-45 | 0.88 | 0.72 | 0.16 | 1.23 |
| SE_basal | Sema5a | 2.11 | 2.94E-49 | 6.29E-45 | 0.43 | 0.17 | 0.26 | 2.47 |
| SE_basal | Mbnl2 | 0.89 | 1.66E-48 | 3.54E-44 | 0.91 | 0.77 | 0.13 | 1.17 |
| SE_basal | Tmem178 | 2.29 | 2.73E-48 | 5.84E-44 | 0.32 | 0.09 | 0.22 | 3.39 |
| SE_basal | Ahnak | 1.34 | 9.96E-48 | 2.13E-43 | 0.63 | 0.33 | 0.30 | 1.91 |
| SE_basal | Fzd10 | 2.33 | 1.09E-47 | 2.33E-43 | 0.34 | 0.11 | 0.23 | 3.04 |
| SE_basal | Tspan18 | 3.07 | 3.67E-47 | 7.83E-43 | 0.19 | 0.03 | 0.16 | 5.56 |
| SE_basal | Malat1 | 0.63 | 4.68E-47 | 1.00E-42 | 1.00 | 1.00 | 0.00 | 1.00 |
| SE_basal | Plxna4 | 3.06 | 1.30E-46 | 2.77E-42 | 0.20 | 0.04 | 0.16 | 5.18 |
| SE_basal | Ehbp1 | 1.33 | 1.50E-46 | 3.21E-42 | 0.69 | 0.47 | 0.22 | 1.46 |
| SE_basal | Npnt | 1.29 | 4.48E-46 | 9.58E-42 | 0.68 | 0.42 | 0.26 | 1.61 |
| SE_basal | Tfap2c | 1.83 | 4.90E-46 | 1.05E-41 | 0.45 | 0.19 | 0.26 | 2.34 |
| SE_basal | Cib2 | 1.85 | 2.08E-45 | 4.44E-41 | 0.40 | 0.15 | 0.25 | 2.62 |
| SE_basal | Bnc1 | 2.63 | 2.38E-45 | 5.09E-41 | 0.16 | 0.02 | 0.14 | 6.67 |
| SE_basal | Cd59a | 1.44 | 6.31E-45 | 1.35E-40 | 0.55 | 0.27 | 0.28 | 2.01 |

**Supplementary Table 4. Marker genes for the annotated cell populations of integrated clustering of E11.5 control and NotchGOF λ-junction ectoderm cells.**

| | | | | | | | | |
|---|---|---|---|---|---|---|---|---|
| SE_basal | Ajuba | 1.33 | 7.17E-45 | 1.53E-40 | 0.65 | 0.40 | 0.25 | 1.63 |
| SE_basal | Tbx3 | 2.31 | 1.78E-44 | 3.80E-40 | 0.25 | 0.06 | 0.19 | 3.92 |
| SE_basal | Pltp | 1.21 | 6.95E-44 | 1.49E-39 | 0.70 | 0.44 | 0.27 | 1.61 |
| SE_basal | Chst15 | 2.16 | 1.11E-42 | 2.36E-38 | 0.33 | 0.12 | 0.21 | 2.84 |
| SE_basal | Ntf5 | 1.38 | 2.28E-42 | 4.88E-38 | 0.56 | 0.29 | 0.27 | 1.93 |
| SE_basal | Ifitm2 | 0.59 | 4.86E-42 | 1.04E-37 | 0.97 | 0.93 | 0.04 | 1.05 |
| SE_basal | Nckap5 | 1.68 | 5.24E-42 | 1.12E-37 | 0.53 | 0.28 | 0.24 | 1.86 |
| SE_basal | Lratd2 | 1.62 | 3.43E-41 | 7.33E-37 | 0.55 | 0.31 | 0.24 | 1.77 |
| SE_basal | Laptm4a | 0.58 | 4.98E-41 | 1.06E-36 | 0.99 | 0.96 | 0.03 | 1.04 |
| SE_basal | Adgrl2 | 1.00 | 5.95E-41 | 1.27E-36 | 0.83 | 0.69 | 0.14 | 1.20 |
| SE_basal | Fam162a | 0.64 | 7.67E-41 | 1.64E-36 | 0.95 | 0.93 | 0.02 | 1.03 |
| SE_basal | Artn | 3.32 | 2.05E-40 | 4.37E-36 | 0.12 | 0.01 | 0.10 | 8.29 |
| SE_basal | Sorcs2 | 2.33 | 2.86E-40 | 6.10E-36 | 0.26 | 0.08 | 0.19 | 3.37 |
| SE_basal | Luzp1 | 0.74 | 5.24E-39 | 1.12E-34 | 0.88 | 0.80 | 0.09 | 1.11 |
| SE_basal | Vcan | 0.70 | 5.59E-39 | 1.19E-34 | 0.91 | 0.73 | 0.18 | 1.25 |
| SE_basal | Sox14 | 2.53 | 8.11E-39 | 1.73E-34 | 0.17 | 0.04 | 0.14 | 4.97 |
| SE_basal | Slc2a3 | 1.14 | 9.70E-39 | 2.07E-34 | 0.58 | 0.33 | 0.26 | 1.79 |
| SE_basal | Nrp2 | 1.72 | 2.07E-38 | 4.42E-34 | 0.37 | 0.15 | 0.22 | 2.49 |
| SE_basal | Foxp1 | 0.74 | 3.72E-38 | 7.95E-34 | 0.93 | 0.86 | 0.07 | 1.09 |
| SE_basal | Ackr3 | 1.55 | 5.67E-38 | 1.21E-33 | 0.57 | 0.36 | 0.22 | 1.60 |
| SE_basal | Arhgdib | 1.71 | 1.17E-37 | 2.49E-33 | 0.42 | 0.19 | 0.23 | 2.24 |
| SE_basal | mt-Co1 | 0.38 | 1.21E-37 | 2.60E-33 | 1.00 | 0.99 | 0.01 | 1.01 |
| SE_basal | Ccdc68 | 1.96 | 2.13E-37 | 4.55E-33 | 0.32 | 0.11 | 0.20 | 2.81 |
| SE_basal | Slc35b3 | 1.64 | 5.03E-37 | 1.07E-32 | 0.47 | 0.24 | 0.23 | 1.96 |
| SE_basal | Ttc28 | 1.24 | 6.73E-37 | 1.44E-32 | 0.63 | 0.43 | 0.21 | 1.49 |
| SE_basal | Parm1 | 1.51 | 9.19E-37 | 1.96E-32 | 0.42 | 0.18 | 0.24 | 2.30 |
| SE_basal | Arf5 | 0.39 | 1.63E-36 | 3.48E-32 | 0.99 | 0.99 | 0.00 | 1.00 |
| SE_basal | Selenow | 0.35 | 3.41E-36 | 7.29E-32 | 1.00 | 0.99 | 0.01 | 1.01 |
| SE_basal | Ednrb | 3.21 | 3.56E-36 | 7.60E-32 | 0.14 | 0.03 | 0.12 | 5.72 |
| SE_basal | Lmo1 | 1.90 | 4.00E-36 | 8.54E-32 | 0.43 | 0.21 | 0.22 | 2.05 |
| SE_basal | Nbl1 | 2.46 | 6.07E-36 | 1.30E-31 | 0.23 | 0.06 | 0.16 | 3.53 |
| SE_basal | Gadd45g | 0.57 | 2.33E-35 | 4.98E-31 | 0.60 | 0.31 | 0.28 | 1.91 |
| SE_basal | Dst | 1.10 | 2.41E-35 | 5.15E-31 | 0.69 | 0.46 | 0.23 | 1.50 |
| SE_basal | Antxr1 | 1.65 | 7.32E-35 | 1.56E-30 | 0.45 | 0.23 | 0.22 | 1.93 |
| SE_basal | Tmem132c | 2.38 | 1.59E-34 | 3.40E-30 | 0.29 | 0.11 | 0.18 | 2.74 |
| SE_basal | Sdc1 | 1.08 | 1.93E-34 | 4.13E-30 | 0.73 | 0.57 | 0.16 | 1.28 |
| SE_basal | Znrf3 | 1.51 | 6.41E-34 | 1.37E-29 | 0.49 | 0.27 | 0.22 | 1.82 |
| SE_basal | Pla2r1 | 1.58 | 7.91E-33 | 1.69E-28 | 0.48 | 0.25 | 0.22 | 1.87 |
| SE_basal | Igfbp5 | 0.48 | 1.03E-32 | 2.19E-28 | 0.99 | 0.96 | 0.03 | 1.03 |
| SE_basal | Rest | 0.75 | 1.19E-32 | 2.53E-28 | 0.84 | 0.66 | 0.17 | 1.26 |
| SE_basal | Dlx1 | 1.75 | 1.88E-32 | 4.01E-28 | 0.32 | 0.13 | 0.20 | 2.56 |
| SE_basal | Kcnq1ot1 | 0.70 | 1.93E-32 | 4.12E-28 | 0.95 | 0.94 | 0.01 | 1.01 |
| SE_basal | Fzd3 | 0.78 | 2.88E-32 | 6.14E-28 | 0.91 | 0.84 | 0.08 | 1.09 |
| SE_basal | Rap2c | 0.90 | 1.67E-31 | 3.58E-27 | 0.78 | 0.65 | 0.12 | 1.19 |
| SE_basal | 2610204G07Rik | 2.99 | 3.29E-31 | 7.03E-27 | 0.12 | 0.02 | 0.10 | 5.90 |
| SE_basal | Pgam1 | 0.37 | 4.31E-31 | 9.21E-27 | 1.00 | 0.99 | 0.01 | 1.01 |
| SE_basal | AY036118 | 1.01 | 7.20E-31 | 1.54E-26 | 0.77 | 0.63 | 0.14 | 1.23 |
| SE_basal | Pard6g | 0.78 | 1.26E-30 | 2.70E-26 | 0.77 | 0.63 | 0.14 | 1.22 |
| SE_basal | Frmd4b | 1.11 | 1.70E-30 | 3.62E-26 | 0.66 | 0.48 | 0.18 | 1.38 |
| SE_basal | P4ha1 | 1.16 | 1.93E-30 | 4.12E-26 | 0.67 | 0.51 | 0.16 | 1.30 |
| SE_basal | Igf2 | 0.58 | 4.01E-30 | 8.56E-26 | 0.91 | 0.80 | 0.11 | 1.14 |
| SE_basal | Cdh6 | 2.01 | 7.79E-30 | 1.66E-25 | 0.29 | 0.11 | 0.18 | 2.62 |
| SE_basal | Dkk4 | 2.28 | 8.77E-30 | 1.87E-25 | 0.11 | 0.02 | 0.09 | 6.22 |
| SE_basal | Btbd11 | 2.83 | 1.84E-29 | 3.93E-25 | 0.13 | 0.03 | 0.10 | 5.16 |
| SE_basal | Pcdh18 | 1.77 | 4.25E-29 | 9.09E-25 | 0.35 | 0.16 | 0.19 | 2.18 |

**Supplementary Table 4. Marker genes for the annotated cell populations of integrated clustering of E11.5 control and NotchGOF λ-junction ectoderm cells.**

| | | | | | | | | |
|---|---|---|---|---|---|---|---|---|
| SE_basal | Filip1l | 1.38 | 7.47E-29 | 1.60E-24 | 0.45 | 0.25 | 0.21 | 1.84 |
| SE_basal | Zfp462 | 0.73 | 8.73E-29 | 1.87E-24 | 0.82 | 0.73 | 0.09 | 1.13 |
| SE_basal | Dusp7 | 1.40 | 1.35E-28 | 2.89E-24 | 0.48 | 0.29 | 0.19 | 1.66 |
| SE_basal | Atxn7l3b | 0.32 | 1.52E-28 | 3.25E-24 | 0.99 | 0.99 | 0.00 | 1.00 |
| SE_basal | Prdx6 | 0.48 | 4.14E-28 | 8.84E-24 | 0.96 | 0.95 | 0.01 | 1.01 |
| SE_basal | Bcam | 0.62 | 9.94E-28 | 2.12E-23 | 0.88 | 0.71 | 0.17 | 1.24 |
| SE_basal | Gdf6 | 2.51 | 1.45E-27 | 3.10E-23 | 0.13 | 0.03 | 0.10 | 4.78 |
| SE_basal | Arhgap42 | 1.36 | 3.04E-27 | 6.50E-23 | 0.45 | 0.26 | 0.20 | 1.76 |
| SE_basal | Otx1 | 1.21 | 3.39E-27 | 7.24E-23 | 0.58 | 0.42 | 0.16 | 1.38 |
| SE_basal | Dag1 | 0.92 | 1.18E-26 | 2.53E-22 | 0.73 | 0.62 | 0.11 | 1.17 |
| SE_basal | Dapl1 | 1.03 | 1.92E-26 | 4.11E-22 | 0.47 | 0.25 | 0.22 | 1.90 |
| SE_basal | Hpca | 2.29 | 2.37E-26 | 5.06E-22 | 0.10 | 0.02 | 0.09 | 5.78 |
| SE_basal | Bgn | 2.81 | 4.56E-26 | 9.75E-22 | 0.12 | 0.02 | 0.09 | 5.04 |
| SE_basal | Tmem123 | 1.05 | 7.42E-26 | 1.58E-21 | 0.60 | 0.43 | 0.16 | 1.38 |
| SE_basal | Mbnl1 | 1.24 | 9.52E-26 | 2.03E-21 | 0.58 | 0.41 | 0.17 | 1.40 |
| SE_basal | Rbbp8 | 0.78 | 1.14E-25 | 2.43E-21 | 0.77 | 0.67 | 0.10 | 1.15 |
| SE_basal | Tgfbi | 2.87 | 1.31E-25 | 2.80E-21 | 0.10 | 0.02 | 0.08 | 5.88 |
| SE_basal | Irf6 | 1.17 | 1.77E-25 | 3.78E-21 | 0.59 | 0.39 | 0.20 | 1.49 |
| SE_basal | Aldh2 | 0.76 | 2.07E-25 | 4.41E-21 | 0.77 | 0.69 | 0.08 | 1.12 |
| SE_basal | Sema3e | 2.62 | 2.51E-25 | 5.37E-21 | 0.11 | 0.02 | 0.09 | 5.30 |
| SE_basal | Nxn | 0.59 | 4.27E-25 | 9.13E-21 | 0.85 | 0.74 | 0.11 | 1.15 |
| SE_basal | Calm2 | 0.33 | 6.21E-25 | 1.33E-20 | 0.99 | 0.99 | 0.01 | 1.01 |
| SE_basal | Ier5 | 0.59 | 9.09E-25 | 1.94E-20 | 0.92 | 0.83 | 0.08 | 1.10 |
| SE_basal | Slc3a2 | 0.84 | 9.92E-25 | 2.12E-20 | 0.78 | 0.66 | 0.12 | 1.18 |
| SE_basal | Mylip | 1.45 | 1.61E-24 | 3.44E-20 | 0.36 | 0.19 | 0.18 | 1.94 |
| SE_basal | Etaa1os | 2.07 | 1.98E-24 | 4.22E-20 | 0.21 | 0.07 | 0.14 | 2.88 |
| SE_basal | Meis2 | 0.40 | 3.65E-24 | 7.80E-20 | 0.98 | 0.95 | 0.03 | 1.03 |
| SE_basal | Blvra | 1.70 | 4.48E-24 | 9.58E-20 | 0.28 | 0.12 | 0.16 | 2.36 |
| SE_basal | mt-Co2 | 0.27 | 7.26E-24 | 1.55E-19 | 1.00 | 1.00 | 0.00 | 1.00 |
| SE_basal | Itm2c | 0.95 | 7.33E-24 | 1.56E-19 | 0.69 | 0.59 | 0.10 | 1.17 |
| SE_basal | Pdpn | 1.07 | 1.11E-23 | 2.38E-19 | 0.53 | 0.35 | 0.19 | 1.53 |
| SE_basal | Mycn | 1.32 | 1.21E-23 | 2.59E-19 | 0.51 | 0.34 | 0.17 | 1.51 |
| SE_basal | Nsg1 | 0.77 | 1.70E-23 | 3.64E-19 | 0.74 | 0.60 | 0.14 | 1.24 |
| SE_basal | Anxa2 | 0.44 | 1.84E-23 | 3.93E-19 | 0.91 | 0.74 | 0.17 | 1.23 |
| SE_basal | Cth | 1.74 | 2.14E-23 | 4.57E-19 | 0.30 | 0.15 | 0.16 | 2.08 |
| SE_basal | Mir100hg | 0.81 | 4.18E-23 | 8.93E-19 | 0.74 | 0.62 | 0.12 | 1.19 |
| SE_basal | Pnrc1 | 0.48 | 4.98E-23 | 1.06E-18 | 0.95 | 0.90 | 0.05 | 1.06 |
| SE_basal | Dnajc1 | 0.97 | 5.44E-23 | 1.16E-18 | 0.72 | 0.65 | 0.08 | 1.12 |
| SE_basal | Trps1 | 1.02 | 1.05E-22 | 2.24E-18 | 0.64 | 0.52 | 0.12 | 1.22 |
| SE_basal | Pla2g4a | 1.64 | 1.38E-22 | 2.94E-18 | 0.23 | 0.09 | 0.14 | 2.61 |
| SE_basal | Cnih2 | 1.27 | 1.70E-22 | 3.62E-18 | 0.30 | 0.14 | 0.16 | 2.14 |
| SE_basal | Afap1l1 | 1.71 | 1.71E-22 | 3.64E-18 | 0.25 | 0.10 | 0.15 | 2.41 |
| SE_basal | Col4a1 | 0.72 | 2.28E-22 | 4.87E-18 | 0.78 | 0.67 | 0.11 | 1.16 |
| SE_basal | Megf6 | 1.64 | 3.01E-22 | 6.43E-18 | 0.33 | 0.17 | 0.16 | 1.99 |
| SE_basal | Fundc2 | 0.71 | 3.30E-22 | 7.04E-18 | 0.76 | 0.69 | 0.07 | 1.10 |
| SE_basal | Ctnnal1 | 1.11 | 4.22E-22 | 9.01E-18 | 0.60 | 0.48 | 0.12 | 1.25 |
| SE_basal | Tmem132a | 0.78 | 6.89E-22 | 1.47E-17 | 0.74 | 0.66 | 0.09 | 1.13 |
| SE_basal | Zfp503 | 0.86 | 1.23E-21 | 2.64E-17 | 0.65 | 0.49 | 0.16 | 1.33 |
| SE_basal | Itm2b | 0.26 | 2.38E-21 | 5.08E-17 | 0.98 | 0.93 | 0.04 | 1.05 |
| SE_basal | Nudt10 | 1.54 | 2.52E-21 | 5.39E-17 | 0.31 | 0.15 | 0.16 | 2.02 |
| SE_basal | mt-Co3 | 0.27 | 3.13E-21 | 6.69E-17 | 1.00 | 1.00 | 0.01 | 1.01 |
| SE_basal | Gstm5 | 0.59 | 4.36E-21 | 9.32E-17 | 0.86 | 0.82 | 0.04 | 1.05 |
| SE_basal | Calml3 | 2.24 | 5.20E-21 | 1.11E-16 | 0.14 | 0.04 | 0.10 | 3.56 |
| SE_basal | Cnr1 | 1.93 | 6.09E-21 | 1.30E-16 | 0.17 | 0.06 | 0.11 | 2.97 |
| SE_basal | Greb1 | 1.51 | 6.15E-21 | 1.31E-16 | 0.33 | 0.18 | 0.16 | 1.89 |

**Supplementary Table 4. Marker genes for the annotated cell populations of integrated clustering of E11.5 control and NotchGOF λ-junction ectoderm cells.**

| | | | | | | | | |
|---|---|---|---|---|---|---|---|---|
| SE_basal | Sfr1 | 0.33 | 8.10E-21 | 1.73E-16 | 0.99 | 0.98 | 0.01 | 1.01 |
| SE_basal | Cpe | 0.82 | 9.05E-21 | 1.93E-16 | 0.61 | 0.45 | 0.16 | 1.34 |
| SE_basal | Efcab1 | 1.63 | 9.99E-21 | 2.13E-16 | 0.21 | 0.08 | 0.13 | 2.58 |
| SE_basal | Atpif1 | 0.26 | 1.28E-20 | 2.74E-16 | 0.99 | 0.99 | 0.00 | 1.00 |
| SE_basal | Klhl13 | 1.01 | 1.91E-20 | 4.09E-16 | 0.54 | 0.39 | 0.15 | 1.39 |
| SE_basal | Sox12 | 0.77 | 2.10E-20 | 4.49E-16 | 0.69 | 0.59 | 0.10 | 1.17 |
| SE_basal | Prps2 | 1.29 | 2.67E-20 | 5.71E-16 | 0.46 | 0.31 | 0.16 | 1.51 |
| SE_basal | Pde10a | 2.18 | 2.80E-20 | 5.97E-16 | 0.13 | 0.04 | 0.09 | 3.69 |
| SE_basal | Mfge8 | 0.73 | 3.09E-20 | 6.61E-16 | 0.68 | 0.55 | 0.13 | 1.25 |
| SE_basal | Sox6 | 1.92 | 3.56E-20 | 7.61E-16 | 0.22 | 0.09 | 0.13 | 2.48 |
| SE_basal | Msx1 | 0.55 | 4.09E-20 | 8.74E-16 | 0.87 | 0.72 | 0.16 | 1.22 |
| SE_basal | Npc2 | 0.46 | 6.76E-20 | 1.44E-15 | 0.94 | 0.92 | 0.02 | 1.02 |
| SE_basal | Smad7 | 0.66 | 8.29E-20 | 1.77E-15 | 0.73 | 0.60 | 0.14 | 1.23 |
| SE_basal | Gnai1 | 1.74 | 8.99E-20 | 1.92E-15 | 0.23 | 0.10 | 0.13 | 2.38 |
| SE_basal | Phlda2 | 2.58 | 9.47E-20 | 2.02E-15 | 0.13 | 0.04 | 0.09 | 3.41 |
| SE_basal | Bzw2 | 0.38 | 1.98E-19 | 4.23E-15 | 0.94 | 0.95 | 0.00 | 1.00 |
| SE_basal | Rxfp1 | 2.19 | 2.23E-19 | 4.77E-15 | 0.14 | 0.04 | 0.10 | 3.31 |
| SE_basal | Serpinb6b | 2.09 | 2.92E-19 | 6.24E-15 | 0.16 | 0.05 | 0.11 | 3.06 |
| SE_basal | Erf | 1.11 | 3.17E-19 | 6.77E-15 | 0.52 | 0.37 | 0.15 | 1.40 |
| SE_basal | Snhg11 | 1.90 | 4.91E-19 | 1.05E-14 | 0.18 | 0.07 | 0.11 | 2.73 |
| SE_basal | Cox6b2 | 0.87 | 5.52E-19 | 1.18E-14 | 0.62 | 0.50 | 0.12 | 1.24 |
| SE_basal | Cd109 | 2.31 | 7.46E-19 | 1.59E-14 | 0.13 | 0.04 | 0.09 | 3.39 |
| SE_basal | Slc39a10 | 0.63 | 9.52E-19 | 2.03E-14 | 0.82 | 0.75 | 0.06 | 1.09 |
| SE_basal | Arhgap22 | 2.21 | 1.01E-18 | 2.15E-14 | 0.12 | 0.04 | 0.09 | 3.49 |
| SE_basal | Fam89a | 1.36 | 1.12E-18 | 2.40E-14 | 0.30 | 0.15 | 0.15 | 2.02 |
| SE_basal | Emp1 | 1.09 | 1.18E-18 | 2.53E-14 | 0.31 | 0.16 | 0.15 | 1.97 |
| SE_basal | Ipmk | 1.35 | 1.26E-18 | 2.69E-14 | 0.37 | 0.23 | 0.15 | 1.65 |
| SE_basal | Lpp | 0.88 | 2.42E-18 | 5.18E-14 | 0.58 | 0.44 | 0.14 | 1.31 |
| SE_basal | Cst3 | 0.43 | 3.68E-18 | 7.85E-14 | 0.95 | 0.93 | 0.02 | 1.02 |
| SE_basal | Sulf2 | 1.09 | 4.38E-18 | 9.35E-14 | 0.47 | 0.32 | 0.15 | 1.47 |
| SE_basal | Arpc2 | 0.33 | 4.44E-18 | 9.48E-14 | 0.96 | 0.96 | 0.01 | 1.01 |
| SE_basal | Abracl | 0.34 | 6.01E-18 | 1.28E-13 | 0.94 | 0.92 | 0.01 | 1.02 |
| SE_basal | Castor1 | 2.23 | 6.15E-18 | 1.31E-13 | 0.14 | 0.04 | 0.09 | 3.11 |
| SE_basal | Cacng4 | 0.92 | 7.72E-18 | 1.65E-13 | 0.49 | 0.33 | 0.16 | 1.48 |
| SE_basal | Gabarapl1 | 1.11 | 1.61E-17 | 3.45E-13 | 0.51 | 0.38 | 0.13 | 1.33 |
| SE_basal | Pkp3 | 1.26 | 4.39E-17 | 9.38E-13 | 0.38 | 0.24 | 0.15 | 1.62 |
| SE_basal | Pcdh7 | 0.69 | 6.70E-17 | 1.43E-12 | 0.72 | 0.65 | 0.08 | 1.12 |
| SE_basal | Dlx3 | 1.05 | 6.87E-17 | 1.47E-12 | 0.43 | 0.27 | 0.16 | 1.58 |
| SE_basal | Ddr1 | 0.76 | 7.04E-17 | 1.50E-12 | 0.59 | 0.44 | 0.15 | 1.33 |
| SE_basal | Gga2 | 0.74 | 8.09E-17 | 1.73E-12 | 0.61 | 0.50 | 0.11 | 1.22 |
| SE_basal | Rtn4ip1 | 1.81 | 9.66E-17 | 2.06E-12 | 0.18 | 0.07 | 0.11 | 2.47 |
| SE_basal | Arid5b | 0.69 | 1.02E-16 | 2.18E-12 | 0.67 | 0.55 | 0.12 | 1.23 |
| SE_basal | Cbr2 | 1.95 | 1.30E-16 | 2.78E-12 | 0.11 | 0.03 | 0.08 | 3.60 |
| SE_basal | Myo1b | 0.57 | 1.39E-16 | 2.97E-12 | 0.72 | 0.62 | 0.10 | 1.17 |
| SE_basal | Nudt11 | 1.15 | 1.49E-16 | 3.19E-12 | 0.41 | 0.27 | 0.14 | 1.53 |
| SE_basal | Mbd2 | 0.56 | 1.55E-16 | 3.30E-12 | 0.70 | 0.60 | 0.09 | 1.15 |
| SE_basal | Card10 | 2.22 | 1.66E-16 | 3.56E-12 | 0.12 | 0.04 | 0.08 | 3.10 |
| SE_basal | Ptpn13 | 0.60 | 1.98E-16 | 4.22E-12 | 0.69 | 0.56 | 0.13 | 1.24 |
| SE_basal | Kitl | 1.68 | 2.06E-16 | 4.41E-12 | 0.25 | 0.13 | 0.13 | 2.02 |
| SE_basal | Gprc5b | 1.52 | 2.08E-16 | 4.44E-12 | 0.26 | 0.13 | 0.13 | 1.96 |
| SE_basal | Pxn | 1.48 | 3.16E-16 | 6.76E-12 | 0.29 | 0.16 | 0.13 | 1.82 |
| SE_basal | Dkk2 | 1.80 | 3.18E-16 | 6.79E-12 | 0.13 | 0.04 | 0.09 | 3.05 |
| SE_basal | Gng11 | 1.59 | 4.13E-16 | 8.83E-12 | 0.22 | 0.11 | 0.12 | 2.13 |
| SE_basal | Abca1 | 1.56 | 5.04E-16 | 1.08E-11 | 0.23 | 0.11 | 0.12 | 2.13 |
| SE_basal | Lbh | 0.97 | 5.99E-16 | 1.28E-11 | 0.41 | 0.25 | 0.15 | 1.59 |

**Supplementary Table 4. Marker genes for the annotated cell populations of integrated clustering of E11.5 control and NotchGOF λ-junction ectoderm cells.**

| | | | | | | | | |
|---|---|---|---|---|---|---|---|---|
| SE_basal | Ctsb | 0.66 | 6.21E-16 | 1.33E-11 | 0.71 | 0.64 | 0.07 | 1.11 |
| SE_basal | Pcdh9 | 1.41 | 8.34E-16 | 1.78E-11 | 0.31 | 0.17 | 0.13 | 1.77 |
| SE_basal | Ptn | 0.50 | 8.93E-16 | 1.91E-11 | 0.88 | 0.86 | 0.02 | 1.02 |
| SE_basal | Krt5 | 0.74 | 1.81E-15 | 3.86E-11 | 0.49 | 0.32 | 0.17 | 1.52 |
| SE_basal | Lars2 | 0.71 | 1.88E-15 | 4.01E-11 | 0.64 | 0.51 | 0.13 | 1.25 |
| SE_basal | Pcyt1b | 1.46 | 2.46E-15 | 5.25E-11 | 0.23 | 0.11 | 0.12 | 2.09 |
| SE_basal | Serpinb6a | 0.78 | 3.00E-15 | 6.41E-11 | 0.57 | 0.47 | 0.11 | 1.22 |
| SE_basal | Eef2k | 1.09 | 3.57E-15 | 7.63E-11 | 0.30 | 0.17 | 0.14 | 1.82 |
| SE_basal | Susd4 | 1.16 | 4.00E-15 | 8.54E-11 | 0.42 | 0.29 | 0.13 | 1.44 |
| SE_basal | Rb1 | 0.94 | 4.33E-15 | 9.26E-11 | 0.42 | 0.28 | 0.15 | 1.53 |
| SE_basal | Purb | 0.43 | 4.94E-15 | 1.05E-10 | 0.91 | 0.89 | 0.02 | 1.02 |
| SE_basal | Rapgefl1 | 1.73 | 5.18E-15 | 1.11E-10 | 0.16 | 0.06 | 0.10 | 2.54 |
| SE_basal | Sh3bp5 | 0.87 | 5.78E-15 | 1.23E-10 | 0.50 | 0.37 | 0.13 | 1.35 |
| SE_basal | Osbpl8 | 1.03 | 7.18E-15 | 1.53E-10 | 0.39 | 0.26 | 0.14 | 1.53 |
| SE_basal | Slc1a3 | 1.29 | 8.58E-15 | 1.83E-10 | 0.32 | 0.19 | 0.13 | 1.70 |
| SE_basal | Rab38 | 1.01 | 1.48E-14 | 3.16E-10 | 0.33 | 0.19 | 0.14 | 1.73 |
| SE_basal | Gpc2 | 0.90 | 1.52E-14 | 3.26E-10 | 0.49 | 0.35 | 0.14 | 1.39 |
| SE_basal | Apbb2 | 1.23 | 1.53E-14 | 3.28E-10 | 0.39 | 0.27 | 0.12 | 1.45 |
| SE_basal | Tes | 0.65 | 1.67E-14 | 3.57E-10 | 0.69 | 0.61 | 0.08 | 1.13 |
| SE_basal | Dennd2c | 1.43 | 2.14E-14 | 4.58E-10 | 0.20 | 0.09 | 0.11 | 2.21 |
| SE_basal | Cited2 | 0.58 | 2.42E-14 | 5.18E-10 | 0.77 | 0.68 | 0.10 | 1.14 |
| SE_basal | Fjx1 | 0.53 | 3.04E-14 | 6.50E-10 | 0.87 | 0.79 | 0.08 | 1.11 |
| SE_basal | Hmgn3 | 0.52 | 3.25E-14 | 6.95E-10 | 0.76 | 0.71 | 0.05 | 1.07 |
| SE_basal | Man1a | 2.03 | 5.22E-14 | 1.11E-09 | 0.12 | 0.04 | 0.08 | 2.84 |
| SE_basal | Fry | 1.29 | 6.74E-14 | 1.44E-09 | 0.25 | 0.13 | 0.12 | 1.92 |
| SE_basal | Sema3c | 1.37 | 6.78E-14 | 1.45E-09 | 0.37 | 0.24 | 0.13 | 1.53 |
| SE_basal | Rarg | 1.10 | 6.96E-14 | 1.49E-09 | 0.40 | 0.28 | 0.13 | 1.45 |
| SE_basal | Efnb2 | 0.65 | 8.32E-14 | 1.78E-09 | 0.68 | 0.59 | 0.09 | 1.15 |
| SE_basal | Rftn1 | 2.17 | 9.51E-14 | 2.03E-09 | 0.11 | 0.04 | 0.08 | 2.92 |
| SE_basal | Bri3 | 0.29 | 1.19E-13 | 2.55E-09 | 0.97 | 0.97 | 0.00 | 1.00 |
| SE_basal | Gpx8 | 0.62 | 1.65E-13 | 3.52E-09 | 0.71 | 0.65 | 0.06 | 1.09 |
| SE_basal | Ccdc126 | 1.56 | 1.86E-13 | 3.98E-09 | 0.18 | 0.08 | 0.10 | 2.22 |
| SE_basal | Irx2 | 1.85 | 2.04E-13 | 4.35E-09 | 0.13 | 0.05 | 0.08 | 2.76 |
| SE_basal | Tgif1 | 0.53 | 2.76E-13 | 5.89E-09 | 0.75 | 0.68 | 0.07 | 1.10 |
| SE_basal | Ucp2 | 0.53 | 3.32E-13 | 7.09E-09 | 0.58 | 0.47 | 0.12 | 1.25 |
| SE_basal | Helb | 1.28 | 3.47E-13 | 7.40E-09 | 0.24 | 0.13 | 0.11 | 1.90 |
| SE_basal | Ldb1 | 0.57 | 3.53E-13 | 7.53E-09 | 0.70 | 0.64 | 0.06 | 1.09 |
| SE_basal | Atp2c2 | 1.84 | 3.98E-13 | 8.50E-09 | 0.13 | 0.05 | 0.08 | 2.70 |
| SE_basal | Gm16141 | 1.89 | 4.23E-13 | 9.04E-09 | 0.13 | 0.05 | 0.08 | 2.61 |
| SE_basal | Ddit4 | 0.74 | 5.85E-13 | 1.25E-08 | 0.54 | 0.43 | 0.11 | 1.26 |
| SE_basal | Trp53inp1 | 1.06 | 9.18E-13 | 1.96E-08 | 0.38 | 0.25 | 0.13 | 1.51 |
| SE_basal | Arhgap24 | 1.70 | 9.62E-13 | 2.06E-08 | 0.16 | 0.07 | 0.09 | 2.33 |
| SE_basal | Adamts18 | 0.91 | 9.96E-13 | 2.13E-08 | 0.39 | 0.26 | 0.14 | 1.53 |
| SE_basal | Cep164 | 0.57 | 1.01E-12 | 2.16E-08 | 0.71 | 0.66 | 0.05 | 1.08 |
| SE_basal | Gpx1 | 0.30 | 1.39E-12 | 2.97E-08 | 0.94 | 0.95 | -0.01 | 0.99 |
| SE_basal | Mast4 | 0.76 | 1.42E-12 | 3.03E-08 | 0.58 | 0.49 | 0.09 | 1.18 |
| SE_basal | Rab12 | 0.76 | 1.80E-12 | 3.84E-08 | 0.55 | 0.46 | 0.10 | 1.21 |
| SE_basal | Tdrp | 1.54 | 1.85E-12 | 3.96E-08 | 0.19 | 0.09 | 0.10 | 2.10 |
| SE_basal | Itpripl2 | 2.02 | 2.34E-12 | 4.99E-08 | 0.11 | 0.04 | 0.07 | 2.80 |
| SE_basal | Hmga2 | 0.38 | 2.63E-12 | 5.61E-08 | 0.95 | 0.92 | 0.03 | 1.03 |
| SE_basal | Flna | 0.54 | 2.63E-12 | 5.63E-08 | 0.74 | 0.69 | 0.05 | 1.08 |
| SE_basal | Mcee | 0.58 | 2.96E-12 | 6.33E-08 | 0.64 | 0.55 | 0.08 | 1.15 |
| SE_basal | Tulp4 | 0.44 | 3.11E-12 | 6.65E-08 | 0.80 | 0.74 | 0.06 | 1.08 |
| SE_basal | Top2b | 0.31 | 3.60E-12 | 7.69E-08 | 0.92 | 0.93 | -0.01 | 0.99 |
| SE_basal | Gimap9 | 1.22 | 4.08E-12 | 8.71E-08 | 0.23 | 0.12 | 0.11 | 1.87 |

**Supplementary Table 4. Marker genes for the annotated cell populations of integrated clustering of E11.5 control and NotchGOF λ-junction ectoderm cells.**

| | | | | | | | | |
|---|---|---|---|---|---|---|---|---|
| SE_basal | Ank3 | 0.44 | 4.08E-12 | 8.72E-08 | 0.73 | 0.63 | 0.10 | 1.16 |
| SE_basal | Apoe | 0.77 | 4.17E-12 | 8.90E-08 | 0.59 | 0.49 | 0.10 | 1.20 |
| SE_basal | Fbln1 | 0.59 | 4.18E-12 | 8.94E-08 | 0.70 | 0.65 | 0.05 | 1.08 |
| SE_basal | Kctd1 | 0.61 | 4.50E-12 | 9.60E-08 | 0.61 | 0.49 | 0.12 | 1.25 |
| SE_basal | Sema3d | 0.67 | 4.52E-12 | 9.66E-08 | 0.53 | 0.41 | 0.12 | 1.30 |
| SE_basal | Zfp57 | 1.34 | 5.41E-12 | 1.15E-07 | 0.27 | 0.16 | 0.11 | 1.67 |
| SE_basal | Avpi1 | 0.90 | 5.69E-12 | 1.21E-07 | 0.45 | 0.35 | 0.10 | 1.28 |
| SE_basal | Rap2b | 0.62 | 5.74E-12 | 1.23E-07 | 0.60 | 0.53 | 0.08 | 1.14 |
| SE_basal | Vwa1 | 1.20 | 7.35E-12 | 1.57E-07 | 0.21 | 0.10 | 0.10 | 1.99 |
| SE_basal | Sema4g | 1.39 | 7.37E-12 | 1.57E-07 | 0.24 | 0.13 | 0.11 | 1.80 |
| SE_basal | Mmp2 | 1.38 | 7.38E-12 | 1.58E-07 | 0.21 | 0.11 | 0.10 | 1.93 |
| SE_basal | Tpm2 | 1.23 | 9.81E-12 | 2.10E-07 | 0.29 | 0.18 | 0.11 | 1.63 |
| SE_basal | Yap1 | 0.55 | 1.61E-11 | 3.44E-07 | 0.67 | 0.61 | 0.06 | 1.11 |
| SE_basal | Scmh1 | 0.95 | 1.71E-11 | 3.65E-07 | 0.41 | 0.30 | 0.11 | 1.36 |
| SE_basal | 6330403K07Rik | 0.37 | 2.11E-11 | 4.51E-07 | 0.87 | 0.85 | 0.02 | 1.02 |
| SE_basal | Ahcyl2 | 0.76 | 2.31E-11 | 4.93E-07 | 0.47 | 0.36 | 0.10 | 1.28 |
| SE_basal | Ppp1r3b | 1.91 | 2.82E-11 | 6.03E-07 | 0.12 | 0.05 | 0.07 | 2.57 |
| SE_basal | Smad5 | 0.73 | 3.12E-11 | 6.67E-07 | 0.55 | 0.47 | 0.08 | 1.18 |
| SE_basal | Fblim1 | 0.72 | 3.20E-11 | 6.84E-07 | 0.52 | 0.40 | 0.12 | 1.29 |
| SE_basal | Col5a1 | 1.16 | 3.55E-11 | 7.58E-07 | 0.28 | 0.18 | 0.11 | 1.60 |
| SE_basal | Slc15a2 | 1.26 | 3.60E-11 | 7.69E-07 | 0.25 | 0.14 | 0.11 | 1.78 |
| SE_basal | Cisd1 | 0.37 | 3.81E-11 | 8.14E-07 | 0.81 | 0.79 | 0.02 | 1.03 |
| SE_basal | Crppa | 1.51 | 4.12E-11 | 8.80E-07 | 0.14 | 0.06 | 0.08 | 2.31 |
| SE_basal | Ogt | 0.47 | 4.38E-11 | 9.36E-07 | 0.83 | 0.81 | 0.02 | 1.02 |
| SE_basal | Acaa2 | 0.52 | 5.03E-11 | 1.07E-06 | 0.76 | 0.71 | 0.05 | 1.07 |
| SE_basal | Khdrbs1 | 0.27 | 5.16E-11 | 1.10E-06 | 0.94 | 0.94 | 0.00 | 1.00 |
| SE_basal | Nt5c | 0.52 | 5.31E-11 | 1.13E-06 | 0.71 | 0.66 | 0.04 | 1.06 |
| SE_basal | Smarcad1 | 0.43 | 5.86E-11 | 1.25E-06 | 0.77 | 0.77 | 0.01 | 1.01 |
| SE_basal | Plxdc2 | 0.78 | 5.90E-11 | 1.26E-06 | 0.45 | 0.34 | 0.11 | 1.33 |
| SE_basal | Ccdc50 | 0.40 | 7.23E-11 | 1.54E-06 | 0.78 | 0.76 | 0.02 | 1.02 |
| SE_basal | Dgkh | 1.78 | 8.46E-11 | 1.81E-06 | 0.11 | 0.04 | 0.07 | 2.62 |
| SE_basal | Cox7a2l | 0.28 | 8.82E-11 | 1.88E-06 | 0.98 | 0.98 | 0.00 | 1.00 |
| SE_basal | Sytl1 | 1.83 | 8.89E-11 | 1.90E-06 | 0.10 | 0.04 | 0.06 | 2.78 |
| SE_basal | Mllt10 | 0.43 | 1.04E-10 | 2.22E-06 | 0.77 | 0.72 | 0.05 | 1.07 |
| SE_basal | Nebl | 0.87 | 1.04E-10 | 2.22E-06 | 0.50 | 0.39 | 0.11 | 1.28 |
| SE_basal | Pdcd4 | 0.33 | 1.22E-10 | 2.60E-06 | 0.84 | 0.78 | 0.06 | 1.08 |
| SE_basal | Slc16a13 | 1.25 | 1.23E-10 | 2.62E-06 | 0.21 | 0.12 | 0.10 | 1.86 |
| SE_basal | Ctdspl | 0.64 | 1.28E-10 | 2.74E-06 | 0.56 | 0.48 | 0.08 | 1.17 |
| SE_basal | Mtf2 | 0.38 | 1.33E-10 | 2.85E-06 | 0.80 | 0.78 | 0.02 | 1.03 |
| SE_basal | Adamts17 | 1.46 | 1.93E-10 | 4.12E-06 | 0.20 | 0.11 | 0.09 | 1.86 |
| SE_basal | 1810058I24Rik | 0.27 | 2.13E-10 | 4.56E-06 | 0.94 | 0.93 | 0.00 | 1.00 |
| SE_basal | Ctr9 | 0.48 | 2.29E-10 | 4.90E-06 | 0.71 | 0.68 | 0.03 | 1.04 |
| SE_basal | Xist | 0.43 | 2.44E-10 | 5.22E-06 | 0.82 | 0.92 | -0.10 | 0.89 |
| SE_basal | Tasor2 | 0.67 | 3.13E-10 | 6.69E-06 | 0.54 | 0.44 | 0.09 | 1.21 |
| SE_basal | Osbpl3 | 1.34 | 3.34E-10 | 7.12E-06 | 0.25 | 0.15 | 0.10 | 1.66 |
| SE_basal | Cachd1 | 0.90 | 3.34E-10 | 7.13E-06 | 0.43 | 0.33 | 0.11 | 1.33 |
| SE_basal | Cers6 | 1.11 | 3.43E-10 | 7.32E-06 | 0.33 | 0.23 | 0.10 | 1.46 |
| SE_basal | Amotl1 | 0.37 | 3.47E-10 | 7.41E-06 | 0.77 | 0.69 | 0.08 | 1.12 |
| SE_basal | Hunk | 0.54 | 4.22E-10 | 9.00E-06 | 0.64 | 0.58 | 0.06 | 1.10 |
| SE_basal | Ube2g1 | 0.57 | 4.98E-10 | 1.06E-05 | 0.61 | 0.56 | 0.05 | 1.09 |
| SE_basal | Fzd10os | 1.18 | 6.88E-10 | 1.47E-05 | 0.24 | 0.14 | 0.10 | 1.69 |
| SE_basal | Ssbp2 | 0.47 | 7.18E-10 | 1.53E-05 | 0.72 | 0.69 | 0.03 | 1.04 |
| SE_basal | Cited4 | 1.51 | 7.77E-10 | 1.66E-05 | 0.12 | 0.05 | 0.07 | 2.35 |
| SE_basal | Pid1 | 1.35 | 7.91E-10 | 1.69E-05 | 0.13 | 0.06 | 0.07 | 2.31 |
| SE_basal | Sh3bgrl | 0.37 | 9.55E-10 | 2.04E-05 | 0.78 | 0.75 | 0.02 | 1.03 |

**Supplementary Table 4. Marker genes for the annotated cell populations of integrated clustering of E11.5 control and NotchGOF λ-junction ectoderm cells.**

| | | | | | | | | |
|---|---|---|---|---|---|---|---|---|
| SE_basal | Spats2l | 0.63 | 9.81E-10 | 2.10E-05 | 0.45 | 0.33 | 0.12 | 1.38 |
| SE_basal | Fat1 | 0.51 | 1.00E-09 | 2.14E-05 | 0.70 | 0.64 | 0.06 | 1.10 |
| SE_basal | Gja1 | 0.34 | 1.04E-09 | 2.23E-05 | 0.78 | 0.70 | 0.08 | 1.12 |
| SE_basal | Cpxm1 | 1.01 | 1.07E-09 | 2.28E-05 | 0.36 | 0.26 | 0.10 | 1.39 |
| SE_basal | Acyp1 | 0.61 | 1.13E-09 | 2.42E-05 | 0.58 | 0.49 | 0.09 | 1.17 |
| SE_basal | Zbtb34 | 1.38 | 1.15E-09 | 2.46E-05 | 0.19 | 0.10 | 0.09 | 1.85 |
| SE_basal | Pnisr | 0.30 | 1.16E-09 | 2.47E-05 | 0.93 | 0.93 | 0.00 | 1.00 |
| SE_basal | Jak1 | 0.57 | 1.19E-09 | 2.55E-05 | 0.61 | 0.55 | 0.05 | 1.10 |
| SE_basal | Trim32 | 0.86 | 1.22E-09 | 2.62E-05 | 0.45 | 0.37 | 0.08 | 1.23 |
| SE_basal | Klk8 | 0.60 | 1.51E-09 | 3.22E-05 | 0.63 | 0.56 | 0.07 | 1.13 |
| SE_basal | D16Ertd472e | 0.93 | 1.57E-09 | 3.36E-05 | 0.33 | 0.23 | 0.10 | 1.44 |
| SE_basal | Smarcc1 | 0.26 | 1.67E-09 | 3.56E-05 | 0.97 | 0.97 | -0.01 | 0.99 |
| SE_basal | Slc2a1 | 0.57 | 1.87E-09 | 4.00E-05 | 0.72 | 0.69 | 0.04 | 1.05 |
| SE_basal | Pmaip1 | 1.30 | 2.73E-09 | 5.84E-05 | 0.17 | 0.09 | 0.08 | 1.93 |
| SE_basal | Nox4 | 1.61 | 3.09E-09 | 6.60E-05 | 0.16 | 0.08 | 0.08 | 1.98 |
| SE_basal | Dnase2a | 0.66 | 3.15E-09 | 6.72E-05 | 0.52 | 0.43 | 0.08 | 1.19 |
| SE_basal | Phlda3 | 0.36 | 3.19E-09 | 6.81E-05 | 0.60 | 0.49 | 0.11 | 1.22 |
| SE_basal | Trp53 | 0.33 | 3.30E-09 | 7.05E-05 | 0.89 | 0.89 | 0.00 | 1.00 |
| SE_basal | Erich1 | 0.73 | 3.44E-09 | 7.34E-05 | 0.48 | 0.39 | 0.09 | 1.22 |
| SE_basal | Dzip1 | 0.99 | 4.16E-09 | 8.89E-05 | 0.30 | 0.20 | 0.10 | 1.48 |
| SE_basal | Stxbp4 | 0.82 | 4.33E-09 | 9.24E-05 | 0.41 | 0.33 | 0.09 | 1.27 |
| SE_basal | Cox4i2 | 0.95 | 4.37E-09 | 9.33E-05 | 0.55 | 0.48 | 0.07 | 1.14 |
| SE_basal | Slc39a6 | 0.59 | 4.63E-09 | 9.90E-05 | 0.55 | 0.48 | 0.07 | 1.15 |
| SE_basal | Cldn1 | 0.77 | 5.55E-09 | 1.19E-04 | 0.18 | 0.10 | 0.09 | 1.91 |
| SE_basal | Dhrs3 | 1.21 | 5.88E-09 | 1.26E-04 | 0.30 | 0.20 | 0.10 | 1.47 |
| SE_basal | Prdx5 | 0.36 | 5.93E-09 | 1.27E-04 | 0.81 | 0.81 | 0.00 | 1.00 |
| SE_basal | Bmp7 | 0.77 | 7.07E-09 | 1.51E-04 | 0.49 | 0.40 | 0.09 | 1.22 |
| SE_basal | Mxd4 | 0.67 | 9.10E-09 | 1.94E-04 | 0.51 | 0.41 | 0.10 | 1.23 |
| SE_basal | St6gal2 | 1.47 | 1.02E-08 | 2.19E-04 | 0.14 | 0.07 | 0.07 | 2.08 |
| SE_basal | L3hypdh | 1.22 | 1.04E-08 | 2.23E-04 | 0.22 | 0.14 | 0.09 | 1.64 |
| SE_basal | Arf4 | 0.26 | 1.15E-08 | 2.46E-04 | 0.95 | 0.94 | 0.00 | 1.00 |
| SE_basal | Maco1 | 0.39 | 1.23E-08 | 2.62E-04 | 0.76 | 0.72 | 0.03 | 1.05 |
| SE_basal | Srpk2 | 0.48 | 1.30E-08 | 2.77E-04 | 0.72 | 0.69 | 0.02 | 1.03 |
| SE_basal | Heph | 1.03 | 1.47E-08 | 3.15E-04 | 0.35 | 0.26 | 0.09 | 1.34 |
| SE_basal | Prickle1 | 1.34 | 1.53E-08 | 3.27E-04 | 0.13 | 0.06 | 0.07 | 2.15 |
| SE_basal | Slc25a17 | 0.36 | 1.63E-08 | 3.49E-04 | 0.75 | 0.71 | 0.04 | 1.06 |
| SE_basal | Rab18 | 0.32 | 1.67E-08 | 3.56E-04 | 0.80 | 0.78 | 0.02 | 1.03 |
| SE_basal | Rere | 0.46 | 1.70E-08 | 3.64E-04 | 0.69 | 0.64 | 0.05 | 1.08 |
| SE_basal | Dlg5 | 0.77 | 1.71E-08 | 3.66E-04 | 0.46 | 0.38 | 0.07 | 1.19 |
| SE_basal | Bdnf | 0.97 | 1.96E-08 | 4.19E-04 | 0.27 | 0.18 | 0.10 | 1.54 |
| SE_basal | Ilrun | 0.61 | 2.14E-08 | 4.58E-04 | 0.54 | 0.47 | 0.07 | 1.15 |
| SE_basal | Fhit | 1.51 | 2.16E-08 | 4.61E-04 | 0.14 | 0.07 | 0.07 | 1.99 |
| SE_basal | Taf4b | 1.77 | 2.36E-08 | 5.04E-04 | 0.10 | 0.04 | 0.06 | 2.32 |
| SE_basal | Tanc2 | 0.83 | 2.36E-08 | 5.05E-04 | 0.31 | 0.22 | 0.10 | 1.44 |
| SE_basal | Eef1e1 | 0.41 | 2.42E-08 | 5.17E-04 | 0.69 | 0.68 | 0.01 | 1.02 |
| SE_basal | Ddx6 | 0.31 | 2.54E-08 | 5.42E-04 | 0.87 | 0.86 | 0.00 | 1.00 |
| SE_basal | Kit | 1.11 | 2.70E-08 | 5.76E-04 | 0.26 | 0.17 | 0.09 | 1.52 |
| SE_basal | Rac3 | 0.66 | 2.72E-08 | 5.80E-04 | 0.49 | 0.41 | 0.08 | 1.19 |
| SE_basal | Atp6v1a | 0.60 | 2.75E-08 | 5.88E-04 | 0.58 | 0.54 | 0.04 | 1.08 |
| SE_basal | Cfap53 | 1.77 | 3.03E-08 | 6.47E-04 | 0.11 | 0.05 | 0.06 | 2.25 |
| SE_basal | Eif2s3y | 1.43 | 3.47E-08 | 7.40E-04 | 0.10 | 0.04 | 0.06 | 2.37 |
| SE_basal | Tnrc18 | 0.61 | 3.73E-08 | 7.96E-04 | 0.62 | 0.56 | 0.06 | 1.11 |
| SE_basal | Stox2 | 0.46 | 3.86E-08 | 8.25E-04 | 0.73 | 0.69 | 0.04 | 1.05 |
| SE_basal | Ccnd3 | 0.36 | 4.23E-08 | 9.03E-04 | 0.83 | 0.80 | 0.03 | 1.04 |
| SE_basal | Frem2 | 0.43 | 4.32E-08 | 9.24E-04 | 0.69 | 0.62 | 0.07 | 1.11 |

**Supplementary Table 4. Marker genes for the annotated cell populations of integrated clustering of E11.5 control and NotchGOF λ-junction ectoderm cells.**

| | | | | | | | | |
|---|---|---|---|---|---|---|---|---|
| SE_basal | Dusp9 | 0.96 | 4.87E-08 | 1.04E-03 | 0.32 | 0.23 | 0.09 | 1.38 |
| SE_basal | Nell2 | 1.02 | 4.95E-08 | 1.06E-03 | 0.19 | 0.11 | 0.08 | 1.72 |
| SE_basal | Commd3 | 0.31 | 5.24E-08 | 1.12E-03 | 0.82 | 0.83 | -0.01 | 0.99 |
| SE_basal | Enah | 0.33 | 5.74E-08 | 1.23E-03 | 0.88 | 0.87 | 0.01 | 1.01 |
| SE_basal | Jarid2 | 0.52 | 5.76E-08 | 1.23E-03 | 0.59 | 0.55 | 0.04 | 1.08 |
| SE_basal | Prkar2b | 0.89 | 6.18E-08 | 1.32E-03 | 0.38 | 0.30 | 0.08 | 1.27 |
| SE_basal | Slc26a2 | 1.04 | 6.24E-08 | 1.33E-03 | 0.25 | 0.16 | 0.09 | 1.52 |
| SE_basal | Ccdc102a | 0.89 | 7.89E-08 | 1.68E-03 | 0.31 | 0.23 | 0.09 | 1.38 |
| SE_basal | Flywch2 | 0.58 | 8.09E-08 | 1.73E-03 | 0.55 | 0.49 | 0.07 | 1.14 |
| SE_basal | Fryl | 0.75 | 8.23E-08 | 1.76E-03 | 0.46 | 0.38 | 0.07 | 1.19 |
| SE_basal | Zfp90 | 0.75 | 8.52E-08 | 1.82E-03 | 0.40 | 0.32 | 0.08 | 1.26 |
| SE_basal | Cntnap2 | 1.25 | 8.88E-08 | 1.90E-03 | 0.22 | 0.14 | 0.08 | 1.55 |
| SE_basal | Large2 | 1.02 | 9.53E-08 | 2.04E-03 | 0.31 | 0.23 | 0.08 | 1.36 |
| SE_basal | Hbb-y | 0.69 | 1.02E-07 | 2.18E-03 | 0.98 | 0.98 | 0.00 | 1.00 |
| SE_basal | Dclre1c | 1.12 | 1.03E-07 | 2.20E-03 | 0.17 | 0.09 | 0.08 | 1.81 |
| SE_basal | Lpin2 | 0.53 | 1.03E-07 | 2.21E-03 | 0.53 | 0.46 | 0.07 | 1.15 |
| SE_basal | Znrf1 | 0.45 | 1.06E-07 | 2.27E-03 | 0.59 | 0.56 | 0.03 | 1.05 |
| SE_basal | Glul | 0.62 | 1.07E-07 | 2.28E-03 | 0.52 | 0.46 | 0.06 | 1.13 |
| SE_basal | Hs2st1 | 0.50 | 1.08E-07 | 2.31E-03 | 0.57 | 0.53 | 0.05 | 1.09 |
| SE_basal | Robo1 | 0.65 | 1.11E-07 | 2.37E-03 | 0.47 | 0.40 | 0.08 | 1.19 |
| SE_basal | Comtd1 | 1.07 | 1.13E-07 | 2.42E-03 | 0.24 | 0.16 | 0.09 | 1.54 |
| SE_basal | Ston2 | 0.42 | 1.15E-07 | 2.46E-03 | 0.67 | 0.60 | 0.07 | 1.11 |
| SE_basal | Zfp644 | 0.29 | 1.19E-07 | 2.54E-03 | 0.85 | 0.84 | 0.01 | 1.01 |
| SE_basal | Barx2 | 1.05 | 1.23E-07 | 2.63E-03 | 0.24 | 0.16 | 0.09 | 1.56 |
| SE_basal | Usp3 | 0.40 | 1.41E-07 | 3.01E-03 | 0.72 | 0.71 | 0.00 | 1.01 |
| SE_basal | Pakap | 1.38 | 1.51E-07 | 3.23E-03 | 0.12 | 0.06 | 0.06 | 2.00 |
| SE_basal | Ppic | 0.54 | 1.67E-07 | 3.56E-03 | 0.62 | 0.59 | 0.03 | 1.05 |
| SE_basal | Gjb3 | 0.92 | 1.69E-07 | 3.61E-03 | 0.22 | 0.14 | 0.08 | 1.59 |
| SE_basal | Smo | 0.38 | 1.81E-07 | 3.87E-03 | 0.72 | 0.69 | 0.03 | 1.04 |
| SE_basal | Map4k5 | 0.76 | 1.99E-07 | 4.25E-03 | 0.45 | 0.37 | 0.07 | 1.19 |
| SE_basal | Cdc42ep3 | 0.78 | 2.12E-07 | 4.53E-03 | 0.26 | 0.17 | 0.09 | 1.52 |
| SE_basal | Gjb6 | 1.71 | 2.20E-07 | 4.69E-03 | 0.12 | 0.06 | 0.06 | 2.00 |
| SE_basal | Armcx1 | 0.72 | 2.30E-07 | 4.91E-03 | 0.43 | 0.36 | 0.07 | 1.20 |
| SE_basal | Ptpn14 | 0.81 | 2.32E-07 | 4.96E-03 | 0.31 | 0.22 | 0.09 | 1.41 |
| SE_basal | Palmd | 1.00 | 2.33E-07 | 4.98E-03 | 0.11 | 0.05 | 0.06 | 2.16 |
| SE_basal | Ctso | 1.28 | 2.34E-07 | 4.99E-03 | 0.20 | 0.13 | 0.08 | 1.61 |
| SE_basal | Tmem44 | 1.13 | 2.46E-07 | 5.25E-03 | 0.21 | 0.13 | 0.08 | 1.59 |
| SE_basal | Map4k4 | 0.29 | 2.69E-07 | 5.75E-03 | 0.89 | 0.89 | 0.00 | 1.00 |
| SE_basal | Ercc5 | 0.92 | 2.75E-07 | 5.87E-03 | 0.38 | 0.30 | 0.07 | 1.24 |
| SE_basal | Socs7 | 0.81 | 2.78E-07 | 5.94E-03 | 0.37 | 0.29 | 0.07 | 1.25 |
| SE_basal | 2610307P16Rik | 1.02 | 3.02E-07 | 6.45E-03 | 0.28 | 0.19 | 0.08 | 1.44 |
| SE_basal | Ttc7 | 1.40 | 3.07E-07 | 6.57E-03 | 0.12 | 0.06 | 0.06 | 1.97 |
| SE_basal | Arl4a | 1.21 | 3.33E-07 | 7.11E-03 | 0.23 | 0.16 | 0.08 | 1.50 |
| SE_basal | Phip | 0.28 | 3.41E-07 | 7.28E-03 | 0.88 | 0.88 | 0.00 | 1.00 |
| SE_basal | Shf | 0.84 | 3.52E-07 | 7.52E-03 | 0.28 | 0.20 | 0.09 | 1.43 |
| SE_basal | Ptprd | 0.53 | 3.56E-07 | 7.61E-03 | 0.61 | 0.56 | 0.06 | 1.10 |
| SE_basal | Wnk2 | 1.12 | 3.66E-07 | 7.81E-03 | 0.18 | 0.11 | 0.07 | 1.68 |
| SE_basal | Chd9 | 0.66 | 4.02E-07 | 8.58E-03 | 0.54 | 0.50 | 0.04 | 1.08 |
| SE_basal | Slc25a27 | 0.54 | 4.08E-07 | 8.72E-03 | 0.59 | 0.54 | 0.05 | 1.09 |
| SE_basal | Sept10 | 0.68 | 4.17E-07 | 8.91E-03 | 0.43 | 0.36 | 0.07 | 1.19 |
| SE_basal | Bnip3 | 0.47 | 5.28E-07 | 1.13E-02 | 0.53 | 0.45 | 0.08 | 1.19 |
| SE_basal | Qk | 0.26 | 5.55E-07 | 1.18E-02 | 0.88 | 0.88 | 0.00 | 1.00 |
| SE_basal | Tro | 0.76 | 5.57E-07 | 1.19E-02 | 0.33 | 0.25 | 0.08 | 1.33 |
| SE_basal | Pygl | 1.02 | 5.62E-07 | 1.20E-02 | 0.25 | 0.17 | 0.08 | 1.46 |
| SE_basal | Hadha | 0.35 | 5.89E-07 | 1.26E-02 | 0.76 | 0.75 | 0.02 | 1.02 |

**Supplementary Table 4. Marker genes for the annotated cell populations of integrated clustering of E11.5 control and NotchGOF λ-junction ectoderm cells.**

| | | | | | | | | |
|---|---|---|---|---|---|---|---|---|
| SE_basal | Gosr2 | 0.81 | 6.02E-07 | 1.29E-02 | 0.40 | 0.33 | 0.07 | 1.21 |
| SE_basal | Cep85l | 1.57 | 6.39E-07 | 1.36E-02 | 0.10 | 0.05 | 0.05 | 2.13 |
| SE_basal | Lratd1 | 0.87 | 7.21E-07 | 1.54E-02 | 0.20 | 0.12 | 0.08 | 1.62 |
| SE_basal | Zfp950 | 0.71 | 7.26E-07 | 1.55E-02 | 0.43 | 0.37 | 0.06 | 1.17 |
| SE_basal | Txnrd1 | 0.27 | 7.33E-07 | 1.57E-02 | 0.87 | 0.87 | -0.01 | 0.99 |
| SE_basal | Scoc | 0.52 | 7.36E-07 | 1.57E-02 | 0.58 | 0.56 | 0.02 | 1.04 |
| SE_basal | Eva1a | 1.08 | 7.51E-07 | 1.60E-02 | 0.26 | 0.18 | 0.08 | 1.41 |
| SE_basal | Chd6 | 0.33 | 7.60E-07 | 1.62E-02 | 0.77 | 0.76 | 0.01 | 1.02 |
| SE_basal | Net1 | 0.50 | 8.30E-07 | 1.77E-02 | 0.56 | 0.50 | 0.06 | 1.12 |
| SE_basal | Sord | 0.73 | 8.41E-07 | 1.80E-02 | 0.31 | 0.23 | 0.08 | 1.36 |
| SE_basal | Snap47 | 0.40 | 8.48E-07 | 1.81E-02 | 0.70 | 0.68 | 0.02 | 1.03 |
| SE_basal | Nenf | 0.39 | 9.17E-07 | 1.96E-02 | 0.72 | 0.72 | 0.00 | 1.00 |
| SE_basal | Klhl24 | 0.45 | 9.18E-07 | 1.96E-02 | 0.69 | 0.63 | 0.05 | 1.09 |
| SE_basal | Sowahc | 1.09 | 9.20E-07 | 1.97E-02 | 0.24 | 0.16 | 0.08 | 1.48 |
| SE_basal | Nme4 | 0.32 | 1.05E-06 | 2.24E-02 | 0.72 | 0.73 | -0.01 | 0.99 |
| SE_basal | Macf1 | 0.35 | 1.11E-06 | 2.36E-02 | 0.84 | 0.83 | 0.01 | 1.01 |
| SE_basal | Vwa5a | 0.91 | 1.13E-06 | 2.42E-02 | 0.24 | 0.16 | 0.08 | 1.50 |
| SE_basal | Sik1 | 1.04 | 1.16E-06 | 2.48E-02 | 0.23 | 0.16 | 0.08 | 1.49 |
| SE_basal | Micall1 | 0.99 | 1.30E-06 | 2.78E-02 | 0.19 | 0.12 | 0.07 | 1.62 |
| SE_basal | Ypel3 | 0.37 | 1.55E-06 | 3.31E-02 | 0.63 | 0.57 | 0.06 | 1.11 |
| SE_basal | Frmd6 | 0.65 | 1.59E-06 | 3.39E-02 | 0.31 | 0.23 | 0.09 | 1.39 |
| SE_basal | Srek1ip1 | 0.43 | 1.63E-06 | 3.49E-02 | 0.70 | 0.68 | 0.01 | 1.02 |
| SE_basal | Cnst | 1.04 | 1.65E-06 | 3.52E-02 | 0.21 | 0.14 | 0.07 | 1.52 |
| SE_basal | Eml4 | 0.37 | 1.73E-06 | 3.71E-02 | 0.72 | 0.71 | 0.01 | 1.01 |
| SE_basal | Itga3 | 0.92 | 1.85E-06 | 3.95E-02 | 0.32 | 0.25 | 0.07 | 1.29 |
| SE_basal | Atf7ip | 0.42 | 1.91E-06 | 4.08E-02 | 0.65 | 0.64 | 0.02 | 1.02 |
| SE_basal | Dancr | 0.52 | 2.08E-06 | 4.45E-02 | 0.60 | 0.55 | 0.05 | 1.09 |
| palate | Foxe1 | 4.30 | 0 | 0 | 0.96 | 0.12 | 0.84 | 7.89 |
| palate | Gm12446 | 4.61 | 0 | 0 | 0.86 | 0.06 | 0.80 | 14.05 |
| palate | Barx1 | 4.78 | 3.80E-276 | 8.11E-272 | 0.74 | 0.06 | 0.67 | 11.87 |
| palate | Cgnl1 | 3.88 | 3.01E-208 | 6.42E-204 | 0.80 | 0.13 | 0.68 | 6.38 |
| palate | Krt19 | 3.24 | 3.94E-151 | 8.42E-147 | 0.88 | 0.25 | 0.63 | 3.57 |
| palate | Cryab | 3.77 | 1.57E-146 | 3.34E-142 | 0.71 | 0.14 | 0.57 | 4.95 |
| palate | Foxq1 | 4.61 | 8.12E-146 | 1.73E-141 | 0.31 | 0.01 | 0.30 | 22.07 |
| palate | S100a6 | 3.19 | 8.87E-137 | 1.89E-132 | 0.79 | 0.20 | 0.59 | 3.90 |
| palate | Csrp2 | 2.48 | 6.15E-133 | 1.31E-128 | 1.00 | 0.83 | 0.17 | 1.21 |
| palate | Foxa1 | 4.68 | 2.73E-126 | 5.83E-122 | 0.29 | 0.02 | 0.27 | 19.13 |
| palate | H19 | 2.13 | 2.30E-125 | 4.92E-121 | 1.00 | 0.87 | 0.13 | 1.15 |
| palate | Cdo1 | 2.55 | 3.25E-122 | 6.95E-118 | 0.94 | 0.44 | 0.50 | 2.14 |
| palate | Tbx1 | 3.89 | 6.18E-121 | 1.32E-116 | 0.35 | 0.03 | 0.32 | 12.39 |
| palate | Upk2 | 3.85 | 1.74E-120 | 3.72E-116 | 0.40 | 0.04 | 0.36 | 10.00 |
| palate | Amot | 2.42 | 1.73E-118 | 3.70E-114 | 0.96 | 0.55 | 0.41 | 1.74 |
| palate | Barx2 | 2.51 | 3.42E-113 | 7.31E-109 | 0.66 | 0.13 | 0.53 | 4.98 |
| palate | Upk3bl | 2.60 | 6.80E-113 | 1.45E-108 | 0.83 | 0.27 | 0.56 | 3.11 |
| palate | Krt8 | 1.58 | 4.96E-110 | 1.06E-105 | 1.00 | 0.81 | 0.19 | 1.24 |
| palate | Dmrt2 | 2.90 | 7.77E-109 | 1.66E-104 | 0.49 | 0.07 | 0.42 | 6.86 |
| palate | Nr2f2 | 2.09 | 3.58E-104 | 7.65E-100 | 0.94 | 0.57 | 0.38 | 1.67 |
| palate | Plcd1 | 2.71 | 2.71E-92 | 5.78E-88 | 0.51 | 0.10 | 0.42 | 5.34 |
| palate | Anxa1 | 3.42 | 4.74E-92 | 1.01E-87 | 0.36 | 0.04 | 0.32 | 8.33 |
| palate | Siva1 | 1.17 | 9.97E-91 | 2.13E-86 | 1.00 | 0.97 | 0.03 | 1.03 |
| palate | Atox1 | 1.07 | 7.54E-90 | 1.61E-85 | 1.00 | 0.98 | 0.02 | 1.02 |
| palate | Krt7 | 2.44 | 3.95E-87 | 8.45E-83 | 0.66 | 0.18 | 0.48 | 3.63 |
| palate | Anxa2 | 1.40 | 1.17E-83 | 2.50E-79 | 1.00 | 0.74 | 0.25 | 1.34 |
| palate | Tacstd2 | 1.69 | 5.00E-83 | 1.07E-78 | 0.76 | 0.24 | 0.52 | 3.12 |
| palate | Pon2 | 2.23 | 1.40E-82 | 2.99E-78 | 0.75 | 0.26 | 0.48 | 2.83 |

**Supplementary Table 4. Marker genes for the annotated cell populations of integrated clustering of E11.5 control and NotchGOF λ-junction ectoderm cells.**

| | | | | | | | | |
|---|---|---|---|---|---|---|---|---|
| palate | Glrx | 2.35 | 4.68E-81 | 1.00E-76 | 0.74 | 0.27 | 0.47 | 2.74 |
| palate | Nrk | 2.89 | 1.07E-79 | 2.28E-75 | 0.49 | 0.10 | 0.38 | 4.68 |
| palate | Igf2 | 1.60 | 2.59E-79 | 5.53E-75 | 0.93 | 0.81 | 0.12 | 1.14 |
| palate | Pitx2 | 1.39 | 4.77E-79 | 1.02E-74 | 0.76 | 0.22 | 0.54 | 3.45 |
| palate | Slc39a8 | 2.80 | 1.03E-77 | 2.19E-73 | 0.43 | 0.08 | 0.35 | 5.53 |
| palate | Fzd5 | 3.62 | 1.80E-77 | 3.84E-73 | 0.26 | 0.03 | 0.24 | 9.78 |
| palate | Isoc1 | 1.60 | 4.08E-76 | 8.71E-72 | 0.94 | 0.69 | 0.25 | 1.36 |
| palate | Krt18 | 1.19 | 3.52E-74 | 7.53E-70 | 1.00 | 0.86 | 0.14 | 1.16 |
| palate | S100a10 | 1.27 | 1.29E-69 | 2.75E-65 | 1.00 | 0.73 | 0.26 | 1.36 |
| palate | Hspb2 | 2.54 | 2.38E-69 | 5.08E-65 | 0.47 | 0.11 | 0.37 | 4.45 |
| palate | Tpm1 | 1.07 | 2.58E-69 | 5.52E-65 | 1.00 | 0.93 | 0.07 | 1.07 |
| palate | Lypd2 | 2.29 | 4.13E-65 | 8.81E-61 | 0.52 | 0.14 | 0.38 | 3.80 |
| palate | Arhgap29 | 1.60 | 5.09E-65 | 1.09E-60 | 0.82 | 0.38 | 0.45 | 2.19 |
| palate | Igdcc3 | 1.83 | 5.88E-65 | 1.26E-60 | 0.76 | 0.32 | 0.44 | 2.39 |
| palate | Grhl3 | 1.88 | 8.83E-63 | 1.89E-58 | 0.62 | 0.19 | 0.43 | 3.21 |
| palate | Cldn8 | 2.27 | 4.10E-62 | 8.76E-58 | 0.45 | 0.11 | 0.35 | 4.31 |
| palate | Arl4c | 1.80 | 9.55E-62 | 2.04E-57 | 0.87 | 0.56 | 0.31 | 1.55 |
| palate | Plagl1 | 1.88 | 2.24E-60 | 4.79E-56 | 0.79 | 0.41 | 0.38 | 1.94 |
| palate | Nrtn | 1.92 | 6.06E-60 | 1.29E-55 | 0.72 | 0.30 | 0.42 | 2.43 |
| palate | Dpp6 | 3.83 | 1.76E-59 | 3.77E-55 | 0.15 | 0.01 | 0.14 | 16.33 |
| palate | Slc38a4 | 1.93 | 1.97E-59 | 4.21E-55 | 0.67 | 0.25 | 0.42 | 2.65 |
| palate | Tcf7l2 | 1.40 | 9.86E-59 | 2.11E-54 | 0.93 | 0.72 | 0.21 | 1.29 |
| palate | Adamts18 | 1.73 | 2.74E-57 | 5.85E-53 | 0.66 | 0.25 | 0.42 | 2.71 |
| palate | Pof1b | 3.12 | 1.10E-55 | 2.36E-51 | 0.25 | 0.03 | 0.21 | 7.21 |
| palate | Gask1b | 2.45 | 2.70E-55 | 5.77E-51 | 0.30 | 0.05 | 0.25 | 5.92 |
| palate | Lmod2 | 4.64 | 8.41E-55 | 1.80E-50 | 0.11 | 0.00 | 0.11 | 27.25 |
| palate | Lamc2 | 2.26 | 9.29E-55 | 1.99E-50 | 0.51 | 0.15 | 0.35 | 3.31 |
| palate | Evpl | 2.30 | 1.07E-53 | 2.29E-49 | 0.42 | 0.10 | 0.32 | 4.15 |
| palate | Sytl2 | 4.13 | 1.42E-53 | 3.04E-49 | 0.14 | 0.01 | 0.13 | 15.11 |
| palate | Fgfbp1 | 3.74 | 2.19E-53 | 4.67E-49 | 0.17 | 0.02 | 0.15 | 11.07 |
| palate | Gm16136 | 1.19 | 4.90E-53 | 1.05E-48 | 0.94 | 0.63 | 0.31 | 1.49 |
| palate | Wfdc2 | 1.18 | 9.84E-53 | 2.10E-48 | 0.99 | 0.81 | 0.18 | 1.22 |
| palate | Mt1 | 1.64 | 1.09E-51 | 2.33E-47 | 0.85 | 0.48 | 0.37 | 1.75 |
| palate | Mt2 | 1.74 | 1.98E-51 | 4.24E-47 | 0.65 | 0.25 | 0.40 | 2.60 |
| palate | Isl2 | 2.30 | 7.86E-51 | 1.68E-46 | 0.45 | 0.12 | 0.32 | 3.62 |
| palate | Klf5 | 1.49 | 9.90E-51 | 2.12E-46 | 0.77 | 0.39 | 0.38 | 1.98 |
| palate | Fxyd3 | 1.69 | 7.39E-50 | 1.58E-45 | 0.69 | 0.29 | 0.41 | 2.42 |
| palate | Cald1 | 0.94 | 2.31E-49 | 4.94E-45 | 0.97 | 0.69 | 0.28 | 1.41 |
| palate | Krt15 | 1.77 | 3.60E-49 | 7.70E-45 | 0.47 | 0.14 | 0.34 | 3.50 |
| palate | Ahnak | 1.52 | 6.61E-49 | 1.41E-44 | 0.73 | 0.34 | 0.39 | 2.17 |
| palate | Rab27b | 1.59 | 1.49E-48 | 3.19E-44 | 0.69 | 0.30 | 0.39 | 2.31 |
| palate | Gpc3 | 0.84 | 4.54E-48 | 9.70E-44 | 1.00 | 0.83 | 0.17 | 1.20 |
| palate | Mycl | 1.16 | 7.44E-48 | 1.59E-43 | 0.92 | 0.63 | 0.30 | 1.47 |
| palate | Npnt | 1.49 | 1.73E-47 | 3.69E-43 | 0.80 | 0.43 | 0.37 | 1.87 |
| palate | Fam210b | 2.23 | 9.45E-47 | 2.02E-42 | 0.53 | 0.20 | 0.34 | 2.73 |
| palate | Nudt4 | 1.13 | 1.27E-46 | 2.70E-42 | 0.94 | 0.77 | 0.17 | 1.22 |
| palate | Palm3 | 1.64 | 2.87E-46 | 6.13E-42 | 0.70 | 0.33 | 0.37 | 2.12 |
| palate | Anxa8 | 2.31 | 4.07E-46 | 8.69E-42 | 0.35 | 0.08 | 0.27 | 4.33 |
| palate | Arg1 | 2.84 | 1.11E-45 | 2.37E-41 | 0.24 | 0.04 | 0.20 | 6.05 |
| palate | Cstb | 0.86 | 9.33E-45 | 1.99E-40 | 0.99 | 0.84 | 0.15 | 1.18 |
| palate | Cfc1 | 3.17 | 3.24E-44 | 6.92E-40 | 0.16 | 0.02 | 0.14 | 9.29 |
| palate | Ccnd2 | 0.94 | 1.28E-42 | 2.74E-38 | 0.99 | 0.77 | 0.21 | 1.27 |
| palate | Anxa3 | 2.40 | 2.63E-42 | 5.62E-38 | 0.28 | 0.06 | 0.22 | 4.91 |
| palate | Sfn | 0.66 | 1.23E-41 | 2.63E-37 | 0.99 | 0.68 | 0.31 | 1.45 |
| palate | Mmp11 | 1.49 | 3.33E-41 | 7.11E-37 | 0.73 | 0.37 | 0.35 | 1.95 |

**Supplementary Table 4. Marker genes for the annotated cell populations of integrated clustering of E11.5 control and NotchGOF λ-junction ectoderm cells.**

| | | | | | | | | |
|---|---|---|---|---|---|---|---|---|
| palate | Lcp1 | 3.42 | 5.18E-41 | 1.11E-36 | 0.11 | 0.01 | 0.10 | 13.63 |
| palate | Afap1l1 | 2.17 | 2.56E-40 | 5.46E-36 | 0.37 | 0.10 | 0.27 | 3.60 |
| palate | Tmod3 | 1.27 | 2.84E-39 | 6.06E-35 | 0.87 | 0.65 | 0.22 | 1.34 |
| palate | Ccdc68 | 1.81 | 3.52E-39 | 7.51E-35 | 0.40 | 0.12 | 0.29 | 3.39 |
| palate | Unc5c | 1.15 | 6.25E-39 | 1.33E-34 | 0.69 | 0.31 | 0.38 | 2.23 |
| palate | Cldn23 | 1.93 | 2.99E-38 | 6.40E-34 | 0.42 | 0.14 | 0.28 | 3.04 |
| palate | Shh | 1.96 | 4.42E-38 | 9.43E-34 | 0.25 | 0.05 | 0.20 | 5.08 |
| palate | 2200002D01Rik | 2.52 | 4.71E-38 | 1.01E-33 | 0.26 | 0.06 | 0.21 | 4.73 |
| palate | Dsp | 0.82 | 7.17E-38 | 1.53E-33 | 0.98 | 0.86 | 0.12 | 1.14 |
| palate | Igfbp5 | 0.74 | 1.12E-37 | 2.39E-33 | 1.00 | 0.96 | 0.04 | 1.04 |
| palate | Hspb1 | 1.12 | 2.67E-37 | 5.69E-33 | 0.74 | 0.36 | 0.38 | 2.04 |
| palate | 2610016A17Rik | 2.99 | 1.04E-36 | 2.22E-32 | 0.19 | 0.03 | 0.16 | 6.30 |
| palate | Arid3a | 1.42 | 4.19E-36 | 8.95E-32 | 0.67 | 0.34 | 0.33 | 1.96 |
| palate | Utrn | 0.98 | 4.19E-36 | 8.95E-32 | 0.93 | 0.71 | 0.22 | 1.31 |
| palate | Ralbp1 | 0.84 | 6.78E-36 | 1.45E-31 | 0.96 | 0.82 | 0.14 | 1.17 |
| palate | Ghr | 2.24 | 9.42E-36 | 2.01E-31 | 0.28 | 0.07 | 0.22 | 4.16 |
| palate | Foxc1 | 1.88 | 1.47E-35 | 3.14E-31 | 0.40 | 0.13 | 0.27 | 3.09 |
| palate | Sema3c | 1.47 | 1.54E-35 | 3.28E-31 | 0.56 | 0.24 | 0.33 | 2.38 |
| palate | Zbtb42 | 2.62 | 2.27E-34 | 4.86E-30 | 0.19 | 0.03 | 0.16 | 5.91 |
| palate | Calm1 | 0.42 | 3.84E-34 | 8.20E-30 | 1.00 | 0.99 | 0.01 | 1.01 |
| palate | Fgfr3 | 2.51 | 1.30E-33 | 2.78E-29 | 0.21 | 0.04 | 0.17 | 5.15 |
| palate | Perp | 0.78 | 2.52E-33 | 5.37E-29 | 0.91 | 0.52 | 0.39 | 1.74 |
| palate | Ogfrl1 | 1.85 | 1.32E-32 | 2.81E-28 | 0.44 | 0.17 | 0.28 | 2.65 |
| palate | Mfge8 | 1.27 | 1.91E-32 | 4.07E-28 | 0.81 | 0.54 | 0.27 | 1.49 |
| palate | B230208H11Rik | 2.41 | 2.28E-31 | 4.86E-27 | 0.22 | 0.05 | 0.17 | 4.76 |
| palate | Hebp1 | 1.45 | 5.32E-31 | 1.14E-26 | 0.51 | 0.22 | 0.29 | 2.37 |
| palate | Pmp22 | 3.02 | 8.80E-31 | 1.88E-26 | 0.14 | 0.02 | 0.12 | 7.16 |
| palate | Grb10 | 0.63 | 1.12E-29 | 2.39E-25 | 1.00 | 0.93 | 0.07 | 1.08 |
| palate | Srd5a1 | 1.29 | 1.89E-29 | 4.03E-25 | 0.62 | 0.32 | 0.30 | 1.92 |
| palate | Tinagl1 | 2.05 | 3.10E-29 | 6.63E-25 | 0.29 | 0.08 | 0.21 | 3.54 |
| palate | A730020E08Rik | 2.01 | 3.16E-29 | 6.76E-25 | 0.28 | 0.08 | 0.20 | 3.67 |
| palate | Slc2a3 | 0.93 | 3.85E-29 | 8.23E-25 | 0.65 | 0.34 | 0.32 | 1.94 |
| palate | Aldh3b2 | 2.76 | 4.20E-29 | 8.97E-25 | 0.16 | 0.03 | 0.13 | 5.79 |
| palate | Ppl | 1.21 | 9.75E-29 | 2.08E-24 | 0.42 | 0.15 | 0.27 | 2.77 |
| palate | Cmas | 0.95 | 1.54E-28 | 3.28E-24 | 0.84 | 0.64 | 0.20 | 1.31 |
| palate | Cers4 | 1.36 | 1.68E-28 | 3.58E-24 | 0.66 | 0.36 | 0.30 | 1.82 |
| palate | Col5a2 | 1.51 | 2.02E-28 | 4.31E-24 | 0.41 | 0.15 | 0.26 | 2.78 |
| palate | Alcam | 1.36 | 6.34E-28 | 1.35E-23 | 0.46 | 0.19 | 0.28 | 2.48 |
| palate | Igf2bp3 | 0.67 | 6.37E-28 | 1.36E-23 | 0.95 | 0.86 | 0.09 | 1.10 |
| palate | Fignl2 | 1.91 | 1.15E-27 | 2.45E-23 | 0.25 | 0.07 | 0.19 | 3.78 |
| palate | Arf1 | 0.40 | 1.52E-27 | 3.24E-23 | 0.99 | 0.98 | 0.01 | 1.01 |
| palate | Iqgap1 | 0.87 | 1.54E-27 | 3.29E-23 | 0.85 | 0.67 | 0.17 | 1.26 |
| palate | Cd9 | 0.73 | 1.60E-27 | 3.42E-23 | 0.92 | 0.70 | 0.21 | 1.30 |
| palate | Ppia | 0.26 | 2.50E-27 | 5.33E-23 | 1.00 | 1.00 | 0.00 | 1.00 |
| palate | Myh14 | 1.35 | 2.61E-27 | 5.58E-23 | 0.56 | 0.29 | 0.28 | 1.97 |
| palate | Prom2 | 1.77 | 7.27E-27 | 1.55E-22 | 0.34 | 0.12 | 0.23 | 2.96 |
| palate | Il17re | 1.63 | 1.18E-26 | 2.53E-22 | 0.42 | 0.17 | 0.25 | 2.47 |
| palate | Ppp1r13b | 1.69 | 2.19E-26 | 4.69E-22 | 0.34 | 0.12 | 0.23 | 2.91 |
| palate | Prss22 | 1.24 | 3.46E-26 | 7.39E-22 | 0.51 | 0.23 | 0.28 | 2.24 |
| palate | Platr21 | 1.91 | 1.36E-25 | 2.91E-21 | 0.29 | 0.09 | 0.20 | 3.20 |
| palate | Hsdl2 | 1.30 | 7.56E-25 | 1.62E-20 | 0.59 | 0.33 | 0.26 | 1.78 |
| palate | Alox12 | 2.08 | 8.08E-25 | 1.73E-20 | 0.25 | 0.07 | 0.18 | 3.47 |
| palate | Smtnl2 | 1.38 | 8.40E-25 | 1.79E-20 | 0.51 | 0.25 | 0.26 | 2.08 |
| palate | Pwwp2b | 1.49 | 1.55E-24 | 3.32E-20 | 0.37 | 0.14 | 0.24 | 2.71 |
| palate | Ptpn13 | 0.82 | 2.55E-24 | 5.45E-20 | 0.81 | 0.56 | 0.25 | 1.44 |

**Supplementary Table 4. Marker genes for the annotated cell populations of integrated clustering of E11.5 control and NotchGOF λ-junction ectoderm cells.**

| | | | | | | | | |
|---|---|---|---|---|---|---|---|---|
| palate | Wwc2 | 0.93 | 2.82E-24 | 6.02E-20 | 0.82 | 0.59 | 0.22 | 1.37 |
| palate | Hsd17b14 | 2.90 | 3.26E-24 | 6.97E-20 | 0.14 | 0.02 | 0.11 | 5.67 |
| palate | Pcdh1 | 1.79 | 5.40E-24 | 1.15E-19 | 0.29 | 0.09 | 0.20 | 3.12 |
| palate | Fstl1 | 0.90 | 6.82E-24 | 1.46E-19 | 0.79 | 0.60 | 0.20 | 1.33 |
| palate | Cyb5r3 | 0.52 | 1.09E-23 | 2.32E-19 | 0.97 | 0.93 | 0.04 | 1.04 |
| palate | Foxp1 | 0.58 | 1.37E-23 | 2.94E-19 | 0.96 | 0.86 | 0.10 | 1.11 |
| palate | Nectin4 | 0.83 | 1.61E-23 | 3.43E-19 | 0.80 | 0.52 | 0.28 | 1.54 |
| palate | Esrp2 | 1.23 | 1.84E-23 | 3.92E-19 | 0.52 | 0.25 | 0.27 | 2.11 |
| palate | Ctsh | 1.63 | 2.93E-23 | 6.26E-19 | 0.32 | 0.11 | 0.21 | 2.82 |
| palate | Tek | 2.72 | 4.74E-23 | 1.01E-18 | 0.12 | 0.02 | 0.10 | 6.16 |
| palate | Rassf8 | 2.04 | 4.84E-23 | 1.03E-18 | 0.29 | 0.10 | 0.19 | 2.93 |
| palate | Mpp7 | 0.99 | 5.70E-23 | 1.22E-18 | 0.74 | 0.54 | 0.20 | 1.38 |
| palate | Mansc4 | 2.18 | 8.90E-23 | 1.90E-18 | 0.17 | 0.04 | 0.13 | 4.47 |
| palate | Limk2 | 1.08 | 9.83E-23 | 2.10E-18 | 0.60 | 0.34 | 0.26 | 1.78 |
| palate | Phactr2 | 1.17 | 1.50E-22 | 3.21E-18 | 0.60 | 0.36 | 0.24 | 1.67 |
| palate | Stx19 | 2.33 | 3.11E-22 | 6.64E-18 | 0.14 | 0.03 | 0.12 | 5.11 |
| palate | Cdh1 | 0.76 | 9.74E-22 | 2.08E-17 | 0.83 | 0.65 | 0.18 | 1.28 |
| palate | Chst5 | 2.12 | 1.43E-21 | 3.05E-17 | 0.20 | 0.06 | 0.15 | 3.71 |
| palate | Fam83f | 2.26 | 1.43E-21 | 3.06E-17 | 0.16 | 0.03 | 0.12 | 4.56 |
| palate | Gpc4 | 1.18 | 1.96E-21 | 4.18E-17 | 0.55 | 0.30 | 0.25 | 1.85 |
| palate | Eci1 | 0.91 | 2.38E-21 | 5.09E-17 | 0.75 | 0.52 | 0.23 | 1.44 |
| palate | Shisa4 | 1.24 | 2.42E-21 | 5.16E-17 | 0.48 | 0.23 | 0.25 | 2.10 |
| palate | Rbms1 | 0.62 | 3.58E-21 | 7.64E-17 | 0.93 | 0.84 | 0.08 | 1.10 |
| palate | Idh2 | 0.56 | 4.33E-21 | 9.26E-17 | 0.96 | 0.87 | 0.09 | 1.11 |
| palate | Nr3c1 | 1.24 | 4.69E-21 | 1.00E-16 | 0.51 | 0.26 | 0.25 | 1.95 |
| palate | Elf5 | 1.28 | 7.16E-21 | 1.53E-16 | 0.47 | 0.23 | 0.24 | 2.07 |
| palate | Epb41l5 | 0.92 | 1.46E-20 | 3.13E-16 | 0.73 | 0.51 | 0.22 | 1.44 |
| palate | Cers3 | 2.15 | 2.68E-20 | 5.72E-16 | 0.17 | 0.04 | 0.13 | 4.15 |
| palate | 2310030G06Rik | 1.29 | 2.71E-20 | 5.79E-16 | 0.49 | 0.26 | 0.23 | 1.89 |
| palate | Gjb3 | 1.28 | 2.86E-20 | 6.10E-16 | 0.35 | 0.14 | 0.21 | 2.56 |
| palate | Junb | 1.30 | 3.65E-20 | 7.80E-16 | 0.44 | 0.20 | 0.24 | 2.18 |
| palate | Dstn | 0.35 | 4.19E-20 | 8.96E-16 | 0.99 | 0.98 | 0.01 | 1.01 |
| palate | S100a11 | 0.44 | 4.82E-20 | 1.03E-15 | 1.00 | 0.90 | 0.10 | 1.11 |
| palate | Ldhb | 0.48 | 6.08E-20 | 1.30E-15 | 0.98 | 0.87 | 0.12 | 1.13 |
| palate | Ccnd3 | 0.61 | 6.48E-20 | 1.38E-15 | 0.92 | 0.80 | 0.12 | 1.15 |
| palate | Sh3tc2 | 2.02 | 7.06E-20 | 1.51E-15 | 0.18 | 0.05 | 0.13 | 3.85 |
| palate | Fhl1 | 1.32 | 9.85E-20 | 2.10E-15 | 0.47 | 0.24 | 0.23 | 1.95 |
| palate | Spsb1 | 1.73 | 9.97E-20 | 2.13E-15 | 0.21 | 0.06 | 0.15 | 3.41 |
| palate | Cldn7 | 0.61 | 3.13E-19 | 6.69E-15 | 0.97 | 0.84 | 0.14 | 1.16 |
| palate | Eps8l2 | 1.62 | 3.42E-19 | 7.31E-15 | 0.34 | 0.15 | 0.20 | 2.35 |
| palate | Ier2 | 0.64 | 6.17E-19 | 1.32E-14 | 0.90 | 0.79 | 0.12 | 1.15 |
| palate | Fign | 1.49 | 6.93E-19 | 1.48E-14 | 0.39 | 0.18 | 0.21 | 2.15 |
| palate | Sfr1 | 0.39 | 7.39E-19 | 1.58E-14 | 1.00 | 0.98 | 0.02 | 1.02 |
| palate | Sema3d | 0.82 | 8.38E-19 | 1.79E-14 | 0.67 | 0.41 | 0.26 | 1.65 |
| palate | Rai14 | 1.28 | 9.38E-19 | 2.00E-14 | 0.51 | 0.29 | 0.22 | 1.77 |
| palate | Jph1 | 1.87 | 1.04E-18 | 2.22E-14 | 0.25 | 0.09 | 0.16 | 2.86 |
| palate | Fhl2 | 2.00 | 1.32E-18 | 2.83E-14 | 0.19 | 0.05 | 0.13 | 3.56 |
| palate | Fabp5 | 0.31 | 1.37E-18 | 2.93E-14 | 1.00 | 0.99 | 0.01 | 1.01 |
| palate | Dmrta2 | 0.87 | 1.39E-18 | 2.98E-14 | 0.74 | 0.50 | 0.25 | 1.50 |
| palate | S100a16 | 1.73 | 1.55E-18 | 3.32E-14 | 0.33 | 0.14 | 0.19 | 2.39 |
| palate | Lypd6 | 1.40 | 2.05E-18 | 4.38E-14 | 0.41 | 0.19 | 0.22 | 2.11 |
| palate | B130024G19Rik | 2.09 | 2.35E-18 | 5.01E-14 | 0.16 | 0.04 | 0.12 | 4.08 |
| palate | Lgi2 | 1.11 | 2.39E-18 | 5.11E-14 | 0.52 | 0.28 | 0.24 | 1.83 |
| palate | Fuom | 1.05 | 2.45E-18 | 5.24E-14 | 0.55 | 0.31 | 0.24 | 1.75 |
| palate | F3 | 1.34 | 2.64E-18 | 5.64E-14 | 0.43 | 0.22 | 0.22 | 1.99 |

**Supplementary Table 4. Marker genes for the annotated cell populations of integrated clustering of E11.5 control and NotchGOF λ-junction ectoderm cells.**

| | | | | | | | | |
|---|---|---|---|---|---|---|---|---|
| palate | Spats2l | 0.89 | 2.71E-18 | 5.80E-14 | 0.57 | 0.33 | 0.25 | 1.75 |
| palate | F11r | 0.76 | 3.57E-18 | 7.64E-14 | 0.80 | 0.60 | 0.21 | 1.34 |
| palate | Rflnb | 0.88 | 4.05E-18 | 8.65E-14 | 0.74 | 0.51 | 0.23 | 1.46 |
| palate | Net1 | 0.97 | 4.41E-18 | 9.43E-14 | 0.69 | 0.49 | 0.20 | 1.40 |
| palate | Arhgap24 | 1.60 | 6.01E-18 | 1.28E-13 | 0.22 | 0.07 | 0.15 | 3.16 |
| palate | Mapkapk2 | 0.81 | 7.54E-18 | 1.61E-13 | 0.73 | 0.54 | 0.19 | 1.36 |
| palate | Negr1 | 2.07 | 1.04E-17 | 2.23E-13 | 0.15 | 0.04 | 0.11 | 4.08 |
| palate | Capg | 0.99 | 1.25E-17 | 2.67E-13 | 0.58 | 0.34 | 0.24 | 1.69 |
| palate | Macc1 | 1.20 | 2.48E-17 | 5.29E-13 | 0.42 | 0.21 | 0.21 | 1.99 |
| palate | Dusp9 | 1.43 | 4.92E-17 | 1.05E-12 | 0.43 | 0.23 | 0.21 | 1.90 |
| palate | Scaper | 1.41 | 5.48E-17 | 1.17E-12 | 0.54 | 0.33 | 0.21 | 1.63 |
| palate | Pik3ip1 | 1.01 | 1.01E-16 | 2.17E-12 | 0.40 | 0.18 | 0.21 | 2.18 |
| palate | Arap2 | 1.49 | 1.26E-16 | 2.69E-12 | 0.31 | 0.13 | 0.18 | 2.34 |
| palate | Gca | 1.95 | 1.29E-16 | 2.76E-12 | 0.21 | 0.07 | 0.14 | 2.93 |
| palate | Pfn2 | 0.69 | 1.30E-16 | 2.77E-12 | 0.84 | 0.70 | 0.15 | 1.21 |
| palate | Pdpn | 1.00 | 2.15E-16 | 4.60E-12 | 0.58 | 0.36 | 0.22 | 1.63 |
| palate | Egfl6 | 1.38 | 3.64E-16 | 7.77E-12 | 0.37 | 0.17 | 0.19 | 2.10 |
| palate | Ctdspl | 0.74 | 4.94E-16 | 1.06E-11 | 0.70 | 0.48 | 0.23 | 1.47 |
| palate | Lifr | 1.39 | 4.95E-16 | 1.06E-11 | 0.28 | 0.12 | 0.17 | 2.42 |
| palate | Ptprf | 0.42 | 6.27E-16 | 1.34E-11 | 0.99 | 0.96 | 0.03 | 1.03 |
| palate | Hs6st2 | 0.73 | 8.91E-16 | 1.90E-11 | 0.75 | 0.53 | 0.22 | 1.41 |
| palate | Lmnb1 | 0.47 | 8.95E-16 | 1.91E-11 | 0.99 | 0.93 | 0.06 | 1.07 |
| palate | Egln3 | 1.26 | 1.22E-15 | 2.61E-11 | 0.37 | 0.18 | 0.19 | 2.05 |
| palate | C430049B03Rik | 1.50 | 1.37E-15 | 2.94E-11 | 0.31 | 0.14 | 0.17 | 2.22 |
| palate | Nebl | 0.58 | 2.17E-15 | 4.63E-11 | 0.63 | 0.39 | 0.24 | 1.62 |
| palate | Sox2 | 0.42 | 2.57E-15 | 5.49E-11 | 0.95 | 0.69 | 0.26 | 1.38 |
| palate | Fos | 1.35 | 3.04E-15 | 6.49E-11 | 0.27 | 0.11 | 0.16 | 2.48 |
| palate | Pon3 | 2.34 | 3.05E-15 | 6.51E-11 | 0.11 | 0.02 | 0.08 | 4.61 |
| palate | Rxfp1 | 1.94 | 3.73E-15 | 7.96E-11 | 0.16 | 0.05 | 0.11 | 3.43 |
| palate | Gadd45b | 1.14 | 5.82E-15 | 1.24E-10 | 0.39 | 0.20 | 0.19 | 1.99 |
| palate | Jade3 | 1.23 | 5.91E-15 | 1.26E-10 | 0.34 | 0.16 | 0.18 | 2.17 |
| palate | Dcaf4 | 0.97 | 6.10E-15 | 1.30E-10 | 0.54 | 0.34 | 0.20 | 1.59 |
| palate | Phldb3 | 1.64 | 6.16E-15 | 1.32E-10 | 0.22 | 0.08 | 0.14 | 2.76 |
| palate | Plch2 | 1.52 | 6.70E-15 | 1.43E-10 | 0.25 | 0.10 | 0.15 | 2.50 |
| palate | Dlc1 | 1.55 | 7.20E-15 | 1.54E-10 | 0.22 | 0.08 | 0.14 | 2.79 |
| palate | Lima1 | 0.74 | 7.76E-15 | 1.66E-10 | 0.68 | 0.47 | 0.21 | 1.45 |
| palate | Zswim8 | 0.88 | 8.27E-15 | 1.77E-10 | 0.58 | 0.37 | 0.21 | 1.57 |
| palate | Fosb | 1.51 | 8.29E-15 | 1.77E-10 | 0.11 | 0.02 | 0.08 | 4.61 |
| palate | Myl12a | 0.36 | 8.31E-15 | 1.78E-10 | 0.98 | 0.93 | 0.04 | 1.05 |
| palate | Serpinb6a | 0.74 | 1.66E-14 | 3.55E-10 | 0.68 | 0.47 | 0.21 | 1.45 |
| palate | Wnt4 | 0.32 | 1.72E-14 | 3.68E-10 | 0.94 | 0.70 | 0.24 | 1.35 |
| palate | Dpysl3 | 0.34 | 1.88E-14 | 4.02E-10 | 0.80 | 0.56 | 0.25 | 1.45 |
| palate | Bcl9l | 0.94 | 1.99E-14 | 4.24E-10 | 0.53 | 0.32 | 0.21 | 1.66 |
| palate | Zfp568 | 1.03 | 2.25E-14 | 4.80E-10 | 0.52 | 0.33 | 0.20 | 1.60 |
| palate | Tcf7l1 | 0.75 | 2.40E-14 | 5.13E-10 | 0.77 | 0.56 | 0.20 | 1.36 |
| palate | Ndufa11 | 0.35 | 2.41E-14 | 5.14E-10 | 0.99 | 0.95 | 0.04 | 1.04 |
| palate | Tceal9 | 0.27 | 2.61E-14 | 5.57E-10 | 1.00 | 0.99 | 0.01 | 1.01 |
| palate | Bmp7 | 0.76 | 3.04E-14 | 6.50E-10 | 0.60 | 0.40 | 0.21 | 1.52 |
| palate | Eif1a | 0.69 | 3.76E-14 | 8.03E-10 | 0.82 | 0.68 | 0.14 | 1.20 |
| palate | Pfdn1 | 0.36 | 3.81E-14 | 8.13E-10 | 0.98 | 0.94 | 0.04 | 1.04 |
| palate | Rest | 0.56 | 4.95E-14 | 1.06E-09 | 0.88 | 0.67 | 0.21 | 1.31 |
| palate | Sos1 | 0.83 | 5.15E-14 | 1.10E-09 | 0.59 | 0.39 | 0.20 | 1.51 |
| palate | Crip1 | 0.84 | 5.69E-14 | 1.21E-09 | 0.59 | 0.39 | 0.20 | 1.51 |
| palate | Rbbp7 | 0.51 | 5.75E-14 | 1.23E-09 | 0.93 | 0.85 | 0.08 | 1.10 |
| palate | Cgn | 0.86 | 6.46E-14 | 1.38E-09 | 0.57 | 0.38 | 0.19 | 1.50 |

**Supplementary Table 4. Marker genes for the annotated cell populations of integrated clustering of E11.5 control and NotchGOF λ-junction ectoderm cells.**

| | | | | | | | | |
|---|---|---|---|---|---|---|---|---|
| palate | Stard3nl | 0.79 | 9.30E-14 | 1.99E-09 | 0.70 | 0.53 | 0.17 | 1.33 |
| palate | Stard10 | 0.62 | 9.77E-14 | 2.09E-09 | 0.86 | 0.72 | 0.14 | 1.19 |
| palate | Endod1 | 1.24 | 9.79E-14 | 2.09E-09 | 0.32 | 0.15 | 0.17 | 2.11 |
| palate | Lyrm9 | 1.14 | 1.01E-13 | 2.15E-09 | 0.39 | 0.21 | 0.18 | 1.83 |
| palate | Klf6 | 1.02 | 1.09E-13 | 2.33E-09 | 0.43 | 0.23 | 0.20 | 1.87 |
| palate | Foxp4 | 0.79 | 1.13E-13 | 2.41E-09 | 0.56 | 0.36 | 0.20 | 1.55 |
| palate | Ppp1r14c | 2.17 | 1.24E-13 | 2.64E-09 | 0.14 | 0.04 | 0.10 | 3.41 |
| palate | Parp1 | 0.42 | 1.60E-13 | 3.42E-09 | 0.94 | 0.88 | 0.06 | 1.07 |
| palate | Erbb2 | 0.86 | 1.74E-13 | 3.72E-09 | 0.53 | 0.31 | 0.21 | 1.68 |
| palate | 4930523C07Rik | 1.67 | 1.75E-13 | 3.74E-09 | 0.23 | 0.09 | 0.14 | 2.47 |
| palate | Plet1 | 2.06 | 2.40E-13 | 5.14E-09 | 0.14 | 0.04 | 0.10 | 3.33 |
| palate | Ece1 | 0.77 | 2.58E-13 | 5.50E-09 | 0.65 | 0.45 | 0.20 | 1.44 |
| palate | Acadl | 0.75 | 2.85E-13 | 6.08E-09 | 0.62 | 0.42 | 0.21 | 1.49 |
| palate | Lrrk1 | 0.96 | 3.68E-13 | 7.86E-09 | 0.49 | 0.30 | 0.20 | 1.66 |
| palate | Tjp2 | 0.75 | 4.20E-13 | 8.98E-09 | 0.66 | 0.47 | 0.19 | 1.42 |
| palate | Pdlim1 | 0.32 | 4.31E-13 | 9.21E-09 | 0.84 | 0.64 | 0.21 | 1.32 |
| palate | Sh3bgrl3 | 0.37 | 4.42E-13 | 9.44E-09 | 0.98 | 0.91 | 0.07 | 1.07 |
| palate | Ank3 | 0.59 | 4.53E-13 | 9.68E-09 | 0.79 | 0.63 | 0.17 | 1.27 |
| palate | Fgfr2 | 0.51 | 5.35E-13 | 1.14E-08 | 0.70 | 0.46 | 0.24 | 1.52 |
| palate | Ccn2 | 1.74 | 6.17E-13 | 1.32E-08 | 0.21 | 0.08 | 0.13 | 2.51 |
| palate | Gls | 0.92 | 7.24E-13 | 1.55E-08 | 0.62 | 0.47 | 0.15 | 1.33 |
| palate | Itprid2 | 1.36 | 8.97E-13 | 1.92E-08 | 0.26 | 0.12 | 0.15 | 2.26 |
| palate | Tbrg1 | 0.47 | 9.12E-13 | 1.95E-08 | 0.93 | 0.84 | 0.08 | 1.10 |
| palate | Ywhab | 0.32 | 1.22E-12 | 2.60E-08 | 0.99 | 0.96 | 0.03 | 1.03 |
| palate | Lacc1 | 1.74 | 1.24E-12 | 2.64E-08 | 0.20 | 0.08 | 0.12 | 2.61 |
| palate | Itgb1 | 0.41 | 1.39E-12 | 2.96E-08 | 0.91 | 0.85 | 0.07 | 1.08 |
| palate | Kank1 | 0.88 | 1.51E-12 | 3.23E-08 | 0.47 | 0.28 | 0.19 | 1.68 |
| palate | Mbnl3 | 1.17 | 1.69E-12 | 3.61E-08 | 0.33 | 0.16 | 0.16 | 2.01 |
| palate | Gsto1 | 0.84 | 1.97E-12 | 4.22E-08 | 0.58 | 0.40 | 0.18 | 1.45 |
| palate | Rassf9 | 1.08 | 2.03E-12 | 4.34E-08 | 0.41 | 0.23 | 0.18 | 1.76 |
| palate | Krcc1 | 0.60 | 2.66E-12 | 5.68E-08 | 0.79 | 0.62 | 0.16 | 1.26 |
| palate | Sumo3 | 0.39 | 2.91E-12 | 6.21E-08 | 0.95 | 0.91 | 0.04 | 1.05 |
| palate | Bcam | 0.43 | 3.35E-12 | 7.15E-08 | 0.90 | 0.72 | 0.18 | 1.25 |
| palate | Anp32b | 0.26 | 3.93E-12 | 8.39E-08 | 1.00 | 0.99 | 0.01 | 1.01 |
| palate | Rassf10 | 0.86 | 4.68E-12 | 9.99E-08 | 0.43 | 0.24 | 0.19 | 1.80 |
| palate | Cmtm8 | 0.69 | 5.16E-12 | 1.10E-07 | 0.69 | 0.52 | 0.16 | 1.31 |
| palate | Ppp2r2b | 0.69 | 5.23E-12 | 1.12E-07 | 0.57 | 0.37 | 0.20 | 1.53 |
| palate | Spint2 | 0.29 | 6.46E-12 | 1.38E-07 | 0.99 | 0.97 | 0.02 | 1.02 |
| palate | Aplp2 | 0.30 | 8.34E-12 | 1.78E-07 | 0.95 | 0.92 | 0.03 | 1.04 |
| palate | Ubl4a | 0.68 | 9.67E-12 | 2.07E-07 | 0.72 | 0.57 | 0.15 | 1.26 |
| palate | Vps36 | 0.47 | 9.91E-12 | 2.12E-07 | 0.95 | 0.85 | 0.10 | 1.12 |
| palate | Elf1 | 0.60 | 1.00E-11 | 2.14E-07 | 0.68 | 0.50 | 0.18 | 1.37 |
| palate | Kctd11 | 1.42 | 1.12E-11 | 2.40E-07 | 0.20 | 0.08 | 0.12 | 2.55 |
| palate | Ubald1 | 0.74 | 1.21E-11 | 2.58E-07 | 0.57 | 0.38 | 0.19 | 1.50 |
| palate | 1810058I24Rik | 0.30 | 1.25E-11 | 2.67E-07 | 0.98 | 0.93 | 0.05 | 1.05 |
| palate | Lims1 | 0.68 | 1.27E-11 | 2.71E-07 | 0.68 | 0.52 | 0.16 | 1.31 |
| palate | 4930556M19Rik | 2.04 | 1.30E-11 | 2.77E-07 | 0.11 | 0.03 | 0.08 | 3.63 |
| palate | Myo1c | 1.62 | 1.35E-11 | 2.87E-07 | 0.20 | 0.08 | 0.12 | 2.48 |
| palate | St7 | 0.90 | 1.48E-11 | 3.15E-07 | 0.50 | 0.32 | 0.18 | 1.56 |
| palate | Tmem256 | 0.36 | 1.62E-11 | 3.46E-07 | 0.94 | 0.88 | 0.06 | 1.07 |
| palate | Cystm1 | 0.73 | 1.82E-11 | 3.89E-07 | 0.53 | 0.35 | 0.18 | 1.50 |
| palate | Pdcd10 | 0.50 | 1.92E-11 | 4.11E-07 | 0.81 | 0.70 | 0.11 | 1.16 |
| palate | Sri | 0.49 | 1.94E-11 | 4.14E-07 | 0.82 | 0.71 | 0.11 | 1.15 |
| palate | Hpcal1 | 0.60 | 1.95E-11 | 4.16E-07 | 0.65 | 0.46 | 0.19 | 1.42 |
| palate | Wnt7b | 0.49 | 2.38E-11 | 5.08E-07 | 0.33 | 0.16 | 0.17 | 2.09 |

**Supplementary Table 4. Marker genes for the annotated cell populations of integrated clustering of E11.5 control and NotchGOF λ-junction ectoderm cells.**

| | | | | | | | | |
|---|---|---|---|---|---|---|---|---|
| palate | Amotl1 | 0.47 | 2.38E-11 | 5.08E-07 | 0.83 | 0.69 | 0.14 | 1.20 |
| palate | Akt1 | 0.61 | 2.68E-11 | 5.72E-07 | 0.77 | 0.67 | 0.10 | 1.16 |
| palate | Ptpn14 | 0.92 | 2.74E-11 | 5.86E-07 | 0.40 | 0.22 | 0.18 | 1.79 |
| palate | Adgrg6 | 1.95 | 2.84E-11 | 6.08E-07 | 0.10 | 0.03 | 0.08 | 3.78 |
| palate | Arhgef16 | 1.10 | 2.89E-11 | 6.17E-07 | 0.37 | 0.21 | 0.16 | 1.77 |
| palate | Slc15a2 | 1.10 | 3.71E-11 | 7.93E-07 | 0.29 | 0.14 | 0.15 | 2.06 |
| palate | Etfa | 0.46 | 3.75E-11 | 8.02E-07 | 0.86 | 0.75 | 0.11 | 1.15 |
| palate | Odc1 | 0.45 | 3.98E-11 | 8.50E-07 | 0.92 | 0.83 | 0.09 | 1.11 |
| palate | Sntb1 | 1.63 | 5.35E-11 | 1.14E-06 | 0.17 | 0.07 | 0.11 | 2.62 |
| palate | Ptch1 | 0.68 | 5.48E-11 | 1.17E-06 | 0.53 | 0.35 | 0.19 | 1.53 |
| palate | Lrp2 | 0.98 | 6.51E-11 | 1.39E-06 | 0.38 | 0.21 | 0.16 | 1.76 |
| palate | Fkbp5 | 0.89 | 6.84E-11 | 1.46E-06 | 0.48 | 0.31 | 0.17 | 1.54 |
| palate | Cldn3 | 0.51 | 7.85E-11 | 1.68E-06 | 0.82 | 0.62 | 0.20 | 1.32 |
| palate | Prr15l | 1.21 | 8.14E-11 | 1.74E-06 | 0.28 | 0.14 | 0.14 | 2.02 |
| palate | Myh9 | 0.56 | 9.45E-11 | 2.02E-06 | 0.84 | 0.75 | 0.09 | 1.12 |
| palate | Rnf128 | 1.66 | 1.04E-10 | 2.22E-06 | 0.17 | 0.07 | 0.11 | 2.56 |
| palate | Cdca7 | 0.50 | 1.05E-10 | 2.24E-06 | 0.81 | 0.64 | 0.17 | 1.27 |
| palate | Srrm1 | 0.30 | 1.07E-10 | 2.29E-06 | 0.98 | 0.97 | 0.02 | 1.02 |
| palate | Atp6v1d | 0.55 | 1.14E-10 | 2.43E-06 | 0.79 | 0.67 | 0.12 | 1.18 |
| palate | Gab1 | 0.81 | 1.15E-10 | 2.46E-06 | 0.46 | 0.28 | 0.18 | 1.63 |
| palate | Ctnna1 | 0.38 | 1.37E-10 | 2.94E-06 | 0.93 | 0.87 | 0.06 | 1.07 |
| palate | Vmac | 1.04 | 1.38E-10 | 2.94E-06 | 0.30 | 0.15 | 0.15 | 1.95 |
| palate | Tcf15 | 1.96 | 1.48E-10 | 3.16E-06 | 0.12 | 0.04 | 0.08 | 3.18 |
| palate | Tap1 | 1.62 | 1.53E-10 | 3.27E-06 | 0.17 | 0.07 | 0.11 | 2.56 |
| palate | Hacd2 | 0.80 | 1.55E-10 | 3.30E-06 | 0.55 | 0.39 | 0.16 | 1.41 |
| palate | Hnrnpa3 | 0.26 | 1.64E-10 | 3.50E-06 | 1.00 | 1.00 | 0.01 | 1.01 |
| palate | Cnst | 1.07 | 1.67E-10 | 3.57E-06 | 0.28 | 0.14 | 0.14 | 2.02 |
| palate | Eppk1 | 1.40 | 1.67E-10 | 3.57E-06 | 0.19 | 0.08 | 0.11 | 2.43 |
| palate | Ndufb3 | 0.34 | 2.14E-10 | 4.57E-06 | 0.95 | 0.90 | 0.05 | 1.06 |
| palate | Cldn4 | 0.63 | 2.42E-10 | 5.16E-06 | 0.57 | 0.41 | 0.16 | 1.40 |
| palate | Mgat3 | 1.30 | 2.50E-10 | 5.34E-06 | 0.25 | 0.12 | 0.13 | 2.11 |
| palate | Ugt8a | 1.61 | 2.55E-10 | 5.44E-06 | 0.14 | 0.05 | 0.09 | 2.80 |
| palate | Anapc13 | 0.38 | 2.74E-10 | 5.85E-06 | 0.93 | 0.84 | 0.09 | 1.10 |
| palate | Prrg3 | 1.11 | 2.76E-10 | 5.89E-06 | 0.31 | 0.16 | 0.15 | 1.90 |
| palate | Kcnk1 | 0.51 | 2.80E-10 | 5.99E-06 | 0.72 | 0.58 | 0.14 | 1.25 |
| palate | Fignl1 | 0.74 | 3.24E-10 | 6.92E-06 | 0.61 | 0.47 | 0.14 | 1.29 |
| palate | Anxa5 | 0.47 | 3.33E-10 | 7.12E-06 | 0.79 | 0.62 | 0.17 | 1.28 |
| palate | Jup | 0.58 | 3.55E-10 | 7.58E-06 | 0.67 | 0.51 | 0.16 | 1.32 |
| palate | Snx31 | 1.80 | 3.99E-10 | 8.53E-06 | 0.12 | 0.04 | 0.08 | 3.16 |
| palate | Ranbp1 | 0.26 | 4.01E-10 | 8.56E-06 | 1.00 | 0.99 | 0.02 | 1.02 |
| palate | Id1 | 0.34 | 4.35E-10 | 9.30E-06 | 0.99 | 0.93 | 0.06 | 1.07 |
| palate | Sdc4 | 0.48 | 4.43E-10 | 9.45E-06 | 0.79 | 0.64 | 0.15 | 1.23 |
| palate | Sptlc3 | 1.09 | 4.48E-10 | 9.57E-06 | 0.18 | 0.07 | 0.11 | 2.53 |
| palate | Cbx6 | 0.80 | 4.56E-10 | 9.75E-06 | 0.52 | 0.36 | 0.16 | 1.43 |
| palate | Klc3 | 0.47 | 4.70E-10 | 1.00E-05 | 0.57 | 0.39 | 0.18 | 1.47 |
| palate | Nr2f6 | 0.47 | 5.18E-10 | 1.11E-05 | 0.86 | 0.74 | 0.12 | 1.16 |
| palate | Esrp1 | 0.48 | 6.16E-10 | 1.32E-05 | 0.77 | 0.64 | 0.14 | 1.22 |
| palate | Cxxc4 | 0.74 | 6.32E-10 | 1.35E-05 | 0.40 | 0.25 | 0.16 | 1.64 |
| palate | Tmem109 | 0.86 | 8.06E-10 | 1.72E-05 | 0.41 | 0.25 | 0.16 | 1.62 |
| palate | Cebpa | 1.41 | 9.25E-10 | 1.98E-05 | 0.19 | 0.08 | 0.11 | 2.37 |
| palate | Smarca5 | 0.25 | 9.33E-10 | 1.99E-05 | 0.98 | 0.97 | 0.01 | 1.01 |
| palate | Erbb3 | 0.51 | 9.47E-10 | 2.02E-05 | 0.71 | 0.53 | 0.17 | 1.33 |
| palate | Shisa2 | 1.19 | 9.62E-10 | 2.06E-05 | 0.22 | 0.10 | 0.12 | 2.14 |
| palate | Larp7 | 0.42 | 9.67E-10 | 2.07E-05 | 0.88 | 0.77 | 0.11 | 1.14 |
| palate | Nab2 | 1.04 | 1.10E-09 | 2.34E-05 | 0.34 | 0.20 | 0.15 | 1.74 |

**Supplementary Table 4. Marker genes for the annotated cell populations of integrated clustering of E11.5 control and NotchGOF λ-junction ectoderm cells.**

| | | | | | | | | |
|---|---|---|---|---|---|---|---|---|
| palate | Fermt1 | 0.38 | 1.12E-09 | 2.40E-05 | 0.79 | 0.54 | 0.25 | 1.45 |
| palate | Tln1 | 0.49 | 1.17E-09 | 2.50E-05 | 0.77 | 0.65 | 0.12 | 1.18 |
| palate | Efna1 | 0.58 | 1.50E-09 | 3.20E-05 | 0.57 | 0.39 | 0.18 | 1.46 |
| palate | Slfn9 | 0.85 | 1.74E-09 | 3.71E-05 | 0.40 | 0.24 | 0.16 | 1.66 |
| palate | Ifitm2 | 0.27 | 2.21E-09 | 4.73E-05 | 0.98 | 0.93 | 0.05 | 1.06 |
| palate | Rab15 | 0.59 | 2.39E-09 | 5.11E-05 | 0.60 | 0.41 | 0.19 | 1.45 |
| palate | Lmna | 0.88 | 2.59E-09 | 5.53E-05 | 0.33 | 0.18 | 0.15 | 1.81 |
| palate | Amer1 | 0.51 | 2.65E-09 | 5.65E-05 | 0.72 | 0.58 | 0.14 | 1.25 |
| palate | Usp43 | 1.53 | 2.85E-09 | 6.09E-05 | 0.14 | 0.05 | 0.09 | 2.69 |
| palate | Isl1 | 0.31 | 2.93E-09 | 6.26E-05 | 0.92 | 0.75 | 0.18 | 1.23 |
| palate | Hip1r | 0.96 | 3.00E-09 | 6.42E-05 | 0.36 | 0.21 | 0.15 | 1.71 |
| palate | Slc44a2 | 0.69 | 3.01E-09 | 6.42E-05 | 0.54 | 0.39 | 0.15 | 1.39 |
| palate | Hmg20b | 0.46 | 3.32E-09 | 7.09E-05 | 0.85 | 0.76 | 0.09 | 1.12 |
| palate | Cxcl12 | 1.30 | 3.41E-09 | 7.29E-05 | 0.25 | 0.13 | 0.12 | 1.95 |
| palate | Purb | 0.34 | 4.47E-09 | 9.55E-05 | 0.96 | 0.89 | 0.07 | 1.08 |
| palate | Flywch2 | 0.65 | 4.64E-09 | 9.91E-05 | 0.62 | 0.48 | 0.14 | 1.28 |
| palate | Aqp11 | 1.68 | 4.71E-09 | 1.01E-04 | 0.14 | 0.05 | 0.09 | 2.64 |
| palate | Kras | 0.34 | 5.78E-09 | 1.24E-04 | 0.91 | 0.83 | 0.08 | 1.09 |
| palate | Brwd3 | 0.62 | 5.98E-09 | 1.28E-04 | 0.65 | 0.50 | 0.14 | 1.29 |
| palate | Stx3 | 0.63 | 6.06E-09 | 1.30E-04 | 0.59 | 0.43 | 0.16 | 1.38 |
| palate | Dek | 0.31 | 6.18E-09 | 1.32E-04 | 0.99 | 0.93 | 0.07 | 1.07 |
| palate | Cdc42ep5 | 0.49 | 6.80E-09 | 1.45E-04 | 0.73 | 0.59 | 0.15 | 1.25 |
| palate | Hes1 | 0.35 | 7.60E-09 | 1.62E-04 | 0.86 | 0.72 | 0.13 | 1.19 |
| palate | Lrrfip1 | 0.68 | 7.90E-09 | 1.69E-04 | 0.62 | 0.50 | 0.12 | 1.24 |
| palate | Sh2d4a | 1.54 | 9.04E-09 | 1.93E-04 | 0.12 | 0.04 | 0.08 | 2.81 |
| palate | Arhgap23 | 1.04 | 1.04E-08 | 2.22E-04 | 0.23 | 0.11 | 0.12 | 2.07 |
| palate | Syngr1 | 0.93 | 1.05E-08 | 2.23E-04 | 0.36 | 0.22 | 0.14 | 1.67 |
| palate | Emp1 | 0.46 | 1.09E-08 | 2.32E-04 | 0.31 | 0.17 | 0.15 | 1.86 |
| palate | Tnk1 | 1.08 | 1.10E-08 | 2.35E-04 | 0.25 | 0.13 | 0.12 | 1.95 |
| palate | Bmp8b | 1.83 | 1.15E-08 | 2.45E-04 | 0.10 | 0.03 | 0.07 | 3.09 |
| palate | Ptprm | 1.42 | 1.27E-08 | 2.70E-04 | 0.20 | 0.10 | 0.11 | 2.08 |
| palate | Pdcd4 | 0.31 | 1.33E-08 | 2.83E-04 | 0.91 | 0.78 | 0.13 | 1.17 |
| palate | Col4a1 | 0.39 | 1.34E-08 | 2.86E-04 | 0.80 | 0.68 | 0.12 | 1.18 |
| palate | Bex4 | 0.29 | 1.50E-08 | 3.20E-04 | 0.98 | 0.93 | 0.05 | 1.05 |
| palate | Egr1 | 0.79 | 1.61E-08 | 3.44E-04 | 0.21 | 0.10 | 0.11 | 2.15 |
| palate | Serpini1 | 1.20 | 1.62E-08 | 3.45E-04 | 0.12 | 0.04 | 0.08 | 2.81 |
| palate | S100a1 | 1.48 | 1.63E-08 | 3.49E-04 | 0.15 | 0.06 | 0.09 | 2.44 |
| palate | Aktip | 0.59 | 1.65E-08 | 3.53E-04 | 0.62 | 0.46 | 0.16 | 1.34 |
| palate | Myzap | 1.59 | 1.66E-08 | 3.55E-04 | 0.12 | 0.04 | 0.08 | 2.85 |
| palate | Tmem79 | 1.52 | 1.76E-08 | 3.76E-04 | 0.17 | 0.08 | 0.10 | 2.27 |
| palate | Cntnap2 | 1.06 | 1.95E-08 | 4.16E-04 | 0.26 | 0.14 | 0.12 | 1.85 |
| palate | Ccser1 | 1.23 | 2.03E-08 | 4.33E-04 | 0.17 | 0.07 | 0.10 | 2.33 |
| palate | Rnh1 | 0.36 | 2.15E-08 | 4.58E-04 | 0.90 | 0.77 | 0.13 | 1.16 |
| palate | Tuft1 | 1.14 | 2.26E-08 | 4.82E-04 | 0.17 | 0.07 | 0.10 | 2.33 |
| palate | Dock9 | 0.95 | 3.00E-08 | 6.40E-04 | 0.33 | 0.19 | 0.13 | 1.68 |
| palate | Dap | 0.49 | 3.29E-08 | 7.02E-04 | 0.78 | 0.68 | 0.10 | 1.15 |
| palate | Dag1 | 0.50 | 3.31E-08 | 7.08E-04 | 0.73 | 0.63 | 0.10 | 1.16 |
| palate | H2afj | 0.30 | 4.19E-08 | 8.94E-04 | 0.96 | 0.90 | 0.06 | 1.07 |
| palate | Pkp4 | 0.55 | 4.19E-08 | 8.94E-04 | 0.70 | 0.58 | 0.12 | 1.21 |
| palate | Llgl2 | 0.86 | 4.40E-08 | 9.40E-04 | 0.31 | 0.18 | 0.13 | 1.74 |
| palate | Car13 | 1.76 | 4.51E-08 | 9.64E-04 | 0.12 | 0.04 | 0.07 | 2.66 |
| palate | Inf2 | 1.53 | 4.53E-08 | 9.68E-04 | 0.11 | 0.04 | 0.07 | 2.79 |
| palate | Impdh1 | 0.97 | 4.54E-08 | 9.71E-04 | 0.29 | 0.17 | 0.12 | 1.73 |
| palate | Cldn6 | 0.29 | 4.57E-08 | 9.76E-04 | 0.96 | 0.87 | 0.09 | 1.10 |
| palate | Rab14 | 0.25 | 5.13E-08 | 1.10E-03 | 0.94 | 0.93 | 0.01 | 1.01 |

**Supplementary Table 4. Marker genes for the annotated cell populations of integrated clustering of E11.5 control and NotchGOF λ-junction ectoderm cells.**

| | | | | | | | | |
|---|---|---|---|---|---|---|---|---|
| palate | Mark2 | 0.49 | 5.38E-08 | 1.15E-03 | 0.72 | 0.58 | 0.14 | 1.25 |
| palate | Irf6 | 0.27 | 5.79E-08 | 1.24E-03 | 0.62 | 0.40 | 0.21 | 1.52 |
| palate | Erbin | 0.47 | 5.83E-08 | 1.25E-03 | 0.71 | 0.56 | 0.14 | 1.25 |
| palate | Tmem54 | 0.63 | 6.87E-08 | 1.47E-03 | 0.61 | 0.47 | 0.14 | 1.29 |
| palate | Pcbd1 | 0.88 | 7.37E-08 | 1.57E-03 | 0.36 | 0.23 | 0.13 | 1.58 |
| palate | Lypd3 | 1.02 | 7.63E-08 | 1.63E-03 | 0.21 | 0.10 | 0.11 | 2.06 |
| palate | Morc4 | 0.99 | 7.83E-08 | 1.67E-03 | 0.26 | 0.14 | 0.12 | 1.84 |
| palate | Tfap2c | 0.69 | 7.89E-08 | 1.69E-03 | 0.36 | 0.21 | 0.14 | 1.66 |
| palate | Prkci | 0.59 | 8.03E-08 | 1.72E-03 | 0.57 | 0.43 | 0.14 | 1.33 |
| palate | Nabp1 | 1.52 | 8.06E-08 | 1.72E-03 | 0.14 | 0.06 | 0.08 | 2.41 |
| palate | Xpa | 0.57 | 8.44E-08 | 1.80E-03 | 0.63 | 0.47 | 0.16 | 1.33 |
| palate | Tead1 | 0.37 | 9.03E-08 | 1.93E-03 | 0.86 | 0.76 | 0.10 | 1.12 |
| palate | Coro1c | 0.66 | 9.60E-08 | 2.05E-03 | 0.46 | 0.32 | 0.14 | 1.43 |
| palate | Mafb | 0.71 | 1.00E-07 | 2.15E-03 | 0.28 | 0.15 | 0.13 | 1.84 |
| palate | Msh2 | 0.45 | 1.02E-07 | 2.18E-03 | 0.79 | 0.69 | 0.10 | 1.14 |
| palate | Bcat1 | 1.15 | 1.03E-07 | 2.21E-03 | 0.24 | 0.13 | 0.11 | 1.86 |
| palate | Reep6 | 1.46 | 1.04E-07 | 2.23E-03 | 0.17 | 0.08 | 0.09 | 2.12 |
| palate | Etl4 | 0.43 | 1.11E-07 | 2.37E-03 | 0.71 | 0.57 | 0.14 | 1.24 |
| palate | Tnfaip8 | 0.48 | 1.12E-07 | 2.39E-03 | 0.71 | 0.56 | 0.15 | 1.27 |
| palate | Plpp2 | 0.59 | 1.34E-07 | 2.85E-03 | 0.53 | 0.37 | 0.15 | 1.42 |
| palate | Efna5 | 0.50 | 1.35E-07 | 2.88E-03 | 0.65 | 0.52 | 0.13 | 1.25 |
| palate | Bcl2l11 | 0.48 | 1.45E-07 | 3.09E-03 | 0.53 | 0.38 | 0.15 | 1.40 |
| palate | Nynrin | 0.54 | 1.45E-07 | 3.10E-03 | 0.66 | 0.53 | 0.13 | 1.25 |
| palate | Capn1 | 1.61 | 1.54E-07 | 3.29E-03 | 0.12 | 0.05 | 0.07 | 2.60 |
| palate | Fam3c | 0.55 | 1.56E-07 | 3.33E-03 | 0.59 | 0.47 | 0.12 | 1.27 |
| palate | Adamts15 | 1.53 | 1.66E-07 | 3.54E-03 | 0.11 | 0.04 | 0.07 | 2.66 |
| palate | Synj2 | 1.72 | 1.67E-07 | 3.56E-03 | 0.13 | 0.05 | 0.08 | 2.50 |
| palate | Creb3l2 | 0.57 | 1.96E-07 | 4.19E-03 | 0.47 | 0.32 | 0.15 | 1.48 |
| palate | Shroom4 | 1.32 | 2.00E-07 | 4.27E-03 | 0.14 | 0.06 | 0.08 | 2.37 |
| palate | Ajuba | 0.56 | 2.05E-07 | 4.39E-03 | 0.57 | 0.42 | 0.15 | 1.35 |
| palate | Reep5 | 0.43 | 2.09E-07 | 4.46E-03 | 0.69 | 0.55 | 0.13 | 1.24 |
| palate | Naa38 | 0.38 | 2.10E-07 | 4.49E-03 | 0.85 | 0.76 | 0.10 | 1.13 |
| palate | Arhgap8 | 1.07 | 2.12E-07 | 4.54E-03 | 0.27 | 0.16 | 0.11 | 1.72 |
| palate | Tagln2 | 0.70 | 2.31E-07 | 4.93E-03 | 0.41 | 0.27 | 0.14 | 1.51 |
| palate | Mdc1 | 0.62 | 2.57E-07 | 5.49E-03 | 0.51 | 0.37 | 0.14 | 1.39 |
| palate | Nbeal1 | 0.51 | 2.80E-07 | 5.98E-03 | 0.62 | 0.48 | 0.14 | 1.30 |
| palate | Cyb5r1 | 0.69 | 2.84E-07 | 6.06E-03 | 0.42 | 0.28 | 0.14 | 1.49 |
| palate | Tjp1 | 0.44 | 2.86E-07 | 6.12E-03 | 0.74 | 0.63 | 0.11 | 1.17 |
| palate | Sh3bgrl2 | 1.43 | 2.95E-07 | 6.30E-03 | 0.16 | 0.08 | 0.09 | 2.13 |
| palate | Arhgap18 | 0.82 | 2.97E-07 | 6.35E-03 | 0.33 | 0.20 | 0.13 | 1.65 |
| palate | Epb41l4b | 1.43 | 3.10E-07 | 6.62E-03 | 0.19 | 0.10 | 0.10 | 2.00 |
| palate | Capn2 | 0.62 | 3.15E-07 | 6.72E-03 | 0.54 | 0.39 | 0.15 | 1.37 |
| palate | Zfp711 | 0.76 | 3.34E-07 | 7.13E-03 | 0.33 | 0.19 | 0.13 | 1.68 |
| palate | Fam83h | 1.27 | 3.47E-07 | 7.41E-03 | 0.18 | 0.09 | 0.09 | 2.06 |
| palate | Mprip | 0.36 | 3.55E-07 | 7.59E-03 | 0.80 | 0.73 | 0.07 | 1.10 |
| palate | Tk1 | 0.50 | 3.57E-07 | 7.62E-03 | 0.60 | 0.45 | 0.15 | 1.35 |
| palate | Cryl1 | 0.78 | 3.72E-07 | 7.95E-03 | 0.40 | 0.27 | 0.13 | 1.50 |
| palate | Dnmt1 | 0.38 | 4.00E-07 | 8.54E-03 | 0.83 | 0.73 | 0.10 | 1.14 |
| palate | Fam241a | 1.12 | 4.28E-07 | 9.15E-03 | 0.20 | 0.11 | 0.10 | 1.94 |
| palate | Thrap3 | 0.30 | 4.32E-07 | 9.23E-03 | 0.96 | 0.92 | 0.04 | 1.04 |
| palate | Mme | 0.68 | 4.47E-07 | 9.55E-03 | 0.30 | 0.18 | 0.12 | 1.69 |
| palate | Snx5 | 0.28 | 4.59E-07 | 9.80E-03 | 0.89 | 0.82 | 0.07 | 1.09 |
| palate | Sestd1 | 0.58 | 4.78E-07 | 1.02E-02 | 0.59 | 0.46 | 0.13 | 1.28 |
| palate | Tmed2 | 0.25 | 4.80E-07 | 1.02E-02 | 0.96 | 0.92 | 0.04 | 1.04 |
| palate | Cenpb | 0.39 | 4.96E-07 | 1.06E-02 | 0.80 | 0.70 | 0.10 | 1.15 |

**Supplementary Table 4. Marker genes for the annotated cell populations of integrated clustering of E11.5 control and NotchGOF λ-junction ectoderm cells.**

| | | | | | | | | |
|---|---|---|---|---|---|---|---|---|
| palate | Plagl2 | 0.61 | 5.17E-07 | 1.10E-02 | 0.59 | 0.47 | 0.12 | 1.25 |
| palate | Jarid2 | 0.45 | 5.34E-07 | 1.14E-02 | 0.66 | 0.54 | 0.11 | 1.21 |
| palate | Fzd1 | 0.91 | 5.57E-07 | 1.19E-02 | 0.29 | 0.17 | 0.12 | 1.70 |
| palate | Plcl1 | 1.12 | 6.12E-07 | 1.31E-02 | 0.11 | 0.04 | 0.07 | 2.65 |
| palate | Stk26 | 0.46 | 6.32E-07 | 1.35E-02 | 0.63 | 0.46 | 0.16 | 1.35 |
| palate | Six6 | 0.59 | 6.37E-07 | 1.36E-02 | 0.42 | 0.27 | 0.14 | 1.51 |
| palate | Trim29 | 0.62 | 6.72E-07 | 1.44E-02 | 0.32 | 0.18 | 0.13 | 1.72 |
| palate | 9130017K11Rik | 1.80 | 6.78E-07 | 1.45E-02 | 0.10 | 0.04 | 0.06 | 2.62 |
| palate | Tgm1 | 1.11 | 6.93E-07 | 1.48E-02 | 0.12 | 0.05 | 0.07 | 2.47 |
| palate | Colgalt1 | 0.56 | 7.02E-07 | 1.50E-02 | 0.59 | 0.45 | 0.13 | 1.29 |
| palate | Tbc1d4 | 1.62 | 7.30E-07 | 1.56E-02 | 0.18 | 0.09 | 0.09 | 1.95 |
| palate | Capn6 | 0.43 | 8.02E-07 | 1.71E-02 | 0.59 | 0.44 | 0.15 | 1.34 |
| palate | Ccdc18 | 0.69 | 8.91E-07 | 1.90E-02 | 0.51 | 0.38 | 0.13 | 1.34 |
| palate | Fn1 | 0.38 | 9.21E-07 | 1.97E-02 | 0.70 | 0.58 | 0.12 | 1.20 |
| palate | Mcm5 | 0.38 | 9.42E-07 | 2.01E-02 | 0.77 | 0.63 | 0.15 | 1.23 |
| palate | Timeless | 0.57 | 9.74E-07 | 2.08E-02 | 0.59 | 0.46 | 0.13 | 1.28 |
| palate | Smagp | 1.34 | 9.76E-07 | 2.08E-02 | 0.13 | 0.06 | 0.08 | 2.32 |
| palate | Timp2 | 0.54 | 1.00E-06 | 2.14E-02 | 0.64 | 0.51 | 0.14 | 1.27 |
| palate | Ssna1 | 0.29 | 1.15E-06 | 2.46E-02 | 0.89 | 0.82 | 0.07 | 1.09 |
| palate | Pik3cb | 1.00 | 1.23E-06 | 2.62E-02 | 0.25 | 0.15 | 0.11 | 1.72 |
| palate | Hadh | 0.32 | 1.25E-06 | 2.66E-02 | 0.85 | 0.76 | 0.09 | 1.12 |
| palate | Atic | 0.36 | 1.25E-06 | 2.68E-02 | 0.82 | 0.68 | 0.14 | 1.20 |
| palate | Tcim | 0.89 | 1.27E-06 | 2.70E-02 | 0.39 | 0.26 | 0.12 | 1.46 |
| palate | Shtn1 | 0.58 | 1.28E-06 | 2.73E-02 | 0.31 | 0.20 | 0.12 | 1.59 |
| palate | Elovl6 | 0.49 | 1.29E-06 | 2.75E-02 | 0.64 | 0.52 | 0.12 | 1.24 |
| palate | Hdhd3 | 0.83 | 1.31E-06 | 2.80E-02 | 0.33 | 0.21 | 0.12 | 1.55 |
| palate | Msn | 0.89 | 1.38E-06 | 2.94E-02 | 0.25 | 0.14 | 0.10 | 1.73 |
| palate | Nrarp | 0.29 | 1.55E-06 | 3.32E-02 | 0.77 | 0.60 | 0.18 | 1.29 |
| palate | Spc24 | 0.40 | 1.57E-06 | 3.35E-02 | 0.80 | 0.65 | 0.14 | 1.22 |
| palate | Wsb1 | 0.38 | 1.59E-06 | 3.40E-02 | 0.76 | 0.66 | 0.10 | 1.16 |
| palate | St14 | 0.60 | 1.64E-06 | 3.49E-02 | 0.59 | 0.47 | 0.11 | 1.23 |
| palate | Ifit2 | 1.34 | 1.64E-06 | 3.50E-02 | 0.11 | 0.05 | 0.07 | 2.40 |
| palate | Sh3rf1 | 0.57 | 1.66E-06 | 3.55E-02 | 0.46 | 0.32 | 0.14 | 1.43 |
| palate | Ccdc112 | 0.40 | 1.75E-06 | 3.73E-02 | 0.75 | 0.64 | 0.11 | 1.17 |
| palate | Fzd8 | 0.49 | 1.79E-06 | 3.83E-02 | 0.43 | 0.30 | 0.14 | 1.47 |
| palate | Orc1 | 0.99 | 1.91E-06 | 4.07E-02 | 0.21 | 0.12 | 0.10 | 1.83 |
| palate | Map3k20 | 0.83 | 1.93E-06 | 4.13E-02 | 0.29 | 0.18 | 0.11 | 1.62 |
| palate | Metrnl | 0.75 | 1.96E-06 | 4.19E-02 | 0.29 | 0.17 | 0.12 | 1.68 |
| palate | Pla2g4a | 0.92 | 1.97E-06 | 4.22E-02 | 0.19 | 0.10 | 0.09 | 1.94 |
| palate | Ahdc1 | 0.70 | 1.98E-06 | 4.22E-02 | 0.34 | 0.22 | 0.12 | 1.54 |
| palate | Sh3d19 | 0.52 | 2.18E-06 | 4.65E-02 | 0.48 | 0.34 | 0.15 | 1.43 |
| palate | Fkbp9 | 0.57 | 2.28E-06 | 4.87E-02 | 0.54 | 0.42 | 0.12 | 1.28 |
| fusion | Fgf8 | 3.79 | 5.37E-150 | 1.15E-145 | 0.88 | 0.24 | 0.63 | 3.62 |
| fusion | Cdkn2b | 4.19 | 4.87E-142 | 1.04E-137 | 0.55 | 0.07 | 0.48 | 7.73 |
| fusion | Wnt5a | 2.96 | 1.19E-132 | 2.54E-128 | 0.95 | 0.40 | 0.55 | 2.38 |
| fusion | Gad1 | 3.71 | 6.40E-129 | 1.37E-124 | 0.40 | 0.04 | 0.37 | 11.22 |
| fusion | Cldn1 | 3.75 | 5.18E-126 | 1.11E-121 | 0.54 | 0.08 | 0.46 | 6.97 |
| fusion | Cdkn1a | 3.41 | 3.63E-122 | 7.75E-118 | 0.91 | 0.36 | 0.55 | 2.54 |
| fusion | Itm2b | 2.47 | 9.75E-121 | 2.08E-116 | 1.00 | 0.93 | 0.07 | 1.07 |
| fusion | Runx1 | 2.91 | 1.47E-119 | 3.14E-115 | 0.96 | 0.49 | 0.47 | 1.96 |
| fusion | Gcg | 6.74 | 4.80E-118 | 1.03E-113 | 0.19 | 0.00 | 0.19 | 48.00 |
| fusion | Inhba | 3.84 | 3.46E-114 | 7.39E-110 | 0.48 | 0.07 | 0.41 | 7.24 |
| fusion | Fst | 4.85 | 5.94E-114 | 1.27E-109 | 0.40 | 0.04 | 0.36 | 9.18 |
| fusion | Msx1 | 2.25 | 6.25E-108 | 1.33E-103 | 0.99 | 0.72 | 0.28 | 1.38 |
| fusion | Tgfb1 | 2.81 | 1.67E-105 | 3.57E-101 | 0.64 | 0.14 | 0.50 | 4.64 |

**Supplementary Table 4. Marker genes for the annotated cell populations of integrated clustering of E11.5 control and NotchGOF λ-junction ectoderm cells.**

| | | | | | | | | |
|---|---|---|---|---|---|---|---|---|
| fusion | Cdkn2a | 4.96 | 2.32E-102 | 4.95E-98 | 0.27 | 0.02 | 0.25 | 15.06 |
| fusion | Gadd45g | 3.34 | 2.90E-99 | 6.19E-95 | 0.81 | 0.32 | 0.50 | 2.58 |
| fusion | Fabp5 | 1.53 | 6.22E-97 | 1.33E-92 | 1.00 | 0.99 | 0.01 | 1.01 |
| fusion | Notum | 3.54 | 9.39E-95 | 2.01E-90 | 0.38 | 0.05 | 0.34 | 8.00 |
| fusion | Pkp1 | 3.05 | 1.85E-92 | 3.95E-88 | 0.47 | 0.08 | 0.39 | 5.96 |
| fusion | Bambi | 3.13 | 1.68E-90 | 3.58E-86 | 0.88 | 0.46 | 0.41 | 1.89 |
| fusion | Gja1 | 2.03 | 7.33E-89 | 1.57E-84 | 0.97 | 0.69 | 0.28 | 1.41 |
| fusion | Tgfb2 | 2.67 | 7.62E-89 | 1.63E-84 | 0.94 | 0.56 | 0.38 | 1.67 |
| fusion | Zfp385c | 2.97 | 6.71E-85 | 1.43E-80 | 0.50 | 0.10 | 0.40 | 4.98 |
| fusion | Lmo2 | 2.39 | 1.35E-84 | 2.89E-80 | 0.76 | 0.27 | 0.49 | 2.81 |
| fusion | Sfn | 1.81 | 5.12E-84 | 1.09E-79 | 0.99 | 0.69 | 0.31 | 1.45 |
| fusion | Sp5 | 4.52 | 6.34E-84 | 1.35E-79 | 0.26 | 0.02 | 0.24 | 11.77 |
| fusion | Lratd1 | 3.05 | 9.60E-82 | 2.05E-77 | 0.51 | 0.11 | 0.40 | 4.77 |
| fusion | Lgalsl | 2.60 | 5.70E-80 | 1.22E-75 | 0.84 | 0.45 | 0.40 | 1.89 |
| fusion | Igfbp5 | 1.67 | 2.53E-79 | 5.40E-75 | 1.00 | 0.96 | 0.04 | 1.04 |
| fusion | Dlx2 | 2.12 | 7.10E-78 | 1.52E-73 | 0.82 | 0.32 | 0.51 | 2.60 |
| fusion | Aopep | 1.94 | 5.60E-76 | 1.20E-71 | 0.92 | 0.60 | 0.32 | 1.54 |
| fusion | Atxn1 | 3.35 | 3.15E-75 | 6.72E-71 | 0.45 | 0.09 | 0.36 | 4.91 |
| fusion | Gpr87 | 4.24 | 3.25E-75 | 6.95E-71 | 0.24 | 0.02 | 0.22 | 11.38 |
| fusion | Gal | 5.71 | 1.90E-74 | 4.06E-70 | 0.23 | 0.02 | 0.21 | 11.55 |
| fusion | Foxn3 | 1.73 | 5.89E-74 | 1.26E-69 | 0.93 | 0.70 | 0.22 | 1.31 |
| fusion | Glcci1 | 2.30 | 6.11E-74 | 1.31E-69 | 0.86 | 0.48 | 0.38 | 1.78 |
| fusion | S100a11 | 1.10 | 9.75E-71 | 2.08E-66 | 1.00 | 0.90 | 0.10 | 1.11 |
| fusion | Ambn | 7.85 | 7.27E-69 | 1.55E-64 | 0.11 | 0.00 | 0.11 | 55.00 |
| fusion | Pik3r1 | 1.82 | 1.41E-68 | 3.01E-64 | 0.92 | 0.58 | 0.35 | 1.60 |
| fusion | Adamts9 | 2.74 | 3.30E-66 | 7.05E-62 | 0.64 | 0.24 | 0.41 | 2.74 |
| fusion | Krt14 | 1.97 | 9.29E-66 | 1.98E-61 | 0.88 | 0.44 | 0.44 | 2.00 |
| fusion | Lef1 | 2.21 | 9.99E-66 | 2.13E-61 | 0.77 | 0.36 | 0.41 | 2.16 |
| fusion | Trp63 | 1.51 | 4.35E-65 | 9.29E-61 | 0.93 | 0.49 | 0.44 | 1.91 |
| fusion | Tspan7 | 1.90 | 2.22E-64 | 4.74E-60 | 0.82 | 0.45 | 0.37 | 1.81 |
| fusion | Rb1 | 2.28 | 6.48E-64 | 1.38E-59 | 0.68 | 0.27 | 0.42 | 2.55 |
| fusion | Cd2ap | 1.40 | 8.34E-64 | 1.78E-59 | 0.96 | 0.80 | 0.16 | 1.21 |
| fusion | Tafa2 | 2.71 | 5.57E-60 | 1.19E-55 | 0.45 | 0.11 | 0.34 | 4.14 |
| fusion | Baiap2 | 2.45 | 1.98E-58 | 4.23E-54 | 0.57 | 0.20 | 0.38 | 2.92 |
| fusion | Krt5 | 2.07 | 2.12E-58 | 4.53E-54 | 0.73 | 0.32 | 0.42 | 2.32 |
| fusion | Spock2 | 3.47 | 6.53E-58 | 1.40E-53 | 0.44 | 0.11 | 0.33 | 3.99 |
| fusion | Sox11 | 0.96 | 2.16E-57 | 4.62E-53 | 1.00 | 0.99 | 0.00 | 1.00 |
| fusion | Wnt6 | 1.32 | 3.85E-57 | 8.23E-53 | 0.84 | 0.35 | 0.49 | 2.43 |
| fusion | Clta | 0.89 | 2.22E-56 | 4.75E-52 | 0.99 | 0.98 | 0.01 | 1.01 |
| fusion | Eef1a1 | 0.38 | 9.36E-56 | 2.00E-51 | 1.00 | 1.00 | 0.00 | 1.00 |
| fusion | Diras2 | 2.99 | 7.52E-55 | 1.61E-50 | 0.36 | 0.07 | 0.28 | 4.82 |
| fusion | Epha4 | 1.51 | 1.49E-54 | 3.17E-50 | 0.87 | 0.54 | 0.33 | 1.61 |
| fusion | Krt28 | 6.43 | 2.24E-53 | 4.78E-49 | 0.11 | 0.00 | 0.10 | 26.50 |
| fusion | Tmtc1 | 2.81 | 2.81E-53 | 6.00E-49 | 0.29 | 0.05 | 0.24 | 5.92 |
| fusion | Rfk | 1.61 | 6.03E-52 | 1.29E-47 | 0.85 | 0.50 | 0.35 | 1.69 |
| fusion | Fam49a | 2.80 | 3.27E-51 | 6.99E-47 | 0.37 | 0.09 | 0.28 | 4.34 |
| fusion | Sat1 | 2.25 | 2.50E-50 | 5.34E-46 | 0.64 | 0.28 | 0.36 | 2.26 |
| fusion | Ly6h | 1.80 | 8.47E-50 | 1.81E-45 | 0.80 | 0.48 | 0.32 | 1.67 |
| fusion | Frmd6 | 2.22 | 1.08E-49 | 2.31E-45 | 0.57 | 0.21 | 0.36 | 2.67 |
| fusion | Dcbld1 | 1.85 | 2.53E-49 | 5.39E-45 | 0.72 | 0.38 | 0.35 | 1.93 |
| fusion | Palld | 1.49 | 2.93E-49 | 6.26E-45 | 0.85 | 0.63 | 0.22 | 1.34 |
| fusion | Arid5b | 1.62 | 5.48E-47 | 1.17E-42 | 0.81 | 0.54 | 0.27 | 1.50 |
| fusion | Rpl7 | 0.37 | 6.77E-47 | 1.45E-42 | 1.00 | 1.00 | 0.00 | 1.00 |
| fusion | Rasl11b | 1.93 | 8.42E-47 | 1.80E-42 | 0.62 | 0.26 | 0.37 | 2.42 |
| fusion | Dkk4 | 4.05 | 2.50E-46 | 5.34E-42 | 0.18 | 0.02 | 0.16 | 8.80 |

**Supplementary Table 4. Marker genes for the annotated cell populations of integrated clustering of E11.5 control and NotchGOF λ-junction ectoderm cells.**

| | | | | | | | | |
|---|---|---|---|---|---|---|---|---|
| fusion | Nav2 | 1.22 | 7.41E-46 | 1.58E-41 | 0.90 | 0.69 | 0.21 | 1.30 |
| fusion | 5033430I15Rik | 3.00 | 8.14E-46 | 1.74E-41 | 0.31 | 0.07 | 0.25 | 4.55 |
| fusion | Ldlrad4 | 2.97 | 4.48E-45 | 9.58E-41 | 0.24 | 0.04 | 0.20 | 6.13 |
| fusion | Dlx1 | 1.96 | 3.76E-44 | 8.04E-40 | 0.44 | 0.13 | 0.31 | 3.43 |
| fusion | Lurap1l | 2.04 | 5.18E-43 | 1.11E-38 | 0.58 | 0.24 | 0.34 | 2.38 |
| fusion | Slc16a10 | 4.29 | 1.22E-42 | 2.61E-38 | 0.16 | 0.02 | 0.14 | 8.94 |
| fusion | Pnldc1 | 2.86 | 1.72E-41 | 3.67E-37 | 0.24 | 0.04 | 0.20 | 5.56 |
| fusion | Dsp | 0.99 | 4.23E-41 | 9.04E-37 | 0.98 | 0.86 | 0.11 | 1.13 |
| fusion | Insyn2b | 2.49 | 6.74E-41 | 1.44E-36 | 0.28 | 0.06 | 0.22 | 4.86 |
| fusion | Smad7 | 1.26 | 1.21E-40 | 2.58E-36 | 0.86 | 0.60 | 0.26 | 1.44 |
| fusion | Plce1 | 1.87 | 1.59E-40 | 3.39E-36 | 0.64 | 0.31 | 0.33 | 2.05 |
| fusion | Tagln | 2.90 | 2.47E-40 | 5.29E-36 | 0.23 | 0.04 | 0.19 | 5.68 |
| fusion | Kdm5b | 0.83 | 5.44E-40 | 1.16E-35 | 0.97 | 0.92 | 0.05 | 1.05 |
| fusion | Apcdd1 | 2.27 | 7.30E-40 | 1.56E-35 | 0.41 | 0.13 | 0.29 | 3.27 |
| fusion | Ptprz1 | 2.05 | 7.66E-40 | 1.64E-35 | 0.52 | 0.20 | 0.32 | 2.56 |
| fusion | Ptn | 1.41 | 9.02E-40 | 1.93E-35 | 0.94 | 0.86 | 0.08 | 1.09 |
| fusion | Retreg1 | 1.93 | 9.54E-40 | 2.04E-35 | 0.55 | 0.24 | 0.32 | 2.33 |
| fusion | Npm3 | 0.98 | 1.79E-39 | 3.83E-35 | 0.92 | 0.87 | 0.05 | 1.05 |
| fusion | Ermp1 | 1.45 | 2.43E-39 | 5.18E-35 | 0.73 | 0.41 | 0.32 | 1.77 |
| fusion | Fam89a | 2.04 | 2.43E-39 | 5.18E-35 | 0.44 | 0.15 | 0.30 | 3.06 |
| fusion | Parm1 | 1.88 | 3.62E-39 | 7.74E-35 | 0.50 | 0.19 | 0.31 | 2.64 |
| fusion | Aspscr1 | 1.54 | 4.08E-39 | 8.72E-35 | 0.76 | 0.50 | 0.25 | 1.50 |
| fusion | Amtn | 4.21 | 4.78E-39 | 1.02E-34 | 0.10 | 0.01 | 0.10 | 14.57 |
| fusion | Malat1 | 0.68 | 4.90E-39 | 1.05E-34 | 1.00 | 1.00 | 0.00 | 1.00 |
| fusion | Pmepa1 | 1.84 | 7.69E-39 | 1.64E-34 | 0.66 | 0.35 | 0.30 | 1.87 |
| fusion | Me1 | 2.11 | 9.61E-39 | 2.05E-34 | 0.48 | 0.17 | 0.31 | 2.77 |
| fusion | Mbnl2 | 1.03 | 2.40E-38 | 5.12E-34 | 0.93 | 0.78 | 0.15 | 1.19 |
| fusion | Amotl1 | 1.21 | 3.73E-38 | 7.97E-34 | 0.87 | 0.69 | 0.18 | 1.26 |
| fusion | Erc2 | 2.61 | 5.95E-38 | 1.27E-33 | 0.36 | 0.10 | 0.26 | 3.47 |
| fusion | Tpm1 | 1.06 | 6.56E-38 | 1.40E-33 | 0.97 | 0.93 | 0.04 | 1.05 |
| fusion | Peg3 | 1.21 | 9.59E-38 | 2.05E-33 | 0.93 | 0.82 | 0.11 | 1.14 |
| fusion | Saraf | 1.04 | 9.95E-38 | 2.13E-33 | 0.89 | 0.65 | 0.25 | 1.38 |
| fusion | Cbr3 | 3.81 | 1.30E-37 | 2.77E-33 | 0.13 | 0.01 | 0.11 | 10.42 |
| fusion | Hspb8 | 1.31 | 1.42E-37 | 3.04E-33 | 0.79 | 0.47 | 0.32 | 1.68 |
| fusion | Cux1 | 0.91 | 2.68E-37 | 5.73E-33 | 0.94 | 0.84 | 0.10 | 1.12 |
| fusion | Abracl | 0.79 | 1.14E-36 | 2.44E-32 | 0.96 | 0.92 | 0.04 | 1.04 |
| fusion | Perp | 0.97 | 2.12E-36 | 4.54E-32 | 0.90 | 0.52 | 0.38 | 1.72 |
| fusion | Zfhx3 | 0.74 | 2.31E-36 | 4.94E-32 | 0.99 | 0.92 | 0.07 | 1.08 |
| fusion | Proser2 | 2.02 | 2.50E-36 | 5.34E-32 | 0.42 | 0.14 | 0.28 | 2.99 |
| fusion | Pdcd4 | 1.06 | 3.02E-36 | 6.46E-32 | 0.93 | 0.78 | 0.15 | 1.19 |
| fusion | Nrip1 | 1.09 | 7.82E-36 | 1.67E-31 | 0.94 | 0.81 | 0.13 | 1.16 |
| fusion | Trp53inp1 | 1.82 | 1.12E-35 | 2.39E-31 | 0.55 | 0.24 | 0.31 | 2.27 |
| fusion | Pdgfrl | 1.71 | 2.61E-35 | 5.58E-31 | 0.62 | 0.32 | 0.31 | 1.97 |
| fusion | Pcdh9 | 1.91 | 4.38E-35 | 9.37E-31 | 0.46 | 0.17 | 0.29 | 2.70 |
| fusion | Rps27 | 0.31 | 1.02E-34 | 2.19E-30 | 1.00 | 1.00 | 0.00 | 1.00 |
| fusion | Cbr1 | 1.20 | 1.36E-34 | 2.90E-30 | 0.89 | 0.72 | 0.17 | 1.24 |
| fusion | Zfp503 | 1.58 | 2.72E-34 | 5.81E-30 | 0.75 | 0.49 | 0.25 | 1.51 |
| fusion | H3f3b | 0.39 | 3.03E-34 | 6.48E-30 | 1.00 | 1.00 | 0.00 | 1.00 |
| fusion | Dact1 | 2.52 | 4.04E-34 | 8.64E-30 | 0.39 | 0.13 | 0.26 | 2.94 |
| fusion | Hras | 0.65 | 4.82E-34 | 1.03E-29 | 0.98 | 0.90 | 0.08 | 1.09 |
| fusion | Rpl18a | 0.27 | 5.37E-34 | 1.15E-29 | 1.00 | 1.00 | 0.00 | 1.00 |
| fusion | 6330403K07Rik | 0.95 | 6.70E-34 | 1.43E-29 | 0.93 | 0.85 | 0.09 | 1.10 |
| fusion | Klf4 | 2.15 | 7.07E-34 | 1.51E-29 | 0.46 | 0.18 | 0.28 | 2.54 |
| fusion | Skil | 1.35 | 7.87E-34 | 1.68E-29 | 0.79 | 0.56 | 0.23 | 1.41 |
| fusion | Eya2 | 1.18 | 8.03E-34 | 1.71E-29 | 0.68 | 0.35 | 0.33 | 1.93 |

**Supplementary Table 4. Marker genes for the annotated cell populations of integrated clustering of E11.5 control and NotchGOF λ-junction ectoderm cells.**

| | | | | | | | | |
|---|---|---|---|---|---|---|---|---|
| fusion | Fau | 0.26 | 8.46E-34 | 1.81E-29 | 1.00 | 1.00 | 0.00 | 1.00 |
| fusion | Ptbp3 | 1.26 | 1.12E-33 | 2.40E-29 | 0.83 | 0.68 | 0.14 | 1.21 |
| fusion | Ctnnb1 | 0.59 | 1.64E-33 | 3.50E-29 | 0.99 | 0.99 | 0.01 | 1.01 |
| fusion | Fxyd3 | 1.40 | 2.78E-33 | 5.93E-29 | 0.63 | 0.29 | 0.34 | 2.17 |
| fusion | Cald1 | 1.28 | 7.32E-33 | 1.56E-28 | 0.91 | 0.69 | 0.22 | 1.31 |
| fusion | Gad2 | 1.87 | 7.72E-33 | 1.65E-28 | 0.57 | 0.28 | 0.29 | 2.06 |
| fusion | Itgb5 | 1.10 | 1.23E-32 | 2.62E-28 | 0.78 | 0.48 | 0.30 | 1.61 |
| fusion | Ier5 | 0.87 | 4.15E-32 | 8.87E-28 | 0.95 | 0.84 | 0.12 | 1.14 |
| fusion | Irf6 | 1.23 | 4.84E-32 | 1.03E-27 | 0.72 | 0.40 | 0.33 | 1.82 |
| fusion | Pcdh17 | 1.61 | 5.41E-32 | 1.16E-27 | 0.66 | 0.36 | 0.30 | 1.85 |
| fusion | Vcan | 1.18 | 1.25E-31 | 2.67E-27 | 0.92 | 0.74 | 0.18 | 1.24 |
| fusion | Pnrc1 | 0.74 | 1.29E-31 | 2.75E-27 | 0.97 | 0.90 | 0.06 | 1.07 |
| fusion | Klhl24 | 1.08 | 1.96E-31 | 4.18E-27 | 0.85 | 0.63 | 0.22 | 1.36 |
| fusion | Gtf2i | 0.75 | 2.13E-31 | 4.55E-27 | 0.97 | 0.90 | 0.07 | 1.08 |
| fusion | Acot1 | 1.68 | 2.67E-31 | 5.71E-27 | 0.61 | 0.33 | 0.28 | 1.84 |
| fusion | Ndrg1 | 2.93 | 5.59E-31 | 1.19E-26 | 0.19 | 0.04 | 0.15 | 5.37 |
| fusion | Stac | 3.86 | 8.87E-31 | 1.90E-26 | 0.11 | 0.01 | 0.10 | 9.64 |
| fusion | Myo10 | 1.12 | 1.31E-30 | 2.80E-26 | 0.84 | 0.70 | 0.14 | 1.20 |
| fusion | Calcoco1 | 1.55 | 1.34E-30 | 2.87E-26 | 0.57 | 0.28 | 0.29 | 2.04 |
| fusion | Kcnj2 | 2.13 | 1.45E-30 | 3.10E-26 | 0.39 | 0.14 | 0.25 | 2.78 |
| fusion | Socs2 | 1.19 | 1.89E-30 | 4.03E-26 | 0.89 | 0.78 | 0.12 | 1.15 |
| fusion | Fgf9 | 1.55 | 2.27E-30 | 4.85E-26 | 0.69 | 0.41 | 0.27 | 1.67 |
| fusion | Map1lc3b | 0.65 | 3.41E-30 | 7.28E-26 | 0.95 | 0.93 | 0.02 | 1.02 |
| fusion | Gabarap | 0.59 | 3.93E-30 | 8.39E-26 | 0.99 | 0.97 | 0.02 | 1.02 |
| fusion | Phlda3 | 1.35 | 7.97E-30 | 1.70E-25 | 0.75 | 0.49 | 0.26 | 1.54 |
| fusion | Chd3 | 0.68 | 1.36E-29 | 2.92E-25 | 0.98 | 0.94 | 0.04 | 1.05 |
| fusion | Kdm6b | 1.08 | 3.27E-29 | 6.98E-25 | 0.86 | 0.72 | 0.14 | 1.19 |
| fusion | Ypel3 | 1.04 | 5.00E-29 | 1.07E-24 | 0.78 | 0.56 | 0.22 | 1.39 |
| fusion | Sh3bp5 | 1.52 | 5.14E-29 | 1.10E-24 | 0.63 | 0.37 | 0.25 | 1.68 |
| fusion | Dclre1c | 2.50 | 8.56E-29 | 1.83E-24 | 0.30 | 0.09 | 0.21 | 3.31 |
| fusion | Irx1 | 2.04 | 1.23E-28 | 2.62E-24 | 0.26 | 0.06 | 0.19 | 3.98 |
| fusion | Tiam2 | 3.13 | 1.38E-28 | 2.95E-24 | 0.16 | 0.03 | 0.13 | 5.75 |
| fusion | Syt1 | 1.76 | 1.49E-28 | 3.19E-24 | 0.38 | 0.13 | 0.25 | 2.91 |
| fusion | Tgif1 | 1.05 | 1.92E-28 | 4.11E-24 | 0.83 | 0.68 | 0.16 | 1.23 |
| fusion | Cmip | 1.27 | 9.20E-28 | 1.97E-23 | 0.72 | 0.50 | 0.22 | 1.44 |
| fusion | Cdh6 | 1.95 | 2.41E-27 | 5.16E-23 | 0.35 | 0.12 | 0.23 | 2.97 |
| fusion | Jun | 1.61 | 3.50E-27 | 7.47E-23 | 0.74 | 0.53 | 0.21 | 1.39 |
| fusion | Dkk2 | 3.39 | 4.62E-27 | 9.88E-23 | 0.20 | 0.04 | 0.15 | 4.45 |
| fusion | Ly6g6e | 2.25 | 4.69E-27 | 1.00E-22 | 0.24 | 0.06 | 0.18 | 3.98 |
| fusion | Nxph3 | 2.17 | 5.54E-27 | 1.18E-22 | 0.22 | 0.05 | 0.17 | 4.23 |
| fusion | Kcnmb4os2 | 1.68 | 6.18E-27 | 1.32E-22 | 0.49 | 0.23 | 0.26 | 2.15 |
| fusion | Ndufa4l2 | 2.21 | 7.25E-27 | 1.55E-22 | 0.29 | 0.09 | 0.20 | 3.30 |
| fusion | Npc2 | 0.66 | 1.21E-26 | 2.58E-22 | 0.97 | 0.92 | 0.05 | 1.06 |
| fusion | Pcsk5 | 2.07 | 1.73E-26 | 3.69E-22 | 0.31 | 0.10 | 0.21 | 3.05 |
| fusion | Znrf3 | 1.61 | 2.93E-26 | 6.26E-22 | 0.53 | 0.28 | 0.26 | 1.92 |
| fusion | Dapl1 | 1.73 | 4.52E-26 | 9.65E-22 | 0.53 | 0.26 | 0.28 | 2.08 |
| fusion | Rgs12 | 1.36 | 5.64E-26 | 1.20E-21 | 0.67 | 0.45 | 0.22 | 1.48 |
| fusion | Fnbp1 | 1.89 | 6.67E-26 | 1.42E-21 | 0.44 | 0.20 | 0.24 | 2.20 |
| fusion | Minar2 | 1.83 | 8.30E-26 | 1.77E-21 | 0.38 | 0.15 | 0.23 | 2.57 |
| fusion | Mtss1 | 1.25 | 1.06E-25 | 2.27E-21 | 0.67 | 0.42 | 0.25 | 1.58 |
| fusion | Bnc1 | 2.82 | 1.29E-25 | 2.74E-21 | 0.17 | 0.03 | 0.13 | 5.16 |
| fusion | Csnk1e | 0.72 | 1.63E-25 | 3.47E-21 | 0.94 | 0.87 | 0.07 | 1.08 |
| fusion | Irx2 | 2.44 | 1.63E-25 | 3.49E-21 | 0.20 | 0.05 | 0.15 | 4.35 |
| fusion | Sox9 | 1.50 | 1.96E-25 | 4.18E-21 | 0.78 | 0.57 | 0.21 | 1.37 |
| fusion | Aplp2 | 0.73 | 2.15E-25 | 4.60E-21 | 0.96 | 0.92 | 0.04 | 1.04 |

**Supplementary Table 4. Marker genes for the annotated cell populations of integrated clustering of E11.5 control and NotchGOF λ-junction ectoderm cells.**

| | | | | | | | | |
|---|---|---|---|---|---|---|---|---|
| fusion | Actg1 | 0.41 | 2.99E-25 | 6.39E-21 | 1.00 | 1.00 | 0.00 | 1.00 |
| fusion | Pag1 | 2.50 | 3.39E-25 | 7.23E-21 | 0.20 | 0.05 | 0.15 | 4.26 |
| fusion | Igfbp2 | 0.77 | 6.50E-25 | 1.39E-20 | 1.00 | 0.99 | 0.01 | 1.01 |
| fusion | Gxylt2 | 2.38 | 6.80E-25 | 1.45E-20 | 0.28 | 0.09 | 0.19 | 3.12 |
| fusion | Hs3st1 | 2.57 | 9.95E-25 | 2.13E-20 | 0.23 | 0.06 | 0.17 | 3.67 |
| fusion | Neat1 | 2.23 | 2.05E-24 | 4.38E-20 | 0.25 | 0.07 | 0.18 | 3.53 |
| fusion | Igfbp4 | 1.28 | 2.55E-24 | 5.44E-20 | 0.88 | 0.73 | 0.14 | 1.19 |
| fusion | Gadd45b | 1.69 | 5.22E-24 | 1.11E-19 | 0.44 | 0.19 | 0.24 | 2.25 |
| fusion | Mid1ip1 | 1.86 | 6.16E-24 | 1.32E-19 | 0.59 | 0.37 | 0.22 | 1.60 |
| fusion | Trps1 | 1.29 | 9.01E-24 | 1.92E-19 | 0.70 | 0.53 | 0.18 | 1.33 |
| fusion | Gnpda2 | 1.48 | 9.09E-24 | 1.94E-19 | 0.60 | 0.39 | 0.21 | 1.54 |
| fusion | Sema6a | 1.18 | 1.10E-23 | 2.36E-19 | 0.78 | 0.61 | 0.18 | 1.29 |
| fusion | Prss23 | 2.02 | 1.34E-23 | 2.86E-19 | 0.52 | 0.28 | 0.24 | 1.85 |
| fusion | Med13 | 1.29 | 1.92E-23 | 4.10E-19 | 0.75 | 0.59 | 0.16 | 1.28 |
| fusion | Frmd4b | 1.12 | 2.02E-23 | 4.32E-19 | 0.71 | 0.49 | 0.22 | 1.45 |
| fusion | Eef2 | 0.33 | 2.04E-23 | 4.36E-19 | 1.00 | 1.00 | 0.00 | 1.00 |
| fusion | Sp6 | 1.99 | 2.05E-23 | 4.38E-19 | 0.31 | 0.11 | 0.20 | 2.83 |
| fusion | Fat3 | 1.94 | 2.17E-23 | 4.63E-19 | 0.35 | 0.14 | 0.21 | 2.52 |
| fusion | Arpp21 | 3.08 | 3.26E-23 | 6.97E-19 | 0.13 | 0.02 | 0.11 | 5.78 |
| fusion | Bdnf | 2.05 | 3.53E-23 | 7.54E-19 | 0.40 | 0.17 | 0.22 | 2.30 |
| fusion | Fbxw4 | 1.74 | 3.60E-23 | 7.68E-19 | 0.41 | 0.18 | 0.23 | 2.28 |
| fusion | Rdh10 | 1.95 | 4.34E-23 | 9.26E-19 | 0.53 | 0.29 | 0.24 | 1.83 |
| fusion | Tmem178 | 1.76 | 4.60E-23 | 9.82E-19 | 0.31 | 0.11 | 0.20 | 2.90 |
| fusion | Ube2h | 1.18 | 5.35E-23 | 1.14E-18 | 0.69 | 0.50 | 0.20 | 1.39 |
| fusion | Fhl2 | 2.53 | 6.75E-23 | 1.44E-18 | 0.20 | 0.05 | 0.15 | 3.92 |
| fusion | Mylip | 1.70 | 1.14E-22 | 2.43E-18 | 0.42 | 0.19 | 0.23 | 2.20 |
| fusion | Gpr4 | 2.60 | 1.18E-22 | 2.52E-18 | 0.11 | 0.02 | 0.10 | 6.33 |
| fusion | Trim16 | 2.84 | 1.21E-22 | 2.58E-18 | 0.17 | 0.04 | 0.13 | 4.58 |
| fusion | 1810058I24Rik | 0.62 | 1.49E-22 | 3.19E-18 | 0.95 | 0.93 | 0.02 | 1.02 |
| fusion | Fblim1 | 1.26 | 1.97E-22 | 4.20E-18 | 0.62 | 0.40 | 0.22 | 1.55 |
| fusion | Zfand5 | 0.93 | 2.00E-22 | 4.28E-18 | 0.89 | 0.83 | 0.07 | 1.08 |
| fusion | Scnn1a | 3.02 | 2.64E-22 | 5.65E-18 | 0.12 | 0.02 | 0.10 | 6.21 |
| fusion | Fosl2 | 1.88 | 3.17E-22 | 6.76E-18 | 0.26 | 0.08 | 0.18 | 3.31 |
| fusion | Mir100hg | 0.86 | 3.67E-22 | 7.83E-18 | 0.82 | 0.62 | 0.20 | 1.32 |
| fusion | Robo1 | 1.35 | 4.62E-22 | 9.86E-18 | 0.61 | 0.39 | 0.22 | 1.56 |
| fusion | Dlx3 | 1.41 | 5.13E-22 | 1.10E-17 | 0.51 | 0.27 | 0.24 | 1.89 |
| fusion | Btrc | 1.83 | 1.17E-21 | 2.50E-17 | 0.53 | 0.32 | 0.21 | 1.66 |
| fusion | Chst3 | 2.42 | 1.31E-21 | 2.80E-17 | 0.18 | 0.05 | 0.14 | 4.00 |
| fusion | Ablim1 | 0.86 | 1.73E-21 | 3.70E-17 | 0.76 | 0.54 | 0.22 | 1.41 |
| fusion | Mfsd1 | 1.22 | 1.91E-21 | 4.08E-17 | 0.62 | 0.39 | 0.23 | 1.58 |
| fusion | Serinc5 | 1.91 | 2.04E-21 | 4.35E-17 | 0.31 | 0.11 | 0.19 | 2.73 |
| fusion | Bcl11b | 0.83 | 2.93E-21 | 6.25E-17 | 0.86 | 0.71 | 0.15 | 1.20 |
| fusion | Tmem59 | 0.56 | 3.44E-21 | 7.35E-17 | 0.96 | 0.89 | 0.07 | 1.08 |
| fusion | Lmo4 | 0.62 | 3.72E-21 | 7.95E-17 | 0.95 | 0.91 | 0.04 | 1.04 |
| fusion | Mfap4 | 0.67 | 5.54E-21 | 1.18E-16 | 0.84 | 0.65 | 0.20 | 1.30 |
| fusion | Slc39a14 | 2.22 | 5.77E-21 | 1.23E-16 | 0.22 | 0.07 | 0.16 | 3.34 |
| fusion | Filip1l | 1.58 | 6.72E-21 | 1.44E-16 | 0.49 | 0.26 | 0.23 | 1.89 |
| fusion | Cited4 | 2.23 | 8.13E-21 | 1.74E-16 | 0.19 | 0.05 | 0.14 | 3.84 |
| fusion | Msx2 | 1.12 | 8.53E-21 | 1.82E-16 | 0.63 | 0.39 | 0.24 | 1.61 |
| fusion | Nab1 | 1.43 | 1.02E-20 | 2.18E-16 | 0.46 | 0.23 | 0.23 | 1.99 |
| fusion | Kcnmb4 | 1.24 | 1.22E-20 | 2.61E-16 | 0.60 | 0.39 | 0.21 | 1.54 |
| fusion | Tctex1d1 | 1.59 | 1.50E-20 | 3.20E-16 | 0.28 | 0.10 | 0.18 | 2.93 |
| fusion | Gm49708 | 1.27 | 1.68E-20 | 3.60E-16 | 0.68 | 0.48 | 0.20 | 1.42 |
| fusion | Cox7a2l | 0.42 | 2.11E-20 | 4.51E-16 | 0.99 | 0.98 | 0.01 | 1.01 |
| fusion | Nedd9 | 1.41 | 2.83E-20 | 6.05E-16 | 0.52 | 0.30 | 0.22 | 1.74 |

**Supplementary Table 4. Marker genes for the annotated cell populations of integrated clustering of E11.5 control and NotchGOF λ-junction ectoderm cells.**

| | | | | | | | | |
|---|---|---|---|---|---|---|---|---|
| fusion | Bmp4 | 1.42 | 5.50E-20 | 1.18E-15 | 0.72 | 0.53 | 0.19 | 1.37 |
| fusion | Slc25a17 | 0.72 | 5.81E-20 | 1.24E-15 | 0.85 | 0.70 | 0.14 | 1.20 |
| fusion | Sptlc3 | 2.29 | 6.16E-20 | 1.32E-15 | 0.22 | 0.07 | 0.15 | 3.28 |
| fusion | Col9a3 | 1.67 | 7.70E-20 | 1.64E-15 | 0.34 | 0.14 | 0.20 | 2.44 |
| fusion | Tnfrsf1a | 1.44 | 9.81E-20 | 2.10E-15 | 0.53 | 0.31 | 0.21 | 1.67 |
| fusion | Rel | 1.72 | 1.14E-19 | 2.43E-15 | 0.35 | 0.15 | 0.20 | 2.35 |
| fusion | Ddhd1 | 1.49 | 1.34E-19 | 2.87E-15 | 0.45 | 0.23 | 0.22 | 1.94 |
| fusion | Ppp1r13l | 1.61 | 1.49E-19 | 3.19E-15 | 0.39 | 0.19 | 0.21 | 2.11 |
| fusion | Ddi2 | 1.13 | 2.29E-19 | 4.90E-15 | 0.65 | 0.47 | 0.18 | 1.39 |
| fusion | Notch1 | 0.96 | 4.03E-19 | 8.61E-15 | 0.75 | 0.62 | 0.12 | 1.20 |
| fusion | Ell2 | 1.79 | 4.22E-19 | 9.01E-15 | 0.36 | 0.17 | 0.20 | 2.17 |
| fusion | Itm2c | 0.85 | 4.55E-19 | 9.73E-15 | 0.75 | 0.59 | 0.16 | 1.27 |
| fusion | Kif21a | 0.81 | 5.72E-19 | 1.22E-14 | 0.84 | 0.74 | 0.10 | 1.13 |
| fusion | S100a10 | 0.55 | 5.94E-19 | 1.27E-14 | 0.92 | 0.74 | 0.18 | 1.24 |
| fusion | Kdm5a | 0.74 | 8.22E-19 | 1.76E-14 | 0.88 | 0.80 | 0.08 | 1.10 |
| fusion | Abca1 | 1.76 | 8.82E-19 | 1.88E-14 | 0.29 | 0.11 | 0.18 | 2.63 |
| fusion | Map1lc3a | 0.76 | 1.76E-18 | 3.76E-14 | 0.84 | 0.73 | 0.11 | 1.15 |
| fusion | Mir205hg | 0.94 | 2.07E-18 | 4.43E-14 | 0.68 | 0.43 | 0.25 | 1.57 |
| fusion | Pla2r1 | 1.28 | 2.64E-18 | 5.63E-14 | 0.49 | 0.27 | 0.22 | 1.84 |
| fusion | Txnip | 1.15 | 2.67E-18 | 5.71E-14 | 0.64 | 0.44 | 0.20 | 1.45 |
| fusion | Cd59a | 1.20 | 2.83E-18 | 6.04E-14 | 0.52 | 0.29 | 0.23 | 1.78 |
| fusion | Bcl2l11 | 1.52 | 2.93E-18 | 6.26E-14 | 0.57 | 0.38 | 0.19 | 1.50 |
| fusion | Rere | 0.99 | 3.55E-18 | 7.58E-14 | 0.76 | 0.64 | 0.12 | 1.19 |
| fusion | Pcmtd1 | 0.99 | 4.53E-18 | 9.67E-14 | 0.69 | 0.51 | 0.18 | 1.35 |
| fusion | Nxn | 0.69 | 5.61E-18 | 1.20E-13 | 0.89 | 0.75 | 0.15 | 1.20 |
| fusion | Plp2 | 0.82 | 5.92E-18 | 1.26E-13 | 0.81 | 0.69 | 0.11 | 1.16 |
| fusion | Foxo1 | 1.31 | 8.28E-18 | 1.77E-13 | 0.49 | 0.28 | 0.21 | 1.74 |
| fusion | Tns4 | 2.68 | 8.72E-18 | 1.86E-13 | 0.12 | 0.03 | 0.10 | 4.88 |
| fusion | Wls | 0.54 | 1.04E-17 | 2.23E-13 | 0.95 | 0.79 | 0.15 | 1.19 |
| fusion | Rnf149 | 1.58 | 1.33E-17 | 2.84E-13 | 0.47 | 0.28 | 0.19 | 1.70 |
| fusion | Atp6v1g1 | 0.42 | 1.38E-17 | 2.95E-13 | 0.98 | 0.97 | 0.01 | 1.01 |
| fusion | B9d2 | 1.39 | 1.48E-17 | 3.15E-13 | 0.47 | 0.27 | 0.21 | 1.77 |
| fusion | Eif3f | 0.28 | 1.78E-17 | 3.80E-13 | 1.00 | 0.99 | 0.01 | 1.01 |
| fusion | Wipi1 | 1.34 | 2.38E-17 | 5.08E-13 | 0.50 | 0.30 | 0.21 | 1.70 |
| fusion | Bmpr2 | 0.87 | 2.75E-17 | 5.88E-13 | 0.75 | 0.58 | 0.17 | 1.30 |
| fusion | Cnksr2 | 1.25 | 2.79E-17 | 5.96E-13 | 0.53 | 0.33 | 0.21 | 1.63 |
| fusion | Ogt | 0.56 | 2.85E-17 | 6.10E-13 | 0.91 | 0.80 | 0.11 | 1.14 |
| fusion | Gnas | 0.29 | 6.60E-17 | 1.41E-12 | 1.00 | 1.00 | 0.00 | 1.00 |
| fusion | Cpne8 | 2.04 | 1.01E-16 | 2.16E-12 | 0.22 | 0.08 | 0.15 | 2.91 |
| fusion | Bnip3 | 1.22 | 1.03E-16 | 2.19E-12 | 0.63 | 0.45 | 0.18 | 1.40 |
| fusion | Rab6a | 0.78 | 1.16E-16 | 2.48E-12 | 0.84 | 0.76 | 0.08 | 1.11 |
| fusion | Supt4a | 0.44 | 1.22E-16 | 2.62E-12 | 0.94 | 0.91 | 0.04 | 1.04 |
| fusion | Nectin3 | 0.77 | 1.35E-16 | 2.87E-12 | 0.85 | 0.73 | 0.12 | 1.16 |
| fusion | Prickle1 | 2.22 | 1.59E-16 | 3.40E-12 | 0.19 | 0.06 | 0.13 | 3.19 |
| fusion | Kctd1 | 0.77 | 2.20E-16 | 4.70E-12 | 0.70 | 0.49 | 0.21 | 1.44 |
| fusion | Afdn | 0.57 | 2.34E-16 | 4.99E-12 | 0.90 | 0.86 | 0.04 | 1.05 |
| fusion | Tmem140 | 2.68 | 3.35E-16 | 7.16E-12 | 0.11 | 0.02 | 0.09 | 5.05 |
| fusion | Arl4c | 0.90 | 4.05E-16 | 8.66E-12 | 0.74 | 0.57 | 0.17 | 1.30 |
| fusion | Lamp2 | 0.71 | 4.89E-16 | 1.04E-11 | 0.76 | 0.60 | 0.16 | 1.27 |
| fusion | Duoxa1 | 2.74 | 4.90E-16 | 1.05E-11 | 0.11 | 0.02 | 0.09 | 4.78 |
| fusion | Plec | 1.03 | 5.61E-16 | 1.20E-11 | 0.62 | 0.44 | 0.18 | 1.40 |
| fusion | Enpp2 | 3.15 | 5.77E-16 | 1.23E-11 | 0.13 | 0.03 | 0.10 | 4.17 |
| fusion | Chmp5 | 0.61 | 6.55E-16 | 1.40E-11 | 0.89 | 0.79 | 0.10 | 1.12 |
| fusion | Snai2 | 1.26 | 6.92E-16 | 1.48E-11 | 0.37 | 0.17 | 0.20 | 2.16 |
| fusion | Rhob | 1.00 | 9.02E-16 | 1.93E-11 | 0.66 | 0.47 | 0.18 | 1.39 |

**Supplementary Table 4. Marker genes for the annotated cell populations of integrated clustering of E11.5 control and NotchGOF λ-junction ectoderm cells.**

| | | | | | | | | |
|---|---|---|---|---|---|---|---|---|
| fusion | Pygl | 1.50 | 1.17E-15 | 2.49E-11 | 0.35 | 0.17 | 0.18 | 2.06 |
| fusion | Tnks2 | 0.55 | 1.23E-15 | 2.62E-11 | 0.89 | 0.88 | 0.01 | 1.02 |
| fusion | Mafg | 0.98 | 1.37E-15 | 2.92E-11 | 0.68 | 0.53 | 0.15 | 1.28 |
| fusion | Ppfibp1 | 0.89 | 1.77E-15 | 3.78E-11 | 0.75 | 0.61 | 0.14 | 1.23 |
| fusion | Klhl8 | 1.53 | 1.86E-15 | 3.97E-11 | 0.32 | 0.15 | 0.17 | 2.16 |
| fusion | Nectin4 | 0.71 | 1.88E-15 | 4.01E-11 | 0.73 | 0.52 | 0.21 | 1.39 |
| fusion | Ifngr2 | 1.36 | 1.90E-15 | 4.05E-11 | 0.46 | 0.27 | 0.19 | 1.71 |
| fusion | Cdh3 | 0.75 | 2.04E-15 | 4.36E-11 | 0.73 | 0.56 | 0.16 | 1.29 |
| fusion | Ppp1r12b | 1.31 | 2.20E-15 | 4.69E-11 | 0.33 | 0.15 | 0.18 | 2.15 |
| fusion | Ptk7 | 0.95 | 2.22E-15 | 4.74E-11 | 0.66 | 0.48 | 0.17 | 1.36 |
| fusion | Hs3st6 | 1.94 | 2.25E-15 | 4.80E-11 | 0.26 | 0.11 | 0.15 | 2.44 |
| fusion | Vgll4 | 0.69 | 2.37E-15 | 5.06E-11 | 0.92 | 0.85 | 0.07 | 1.08 |
| fusion | Khdrbs2 | 2.32 | 2.67E-15 | 5.70E-11 | 0.13 | 0.03 | 0.10 | 3.91 |
| fusion | Zfp704 | 0.84 | 3.13E-15 | 6.69E-11 | 0.73 | 0.62 | 0.11 | 1.18 |
| fusion | Ptger4 | 2.93 | 4.74E-15 | 1.01E-10 | 0.11 | 0.02 | 0.08 | 4.61 |
| fusion | Dst | 1.03 | 4.78E-15 | 1.02E-10 | 0.68 | 0.47 | 0.20 | 1.42 |
| fusion | Akirin2 | 0.66 | 5.47E-15 | 1.17E-10 | 0.88 | 0.81 | 0.07 | 1.08 |
| fusion | Cited2 | 0.89 | 7.19E-15 | 1.54E-10 | 0.79 | 0.68 | 0.11 | 1.16 |
| fusion | Bcl9 | 0.72 | 9.61E-15 | 2.05E-10 | 0.79 | 0.68 | 0.11 | 1.16 |
| fusion | Ptpn13 | 0.77 | 1.08E-14 | 2.31E-10 | 0.73 | 0.57 | 0.17 | 1.30 |
| fusion | Axin2 | 1.57 | 1.29E-14 | 2.76E-10 | 0.43 | 0.25 | 0.18 | 1.70 |
| fusion | Mtmr11 | 2.19 | 1.39E-14 | 2.96E-10 | 0.18 | 0.06 | 0.12 | 3.03 |
| fusion | Gm42418 | 0.33 | 1.53E-14 | 3.26E-10 | 1.00 | 1.00 | 0.00 | 1.00 |
| fusion | Sema3d | 1.19 | 1.87E-14 | 4.00E-10 | 0.60 | 0.41 | 0.19 | 1.47 |
| fusion | Vwa5a | 1.47 | 1.89E-14 | 4.04E-10 | 0.33 | 0.16 | 0.17 | 2.10 |
| fusion | Cnn2 | 1.60 | 2.39E-14 | 5.10E-10 | 0.49 | 0.30 | 0.19 | 1.61 |
| fusion | Atxn7l3b | 0.29 | 2.64E-14 | 5.65E-10 | 1.00 | 0.99 | 0.01 | 1.01 |
| fusion | Cst3 | 0.40 | 2.71E-14 | 5.79E-10 | 0.97 | 0.93 | 0.04 | 1.05 |
| fusion | Peli1 | 0.78 | 2.90E-14 | 6.20E-10 | 0.78 | 0.65 | 0.13 | 1.20 |
| fusion | Slc7a11 | 1.43 | 3.11E-14 | 6.64E-10 | 0.29 | 0.12 | 0.17 | 2.34 |
| fusion | Jup | 0.80 | 3.44E-14 | 7.34E-10 | 0.68 | 0.51 | 0.16 | 1.32 |
| fusion | Map4k4 | 0.53 | 3.48E-14 | 7.43E-10 | 0.89 | 0.89 | 0.01 | 1.01 |
| fusion | Plod2 | 1.04 | 3.65E-14 | 7.80E-10 | 0.65 | 0.47 | 0.18 | 1.37 |
| fusion | Igfbp3 | 1.94 | 4.86E-14 | 1.04E-09 | 0.54 | 0.38 | 0.16 | 1.41 |
| fusion | Pgpep1 | 1.26 | 4.92E-14 | 1.05E-09 | 0.51 | 0.34 | 0.17 | 1.51 |
| fusion | Serf1 | 0.52 | 4.98E-14 | 1.06E-09 | 0.94 | 0.93 | 0.01 | 1.01 |
| fusion | Lmo7 | 1.06 | 6.35E-14 | 1.36E-09 | 0.60 | 0.45 | 0.16 | 1.35 |
| fusion | Churc1 | 0.81 | 6.70E-14 | 1.43E-09 | 0.81 | 0.72 | 0.08 | 1.12 |
| fusion | Cdkn1c | 0.90 | 6.93E-14 | 1.48E-09 | 0.91 | 0.88 | 0.03 | 1.04 |
| fusion | Col17a1 | 1.12 | 8.02E-14 | 1.71E-09 | 0.29 | 0.12 | 0.17 | 2.36 |
| fusion | Tmem30b | 1.54 | 1.11E-13 | 2.37E-09 | 0.43 | 0.27 | 0.17 | 1.62 |
| fusion | Palmd | 2.28 | 1.16E-13 | 2.47E-09 | 0.16 | 0.05 | 0.11 | 3.16 |
| fusion | Socs7 | 1.17 | 1.18E-13 | 2.53E-09 | 0.47 | 0.29 | 0.18 | 1.61 |
| fusion | Xpr1 | 1.19 | 1.25E-13 | 2.66E-09 | 0.58 | 0.41 | 0.17 | 1.41 |
| fusion | Sertad4 | 1.10 | 1.25E-13 | 2.67E-09 | 0.49 | 0.32 | 0.18 | 1.55 |
| fusion | Lbh | 1.25 | 2.08E-13 | 4.45E-09 | 0.44 | 0.26 | 0.17 | 1.67 |
| fusion | Kdm1a | 0.38 | 2.64E-13 | 5.64E-09 | 0.97 | 0.94 | 0.03 | 1.04 |
| fusion | Ypel5 | 1.23 | 2.82E-13 | 6.01E-09 | 0.43 | 0.25 | 0.18 | 1.71 |
| fusion | Atp6v0e | 0.62 | 2.85E-13 | 6.09E-09 | 0.82 | 0.77 | 0.04 | 1.06 |
| fusion | Rgs2 | 1.26 | 3.07E-13 | 6.56E-09 | 0.47 | 0.29 | 0.19 | 1.65 |
| fusion | Myl12a | 0.47 | 4.38E-13 | 9.35E-09 | 0.97 | 0.94 | 0.03 | 1.04 |
| fusion | Sulf1 | 0.74 | 4.49E-13 | 9.58E-09 | 0.79 | 0.66 | 0.13 | 1.20 |
| fusion | Fbxo32 | 2.17 | 6.03E-13 | 1.29E-08 | 0.18 | 0.07 | 0.12 | 2.77 |
| fusion | Cd151 | 0.66 | 6.34E-13 | 1.35E-08 | 0.79 | 0.66 | 0.13 | 1.20 |
| fusion | Itgb6 | 1.35 | 6.71E-13 | 1.43E-08 | 0.32 | 0.16 | 0.16 | 2.03 |

**Supplementary Table 4. Marker genes for the annotated cell populations of integrated clustering of E11.5 control and NotchGOF λ-junction ectoderm cells.**

| | | | | | | | | |
|---|---|---|---|---|---|---|---|---|
| fusion | Tfap2a | 0.87 | 6.79E-13 | 1.45E-08 | 0.53 | 0.34 | 0.20 | 1.59 |
| fusion | Lrrk2 | 2.30 | 8.11E-13 | 1.73E-08 | 0.12 | 0.03 | 0.09 | 3.81 |
| fusion | Eppk1 | 2.00 | 8.92E-13 | 1.91E-08 | 0.20 | 0.08 | 0.13 | 2.58 |
| fusion | Spsb1 | 2.48 | 8.95E-13 | 1.91E-08 | 0.18 | 0.06 | 0.11 | 2.79 |
| fusion | Tulp4 | 0.59 | 9.49E-13 | 2.03E-08 | 0.84 | 0.75 | 0.09 | 1.12 |
| fusion | Slc16a7 | 2.44 | 1.27E-12 | 2.72E-08 | 0.14 | 0.04 | 0.10 | 3.20 |
| fusion | Col23a1 | 2.47 | 1.49E-12 | 3.17E-08 | 0.11 | 0.03 | 0.08 | 3.80 |
| fusion | Nsg1 | 0.72 | 1.49E-12 | 3.19E-08 | 0.72 | 0.61 | 0.12 | 1.19 |
| fusion | Col4a2 | 0.78 | 1.60E-12 | 3.42E-08 | 0.66 | 0.50 | 0.17 | 1.33 |
| fusion | Adgrl3 | 1.33 | 1.73E-12 | 3.69E-08 | 0.48 | 0.31 | 0.17 | 1.55 |
| fusion | Rab38 | 1.34 | 1.78E-12 | 3.81E-08 | 0.37 | 0.19 | 0.17 | 1.88 |
| fusion | Inpp5f | 1.43 | 1.90E-12 | 4.06E-08 | 0.42 | 0.26 | 0.16 | 1.63 |
| fusion | Fzd10os | 1.21 | 2.15E-12 | 4.60E-08 | 0.31 | 0.15 | 0.16 | 2.10 |
| fusion | Kidins220 | 1.01 | 2.68E-12 | 5.72E-08 | 0.52 | 0.36 | 0.16 | 1.44 |
| fusion | Usp22 | 0.60 | 2.92E-12 | 6.24E-08 | 0.78 | 0.66 | 0.13 | 1.19 |
| fusion | Cpm | 0.62 | 2.98E-12 | 6.37E-08 | 0.73 | 0.52 | 0.21 | 1.39 |
| fusion | Rb1cc1 | 0.64 | 3.08E-12 | 6.58E-08 | 0.78 | 0.70 | 0.08 | 1.12 |
| fusion | Ttc3 | 0.36 | 3.50E-12 | 7.48E-08 | 1.00 | 0.99 | 0.01 | 1.01 |
| fusion | Lrrc8d | 1.16 | 3.76E-12 | 8.02E-08 | 0.48 | 0.32 | 0.16 | 1.50 |
| fusion | Pdgfc | 0.56 | 3.90E-12 | 8.33E-08 | 0.85 | 0.76 | 0.09 | 1.12 |
| fusion | Kdm7a | 0.96 | 4.03E-12 | 8.61E-08 | 0.57 | 0.41 | 0.16 | 1.39 |
| fusion | Pcmtd2 | 0.83 | 4.20E-12 | 8.98E-08 | 0.57 | 0.41 | 0.16 | 1.39 |
| fusion | Mapk6 | 0.73 | 4.22E-12 | 9.00E-08 | 0.68 | 0.54 | 0.14 | 1.26 |
| fusion | Trp53inp2 | 1.76 | 4.90E-12 | 1.05E-07 | 0.19 | 0.07 | 0.12 | 2.59 |
| fusion | Thsd7b | 2.00 | 5.00E-12 | 1.07E-07 | 0.17 | 0.06 | 0.11 | 2.89 |
| fusion | Ddr1 | 0.76 | 5.15E-12 | 1.10E-07 | 0.61 | 0.45 | 0.16 | 1.36 |
| fusion | Gpr107 | 1.07 | 6.15E-12 | 1.31E-07 | 0.49 | 0.34 | 0.15 | 1.45 |
| fusion | Vezf1 | 0.49 | 7.02E-12 | 1.50E-07 | 0.87 | 0.82 | 0.05 | 1.06 |
| fusion | Cebpb | 1.64 | 8.14E-12 | 1.74E-07 | 0.39 | 0.24 | 0.15 | 1.65 |
| fusion | Adamts18 | 1.24 | 8.25E-12 | 1.76E-07 | 0.43 | 0.26 | 0.17 | 1.63 |
| fusion | Pdlim4 | 0.86 | 9.12E-12 | 1.95E-07 | 0.58 | 0.43 | 0.15 | 1.35 |
| fusion | Klc3 | 0.71 | 1.04E-11 | 2.21E-07 | 0.55 | 0.39 | 0.16 | 1.41 |
| fusion | Fzd6 | 1.06 | 1.04E-11 | 2.23E-07 | 0.48 | 0.32 | 0.16 | 1.50 |
| fusion | D7Ertd443e | 1.56 | 1.12E-11 | 2.40E-07 | 0.22 | 0.09 | 0.13 | 2.38 |
| fusion | Sesn1 | 1.43 | 1.28E-11 | 2.72E-07 | 0.36 | 0.20 | 0.16 | 1.78 |
| fusion | Tns3 | 1.09 | 1.30E-11 | 2.78E-07 | 0.48 | 0.33 | 0.15 | 1.44 |
| fusion | Jag1 | 0.77 | 1.35E-11 | 2.89E-07 | 0.71 | 0.55 | 0.16 | 1.29 |
| fusion | Dync1i1 | 1.43 | 1.56E-11 | 3.33E-07 | 0.26 | 0.12 | 0.14 | 2.17 |
| fusion | Chmp1b | 0.84 | 1.65E-11 | 3.52E-07 | 0.57 | 0.43 | 0.15 | 1.35 |
| fusion | Osbpl8 | 1.15 | 1.68E-11 | 3.59E-07 | 0.43 | 0.26 | 0.16 | 1.62 |
| fusion | Cdca7 | 0.63 | 1.85E-11 | 3.95E-07 | 0.78 | 0.64 | 0.14 | 1.21 |
| fusion | Sh3glb1 | 0.49 | 1.95E-11 | 4.16E-07 | 0.84 | 0.79 | 0.05 | 1.07 |
| fusion | Myo1b | 0.61 | 2.11E-11 | 4.50E-07 | 0.76 | 0.62 | 0.14 | 1.22 |
| fusion | Ddah2 | 0.33 | 2.55E-11 | 5.46E-07 | 0.99 | 0.97 | 0.02 | 1.02 |
| fusion | Ifitm1 | 1.05 | 2.56E-11 | 5.47E-07 | 0.35 | 0.19 | 0.16 | 1.85 |
| fusion | Alpl | 1.55 | 2.70E-11 | 5.77E-07 | 0.31 | 0.16 | 0.15 | 1.89 |
| fusion | Prag1 | 1.17 | 2.83E-11 | 6.04E-07 | 0.45 | 0.29 | 0.16 | 1.56 |
| fusion | Rora | 1.76 | 2.90E-11 | 6.19E-07 | 0.21 | 0.09 | 0.12 | 2.36 |
| fusion | Cpxm1 | 1.10 | 3.18E-11 | 6.79E-07 | 0.42 | 0.26 | 0.16 | 1.61 |
| fusion | Fmnl2 | 1.00 | 3.66E-11 | 7.82E-07 | 0.60 | 0.47 | 0.13 | 1.28 |
| fusion | Bcl11a | 0.59 | 3.80E-11 | 8.13E-07 | 0.86 | 0.80 | 0.07 | 1.09 |
| fusion | Hbp1 | 0.62 | 3.92E-11 | 8.36E-07 | 0.77 | 0.63 | 0.14 | 1.22 |
| fusion | Pax7 | 2.40 | 4.35E-11 | 9.29E-07 | 0.11 | 0.03 | 0.08 | 3.66 |
| fusion | Bckdha | 0.87 | 4.59E-11 | 9.81E-07 | 0.61 | 0.46 | 0.15 | 1.34 |
| fusion | Aff4 | 0.71 | 5.00E-11 | 1.07E-06 | 0.75 | 0.66 | 0.09 | 1.14 |

**Supplementary Table 4. Marker genes for the annotated cell populations of integrated clustering of E11.5 control and NotchGOF λ-junction ectoderm cells.**

| | | | | | | | | |
|---|---|---|---|---|---|---|---|---|
| fusion | Rap2c | 0.68 | 5.23E-11 | 1.12E-06 | 0.75 | 0.66 | 0.09 | 1.13 |
| fusion | Usp46 | 0.65 | 5.26E-11 | 1.12E-06 | 0.78 | 0.72 | 0.06 | 1.09 |
| fusion | Ctsl | 0.44 | 5.34E-11 | 1.14E-06 | 0.92 | 0.86 | 0.06 | 1.07 |
| fusion | Irs2 | 1.00 | 6.21E-11 | 1.33E-06 | 0.57 | 0.44 | 0.13 | 1.30 |
| fusion | Gm15222 | 1.38 | 6.42E-11 | 1.37E-06 | 0.29 | 0.14 | 0.14 | 1.99 |
| fusion | Rsrp1 | 0.40 | 6.87E-11 | 1.47E-06 | 0.94 | 0.92 | 0.03 | 1.03 |
| fusion | Jmy | 0.85 | 7.21E-11 | 1.54E-06 | 0.60 | 0.48 | 0.13 | 1.27 |
| fusion | Zfas1 | 0.48 | 8.68E-11 | 1.85E-06 | 0.89 | 0.80 | 0.09 | 1.11 |
| fusion | Creb3l2 | 1.01 | 9.86E-11 | 2.11E-06 | 0.48 | 0.32 | 0.16 | 1.52 |
| fusion | Ddb1 | 0.44 | 1.49E-10 | 3.18E-06 | 0.92 | 0.89 | 0.03 | 1.03 |
| fusion | Macf1 | 0.57 | 1.52E-10 | 3.24E-06 | 0.88 | 0.83 | 0.05 | 1.06 |
| fusion | Nhsl1 | 0.96 | 1.63E-10 | 3.49E-06 | 0.47 | 0.32 | 0.15 | 1.47 |
| fusion | Lpp | 0.89 | 1.78E-10 | 3.80E-06 | 0.59 | 0.45 | 0.14 | 1.31 |
| fusion | Dnase2a | 0.90 | 1.93E-10 | 4.13E-06 | 0.58 | 0.43 | 0.14 | 1.33 |
| fusion | Nfil3 | 2.00 | 2.10E-10 | 4.49E-06 | 0.19 | 0.08 | 0.11 | 2.34 |
| fusion | Dusp2 | 2.11 | 2.35E-10 | 5.01E-06 | 0.12 | 0.04 | 0.08 | 3.28 |
| fusion | Arl5b | 0.76 | 2.46E-10 | 5.26E-06 | 0.60 | 0.45 | 0.15 | 1.33 |
| fusion | Dlgap4 | 0.64 | 2.50E-10 | 5.35E-06 | 0.79 | 0.69 | 0.10 | 1.14 |
| fusion | Ndufa6 | 0.41 | 2.68E-10 | 5.73E-06 | 0.97 | 0.94 | 0.03 | 1.03 |
| fusion | Satb1 | 0.73 | 3.28E-10 | 7.01E-06 | 0.68 | 0.57 | 0.11 | 1.20 |
| fusion | Crebrf | 1.13 | 3.29E-10 | 7.03E-06 | 0.31 | 0.17 | 0.15 | 1.88 |
| fusion | Usp3 | 0.54 | 3.33E-10 | 7.11E-06 | 0.78 | 0.71 | 0.07 | 1.09 |
| fusion | Frem2 | 0.56 | 3.38E-10 | 7.22E-06 | 0.75 | 0.62 | 0.13 | 1.21 |
| fusion | Slc16a2 | 1.63 | 3.42E-10 | 7.30E-06 | 0.24 | 0.12 | 0.13 | 2.06 |
| fusion | Hspb1 | 0.60 | 3.43E-10 | 7.32E-06 | 0.57 | 0.38 | 0.19 | 1.50 |
| fusion | Fermt1 | 0.56 | 4.10E-10 | 8.75E-06 | 0.73 | 0.55 | 0.19 | 1.34 |
| fusion | Gm4258 | 0.89 | 4.33E-10 | 9.26E-06 | 0.62 | 0.51 | 0.11 | 1.23 |
| fusion | Sgpl1 | 0.80 | 5.53E-10 | 1.18E-05 | 0.55 | 0.41 | 0.14 | 1.34 |
| fusion | Zdhhc20 | 0.66 | 5.83E-10 | 1.25E-05 | 0.71 | 0.61 | 0.11 | 1.17 |
| fusion | Hes1 | 0.74 | 6.84E-10 | 1.46E-05 | 0.79 | 0.73 | 0.06 | 1.09 |
| fusion | Cnpy2 | 0.37 | 6.95E-10 | 1.49E-05 | 0.96 | 0.95 | 0.01 | 1.01 |
| fusion | Pik3ip1 | 0.94 | 6.99E-10 | 1.49E-05 | 0.33 | 0.19 | 0.15 | 1.78 |
| fusion | Btg1 | 0.67 | 7.71E-10 | 1.65E-05 | 0.80 | 0.75 | 0.05 | 1.07 |
| fusion | Dpysl3 | 0.34 | 9.04E-10 | 1.93E-05 | 0.72 | 0.56 | 0.16 | 1.28 |
| fusion | Unc5c | 0.73 | 1.04E-09 | 2.22E-05 | 0.49 | 0.33 | 0.17 | 1.52 |
| fusion | Vwa1 | 1.55 | 1.19E-09 | 2.55E-05 | 0.23 | 0.11 | 0.12 | 2.10 |
| fusion | Mafb | 1.40 | 1.20E-09 | 2.55E-05 | 0.29 | 0.15 | 0.13 | 1.88 |
| fusion | Dok5 | 0.79 | 1.33E-09 | 2.83E-05 | 0.54 | 0.37 | 0.17 | 1.46 |
| fusion | Ago1 | 0.61 | 1.45E-09 | 3.10E-05 | 0.71 | 0.62 | 0.09 | 1.15 |
| fusion | Zfp750 | 1.14 | 1.48E-09 | 3.16E-05 | 0.34 | 0.19 | 0.14 | 1.74 |
| fusion | Dusp23 | 1.41 | 1.48E-09 | 3.16E-05 | 0.23 | 0.11 | 0.12 | 2.08 |
| fusion | Cdc14a | 1.02 | 1.50E-09 | 3.20E-05 | 0.50 | 0.37 | 0.14 | 1.37 |
| fusion | Rest | 0.49 | 1.50E-09 | 3.21E-05 | 0.80 | 0.68 | 0.12 | 1.18 |
| fusion | Dhrs3 | 1.36 | 1.71E-09 | 3.64E-05 | 0.34 | 0.21 | 0.13 | 1.65 |
| fusion | Erich1 | 0.78 | 1.72E-09 | 3.67E-05 | 0.53 | 0.39 | 0.14 | 1.36 |
| fusion | Serinc1 | 0.55 | 1.83E-09 | 3.92E-05 | 0.75 | 0.66 | 0.09 | 1.13 |
| fusion | Prxl2a | 0.65 | 1.84E-09 | 3.93E-05 | 0.68 | 0.58 | 0.10 | 1.16 |
| fusion | Gdi1 | 0.64 | 2.08E-09 | 4.45E-05 | 0.59 | 0.45 | 0.14 | 1.31 |
| fusion | Adam19 | 2.11 | 2.13E-09 | 4.56E-05 | 0.10 | 0.03 | 0.07 | 3.29 |
| fusion | Nradd | 0.66 | 2.29E-09 | 4.89E-05 | 0.61 | 0.49 | 0.12 | 1.24 |
| fusion | Mob4 | 0.44 | 2.37E-09 | 5.06E-05 | 0.85 | 0.81 | 0.04 | 1.05 |
| fusion | Unc119 | 0.54 | 2.38E-09 | 5.08E-05 | 0.84 | 0.76 | 0.08 | 1.11 |
| fusion | Unc5b | 0.72 | 2.53E-09 | 5.39E-05 | 0.64 | 0.53 | 0.11 | 1.22 |
| fusion | Tbx2 | 1.62 | 2.64E-09 | 5.63E-05 | 0.26 | 0.14 | 0.12 | 1.87 |
| fusion | Tax1bp1 | 0.37 | 2.71E-09 | 5.79E-05 | 0.95 | 0.92 | 0.02 | 1.03 |

**Supplementary Table 4. Marker genes for the annotated cell populations of integrated clustering of E11.5 control and NotchGOF λ-junction ectoderm cells.**

| | | | | | | | | |
|---|---|---|---|---|---|---|---|---|
| fusion | Atp6v0b | 0.47 | 2.77E-09 | 5.93E-05 | 0.84 | 0.77 | 0.07 | 1.09 |
| fusion | Baz2b | 0.47 | 2.86E-09 | 6.12E-05 | 0.87 | 0.81 | 0.06 | 1.07 |
| fusion | Abcg1 | 0.75 | 2.90E-09 | 6.19E-05 | 0.44 | 0.28 | 0.15 | 1.53 |
| fusion | Mex3a | 0.38 | 2.90E-09 | 6.19E-05 | 0.93 | 0.87 | 0.06 | 1.07 |
| fusion | Btbd7 | 0.57 | 2.99E-09 | 6.40E-05 | 0.78 | 0.71 | 0.07 | 1.09 |
| fusion | Heph | 0.92 | 3.17E-09 | 6.78E-05 | 0.41 | 0.26 | 0.15 | 1.58 |
| fusion | Map3k20 | 1.30 | 3.21E-09 | 6.86E-05 | 0.31 | 0.18 | 0.13 | 1.71 |
| fusion | Alcam | 1.53 | 3.37E-09 | 7.20E-05 | 0.33 | 0.20 | 0.14 | 1.69 |
| fusion | Atp6v0d1 | 0.65 | 3.88E-09 | 8.28E-05 | 0.65 | 0.56 | 0.09 | 1.17 |
| fusion | Ubald2 | 0.52 | 4.01E-09 | 8.56E-05 | 0.78 | 0.71 | 0.07 | 1.10 |
| fusion | Ptprk | 0.59 | 4.15E-09 | 8.87E-05 | 0.69 | 0.59 | 0.10 | 1.17 |
| fusion | Rrbp1 | 0.46 | 4.21E-09 | 8.99E-05 | 0.93 | 0.89 | 0.03 | 1.04 |
| fusion | Trip12 | 0.53 | 4.22E-09 | 9.00E-05 | 0.77 | 0.71 | 0.06 | 1.09 |
| fusion | Myh9 | 0.56 | 4.45E-09 | 9.50E-05 | 0.81 | 0.75 | 0.06 | 1.08 |
| fusion | Rnaset2a | 0.94 | 4.75E-09 | 1.01E-04 | 0.61 | 0.53 | 0.08 | 1.16 |
| fusion | Ppdpf | 0.27 | 4.97E-09 | 1.06E-04 | 0.98 | 0.97 | 0.01 | 1.01 |
| fusion | Stx18 | 0.67 | 5.81E-09 | 1.24E-04 | 0.54 | 0.41 | 0.13 | 1.31 |
| fusion | Hspa2 | 1.18 | 5.99E-09 | 1.28E-04 | 0.26 | 0.13 | 0.13 | 1.96 |
| fusion | Tanc2 | 1.21 | 6.00E-09 | 1.28E-04 | 0.36 | 0.22 | 0.14 | 1.63 |
| fusion | Nsa2 | 0.26 | 6.03E-09 | 1.29E-04 | 0.97 | 0.97 | 0.00 | 1.00 |
| fusion | Renbp | 1.53 | 6.76E-09 | 1.44E-04 | 0.22 | 0.11 | 0.11 | 2.02 |
| fusion | Ankhd1 | 0.64 | 7.75E-09 | 1.66E-04 | 0.67 | 0.58 | 0.10 | 1.16 |
| fusion | Prrc2b | 0.44 | 7.79E-09 | 1.67E-04 | 0.85 | 0.79 | 0.06 | 1.08 |
| fusion | Itgb4 | 1.23 | 8.00E-09 | 1.71E-04 | 0.24 | 0.12 | 0.12 | 2.01 |
| fusion | Grina | 1.12 | 8.22E-09 | 1.75E-04 | 0.43 | 0.30 | 0.13 | 1.42 |
| fusion | Ulk1 | 0.99 | 8.29E-09 | 1.77E-04 | 0.40 | 0.27 | 0.13 | 1.49 |
| fusion | 2610035D17Rik | 1.87 | 9.39E-09 | 2.01E-04 | 0.13 | 0.05 | 0.08 | 2.72 |
| fusion | Hsdl2 | 1.06 | 9.77E-09 | 2.09E-04 | 0.46 | 0.34 | 0.12 | 1.36 |
| fusion | Ube2b | 0.31 | 9.78E-09 | 2.09E-04 | 0.94 | 0.95 | -0.01 | 0.99 |
| fusion | Dkk3 | 1.32 | 9.95E-09 | 2.13E-04 | 0.28 | 0.16 | 0.12 | 1.77 |
| fusion | Tacc1 | 1.29 | 1.03E-08 | 2.20E-04 | 0.31 | 0.19 | 0.13 | 1.67 |
| fusion | Nfic | 1.31 | 1.18E-08 | 2.51E-04 | 0.28 | 0.16 | 0.12 | 1.75 |
| fusion | Oga | 0.48 | 1.24E-08 | 2.65E-04 | 0.80 | 0.72 | 0.08 | 1.11 |
| fusion | Sbk1 | 0.58 | 1.37E-08 | 2.92E-04 | 0.67 | 0.56 | 0.11 | 1.19 |
| fusion | Klk8 | 0.60 | 1.39E-08 | 2.96E-04 | 0.66 | 0.56 | 0.11 | 1.19 |
| fusion | Mbtd1 | 0.43 | 1.44E-08 | 3.07E-04 | 0.84 | 0.77 | 0.07 | 1.09 |
| fusion | Arl8a | 0.71 | 1.44E-08 | 3.08E-04 | 0.52 | 0.38 | 0.14 | 1.37 |
| fusion | Itga11 | 1.95 | 1.52E-08 | 3.24E-04 | 0.10 | 0.03 | 0.07 | 3.09 |
| fusion | E130102H24Rik | 1.98 | 1.53E-08 | 3.28E-04 | 0.11 | 0.04 | 0.07 | 2.97 |
| fusion | Wnt5b | 1.47 | 1.63E-08 | 3.48E-04 | 0.25 | 0.13 | 0.12 | 1.89 |
| fusion | Sdc4 | 0.58 | 1.71E-08 | 3.65E-04 | 0.73 | 0.65 | 0.08 | 1.12 |
| fusion | Glmp | 0.65 | 1.79E-08 | 3.83E-04 | 0.61 | 0.49 | 0.12 | 1.25 |
| fusion | 2410006H16Rik | 0.43 | 1.81E-08 | 3.86E-04 | 0.90 | 0.86 | 0.04 | 1.05 |
| fusion | Camk2n1 | 0.65 | 1.88E-08 | 4.02E-04 | 0.23 | 0.12 | 0.12 | 2.01 |
| fusion | Mapkbp1 | 1.67 | 1.93E-08 | 4.13E-04 | 0.17 | 0.07 | 0.09 | 2.32 |
| fusion | Pax3 | 1.75 | 1.96E-08 | 4.19E-04 | 0.14 | 0.06 | 0.09 | 2.52 |
| fusion | Tet2 | 0.95 | 1.99E-08 | 4.24E-04 | 0.34 | 0.21 | 0.13 | 1.63 |
| fusion | Meg3 | 0.64 | 2.03E-08 | 4.34E-04 | 0.83 | 0.76 | 0.07 | 1.09 |
| fusion | Wdr26 | 0.53 | 2.38E-08 | 5.09E-04 | 0.81 | 0.75 | 0.06 | 1.08 |
| fusion | Capg | 0.81 | 2.52E-08 | 5.38E-04 | 0.48 | 0.35 | 0.13 | 1.36 |
| fusion | Sbspon | 2.18 | 2.66E-08 | 5.68E-04 | 0.11 | 0.04 | 0.07 | 2.94 |
| fusion | Pkp3 | 0.79 | 2.86E-08 | 6.12E-04 | 0.39 | 0.24 | 0.14 | 1.59 |
| fusion | Pdlim5 | 1.35 | 2.88E-08 | 6.14E-04 | 0.29 | 0.17 | 0.12 | 1.70 |
| fusion | Pak2 | 0.40 | 2.94E-08 | 6.29E-04 | 0.91 | 0.89 | 0.02 | 1.02 |
| fusion | Amn1 | 0.64 | 3.09E-08 | 6.59E-04 | 0.61 | 0.50 | 0.11 | 1.22 |

**Supplementary Table 4. Marker genes for the annotated cell populations of integrated clustering of E11.5 control and NotchGOF λ-junction ectoderm cells.**

| | | | | | | | | |
|---|---|---|---|---|---|---|---|---|
| fusion | Myl9 | 1.63 | 3.10E-08 | 6.63E-04 | 0.31 | 0.19 | 0.12 | 1.65 |
| fusion | Cdc42ep3 | 1.31 | 3.11E-08 | 6.64E-04 | 0.30 | 0.17 | 0.13 | 1.74 |
| fusion | Sesn2 | 1.50 | 3.21E-08 | 6.85E-04 | 0.21 | 0.11 | 0.11 | 2.02 |
| fusion | Spry1 | 1.30 | 3.23E-08 | 6.89E-04 | 0.44 | 0.32 | 0.12 | 1.38 |
| fusion | Kxd1 | 0.49 | 3.24E-08 | 6.91E-04 | 0.73 | 0.63 | 0.10 | 1.16 |
| fusion | Cacna1b | 0.93 | 3.29E-08 | 7.03E-04 | 0.43 | 0.30 | 0.13 | 1.43 |
| fusion | Siva1 | 0.40 | 3.37E-08 | 7.20E-04 | 0.98 | 0.97 | 0.01 | 1.01 |
| fusion | Tmtc2 | 0.94 | 4.03E-08 | 8.62E-04 | 0.31 | 0.18 | 0.13 | 1.74 |
| fusion | Arid3b | 0.66 | 4.08E-08 | 8.71E-04 | 0.55 | 0.41 | 0.14 | 1.35 |
| fusion | Sqstm1 | 0.61 | 4.26E-08 | 9.11E-04 | 0.58 | 0.46 | 0.13 | 1.28 |
| fusion | Qk | 0.38 | 4.29E-08 | 9.16E-04 | 0.91 | 0.87 | 0.03 | 1.04 |
| fusion | Ccdc28a | 1.01 | 4.34E-08 | 9.27E-04 | 0.28 | 0.16 | 0.12 | 1.79 |
| fusion | Mgat5 | 0.84 | 4.54E-08 | 9.69E-04 | 0.43 | 0.29 | 0.14 | 1.48 |
| fusion | Snhg6 | 0.48 | 4.56E-08 | 9.75E-04 | 0.78 | 0.73 | 0.05 | 1.07 |
| fusion | Cd81 | 0.34 | 4.68E-08 | 1.00E-03 | 0.94 | 0.92 | 0.02 | 1.02 |
| fusion | Bmp7 | 0.79 | 4.69E-08 | 1.00E-03 | 0.53 | 0.40 | 0.13 | 1.32 |
| fusion | Slc30a4 | 1.55 | 4.70E-08 | 1.00E-03 | 0.22 | 0.11 | 0.11 | 1.95 |
| fusion | Bag3 | 1.72 | 4.82E-08 | 1.03E-03 | 0.16 | 0.07 | 0.09 | 2.27 |
| fusion | Plin2 | 0.88 | 5.06E-08 | 1.08E-03 | 0.49 | 0.36 | 0.13 | 1.36 |
| fusion | Mtus1 | 0.93 | 6.33E-08 | 1.35E-03 | 0.42 | 0.29 | 0.13 | 1.43 |
| fusion | Rnf167 | 0.63 | 6.50E-08 | 1.39E-03 | 0.58 | 0.47 | 0.12 | 1.25 |
| fusion | Hey1 | 0.47 | 6.57E-08 | 1.40E-03 | 0.86 | 0.74 | 0.12 | 1.16 |
| fusion | Slc12a4 | 1.05 | 6.59E-08 | 1.41E-03 | 0.33 | 0.20 | 0.13 | 1.64 |
| fusion | Ccng2 | 0.78 | 7.12E-08 | 1.52E-03 | 0.51 | 0.39 | 0.12 | 1.31 |
| fusion | Etv3 | 0.90 | 7.90E-08 | 1.69E-03 | 0.41 | 0.29 | 0.13 | 1.45 |
| fusion | Tulp1 | 1.40 | 8.10E-08 | 1.73E-03 | 0.20 | 0.10 | 0.10 | 1.98 |
| fusion | Skp1a | 0.26 | 8.27E-08 | 1.77E-03 | 0.99 | 0.97 | 0.02 | 1.02 |
| fusion | Atp6v1e1 | 0.52 | 8.39E-08 | 1.79E-03 | 0.74 | 0.69 | 0.05 | 1.08 |
| fusion | Rtn4 | 0.29 | 8.50E-08 | 1.82E-03 | 0.98 | 0.96 | 0.03 | 1.03 |
| fusion | Zkscan3 | 0.45 | 9.50E-08 | 2.03E-03 | 0.77 | 0.67 | 0.10 | 1.15 |
| fusion | Tiam1 | 0.69 | 9.68E-08 | 2.07E-03 | 0.57 | 0.45 | 0.12 | 1.26 |
| fusion | A430035B10Rik | 1.59 | 9.72E-08 | 2.08E-03 | 0.11 | 0.04 | 0.07 | 2.79 |
| fusion | Filip1 | 1.97 | 1.01E-07 | 2.16E-03 | 0.16 | 0.07 | 0.09 | 2.21 |
| fusion | Gng12 | 1.03 | 1.02E-07 | 2.18E-03 | 0.35 | 0.23 | 0.12 | 1.52 |
| fusion | Kremen1 | 1.07 | 1.04E-07 | 2.23E-03 | 0.44 | 0.31 | 0.12 | 1.39 |
| fusion | Elk3 | 1.10 | 1.05E-07 | 2.24E-03 | 0.31 | 0.19 | 0.12 | 1.64 |
| fusion | C1qtnf12 | 1.36 | 1.06E-07 | 2.26E-03 | 0.34 | 0.22 | 0.11 | 1.50 |
| fusion | Clip3 | 0.73 | 1.07E-07 | 2.29E-03 | 0.55 | 0.46 | 0.10 | 1.21 |
| fusion | Smarce1 | 0.26 | 1.34E-07 | 2.85E-03 | 0.98 | 0.97 | 0.00 | 1.00 |
| fusion | Rnf24 | 0.79 | 1.34E-07 | 2.86E-03 | 0.53 | 0.41 | 0.13 | 1.32 |
| fusion | Pura | 0.51 | 1.42E-07 | 3.04E-03 | 0.72 | 0.64 | 0.08 | 1.13 |
| fusion | Ppp1r36 | 1.06 | 1.47E-07 | 3.13E-03 | 0.28 | 0.16 | 0.12 | 1.77 |
| fusion | Fgd6 | 1.00 | 1.55E-07 | 3.32E-03 | 0.34 | 0.22 | 0.12 | 1.57 |
| fusion | Zmiz1 | 0.62 | 1.62E-07 | 3.46E-03 | 0.70 | 0.62 | 0.08 | 1.12 |
| fusion | Tmem184b | 1.14 | 1.68E-07 | 3.58E-03 | 0.32 | 0.20 | 0.11 | 1.56 |
| fusion | Lpin2 | 0.78 | 1.74E-07 | 3.71E-03 | 0.55 | 0.46 | 0.09 | 1.19 |
| fusion | Sox14 | 1.50 | 1.74E-07 | 3.72E-03 | 0.12 | 0.05 | 0.08 | 2.60 |
| fusion | Ehbp1 | 0.68 | 1.77E-07 | 3.78E-03 | 0.60 | 0.49 | 0.11 | 1.23 |
| fusion | Rgl1 | 1.69 | 2.00E-07 | 4.28E-03 | 0.14 | 0.06 | 0.08 | 2.36 |
| fusion | Mxd4 | 0.62 | 2.04E-07 | 4.35E-03 | 0.54 | 0.41 | 0.13 | 1.31 |
| fusion | Ass1 | 1.10 | 2.20E-07 | 4.71E-03 | 0.22 | 0.12 | 0.11 | 1.91 |
| fusion | Mbd2 | 0.48 | 2.39E-07 | 5.11E-03 | 0.69 | 0.61 | 0.08 | 1.12 |
| fusion | Bach1 | 0.69 | 2.40E-07 | 5.13E-03 | 0.53 | 0.42 | 0.11 | 1.26 |
| fusion | Nfe2l3 | 1.22 | 2.43E-07 | 5.19E-03 | 0.24 | 0.14 | 0.11 | 1.80 |
| fusion | Kcnj15 | 1.60 | 2.62E-07 | 5.59E-03 | 0.13 | 0.06 | 0.08 | 2.42 |

**Supplementary Table 4. Marker genes for the annotated cell populations of integrated clustering of E11.5 control and NotchGOF λ-junction ectoderm cells.**

| | | | | | | | | |
|---|---|---|---|---|---|---|---|---|
| fusion | Macrod2 | 1.22 | 2.70E-07 | 5.76E-03 | 0.46 | 0.35 | 0.11 | 1.32 |
| fusion | Ahr | 1.52 | 2.73E-07 | 5.82E-03 | 0.28 | 0.18 | 0.11 | 1.61 |
| fusion | Lrp1 | 0.71 | 2.77E-07 | 5.92E-03 | 0.54 | 0.43 | 0.11 | 1.25 |
| fusion | Txndc17 | 0.31 | 2.77E-07 | 5.93E-03 | 0.95 | 0.95 | 0.00 | 1.00 |
| fusion | F11r | 0.49 | 2.82E-07 | 6.02E-03 | 0.69 | 0.61 | 0.09 | 1.14 |
| fusion | Htra1 | 1.69 | 3.02E-07 | 6.44E-03 | 0.14 | 0.06 | 0.08 | 2.31 |
| fusion | Fzd3 | 0.29 | 3.36E-07 | 7.17E-03 | 0.89 | 0.84 | 0.05 | 1.05 |
| fusion | Slc25a27 | 0.55 | 3.46E-07 | 7.39E-03 | 0.64 | 0.54 | 0.09 | 1.17 |
| fusion | Trappc6a | 0.47 | 3.47E-07 | 7.42E-03 | 0.72 | 0.66 | 0.06 | 1.09 |
| fusion | Plekhg3 | 1.08 | 3.48E-07 | 7.43E-03 | 0.38 | 0.27 | 0.12 | 1.43 |
| fusion | Dusp11 | 0.46 | 3.53E-07 | 7.55E-03 | 0.73 | 0.66 | 0.07 | 1.10 |
| fusion | Herc3 | 1.48 | 3.57E-07 | 7.63E-03 | 0.24 | 0.14 | 0.10 | 1.76 |
| fusion | Sinhcaf | 0.27 | 3.68E-07 | 7.86E-03 | 0.95 | 0.94 | 0.01 | 1.01 |
| fusion | Rara | 0.73 | 3.74E-07 | 7.99E-03 | 0.55 | 0.45 | 0.10 | 1.22 |
| fusion | Nfat5 | 0.43 | 4.10E-07 | 8.75E-03 | 0.72 | 0.63 | 0.09 | 1.14 |
| fusion | Gpbp1l1 | 0.65 | 4.14E-07 | 8.85E-03 | 0.56 | 0.45 | 0.11 | 1.25 |
| fusion | Nkd1 | 1.50 | 4.23E-07 | 9.04E-03 | 0.22 | 0.12 | 0.10 | 1.82 |
| fusion | Tspan5 | 0.93 | 4.24E-07 | 9.05E-03 | 0.44 | 0.34 | 0.10 | 1.30 |
| fusion | Znrf1 | 0.48 | 4.42E-07 | 9.44E-03 | 0.66 | 0.56 | 0.10 | 1.17 |
| fusion | Kmt2e | 0.38 | 4.43E-07 | 9.47E-03 | 0.88 | 0.86 | 0.02 | 1.02 |
| fusion | Anxa4 | 1.08 | 4.49E-07 | 9.59E-03 | 0.27 | 0.16 | 0.11 | 1.72 |
| fusion | Dok1 | 1.51 | 4.57E-07 | 9.76E-03 | 0.20 | 0.11 | 0.10 | 1.91 |
| fusion | Cpeb4 | 1.53 | 4.60E-07 | 9.83E-03 | 0.15 | 0.07 | 0.08 | 2.19 |
| fusion | Trappc2l | 0.48 | 4.77E-07 | 1.02E-02 | 0.68 | 0.61 | 0.07 | 1.12 |
| fusion | Mylk | 1.17 | 4.82E-07 | 1.03E-02 | 0.31 | 0.20 | 0.11 | 1.55 |
| fusion | Rap2b | 0.71 | 4.89E-07 | 1.04E-02 | 0.61 | 0.53 | 0.08 | 1.15 |
| fusion | Cuedc1 | 0.63 | 5.34E-07 | 1.14E-02 | 0.61 | 0.51 | 0.10 | 1.20 |
| fusion | Ubxn6 | 0.63 | 5.64E-07 | 1.21E-02 | 0.59 | 0.49 | 0.10 | 1.20 |
| fusion | Neil2 | 1.80 | 6.17E-07 | 1.32E-02 | 0.10 | 0.04 | 0.06 | 2.68 |
| fusion | Ankrd12 | 0.66 | 6.89E-07 | 1.47E-02 | 0.53 | 0.42 | 0.11 | 1.26 |
| fusion | Arhgef19 | 0.68 | 6.98E-07 | 1.49E-02 | 0.58 | 0.49 | 0.10 | 1.20 |
| fusion | Lhfp | 0.86 | 7.07E-07 | 1.51E-02 | 0.44 | 0.31 | 0.13 | 1.42 |
| fusion | Akap9 | 0.31 | 7.39E-07 | 1.58E-02 | 0.90 | 0.87 | 0.03 | 1.03 |
| fusion | Arhgap42 | 0.84 | 7.46E-07 | 1.59E-02 | 0.39 | 0.27 | 0.12 | 1.43 |
| fusion | Bcam | 0.46 | 7.72E-07 | 1.65E-02 | 0.82 | 0.72 | 0.10 | 1.14 |
| fusion | Crlf1 | 1.42 | 7.91E-07 | 1.69E-02 | 0.22 | 0.13 | 0.10 | 1.78 |
| fusion | Hook3 | 0.54 | 8.06E-07 | 1.72E-02 | 0.64 | 0.53 | 0.11 | 1.20 |
| fusion | Eps15 | 0.78 | 8.38E-07 | 1.79E-02 | 0.44 | 0.32 | 0.11 | 1.34 |
| fusion | Metap2 | 0.28 | 8.59E-07 | 1.84E-02 | 0.98 | 0.97 | 0.01 | 1.01 |
| fusion | Tcf4 | 0.43 | 8.97E-07 | 1.92E-02 | 0.91 | 0.89 | 0.03 | 1.03 |
| fusion | Ddx17 | 0.36 | 9.06E-07 | 1.94E-02 | 0.89 | 0.87 | 0.02 | 1.02 |
| fusion | Rps27rt | 0.40 | 9.55E-07 | 2.04E-02 | 0.84 | 0.78 | 0.06 | 1.08 |
| fusion | Rasgef1b | 1.22 | 9.62E-07 | 2.06E-02 | 0.18 | 0.09 | 0.09 | 1.98 |
| fusion | Sppl3 | 0.57 | 9.91E-07 | 2.12E-02 | 0.71 | 0.64 | 0.06 | 1.10 |
| fusion | Sema6c | 1.30 | 1.02E-06 | 2.19E-02 | 0.22 | 0.13 | 0.10 | 1.79 |
| fusion | Castor1 | 1.52 | 1.03E-06 | 2.19E-02 | 0.12 | 0.05 | 0.07 | 2.44 |
| fusion | Elf1 | 0.54 | 1.07E-06 | 2.30E-02 | 0.61 | 0.50 | 0.11 | 1.22 |
| fusion | Gpcpd1 | 1.02 | 1.12E-06 | 2.39E-02 | 0.32 | 0.21 | 0.11 | 1.53 |
| fusion | Gpr153 | 1.50 | 1.13E-06 | 2.42E-02 | 0.11 | 0.04 | 0.07 | 2.59 |
| fusion | Sik1 | 1.03 | 1.14E-06 | 2.44E-02 | 0.27 | 0.16 | 0.11 | 1.69 |
| fusion | Nfatc4 | 0.89 | 1.16E-06 | 2.47E-02 | 0.44 | 0.34 | 0.10 | 1.30 |
| fusion | Mrps33 | 0.27 | 1.19E-06 | 2.53E-02 | 0.95 | 0.94 | 0.01 | 1.01 |
| fusion | Glul | 0.64 | 1.19E-06 | 2.54E-02 | 0.57 | 0.46 | 0.11 | 1.23 |
| fusion | Rflnb | 0.64 | 1.34E-06 | 2.86E-02 | 0.62 | 0.52 | 0.10 | 1.20 |
| fusion | Kansl1l | 1.04 | 1.36E-06 | 2.91E-02 | 0.31 | 0.20 | 0.11 | 1.56 |

**Supplementary Table 4. Marker genes for the annotated cell populations of integrated clustering of E11.5 control and NotchGOF λ-junction ectoderm cells.**

| | | | | | | | | |
|---|---|---|---|---|---|---|---|---|
| fusion | Pink1 | 0.87 | 1.42E-06 | 3.03E-02 | 0.36 | 0.25 | 0.11 | 1.44 |
| fusion | Efna1 | 0.56 | 1.45E-06 | 3.10E-02 | 0.52 | 0.40 | 0.12 | 1.30 |
| fusion | Smad5 | 0.63 | 1.61E-06 | 3.44E-02 | 0.57 | 0.47 | 0.10 | 1.22 |
| fusion | Ctnnd1 | 0.63 | 1.62E-06 | 3.45E-02 | 0.60 | 0.52 | 0.08 | 1.16 |
| fusion | Cdk19 | 0.84 | 1.64E-06 | 3.50E-02 | 0.40 | 0.30 | 0.11 | 1.36 |
| fusion | Ebpl | 0.54 | 1.76E-06 | 3.75E-02 | 0.64 | 0.55 | 0.09 | 1.17 |
| fusion | Heca | 0.77 | 1.77E-06 | 3.79E-02 | 0.38 | 0.26 | 0.12 | 1.46 |
| fusion | Kdm3a | 0.65 | 1.80E-06 | 3.85E-02 | 0.56 | 0.47 | 0.09 | 1.20 |
| fusion | Cttnbp2nl | 0.77 | 1.87E-06 | 4.00E-02 | 0.51 | 0.43 | 0.08 | 1.19 |
| fusion | Stim1 | 1.03 | 1.88E-06 | 4.01E-02 | 0.28 | 0.17 | 0.11 | 1.64 |
| fusion | Rbm5 | 0.37 | 1.90E-06 | 4.06E-02 | 0.84 | 0.81 | 0.03 | 1.04 |
| fusion | Lrrc58 | 0.47 | 1.91E-06 | 4.07E-02 | 0.68 | 0.61 | 0.07 | 1.11 |
| fusion | Cir1 | 0.49 | 2.00E-06 | 4.28E-02 | 0.71 | 0.64 | 0.07 | 1.11 |
| fusion | Hmox1 | 1.26 | 2.01E-06 | 4.29E-02 | 0.17 | 0.09 | 0.08 | 1.99 |
| fusion | Pard6g | 0.53 | 2.03E-06 | 4.34E-02 | 0.69 | 0.65 | 0.05 | 1.07 |
| fusion | H2-T23 | 1.15 | 2.09E-06 | 4.47E-02 | 0.25 | 0.15 | 0.10 | 1.69 |
| fusion | Numb | 1.03 | 2.15E-06 | 4.59E-02 | 0.42 | 0.32 | 0.10 | 1.31 |
| fusion | Gabarapl2 | 0.34 | 2.19E-06 | 4.67E-02 | 0.88 | 0.87 | 0.01 | 1.01 |
| fusion | Zfp637 | 0.48 | 2.27E-06 | 4.85E-02 | 0.71 | 0.65 | 0.06 | 1.09 |
| fusion | 4930402H24Rik | 1.01 | 2.30E-06 | 4.91E-02 | 0.26 | 0.16 | 0.10 | 1.65 |
| OFN | Lhx2 | 10.06 | 0 | 0 | 0.93 | 0.02 | 0.91 | 38.71 |
| OFN | Nhlh2 | 11.10 | 0 | 0 | 0.87 | 0.00 | 0.86 | 216.25 |
| OFN | Igfbpl1 | 9.51 | 0 | 0 | 0.85 | 0.01 | 0.84 | 71.00 |
| OFN | Ppp1r14a | 7.64 | 0 | 0 | 0.88 | 0.04 | 0.84 | 20.88 |
| OFN | Ebf2 | 9.72 | 0 | 0 | 0.83 | 0.00 | 0.83 | 208.00 |
| OFN | Neurod1 | 11.59 | 0 | 0 | 0.83 | 0.00 | 0.82 | 275.33 |
| OFN | Insm1 | 7.47 | 0 | 0 | 0.85 | 0.02 | 0.82 | 36.74 |
| OFN | Elavl3 | 9.48 | 0 | 0 | 0.83 | 0.01 | 0.82 | 64.00 |
| OFN | Srrm4 | 9.29 | 0 | 0 | 0.79 | 0.01 | 0.78 | 112.43 |
| OFN | Neurog1 | 8.96 | 0 | 0 | 0.74 | 0.01 | 0.73 | 147.00 |
| OFN | Nhlh1 | 11.32 | 0 | 0 | 0.70 | 0.00 | 0.70 | 703.00 |
| OFN | Gng8 | 8.70 | 0 | 0 | 0.66 | 0.01 | 0.65 | 54.83 |
| OFN | Ebf3 | 7.79 | 0 | 0 | 0.66 | 0.02 | 0.64 | 32.90 |
| OFN | Sowaha | 10.18 | 0 | 0 | 0.63 | 0.00 | 0.63 | 626.00 |
| OFN | Btbd17 | 7.71 | 0 | 0 | 0.63 | 0.02 | 0.62 | 39.50 |
| OFN | Miat | 6.47 | 0 | 0 | 0.59 | 0.01 | 0.58 | 54.00 |
| OFN | St18 | 10.57 | 0 | 0 | 0.55 | 0.00 | 0.55 | 548.00 |
| OFN | Cdhr1 | 8.38 | 0 | 0 | 0.56 | 0.01 | 0.55 | 61.67 |
| OFN | Ina | 10.42 | 0 | 0 | 0.54 | 0.01 | 0.53 | 76.43 |
| OFN | Slc17a6 | 11.09 | 0 | 0 | 0.50 | 0.00 | 0.50 | #NA |
| OFN | Olig2 | 11.15 | 0 | 0 | 0.48 | 0.00 | 0.48 | 477.00 |
| OFN | Celf3 | 8.37 | 0 | 0 | 0.46 | 0.00 | 0.46 | 152.67 |
| OFN | Atp8b4 | 10.14 | 0 | 0 | 0.43 | 0.00 | 0.43 | 216.00 |
| OFN | Myt1l | 10.05 | 0 | 0 | 0.43 | 0.00 | 0.42 | 142.00 |
| OFN | Wdr66 | 6.38 | 2.39E-277 | 5.11E-273 | 0.72 | 0.04 | 0.69 | 20.66 |
| OFN | Elavl4 | 7.98 | 2.94E-268 | 6.27E-264 | 0.91 | 0.08 | 0.83 | 11.67 |
| OFN | Cbfa2t3 | 7.17 | 1.47E-267 | 3.14E-263 | 0.42 | 0.01 | 0.41 | 83.80 |
| OFN | Tubb3 | 8.09 | 4.71E-261 | 1.01E-256 | 0.99 | 0.11 | 0.89 | 9.29 |
| OFN | Ttc9b | 7.86 | 8.30E-258 | 1.77E-253 | 0.41 | 0.01 | 0.40 | 81.20 |
| OFN | Rbm24 | 6.56 | 1.14E-257 | 2.44E-253 | 0.52 | 0.01 | 0.50 | 36.86 |
| OFN | Dscaml1 | 7.97 | 6.66E-248 | 1.42E-243 | 0.34 | 0.00 | 0.33 | 167.50 |
| OFN | Kif19a | 6.20 | 5.64E-247 | 1.20E-242 | 0.59 | 0.02 | 0.56 | 25.52 |
| OFN | Nxph4 | 8.10 | 1.76E-239 | 3.76E-235 | 0.34 | 0.00 | 0.33 | 167.50 |
| OFN | Akap6 | 8.99 | 3.70E-233 | 7.91E-229 | 0.32 | 0.00 | 0.32 | 161.50 |
| OFN | Syp | 5.95 | 3.17E-232 | 6.78E-228 | 0.62 | 0.03 | 0.59 | 19.97 |

**Supplementary Table 4. Marker genes for the annotated cell populations of integrated clustering of E11.5 control and NotchGOF λ-junction ectoderm cells.**

| | | | | | | | | |
|---|---|---|---|---|---|---|---|---|
| OFN | Emx2 | 6.68 | 7.90E-228 | 1.69E-223 | 0.50 | 0.02 | 0.48 | 31.06 |
| OFN | Map6 | 7.91 | 3.80E-226 | 8.11E-222 | 0.36 | 0.00 | 0.35 | 88.75 |
| OFN | Ndrg4 | 6.21 | 3.67E-224 | 7.84E-220 | 0.54 | 0.02 | 0.51 | 25.48 |
| OFN | Syt13 | 5.97 | 1.43E-222 | 3.06E-218 | 0.55 | 0.02 | 0.53 | 23.83 |
| OFN | Dcc | 6.29 | 4.20E-217 | 8.98E-213 | 0.50 | 0.02 | 0.48 | 26.47 |
| OFN | Igsf21 | 8.34 | 8.40E-201 | 1.79E-196 | 0.28 | 0.00 | 0.28 | 142.00 |
| OFN | Slc17a7 | 8.67 | 1.23E-195 | 2.62E-191 | 0.24 | 0.00 | 0.24 | #NA |
| OFN | Scg3 | 6.43 | 6.28E-191 | 1.34E-186 | 0.43 | 0.01 | 0.41 | 30.43 |
| OFN | Rgs20 | 7.78 | 1.05E-190 | 2.24E-186 | 0.29 | 0.00 | 0.29 | 96.67 |
| OFN | Nrn1 | 10.64 | 2.74E-190 | 5.85E-186 | 0.23 | 0.00 | 0.23 | #NA |
| OFN | Mfng | 5.85 | 6.54E-190 | 1.40E-185 | 0.79 | 0.09 | 0.71 | 9.13 |
| OFN | Smim18 | 8.98 | 8.79E-185 | 1.88E-180 | 0.23 | 0.00 | 0.23 | #NA |
| OFN | Olig1 | 7.94 | 3.81E-184 | 8.14E-180 | 0.31 | 0.01 | 0.31 | 62.00 |
| OFN | Inka2 | 6.18 | 1.15E-181 | 2.45E-177 | 0.37 | 0.01 | 0.36 | 34.00 |
| OFN | Tagln3 | 6.22 | 1.53E-181 | 3.27E-177 | 0.77 | 0.08 | 0.69 | 9.37 |
| OFN | Myt1 | 8.14 | 1.20E-180 | 2.57E-176 | 0.24 | 0.00 | 0.24 | 239.00 |
| OFN | 5730403I07Rik | 8.97 | 4.22E-180 | 9.02E-176 | 0.23 | 0.00 | 0.23 | 232.00 |
| OFN | Abcd2 | 6.17 | 1.26E-179 | 2.70E-175 | 0.35 | 0.01 | 0.34 | 43.50 |
| OFN | Clvs1 | 7.54 | 1.32E-177 | 2.83E-173 | 0.29 | 0.00 | 0.29 | 72.50 |
| OFN | Actl6b | 9.11 | 2.90E-174 | 6.19E-170 | 0.22 | 0.00 | 0.22 | 219.00 |
| OFN | Fstl5 | 7.50 | 3.16E-173 | 6.75E-169 | 0.37 | 0.01 | 0.36 | 31.17 |
| OFN | Chgb | 8.90 | 4.30E-173 | 9.19E-169 | 0.25 | 0.00 | 0.24 | 122.50 |
| OFN | Chga | 6.76 | 4.50E-170 | 9.60E-166 | 0.54 | 0.04 | 0.50 | 15.29 |
| OFN | Zeb1 | 5.77 | 1.26E-169 | 2.69E-165 | 0.42 | 0.02 | 0.40 | 24.65 |
| OFN | Dcx | 5.44 | 2.38E-169 | 5.09E-165 | 0.66 | 0.06 | 0.60 | 10.97 |
| OFN | Rgs4 | 9.56 | 7.74E-168 | 1.65E-163 | 0.24 | 0.00 | 0.24 | 119.50 |
| OFN | Runx1t1 | 5.20 | 6.57E-164 | 1.40E-159 | 0.53 | 0.04 | 0.49 | 15.11 |
| OFN | Pirt | 9.33 | 3.50E-163 | 7.47E-159 | 0.20 | 0.00 | 0.20 | #NA |
| OFN | Tuba4a | 5.78 | 1.00E-160 | 2.14E-156 | 0.63 | 0.06 | 0.57 | 10.98 |
| OFN | Gap43 | 7.01 | 3.60E-160 | 7.68E-156 | 0.61 | 0.05 | 0.55 | 11.65 |
| OFN | Pcsk2 | 8.24 | 2.66E-157 | 5.67E-153 | 0.23 | 0.00 | 0.22 | 113.00 |
| OFN | Stmn4 | 7.12 | 2.56E-153 | 5.46E-149 | 0.29 | 0.01 | 0.28 | 41.43 |
| OFN | A230051N06Rik | 8.62 | 1.28E-152 | 2.73E-148 | 0.19 | 0.00 | 0.19 | 194.00 |
| OFN | Camk2n1 | 5.30 | 6.23E-150 | 1.33E-145 | 0.74 | 0.10 | 0.64 | 7.58 |
| OFN | Ebf1 | 5.23 | 6.43E-149 | 1.37E-144 | 0.78 | 0.11 | 0.67 | 6.85 |
| OFN | Gng3 | 6.26 | 1.56E-148 | 3.34E-144 | 0.57 | 0.05 | 0.52 | 11.59 |
| OFN | Ppp1r17 | 11.47 | 4.51E-147 | 9.63E-143 | 0.17 | 0.00 | 0.17 | #NA |
| OFN | Gfy | 9.49 | 5.58E-147 | 1.19E-142 | 0.18 | 0.00 | 0.18 | #NA |
| OFN | Gm26871 | 8.94 | 6.02E-147 | 1.29E-142 | 0.18 | 0.00 | 0.18 | #NA |
| OFN | Chrna3 | 9.54 | 6.02E-147 | 1.29E-142 | 0.18 | 0.00 | 0.18 | #NA |
| OFN | Tmem163 | 6.36 | 1.06E-145 | 2.25E-141 | 0.32 | 0.01 | 0.31 | 29.36 |
| OFN | Coro2a | 5.14 | 2.78E-145 | 5.95E-141 | 0.54 | 0.04 | 0.50 | 12.32 |
| OFN | Fezf1 | 5.02 | 4.71E-145 | 1.01E-140 | 0.67 | 0.08 | 0.59 | 8.87 |
| OFN | Pnmal1 | 7.78 | 1.23E-142 | 2.63E-138 | 0.21 | 0.00 | 0.21 | 106.50 |
| OFN | Cacna2d1 | 5.67 | 1.59E-141 | 3.39E-137 | 0.53 | 0.04 | 0.49 | 12.02 |
| OFN | Lamp5 | 7.99 | 9.87E-138 | 2.11E-133 | 0.19 | 0.00 | 0.19 | 187.00 |
| OFN | Syt4 | 5.03 | 1.92E-135 | 4.10E-131 | 0.39 | 0.02 | 0.37 | 18.76 |
| OFN | Srrm3 | 6.23 | 6.58E-135 | 1.41E-130 | 0.29 | 0.01 | 0.28 | 32.22 |
| OFN | Art5 | 6.99 | 7.53E-134 | 1.61E-129 | 0.21 | 0.00 | 0.20 | 103.00 |
| OFN | Rtn1 | 5.96 | 3.06E-133 | 6.54E-129 | 0.92 | 0.22 | 0.70 | 4.14 |
| OFN | Thsd7b | 4.49 | 3.65E-130 | 7.80E-126 | 0.52 | 0.05 | 0.48 | 11.62 |
| OFN | Dll3 | 5.37 | 3.38E-129 | 7.22E-125 | 0.30 | 0.01 | 0.29 | 29.70 |
| OFN | Rgs16 | 5.69 | 1.02E-128 | 2.19E-124 | 0.28 | 0.01 | 0.27 | 34.63 |
| OFN | Foxo6 | 5.41 | 3.03E-127 | 6.46E-123 | 0.47 | 0.04 | 0.43 | 12.92 |
| OFN | Gdap1l1 | 5.53 | 6.97E-127 | 1.49E-122 | 0.50 | 0.04 | 0.46 | 11.43 |

**Supplementary Table 4. Marker genes for the annotated cell populations of integrated clustering of E11.5 control and NotchGOF λ-junction ectoderm cells.**

| | | | | | | | | |
|---|---|---|---|---|---|---|---|---|
| OFN | Tmprss6 | 8.21 | 2.81E-126 | 6.01E-122 | 0.17 | 0.00 | 0.17 | 168.00 |
| OFN | Rd3 | 9.94 | 2.23E-125 | 4.77E-121 | 0.16 | 0.00 | 0.16 | #NA |
| OFN | Gda | 7.62 | 8.23E-122 | 1.76E-117 | 0.17 | 0.00 | 0.17 | 168.00 |
| OFN | Clmp | 4.91 | 3.44E-121 | 7.35E-117 | 0.60 | 0.07 | 0.53 | 8.11 |
| OFN | Gm27197 | 9.13 | 4.68E-120 | 9.99E-116 | 0.14 | 0.00 | 0.14 | #NA |
| OFN | Gm13589 | 6.83 | 4.14E-119 | 8.84E-115 | 0.20 | 0.00 | 0.20 | 66.67 |
| OFN | Islr2 | 8.18 | 4.66E-119 | 9.95E-115 | 0.17 | 0.00 | 0.17 | 87.00 |
| OFN | Nol4 | 6.10 | 1.02E-118 | 2.19E-114 | 0.27 | 0.01 | 0.26 | 29.44 |
| OFN | Cend1 | 6.14 | 4.82E-118 | 1.03E-113 | 0.24 | 0.01 | 0.23 | 39.83 |
| OFN | Gfra1 | 5.15 | 3.14E-117 | 6.70E-113 | 0.38 | 0.02 | 0.36 | 15.88 |
| OFN | Vwa5b2 | 6.73 | 8.89E-117 | 1.90E-112 | 0.18 | 0.00 | 0.18 | 90.50 |
| OFN | Stmn3 | 5.75 | 2.03E-116 | 4.34E-112 | 0.65 | 0.10 | 0.56 | 6.79 |
| OFN | Sct | 7.02 | 3.93E-114 | 8.39E-110 | 0.45 | 0.04 | 0.41 | 11.71 |
| OFN | Scrt1 | 6.18 | 4.01E-113 | 8.56E-109 | 0.25 | 0.01 | 0.24 | 30.63 |
| OFN | Brsk2 | 4.57 | 4.51E-112 | 9.64E-108 | 0.43 | 0.04 | 0.40 | 12.34 |
| OFN | Dner | 5.25 | 1.42E-111 | 3.04E-107 | 0.27 | 0.01 | 0.26 | 26.50 |
| OFN | Syt2 | 6.36 | 2.98E-111 | 6.37E-107 | 0.23 | 0.01 | 0.22 | 37.67 |
| OFN | Shisal1 | 5.07 | 4.90E-111 | 1.05E-106 | 0.36 | 0.02 | 0.33 | 16.14 |
| OFN | Mast1 | 5.09 | 2.11E-109 | 4.52E-105 | 0.28 | 0.01 | 0.27 | 23.08 |
| OFN | Stmn2 | 6.00 | 2.07E-108 | 4.42E-104 | 0.79 | 0.18 | 0.62 | 4.46 |
| OFN | Col16a1 | 6.75 | 4.66E-107 | 9.95E-103 | 0.25 | 0.01 | 0.24 | 28.00 |
| OFN | Soga3 | 4.36 | 5.74E-106 | 1.23E-101 | 0.74 | 0.15 | 0.59 | 4.95 |
| OFN | Myo7a | 5.27 | 1.18E-105 | 2.53E-101 | 0.27 | 0.01 | 0.26 | 22.58 |
| OFN | Jakmip2 | 4.57 | 1.54E-105 | 3.29E-101 | 0.47 | 0.05 | 0.42 | 10.11 |
| OFN | Nrxn1 | 4.82 | 1.78E-104 | 3.81E-100 | 0.53 | 0.06 | 0.47 | 8.27 |
| OFN | Ckmt1 | 5.27 | 6.12E-104 | 1.31E-99 | 0.67 | 0.12 | 0.55 | 5.64 |
| OFN | Gm31819 | 8.78 | 7.11E-104 | 1.52E-99 | 0.12 | 0.00 | 0.12 | #NA |
| OFN | Stxbp1 | 4.83 | 1.01E-103 | 2.16E-99 | 0.35 | 0.02 | 0.33 | 15.13 |
| OFN | Cotl1 | 4.59 | 1.61E-103 | 3.43E-99 | 0.98 | 0.47 | 0.51 | 2.08 |
| OFN | Lingo2 | 6.14 | 1.36E-100 | 2.91E-96 | 0.23 | 0.01 | 0.22 | 29.00 |
| OFN | Spock1 | 6.38 | 9.06E-100 | 1.94E-95 | 0.21 | 0.01 | 0.20 | 34.33 |
| OFN | Rassf4 | 5.78 | 1.35E-99 | 2.89E-95 | 0.27 | 0.01 | 0.26 | 20.85 |
| OFN | Lrrc10b | 6.45 | 4.94E-99 | 1.06E-94 | 0.18 | 0.00 | 0.18 | 45.25 |
| OFN | Sez6l2 | 6.36 | 7.20E-99 | 1.54E-94 | 0.19 | 0.00 | 0.18 | 46.75 |
| OFN | Map1a | 4.41 | 1.35E-98 | 2.88E-94 | 0.45 | 0.05 | 0.40 | 9.89 |
| OFN | Ntrk1 | 9.46 | 1.74E-98 | 3.72E-94 | 0.12 | 0.00 | 0.12 | #NA |
| OFN | Crmp1 | 4.73 | 5.65E-98 | 1.21E-93 | 0.68 | 0.13 | 0.55 | 5.25 |
| OFN | Tesc | 7.47 | 7.75E-98 | 1.65E-93 | 0.28 | 0.02 | 0.27 | 18.93 |
| OFN | Cplx2 | 4.85 | 3.90E-97 | 8.34E-93 | 0.77 | 0.19 | 0.58 | 4.15 |
| OFN | Nsg2 | 5.40 | 6.29E-97 | 1.34E-92 | 0.35 | 0.03 | 0.32 | 13.92 |
| OFN | Rnf112 | 5.62 | 6.41E-97 | 1.37E-92 | 0.20 | 0.01 | 0.19 | 33.33 |
| OFN | Mreg | 5.26 | 7.16E-97 | 1.53E-92 | 0.34 | 0.02 | 0.32 | 14.25 |
| OFN | Atp1a3 | 7.65 | 2.18E-95 | 4.65E-91 | 0.14 | 0.00 | 0.13 | 135.00 |
| OFN | Gm17322 | 7.73 | 1.01E-94 | 2.16E-90 | 0.17 | 0.00 | 0.17 | 56.00 |
| OFN | Asic1 | 5.84 | 2.52E-94 | 5.39E-90 | 0.19 | 0.01 | 0.18 | 37.40 |
| OFN | Nrxn2 | 5.14 | 2.84E-94 | 6.08E-90 | 0.32 | 0.02 | 0.30 | 14.68 |
| OFN | Ccdc184 | 8.73 | 1.73E-93 | 3.69E-89 | 0.12 | 0.00 | 0.12 | 123.00 |
| OFN | Sgpp2 | 4.75 | 2.20E-93 | 4.69E-89 | 0.31 | 0.02 | 0.29 | 15.50 |
| OFN | Kcnk10 | 9.12 | 4.25E-93 | 9.09E-89 | 0.11 | 0.00 | 0.11 | #NA |
| OFN | Tmsb4x | 2.63 | 1.64E-92 | 3.49E-88 | 1.00 | 1.00 | 0.00 | 1.00 |
| OFN | Cdyl2 | 5.61 | 3.01E-92 | 6.42E-88 | 0.35 | 0.03 | 0.32 | 12.43 |
| OFN | Gadd45a | 4.72 | 3.56E-92 | 7.61E-88 | 0.76 | 0.19 | 0.56 | 3.95 |
| OFN | Scn9a | 4.58 | 1.24E-91 | 2.65E-87 | 0.37 | 0.03 | 0.34 | 11.33 |
| OFN | Serpini1 | 4.28 | 1.09E-89 | 2.32E-85 | 0.38 | 0.04 | 0.35 | 10.89 |
| OFN | Gm3234 | 6.30 | 1.44E-89 | 3.08E-85 | 0.17 | 0.00 | 0.16 | 42.00 |

**Supplementary Table 4. Marker genes for the annotated cell populations of integrated clustering of E11.5 control and NotchGOF λ-junction ectoderm cells.**

| | | | | | | | | |
|---|---|---|---|---|---|---|---|---|
| OFN | Cerkl | 6.48 | 1.49E-89 | 3.18E-85 | 0.14 | 0.00 | 0.14 | 71.00 |
| OFN | Cpeb1 | 8.63 | 3.70E-88 | 7.91E-84 | 0.12 | 0.00 | 0.12 | 116.00 |
| OFN | Uncx | 4.93 | 4.70E-88 | 1.00E-83 | 0.53 | 0.08 | 0.45 | 6.61 |
| OFN | Lhx3 | 8.16 | 1.03E-87 | 2.21E-83 | 0.10 | 0.00 | 0.10 | #NA |
| OFN | Chrna9 | 8.57 | 1.03E-87 | 2.21E-83 | 0.10 | 0.00 | 0.10 | #NA |
| OFN | Neurod4 | 9.82 | 1.03E-87 | 2.21E-83 | 0.10 | 0.00 | 0.10 | #NA |
| OFN | Apba1 | 8.67 | 1.10E-87 | 2.34E-83 | 0.11 | 0.00 | 0.11 | #NA |
| OFN | Ccdc187 | 8.56 | 1.11E-87 | 2.37E-83 | 0.11 | 0.00 | 0.11 | #NA |
| OFN | Eef1a2 | 8.33 | 1.12E-87 | 2.40E-83 | 0.11 | 0.00 | 0.11 | #NA |
| OFN | Emx2os | 5.64 | 1.55E-87 | 3.32E-83 | 0.18 | 0.01 | 0.18 | 36.20 |
| OFN | Rundc3a | 4.40 | 5.95E-86 | 1.27E-81 | 0.52 | 0.08 | 0.45 | 6.79 |
| OFN | Pcdh8 | 5.13 | 8.31E-86 | 1.78E-81 | 0.32 | 0.02 | 0.29 | 13.17 |
| OFN | Cadps | 4.94 | 2.02E-85 | 4.31E-81 | 0.30 | 0.02 | 0.28 | 14.14 |
| OFN | Map1b | 4.85 | 3.86E-85 | 8.25E-81 | 0.95 | 0.56 | 0.39 | 1.71 |
| OFN | Fam110a | 4.09 | 1.29E-84 | 2.75E-80 | 0.83 | 0.28 | 0.56 | 3.03 |
| OFN | Dusp4 | 4.25 | 4.24E-84 | 9.06E-80 | 0.48 | 0.06 | 0.41 | 7.45 |
| OFN | Apc2 | 5.32 | 1.07E-83 | 2.29E-79 | 0.25 | 0.01 | 0.24 | 18.00 |
| OFN | 4930578M01Rik | 4.63 | 4.27E-83 | 9.12E-79 | 0.38 | 0.04 | 0.34 | 9.77 |
| OFN | Trim66 | 8.33 | 2.57E-82 | 5.50E-78 | 0.10 | 0.00 | 0.10 | #NA |
| OFN | Mapk10 | 5.82 | 3.41E-82 | 7.28E-78 | 0.19 | 0.01 | 0.19 | 27.71 |
| OFN | Stmn1 | 1.64 | 1.07E-81 | 2.28E-77 | 1.00 | 1.00 | 0.00 | 1.00 |
| OFN | Serping1 | 4.55 | 1.19E-81 | 2.55E-77 | 0.40 | 0.04 | 0.36 | 9.09 |
| OFN | Grasp | 4.46 | 5.62E-81 | 1.20E-76 | 0.68 | 0.17 | 0.52 | 4.05 |
| OFN | Plk3 | 5.53 | 8.72E-81 | 1.86E-76 | 0.24 | 0.01 | 0.23 | 18.38 |
| OFN | Rab6b | 5.61 | 1.39E-80 | 2.96E-76 | 0.34 | 0.03 | 0.31 | 10.69 |
| OFN | Tuba1a | 3.37 | 1.18E-78 | 2.53E-74 | 0.99 | 0.98 | 0.02 | 1.02 |
| OFN | Gnas | 1.63 | 2.73E-78 | 5.84E-74 | 1.00 | 1.00 | 0.00 | 1.00 |
| OFN | Nacad | 5.70 | 8.14E-78 | 1.74E-73 | 0.16 | 0.00 | 0.15 | 38.75 |
| OFN | Gm14226 | 7.38 | 9.59E-78 | 2.05E-73 | 0.12 | 0.00 | 0.12 | 61.50 |
| OFN | Basp1 | 2.35 | 3.57E-77 | 7.63E-73 | 0.99 | 0.97 | 0.03 | 1.03 |
| OFN | Cntn1 | 5.41 | 1.60E-76 | 3.42E-72 | 0.22 | 0.01 | 0.21 | 19.91 |
| OFN | Dkkl1 | 5.16 | 7.58E-76 | 1.62E-71 | 0.20 | 0.01 | 0.19 | 22.22 |
| OFN | Galnt17 | 4.93 | 1.04E-75 | 2.22E-71 | 0.21 | 0.01 | 0.20 | 20.60 |
| OFN | Rem2 | 4.47 | 3.50E-75 | 7.48E-71 | 0.50 | 0.08 | 0.42 | 6.29 |
| OFN | Mapt | 5.45 | 3.55E-74 | 7.58E-70 | 0.33 | 0.03 | 0.30 | 10.28 |
| OFN | Gdap1 | 4.19 | 4.24E-73 | 9.05E-69 | 0.38 | 0.05 | 0.34 | 8.47 |
| OFN | Coro1a | 4.83 | 6.29E-71 | 1.34E-66 | 0.36 | 0.04 | 0.32 | 8.80 |
| OFN | Rap1gap2 | 4.84 | 2.81E-70 | 5.99E-66 | 0.26 | 0.02 | 0.24 | 13.58 |
| OFN | Cadm3 | 4.70 | 3.28E-70 | 7.01E-66 | 0.27 | 0.02 | 0.25 | 12.32 |
| OFN | Arhgdig | 5.92 | 1.22E-69 | 2.60E-65 | 0.66 | 0.18 | 0.48 | 3.66 |
| OFN | Kcnh3 | 6.94 | 2.65E-69 | 5.67E-65 | 0.10 | 0.00 | 0.10 | 103.00 |
| OFN | Ada | 4.48 | 2.66E-69 | 5.69E-65 | 0.40 | 0.05 | 0.35 | 7.55 |
| OFN | Gm16364 | 7.26 | 1.23E-67 | 2.62E-63 | 0.11 | 0.00 | 0.11 | 55.00 |
| OFN | Prnp | 3.51 | 2.19E-67 | 4.68E-63 | 0.66 | 0.17 | 0.49 | 3.85 |
| OFN | Grik2 | 5.25 | 7.03E-67 | 1.50E-62 | 0.17 | 0.01 | 0.17 | 21.75 |
| OFN | Trp53i11 | 3.19 | 8.78E-67 | 1.88E-62 | 0.86 | 0.41 | 0.45 | 2.11 |
| OFN | Kif5a | 4.49 | 1.56E-66 | 3.34E-62 | 0.37 | 0.05 | 0.32 | 7.83 |
| OFN | Selenom | 4.71 | 7.00E-66 | 1.50E-61 | 0.83 | 0.39 | 0.44 | 2.15 |
| OFN | Nefl | 7.77 | 3.57E-65 | 7.64E-61 | 0.14 | 0.00 | 0.14 | 35.50 |
| OFN | Fam169b | 6.61 | 1.30E-64 | 2.78E-60 | 0.11 | 0.00 | 0.11 | 55.00 |
| OFN | Svop | 6.36 | 4.59E-64 | 9.81E-60 | 0.12 | 0.00 | 0.11 | 58.00 |
| OFN | Hes6 | 4.28 | 4.80E-64 | 1.03E-59 | 0.87 | 0.52 | 0.35 | 1.67 |
| OFN | Fam57b | 5.02 | 1.99E-63 | 4.25E-59 | 0.21 | 0.01 | 0.20 | 15.21 |
| OFN | Kcnmb2 | 4.87 | 3.36E-63 | 7.18E-59 | 0.32 | 0.04 | 0.28 | 9.03 |
| OFN | Map2 | 3.67 | 4.70E-63 | 1.00E-58 | 0.71 | 0.24 | 0.47 | 2.98 |

**Supplementary Table 4. Marker genes for the annotated cell populations of integrated clustering of E11.5 control and NotchGOF λ-junction ectoderm cells.**

| | | | | | | | | |
|---|---|---|---|---|---|---|---|---|
| OFN | L1cam | 5.30 | 5.61E-62 | 1.20E-57 | 0.20 | 0.01 | 0.19 | 16.67 |
| OFN | D930020B18Rik | 4.39 | 6.31E-62 | 1.35E-57 | 0.37 | 0.05 | 0.32 | 7.36 |
| OFN | Snca | 4.36 | 9.16E-62 | 1.96E-57 | 0.53 | 0.11 | 0.42 | 4.77 |
| OFN | Ptms | 1.71 | 6.55E-61 | 1.40E-56 | 0.95 | 0.98 | -0.03 | 0.97 |
| OFN | Cnrip1 | 4.82 | 1.44E-60 | 3.07E-56 | 0.25 | 0.02 | 0.23 | 11.45 |
| OFN | Fam78b | 5.00 | 2.16E-60 | 4.61E-56 | 0.23 | 0.02 | 0.21 | 13.29 |
| OFN | Actb | 1.00 | 9.41E-60 | 2.01E-55 | 1.00 | 1.00 | 0.00 | 1.00 |
| OFN | Scg2 | 5.70 | 2.18E-59 | 4.65E-55 | 0.16 | 0.01 | 0.15 | 20.13 |
| OFN | Tubb2b | 2.14 | 3.48E-59 | 7.43E-55 | 0.98 | 0.92 | 0.06 | 1.06 |
| OFN | Fam219a | 4.40 | 4.20E-59 | 8.97E-55 | 0.23 | 0.02 | 0.21 | 12.56 |
| OFN | a | 5.18 | 1.90E-57 | 4.05E-53 | 0.19 | 0.01 | 0.18 | 14.92 |
| OFN | Eif4e3 | 3.14 | 4.60E-57 | 9.82E-53 | 0.66 | 0.21 | 0.45 | 3.21 |
| OFN | Pde2a | 6.47 | 4.94E-57 | 1.05E-52 | 0.10 | 0.00 | 0.10 | 51.50 |
| OFN | Dpysl3 | 2.73 | 9.41E-57 | 2.01E-52 | 0.90 | 0.56 | 0.35 | 1.62 |
| OFN | Prex1 | 3.43 | 1.05E-56 | 2.23E-52 | 0.50 | 0.11 | 0.39 | 4.61 |
| OFN | Frrs1l | 5.22 | 1.07E-56 | 2.28E-52 | 0.15 | 0.01 | 0.14 | 21.14 |
| OFN | Atp1b1 | 4.15 | 1.50E-56 | 3.20E-52 | 0.54 | 0.13 | 0.41 | 4.04 |
| OFN | Btg2 | 3.28 | 3.75E-56 | 8.00E-52 | 0.76 | 0.30 | 0.46 | 2.54 |
| OFN | Atf5 | 4.46 | 1.20E-55 | 2.56E-51 | 0.65 | 0.21 | 0.44 | 3.12 |
| OFN | Nrsn1 | 5.07 | 2.24E-55 | 4.77E-51 | 0.15 | 0.01 | 0.14 | 21.14 |
| OFN | Scn3a | 4.86 | 5.65E-55 | 1.21E-50 | 0.22 | 0.02 | 0.20 | 12.17 |
| OFN | P2rx3 | 5.36 | 2.00E-54 | 4.28E-50 | 0.14 | 0.01 | 0.13 | 22.50 |
| OFN | Brinp2 | 4.37 | 4.84E-54 | 1.03E-49 | 0.21 | 0.02 | 0.19 | 12.88 |
| OFN | Psip1 | 1.28 | 5.84E-54 | 1.25E-49 | 0.98 | 0.98 | 0.00 | 1.00 |
| OFN | Mllt11 | 4.07 | 3.67E-53 | 7.84E-49 | 0.75 | 0.32 | 0.43 | 2.34 |
| OFN | Ptma | 0.87 | 7.37E-53 | 1.57E-48 | 1.00 | 1.00 | 0.00 | 1.00 |
| OFN | Ckb | 3.00 | 1.92E-52 | 4.10E-48 | 0.77 | 0.34 | 0.44 | 2.30 |
| OFN | Uchl1 | 3.04 | 3.42E-52 | 7.31E-48 | 0.90 | 0.64 | 0.26 | 1.40 |
| OFN | Gprin1 | 4.02 | 4.13E-52 | 8.82E-48 | 0.27 | 0.03 | 0.24 | 9.14 |
| OFN | Pou2f2 | 5.94 | 5.77E-52 | 1.23E-47 | 0.15 | 0.01 | 0.14 | 18.50 |
| OFN | Rnf182 | 4.17 | 8.62E-52 | 1.84E-47 | 0.59 | 0.18 | 0.41 | 3.26 |
| OFN | Tppp3 | 6.42 | 1.04E-51 | 2.22E-47 | 0.17 | 0.01 | 0.16 | 15.27 |
| OFN | Phactr1 | 4.61 | 1.07E-51 | 2.29E-47 | 0.25 | 0.03 | 0.23 | 9.33 |
| OFN | Shisa7 | 4.14 | 1.33E-51 | 2.85E-47 | 0.23 | 0.02 | 0.21 | 10.76 |
| OFN | Ripor2 | 5.09 | 1.40E-51 | 2.99E-47 | 0.14 | 0.01 | 0.13 | 22.50 |
| OFN | Flrt1 | 4.19 | 1.87E-51 | 3.99E-47 | 0.25 | 0.03 | 0.22 | 9.80 |
| OFN | Cntn2 | 4.17 | 4.64E-51 | 9.92E-47 | 0.36 | 0.06 | 0.30 | 6.12 |
| OFN | Pnma2 | 4.30 | 5.56E-51 | 1.19E-46 | 0.25 | 0.03 | 0.23 | 9.33 |
| OFN | Actg1 | 1.04 | 5.84E-51 | 1.25E-46 | 0.99 | 1.00 | -0.01 | 0.99 |
| OFN | Snap25 | 4.63 | 3.35E-50 | 7.16E-46 | 0.23 | 0.02 | 0.21 | 10.09 |
| OFN | Nnat | 2.82 | 5.22E-50 | 1.11E-45 | 0.96 | 0.95 | 0.00 | 1.00 |
| OFN | Riiad1 | 4.88 | 6.02E-50 | 1.29E-45 | 0.16 | 0.01 | 0.15 | 17.22 |
| OFN | Ssc5d | 4.76 | 6.33E-49 | 1.35E-44 | 0.16 | 0.01 | 0.15 | 16.10 |
| OFN | Necab3 | 6.01 | 1.24E-48 | 2.64E-44 | 0.15 | 0.01 | 0.14 | 18.50 |
| OFN | Dysf | 5.15 | 3.73E-48 | 7.97E-44 | 0.12 | 0.00 | 0.11 | 29.00 |
| OFN | Aplp1 | 3.49 | 4.78E-48 | 1.02E-43 | 0.65 | 0.23 | 0.41 | 2.78 |
| OFN | Rab3a | 3.82 | 5.42E-48 | 1.16E-43 | 0.44 | 0.09 | 0.35 | 4.67 |
| OFN | Nova2 | 4.02 | 8.38E-48 | 1.79E-43 | 0.27 | 0.03 | 0.24 | 7.97 |
| OFN | Ucp2 | 3.02 | 5.55E-47 | 1.19E-42 | 0.83 | 0.47 | 0.36 | 1.78 |
| OFN | Pcsk1n | 6.10 | 8.50E-47 | 1.81E-42 | 0.20 | 0.02 | 0.18 | 11.11 |
| OFN | Scg5 | 2.84 | 4.25E-46 | 9.09E-42 | 0.61 | 0.19 | 0.41 | 3.12 |
| OFN | Eid1 | 1.33 | 1.40E-45 | 3.00E-41 | 0.94 | 0.97 | -0.04 | 0.96 |
| OFN | Mfap4 | 2.47 | 1.85E-45 | 3.95E-41 | 0.87 | 0.65 | 0.21 | 1.33 |
| OFN | B630019K06Rik | 3.61 | 2.39E-45 | 5.11E-41 | 0.41 | 0.09 | 0.33 | 4.80 |
| OFN | Adgrb2 | 3.89 | 2.51E-45 | 5.37E-41 | 0.21 | 0.02 | 0.19 | 10.14 |

**Supplementary Table 4. Marker genes for the annotated cell populations of integrated clustering of E11.5 control and NotchGOF λ-junction ectoderm cells.**

| | | | | | | | | |
|---|---|---|---|---|---|---|---|---|
| OFN | Eya2 | 2.54 | 4.45E-45 | 9.51E-41 | 0.78 | 0.36 | 0.42 | 2.18 |
| OFN | Tnik | 3.14 | 4.97E-45 | 1.06E-40 | 0.43 | 0.10 | 0.34 | 4.55 |
| OFN | Kif1a | 3.75 | 5.30E-45 | 1.13E-40 | 0.40 | 0.08 | 0.32 | 4.88 |
| OFN | Plk4 | 2.98 | 5.68E-45 | 1.21E-40 | 0.80 | 0.51 | 0.30 | 1.58 |
| OFN | 9330159F19Rik | 3.47 | 7.40E-45 | 1.58E-40 | 0.28 | 0.04 | 0.25 | 7.28 |
| OFN | Hecw1 | 3.74 | 8.69E-45 | 1.86E-40 | 0.24 | 0.03 | 0.21 | 8.54 |
| OFN | Ap3b2 | 3.62 | 2.57E-44 | 5.50E-40 | 0.36 | 0.07 | 0.29 | 5.39 |
| OFN | Spsb4 | 3.12 | 3.61E-44 | 7.71E-40 | 0.48 | 0.12 | 0.36 | 3.97 |
| OFN | Rgs11 | 3.79 | 4.00E-44 | 8.54E-40 | 0.25 | 0.03 | 0.22 | 7.88 |
| OFN | Cep152 | 2.94 | 4.17E-44 | 8.90E-40 | 0.57 | 0.19 | 0.39 | 3.10 |
| OFN | Gnrh1 | 9.24 | 4.67E-44 | 9.97E-40 | 0.14 | 0.01 | 0.13 | 16.88 |
| OFN | Klf7 | 3.53 | 7.85E-44 | 1.68E-39 | 0.68 | 0.29 | 0.38 | 2.30 |
| OFN | Mapk8ip2 | 3.43 | 1.08E-43 | 2.31E-39 | 0.43 | 0.10 | 0.33 | 4.48 |
| OFN | Fgd4 | 2.82 | 1.21E-43 | 2.58E-39 | 0.58 | 0.18 | 0.40 | 3.23 |
| OFN | Sh2d3c | 3.54 | 2.43E-43 | 5.20E-39 | 0.28 | 0.04 | 0.24 | 7.10 |
| OFN | 1500004A13Rik | 3.85 | 2.80E-43 | 5.99E-39 | 0.39 | 0.08 | 0.31 | 4.84 |
| OFN | Sult4a1 | 3.59 | 3.23E-43 | 6.90E-39 | 0.37 | 0.07 | 0.30 | 5.11 |
| OFN | Parp6 | 3.21 | 3.51E-43 | 7.51E-39 | 0.46 | 0.11 | 0.35 | 4.09 |
| OFN | A330009N23Rik | 4.32 | 4.57E-43 | 9.76E-39 | 0.27 | 0.04 | 0.23 | 7.36 |
| OFN | Add2 | 3.58 | 6.20E-43 | 1.32E-38 | 0.31 | 0.05 | 0.26 | 6.20 |
| OFN | Elavl2 | 2.65 | 8.11E-43 | 1.73E-38 | 0.76 | 0.40 | 0.35 | 1.88 |
| OFN | Cyth2 | 2.10 | 1.18E-42 | 2.51E-38 | 0.85 | 0.64 | 0.21 | 1.33 |
| OFN | Dusp8 | 4.09 | 4.98E-42 | 1.06E-37 | 0.28 | 0.04 | 0.24 | 6.60 |
| OFN | Gm2694 | 3.68 | 7.57E-42 | 1.62E-37 | 0.34 | 0.06 | 0.28 | 5.43 |
| OFN | Ackr1 | 5.48 | 1.00E-41 | 2.14E-37 | 0.10 | 0.00 | 0.10 | 25.75 |
| OFN | Inka1 | 3.18 | 3.79E-41 | 8.10E-37 | 0.36 | 0.07 | 0.29 | 5.16 |
| OFN | Trim36 | 3.62 | 7.04E-41 | 1.50E-36 | 0.30 | 0.05 | 0.25 | 6.19 |
| OFN | Ddx5 | 0.69 | 2.30E-40 | 4.90E-36 | 0.98 | 1.00 | -0.02 | 0.98 |
| OFN | Trp53inp2 | 3.00 | 2.96E-40 | 6.32E-36 | 0.36 | 0.07 | 0.29 | 5.16 |
| OFN | Dll1 | 4.59 | 3.46E-40 | 7.40E-36 | 0.13 | 0.01 | 0.12 | 16.13 |
| OFN | Sox11 | 1.06 | 5.92E-40 | 1.27E-35 | 0.94 | 1.00 | -0.06 | 0.94 |
| OFN | Ndufaf3 | 2.70 | 7.66E-40 | 1.64E-35 | 0.74 | 0.41 | 0.32 | 1.79 |
| OFN | Tmcc2 | 2.97 | 2.79E-39 | 5.95E-35 | 0.32 | 0.06 | 0.26 | 5.64 |
| OFN | Cyfip2 | 3.34 | 5.89E-39 | 1.26E-34 | 0.51 | 0.16 | 0.35 | 3.11 |
| OFN | Elmo1 | 4.30 | 7.55E-39 | 1.61E-34 | 0.22 | 0.03 | 0.19 | 8.11 |
| OFN | Cnih2 | 3.67 | 8.90E-39 | 1.90E-34 | 0.49 | 0.15 | 0.34 | 3.36 |
| OFN | Sh3bgrl | 1.82 | 1.27E-38 | 2.71E-34 | 0.87 | 0.75 | 0.12 | 1.16 |
| OFN | Prdx2 | 0.75 | 1.63E-38 | 3.49E-34 | 1.00 | 1.00 | 0.00 | 1.00 |
| OFN | Col9a2 | 4.40 | 1.71E-38 | 3.66E-34 | 0.45 | 0.12 | 0.32 | 3.62 |
| OFN | C530008M17Rik | 2.47 | 2.16E-38 | 4.61E-34 | 0.74 | 0.39 | 0.34 | 1.88 |
| OFN | Unc13a | 3.91 | 5.09E-38 | 1.09E-33 | 0.19 | 0.02 | 0.17 | 9.24 |
| OFN | Nol4l | 2.87 | 5.91E-38 | 1.26E-33 | 0.50 | 0.15 | 0.35 | 3.34 |
| OFN | Ubb | 0.70 | 8.91E-38 | 1.90E-33 | 0.99 | 1.00 | -0.01 | 0.99 |
| OFN | Npr2 | 4.21 | 1.25E-37 | 2.67E-33 | 0.14 | 0.01 | 0.13 | 12.91 |
| OFN | Efna3 | 4.33 | 1.32E-37 | 2.82E-33 | 0.37 | 0.09 | 0.29 | 4.35 |
| OFN | Cfl1 | 0.70 | 2.10E-37 | 4.49E-33 | 0.99 | 1.00 | -0.01 | 0.99 |
| OFN | Kdm7a | 2.21 | 2.24E-37 | 4.79E-33 | 0.76 | 0.41 | 0.35 | 1.85 |
| OFN | Tubb5 | 0.84 | 3.66E-37 | 7.81E-33 | 1.00 | 1.00 | 0.00 | 1.00 |
| OFN | Robo2 | 2.76 | 3.82E-37 | 8.16E-33 | 0.60 | 0.23 | 0.37 | 2.59 |
| OFN | Ache | 5.47 | 6.30E-37 | 1.35E-32 | 0.11 | 0.01 | 0.10 | 18.33 |
| OFN | Nkain4 | 4.38 | 7.75E-37 | 1.66E-32 | 0.23 | 0.03 | 0.20 | 7.29 |
| OFN | Tox3 | 2.72 | 7.85E-37 | 1.68E-32 | 0.67 | 0.30 | 0.37 | 2.22 |
| OFN | Celf6 | 5.42 | 1.59E-36 | 3.40E-32 | 0.12 | 0.01 | 0.11 | 16.57 |
| OFN | Akr1b8 | 3.54 | 1.64E-36 | 3.50E-32 | 0.21 | 0.03 | 0.18 | 8.24 |
| OFN | Dpysl4 | 3.09 | 2.96E-36 | 6.32E-32 | 0.41 | 0.10 | 0.30 | 3.98 |

**Supplementary Table 4. Marker genes for the annotated cell populations of integrated clustering of E11.5 control and NotchGOF λ-junction ectoderm cells.**

| | | | | | | | | |
|---|---|---|---|---|---|---|---|---|
| OFN | Pcbp4 | 1.34 | 4.28E-36 | 9.15E-32 | 0.91 | 0.91 | 0.00 | 1.00 |
| OFN | Kcne1l | 4.24 | 8.03E-36 | 1.72E-31 | 0.26 | 0.04 | 0.22 | 6.14 |
| OFN | Lzts1 | 3.49 | 9.52E-36 | 2.03E-31 | 0.21 | 0.03 | 0.18 | 7.92 |
| OFN | Cemip2 | 2.32 | 1.10E-35 | 2.35E-31 | 0.79 | 0.60 | 0.19 | 1.31 |
| OFN | Numbl | 2.91 | 1.39E-35 | 2.96E-31 | 0.53 | 0.18 | 0.35 | 2.94 |
| OFN | AC109619.1 | 4.85 | 5.91E-35 | 1.26E-30 | 0.11 | 0.01 | 0.10 | 15.71 |
| OFN | Celsr3 | 4.50 | 1.18E-34 | 2.51E-30 | 0.12 | 0.01 | 0.11 | 13.67 |
| OFN | Gse1 | 2.50 | 1.59E-34 | 3.39E-30 | 0.61 | 0.26 | 0.35 | 2.35 |
| OFN | Rmst | 3.71 | 3.02E-34 | 6.45E-30 | 0.25 | 0.04 | 0.21 | 6.28 |
| OFN | Rgmb | 3.16 | 5.53E-34 | 1.18E-29 | 0.46 | 0.14 | 0.32 | 3.37 |
| OFN | Fam43a | 4.78 | 7.53E-34 | 1.61E-29 | 0.12 | 0.01 | 0.11 | 13.67 |
| OFN | Rab3b | 4.22 | 1.13E-33 | 2.41E-29 | 0.28 | 0.05 | 0.23 | 5.26 |
| OFN | Pnmal2 | 3.58 | 1.15E-33 | 2.45E-29 | 0.27 | 0.05 | 0.22 | 5.53 |
| OFN | Baz1a | 2.82 | 1.73E-33 | 3.69E-29 | 0.80 | 0.58 | 0.22 | 1.38 |
| OFN | Kif5c | 2.39 | 2.51E-33 | 5.36E-29 | 0.81 | 0.59 | 0.23 | 1.39 |
| OFN | Calm1 | 1.38 | 7.00E-33 | 1.50E-28 | 0.95 | 0.99 | -0.04 | 0.96 |
| OFN | Trim62 | 3.09 | 1.25E-32 | 2.67E-28 | 0.37 | 0.09 | 0.28 | 3.98 |
| OFN | Ppp1r14b | 0.78 | 1.54E-32 | 3.28E-28 | 0.96 | 0.99 | -0.03 | 0.97 |
| OFN | Celf5 | 4.20 | 4.22E-32 | 9.02E-28 | 0.16 | 0.02 | 0.14 | 8.94 |
| OFN | Ppp2r5b | 2.85 | 4.58E-32 | 9.78E-28 | 0.30 | 0.06 | 0.24 | 4.89 |
| OFN | Pclo | 2.77 | 8.68E-32 | 1.86E-27 | 0.60 | 0.28 | 0.32 | 2.16 |
| OFN | Fyn | 2.26 | 1.09E-31 | 2.34E-27 | 0.73 | 0.40 | 0.33 | 1.84 |
| OFN | Stil | 2.90 | 1.53E-31 | 3.26E-27 | 0.48 | 0.17 | 0.31 | 2.84 |
| OFN | Nat14 | 3.21 | 1.98E-31 | 4.23E-27 | 0.26 | 0.05 | 0.21 | 5.49 |
| OFN | Atl1 | 2.83 | 1.99E-31 | 4.24E-27 | 0.31 | 0.07 | 0.24 | 4.70 |
| OFN | Amotl2 | 2.32 | 2.54E-31 | 5.42E-27 | 0.68 | 0.36 | 0.32 | 1.88 |
| OFN | Trnp1 | 4.29 | 3.12E-31 | 6.67E-27 | 0.19 | 0.03 | 0.16 | 7.48 |
| OFN | Ube2s | 1.43 | 3.58E-31 | 7.66E-27 | 0.95 | 0.96 | -0.01 | 0.99 |
| OFN | Marcks | 0.71 | 4.40E-31 | 9.40E-27 | 0.95 | 0.99 | -0.04 | 0.96 |
| OFN | E130114P18Rik | 3.77 | 8.63E-31 | 1.84E-26 | 0.26 | 0.05 | 0.21 | 5.49 |
| OFN | Hspa4l | 2.45 | 1.06E-30 | 2.27E-26 | 0.58 | 0.24 | 0.34 | 2.38 |
| OFN | Fam131a | 3.82 | 1.41E-30 | 3.02E-26 | 0.17 | 0.02 | 0.15 | 8.40 |
| OFN | Mturn | 2.99 | 1.85E-30 | 3.96E-26 | 0.31 | 0.07 | 0.24 | 4.49 |
| OFN | Calm2 | 0.91 | 1.96E-30 | 4.18E-26 | 0.95 | 0.99 | -0.04 | 0.96 |
| OFN | Bcl2l1 | 2.07 | 2.25E-30 | 4.80E-26 | 0.74 | 0.45 | 0.29 | 1.64 |
| OFN | Gabarapl2 | 1.32 | 2.98E-30 | 6.36E-26 | 0.89 | 0.87 | 0.02 | 1.02 |
| OFN | 1110017D15Rik | 4.16 | 3.19E-30 | 6.82E-26 | 0.14 | 0.01 | 0.12 | 10.38 |
| OFN | Dynll1 | 0.82 | 4.02E-30 | 8.58E-26 | 0.99 | 1.00 | 0.00 | 1.00 |
| OFN | Dpysl5 | 2.21 | 4.20E-30 | 8.98E-26 | 0.69 | 0.38 | 0.31 | 1.84 |
| OFN | Ccser2 | 2.24 | 5.76E-30 | 1.23E-25 | 0.62 | 0.29 | 0.33 | 2.11 |
| OFN | Dscam | 4.16 | 9.43E-30 | 2.01E-25 | 0.10 | 0.01 | 0.10 | 14.71 |
| OFN | Ywhah | 1.58 | 1.44E-29 | 3.07E-25 | 0.89 | 0.80 | 0.09 | 1.11 |
| OFN | Adgrg1 | 2.51 | 2.24E-29 | 4.79E-25 | 0.42 | 0.13 | 0.29 | 3.33 |
| OFN | Afap1 | 2.65 | 2.97E-29 | 6.35E-25 | 0.53 | 0.21 | 0.32 | 2.57 |
| OFN | Sept3 | 2.40 | 3.50E-29 | 7.49E-25 | 0.61 | 0.30 | 0.31 | 2.03 |
| OFN | Jpt1 | 1.08 | 7.18E-29 | 1.53E-24 | 0.94 | 0.97 | -0.03 | 0.97 |
| OFN | Mdm1 | 2.78 | 7.20E-29 | 1.54E-24 | 0.60 | 0.29 | 0.31 | 2.05 |
| OFN | Cep170 | 2.09 | 8.20E-29 | 1.75E-24 | 0.68 | 0.39 | 0.30 | 1.77 |
| OFN | Stx1a | 2.93 | 8.50E-29 | 1.82E-24 | 0.30 | 0.07 | 0.23 | 4.39 |
| OFN | Dbn1 | 1.41 | 9.60E-29 | 2.05E-24 | 0.84 | 0.87 | -0.03 | 0.96 |
| OFN | Tmsb10 | 0.98 | 1.09E-28 | 2.33E-24 | 1.00 | 1.00 | 0.00 | 1.00 |
| OFN | Trim24 | 2.06 | 1.13E-28 | 2.41E-24 | 0.77 | 0.60 | 0.17 | 1.29 |
| OFN | Rufy3 | 3.61 | 1.32E-28 | 2.82E-24 | 0.50 | 0.20 | 0.30 | 2.49 |
| OFN | Nrp1 | 3.69 | 1.54E-28 | 3.29E-24 | 0.23 | 0.04 | 0.19 | 5.79 |
| OFN | Casp3 | 1.89 | 1.62E-28 | 3.47E-24 | 0.83 | 0.70 | 0.13 | 1.19 |

**Supplementary Table 4. Marker genes for the annotated cell populations of integrated clustering of E11.5 control and NotchGOF λ-junction ectoderm cells.**

| | | | | | | | | |
|---|---|---|---|---|---|---|---|---|
| OFN | Dpysl2 | 1.84 | 1.70E-28 | 3.63E-24 | 0.80 | 0.63 | 0.18 | 1.28 |
| OFN | Cmpk1 | 1.51 | 2.64E-28 | 5.64E-24 | 0.83 | 0.72 | 0.11 | 1.15 |
| OFN | Shox2 | 5.72 | 2.93E-28 | 6.26E-24 | 0.14 | 0.02 | 0.13 | 8.88 |
| OFN | Smoc1 | 3.77 | 3.86E-28 | 8.23E-24 | 0.12 | 0.01 | 0.11 | 11.60 |
| OFN | Snap91 | 3.11 | 4.85E-28 | 1.04E-23 | 0.22 | 0.04 | 0.18 | 5.92 |
| OFN | Tpd52 | 1.48 | 5.50E-28 | 1.17E-23 | 0.88 | 0.81 | 0.07 | 1.09 |
| OFN | Col26a1 | 3.01 | 6.50E-28 | 1.39E-23 | 0.32 | 0.08 | 0.24 | 4.04 |
| OFN | Gpx2 | 4.51 | 9.87E-28 | 2.11E-23 | 0.10 | 0.01 | 0.10 | 12.88 |
| OFN | Myo1b | 1.55 | 1.42E-27 | 3.04E-23 | 0.81 | 0.62 | 0.18 | 1.30 |
| OFN | Klc3 | 2.70 | 1.56E-27 | 3.33E-23 | 0.68 | 0.39 | 0.29 | 1.75 |
| OFN | Neurl1a | 2.78 | 2.53E-27 | 5.40E-23 | 0.27 | 0.06 | 0.21 | 4.75 |
| OFN | Bcl7a | 1.51 | 4.92E-27 | 1.05E-22 | 0.83 | 0.77 | 0.06 | 1.08 |
| OFN | Zc3h12c | 2.60 | 9.38E-27 | 2.00E-22 | 0.37 | 0.11 | 0.26 | 3.40 |
| OFN | Tomm34 | 1.94 | 1.04E-26 | 2.23E-22 | 0.69 | 0.43 | 0.26 | 1.62 |
| OFN | H3f3a | 0.40 | 1.08E-26 | 2.31E-22 | 1.00 | 1.00 | 0.00 | 1.00 |
| OFN | Nlk | 2.26 | 1.11E-26 | 2.38E-22 | 0.66 | 0.40 | 0.26 | 1.66 |
| OFN | Dnaja1 | 0.67 | 1.31E-26 | 2.79E-22 | 0.94 | 0.99 | -0.05 | 0.95 |
| OFN | Hid1 | 3.69 | 1.37E-26 | 2.92E-22 | 0.17 | 0.02 | 0.15 | 7.30 |
| OFN | Ccsap | 2.98 | 1.46E-26 | 3.12E-22 | 0.32 | 0.08 | 0.24 | 3.89 |
| OFN | Rab39b | 2.57 | 1.82E-26 | 3.89E-22 | 0.41 | 0.13 | 0.28 | 3.22 |
| OFN | Smarcd2 | 2.40 | 1.87E-26 | 3.99E-22 | 0.65 | 0.35 | 0.29 | 1.84 |
| OFN | Nrg1 | 2.29 | 2.06E-26 | 4.40E-22 | 0.57 | 0.26 | 0.32 | 2.22 |
| OFN | Ank2 | 3.19 | 3.26E-26 | 6.96E-22 | 0.59 | 0.31 | 0.28 | 1.91 |
| OFN | Nefm | 5.19 | 4.06E-26 | 8.66E-22 | 0.17 | 0.02 | 0.14 | 7.00 |
| OFN | Ttll7 | 3.26 | 4.93E-26 | 1.05E-21 | 0.26 | 0.06 | 0.20 | 4.61 |
| OFN | Ssbp3 | 1.17 | 6.09E-26 | 1.30E-21 | 0.86 | 0.89 | -0.03 | 0.96 |
| OFN | Selenok | 1.35 | 1.06E-25 | 2.27E-21 | 0.88 | 0.90 | -0.02 | 0.97 |
| OFN | Map9 | 2.29 | 1.42E-25 | 3.04E-21 | 0.56 | 0.26 | 0.30 | 2.15 |
| OFN | Rbfox3 | 3.85 | 1.80E-25 | 3.85E-21 | 0.12 | 0.01 | 0.11 | 10.55 |
| OFN | Ralgds | 2.20 | 2.68E-25 | 5.72E-21 | 0.54 | 0.23 | 0.31 | 2.32 |
| OFN | Ccp110 | 2.47 | 3.24E-25 | 6.92E-21 | 0.63 | 0.34 | 0.28 | 1.82 |
| OFN | Map1lc3b | 1.15 | 3.47E-25 | 7.42E-21 | 0.89 | 0.94 | -0.05 | 0.95 |
| OFN | Eif1 | 0.51 | 4.06E-25 | 8.67E-21 | 0.99 | 1.00 | -0.01 | 0.99 |
| OFN | Ell3 | 2.93 | 7.88E-25 | 1.68E-20 | 0.36 | 0.11 | 0.25 | 3.38 |
| OFN | Gdi1 | 2.17 | 8.78E-25 | 1.88E-20 | 0.70 | 0.45 | 0.25 | 1.56 |
| OFN | Tle1 | 1.91 | 9.21E-25 | 1.97E-20 | 0.77 | 0.54 | 0.23 | 1.43 |
| OFN | Pex5l | 3.77 | 1.19E-24 | 2.55E-20 | 0.15 | 0.02 | 0.13 | 7.40 |
| OFN | Rcor2 | 1.88 | 1.44E-24 | 3.07E-20 | 0.70 | 0.49 | 0.22 | 1.44 |
| OFN | Gm5468 | 4.26 | 1.47E-24 | 3.14E-20 | 0.10 | 0.01 | 0.09 | 11.44 |
| OFN | Kdm4b | 2.35 | 1.64E-24 | 3.50E-20 | 0.45 | 0.16 | 0.29 | 2.80 |
| OFN | Akna | 2.97 | 2.67E-24 | 5.71E-20 | 0.27 | 0.06 | 0.21 | 4.42 |
| OFN | Prkar1a | 1.36 | 2.80E-24 | 5.99E-20 | 0.88 | 0.85 | 0.03 | 1.04 |
| OFN | Cxcr4 | 2.48 | 3.14E-24 | 6.70E-20 | 0.46 | 0.17 | 0.29 | 2.65 |
| OFN | Amer2 | 2.28 | 3.45E-24 | 7.38E-20 | 0.43 | 0.15 | 0.28 | 2.92 |
| OFN | Kcnma1 | 4.68 | 3.84E-24 | 8.21E-20 | 0.12 | 0.01 | 0.10 | 9.67 |
| OFN | Kdm1a | 1.02 | 4.14E-24 | 8.85E-20 | 0.89 | 0.94 | -0.05 | 0.95 |
| OFN | Thsd7a | 4.99 | 5.03E-24 | 1.07E-19 | 0.12 | 0.01 | 0.10 | 9.67 |
| OFN | Syt11 | 2.56 | 5.37E-24 | 1.15E-19 | 0.61 | 0.34 | 0.28 | 1.82 |
| OFN | Dusp26 | 3.78 | 5.46E-24 | 1.17E-19 | 0.16 | 0.02 | 0.13 | 7.05 |
| OFN | Mbd2 | 1.76 | 5.79E-24 | 1.24E-19 | 0.79 | 0.61 | 0.18 | 1.29 |
| OFN | Wdr47 | 2.23 | 5.98E-24 | 1.28E-19 | 0.39 | 0.12 | 0.27 | 3.20 |
| OFN | Gng2 | 1.96 | 7.28E-24 | 1.56E-19 | 0.68 | 0.43 | 0.26 | 1.60 |
| OFN | Ncam1 | 3.45 | 1.32E-23 | 2.81E-19 | 0.60 | 0.33 | 0.27 | 1.83 |
| OFN | Ftl1 | 0.49 | 1.97E-23 | 4.21E-19 | 1.00 | 1.00 | 0.00 | 1.00 |
| OFN | Mxi1 | 1.75 | 2.00E-23 | 4.27E-19 | 0.77 | 0.57 | 0.20 | 1.35 |

**Supplementary Table 4. Marker genes for the annotated cell populations of integrated clustering of E11.5 control and NotchGOF λ-junction ectoderm cells.**

| | | | | | | | | |
|---|---|---|---|---|---|---|---|---|
| OFN | Aff3 | 2.93 | 2.15E-23 | 4.59E-19 | 0.28 | 0.07 | 0.21 | 3.96 |
| OFN | Rhbdl3 | 3.63 | 3.61E-23 | 7.72E-19 | 0.12 | 0.01 | 0.10 | 8.92 |
| OFN | Olfm1 | 2.27 | 5.33E-23 | 1.14E-18 | 0.60 | 0.32 | 0.28 | 1.86 |
| OFN | Dubr | 3.11 | 1.05E-22 | 2.24E-18 | 0.18 | 0.03 | 0.15 | 5.84 |
| OFN | Hspa12a | 3.23 | 1.15E-22 | 2.46E-18 | 0.17 | 0.03 | 0.15 | 6.00 |
| OFN | Lsamp | 4.35 | 1.63E-22 | 3.48E-18 | 0.12 | 0.01 | 0.10 | 8.92 |
| OFN | Smarca5 | 0.75 | 1.97E-22 | 4.20E-18 | 0.94 | 0.97 | -0.04 | 0.96 |
| OFN | Shtn1 | 2.97 | 2.26E-22 | 4.83E-18 | 0.46 | 0.19 | 0.26 | 2.36 |
| OFN | Bex2 | 1.07 | 2.30E-22 | 4.92E-18 | 0.92 | 0.88 | 0.04 | 1.05 |
| OFN | Cdk5r1 | 2.98 | 2.38E-22 | 5.09E-18 | 0.26 | 0.06 | 0.20 | 4.10 |
| OFN | Tsg101 | 1.19 | 2.46E-22 | 5.25E-18 | 0.87 | 0.86 | 0.01 | 1.01 |
| OFN | Atp6v1g1 | 0.90 | 5.90E-22 | 1.26E-17 | 0.91 | 0.97 | -0.06 | 0.94 |
| OFN | Tmcc3 | 2.62 | 9.81E-22 | 2.10E-17 | 0.33 | 0.10 | 0.23 | 3.26 |
| OFN | Lrp11 | 2.65 | 1.16E-21 | 2.48E-17 | 0.27 | 0.07 | 0.20 | 3.93 |
| OFN | Adam19 | 2.78 | 1.20E-21 | 2.56E-17 | 0.17 | 0.03 | 0.14 | 5.80 |
| OFN | Ddah2 | 0.94 | 1.53E-21 | 3.26E-17 | 0.94 | 0.98 | -0.04 | 0.96 |
| OFN | Ctnna2 | 2.81 | 1.71E-21 | 3.64E-17 | 0.27 | 0.07 | 0.20 | 3.82 |
| OFN | Lcorl | 2.11 | 1.78E-21 | 3.80E-17 | 0.65 | 0.40 | 0.25 | 1.62 |
| OFN | Cpeb4 | 2.74 | 2.64E-21 | 5.64E-17 | 0.27 | 0.07 | 0.20 | 3.96 |
| OFN | Slc43a2 | 2.64 | 2.73E-21 | 5.84E-17 | 0.25 | 0.06 | 0.19 | 4.06 |
| OFN | Trim9 | 3.80 | 2.81E-21 | 6.00E-17 | 0.11 | 0.01 | 0.10 | 9.17 |
| OFN | Spock2 | 2.10 | 2.82E-21 | 6.02E-17 | 0.37 | 0.12 | 0.25 | 3.07 |
| OFN | Rbfox2 | 1.43 | 2.86E-21 | 6.10E-17 | 0.79 | 0.71 | 0.08 | 1.12 |
| OFN | Pkig | 1.33 | 4.56E-21 | 9.74E-17 | 0.87 | 0.75 | 0.12 | 1.15 |
| OFN | Cadm1 | 1.44 | 5.30E-21 | 1.13E-16 | 0.80 | 0.71 | 0.09 | 1.12 |
| OFN | Arpc5 | 1.27 | 7.83E-21 | 1.67E-16 | 0.86 | 0.82 | 0.03 | 1.04 |
| OFN | Tenm1 | 3.46 | 8.61E-21 | 1.84E-16 | 0.14 | 0.02 | 0.12 | 6.76 |
| OFN | Kalrn | 2.33 | 8.65E-21 | 1.85E-16 | 0.43 | 0.17 | 0.26 | 2.49 |
| OFN | Csrnp3 | 3.10 | 8.67E-21 | 1.85E-16 | 0.14 | 0.02 | 0.12 | 7.11 |
| OFN | Dctn3 | 1.27 | 8.88E-21 | 1.90E-16 | 0.82 | 0.81 | 0.01 | 1.01 |
| OFN | Selenow | 0.79 | 1.11E-20 | 2.36E-16 | 0.96 | 0.99 | -0.04 | 0.96 |
| OFN | Caly | 3.67 | 1.48E-20 | 3.15E-16 | 0.11 | 0.01 | 0.10 | 8.46 |
| OFN | Snta1 | 3.05 | 1.59E-20 | 3.39E-16 | 0.16 | 0.03 | 0.13 | 5.75 |
| OFN | Pdrg1 | 2.04 | 1.59E-20 | 3.39E-16 | 0.76 | 0.57 | 0.18 | 1.32 |
| OFN | Calm3 | 1.27 | 1.73E-20 | 3.69E-16 | 0.87 | 0.84 | 0.03 | 1.04 |
| OFN | Eif4g2 | 0.51 | 2.70E-20 | 5.77E-16 | 0.94 | 1.00 | -0.06 | 0.94 |
| OFN | Spns2 | 3.78 | 2.83E-20 | 6.05E-16 | 0.10 | 0.01 | 0.09 | 8.58 |
| OFN | Rnf5 | 1.56 | 3.19E-20 | 6.81E-16 | 0.81 | 0.70 | 0.12 | 1.17 |
| OFN | Acap3 | 2.62 | 3.53E-20 | 7.54E-16 | 0.23 | 0.06 | 0.18 | 4.22 |
| OFN | Stk33 | 4.42 | 3.85E-20 | 8.21E-16 | 0.11 | 0.01 | 0.10 | 8.46 |
| OFN | Hnrnpa2b1 | 0.52 | 4.53E-20 | 9.67E-16 | 0.98 | 1.00 | -0.02 | 0.98 |
| OFN | Gm13648 | 2.89 | 4.83E-20 | 1.03E-15 | 0.32 | 0.10 | 0.22 | 3.13 |
| OFN | Kif5b | 0.79 | 7.15E-20 | 1.53E-15 | 0.91 | 0.95 | -0.04 | 0.96 |
| OFN | Gadd45g | 1.31 | 7.24E-20 | 1.55E-15 | 0.65 | 0.33 | 0.32 | 1.95 |
| OFN | Mybl1 | 3.64 | 1.07E-19 | 2.29E-15 | 0.25 | 0.07 | 0.19 | 3.82 |
| OFN | Atp8a2 | 3.69 | 1.11E-19 | 2.36E-15 | 0.12 | 0.02 | 0.11 | 7.24 |
| OFN | Shb | 2.19 | 1.12E-19 | 2.39E-15 | 0.52 | 0.25 | 0.26 | 2.05 |
| OFN | Crabp2 | 0.92 | 1.36E-19 | 2.90E-15 | 0.96 | 0.85 | 0.12 | 1.14 |
| OFN | Csrp1 | 2.54 | 1.37E-19 | 2.93E-15 | 0.42 | 0.17 | 0.25 | 2.46 |
| OFN | Napb | 2.66 | 1.40E-19 | 2.98E-15 | 0.24 | 0.06 | 0.18 | 4.05 |
| OFN | Enho | 1.91 | 2.03E-19 | 4.33E-15 | 0.59 | 0.33 | 0.26 | 1.78 |
| OFN | Srgap1 | 1.83 | 2.31E-19 | 4.94E-15 | 0.64 | 0.40 | 0.24 | 1.60 |
| OFN | Mical2 | 2.70 | 2.64E-19 | 5.64E-15 | 0.22 | 0.05 | 0.17 | 4.21 |
| OFN | Riok3 | 1.64 | 2.80E-19 | 5.98E-15 | 0.68 | 0.47 | 0.21 | 1.44 |
| OFN | Chn1 | 2.30 | 4.51E-19 | 9.63E-15 | 0.52 | 0.27 | 0.25 | 1.94 |

**Supplementary Table 4. Marker genes for the annotated cell populations of integrated clustering of E11.5 control and NotchGOF λ-junction ectoderm cells.**

| | | | | | | | | |
|---|---|---|---|---|---|---|---|---|
| OFN | 2310040G24Rik | 3.95 | 5.45E-19 | 1.16E-14 | 0.17 | 0.03 | 0.14 | 5.25 |
| OFN | Crtc3 | 2.47 | 9.66E-19 | 2.06E-14 | 0.40 | 0.17 | 0.23 | 2.41 |
| OFN | Dynlrb1 | 0.96 | 1.03E-18 | 2.19E-14 | 0.91 | 0.94 | -0.03 | 0.97 |
| OFN | Carhsp1 | 1.67 | 1.14E-18 | 2.44E-14 | 0.73 | 0.58 | 0.15 | 1.26 |
| OFN | Cystm1 | 2.53 | 1.16E-18 | 2.48E-14 | 0.58 | 0.35 | 0.23 | 1.64 |
| OFN | Usp29 | 2.04 | 1.32E-18 | 2.81E-14 | 0.41 | 0.16 | 0.25 | 2.54 |
| OFN | Klf13 | 1.80 | 1.41E-18 | 3.01E-14 | 0.63 | 0.41 | 0.22 | 1.54 |
| OFN | Gdf11 | 1.97 | 1.42E-18 | 3.02E-14 | 0.51 | 0.26 | 0.25 | 1.97 |
| OFN | Chrnb2 | 3.26 | 1.47E-18 | 3.14E-14 | 0.12 | 0.02 | 0.10 | 7.25 |
| OFN | Cox6a1 | 0.76 | 1.52E-18 | 3.25E-14 | 0.93 | 0.95 | -0.02 | 0.97 |
| OFN | Map3k13 | 4.01 | 1.85E-18 | 3.96E-14 | 0.13 | 0.02 | 0.11 | 6.45 |
| OFN | Ddit4 | 1.86 | 2.08E-18 | 4.44E-14 | 0.68 | 0.43 | 0.25 | 1.58 |
| OFN | Fam124a | 3.11 | 2.58E-18 | 5.50E-14 | 0.15 | 0.03 | 0.12 | 5.69 |
| OFN | Nbea | 1.80 | 2.68E-18 | 5.73E-14 | 0.60 | 0.36 | 0.24 | 1.65 |
| OFN | Tekt2 | 3.67 | 3.04E-18 | 6.49E-14 | 0.13 | 0.02 | 0.11 | 6.45 |
| OFN | Kcnh8 | 2.40 | 3.20E-18 | 6.84E-14 | 0.37 | 0.14 | 0.23 | 2.69 |
| OFN | Enc1 | 2.58 | 3.23E-18 | 6.89E-14 | 0.49 | 0.25 | 0.24 | 1.98 |
| OFN | Rnf165 | 1.94 | 3.26E-18 | 6.97E-14 | 0.52 | 0.27 | 0.25 | 1.93 |
| OFN | Gjb5 | 3.80 | 4.51E-18 | 9.62E-14 | 0.12 | 0.02 | 0.11 | 6.83 |
| OFN | Smpd3 | 2.85 | 4.53E-18 | 9.68E-14 | 0.16 | 0.03 | 0.13 | 5.54 |
| OFN | Cdkn2d | 2.26 | 4.65E-18 | 9.94E-14 | 0.55 | 0.32 | 0.23 | 1.73 |
| OFN | Apbb1 | 2.12 | 5.20E-18 | 1.11E-13 | 0.57 | 0.34 | 0.23 | 1.67 |
| OFN | Fkbp1a | 0.84 | 5.63E-18 | 1.20E-13 | 0.90 | 0.94 | -0.05 | 0.95 |
| OFN | Tmem198 | 4.02 | 6.52E-18 | 1.39E-13 | 0.11 | 0.02 | 0.10 | 7.33 |
| OFN | Tnfrsf21 | 2.45 | 6.63E-18 | 1.42E-13 | 0.30 | 0.09 | 0.20 | 3.16 |
| OFN | Fdps | 1.14 | 8.35E-18 | 1.78E-13 | 0.85 | 0.82 | 0.04 | 1.04 |
| OFN | Ube2b | 0.84 | 9.80E-18 | 2.09E-13 | 0.90 | 0.95 | -0.05 | 0.94 |
| OFN | B3gnt4 | 2.79 | 1.02E-17 | 2.18E-13 | 0.17 | 0.03 | 0.14 | 5.09 |
| OFN | Fbxl16 | 3.94 | 1.07E-17 | 2.29E-13 | 0.14 | 0.02 | 0.11 | 5.87 |
| OFN | Enox1 | 3.07 | 1.68E-17 | 3.59E-13 | 0.20 | 0.05 | 0.15 | 4.26 |
| OFN | Ctxn1 | 1.46 | 1.71E-17 | 3.65E-13 | 0.81 | 0.79 | 0.02 | 1.02 |
| OFN | Atp6v0b | 1.28 | 2.22E-17 | 4.74E-13 | 0.80 | 0.77 | 0.03 | 1.04 |
| OFN | Gabarap | 0.77 | 2.57E-17 | 5.49E-13 | 0.91 | 0.97 | -0.06 | 0.94 |
| OFN | Stx7 | 2.78 | 2.67E-17 | 5.70E-13 | 0.48 | 0.24 | 0.24 | 1.97 |
| OFN | Rala | 0.98 | 2.72E-17 | 5.80E-13 | 0.85 | 0.84 | 0.00 | 1.00 |
| OFN | Mospd1 | 2.13 | 2.90E-17 | 6.18E-13 | 0.50 | 0.25 | 0.25 | 2.00 |
| OFN | Syngr3 | 3.38 | 3.18E-17 | 6.79E-13 | 0.12 | 0.02 | 0.10 | 6.82 |
| OFN | Ywhag | 1.63 | 3.46E-17 | 7.40E-13 | 0.73 | 0.63 | 0.10 | 1.17 |
| OFN | Dync1i2 | 0.94 | 4.13E-17 | 8.81E-13 | 0.85 | 0.88 | -0.03 | 0.96 |
| OFN | Atp8a1 | 2.64 | 4.32E-17 | 9.23E-13 | 0.30 | 0.10 | 0.20 | 2.97 |
| OFN | Mpc1 | 1.80 | 4.40E-17 | 9.41E-13 | 0.67 | 0.48 | 0.20 | 1.41 |
| OFN | Mapk8ip1 | 2.12 | 4.86E-17 | 1.04E-12 | 0.58 | 0.36 | 0.22 | 1.62 |
| OFN | Tmem176b | 1.48 | 5.24E-17 | 1.12E-12 | 0.74 | 0.60 | 0.13 | 1.22 |
| OFN | Samd14 | 1.99 | 8.58E-17 | 1.83E-12 | 0.50 | 0.26 | 0.24 | 1.91 |
| OFN | Cep120 | 1.90 | 8.87E-17 | 1.89E-12 | 0.47 | 0.22 | 0.24 | 2.09 |
| OFN | B230216N24Rik | 3.61 | 1.05E-16 | 2.24E-12 | 0.10 | 0.01 | 0.09 | 7.36 |
| OFN | Ccdc88a | 1.15 | 1.30E-16 | 2.78E-12 | 0.77 | 0.70 | 0.07 | 1.11 |
| OFN | Rtl8a | 2.35 | 1.45E-16 | 3.10E-12 | 0.54 | 0.30 | 0.24 | 1.79 |
| OFN | Racgap1 | 1.50 | 1.48E-16 | 3.17E-12 | 0.72 | 0.51 | 0.21 | 1.41 |
| OFN | Cox7a2 | 0.51 | 1.62E-16 | 3.46E-12 | 0.94 | 0.99 | -0.05 | 0.95 |
| OFN | Dclk1 | 2.52 | 1.63E-16 | 3.48E-12 | 0.41 | 0.18 | 0.22 | 2.21 |
| OFN | Cxxc4 | 2.58 | 1.85E-16 | 3.96E-12 | 0.47 | 0.25 | 0.22 | 1.88 |
| OFN | Aldh1b1 | 3.03 | 2.27E-16 | 4.85E-12 | 0.12 | 0.02 | 0.10 | 6.44 |
| OFN | Actr3b | 3.16 | 2.31E-16 | 4.92E-12 | 0.14 | 0.03 | 0.12 | 5.26 |
| OFN | Gabrb3 | 2.94 | 2.87E-16 | 6.14E-12 | 0.11 | 0.02 | 0.09 | 6.88 |

**Supplementary Table 4. Marker genes for the annotated cell populations of integrated clustering of E11.5 control and NotchGOF λ-junction ectoderm cells.**

| | | | | | | | | |
|---|---|---|---|---|---|---|---|---|
| OFN | Gnao1 | 2.46 | 3.46E-16 | 7.40E-12 | 0.39 | 0.18 | 0.22 | 2.25 |
| OFN | Snx32 | 3.16 | 3.86E-16 | 8.24E-12 | 0.12 | 0.02 | 0.10 | 5.86 |
| OFN | Ap3m2 | 2.58 | 3.96E-16 | 8.46E-12 | 0.19 | 0.05 | 0.14 | 4.16 |
| OFN | Shd | 2.81 | 4.38E-16 | 9.35E-12 | 0.27 | 0.09 | 0.18 | 3.12 |
| OFN | Sept5 | 2.68 | 4.52E-16 | 9.66E-12 | 0.18 | 0.04 | 0.14 | 4.21 |
| OFN | Tdrkh | 2.40 | 5.33E-16 | 1.14E-11 | 0.28 | 0.09 | 0.18 | 2.98 |
| OFN | Mapk8 | 1.49 | 5.98E-16 | 1.28E-11 | 0.57 | 0.34 | 0.24 | 1.71 |
| OFN | Palm | 1.93 | 7.83E-16 | 1.67E-11 | 0.61 | 0.41 | 0.20 | 1.49 |
| OFN | Ypel3 | 1.65 | 8.90E-16 | 1.90E-11 | 0.72 | 0.57 | 0.15 | 1.26 |
| OFN | Lpcat2 | 2.48 | 9.18E-16 | 1.96E-11 | 0.21 | 0.05 | 0.15 | 3.81 |
| OFN | Rsph9 | 2.39 | 1.06E-15 | 2.25E-11 | 0.31 | 0.11 | 0.20 | 2.82 |
| OFN | Hdac2 | 0.93 | 1.09E-15 | 2.33E-11 | 0.85 | 0.89 | -0.03 | 0.96 |
| OFN | Pth1r | 2.47 | 1.56E-15 | 3.33E-11 | 0.20 | 0.05 | 0.15 | 3.85 |
| OFN | Mgst3 | 2.43 | 1.61E-15 | 3.45E-11 | 0.65 | 0.49 | 0.16 | 1.32 |
| OFN | Atp5f1 | 0.48 | 2.86E-15 | 6.12E-11 | 0.95 | 0.99 | -0.04 | 0.96 |
| OFN | Akap17b | 2.48 | 2.96E-15 | 6.32E-11 | 0.35 | 0.14 | 0.21 | 2.47 |
| OFN | Ccdc96 | 2.72 | 3.62E-15 | 7.73E-11 | 0.16 | 0.04 | 0.13 | 4.47 |
| OFN | Gm1673 | 0.81 | 3.63E-15 | 7.75E-11 | 0.88 | 0.93 | -0.05 | 0.94 |
| OFN | Gdpd1 | 1.89 | 3.83E-15 | 8.18E-11 | 0.61 | 0.40 | 0.21 | 1.53 |
| OFN | Hpcal1 | 1.71 | 4.60E-15 | 9.83E-11 | 0.65 | 0.46 | 0.19 | 1.41 |
| OFN | Draxin | 3.30 | 5.06E-15 | 1.08E-10 | 0.10 | 0.02 | 0.09 | 6.44 |
| OFN | 1700088E04Rik | 2.49 | 5.55E-15 | 1.19E-10 | 0.24 | 0.07 | 0.17 | 3.23 |
| OFN | Zfp386 | 1.78 | 6.85E-15 | 1.46E-10 | 0.65 | 0.44 | 0.21 | 1.48 |
| OFN | Nfasc | 2.59 | 7.41E-15 | 1.58E-10 | 0.27 | 0.09 | 0.18 | 2.91 |
| OFN | Maml3 | 1.46 | 7.86E-15 | 1.68E-10 | 0.61 | 0.40 | 0.21 | 1.53 |
| OFN | Tecpr1 | 1.84 | 7.97E-15 | 1.70E-10 | 0.36 | 0.15 | 0.21 | 2.44 |
| OFN | C2cd2l | 2.76 | 8.08E-15 | 1.73E-10 | 0.18 | 0.05 | 0.14 | 4.02 |
| OFN | Eif4a1 | 0.44 | 8.98E-15 | 1.92E-10 | 0.98 | 1.00 | -0.02 | 0.98 |
| OFN | Sdk1 | 2.92 | 9.03E-15 | 1.93E-10 | 0.26 | 0.09 | 0.17 | 3.00 |
| OFN | Cecr2 | 1.50 | 9.52E-15 | 2.03E-10 | 0.60 | 0.38 | 0.22 | 1.59 |
| OFN | Pcolce2 | 3.79 | 1.04E-14 | 2.21E-10 | 0.21 | 0.06 | 0.15 | 3.44 |
| OFN | Rab3c | 3.20 | 1.32E-14 | 2.81E-10 | 0.17 | 0.04 | 0.13 | 4.10 |
| OFN | Onecut2 | 3.71 | 1.41E-14 | 3.02E-10 | 0.16 | 0.04 | 0.12 | 4.24 |
| OFN | Art1 | 3.79 | 1.43E-14 | 3.05E-10 | 0.12 | 0.02 | 0.10 | 5.35 |
| OFN | Tfeb | 2.82 | 1.57E-14 | 3.35E-10 | 0.16 | 0.04 | 0.12 | 4.35 |
| OFN | Plppr2 | 2.74 | 1.75E-14 | 3.75E-10 | 0.18 | 0.05 | 0.14 | 3.93 |
| OFN | BC031181 | 1.19 | 1.88E-14 | 4.01E-10 | 0.79 | 0.78 | 0.02 | 1.02 |
| OFN | Vim | 2.08 | 1.91E-14 | 4.08E-10 | 0.77 | 0.72 | 0.05 | 1.07 |
| OFN | Dnajb6 | 0.98 | 2.33E-14 | 4.97E-10 | 0.87 | 0.90 | -0.02 | 0.97 |
| OFN | Arhgap33 | 2.00 | 2.55E-14 | 5.46E-10 | 0.32 | 0.12 | 0.19 | 2.57 |
| OFN | Traf4 | 1.49 | 3.06E-14 | 6.53E-10 | 0.67 | 0.51 | 0.16 | 1.32 |
| OFN | 1110008P14Rik | 1.80 | 3.13E-14 | 6.68E-10 | 0.57 | 0.36 | 0.21 | 1.59 |
| OFN | Ppp3ca | 1.45 | 3.37E-14 | 7.19E-10 | 0.67 | 0.54 | 0.13 | 1.24 |
| OFN | Plekho1 | 1.47 | 3.37E-14 | 7.20E-10 | 0.56 | 0.35 | 0.20 | 1.57 |
| OFN | Dbndd1 | 2.33 | 3.60E-14 | 7.69E-10 | 0.26 | 0.09 | 0.17 | 2.93 |
| OFN | Slc43a1 | 2.09 | 3.72E-14 | 7.95E-10 | 0.28 | 0.10 | 0.18 | 2.81 |
| OFN | Stx2 | 2.02 | 3.76E-14 | 8.03E-10 | 0.36 | 0.16 | 0.20 | 2.23 |
| OFN | Nmnat2 | 4.19 | 3.97E-14 | 8.47E-10 | 0.10 | 0.02 | 0.09 | 6.06 |
| OFN | Scube1 | 2.78 | 4.91E-14 | 1.05E-09 | 0.18 | 0.05 | 0.13 | 3.85 |
| OFN | Ccdc28b | 1.72 | 5.07E-14 | 1.08E-09 | 0.56 | 0.35 | 0.21 | 1.61 |
| OFN | Rapgef5 | 2.37 | 5.30E-14 | 1.13E-09 | 0.37 | 0.17 | 0.20 | 2.22 |
| OFN | Dda1 | 1.49 | 5.63E-14 | 1.20E-09 | 0.70 | 0.56 | 0.15 | 1.26 |
| OFN | Klc1 | 1.98 | 6.13E-14 | 1.31E-09 | 0.65 | 0.49 | 0.16 | 1.32 |
| OFN | Tmeff1 | 1.25 | 6.32E-14 | 1.35E-09 | 0.76 | 0.68 | 0.07 | 1.11 |
| OFN | Dram2 | 1.72 | 6.46E-14 | 1.38E-09 | 0.40 | 0.18 | 0.22 | 2.23 |

**Supplementary Table 4. Marker genes for the annotated cell populations of integrated clustering of E11.5 control and NotchGOF λ-junction ectoderm cells.**

| | | | | | | | | |
|---|---|---|---|---|---|---|---|---|
| OFN | Ciapin1 | 1.22 | 6.73E-14 | 1.44E-09 | 0.78 | 0.69 | 0.09 | 1.14 |
| OFN | Acsl1 | 2.30 | 1.13E-13 | 2.41E-09 | 0.21 | 0.06 | 0.15 | 3.33 |
| OFN | Gamt | 3.36 | 1.16E-13 | 2.47E-09 | 0.24 | 0.08 | 0.16 | 3.03 |
| OFN | Agtpbp1 | 2.28 | 1.16E-13 | 2.48E-09 | 0.28 | 0.10 | 0.17 | 2.69 |
| OFN | Apba2 | 2.37 | 1.19E-13 | 2.54E-09 | 0.32 | 0.13 | 0.19 | 2.43 |
| OFN | Cfap36 | 1.59 | 1.27E-13 | 2.72E-09 | 0.69 | 0.49 | 0.20 | 1.40 |
| OFN | Plcxd3 | 3.76 | 1.39E-13 | 2.97E-09 | 0.11 | 0.02 | 0.09 | 5.50 |
| OFN | Hsp90aa1 | 0.45 | 1.47E-13 | 3.14E-09 | 0.97 | 1.00 | -0.03 | 0.97 |
| OFN | Nav1 | 1.35 | 1.47E-13 | 3.14E-09 | 0.67 | 0.51 | 0.16 | 1.31 |
| OFN | Mdh2 | 0.83 | 1.60E-13 | 3.41E-09 | 0.88 | 0.94 | -0.06 | 0.94 |
| OFN | Mex3a | 0.91 | 1.69E-13 | 3.62E-09 | 0.83 | 0.88 | -0.04 | 0.95 |
| OFN | Zmiz1 | 1.32 | 1.82E-13 | 3.89E-09 | 0.73 | 0.62 | 0.11 | 1.17 |
| OFN | Hist3h2ba | 1.59 | 2.00E-13 | 4.28E-09 | 0.85 | 0.83 | 0.02 | 1.03 |
| OFN | A330084C13Rik | 2.75 | 2.11E-13 | 4.50E-09 | 0.11 | 0.02 | 0.09 | 5.50 |
| OFN | Pak7 | 2.77 | 2.12E-13 | 4.54E-09 | 0.15 | 0.03 | 0.11 | 4.35 |
| OFN | Fam89b | 1.17 | 2.18E-13 | 4.66E-09 | 0.80 | 0.75 | 0.05 | 1.06 |
| OFN | Arl8a | 1.95 | 2.28E-13 | 4.86E-09 | 0.56 | 0.38 | 0.18 | 1.46 |
| OFN | Arfgef3 | 3.19 | 2.41E-13 | 5.15E-09 | 0.20 | 0.06 | 0.14 | 3.33 |
| OFN | Ttll1 | 2.21 | 2.93E-13 | 6.26E-09 | 0.39 | 0.19 | 0.21 | 2.13 |
| OFN | 2810455O05Rik | 3.31 | 3.14E-13 | 6.71E-09 | 0.10 | 0.02 | 0.09 | 5.72 |
| OFN | Slc26a10 | 2.44 | 3.39E-13 | 7.25E-09 | 0.14 | 0.03 | 0.11 | 4.66 |
| OFN | Rnasek | 1.06 | 3.43E-13 | 7.34E-09 | 0.77 | 0.73 | 0.04 | 1.06 |
| OFN | Cntfr | 2.33 | 3.58E-13 | 7.65E-09 | 0.33 | 0.14 | 0.19 | 2.32 |
| OFN | Grcc10 | 0.83 | 5.30E-13 | 1.13E-08 | 0.92 | 0.98 | -0.05 | 0.95 |
| OFN | Srp14 | 0.74 | 5.67E-13 | 1.21E-08 | 0.90 | 0.95 | -0.05 | 0.95 |
| OFN | Jarid2 | 1.33 | 6.22E-13 | 1.33E-08 | 0.67 | 0.55 | 0.12 | 1.23 |
| OFN | Phyhipl | 2.14 | 6.65E-13 | 1.42E-08 | 0.27 | 0.10 | 0.17 | 2.74 |
| OFN | Dmxl2 | 2.25 | 7.67E-13 | 1.64E-08 | 0.23 | 0.08 | 0.16 | 3.01 |
| OFN | Map3k12 | 1.68 | 7.90E-13 | 1.69E-08 | 0.36 | 0.16 | 0.20 | 2.25 |
| OFN | Ubald2 | 1.11 | 9.14E-13 | 1.95E-08 | 0.78 | 0.71 | 0.07 | 1.10 |
| OFN | Mapre2 | 2.24 | 9.46E-13 | 2.02E-08 | 0.45 | 0.23 | 0.21 | 1.93 |
| OFN | Gal | 0.98 | 1.02E-12 | 2.18E-08 | 0.14 | 0.03 | 0.11 | 4.50 |
| OFN | Gm42372 | 2.60 | 1.04E-12 | 2.22E-08 | 0.16 | 0.04 | 0.12 | 3.97 |
| OFN | Plekha6 | 2.91 | 1.06E-12 | 2.27E-08 | 0.16 | 0.04 | 0.12 | 3.83 |
| OFN | Astn1 | 2.58 | 1.07E-12 | 2.28E-08 | 0.16 | 0.04 | 0.12 | 3.83 |
| OFN | Tanc2 | 1.59 | 1.08E-12 | 2.31E-08 | 0.43 | 0.22 | 0.21 | 1.96 |
| OFN | Cep250 | 1.74 | 1.11E-12 | 2.36E-08 | 0.44 | 0.24 | 0.20 | 1.85 |
| OFN | Gm10544 | 2.87 | 1.16E-12 | 2.47E-08 | 0.15 | 0.04 | 0.11 | 4.11 |
| OFN | Fscn1 | 0.80 | 1.17E-12 | 2.49E-08 | 0.83 | 0.86 | -0.04 | 0.96 |
| OFN | Psmc5 | 0.71 | 1.28E-12 | 2.72E-08 | 0.85 | 0.90 | -0.04 | 0.95 |
| OFN | Gck | 2.73 | 1.31E-12 | 2.79E-08 | 0.16 | 0.04 | 0.12 | 3.83 |
| OFN | Khdrbs1 | 0.61 | 1.43E-12 | 3.05E-08 | 0.89 | 0.95 | -0.06 | 0.94 |
| OFN | Dctn1 | 1.59 | 1.53E-12 | 3.26E-08 | 0.65 | 0.51 | 0.13 | 1.26 |
| OFN | Ap2m1 | 0.86 | 1.73E-12 | 3.69E-08 | 0.83 | 0.83 | 0.00 | 1.00 |
| OFN | Zfp428 | 1.20 | 1.78E-12 | 3.80E-08 | 0.75 | 0.67 | 0.08 | 1.11 |
| OFN | Kif3c | 1.97 | 2.61E-12 | 5.57E-08 | 0.28 | 0.11 | 0.17 | 2.54 |
| OFN | Gpm6b | 2.61 | 2.88E-12 | 6.14E-08 | 0.25 | 0.09 | 0.16 | 2.68 |
| OFN | Ptpn6 | 2.73 | 3.13E-12 | 6.69E-08 | 0.23 | 0.08 | 0.15 | 2.97 |
| OFN | Cadps2 | 2.38 | 3.25E-12 | 6.95E-08 | 0.31 | 0.13 | 0.18 | 2.31 |
| OFN | Dact1 | 1.48 | 3.58E-12 | 7.65E-08 | 0.34 | 0.14 | 0.19 | 2.38 |
| OFN | Atp6v1e1 | 1.11 | 4.14E-12 | 8.84E-08 | 0.74 | 0.69 | 0.05 | 1.08 |
| OFN | Unc45a | 1.85 | 4.70E-12 | 1.00E-07 | 0.48 | 0.28 | 0.20 | 1.73 |
| OFN | Kctd13 | 1.72 | 5.81E-12 | 1.24E-07 | 0.34 | 0.16 | 0.19 | 2.19 |
| OFN | Mien1 | 1.12 | 6.79E-12 | 1.45E-07 | 0.75 | 0.68 | 0.07 | 1.10 |
| OFN | Fam20c | 2.24 | 7.87E-12 | 1.68E-07 | 0.23 | 0.08 | 0.15 | 2.90 |

**Supplementary Table 4. Marker genes for the annotated cell populations of integrated clustering of E11.5 control and NotchGOF λ-junction ectoderm cells.**

| | | | | | | | | |
|---|---|---|---|---|---|---|---|---|
| OFN | Zcrb1 | 0.80 | 8.18E-12 | 1.75E-07 | 0.85 | 0.90 | -0.05 | 0.95 |
| OFN | Rtn4 | 0.74 | 8.37E-12 | 1.79E-07 | 0.83 | 0.96 | -0.13 | 0.86 |
| OFN | Kcnn2 | 2.73 | 8.42E-12 | 1.80E-07 | 0.12 | 0.03 | 0.09 | 4.64 |
| OFN | Gm8797 | 0.81 | 8.69E-12 | 1.86E-07 | 0.88 | 0.93 | -0.04 | 0.96 |
| OFN | Lman2l | 1.58 | 9.56E-12 | 2.04E-07 | 0.55 | 0.36 | 0.19 | 1.54 |
| OFN | Foxg1 | 1.30 | 1.02E-11 | 2.18E-07 | 0.65 | 0.54 | 0.11 | 1.20 |
| OFN | Zbtb18 | 1.93 | 1.09E-11 | 2.33E-07 | 0.41 | 0.23 | 0.19 | 1.83 |
| OFN | Hpca | 3.82 | 1.12E-11 | 2.39E-07 | 0.12 | 0.03 | 0.09 | 4.64 |
| OFN | Crip2 | 1.93 | 1.15E-11 | 2.46E-07 | 0.61 | 0.42 | 0.20 | 1.47 |
| OFN | Evl | 1.16 | 1.17E-11 | 2.49E-07 | 0.78 | 0.73 | 0.05 | 1.07 |
| OFN | Mcf2l | 2.02 | 1.28E-11 | 2.74E-07 | 0.30 | 0.13 | 0.17 | 2.34 |
| OFN | Jund | 0.79 | 1.33E-11 | 2.84E-07 | 0.87 | 0.93 | -0.06 | 0.94 |
| OFN | Cog1 | 1.97 | 1.34E-11 | 2.85E-07 | 0.32 | 0.14 | 0.18 | 2.31 |
| OFN | Sertad2 | 1.26 | 1.42E-11 | 3.04E-07 | 0.66 | 0.52 | 0.14 | 1.26 |
| OFN | Stat3 | 2.11 | 1.71E-11 | 3.66E-07 | 0.25 | 0.09 | 0.15 | 2.69 |
| OFN | Katnal1 | 2.04 | 1.73E-11 | 3.69E-07 | 0.32 | 0.14 | 0.18 | 2.32 |
| OFN | Znrf1 | 1.29 | 1.83E-11 | 3.91E-07 | 0.69 | 0.56 | 0.13 | 1.23 |
| OFN | Oaf | 2.06 | 1.85E-11 | 3.95E-07 | 0.31 | 0.14 | 0.17 | 2.18 |
| OFN | Ksr2 | 2.73 | 1.86E-11 | 3.98E-07 | 0.16 | 0.04 | 0.11 | 3.69 |
| OFN | Tbcb | 0.82 | 2.11E-11 | 4.51E-07 | 0.82 | 0.85 | -0.03 | 0.96 |
| OFN | Hecw2 | 2.70 | 2.35E-11 | 5.02E-07 | 0.14 | 0.03 | 0.10 | 3.97 |
| OFN | Dusp1 | 1.83 | 2.45E-11 | 5.23E-07 | 0.43 | 0.24 | 0.19 | 1.80 |
| OFN | Nrp2 | 2.16 | 2.60E-11 | 5.56E-07 | 0.36 | 0.17 | 0.19 | 2.09 |
| OFN | E2f1 | 1.74 | 2.69E-11 | 5.75E-07 | 0.39 | 0.21 | 0.19 | 1.89 |
| OFN | Cspg5 | 2.95 | 2.76E-11 | 5.90E-07 | 0.14 | 0.04 | 0.11 | 3.84 |
| OFN | Efr3b | 2.51 | 2.80E-11 | 5.98E-07 | 0.19 | 0.06 | 0.13 | 3.08 |
| OFN | Psmc2 | 0.86 | 4.03E-11 | 8.61E-07 | 0.83 | 0.87 | -0.04 | 0.95 |
| OFN | Bmerb1 | 3.03 | 4.05E-11 | 8.64E-07 | 0.13 | 0.03 | 0.10 | 4.16 |
| OFN | Ndufb1-ps | 0.74 | 4.06E-11 | 8.68E-07 | 0.89 | 0.93 | -0.04 | 0.95 |
| OFN | Tgfbr1 | 1.65 | 4.37E-11 | 9.33E-07 | 0.68 | 0.61 | 0.07 | 1.12 |
| OFN | C9orf72 | 2.41 | 4.39E-11 | 9.38E-07 | 0.28 | 0.12 | 0.16 | 2.37 |
| OFN | Saraf | 1.50 | 4.74E-11 | 1.01E-06 | 0.72 | 0.66 | 0.05 | 1.08 |
| OFN | Pgm2l1 | 1.54 | 5.58E-11 | 1.19E-06 | 0.41 | 0.22 | 0.19 | 1.86 |
| OFN | Rbm18 | 1.33 | 5.61E-11 | 1.20E-06 | 0.64 | 0.49 | 0.15 | 1.29 |
| OFN | Ywhaq | 0.41 | 5.97E-11 | 1.27E-06 | 0.92 | 0.99 | -0.07 | 0.93 |
| OFN | Dynlt1f | 1.02 | 6.42E-11 | 1.37E-06 | 0.79 | 0.75 | 0.05 | 1.06 |
| OFN | Smchd1 | 1.82 | 6.43E-11 | 1.37E-06 | 0.79 | 0.84 | -0.04 | 0.95 |
| OFN | Ttc39b | 2.43 | 6.88E-11 | 1.47E-06 | 0.19 | 0.06 | 0.13 | 3.12 |
| OFN | Dync1li1 | 1.37 | 6.98E-11 | 1.49E-06 | 0.64 | 0.50 | 0.14 | 1.27 |
| OFN | Ciao2b | 1.09 | 8.21E-11 | 1.75E-06 | 0.76 | 0.74 | 0.02 | 1.02 |
| OFN | Gulo | 3.22 | 8.92E-11 | 1.91E-06 | 0.10 | 0.02 | 0.08 | 4.90 |
| OFN | 0610010F05Rik | 1.42 | 8.94E-11 | 1.91E-06 | 0.52 | 0.33 | 0.18 | 1.54 |
| OFN | Mkrn1 | 1.10 | 1.02E-10 | 2.17E-06 | 0.74 | 0.65 | 0.09 | 1.13 |
| OFN | Dact3 | 2.01 | 1.08E-10 | 2.30E-06 | 0.28 | 0.12 | 0.16 | 2.39 |
| OFN | Lmo1 | 2.13 | 1.09E-10 | 2.32E-06 | 0.42 | 0.23 | 0.19 | 1.85 |
| OFN | Pdlim7 | 1.03 | 1.14E-10 | 2.43E-06 | 0.73 | 0.62 | 0.11 | 1.17 |
| OFN | Cep63 | 1.38 | 1.18E-10 | 2.51E-06 | 0.52 | 0.35 | 0.18 | 1.50 |
| OFN | Pcdh20 | 2.55 | 1.34E-10 | 2.87E-06 | 0.12 | 0.03 | 0.09 | 4.30 |
| OFN | Cby1 | 1.41 | 1.38E-10 | 2.95E-06 | 0.63 | 0.46 | 0.17 | 1.38 |
| OFN | Eml1 | 1.84 | 1.74E-10 | 3.72E-06 | 0.34 | 0.17 | 0.17 | 2.01 |
| OFN | Csnk1e | 0.85 | 1.76E-10 | 3.75E-06 | 0.83 | 0.88 | -0.04 | 0.95 |
| OFN | Atp5md | 0.53 | 1.96E-10 | 4.18E-06 | 0.92 | 0.98 | -0.07 | 0.93 |
| OFN | Ppp1r11 | 0.91 | 1.97E-10 | 4.21E-06 | 0.82 | 0.82 | 0.00 | 1.00 |
| OFN | Tmcc1 | 1.22 | 2.10E-10 | 4.48E-06 | 0.62 | 0.46 | 0.16 | 1.35 |
| OFN | Rnf32 | 2.40 | 2.92E-10 | 6.23E-06 | 0.13 | 0.03 | 0.10 | 3.91 |

**Supplementary Table 4. Marker genes for the annotated cell populations of integrated clustering of E11.5 control and NotchGOF λ-junction ectoderm cells.**

| | | | | | | | | |
|---|---|---|---|---|---|---|---|---|
| OFN | Rims2 | 2.04 | 2.93E-10 | 6.27E-06 | 0.38 | 0.21 | 0.18 | 1.86 |
| OFN | Gm5860 | 3.43 | 3.46E-10 | 7.40E-06 | 0.12 | 0.03 | 0.09 | 3.97 |
| OFN | H3f3b | 0.28 | 3.56E-10 | 7.61E-06 | 1.00 | 1.00 | 0.00 | 1.00 |
| OFN | Hspa8 | 0.34 | 3.58E-10 | 7.64E-06 | 0.98 | 1.00 | -0.02 | 0.98 |
| OFN | Ndufa4l2 | 2.36 | 3.71E-10 | 7.92E-06 | 0.24 | 0.10 | 0.14 | 2.52 |
| OFN | Chd7 | 1.02 | 4.10E-10 | 8.76E-06 | 0.75 | 0.79 | -0.04 | 0.95 |
| OFN | Tspyl4 | 2.03 | 4.10E-10 | 8.77E-06 | 0.40 | 0.24 | 0.16 | 1.69 |
| OFN | Smarcc2 | 1.08 | 4.56E-10 | 9.74E-06 | 0.71 | 0.62 | 0.09 | 1.15 |
| OFN | Syt1 | 2.10 | 4.81E-10 | 1.03E-05 | 0.31 | 0.14 | 0.17 | 2.23 |
| OFN | Gnb4 | 2.63 | 4.92E-10 | 1.05E-05 | 0.12 | 0.03 | 0.09 | 3.97 |
| OFN | Cdc42ep3 | 1.72 | 5.01E-10 | 1.07E-05 | 0.35 | 0.17 | 0.18 | 2.02 |
| OFN | Tubb2a | 1.83 | 5.94E-10 | 1.27E-05 | 0.52 | 0.36 | 0.17 | 1.46 |
| OFN | Chd4 | 0.68 | 6.44E-10 | 1.37E-05 | 0.88 | 0.97 | -0.09 | 0.91 |
| OFN | Akirin1 | 1.38 | 6.80E-10 | 1.45E-05 | 0.66 | 0.58 | 0.08 | 1.14 |
| OFN | Trim35 | 1.07 | 6.85E-10 | 1.46E-05 | 0.68 | 0.55 | 0.13 | 1.23 |
| OFN | Dab2ip | 1.33 | 6.88E-10 | 1.47E-05 | 0.63 | 0.53 | 0.10 | 1.19 |
| OFN | Fam122b | 1.88 | 7.20E-10 | 1.54E-05 | 0.23 | 0.09 | 0.14 | 2.63 |
| OFN | Cep97 | 2.01 | 7.93E-10 | 1.69E-05 | 0.39 | 0.23 | 0.16 | 1.71 |
| OFN | Mtcl1 | 1.11 | 8.15E-10 | 1.74E-05 | 0.50 | 0.31 | 0.19 | 1.63 |
| OFN | Npdc1 | 1.61 | 8.84E-10 | 1.89E-05 | 0.49 | 0.32 | 0.17 | 1.51 |
| OFN | Kctd10 | 1.69 | 9.06E-10 | 1.94E-05 | 0.43 | 0.25 | 0.18 | 1.71 |
| OFN | Phf21b | 2.49 | 9.30E-10 | 1.99E-05 | 0.19 | 0.07 | 0.13 | 2.85 |
| OFN | Prmt2 | 1.61 | 1.07E-09 | 2.29E-05 | 0.50 | 0.35 | 0.15 | 1.42 |
| OFN | Bpgm | 1.35 | 1.11E-09 | 2.37E-05 | 0.57 | 0.42 | 0.15 | 1.35 |
| OFN | Tcta | 2.16 | 1.12E-09 | 2.40E-05 | 0.23 | 0.09 | 0.14 | 2.54 |
| OFN | Gm43260 | 2.14 | 1.12E-09 | 2.40E-05 | 0.16 | 0.05 | 0.11 | 3.30 |
| OFN | Fam171b | 2.08 | 1.17E-09 | 2.51E-05 | 0.28 | 0.13 | 0.15 | 2.18 |
| OFN | Cycs | 0.77 | 1.18E-09 | 2.52E-05 | 0.87 | 0.93 | -0.06 | 0.93 |
| OFN | Klhl29 | 2.00 | 1.27E-09 | 2.72E-05 | 0.14 | 0.04 | 0.10 | 3.55 |
| OFN | Mapre3 | 2.28 | 1.53E-09 | 3.27E-05 | 0.28 | 0.13 | 0.15 | 2.20 |
| OFN | Ywhae | 0.37 | 1.54E-09 | 3.29E-05 | 0.96 | 1.00 | -0.04 | 0.96 |
| OFN | Arpc3 | 0.67 | 1.58E-09 | 3.37E-05 | 0.87 | 0.92 | -0.05 | 0.95 |
| OFN | Tmem41a | 1.48 | 1.62E-09 | 3.45E-05 | 0.34 | 0.17 | 0.17 | 2.03 |
| OFN | Dennd1a | 1.40 | 1.78E-09 | 3.81E-05 | 0.37 | 0.20 | 0.17 | 1.85 |
| OFN | Arl6 | 1.68 | 2.06E-09 | 4.39E-05 | 0.54 | 0.38 | 0.16 | 1.42 |
| OFN | Tshz1 | 1.71 | 2.06E-09 | 4.40E-05 | 0.59 | 0.48 | 0.11 | 1.23 |
| OFN | Zbtb20 | 2.19 | 2.07E-09 | 4.43E-05 | 0.45 | 0.30 | 0.15 | 1.49 |
| OFN | Clcn2 | 1.66 | 2.16E-09 | 4.62E-05 | 0.21 | 0.08 | 0.13 | 2.75 |
| OFN | Vamp2 | 1.81 | 2.39E-09 | 5.10E-05 | 0.47 | 0.32 | 0.15 | 1.46 |
| OFN | Csnk1g3 | 1.09 | 2.58E-09 | 5.51E-05 | 0.75 | 0.67 | 0.08 | 1.12 |
| OFN | H13 | 0.93 | 2.86E-09 | 6.11E-05 | 0.72 | 0.65 | 0.08 | 1.12 |
| OFN | Olfm2 | 1.81 | 3.06E-09 | 6.53E-05 | 0.34 | 0.18 | 0.17 | 1.95 |
| OFN | Actr10 | 1.00 | 3.11E-09 | 6.64E-05 | 0.73 | 0.69 | 0.04 | 1.05 |
| OFN | Pik3r3 | 1.96 | 3.16E-09 | 6.76E-05 | 0.37 | 0.20 | 0.17 | 1.85 |
| OFN | Atp6v0d1 | 1.15 | 3.27E-09 | 6.99E-05 | 0.70 | 0.56 | 0.14 | 1.25 |
| OFN | Peli1 | 0.94 | 3.30E-09 | 7.04E-05 | 0.72 | 0.66 | 0.06 | 1.09 |
| OFN | Chchd2 | 0.28 | 3.33E-09 | 7.12E-05 | 0.98 | 1.00 | -0.02 | 0.98 |
| OFN | Ahi1 | 1.35 | 3.36E-09 | 7.17E-05 | 0.51 | 0.33 | 0.18 | 1.53 |
| OFN | Cacna1h | 2.32 | 3.37E-09 | 7.19E-05 | 0.14 | 0.04 | 0.10 | 3.38 |
| OFN | 2700097O09Rik | 1.66 | 3.52E-09 | 7.52E-05 | 0.48 | 0.33 | 0.15 | 1.45 |
| OFN | Kras | 0.86 | 3.67E-09 | 7.83E-05 | 0.81 | 0.84 | -0.03 | 0.97 |
| OFN | 3110039M20Rik | 1.97 | 3.86E-09 | 8.25E-05 | 0.45 | 0.30 | 0.15 | 1.50 |
| OFN | Twf1 | 1.85 | 5.14E-09 | 1.10E-04 | 0.63 | 0.54 | 0.08 | 1.15 |
| OFN | Trappc2l | 1.34 | 5.30E-09 | 1.13E-04 | 0.68 | 0.61 | 0.07 | 1.12 |
| OFN | Pdap1 | 0.52 | 5.38E-09 | 1.15E-04 | 0.93 | 0.99 | -0.06 | 0.94 |

**Supplementary Table 4. Marker genes for the annotated cell populations of integrated clustering of E11.5 control and NotchGOF λ-junction ectoderm cells.**

| | | | | | | | | |
|---|---|---|---|---|---|---|---|---|
| OFN | Phyh | 1.37 | 5.58E-09 | 1.19E-04 | 0.49 | 0.33 | 0.16 | 1.48 |
| OFN | Gfod1 | 2.60 | 5.84E-09 | 1.25E-04 | 0.11 | 0.03 | 0.08 | 3.93 |
| OFN | Mapk3 | 1.32 | 5.99E-09 | 1.28E-04 | 0.61 | 0.48 | 0.12 | 1.25 |
| OFN | Tle4 | 1.33 | 6.42E-09 | 1.37E-04 | 0.63 | 0.54 | 0.09 | 1.17 |
| OFN | Chchd6 | 1.35 | 6.48E-09 | 1.38E-04 | 0.65 | 0.53 | 0.13 | 1.24 |
| OFN | BC030867 | 2.20 | 6.88E-09 | 1.47E-04 | 0.24 | 0.10 | 0.14 | 2.30 |
| OFN | Pou3f1 | 2.24 | 7.07E-09 | 1.51E-04 | 0.14 | 0.04 | 0.10 | 3.38 |
| OFN | Peg13 | 2.13 | 7.67E-09 | 1.64E-04 | 0.16 | 0.05 | 0.11 | 2.98 |
| OFN | Arhgef2 | 1.27 | 8.27E-09 | 1.77E-04 | 0.53 | 0.39 | 0.14 | 1.36 |
| OFN | Mtch1 | 0.57 | 8.35E-09 | 1.78E-04 | 0.94 | 0.97 | -0.04 | 0.96 |
| OFN | Atf4 | 0.99 | 8.96E-09 | 1.91E-04 | 0.74 | 0.68 | 0.07 | 1.10 |
| OFN | Snx11 | 1.78 | 9.65E-09 | 2.06E-04 | 0.31 | 0.15 | 0.16 | 2.01 |
| OFN | Gpr85 | 2.47 | 9.85E-09 | 2.10E-04 | 0.11 | 0.03 | 0.08 | 3.79 |
| OFN | Cxxc5 | 1.30 | 1.10E-08 | 2.35E-04 | 0.67 | 0.59 | 0.09 | 1.15 |
| OFN | Ndrg3 | 1.85 | 1.12E-08 | 2.40E-04 | 0.39 | 0.22 | 0.17 | 1.74 |
| OFN | Pbx3 | 1.17 | 1.18E-08 | 2.51E-04 | 0.68 | 0.59 | 0.09 | 1.16 |
| OFN | Mterf2 | 1.55 | 1.20E-08 | 2.56E-04 | 0.39 | 0.22 | 0.17 | 1.74 |
| OFN | Ndufa11 | 0.71 | 1.20E-08 | 2.56E-04 | 0.92 | 0.95 | -0.03 | 0.96 |
| OFN | Sdccag8 | 2.02 | 1.26E-08 | 2.69E-04 | 0.28 | 0.14 | 0.15 | 2.10 |
| OFN | Chfr | 1.24 | 1.29E-08 | 2.75E-04 | 0.48 | 0.31 | 0.17 | 1.55 |
| OFN | Lrrn3 | 1.60 | 1.44E-08 | 3.07E-04 | 0.30 | 0.15 | 0.15 | 2.05 |
| OFN | Shisa4 | 1.39 | 1.45E-08 | 3.10E-04 | 0.40 | 0.24 | 0.16 | 1.69 |
| OFN | Vps28 | 1.19 | 1.47E-08 | 3.14E-04 | 0.70 | 0.62 | 0.09 | 1.14 |
| OFN | Podxl2 | 1.49 | 1.53E-08 | 3.27E-04 | 0.52 | 0.38 | 0.14 | 1.37 |
| OFN | Gsk3b | 0.77 | 1.58E-08 | 3.38E-04 | 0.83 | 0.91 | -0.07 | 0.92 |
| OFN | Ankrd13b | 1.70 | 1.60E-08 | 3.41E-04 | 0.34 | 0.18 | 0.15 | 1.85 |
| OFN | H2afy | 0.42 | 1.60E-08 | 3.43E-04 | 0.92 | 0.98 | -0.07 | 0.93 |
| OFN | Clip3 | 0.97 | 1.63E-08 | 3.48E-04 | 0.59 | 0.46 | 0.14 | 1.30 |
| OFN | Dennd2a | 1.51 | 1.66E-08 | 3.54E-04 | 0.41 | 0.24 | 0.17 | 1.72 |
| OFN | Gde1 | 1.23 | 1.77E-08 | 3.79E-04 | 0.61 | 0.50 | 0.11 | 1.21 |
| OFN | Asxl3 | 1.55 | 1.99E-08 | 4.24E-04 | 0.47 | 0.32 | 0.16 | 1.50 |
| OFN | Mob3b | 1.70 | 2.01E-08 | 4.30E-04 | 0.23 | 0.10 | 0.13 | 2.32 |
| OFN | Plppr3 | 1.18 | 2.05E-08 | 4.39E-04 | 0.48 | 0.31 | 0.16 | 1.52 |
| OFN | Kif3a | 1.14 | 2.06E-08 | 4.41E-04 | 0.68 | 0.59 | 0.10 | 1.16 |
| OFN | Dusp18 | 1.74 | 2.19E-08 | 4.67E-04 | 0.24 | 0.11 | 0.13 | 2.28 |
| OFN | Cpe | 1.20 | 2.21E-08 | 4.72E-04 | 0.59 | 0.47 | 0.13 | 1.27 |
| OFN | Fam204a | 1.08 | 2.23E-08 | 4.76E-04 | 0.72 | 0.63 | 0.09 | 1.14 |
| OFN | Lrrc75a | 1.58 | 2.34E-08 | 4.99E-04 | 0.36 | 0.19 | 0.16 | 1.83 |
| OFN | Trappc4 | 0.87 | 2.87E-08 | 6.13E-04 | 0.80 | 0.81 | -0.01 | 0.98 |
| OFN | Idh1 | 1.10 | 3.18E-08 | 6.79E-04 | 0.65 | 0.55 | 0.10 | 1.18 |
| OFN | Gm49066 | 1.80 | 3.21E-08 | 6.85E-04 | 0.18 | 0.07 | 0.11 | 2.66 |
| OFN | Susd6 | 1.59 | 3.38E-08 | 7.23E-04 | 0.37 | 0.21 | 0.16 | 1.74 |
| OFN | Baz2b | 0.91 | 3.58E-08 | 7.65E-04 | 0.79 | 0.81 | -0.02 | 0.98 |
| OFN | Pkn1 | 1.45 | 3.69E-08 | 7.88E-04 | 0.44 | 0.29 | 0.15 | 1.54 |
| OFN | Slc27a2 | 2.11 | 3.79E-08 | 8.10E-04 | 0.16 | 0.06 | 0.10 | 2.82 |
| OFN | Pdxp | 1.29 | 3.81E-08 | 8.15E-04 | 0.50 | 0.34 | 0.16 | 1.47 |
| OFN | Gdap2 | 1.74 | 3.97E-08 | 8.47E-04 | 0.23 | 0.10 | 0.13 | 2.31 |
| OFN | Ncoa1 | 1.92 | 4.18E-08 | 8.93E-04 | 0.36 | 0.21 | 0.15 | 1.70 |
| OFN | Psmb7 | 0.61 | 4.94E-08 | 1.05E-03 | 0.86 | 0.91 | -0.05 | 0.94 |
| OFN | Ywhab | 0.51 | 5.00E-08 | 1.07E-03 | 0.91 | 0.97 | -0.06 | 0.94 |
| OFN | Tmem131l | 1.03 | 5.46E-08 | 1.17E-03 | 0.59 | 0.48 | 0.11 | 1.22 |
| OFN | Rc3h2 | 1.11 | 5.59E-08 | 1.19E-03 | 0.66 | 0.56 | 0.10 | 1.18 |
| OFN | Slco3a1 | 2.36 | 5.62E-08 | 1.20E-03 | 0.14 | 0.04 | 0.09 | 3.14 |
| OFN | 2900093K20Rik | 1.82 | 6.20E-08 | 1.32E-03 | 0.23 | 0.10 | 0.13 | 2.23 |
| OFN | Sh3gl3 | 1.91 | 7.07E-08 | 1.51E-03 | 0.21 | 0.09 | 0.12 | 2.39 |

**Supplementary Table 4. Marker genes for the annotated cell populations of integrated clustering of E11.5 control and NotchGOF λ-junction ectoderm cells.**

| | | | | | | | | |
|---|---|---|---|---|---|---|---|---|
| OFN | Magee1 | 2.01 | 7.73E-08 | 1.65E-03 | 0.18 | 0.07 | 0.11 | 2.62 |
| OFN | Apip | 1.23 | 7.81E-08 | 1.67E-03 | 0.56 | 0.40 | 0.16 | 1.40 |
| OFN | Khdrbs2 | 1.79 | 7.83E-08 | 1.67E-03 | 0.12 | 0.04 | 0.09 | 3.32 |
| OFN | Esyt1 | 1.90 | 7.83E-08 | 1.67E-03 | 0.19 | 0.08 | 0.12 | 2.49 |
| OFN | Zcchc18 | 1.40 | 7.91E-08 | 1.69E-03 | 0.57 | 0.44 | 0.13 | 1.29 |
| OFN | Ndn | 1.63 | 7.96E-08 | 1.70E-03 | 0.67 | 0.61 | 0.06 | 1.10 |
| OFN | Arpc1a | 0.65 | 8.16E-08 | 1.74E-03 | 0.87 | 0.91 | -0.05 | 0.95 |
| OFN | Bach2 | 1.41 | 9.28E-08 | 1.98E-03 | 0.40 | 0.24 | 0.16 | 1.65 |
| OFN | Pak3 | 1.48 | 9.85E-08 | 2.10E-03 | 0.45 | 0.30 | 0.14 | 1.47 |
| OFN | Ankrd13a | 1.53 | 1.01E-07 | 2.16E-03 | 0.41 | 0.27 | 0.15 | 1.55 |
| OFN | Atp6v1h | 0.99 | 1.01E-07 | 2.16E-03 | 0.54 | 0.38 | 0.16 | 1.43 |
| OFN | Ufm1 | 0.97 | 1.01E-07 | 2.16E-03 | 0.72 | 0.68 | 0.04 | 1.06 |
| OFN | Rnd2 | 2.16 | 1.04E-07 | 2.23E-03 | 0.24 | 0.11 | 0.13 | 2.19 |
| OFN | Rtl8b | 1.88 | 1.06E-07 | 2.26E-03 | 0.43 | 0.28 | 0.15 | 1.52 |
| OFN | Igf1r | 0.79 | 1.06E-07 | 2.27E-03 | 0.78 | 0.83 | -0.05 | 0.94 |
| OFN | Ttc5 | 0.92 | 1.07E-07 | 2.28E-03 | 0.69 | 0.63 | 0.06 | 1.10 |
| OFN | Ssh2 | 1.19 | 1.09E-07 | 2.32E-03 | 0.43 | 0.28 | 0.15 | 1.54 |
| OFN | Lrrc42 | 1.25 | 1.11E-07 | 2.38E-03 | 0.70 | 0.71 | -0.01 | 0.98 |
| OFN | Capn10 | 1.23 | 1.17E-07 | 2.49E-03 | 0.37 | 0.22 | 0.16 | 1.71 |
| OFN | Sgip1 | 2.34 | 1.18E-07 | 2.52E-03 | 0.10 | 0.03 | 0.07 | 3.55 |
| OFN | Klf12 | 2.40 | 1.23E-07 | 2.63E-03 | 0.23 | 0.10 | 0.12 | 2.22 |
| OFN | Msi2 | 0.75 | 1.28E-07 | 2.72E-03 | 0.77 | 0.75 | 0.01 | 1.02 |
| OFN | Chmp5 | 0.70 | 1.29E-07 | 2.76E-03 | 0.79 | 0.80 | -0.01 | 0.99 |
| OFN | Prkx | 1.79 | 1.33E-07 | 2.85E-03 | 0.33 | 0.19 | 0.14 | 1.77 |
| OFN | Tmem151b | 1.81 | 1.36E-07 | 2.91E-03 | 0.19 | 0.08 | 0.11 | 2.49 |
| OFN | Rtn2 | 1.67 | 1.43E-07 | 3.05E-03 | 0.33 | 0.18 | 0.15 | 1.81 |
| OFN | Katnal2 | 1.98 | 1.56E-07 | 3.33E-03 | 0.17 | 0.07 | 0.11 | 2.60 |
| OFN | Tmod2 | 1.98 | 1.63E-07 | 3.48E-03 | 0.25 | 0.13 | 0.13 | 2.02 |
| OFN | Sv2a | 2.59 | 1.65E-07 | 3.52E-03 | 0.11 | 0.03 | 0.08 | 3.44 |
| OFN | Smap2 | 1.49 | 1.87E-07 | 4.00E-03 | 0.42 | 0.27 | 0.15 | 1.55 |
| OFN | Bbip1 | 1.00 | 1.90E-07 | 4.05E-03 | 0.68 | 0.61 | 0.07 | 1.11 |
| OFN | Ube2m | 0.50 | 1.92E-07 | 4.11E-03 | 0.88 | 0.95 | -0.06 | 0.93 |
| OFN | Malat1 | 0.73 | 2.07E-07 | 4.42E-03 | 0.93 | 1.00 | -0.07 | 0.93 |
| OFN | Bmpr1b | 1.83 | 2.08E-07 | 4.44E-03 | 0.19 | 0.08 | 0.11 | 2.46 |
| OFN | Rhobtb2 | 2.14 | 2.11E-07 | 4.52E-03 | 0.19 | 0.08 | 0.11 | 2.46 |
| OFN | Ift22 | 1.20 | 2.13E-07 | 4.54E-03 | 0.50 | 0.38 | 0.12 | 1.32 |
| OFN | Kif1b | 1.26 | 2.13E-07 | 4.56E-03 | 0.68 | 0.65 | 0.03 | 1.04 |
| OFN | Shc2 | 2.58 | 2.19E-07 | 4.67E-03 | 0.18 | 0.07 | 0.11 | 2.45 |
| OFN | Kcnh2 | 2.10 | 2.24E-07 | 4.78E-03 | 0.14 | 0.05 | 0.09 | 2.84 |
| OFN | 4930452B06Rik | 2.35 | 2.28E-07 | 4.88E-03 | 0.28 | 0.15 | 0.14 | 1.91 |
| OFN | Nedd4l | 1.47 | 2.29E-07 | 4.89E-03 | 0.39 | 0.25 | 0.14 | 1.56 |
| OFN | Gm36839 | 2.16 | 2.33E-07 | 4.98E-03 | 0.12 | 0.04 | 0.08 | 3.15 |
| OFN | Gm4419 | 2.49 | 2.36E-07 | 5.04E-03 | 0.10 | 0.03 | 0.07 | 3.43 |
| OFN | Nelfcd | 0.89 | 2.37E-07 | 5.07E-03 | 0.65 | 0.58 | 0.07 | 1.13 |
| OFN | Epb41 | 1.01 | 2.48E-07 | 5.29E-03 | 0.74 | 0.75 | -0.01 | 0.98 |
| OFN | Ccpg1 | 1.27 | 2.55E-07 | 5.46E-03 | 0.40 | 0.24 | 0.16 | 1.67 |
| OFN | Clstn2 | 2.45 | 2.72E-07 | 5.82E-03 | 0.10 | 0.03 | 0.07 | 3.43 |
| OFN | Rac3 | 1.58 | 2.73E-07 | 5.83E-03 | 0.51 | 0.41 | 0.10 | 1.24 |
| OFN | 4930402H24Rik | 1.40 | 2.80E-07 | 5.98E-03 | 0.30 | 0.16 | 0.14 | 1.89 |
| OFN | Cenpj | 1.31 | 2.82E-07 | 6.03E-03 | 0.47 | 0.34 | 0.13 | 1.39 |
| OFN | Sbk1 | 1.08 | 2.83E-07 | 6.04E-03 | 0.62 | 0.57 | 0.05 | 1.09 |
| OFN | Rab3d | 2.36 | 2.87E-07 | 6.13E-03 | 0.13 | 0.04 | 0.09 | 3.00 |
| OFN | Btbd10 | 1.58 | 2.94E-07 | 6.29E-03 | 0.41 | 0.27 | 0.14 | 1.53 |
| OFN | Pim1 | 1.49 | 3.06E-07 | 6.53E-03 | 0.57 | 0.45 | 0.12 | 1.26 |
| OFN | Pim3 | 2.05 | 3.19E-07 | 6.82E-03 | 0.52 | 0.41 | 0.11 | 1.27 |

**Supplementary Table 4. Marker genes for the annotated cell populations of integrated clustering of E11.5 control and NotchGOF λ-junction ectoderm cells.**

| | | | | | | | | |
|---|---|---|---|---|---|---|---|---|
| OFN | Prelid3b | 1.02 | 3.22E-07 | 6.89E-03 | 0.67 | 0.59 | 0.08 | 1.14 |
| OFN | Ttbk2 | 1.47 | 3.29E-07 | 7.03E-03 | 0.25 | 0.12 | 0.13 | 2.07 |
| OFN | Ror2 | 1.90 | 3.32E-07 | 7.09E-03 | 0.15 | 0.05 | 0.09 | 2.74 |
| OFN | Ndufb10 | 0.74 | 3.38E-07 | 7.21E-03 | 0.86 | 0.90 | -0.04 | 0.96 |
| OFN | Mindy3 | 1.30 | 3.40E-07 | 7.26E-03 | 0.39 | 0.25 | 0.14 | 1.58 |
| OFN | Nkiras1 | 2.15 | 3.68E-07 | 7.86E-03 | 0.25 | 0.12 | 0.13 | 2.03 |
| OFN | Rasgef1b | 1.92 | 3.86E-07 | 8.24E-03 | 0.21 | 0.10 | 0.12 | 2.24 |
| OFN | 4833439L19Rik | 1.38 | 4.13E-07 | 8.82E-03 | 0.55 | 0.46 | 0.09 | 1.19 |
| OFN | Trappc6b | 0.98 | 4.28E-07 | 9.15E-03 | 0.63 | 0.53 | 0.10 | 1.18 |
| OFN | Smarce1 | 0.34 | 4.29E-07 | 9.17E-03 | 0.91 | 0.98 | -0.06 | 0.93 |
| OFN | Coq10b | 1.33 | 4.40E-07 | 9.41E-03 | 0.32 | 0.18 | 0.14 | 1.79 |
| OFN | Sel1l3 | 1.92 | 4.69E-07 | 1.00E-02 | 0.17 | 0.07 | 0.10 | 2.49 |
| OFN | Mab21l2 | 2.51 | 5.07E-07 | 1.08E-02 | 0.15 | 0.06 | 0.09 | 2.64 |
| OFN | Med22 | 1.57 | 5.39E-07 | 1.15E-02 | 0.43 | 0.28 | 0.14 | 1.51 |
| OFN | Scrn1 | 2.38 | 5.41E-07 | 1.16E-02 | 0.24 | 0.12 | 0.12 | 1.98 |
| OFN | Sema6c | 1.33 | 5.77E-07 | 1.23E-02 | 0.26 | 0.13 | 0.13 | 2.05 |
| OFN | Ccdc34 | 0.69 | 6.21E-07 | 1.33E-02 | 0.85 | 0.86 | -0.02 | 0.98 |
| OFN | Ift43 | 1.35 | 6.40E-07 | 1.37E-02 | 0.72 | 0.67 | 0.05 | 1.08 |
| OFN | Lmna | 1.82 | 6.41E-07 | 1.37E-02 | 0.32 | 0.19 | 0.14 | 1.72 |
| OFN | Svip | 1.61 | 6.84E-07 | 1.46E-02 | 0.40 | 0.26 | 0.14 | 1.54 |
| OFN | Mex3b | 0.90 | 6.86E-07 | 1.46E-02 | 0.59 | 0.47 | 0.12 | 1.26 |
| OFN | Nabp1 | 1.89 | 6.90E-07 | 1.47E-02 | 0.16 | 0.06 | 0.10 | 2.63 |
| OFN | Wsb2 | 1.29 | 7.02E-07 | 1.50E-02 | 0.47 | 0.33 | 0.13 | 1.40 |
| OFN | Pacsin1 | 1.30 | 7.51E-07 | 1.60E-02 | 0.31 | 0.17 | 0.14 | 1.82 |
| OFN | Tmem108 | 1.09 | 7.53E-07 | 1.61E-02 | 0.46 | 0.30 | 0.16 | 1.53 |
| OFN | Atg4a | 2.40 | 7.72E-07 | 1.65E-02 | 0.16 | 0.06 | 0.10 | 2.63 |
| OFN | Rnf14 | 1.20 | 8.01E-07 | 1.71E-02 | 0.47 | 0.34 | 0.13 | 1.39 |
| OFN | Armh4 | 2.23 | 8.16E-07 | 1.74E-02 | 0.30 | 0.18 | 0.12 | 1.66 |
| OFN | Rassf3 | 1.79 | 8.17E-07 | 1.75E-02 | 0.25 | 0.13 | 0.12 | 1.96 |
| OFN | Vbp1 | 0.82 | 8.30E-07 | 1.77E-02 | 0.73 | 0.68 | 0.04 | 1.07 |
| OFN | March8 | 1.32 | 8.49E-07 | 1.81E-02 | 0.39 | 0.25 | 0.14 | 1.57 |
| OFN | Psmd6 | 0.68 | 9.10E-07 | 1.94E-02 | 0.85 | 0.86 | -0.01 | 0.99 |
| OFN | Gnb1 | 0.52 | 9.22E-07 | 1.97E-02 | 0.88 | 0.97 | -0.09 | 0.91 |
| OFN | Akr1a1 | 0.57 | 9.36E-07 | 2.00E-02 | 0.88 | 0.97 | -0.08 | 0.92 |
| OFN | Ypel5 | 1.09 | 9.71E-07 | 2.08E-02 | 0.42 | 0.26 | 0.16 | 1.64 |
| OFN | Lrrc49 | 1.24 | 1.05E-06 | 2.24E-02 | 0.32 | 0.18 | 0.14 | 1.76 |
| OFN | Morn4 | 1.51 | 1.09E-06 | 2.33E-02 | 0.29 | 0.16 | 0.13 | 1.85 |
| OFN | Ubl5 | 0.43 | 1.09E-06 | 2.34E-02 | 0.93 | 0.99 | -0.06 | 0.94 |
| OFN | Spa17 | 1.70 | 1.10E-06 | 2.35E-02 | 0.29 | 0.16 | 0.13 | 1.85 |
| OFN | Mtmr6 | 2.18 | 1.11E-06 | 2.37E-02 | 0.24 | 0.12 | 0.12 | 1.98 |
| OFN | Asb1 | 2.56 | 1.20E-06 | 2.57E-02 | 0.14 | 0.05 | 0.09 | 2.76 |
| OFN | Hbb-y | 2.36 | 1.26E-06 | 2.69E-02 | 0.94 | 0.98 | -0.04 | 0.96 |
| OFN | Map2k1 | 1.62 | 1.28E-06 | 2.73E-02 | 0.50 | 0.40 | 0.10 | 1.24 |
| OFN | Slco5a1 | 1.58 | 1.30E-06 | 2.78E-02 | 0.30 | 0.17 | 0.13 | 1.78 |
| OFN | Glipr2 | 2.27 | 1.35E-06 | 2.88E-02 | 0.17 | 0.07 | 0.10 | 2.40 |
| OFN | Zfp667 | 1.68 | 1.36E-06 | 2.90E-02 | 0.27 | 0.14 | 0.13 | 1.88 |
| OFN | Actr1a | 1.16 | 1.39E-06 | 2.98E-02 | 0.56 | 0.44 | 0.11 | 1.26 |
| OFN | Pfdn4 | 0.91 | 1.40E-06 | 2.98E-02 | 0.79 | 0.78 | 0.02 | 1.02 |
| OFN | 1110006O24Rik | 2.94 | 1.42E-06 | 3.04E-02 | 0.12 | 0.04 | 0.08 | 2.97 |
| OFN | Gnaz | 2.30 | 1.46E-06 | 3.11E-02 | 0.14 | 0.05 | 0.09 | 2.63 |
| OFN | Garnl3 | 2.15 | 1.52E-06 | 3.24E-02 | 0.11 | 0.04 | 0.08 | 3.14 |
| OFN | Nab1 | 1.16 | 1.57E-06 | 3.35E-02 | 0.39 | 0.24 | 0.15 | 1.61 |
| OFN | Ahsa2 | 0.87 | 1.61E-06 | 3.45E-02 | 0.68 | 0.63 | 0.04 | 1.07 |
| OFN | Taf13 | 1.13 | 1.65E-06 | 3.53E-02 | 0.60 | 0.49 | 0.11 | 1.22 |
| OFN | Synj1 | 1.78 | 1.70E-06 | 3.64E-02 | 0.23 | 0.12 | 0.12 | 1.98 |

**Supplementary Table 4. Marker genes for the annotated cell populations of integrated clustering of E11.5 control and NotchGOF λ-junction ectoderm cells.**

| | | | | | | | | |
|---|---|---|---|---|---|---|---|---|
| OFN | Six1 | 0.83 | 1.73E-06 | 3.70E-02 | 0.74 | 0.69 | 0.05 | 1.08 |
| OFN | Purg | 1.80 | 1.83E-06 | 3.92E-02 | 0.40 | 0.27 | 0.13 | 1.47 |
| OFN | Nin | 1.31 | 1.86E-06 | 3.98E-02 | 0.41 | 0.28 | 0.13 | 1.48 |
| OFN | Fam227a | 2.01 | 1.87E-06 | 4.00E-02 | 0.14 | 0.05 | 0.09 | 2.63 |
| OFN | Prtn3 | 1.59 | 1.88E-06 | 4.01E-02 | 0.22 | 0.10 | 0.12 | 2.11 |
| OFN | Cib2 | 1.41 | 1.94E-06 | 4.15E-02 | 0.31 | 0.18 | 0.13 | 1.76 |
| OFN | Frs3 | 3.14 | 2.14E-06 | 4.58E-02 | 0.12 | 0.04 | 0.08 | 2.97 |
| OFN | Wscd1 | 2.27 | 2.34E-06 | 5.00E-02 | 0.19 | 0.08 | 0.10 | 2.23 |

**Supplementary Table 5. Differentially expressed genes in the mutant λ-junction ectoderm cells.**

| gene | avg_log2FC | p_val | p_val_adj | pct.1 | pct.2 | pct1-pct2 | pct1/pct2 |
|---|---|---|---|---|---|---|---|
| Uncx | 6.61 | 4.96E-44 | 1.03E-39 | 0.11 | 0 | 0.11 | 112 |
| Hes5 | 5.81 | 5.31E-262 | 1.11E-257 | 0.56 | 0.04 | 0.52 | 14.61 |
| Oxtr | 5.51 | 4.38E-166 | 9.11E-162 | 0.38 | 0.01 | 0.37 | 29.38 |
| Heyl | 5.26 | 6.57E-127 | 1.37E-122 | 0.31 | 0.01 | 0.3 | 25.83 |
| Sox1 | 5.11 | 6.28E-25 | 1.31E-20 | 0.06 | 0 | 0.06 | 64 |
| Cdkl1 | 4.65 | 2.73E-48 | 5.68E-44 | 0.14 | 0.01 | 0.13 | 15.44 |
| Rhcg | 4.62 | 8.28E-21 | 1.72E-16 | 0.06 | 0 | 0.06 | 15 |
| Ap3b2 | 4.29 | 8.27E-33 | 1.72E-28 | 0.09 | 0 | 0.09 | 22.75 |
| Mir9-3hg | 4.16 | 3.72E-49 | 7.74E-45 | 0.15 | 0.01 | 0.14 | 14.5 |
| Tctex1d1 | 4 | 1.84E-45 | 3.82E-41 | 0.15 | 0.02 | 0.13 | 8.71 |
| Stmn3 | 3.99 | 2.93E-40 | 6.10E-36 | 0.13 | 0.01 | 0.11 | 10.42 |
| Jph4 | 3.96 | 3.28E-92 | 6.82E-88 | 0.27 | 0.03 | 0.24 | 8.84 |
| AW011738 | 3.49 | 6.19E-151 | 1.29E-146 | 0.41 | 0.05 | 0.36 | 8.72 |
| Pkp1 | 3.49 | 7.76E-44 | 1.61E-39 | 0.14 | 0.02 | 0.13 | 8.94 |
| Kazald1 | 3.45 | 1.09E-35 | 2.28E-31 | 0.12 | 0.01 | 0.1 | 9.58 |
| Tnfaip2 | 3.45 | 3.17E-213 | 6.59E-209 | 0.58 | 0.13 | 0.45 | 4.47 |
| Rab3c | 3.26 | 2.70E-17 | 5.61E-13 | 0.05 | 0 | 0.05 | 13 |
| Nipal1 | 3.12 | 4.17E-18 | 8.66E-14 | 0.05 | 0 | 0.05 | 13.25 |
| Dipk1c | 3.1 | 3.13E-16 | 6.52E-12 | 0.05 | 0.01 | 0.05 | 10.2 |
| Rhov | 3.03 | 1.96E-114 | 4.07E-110 | 0.32 | 0.03 | 0.29 | 12.64 |
| Rims4 | 2.67 | 2.52E-19 | 5.25E-15 | 0.07 | 0.01 | 0.06 | 6 |
| Nrarp | 2.64 | 0 | 0 | 0.77 | 0.23 | 0.54 | 3.39 |
| St8sia6 | 2.59 | 1.44E-48 | 3.00E-44 | 0.22 | 0.06 | 0.16 | 3.82 |
| Zfp750 | 2.56 | 7.36E-72 | 1.53E-67 | 0.27 | 0.05 | 0.22 | 5.15 |
| Six6 | 2.55 | 1.55E-122 | 3.23E-118 | 0.37 | 0.05 | 0.32 | 7.44 |
| Stmn2 | 2.55 | 1.07E-49 | 2.22E-45 | 0.23 | 0.06 | 0.17 | 3.67 |
| C130071C03Rik | 2.45 | 3.14E-62 | 6.53E-58 | 0.26 | 0.06 | 0.2 | 4.19 |
| Sgms2 | 2.43 | 1.11E-16 | 2.30E-12 | 0.08 | 0.02 | 0.06 | 4.22 |
| Pof1b | 2.39 | 6.94E-15 | 1.44E-10 | 0.06 | 0.01 | 0.05 | 4.5 |
| Cpm | 2.32 | 8.73E-175 | 1.82E-170 | 0.66 | 0.26 | 0.4 | 2.56 |
| Rcsd1 | 2.27 | 6.94E-51 | 1.44E-46 | 0.22 | 0.05 | 0.17 | 4.13 |
| Kcnj15 | 2.26 | 3.05E-16 | 6.33E-12 | 0.08 | 0.02 | 0.06 | 3.9 |
| Cbln2 | 2.25 | 1.81E-43 | 3.77E-39 | 0.19 | 0.04 | 0.14 | 4.51 |
| Gsta3 | 2.23 | 8.18E-17 | 1.70E-12 | 0.08 | 0.02 | 0.06 | 4.28 |
| Ncam1 | 2.23 | 2.45E-92 | 5.10E-88 | 0.43 | 0.15 | 0.28 | 2.82 |
| Ttyh1 | 2.23 | 1.53E-27 | 3.18E-23 | 0.12 | 0.03 | 0.09 | 4.58 |
| Nav3 | 2.18 | 7.50E-115 | 1.56E-110 | 0.48 | 0.15 | 0.33 | 3.22 |
| Cndp1 | 2.13 | 1.61E-10 | 3.34E-06 | 0.05 | 0.02 | 0.04 | 3.6 |
| D7Ertd443e | 2.13 | 2.67E-22 | 5.54E-18 | 0.13 | 0.04 | 0.09 | 3.12 |
| Kcnb1 | 2.13 | 9.99E-22 | 2.08E-17 | 0.12 | 0.03 | 0.08 | 3.38 |
| C230057A21Rik | 2.1 | 3.71E-11 | 7.73E-07 | 0.05 | 0.01 | 0.04 | 3.79 |
| Fgf15 | 2.1 | 1.53E-28 | 3.17E-24 | 0.13 | 0.03 | 0.1 | 4.19 |
| Faah | 2.09 | 1.27E-62 | 2.63E-58 | 0.29 | 0.08 | 0.21 | 3.54 |
| Fzd9 | 2.08 | 1.97E-11 | 4.10E-07 | 0.07 | 0.02 | 0.05 | 2.92 |
| Prkg1 | 2.06 | 2.42E-65 | 5.02E-61 | 0.33 | 0.11 | 0.22 | 2.98 |
| Crb2 | 2.05 | 1.99E-22 | 4.14E-18 | 0.11 | 0.03 | 0.08 | 3.68 |
| Jag1 | 2.01 | 6.32E-180 | 1.32E-175 | 0.67 | 0.27 | 0.4 | 2.52 |
| Cgnl1 | 1.97 | 4.36E-23 | 9.06E-19 | 0.21 | 0.1 | 0.11 | 2.04 |

**Supplementary Table 5. Differentially expressed genes in the mutant λ-junction ectoderm cells.**

| | | | | | | | |
|---|---|---|---|---|---|---|---|
| Filip1 | 1.97 | 2.04E-21 | 4.24E-17 | 0.1 | 0.03 | 0.08 | 4.08 |
| Gm266 | 1.94 | 1.43E-115 | 2.97E-111 | 0.64 | 0.34 | 0.3 | 1.87 |
| Plet1 | 1.93 | 5.07E-13 | 1.05E-08 | 0.06 | 0.02 | 0.05 | 4.07 |
| Grhl1 | 1.92 | 5.84E-21 | 1.22E-16 | 0.11 | 0.03 | 0.08 | 3.55 |
| Nectin4 | 1.9 | 3.65E-175 | 7.59E-171 | 0.65 | 0.23 | 0.42 | 2.86 |
| Slfn2 | 1.89 | 4.57E-11 | 9.50E-07 | 0.06 | 0.02 | 0.04 | 3.47 |
| Itih5 | 1.89 | 3.35E-16 | 6.97E-12 | 0.11 | 0.04 | 0.07 | 2.79 |
| Nrcam | 1.87 | 4.70E-42 | 9.79E-38 | 0.24 | 0.08 | 0.16 | 3.05 |
| Enpp2 | 1.86 | 4.74E-07 | 9.85E-03 | 0.06 | 0.03 | 0.03 | 2.18 |
| Gdap1l1 | 1.85 | 3.55E-11 | 7.37E-07 | 0.06 | 0.02 | 0.04 | 3.67 |
| Fxyd7 | 1.84 | 9.65E-19 | 2.01E-14 | 0.1 | 0.03 | 0.07 | 3.54 |
| Gm26793 | 1.84 | 3.26E-17 | 6.78E-13 | 0.09 | 0.03 | 0.07 | 3.54 |
| Notch1 | 1.81 | 4.16E-189 | 8.64E-185 | 0.73 | 0.32 | 0.42 | 2.31 |
| Tmem37 | 1.79 | 4.38E-13 | 9.11E-09 | 0.08 | 0.03 | 0.05 | 2.77 |
| Hey1 | 1.79 | 6.92E-286 | 1.44E-281 | 0.89 | 0.44 | 0.46 | 2.05 |
| Gas7 | 1.78 | 4.71E-21 | 9.80E-17 | 0.13 | 0.05 | 0.09 | 2.91 |
| Qpct | 1.78 | 3.71E-12 | 7.71E-08 | 0.07 | 0.02 | 0.05 | 3 |
| Soga3 | 1.76 | 1.83E-27 | 3.81E-23 | 0.17 | 0.06 | 0.11 | 2.81 |
| Cox4i2 | 1.75 | 8.51E-110 | 1.77E-105 | 0.58 | 0.28 | 0.3 | 2.1 |
| Igdcc3 | 1.71 | 2.31E-71 | 4.81E-67 | 0.43 | 0.18 | 0.24 | 2.34 |
| Plce1 | 1.71 | 2.51E-63 | 5.22E-59 | 0.42 | 0.19 | 0.23 | 2.23 |
| Cdh4 | 1.7 | 9.28E-17 | 1.93E-12 | 0.13 | 0.05 | 0.07 | 2.42 |
| Endod1 | 1.7 | 1.07E-28 | 2.23E-24 | 0.2 | 0.08 | 0.12 | 2.52 |
| Smagp | 1.7 | 1.66E-11 | 3.46E-07 | 0.07 | 0.03 | 0.05 | 2.81 |
| Pon2 | 1.68 | 1.94E-68 | 4.04E-64 | 0.37 | 0.13 | 0.24 | 2.83 |
| St6galnac5 | 1.66 | 1.40E-08 | 2.91E-04 | 0.06 | 0.02 | 0.04 | 2.57 |
| Gnao1 | 1.66 | 5.32E-32 | 1.11E-27 | 0.21 | 0.08 | 0.13 | 2.66 |
| Ccdc160 | 1.65 | 3.46E-15 | 7.20E-11 | 0.09 | 0.03 | 0.06 | 3.27 |
| Rundc3a | 1.65 | 4.88E-17 | 1.02E-12 | 0.09 | 0.03 | 0.07 | 3.36 |
| Lama1 | 1.64 | 2.61E-21 | 5.42E-17 | 0.14 | 0.05 | 0.09 | 2.78 |
| Dab1 | 1.64 | 1.19E-105 | 2.48E-101 | 0.58 | 0.27 | 0.31 | 2.15 |
| Gm35040 | 1.63 | 1.43E-13 | 2.97E-09 | 0.08 | 0.02 | 0.05 | 3.26 |
| Maneal | 1.61 | 4.98E-12 | 1.04E-07 | 0.06 | 0.02 | 0.04 | 3.93 |
| Cldn10 | 1.6 | 1.35E-11 | 2.80E-07 | 0.1 | 0.04 | 0.06 | 2.34 |
| Lrp11 | 1.58 | 6.36E-11 | 1.32E-06 | 0.08 | 0.03 | 0.05 | 2.45 |
| Slit3 | 1.57 | 7.38E-15 | 1.54E-10 | 0.12 | 0.05 | 0.07 | 2.39 |
| Pld1 | 1.56 | 1.80E-09 | 3.74E-05 | 0.05 | 0.02 | 0.04 | 3.4 |
| Cd83 | 1.55 | 3.04E-08 | 6.32E-04 | 0.05 | 0.02 | 0.03 | 2.7 |
| 4930426D05Rik | 1.54 | 5.94E-21 | 1.24E-16 | 0.15 | 0.06 | 0.09 | 2.59 |
| Mapk8ip2 | 1.53 | 8.81E-18 | 1.83E-13 | 0.11 | 0.04 | 0.07 | 2.85 |
| Exoc3l4 | 1.53 | 5.89E-31 | 1.22E-26 | 0.22 | 0.09 | 0.13 | 2.52 |
| Socs2 | 1.52 | 8.67E-179 | 1.80E-174 | 0.86 | 0.6 | 0.26 | 1.43 |
| Slc22a23 | 1.52 | 6.36E-59 | 1.32E-54 | 0.37 | 0.15 | 0.22 | 2.46 |
| Rasgrp2 | 1.5 | 1.25E-10 | 2.59E-06 | 0.06 | 0.02 | 0.04 | 3 |
| Tgm1 | 1.5 | 3.03E-10 | 6.30E-06 | 0.07 | 0.02 | 0.04 | 2.79 |
| Phf21b | 1.48 | 4.03E-12 | 8.38E-08 | 0.09 | 0.03 | 0.05 | 2.66 |
| 2600006K01Rik | 1.47 | 5.97E-08 | 1.24E-03 | 0.05 | 0.02 | 0.03 | 2.78 |
| Gucy1b1 | 1.46 | 5.11E-12 | 1.06E-07 | 0.09 | 0.04 | 0.05 | 2.38 |
| Prss12 | 1.45 | 7.76E-09 | 1.61E-04 | 0.07 | 0.03 | 0.04 | 2.44 |

**Supplementary Table 5. Differentially expressed genes in the mutant λ-junction ectoderm cells.**

| | | | | | | | |
|---|---|---|---|---|---|---|---|
| Ephb1 | 1.45 | 1.33E-71 | 2.78E-67 | 0.45 | 0.2 | 0.25 | 2.25 |
| Adgrb3 | 1.44 | 4.02E-22 | 8.37E-18 | 0.15 | 0.05 | 0.09 | 2.79 |
| Zmat4 | 1.44 | 3.91E-21 | 8.13E-17 | 0.17 | 0.07 | 0.1 | 2.32 |
| Tmem200a | 1.44 | 1.21E-19 | 2.51E-15 | 0.14 | 0.05 | 0.09 | 2.62 |
| Pcdh1 | 1.43 | 3.20E-17 | 6.66E-13 | 0.13 | 0.05 | 0.08 | 2.55 |
| Alpl | 1.41 | 1.11E-18 | 2.31E-14 | 0.2 | 0.11 | 0.1 | 1.91 |
| Myo5c | 1.41 | 5.75E-25 | 1.20E-20 | 0.18 | 0.07 | 0.11 | 2.53 |
| Cyp4v3 | 1.4 | 3.14E-15 | 6.54E-11 | 0.12 | 0.05 | 0.07 | 2.5 |
| Fzd4 | 1.4 | 1.44E-28 | 3.00E-24 | 0.25 | 0.12 | 0.13 | 2.09 |
| Dmbx1 | 1.39 | 9.30E-30 | 1.94E-25 | 0.2 | 0.08 | 0.13 | 2.6 |
| Lypd2 | 1.39 | 2.76E-21 | 5.74E-17 | 0.19 | 0.08 | 0.1 | 2.2 |
| Gucy1a1 | 1.38 | 3.68E-37 | 7.64E-33 | 0.25 | 0.1 | 0.15 | 2.62 |
| Camk2n2 | 1.38 | 9.79E-13 | 2.04E-08 | 0.1 | 0.04 | 0.06 | 2.34 |
| Tex15 | 1.38 | 4.70E-35 | 9.77E-31 | 0.31 | 0.16 | 0.16 | 2.01 |
| Nebl | 1.37 | 1.95E-84 | 4.06E-80 | 0.48 | 0.19 | 0.29 | 2.57 |
| Adamts15 | 1.35 | 2.90E-08 | 6.03E-04 | 0.06 | 0.02 | 0.04 | 2.57 |
| Tox2 | 1.35 | 2.70E-23 | 5.61E-19 | 0.15 | 0.05 | 0.1 | 2.98 |
| Adi1 | 1.35 | 2.03E-65 | 4.23E-61 | 0.54 | 0.31 | 0.22 | 1.71 |
| Hey2 | 1.34 | 2.53E-14 | 5.26E-10 | 0.12 | 0.05 | 0.07 | 2.36 |
| Rbp1 | 1.34 | 1.83E-156 | 3.81E-152 | 0.97 | 0.88 | 0.09 | 1.1 |
| Aqp3 | 1.33 | 3.94E-08 | 8.19E-04 | 0.05 | 0.02 | 0.03 | 2.83 |
| Glrx | 1.32 | 1.23E-48 | 2.56E-44 | 0.36 | 0.17 | 0.19 | 2.14 |
| Acsl6 | 1.32 | 9.19E-10 | 1.91E-05 | 0.06 | 0.02 | 0.04 | 2.74 |
| Fxyd1 | 1.32 | 7.32E-09 | 1.52E-04 | 0.06 | 0.02 | 0.04 | 2.68 |
| B4galt5 | 1.32 | 1.58E-14 | 3.29E-10 | 0.15 | 0.07 | 0.08 | 2.07 |
| Gsap | 1.31 | 5.92E-17 | 1.23E-12 | 0.14 | 0.06 | 0.08 | 2.4 |
| 0610009E02Rik | 1.3 | 4.75E-10 | 9.88E-06 | 0.09 | 0.04 | 0.05 | 2.17 |
| Stbd1 | 1.3 | 1.93E-15 | 4.02E-11 | 0.17 | 0.09 | 0.08 | 1.97 |
| Fam189a2 | 1.29 | 4.08E-11 | 8.49E-07 | 0.1 | 0.04 | 0.06 | 2.25 |
| Plagl1 | 1.29 | 3.13E-70 | 6.51E-66 | 0.53 | 0.27 | 0.26 | 1.95 |
| Fhod3 | 1.27 | 2.58E-07 | 5.37E-03 | 0.09 | 0.05 | 0.04 | 1.83 |
| Pde1b | 1.27 | 6.38E-11 | 1.33E-06 | 0.09 | 0.04 | 0.05 | 2.39 |
| Notch3 | 1.27 | 3.80E-96 | 7.89E-92 | 0.63 | 0.35 | 0.28 | 1.79 |
| Upk2 | 1.27 | 2.89E-08 | 6.01E-04 | 0.08 | 0.03 | 0.04 | 2.21 |
| Klhl14 | 1.26 | 2.81E-08 | 5.84E-04 | 0.06 | 0.03 | 0.04 | 2.46 |
| Tmprss13 | 1.26 | 1.46E-10 | 3.04E-06 | 0.08 | 0.03 | 0.05 | 2.53 |
| Kbtbd11 | 1.26 | 9.30E-08 | 1.93E-03 | 0.06 | 0.03 | 0.04 | 2.38 |
| Elavl2 | 1.25 | 6.28E-54 | 1.31E-49 | 0.46 | 0.24 | 0.22 | 1.93 |
| Cadps2 | 1.25 | 2.58E-17 | 5.37E-13 | 0.15 | 0.07 | 0.08 | 2.24 |
| Prtg | 1.24 | 3.36E-27 | 7.00E-23 | 0.26 | 0.13 | 0.13 | 2 |
| Smtnl2 | 1.24 | 1.17E-38 | 2.43E-34 | 0.31 | 0.14 | 0.17 | 2.27 |
| Gca | 1.23 | 4.63E-11 | 9.63E-07 | 0.1 | 0.04 | 0.05 | 2.32 |
| Trpc1 | 1.23 | 2.27E-35 | 4.72E-31 | 0.29 | 0.13 | 0.16 | 2.22 |
| Foxc1 | 1.22 | 3.41E-23 | 7.09E-19 | 0.22 | 0.1 | 0.11 | 2.1 |
| Tmem229b | 1.22 | 8.55E-10 | 1.78E-05 | 0.08 | 0.03 | 0.05 | 2.45 |
| Itga6 | 1.22 | 2.78E-119 | 5.77E-115 | 0.82 | 0.62 | 0.2 | 1.32 |
| Pde1c | 1.22 | 1.51E-13 | 3.14E-09 | 0.13 | 0.07 | 0.07 | 2.06 |
| Gdap1 | 1.21 | 1.40E-08 | 2.92E-04 | 0.05 | 0.02 | 0.04 | 2.84 |
| Bicdl2 | 1.21 | 1.09E-06 | 2.27E-02 | 0.07 | 0.03 | 0.03 | 2.03 |

**Supplementary Table 5. Differentially expressed genes in the mutant λ-junction ectoderm cells.**

| | | | | | | | |
|---|---|---|---|---|---|---|---|
| Dtx4 | 1.21 | 1.74E-10 | 3.62E-06 | 0.08 | 0.03 | 0.05 | 2.47 |
| Mapk12 | 1.21 | 8.77E-41 | 1.82E-36 | 0.34 | 0.16 | 0.18 | 2.12 |
| Gpc1 | 1.2 | 8.31E-62 | 1.73E-57 | 0.56 | 0.35 | 0.21 | 1.59 |
| Emid1 | 1.18 | 4.67E-26 | 9.70E-22 | 0.21 | 0.09 | 0.12 | 2.32 |
| Moxd1 | 1.18 | 2.27E-08 | 4.73E-04 | 0.07 | 0.03 | 0.04 | 2.21 |
| D030055H07Rik | 1.17 | 1.57E-09 | 3.26E-05 | 0.08 | 0.03 | 0.04 | 2.33 |
| Syt1 | 1.17 | 1.51E-13 | 3.14E-09 | 0.15 | 0.08 | 0.07 | 1.94 |
| Ahr | 1.16 | 3.66E-24 | 7.61E-20 | 0.22 | 0.1 | 0.12 | 2.2 |
| Cldn23 | 1.15 | 8.51E-19 | 1.77E-14 | 0.18 | 0.08 | 0.09 | 2.16 |
| Snx31 | 1.15 | 2.16E-06 | 4.49E-02 | 0.05 | 0.02 | 0.03 | 2.32 |
| Rtn2 | 1.15 | 4.16E-20 | 8.65E-16 | 0.2 | 0.1 | 0.1 | 2.06 |
| Col9a2 | 1.15 | 1.63E-11 | 3.40E-07 | 0.15 | 0.08 | 0.07 | 1.81 |
| Ano1 | 1.14 | 1.16E-58 | 2.42E-54 | 0.52 | 0.3 | 0.23 | 1.78 |
| Ppp2r2b | 1.14 | 2.37E-53 | 4.93E-49 | 0.43 | 0.21 | 0.22 | 2.07 |
| Scg5 | 1.14 | 3.61E-23 | 7.51E-19 | 0.22 | 0.1 | 0.11 | 2.11 |
| Ppl | 1.13 | 6.22E-25 | 1.29E-20 | 0.2 | 0.08 | 0.12 | 2.38 |
| Tbc1d30 | 1.13 | 3.33E-10 | 6.93E-06 | 0.12 | 0.07 | 0.06 | 1.88 |
| Frmd4a | 1.13 | 3.56E-68 | 7.40E-64 | 0.59 | 0.36 | 0.23 | 1.65 |
| Rab11fip1 | 1.12 | 7.80E-38 | 1.62E-33 | 0.36 | 0.19 | 0.17 | 1.9 |
| Crym | 1.12 | 1.20E-27 | 2.50E-23 | 0.33 | 0.19 | 0.14 | 1.76 |
| Rims2 | 1.11 | 8.43E-22 | 1.75E-17 | 0.23 | 0.12 | 0.11 | 1.95 |
| Ermn | 1.1 | 2.25E-23 | 4.67E-19 | 0.21 | 0.1 | 0.12 | 2.2 |
| Sh2d4a | 1.1 | 7.92E-10 | 1.65E-05 | 0.06 | 0.02 | 0.04 | 2.95 |
| Id4 | 1.1 | 4.39E-53 | 9.12E-49 | 0.64 | 0.45 | 0.18 | 1.4 |
| Cobl | 1.09 | 1.22E-18 | 2.54E-14 | 0.2 | 0.1 | 0.1 | 1.96 |
| Dach1 | 1.09 | 4.05E-69 | 8.42E-65 | 0.57 | 0.31 | 0.26 | 1.84 |
| Zdhhc2 | 1.09 | 3.29E-31 | 6.85E-27 | 0.41 | 0.26 | 0.15 | 1.58 |
| Vim | 1.09 | 1.42E-82 | 2.96E-78 | 0.79 | 0.61 | 0.17 | 1.28 |
| S100a16 | 1.08 | 9.05E-19 | 1.88E-14 | 0.18 | 0.08 | 0.1 | 2.16 |
| Slitrk5 | 1.08 | 1.49E-07 | 3.10E-03 | 0.09 | 0.05 | 0.04 | 1.87 |
| Cebpa | 1.07 | 6.38E-09 | 1.33E-04 | 0.09 | 0.05 | 0.05 | 2 |
| Hlf | 1.07 | 2.79E-07 | 5.81E-03 | 0.07 | 0.03 | 0.04 | 2.17 |
| Cxcl12 | 1.07 | 4.46E-21 | 9.27E-17 | 0.19 | 0.08 | 0.1 | 2.23 |
| Stxbp6 | 1.07 | 2.11E-24 | 4.39E-20 | 0.24 | 0.11 | 0.12 | 2.08 |
| Phlpp1 | 1.07 | 9.00E-28 | 1.87E-23 | 0.31 | 0.17 | 0.14 | 1.85 |
| Lhfpl4 | 1.06 | 1.96E-10 | 4.07E-06 | 0.06 | 0.02 | 0.04 | 2.91 |
| Rassf10 | 1.06 | 1.06E-25 | 2.20E-21 | 0.28 | 0.15 | 0.13 | 1.89 |
| Pclo | 1.05 | 5.64E-29 | 1.17E-24 | 0.3 | 0.16 | 0.15 | 1.91 |
| Gm19938 | 1.05 | 9.24E-13 | 1.92E-08 | 0.11 | 0.05 | 0.06 | 2.29 |
| Nrk | 1.05 | 5.49E-11 | 1.14E-06 | 0.16 | 0.1 | 0.07 | 1.71 |
| Synm | 1.05 | 1.58E-07 | 3.28E-03 | 0.08 | 0.04 | 0.04 | 2.05 |
| Chst2 | 1.05 | 6.10E-16 | 1.27E-11 | 0.2 | 0.11 | 0.09 | 1.83 |
| Grhl3 | 1.05 | 3.98E-22 | 8.28E-18 | 0.26 | 0.14 | 0.12 | 1.88 |
| Prom2 | 1.04 | 1.32E-12 | 2.75E-08 | 0.15 | 0.08 | 0.07 | 1.95 |
| Bcl2l14 | 1.04 | 1.39E-06 | 2.90E-02 | 0.06 | 0.03 | 0.03 | 2.1 |
| Scrn1 | 1.03 | 2.92E-14 | 6.07E-10 | 0.14 | 0.07 | 0.07 | 2.12 |
| Itgb8 | 1.03 | 8.08E-21 | 1.68E-16 | 0.24 | 0.13 | 0.11 | 1.87 |
| Cited4 | 1.03 | 1.77E-06 | 3.67E-02 | 0.07 | 0.03 | 0.03 | 1.97 |
| A330048O09Rik | 1.03 | 4.05E-10 | 8.42E-06 | 0.08 | 0.03 | 0.05 | 2.5 |

**Supplementary Table 5. Differentially expressed genes in the mutant λ-junction ectoderm cells.**

| | | | | | | | |
|---|---|---|---|---|---|---|---|
| Sox3 | 1.02 | 5.93E-23 | 1.23E-18 | 0.19 | 0.08 | 0.11 | 2.35 |
| Crmp1 | 1.02 | 7.64E-09 | 1.59E-04 | 0.15 | 0.09 | 0.06 | 1.62 |
| Eva1a | 1.02 | 2.80E-17 | 5.83E-13 | 0.24 | 0.14 | 0.1 | 1.7 |
| Egfl6 | 1.02 | 7.90E-22 | 1.64E-17 | 0.22 | 0.11 | 0.11 | 2.07 |
| Gpr19 | 1.02 | 4.63E-09 | 9.63E-05 | 0.09 | 0.05 | 0.05 | 2.04 |
| Eaf2 | 1.01 | 5.44E-07 | 1.13E-02 | 0.1 | 0.06 | 0.04 | 1.75 |
| Npas3 | 1.01 | 2.84E-12 | 5.90E-08 | 0.14 | 0.07 | 0.07 | 1.94 |
| Mt1 | 1.01 | 3.12E-39 | 6.50E-35 | 0.56 | 0.39 | 0.18 | 1.45 |
| Gstm1 | 1.01 | 9.70E-14 | 2.02E-09 | 0.39 | 0.3 | 0.09 | 1.29 |
| Myh14 | 1 | 4.92E-32 | 1.02E-27 | 0.35 | 0.19 | 0.16 | 1.82 |
| Snca | 1 | 7.58E-11 | 1.58E-06 | 0.13 | 0.07 | 0.06 | 1.92 |
| Trmt9b | 1 | 1.46E-15 | 3.04E-11 | 0.21 | 0.12 | 0.09 | 1.78 |
| Nr2f1 | 1 | 2.11E-22 | 4.38E-18 | 0.29 | 0.17 | 0.12 | 1.73 |
| Foxp2 | 1 | 1.20E-23 | 2.50E-19 | 0.29 | 0.16 | 0.13 | 1.79 |
| Kazn | 1 | 2.26E-33 | 4.70E-29 | 0.36 | 0.2 | 0.16 | 1.81 |
| 4930523C07Rik | 0.99 | 3.36E-12 | 6.99E-08 | 0.12 | 0.06 | 0.07 | 2.12 |
| Fbn2 | 0.99 | 5.50E-51 | 1.14E-46 | 0.57 | 0.37 | 0.21 | 1.56 |
| Tmem79 | 0.98 | 7.65E-07 | 1.59E-02 | 0.09 | 0.05 | 0.04 | 1.78 |
| Plcd1 | 0.98 | 2.64E-11 | 5.49E-07 | 0.14 | 0.08 | 0.07 | 1.85 |
| Carmil3 | 0.98 | 9.69E-11 | 2.02E-06 | 0.11 | 0.05 | 0.06 | 2.14 |
| Arg1 | 0.98 | 1.28E-07 | 2.67E-03 | 0.07 | 0.03 | 0.04 | 2.19 |
| Ctsh | 0.98 | 3.31E-11 | 6.88E-07 | 0.15 | 0.08 | 0.07 | 1.81 |
| Cryab | 0.98 | 4.73E-18 | 9.83E-14 | 0.21 | 0.11 | 0.1 | 1.91 |
| Capn5 | 0.97 | 3.17E-07 | 6.59E-03 | 0.07 | 0.04 | 0.04 | 2.03 |
| Exph5 | 0.97 | 1.23E-15 | 2.55E-11 | 0.17 | 0.09 | 0.09 | 1.99 |
| Atl1 | 0.97 | 1.84E-09 | 3.82E-05 | 0.08 | 0.03 | 0.05 | 2.32 |
| Mylk | 0.97 | 2.08E-18 | 4.34E-14 | 0.24 | 0.13 | 0.11 | 1.78 |
| Fezf1 | 0.97 | 1.72E-09 | 3.58E-05 | 0.09 | 0.04 | 0.05 | 2.2 |
| Ric3 | 0.97 | 4.69E-11 | 9.75E-07 | 0.13 | 0.07 | 0.06 | 1.95 |
| Med12l | 0.96 | 1.20E-07 | 2.49E-03 | 0.1 | 0.06 | 0.05 | 1.8 |
| Sorbs2 | 0.96 | 7.18E-28 | 1.49E-23 | 0.44 | 0.31 | 0.14 | 1.44 |
| Spry4 | 0.96 | 8.69E-11 | 1.81E-06 | 0.13 | 0.07 | 0.06 | 1.9 |
| Ptprz1 | 0.96 | 8.64E-18 | 1.80E-13 | 0.25 | 0.14 | 0.11 | 1.75 |
| Ggt7 | 0.96 | 1.89E-07 | 3.94E-03 | 0.07 | 0.03 | 0.04 | 2.12 |
| Pkdcc | 0.96 | 1.74E-29 | 3.62E-25 | 0.39 | 0.24 | 0.15 | 1.62 |
| Notch2 | 0.95 | 2.17E-22 | 4.51E-18 | 0.37 | 0.24 | 0.13 | 1.52 |
| Gcnt2 | 0.95 | 1.37E-09 | 2.84E-05 | 0.1 | 0.05 | 0.05 | 2.09 |
| Ly6g6e | 0.95 | 4.37E-07 | 9.10E-03 | 0.08 | 0.04 | 0.04 | 1.95 |
| Proser2 | 0.95 | 9.29E-12 | 1.93E-07 | 0.18 | 0.1 | 0.07 | 1.7 |
| Usp43 | 0.95 | 1.30E-08 | 2.70E-04 | 0.07 | 0.03 | 0.04 | 2.43 |
| Hes1 | 0.95 | 3.44E-86 | 7.15E-82 | 0.81 | 0.6 | 0.2 | 1.33 |
| Fam13c | 0.95 | 1.54E-06 | 3.20E-02 | 0.09 | 0.05 | 0.04 | 1.8 |
| Rgcc | 0.95 | 1.20E-07 | 2.50E-03 | 0.13 | 0.08 | 0.05 | 1.61 |
| Pea15a | 0.95 | 2.18E-34 | 4.54E-30 | 0.4 | 0.23 | 0.17 | 1.73 |
| Hspb2 | 0.94 | 1.32E-16 | 2.75E-12 | 0.16 | 0.07 | 0.09 | 2.16 |
| Slc15a2 | 0.94 | 1.68E-12 | 3.50E-08 | 0.17 | 0.09 | 0.07 | 1.79 |
| Barx1 | 0.94 | 1.37E-13 | 2.85E-09 | 0.16 | 0.08 | 0.08 | 1.93 |
| Ptges | 0.94 | 3.70E-08 | 7.69E-04 | 0.11 | 0.06 | 0.05 | 1.8 |
| Nup210 | 0.93 | 1.87E-13 | 3.89E-09 | 0.21 | 0.12 | 0.08 | 1.68 |

**Supplementary Table 5. Differentially expressed genes in the mutant λ-junction ectoderm cells.**

| | | | | | | | |
|---|---|---|---|---|---|---|---|
| Crip2 | 0.93 | 9.93E-36 | 2.07E-31 | 0.45 | 0.29 | 0.17 | 1.58 |
| Cebpb | 0.93 | 5.48E-16 | 1.14E-11 | 0.28 | 0.18 | 0.1 | 1.58 |
| A830082K12Rik | 0.93 | 5.72E-07 | 1.19E-02 | 0.08 | 0.04 | 0.04 | 1.91 |
| Dsc2 | 0.91 | 6.36E-22 | 1.32E-17 | 0.26 | 0.14 | 0.12 | 1.86 |
| Eppk1 | 0.9 | 2.25E-07 | 4.69E-03 | 0.1 | 0.06 | 0.04 | 1.78 |
| Mapk11 | 0.9 | 1.26E-11 | 2.62E-07 | 0.14 | 0.07 | 0.07 | 1.9 |
| Muc1 | 0.9 | 4.84E-08 | 1.01E-03 | 0.09 | 0.05 | 0.04 | 1.98 |
| Cdk6 | 0.87 | 1.04E-44 | 2.15E-40 | 0.59 | 0.4 | 0.18 | 1.46 |
| Evpl | 0.87 | 7.02E-09 | 1.46E-04 | 0.13 | 0.08 | 0.06 | 1.72 |
| Stox1 | 0.87 | 2.38E-08 | 4.96E-04 | 0.09 | 0.04 | 0.05 | 2.02 |
| Gm42047 | 0.87 | 4.89E-09 | 1.02E-04 | 0.12 | 0.07 | 0.05 | 1.82 |
| Paqr4 | 0.87 | 2.37E-08 | 4.93E-04 | 0.09 | 0.05 | 0.05 | 1.98 |
| Srgap3 | 0.87 | 2.08E-23 | 4.33E-19 | 0.26 | 0.14 | 0.13 | 1.91 |
| Klf4 | 0.87 | 1.79E-17 | 3.72E-13 | 0.24 | 0.14 | 0.1 | 1.75 |
| Lurap1l | 0.86 | 4.84E-16 | 1.01E-11 | 0.29 | 0.19 | 0.1 | 1.54 |
| Egfr | 0.86 | 2.21E-16 | 4.59E-12 | 0.26 | 0.16 | 0.1 | 1.65 |
| Ppfibp1 | 0.86 | 7.66E-52 | 1.59E-47 | 0.67 | 0.5 | 0.17 | 1.35 |
| Fjx1 | 0.86 | 4.73E-86 | 9.84E-82 | 0.84 | 0.64 | 0.2 | 1.3 |
| Cgn | 0.86 | 8.21E-31 | 1.71E-26 | 0.42 | 0.26 | 0.16 | 1.61 |
| P2rx2 | 0.85 | 3.40E-10 | 7.08E-06 | 0.09 | 0.04 | 0.05 | 2.24 |
| Mn1 | 0.85 | 9.93E-16 | 2.06E-11 | 0.22 | 0.13 | 0.1 | 1.75 |
| Thrb | 0.85 | 7.43E-12 | 1.55E-07 | 0.18 | 0.1 | 0.07 | 1.71 |
| Ppp1r1a | 0.85 | 6.47E-24 | 1.35E-19 | 0.35 | 0.22 | 0.13 | 1.6 |
| Cldn3 | 0.84 | 7.69E-58 | 1.60E-53 | 0.68 | 0.45 | 0.23 | 1.52 |
| Kremen1 | 0.84 | 6.54E-24 | 1.36E-19 | 0.35 | 0.22 | 0.14 | 1.63 |
| Cd82 | 0.84 | 4.15E-09 | 8.64E-05 | 0.15 | 0.09 | 0.06 | 1.67 |
| Dlk1 | 0.84 | 1.15E-113 | 2.39E-109 | 0.74 | 0.45 | 0.3 | 1.67 |
| Hspa4l | 0.84 | 9.43E-17 | 1.96E-12 | 0.27 | 0.17 | 0.1 | 1.62 |
| Tbc1d9 | 0.84 | 3.60E-09 | 7.49E-05 | 0.14 | 0.09 | 0.06 | 1.69 |
| Pwwp3b | 0.83 | 1.43E-18 | 2.97E-14 | 0.35 | 0.23 | 0.11 | 1.48 |
| Parp9 | 0.83 | 7.95E-07 | 1.65E-02 | 0.08 | 0.05 | 0.04 | 1.83 |
| Trpm3 | 0.83 | 1.90E-12 | 3.96E-08 | 0.19 | 0.11 | 0.08 | 1.75 |
| Tshz3 | 0.83 | 3.13E-12 | 6.52E-08 | 0.15 | 0.08 | 0.07 | 1.88 |
| Pcdh17 | 0.83 | 5.85E-30 | 1.22E-25 | 0.41 | 0.25 | 0.16 | 1.65 |
| Pls1 | 0.83 | 1.12E-21 | 2.33E-17 | 0.33 | 0.2 | 0.13 | 1.64 |
| Nfe2l3 | 0.82 | 6.94E-13 | 1.44E-08 | 0.15 | 0.08 | 0.07 | 1.9 |
| Prrx1 | 0.82 | 1.56E-29 | 3.24E-25 | 0.33 | 0.17 | 0.16 | 1.92 |
| Pax6 | 0.82 | 7.00E-46 | 1.46E-41 | 0.55 | 0.33 | 0.22 | 1.65 |
| Cd9 | 0.82 | 2.64E-64 | 5.49E-60 | 0.75 | 0.55 | 0.2 | 1.36 |
| Baiap2 | 0.82 | 1.16E-08 | 2.42E-04 | 0.24 | 0.18 | 0.07 | 1.38 |
| Tspyl5 | 0.81 | 1.94E-06 | 4.03E-02 | 0.09 | 0.05 | 0.04 | 1.72 |
| Pdgfrl | 0.81 | 1.33E-20 | 2.76E-16 | 0.37 | 0.24 | 0.12 | 1.51 |
| Irf6 | 0.81 | 1.15E-16 | 2.39E-12 | 0.44 | 0.34 | 0.1 | 1.29 |
| Prr5 | 0.81 | 4.37E-14 | 9.10E-10 | 0.2 | 0.11 | 0.09 | 1.75 |
| G0s2 | 0.81 | 1.39E-06 | 2.90E-02 | 0.11 | 0.07 | 0.04 | 1.65 |
| Ntm | 0.8 | 1.89E-13 | 3.94E-09 | 0.26 | 0.17 | 0.09 | 1.53 |
| Fhdc1 | 0.8 | 2.16E-10 | 4.49E-06 | 0.14 | 0.08 | 0.06 | 1.78 |
| Iqgap2 | 0.8 | 7.22E-25 | 1.50E-20 | 0.39 | 0.25 | 0.14 | 1.57 |
| Palm3 | 0.8 | 2.22E-22 | 4.61E-18 | 0.38 | 0.25 | 0.13 | 1.51 |

**Supplementary Table 5. Differentially expressed genes in the mutant λ-junction ectoderm cells.**

| | | | | | | | |
|---|---|---|---|---|---|---|---|
| Cdh2 | 0.8 | 5.36E-40 | 1.11E-35 | 0.64 | 0.48 | 0.16 | 1.34 |
| Mdfic | 0.8 | 2.80E-13 | 5.82E-09 | 0.24 | 0.15 | 0.09 | 1.57 |
| Kcnh8 | 0.8 | 4.59E-12 | 9.54E-08 | 0.15 | 0.08 | 0.07 | 1.89 |
| Fam181b | 0.79 | 7.56E-08 | 1.57E-03 | 0.14 | 0.09 | 0.05 | 1.6 |
| Ust | 0.79 | 1.64E-10 | 3.42E-06 | 0.15 | 0.09 | 0.07 | 1.73 |
| Nectin3 | 0.79 | 4.27E-75 | 8.88E-71 | 0.79 | 0.58 | 0.2 | 1.35 |
| Isl2 | 0.79 | 8.21E-08 | 1.71E-03 | 0.17 | 0.12 | 0.06 | 1.47 |
| Slitrk6 | 0.79 | 8.10E-17 | 1.69E-12 | 0.24 | 0.14 | 0.1 | 1.77 |
| Tspan12 | 0.78 | 1.93E-34 | 4.01E-30 | 0.5 | 0.33 | 0.17 | 1.52 |
| Mt2 | 0.78 | 5.73E-23 | 1.19E-18 | 0.31 | 0.18 | 0.13 | 1.74 |
| Usp25 | 0.78 | 2.29E-50 | 4.75E-46 | 0.71 | 0.55 | 0.16 | 1.29 |
| Tyro3 | 0.78 | 1.80E-12 | 3.74E-08 | 0.26 | 0.17 | 0.08 | 1.49 |
| Tst | 0.78 | 3.37E-17 | 7.02E-13 | 0.29 | 0.18 | 0.11 | 1.61 |
| Pmm1 | 0.78 | 1.03E-24 | 2.15E-20 | 0.39 | 0.25 | 0.14 | 1.58 |
| Rhbdf1 | 0.78 | 4.32E-12 | 8.98E-08 | 0.23 | 0.15 | 0.08 | 1.57 |
| Rab15 | 0.78 | 1.82E-33 | 3.79E-29 | 0.46 | 0.29 | 0.17 | 1.6 |
| Gm48435 | 0.78 | 1.64E-07 | 3.41E-03 | 0.12 | 0.07 | 0.05 | 1.69 |
| Cdc14a | 0.78 | 6.01E-20 | 1.25E-15 | 0.41 | 0.29 | 0.12 | 1.43 |
| Gja1 | 0.78 | 6.24E-15 | 1.30E-10 | 0.73 | 0.66 | 0.08 | 1.12 |
| Tceal3 | 0.78 | 6.80E-18 | 1.41E-13 | 0.2 | 0.1 | 0.1 | 2.01 |
| Pik3r1 | 0.77 | 1.37E-29 | 2.85E-25 | 0.63 | 0.49 | 0.14 | 1.27 |
| Cacna1b | 0.77 | 3.46E-14 | 7.19E-10 | 0.32 | 0.23 | 0.1 | 1.43 |
| Nhs | 0.77 | 3.07E-26 | 6.38E-22 | 0.48 | 0.34 | 0.14 | 1.42 |
| Bspry | 0.77 | 5.37E-18 | 1.12E-13 | 0.29 | 0.18 | 0.11 | 1.62 |
| B230118H07Rik | 0.77 | 4.02E-37 | 8.36E-33 | 0.62 | 0.47 | 0.15 | 1.31 |
| Sox9 | 0.77 | 2.27E-64 | 4.73E-60 | 0.69 | 0.43 | 0.27 | 1.63 |
| Dsp | 0.77 | 6.62E-75 | 1.38E-70 | 0.9 | 0.75 | 0.15 | 1.2 |
| Maf | 0.76 | 2.75E-16 | 5.73E-12 | 0.17 | 0.08 | 0.09 | 2.1 |
| Slc9a3r1 | 0.76 | 1.30E-61 | 2.71E-57 | 0.79 | 0.61 | 0.18 | 1.29 |
| Dcbld1 | 0.76 | 3.78E-20 | 7.86E-16 | 0.44 | 0.32 | 0.12 | 1.39 |
| Pkia | 0.75 | 1.12E-11 | 2.33E-07 | 0.24 | 0.16 | 0.08 | 1.52 |
| Fbn1 | 0.75 | 1.44E-13 | 3.00E-09 | 0.26 | 0.17 | 0.09 | 1.54 |
| Pde5a | 0.75 | 3.02E-15 | 6.29E-11 | 0.3 | 0.2 | 0.1 | 1.5 |
| Lrrk1 | 0.74 | 1.04E-20 | 2.15E-16 | 0.35 | 0.22 | 0.13 | 1.58 |
| Shisa2 | 0.74 | 4.17E-08 | 8.68E-04 | 0.13 | 0.08 | 0.05 | 1.67 |
| Fgf13 | 0.74 | 2.81E-14 | 5.85E-10 | 0.21 | 0.12 | 0.09 | 1.74 |
| Mfsd13a | 0.74 | 1.54E-08 | 3.19E-04 | 0.1 | 0.06 | 0.05 | 1.89 |
| Nes | 0.73 | 2.12E-31 | 4.40E-27 | 0.55 | 0.41 | 0.14 | 1.34 |
| Acsf2 | 0.73 | 1.56E-07 | 3.24E-03 | 0.15 | 0.1 | 0.05 | 1.55 |
| Plch1 | 0.73 | 5.85E-20 | 1.22E-15 | 0.37 | 0.24 | 0.13 | 1.53 |
| Shisa3 | 0.73 | 2.89E-09 | 6.02E-05 | 0.05 | 0.02 | 0.04 | 3.06 |
| Sh3gl2 | 0.73 | 4.40E-07 | 9.15E-03 | 0.14 | 0.09 | 0.05 | 1.54 |
| Dmrta1 | 0.72 | 2.06E-07 | 4.29E-03 | 0.1 | 0.05 | 0.04 | 1.81 |
| Mboat1 | 0.72 | 1.51E-08 | 3.15E-04 | 0.12 | 0.07 | 0.05 | 1.76 |
| B830012L14Rik | 0.72 | 5.63E-08 | 1.17E-03 | 0.11 | 0.06 | 0.05 | 1.84 |
| Mgst1 | 0.72 | 2.15E-44 | 4.47E-40 | 0.73 | 0.57 | 0.16 | 1.28 |
| Lypd3 | 0.72 | 6.40E-08 | 1.33E-03 | 0.12 | 0.07 | 0.05 | 1.74 |
| Rpia | 0.72 | 4.19E-23 | 8.72E-19 | 0.4 | 0.27 | 0.14 | 1.52 |
| Erbb3 | 0.72 | 9.60E-39 | 2.00E-34 | 0.59 | 0.4 | 0.19 | 1.48 |

**Supplementary Table 5. Differentially expressed genes in the mutant λ-junction ectoderm cells.**

| | | | | | | | |
|---|---|---|---|---|---|---|---|
| Cmtm8 | 0.72 | 8.31E-34 | 1.73E-29 | 0.57 | 0.41 | 0.16 | 1.38 |
| Fgf10 | 0.71 | 3.24E-08 | 6.73E-04 | 0.15 | 0.09 | 0.06 | 1.6 |
| 2900026A02Rik | 0.71 | 1.63E-22 | 3.40E-18 | 0.39 | 0.25 | 0.14 | 1.55 |
| C330021F23Rik | 0.71 | 4.07E-09 | 8.47E-05 | 0.12 | 0.07 | 0.06 | 1.83 |
| Mme | 0.71 | 9.11E-09 | 1.89E-04 | 0.26 | 0.19 | 0.07 | 1.36 |
| Adgrl3 | 0.71 | 1.20E-13 | 2.49E-09 | 0.37 | 0.27 | 0.1 | 1.38 |
| Nuak1 | 0.71 | 2.94E-16 | 6.11E-12 | 0.31 | 0.2 | 0.11 | 1.54 |
| Amot | 0.71 | 5.09E-23 | 1.06E-18 | 0.6 | 0.48 | 0.13 | 1.26 |
| Tmem159 | 0.7 | 2.59E-19 | 5.38E-15 | 0.45 | 0.33 | 0.12 | 1.36 |
| Apc | 0.7 | 2.90E-48 | 6.04E-44 | 0.79 | 0.66 | 0.13 | 1.2 |
| Nrtn | 0.7 | 7.29E-14 | 1.52E-09 | 0.35 | 0.25 | 0.1 | 1.39 |
| Rad18 | 0.7 | 1.43E-24 | 2.98E-20 | 0.46 | 0.33 | 0.14 | 1.41 |
| Hacd4 | 0.7 | 4.47E-07 | 9.30E-03 | 0.15 | 0.1 | 0.05 | 1.54 |
| Pcdh11x | 0.7 | 1.53E-10 | 3.18E-06 | 0.16 | 0.09 | 0.07 | 1.75 |
| Sdc2 | 0.7 | 7.86E-18 | 1.63E-13 | 0.36 | 0.24 | 0.11 | 1.47 |
| Dmrta2 | 0.7 | 2.23E-34 | 4.63E-30 | 0.55 | 0.37 | 0.18 | 1.48 |
| Krt8 | 0.7 | 9.64E-74 | 2.01E-69 | 0.86 | 0.73 | 0.13 | 1.18 |
| Slc2a3 | 0.69 | 1.81E-29 | 3.76E-25 | 0.4 | 0.24 | 0.16 | 1.69 |
| Gm21887 | 0.69 | 6.34E-08 | 1.32E-03 | 0.16 | 0.11 | 0.06 | 1.53 |
| Dgka | 0.69 | 1.39E-09 | 2.89E-05 | 0.17 | 0.11 | 0.07 | 1.62 |
| Mef2c | 0.69 | 1.13E-07 | 2.35E-03 | 0.16 | 0.1 | 0.06 | 1.55 |
| Kcnk1 | 0.69 | 2.27E-35 | 4.71E-31 | 0.62 | 0.46 | 0.15 | 1.33 |
| Slc38a4 | 0.68 | 9.37E-12 | 1.95E-07 | 0.32 | 0.23 | 0.09 | 1.37 |
| Poglut3 | 0.68 | 6.90E-09 | 1.44E-04 | 0.2 | 0.14 | 0.07 | 1.48 |
| Khdrbs3 | 0.68 | 4.63E-70 | 9.62E-66 | 0.92 | 0.8 | 0.11 | 1.14 |
| Prkch | 0.68 | 5.30E-10 | 1.10E-05 | 0.22 | 0.15 | 0.07 | 1.5 |
| Ppp1r13l | 0.68 | 3.31E-09 | 6.88E-05 | 0.22 | 0.15 | 0.07 | 1.47 |
| Epn3 | 0.67 | 5.32E-07 | 1.11E-02 | 0.19 | 0.13 | 0.06 | 1.41 |
| Cyp39a1 | 0.67 | 2.18E-10 | 4.54E-06 | 0.26 | 0.18 | 0.08 | 1.42 |
| Slc7a5 | 0.67 | 8.25E-08 | 1.71E-03 | 0.09 | 0.05 | 0.04 | 1.88 |
| Il17re | 0.67 | 3.04E-10 | 6.32E-06 | 0.21 | 0.13 | 0.07 | 1.55 |
| Plekhh1 | 0.67 | 7.95E-09 | 1.65E-04 | 0.17 | 0.11 | 0.06 | 1.57 |
| Wfdc2 | 0.67 | 1.05E-60 | 2.18E-56 | 0.86 | 0.72 | 0.14 | 1.2 |
| Tom1 | 0.67 | 3.64E-08 | 7.58E-04 | 0.16 | 0.1 | 0.06 | 1.57 |
| Lamc2 | 0.67 | 5.77E-09 | 1.20E-04 | 0.19 | 0.12 | 0.06 | 1.52 |
| Lrrc8b | 0.66 | 2.55E-10 | 5.31E-06 | 0.23 | 0.16 | 0.08 | 1.49 |
| Casp7 | 0.66 | 3.92E-09 | 8.15E-05 | 0.19 | 0.12 | 0.07 | 1.54 |
| Syt7 | 0.66 | 5.45E-09 | 1.13E-04 | 0.16 | 0.09 | 0.06 | 1.65 |
| Igf2r | 0.66 | 2.75E-32 | 5.72E-28 | 0.62 | 0.47 | 0.15 | 1.32 |
| Slc35f1 | 0.66 | 9.72E-07 | 2.02E-02 | 0.12 | 0.07 | 0.05 | 1.62 |
| Me1 | 0.66 | 6.70E-09 | 1.39E-04 | 0.23 | 0.16 | 0.07 | 1.42 |
| Jup | 0.66 | 3.68E-27 | 7.65E-23 | 0.54 | 0.4 | 0.14 | 1.35 |
| Lmo7 | 0.65 | 2.02E-22 | 4.19E-18 | 0.48 | 0.34 | 0.14 | 1.43 |
| Erbb2 | 0.65 | 1.74E-13 | 3.61E-09 | 0.36 | 0.27 | 0.1 | 1.36 |
| Rprm | 0.65 | 9.67E-20 | 2.01E-15 | 0.42 | 0.29 | 0.14 | 1.47 |
| Xylt1 | 0.65 | 1.75E-06 | 3.64E-02 | 0.12 | 0.07 | 0.05 | 1.61 |
| Prrg3 | 0.65 | 2.23E-10 | 4.64E-06 | 0.2 | 0.12 | 0.07 | 1.59 |
| Nfix | 0.64 | 5.41E-10 | 1.13E-05 | 0.13 | 0.07 | 0.06 | 1.81 |
| Capg | 0.64 | 2.38E-15 | 4.96E-11 | 0.39 | 0.28 | 0.11 | 1.38 |

**Supplementary Table 5. Differentially expressed genes in the mutant λ-junction ectoderm cells.**

| | | | | | | | |
|---|---|---|---|---|---|---|---|
| Tspan33 | 0.64 | 2.42E-07 | 5.03E-03 | 0.15 | 0.1 | 0.05 | 1.53 |
| B3gnt7 | 0.64 | 1.56E-07 | 3.24E-03 | 0.13 | 0.08 | 0.05 | 1.65 |
| Epb41l2 | 0.64 | 2.76E-22 | 5.74E-18 | 0.56 | 0.43 | 0.13 | 1.31 |
| Elf5 | 0.63 | 9.25E-20 | 1.92E-15 | 0.28 | 0.16 | 0.12 | 1.75 |
| Mtfp1 | 0.63 | 6.84E-07 | 1.42E-02 | 0.1 | 0.06 | 0.04 | 1.75 |
| Gm16136 | 0.63 | 4.53E-24 | 9.43E-20 | 0.68 | 0.59 | 0.1 | 1.17 |
| C1qtnf12 | 0.63 | 1.08E-06 | 2.24E-02 | 0.25 | 0.19 | 0.06 | 1.31 |
| Clcc1 | 0.62 | 4.35E-21 | 9.05E-17 | 0.51 | 0.38 | 0.13 | 1.34 |
| Map7d2 | 0.62 | 6.81E-13 | 1.42E-08 | 0.23 | 0.15 | 0.09 | 1.61 |
| Arhgap27os2 | 0.62 | 4.17E-12 | 8.68E-08 | 0.25 | 0.16 | 0.09 | 1.55 |
| Dtx3l | 0.62 | 8.24E-09 | 1.71E-04 | 0.12 | 0.07 | 0.05 | 1.78 |
| Rilpl2 | 0.62 | 2.22E-11 | 4.61E-07 | 0.28 | 0.2 | 0.09 | 1.44 |
| Ccdc85c | 0.62 | 8.13E-13 | 1.69E-08 | 0.23 | 0.15 | 0.09 | 1.58 |
| Ltbr | 0.62 | 3.38E-12 | 7.03E-08 | 0.25 | 0.16 | 0.09 | 1.53 |
| Pnck | 0.62 | 9.11E-09 | 1.89E-04 | 0.19 | 0.13 | 0.06 | 1.51 |
| Prr36 | 0.61 | 4.17E-11 | 8.67E-07 | 0.28 | 0.19 | 0.09 | 1.45 |
| Sema4d | 0.61 | 1.70E-06 | 3.54E-02 | 0.14 | 0.09 | 0.05 | 1.52 |
| Pygl | 0.61 | 2.81E-08 | 5.84E-04 | 0.2 | 0.14 | 0.07 | 1.48 |
| Tmem121 | 0.61 | 8.71E-13 | 1.81E-08 | 0.22 | 0.14 | 0.08 | 1.59 |
| B3galnt1 | 0.61 | 6.72E-09 | 1.40E-04 | 0.19 | 0.12 | 0.07 | 1.52 |
| Gnptg | 0.61 | 1.96E-12 | 4.07E-08 | 0.32 | 0.23 | 0.09 | 1.4 |
| Sort1 | 0.61 | 6.46E-12 | 1.34E-07 | 0.35 | 0.26 | 0.09 | 1.36 |
| Camsap3 | 0.61 | 1.64E-15 | 3.42E-11 | 0.3 | 0.2 | 0.11 | 1.54 |
| Igsf9b | 0.6 | 9.69E-08 | 2.02E-03 | 0.18 | 0.12 | 0.06 | 1.49 |
| Arl4c | 0.6 | 9.40E-23 | 1.96E-18 | 0.61 | 0.48 | 0.13 | 1.28 |
| Pawr | 0.6 | 3.44E-19 | 7.16E-15 | 0.41 | 0.28 | 0.13 | 1.45 |
| Shisa4 | 0.59 | 5.90E-13 | 1.23E-08 | 0.28 | 0.19 | 0.09 | 1.5 |
| Sox13 | 0.59 | 6.49E-09 | 1.35E-04 | 0.23 | 0.16 | 0.07 | 1.46 |
| Syngr1 | 0.59 | 4.09E-11 | 8.50E-07 | 0.24 | 0.16 | 0.08 | 1.52 |
| Rhou | 0.59 | 1.12E-22 | 2.33E-18 | 0.53 | 0.4 | 0.13 | 1.33 |
| Kctd6 | 0.59 | 2.53E-12 | 5.26E-08 | 0.27 | 0.18 | 0.09 | 1.5 |
| Acaa2 | 0.58 | 3.14E-42 | 6.52E-38 | 0.77 | 0.6 | 0.17 | 1.28 |
| Olfm1 | 0.58 | 3.10E-11 | 6.46E-07 | 0.35 | 0.26 | 0.09 | 1.35 |
| Vill | 0.58 | 9.38E-07 | 1.95E-02 | 0.18 | 0.13 | 0.05 | 1.43 |
| Ocln | 0.58 | 4.35E-13 | 9.04E-09 | 0.32 | 0.23 | 0.1 | 1.43 |
| Grtp1 | 0.57 | 1.26E-09 | 2.63E-05 | 0.26 | 0.18 | 0.08 | 1.42 |
| Sfrp2 | 0.57 | 2.86E-17 | 5.94E-13 | 0.56 | 0.45 | 0.11 | 1.25 |
| Rassf9 | 0.57 | 1.11E-10 | 2.30E-06 | 0.28 | 0.19 | 0.08 | 1.43 |
| Gstt3 | 0.57 | 3.80E-08 | 7.90E-04 | 0.14 | 0.09 | 0.05 | 1.63 |
| Pck2 | 0.56 | 1.60E-07 | 3.33E-03 | 0.2 | 0.14 | 0.06 | 1.42 |
| Pkp4 | 0.56 | 3.57E-29 | 7.42E-25 | 0.63 | 0.49 | 0.14 | 1.29 |
| Cttnbp2 | 0.56 | 1.56E-06 | 3.24E-02 | 0.14 | 0.09 | 0.05 | 1.52 |
| Mtss1 | 0.55 | 7.46E-15 | 1.55E-10 | 0.47 | 0.36 | 0.11 | 1.3 |
| Mgat5 | 0.55 | 4.07E-12 | 8.47E-08 | 0.33 | 0.24 | 0.1 | 1.4 |
| Nectin1 | 0.55 | 4.07E-15 | 8.47E-11 | 0.4 | 0.29 | 0.11 | 1.39 |
| Echdc1 | 0.55 | 1.16E-14 | 2.41E-10 | 0.4 | 0.29 | 0.11 | 1.36 |
| Necab2 | 0.54 | 1.74E-16 | 3.63E-12 | 0.32 | 0.21 | 0.11 | 1.54 |
| Cux2 | 0.54 | 1.21E-06 | 2.51E-02 | 0.18 | 0.12 | 0.06 | 1.45 |
| Id3 | 0.54 | 3.16E-46 | 6.58E-42 | 0.98 | 0.96 | 0.02 | 1.03 |

**Supplementary Table 5. Differentially expressed genes in the mutant λ-junction ectoderm cells.**

| | | | | | | | |
|---|---|---|---|---|---|---|---|
| 2310030G06Rik | 0.54 | 8.59E-11 | 1.79E-06 | 0.3 | 0.21 | 0.09 | 1.41 |
| Nuak2 | 0.54 | 1.02E-07 | 2.11E-03 | 0.15 | 0.1 | 0.06 | 1.58 |
| Gnai2 | 0.54 | 9.30E-95 | 1.93E-90 | 0.98 | 0.94 | 0.04 | 1.04 |
| Rian | 0.54 | 5.54E-26 | 1.15E-21 | 0.51 | 0.35 | 0.16 | 1.44 |
| Sptbn2 | 0.53 | 2.93E-11 | 6.09E-07 | 0.33 | 0.24 | 0.09 | 1.36 |
| Frk | 0.53 | 1.13E-06 | 2.35E-02 | 0.17 | 0.12 | 0.05 | 1.45 |
| Rasef | 0.53 | 2.11E-08 | 4.39E-04 | 0.17 | 0.11 | 0.06 | 1.57 |
| Wwc1 | 0.53 | 8.29E-24 | 1.72E-19 | 0.61 | 0.48 | 0.13 | 1.28 |
| Mapk8ip1 | 0.53 | 5.64E-16 | 1.17E-11 | 0.39 | 0.28 | 0.11 | 1.39 |
| Phyhipl | 0.53 | 1.14E-06 | 2.38E-02 | 0.11 | 0.07 | 0.04 | 1.67 |
| Sorl1 | 0.53 | 7.03E-14 | 1.46E-09 | 0.37 | 0.27 | 0.1 | 1.37 |
| Irs2 | 0.53 | 2.15E-15 | 4.47E-11 | 0.49 | 0.38 | 0.11 | 1.29 |
| Fzd7 | 0.52 | 1.33E-11 | 2.76E-07 | 0.38 | 0.3 | 0.09 | 1.29 |
| St14 | 0.52 | 2.61E-18 | 5.42E-14 | 0.5 | 0.38 | 0.12 | 1.33 |
| Samd5 | 0.51 | 2.68E-23 | 5.57E-19 | 0.72 | 0.62 | 0.1 | 1.16 |
| Sall2 | 0.51 | 1.18E-15 | 2.46E-11 | 0.52 | 0.42 | 0.1 | 1.24 |
| Amfr | 0.51 | 9.80E-16 | 2.04E-11 | 0.45 | 0.33 | 0.11 | 1.34 |
| Celsr1 | 0.51 | 1.49E-11 | 3.11E-07 | 0.38 | 0.29 | 0.09 | 1.31 |
| Fam126a | 0.51 | 7.16E-08 | 1.49E-03 | 0.31 | 0.24 | 0.07 | 1.28 |
| Ces1d | 0.51 | 8.05E-09 | 1.67E-04 | 0.19 | 0.13 | 0.07 | 1.53 |
| Carmil1 | 0.51 | 6.46E-08 | 1.34E-03 | 0.25 | 0.18 | 0.07 | 1.39 |
| Hspb8 | 0.5 | 1.31E-07 | 2.73E-03 | 0.5 | 0.45 | 0.05 | 1.12 |
| Aplp2 | 0.5 | 2.78E-53 | 5.79E-49 | 0.92 | 0.79 | 0.12 | 1.15 |
| Sfn | 0.5 | 6.47E-16 | 1.35E-11 | 0.73 | 0.66 | 0.07 | 1.1 |
| Snn | 0.5 | 7.64E-16 | 1.59E-11 | 0.63 | 0.54 | 0.09 | 1.16 |
| Marveld2 | 0.5 | 1.98E-06 | 4.12E-02 | 0.24 | 0.18 | 0.06 | 1.33 |
| Otx2 | 0.5 | 3.45E-32 | 7.17E-28 | 0.44 | 0.25 | 0.19 | 1.74 |
| Nckap5 | 0.5 | 1.29E-13 | 2.68E-09 | 0.34 | 0.23 | 0.11 | 1.48 |
| Celf4 | 0.49 | 2.35E-13 | 4.89E-09 | 0.33 | 0.23 | 0.1 | 1.44 |
| Unc5c | 0.49 | 1.41E-17 | 2.93E-13 | 0.38 | 0.26 | 0.12 | 1.45 |
| Airn | 0.49 | 1.81E-10 | 3.76E-06 | 0.32 | 0.24 | 0.08 | 1.34 |
| Cdh1 | 0.49 | 5.62E-28 | 1.17E-23 | 0.68 | 0.54 | 0.14 | 1.26 |
| mt-Nd6 | 0.49 | 3.90E-13 | 8.12E-09 | 0.51 | 0.42 | 0.09 | 1.22 |
| Snhg14 | 0.49 | 6.68E-08 | 1.39E-03 | 0.27 | 0.2 | 0.07 | 1.35 |
| Plod2 | 0.49 | 2.90E-18 | 6.03E-14 | 0.53 | 0.41 | 0.12 | 1.31 |
| Nsun4 | 0.49 | 4.24E-09 | 8.82E-05 | 0.23 | 0.16 | 0.07 | 1.45 |
| H2-K1 | 0.49 | 4.80E-11 | 9.99E-07 | 0.27 | 0.19 | 0.09 | 1.46 |
| Ccdc88c | 0.49 | 2.07E-20 | 4.31E-16 | 0.58 | 0.45 | 0.13 | 1.3 |
| Itgb6 | 0.49 | 1.33E-06 | 2.77E-02 | 0.18 | 0.13 | 0.05 | 1.43 |
| Nab1 | 0.48 | 9.63E-09 | 2.00E-04 | 0.26 | 0.19 | 0.07 | 1.37 |
| Zbtb18 | 0.48 | 1.52E-06 | 3.17E-02 | 0.23 | 0.17 | 0.06 | 1.33 |
| Nr2f2 | 0.48 | 1.46E-16 | 3.03E-12 | 0.63 | 0.52 | 0.11 | 1.22 |
| Mpzl2 | 0.48 | 8.38E-26 | 1.74E-21 | 0.75 | 0.63 | 0.11 | 1.18 |
| Plpp2 | 0.48 | 7.13E-11 | 1.48E-06 | 0.41 | 0.32 | 0.09 | 1.28 |
| Clic6 | 0.48 | 2.08E-07 | 4.32E-03 | 0.23 | 0.16 | 0.06 | 1.38 |
| Dhcr7 | 0.48 | 1.69E-10 | 3.52E-06 | 0.37 | 0.28 | 0.09 | 1.31 |
| Cad | 0.48 | 1.19E-08 | 2.47E-04 | 0.39 | 0.31 | 0.07 | 1.24 |
| Rnf165 | 0.47 | 5.22E-10 | 1.09E-05 | 0.29 | 0.21 | 0.08 | 1.38 |
| Esrp2 | 0.47 | 4.78E-10 | 9.94E-06 | 0.28 | 0.2 | 0.08 | 1.41 |

**Supplementary Table 5. Differentially expressed genes in the mutant λ-junction ectoderm cells.**

| | | | | | | | |
|---|---|---|---|---|---|---|---|
| Nck2 | 0.47 | 2.28E-07 | 4.73E-03 | 0.3 | 0.23 | 0.07 | 1.31 |
| Tekt1 | 0.47 | 8.97E-07 | 1.87E-02 | 0.13 | 0.08 | 0.05 | 1.59 |
| Tmem238 | 0.47 | 3.15E-13 | 6.54E-09 | 0.43 | 0.33 | 0.1 | 1.32 |
| H2-D1 | 0.47 | 1.38E-12 | 2.86E-08 | 0.36 | 0.26 | 0.1 | 1.39 |
| Bcl2l11 | 0.47 | 9.42E-07 | 1.96E-02 | 0.42 | 0.36 | 0.06 | 1.17 |
| Eya1 | 0.46 | 3.75E-31 | 7.80E-27 | 0.65 | 0.47 | 0.17 | 1.36 |
| Syngr2 | 0.46 | 3.24E-08 | 6.75E-04 | 0.29 | 0.22 | 0.07 | 1.32 |
| Plxnb2 | 0.46 | 1.08E-18 | 2.24E-14 | 0.64 | 0.52 | 0.12 | 1.23 |
| Med13l | 0.46 | 2.22E-15 | 4.62E-11 | 0.56 | 0.46 | 0.1 | 1.22 |
| Plekhg1 | 0.46 | 2.82E-07 | 5.88E-03 | 0.31 | 0.25 | 0.07 | 1.26 |
| Epb41l4a | 0.46 | 7.06E-07 | 1.47E-02 | 0.22 | 0.16 | 0.06 | 1.39 |
| Gipc2 | 0.46 | 1.01E-10 | 2.11E-06 | 0.19 | 0.11 | 0.07 | 1.65 |
| Ctsc | 0.46 | 3.21E-08 | 6.67E-04 | 0.22 | 0.16 | 0.07 | 1.42 |
| Rhoc | 0.46 | 6.03E-30 | 1.26E-25 | 0.73 | 0.58 | 0.15 | 1.25 |
| Amt | 0.46 | 5.68E-07 | 1.18E-02 | 0.26 | 0.19 | 0.06 | 1.34 |
| Por | 0.46 | 2.50E-12 | 5.20E-08 | 0.46 | 0.37 | 0.1 | 1.26 |
| Acot7 | 0.46 | 3.80E-14 | 7.90E-10 | 0.44 | 0.34 | 0.1 | 1.31 |
| Rab25 | 0.46 | 1.42E-26 | 2.95E-22 | 0.66 | 0.52 | 0.14 | 1.27 |
| Cndp2 | 0.45 | 9.33E-15 | 1.94E-10 | 0.53 | 0.42 | 0.11 | 1.25 |
| Ripply3 | 0.45 | 2.13E-08 | 4.43E-04 | 0.2 | 0.13 | 0.07 | 1.49 |
| Cldn6 | 0.45 | 2.32E-41 | 4.83E-37 | 0.89 | 0.74 | 0.14 | 1.19 |
| Tacstd2 | 0.45 | 6.33E-19 | 1.32E-14 | 0.31 | 0.19 | 0.12 | 1.64 |
| Gm10282 | 0.45 | 1.19E-06 | 2.48E-02 | 0.19 | 0.14 | 0.06 | 1.42 |
| Cyb5r3 | 0.44 | 2.30E-45 | 4.78E-41 | 0.94 | 0.86 | 0.07 | 1.08 |
| Lrrfip1 | 0.44 | 5.47E-16 | 1.14E-11 | 0.54 | 0.43 | 0.11 | 1.27 |
| Tgfb2 | 0.44 | 1.39E-25 | 2.90E-21 | 0.63 | 0.47 | 0.16 | 1.34 |
| Arhgap32 | 0.44 | 6.90E-07 | 1.44E-02 | 0.27 | 0.21 | 0.06 | 1.29 |
| Dsg2 | 0.44 | 6.33E-20 | 1.32E-15 | 0.51 | 0.37 | 0.14 | 1.38 |
| Fkbp5 | 0.44 | 2.11E-07 | 4.39E-03 | 0.37 | 0.29 | 0.07 | 1.24 |
| Atp11b | 0.44 | 2.36E-14 | 4.90E-10 | 0.55 | 0.45 | 0.1 | 1.22 |
| Tiam1 | 0.44 | 1.64E-08 | 3.41E-04 | 0.48 | 0.41 | 0.07 | 1.17 |
| Tjp2 | 0.44 | 9.59E-13 | 1.99E-08 | 0.51 | 0.42 | 0.09 | 1.22 |
| Fndc3b | 0.44 | 5.44E-11 | 1.13E-06 | 0.35 | 0.26 | 0.09 | 1.35 |
| Cbarp | 0.44 | 6.30E-08 | 1.31E-03 | 0.23 | 0.16 | 0.07 | 1.43 |
| Wdfy2 | 0.44 | 3.89E-07 | 8.09E-03 | 0.22 | 0.16 | 0.06 | 1.39 |
| Cep89 | 0.44 | 1.41E-09 | 2.93E-05 | 0.38 | 0.3 | 0.08 | 1.27 |
| Eml5 | 0.44 | 7.74E-07 | 1.61E-02 | 0.25 | 0.18 | 0.07 | 1.36 |
| Cyb5a | 0.43 | 1.89E-55 | 3.93E-51 | 0.97 | 0.94 | 0.03 | 1.03 |
| Irf2 | 0.43 | 4.42E-16 | 9.19E-12 | 0.46 | 0.33 | 0.12 | 1.37 |
| Lrrn3 | 0.43 | 6.48E-07 | 1.35E-02 | 0.16 | 0.1 | 0.05 | 1.5 |
| Acap2 | 0.43 | 2.79E-12 | 5.81E-08 | 0.49 | 0.38 | 0.1 | 1.27 |
| Bmp4 | 0.43 | 8.46E-17 | 1.76E-12 | 0.59 | 0.47 | 0.12 | 1.26 |
| Thbs3 | 0.43 | 1.61E-06 | 3.34E-02 | 0.16 | 0.11 | 0.05 | 1.47 |
| Vcan | 0.43 | 3.28E-12 | 6.82E-08 | 0.76 | 0.7 | 0.06 | 1.08 |
| Ablim1 | 0.43 | 7.32E-10 | 1.52E-05 | 0.56 | 0.49 | 0.07 | 1.14 |
| Cystm1 | 0.43 | 2.23E-11 | 4.64E-07 | 0.36 | 0.26 | 0.1 | 1.37 |
| Tacc2 | 0.43 | 1.80E-18 | 3.74E-14 | 0.58 | 0.46 | 0.11 | 1.24 |
| Ccdc120 | 0.43 | 1.75E-06 | 3.64E-02 | 0.2 | 0.14 | 0.05 | 1.38 |
| Cers4 | 0.42 | 4.82E-11 | 1.00E-06 | 0.4 | 0.3 | 0.1 | 1.32 |

**Supplementary Table 5. Differentially expressed genes in the mutant λ-junction ectoderm cells.**

| | | | | | | | |
|---|---|---|---|---|---|---|---|
| Pacsin3 | 0.42 | 2.69E-09 | 5.59E-05 | 0.35 | 0.27 | 0.08 | 1.31 |
| Gjb2 | 0.42 | 5.17E-11 | 1.08E-06 | 0.2 | 0.12 | 0.08 | 1.66 |
| Ier5l | 0.42 | 2.37E-17 | 4.93E-13 | 0.7 | 0.59 | 0.11 | 1.18 |
| Unc13b | 0.42 | 3.25E-07 | 6.76E-03 | 0.26 | 0.19 | 0.06 | 1.33 |
| Cyb561 | 0.42 | 2.72E-07 | 5.66E-03 | 0.21 | 0.15 | 0.06 | 1.42 |
| App | 0.42 | 2.99E-29 | 6.22E-25 | 0.85 | 0.74 | 0.11 | 1.15 |
| Hdac4 | 0.42 | 2.32E-06 | 4.82E-02 | 0.29 | 0.23 | 0.06 | 1.27 |
| Ppp1r9a | 0.42 | 1.18E-12 | 2.45E-08 | 0.54 | 0.44 | 0.1 | 1.22 |
| Krt18 | 0.42 | 1.76E-38 | 3.67E-34 | 0.9 | 0.81 | 0.09 | 1.11 |
| Rbpms | 0.42 | 7.50E-13 | 1.56E-08 | 0.55 | 0.45 | 0.09 | 1.21 |
| Bex2 | 0.41 | 1.03E-33 | 2.15E-29 | 0.9 | 0.82 | 0.08 | 1.1 |
| Mapk13 | 0.41 | 1.09E-20 | 2.26E-16 | 0.64 | 0.51 | 0.13 | 1.24 |
| Smim1 | 0.41 | 1.13E-09 | 2.35E-05 | 0.28 | 0.2 | 0.08 | 1.41 |
| Ppp1r15b | 0.41 | 1.88E-06 | 3.91E-02 | 0.33 | 0.27 | 0.06 | 1.24 |
| Sapcd2 | 0.41 | 2.03E-09 | 4.23E-05 | 0.32 | 0.25 | 0.08 | 1.32 |
| Fndc3c1 | 0.41 | 2.50E-09 | 5.20E-05 | 0.37 | 0.29 | 0.08 | 1.28 |
| Efna1 | 0.41 | 1.00E-08 | 2.08E-04 | 0.42 | 0.34 | 0.08 | 1.24 |
| Nudt4 | 0.41 | 4.17E-24 | 8.68E-20 | 0.81 | 0.73 | 0.08 | 1.11 |
| Klhl23 | 0.41 | 4.22E-11 | 8.77E-07 | 0.55 | 0.47 | 0.09 | 1.18 |
| Sox21 | 0.41 | 1.37E-07 | 2.86E-03 | 0.14 | 0.09 | 0.05 | 1.6 |
| Lrp2 | 0.4 | 1.04E-07 | 2.17E-03 | 0.23 | 0.17 | 0.07 | 1.4 |
| Mocs2 | 0.4 | 1.38E-16 | 2.88E-12 | 0.58 | 0.47 | 0.11 | 1.23 |
| Rab27b | 0.4 | 1.55E-06 | 3.21E-02 | 0.34 | 0.27 | 0.07 | 1.25 |
| Tspan13 | 0.4 | 1.40E-24 | 2.91E-20 | 0.76 | 0.66 | 0.11 | 1.16 |
| Ephb2 | 0.4 | 5.08E-11 | 1.06E-06 | 0.45 | 0.36 | 0.1 | 1.27 |
| Hebp2 | 0.39 | 7.70E-08 | 1.60E-03 | 0.35 | 0.27 | 0.08 | 1.28 |
| Mpp7 | 0.39 | 4.68E-13 | 9.74E-09 | 0.57 | 0.48 | 0.09 | 1.2 |
| Unc5b | 0.39 | 1.84E-13 | 3.82E-09 | 0.56 | 0.46 | 0.11 | 1.23 |
| Slc12a2 | 0.39 | 1.73E-26 | 3.60E-22 | 0.73 | 0.6 | 0.14 | 1.23 |
| Gm26917 | 0.39 | 1.76E-14 | 3.65E-10 | 0.66 | 0.56 | 0.1 | 1.18 |
| Map4k5 | 0.39 | 1.25E-08 | 2.61E-04 | 0.41 | 0.34 | 0.08 | 1.23 |
| Mest | 0.39 | 4.64E-40 | 9.65E-36 | 0.93 | 0.86 | 0.07 | 1.08 |
| Plpp3 | 0.39 | 7.78E-13 | 1.62E-08 | 0.51 | 0.39 | 0.12 | 1.3 |
| Sel1l | 0.39 | 4.51E-08 | 9.38E-04 | 0.43 | 0.36 | 0.08 | 1.21 |
| Map7 | 0.38 | 2.73E-11 | 5.68E-07 | 0.44 | 0.34 | 0.09 | 1.27 |
| Spint1 | 0.38 | 4.92E-10 | 1.02E-05 | 0.42 | 0.32 | 0.09 | 1.28 |
| Tsix | 0.38 | 1.93E-11 | 4.02E-07 | 0.42 | 0.31 | 0.11 | 1.34 |
| Sdc4 | 0.38 | 5.98E-14 | 1.24E-09 | 0.66 | 0.57 | 0.1 | 1.17 |
| Macc1 | 0.38 | 1.22E-06 | 2.54E-02 | 0.24 | 0.18 | 0.06 | 1.35 |
| Hist1h2ap | 0.38 | 1.90E-07 | 3.96E-03 | 0.74 | 0.69 | 0.04 | 1.06 |
| Lypd6 | 0.38 | 6.94E-08 | 1.44E-03 | 0.24 | 0.17 | 0.07 | 1.42 |
| Wrb | 0.38 | 2.34E-06 | 4.88E-02 | 0.3 | 0.24 | 0.06 | 1.26 |
| Xist | 0.38 | 4.57E-46 | 9.50E-42 | 0.95 | 0.68 | 0.27 | 1.4 |
| Mtss2 | 0.38 | 1.93E-07 | 4.01E-03 | 0.42 | 0.35 | 0.07 | 1.21 |
| Phactr2 | 0.38 | 3.46E-08 | 7.20E-04 | 0.42 | 0.35 | 0.07 | 1.21 |
| Tcim | 0.38 | 2.53E-08 | 5.27E-04 | 0.32 | 0.24 | 0.08 | 1.31 |
| Arhgap18 | 0.37 | 8.80E-07 | 1.83E-02 | 0.22 | 0.16 | 0.06 | 1.38 |
| Cenpf | 0.37 | 2.53E-12 | 5.26E-08 | 0.72 | 0.63 | 0.09 | 1.14 |
| Etv5 | 0.37 | 1.01E-08 | 2.09E-04 | 0.29 | 0.21 | 0.08 | 1.37 |

**Supplementary Table 5. Differentially expressed genes in the mutant λ-junction ectoderm cells.**

| | | | | | | | |
|---|---|---|---|---|---|---|---|
| Atl2 | 0.37 | 3.05E-08 | 6.34E-04 | 0.33 | 0.25 | 0.08 | 1.31 |
| Knl1 | 0.37 | 1.45E-11 | 3.02E-07 | 0.52 | 0.43 | 0.1 | 1.23 |
| Epb41l3 | 0.37 | 7.77E-10 | 1.62E-05 | 0.53 | 0.45 | 0.08 | 1.18 |
| Tmem106c | 0.37 | 6.76E-07 | 1.41E-02 | 0.31 | 0.24 | 0.07 | 1.28 |
| Msmo1 | 0.37 | 1.73E-10 | 3.59E-06 | 0.52 | 0.44 | 0.09 | 1.2 |
| Bag2 | 0.37 | 5.32E-12 | 1.11E-07 | 0.39 | 0.3 | 0.1 | 1.34 |
| Ide | 0.37 | 1.95E-12 | 4.05E-08 | 0.49 | 0.39 | 0.1 | 1.26 |
| Crip1 | 0.37 | 8.81E-13 | 1.83E-08 | 0.44 | 0.34 | 0.11 | 1.31 |
| Shmt1 | 0.37 | 1.04E-06 | 2.15E-02 | 0.39 | 0.32 | 0.07 | 1.22 |
| Spata24 | 0.37 | 5.26E-08 | 1.09E-03 | 0.33 | 0.25 | 0.07 | 1.29 |
| Reep3 | 0.37 | 8.84E-13 | 1.84E-08 | 0.66 | 0.58 | 0.09 | 1.15 |
| Gas2l3 | 0.36 | 2.56E-07 | 5.32E-03 | 0.27 | 0.2 | 0.07 | 1.34 |
| E2f5 | 0.36 | 9.01E-11 | 1.87E-06 | 0.48 | 0.39 | 0.09 | 1.23 |
| Anxa5 | 0.36 | 1.13E-14 | 2.36E-10 | 0.67 | 0.57 | 0.09 | 1.16 |
| Ppm1l | 0.36 | 3.16E-07 | 6.58E-03 | 0.44 | 0.37 | 0.07 | 1.18 |
| Cldn7 | 0.36 | 3.30E-31 | 6.86E-27 | 0.86 | 0.72 | 0.14 | 1.2 |
| Cit | 0.36 | 6.10E-09 | 1.27E-04 | 0.44 | 0.36 | 0.08 | 1.22 |
| Myo5b | 0.36 | 9.97E-07 | 2.07E-02 | 0.16 | 0.11 | 0.05 | 1.48 |
| Cited2 | 0.35 | 2.44E-17 | 5.07E-13 | 0.71 | 0.61 | 0.11 | 1.18 |
| Clu | 0.35 | 1.02E-09 | 2.12E-05 | 0.38 | 0.3 | 0.09 | 1.29 |
| Sema3f | 0.35 | 5.25E-07 | 1.09E-02 | 0.41 | 0.34 | 0.07 | 1.2 |
| Hmgn2 | 0.35 | 5.70E-58 | 1.19E-53 | 0.98 | 0.97 | 0.01 | 1.01 |
| Grhpr | 0.35 | 2.29E-09 | 4.76E-05 | 0.53 | 0.44 | 0.09 | 1.2 |
| Sfrp1 | 0.35 | 2.70E-18 | 5.62E-14 | 0.58 | 0.46 | 0.12 | 1.27 |
| Cenpb | 0.35 | 8.35E-17 | 1.74E-12 | 0.73 | 0.64 | 0.09 | 1.15 |
| Fgfr1 | 0.35 | 2.57E-08 | 5.35E-04 | 0.46 | 0.39 | 0.07 | 1.18 |
| Sh3glb2 | 0.35 | 1.19E-07 | 2.47E-03 | 0.42 | 0.35 | 0.07 | 1.21 |
| Rfk | 0.35 | 3.50E-10 | 7.27E-06 | 0.55 | 0.47 | 0.09 | 1.19 |
| Sod1 | 0.35 | 8.58E-19 | 1.79E-14 | 0.8 | 0.71 | 0.09 | 1.13 |
| Ccdc115 | 0.35 | 4.51E-14 | 9.38E-10 | 0.63 | 0.52 | 0.11 | 1.21 |
| Nup188 | 0.35 | 6.59E-08 | 1.37E-03 | 0.44 | 0.36 | 0.07 | 1.2 |
| Clstn1 | 0.35 | 2.93E-11 | 6.09E-07 | 0.57 | 0.48 | 0.09 | 1.18 |
| Ptk7 | 0.35 | 1.59E-08 | 3.31E-04 | 0.5 | 0.43 | 0.07 | 1.16 |
| Msi1 | 0.34 | 3.44E-10 | 7.16E-06 | 0.5 | 0.42 | 0.08 | 1.2 |
| Pla2g12a | 0.34 | 3.23E-14 | 6.72E-10 | 0.65 | 0.54 | 0.1 | 1.19 |
| Strn | 0.34 | 1.46E-06 | 3.03E-02 | 0.25 | 0.18 | 0.06 | 1.33 |
| Timeless | 0.34 | 1.00E-09 | 2.09E-05 | 0.51 | 0.42 | 0.09 | 1.21 |
| Nrm | 0.34 | 4.65E-09 | 9.68E-05 | 0.41 | 0.33 | 0.08 | 1.25 |
| Igfbp3 | 0.34 | 4.39E-26 | 9.13E-22 | 0.43 | 0.27 | 0.15 | 1.56 |
| Cdt1 | 0.34 | 2.33E-07 | 4.84E-03 | 0.44 | 0.37 | 0.07 | 1.19 |
| Pcbd1 | 0.34 | 4.45E-10 | 9.26E-06 | 0.27 | 0.19 | 0.08 | 1.44 |
| Lin28b | 0.34 | 8.33E-07 | 1.73E-02 | 0.3 | 0.23 | 0.07 | 1.28 |
| Naf1 | 0.34 | 1.48E-11 | 3.08E-07 | 0.5 | 0.4 | 0.1 | 1.24 |
| Rcn1 | 0.34 | 4.24E-24 | 8.81E-20 | 0.86 | 0.78 | 0.09 | 1.11 |
| Ctnnd2 | 0.34 | 9.61E-10 | 2.00E-05 | 0.47 | 0.37 | 0.1 | 1.26 |
| Tkt | 0.34 | 8.55E-28 | 1.78E-23 | 0.9 | 0.83 | 0.07 | 1.08 |
| Wwtr1 | 0.34 | 5.21E-09 | 1.08E-04 | 0.36 | 0.28 | 0.08 | 1.29 |
| Cmtm7 | 0.34 | 5.56E-08 | 1.16E-03 | 0.42 | 0.34 | 0.07 | 1.22 |
| Eml1 | 0.34 | 1.24E-06 | 2.58E-02 | 0.2 | 0.14 | 0.06 | 1.39 |

**Supplementary Table 5. Differentially expressed genes in the mutant λ-junction ectoderm cells.**

| | | | | | | | |
|---|---|---|---|---|---|---|---|
| Cdc42ep5 | 0.34 | 2.57E-15 | 5.35E-11 | 0.64 | 0.53 | 0.11 | 1.21 |
| Atox1 | 0.34 | 1.18E-17 | 2.45E-13 | 0.98 | 0.97 | 0.01 | 1.01 |
| Mtln | 0.33 | 2.21E-11 | 4.60E-07 | 0.57 | 0.48 | 0.09 | 1.19 |
| Hist1h4i | 0.33 | 2.63E-08 | 5.48E-04 | 0.47 | 0.39 | 0.08 | 1.2 |
| Ndrg2 | 0.33 | 1.51E-08 | 3.14E-04 | 0.37 | 0.29 | 0.08 | 1.29 |
| Xpr1 | 0.33 | 3.73E-10 | 7.75E-06 | 0.43 | 0.35 | 0.09 | 1.25 |
| Tfrc | 0.33 | 1.01E-09 | 2.11E-05 | 0.52 | 0.44 | 0.09 | 1.2 |
| Slc3a2 | 0.33 | 1.40E-08 | 2.90E-04 | 0.68 | 0.62 | 0.06 | 1.09 |
| Ipo5 | 0.33 | 2.37E-13 | 4.94E-09 | 0.73 | 0.65 | 0.08 | 1.12 |
| Prc1 | 0.33 | 2.51E-07 | 5.22E-03 | 0.57 | 0.5 | 0.07 | 1.14 |
| Slfn9 | 0.33 | 7.24E-10 | 1.51E-05 | 0.29 | 0.21 | 0.08 | 1.41 |
| Wdr76 | 0.33 | 1.80E-07 | 3.75E-03 | 0.39 | 0.31 | 0.08 | 1.25 |
| Dut | 0.33 | 4.62E-24 | 9.61E-20 | 0.92 | 0.88 | 0.04 | 1.04 |
| Alg13 | 0.33 | 1.79E-06 | 3.72E-02 | 0.29 | 0.23 | 0.06 | 1.27 |
| Idi1 | 0.33 | 1.49E-10 | 3.11E-06 | 0.63 | 0.55 | 0.09 | 1.16 |
| Hells | 0.33 | 5.01E-14 | 1.04E-09 | 0.76 | 0.69 | 0.08 | 1.11 |
| Atp1a1 | 0.33 | 7.47E-14 | 1.55E-09 | 0.69 | 0.6 | 0.1 | 1.16 |
| Camk1 | 0.33 | 9.49E-09 | 1.97E-04 | 0.44 | 0.36 | 0.09 | 1.24 |
| Hacd2 | 0.32 | 3.57E-08 | 7.41E-04 | 0.42 | 0.34 | 0.08 | 1.23 |
| Bend7 | 0.32 | 9.25E-07 | 1.92E-02 | 0.27 | 0.2 | 0.07 | 1.33 |
| Pitpnc1 | 0.32 | 1.65E-07 | 3.43E-03 | 0.46 | 0.4 | 0.07 | 1.17 |
| Mki67 | 0.32 | 2.96E-13 | 6.15E-09 | 0.69 | 0.6 | 0.1 | 1.16 |
| Knstrn | 0.32 | 4.59E-10 | 9.55E-06 | 0.63 | 0.55 | 0.08 | 1.14 |
| Tnfrsf12a | 0.32 | 5.51E-07 | 1.15E-02 | 0.24 | 0.17 | 0.06 | 1.37 |
| Tubb2b | 0.32 | 1.40E-13 | 2.91E-09 | 0.92 | 0.9 | 0.02 | 1.03 |
| Cltb | 0.32 | 1.53E-06 | 3.19E-02 | 0.34 | 0.27 | 0.07 | 1.25 |
| Prkd3 | 0.32 | 4.08E-10 | 8.48E-06 | 0.59 | 0.5 | 0.08 | 1.17 |
| Ppp1r2 | 0.32 | 8.61E-16 | 1.79E-11 | 0.78 | 0.7 | 0.08 | 1.12 |
| Klf13 | 0.32 | 5.65E-07 | 1.17E-02 | 0.43 | 0.36 | 0.07 | 1.21 |
| Dhrs4 | 0.32 | 9.31E-13 | 1.94E-08 | 0.72 | 0.67 | 0.05 | 1.08 |
| Fdps | 0.31 | 1.36E-09 | 2.82E-05 | 0.82 | 0.79 | 0.03 | 1.04 |
| Vars | 0.31 | 6.87E-11 | 1.43E-06 | 0.62 | 0.53 | 0.09 | 1.16 |
| Ckap4 | 0.31 | 1.26E-16 | 2.62E-12 | 0.79 | 0.71 | 0.08 | 1.12 |
| Actn1 | 0.31 | 4.14E-11 | 8.60E-07 | 0.62 | 0.53 | 0.1 | 1.18 |
| Nynrin | 0.31 | 5.67E-09 | 1.18E-04 | 0.57 | 0.49 | 0.08 | 1.16 |
| Sh3rf1 | 0.31 | 3.14E-08 | 6.53E-04 | 0.36 | 0.27 | 0.08 | 1.3 |
| Cldn4 | 0.31 | 3.30E-07 | 6.87E-03 | 0.43 | 0.35 | 0.08 | 1.22 |
| Ccna2 | 0.31 | 1.27E-09 | 2.63E-05 | 0.61 | 0.53 | 0.08 | 1.16 |
| Sypl | 0.31 | 7.51E-12 | 1.56E-07 | 0.62 | 0.53 | 0.09 | 1.17 |
| Dtl | 0.31 | 2.67E-10 | 5.54E-06 | 0.59 | 0.5 | 0.09 | 1.18 |
| Lad1 | 0.31 | 2.92E-08 | 6.07E-04 | 0.37 | 0.29 | 0.08 | 1.29 |
| St7 | 0.31 | 1.86E-06 | 3.86E-02 | 0.35 | 0.29 | 0.07 | 1.24 |
| Plp2 | 0.31 | 1.73E-13 | 3.59E-09 | 0.7 | 0.61 | 0.09 | 1.15 |
| Yes1 | 0.31 | 1.07E-06 | 2.23E-02 | 0.35 | 0.29 | 0.07 | 1.24 |
| Lbr | 0.31 | 8.35E-10 | 1.74E-05 | 0.56 | 0.48 | 0.08 | 1.17 |
| Mogs | 0.31 | 5.97E-07 | 1.24E-02 | 0.39 | 0.33 | 0.07 | 1.2 |
| Racgap1 | 0.31 | 8.51E-10 | 1.77E-05 | 0.55 | 0.47 | 0.08 | 1.17 |
| Aig1 | 0.31 | 3.77E-14 | 7.84E-10 | 0.56 | 0.45 | 0.11 | 1.25 |
| Cdkn1b | 0.3 | 3.38E-07 | 7.03E-03 | 0.46 | 0.39 | 0.07 | 1.18 |

**Supplementary Table 5. Differentially expressed genes in the mutant λ-junction ectoderm cells.**

| | | | | | | | |
|---|---|---|---|---|---|---|---|
| Wee1 | 0.3 | 2.15E-09 | 4.46E-05 | 0.48 | 0.39 | 0.09 | 1.22 |
| Tmem189 | 0.3 | 1.63E-07 | 3.40E-03 | 0.44 | 0.37 | 0.07 | 1.19 |
| Ext2 | 0.3 | 3.46E-07 | 7.19E-03 | 0.47 | 0.39 | 0.08 | 1.2 |
| Cd3eap | 0.3 | 5.33E-09 | 1.11E-04 | 0.63 | 0.56 | 0.07 | 1.13 |
| Cdca8 | 0.3 | 1.16E-11 | 2.41E-07 | 0.74 | 0.66 | 0.08 | 1.12 |
| Litaf | 0.3 | 2.38E-07 | 4.94E-03 | 0.51 | 0.43 | 0.07 | 1.17 |
| Nek6 | 0.3 | 1.82E-06 | 3.78E-02 | 0.34 | 0.27 | 0.07 | 1.24 |
| Hmgn5 | 0.3 | 5.09E-15 | 1.06E-10 | 0.76 | 0.66 | 0.1 | 1.15 |
| Tsen15 | 0.3 | 4.19E-10 | 8.71E-06 | 0.62 | 0.55 | 0.08 | 1.14 |
| Bex4 | 0.3 | 3.40E-30 | 7.07E-26 | 0.94 | 0.89 | 0.05 | 1.06 |
| Iqgap1 | 0.3 | 3.25E-12 | 6.77E-08 | 0.67 | 0.58 | 0.09 | 1.16 |
| Tor2a | 0.3 | 1.03E-07 | 2.14E-03 | 0.44 | 0.37 | 0.07 | 1.2 |
| Rad51ap1 | 0.3 | 1.28E-07 | 2.66E-03 | 0.46 | 0.37 | 0.08 | 1.22 |
| Cpne3 | 0.29 | 9.73E-10 | 2.02E-05 | 0.66 | 0.59 | 0.07 | 1.11 |
| Plxnb1 | 0.29 | 1.64E-08 | 3.41E-04 | 0.45 | 0.37 | 0.08 | 1.21 |
| Cenpm | 0.29 | 1.77E-09 | 3.68E-05 | 0.56 | 0.48 | 0.08 | 1.17 |
| Epcam | 0.29 | 6.17E-33 | 1.28E-28 | 0.98 | 0.94 | 0.04 | 1.04 |
| Myo10 | 0.29 | 3.32E-13 | 6.91E-09 | 0.72 | 0.62 | 0.09 | 1.15 |
| Aktip | 0.29 | 5.57E-07 | 1.16E-02 | 0.48 | 0.41 | 0.07 | 1.17 |
| Gipc1 | 0.29 | 4.57E-15 | 9.50E-11 | 0.78 | 0.7 | 0.09 | 1.12 |
| Kif15 | 0.29 | 4.16E-11 | 8.66E-07 | 0.63 | 0.54 | 0.09 | 1.18 |
| Tubb4b | 0.29 | 3.59E-16 | 7.47E-12 | 0.84 | 0.77 | 0.07 | 1.09 |
| Meg3 | 0.29 | 8.43E-36 | 1.75E-31 | 0.81 | 0.63 | 0.18 | 1.29 |
| E2f8 | 0.29 | 2.04E-08 | 4.23E-04 | 0.29 | 0.22 | 0.08 | 1.36 |
| Cxxc5 | 0.29 | 1.86E-08 | 3.86E-04 | 0.6 | 0.52 | 0.08 | 1.15 |
| Mark2 | 0.29 | 6.27E-10 | 1.30E-05 | 0.61 | 0.54 | 0.07 | 1.14 |
| Srd5a1 | 0.29 | 7.08E-07 | 1.47E-02 | 0.36 | 0.29 | 0.07 | 1.23 |
| Eci1 | 0.29 | 4.31E-09 | 8.96E-05 | 0.57 | 0.5 | 0.08 | 1.15 |
| Cmas | 0.28 | 9.32E-09 | 1.94E-04 | 0.66 | 0.6 | 0.07 | 1.11 |
| P3h4 | 0.28 | 8.50E-07 | 1.77E-02 | 0.34 | 0.27 | 0.07 | 1.27 |
| Cmss1 | 0.28 | 1.13E-09 | 2.35E-05 | 0.5 | 0.4 | 0.1 | 1.25 |
| Sox2 | 0.28 | 3.43E-28 | 7.13E-24 | 0.76 | 0.57 | 0.19 | 1.33 |
| Fxyd6 | 0.28 | 9.94E-11 | 2.07E-06 | 0.61 | 0.52 | 0.09 | 1.18 |
| Clspn | 0.28 | 3.25E-11 | 6.76E-07 | 0.58 | 0.46 | 0.11 | 1.24 |
| Tk1 | 0.28 | 4.36E-09 | 9.06E-05 | 0.51 | 0.42 | 0.09 | 1.22 |
| 2310009B15Rik | 0.28 | 1.86E-08 | 3.87E-04 | 0.55 | 0.47 | 0.08 | 1.18 |
| Ezr | 0.28 | 5.07E-23 | 1.05E-18 | 0.92 | 0.85 | 0.06 | 1.08 |
| Incenp | 0.28 | 4.77E-09 | 9.91E-05 | 0.59 | 0.5 | 0.1 | 1.2 |
| Cab39 | 0.28 | 1.78E-10 | 3.70E-06 | 0.68 | 0.61 | 0.07 | 1.11 |
| Kif23 | 0.28 | 3.42E-08 | 7.11E-04 | 0.47 | 0.38 | 0.09 | 1.22 |
| Ptpn13 | 0.28 | 1.47E-07 | 3.06E-03 | 0.59 | 0.51 | 0.08 | 1.15 |
| Cyp51 | 0.28 | 2.66E-09 | 5.53E-05 | 0.63 | 0.55 | 0.08 | 1.15 |
| Nr2e1 | 0.28 | 5.63E-08 | 1.17E-03 | 0.19 | 0.13 | 0.06 | 1.49 |
| Camk2d | 0.28 | 1.99E-06 | 4.14E-02 | 0.27 | 0.2 | 0.06 | 1.32 |
| Pimreg | 0.28 | 9.81E-07 | 2.04E-02 | 0.52 | 0.45 | 0.07 | 1.15 |
| Epb41l5 | 0.28 | 7.52E-08 | 1.56E-03 | 0.54 | 0.46 | 0.08 | 1.17 |
| Ston2 | 0.28 | 3.66E-09 | 7.62E-05 | 0.61 | 0.54 | 0.08 | 1.14 |
| Tom1l1 | 0.28 | 2.52E-10 | 5.25E-06 | 0.5 | 0.4 | 0.1 | 1.25 |
| Pcbd2 | 0.27 | 1.81E-08 | 3.77E-04 | 0.48 | 0.4 | 0.08 | 1.2 |

**Supplementary Table 5. Differentially expressed genes in the mutant λ-junction ectoderm cells.**

| | | | | | | | |
|---|---|---|---|---|---|---|---|
| Nav1 | 0.27 | 1.83E-09 | 3.81E-05 | 0.54 | 0.46 | 0.09 | 1.19 |
| Txndc5 | 0.27 | 1.37E-08 | 2.84E-04 | 0.65 | 0.57 | 0.07 | 1.13 |
| Ssbp4 | 0.27 | 4.08E-08 | 8.48E-04 | 0.54 | 0.46 | 0.08 | 1.18 |
| Ddx39 | 0.27 | 1.18E-12 | 2.46E-08 | 0.78 | 0.72 | 0.06 | 1.08 |
| Mcm3 | 0.27 | 5.14E-11 | 1.07E-06 | 0.74 | 0.66 | 0.08 | 1.12 |
| Ptbp3 | 0.27 | 1.32E-12 | 2.76E-08 | 0.71 | 0.62 | 0.09 | 1.14 |
| Sp8 | 0.27 | 7.71E-18 | 1.60E-13 | 0.56 | 0.4 | 0.16 | 1.4 |
| Mcm10 | 0.27 | 2.12E-06 | 4.40E-02 | 0.33 | 0.27 | 0.06 | 1.24 |
| Syce2 | 0.27 | 1.12E-09 | 2.33E-05 | 0.59 | 0.5 | 0.08 | 1.17 |
| Fzd2 | 0.27 | 8.32E-16 | 1.73E-11 | 0.83 | 0.74 | 0.09 | 1.12 |
| Pak4 | 0.27 | 2.32E-07 | 4.83E-03 | 0.53 | 0.46 | 0.07 | 1.15 |
| Col4a1 | 0.26 | 4.60E-13 | 9.58E-09 | 0.71 | 0.6 | 0.11 | 1.18 |
| Nr2f6 | 0.26 | 2.72E-13 | 5.65E-09 | 0.76 | 0.69 | 0.07 | 1.1 |
| Erg28 | 0.26 | 2.31E-08 | 4.81E-04 | 0.71 | 0.65 | 0.05 | 1.08 |
| Mrps28 | 0.26 | 6.12E-11 | 1.27E-06 | 0.7 | 0.62 | 0.08 | 1.12 |
| Rnh1 | 0.26 | 4.24E-16 | 8.81E-12 | 0.8 | 0.71 | 0.09 | 1.13 |
| Satb1 | 0.26 | 8.74E-08 | 1.82E-03 | 0.59 | 0.52 | 0.07 | 1.13 |
| Krcc1 | 0.26 | 3.95E-11 | 8.21E-07 | 0.67 | 0.57 | 0.1 | 1.17 |
| Ncaph2 | 0.26 | 9.05E-08 | 1.88E-03 | 0.64 | 0.58 | 0.06 | 1.09 |
| Rrp15 | 0.26 | 1.74E-10 | 3.62E-06 | 0.71 | 0.64 | 0.07 | 1.1 |
| Hibadh | 0.26 | 3.07E-08 | 6.39E-04 | 0.64 | 0.57 | 0.07 | 1.12 |
| Dhfr | 0.26 | 5.74E-11 | 1.19E-06 | 0.64 | 0.54 | 0.1 | 1.19 |
| Tmx2 | 0.26 | 4.61E-08 | 9.58E-04 | 0.6 | 0.52 | 0.08 | 1.14 |
| Clip1 | 0.26 | 2.08E-07 | 4.33E-03 | 0.55 | 0.48 | 0.07 | 1.15 |
| Arl8b | 0.26 | 8.36E-08 | 1.74E-03 | 0.55 | 0.48 | 0.07 | 1.15 |
| Scaf4 | 0.26 | 1.95E-07 | 4.06E-03 | 0.44 | 0.36 | 0.08 | 1.22 |
| Rad23a | 0.26 | 1.96E-07 | 4.07E-03 | 0.53 | 0.45 | 0.08 | 1.17 |
| Fzd3 | 0.26 | 1.58E-13 | 3.29E-09 | 0.84 | 0.74 | 0.1 | 1.13 |
| Oard1 | 0.25 | 3.38E-12 | 7.02E-08 | 0.75 | 0.67 | 0.07 | 1.11 |
| Acadl | 0.25 | 4.11E-08 | 8.55E-04 | 0.47 | 0.39 | 0.09 | 1.22 |
| Zfhx2 | 0.25 | 2.20E-06 | 4.57E-02 | 0.5 | 0.43 | 0.07 | 1.16 |
| Pigyl | 0.25 | 3.18E-07 | 6.62E-03 | 0.51 | 0.44 | 0.07 | 1.15 |
| H2afx | 0.25 | 8.62E-14 | 1.79E-09 | 0.91 | 0.89 | 0.02 | 1.02 |
| Pbx3 | 0.25 | 2.06E-09 | 4.27E-05 | 0.61 | 0.53 | 0.08 | 1.15 |
| Il17rd | 0.25 | 3.36E-08 | 7.00E-04 | 0.53 | 0.44 | 0.09 | 1.19 |
| Tmed1 | 0.25 | 2.59E-08 | 5.38E-04 | 0.68 | 0.6 | 0.07 | 1.12 |
| Map1lc3a | -0.25 | 1.77E-07 | 3.68E-03 | 0.73 | 0.75 | -0.03 | 0.97 |
| Basp1 | -0.25 | 9.26E-23 | 1.93E-18 | 0.96 | 0.95 | 0 | 1 |
| Btg1 | -0.25 | 1.22E-06 | 2.54E-02 | 0.74 | 0.75 | -0.01 | 0.99 |
| Rbis | -0.25 | 5.17E-11 | 1.08E-06 | 0.82 | 0.84 | -0.02 | 0.97 |
| Selenow | -0.26 | 1.45E-35 | 3.01E-31 | 0.98 | 0.99 | 0 | 1 |
| Rbpj | -0.26 | 7.64E-07 | 1.59E-02 | 0.65 | 0.68 | -0.03 | 0.95 |
| Ubc | -0.27 | 8.22E-22 | 1.71E-17 | 0.97 | 0.97 | 0 | 1 |
| Tceal8 | -0.27 | 1.01E-11 | 2.10E-07 | 0.85 | 0.86 | -0.02 | 0.98 |
| Rnd3 | -0.27 | 6.33E-09 | 1.32E-04 | 0.72 | 0.75 | -0.03 | 0.96 |
| Limd2 | -0.27 | 1.24E-09 | 2.58E-05 | 0.77 | 0.8 | -0.03 | 0.97 |
| Trappc4 | -0.27 | 1.39E-11 | 2.89E-07 | 0.8 | 0.81 | -0.01 | 0.99 |
| Rb1cc1 | -0.27 | 2.87E-07 | 5.97E-03 | 0.67 | 0.71 | -0.03 | 0.95 |
| Chmp2b | -0.27 | 1.22E-08 | 2.53E-04 | 0.74 | 0.76 | -0.02 | 0.97 |

**Supplementary Table 5. Differentially expressed genes in the mutant λ-junction ectoderm cells.**

| | | | | | | | |
|---|---|---|---|---|---|---|---|
| Chd3 | -0.28 | 5.66E-18 | 1.18E-13 | 0.93 | 0.95 | -0.02 | 0.98 |
| Phlda3 | -0.28 | 6.71E-13 | 1.40E-08 | 0.49 | 0.59 | -0.1 | 0.83 |
| Commd3 | -0.28 | 5.04E-13 | 1.05E-08 | 0.82 | 0.83 | -0.01 | 0.99 |
| Igfbp4 | -0.28 | 2.53E-09 | 5.27E-05 | 0.73 | 0.77 | -0.04 | 0.94 |
| Khdrbs1 | -0.28 | 4.35E-32 | 9.04E-28 | 0.93 | 0.94 | -0.01 | 0.99 |
| Ip6k2 | -0.29 | 1.33E-06 | 2.78E-02 | 0.48 | 0.54 | -0.06 | 0.89 |
| Cd151 | -0.29 | 6.34E-07 | 1.32E-02 | 0.62 | 0.67 | -0.04 | 0.94 |
| Serpinh1 | -0.29 | 9.86E-26 | 2.05E-21 | 0.94 | 0.92 | 0.02 | 1.02 |
| Pnrc1 | -0.29 | 4.77E-22 | 9.91E-18 | 0.89 | 0.91 | -0.03 | 0.97 |
| Gm42418 | -0.29 | 1.60E-12 | 3.33E-08 | 1 | 0.99 | 0 | 1 |
| Gnas | -0.29 | 8.07E-47 | 1.68E-42 | 1 | 0.99 | 0 | 1 |
| Clta | -0.29 | 1.00E-33 | 2.08E-29 | 0.97 | 0.97 | 0 | 1 |
| Cbr1 | -0.3 | 1.21E-10 | 2.52E-06 | 0.72 | 0.76 | -0.04 | 0.95 |
| Ppp2r2d | -0.3 | 1.48E-06 | 3.08E-02 | 0.61 | 0.64 | -0.03 | 0.95 |
| Skp1a | -0.3 | 2.21E-30 | 4.60E-26 | 0.97 | 0.97 | 0 | 1 |
| Aldh2 | -0.3 | 1.39E-07 | 2.88E-03 | 0.7 | 0.71 | -0.01 | 0.99 |
| Snap47 | -0.3 | 1.52E-08 | 3.16E-04 | 0.67 | 0.72 | -0.05 | 0.93 |
| Rbfox2 | -0.3 | 9.74E-09 | 2.03E-04 | 0.69 | 0.72 | -0.03 | 0.96 |
| Hdac2 | -0.3 | 2.94E-16 | 6.11E-12 | 0.87 | 0.88 | -0.01 | 0.99 |
| Nxn | -0.31 | 3.08E-15 | 6.41E-11 | 0.74 | 0.79 | -0.05 | 0.94 |
| Atad2b | -0.31 | 1.49E-06 | 3.09E-02 | 0.6 | 0.65 | -0.05 | 0.92 |
| Ifitm2 | -0.31 | 5.94E-19 | 1.23E-14 | 0.92 | 0.91 | 0.01 | 1.01 |
| Smad5 | -0.31 | 1.13E-07 | 2.35E-03 | 0.47 | 0.53 | -0.06 | 0.89 |
| Myo1b | -0.32 | 3.82E-07 | 7.95E-03 | 0.61 | 0.65 | -0.04 | 0.94 |
| Tle5 | -0.32 | 4.51E-30 | 9.38E-26 | 0.95 | 0.97 | -0.02 | 0.98 |
| Skil | -0.32 | 6.15E-08 | 1.28E-03 | 0.56 | 0.62 | -0.06 | 0.9 |
| Gm7324 | -0.33 | 1.94E-07 | 4.04E-03 | 0.57 | 0.62 | -0.05 | 0.92 |
| Lsm14b | -0.33 | 7.03E-07 | 1.46E-02 | 0.42 | 0.48 | -0.06 | 0.87 |
| Mat2b | -0.33 | 1.11E-08 | 2.31E-04 | 0.58 | 0.64 | -0.06 | 0.91 |
| Nfib | -0.33 | 5.70E-12 | 1.19E-07 | 0.76 | 0.78 | -0.02 | 0.97 |
| Tmsb15b2 | -0.33 | 2.12E-06 | 4.41E-02 | 0.5 | 0.55 | -0.05 | 0.9 |
| Ube2e3 | -0.33 | 1.38E-12 | 2.88E-08 | 0.89 | 0.9 | 0 | 1 |
| Bcl11b | -0.34 | 2.04E-10 | 4.24E-06 | 0.69 | 0.75 | -0.05 | 0.93 |
| Rps27rt | -0.34 | 3.04E-17 | 6.33E-13 | 0.77 | 0.82 | -0.05 | 0.93 |
| Hbb-y | -0.35 | 1.94E-66 | 4.04E-62 | 0.97 | 0.98 | -0.01 | 0.99 |
| Zfhx3 | -0.35 | 1.96E-15 | 4.08E-11 | 0.92 | 0.9 | 0.02 | 1.02 |
| Gm8797 | -0.35 | 3.33E-31 | 6.92E-27 | 0.91 | 0.92 | -0.01 | 0.99 |
| Apoe | -0.36 | 1.09E-07 | 2.27E-03 | 0.48 | 0.54 | -0.06 | 0.89 |
| Znrf1 | -0.36 | 5.54E-07 | 1.15E-02 | 0.55 | 0.59 | -0.04 | 0.93 |
| Ccnd2 | -0.36 | 1.98E-21 | 4.13E-17 | 0.8 | 0.85 | -0.06 | 0.93 |
| Grb10 | -0.36 | 1.22E-31 | 2.54E-27 | 0.93 | 0.93 | 0 | 1 |
| Cmip | -0.36 | 1.73E-09 | 3.60E-05 | 0.49 | 0.56 | -0.07 | 0.88 |
| Luzp1 | -0.37 | 6.26E-17 | 1.30E-12 | 0.8 | 0.82 | -0.02 | 0.98 |
| Dlx3 | -0.37 | 2.05E-06 | 4.27E-02 | 0.26 | 0.32 | -0.07 | 0.8 |
| Sox11 | -0.37 | 8.27E-41 | 1.72E-36 | 0.98 | 0.98 | 0 | 1 |
| Hba-x | -0.37 | 4.64E-41 | 9.65E-37 | 0.86 | 0.91 | -0.05 | 0.95 |
| Smim20 | -0.38 | 1.01E-08 | 2.11E-04 | 0.5 | 0.57 | -0.07 | 0.88 |
| Malat1 | -0.38 | 4.00E-33 | 8.33E-29 | 0.99 | 0.98 | 0.01 | 1.01 |
| Smarca2 | -0.38 | 1.25E-09 | 2.60E-05 | 0.58 | 0.63 | -0.06 | 0.91 |

**Supplementary Table 5. Differentially expressed genes in the mutant λ-junction ectoderm cells.**

| | | | | | | | |
|---|---|---|---|---|---|---|---|
| Emb | -0.38 | 4.21E-07 | 8.75E-03 | 0.71 | 0.73 | -0.02 | 0.97 |
| Meis2 | -0.38 | 5.06E-28 | 1.05E-23 | 0.96 | 0.95 | 0.01 | 1.01 |
| Serinc2 | -0.38 | 1.16E-08 | 2.41E-04 | 0.54 | 0.6 | -0.06 | 0.9 |
| Ndufaf4 | -0.38 | 6.22E-08 | 1.29E-03 | 0.57 | 0.62 | -0.05 | 0.92 |
| Bcam | -0.38 | 5.23E-13 | 1.09E-08 | 0.7 | 0.73 | -0.03 | 0.96 |
| Ube2g1 | -0.38 | 1.06E-08 | 2.20E-04 | 0.56 | 0.61 | -0.05 | 0.91 |
| Mfge8 | -0.38 | 7.78E-07 | 1.62E-02 | 0.54 | 0.59 | -0.05 | 0.92 |
| Tjap1 | -0.38 | 1.49E-06 | 3.11E-02 | 0.34 | 0.4 | -0.06 | 0.84 |
| Flywch2 | -0.39 | 1.88E-10 | 3.91E-06 | 0.48 | 0.56 | -0.08 | 0.86 |
| Hba-a2 | -0.39 | 3.48E-12 | 7.24E-08 | 0.64 | 0.68 | -0.04 | 0.94 |
| Ypel1 | -0.39 | 6.97E-11 | 1.45E-06 | 0.54 | 0.62 | -0.07 | 0.88 |
| Mex3b | -0.39 | 3.64E-12 | 7.57E-08 | 0.47 | 0.55 | -0.08 | 0.85 |
| Tulp4 | -0.39 | 1.17E-14 | 2.43E-10 | 0.71 | 0.77 | -0.06 | 0.92 |
| Dok5 | -0.4 | 4.29E-10 | 8.91E-06 | 0.35 | 0.44 | -0.09 | 0.79 |
| Eid1 | -0.4 | 3.24E-70 | 6.75E-66 | 0.96 | 0.97 | -0.01 | 0.99 |
| Bcl11a | -0.4 | 2.78E-21 | 5.79E-17 | 0.78 | 0.81 | -0.03 | 0.96 |
| Fam162a | -0.4 | 3.50E-24 | 7.29E-20 | 0.93 | 0.93 | 0 | 1 |
| Plekho1 | -0.4 | 4.90E-08 | 1.02E-03 | 0.37 | 0.44 | -0.07 | 0.84 |
| Gpc3 | -0.4 | 3.46E-11 | 7.19E-07 | 0.83 | 0.82 | 0.01 | 1.01 |
| 1500009L16Rik | -0.4 | 1.86E-06 | 3.87E-02 | 0.53 | 0.57 | -0.04 | 0.93 |
| Chfr | -0.4 | 2.28E-06 | 4.73E-02 | 0.28 | 0.34 | -0.06 | 0.83 |
| Ywhah | -0.4 | 1.11E-24 | 2.31E-20 | 0.79 | 0.83 | -0.04 | 0.95 |
| Fermt1 | -0.41 | 5.78E-11 | 1.20E-06 | 0.53 | 0.61 | -0.08 | 0.87 |
| Parm1 | -0.41 | 1.46E-08 | 3.05E-04 | 0.18 | 0.25 | -0.07 | 0.72 |
| Lmo4 | -0.41 | 2.03E-24 | 4.22E-20 | 0.9 | 0.93 | -0.03 | 0.97 |
| Glrx2 | -0.41 | 1.25E-17 | 2.59E-13 | 0.68 | 0.73 | -0.05 | 0.93 |
| Nt5c | -0.41 | 1.31E-14 | 2.73E-10 | 0.64 | 0.71 | -0.06 | 0.91 |
| Azin1 | -0.42 | 3.59E-12 | 7.47E-08 | 0.55 | 0.61 | -0.06 | 0.9 |
| Serpinb6a | -0.42 | 1.17E-08 | 2.44E-04 | 0.45 | 0.51 | -0.06 | 0.88 |
| Casp6 | -0.42 | 1.08E-10 | 2.25E-06 | 0.49 | 0.57 | -0.08 | 0.87 |
| Lpp | -0.42 | 1.82E-09 | 3.79E-05 | 0.46 | 0.53 | -0.07 | 0.86 |
| Tert | -0.42 | 1.57E-06 | 3.26E-02 | 0.1 | 0.14 | -0.05 | 0.67 |
| Chp1 | -0.42 | 1.04E-07 | 2.16E-03 | 0.35 | 0.41 | -0.07 | 0.84 |
| Calcoco1 | -0.42 | 1.46E-06 | 3.03E-02 | 0.28 | 0.34 | -0.06 | 0.81 |
| Ddit4 | -0.43 | 1.03E-07 | 2.14E-03 | 0.41 | 0.48 | -0.06 | 0.87 |
| Fblim1 | -0.43 | 1.02E-10 | 2.12E-06 | 0.4 | 0.48 | -0.08 | 0.83 |
| Psph | -0.44 | 2.08E-06 | 4.32E-02 | 0.31 | 0.36 | -0.06 | 0.84 |
| Cox6b2 | -0.44 | 2.00E-09 | 4.15E-05 | 0.51 | 0.58 | -0.06 | 0.89 |
| Ociad2 | -0.44 | 1.68E-06 | 3.50E-02 | 0.28 | 0.34 | -0.06 | 0.82 |
| Dlg5 | -0.44 | 4.24E-07 | 8.83E-03 | 0.39 | 0.45 | -0.06 | 0.86 |
| Rac3 | -0.45 | 3.70E-07 | 7.69E-03 | 0.39 | 0.45 | -0.06 | 0.86 |
| Erich1 | -0.45 | 2.03E-06 | 4.21E-02 | 0.37 | 0.43 | -0.06 | 0.87 |
| 6330403K07Rik | -0.46 | 5.67E-31 | 1.18E-26 | 0.84 | 0.87 | -0.03 | 0.96 |
| Tle4 | -0.47 | 6.16E-08 | 1.28E-03 | 0.53 | 0.57 | -0.05 | 0.92 |
| Hmgn3 | -0.47 | 6.06E-20 | 1.26E-15 | 0.7 | 0.76 | -0.06 | 0.92 |
| Dusp7 | -0.47 | 4.78E-09 | 9.95E-05 | 0.32 | 0.4 | -0.07 | 0.82 |
| Atxn10 | -0.47 | 1.85E-28 | 3.85E-24 | 0.89 | 0.91 | -0.02 | 0.98 |
| Pdlim1 | -0.47 | 5.79E-19 | 1.20E-14 | 0.63 | 0.71 | -0.08 | 0.88 |
| Amotl2 | -0.47 | 5.39E-09 | 1.12E-04 | 0.35 | 0.42 | -0.07 | 0.83 |

**Supplementary Table 5. Differentially expressed genes in the mutant λ-junction ectoderm cells.**

| | | | | | | | |
|---|---|---|---|---|---|---|---|
| Wls | -0.47 | 7.15E-24 | 1.49E-19 | 0.78 | 0.81 | -0.03 | 0.97 |
| Lbh | -0.48 | 9.16E-09 | 1.91E-04 | 0.25 | 0.32 | -0.07 | 0.77 |
| Glul | -0.48 | 6.86E-16 | 1.43E-11 | 0.45 | 0.55 | -0.11 | 0.81 |
| Nkd1 | -0.49 | 1.30E-06 | 2.71E-02 | 0.15 | 0.2 | -0.05 | 0.73 |
| Pkp3 | -0.49 | 2.38E-07 | 4.95E-03 | 0.23 | 0.29 | -0.06 | 0.78 |
| Fth1 | -0.49 | 2.70E-75 | 5.61E-71 | 0.99 | 0.99 | 0 | 1 |
| Twist1 | -0.49 | 8.32E-12 | 1.73E-07 | 0.28 | 0.37 | -0.09 | 0.75 |
| Fmnl2 | -0.5 | 2.03E-12 | 4.21E-08 | 0.44 | 0.52 | -0.08 | 0.85 |
| Zfp57 | -0.5 | 2.44E-08 | 5.07E-04 | 0.18 | 0.25 | -0.07 | 0.73 |
| Nrep | -0.5 | 1.11E-27 | 2.30E-23 | 0.79 | 0.84 | -0.05 | 0.94 |
| Ctsl | -0.51 | 2.81E-19 | 5.85E-15 | 0.84 | 0.81 | 0.03 | 1.03 |
| Rb1 | -0.51 | 1.42E-10 | 2.95E-06 | 0.25 | 0.34 | -0.09 | 0.74 |
| Itgb5 | -0.51 | 7.46E-14 | 1.55E-09 | 0.49 | 0.58 | -0.09 | 0.85 |
| Txndc17 | -0.51 | 4.88E-30 | 1.02E-25 | 0.93 | 0.93 | 0 | 1 |
| P4ha1 | -0.51 | 1.88E-14 | 3.92E-10 | 0.5 | 0.58 | -0.09 | 0.85 |
| Msx2 | -0.52 | 5.18E-09 | 1.08E-04 | 0.41 | 0.48 | -0.07 | 0.85 |
| 1600020E01Rik | -0.52 | 1.10E-06 | 2.29E-02 | 0.2 | 0.25 | -0.06 | 0.77 |
| Ccser2 | -0.52 | 1.64E-06 | 3.40E-02 | 0.28 | 0.34 | -0.06 | 0.83 |
| Enho | -0.52 | 2.89E-11 | 6.01E-07 | 0.32 | 0.41 | -0.08 | 0.79 |
| Cotl1 | -0.52 | 1.06E-13 | 2.21E-09 | 0.44 | 0.53 | -0.09 | 0.83 |
| Thbs1 | -0.52 | 5.46E-09 | 1.14E-04 | 0.2 | 0.27 | -0.07 | 0.74 |
| Afap1l1 | -0.53 | 5.12E-07 | 1.06E-02 | 0.1 | 0.15 | -0.05 | 0.68 |
| Tbc1d10b | -0.54 | 6.49E-09 | 1.35E-04 | 0.29 | 0.36 | -0.07 | 0.8 |
| Id2 | -0.54 | 4.57E-35 | 9.51E-31 | 0.82 | 0.88 | -0.06 | 0.93 |
| Sema5a | -0.54 | 9.47E-11 | 1.97E-06 | 0.21 | 0.29 | -0.08 | 0.72 |
| Rtl8a | -0.54 | 1.01E-08 | 2.11E-04 | 0.29 | 0.37 | -0.08 | 0.8 |
| Smad7 | -0.54 | 7.32E-25 | 1.52E-20 | 0.59 | 0.69 | -0.1 | 0.85 |
| Tbc1d1 | -0.55 | 2.82E-09 | 5.86E-05 | 0.41 | 0.47 | -0.06 | 0.87 |
| Kit | -0.55 | 2.69E-08 | 5.59E-04 | 0.15 | 0.22 | -0.06 | 0.7 |
| Limch1 | -0.55 | 7.15E-11 | 1.49E-06 | 0.37 | 0.45 | -0.08 | 0.83 |
| Susd4 | -0.55 | 1.38E-11 | 2.87E-07 | 0.26 | 0.35 | -0.09 | 0.76 |
| Avpi1 | -0.55 | 1.28E-11 | 2.67E-07 | 0.35 | 0.43 | -0.08 | 0.81 |
| Chst15 | -0.55 | 3.94E-07 | 8.18E-03 | 0.14 | 0.19 | -0.05 | 0.72 |
| Eef2k | -0.55 | 8.10E-08 | 1.69E-03 | 0.17 | 0.24 | -0.06 | 0.73 |
| Ackr3 | -0.56 | 3.44E-15 | 7.15E-11 | 0.36 | 0.45 | -0.1 | 0.78 |
| Otx1 | -0.56 | 8.57E-12 | 1.78E-07 | 0.4 | 0.47 | -0.07 | 0.85 |
| Dzip1 | -0.56 | 7.79E-07 | 1.62E-02 | 0.22 | 0.28 | -0.06 | 0.79 |
| Slc25a33 | -0.56 | 1.08E-06 | 2.24E-02 | 0.2 | 0.25 | -0.06 | 0.78 |
| Ttc23 | -0.56 | 3.74E-10 | 7.78E-06 | 0.29 | 0.37 | -0.08 | 0.79 |
| Cnn2 | -0.56 | 1.27E-08 | 2.64E-04 | 0.35 | 0.42 | -0.07 | 0.84 |
| Gm10320 | -0.56 | 2.30E-19 | 4.79E-15 | 0.32 | 0.46 | -0.13 | 0.71 |
| Hspb1 | -0.57 | 3.66E-08 | 7.61E-04 | 0.37 | 0.43 | -0.07 | 0.84 |
| Cib2 | -0.57 | 8.14E-13 | 1.69E-08 | 0.16 | 0.25 | -0.09 | 0.66 |
| Pltp | -0.57 | 5.83E-16 | 1.21E-11 | 0.43 | 0.52 | -0.1 | 0.81 |
| Bmi1 | -0.57 | 4.05E-13 | 8.42E-09 | 0.37 | 0.46 | -0.09 | 0.81 |
| Ldhb | -0.58 | 5.80E-62 | 1.21E-57 | 0.85 | 0.9 | -0.05 | 0.94 |
| Tbx3 | -0.58 | 1.73E-09 | 3.61E-05 | 0.12 | 0.19 | -0.07 | 0.65 |
| Sh3bgrl | -0.59 | 3.45E-44 | 7.18E-40 | 0.72 | 0.83 | -0.11 | 0.86 |
| Gpc2 | -0.59 | 5.36E-13 | 1.11E-08 | 0.34 | 0.43 | -0.09 | 0.79 |

**Supplementary Table 5. Differentially expressed genes in the mutant λ-junction ectoderm cells.**

| | | | | | | | |
|---|---|---|---|---|---|---|---|
| Ets2 | -0.59 | 4.44E-10 | 9.23E-06 | 0.24 | 0.32 | -0.08 | 0.76 |
| Nampt | -0.59 | 5.16E-13 | 1.07E-08 | 0.32 | 0.41 | -0.09 | 0.77 |
| Pdpn | -0.6 | 3.41E-14 | 7.09E-10 | 0.33 | 0.43 | -0.1 | 0.77 |
| Snhg9 | -0.6 | 6.44E-08 | 1.34E-03 | 0.2 | 0.27 | -0.07 | 0.75 |
| Tox3 | -0.6 | 6.83E-07 | 1.42E-02 | 0.28 | 0.34 | -0.06 | 0.83 |
| Prkx | -0.61 | 5.88E-08 | 1.22E-03 | 0.17 | 0.23 | -0.06 | 0.73 |
| Npnt | -0.61 | 5.07E-22 | 1.05E-17 | 0.4 | 0.53 | -0.12 | 0.76 |
| Man2a1 | -0.61 | 2.51E-10 | 5.22E-06 | 0.23 | 0.31 | -0.08 | 0.74 |
| Itga3 | -0.61 | 2.55E-08 | 5.30E-04 | 0.22 | 0.29 | -0.07 | 0.77 |
| Slc50a1 | -0.61 | 4.71E-07 | 9.79E-03 | 0.23 | 0.29 | -0.06 | 0.8 |
| Slc16a13 | -0.61 | 1.26E-06 | 2.61E-02 | 0.11 | 0.16 | -0.05 | 0.69 |
| Dcxr | -0.62 | 9.01E-09 | 1.87E-04 | 0.14 | 0.2 | -0.06 | 0.69 |
| Tpm2 | -0.62 | 2.84E-09 | 5.90E-05 | 0.21 | 0.28 | -0.07 | 0.75 |
| Fam171b | -0.62 | 1.29E-07 | 2.68E-03 | 0.14 | 0.19 | -0.06 | 0.71 |
| Zfp503 | -0.63 | 4.26E-35 | 8.86E-31 | 0.46 | 0.61 | -0.15 | 0.75 |
| Nnat | -0.63 | 2.92E-36 | 6.08E-32 | 0.93 | 0.94 | -0.01 | 0.99 |
| Rbbp8 | -0.63 | 9.69E-33 | 2.02E-28 | 0.64 | 0.75 | -0.1 | 0.86 |
| Krt14 | -0.64 | 1.62E-31 | 3.37E-27 | 0.41 | 0.58 | -0.17 | 0.71 |
| Lef1 | -0.64 | 2.89E-26 | 6.00E-22 | 0.35 | 0.51 | -0.16 | 0.69 |
| Vwa5a | -0.64 | 4.30E-09 | 8.95E-05 | 0.14 | 0.21 | -0.07 | 0.69 |
| Ptpn14 | -0.64 | 6.27E-13 | 1.30E-08 | 0.21 | 0.3 | -0.09 | 0.7 |
| 1110008P14Rik | -0.64 | 5.53E-13 | 1.15E-08 | 0.35 | 0.43 | -0.09 | 0.8 |
| Dpysl3 | -0.64 | 3.00E-18 | 6.24E-14 | 0.52 | 0.63 | -0.11 | 0.83 |
| Nrp2 | -0.65 | 2.78E-12 | 5.78E-08 | 0.18 | 0.26 | -0.08 | 0.68 |
| Capn6 | -0.65 | 2.59E-21 | 5.38E-17 | 0.44 | 0.56 | -0.12 | 0.79 |
| Sncaip | -0.65 | 9.97E-08 | 2.07E-03 | 0.16 | 0.22 | -0.06 | 0.72 |
| Tmem47 | -0.65 | 4.98E-12 | 1.04E-07 | 0.42 | 0.5 | -0.08 | 0.84 |
| Rab38 | -0.66 | 3.52E-07 | 7.33E-03 | 0.18 | 0.24 | -0.06 | 0.76 |
| Adamts18 | -0.66 | 2.87E-17 | 5.97E-13 | 0.22 | 0.33 | -0.11 | 0.66 |
| Mycn | -0.66 | 2.80E-16 | 5.82E-12 | 0.34 | 0.45 | -0.1 | 0.77 |
| Trp53inp1 | -0.66 | 1.91E-18 | 3.97E-14 | 0.22 | 0.34 | -0.12 | 0.65 |
| Pdgfa | -0.66 | 1.35E-71 | 2.80E-67 | 0.92 | 0.95 | -0.03 | 0.97 |
| Ralgapa2 | -0.67 | 8.79E-08 | 1.83E-03 | 0.18 | 0.24 | -0.06 | 0.75 |
| Plk2 | -0.67 | 3.22E-15 | 6.71E-11 | 0.35 | 0.45 | -0.1 | 0.78 |
| Rest | -0.67 | 5.18E-55 | 1.08E-50 | 0.67 | 0.79 | -0.13 | 0.84 |
| Gm49708 | -0.68 | 8.65E-08 | 1.80E-03 | 0.51 | 0.53 | -0.02 | 0.96 |
| Cpe | -0.68 | 3.15E-21 | 6.56E-17 | 0.42 | 0.53 | -0.11 | 0.79 |
| Ntf5 | -0.68 | 2.04E-24 | 4.24E-20 | 0.27 | 0.41 | -0.14 | 0.66 |
| Cd24a | -0.68 | 8.35E-131 | 1.74E-126 | 0.95 | 0.97 | -0.02 | 0.98 |
| Tinagl1 | -0.68 | 1.03E-06 | 2.15E-02 | 0.07 | 0.12 | -0.04 | 0.63 |
| D16Ertd472e | -0.7 | 3.08E-18 | 6.41E-14 | 0.22 | 0.33 | -0.11 | 0.67 |
| Pla2g4a | -0.7 | 4.98E-08 | 1.04E-03 | 0.12 | 0.17 | -0.06 | 0.67 |
| Lhfp | -0.7 | 5.87E-12 | 1.22E-07 | 0.29 | 0.37 | -0.08 | 0.79 |
| Copz2 | -0.7 | 1.77E-14 | 3.67E-10 | 0.23 | 0.33 | -0.1 | 0.7 |
| Hbb-bh1 | -0.7 | 1.50E-31 | 3.13E-27 | 0.34 | 0.49 | -0.15 | 0.69 |
| Tfap2a | -0.71 | 1.51E-11 | 3.14E-07 | 0.33 | 0.41 | -0.08 | 0.81 |
| Pdgfc | -0.71 | 7.19E-52 | 1.49E-47 | 0.73 | 0.81 | -0.09 | 0.9 |
| Sox12 | -0.71 | 3.52E-39 | 7.32E-35 | 0.58 | 0.7 | -0.13 | 0.82 |
| Samd1 | -0.72 | 4.84E-20 | 1.01E-15 | 0.3 | 0.42 | -0.12 | 0.72 |

**Supplementary Table 5. Differentially expressed genes in the mutant λ-junction ectoderm cells.**

| | | | | | | | |
|---|---|---|---|---|---|---|---|
| Homer2 | -0.72 | 1.13E-09 | 2.35E-05 | 0.19 | 0.26 | -0.07 | 0.73 |
| Gstk1 | -0.73 | 1.43E-08 | 2.97E-04 | 0.14 | 0.2 | -0.06 | 0.69 |
| Map1b | -0.73 | 4.20E-29 | 8.74E-25 | 0.52 | 0.65 | -0.13 | 0.8 |
| Cdk5r1 | -0.73 | 5.79E-07 | 1.20E-02 | 0.06 | 0.1 | -0.04 | 0.6 |
| Pla2r1 | -0.74 | 3.39E-18 | 7.04E-14 | 0.23 | 0.34 | -0.11 | 0.68 |
| Uchl1 | -0.74 | 4.70E-29 | 9.78E-25 | 0.59 | 0.68 | -0.09 | 0.87 |
| Mfap4 | -0.74 | 2.02E-25 | 4.19E-21 | 0.65 | 0.71 | -0.06 | 0.91 |
| Tnik | -0.74 | 6.56E-09 | 1.36E-04 | 0.1 | 0.16 | -0.06 | 0.65 |
| Vwa1 | -0.74 | 1.06E-11 | 2.20E-07 | 0.09 | 0.16 | -0.07 | 0.58 |
| Ntn3 | -0.75 | 2.64E-07 | 5.48E-03 | 0.07 | 0.11 | -0.04 | 0.62 |
| Tmem178 | -0.75 | 2.06E-08 | 4.28E-04 | 0.1 | 0.15 | -0.05 | 0.64 |
| Col5a2 | -0.75 | 1.69E-10 | 3.51E-06 | 0.2 | 0.27 | -0.07 | 0.73 |
| Fry | -0.75 | 1.24E-09 | 2.58E-05 | 0.11 | 0.17 | -0.06 | 0.65 |
| Plin3 | -0.75 | 1.21E-35 | 2.51E-31 | 0.5 | 0.64 | -0.14 | 0.78 |
| Asb7 | -0.77 | 5.41E-08 | 1.13E-03 | 0.23 | 0.3 | -0.06 | 0.79 |
| Trp63 | -0.77 | 5.95E-39 | 1.24E-34 | 0.45 | 0.62 | -0.17 | 0.73 |
| Ccng1 | -0.77 | 4.34E-18 | 9.02E-14 | 0.2 | 0.31 | -0.11 | 0.65 |
| Minar2 | -0.78 | 2.24E-12 | 4.66E-08 | 0.13 | 0.21 | -0.08 | 0.64 |
| Cd59a | -0.78 | 4.58E-15 | 9.53E-11 | 0.27 | 0.37 | -0.1 | 0.72 |
| St3gal1 | -0.79 | 7.23E-07 | 1.50E-02 | 0.04 | 0.07 | -0.03 | 0.55 |
| Mir100hg | -0.79 | 3.39E-46 | 7.05E-42 | 0.6 | 0.73 | -0.13 | 0.82 |
| Htra1 | -0.8 | 7.95E-09 | 1.65E-04 | 0.05 | 0.09 | -0.04 | 0.54 |
| Arhgef28 | -0.8 | 3.22E-10 | 6.69E-06 | 0.13 | 0.19 | -0.07 | 0.65 |
| Nell2 | -0.8 | 3.55E-08 | 7.38E-04 | 0.09 | 0.15 | -0.05 | 0.64 |
| Col17a1 | -0.8 | 7.42E-16 | 1.54E-11 | 0.1 | 0.18 | -0.08 | 0.54 |
| Insyn2b | -0.8 | 2.13E-07 | 4.42E-03 | 0.06 | 0.1 | -0.04 | 0.59 |
| Igf1 | -0.8 | 1.74E-10 | 3.63E-06 | 0.11 | 0.17 | -0.06 | 0.63 |
| Pax7 | -0.81 | 3.37E-08 | 7.01E-04 | 0.06 | 0.1 | -0.04 | 0.57 |
| Fam83b | -0.81 | 2.14E-06 | 4.45E-02 | 0.06 | 0.09 | -0.04 | 0.61 |
| Trp53inp2 | -0.81 | 1.39E-09 | 2.89E-05 | 0.06 | 0.11 | -0.05 | 0.55 |
| Cobll1 | -0.81 | 4.94E-10 | 1.03E-05 | 0.08 | 0.14 | -0.06 | 0.59 |
| Gng11 | -0.81 | 1.86E-08 | 3.88E-04 | 0.13 | 0.19 | -0.06 | 0.68 |
| Igfbp2 | -0.82 | 6.58E-125 | 1.37E-120 | 0.97 | 0.96 | 0.01 | 1.01 |
| Met | -0.82 | 6.53E-10 | 1.36E-05 | 0.09 | 0.14 | -0.06 | 0.61 |
| Mir205hg | -0.83 | 1.02E-27 | 2.11E-23 | 0.38 | 0.52 | -0.14 | 0.73 |
| Ajuba | -0.83 | 6.20E-29 | 1.29E-24 | 0.39 | 0.52 | -0.13 | 0.75 |
| Ccdc68 | -0.83 | 3.61E-08 | 7.50E-04 | 0.12 | 0.18 | -0.06 | 0.68 |
| Dusp4 | -0.84 | 2.52E-08 | 5.24E-04 | 0.05 | 0.1 | -0.04 | 0.56 |
| Cald1 | -0.85 | 3.14E-56 | 6.52E-52 | 0.69 | 0.8 | -0.11 | 0.86 |
| Chst11 | -0.85 | 4.06E-14 | 8.45E-10 | 0.13 | 0.22 | -0.08 | 0.62 |
| Cacna1d | -0.85 | 7.31E-07 | 1.52E-02 | 0.06 | 0.1 | -0.04 | 0.61 |
| Snhg11 | -0.85 | 1.93E-09 | 4.01E-05 | 0.07 | 0.12 | -0.05 | 0.59 |
| Car14 | -0.86 | 1.31E-24 | 2.73E-20 | 0.23 | 0.36 | -0.13 | 0.64 |
| Prtn3 | -0.86 | 1.82E-10 | 3.78E-06 | 0.09 | 0.15 | -0.06 | 0.6 |
| Olfm2 | -0.86 | 1.38E-15 | 2.86E-11 | 0.14 | 0.23 | -0.09 | 0.61 |
| Gnai1 | -0.86 | 1.23E-12 | 2.55E-08 | 0.1 | 0.17 | -0.07 | 0.58 |
| Cth | -0.86 | 6.30E-12 | 1.31E-07 | 0.16 | 0.23 | -0.08 | 0.67 |
| Blvra | -0.86 | 9.90E-16 | 2.06E-11 | 0.12 | 0.21 | -0.09 | 0.59 |
| Arhgap24 | -0.87 | 2.09E-07 | 4.35E-03 | 0.08 | 0.12 | -0.05 | 0.64 |

**Supplementary Table 5. Differentially expressed genes in the mutant λ-junction ectoderm cells.**

| | | | | | | | |
|---|---|---|---|---|---|---|---|
| Klhl8 | -0.87 | 1.56E-12 | 3.23E-08 | 0.13 | 0.21 | -0.08 | 0.63 |
| Dapl1 | -0.87 | 3.56E-18 | 7.41E-14 | 0.24 | 0.35 | -0.11 | 0.68 |
| Ctsf | -0.88 | 2.30E-08 | 4.79E-04 | 0.1 | 0.16 | -0.05 | 0.65 |
| Slc7a10 | -0.88 | 3.87E-08 | 8.06E-04 | 0.04 | 0.07 | -0.04 | 0.5 |
| Jdp2 | -0.88 | 1.72E-09 | 3.58E-05 | 0.08 | 0.14 | -0.05 | 0.6 |
| Kctd12 | -0.88 | 1.16E-06 | 2.40E-02 | 0.13 | 0.18 | -0.05 | 0.72 |
| Pax3 | -0.88 | 1.72E-14 | 3.57E-10 | 0.11 | 0.18 | -0.08 | 0.58 |
| Spats2l | -0.89 | 2.23E-29 | 4.65E-25 | 0.3 | 0.45 | -0.15 | 0.67 |
| Nog | -0.89 | 3.03E-11 | 6.31E-07 | 0.1 | 0.17 | -0.07 | 0.61 |
| Cnih2 | -0.9 | 3.83E-07 | 7.96E-03 | 0.16 | 0.21 | -0.05 | 0.75 |
| Hes6 | -0.91 | 5.36E-18 | 1.12E-13 | 0.48 | 0.56 | -0.08 | 0.85 |
| Bach2 | -0.91 | 1.71E-18 | 3.56E-14 | 0.23 | 0.34 | -0.11 | 0.69 |
| Fgf9 | -0.91 | 4.57E-25 | 9.51E-21 | 0.37 | 0.48 | -0.11 | 0.77 |
| Lgals1 | -0.92 | 2.27E-29 | 4.71E-25 | 0.38 | 0.53 | -0.15 | 0.72 |
| Gdf6 | -0.92 | 2.33E-07 | 4.85E-03 | 0.04 | 0.08 | -0.04 | 0.55 |
| Gadd45g | -0.92 | 2.10E-90 | 4.36E-86 | 0.24 | 0.53 | -0.3 | 0.44 |
| Mxd4 | -0.93 | 1.67E-36 | 3.47E-32 | 0.38 | 0.54 | -0.16 | 0.71 |
| Ddhd2 | -0.93 | 2.51E-28 | 5.22E-24 | 0.24 | 0.38 | -0.14 | 0.63 |
| Sertad1 | -0.93 | 4.94E-14 | 1.03E-09 | 0.11 | 0.19 | -0.08 | 0.59 |
| Arhgdib | -0.94 | 2.51E-24 | 5.21E-20 | 0.22 | 0.35 | -0.13 | 0.64 |
| Plbd1 | -0.94 | 1.19E-07 | 2.48E-03 | 0.03 | 0.06 | -0.03 | 0.47 |
| Adamts5 | -0.95 | 1.32E-06 | 2.74E-02 | 0.05 | 0.09 | -0.04 | 0.59 |
| Bnc2 | -0.95 | 3.08E-10 | 6.40E-06 | 0.08 | 0.14 | -0.06 | 0.59 |
| Gse1 | -0.96 | 4.26E-22 | 8.86E-18 | 0.23 | 0.35 | -0.12 | 0.66 |
| Irx5 | -0.96 | 1.50E-10 | 3.11E-06 | 0.25 | 0.31 | -0.06 | 0.8 |
| Emp3 | -0.96 | 7.57E-07 | 1.57E-02 | 0.07 | 0.11 | -0.04 | 0.63 |
| Rapgefl1 | -0.96 | 1.03E-07 | 2.14E-03 | 0.06 | 0.11 | -0.04 | 0.59 |
| Ifitm3 | -0.98 | 1.20E-27 | 2.50E-23 | 0.39 | 0.53 | -0.13 | 0.75 |
| Pik3r3 | -0.98 | 9.40E-14 | 1.96E-09 | 0.19 | 0.27 | -0.08 | 0.7 |
| Plxna2 | -0.98 | 2.45E-17 | 5.09E-13 | 0.24 | 0.35 | -0.1 | 0.71 |
| Selenbp1 | -0.98 | 6.28E-10 | 1.31E-05 | 0.05 | 0.09 | -0.05 | 0.51 |
| Piezo2 | -0.98 | 2.02E-17 | 4.21E-13 | 0.17 | 0.26 | -0.1 | 0.63 |
| Galnt18 | -1 | 1.89E-10 | 3.93E-06 | 0.08 | 0.13 | -0.06 | 0.58 |
| Ripk3 | -1 | 1.10E-06 | 2.28E-02 | 0.05 | 0.09 | -0.04 | 0.59 |
| Ogfrl1 | -1 | 1.68E-27 | 3.49E-23 | 0.17 | 0.3 | -0.13 | 0.56 |
| Ppp1r3b | -1 | 3.80E-12 | 7.90E-08 | 0.04 | 0.09 | -0.05 | 0.45 |
| Fzd10 | -1 | 4.18E-08 | 8.69E-04 | 0.13 | 0.18 | -0.06 | 0.7 |
| Adamts17 | -1.01 | 6.18E-16 | 1.28E-11 | 0.09 | 0.17 | -0.08 | 0.54 |
| Itgav | -1.03 | 1.19E-24 | 2.48E-20 | 0.19 | 0.31 | -0.13 | 0.6 |
| Ttll7 | -1.03 | 2.97E-10 | 6.17E-06 | 0.05 | 0.1 | -0.05 | 0.51 |
| Nbl1 | -1.03 | 1.22E-25 | 2.55E-21 | 0.11 | 0.23 | -0.11 | 0.5 |
| Sema3a | -1.03 | 7.05E-19 | 1.47E-14 | 0.45 | 0.54 | -0.09 | 0.83 |
| Crlf1 | -1.03 | 6.70E-22 | 1.39E-17 | 0.17 | 0.29 | -0.12 | 0.6 |
| Cdkn1c | -1.04 | 1.87E-151 | 3.90E-147 | 0.85 | 0.96 | -0.11 | 0.89 |
| Dmtn | -1.04 | 2.40E-08 | 4.99E-04 | 0.08 | 0.13 | -0.05 | 0.62 |
| Runx2 | -1.05 | 1.11E-06 | 2.31E-02 | 0.05 | 0.09 | -0.04 | 0.6 |
| Bnip3 | -1.06 | 6.60E-63 | 1.37E-58 | 0.39 | 0.61 | -0.22 | 0.65 |
| Adamts19 | -1.06 | 1.82E-14 | 3.79E-10 | 0.09 | 0.17 | -0.07 | 0.56 |
| Dnm3 | -1.06 | 2.59E-09 | 5.38E-05 | 0.04 | 0.08 | -0.04 | 0.47 |

**Supplementary Table 5. Differentially expressed genes in the mutant λ-junction ectoderm cells.**

| | | | | | | | |
|---|---|---|---|---|---|---|---|
| Pde10a | -1.06 | 2.23E-08 | 4.65E-04 | 0.03 | 0.07 | -0.04 | 0.49 |
| Sord | -1.07 | 3.89E-20 | 8.09E-16 | 0.21 | 0.32 | -0.11 | 0.65 |
| Lmod1 | -1.07 | 5.54E-07 | 1.15E-02 | 0.05 | 0.08 | -0.04 | 0.57 |
| Col1a1 | -1.1 | 2.46E-08 | 5.12E-04 | 0.06 | 0.1 | -0.04 | 0.56 |
| Tanc2 | -1.12 | 1.05E-21 | 2.17E-17 | 0.19 | 0.3 | -0.11 | 0.62 |
| Arl4a | -1.12 | 2.55E-13 | 5.30E-09 | 0.14 | 0.22 | -0.08 | 0.64 |
| Ucp2 | -1.13 | 3.46E-59 | 7.20E-55 | 0.4 | 0.6 | -0.2 | 0.66 |
| Efcab1 | -1.14 | 6.77E-15 | 1.41E-10 | 0.07 | 0.14 | -0.07 | 0.52 |
| Cyp26b1 | -1.14 | 3.66E-07 | 7.60E-03 | 0.03 | 0.07 | -0.03 | 0.52 |
| Pcyt1b | -1.15 | 5.19E-15 | 1.08E-10 | 0.1 | 0.18 | -0.08 | 0.56 |
| Nrxn3 | -1.15 | 4.32E-12 | 8.98E-08 | 0.08 | 0.14 | -0.06 | 0.56 |
| Irx3 | -1.15 | 1.21E-10 | 2.51E-06 | 0.22 | 0.28 | -0.06 | 0.78 |
| Rbms3 | -1.15 | 1.40E-23 | 2.91E-19 | 0.17 | 0.28 | -0.11 | 0.6 |
| Snai2 | -1.16 | 1.19E-38 | 2.47E-34 | 0.19 | 0.35 | -0.16 | 0.54 |
| Stra6 | -1.17 | 5.83E-24 | 1.21E-19 | 0.25 | 0.36 | -0.11 | 0.68 |
| Adarb2 | -1.18 | 8.18E-07 | 1.70E-02 | 0.03 | 0.07 | -0.03 | 0.52 |
| Insl6 | -1.19 | 6.60E-08 | 1.37E-03 | 0.04 | 0.07 | -0.04 | 0.53 |
| Gxylt2 | -1.19 | 7.41E-25 | 1.54E-20 | 0.1 | 0.2 | -0.1 | 0.48 |
| Egr1 | -1.19 | 1.46E-08 | 3.04E-04 | 0.09 | 0.14 | -0.05 | 0.64 |
| Ankrd29 | -1.19 | 3.44E-07 | 7.15E-03 | 0.03 | 0.06 | -0.03 | 0.5 |
| Klhl29 | -1.19 | 5.10E-07 | 1.06E-02 | 0.03 | 0.07 | -0.03 | 0.52 |
| Ptch2 | -1.2 | 1.01E-06 | 2.09E-02 | 0.05 | 0.09 | -0.04 | 0.6 |
| Fgfr2 | -1.2 | 7.68E-57 | 1.60E-52 | 0.42 | 0.6 | -0.18 | 0.7 |
| Cpne8 | -1.21 | 1.62E-11 | 3.37E-07 | 0.07 | 0.13 | -0.06 | 0.56 |
| Meis1 | -1.22 | 1.67E-101 | 3.48E-97 | 0.52 | 0.73 | -0.21 | 0.71 |
| Timp3 | -1.23 | 4.05E-12 | 8.43E-08 | 0.07 | 0.13 | -0.06 | 0.54 |
| Aldh1a3 | -1.23 | 3.40E-30 | 7.07E-26 | 0.56 | 0.63 | -0.08 | 0.88 |
| Myl9 | -1.23 | 1.80E-19 | 3.75E-15 | 0.19 | 0.29 | -0.11 | 0.64 |
| Btg2 | -1.24 | 7.47E-15 | 1.55E-10 | 0.26 | 0.35 | -0.09 | 0.75 |
| Smad9 | -1.25 | 1.35E-09 | 2.82E-05 | 0.03 | 0.07 | -0.04 | 0.44 |
| Ppargc1a | -1.26 | 5.73E-11 | 1.19E-06 | 0.04 | 0.08 | -0.04 | 0.46 |
| Zfp365 | -1.26 | 9.38E-08 | 1.95E-03 | 0.03 | 0.06 | -0.03 | 0.48 |
| Rasgef1b | -1.26 | 1.15E-14 | 2.40E-10 | 0.07 | 0.14 | -0.07 | 0.51 |
| Eno3 | -1.27 | 1.90E-19 | 3.95E-15 | 0.19 | 0.29 | -0.1 | 0.65 |
| Gm32585 | -1.27 | 1.26E-43 | 2.63E-39 | 0.16 | 0.33 | -0.17 | 0.48 |
| 1700001K23Rik | -1.28 | 4.24E-08 | 8.81E-04 | 0.03 | 0.06 | -0.03 | 0.48 |
| Nt5e | -1.29 | 6.74E-13 | 1.40E-08 | 0.02 | 0.06 | -0.04 | 0.33 |
| Pou3f1 | -1.31 | 4.19E-07 | 8.72E-03 | 0.03 | 0.06 | -0.03 | 0.5 |
| Gm16141 | -1.31 | 1.06E-17 | 2.21E-13 | 0.04 | 0.1 | -0.06 | 0.38 |
| Pfkp | -1.32 | 5.90E-09 | 1.23E-04 | 0.04 | 0.07 | -0.04 | 0.48 |
| Crnde | -1.33 | 1.07E-19 | 2.23E-15 | 0.09 | 0.17 | -0.08 | 0.5 |
| Tent5a | -1.37 | 1.08E-12 | 2.24E-08 | 0.05 | 0.1 | -0.05 | 0.48 |
| Maob | -1.37 | 8.60E-13 | 1.79E-08 | 0.04 | 0.09 | -0.05 | 0.45 |
| Gap43 | -1.37 | 9.20E-16 | 1.91E-11 | 0.09 | 0.16 | -0.07 | 0.54 |
| Fam222a | -1.37 | 1.35E-10 | 2.82E-06 | 0.03 | 0.06 | -0.04 | 0.4 |
| Tmem59l | -1.37 | 3.94E-22 | 8.19E-18 | 0.08 | 0.18 | -0.09 | 0.47 |
| Rgs6 | -1.39 | 2.42E-08 | 5.02E-04 | 0.02 | 0.05 | -0.03 | 0.42 |
| Tubg2 | -1.39 | 1.31E-06 | 2.72E-02 | 0.03 | 0.07 | -0.03 | 0.52 |
| Tnfrsf19 | -1.39 | 3.25E-60 | 6.75E-56 | 0.18 | 0.39 | -0.21 | 0.47 |

**Supplementary Table 5. Differentially expressed genes in the mutant λ-junction ectoderm cells.**

| | | | | | | | |
|---|---|---|---|---|---|---|---|
| Cthrc1 | -1.4 | 1.89E-20 | 3.94E-16 | 0.1 | 0.19 | -0.09 | 0.52 |
| Wnt9b | -1.4 | 1.53E-34 | 3.19E-30 | 0.08 | 0.2 | -0.12 | 0.39 |
| Eya2 | -1.41 | 3.51E-86 | 7.31E-82 | 0.27 | 0.54 | -0.27 | 0.5 |
| Pid1 | -1.42 | 3.13E-18 | 6.51E-14 | 0.06 | 0.13 | -0.07 | 0.46 |
| Lmo1 | -1.43 | 3.48E-25 | 7.23E-21 | 0.18 | 0.3 | -0.12 | 0.61 |
| Hpse2 | -1.44 | 2.76E-09 | 5.74E-05 | 0.02 | 0.06 | -0.03 | 0.42 |
| Camk4 | -1.45 | 3.61E-14 | 7.52E-10 | 0.03 | 0.08 | -0.05 | 0.37 |
| Dclre1c | -1.46 | 3.72E-19 | 7.73E-15 | 0.08 | 0.16 | -0.08 | 0.48 |
| Jag2 | -1.46 | 3.77E-81 | 7.85E-77 | 0.19 | 0.43 | -0.25 | 0.43 |
| Etaa1os | -1.46 | 5.39E-24 | 1.12E-19 | 0.06 | 0.15 | -0.09 | 0.42 |
| 8430419K02Rik | -1.47 | 7.98E-10 | 1.66E-05 | 0.02 | 0.05 | -0.03 | 0.37 |
| Ptger4 | -1.48 | 3.49E-10 | 7.25E-06 | 0.02 | 0.05 | -0.03 | 0.36 |
| Tspan18 | -1.48 | 2.24E-11 | 4.66E-07 | 0.04 | 0.08 | -0.04 | 0.45 |
| Sgk3 | -1.5 | 1.17E-37 | 2.43E-33 | 0.1 | 0.23 | -0.13 | 0.41 |
| Gm2694 | -1.51 | 1.04E-15 | 2.16E-11 | 0.04 | 0.1 | -0.06 | 0.41 |
| Wnt6 | -1.52 | 6.19E-68 | 1.29E-63 | 0.3 | 0.52 | -0.22 | 0.58 |
| Tfap2b | -1.53 | 7.55E-33 | 1.57E-28 | 0.17 | 0.3 | -0.13 | 0.56 |
| Spock3 | -1.54 | 1.82E-08 | 3.80E-04 | 0.04 | 0.07 | -0.04 | 0.49 |
| Cxcl14 | -1.55 | 1.33E-57 | 2.78E-53 | 0.16 | 0.35 | -0.2 | 0.44 |
| Jakmip2 | -1.56 | 1.68E-14 | 3.49E-10 | 0.04 | 0.1 | -0.06 | 0.43 |
| Has3 | -1.57 | 1.03E-12 | 2.15E-08 | 0.02 | 0.06 | -0.04 | 0.34 |
| Arhgap22 | -1.58 | 7.97E-20 | 1.66E-15 | 0.03 | 0.08 | -0.06 | 0.3 |
| Tubb3 | -1.6 | 2.26E-19 | 4.70E-15 | 0.09 | 0.18 | -0.09 | 0.51 |
| Lgr6 | -1.63 | 2.28E-11 | 4.75E-07 | 0.02 | 0.06 | -0.04 | 0.36 |
| Trp73 | -1.64 | 1.40E-12 | 2.91E-08 | 0.02 | 0.05 | -0.04 | 0.32 |
| 2610204G07Rik | -1.65 | 7.20E-13 | 1.50E-08 | 0.02 | 0.06 | -0.04 | 0.33 |
| Lmcd1 | -1.65 | 1.75E-17 | 3.64E-13 | 0.05 | 0.12 | -0.07 | 0.43 |
| Tnnt1 | -1.67 | 8.77E-19 | 1.82E-14 | 0.06 | 0.14 | -0.07 | 0.46 |
| Rnase4 | -1.67 | 3.20E-37 | 6.65E-33 | 0.08 | 0.2 | -0.12 | 0.38 |
| Sp5 | -1.67 | 2.84E-07 | 5.91E-03 | 0.03 | 0.06 | -0.03 | 0.51 |
| Ebf1 | -1.71 | 4.29E-33 | 8.92E-29 | 0.12 | 0.24 | -0.13 | 0.48 |
| Kremen2 | -1.71 | 3.01E-46 | 6.27E-42 | 0.13 | 0.29 | -0.16 | 0.45 |
| Dcx | -1.71 | 2.72E-13 | 5.67E-09 | 0.04 | 0.09 | -0.05 | 0.43 |
| Lfng | -1.71 | 1.69E-17 | 3.51E-13 | 0.03 | 0.08 | -0.05 | 0.34 |
| Fgf17 | -1.72 | 7.41E-21 | 1.54E-16 | 0.13 | 0.23 | -0.1 | 0.57 |
| Sema3c | -1.74 | 5.80E-74 | 1.21E-69 | 0.17 | 0.4 | -0.23 | 0.44 |
| Ror1 | -1.75 | 2.78E-12 | 5.77E-08 | 0.03 | 0.07 | -0.04 | 0.38 |
| Kitl | -1.79 | 8.05E-40 | 1.68E-35 | 0.1 | 0.23 | -0.13 | 0.42 |
| Phlda2 | -1.79 | 3.18E-16 | 6.62E-12 | 0.03 | 0.09 | -0.06 | 0.38 |
| Gsn | -1.8 | 1.85E-29 | 3.84E-25 | 0.06 | 0.16 | -0.1 | 0.39 |
| Sox14 | -1.86 | 9.28E-24 | 1.93E-19 | 0.03 | 0.1 | -0.07 | 0.31 |
| Bmp2 | -1.86 | 3.15E-07 | 6.56E-03 | 0.03 | 0.06 | -0.03 | 0.48 |
| Wnt7b | -1.89 | 1.67E-43 | 3.48E-39 | 0.12 | 0.27 | -0.15 | 0.45 |
| Cnr1 | -1.9 | 6.89E-23 | 1.43E-18 | 0.05 | 0.13 | -0.08 | 0.39 |
| Tspan2 | -1.91 | 9.17E-25 | 1.91E-20 | 0.04 | 0.11 | -0.07 | 0.32 |
| Pcp4l1 | -1.92 | 4.92E-40 | 1.02E-35 | 0.04 | 0.15 | -0.11 | 0.28 |
| Man1a | -1.94 | 1.34E-24 | 2.79E-20 | 0.03 | 0.1 | -0.07 | 0.32 |
| Wnt3 | -1.95 | 1.38E-53 | 2.86E-49 | 0.07 | 0.22 | -0.15 | 0.32 |
| Lgals7 | -2.01 | 1.82E-25 | 3.78E-21 | 0.03 | 0.1 | -0.07 | 0.29 |

**Supplementary Table 5. Differentially expressed genes in the mutant λ-junction ectoderm cells.**

| | | | | | | | |
|---|---|---|---|---|---|---|---|
| Pthlh | -2.09 | 2.31E-18 | 4.80E-14 | 0.03 | 0.09 | -0.06 | 0.37 |
| Thsd7b | -2.14 | 2.89E-15 | 6.00E-11 | 0.03 | 0.07 | -0.05 | 0.35 |
| Brinp1 | -2.16 | 7.58E-19 | 1.58E-14 | 0.02 | 0.07 | -0.05 | 0.26 |
| Scn3a | -2.18 | 1.58E-19 | 3.29E-15 | 0.01 | 0.05 | -0.04 | 0.2 |
| Syndig1 | -2.19 | 2.59E-20 | 5.39E-16 | 0.03 | 0.09 | -0.06 | 0.33 |
| Scn9a | -2.24 | 8.38E-13 | 1.74E-08 | 0.02 | 0.05 | -0.04 | 0.31 |
| Dkk4 | -2.29 | 5.17E-14 | 1.07E-09 | 0.02 | 0.05 | -0.04 | 0.28 |
| Dcn | -2.32 | 1.79E-17 | 3.72E-13 | 0.02 | 0.06 | -0.04 | 0.26 |
| Bmerb1 | -2.33 | 1.99E-22 | 4.14E-18 | 0.02 | 0.08 | -0.06 | 0.26 |
| Gria2 | -2.34 | 1.23E-35 | 2.55E-31 | 0.05 | 0.15 | -0.1 | 0.31 |
| Pcp4 | -2.35 | 1.64E-21 | 3.42E-17 | 0.03 | 0.09 | -0.06 | 0.31 |
| Edaradd | -2.37 | 3.59E-62 | 7.47E-58 | 0.04 | 0.18 | -0.14 | 0.21 |
| Tac2 | -2.41 | 2.73E-11 | 5.68E-07 | 0.03 | 0.06 | -0.04 | 0.4 |
| Rxfp1 | -2.41 | 1.58E-25 | 3.30E-21 | 0.03 | 0.1 | -0.07 | 0.29 |
| Tagln | -2.43 | 9.08E-15 | 1.89E-10 | 0.04 | 0.09 | -0.06 | 0.41 |
| Sostdc1 | -2.54 | 4.23E-74 | 8.80E-70 | 0.2 | 0.41 | -0.21 | 0.49 |
| Unc13c | -2.55 | 1.44E-55 | 2.99E-51 | 0.03 | 0.15 | -0.12 | 0.19 |
| Mgp | -2.63 | 1.47E-30 | 3.07E-26 | 0.04 | 0.13 | -0.09 | 0.33 |
| Krt15 | -2.63 | 5.48E-94 | 1.14E-89 | 0.08 | 0.31 | -0.22 | 0.27 |
| Bnc1 | -2.65 | 5.51E-37 | 1.14E-32 | 0.02 | 0.09 | -0.08 | 0.16 |
| Cbr2 | -2.7 | 5.06E-33 | 1.05E-28 | 0.02 | 0.09 | -0.07 | 0.19 |
| Tmeff2 | -2.76 | 2.82E-29 | 5.87E-25 | 0.02 | 0.09 | -0.07 | 0.24 |
| Lmx1b | -2.77 | 5.61E-26 | 1.17E-21 | 0.01 | 0.06 | -0.05 | 0.15 |
| Npr3 | -2.91 | 4.64E-48 | 9.64E-44 | 0.02 | 0.12 | -0.1 | 0.16 |
| C130021I20Rik | -3.36 | 4.18E-36 | 8.70E-32 | 0.01 | 0.07 | -0.06 | 0.1 |
| Ascl1 | -4.06 | 2.80E-28 | 5.82E-24 | 0.01 | 0.06 | -0.05 | 0.12 |
| Dlk2 | -4.16 | 2.57E-40 | 5.35E-36 | 0.01 | 0.07 | -0.07 | 0.08 |
| Wnt10a | -4.64 | 6.41E-54 | 1.33E-49 | 0 | 0.08 | -0.08 | 0.04 |
| Insm1 | -5.04 | 3.00E-37 | 6.25E-33 | 0 | 0.06 | -0.06 | 0.05 |
| Rprml | -5.11 | 4.32E-86 | 8.99E-82 | 0.01 | 0.14 | -0.13 | 0.05 |
| Irx4 | -5.87 | 7.92E-44 | 1.65E-39 | 0 | 0.06 | -0.06 | 0.02 |
| Uty | -7.85 | 8.05E-73 | 1.67E-68 | 0 | 0.1 | -0.1 | 0 |
| Kdm5d | -7.87 | 1.37E-75 | 2.84E-71 | 0 | 0.1 | -0.1 | 0 |
| Ddx3y | -8.58 | 8.95E-111 | 1.86E-106 | 0 | 0.15 | -0.15 | 0 |
| Eif2s3y | -8.84 | 3.20E-123 | 6.65E-119 | 0 | 0.16 | -0.16 | 0 |

**Supplementary Table 6. Differentially expressed genes in the mutant λ-junction mesenchymal cells.**

| gene | avg_log2FC | p_val | p_val_adj | pct.1 | pct.2 | pct1-pct2 | pct1/pct2 |
|---|---|---|---|---|---|---|---|
| Hes5 | 5.68 | 1.83E-94 | 3.80E-90 | 0.06 | 0 | 0.06 | 62 |
| Ebf2 | 2.25 | 3.45E-47 | 7.18E-43 | 0.07 | 0.02 | 0.05 | 3.43 |
| Irx1 | 2.1 | 3.60E-54 | 7.49E-50 | 0.08 | 0.03 | 0.06 | 3.36 |
| Nefl | 1.67 | 4.77E-22 | 9.92E-18 | 0.06 | 0.03 | 0.03 | 2.32 |
| Dsp | 1.65 | 2.36E-26 | 4.91E-22 | 0.06 | 0.02 | 0.04 | 2.75 |
| Penk | 1.56 | 2.89E-17 | 6.02E-13 | 0.14 | 0.09 | 0.04 | 1.47 |
| Ebf1 | 1.55 | 1.85E-162 | 3.85E-158 | 0.46 | 0.25 | 0.2 | 1.79 |
| Aldh1a2 | 1.55 | 8.33E-58 | 1.73E-53 | 0.13 | 0.06 | 0.08 | 2.42 |
| Slc9a3r1 | 1.53 | 7.52E-32 | 1.56E-27 | 0.09 | 0.04 | 0.05 | 2.23 |
| Igfbp5 | 1.48 | 9.11E-81 | 1.89E-76 | 0.3 | 0.18 | 0.13 | 1.72 |
| Barx1 | 1.47 | 6.12E-132 | 1.27E-127 | 0.36 | 0.18 | 0.18 | 1.98 |
| Tbx22 | 1.47 | 6.70E-101 | 1.39E-96 | 0.22 | 0.09 | 0.13 | 2.41 |
| Cobl | 1.45 | 3.80E-22 | 7.91E-18 | 0.06 | 0.03 | 0.03 | 2.32 |
| Tfap2b | 1.45 | 2.76E-16 | 5.74E-12 | 0.07 | 0.04 | 0.03 | 1.82 |
| Foxd2os | 1.38 | 6.29E-88 | 1.31E-83 | 0.27 | 0.14 | 0.13 | 1.91 |
| Foxd2 | 1.28 | 3.10E-63 | 6.45E-59 | 0.21 | 0.11 | 0.1 | 1.95 |
| Cldn11 | 1.27 | 4.51E-65 | 9.37E-61 | 0.2 | 0.1 | 0.1 | 2.05 |
| Cdc42ep3 | 1.24 | 4.95E-45 | 1.03E-40 | 0.17 | 0.09 | 0.08 | 1.88 |
| Itm2a | 1.23 | 7.76E-196 | 1.61E-191 | 0.68 | 0.49 | 0.19 | 1.38 |
| Dlx2 | 1.21 | 1.12E-25 | 2.34E-21 | 0.08 | 0.04 | 0.04 | 2.08 |
| Foxc2 | 1.21 | 8.72E-63 | 1.81E-58 | 0.2 | 0.1 | 0.1 | 1.97 |
| Lhx6 | 1.19 | 1.67E-37 | 3.47E-33 | 0.11 | 0.05 | 0.06 | 2.16 |
| Sfn | 1.16 | 4.64E-24 | 9.64E-20 | 0.06 | 0.03 | 0.04 | 2.29 |
| Foxc1 | 1.15 | 1.00E-135 | 2.09E-131 | 0.44 | 0.26 | 0.18 | 1.71 |
| Pkdcc | 1.14 | 1.01E-109 | 2.10E-105 | 0.44 | 0.28 | 0.16 | 1.58 |
| Scx | 1.13 | 1.27E-76 | 2.64E-72 | 0.28 | 0.15 | 0.12 | 1.81 |
| Osr1 | 1.11 | 8.43E-28 | 1.75E-23 | 0.12 | 0.07 | 0.05 | 1.78 |
| Hic1 | 1.1 | 1.94E-13 | 4.04E-09 | 0.07 | 0.04 | 0.03 | 1.66 |
| Nrp1 | 1.08 | 1.54E-59 | 3.21E-55 | 0.31 | 0.2 | 0.11 | 1.55 |
| Slc1a3 | 1.08 | 4.60E-70 | 9.56E-66 | 0.31 | 0.19 | 0.12 | 1.63 |
| Cthrc1 | 1.08 | 1.64E-48 | 3.41E-44 | 0.28 | 0.19 | 0.09 | 1.5 |
| Islr | 1.05 | 5.25E-29 | 1.09E-24 | 0.14 | 0.08 | 0.06 | 1.7 |
| Cdh15 | 1.03 | 7.46E-16 | 1.55E-11 | 0.07 | 0.04 | 0.03 | 1.86 |
| Cpm | 1.02 | 2.34E-20 | 4.87E-16 | 0.09 | 0.05 | 0.04 | 1.83 |
| Col9a1 | 1.02 | 5.28E-106 | 1.10E-101 | 0.46 | 0.3 | 0.16 | 1.54 |
| Pcdh11x | 1 | 2.44E-33 | 5.08E-29 | 0.14 | 0.08 | 0.06 | 1.82 |
| Hist3h2ba | 0.99 | 1.83E-18 | 3.81E-14 | 0.12 | 0.08 | 0.04 | 1.53 |
| Asb4 | 0.98 | 3.39E-90 | 7.06E-86 | 0.46 | 0.31 | 0.14 | 1.46 |
| Krt8 | 0.97 | 7.45E-18 | 1.55E-13 | 0.1 | 0.06 | 0.04 | 1.67 |
| Ebf3 | 0.95 | 1.05E-31 | 2.18E-27 | 0.25 | 0.18 | 0.07 | 1.4 |
| Thbs1 | 0.94 | 2.26E-11 | 4.70E-07 | 0.11 | 0.07 | 0.03 | 1.42 |
| Foxp2 | 0.93 | 2.28E-109 | 4.75E-105 | 0.5 | 0.33 | 0.17 | 1.52 |
| Nr2f1 | 0.91 | 1.35E-111 | 2.80E-107 | 0.48 | 0.31 | 0.17 | 1.55 |
| Spint2 | 0.91 | 9.93E-22 | 2.06E-17 | 0.13 | 0.08 | 0.05 | 1.59 |
| Erbb3 | 0.88 | 3.08E-18 | 6.40E-14 | 0.08 | 0.05 | 0.04 | 1.79 |
| AI314278 | 0.88 | 4.51E-15 | 9.38E-11 | 0.09 | 0.05 | 0.03 | 1.63 |
| Mgp | 0.88 | 1.76E-19 | 3.66E-15 | 0.1 | 0.06 | 0.04 | 1.68 |

**Supplementary Table 6. Differentially expressed genes in the mutant λ-junction mesenchymal cells.**

| | | | | | | | |
|---|---|---|---|---|---|---|---|
| Cldn7 | 0.88 | 1.32E-12 | 2.75E-08 | 0.06 | 0.04 | 0.03 | 1.72 |
| Shisa2 | 0.86 | 1.63E-22 | 3.39E-18 | 0.15 | 0.09 | 0.05 | 1.57 |
| Dab1 | 0.86 | 1.67E-14 | 3.47E-10 | 0.07 | 0.04 | 0.03 | 1.78 |
| Fzd4 | 0.85 | 2.80E-63 | 5.83E-59 | 0.34 | 0.22 | 0.12 | 1.54 |
| Prkg2 | 0.85 | 4.41E-13 | 9.16E-09 | 0.08 | 0.05 | 0.03 | 1.6 |
| Lypd6 | 0.85 | 2.41E-15 | 5.02E-11 | 0.13 | 0.09 | 0.04 | 1.44 |
| Epcam | 0.84 | 1.97E-25 | 4.09E-21 | 0.17 | 0.11 | 0.06 | 1.54 |
| Frem2 | 0.84 | 2.24E-12 | 4.65E-08 | 0.08 | 0.05 | 0.03 | 1.56 |
| Sfrp2 | 0.84 | 4.29E-40 | 8.93E-36 | 0.7 | 0.68 | 0.02 | 1.03 |
| Egfr | 0.83 | 3.04E-13 | 6.32E-09 | 0.08 | 0.05 | 0.03 | 1.6 |
| Sox9 | 0.83 | 9.66E-173 | 2.01E-168 | 0.73 | 0.54 | 0.19 | 1.35 |
| Cdh3 | 0.81 | 3.39E-17 | 7.05E-13 | 0.13 | 0.09 | 0.04 | 1.49 |
| Bicd1 | 0.8 | 7.31E-08 | 1.52E-03 | 0.06 | 0.04 | 0.02 | 1.46 |
| Rhoj | 0.8 | 1.24E-24 | 2.58E-20 | 0.18 | 0.12 | 0.06 | 1.51 |
| Ngfr | 0.79 | 3.02E-09 | 6.27E-05 | 0.07 | 0.05 | 0.02 | 1.51 |
| Nr2f2 | 0.79 | 1.69E-56 | 3.51E-52 | 0.36 | 0.25 | 0.11 | 1.45 |
| Gm266 | 0.78 | 5.87E-45 | 1.22E-40 | 0.3 | 0.2 | 0.1 | 1.49 |
| A830082K12Rik | 0.78 | 1.80E-14 | 3.74E-10 | 0.1 | 0.07 | 0.04 | 1.53 |
| Pltp | 0.78 | 1.44E-07 | 2.99E-03 | 0.07 | 0.05 | 0.02 | 1.4 |
| Notch1 | 0.78 | 3.95E-22 | 8.22E-18 | 0.15 | 0.1 | 0.05 | 1.55 |
| Tgfbi | 0.78 | 1.66E-53 | 3.45E-49 | 0.44 | 0.34 | 0.1 | 1.29 |
| Pcp2 | 0.78 | 8.81E-11 | 1.83E-06 | 0.08 | 0.05 | 0.03 | 1.52 |
| Tmem37 | 0.77 | 2.58E-16 | 5.36E-12 | 0.1 | 0.07 | 0.04 | 1.58 |
| Krt18 | 0.76 | 6.91E-22 | 1.44E-17 | 0.14 | 0.09 | 0.05 | 1.58 |
| Sned1 | 0.76 | 7.50E-27 | 1.56E-22 | 0.21 | 0.14 | 0.07 | 1.46 |
| Adra2a | 0.75 | 4.74E-11 | 9.86E-07 | 0.06 | 0.04 | 0.02 | 1.63 |
| Col2a1 | 0.73 | 6.24E-123 | 1.30E-118 | 0.71 | 0.58 | 0.13 | 1.22 |
| Mecom | 0.73 | 1.11E-88 | 2.31E-84 | 0.49 | 0.34 | 0.16 | 1.46 |
| Tmem238 | 0.73 | 3.60E-09 | 7.49E-05 | 0.09 | 0.06 | 0.03 | 1.41 |
| Ak1 | 0.72 | 1.93E-11 | 4.02E-07 | 0.11 | 0.08 | 0.03 | 1.43 |
| Pon2 | 0.71 | 2.61E-12 | 5.44E-08 | 0.12 | 0.08 | 0.04 | 1.43 |
| Crym | 0.71 | 4.27E-93 | 8.87E-89 | 0.66 | 0.52 | 0.14 | 1.27 |
| Mapk12 | 0.7 | 5.29E-09 | 1.10E-04 | 0.06 | 0.04 | 0.02 | 1.52 |
| Ptx3 | 0.7 | 5.86E-14 | 1.22E-09 | 0.17 | 0.13 | 0.04 | 1.32 |
| Stc1 | 0.69 | 1.86E-07 | 3.86E-03 | 0.08 | 0.05 | 0.02 | 1.41 |
| Sec24d | 0.69 | 1.03E-12 | 2.14E-08 | 0.11 | 0.08 | 0.03 | 1.43 |
| Morc4 | 0.69 | 3.39E-18 | 7.06E-14 | 0.18 | 0.13 | 0.05 | 1.4 |
| Eya2 | 0.68 | 1.39E-21 | 2.89E-17 | 0.23 | 0.17 | 0.06 | 1.34 |
| Six3 | 0.68 | 3.85E-07 | 8.00E-03 | 0.05 | 0.03 | 0.02 | 1.52 |
| Zfp536 | 0.68 | 5.06E-08 | 1.05E-03 | 0.1 | 0.07 | 0.03 | 1.34 |
| Synpo2 | 0.68 | 1.33E-14 | 2.77E-10 | 0.12 | 0.08 | 0.04 | 1.49 |
| Apoe | 0.67 | 5.18E-35 | 1.08E-30 | 0.46 | 0.39 | 0.07 | 1.19 |
| Trf | 0.67 | 1.25E-06 | 2.60E-02 | 0.06 | 0.04 | 0.02 | 1.44 |
| Cxcl12 | 0.66 | 2.41E-16 | 5.02E-12 | 0.22 | 0.17 | 0.05 | 1.29 |
| Shox2 | 0.65 | 2.43E-16 | 5.06E-12 | 0.2 | 0.15 | 0.05 | 1.34 |
| Cited1 | 0.65 | 3.76E-13 | 7.82E-09 | 0.18 | 0.14 | 0.04 | 1.3 |
| Lrig3 | 0.65 | 7.06E-21 | 1.47E-16 | 0.25 | 0.19 | 0.06 | 1.31 |
| Adcy4 | 0.65 | 1.30E-09 | 2.71E-05 | 0.09 | 0.06 | 0.03 | 1.44 |

**Supplementary Table 6. Differentially expressed genes in the mutant λ-junction mesenchymal cells.**

| | | | | | | | |
|---|---|---|---|---|---|---|---|
| Tcf15 | 0.65 | 9.76E-37 | 2.03E-32 | 0.41 | 0.32 | 0.09 | 1.28 |
| Foxd1 | 0.64 | 1.04E-25 | 2.16E-21 | 0.18 | 0.12 | 0.06 | 1.53 |
| Mdfic | 0.64 | 2.08E-20 | 4.33E-16 | 0.27 | 0.21 | 0.06 | 1.28 |
| Dmd | 0.64 | 1.32E-06 | 2.76E-02 | 0.08 | 0.06 | 0.02 | 1.33 |
| Npnt | 0.64 | 2.95E-18 | 6.14E-14 | 0.21 | 0.16 | 0.05 | 1.33 |
| Loxl1 | 0.63 | 3.11E-07 | 6.47E-03 | 0.1 | 0.08 | 0.02 | 1.31 |
| Gm21887 | 0.63 | 6.42E-14 | 1.33E-09 | 0.16 | 0.12 | 0.04 | 1.34 |
| B4galt4 | 0.63 | 2.44E-18 | 5.07E-14 | 0.25 | 0.19 | 0.05 | 1.27 |
| Foxp1 | 0.63 | 4.20E-87 | 8.73E-83 | 0.61 | 0.46 | 0.15 | 1.32 |
| Kcnk1 | 0.62 | 9.72E-07 | 2.02E-02 | 0.08 | 0.06 | 0.02 | 1.37 |
| Dlk1 | 0.6 | 1.56E-55 | 3.24E-51 | 0.43 | 0.31 | 0.12 | 1.39 |
| Amt | 0.6 | 2.63E-12 | 5.48E-08 | 0.15 | 0.11 | 0.04 | 1.35 |
| Adamts9 | 0.6 | 1.73E-09 | 3.60E-05 | 0.13 | 0.1 | 0.03 | 1.3 |
| Gm13429 | 0.6 | 9.67E-09 | 2.01E-04 | 0.09 | 0.06 | 0.03 | 1.41 |
| Flrt2 | 0.59 | 5.14E-73 | 1.07E-68 | 0.6 | 0.48 | 0.12 | 1.25 |
| Col9a2 | 0.59 | 1.25E-09 | 2.60E-05 | 0.12 | 0.09 | 0.03 | 1.34 |
| Pcdh7 | 0.59 | 2.07E-45 | 4.31E-41 | 0.39 | 0.28 | 0.11 | 1.38 |
| Gsc | 0.59 | 7.95E-62 | 1.65E-57 | 0.46 | 0.33 | 0.14 | 1.42 |
| Igdcc3 | 0.58 | 9.73E-16 | 2.02E-11 | 0.25 | 0.2 | 0.05 | 1.24 |
| Map1b | 0.58 | 4.96E-49 | 1.03E-44 | 0.71 | 0.64 | 0.06 | 1.1 |
| Six2 | 0.57 | 1.82E-14 | 3.78E-10 | 0.47 | 0.43 | 0.03 | 1.08 |
| Rragd | 0.57 | 3.07E-13 | 6.39E-09 | 0.18 | 0.14 | 0.04 | 1.32 |
| Prrx1 | 0.57 | 3.22E-147 | 6.70E-143 | 0.89 | 0.8 | 0.1 | 1.12 |
| Slitrk5 | 0.57 | 2.46E-19 | 5.12E-15 | 0.23 | 0.18 | 0.06 | 1.33 |
| Dusp6 | 0.57 | 6.05E-43 | 1.26E-38 | 0.44 | 0.35 | 0.1 | 1.28 |
| Dtna | 0.57 | 3.41E-08 | 7.10E-04 | 0.1 | 0.07 | 0.03 | 1.36 |
| Spred3 | 0.56 | 8.19E-07 | 1.70E-02 | 0.1 | 0.07 | 0.02 | 1.3 |
| Ntng1 | 0.56 | 3.13E-08 | 6.50E-04 | 0.17 | 0.14 | 0.03 | 1.22 |
| E130114P18Rik | 0.56 | 3.19E-35 | 6.63E-31 | 0.34 | 0.26 | 0.09 | 1.34 |
| Lhfp | 0.55 | 5.24E-13 | 1.09E-08 | 0.24 | 0.2 | 0.04 | 1.21 |
| Prss23 | 0.55 | 3.23E-35 | 6.72E-31 | 0.38 | 0.29 | 0.09 | 1.32 |
| 2700033N17Rik | 0.55 | 2.46E-08 | 5.12E-04 | 0.1 | 0.08 | 0.03 | 1.35 |
| Hs6st2 | 0.55 | 3.44E-19 | 7.15E-15 | 0.25 | 0.19 | 0.06 | 1.3 |
| Itga4 | 0.54 | 3.50E-13 | 7.28E-09 | 0.27 | 0.22 | 0.05 | 1.2 |
| Spry2 | 0.53 | 1.90E-17 | 3.95E-13 | 0.34 | 0.29 | 0.05 | 1.17 |
| Foxf1 | 0.53 | 2.52E-27 | 5.25E-23 | 0.29 | 0.22 | 0.08 | 1.36 |
| Lgr4 | 0.53 | 1.34E-10 | 2.78E-06 | 0.17 | 0.14 | 0.04 | 1.27 |
| Klk8 | 0.52 | 3.24E-07 | 6.74E-03 | 0.1 | 0.08 | 0.02 | 1.31 |
| Meis2 | 0.52 | 9.67E-60 | 2.01E-55 | 0.67 | 0.58 | 0.09 | 1.15 |
| Mafb | 0.52 | 1.17E-11 | 2.43E-07 | 0.27 | 0.23 | 0.04 | 1.19 |
| Ezr | 0.52 | 4.11E-08 | 8.56E-04 | 0.12 | 0.09 | 0.03 | 1.3 |
| Slitrk4 | 0.52 | 7.75E-07 | 1.61E-02 | 0.08 | 0.06 | 0.02 | 1.36 |
| Arhgap31 | 0.5 | 3.10E-13 | 6.44E-09 | 0.25 | 0.2 | 0.04 | 1.22 |
| Tox | 0.5 | 9.75E-19 | 2.03E-14 | 0.31 | 0.26 | 0.06 | 1.23 |
| Hmcn1 | 0.5 | 4.20E-42 | 8.74E-38 | 0.7 | 0.64 | 0.06 | 1.09 |
| Wfdc2 | 0.5 | 3.02E-23 | 6.29E-19 | 0.41 | 0.34 | 0.06 | 1.18 |
| Stxbp6 | 0.49 | 2.87E-10 | 5.97E-06 | 0.14 | 0.11 | 0.03 | 1.31 |
| Angptl1 | 0.49 | 3.46E-29 | 7.19E-25 | 0.35 | 0.27 | 0.08 | 1.31 |

**Supplementary Table 6. Differentially expressed genes in the mutant λ-junction mesenchymal cells.**

| | | | | | | | |
|---|---|---|---|---|---|---|---|
| Etv5 | 0.49 | 1.84E-30 | 3.82E-26 | 0.4 | 0.32 | 0.08 | 1.25 |
| Lrrc8d | 0.48 | 9.88E-08 | 2.06E-03 | 0.18 | 0.15 | 0.03 | 1.2 |
| Isl2 | 0.48 | 4.56E-09 | 9.49E-05 | 0.15 | 0.12 | 0.03 | 1.28 |
| Lin28b | 0.48 | 1.53E-10 | 3.19E-06 | 0.2 | 0.17 | 0.04 | 1.23 |
| Selenop | 0.47 | 1.36E-17 | 2.83E-13 | 0.25 | 0.19 | 0.06 | 1.29 |
| 2610002M06Rik | 0.47 | 3.33E-15 | 6.92E-11 | 0.29 | 0.25 | 0.05 | 1.2 |
| Nrk | 0.47 | 3.47E-09 | 7.22E-05 | 0.19 | 0.15 | 0.03 | 1.22 |
| Mgst1 | 0.46 | 4.16E-09 | 8.65E-05 | 0.31 | 0.28 | 0.03 | 1.11 |
| Spry1 | 0.46 | 3.00E-18 | 6.23E-14 | 0.31 | 0.26 | 0.06 | 1.22 |
| Map2 | 0.46 | 6.70E-08 | 1.39E-03 | 0.23 | 0.2 | 0.03 | 1.14 |
| Cygb | 0.45 | 3.60E-08 | 7.48E-04 | 0.13 | 0.1 | 0.03 | 1.28 |
| Etv4 | 0.45 | 6.38E-28 | 1.33E-23 | 0.46 | 0.39 | 0.07 | 1.17 |
| Asphd2 | 0.44 | 3.02E-08 | 6.27E-04 | 0.14 | 0.11 | 0.03 | 1.27 |
| Peg3 | 0.44 | 1.33E-89 | 2.76E-85 | 0.89 | 0.83 | 0.06 | 1.08 |
| Ccnd2 | 0.44 | 5.62E-90 | 1.17E-85 | 0.94 | 0.91 | 0.03 | 1.03 |
| Dkk2 | 0.44 | 6.17E-07 | 1.28E-02 | 0.36 | 0.34 | 0.02 | 1.07 |
| Mab21l1 | 0.44 | 3.65E-10 | 7.60E-06 | 0.24 | 0.2 | 0.04 | 1.2 |
| Bex4 | 0.43 | 1.85E-37 | 3.86E-33 | 0.6 | 0.54 | 0.07 | 1.13 |
| Gmds | 0.43 | 9.76E-16 | 2.03E-11 | 0.41 | 0.37 | 0.04 | 1.11 |
| Phlda1 | 0.42 | 1.60E-15 | 3.33E-11 | 0.46 | 0.42 | 0.04 | 1.11 |
| Slit2 | 0.42 | 2.48E-31 | 5.16E-27 | 0.43 | 0.34 | 0.09 | 1.26 |
| Rgs2 | 0.42 | 4.28E-15 | 8.91E-11 | 0.38 | 0.34 | 0.05 | 1.14 |
| Rprm | 0.41 | 3.80E-08 | 7.90E-04 | 0.11 | 0.08 | 0.03 | 1.34 |
| Cdh2 | 0.41 | 9.38E-12 | 1.95E-07 | 0.41 | 0.37 | 0.04 | 1.11 |
| Gm42047 | 0.4 | 1.05E-11 | 2.17E-07 | 0.18 | 0.13 | 0.04 | 1.32 |
| Fendrr | 0.4 | 1.96E-07 | 4.08E-03 | 0.15 | 0.12 | 0.03 | 1.25 |
| Ier2 | 0.4 | 4.68E-82 | 9.72E-78 | 0.83 | 0.77 | 0.06 | 1.08 |
| Foxp4 | 0.4 | 2.61E-16 | 5.42E-12 | 0.48 | 0.45 | 0.03 | 1.07 |
| Plpp3 | 0.4 | 1.46E-35 | 3.04E-31 | 0.78 | 0.76 | 0.02 | 1.03 |
| Frzb | 0.39 | 1.66E-35 | 3.46E-31 | 0.53 | 0.44 | 0.09 | 1.21 |
| Bex1 | 0.39 | 1.93E-28 | 4.01E-24 | 0.52 | 0.45 | 0.07 | 1.16 |
| Nhs | 0.39 | 6.47E-14 | 1.35E-09 | 0.34 | 0.3 | 0.05 | 1.16 |
| Hsd11b2 | 0.39 | 3.29E-07 | 6.85E-03 | 0.15 | 0.12 | 0.03 | 1.25 |
| Mov10 | 0.39 | 3.68E-08 | 7.65E-04 | 0.21 | 0.17 | 0.03 | 1.18 |
| Cdk6 | 0.38 | 1.17E-09 | 2.44E-05 | 0.31 | 0.28 | 0.04 | 1.13 |
| Cdkn1c | 0.38 | 1.15E-73 | 2.39E-69 | 0.99 | 0.99 | 0.01 | 1.01 |
| Sobp | 0.37 | 6.14E-19 | 1.28E-14 | 0.42 | 0.37 | 0.05 | 1.15 |
| Eya4 | 0.37 | 3.31E-17 | 6.88E-13 | 0.36 | 0.3 | 0.06 | 1.19 |
| App | 0.36 | 3.14E-53 | 6.54E-49 | 0.83 | 0.78 | 0.05 | 1.06 |
| Rbp1 | 0.36 | 1.26E-71 | 2.61E-67 | 0.96 | 0.94 | 0.02 | 1.02 |
| Col25a1 | 0.36 | 5.01E-09 | 1.04E-04 | 0.28 | 0.24 | 0.04 | 1.17 |
| Fli1 | 0.36 | 1.63E-20 | 3.39E-16 | 0.51 | 0.46 | 0.05 | 1.1 |
| Traf4 | 0.35 | 3.65E-39 | 7.58E-35 | 0.71 | 0.66 | 0.05 | 1.08 |
| Inka1 | 0.35 | 2.11E-31 | 4.38E-27 | 0.66 | 0.61 | 0.05 | 1.08 |
| Epha3 | 0.35 | 2.02E-26 | 4.19E-22 | 0.52 | 0.45 | 0.07 | 1.16 |
| Mn1 | 0.34 | 7.44E-10 | 1.55E-05 | 0.36 | 0.32 | 0.04 | 1.11 |
| Gkap1 | 0.34 | 1.51E-07 | 3.14E-03 | 0.37 | 0.35 | 0.02 | 1.07 |
| Slc24a5 | 0.34 | 8.04E-09 | 1.67E-04 | 0.41 | 0.39 | 0.02 | 1.06 |

**Supplementary Table 6. Differentially expressed genes in the mutant λ-junction mesenchymal cells.**

| | | | | | | | |
|---|---|---|---|---|---|---|---|
| Ccnd1 | 0.33 | 6.36E-58 | 1.32E-53 | 0.89 | 0.84 | 0.05 | 1.06 |
| Rab3il1 | 0.33 | 1.77E-08 | 3.68E-04 | 0.37 | 0.35 | 0.03 | 1.07 |
| Usp29 | 0.33 | 1.51E-07 | 3.13E-03 | 0.25 | 0.21 | 0.03 | 1.15 |
| Hmga1 | 0.32 | 1.33E-17 | 2.76E-13 | 0.51 | 0.47 | 0.04 | 1.09 |
| Fndc3c1 | 0.31 | 4.16E-18 | 8.65E-14 | 0.57 | 0.53 | 0.04 | 1.07 |
| Il11ra1 | 0.3 | 6.72E-28 | 1.40E-23 | 0.8 | 0.78 | 0.02 | 1.03 |
| Mrrf | 0.3 | 2.53E-09 | 5.27E-05 | 0.37 | 0.33 | 0.04 | 1.1 |
| Pbx3 | 0.3 | 3.79E-19 | 7.89E-15 | 0.55 | 0.5 | 0.05 | 1.09 |
| Kctd1 | 0.3 | 2.38E-14 | 4.95E-10 | 0.5 | 0.47 | 0.04 | 1.08 |
| Xist | 0.3 | 1.28E-75 | 2.67E-71 | 1 | 0.77 | 0.23 | 1.3 |
| Cdc42se1 | 0.29 | 3.51E-09 | 7.30E-05 | 0.38 | 0.35 | 0.03 | 1.09 |
| Bcl2l11 | 0.29 | 6.86E-09 | 1.43E-04 | 0.36 | 0.33 | 0.03 | 1.1 |
| Nectin2 | 0.29 | 1.09E-08 | 2.26E-04 | 0.47 | 0.44 | 0.03 | 1.06 |
| Fkbp11 | 0.29 | 1.69E-06 | 3.51E-02 | 0.36 | 0.34 | 0.02 | 1.06 |
| Osr2 | 0.29 | 9.30E-09 | 1.93E-04 | 0.13 | 0.1 | 0.03 | 1.3 |
| Nrip1 | 0.28 | 9.04E-27 | 1.88E-22 | 0.77 | 0.74 | 0.03 | 1.04 |
| Amot | 0.28 | 2.23E-07 | 4.64E-03 | 0.48 | 0.46 | 0.01 | 1.03 |
| Cited2 | 0.27 | 5.75E-17 | 1.20E-12 | 0.65 | 0.62 | 0.03 | 1.06 |
| Ssbp4 | 0.27 | 3.43E-26 | 7.14E-22 | 0.74 | 0.71 | 0.03 | 1.04 |
| Igfbp2 | 0.27 | 1.98E-27 | 4.11E-23 | 0.75 | 0.69 | 0.06 | 1.09 |
| Ier5l | 0.26 | 6.32E-22 | 1.31E-17 | 0.72 | 0.69 | 0.03 | 1.04 |
| Hmgn5 | 0.26 | 1.27E-15 | 2.65E-11 | 0.72 | 0.7 | 0.02 | 1.03 |
| H19 | 0.26 | 1.63E-18 | 3.40E-14 | 0.84 | 0.83 | 0.02 | 1.02 |
| Rps6ka6 | 0.25 | 3.99E-07 | 8.31E-03 | 0.47 | 0.45 | 0.02 | 1.04 |
| Cib2 | -0.25 | 6.65E-16 | 1.38E-11 | 0.26 | 0.33 | -0.07 | 0.78 |
| Prkdc | -0.25 | 7.44E-10 | 1.55E-05 | 0.18 | 0.23 | -0.05 | 0.79 |
| Snhg11 | -0.25 | 1.13E-07 | 2.36E-03 | 0.1 | 0.13 | -0.03 | 0.76 |
| Fsd1l | -0.25 | 1.90E-11 | 3.95E-07 | 0.26 | 0.32 | -0.06 | 0.82 |
| Cyp39a1 | -0.25 | 9.86E-08 | 2.05E-03 | 0.11 | 0.15 | -0.03 | 0.77 |
| Mettl7a1 | -0.25 | 2.25E-09 | 4.68E-05 | 0.12 | 0.16 | -0.04 | 0.76 |
| Btaf1 | -0.25 | 6.41E-14 | 1.33E-09 | 0.33 | 0.4 | -0.07 | 0.83 |
| Dmpk | -0.25 | 1.75E-06 | 3.65E-02 | 0.08 | 0.11 | -0.03 | 0.76 |
| Rictor | -0.25 | 2.62E-14 | 5.46E-10 | 0.35 | 0.41 | -0.07 | 0.83 |
| Pex5l | -0.25 | 7.72E-11 | 1.60E-06 | 0.1 | 0.14 | -0.04 | 0.72 |
| Cd47 | -0.25 | 2.10E-13 | 4.36E-09 | 0.34 | 0.41 | -0.07 | 0.83 |
| Dapk1 | -0.25 | 8.95E-14 | 1.86E-09 | 0.24 | 0.3 | -0.06 | 0.79 |
| Slx4 | -0.25 | 3.46E-07 | 7.20E-03 | 0.12 | 0.15 | -0.03 | 0.79 |
| Sirt4 | -0.25 | 1.36E-06 | 2.83E-02 | 0.11 | 0.13 | -0.03 | 0.79 |
| Kat6b | -0.25 | 5.65E-14 | 1.18E-09 | 0.47 | 0.54 | -0.07 | 0.87 |
| Nfatc4 | -0.25 | 2.04E-25 | 4.25E-21 | 0.61 | 0.7 | -0.09 | 0.88 |
| Eml1 | -0.25 | 4.51E-08 | 9.38E-04 | 0.15 | 0.19 | -0.04 | 0.8 |
| Fanca | -0.26 | 4.81E-07 | 1.00E-02 | 0.1 | 0.13 | -0.03 | 0.77 |
| Dpf1 | -0.26 | 2.95E-10 | 6.13E-06 | 0.18 | 0.22 | -0.05 | 0.78 |
| Smad7 | -0.26 | 4.50E-37 | 9.35E-33 | 0.72 | 0.81 | -0.1 | 0.88 |
| Stx2 | -0.26 | 7.21E-13 | 1.50E-08 | 0.18 | 0.24 | -0.06 | 0.77 |
| Bcl9l | -0.26 | 1.28E-14 | 2.67E-10 | 0.31 | 0.39 | -0.07 | 0.81 |
| Klhl11 | -0.26 | 6.77E-10 | 1.41E-05 | 0.17 | 0.21 | -0.05 | 0.79 |
| Nradd | -0.26 | 3.93E-20 | 8.17E-16 | 0.52 | 0.61 | -0.09 | 0.86 |

**Supplementary Table 6. Differentially expressed genes in the mutant λ-junction mesenchymal cells.**

| | | | | | | | |
|---|---|---|---|---|---|---|---|
| Jrkl | -0.26 | 2.80E-09 | 5.83E-05 | 0.14 | 0.18 | -0.04 | 0.77 |
| Rnase4 | -0.26 | 5.26E-07 | 1.09E-02 | 0.07 | 0.1 | -0.03 | 0.74 |
| Cdkl3 | -0.26 | 4.25E-08 | 8.85E-04 | 0.12 | 0.16 | -0.04 | 0.78 |
| Susd6 | -0.26 | 1.29E-08 | 2.68E-04 | 0.17 | 0.21 | -0.04 | 0.81 |
| Ifit2 | -0.26 | 1.33E-07 | 2.77E-03 | 0.09 | 0.12 | -0.03 | 0.76 |
| Tnks1bp1 | -0.26 | 1.93E-08 | 4.02E-04 | 0.17 | 0.21 | -0.04 | 0.81 |
| A730063M14Rik | -0.26 | 4.35E-08 | 9.05E-04 | 0.12 | 0.16 | -0.04 | 0.78 |
| Tubb2a | -0.26 | 1.17E-11 | 2.44E-07 | 0.23 | 0.28 | -0.06 | 0.8 |
| Ncam1 | -0.26 | 4.39E-15 | 9.14E-11 | 0.36 | 0.43 | -0.07 | 0.83 |
| Csrp1 | -0.26 | 2.06E-15 | 4.28E-11 | 0.35 | 0.42 | -0.08 | 0.82 |
| Pdzd4 | -0.26 | 5.70E-08 | 1.18E-03 | 0.15 | 0.18 | -0.04 | 0.8 |
| Kif3c | -0.26 | 1.25E-08 | 2.59E-04 | 0.11 | 0.14 | -0.03 | 0.76 |
| Kntc1 | -0.27 | 2.04E-08 | 4.25E-04 | 0.19 | 0.23 | -0.04 | 0.82 |
| Clmp | -0.27 | 9.30E-48 | 1.93E-43 | 0.82 | 0.89 | -0.07 | 0.92 |
| Ninl | -0.27 | 4.06E-10 | 8.44E-06 | 0.19 | 0.24 | -0.05 | 0.8 |
| Osbpl8 | -0.27 | 2.75E-09 | 5.72E-05 | 0.2 | 0.24 | -0.05 | 0.81 |
| AC149090.1 | -0.27 | 7.26E-12 | 1.51E-07 | 0.21 | 0.27 | -0.06 | 0.79 |
| Smad6 | -0.27 | 8.51E-13 | 1.77E-08 | 0.23 | 0.28 | -0.06 | 0.8 |
| Fbxl7 | -0.27 | 4.87E-15 | 1.01E-10 | 0.5 | 0.57 | -0.07 | 0.87 |
| 5330434G04Rik | -0.27 | 2.61E-07 | 5.43E-03 | 0.1 | 0.13 | -0.03 | 0.77 |
| Syt11 | -0.27 | 2.43E-14 | 5.05E-10 | 0.28 | 0.34 | -0.07 | 0.8 |
| Gpr161 | -0.27 | 2.33E-07 | 4.85E-03 | 0.1 | 0.13 | -0.03 | 0.77 |
| Pakap.1 | -0.27 | 2.73E-22 | 5.67E-18 | 0.43 | 0.52 | -0.09 | 0.82 |
| Mapk1ip1 | -0.27 | 1.76E-06 | 3.67E-02 | 0.08 | 0.11 | -0.03 | 0.77 |
| Zfp982 | -0.27 | 5.36E-07 | 1.11E-02 | 0.1 | 0.13 | -0.03 | 0.78 |
| Rps6ka3 | -0.27 | 2.18E-08 | 4.53E-04 | 0.46 | 0.51 | -0.05 | 0.91 |
| Zfp992 | -0.27 | 1.15E-07 | 2.39E-03 | 0.11 | 0.15 | -0.03 | 0.78 |
| Kcne1l | -0.27 | 3.68E-11 | 7.66E-07 | 0.14 | 0.18 | -0.05 | 0.75 |
| Dnah7a | -0.27 | 2.18E-12 | 4.53E-08 | 0.13 | 0.18 | -0.05 | 0.73 |
| Nfix | -0.27 | 1.51E-08 | 3.13E-04 | 0.15 | 0.19 | -0.04 | 0.79 |
| Paqr3 | -0.27 | 8.04E-07 | 1.67E-02 | 0.07 | 0.1 | -0.02 | 0.75 |
| Hba-x | -0.27 | 2.72E-136 | 5.66E-132 | 0.85 | 0.93 | -0.08 | 0.92 |
| Ttpa | -0.27 | 1.97E-06 | 4.10E-02 | 0.08 | 0.11 | -0.03 | 0.76 |
| Ldha | -0.27 | 9.68E-67 | 2.01E-62 | 0.97 | 0.98 | -0.02 | 0.98 |
| Msrb2 | -0.27 | 3.60E-15 | 7.49E-11 | 0.33 | 0.4 | -0.07 | 0.82 |
| Rnd3 | -0.28 | 2.06E-28 | 4.28E-24 | 0.61 | 0.7 | -0.09 | 0.87 |
| Gli3 | -0.28 | 1.97E-28 | 4.09E-24 | 0.59 | 0.69 | -0.1 | 0.86 |
| Eya1 | -0.28 | 6.68E-20 | 1.39E-15 | 0.46 | 0.54 | -0.08 | 0.85 |
| Prrg1 | -0.28 | 1.52E-10 | 3.16E-06 | 0.21 | 0.26 | -0.05 | 0.8 |
| Ccdc18 | -0.28 | 6.26E-11 | 1.30E-06 | 0.26 | 0.31 | -0.05 | 0.83 |
| Fam174b | -0.28 | 2.12E-06 | 4.40E-02 | 0.1 | 0.13 | -0.03 | 0.79 |
| Rab30 | -0.28 | 6.30E-10 | 1.31E-05 | 0.15 | 0.2 | -0.04 | 0.78 |
| Efr3b | -0.28 | 1.81E-07 | 3.76E-03 | 0.09 | 0.12 | -0.03 | 0.76 |
| Bnip3 | -0.28 | 4.45E-14 | 9.26E-10 | 0.37 | 0.44 | -0.07 | 0.85 |
| Scd1 | -0.28 | 1.45E-09 | 3.02E-05 | 0.2 | 0.25 | -0.05 | 0.81 |
| Plekha5 | -0.28 | 5.75E-20 | 1.20E-15 | 0.46 | 0.55 | -0.08 | 0.85 |
| Hist1h2be | -0.28 | 1.23E-09 | 2.55E-05 | 0.28 | 0.32 | -0.05 | 0.85 |
| Kcnmb4 | -0.28 | 1.12E-16 | 2.33E-12 | 0.28 | 0.35 | -0.08 | 0.79 |

**Supplementary Table 6. Differentially expressed genes in the mutant λ-junction mesenchymal cells.**

| | | | | | | | |
|---|---|---|---|---|---|---|---|
| Enpp2 | -0.28 | 6.28E-10 | 1.31E-05 | 0.12 | 0.16 | -0.04 | 0.76 |
| Plpp1 | -0.28 | 2.29E-07 | 4.77E-03 | 0.14 | 0.17 | -0.04 | 0.8 |
| Usp2 | -0.29 | 8.36E-09 | 1.74E-04 | 0.14 | 0.18 | -0.04 | 0.78 |
| S100a10 | -0.29 | 6.01E-12 | 1.25E-07 | 0.47 | 0.53 | -0.06 | 0.89 |
| Tapbp | -0.29 | 1.30E-06 | 2.70E-02 | 0.06 | 0.08 | -0.02 | 0.71 |
| Gxylt2 | -0.29 | 2.67E-14 | 5.56E-10 | 0.28 | 0.35 | -0.07 | 0.81 |
| Car13 | -0.29 | 2.94E-08 | 6.11E-04 | 0.1 | 0.13 | -0.03 | 0.76 |
| Megf9 | -0.29 | 1.23E-09 | 2.57E-05 | 0.17 | 0.21 | -0.04 | 0.79 |
| Fbln5 | -0.29 | 3.27E-13 | 6.80E-09 | 0.15 | 0.2 | -0.05 | 0.75 |
| Rttn | -0.29 | 5.41E-08 | 1.13E-03 | 0.15 | 0.19 | -0.04 | 0.8 |
| Tuba1a | -0.29 | 6.11E-109 | 1.27E-104 | 0.99 | 1 | 0 | 1 |
| Npl | -0.29 | 1.84E-06 | 3.82E-02 | 0.09 | 0.12 | -0.03 | 0.78 |
| Ccdc82 | -0.29 | 2.63E-08 | 5.47E-04 | 0.1 | 0.13 | -0.03 | 0.76 |
| Cox6b2 | -0.29 | 2.98E-20 | 6.20E-16 | 0.38 | 0.47 | -0.09 | 0.81 |
| Hist1h1a | -0.29 | 1.51E-08 | 3.14E-04 | 0.42 | 0.46 | -0.04 | 0.91 |
| Slc25a37 | -0.29 | 1.38E-10 | 2.87E-06 | 0.17 | 0.22 | -0.05 | 0.78 |
| Aopep | -0.29 | 5.53E-32 | 1.15E-27 | 0.6 | 0.69 | -0.09 | 0.87 |
| Mlf1 | -0.29 | 1.20E-10 | 2.49E-06 | 0.22 | 0.27 | -0.05 | 0.81 |
| Hist1h1b | -0.29 | 9.04E-08 | 1.88E-03 | 0.64 | 0.68 | -0.03 | 0.95 |
| Ccng2 | -0.29 | 3.81E-21 | 7.93E-17 | 0.41 | 0.5 | -0.09 | 0.83 |
| Smug1 | -0.29 | 2.90E-07 | 6.03E-03 | 0.1 | 0.13 | -0.03 | 0.77 |
| Camk2n1 | -0.29 | 2.60E-07 | 5.41E-03 | 0.2 | 0.24 | -0.04 | 0.83 |
| Edil3 | -0.3 | 3.27E-08 | 6.81E-04 | 0.21 | 0.25 | -0.04 | 0.83 |
| Dnajc12 | -0.3 | 1.06E-10 | 2.20E-06 | 0.16 | 0.2 | -0.05 | 0.78 |
| Net1 | -0.3 | 1.32E-23 | 2.75E-19 | 0.58 | 0.67 | -0.09 | 0.87 |
| Cdon | -0.3 | 5.07E-11 | 1.06E-06 | 0.2 | 0.25 | -0.05 | 0.8 |
| Samd8 | -0.3 | 8.34E-12 | 1.74E-07 | 0.2 | 0.25 | -0.05 | 0.79 |
| Sgk1 | -0.3 | 4.24E-16 | 8.82E-12 | 0.23 | 0.3 | -0.07 | 0.77 |
| Gm3839 | -0.3 | 3.55E-08 | 7.38E-04 | 0.14 | 0.17 | -0.04 | 0.79 |
| Fmnl2 | -0.3 | 1.90E-22 | 3.96E-18 | 0.4 | 0.5 | -0.09 | 0.81 |
| Amer2 | -0.3 | 7.78E-07 | 1.62E-02 | 0.09 | 0.11 | -0.03 | 0.77 |
| Fbln2 | -0.3 | 1.47E-26 | 3.05E-22 | 0.64 | 0.72 | -0.08 | 0.89 |
| Zbtb8a | -0.3 | 6.07E-07 | 1.26E-02 | 0.06 | 0.08 | -0.02 | 0.73 |
| Slc2a4 | -0.3 | 2.95E-13 | 6.13E-09 | 0.17 | 0.22 | -0.05 | 0.76 |
| Me1 | -0.3 | 7.72E-16 | 1.61E-11 | 0.25 | 0.31 | -0.07 | 0.78 |
| Rassf3 | -0.3 | 5.98E-20 | 1.24E-15 | 0.41 | 0.5 | -0.09 | 0.82 |
| Prnp | -0.3 | 1.88E-11 | 3.91E-07 | 0.15 | 0.2 | -0.05 | 0.77 |
| Slc17a5 | -0.3 | 1.49E-11 | 3.10E-07 | 0.16 | 0.21 | -0.05 | 0.77 |
| Daam2 | -0.3 | 2.36E-22 | 4.92E-18 | 0.36 | 0.45 | -0.09 | 0.8 |
| Rnf128 | -0.3 | 8.45E-12 | 1.76E-07 | 0.15 | 0.2 | -0.05 | 0.76 |
| D330041H03Rik | -0.31 | 1.79E-07 | 3.73E-03 | 0.08 | 0.11 | -0.03 | 0.74 |
| Pth1r | -0.31 | 5.64E-45 | 1.17E-40 | 0.72 | 0.82 | -0.1 | 0.88 |
| Lgals1 | -0.31 | 8.73E-81 | 1.82E-76 | 0.98 | 0.99 | -0.01 | 0.99 |
| Vcam1 | -0.31 | 1.11E-29 | 2.31E-25 | 0.54 | 0.64 | -0.1 | 0.85 |
| Ubr5 | -0.31 | 9.54E-29 | 1.98E-24 | 0.65 | 0.73 | -0.08 | 0.89 |
| Celf4 | -0.31 | 8.98E-11 | 1.87E-06 | 0.19 | 0.24 | -0.05 | 0.8 |
| Efnb1 | -0.31 | 1.41E-33 | 2.94E-29 | 0.6 | 0.7 | -0.1 | 0.86 |
| Tead1 | -0.31 | 5.63E-30 | 1.17E-25 | 0.54 | 0.64 | -0.1 | 0.84 |

**Supplementary Table 6. Differentially expressed genes in the mutant λ-junction mesenchymal cells.**

| | | | | | | | |
|---|---|---|---|---|---|---|---|
| Prtn3 | -0.31 | 3.41E-12 | 7.09E-08 | 0.13 | 0.18 | -0.05 | 0.73 |
| Nid1 | -0.31 | 5.45E-29 | 1.13E-24 | 0.39 | 0.5 | -0.11 | 0.79 |
| Rnf26 | -0.31 | 1.88E-18 | 3.91E-14 | 0.34 | 0.42 | -0.08 | 0.82 |
| Slc38a6 | -0.32 | 9.09E-11 | 1.89E-06 | 0.14 | 0.18 | -0.04 | 0.76 |
| Hacd4 | -0.32 | 2.34E-17 | 4.86E-13 | 0.27 | 0.34 | -0.07 | 0.78 |
| Hba-a2 | -0.32 | 7.56E-43 | 1.57E-38 | 0.58 | 0.7 | -0.11 | 0.84 |
| Bcas3 | -0.32 | 6.03E-14 | 1.25E-09 | 0.21 | 0.27 | -0.06 | 0.78 |
| Epb41l5 | -0.32 | 3.14E-12 | 6.53E-08 | 0.15 | 0.2 | -0.05 | 0.76 |
| Asns | -0.32 | 3.31E-25 | 6.88E-21 | 0.43 | 0.52 | -0.09 | 0.82 |
| A430105J06Rik | -0.32 | 1.93E-06 | 4.01E-02 | 0.05 | 0.06 | -0.02 | 0.7 |
| Hist1h2ab | -0.32 | 1.09E-11 | 2.28E-07 | 0.37 | 0.42 | -0.05 | 0.88 |
| Ucp2 | -0.32 | 6.96E-15 | 1.45E-10 | 0.23 | 0.29 | -0.06 | 0.78 |
| Cep128 | -0.32 | 8.01E-11 | 1.67E-06 | 0.15 | 0.2 | -0.04 | 0.77 |
| Zfp974 | -0.32 | 1.58E-07 | 3.29E-03 | 0.07 | 0.1 | -0.03 | 0.73 |
| Lyst | -0.32 | 4.17E-09 | 8.68E-05 | 0.11 | 0.15 | -0.04 | 0.76 |
| Gm5577 | -0.32 | 5.17E-08 | 1.07E-03 | 0.06 | 0.09 | -0.03 | 0.72 |
| Cd99l2 | -0.33 | 4.71E-17 | 9.79E-13 | 0.25 | 0.32 | -0.07 | 0.78 |
| Rasl10b | -0.33 | 1.62E-06 | 3.38E-02 | 0.05 | 0.06 | -0.02 | 0.7 |
| Eno1b | -0.33 | 1.07E-20 | 2.23E-16 | 0.28 | 0.37 | -0.08 | 0.77 |
| Pdgfrl | -0.33 | 4.75E-13 | 9.88E-09 | 0.12 | 0.17 | -0.05 | 0.72 |
| Sgms1 | -0.33 | 1.24E-10 | 2.58E-06 | 0.14 | 0.19 | -0.04 | 0.76 |
| 2810429I04Rik | -0.33 | 7.62E-10 | 1.59E-05 | 0.14 | 0.18 | -0.04 | 0.77 |
| Stradb | -0.33 | 9.66E-15 | 2.01E-10 | 0.16 | 0.22 | -0.06 | 0.74 |
| Pip4k2a | -0.33 | 8.93E-08 | 1.86E-03 | 0.08 | 0.11 | -0.03 | 0.75 |
| Gpc2 | -0.33 | 2.22E-26 | 4.62E-22 | 0.38 | 0.48 | -0.1 | 0.79 |
| Gse1 | -0.33 | 2.42E-17 | 5.03E-13 | 0.28 | 0.35 | -0.07 | 0.79 |
| Hbegf | -0.33 | 2.92E-07 | 6.08E-03 | 0.07 | 0.1 | -0.03 | 0.74 |
| Rassf9 | -0.33 | 3.28E-12 | 6.81E-08 | 0.13 | 0.17 | -0.05 | 0.74 |
| Vim | -0.33 | 2.01E-55 | 4.18E-51 | 0.89 | 0.92 | -0.03 | 0.96 |
| Bcl3 | -0.33 | 4.12E-07 | 8.57E-03 | 0.06 | 0.08 | -0.02 | 0.71 |
| St8sia4 | -0.34 | 5.84E-08 | 1.22E-03 | 0.09 | 0.12 | -0.03 | 0.75 |
| Agtpbp1 | -0.34 | 1.18E-14 | 2.44E-10 | 0.16 | 0.22 | -0.06 | 0.75 |
| Ret | -0.34 | 1.79E-06 | 3.72E-02 | 0.07 | 0.1 | -0.02 | 0.75 |
| Wdr47 | -0.34 | 7.07E-07 | 1.47E-02 | 0.07 | 0.1 | -0.03 | 0.74 |
| Lnpep | -0.34 | 4.93E-18 | 1.03E-13 | 0.27 | 0.34 | -0.07 | 0.79 |
| AI413582 | -0.34 | 7.58E-07 | 1.58E-02 | 0.06 | 0.09 | -0.02 | 0.74 |
| Atoh8 | -0.34 | 1.88E-08 | 3.90E-04 | 0.1 | 0.14 | -0.03 | 0.76 |
| Pex11a | -0.34 | 4.75E-08 | 9.89E-04 | 0.1 | 0.13 | -0.03 | 0.76 |
| Reck | -0.34 | 5.69E-20 | 1.18E-15 | 0.28 | 0.36 | -0.08 | 0.78 |
| Pax7 | -0.34 | 3.77E-24 | 7.84E-20 | 0.27 | 0.36 | -0.09 | 0.76 |
| Mmp23 | -0.34 | 8.43E-07 | 1.75E-02 | 0.06 | 0.08 | -0.02 | 0.73 |
| B230354K17Rik | -0.34 | 7.59E-22 | 1.58E-17 | 0.31 | 0.4 | -0.09 | 0.78 |
| Ttll7 | -0.34 | 3.19E-07 | 6.64E-03 | 0.07 | 0.09 | -0.02 | 0.74 |
| Cox4i2 | -0.34 | 2.17E-15 | 4.51E-11 | 0.18 | 0.24 | -0.06 | 0.75 |
| Ppp1r14a | -0.34 | 1.71E-06 | 3.57E-02 | 0.06 | 0.08 | -0.02 | 0.73 |
| Hist1h4d | -0.34 | 1.60E-16 | 3.32E-12 | 0.44 | 0.5 | -0.06 | 0.88 |
| Pxylp1 | -0.35 | 1.24E-09 | 2.57E-05 | 0.11 | 0.15 | -0.04 | 0.76 |
| 2610008E11Rik | -0.35 | 1.27E-26 | 2.65E-22 | 0.39 | 0.49 | -0.1 | 0.79 |

**Supplementary Table 6. Differentially expressed genes in the mutant λ-junction mesenchymal cells.**

| | | | | | | | |
|---|---|---|---|---|---|---|---|
| Rai14 | -0.35 | 1.16E-65 | 2.41E-61 | 0.74 | 0.84 | -0.1 | 0.88 |
| Tcf12 | -0.35 | 5.37E-112 | 1.12E-107 | 0.93 | 0.97 | -0.04 | 0.96 |
| Zgrf1 | -0.35 | 1.72E-14 | 3.58E-10 | 0.18 | 0.23 | -0.06 | 0.76 |
| Ackr3 | -0.35 | 3.53E-32 | 7.33E-28 | 0.37 | 0.48 | -0.11 | 0.77 |
| Tle2 | -0.35 | 2.83E-13 | 5.88E-09 | 0.12 | 0.17 | -0.05 | 0.73 |
| Gli2 | -0.35 | 6.19E-21 | 1.29E-16 | 0.25 | 0.33 | -0.08 | 0.76 |
| Gpr45 | -0.35 | 6.59E-17 | 1.37E-12 | 0.19 | 0.25 | -0.06 | 0.75 |
| Bambi | -0.35 | 1.47E-60 | 3.05E-56 | 0.65 | 0.77 | -0.12 | 0.84 |
| Tspo | -0.35 | 3.27E-20 | 6.81E-16 | 0.29 | 0.37 | -0.08 | 0.79 |
| Pfkfb4 | -0.36 | 1.08E-08 | 2.24E-04 | 0.08 | 0.11 | -0.03 | 0.73 |
| Trp53i11 | -0.36 | 6.78E-14 | 1.41E-09 | 0.14 | 0.19 | -0.05 | 0.73 |
| Cacnb3 | -0.36 | 2.16E-18 | 4.50E-14 | 0.19 | 0.25 | -0.07 | 0.74 |
| Nfil3 | -0.36 | 3.64E-11 | 7.57E-07 | 0.13 | 0.17 | -0.04 | 0.76 |
| Gata3 | -0.36 | 5.54E-44 | 1.15E-39 | 0.55 | 0.67 | -0.12 | 0.82 |
| Cux2 | -0.36 | 5.46E-07 | 1.13E-02 | 0.07 | 0.1 | -0.03 | 0.74 |
| Mapkbp1 | -0.36 | 3.00E-08 | 6.25E-04 | 0.09 | 0.12 | -0.03 | 0.74 |
| Chsy3 | -0.36 | 5.30E-07 | 1.10E-02 | 0.06 | 0.08 | -0.02 | 0.71 |
| Dkk3 | -0.36 | 1.21E-11 | 2.52E-07 | 0.09 | 0.13 | -0.04 | 0.71 |
| Parvb | -0.36 | 2.19E-09 | 4.55E-05 | 0.12 | 0.16 | -0.04 | 0.77 |
| Zadh2 | -0.36 | 1.03E-07 | 2.15E-03 | 0.08 | 0.11 | -0.03 | 0.74 |
| Nrp2 | -0.36 | 4.07E-13 | 8.47E-09 | 0.41 | 0.47 | -0.05 | 0.89 |
| Tle1 | -0.37 | 1.83E-36 | 3.80E-32 | 0.5 | 0.61 | -0.11 | 0.82 |
| Lsamp | -0.37 | 5.99E-07 | 1.25E-02 | 0.05 | 0.07 | -0.02 | 0.71 |
| Mon1a | -0.37 | 2.34E-06 | 4.86E-02 | 0.05 | 0.07 | -0.02 | 0.73 |
| Crip1 | -0.37 | 1.54E-16 | 3.21E-12 | 0.35 | 0.42 | -0.07 | 0.84 |
| Ptprz1 | -0.37 | 3.50E-07 | 7.28E-03 | 0.06 | 0.09 | -0.02 | 0.73 |
| Eepd1 | -0.37 | 9.97E-13 | 2.07E-08 | 0.13 | 0.17 | -0.05 | 0.73 |
| Fhod3 | -0.38 | 6.80E-12 | 1.41E-07 | 0.08 | 0.12 | -0.04 | 0.69 |
| Ppp1r12b | -0.38 | 3.79E-09 | 7.88E-05 | 0.08 | 0.12 | -0.03 | 0.72 |
| Gch1 | -0.38 | 1.10E-11 | 2.30E-07 | 0.07 | 0.1 | -0.03 | 0.66 |
| Emb | -0.38 | 2.97E-23 | 6.18E-19 | 0.27 | 0.36 | -0.09 | 0.76 |
| Rgs5 | -0.38 | 3.84E-12 | 7.99E-08 | 0.09 | 0.13 | -0.04 | 0.7 |
| Fat4 | -0.38 | 8.45E-24 | 1.76E-19 | 0.3 | 0.39 | -0.09 | 0.77 |
| Cyp2d22 | -0.38 | 5.99E-08 | 1.25E-03 | 0.05 | 0.07 | -0.02 | 0.68 |
| Enc1 | -0.38 | 4.49E-16 | 9.33E-12 | 0.18 | 0.24 | -0.06 | 0.76 |
| Tshz3 | -0.38 | 1.51E-09 | 3.14E-05 | 0.09 | 0.12 | -0.03 | 0.73 |
| Ptgr1 | -0.38 | 2.09E-24 | 4.34E-20 | 0.26 | 0.35 | -0.09 | 0.75 |
| Kcp | -0.38 | 2.12E-14 | 4.42E-10 | 0.12 | 0.17 | -0.05 | 0.71 |
| Fam241a | -0.38 | 2.24E-07 | 4.66E-03 | 0.07 | 0.1 | -0.03 | 0.73 |
| Celf2 | -0.38 | 8.47E-10 | 1.76E-05 | 0.13 | 0.17 | -0.04 | 0.77 |
| Crabp1 | -0.39 | 3.67E-119 | 7.63E-115 | 0.98 | 0.99 | -0.01 | 0.99 |
| Cxxc4 | -0.39 | 1.04E-15 | 2.17E-11 | 0.3 | 0.37 | -0.07 | 0.82 |
| Tppp3 | -0.39 | 7.01E-07 | 1.46E-02 | 0.04 | 0.06 | -0.02 | 0.68 |
| Sv2a | -0.39 | 7.57E-07 | 1.57E-02 | 0.08 | 0.11 | -0.03 | 0.76 |
| Gm10561 | -0.39 | 1.98E-06 | 4.11E-02 | 0.06 | 0.08 | -0.02 | 0.73 |
| Acsl4 | -0.39 | 8.18E-10 | 1.70E-05 | 0.12 | 0.16 | -0.04 | 0.77 |
| Fgf10 | -0.39 | 8.31E-08 | 1.73E-03 | 0.1 | 0.13 | -0.03 | 0.77 |
| Rora | -0.39 | 1.38E-18 | 2.87E-14 | 0.19 | 0.25 | -0.07 | 0.74 |

**Supplementary Table 6. Differentially expressed genes in the mutant λ-junction mesenchymal cells.**

| | | | | | | | |
|---|---|---|---|---|---|---|---|
| Mir100hg | -0.39 | 1.47E-77 | 3.05E-73 | 0.81 | 0.9 | -0.09 | 0.9 |
| Tango2 | -0.4 | 7.76E-12 | 1.61E-07 | 0.1 | 0.14 | -0.04 | 0.72 |
| Entpd1 | -0.4 | 9.02E-09 | 1.88E-04 | 0.06 | 0.09 | -0.03 | 0.69 |
| Phactr3 | -0.4 | 1.93E-12 | 4.01E-08 | 0.09 | 0.12 | -0.04 | 0.69 |
| Anxa7 | -0.4 | 7.49E-25 | 1.56E-20 | 0.34 | 0.43 | -0.09 | 0.79 |
| Atp10a | -0.4 | 2.16E-07 | 4.49E-03 | 0.04 | 0.06 | -0.02 | 0.67 |
| Msl3 | -0.4 | 2.85E-33 | 5.92E-29 | 0.37 | 0.48 | -0.11 | 0.77 |
| Eln | -0.4 | 1.34E-09 | 2.78E-05 | 0.09 | 0.12 | -0.03 | 0.73 |
| Ago4 | -0.4 | 7.04E-09 | 1.46E-04 | 0.06 | 0.09 | -0.03 | 0.7 |
| Mpped2 | -0.4 | 2.45E-96 | 5.10E-92 | 0.9 | 0.96 | -0.05 | 0.95 |
| Unc5b | -0.4 | 1.93E-33 | 4.01E-29 | 0.32 | 0.42 | -0.11 | 0.75 |
| Stbd1 | -0.4 | 1.43E-09 | 2.98E-05 | 0.08 | 0.12 | -0.03 | 0.72 |
| Psd3 | -0.4 | 1.39E-24 | 2.89E-20 | 0.22 | 0.3 | -0.08 | 0.73 |
| Slc29a1 | -0.4 | 7.57E-08 | 1.57E-03 | 0.07 | 0.1 | -0.03 | 0.73 |
| Kremen1 | -0.4 | 1.32E-08 | 2.74E-04 | 0.08 | 0.11 | -0.03 | 0.74 |
| Fos | -0.4 | 6.82E-10 | 1.42E-05 | 0.1 | 0.14 | -0.04 | 0.73 |
| Cend1 | -0.4 | 3.67E-08 | 7.62E-04 | 0.07 | 0.09 | -0.03 | 0.71 |
| Hccs | -0.41 | 1.03E-26 | 2.15E-22 | 0.3 | 0.39 | -0.09 | 0.77 |
| Slc40a1 | -0.41 | 7.85E-07 | 1.63E-02 | 0.04 | 0.06 | -0.02 | 0.69 |
| Bbs9 | -0.41 | 5.93E-09 | 1.23E-04 | 0.08 | 0.11 | -0.03 | 0.73 |
| Gm28578 | -0.41 | 9.77E-08 | 2.03E-03 | 0.04 | 0.07 | -0.02 | 0.68 |
| Ebf4 | -0.41 | 6.20E-08 | 1.29E-03 | 0.03 | 0.05 | -0.02 | 0.63 |
| Fcgrt | -0.41 | 2.75E-18 | 5.72E-14 | 0.16 | 0.22 | -0.06 | 0.72 |
| Zfand4 | -0.41 | 6.52E-12 | 1.36E-07 | 0.08 | 0.11 | -0.04 | 0.68 |
| Hecw2 | -0.41 | 1.12E-10 | 2.33E-06 | 0.06 | 0.09 | -0.03 | 0.67 |
| Mtcp1 | -0.42 | 3.48E-07 | 7.25E-03 | 0.06 | 0.08 | -0.02 | 0.73 |
| Tsga10 | -0.42 | 3.00E-14 | 6.25E-10 | 0.09 | 0.13 | -0.04 | 0.68 |
| Plcb1 | -0.42 | 8.31E-11 | 1.73E-06 | 0.1 | 0.14 | -0.04 | 0.73 |
| Nqo1 | -0.42 | 3.92E-12 | 8.16E-08 | 0.1 | 0.14 | -0.04 | 0.71 |
| Hspb1 | -0.42 | 3.74E-09 | 7.79E-05 | 0.06 | 0.09 | -0.03 | 0.69 |
| Cdh6 | -0.42 | 2.72E-35 | 5.66E-31 | 0.3 | 0.4 | -0.11 | 0.74 |
| Emilin2 | -0.42 | 1.90E-32 | 3.96E-28 | 0.28 | 0.38 | -0.1 | 0.73 |
| Pcsk5 | -0.43 | 9.13E-44 | 1.90E-39 | 0.41 | 0.53 | -0.12 | 0.77 |
| Gm26930 | -0.43 | 6.64E-10 | 1.38E-05 | 0.07 | 0.1 | -0.03 | 0.69 |
| Twist1 | -0.43 | 5.46E-151 | 1.14E-146 | 0.96 | 0.99 | -0.03 | 0.97 |
| Id2 | -0.43 | 2.18E-120 | 4.53E-116 | 0.88 | 0.95 | -0.07 | 0.92 |
| Rspo2 | -0.43 | 2.64E-47 | 5.48E-43 | 0.44 | 0.57 | -0.13 | 0.78 |
| Tubb2b | -0.43 | 3.25E-33 | 6.77E-29 | 0.56 | 0.64 | -0.08 | 0.87 |
| Tnfaip6 | -0.43 | 4.83E-56 | 1.00E-51 | 0.59 | 0.7 | -0.11 | 0.84 |
| Tiam2 | -0.44 | 9.54E-14 | 1.99E-09 | 0.1 | 0.14 | -0.04 | 0.69 |
| Ankrd44 | -0.44 | 1.22E-35 | 2.53E-31 | 0.29 | 0.4 | -0.11 | 0.73 |
| Cd1d1 | -0.44 | 1.57E-19 | 3.27E-15 | 0.15 | 0.22 | -0.06 | 0.71 |
| Meis1 | -0.44 | 1.70E-24 | 3.55E-20 | 0.4 | 0.48 | -0.08 | 0.84 |
| Tmem51 | -0.44 | 2.35E-09 | 4.89E-05 | 0.07 | 0.1 | -0.03 | 0.7 |
| Phldb2 | -0.44 | 8.56E-43 | 1.78E-38 | 0.61 | 0.71 | -0.1 | 0.86 |
| Irx3 | -0.44 | 2.89E-15 | 6.01E-11 | 0.27 | 0.33 | -0.06 | 0.82 |
| Pakap | -0.44 | 2.90E-15 | 6.03E-11 | 0.12 | 0.17 | -0.05 | 0.71 |
| Fibin | -0.44 | 7.77E-11 | 1.62E-06 | 0.15 | 0.19 | -0.04 | 0.78 |

**Supplementary Table 6. Differentially expressed genes in the mutant λ-junction mesenchymal cells.**

| | | | | | | | |
|---|---|---|---|---|---|---|---|
| Igsf10 | -0.44 | 1.74E-16 | 3.61E-12 | 0.25 | 0.32 | -0.07 | 0.79 |
| Igf1 | -0.45 | 2.58E-47 | 5.38E-43 | 0.57 | 0.68 | -0.11 | 0.84 |
| Kcnb2 | -0.45 | 9.87E-10 | 2.05E-05 | 0.06 | 0.08 | -0.03 | 0.67 |
| Xaf1 | -0.45 | 2.03E-11 | 4.23E-07 | 0.04 | 0.07 | -0.03 | 0.61 |
| Chst3 | -0.45 | 8.57E-09 | 1.78E-04 | 0.06 | 0.08 | -0.03 | 0.7 |
| Ddit4l | -0.45 | 3.18E-21 | 6.61E-17 | 0.24 | 0.31 | -0.07 | 0.76 |
| Gm3830 | -0.45 | 8.96E-11 | 1.86E-06 | 0.07 | 0.1 | -0.03 | 0.69 |
| G430095P16Rik | -0.46 | 3.32E-11 | 6.91E-07 | 0.1 | 0.14 | -0.04 | 0.73 |
| C430049B03Rik | -0.46 | 3.46E-37 | 7.19E-33 | 0.37 | 0.48 | -0.11 | 0.77 |
| Arhgdib | -0.46 | 2.56E-58 | 5.33E-54 | 0.53 | 0.66 | -0.13 | 0.8 |
| Pam | -0.46 | 3.03E-13 | 6.30E-09 | 0.14 | 0.18 | -0.05 | 0.74 |
| Tril | -0.46 | 2.79E-72 | 5.81E-68 | 0.59 | 0.73 | -0.13 | 0.82 |
| Ccn4 | -0.46 | 4.08E-08 | 8.48E-04 | 0.04 | 0.06 | -0.02 | 0.65 |
| Chst2 | -0.46 | 3.73E-26 | 7.77E-22 | 0.18 | 0.26 | -0.08 | 0.7 |
| Tcf7 | -0.46 | 5.93E-23 | 1.23E-18 | 0.14 | 0.2 | -0.07 | 0.67 |
| S1pr3 | -0.47 | 1.94E-33 | 4.04E-29 | 0.32 | 0.42 | -0.1 | 0.76 |
| Emid1 | -0.47 | 6.72E-08 | 1.40E-03 | 0.06 | 0.08 | -0.02 | 0.7 |
| Prickle1 | -0.47 | 1.54E-28 | 3.21E-24 | 0.24 | 0.33 | -0.09 | 0.73 |
| Stard8 | -0.47 | 1.36E-10 | 2.82E-06 | 0.08 | 0.11 | -0.03 | 0.71 |
| Stxbp2 | -0.47 | 1.69E-10 | 3.51E-06 | 0.04 | 0.07 | -0.03 | 0.61 |
| Cacnb2 | -0.47 | 6.88E-09 | 1.43E-04 | 0.06 | 0.08 | -0.03 | 0.69 |
| Gfra1 | -0.47 | 8.55E-08 | 1.78E-03 | 0.04 | 0.06 | -0.02 | 0.66 |
| Elmod2 | -0.47 | 5.93E-27 | 1.23E-22 | 0.2 | 0.28 | -0.08 | 0.71 |
| Hist1h2bg | -0.47 | 1.98E-12 | 4.11E-08 | 0.08 | 0.12 | -0.04 | 0.68 |
| Ntm | -0.47 | 9.17E-09 | 1.91E-04 | 0.06 | 0.08 | -0.03 | 0.68 |
| Alx4 | -0.48 | 7.03E-62 | 1.46E-57 | 0.49 | 0.63 | -0.14 | 0.78 |
| 2900060N12Rik | -0.48 | 4.76E-10 | 9.89E-06 | 0.05 | 0.08 | -0.03 | 0.65 |
| Minar2 | -0.48 | 2.88E-09 | 5.98E-05 | 0.07 | 0.1 | -0.03 | 0.71 |
| Nrgn | -0.48 | 1.14E-07 | 2.38E-03 | 0.04 | 0.07 | -0.02 | 0.68 |
| Mogat2 | -0.48 | 3.82E-22 | 7.95E-18 | 0.12 | 0.19 | -0.06 | 0.66 |
| B3galt1 | -0.48 | 5.46E-10 | 1.14E-05 | 0.05 | 0.08 | -0.03 | 0.65 |
| Kif26b | -0.48 | 8.49E-67 | 1.77E-62 | 0.47 | 0.62 | -0.15 | 0.76 |
| Mab21l2 | -0.48 | 7.82E-59 | 1.63E-54 | 0.61 | 0.72 | -0.11 | 0.84 |
| Slc16a10 | -0.48 | 3.00E-19 | 6.24E-15 | 0.09 | 0.14 | -0.05 | 0.64 |
| Anxa11 | -0.49 | 2.00E-14 | 4.17E-10 | 0.15 | 0.2 | -0.05 | 0.75 |
| Robo1 | -0.49 | 2.48E-54 | 5.16E-50 | 0.45 | 0.57 | -0.13 | 0.78 |
| Ankrd12 | -0.49 | 4.96E-58 | 1.03E-53 | 0.51 | 0.64 | -0.13 | 0.8 |
| Vamp5 | -0.49 | 1.05E-06 | 2.18E-02 | 0.03 | 0.05 | -0.02 | 0.65 |
| Hs3st6 | -0.49 | 1.25E-14 | 2.60E-10 | 0.06 | 0.1 | -0.04 | 0.63 |
| C630043F03Rik | -0.49 | 2.21E-20 | 4.60E-16 | 0.14 | 0.2 | -0.06 | 0.68 |
| Tmem26 | -0.49 | 2.76E-30 | 5.73E-26 | 0.29 | 0.38 | -0.1 | 0.75 |
| Slc16a3 | -0.49 | 2.14E-35 | 4.46E-31 | 0.25 | 0.35 | -0.1 | 0.71 |
| St6galnac5 | -0.49 | 5.91E-11 | 1.23E-06 | 0.04 | 0.07 | -0.03 | 0.62 |
| Snai2 | -0.49 | 3.13E-57 | 6.50E-53 | 0.52 | 0.64 | -0.12 | 0.81 |
| Tmem100 | -0.5 | 3.20E-21 | 6.66E-17 | 0.15 | 0.22 | -0.06 | 0.7 |
| Col16a1 | -0.5 | 6.95E-10 | 1.45E-05 | 0.05 | 0.08 | -0.03 | 0.65 |
| Cd34 | -0.5 | 5.66E-19 | 1.18E-14 | 0.12 | 0.18 | -0.06 | 0.68 |
| Cacna1c | -0.5 | 3.92E-17 | 8.15E-13 | 0.1 | 0.15 | -0.05 | 0.67 |

**Supplementary Table 6. Differentially expressed genes in the mutant λ-junction mesenchymal cells.**

| | | | | | | | |
|---|---|---|---|---|---|---|---|
| Smad9 | -0.5 | 4.50E-10 | 9.37E-06 | 0.05 | 0.08 | -0.03 | 0.66 |
| Col5a2 | -0.5 | 2.09E-73 | 4.35E-69 | 0.61 | 0.73 | -0.12 | 0.83 |
| Spock3 | -0.5 | 8.37E-22 | 1.74E-17 | 0.14 | 0.2 | -0.06 | 0.68 |
| Bcl11b | -0.5 | 1.50E-66 | 3.11E-62 | 0.52 | 0.66 | -0.14 | 0.78 |
| Plekhg1 | -0.51 | 1.63E-12 | 3.39E-08 | 0.07 | 0.1 | -0.04 | 0.66 |
| Adgre5 | -0.51 | 4.35E-25 | 9.06E-21 | 0.12 | 0.19 | -0.07 | 0.64 |
| Akr1b8 | -0.51 | 2.50E-12 | 5.20E-08 | 0.08 | 0.11 | -0.04 | 0.68 |
| Cacna1g | -0.51 | 3.71E-56 | 7.71E-52 | 0.37 | 0.51 | -0.14 | 0.73 |
| C030014I23Rik | -0.51 | 1.38E-08 | 2.87E-04 | 0.03 | 0.06 | -0.02 | 0.62 |
| Tspan2 | -0.51 | 3.76E-16 | 7.82E-12 | 0.09 | 0.14 | -0.05 | 0.66 |
| Trps1 | -0.51 | 7.66E-11 | 1.59E-06 | 0.47 | 0.51 | -0.04 | 0.92 |
| Pdlim2 | -0.51 | 4.56E-23 | 9.49E-19 | 0.17 | 0.24 | -0.07 | 0.7 |
| Emp2 | -0.51 | 7.34E-14 | 1.53E-09 | 0.07 | 0.11 | -0.04 | 0.65 |
| Gap43 | -0.51 | 6.61E-63 | 1.37E-58 | 0.47 | 0.61 | -0.14 | 0.77 |
| Fam181b | -0.52 | 3.79E-13 | 7.88E-09 | 0.09 | 0.13 | -0.04 | 0.69 |
| Msx2 | -0.52 | 1.94E-90 | 4.04E-86 | 0.36 | 0.55 | -0.19 | 0.65 |
| Baiap2l2 | -0.52 | 2.75E-22 | 5.73E-18 | 0.12 | 0.18 | -0.06 | 0.65 |
| Dclk2 | -0.52 | 2.54E-18 | 5.27E-14 | 0.11 | 0.16 | -0.05 | 0.67 |
| Sipa1l2 | -0.52 | 3.05E-19 | 6.35E-15 | 0.12 | 0.17 | -0.06 | 0.68 |
| 2810405F17Rik | -0.52 | 5.79E-07 | 1.21E-02 | 0.03 | 0.05 | -0.02 | 0.66 |
| Ankrd29 | -0.52 | 2.35E-16 | 4.89E-12 | 0.09 | 0.14 | -0.05 | 0.67 |
| Nr3c1 | -0.52 | 2.06E-13 | 4.28E-09 | 0.1 | 0.15 | -0.04 | 0.71 |
| Dpysl3 | -0.52 | 1.42E-54 | 2.95E-50 | 0.36 | 0.5 | -0.14 | 0.73 |
| Serpine2 | -0.52 | 2.29E-120 | 4.77E-116 | 0.78 | 0.88 | -0.11 | 0.88 |
| Hist1h2an | -0.53 | 1.26E-27 | 2.62E-23 | 0.29 | 0.37 | -0.08 | 0.78 |
| Filip1l | -0.53 | 8.72E-30 | 1.81E-25 | 0.21 | 0.3 | -0.09 | 0.7 |
| Sh3gl3 | -0.53 | 5.89E-42 | 1.23E-37 | 0.37 | 0.48 | -0.11 | 0.77 |
| Armh4 | -0.53 | 6.94E-19 | 1.44E-14 | 0.12 | 0.17 | -0.06 | 0.68 |
| Pcdhgb6 | -0.53 | 3.02E-07 | 6.27E-03 | 0.03 | 0.05 | -0.02 | 0.64 |
| Sema5a | -0.54 | 7.98E-52 | 1.66E-47 | 0.39 | 0.52 | -0.13 | 0.75 |
| Olfml2b | -0.54 | 4.04E-13 | 8.39E-09 | 0.05 | 0.08 | -0.03 | 0.61 |
| Kif5a | -0.54 | 1.88E-07 | 3.91E-03 | 0.04 | 0.06 | -0.02 | 0.66 |
| Scn3a | -0.54 | 5.34E-11 | 1.11E-06 | 0.06 | 0.09 | -0.03 | 0.65 |
| Bmerb1 | -0.54 | 7.05E-16 | 1.47E-11 | 0.09 | 0.14 | -0.05 | 0.67 |
| Slitrk6 | -0.55 | 4.89E-46 | 1.02E-41 | 0.33 | 0.45 | -0.12 | 0.74 |
| Itga8 | -0.55 | 8.74E-38 | 1.82E-33 | 0.21 | 0.31 | -0.1 | 0.67 |
| A830018L16Rik | -0.55 | 9.87E-17 | 2.05E-12 | 0.13 | 0.19 | -0.05 | 0.72 |
| Erc2 | -0.55 | 3.32E-11 | 6.91E-07 | 0.04 | 0.07 | -0.03 | 0.62 |
| Prdm16os | -0.55 | 1.54E-21 | 3.20E-17 | 0.13 | 0.2 | -0.06 | 0.68 |
| Stx18 | -0.55 | 1.30E-77 | 2.71E-73 | 0.49 | 0.64 | -0.15 | 0.76 |
| Ctnna2 | -0.55 | 1.12E-06 | 2.32E-02 | 0.04 | 0.05 | -0.02 | 0.66 |
| Tmem169 | -0.55 | 2.32E-08 | 4.83E-04 | 0.03 | 0.05 | -0.02 | 0.62 |
| Snhg9 | -0.55 | 4.47E-31 | 9.30E-27 | 0.17 | 0.25 | -0.09 | 0.66 |
| Galnt5 | -0.56 | 3.97E-09 | 8.26E-05 | 0.05 | 0.07 | -0.02 | 0.66 |
| S100a6 | -0.56 | 1.84E-18 | 3.83E-14 | 0.12 | 0.18 | -0.05 | 0.69 |
| Pde3a | -0.56 | 1.82E-26 | 3.78E-22 | 0.14 | 0.21 | -0.07 | 0.66 |
| Gm973 | -0.56 | 2.67E-09 | 5.55E-05 | 0.04 | 0.06 | -0.02 | 0.61 |
| Pax3 | -0.56 | 2.56E-80 | 5.33E-76 | 0.53 | 0.68 | -0.15 | 0.78 |

**Supplementary Table 6. Differentially expressed genes in the mutant λ-junction mesenchymal cells.**

| | | | | | | | |
|---|---|---|---|---|---|---|---|
| Adamts19 | -0.57 | 7.78E-48 | 1.62E-43 | 0.25 | 0.37 | -0.12 | 0.67 |
| Thy1 | -0.57 | 1.13E-08 | 2.35E-04 | 0.04 | 0.06 | -0.02 | 0.63 |
| Srpx | -0.57 | 3.91E-09 | 8.14E-05 | 0.03 | 0.06 | -0.02 | 0.62 |
| Jakmip2 | -0.58 | 5.65E-25 | 1.17E-20 | 0.15 | 0.22 | -0.07 | 0.68 |
| Cacng4 | -0.58 | 1.29E-37 | 2.69E-33 | 0.2 | 0.3 | -0.1 | 0.67 |
| Hbb-bh1 | -0.58 | 2.31E-104 | 4.80E-100 | 0.3 | 0.48 | -0.18 | 0.62 |
| Adamts20 | -0.58 | 1.16E-35 | 2.42E-31 | 0.15 | 0.24 | -0.09 | 0.63 |
| Nbl1 | -0.59 | 1.24E-122 | 2.57E-118 | 0.56 | 0.74 | -0.18 | 0.76 |
| Dpep1 | -0.59 | 1.37E-40 | 2.86E-36 | 0.26 | 0.36 | -0.11 | 0.7 |
| Syne1 | -0.59 | 3.26E-33 | 6.79E-29 | 0.13 | 0.21 | -0.08 | 0.61 |
| Clgn | -0.59 | 1.42E-13 | 2.94E-09 | 0.05 | 0.08 | -0.03 | 0.6 |
| Sgcd | -0.59 | 7.13E-42 | 1.48E-37 | 0.25 | 0.36 | -0.11 | 0.69 |
| Prkcq | -0.59 | 2.06E-08 | 4.29E-04 | 0.03 | 0.05 | -0.02 | 0.63 |
| Zfhx3 | -0.6 | 1.29E-63 | 2.68E-59 | 0.79 | 0.85 | -0.07 | 0.92 |
| Spink2 | -0.6 | 4.47E-10 | 9.30E-06 | 0.04 | 0.06 | -0.02 | 0.61 |
| Ednra | -0.62 | 5.83E-69 | 1.21E-64 | 0.39 | 0.53 | -0.15 | 0.73 |
| Gm15222 | -0.62 | 5.40E-25 | 1.12E-20 | 0.09 | 0.14 | -0.06 | 0.59 |
| Lix1 | -0.62 | 3.99E-158 | 8.30E-154 | 0.78 | 0.89 | -0.11 | 0.87 |
| Gas1 | -0.62 | 1.32E-154 | 2.74E-150 | 0.72 | 0.86 | -0.13 | 0.85 |
| Runx1 | -0.62 | 7.60E-12 | 1.58E-07 | 0.06 | 0.09 | -0.03 | 0.66 |
| Ptn | -0.62 | 9.21E-194 | 1.92E-189 | 0.97 | 0.99 | -0.02 | 0.98 |
| Kcnk2 | -0.63 | 1.03E-41 | 2.14E-37 | 0.24 | 0.35 | -0.11 | 0.69 |
| Cd109 | -0.63 | 8.22E-10 | 1.71E-05 | 0.03 | 0.06 | -0.02 | 0.61 |
| Ankrd45 | -0.64 | 6.16E-13 | 1.28E-08 | 0.05 | 0.09 | -0.03 | 0.62 |
| Mc4r | -0.64 | 1.01E-14 | 2.10E-10 | 0.05 | 0.09 | -0.03 | 0.61 |
| Olfm2 | -0.64 | 1.25E-18 | 2.61E-14 | 0.07 | 0.11 | -0.04 | 0.6 |
| Ahnak | -0.65 | 2.79E-79 | 5.81E-75 | 0.43 | 0.58 | -0.15 | 0.74 |
| S1pr1 | -0.65 | 5.55E-10 | 1.15E-05 | 0.04 | 0.06 | -0.02 | 0.62 |
| Mylk | -0.65 | 4.13E-37 | 8.58E-33 | 0.12 | 0.21 | -0.08 | 0.59 |
| Gm48898 | -0.66 | 7.16E-07 | 1.49E-02 | 0.03 | 0.05 | -0.02 | 0.67 |
| Cachd1 | -0.67 | 6.92E-100 | 1.44E-95 | 0.52 | 0.68 | -0.16 | 0.77 |
| Dnm3 | -0.67 | 5.36E-36 | 1.11E-31 | 0.17 | 0.26 | -0.09 | 0.66 |
| Crnde | -0.68 | 4.26E-27 | 8.87E-23 | 0.14 | 0.21 | -0.07 | 0.66 |
| Zfp503 | -0.68 | 3.12E-47 | 6.49E-43 | 0.3 | 0.41 | -0.11 | 0.73 |
| Zeb2os | -0.68 | 6.74E-14 | 1.40E-09 | 0.04 | 0.07 | -0.03 | 0.57 |
| Mgarp | -0.69 | 2.32E-16 | 4.82E-12 | 0.05 | 0.08 | -0.04 | 0.57 |
| Plxna2 | -0.7 | 7.38E-30 | 1.54E-25 | 0.19 | 0.27 | -0.08 | 0.7 |
| Arhgap6 | -0.7 | 3.93E-18 | 8.17E-14 | 0.06 | 0.1 | -0.04 | 0.59 |
| 1110017D15Rik | -0.7 | 8.61E-21 | 1.79E-16 | 0.08 | 0.13 | -0.05 | 0.62 |
| Pcdh9 | -0.7 | 5.51E-15 | 1.15E-10 | 0.08 | 0.12 | -0.04 | 0.66 |
| Sertad4 | -0.71 | 2.79E-80 | 5.80E-76 | 0.33 | 0.48 | -0.15 | 0.68 |
| a | -0.71 | 4.42E-34 | 9.20E-30 | 0.14 | 0.22 | -0.08 | 0.63 |
| Gadd45b | -0.72 | 1.93E-26 | 4.02E-22 | 0.1 | 0.16 | -0.06 | 0.61 |
| Svep1 | -0.72 | 3.19E-29 | 6.64E-25 | 0.08 | 0.15 | -0.06 | 0.57 |
| Plk2 | -0.72 | 2.94E-75 | 6.12E-71 | 0.3 | 0.45 | -0.15 | 0.67 |
| Msx1 | -0.73 | 5.65E-207 | 1.17E-202 | 0.63 | 0.85 | -0.22 | 0.75 |
| Bdnf | -0.74 | 1.63E-54 | 3.38E-50 | 0.28 | 0.4 | -0.13 | 0.69 |
| Cacna1d | -0.75 | 2.40E-68 | 4.99E-64 | 0.24 | 0.38 | -0.14 | 0.63 |

**Supplementary Table 6. Differentially expressed genes in the mutant λ-junction mesenchymal cells.**

| | | | | | | | |
|---|---|---|---|---|---|---|---|
| H2-Ab1 | -0.75 | 6.88E-17 | 1.43E-12 | 0.05 | 0.08 | -0.04 | 0.57 |
| Sdk1 | -0.75 | 3.36E-19 | 6.99E-15 | 0.06 | 0.11 | -0.04 | 0.59 |
| Samd1 | -0.75 | 7.52E-117 | 1.56E-112 | 0.34 | 0.54 | -0.2 | 0.64 |
| Gm49708 | -0.75 | 4.39E-160 | 9.12E-156 | 0.62 | 0.78 | -0.17 | 0.79 |
| Pmaip1 | -0.76 | 4.23E-69 | 8.80E-65 | 0.23 | 0.38 | -0.14 | 0.62 |
| L1cam | -0.76 | 4.68E-30 | 9.73E-26 | 0.07 | 0.14 | -0.06 | 0.54 |
| Tmem132c | -0.76 | 1.51E-84 | 3.14E-80 | 0.32 | 0.48 | -0.16 | 0.67 |
| Atp1b1 | -0.77 | 1.77E-14 | 3.68E-10 | 0.06 | 0.09 | -0.04 | 0.62 |
| Fzd10os | -0.77 | 5.87E-53 | 1.22E-48 | 0.17 | 0.28 | -0.11 | 0.6 |
| Hpse2 | -0.78 | 1.17E-61 | 2.42E-57 | 0.17 | 0.3 | -0.13 | 0.58 |
| Zeb2 | -0.79 | 8.52E-76 | 1.77E-71 | 0.42 | 0.55 | -0.13 | 0.76 |
| Kcnb1 | -0.8 | 8.54E-17 | 1.78E-12 | 0.04 | 0.07 | -0.03 | 0.54 |
| Gm10320 | -0.8 | 6.40E-149 | 1.33E-144 | 0.3 | 0.53 | -0.23 | 0.56 |
| Nkd2 | -0.81 | 3.75E-37 | 7.80E-33 | 0.13 | 0.21 | -0.08 | 0.61 |
| Irx5 | -0.81 | 4.10E-38 | 8.52E-34 | 0.26 | 0.35 | -0.09 | 0.74 |
| Tnfrsf19 | -0.81 | 1.01E-31 | 2.10E-27 | 0.09 | 0.16 | -0.07 | 0.58 |
| Rtn1 | -0.81 | 1.77E-52 | 3.68E-48 | 0.22 | 0.33 | -0.11 | 0.66 |
| Nts | -0.82 | 1.02E-10 | 2.12E-06 | 0.06 | 0.09 | -0.03 | 0.68 |
| Gm9866 | -0.82 | 2.23E-30 | 4.64E-26 | 0.07 | 0.13 | -0.06 | 0.53 |
| Wnt5a | -0.82 | 8.96E-231 | 1.86E-226 | 0.64 | 0.84 | -0.2 | 0.76 |
| Pcdh10 | -0.83 | 8.05E-24 | 1.67E-19 | 0.09 | 0.15 | -0.06 | 0.62 |
| Rgs4 | -0.83 | 1.02E-15 | 2.12E-11 | 0.03 | 0.06 | -0.03 | 0.52 |
| Sdk2 | -0.85 | 6.42E-19 | 1.34E-14 | 0.05 | 0.08 | -0.04 | 0.55 |
| Nkd1 | -0.85 | 1.42E-125 | 2.94E-121 | 0.36 | 0.55 | -0.2 | 0.65 |
| Igfbp7 | -0.87 | 6.10E-138 | 1.27E-133 | 0.25 | 0.46 | -0.21 | 0.54 |
| Cntfr | -0.87 | 1.03E-60 | 2.13E-56 | 0.2 | 0.32 | -0.12 | 0.62 |
| Egfl6 | -0.88 | 1.32E-24 | 2.74E-20 | 0.06 | 0.11 | -0.05 | 0.54 |
| Cxcl13 | -0.88 | 1.13E-13 | 2.36E-09 | 0.03 | 0.05 | -0.03 | 0.51 |
| Runx2 | -0.88 | 1.19E-50 | 2.47E-46 | 0.16 | 0.26 | -0.1 | 0.61 |
| Klf4 | -0.89 | 1.10E-55 | 2.29E-51 | 0.16 | 0.28 | -0.11 | 0.59 |
| Dync1i1 | -0.9 | 3.41E-15 | 7.09E-11 | 0.03 | 0.06 | -0.03 | 0.52 |
| Ntrk2 | -0.9 | 1.55E-16 | 3.22E-12 | 0.03 | 0.06 | -0.03 | 0.48 |
| Hhip | -0.91 | 4.29E-20 | 8.92E-16 | 0.04 | 0.08 | -0.04 | 0.51 |
| Angpt1 | -0.91 | 1.27E-47 | 2.65E-43 | 0.14 | 0.24 | -0.1 | 0.59 |
| Bmp4 | -0.91 | 1.83E-130 | 3.81E-126 | 0.45 | 0.63 | -0.18 | 0.72 |
| Col7a1 | -0.92 | 3.95E-13 | 8.21E-09 | 0.03 | 0.05 | -0.02 | 0.52 |
| AW551984 | -0.92 | 7.25E-46 | 1.51E-41 | 0.12 | 0.21 | -0.09 | 0.56 |
| Rspo1 | -0.93 | 4.55E-28 | 9.46E-24 | 0.07 | 0.12 | -0.06 | 0.55 |
| Col6a1 | -0.94 | 8.92E-55 | 1.85E-50 | 0.14 | 0.25 | -0.11 | 0.57 |
| Ccbe1 | -0.94 | 1.20E-32 | 2.49E-28 | 0.05 | 0.1 | -0.06 | 0.46 |
| Lrrc17 | -0.94 | 1.07E-36 | 2.23E-32 | 0.07 | 0.14 | -0.07 | 0.51 |
| Kctd12 | -0.95 | 6.01E-80 | 1.25E-75 | 0.28 | 0.43 | -0.15 | 0.66 |
| Gm30564 | -0.95 | 2.91E-46 | 6.06E-42 | 0.09 | 0.18 | -0.09 | 0.51 |
| Rmst | -0.95 | 6.34E-29 | 1.32E-24 | 0.06 | 0.12 | -0.06 | 0.53 |
| Pid1 | -0.96 | 5.58E-84 | 1.16E-79 | 0.17 | 0.32 | -0.15 | 0.54 |
| Myorg | -0.97 | 3.02E-31 | 6.29E-27 | 0.06 | 0.12 | -0.06 | 0.52 |
| Lef1 | -0.98 | 4.07E-139 | 8.47E-135 | 0.41 | 0.61 | -0.2 | 0.67 |
| Tubb3 | -0.98 | 1.26E-60 | 2.62E-56 | 0.14 | 0.25 | -0.11 | 0.56 |

**Supplementary Table 6. Differentially expressed genes in the mutant λ-junction mesenchymal cells.**

| | | | | | | | |
|---|---|---|---|---|---|---|---|
| Col6a2 | -0.98 | 1.08E-51 | 2.25E-47 | 0.15 | 0.25 | -0.1 | 0.59 |
| Ifitm1 | -0.99 | 3.17E-64 | 6.59E-60 | 0.14 | 0.25 | -0.12 | 0.54 |
| Fzd10 | -1.01 | 8.96E-50 | 1.86E-45 | 0.09 | 0.18 | -0.09 | 0.51 |
| Thrb | -1.01 | 5.04E-19 | 1.05E-14 | 0.03 | 0.07 | -0.03 | 0.49 |
| Axin2 | -1.04 | 1.09E-133 | 2.27E-129 | 0.28 | 0.48 | -0.2 | 0.59 |
| Zar1 | -1.05 | 1.29E-20 | 2.68E-16 | 0.03 | 0.06 | -0.03 | 0.44 |
| Bmp5 | -1.06 | 6.01E-22 | 1.25E-17 | 0.02 | 0.06 | -0.03 | 0.42 |
| Plekha2 | -1.09 | 3.35E-26 | 6.96E-22 | 0.03 | 0.08 | -0.04 | 0.44 |
| Tll1 | -1.1 | 6.03E-15 | 1.25E-10 | 0.03 | 0.05 | -0.03 | 0.49 |
| Fam162b | -1.19 | 2.83E-29 | 5.89E-25 | 0.04 | 0.08 | -0.05 | 0.44 |
| Pgm5 | -1.2 | 9.53E-29 | 1.98E-24 | 0.03 | 0.08 | -0.04 | 0.42 |
| Col1a1 | -1.21 | 1.40E-148 | 2.90E-144 | 0.31 | 0.51 | -0.2 | 0.61 |
| Dlx5 | -1.21 | 5.43E-69 | 1.13E-64 | 0.11 | 0.23 | -0.11 | 0.5 |
| Dlg2 | -1.22 | 2.27E-50 | 4.72E-46 | 0.05 | 0.13 | -0.08 | 0.42 |
| Emp1 | -1.25 | 9.65E-44 | 2.01E-39 | 0.07 | 0.14 | -0.07 | 0.48 |
| Ndnf | -1.28 | 1.37E-107 | 2.86E-103 | 0.2 | 0.37 | -0.16 | 0.56 |
| Dlx6 | -1.29 | 8.31E-36 | 1.73E-31 | 0.03 | 0.08 | -0.05 | 0.38 |
| Sp5 | -1.3 | 3.23E-21 | 6.72E-17 | 0.03 | 0.06 | -0.03 | 0.43 |
| Glis1 | -1.34 | 8.21E-26 | 1.71E-21 | 0.02 | 0.06 | -0.04 | 0.39 |
| Robo2 | -1.35 | 3.40E-68 | 7.07E-64 | 0.13 | 0.25 | -0.11 | 0.53 |
| Cyp26b1 | -1.35 | 1.95E-77 | 4.05E-73 | 0.1 | 0.22 | -0.12 | 0.47 |
| Tlx2 | -1.36 | 1.32E-23 | 2.75E-19 | 0.02 | 0.06 | -0.03 | 0.4 |
| Wif1 | -1.47 | 1.80E-66 | 3.74E-62 | 0.07 | 0.16 | -0.1 | 0.41 |
| Cpxm2 | -1.49 | 1.66E-41 | 3.45E-37 | 0.03 | 0.09 | -0.06 | 0.37 |
| Anxa2 | -1.53 | 8.21E-142 | 1.71E-137 | 0.2 | 0.39 | -0.19 | 0.51 |
| Apcdd1 | -1.53 | 5.92E-129 | 1.23E-124 | 0.16 | 0.34 | -0.17 | 0.49 |
| Cacna2d3 | -1.62 | 3.33E-53 | 6.92E-49 | 0.04 | 0.11 | -0.07 | 0.35 |
| Calb2 | -1.71 | 5.81E-35 | 1.21E-30 | 0.04 | 0.1 | -0.06 | 0.44 |
| Epha8 | -1.85 | 5.24E-49 | 1.09E-44 | 0.02 | 0.08 | -0.06 | 0.29 |
| Hey2 | -1.91 | 2.16E-44 | 4.48E-40 | 0.04 | 0.1 | -0.06 | 0.38 |
| Cxcl14 | -2.04 | 5.56E-90 | 1.16E-85 | 0.08 | 0.19 | -0.12 | 0.39 |
| Smoc2 | -2.43 | 1.78E-49 | 3.70E-45 | 0.02 | 0.07 | -0.05 | 0.23 |
| Pax1 | -2.66 | 6.65E-69 | 1.38E-64 | 0.01 | 0.08 | -0.06 | 0.17 |
| Stmn2 | -2.82 | 2.91E-56 | 6.06E-52 | 0.03 | 0.1 | -0.07 | 0.33 |
| Uty | -8.63 | 3.23E-139 | 6.72E-135 | 0 | 0.09 | -0.09 | 0 |
| Kdm5d | -8.86 | 1.03E-157 | 2.14E-153 | 0 | 0.11 | -0.11 | 0 |
| Eif2s3y | -9.71 | 2.77E-256 | 5.77E-252 | 0 | 0.17 | -0.17 | 0 |
| Ddx3y | -9.75 | 1.09E-253 | 2.27E-249 | 0 | 0.17 | -0.17 | 0 |

**Supplementary Table 7. Enriched terms detected using Enrichr for differentially expressed genes in the mutant λ-junction ectoderm or mesenchymal cells.**
**All Ect GO BP 2025**

| Term | Overlap | P-value | Adjusted P-value | Odds Ratio | Combined Score | Genes |
|---|---|---|---|---|---|---|
| Generation of Neurons (GO:0048699) | 51/192 | 8.22E-20 | 2.89E-16 | 5.82 | 255.76 | FZD10;CELSR1;SOX21;SH3RF1;SOX2;SOX3;SOX1;NRCAM;MEF2C;WNT9B;SOX14;SOX11;NAV1;SOX12;EML1;RUNX2;TUBB2B;MAP1B;DDIT4;DCX;LMX1B;RTN2;SEMA3A;ASCL1;EGFR;WNT6;RACGAP1;ERBB3;ERBB2;MARK2;WNT3;TGFB2;WNT10A;FZD2;IRX5;FZD4;FZD7;IRX3;WNT7B;IRX4;FZD9;CDK6;ID2;ID4;ID3;PLXNB2;INSM1;FGF13;FGFR1;ADGRL3;CDK5R1 |
| Neuron Differentiation (GO:0030182) | 40/181 | 5.84E-13 | 1.03E-09 | 4.52 | 127.37 | RTN2;FZD10;ASCL1;SOX21;EGFR;SOX2;SOX3;WNT6;SOX1;BRINP1;ERBB3;ERBB2;OTX2;WNT3;MEF2C;TGFB2;WNT10A;FZD2;IRX5;FZD4;FZD7;IRX3;WNT7B;IRX4;FZD9;PBX3;WNT9B;SOX14;SOX11;PAX6;SOX12;RUNX2;ISL2;ID2;DDIT4;ID4;ID3;LMX1B;INSM1;CDK5R1 |
| Positive Regulation of Cell Differentiation (GO:0045597) | 53/297 | 1.03E-12 | 1.20E-09 | 3.48 | 96.12 | APP;BTG1;GJA1;PPP1R13L;NRCAM;SOX9;MEF2C;PRKCH;IGFBP3;SOX11;TENT5A;NPNT;RUNX2;TIAM1;OLFM1;SFRP1;PCP4;SFRP2;WDFY2;CRB2;FBN2;FOXC1;CEBPB;PKDCC;LAMA1;LEF1;TACSTD2;TWIST1;ASCL1;GPC1;PDPN;GCNT2;WNT3;WWTR1;TGFB2;ZFHX3;WNT7B;IGF1;GDF6;SMARCA2;SMAD5;MAPK12;COL1A1;BMP4;HEYL;MAPK11;BMP2;REST;RGCC;ID2;CAMK1;INSM1;FGFR1 |
| Regulation of Cell Migration (GO:0030334) | 71/479 | 1.98E-12 | 1.74E-09 | 2.81 | 75.68 | SEMA5A;CITED2;CLSTN1;LAMC2;MYLK;AMOT;DACH1;FGF9;DPYSL3;ENPP2;CTSH;CPNE3;EPHB2;GLUL;LIMCH1;LMO4;NOG;RHOC;TIAM1;CLDN4;TMEFF2;SFRP1;CLDN3;SFRP2;NAV3;KIT;ITGA6;SGK3;BEX4;FBN2;HDAC2;CD151;NOTCH1;SEMA3C;SEMA3A;LEF1;PDGFA;PIK3R3;TWIST1;PRKX;SEMA3F;THBS1;EGFR;NKD1;MGAT5;PDGFC;PDPN;GCNT2;PLXNA2;CARMIL1;JAG1;SEMA4D;FZD4;IGF1;PLET1;SORL1;SMAD7;COL1A1;CAMSAP3;FGF17;BMP2;CXCL12;DLG5;SNAI2;PLXNB2;PLXNB1;EPPK1;AMOTL2;LGR6;FGF13;FGF10 |
| Notch Signaling Pathway (GO:0007219) | 17/42 | 1.05E-10 | 7.37E-08 | 10.70 | 245.77 | NOTCH2;JAG2;NOTCH3;NOTCH1;JAG1;DTX3L;NRARP;SORBS2;ASCL1;DTX4;RBPJ;HEYL;BMP2;HEY1;HEY2;SNAI2;HES5 |
| Negative Regulation of Transcription by RNA Polymerase II (GO:0000122) | 88/732 | 4.65E-10 | 2.34E-07 | 2.21 | 47.49 | RB1;APP;THRB;CITED2;WWC1;CHD3;NR2E1;RBPJ;SOX21;BACH2;ETS2;AMOT;SOX2;SOX3;DACH1;SOX1;HEY1;HEY2;SOX9;LRRFIP1;TLE4;MEF2C;MSX2;KCNH8;NOG;SOX13;SOX14;SOX11;PAX6;SOX12;PARP9;ETV5;TLE5;EID1;RIPPLY3;MAF;IRF2;EZR;JDP2;HDAC4;CEBPA;CEBPB;HDAC2;FOXC1;NFIX;NOTCH1;SATB1;LEF1;TWIST1;ASCL1;CXXC5;CUX2;PLAGL1;SAMD1;CRYM;SKIL;HES5;E2F8;SNCA;ZBTB18;TFAP2A;TFAP2B;ZFHX3;SEMA4D;BCL11A;IRX3;NR2F1;NR2F2;KLF4;SMARCA2;MEIS2;TBX3;NR2F6;SMAD7;BMP4;EFNA1;BMP2;REST;NFIB;ID2;ID4;SNAI2;ID3;AMOTL2;INSM1;FRK;FGFR2;MXD4 |

**Supplementary Table 7. Enriched terms detected using Enrichr for differentially expressed genes in the mutant λ-junction ectoderm or mesenchymal cells.**
**All Ect GO BP 2025**

| | | | | | | |
|---|---|---|---|---|---|---|
| Positive Regulation of Cell Population Proliferation (GO:0008284) | 66/484 | 4.66E-10 | 2.34E-07 | 2.54 | 54.48 | PID1;BNC1;IRS2;LAMC2;RBPJ;GJA1;HMGN5;CCND2;FGF9;NAMPT;CTSH;ITGAV;SOX9;MEF2C;PRKCH;SOX11;TIAM1;SFRP1;SFRP2;SAPCD2;KIT;PRTN3;TSPYL5;HDAC4;HDAC2;NOTCH1;LEF1;PDGFA;PIK3R1;THBS1;EGFR;PTHLH;GNAI2;TERT;RASSF10;RAB25;SERTAD1;ERBB3;EPCAM;PDGFC;ERBB2;GCNT2;HES5;WWTR1;TFAP2B;TGFB2;FZD9;IGF1;SMARCA2;MEIS2;TBX3;BMP4;FGF17;BMP2;MEIS1;CDK6;PRC1;ID2;INSM1;FGF13;NES;SLC25A33;CRLF1;FGFR2;FGF10;FGFR1 |
| Negative Regulation of Macromolecule Biosynthetic Process (GO:0010558) | 37/202 | 1.29E-09 | 5.65E-07 | 3.56 | 72.89 | RB1;NOTCH2;DDX3Y;APP;HDAC2;NOTCH1;CITED2;DACH1;TERT;HEY2;GRB10;ITGB8;APOE;SH3GL2;STOX1;MEF2C;TGFB2;SOX11;IGF1;PARP9;KLF4;SMAD5;TLE5;BMP4;SLC7A5;OLFM1;SFRP1;OCLN;TMEFF2;BMP2;REST;CLDN3;SFRP2;MYCN;ID2;FGF13;RBMS3 |
| Cardiac Epithelial to Mesenchymal Transition (GO:0060317) | 11/19 | 1.83E-09 | 7.15E-07 | 21.54 | 433.32 | BMP4;HEYL;OLFM1;TGFB2;BMP2;NOTCH1;HEY1;HEY2;SNAI2;RBPJ;TBX3 |
| Regulation of Neuron Differentiation (GO:0045664) | 21/76 | 2.60E-09 | 9.15E-07 | 6.02 | 118.92 | APP;MEF2C;ZFHX3;ZFHX2;SOX11;NREP;ASCL1;SOX2;SOX3;TIAM1;HEYL;PCP4;BMP2;REST;HEY1;ID2;NRCAM;HES5;YWHAH;CDK5R1;FGFR1 |

**Supplementary Table 7. Enriched terms detected using Enrichr for differentially expressed genes in the mutant λ-junction ectoderm or mesenchymal cells.**
**All Ect GO CC 2025**

| Term | Overlap | P-value | Adjusted P-value | Odds Ratio | Combined Score | Genes |
|---|---|---|---|---|---|---|
| Cell-Cell Junction (GO:0005911) | 49/316 | 1.18E-09 | 3.45E-07 | 2.93 | 60.19 | CTNND2;AQP3;TMEM47;GJA1;CDH4;POF1B;CDH2;ACTN1;KAZN;FRMD4A;TIAM1;CLDN10;CLDN4;CLDN3;SAPCD2;PKP1;PKP4;DSG2;PKP3;EZR;PLPP3;AJUBA;CRB2;COL17A1;NOTCH1;LUZP1;CNN2;PDLIM1;CCDC85C;MARVELD2;EPB41L3;STXBP6;PCDH1;PAK4;JAG1;JUP;BAIAP2;MPP7;SMAD7;GJB2;OCLN;KRT18;PTK7;DLG5;NECTIN4;NECTIN3;ADGRL3;TJP2;NECTIN1 |
| Apical Junction Complex (GO:0043296) | 25/107 | 3.77E-09 | 5.31E-07 | 4.81 | 93.36 | CRB2;AMOT;POF1B;CCDC85C;MARVELD2;EPCAM;CLDN23;CGNL1;JUP;FRMD4A;CGN;MPP7;CLDN6;CAMSAP3;CLDN10;CLDN4;OCLN;CLDN3;SAPCD2;CLDN7;AMOTL2;STRN;EPPK1;NECTIN3;TJP2 |
| Polymeric Cytoskeletal Fiber (GO:0099513) | 44/280 | 5.46E-09 | 5.31E-07 | 2.97 | 56.44 | BEX4;RTN2;SYNM;LUZP1;LMOD1;IQGAP1;IQGAP2;RCSD1;EVPL;PPL;KIF15;AMOT;BCL2L11;POF1B;BAG2;TUBB3;COTL1;MARK2;PLS1;DMTN;ACTN1;TPM2;KRT8;PAWR;COBL;KIF23;TUBG2;FHDC1;TUBB4B;EML1;DNM3;TUBB2B;CLIP1;MYO1B;KRT18;MAP1B;KRT14;MAP7D2;PKP1;EPPK1;VIM;EZR;FGF13;NES |
| Tight Junction (GO:0070160) | 21/96 | 2.19E-07 | 1.60E-05 | 4.41 | 67.58 | LUZP1;FRMD4A;CGN;MPP7;AMOT;CLDN6;CLDN10;GJA1;CLDN4;OCLN;CLDN3;POF1B;MARVELD2;EPCAM;CLDN23;CLDN7;AMOTL2;EPPK1;STRN;CGNL1;TJP2 |
| Adherens Junction (GO:0005912) | 27/151 | 3.56E-07 | 1.76E-05 | 3.44 | 51.02 | NOTCH1;CTNND2;PDLIM1;CDH4;CDH2;POF1B;CCDC85C;STXBP6;PAK4;JAG1;JUP;FRMD4A;BAIAP2;MPP7;SMAD7;CAMSAP3;KRT18;DLG5;PKP1;PKP4;PKP3;EZR;PLPP3;NECTIN4;AJUBA;NECTIN3;NECTIN1 |
| Cytoskeleton (GO:0005856) | 76/694 | 3.61E-07 | 1.76E-05 | 1.97 | 29.23 | LAD1;SYNM;LMOD1;HSPB1;PPP1R9A;CLU;MYLK;LIMD2;EPB41L4A;PCLO;TUBB3;BASP1;ARHGDIB;TEKT1;BMERB1;LRRFIP1;GAS2L3;LIMCH1;TPM2;KRT8;PAWR;NAV1;TUBG2;EML1;MTSS1;MTSS2;TUBB2B;CLIP1;CDC42EP5;INCENP;MAP7D2;DCX;PKP4;TNIK;EZR;LUZP1;KLHL14;CDCA8;IQGAP1;IQGAP2;EVPL;PPL;CNN2;PDLIM1;EPB41L5;ABLIM1;NUAK1;CALD1;EPB41L2;EPB41L3;MAP7;VILL;MPZL2;SNCA;DMTN;GSN;JUP;GRHL3;SORBS2;TUBB4B;BAIAP2;KLF4;SMARCA2;ARHGAP32;MYO1B;KRT18;PRC1;MYO5B;KRT15;MYO5C;KRT14;CCSER2;FILIP1;EPPK1;VIM;NES |
| Bicellular Tight Junction (GO:0005923) | 18/81 | 1.24E-06 | 5.19E-05 | 4.49 | 61.03 | FRMD4A;CGN;MPP7;CLDN6;CLDN10;CLDN4;OCLN;CLDN3;POF1B;MARVELD2;EPCAM;CLDN23;CLDN7;AMOTL2;EPPK1;STRN;CGNL1;TJP2 |

**Supplementary Table 7. Enriched terms detected using Enrichr for differentially expressed genes in the mutant λ-junction ectoderm or mesenchymal cells.**
**All Ect GO CC 2025**

| | | | | | | |
|---|---|---|---|---|---|---|
| Collagen-Containing Extracellular Matrix (GO:0062023) | 47/383 | 3.15E-06 | 1.05E-04 | 2.22 | 28.12 | COL17A1;FBN2;ITIH5;TINAGL1;CD151;LAMA1;SDC2;HTRA1;PLOD2;LAMC2;CLU;THBS1;THBS3;LGALS1;CDH2;CTSL;GPC1;SERPINH1;CTSH;TIMP3;NCAM1;APOE;CTSF;CTSC;CTHRC1;TGFB2;ANXA5;NPNT;KAZALD1;DCN;COL1A1;MFAP4;BCAM;SFRP1;VCAN;SFRP2;CXCL12;COL4A1;COL5A2;MGP;VWA1;PRTN3;COL9A2;MFGE8;FGFR2;FBN1;FGF10 |
| Actin Filament (GO:0005884) | 17/78 | 3.25E-06 | 1.05E-04 | 4.38 | 55.29 | DMTN;LUZP1;ACTN1;TPM2;LMOD1;COBL;PAWR;IQGAP1;RCSD1;FHDC1;AMOT;MYO1B;POF1B;COTL1;EZR;MARK2;PLS1 |
| Actin Cytoskeleton (GO:0015629) | 43/361 | 1.72E-05 | 5.01E-04 | 2.14 | 23.49 | LAD1;KLHL14;LUZP1;LMOD1;IQGAP1;IQGAP2;RCSD1;PPP1R9A;MYLK;LIMD2;AMOT;CNN2;ABLIM1;POF1B;CDH2;CALD1;COTL1;VILL;MARK2;SPTBN2;PLS1;SNCA;GAS2L3;DMTN;GSN;LIMCH1;ACTN1;TPM2;PAWR;COBL;GRHL3;SORBS2;FHDC1;BAIAP2;MTSS1;MTSS2;ARHGAP32;MYO1B;MYO5B;MYO5C;FILIP1;AMOTL2;EZR |

**Supplementary Table 7. Enriched terms detected using Enrichr for differentially expressed genes in the mutant λ-junction ectoderm or mesenchymal cells.**
**All Ect GO MF 2025**

| Term | Overlap | P-value | Adjusted P-value | Odds Ratio | Combined Score | Genes |
|---|---|---|---|---|---|---|
| Sequence-Specific Double-Stranded DNA Binding (GO:1990837) | 79/706 | 8.62E-08 | 5.66E-05 | 2.02 | 32.91 | THRB;BNC1;MCM10;AHR;CHD3;NR2E1;BACH2;ETS2;SOX2;DMBX1;SOX3;SIX6;HEY1;BASP1;HEY2;SOX9;HES6;MEF2C;KLF13;MSX2;KCNH8;SOX13;SOX14;PAX3;SOX11;PAX6;SOX12;POU3F1;RUNX2;ETV5;DMRTA2;ISL2;MAF;MYCN;PAX7;DMRTA1;ELF5;IRF2;LMX1B;IRF6;JDP2;HDAC4;CEBPA;CEBPB;FOXC1;NFIX;DLX3;HLF;LEF1;ASCL1;CENPB;CUX2;OTX2;OTX1;HES5;E2F8;SNCA;ZBTB18;TFAP2A;EGR1;TFAP2B;ZFHX3;IRX5;BCL11B;IRX3;NR2F1;UNCX;KLF4;GRHL1;SMARCA2;SMAD5;MEIS2;TBX3;NR2F6;LIN28B;REST;NFIB;SNAI2;SP8 |
| Sequence-Specific DNA Binding (GO:0043565) | 77/718 | 6.92E-07 | 2.27E-04 | 1.92 | 27.29 | CALCOCO1;THRB;AHR;NR2E1;RBPJ;BACH2;ETS2;SOX2;DMBX1;SOX3;SIX6;HEY1;HEY2;SOX9;HES6;MEF2C;KLF13;MSX2;KCNH8;SOX13;SOX14;PAX3;PAX6;SOX12;POU3F1;FOXP2;RUNX2;ETV5;DMRTA2;ISL2;MAF;MYCN;PAX7;DMRTA1;ELF5;IRF2;LMX1B;IRF6;JDP2;CEBPB;FOXC1;NFIX;DLX3;HLF;SATB1;LEF1;ASCL1;CXXC5;CENPB;CUX2;TERT;OTX2;OTX1;PPARGC1A;HES5;E2F8;ZBTB18;TFAP2A;EGR1;TFAP2B;IRX5;BCL11B;IRX3;NR2F1;UNCX;GRHL3;NR2F2;KLF4;GRHL1;SMAD5;MEIS2;TBX3;NR2F6;LIN28B;NFIB;SNAI2;SP8 |
| Cadherin Binding (GO:0045296) | 42/317 | 1.47E-06 | 3.22E-04 | 2.42 | 32.51 | LAD1;NOTCH3;CTNND2;ARHGAP18;SLC3A2;IQGAP1;EVPL;PPL;EGFR;CNN2;PDLIM1;CDH4;CDH2;EPCAM;CALD1;PPP1R13L;COBLL1;STXBP6;LRRFIP1;MARK2;SPTBN2;PAK4;SH3GLB2;JUP;CGN;BAIAP2;MPP7;PPFIBP1;MYO1B;KRT18;FMNL2;GIPC1;CHMP2B;PLIN3;PKP1;PKP4;ITGA6;PKP3;EZR;PFKP;CDK5R1;TJP2 |
| Insulin-Like Growth Factor Binding (GO:0005520) | 7/15 | 1.22E-05 | 1.89E-03 | 13.66 | 154.56 | IGFBP4;IGFBP3;IGFBP2;ITGA6;ITGAV;LRP2;IGF2R |
| Double-Stranded DNA Binding (GO:0003690) | 67/651 | 1.44E-05 | 1.89E-03 | 1.83 | 20.38 | THRB;AHR;NR2E1;BACH2;ETS2;DMBX1;SOX3;SIX6;HEY1;HEY2;SOX9;HES6;MEF2C;KLF13;MSX2;KCNH8;SOX14;PAX3;PAX6;SOX12;POU3F1;RUNX2;ETV5;DMRTA2;ISL2;MAF;MYCN;PAX7;DMRTA1;ELF5;IRF2;LMX1B;IRF6;JDP2;CEBPB;NFIX;DLX3;HLF;SATB1;LEF1;ASCL1;EGFR;RAD51AP1;CUX2;OTX2;OTX1;HES5;E2F8;ZBTB18;TFAP2A;EGR1;TFAP2B;IRX5;BCL11B;IRX3;NR2F1;UNCX;KLF4;GRHL1;SMAD5;MEIS2;TBX3;NR2F6;LIN28B;NFIB;SNAI2;SP8 |
| DNA-binding Transcription Activator Activity, RNA Polymerase II-specific (GO:0001228) | 39/318 | 2.15E-05 | 2.36E-03 | 2.21 | 23.75 | NOTCH2;CEBPA;FOXC1;NOTCH1;HLF;LEF1;RBPJ;SOX21;LITAF;SOX2;SOX3;SOX1;SALL2;PLAGL1;OTX2;SOX9;OTX1;TFAP2A;EGR1;TFAP2B;MEF2C;PRRX1;KCNH8;IRX3;IRX4;SOX14;SOX11;PAX6;GRHL3;SOX12;GRHL1;KLF4;MEIS2;ETV5;TBX3;MEIS1;MYCN;NFIB;IRF2 |
| Insulin-Like Growth Factor I Binding (GO:0031994) | 6/12 | 3.26E-05 | 2.72E-03 | 15.60 | 161.17 | IGFBP4;IGFBP3;IGFBP2;ITGA6;ITGAV;LRP2 |
| Cadherin Binding Involved in Cell-Cell Adhesion (GO:0098641) | 7/17 | 3.31E-05 | 2.72E-03 | 10.93 | 112.72 | PDLIM1;KRT18;EPCAM;PKP3;STXBP6;BAIAP2;PAK4 |

**Supplementary Table 7. Enriched terms detected using Enrichr for differentially expressed genes in the mutant λ-junction ectoderm or mesenchymal cells.**
**All Ect GO MF 2025**

| | | | | | | |
|---|---|---|---|---|---|---|
| Cis-Regulatory Region Sequence-Specific DNA Binding (GO:0000987) | 94/1035 | 4.80E-05 | 3.50E-03 | 1.60 | 15.89 | CALCOCO1;THRB;NR2E1;RBPJ;LITAF;SOX21;ETS2;BACH2;SOX2;SOX3;SIX6;DACH1;SOX1;HEY1;HEY2;UTY;LRRFIP1;SOX9;FEZF1;HES6;MEF2C;KLF13;KCNH8;SOX13;EBF1;SOX14;PAX3;SOX11;PAX6;SOX12;POU3F1;FOXP2;RUNX2;DMRTA2;ISL2;MAF;MYCN;PAX7;DMRTA1;IRF2;IRF6;JDP2;NOTCH2;HDAC4;CEBPA;FOXC1;CEBPB;DLX3;NFIX;NOTCH1;HLF;SATB1;LEF1;TWIST1;ASCL1;NTN3;MUC1;PLAGL1;OTX2;OTX1;E2F5;SKIL;HES5;E2F8;TFAP2A;EGR1;TFAP2B;ZFHX3;ZFHX2;PRRX1;IRX5;BCL11B;BCL11A;IRX3;IRX4;NR2F1;SMAD9;GRHL3;NR2F2;KLF4;GRHL1;SMAD5;MEIS2;TBX3;NR2F6;HEYL;REST;NFIB;NFE2L3;SP5;SNAI2;SP8;INSM1;MXD4 |
| DNA-binding Transcription Repressor Activity, RNA Polymerase II-specific (GO:0001227) | 28/210 | 7.08E-05 | 4.65E-03 | 2.42 | 23.14 | CEBPB;SATB1;ASCL1;ETS2;DACH1;HEY1;PLAGL1;HEY2;LRRFIP1;SKIL;HES5;E2F8;TFAP2A;ZBTB18;ZFHX3;MSX2;BCL11A;KCNH8;IRX3;SOX13;NR2F1;FOXP2;TBX3;HEYL;REST;SNAI2;INSM1;JDP2 |

**Supplementary Table 7. Enriched terms detected using Enrichr for differentially expressed genes in the mutant λ-junction ectoderm or mesenchymal cells. All Ectoderm Diseases 2025**

| Term | Overlap | P-value | Adjusted P-value | Odds Ratio | Combined Score | Genes |
|---|---|---|---|---|---|---|
| OROFACIAL CLEFT | 21/104 | 9.00E-07 | 1.27E-04 | 3.98 | 55.41 | SEMA4D;DGKA;NOG;KRT8;WNT9B;RASSF9;TANC2;DLK1;GADD45G;CUX2;KRT18;CMSS1;ERBB3;PAX7;IRF2;RPRML;IRF6;WNT3;CBLN2;FGF10;FGFR1 |
| LIP DISEASE | 21/104 | 9.00E-07 | 1.27E-04 | 3.98 | 55.41 | SEMA4D;DGKA;NOG;KRT8;WNT9B;RASSF9;TANC2;DLK1;GADD45G;CUX2;KRT18;CMSS1;ERBB3;PAX7;IRF2;RPRML;IRF6;WNT3;CBLN2;FGF10;FGFR1 |
| CLEFT LIP | 21/104 | 9.00E-07 | 1.27E-04 | 3.98 | 55.41 | SEMA4D;DGKA;NOG;KRT8;WNT9B;RASSF9;TANC2;DLK1;GADD45G;CUX2;KRT18;CMSS1;ERBB3;PAX7;IRF2;RPRML;IRF6;WNT3;CBLN2;FGF10;FGFR1 |
| PHYSICAL DISORDER | 26/185 | 5.20E-05 | 4.56E-03 | 2.57 | 25.38 | PKDCC;HIBADH;DGKA;CUX2;ERBB3;RPRML;WNT3;ZFHX3;SEMA4D;EYA1;NOG;KRT8;WNT9B;RASSF9;TANC2;DLK1;GADD45G;KRT18;CMSS1;PAX7;IRF2;SCG5;IRF6;CBLN2;FGF10;FGFR1 |
| MOUTH DISEASE | 40/343 | 5.38E-05 | 4.56E-03 | 2.09 | 20.51 | DGKA;TMEM59L;PITPNC1;NPAS3;THBS3;WNT6;MUC1;CUX2;CCND2;TERT;ERBB3;RPRML;ITGB6;WNT3;WNT10A;ZFHX3;SEMA4D;NOG;KRT8;PBX3;WNT9B;RASSF9;TUBG2;SLC50A1;TANC2;DLK1;GADD45G;EFNA1;CMSS1;KRT18;PPP1R3B;PAX7;DMRTA1;IRF2;IRF6;RHCG;CRLF1;CBLN2;FGFR1;FGF10 |

**Supplementary Table 7. Enriched terms detected using Enrichr for differentially expressed genes in the mutant λ-junction ectoderm or mesenchymal cells. All Ectoderm Diseases 2025**

| | | | | | | |
|---|---|---|---|---|---|---|
| NERVOUS SYSTEM DISEASE | 184/2371 | 1.85E-04 | 1.30E-02 | 1.36 | 11.70 | IFITM3;APP;CMAS;IFITM2;NAB1;STMN3;ENO3;CLU;CMIP;RIMS2;CDH4;LGALS1;CCND2;TUBB3;DPYSL3;SCN9A;GRB10;PLCE1;NRTN;LMO1;MINAR2;UNC13C;SLC15A2;UNC5B;ACTN1;TPM2;FNDC3B;KAZN;FRMD4A;LMO7;POU3F1;MAPK8IP2;PDGFRL;TANC2;RUNX2;ISL2;BCAM;MAF;ADGRB3;COL4A1;TMEM79;PKP4;ROR1;PRTN3;LMX1B;EMID1;XPR1;CD151;CTTNBP2;ARHGEF28;TACSTD2;HTRA1;CDCA8;TWIST1;IQGAP1;NDRG2;LPP;MED12L;NPAS3;CNN2;ADAMTS15;ZFP57;TERT;ADAMTS19;CALD1;OARD1;ADAMTS18;COBLL1;EPB41L3;ADAMTS17;PLXNA2;TSPAN2;APOE;PPARGC1A;SCN3A;SKIL;PLTP;WNT3;TRPM3;KLHL29;FDPS;TFAP2B;ZFHX3;GSTM1;YES1;AFAP1L1;RBPMS;BCL11B;FZD4;GRTP1;NCAPH2;FZD7;RAB27B;SMARCA2;GNAO1;EFNA1;ARHGAP32;BMP2;GJB2;LIN28B;DAB1;FXYD1;COL5A2;FXYD7;PLCH1;FBN1;CDKN1C;CLIC6;NRP2;ITGB5;BNC2;NRARP;SLC2A3;IDE;SIX6;CASP7;PCLO;ALDH2;CASP6;RAC3;TIMP3;TMED1;RHBDF1;PCYT1B;PLA2G12A;DCXR;WNT9B;KREMEN1;COPZ2;COBL;TEX15;RHOC;ETV5;MTSS2;ARL4C;DNM3;SLC7A5;TOX3;DMRTA1;FAM181B;SGK3;MET;GAS7;SAMD5;FOXC1;CAMK2D;SEMA3C;CCDC68;TULP4;SEMA3F;EGFR;PTHLH;NTN3;GNAI2;ATAD2B;JAKMIP2;THBS3;PRTG;ZNRF1;MUC1;SERTAD1;RPRML;RIC3;DUT;PRRX1;EYA2;AMFR;IRX4;UNCX;SLC50A1;SNCAIP;CHFR;REST;ERG28;MEIS1;CMSS1;FMNL2;PRC1;TSPAN18;FILIP1;TACC2;CYB561;FRK;ADGRL3 |
| BREAST CARCINOMA | 41/392 | 4.59E-04 | 2.78E-02 | 1.84 | 14.16 | CLIC6;RAB3C;PID1;HLF;FRY;PTHLH;BCL2L11;CUX2;TERT;CDH2;GCNT2;RBBP8;EMB;POGLUT3;WNT3;EXPH5;TFAP2A;GSTM1;LMO4;WNT9B;EBF1;KREMEN1;TENT5A;KLF4;TBX3;NR2F6;TOM1L1;EXT2;TOX3;CMSS1;INCENP;PRC1;DMRTA1;CCDC88C;ECHDC1;ITGA6;LGR6;CNIH2;EMID1;FGFR2;SSBP4 |
| GASTRIC ADENOCARCINOMA | 5/13 | 6.87E-04 | 3.64E-02 | 9.74 | 70.95 | EFNA1;FDPS;MUC1;SLC50A1;THBS3 |

**Supplementary Table 7. Enriched terms detected using Enrichr for differentially expressed genes in the mutant λ-junction ectoderm or mesenchymal cells. All Ectoderm Diseases 2025**

| | | | | | | |
|---|---|---|---|---|---|---|
| CENTRAL NERVOUS SYSTEM DISEASE | 98/1178 | 7.74E-04 | 3.65E-02 | 1.45 | 10.35 | IFITM3;IFITM2;CMAS;NAB1;STMN3;ENO3;CLU;CMIP;RIMS2;CDH4;CCND2;SCN9A;PLCE1;LMO1;UNC13C;UNC5B;ACTN1;TPM2;KAZN;FRMD4A;TANC2;ISL2;BCAM;MAF;ADGRB3;COL4A1;TMEM79;PKP4;ROR1;LMX1B;XPR1;TACSTD2;HTRA1;TWIST1;NDRG2;MED12L;NPAS3;TERT;OARD1;EPB41L3;PLXNA2;TSPAN2;APOE;PPARGC1A;SKIL;TRPM3;KLHL29;FDPS;TFAP2B;ZFHX3;GSTM1;YES1;BCL11B;GRTP1;FZD7;SMARCA2;GNAO1;ARHGAP32;DAB1;PLCH1;FBN1;CDKN1C;CLIC6;NRARP;SLC2A3;IDE;ALDH2;RAC3;TMED1;RHBDF1;PCYT1B;DCXR;WNT9B;COPZ2;ARL4C;TOX3;DMRTA1;FAM181B;SGK3;SAMD5;CAMK2D;EGFR;ATAD2B;THBS3;MUC1;RPRML;DUT;PRRX1;AMFR;IRX4;UNCX;SLC50A1;SNCAIP;CHFR;MEIS1;PRC1;FILIP1;CYB561 |

**Supplementary Table 7. Enriched terms detected using Enrichr for differentially expressed genes in the mutant λ-junction ectoderm or mesenchymal cells.**
**Ect Up GO BP**

| Term | Overlap | P-value | Adjusted P-value | Odds Ratio | Combined Score | Genes |
|---|---|---|---|---|---|---|
| Generation of Neurons (GO:0048699) | 29/192 | 2.96E-10 | 8.36E-07 | 4.58 | 100.45 | RTN2;CELSR1;SOX21;SH3RF1;EGFR;SOX2;SOX3;SOX1;RACGAP1;ERBB3;ERBB2;NRCAM;MARK2;MEF2C;TGFB2;FZD2;FZD4;FZD7;FZD9;NAV1;EML1;TUBB2B;CDK6;ID4;ID3;PLXNB2;FGF13;FGFR1;ADGRL3 |
| Epithelial Cell Development (GO:0002064) | 10/25 | 1.31E-08 | 1.84E-05 | 16.86 | 306.00 | BMP4;CDK6;CLDN3;CDH2;POF1B;RAB25;RILPL2;TMEM79;PAX6;EXPH5 |
| Notch Signaling Pathway (GO:0007219) | 12/42 | 3.72E-08 | 3.50E-05 | 10.13 | 173.34 | NOTCH2;NOTCH3;HEYL;NOTCH1;JAG1;HEY1;DTX3L;HEY2;NRARP;SORBS2;DTX4;HES5 |
| Pulmonary Valve Morphogenesis (GO:0003184) | 8/19 | 2.41E-07 | 1.71E-04 | 18.34 | 279.50 | NOTCH2;BMP4;HEYL;TGFB2;JAG1;NOTCH1;HEY1;HEY2 |
| Intracellular Signaling Cassette (GO:0141124) | 35/354 | 3.55E-07 | 1.83E-04 | 2.82 | 41.93 | APP;CEBPA;CEBPB;GUCY1B1;PDE1C;PDE1B;CLSTN1;ARHGAP18;PIK3R1;LRP2;PPP1R9A;EGFR;NUDT4;ERBB3;ERBB2;PLCE1;NRTN;WWTR1;MEF2C;GUCY1A1;RBPMS;AMFR;MAPK8IP2;RUNDC3A;MAPK12;TIAM1;ARHGAP32;MAF;CCDC88C;TYRO3;NFE2L3;PDE5A;FGF13;PLCD1;FGFR1 |
| Atrioventricular Valve Morphogenesis (GO:0003181) | 8/20 | 3.89E-07 | 1.83E-04 | 16.81 | 248.18 | EFNA1;HEYL;OLFM1;TGFB2;NOTCH1;HEY1;HEY2;SLIT3 |
| Positive Regulation of Cell Differentiation (GO:0045597) | 31/297 | 5.13E-07 | 2.07E-04 | 2.99 | 43.31 | CRB2;FBN2;APP;FOXC1;CEBPB;PKDCC;LAMA1;TACSTD2;GJA1;PPP1R13L;GPC1;GCNT2;NRCAM;SOX9;WWTR1;MEF2C;TGFB2;PRKCH;IGFBP3;MAPK12;BMP4;TIAM1;HEYL;OLFM1;MAPK11;SFRP1;SFRP2;RGCC;CAMK1;WDFY2;FGFR1 |
| Pulmonary Valve Development (GO:0003177) | 8/21 | 6.06E-07 | 2.13E-04 | 15.52 | 222.17 | NOTCH2;BMP4;HEYL;TGFB2;JAG1;NOTCH1;HEY1;HEY2 |
| Nervous System Development (GO:0007399) | 42/480 | 6.93E-07 | 2.13E-04 | 2.48 | 35.11 | NOTCH2;HDAC4;APP;STMN3;CRMP1;MSI1;NR2E1;CELSR1;ZDHHC2;DMBX1;SOX3;CDH2;PTPRZ1;ERBB3;SLITRK6;NRCAM;NRTN;PCDH1;EPHB2;EPHB1;SH3GL2;HES5;MEF2C;JAG1;APLP2;FZD9;NR2F1;PAX6;NAV1;NR2F2;NR2F6;OLFM1;VCAN;NAV3;TYRO3;PLXNB2;PLXNB1;SP8;FGF13;NES;ADGRL3;FGF10 |
| Apical Junction Assembly (GO:0043297) | 12/54 | 7.54E-07 | 2.13E-04 | 7.23 | 101.96 | CLDN6;CLDN10;CLDN4;OCLN;CLDN3;POF1B;MARVELD2;CLDN23;CLDN7;STRN;RHOC;MPP7 |

**Supplementary Table 7. Enriched terms detected using Enrichr for differentially expressed genes in the mutant λ-junction ectoderm or mesenchymal cells. Ect Up GO CC 2025**

| Term | Overlap | P-value | Adjusted P-value | Odds Ratio | Combined Score | Genes |
|---|---|---|---|---|---|---|
| Apical Junction Complex (GO:0043296) | 24/107 | 1.92E-12 | 4.52E-10 | 7.42 | 200.21 | CRB2;JUP;FRMD4A;CGN;MPP7;AMOT;CLDN6;CAMSAP3;CLDN10;CLDN4;OCLN;CLDN3;POF1B;SAPCD2;CCDC85C;MARVELD2;EPCAM;CLDN7;CLDN23;STRN;EPPK1;NECTIN3;CGNL1;TJP2 |
| Cell-Cell Junction (GO:0005911) | 40/316 | 3.76E-11 | 4.42E-09 | 3.76 | 90.33 | CRB2;NOTCH1;CTNND2;AQP3;GJA1;CDH4;POF1B;CDH2;CCDC85C;MARVELD2;EPB41L3;STXBP6;PCDH1;PAK4;JAG1;JUP;ACTN1;KAZN;FRMD4A;BAIAP2;MPP7;TIAM1;CLDN10;CLDN4;GJB2;OCLN;CLDN3;KRT18;SAPCD2;PTK7;PKP1;PKP4;DSG2;EZR;PLPP3;NECTIN4;NECTIN3;ADGRL3;TJP2;NECTIN1 |
| Tight Junction (GO:0070160) | 19/96 | 3.68E-09 | 2.88E-07 | 6.29 | 122.21 | FRMD4A;CGN;MPP7;AMOT;CLDN6;CLDN10;GJA1;CLDN4;OCLN;CLDN3;POF1B;MARVELD2;EPCAM;CLDN23;CLDN7;EPPK1;STRN;CGNL1;TJP2 |
| Bicellular Tight Junction (GO:0005923) | 17/81 | 9.60E-09 | 5.60E-07 | 6.76 | 124.81 | FRMD4A;CGN;MPP7;CLDN6;CLDN10;CLDN4;OCLN;CLDN3;POF1B;MARVELD2;EPCAM;CLDN23;CLDN7;EPPK1;STRN;CGNL1;TJP2 |
| Polymeric Cytoskeletal Fiber (GO:0099513) | 33/280 | 1.19E-08 | 5.60E-07 | 3.44 | 62.79 | BEX4;RTN2;SYNM;IQGAP1;IQGAP2;RCSD1;EVPL;PPL;KIF15;AMOT;BCL2L11;POF1B;BAG2;MARK2;PLS1;ACTN1;KRT8;PAWR;COBL;KIF23;FHDC1;TUBB4B;EML1;TUBB2B;CLIP1;KRT18;MAP7D2;PKP1;EPPK1;VIM;EZR;FGF13;NES |
| Adherens Junction (GO:0005912) | 22/151 | 8.00E-08 | 3.13E-06 | 4.35 | 71.17 | NOTCH1;JAG1;JUP;CTNND2;FRMD4A;BAIAP2;MPP7;CAMSAP3;CDH4;KRT18;CDH2;POF1B;CCDC85C;PKP1;PKP4;STXBP6;EZR;PLPP3;NECTIN4;NECTIN3;PAK4;NECTIN1 |
| Cytoskeleton (GO:0005856) | 53/694 | 1.73E-06 | 5.82E-05 | 2.14 | 28.44 | LAD1;SYNM;PPP1R9A;CLU;MYLK;EPB41L4A;PCLO;TEKT1;LRRFIP1;GAS2L3;KRT8;PAWR;NAV1;EML1;MTSS1;MTSS2;TUBB2B;CLIP1;CDC42EP5;INCENP;MAP7D2;PKP4;EZR;KLHL14;CDCA8;IQGAP1;IQGAP2;EVPL;PPL;EPB41L5;ABLIM1;NUAK1;EPB41L2;EPB41L3;MAP7;VILL;MPZL2;SNCA;JUP;GRHL3;SORBS2;TUBB4B;BAIAP2;KLF4;ARHGAP32;KRT18;PRC1;MYO5B;MYO5C;FILIP1;EPPK1;VIM;NES |
| Desmosome (GO:0030057) | 7/22 | 1.25E-05 | 3.68E-04 | 11.75 | 132.65 | JUP;CDH2;POF1B;KAZN;PKP1;DSG2;PKP4 |
| Actin Cytoskeleton (GO:0015629) | 31/361 | 2.84E-05 | 7.41E-04 | 2.40 | 25.15 | LAD1;KLHL14;IQGAP1;IQGAP2;RCSD1;PPP1R9A;MYLK;AMOT;ABLIM1;POF1B;CDH2;VILL;MARK2;SPTBN2;PLS1;SNCA;GAS2L3;ACTN1;PAWR;COBL;GRHL3;SORBS2;FHDC1;BAIAP2;MTSS1;MTSS2;ARHGAP32;MYO5B;MYO5C;FILIP1;EZR |
| Cell-Cell Contact Zone (GO:0044291) | 9/48 | 7.72E-05 | 1.70E-03 | 5.82 | 55.11 | TIAM1;GJA1;CDH2;JUP;FXYD1;DSG2;PKP4;FGF13;TJP2 |

**Supplementary Table 7. Enriched terms detected using Enrichr for differentially expressed genes in the mutant λ-junction ectoderm or mesenchymal cells.**
**Ect Up GO MF 2025**

| Term | Overlap | P-value | Adjusted P-value | Odds Ratio | Combined Score | Genes |
|---|---|---|---|---|---|---|
| Cadherin Binding (GO:0045296) | 31/317 | 2.07E-06 | 1.04E-03 | 2.78 | 36.36 | LAD1;NOTCH3;CTNND2;ARHGAP18;SLC3A2;IQGAP1;EVPL;PPL;EGFR;CDH4;CDH2;EPCAM;PPP1R13L;STXBP6;LRRFIP1;MARK2;SPTBN2;PAK4;SH3GLB2;JUP;CGN;BAIAP2;MPP7;PPFIBP1;KRT18;GIPC1;PKP1;PKP4;ITGA6;EZR;TJP2 |
| Wnt Receptor Activity (GO:0042813) | 5/14 | 1.25E-04 | 2.39E-02 | 13.96 | 125.44 | FZD2;FZD4;FZD7;FZD9;TSPAN12 |
| Kinesin Binding (GO:0019894) | 7/31 | 1.43E-04 | 2.39E-02 | 7.34 | 65.00 | PRC1;NEK6;CLSTN1;MAP7D2;MAPK8IP2;SNCA;MAPK8IP1 |
| Cadherin Binding Involved in Cell-Cell Adhesion (GO:0098641) | 5/17 | 3.51E-04 | 3.27E-02 | 10.47 | 83.26 | KRT18;EPCAM;STXBP6;BAIAP2;PAK4 |
| MAP-kinase Scaffold Activity (GO:0005078) | 4/10 | 3.80E-04 | 3.27E-02 | 16.73 | 131.76 | IQGAP1;MAPK8IP2;SH3RF1;MAPK8IP1 |
| Tubulin Binding (GO:0015631) | 26/326 | 3.92E-04 | 3.27E-02 | 2.21 | 17.30 | BEX4;STMN2;STMN3;KIF15;GJA1;BCL2L11;RACGAP1;SNCA;GAS2L3;UNC5C;KIF23;KNL1;CGN;FHDC1;EML1;ARL4C;EML5;CLIP1;NAV3;PRC1;REEP3;CCDC88C;MAP7D2;EZR;FGF13;ARL8B |

**Supplementary Table 7. Enriched terms detected using Enrichr for differentially expressed genes in the mutant λ-junction ectoderm or mesenchymal cells. Ect Up Diseaaes 2025**

| Term | Overlap | P-value | Adjusted P-value | Odds Ratio | Combined Score | Genes |
|---|---|---|---|---|---|---|
| OROFACIAL CLEFT | 14/104 | 4.48E-05 | 5.73E-03 | 3.94 | 39.44 | SEMA4D;DGKA;KRT8;RASSF9;DLK1;CUX2;KRT18;CMSS1;ERBB3;IRF2;IRF6;CBLN2;FGF10;FGFR1 |
| LIP DISEASE | 14/104 | 4.48E-05 | 5.73E-03 | 3.94 | 39.44 | SEMA4D;DGKA;KRT8;RASSF9;DLK1;CUX2;KRT18;CMSS1;ERBB3;IRF2;IRF6;CBLN2;FGF10;FGFR1 |
| CLEFT LIP | 14/104 | 4.48E-05 | 5.73E-03 | 3.94 | 39.44 | SEMA4D;DGKA;KRT8;RASSF9;DLK1;CUX2;KRT18;CMSS1;ERBB3;IRF2;IRF6;CBLN2;FGF10;FGFR1 |
| PHYSICAL DISORDER | 18/185 | 2.96E-04 | 2.84E-02 | 2.73 | 22.20 | PKDCC;HIBADH;SEMA4D;EYA1;DGKA;KRT8;RASSF9;DLK1;CUX2;KRT18;CMSS1;ERBB3;IRF2;SCG5;IRF6;CBLN2;FGF10;FGFR1 |

**Supplementary Table 7. Enriched terms detected using Enrichr for differentially expressed genes in the mutant λ-junction ectoderm or mesenchymal cells.**
**Ect Down GO BP 2025**

| Term | Overlap | P-value | Adjusted P-value | Odds Ratio | Combined Score | Genes |
|---|---|---|---|---|---|---|
| Generation of Neurons (GO:0048699) | 22/192 | 2.83E-10 | 6.21E-07 | 6.00 | 131.98 | WNT10A;IRX5;SEMA3A;IRX3;WNT7B;IRX4;WNT9B;SOX14;SOX11;FZD10;SOX12;ASCL1;RUNX2;WNT6;MAP1B;ID2;DDIT4;DCX;LMX1B;INSM1;WNT3;CDK5R1 |
| Neuron Differentiation (GO:0030182) | 20/181 | 3.49E-09 | 3.83E-06 | 5.74 | 111.73 | WNT10A;IRX5;IRX3;WNT7B;IRX4;WNT9B;SOX14;SOX11;FZD10;SOX12;ASCL1;RUNX2;WNT6;BRINP1;ID2;DDIT4;LMX1B;INSM1;WNT3;CDK5R1 |
| Regulation of Cell Migration (GO:0030334) | 33/479 | 8.03E-09 | 5.88E-06 | 3.47 | 64.77 | SEMA5A;HDAC2;CD151;SEMA3C;SEMA3A;LEF1;PDGFA;TWIST1;PIK3R3;PRKX;THBS1;NKD1;FGF9;DPYSL3;PDGFC;PDPN;PLXNA2;GLUL;LIMCH1;LMO4;NOG;IGF1;SMAD7;COL1A1;FGF17;TMEFF2;BMP2;DLG5;KIT;SNAI2;SGK3;AMOTL2;LGR6 |
| Regulation of Osteoblast Differentiation (GO:0045667) | 13/92 | 1.09E-07 | 5.27E-05 | 7.51 | 120.34 | WNT7B;NOG;TWIST1;SOX11;TENT5A;IGF1;NPNT;SMAD5;RUNX2;REST;SNAI2;WNT3;FGFR2 |
| Positive Regulation of Cell Population Proliferation (GO:0008284) | 31/484 | 1.20E-07 | 5.27E-05 | 3.20 | 50.94 | HDAC2;PID1;BNC1;LEF1;PDGFA;RBPJ;THBS1;PTHLH;CCND2;TERT;FGF9;SERTAD1;NAMPT;PDGFC;ITGAV;TFAP2B;SOX11;IGF1;SMARCA2;MEIS2;TBX3;FGF17;BMP2;MEIS1;ID2;KIT;PRTN3;INSM1;SLC25A33;CRLF1;FGFR2 |
| Regulation of Cell Differentiation (GO:0045595) | 18/201 | 5.26E-07 | 1.92E-04 | 4.52 | 65.30 | TFAP2A;TFAP2B;ZFHX3;HDAC2;LEF1;NREP;CHD3;SMARCA2;RUNX2;PTHLH;SMAD7;REST;ID2;SNAI2;INSM1;FGFR2;CDK5R1;YWHAH |
| Positive Regulation of Cell Differentiation (GO:0045597) | 22/297 | 7.85E-07 | 2.46E-04 | 3.69 | 51.88 | ZFHX3;BTG1;WNT7B;LEF1;TWIST1;SOX11;IGF1;TENT5A;GDF6;ASCL1;NPNT;SMARCA2;SMAD5;RUNX2;COL1A1;PCP4;BMP2;REST;ID2;PDPN;INSM1;WNT3 |
| Negative Regulation of Transcription by RNA Polymerase II (GO:0000122) | 38/732 | 9.60E-07 | 2.63E-04 | 2.57 | 35.61 | RB1;HDAC2;LEF1;TWIST1;CHD3;RBPJ;ASCL1;BACH2;ETS2;SAMD1;SKIL;TLE4;TFAP2A;TFAP2B;ZFHX3;MSX2;BCL11A;IRX3;NOG;SOX14;SOX11;SOX12;SMARCA2;MEIS2;TBX3;TLE5;SMAD7;EID1;BMP2;REST;NFIB;ID2;SNAI2;AMOTL2;INSM1;JDP2;FGFR2;MXD4 |
| Negative Regulation of Cell Differentiation (GO:0045596) | 18/220 | 1.95E-06 | 4.77E-04 | 4.09 | 53.73 | RB1;ZFHX3;NOG;TWIST1;ASCL1;SMARCA2;MEIS2;PTHLH;TBX3;SMAD7;BMP2;MEIS1;REST;ID2;COL5A2;SNAI2;SOSTDC1;ITGAV |
| Positive Regulation of DNA-templated Transcription (GO:0045893) | 54/1274 | 2.81E-06 | 6.17E-04 | 2.10 | 26.88 | CDKN1C;CALCOCO1;PID1;BNC1;CHD3;RBPJ;ETS2;NAMPT;YWHAH;LMO1;HES6;SOX14;PAX3;SOX11;SOX12;POU3F1;RUNX2;DCN;TOX3;MYCN;LMX1B;HDAC2;DLX3;LEF1;TWIST1;ASCL1;TERT;HAS3;APOE;OTX1;PPARGC1A;TFAP2A;EGR1;TFAP2B;ZFHX3;BCL11B;BCL11A;IRX3;IRX4;SMAD9;IGF1;GDF6;SMARCA2;SMAD5;MEIS2;TBX3;COL1A1;BMP2;REST;MEIS1;NFIB;ID2;CAMK4;FGFR2 |

**Supplementary Table 7. Enriched terms detected using Enrichr for differentially expressed genes in the mutant λ-junction ectoderm or mesenchymal cells.**
**Ect Down GO CC 2025**

| Term | Overlap | P-value | Adjusted P-value | Odds Ratio | Combined Score | Genes |
|---|---|---|---|---|---|---|
| Collagen-Containing Extracellular Matrix (GO:0062023) | 23/383 | 1.49E-05 | 3.00E-03 | 2.94 | 32.69 | COL17A1;TINAGL1;CD151;HTRA1;NPNT;THBS1;DCN;COL1A1;MFAP4;BCAM;LGALS1;CTSL;COL5A2;MGP;SERPINH1;TIMP3;VWA1;PRTN3;APOE;CTSF;MFGE8;FGFR2;CTHRC1 |
| Filopodium Membrane (GO:0031527) | 4/11 | 6.75E-05 | 5.29E-03 | 25.63 | 246.12 | GAP43;ITGA3;PDPN;ITGAV |
| Endoplasmic Reticulum Lumen (GO:0005788) | 18/289 | 7.90E-05 | 5.29E-03 | 3.04 | 28.68 | COL17A1;IGFBP4;WNT7B;PDGFA;THBS1;COL1A1;WNT6;ADAMTS5;LGALS1;P4HA1;PDGFC;COL5A2;SERPINH1;GPC3;VWA1;APOE;MFGE8;WNT3 |

**Supplementary Table 7. Enriched terms detected using Enrichr for differentially expressed genes in the mutant λ-junction ectoderm or mesenchymal cells.**
**Ect Down GO MF**

| Term | Overlap | P-value | Adjusted P-value | Odds Ratio | Combined Score | Genes |
|---|---|---|---|---|---|---|
| Sequence-Specific Double-Stranded DNA Binding (GO:1990837) | 35/706 | 7.19E-06 | 2.90E-03 | 2.43 | 28.81 | DLX3;BNC1;LEF1;CHD3;ASCL1;BACH2;ETS2;BASP1;OTX1;TFAP2A;HES6;EGR1;TFAP2B;ZFHX3;IRX5;BCL11B;MSX2;IRX3;SOX14;PAX3;SOX11;SOX12;POU3F1;SMARCA2;SMAD5;MEIS2;TBX3;RUNX2;REST;MYCN;PAX7;NFIB;SNAI2;LMX1B;JDP2 |
| Cis-Regulatory Region Sequence-Specific DNA Binding (GO:0000987) | 42/1035 | 1.02E-04 | 1.54E-02 | 1.97 | 18.12 | CALCOCO1;DLX3;LEF1;TWIST1;RBPJ;ASCL1;BACH2;ETS2;NTN3;UTY;OTX1;SKIL;TFAP2A;HES6;EGR1;TFAP2B;ZFHX3;IRX5;BCL11B;BCL11A;IRX3;IRX4;EBF1;SOX14;PAX3;SMAD9;SOX11;SOX12;POU3F1;SMAD5;MEIS2;TBX3;RUNX2;REST;MYCN;PAX7;NFIB;SNAI2;SP5;INSM1;JDP2;MXD4 |
| Sequence-Specific DNA Binding (GO:0043565) | 32/718 | 1.35E-04 | 1.54E-02 | 2.16 | 19.23 | CALCOCO1;DLX3;LEF1;RBPJ;ASCL1;BACH2;ETS2;TERT;OTX1;PPARGC1A;TFAP2A;HES6;EGR1;TFAP2B;IRX5;BCL11B;MSX2;IRX3;SOX14;PAX3;SOX12;POU3F1;SMAD5;MEIS2;TBX3;RUNX2;MYCN;PAX7;NFIB;SNAI2;LMX1B;JDP2 |
| RNA Polymerase II Cis-Regulatory Region Sequence-Specific DNA Binding (GO:0000978) | 42/1054 | 1.52E-04 | 1.54E-02 | 1.93 | 17.00 | CALCOCO1;DLX3;LEF1;TWIST1;RBPJ;ASCL1;BACH2;ETS2;NTN3;UTY;OTX1;SKIL;TFAP2A;HES6;EGR1;TFAP2B;ZFHX3;IRX5;BCL11B;BCL11A;IRX3;IRX4;EBF1;SOX14;PAX3;SMAD9;SOX11;SOX12;POU3F1;SMAD5;MEIS2;TBX3;RUNX2;REST;MYCN;PAX7;NFIB;SNAI2;SP5;INSM1;JDP2;MXD4 |
| DNA-binding Transcription Factor Binding (GO:0140297) | 17/294 | 2.99E-04 | 1.99E-02 | 2.80 | 22.71 | RB1;LMO1;HES6;HDAC2;RBFOX2;BCL11A;LMO4;TWIST1;HSPB1;RBPJ;ASCL1;ETS2;TBX3;REST;ID2;RBBP8;PPARGC1A |
| Receptor Ligand Activity (GO:0048018) | 20/380 | 3.13E-04 | 1.99E-02 | 2.54 | 20.49 | JAG2;SEMA5A;WNT10A;SEMA3C;WNT7B;SEMA3A;WNT9B;PDGFA;IGF1;GDF6;PTHLH;FGF17;WNT6;INSL6;BMP2;LGALS1;FGF9;APOE;CRLF1;WNT3 |
| RNA Polymerase II Transcription Regulatory Region Sequence-Specific DNA Binding (GO:0000977) | 46/1236 | 3.44E-04 | 1.99E-02 | 1.80 | 14.37 | RB1;CALCOCO1;DLX3;LEF1;TWIST1;RBPJ;ASCL1;BACH2;ETS2;NTN3;ZFP57;UTY;OTX1;SKIL;TFAP2A;HES6;EGR1;TFAP2B;ZFHX3;IRX5;BCL11B;MSX2;BCL11A;IRX3;IRX4;EBF1;SOX14;PAX3;SMAD9;SOX11;SOX12;POU3F1;SMAD5;MEIS2;TBX3;RUNX2;REST;MYCN;PAX7;NFIB;SNAI2;SP5;INSM1;LMX1B;JDP2;MXD4 |
| Double-Stranded DNA Binding (GO:0003690) | 28/651 | 6.13E-04 | 3.10E-02 | 2.07 | 15.28 | DLX3;LEF1;ASCL1;BACH2;ETS2;OTX1;TFAP2A;HES6;EGR1;TFAP2B;IRX5;BCL11B;MSX2;IRX3;SOX14;PAX3;SOX12;POU3F1;SMAD5;MEIS2;TBX3;RUNX2;MYCN;PAX7;NFIB;SNAI2;LMX1B;JDP2 |
| DNA-binding Transcription Repressor Activity, RNA Polymerase II-specific (GO:0001227) | 13/210 | 8.02E-04 | 3.60E-02 | 2.99 | 21.33 | TFAP2A;ZFHX3;MSX2;BCL11A;IRX3;ASCL1;ETS2;TBX3;REST;SNAI2;INSM1;SKIL;JDP2 |
| Frizzled Binding (GO:0005109) | 5/35 | 9.49E-04 | 3.83E-02 | 7.48 | 52.08 | WNT6;WNT10A;WNT7B;WNT9B;WNT3 |

**Supplementary Table 7. Enriched terms detected using Enrichr for differentially expressed genes in the mutant λ-junction ectoderm or mesenchymal cells.**
**All Mes GO BP 2025**

| Term | Overlap | P-value | Adjusted P-value | Odds Ratio | Combined Score | Genes |
|---|---|---|---|---|---|---|
| Regulation of Canonical Wnt Signaling Pathway (GO:0060828) | 32/216 | 7.05E-12 | 1.99E-08 | 4.91 | 126.04 | SEMA5A;APP;AMER2;PRICKLE1;LYPD6;GLI3;EGFR;NKD1;NKD2;FRZB;UBR5;RSPO2;SOX9;APOE;RSPO1;TLE2;TLE1;IGFBP2;WNT5A;KREMEN1;AXIN2;DKK2;DKK3;COL1A1;ZEB2;SFRP2;DAAM2;BAMBI;SNAI2;CDH15;RECK;LGR4 |
| Regulation of Endothelial Cell Migration (GO:0010594) | 20/88 | 2.41E-11 | 3.40E-08 | 8.21 | 200.63 | NRP1;BCAS3;FOXC2;NRP2;ANGPT1;WNT5A;STC1;EMP2;GATA3;PTN;NR2F2;THBS1;FOXP1;SMOC2;RHOJ;ATOH8;APOE;SLIT2;PLPP3;ADAMTS9 |
| Negative Regulation of Wnt Signaling Pathway (GO:0030178) | 25/146 | 5.94E-11 | 5.58E-08 | 5.79 | 136.35 | APP;AMER2;CITED1;PRICKLE1;APCDD1;CXXC4;GLI3;NKD1;NKD2;FRZB;WIF1;SOX9;APOE;TLE2;TLE1;IGFBP2;WNT5A;KREMEN1;SHISA2;AXIN2;DKK2;HIC1;DKK3;SFRP2;SNAI2 |
| Regulation of Cell Migration (GO:0030334) | 48/479 | 9.65E-11 | 6.81E-08 | 3.18 | 73.28 | RET;SEMA5A;BEX4;IFITM1;NOTCH1;SERPINE2;CITED2;SYNPO2;LEF1;STC1;TWIST1;RORA;THY1;PTN;THBS1;EGFR;NKD1;MYLK;ROBO1;AMOT;DPYSL3;PLXNA2;ENPP2;EMILIN2;RICTOR;CTNNA2;SLIT2;NGFR;IGFBP5;FZD4;ANKRD44;EMP2;IGF1;ADRA2A;SMAD7;COL1A1;IGSF10;CXCL12;SFRP2;DAAM2;RHOJ;SNAI2;DPEP1;SPRY2;SGK1;RECK;HBEGF;FGF10 |
| Positive Regulation of DNA-templated Transcription (GO:0045893) | 90/1274 | 2.42E-10 | 1.37E-07 | 2.22 | 49.23 | CDKN1C;RET;APP;PID1;THRB;CITED1;CITED2;SCX;RORA;NR3C1;GLI3;GLI2;RPS6KA3;HMGN5;MECOM;HEY2;SOX9;TEAD1;IER2;BCAS3;OSR2;FOXD2;TCF15;ANXA2;PEG3;OSR1;TCF12;WNT5A;PAX3;MSL3;EBF3;FOS;EBF4;ETV4;RUNX2;ETV5;RUNX1;ISL2;ZEB2;SFRP2;MDFIC;KAT6B;PLCB1;TOX;PLPP3;FOXC2;DLX2;FOXC1;NFIX;NOTCH1;PRKDC;DLX5;LEF1;TWIST1;GATA3;EGFR;GLIS1;NFIL3;EPCAM;NRIP1;ATOH8;APOE;HES5;S100A10;BCL9L;NGFR;TFAP2B;ZFHX3;PRRX1;BCL11B;FZD4;FOXF1;IRX3;HMGA1;SMAD9;NR2F2;IGF1;KLF4;MEIS2;NFATC4;BMP5;COL1A1;BMP4;MEIS1;MAFB;BAMBI;TLX2;ID2;BCL3;SSBP4 |
| Negative Regulation of DNA-templated Transcription (GO:0045892) | 76/1006 | 3.57E-10 | 1.68E-07 | 2.37 | 51.57 | CDKN1C;APP;THRB;CITED1;CITED2;NR3C1;GLI3;GLI2;AMOT;MECOM;CCND1;TRPS1;HEY2;SIX3;SOX9;OSR2;TLE2;TLE1;FOXD1;PEG3;MSX2;OSR1;USP2;WNT5A;KCTD1;HIC1;FOXP2;RUNX2;ETV5;DKK3;FOXP1;RUNX1;ZEB2;SFRP2;MDFIC;KAT6B;PLCB1;EZR;FOXC2;ZBTB8A;FOXC1;NFIX;NOTCH1;TSHZ3;SHOX2;LEF1;TWIST1;GATA3;PRICKLE1;GLIS1;BTAF1;CUX2;NFIL3;TBX22;NRIP1;ATOH8;SAMD1;CRYM;HES5;IRX1;TFAP2B;PRNP;ZFHX3;IRX3;NR2F1;HMGA1;NR2F2;KLF4;MEIS2;NFATC4;SMAD7;BMP4;MAFB;ID2;BCL3;SNAI2 |

**Supplementary Table 7. Enriched terms detected using Enrichr for differentially expressed genes in the mutant λ-junction ectoderm or mesenchymal cells.**
**All Mes GO BP 2025**

| | | | | | | |
|---|---|---|---|---|---|---|
| Axonogenesis (GO:0007409) | 28/200 | 5.56E-10 | 2.10E-07 | 4.57 | 97.39 | RET;ROBO2;SEMA5A;APP;NRP1;NRP2;NOTCH1;GLI2;EFNB1;TUBB3;SLITRK4;KIF5A;SLITRK6;SLITRK5;EMB;CTNNA2;SLIT2;BCL11B;EPHA8;BDNF;WNT5A;L1CAM;ISL2;CXCL12;MAP1B;S100A6;PRKCQ;EPHA3 |
| Positive Regulation of Cell Projection Organization (GO:0031346) | 21/115 | 5.96E-10 | 2.10E-07 | 6.23 | 132.40 | RET;ROBO2;CDKL3;NTRK2;BDNF;FZD4;ANKRD44;WNT5A;STMN2;NDNF;PTN;ROBO1;TIAM2;CUX2;DPYSL3;GPC2;DMD;APOE;SLIT2;TOX;EPHA3 |
| Negative Regulation of Cell Differentiation (GO:0045596) | 29/220 | 1.16E-09 | 3.63E-07 | 4.26 | 87.76 | NOTCH1;CITED1;TNFAIP6;TWIST1;RORA;GATA3;EGFR;SPRED3;CSRP1;SIX3;SOX9;HES5;ZFHX3;OSR1;WNT5A;SMAD6;MEIS2;BMP5;SMAD7;BMP4;MEIS1;SFRP2;CDK6;BAMBI;ID2;COL5A2;SNAI2;SPRY2;SPRY1 |
| Skeletal System Development (GO:0001501) | 24/160 | 2.33E-09 | 6.58E-07 | 4.93 | 98.08 | DLX2;PKDCC;DLX5;HHIP;DLX6;WNT5A;SHOX2;STC1;IGF1;PTH1R;PCSK5;BMP5;GLI2;PAX1;COL1A1;RPS6KA3;COL2A1;FRZB;TRPS1;ALX4;TLL1;SOX9;COL9A2;HES5 |

**Supplementary Table 7. Enriched terms detected using Enrichr for differentially expressed genes in the mutant λ-junction ectoderm or mesenchymal cells.**
**All Mes GO CC 2025**

| Term | Overlap | P-value | Adjusted P-value | Odds Ratio | Combined Score | Genes |
|---|---|---|---|---|---|---|
| Collagen-Containing Extracellular Matrix (GO:0062023) | 47/383 | 1.13E-13 | 2.47E-11 | 4.01 | 119.41 | SRPX;COL16A1;SERPINE2;NID1;FBLN2;THBS1;LOXL1;FBLN5;LGALS1;CDH2;ANXA7;EMILIN2;IGFBP7;NCAM1;HMCN1;APOE;ADAMTS9;EDIL3;CDON;S100A10;CTHRC1;ANXA2;ANGPT1;COL25A1;WNT5A;ANXA11;L1CAM;NPNT;ADAMTS20;COL1A1;SMOC2;SFRP2;COL2A1;CXCL12;COL6A2;COL7A1;COL5A2;COL6A1;MGP;COL9A1;S100A6;PRTN3;COL9A2;TGFBI;FREM2;ANGPTL1;FGF10 |
| Adherens Junction (GO:0005912) | 16/151 | 9.43E-05 | 1.03E-02 | 3.28 | 30.37 | NOTCH1;ANXA2;SMAD7;CDH6;PDLIM2;KRT18;CDH2;TNKS1BP1;PGM5;HMCN1;STXBP6;CDH15;CTNNA2;EZR;PLPP3;NECTIN2 |
| Endoplasmic Reticulum Lumen (GO:0005788) | 23/289 | 2.74E-04 | 1.66E-02 | 2.40 | 19.67 | APP;COL16A1;IGFBP5;BDNF;COL25A1;ANKRD44;WNT5A;PRSS23;THBS1;COL1A1;BMP4;LGALS1;COL2A1;CDH2;COL6A2;COL7A1;PENK;COL5A2;COL6A1;COL9A1;IGFBP7;APOE;COL9A2 |
| Voltage-Gated Calcium Channel Complex (GO:0005891) | 7/39 | 3.90E-04 | 1.66E-02 | 6.00 | 47.10 | CACNB2;CACNB3;CACNA2D3;CACHD1;CACNA1D;CACNA1C;CACNA1G |
| beta-catenin-TCF Complex (GO:1990907) | 4/11 | 4.23E-04 | 1.66E-02 | 15.63 | 121.40 | TLE1;LEF1;TCF7;BCL9L |
| Neuron Projection (GO:0043005) | 36/560 | 4.56E-04 | 1.66E-02 | 1.92 | 14.74 | RET;APP;NRP1;NRP2;RTN1;STMN2;SLC1A3;RORA;THY1;CALB2;CDH2;TUBB3;SV2A;PENK;NEFL;NCAM1;EMB;APOE;CTNNA2;CTHRC1;NTRK2;PRNP;FZD4;KCNB1;BDNF;EPHA8;PLK2;KCNB2;BNIP3;COBL;L1CAM;CLDN11;MAP1B;VIM;KCNK2;EPHA3 |
| Vesicle (GO:0031982) | 18/230 | 1.47E-03 | 4.60E-02 | 2.34 | 15.29 | SERPINE2;ANXA2;AHNAK;CEND1;ANXA11;SLC2A4;ATP1B1;FBLN2;EGFR;BMP5;BAIAP2L2;TUBB2A;COL6A2;ANXA7;APOE;EZR;EDIL3;DYNC1I1 |
| Axon (GO:0030424) | 17/217 | 1.95E-03 | 4.95E-02 | 2.34 | 14.64 | RET;APP;NRP1;NTRK2;NRP2;KCNB1;BDNF;COBL;RORA;L1CAM;CLDN11;TUBB3;MAP1B;NEFL;EMB;CTNNA2;VIM |
| Calcium Channel Complex (GO:0034704) | 7/51 | 2.04E-03 | 4.95E-02 | 4.36 | 27.03 | CACNB2;CACNB3;CACNA2D3;CACHD1;CACNA1D;CACNA1C;CACNA1G |

**Supplementary Table 7. Enriched terms detected using Enrichr for differentially expressed genes in the mutant λ-junction ectoderm or mesenchymal cells.**
**Mes Up GO CC 2025**

| Term | Overlap | P-value | Adjusted P-value | Odds Ratio | Combined Score | Genes |
|---|---|---|---|---|---|---|
| Collagen-Containing Extracellular Matrix (GO:0062023) | 18/383 | 1.88E-07 | 2.53E-05 | 4.861414366 | 75.29526061 | COL25A1;NPNT;THBS1;LOXL1;SFRP2;COL2A1;CXCL12;CDH2;MGP;COL9A1;HMCN1;APOE;COL9A2;TGFBI;ADAMTS9;FREM2;ANGPTL1;CTHRC1 |
| Adherens Junction (GO:0005912) | 9/151 | 3.83E-05 | 2.58E-03 | 6.043586763 | 61.4674282 | NOTCH1;KRT18;CDH2;HMCN1;STXBP6;CDH15;EZR;PLPP3;NECTIN2 |
| Nucleus (GO:0005634) | 77/4879 | 1.11E-04 | 4.98E-03 | 1.740957874 | 15.85687217 | CDKN1C;APP;CITED1;CITED2;SCX;SOBP;PRSS23;RGS2;CCND2;HMGN5;MECOM;CCND1;SIX2;SIX3;SOX9;PHLDA1;IER2;OSR2;FOXD1;MORC4;TCF15;PEG3;OSR1;KRT8;KCTD1;EBF3;FOXP4;ETV4;HIC1;FOXP2;ETV5;FOXP1;MN1;ISL2;MDFIC;TRAF4;RRAGD;RBP1;TOX;BEX4;FOXC2;FOXC1;NOTCH1;SYNPO2;SHOX2;AK1;STC1;BARX1;EGFR;CYGB;TBX22;GSC;NRIP1;DMD;LRIG3;APOE;PLTP;HES5;IRX1;TFAP2B;PRRX1;FOXF1;EYA2;EYA4;NR2F1;HMGA1;NR2F2;USP29;MAPK12;MOV10;LIN28B;MAFB;CDK6;LHX6;SPRY2;SSBP4;INKA1 |
| Endoplasmic Reticulum Lumen (GO:0005788) | 11/289 | 3.08E-04 | 1.04E-02 | 3.78362604 | 30.59126905 | APP;COL2A1;CDH2;IGFBP5;COL25A1;PENK;COL9A1;APOE;COL9A2;PRSS23;THBS1 |
| Cell-Cell Junction (GO:0005911) | 11/316 | 6.51E-04 | 1.71E-02 | 3.443908711 | 25.26846197 | CLDN11;KRT18;NOTCH1;FLRT2;CDH2;HMCN1;STXBP6;CDH15;EZR;PLPP3;NECTIN2 |
| Intracellular Membrane-Bounded Organelle (GO:0043231) | 82/5597 | 7.58E-04 | 1.71E-02 | 1.595293765 | 11.46198008 | CDKN1C;APP;CITED1;CITED2;SCX;SPINT2;SOBP;PRSS23;RGS2;CCND2;HMGN5;MECOM;CCND1;SIX2;SIX3;SOX9;PHLDA1;ADAMTS9;IER2;OSR2;FOXD1;MORC4;TCF15;PEG3;OSR1;KRT8;KCTD1;EBF3;FOXP4;ETV4;HIC1;FOXP2;ETV5;FOXP1;MN1;ISL2;MDFIC;TRAF4;RRAGD;RBP1;TOX;BEX4;FOXC2;FOXC1;NOTCH1;SYNPO2;SHOX2;AK1;STC1;BARX1;EGFR;CYGB;HSD11B2;TBX22;GSC;NRIP1;DMD;LRIG3;APOE;PLTP;HES5;IRX1;TFAP2B;PRRX1;FOXF1;EYA2;EYA4;NR2F1;HMGA1;NR2F2;USP29;MAPK12;MOV10;LIN28B;MAFB;CDK6;LHX6;SPRY2;SEC24D;KCNK1;SSBP4;INKA1 |
| High-Density Lipoprotein Particle (GO:0034364) | 3/19 | 1.05E-03 | 2.02E-02 | 17.484375 | 120.0024869 | APP;APOE;PLTP |
| Platelet Alpha Granule (GO:0031091) | 5/90 | 2.87E-03 | 4.84E-02 | 5.518207283 | 32.29375759 | APP;ISLR;PCDH7;SELENOP;THBS1 |
| Polymeric Cytoskeletal Fiber (GO:0099513) | 9/280 | 3.40E-03 | 4.84E-02 | 3.145953498 | 17.88515193 | BEX4;KRT18;BCL2L11;MAP1B;KRT8;NEFL;COBL;EZR;AMOT |
| Cell-Substrate Junction (GO:0030055) | 11/395 | 3.78E-03 | 4.84E-02 | 2.724303002 | 15.19982911 | NRP1;FLRT2;CDH2;ITGA4;SYNPO2;HMGA1;DMD;HMCN1;EZR;EGFR;NECTIN2 |

**Supplementary Table 7. Enriched terms detected using Enrichr for differentially expressed genes in the mutant λ-junction ectoderm or mesenchymal cells. Mes Up Diseases 2025**

| Term | Overlap | P-value | Adjusted P-value | Odds Ratio | Combined Score | Genes |
|---|---|---|---|---|---|---|
| OROFACIAL CLEFT | 7/104 | 1.32E-04 | 1.01E-02 | 6.83 | 61.00 | MAFB;KRT18;ERBB3;KRT8;SPRY2;EPHA3;DLK1 |
| LIP DISEASE | 7/104 | 1.32E-04 | 1.01E-02 | 6.83 | 61.00 | MAFB;KRT18;ERBB3;KRT8;SPRY2;EPHA3;DLK1 |
| CLEFT LIP | 7/104 | 1.32E-04 | 1.01E-02 | 6.83 | 61.00 | MAFB;KRT18;ERBB3;KRT8;SPRY2;EPHA3;DLK1 |
| GLAUCOMA | 21/799 | 1.52E-04 | 1.01E-02 | 2.64 | 23.25 | APP;FOXC2;TFAP2B;SNED1;FOXC1;FZD4;FOXF1;EYA2;OSR1;AK1;FOXP4;PRSS23;FLI1;LOXL1;ETV5;MOV10;MECOM;CCND1;GMDS;SIX2;SIX3 |
| EYE DISEASE | 26/1112 | 1.63E-04 | 1.01E-02 | 2.37 | 20.66 | APP;FOXC2;SNED1;FOXC1;AK1;PRSS23;LOXL1;MECOM;CCND1;GMDS;SIX2;SIX3;APOE;PLTP;ADAMTS9;TFAP2B;FZD4;FOXF1;EYA2;OSR1;FOXP4;FLI1;ETV5;MOV10;ALDH1A2;NECTIN2 |
| SENSORY SYSTEM DISEASE | 28/1272 | 2.45E-04 | 1.26E-02 | 2.23 | 18.55 | APP;FOXC2;SNED1;FOXC1;AK1;PRSS23;LOXL1;MECOM;CCND1;GMDS;SIX2;SIX3;APOE;PLTP;ADAMTS9;TFAP2B;FZD4;FOXF1;EYA2;OSR1;EYA4;COBL;FOXP4;FLI1;ETV5;MOV10;ALDH1A2;NECTIN2 |
| OPEN-ANGLE GLAUCOMA | 20/782 | 3.12E-04 | 1.37E-02 | 2.56 | 20.67 | APP;FOXC2;TFAP2B;SNED1;FOXC1;FZD4;FOXF1;EYA2;OSR1;FOXP4;PRSS23;FLI1;LOXL1;ETV5;MOV10;MECOM;CCND1;GMDS;SIX2;SIX3 |
| NERVOUS SYSTEM DISEASE | 43/2371 | 3.86E-04 | 1.49E-02 | 1.87 | 14.73 | CDC42SE1;CDKN1C;APP;FOXC2;SNED1;FOXC1;CITED1;AK1;PRSS23;EGFR;LOXL1;CCND2;MECOM;CCND1;GMDS;SIX2;SIX3;NRIP1;APOE;PLTP;ADAMTS9;TFAP2B;PRRX1;FZD4;FOXF1;EYA2;OSR1;PCDH7;EYA4;COBL;EBF3;FOXP4;FLI1;ETV5;CLDN11;MOV10;ISL2;LIN28B;DAB1;ALDH1A2;CDH15;SLC24A5;NECTIN2 |
| PLANTAR FASCIAL FIBROMATOSIS | 8/166 | 4.42E-04 | 1.51E-02 | 4.80 | 37.08 | MAFB;EBF1;SCX;EBF2;FOXP4;ETV4;USP29;SSBP4 |
| PHYSICAL DISORDER | 8/185 | 9.00E-04 | 2.77E-02 | 4.28 | 30.02 | PKDCC;MAFB;KRT18;ERBB3;KRT8;SPRY2;EPHA3;DLK1 |

**Supplementary Table 7. Enriched terms detected using Enrichr for differentially expressed genes in the mutant λ-junction ectoderm or mesenchymal cells.**
**Mes Down GO BP 2025**

| Term | Overlap | P-value | Adjusted P-value | Odds Ratio | Combined Score | Genes |
|---|---|---|---|---|---|---|
| Regulation of Canonical Wnt Signaling Pathway (GO:0060828) | 21/216 | 1.02E-07 | 1.20E-04 | 4.40 | 70.78 | SEMA5A;AMER2;TLE2;TLE1;WNT5A;KREMEN1;PRICKLE1;AXIN2;GLI3;NKD1;DKK3;NKD2;COL1A1;ZEB2;DAAM2;BAMBI;UBR5;RSPO2;SNAI2;RSPO1;RECK |
| Nervous System Development (GO:0007399) | 33/480 | 1.34E-07 | 1.20E-04 | 3.05 | 48.31 | SEMA5A;ROBO2;CNTFR;SH3GL3;DLX5;DLX6;JRKL;PTN;NRGN;ROBO1;RPS6KA3;CYP26B1;PTPRZ1;DPF1;ENC1;PLXNA2;SLITRK6;ST8SIA4;CDON;PRNP;BDNF;LSAMP;PAX3;GFRA1;L1CAM;DNM3;SDK1;ZEB2;PAX7;MAB21L2;FAT4;SDK2;FGF10 |
| Positive Regulation of Cell Projection Organization (GO:0031346) | 15/115 | 1.57E-07 | 1.20E-04 | 6.08 | 95.23 | RET;ROBO2;CDKL3;NTRK2;BDNF;ANKRD44;WNT5A;STMN2;NDNF;PTN;ROBO1;TIAM2;CUX2;DPYSL3;GPC2 |
| Axon Guidance (GO:0007411) | 17/152 | 2.29E-07 | 1.26E-04 | 5.11 | 78.18 | SEMA5A;RET;ROBO2;NRP2;EPHA8;UNC5B;BDNF;WNT5A;L1CAM;GLI2;EFNB1;GAP43;TUBB3;KIF5A;PRKCQ;NCAM1;EMB |
| Aortic Valve Morphogenesis (GO:0003180) | 9/38 | 2.74E-07 | 1.26E-04 | 12.46 | 188.31 | ROBO2;BMP4;HEY2;TWIST1;SNAI2;GATA3;AXIN2;SMAD6;ROBO1 |
| Aortic Valve Development (GO:0003176) | 9/41 | 5.52E-07 | 2.11E-04 | 11.29 | 162.73 | ROBO2;BMP4;HEY2;TWIST1;SNAI2;GATA3;AXIN2;SMAD6;ROBO1 |
| Positive Regulation of Neuron Projection Development (GO:0010976) | 13/99 | 9.72E-07 | 3.19E-04 | 6.10 | 84.49 | RET;NTRK2;BDNF;ANKRD44;WNT5A;STMN2;NDNF;PTN;BMP5;TUBB2B;CUX2;DPYSL3;GPC2 |
| Regulation of Cell Migration (GO:0030334) | 31/479 | 1.18E-06 | 3.38E-04 | 2.85 | 38.89 | RET;SEMA5A;IFITM1;SERPINE2;LEF1;TWIST1;RORA;THY1;PTN;NKD1;MYLK;ROBO1;DPYSL3;PLXNA2;ENPP2;EMILIN2;RICTOR;CTNNA2;ANKRD44;EMP2;IGF1;SMAD7;COL1A1;IGSF10;DAAM2;SNAI2;DPEP1;SGK1;RECK;HBEGF;FGF10 |
| Axonogenesis (GO:0007409) | 18/200 | 2.55E-06 | 6.49E-04 | 4.02 | 51.72 | RET;ROBO2;SEMA5A;NRP2;BCL11B;EPHA8;BDNF;WNT5A;L1CAM;GLI2;EFNB1;TUBB3;KIF5A;S100A6;SLITRK6;PRKCQ;EMB;CTNNA2 |
| Negative Regulation of Wnt Signaling Pathway (GO:0030178) | 15/146 | 3.44E-06 | 7.90E-04 | 4.63 | 58.26 | AMER2;TLE2;TLE1;WNT5A;KREMEN1;AXIN2;PRICKLE1;APCDD1;CXXC4;GLI3;NKD1;DKK3;NKD2;WIF1;SNAI2 |

**Supplementary Table 7. Enriched terms detected using Enrichr for differentially expressed genes in the mutant λ-junction ectoderm or mesenchymal cells.**
**Mes Down GO CC 2025**

| Term | Overlap | P-value | Adjusted P-value | Odds Ratio | Combined Score | Genes |
|---|---|---|---|---|---|---|
| Collagen-Containing Extracellular Matrix (GO:0062023) | 29/383 | 1.04E-07 | 1.89E-05 | 3.37 | 54.26 | SRPX;COL16A1;SERPINE2;NID1;FBLN2;FBLN5;LGALS1;ANXA7;EMILIN2;IGFBP7;NCAM1;EDIL3;CDON;S100A10;ANXA2;ANGPT1;WNT5A;ANXA11;L1CAM;ADAMTS20;COL1A1;SMOC2;COL6A2;COL7A1;COL5A2;COL6A1;S100A6;PRTN3;FGF10 |
| Voltage-Gated Calcium Channel Complex (GO:0005891) | 7/39 | 4.17E-05 | 3.79E-03 | 8.75 | 88.22 | CACNB2;CACNB3;CACNA2D3;CACHD1;CACNA1D;CACNA1C;CACNA1G |
| beta-catenin-TCF Complex (GO:1990907) | 4/11 | 1.06E-04 | 6.41E-03 | 22.74 | 208.18 | TLE1;LEF1;TCF7;BCL9L |
| Calcium Channel Complex (GO:0034704) | 7/51 | 2.43E-04 | 1.11E-02 | 6.36 | 52.91 | CACNB2;CACNB3;CACNA2D3;CACHD1;CACNA1D;CACNA1C;CACNA1G |
| Vesicle (GO:0031982) | 15/230 | 6.29E-04 | 2.29E-02 | 2.81 | 20.71 | SERPINE2;ANXA2;AHNAK;CEND1;ANXA11;SLC2A4;ATP1B1;FBLN2;BMP5;BAIAP2L2;TUBB2A;COL6A2;ANXA7;EDIL3;DYNC1I1 |
